# ROS Impair Mitophagy via PARylation of PINK1

**DOI:** 10.64898/2026.06.18.733102

**Authors:** Liangliang Gao, He-Ling Wang, Xuxu Zhuang, Dade Rong, Xiang-Zheng Gao, Liming Xie, Zhuo Wang, Mingzhu Tang, Yiguan Chen, Yichi Zhang, Alexander D. Carlsson, Liming Wang, Guang Lu, Jia-Hong Lu, Evandro F. Fang, Han-Ming Shen

## Abstract

Mitophagy is the process of selective autophagic clearance of damaged mitochondria and is closely implicated in neurodegenerative disease. PTEN-induced kinase 1 (PINK1) and a RBR E3 ubiquitin-protein ligase (Parkin) constitute a positive feedback loop in mitophagy initiation. It is known that reactive oxygen species (ROS) modulate mitophagy, while the exact regulatory mechanism remains largely elusive. Here, we found that exogenously applied ROS effectively block mitophagy induced by acute mitochondrial damage agents, which could be reversed by antioxidants. Mechanistically, ROS activate poly(ADP-ribose) polymerase 1 (PARP1), and suppression of PARP1 eliminates the inhibitory effect of ROS on mitophagy. Notably, PARP1 directly interacts with PINK1 and mediates its PARylation at residue E417, thereby negatively regulating PINK1 function. Collectively, our study identifies PARylation as a new form of post-translational modification of PINK1 and reveals a novel mechanism underlying the regulatory role of ROS in mitophagy by PARP1 activation and PARylation of PINK1.

**In brief:** Gao et al. demonstrate that exogenous ROS inhibit mitophagy. Mechanistically, ROS activate PARP1, which mediates PARylation of PINK1, a central regulator of mitophagy, leading to its functional impairment. This study reveals a novel regulatory mechanism of ROS on mitophagy through PARP1 activation and identifies PARylation as a novel form of post-translational modification of PINK1.

**Highlights:**

- ROS block PINK1-Parkin-mediated mitophagy.
- ROS activate PARP1.
- PARP1 suppression eliminates the inhibitory effect of ROS on mitophagy.
- PARylation of PINK1 by PARP1 impairs its activity and mitophagy.

## INTRODUCTION

Selective autophagy of damaged mitochondria, termed mitophagy, serves as an essential mitochondrial quality control mechanism [1–3]. This evolutionarily conserved process enables timely removal of damaged or superfluous mitochondria through lysosomal degradation, thereby maintaining the physiological homeostasis of mitochondria [4–7]. Compromised mitophagy contributes to multiple pathological conditions, especially neurodegenerative disorders [8–11]. Recent advances have identified multiple molecular pathways driving mitophagy, one of the most comprehensively characterized is PINK1 (PTEN-induced kinase 1)-Parkin (a RBR E3 ubiquitin-protein ligase)-mediated mitophagy [12–15]. Briefly, in healthy mitochondria, PINK1, once synthesized in the cytosol, imports into mitochondria via the TOM and TIM complexes, followed by presenilin-associated rhomboid-like protease (PARL)-mediated cleavage and proteasomal degradation [16, 17]. Upon mitochondrial damage with mitochondrial depolarization, PINK1 accumulates and stabilizes on the outer membrane of mitochondria (OMM) and its kinase activity is rapidly activated through dimerization and auto-phosphorylation [18–20]. The activated PINK1 then phosphorylates two important substrates: ubiquitin (Ub) and Parkin, both at Ser65 and this process constitutes a feedforward loop in the initiation stage of mitophagy [21–25].

Reactive oxygen species (ROS) are unstable oxygen derivatives comprising three primary forms: superoxide anion (O_2_^•-^), hydrogen peroxide (H_2_O_2_), and hydroxyl radicals (•OH) [26–28]. These molecules serve as critical signaling molecules modulating multiple biological processes [29, 30], while excessive ROS induce oxidative damage to cellular components including DNA, lipids, and proteins [31–33]. Current research reveals a reciprocal relationship between mitophagy and ROS: treatment with mitochondrial damage agents induces mitophagy, which enhances ROS production [34, 35], while higher levels of ROS are capable of initiating mitophagy [36, 37] and sustaining its progression following Parkin translocation [38]. Reciprocally, effective mitophagy helps reduce mitochondrial ROS production [39]. At present, it has been reported that ROS regulate mitophagy through several key signaling pathways such as NF-κB, mTOR, and SIRT1 [40], the exact role of ROS in the PINK1-Parkin pathway remains to be further elucidated.

One of the consequences of higher levels of ROS and oxidative stress is to induce DNA damage such as double strand breaks, leading to the activation of poly(ADP-ribose) polymerase 1 (PARP1) for the purpose of DNA damage repair [41, 42]. Activated PARP1 catalyzes the polymerization of the ADP-ribose units originating from NAD^+^, resulting in the conjugation of PAR polymers onto specific target proteins, a process known as poly(ADP-ribosyl)ation (PARylation) [43–45]. At present, there is evidence demonstrated that PARP1 modulates mitophagy by inhibiting neuroprotective factor SIRT1 [46], the direct regulatory mechanisms of PARP1 in PINK1-Parkin-mediated mitophagy are poorly characterized.

Herein, we demonstrated that high levels of ROS and oxidative stress suppress mitophagy via activation of PARP1, leading to PARylation of PINK1, a process causing impairment of PINK1 function, thereby abrogating the mitophagy process. Our findings thus provide a new insight for the regulatory role of ROS in mitophagy and provide novel therapeutic directions for mitophagy-related diseases through targeting the ROS-PARP1 pathway.

## RESULTS

### ROS inhibit PINK1-Parkin-mediated mitophagy induced by acute mitochondrial damage agents

At present, the exact role of ROS and oxidative stress in mitophagy remains controversial and inconclusive [47]. To further elucidate the role of ROS in PINK1-Parkin-mediated mitophagy, we utilized two prooxidants: oxidopamine hydrochloride (6-OHDA) and H_2_O_2_. 6-OHDA is a compound that generates ROS through auto-oxidation and is commonly used to establish Parkinson’s disease models in mice [48, 49]. H_2_O_2_ itself is a type of ROS. YFP-Parkin-HeLa cells were treated with the mitophagy inducer CCCP or Oligomycin/Antimycin A (O/A), each combined with different concentrations of either 6-OHDA or H_2_O_2_. Strikingly, both 6-OHDA and H_2_O_2_ suppressed CCCP or O/A induced mitophagy measured by increased mitochondrial protein degradation in a dose-dependent manner (Figure 1A and S1A). The inhibitory effect of 6-OHDA and H_2_O_2_ on mitophagy was also found to be time-dependent (Figure 1B and S1B). To further corroborate the effects of ROS on mitophagy, mitochondrial targeted Keima (mt-Keima) assay was utilized to quantify mitophagy, following established protocols [50–52]. We observed that both CCCP- and O/A-induced mitophagy was completely blocked with the addition of 6-OHDA (Figure 1C-1D). Similar results were found upon H_2_O_2_ treatment (Figure S1C-S1D). Collectively, these data establish ROS as a potent negative regulator of PINK1-Parkin-mediated mitophagy.

**Figure 1.**
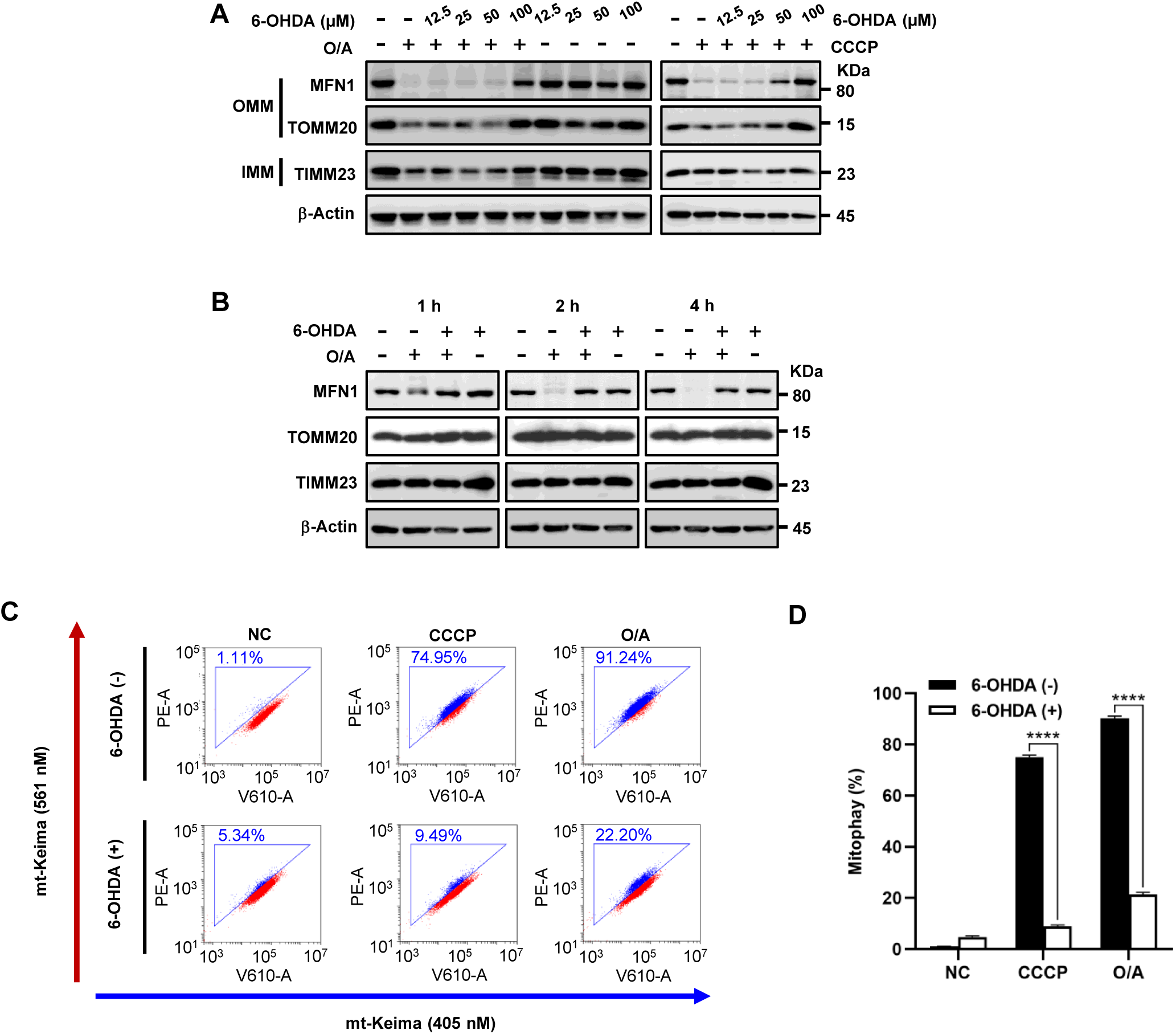
ROS inhibit mitophagy induced by acute mitochondrial damage agents. (A) The effect of different concentrations of 6-OHDA on CCCP or O/A induced mitochondrial protein degradation. YFP-Parkin-HeLa cells were treated with CCCP (20 µM) or O/A (1 µM/1 µM) combined with different concentrations of 6-OHDA for 6 h. Mitochondrial protein degradation was detected by immunoblotting. (B) The effect of 6-OHDA on O/A induced mitochondrial protein degradation over time. YFP-Parkin-HeLa cells were treated with or without O/A (1 µM/1 µM), 6-OHDA (100 µM) or in combination, and harvested at the indicated time points for immunoblotting. (C) mt-Keima-HeLa cells were treated with CCCP (20 µM) or O/A (1 µM/1 µM) combined with 6-OHDA (100 µM) for 4 h, followed by flow cytometry analysis at 405 nm (neutral pH) and 561 nm (acidic pH) excitation. Cells gated with increased 561 nm excitation represent populations undergoing mitophagy. (D) Mitophagy level was quantified in mt-Keima-HeLa cells following treatment as described in (C). bars, ± SD of 3 independent experiments; ****, P < 0.0001 (Two-way ANOVA test).

### ROS inhibit PINK1-Parkin-mediated mitophagy via blockage of PINK1 activation and Parkin recruitment

Having demonstrated the capacity of ROS to suppress mitophagy, we next set to examine the underlying mechanisms by testing the effect of ROS on PINK1 activation and Parkin recruitment to mitochondria, the two key events at the early stage of mitophagy. The level of pSer65-Ub is a well-recognized marker of PINK1 activity [53–55]. Immunoblotting results revealed that 6-OHDA markedly reduced CCCP or O/A-induced stabilization of full-length PINK1 and pSer65-Ub level (Figure 2A). Similar inhibitory effects were observed in embryonic kidney (HEK293T) and neuronal (SH-SY5Y) cell models, both with endogenous expression of Parkin (Figure 2B-2C). Live-cell imaging of YFP-Parkin-HeLa cells showed that 6-OHDA significantly blocked Parkin mitochondrial recruitment in cells treated with CCCP or O/A (Figure 2D). Notably, H_2_O_2_ had similar effect on PINK1 activation and Parkin recruitment (Figure 2E-2F and S1E-S1F). Taken together, these data suggest that ROS suppress mitophagy by impairing PINK1 activation and Parkin mitochondrial recruitment.

**Figure 2.**
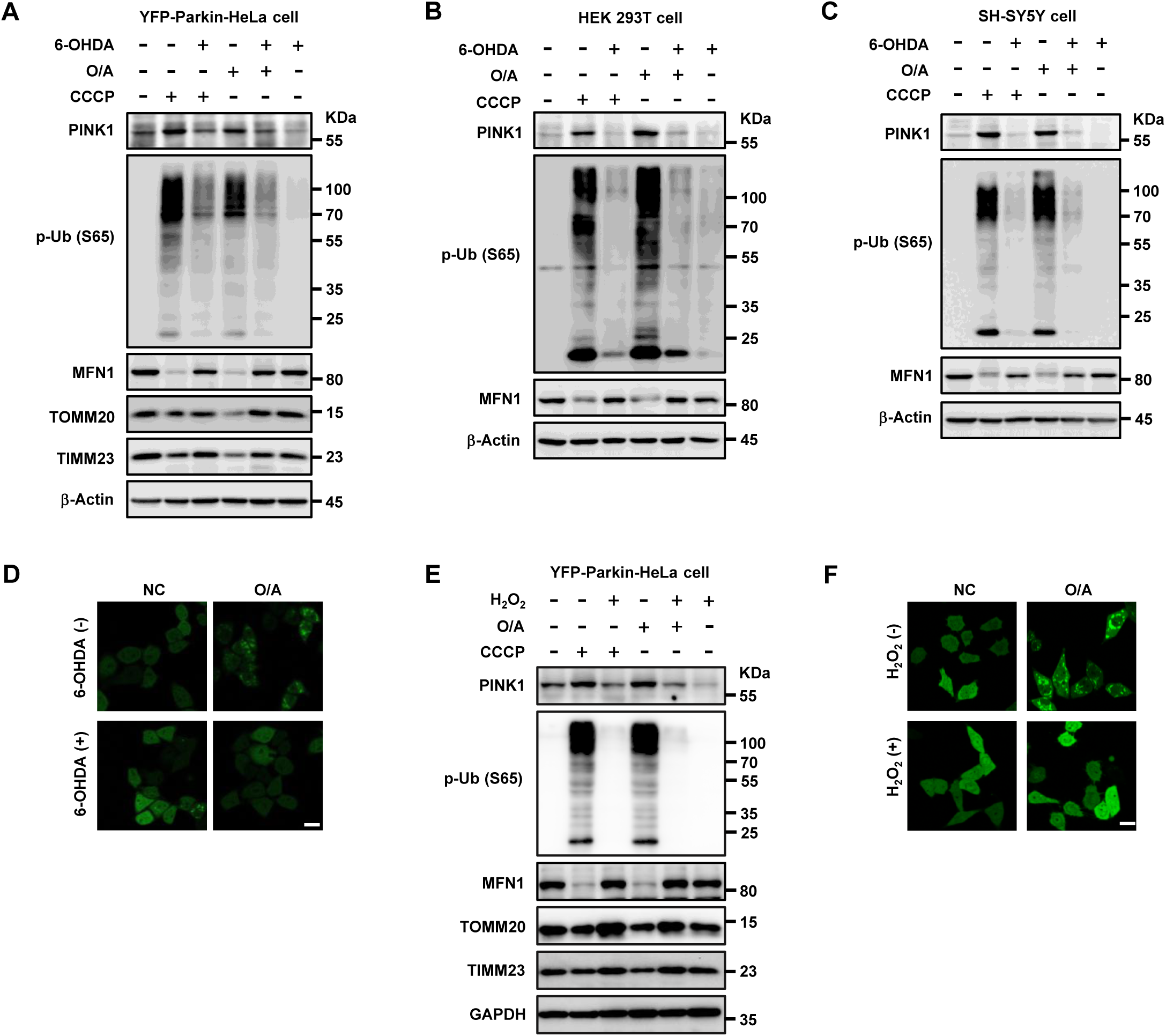
ROS inhibit mitophagy via blockage of PINK1 activation and Parkin recruitment. (A) and (E) YFP-Parkin-HeLa cells were treated with CCCP (20 µM) or O/A (1 µM/1 µM) combined with 6-OHDA (100 µM) (A) or H_2_O_2_ (200 µM) (E) for 6 h and harvested for immunoblotting analysis with the indicated antibodies. (B-C) HEK 293T cells (B) and SH-SY5Y cells (C) were treated with CCCP (20 µM) or O/A (1 µM/1 µM) combined with 6-OHDA (100 µM) for 8 h and harvested for immunoblotting analysis with the indicated antibodies. (D) and (F) YFP-Parkin-HeLa cells were treated with CCCP (20 µM) or O/A (1 µM/1 µM) combined with 6-OHDA (100 µM) (D) or H_2_O_2_ (200 µM) (F) for 1 h and imaged under confocal microscope. Scale bar: 20 μm.

To further understand the mechanisms underlying the inhibitor effect of ROS on PINK1, we tested whether ROS affect PINK1 transcription or translation upon mitochondrial damage. Quantitative RT-PCR analyses revealed no significant differences in PINK1 mRNA levels with or without treatment with H_2_O_2_ (Figure S2A). Under basal conditions, PINK1 undergoes rapid N-end rule-mediated proteasomal degradation [56] and the cleaved PINK1 can be stabilized by proteasome inhibitor [16]. MG132 is a commonly used proteasome inhibitor that specifically binds to the β5 subunit of the 20S proteasome and block its proteolytic activity [57]. Pretreatment with MG132, followed by the addition of O/A together with either 6-OHDA or H_2_O_2_, resulted in the detection of stabilized cleaved PINK1 (Figure S2B-S2C, lane #5), indicating that ROS did not alter the synthesis of the full-length PINK1. While O/A-triggered mitochondrial depolarization, ROS-elevating treatments failed to restore the collapsed mitochondrial membrane potential (MMP) caused by O/A (Figure S2D-S2E), indicating that the effect of ROS on PINK1 and Parkin is independent of MMP. Thus, our data exclude the possibility that the inhibitory effect of ROS on PINK1 activation is acting via inhibition of PINK1 transcription and translation or by restoration of MMP.

#### Antioxidants abolish the inhibitory effect of ROS on PINK1-Parkin-mediated mitophagy

To further establish the inhibitory role of ROS in mitophagy, here we utilized N-acetylcysteine (NAC), a potent ROS scavenger. NAC treatment reversed ROS-induced inhibition of mitophagy: it effectively abolished the inhibitory effects of either 6-OHDA or H_2_O_2_ on PINK1 and pSer65-Ub and restored mitochondrial protein degradation (Figure 3A and S3A) and Parkin mitochondrial recruitment (Figure 3B and S3B). Mitophagy restoration by NAC was also quantified via mt-Keima assay (Figure 3C-3D and S3C-S3D). Similar results were also found in HEK293T and SH-SY5Y cells (Figure 3E-3F and S3E-S3F), suggesting that the effect of NAC is not cell-type specific.

**Figure 3.**
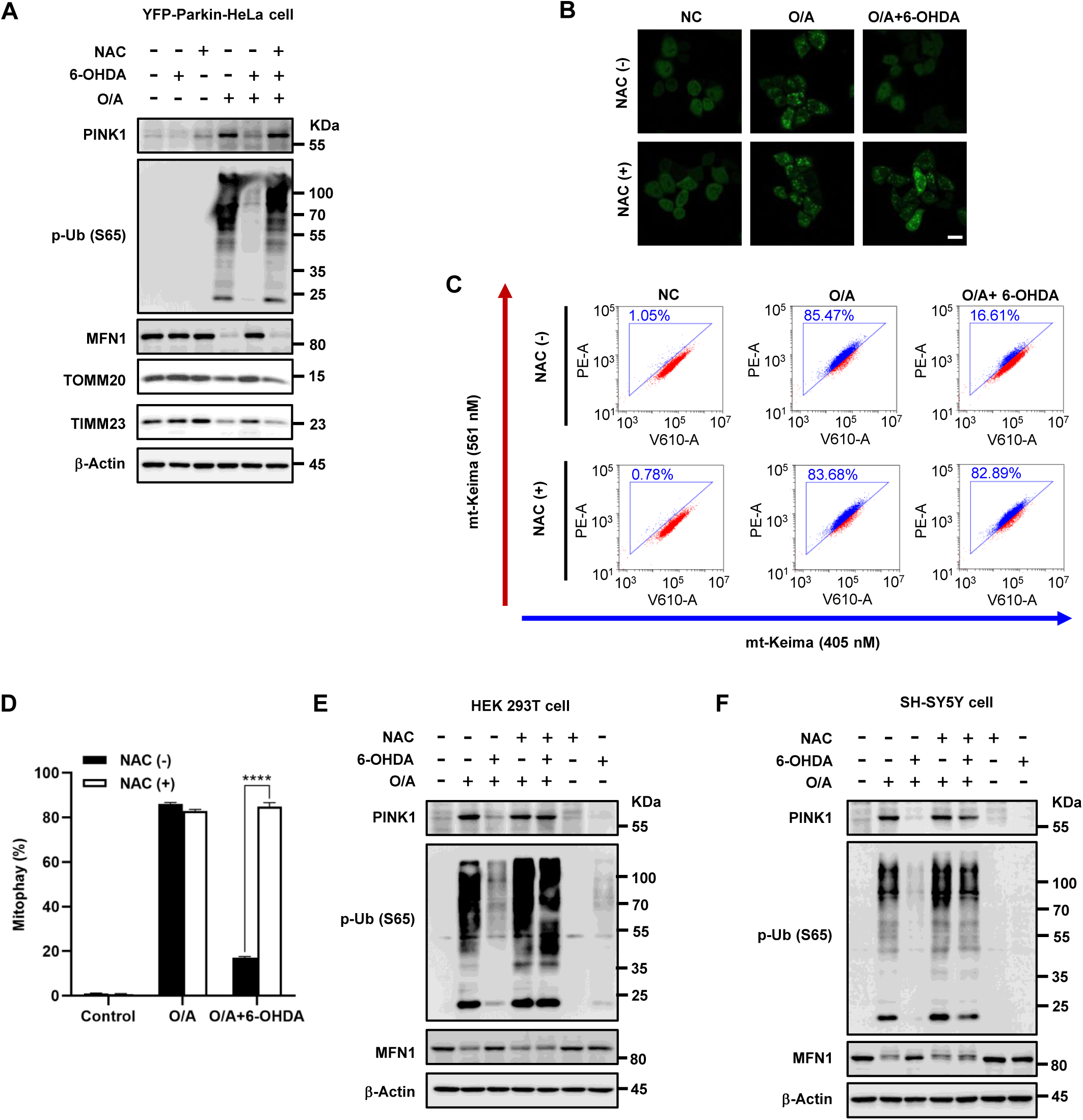
Antioxidants abolish the inhibitory effect of ROS on mitophagy. (A) YFP-Parkin-HeLa cells were pre-treated with NAC (5 mM) for 1.5 h and then treated with CCCP (20 µM) or O/A (1 µM/1 µM) combined with 6-OHDA (100 µM) for 6 h. Cells were harvested for immunoblotting analysis with the indicated antibodies. (B) YFP-Parkin-HeLa cells were treated with O/A (1 µM/1 µM) and 6-OHDA (100 µM) in the presence or absence of NAC (5 mM) for 1 h and imaged under confocal microscope. Scale bar: 20 μm. (C) mt-Keima-HeLa cells were pre-treated with NAC for 1.5 h and then treated with O/A (1 µM/1 µM) and 6-OHDA (100 µM) for 4 h, followed by flow cytometry analysis at 405 nm (neutral pH) and 561 nm (acidic pH) excitation. (D) Mitophagy level was quantified in mt-Keima-HeLa cells following treatment as described in (C). bars, ± SD of 3 independent experiments; ****, P < 0.0001 (Two-way ANOVA test). (E-F) HEK 293T cells (E) and SH-SY5Y cells (F) were pre-treated with different concentrations of NAC for 1.5 h and then treated with O/A (1 µM/1 µM) and 6-OHDA (100 µM) for 8 h. Cells were harvested for immunoblotting analysis with the indicated antibodies.

### ROS activate PARP1 and deplete intracellular ATP and NAD^+^

It is well known that one of the main consequences of ROS and oxidative stress is DNA damage such as DNA double strand breaks, leading to PARP1 activation, protein PARylation and the depletion of intracellular ATP and NAD^+^, as part of the DNA damage response (Figure 4A) [58, 59]. Consistently, we observed that both 6-OHDA and H_2_O_2_ treatment increased global PARylation levels, in a process dependent on PARP1 activation as evidenced by complete PARylation ablation in PARP1-knockout (KO) cells (Figure 4B and S4A). In contrast, O/A treatment alone did not induce PARP1 activation (Figure 4B and S4A). As expected, PARP1 activation caused rapid depletion of intracellular ATP and NAD⁺ (Figure 4C-4D and S4B-S4C). Olaparib is a specific inhibitor of PARP1 and inhibits the enzyme activity of PARP1 by competitively combining with its NAD^+^ binding site [60]. The depletion of ATP and NAD^+^ caused by ROS can be partially reversed by Olaparib treatment (Figure 4E-4F and S4D-S4E). The above data thus suggest that PARP1 activation is an important biological event upon oxidative stress.

**Figure 4.**
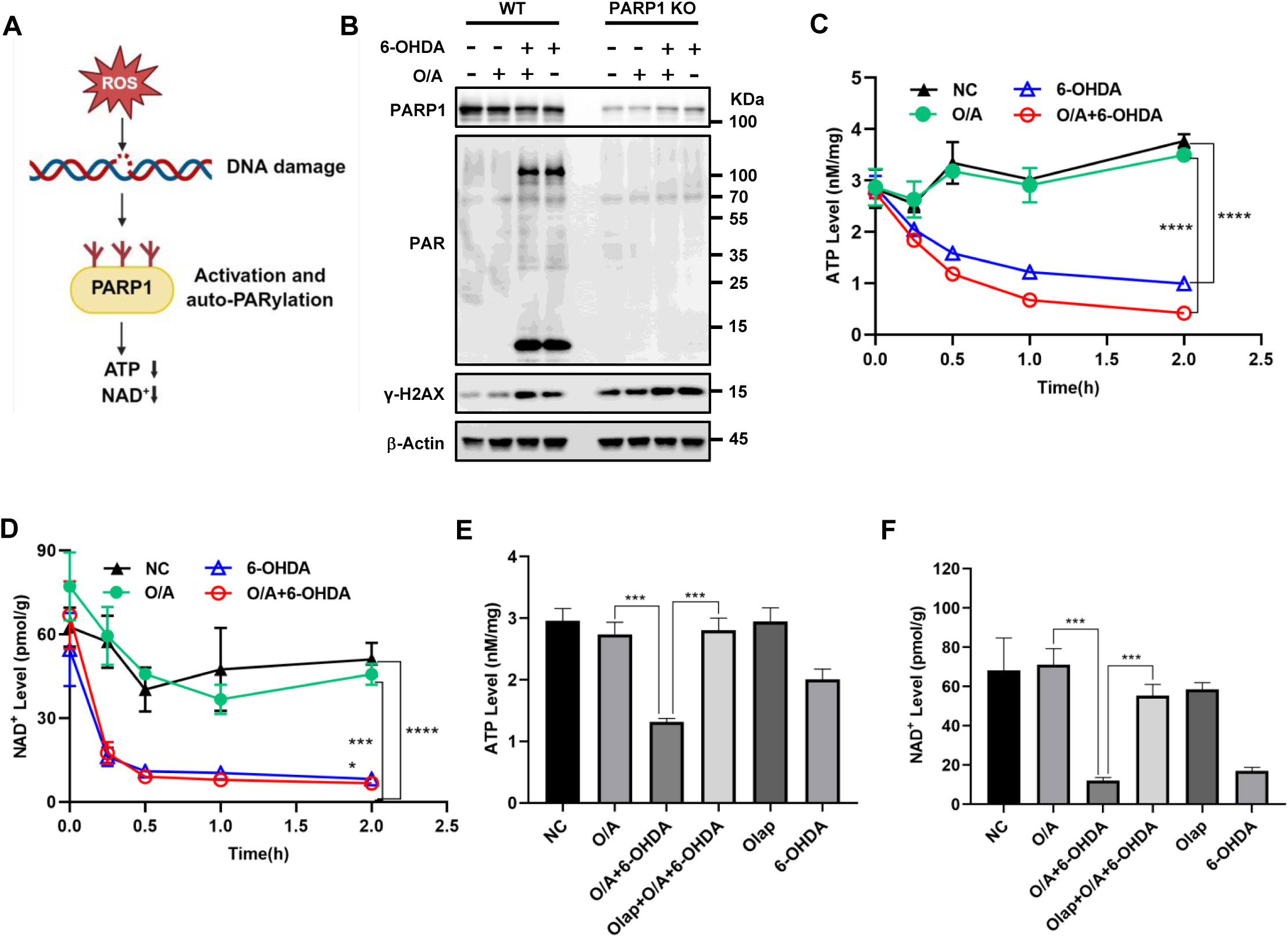
ROS activate PARP1 and deplete intracellular ATP and NAD^+^. (A) PARP1 activation induced by DNA damage depletes intracellular ATP and NAD^+^. (B) Wild-type (WT) and PARP1 knockout (KO) YFP-Parkin-HeLa cells were treated with or without O/A (1 µM/1 µM), 6-OHDA (100 µM) or in combination for 10 min and harvested for immunoblotting. (C) Intracellular ATP levels response to 6-OHDA treatment over time. YFP-Parkin-HeLa cells treated with or without O/A (1 µM/1 µM), 6-OHDA (100 µM) or in combination for different times. ATP levels were measured via luciferase assay. bars, ± SD of 3 independent experiments; ****, P < 0.0001 (Two-way ANOVA test). (D) Intracellular NAD^+^ levels response to 6-OHDA treatment over time. YFP-Parkin-HeLa cells were treated as in (C). NAD^+^ levels were quantified via WST-8. Statistical analysis was performed as in (C). (E-F) Intracellular ATP and NAD^+^ levels response to Olaparib. Cells were pre-treated by Olaparib for 1.5 h and then treated with O/A (1 µM/1 µM) and 6-OHDA (100 µM) for 2 h. bars, ± SD of 3 independent experiments; ns, no significance, ***, P < 0.001; ****, P < 0.0001 (Student’s t-test).

### Suppression of PARP1 abolishes the inhibitory effect of ROS on PINK1-Parkin-mediated mitophagy

Based on the results that ROS activate PARP1, we next explored the potential role of PARP1 activation in ROS-mediated mitophagy inhibition. Temporal correlation analyses revealed that PARP1 activation induced by prooxidant treatment (6-OHDA or H_2_O_2_), as evidenced by increased PARylation, paralleled the concomitant inhibition of mitophagy (Figure 5A and S5A). Similarly, the inhibitory effect of prooxidant treatment on mitophagy correlates with the increase of PARylation levels in a dose-dependent manner (Figure 5B and S5B). These data suggest that the elevated PARylation levels induced by activation of PARP1 might be functionally implicated in ROS-mediated mitophagy inhibition. To test this notion, we utilized pharmacologic PARP1 inhibitors Olaparib, and found that Olaparib effectively reversed the inhibitory effect of ROS on mitophagy, restoring PINK1 and pSer65-Ub levels (Figure 5C and S5C-S5G). Consistent results were obtained with the mt-Keima assay (Figure 5D-5E and S5H-S5I) and Parkin recruitment (Figure 5F and S5J). Moreover, genetic deletion of PARP1 via siRNA (knockdown) or CRISPR-KO fully abolished the blockage of ROS on mitophagy (Figure 5G-5H and S5K-S5L). Of note, as shown in Figure 5G and S5K, we employed two distinct PARP1 siRNA to generate two PARP1-knockdown (KD) cell lines. Compared with PARP1-KD-1, PARP1-KD-2 achieved a more efficient knockdown, as shown by a greater reduction in PARP1 activation. Consistent with these differential effects on PARP1, PARP1-KD-2 showed much significant effect on the ROS-induced inhibition of mitophagy compared to PARP1-KD-1 cell line (Figure 5G and S5K). Taken together, these data suggest that the activation of PARP1 induced by ROS is implicated in the inhibitory effect of ROS on mitophagy.

**Figure 5.**
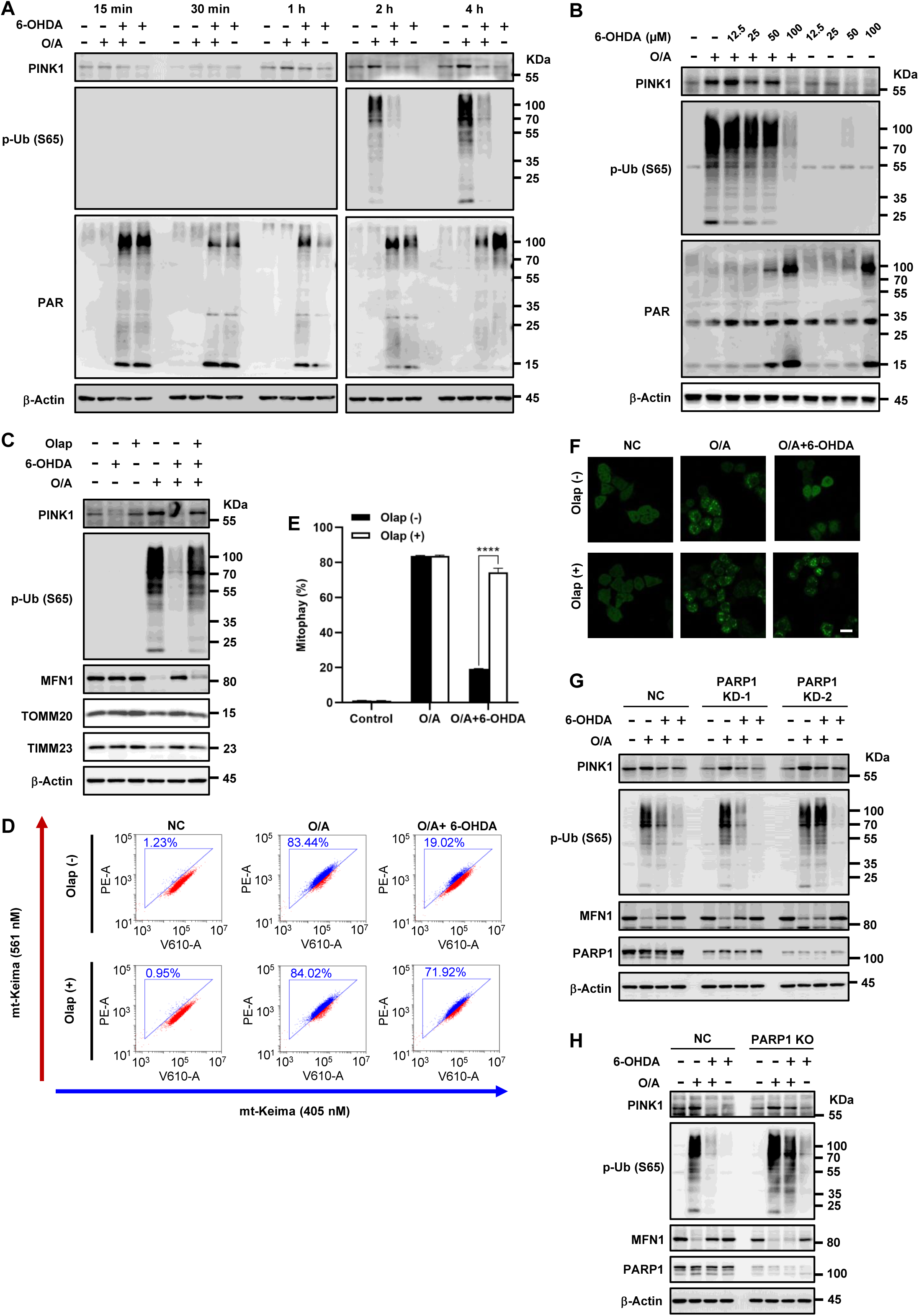
Suppression of PARP1 abolish the inhibitory effect of ROS on mitophagy. (A) YFP-Parkin-HeLa cells were treated with or without O/A (1 µM/1 µM), 6-OHDA (100 µM) or in combination, and harvested at the indicated time points for immunoblotting. (B) YFP-Parkin-HeLa cells were treated with O/A (1 µM/1 µM) and different concentrations of 6-OHDA. Cells were harvested for immunoblotting analysis with the indicated antibodies. (C) YFP-Parkin-HeLa cells were pre-treated with Olaparib (1 µM) for 1.5 h and then treated with O/A (1 µM/1 µM) and 6-OHDA (100 µM) for 6 h. Cells were harvested for immunoblotting analysis with the indicated antibodies. (D) mt-Keima-HeLa cells were treated with O/A (1 µM/1 µM) and 6-OHDA (100 µM) in the presence or absence of Olaparib (1 µM) for 4 h, followed by flow cytometry analysis at 405 nm (neutral pH) and 561 nm (acidic pH) excitation. (E) Mitophagy level was quantified in mt-Keima-HeLa cells following treatment as described in (D). bars, ± SD of 3 independent experiments; ****, P < 0.0001 (Two-way ANOVA test). (F) YFP-Parkin-HeLa cells were treated as in (D) for 1 h and imaged under confocal microscope. Scale bar: 20 μm. (G) WT and PARP1 KD YFP-Parkin-HeLa cells were treated with or without O/A (1 µM/1 µM), 6-OHDA (100 µM) or in combination for 6 h and harvested for immunoblotting. (H) WT and PARP1 KO YFP-Parkin-HeLa cells were treated as in (G) and harvested for immunoblotting.

### PARylation of PINK1 at damaged mitochondria impairs its function

Having established the critical role of PARP1 in ROS-mediated mitophagy inhibition, we next attempted to explore the molecular mechanisms underlying the effect of PARP1 on mitophagy. Based on the facts that PARP1-mediated protein PARylation regulates various cellular processes, as a form of post-translational modifications (PTMs) [61, 62], we speculated that ROS could inhibit mitophagy via inducing the PARylation of some key components of the mitophagy machinery. Subcellular fractionation revealed that both 6-OHDA and H_2_O_2_ treatment significantly elevated mitochondrial PARylation levels (Figure 6A and S6A). Given the well-known fact that PINK1 accumulates on outer mitochondrial membrane (OMM) during mitophagy initiation, we postulated that PINK1 might undergo this form of PTM. To test this possibility, we pulled down PINK1 protein in YFP-Parkin-HeLa cells stably expressing PINK1-MYC for mass spectrometry (MS) and found PARP1 as an interacting protein of PINK1 (Table S1). Consistently, co-IP experiments also showed an interaction between PINK1 and PARP1 (Figure 6B and S6B). Moreover, PARylation of PINK1 was detected by anti-PAR antibody upon either 6-OHDA or H_2_O_2_ treatment, and this modification was fully eliminated by PARP1 inhibitor Olaparib (Figure 6B and S6B). Thus, data from this part of our study suggest the possibility that PINK1 may subject to PARylation by PARP1.

**Figure 6.**
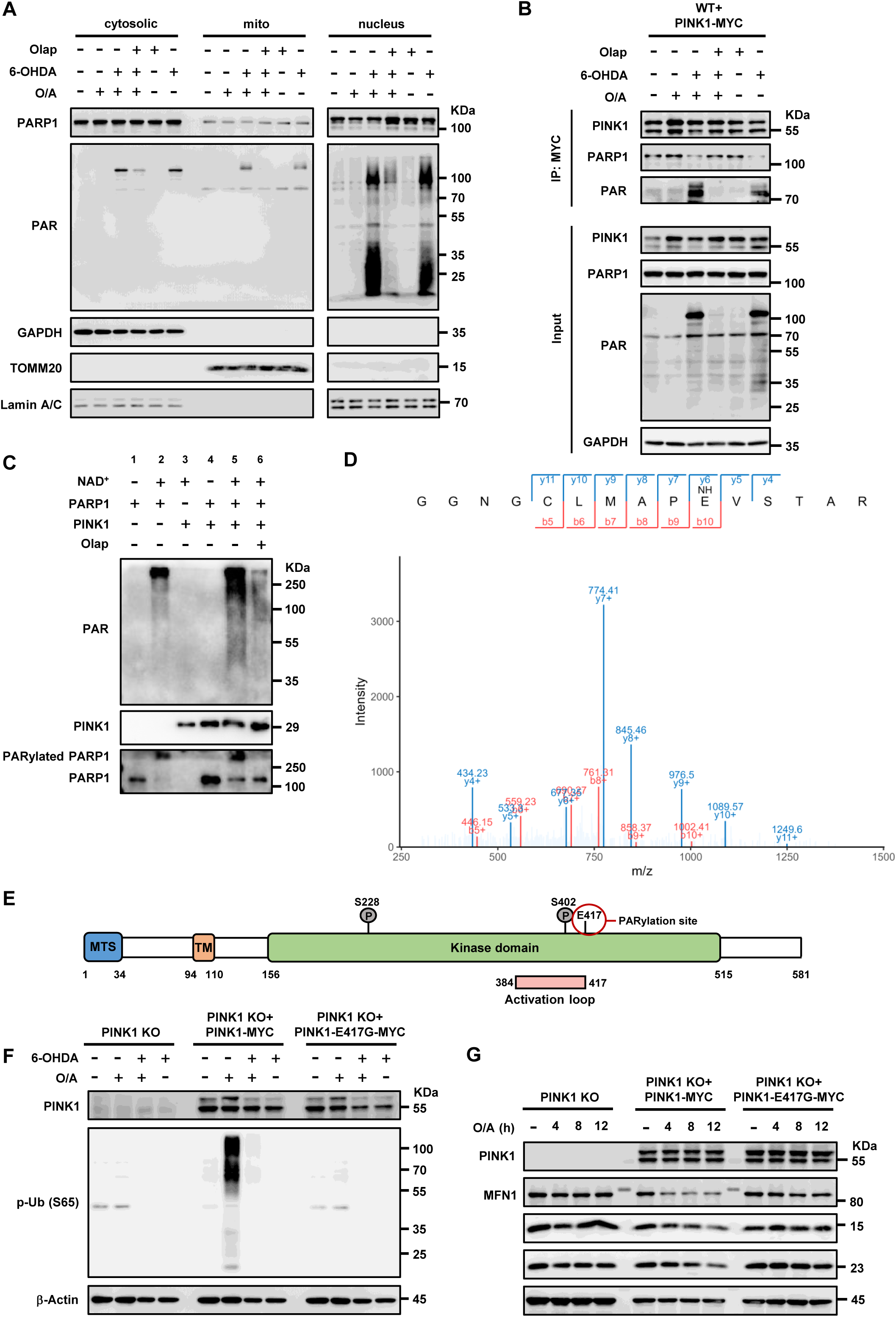
PARylation of PINK1 at damaged mitochondria impairs its function. (A) YFP-Parkin-HeLa cells were pre-treated with Olaparib (1 µM) for 1.5 h and then treated with O/A (1 µM/1 µM) and 6-OHDA (100 µM) for 10 min. Subcellular fractionation was then performed to isolate the mitochondrial, nucleus, and cytosolic fractions. TOMM20, Lamin A/C and GAPDH were used as mitochondrial, nucleus and cytosolic markers, respectively. (B) YFP-Parkin-HeLa cells stably expressing PINK1-MYC pre-treated with Olaparib (1 µM) for 1.5 h and then treated with O/A (1 µM/1 µM) and 6-OHDA (100 µM) for 10 min. Cells were lysed with IP lysis buffer. And the cell lysates were subjected to MYC IP and analyzed by immunoblotting. (C) *In vitro* PARylation assays were performed with the indicated purified recombinant proteins and analyzed by immunoblotting. (D) Mass spectrometry analysis reveals the PARylation site of PINK1 on E417. (E) The domain structure of PINK1. PINK1 contains mitochondrial targeting sequence (MTS), transmembrane domain (TM), and kinase domain. The activation loop of PINK1 comprises amino acids 384-417. (F) PINK1-MYC and PINK1-E417G-MYC were reconstituted in PINK1 KO YFP-Parkin-HeLa cells. Cells were treated with or without O/A (1 µM/1 µM), 6-OHDA (100 µM) or in combination for 6 h and harvested for immunoblotting.

To prove this notion, we then performed *in vitro* PARylation assay to test whether PINK1 was directly PARylated by PARP1. First, PARP1 was auto-PARylated with the presence of NAD^+^ in the reaction (Figure 6C, lane #1 and #2). Second, PARylated PINK1 was detected with the presence of purified PINK1 protein into the reaction (Figure 6C, lane #5). Finally, PARylated PINK1 was almost completely eliminated by PARP1 inhibitor Olaparib (Figure 6C, lane #6).

With the above-mentioned biochemical data showing PARP1-meidated PARylation of PINK1, we attempted to identify the PARylation site, using MS for the analysis of Asp and Glu-ADP-ribosylation according to established methods [63, 64]. Our data identified E417 as a PARylation site on PINK1 (Figure 6D), which resides within its kinase activation loop (Figure 6E).

Moreover, mutating E417 to either glycine (G) or glutamine (Q) impaired PINK1 kinase activity (Figure 6F and S6C–S6E). Since PINK1 activity is essential for mitophagy, we next examined whether this mutation also affects mitochondrial protein degradation. Consistently, the E417 mutation abolished the degradation of mitochondrial proteins induced by O/A (Figure 6G). These data suggest that PARP1-mediated modification of this residue impairs PINK1 function and inhibits mitophagy.

Collectively, these findings reveal a novel mechanism underlying the regulatory role of ROS in mitophagy: ROS-mediated activation of PARP1 directly PARylates PINK1, impairs its kinase activity and thereby inhibiting mitophagy.

## DISCUSSION

In this study, we identified a novel regulatory mechanism of ROS on PINK1-Parkin-mediated mitophagy. Initially, we observed that ROS significantly inhibited mitophagy triggered by acute mitochondrial damage agents (Figure 1-2). This inhibitory effect was specifically reversed by antioxidant (Figure 3). Mechanistic investigations revealed that ROS induced DNA damage response marked by PARP1 activation and elevated PARylation levels, while concomitantly depleting intracellular ATP and NAD^+^ (Figure 4). Furthermore, genetic deletion (KD/KO) or pharmacological inhibition of PARP1 completely abrogated the effect of ROS on mitophagy, establishing the critical role of PARP1 in this process (Figure 5). Through biochemical analyses, we demonstrated that PARP1 directly PARylates PINK1 at residue E417, thereby impairing its functional activity (Figure 6). Collectively, our findings reveal that ROS modulate PINK1-Parkin-mediated mitophagy through PARP1-mediated PARylation of PINK1 (Figure 7).

**Figure 7.**
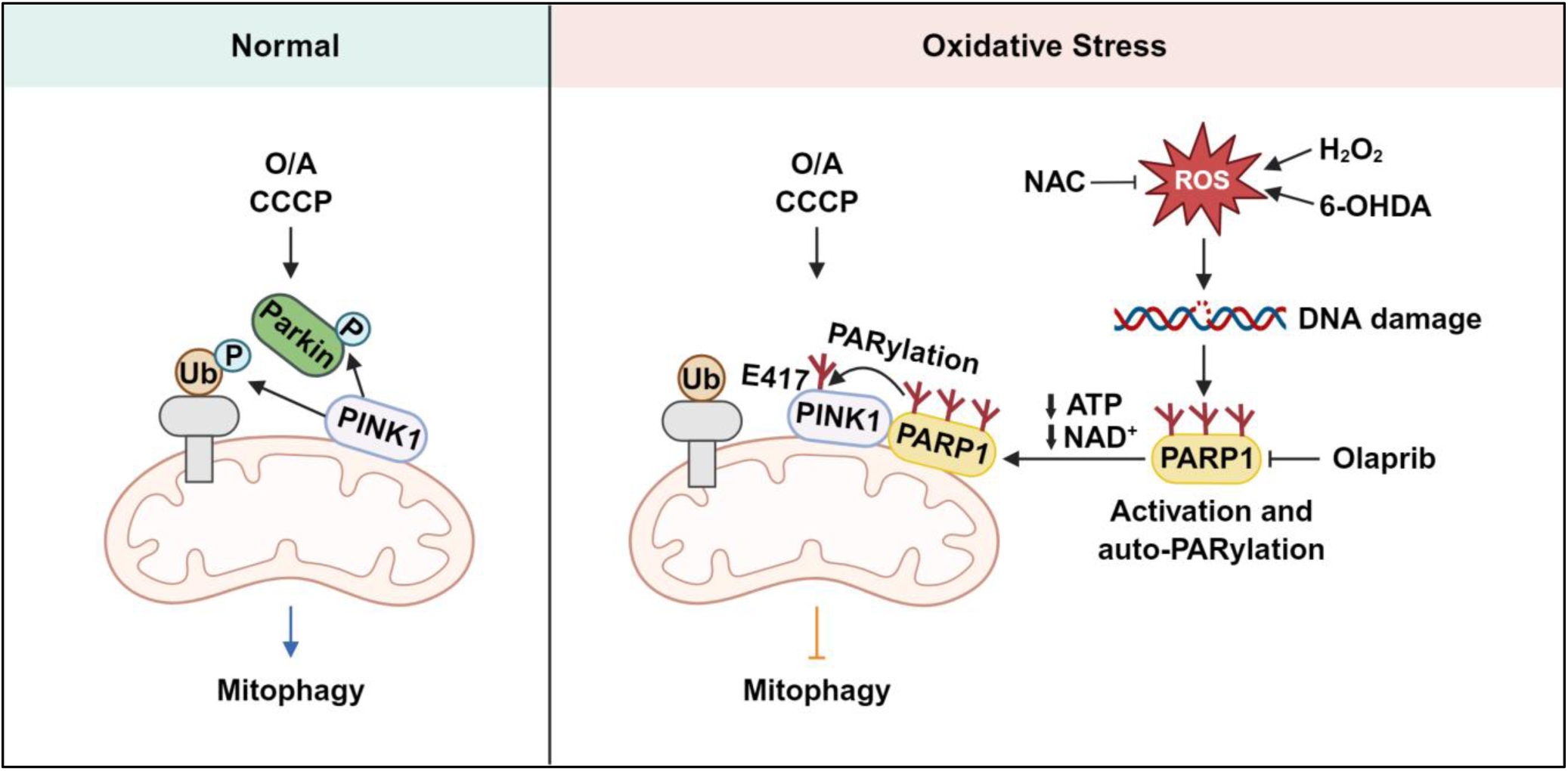
Schematic diagram for the regulatory role of ROS in mitophagy. ROS-induced DNA damage activates PARP1, contributes to PARylation and ATP/NAD^+^ reduction. PARP1 mediates PARylation of PINK1, impairing its function and further inhibiting mitophagy.

ROS play multifaceted roles in cellular homeostasis and redox regulation [65]. Several research has demonstrated that both 6-OHDA and H_2_O_2_ can exert differential regulatory effects on mitophagy, while some of these findings align with our observations, others appear contradictory. For 6-OHDA, some studies reported that it induces excessive Drp1-dependent mitochondrial fission, generating fragmented, depolarized mitochondria that are subsequently cleared by mitophagy [66, 67]. Notably, in those studies, a marked alteration of mitochondrial membrane potential in SH-SY5Y cells required treatment with 50 μM 6-OHDA for more than 9 hours, whereas our experiments employed different treatment conditions (100 μM, 6 h). Moreover, the effect of 6-OHDA specifically on PINK1-Parkin-mediated mitophagy was not directly examined. In contrast, other work demonstrated that 6-OHDA inhibits mitophagy through downregulating the level of key proteins such as PINK1 and Parkin [68, 69], which is consistent with our results. Similarly, studies using H_2_O_2_ as a direct ROS source have yielded conflicting results. For instance, Frank et al. proposed that mild oxidative stress can induce mitophagy through mitochondrial fission-dependent mechanisms [36]. Several key distinctions exist between their study and ours: First, they assessed general mitophagy levels, whereas we focused specifically on the PINK1-Parkin-dependent pathway. Second, they utilized HeLa cells, which lack endogenous Parkin, while our experiments employed HeLa cells stably overexpressing Parkin, a well-established model for investigating PINK1-Parkin-mediated mitophagy, as well as two other cells with endogenous Parkin. Furthermore, there was a substantial difference in H_2_O_2_ treatment conditions: suggesting that varying ROS concentrations may exert divergent effects on mitophagy. On the other hand, several earlier studies have discovered similar inhibitory effect of ROS on mitophagy, which are consistent with our results, while different underlying mechanisms were proposed. For instance, Zhang et al. reported that H_2_O_2_ inhibits mitophagy through reducing the mRNA levels of PINK1 and Parkin [70], while in our hands, both transcriptional and translational level of PINK1 were not affected upon prooxidant treatment (Figure S2A-S2C). This mechanistic variation may stem from differences in cellular models. Zhang et al. used retinal pigment epithelial (RPE) cells, which exhibit high basal PINK1 protein levels, reflecting their elevated energy demand and adaptive response to persistent oxidative stress [71, 72]. In contrast, PINK1 protein is barely detectable under baseline conditions in our model. Additionally, another study demonstrated that H_2_O_2_ can inhibit the ubiquitin kinase activity of PINK1, identifying Cys166 as a reversible oxidation site in human PINK1 that enables ROS to regulate its activity [53]. However, in our study, the ROS generated by 6-OHDA and H_2_O_2_ appear to inhibit PINK1 predominantly through PARP1 activation and subsequent PARylation, as supported by our PARP1 knockdown and inhibitor experiments. While we cannot exclude the possibility that direct oxidation of Cys166 occurs in parallel, the strong dependence of mitophagy inhibition on PARP1 indicates that PARylation is a major mechanism underlying ROS-mediated PINK1 suppression in our model. Taken together, these findings suggest that ROS may exert pleiotropic roles in mitophagy regulation, depending on their concentration, the cellular context, and specific signaling pathway engaged. This further highlights the importance of targeting ROS as a potential strategy for modulating mitophagy.

PTMs constitute a class of chemical alterations to amino acid side chains that occur after protein biosynthesis [73, 74]. These modifications play pivotal roles in cellular processes due to their ability to modulate protein activity, interaction networks, and subcellular localization [75, 76]. Well-characterized PTMs include phosphorylation, ubiquitination, methylation, acetylation, and O-GlcNAcylation [77, 78]. In addition, some novel PTMs such as PARylation, crotonylation, malonylation, succinylation, and lactylation have been proposed in recent years [79]. PINK1 is a mitochondrial serine/threonine kinase involve in mitochondrial quality control, undergoes multiple PTM-mediated regulation. Under normal conditions, the ubiquitination of cleaved PINK1 contributes to proteasomal degradation, maintaining low level of PINK1 [80, 81]. When mitochondria depolarize, PINK1 accumulates on the OMM where it undergoes autophosphorylation, a critical activation step that initiates mitophagy by recruiting downstream effectors [82, 83]. Beyond these canonical modifications, recent work from our lab has shown that PINK1 activity is modulated by O-GlcNAcylation, a process with important implication in PINK1 activity [84]. Another study has identified that PINK1 undergoes SUMOylation, and the removing of this modification markedly decreases phosphorylated ubiquitin levels and consequently impairs mitophagy. In this study, we identified PARylation as a previously unrecognized PTM governing PINK1 function (Figure 6). We demonstrated that ROS-induced activation of PARP1 directly mediated PINK1 PARylation, disrupting its activity and impairing mitophagy (Figure 7).

PARP1 is an enzyme essential for numerous cellular functions, including DNA repair, transcription regulation, and chromatin remodeling [85, 86]. While it is well known that PAPR1 is primarily a nuclear protein, the mitochondrial presence of PARP1 remains contentious. Current literatures present conflicting evidence, with some studies demonstrating mitochondrial PARP1 localization [87–89], while others refute this observation [90]. Our subcellular fractionation analyses revealed significant PARP1 accumulation and PARylation proteins on mitochondria (Figure 6A and S6A). The origin of PARP1 within mitochondria remains controversial. Some studies indicates that PARP1 may localize directly to mitochondria [88], whereas others suggest that it translocates from the nucleus to mitochondria via interactions with mitochondrial proteins [87, 91, 92]. The latter perspective is further supported by the identification of mitochondrial-nuclear membrane contact sites, which play a vital role in mediating their communication [93–95]. In this study, we investigated the role of PARP1 in PINK1-Parkin-mediated mitophagy. Previous reports suggest that PARP1 modulates PINK1-Parkin-mediated mitophagy through suppression of the NAD^+^- SIRT1-PGC-1α axis [46], whereas 6-OHDA and H_2_O_2_ showed no significant impact on this pathway in our model (data not shown). Our study proposed a novel mechanism for the direct regulatory role of PARP1 in mitophagy: Co-IP and MS analyses confirmed the physical interaction between PARP1 and PINK1 (Figure 6B and S6B, Table S1). Under oxidative stress conditions, PARP1 directly PARylated PINK1, impairing its activity and subsequent recruitment of Parkin to depolarized mitochondria. Genetic suppression of PARP1 or pharmacological inhibition of its activity restored PINK1 functionality, rescuing mitophagy in stressed cells (Figure 5 and S5). These findings collectively identify PARP1 as a critical mediator that couples oxidative stress signaling to mitochondrial quality control through direct modification of PINK1.

Following the MS identification of E417 as the PINK1 PARylation site, we anticipated that mutating this residue would abolish PINK1 PARylation in cells co-treated with O/A and either 6-OHDA or H_2_O_2_, thereby restoring PINK1 function and rescuing mitophagy. However, contrary to our expectations, mutation of this residue (E417) resulted in a complete loss of PINK1 kinase activity (Figure 6F and S6C). To rule out the possibility that this effect was specific to the glycine (G) substitution, we also generated a glutamine (Q) mutant, an amino acid structurally similar to glutamic acid (E), the E417Q PINK1 mutant exhibited the same loss of kinase activity as E417G (Figure S6D-S6E). It is reported that E417 is located in the kinase activation loop of PINK1 [96, 97], evolutionarily conserved [98], and has been identified as a Parkinson’s disease-related mutation site [99]. Several studies highlighted its critical importance for PINK1 function and kinase activity. For instance, the E417G mutation has been shown to impair PINK1 phosphorylation and Parkin recruitment [100], and other studies demonstrated that this mutation impairs PINK1 kinase activity [98, 101]. These findings align with our results and support the conclusion that PARylation modification likely disrupts the function of this critical residue, consequently impairing PINK1 activity. Moreover, structural data reveal that E417 is exposed on the surface of PINK1 [102], further supporting the possibility that it serves as a direct PARylation target.

In summary, we identify a novel mechanism by which ROS suppress PINK1-Parkin-mediated mitophagy through PARP1-driven PARylation of PINK1. Data from this study thus demonstrate the possibility of targeting ROS-PARP1 pathway through genetic and pharmacological methods to enhance mitophagy as a novel therapeutic strategy for pathological condition with defective mitophagy such as neurodegenerative disorders.

### Limitations of the study

While this work provides novel insights into ROS-mediated mitophagy regulation, there are several limitations. First, the proteolytic mechanism responsible for PARylated PINK1 clearance remains uncharacterized. Second, the source of PARP1 responsible for PARylation of PINK remains elusive. It might come from nuclear-localized PARP1 that translocate to mitochondria and mediate mitochondrial protein PARylation, or it might be part of the mitochondrial PARP1 responsible for the observed PINK1 PARylation. Finally, the biological functions of PARP1-mediated mitochondrial PARylation have yet to be established. Employing *in vivo* models to assess the therapeutic potential of modulating ROS-PARP1 for neurodegenerative diseases warrant further investigations,

## RESOURCE AVAILABILITY

### Lead contact

Requests for further information and resources should be directed to and will be fulfilled by the lead contact, Han-Ming Shen.

### Materials availability

All unique/stable reagents generated in this study are available from the lead contact with a completed materials transfer agreement.

### Data and code availability

- This paper does not report original code.
- Any additional information required to reanalyze the data reported in this paper is available from the lead contact upon request

## ACKNOWLEDGMENTS

We thank the members of the Shen Lab for their valuable discussions. L.G., D.R., X.G., L.X., M.T., Y.C., and Y.Z. were supported by research scholarship/assistantship from the University of Macau. This project was supported by the following research grants: Science and Technology Development Fund, Macau SAR (FDCT0031/2021/A1, FDCT0081/2022/AMJ, FDCT0036/2024/RIB1) and University of Macau (CPG2024-0035-FHS, CPG2025-0031-FHS) to H.S., and FDCT0052/2025/AIJ to E.F.F. and H.S.

## AUTHOR CONTRIBUTIONS

Conceptualization, H.S. and L.G.; Data curation, L.G., H.W., X.Z., D.R., and X.G.; Formal analysis, L.G., G.L. and H.S.; Funding acquisition, H.S.; Investigation, L.G., H.W., X.Z., D.R., X.G., L.X., Z.W., M.T., Y.C., Y.Z., A.D.C., G.L., and H.S.; Methodology, L.G., H.W., X.Z., D.R., X.G., L.X., Z.W., M.T., Y.C., Y.Z., L.W., G.L., J.L. and E.F.F.; Project administration, L.G. and H.S.; Resources, H.S.; Supervision, H.S.; Validation, L.G.; Writing – original draft, L.G.; Writing – review & editing, L.G., J.L., E.F.F. and H.S.

## DECLARATION OF INTERESTS

EFF is a co-owner of Fang-S Consultation AS (Organization number 931 410 717) and NO-Age AS (Organization number 933 219 127); he has an MTA with LMITO Therapeutics Inc (South Korea), a CRADA arrangement with ChromaDex (USA), a commercialization agreement with Molecule AG/VITADAO, an MTAs with GeneHarbor (Hong Kong) Biotechnologies Limited, and a data license option agreement with Hong Kong Longevity Science Laboratory (Hong Kong); he is a consultant to MindRank AI (China), NYO3 (Norway), AgeLab (Vitality Nordic AS, Norway), and Hong Kong Longevity Science Laboratory (Hong Kong).

All other authors declare no conflict of interests.

## DECLARATION OF GENERATIVE AI AND AI-ASSISTED TECHNOLOGIES

During the preparation of this work, the author(s) used Deepseek in order to correct grammatical errors. After using this tool or service, the author(s) reviewed and edited the content as needed and take(s) full responsibility for the content of the publication.

## STAR★METHODS

### KEY RESOURCES TABLE

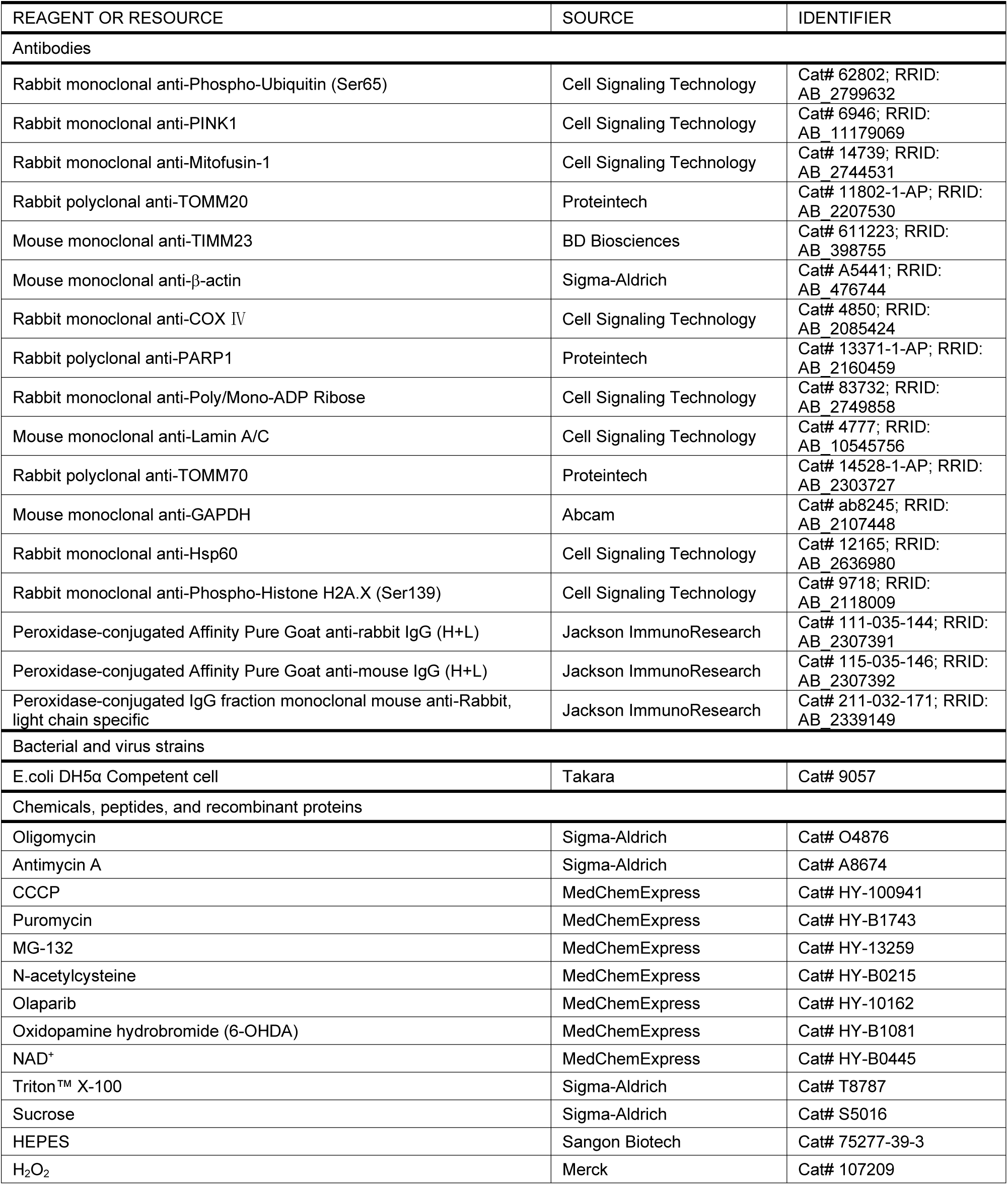
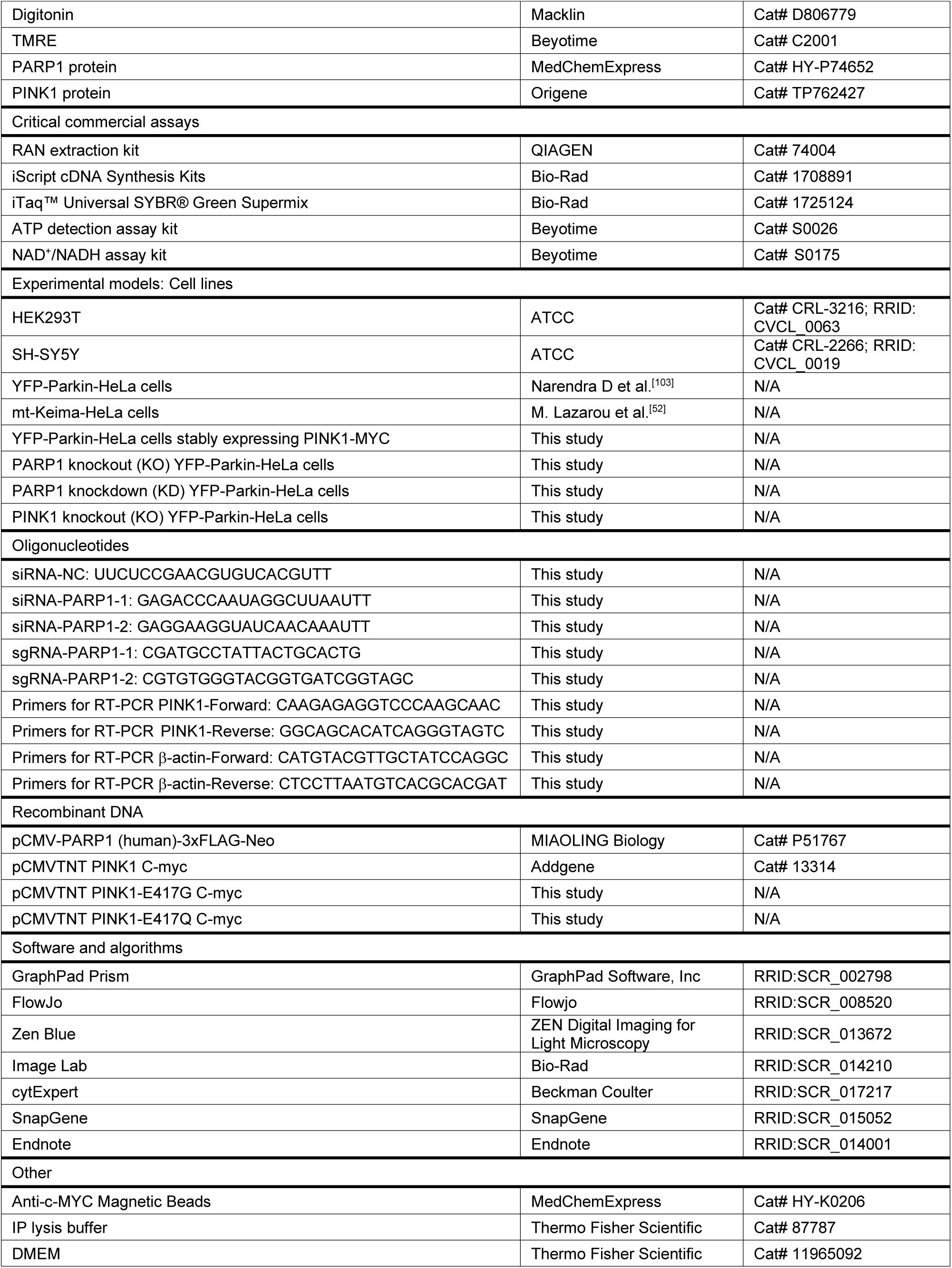
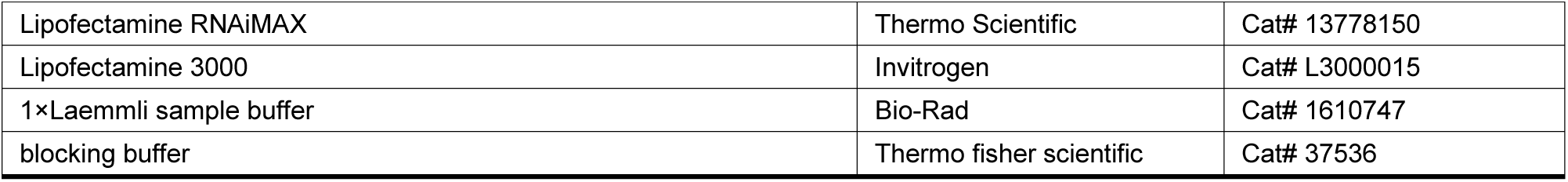

### EXPERIMENTAL MODEL AND STUDY PARTICIPANT DETAILS

Hela, HEK293T, and SH-SY5Y cells were cultured in DMEM with 10% fetal bovine serum and 1% Penicillin-Streptomycin at 37°C incubator with 5% CO2.

HeLa cells stably expressing YFP-Parkin (YFP-Parkin-HeLa cells) and mitoKeima (mt-Keima-HeLa cells) were kind gifts from Dr. Richard Youle (National Institute of Neurological Disorders and Stroke, NIH, USA). YFP-Parkin-HeLa cells stably expressing PINK1-MYC were kind gifts from Dr. Guang Lu (Sun Yat-sen University, Guangzhou, China). PINK1 knockout (KO) YFP-Parkin-HeLa cells were kind gifts from Dr. Liming Wang (Hunan University, Changsha, China).

### METHOD DETAILS

#### PARP1 KO cell generation by CRISPR-Cas9

CRISPR-Cas9-mediated PARP1 KO was performed using the lentiCRISPR v2 system. Lentiviral particles were generated by co-transfecting HEK293T cells with lentiCRISPRv2-sgRNA, psPAX2, and pMD2.G using polyethylenimine (PEI) at 3:2:1 molar ratio. Supernatants containing lentivirus were collected and transduced to YFP-Parkin-HeLa cells. The transfected cells were reseeded to a new dish after 48 h and selected by 2 µg/mL puromycin for another 48 h. Subsequently, the cells were subcultured at a ratio of 1:2000 into selective medium with 2 µg/mL puromycin, culturing for 2 weeks to obtain single cell cloning. PARP1 KO effect was verified by immunoblotting. The LentiCRISPR v2-Puro-PARP1-sgRNA plasmid was purchased from QINGKE.

#### Immunoblotting analysis

After indicated treatments, cells were washed with ice cold PBS and lysed using lysis buffer (62.5 mM Tris, 20% glycerol, 2% SDS, phosphatase and protease inhibitors, 1 mM dithiothreitol (DTT)). Equal amounts of protein were loaded onto SDS-PAGE gels to separate and transferred to polyvinylidene difluoride (PVDF) membranes. The membranes were blocked with blocking buffer (Thermo fisher scientific, 37536) for 30 min at room temperature, then incubated with appropriate primary and secondary antibodies. Immobilon Forte Western HRP substrate (Millipore, WBLUF0500) was used to enhance the chemiluminescence signal, and protein signal was detected by the ChemiDoc™ Touch Imaging System. Immunoblotting results were analyzed with Image Lab software.

#### RNA extraction and quantitative RT-PCR

The total RNAs were isolated from the treated YFP-Parkin-HeLa cells using RAN extraction kit (QIAGEN, 74004) following the manufacturer’s protocol. cDNAs were synthesized using iScript cDNA Synthesis Kits (Bio-Rad, 1708891). The reaction system of RT-qPCR was prepared according to the instruction of iTaq™ Universal SYBR® Green Supermix (Bio-Rad, 1725124). Each sample had three replicates. RT-qPCR was performed on Bio-Rad CFX96 Touch Real-Time PCR Detection System. The PCR primers were synthesized by QINGKE.

#### ATP and NAD^+^/NADH quantitation

The intracellular ATP and NAD^+^ levels were measured by ATP detection assay kit (Beyotime, S0026) and NAD^+^/NADH assay kit (Beyotime, S0175) following the provided protocols. Each biological group was analyzed with three technical replicates. The absorbance was measured by VICTOR x3 multilabel plate reader.

#### Confocal microscopy

The appropriate density of cells was plated onto glass bottom cell culture dishes (NEST, 801002). After indicated treatments, cells were imaged by Carl Zeiss LSM710 Confocal, using 63× oil lens, appropriate excitation wavelength and laser Power. The images could be analyzed by ZEN Microscopy Software.

#### Mitochondrial membrane potential assay

After treating cells with indicated drugs for 1 h, the medium was exchanged for TMRE work solution (Beyotime, C2001) and incubated at 37°C for 20 min. Subsequently, the cells were washed with prewarmed PBS for three times and digested by trypsin for collection. TMRE was determined at Ex/Em wavelengths of 550nm/575nm. Fluorescence intensity of cells was detected by CytoFLEX flow cytometer. Each biological group was analyzed with three technical replicates, collecting at least 10,000 events per replicate. Quantification analysis was performed by FlowJo software.

#### Immunoprecipitation

After indicated treatment, cells were washed by ice cold PBS for two times and lysed in IP lysis buffer (Thermo Scientific, 87787) containing phosphatase inhibitor and proteinase inhibitor on ice for 10 min. Lysates were centrifuged at 14,000 g for 10 min at 4°C to collect supernatants. The collected supernatant was mixed with 20 µL Anti-c-MYC Magnetic Beads (MCE, HY-K0206) and gently incubated under constant rotation (15 rpm) at 4°C for 16 h. Beads were subsequently washed with PBST (1×PBS + 0.5% Tween-20, pH 7.4) for five times. Immunoprecipitants were eluted from beads by boiling in 1×Laemmli sample buffer (Bio-Rad, #1610747) at 95°C for 5 min and analyzed with immunoblotting.

#### Subcellular fractionation

YFP-Parkin-HeLa cells were seeded to 10 cm cell culture dish and cultured to approximately 90% confluency. After the designed treatment, cells were pelleted at 300 g for 5 min at 4°C, resuspended in 800 µL Digitonin buffer (150 mM NaCl, 50 mM HEPES, 50 ug/mL digitonin, protease and phosphatase inhibitors) and incubated under constant rotation at 4°C for 10 min. Lysate was centrifuged at 2,000 g for 10 min at 4°C to separate cytosolic fractions (Supernatants) and other cellular components (pellet). Supernatants (cytosolic) were transferred to new tubes and centrifuged at 20,000 g for 20 min at 4°C to clear other cellular components. The centrifugation was performed three times with the supernatant being collected after each spin to obtain final cytosolic fraction. To separate the mitochondrial and nuclear fractions, the pellet from the previous step was washed with ice cold PBS for three times to remove the Digitonin buffer containing cytosolic fractions and centrifuged at 2,000 g each time for 5 min at 4°C. Subsequently, 800 µL NP-40 buffer (150 mM NaCl, 50 mM HEPES, 1% NP-40, protease and phosphatase inhibitors) was added to resuspend the pellet and incubated under constant rotation at 4°C for 30 min. After centrifuging the samples at 7,000 g for 10 minutes at 4°C, the supernatants containing the mitochondria were collected. The centrifugation was performed three times with the supernatant being collected after each spin to obtain final mitochondrial fraction. The pellet was then washed with ice cold PBS for three times to remove the NP-40 buffer containing mitochondrial fractions, and the remaining pellet was the nuclear.

#### *In vitro* PARylation assay

The indicated amount of purified PARP1 protein (MCE, HY-P74652) and PINK1 protein (origene, TP762427) were incubated in reaction buffer (50 mM Tris-HCl pH 7.5, 4 mM MgCl_2_, 20 mM NaCl and 250 μM fresh DTT) with 1 mM NAD^+^ and 400 ng sheared salmon sperm DNA at 25°C for 2 h. PARylation reactions were stopped by adding 1×Laemmli sample buffer and boiling at 95°C for 10 min. The reaction productions were analyzed by immunoblotting.

#### Mass spectrometry analysis of PINK1 PARylation

An *in vitro* PARylation assay was conducted following the protocols described above. The resulting reaction products were dissolved in 50 μL of 0.5 M NH_2_OH solution and incubated at 37°C for 2-4 hours. After desalting, the samples were digested with trypsin and subsequently subjected to mass spectrometry (MS) analysis. The tryptic peptides were dissolved in solvent A and directly loaded onto a homemade reversed-phase analytical column (25 cm in length, 100 μm i.d.). The mobile phase consisted of solvent A (0.1% formic acid in water with 2% acetonitrile) and solvent B (0.1% formic acid in acetonitrile). Peptides were separated with the following gradient: 0-14 min, 6%-24% B; 14-16 min, 24%-35% B; 16-18 min, 35%-80% B; 18-20 min, 80% B, at a constant flow rate of 500 nL/min on a NanoElute UHPLC system (Bruker Daltonics). The eluted peptides were ionized via a capillary source and analyzed using a timsTOF Pro 2 mass spectrometer. The electrospray voltage was set to 1.75 kV. Both precursor and fragment ions were detected with the TOF detector. Data were acquired in data-independent acquisition parallel accumulation serial fragmentation (dia-PASEF) mode. The full MS scan range was set from 300 to 1500 m/z. Each full MS scan cycle was followed by 20 PASEF MS/MS scans. The MS/MS scan range was set from 400 to 850 m/z, with an isolation window of 7 m/z. All MS analyses were performed by Hangzhou Jingjie PTM BIO.

#### Data processing for PINK1 PARylation

MS analysis of Asp- and Glu-ADP-ribosylation was conducted following an established protocol [63]. The DIA data were processed using Spectronaut (v.18) software. Tandem mass spectra were searched against the Homo_sapiens_9606_SP_20231220.fasta (20,429 entries) concatenated with a reverse decoy database. Trypsin/P was specified as the cleavage enzyme, allowing up to four missing cleavages. Carbamidomethyl of cysteine was set as a fixed modification. Acetylation of protein N-terminal, oxidation of methionine, and ADP-ribosylation were specified as variable modifications. False discovery rate (FDR) of protein, peptide and PSM was adjusted to < 1%.

### QUANTIFICATION AND STATISTICAL ANALYSIS

All data were statistically analyzed using GraphPad Prism 8. Student’s t-test was used to compare the differences between two groups. Two-way ANOVA was applied to compare multiple groups based on two factors. Data are presented as means ± SD from at least three independent experiments. A p-value < 0.05 was considered statistically significant.

