## Supplementary material for "ROS Impair Mitophagy via PARylation of PINK1": Suppl Figure S1-S6-Suppl table

#### Supplemental Figures and Legends

**Figure S1, related to Figure 1: ROS inhibit mitophagy induced by acute mitochondrial damage agents.**

**Figure S2, related to Figure 2: The inhibitory effect of ROS on mitophagy is not due to its effect on PINK1 transcription and translation level or mitochondrial membrane potential.**

**Figure S3, related to Figure 3: Antioxidants abolish the inhibitory effect of ROS on mitophagy.**

**Figure S4, related to Figure 4: ROS activate PARP1 and deplete intracellular ATP and $NAD^+$.**

**Figure S5, related to Figure 5: Suppression of PARP1 abolish the inhibitory effect of ROS on mitophagy.**

**Figure S6, related to Figure 6: PARylation of PINK1 at damaged mitochondria impairs its function.**

**Figure S1**

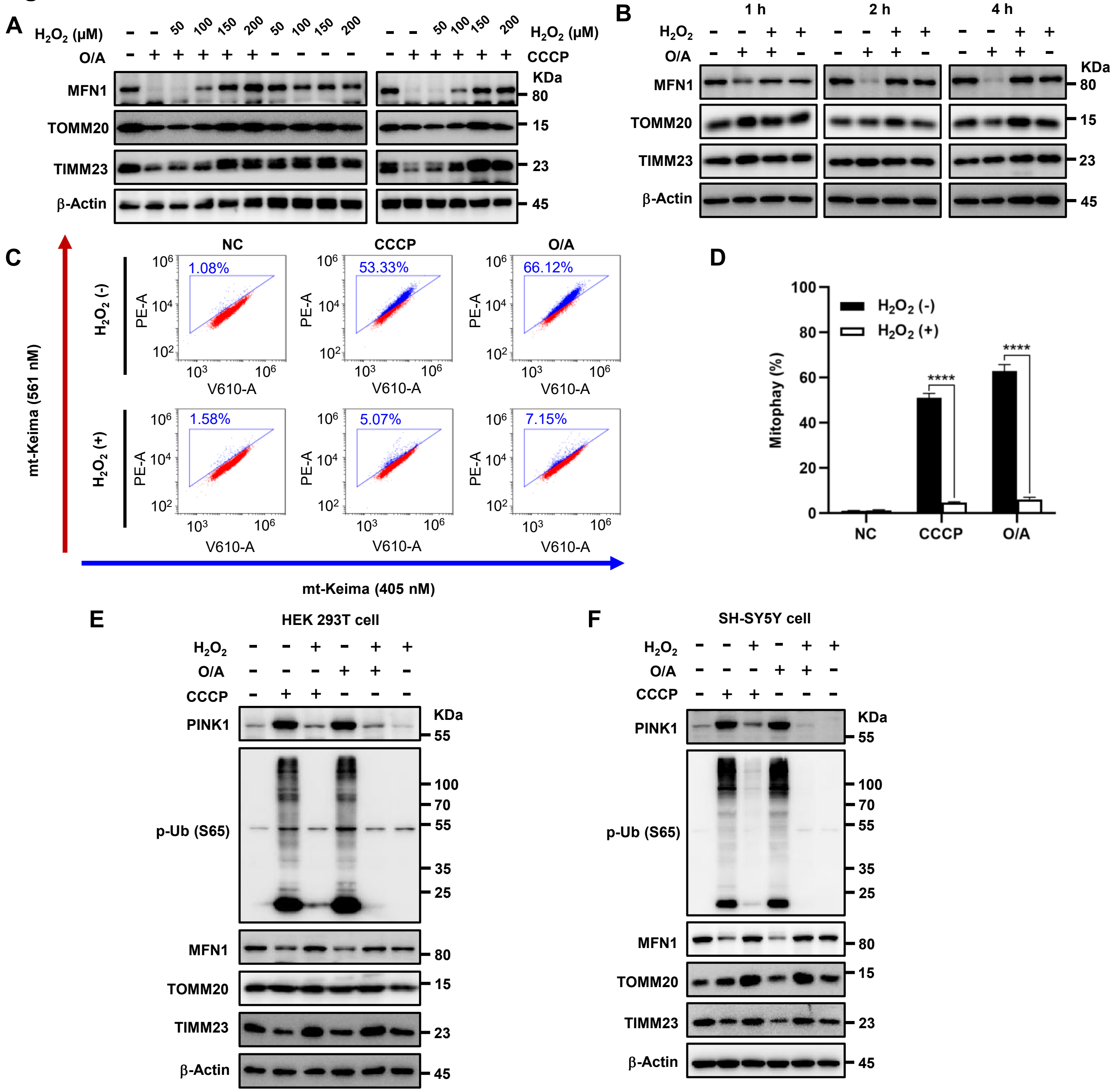

**Figure S1. ROS inhibit mitophagy induced by acute mitochondrial damage agents. Related to Figure 1.**

(A) The effect of different concentrations of $H_2O_2$ on CCCP or O/A induced mitochondrial protein degradation. YFP-Parkin-HeLa cells were treated with CCCP (20 µM) or O/A (1 µM/1 µM) combined with different concentrations of $H_2O_2$ for 6 h. Mitochondrial protein degradation was detected by immunoblotting. (B) The effect of $H_2O_2$ on O/A induced mitochondrial protein degradation over time. YFP-Parkin-HeLa cells were treated with or without O/A (1 µM/1 µM), $H_2O_2$ (200 µM) or in combination, and harvested at the indicated time points for immunoblotting. (C) mt-Keima-HeLa cells were treated with CCCP (20 µM) or O/A (1 µM/1 µM) combined with $H_2O_2$ (200 µM) for 4 h, followed by flow cytometry analysis at 405 nm (neutral pH) and 561 nm (acidic pH) excitation. Cells gated with increased 561 nm excitation represent populations undergoing mitophagy. (D) Mitophagy level was quantified in mt-Keima-HeLa cells following treatment as described in (C). bars, ± SD of 3 independent experiments; ****, $P < 0.0001$ (Two-way ANOVA test). (E-F) HEK 293T cells (E) and SH-SY5Y cells (F) were treated with CCCP (20 µM) or O/A (1 µM/1 µM) combined $H_2O_2$ (200 µM) for 6 h and harvested for immunoblotting analysis with the indicated antibodies.

#### Figure S2

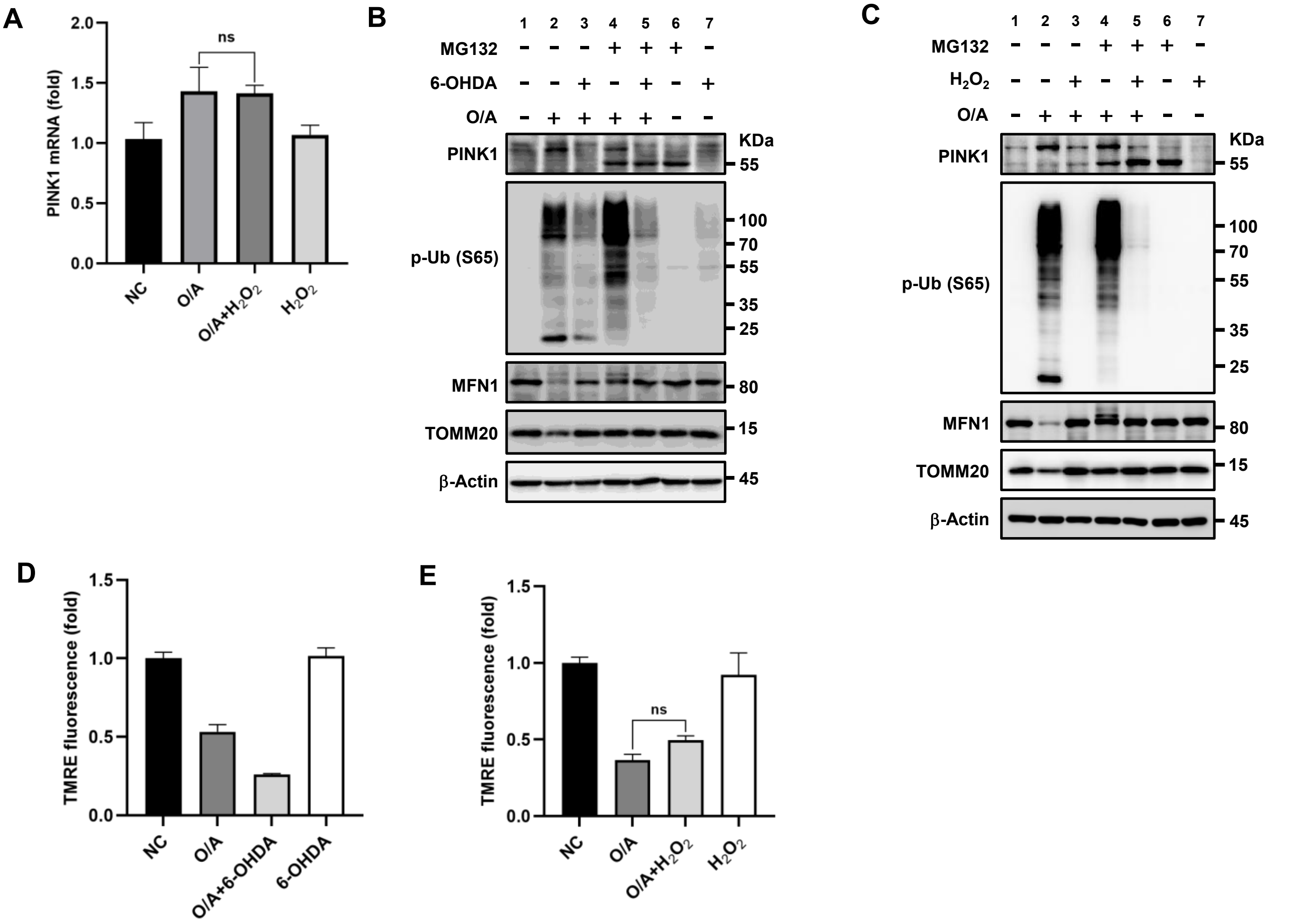

**Figure S2. The inhibitory effect of ROS on mitophagy is not due to its effect on PINK1 transcription and translation level or mitochondrial membrane potential. Related to Figure 2.**

(A) mRNA levels of PINK1. YFP-Parkin-HeLa cells were treated with O/A (1 μM/1 μM) combined with $H_2O_2$ (200 μM) for 2 h, RNA was extracted from each sample and quantified by real-time PCR. (B-C) YFP-Parkin-HeLa cells were pre-treated with MG132 (5 μM) for 1.5 h and then treated with O/A (1 μM/1 μM) combined with 6-OHDA (100 μM) (B) or $H_2O_2$ (200 μM) (C) for 6 h. Cells were harvested for immunoblotting analysis with the indicated antibodies. (D-E) YFP-Parkin-HeLa cells were treated with O/A (1 μM/1 μM) combined with 6-OHDA (100 μM) (D) or $H_2O_2$ (200 μM) (E) for 1 h. Mitochondrial membrane potential was measured using TMRE reagent by flow cytometry. bars, ± SD of 3 independent experiments; ns, no significance, $p>0.05$ (Student's t-test).

**Figure S3**

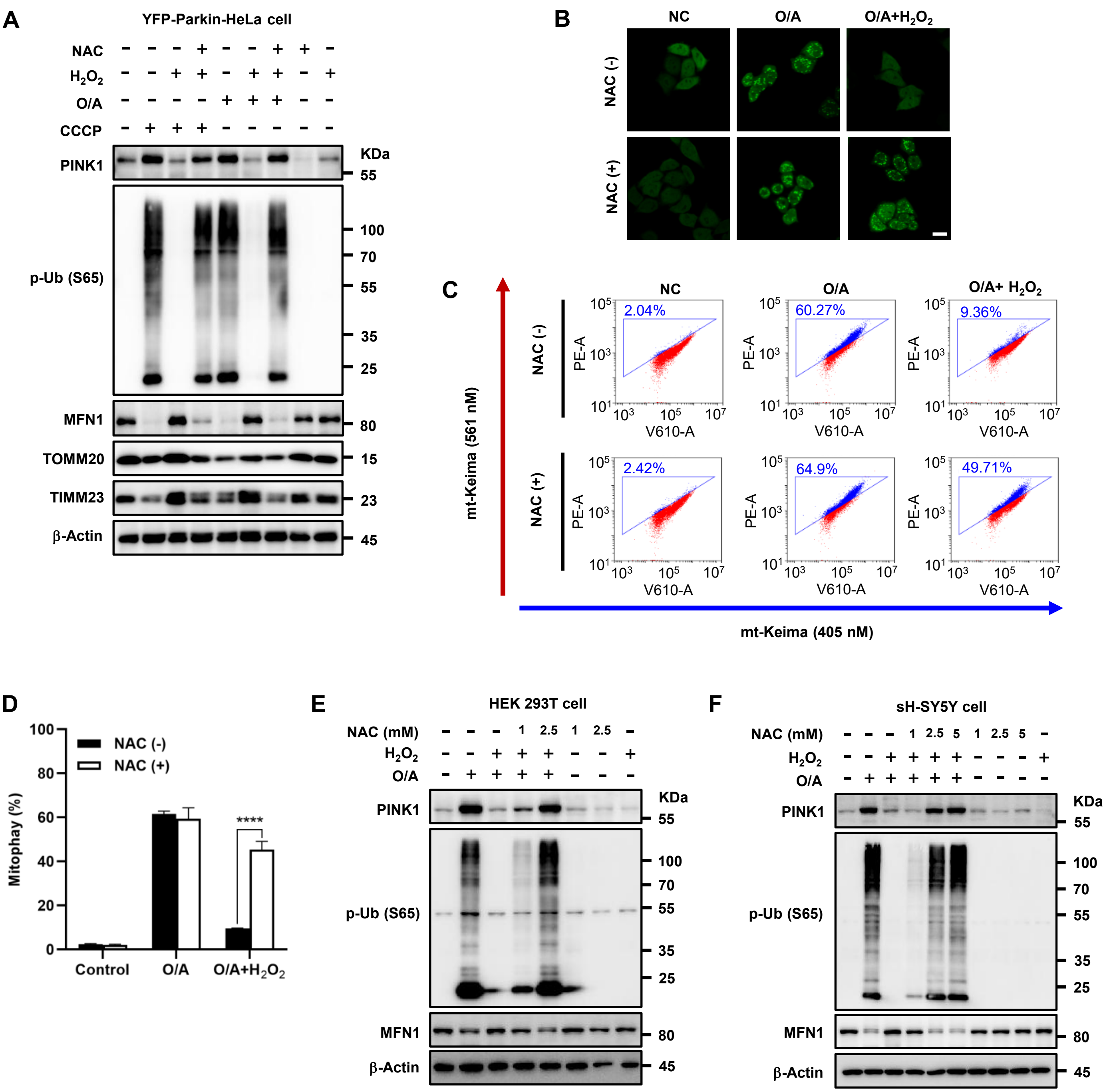

**Figure S3. Antioxidants abolish the inhibitory effect of ROS on mitophagy. Related to Figure 3.**

(A) YFP-Parkin-HeLa cells were pre-treated with NAC (5 mM) for 1.5 h and then treated with CCCP (20 μM) or O/A (1 μM/1 μM) combined with $H_2O_2$ (200 μM) for 6 h. Cells were harvested for immunoblotting analysis with the indicated antibodies. (B) YFP-Parkin-HeLa cells were treated with O/A (1 μM/1 μM) and $H_2O_2$ (200 μM) in the presence or absence of NAC (5 mM) for 1 h and imaged under confocal microscope. Scale bar: 20 μm. (C) mt-Keima-HeLa cells were pre-treated with NAC for 1.5 h and then treated with O/A (1 μM/1 μM) and $H_2O_2$ (200 μM) for 4 h, followed by flow cytometry analysis at 405 nm (neutral pH) and 561 nm (acidic pH) excitation. (D) Mitophagy level was quantified in mt-Keima-HeLa cells following treatment as described in (C). bars, ± SD of 3 independent experiments; ****, $P < 0.0001$ (Two-way ANOVA test). (E-F) HEK 293T cells (E) and SH-SY5Y cells (F) were pre-treated with different concentrations of NAC for 1.5 h and then treated with O/A (1 μM/1 μM) and $H_2O_2$ (200 μM) for 6 h. Cells were harvested for immunoblotting analysis with the indicated antibodies.

#### Figure S4

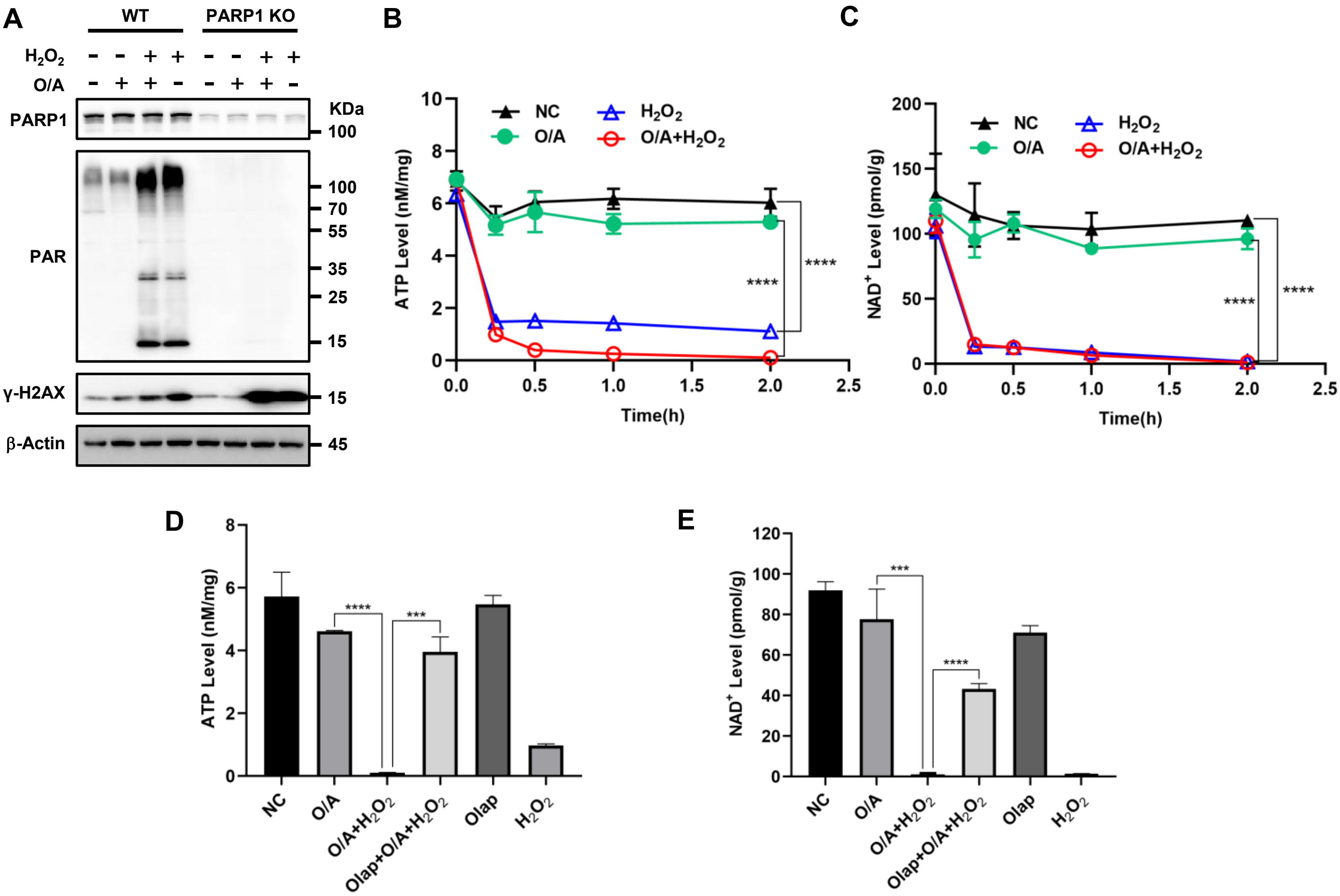

**Figure S4. ROS activate PARP1 and deplete intracellular ATP and NAD$^+$. Related to Figure 4.**

(A) Wild-type (WT) and PARP1 knockout (KO) YFP-Parkin-HeLa cells were treated with or without O/A (1 μM/1 μM), $H_2O_2$ (200 μM) or in combination for 30 min and harvested for immunoblotting. (B) Intracellular ATP levels response to $H_2O_2$ treatment over time. YFP-Parkin-HeLa cells treated with or without O/A (1 μM/1 μM), $H_2O_2$ (200 μM) or in combination for different times. ATP levels were measured via luciferase assay. bars, ± SD of 3 independent experiments; ****, $P < 0.0001$ (Two-way ANOVA test). (C) Intracellular NAD$^+$ levels response to $H_2O_2$ treatment over time. YFP-Parkin-HeLa cells were treated as in (B). NAD$^+$ levels were quantified via WST-8. Statistical analysis was performed as in (B). (D-E) Intracellular ATP and NAD$^+$ levels response to Olaparib. Cells were pre-treated by Olaparib for 1.5 h and then treated with O/A (1 μM/1 μM) and $H_2O_2$ (200 μM) for 2 h. bars, ± SD of 3 independent experiments; ns, no significance, ***, $P < 0.001$; ****, $P < 0.0001$ (Student's t-test).

### Figure S5

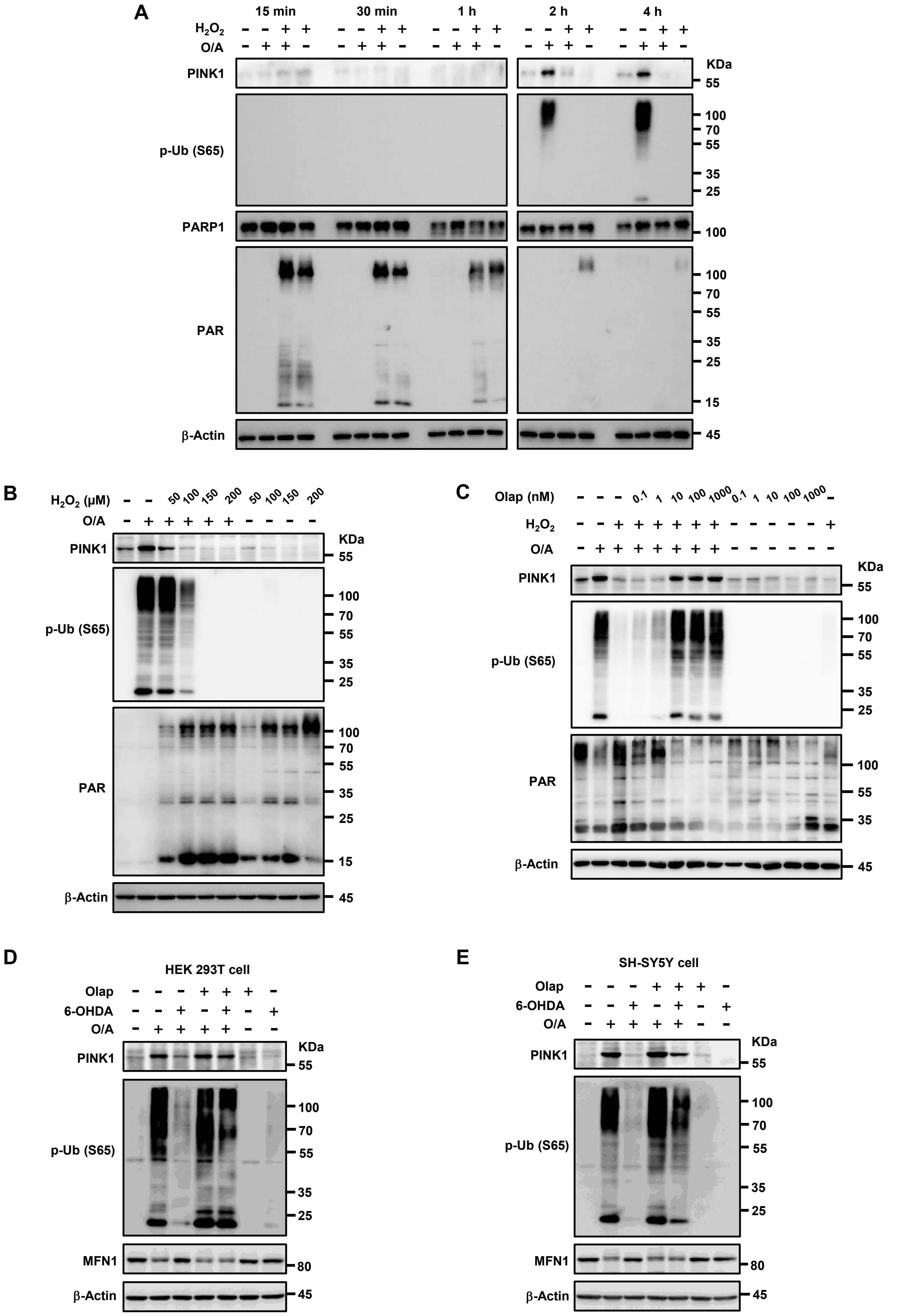
A
15 min
30 min
1 h
2 h
4 h
H2O2
O/A
PINK1
p-Ub (S65)
PARP1
PAR
β-Actin
KDa
55
100
70
55
35
25
100
100
70
55
35
25
15
45
B
H2O2 (µM)
50 100 150 200 50 100 150 200
O/A
PINK1
p-Ub (S65)
PAR
β-Actin
KDa
55
100
70
55
35
25
100
70
55
35
25
15
45
C
Olap (nM)
0.1 1 10 100 1000 0.1 1 10 100 1000
H2O2
O/A
PINK1
p-Ub (S65)
PAR
β-Actin
KDa
55
100
70
55
35
25
100
55
35
45
D
HEK 293T cell
Olap
6-OHDA
O/A
PINK1
p-Ub (S65)
MFN1
β-Actin
KDa
55
100
70
55
35
25
80
45
E
SH-SY5Y cell
Olap
6-OHDA
O/A
PINK1
p-Ub (S65)
MFN1
β-Actin
KDa
55
100
70
55
35
25
80
45

#### Figure S5

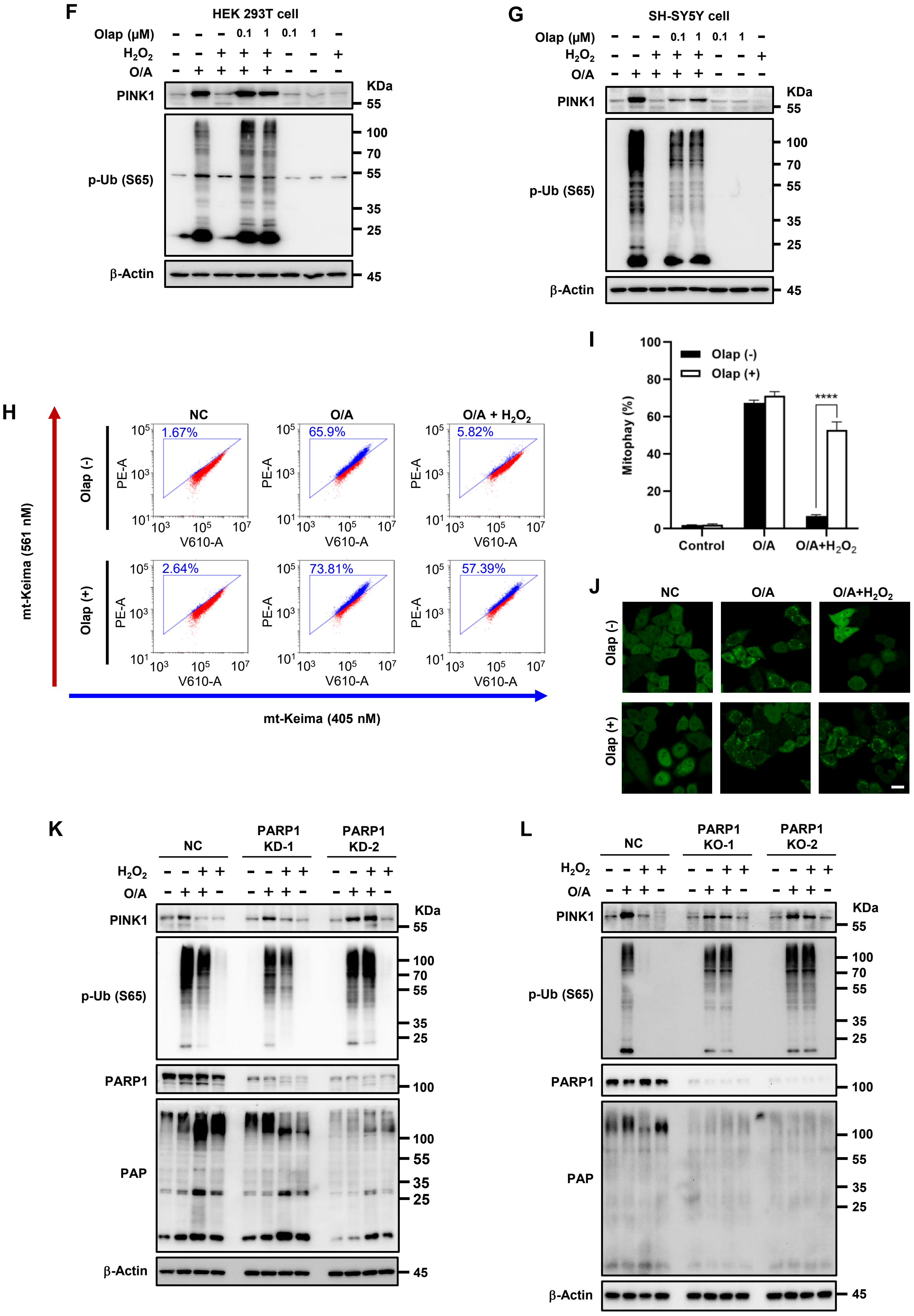

F
HEK 293T cell
Olap (μM)
0.1
1
0.1
1
H2O2
O/A
KDa
PINK1
55
p-Ub (S65)
100
70
55
35
25
β-Actin
45
G
SH-SY5Y cell
Olap (μM)
H2O2
O/A
PINK1
p-Ub (S65)
β-Actin
H
mt-Keima (561 nM)
NC
O/A
O/A + H2O2
Olap (-)
Olap (+)
PE-A
V610-A
1.67%
65.9%
5.82%
2.64%
73.81%
57.39%
mt-Keima (405 nM)
I
Mitophay (%)
Olap (-)
Olap (+)
****
Control
O/A
O/A+H2O2
J
NC
O/A
O/A+H2O2
Olap (-)
Olap (+)
K
NC
PARP1 KD-1
PARP1 KD-2
H2O2
O/A
PINK1
p-Ub (S65)
PARP1
PAP
β-Actin
L
NC
PARP1 KO-1
PARP1 KO-2
H2O2
O/A
PINK1
p-Ub (S65)
PARP1
PAP
β-Actin

**Figure S5. Suppression of PARP1 abolish the inhibitory effect of ROS on mitophagy. Related to Figure 5.**

(A) YFP-Parkin-HeLa cells were treated with or without O/A (1 μM/1 μM), $H_2O_2$ (200 μM) or in combination, and harvested at the indicated time points for immunoblotting. (B) YFP-Parkin-HeLa cells were treated with O/A (1 μM/1 μM) and different concentrations of $H_2O_2$. Cells were harvested for immunoblotting analysis with the indicated antibodies. (C) YFP-Parkin-HeLa cells were pre-treated with different concentrations of Olaparib for 1.5 h and then treated with O/A (1 μM/1 μM) and $H_2O_2$ (200 μM) for 6 h. Cells were harvested for immunoblotting analysis with the indicated antibodies. (D-E) HEK 293T cells (D) and SH-SY5Y cells (E) were pre-treated with different concentrations of Olaparib for 1.5 h and then treated with O/A (1 μM/1 μM) and 6-OHDA (100 μM) for 8 h. Cells were harvested for immunoblotting analysis with the indicated antibodies. (F-G) HEK 293T cells (F) and SH-SY5Y cells (G) were pre-treated with different concentrations of Olaparib for 1.5 h and then treated with O/A (1 μM/1 μM) and $H_2O_2$ (200 μM) for 6 h. Cells were harvested for immunoblotting analysis with the indicated antibodies. (H) mt-Keima-HeLa cells were treated with O/A (1 μM/1 μM) and $H_2O_2$ (200 μM) in the presence or absence of Olaparib (1 μM) for 4 h, followed by flow cytometry analysis at 405 nm (neutral pH) and 561 nm (acidic pH) excitation. (I) Mitophagy level was quantified in mt-Keima-HeLa cells following treatment as described in (H). bars, ± SD of 3 independent experiments; ****, $P < 0.0001$ (Two-way ANOVA test). (J) YFP-Parkin-HeLa cells were treated as in (H) for 1 h and imaged under confocal microscope. Scale bar: 20 μm. (K) WT and PARP1 KD YFP-Parkin-HeLa cells were treated with or without O/A (1 μM/1 μM), $H_2O_2$ (200 μM) or in combination for 6 h and harvested for immunoblotting. (L) WT and PARP1 KO YFP-Parkin-HeLa cells were treated as in (K) and harvested for immunoblotting.

**Figure S6**

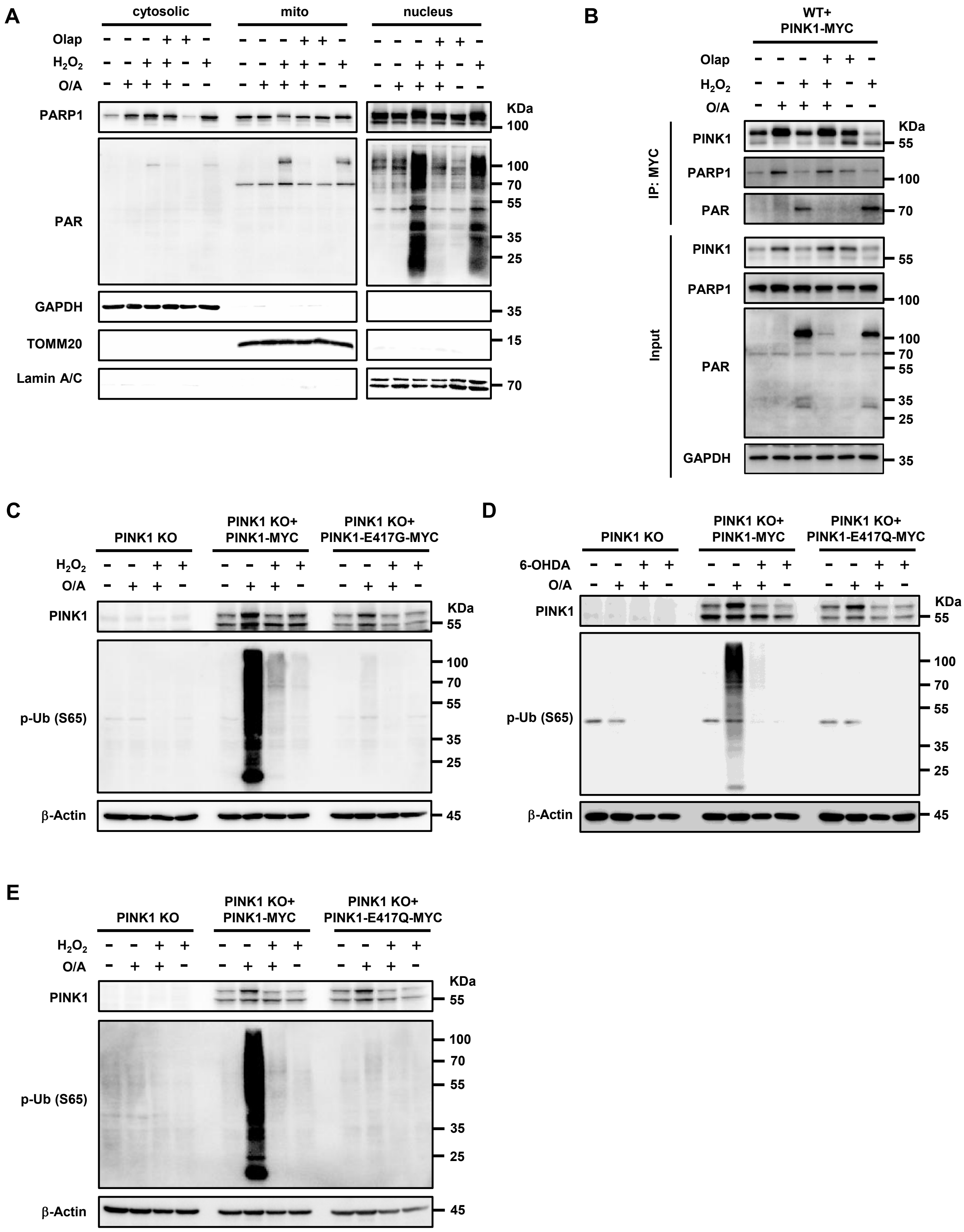

A
cytosolic
mito
nucleus
Olap
$H_2O_2$
O/A
PARP1
PAR
GAPDH
TOMM20
Lamin A/C
KDa
100
70
55
35
25
15
B
WT+
PINK1-MYC
IP: MYC
Input
PINK1
PARP1
PAR
GAPDH
C
PINK1 KO
PINK1 KO+
PINK1-MYC
PINK1-E417G-MYC
p-Ub (S65)
β-Actin
45
D
6-OHDA
PINK1-E417Q-MYC
E

**Figure S6. PARylation of PINK1 at damaged mitochondria impairs its function. Related to Figure 6.**

(A) YFP-Parkin-HeLa cells were pre-treated with Olaparib (1 μM) for 1.5 h and then treated with O/A (1 μM/1 μM) and $H_2O_2$ (200 μM) for 30 min. Subcellular fractionation was then performed to isolate the mitochondrial, nucleus, and cytosolic fractions. TOMM20, Lamin A/C and GAPDH were used as mitochondrial, nucleus and cytosolic markers, respectively. (B) YFP-Parkin-HeLa cells stably expressing PINK1-MYC pre-treated with Olaparib (1 μM) for 1.5 h and then treated with O/A (1 μM/1 μM) and $H_2O_2$ (200 μM) for 30 min. Cells were lysed with IP lysis buffer. And the cell lysates were subjected to MYC IP and analyzed by immunoblotting. (C) PINK1-MYC and PINK1-E417Q-MYC were reconstituted in PINK1 KO YFP-Parkin-HeLa cells. Cells were treated with or without O/A (1 μM/1 μM), 6-OHDA (100 μM) or in combination for 6 h and harvested for immunoblotting. (D-E) PINK1 KO YFP-Parkin HeLa cells were reconstituted with PINK1-MYC or the indicated E417 mutant (E417G in D, E417Q in E). Cells were treated with or without O/A (1 μM/1 μM), $H_2O_2$ (200 μM) or in combination for 6 h and harvested for immunoblotting.

**Table S1**

| Header line description of table 1 | |
|---|---|
| **Peptides** | Number of peptides which spectrum hit |
| **Unique peptides** | Number of unique peptides which spectrums hit |
| **Identified proteins** | Number of proteins detected by spectrum search analysis |

| Header line description of worksheet "Protein_ident" | |
|---|---|
| **Protein accession** | Protein accession number of database used for search |
| **Protein description** | Protein functional description |
| **Gene name** | Name of the gene associated to the protein sequence |
| **MW [kDa]** | Protein molecular weight, unit [kDa] |
| **Coverage [%]** | Percent of identified peptide sequence covering in protein sequence |
| **Peptides** | Number of peptides which spectrums hit |
| **Unique peptides** | Number of unique peptides which spectrums hit |

| Header line description of worksheet "Peptide_ident" | |
|---|---|
| **Sequence** | Identified peptide amino acid sequence |
| **Length** | Identified peptide amino acid length |
| **Protein accession** | Protein accession number of database used for search |
| **Protein description** | Protein functional description |
| **Gene name** | Name of the gene associated to the protein sequence |
| **MW [kDa]** | Protein molecular weight, unit [kDa] |
| **Charge** | The charge-state of the peptide |
| **Score** | Andromeda score for the best associated MS/MS spectrum |
| **Unique [yes/no]** | The peptide is or no unique to a single protein group |
| **PEP** | The maximal posterior error probability for peptides |
| **m/z** | The recalibrated mass-over-charge value of the peptide |

**Table 1. Spectrum database search analysis summary**

| Peptides | Unique peptides | Identified proteins |
|---|---|---|
| 35080 | 33244 | 4414 |

| Protein accession | Protein description | Gene name | MW [kDa] | Coverage [%] | Peptides | Unique peptides | Intensity PINK1 |
|---|---|---|---|---|---|---|---|
| A0A0B4J2F | Putative serine/threonine-protein kinase S | SIK1B | 84.931 | 1.4 | 1 | 1 | 72.787872 |
| A0AV96 | RNA-binding protein 47 OS=Homo sapie | RBM47 | 64.1 | 25.5 | 16 | 15 | 3003.6541 |
| A0FGR8 | Extended synaptotagmin-2 OS=Homo sap | ESYT2 | 102.358 | 17.9 | 14 | 14 | 262.78146 |
| A0JLT2 | Mediator of RNA polymerase II transcript | MED19 | 26.274 | 8.2 | 2 | 2 | 54.077255 |
| A0JNW5 | Bridge-like lipid transfer protein family m | BLTP3B | 164.199 | 0.6 | 1 | 1 | 448.94446 |
| A1A4S6 | Rho GTPase-activating protein 10 OS=Hc | ARHGAP1( | 89.375 | 1.3 | 1 | 1 | 61.345829 |
| A1X283 | SH3 and PX domain-containing protein 2I | SH3PXD2B | 101.58 | 3.1 | 2 | 2 | 180.01343 |
| A2RRP1 | NBAS subunit of NRZ tethering complex | NBAS | 268.572 | 0.3 | 1 | 1 | 643.17908 |
| A2RUB6 | Coiled-coil domain-containing protein 66 | CCDC66 | 109.412 | 3.8 | 3 | 3 | 37.851402 |
| A2VDJ0 | Transmembrane protein 131-like OS=Hor | TMEM131I | 179.34 | 2.1 | 3 | 3 | 46.304497 |
| A3KMH1 | von Willebrand factor A domain-containii | VWA8 | 214.825 | 1.6 | 3 | 3 | 65.731415 |
| A4D1E9 | GTP-binding protein 10 OS=Homo sapier | GTPBP10 | 42.933 | 43.4 | 13 | 13 | 350.74881 |
| A5A3E0 | POTE ankyrin domain family member F ( | POTEF | 121.445 | 0.7 | 1 | 0 | 1578.1166 |
| A5D8V6 | Vacuolar protein sorting-associated protei | VPS37C | 38.659 | 3.4 | 1 | 1 | 58.857521 |
| A5YKK6 | CCR4-NOT transcription complex subuni | CNOT1 | 266.939 | 27.5 | 60 | 60 | 789.10925 |
| A6NHR9 | Structural maintenance of chromosomes fl | SMCHD1 | 226.374 | 15.8 | 26 | 26 | 133.26465 |
| A7E2V4 | Zinc finger SWIM domain-containing pro | ZSWIM8 | 197.298 | 4.2 | 7 | 7 | 115.85718 |
| A8MT69 | Centromere protein X OS=Homo sapiens | CENPX | 8.96 | 21 | 2 | 2 | 296.77283 |
| A8MV65 | Transcription cofactor vestigial-like prote | VGLL3 | 36.009 | 14.1 | 4 | 4 | 284.54199 |
| A9QM74 | Importin subunit alpha-8 OS=Homo sapie | KPNA7 | 56.938 | 2.1 | 1 | 1 | 4740.5049 |
| B2RTY4 | Unconventional myosin-IXa OS=Homo sa | MYO9A | 292.706 | 4 | 9 | 9 | 115.24463 |
| C9JLW8 | Mapk-regulated corepressor-interacting pı | MCRIP1 | 10.921 | 32 | 3 | 3 | 493.46481 |
| L0R8F8 | Mitochondrial ribosome and complex I as | MIEF1 | 8.445 | 11.4 | 1 | 1 | 487.30414 |
| O00116 | Alkyldihydroxyacetonephosphate synthas | AGPS | 72.912 | 9.1 | 5 | 5 | 57.092434 |
| O00139 | Kinesin-like protein KIF2A OS=Homo sa | KIF2A | 79.955 | 31 | 21 | 19 | 1597.8885 |
| O00142 | Thymidine kinase 2, mitochondrial OS=H | TK2 | 31.005 | 8.3 | 2 | 2 | 45.412766 |
| O00148 | ATP-dependent RNA helicase DDX39A ( | DDX39A | 49.13 | 15 | 6 | 2 | 726.01782 |
| O00159 | Unconventional myosin-Ic OS=Homo sap | MYO1C | 121.682 | 20.8 | 21 | 21 | 542.64252 |
| O00165 | HCLS1-associated protein X-1 OS=Homc | HAX1 | 31.621 | 7.9 | 2 | 2 | 123.64433 |
| O00170 | AH receptor-interacting protein OS=Hom | AIP | 37.665 | 4.2 | 1 | 1 | 93.981895 |
| O00186 | Syntaxin-binding protein 3 OS=Homo sap | STXBP3 | 67.765 | 2.5 | 1 | 1 | 30.573503 |
| O00187 | Mannan-binding lectin serine protease 2 ( | MASP2 | 75.703 | 1.9 | 1 | 1 | 34.54718 |
| O00203 | AP-3 complex subunit beta-1 OS=Homo s | AP3B1 | 121.321 | 14.1 | 14 | 11 | 129.97688 |

| | | | | | | | |
|---|---|---|---|---|---|---|---|
| O00217 | NADH dehydrogenase [ubiquinone] iron- | NDUFS8 | 23.706 | 11 | 2 | 2 | 138.03517 |
| O00221 | NF-kappa-B inhibitor epsilon OS=Homo : | NFKBIE | 52.864 | 2.4 | 1 | 1 | 81.267899 |
| O00231 | 26S proteasome non-ATPase regulatory sı | PSMD11 | 47.464 | 8.3 | 3 | 3 | 35.001183 |
| O00233 | 26S proteasome non-ATPase regulatory sı | PSMD9 | 24.682 | 5.4 | 1 | 1 | 45.532452 |
| O00257 | E3 SUMO-protein ligase CBX4 OS=Hom | CBX4 | 61.368 | 23.9 | 12 | 12 | 481.25629 |
| O00267 | Transcription elongation factor SPT5 OS= | SUPT5H | 121 | 14.8 | 13 | 13 | 206.06709 |
| O00287 | Regulatory factor X-associated protein OS | RFXAP | 28.233 | 9.6 | 2 | 2 | 42.433002 |
| O00299 | Chloride intracellular channel protein 1 O | CLIC1 | 26.923 | 5 | 1 | 1 | 76.704803 |
| O00303 | Eukaryotic translation initiation factor 3 sı | EIF3F | 37.564 | 37 | 9 | 9 | 2518.2542 |
| O00308 | NEDD4-like E3 ubiquitin-protein ligase V | WWP2 | 98.913 | 9.3 | 7 | 5 | 163.5528 |
| O00311 | Cell division cycle 7-related protein kinas | CDC7 | 63.889 | 19.5 | 10 | 10 | 350.92352 |
| O00327 | Basic helix-loop-helix ARNT-like protein | BMAL1 | 68.762 | 8.1 | 4 | 4 | 88.018524 |
| O00391 | Sulfhydryl oxidase 1 OS=Homo sapiens C | QSOX1 | 82.578 | 4.3 | 3 | 3 | 57.703682 |
| O00410 | Importin-5 OS=Homo sapiens OX=9606 ( | IPO5 | 123.63 | 17 | 16 | 15 | 203.69476 |
| O00411 | DNA-directed RNA polymerase, mitochoı | POLRMT | 138.621 | 31.3 | 33 | 33 | 1166.0115 |
| O00418 | Eukaryotic elongation factor 2 kinase OS= | EEF2K | 82.145 | 4 | 2 | 2 | 48.086136 |
| O00422 | Histone deacetylase complex subunit SAP | SAP18 | 17.562 | 26.8 | 5 | 5 | 131.25879 |
| O00423 | Echinoderm microtubule-associated prote | EML1 | 89.861 | 1.2 | 1 | 1 | 54.390907 |
| O00425 | Insulin-like growth factor 2 mRNA-bindir | IGF2BP3 | 63.706 | 38.2 | 22 | 19 | 17522.447 |
| O00442 | RNA 3'-terminal phosphate cyclase OS=H | RTCA | 39.337 | 21.6 | 6 | 6 | 510.60327 |
| O00443 | Phosphatidylinositol 4-phosphate 3-kinase | PIK3C2A | 190.68 | 3 | 4 | 4 | 43.592625 |
| O00461 | Golgi integral membrane protein 4 OS=H | GOLIM4 | 81.881 | 3.9 | 2 | 2 | 29.468327 |
| O00463 | TNF receptor-associated factor 5 OS=Hoı | TRAF5 | 64.406 | 2 | 1 | 1 | 31.425241 |
| O00468 | Agrin OS=Homo sapiens OX=9606 GN=. | AGRN | 217.32 | 1.2 | 2 | 2 | 60.406685 |
| O00483 | Cytochrome c oxidase subunit NDUFA4 ( | NDUFA4 | 9.37 | 37 | 3 | 3 | 966.70715 |
| O00487 | 26S proteasome non-ATPase regulatory sı | PSMD14 | 34.578 | 4.2 | 1 | 1 | 184.24059 |
| O00488 | Zinc finger protein 593 OS=Homo sapien: | ZNF593 | 15.2 | 17.2 | 2 | 2 | 790.42401 |
| O00505 | Importin subunit alpha-4 OS=Homo sapie | KPNA3 | 57.811 | 17.5 | 9 | 5 | 580.82813 |
| O00522 | Krev interaction trapped protein 1 OS=Hc | KRIT1 | 84.349 | 3.3 | 2 | 2 | 42.405281 |
| O00541 | Pescadillo homolog OS=Homo sapiens O. | PES1 | 68.003 | 26.2 | 19 | 19 | 1981.8521 |
| O00560 | Syntenin-1 OS=Homo sapiens OX=9606 ( | SDCBP | 32.445 | 20.5 | 4 | 4 | 788.46472 |
| O00566 | U3 small nucleolar ribonucleoprotein prot | MPHOSPH | 78.864 | 16.6 | 9 | 9 | 167.91574 |
| O00567 | Nucleolar protein 56 OS=Homo sapiens C | NOP56 | 66.05 | 24.7 | 14 | 14 | 705.43976 |
| O00571 | ATP-dependent RNA helicase DDX3X O | DDX3X | 73.244 | 48.9 | 34 | 8 | 34354.242 |
| O00622 | CCN family member 1 OS=Homo sapiens | CCN1 | 42.027 | 27.3 | 10 | 10 | 300.72433 |

| | | | | | | | |
|---|---|---|---|---|---|---|---|
| O00625 | Pirin OS=Homo sapiens OX=9606 GN=P | PIR | 32.114 | 9.7 | 3 | 3 | 95.964058 |
| O00629 | Importin subunit alpha-3 OS=Homo sapie | KPNA4 | 57.887 | 6.1 | 3 | 3 | 262.11615 |
| O00712 | Nuclear factor 1 B-type OS=Homo sapien | NFIB | 47.442 | 32.1 | 12 | 12 | 1065.3621 |
| O00716 | Transcription factor E2F3 OS=Homo sapi | E2F3 | 49.162 | 14.8 | 6 | 4 | 126.54906 |
| O00743 | Serine/threonine-protein phosphatase 6 ca | PPP6C | 35.145 | 20.3 | 6 | 6 | 394.9483 |
| O00750 | Phosphatidylinositol 4-phosphate 3-kinase | PIK3C2B | 184.768 | 5.5 | 8 | 8 | 206.0387 |
| O00767 | Stearoyl-CoA desaturase OS=Homo sapie | SCD | 41.523 | 7 | 2 | 2 | 237.3904 |
| O14519 | Cyclin-dependent kinase 2-associated pro | CDK2AP1 | 12.365 | 15.7 | 2 | 2 | 83.861992 |
| O14556 | Glyceraldehyde-3-phosphate dehydrogena | GAPDHS | 44.502 | 2.7 | 1 | 1 | 395.08673 |
| O14561 | Acyl carrier protein, mitochondrial OS=H | NDUFAB1 | 17.418 | 14.7 | 2 | 2 | 257.28241 |
| O14578 | Citron Rho-interacting kinase OS=Homo : | CIT | 231.431 | 10.8 | 20 | 20 | 167.21295 |
| O14579 | Coatomer subunit epsilon OS=Homo sapi | COPE | 34.483 | 8.1 | 2 | 2 | 163.99704 |
| O14617 | AP-3 complex subunit delta-1 OS=Homo | AP3D1 | 130.158 | 13.6 | 14 | 14 | 238.13699 |
| O14639 | Actin-binding LIM protein 1 OS=Homo s | ABLIM1 | 87.688 | 8.5 | 6 | 6 | 140.46024 |
| O14640 | Segment polarity protein dishevelled hom | DVL1 | 75.187 | 10.5 | 6 | 1 | 226.89986 |
| O14641 | Segment polarity protein dishevelled hom | DVL2 | 78.948 | 3.7 | 2 | 2 | 110.0701 |
| O14646 | Chromodomain-helicase-DNA-binding pr | CHD1 | 196.688 | 16.8 | 29 | 23 | 605.99036 |
| O14647 | Chromodomain-helicase-DNA-binding pr | CHD2 | 211.344 | 5.7 | 10 | 10 | 165.32986 |
| O14681 | Etoposide-induced protein 2.4 homolog O | EI24 | 38.965 | 5.6 | 2 | 2 | 91.26899 |
| O14713 | Integrin beta-1-binding protein 1 OS=Hor | ITGB1BP1 | 21.783 | 6.5 | 1 | 1 | 90.875504 |
| O14730 | Serine/threonine-protein kinase RIO3 OS= | RIOK3 | 59.094 | 6 | 3 | 3 | 163.28033 |
| O14733 | Dual specificity mitogen-activated protein | MAP2K7 | 47.485 | 20.8 | 8 | 8 | 206.29387 |
| O14734 | Acyl-coenzyme A thioesterase 8 OS=Hon | ACOT8 | 35.915 | 2.8 | 1 | 1 | 110.47009 |
| O14737 | Programmed cell death protein 5 OS=Hor | PDCD5 | 14.286 | 8.8 | 1 | 1 | 10.914261 |
| O14744 | Protein arginine N-methyltransferase 5 OS | PRMT5 | 72.684 | 24.2 | 13 | 13 | 826.03394 |
| O14757 | Serine/threonine-protein kinase Chk1 OS= | CHEK1 | 54.434 | 15.1 | 6 | 6 | 119.66503 |
| O14776 | Transcription elongation regulator 1 OS=I | TCERG1 | 123.902 | 9.7 | 10 | 10 | 178.3421 |
| O14777 | Kinetochore protein NDC80 homolog OS | NDC80 | 73.913 | 5.1 | 3 | 3 | 158.65961 |
| O14795 | Protein unc-13 homolog B OS=Homo sap | UNC13B | 180.679 | 8.1 | 11 | 7 | 115.48705 |
| O14802 | DNA-directed RNA polymerase III subun | POLR3A | 155.642 | 16.8 | 21 | 21 | 229.07181 |
| O14818 | Proteasome subunit alpha type-7 OS=Hon | PSMA7 | 27.887 | 31 | 5 | 2 | 299.26218 |
| O14828 | Secretory carrier-associated membrane pr | SCAMP3 | 38.287 | 13 | 3 | 3 | 185.73904 |
| O14874 | [3-methyl-2-oxobutanoate dehydrogenase | BCKDK | 46.361 | 5.3 | 2 | 2 | 70.439285 |
| O14879 | Interferon-induced protein with tetratricop | IFIT3 | 55.985 | 17.8 | 6 | 6 | 117.36469 |
| O14880 | Glutathione S-transferase 3, mitochondria | MGST3 | 16.517 | 17.8 | 2 | 2 | 1058.3557 |

| | | | | | | | |
|---|---|---|---|---|---|---|---|
| O14893 | Gem-associated protein 2 OS=Homo sapi | GEMIN2 | 31.586 | 13.9 | 4 | 4 | 504.10358 |
| O14901 | Krueppel-like factor 11 OS=Homo sapien | KLF11 | 55.14 | 7.2 | 3 | 3 | 72.152969 |
| O14908 | PDZ domain-containing protein GIPC1 O | GIPC1 | 36.05 | 11.7 | 3 | 2 | 67.551575 |
| O14925 | Mitochondrial import inner membrane tra | TIMM23 | 21.944 | 5.3 | 1 | 1 | 87.812599 |
| O14929 | Histone acetyltransferase type B catalytic | HAT1 | 49.541 | 9.5 | 4 | 4 | 164.12437 |
| O14936 | Peripheral plasma membrane protein CAS | CASK | 105.123 | 1 | 1 | 1 | 130.49898 |
| O14949 | Cytochrome b-c1 complex subunit 8 OS= | UQCRQ | 9.907 | 22 | 2 | 2 | 411.22488 |
| O14950 | Myosin regulatory light chain 12B OS=H | MYL12B | 19.78 | 36.6 | 6 | 2 | 6202.6323 |
| O14964 | Hepatocyte growth factor-regulated tyrosi | HGS | 86.193 | 1.5 | 1 | 1 | 31.619755 |
| O14965 | Aurora kinase A OS=Homo sapiens OX= | AURKA | 45.824 | 41.2 | 15 | 14 | 3173.666 |
| O14966 | Ras-related protein Rab-7L1 OS=Homo s | RAB29 | 23.156 | 21.2 | 4 | 3 | 123.59475 |
| O14974 | Protein phosphatase 1 regulatory subunit | PPP1R12A | 115.281 | 17.1 | 15 | 14 | 357.19305 |
| O14975 | Long-chain fatty acid transport protein 2 ( | SLC27A2 | 70.313 | 5.3 | 3 | 3 | 67.069031 |
| O14979 | Heterogeneous nuclear ribonucleoprotein | HNRNPDL | 46.438 | 22.6 | 9 | 7 | 21918.213 |
| O14980 | Exportin-1 OS=Homo sapiens OX=9606 ( | XPO1 | 123.386 | 6.4 | 6 | 6 | 104.67975 |
| O15013 | Rho guanine nucleotide exchange factor 1 | ARHGEF1C | 151.612 | 3.1 | 2 | 2 | 17.85903 |
| O15014 | Zinc finger protein 609 OS=Homo sapien | ZNF609 | 151.191 | 14.8 | 18 | 17 | 647.19702 |
| O15021 | Microtubule-associated serine/threonine-p | MAST4 | 284.098 | 1 | 2 | 1 | 101.70504 |
| O15027 | Protein transport protein Sec16A OS=Hor | SEC16A | 251.894 | 25.8 | 48 | 48 | 2929.3027 |
| O15042 | U2 snRNP-associated SURP motif-contai | U2SURP | 118.292 | 24.4 | 25 | 25 | 5463.8716 |
| O15049 | NEDD4-binding protein 3 OS=Homo sap | N4BP3 | 60.47 | 4.2 | 2 | 2 | 56.806171 |
| O15054 | Lysine-specific demethylase 6B OS=Hom | KDM6B | 176.633 | 3.6 | 5 | 5 | 179.85426 |
| O15061 | Synemin OS=Homo sapiens OX=9606 GI | SYNM | 172.868 | 2 | 3 | 3 | 61.295868 |
| O15067 | Phosphoribosylformylglycinamidine synth | PFAS | 144.735 | 3.4 | 4 | 4 | 50.774788 |
| O15078 | Centrosomal protein of 290 kDa OS=Hon | CEP290 | 290.386 | 0.6 | 2 | 2 | 108.67847 |
| O15084 | Serine/threonine-protein phosphatase 6 re | ANKRD28 | 112.966 | 25.3 | 18 | 16 | 603.93823 |
| O15116 | U6 snRNA-associated Sm-like protein LS | LSM1 | 15.18 | 10.5 | 1 | 1 | 679.38312 |
| O15119 | T-box transcription factor TBX3 OS=Hor | TBX3 | 79.389 | 4.8 | 3 | 3 | 54.501186 |
| O15131 | Importin subunit alpha-6 OS=Homo sapie | KPNA5 | 60.667 | 2.8 | 1 | 1 | 103.18683 |
| O15143 | Actin-related protein 2/3 complex subunit | ARPC1B | 40.95 | 6.2 | 2 | 2 | 114.36604 |
| O15144 | Actin-related protein 2/3 complex subunit | ARPC2 | 34.334 | 6 | 2 | 2 | 112.33884 |
| O15145 | Actin-related protein 2/3 complex subunit | ARPC3 | 20.547 | 3.9 | 1 | 1 | 103.04248 |
| O15156 | Zinc finger and BTB domain-containing p | ZBTB7B | 58.027 | 6.9 | 2 | 2 | 45.946289 |
| O15160 | DNA-directed RNA polymerases I and III | POLR1C | 39.25 | 23.7 | 7 | 7 | 380.10147 |
| O15169 | Axin-1 OS=Homo sapiens OX=9606 GN= | AXIN1 | 95.635 | 10.6 | 9 | 9 | 155.32581 |

| | | | | | | | |
|---|---|---|---|---|---|---|---|
| O15182 | Centrin-3 OS=Homo sapiens OX=9606 G | CETN3 | 19.55 | 7.2 | 1 | 1 | 78.202286 |
| O15212 | Prefoldin subunit 6 OS=Homo sapiens OX | PFDN6 | 14.583 | 29.5 | 3 | 3 | 80.122993 |
| O15213 | WD repeat-containing protein 46 OS=Hoi | WDR46 | 68.072 | 15.1 | 8 | 8 | 190.62411 |
| O15226 | NF-kappa-B-repressing factor OS=Homo | NKRF | 77.673 | 31.2 | 22 | 22 | 838.97485 |
| O15228 | Dihydroxyacetone phosphate acyltransfera | GNPAT | 77.188 | 5.3 | 3 | 3 | 61.361561 |
| O15230 | Laminin subunit alpha-5 OS=Homo sapiei | LAMA5 | 399.738 | 0.4 | 1 | 1 | 40.512444 |
| O15234 | Protein CASC3 OS=Homo sapiens OX=9 | CASC3 | 76.278 | 29.2 | 20 | 20 | 7343.6597 |
| O15258 | Protein RER1 OS=Homo sapiens OX=96( | RER1 | 22.958 | 15.3 | 3 | 3 | 173.85899 |
| O15265 | Ataxin-7 OS=Homo sapiens OX=9606 GI | ATXN7 | 95.452 | 1 | 1 | 1 | 53.720921 |
| O15269 | Serine palmitoyltransferase 1 OS=Homo s | SPTLC1 | 52.744 | 12.9 | 6 | 6 | 240.78874 |
| O15270 | Serine palmitoyltransferase 2 OS=Homo s | SPTLC2 | 62.925 | 8 | 4 | 4 | 91.228508 |
| O15294 | UDP-N-acetylglucosamine--peptide N-ac | OGT | 116.925 | 16.1 | 17 | 17 | 686.80798 |
| O15304 | Apoptosis regulatory protein Siva OS=Ho | SIVA1 | 18.695 | 17.7 | 3 | 3 | 210.38765 |
| O15318 | DNA-directed RNA polymerase III subun | POLR3G | 25.915 | 10.8 | 2 | 2 | 94.155411 |
| O15344 | E3 ubiquitin-protein ligase Midline-1 OS= | MID1 | 75.251 | 10 | 6 | 6 | 74.589355 |
| O15355 | Protein phosphatase 1G OS=Homo sapien | PPM1G | 59.272 | 12.3 | 5 | 5 | 112.02061 |
| O15357 | Phosphatidylinositol 3,4,5-trisphosphate 5 | INPPL1 | 138.599 | 3 | 3 | 3 | 67.16581 |
| O15360 | Fanconi anemia group A protein OS=Hon | FANCA | 162.776 | 2 | 3 | 3 | 36.254333 |
| O15371 | Eukaryotic translation initiation factor 3 s | EIF3D | 63.973 | 24.6 | 12 | 12 | 2693.0776 |
| O15372 | Eukaryotic translation initiation factor 3 s | EIF3H | 39.931 | 43.8 | 14 | 14 | 1641.0835 |
| O15379 | Histone deacetylase 3 OS=Homo sapiens | HDAC3 | 48.848 | 3.7 | 2 | 2 | 59.936623 |
| O15381 | Nuclear valosin-containing protein-like O | NVL | 95.051 | 13.3 | 9 | 9 | 140.15344 |
| O15391 | Transcription factor YY2 OS=Homo sapi | YY2 | 41.348 | 4.3 | 1 | 1 | 258.4491 |
| O15392 | Baculoviral IAP repeat-containing protein | BIRC5 | 16.389 | 16.9 | 3 | 3 | 64.738991 |
| O15397 | Importin-8 OS=Homo sapiens OX=9606 ( | IPO8 | 119.939 | 4.5 | 4 | 4 | 46.099731 |
| O15427 | Monocarboxylate transporter 4 OS=Homo | SLC16A3 | 49.47 | 17.6 | 8 | 8 | 551.04517 |
| O15440 | ATP-binding cassette sub-family C memb | ABCC5 | 160.661 | 2.6 | 4 | 3 | 82.205269 |
| O15446 | DNA-directed RNA polymerase I subunit | POLR1G | 54.986 | 10 | 4 | 4 | 372.42279 |
| O15460 | Prolyl 4-hydroxylase subunit alpha-2 OS= | P4HA2 | 60.902 | 6.7 | 3 | 3 | 47.85107 |
| O15479 | Melanoma-associated antigen B2 OS=Hoi | MAGEB2 | 35.278 | 32 | 11 | 11 | 2080.1899 |
| O15504 | Nucleoporin NUP42 OS=Homo sapiens C | NUP42 | 44.872 | 11.3 | 4 | 4 | 275.86707 |
| O15511 | Actin-related protein 2/3 complex subunit | ARPC5 | 16.321 | 16.6 | 2 | 2 | 83.107735 |
| O15516 | Circadian locomoter output cycles protein | CLOCK | 95.304 | 4.4 | 3 | 3 | 38.639114 |
| O15523 | ATP-dependent RNA helicase DDX3Y O | DDX3Y | 73.154 | 1.4 | 1 | 1 | 52.053806 |
| O15533 | Tapasin OS=Homo sapiens OX=9606 GN | TAPBP | 47.571 | 6.9 | 3 | 3 | 76.606346 |

| | | | | | | | |
|---|---|---|---|---|---|---|---|
| O15541 | E3 ubiquitin-protein ligase RNF113A OS= | RNF113A | 38.788 | 29.7 | 9 | 8 | 301.30429 |
| O43143 | ATP-dependent RNA helicase DHX15 OS | DHX15 | 90.933 | 31.1 | 27 | 27 | 5575.4014 |
| O43148 | mRNA cap guanine-N7 methyltransferase | RNMT | 54.845 | 22.1 | 12 | 12 | 1121.4702 |
| O43159 | Ribosomal RNA-processing protein 8 OS= | RRP8 | 50.715 | 5 | 2 | 2 | 118.79513 |
| O43164 | E3 ubiquitin-protein ligase Praja-2 OS=H | PJA2 | 78.215 | 16.1 | 8 | 8 | 132.09665 |
| O43167 | Zinc finger and BTB domain-containing p | ZBTB24 | 78.283 | 9 | 6 | 6 | 133.72115 |
| O43172 | U4/U6 small nuclear ribonucleoprotein Pr | PRPF4 | 58.45 | 40 | 18 | 18 | 905.25159 |
| O43175 | D-3-phosphoglycerate dehydrogenase OS= | PHGDH | 56.651 | 20.3 | 8 | 8 | 234.24869 |
| O43242 | 26S proteasome non-ATPase regulatory su | PSMD3 | 60.978 | 27.7 | 12 | 12 | 350.41208 |
| O43248 | Homeobox protein Hox-C11 OS=Homo s | HOXC11 | 33.748 | 21.7 | 6 | 4 | 286.39645 |
| O43251 | RNA binding protein fox-1 homolog 2 OS | RBFOX2 | 41.374 | 24.9 | 9 | 1 | 1715.6344 |
| O43290 | U4/U6.U5 tri-snRNP-associated protein 1 | SART1 | 90.255 | 26.1 | 18 | 18 | 435.4859 |
| O43293 | Death-associated protein kinase 3 OS=Ho | DAPK3 | 52.536 | 10.6 | 4 | 4 | 117.393 |
| O43296 | Zinc finger protein 264 OS=Homo sapiens | ZNF264 | 70.587 | 3.8 | 2 | 2 | 36.574875 |
| O43298 | Zinc finger and BTB domain-containing p | ZBTB43 | 52.631 | 15.2 | 6 | 6 | 548.05542 |
| O43310 | CBP80/20-dependent translation initiation | CTIF | 67.587 | 8.5 | 5 | 5 | 131.87411 |
| O43314 | Inositol hexakisphosphate and diphosphoi | PPIP5K2 | 140.407 | 1.9 | 2 | 2 | 54.311771 |
| O43324 | Eukaryotic translation elongation factor 1 | EEF1E1 | 19.811 | 23 | 4 | 4 | 274.10611 |
| O43353 | Receptor-interacting serine/threonine-prot | RIPK2 | 61.195 | 19.3 | 9 | 9 | 329.86313 |
| O43379 | WD repeat-containing protein 62 OS=Hom | WDR62 | 165.954 | 3.9 | 5 | 5 | 61.055668 |
| O43390 | Heterogeneous nuclear ribonucleoprotein | HNRNPR | 70.944 | 38.9 | 24 | 17 | 9561.2021 |
| O43395 | U4/U6 small nuclear ribonucleoprotein Pr | PRPF3 | 77.529 | 25 | 18 | 18 | 882.05511 |
| O43422 | 52 kDa repressor of the inhibitor of the pr | THAP12 | 87.704 | 40.2 | 26 | 26 | 1257.5909 |
| O43426 | Synaptojanin-1 OS=Homo sapiens OX=96 | SYNJ1 | 173.104 | 0.6 | 1 | 1 | 56.859047 |
| O43432 | Eukaryotic translation initiation factor 4 g | EIF4G3 | 176.653 | 30.7 | 53 | 49 | 2619.5684 |
| O43435 | T-box transcription factor TBX1 OS=Hom | TBX1 | 43.133 | 5 | 2 | 2 | 48.782436 |
| O43439 | Protein CBFA2T2 OS=Homo sapiens OX | CBFA2T2 | 67.133 | 1.8 | 1 | 1 | 113.06348 |
| O43447 | Peptidyl-prolyl cis-trans isomerase H OS= | PPIH | 19.209 | 28.2 | 5 | 5 | 532.80804 |
| O43464 | Serine protease HTRA2, mitochondrial O | HTRA2 | 48.841 | 7.2 | 3 | 3 | 77.493149 |
| O43474 | Krueppel-like factor 4 OS=Homo sapiens | KLF4 | 54.671 | 5.3 | 2 | 2 | 476.8226 |
| O43491 | Band 4.1-like protein 2 OS=Homo sapiens | EPB41L2 | 112.589 | 1.9 | 2 | 2 | 31.213449 |
| O43524 | Forkhead box protein O3 OS=Homo sapie | FOXO3 | 71.277 | 5.6 | 3 | 2 | 143.80872 |
| O43572 | A-kinase anchor protein 10, mitochondria | AKAP10 | 73.819 | 1.4 | 1 | 1 | 156.70978 |
| O43583 | Density-regulated protein OS=Homo sapi | DENR | 22.093 | 20.7 | 4 | 4 | 140.78357 |
| O43592 | Exportin-T OS=Homo sapiens OX=9606 | XPOT | 109.965 | 4.8 | 4 | 4 | 72.173996 |

| | | | | | | | |
|---|---|---|---|---|---|---|---|
| O43615 | Mitochondrial import inner membrane tra | TIMM44 | 51.356 | 2.2 | 1 | 1 | 32.213516 |
| O43633 | Charged multivesicular body protein 2a O | CHMP2A | 25.104 | 19.8 | 5 | 5 | 211.40138 |
| O43660 | Pleiotropic regulator 1 OS=Homo sapiens | PLRG1 | 57.194 | 26.7 | 12 | 12 | 214.75516 |
| O43663 | Protein regulator of cytokinesis 1 OS=Hoi | PRC1 | 71.607 | 39.7 | 25 | 25 | 2799.6812 |
| O43670 | BUB3-interacting and GLEBS motif-cont | ZNF207 | 50.751 | 13.4 | 6 | 6 | 2072.2971 |
| O43674 | NADH dehydrogenase [ubiquinone] 1 bet | NDUFB5 | 21.751 | 12.7 | 2 | 2 | 188.16373 |
| O43678 | NADH dehydrogenase [ubiquinone] 1 alp | NDUFA2 | 10.922 | 10.1 | 1 | 1 | 28.364439 |
| O43681 | ATPase GET3 OS=Homo sapiens OX=96 | GET3 | 38.793 | 2.9 | 1 | 1 | 58.666294 |
| O43683 | Mitotic checkpoint serine/threonine-prote | BUB1 | 122.376 | 1.5 | 1 | 1 | 22.328293 |
| O43684 | Mitotic checkpoint protein BUB3 OS=Ho | BUB3 | 37.155 | 50.3 | 12 | 12 | 3148.7478 |
| O43707 | Alpha-actinin-4 OS=Homo sapiens OX=9 | ACTN4 | 104.855 | 8.2 | 6 | 6 | 153.96423 |
| O43709 | Probable 18S rRNA (guanine-N(7))-meth | BUD23 | 31.881 | 19.9 | 4 | 4 | 327.6951 |
| O43711 | T-cell leukemia homeobox protein 3 OS= | TLX3 | 31.868 | 2.7 | 1 | 1 | 143.20013 |
| O43719 | HIV Tat-specific factor 1 OS=Homo sapi | HTATSF1 | 85.853 | 2.8 | 2 | 2 | 79.613258 |
| O43734 | E3 ubiquitin ligase TRAF3IP2 OS=Homo | TRAF3IP2 | 64.667 | 4.7 | 4 | 4 | 140.59052 |
| O43739 | Cytohesin-3 OS=Homo sapiens OX=9606 | CYTH3 | 46.349 | 5 | 2 | 2 | 37.394775 |
| O43747 | AP-1 complex subunit gamma-1 OS=Hon | AP1G1 | 91.352 | 4.9 | 4 | 4 | 42.176903 |
| O43776 | Asparagine--tRNA ligase, cytoplasmic OS | NARS1 | 62.943 | 15.5 | 8 | 8 | 329.26175 |
| O43781 | Dual specificity tyrosine-phosphorylation- | DYRK3 | 65.714 | 4.1 | 2 | 2 | 39.570259 |
| O43791 | Speckle-type POZ protein OS=Homo sapi | SPOP | 42.133 | 28.9 | 8 | 7 | 301.52734 |
| O43795 | Unconventional myosin-Ib OS=Homo sap | MYO1B | 131.986 | 16.9 | 18 | 15 | 350.63068 |
| O43808 | Peroxisomal membrane protein PMP34 O | SLC25A17 | 34.567 | 20.2 | 6 | 6 | 200.23184 |
| O43809 | Cleavage and polyadenylation specificity | NUDT21 | 26.228 | 30.4 | 8 | 8 | 2874.6443 |
| O43813 | Glutathione S-transferase LANCL1 OS=H | LANCL1 | 45.284 | 9.8 | 4 | 4 | 310.29504 |
| O43818 | U3 small nucleolar RNA-interacting prote | RRP9 | 51.841 | 13.3 | 7 | 7 | 288.70029 |
| O43819 | Protein SCO2 homolog, mitochondrial OS | SCO2 | 29.81 | 7.1 | 2 | 2 | 98.716774 |
| O43823 | A-kinase anchor protein 8 OS=Homo sapi | AKAP8 | 76.109 | 24.3 | 15 | 15 | 4396.4912 |
| O43837 | Isocitrate dehydrogenase [NAD] subunit b | IDH3B | 42.184 | 6 | 2 | 2 | 45.487366 |
| O43847 | Nardilysin OS=Homo sapiens OX=9606 ( | NRDC | 131.701 | 5 | 4 | 4 | 49.890175 |
| O43852 | Calumenin OS=Homo sapiens OX=9606 ( | CALU | 37.107 | 46.3 | 14 | 14 | 8580.7705 |
| O43865 | S-adenosylhomocysteine hydrolase-like pi | AHCYL1 | 58.952 | 11.1 | 5 | 2 | 116.96698 |
| O43896 | Kinesin-like protein KIF1C OS=Homo sa | KIF1C | 122.947 | 34.3 | 34 | 34 | 1552.0354 |
| O43900 | Prickle planar cell polarity protein 3 OS=I | PRICKLE3 | 68.61 | 8.6 | 5 | 5 | 121.00668 |
| O43909 | Exostosin-like 3 OS=Homo sapiens OX=9 | EXTL3 | 104.75 | 2.9 | 3 | 3 | 56.407318 |
| O43913 | Origin recognition complex subunit 5 OS= | ORC5 | 50.284 | 2.1 | 1 | 1 | 87.115799 |

| | | | | | | | |
|---|---|---|---|---|---|---|---|
| O43920 | NADH dehydrogenase [ubiquinone] iron- | NDUFS5 | 12.518 | 18.9 | 2 | 2 | 43.0177 |
| O43929 | Origin recognition complex subunit 4 OS= | ORC4 | 50.378 | 9.2 | 3 | 3 | 74.981133 |
| O43933 | Peroxisomal ATPase PEX1 OS=Homo sa | PEX1 | 142.868 | 19.3 | 22 | 22 | 626.0434 |
| O60216 | Double-strand-break repair protein rad21 | RAD21 | 71.69 | 17.3 | 9 | 8 | 120.82719 |
| O60231 | Pre-mRNA-splicing factor ATP-depender | DHX16 | 119.264 | 14.8 | 15 | 14 | 1120.6477 |
| O60232 | Protein ZNRD2 OS=Homo sapiens OX=9 | ZNRD2 | 21.475 | 4 | 1 | 1 | 104.82904 |
| O60244 | Mediator of RNA polymerase II transcript | MED14 | 160.607 | 10.4 | 14 | 14 | 111.37226 |
| O60260 | E3 ubiquitin-protein ligase parkin OS=Ho | PRKN | 51.641 | 35.3 | 13 | 13 | 941.85168 |
| O60264 | SWI/SNF-related matrix-associated actin- | SMARCA5 | 121.906 | 20.2 | 21 | 12 | 528.45886 |
| O60281 | Zinc finger protein 292 OS=Homo sapien: | ZNF292 | 304.816 | 0.3 | 1 | 1 | 134.2377 |
| O60292 | Signal-induced proliferation-associated 1- | SIPA1L3 | 194.61 | 2.1 | 3 | 3 | 71.741035 |
| O60293 | Zinc finger C3H1 domain-containing prot | ZFC3H1 | 226.356 | 10.4 | 19 | 19 | 114.76003 |
| O60306 | RNA helicase aquarius OS=Homo sapiens | AQR | 171.296 | 13.7 | 19 | 19 | 246.8744 |
| O60307 | Microtubule-associated serine/threonine-p | MAST3 | 143.138 | 3.3 | 3 | 3 | 108.30832 |
| O60313 | Dynamin-like 120 kDa protein, mitochonc | OPA1 | 111.631 | 8.6 | 7 | 7 | 99.830139 |
| O60318 | Germinal-center associated nuclear protei | MCM3AP | 218.405 | 3.6 | 6 | 6 | 60.046291 |
| O60333 | Kinesin-like protein KIF1B OS=Homo sa | KIF1B | 204.477 | 11 | 21 | 13 | 2486.6545 |
| O60341 | Lysine-specific histone demethylase 1A O | KDM1A | 92.903 | 6.1 | 5 | 5 | 61.766747 |
| O60343 | TBC1 domain family member 4 OS=Hom | TBC1D4 | 146.564 | 7 | 7 | 7 | 61.648891 |
| O60427 | Acyl-CoA (8-3)-desaturase OS=Homo sap | FADS1 | 51.965 | 4.7 | 2 | 2 | 101.68407 |
| O60479 | Homeobox protein DLX-3 OS=Homo sap | DLX3 | 31.739 | 17.4 | 4 | 4 | 153.61269 |
| O60488 | Long-chain-fatty-acid--CoA ligase 4 OS=] | ACSL4 | 79.189 | 5.9 | 4 | 4 | 65.598473 |
| O60493 | Sorting nexin-3 OS=Homo sapiens OX=9 | SNX3 | 18.763 | 13.6 | 2 | 2 | 50.750443 |
| O60499 | Syntaxin-10 OS=Homo sapiens OX=9606 | STX10 | 28.115 | 17.3 | 3 | 3 | 44.361397 |
| O60502 | Protein O-GlcNAcase OS=Homo sapiens | OGA | 102.916 | 2.2 | 1 | 1 | 45.796143 |
| O60504 | Vinexin OS=Homo sapiens OX=9606 GN | SORBS3 | 75.342 | 5.5 | 3 | 3 | 176.43298 |
| O60506 | Heterogeneous nuclear ribonucleoprotein | SYNCRIP | 69.603 | 22.8 | 14 | 14 | 9804.5068 |
| O60508 | Pre-mRNA-processing factor 17 OS=Hon | CDC40 | 65.522 | 35.2 | 16 | 16 | 239.78566 |
| O60524 | Ribosome quality control complex subuni | NEMF | 122.955 | 19.5 | 20 | 20 | 679.63605 |
| O60547 | GDP-mannose 4,6 dehydratase OS=Homo | GMDS | 41.95 | 12.9 | 4 | 4 | 100.02417 |
| O60551 | Glycylpeptide N-tetradecanoyltransferase | NMT2 | 56.981 | 2.2 | 1 | 1 | 98.247017 |
| O60563 | Cyclin-T1 OS=Homo sapiens OX=9606 ( | CCNT1 | 80.685 | 11.8 | 7 | 5 | 167.09714 |
| O60568 | Multifunctional procollagen lysine hydrox | PLOD3 | 84.786 | 8.3 | 6 | 6 | 111.53621 |
| O60573 | Eukaryotic translation initiation factor 4E | EIF4E2 | 28.362 | 37.1 | 8 | 8 | 1193.6563 |
| O60645 | Exocyst complex component 3 OS=Homo | EXOC3 | 85.568 | 4 | 3 | 3 | 46.253174 |

| | | | | | | | |
|---|---|---|---|---|---|---|---|
| O60674 | Tyrosine-protein kinase JAK2 OS=Homo | JAK2 | 130.674 | 0.8 | 1 | 1 | 21.490927 |
| O60678 | Protein arginine N-methyltransferase 3 OS | PRMT3 | 59.904 | 13.9 | 7 | 7 | 112.8755 |
| O60684 | Importin subunit alpha-7 OS=Homo sapie | KPNA6 | 60.03 | 14.7 | 7 | 2 | 343.88986 |
| O60701 | UDP-glucose 6-dehydrogenase OS=Homo | UGDH | 55.025 | 15 | 6 | 6 | 95.130585 |
| O60716 | Catenin delta-1 OS=Homo sapiens OX=96 | CTNND1 | 108.171 | 24 | 21 | 21 | 1188.8817 |
| O60749 | Sorting nexin-2 OS=Homo sapiens OX=9 | SNX2 | 58.472 | 2.1 | 1 | 1 | 54.88567 |
| O60762 | Dolichol-phosphate mannosyltransferase s | DPM1 | 29.635 | 36.2 | 8 | 8 | 1526.5944 |
| O60783 | Small ribosomal subunit protein uS14m O | MRPS14 | 15.139 | 28.9 | 3 | 3 | 166.72768 |
| O60814 | Histone H2B type 1-K OS=Homo sapiens | H2BC12 | 13.891 | 7.9 | 2 | 2 | 3156.9365 |
| O60825 | 6-phosphofructo-2-kinase/fructose-2,6-bis | PFKFB2 | 58.477 | 30.3 | 14 | 10 | 134.16107 |
| O60828 | Polyglutamine-binding protein 1 OS=Hon | PQBP1 | 30.473 | 12.8 | 3 | 3 | 266.96246 |
| O60830 | Mitochondrial import inner membrane tra | TIMM17B | 18.274 | 25.6 | 3 | 3 | 105.34328 |
| O60832 | H/ACA ribonucleoprotein complex subun | DKC1 | 57.675 | 40.9 | 22 | 22 | 1110.182 |
| O60841 | Eukaryotic translation initiation factor 5B | EIF5B | 138.827 | 31.8 | 34 | 34 | 1659.1633 |
| O60870 | DNA/RNA-binding protein KIN17 OS=H | KIN | 45.374 | 33.6 | 13 | 13 | 549.34436 |
| O60884 | DnaJ homolog subfamily A member 2 OS | DNAJA2 | 45.746 | 19.4 | 8 | 8 | 893.26776 |
| O60888 | Protein CutA OS=Homo sapiens OX=960 | CUTA | 19.117 | 7.8 | 1 | 1 | 405.42188 |
| O60942 | mRNA-capping enzyme OS=Homo sapier | RNGTT | 68.557 | 10.4 | 5 | 5 | 66.646706 |
| O75030 | Microphthalmia-associated transcription f | MITF | 58.796 | 7.2 | 4 | 4 | 112.95355 |
| O75044 | SLIT-ROBO Rho GTPase-activating prot | SRGAP2 | 120.871 | 19 | 19 | 10 | 281.84985 |
| O75063 | Glycosaminoglycan xylosylkinase OS=Hc | FAM20B | 46.433 | 37.9 | 12 | 12 | 728.85522 |
| O75083 | WD repeat-containing protein 1 OS=Hom | WDR1 | 66.194 | 7.9 | 3 | 3 | 83.442619 |
| O75106 | Retina-specific copper amine oxidase OS= | AOC2 | 83.674 | 1.2 | 1 | 1 | 200.16373 |
| O75116 | Rho-associated protein kinase 2 OS=Hom | ROCK2 | 160.9 | 9.7 | 12 | 9 | 125.37067 |
| O75122 | CLIP-associating protein 2 OS=Homo sap | CLASP2 | 141.064 | 32 | 38 | 38 | 1413.025 |
| O75127 | Pentatricopeptide repeat-containing protei | PTCD1 | 78.856 | 34.3 | 21 | 21 | 1921.6741 |
| O75128 | Protein cordon-bleu OS=Homo sapiens O | COBL | 135.617 | 2.2 | 2 | 2 | 45.666782 |
| O75131 | Copine-3 OS=Homo sapiens OX=9606 G | CPNE3 | 60.131 | 14.3 | 7 | 7 | 84.222183 |
| O75147 | Obscurin-like protein 1 OS=Homo sapien | OBSL1 | 206.947 | 17.8 | 29 | 29 | 1035.2683 |
| O75150 | E3 ubiquitin-protein ligase BRE1B OS=H | RNF40 | 113.679 | 4.2 | 3 | 2 | 41.822479 |
| O75151 | Lysine-specific demethylase PHF2 OS=H | PHF2 | 120.775 | 5.7 | 5 | 5 | 108.84593 |
| O75152 | Zinc finger CCCH domain-containing pro | ZC3H11A | 89.132 | 40.4 | 31 | 23 | 2371.9053 |
| O75161 | Nephrocystin-4 OS=Homo sapiens OX=9 | NPHP4 | 157.599 | 0.6 | 1 | 1 | 22.80262 |
| O75165 | DnaJ homolog subfamily C member 13 O | DNAJC13 | 254.415 | 27.7 | 60 | 60 | 1173.0485 |
| O75170 | Serine/threonine-protein phosphatase 6 re | PPP6R2 | 104.942 | 2.5 | 2 | 2 | 52.610641 |

| | | | | | | | |
|---|---|---|---|---|---|---|---|
| O75175 | CCR4-NOT transcription complex subuni | CNOT3 | 81.872 | 10.8 | 9 | 9 | 253.0007 |
| O75177 | Calcium-responsive transactivator OS=Hc | SS18L1 | 42.991 | 9.3 | 4 | 4 | 122.88813 |
| O75179 | Ankyrin repeat domain-containing protein | ANKRD17 | 274.258 | 11.9 | 27 | 27 | 631.39429 |
| O75190 | DnaJ homolog subfamily B member 6 OS | DNAJB6 | 36.087 | 22.7 | 7 | 5 | 818.82697 |
| O75251 | NADH dehydrogenase [ubiquinone] iron- | NDUFS7 | 23.564 | 15.5 | 3 | 3 | 255.34532 |
| O75306 | NADH dehydrogenase [ubiquinone] iron- | NDUFS2 | 52.546 | 13.8 | 6 | 6 | 107.05428 |
| O75317 | Ubiquitin carboxyl-terminal hydrolase 12 | USP12 | 42.858 | 2.2 | 1 | 1 | 61.808121 |
| O75319 | RNA/RNP complex-1-interacting phospha | DUSP11 | 43.711 | 5.3 | 2 | 2 | 54.463535 |
| O75330 | Hyaluronan mediated motility receptor OS | HMMR | 84.101 | 43 | 33 | 33 | 1856.7297 |
| O75340 | Programmed cell death protein 6 OS=Hor | PDCD6 | 21.869 | 23.6 | 4 | 4 | 267.83899 |
| O75348 | V-type proton ATPase subunit G 1 OS=H | ATP6V1G1 | 13.758 | 9.3 | 1 | 1 | 74.500687 |
| O75362 | Zinc finger protein 217 OS=Homo sapiens | ZNF217 | 115.272 | 2.8 | 2 | 2 | 49.953156 |
| O75367 | Core histone macro-H2A.1 OS=Homo sap | MACROH2 | 39.184 | 25.5 | 7 | 6 | 282.83301 |
| O75369 | Filamin-B OS=Homo sapiens OX=9606 ( | FLNB | 278.165 | 4.8 | 9 | 9 | 46.519947 |
| O75382 | Tripartite motif-containing protein 3 OS=] | TRIM3 | 80.83 | 27.8 | 17 | 16 | 596.6355 |
| O75386 | Tubby-related protein 3 OS=Homo sapien | TULP3 | 49.643 | 13.3 | 5 | 5 | 158.77394 |
| O75390 | Citrate synthase, mitochondrial OS=Homo | CS | 51.713 | 2.4 | 1 | 1 | 86.279228 |
| O75396 | Vesicle-trafficking protein SEC22b OS=F | SEC22B | 24.741 | 20.5 | 4 | 4 | 151.00658 |
| O75400 | Pre-mRNA-processing factor 40 homolog | PRPF40A | 108.805 | 8.5 | 8 | 7 | 207.347 |
| O75420 | GRB10-interacting GYF protein 1 OS=Hc | GIGYF1 | 114.602 | 15.7 | 13 | 13 | 252.41049 |
| O75436 | Vacuolar protein sorting-associated protei | VPS26A | 38.17 | 2.4 | 1 | 1 | 62.005417 |
| O75438 | NADH dehydrogenase [ubiquinone] 1 bet | NDUFB1 | 6.962 | 13.8 | 1 | 1 | 28.136671 |
| O75439 | Mitochondrial-processing peptidase subur | PMPCB | 54.367 | 16 | 8 | 8 | 175.17526 |
| O75446 | Histone deacetylase complex subunit SAF | SAP30 | 23.307 | 4.1 | 1 | 1 | 26.148865 |
| O75449 | Katanin p60 ATPase-containing subunit A | KATNA1 | 55.965 | 15.5 | 7 | 6 | 187.5939 |
| O75475 | PC4 and SFRS1-interacting protein OS=F | PSIP1 | 60.104 | 28.3 | 14 | 13 | 2016.0177 |
| O75477 | Erlin-1 OS=Homo sapiens OX=9606 GN= | ERLIN1 | 39.172 | 15.2 | 4 | 2 | 82.489845 |
| O75489 | NADH dehydrogenase [ubiquinone] iron- | NDUFS3 | 30.242 | 9.1 | 2 | 2 | 176.62102 |
| O75494 | Serine/arginine-rich splicing factor 10 OS | SRSF10 | 31.301 | 19.8 | 5 | 3 | 1716.832 |
| O75496 | Geminin OS=Homo sapiens OX=9606 GN | GMNN | 23.566 | 13.4 | 2 | 2 | 87.374321 |
| O75521 | Enoyl-CoA delta isomerase 2 OS=Homo s | ECI2 | 43.586 | 2.3 | 1 | 1 | 14.014915 |
| O75528 | Transcriptional adapter 3 OS=Homo sapie | TADA3 | 48.903 | 10.6 | 4 | 4 | 125.60054 |
| O75529 | TAF5-like RNA polymerase II p300/CBP | TAF5L | 66.156 | 9.5 | 5 | 5 | 84.499237 |
| O75530 | Polycomb protein EED OS=Homo sapien | EED | 50.198 | 48.3 | 20 | 20 | 1417.0485 |
| O75531 | Barrier-to-autointegration factor OS=Hon | BANF1 | 10.059 | 58.4 | 6 | 6 | 911.53558 |

| | | | | | | | |
|---|---|---|---|---|---|---|---|
| O75533 | Splicing factor 3B subunit 1 OS=Homo sa | SF3B1 | 145.831 | 25.5 | 29 | 29 | 736.49188 |
| O75534 | Cold shock domain-containing protein E1 | CSDE1 | 88.885 | 38.2 | 33 | 33 | 5910.8945 |
| O75569 | Interferon-inducible double-stranded RNA | PRKRA | 34.405 | 44.4 | 10 | 10 | 1608.7642 |
| O75581 | Low-density lipoprotein receptor-related p | LRP6 | 180.43 | 6.5 | 9 | 6 | 126.84742 |
| O75592 | E3 ubiquitin-protein ligase MYCBP2 OS= | MYCBP2 | 513.637 | 15.9 | 63 | 63 | 340.40979 |
| O75607 | Nucleoplasmin-3 OS=Homo sapiens OX= | NPM3 | 19.344 | 26.4 | 3 | 3 | 206.66229 |
| O75616 | GTPase Era, mitochondrial OS=Homo sap | ERAL1 | 48.351 | 27.9 | 11 | 11 | 1240.5219 |
| O75643 | U5 small nuclear ribonucleoprotein 200 k | SNRNP200 | 244.508 | 26.5 | 54 | 54 | 790.04681 |
| O75663 | TIP41-like protein OS=Homo sapiens OX | TIPRL | 31.444 | 4 | 1 | 1 | 49.531811 |
| O75665 | Centriole and centriolar satellite protein C | OFD1 | 116.672 | 1.4 | 1 | 1 | 59.876427 |
| O75676 | Ribosomal protein S6 kinase alpha-4 OS= | RPS6KA4 | 85.606 | 14.1 | 9 | 8 | 183.11514 |
| O75691 | Small subunit processome component 20 l | UTP20 | 318.386 | 12.4 | 32 | 32 | 371.67346 |
| O75694 | Nuclear pore complex protein Nup155 OS | NUP155 | 155.2 | 18.4 | 19 | 19 | 280.67889 |
| O75717 | WD repeat and HMG-box DNA-binding p | WDHD1 | 125.968 | 0.8 | 1 | 1 | 14.154386 |
| O75718 | Cartilage-associated protein OS=Homo sa | CRTAP | 46.562 | 8 | 3 | 3 | 55.528923 |
| O75746 | Electrogenic aspartate/glutamate antiporte | SLC25A12 | 74.762 | 15.9 | 11 | 5 | 736.67841 |
| O75808 | Calpain-15 OS=Homo sapiens OX=9606 | CAPN15 | 117.315 | 3.9 | 3 | 3 | 55.831928 |
| O75815 | Breast cancer anti-estrogen resistance pro | BCAR3 | 92.566 | 4.1 | 3 | 3 | 48.843124 |
| O75817 | Ribonuclease P protein subunit p20 OS=H | POP7 | 15.651 | 23.6 | 3 | 3 | 1354.3929 |
| O75818 | Ribonuclease P protein subunit p40 OS=H | RPP40 | 41.834 | 22.3 | 6 | 6 | 144.38649 |
| O75821 | Eukaryotic translation initiation factor 3 su | EIF3G | 35.612 | 34.1 | 8 | 8 | 1269.3926 |
| O75822 | Eukaryotic translation initiation factor 3 su | EIF3J | 29.063 | 8.9 | 2 | 2 | 239.11575 |
| O75909 | Cyclin-K OS=Homo sapiens OX=9606 G | CCNK | 64.24 | 6.6 | 3 | 3 | 141.48538 |
| O75915 | PRA1 family protein 3 OS=Homo sapiens | ARL6IP5 | 21.615 | 5.9 | 1 | 1 | 35.847301 |
| O75925 | E3 SUMO-protein ligase PIAS1 OS=Hom | PIAS1 | 71.836 | 12.4 | 7 | 6 | 167.86725 |
| O75934 | Pre-mRNA-splicing factor SPF27 OS=Ho | BCAS2 | 26.132 | 30.7 | 8 | 8 | 1197.3436 |
| O75940 | Survival of motor neuron-related-splicing | SMNDC1 | 26.712 | 27.7 | 8 | 8 | 587.80316 |
| O75955 | Flotillin-1 OS=Homo sapiens OX=9606 G | FLOT1 | 47.356 | 30.4 | 10 | 10 | 181.31299 |
| O75962 | Triple functional domain protein OS=Hom | TRIO | 346.901 | 2.6 | 8 | 6 | 61.872677 |
| O76003 | Glutaredoxin-3 OS=Homo sapiens OX=96 | GLRX3 | 37.433 | 6.3 | 2 | 2 | 39.280079 |
| O76021 | Ribosomal L1 domain-containing protein | RSL1D1 | 54.973 | 20.2 | 12 | 12 | 1485.0468 |
| O76031 | ATP-dependent Clp protease ATP-binding | CLPX | 69.224 | 16.9 | 9 | 9 | 115.58659 |
| O76039 | Cyclin-dependent kinase-like 5 OS=Homo | CDKL5 | 107.52 | 13.3 | 10 | 10 | 149.70531 |
| O76071 | Probable cytosolic iron-sulfur protein asse | CIAO1 | 37.841 | 22.1 | 7 | 7 | 392.50293 |
| O76094 | Signal recognition particle subunit SRP72 | SRP72 | 74.607 | 26.7 | 16 | 16 | 469.13312 |

| | | | | | | | |
|---|---|---|---|---|---|---|---|
| O94761 | ATP-dependent DNA helicase Q4 OS=Hc | RECQL4 | 133.068 | 12.4 | 12 | 12 | 314.06897 |
| O94762 | ATP-dependent DNA helicase Q5 OS=Hc | RECQL5 | 108.859 | 17.5 | 16 | 16 | 226.67255 |
| O94763 | Unconventional prefoldin RPB5 interacto | URI1 | 59.833 | 14 | 5 | 5 | 95.427292 |
| O94766 | Galactosylgalactosylxylosylprotein 3-beta | B3GAT3 | 37.122 | 6.9 | 2 | 2 | 150.44864 |
| O94776 | Metastasis-associated protein MTA2 OS= | MTA2 | 75.024 | 19.5 | 12 | 10 | 370.00455 |
| O94782 | Ubiquitin carboxyl-terminal hydrolase 1 C | USP1 | 88.208 | 3.1 | 2 | 2 | 30.565868 |
| O94804 | Serine/threonine-protein kinase 10 OS=Hc | STK10 | 112.136 | 2.3 | 2 | 1 | 20.64142 |
| O94806 | Serine/threonine-protein kinase D3 OS=H | PRKD3 | 100.471 | 13.9 | 10 | 7 | 346.5206 |
| O94822 | E3 ubiquitin-protein ligase listerin OS=Hc | LTN1 | 200.553 | 1.8 | 3 | 3 | 48.155922 |
| O94826 | Mitochondrial import receptor subunit TC | TOMM70 | 67.455 | 1.2 | 1 | 1 | 25.951998 |
| O94842 | TOX high mobility group box family men | TOX4 | 66.195 | 12.6 | 7 | 5 | 1194.4005 |
| O94851 | [F-actin]-monooxygenase MICAL2 OS=H | MICAL2 | 219.062 | 8 | 13 | 11 | 236.19673 |
| O94864 | STAGA complex 65 subunit gamma OS= | SUPT7L | 46.193 | 5.8 | 2 | 2 | 139.39529 |
| O94874 | E3 UFM1-protein ligase 1 OS=Homo sap | UFL1 | 89.596 | 5.2 | 3 | 3 | 48.404762 |
| O94875 | Sorbin and SH3 domain-containing protei | SORBS2 | 124.108 | 10.8 | 11 | 11 | 291.70914 |
| O94880 | PHD finger protein 14 OS=Homo sapiens | PHF14 | 106.987 | 5 | 4 | 4 | 56.063881 |
| O94887 | FERM, ARHGEF and pleckstrin domain-c | FARP2 | 119.889 | 19.6 | 17 | 17 | 1024.7678 |
| O94905 | Erlin-2 OS=Homo sapiens OX=9606 GN= | ERLIN2 | 37.84 | 8 | 2 | 2 | 33.571159 |
| O94906 | Pre-mRNA-processing factor 6 OS=Homo | PRPF6 | 106.925 | 14.6 | 14 | 14 | 271.97235 |
| O94913 | Pre-mRNA cleavage complex 2 protein Pc | PCF11 | 173.051 | 11.4 | 17 | 17 | 350.00705 |
| O94916 | Nuclear factor of activated T-cells 5 OS=l | NFAT5 | 165.764 | 4.4 | 5 | 5 | 63.016834 |
| O94923 | D-glucuronyl C5-epimerase OS=Homo sa | GLCE | 70.102 | 4.1 | 2 | 2 | 52.371605 |
| O94925 | Glutaminase kidney isoform, mitochondri | GLS | 73.462 | 4.3 | 2 | 2 | 19.698631 |
| O94927 | HAUS augmin-like complex subunit 5 OS | HAUS5 | 71.683 | 10.7 | 6 | 6 | 118.76665 |
| O94955 | Rho-related BTB domain-containing prote | RHOBTB3 | 69.414 | 6.1 | 3 | 3 | 72.78479 |
| O94964 | Protein SOGA1 OS=Homo sapiens OX=9 | SOGA1 | 159.76 | 26.1 | 34 | 34 | 461.19833 |
| O94967 | WD repeat-containing protein 47 OS=Hoi | WDR47 | 101.95 | 1.5 | 1 | 1 | 10.491266 |
| O94973 | AP-2 complex subunit alpha-2 OS=Homo | AP2A2 | 103.961 | 16.3 | 14 | 14 | 602.36859 |
| O94986 | Centrosomal protein of 152 kDa OS=Hon | CEP152 | 195.627 | 23.8 | 40 | 40 | 793.50281 |
| O94992 | Protein HEXIM1 OS=Homo sapiens OX= | HEXIM1 | 40.624 | 11.1 | 4 | 4 | 242.86728 |
| O95049 | Tight junction protein ZO-3 OS=Homo sa | TJP3 | 101.397 | 11.5 | 10 | 10 | 119.36637 |
| O95059 | Ribonuclease P protein subunit p14 OS=H | RPP14 | 13.694 | 24.2 | 3 | 3 | 163.37265 |
| O95070 | Protein YIF1A OS=Homo sapiens OX=9( | YIF1A | 32.012 | 9.2 | 2 | 2 | 92.62709 |
| O95071 | E3 ubiquitin-protein ligase UBR5 OS=Ho | UBR5 | 309.353 | 3.6 | 9 | 9 | 77.435364 |
| O95139 | NADH dehydrogenase [ubiquinone] 1 bet | NDUFB6 | 15.49 | 5.5 | 1 | 1 | 56.371101 |

| | | | | | | | |
|---|---|---|---|---|---|---|---|
| O95147 | Dual specificity protein phosphatase 14 O | DUSP14 | 22.255 | 5.6 | 1 | 1 | 31.735893 |
| O95159 | Zinc finger protein-like 1 OS=Homo sapie | ZFPL1 | 34.114 | 8.7 | 3 | 3 | 152.45604 |
| O95163 | Elongator complex protein 1 OS=Homo sa | ELP1 | 150.255 | 19.6 | 23 | 23 | 514.24304 |
| O95168 | NADH dehydrogenase [ubiquinone] 1 bet | NDUFB4 | 15.209 | 23.3 | 3 | 3 | 351.28473 |
| O95202 | Mitochondrial proton/calcium exchanger | LETM1 | 83.355 | 2 | 1 | 1 | 38.389893 |
| O95218 | Zinc finger Ran-binding domain-containir | ZRANB2 | 37.405 | 32.7 | 11 | 11 | 609.00342 |
| O95229 | ZW10 interactor OS=Homo sapiens OX= | ZWINT | 31.294 | 4.3 | 1 | 1 | 74.724808 |
| O95232 | Luc7-like protein 3 OS=Homo sapiens OX | LUC7L3 | 51.467 | 18.3 | 7 | 7 | 1355.6721 |
| O95235 | Kinesin-like protein KIF20A OS=Homo s | KIF20A | 100.278 | 20.7 | 16 | 16 | 117.01137 |
| O95239 | Chromosome-associated kinesin KIF4A C | KIF4A | 139.882 | 24.6 | 25 | 16 | 546.64392 |
| O95243 | Methyl-CpG-binding domain protein 4 OS | MBD4 | 66.051 | 31.4 | 18 | 18 | 321.05969 |
| O95248 | Myotubularin-related protein 5 OS=Homo | SBF1 | 208.444 | 11.9 | 21 | 19 | 167.435 |
| O95251 | Histone acetyltransferase KAT7 OS=Hom | KAT7 | 70.643 | 23.7 | 15 | 14 | 353.03256 |
| O95255 | ATP-binding cassette sub-family C memb | ABCC6 | 164.907 | 0.5 | 1 | 1 | 733.05389 |
| O95263 | High affinity cAMP-specific and IBMX-ir | PDE8B | 98.979 | 1.4 | 1 | 1 | 42.556061 |
| O95271 | Poly [ADP-ribose] polymerase tankyrase- | TNKS | 142.04 | 23.2 | 27 | 20 | 351.7691 |
| O95279 | Potassium channel subfamily K member 5 | KCNK5 | 55.13 | 1.4 | 1 | 1 | 124.07233 |
| O95292 | Vesicle-associated membrane protein-asso | VAPB | 27.229 | 21 | 4 | 4 | 130.68517 |
| O95295 | SNARE-associated protein Snapin OS=Ho | SNAPIN | 14.874 | 26.5 | 3 | 3 | 39.620674 |
| O95299 | NADH dehydrogenase [ubiquinone] 1 alp | NDUFA10 | 40.751 | 32.7 | 11 | 11 | 630.27472 |
| O95347 | Structural maintenance of chromosomes p | SMC2 | 135.657 | 22.3 | 23 | 23 | 352.07336 |
| O95363 | Phenylalanine--tRNA ligase, mitochondria | FARS2 | 52.357 | 31.7 | 13 | 13 | 794.78546 |
| O95365 | Zinc finger and BTB domain-containing p | ZBTB7A | 61.44 | 5.8 | 2 | 2 | 151.24033 |
| O95373 | Importin-7 OS=Homo sapiens OX=9606 ( | IPO7 | 119.517 | 12 | 12 | 11 | 400.8667 |
| O95391 | Pre-mRNA-splicing factor SLU7 OS=Hon | SLU7 | 68.387 | 33.1 | 19 | 19 | 477.18692 |
| O95400 | CD2 antigen cytoplasmic tail-binding prot | CD2BP2 | 37.647 | 16.1 | 5 | 5 | 119.68014 |
| O95405 | Zinc finger FYVE domain-containing prot | ZFYVE9 | 156.403 | 3.2 | 4 | 4 | 55.642525 |
| O95409 | Zinc finger protein ZIC 2 OS=Homo sapie | ZIC2 | 55.006 | 5.6 | 3 | 2 | 391.54303 |
| O95425 | Supervillin OS=Homo sapiens OX=9606 | SVIL | 247.747 | 1.9 | 3 | 3 | 36.72522 |
| O95429 | BAG family molecular chaperone regulato | BAG4 | 49.595 | 13.8 | 7 | 7 | 467.63397 |
| O95433 | Activator of 90 kDa heat shock protein A | AHSA1 | 38.275 | 10.4 | 3 | 3 | 68.695702 |
| O95453 | Poly(A)-specific ribonuclease PARN OS= | PARN | 73.451 | 21.8 | 14 | 14 | 408.63287 |
| O95470 | Sphingosine-1-phosphate lyase 1 OS=Hon | SGPL1 | 63.524 | 17.1 | 9 | 9 | 337.44281 |
| O95486 | Protein transport protein Sec24A OS=Hon | SEC24A | 119.75 | 2.5 | 2 | 2 | 33.211227 |
| O95487 | Protein transport protein Sec24B OS=Hon | SEC24B | 137.418 | 5.4 | 6 | 6 | 80.19677 |

| | | | | | | | |
|---|---|---|---|---|---|---|---|
| O95497 | Pantetheinase OS=Homo sapiens OX=960 | VNN1 | 57.012 | 1.8 | 1 | 1 | 266.98541 |
| O95503 | Chromobox protein homolog 6 OS=Homo | CBX6 | 43.899 | 11.9 | 5 | 3 | 99.006874 |
| O95544 | NAD kinase OS=Homo sapiens OX=9606 | NADK | 49.229 | 16.1 | 6 | 6 | 166.41624 |
| O95551 | Tyrosyl-DNA phosphodiesterase 2 OS=H | TDP2 | 40.93 | 7.7 | 3 | 3 | 81.494598 |
| O95563 | Mitochondrial pyruvate carrier 2 OS=Hor | MPC2 | 14.279 | 9.4 | 1 | 1 | 51.730103 |
| O95571 | Persulfide dioxygenase ETHE1, mitochon | ETHE1 | 27.873 | 17.7 | 4 | 4 | 161.2576 |
| O95573 | Fatty acid CoA ligase Acsl3 OS=Homo sa | ACSL3 | 80.421 | 23.1 | 13 | 12 | 281.21738 |
| O95602 | DNA-directed RNA polymerase I subunit | POLR1A | 194.812 | 4.6 | 6 | 6 | 124.55865 |
| O95613 | Pericentrin OS=Homo sapiens OX=9606 | PCNT | 378.038 | 15 | 43 | 43 | 145.55287 |
| O95619 | YEATS domain-containing protein 4 OS= | YEATS4 | 26.5 | 24.7 | 5 | 5 | 176.33052 |
| O95630 | STAM-binding protein OS=Homo sapiens | STAMBP | 48.077 | 2.8 | 1 | 1 | 37.786404 |
| O95633 | Follistatin-related protein 3 OS=Homo sa | FSTL3 | 27.664 | 23.6 | 5 | 5 | 134.63425 |
| O95639 | Cleavage and polyadenylation specificity | CPSF4 | 30.255 | 16.4 | 4 | 3 | 122.91219 |
| O95644 | Nuclear factor of activated T-cells, cytopl | NFATC1 | 101.243 | 4.9 | 4 | 4 | 76.773857 |
| O95672 | Endothelin-converting enzyme-like 1 OS= | ECEL1 | 87.792 | 13.5 | 9 | 9 | 142.14519 |
| O95677 | Eyes absent homolog 4 OS=Homo sapiens | EYA4 | 69.506 | 3.8 | 2 | 1 | 36.614304 |
| O95684 | Centrosomal protein 43 OS=Homo sapien | CEP43 | 43.065 | 18.3 | 6 | 6 | 131.51195 |
| O95707 | Ribonuclease P protein subunit p29 OS=H | POP4 | 25.425 | 18.6 | 4 | 4 | 535.01868 |
| O95714 | E3 ubiquitin-protein ligase HERC2 OS=H | HERC2 | 527.228 | 6.7 | 27 | 21 | 118.02499 |
| O95751 | Protein LDOC1 OS=Homo sapiens OX=9 | LDOC1 | 16.969 | 9.6 | 1 | 1 | 120.60863 |
| O95758 | Polypyrimidine tract-binding protein 3 OS | PTBP3 | 59.69 | 15.8 | 8 | 6 | 6570.2246 |
| O95782 | AP-2 complex subunit alpha-1 OS=Homo | AP2A1 | 107.546 | 34 | 30 | 23 | 1316.064 |
| O95785 | Protein Wiz OS=Homo sapiens OX=9606 | WIZ | 178.675 | 7.1 | 9 | 9 | 167.83615 |
| O95793 | Double-stranded RNA-binding protein Sta | STAU1 | 63.183 | 49.2 | 25 | 25 | 1991.9951 |
| O95801 | Tetratricopeptide repeat protein 4 OS=Ho | TTC4 | 44.679 | 17.1 | 6 | 6 | 163.0817 |
| O95816 | BAG family molecular chaperone regulato | BAG2 | 23.772 | 53.1 | 10 | 10 | 2469.0276 |
| O95817 | BAG family molecular chaperone regulato | BAG3 | 61.595 | 29.2 | 14 | 14 | 609.95526 |
| O95819 | Mitogen-activated protein kinase kinase k | MAP4K4 | 142.101 | 16.7 | 18 | 12 | 421.53836 |
| O95831 | Apoptosis-inducing factor 1, mitochondri | AIFM1 | 66.901 | 21.7 | 10 | 10 | 403.56525 |
| O95835 | Serine/threonine-protein kinase LATS1 O | LATS1 | 126.87 | 1.9 | 2 | 2 | 39.843567 |
| O95863 | Zinc finger protein SNAI1 OS=Homo sap | SNAI1 | 29.084 | 8.7 | 2 | 2 | 110.10359 |
| O95870 | Phosphatidylserine lipase ABHD16A OS= | ABHD16A | 63.244 | 3.8 | 2 | 2 | 160.95856 |
| O95900 | Pseudouridylate synthase TRUB2, mitoch | TRUB2 | 36.695 | 45.3 | 15 | 15 | 1736.6027 |
| O95905 | Protein ecdysoneless homolog OS=Homo | ECD | 72.759 | 3.4 | 2 | 2 | 123.67223 |
| O95926 | Pre-mRNA-splicing factor SYF2 OS=Hor | SYF2 | 28.723 | 4.5 | 1 | 1 | 104.35349 |

| | | | | | | | |
|---|---|---|---|---|---|---|---|
| O95983 | Methyl-CpG-binding domain protein 3 OS | MBD3 | 32.845 | 12.4 | 3 | 3 | 35.04789 |
| O95985 | DNA topoisomerase 3-beta-1 OS=Homo s | TOP3B | 96.662 | 35.2 | 28 | 28 | 3765.7368 |
| O95995 | Dynein regulatory complex subunit 4 OS= | GAS8 | 56.356 | 10.5 | 4 | 4 | 79.932281 |
| O95997 | Securin OS=Homo sapiens OX=9606 GN | PTTG1 | 22.025 | 17.8 | 3 | 1 | 86.434769 |
| O95999 | B-cell lymphoma/leukemia 10 OS=Homo | BCL10 | 26.252 | 14.2 | 3 | 3 | 94.291855 |
| O96000 | NADH dehydrogenase [ubiquinone] 1 bet | NDUFB10 | 20.777 | 19.2 | 3 | 3 | 135.22192 |
| O96008 | Mitochondrial import receptor subunit TC | TOMM40 | 37.894 | 10.2 | 3 | 3 | 82.454575 |
| O96011 | Peroxisomal membrane protein 11B OS=I | PEX11B | 28.432 | 16.6 | 4 | 4 | 89.233078 |
| O96013 | Serine/threonine-protein kinase PAK 4 OS | PAK4 | 64.073 | 2.5 | 1 | 1 | 65.690964 |
| O96019 | Actin-like protein 6A OS=Homo sapiens ( | ACTL6A | 47.461 | 34 | 11 | 9 | 1053.2239 |
| O96020 | G1/S-specific cyclin-E2 OS=Homo sapier | CCNE2 | 46.758 | 3 | 1 | 1 | 61.69743 |
| O96028 | Histone-lysine N-methyltransferase NSD2 | NSD2 | 152.259 | 7.4 | 8 | 8 | 161.45676 |
| P00338 | L-lactate dehydrogenase A chain OS=Hor | LDHA | 36.689 | 19.3 | 6 | 5 | 137.75804 |
| P00367 | Glutamate dehydrogenase 1, mitochondria | GLUD1 | 61.398 | 8.6 | 5 | 5 | 110.90511 |
| P00390 | Glutathione reductase, mitochondrial OS= | GSR | 56.258 | 2.3 | 1 | 1 | 108.74495 |
| P00403 | Cytochrome c oxidase subunit 2 OS=Hom | MT-CO2 | 25.566 | 4.4 | 1 | 1 | 385.00342 |
| P00450 | Ceruloplasmin OS=Homo sapiens OX=96 | CP | 122.206 | 5.6 | 6 | 6 | 839.28424 |
| P00488 | Coagulation factor XIII A chain OS=Hom | F13A1 | 83.269 | 2.2 | 1 | 1 | 149.52461 |
| P00492 | Hypoxanthine-guanine phosphoribosyltrai | HPRT1 | 24.58 | 18.3 | 3 | 3 | 187.24989 |
| P00533 | Epidermal growth factor receptor OS=Ho | EGFR | 134.278 | 1 | 1 | 1 | 13.374226 |
| P00558 | Phosphoglycerate kinase 1 OS=Homo sap | PGK1 | 44.615 | 2.4 | 1 | 1 | 75.74086 |
| P00734 | Prothrombin OS=Homo sapiens OX=960( | F2 | 70.037 | 7.7 | 4 | 4 | 363.41177 |
| P00738 | Haptoglobin OS=Homo sapiens OX=960( | HP | 45.206 | 8.1 | 4 | 1 | 852.42755 |
| P00747 | Plasminogen OS=Homo sapiens OX=960 | PLG | 90.569 | 6.3 | 6 | 4 | 1687.573 |
| P00846 | ATP synthase subunit a OS=Homo sapien | MT-ATP6 | 24.818 | 4.4 | 1 | 1 | 134.57587 |
| P00966 | Argininosuccinate synthase OS=Homo saj | ASS1 | 46.531 | 19.2 | 9 | 9 | 116.63448 |
| P00973 | 2'-5'-oligoadenylate synthase 1 OS=Homo | OAS1 | 46.029 | 3.2 | 1 | 1 | 61.820007 |
| P01008 | Antithrombin-III OS=Homo sapiens OX= | SERPINC1 | 52.603 | 5.8 | 3 | 3 | 569.08575 |
| P01011 | Alpha-1-antichymotrypsin OS=Homo sap | SERPINA3 | 47.651 | 2.8 | 1 | 1 | 50.523445 |
| P01023 | Alpha-2-macroglobulin OS=Homo sapien | A2M | 163.291 | 1.3 | 2 | 1 | 260.28983 |
| P01024 | Complement C3 OS=Homo sapiens OX=9 | C3 | 187.149 | 4.4 | 7 | 7 | 424.86438 |
| P01042 | Kininogen-1 OS=Homo sapiens OX=960( | KNG1 | 71.958 | 1.4 | 1 | 1 | 99.45343 |
| P01100 | Protein c-Fos OS=Homo sapiens OX=960 | FOS | 40.696 | 3.7 | 1 | 1 | 51.03791 |
| P01106 | Myc proto-oncogene protein OS=Homo sa | MYC | 50.566 | 13.9 | 6 | 5 | 648.30664 |
| P01111 | GTPase NRas OS=Homo sapiens OX=96 | NRAS | 21.23 | 12.2 | 2 | 1 | 138.43034 |

| | | | | | | | |
|---|---|---|---|---|---|---|---|
| P01130 | Low-density lipoprotein receptor OS=Hor | LDLR | 95.377 | 5.3 | 4 | 4 | 114.65878 |
| P01344 | Insulin-like growth factor II OS=Homo sa | IGF2 | 20.141 | 5 | 1 | 1 | 1309.2108 |
| P01591 | Immunoglobulin J chain OS=Homo sapier | JCHAIN | 18.099 | 6.3 | 1 | 1 | 192.97881 |
| P01859 | Immunoglobulin heavy constant gamma 2 | IGHG2 | 43.806 | 2.3 | 1 | 1 | 2575.3381 |
| P01889 | HLA class I histocompatibility antigen, B | HLA-B | 40.461 | 14.6 | 4 | 0 | 108.31017 |
| P02042 | Hemoglobin subunit delta OS=Homo sapi | HBD | 16.056 | 6.1 | 1 | 1 | 852.44843 |
| P02100 | Hemoglobin subunit epsilon OS=Homo sa | HBE1 | 16.203 | 12.9 | 2 | 1 | 1426.7236 |
| P02452 | Collagen alpha-1(I) chain OS=Homo sapi | COL1A1 | 138.912 | 6.5 | 9 | 9 | 1208.7498 |
| P02461 | Collagen alpha-1(III) chain OS=Homo sa | COL3A1 | 138.565 | 4.9 | 6 | 6 | 2446.9951 |
| P02462 | Collagen alpha-1(IV) chain OS=Homo sa | COL4A1 | 160.611 | 1.8 | 2 | 1 | 144.91388 |
| P02533 | Keratin, type I cytoskeletal 14 OS=Homo | KRT14 | 51.562 | 20.1 | 9 | 2 | 833.17615 |
| P02538 | Keratin, type II cytoskeletal 6A OS=Hom | KRT6A | 60.045 | 32.3 | 18 | 0 | 1567.8706 |
| P02545 | Prelamin-A/C OS=Homo sapiens OX=96 | LMNA | 74.14 | 41.7 | 30 | 28 | 939.05554 |
| P02656 | Apolipoprotein C-III OS=Homo sapiens ( | APOC3 | 10.853 | 19.2 | 2 | 2 | 47.666374 |
| P02671 | Fibrinogen alpha chain OS=Homo sapiens | FGA | 94.974 | 1.3 | 3 | 3 | 1410.8553 |
| P02675 | Fibrinogen beta chain OS=Homo sapiens | FGB | 55.929 | 17.9 | 10 | 10 | 6538.1353 |
| P02679 | Fibrinogen gamma chain OS=Homo sapie | FGG | 51.512 | 11.3 | 5 | 5 | 2797.4717 |
| P02749 | Beta-2-glycoprotein 1 OS=Homo sapiens | APOH | 38.299 | 2.6 | 1 | 1 | 471.22534 |
| P02751 | Fibronectin OS=Homo sapiens OX=9606 | FN1 | 272.321 | 13.5 | 28 | 28 | 1426.1769 |
| P02760 | Protein AMBP OS=Homo sapiens OX=96 | AMBP | 39 | 2 | 1 | 1 | 1747.7454 |
| P02768 | Albumin OS=Homo sapiens OX=9606 GI | ALB | 69.367 | 15.3 | 13 | 13 | 48266.438 |
| P02786 | Transferrin receptor protein 1 OS=Homo | TFRC | 84.872 | 13.9 | 9 | 9 | 117.68515 |
| P02788 | Lactotransferrin OS=Homo sapiens OX=9 | LTF | 78.182 | 3 | 2 | 2 | 3754.7024 |
| P02790 | Hemopexin OS=Homo sapiens OX=9606 | HPX | 51.677 | 1.7 | 1 | 1 | 215.61758 |
| P02792 | Ferritin light chain OS=Homo sapiens OX | FTL | 20.02 | 9.1 | 1 | 1 | 17.392036 |
| P02794 | Ferritin heavy chain OS=Homo sapiens O | FTH1 | 21.226 | 13.7 | 2 | 2 | 103.27927 |
| P02795 | Metallothionein-2 OS=Homo sapiens OX | MT2A | 6.043 | 13.1 | 1 | 1 | 245.603 |
| P03928 | ATP synthase protein 8 OS=Homo sapien | MT-ATP8 | 7.992 | 48.5 | 3 | 3 | 396.15176 |
| P04003 | C4b-binding protein alpha chain OS=Hon | C4BPA | 67.034 | 1.8 | 1 | 1 | 32.760994 |
| P04004 | Vitronectin OS=Homo sapiens OX=9606 | VTN | 54.306 | 6.7 | 3 | 3 | 186.49768 |
| P04040 | Catalase OS=Homo sapiens OX=9606 GN | CAT | 59.757 | 2.8 | 1 | 1 | 116.56654 |
| P04049 | RAF proto-oncogene serine/threonine-pro | RAF1 | 73.052 | 6.9 | 4 | 3 | 133.94733 |
| P04075 | Fructose-bisphosphate aldolase A OS=Ho | ALDOA | 39.421 | 16.2 | 5 | 5 | 542.09021 |
| P04080 | Cystatin-B OS=Homo sapiens OX=9606 ( | CSTB | 11.14 | 33.7 | 3 | 3 | 219.69165 |
| P04114 | Apolipoprotein B-100 OS=Homo sapiens | APOB | 515.605 | 0.4 | 2 | 2 | 150.88519 |

| | | | | | | | |
|---|---|---|---|---|---|---|---|
| P04150 | Glucocorticoid receptor OS=Homo sapier | NR3C1 | 85.66 | 5.1 | 3 | 3 | 53.353199 |
| P04156 | Major prion protein OS=Homo sapiens O. | PRNP | 27.662 | 9.1 | 2 | 2 | 68.526573 |
| P04183 | Thymidine kinase, cytosolic OS=Homo sa | TK1 | 25.469 | 47.9 | 9 | 9 | 277.98441 |
| P04264 | Keratin, type II cytoskeletal 1 OS=Homo | KRT1 | 66.039 | 27 | 19 | 16 | 2319.4541 |
| P04406 | Glyceraldehyde-3-phosphate dehydrogena | GAPDH | 36.054 | 31.3 | 8 | 8 | 3457.2903 |
| P04439 | HLA class I histocompatibility antigen, A | HLA-A | 40.841 | 3.8 | 1 | 1 | 739.61365 |
| P04632 | Calpain small subunit 1 OS=Homo sapien | CAPNS1 | 28.316 | 4.9 | 1 | 1 | 152.15987 |
| P04792 | Heat shock protein beta-1 OS=Homo sapi | HSPB1 | 22.783 | 41 | 8 | 8 | 2090.0247 |
| P04843 | Dolichyl-diphosphooligosaccharide--prote | RPN1 | 68.57 | 36.4 | 20 | 20 | 811.52533 |
| P04844 | Dolichyl-diphosphooligosaccharide--prote | RPN2 | 69.284 | 7.1 | 3 | 3 | 88.13662 |
| P04899 | Guanine nucleotide-binding protein G(i) s | GNAI2 | 40.451 | 12.7 | 4 | 2 | 224.32397 |
| P05019 | Insulin-like growth factor I OS=Homo sap | IGF1 | 21.842 | 21.5 | 4 | 4 | 178.86647 |
| P05023 | Sodium/potassium-transporting ATPase su | ATP1A1 | 112.897 | 17.6 | 15 | 9 | 332.52548 |
| P05026 | Sodium/potassium-transporting ATPase su | ATP1B1 | 35.062 | 4.3 | 1 | 1 | 77.949692 |
| P05067 | Amyloid-beta precursor protein OS=Hom | APP | 86.944 | 4.7 | 3 | 2 | 95.904755 |
| P05114 | Non-histone chromosomal protein HMG-1 | HMGN1 | 10.659 | 13 | 1 | 1 | 44.161098 |
| P05141 | ADP/ATP translocase 2 OS=Homo sapier | SLC25A5 | 32.853 | 45.3 | 15 | 6 | 12626.752 |
| P05156 | Complement factor I OS=Homo sapiens C | CFI | 65.751 | 3.4 | 2 | 2 | 2305.8389 |
| P05187 | Alkaline phosphatase, placental type OS= | ALPP | 57.954 | 17 | 8 | 1 | 102.18729 |
| P05198 | Eukaryotic translation initiation factor 2 su | EIF2S1 | 36.113 | 56.2 | 22 | 22 | 4001.2219 |
| P05204 | Non-histone chromosomal protein HMG-1 | HMGN2 | 9.393 | 13.3 | 1 | 1 | 21.88023 |
| P05362 | Intercellular adhesion molecule 1 OS=Ho | ICAM1 | 57.826 | 13.2 | 5 | 5 | 128.03687 |
| P05387 | Large ribosomal subunit protein P2 OS=H | RPLP2 | 11.665 | 50.4 | 8 | 8 | 19895.213 |
| P05388 | Large ribosomal subunit protein uL10 OS | RPLP0 | 34.274 | 38.8 | 9 | 2 | 22452.035 |
| P05423 | DNA-directed RNA polymerase III subun | POLR3D | 44.396 | 6.3 | 2 | 2 | 73.276703 |
| P05455 | Lupus La protein OS=Homo sapiens OX= | SSB | 46.838 | 42.2 | 19 | 19 | 2486.9829 |
| P05549 | Transcription factor AP-2-alpha OS=Hom | TFAP2A | 48.063 | 30 | 13 | 5 | 2949.3662 |
| P05783 | Keratin, type I cytoskeletal 18 OS=Homo | KRT18 | 48.058 | 49.3 | 22 | 20 | 3241.9009 |
| P05787 | Keratin, type II cytoskeletal 8 OS=Homo | KRT8 | 53.705 | 30.4 | 15 | 14 | 3901.6812 |
| P05976 | Myosin light chain 1/3, skeletal muscle isc | MYL1 | 21.146 | 6.2 | 1 | 1 | 638.09369 |
| P06312 | Immunoglobulin kappa variable 4-1 OS=H | IGKV4-1 | 13.381 | 14.9 | 2 | 2 | 4194.6064 |
| P06396 | Gelsolin OS=Homo sapiens OX=9606 GN | GSN | 85.698 | 5 | 2 | 2 | 55.03997 |
| P06493 | Cyclin-dependent kinase 1 OS=Homo sap | CDK1 | 34.096 | 28.6 | 8 | 8 | 435.55801 |
| P06576 | ATP synthase subunit beta, mitochondrial | ATP5F1B | 56.56 | 26.5 | 10 | 10 | 270.8082 |
| P06703 | Protein S100-A6 OS=Homo sapiens OX= | S100A6 | 10.18 | 16.7 | 2 | 2 | 586.33307 |

| | | | | | | | |
|---|---|---|---|---|---|---|---|
| P06730 | Eukaryotic translation initiation factor 4E | EIF4E | 25.098 | 31.3 | 7 | 7 | 5568.3521 |
| P06733 | Alpha-enolase OS=Homo sapiens OX=96 | ENO1 | 47.169 | 23.3 | 9 | 9 | 494.48764 |
| P06744 | Glucose-6-phosphate isomerase OS=Hom | GPI | 63.148 | 2.7 | 1 | 1 | 29.661901 |
| P06746 | DNA polymerase beta OS=Homo sapiens | POLB | 38.178 | 18.2 | 6 | 6 | 141.98044 |
| P06748 | Nucleophosmin OS=Homo sapiens OX=9 | NPM1 | 32.576 | 25.9 | 7 | 7 | 3062.4429 |
| P06753 | Tropomyosin alpha-3 chain OS=Homo sa | TPM3 | 32.95 | 8.4 | 3 | 3 | 1150.7344 |
| P06899 | Histone H2B type 1-J OS=Homo sapiens | H2BC11 | 13.905 | 23.8 | 4 | 0 | 19373.402 |
| P07195 | L-lactate dehydrogenase B chain OS=Hor | LDHB | 36.639 | 10.2 | 3 | 3 | 87.77533 |
| P07237 | Protein disulfide-isomerase OS=Homo sa | P4HB | 57.117 | 22.6 | 11 | 11 | 203.59227 |
| P07305 | Histone H1.0 OS=Homo sapiens OX=960 | H1-0 | 20.863 | 11.9 | 2 | 2 | 228.85147 |
| P07355 | Annexin A2 OS=Homo sapiens OX=9606 | ANXA2 | 38.605 | 40.1 | 13 | 3 | 2103.0835 |
| P07358 | Complement component C8 beta chain OS | C8B | 66.948 | 1.9 | 1 | 1 | 34.16618 |
| P07437 | Tubulin beta chain OS=Homo sapiens OX | TUBB | 49.671 | 12.2 | 3 | 3 | 12974.146 |
| P07477 | Serine protease 1 OS=Homo sapiens OX= | PRSS1 | 26.559 | 3.2 | 1 | 1 | 10679.297 |
| P07711 | Procathepsin L OS=Homo sapiens OX=9 | CTSL | 37.565 | 8.4 | 2 | 2 | 172.18993 |
| P07737 | Profilin-1 OS=Homo sapiens OX=9606 G | PFN1 | 15.055 | 53.6 | 6 | 6 | 146.93076 |
| P07741 | Adenine phosphoribosyltransferase OS=H | APRT | 19.608 | 12.8 | 2 | 2 | 128.33023 |
| P07814 | Bifunctional glutamate/proline--tRNA liga | EPRS1 | 170.591 | 22.5 | 29 | 29 | 344.63422 |
| P07900 | Heat shock protein HSP 90-alpha OS=Ho | HSP90AA1 | 84.66 | 23.6 | 17 | 12 | 829.23511 |
| P07910 | Heterogeneous nuclear ribonucleoproteins | HNRNPC | 33.67 | 38.2 | 14 | 6 | 9136.5938 |
| P07919 | Cytochrome b-c1 complex subunit 6, mito | UQCRH | 10.739 | 38.5 | 4 | 3 | 637.08356 |
| P07942 | Laminin subunit beta-1 OS=Homo sapiens | LAMB1 | 198.039 | 2.4 | 4 | 4 | 109.79915 |
| P07947 | Tyrosine-protein kinase Yes OS=Homo sa | YES1 | 60.802 | 11 | 6 | 1 | 214.72697 |
| P07948 | Tyrosine-protein kinase Lyn OS=Homo sa | LYN | 58.575 | 25.8 | 11 | 10 | 188.41225 |
| P07951 | Tropomyosin beta chain OS=Homo sapien | TPM2 | 32.851 | 36.3 | 13 | 4 | 2644.4146 |
| P07992 | DNA excision repair protein ERCC-1 OS= | ERCC1 | 32.563 | 11.1 | 3 | 3 | 47.297306 |
| P07996 | Thrombospondin-1 OS=Homo sapiens OX | THBS1 | 129.383 | 10.5 | 11 | 11 | 238.65141 |
| P08047 | Transcription factor Sp1 OS=Homo sapie | SP1 | 80.693 | 3.7 | 4 | 4 | 94.847458 |
| P08123 | Collagen alpha-2(I) chain OS=Homo sapi | COL1A2 | 129.314 | 1.4 | 1 | 1 | 243.10533 |
| P08134 | Rho-related GTP-binding protein RhoC O | RHOC | 22.007 | 28 | 5 | 1 | 172.54149 |
| P08174 | Complement decay-accelerating factor OS | CD55 | 41.401 | 2.1 | 1 | 1 | 91.762695 |
| P08195 | Amino acid transporter heavy chain SLC3 | SLC3A2 | 67.994 | 8.7 | 5 | 5 | 137.05179 |
| P08237 | ATP-dependent 6-phosphofructokinase, m | PFKM | 85.183 | 27.1 | 19 | 17 | 623.48822 |
| P08238 | Heat shock protein HSP 90-beta OS=Hom | HSP90AB1 | 83.265 | 34.9 | 26 | 10 | 1822.5649 |
| P08240 | Signal recognition particle receptor subun | SRPRA | 69.812 | 24.5 | 14 | 14 | 461.63348 |

| | | | | | | | |
|---|---|---|---|---|---|---|---|
| P08243 | Asparagine synthetase [glutamine-hydroly | ASNS | 64.37 | 23.5 | 10 | 10 | 167.56171 |
| P08559 | Pyruvate dehydrogenase E1 component su | PDHA1 | 43.296 | 19.5 | 7 | 6 | 215.46101 |
| P08572 | Collagen alpha-2(IV) chain OS=Homo sa | COL4A2 | 167.554 | 2.2 | 3 | 3 | 80.661049 |
| P08574 | Cytochrome c1, heme protein, mitochondı | CYC1 | 35.422 | 21.2 | 7 | 7 | 705.65869 |
| P08579 | U2 small nuclear ribonucleoprotein B'' OS | SNRPB2 | 25.487 | 23.1 | 5 | 4 | 706.34351 |
| P08621 | U1 small nuclear ribonucleoprotein 70 kD | SNRNP70 | 51.557 | 29.1 | 15 | 15 | 5051.1362 |
| P08651 | Nuclear factor 1 C-type OS=Homo sapien | NFIC | 55.675 | 36.6 | 15 | 15 | 1641.3596 |
| P08670 | Vimentin OS=Homo sapiens OX=9606 G | VIM | 53.652 | 53.9 | 25 | 21 | 4799.5474 |
| P08708 | Small ribosomal subunit protein eS17 OS= | RPS17 | 15.551 | 57.8 | 10 | 10 | 13628.079 |
| P08729 | Keratin, type II cytoskeletal 7 OS=Homo | KRT7 | 51.386 | 46.9 | 22 | 17 | 2016.174 |
| P08779 | Keratin, type I cytoskeletal 16 OS=Homo | KRT16 | 51.268 | 5.7 | 2 | 2 | 46.736332 |
| P09012 | U1 small nuclear ribonucleoprotein A OS= | SNRPA | 31.28 | 24.8 | 7 | 7 | 1736.8352 |
| P09038 | Fibroblast growth factor 2 OS=Homo sapi | FGF2 | 30.77 | 13.5 | 4 | 4 | 213.10463 |
| P09132 | Signal recognition particle 19 kDa protein | SRP19 | 16.156 | 18.1 | 2 | 2 | 147.2774 |
| P09234 | U1 small nuclear ribonucleoprotein C OS= | SNRPC | 17.394 | 7.5 | 1 | 1 | 1063.0861 |
| P09382 | Galectin-1 OS=Homo sapiens OX=9606 ( | LGALS1 | 14.716 | 46.7 | 6 | 6 | 1543.483 |
| P09429 | High mobility group protein B1 OS=Hom | HMGB1 | 24.894 | 36.3 | 7 | 2 | 206.95856 |
| P09493 | Tropomyosin alpha-1 chain OS=Homo sa | TPM1 | 32.709 | 9.9 | 3 | 3 | 653.16858 |
| P09496 | Clathrin light chain A OS=Homo sapiens | CLTA | 27.077 | 9.7 | 3 | 3 | 289.10886 |
| P09497 | Clathrin light chain B OS=Homo sapiens ( | CLTB | 25.191 | 11.8 | 3 | 3 | 232.31592 |
| P09543 | 2',3'-cyclic-nucleotide 3'-phosphodiesteras | CNP | 47.579 | 29.2 | 12 | 12 | 407.1933 |
| P09601 | Heme oxygenase 1 OS=Homo sapiens OX | HMOX1 | 32.819 | 8.3 | 2 | 2 | 81.493294 |
| P09629 | Homeobox protein Hox-B7 OS=Homo sa | HOXB7 | 24.015 | 10.1 | 2 | 2 | 121.60526 |
| P09651 | Heterogeneous nuclear ribonucleoprotein | HNRNPA1 | 38.747 | 54.3 | 18 | 2 | 33608.66 |
| P09661 | U2 small nuclear ribonucleoprotein A' OS | SNRPA1 | 28.416 | 29.4 | 7 | 7 | 1710.1064 |
| P09669 | Cytochrome c oxidase subunit 6C OS=Ho | COX6C | 8.782 | 33.3 | 4 | 4 | 112.58765 |
| P09874 | Poly [ADP-ribose] polymerase 1 OS=Hor | PARP1 | 113.084 | 36.2 | 33 | 33 | 2277.4238 |
| P09884 | DNA polymerase alpha catalytic subunit ( | POLA1 | 165.913 | 4.5 | 6 | 6 | 195.00795 |
| P09913 | Interferon-induced protein with tetratricop | IFIT2 | 54.633 | 4 | 2 | 2 | 30.965153 |
| P09914 | Interferon-induced protein with tetratricop | IFIT1 | 55.361 | 7.5 | 3 | 3 | 93.724861 |
| P0C0S5 | Histone H2A.Z OS=Homo sapiens OX=9 | H2AZ1 | 13.553 | 13.3 | 2 | 1 | 32036.408 |
| P0C1Z6 | TCF3 fusion partner OS=Homo sapiens C | TFPT | 28.279 | 5.9 | 1 | 1 | 85.29229 |
| P0C2W1 | F-box/SPRY domain-containing protein 1 | FBXO45 | 30.633 | 10.5 | 3 | 3 | 71.783257 |
| P0CW18 | Serine protease 56 OS=Homo sapiens OX | PRSS56 | 64.598 | 22.7 | 13 | 13 | 513.91058 |
| P0DI83 | Ras-related protein Rab-34, isoform NAR | RAB34 | 21.119 | 17.2 | 2 | 2 | 92.71714 |

| | | | | | | | |
|---|---|---|---|---|---|---|---|
| P0DMV8 | Heat shock 70 kDa protein 1A OS=Homo | HSPA1A | 70.053 | 37.4 | 21 | 11 | 12054.264 |
| P0DP23 | Calmodulin-1 OS=Homo sapiens OX=96( | CALM1 | 16.838 | 24.2 | 4 | 4 | 1221.1339 |
| P0DP91 | Chimeric ERCC6-PGBD3 protein OS=Hc | ERCC6 | 119.487 | 21.1 | 20 | 0 | 613.96228 |
| P0DPB6 | DNA-directed RNA polymerases I and III | POLR1D | 15.238 | 18 | 2 | 2 | 174.4043 |
| P10074 | Telomere zinc finger-associated protein O | ZBTB48 | 77.054 | 4.7 | 2 | 2 | 55.365646 |
| P10114 | Ras-related protein Rap-2a OS=Homo sap | RAP2A | 20.616 | 11.5 | 2 | 2 | 48.195629 |
| P10124 | Serglycin OS=Homo sapiens OX=9606 G | SRGN | 17.652 | 8.2 | 1 | 1 | 82.467056 |
| P10155 | RNA-binding protein RO60 OS=Homo sa | RO60 | 60.671 | 15.2 | 7 | 7 | 237.95691 |
| P10321 | HLA class I histocompatibility antigen, C | HLA-C | 40.649 | 3.3 | 1 | 1 | 221.47856 |
| P10398 | Serine/threonine-protein kinase A-Raf OS | ARAF | 67.586 | 23.9 | 13 | 7 | 209.36362 |
| P10412 | Histone H1.4 OS=Homo sapiens OX=960 | H1-4 | 21.866 | 22.8 | 7 | 1 | 19145.783 |
| P10515 | Dihydrolipoyllysine-residue acetyltransfer | DLAT | 68.997 | 12.5 | 7 | 7 | 85.031433 |
| P10589 | COUP transcription factor 1 OS=Homo sa | NR2F1 | 46.156 | 6.6 | 3 | 1 | 109.69504 |
| P10599 | Thioredoxin OS=Homo sapiens OX=960( | TXN | 11.738 | 42.9 | 5 | 5 | 1635.1112 |
| P10620 | Microsomal glutathione S-transferase 1 O | MGST1 | 17.599 | 16.1 | 2 | 2 | 186.45874 |
| P10809 | 60 kDa heat shock protein, mitochondrial | HSPD1 | 61.055 | 37.7 | 17 | 17 | 946.98767 |
| P10909 | Clusterin OS=Homo sapiens OX=9606 G | CLU | 52.495 | 5.1 | 1 | 1 | 13.552837 |
| P11021 | Endoplasmic reticulum chaperone BiP OS | HSPA5 | 72.333 | 37.9 | 23 | 23 | 3244.5837 |
| P11047 | Laminin subunit gamma-1 OS=Homo sap | LAMC1 | 177.603 | 4.6 | 6 | 6 | 133.85225 |
| P11142 | Heat shock cognate 71 kDa protein OS=H | HSPA8 | 70.899 | 41.5 | 22 | 17 | 14293.025 |
| P11166 | Solute carrier family 2, facilitated glucose | SLC2A1 | 54.084 | 1.8 | 1 | 1 | 32.902245 |
| P11171 | Protein 4.1 OS=Homo sapiens OX=9606 | EPB41 | 97.017 | 4.9 | 3 | 2 | 39.582546 |
| P11172 | Uridine 5'-monophosphate synthase OS=H | UMPS | 52.222 | 9.8 | 4 | 4 | 67.118179 |
| P11177 | Pyruvate dehydrogenase E1 component su | PDHB | 39.234 | 6.7 | 2 | 2 | 78.621826 |
| P11182 | Lipoamide acyltransferase component of b | DBT | 53.488 | 10.4 | 5 | 5 | 103.72279 |
| P11233 | Ras-related protein Ral-A OS=Homo sapi | RALA | 23.567 | 10.2 | 2 | 1 | 46.968346 |
| P11274 | Breakpoint cluster region protein OS=Ho | BCR | 142.82 | 7.5 | 8 | 7 | 144.38144 |
| P11308 | Transcriptional regulator ERG OS=Homo | ERG | 53.838 | 7.1 | 2 | 2 | 67.898651 |
| P11310 | Medium-chain specific acyl-CoA dehydro | ACADM | 46.589 | 11.4 | 3 | 3 | 130.79568 |
| P11387 | DNA topoisomerase 1 OS=Homo sapiens | TOP1 | 90.726 | 36.7 | 31 | 26 | 2178.5078 |
| P11388 | DNA topoisomerase 2-alpha OS=Homo sa | TOP2A | 174.386 | 23.4 | 37 | 37 | 1819.3597 |
| P11413 | Glucose-6-phosphate 1-dehydrogenase OS | G6PD | 59.257 | 15.5 | 8 | 8 | 161.44363 |
| P11441 | Ubiquitin-like protein 4A OS=Homo sapi | UBL4A | 17.777 | 11.5 | 2 | 2 | 70.108566 |
| P11532 | Dystrophin OS=Homo sapiens OX=9606 | DMD | 426.75 | 3.5 | 12 | 10 | 650.55688 |
| P11586 | C-1-tetrahydrofolate synthase, cytoplasmi | MTHFD1 | 101.532 | 8.9 | 7 | 7 | 95.618729 |

| | | | | | | | |
|---|---|---|---|---|---|---|---|
| P11717 | Cation-independent mannose-6-phosphate | IGF2R | 274.375 | 7.7 | 17 | 17 | 102.43893 |
| P11802 | Cyclin-dependent kinase 4 OS=Homo sap | CDK4 | 33.73 | 20.5 | 5 | 5 | 126.57913 |
| P11908 | Ribose-phosphate pyrophosphokinase 2 C | PRPS2 | 34.77 | 7.9 | 2 | 2 | 53.315884 |
| P11940 | Polyadenylate-binding protein 1 OS=Hon | PABPC1 | 70.671 | 42.5 | 30 | 9 | 93174.906 |
| P12004 | Proliferating cell nuclear antigen OS=Hor | PCNA | 28.769 | 23.8 | 6 | 6 | 212.74901 |
| P12235 | ADP/ATP translocase 1 OS=Homo sapier | SLC25A4 | 33.065 | 21.1 | 6 | 4 | 894.16522 |
| P12236 | ADP/ATP translocase 3 OS=Homo sapier | SLC25A6 | 32.867 | 10.4 | 4 | 4 | 193.53969 |
| P12268 | Inosine-5'-monophosphate dehydrogenase | IMPDH2 | 55.805 | 35.4 | 15 | 14 | 665.79968 |
| P12270 | Nucleoprotein TPR OS=Homo sapiens O | TPR | 267.293 | 5.7 | 12 | 12 | 94.396515 |
| P12277 | Creatine kinase B-type OS=Homo sapiens | CKB | 42.645 | 9.4 | 3 | 3 | 113.10501 |
| P12532 | Creatine kinase U-type, mitochondrial OS | CKMT1A | 47.037 | 12.7 | 5 | 4 | 410.30496 |
| P12814 | Alpha-actinin-1 OS=Homo sapiens OX=9 | ACTN1 | 103.058 | 19.1 | 15 | 5 | 268.74573 |
| P12882 | Myosin-1 OS=Homo sapiens OX=9606 G | MYH1 | 223.146 | 2.6 | 4 | 1 | 153.90625 |
| P12956 | X-ray repair cross-complementing protein | XRCC6 | 69.844 | 42 | 26 | 26 | 13746.142 |
| P13010 | X-ray repair cross-complementing protein | XRCC5 | 82.705 | 21 | 19 | 19 | 10877.92 |
| P13073 | Cytochrome c oxidase subunit 4 isoform 1 | COX4I1 | 19.577 | 19.5 | 3 | 3 | 107.09291 |
| P13196 | 5-aminolevulinate synthase, non-specific, | ALAS1 | 70.581 | 7.2 | 4 | 4 | 61.441128 |
| P13378 | Homeobox protein Hox-D8 OS=Homo sa | HOXD8 | 31.911 | 3.4 | 1 | 0 | 125.03669 |
| P13489 | Ribonuclease inhibitor OS=Homo sapiens | RNH1 | 49.974 | 18.7 | 7 | 7 | 137.26704 |
| P13497 | Bone morphogenetic protein 1 OS=Homo | BMP1 | 111.249 | 4.3 | 4 | 4 | 38.616673 |
| P13498 | Cytochrome b-245 light chain OS=Homo | CYBA | 21.013 | 5.1 | 1 | 1 | 106.38649 |
| P13639 | Elongation factor 2 OS=Homo sapiens O | EEF2 | 95.339 | 31 | 27 | 27 | 1164.1759 |
| P13645 | Keratin, type I cytoskeletal 10 OS=Homo | KRT10 | 58.828 | 21.9 | 14 | 12 | 1446.4189 |
| P13647 | Keratin, type II cytoskeletal 5 OS=Homo | KRT5 | 62.379 | 13.9 | 7 | 7 | 123.27458 |
| P13674 | Prolyl 4-hydroxylase subunit alpha-1 OS= | P4HA1 | 61.05 | 21.3 | 9 | 9 | 179.46519 |
| P13682 | Zinc finger protein 35 OS=Homo sapiens | ZNF35 | 59.09 | 3.8 | 2 | 2 | 27.306412 |
| P13716 | Delta-aminolevulinic acid dehydratase OS | ALAD | 36.295 | 9.1 | 3 | 3 | 56.269592 |
| P13796 | Plastin-2 OS=Homo sapiens OX=9606 G | LCP1 | 70.289 | 1.3 | 1 | 1 | 17.623938 |
| P13797 | Plastin-3 OS=Homo sapiens OX=9606 G | PLS3 | 70.812 | 8.6 | 5 | 2 | 87.386627 |
| P13807 | Glycogen [starch] synthase, muscle OS=H | GYS1 | 83.786 | 17.9 | 12 | 11 | 932.14624 |
| P13861 | cAMP-dependent protein kinase type II-al | PRKAR2A | 45.519 | 31.2 | 11 | 9 | 979.46497 |
| P13984 | General transcription factor IIF subunit 2 | GTF2F2 | 28.381 | 4.4 | 1 | 1 | 23.622341 |
| P13987 | CD59 glycoprotein OS=Homo sapiens O | CD59 | 14.178 | 7.8 | 1 | 1 | 26.828384 |
| P13994 | Probable splicing factor YJU2B OS=Hon | YJU2B | 44.803 | 12.9 | 5 | 5 | 60.428982 |
| P13995 | Bifunctional methylenetetrahydrofolate de | MTHFD2 | 37.896 | 7.7 | 2 | 2 | 135.82262 |

| | | | | | | | |
|---|---|---|---|---|---|---|---|
| P14210 | Hepatocyte growth factor OS=Homo sapi | HGF | 83.134 | 1.1 | 1 | 1 | 70.239464 |
| P14324 | Farnesyl pyrophosphate synthase OS=Hoı | FDPS | 48.276 | 7.2 | 3 | 3 | 51.842743 |
| P14373 | Zinc finger protein RFP OS=Homo sapier | TRIM27 | 58.49 | 8.8 | 4 | 3 | 76.871849 |
| P14406 | Cytochrome c oxidase subunit 7A2, mitoc | COX7A2 | 9.396 | 15.7 | 1 | 1 | 225.09439 |
| P14543 | Nidogen-1 OS=Homo sapiens OX=9606 ( | NID1 | 136.378 | 2.8 | 3 | 2 | 147.78915 |
| P14618 | Pyruvate kinase PKM OS=Homo sapiens | PKM | 57.937 | 42.4 | 18 | 18 | 476.77835 |
| P14625 | Endoplasmin OS=Homo sapiens OX=960 | HSP90B1 | 92.469 | 11 | 8 | 8 | 146.91187 |
| P14635 | G2/mitotic-specific cyclin-B1 OS=Homo | CCNB1 | 48.338 | 14.1 | 6 | 5 | 220.10831 |
| P14649 | Myosin light chain 6B OS=Homo sapiens | MYL6B | 22.764 | 33.2 | 7 | 5 | 5547.3125 |
| P14678 | Small nuclear ribonucleoprotein-associate | SNRPB | 24.611 | 24.6 | 6 | 1 | 1483.1577 |
| P14735 | Insulin-degrading enzyme OS=Homo sapi | IDE | 117.969 | 3.9 | 4 | 4 | 122.75185 |
| P14854 | Cytochrome c oxidase subunit 6B1 OS=H | COX6B1 | 10.193 | 14 | 1 | 1 | 99.343224 |
| P14866 | Heterogeneous nuclear ribonucleoprotein | HNRNPL | 64.133 | 41.3 | 18 | 18 | 10853.634 |
| P14868 | Aspartate--tRNA ligase, cytoplasmic OS= | DARS1 | 57.137 | 34.1 | 17 | 17 | 238.61536 |
| P14921 | Protein C-ets-1 OS=Homo sapiens OX=9( | ETS1 | 50.408 | 4.5 | 2 | 2 | 108.25716 |
| P14923 | Junction plakoglobin OS=Homo sapiens ( | JUP | 81.745 | 11.9 | 6 | 6 | 80.8144 |
| P15121 | Aldo-keto reductase family 1 member B1 | AKR1B1 | 35.854 | 3.2 | 1 | 1 | 25.259211 |
| P15170 | Eukaryotic peptide chain release factor G | GSPT1 | 55.756 | 20.4 | 10 | 2 | 678.27264 |
| P15291 | Beta-1,4-galactosyltransferase 1 OS=Hon | B4GALT1 | 43.921 | 2.8 | 1 | 1 | 45.943382 |
| P15311 | Ezrin OS=Homo sapiens OX=9606 GN=I | EZR | 69.413 | 9.9 | 5 | 5 | 87.587593 |
| P15408 | Fos-related antigen 2 OS=Homo sapiens ( | FOSL2 | 35.194 | 7.4 | 2 | 2 | 83.111748 |
| P15531 | Nucleoside diphosphate kinase A OS=Hoı | NME1 | 17.149 | 15.8 | 2 | 2 | 84.433762 |
| P15559 | NAD(P)H dehydrogenase [quinone] 1 OS | NQO1 | 30.868 | 4.7 | 1 | 1 | 256.41034 |
| P15622 | Zinc finger protein 250 OS=Homo sapien: | ZNF250 | 63.468 | 10 | 6 | 6 | 86.589874 |
| P15880 | Small ribosomal subunit protein uS5 OS= | RPS2 | 31.325 | 34.5 | 10 | 10 | 32088.123 |
| P15924 | Desmoplakin OS=Homo sapiens OX=960 | DSP | 331.774 | 36.2 | 112 | 112 | 2452.5974 |
| P15927 | Replication protein A 32 kDa subunit OS= | RPA2 | 29.247 | 10.4 | 3 | 3 | 304.78125 |
| P16035 | Metalloproteinase inhibitor 2 OS=Homo s | TIMP2 | 24.4 | 6.4 | 1 | 1 | 64.217613 |
| P16070 | CD44 antigen OS=Homo sapiens OX=96( | CD44 | 81.538 | 5.4 | 4 | 4 | 238.61237 |
| P16220 | Cyclic AMP-responsive element-binding ] | CREB1 | 35.137 | 7 | 2 | 1 | 194.1266 |
| P16383 | Intron Large complex component GCFC2 | GCFC2 | 89.385 | 6.4 | 5 | 5 | 45.852638 |
| P16401 | Histone H1.5 OS=Homo sapiens OX=960 | H1-5 | 22.581 | 22.1 | 6 | 6 | 6933.4321 |
| P16403 | Histone H1.2 OS=Homo sapiens OX=960 | H1-2 | 21.365 | 11.7 | 2 | 2 | 2508.2405 |
| P16435 | NADPH--cytochrome P450 reductase OS | POR | 76.69 | 3.4 | 2 | 2 | 49.030476 |
| P16615 | Sarcoplasmic/endoplasmic reticulum calci | ATP2A2 | 114.757 | 18.4 | 15 | 9 | 247.91585 |

| | | | | | | | |
|---|---|---|---|---|---|---|---|
| P16989 | Y-box-binding protein 3 OS=Homo sapie | YBX3 | 40.09 | 22.6 | 12 | 9 | 12537.684 |
| P17020 | Zinc finger protein 16 OS=Homo sapiens | ZNF16 | 76.472 | 17.3 | 11 | 9 | 283.57697 |
| P17028 | Zinc finger protein 24 OS=Homo sapiens | ZNF24 | 42.156 | 25.3 | 9 | 9 | 442.00775 |
| P17036 | Zinc finger protein 3 OS=Homo sapiens C | ZNF3 | 50.916 | 5.8 | 3 | 3 | 193.93105 |
| P17081 | Rho-related GTP-binding protein RhoQ C | RHOQ | 22.66 | 4.4 | 1 | 1 | 41.656784 |
| P17096 | High mobility group protein HMG-I/HMC | HMGA1 | 11.677 | 23.4 | 3 | 3 | 1858.5536 |
| P17152 | Transmembrane protein 11, mitochondrial | TMEM11 | 21.541 | 8.3 | 1 | 1 | 103.5674 |
| P17275 | Transcription factor JunB OS=Homo sapi | JUNB | 35.88 | 16.7 | 5 | 5 | 392.14078 |
| P17480 | Nucleolar transcription factor 1 OS=Hom | UBTF | 89.406 | 25.5 | 20 | 20 | 1070.2356 |
| P17509 | Homeobox protein Hox-B6 OS=Homo sa | HOXB6 | 25.433 | 6.2 | 1 | 1 | 69.011528 |
| P17544 | Cyclic AMP-dependent transcription facto | ATF7 | 51.757 | 5.8 | 2 | 2 | 72.864731 |
| P17612 | cAMP-dependent protein kinase catalytic | PRKACA | 40.59 | 20.8 | 8 | 7 | 414.52573 |
| P17655 | Calpain-2 catalytic subunit OS=Homo sap | CAPN2 | 79.996 | 9.1 | 6 | 6 | 109.47121 |
| P17676 | CCAAT/enhancer-binding protein beta OS | CEBPB | 36.106 | 22.6 | 6 | 6 | 388.72766 |
| P17706 | Tyrosine-protein phosphatase non-recepto | PTPN2 | 48.474 | 8.7 | 3 | 3 | 113.97708 |
| P17812 | CTP synthase 1 OS=Homo sapiens OX=9 | CTPS1 | 66.691 | 21.3 | 13 | 12 | 359.67929 |
| P17844 | Probable ATP-dependent RNA helicase D | DDX5 | 69.149 | 52.3 | 38 | 29 | 49783.91 |
| P17858 | ATP-dependent 6-phosphofructokinase, li | PFKL | 85.019 | 15.8 | 11 | 11 | 413.27136 |
| P17931 | Galectin-3 OS=Homo sapiens OX=9606 ( | LGALS3 | 26.153 | 22.8 | 5 | 5 | 822.24146 |
| P17936 | Insulin-like growth factor-binding protein | IGFBP3 | 31.675 | 8.9 | 3 | 3 | 512.76636 |
| P17980 | 26S proteasome regulatory subunit 6A OS | PSMC3 | 49.204 | 16.2 | 6 | 6 | 107.6725 |
| P17987 | T-complex protein 1 subunit alpha OS=Ho | TCP1 | 60.344 | 37.2 | 19 | 19 | 353.90701 |
| P18031 | Tyrosine-protein phosphatase non-recepto | PTPN1 | 49.967 | 25.3 | 9 | 9 | 220.73701 |
| P18074 | General transcription and DNA repair fact | ERCC2 | 86.91 | 15.9 | 11 | 11 | 148.73256 |
| P18077 | Large ribosomal subunit protein eL33 OS | RPL35A | 12.538 | 36.4 | 5 | 5 | 5617.9126 |
| P18085 | ADP-ribosylation factor 4 OS=Homo sapi | ARF4 | 20.511 | 31.7 | 5 | 5 | 259.30667 |
| P18124 | Large ribosomal subunit protein uL30 OS | RPL7 | 29.226 | 50.4 | 16 | 16 | 15442.168 |
| P18583 | Protein SON OS=Homo sapiens OX=960 | SON | 263.83 | 9.3 | 17 | 17 | 289.85965 |
| P18615 | Negative elongation factor E OS=Homo s | NELFE | 43.24 | 21.8 | 7 | 7 | 1131.1381 |
| P18621 | Large ribosomal subunit protein uL22 OS | RPL17 | 21.398 | 27.7 | 5 | 5 | 23762.465 |
| P18754 | Regulator of chromosome condensation C | RCC1 | 44.97 | 19.5 | 5 | 5 | 118.05608 |
| P18850 | Cyclic AMP-dependent transcription facto | ATF6 | 74.585 | 9.9 | 6 | 6 | 157.42535 |
| P18887 | DNA repair protein XRCC1 OS=Homo sa | XRCC1 | 69.498 | 21.8 | 13 | 13 | 1117.1071 |
| P19338 | Nucleolin OS=Homo sapiens OX=9606 C | NCL | 76.615 | 28.3 | 24 | 24 | 20311.473 |
| P19367 | Hexokinase-1 OS=Homo sapiens OX=96( | HK1 | 102.486 | 1.3 | 1 | 1 | 56.894119 |

| | | | | | | | |
|---|---|---|---|---|---|---|---|
| P19387 | DNA-directed RNA polymerase II subuni | POLR2C | 31.442 | 12.4 | 3 | 3 | 84.941246 |
| P19388 | DNA-directed RNA polymerases I, II, anc | POLR2E | 24.552 | 28.1 | 6 | 6 | 217.34723 |
| P19404 | NADH dehydrogenase [ubiquinone] flavo | NDUFV2 | 27.392 | 5.2 | 1 | 1 | 54.317524 |
| P19447 | General transcription and DNA repair fact | ERCC3 | 89.278 | 33.4 | 20 | 20 | 395.48685 |
| P19474 | E3 ubiquitin-protein ligase TRIM21 OS=I | TRIM21 | 54.17 | 50.1 | 23 | 23 | 15129.873 |
| P19484 | Transcription factor EB OS=Homo sapier | TFEB | 52.865 | 6.5 | 2 | 2 | 43.958115 |
| P19525 | Interferon-induced, double-stranded RNA | EIF2AK2 | 62.095 | 30.5 | 19 | 19 | 4390.0273 |
| P19532 | Transcription factor E3 OS=Homo sapien | TFE3 | 61.522 | 13.7 | 8 | 6 | 305.39554 |
| P19623 | Spermidine synthase OS=Homo sapiens C | SRM | 33.825 | 7 | 2 | 2 | 81.761246 |
| P19784 | Casein kinase II subunit alpha' OS=Homo | CSNK2A2 | 41.214 | 19.1 | 6 | 6 | 213.45834 |
| P19823 | Inter-alpha-trypsin inhibitor heavy chain I | ITIH2 | 106.464 | 1.2 | 1 | 1 | 57.00037 |
| P20020 | Plasma membrane calcium-transporting A | ATP2B1 | 134.685 | 2 | 2 | 1 | 49.530945 |
| P20042 | Eukaryotic translation initiation factor 2 s | EIF2S2 | 38.389 | 41.4 | 19 | 19 | 3295.8254 |
| P20073 | Annexin A7 OS=Homo sapiens OX=9606 | ANXA7 | 52.74 | 18 | 8 | 8 | 131.50061 |
| P20226 | TATA-box-binding protein OS=Homo sa | TBP | 37.699 | 3.2 | 1 | 1 | 92.214691 |
| P20248 | Cyclin-A2 OS=Homo sapiens OX=9606 ( | CCNA2 | 48.551 | 10.2 | 3 | 3 | 49.490906 |
| P20290 | Transcription factor BTF3 OS=Homo sap | BTF3 | 22.168 | 39.8 | 5 | 5 | 171.40453 |
| P20339 | Ras-related protein Rab-5A OS=Homo sa | RAB5A | 23.659 | 16.3 | 3 | 1 | 61.787468 |
| P20340 | Ras-related protein Rab-6A OS=Homo sa | RAB6A | 23.593 | 12.5 | 2 | 1 | 37.982376 |
| P20585 | DNA mismatch repair protein Msh3 OS=I | MSH3 | 127.412 | 17.7 | 18 | 18 | 351.23413 |
| P20618 | Proteasome subunit beta type-1 OS=Hom | PSMB1 | 26.49 | 10 | 2 | 2 | 121.67102 |
| P20674 | Cytochrome c oxidase subunit 5A, mitoch | COX5A | 16.763 | 6 | 1 | 1 | 123.26364 |
| P20700 | Lamin-B1 OS=Homo sapiens OX=9606 ( | LMNB1 | 66.409 | 23 | 13 | 13 | 279.0943 |
| P20719 | Homeobox protein Hox-A5 OS=Homo sa | HOXA5 | 29.346 | 35.2 | 7 | 5 | 257.03058 |
| P20742 | Pregnancy zone protein OS=Homo sapien | PZP | 163.863 | 0.7 | 1 | 1 | 114.8447 |
| P20749 | B-cell lymphoma 3 protein OS=Homo sap | BCL3 | 47.584 | 12.8 | 5 | 5 | 81.919441 |
| P20839 | Inosine-5'-monophosphate dehydrogenase | IMPDH1 | 55.406 | 12.6 | 5 | 5 | 159.12003 |
| P20908 | Collagen alpha-1(V) chain OS=Homo sap | COL5A1 | 183.56 | 2.3 | 4 | 4 | 204.47888 |
| P21127 | Cyclin-dependent kinase 11B OS=Homo | CDK11B | 92.62 | 14.2 | 11 | 11 | 143.28191 |
| P21266 | Glutathione S-transferase Mu 3 OS=Hom | GSTM3 | 26.56 | 11.1 | 2 | 2 | 77.13472 |
| P21291 | Cysteine and glycine-rich protein 1 OS=H | CSRP1 | 20.568 | 35.2 | 6 | 6 | 1027.9768 |
| P21333 | Filamin-A OS=Homo sapiens OX=9606 ( | FLNA | 280.739 | 14.1 | 30 | 30 | 495.93179 |
| P21359 | Neurofibromin OS=Homo sapiens OX=9( | NF1 | 319.373 | 1.5 | 4 | 4 | 37.547344 |
| P21399 | Cytoplasmic aconitate hydratase OS=Hon | ACO1 | 98.399 | 8.3 | 7 | 7 | 140.9727 |
| P21580 | Tumor necrosis factor alpha-induced prot | TNFAIP3 | 89.614 | 27.7 | 20 | 20 | 248.35558 |

| | | | | | | | |
|---|---|---|---|---|---|---|---|
| P21675 | Transcription initiation factor TFIID subu | TAF1 | 214.714 | 0.6 | 1 | 1 | 17.75909 |
| P21796 | Voltage-dependent anion-selective channe | VDAC1 | 30.773 | 16.3 | 4 | 4 | 93.363205 |
| P21912 | Succinate dehydrogenase [ubiquinone] irc | SDHB | 31.63 | 15.7 | 4 | 4 | 149.79283 |
| P21964 | Catechol O-methyltransferase OS=Homo | COMT | 30.038 | 5.9 | 1 | 1 | 68.755692 |
| P21980 | Protein-glutamine gamma-glutamyltransfe | TGM2 | 77.329 | 16.4 | 10 | 8 | 331.53229 |
| P22061 | Protein-L-isoaspartate(D-aspartate) O-me | PCMT1 | 24.637 | 39.2 | 10 | 10 | 2452.7976 |
| P22087 | rRNA 2'-O-methyltransferase fibrillarin O | FBL | 33.785 | 38.9 | 11 | 9 | 2347.2166 |
| P22102 | Trifunctional purine biosynthetic protein a | GART | 107.768 | 9 | 8 | 8 | 130.32004 |
| P22234 | Bifunctional phosphoribosylaminoimidazo | PAICS | 47.08 | 5.9 | 2 | 2 | 83.77713 |
| P22314 | Ubiquitin-like modifier-activating enzyme | UBA1 | 117.849 | 6.1 | 5 | 5 | 71.06179 |
| P22392 | Nucleoside diphosphate kinase B OS=Ho | NME2 | 17.299 | 38.2 | 5 | 0 | 167.40457 |
| P22415 | Upstream stimulatory factor 1 OS=Homo | USF1 | 33.539 | 3.2 | 1 | 1 | 44.685394 |
| P22626 | Heterogeneous nuclear ribonucleoproteins | HNRNPA2I | 37.43 | 53.8 | 17 | 17 | 44988.754 |
| P22695 | Cytochrome b-c1 complex subunit 2, mitc | UQCRC2 | 48.443 | 18.8 | 6 | 6 | 163.88593 |
| P22830 | Ferrochelatase, mitochondrial OS=Homo | FECH | 47.863 | 5.7 | 2 | 2 | 80.337204 |
| P23025 | DNA repair protein complementing XP-A | XPA | 31.368 | 5.5 | 1 | 1 | 169.59409 |
| P23142 | Fibulin-1 OS=Homo sapiens OX=9606 G | FBLN1 | 77.215 | 13.4 | 7 | 7 | 274.84235 |
| P23193 | Transcription elongation factor A protein | TCEA1 | 33.97 | 18.6 | 5 | 3 | 99.863602 |
| P23246 | Splicing factor, proline- and glutamine-ric | SFPQ | 76.15 | 42.4 | 30 | 30 | 38266.578 |
| P23258 | Tubulin gamma-1 chain OS=Homo sapier | TUBG1 | 51.17 | 24.2 | 12 | 3 | 1663.6588 |
| P23284 | Peptidyl-prolyl cis-trans isomerase B OS= | PPIB | 23.743 | 31.5 | 6 | 6 | 641.40088 |
| P23368 | NAD-dependent malic enzyme, mitochon | ME2 | 65.444 | 6.3 | 3 | 3 | 73.913383 |
| P23381 | Tryptophan--tRNA ligase, cytoplasmic OS | WARS1 | 53.166 | 2.3 | 1 | 1 | 18.005459 |
| P23396 | Small ribosomal subunit protein uS3 OS= | RPS3 | 26.689 | 79 | 19 | 19 | 45118.746 |
| P23458 | Tyrosine-protein kinase JAK1 OS=Homo | JAK1 | 133.278 | 28.6 | 30 | 29 | 349.46771 |
| P23497 | Nuclear autoantigen Sp-100 OS=Homo sa | SP100 | 100.417 | 4.7 | 4 | 3 | 136.82607 |
| P23526 | Adenosylhomocysteinase OS=Homo sapi | AHCY | 47.717 | 10 | 4 | 4 | 72.487305 |
| P23528 | Cofilin-1 OS=Homo sapiens OX=9606 G | CFL1 | 18.503 | 44 | 8 | 7 | 1218.5891 |
| P23588 | Eukaryotic translation initiation factor 4B | EIF4B | 69.152 | 23.9 | 15 | 15 | 864.05945 |
| P23769 | Endothelial transcription factor GATA-2 ( | GATA2 | 50.5 | 6.9 | 3 | 2 | 217.55962 |
| P23919 | Thymidylate kinase OS=Homo sapiens O | DTYMK | 23.82 | 10.4 | 2 | 2 | 83.274094 |
| P23921 | Ribonucleoside-diphosphate reductase lar | RRM1 | 90.071 | 16 | 12 | 12 | 219.06294 |
| P24390 | ER lumen protein-retaining receptor 1 OS | KDELR1 | 24.543 | 5.2 | 1 | 1 | 31.808434 |
| P24534 | Elongation factor 1-beta OS=Homo sapie | EEF1B2 | 24.764 | 24 | 5 | 5 | 16398.367 |
| P24539 | ATP synthase F(0) complex subunit B1, n | ATP5PB | 28.909 | 34.8 | 8 | 8 | 465.37589 |

| | | | | | | | |
|---|---|---|---|---|---|---|---|
| P24593 | Insulin-like growth factor-binding protein | IGFBP5 | 30.571 | 3.7 | 1 | 1 | 104.49757 |
| P24666 | Low molecular weight phosphotyrosine pr | ACP1 | 18.043 | 34.2 | 5 | 5 | 761.24451 |
| P24752 | Acetyl-CoA acetyltransferase, mitochondr | ACAT1 | 45.2 | 3 | 1 | 1 | 82.661804 |
| P24844 | Myosin regulatory light polypeptide 9 OS | MYL9 | 19.828 | 6.4 | 1 | 1 | 38.418186 |
| P24863 | Cyclin-C OS=Homo sapiens OX=9606 G | CCNC | 33.243 | 7.4 | 2 | 2 | 101.02803 |
| P24928 | DNA-directed RNA polymerase II subuni | POLR2A | 217.177 | 18.2 | 32 | 32 | 345.36649 |
| P24941 | Cyclin-dependent kinase 2 OS=Homo sap | CDK2 | 33.93 | 25.8 | 7 | 5 | 372.70258 |
| P25054 | Adenomatous polyposis coli protein OS=I | APC | 311.646 | 10.8 | 26 | 25 | 204.8381 |
| P25205 | DNA replication licensing factor MCM3 ( | MCM3 | 90.981 | 33.9 | 26 | 26 | 659.57831 |
| P25325 | 3-mercaptopyruvate sulfurtransferase OS= | MPST | 33.179 | 8.1 | 1 | 1 | 104.01199 |
| P25391 | Laminin subunit alpha-1 OS=Homo sapie | LAMA1 | 337.084 | 12.3 | 32 | 32 | 283.25787 |
| P25398 | Small ribosomal subunit protein eS12 OS | RPS12 | 14.515 | 54.5 | 8 | 8 | 17968.346 |
| P25445 | Tumor necrosis factor receptor superfamil | FAS | 37.733 | 13.4 | 5 | 5 | 165.99965 |
| P25490 | Transcriptional repressor protein YY1 OS | YY1 | 44.713 | 27.5 | 13 | 4 | 835.27844 |
| P25685 | DnaJ homolog subfamily B member 1 OS | DNAJB1 | 38.045 | 17.1 | 5 | 4 | 237.15221 |
| P25705 | ATP synthase subunit alpha, mitochondria | ATP5F1A | 59.751 | 33.1 | 17 | 17 | 675.82062 |
| P25786 | Proteasome subunit alpha type-1 OS=Hon | PSMA1 | 29.556 | 12.2 | 2 | 2 | 45.261818 |
| P25789 | Proteasome subunit alpha type-4 OS=Hon | PSMA4 | 29.484 | 16.5 | 4 | 4 | 161.77715 |
| P25963 | NF-kappa-B inhibitor alpha OS=Homo sa | NFKBIA | 35.609 | 2.5 | 1 | 1 | 118.77834 |
| P26038 | Moesin OS=Homo sapiens OX=9606 GN | MSN | 67.821 | 15.1 | 9 | 9 | 202.14964 |
| P26045 | Tyrosine-protein phosphatase non-recepto | PTPN3 | 103.99 | 12.7 | 11 | 11 | 103.91849 |
| P26196 | Probable ATP-dependent RNA helicase D | DDX6 | 54.417 | 36.6 | 18 | 18 | 16762.041 |
| P26358 | DNA (cytosine-5)-methyltransferase 1 OS | DNMT1 | 183.166 | 22.3 | 33 | 33 | 331.966 |
| P26367 | Paired box protein Pax-6 OS=Homo sapie | PAX6 | 46.684 | 3.1 | 1 | 1 | 39.689548 |
| P26368 | Splicing factor U2AF 65 kDa subunit OS= | U2AF2 | 53.501 | 31.8 | 10 | 10 | 5230.8555 |
| P26373 | Large ribosomal subunit protein eL13 OS | RPL13 | 24.262 | 48.3 | 12 | 12 | 19165.813 |
| P26447 | Protein S100-A4 OS=Homo sapiens OX= | S100A4 | 11.729 | 28.7 | 3 | 3 | 372.20404 |
| P26572 | Alpha-1,3-mannosyl-glycoprotein 2-beta-I | MGAT1 | 50.879 | 8.3 | 3 | 3 | 68.566742 |
| P26583 | High mobility group protein B2 OS=Hom | HMGB2 | 24.034 | 6.2 | 1 | 1 | 84.105812 |
| P26599 | Polypyrimidine tract-binding protein 1 OS | PTBP1 | 59.634 | 20.8 | 8 | 8 | 14845.415 |
| P26639 | Threonine--tRNA ligase 1, cytoplasmic O | TARS1 | 83.436 | 15.2 | 10 | 10 | 372.90277 |
| P26640 | Valine--tRNA ligase OS=Homo sapiens C | VARS1 | 140.477 | 36.6 | 45 | 45 | 2370.4014 |
| P26641 | Elongation factor 1-gamma OS=Homo sa | EEF1G | 50.119 | 45.3 | 21 | 21 | 17204.01 |
| P27105 | Stomatin OS=Homo sapiens OX=9606 G | STOM | 31.731 | 10.1 | 3 | 3 | 70.798943 |
| P27169 | Serum paraoxonase/arylesterase 1 OS=Hc | PON1 | 39.732 | 3.1 | 1 | 1 | 226.63841 |

| | | | | | | | |
|---|---|---|---|---|---|---|---|
| P27348 | 14-3-3 protein theta OS=Homo sapiens O | YWHAQ | 27.765 | 16.7 | 5 | 5 | 1656.3788 |
| P27448 | MAP/microtubule affinity-regulating kina | MARK3 | 84.429 | 30.1 | 19 | 12 | 1372.2053 |
| P27540 | Aryl hydrocarbon receptor nuclear translo | ARNT | 86.637 | 13.4 | 7 | 7 | 96.362778 |
| P27635 | Large ribosomal subunit protein uL16 OS | RPL10 | 24.577 | 33.2 | 8 | 4 | 22421.914 |
| P27694 | Replication protein A 70 kDa DNA-bindi | RPA1 | 68.139 | 24.8 | 16 | 16 | 304.55997 |
| P27695 | DNA-(apurinic or apyrimidinic site) endo | APEX1 | 35.555 | 7.2 | 2 | 2 | 118.16006 |
| P27708 | CAD protein OS=Homo sapiens OX=960 | CAD | 242.984 | 16.5 | 32 | 32 | 258.65408 |
| P27797 | Calreticulin OS=Homo sapiens OX=9606 | CALR | 48.142 | 10.8 | 4 | 4 | 72.882889 |
| P27815 | cAMP-specific 3',5'-cyclic phosphodiester | PDE4A | 98.144 | 1.1 | 1 | 1 | 15.370816 |
| P27816 | Microtubule-associated protein 4 OS=Hon | MAP4 | 121.005 | 33.2 | 33 | 33 | 2497.6895 |
| P27824 | Calnexin OS=Homo sapiens OX=9606 G | CANX | 67.569 | 9 | 5 | 5 | 118.196 |
| P28066 | Proteasome subunit alpha type-5 OS=Hon | PSMA5 | 26.412 | 18.7 | 3 | 3 | 86.165855 |
| P28070 | Proteasome subunit beta type-4 OS=Hom | PSMB4 | 29.205 | 17.4 | 3 | 3 | 151.92323 |
| P28074 | Proteasome subunit beta type-5 OS=Hom | PSMB5 | 28.481 | 3.8 | 1 | 1 | 52.546444 |
| P28288 | ATP-binding cassette sub-family D memb | ABCD3 | 75.476 | 21.5 | 14 | 14 | 687.89777 |
| P28289 | Tropomodulin-1 OS=Homo sapiens OX=! | TMOD1 | 40.57 | 8.9 | 3 | 3 | 61.507896 |
| P28331 | NADH-ubiquinone oxidoreductase 75 kD | NDUFS1 | 79.468 | 11.6 | 7 | 7 | 132.3175 |
| P28340 | DNA polymerase delta catalytic subunit C | POLD1 | 123.632 | 24.6 | 23 | 23 | 337.03799 |
| P28347 | Transcriptional enhancer factor TEF-1 OS | TEAD1 | 47.946 | 5.2 | 2 | 2 | 86.040192 |
| P28370 | Probable global transcription activator SN | SMARCA1 | 122.605 | 11.6 | 12 | 12 | 185.44142 |
| P28482 | Mitogen-activated protein kinase 1 OS=H | MAPK1 | 41.39 | 23.9 | 9 | 5 | 384.97351 |
| P28702 | Retinoic acid receptor RXR-beta OS=Hor | RXRB | 56.922 | 5.6 | 3 | 3 | 81.676422 |
| P28715 | DNA excision repair protein ERCC-5 OS= | ERCC5 | 133.109 | 16.9 | 18 | 18 | 321.79181 |
| P28799 | Progranulin OS=Homo sapiens OX=9606 | GRN | 63.545 | 5.2 | 3 | 3 | 195.18388 |
| P29083 | General transcription factor IIE subunit 1 | GTF2E1 | 49.453 | 8.9 | 3 | 3 | 78.270683 |
| P29084 | Transcription initiation factor IIE subunit | GTF2E2 | 33.044 | 16.2 | 4 | 4 | 188.82491 |
| P29144 | Tripeptidyl-peptidase 2 OS=Homo sapien | TPP2 | 138.35 | 2.9 | 3 | 3 | 29.048635 |
| P29279 | CCN family member 2 OS=Homo sapiens | CCN2 | 38.092 | 2.6 | 1 | 1 | 33.579464 |
| P29353 | SHC-transforming protein 1 OS=Homo sa | SHC1 | 62.823 | 6.5 | 3 | 3 | 87.858795 |
| P29372 | DNA-3-methyladenine glycosylase OS=H | MPG | 32.869 | 54.4 | 12 | 12 | 617.89093 |
| P29375 | Lysine-specific demethylase 5A OS=Hom | KDM5A | 192.096 | 2.7 | 3 | 3 | 33.988106 |
| P29401 | Transketolase OS=Homo sapiens OX=96( | TKT | 67.878 | 1.4 | 1 | 1 | 41.690086 |
| P29558 | RNA-binding motif, single-stranded-inter | RBMS1 | 44.506 | 24.4 | 8 | 4 | 4581.5625 |
| P29597 | Non-receptor tyrosine-protein kinase TYK | TYK2 | 133.65 | 8.6 | 10 | 10 | 147.00526 |
| P29692 | Elongation factor 1-delta OS=Homo sapie | EEF1D | 31.122 | 53 | 14 | 12 | 10975.504 |

| | | | | | | | |
|---|---|---|---|---|---|---|---|
| P29966 | Myristoylated alanine-rich C-kinase subst | MARCKS | 31.555 | 5.4 | 1 | 1 | 50.301857 |
| P30041 | Peroxiredoxin-6 OS=Homo sapiens OX=9 | PRDX6 | 25.035 | 29.9 | 7 | 7 | 276.40277 |
| P30044 | Peroxiredoxin-5, mitochondrial OS=Hom | PRDX5 | 22.087 | 5.1 | 1 | 1 | 107.51529 |
| P30048 | Thioredoxin-dependent peroxide reductas | PRDX3 | 27.693 | 30.5 | 5 | 5 | 213.25687 |
| P30049 | ATP synthase subunit delta, mitochondria | ATP5F1D | 17.49 | 22.6 | 3 | 3 | 113.02562 |
| P30050 | Large ribosomal subunit protein uL11 OS | RPL12 | 17.819 | 52.1 | 7 | 7 | 18715.58 |
| P30101 | Protein disulfide-isomerase A3 OS=Homo | PDIA3 | 56.783 | 19.8 | 9 | 9 | 102.87015 |
| P30153 | Serine/threonine-protein phosphatase 2A ( | PPP2R1A | 65.309 | 32.6 | 18 | 13 | 1160.73 |
| P30154 | Serine/threonine-protein phosphatase 2A ( | PPP2R1B | 66.214 | 6.8 | 3 | 3 | 71.225174 |
| P30260 | Cell division cycle protein 27 homolog OS | CDC27 | 91.868 | 12.7 | 10 | 10 | 254.0322 |
| P30291 | Wee1-like protein kinase OS=Homo sapie | WEE1 | 71.598 | 18.7 | 12 | 12 | 364.63922 |
| P30414 | NK-tumor recognition protein OS=Homo | NKTR | 165.677 | 3.5 | 4 | 4 | 120.76672 |
| P30530 | Tyrosine-protein kinase receptor UFO OS | AXL | 98.337 | 8.2 | 6 | 5 | 92.916794 |
| P30536 | Translocator protein OS=Homo sapiens C | TSPO | 18.828 | 4.7 | 1 | 1 | 389.51556 |
| P30622 | CAP-Gly domain-containing linker protei | CLIP1 | 162.247 | 1 | 1 | 1 | 54.183208 |
| P30626 | Sorcin OS=Homo sapiens OX=9606 GN= | SRI | 21.677 | 16.2 | 3 | 3 | 46.245529 |
| P30876 | DNA-directed RNA polymerase II subuni | POLR2B | 133.897 | 10.6 | 12 | 12 | 118.86954 |
| P31040 | Succinate dehydrogenase [ubiquinone] fla | SDHA | 72.692 | 20 | 11 | 11 | 203.53967 |
| P31153 | S-adenosylmethionine synthase isoform ty | MAT2A | 43.661 | 28.1 | 9 | 7 | 347.63251 |
| P31274 | Homeobox protein Hox-C9 OS=Homo sa | HOXC9 | 29.248 | 16.5 | 3 | 3 | 168.07472 |
| P31327 | Carbamoyl-phosphate synthase [ammonia | CPS1 | 164.94 | 25.3 | 34 | 33 | 584.98199 |
| P31350 | Ribonucleoside-diphosphate reductase sul | RRM2 | 44.878 | 5.7 | 2 | 2 | 98.579239 |
| P31431 | Syndecan-4 OS=Homo sapiens OX=9606 | SDC4 | 21.642 | 24.7 | 4 | 4 | 479.13989 |
| P31483 | Cytotoxic granule associated RNA bindin | TIA1 | 42.964 | 14.2 | 5 | 2 | 1457.1317 |
| P31689 | DnaJ homolog subfamily A member 1 OS | DNAJA1 | 44.869 | 33 | 12 | 12 | 2702.9854 |
| P31749 | RAC-alpha serine/threonine-protein kinas | AKT1 | 55.687 | 1.7 | 1 | 1 | 140.45848 |
| P31930 | Cytochrome b-c1 complex subunit 1, mito | UQCRC1 | 52.646 | 11.2 | 4 | 4 | 64.037392 |
| P31942 | Heterogeneous nuclear ribonucleoprotein | HNRNPH3 | 36.927 | 39.6 | 10 | 10 | 11799.419 |
| P31943 | Heterogeneous nuclear ribonucleoprotein | HNRNPH1 | 49.23 | 35.4 | 15 | 8 | 48656.043 |
| P31946 | 14-3-3 protein beta/alpha OS=Homo sapi | YWHAB | 28.083 | 27.2 | 7 | 2 | 447.82309 |
| P31947 | 14-3-3 protein sigma OS=Homo sapiens ( | SFN | 27.775 | 8.9 | 2 | 2 | 102.83179 |
| P31948 | Stress-induced-phosphoprotein 1 OS=Ho | STIP1 | 62.64 | 5.3 | 3 | 3 | 56.545574 |
| P32121 | Beta-arrestin-2 OS=Homo sapiens OX=9( | ARRB2 | 46.106 | 12.2 | 5 | 5 | 111.45586 |
| P32242 | Homeobox protein OTX1 OS=Homo sapi | OTX1 | 37.328 | 11.3 | 4 | 2 | 446.2345 |
| P32320 | Cytidine deaminase OS=Homo sapiens O | CDA | 16.185 | 15.1 | 1 | 1 | 177.34071 |

| | | | | | | | |
|---|---|---|---|---|---|---|---|
| P32322 | Pyrroline-5-carboxylate reductase 1, mito | PYCR1 | 33.361 | 10.7 | 3 | 2 | 98.167946 |
| P32754 | 4-hydroxyphenylpyruvate dioxygenase OS | HPD | 44.935 | 15.5 | 6 | 6 | 77.893837 |
| P32780 | General transcription factor IIH subunit 1 | GTF2H1 | 62.032 | 19.2 | 10 | 10 | 308.59277 |
| P32969 | Large ribosomal subunit protein uL6 OS= | RPL9 | 21.864 | 51 | 9 | 9 | 12966.938 |
| P32970 | CD70 antigen OS=Homo sapiens OX=96 | CD70 | 21.119 | 17.6 | 3 | 3 | 103.3069 |
| P33121 | Long-chain-fatty-acid--CoA ligase 1 OS= | ACSL1 | 77.944 | 2.9 | 2 | 1 | 54.41468 |
| P33176 | Kinesin-1 heavy chain OS=Homo sapiens | KIF5B | 109.685 | 29.4 | 24 | 17 | 402.17755 |
| P33240 | Cleavage stimulation factor subunit 2 OS= | CSTF2 | 60.96 | 3.6 | 2 | 1 | 100.15247 |
| P33316 | Deoxyuridine 5'-triphosphate nucleotidoh | DUT | 26.564 | 23.4 | 5 | 5 | 161.32388 |
| P33897 | ATP-binding cassette sub-family D memb | ABCD1 | 82.937 | 12.3 | 8 | 7 | 101.12115 |
| P33981 | Dual specificity protein kinase TTK OS=I | TTK | 97.073 | 12.3 | 8 | 8 | 71.883575 |
| P33991 | DNA replication licensing factor MCM4 ( | MCM4 | 96.558 | 11.5 | 8 | 8 | 124.38725 |
| P33992 | DNA replication licensing factor MCM5 ( | MCM5 | 82.286 | 40.5 | 28 | 28 | 832.08185 |
| P33993 | DNA replication licensing factor MCM7 ( | MCM7 | 81.308 | 23.8 | 15 | 15 | 201.26106 |
| P34096 | Ribonuclease 4 OS=Homo sapiens OX=9 | RNASE4 | 16.841 | 7.5 | 1 | 1 | 82.426262 |
| P34741 | Syndecan-2 OS=Homo sapiens OX=9606 | SDC2 | 22.16 | 10.9 | 2 | 2 | 192.95973 |
| P34896 | Serine hydroxymethyltransferase, cytosoli | SHMT1 | 53.083 | 2.9 | 1 | 1 | 55.571369 |
| P34897 | Serine hydroxymethyltransferase, mitochc | SHMT2 | 55.993 | 18.1 | 7 | 7 | 113.2317 |
| P34932 | Heat shock 70 kDa protein 4 OS=Homo s | HSPA4 | 94.331 | 9.4 | 6 | 6 | 131.89793 |
| P34947 | G protein-coupled receptor kinase 5 OS=I | GRK5 | 67.787 | 8.6 | 4 | 2 | 51.988438 |
| P35030 | Trypsin-3 OS=Homo sapiens OX=9606 C | PRSS3 | 32.529 | 4.3 | 1 | 1 | 6086.2832 |
| P35052 | Glypican-1 OS=Homo sapiens OX=9606 | GPC1 | 61.681 | 9 | 4 | 4 | 80.011322 |
| P35080 | Profilin-2 OS=Homo sapiens OX=9606 G | PFN2 | 15.047 | 36.4 | 3 | 3 | 74.410294 |
| P35221 | Catenin alpha-1 OS=Homo sapiens OX=9 | CTNNA1 | 100.072 | 12.5 | 10 | 8 | 201.69418 |
| P35222 | Catenin beta-1 OS=Homo sapiens OX=96 | CTNNB1 | 85.497 | 14.5 | 9 | 7 | 97.932297 |
| P35226 | Polycomb complex protein BMI-1 OS=H | BMI1 | 36.949 | 7.7 | 2 | 2 | 39.360149 |
| P35232 | Prohibitin 1 OS=Homo sapiens OX=9606 | PHB1 | 29.805 | 4 | 1 | 1 | 38.379505 |
| P35237 | Serpin B6 OS=Homo sapiens OX=9606 C | SERPINB6 | 42.622 | 2.4 | 1 | 1 | 41.914886 |
| P35240 | Merlin OS=Homo sapiens OX=9606 GN= | NF2 | 69.691 | 13.8 | 8 | 8 | 135.29837 |
| P35241 | Radixin OS=Homo sapiens OX=9606 GN | RDX | 68.564 | 18 | 12 | 5 | 660.29236 |
| P35244 | Replication protein A 14 kDa subunit OS= | RPA3 | 13.569 | 18.2 | 2 | 2 | 181.31676 |
| P35249 | Replication factor C subunit 4 OS=Homo | RFC4 | 39.682 | 37.7 | 13 | 13 | 1142.8928 |
| P35250 | Replication factor C subunit 2 OS=Homo | RFC2 | 39.158 | 36.2 | 12 | 12 | 1350.9457 |
| P35251 | Replication factor C subunit 1 OS=Homo | RFC1 | 128.255 | 27.5 | 32 | 32 | 689.69678 |
| P35268 | Large ribosomal subunit protein eL22 OS | RPL22 | 14.787 | 18.8 | 2 | 2 | 37695.586 |

| | | | | | | | |
|---|---|---|---|---|---|---|---|
| P35269 | General transcription factor IIF subunit 1 ( | GTF2F1 | 58.241 | 7.7 | 3 | 3 | 38.337353 |
| P35520 | Cystathionine beta-synthase OS=Homo sa | CBS | 60.587 | 4 | 2 | 2 | 93.068214 |
| P35527 | Keratin, type I cytoskeletal 9 OS=Homo s | KRT9 | 62.065 | 20.5 | 13 | 13 | 571.88428 |
| P35555 | Fibrillin-1 OS=Homo sapiens OX=9606 ( | FBN1 | 312.298 | 1.6 | 4 | 4 | 62.811111 |
| P35579 | Myosin-9 OS=Homo sapiens OX=9606 G | MYH9 | 226.533 | 42.3 | 91 | 72 | 7884.6797 |
| P35580 | Myosin-10 OS=Homo sapiens OX=9606 ( | MYH10 | 229 | 28.4 | 53 | 49 | 1095.4663 |
| P35606 | Coatomer subunit beta' OS=Homo sapiens | COPB2 | 102.488 | 16 | 15 | 15 | 178.10623 |
| P35611 | Alpha-adducin OS=Homo sapiens OX=96 | ADD1 | 80.956 | 7.9 | 5 | 5 | 179.72697 |
| P35613 | Basigin OS=Homo sapiens OX=9606 GN | BSG | 42.201 | 8.6 | 3 | 3 | 306.23935 |
| P35625 | Metalloproteinase inhibitor 3 OS=Homo s | TIMP3 | 24.145 | 19 | 4 | 4 | 222.74734 |
| P35637 | RNA-binding protein FUS OS=Homo sap | FUS | 53.426 | 16.3 | 9 | 7 | 37557.492 |
| P35658 | Nuclear pore complex protein Nup214 OS | NUP214 | 213.62 | 2 | 3 | 3 | 32.838593 |
| P35659 | Protein DEK OS=Homo sapiens OX=960 | DEK | 42.675 | 28 | 10 | 10 | 1048.844 |
| P35713 | Transcription factor SOX-18 OS=Homo s | SOX18 | 40.892 | 10.7 | 3 | 3 | 125.55516 |
| P35869 | Aryl hydrocarbon receptor OS=Homo sap | AHR | 96.148 | 11.9 | 8 | 8 | 141.59286 |
| P35908 | Keratin, type II cytoskeletal 2 epidermal ( | KRT2 | 65.433 | 23.2 | 13 | 12 | 522.30505 |
| P35998 | 26S proteasome regulatory subunit 7 OS= | PSMC2 | 48.634 | 13.9 | 5 | 5 | 97.682472 |
| P36402 | Transcription factor 7 OS=Homo sapiens | TCF7 | 41.642 | 2.9 | 1 | 1 | 63.42453 |
| P36542 | ATP synthase subunit gamma, mitochondr | ATP5F1C | 32.997 | 24.8 | 8 | 8 | 1959.4368 |
| P36578 | Large ribosomal subunit protein uL4 OS= | RPL4 | 47.698 | 40.5 | 17 | 17 | 36442.141 |
| P36776 | Lon protease homolog, mitochondrial OS= | LONP1 | 106.49 | 5 | 4 | 4 | 26.252106 |
| P36873 | Serine/threonine-protein phosphatase PP1 | PPP1CC | 36.984 | 37.5 | 12 | 3 | 2344.5234 |
| P36897 | TGF-beta receptor type-1 OS=Homo sapi | TGFBR1 | 55.96 | 3.8 | 2 | 1 | 55.81213 |
| P36954 | DNA-directed RNA polymerase II subuni | POLR2I | 14.524 | 25.6 | 3 | 3 | 91.134842 |
| P36956 | Sterol regulatory element-binding protein | SREBF1 | 121.676 | 3.2 | 3 | 3 | 41.164654 |
| P36957 | Dihydrolipoyllysine-residue succinyltrans | DLST | 48.756 | 4.4 | 2 | 2 | 56.975872 |
| P37108 | Signal recognition particle 14 kDa protein | SRP14 | 14.57 | 68.4 | 8 | 8 | 2621.1453 |
| P37198 | Nuclear pore glycoprotein p62 OS=Homo | NUP62 | 53.255 | 3.8 | 2 | 2 | 52.041664 |
| P37235 | Hippocalcin-like protein 1 OS=Homo sap | HPCAL1 | 22.314 | 10.4 | 2 | 1 | 44.982388 |
| P37268 | Squalene synthase OS=Homo sapiens OX | FDFT1 | 48.116 | 7.9 | 3 | 3 | 41.470673 |
| P37802 | Transgelin-2 OS=Homo sapiens OX=960( | TAGLN2 | 22.392 | 18.6 | 3 | 3 | 64.969559 |
| P38117 | Electron transfer flavoprotein subunit beta | ETFB | 27.844 | 4.7 | 1 | 1 | 108.03108 |
| P38159 | RNA-binding motif protein, X chromosom | RBMX | 42.332 | 33.2 | 15 | 3 | 9048.7363 |
| P38398 | Breast cancer type 1 susceptibility protein | BRCA1 | 207.721 | 22.2 | 38 | 38 | 623.12738 |
| P38432 | Coilin OS=Homo sapiens OX=9606 GN= | COIL | 62.609 | 6.6 | 3 | 3 | 108.49046 |

| | | | | | | | |
|---|---|---|---|---|---|---|---|
| P38435 | Vitamin K-dependent gamma-carboxylase | GGCX | 87.561 | 1.1 | 1 | 1 | 24.7803 |
| P38606 | V-type proton ATPase catalytic subunit A | ATP6V1A | 68.305 | 6 | 3 | 3 | 50.210793 |
| P38646 | Stress-70 protein, mitochondrial OS=Hom | HSPA9 | 73.681 | 41.2 | 27 | 27 | 5014.3682 |
| P38919 | Eukaryotic initiation factor 4A-III OS=Hc | EIF4A3 | 46.872 | 46.5 | 19 | 19 | 10571.905 |
| P38935 | DNA-binding protein SMUBP-2 OS=Hor | IGHMBP2 | 109.15 | 23 | 22 | 22 | 536.85059 |
| P39019 | Small ribosomal subunit protein eS19 OS= | RPS19 | 16.061 | 40.7 | 9 | 9 | 22581.291 |
| P39023 | Large ribosomal subunit protein uL3 OS= | RPL3 | 46.109 | 40.4 | 19 | 17 | 25793.85 |
| P39060 | Collagen alpha-1(XVIII) chain OS=Homc | COL18A1 | 178.188 | 0.5 | 1 | 1 | 66.626709 |
| P39656 | Dolichyl-diphosphooligosaccharide--prote | DDOST | 50.801 | 19.7 | 8 | 8 | 211.90428 |
| P39687 | Acidic leucine-rich nuclear phosphoprotei | ANP32A | 28.586 | 4.4 | 1 | 0 | 20.973162 |
| P39748 | Flap endonuclease 1 OS=Homo sapiens C | FEN1 | 42.593 | 2.9 | 1 | 1 | 91.778282 |
| P39880 | Homeobox protein cut-like 1 OS=Homo s | CUX1 | 164.188 | 10.4 | 13 | 5 | 115.28799 |
| P40222 | Alpha-taxilin OS=Homo sapiens OX=960 | TXLNA | 61.892 | 20.7 | 12 | 11 | 429.74078 |
| P40227 | T-complex protein 1 subunit zeta OS=Hoi | CCT6A | 58.025 | 12.8 | 7 | 4 | 145.13525 |
| P40424 | Pre-B-cell leukemia transcription factor 1 | PBX1 | 46.626 | 8.4 | 3 | 1 | 54.885681 |
| P40429 | Large ribosomal subunit protein uL13 OS | RPL13A | 23.578 | 37.9 | 8 | 5 | 20387.814 |
| P40616 | ADP-ribosylation factor-like protein 1 OS | ARL1 | 20.418 | 26 | 4 | 4 | 157.12207 |
| P40692 | DNA mismatch repair protein Mlh1 OS=I | MLH1 | 84.601 | 22.8 | 16 | 16 | 243.97737 |
| P40763 | Signal transducer and activator of transcri | STAT3 | 88.068 | 7 | 5 | 5 | 72.353394 |
| P40818 | Ubiquitin carboxyl-terminal hydrolase 8 C | USP8 | 127.524 | 2.1 | 2 | 2 | 40.070091 |
| P40926 | Malate dehydrogenase, mitochondrial OS= | MDH2 | 35.504 | 6.8 | 2 | 2 | 60.264549 |
| P40937 | Replication factor C subunit 5 OS=Homo | RFC5 | 38.497 | 40 | 13 | 13 | 1075.9056 |
| P40938 | Replication factor C subunit 3 OS=Homo | RFC3 | 40.557 | 41.3 | 14 | 14 | 1082.8132 |
| P40939 | Trifunctional enzyme subunit alpha, mitoc | HADHA | 83 | 22.3 | 16 | 16 | 521.08002 |
| P41091 | Eukaryotic translation initiation factor 2 s | EIF2S3 | 51.11 | 39.2 | 17 | 4 | 4123.9956 |
| P41134 | DNA-binding protein inhibitor ID-1 OS=I | ID1 | 16.133 | 11.6 | 1 | 1 | 35.396069 |
| P41161 | ETS translocation variant 5 OS=Homo sa | ETV5 | 57.838 | 5.5 | 3 | 2 | 222.63298 |
| P41182 | B-cell lymphoma 6 protein OS=Homo sap | BCL6 | 78.847 | 16.7 | 9 | 8 | 147.85368 |
| P41208 | Centrin-2 OS=Homo sapiens OX=9606 G | CETN2 | 19.739 | 5.2 | 1 | 1 | 44.792374 |
| P41214 | Eukaryotic translation initiation factor 2D | EIF2D | 64.707 | 14.2 | 7 | 7 | 222.53252 |
| P41223 | Protein BUD31 homolog OS=Homo sapie | BUD31 | 17 | 20.8 | 3 | 3 | 83.29686 |
| P41227 | N-alpha-acetyltransferase 10 OS=Homo s | NAA10 | 26.459 | 24.7 | 5 | 3 | 86.738678 |
| P41250 | Glycine--tRNA ligase OS=Homo sapiens | GARS1 | 83.166 | 9.9 | 7 | 7 | 114.79602 |
| P41252 | Isoleucine--tRNA ligase, cytoplasmic OS= | IARS1 | 144.499 | 15.2 | 19 | 19 | 188.87602 |
| P41743 | Protein kinase C iota type OS=Homo sapi | PRKCI | 68.263 | 22.7 | 10 | 9 | 660.48627 |

| | | | | | | | |
|---|---|---|---|---|---|---|---|
| P42025 | Beta-centractin OS=Homo sapiens OX=96 | ACTR1B | 42.294 | 4 | 1 | 1 | 318.29428 |
| P42126 | Enoyl-CoA delta isomerase 1, mitochondr | ECI1 | 32.816 | 9.3 | 2 | 2 | 158.65752 |
| P42166 | Lamina-associated polypeptide 2, isoform | TMPO | 75.493 | 32 | 19 | 13 | 2374.6814 |
| P42167 | Lamina-associated polypeptide 2, isoform | TMPO | 50.671 | 14.1 | 5 | 5 | 866.82477 |
| P42224 | Signal transducer and activator of transcri | STAT1 | 87.335 | 1.3 | 1 | 1 | 34.233913 |
| P42226 | Signal transducer and activator of transcri | STAT6 | 94.136 | 2.2 | 2 | 2 | 54.816944 |
| P42285 | Exosome RNA helicase MTR4 OS=Homo | MTREX | 117.805 | 29.7 | 28 | 28 | 686.31342 |
| P42566 | Epidermal growth factor receptor substrat | EPS15 | 98.656 | 9.4 | 7 | 7 | 94.73378 |
| P42568 | Protein AF-9 OS=Homo sapiens OX=960 | MLLT3 | 63.352 | 2.8 | 1 | 1 | 108.90061 |
| P42677 | Small ribosomal subunit protein eS27 OS= | RPS27 | 9.462 | 14.3 | 2 | 2 | 10859.528 |
| P42679 | Megakaryocyte-associated tyrosine-protei | MATK | 56.469 | 10.3 | 5 | 4 | 115.32308 |
| P42680 | Tyrosine-protein kinase Tec OS=Homo sa | TEC | 73.582 | 7.9 | 5 | 5 | 79.890091 |
| P42694 | Probable helicase with zinc finger domain | HELZ | 218.971 | 12.7 | 23 | 23 | 212.01915 |
| P42695 | Condensin-2 complex subunit D3 OS=Ho | NCAPD3 | 168.891 | 6.9 | 10 | 10 | 157.06071 |
| P42704 | Leucine-rich PPR motif-containing protei | LRPPRC | 157.906 | 38.5 | 57 | 57 | 2515.1025 |
| P42766 | Large ribosomal subunit protein uL29 OS | RPL35 | 14.552 | 32.5 | 7 | 7 | 19517.777 |
| P42771 | Cyclin-dependent kinase inhibitor 2A OS= | CDKN2A | 16.533 | 15.4 | 2 | 1 | 215.54439 |
| P43034 | Platelet-activating factor acetylhydrolase I | PAFAH1B1 | 46.638 | 3.7 | 1 | 1 | 70.59343 |
| P43243 | Matrin-3 OS=Homo sapiens OX=9606 GI | MATR3 | 94.624 | 33.1 | 24 | 24 | 7656.7754 |
| P43246 | DNA mismatch repair protein Msh2 OS=I | MSH2 | 104.744 | 18.2 | 17 | 17 | 371.5838 |
| P43304 | Glycerol-3-phosphate dehydrogenase, mit | GPD2 | 80.853 | 1.5 | 1 | 1 | 40.974022 |
| P43307 | Translocon-associated protein subunit alp | SSR1 | 32.236 | 21.7 | 5 | 5 | 314.27225 |
| P43353 | Aldehyde dehydrogenase family 3 membe | ALDH3B1 | 51.84 | 4.3 | 2 | 1 | 488.78406 |
| P43354 | Nuclear receptor subfamily 4 group A me | NR4A2 | 66.591 | 8.5 | 3 | 2 | 115.23551 |
| P43355 | Melanoma-associated antigen 1 OS=Hom | MAGEA1 | 34.343 | 11 | 3 | 3 | 94.172722 |
| P43361 | Melanoma-associated antigen 8 OS=Hom | MAGEA8 | 35.215 | 13.5 | 3 | 1 | 140.95755 |
| P43378 | Tyrosine-protein phosphatase non-recepto | PTPN9 | 68.021 | 1.5 | 1 | 1 | 19.749256 |
| P43487 | Ran-specific GTPase-activating protein O | RANBP1 | 23.311 | 10.9 | 2 | 2 | 76.254997 |
| P43490 | Nicotinamide phosphoribosyltransferase ( | NAMPT | 55.522 | 4.3 | 2 | 2 | 35.296955 |
| P43686 | 26S proteasome regulatory subunit 6B OS | PSMC4 | 47.367 | 6 | 2 | 2 | 39.436718 |
| P43897 | Elongation factor Ts, mitochondrial OS=H | TSFM | 35.391 | 20.9 | 6 | 6 | 202.19406 |
| P45880 | Voltage-dependent anion-selective channe | VDAC2 | 31.567 | 15.6 | 4 | 4 | 310.77942 |
| P45973 | Chromobox protein homolog 5 OS=Homo | CBX5 | 22.225 | 39.3 | 6 | 5 | 366.18613 |
| P45984 | Mitogen-activated protein kinase 9 OS=H | MAPK9 | 48.14 | 5.9 | 2 | 1 | 114.61984 |
| P46013 | Proliferation marker protein Ki-67 OS=Ho | MKI67 | 358.694 | 25 | 58 | 58 | 729.33545 |

| | | | | | | | |
|---|---|---|---|---|---|---|---|
| P46019 | Phosphorylase b kinase regulatory subunit | PHKA2 | 138.409 | 0.7 | 1 | 1 | 55.471851 |
| P46060 | Ran GTPase-activating protein 1 OS=Hor | RANGAP1 | 63.542 | 23.2 | 12 | 12 | 179.11548 |
| P46063 | ATP-dependent DNA helicase Q1 OS=Hc | RECQL | 73.458 | 32.4 | 18 | 18 | 1641.2229 |
| P46087 | Probable 28S rRNA (cytosine(4447)-C(5) | NOP2 | 89.302 | 27.1 | 20 | 20 | 1239.6056 |
| P46109 | Crk-like protein OS=Homo sapiens OX=9 | CRKL | 33.777 | 4 | 1 | 1 | 197.53952 |
| P46199 | Translation initiation factor IF-2, mitocho | MTIF2 | 81.318 | 3 | 2 | 2 | 37.560806 |
| P46379 | Large proline-rich protein BAG6 OS=Hor | BAG6 | 119.409 | 0.9 | 1 | 1 | 129.29494 |
| P46459 | Vesicle-fusing ATPase OS=Homo sapiens | NSF | 82.595 | 1.7 | 1 | 1 | 36.333908 |
| P46734 | Dual specificity mitogen-activated protein | MAP2K3 | 39.319 | 22.8 | 7 | 7 | 244.56908 |
| P46736 | Lys-63-specific deubiquitinase BRCC36 ( | BRCC3 | 36.072 | 38.6 | 9 | 9 | 434.13931 |
| P46776 | Large ribosomal subunit protein uL15 OS | RPL27A | 16.562 | 28.4 | 5 | 5 | 43742.797 |
| P46777 | Large ribosomal subunit protein uL18 OS | RPL5 | 34.363 | 45.8 | 16 | 16 | 6507.3677 |
| P46778 | Large ribosomal subunit protein eL21 OS | RPL21 | 18.565 | 27.5 | 6 | 6 | 6523.1099 |
| P46779 | Large ribosomal subunit protein eL28 OS | RPL28 | 15.748 | 47.4 | 9 | 9 | 12509.563 |
| P46781 | Small ribosomal subunit protein uS4 OS= | RPS9 | 22.592 | 54.1 | 16 | 16 | 48314.73 |
| P46782 | Small ribosomal subunit protein uS7 OS= | RPS5 | 22.877 | 36.3 | 9 | 9 | 18954.465 |
| P46783 | Small ribosomal subunit protein eS10 OS | RPS10 | 18.898 | 52.1 | 9 | 4 | 10271.71 |
| P46939 | Utrophin OS=Homo sapiens OX=9606 G | UTRN | 394.467 | 0.3 | 1 | 1 | 29.030697 |
| P46940 | Ras GTPase-activating-like protein IQGA | IQGAP1 | 189.252 | 23.3 | 37 | 35 | 641.28259 |
| P46976 | Glycogenin-1 OS=Homo sapiens OX=96( | GYG1 | 39.384 | 14.9 | 5 | 5 | 595.00934 |
| P46977 | Dolichyl-diphosphooligosaccharide--prote | STT3A | 80.53 | 5.5 | 4 | 4 | 70.654976 |
| P47736 | Rap1 GTPase-activating protein 1 OS=Hc | RAP1GAP | 73.362 | 1.4 | 1 | 1 | 68.840874 |
| P47755 | F-actin-capping protein subunit alpha-2 O | CAPZA2 | 32.95 | 33.2 | 8 | 6 | 1746.1682 |
| P47756 | F-actin-capping protein subunit beta OS=l | CAPZB | 30.629 | 51.5 | 13 | 13 | 3708.4688 |
| P47895 | Retinaldehyde dehydrogenase 3 OS=Hom | ALDH1A3 | 56.109 | 4.9 | 2 | 2 | 21.652819 |
| P47897 | Glutamine--tRNA ligase OS=Homo sapie | QARS1 | 87.799 | 30.1 | 19 | 19 | 551.81866 |
| P47914 | Large ribosomal subunit protein eL29 OS | RPL29 | 17.753 | 14.5 | 2 | 2 | 20655.871 |
| P47974 | mRNA decay activator protein ZFP36L2 ( | ZFP36L2 | 51.063 | 16 | 6 | 6 | 787.90918 |
| P47985 | Cytochrome b-c1 complex subunit Rieske | UQCRFS1 | 29.668 | 10.9 | 3 | 1 | 64.560562 |
| P48047 | ATP synthase subunit O, mitochondrial O | ATP5PO | 23.278 | 34.3 | 7 | 7 | 594.40625 |
| P48059 | LIM and senescent cell antigen-like-conta | LIMS1 | 37.252 | 13.8 | 4 | 2 | 116.18919 |
| P48200 | Iron-responsive element-binding protein 2 | IREB2 | 105.059 | 13.7 | 11 | 11 | 154.58334 |
| P48382 | DNA-binding protein RFX5 OS=Homo sa | RFX5 | 65.323 | 2.4 | 1 | 1 | 17.611843 |
| P48444 | Coatomer subunit delta OS=Homo sapien | ARCN1 | 57.211 | 28.6 | 14 | 14 | 178.05908 |
| P48507 | Glutamate--cysteine ligase regulatory subu | GCLM | 30.728 | 8.8 | 2 | 2 | 210.76523 |

| | | | | | | | |
|---|---|---|---|---|---|---|---|
| P48634 | Protein PRRC2A OS=Homo sapiens OX= | PRRC2A | 228.864 | 33.3 | 59 | 59 | 4949.853 |
| P48643 | T-complex protein 1 subunit epsilon OS=] | CCT5 | 59.672 | 22.6 | 12 | 12 | 235.67 |
| P48651 | Phosphatidylserine synthase 1 OS=Homo | PTDSS1 | 55.528 | 14 | 6 | 6 | 502.74533 |
| P48681 | Nestin OS=Homo sapiens OX=9606 GN= | NES | 177.439 | 6.1 | 9 | 9 | 100.87017 |
| P48729 | Casein kinase I isoform alpha OS=Homo : | CSNK1A1 | 38.915 | 25.2 | 8 | 3 | 957.13074 |
| P48730 | Casein kinase I isoform delta OS=Homo s | CSNK1D | 47.331 | 6.3 | 2 | 2 | 1217.5308 |
| P48740 | Mannan-binding lectin serine protease 1 C | MASP1 | 79.247 | 2 | 1 | 1 | 109.47717 |
| P48960 | Adhesion G protein-coupled receptor E5 ( | ADGRE5 | 91.87 | 1.1 | 1 | 1 | 37.785538 |
| P49005 | DNA polymerase delta subunit 2 OS=Hor | POLD2 | 51.29 | 8.1 | 3 | 3 | 34.466866 |
| P49137 | MAP kinase-activated protein kinase 2 OS | MAPKAPK | 45.568 | 3 | 1 | 1 | 27.992748 |
| P49189 | 4-trimethylaminobutyraldehyde dehydrog | ALDH9A1 | 53.802 | 2.4 | 1 | 1 | 28.978949 |
| P49207 | Large ribosomal subunit protein eL34 OS | RPL34 | 13.293 | 36.8 | 6 | 6 | 12658.188 |
| P49327 | Fatty acid synthase OS=Homo sapiens OX | FASN | 273.427 | 20.4 | 41 | 41 | 332.73938 |
| P49336 | Cyclin-dependent kinase 8 OS=Homo sap | CDK8 | 53.284 | 6.5 | 3 | 1 | 100.63219 |
| P49368 | T-complex protein 1 subunit gamma OS=] | CCT3 | 60.534 | 38.2 | 19 | 19 | 323.71432 |
| P49407 | Beta-arrestin-1 OS=Homo sapiens OX=96 | ARRB1 | 47.066 | 9.6 | 4 | 4 | 81.063072 |
| P49411 | Elongation factor Tu, mitochondrial OS=I | TUFM | 49.875 | 47.9 | 21 | 21 | 2871.3872 |
| P49427 | Ubiquitin-conjugating enzyme E2 R1 OS= | CDC34 | 26.737 | 12.7 | 3 | 3 | 89.987839 |
| P49450 | Histone H3-like centromeric protein A OS | CENPA | 15.991 | 5 | 1 | 1 | 117.1261 |
| P49454 | Centromere protein F OS=Homo sapiens ( | CENPF | 357.528 | 4.5 | 12 | 12 | 101.29215 |
| P49458 | Signal recognition particle 9 kDa protein ( | SRP9 | 10.112 | 53.5 | 5 | 5 | 7696.4248 |
| P49588 | Alanine--tRNA ligase, cytoplasmic OS=H | AARS1 | 106.811 | 24.2 | 19 | 19 | 336.4743 |
| P49589 | Cysteine--tRNA ligase, cytoplasmic OS=I | CARS1 | 85.474 | 28.2 | 19 | 19 | 696.62244 |
| P49590 | Histidine--tRNA ligase, mitochondrial OS | HARS2 | 56.889 | 23.3 | 11 | 7 | 144.59654 |
| P49643 | DNA primase large subunit OS=Homo sa | PRIM2 | 58.806 | 12 | 5 | 5 | 91.96859 |
| P49674 | Casein kinase I isoform epsilon OS=Hom | CSNK1E | 47.316 | 33.7 | 13 | 5 | 739.81213 |
| P49703 | ADP-ribosylation factor-like protein 4D C | ARL4D | 22.156 | 7.5 | 1 | 1 | 48.174187 |
| P49711 | Transcriptional repressor CTCF OS=Hom | CTCF | 82.786 | 8.7 | 7 | 6 | 128.91174 |
| P49716 | CCAAT/enhancer-binding protein delta O | CEBPD | 28.468 | 9.7 | 2 | 2 | 68.389732 |
| P49721 | Proteasome subunit beta type-2 OS=Hom | PSMB2 | 22.837 | 5 | 1 | 1 | 90.728859 |
| P49747 | Cartilage oligomeric matrix protein OS=H | COMP | 82.861 | 1.5 | 1 | 1 | 41.998047 |
| P49748 | Very long-chain specific acyl-CoA dehyd | ACADVL | 70.391 | 2.3 | 1 | 1 | 30.817316 |
| P49755 | Transmembrane emp24 domain-containin | TMED10 | 24.976 | 14.6 | 3 | 3 | 115.29362 |
| P49756 | RNA-binding protein 25 OS=Homo sapie | RBM25 | 100.186 | 12.3 | 10 | 10 | 192.3595 |
| P49757 | Protein numb homolog OS=Homo sapiens | NUMB | 70.804 | 21.5 | 13 | 10 | 350.24368 |

| | | | | | | | |
|---|---|---|---|---|---|---|---|
| P49759 | Dual specificity protein kinase CLK1 OS= | CLK1 | 57.291 | 16.9 | 8 | 7 | 149.27399 |
| P49760 | Dual specificity protein kinase CLK2 OS= | CLK2 | 60.091 | 15 | 6 | 6 | 190.50871 |
| P49761 | Dual specificity protein kinase CLK3 OS= | CLK3 | 73.516 | 21.5 | 10 | 10 | 758.0769 |
| P49770 | Translation initiation factor eIF-2B subun | EIF2B2 | 38.99 | 15.1 | 4 | 4 | 88.12944 |
| P49790 | Nuclear pore complex protein Nup153 OS | NUP153 | 153.939 | 15.3 | 19 | 19 | 205.75299 |
| P49792 | E3 SUMO-protein ligase RanBP2 OS=Hc | RANBP2 | 358.2 | 9.4 | 24 | 15 | 180.91086 |
| P49796 | Regulator of G-protein signaling 3 OS=H | RGS3 | 132.336 | 4.6 | 5 | 5 | 58.644711 |
| P49821 | NADH dehydrogenase [ubiquinone] flavo | NDUFV1 | 50.818 | 4.3 | 2 | 2 | 75.686417 |
| P49841 | Glycogen synthase kinase-3 beta OS=Hon | GSK3B | 46.745 | 15.2 | 7 | 3 | 290.8757 |
| P49915 | GMP synthase [glutamine-hydrolyzing] O | GMPS | 76.716 | 11.4 | 7 | 7 | 126.11019 |
| P49916 | DNA ligase 3 OS=Homo sapiens OX=96( | LIG3 | 112.907 | 38.5 | 37 | 37 | 1987.4152 |
| P49959 | Double-strand break repair protein MRE1 | MRE11 | 80.594 | 16.2 | 11 | 11 | 188.31784 |
| P50213 | Isocitrate dehydrogenase [NAD] subunit a | IDH3A | 39.592 | 6 | 2 | 2 | 71.446426 |
| P50221 | Homeobox protein MOX-1 OS=Homo sa | MEOX1 | 27.997 | 3.9 | 1 | 1 | 37.751915 |
| P50238 | Cysteine-rich protein 1 OS=Homo sapiens | CRIP1 | 8.533 | 71.4 | 6 | 6 | 657.33221 |
| P50402 | Emerin OS=Homo sapiens OX=9606 GN | EMD | 28.994 | 34.6 | 10 | 10 | 1665.3192 |
| P50416 | Carnitine O-palmitoyltransferase 1, liver i | CPT1A | 88.368 | 27.3 | 20 | 19 | 759.04303 |
| P50454 | Serpin H1 OS=Homo sapiens OX=9606 ( | SERPINH1 | 46.441 | 18.2 | 6 | 6 | 205.52525 |
| P50542 | Peroxisomal targeting signal 1 receptor O | PEX5 | 70.865 | 21.3 | 11 | 11 | 133.55675 |
| P50548 | ETS domain-containing transcription fact | ERF | 58.703 | 8 | 3 | 3 | 116.04021 |
| P50579 | Methionine aminopeptidase 2 OS=Homo | METAP2 | 52.892 | 15.5 | 7 | 7 | 693.79675 |
| P50613 | Cyclin-dependent kinase 7 OS=Homo sap | CDK7 | 39.039 | 27.5 | 7 | 7 | 226.49788 |
| P50616 | Protein Tob1 OS=Homo sapiens OX=960 | TOB1 | 38.155 | 4.6 | 1 | 1 | 22.317057 |
| P50748 | Kinetochore-associated protein 1 OS=Hor | KNTC1 | 250.749 | 1.1 | 1 | 1 | 72.624908 |
| P50750 | Cyclin-dependent kinase 9 OS=Homo sap | CDK9 | 42.778 | 16.4 | 5 | 5 | 160.73418 |
| P50914 | Large ribosomal subunit protein eL14 OS | RPL14 | 23.432 | 24.2 | 5 | 5 | 12485.94 |
| P50990 | T-complex protein 1 subunit theta OS=Hc | CCT8 | 59.621 | 33.2 | 17 | 17 | 213.80051 |
| P50991 | T-complex protein 1 subunit delta OS=Hc | CCT4 | 57.925 | 28.9 | 13 | 13 | 352.45877 |
| P51114 | RNA-binding protein FXR1 OS=Homo sa | FXR1 | 69.721 | 45.1 | 24 | 24 | 26776.736 |
| P51116 | RNA-binding protein FXR2 OS=Homo sa | FXR2 | 74.224 | 41.6 | 23 | 21 | 26858.475 |
| P51148 | Ras-related protein Rab-5C OS=Homo sa | RAB5C | 23.483 | 12 | 2 | 2 | 48.534828 |
| P51149 | Ras-related protein Rab-7a OS=Homo sap | RAB7A | 23.49 | 6.8 | 1 | 1 | 114.00082 |
| P51151 | Ras-related protein Rab-9A OS=Homo sa | RAB9A | 22.838 | 26.4 | 4 | 4 | 124.52281 |
| P51153 | Ras-related protein Rab-13 OS=Homo sap | RAB13 | 22.775 | 33.5 | 6 | 5 | 609.70801 |
| P51398 | Small ribosomal subunit protein mS29 OS | DAP3 | 45.567 | 34.4 | 14 | 14 | 2272.0715 |

| | | | | | | | |
|---|---|---|---|---|---|---|---|
| P51530 | DNA replication ATP-dependent helicase | DNA2 | 120.415 | 6.1 | 6 | 6 | 680.44714 |
| P51531 | Probable global transcription activator SN | SMARCA2 | 181.28 | 6.3 | 8 | 8 | 165.41139 |
| P51532 | Transcription activator BRG1 OS=Homo | SMARCA4 | 184.646 | 21.8 | 34 | 21 | 1384.4406 |
| P51553 | Isocitrate dehydrogenase [NAD] subunit g | IDH3G | 42.795 | 5.1 | 1 | 1 | 25.763578 |
| P51570 | Galactokinase OS=Homo sapiens OX=96 | GALK1 | 42.273 | 12.2 | 4 | 4 | 58.224236 |
| P51571 | Translocon-associated protein subunit del | SSR4 | 18.999 | 19.7 | 3 | 3 | 339.69766 |
| P51572 | B-cell receptor-associated protein 31 OS= | BCAP31 | 27.992 | 8.1 | 2 | 2 | 124.55116 |
| P51608 | Methyl-CpG-binding protein 2 OS=Homo | MECP2 | 52.441 | 20.4 | 8 | 8 | 114.21458 |
| P51610 | Host cell factor 1 OS=Homo sapiens OX= | HCFC1 | 208.733 | 7.3 | 12 | 12 | 131.20633 |
| P51617 | Interleukin-1 receptor-associated kinase 1 | IRAK1 | 76.537 | 11.4 | 7 | 7 | 83.748436 |
| P51648 | Aldehyde dehydrogenase family 3 membe | ALDH3A2 | 54.848 | 19.8 | 8 | 8 | 405.93399 |
| P51659 | Peroxisomal multifunctional enzyme type | HSD17B4 | 79.687 | 1.6 | 1 | 1 | 25.5301 |
| P51798 | H(+)/Cl(-) exchange transporter 7 OS=Ho | CLCN7 | 88.68 | 1.4 | 1 | 1 | 69.730347 |
| P51808 | Dynein light chain Tctex-type 3 OS=Hom | DYNLT3 | 13.062 | 14.7 | 1 | 1 | 40.919525 |
| P51812 | Ribosomal protein S6 kinase alpha-3 OS= | RPS6KA3 | 83.736 | 29.6 | 18 | 11 | 262.89331 |
| P51825 | AF4/FMR2 family member 1 OS=Homo s | AFF1 | 131.422 | 6.5 | 8 | 8 | 88.163742 |
| P51946 | Cyclin-H OS=Homo sapiens OX=9606 G | CCNH | 37.644 | 13.3 | 6 | 6 | 103.51676 |
| P51948 | CDK-activating kinase assembly factor M | MNAT1 | 35.823 | 6.8 | 2 | 2 | 78.795074 |
| P51955 | Serine/threonine-protein kinase Nek2 OS= | NEK2 | 51.764 | 22 | 9 | 9 | 146.50864 |
| P51957 | Serine/threonine-protein kinase Nek4 OS= | NEK4 | 94.598 | 2.9 | 2 | 2 | 44.327797 |
| P51959 | Cyclin-G1 OS=Homo sapiens OX=9606 ( | CCNG1 | 34.074 | 21.7 | 5 | 5 | 71.839111 |
| P51965 | Ubiquitin-conjugating enzyme E2 E1 OS= | UBE2E1 | 21.405 | 5.7 | 1 | 1 | 54.196674 |
| P51970 | NADH dehydrogenase [ubiquinone] 1 alp | NDUFA8 | 20.106 | 31.4 | 4 | 4 | 36.634705 |
| P51991 | Heterogeneous nuclear ribonucleoprotein | HNRNPA3 | 39.595 | 42.9 | 14 | 14 | 14836.013 |
| P52272 | Heterogeneous nuclear ribonucleoprotein | HNRNPM | 77.516 | 51.1 | 33 | 33 | 11698.647 |
| P52292 | Importin subunit alpha-1 OS=Homo sapie | KPNA2 | 57.862 | 25.1 | 15 | 15 | 3898.5046 |
| P52294 | Importin subunit alpha-5 OS=Homo sapie | KPNA1 | 60.222 | 4.1 | 2 | 2 | 428.54169 |
| P52298 | Nuclear cap-binding protein subunit 2 OS | NCBP2 | 18.002 | 25.6 | 4 | 3 | 4973.0952 |
| P52434 | DNA-directed RNA polymerases I, II, and | POLR2H | 17.144 | 22 | 3 | 3 | 119.32385 |
| P52564 | Dual specificity mitogen-activated protein | MAP2K6 | 37.493 | 9.9 | 3 | 2 | 95.433197 |
| P52565 | Rho GDP-dissociation inhibitor 1 OS=Ho | ARHGDIA | 23.208 | 5.4 | 1 | 1 | 61.651733 |
| P52597 | Heterogeneous nuclear ribonucleoprotein | HNRNPF | 45.672 | 28.4 | 11 | 11 | 16257.59 |
| P52701 | DNA mismatch repair protein Msh6 OS=I | MSH6 | 152.786 | 22.3 | 26 | 26 | 367.58496 |
| P52732 | Kinesin-like protein KIF11 OS=Homo sa | KIF11 | 119.159 | 4.4 | 4 | 4 | 58.370892 |
| P52756 | RNA-binding protein 5 OS=Homo sapien | RBM5 | 92.154 | 22.2 | 20 | 20 | 1131.9879 |

| | | | | | | | |
|---|---|---|---|---|---|---|---|
| P52815 | Large ribosomal subunit protein bL12m C | MRPL12 | 21.349 | 23.2 | 5 | 5 | 1667.6832 |
| P52907 | F-actin-capping protein subunit alpha-1 O | CAPZA1 | 32.923 | 32.2 | 7 | 7 | 1335.9631 |
| P52948 | Nuclear pore complex protein Nup98-Nup | NUP98 | 197.58 | 7.4 | 13 | 13 | 203.03062 |
| P52952 | Homeobox protein Nkx-2.5 OS=Homo sa | NKX2-5 | 34.918 | 17.6 | 5 | 4 | 379.93625 |
| P53007 | Tricarboxylate transport protein, mitochoı | SLC25A1 | 34.013 | 25.1 | 8 | 8 | 1761.6406 |
| P53041 | Serine/threonine-protein phosphatase 5 OS | PPP5C | 56.879 | 4.6 | 2 | 2 | 49.873528 |
| P53350 | Serine/threonine-protein kinase PLK1 OS | PLK1 | 68.255 | 30.8 | 18 | 18 | 1119.323 |
| P53355 | Death-associated protein kinase 1 OS=Ho | DAPK1 | 160.047 | 0.6 | 1 | 1 | 110.41029 |
| P53396 | ATP-citrate synthase OS=Homo sapiens ( | ACLY | 120.84 | 8.7 | 8 | 8 | 103.09123 |
| P53539 | Protein FosB OS=Homo sapiens OX=960 | FOSB | 35.928 | 7.4 | 2 | 2 | 125.55348 |
| P53582 | Methionine aminopeptidase 1 OS=Homo | METAP1 | 43.216 | 22.3 | 5 | 5 | 93.472855 |
| P53611 | Geranylgeranyl transferase type-2 subunit | RABGGTB | 36.925 | 3.3 | 1 | 1 | 89.291153 |
| P53618 | Coatomer subunit beta OS=Homo sapiens | COPB1 | 107.143 | 20.9 | 16 | 16 | 285.44037 |
| P53621 | Coatomer subunit alpha OS=Homo sapien | COPA | 138.346 | 37.9 | 38 | 38 | 481.80746 |
| P53634 | Dipeptidyl peptidase 1 OS=Homo sapiens | CTSC | 51.854 | 2.6 | 1 | 1 | 43.862865 |
| P53675 | Clathrin heavy chain 2 OS=Homo sapiens | CLTCL1 | 187.03 | 8.6 | 12 | 2 | 633.62646 |
| P53680 | AP-2 complex subunit sigma OS=Homo s | AP2S1 | 17.018 | 26.1 | 4 | 4 | 1421.3364 |
| P53701 | Holocytochrome c-type synthase OS=Hor | HCCS | 30.602 | 8.6 | 2 | 2 | 58.167114 |
| P53803 | DNA-directed RNA polymerases I, II, anc | POLR2K | 7.005 | 12.1 | 1 | 1 | 120.92618 |
| P53804 | E3 ubiquitin-protein ligase TTC3 OS=Ho | TTC3 | 229.869 | 3.7 | 6 | 6 | 73.012329 |
| P53814 | Smoothelin OS=Homo sapiens OX=9606 | SMTN | 99.059 | 6.2 | 5 | 5 | 58.833538 |
| P53985 | Monocarboxylate transporter 1 OS=Homo | SLC16A1 | 53.945 | 10.2 | 4 | 4 | 322.23273 |
| P53990 | IST1 homolog OS=Homo sapiens OX=96 | IST1 | 39.751 | 4.9 | 2 | 2 | 67.133453 |
| P53992 | Protein transport protein Sec24C OS=Hor | SEC24C | 118.325 | 14.9 | 12 | 12 | 303.58386 |
| P53999 | Activated RNA polymerase II transcriptio | SUB1 | 14.396 | 29.9 | 3 | 3 | 235.41528 |
| P54098 | DNA polymerase subunit gamma-1 OS=H | POLG | 139.563 | 2 | 2 | 2 | 57.811493 |
| P54105 | Methylosome subunit pICln OS=Homo sa | CLNS1A | 26.216 | 11 | 2 | 2 | 200.60416 |
| P54132 | RecQ-like DNA helicase BLM OS=Homo | BLM | 159.001 | 20.1 | 24 | 24 | 287.08389 |
| P54136 | Arginine--tRNA ligase, cytoplasmic OS=I | RARS1 | 75.379 | 32.4 | 20 | 20 | 374.63193 |
| P54259 | Atrophin-1 OS=Homo sapiens OX=9606 | ATN1 | 125.415 | 1 | 1 | 1 | 41.195972 |
| P54274 | Telomeric repeat-binding factor 1 OS=Ho | TERF1 | 50.246 | 23.5 | 9 | 9 | 236.68327 |
| P54277 | PMS1 protein homolog 1 OS=Homo sapi | PMS1 | 105.831 | 2.3 | 2 | 2 | 40.675808 |
| P54278 | Mismatch repair endonuclease PMS2 OS= | PMS2 | 95.797 | 12.2 | 8 | 8 | 108.13372 |
| P54284 | Voltage-dependent L-type calcium channe | CACNB3 | 54.532 | 4.5 | 2 | 1 | 78.490356 |
| P54578 | Ubiquitin carboxyl-terminal hydrolase 14 | USP14 | 56.07 | 1.6 | 1 | 1 | 19.858801 |

| | | | | | | | |
|---|---|---|---|---|---|---|---|
| P54619 | 5'-AMP-activated protein kinase subunit g | PRKAG1 | 37.58 | 15.7 | 5 | 4 | 100.08071 |
| P54646 | 5'-AMP-activated protein kinase catalytic | PRKAA2 | 62.32 | 5.1 | 2 | 1 | 69.707825 |
| P54709 | Sodium/potassium-transporting ATPase su | ATP1B3 | 31.513 | 3.9 | 1 | 1 | 49.561691 |
| P54727 | UV excision repair protein RAD23 homol | RAD23B | 43.172 | 7.6 | 3 | 3 | 91.849251 |
| P54886 | Delta-1-pyrroline-5-carboxylate synthase | ALDH18A1 | 87.303 | 8.1 | 6 | 6 | 116.9916 |
| P55010 | Eukaryotic translation initiation factor 5 C | EIF5 | 49.223 | 2.3 | 1 | 1 | 143.72462 |
| P55055 | Oxysterols receptor LXR-beta OS=Homo | NR1H2 | 50.974 | 2.4 | 1 | 1 | 105.42067 |
| P55060 | Exportin-2 OS=Homo sapiens OX=9606 ( | CSE1L | 110.417 | 13.3 | 13 | 13 | 260.50668 |
| P55072 | Transitional endoplasmic reticulum ATPa | VCP | 89.322 | 20.5 | 14 | 14 | 167.86357 |
| P55081 | Microfibrillar-associated protein 1 OS=Ho | MFAP1 | 51.959 | 11.6 | 5 | 5 | 226.16472 |
| P55084 | Trifunctional enzyme subunit beta, mitoch | HADHB | 51.295 | 24.9 | 12 | 12 | 409.28378 |
| P55160 | Nck-associated protein 1-like OS=Homo s | NCKAP1L | 128.154 | 1.6 | 2 | 2 | 43.190464 |
| P55196 | Afadin OS=Homo sapiens OX=9606 GN= | AFDN | 206.805 | 2.2 | 3 | 3 | 81.996964 |
| P55199 | RNA polymerase II elongation factor ELL | ELL | 68.265 | 8.5 | 5 | 5 | 48.518837 |
| P55201 | Peregrin OS=Homo sapiens OX=9606 GN | BRPF1 | 137.499 | 3.5 | 5 | 5 | 125.24071 |
| P55209 | Nucleosome assembly protein 1-like 1 OS | NAP1L1 | 45.375 | 5.6 | 3 | 2 | 1008.4865 |
| P55265 | Double-stranded RNA-specific adenosine | ADAR | 136.067 | 32.2 | 36 | 36 | 1870.3094 |
| P55268 | Laminin subunit beta-2 OS=Homo sapiens | LAMB2 | 195.982 | 1.3 | 2 | 2 | 50.231518 |
| P55317 | Hepatocyte nuclear factor 3-alpha OS=Ho | FOXA1 | 49.148 | 2.3 | 1 | 1 | 100.70558 |
| P55735 | Protein SEC13 homolog OS=Homo sapie | SEC13 | 35.541 | 34.8 | 9 | 9 | 1494.4579 |
| P55769 | NHP2-like protein 1 OS=Homo sapiens C | SNU13 | 14.174 | 41.4 | 5 | 5 | 1170.6178 |
| P55786 | Puromycin-sensitive aminopeptidase OS= | NPEPPS | 103.277 | 1 | 1 | 1 | 22.307293 |
| P55795 | Heterogeneous nuclear ribonucleoprotein | HNRNPH2 | 49.264 | 18.5 | 5 | 5 | 5522.1396 |
| P55884 | Eukaryotic translation initiation factor 3 su | EIF3B | 92.482 | 33 | 26 | 26 | 2015.0099 |
| P56134 | ATP synthase subunit f, mitochondrial OS | ATP5MF | 10.918 | 27.7 | 2 | 2 | 372.30017 |
| P56177 | Homeobox protein DLX-1 OS=Homo sap | DLX1 | 27.32 | 29.8 | 6 | 4 | 190.08366 |
| P56178 | Homeobox protein DLX-5 OS=Homo sap | DLX5 | 31.54 | 7.6 | 2 | 2 | 316.43436 |
| P56182 | Ribosomal RNA processing protein 1 hon | RRP1 | 52.84 | 26.2 | 13 | 13 | 590.13672 |
| P56192 | Methionine--tRNA ligase, cytoplasmic OS | MARS1 | 101.116 | 15.1 | 11 | 11 | 183.99355 |
| P56199 | Integrin alpha-1 OS=Homo sapiens OX=9 | ITGA1 | 130.848 | 0.7 | 1 | 1 | 67.751099 |
| P56270 | Myc-associated zinc finger protein OS=H | MAZ | 48.608 | 16.6 | 7 | 7 | 188.01015 |
| P56381 | ATP synthase subunit epsilon, mitochondr | ATP5F1E | 5.78 | 15.7 | 1 | 1 | 179.26093 |
| P56524 | Histone deacetylase 4 OS=Homo sapiens | HDAC4 | 119.04 | 0.8 | 1 | 1 | 33.411053 |
| P56537 | Eukaryotic translation initiation factor 6 C | EIF6 | 26.6 | 36.7 | 6 | 6 | 688.2171 |
| P56545 | C-terminal-binding protein 2 OS=Homo s | CTBP2 | 48.945 | 15.3 | 7 | 4 | 244.97121 |

| | | | | | | | |
|---|---|---|---|---|---|---|---|
| P56937 | 3-keto-steroid reductase/17-beta-hydroxys | HSD17B7 | 38.207 | 3.5 | 1 | 1 | 44.32972 |
| P56945 | Breast cancer anti-estrogen resistance pro | BCAR1 | 93.372 | 2.5 | 2 | 2 | 70.304855 |
| P57088 | Transmembrane protein 33 OS=Homo sap | TMEM33 | 27.979 | 33.6 | 10 | 10 | 806.72314 |
| P57678 | Gem-associated protein 4 OS=Homo sapi | GEMIN4 | 120.038 | 28.4 | 29 | 29 | 4014.3164 |
| P57682 | Krueppel-like factor 3 OS=Homo sapiens | KLF3 | 38.829 | 3.5 | 1 | 1 | 54.583584 |
| P57721 | Poly(rC)-binding protein 3 OS=Homo sap | PCBP3 | 39.466 | 3.2 | 1 | 1 | 4037.5339 |
| P57740 | Nuclear pore complex protein Nup107 OS | NUP107 | 106.375 | 7.6 | 7 | 7 | 160.60133 |
| P57772 | Selenocysteine-specific elongation factor | EEFSEC | 65.305 | 20 | 10 | 10 | 187.83759 |
| P58012 | Forkhead box protein L2 OS=Homo sapie | FOXL2 | 38.773 | 15.2 | 3 | 3 | 83.531357 |
| P58107 | Epiplakin OS=Homo sapiens OX=9606 G | EPPK1 | 555.659 | 2.9 | 2 | 2 | 30.305223 |
| P59998 | Actin-related protein 2/3 complex subunit | ARPC4 | 19.668 | 16.1 | 3 | 3 | 66.704224 |
| P60059 | Protein transport protein Sec61 subunit ga | SEC61G | 7.742 | 19.1 | 1 | 1 | 74.33139 |
| P60174 | Triosephosphate isomerase OS=Homo sap | TPI1 | 26.67 | 5.2 | 1 | 1 | 40.052074 |
| P60228 | Eukaryotic translation initiation factor 3 s | EIF3E | 52.221 | 29.2 | 14 | 14 | 2400.4099 |
| P60468 | Protein transport protein Sec61 subunit be | SEC61B | 9.975 | 11.5 | 1 | 1 | 54.549507 |
| P60510 | Serine/threonine-protein phosphatase 4 ca | PPP4C | 35.08 | 7.8 | 2 | 2 | 70.903442 |
| P60660 | Myosin light polypeptide 6 OS=Homo sap | MYL6 | 16.931 | 22.5 | 3 | 3 | 4423.4058 |
| P60709 | Actin, cytoplasmic 1 OS=Homo sapiens C | ACTB | 41.737 | 44.3 | 14 | 5 | 24091.025 |
| P60842 | Eukaryotic initiation factor 4A-I OS=Hon | EIF4A1 | 46.154 | 23.6 | 9 | 9 | 1459.6696 |
| P60866 | Small ribosomal subunit protein uS10 OS | RPS20 | 13.373 | 26.1 | 4 | 4 | 24314.855 |
| P60891 | Ribose-phosphate pyrophosphokinase 1 O | PRPS1 | 34.835 | 29.2 | 7 | 3 | 1285.9843 |
| P60900 | Proteasome subunit alpha type-6 OS=Hon | PSMA6 | 27.4 | 5.3 | 1 | 1 | 62.873459 |
| P60903 | Protein S100-A10 OS=Homo sapiens OX | S100A10 | 11.204 | 27.8 | 2 | 2 | 268.39688 |
| P60953 | Cell division control protein 42 homolog | CDC42 | 21.259 | 23.6 | 4 | 4 | 132.87692 |
| P60981 | Destrin OS=Homo sapiens OX=9606 GN | DSTN | 18.506 | 24.2 | 4 | 4 | 415.4292 |
| P61009 | Signal peptidase complex subunit 3 OS=H | SPCS3 | 20.314 | 3.9 | 1 | 1 | 65.015831 |
| P61011 | Signal recognition particle subunit SRP54 | SRP54 | 55.705 | 5 | 2 | 2 | 38.682983 |
| P61019 | Ras-related protein Rab-2A OS=Homo sa | RAB2A | 23.546 | 12.7 | 2 | 1 | 47.230713 |
| P61024 | Cyclin-dependent kinases regulatory subu | CKS1B | 9.661 | 12.7 | 1 | 1 | 25.387562 |
| P61073 | C-X-C chemokine receptor type 4 OS=Hc | CXCR4 | 39.746 | 3.4 | 1 | 1 | 141.42879 |
| P61077 | Ubiquitin-conjugating enzyme E2 D3 OS | UBE2D3 | 16.688 | 7.5 | 1 | 1 | 149.37094 |
| P61088 | Ubiquitin-conjugating enzyme E2 N OS=l | UBE2N | 17.138 | 12.5 | 2 | 2 | 39.785759 |
| P61106 | Ras-related protein Rab-14 OS=Homo sap | RAB14 | 23.897 | 18.1 | 3 | 3 | 59.150841 |
| P61158 | Actin-related protein 3 OS=Homo sapiens | ACTR3 | 47.372 | 13.9 | 5 | 5 | 73.614258 |
| P61160 | Actin-related protein 2 OS=Homo sapiens | ACTR2 | 44.761 | 12.9 | 4 | 4 | 159.70114 |

| | | | | | | | |
|---|---|---|---|---|---|---|---|
| P61204 | ADP-ribosylation factor 3 OS=Homo sapi | ARF3 | 20.601 | 42 | 6 | 3 | 1062.6793 |
| P61221 | ATP-binding cassette sub-family E memb | ABCE1 | 67.315 | 32.1 | 18 | 18 | 1419.9631 |
| P61244 | Protein max OS=Homo sapiens OX=9606 | MAX | 18.275 | 24.4 | 4 | 4 | 480.38168 |
| P61247 | Small ribosomal subunit protein eS1 OS= | RPS3A | 29.945 | 54.9 | 17 | 17 | 32712.762 |
| P61254 | Large ribosomal subunit protein uL24 OS | RPL26 | 17.259 | 45.5 | 10 | 2 | 31768.191 |
| P61289 | Proteasome activator complex subunit 3 C | PSME3 | 29.507 | 21.3 | 5 | 5 | 101.68883 |
| P61313 | Large ribosomal subunit protein eL15 OS | RPL15 | 24.147 | 44.6 | 14 | 14 | 15776.799 |
| P61326 | Protein mago nashi homolog OS=Homo s | MAGOH | 17.164 | 51.4 | 7 | 7 | 6378.4521 |
| P61353 | Large ribosomal subunit protein eL27 OS | RPL27 | 15.798 | 48.5 | 7 | 7 | 9560.1201 |
| P61371 | Insulin gene enhancer protein ISL-1 OS=I | ISL1 | 39.036 | 2.6 | 1 | 1 | 38.848701 |
| P61513 | Large ribosomal subunit protein eL43 OS | RPL37A | 10.276 | 31.5 | 4 | 3 | 4893.7012 |
| P61586 | Transforming protein RhoA OS=Homo sa | RHOA | 21.769 | 6.2 | 1 | 1 | 9.4949217 |
| P61587 | Rho-related GTP-binding protein RhoE O | RND3 | 27.369 | 13.1 | 3 | 3 | 42.88821 |
| P61619 | Protein transport protein Sec61 subunit al | SEC61A1 | 52.265 | 7.6 | 4 | 2 | 242.78712 |
| P61758 | Prefoldin subunit 3 OS=Homo sapiens OX | VBP1 | 22.626 | 16.8 | 3 | 3 | 50.11438 |
| P61769 | Beta-2-microglobulin OS=Homo sapiens ( | B2M | 13.715 | 8.4 | 1 | 1 | 58.320831 |
| P61803 | Dolichyl-diphosphooligosaccharide--prote | DAD1 | 12.497 | 10.6 | 1 | 1 | 64.877243 |
| P61927 | Large ribosomal subunit protein eL37 OS | RPL37 | 11.078 | 7.2 | 1 | 1 | 2931.5134 |
| P61960 | Ubiquitin-fold modifier 1 OS=Homo sapi | UFM1 | 9.118 | 17.6 | 1 | 1 | 249.71428 |
| P61962 | DDB1- and CUL4-associated factor 7 OS | DCAF7 | 38.927 | 35.7 | 11 | 11 | 427.3111 |
| P61964 | WD repeat-containing protein 5 OS=Hom | WDR5 | 36.589 | 34.7 | 9 | 8 | 267.76886 |
| P61968 | LIM domain transcription factor LMO4 C | LMO4 | 17.994 | 10.3 | 2 | 2 | 71.369202 |
| P61978 | Heterogeneous nuclear ribonucleoprotein | HNRNPK | 50.977 | 36.3 | 18 | 18 | 33521.348 |
| P61981 | 14-3-3 protein gamma OS=Homo sapiens | YWHAG | 28.303 | 28.3 | 7 | 6 | 254.97478 |
| P62072 | Mitochondrial import inner membrane tra | TIMM10 | 10.333 | 22.2 | 2 | 2 | 92.390739 |
| P62081 | Small ribosomal subunit protein eS7 OS= | RPS7 | 22.127 | 30.9 | 4 | 4 | 18882.838 |
| P62136 | Serine/threonine-protein phosphatase PP1 | PPP1CA | 37.513 | 12.7 | 4 | 4 | 1099.0635 |
| P62140 | Serine/threonine-protein phosphatase PP1 | PPP1CB | 37.187 | 18.3 | 5 | 5 | 601.14264 |
| P62166 | Neuronal calcium sensor 1 OS=Homo sap | NCS1 | 21.879 | 5.8 | 1 | 1 | 97.201035 |
| P62191 | 26S proteasome regulatory subunit 4 OS= | PSMC1 | 49.185 | 4.8 | 2 | 2 | 81.916214 |
| P62195 | 26S proteasome regulatory subunit 8 OS= | PSMC5 | 45.626 | 9.1 | 3 | 3 | 104.80038 |
| P62241 | Small ribosomal subunit protein eS8 OS= | RPS8 | 24.206 | 51 | 11 | 11 | 37490.398 |
| P62244 | Small ribosomal subunit protein uS8 OS= | RPS15A | 14.84 | 60.8 | 8 | 8 | 42813.828 |
| P62249 | Small ribosomal subunit protein uS9 OS= | RPS16 | 16.446 | 50.7 | 10 | 10 | 32234.336 |
| P62258 | 14-3-3 protein epsilon OS=Homo sapiens | YWHAE | 29.174 | 37.6 | 9 | 9 | 570.79572 |

| | | | | | | | |
|---|---|---|---|---|---|---|---|
| P62263 | Small ribosomal subunit protein uS11 OS= | RPS14 | 16.273 | 39.1 | 7 | 7 | 46829.445 |
| P62266 | Small ribosomal subunit protein uS12 OS= | RPS23 | 15.808 | 30.1 | 6 | 6 | 26309.459 |
| P62269 | Small ribosomal subunit protein uS13 OS= | RPS18 | 17.719 | 54.6 | 12 | 12 | 40422.328 |
| P62273 | Small ribosomal subunit protein uS14 OS= | RPS29 | 6.677 | 33.9 | 3 | 3 | 2393.4104 |
| P62277 | Small ribosomal subunit protein uS15 OS= | RPS13 | 17.223 | 45.7 | 9 | 9 | 45297.113 |
| P62280 | Small ribosomal subunit protein uS17 OS= | RPS11 | 18.431 | 57 | 14 | 14 | 25328.166 |
| P62304 | Small nuclear ribonucleoprotein E OS=Ho | SNRPE | 10.804 | 38 | 3 | 3 | 986.54956 |
| P62306 | Small nuclear ribonucleoprotein F OS=Ho | SNRPF | 9.726 | 50 | 6 | 6 | 935.52136 |
| P62310 | U6 snRNA-associated Sm-like protein LS | LSM3 | 11.846 | 11.8 | 1 | 1 | 1702.0432 |
| P62312 | U6 snRNA-associated Sm-like protein LS | LSM6 | 9.128 | 37.5 | 4 | 4 | 517.31097 |
| P62314 | Small nuclear ribonucleoprotein Sm D1 O | SNRPD1 | 13.282 | 37.8 | 4 | 4 | 916.82648 |
| P62316 | Small nuclear ribonucleoprotein Sm D2 O | SNRPD2 | 13.527 | 45.8 | 6 | 6 | 2932.4265 |
| P62318 | Small nuclear ribonucleoprotein Sm D3 O | SNRPD3 | 13.917 | 31.7 | 3 | 3 | 2783.5417 |
| P62330 | ADP-ribosylation factor 6 OS=Homo sapi | ARF6 | 20.083 | 14.9 | 2 | 2 | 204.26132 |
| P62333 | 26S proteasome regulatory subunit 10B O | PSMC6 | 44.173 | 6.2 | 2 | 2 | 50.026379 |
| P62424 | Large ribosomal subunit protein eL8 OS= | RPL7A | 29.996 | 40.6 | 16 | 16 | 24957.053 |
| P62491 | Ras-related protein Rab-11A OS=Homo s | RAB11A | 24.394 | 10.2 | 2 | 2 | 57.162052 |
| P62495 | Eukaryotic peptide chain release factor su | ETF1 | 49.031 | 15.3 | 6 | 6 | 263.23041 |
| P62633 | CCHC-type zinc finger nucleic acid bindi | CNBP | 19.463 | 44.6 | 9 | 9 | 9854.9971 |
| P62701 | Small ribosomal subunit protein eS4, X is | RPS4X | 29.598 | 43.7 | 13 | 4 | 37287.574 |
| P62714 | Serine/threonine-protein phosphatase 2A ( | PPP2CB | 35.576 | 15.5 | 4 | 1 | 481.7287 |
| P62736 | Actin, aortic smooth muscle OS=Homo sa | ACTA2 | 42.009 | 3.4 | 1 | 0 | 1553.5259 |
| P62750 | Large ribosomal subunit protein uL23 OS= | RPL23A | 17.696 | 44.2 | 10 | 10 | 23955.559 |
| P62753 | Small ribosomal subunit protein eS6 OS= | RPS6 | 28.681 | 29.3 | 8 | 8 | 33764.984 |
| P62805 | Histone H4 OS=Homo sapiens OX=9606 | H4C1 | 11.368 | 51.5 | 6 | 6 | 31553.311 |
| P62826 | GTP-binding nuclear protein Ran OS=Ho | RAN | 24.424 | 46.3 | 13 | 13 | 2362.2615 |
| P62829 | Large ribosomal subunit protein uL14 OS= | RPL23 | 14.866 | 57.9 | 7 | 7 | 14258.029 |
| P62834 | Ras-related protein Rap-1A OS=Homo sa | RAP1A | 20.988 | 26.6 | 4 | 1 | 182.00415 |
| P62841 | Small ribosomal subunit protein uS19 OS= | RPS15 | 17.041 | 39.3 | 7 | 7 | 3757.8672 |
| P62847 | Small ribosomal subunit protein eS24 OS= | RPS24 | 15.424 | 26.3 | 4 | 4 | 4341.1035 |
| P62851 | Small ribosomal subunit protein eS25 OS= | RPS25 | 13.743 | 36 | 6 | 6 | 43879.664 |
| P62854 | Small ribosomal subunit protein eS26 OS= | RPS26 | 13.016 | 53.9 | 7 | 1 | 3167.3176 |
| P62857 | Small ribosomal subunit protein eS28 OS= | RPS28 | 7.842 | 46.4 | 3 | 3 | 6058.1177 |
| P62861 | Ubiquitin-like FUBI-ribosomal protein eS | FAU | 14.39 | 8.3 | 2 | 2 | 34311.328 |
| P62873 | Guanine nucleotide-binding protein G(I)/( | GNB1 | 37.377 | 3.5 | 1 | 1 | 47.157677 |

| | | | | | | | |
|---|---|---|---|---|---|---|---|
| P62875 | DNA-directed RNA polymerases I, II, and | POLR2L | 7.646 | 29.9 | 2 | 2 | 286.24518 |
| P62877 | E3 ubiquitin-protein ligase RBX1 OS=Ho | RBX1 | 12.274 | 24.1 | 2 | 2 | 420.69678 |
| P62879 | Guanine nucleotide-binding protein G(I)/( | GNB2 | 37.332 | 12.1 | 4 | 0 | 291.15869 |
| P62888 | Large ribosomal subunit protein eL30 OS | RPL30 | 12.785 | 53 | 7 | 7 | 12278.539 |
| P62891 | Large ribosomal subunit protein eL39 OS | RPL39 | 6.407 | 19.6 | 2 | 2 | 1309.5938 |
| P62899 | Large ribosomal subunit protein eL31 OS | RPL31 | 14.463 | 45.6 | 6 | 6 | 27984.217 |
| P62906 | Large ribosomal subunit protein uL1 OS= | RPL10A | 24.832 | 49.8 | 14 | 14 | 6695.4683 |
| P62910 | Large ribosomal subunit protein eL32 OS | RPL32 | 15.86 | 46.7 | 8 | 8 | 14671.523 |
| P62913 | Large ribosomal subunit protein uL5 OS= | RPL11 | 20.253 | 47.2 | 8 | 8 | 25485.111 |
| P62917 | Large ribosomal subunit protein uL2 OS= | RPL8 | 28.025 | 55.6 | 12 | 12 | 22961.891 |
| P62937 | Peptidyl-prolyl cis-trans isomerase A OS= | PPIA | 18.013 | 32.7 | 5 | 4 | 1297.5355 |
| P62979 | Ubiquitin-ribosomal protein eS31 fusion p | RPS27A | 17.965 | 41.7 | 6 | 3 | 5684.6631 |
| P62987 | Ubiquitin-ribosomal protein eL40 fusion p | UBA52 | 14.729 | 7 | 1 | 1 | 142.18269 |
| P62993 | Growth factor receptor-bound protein 2 O | GRB2 | 25.207 | 40.6 | 9 | 9 | 571.98486 |
| P62995 | Transformer-2 protein homolog beta OS= | TRA2B | 33.666 | 33.3 | 8 | 6 | 3148.0508 |
| P63000 | Ras-related C3 botulinum toxin substrate | RAC1 | 21.451 | 27.1 | 5 | 1 | 354.9447 |
| P63010 | AP-2 complex subunit beta OS=Homo sap | AP2B1 | 104.553 | 33.5 | 29 | 15 | 1275.2606 |
| P63092 | Guanine nucleotide-binding protein G(s) s | GNAS | 45.665 | 12.9 | 5 | 5 | 276.09152 |
| P63098 | Calcineurin subunit B type 1 OS=Homo s | PPP3R1 | 19.3 | 5.9 | 1 | 1 | 68.276688 |
| P63104 | 14-3-3 protein zeta/delta OS=Homo sapie | YWHAZ | 27.746 | 21.2 | 5 | 5 | 497.74136 |
| P63151 | Serine/threonine-protein phosphatase 2A : | PPP2R2A | 51.693 | 19.9 | 8 | 4 | 266.83914 |
| P63167 | Dynein light chain 1, cytoplasmic OS=Ho | DYNLL1 | 10.366 | 50.6 | 4 | 1 | 4869.4722 |
| P63173 | Large ribosomal subunit protein eL38 OS | RPL38 | 8.218 | 54.3 | 6 | 6 | 4912.4067 |
| P63208 | S-phase kinase-associated protein 1 OS=H | SKP1 | 18.658 | 44.2 | 8 | 8 | 771.99554 |
| P63220 | Small ribosomal subunit protein eS21 OS | RPS21 | 9.112 | 39.8 | 3 | 3 | 3908.7461 |
| P63241 | Eukaryotic translation initiation factor 5A | EIF5A | 16.833 | 20.8 | 4 | 1 | 1167.3827 |
| P63244 | Small ribosomal subunit protein RACK1 ( | RACK1 | 35.077 | 51.1 | 14 | 14 | 6498.9126 |
| P63272 | Transcription elongation factor SPT4 OS= | SUPT4H1 | 13.194 | 21.4 | 2 | 2 | 273.37875 |
| P63279 | SUMO-conjugating enzyme UBC9 OS=H | UBE2I | 18.007 | 7.6 | 1 | 1 | 56.6329 |
| P67775 | Serine/threonine-protein phosphatase 2A ( | PPP2CA | 35.595 | 2.6 | 1 | 1 | 130.44347 |
| P67809 | Y-box-binding protein 1 OS=Homo sapie | YBX1 | 35.925 | 30.6 | 8 | 8 | 15934.13 |
| P67812 | Signal peptidase complex catalytic subuni | SEC11A | 20.626 | 3.9 | 1 | 1 | 70.412903 |
| P67870 | Casein kinase II subunit beta OS=Homo s | CSNK2B | 24.943 | 11.6 | 2 | 2 | 214.19016 |
| P67936 | Tropomyosin alpha-4 chain OS=Homo sa | TPM4 | 28.522 | 14.5 | 4 | 4 | 297.33334 |
| P68036 | Ubiquitin-conjugating enzyme E2 L3 OS= | UBE2L3 | 17.862 | 20.8 | 3 | 1 | 88.724182 |

| | | | | | | | |
|---|---|---|---|---|---|---|---|
| P68104 | Elongation factor 1-alpha 1 OS=Homo saj | EEF1A1 | 50.141 | 35.1 | 14 | 7 | 18942.715 |
| P68363 | Tubulin alpha-1B chain OS=Homo sapien | TUBA1B | 50.152 | 32.6 | 14 | 0 | 7640.3438 |
| P68366 | Tubulin alpha-4A chain OS=Homo sapier | TUBA4A | 49.925 | 4.2 | 2 | 2 | 1280.632 |
| P68371 | Tubulin beta-4B chain OS=Homo sapiens | TUBB4B | 49.831 | 48.1 | 18 | 1 | 15648.046 |
| P68400 | Casein kinase II subunit alpha OS=Homo | CSNK2A1 | 45.144 | 10.5 | 4 | 1 | 75.425301 |
| P68431 | Histone H3.1 OS=Homo sapiens OX=960 | H3C1 | 15.405 | 5.1 | 1 | 1 | 256.10181 |
| P69905 | Hemoglobin subunit alpha OS=Homo sap | HBA1 | 15.258 | 36.6 | 4 | 4 | 757.93323 |
| P78316 | Nucleolar protein 14 OS=Homo sapiens C | NOP14 | 97.669 | 32.7 | 27 | 27 | 1058.0149 |
| P78317 | E3 ubiquitin-protein ligase RNF4 OS=Ho | RNF4 | 21.319 | 10 | 2 | 2 | 221.40536 |
| P78318 | Immunoglobulin-binding protein 1 OS=H | IGBP1 | 39.222 | 9.7 | 3 | 3 | 125.75694 |
| P78332 | RNA-binding protein 6 OS=Homo sapien | RBM6 | 128.644 | 42.7 | 49 | 48 | 7500.2466 |
| P78337 | Pituitary homeobox 1 OS=Homo sapiens | PITX1 | 34.129 | 23.6 | 7 | 6 | 1259.5559 |
| P78344 | Eukaryotic translation initiation factor 4 g | EIF4G2 | 102.362 | 28.4 | 27 | 27 | 788.07483 |
| P78345 | Ribonuclease P protein subunit p38 OS=H | RPP38 | 31.835 | 50.9 | 15 | 15 | 579.20288 |
| P78346 | Ribonuclease P protein subunit p30 OS=H | RPP30 | 29.322 | 56.7 | 12 | 12 | 387.37344 |
| P78347 | General transcription factor II-I OS=Hom | GTF2I | 112.417 | 16.9 | 16 | 16 | 284.41461 |
| P78362 | SRSF protein kinase 2 OS=Homo sapiens | SRPK2 | 77.527 | 22.1 | 13 | 11 | 2718.9678 |
| P78368 | Casein kinase I isoform gamma-2 OS=Ho | CSNK1G2 | 47.458 | 9.6 | 3 | 3 | 80.207619 |
| P78371 | T-complex protein 1 subunit beta OS=Ho | CCT2 | 57.489 | 31.2 | 13 | 13 | 263.97849 |
| P78406 | mRNA export factor RAE1 OS=Homo saj | RAE1 | 40.969 | 22.3 | 6 | 6 | 459.46231 |
| P78411 | Iroquois-class homeodomain protein IRX- | IRX5 | 50.361 | 7 | 3 | 3 | 114.91071 |
| P78524 | DENN domain-containing protein 2B OS= | DENND2B | 126.485 | 3.3 | 3 | 3 | 144.9427 |
| P78527 | DNA-dependent protein kinase catalytic s | PRKDC | 469.089 | 19.1 | 73 | 73 | 376.74475 |
| P78549 | Endonuclease III-like protein 1 OS=Homo | NTHL1 | 34.39 | 18.6 | 5 | 5 | 360.26563 |
| P78563 | Double-stranded RNA-specific editase 1 C | ADARB1 | 80.764 | 20.2 | 14 | 14 | 362.32095 |
| P81605 | Dermcidin OS=Homo sapiens OX=9606 ( | DCD | 11.284 | 10 | 1 | 1 | 37.533024 |
| P82650 | Small ribosomal subunit protein mS22 OS | MRPS22 | 41.281 | 40 | 16 | 16 | 2095.5105 |
| P82914 | Small ribosomal subunit protein uS15m O | MRPS15 | 29.843 | 31.9 | 10 | 10 | 505.82904 |
| P82930 | Small ribosomal subunit protein mS34 OS | MRPS34 | 25.651 | 39.4 | 8 | 8 | 1181.3978 |
| P82932 | Small ribosomal subunit protein bS6m OS | MRPS6 | 14.227 | 32.8 | 4 | 4 | 1139.0546 |
| P83111 | Serine beta-lactamase-like protein LACTI | LACTB | 60.694 | 25.6 | 13 | 13 | 1753.4504 |
| P83369 | U7 snRNA-associated Sm-like protein LS | LSM11 | 39.5 | 11.4 | 4 | 4 | 157.59525 |
| P83731 | Large ribosomal subunit protein eL24 OS | RPL24 | 17.779 | 39.5 | 8 | 8 | 17036.988 |
| P83876 | Thioredoxin-like protein 4A OS=Homo sa | TXNL4A | 16.787 | 16.2 | 2 | 2 | 97.751266 |
| P83881 | Large ribosomal subunit protein eL42 OS | RPL36A | 12.441 | 41.5 | 8 | 2 | 10451.488 |

| | | | | | | | |
|---|---|---|---|---|---|---|---|
| P83916 | Chromobox protein homolog 1 OS=Homo | CBX1 | 21.418 | 12.4 | 2 | 2 | 61.528633 |
| P84022 | Mothers against decapentaplegic homolog | SMAD3 | 48.081 | 9.9 | 4 | 4 | 255.76833 |
| P84085 | ADP-ribosylation factor 5 OS=Homo sapi | ARF5 | 20.53 | 30 | 5 | 5 | 115.56999 |
| P84090 | Enhancer of rudimentary homolog OS=Ho | ERH | 12.259 | 27.9 | 3 | 3 | 3484.2822 |
| P84095 | Rho-related GTP-binding protein RhoG O | RHOG | 21.309 | 11.5 | 2 | 2 | 248.87396 |
| P84098 | Large ribosomal subunit protein eL19 OS | RPL19 | 23.466 | 27.6 | 7 | 7 | 18222.188 |
| P84103 | Serine/arginine-rich splicing factor 3 OS= | SRSF3 | 19.33 | 32.3 | 6 | 5 | 16714.463 |
| P85037 | Forkhead box protein K1 OS=Homo sapie | FOXK1 | 75.458 | 15.1 | 10 | 10 | 279.03714 |
| P85299 | Proline-rich protein 5 OS=Homo sapiens | PRR5 | 42.754 | 3.9 | 1 | 1 | 21.787712 |
| P98082 | Disabled homolog 2 OS=Homo sapiens O | DAB2 | 82.448 | 1.7 | 1 | 1 | 62.632092 |
| P98095 | Fibulin-2 OS=Homo sapiens OX=9606 G | FBLN2 | 126.573 | 1.8 | 2 | 2 | 56.402153 |
| P98160 | Basement membrane-specific heparan sul | HSPG2 | 468.83 | 1.2 | 4 | 4 | 117.76743 |
| P98164 | Low-density lipoprotein receptor-related p | LRP2 | 521.958 | 0.6 | 1 | 1 | 8.7202435 |
| P98171 | Rho GTPase-activating protein 4 OS=Hor | ARHGAP4 | 105.026 | 8.4 | 6 | 6 | 103.40919 |
| P98175 | RNA-binding protein 10 OS=Homo sapie | RBM10 | 103.533 | 22.9 | 23 | 20 | 3209.2466 |
| P98179 | RNA-binding protein 3 OS=Homo sapiens | RBM3 | 17.171 | 24.2 | 3 | 3 | 786.8941 |
| P99999 | Cytochrome c OS=Homo sapiens OX=960 | CYCS | 11.749 | 10.5 | 1 | 1 | 160.53041 |
| Q00059 | Transcription factor A, mitochondrial OS= | TFAM | 29.097 | 28.9 | 10 | 10 | 3993.4502 |
| Q00325 | Solute carrier family 25 member 3 OS=Ho | SLC25A3 | 40.095 | 19.1 | 8 | 8 | 3128.168 |
| Q00341 | Vigilin OS=Homo sapiens OX=9606 GN= | HDLBP | 141.456 | 30.4 | 34 | 34 | 744.84991 |
| Q00403 | Transcription initiation factor IIB OS=Ho | GTF2B | 34.833 | 32.9 | 10 | 10 | 615.40149 |
| Q00444 | Homeobox protein Hox-C5 OS=Homo sa | HOXC5 | 24.977 | 15.3 | 2 | 2 | 248.97357 |
| Q00534 | Cyclin-dependent kinase 6 OS=Homo sap | CDK6 | 36.939 | 2.8 | 1 | 1 | 59.461342 |
| Q00535 | Cyclin-dependent kinase 5 OS=Homo sap | CDK5 | 33.305 | 19.5 | 5 | 5 | 138.10512 |
| Q00537 | Cyclin-dependent kinase 17 OS=Homo sa | CDK17 | 59.583 | 18 | 9 | 5 | 474.80371 |
| Q00577 | Transcriptional activator protein Pur-alpha | PURA | 34.911 | 24.8 | 9 | 9 | 4810.5684 |
| Q00587 | Cdc42 effector protein 1 OS=Homo sapie | CDC42EP1 | 40.295 | 9.7 | 3 | 3 | 237.69701 |
| Q00610 | Clathrin heavy chain 1 OS=Homo sapiens | CLTC | 191.615 | 25.2 | 36 | 36 | 1007.999 |
| Q00653 | Nuclear factor NF-kappa-B p100 subunit | NFKB2 | 96.75 | 18.8 | 15 | 15 | 495.40744 |
| Q00839 | Heterogeneous nuclear ribonucleoprotein | HNRNPU | 90.585 | 26.4 | 23 | 23 | 23043.859 |
| Q01081 | Splicing factor U2AF 35 kDa subunit OS= | U2AF1 | 27.873 | 22.5 | 7 | 4 | 3563.4473 |
| Q01082 | Spectrin beta chain, non-erythrocytic 1 OS | SPTBN1 | 274.61 | 15.7 | 31 | 29 | 175.0649 |
| Q01085 | Nucleolysin TIAR OS=Homo sapiens OX | TIAL1 | 41.591 | 12.3 | 3 | 3 | 620.32642 |
| Q01105 | Protein SET OS=Homo sapiens OX=9606 | SET | 33.489 | 7.2 | 2 | 1 | 81.015747 |
| Q01130 | Serine/arginine-rich splicing factor 2 OS= | SRSF2 | 25.477 | 3.6 | 1 | 1 | 1333.4639 |

| | | | | | | | |
|---|---|---|---|---|---|---|---|
| Q01167 | Forkhead box protein K2 OS=Homo sapie | FOXK2 | 69.062 | 3.9 | 2 | 2 | 37.947277 |
| Q01196 | Runt-related transcription factor 1 OS=Hc | RUNX1 | 48.737 | 4.9 | 2 | 2 | 77.601028 |
| Q01518 | Adenylyl cyclase-associated protein 1 OS | CAP1 | 51.902 | 9.7 | 4 | 4 | 57.735397 |
| Q01628 | Interferon-induced transmembrane proteir | IFITM3 | 14.633 | 30.8 | 2 | 1 | 44.546654 |
| Q01650 | Large neutral amino acids transporter sma | SLC7A5 | 55.011 | 4.5 | 2 | 2 | 58.101158 |
| Q01658 | Protein Dr1 OS=Homo sapiens OX=9606 | DR1 | 19.444 | 7.4 | 1 | 1 | 189.32526 |
| Q01664 | Transcription factor AP-4 OS=Homo sapi | TFAP4 | 38.726 | 14.2 | 5 | 5 | 183.94139 |
| Q01780 | Exosome complex component 10 OS=Hoi | EXOSC10 | 100.832 | 27.1 | 22 | 22 | 402.0549 |
| Q01804 | OTU domain-containing protein 4 OS=Hc | OTUD4 | 124.045 | 17.7 | 18 | 18 | 644.03479 |
| Q01813 | ATP-dependent 6-phosphofructokinase, p | PFKP | 85.597 | 26.4 | 20 | 20 | 1405.1179 |
| Q01831 | DNA repair protein complementing XP-C | XPC | 105.953 | 4.7 | 4 | 4 | 70.879166 |
| Q01844 | RNA-binding protein EWS OS=Homo saj | EWSR1 | 68.479 | 9 | 5 | 5 | 11703.937 |
| Q01955 | Collagen alpha-3(IV) chain OS=Homo saj | COL4A3 | 161.813 | 0.6 | 1 | 1 | 847.05524 |
| Q01959 | Sodium-dependent dopamine transporter ( | SLC6A3 | 68.495 | 3.5 | 1 | 1 | 81.885597 |
| Q01968 | Inositol polyphosphate 5-phosphatase OC | OCRL | 104.205 | 9.1 | 7 | 7 | 106.20927 |
| Q01970 | 1-phosphatidylinositol 4,5-bisphosphate p | PLCB3 | 138.799 | 2 | 2 | 2 | 26.429909 |
| Q02040 | A-kinase anchor protein 17A OS=Homo s | AKAP17A | 80.736 | 8.8 | 6 | 6 | 148.3862 |
| Q02086 | Transcription factor Sp2 OS=Homo sapie | SP2 | 64.901 | 2.6 | 1 | 1 | 43.70882 |
| Q02218 | 2-oxoglutarate dehydrogenase complex cc | OGDH | 115.936 | 8 | 8 | 3 | 79.595047 |
| Q02224 | Centromere-associated protein E OS=Hor | CENPE | 316.416 | 4.7 | 11 | 11 | 123.5271 |
| Q02241 | Kinesin-like protein KIF23 OS=Homo saj | KIF23 | 110.059 | 39 | 33 | 33 | 2590.5974 |
| Q02487 | Desmocollin-2 OS=Homo sapiens OX=96 | DSC2 | 99.962 | 1.4 | 1 | 1 | 73.66407 |
| Q02535 | DNA-binding protein inhibitor ID-3 OS=I | ID3 | 13 | 20.2 | 2 | 2 | 45.171494 |
| Q02543 | Large ribosomal subunit protein eL20 OS | RPL18A | 20.763 | 52.8 | 10 | 10 | 23636.828 |
| Q02750 | Dual specificity mitogen-activated protein | MAP2K1 | 43.44 | 2.8 | 1 | 1 | 42.064445 |
| Q02790 | Peptidyl-prolyl cis-trans isomerase FKBP | FKBP4 | 51.805 | 3.1 | 1 | 1 | 53.986847 |
| Q02809 | Procollagen-lysine,2-oxoglutarate 5-dioxy | PLOD1 | 83.551 | 19.7 | 12 | 12 | 171.12914 |
| Q02878 | Large ribosomal subunit protein eL6 OS= | RPL6 | 32.728 | 34 | 12 | 12 | 28784.947 |
| Q02880 | DNA topoisomerase 2-beta OS=Homo saj | TOP2B | 183.268 | 31.4 | 49 | 35 | 2538.467 |
| Q02930 | Cyclic AMP-responsive element-binding j | CREB5 | 56.919 | 1.8 | 1 | 1 | 38.853016 |
| Q02978 | Mitochondrial 2-oxoglutarate/malate carri | SLC25A11 | 34.062 | 35.4 | 10 | 10 | 329.54276 |
| Q03001 | Dystonin OS=Homo sapiens OX=9606 G | DST | 860.662 | 21.5 | 153 | 153 | 910.96655 |
| Q03111 | Protein ENL OS=Homo sapiens OX=960 | MLLT1 | 62.057 | 15.2 | 7 | 7 | 220.71989 |
| Q03112 | Histone-lysine N-methyltransferase MEC | MECOM | 138.136 | 2.6 | 2 | 2 | 42.797886 |
| Q03164 | Histone-lysine N-methyltransferase 2A OS | KMT2A | 431.765 | 2.1 | 5 | 5 | 69.01712 |

| | | | | | | | |
|---|---|---|---|---|---|---|---|
| Q03252 | Lamin-B2 OS=Homo sapiens OX=9606 ( | LMNB2 | 69.949 | 14.2 | 8 | 8 | 94.544228 |
| Q03468 | DNA excision repair protein ERCC-6 OS= | ERCC6 | 168.417 | 14.1 | 19 | 19 | 386.99063 |
| Q03519 | Antigen peptide transporter 2 OS=Homo s | TAP2 | 75.664 | 1.2 | 1 | 1 | 92.752022 |
| Q04206 | Transcription factor p65 OS=Homo sapier | RELA | 60.22 | 7.4 | 4 | 4 | 90.590385 |
| Q04637 | Eukaryotic translation initiation factor 4 g | EIF4G1 | 175.492 | 28.1 | 49 | 49 | 7987.0029 |
| Q04721 | Neurogenic locus notch homolog protein 2 | NOTCH2 | 265.405 | 3.8 | 8 | 8 | 88.058533 |
| Q04726 | Transducin-like enhancer protein 3 OS=H | TLE3 | 83.417 | 1.9 | 1 | 1 | 40.29557 |
| Q04837 | Single-stranded DNA-binding protein, mi | SSBP1 | 17.26 | 40.5 | 5 | 5 | 231.73 |
| Q04864 | Proto-oncogene c-Rel OS=Homo sapiens | REL | 68.52 | 21 | 11 | 11 | 447.746 |
| Q04917 | 14-3-3 protein eta OS=Homo sapiens OX= | YWHAH | 28.219 | 10.6 | 2 | 2 | 224.28885 |
| Q05397 | Focal adhesion kinase 1 OS=Homo sapier | PTK2 | 119.234 | 10.6 | 10 | 9 | 132.86755 |
| Q05519 | Serine/arginine-rich splicing factor 11 OS | SRSF11 | 53.543 | 16.5 | 7 | 7 | 1190.4824 |
| Q05823 | 2-5A-dependent ribonuclease OS=Homo s | RNASEL | 83.533 | 1.5 | 1 | 1 | 43.709747 |
| Q05932 | Folylpolyglutamate synthase, mitochondri | FPGS | 64.61 | 3.4 | 2 | 2 | 55.489597 |
| Q06033 | Inter-alpha-trypsin inhibitor heavy chain H | ITIH3 | 99.85 | 2.5 | 2 | 2 | 218.81114 |
| Q06124 | Tyrosine-protein phosphatase non-recepto | PTPN11 | 68.011 | 2.5 | 2 | 2 | 36.300629 |
| Q06210 | Glutamine--fructose-6-phosphate aminotra | GFPT1 | 78.807 | 14.7 | 8 | 8 | 121.03413 |
| Q06265 | Exosome complex component RRP45 OS | EXOSC9 | 48.95 | 17.5 | 7 | 7 | 730.36438 |
| Q06330 | Recombining binding protein suppressor o | RBPJ | 55.638 | 29.8 | 14 | 13 | 419.65692 |
| Q06481 | Amyloid beta precursor like protein 2 OS= | APLP2 | 86.956 | 2.9 | 2 | 2 | 48.268078 |
| Q06587 | E3 ubiquitin-protein ligase RING1 OS=H | RING1 | 42.43 | 8.4 | 3 | 3 | 159.76021 |
| Q06787 | Fragile X messenger ribonucleoprotein 1 ( | FMR1 | 71.175 | 33.2 | 19 | 19 | 9906.1201 |
| Q06830 | Peroxiredoxin-1 OS=Homo sapiens OX=9 | PRDX1 | 22.111 | 68.8 | 12 | 10 | 896.07355 |
| Q07020 | Large ribosomal subunit protein eL18 OS= | RPL18 | 21.635 | 46.3 | 9 | 9 | 30488.342 |
| Q07021 | Complement component 1 Q subcompone | C1QBP | 31.363 | 22 | 4 | 4 | 724.98596 |
| Q07065 | Cytoskeleton-associated protein 4 OS=Ho | CKAP4 | 66.023 | 38.4 | 21 | 21 | 1655.5364 |
| Q07157 | Tight junction protein ZO-1 OS=Homo sa | TJP1 | 195.46 | 32.7 | 49 | 48 | 1175.0083 |
| Q07352 | mRNA decay activator protein ZFP36L1 ( | ZFP36L1 | 36.314 | 18.6 | 6 | 5 | 290.22415 |
| Q07666 | KH domain-containing, RNA-binding, sig | KHDRBS1 | 48.228 | 22.8 | 12 | 8 | 40744.016 |
| Q07866 | Kinesin light chain 1 OS=Homo sapiens ( | KLC1 | 65.31 | 3.8 | 2 | 1 | 40.952824 |
| Q07955 | Serine/arginine-rich splicing factor 1 OS= | SRSF1 | 27.745 | 44 | 13 | 12 | 9953.7676 |
| Q08043 | Alpha-actinin-3 OS=Homo sapiens OX=9 | ACTN3 | 103.242 | 0.9 | 1 | 1 | 1808.8347 |
| Q08050 | Forkhead box protein M1 OS=Homo sapi | FOXM1 | 84.283 | 8.8 | 7 | 7 | 124.75815 |
| Q08170 | Serine/arginine-rich splicing factor 4 OS= | SRSF4 | 56.679 | 9.1 | 4 | 4 | 279.79047 |
| Q08209 | Protein phosphatase 3 catalytic subunit al | PPP3CA | 58.688 | 14 | 6 | 4 | 511.18985 |

| | | | | | | | |
|---|---|---|---|---|---|---|---|
| Q08211 | ATP-dependent RNA helicase A OS=Hor | DHX9 | 140.959 | 32.5 | 41 | 41 | 9341.3496 |
| Q08378 | Golgin subfamily A member 3 OS=Homo | GOLGA3 | 167.355 | 0.5 | 1 | 1 | 147.90251 |
| Q08379 | Golgin subfamily A member 2 OS=Homo | GOLGA2 | 113.087 | 24.7 | 20 | 20 | 327.17657 |
| Q08380 | Galectin-3-binding protein OS=Homo sap | LGALS3BP | 65.331 | 6 | 3 | 3 | 109.39732 |
| Q08945 | FACT complex subunit SSRP1 OS=Homo | SSRP1 | 81.075 | 21.2 | 13 | 13 | 240.42911 |
| Q08AD1 | Calmodulin-regulated spectrin-associated | CAMSAP2 | 168.089 | 21.3 | 26 | 26 | 439.19641 |
| Q08AE8 | Protein spire homolog 1 OS=Homo sapier | SPIRE1 | 85.544 | 1.5 | 1 | 1 | 22.719164 |
| Q08AG7 | Mitotic-spindle organizing protein 1 OS=] | MZT1 | 8.479 | 13.4 | 1 | 1 | 856.58191 |
| Q08J23 | RNA cytosine C(5)-methyltransferase NS | NSUN2 | 86.471 | 39 | 32 | 32 | 4893.2363 |
| Q09028 | Histone-binding protein RBBP4 OS=Hon | RBBP4 | 47.656 | 16.7 | 8 | 3 | 978.5799 |
| Q09161 | Nuclear cap-binding protein subunit 1 OS | NCBP1 | 91.84 | 24.2 | 17 | 17 | 11274.687 |
| Q09472 | Histone acetyltransferase p300 OS=Homo | EP300 | 264.161 | 1.1 | 2 | 2 | 56.339035 |
| Q09666 | Neuroblast differentiation-associated prot | AHNAK | 629.102 | 24.8 | 79 | 79 | 606.45654 |
| Q0PNE2 | Elongator complex protein 6 OS=Homo s | ELP6 | 29.794 | 10.9 | 3 | 3 | 43.34428 |
| Q0VG06 | Fanconi anemia core complex-associated | FAAP100 | 93.434 | 1.2 | 1 | 1 | 43.661808 |
| Q10471 | Polypeptide N-acetylgalactosaminyltransf | GALNT2 | 64.733 | 28 | 15 | 15 | 350.65384 |
| Q10472 | Polypeptide N-acetylgalactosaminyltransf | GALNT1 | 64.219 | 5.2 | 3 | 1 | 44.921791 |
| Q10567 | AP-1 complex subunit beta-1 OS=Homo s | AP1B1 | 104.607 | 2.8 | 2 | 2 | 39.244732 |
| Q10570 | Cleavage and polyadenylation specificity | CPSF1 | 160.884 | 9.8 | 13 | 13 | 333.29575 |
| Q12769 | Nuclear pore complex protein Nup160 OS | NUP160 | 162.122 | 12.5 | 16 | 16 | 190.43576 |
| Q12772 | Sterol regulatory element-binding protein | SREBF2 | 123.688 | 6.6 | 7 | 7 | 152.65891 |
| Q12774 | Rho guanine nucleotide exchange factor 5 | ARHGEF5 | 176.8 | 11.8 | 16 | 14 | 214.6349 |
| Q12778 | Forkhead box protein O1 OS=Homo sapie | FOXO1 | 69.662 | 12.1 | 7 | 4 | 304.1972 |
| Q12788 | Transducin beta-like protein 3 OS=Homo | TBL3 | 89.035 | 23.1 | 14 | 14 | 356.55209 |
| Q12789 | General transcription factor 3C polypeptic | GTF3C1 | 238.875 | 16.5 | 33 | 33 | 341.50244 |
| Q12792 | Twinfilin-1 OS=Homo sapiens OX=9606 | TWF1 | 40.283 | 22.3 | 7 | 7 | 110.15695 |
| Q12797 | Aspartyl/asparaginyl beta-hydroxylase OS | ASPH | 85.863 | 29.6 | 25 | 25 | 4770.9707 |
| Q12800 | Alpha-globin transcription factor CP2 OS | TFCP2 | 57.256 | 14.1 | 6 | 4 | 539.10583 |
| Q12802 | A-kinase anchor protein 13 OS=Homo sap | AKAP13 | 307.551 | 13 | 30 | 30 | 285.04333 |
| Q12805 | EGF-containing fibulin-like extracellular | EFEMP1 | 54.641 | 20.5 | 9 | 9 | 280.54837 |
| Q12815 | Tastin OS=Homo sapiens OX=9606 GN= | TROAP | 83.858 | 24.3 | 17 | 17 | 646.92676 |
| Q12824 | SWI/SNF-related matrix-associated actin- | SMARCB1 | 44.142 | 8.3 | 3 | 3 | 124.23424 |
| Q12830 | Nucleosome-remodeling factor subunit BI | BPTF | 338.263 | 4.2 | 11 | 11 | 758.81982 |
| Q12834 | Cell division cycle protein 20 homolog OS | CDC20 | 54.723 | 28.1 | 11 | 11 | 716.45007 |
| Q12849 | G-rich sequence factor 1 OS=Homo sapie | GRSF1 | 53.127 | 29.6 | 14 | 14 | 1729.498 |

| | | | | | | | |
|---|---|---|---|---|---|---|---|
| Q12857 | Nuclear factor 1 A-type OS=Homo sapien | NFIA | 55.945 | 31.8 | 12 | 8 | 1902.4587 |
| Q12873 | Chromodomain-helicase-DNA-binding pr | CHD3 | 226.592 | 13.2 | 24 | 15 | 401.75879 |
| Q12874 | Splicing factor 3A subunit 3 OS=Homo sa | SF3A3 | 58.849 | 17.2 | 9 | 9 | 515.04407 |
| Q12899 | Tripartite motif-containing protein 26 OS= | TRIM26 | 62.166 | 29.5 | 15 | 15 | 1053.5294 |
| Q12904 | Aminoacyl tRNA synthase complex-intera | AIMP1 | 34.353 | 26.3 | 6 | 6 | 268.14282 |
| Q12905 | Interleukin enhancer-binding factor 2 OS= | ILF2 | 43.063 | 35.1 | 13 | 13 | 12687.748 |
| Q12906 | Interleukin enhancer-binding factor 3 OS= | ILF3 | 95.339 | 39.4 | 31 | 27 | 13709.942 |
| Q12931 | Heat shock protein 75 kDa, mitochondrial | TRAP1 | 80.11 | 7.4 | 4 | 4 | 1073.4248 |
| Q12933 | TNF receptor-associated factor 2 OS=Hor | TRAF2 | 55.86 | 7.2 | 3 | 3 | 116.73358 |
| Q12947 | Forkhead box protein F2 OS=Homo sapie | FOXF2 | 45.994 | 9.7 | 4 | 3 | 688.20856 |
| Q12948 | Forkhead box protein C1 OS=Homo sapie | FOXC1 | 56.789 | 13.9 | 6 | 6 | 929.10791 |
| Q12959 | Disks large homolog 1 OS=Homo sapiens | DLG1 | 100.456 | 12.3 | 9 | 7 | 183.08994 |
| Q12965 | Unconventional myosin-Ie OS=Homo sap | MYO1E | 127.062 | 33.5 | 33 | 29 | 1364.7656 |
| Q12968 | Nuclear factor of activated T-cells, cytopl | NFATC3 | 115.594 | 21.8 | 18 | 18 | 1212.7947 |
| Q12979 | Active breakpoint cluster region-related p | ABR | 97.598 | 1.5 | 1 | 1 | 23.302753 |
| Q12980 | GATOR1 complex protein NPRL3 OS=H | NPRL3 | 63.605 | 2.3 | 1 | 1 | 32.662529 |
| Q12986 | Transcriptional repressor NF-X1 OS=Hor | NFX1 | 124.395 | 49.7 | 47 | 47 | 1628.0795 |
| Q12996 | Cleavage stimulation factor subunit 3 OS= | CSTF3 | 82.922 | 7.7 | 5 | 5 | 138.31841 |
| Q13011 | Delta(3,5)-Delta(2,4)-dienoyl-CoA isome | ECH1 | 35.817 | 4.3 | 1 | 1 | 64.23114 |
| Q13042 | Cell division cycle protein 16 homolog O | CDC16 | 71.656 | 12.1 | 7 | 7 | 187.23877 |
| Q13045 | Protein flightless-1 homolog OS=Homo sa | FLII | 144.752 | 16.5 | 17 | 17 | 555.42419 |
| Q13049 | E3 ubiquitin-protein ligase TRIM32 OS=I | TRIM32 | 71.989 | 6.1 | 3 | 3 | 56.987751 |
| Q13064 | Probable E3 ubiquitin-protein ligase mak | MKRN3 | 55.645 | 4.3 | 2 | 2 | 45.01487 |
| Q13084 | Large ribosomal subunit protein bL28m C | MRPL28 | 30.157 | 25.8 | 8 | 8 | 234.82753 |
| Q13085 | Acetyl-CoA carboxylase 1 OS=Homo sap | ACACA | 265.554 | 3.4 | 7 | 5 | 78.850616 |
| Q13098 | COP9 signalosome complex subunit 1 OS | GPS1 | 55.537 | 4.3 | 2 | 2 | 65.064331 |
| Q13111 | Chromatin assembly factor 1 subunit A O | CHAF1A | 106.91 | 8.5 | 7 | 7 | 256.91345 |
| Q13112 | Chromatin assembly factor 1 subunit B O | CHAF1B | 61.493 | 15.7 | 8 | 8 | 463.41034 |
| Q13118 | Krueppel-like factor 10 OS=Homo sapien | KLF10 | 52.556 | 20 | 10 | 10 | 267.02563 |
| Q13123 | Protein Red OS=Homo sapiens OX=9606 | IK | 65.603 | 28.5 | 17 | 17 | 910.14795 |
| Q13131 | 5'-AMP-activated protein kinase catalytic | PRKAA1 | 64.01 | 11.8 | 6 | 6 | 165.24246 |
| Q13136 | Liprin-alpha-1 OS=Homo sapiens OX=96 | PPFIA1 | 135.779 | 17.6 | 18 | 13 | 270.19174 |
| Q13137 | Calcium-binding and coiled-coil domain-c | CALCOCO | 52.254 | 14.6 | 5 | 5 | 162.25336 |
| Q13144 | Translation initiation factor eIF-2B subun | EIF2B5 | 80.38 | 7.5 | 5 | 5 | 91.545547 |
| Q13148 | TAR DNA-binding protein 43 OS=Homo | TARDBP | 44.74 | 14.5 | 7 | 7 | 1259.825 |

| | | | | | | | |
|---|---|---|---|---|---|---|---|
| Q13151 | Heterogeneous nuclear ribonucleoprotein | HNRNPA0 | 30.841 | 26.6 | 5 | 5 | 5631.8447 |
| Q13155 | Aminoacyl tRNA synthase complex-intera | AIMP2 | 35.349 | 12.2 | 4 | 4 | 414.24194 |
| Q13162 | Peroxiredoxin-4 OS=Homo sapiens OX=9 | PRDX4 | 30.54 | 17.7 | 4 | 4 | 129.12794 |
| Q13185 | Chromobox protein homolog 3 OS=Homo | CBX3 | 20.812 | 27.3 | 6 | 6 | 331.02258 |
| Q13188 | Serine/threonine-protein kinase 3 OS=Ho | STK3 | 56.302 | 14.5 | 6 | 2 | 195.68385 |
| Q13190 | Syntaxin-5 OS=Homo sapiens OX=9606 | STX5 | 39.673 | 5.9 | 2 | 2 | 40.253407 |
| Q13191 | E3 ubiquitin-protein ligase CBL-B OS=H | CBLB | 109.45 | 1.2 | 1 | 1 | 42.717102 |
| Q13200 | 26S proteasome non-ATPase regulatory s | PSMD2 | 100.2 | 6.4 | 4 | 4 | 51.969391 |
| Q13206 | Probable ATP-dependent RNA helicase D | DDX10 | 100.888 | 19 | 15 | 15 | 98.143463 |
| Q13214 | Semaphorin-3B OS=Homo sapiens OX=9 | SEMA3B | 83.122 | 19.6 | 11 | 11 | 196.11528 |
| Q13242 | Serine/arginine-rich splicing factor 9 OS= | SRSF9 | 25.543 | 44.3 | 10 | 10 | 4044.6538 |
| Q13243 | Serine/arginine-rich splicing factor 5 OS= | SRSF5 | 31.264 | 24.3 | 7 | 7 | 2347.5762 |
| Q13247 | Serine/arginine-rich splicing factor 6 OS= | SRSF6 | 39.587 | 33.4 | 12 | 7 | 6629.396 |
| Q13263 | Transcription intermediary factor 1-beta C | TRIM28 | 88.55 | 21 | 15 | 15 | 753.60669 |
| Q13275 | Semaphorin-3F OS=Homo sapiens OX=9 | SEMA3F | 88.382 | 5.4 | 4 | 4 | 100.64632 |
| Q13283 | Ras GTPase-activating protein-binding pr | G3BP1 | 52.165 | 53.6 | 25 | 25 | 34460.523 |
| Q13309 | S-phase kinase-associated protein 2 OS=H | SKP2 | 47.761 | 2.4 | 1 | 1 | 70.377892 |
| Q13310 | Polyadenylate-binding protein 4 OS=Hom | PABPC4 | 70.783 | 39.1 | 23 | 19 | 61801.996 |
| Q13322 | Growth factor receptor-bound protein 10 | GRB10 | 67.232 | 1.7 | 1 | 1 | 54.99472 |
| Q13325 | Interferon-induced protein with tetratricop | IFIT5 | 55.847 | 2.1 | 1 | 1 | 71.38649 |
| Q13347 | Eukaryotic translation initiation factor 3 s | EIF3I | 36.502 | 41.8 | 12 | 12 | 876.95502 |
| Q13356 | RING-type E3 ubiquitin-protein ligase PP | PPIL2 | 58.824 | 35.2 | 17 | 17 | 542.8974 |
| Q13362 | Serine/threonine-protein phosphatase 2A | PPP2R5C | 61.061 | 5.5 | 3 | 1 | 37.939583 |
| Q13370 | cGMP-inhibited 3',5'-cyclic phosphodieste | PDE3B | 124.334 | 0.6 | 1 | 1 | 45.275581 |
| Q13371 | Phosducin-like protein OS=Homo sapiens | PDCL | 34.282 | 3 | 1 | 1 | 46.303093 |
| Q13393 | Phospholipase D1 OS=Homo sapiens OX | PLD1 | 124.185 | 4.2 | 4 | 4 | 56.995251 |
| Q13395 | Probable methyltransferase TARBP1 OS= | TARBP1 | 181.675 | 7.7 | 9 | 9 | 119.03873 |
| Q13404 | Ubiquitin-conjugating enzyme E2 variant | UBE2V1 | 16.495 | 6.8 | 1 | 1 | 27.144182 |
| Q13405 | Large ribosomal subunit protein mL49 OS | MRPL49 | 19.199 | 43.4 | 8 | 8 | 1019.0402 |
| Q13409 | Cytoplasmic dynein 1 intermediate chain 2 | DYNC1I2 | 71.457 | 2.2 | 1 | 1 | 37.901634 |
| Q13415 | Origin recognition complex subunit 1 OS= | ORC1 | 97.35 | 12.1 | 10 | 10 | 96.718918 |
| Q13416 | Origin recognition complex subunit 2 OS= | ORC2 | 65.972 | 6.9 | 4 | 4 | 103.92638 |
| Q13418 | Integrin-linked protein kinase OS=Homo | ILK | 51.42 | 12.6 | 5 | 5 | 109.26837 |
| Q13425 | Beta-2-syntrophin OS=Homo sapiens OX | SNTB2 | 57.95 | 9.3 | 5 | 5 | 76.432892 |
| Q13427 | Peptidyl-prolyl cis-trans isomerase G OS= | PPIG | 88.618 | 7.6 | 5 | 5 | 162.42874 |

| | | | | | | | |
|---|---|---|---|---|---|---|---|
| Q13428 | Treacle protein OS=Homo sapiens OX=9( | TCOF1 | 152.107 | 11.7 | 16 | 16 | 690.63245 |
| Q13435 | Splicing factor 3B subunit 2 OS=Homo sa | SF3B2 | 100.228 | 30.6 | 25 | 25 | 2306.4492 |
| Q13451 | Peptidyl-prolyl cis-trans isomerase FKBP: | FKBP5 | 51.213 | 4.2 | 2 | 2 | 77.207596 |
| Q13459 | Unconventional myosin-IXb OS=Homo sa | MYO9B | 243.401 | 17.6 | 34 | 31 | 353.83804 |
| Q13461 | Forkhead box protein E3 OS=Homo sapie | FOXE3 | 33.235 | 3.4 | 1 | 1 | 483.2319 |
| Q13469 | Nuclear factor of activated T-cells, cytopl | NFATC2 | 100.146 | 3.4 | 3 | 3 | 51.537266 |
| Q13470 | Non-receptor tyrosine-protein kinase TNK | TNK1 | 72.468 | 17.7 | 10 | 10 | 173.1555 |
| Q13472 | DNA topoisomerase 3-alpha OS=Homo sa | TOP3A | 112.373 | 17.1 | 14 | 13 | 840.49677 |
| Q13501 | Sequestosome-1 OS=Homo sapiens OX=9 | SQSTM1 | 47.688 | 20.9 | 8 | 8 | 954.66302 |
| Q13503 | Mediator of RNA polymerase II transcript | MED21 | 15.565 | 9.7 | 1 | 1 | 121.62696 |
| Q13505 | Metaxin-1 OS=Homo sapiens OX=9606 ( | MTX1 | 51.463 | 1.9 | 1 | 1 | 72.963394 |
| Q13509 | Tubulin beta-3 chain OS=Homo sapiens C | TUBB3 | 50.433 | 11.6 | 4 | 4 | 213.76791 |
| Q13523 | Serine/threonine-protein kinase PRP4 hon | PRPF4B | 116.987 | 7.8 | 7 | 7 | 129.7245 |
| Q13546 | Receptor-interacting serine/threonine-prot | RIPK1 | 75.931 | 12.1 | 8 | 8 | 126.57759 |
| Q13547 | Histone deacetylase 1 OS=Homo sapiens | HDAC1 | 55.103 | 28.6 | 12 | 5 | 455.09778 |
| Q13554 | Calcium/calmodulin-dependent protein ki | CAMK2B | 72.678 | 9.3 | 6 | 0 | 319.39005 |
| Q13555 | Calcium/calmodulin-dependent protein ki | CAMK2G | 62.608 | 7.3 | 4 | 4 | 212.29594 |
| Q13557 | Calcium/calmodulin-dependent protein ki | CAMK2D | 56.37 | 8.4 | 4 | 4 | 59.147324 |
| Q13569 | G/T mismatch-specific thymine DNA glyc | TDG | 46.054 | 8 | 3 | 3 | 572.11389 |
| Q13573 | SNW domain-containing protein 1 OS=H | SNW1 | 61.495 | 30.2 | 16 | 16 | 1533.5874 |
| Q13574 | Diacylglycerol kinase zeta OS=Homo sap | DGKZ | 103.981 | 5.3 | 4 | 2 | 71.440765 |
| Q13595 | Transformer-2 protein homolog alpha OS= | TRA2A | 32.689 | 26.2 | 6 | 6 | 1651.3876 |
| Q13601 | KRR1 small subunit processome compone | KRR1 | 43.665 | 21 | 7 | 7 | 535.73773 |
| Q13608 | Peroxisomal ATPase PEX6 OS=Homo sa | PEX6 | 104.062 | 17.6 | 14 | 14 | 540.94373 |
| Q13610 | Periodic tryptophan protein 1 homolog OS | PWP1 | 55.828 | 32.5 | 13 | 13 | 1081.507 |
| Q13613 | Myotubularin-related protein 1 OS=Homo | MTMR1 | 74.678 | 9 | 6 | 5 | 130.42033 |
| Q13614 | Myotubularin-related protein 2 OS=Homo | MTMR2 | 73.382 | 8.1 | 5 | 5 | 136.73807 |
| Q13615 | Myotubularin-related protein 3 OS=Homo | MTMR3 | 133.619 | 1.1 | 1 | 1 | 79.190094 |
| Q13616 | Cullin-1 OS=Homo sapiens OX=9606 GN | CUL1 | 89.679 | 8.6 | 6 | 6 | 60.44125 |
| Q13617 | Cullin-2 OS=Homo sapiens OX=9606 GN | CUL2 | 86.984 | 2.6 | 2 | 2 | 73.175705 |
| Q13618 | Cullin-3 OS=Homo sapiens OX=9606 GN | CUL3 | 88.931 | 10.7 | 7 | 7 | 96.611404 |
| Q13619 | Cullin-4A OS=Homo sapiens OX=9606 ( | CUL4A | 87.681 | 17.4 | 12 | 8 | 101.00047 |
| Q13625 | Apoptosis-stimulating of p53 protein 2 OS | TP53BP2 | 125.617 | 2.2 | 2 | 2 | 34.699284 |
| Q13627 | Dual specificity tyrosine-phosphorylation- | DYRK1A | 85.584 | 17.2 | 13 | 7 | 418.44122 |
| Q13643 | Four and a half LIM domains protein 3 OS | FHL3 | 31.192 | 22.1 | 7 | 7 | 338.41479 |

| | | | | | | | |
|---|---|---|---|---|---|---|---|
| Q13671 | Ras and Rab interactor 1 OS=Homo sapie | RIN1 | 84.099 | 7.3 | 5 | 5 | 66.262749 |
| Q13685 | Angio-associated migratory cell protein O | AAMP | 46.751 | 15.2 | 5 | 5 | 63.829346 |
| Q13724 | Mannosyl-oligosaccharide glucosidase OS | MOGS | 91.918 | 19.7 | 15 | 15 | 582.01147 |
| Q13751 | Laminin subunit beta-3 OS=Homo sapiens | LAMB3 | 129.573 | 13.9 | 13 | 13 | 206.75377 |
| Q13769 | THO complex subunit 5 homolog OS=Ho | THOC5 | 78.508 | 10.4 | 7 | 7 | 174.1505 |
| Q13772 | Nuclear receptor coactivator 4 OS=Homo | NCOA4 | 69.727 | 4.7 | 3 | 3 | 77.31601 |
| Q13795 | ADP-ribosylation factor-related protein 1 | ARFRP1 | 22.615 | 8 | 1 | 1 | 18.217407 |
| Q13813 | Spectrin alpha chain, non-erythrocytic 1 C | SPTAN1 | 284.54 | 18.6 | 41 | 41 | 145.97418 |
| Q13823 | Nucleolar GTP-binding protein 2 OS=Hor | GNL2 | 83.655 | 31.7 | 28 | 28 | 1194.5845 |
| Q13838 | Spliceosome RNA helicase DDX39B OS= | DDX39B | 48.992 | 7.7 | 3 | 3 | 58.3871 |
| Q13868 | Exosome complex component RRP4 OS= | EXOSC2 | 32.789 | 23.9 | 5 | 5 | 611.23877 |
| Q13885 | Tubulin beta-2A chain OS=Homo sapiens | TUBB2A | 49.907 | 14.2 | 4 | 1 | 12037.79 |
| Q13887 | Krueppel-like factor 5 OS=Homo sapiens | KLF5 | 50.792 | 3.9 | 2 | 1 | 252.85934 |
| Q13888 | General transcription factor IIH subunit 2 | GTF2H2 | 44.419 | 26.1 | 9 | 1 | 263.60016 |
| Q13889 | General transcription factor IIH subunit 3 | GTF2H3 | 34.379 | 9.4 | 3 | 3 | 100.86503 |
| Q13895 | Bystin OS=Homo sapiens OX=9606 GN= | BYSL | 49.602 | 42.1 | 16 | 16 | 3866.8062 |
| Q13907 | Isopentenyl-diphosphate Delta-isomerase | IDI1 | 26.32 | 11 | 3 | 3 | 109.09793 |
| Q13951 | Core-binding factor subunit beta OS=Hon | CBFB | 21.509 | 18.1 | 3 | 3 | 33.914127 |
| Q14004 | Cyclin-dependent kinase 13 OS=Homo sa | CDK13 | 164.924 | 5.2 | 6 | 6 | 122.66322 |
| Q14008 | Cytoskeleton-associated protein 5 OS=Ho | CKAP5 | 225.495 | 29.9 | 56 | 56 | 932.11639 |
| Q14011 | Cold-inducible RNA-binding protein OS= | CIRBP | 18.648 | 20.9 | 3 | 3 | 2370.2236 |
| Q14103 | Heterogeneous nuclear ribonucleoprotein | HNRNPD | 38.435 | 24.2 | 10 | 10 | 14528.584 |
| Q14106 | Protein Tob2 OS=Homo sapiens OX=960 | TOB2 | 36.633 | 12.5 | 3 | 3 | 52.669163 |
| Q14112 | Nidogen-2 OS=Homo sapiens OX=9606 ( | NID2 | 151.254 | 0.7 | 1 | 1 | 49.29488 |
| Q14114 | Low-density lipoprotein receptor-related p | LRP8 | 105.634 | 5.3 | 5 | 5 | 108.71551 |
| Q14119 | Vascular endothelial zinc finger 1 OS=Ho | VEZF1 | 56.932 | 4 | 2 | 2 | 141.13522 |
| Q14126 | Desmoglein-2 OS=Homo sapiens OX=96( | DSG2 | 122.294 | 3.2 | 3 | 3 | 29.804777 |
| Q14134 | Tripartite motif-containing protein 29 OS= | TRIM29 | 65.835 | 30.6 | 16 | 16 | 519.96741 |
| Q14137 | Ribosome biogenesis protein BOP1 OS=H | BOP1 | 83.63 | 16.9 | 13 | 13 | 357.59305 |
| Q14141 | Septin-6 OS=Homo sapiens OX=9606 GN | SEPTIN6 | 49.717 | 6 | 2 | 1 | 66.3508 |
| Q14142 | Tripartite motif-containing protein 14 OS= | TRIM14 | 49.773 | 31 | 12 | 12 | 548.27869 |
| Q14145 | Kelch-like ECH-associated protein 1 OS= | KEAP1 | 69.667 | 30.1 | 18 | 18 | 279.28925 |
| Q14147 | Probable ATP-dependent RNA helicase D | DHX34 | 128.12 | 17.1 | 16 | 16 | 156.30405 |
| Q14151 | Scaffold attachment factor B2 OS=Homo | SAFB2 | 107.474 | 27.3 | 27 | 19 | 1368.6062 |
| Q14152 | Eukaryotic translation initiation factor 3 s | EIF3A | 166.57 | 39.1 | 58 | 58 | 2707.3113 |

| | | | | | | | |
|---|---|---|---|---|---|---|---|
| Q14153 | Protein FAM53B OS=Homo sapiens OX= | FAM53B | 45.768 | 9.2 | 3 | 3 | 179.18898 |
| Q14157 | Ubiquitin-associated protein 2-like OS=H | UBAP2L | 114.535 | 19.2 | 16 | 16 | 4420.5195 |
| Q14160 | Protein scribble homolog OS=Homo sapie | SCRIB | 174.915 | 1.4 | 2 | 1 | 46.456276 |
| Q14181 | DNA polymerase alpha subunit B OS=Ho | POLA2 | 65.948 | 7.9 | 4 | 4 | 86.363052 |
| Q14186 | Transcription factor Dp-1 OS=Homo sapi | TFDP1 | 45.071 | 10 | 2 | 2 | 134.45522 |
| Q14191 | Bifunctional 3'-5' exonuclease/ATP-deper | WRN | 162.461 | 16.1 | 20 | 20 | 155.31929 |
| Q14192 | Four and a half LIM domains protein 2 O: | FHL2 | 32.193 | 39.1 | 10 | 10 | 951.42267 |
| Q14194 | Dihydropyrimidinase-related protein 1 OS | CRMP1 | 62.184 | 2.3 | 1 | 1 | 35.005371 |
| Q14197 | Large ribosomal subunit protein mL62 OS | MRPL58 | 23.631 | 31.6 | 6 | 6 | 112.67745 |
| Q14203 | Dynactin subunit 1 OS=Homo sapiens OX | DCTN1 | 141.695 | 1.5 | 2 | 2 | 45.723495 |
| Q14204 | Cytoplasmic dynein 1 heavy chain 1 OS=I | DYNC1H1 | 532.408 | 2.4 | 11 | 11 | 121.21152 |
| Q14232 | Translation initiation factor eIF-2B subun | EIF2B1 | 33.713 | 33.1 | 9 | 9 | 222.64825 |
| Q14240 | Eukaryotic initiation factor 4A-II OS=Hor | EIF4A2 | 46.403 | 52.3 | 20 | 9 | 6194.9551 |
| Q14241 | Elongin-A OS=Homo sapiens OX=9606 ( | ELOA | 89.909 | 18.8 | 16 | 16 | 645.85773 |
| Q14244 | Ensconsin OS=Homo sapiens OX=9606 C | MAP7 | 84.052 | 23.9 | 17 | 17 | 1041.6025 |
| Q14247 | Src substrate cortactin OS=Homo sapiens | CTTN | 61.587 | 30.2 | 17 | 17 | 6214.1812 |
| Q14249 | Endonuclease G, mitochondrial OS=Hom | ENDOG | 32.621 | 18.5 | 5 | 5 | 186.25804 |
| Q14254 | Flotillin-2 OS=Homo sapiens OX=9606 C | FLOT2 | 47.065 | 15.7 | 6 | 6 | 122.4464 |
| Q14257 | Reticulocalbin-2 OS=Homo sapiens OX=! | RCN2 | 36.877 | 15.8 | 5 | 5 | 920.72479 |
| Q14258 | E3 ubiquitin/ISG15 ligase TRIM25 OS=F | TRIM25 | 70.974 | 38.7 | 24 | 24 | 4362.5381 |
| Q14296 | Fas-activated serine/threonine kinase OS= | FASTK | 61.105 | 10 | 5 | 5 | 269.60107 |
| Q14318 | Peptidyl-prolyl cis-trans isomerase FKBP: | FKBP8 | 44.562 | 9.7 | 3 | 3 | 93.692024 |
| Q14331 | Protein FRG1 OS=Homo sapiens OX=96( | FRG1 | 29.173 | 32.2 | 8 | 8 | 1309.5605 |
| Q14444 | Caprin-1 OS=Homo sapiens OX=9606 GI | CAPRIN1 | 78.367 | 19.2 | 16 | 16 | 50911.449 |
| Q14451 | Growth factor receptor-bound protein 7 O | GRB7 | 59.682 | 3.9 | 2 | 2 | 288.9231 |
| Q14493 | Histone RNA hairpin-binding protein OS= | SLBP | 31.286 | 19.3 | 5 | 5 | 749.53174 |
| Q14498 | RNA-binding protein 39 OS=Homo sapie | RBM39 | 59.38 | 29.1 | 13 | 11 | 2763.0422 |
| Q14517 | Protocadherin Fat 1 OS=Homo sapiens O. | FAT1 | 506.273 | 12.8 | 51 | 51 | 389.95026 |
| Q14527 | Helicase-like transcription factor OS=Hor | HLTF | 113.929 | 25.7 | 25 | 25 | 962.23438 |
| Q14558 | Phosphoribosyl pyrophosphate synthase-a | PRPSAP1 | 39.394 | 17.4 | 4 | 4 | 65.933327 |
| Q14562 | ATP-dependent RNA helicase DHX8 OS= | DHX8 | 139.315 | 16.5 | 17 | 17 | 257.37509 |
| Q14573 | Inositol 1,4,5-trisphosphate receptor type | ITPR3 | 304.107 | 2.4 | 6 | 6 | 67.717285 |
| Q14587 | Zinc finger protein 268 OS=Homo sapien: | ZNF268 | 108.374 | 3.8 | 3 | 1 | 273.905 |
| Q14592 | Zinc finger protein 460 OS=Homo sapien: | ZNF460 | 63.665 | 10 | 5 | 3 | 112.90022 |
| Q14624 | Inter-alpha-trypsin inhibitor heavy chain I | ITIH4 | 103.358 | 3.9 | 4 | 4 | 438.83463 |

| | | | | | | | |
|---|---|---|---|---|---|---|---|
| Q14643 | Inositol 1,4,5-trisphosphate receptor type | ITPR1 | 313.93 | 2.2 | 6 | 4 | 54.576637 |
| Q14644 | Ras GTPase-activating protein 3 OS=Hon | RASA3 | 95.699 | 6.7 | 6 | 6 | 103.32102 |
| Q14669 | E3 ubiquitin-protein ligase TRIP12 OS=H | TRIP12 | 220.434 | 11.4 | 20 | 20 | 206.03215 |
| Q14671 | Pumilio homolog 1 OS=Homo sapiens OX | PUM1 | 126.474 | 20.9 | 24 | 20 | 2833.4031 |
| Q14674 | Separin OS=Homo sapiens OX=9606 GN | ESPL1 | 233.176 | 14.3 | 24 | 24 | 263.56592 |
| Q14676 | Mediator of DNA damage checkpoint pro | MDC1 | 226.667 | 6 | 11 | 11 | 92.977913 |
| Q14677 | Clathrin interactor 1 OS=Homo sapiens O | CLINT1 | 68.26 | 14.7 | 7 | 7 | 82.052345 |
| Q14678 | KN motif and ankyrin repeat domain-cont | KANK1 | 147.29 | 4.3 | 5 | 5 | 64.947891 |
| Q14679 | Tubulin monoglutamylase TTLL4 OS=Ho | TTLL4 | 133.379 | 9 | 10 | 10 | 155.63188 |
| Q14680 | Maternal embryonic leucine zipper kinase | MELK | 74.643 | 23.5 | 16 | 16 | 355.31 |
| Q14683 | Structural maintenance of chromosomes p | SMC1A | 143.233 | 19.5 | 23 | 23 | 276.466 |
| Q14684 | Ribosomal RNA processing protein 1 hon | RRP1B | 84.428 | 21.9 | 16 | 16 | 540.63458 |
| Q14690 | Protein RRP5 homolog OS=Homo sapien | PDCD11 | 208.701 | 19.9 | 33 | 33 | 291.75131 |
| Q14692 | Ribosome biogenesis protein BMS1 homo | BMS1 | 145.808 | 26.6 | 31 | 31 | 1397.1115 |
| Q14693 | Phosphatidate phosphatase LPIN1 OS=Ho | LPIN1 | 98.665 | 3.8 | 3 | 3 | 73.210442 |
| Q14694 | Ubiquitin carboxyl-terminal hydrolase 10 | USP10 | 87.134 | 36 | 33 | 33 | 30422.428 |
| Q14697 | Neutral alpha-glucosidase AB OS=Homo | GANAB | 106.874 | 0.7 | 1 | 1 | 16.670938 |
| Q14703 | Membrane-bound transcription factor site | MBTPS1 | 117.749 | 1 | 1 | 1 | 49.341988 |
| Q14739 | Delta(14)-sterol reductase LBR OS=Hom | LBR | 70.704 | 9.6 | 7 | 7 | 930.66846 |
| Q14764 | Major vault protein OS=Homo sapiens O | MVP | 99.327 | 20.3 | 17 | 17 | 288.86832 |
| Q14781 | Chromobox protein homolog 2 OS=Homo | CBX2 | 56.081 | 12 | 6 | 6 | 114.44533 |
| Q14789 | Golgin subfamily B member 1 OS=Homo | GOLGB1 | 376.02 | 0.7 | 2 | 2 | 40.648979 |
| Q14807 | Kinesin-like protein KIF22 OS=Homo sa | KIF22 | 73.263 | 18.2 | 10 | 10 | 122.62528 |
| Q14839 | Chromodomain-helicase-DNA-binding pr | CHD4 | 218.005 | 12.9 | 22 | 18 | 315.19232 |
| Q14847 | LIM and SH3 domain protein 1 OS=Hom | LASP1 | 29.718 | 35.2 | 9 | 9 | 480.19928 |
| Q14938 | Nuclear factor 1 X-type OS=Homo sapien | NFIX | 55.099 | 14.7 | 7 | 7 | 414.71826 |
| Q14964 | Ras-related protein Rab-39A OS=Homo s | RAB39A | 25.007 | 4.1 | 1 | 1 | 36.933662 |
| Q14966 | Zinc finger protein 638 OS=Homo sapien | ZNF638 | 220.625 | 16.4 | 29 | 29 | 368.4267 |
| Q14974 | Importin subunit beta-1 OS=Homo sapien | KPNB1 | 97.171 | 16.1 | 12 | 12 | 359.5744 |
| Q14978 | Nucleolar and coiled-body phosphoprotei | NOLC1 | 73.604 | 12.3 | 7 | 7 | 129.06097 |
| Q14980 | Nuclear mitotic apparatus protein 1 OS=H | NUMA1 | 238.26 | 8.9 | 16 | 16 | 81.697586 |
| Q14995 | Nuclear receptor subfamily 1 group D me | NR1D2 | 64.644 | 5.2 | 3 | 3 | 62.908115 |
| Q14997 | Proteasome activator complex subunit 4 C | PSME4 | 211.334 | 0.8 | 1 | 1 | 58.44529 |
| Q14999 | Cullin-7 OS=Homo sapiens OX=9606 GN | CUL7 | 191.161 | 1.5 | 2 | 2 | 37.390717 |
| Q149N8 | E3 ubiquitin-protein ligase SHPRH OS=H | SHPRH | 193.08 | 4.1 | 6 | 6 | 66.107773 |

| | | | | | | | |
|---|---|---|---|---|---|---|---|
| Q14C86 | GTPase-activating protein and VPS9 dom | GAPVD1 | 164.98 | 0.8 | 1 | 1 | 47.152725 |
| Q14CB8 | Rho GTPase-activating protein 19 OS=Hc | ARHGAP19 | 55.757 | 12.3 | 5 | 5 | 81.755096 |
| Q14CW9 | Ataxin-7-like protein 3 OS=Homo sapiens | ATXN7L3 | 38.651 | 4.6 | 1 | 1 | 25.136385 |
| Q14CZ7 | FAST kinase domain-containing protein 3 | FASTKD3 | 75.69 | 11.5 | 8 | 8 | 145.25859 |
| Q15003 | Condensin complex subunit 2 OS=Homo | NCAPH | 82.564 | 1.6 | 1 | 1 | 83.481232 |
| Q15004 | PCNA-associated factor OS=Homo sapiei | PCLAF | 11.987 | 45 | 4 | 4 | 33.915344 |
| Q15005 | Signal peptidase complex subunit 2 OS=F | SPCS2 | 25.003 | 3.5 | 1 | 1 | 86.00515 |
| Q15007 | Pre-mRNA-splicing regulator WTAP OS= | WTAP | 44.244 | 11.9 | 4 | 4 | 165.33482 |
| Q15008 | 26S proteasome non-ATPase regulatory s | PSMD6 | 45.532 | 3.9 | 1 | 1 | 20.73295 |
| Q15014 | Mortality factor 4-like protein 2 OS=Hom | MORF4L2 | 32.308 | 48.3 | 12 | 10 | 1096.686 |
| Q15019 | Septin-2 OS=Homo sapiens OX=9606 GN | SEPTIN2 | 41.488 | 15.5 | 4 | 4 | 75.031303 |
| Q15020 | Squamous cell carcinoma antigen recogni | SART3 | 109.935 | 24.3 | 22 | 22 | 366.99374 |
| Q15021 | Condensin complex subunit 1 OS=Homo | NCAPD2 | 157.183 | 7 | 9 | 9 | 92.050674 |
| Q15022 | Polycomb protein SUZ12 OS=Homo sapi | SUZ12 | 83.055 | 11.8 | 9 | 9 | 263.47595 |
| Q15024 | Exosome complex component RRP42 OS | EXOSC7 | 31.822 | 25.8 | 6 | 6 | 727.7984 |
| Q15025 | TNFAIP3-interacting protein 1 OS=Homo | TNIP1 | 71.865 | 14.6 | 8 | 8 | 208.03699 |
| Q15029 | 116 kDa U5 small nuclear ribonucleoprot | EFTUD2 | 109.436 | 24.5 | 22 | 22 | 566.77295 |
| Q15046 | Lysine--tRNA ligase OS=Homo sapiens C | KARS1 | 68.049 | 18.1 | 11 | 11 | 522.29413 |
| Q15050 | Ribosome biogenesis regulatory protein h | RRS1 | 41.194 | 25.5 | 10 | 10 | 286.27271 |
| Q15054 | DNA polymerase delta subunit 3 OS=Hor | POLD3 | 51.401 | 10.5 | 4 | 4 | 122.16936 |
| Q15056 | Eukaryotic translation initiation factor 4H | EIF4H | 27.386 | 5.2 | 1 | 1 | 162.17163 |
| Q15058 | Kinesin-like protein KIF14 OS=Homo sap | KIF14 | 186.492 | 27.3 | 42 | 42 | 805.28442 |
| Q15061 | WD repeat-containing protein 43 OS=Hor | WDR43 | 74.891 | 19.8 | 12 | 12 | 360.2233 |
| Q15063 | Periostin OS=Homo sapiens OX=9606 GI | POSTN | 93.314 | 1.6 | 1 | 1 | 66.859962 |
| Q15070 | Mitochondrial inner membrane protein O) | OXA1L | 48.548 | 13.8 | 6 | 6 | 122.6077 |
| Q15072 | Zinc finger protein OZF OS=Homo sapiei | ZNF146 | 33.308 | 9.9 | 3 | 3 | 138.32523 |
| Q15084 | Protein disulfide-isomerase A6 OS=Homo | PDIA6 | 48.122 | 15.5 | 5 | 5 | 172.19055 |
| Q15118 | [Pyruvate dehydrogenase (acetyl-transferr | PDK1 | 49.245 | 2.5 | 1 | 1 | 26.056808 |
| Q15119 | [Pyruvate dehydrogenase (acetyl-transferr | PDK2 | 46.154 | 3.2 | 1 | 1 | 41.829853 |
| Q15120 | [Pyruvate dehydrogenase (acetyl-transferr | PDK3 | 46.939 | 23.4 | 8 | 7 | 199.87315 |
| Q15125 | 3-beta-hydroxysteroid-Delta(8),Delta(7)-i | EBP | 26.353 | 9.6 | 2 | 2 | 184.54343 |
| Q15149 | Plectin OS=Homo sapiens OX=9606 GN= | PLEC | 531.791 | 42 | 195 | 190 | 2029.2117 |
| Q15154 | Pericentriolar material 1 protein OS=Hom | PCM1 | 228.545 | 9 | 16 | 16 | 152.92316 |
| Q15172 | Serine/threonine-protein phosphatase 2A : | PPP2R5A | 56.194 | 14.8 | 6 | 6 | 95.200012 |
| Q15185 | Prostaglandin E synthase 3 OS=Homo sap | PTGES3 | 18.698 | 26.2 | 4 | 4 | 255.25682 |

| | | | | | | | |
|---|---|---|---|---|---|---|---|
| Q15208 | Serine/threonine-protein kinase 38 OS=H | STK38 | 54.191 | 2.4 | 1 | 1 | 92.261909 |
| Q15233 | Non-POU domain-containing octamer-bin | NONO | 54.232 | 39.7 | 22 | 21 | 42770.648 |
| Q15262 | Receptor-type tyrosine-protein phosphatas | PTPRK | 162.103 | 3.1 | 3 | 3 | 50.772671 |
| Q15269 | Periodic tryptophan protein 2 homolog O | PWP2 | 102.453 | 11.4 | 10 | 10 | 274.02011 |
| Q15286 | Ras-related protein Rab-35 OS=Homo sa | RAB35 | 23.026 | 32.3 | 5 | 4 | 411.26352 |
| Q15287 | RNA-binding protein with serine-rich don | RNPS1 | 34.209 | 18.4 | 4 | 4 | 786.24384 |
| Q15291 | Retinoblastoma-binding protein 5 OS=Ho | RBBP5 | 59.153 | 2 | 1 | 1 | 54.892895 |
| Q15293 | Reticulocalbin-1 OS=Homo sapiens OX= | RCN1 | 38.89 | 42.3 | 14 | 14 | 10724.358 |
| Q15334 | Lethal(2) giant larvae protein homolog 1 ( | LLGL1 | 115.419 | 11.7 | 9 | 9 | 193.16623 |
| Q15345 | Leucine-rich repeat-containing protein 41 | LRRC41 | 88.65 | 7.4 | 5 | 5 | 134.49745 |
| Q15361 | Transcription termination factor 1 OS=Hc | TTF1 | 103.052 | 1.4 | 1 | 1 | 38.54916 |
| Q15365 | Poly(rC)-binding protein 1 OS=Homo sap | PCBP1 | 37.498 | 44.7 | 10 | 10 | 6060.5703 |
| Q15366 | Poly(rC)-binding protein 2 OS=Homo sap | PCBP2 | 38.581 | 42.5 | 12 | 8 | 13593.577 |
| Q15369 | Elongin-C OS=Homo sapiens OX=9606 ( | ELOC | 12.474 | 47.3 | 5 | 5 | 875.21252 |
| Q15370 | Elongin-B OS=Homo sapiens OX=9606 ( | ELOB | 13.133 | 52.5 | 7 | 7 | 1022.7741 |
| Q15382 | GTP-binding protein Rheb OS=Homo sap | RHEB | 20.498 | 20.1 | 3 | 3 | 123.08765 |
| Q15386 | Ubiquitin-protein ligase E3C OS=Homo s | UBE3C | 123.923 | 2.1 | 2 | 2 | 76.933716 |
| Q15388 | Mitochondrial import receptor subunit TC | TOMM20 | 16.298 | 18.6 | 3 | 3 | 1075.734 |
| Q15392 | Delta(24)-sterol reductase OS=Homo sapi | DHCR24 | 60.102 | 12.6 | 7 | 7 | 432.70554 |
| Q15393 | Splicing factor 3B subunit 3 OS=Homo sa | SF3B3 | 135.578 | 30.2 | 31 | 31 | 1633.4279 |
| Q15397 | Pumilio homolog 3 OS=Homo sapiens OX | PUM3 | 73.585 | 15 | 11 | 11 | 169.5078 |
| Q15418 | Ribosomal protein S6 kinase alpha-1 OS= | RPS6KA1 | 82.723 | 1.5 | 1 | 1 | 49.312988 |
| Q15424 | Scaffold attachment factor B1 OS=Homo | SAFB | 102.642 | 17.7 | 14 | 14 | 1422.4071 |
| Q15427 | Splicing factor 3B subunit 4 OS=Homo sa | SF3B4 | 44.386 | 6.4 | 2 | 2 | 315.1268 |
| Q15428 | Splicing factor 3A subunit 2 OS=Homo sa | SF3A2 | 49.256 | 25.4 | 10 | 10 | 266.58878 |
| Q15436 | Protein transport protein Sec23A OS=Hor | SEC23A | 86.161 | 14.1 | 9 | 7 | 182.73309 |
| Q15437 | Protein transport protein Sec23B OS=Hor | SEC23B | 86.48 | 21.4 | 14 | 14 | 204.47249 |
| Q15459 | Splicing factor 3A subunit 1 OS=Homo sa | SF3A1 | 88.887 | 18.5 | 13 | 13 | 613.36212 |
| Q15464 | SH2 domain-containing adapter protein B | SHB | 55.043 | 12 | 6 | 6 | 141.70113 |
| Q15468 | SCL-interrupting locus protein OS=Homo | STIL | 142.955 | 1 | 1 | 1 | 16.12764 |
| Q15475 | Homeobox protein SIX1 OS=Homo sapie | SIX1 | 32.211 | 4.6 | 1 | 1 | 31.485125 |
| Q15477 | Superkiller complex protein 2 OS=Homo | SKIC2 | 137.756 | 20.7 | 23 | 23 | 342.6745 |
| Q15532 | Protein SSXT OS=Homo sapiens OX=96 | SS18 | 45.93 | 6.2 | 3 | 3 | 176.53912 |
| Q15545 | Transcription initiation factor TFIID subu | TAF7 | 40.26 | 9.7 | 3 | 3 | 57.186058 |
| Q15554 | Telomeric repeat-binding factor 2 OS=Ho | TERF2 | 59.594 | 28.2 | 13 | 13 | 219.49901 |

| | | | | | | | |
|---|---|---|---|---|---|---|---|
| Q15560 | Transcription elongation factor A protein | TCEA2 | 33.601 | 3.7 | 1 | 1 | 20.322367 |
| Q15561 | Transcriptional enhancer factor TEF-3 OS | TEAD4 | 48.329 | 2.3 | 1 | 1 | 33.092537 |
| Q15583 | Homeobox protein TGIF1 OS=Homo sap | TGIF1 | 43.013 | 4.2 | 2 | 2 | 34.353718 |
| Q15599 | Na(+)/H(+) exchange regulatory cofactor | NHERF2 | 37.414 | 3.3 | 1 | 1 | 33.755703 |
| Q15629 | Translocating chain-associated membrane | TRAM1 | 43.072 | 8.6 | 3 | 3 | 341.59338 |
| Q15631 | Translin OS=Homo sapiens OX=9606 GN | TSN | 26.183 | 30.3 | 6 | 6 | 205.2093 |
| Q15633 | RISC-loading complex subunit TARBP2 ( | TARBP2 | 39.04 | 31.7 | 11 | 11 | 1454.3219 |
| Q15637 | Splicing factor 1 OS=Homo sapiens OX= | SF1 | 68.33 | 24.7 | 13 | 13 | 6273.8906 |
| Q15642 | Cdc42-interacting protein 4 OS=Homo sa | TRIP10 | 68.353 | 2.2 | 1 | 1 | 82.311722 |
| Q15645 | Pachytene checkpoint protein 2 homolog ( | TRIP13 | 48.551 | 28.5 | 10 | 10 | 196.30386 |
| Q15646 | 2'-5'-oligoadenylate synthase-like protein ( | OASL | 59.227 | 26.8 | 13 | 13 | 1176.7086 |
| Q15648 | Mediator of RNA polymerase II transcript | MED1 | 168.479 | 9 | 11 | 11 | 177.39403 |
| Q15649 | Zinc finger HIT domain-containing protei | ZNHIT3 | 17.608 | 34.8 | 5 | 5 | 245.06575 |
| Q15650 | Activating signal cointegrator 1 OS=Hom | TRIP4 | 66.147 | 35.8 | 21 | 21 | 5468.249 |
| Q15652 | Probable JmjC domain-containing histone | JMJD1C | 284.525 | 2.6 | 6 | 6 | 45.937489 |
| Q15654 | Thyroid receptor-interacting protein 6 OS | TRIP6 | 50.288 | 21 | 7 | 7 | 774.78729 |
| Q15678 | Tyrosine-protein phosphatase non-recepto | PTPN14 | 135.262 | 0.8 | 1 | 1 | 39.543522 |
| Q15691 | Microtubule-associated protein RP/EB fa | MAPRE1 | 30 | 10.4 | 2 | 2 | 31.126205 |
| Q15699 | ALX homeobox protein 1 OS=Homo sapi | ALX1 | 36.962 | 10.1 | 3 | 3 | 154.01758 |
| Q15717 | ELAV-like protein 1 OS=Homo sapiens ( | ELAVL1 | 36.092 | 33.1 | 10 | 10 | 29927.375 |
| Q15746 | Myosin light chain kinase, smooth muscle | MYLK | 210.716 | 0.4 | 1 | 1 | 74.155617 |
| Q15776 | Zinc finger protein with KRAB and SCAN | ZKSCAN8 | 65.816 | 6.1 | 3 | 3 | 69.493309 |
| Q15796 | Mothers against decapentaplegic homolog | SMAD2 | 52.307 | 16.7 | 7 | 3 | 512.45947 |
| Q15797 | Mothers against decapentaplegic homolog | SMAD1 | 52.261 | 2.8 | 1 | 1 | 28.284817 |
| Q15834 | Coiled-coil domain-containing protein 85 | CCDC85B | 22.091 | 10.4 | 2 | 2 | 50.656342 |
| Q15906 | Vacuolar protein sorting-associated protei | VPS72 | 40.595 | 15.4 | 6 | 6 | 198.39807 |
| Q15910 | Histone-lysine N-methyltransferase EZH2 | EZH2 | 85.364 | 15.7 | 9 | 7 | 232.95009 |
| Q16181 | Septin-7 OS=Homo sapiens OX=9606 GN | SEPTIN7 | 50.68 | 5 | 2 | 2 | 68.262459 |
| Q16186 | Proteasomal ubiquitin receptor ADRM1 ( | ADRM1 | 42.154 | 3.9 | 1 | 1 | 32.41935 |
| Q16204 | Coiled-coil domain-containing protein 6 ( | CCDC6 | 53.291 | 3 | 1 | 1 | 27.893858 |
| Q16206 | Ecto-NOX disulfide-thiol exchanger 2 OS | ENOX2 | 70.083 | 22.6 | 13 | 13 | 283.07529 |
| Q16342 | Programmed cell death protein 2 OS=Hor | PDCD2 | 38.593 | 15.4 | 5 | 5 | 188.53442 |
| Q16363 | Laminin subunit alpha-4 OS=Homo sapie | LAMA4 | 202.525 | 0.8 | 1 | 1 | 76.59967 |
| Q16394 | Exostosin-1 OS=Homo sapiens OX=9606 | EXT1 | 86.255 | 9.2 | 5 | 5 | 205.58606 |
| Q16401 | 26S proteasome non-ATPase regulatory s | PSMD5 | 56.196 | 1.8 | 1 | 1 | 47.140244 |

| | | | | | | | |
|---|---|---|---|---|---|---|---|
| Q16513 | Serine/threonine-protein kinase N2 OS=H | PKN2 | 112.035 | 22.8 | 19 | 19 | 234.20222 |
| Q16527 | Cysteine and glycine-rich protein 2 OS=H | CSRP2 | 20.954 | 46.6 | 7 | 7 | 218.24706 |
| Q16531 | DNA damage-binding protein 1 OS=Hom | DDB1 | 126.968 | 21.3 | 25 | 25 | 327.40051 |
| Q16537 | Serine/threonine-protein phosphatase 2A : | PPP2R5E | 54.7 | 25.9 | 11 | 8 | 519.56372 |
| Q16540 | Large ribosomal subunit protein uL23m C | MRPL23 | 17.782 | 34 | 5 | 5 | 924.6756 |
| Q16543 | Hsp90 co-chaperone Cdc37 OS=Homo sa | CDC37 | 44.469 | 21.7 | 7 | 7 | 979.51953 |
| Q16576 | Histone-binding protein RBBP7 OS=Hom | RBBP7 | 47.821 | 12.5 | 3 | 3 | 402.11331 |
| Q16594 | Transcription initiation factor TFIID subu | TAF9 | 28.974 | 9.5 | 2 | 1 | 67.489403 |
| Q16600 | Zinc finger protein 239 OS=Homo sapien: | ZNF239 | 51.591 | 18.1 | 7 | 7 | 373.44217 |
| Q16610 | Extracellular matrix protein 1 OS=Homo : | ECM1 | 60.675 | 6.5 | 4 | 4 | 3223.2888 |
| Q16629 | Serine/arginine-rich splicing factor 7 OS= | SRSF7 | 27.367 | 18.9 | 5 | 5 | 4227.2456 |
| Q16630 | Cleavage and polyadenylation specificity | CPSF6 | 59.21 | 15.1 | 8 | 8 | 2030.2123 |
| Q16637 | Survival motor neuron protein OS=Homo | SMN1 | 31.849 | 22.8 | 6 | 6 | 2595.5103 |
| Q16643 | Drebrin OS=Homo sapiens OX=9606 GN | DBN1 | 71.43 | 12.2 | 8 | 8 | 206.24995 |
| Q16656 | Nuclear respiratory factor 1 OS=Homo sa | NRF1 | 53.542 | 2.4 | 1 | 1 | 226.67139 |
| Q16658 | Fascin OS=Homo sapiens OX=9606 GN= | FSCN1 | 54.53 | 10.1 | 5 | 5 | 112.73743 |
| Q16659 | Mitogen-activated protein kinase 6 OS=H | MAPK6 | 82.681 | 3.1 | 2 | 1 | 138.55998 |
| Q16666 | Gamma-interferon-inducible protein 16 O | IFI16 | 88.256 | 31 | 24 | 21 | 2825.2336 |
| Q16670 | Zinc finger and SCAN domain-containing | ZSCAN26 | 55.254 | 6.3 | 3 | 3 | 119.69373 |
| Q16678 | Cytochrome P450 1B1 OS=Homo sapiens | CYP1B1 | 60.846 | 35.9 | 19 | 19 | 3340.7976 |
| Q16698 | 2,4-dienoyl-CoA reductase [(3E)-enoyl-C | DECR1 | 36.068 | 3.9 | 1 | 1 | 68.771439 |
| Q16706 | Alpha-mannosidase 2 OS=Homo sapiens | MAN2A1 | 131.141 | 11.5 | 12 | 12 | 154.76233 |
| Q16762 | Thiosulfate sulfurtransferase OS=Homo s: | TST | 33.429 | 21.5 | 5 | 5 | 118.49113 |
| Q16763 | Ubiquitin-conjugating enzyme E2 S OS=I | UBE2S | 23.846 | 15.8 | 3 | 3 | 115.38255 |
| Q16795 | NADH dehydrogenase [ubiquinone] 1 alp | NDUFA9 | 42.51 | 44.6 | 16 | 16 | 774.03644 |
| Q16831 | Uridine phosphorylase 1 OS=Homo sapie | UPP1 | 33.935 | 11 | 3 | 3 | 30.76766 |
| Q16850 | Lanosterol 14-alpha demethylase OS=Hor | CYP51A1 | 57.279 | 2.9 | 1 | 1 | 57.804737 |
| Q16875 | 6-phosphofructo-2-kinase/fructose-2,6-bis | PFKFB3 | 59.61 | 10.4 | 4 | 4 | 109.43094 |
| Q16891 | MICOS complex subunit MIC60 OS=Hor | IMMT | 83.678 | 14.5 | 10 | 10 | 131.38612 |
| Q17RS7 | Flap endonuclease GEN homolog 1 OS=H | GEN1 | 102.885 | 5.7 | 4 | 4 | 96.491913 |
| Q17RW2 | Collagen alpha-1(XXIV) chain OS=Homo | COL24A1 | 175.497 | 1.1 | 1 | 1 | 100.8071 |
| Q17RY0 | Cytoplasmic polyadenylation element-bin | CPEB4 | 80.152 | 13.3 | 10 | 10 | 1380.0587 |
| Q27J81 | Inverted formin-2 OS=Homo sapiens OX= | INF2 | 135.625 | 3 | 3 | 3 | 76.762779 |
| Q29RF7 | Sister chromatid cohesion protein PDS5 h | PDS5A | 150.831 | 7.3 | 8 | 7 | 91.587639 |
| Q2KHR2 | DNA-binding protein RFX7 OS=Homo sa | RFX7 | 146.896 | 11.2 | 12 | 12 | 138.08542 |

| | | | | | | | |
|---|---|---|---|---|---|---|---|
| Q2M1P5 | Kinesin-like protein KIF7 OS=Homo sapi | KIF7 | 150.588 | 6.3 | 8 | 8 | 119.32655 |
| Q2M2I8 | AP2-associated protein kinase 1 OS=Hon | AAK1 | 103.885 | 7 | 5 | 5 | 81.391235 |
| Q2M389 | WASH complex subunit 4 OS=Homo sap | WASHC4 | 136.404 | 2.6 | 3 | 3 | 56.295147 |
| Q2NKJ3 | CST complex subunit CTC1 OS=Homo sa | CTC1 | 134.609 | 2 | 2 | 2 | 68.518242 |
| Q2T9J0 | Peroxisomal leader peptide-processing pro | TYSND1 | 59.309 | 2.5 | 1 | 1 | 31.318047 |
| Q2TAK8 | PWWP domain-containing DNA repair fa | PWWP3A | 78.637 | 5.8 | 3 | 3 | 74.978394 |
| Q2TAY7 | WD40 repeat-containing protein SMU1 O | SMU1 | 57.544 | 41.7 | 17 | 17 | 1923.454 |
| Q2TBE0 | CWF19-like protein 2 OS=Homo sapiens | CWF19L2 | 103.787 | 12.6 | 11 | 11 | 142.70602 |
| Q2VPK5 | Cytoplasmic tRNA 2-thiolation protein 2 ( | CTU2 | 56.108 | 10.7 | 5 | 5 | 141.16116 |
| Q32MZ4 | Leucine-rich repeat flightless-interacting p | LRRFIP1 | 89.254 | 14.9 | 10 | 10 | 181.52689 |
| Q32P41 | tRNA (guanine(37)-N1)-methyltransferase | TRMT5 | 58.257 | 1.8 | 1 | 1 | 260.30194 |
| Q32P44 | Echinoderm microtubule-associated prote | EML3 | 95.197 | 14.5 | 11 | 11 | 261.50684 |
| Q38SD2 | Leucine-rich repeat serine/threonine-prote | LRRK1 | 225.393 | 5 | 9 | 9 | 65.125305 |
| Q3B7T1 | Erythroid differentiation-related factor 1 ( | EDRF1 | 138.529 | 2.5 | 3 | 3 | 48.9837 |
| Q3KR37 | Protein Aster-B OS=Homo sapiens OX=9 | GRAMD1B | 85.401 | 1.1 | 1 | 1 | 98.724335 |
| Q3L8U1 | Chromodomain-helicase-DNA-binding pr | CHD9 | 326.022 | 10.4 | 28 | 16 | 426.95392 |
| Q3MHD2 | Protein LSM12 OS=Homo sapiens OX=9 | LSM12 | 21.701 | 45.1 | 6 | 6 | 27389.285 |
| Q3V6T2 | Girdin OS=Homo sapiens OX=9606 GN= | CCDC88A | 216.042 | 1.2 | 2 | 2 | 31.978966 |
| Q3ZCM7 | Tubulin beta-8 chain OS=Homo sapiens ( | TUBB8 | 49.776 | 7.2 | 1 | 1 | 208.63918 |
| Q3ZCQ8 | Mitochondrial import inner membrane tra | TIMM50 | 39.647 | 11.9 | 4 | 4 | 293.18243 |
| Q49A26 | Cytokine-like nuclear factor N-PAC OS=I | GLYR1 | 60.548 | 22.6 | 11 | 11 | 378.69278 |
| Q49A88 | Coiled-coil domain-containing protein 14 | CCDC14 | 106.302 | 6 | 4 | 4 | 59.43116 |
| Q49AM1 | Transcription termination factor 2, mitoch | MTERF2 | 44.414 | 2.3 | 1 | 1 | 38.977825 |
| Q4AC94 | C2 domain-containing protein 3 OS=Hom | C2CD3 | 260.389 | 4.1 | 8 | 8 | 82.103973 |
| Q4G0J3 | La-related protein 7 OS=Homo sapiens O | LARP7 | 66.899 | 10 | 5 | 5 | 152.02365 |
| Q4KMP7 | TBC1 domain family member 10B OS=H | TBC1D10B | 87.199 | 4.3 | 3 | 3 | 62.602444 |
| Q4VC44 | FLYWCH-type zinc finger-containing pro | FLYWCH1 | 80.109 | 1.3 | 1 | 1 | 760.24231 |
| Q504Q3 | PAN2-PAN3 deadenylation complex cata | PAN2 | 135.368 | 15.1 | 17 | 17 | 195.0627 |
| Q52LJ0 | Protein FAM98B OS=Homo sapiens OX= | FAM98B | 45.548 | 17.8 | 6 | 6 | 343.65997 |
| Q53EL6 | Programmed cell death protein 4 OS=Hor | PDCD4 | 51.736 | 11.5 | 5 | 5 | 333.12885 |
| Q53EP0 | Fibronectin type III domain-containing pro | FNDC3B | 132.889 | 16.5 | 17 | 17 | 668.01227 |
| Q53ET0 | CREB-regulated transcription coactivator | CRTC2 | 73.302 | 4.9 | 3 | 3 | 114.98605 |
| Q53EZ4 | Centrosomal protein of 55 kDa OS=Homo | CEP55 | 54.179 | 39.4 | 18 | 18 | 990.22784 |
| Q53F19 | Nuclear cap-binding protein subunit 3 OS | NCBP3 | 70.593 | 25.6 | 15 | 15 | 3483.0796 |
| Q53G59 | Kelch-like protein 12 OS=Homo sapiens ( | KLHL12 | 63.278 | 25.9 | 12 | 12 | 350.59229 |

| | | | | | | |
|---|---|---|---|---|---|---|
| Q53GA4 | Pleckstrin homology-like domain family A | PHLDA2 | 17.093 | 5.9 | 1 | 1 | 568.03857 |
| Q53GQ0 | Very-long-chain 3-oxoacyl-CoA reductase | HSD17B12 | 34.325 | 19.6 | 5 | 5 | 104.50337 |
| Q53GS7 | mRNA export factor GLE1 OS=Homo sap | GLE1 | 79.837 | 23.5 | 16 | 16 | 464.84244 |
| Q53GS9 | Ubiquitin carboxyl-terminal hydrolase 39 | USP39 | 65.381 | 13.3 | 7 | 7 | 297.03931 |
| Q53H96 | Pyrroline-5-carboxylate reductase 3 OS=H | PYCR3 | 28.664 | 3.6 | 1 | 1 | 145.48396 |
| Q53HL2 | Borealin OS=Homo sapiens OX=9606 GN | CDCA8 | 31.323 | 22.5 | 6 | 6 | 309.29395 |
| Q53R41 | FAST kinase domain-containing protein 1 | FASTKD1 | 97.411 | 2.7 | 2 | 2 | 39.454475 |
| Q53T94 | TATA box-binding protein-associated fac | TAF1B | 68.832 | 3.1 | 2 | 2 | 38.324799 |
| Q562E7 | WD repeat-containing protein 81 OS=Hon | WDR81 | 211.697 | 2.3 | 4 | 4 | 80.119133 |
| Q562F6 | Shugoshin 2 OS=Homo sapiens OX=9606 | SGO2 | 144.74 | 8.5 | 11 | 11 | 136.04065 |
| Q56NI9 | N-acetyltransferase ESCO2 OS=Homo sa | ESCO2 | 68.307 | 19.3 | 10 | 10 | 210.81056 |
| Q58A45 | PAN2-PAN3 deadenylation complex subu | PAN3 | 95.613 | 28 | 21 | 21 | 573.80292 |
| Q58EX7 | Puratrophin-1 OS=Homo sapiens OX=960 | PLEKHG4 | 130.803 | 2.9 | 3 | 3 | 37.073906 |
| Q5BJF2 | Sigma intracellular receptor 2 OS=Homo | TMEM97 | 20.848 | 4.5 | 1 | 1 | 133.45387 |
| Q5BJF6 | Outer dense fiber protein 2 OS=Homo sap | ODF2 | 95.402 | 19.2 | 14 | 14 | 120.90117 |
| Q5BJH7 | Protein YIF1B OS=Homo sapiens OX=96 | YIF1B | 34.436 | 4.1 | 1 | 1 | 45.424168 |
| Q5BKZ1 | DBIRD complex subunit ZNF326 OS=Ho | ZNF326 | 65.654 | 31.8 | 15 | 15 | 675.97595 |
| Q5C9Z4 | Nucleolar MIF4G domain-containing prot | NOM1 | 96.258 | 14.9 | 12 | 12 | 308.29587 |
| Q5D0E6 | DALR anticodon-binding domain-contain | DALRD3 | 59.363 | 5 | 3 | 3 | 66.261864 |
| Q5D1E8 | Endoribonuclease ZC3H12A OS=Homo s | ZC3H12A | 65.7 | 7 | 3 | 3 | 64.095901 |
| Q5EBL8 | PDZ domain-containing protein 11 OS=H | PDZD11 | 16.132 | 15.7 | 2 | 2 | 132.79538 |
| Q5F1R6 | DnaJ homolog subfamily C member 21 O | DNAJC21 | 62.028 | 17.3 | 10 | 10 | 724.59662 |
| Q5FBB7 | Shugoshin 1 OS=Homo sapiens OX=9606 | SGO1 | 64.19 | 1.8 | 1 | 1 | 29.928768 |
| Q5H9R7 | Serine/threonine-protein phosphatase 6 re | PPP6R3 | 97.67 | 23.6 | 17 | 17 | 177.22855 |
| Q5HYI8 | Rab-like protein 3 OS=Homo sapiens OX | RABL3 | 26.423 | 4.7 | 1 | 1 | 170.46077 |
| Q5HYJ3 | Protein FAM76B OS=Homo sapiens OX= | FAM76B | 38.708 | 24.5 | 7 | 6 | 268.69415 |
| Q5JPH6 | Probable glutamate--tRNA ligase, mitoch | EARS2 | 58.689 | 9.4 | 4 | 4 | 67.461128 |
| Q5JRA6 | Transport and Golgi organization protein | MIA3 | 213.703 | 0.6 | 1 | 1 | 59.476746 |
| Q5JST6 | EF-hand domain-containing family memb | EFHC2 | 87.397 | 1.2 | 1 | 1 | 86.918556 |
| Q5JTC6 | APC membrane recruitment protein 1 OS= | AMER1 | 124.03 | 1 | 1 | 1 | 49.251427 |
| Q5JTH9 | RRP12-like protein OS=Homo sapiens OX | RRP12 | 143.703 | 23.6 | 29 | 29 | 1358.1719 |
| Q5JTV8 | Torsin-1A-interacting protein 1 OS=Hom | TOR1AIP1 | 66.249 | 4.5 | 2 | 2 | 19.381477 |
| Q5JTW2 | Centrosomal protein of 78 kDa OS=Homo | CEP78 | 76.397 | 12.5 | 9 | 9 | 225.97067 |
| Q5JTZ9 | Alanine--tRNA ligase, mitochondrial OS= | AARS2 | 107.341 | 9.2 | 8 | 8 | 72.708809 |
| Q5JVF3 | PCI domain-containing protein 2 OS=Hor | PCID2 | 46.03 | 38.6 | 13 | 13 | 372.8512 |

| | | | | | | | |
|---|---|---|---|---|---|---|---|
| Q5JVS0 | Intracellular hyaluronan-binding protein 4 | HABP4 | 45.786 | 1.9 | 1 | 1 | 308.89783 |
| Q5JXC2 | Migration and invasion-inhibitory protein | MIIP | 42.825 | 3.4 | 1 | 1 | 51.719551 |
| Q5K651 | Sterile alpha motif domain-containing pro | SAMD9 | 184.282 | 0.5 | 1 | 1 | 135.03137 |
| Q5M775 | Cytospin-B OS=Homo sapiens OX=9606 | SPECC1 | 118.585 | 2.1 | 2 | 2 | 48.744186 |
| Q5MIZ7 | Serine/threonine-protein phosphatase 4 re | PPP4R3B | 97.459 | 2.8 | 2 | 2 | 35.652267 |
| Q5PRF9 | Protein Smaug homolog 2 OS=Homo sapi | SAMD4B | 75.484 | 13.4 | 7 | 7 | 344.30563 |
| Q5QJE6 | Deoxynucleotidyltransferase terminal-inte | DNTTIP2 | 84.47 | 26.1 | 17 | 17 | 1371.3706 |
| Q5QP82 | DDB1- and CUL4-associated factor 10 O | DCAF10 | 60.583 | 8.6 | 4 | 4 | 48.052322 |
| Q5RKV6 | Exosome complex component MTR3 OS= | EXOSC6 | 28.236 | 28.3 | 7 | 7 | 523.82916 |
| Q5SNT2 | Transmembrane protein 201 OS=Homo sa | TMEM201 | 72.236 | 6.6 | 4 | 4 | 64.488083 |
| Q5SRE5 | Nucleoporin NUP188 OS=Homo sapiens | NUP188 | 196.043 | 4.1 | 6 | 6 | 79.219048 |
| Q5SSJ5 | Heterochromatin protein 1-binding protei | HP1BP3 | 61.207 | 13 | 7 | 7 | 146.71135 |
| Q5ST30 | Valine--tRNA ligase, mitochondrial OS=H | VARS2 | 118.49 | 1.8 | 2 | 2 | 60.158203 |
| Q5SW79 | Centrosomal protein of 170 kDa OS=Hon | CEP170 | 175.293 | 21.3 | 28 | 22 | 645.55066 |
| Q5SY16 | Polynucleotide 5'-hydroxyl-kinase NOL9 | NOL9 | 79.323 | 5 | 3 | 3 | 39.199142 |
| Q5SZL2 | Centrosomal protein of 85 kDa-like OS=H | CEP85L | 91.809 | 8.8 | 7 | 7 | 79.643044 |
| Q5T0N5 | Formin-binding protein 1-like OS=Homo | FNBP1L | 70.066 | 10.7 | 6 | 6 | 174.95317 |
| Q5T200 | Zinc finger CCCH domain-containing pro | ZC3H13 | 196.636 | 3.8 | 6 | 6 | 151.79877 |
| Q5T280 | Putative methyltransferase C9orf114 OS= | SPOUT1 | 42.009 | 26.3 | 10 | 10 | 827.01654 |
| Q5T2T1 | MAGUK p55 subfamily member 7 OS=H | MPP7 | 65.524 | 29 | 13 | 13 | 383.24258 |
| Q5T440 | Putative transferase CAF17, mitochondria | IBA57 | 38.155 | 10.1 | 3 | 3 | 129.20619 |
| Q5T5P2 | Sickle tail protein homolog OS=Homo sa | KIAA1217 | 214.116 | 3.3 | 6 | 6 | 55.381905 |
| Q5T5U3 | Rho GTPase-activating protein 21 OS=Hc | ARHGAP21 | 217.463 | 10.3 | 18 | 18 | 102.01345 |
| Q5T5X7 | BEN domain-containing protein 3 OS=Hc | BEND3 | 94.475 | 1.8 | 1 | 1 | 48.834473 |
| Q5T5Y3 | Calmodulin-regulated spectrin-associated | CAMSAP1 | 177.972 | 16 | 21 | 21 | 317.18823 |
| Q5T6F0 | DDB1- and CUL4-associated factor 12 O | DCAF12 | 50.518 | 6.4 | 3 | 3 | 88.961731 |
| Q5T6F2 | Ubiquitin-associated protein 2 OS=Homo | UBAP2 | 117.116 | 13.6 | 12 | 12 | 3833.457 |
| Q5T6S3 | PHD finger protein 19 OS=Homo sapiens | PHF19 | 65.591 | 24 | 12 | 12 | 169.81586 |
| Q5T749 | Keratinocyte proline-rich protein OS=Hor | KPRP | 64.136 | 1.6 | 1 | 1 | 124.45901 |
| Q5T7B8 | Kinesin-like protein KIF24 OS=Homo sa | KIF24 | 151.904 | 1.2 | 1 | 1 | 55.531357 |
| Q5T7V8 | RAB6-interacting golgin OS=Homo sapie | GORAB | 42.266 | 2.4 | 1 | 1 | 90.530594 |
| Q5T7W7 | Thiosulfate sulfurtransferase/rhodanese-li | TSTD2 | 58.263 | 3.3 | 2 | 2 | 41.230927 |
| Q5T9A4 | ATPase family AAA domain-containing p | ATAD3B | 72.573 | 12.5 | 7 | 4 | 706.43298 |
| Q5TAQ9 | DDB1- and CUL4-associated factor 8 OS | DCAF8 | 66.853 | 6 | 3 | 3 | 27.861443 |
| Q5TAX3 | Terminal uridylyltransferase 4 OS=Homo | TUT4 | 185.166 | 10 | 17 | 17 | 191.57019 |

| | | | | | | | |
|---|---|---|---|---|---|---|---|
| Q5TB30 | DEP domain-containing protein 1A OS=H | DEPDC1 | 92.96 | 10.5 | 8 | 8 | 115.35883 |
| Q5TC82 | Roquin-1 OS=Homo sapiens OX=9606 G | RC3H1 | 125.737 | 15.3 | 13 | 13 | 989.9104 |
| Q5TCZ1 | SH3 and PX domain-containing protein 2. | SH3PXD2A | 125.29 | 10.4 | 11 | 11 | 192.70137 |
| Q5TGY3 | Transcription factor Gibbin OS=Homo sa | AHDC1 | 168.35 | 9.1 | 12 | 12 | 155.98004 |
| Q5TKA1 | Protein lin-9 homolog OS=Homo sapiens | LIN9 | 61.946 | 3.5 | 2 | 2 | 28.813957 |
| Q5U5Q3 | RNA-binding E3 ubiquitin-protein ligase | MEX3C | 69.367 | 1.7 | 1 | 1 | 87.997757 |
| Q5UIP0 | Telomere-associated protein RIF1 OS=Hc | RIF1 | 274.466 | 8.9 | 16 | 16 | 163.70918 |
| Q5VSY0 | G kinase-anchoring protein 1 OS=Homo s | GKAP1 | 42.079 | 17.8 | 6 | 6 | 135.63908 |
| Q5VT06 | Centrosome-associated protein 350 OS=H | CEP350 | 350.931 | 3.9 | 10 | 10 | 102.45839 |
| Q5VT25 | Serine/threonine-protein kinase MRCK al | CDC42BPA | 197.307 | 9.1 | 13 | 13 | 130.93588 |
| Q5VT52 | Regulation of nuclear pre-mRNA domain- | RPRD2 | 156.02 | 1.8 | 2 | 2 | 37.761612 |
| Q5VTB9 | E3 ubiquitin-protein ligase RNF220 OS=I | RNF220 | 62.766 | 8.1 | 4 | 4 | 601.51184 |
| Q5VU43 | Myomegalin OS=Homo sapiens OX=960( | PDE4DIP | 265.103 | 3 | 7 | 7 | 99.013405 |
| Q5VUA4 | Zinc finger protein 318 OS=Homo sapien: | ZNF318 | 251.113 | 11.9 | 25 | 25 | 242.59604 |
| Q5VV42 | Threonylcarbamoyladenosine tRNA meth | CDKAL1 | 65.112 | 15.4 | 8 | 8 | 191.92035 |
| Q5VW36 | Focadhesin OS=Homo sapiens OX=9606 | FOCAD | 200.059 | 10.2 | 18 | 18 | 199.82976 |
| Q5VWN6 | Protein TASOR 2 OS=Homo sapiens OX: | TASOR2 | 268.843 | 1.9 | 4 | 4 | 51.359722 |
| Q5VWQ8 | Disabled homolog 2-interacting protein O | DAB2IP | 131.626 | 6.2 | 6 | 6 | 116.38246 |
| Q5VWZ2 | Lysophospholipase-like protein 1 OS=Ho: | LYPLAL1 | 26.317 | 4.2 | 1 | 1 | 69.911751 |
| Q5VYK3 | Proteasome adapter and scaffold protein E | ECPAS | 204.291 | 0.5 | 1 | 1 | 24.069988 |
| Q5VYS8 | Terminal uridylyltransferase 7 OS=Homo | TUT7 | 171.23 | 18.1 | 25 | 23 | 200.86125 |
| Q5VZ89 | DENN domain-containing protein 4C OS= | DENND4C | 212.711 | 1.5 | 2 | 2 | 46.233192 |
| Q5VZE5 | N-alpha-acetyltransferase 35, NatC auxili: | NAA35 | 83.639 | 4.1 | 3 | 3 | 58.360386 |
| Q5VZF2 | Muscleblind-like protein 2 OS=Homo sap | MBNL2 | 40.518 | 16.9 | 4 | 4 | 278.29688 |
| Q5VZK9 | F-actin-uncapping protein LRRC16A OS= | CARMIL1 | 151.557 | 5.7 | 6 | 6 | 79.903717 |
| Q5VZM2 | Ras-related GTP-binding protein B OS=H | RRAGB | 43.251 | 5.3 | 2 | 2 | 42.098419 |
| Q5W0B1 | ORC ubiquitin ligase 1 OS=Homo sapiens | OBI1 | 81.117 | 17.5 | 11 | 11 | 173.16751 |
| Q5XG87 | Terminal nucleotidyltransferase 4A OS=F | TENT4A | 84.651 | 8.7 | 6 | 6 | 161.69556 |
| Q63HN8 | E3 ubiquitin-protein ligase RNF213 OS=I | RNF213 | 591.408 | 6.3 | 29 | 29 | 142.42488 |
| Q63HR2 | Tensin-2 OS=Homo sapiens OX=9606 GI | TNS2 | 152.581 | 11.9 | 14 | 14 | 118.69424 |
| Q63ZY3 | KN motif and ankyrin repeat domain-cont | KANK2 | 91.175 | 7.4 | 6 | 6 | 152.3541 |
| Q64LD2 | WD repeat-containing protein 25 OS=Hoi | WDR25 | 60.162 | 11.2 | 6 | 6 | 114.69925 |
| Q659A1 | Little elongation complex subunit 2 OS=F | ICE2 | 110.012 | 2 | 2 | 2 | 52.964165 |
| Q659C4 | La-related protein 1B OS=Homo sapiens ( | LARP1B | 105.323 | 8.4 | 7 | 7 | 224.27631 |
| Q66K74 | Microtubule-associated protein 1S OS=H( | MAP1S | 112.212 | 7 | 6 | 6 | 122.45975 |

| | | | | | | | |
|---|---|---|---|---|---|---|---|
| Q66PJ3 | ADP-ribosylation factor-like protein 6-int | ARL6IP4 | 26.375 | 21.5 | 4 | 4 | 569.02045 |
| Q68CP9 | AT-rich interactive domain-containing pro | ARID2 | 197.391 | 1.1 | 2 | 2 | 51.345188 |
| Q68CQ4 | U3 small nucleolar RNA-associated prote | UTP25 | 87.056 | 27.2 | 18 | 18 | 200.86786 |
| Q68CZ1 | Protein fantom OS=Homo sapiens OX=96 | RPGRIP1L | 151.202 | 5.5 | 6 | 6 | 66.495056 |
| Q68CZ2 | Tensin-3 OS=Homo sapiens OX=9606 Gl | TNS3 | 155.266 | 6.2 | 7 | 7 | 100.70272 |
| Q68CZ6 | HAUS augmin-like complex subunit 3 OS | HAUS3 | 69.651 | 11.8 | 6 | 6 | 40.770103 |
| Q68D10 | Protein SPT2 homolog OS=Homo sapiens | SPTY2D1 | 75.599 | 34.3 | 24 | 24 | 551.86597 |
| Q68E01 | Integrator complex subunit 3 OS=Homo s | INTS3 | 118.07 | 7.2 | 7 | 7 | 63.550758 |
| Q69YN4 | Protein virilizer homolog OS=Homo sapie | VIRMA | 202.025 | 6.5 | 12 | 12 | 135.51967 |
| Q69YQ0 | Cytospin-A OS=Homo sapiens OX=9606 | SPECC1L | 124.544 | 2.4 | 2 | 2 | 25.591103 |
| Q6DN12 | Multiple C2 and transmembrane domain-c | MCTP2 | 99.596 | 5.4 | 4 | 4 | 23.942688 |
| Q6DN90 | IQ motif and SEC7 domain-containing pro | IQSEC1 | 108.315 | 21.8 | 19 | 15 | 410.91492 |
| Q6FI81 | Anamorsin OS=Homo sapiens OX=9606 | CIAPIN1 | 33.583 | 51 | 13 | 13 | 447.68326 |
| Q6FIF0 | AN1-type zinc finger protein 6 OS=Homo | ZFAND6 | 22.556 | 4.3 | 1 | 1 | 73.667229 |
| Q6GQQ9 | OTU domain-containing protein 7B OS=H | OTUD7B | 92.526 | 11.5 | 8 | 7 | 124.11989 |
| Q6I9Y2 | THO complex subunit 7 homolog OS=Ho | THOC7 | 23.743 | 27.9 | 5 | 5 | 135.72929 |
| Q6IA86 | Elongator complex protein 2 OS=Homo s | ELP2 | 92.501 | 1.6 | 1 | 1 | 63.285774 |
| Q6IAA8 | Ragulator complex protein LAMTOR1 O | LAMTOR1 | 17.745 | 6.8 | 1 | 1 | 105.11781 |
| Q6IAN0 | Dehydrogenase/reductase SDR family me | DHRS7B | 35.12 | 25.5 | 7 | 7 | 132.26964 |
| Q6IBW4 | Condensin-2 complex subunit H2 OS=Ho | NCAPH2 | 68.227 | 11.4 | 7 | 7 | 113.25822 |
| Q6IE81 | Protein Jade-1 OS=Homo sapiens OX=96 | JADE1 | 95.534 | 1.1 | 1 | 0 | 53.082371 |
| Q6IEG0 | U11/U12 small nuclear ribonucleoprotein | SNRNP48 | 39.966 | 6.8 | 2 | 2 | 54.069992 |
| Q6IMN6 | Caprin-2 OS=Homo sapiens OX=9606 Gl | CAPRIN2 | 125.925 | 5.9 | 5 | 5 | 57.587513 |
| Q6IN84 | rRNA methyltransferase 1, mitochondrial | MRM1 | 38.639 | 25.5 | 9 | 9 | 675.77576 |
| Q6IPM2 | IQ domain-containing protein E OS=Hom | IQCE | 77.298 | 1.4 | 1 | 1 | 40.585186 |
| Q6IQ16 | Speckle-type POZ protein-like OS=Homo | SPOPL | 44.648 | 8.4 | 3 | 3 | 50.253639 |
| Q6IQ19 | Centriole, cilia and spindle-associated pro | CCSAP | 30.217 | 15.9 | 4 | 4 | 168.06602 |
| Q6IQ23 | Pleckstrin homology domain-containing fa | PLEKHA7 | 127.135 | 1.9 | 2 | 2 | 88.499939 |
| Q6IQ49 | Splicing regulator SDE2 OS=Homo sapie | SDE2 | 49.742 | 17.5 | 7 | 7 | 274.07294 |
| Q6KC79 | Nipped-B-like protein OS=Homo sapiens | NIPBL | 316.052 | 5.1 | 13 | 13 | 92.740326 |
| Q6L8Q7 | 2',5'-phosphodiesterase 12 OS=Homo sap | PDE12 | 67.352 | 2 | 1 | 1 | 50.362011 |
| Q6MZQ0 | Proline-rich protein 5-like OS=Homo sapi | PRR5L | 40.836 | 7.1 | 2 | 2 | 17.574112 |
| Q6N022 | Teneurin-4 OS=Homo sapiens OX=9606 | TENM4 | 307.957 | 0.4 | 1 | 1 | 22.745352 |
| Q6NS38 | DNA oxidative demethylase ALKBH2 OS | ALKBH2 | 29.323 | 17.2 | 4 | 4 | 234.75987 |
| Q6NSJ5 | Volume-regulated anion channel subunit I | LRRC8E | 90.248 | 6.7 | 4 | 4 | 77.526772 |

| | | | | | | | |
|---|---|---|---|---|---|---|---|
| Q6NTE8 | MRN complex-interacting protein OS=Hc | MRNIP | 37.744 | 17.8 | 5 | 5 | 165.3838 |
| Q6NUK1 | Mitochondrial adenyl nucleotide antiporte | SLC25A24 | 53.355 | 15.1 | 6 | 6 | 103.05676 |
| Q6NUQ4 | Transmembrane protein 214 OS=Homo sa | TMEM214 | 77.151 | 11.3 | 7 | 7 | 270.9921 |
| Q6NYC1 | Bifunctional arginine demethylase and lys | JMJD6 | 46.462 | 7.2 | 2 | 2 | 42.5994 |
| Q6NYC8 | Phostensin OS=Homo sapiens OX=9606 ( | PPP1R18 | 67.943 | 4.6 | 3 | 3 | 59.019501 |
| Q6NZI2 | Caveolae-associated protein 1 OS=Homo | CAVIN1 | 43.477 | 9.5 | 4 | 4 | 195.46132 |
| Q6NZY4 | Zinc finger CCHC domain-containing pro | ZCCHC8 | 78.578 | 16.8 | 9 | 9 | 315.97842 |
| Q6P087 | Mitochondrial mRNA pseudouridine syntl | RPUSD3 | 38.461 | 31.6 | 9 | 9 | 987.52386 |
| Q6P0Q8 | Microtubule-associated serine/threonine-p | MAST2 | 196.436 | 11.9 | 20 | 17 | 191.67061 |
| Q6P158 | Putative ATP-dependent RNA helicase D | DHX57 | 155.605 | 29 | 38 | 35 | 4202.7607 |
| Q6P1J9 | Parafibromin OS=Homo sapiens OX=960 | CDC73 | 60.577 | 43.9 | 22 | 22 | 567.05957 |
| Q6P1K2 | Polyamine-modulated factor 1 OS=Homo | PMF1 | 23.34 | 10.7 | 2 | 2 | 67.155182 |
| Q6P1L5 | Protein FAM117B OS=Homo sapiens OX | FAM117B | 61.968 | 6.1 | 3 | 3 | 13.928834 |
| Q6P1M0 | Long-chain fatty acid transport protein 4 ( | SLC27A4 | 72.065 | 14.6 | 8 | 8 | 153.60776 |
| Q6P1M3 | LLGL scribble cell polarity complex comj | LLGL2 | 113.448 | 18.1 | 17 | 17 | 296.71225 |
| Q6P1N0 | Coiled-coil and C2 domain-containing prc | CC2D1A | 104.063 | 13.5 | 13 | 13 | 589.12909 |
| Q6P1Q0 | LETM1 domain-containing protein 1 OS= | LETMD1 | 41.791 | 21.4 | 7 | 7 | 323.03189 |
| Q6P1X5 | Transcription initiation factor TFIID subu | TAF2 | 136.972 | 16.8 | 18 | 18 | 324.05634 |
| Q6P2C8 | Mediator of RNA polymerase II transcript | MED27 | 35.432 | 20.6 | 5 | 5 | 75.684937 |
| Q6P2E9 | Enhancer of mRNA-decapping protein 4 ( | EDC4 | 151.662 | 33.5 | 34 | 34 | 3537.8545 |
| Q6P2H3 | Centrosomal protein of 85 kDa OS=Homc | CEP85 | 85.64 | 20.6 | 14 | 14 | 593.0437 |
| Q6P2Q9 | Pre-mRNA-processing-splicing factor 8 C | PRPF8 | 273.601 | 24.6 | 54 | 54 | 546.3335 |
| Q6P4A7 | Sideroflexin-4 OS=Homo sapiens OX=96 | SFXN4 | 37.999 | 19.9 | 6 | 6 | 154.02156 |
| Q6P5Z2 | Serine/threonine-protein kinase N3 OS=H | PKN3 | 99.421 | 16.6 | 13 | 13 | 409.69583 |
| Q6P6C2 | RNA demethylase ALKBH5 OS=Homo s | ALKBH5 | 44.256 | 28.9 | 10 | 10 | 830.27466 |
| Q6P9F0 | Coiled-coil domain-containing protein 62 | CCDC62 | 77.749 | 1 | 1 | 1 | 27.877399 |
| Q6PD62 | RNA polymerase-associated protein CTR | CTR9 | 133.503 | 4.9 | 5 | 5 | 90.483704 |
| Q6PF05 | Tetratricopeptide repeat protein 23-like O | TTC23L | 40.837 | 2.2 | 1 | 1 | 181.12907 |
| Q6PGN9 | Proline/serine-rich coiled-coil protein 1 O | PSRC1 | 38.797 | 25.9 | 8 | 8 | 493.60715 |
| Q6PGP7 | Superkiller complex protein 3 OS=Homo | SKIC3 | 175.487 | 17.7 | 23 | 23 | 221.07996 |
| Q6PI48 | Aspartate--tRNA ligase, mitochondrial OS | DARS2 | 73.563 | 1.9 | 1 | 1 | 66.018799 |
| Q6PID8 | Kelch domain-containing protein 10 OS=l | KLHDC10 | 49.098 | 3.8 | 2 | 2 | 39.112915 |
| Q6PIU2 | Neutral cholesterol ester hydrolase 1 OS= | NCEH1 | 45.808 | 5.9 | 2 | 2 | 101.05383 |
| Q6PIY7 | Poly(A) RNA polymerase GLD2 OS=Hor | TENT2 | 56.028 | 11.6 | 5 | 5 | 144.12822 |
| Q6PJF5 | Inactive rhomboid protein 2 OS=Homo sa | RHBDF2 | 96.687 | 2.1 | 2 | 2 | 66.682846 |

| | | | | | | | |
|---|---|---|---|---|---|---|---|
| Q6PJG6 | BRCA1-associated ATM activator 1 OS= | BRAT1 | 88.12 | 4.9 | 3 | 3 | 42.077381 |
| Q6PJI9 | GATOR2 complex protein WDR59 OS=F | WDR59 | 109.794 | 9.5 | 7 | 7 | 145.9801 |
| Q6PJP8 | DNA cross-link repair 1A protein OS=Ho | DCLRE1A | 116.4 | 2.9 | 3 | 3 | 51.216991 |
| Q6PJT7 | Zinc finger CCCH domain-containing pro | ZC3H14 | 82.876 | 11.7 | 8 | 8 | 96.918701 |
| Q6PJW8 | Consortin OS=Homo sapiens OX=9606 C | CNST | 79.597 | 12.1 | 9 | 9 | 647.99689 |
| Q6PKG0 | La-related protein 1 OS=Homo sapiens O | LARP1 | 123.511 | 33.6 | 41 | 36 | 29616.965 |
| Q6PL18 | ATPase family AAA domain-containing p | ATAD2 | 158.555 | 1.7 | 2 | 1 | 49.601563 |
| Q6Q0C0 | E3 ubiquitin-protein ligase TRAF7 OS=H | TRAF7 | 74.61 | 31.6 | 18 | 18 | 281.29285 |
| Q6R327 | Rapamycin-insensitive companion of mT( | RICTOR | 192.218 | 19.4 | 32 | 32 | 290.85324 |
| Q6RFH5 | WD repeat-containing protein 74 OS=Ho | WDR74 | 42.442 | 39.2 | 14 | 14 | 516.16174 |
| Q6SJ93 | Serine protease FAM111B OS=Homo sap | FAM111B | 84.675 | 22.5 | 15 | 15 | 318.83643 |
| Q6SPF0 | Sterile alpha motif domain-containing pro | SAMD1 | 56.052 | 5.6 | 3 | 3 | 93.730423 |
| Q6T4R5 | Actin remodeling regulator NHS OS=Hor | NHS | 179.136 | 14.1 | 18 | 18 | 288.42435 |
| Q6UB35 | Monofunctional C1-tetrahydrofolate synth | MTHFD1L | 105.79 | 16.9 | 14 | 14 | 313.1889 |
| Q6UB99 | Ankyrin repeat domain-containing protein | ANKRD11 | 297.914 | 2.8 | 7 | 7 | 52.548347 |
| Q6ULP2 | Aftiphilin OS=Homo sapiens OX=9606 G | AFTPH | 102.113 | 2.8 | 2 | 2 | 20.034554 |
| Q6UN15 | Pre-mRNA 3'-end-processing factor FIP1 | FIP1L1 | 66.527 | 10.4 | 6 | 6 | 917.65692 |
| Q6UUV7 | CREB-regulated transcription coactivator | CRTC3 | 66.96 | 1.8 | 1 | 1 | 81.004471 |
| Q6UWP7 | Lysocardiolipin acyltransferase 1 OS=Ho | LCLAT1 | 48.921 | 14 | 6 | 6 | 278.9187 |
| Q6UWZ7 | BRCA1-A complex subunit Abraxas 1 OS | ABRAXAS | 46.663 | 29.1 | 10 | 10 | 478.30627 |
| Q6UX04 | Spliceosome-associated protein CWC27 h | CWC27 | 53.848 | 2.3 | 1 | 1 | 34.575386 |
| Q6UXN9 | WD repeat-containing protein 82 OS=Ho | WDR82 | 35.08 | 47.6 | 14 | 14 | 1495.5228 |
| Q6VY07 | Phosphofurin acidic cluster sorting protei | PACS1 | 104.899 | 1.6 | 1 | 1 | 50.018646 |
| Q6W2J9 | BCL-6 corepressor OS=Homo sapiens OX | BCOR | 192.189 | 5.4 | 7 | 6 | 202.28496 |
| Q6WCQ1 | Myosin phosphatase Rho-interacting prot | MPRIP | 116.534 | 3.5 | 3 | 3 | 31.472345 |
| Q6WKZ4 | Rab11 family-interacting protein 1 OS=H | RAB11FIP1 | 137.168 | 2.7 | 3 | 1 | 103.50793 |
| Q6XE24 | RNA-binding motif, single-stranded-inter | RBMS3 | 47.84 | 3.4 | 1 | 1 | 89.121193 |
| Q6XZF7 | Dynamin-binding protein OS=Homo sapi | DNMBP | 177.348 | 2.7 | 3 | 3 | 63.400246 |
| Q6Y7W6 | GRB10-interacting GYF protein 2 OS=H | GIGYF2 | 150.07 | 17.2 | 21 | 21 | 2259.4956 |
| Q6YN16 | Hydroxysteroid dehydrogenase-like protei | HSDL2 | 45.395 | 12.7 | 4 | 4 | 97.961296 |
| Q6ZMS7 | Zinc finger protein 783 OS=Homo sapien | ZNF783 | 61.015 | 3.3 | 1 | 1 | 6.1375723 |
| Q6ZN04 | RNA-binding protein MEX3B OS=Homo | MEX3B | 58.833 | 11.4 | 6 | 4 | 166.40533 |
| Q6ZN18 | Zinc finger protein AEBP2 OS=Homo sap | AEBP2 | 54.467 | 7.5 | 3 | 3 | 100.1881 |
| Q6ZN55 | Zinc finger protein 574 OS=Homo sapien | ZNF574 | 98.901 | 24 | 19 | 19 | 929.90179 |
| Q6ZNA4 | E3 ubiquitin-protein ligase Arkadia OS=H | RNF111 | 108.862 | 3.4 | 3 | 3 | 72.156029 |

| | | | | | | | |
|---|---|---|---|---|---|---|---|
| Q6ZQX7 | Protein LIAT1 OS=Homo sapiens OX=9( | LIAT1 | 49.656 | 22.1 | 1 | 1 | 76.807053 |
| Q6ZRP7 | Sulfhydryl oxidase 2 OS=Homo sapiens C | QSOX2 | 77.529 | 5.6 | 3 | 3 | 40.548557 |
| Q6ZRV2 | Protein FAM83H OS=Homo sapiens OX= | FAM83H | 127.123 | 12.6 | 11 | 11 | 148.6142 |
| Q6ZRY4 | RNA-binding protein with multiple splicir | RBPMS2 | 22.497 | 17.7 | 3 | 3 | 345.55328 |
| Q6ZSY5 | Protein phosphatase 1 regulatory subunit 3 | PPP1R3F | 82.798 | 3.9 | 1 | 1 | 49.937653 |
| Q6ZSZ5 | Rho guanine nucleotide exchange factor 1 | ARHGEF18 | 151.643 | 1.3 | 2 | 2 | 82.220825 |
| Q6ZUT1 | Uncharacterized protein NKAPD1 OS=H | NKAPD1 | 34.11 | 3.8 | 1 | 1 | 89.572945 |
| Q6ZW31 | Rho GTPase-activating protein SYDE1 O | SYDE1 | 79.793 | 27.6 | 18 | 18 | 886.95172 |
| Q6ZWJ1 | Syntaxin-binding protein 4 OS=Homo sap | STXBP4 | 61.662 | 2.4 | 1 | 1 | 40.383606 |
| Q6ZXV5 | Protein O-mannosyl-transferase TMTC3 ( | TMTC3 | 104.009 | 3.9 | 3 | 3 | 131.87477 |
| Q6ZYL4 | General transcription factor IIH subunit 5 | GTF2H5 | 8.054 | 56.3 | 3 | 3 | 147.46629 |
| Q709F0 | Acyl-CoA dehydrogenase family member | ACAD11 | 87.265 | 10.5 | 8 | 8 | 202.80202 |
| Q70CQ4 | Ubiquitin carboxyl-terminal hydrolase 31 | USP31 | 146.652 | 4.1 | 5 | 5 | 82.108292 |
| Q70E73 | Ras-associated and pleckstrin homology d | RAPH1 | 135.256 | 1.9 | 2 | 2 | 62.207359 |
| Q70EK8 | Inactive ubiquitin carboxyl-terminal hydro | USP53 | 120.807 | 4.6 | 4 | 4 | 43.201759 |
| Q70EL1 | Inactive ubiquitin carboxyl-terminal hydro | USP54 | 187.389 | 3.7 | 5 | 5 | 61.570862 |
| Q70EL4 | Ubiquitin carboxyl-terminal hydrolase 43 | USP43 | 122.81 | 8.5 | 7 | 7 | 70.617706 |
| Q70SY1 | Cyclic AMP-responsive element-binding p | CREB3L2 | 57.415 | 4 | 2 | 1 | 51.993599 |
| Q70UQ0 | Inhibitor of nuclear factor kappa-B kinase | IKBIP | 39.309 | 6 | 2 | 2 | 79.119644 |
| Q712K3 | Ubiquitin-conjugating enzyme E2 R2 OS= | UBE2R2 | 27.166 | 9.2 | 2 | 2 | 140.94334 |
| Q71F23 | Centromere protein U OS=Homo sapiens | CENPU | 47.522 | 11.7 | 4 | 4 | 86.231873 |
| Q71RC2 | La-related protein 4 OS=Homo sapiens O | LARP4 | 80.596 | 35.8 | 23 | 23 | 14701.138 |
| Q71UM5 | Ribosomal protein eS27-like OS=Homo s | RPS27L | 9.478 | 39.3 | 4 | 2 | 8488.5352 |
| Q765P7 | Protein MTSS 2 OS=Homo sapiens OX=9 | MTSS2 | 79.93 | 9.9 | 6 | 6 | 55.645992 |
| Q76FK4 | Nucleolar protein 8 OS=Homo sapiens O | NOL8 | 131.617 | 5.1 | 6 | 6 | 56.492611 |
| Q76M96 | Coiled-coil domain-containing protein 80 | CCDC80 | 108.174 | 12.2 | 11 | 11 | 154.49776 |
| Q76N32 | Centrosomal protein of 68 kDa OS=Homo | CEP68 | 81.103 | 1.3 | 1 | 1 | 62.726273 |
| Q76N89 | E3 ubiquitin-protein ligase HECW1 OS=H | HECW1 | 179.554 | 3.4 | 5 | 4 | 78.314903 |
| Q7KZ85 | Transcription elongation factor SPT6 OS= | SUPT6H | 199.073 | 0.7 | 1 | 1 | 38.241577 |
| Q7KZF4 | Staphylococcal nuclease domain-containin | SND1 | 101.997 | 30.2 | 26 | 26 | 509.05881 |
| Q7KZI7 | Serine/threonine-protein kinase MARK2 ( | MARK2 | 87.911 | 29.6 | 20 | 20 | 1470.8813 |
| Q7KZN9 | Cytochrome c oxidase assembly protein C | COX15 | 46.031 | 4.9 | 2 | 2 | 81.387466 |
| Q7L014 | Probable ATP-dependent RNA helicase D | DDX46 | 117.363 | 20.5 | 21 | 21 | 330.92313 |
| Q7L0Y3 | tRNA methyltransferase 10 homolog C OS | TRMT10C | 47.347 | 41.7 | 17 | 17 | 777.92053 |
| Q7L1Q6 | eIF5-mimic protein 2 OS=Homo sapiens ( | BZW1 | 48.044 | 9.5 | 4 | 4 | 120.62996 |

| | | | | | | | |
|---|---|---|---|---|---|---|---|
| Q7L1W4 | Volume-regulated anion channel subunit I | LRRC8D | 98.202 | 10 | 7 | 7 | 81.600479 |
| Q7L2E3 | ATP-dependent RNA helicase DHX30 OS | DHX30 | 133.939 | 51.5 | 59 | 59 | 12430.425 |
| Q7L2H7 | Eukaryotic translation initiation factor 3 s | EIF3M | 42.503 | 31.3 | 10 | 10 | 2117.24 |
| Q7L2J0 | 7SK snRNA methylphosphate capping en: | MEPCE | 74.356 | 12.2 | 8 | 8 | 502.89316 |
| Q7L2Z9 | Centromere protein Q OS=Homo sapiens | CENPQ | 30.596 | 9.3 | 2 | 2 | 37.863308 |
| Q7L4I2 | Arginine/serine-rich coiled-coil protein 2 | RSRC2 | 50.56 | 13.6 | 6 | 6 | 368.45477 |
| Q7L576 | Cytoplasmic FMR1-interacting protein 1 ( | CYFIP1 | 145.183 | 17.3 | 20 | 13 | 420.35498 |
| Q7L590 | Protein MCM10 homolog OS=Homo sapi | MCM10 | 98.184 | 18.4 | 15 | 15 | 330.31912 |
| Q7L592 | Protein arginine methyltransferase NDUF. | NDUFAF7 | 49.238 | 2.5 | 1 | 1 | 38.651127 |
| Q7L5Y9 | E3 ubiquitin-protein transferase MAEA O | MAEA | 45.288 | 4 | 2 | 2 | 38.853817 |
| Q7L622 | G2/M phase-specific E3 ubiquitin-protein | G2E3 | 80.505 | 4.5 | 3 | 3 | 74.674507 |
| Q7L7X3 | Serine/threonine-protein kinase TAO1 OS | TAOK1 | 116.071 | 9.2 | 9 | 7 | 130.77939 |
| Q7L804 | Rab11 family-interacting protein 2 OS=H | RAB11FIP2 | 58.28 | 9.4 | 4 | 4 | 88.777527 |
| Q7L8L6 | FAST kinase domain-containing protein 5 | FASTKD5 | 86.575 | 8.6 | 6 | 6 | 95.031281 |
| Q7LBR1 | Charged multivesicular body protein 1b O | CHMP1B | 22.11 | 4 | 1 | 1 | 288.18726 |
| Q7LGA3 | Heparan sulfate 2-O-sulfotransferase 1 OS | HS2ST1 | 41.882 | 18.8 | 7 | 7 | 532.90417 |
| Q7LGC8 | Carbohydrate sulfotransferase 3 OS=Hom | CHST3 | 54.707 | 5.6 | 2 | 2 | 68.033928 |
| Q7RTP6 | [F-actin]-monooxygenase MICAL3 OS=H | MICAL3 | 224.295 | 1.5 | 3 | 3 | 64.188004 |
| Q7RTV0 | PHD finger-like domain-containing protei | PHF5A | 12.406 | 22.7 | 3 | 3 | 399.43634 |
| Q7Z2E3 | Aprataxin OS=Homo sapiens OX=9606 C | APTX | 40.741 | 24.2 | 8 | 8 | 404.38354 |
| Q7Z2W4 | Zinc finger CCCH-type antiviral protein 1 | ZC3HAV1 | 101.432 | 47.5 | 47 | 46 | 9602.7861 |
| Q7Z2Z1 | Treslin OS=Homo sapiens OX=9606 GN= | TICRR | 210.857 | 9.1 | 14 | 14 | 229.15936 |
| Q7Z333 | Probable helicase senataxin OS=Homo sa | SETX | 302.88 | 10.4 | 27 | 27 | 253.07936 |
| Q7Z3B4 | Nucleoporin p54 OS=Homo sapiens OX= | NUP54 | 55.436 | 4.3 | 2 | 2 | 52.87365 |
| Q7Z3I7 | Zinc finger protein 572 OS=Homo sapien: | ZNF572 | 61.239 | 5.7 | 3 | 3 | 56.337009 |
| Q7Z3K3 | Pogo transposable element with ZNF dom | POGZ | 155.344 | 6.3 | 7 | 7 | 100.68993 |
| Q7Z3K6 | Mesoderm induction early response protei | MIER3 | 61.437 | 3.5 | 2 | 2 | 32.218742 |
| Q7Z3T8 | Zinc finger FYVE domain-containing pro | ZFYVE16 | 168.904 | 2.3 | 3 | 3 | 43.16013 |
| Q7Z401 | C-myc promoter-binding protein OS=Hon | DENND4A | 209.245 | 4.3 | 6 | 6 | 91.738457 |
| Q7Z403 | Transmembrane channel-like protein 6 OS | TMC6 | 90.046 | 2 | 1 | 1 | 212.14302 |
| Q7Z406 | Myosin-14 OS=Homo sapiens OX=9606 ( | MYH14 | 227.871 | 0.4 | 1 | 1 | 382.8136 |
| Q7Z417 | FMR1-interacting protein NUFIP2 OS=H | NUFIP2 | 76.122 | 37.7 | 25 | 25 | 32621.734 |
| Q7Z460 | CLIP-associating protein 1 OS=Homo sap | CLASP1 | 169.451 | 33.4 | 49 | 43 | 1390.891 |
| Q7Z478 | ATP-dependent RNA helicase DHX29 OS | DHX29 | 155.236 | 23.8 | 29 | 29 | 790.42169 |
| Q7Z4H7 | HAUS augmin-like complex subunit 6 OS | HAUS6 | 108.622 | 12.5 | 11 | 11 | 257.64435 |

| | | | | | | | |
|---|---|---|---|---|---|---|---|
| Q7Z4S6 | Kinesin-like protein KIF21A OS=Homo s | KIF21A | 187.18 | 2.7 | 4 | 3 | 56.454933 |
| Q7Z569 | BRCA1-associated protein OS=Homo sap | BRAP | 67.305 | 18.2 | 10 | 10 | 139.44037 |
| Q7Z5J4 | Retinoic acid-induced protein 1 OS=Hom | RAI1 | 203.353 | 1.8 | 3 | 3 | 103.20126 |
| Q7Z5K2 | Wings apart-like protein homolog OS=Ho | WAPL | 132.946 | 7.1 | 8 | 8 | 88.84568 |
| Q7Z5Q1 | Cytoplasmic polyadenylation element-bin | CPEB2 | 64.945 | 18 | 10 | 4 | 1134.9935 |
| Q7Z5W3 | RNA 5'-monophosphate methyltransferase | BCDIN3D | 33.201 | 4.1 | 1 | 1 | 40.213207 |
| Q7Z628 | Neuroepithelial cell-transforming gene 1 p | NET1 | 67.741 | 7.2 | 4 | 3 | 131.4543 |
| Q7Z6B7 | SLIT-ROBO Rho GTPase-activating prot | SRGAP1 | 124.265 | 29.3 | 27 | 24 | 454.67175 |
| Q7Z6E9 | E3 ubiquitin-protein ligase RBBP6 OS=H | RBBP6 | 201.565 | 13.8 | 20 | 20 | 267.44568 |
| Q7Z6M4 | Transcription termination factor 4, mitoch | MTERF4 | 43.958 | 23.4 | 8 | 8 | 100.48688 |
| Q7Z6Z7 | E3 ubiquitin-protein ligase HUWE1 OS=I | HUWE1 | 481.891 | 1.6 | 5 | 5 | 28.735758 |
| Q7Z739 | YTH domain-containing family protein 3 | YTHDF3 | 63.862 | 31.1 | 19 | 11 | 4993.1401 |
| Q7Z7C8 | Transcription initiation factor TFIID subu | TAF8 | 34.263 | 5.2 | 1 | 1 | 176.39291 |
| Q7Z7H8 | Large ribosomal subunit protein uL10m C | MRPL10 | 29.283 | 18.4 | 4 | 4 | 277.73953 |
| Q7Z7K6 | Centromere protein V OS=Homo sapiens | CENPV | 29.946 | 26.2 | 6 | 6 | 441.8985 |
| Q86SF2 | N-acetylgalactosaminyltransferase 7 OS=I | GALNT7 | 75.39 | 4.9 | 3 | 3 | 58.647209 |
| Q86SQ0 | Pleckstrin homology-like domain family E | PHLDB2 | 142.159 | 16.9 | 19 | 19 | 166.96964 |
| Q86T03 | Type 1 phosphatidylinositol 4,5-bisphospl | PIP4P1 | 29.47 | 4 | 1 | 1 | 35.577995 |
| Q86T24 | Transcriptional regulator Kaiso OS=Homo | ZBTB33 | 74.485 | 1.9 | 1 | 1 | 44.84301 |
| Q86T82 | Ubiquitin carboxyl-terminal hydrolase 37 | USP37 | 110.17 | 1.7 | 2 | 2 | 8.8726416 |
| Q86TB9 | Protein PAT1 homolog 1 OS=Homo sapie | PATL1 | 86.851 | 24 | 18 | 18 | 954.70392 |
| Q86TC9 | Myopalladin OS=Homo sapiens OX=960 | MYPN | 145.258 | 6.3 | 8 | 8 | 120.3927 |
| Q86TG7 | Retrotransposon-derived protein PEG10 C | PEG10 | 80.173 | 30.9 | 17 | 17 | 3092.6995 |
| Q86TJ2 | Transcriptional adapter 2-beta OS=Homo | TADA2B | 48.471 | 10.5 | 4 | 4 | 80.841377 |
| Q86U06 | Probable RNA-binding protein 23 OS=Ho | RBM23 | 48.731 | 15.3 | 6 | 6 | 140.59648 |
| Q86U42 | Polyadenylate-binding protein 2 OS=Hom | PABPN1 | 32.75 | 9.5 | 3 | 3 | 1102.3397 |
| Q86U70 | LIM domain-binding protein 1 OS=Homo | LDB1 | 46.533 | 10.9 | 4 | 3 | 166.96178 |
| Q86U86 | Protein polybromo-1 OS=Homo sapiens C | PBRM1 | 192.948 | 2.1 | 4 | 4 | 62.615463 |
| Q86U90 | Threonylcarbamoyl-AMP synthase OS=H | YRDC | 29.328 | 12.5 | 3 | 3 | 59.181358 |
| Q86UE4 | Protein LYRIC OS=Homo sapiens OX=9 | MTDH | 63.837 | 23 | 12 | 12 | 3175.2466 |
| Q86UK7 | E3 ubiquitin-protein ligase ZNF598 OS=H | ZNF598 | 98.638 | 30.9 | 22 | 22 | 924.32898 |
| Q86UL3 | Glycerol-3-phosphate acyltransferase 4 O | GPAT4 | 52.072 | 2.6 | 1 | 1 | 78.331993 |
| Q86UP2 | Kinectin OS=Homo sapiens OX=9606 GN | KTN1 | 156.276 | 42 | 59 | 59 | 2060.6042 |
| Q86UQ0 | Zinc finger protein 589 OS=Homo sapien | ZNF589 | 41.19 | 4.7 | 2 | 2 | 113.03429 |
| Q86US8 | Telomerase-binding protein EST1A OS=H | SMG6 | 160.462 | 22.7 | 32 | 32 | 1168.0341 |

| | | | | | | | |
|---|---|---|---|---|---|---|---|
| Q86UT8 | Centrosomal AT-AC splicing factor OS=I | CENATAC | 37.975 | 7.8 | 3 | 3 | 97.431007 |
| Q86UU1 | Pleckstrin homology-like domain family E | PHLDB1 | 151.162 | 2.7 | 3 | 3 | 64.320435 |
| Q86UW6 | NEDD4-binding protein 2 OS=Homo sap | N4BP2 | 198.802 | 8 | 12 | 12 | 93.471008 |
| Q86UW9 | Probable E3 ubiquitin-protein ligase DTX | DTX2 | 67.247 | 2.9 | 2 | 2 | 42.531551 |
| Q86V48 | Leucine zipper protein 1 OS=Homo sapie | LUZP1 | 120.275 | 25.9 | 23 | 23 | 220.74811 |
| Q86V81 | THO complex subunit 4 OS=Homo sapie | ALYREF | 26.888 | 38.9 | 9 | 9 | 5845.1914 |
| Q86VI3 | Ras GTPase-activating-like protein IQGA | IQGAP3 | 184.699 | 6.1 | 8 | 8 | 301.81924 |
| Q86VM9 | Zinc finger CCCH domain-containing pro | ZC3H18 | 106.379 | 12.1 | 10 | 10 | 318.08569 |
| Q86VP1 | Tax1-binding protein 1 OS=Homo sapien | TAX1BP1 | 90.878 | 11.3 | 9 | 9 | 99.81752 |
| Q86VP6 | Cullin-associated NEDD8-dissociated pro | CAND1 | 136.376 | 0.9 | 1 | 1 | 61.964386 |
| Q86VR2 | Reticulophagy regulator 3 OS=Homo sapi | RETREG3 | 51.397 | 5.2 | 2 | 2 | 83.358292 |
| Q86VS8 | Protein Hook homolog 3 OS=Homo sapie | HOOK3 | 83.126 | 4.9 | 3 | 3 | 49.627312 |
| Q86W42 | THO complex subunit 6 homolog OS=Ho | THOC6 | 37.535 | 29 | 8 | 8 | 184.34436 |
| Q86W50 | RNA N6-adenosine-methyltransferase MI | METTL16 | 63.622 | 14.9 | 8 | 8 | 149.49318 |
| Q86W56 | Poly(ADP-ribose) glycohydrolase OS=Ho | PARG | 111.111 | 17 | 15 | 15 | 157.61781 |
| Q86W92 | Liprin-beta-1 OS=Homo sapiens OX=960 | PPFIBP1 | 114.025 | 37 | 34 | 32 | 1313.9567 |
| Q86WA8 | Lon protease homolog 2, peroxisomal OS | LONP2 | 94.617 | 20.9 | 16 | 16 | 1600.5343 |
| Q86WB0 | Zinc finger C3HC-type protein 1 OS=Hor | ZC3HC1 | 55.262 | 2 | 1 | 1 | 8.6109056 |
| Q86WP2 | Vasculin OS=Homo sapiens OX=9606 GI | GPBP1 | 53.34 | 7.2 | 3 | 3 | 305.27161 |
| Q86WX3 | Active regulator of SIRT1 OS=Homo sap | RPS19BP1 | 15.434 | 9.6 | 1 | 1 | 166.34123 |
| Q86X02 | Cerebellar degeneration-related protein 2- | CDR2L | 53.011 | 12.5 | 6 | 6 | 127.69696 |
| Q86X52 | Chondroitin sulfate synthase 1 OS=Homo | CHSY1 | 91.785 | 21.8 | 17 | 14 | 614.18396 |
| Q86X55 | Histone-arginine methyltransferase CARM | CARM1 | 65.854 | 9.2 | 5 | 5 | 68.972496 |
| Q86X67 | NAD(P)H pyrophosphatase NUDT13, mi | NUDT13 | 39.689 | 2.3 | 1 | 1 | 87.925529 |
| Q86X95 | Corepressor interacting with RBPJ 1 OS= | CIR1 | 52.314 | 10.9 | 6 | 6 | 230.58263 |
| Q86XI2 | Condensin-2 complex subunit G2 OS=Ho | NCAPG2 | 130.961 | 8.9 | 11 | 11 | 171.40416 |
| Q86XJ1 | GAS2-like protein 3 OS=Homo sapiens C | GAS2L3 | 75.215 | 9.5 | 6 | 6 | 152.48361 |
| Q86XK2 | F-box only protein 11 OS=Homo sapiens | FBXO11 | 103.585 | 7.2 | 6 | 6 | 120.43185 |
| Q86XL3 | Ankyrin repeat and LEM domain-containi | ANKLE2 | 104.115 | 4.8 | 4 | 4 | 54.042221 |
| Q86XN8 | RNA-binding protein MEX3D OS=Homo | MEX3D | 64.883 | 9.1 | 6 | 6 | 264.45572 |
| Q86XP3 | ATP-dependent RNA helicase DDX42 OS | DDX42 | 102.976 | 13.5 | 11 | 11 | 166.11391 |
| Q86XR8 | Centrosomal protein of 57 kDa OS=Homo | CEP57 | 57.089 | 8.4 | 4 | 4 | 69.60257 |
| Q86Y07 | Serine/threonine-protein kinase VRK2 OS | VRK2 | 58.141 | 25.2 | 11 | 11 | 647.85541 |
| Q86Y38 | Xylosyltransferase 1 OS=Homo sapiens C | XYLT1 | 107.57 | 0.7 | 1 | 1 | 73.924866 |
| Q86Y39 | NADH dehydrogenase [ubiquinone] 1 alp | NDUFA11 | 14.853 | 17 | 2 | 2 | 86.195831 |

| | | | | | | |
|---|---|---|---|---|---|---|
| Q86Y91 | Kinesin-like protein KIF18B OS=Homo s | KIF18B | 93.011 | 9.6 | 7 | 7 | 116.66246 |
| Q86YD1 | Prostate tumor-overexpressed gene 1 prot | PTOV1 | 46.869 | 4.8 | 2 | 1 | 63.22369 |
| Q86YP4 | Transcriptional repressor p66-alpha OS=H | GATAD2A | 68.063 | 24.5 | 12 | 11 | 197.26543 |
| Q86YT6 | E3 ubiquitin-protein ligase MIB1 OS=Ho | MIB1 | 110.136 | 12.8 | 11 | 10 | 143.04575 |
| Q86YV5 | Inactive tyrosine-protein kinase PRAG1 C | PRAG1 | 149.624 | 0.7 | 1 | 1 | 45.820568 |
| Q86YV9 | BLOC-2 complex member HPS6 OS=Hon | HPS6 | 82.975 | 8 | 5 | 5 | 83.260162 |
| Q86YW0 | 1-phosphatidylinositol 4,5-bisphosphate p | PLCZ1 | 70.412 | 1.2 | 1 | 1 | 410.77817 |
| Q86YZ3 | Hornerin OS=Homo sapiens OX=9606 GI | HRNR | 282.39 | 1.7 | 1 | 1 | 99.295937 |
| Q8IU60 | m7GpppN-mRNA hydrolase OS=Homo s | DCP2 | 48.424 | 13.3 | 6 | 6 | 418.95712 |
| Q8IUC4 | Rhophilin-2 OS=Homo sapiens OX=9606 | RHPN2 | 76.994 | 3.4 | 2 | 2 | 59.224968 |
| Q8IUD2 | ELKS/Rab6-interacting/CAST family mei | ERC1 | 128.087 | 23.9 | 27 | 20 | 341.13916 |
| Q8IUE6 | Histone H2A type 2-B OS=Homo sapiens | H2AC21 | 13.996 | 5.4 | 1 | 1 | 851.16815 |
| Q8IUF8 | Ribosomal oxygenase 2 OS=Homo sapien | RIOX2 | 52.801 | 20.4 | 9 | 9 | 528.05151 |
| Q8IUH3 | RNA-binding protein 45 OS=Homo sapie | RBM45 | 53.503 | 3.4 | 2 | 2 | 250.57257 |
| Q8IUR6 | CREB3 regulatory factor OS=Homo sapie | CREBRF | 72.149 | 2.2 | 1 | 1 | 91.863899 |
| Q8IUR7 | Armadillo repeat-containing protein 8 OS | ARMC8 | 75.51 | 4.8 | 3 | 3 | 46.430305 |
| Q8IUX4 | DNA dC->dU-editing enzyme APOBEC-3 | APOBEC3F | 45.021 | 18.8 | 6 | 4 | 531.50537 |
| Q8IV48 | 3'-5' exoribonuclease 1 OS=Homo sapiens | ERI1 | 40.064 | 26.9 | 7 | 7 | 585.38232 |
| Q8IVF7 | Formin-like protein 3 OS=Homo sapiens ( | FMNL3 | 117.213 | 4.3 | 4 | 4 | 34.678204 |
| Q8IVH8 | Mitogen-activated protein kinase kinase k | MAP4K3 | 101.316 | 11.6 | 9 | 9 | 244.21849 |
| Q8IVL1 | Neuron navigator 2 OS=Homo sapiens OX | NAV2 | 268.168 | 4.5 | 9 | 9 | 76.406136 |
| Q8IVM0 | Coiled-coil domain-containing protein 50 | CCDC50 | 35.822 | 3.6 | 1 | 1 | 68.840179 |
| Q8IVS2 | Malonyl-CoA-acyl carrier protein transacy | MCAT | 42.962 | 32.8 | 10 | 10 | 1117.946 |
| Q8IVT2 | Mitotic interactor and substrate of PLK1 ( | MISP | 75.357 | 14.9 | 9 | 9 | 213.56613 |
| Q8IVT5 | Kinase suppressor of Ras 1 OS=Homo sap | KSR1 | 102.16 | 10.1 | 9 | 9 | 124.16637 |
| Q8IWA0 | WD repeat-containing protein 75 OS=Hon | WDR75 | 94.499 | 15.8 | 11 | 11 | 292.94531 |
| Q8IWC1 | MAP7 domain-containing protein 3 OS=H | MAP7D3 | 98.43 | 30.8 | 25 | 25 | 936.11414 |
| Q8IWI9 | MAX gene-associated protein OS=Homo | MGA | 336.16 | 1.3 | 4 | 4 | 312.82675 |
| Q8IWK6 | Adhesion G protein-coupled receptor A3 ( | ADGRA3 | 146.151 | 0.5 | 1 | 1 | 172.93562 |
| Q8IWS0 | PHD finger protein 6 OS=Homo sapiens ( | PHF6 | 41.291 | 28.5 | 10 | 10 | 2128.4153 |
| Q8IWT6 | Volume-regulated anion channel subunit I | LRRC8A | 94.199 | 12 | 9 | 7 | 126.01548 |
| Q8IWV8 | E3 ubiquitin-protein ligase UBR2 OS=Ho | UBR2 | 200.539 | 0.7 | 1 | 1 | 72.962883 |
| Q8IWW6 | Rho GTPase-activating protein 12 OS=Ho | ARHGAP12 | 96.255 | 7.4 | 6 | 6 | 74.723648 |
| Q8IWX8 | Calcium homeostasis endoplasmic reticulu | CHERP | 103.703 | 20.2 | 18 | 18 | 3391.7466 |
| Q8IWZ3 | Ankyrin repeat and KH domain-containing | ANKHD1 | 269.458 | 19.3 | 42 | 30 | 1361.0809 |

| | | | | | | | |
|---|---|---|---|---|---|---|---|
| Q8IWZ8 | SURP and G-patch domain-containing pro | SUGP1 | 72.471 | 10.1 | 5 | 5 | 66.563301 |
| Q8IX01 | SURP and G-patch domain-containing pro | SUGP2 | 120.208 | 22.6 | 24 | 24 | 523.75916 |
| Q8IX12 | Cell division cycle and apoptosis regulato | CCAR1 | 132.822 | 11.1 | 13 | 13 | 644.23798 |
| Q8IX90 | Spindle and kinetochore-associated protei | SKA3 | 46.36 | 8.3 | 3 | 3 | 61.895004 |
| Q8IXB1 | DnaJ homolog subfamily C member 10 O | DNAJC10 | 91.08 | 11 | 8 | 8 | 86.452179 |
| Q8IXH7 | Negative elongation factor C/D OS=Hom | NELFCD | 66.247 | 17.8 | 8 | 8 | 172.76102 |
| Q8IXI2 | Mitochondrial Rho GTPase 1 OS=Homo : | RHOT1 | 70.785 | 9.4 | 5 | 5 | 99.838699 |
| Q8IXK0 | Polyhomeotic-like protein 2 OS=Homo sa | PHC2 | 90.714 | 6.4 | 5 | 4 | 124.11315 |
| Q8IXM3 | Large ribosomal subunit protein mL41 OS | MRPL41 | 15.383 | 27.7 | 4 | 4 | 760.22687 |
| Q8IXQ6 | Protein mono-ADP-ribosyltransferase PA | PARP9 | 96.344 | 17.4 | 14 | 14 | 186.26648 |
| Q8IXW5 | Putative RNA polymerase II subunit B1 C | RPAP2 | 69.509 | 18.8 | 9 | 9 | 79.245659 |
| Q8IXZ2 | Zinc finger CCCH domain-containing pro | ZC3H3 | 101.942 | 2.5 | 2 | 2 | 58.739597 |
| Q8IY17 | Patatin-like phospholipase domain-contai | PNPLA6 | 150.954 | 19.6 | 25 | 23 | 365.80737 |
| Q8IY18 | Structural maintenance of chromosomes p | SMC5 | 128.807 | 23.6 | 24 | 24 | 293.52661 |
| Q8IY21 | Probable ATP-dependent RNA helicase D | DDX60 | 197.854 | 19.3 | 29 | 29 | 188.60945 |
| Q8IY31 | Intraflagellar transport protein 20 homolo | IFT20 | 15.281 | 10.6 | 1 | 1 | 60.830311 |
| Q8IY37 | Probable ATP-dependent RNA helicase D | DHX37 | 129.546 | 31.5 | 36 | 36 | 1147.7922 |
| Q8IY81 | pre-rRNA 2'-O-ribose RNA methyltransfe | FTSJ3 | 96.559 | 19.5 | 16 | 16 | 382.81174 |
| Q8IY82 | Dynein regulatory complex subunit 7 OS= | DRC7 | 103.498 | 0.8 | 1 | 1 | 88.495987 |
| Q8IY92 | Structure-specific endonuclease subunit S | SLX4 | 200.012 | 3.2 | 6 | 6 | 95.199211 |
| Q8IYA6 | Cytoskeleton-associated protein 2-like OS | CKAP2L | 83.587 | 17.6 | 13 | 13 | 193.59726 |
| Q8IYB3 | Serine/arginine repetitive matrix protein 1 | SRRM1 | 102.336 | 12.3 | 10 | 10 | 382.04044 |
| Q8IYB7 | DIS3-like exonuclease 2 OS=Homo sapie | DIS3L2 | 99.28 | 24.9 | 21 | 21 | 246.78368 |
| Q8IYB8 | ATP-dependent RNA helicase SUPV3L1, | SUPV3L1 | 87.992 | 23.5 | 17 | 17 | 618.28296 |
| Q8IYD1 | Eukaryotic peptide chain release factor G | GSPT2 | 68.884 | 2.7 | 1 | 1 | 22.109385 |
| Q8IYD8 | Fanconi anemia group M protein OS=Hor | FANCM | 232.192 | 8.2 | 14 | 14 | 130.24388 |
| Q8IYM2 | Ribonuclease SLFN12 OS=Homo sapiens | SLFN12 | 66.973 | 6.7 | 4 | 3 | 64.907295 |
| Q8IYT2 | Cap-specific mRNA (nucleoside-2'-O-)-m | CMTR2 | 88.12 | 17 | 12 | 12 | 300.76978 |
| Q8IYT4 | Katanin p60 ATPase-containing subunit A | KATNAL2 | 61.253 | 2.2 | 1 | 1 | 76.207802 |
| Q8IYU8 | Calcium uptake protein 2, mitochondrial ( | MICU2 | 49.667 | 17.7 | 6 | 6 | 295.11224 |
| Q8IYW5 | E3 ubiquitin-protein ligase RNF168 OS=I | RNF168 | 65.02 | 24.7 | 17 | 17 | 1254.9626 |
| Q8IZ52 | Chondroitin sulfate synthase 2 OS=Homo | CHPF | 85.468 | 16.4 | 12 | 12 | 256.11649 |
| Q8IZ69 | tRNA (uracil-5-)-methyltransferase homol | TRMT2A | 68.726 | 33 | 20 | 20 | 1115.1294 |
| Q8IZE3 | Protein-associating with the carboxyl-term | SCYL3 | 82.858 | 1.6 | 1 | 1 | 28.900433 |
| Q8IZH2 | 5'-3' exoribonuclease 1 OS=Homo sapiens | XRN1 | 194.108 | 24.3 | 39 | 39 | 1031.5305 |

| | | | | | | | |
|---|---|---|---|---|---|---|---|
| Q8IZL8 | Proline-, glutamic acid- and leucine-rich p | PELP1 | 119.7 | 4.9 | 5 | 5 | 149.78931 |
| Q8IZP0 | Abl interactor 1 OS=Homo sapiens OX=9 | ABI1 | 55.081 | 5.3 | 2 | 2 | 219.9928 |
| Q8IZT6 | Abnormal spindle-like microcephaly-asso | ASPM | 409.801 | 10.1 | 31 | 31 | 138.03088 |
| Q8N0U8 | Vitamin K epoxide reductase complex sul | VKORC1L1 | 19.836 | 5.7 | 1 | 1 | 56.782089 |
| Q8N0V3 | Putative ribosome-binding factor A, mitoc | RBFA | 38.36 | 9.3 | 3 | 3 | 149.51585 |
| Q8N0X7 | Spartin OS=Homo sapiens OX=9606 GN= | SPART | 72.834 | 6.3 | 3 | 3 | 47.967354 |
| Q8N0Y2 | Zinc finger protein 444 OS=Homo sapien: | ZNF444 | 35.204 | 5.8 | 2 | 2 | 30.503735 |
| Q8N0Z3 | Spindle and centriole-associated protein 1 | SPICE1 | 96.265 | 5.4 | 4 | 4 | 53.399601 |
| Q8N108 | Mesoderm induction early response protei | MIER1 | 57.984 | 7 | 3 | 3 | 66.894577 |
| Q8N122 | Regulatory-associated protein of mTOR C | RPTOR | 149.038 | 7.1 | 8 | 8 | 143.21016 |
| Q8N157 | Jouberin OS=Homo sapiens OX=9606 GN | AHI1 | 137.116 | 4.1 | 4 | 4 | 55.587471 |
| Q8N163 | Cell cycle and apoptosis regulator protein | CCAR2 | 102.902 | 21.1 | 17 | 17 | 294.40125 |
| Q8N1B3 | Cyclin-Q OS=Homo sapiens OX=9606 G | CCNQ | 28.369 | 10.9 | 2 | 2 | 78.584961 |
| Q8N1E6 | F-box/LRR-repeat protein 14 OS=Homo s | FBXL14 | 45.887 | 2.6 | 1 | 1 | 105.04183 |
| Q8N1F7 | Nuclear pore complex protein Nup93 OS= | NUP93 | 93.489 | 26.3 | 19 | 19 | 307.75677 |
| Q8N1G0 | Zinc finger protein 687 OS=Homo sapien: | ZNF687 | 129.529 | 2.1 | 2 | 2 | 71.911095 |
| Q8N1G1 | RNA exonuclease 1 homolog OS=Homo s | REXO1 | 131.51 | 2.9 | 3 | 3 | 99.177399 |
| Q8N1G2 | Cap-specific mRNA (nucleoside-2'-O-)-m | CMTR1 | 95.321 | 32.9 | 27 | 27 | 576.29224 |
| Q8N1W1 | Rho guanine nucleotide exchange factor 2 | ARHGEF28 | 191.891 | 2.5 | 4 | 4 | 29.227211 |
| Q8N2A8 | Mitochondrial cardiolipin hydrolase OS=I | PLD6 | 28.273 | 11.5 | 3 | 3 | 81.704605 |
| Q8N2K0 | Lysophosphatidylserine lipase ABHD12 ( | ABHD12 | 45.097 | 7.5 | 3 | 3 | 96.575478 |
| Q8N2M8 | CLK4-associating serine/arginine rich pro | CLASRP | 77.162 | 2.1 | 1 | 1 | 66.588058 |
| Q8N2S1 | Latent-transforming growth factor beta-bi | LTBP4 | 173.436 | 2 | 3 | 3 | 62.719971 |
| Q8N2Z9 | Centromere protein S OS=Homo sapiens ( | CENPS | 15.893 | 25.4 | 3 | 3 | 201.01169 |
| Q8N3C0 | Activating signal cointegrator 1 complex s | ASCC3 | 251.461 | 47 | 98 | 98 | 9000.4043 |
| Q8N3E9 | 1-phosphatidylinositol 4,5-bisphosphate p | PLCD3 | 89.259 | 6.8 | 5 | 5 | 92.842392 |
| Q8N3F8 | MICAL-like protein 1 OS=Homo sapiens | MICALL1 | 93.442 | 1.6 | 1 | 1 | 21.50309 |
| Q8N3R9 | Protein PALS1 OS=Homo sapiens OX=9( | PALS1 | 77.294 | 2.7 | 2 | 2 | 68.154213 |
| Q8N3U4 | Cohesin subunit SA-2 OS=Homo sapiens | STAG2 | 141.327 | 3.9 | 5 | 2 | 60.8251 |
| Q8N441 | Fibroblast growth factor receptor-like 1 O | FGFRL1 | 54.537 | 9.1 | 4 | 4 | 96.933083 |
| Q8N4C8 | Misshapen-like kinase 1 OS=Homo sapier | MINK1 | 149.822 | 4.4 | 4 | 4 | 47.032265 |
| Q8N4J0 | Carnosine N-methyltransferase OS=Homo | CARNMT1 | 47.186 | 5.1 | 2 | 2 | 72.010635 |
| Q8N4Q1 | Mitochondrial intermembrane space impo | CHCHD4 | 15.997 | 6.3 | 1 | 1 | 61.069 |
| Q8N511 | Transmembrane protein 199 OS=Homo sa | TMEM199 | 23.13 | 10.6 | 2 | 2 | 43.145668 |
| Q8N554 | Zinc finger protein 276 OS=Homo sapien: | ZNF276 | 67.219 | 2 | 1 | 1 | 25.965481 |

| | | | | | | | |
|---|---|---|---|---|---|---|---|
| Q8N556 | Actin filament-associated protein 1 OS=H | AFAP1 | 80.725 | 6.2 | 5 | 4 | 149.45728 |
| Q8N568 | Serine/threonine-protein kinase DCLK2 C | DCLK2 | 83.606 | 3 | 2 | 2 | 21.840601 |
| Q8N5A5 | Zinc finger CCCH-type with G patch dom | ZGPAT | 57.359 | 30.9 | 13 | 13 | 302.15427 |
| Q8N5F7 | NF-kappa-B-activating protein OS=Homo | NKAP | 47.139 | 15.7 | 5 | 4 | 212.19199 |
| Q8N5G0 | Small integral membrane protein 20 OS=H | SMIM20 | 7.703 | 20.9 | 1 | 1 | 100.07994 |
| Q8N5G2 | Macoilin OS=Homo sapiens OX=9606 GN | MACO1 | 76.178 | 19.4 | 10 | 10 | 243.66518 |
| Q8N5I2 | Arrestin domain-containing protein 1 OS= | ARRDC1 | 45.982 | 21.7 | 7 | 7 | 235.55405 |
| Q8N5K1 | CDGSH iron-sulfur domain-containing pr | CISD2 | 15.279 | 10.4 | 1 | 1 | 148.75725 |
| Q8N5P1 | Zinc finger CCCH domain-containing pro | ZC3H8 | 33.576 | 24.7 | 8 | 8 | 708.63525 |
| Q8N5U6 | E3 ubiquitin-protein ligase RNF10 OS=H | RNF10 | 89.927 | 12.9 | 8 | 8 | 295.32797 |
| Q8N5Y2 | Male-specific lethal 3 homolog OS=Homo | MSL3 | 59.824 | 7.9 | 4 | 4 | 87.038132 |
| Q8N680 | Zinc finger and BTB domain-containing p | ZBTB2 | 57.337 | 1.8 | 1 | 1 | 48.491669 |
| Q8N684 | Cleavage and polyadenylation specificity | CPSF7 | 52.051 | 26.8 | 12 | 12 | 4029.2654 |
| Q8N6G5 | Chondroitin sulfate N-acetylgalactosamin | CSGALNA( | 62.573 | 2 | 1 | 1 | 54.546432 |
| Q8N6M0 | Deubiquitinase OTUD6B OS=Homo sapi | OTUD6B | 33.813 | 4.4 | 1 | 1 | 70.007317 |
| Q8N6R0 | eEF1A lysine and N-terminal methyltrans | METTL13 | 78.768 | 29.2 | 16 | 16 | 448.17902 |
| Q8N6T3 | ADP-ribosylation factor GTPase-activatin | ARFGAP1 | 44.668 | 23.4 | 7 | 7 | 418.91266 |
| Q8N6T7 | NAD-dependent protein deacylase sirtuin- | SIRT6 | 39.119 | 29.3 | 9 | 9 | 183.65976 |
| Q8N720 | Zinc finger protein 655 OS=Homo sapiens | ZNF655 | 57.407 | 13.6 | 6 | 6 | 140.05418 |
| Q8N726 | Tumor suppressor ARF OS=Homo sapien | CDKN2A | 13.904 | 12.1 | 2 | 2 | 384.4231 |
| Q8N7H5 | RNA polymerase II-associated factor 1 ho | PAF1 | 59.976 | 9.4 | 4 | 4 | 64.439682 |
| Q8N884 | Cyclic GMP-AMP synthase OS=Homo sa | CGAS | 58.815 | 31.2 | 15 | 15 | 804.49792 |
| Q8N983 | Large ribosomal subunit protein mL43 OS | MRPL43 | 23.431 | 23.3 | 5 | 5 | 670.30396 |
| Q8N999 | RNA ligase 1 OS=Homo sapiens OX=960 | RLIG1 | 37.491 | 3.1 | 1 | 1 | 134.2408 |
| Q8N9N2 | Activating signal cointegrator 1 complex s | ASCC1 | 45.51 | 25.5 | 11 | 11 | 3996.0623 |
| Q8N9N5 | Protein BANP OS=Homo sapiens OX=96 | BANP | 56.495 | 5.2 | 2 | 2 | 48.977554 |
| Q8N9Q2 | Protein SREK1IP1 OS=Homo sapiens OX | SREK1IP1 | 18.177 | 16.1 | 3 | 3 | 97.632339 |
| Q8N9R8 | Protein SCAI OS=Homo sapiens OX=960 | SCAI | 70.4 | 8.4 | 5 | 5 | 84.832054 |
| Q8NAT1 | Protein O-linked-mannose beta-1,4-N-ace | POMGNT2 | 66.616 | 7.1 | 4 | 4 | 69.705292 |
| Q8NAV1 | Pre-mRNA-splicing factor 38A OS=Homo | PRPF38A | 37.477 | 22.4 | 7 | 7 | 180.76004 |
| Q8NB16 | Mixed lineage kinase domain-like protein | MLKL | 54.48 | 7.9 | 4 | 4 | 74.229416 |
| Q8NB78 | Lysine-specific histone demethylase 2 OS | KDM1B | 92.099 | 5.5 | 4 | 4 | 45.711533 |
| Q8NB90 | ATPase family gene 2 protein homolog A | AFG2A | 97.905 | 20.2 | 16 | 15 | 701.00555 |
| Q8NBF1 | Zinc finger protein GLIS1 OS=Homo sap | GLIS1 | 65.977 | 2.6 | 1 | 1 | 41.981857 |
| Q8NBJ4 | Golgi membrane protein 1 OS=Homo sap | GOLM1 | 45.334 | 5.7 | 2 | 2 | 1029.8292 |

| | | | | | | | |
|---|---|---|---|---|---|---|---|
| Q8NBJ5 | Procollagen galactosyltransferase 1 OS=H | COLGALT1 | 71.637 | 11.4 | 6 | 6 | 176.50554 |
| Q8NBM4 | Ubiquitin-associated domain-containing p | UBAC2 | 38.964 | 12.2 | 4 | 4 | 163.44849 |
| Q8NBN7 | Retinol dehydrogenase 13 OS=Homo sapi | RDH13 | 35.933 | 3.9 | 1 | 1 | 45.980827 |
| Q8NBQ5 | Estradiol 17-beta-dehydrogenase 11 OS=I | HSD17B11 | 32.964 | 16.7 | 5 | 5 | 137.03737 |
| Q8NBS9 | Thioredoxin domain-containing protein 5 | TXNDC5 | 47.629 | 4.6 | 2 | 2 | 39.728901 |
| Q8NBT0 | POC1 centriolar protein homolog A OS=I | POC1A | 45.009 | 16 | 6 | 6 | 116.19524 |
| Q8NBU5 | Outer mitochondrial transmembrane helix | ATAD1 | 40.745 | 14.7 | 5 | 5 | 64.66259 |
| Q8NBZ0 | INO80 complex subunit E OS=Homo sap | INO80E | 26.479 | 4.9 | 1 | 1 | 25.580057 |
| Q8NC51 | SERPINE1 mRNA-binding protein 1 OS= | SERBP1 | 44.966 | 44.6 | 24 | 24 | 9355.9961 |
| Q8NC56 | LEM domain-containing protein 2 OS=Ho | LEMD2 | 56.976 | 15.1 | 7 | 7 | 205.44998 |
| Q8NC60 | Nitric oxide-associated protein 1 OS=Hor | NOA1 | 78.458 | 26.4 | 15 | 15 | 723.4928 |
| Q8NCA5 | Protein FAM98A OS=Homo sapiens OX= | FAM98A | 55.273 | 29.3 | 16 | 13 | 31519.137 |
| Q8NCD3 | Holliday junction recognition protein OS= | HJURP | 83.54 | 4.7 | 3 | 3 | 210.06226 |
| Q8NCE0 | tRNA-splicing endonuclease subunit Sen2 | TSEN2 | 53.247 | 3.9 | 2 | 2 | 44.426682 |
| Q8NCE2 | Myotubularin-related protein 14 OS=Hom | MTMR14 | 72.204 | 1.2 | 1 | 1 | 764.10382 |
| Q8NCL4 | Polypeptide N-acetylgalactosaminyltransf | GALNT6 | 71.159 | 7.6 | 4 | 4 | 121.95436 |
| Q8NCN4 | E3 ubiquitin-protein ligase RNF169 OS=I | RNF169 | 77.195 | 11.4 | 6 | 6 | 134.50923 |
| Q8NCP5 | Zinc finger and BTB domain-containing p | ZBTB44 | 63.848 | 1.9 | 1 | 1 | 22.10239 |
| Q8ND04 | Nonsense-mediated mRNA decay factor S | SMG8 | 109.684 | 4.8 | 4 | 4 | 69.345825 |
| Q8ND56 | Protein LSM14 homolog A OS=Homo sa | LSM14A | 50.53 | 18.6 | 10 | 9 | 2330.8706 |
| Q8NDD1 | Uncharacterized protein C1orf131 OS=Ho | C1orf131 | 32.611 | 26.6 | 6 | 6 | 352.02863 |
| Q8NDF8 | Terminal nucleotidyltransferase 4B OS=H | TENT4B | 63.267 | 6.8 | 3 | 3 | 133.58365 |
| Q8NDT2 | Putative RNA-binding protein 15B OS=H | RBM15B | 97.206 | 10.1 | 9 | 9 | 104.50991 |
| Q8NDV7 | Trinucleotide repeat-containing gene 6A p | TNRC6A | 210.298 | 19.8 | 31 | 30 | 1169.8326 |
| Q8NDZ4 | Divergent protein kinase domain 2A OS= | DIPK2A | 49.482 | 23.7 | 9 | 9 | 209.60776 |
| Q8NE35 | Cytoplasmic polyadenylation element-bin | CPEB3 | 76.015 | 5.9 | 3 | 3 | 93.746475 |
| Q8NE71 | ATP-binding cassette sub-family F memb | ABCF1 | 95.926 | 35.9 | 28 | 28 | 3058.3203 |
| Q8NE86 | Calcium uniporter protein, mitochondrial | MCU | 39.867 | 6.6 | 2 | 2 | 110.17256 |
| Q8NEJ9 | Neuroguidin OS=Homo sapiens OX=9606 | NGDN | 35.895 | 24.1 | 6 | 6 | 170.81825 |
| Q8NEL9 | Phospholipase DDHD1 OS=Homo sapien | DDHD1 | 100.436 | 4.1 | 4 | 4 | 55.862759 |
| Q8NEM0 | Microcephalin OS=Homo sapiens OX=96 | MCPH1 | 92.85 | 9 | 6 | 6 | 44.68716 |
| Q8NEM7 | Transcription factor SPT20 homolog OS= | SUPT20H | 85.79 | 3.7 | 2 | 2 | 26.5975 |
| Q8NEW0 | Zinc transporter 7 OS=Homo sapiens OX= | SLC30A7 | 41.626 | 5.6 | 2 | 2 | 125.67345 |
| Q8NEY1 | Neuron navigator 1 OS=Homo sapiens OX | NAV1 | 202.472 | 23.1 | 36 | 36 | 395.07687 |
| Q8NEY8 | Periphilin-1 OS=Homo sapiens OX=9606 | PPHLN1 | 52.738 | 29.7 | 14 | 14 | 679.14581 |

| | | | | | | | |
|---|---|---|---|---|---|---|---|
| Q8NEZ5 | F-box only protein 22 OS=Homo sapiens | FBXO22 | 44.509 | 8.7 | 3 | 3 | 56.132542 |
| Q8NF37 | Lysophosphatidylcholine acyltransferase 1 | LPCAT1 | 59.152 | 9.2 | 5 | 5 | 96.713486 |
| Q8NF91 | Nesprin-1 OS=Homo sapiens OX=9606 C | SYNE1 | 1011.086 | 0.2 | 2 | 2 | 124.08189 |
| Q8NFF5 | FAD synthase OS=Homo sapiens OX=96 | FLAD1 | 65.266 | 6.8 | 4 | 4 | 62.043827 |
| Q8NFG4 | Folliculin OS=Homo sapiens OX=9606 G | FLCN | 64.474 | 2.1 | 1 | 1 | 21.469122 |
| Q8NFH3 | Nucleoporin Nup43 OS=Homo sapiens O | NUP43 | 42.151 | 4.7 | 2 | 2 | 81.043884 |
| Q8NFH4 | Nucleoporin Nup37 OS=Homo sapiens O | NUP37 | 36.708 | 3.7 | 1 | 1 | 181.43198 |
| Q8NFH5 | Nucleoporin NUP35 OS=Homo sapiens C | NUP35 | 34.774 | 12.3 | 3 | 3 | 52.01754 |
| Q8NFJ5 | Retinoic acid-induced protein 3 OS=Hom | GPRC5A | 40.252 | 6.2 | 2 | 2 | 305.9371 |
| Q8NFT6 | Protein DBF4 homolog B OS=Homo sapi | DBF4B | 67.243 | 6.7 | 3 | 3 | 43.627861 |
| Q8NFW8 | N-acylneuraminate cytidylyltransferase O | CMAS | 48.38 | 31.6 | 13 | 13 | 3022.021 |
| Q8NFZ5 | TNFAIP3-interacting protein 2 OS=Homo | TNIP2 | 48.7 | 9.6 | 4 | 4 | 76.616661 |
| Q8NG31 | Kinetochore scaffold 1 OS=Homo sapiens | KNL1 | 265.392 | 1.8 | 4 | 4 | 31.539694 |
| Q8NHM5 | Lysine-specific demethylase 2B OS=Hom | KDM2B | 152.615 | 4.1 | 5 | 5 | 106.98627 |
| Q8NHQ9 | ATP-dependent RNA helicase DDX55 OS | DDX55 | 68.547 | 26.2 | 17 | 17 | 851.31842 |
| Q8NHV4 | Protein NEDD1 OS=Homo sapiens OX=9 | NEDD1 | 71.967 | 21.5 | 12 | 12 | 582.12933 |
| Q8NHZ8 | Anaphase-promoting complex subunit CD | CDC26 | 9.777 | 12.9 | 1 | 1 | 343.44174 |
| Q8NI08 | Nuclear receptor coactivator 7 OS=Homo | NCOA7 | 106.162 | 6.8 | 5 | 5 | 117.85795 |
| Q8NI27 | THO complex subunit 2 OS=Homo sapien | THOC2 | 182.775 | 14.5 | 20 | 20 | 434.5636 |
| Q8NI36 | WD repeat-containing protein 36 OS=Hon | WDR36 | 105.323 | 23.6 | 20 | 20 | 327.68478 |
| Q8NI37 | Protein phosphatase PTC7 homolog OS=I | PPTC7 | 32.646 | 4.9 | 1 | 1 | 81.387169 |
| Q8NI60 | Atypical kinase COQ8A, mitochondrial O | COQ8A | 71.951 | 16.1 | 8 | 8 | 115.87402 |
| Q8NI77 | Kinesin-like protein KIF18A OS=Homo s | KIF18A | 102.281 | 16.9 | 14 | 13 | 294.43106 |
| Q8TA86 | Retinitis pigmentosa 9 protein OS=Homo | RP9 | 26.107 | 26.7 | 5 | 5 | 229.75523 |
| Q8TAA9 | Vang-like protein 1 OS=Homo sapiens O | VANGL1 | 59.975 | 4.8 | 2 | 2 | 46.99807 |
| Q8TAD8 | Smad nuclear-interacting protein 1 OS=H | SNIP1 | 45.778 | 22.2 | 7 | 7 | 482.64056 |
| Q8TAE8 | Large ribosomal subunit protein mL64 OS | GADD45GI | 25.384 | 19.8 | 4 | 4 | 352.63724 |
| Q8TAF3 | WD repeat-containing protein 48 OS=Hon | WDR48 | 76.211 | 18 | 10 | 10 | 130.43652 |
| Q8TAP8 | Protein phosphatase 1 regulatory subunit 3 | PPP1R35 | 27.954 | 17 | 4 | 4 | 123.30681 |
| Q8TAP9 | M-phase-specific PLK1-interacting protei | MPLKIP | 19.147 | 14 | 2 | 2 | 35.288559 |
| Q8TAQ2 | SWI/SNF complex subunit SMARCC2 O | SMARCC2 | 132.88 | 4.1 | 4 | 4 | 69.656708 |
| Q8TAT5 | Endonuclease 8-like 3 OS=Homo sapiens | NEIL3 | 67.77 | 1.7 | 1 | 1 | 83.548462 |
| Q8TAT6 | Nuclear protein localization protein 4 hon | NPLOC4 | 68.121 | 3.9 | 2 | 2 | 57.997742 |
| Q8TB45 | DEP domain-containing mTOR-interactin | DEPTOR | 46.294 | 7.3 | 1 | 1 | 238.45337 |
| Q8TB61 | Adenosine 3'-phospho 5'-phosphosulfate t | SLC35B2 | 47.515 | 8.1 | 3 | 3 | 157.84811 |

| | | | | | | | |
|---|---|---|---|---|---|---|---|
| Q8TB72 | Pumilio homolog 2 OS=Homo sapiens OX | PUM2 | 114.216 | 16.9 | 16 | 16 | 1157.9941 |
| Q8TBC3 | SH3KBP1-binding protein 1 OS=Homo sa | SHKBP1 | 76.344 | 2.3 | 1 | 1 | 31.00292 |
| Q8TBE0 | Bromo adjacent homology domain-contai | BAHD1 | 84.653 | 3.1 | 2 | 2 | 33.417961 |
| Q8TBM8 | DnaJ homolog subfamily B member 14 O | DNAJB14 | 42.516 | 15.6 | 5 | 5 | 124.9131 |
| Q8TBX8 | Phosphatidylinositol 5-phosphate 4-kinase | PIP4K2C | 47.3 | 12.8 | 5 | 5 | 80.491356 |
| Q8TC07 | TBC1 domain family member 15 OS=Hon | TBC1D15 | 79.491 | 9.1 | 5 | 5 | 57.693314 |
| Q8TC92 | Ecto-NOX disulfide-thiol exchanger 1 OS | ENOX1 | 73.349 | 20.4 | 14 | 11 | 1306.1829 |
| Q8TCJ2 | Dolichyl-diphosphooligosaccharide--prote | STT3B | 93.675 | 11.7 | 10 | 9 | 340.9086 |
| Q8TCS8 | Polyribonucleotide nucleotidyltransferase | PNPT1 | 85.951 | 25.2 | 18 | 18 | 621.61694 |
| Q8TCU4 | Centrosome-associated protein ALMS1 O | ALMS1 | 461.063 | 1.3 | 5 | 5 | 43.151569 |
| Q8TCY9 | Up-regulator of cell proliferation OS=Hon | URGCP | 104.988 | 12 | 11 | 11 | 165.67017 |
| Q8TD17 | Zinc finger protein 398 OS=Homo sapiens | ZNF398 | 71.312 | 1.2 | 1 | 1 | 29.199919 |
| Q8TD19 | Serine/threonine-protein kinase Nek9 OS= | NEK9 | 107.169 | 7.4 | 6 | 6 | 65.599022 |
| Q8TDB6 | E3 ubiquitin-protein ligase DTX3L OS=H | DTX3L | 83.555 | 27.2 | 18 | 18 | 932.03723 |
| Q8TDD1 | ATP-dependent RNA helicase DDX54 OS | DDX54 | 98.595 | 20.3 | 16 | 16 | 514.75732 |
| Q8TDM6 | Disks large homolog 5 OS=Homo sapiens | DLG5 | 213.868 | 13 | 23 | 23 | 154.67992 |
| Q8TDR0 | TRAF3-interacting protein 1 OS=Homo s | TRAF3IP1 | 78.632 | 1.4 | 1 | 1 | 62.550354 |
| Q8TDX7 | Serine/threonine-protein kinase Nek7 OS= | NEK7 | 34.551 | 26.8 | 8 | 7 | 138.91656 |
| Q8TDY2 | RB1-inducible coiled-coil protein 1 OS=H | RB1CC1 | 183.091 | 0.6 | 1 | 1 | 28.338139 |
| Q8TE02 | Elongator complex protein 5 OS=Homo sa | ELP5 | 34.842 | 4.4 | 1 | 1 | 72.818398 |
| Q8TED0 | U3 small nucleolar RNA-associated prote | UTP15 | 58.416 | 25.1 | 12 | 12 | 311.32431 |
| Q8TED1 | Probable glutathione peroxidase 8 OS=Ho | GPX8 | 23.882 | 24.4 | 5 | 5 | 525.76831 |
| Q8TEP8 | Centrosomal protein of 192 kDa OS=Hon | CEP192 | 279.112 | 1.1 | 2 | 2 | 43.962921 |
| Q8TEQ6 | Gem-associated protein 5 OS=Homo sapi | GEMIN5 | 168.59 | 33.9 | 46 | 46 | 2502.7004 |
| Q8TER5 | Rho guanine nucleotide exchange factor 4 | ARHGEF40 | 164.658 | 14.4 | 20 | 20 | 315.41241 |
| Q8TEU7 | Rap guanine nucleotide exchange factor 6 | RAPGEF6 | 179.423 | 5.5 | 7 | 7 | 262.52945 |
| Q8TEW0 | Partitioning defective 3 homolog OS=Hon | PARD3 | 151.424 | 27.4 | 34 | 33 | 548.83392 |
| Q8TEX9 | Importin-4 OS=Homo sapiens OX=9606 ( | IPO4 | 118.716 | 6.1 | 7 | 7 | 100.60805 |
| Q8TF01 | Arginine/serine-rich protein PNISR OS=H | PNISR | 92.577 | 2.6 | 2 | 2 | 61.073833 |
| Q8TF40 | Folliculin-interacting protein 1 OS=Homo | FNIP1 | 130.556 | 2 | 2 | 2 | 64.195869 |
| Q8TF42 | Ubiquitin-associated and SH3 domain-con | UBASH3B | 72.696 | 10.8 | 6 | 6 | 108.06075 |
| Q8TF46 | DIS3-like exonuclease 1 OS=Homo sapie | DIS3L | 120.788 | 21.4 | 20 | 20 | 376.03516 |
| Q8TF50 | Zinc finger protein 526 OS=Homo sapiens | ZNF526 | 73.622 | 2.5 | 2 | 2 | 75.083931 |
| Q8TF63 | Dendritic cell nuclear protein 1 OS=Homo | DCANP1 | 26.704 | 3.3 | 1 | 1 | 130.68881 |
| Q8TF72 | Protein Shroom3 OS=Homo sapiens OX= | SHROOM3 | 216.858 | 10.2 | 15 | 15 | 199.41933 |

| | | | | | | | |
|---|---|---|---|---|---|---|---|
| Q8TF76 | Serine/threonine-protein kinase haspin OS | HASPIN | 88.496 | 1.5 | 1 | 1 | 24.716591 |
| Q8WTP8 | Apoptosis-enhancing nuclease OS=Homo | AEN | 36.351 | 13.5 | 4 | 4 | 76.504311 |
| Q8WTT2 | Nucleolar complex protein 3 homolog OS | NOC3L | 92.548 | 13 | 9 | 9 | 141.27454 |
| Q8WU90 | Zinc finger CCCH domain-containing pro | ZC3H15 | 48.603 | 24.6 | 11 | 11 | 813.58777 |
| Q8WUA4 | General transcription factor 3C polypeptic | GTF3C2 | 100.68 | 20.7 | 15 | 15 | 690.11731 |
| Q8WUB8 | PHD finger protein 10 OS=Homo sapiens | PHF10 | 56.051 | 12 | 6 | 6 | 128.21252 |
| Q8WUF5 | RelA-associated inhibitor OS=Homo sapi | PPP1R13L | 89.092 | 2.8 | 2 | 2 | 47.009159 |
| Q8WUK0 | Phosphatidylglycerophosphatase and prot | PTPMT1 | 22.844 | 33.3 | 7 | 7 | 162.18044 |
| Q8WUM0 | Nuclear pore complex protein Nup133 OS | NUP133 | 128.98 | 8 | 9 | 9 | 148.93181 |
| Q8WUM4 | Programmed cell death 6-interacting prote | PDCD6IP | 96.024 | 21.2 | 17 | 17 | 336.95709 |
| Q8WUQ7 | Splicing factor Cactin OS=Homo sapiens | CACTIN | 88.702 | 10.9 | 7 | 7 | 226.61308 |
| Q8WUU5 | GATA zinc finger domain-containing pro | GATAD1 | 28.691 | 4.8 | 1 | 1 | 71.46418 |
| Q8WUW1 | Protein BRICK1 OS=Homo sapiens OX= | BRK1 | 8.745 | 54.7 | 4 | 4 | 206.89452 |
| Q8WV16 | DDB1- and CUL4-associated factor 4 OS | DCAF4 | 55.694 | 8.9 | 4 | 4 | 97.117981 |
| Q8WV22 | Non-structural maintenance of chromoson | NSMCE1 | 30.856 | 12.4 | 3 | 3 | 210.69994 |
| Q8WV41 | Sorting nexin-33 OS=Homo sapiens OX= | SNX33 | 65.265 | 12 | 6 | 6 | 200.33078 |
| Q8WV44 | E3 ubiquitin-protein ligase TRIM41 OS=I | TRIM41 | 71.67 | 21.3 | 13 | 13 | 545.39258 |
| Q8WV92 | MIT domain-containing protein 1 OS=Ho | MITD1 | 29.315 | 32.9 | 6 | 6 | 148.37311 |
| Q8WVB6 | Chromosome transmission fidelity protein | CHTF18 | 107.383 | 12.8 | 9 | 9 | 131.94116 |
| Q8WVC0 | RNA polymerase-associated protein LEO | LEO1 | 75.405 | 3.9 | 2 | 2 | 58.865372 |
| Q8WVD3 | E3 ubiquitin-protein ligase RNF138 OS=I | RNF138 | 28.193 | 23.3 | 6 | 6 | 357.18332 |
| Q8WVK2 | U4/U6.U5 small nuclear ribonucleoprotei | SNRNP27 | 18.861 | 29.7 | 4 | 4 | 87.912735 |
| Q8WVK7 | Spindle and kinetochore-associated protei | SKA2 | 14.189 | 9.1 | 1 | 1 | 107.32784 |
| Q8WVM0 | Dimethyladenosine transferase 1, mitocho | TFB1M | 39.543 | 44.2 | 13 | 13 | 1033.4548 |
| Q8WVS4 | Cytoplasmic dynein 2 intermediate chain | DYNC2I1 | 122.571 | 2.1 | 2 | 2 | 72.58699 |
| Q8WVV9 | Heterogeneous nuclear ribonucleoprotein | HNRNPLL | 60.084 | 30.1 | 14 | 14 | 1708.059 |
| Q8WVX9 | Fatty acyl-CoA reductase 1 OS=Homo saj | FAR1 | 59.357 | 17.7 | 8 | 8 | 462.9104 |
| Q8WW12 | PEST proteolytic signal-containing nuclea | PCNP | 18.925 | 27 | 4 | 4 | 64.464607 |
| Q8WWC4 | m-AAA protease-interacting protein 1, mi | MAIP1 | 32.545 | 18.2 | 5 | 5 | 263.74045 |
| Q8WWH5 | Pseudouridylate synthase TRUB1 OS=Ho | TRUB1 | 37.253 | 17.2 | 5 | 5 | 95.907211 |
| Q8WWI1 | LIM domain only protein 7 OS=Homo saj | LMO7 | 192.696 | 13.4 | 20 | 20 | 295.56311 |
| Q8WWK9 | Cytoskeleton-associated protein 2 OS=Ho | CKAP2 | 76.987 | 25.5 | 15 | 15 | 479.36902 |
| Q8WWM7 | Ataxin-2-like protein OS=Homo sapiens ( | ATXN2L | 113.374 | 29.2 | 37 | 37 | 43492.004 |
| Q8WWN8 | Arf-GAP with Rho-GAP domain, ANK re | ARAP3 | 169.845 | 1.9 | 3 | 3 | 56.155884 |
| Q8WWQ0 | PH-interacting protein OS=Homo sapiens | PHIP | 206.69 | 5.2 | 8 | 7 | 71.72876 |

| | | | | | | | |
|---|---|---|---|---|---|---|---|
| Q8WWY3 | U4/U6 small nuclear ribonucleoprotein Pr | PRPF31 | 55.457 | 22.6 | 10 | 10 | 603.71875 |
| Q8WX92 | Negative elongation factor B OS=Homo s | NELFB | 65.698 | 22.1 | 12 | 12 | 204.19934 |
| Q8WX93 | Palladin OS=Homo sapiens OX=9606 GN | PALLD | 150.564 | 14.8 | 18 | 17 | 206.85524 |
| Q8WXA9 | Splicing regulatory glutamine/lysine-rich | SREK1 | 59.381 | 11 | 4 | 4 | 88.713966 |
| Q8WXD5 | Gem-associated protein 6 OS=Homo sapi | GEMIN6 | 18.825 | 35.3 | 6 | 6 | 1258.5157 |
| Q8WXE9 | Stonin-2 OS=Homo sapiens OX=9606 GN | STON2 | 101.166 | 2.2 | 2 | 2 | 47.158844 |
| Q8WXF1 | Paraspeckle component 1 OS=Homo sapi | PSPC1 | 58.744 | 24.9 | 13 | 13 | 1650.2097 |
| Q8WXG6 | MAP kinase-activating death domain prot | MADD | 183.304 | 0.5 | 1 | 1 | 28.409859 |
| Q8WXI9 | Transcriptional repressor p66-beta OS=H | GATAD2B | 65.261 | 11.5 | 6 | 6 | 158.0103 |
| Q8WXX5 | DnaJ homolog subfamily C member 9 OS | DNAJC9 | 29.91 | 30 | 9 | 9 | 1659.6721 |
| Q8WY36 | HMG box transcription factor BBX OS=H | BBX | 105.131 | 18.9 | 15 | 15 | 811.55109 |
| Q8WY91 | Peroxynitrite isomerase THAP4 OS=Hom | THAP4 | 62.89 | 15.6 | 7 | 7 | 102.54092 |
| Q8WYH8 | Inhibitor of growth protein 5 OS=Homo s | ING5 | 27.751 | 15.4 | 4 | 3 | 105.56415 |
| Q8WYP5 | Protein ELYS OS=Homo sapiens OX=96 | AHCTF1 | 252.499 | 12.6 | 22 | 22 | 131.21187 |
| Q8WYQ5 | Microprocessor complex subunit DGCR8 | DGCR8 | 86.046 | 28.3 | 20 | 20 | 540.34998 |
| Q8WZA1 | Protein O-linked-mannose beta-1,2-N-ace | POMGNT1 | 75.252 | 3.9 | 2 | 2 | 46.176559 |
| Q92482 | Aquaporin-3 OS=Homo sapiens OX=960 | AQP3 | 31.544 | 2.7 | 1 | 1 | 22.467327 |
| Q92499 | ATP-dependent RNA helicase DDX1 OS= | DDX1 | 82.433 | 47.8 | 33 | 33 | 43259.91 |
| Q92504 | Zinc transporter SLC39A7 OS=Homo sap | SLC39A7 | 50.119 | 7.7 | 3 | 3 | 705.71021 |
| Q92520 | Protein FAM3C OS=Homo sapiens OX=9 | FAM3C | 24.681 | 19.4 | 4 | 4 | 100.73126 |
| Q92522 | Histone H1.10 OS=Homo sapiens OX=96 | H1-10 | 22.488 | 23 | 6 | 6 | 2263.749 |
| Q92530 | Proteasome inhibitor PI31 subunit OS=H | PSMF1 | 29.817 | 4.4 | 1 | 1 | 71.301414 |
| Q92538 | Golgi-specific brefeldin A-resistance guar | GBF1 | 206.446 | 0.8 | 1 | 1 | 357.61292 |
| Q92540 | Nonsense-mediated mRNA decay factor S | SMG7 | 127.282 | 18.2 | 19 | 19 | 418.61423 |
| Q92541 | RNA polymerase-associated protein RTF | RTF1 | 80.314 | 3.2 | 2 | 2 | 55.219776 |
| Q92547 | DNA topoisomerase 2-binding protein 1 ( | TOPBP1 | 170.679 | 26.3 | 33 | 33 | 389.2272 |
| Q92552 | Small ribosomal subunit protein mS27 OS | MRPS27 | 47.612 | 42 | 19 | 19 | 1595.4738 |
| Q92570 | Nuclear receptor subfamily 4 group A me | NR4A3 | 68.23 | 7.8 | 6 | 6 | 730.38287 |
| Q92598 | Heat shock protein 105 kDa OS=Homo sa | HSPH1 | 96.865 | 24.6 | 18 | 16 | 473.1427 |
| Q92600 | CCR4-NOT transcription complex subuni | CNOT9 | 33.632 | 38.5 | 10 | 10 | 278.28699 |
| Q92604 | Acyl-CoA:lysophosphatidylglycerol acylt | LPGAT1 | 43.09 | 5.7 | 2 | 2 | 70.099663 |
| Q92613 | Protein Jade-3 OS=Homo sapiens OX=96 | JADE3 | 93.809 | 12.8 | 11 | 10 | 278.73169 |
| Q92614 | Unconventional myosin-XVIIIa OS=Hom | MYO18A | 233.116 | 13.2 | 23 | 23 | 191.60265 |
| Q92615 | La-related protein 4B OS=Homo sapiens ( | LARP4B | 80.552 | 28.6 | 22 | 22 | 5397.0684 |
| Q92620 | Pre-mRNA-splicing factor ATP-dependen | DHX38 | 140.503 | 11.3 | 12 | 12 | 165.61238 |

| | | | | | | | |
|---|---|---|---|---|---|---|---|
| Q92621 | Nuclear pore complex protein Nup205 OS | NUP205 | 227.922 | 6.2 | 10 | 10 | 93.868095 |
| Q92626 | Peroxidasin homolog OS=Homo sapiens ( | PXDN | 165.275 | 4.9 | 7 | 7 | 187.77556 |
| Q92665 | Small ribosomal subunit protein mS31 OS | MRPS31 | 45.319 | 13.2 | 5 | 5 | 436.37125 |
| Q92667 | A-kinase anchor protein 1, mitochondrial | AKAP1 | 97.342 | 29.1 | 23 | 23 | 1343.4834 |
| Q92674 | Centromere protein I OS=Homo sapiens ( | CENPI | 86.721 | 1.3 | 1 | 1 | 24.315479 |
| Q92685 | Dol-P-Man:Man(5)GlcNAc(2)-PP-Dol al | ALG3 | 50.127 | 4.1 | 2 | 2 | 107.62534 |
| Q92688 | Acidic leucine-rich nuclear phosphoprotei | ANP32B | 28.788 | 13.5 | 3 | 2 | 286.80832 |
| Q92698 | DNA repair and recombination protein R/ | RAD54L | 84.352 | 1.2 | 1 | 1 | 47.287827 |
| Q92738 | USP6 N-terminal-like protein OS=Homo | USP6NL | 94.105 | 4.6 | 3 | 3 | 55.333744 |
| Q92754 | Transcription factor AP-2 gamma OS=Ho | TFAP2C | 49.177 | 15.6 | 5 | 5 | 1074.1003 |
| Q92759 | General transcription factor IIH subunit 4 | GTF2H4 | 52.187 | 19.9 | 7 | 7 | 147.3914 |
| Q92766 | Ras-responsive element-binding protein 1 | RREB1 | 181.421 | 1.8 | 3 | 3 | 111.73865 |
| Q92769 | Histone deacetylase 2 OS=Homo sapiens | HDAC2 | 55.365 | 12.5 | 6 | 6 | 320.93323 |
| Q92785 | Zinc finger protein ubi-d4 OS=Homo sapi | DPF2 | 44.156 | 12.8 | 4 | 4 | 97.794357 |
| Q92791 | Endoplasmic reticulum protein SC65 OS= | P3H4 | 50.382 | 2.5 | 1 | 1 | 32.235851 |
| Q92804 | TATA-binding protein-associated factor 2 | TAF15 | 61.83 | 18.2 | 12 | 12 | 7796.7402 |
| Q92830 | Histone acetyltransferase KAT2A OS=Hc | KAT2A | 93.926 | 3.1 | 3 | 1 | 103.43522 |
| Q92841 | Probable ATP-dependent RNA helicase D | DDX17 | 80.273 | 37.4 | 27 | 27 | 11374.445 |
| Q92845 | Kinesin-associated protein 3 OS=Homo sa | KIFAP3 | 91.206 | 4.2 | 3 | 3 | 49.515209 |
| Q92878 | DNA repair protein RAD50 OS=Homo sa | RAD50 | 153.893 | 8.5 | 11 | 11 | 152.89693 |
| Q92879 | CUGBP Elav-like family member 1 OS=H | CELF1 | 52.064 | 18.5 | 9 | 8 | 2695.0715 |
| Q92888 | Rho guanine nucleotide exchange factor 1 | ARHGEF1 | 102.436 | 5.7 | 4 | 4 | 98.374809 |
| Q92889 | DNA repair endonuclease XPF OS=Homo | ERCC4 | 104.486 | 7.1 | 6 | 6 | 150.9436 |
| Q92890 | Ubiquitin recognition factor in ER-associa | UFD1 | 34.501 | 3.9 | 1 | 1 | 57.341042 |
| Q92896 | Golgi apparatus protein 1 OS=Homo sapi | GLG1 | 134.552 | 12.7 | 13 | 13 | 172.89018 |
| Q92900 | Regulator of nonsense transcripts 1 OS=H | UPF1 | 124.346 | 42.5 | 45 | 45 | 18727.662 |
| Q92908 | Transcription factor GATA-6 OS=Homo | GATA6 | 60.033 | 4.2 | 2 | 2 | 444.18665 |
| Q92917 | G-patch domain and KOW motifs-contain | GPKOW | 52.229 | 5.3 | 2 | 2 | 38.185394 |
| Q92922 | SWI/SNF complex subunit SMARCC1 O | SMARCC1 | 122.868 | 13.8 | 13 | 9 | 326.63287 |
| Q92925 | SWI/SNF-related matrix-associated actin- | SMARCD2 | 58.921 | 2.4 | 1 | 1 | 41.477142 |
| Q92945 | Far upstream element-binding protein 2 O | KHSRP | 73.116 | 29.4 | 21 | 21 | 3935.6758 |
| Q92947 | Glutaryl-CoA dehydrogenase, mitochondr | GCDH | 48.128 | 4.3 | 2 | 2 | 152.49937 |
| Q92954 | Proteoglycan 4 OS=Homo sapiens OX=9( | PRG4 | 151.062 | 9.7 | 1 | 1 | 250.59146 |
| Q92966 | snRNA-activating protein complex subuni | SNAPC3 | 46.753 | 2.7 | 1 | 1 | 51.059517 |
| Q92973 | Transportin-1 OS=Homo sapiens OX=96( | TNPO1 | 102.356 | 11.6 | 9 | 8 | 180.30109 |

| | | | | | | | |
|---|---|---|---|---|---|---|---|
| Q92974 | Rho guanine nucleotide exchange factor 2 | ARHGEF2 | 111.543 | 25.8 | 24 | 24 | 1375.9375 |
| Q92979 | Ribosomal RNA small subunit methyltran | EMG1 | 26.721 | 43 | 9 | 9 | 3482.9968 |
| Q92989 | Polyribonucleotide 5'-hydroxyl-kinase Clp | CLP1 | 47.646 | 11.5 | 5 | 5 | 230.11165 |
| Q92993 | Histone acetyltransferase KAT5 OS=Hom | KAT5 | 58.583 | 4.9 | 2 | 2 | 72.146797 |
| Q92994 | Transcription factor IIIB 90 kDa subunit ( | BRF1 | 73.841 | 1.5 | 1 | 1 | 41.724312 |
| Q92997 | Segment polarity protein dishevelled hom | DVL3 | 78.055 | 6.6 | 4 | 4 | 86.130959 |
| Q93008 | Probable ubiquitin carboxyl-terminal hydr | USP9X | 290.463 | 6.5 | 16 | 7 | 103.56091 |
| Q93009 | Ubiquitin carboxyl-terminal hydrolase 7 C | USP7 | 128.303 | 26.2 | 27 | 27 | 334.56564 |
| Q93034 | Cullin-5 OS=Homo sapiens OX=9606 GN | CUL5 | 90.956 | 6.7 | 5 | 5 | 78.849548 |
| Q93050 | V-type proton ATPase 116 kDa subunit a | ATP6V0A1 | 96.413 | 2.2 | 2 | 1 | 23.79635 |
| Q93052 | Lipoma-preferred partner OS=Homo sapi | LPP | 65.747 | 5.7 | 3 | 3 | 187.89749 |
| Q93062 | RNA-binding protein with multiple splicir | RBPMS | 21.802 | 17.9 | 3 | 3 | 682.21216 |
| Q93063 | Exostosin-2 OS=Homo sapiens OX=9606 | EXT2 | 82.255 | 13.1 | 8 | 8 | 209.53966 |
| Q93074 | Mediator of RNA polymerase II transcript | MED12 | 243.082 | 3.7 | 7 | 6 | 83.554146 |
| Q93096 | Protein tyrosine phosphatase type IVA 1 ( | PTP4A1 | 19.816 | 22 | 4 | 2 | 116.38689 |
| Q969E8 | Pre-rRNA-processing protein TSR2 homo | TSR2 | 20.895 | 6.8 | 1 | 1 | 52.100124 |
| Q969F9 | BLOC-2 complex member HPS3 OS=Hor | HPS3 | 113.736 | 3.4 | 3 | 3 | 50.32478 |
| Q969G3 | SWI/SNF-related matrix-associated actin- | SMARCE1 | 46.65 | 3.6 | 1 | 1 | 49.219536 |
| Q969H6 | Ribonuclease P/MRP protein subunit POF | POP5 | 18.82 | 16.6 | 2 | 2 | 60.916801 |
| Q969J2 | Zinc finger protein with KRAB and SCAN | ZKSCAN4 | 61.58 | 11 | 4 | 4 | 64.165916 |
| Q969K3 | E3 ubiquitin-protein ligase RNF34 OS=H | RNF34 | 41.641 | 6.5 | 2 | 2 | 21.34243 |
| Q969L4 | U7 snRNA-associated Sm-like protein LS | LSM10 | 14.081 | 16.3 | 2 | 2 | 99.254761 |
| Q969M3 | Protein YIPF5 OS=Homo sapiens OX=96 | YIPF5 | 27.99 | 8.2 | 2 | 2 | 324.46454 |
| Q969N2 | GPI transamidase component PIG-T OS= | PIGT | 65.7 | 2.2 | 1 | 1 | 63.462635 |
| Q969P6 | DNA topoisomerase I, mitochondrial OS= | TOP1MT | 69.872 | 7.8 | 4 | 4 | 66.644882 |
| Q969Q6 | Serine/threonine-protein phosphatase 2A r | PPP2R3C | 53.316 | 2.4 | 1 | 1 | 51.957493 |
| Q969R5 | Lethal(3)malignant brain tumor-like prote | L3MBTL2 | 79.111 | 11.9 | 7 | 6 | 177.2991 |
| Q969S3 | Cytoplasmic 60S subunit biogenesis facto | ZNF622 | 54.273 | 27.9 | 12 | 12 | 887.64087 |
| Q969T9 | WW domain-binding protein 2 OS=Homo | WBP2 | 28.087 | 7.7 | 2 | 2 | 91.042404 |
| Q969V3 | BOS complex subunit NCLN OS=Homo s | NCLN | 62.975 | 6.4 | 3 | 3 | 64.201324 |
| Q969V4 | Tektin-1 OS=Homo sapiens OX=9606 GN | TEKT1 | 48.284 | 2.9 | 1 | 1 | 212.87248 |
| Q969V5 | Mitochondrial ubiquitin ligase activator o | MUL1 | 39.8 | 2.3 | 1 | 1 | 193.52121 |
| Q969V6 | Myocardin-related transcription factor A ( | MRTFA | 98.919 | 6.9 | 4 | 4 | 48.775131 |
| Q969X5 | Endoplasmic reticulum-Golgi intermediate | ERGIC1 | 32.593 | 14.8 | 4 | 4 | 91.889481 |
| Q969X6 | U3 small nucleolar RNA-associated prote | UTP4 | 76.891 | 19.7 | 10 | 10 | 617.05273 |

| | | | | | | | |
|---|---|---|---|---|---|---|---|
| Q969Z0 | FAST kinase domain-containing protein 4 | TBRG4 | 70.739 | 3.5 | 2 | 2 | 60.955299 |
| Q969Z3 | Mitochondrial amidoxime reducing comp | MTARC2 | 38.024 | 11.9 | 3 | 2 | 158.27295 |
| Q96A26 | Protein FAM162A OS=Homo sapiens OX | FAM162A | 17.343 | 7.1 | 1 | 1 | 51.171894 |
| Q96A33 | PAT complex subunit CCDC47 OS=Hom | CCDC47 | 55.874 | 18.2 | 10 | 10 | 530.78394 |
| Q96A65 | Exocyst complex component 4 OS=Homo | EXOC4 | 110.499 | 7.1 | 6 | 6 | 57.973305 |
| Q96AC1 | Fermitin family homolog 2 OS=Homo sap | FERMT2 | 77.861 | 3.7 | 2 | 2 | 54.535301 |
| Q96AE4 | Far upstream element-binding protein 1 O | FUBP1 | 67.561 | 19.7 | 13 | 9 | 1301.8813 |
| Q96AE7 | Tetratricopeptide repeat protein 17 OS=H | TTC17 | 129.559 | 7.3 | 7 | 7 | 105.2121 |
| Q96AG4 | Leucine-rich repeat-containing protein 59 | LRRC59 | 34.931 | 36.2 | 10 | 10 | 6111.8774 |
| Q96AH0 | SOSS complex subunit B2 OS=Homo sap | NABP1 | 22.424 | 8.8 | 2 | 1 | 179.59003 |
| Q96AP4 | Zinc finger-containing ubiquitin peptidase | ZUP1 | 65.959 | 9.5 | 5 | 5 | 140.00162 |
| Q96AQ6 | Pre-B-cell leukemia transcription factor-ir | PBXIP1 | 80.644 | 4.9 | 3 | 3 | 55.012371 |
| Q96AY2 | Crossover junction endonuclease EME1 C | EME1 | 63.252 | 6.3 | 3 | 3 | 126.44652 |
| Q96B01 | RAD51-associated protein 1 OS=Homo sa | RAD51AP1 | 38.458 | 15.3 | 4 | 4 | 91.589783 |
| Q96B26 | Exosome complex component RRP43 OS | EXOSC8 | 30.04 | 23.2 | 7 | 7 | 432.57568 |
| Q96B36 | Proline-rich AKT1 substrate 1 OS=Homo | AKT1S1 | 27.384 | 5.1 | 1 | 1 | 65.843582 |
| Q96BH1 | E3 ubiquitin-protein ligase RNF25 OS=H | RNF25 | 51.219 | 4.4 | 2 | 2 | 85.007996 |
| Q96BK5 | PIN2/TERF1-interacting telomerase inhib | PINX1 | 37.035 | 4.3 | 1 | 1 | 158.44949 |
| Q96BM9 | ADP-ribosylation factor-like protein 8A C | ARL8A | 21.416 | 4.8 | 1 | 1 | 38.629311 |
| Q96BN2 | Transcriptional adapter 1 OS=Homo sapie | TADA1 | 37.382 | 6 | 2 | 2 | 87.879639 |
| Q96BN8 | Ubiquitin thioesterase otulin OS=Homo sa | OTULIN | 40.263 | 6 | 2 | 2 | 58.733768 |
| Q96BP3 | Peptidylprolyl isomerase domain and WD | PPWD1 | 73.575 | 1.2 | 1 | 1 | 37.985569 |
| Q96BR5 | Cytochrome c oxidase assembly factor 7 C | COA7 | 25.71 | 5.6 | 1 | 1 | 131.49524 |
| Q96BY9 | Store-operated calcium entry-associated re | SARAF | 36.976 | 16.5 | 5 | 5 | 147.8718 |
| Q96C00 | Zinc finger and BTB domain-containing p | ZBTB9 | 50.603 | 2.3 | 1 | 1 | 33.971436 |
| Q96C36 | Pyrroline-5-carboxylate reductase 2 OS=H | PYCR2 | 33.638 | 2.8 | 1 | 1 | 72.957367 |
| Q96C92 | Endosome-associated-trafficking regulato | ENTR1 | 47.962 | 16.1 | 6 | 6 | 241.94386 |
| Q96CB9 | 5-methylcytosine rRNA methyltransferase | NSUN4 | 43.089 | 17.4 | 6 | 6 | 342.88318 |
| Q96CF2 | Charged multivesicular body protein 4c O | CHMP4C | 26.411 | 6.4 | 1 | 1 | 78.954765 |
| Q96CM3 | Pseudouridylate synthase RPUSD4, mitoc | RPUSD4 | 42.206 | 31 | 13 | 13 | 1223.0513 |
| Q96CP6 | Protein Aster-A OS=Homo sapiens OX=9 | GRAMD1A | 80.681 | 4.7 | 3 | 2 | 70.723076 |
| Q96CP7 | TLC domain-containing protein 1 OS=Ho | TLCD1 | 28.548 | 4 | 1 | 1 | 67.470345 |
| Q96CS3 | FAS-associated factor 2 OS=Homo sapier | FAF2 | 52.624 | 2.7 | 1 | 1 | 61.795914 |
| Q96CS7 | Pleckstrin homology domain-containing fa | PLEKHB2 | 24.736 | 5 | 1 | 1 | 53.623997 |
| Q96CT7 | Coiled-coil domain-containing protein 12 | CCDC124 | 25.836 | 23.8 | 5 | 5 | 232.14188 |

| | | | | | | | |
|---|---|---|---|---|---|---|---|
| Q96CU9 | FAD-dependent oxidoreductase domain-c | FOXRED1 | 53.812 | 17.5 | 8 | 8 | 239.08211 |
| Q96CW1 | AP-2 complex subunit mu OS=Homo sapi | AP2M1 | 49.655 | 25.3 | 12 | 12 | 1583.7415 |
| Q96CW5 | Gamma-tubulin complex component 3 OS | TUBGCP3 | 103.572 | 36.6 | 31 | 31 | 1784.4047 |
| Q96D15 | Reticulocalbin-3 OS=Homo sapiens OX= | RCN3 | 37.493 | 49.7 | 10 | 10 | 11194.377 |
| Q96D46 | 60S ribosomal export protein NMD3 OS= | NMD3 | 57.604 | 23.1 | 12 | 12 | 293.53571 |
| Q96D53 | Atypical kinase COQ8B, mitochondrial O | COQ8B | 60.07 | 1.8 | 1 | 1 | 39.762909 |
| Q96D71 | RalBP1-associated Eps domain-containing | REPS1 | 86.663 | 1.4 | 1 | 1 | 62.31461 |
| Q96DA6 | Mitochondrial import inner membrane tra | DNAJC19 | 12.499 | 34.5 | 4 | 4 | 110.02787 |
| Q96DE5 | Anaphase-promoting complex subunit 16 | ANAPC16 | 11.668 | 37.3 | 3 | 3 | 78.365852 |
| Q96DF8 | Splicing factor ESS-2 homolog OS=Homo | ESS2 | 52.569 | 23.5 | 8 | 8 | 128.79701 |
| Q96DH6 | RNA-binding protein Musashi homolog 2 | MSI2 | 35.197 | 25 | 7 | 5 | 2636.1438 |
| Q96DI7 | U5 small nuclear ribonucleoprotein 40 kD | SNRNP40 | 39.311 | 22.4 | 6 | 6 | 375.54486 |
| Q96DP5 | Methionyl-tRNA formyltransferase, mitoc | MTFMT | 43.833 | 23.4 | 8 | 8 | 214.20293 |
| Q96DT7 | Zinc finger and BTB domain-containing p | ZBTB10 | 94.894 | 10.2 | 8 | 8 | 160.38214 |
| Q96DU7 | Inositol-trisphosphate 3-kinase C OS=Hom | ITPKC | 75.208 | 17.7 | 11 | 10 | 177.99272 |
| Q96DV4 | Large ribosomal subunit protein mL38 OS | MRPL38 | 44.597 | 23.7 | 9 | 9 | 224.75105 |
| Q96E11 | Ribosome-recycling factor, mitochondrial | MRRF | 29.278 | 4.6 | 1 | 1 | 80.815918 |
| Q96E22 | Dehydrodolichyl diphosphate synthase co | NUS1 | 33.225 | 8.2 | 2 | 2 | 89.580528 |
| Q96E29 | Transcription termination factor 3, mitoch | MTERF3 | 47.972 | 33.1 | 15 | 15 | 1492.9701 |
| Q96EC8 | Protein YIPF6 OS=Homo sapiens OX=96 | YIPF6 | 26.256 | 8.5 | 2 | 2 | 81.546478 |
| Q96EE3 | Nucleoporin SEH1 OS=Homo sapiens OX | SEH1L | 39.649 | 33.3 | 9 | 9 | 287.06418 |
| Q96EH3 | Mitochondrial assembly of ribosomal larg | MALSU1 | 26.171 | 11.1 | 2 | 2 | 508.10199 |
| Q96EK4 | THAP domain-containing protein 11 OS= | THAP11 | 34.455 | 11.1 | 4 | 4 | 211.6992 |
| Q96EK5 | KIF-binding protein OS=Homo sapiens O | KIFBP | 71.814 | 3.1 | 2 | 2 | 68.143143 |
| Q96EL3 | Large ribosomal subunit protein mL53 OS | MRPL53 | 12.107 | 42.9 | 4 | 4 | 529.36053 |
| Q96EN8 | Molybdenum cofactor sulfurase OS=Hom | MOCOS | 98.12 | 6.2 | 5 | 5 | 125.45483 |
| Q96EP5 | DAZ-associated protein 1 OS=Homo sapi | DAZAP1 | 43.384 | 19.4 | 7 | 7 | 2417.645 |
| Q96ES7 | SAGA-associated factor 29 OS=Homo sap | SGF29 | 33.238 | 7.2 | 2 | 2 | 48.138733 |
| Q96EU6 | Ribosomal RNA processing protein 36 ho | RRP36 | 29.824 | 24.7 | 8 | 7 | 330.78455 |
| Q96EX3 | Cytoplasmic dynein 2 intermediate chain 2 | DYNC2I2 | 57.801 | 13.2 | 6 | 6 | 98.251656 |
| Q96EY1 | DnaJ homolog subfamily A member 3, mi | DNAJA3 | 52.489 | 26.9 | 13 | 13 | 2764.1067 |
| Q96EY4 | Translation machinery-associated protein | TMA16 | 23.865 | 12.8 | 5 | 5 | 569.60559 |
| Q96EY5 | Multivesicular body subunit 12A OS=Hom | MVB12A | 28.784 | 5.5 | 1 | 1 | 90.720062 |
| Q96EY7 | Small ribosomal subunit protein mS39 OS | PTCD3 | 78.55 | 19.4 | 12 | 12 | 482.54834 |
| Q96EZ8 | Microspherule protein 1 OS=Homo sapien | MCRS1 | 51.804 | 19 | 7 | 7 | 66.932938 |

| | | | | | | | |
|---|---|---|---|---|---|---|---|
| Q96F44 | E3 ubiquitin-protein ligase TRIM11 OS=l | TRIM11 | 52.775 | 30.8 | 13 | 12 | 505.05133 |
| Q96F86 | Enhancer of mRNA-decapping protein 3 ( | EDC3 | 56.078 | 45.3 | 19 | 19 | 1255.3489 |
| Q96FF9 | Sororin OS=Homo sapiens OX=9606 GN | CDCA5 | 27.601 | 11.5 | 2 | 2 | 83.323036 |
| Q96FS4 | Signal-induced proliferation-associated pr | SIPA1 | 112.15 | 0.7 | 1 | 1 | 139.94322 |
| Q96FV9 | THO complex subunit 1 OS=Homo sapier | THOC1 | 75.667 | 20.4 | 12 | 12 | 291.43195 |
| Q96FW1 | Ubiquitin thioesterase OTUB1 OS=Homo | OTUB1 | 31.285 | 19.2 | 5 | 5 | 141.32127 |
| Q96FX7 | tRNA (adenine(58)-N(1))-methyltransfera | TRMT61A | 31.382 | 34.3 | 7 | 7 | 337.48785 |
| Q96FZ2 | Abasic site processing protein HMCES O | HMCES | 40.575 | 11 | 3 | 3 | 45.014355 |
| Q96G21 | U3 small nucleolar ribonucleoprotein prot | IMP4 | 33.757 | 19.2 | 6 | 6 | 257.52237 |
| Q96G25 | Mediator of RNA polymerase II transcript | MED8 | 29.08 | 8.6 | 2 | 2 | 44.354614 |
| Q96G46 | tRNA-dihydrouridine(47) synthase [NAD | DUS3L | 72.594 | 18.5 | 10 | 10 | 246.8757 |
| Q96GD4 | Aurora kinase B OS=Homo sapiens OX=9 | AURKB | 39.311 | 16.3 | 5 | 5 | 222.16835 |
| Q96GE4 | Centrosomal protein of 95 kDa OS=Homo | CEP95 | 95.297 | 2.6 | 2 | 2 | 33.989075 |
| Q96GJ1 | tRNA (uracil-5-)-methyltransferase homol | TRMT2B | 56.477 | 9.1 | 4 | 4 | 40.392994 |
| Q96GM5 | SWI/SNF-related matrix-associated actin- | SMARCD1 | 58.233 | 2.5 | 1 | 1 | 43.576244 |
| Q96GM8 | Target of EGR1 protein 1 OS=Homo sapi | TOE1 | 56.548 | 15.9 | 8 | 8 | 982.84094 |
| Q96GQ7 | Probable ATP-dependent RNA helicase D | DDX27 | 89.836 | 21.9 | 16 | 16 | 330.00851 |
| Q96GW9 | Methionine--tRNA ligase, mitochondrial ( | MARS2 | 66.591 | 2.7 | 1 | 1 | 58.944374 |
| Q96GX5 | Serine/threonine-protein kinase greatwall | MASTL | 97.32 | 10.2 | 7 | 7 | 104.20756 |
| Q96H55 | Unconventional myosin-XIX OS=Homo s | MYO19 | 109.136 | 2.3 | 2 | 2 | 49.69413 |
| Q96HA1 | Nuclear envelope pore membrane protein | POM121 | 127.721 | 3.6 | 4 | 1 | 90.215752 |
| Q96HC4 | PDZ and LIM domain protein 5 OS=Hom | PDLIM5 | 63.946 | 17.6 | 9 | 9 | 96.119293 |
| Q96HJ3 | Coiled-coil domain-containing protein 34 | CCDC34 | 43.226 | 10.5 | 4 | 4 | 230.30931 |
| Q96HR3 | Mediator of RNA polymerase II transcript | MED30 | 20.278 | 6.7 | 1 | 1 | 107.80075 |
| Q96HR8 | H/ACA ribonucleoprotein complex non-co | NAF1 | 53.717 | 6.1 | 3 | 3 | 318.1088 |
| Q96HS1 | Serine/threonine-protein phosphatase PGA | PGAM5 | 32.005 | 48.4 | 15 | 15 | 4430.5264 |
| Q96HW7 | Integrator complex subunit 4 OS=Homo s | INTS4 | 108.171 | 1.3 | 1 | 1 | 45.660522 |
| Q96HY7 | 2-oxoadipate dehydrogenase complex con | DHTKD1 | 103.077 | 6.2 | 4 | 4 | 183.88483 |
| Q96I24 | Far upstream element-binding protein 3 O | FUBP3 | 61.641 | 44.6 | 23 | 23 | 5280.0625 |
| Q96I25 | Splicing factor 45 OS=Homo sapiens OX= | RBM17 | 44.962 | 40.1 | 15 | 15 | 2291.7473 |
| Q96I51 | RCC1-like G exchanging factor-like prote | RCC1L | 49.898 | 27.8 | 13 | 13 | 996.90912 |
| Q96I59 | Probable asparagine--tRNA ligase, mitoch | NARS2 | 54.091 | 4.4 | 2 | 2 | 34.701431 |
| Q96IF1 | LIM domain-containing protein ajuba OS= | AJUBA | 56.934 | 17.8 | 8 | 8 | 159.52155 |
| Q96II8 | DISP complex protein LRCH3 OS=Homo | LRCH3 | 86.084 | 8.2 | 5 | 4 | 96.151436 |
| Q96IJ6 | Mannose-1-phosphate guanyltransferase a | GMPPA | 46.292 | 5 | 2 | 2 | 41.88176 |

| | | | | | | | |
|---|---|---|---|---|---|---|---|
| Q96IK5 | Germ cell-less protein-like 1 OS=Homo s | GMCL1 | 58.686 | 10.9 | 5 | 1 | 151.43523 |
| Q96IP4 | Terminal nucleotidyltransferase 5A OS=H | TENT5A | 49.667 | 21.3 | 8 | 8 | 236.09579 |
| Q96IQ9 | Zinc finger protein 414 OS=Homo sapien | ZNF414 | 32.783 | 3.8 | 1 | 1 | 61.799587 |
| Q96IU2 | Zinc finger BED domain-containing prote | ZBED3 | 25.133 | 4.7 | 1 | 1 | 114.09644 |
| Q96IU4 | Putative protein-lysine deacylase ABHD1 | ABHD14B | 22.346 | 6.7 | 1 | 1 | 72.839073 |
| Q96IX5 | ATP synthase membrane subunit K, mitoc | ATP5MK | 6.458 | 44.8 | 3 | 3 | 311.23715 |
| Q96IZ0 | PRKC apoptosis WT1 regulator protein C | PAWR | 36.568 | 11.5 | 3 | 3 | 118.32991 |
| Q96IZ5 | RNA-binding protein 41 OS=Homo sapie | RBM41 | 47.1 | 27.6 | 12 | 12 | 238.88252 |
| Q96IZ7 | Serine/Arginine-related protein 53 OS=H | RSRC1 | 38.678 | 6.6 | 2 | 2 | 64.726044 |
| Q96J01 | THO complex subunit 3 OS=Homo sapie | THOC3 | 38.772 | 35.9 | 9 | 9 | 253.7901 |
| Q96J02 | E3 ubiquitin-protein ligase Itchy homolog | ITCH | 102.803 | 3.4 | 3 | 3 | 35.190327 |
| Q96J84 | Kin of IRRE-like protein 1 OS=Homo sap | KIRREL1 | 83.536 | 7.7 | 5 | 5 | 64.006111 |
| Q96J92 | Serine/threonine-protein kinase WNK4 O | WNK4 | 134.74 | 0.8 | 1 | 1 | 41.261593 |
| Q96JC1 | Vam6/Vps39-like protein OS=Homo sapi | VPS39 | 101.81 | 22.9 | 18 | 18 | 321.93097 |
| Q96JM3 | Chromosome alignment-maintaining phos | CHAMP1 | 89.1 | 20.6 | 16 | 16 | 248.1243 |
| Q96JP5 | E3 ubiquitin-protein ligase ZFP91 OS=H | ZFP91 | 63.446 | 18.6 | 11 | 11 | 157.52823 |
| Q96JY6 | PDZ and LIM domain protein 2 OS=Hom | PDLIM2 | 37.459 | 3.1 | 1 | 1 | 54.576519 |
| Q96K58 | Zinc finger protein 668 OS=Homo sapien | ZNF668 | 67.905 | 13.6 | 8 | 8 | 183.48048 |
| Q96KA5 | Lipid scramblase CLPTM1L OS=Homo s | CLPTM1L | 62.23 | 6.1 | 2 | 2 | 45.420685 |
| Q96KB5 | Lymphokine-activated killer T-cell-origin | PBK | 36.086 | 8.4 | 2 | 2 | 55.899147 |
| Q96KC8 | DnaJ homolog subfamily C member 1 OS | DNAJC1 | 63.883 | 4.3 | 2 | 2 | 81.779297 |
| Q96KG9 | N-terminal kinase-like protein OS=Homo | SCYL1 | 89.632 | 18.4 | 13 | 13 | 534.50867 |
| Q96KM6 | Zinc finger protein 512B OS=Homo sapie | ZNF512B | 97.265 | 8.3 | 6 | 6 | 232.1507 |
| Q96KP1 | Exocyst complex component 2 OS=Homo | EXOC2 | 104.067 | 3.5 | 3 | 3 | 48.736179 |
| Q96KQ4 | Apoptosis-stimulating of p53 protein 1 OS | PPP1R13B | 119.566 | 0.8 | 1 | 1 | 124.93748 |
| Q96KQ7 | Histone-lysine N-methyltransferase EHM | EHMT2 | 132.371 | 7.6 | 8 | 8 | 107.69919 |
| Q96L34 | MAP/microtubule affinity-regulating kina | MARK4 | 82.52 | 3.5 | 2 | 2 | 52.106388 |
| Q96L50 | Leucine-rich repeat protein 1 OS=Homo s | LRR1 | 46.723 | 3.6 | 1 | 1 | 31.864265 |
| Q96L58 | Beta-1,3-galactosyltransferase 6 OS=Hom | B3GALT6 | 37.138 | 16.1 | 5 | 5 | 227.39133 |
| Q96L73 | Histone-lysine N-methyltransferase, H3 ly | NSD1 | 296.652 | 3.2 | 7 | 7 | 69.972664 |
| Q96L92 | Sorting nexin-27 OS=Homo sapiens OX= | SNX27 | 61.266 | 14 | 7 | 7 | 209.11865 |
| Q96LD4 | E3 ubiquitin-protein ligase TRIM47 OS=I | TRIM47 | 69.533 | 11.9 | 6 | 6 | 131.37238 |
| Q96LI5 | CCR4-NOT transcription complex subuni | CNOT6L | 63.002 | 8.1 | 4 | 1 | 118.971 |
| Q96LL9 | DnaJ homolog subfamily C member 30, m | DNAJC30 | 25.962 | 4 | 1 | 1 | 149.57059 |
| Q96LT9 | RNA-binding region-containing protein 3 | RNPC3 | 58.576 | 11.4 | 6 | 6 | 133.49878 |

| | | | | | | | |
|---|---|---|---|---|---|---|---|
| Q96M94 | Kelch-like protein 15 OS=Homo sapiens ( | KLHL15 | 69.775 | 5.6 | 3 | 3 | 44.407539 |
| Q96MF7 | E3 SUMO-protein ligase NSE2 OS=Hom | NSMCE2 | 27.933 | 25.1 | 6 | 6 | 157.80849 |
| Q96MG7 | Non-structural maintenance of chromoson | NSMCE3 | 34.308 | 11.8 | 5 | 5 | 164.74651 |
| Q96MH2 | Protein HEXIM2 OS=Homo sapiens OX= | HEXIM2 | 32.419 | 23.1 | 6 | 6 | 273.55383 |
| Q96MT8 | Centrosomal protein of 63 kDa OS=Homo | CEP63 | 81.344 | 17.5 | 12 | 12 | 284.24573 |
| Q96MU7 | YTH domain-containing protein 1 OS=Ho | YTHDC1 | 84.701 | 14.2 | 11 | 11 | 473.15228 |
| Q96N66 | Lysophospholipid acyltransferase 7 OS=H | MBOAT7 | 52.765 | 5.1 | 2 | 2 | 81.614975 |
| Q96N67 | Dedicator of cytokinesis protein 7 OS=Ho | DOCK7 | 242.561 | 12.9 | 24 | 22 | 193.47775 |
| Q96NB3 | Zinc finger protein 830 OS=Homo sapiens | ZNF830 | 41.999 | 11.6 | 3 | 3 | 29.30097 |
| Q96NC0 | Zinc finger matrin-type protein 2 OS=Hor | ZMAT2 | 23.612 | 13.6 | 3 | 3 | 77.422989 |
| Q96NE9 | FERM domain-containing protein 6 OS=H | FRMD6 | 72.044 | 14.6 | 7 | 7 | 126.64713 |
| Q96NW4 | Ankyrin repeat domain-containing protein | ANKRD27 | 116.985 | 2.5 | 2 | 2 | 47.49202 |
| Q96NY9 | Crossover junction endonuclease MUS81 | MUS81 | 61.173 | 16.3 | 8 | 8 | 265.84836 |
| Q96P11 | 28S rRNA (cytosine-C(5))-methyltransfer | NSUN5 | 46.692 | 45.2 | 18 | 12 | 1500.4348 |
| Q96P47 | Arf-GAP with GTPase, ANK repeat and P | AGAP3 | 95.045 | 8 | 6 | 5 | 860.14233 |
| Q96P70 | Importin-9 OS=Homo sapiens OX=9606 ( | IPO9 | 115.963 | 1.1 | 1 | 1 | 56.79343 |
| Q96PE3 | Inositol polyphosphate-4-phosphatase typ | INPP4A | 109.956 | 5.6 | 5 | 5 | 235.84099 |
| Q96PK6 | RNA-binding protein 14 OS=Homo sapie | RBM14 | 69.492 | 34.5 | 28 | 28 | 12897.497 |
| Q96PM9 | Zinc finger protein 385A OS=Homo sapie | ZNF385A | 40.454 | 38.1 | 11 | 11 | 171.07274 |
| Q96PU4 | E3 ubiquitin-protein ligase UHRF2 OS=H | UHRF2 | 89.986 | 11.2 | 7 | 6 | 725.12274 |
| Q96PU5 | E3 ubiquitin-protein ligase NEDD4-like C | NEDD4L | 111.932 | 1 | 1 | 1 | 31.571447 |
| Q96PU8 | KH domain-containing RNA-binding prot | QKI | 37.671 | 27.6 | 8 | 8 | 495.56412 |
| Q96PV6 | Leukocyte receptor cluster member 8 OS= | LENG8 | 88.158 | 20.2 | 16 | 16 | 657.29199 |
| Q96PV7 | Protein FAM193B OS=Homo sapiens OX | FAM193B | 96.543 | 11.2 | 9 | 9 | 122.25578 |
| Q96PY5 | Formin-like protein 2 OS=Homo sapiens ( | FMNL2 | 123.321 | 15 | 15 | 9 | 108.21319 |
| Q96PZ0 | Pseudouridylate synthase 7 homolog OS= | PUS7 | 75.036 | 31.5 | 17 | 17 | 322.1377 |
| Q96Q15 | Serine/threonine-protein kinase SMG1 OS | SMG1 | 410.501 | 1.8 | 5 | 5 | 105.48431 |
| Q96Q45 | Transmembrane protein 237 OS=Homo sa | TMEM237 | 45.526 | 4.2 | 1 | 1 | 85.213783 |
| Q96Q89 | Kinesin-like protein KIF20B OS=Homo s | KIF20B | 210.63 | 2.1 | 3 | 3 | 38.195038 |
| Q96QC0 | Serine/threonine-protein phosphatase 1 re | PPP1R10 | 99.058 | 23.6 | 21 | 21 | 1515.7949 |
| Q96QD9 | UAP56-interacting factor OS=Homo sapie | FYTTD1 | 35.819 | 9.7 | 3 | 3 | 44.784874 |
| Q96QE3 | ATPase family AAA domain-containing p | ATAD5 | 207.571 | 4.2 | 6 | 6 | 79.255295 |
| Q96QE5 | Transcription elongation factor, mitochon | TEFM | 41.677 | 28.3 | 11 | 11 | 520.33368 |
| Q96QK1 | Vacuolar protein sorting-associated protei | VPS35 | 91.708 | 4.9 | 4 | 4 | 139.32619 |
| Q96QR8 | Transcriptional activator protein Pur-beta | PURB | 33.241 | 26.9 | 10 | 8 | 1036.847 |

| | | | | | | | |
|---|---|---|---|---|---|---|---|
| Q96R06 | Sperm-associated antigen 5 OS=Homo sa| | SPAG5 | 134.422 | 41.3 | 45 | 45 | 4769.9043 |
| Q96RD7 | Pannexin-1 OS=Homo sapiens OX=9606 | PANX1 | 48.051 | 4.9 | 2 | 2 | 96.265007 |
| Q96RE7 | Nucleus accumbens-associated protein 1 ( | NACC1 | 57.259 | 20.5 | 9 | 8 | 181.70708 |
| Q96RF0 | Sorting nexin-18 OS=Homo sapiens OX= | SNX18 | 68.895 | 1.9 | 1 | 1 | 141.23613 |
| Q96RG2 | PAS domain-containing serine/threonine-| | PASK | 142.93 | 5.1 | 6 | 6 | 82.123161 |
| Q96RL1 | BRCA1-A complex subunit RAP80 OS=H | UIMC1 | 79.728 | 16.1 | 13 | 13 | 663.94897 |
| Q96RL7 | Intermembrane lipid transfer protein VPS | VPS13A | 360.277 | 0.6 | 2 | 2 | 738.45123 |
| Q96RP9 | Elongation factor G, mitochondrial OS=H | GFM1 | 83.472 | 18.4 | 12 | 12 | 152.6555 |
| Q96RR1 | Twinkle mtDNA helicase OS=Homo sapi | TWNK | 77.155 | 22.8 | 13 | 13 | 638.75696 |
| Q96RS0 | Trimethylguanosine synthase OS=Homo s | TGS1 | 96.62 | 6 | 5 | 5 | 98.017586 |
| Q96RS6 | NudC domain-containing protein 1 OS=H | NUDCD1 | 66.756 | 1.9 | 1 | 1 | 59.403831 |
| Q96RT1 | Erbin OS=Homo sapiens OX=9606 GN=I | ERBIN | 158.299 | 4.3 | 6 | 6 | 58.066685 |
| Q96RT7 | Gamma-tubulin complex component 6 OS | TUBGCP6 | 200.499 | 17.5 | 23 | 23 | 877.61047 |
| Q96RT8 | Gamma-tubulin complex component 5 OS | TUBGCP5 | 118.321 | 13.4 | 13 | 13 | 269.44196 |
| Q96S15 | GATOR2 complex protein WDR24 OS=H | WDR24 | 88.207 | 3.3 | 2 | 2 | 49.443657 |
| Q96SB4 | SRSF protein kinase 1 OS=Homo sapiens | SRPK1 | 74.325 | 21.1 | 12 | 12 | 3277.7317 |
| Q96SB8 | Structural maintenance of chromosomes p | SMC6 | 126.326 | 8.4 | 8 | 8 | 172.62582 |
| Q96SI9 | Spermatid perinuclear RNA-binding prote | STRBP | 73.653 | 24.7 | 16 | 16 | 649.95178 |
| Q96SN8 | CDK5 regulatory subunit-associated prote | CDK5RAP2 | 215.038 | 16.3 | 28 | 28 | 389.89478 |
| Q96ST2 | Protein IWS1 homolog OS=Homo sapiens | IWS1 | 91.955 | 5.1 | 4 | 4 | 73.289253 |
| Q96ST8 | Centrosomal protein of 89 kDa OS=Homo | CEP89 | 89.59 | 1.4 | 1 | 1 | 40.617645 |
| Q96SZ5 | 2-aminoethanethiol dioxygenase OS=Hon | ADO | 29.751 | 13 | 4 | 4 | 88.728874 |
| Q96SZ6 | Mitochondrial tRNA methylthiotransferas | CDK5RAP1 | 67.689 | 9.2 | 5 | 5 | 194.14058 |
| Q96T21 | Selenocysteine insertion sequence-binding | SECISBP2 | 95.462 | 32 | 23 | 23 | 512.42896 |
| Q96T37 | RNA-binding protein 15 OS=Homo sapie | RBM15 | 107.189 | 23.7 | 19 | 19 | 288.76596 |
| Q96T58 | Msx2-interacting protein OS=Homo sapie | SPEN | 402.249 | 4.2 | 13 | 13 | 123.26841 |
| Q96T60 | Bifunctional polynucleotide phosphatase/l | PNKP | 57.077 | 17.5 | 8 | 8 | 193.03737 |
| Q96T88 | E3 ubiquitin-protein ligase UHRF1 OS=H | UHRF1 | 89.814 | 38.5 | 26 | 26 | 1479.4244 |
| Q96TA2 | ATP-dependent zinc metalloprotease YM | YME1L1 | 86.456 | 22.4 | 14 | 14 | 724.99646 |
| Q96TC7 | Regulator of microtubule dynamics protei | RMDN3 | 52.119 | 10.2 | 4 | 4 | 257.93954 |
| Q99081 | Transcription factor 12 OS=Homo sapiens | TCF12 | 72.965 | 3.7 | 2 | 1 | 49.991127 |
| Q99417 | c-Myc-binding protein OS=Homo sapiens | MYCBP | 11.967 | 48.5 | 5 | 5 | 224.38641 |
| Q99426 | Tubulin-folding cofactor B OS=Homo sa| | TBCB | 27.326 | 4.5 | 1 | 1 | 33.952263 |
| Q99439 | Calponin-2 OS=Homo sapiens OX=9606 | CNN2 | 33.698 | 26.2 | 7 | 6 | 7151.3149 |
| Q99459 | Cell division cycle 5-like protein OS=Hor | CDC5L | 92.251 | 33.2 | 23 | 23 | 1769.3872 |

| | | | | | | |
|---|---|---|---|---|---|---|
| Q99471 | Prefoldin subunit 5 OS=Homo sapiens OX | PFDN5 | 17.329 | 20.8 | 3 | 3 | 27.473618 |
| Q99496 | E3 ubiquitin-protein ligase RING2 OS=H | RNF2 | 37.656 | 27.1 | 7 | 6 | 219.39944 |
| Q99497 | Parkinson disease protein 7 OS=Homo sa | PARK7 | 19.892 | 15.3 | 2 | 2 | 69.988411 |
| Q99501 | GAS2-like protein 1 OS=Homo sapiens C | GAS2L1 | 72.717 | 14 | 9 | 9 | 155.70195 |
| Q99543 | DnaJ homolog subfamily C member 2 OS | DNAJC2 | 71.997 | 20.6 | 11 | 11 | 288.76337 |
| Q99547 | M-phase phosphoprotein 6 OS=Homo sap | MPHOSPH | 19.024 | 11.9 | 2 | 2 | 65.58654 |
| Q99549 | M-phase phosphoprotein 8 OS=Homo sap | MPHOSPH | 97.183 | 5.8 | 4 | 4 | 118.11886 |
| Q99551 | Transcription termination factor 1, mitoch | MTERF1 | 45.779 | 23.8 | 7 | 7 | 130.53302 |
| Q99567 | Nuclear pore complex protein Nup88 OS= | NUP88 | 83.542 | 5.8 | 4 | 4 | 92.057472 |
| Q99569 | Plakophilin-4 OS=Homo sapiens OX=960 | PKP4 | 131.869 | 20.9 | 19 | 18 | 340.93567 |
| Q99570 | Phosphoinositide 3-kinase regulatory subu | PIK3R4 | 153.104 | 8.3 | 10 | 10 | 99.117432 |
| Q99571 | P2X purinoceptor 4 OS=Homo sapiens O | P2RX4 | 43.37 | 8.2 | 3 | 3 | 120.56267 |
| Q99575 | Ribonucleases P/MRP protein subunit PO | POP1 | 114.709 | 25.5 | 23 | 23 | 548.25409 |
| Q99590 | Protein SCAF11 OS=Homo sapiens OX= | SCAF11 | 164.653 | 13.7 | 19 | 19 | 539.72278 |
| Q99598 | Translin-associated protein X OS=Homo | TSNAX | 33.113 | 14.5 | 4 | 4 | 99.669884 |
| Q99615 | DnaJ homolog subfamily C member 7 OS | DNAJC7 | 56.441 | 26.1 | 12 | 12 | 284.79095 |
| Q99622 | Protein C10 OS=Homo sapiens OX=9606 | C12orf57 | 13.179 | 11.9 | 1 | 1 | 116.80617 |
| Q99623 | Prohibitin-2 OS=Homo sapiens OX=9606 | PHB2 | 33.297 | 34.1 | 9 | 9 | 194.43251 |
| Q99653 | Calcineurin B homologous protein 1 OS= | CHP1 | 22.457 | 14.9 | 3 | 3 | 119.681 |
| Q99661 | Kinesin-like protein KIF2C OS=Homo sa | KIF2C | 81.313 | 29.1 | 20 | 19 | 1044.1201 |
| Q99700 | Ataxin-2 OS=Homo sapiens OX=9606 G | ATXN2 | 140.283 | 23.5 | 29 | 28 | 6206.6865 |
| Q99707 | Methionine synthase OS=Homo sapiens C | MTR | 140.528 | 0.9 | 1 | 1 | 39.039936 |
| Q99708 | DNA endonuclease RBBP8 OS=Homo sa | RBBP8 | 101.942 | 2.5 | 2 | 2 | 43.309357 |
| Q99714 | 3-hydroxyacyl-CoA dehydrogenase type- | HSD17B10 | 26.924 | 21.1 | 4 | 4 | 276.43665 |
| Q99715 | Collagen alpha-1(XII) chain OS=Homo sa | COL12A1 | 333.147 | 3.9 | 10 | 10 | 154.686 |
| Q99728 | BRCA1-associated RING domain protein | BARD1 | 86.648 | 12.9 | 9 | 9 | 623.45691 |
| Q99729 | Heterogeneous nuclear ribonucleoprotein | HNRNPAB | 36.225 | 16 | 6 | 6 | 3356.6411 |
| Q99733 | Nucleosome assembly protein 1-like 4 OS | NAP1L4 | 42.824 | 7.7 | 3 | 3 | 117.58234 |
| Q99741 | Cell division control protein 6 homolog O | CDC6 | 62.721 | 26.6 | 13 | 13 | 616.5036 |
| Q99755 | Phosphatidylinositol 4-phosphate 5-kinas | PIP5K1A | 62.634 | 35.6 | 18 | 4 | 960.61884 |
| Q99816 | Tumor susceptibility gene 101 protein OS | TSG101 | 43.945 | 10.3 | 4 | 4 | 113.82841 |
| Q99832 | T-complex protein 1 subunit eta OS=Hom | CCT7 | 59.367 | 18.6 | 9 | 9 | 242.39619 |
| Q99848 | Probable rRNA-processing protein EBP2 | EBNA1BP2 | 34.852 | 34 | 12 | 12 | 696.93512 |
| Q99871 | HAUS augmin-like complex subunit 7 OS | HAUS7 | 39.795 | 15.4 | 4 | 4 | 136.08099 |
| Q99873 | Protein arginine N-methyltransferase 1 O | PRMT1 | 42.462 | 35.8 | 14 | 11 | 895.50531 |

| | | | | | | | |
|---|---|---|---|---|---|---|---|
| Q99933 | BAG family molecular chaperone regulat | BAG1 | 38.779 | 9.3 | 3 | 3 | 239.19514 |
| Q99941 | Cyclic AMP-dependent transcription fact | ATF6B | 76.71 | 3.3 | 2 | 2 | 83.310822 |
| Q99942 | E3 ubiquitin-protein ligase RNF5 OS=Ho | RNF5 | 19.882 | 8.3 | 1 | 1 | 13.853078 |
| Q99958 | Forkhead box protein C2 OS=Homo sapie | FOXC2 | 53.72 | 23.4 | 11 | 9 | 3370.8772 |
| Q99959 | Plakophilin-2 OS=Homo sapiens OX=96( | PKP2 | 97.415 | 15.7 | 12 | 12 | 279.23218 |
| Q99961 | Endophilin-A2 OS=Homo sapiens OX=96 | SH3GL1 | 41.49 | 12 | 4 | 2 | 140.87202 |
| Q99967 | Cbp/p300-interacting transactivator 2 OS= | CITED2 | 28.497 | 4.1 | 1 | 1 | 186.13527 |
| Q99985 | Semaphorin-3C OS=Homo sapiens OX=9 | SEMA3C | 85.208 | 13.8 | 9 | 9 | 138.74695 |
| Q99996 | A-kinase anchor protein 9 OS=Homo sapi | AKAP9 | 452.988 | 24.8 | 88 | 88 | 405.81427 |
| Q9BPX3 | Condensin complex subunit 3 OS=Homo | NCAPG | 114.335 | 3.2 | 3 | 3 | 33.184933 |
| Q9BPX6 | Calcium uptake protein 1, mitochondrial ( | MICU1 | 54.352 | 12.2 | 8 | 8 | 239.34065 |
| Q9BPZ3 | Polyadenylate-binding protein-interacting | PAIP2 | 14.985 | 21.3 | 2 | 2 | 294.17084 |
| Q9BPZ7 | Target of rapamycin complex 2 subunit M | MAPKAP1 | 59.123 | 15.9 | 6 | 6 | 130.11275 |
| Q9BQ39 | ATP-dependent RNA helicase DDX50 OS | DDX50 | 82.566 | 19.8 | 15 | 15 | 431.69937 |
| Q9BQ52 | Zinc phosphodiesterase ELAC protein 2 ( | ELAC2 | 92.219 | 19 | 12 | 12 | 269.08337 |
| Q9BQ61 | Telomerase RNA component interacting I | TRIR | 18.42 | 21.6 | 5 | 5 | 286.65045 |
| Q9BQ67 | Glutamate-rich WD repeat-containing pro | GRWD1 | 49.42 | 39.7 | 11 | 11 | 2294.697 |
| Q9BQ70 | Ribosome quality control complex subuni | TCF25 | 76.667 | 17.3 | 11 | 11 | 274.49066 |
| Q9BQ95 | Evolutionarily conserved signaling interm | ECSIT | 49.149 | 7.4 | 2 | 2 | 26.847092 |
| Q9BQA1 | Methylosome protein WDR77 OS=Homo | WDR77 | 36.725 | 33 | 8 | 8 | 1262.6497 |
| Q9BQB6 | Vitamin K epoxide reductase complex sul | VKORC1 | 18.235 | 14.7 | 2 | 2 | 328.49213 |
| Q9BQE3 | Tubulin alpha-1C chain OS=Homo sapien | TUBA1C | 49.896 | 6.7 | 2 | 1 | 687.78882 |
| Q9BQE4 | Selenoprotein S OS=Homo sapiens OX=9 | SELENOS | 21.163 | 22.8 | 4 | 4 | 124.65519 |
| Q9BQE5 | Apolipoprotein L2 OS=Homo sapiens OX | APOL2 | 37.079 | 5.3 | 2 | 2 | 41.35611 |
| Q9BQE9 | B-cell CLL/lymphoma 7 protein family m | BCL7B | 22.196 | 10.9 | 3 | 1 | 415.10254 |
| Q9BQG0 | Myb-binding protein 1A OS=Homo sapie | MYBBP1A | 148.855 | 23.3 | 28 | 28 | 566.32715 |
| Q9BQP7 | Mitochondrial genome maintenance exon | MGME1 | 39.421 | 13.4 | 4 | 4 | 58.817562 |
| Q9BRA0 | N-alpha-acetyltransferase 38, NatC auxili | NAA38 | 13.515 | 24 | 2 | 2 | 116.73663 |
| Q9BRD0 | BUD13 homolog OS=Homo sapiens OX= | BUD13 | 70.521 | 15.2 | 9 | 9 | 165.60776 |
| Q9BRJ7 | Tudor-interacting repair regulator protein | NUDT16L1 | 23.339 | 55.5 | 11 | 10 | 1377.1887 |
| Q9BRK4 | Leucine zipper putative tumor suppressor | LZTS2 | 72.76 | 3.6 | 2 | 2 | 74.956818 |
| Q9BRK5 | 45 kDa calcium-binding protein OS=Hom | SDF4 | 41.807 | 24 | 7 | 7 | 168.13451 |
| Q9BRP8 | Partner of Y14 and mago OS=Homo sapie | PYM1 | 22.656 | 16.2 | 3 | 3 | 251.16457 |
| Q9BRS2 | Serine/threonine-protein kinase RIO1 OS= | RIOK1 | 65.583 | 21.1 | 11 | 11 | 469.76614 |
| Q9BRU9 | rRNA-processing protein UTP23 homolog | UTP23 | 28.403 | 4.4 | 1 | 1 | 256.17654 |

| | | | | | | | |
|---|---|---|---|---|---|---|---|
| Q9BRX2 | Protein pelota homolog OS=Homo sapien | PELO | 43.36 | 34.3 | 12 | 12 | 682.75122 |
| Q9BRZ2 | E3 ubiquitin-protein ligase TRIM56 OS=l | TRIM56 | 81.488 | 42.5 | 27 | 27 | 1870.2769 |
| Q9BS26 | Endoplasmic reticulum resident protein 4 | ERP44 | 46.972 | 2.2 | 1 | 1 | 40.897308 |
| Q9BS34 | Zinc finger protein 670 OS=Homo sapien | ZNF670 | 44.604 | 9.8 | 4 | 4 | 105.65557 |
| Q9BSD7 | Cancer-related nucleoside-triphosphatase | NTPCR | 20.714 | 31.1 | 5 | 5 | 696.22797 |
| Q9BSE5 | Guanidino acid hydrolase, mitochondrial ( | AGMAT | 37.661 | 5.7 | 2 | 2 | 51.041763 |
| Q9BSH4 | Translational activator of cytochrome c o | TACO1 | 32.478 | 18.5 | 5 | 5 | 128.67392 |
| Q9BSJ2 | Gamma-tubulin complex component 2 OS | TUBGCP2 | 102.535 | 38.6 | 30 | 30 | 1316.9944 |
| Q9BSJ6 | Protein PIMREG OS=Homo sapiens OX= | PIMREG | 27.48 | 36.7 | 7 | 7 | 185.46756 |
| Q9BSJ8 | Extended synaptotagmin-1 OS=Homo sap | ESYT1 | 122.857 | 4.8 | 5 | 5 | 69.033546 |
| Q9BSK2 | Solute carrier family 25 member 33 OS=H | SLC25A33 | 35.376 | 8.1 | 2 | 1 | 78.574249 |
| Q9BSM1 | Polycomb group RING finger protein 1 O | PCGF1 | 30.347 | 7.3 | 2 | 2 | 65.760132 |
| Q9BSU1 | Phagosome assembly factor 1 OS=Homo | PHAF1 | 47.525 | 2.1 | 1 | 1 | 95.32637 |
| Q9BT22 | Chitobiosyldiphosphodolichol beta-mann | ALG1 | 52.519 | 2.6 | 1 | 1 | 42.51009 |
| Q9BT25 | HAUS augmin-like complex subunit 8 OS | HAUS8 | 44.858 | 12.4 | 5 | 5 | 230.31305 |
| Q9BT40 | Inositol polyphosphate 5-phosphatase K ( | INPP5K | 51.091 | 15.4 | 7 | 7 | 260.79715 |
| Q9BT78 | COP9 signalosome complex subunit 4 OS | COPS4 | 46.269 | 9.9 | 3 | 3 | 87.291313 |
| Q9BT92 | Trichoplein keratin filament-binding prot | TCHP | 61.072 | 14.9 | 7 | 7 | 93.716873 |
| Q9BTC0 | Death-inducer obliterator 1 OS=Homo sa | DIDO1 | 243.873 | 15.4 | 30 | 30 | 515.0412 |
| Q9BTD8 | RNA-binding protein 42 OS=Homo sapie | RBM42 | 50.415 | 5.8 | 3 | 3 | 387.08981 |
| Q9BTE3 | Mini-chromosome maintenance complex- | MCMBP | 72.981 | 17.9 | 11 | 11 | 136.75301 |
| Q9BTE6 | Alanyl-tRNA editing protein Aarsd1 OS= | AARSD1 | 45.48 | 8.5 | 3 | 3 | 102.94748 |
| Q9BTE7 | DCN1-like protein 5 OS=Homo sapiens ( | DCUN1D5 | 27.509 | 32.9 | 7 | 6 | 109.24889 |
| Q9BTL3 | RNA guanine-N7 methyltransferase activ | RAMAC | 14.382 | 15.3 | 2 | 1 | 161.25313 |
| Q9BTP7 | Fanconi anemia core complex-associated | FAAP24 | 23.897 | 5.1 | 1 | 1 | 58.151119 |
| Q9BTT4 | Mediator of RNA polymerase II transcrip | MED10 | 15.688 | 6.7 | 1 | 1 | 176.13095 |
| Q9BTV4 | Transmembrane protein 43 OS=Homo sap | TMEM43 | 44.876 | 6 | 2 | 2 | 49.931618 |
| Q9BTX1 | Nucleoporin NDC1 OS=Homo sapiens O | NDC1 | 76.305 | 6.1 | 3 | 3 | 78.006477 |
| Q9BU19 | Zinc finger protein 692 OS=Homo sapien | ZNF692 | 56.969 | 14.3 | 6 | 6 | 120.58157 |
| Q9BU61 | NADH dehydrogenase [ubiquinone] 1 alp | NDUFAF3 | 20.351 | 7.6 | 1 | 1 | 19.619925 |
| Q9BU76 | Multiple myeloma tumor-associated prote | MMTAG2 | 29.412 | 30.4 | 8 | 8 | 1610.8859 |
| Q9BU89 | Deoxyhypusine hydroxylase OS=Homo sa | DOHH | 32.905 | 13.2 | 3 | 3 | 135.26625 |
| Q9BUB7 | Transmembrane protein 70, mitochondrial | TMEM70 | 28.97 | 4.6 | 1 | 1 | 86.615509 |
| Q9BUE6 | Iron-sulfur cluster assembly 1 homolog, m | ISCA1 | 14.18 | 19.4 | 2 | 2 | 104.78099 |
| Q9BUF5 | Tubulin beta-6 chain OS=Homo sapiens ( | TUBB6 | 49.858 | 21.5 | 9 | 4 | 1386.7822 |

| | | | | | | | |
|---|---|---|---|---|---|---|---|
| Q9BUH8 | Brain-enriched guanylate kinase-associate | BEGAIN | 64.804 | 23.9 | 15 | 15 | 290.54767 |
| Q9BUI4 | DNA-directed RNA polymerase III subun | POLR3C | 60.612 | 12.7 | 6 | 6 | 724.42126 |
| Q9BUJ2 | Heterogeneous nuclear ribonucleoprotein | HNRNPUL | 95.739 | 29.2 | 25 | 25 | 9679.5449 |
| Q9BUL5 | PHD finger protein 23 OS=Homo sapiens | PHF23 | 43.819 | 3.2 | 1 | 1 | 44.065151 |
| Q9BUL8 | Programmed cell death protein 10 OS=Hc | PDCD10 | 24.702 | 25.9 | 5 | 5 | 72.35437 |
| Q9BUL9 | Ribonuclease P protein subunit p25 OS=I | RPP25 | 20.633 | 33.7 | 6 | 6 | 250.19069 |
| Q9BUN8 | Derlin-1 OS=Homo sapiens OX=9606 GN | DERL1 | 28.801 | 7.6 | 2 | 2 | 291.88724 |
| Q9BUP0 | EF-hand domain-containing protein D1 O | EFHD1 | 26.928 | 22.2 | 4 | 3 | 134.83322 |
| Q9BUQ8 | Probable ATP-dependent RNA helicase D | DDX23 | 95.583 | 27 | 22 | 22 | 738.08777 |
| Q9BUR4 | Telomerase Cajal body protein 1 OS=Hor | WRAP53 | 59.31 | 4.9 | 2 | 2 | 146.93617 |
| Q9BUZ4 | TNF receptor-associated factor 4 OS=Ho | TRAF4 | 53.543 | 39.6 | 17 | 17 | 722.30188 |
| Q9BV38 | WD repeat-containing protein 18 OS=Ho | WDR18 | 47.406 | 8.1 | 3 | 3 | 73.144913 |
| Q9BV44 | tRNA (guanine(6)-N2)-methyltransferase | THUMPD3 | 57.003 | 8.9 | 4 | 4 | 76.882416 |
| Q9BV57 | Acireductone dioxygenase OS=Homo sap | ADI1 | 21.499 | 53.1 | 8 | 8 | 314.97351 |
| Q9BV68 | E3 ubiquitin-protein ligase RNF126 OS=I | RNF126 | 33.862 | 6.8 | 2 | 2 | 123.51991 |
| Q9BVA0 | Katanin p80 WD40 repeat-containing sub | KATNB1 | 72.334 | 6.4 | 4 | 4 | 45.098602 |
| Q9BVC3 | Sister chromatid cohesion protein DCC1 ( | DSCC1 | 44.826 | 10.9 | 4 | 4 | 46.84528 |
| Q9BVC4 | Target of rapamycin complex subunit LST | MLST8 | 35.877 | 14.4 | 4 | 4 | 68.921013 |
| Q9BVG8 | Kinesin-like protein KIFC3 OS=Homo sa | KIFC3 | 92.775 | 20.9 | 15 | 15 | 295.66779 |
| Q9BVI4 | Nucleolar complex protein 4 homolog OS | NOC4L | 58.468 | 37.8 | 19 | 19 | 846.73651 |
| Q9BVJ6 | U3 small nucleolar RNA-associated prote | UTP14A | 87.978 | 32 | 24 | 16 | 643.38202 |
| Q9BVJ7 | Dual specificity protein phosphatase 23 O | DUSP23 | 16.589 | 44.7 | 6 | 6 | 159.58115 |
| Q9BVK6 | Transmembrane emp24 domain-containin | TMED9 | 27.278 | 4.3 | 1 | 1 | 69.686974 |
| Q9BVK8 | BOS complex subunit TMEM147 OS=Hc | TMEM147 | 25.261 | 4 | 1 | 1 | 83.430038 |
| Q9BVP2 | Guanine nucleotide-binding protein-like 3 | GNL3 | 61.994 | 33.7 | 17 | 17 | 2539.8726 |
| Q9BVQ7 | ATPase family gene 2 protein homolog B | AFG2B | 80.711 | 13.8 | 10 | 10 | 296.19437 |
| Q9BVS4 | Serine/threonine-protein kinase RIO2 OS | RIOK2 | 63.284 | 30.1 | 13 | 13 | 1094.6527 |
| Q9BVS5 | tRNA (adenine(58)-N(1))-methyltransfera | TRMT61B | 52.966 | 15.1 | 6 | 6 | 173.98126 |
| Q9BVV7 | Mitochondrial import inner membrane tra | TIMM21 | 28.203 | 22.2 | 5 | 5 | 231.27632 |
| Q9BW19 | Kinesin-like protein KIFC1 OS=Homo sa | KIFC1 | 73.748 | 34.3 | 21 | 21 | 587.37189 |
| Q9BW27 | Nuclear pore complex protein Nup85 OS | NUP85 | 75.02 | 3.5 | 2 | 2 | 66.168274 |
| Q9BW66 | Cyclin-dependent kinase 2-interacting pro | CINP | 24.324 | 22.6 | 4 | 4 | 118.03702 |
| Q9BW85 | Splicing factor YJU2 OS=Homo sapiens ( | YJU2 | 37.086 | 4 | 1 | 1 | 35.451515 |
| Q9BW92 | Threonine--tRNA ligase, mitochondrial O | TARS2 | 81.037 | 13.9 | 10 | 10 | 407.66937 |
| Q9BWE0 | Replication initiator 1 OS=Homo sapiens | REPIN1 | 63.575 | 18.7 | 10 | 10 | 279.61441 |

| | | | | | | | |
|---|---|---|---|---|---|---|---|
| Q9BWF2 | E3 ubiquitin-protein ligase TRAIP OS=H | TRAIP | 53.295 | 5.5 | 2 | 2 | 97.803619 |
| Q9BWF3 | RNA-binding protein 4 OS=Homo sapien | RBM4 | 40.314 | 36.5 | 14 | 1 | 7383.978 |
| Q9BWG6 | Sodium channel modifier 1 OS=Homo sa | SCNM1 | 25.95 | 31.7 | 6 | 6 | 415.89343 |
| Q9BWJ5 | Splicing factor 3B subunit 5 OS=Homo s | SF3B5 | 10.136 | 32.6 | 2 | 2 | 129.32947 |
| Q9BWM7 | Sideroflexin-3 OS=Homo sapiens OX=96 | SFXN3 | 35.504 | 17.8 | 4 | 4 | 194.14049 |
| Q9BWT3 | Poly(A) polymerase gamma OS=Homo sa | PAPOLG | 82.803 | 10.3 | 6 | 5 | 98.201927 |
| Q9BX10 | GTP-binding protein 2 OS=Homo sapiens | GTPBP2 | 65.769 | 17.3 | 10 | 10 | 289.44592 |
| Q9BX40 | Protein LSM14 homolog B OS=Homo sa | LSM14B | 42.071 | 13 | 6 | 6 | 320.11298 |
| Q9BX63 | Fanconi anemia group J protein OS=Hom | BRIP1 | 140.868 | 10.2 | 10 | 10 | 94.731064 |
| Q9BX70 | BTB/POZ domain-containing protein 2 O | BTBD2 | 55.932 | 2.3 | 1 | 1 | 87.778702 |
| Q9BXF6 | Rab11 family-interacting protein 5 OS=H | RAB11FIP5 | 70.416 | 8.9 | 4 | 4 | 28.689493 |
| Q9BXI6 | TBC1 domain family member 10A OS=H | TBC1D10A | 57.118 | 5.9 | 3 | 2 | 72.820236 |
| Q9BXJ9 | N-alpha-acetyltransferase 15, NatA auxili | NAA15 | 101.273 | 7 | 6 | 5 | 68.30191 |
| Q9BXM7 | Serine/threonine-protein kinase PINK1, m | PINK1 | 62.77 | 35.6 | 19 | 19 | 2754.709 |
| Q9BXP5 | Serrate RNA effector molecule homolog ( | SRRT | 100.667 | 32.2 | 32 | 32 | 2211.5769 |
| Q9BXS4 | Transmembrane protein 59 OS=Homo sa | TMEM59 | 36.224 | 2.8 | 1 | 1 | 31.068098 |
| Q9BXS5 | AP-1 complex subunit mu-1 OS=Homo s | AP1M1 | 48.587 | 15.6 | 6 | 4 | 176.7955 |
| Q9BXS6 | Nucleolar and spindle-associated protein | NUSAP1 | 49.452 | 45.4 | 19 | 19 | 1072.0165 |
| Q9BY12 | S phase cyclin A-associated protein in the | SCAPER | 158.288 | 22.3 | 30 | 30 | 632.54114 |
| Q9BY42 | Replication termination factor 2 OS=Hom | RTF2 | 33.887 | 14.1 | 4 | 4 | 54.864353 |
| Q9BY43 | Charged multivesicular body protein 4a O | CHMP4A | 25.099 | 9.5 | 2 | 2 | 95.541397 |
| Q9BY44 | Eukaryotic translation initiation factor 2A | EIF2A | 64.991 | 42.9 | 19 | 19 | 1172.5833 |
| Q9BY49 | Peroxisomal trans-2-enoyl-CoA reductase | PECR | 32.545 | 4.6 | 1 | 1 | 71.565041 |
| Q9BY77 | Polymerase delta-interacting protein 3 OS | POLDIP3 | 46.09 | 43.9 | 17 | 17 | 3144.5779 |
| Q9BYC5 | Alpha-(1,6)-fucosyltransferase OS=Homo | FUT8 | 66.516 | 2.8 | 2 | 2 | 80.264709 |
| Q9BYG3 | MKI67 FHA domain-interacting nucleola | NIFK | 34.223 | 35.5 | 12 | 12 | 1644.0405 |
| Q9BYG5 | Partitioning defective 6 homolog beta OS | PARD6B | 41.183 | 6.7 | 2 | 2 | 148.14375 |
| Q9BYJ4 | E3 ubiquitin-protein ligase TRIM34 OS=l | TRIM34 | 56.864 | 5.9 | 3 | 1 | 65.49324 |
| Q9BYJ9 | YTH domain-containing family protein 1 | YTHDF1 | 60.874 | 19.7 | 11 | 10 | 1550.7214 |
| Q9BYK8 | Helicase with zinc finger domain 2 OS=H | HELZ2 | 294.651 | 26.8 | 63 | 63 | 546.31555 |
| Q9BYW2 | Histone-lysine N-methyltransferase SETD | SETD2 | 287.597 | 12.6 | 28 | 28 | 174.80467 |
| Q9BZ29 | Dedicator of cytokinesis protein 9 OS=Hc | DOCK9 | 236.446 | 1.4 | 2 | 2 | 35.196198 |
| Q9BZ95 | Histone-lysine N-methyltransferase NSD3 | NSD3 | 161.613 | 7 | 8 | 8 | 160.31085 |
| Q9BZE4 | GTP-binding protein 4 OS=Homo sapiens | GTPBP4 | 73.965 | 45.3 | 28 | 28 | 3128.4294 |
| Q9BZF3 | Oxysterol-binding protein-related protein | OSBPL6 | 106.306 | 3.2 | 3 | 3 | 67.329514 |

| | | | | | | | |
|---|---|---|---|---|---|---|---|
| Q9BZF9 | Uveal autoantigen with coiled-coil domai | UACA | 162.506 | 3 | 4 | 4 | 66.696053 |
| Q9BZG1 | Ras-related protein Rab-34 OS=Homo sa | RAB34 | 29.045 | 30.5 | 7 | 7 | 322.83582 |
| Q9BZH6 | WD repeat-containing protein 11 OS=Ho | WDR11 | 136.685 | 30.6 | 33 | 33 | 2843.3923 |
| Q9BZI7 | Regulator of nonsense transcripts 3B OS= | UPF3B | 57.763 | 40.6 | 20 | 20 | 3280.5947 |
| Q9BZJ0 | Crooked neck-like protein 1 OS=Homo sa | CRNKL1 | 100.452 | 7.9 | 6 | 6 | 136.73917 |
| Q9BZK7 | F-box-like/WD repeat-containing protein | TBL1XR1 | 55.595 | 13.6 | 5 | 3 | 114.99377 |
| Q9BZL6 | Serine/threonine-protein kinase D2 OS=H | PRKD2 | 96.723 | 1.1 | 1 | 1 | 174.04076 |
| Q9BZR9 | E3 ubiquitin-protein ligase TRIM8 OS=H | TRIM8 | 61.489 | 28.1 | 13 | 13 | 428.61264 |
| Q9BZX2 | Uridine-cytidine kinase 2 OS=Homo sapi | UCK2 | 29.3 | 16.5 | 4 | 3 | 497.56726 |
| Q9BZZ5 | Apoptosis inhibitor 5 OS=Homo sapiens ( | API5 | 59.005 | 10.7 | 5 | 5 | 82.887154 |
| Q9C029 | E3 ubiquitin-protein ligase TRIM7 OS=H | TRIM7 | 56.631 | 18.2 | 9 | 9 | 169.35986 |
| Q9C086 | INO80 complex subunit B OS=Homo sap | INO80B | 38.638 | 12.9 | 4 | 4 | 197.85982 |
| Q9C0B0 | RING finger protein unkempt homolog O | UNK | 88.085 | 16.8 | 15 | 14 | 717.59491 |
| Q9C0B5 | Palmitoyltransferase ZDHHC5 OS=Homo | ZDHHC5 | 77.545 | 2.2 | 1 | 1 | 41.788837 |
| Q9C0C2 | 182 kDa tankyrase-1-binding protein OS= | TNKS1BP1 | 181.796 | 24.9 | 34 | 34 | 746.28113 |
| Q9C0C9 | (E3-independent) E2 ubiquitin-conjugatin | UBE2O | 141.294 | 1.9 | 2 | 2 | 39.372383 |
| Q9C0D5 | Protein TANC1 OS=Homo sapiens OX=9 | TANC1 | 202.22 | 12.4 | 20 | 20 | 153.2477 |
| Q9C0D6 | FH2 domain-containing protein 1 OS=Ho | FHDC1 | 124.762 | 3.7 | 3 | 3 | 46.525257 |
| Q9C0F1 | Centrosomal protein of 44 kDa OS=Homo | CEP44 | 44.14 | 18.2 | 6 | 6 | 248.0439 |
| Q9C0F3 | Zinc finger protein 436 OS=Homo sapien | ZNF436 | 54.277 | 4.7 | 2 | 2 | 34.591385 |
| Q9C0J8 | pre-mRNA 3' end processing protein WD | WDR33 | 145.892 | 13.7 | 17 | 17 | 351.25015 |
| Q9GZL7 | Ribosome biogenesis protein WDR12 OS | WDR12 | 47.708 | 19.1 | 7 | 7 | 279.14435 |
| Q9GZM8 | Nuclear distribution protein nudE-like 1 C | NDEL1 | 38.375 | 3.2 | 1 | 1 | 42.324947 |
| Q9GZP9 | Derlin-2 OS=Homo sapiens OX=9606 GN | DERL2 | 27.568 | 4.6 | 1 | 1 | 46.185692 |
| Q9GZR2 | RNA exonuclease 4 OS=Homo sapiens O | REXO4 | 46.672 | 23 | 8 | 8 | 188.34827 |
| Q9GZS1 | DNA-directed RNA polymerase I subunit | POLR1E | 47.261 | 25.3 | 9 | 9 | 262.97366 |
| Q9GZS3 | Superkiller complex protein 8 OS=Homo | SKIC8 | 33.581 | 17.4 | 4 | 4 | 251.11247 |
| Q9GZT3 | SRA stem-loop-interacting RNA-binding | SLIRP | 12.35 | 69.7 | 8 | 8 | 1229.0654 |
| Q9GZT9 | Egl nine homolog 1 OS=Homo sapiens O | EGLN1 | 46.021 | 21.4 | 7 | 7 | 247.89345 |
| Q9GZV5 | WW domain-containing transcription regu | WWTR1 | 44.102 | 2 | 1 | 1 | 59.705921 |
| Q9GZZ6 | Neuronal acetylcholine receptor subunit a | CHRNA10 | 49.705 | 2 | 1 | 1 | 45.039288 |
| Q9GZZ9 | Ubiquitin-like modifier-activating enzyme | UBA5 | 44.864 | 3.2 | 1 | 1 | 76.914215 |
| Q9H000 | E3 ubiquitin-protein ligase makorin-2 OS | MKRN2 | 46.941 | 27.6 | 12 | 12 | 1071.9725 |
| Q9H040 | DNA-dependent metalloprotease SPRTN | SPRTN | 55.134 | 3.1 | 1 | 1 | 68.155708 |
| Q9H074 | Polyadenylate-binding protein-interacting | PAIP1 | 53.525 | 24.8 | 12 | 12 | 3431.3533 |

| | | | | | | | |
|---|---|---|---|---|---|---|---|
| Q9H078 | Mitochondrial disaggregase OS=Homo sa | CLPB | 78.729 | 18.7 | 13 | 13 | 408.33295 |
| Q9H079 | KATNB1-like protein 1 OS=Homo sapier | KATNBL1 | 34.767 | 4.3 | 1 | 1 | 29.446489 |
| Q9H089 | Large subunit GTPase 1 homolog OS=Ho | LSG1 | 75.226 | 33.9 | 22 | 22 | 3779.4563 |
| Q9H093 | NUAK family SNF1-like kinase 2 OS=Hc | NUAK2 | 69.613 | 5.7 | 3 | 3 | 53.027363 |
| Q9H0A0 | RNA cytidine acetyltransferase OS=Homo | NAT10 | 115.73 | 24.7 | 26 | 26 | 718.19617 |
| Q9H0B6 | Kinesin light chain 2 OS=Homo sapiens ( | KLC2 | 68.935 | 14.3 | 8 | 5 | 100.30834 |
| Q9H0D6 | 5'-3' exoribonuclease 2 OS=Homo sapiens | XRN2 | 108.583 | 27.1 | 24 | 24 | 1206.4532 |
| Q9H0G5 | Nuclear speckle splicing regulatory protei | NSRP1 | 66.391 | 20.8 | 11 | 11 | 657.20282 |
| Q9H0H5 | Rac GTPase-activating protein 1 OS=Hor | RACGAP1 | 71.027 | 36.4 | 22 | 22 | 3673.0847 |
| Q9H0I2 | Enkurin domain-containing protein 1 OS= | ENKD1 | 38.76 | 6.1 | 2 | 2 | 110.98527 |
| Q9H0J9 | Protein mono-ADP-ribosyltransferase PA | PARP12 | 79.064 | 25.2 | 16 | 16 | 1758.0239 |
| Q9H0K1 | Serine/threonine-protein kinase SIK2 OS= | SIK2 | 103.916 | 4.8 | 4 | 4 | 49.941212 |
| Q9H0K6 | Pseudouridylate synthase PUS7L OS=Ho | PUS7L | 80.701 | 1.3 | 1 | 1 | 56.089165 |
| Q9H0L4 | Cleavage stimulation factor subunit 2 tau | CSTF2T | 64.437 | 1.6 | 1 | 1 | 49.485298 |
| Q9H0M0 | NEDD4-like E3 ubiquitin-protein ligase V | WWP1 | 105.202 | 2.6 | 2 | 2 | 51.773106 |
| Q9H0S4 | Probable ATP-dependent RNA helicase D | DDX47 | 50.647 | 37.1 | 15 | 15 | 2380.0044 |
| Q9H0U3 | Magnesium transporter protein 1 OS=Hor | MAGT1 | 38.037 | 17 | 6 | 6 | 520.86877 |
| Q9H0U6 | Large ribosomal subunit protein uL18m C | MRPL18 | 20.577 | 17.8 | 3 | 3 | 137.1019 |
| Q9H0W8 | Nonsense-mediated mRNA decay factor S | SMG9 | 57.651 | 3.5 | 1 | 1 | 104.26354 |
| Q9H0Z9 | RNA-binding protein 38 OS=Homo sapie | RBM38 | 25.498 | 35.6 | 7 | 3 | 95.941414 |
| Q9H147 | Deoxynucleotidyltransferase terminal-inte | DNTTIP1 | 37.014 | 27.1 | 9 | 9 | 348.91373 |
| Q9H1A4 | Anaphase-promoting complex subunit 1 C | ANAPC1 | 216.5 | 11.4 | 19 | 19 | 149.95166 |
| Q9H1B5 | Xylosyltransferase 2 OS=Homo sapiens C | XYLT2 | 96.768 | 2.5 | 2 | 2 | 57.064781 |
| Q9H1D9 | DNA-directed RNA polymerase III subun | POLR3F | 35.684 | 14.2 | 4 | 4 | 66.63282 |
| Q9H1I8 | Activating signal cointegrator 1 complex s | ASCC2 | 86.361 | 18 | 14 | 14 | 2870.3958 |
| Q9H1J1 | Regulator of nonsense transcripts 3A OS= | UPF3A | 54.697 | 16.6 | 7 | 7 | 273.64542 |
| Q9H1K1 | Iron-sulfur cluster assembly enzyme ISCU | ISCU | 17.999 | 10.2 | 1 | 1 | 68.946777 |
| Q9H223 | EH domain-containing protein 4 OS=Hon | EHD4 | 61.176 | 12 | 5 | 5 | 73.887383 |
| Q9H254 | Spectrin beta chain, non-erythrocytic 4 OS | SPTBN4 | 288.986 | 0.7 | 2 | 2 | 263.92499 |
| Q9H269 | Vacuolar protein sorting-associated protei | VPS16 | 94.694 | 8.8 | 5 | 5 | 68.086662 |
| Q9H270 | Vacuolar protein sorting-associated protei | VPS11 | 107.837 | 8.8 | 8 | 8 | 130.37022 |
| Q9H2D1 | Solute carrier family 25 member 32 OS=H | SLC25A32 | 35.408 | 11.4 | 3 | 3 | 65.038628 |
| Q9H2D6 | TRIO and F-actin-binding protein OS=Hc | TRIOBP | 261.376 | 1 | 2 | 2 | 48.453529 |
| Q9H2G4 | Testis-specific Y-encoded-like protein 2 ( | TSPYL2 | 79.436 | 2.9 | 2 | 2 | 117.9546 |
| Q9H2H8 | Peptidyl-prolyl cis-trans isomerase-like 3 | PPIL3 | 18.155 | 46 | 7 | 7 | 425.20898 |

| | | | | | | | |
|---|---|---|---|---|---|---|---|
| Q9H2H9 | Sodium-coupled neutral amino acid symp | SLC38A1 | 54.048 | 2.3 | 1 | 1 | 57.400486 |
| Q9H2K2 | Poly [ADP-ribose] polymerase tankyrase- | TNKS2 | 126.918 | 3.9 | 4 | 4 | 73.644913 |
| Q9H2K8 | Serine/threonine-protein kinase TAO3 OS | TAOK3 | 105.406 | 0.9 | 1 | 1 | 70.943237 |
| Q9H2P0 | Activity-dependent neuroprotector homeo | ADNP | 123.563 | 12.9 | 13 | 13 | 166.93852 |
| Q9H2U1 | ATP-dependent DNA/RNA helicase DHX | DHX36 | 114.761 | 35.3 | 33 | 33 | 4617.2178 |
| Q9H2U2 | Inorganic pyrophosphatase 2, mitochondri | PPA2 | 37.921 | 6.6 | 2 | 2 | 336.5636 |
| Q9H2Y7 | Zinc finger protein 106 OS=Homo sapiens | ZNF106 | 208.883 | 13.1 | 18 | 18 | 239.40108 |
| Q9H300 | Presenilin-associated rhomboid-like prote | PARL | 42.191 | 8.2 | 3 | 3 | 118.83726 |
| Q9H307 | Pinin OS=Homo sapiens OX=9606 GN=P | PNN | 81.628 | 15.9 | 10 | 10 | 450.98849 |
| Q9H329 | Band 4.1-like protein 4B OS=Homo sapie | EPB41L4B | 99.713 | 7.7 | 6 | 6 | 267.37131 |
| Q9H3F6 | BTB/POZ domain-containing adapter for | KCTD10 | 35.433 | 17.6 | 5 | 4 | 85.012749 |
| Q9H3H3 | UPF0696 protein C11orf68 OS=Homo sa | C11orf68 | 31.431 | 2.7 | 1 | 1 | 32.201023 |
| Q9H3K6 | BolA-like protein 2 OS=Homo sapiens O | BOLA2 | 10.117 | 43 | 3 | 3 | 388.15262 |
| Q9H3L0 | Cobalamin trafficking protein CblD OS=H | MMADHC | 32.941 | 3 | 1 | 1 | 78.246964 |
| Q9H3P2 | Negative elongation factor A OS=Homo s | NELFA | 57.277 | 18.4 | 8 | 8 | 115.69012 |
| Q9H3P7 | Golgi resident protein GCP60 OS=Homo | ACBD3 | 60.594 | 5.7 | 2 | 2 | 42.037815 |
| Q9H3Q1 | Cdc42 effector protein 4 OS=Homo sapie | CDC42EP4 | 37.98 | 5.9 | 2 | 2 | 95.438988 |
| Q9H3R5 | Centromere protein H OS=Homo sapiens | CENPH | 28.481 | 4.5 | 1 | 1 | 23.838936 |
| Q9H3S7 | Tyrosine-protein phosphatase non-recepto | PTPN23 | 178.974 | 6.9 | 10 | 10 | 89.817543 |
| Q9H410 | Kinetochore-associated protein DSN1 hor | DSN1 | 40.068 | 3.1 | 1 | 1 | 123.2628 |
| Q9H444 | Charged multivesicular body protein 4b O | CHMP4B | 24.951 | 23.2 | 5 | 3 | 272.20236 |
| Q9H4A5 | Golgi phosphoprotein 3-like OS=Homo sa | GOLPH3L | 32.768 | 10.9 | 1 | 1 | 21.106665 |
| Q9H4A6 | Golgi phosphoprotein 3 OS=Homo sapien | GOLPH3 | 33.811 | 8.1 | 2 | 2 | 44.748402 |
| Q9H4B6 | Protein salvador homolog 1 OS=Homo sa | SAV1 | 44.634 | 11.2 | 4 | 4 | 154.23068 |
| Q9H4B7 | Tubulin beta-1 chain OS=Homo sapiens O | TUBB1 | 50.327 | 3.1 | 1 | 1 | 65.556389 |
| Q9H4F8 | SPARC-related modular calcium-binding | SMOC1 | 48.163 | 15.2 | 6 | 6 | 84.700058 |
| Q9H4H8 | Protein FAM83D OS=Homo sapiens OX= | FAM83D | 64.425 | 29.7 | 17 | 17 | 1823.0471 |
| Q9H4K7 | Mitochondrial ribosome-associated GTPa | MTG2 | 43.956 | 18 | 7 | 7 | 73.227188 |
| Q9H4L4 | Sentrin-specific protease 3 OS=Homo sap | SENP3 | 65.01 | 10.3 | 5 | 5 | 151.00368 |
| Q9H4L5 | Oxysterol-binding protein-related protein | OSBPL3 | 101.224 | 34.6 | 24 | 22 | 1704.7166 |
| Q9H4M7 | Pleckstrin homology domain-containing fa | PLEKHA4 | 85.401 | 1.8 | 1 | 1 | 69.622849 |
| Q9H4M9 | EH domain-containing protein 1 OS=Hon | EHD1 | 60.627 | 7.7 | 4 | 1 | 197.0094 |
| Q9H4P4 | E3 ubiquitin-protein ligase NRDP1 OS=H | RNF41 | 35.906 | 18 | 5 | 5 | 290.14941 |
| Q9H4Z2 | Zinc finger protein 335 OS=Homo sapiens | ZNF335 | 144.893 | 4.7 | 6 | 6 | 100.58237 |
| Q9H583 | HEAT repeat-containing protein 1 OS=Ho | HEATR1 | 242.371 | 8.5 | 15 | 15 | 289.96649 |

| | | | | | | | |
|---|---|---|---|---|---|---|---|
| Q9H5H4 | Zinc finger protein 768 OS=Homo sapien: | ZNF768 | 60.229 | 13.1 | 7 | 7 | 176.75711 |
| Q9H5P4 | PDZ domain-containing protein 7 OS=Ho | PDZD7 | 111.752 | 0.7 | 1 | 1 | 130.79745 |
| Q9H5Q4 | Dimethyladenosine transferase 2, mitocho | TFB2M | 45.349 | 41.9 | 15 | 15 | 604.47876 |
| Q9H5U6 | rRNA N6-adenosine-methyltransferase ZC | ZCCHC4 | 59.01 | 18.9 | 10 | 10 | 376.58572 |
| Q9H5V9 | STING ER exit protein OS=Homo sapien | STEEP1 | 25.625 | 21.2 | 4 | 4 | 270.15543 |
| Q9H5Z1 | Probable ATP-dependent RNA helicase D | DHX35 | 78.911 | 2.7 | 2 | 2 | 46.079384 |
| Q9H633 | Ribonuclease P protein subunit p21 OS=H | RPP21 | 17.571 | 21.4 | 3 | 3 | 175.96274 |
| Q9H668 | CST complex subunit STN1 OS=Homo sa | STN1 | 42.12 | 5.7 | 2 | 2 | 42.81699 |
| Q9H694 | Protein bicaudal C homolog 1 OS=Homo | BICC1 | 104.844 | 12.3 | 10 | 10 | 232.21796 |
| Q9H6D7 | HAUS augmin-like complex subunit 4 OS | HAUS4 | 42.4 | 14.3 | 4 | 4 | 80.543228 |
| Q9H6H4 | Receptor expression-enhancing protein 4 ( | REEP4 | 29.396 | 26.1 | 7 | 7 | 735.00439 |
| Q9H6Q4 | Cytosolic iron-sulfur assembly component | CIAO3 | 53.021 | 2.3 | 1 | 1 | 43.359951 |
| Q9H6R0 | ATP-dependent RNA helicase DHX33 OS | DHX33 | 78.875 | 22.3 | 15 | 15 | 193.24318 |
| Q9H6R4 | Nucleolar protein 6 OS=Homo sapiens OX | NOL6 | 127.594 | 16.9 | 17 | 17 | 323.01913 |
| Q9H6R7 | WD repeat and coiled-coil-containing pro | WDCP | 79.136 | 1.5 | 1 | 1 | 70.095245 |
| Q9H6S0 | 3'-5' RNA helicase YTHDC2 OS=Homo s | YTHDC2 | 160.249 | 37.8 | 47 | 47 | 2436.0815 |
| Q9H6S1 | 5-azacytidine-induced protein 2 OS=Hom | AZI2 | 44.935 | 16.3 | 6 | 6 | 135.6951 |
| Q9H6T0 | Epithelial splicing regulatory protein 2 OS | ESRP2 | 78.402 | 21.9 | 14 | 13 | 850.18524 |
| Q9H6T3 | RNA polymerase II-associated protein 3 ( | RPAP3 | 75.719 | 14 | 8 | 8 | 118.8038 |
| Q9H6U6 | BCAS3 microtubule associated cell migra | BCAS3 | 101.237 | 3.2 | 3 | 3 | 66.090881 |
| Q9H6W3 | Ribosomal oxygenase 1 OS=Homo sapien | RIOX1 | 71.086 | 14.4 | 7 | 7 | 144.88237 |
| Q9H707 | Zinc finger protein 552 OS=Homo sapien: | ZNF552 | 46.199 | 26.8 | 10 | 7 | 314.44238 |
| Q9H773 | dCTP pyrophosphatase 1 OS=Homo sapie | DCTPP1 | 18.681 | 21.2 | 3 | 3 | 95.964767 |
| Q9H7B2 | Ribosome production factor 2 homolog O | RPF2 | 35.583 | 21.6 | 7 | 7 | 217.61455 |
| Q9H7B4 | Histone-lysine N-methyltransferase SMYI | SMYD3 | 49.098 | 8.4 | 3 | 3 | 39.165314 |
| Q9H7D7 | WD repeat-containing protein 26 OS=Hoi | WDR26 | 72.124 | 3.9 | 3 | 3 | 67.55307 |
| Q9H7E2 | Tudor domain-containing protein 3 OS=H | TDRD3 | 73.185 | 40.9 | 24 | 24 | 3526.3306 |
| Q9H7E9 | UPF0488 protein C8orf33 OS=Homo sap | C8orf33 | 24.993 | 35.8 | 7 | 7 | 1634.3409 |
| Q9H7H0 | Methyltransferase-like protein 17, mitoch | METTL17 | 50.734 | 12.3 | 6 | 6 | 137.56525 |
| Q9H7N4 | Splicing factor, arginine/serine-rich 19 OS | SCAF1 | 139.27 | 6.1 | 6 | 6 | 47.291634 |
| Q9H7T9 | Aurora kinase A- and ninein-interacting p | AUNIP | 40.253 | 24.1 | 8 | 8 | 242.22659 |
| Q9H7Z3 | Nuclear exosome regulator NRDE2 OS=H | NRDE2 | 132.673 | 4.3 | 3 | 3 | 79.622925 |
| Q9H7Z6 | Histone acetyltransferase KAT8 OS=Hon | KAT8 | 52.403 | 4.4 | 2 | 2 | 61.614658 |
| Q9H814 | Phosphorylated adapter RNA export prote | PHAX | 44.403 | 44.9 | 15 | 15 | 1552.9596 |
| Q9H825 | tRNA N(3)-methylcytidine methyltransfer | METTL8 | 33.387 | 15.8 | 4 | 4 | 242.15837 |

| | | | | | | | |
|---|---|---|---|---|---|---|---|
| Q9H840 | Gem-associated protein 7 OS=Homo sapi | GEMIN7 | 14.537 | 53.4 | 4 | 4 | 269.78598 |
| Q9H845 | Complex I assembly factor ACAD9, mito | ACAD9 | 68.761 | 9.7 | 5 | 5 | 67.464478 |
| Q9H869 | YY1-associated protein 1 OS=Homo sapi | YY1AP1 | 87.945 | 1.5 | 1 | 1 | 109.55231 |
| Q9H8H2 | Probable ATP-dependent RNA helicase D | DDX31 | 94.088 | 18.7 | 13 | 13 | 179.9093 |
| Q9H8M2 | Bromodomain-containing protein 9 OS=H | BRD9 | 67.001 | 2.2 | 1 | 1 | 41.73431 |
| Q9H8V3 | Protein ECT2 OS=Homo sapiens OX=96 | ECT2 | 103.505 | 30.9 | 26 | 26 | 701.19495 |
| Q9H8X2 | Inositol-pentakisphosphate 2-kinase OS=H | IPPK | 56.017 | 12.6 | 6 | 6 | 195.95427 |
| Q9H8Y5 | tRNA endonuclease ANKZF1 OS=Homo | ANKZF1 | 80.927 | 27.3 | 18 | 18 | 956.91199 |
| Q9H910 | Jupiter microtubule associated homolog 2 | JPT2 | 20.064 | 14.7 | 2 | 2 | 54.24543 |
| Q9H936 | Mitochondrial glutamate carrier 1 OS=Ho | SLC25A22 | 34.471 | 19.8 | 6 | 3 | 268.8652 |
| Q9H944 | Mediator of RNA polymerase II transcript | MED20 | 23.222 | 17 | 3 | 3 | 54.358467 |
| Q9H992 | E3 ubiquitin-protein ligase MARCHF7 O | MARCHF7 | 78.051 | 5.1 | 3 | 3 | 37.522675 |
| Q9H9A5 | CCR4-NOT transcription complex subuni | CNOT10 | 82.31 | 26.6 | 17 | 17 | 605.01392 |
| Q9H9A7 | RecQ-mediated genome instability protei | RMI1 | 70.145 | 11 | 6 | 6 | 119.12876 |
| Q9H9B1 | Histone-lysine N-methyltransferase EHM | EHMT1 | 141.467 | 8.1 | 8 | 7 | 110.37794 |
| Q9H9B4 | Sideroflexin-1 OS=Homo sapiens OX=96 | SFXN1 | 35.62 | 17.7 | 5 | 5 | 110.26001 |
| Q9H9G7 | Protein argonaute-3 OS=Homo sapiens O | AGO3 | 97.361 | 26.7 | 18 | 15 | 862.52356 |
| Q9H9H4 | Vacuolar protein sorting-associated protei | VPS37B | 31.308 | 3.9 | 1 | 1 | 25.893494 |
| Q9H9J2 | Large ribosomal subunit protein mL44 OS | MRPL44 | 37.536 | 22 | 7 | 7 | 778.70953 |
| Q9H9L4 | KAT8 regulatory NSL complex subunit 2 | KANSL2 | 55.043 | 5.5 | 3 | 3 | 67.26226 |
| Q9H9P5 | Putative E3 ubiquitin-protein ligase UNK | UNKL | 73.828 | 8.8 | 5 | 5 | 66.121651 |
| Q9H9P8 | L-2-hydroxyglutarate dehydrogenase, mit | L2HGDH | 50.317 | 2.6 | 1 | 1 | 46.893871 |
| Q9H9Q4 | Non-homologous end-joining factor 1 OS | NHEJ1 | 33.338 | 3 | 1 | 1 | 89.262764 |
| Q9H9T3 | Elongator complex protein 3 OS=Homo s | ELP3 | 62.259 | 30.7 | 16 | 16 | 579.49615 |
| Q9H9Y6 | DNA-directed RNA polymerase I subunit | POLR1B | 128.23 | 19.6 | 22 | 22 | 246.04504 |
| Q9HA38 | Zinc finger matrin-type protein 3 OS=Hor | ZMAT3 | 32.06 | 3.8 | 1 | 1 | 63.382492 |
| Q9HA64 | Ketosamine-3-kinase OS=Homo sapiens ( | FN3KRP | 34.413 | 10.4 | 2 | 2 | 46.378788 |
| Q9HAF1 | Chromatin modification-related protein M | MEAF6 | 21.635 | 12 | 3 | 3 | 310.67404 |
| Q9HAN9 | Nicotinamide/nicotinic acid mononucleoti | NMNAT1 | 31.933 | 41.6 | 11 | 11 | 1069.8816 |
| Q9HAQ2 | Kinesin-like protein KIF9 OS=Homo sapi | KIF9 | 89.987 | 1.4 | 1 | 1 | 541.35675 |
| Q9HAS0 | Protein Njmu-R1 OS=Homo sapiens OX= | C17orf75 | 44.622 | 33.3 | 12 | 12 | 824.66791 |
| Q9HAU0 | Pleckstrin homology domain-containing fa | PLEKHA5 | 127.464 | 14 | 14 | 14 | 190.49486 |
| Q9HAU4 | E3 ubiquitin-protein ligase SMURF2 OS= | SMURF2 | 86.196 | 10.3 | 7 | 4 | 115.2384 |
| Q9HAU5 | Regulator of nonsense transcripts 2 OS=H | UPF2 | 147.811 | 31.4 | 44 | 44 | 3170.0015 |
| Q9HAV0 | Guanine nucleotide-binding protein subun | GNB4 | 37.568 | 5.6 | 2 | 2 | 82.296158 |

| | | | | | | | |
|---|---|---|---|---|---|---|---|
| Q9HAV4 | Exportin-5 OS=Homo sapiens OX=9606 ( | XPO5 | 136.312 | 1.7 | 2 | 2 | 38.318657 |
| Q9HAW4 | Claspin OS=Homo sapiens OX=9606 GN | CLSPN | 151.094 | 8.1 | 10 | 10 | 110.88468 |
| Q9HB07 | MYG1 exonuclease OS=Homo sapiens O | MYG1 | 42.478 | 17.6 | 6 | 6 | 343.36447 |
| Q9HB19 | Pleckstrin homology domain-containing fa | PLEKHA2 | 47.256 | 9.2 | 4 | 4 | 76.479012 |
| Q9HB21 | Pleckstrin homology domain-containing fa | PLEKHA1 | 45.554 | 2.7 | 1 | 1 | 28.723467 |
| Q9HB58 | Sp110 nuclear body protein OS=Homo sa | SP110 | 78.396 | 11.9 | 7 | 7 | 115.85871 |
| Q9HB71 | Calcyclin-binding protein OS=Homo sapi | CACYBP | 26.21 | 27.6 | 6 | 6 | 283.08853 |
| Q9HBD1 | Roquin-2 OS=Homo sapiens OX=9606 G | RC3H2 | 131.669 | 17.8 | 20 | 13 | 1524.5211 |
| Q9HBE1 | POZ-, AT hook-, and zinc finger-containi | PATZ1 | 74.06 | 11.9 | 7 | 6 | 549.20276 |
| Q9HBH0 | Rho-related GTP-binding protein RhoF O | RHOF | 23.626 | 28 | 5 | 5 | 163.34534 |
| Q9HBH1 | Peptide deformylase, mitochondrial OS=H | PDF | 27.014 | 19.8 | 5 | 5 | 303.8074 |
| Q9HBL0 | Tensin-1 OS=Homo sapiens OX=9606 GI | TNS1 | 197.023 | 7.3 | 10 | 9 | 127.46959 |
| Q9HC07 | Putative divalent cation/proton antiporter | TMEM165 | 34.906 | 3.4 | 1 | 1 | 79.222252 |
| Q9HC21 | Mitochondrial thiamine pyrophosphate ca | SLC25A19 | 35.512 | 26.9 | 9 | 9 | 278.11111 |
| Q9HC35 | Echinoderm microtubule-associated prote | EML4 | 108.917 | 6.7 | 5 | 5 | 54.968777 |
| Q9HC36 | rRNA methyltransferase 3, mitochondrial | MRM3 | 47.021 | 25 | 11 | 11 | 948.48468 |
| Q9HC52 | Chromobox protein homolog 8 OS=Homo | CBX8 | 43.396 | 23.1 | 9 | 9 | 826.49823 |
| Q9HC62 | Sentrin-specific protease 2 OS=Homo sap | SENP2 | 67.855 | 4.8 | 2 | 2 | 48.323601 |
| Q9HC77 | Centromere protein J OS=Homo sapiens ( | CENPJ | 153.001 | 5.8 | 7 | 7 | 85.784805 |
| Q9HC98 | Serine/threonine-protein kinase Nek6 OS= | NEK6 | 35.715 | 8.6 | 3 | 3 | 60.043701 |
| Q9HCC0 | Methylcrotonoyl-CoA carboxylase beta ch | MCCC2 | 61.334 | 13.7 | 6 | 6 | 58.680721 |
| Q9HCD5 | Nuclear receptor coactivator 5 OS=Homo | NCOA5 | 65.537 | 24.2 | 11 | 11 | 426.69284 |
| Q9HCD6 | Protein TANC2 OS=Homo sapiens OX=9 | TANC2 | 219.65 | 11.4 | 19 | 19 | 154.13518 |
| Q9HCE1 | Helicase MOV-10 OS=Homo sapiens OX | MOV10 | 113.672 | 42.5 | 40 | 40 | 10313.957 |
| Q9HCE7 | E3 ubiquitin-protein ligase SMURF1 OS= | SMURF1 | 86.114 | 2.8 | 2 | 2 | 69.782585 |
| Q9HCG8 | Pre-mRNA-splicing factor CWC22 homol | CWC22 | 105.467 | 3.3 | 3 | 3 | 52.473747 |
| Q9HCH0 | Nck-associated protein 5-like OS=Homo s | NCKAP5L | 139.462 | 0.6 | 1 | 1 | 54.072746 |
| Q9HCI7 | E3 ubiquitin-protein ligase MSL2 OS=Ho | MSL2 | 62.541 | 2.3 | 1 | 1 | 20.37252 |
| Q9HCJ0 | Trinucleotide repeat-containing gene 6C p | TNRC6C | 175.965 | 8 | 10 | 10 | 87.634491 |
| Q9HCK5 | Protein argonaute-4 OS=Homo sapiens O | AGO4 | 97.097 | 1.5 | 1 | 1 | 25.587517 |
| Q9HCK8 | Chromodomain-helicase-DNA-binding pr | CHD8 | 290.52 | 2 | 4 | 4 | 36.30476 |
| Q9HCL2 | Glycerol-3-phosphate acyltransferase 1, m | GPAM | 93.795 | 12.2 | 9 | 9 | 159.56613 |
| Q9HCM4 | Band 4.1-like protein 5 OS=Homo sapiens | EPB41L5 | 81.857 | 15.6 | 10 | 8 | 439.71002 |
| Q9HCP0 | Casein kinase I isoform gamma-1 OS=Ho | CSNK1G1 | 48.512 | 5.2 | 2 | 2 | 28.936239 |
| Q9HCS7 | Pre-mRNA-splicing factor SYF1 OS=Hor | XAB2 | 100.01 | 20.4 | 15 | 15 | 295.11438 |

| | | | | | | | |
|---|---|---|---|---|---|---|---|
| Q9HCU5 | Prolactin regulatory element-binding prot | PREB | 45.469 | 17.5 | 6 | 6 | 294.71503 |
| Q9HD42 | Charged multivesicular body protein 1a O | CHMP1A | 21.703 | 21.4 | 6 | 6 | 511.02606 |
| Q9HD67 | Unconventional myosin-X OS=Homo sap | MYO10 | 237.347 | 8.3 | 15 | 15 | 143.13531 |
| Q9HDC5 | Junctophilin-1 OS=Homo sapiens OX=96 | JPH1 | 71.686 | 15.3 | 10 | 10 | 176.98724 |
| Q9NNW5 | tRNA (34-2'-O)-methyltransferase regulat | WDR6 | 121.725 | 23.6 | 25 | 25 | 1721.9045 |
| Q9NP50 | SIN3-HDAC complex-associated factor C | SINHCAF | 24.853 | 11.8 | 3 | 3 | 93.74556 |
| Q9NP61 | ADP-ribosylation factor GTPase-activatin | ARFGAP3 | 56.929 | 6.8 | 3 | 3 | 52.505001 |
| Q9NP64 | Zinc finger CCHC domain-containing pro | ZCCHC17 | 27.57 | 28.6 | 6 | 6 | 770.23267 |
| Q9NP66 | High mobility group protein 20A OS=Ho | HMG20A | 40.144 | 2.6 | 1 | 1 | 109.97334 |
| Q9NP72 | Ras-related protein Rab-18 OS=Homo sa | RAB18 | 22.978 | 26.2 | 5 | 5 | 325.35211 |
| Q9NP73 | Putative bifunctional UDP-N-acetylgluco | ALG13 | 126.057 | 6.4 | 7 | 7 | 83.800133 |
| Q9NP77 | RNA polymerase II subunit A C-terminal | SSU72 | 22.575 | 6.2 | 1 | 1 | 63.723423 |
| Q9NP79 | Vacuolar protein sorting-associated protei | VTA1 | 33.88 | 15 | 6 | 6 | 140.90923 |
| Q9NP81 | Serine--tRNA ligase, mitochondrial OS=H | SARS2 | 58.283 | 22.8 | 11 | 11 | 452.06824 |
| Q9NP87 | DNA-directed DNA/RNA polymerase mu | POLM | 54.816 | 7.9 | 3 | 3 | 79.761902 |
| Q9NP97 | Dynein light chain roadblock-type 1 OS=I | DYNLRB1 | 10.922 | 29.2 | 2 | 1 | 114.38964 |
| Q9NPA8 | Transcription and mRNA export factor EN | ENY2 | 11.529 | 8.9 | 1 | 1 | 233.82394 |
| Q9NPC8 | Homeobox protein SIX2 OS=Homo sapie | SIX2 | 32.287 | 14.4 | 4 | 2 | 99.534637 |
| Q9NPD3 | Exosome complex component RRP41 OS | EXOSC4 | 26.383 | 24.9 | 5 | 5 | 463.17154 |
| Q9NPE2 | Neugrin OS=Homo sapiens OX=9606 GN | NGRN | 32.408 | 48.1 | 11 | 11 | 1406.8707 |
| Q9NPE3 | H/ACA ribonucleoprotein complex subun | NOP10 | 7.706 | 57.8 | 3 | 3 | 435.27655 |
| Q9NPF0 | CD320 antigen OS=Homo sapiens OX=9 | CD320 | 28.992 | 3.9 | 1 | 1 | 16.636059 |
| Q9NPF5 | DNA methyltransferase 1-associated prot | DMAP1 | 52.993 | 21 | 9 | 9 | 549.10553 |
| Q9NPI1 | Bromodomain-containing protein 7 OS=H | BRD7 | 74.139 | 1.5 | 1 | 1 | 111.59661 |
| Q9NPI6 | mRNA-decapping enzyme 1A OS=Homo | DCP1A | 63.279 | 18.9 | 9 | 9 | 741.10223 |
| Q9NPJ6 | Mediator of RNA polymerase II transcript | MED4 | 29.746 | 16.3 | 4 | 4 | 90.082245 |
| Q9NPL8 | Complex I assembly factor TIMMDC1, m | TIMMDC1 | 32.178 | 7 | 2 | 2 | 100.35222 |
| Q9NQ29 | Putative RNA-binding protein Luc7-like 1 | LUC7L | 43.728 | 34 | 13 | 7 | 3417.5361 |
| Q9NQ55 | Suppressor of SWI4 1 homolog OS=Hom | PPAN | 53.194 | 18.4 | 8 | 8 | 337.7652 |
| Q9NQG5 | Regulation of nuclear pre-mRNA domain- | RPRD1B | 36.9 | 10.7 | 3 | 3 | 32.908348 |
| Q9NQP4 | Prefoldin subunit 4 OS=Homo sapiens OX | PFDN4 | 15.315 | 11.2 | 1 | 1 | 20.151911 |
| Q9NQS7 | Inner centromere protein OS=Homo sapie | INCENP | 105.429 | 10.8 | 9 | 9 | 149.04048 |
| Q9NQT4 | Exosome complex component RRP46 OS | EXOSC5 | 25.249 | 19.1 | 4 | 4 | 472.06396 |
| Q9NQT5 | Exosome complex component RRP40 OS | EXOSC3 | 29.573 | 27.3 | 5 | 5 | 369.25183 |
| Q9NQT8 | Kinesin-like protein KIF13B OS=Homo s | KIF13B | 202.789 | 5.6 | 9 | 9 | 49.900291 |

| | | | | | | | |
|---|---|---|---|---|---|---|---|
| Q9NQW6 | Anillin OS=Homo sapiens OX=9606 GN= | ANLN | 124.199 | 20 | 20 | 20 | 226.77666 |
| Q9NQZ2 | Something about silencing protein 10 OS= | UTP3 | 54.558 | 8.8 | 5 | 5 | 193.50618 |
| Q9NR12 | PDZ and LIM domain protein 7 OS=Hom | PDLIM7 | 49.845 | 29.1 | 11 | 11 | 827.99603 |
| Q9NR30 | Nucleolar RNA helicase 2 OS=Homo sap | DDX21 | 87.345 | 35 | 29 | 27 | 3315.1838 |
| Q9NR31 | GTP-binding protein SAR1a OS=Homo s | SAR1A | 22.367 | 11.1 | 2 | 1 | 72.135742 |
| Q9NR50 | Translation initiation factor eIF-2B subun | EIF2B3 | 50.241 | 11.7 | 4 | 4 | 66.909157 |
| Q9NR56 | Muscleblind-like protein 1 OS=Homo sap | MBNL1 | 41.818 | 18.8 | 7 | 2 | 1385.8019 |
| Q9NR81 | Rho guanine nucleotide exchange factor 3 | ARHGEF3 | 59.783 | 7.4 | 4 | 4 | 62.564011 |
| Q9NR97 | Toll-like receptor 8 OS=Homo sapiens O. | TLR8 | 119.829 | 0.7 | 1 | 1 | 788.54034 |
| Q9NRA8 | Eukaryotic translation initiation factor 4E | EIF4ENIF1 | 108.201 | 18.8 | 13 | 13 | 273.58401 |
| Q9NRC1 | Suppressor of tumorigenicity 7 protein OS | ST7 | 67.167 | 11.5 | 7 | 3 | 135.39798 |
| Q9NRC8 | NAD-dependent protein deacetylase sirtui | SIRT7 | 44.899 | 15.5 | 5 | 5 | 250.90042 |
| Q9NRG0 | Chromatin accessibility complex protein 1 | CHRAC1 | 14.711 | 6.1 | 1 | 1 | 122.99395 |
| Q9NRG9 | Aladin OS=Homo sapiens OX=9606 GN= | AAAS | 59.575 | 8.2 | 4 | 4 | 200.69527 |
| Q9NRL2 | Bromodomain adjacent to zinc finger dom | BAZ1A | 178.703 | 16.8 | 22 | 22 | 217.82166 |
| Q9NRM7 | Serine/threonine-protein kinase LATS2 O | LATS2 | 120.136 | 4.9 | 4 | 3 | 104.671 |
| Q9NRP0 | Oligosaccharyltransferase complex subun | OSTC | 16.83 | 8.1 | 1 | 1 | 90.693085 |
| Q9NRR4 | Ribonuclease 3 OS=Homo sapiens OX=9 | DROSHA | 159.316 | 18.1 | 23 | 23 | 720.57654 |
| Q9NRR8 | CDC42 small effector protein 1 OS=Hom | CDC42SE1 | 8.926 | 15.2 | 1 | 1 | 38.355103 |
| Q9NRW3 | DNA dC->dU-editing enzyme APOBEC-3 | APOBEC3C | 22.826 | 17.4 | 3 | 3 | 759.71088 |
| Q9NRX1 | RNA-binding protein PNO1 OS=Homo sa | PNO1 | 27.925 | 25.8 | 7 | 7 | 2014.3982 |
| Q9NRZ9 | Lymphoid-specific helicase OS=Homo sa | HELLS | 97.074 | 23 | 18 | 18 | 313.35541 |
| Q9NS15 | Latent-transforming growth factor beta-bi | LTBP3 | 139.36 | 0.8 | 1 | 1 | 89.908417 |
| Q9NS23 | Ras association domain-containing protein | RASSF1 | 39.22 | 2 | 1 | 1 | 57.598526 |
| Q9NS37 | CREB/ATF bZIP transcription factor OS= | CREBZF | 37.134 | 3.1 | 1 | 1 | 11.915234 |
| Q9NS69 | Mitochondrial import receptor subunit TC | TOMM22 | 15.522 | 16.2 | 2 | 2 | 145.85477 |
| Q9NS86 | LanC-like protein 2 OS=Homo sapiens O. | LANCL2 | 50.855 | 24 | 10 | 10 | 281.6691 |
| Q9NS91 | E3 ubiquitin-protein ligase RAD18 OS=H | RAD18 | 56.223 | 6.3 | 3 | 3 | 55.399094 |
| Q9NSD9 | Phenylalanine--tRNA ligase beta subunit ( | FARSB | 66.116 | 7.5 | 4 | 4 | 39.423538 |
| Q9NSE4 | Isoleucine--tRNA ligase, mitochondrial O | IARS2 | 113.792 | 3.3 | 3 | 3 | 60.332226 |
| Q9NSP4 | Centromere protein M OS=Homo sapiens | CENPM | 19.737 | 5.6 | 1 | 1 | 50.136795 |
| Q9NSV4 | Protein diaphanous homolog 3 OS=Homo | DIAPH3 | 136.926 | 10 | 11 | 11 | 87.901489 |
| Q9NSY1 | BMP-2-inducible protein kinase OS=Hom | BMP2K | 129.172 | 2.2 | 2 | 2 | 78.741852 |
| Q9NT62 | Ubiquitin-like-conjugating enzyme ATG3 | ATG3 | 35.865 | 7.3 | 2 | 2 | 48.960197 |
| Q9NTI5 | Sister chromatid cohesion protein PDS5 h | PDS5B | 164.668 | 6 | 8 | 8 | 116.33061 |

| | | | | | | | |
|---|---|---|---|---|---|---|---|
| Q9NTJ3 | Structural maintenance of chromosomes p | SMC4 | 147.182 | 22.5 | 27 | 27 | 428.04919 |
| Q9NTJ5 | Phosphatidylinositol-3-phosphatase SAC1 | SACM1L | 66.967 | 4.9 | 3 | 3 | 27.139763 |
| Q9NTK5 | Obg-like ATPase 1 OS=Homo sapiens OX | OLA1 | 44.744 | 5.6 | 2 | 2 | 79.754272 |
| Q9NTW7 | Zinc finger protein 64 OS=Homo sapiens | ZFP64 | 72.217 | 4.8 | 2 | 2 | 45.250538 |
| Q9NUD5 | Zinc finger CCHC domain-containing pro | ZCCHC3 | 43.548 | 27.8 | 12 | 12 | 883.93921 |
| Q9NUI1 | Peroxisomal 2,4-dienoyl-CoA reductase [( | DECR2 | 30.778 | 39 | 9 | 9 | 267.45447 |
| Q9NUK0 | Muscleblind-like protein 3 OS=Homo sap | MBNL3 | 38.532 | 10.5 | 4 | 4 | 593.6698 |
| Q9NUL3 | Double-stranded RNA-binding protein Sta | STAU2 | 62.609 | 44.7 | 28 | 26 | 2793.4922 |
| Q9NUL5 | Shiftless antiviral inhibitor of ribosomal fi | SHFL | 33.111 | 14.1 | 5 | 5 | 487.39053 |
| Q9NUL7 | Probable ATP-dependent RNA helicase D | DDX28 | 59.582 | 34.3 | 17 | 17 | 1689.8258 |
| Q9NUM4 | Transmembrane protein 106B OS=Homo | TMEM106B | 31.128 | 8.4 | 2 | 2 | 159.93643 |
| Q9NUP9 | Protein lin-7 homolog C OS=Homo sapie | LIN7C | 21.834 | 39.1 | 6 | 4 | 380.45724 |
| Q9NUQ2 | 1-acyl-sn-glycerol-3-phosphate acyltransf | AGPAT5 | 42.073 | 6.9 | 2 | 2 | 35.10334 |
| Q9NUQ3 | Gamma-taxilin OS=Homo sapiens OX=9( | TXLNG | 60.586 | 20.5 | 13 | 13 | 266.53049 |
| Q9NUU7 | ATP-dependent RNA helicase DDX19A ( | DDX19A | 53.976 | 9 | 3 | 3 | 53.573902 |
| Q9NUW8 | Tyrosyl-DNA phosphodiesterase 1 OS=H | TDP1 | 68.421 | 7.7 | 4 | 4 | 81.473862 |
| Q9NV06 | DDB1- and CUL4-associated factor 13 O | DCAF13 | 51.403 | 34.4 | 16 | 16 | 470.25638 |
| Q9NV31 | U3 small nucleolar ribonucleoprotein prot | IMP3 | 21.851 | 30.4 | 4 | 4 | 455.31027 |
| Q9NV56 | MRG/MORF4L-binding protein OS=Hon | MRGBP | 22.418 | 13.7 | 2 | 2 | 68.825333 |
| Q9NVC6 | Mediator of RNA polymerase II transcript | MED17 | 72.891 | 14.1 | 8 | 8 | 94.150459 |
| Q9NVF7 | F-box only protein 28 OS=Homo sapiens | FBXO28 | 41.15 | 14.7 | 5 | 5 | 104.96358 |
| Q9NVH0 | Exonuclease 3'-5' domain-containing prot | EXD2 | 70.353 | 33.7 | 19 | 19 | 410.1716 |
| Q9NVH1 | DnaJ homolog subfamily C member 11 O | DNAJC11 | 63.279 | 13.4 | 6 | 6 | 94.088669 |
| Q9NVI7 | ATPase family AAA domain-containing p | ATAD3A | 71.37 | 28.5 | 18 | 4 | 1891.1804 |
| Q9NVJ2 | ADP-ribosylation factor-like protein 8B C | ARL8B | 21.54 | 32.3 | 5 | 1 | 233.53093 |
| Q9NVM9 | Integrator complex subunit 13 OS=Homo | INTS13 | 80.226 | 11.2 | 7 | 7 | 74.623764 |
| Q9NVN8 | Guanine nucleotide-binding protein-like 3 | GNL3L | 65.574 | 36.9 | 20 | 20 | 1425.4446 |
| Q9NVP1 | ATP-dependent RNA helicase DDX18 OS | DDX18 | 75.407 | 38.4 | 22 | 22 | 825.28625 |
| Q9NVU0 | DNA-directed RNA polymerase III subun | POLR3E | 79.899 | 7.3 | 5 | 5 | 90.050812 |
| Q9NVU7 | Protein SDA1 homolog OS=Homo sapien | SDAD1 | 79.872 | 9.6 | 6 | 6 | 145.5193 |
| Q9NVV4 | Poly(A) RNA polymerase, mitochondrial | MTPAP | 66.172 | 23 | 13 | 13 | 2071.5383 |
| Q9NVX0 | HAUS augmin-like complex subunit 2 OS | HAUS2 | 26.934 | 22.6 | 5 | 5 | 143.74364 |
| Q9NVX2 | Notchless protein homolog 1 OS=Homo s | NLE1 | 53.321 | 15.1 | 5 | 5 | 160.06129 |
| Q9NW08 | DNA-directed RNA polymerase III subun | POLR3B | 127.785 | 7.9 | 9 | 9 | 176.94701 |
| Q9NW13 | RNA-binding protein 28 OS=Homo sapie | RBM28 | 85.738 | 28.5 | 21 | 21 | 449.47113 |

| | | | | | | | |
|---|---|---|---|---|---|---|---|
| Q9NW64 | Pre-mRNA-splicing factor RBM22 OS=H | RBM22 | 46.896 | 16.7 | 7 | 7 | 885.79419 |
| Q9NW75 | G patch domain-containing protein 2 OS= | GPATCH2 | 58.944 | 34.5 | 14 | 14 | 1069.2822 |
| Q9NW81 | Distal membrane-arm assembly complex | DMAC2 | 29.268 | 21.8 | 5 | 5 | 237.83432 |
| Q9NWF9 | E3 ubiquitin-protein ligase RNF216 OS=I | RNF216 | 99.406 | 17.9 | 15 | 15 | 174.77571 |
| Q9NWH9 | SAFB-like transcription modulator OS=H | SLTM | 117.149 | 10.1 | 10 | 10 | 163.09619 |
| Q9NWK9 | Box C/D snoRNA protein 1 OS=Homo sa | ZNHIT6 | 53.919 | 30.6 | 13 | 13 | 184.50763 |
| Q9NWQ4 | G patch domain-containing protein 2-like | GPATCH2I | 54.26 | 30.7 | 11 | 11 | 185.9612 |
| Q9NWR8 | Calcium uniporter regulatory subunit MC | MCUB | 39.083 | 3.9 | 1 | 1 | 18.963221 |
| Q9NWS0 | PIH1 domain-containing protein 1 OS=H | PIH1D1 | 32.363 | 3.4 | 1 | 1 | 104.3045 |
| Q9NWT1 | p21-activated protein kinase-interacting p | PAK1IP1 | 43.964 | 34.2 | 11 | 11 | 813.37396 |
| Q9NWT8 | Small ribosomal subunit protein mS38 OS | AURKAIP1 | 22.355 | 11.6 | 2 | 2 | 549.02832 |
| Q9NWU5 | Large ribosomal subunit protein uL22m C | MRPL22 | 23.641 | 34.5 | 7 | 7 | 1102.1149 |
| Q9NWV8 | BRISC and BRCA1-A complex member 1 | BABAM1 | 36.561 | 15.2 | 5 | 5 | 202.49861 |
| Q9NWZ5 | Uridine-cytidine kinase-like 1 OS=Homo | UCKL1 | 61.142 | 2 | 1 | 1 | 73.975525 |
| Q9NWZ8 | Gem-associated protein 8 OS=Homo sapi | GEMIN8 | 28.637 | 29.3 | 8 | 8 | 1164.855 |
| Q9NX04 | AFG2-interacting ribosome maturation fa | AIRIM | 23.373 | 14.3 | 3 | 3 | 133.73868 |
| Q9NX07 | tRNA selenocysteine 1-associated protein | TRNAU1AI | 32.499 | 4.9 | 1 | 1 | 60.481647 |
| Q9NX09 | DNA damage-inducible transcript 4 prote | DDIT4 | 25.371 | 15.9 | 3 | 3 | 49.734772 |
| Q9NX24 | H/ACA ribonucleoprotein complex subun | NHP2 | 17.202 | 5.9 | 1 | 1 | 503.64636 |
| Q9NX46 | ADP-ribosylhydrolase ARH3 OS=Homo | ADPRS | 38.947 | 3.9 | 1 | 1 | 37.690998 |
| Q9NX57 | Ras-related protein Rab-20 OS=Homo sa | RAB20 | 26.278 | 24.8 | 5 | 5 | 153.34439 |
| Q9NX58 | Cell growth-regulating nucleolar protein | LYAR | 43.634 | 23 | 8 | 8 | 765.12921 |
| Q9NX63 | MICOS complex subunit MIC19 OS=Hor | CHCHD3 | 26.153 | 15.4 | 4 | 4 | 152.04272 |
| Q9NX74 | tRNA-dihydrouridine(20) synthase [NAD | DUS2 | 55.051 | 26.2 | 12 | 12 | 291.75378 |
| Q9NX95 | Syntabulin OS=Homo sapiens OX=9606 | SYBU | 72.389 | 7.2 | 4 | 3 | 69.269211 |
| Q9NXC5 | GATOR2 complex protein MIOS OS=Ho | MIOS | 98.585 | 9.7 | 7 | 7 | 98.991524 |
| Q9NXE4 | Sphingomyelin phosphodiesterase 4 OS=I | SMPD4 | 97.81 | 1 | 1 | 1 | 62.61726 |
| Q9NXE8 | Pre-mRNA-splicing factor CWC25 homol | CWC25 | 49.648 | 32.9 | 17 | 17 | 1108.7443 |
| Q9NXH9 | tRNA (guanine(26)-N(2))-dimethyltransfe | TRMT1 | 72.235 | 23.8 | 15 | 15 | 349.92065 |
| Q9NXL9 | DNA helicase MCM9 OS=Homo sapiens | MCM9 | 127.313 | 4.2 | 4 | 4 | 63.453934 |
| Q9NXR5 | Ankyrin repeat domain-containing protein | ANKRD10 | 44.768 | 7.4 | 3 | 3 | 53.062542 |
| Q9NXR7 | BRISC and BRCA1-A complex member 2 | BABAM2 | 43.552 | 14.4 | 6 | 6 | 665.62415 |
| Q9NXS2 | Glutaminyl-peptide cyclotransferase-like | QPCTL | 42.925 | 19.9 | 9 | 9 | 591.20886 |
| Q9NXV2 | BTB/POZ domain-containing protein KC | KCTD5 | 26.093 | 20.5 | 4 | 4 | 123.47267 |
| Q9NXV6 | CDKN2A-interacting protein OS=Homo s | CDKN2AIP | 61.125 | 32.2 | 16 | 16 | 741.17462 |

| | | | | | | | |
|---|---|---|---|---|---|---|---|
| Q9NXW2 | DnaJ homolog subfamily B member 12 O | DNAJB12 | 41.86 | 25.9 | 9 | 9 | 332.54373 |
| Q9NXX6 | Non-structural maintenance of chromoson | NSMCE4A | 44.302 | 16.4 | 6 | 5 | 219.88062 |
| Q9NY12 | H/ACA ribonucleoprotein complex subun | GAR1 | 22.348 | 17.5 | 4 | 4 | 1939.2329 |
| Q9NY61 | Protein AATF OS=Homo sapiens OX=96 | AATF | 63.133 | 8.6 | 4 | 4 | 81.27066 |
| Q9NY93 | Probable ATP-dependent RNA helicase D | DDX56 | 61.59 | 31.1 | 17 | 17 | 1208.7915 |
| Q9NYA1 | Sphingosine kinase 1 OS=Homo sapiens ( | SPHK1 | 42.518 | 2.1 | 1 | 1 | 40.338108 |
| Q9NYB0 | Telomeric repeat-binding factor 2-interact | TERF2IP | 44.26 | 13.8 | 5 | 5 | 88.506531 |
| Q9NYB9 | Abl interactor 2 OS=Homo sapiens OX=9 | ABI2 | 55.664 | 13.3 | 7 | 4 | 166.21507 |
| Q9NYD6 | Homeobox protein Hox-C10 OS=Homo s | HOXC10 | 38.073 | 51.2 | 14 | 12 | 1372.476 |
| Q9NYF8 | Bcl-2-associated transcription factor 1 OS | BCLAF1 | 106.123 | 14.5 | 15 | 15 | 1525.2242 |
| Q9NYH9 | U3 small nucleolar RNA-associated prote | UTP6 | 70.194 | 16.2 | 10 | 10 | 251.455 |
| Q9NYK5 | Large ribosomal subunit protein mL39 OS | MRPL39 | 38.712 | 45 | 16 | 16 | 1603.297 |
| Q9NYL2 | Mitogen-activated protein kinase kinase k | MAP3K20 | 91.156 | 23.4 | 17 | 17 | 611.3136 |
| Q9NYP7 | Elongation of very long chain fatty acids p | ELOVL5 | 35.294 | 4.3 | 1 | 1 | 122.57705 |
| Q9NYS0 | NF-kappa-B inhibitor-interacting Ras-like | NKIRAS1 | 21.643 | 10.4 | 2 | 2 | 75.532516 |
| Q9NYV4 | Cyclin-dependent kinase 12 OS=Homo sa | CDK12 | 164.155 | 9.9 | 11 | 9 | 181.95222 |
| Q9NYW8 | RB-associated KRAB zinc finger protein ( | RBAK | 82.995 | 2.5 | 2 | 2 | 70.399361 |
| Q9NYY3 | Serine/threonine-protein kinase PLK2 OS | PLK2 | 78.237 | 12.4 | 7 | 6 | 103.44685 |
| Q9NYY8 | FAST kinase domain-containing protein 2 | FASTKD2 | 81.463 | 22.1 | 14 | 14 | 1257.4161 |
| Q9NYZ3 | G2 and S phase-expressed protein 1 OS=I | GTSE1 | 76.645 | 13.9 | 8 | 8 | 174.83371 |
| Q9NZ01 | Very-long-chain enoyl-CoA reductase OS | TECR | 36.035 | 32.5 | 11 | 11 | 2021.9929 |
| Q9NZ45 | CDGSH iron-sulfur domain-containing pr | CISD1 | 12.2 | 34.3 | 3 | 3 | 111.34715 |
| Q9NZB2 | Constitutive coactivator of PPAR-gamma- | FAM120A | 121.888 | 45.1 | 42 | 38 | 8990.8887 |
| Q9NZC9 | SWI/SNF-related matrix-associated actin- | SMARCAL | 105.938 | 16.9 | 14 | 14 | 254.7572 |
| Q9NZE8 | Large ribosomal subunit protein bL35m C | MRPL35 | 21.515 | 9.6 | 2 | 2 | 207.53896 |
| Q9NZI7 | Upstream-binding protein 1 OS=Homo sa | UBP1 | 60.492 | 13 | 6 | 6 | 346.35556 |
| Q9NZI8 | Insulin-like growth factor 2 mRNA-bindir | IGF2BP1 | 63.481 | 33.4 | 20 | 20 | 12785.201 |
| Q9NZJ0 | Denticleless protein homolog OS=Homo s | DTL | 79.468 | 11.4 | 7 | 7 | 147.99689 |
| Q9NZJ7 | Mitochondrial carrier homolog 1 OS=Hor | MTCH1 | 41.545 | 19.5 | 7 | 7 | 257.35284 |
| Q9NZL4 | Hsp70-binding protein 1 OS=Homo sapie | HSPBP1 | 39.303 | 11.4 | 4 | 4 | 96.535789 |
| Q9NZM1 | Myoferlin OS=Homo sapiens OX=9606 C | MYOF | 234.709 | 2.3 | 4 | 4 | 48.713352 |
| Q9NZM5 | Ribosome biogenesis protein NOP53 OS= | NOP53 | 54.39 | 13 | 6 | 6 | 374.2326 |
| Q9NZN4 | EH domain-containing protein 2 OS=Hon | EHD2 | 61.162 | 10.9 | 4 | 4 | 65.901169 |
| Q9NZN5 | Rho guanine nucleotide exchange factor 1 | ARHGEF12 | 173.233 | 0.6 | 1 | 1 | 51.211185 |
| Q9NZN8 | CCR4-NOT transcription complex subuni | CNOT2 | 59.738 | 23 | 11 | 11 | 321.53802 |

| | | | | | | | |
|---|---|---|---|---|---|---|---|
| Q9P021 | Cysteine-rich PDZ-binding protein OS=H | CRIPT | 11.216 | 20.8 | 3 | 3 | 291.97449 |
| Q9P035 | Very-long-chain (3R)-3-hydroxyacyl-CoA | HACD3 | 43.16 | 23.2 | 9 | 9 | 929.16223 |
| Q9P0B6 | Coiled-coil domain-containing protein 16 | CCDC167 | 11.46 | 15.5 | 1 | 1 | 33.324852 |
| Q9P0J0 | NADH dehydrogenase [ubiquinone] 1 alp | NDUFA13 | 16.699 | 31.2 | 4 | 4 | 125.20432 |
| Q9P0J7 | E3 ubiquitin-protein ligase KCMF1 OS=H | KCMF1 | 41.946 | 6.3 | 2 | 2 | 23.029144 |
| Q9P0L0 | Vesicle-associated membrane protein-ass | VAPA | 27.894 | 37.8 | 10 | 9 | 552.08276 |
| Q9P0L2 | Serine/threonine-protein kinase MARK1 ( | MARK1 | 89.003 | 25.5 | 16 | 16 | 554.17047 |
| Q9P0S9 | Transmembrane protein 14C OS=Homo s | TMEM14C | 11.565 | 8.9 | 1 | 1 | 46.069748 |
| Q9P0T4 | Zinc finger protein 581 OS=Homo sapien | ZNF581 | 22.011 | 19.3 | 4 | 4 | 130.88196 |
| Q9P0U3 | Sentrin-specific protease 1 OS=Homo sap | SENP1 | 73.481 | 2.6 | 2 | 2 | 61.7645 |
| Q9P0V3 | SH3 domain-binding protein 4 OS=Homo | SH3BP4 | 107.496 | 18.3 | 15 | 15 | 167.27524 |
| Q9P0V9 | Septin-10 OS=Homo sapiens OX=9606 G | SEPTIN10 | 52.593 | 5.7 | 2 | 2 | 37.736042 |
| Q9P1Y5 | Calmodulin-regulated spectrin-associated | CAMSAP3 | 134.75 | 13.9 | 14 | 14 | 215.57132 |
| Q9P1Z2 | Calcium-binding and coiled-coil domain-c | CALCOCO | 77.337 | 3.3 | 2 | 2 | 43.659668 |
| Q9P209 | Centrosomal protein of 72 kDa OS=Hom | CEP72 | 71.718 | 14.7 | 7 | 7 | 142.93285 |
| Q9P253 | Vacuolar protein sorting-associated prote | VPS18 | 110.186 | 1.5 | 1 | 1 | 68.132004 |
| Q9P258 | Protein RCC2 OS=Homo sapiens OX=96 | RCC2 | 56.085 | 41 | 18 | 18 | 3674.1094 |
| Q9P270 | SLAIN motif-containing protein 2 OS=H | SLAIN2 | 62.544 | 22.7 | 11 | 11 | 696.08636 |
| Q9P275 | Ubiquitin carboxyl-terminal hydrolase 36 | USP36 | 122.908 | 11.8 | 12 | 12 | 129.9603 |
| Q9P287 | BRCA2 and CDKN1A-interacting protein | BCCIP | 35.98 | 15.9 | 4 | 4 | 65.298561 |
| Q9P289 | Serine/threonine-protein kinase 26 OS=H | STK26 | 46.529 | 5.8 | 2 | 2 | 72.092003 |
| Q9P2D0 | Inhibitor of Bruton tyrosine kinase OS=H | IBTK | 150.528 | 20.5 | 25 | 25 | 296.97668 |
| Q9P2D1 | Chromodomain-helicase-DNA-binding pr | CHD7 | 335.928 | 10 | 25 | 24 | 501.80197 |
| Q9P2D3 | HEAT repeat-containing protein 5B OS=I | HEATR5B | 224.303 | 2.8 | 5 | 5 | 54.753094 |
| Q9P2E3 | NFX1-type zinc finger-containing protein | ZNFX1 | 220.227 | 24.2 | 41 | 41 | 754.38153 |
| Q9P2E5 | Chondroitin sulfate glucuronyltransferase | CHPF2 | 85.949 | 23.4 | 17 | 17 | 491.61551 |
| Q9P2E9 | Ribosome-binding protein 1 OS=Homo sa | RRBP1 | 152.457 | 32.6 | 48 | 47 | 3080.4192 |
| Q9P2G4 | Microtubule-associated protein 10 OS=H | MAP10 | 100.345 | 1.8 | 1 | 1 | 228.13339 |
| Q9P2G9 | Kelch-like protein 8 OS=Homo sapiens O | KLHL8 | 68.803 | 1.8 | 1 | 1 | 61.689079 |
| Q9P2I0 | Cleavage and polyadenylation specificity | CPSF2 | 88.487 | 18 | 12 | 12 | 222.13107 |
| Q9P2J5 | Leucine--tRNA ligase, cytoplasmic OS=H | LARS1 | 134.467 | 7.5 | 8 | 8 | 99.56321 |
| Q9P2L0 | WD repeat-containing protein 35 OS=Ho | WDR35 | 133.548 | 1.3 | 1 | 1 | 9.2653837 |
| Q9P2N7 | Kelch-like protein 13 OS=Homo sapiens ( | KLHL13 | 73.868 | 9.3 | 5 | 2 | 89.400932 |
| Q9P2R6 | Arginine-glutamic acid dipeptide repeats | RERE | 172.424 | 2 | 3 | 3 | 103.24677 |
| Q9P2S5 | WD repeat-containing protein WRAP73 ( | WRAP73 | 51.589 | 33.3 | 13 | 13 | 434.84521 |

| | | | | | | | |
|---|---|---|---|---|---|---|---|
| Q9P2W1 | Homologous-pairing protein 2 homolog C | PSMC3IP | 24.907 | 6 | 1 | 1 | 67.801033 |
| Q9P2X0 | Dolichol-phosphate mannosyltransferase s | DPM3 | 10.094 | 10.9 | 1 | 1 | 161.56328 |
| Q9UBB4 | Ataxin-10 OS=Homo sapiens OX=9606 C | ATXN10 | 53.489 | 32.2 | 14 | 14 | 2057.5183 |
| Q9UBB5 | Methyl-CpG-binding domain protein 2 OS | MBD2 | 43.255 | 13.4 | 4 | 4 | 67.785362 |
| Q9UBB6 | Neurochondrin OS=Homo sapiens OX=96 | NCDN | 78.865 | 3.7 | 2 | 2 | 50.590988 |
| Q9UBB9 | Tuftelin-interacting protein 11 OS=Homo | TFIP11 | 96.82 | 27.2 | 20 | 20 | 325.37088 |
| Q9UBC2 | Epidermal growth factor receptor substrat | EPS15L1 | 94.256 | 5.6 | 4 | 4 | 58.432526 |
| Q9UBD5 | Origin recognition complex subunit 3 OS= | ORC3 | 82.255 | 9.8 | 6 | 6 | 66.219994 |
| Q9UBF2 | Coatomer subunit gamma-2 OS=Homo sa | COPG2 | 97.623 | 1.3 | 1 | 1 | 60.058216 |
| Q9UBF6 | RING-box protein 2 OS=Homo sapiens O | RNF7 | 12.684 | 10.6 | 1 | 1 | 46.58849 |
| Q9UBI6 | Guanine nucleotide-binding protein G(I)/( | GNG12 | 8.007 | 19.4 | 1 | 1 | 29.222366 |
| Q9UBK9 | Protein UXT OS=Homo sapiens OX=960 | UXT | 18.247 | 7 | 1 | 1 | 89.547577 |
| Q9UBL3 | Set1/Ash2 histone methyltransferase comp | ASH2L | 68.723 | 1.3 | 1 | 1 | 70.916626 |
| Q9UBM7 | 7-dehydrocholesterol reductase OS=Homo | DHCR7 | 54.49 | 4.4 | 2 | 2 | 40.449638 |
| Q9UBP0 | Spastin OS=Homo sapiens OX=9606 GN | SPAST | 67.198 | 17.7 | 9 | 9 | 195.38412 |
| Q9UBQ0 | Vacuolar protein sorting-associated protei | VPS29 | 20.506 | 5.5 | 1 | 1 | 26.577213 |
| Q9UBQ5 | Eukaryotic translation initiation factor 3 s | EIF3K | 25.06 | 24.8 | 4 | 4 | 1936.0159 |
| Q9UBQ6 | Exostosin-like 2 OS=Homo sapiens OX=9 | EXTL2 | 37.466 | 3.3 | 1 | 1 | 44.182411 |
| Q9UBR2 | Cathepsin Z OS=Homo sapiens OX=9606 | CTSZ | 33.868 | 16.2 | 4 | 4 | 411.90573 |
| Q9UBR4 | LIM/homeobox protein Lhx3 OS=Homo s | LHX3 | 43.358 | 4.3 | 2 | 1 | 98.060051 |
| Q9UBS0 | Ribosomal protein S6 kinase beta-2 OS=H | RPS6KB2 | 53.456 | 3.3 | 1 | 1 | 65.715431 |
| Q9UBS4 | DnaJ homolog subfamily B member 11 O | DNAJB11 | 40.514 | 8.7 | 3 | 3 | 421.51086 |
| Q9UBT6 | DNA polymerase kappa OS=Homo sapier | POLK | 98.809 | 8.4 | 6 | 6 | 145.5871 |
| Q9UBU7 | Protein DBF4 homolog A OS=Homo sapi | DBF4 | 76.858 | 15 | 9 | 9 | 141.04451 |
| Q9UBU8 | Mortality factor 4-like protein 1 OS=Hom | MORF4L1 | 41.474 | 29.6 | 7 | 7 | 291.90741 |
| Q9UBU9 | Nuclear RNA export factor 1 OS=Homo s | NXF1 | 70.183 | 26.5 | 18 | 18 | 922.86707 |
| Q9UBV7 | Beta-1,4-galactosyltransferase 7 OS=Hom | B4GALT7 | 37.406 | 5.8 | 2 | 2 | 61.101116 |
| Q9UBV8 | Peflin OS=Homo sapiens OX=9606 GN= | PEF1 | 30.381 | 18.7 | 5 | 5 | 229.78204 |
| Q9UBW8 | COP9 signalosome complex subunit 7a O | COPS7A | 30.277 | 8 | 1 | 1 | 31.576151 |
| Q9UBX3 | Mitochondrial dicarboxylate carrier OS=H | SLC25A10 | 31.283 | 24.4 | 6 | 6 | 1164.1104 |
| Q9UBX5 | Fibulin-5 OS=Homo sapiens OX=9606 G | FBLN5 | 50.181 | 4.7 | 2 | 2 | 45.232098 |
| Q9UBZ4 | DNA-(apurinic or apyrimidinic site) endo | APEX2 | 57.401 | 14.3 | 6 | 6 | 238.12955 |
| Q9UDW3 | Zinc finger matrin-type protein 5 OS=Hor | ZMAT5 | 19.971 | 7.6 | 1 | 1 | 64.000694 |
| Q9UDY2 | Tight junction protein ZO-2 OS=Homo sa | TJP2 | 133.959 | 29.8 | 33 | 33 | 1764.8285 |
| Q9UDY4 | DnaJ homolog subfamily B member 4 OS | DNAJB4 | 37.807 | 16 | 5 | 5 | 252.5786 |

| | | | | | | | |
|---|---|---|---|---|---|---|---|
| Q9UDY8 | Mucosa-associated lymphoid tissue lymph | MALT1 | 92.273 | 20.6 | 15 | 15 | 207.72441 |
| Q9UEG4 | Zinc finger protein 629 OS=Homo sapiens | ZNF629 | 96.621 | 8.6 | 7 | 7 | 108.84198 |
| Q9UER7 | Death domain-associated protein 6 OS=H | DAXX | 81.373 | 10.4 | 6 | 6 | 123.70121 |
| Q9UET6 | tRNA (cytidine(32)/guanosine(34)-2'-O)-r | FTSJ1 | 36.08 | 12.5 | 4 | 4 | 1048.6079 |
| Q9UEU0 | Vesicle transport through interaction with | VTI1B | 26.689 | 5.2 | 1 | 1 | 34.654243 |
| Q9UEY8 | Gamma-adducin OS=Homo sapiens OX=9 | ADD3 | 79.155 | 18 | 11 | 11 | 221.14845 |
| Q9UFC0 | Leucine-rich repeat and WD repeat-contai | LRWD1 | 70.862 | 4.2 | 2 | 2 | 63.111053 |
| Q9UFF9 | CCR4-NOT transcription complex subuni | CNOT8 | 33.541 | 19.2 | 5 | 4 | 144.14563 |
| Q9UFW8 | CGG triplet repeat-binding protein 1 OS= | CGGBP1 | 18.821 | 42.5 | 8 | 8 | 154.53738 |
| Q9UG56 | Phosphatidylserine decarboxylase proenzy | PISD | 46.672 | 8.3 | 3 | 3 | 80.474113 |
| Q9UGJ1 | Gamma-tubulin complex component 4 OS | TUBGCP4 | 76.09 | 27.1 | 13 | 13 | 473.64484 |
| Q9UGL1 | Lysine-specific demethylase 5B OS=Hom | KDM5B | 175.658 | 0.7 | 1 | 1 | 46.787979 |
| Q9UGN5 | Poly [ADP-ribose] polymerase 2 OS=Hor | PARP2 | 66.206 | 21.1 | 12 | 12 | 521.28247 |
| Q9UGP4 | LIM domain-containing protein 1 OS=Ho | LIMD1 | 72.191 | 29.6 | 16 | 16 | 383.66397 |
| Q9UGP8 | Translocation protein SEC63 homolog OS | SEC63 | 87.997 | 11.1 | 7 | 7 | 116.10608 |
| Q9UGR2 | Zinc finger CCCH domain-containing pro | ZC3H7B | 109.858 | 34.4 | 31 | 30 | 1092.3661 |
| Q9UGU0 | Transcription factor 20 OS=Homo sapiens | TCF20 | 211.772 | 4 | 6 | 6 | 98.114494 |
| Q9UGY1 | Nucleolar protein 12 OS=Homo sapiens C | NOL12 | 24.663 | 20.7 | 4 | 4 | 861.87665 |
| Q9UH17 | DNA dC->dU-editing enzyme APOBEC-3 | APOBEC3B | 45.925 | 24.9 | 9 | 3 | 828.99249 |
| Q9UH99 | SUN domain-containing protein 2 OS=Hc | SUN2 | 80.311 | 14.8 | 9 | 9 | 189.80182 |
| Q9UHB6 | LIM domain and actin-binding protein 1 C | LIMA1 | 85.226 | 26 | 19 | 18 | 1137.2703 |
| Q9UHB7 | AF4/FMR2 family member 4 OS=Homo s | AFF4 | 127.46 | 10.1 | 11 | 11 | 235.16678 |
| Q9UHB9 | Signal recognition particle subunit SRP68 | SRP68 | 70.73 | 28.1 | 17 | 17 | 743.60083 |
| Q9UHC7 | E3 ubiquitin-protein ligase makorin-1 OS= | MKRN1 | 53.35 | 19.9 | 9 | 6 | 1742.8945 |
| Q9UHD2 | Serine/threonine-protein kinase TBK1 OS | TBK1 | 83.643 | 22.9 | 15 | 15 | 348.37228 |
| Q9UHD8 | Septin-9 OS=Homo sapiens OX=9606 GN | SEPTIN9 | 65.402 | 21.5 | 13 | 13 | 188.6927 |
| Q9UHD9 | Ubiquilin-2 OS=Homo sapiens OX=9606 | UBQLN2 | 65.697 | 6.2 | 3 | 2 | 31.54365 |
| Q9UHI6 | Probable ATP-dependent RNA helicase D | DDX20 | 92.241 | 31.6 | 23 | 23 | 3927.8254 |
| Q9UHK0 | FMR1-interacting protein NUFIP1 OS=H | NUFIP1 | 56.3 | 17.4 | 8 | 8 | 359.23883 |
| Q9UHN1 | DNA polymerase subunit gamma-2, mitoc | POLG2 | 54.912 | 2.5 | 1 | 1 | 106.30674 |
| Q9UHR4 | Brain-specific angiogenesis inhibitor 1-as | BAIAP2L1 | 56.883 | 24.5 | 12 | 12 | 490.1853 |
| Q9UHR5 | SAP30-binding protein OS=Homo sapiens | SAP30BP | 33.871 | 12.3 | 3 | 3 | 181.66597 |
| Q9UHV9 | Prefoldin subunit 2 OS=Homo sapiens OX | PFDN2 | 16.648 | 29.9 | 4 | 4 | 170.50902 |
| Q9UHX1 | Poly(U)-binding-splicing factor PUF60 O | PUF60 | 59.876 | 25 | 14 | 14 | 1677.7189 |
| Q9UI09 | NADH dehydrogenase [ubiquinone] 1 alp | NDUFA12 | 17.115 | 8.3 | 1 | 1 | 40.60202 |

| | | | | | | | |
|---|---|---|---|---|---|---|---|
| Q9UI10 | Translation initiation factor eIF-2B subun | EIF2B4 | 57.558 | 18.9 | 8 | 8 | 222.92348 |
| Q9UI30 | Multifunctional methyltransferase subunit | TRMT112 | 14.2 | 49.6 | 5 | 5 | 942.90631 |
| Q9UI43 | rRNA methyltransferase 2, mitochondrial | MRM2 | 27.424 | 17.5 | 4 | 4 | 177.85396 |
| Q9UID6 | Zinc finger protein 639 OS=Homo sapien: | ZNF639 | 56.055 | 13.8 | 6 | 6 | 156.89316 |
| Q9UIF9 | Bromodomain adjacent to zinc finger don | BAZ2A | 211.198 | 0.5 | 1 | 1 | 40.930687 |
| Q9UIG0 | Tyrosine-protein kinase BAZ1B OS=Hon | BAZ1B | 170.903 | 13.8 | 16 | 16 | 180.5224 |
| Q9UII4 | E3 ISG15--protein ligase HERC5 OS=Ho | HERC5 | 116.853 | 23.4 | 23 | 23 | 799.56122 |
| Q9UIJ7 | GTP:AMP phosphotransferase AK3, mito | AK3 | 25.566 | 3.1 | 1 | 1 | 150.30086 |
| Q9UIV1 | CCR4-NOT transcription complex subuni | CNOT7 | 32.746 | 13 | 3 | 3 | 182.56346 |
| Q9UJ70 | N-acetyl-D-glucosamine kinase OS=Hom | NAGK | 37.376 | 7 | 2 | 2 | 53.491653 |
| Q9UJA3 | DNA helicase MCM8 OS=Homo sapiens | MCM8 | 93.697 | 20.1 | 13 | 13 | 128.07088 |
| Q9UJA5 | tRNA (adenine(58)-N(1))-methyltransfera | TRMT6 | 55.8 | 27 | 15 | 15 | 497.75067 |
| Q9UJC3 | Protein Hook homolog 1 OS=Homo sapie | HOOK1 | 84.648 | 1.4 | 1 | 1 | 54.422089 |
| Q9UJF2 | Ras GTPase-activating protein nGAP OS= | RASAL2 | 128.558 | 23.7 | 22 | 20 | 284.63754 |
| Q9UJS0 | Electrogenic aspartate/glutamate antiporte | SLC25A13 | 74.176 | 27.6 | 15 | 15 | 580.36218 |
| Q9UJT1 | Tubulin delta chain OS=Homo sapiens O: | TUBD1 | 51.035 | 5.3 | 2 | 2 | 37.38903 |
| Q9UJU6 | Drebrin-like protein OS=Homo sapiens O | DBNL | 48.208 | 6.7 | 2 | 2 | 61.978058 |
| Q9UJV9 | Probable ATP-dependent RNA helicase D | DDX41 | 69.838 | 45.5 | 25 | 25 | 560.37891 |
| Q9UJW0 | Dynactin subunit 4 OS=Homo sapiens OX | DCTN4 | 52.338 | 19.6 | 8 | 8 | 103.24332 |
| Q9UJX2 | Cell division cycle protein 23 homolog O: | CDC23 | 68.835 | 10.7 | 6 | 6 | 90.77961 |
| Q9UJX3 | Anaphase-promoting complex subunit 7 C | ANAPC7 | 63.134 | 23.9 | 13 | 13 | 227.7841 |
| Q9UJX4 | Anaphase-promoting complex subunit 5 C | ANAPC5 | 85.077 | 9.7 | 6 | 6 | 117.22728 |
| Q9UJX5 | Anaphase-promoting complex subunit 4 C | ANAPC4 | 92.117 | 12.5 | 8 | 8 | 87.314461 |
| Q9UJY1 | Heat shock protein beta-8 OS=Homo sapi | HSPB8 | 21.605 | 30.6 | 5 | 5 | 238.63644 |
| Q9UJZ1 | Stomatin-like protein 2, mitochondrial OS | STOML2 | 38.535 | 19.4 | 5 | 5 | 179.02242 |
| Q9UK41 | Vacuolar protein sorting-associated protei | VPS28 | 25.426 | 4.1 | 1 | 1 | 144.25479 |
| Q9UK45 | U6 snRNA-associated Sm-like protein LS | LSM7 | 11.603 | 16.5 | 3 | 3 | 198.16336 |
| Q9UK58 | Cyclin-L1 OS=Homo sapiens OX=9606 C | CCNL1 | 59.634 | 1.7 | 1 | 1 | 16.004593 |
| Q9UK59 | Lariat debranching enzyme OS=Homo sa | DBR1 | 61.555 | 11.4 | 6 | 6 | 353.94141 |
| Q9UK61 | Protein TASOR OS=Homo sapiens OX=9 | TASOR | 189.032 | 14.6 | 22 | 22 | 211.84073 |
| Q9UK73 | Protein fem-1 homolog B OS=Homo sapi | FEM1B | 70.265 | 1.8 | 1 | 1 | 20.844074 |
| Q9UK80 | Ubiquitin carboxyl-terminal hydrolase 21 | USP21 | 62.656 | 1.4 | 1 | 1 | 27.346609 |
| Q9UK99 | F-box only protein 3 OS=Homo sapiens C | FBXO3 | 54.561 | 6.8 | 3 | 3 | 66.977493 |
| Q9UKA9 | Polypyrimidine tract-binding protein 2 OS | PTBP2 | 57.491 | 3.4 | 2 | 2 | 8018.7017 |
| Q9UKB1 | F-box/WD repeat-containing protein 11 O | FBXW11 | 62.091 | 19.4 | 10 | 4 | 293.3222 |

| | | | | | | | |
|---|---|---|---|---|---|---|---|
| Q9UKF6 | Cleavage and polyadenylation specificity | CPSF3 | 77.486 | 13.2 | 9 | 9 | 167.4463 |
| Q9UKI2 | Cdc42 effector protein 3 OS=Homo sapie | CDC42EP3 | 27.679 | 3.9 | 1 | 1 | 46.509827 |
| Q9UKK3 | Protein mono-ADP-ribosyltransferase PA | PARP4 | 192.595 | 12.2 | 19 | 19 | 255.81245 |
| Q9UKK9 | ADP-sugar pyrophosphatase OS=Homo s | NUDT5 | 24.328 | 23.7 | 4 | 4 | 82.849724 |
| Q9UKL0 | REST corepressor 1 OS=Homo sapiens O | RCOR1 | 53.328 | 5.4 | 2 | 2 | 44.151615 |
| Q9UKM7 | Endoplasmic reticulum mannosyl-oligosa | MAN1B1 | 79.58 | 3.3 | 2 | 2 | 36.58765 |
| Q9UKM9 | RNA-binding protein Raly OS=Homo sap | RALY | 32.464 | 24.8 | 8 | 8 | 778.20978 |
| Q9UKN8 | General transcription factor 3C polypeptic | GTF3C4 | 91.983 | 26.5 | 20 | 20 | 362.40118 |
| Q9UKS6 | Protein kinase C and casein kinase substra | PACSIN3 | 48.487 | 28.1 | 12 | 12 | 818.13403 |
| Q9UKT4 | F-box only protein 5 OS=Homo sapiens C | FBXO5 | 50.147 | 7.8 | 3 | 3 | 68.946274 |
| Q9UKV3 | Apoptotic chromatin condensation induce | ACIN1 | 151.862 | 10.7 | 15 | 15 | 278.80017 |
| Q9UKV8 | Protein argonaute-2 OS=Homo sapiens O | AGO2 | 97.209 | 41.4 | 30 | 30 | 3588.998 |
| Q9UKX2 | Myosin-2 OS=Homo sapiens OX=9606 G | MYH2 | 223.045 | 0.5 | 1 | 1 | 142.01222 |
| Q9UKX7 | Nuclear pore complex protein Nup50 OS= | NUP50 | 50.145 | 14.7 | 5 | 5 | 84.647942 |
| Q9UKZ9 | Procollagen C-endopeptidase enhancer 2 ( | PCOLCE2 | 45.717 | 14 | 5 | 5 | 71.757919 |
| Q9UL15 | BAG family molecular chaperone regulato | BAG5 | 51.2 | 15.4 | 5 | 5 | 78.334259 |
| Q9UL18 | Protein argonaute-1 OS=Homo sapiens O | AGO1 | 97.215 | 48.7 | 33 | 17 | 2586.0093 |
| Q9UL25 | Ras-related protein Rab-21 OS=Homo sap | RAB21 | 24.348 | 16.9 | 3 | 3 | 54.105183 |
| Q9UL33 | Trafficking protein particle complex subu | TRAPPC2L | 16.146 | 22.1 | 3 | 3 | 93.395386 |
| Q9UL40 | Zinc finger protein 346 OS=Homo sapien | ZNF346 | 32.933 | 23.5 | 6 | 6 | 366.91348 |
| Q9UL54 | Serine/threonine-protein kinase TAO2 OS | TAOK2 | 138.252 | 6 | 7 | 7 | 235.0593 |
| Q9ULA0 | Aspartyl aminopeptidase OS=Homo sapie | DNPEP | 53.411 | 3.9 | 2 | 2 | 29.367918 |
| Q9ULC4 | Malignant T-cell-amplified sequence 1 OS | MCTS1 | 20.556 | 19.3 | 3 | 1 | 89.555283 |
| Q9ULD2 | Microtubule-associated tumor suppressor | MTUS1 | 141.397 | 25 | 28 | 28 | 989.40552 |
| Q9ULD4 | Bromodomain and PHD finger-containing | BRPF3 | 135.745 | 17.4 | 21 | 18 | 322.30225 |
| Q9ULG6 | Cell cycle progression protein 1 OS=Hom | CCPG1 | 87.341 | 13.6 | 10 | 10 | 233.29863 |
| Q9ULJ3 | Zinc finger and BTB domain-containing p | ZBTB21 | 118.871 | 14.9 | 14 | 14 | 134.99112 |
| Q9ULK4 | Mediator of RNA polymerase II transcript | MED23 | 156.474 | 2.3 | 3 | 3 | 26.86578 |
| Q9ULM6 | CCR4-NOT transcription complex subuni | CNOT6 | 63.307 | 1.8 | 1 | 1 | 46.703693 |
| Q9ULP9 | TBC1 domain family member 24 OS=Hor | TBC1D24 | 62.92 | 2.7 | 1 | 1 | 45.378708 |
| Q9ULR0 | Pre-mRNA-splicing factor ISY1 homolog | ISY1 | 32.993 | 4.6 | 1 | 1 | 159.95453 |
| Q9ULT8 | E3 ubiquitin-protein ligase HECTD1 OS= | HECTD1 | 289.369 | 2 | 5 | 5 | 74.535568 |
| Q9ULV3 | Cip1-interacting zinc finger protein OS=H | CIZ1 | 100.046 | 4.8 | 4 | 4 | 72.739288 |
| Q9ULV4 | Coronin-1C OS=Homo sapiens OX=9606 | CORO1C | 53.25 | 13.9 | 7 | 6 | 228.13295 |
| Q9ULW0 | Targeting protein for Xklp2 OS=Homo sa | TPX2 | 85.654 | 42 | 31 | 31 | 1521.0833 |

| | | | | | | | |
|---|---|---|---|---|---|---|---|
| Q9ULW3 | Activator of basal transcription 1 OS=Hoı | ABT1 | 31.079 | 19.5 | 5 | 5 | 183.83636 |
| Q9ULW5 | Ras-related protein Rab-26 OS=Homo saı | RAB26 | 27.9 | 8.6 | 2 | 2 | 53.998451 |
| Q9ULX3 | RNA-binding protein NOB1 OS=Homo sa | NOB1 | 46.675 | 28.9 | 11 | 11 | 2793.73 |
| Q9ULX6 | A-kinase anchor protein 8-like OS=Homo | AKAP8L | 71.64 | 19.7 | 11 | 11 | 1738.434 |
| Q9ULX9 | Transcription factor MafF OS=Homo sapi | MAFF | 17.761 | 9.8 | 1 | 1 | 27.560879 |
| Q9UM00 | Calcium load-activated calcium channel C | TMCO1 | 27.08 | 13.4 | 3 | 3 | 529.68005 |
| Q9UM11 | Fizzy-related protein homolog OS=Homo | FZR1 | 55.18 | 3.2 | 1 | 1 | 106.2442 |
| Q9UM54 | Unconventional myosin-VI OS=Homo saı | MYO6 | 149.692 | 10 | 12 | 12 | 109.45948 |
| Q9UMS4 | Pre-mRNA-processing factor 19 OS=Hon | PRPF19 | 55.181 | 37.1 | 14 | 14 | 2239.1865 |
| Q9UMY1 | Nucleolar protein 7 OS=Homo sapiens O2 | NOL7 | 29.427 | 24.1 | 7 | 7 | 652.52856 |
| Q9UMZ2 | Synergin gamma OS=Homo sapiens OX= | SYNRG | 140.655 | 3 | 3 | 3 | 50.570938 |
| Q9UN37 | Vacuolar protein sorting-associated protei | VPS4A | 48.898 | 26.1 | 11 | 10 | 434.76678 |
| Q9UN81 | LINE-1 retrotransposable element ORF1 ı | L1RE1 | 40.056 | 26.6 | 8 | 8 | 237.03322 |
| Q9UN86 | Ras GTPase-activating protein-binding pr | G3BP2 | 54.122 | 50.6 | 23 | 20 | 21660.574 |
| Q9UNE7 | E3 ubiquitin-protein ligase CHIP OS=Hoı | STUB1 | 34.857 | 21.1 | 6 | 6 | 144.64372 |
| Q9UNF0 | Protein kinase C and casein kinase substra | PACSIN2 | 55.739 | 2.5 | 1 | 1 | 24.24172 |
| Q9UNF1 | Melanoma-associated antigen D2 OS=Ho: | MAGED2 | 64.955 | 8.6 | 5 | 4 | 155.46188 |
| Q9UNK4 | Group IID secretory phospholipase A2 OS | PLA2G2D | 16.547 | 9 | 1 | 1 | 27.191916 |
| Q9UNL4 | Inhibitor of growth protein 4 OS=Homo s | ING4 | 28.531 | 7.2 | 2 | 2 | 57.333225 |
| Q9UNN5 | FAS-associated factor 1 OS=Homo sapier | FAF1 | 73.955 | 3.7 | 2 | 2 | 21.850788 |
| Q9UNP9 | Peptidyl-prolyl cis-trans isomerase E OS= | PPIE | 33.431 | 20.3 | 5 | 5 | 124.2515 |
| Q9UNQ2 | Probable dimethyladenosine transferase C | DIMT1 | 35.237 | 48.2 | 15 | 15 | 3118.9377 |
| Q9UNX4 | WD repeat-containing protein 3 OS=Hom | WDR3 | 106.099 | 16.1 | 14 | 14 | 337.0463 |
| Q9UNY4 | Transcription termination factor 2 OS=Hc | TTF2 | 129.589 | 1.7 | 2 | 2 | 58.726746 |
| Q9UPN3 | Microtubule-actin cross-linking factor 1, i | MACF1 | 838.308 | 6.3 | 41 | 17 | 600.61346 |
| Q9UPN4 | Centrosomal protein of 131 kDa OS=Hon | CEP131 | 122.149 | 19.7 | 19 | 19 | 165.69534 |
| Q9UPN6 | SR-related and CTD-associated factor 8 C | SCAF8 | 140.52 | 8.8 | 10 | 9 | 393.84985 |
| Q9UPN7 | Serine/threonine-protein phosphatase 6 re | PPP6R1 | 96.724 | 9.9 | 10 | 10 | 652.09106 |
| Q9UPP1 | Histone lysine demethylase PHF8 OS=Ho | PHF8 | 117.865 | 17.5 | 17 | 14 | 558.13214 |
| Q9UPQ0 | LIM and calponin homology domains-con | LIMCH1 | 121.868 | 31.2 | 26 | 26 | 247.7702 |
| Q9UPQ3 | Arf-GAP with GTPase, ANK repeat and I | AGAP1 | 94.471 | 5.3 | 4 | 4 | 175.75307 |
| Q9UPQ9 | Trinucleotide repeat-containing gene 6B p | TNRC6B | 194.003 | 20.6 | 33 | 33 | 1405.5968 |
| Q9UPR3 | Nonsense-mediated mRNA decay factor S | SMG5 | 113.928 | 10 | 11 | 11 | 370.50253 |
| Q9UPS8 | Ankyrin repeat domain-containing protein | ANKRD26 | 196.411 | 5.9 | 10 | 9 | 85.970032 |
| Q9UPT8 | Zinc finger CCCH domain-containing pro | ZC3H4 | 140.257 | 14.7 | 19 | 19 | 976.57147 |

| | | | | | | | |
|---|---|---|---|---|---|---|---|
| Q9UPT9 | Ubiquitin carboxyl-terminal hydrolase 22 | USP22 | 59.961 | 1.9 | 1 | 1 | 131.67276 |
| Q9UPU9 | Protein Smaug homolog 1 OS=Homo sapi | SAMD4A | 79.415 | 20.2 | 12 | 12 | 211.37599 |
| Q9UPV0 | Centrosomal protein of 164 kDa OS=Hon | CEP164 | 164.314 | 2.3 | 3 | 3 | 52.259838 |
| Q9UPW5 | Cytosolic carboxypeptidase 1 OS=Homo : | AGTPBP1 | 138.449 | 1.5 | 2 | 2 | 59.771584 |
| Q9UPY3 | Endoribonuclease Dicer OS=Homo sapier | DICER1 | 218.682 | 14.5 | 24 | 24 | 242.16867 |
| Q9UQ13 | Leucine-rich repeat protein SHOC-2 OS= | SHOC2 | 64.889 | 1.4 | 1 | 1 | 66.723526 |
| Q9UQ35 | Serine/arginine repetitive matrix protein 2 | SRRM2 | 299.616 | 12.6 | 30 | 30 | 948.56036 |
| Q9UQ80 | Proliferation-associated protein 2G4 OS= | PA2G4 | 43.787 | 19 | 7 | 7 | 252.83919 |
| Q9UQ84 | Exonuclease 1 OS=Homo sapiens OX=96 | EXO1 | 94.104 | 20.6 | 15 | 15 | 252.99347 |
| Q9UQB8 | Brain-specific angiogenesis inhibitor 1-as: | BAIAP2 | 60.868 | 33 | 16 | 16 | 427.75372 |
| Q9UQE7 | Structural maintenance of chromosomes p | SMC3 | 141.542 | 19.3 | 20 | 20 | 210.60541 |
| Q9UQN3 | Charged multivesicular body protein 2b C | CHMP2B | 23.907 | 13.6 | 3 | 3 | 115.31223 |
| Q9UQR0 | Sex comb on midleg-like protein 2 OS=H | SCML2 | 77.257 | 12 | 6 | 6 | 118.75664 |
| Q9UQR1 | Zinc finger protein 148 OS=Homo sapien: | ZNF148 | 88.977 | 15.2 | 11 | 11 | 236.3299 |
| Q9Y221 | 60S ribosome subunit biogenesis protein N | NIP7 | 20.463 | 23.9 | 3 | 3 | 111.59689 |
| Q9Y223 | Bifunctional UDP-N-acetylglucosamine 2 | GNE | 79.275 | 23 | 14 | 14 | 212.96069 |
| Q9Y224 | RNA transcription, translation and transpc | RTRAF | 28.069 | 50.8 | 13 | 13 | 19387.221 |
| Q9Y230 | RuvB-like 2 OS=Homo sapiens OX=960€ | RUVBL2 | 51.157 | 36.9 | 17 | 17 | 768.43402 |
| Q9Y232 | Chromodomain Y-like protein OS=Homo | CDYL | 66.482 | 2.5 | 1 | 1 | 63.707306 |
| Q9Y234 | Lipoyltransferase 1, mitochondrial OS=H | LIPT1 | 42.48 | 4 | 1 | 1 | 70.794174 |
| Q9Y251 | Heparanase OS=Homo sapiens OX=9606 | HPSE | 61.149 | 20.1 | 11 | 11 | 495.21005 |
| Q9Y253 | DNA polymerase eta OS=Homo sapiens ( | POLH | 78.414 | 6.3 | 4 | 4 | 63.11182 |
| Q9Y262 | Eukaryotic translation initiation factor 3 s | EIF3L | 66.727 | 24.5 | 17 | 17 | 1718.163 |
| Q9Y265 | RuvB-like 1 OS=Homo sapiens OX=960€ | RUVBL1 | 50.228 | 30.9 | 11 | 11 | 490.08875 |
| Q9Y266 | Nuclear migration protein nudC OS=Hom | NUDC | 38.243 | 24.8 | 7 | 7 | 135.22112 |
| Q9Y277 | Voltage-dependent anion-selective channe | VDAC3 | 30.659 | 4.6 | 1 | 1 | 67.860046 |
| Q9Y283 | Inversin OS=Homo sapiens OX=9606 GN | INVS | 117.827 | 3 | 3 | 3 | 85.047203 |
| Q9Y285 | Phenylalanine--tRNA ligase alpha subunit | FARSA | 57.564 | 23.4 | 10 | 10 | 378.59402 |
| Q9Y294 | Histone chaperone ASF1A OS=Homo sap | ASF1A | 22.969 | 10.8 | 2 | 2 | 698.46753 |
| Q9Y295 | Developmentally-regulated GTP-binding ] | DRG1 | 40.543 | 47.1 | 15 | 15 | 1397.9814 |
| Q9Y297 | F-box/WD repeat-containing protein 1A ( | BTRC | 68.867 | 3.3 | 2 | 2 | 43.643112 |
| Q9Y2A7 | Nck-associated protein 1 OS=Homo sapie | NCKAP1 | 128.79 | 20.5 | 22 | 21 | 484.62219 |
| Q9Y2C4 | Nuclease EXOG, mitochondrial OS=Hom | EXOG | 41.085 | 9.2 | 3 | 3 | 33.44305 |
| Q9Y2D2 | UDP-N-acetylglucosamine transporter OS | SLC35A3 | 35.985 | 4.6 | 1 | 1 | 50.574108 |
| Q9Y2D8 | Afadin- and alpha-actinin-binding protein | SSX2IP | 71.237 | 26.7 | 17 | 17 | 901.83203 |

| | | | | | | | |
|---|---|---|---|---|---|---|---|
| Q9Y2F5 | Little elongation complex subunit 1 OS=H | ICE1 | 247.891 | 3.2 | 6 | 6 | 60.073235 |
| Q9Y2F9 | BTB/POZ domain-containing protein 3 O | BTBD3 | 58.42 | 2.5 | 1 | 1 | 81.048065 |
| Q9Y2G8 | DnaJ homolog subfamily C member 16 O | DNAJC16 | 90.592 | 16.6 | 12 | 12 | 419.43765 |
| Q9Y2G9 | Protein strawberry notch homolog 2 OS=I | SBNO2 | 150.276 | 1.5 | 2 | 1 | 32.775372 |
| Q9Y2I8 | WD repeat-containing protein 37 OS=Hoi | WDR37 | 54.666 | 4.9 | 1 | 1 | 22.719753 |
| Q9Y2K2 | Serine/threonine-protein kinase SIK3 OS= | SIK3 | 144.852 | 14 | 15 | 15 | 190.71304 |
| Q9Y2K5 | R3H domain-containing protein 2 OS=Ho | R3HDM2 | 106.999 | 6.5 | 6 | 6 | 407.22693 |
| Q9Y2K7 | Lysine-specific demethylase 2A OS=Hom | KDM2A | 132.793 | 10.6 | 11 | 11 | 150.41757 |
| Q9Y2L1 | Exosome complex exonuclease RRP44 O: | DIS3 | 109.003 | 12.1 | 10 | 10 | 111.55564 |
| Q9Y2P8 | RNA 3'-terminal phosphate cyclase-like p | RCL1 | 40.843 | 30 | 11 | 11 | 896.7019 |
| Q9Y2Q3 | Glutathione S-transferase kappa 1 OS=Hc | GSTK1 | 25.497 | 6.2 | 1 | 1 | 84.45639 |
| Q9Y2Q5 | Ragulator complex protein LAMTOR2 O: | LAMTOR2 | 13.508 | 8 | 1 | 1 | 65.160797 |
| Q9Y2R4 | Probable ATP-dependent RNA helicase D | DDX52 | 67.467 | 25.5 | 13 | 13 | 405.03391 |
| Q9Y2S7 | Polymerase delta-interacting protein 2 OS | POLDIP2 | 42.034 | 19 | 6 | 6 | 95.193848 |
| Q9Y2T2 | AP-3 complex subunit mu-1 OS=Homo sa | AP3M1 | 46.94 | 11 | 4 | 2 | 234.15985 |
| Q9Y2T3 | Guanine deaminase OS=Homo sapiens O: | GDA | 51.004 | 5.5 | 2 | 2 | 61.289764 |
| Q9Y2U8 | Inner nuclear membrane protein Man1 OS | LEMD3 | 99.998 | 7.6 | 6 | 6 | 49.81414 |
| Q9Y2W1 | Thyroid hormone receptor-associated prot | THRAP3 | 108.666 | 26.2 | 27 | 26 | 5964.373 |
| Q9Y2W2 | WW domain-binding protein 11 OS=Hom | WBP11 | 69.998 | 8.4 | 5 | 5 | 106.59758 |
| Q9Y2X0 | Mediator of RNA polymerase II transcript | MED16 | 96.794 | 3.4 | 3 | 3 | 46.432156 |
| Q9Y2X3 | Nucleolar protein 58 OS=Homo sapiens C | NOP58 | 59.579 | 31.4 | 17 | 17 | 1561.005 |
| Q9Y2X9 | Zinc finger protein 281 OS=Homo sapien: | ZNF281 | 96.915 | 22.5 | 17 | 17 | 1561.7046 |
| Q9Y2Y1 | DNA-directed RNA polymerase III subun | POLR3K | 12.321 | 18.5 | 2 | 2 | 146.11148 |
| Q9Y2Y9 | Krueppel-like factor 13 OS=Homo sapien | KLF13 | 31.18 | 16.7 | 4 | 4 | 114.76035 |
| Q9Y2Z2 | Protein MTO1 homolog, mitochondrial O | MTO1 | 79.964 | 2.1 | 1 | 1 | 42.537895 |
| Q9Y2Z4 | Tyrosine--tRNA ligase, mitochondrial OS | YARS2 | 53.2 | 30.8 | 11 | 11 | 887.07843 |
| Q9Y314 | Nitric oxide synthase-interacting protein ( | NOSIP | 33.172 | 32.2 | 8 | 8 | 402.41891 |
| Q9Y316 | Protein MEMO1 OS=Homo sapiens OX= | MEMO1 | 33.734 | 4.7 | 1 | 1 | 138.45058 |
| Q9Y320 | Thioredoxin-related transmembrane prote | TMX2 | 34.038 | 3.4 | 1 | 1 | 37.025116 |
| Q9Y324 | rRNA-processing protein FCF1 homolog ( | FCF1 | 23.37 | 14.6 | 3 | 3 | 199.16388 |
| Q9Y333 | U6 snRNA-associated Sm-like protein LS | LSM2 | 10.835 | 48.4 | 4 | 4 | 172.6825 |
| Q9Y383 | Putative RNA-binding protein Luc7-like 2 | LUC7L2 | 46.514 | 25.5 | 12 | 12 | 5323.2563 |
| Q9Y388 | RNA-binding motif protein, X-linked 2 O | RBMX2 | 37.336 | 21.7 | 7 | 7 | 407.16214 |
| Q9Y399 | Small ribosomal subunit protein uS2m OS | MRPS2 | 33.25 | 30.1 | 10 | 10 | 881.01825 |
| Q9Y3A2 | Probable U3 small nucleolar RNA-associa | UTP11 | 30.447 | 21.3 | 6 | 6 | 106.77719 |

| | | | | | | | |
|---|---|---|---|---|---|---|---|
| Q9Y3A4 | Ribosomal RNA-processing protein 7 hon | RRP7A | 32.335 | 21.4 | 5 | 5 | 789.70178 |
| Q9Y3B2 | Exosome complex component CSL4 OS= | EXOSC1 | 21.452 | 26.2 | 5 | 5 | 343.9306 |
| Q9Y3B4 | Splicing factor 3B subunit 6 OS=Homo sa | SF3B6 | 14.585 | 48 | 5 | 5 | 813.10938 |
| Q9Y3B7 | Large ribosomal subunit protein uL11m C | MRPL11 | 20.684 | 49.5 | 10 | 10 | 1602.2537 |
| Q9Y3C6 | Peptidyl-prolyl cis-trans isomerase-like 1 | PPIL1 | 18.237 | 25.3 | 4 | 4 | 1484.7048 |
| Q9Y3C7 | Mediator of RNA polymerase II transcript | MED31 | 15.806 | 10.7 | 1 | 1 | 217.56667 |
| Q9Y3C8 | Ubiquitin-fold modifier-conjugating enzy | UFC1 | 19.459 | 6 | 1 | 1 | 53.00975 |
| Q9Y3D3 | Small ribosomal subunit protein bS16m O | MRPS16 | 15.345 | 22.6 | 4 | 4 | 660.2746 |
| Q9Y3D6 | Mitochondrial fission 1 protein OS=Homo | FIS1 | 16.938 | 7.9 | 1 | 1 | 46.833195 |
| Q9Y3D7 | Mitochondrial import inner membrane tra | PAM16 | 13.825 | 14.4 | 2 | 2 | 217.20103 |
| Q9Y3D8 | Adenylate kinase isoenzyme 6 OS=Homo | AK6 | 20.062 | 10.5 | 2 | 2 | 172.7466 |
| Q9Y3D9 | Small ribosomal subunit protein mS23 OS | MRPS23 | 21.771 | 54.7 | 10 | 10 | 1393.7603 |
| Q9Y3E5 | Peptidyl-tRNA hydrolase 2, mitochondria | PTRH2 | 19.194 | 30.2 | 4 | 4 | 67.158638 |
| Q9Y3E7 | Charged multivesicular body protein 3 OS | CHMP3 | 25.074 | 14.4 | 3 | 3 | 198.05791 |
| Q9Y3F4 | Serine-threonine kinase receptor-associate | STRAP | 38.439 | 56.9 | 18 | 18 | 3501.6694 |
| Q9Y3I0 | RNA-splicing ligase RtcB homolog OS=H | RTCB | 55.211 | 50.7 | 25 | 25 | 30424.396 |
| Q9Y3I1 | F-box only protein 7 OS=Homo sapiens C | FBXO7 | 58.503 | 2.1 | 1 | 1 | 61.159725 |
| Q9Y3M8 | StAR-related lipid transfer protein 13 OS= | STARD13 | 124.968 | 2.2 | 2 | 2 | 33.599052 |
| Q9Y3S2 | Zinc finger protein 330 OS=Homo sapiens | ZNF330 | 36.202 | 13.4 | 4 | 4 | 215.30016 |
| Q9Y3T9 | Nucleolar complex protein 2 homolog OS | NOC2L | 84.92 | 11.6 | 9 | 9 | 160.50174 |
| Q9Y3U8 | Large ribosomal subunit protein eL36 OS | RPL36 | 12.254 | 30.5 | 5 | 5 | 15053.912 |
| Q9Y3Y2 | Chromatin target of PRMT1 protein OS=I | CHTOP | 26.397 | 17.7 | 5 | 5 | 766.08728 |
| Q9Y3Z3 | Deoxynucleoside triphosphate triphosphol | SAMHD1 | 72.201 | 6.9 | 4 | 4 | 89.463654 |
| Q9Y446 | Plakophilin-3 OS=Homo sapiens OX=960 | PKP3 | 87.083 | 34.8 | 25 | 25 | 1749.3009 |
| Q9Y448 | Small kinetochore-associated protein OS= | KNSTRN | 35.439 | 28.5 | 12 | 12 | 5007.4951 |
| Q9Y450 | HBS1-like protein OS=Homo sapiens OX | HBS1L | 75.473 | 43 | 24 | 24 | 3046.7292 |
| Q9Y483 | Metal-response element-binding transcrip | MTF2 | 67.09 | 5.6 | 3 | 3 | 71.375458 |
| Q9Y490 | Talin-1 OS=Homo sapiens OX=9606 GN | TLN1 | 269.768 | 5 | 9 | 8 | 1457.6813 |
| Q9Y496 | Kinesin-like protein KIF3A OS=Homo sa | KIF3A | 80.041 | 8.3 | 5 | 5 | 92.968529 |
| Q9Y4A5 | Transformation/transcription domain-asso | TRRAP | 437.601 | 7.6 | 28 | 28 | 140.72897 |
| Q9Y4B5 | Microtubule cross-linking factor 1 OS=Ho | MTCL1 | 209.526 | 47.2 | 87 | 82 | 9198.2285 |
| Q9Y4C2 | TRPM8 channel-associated factor 1 OS=I | TCAF1 | 102.126 | 19.1 | 17 | 17 | 396.41675 |
| Q9Y4C4 | Malignant fibrous histiocytoma-amplified | MFHAS1 | 116.95 | 1.4 | 1 | 1 | 15.740252 |
| Q9Y4C8 | Probable RNA-binding protein 19 OS=Ho | RBM19 | 107.332 | 14 | 11 | 11 | 265.90186 |
| Q9Y4D8 | Probable E3 ubiquitin-protein ligase HEC | HECTD4 | 439.345 | 0.6 | 2 | 2 | 41.350872 |

| | | | | | | | |
|---|---|---|---|---|---|---|---|
| Q9Y4E5 | E3 SUMO-protein ligase ZNF451 OS=Hc | ZNF451 | 121.485 | 3.8 | 4 | 4 | 93.727417 |
| Q9Y4E8 | Ubiquitin carboxyl-terminal hydrolase 15 | USP15 | 112.42 | 8.5 | 8 | 7 | 62.960308 |
| Q9Y4F1 | FERM, ARHGEF and pleckstrin domain-c | FARP1 | 118.633 | 19.4 | 19 | 19 | 378.04123 |
| Q9Y4F3 | Meiosis regulator and mRNA stability fac | MARF1 | 192.86 | 11.3 | 17 | 17 | 215.04492 |
| Q9Y4G8 | Rap guanine nucleotide exchange factor 2 | RAPGEF2 | 167.417 | 6.8 | 9 | 5 | 84.734955 |
| Q9Y4H2 | Insulin receptor substrate 2 OS=Homo sa | IRS2 | 137.335 | 14.6 | 16 | 16 | 481.87827 |
| Q9Y4I1 | Unconventional myosin-Va OS=Homo sa | MYO5A | 215.405 | 0.6 | 1 | 1 | 63.395981 |
| Q9Y4K0 | Lysyl oxidase homolog 2 OS=Homo sapie | LOXL2 | 86.725 | 8.1 | 6 | 6 | 292.27466 |
| Q9Y4K4 | Mitogen-activated protein kinase kinase k | MAP4K5 | 95.024 | 1.5 | 1 | 1 | 85.87471 |
| Q9Y4R8 | Telomere length regulation protein TEL2 | TELO2 | 91.748 | 1.4 | 1 | 1 | 49.962288 |
| Q9Y4W2 | Ribosomal biogenesis protein LAS1L OS= | LAS1L | 83.065 | 11.6 | 8 | 8 | 91.58976 |
| Q9Y4W6 | AFG3-like protein 2 OS=Homo sapiens C | AFG3L2 | 88.584 | 12 | 11 | 11 | 277.37869 |
| Q9Y4X5 | E3 ubiquitin-protein ligase ARIH1 OS=H | ARIH1 | 64.118 | 11 | 6 | 6 | 137.35878 |
| Q9Y4Z0 | U6 snRNA-associated Sm-like protein LS | LSM4 | 15.35 | 20.9 | 3 | 3 | 672.07281 |
| Q9Y508 | E3 ubiquitin-protein ligase RNF114 OS=I | RNF114 | 25.695 | 26.8 | 6 | 6 | 418.72012 |
| Q9Y512 | Sorting and assembly machinery compone | SAMM50 | 51.977 | 9.6 | 4 | 4 | 61.532116 |
| Q9Y520 | Protein PRRC2C OS=Homo sapiens OX= | PRRC2C | 316.912 | 30.1 | 87 | 87 | 18660.971 |
| Q9Y570 | Protein phosphatase methylesterase 1 OS= | PPME1 | 42.316 | 3.4 | 1 | 1 | 41.892006 |
| Q9Y580 | RNA-binding protein 7 OS=Homo sapien | RBM7 | 30.504 | 19.5 | 5 | 5 | 518.49097 |
| Q9Y5A9 | YTH domain-containing family protein 2 | YTHDF2 | 62.335 | 26.1 | 12 | 12 | 3648.9617 |
| Q9Y5B6 | PAX3- and PAX7-binding protein 1 OS=I | PAXBP1 | 104.804 | 15.6 | 12 | 12 | 191.12454 |
| Q9Y5B8 | Nucleoside diphosphate kinase 7 OS=Hor | NME7 | 42.492 | 14.9 | 4 | 4 | 191.22055 |
| Q9Y5B9 | FACT complex subunit SPT16 OS=Homo | SUPT16H | 119.914 | 12.5 | 11 | 11 | 509.21741 |
| Q9Y5J1 | U3 small nucleolar RNA-associated prote | UTP18 | 62.004 | 30.9 | 15 | 15 | 865.8064 |
| Q9Y5K5 | Ubiquitin carboxyl-terminal hydrolase iso | UCHL5 | 37.607 | 7 | 2 | 2 | 40.283669 |
| Q9Y5K6 | CD2-associated protein OS=Homo sapien | CD2AP | 71.452 | 5.2 | 3 | 3 | 32.703682 |
| Q9Y5L4 | Mitochondrial import inner membrane tra | TIMM13 | 10.501 | 36.8 | 3 | 3 | 212.26642 |
| Q9Y5P8 | Serine/threonine-protein phosphatase 2A r | PPP2R3B | 65.061 | 3.3 | 2 | 2 | 41.8843 |
| Q9Y5Q8 | General transcription factor 3C polypeptic | GTF3C5 | 59.571 | 22 | 11 | 11 | 397.59192 |
| Q9Y5Q9 | General transcription factor 3C polypeptic | GTF3C3 | 101.273 | 16 | 13 | 13 | 361.32233 |
| Q9Y5S2 | Serine/threonine-protein kinase MRCK be | CDC42BPB | 194.316 | 13.6 | 21 | 18 | 103.73552 |
| Q9Y5S9 | RNA-binding protein 8A OS=Homo sapie | RBM8A | 19.889 | 15.5 | 3 | 3 | 4179.5986 |
| Q9Y5T4 | DnaJ homolog subfamily C member 15 O | DNAJC15 | 16.383 | 38.7 | 5 | 5 | 123.99723 |
| Q9Y5T5 | Ubiquitin carboxyl-terminal hydrolase 16 | USP16 | 93.571 | 10.1 | 8 | 8 | 290.48877 |
| Q9Y5U2 | U5 small nuclear ribonucleoprotein TSSC | TSSC4 | 34.326 | 10 | 3 | 3 | 29.800848 |

| | | | | | | | |
|---|---|---|---|---|---|---|---|
| Q9Y5W9 | Sorting nexin-11 OS=Homo sapiens OX= | SNX11 | 30.434 | 4.8 | 1 | 1 | 53.264091 |
| Q9Y5X1 | Sorting nexin-9 OS=Homo sapiens OX=9 | SNX9 | 66.592 | 23.5 | 12 | 12 | 3875.3257 |
| Q9Y5X2 | Sorting nexin-8 OS=Homo sapiens OX=9 | SNX8 | 52.57 | 7.5 | 3 | 3 | 46.887127 |
| Q9Y5Y2 | Cytosolic Fe-S cluster assembly factor NU | NUBP2 | 28.826 | 14 | 3 | 3 | 204.84581 |
| Q9Y5Y5 | Peroxisomal membrane protein PEX16 O: | PEX16 | 38.629 | 18.2 | 6 | 6 | 213.6842 |
| Q9Y605 | MORF4 family-associated protein 1 OS=I | MRFAP1 | 14.65 | 23.6 | 3 | 2 | 295.07242 |
| Q9Y606 | Pseudouridylate synthase 1 homolog OS= | PUS1 | 47.471 | 30.4 | 12 | 12 | 575.7525 |
| Q9Y608 | Leucine-rich repeat flightless-interacting p | LRRFIP2 | 82.172 | 12.8 | 9 | 8 | 308.52966 |
| Q9Y613 | FH1/FH2 domain-containing protein 1 OS | FHOD1 | 126.552 | 11.4 | 11 | 11 | 179.28038 |
| Q9Y619 | Mitochondrial ornithine transporter 1 OS= | SLC25A15 | 32.737 | 6.6 | 2 | 2 | 23.716801 |
| Q9Y620 | DNA repair and recombination protein RA | RAD54B | 102.967 | 4.9 | 4 | 4 | 64.352013 |
| Q9Y657 | Spindlin-1 OS=Homo sapiens OX=9606 ( | SPIN1 | 29.601 | 4.6 | 1 | 1 | 77.379158 |
| Q9Y676 | Small ribosomal subunit protein mS40 OS | MRPS18B | 29.396 | 26 | 6 | 6 | 1197.6055 |
| Q9Y678 | Coatomer subunit gamma-1 OS=Homo sa | COPG1 | 97.719 | 8.2 | 7 | 6 | 219.02357 |
| Q9Y679 | Lipid droplet-regulating VLDL assembly | AUP1 | 45.787 | 8.8 | 3 | 3 | 86.565048 |
| Q9Y697 | Cysteine desulfurase OS=Homo sapiens C | NFS1 | 50.196 | 10.9 | 4 | 4 | 87.759285 |
| Q9Y6A4 | Cilia- and flagella-associated protein 20 C | CFAP20 | 22.775 | 36.8 | 7 | 7 | 539.91284 |
| Q9Y6C9 | Mitochondrial carrier homolog 2 OS=Hor | MTCH2 | 33.331 | 11.9 | 3 | 3 | 107.63613 |
| Q9Y6E0 | Serine/threonine-protein kinase 24 OS=H | STK24 | 49.308 | 8.6 | 3 | 1 | 159.42813 |
| Q9Y6E2 | eIF5-mimic protein 1 OS=Homo sapiens ( | BZW2 | 48.163 | 4.5 | 2 | 2 | 73.143951 |
| Q9Y6G3 | Large ribosomal subunit protein mL42 OS | MRPL42 | 16.662 | 14.8 | 1 | 1 | 87.163681 |
| Q9Y6G9 | Cytoplasmic dynein 1 light intermediate c | DYNC1LI1 | 56.58 | 5 | 2 | 2 | 34.021744 |
| Q9Y6J9 | TAF6-like RNA polymerase II p300/CBP | TAF6L | 67.815 | 18.6 | 9 | 9 | 114.15262 |
| Q9Y6K0 | Choline/ethanolaminephosphotransferase | CEPT1 | 46.554 | 2.6 | 1 | 1 | 563.38055 |
| Q9Y6K5 | 2'-5'-oligoadenylate synthase 3 OS=Homo | OAS3 | 121.171 | 30.2 | 30 | 30 | 1185.744 |
| Q9Y6M0 | Testisin OS=Homo sapiens OX=9606 GN | PRSS21 | 34.884 | 16.2 | 4 | 4 | 88.72625 |
| Q9Y6M1 | Insulin-like growth factor 2 mRNA-bindir | IGF2BP2 | 66.122 | 31.2 | 18 | 18 | 3142.4927 |
| Q9Y6M4 | Casein kinase I isoform gamma-3 OS=Ho | CSNK1G3 | 51.389 | 24.4 | 11 | 2 | 383.59894 |
| Q9Y6M9 | NADH dehydrogenase [ubiquinone] 1 bet | NDUFB9 | 21.831 | 14 | 3 | 3 | 127.42625 |
| Q9Y6N5 | Sulfide:quinone oxidoreductase, mitochor | SQOR | 49.961 | 29.1 | 11 | 11 | 397.77448 |
| Q9Y6R0 | Numb-like protein OS=Homo sapiens OX | NUMBL | 64.892 | 15.6 | 8 | 8 | 412.25266 |
| Q9Y6W3 | Calpain-7 OS=Homo sapiens OX=9606 C | CAPN7 | 92.653 | 3.9 | 3 | 3 | 243.80739 |
| Q9Y6X2 | E3 SUMO-protein ligase PIAS3 OS=Hon | PIAS3 | 68.018 | 5.4 | 3 | 2 | 49.927475 |
| Q9Y6X9 | ATPase MORC2 OS=Homo sapiens OX= | MORC2 | 117.824 | 17.2 | 18 | 18 | 311.14029 |
| Q9Y6Y1 | Calmodulin-binding transcription activato | CAMTA1 | 183.672 | 1 | 2 | 2 | 136.18073 |

| | | | | | | | |
|---|---|---|---|---|---|---|---|
| Q9Y6Y8 | SEC23-interacting protein OS=Homo sap | SEC23IP | 111.077 | 1 | 1 | 1 | 21.832525 |
| A0A024RB | Diphosphoinositol polyphosphate phosph | NUDT4B | 20.435 | 7.7 | 1 | 1 | 135.26849 |
| A0A8I5KQ | Small ribosomal subunit protein uS2B OS | RPSA2 | 32.91 | 22.7 | 7 | 7 | 7981.5933 |
| A0PJW6 | Transmembrane protein 223 OS=Homo sa | TMEM223 | 22.049 | 9.4 | 2 | 2 | 69.186356 |
| A1L020 | RNA-binding protein MEX3A OS=Homo | MEX3A | 54.174 | 7.7 | 3 | 3 | 99.624939 |
| A6NHQ4 | Elongin BC and Polycomb repressive con | EPOP | 39.322 | 6.6 | 2 | 2 | 90.567818 |
| A6NKF1 | SAC3 domain-containing protein 1 OS=H | SAC3D1 | 43.554 | 5.9 | 2 | 2 | 63.839439 |
| A6ZKI3 | Retrotransposon Gag-like protein 8C OS= | RTL8C | 13.172 | 18.6 | 2 | 1 | 94.473579 |
| A8K855 | EF-hand calcium-binding domain-contain | EFCAB7 | 71.982 | 5.4 | 3 | 3 | 57.794617 |
| A8MPP1 | Putative ATP-dependent RNA helicase D | DDX11L8 | 101.811 | 2.9 | 2 | 1 | 25.707617 |
| A8MV81 | HIG1 domain family member 1C OS=Ho | HIGD1C | 11.079 | 9.3 | 1 | 1 | 190.32448 |
| E9PAV3 | Nascent polypeptide-associated complex s | NACA | 205.422 | 2.6 | 4 | 3 | 857.86047 |
| G9CGD6 | CNK3/IPCEF1 fusion protein OS=Homo | CNK3/IPCE | 100.365 | 2.7 | 2 | 2 | 36.555618 |
| O00178 | GTP-binding protein 1 OS=Homo sapiens | GTPBP1 | 72.454 | 27.1 | 17 | 17 | 1027.9886 |
| O00458 | Interferon-related developmental regulato | IFRD1 | 50.269 | 15.1 | 7 | 7 | 347.33078 |
| O14545 | TRAF-type zinc finger domain-containing | TRAFD1 | 64.842 | 13.9 | 8 | 8 | 186.50784 |
| O14602 | Eukaryotic translation initiation factor 1A | EIF1AY | 16.443 | 6.9 | 1 | 1 | 292.63483 |
| O15063 | Granule associated Rac and RHOG effect | GARRE1 | 116.02 | 6.5 | 6 | 6 | 80.407089 |
| O15235 | Small ribosomal subunit protein uS12m O | MRPS12 | 15.173 | 44.9 | 6 | 6 | 672.45868 |
| O15370 | Transcription factor SOX-12 OS=Homo s | SOX12 | 34.122 | 8.9 | 3 | 2 | 90.085068 |
| O43414 | ERI1 exoribonuclease 3 OS=Homo sapier | ERI3 | 37.239 | 12.2 | 4 | 4 | 426.12531 |
| O75153 | Clustered mitochondria protein homolog ( | CLUH | 146.67 | 31.2 | 34 | 33 | 2420.3796 |
| O75352 | Mannose-P-dolichol utilization defect 1 p | MPDU1 | 26.638 | 4 | 1 | 1 | 109.8343 |
| O75380 | NADH dehydrogenase [ubiquinone] iron- | NDUFS6 | 13.712 | 18.5 | 2 | 2 | 224.79366 |
| O75394 | Large ribosomal subunit protein bL33m C | MRPL33 | 7.62 | 21.5 | 1 | 1 | 154.57419 |
| O75820 | Zinc finger protein 189 OS=Homo sapien | ZNF189 | 72.976 | 4.5 | 2 | 2 | 95.094666 |
| O75947 | ATP synthase subunit d, mitochondrial O | ATP5PD | 18.492 | 19.9 | 4 | 4 | 274.31171 |
| O75964 | ATP synthase subunit g, mitochondrial O | ATP5MG | 11.429 | 44.7 | 4 | 4 | 305.25851 |
| O95084 | Serine protease 23 OS=Homo sapiens OX | PRSS23 | 43.002 | 21.7 | 7 | 7 | 324.00974 |
| O95625 | Zinc finger and BTB domain-containing p | ZBTB11 | 119.384 | 5.9 | 6 | 6 | 154.15376 |
| O95685 | Protein phosphatase 1 regulatory subunit | PPP1R3D | 32.56 | 9 | 2 | 2 | 52.33313 |
| O95757 | Heat shock 70 kDa protein 4L OS=Homo | HSPA4L | 94.513 | 4.2 | 3 | 3 | 57.52692 |
| P09001 | Large ribosomal subunit protein uL3m O | MRPL3 | 38.633 | 32.2 | 11 | 11 | 869.21979 |
| P0DOX5 | Immunoglobulin gamma-1 heavy chain O | -- | 49.329 | 2.7 | 1 | 1 | 218.8822 |
| P16260 | Solute carrier family 25 member 16 OS=H | SLC25A16 | 36.224 | 6.3 | 2 | 2 | 14.840598 |

| | | | | | | | |
|---|---|---|---|---|---|---|---|
| P17029 | Zinc finger protein with KRAB and SCAN | ZKSCAN1 | 63.631 | 6.9 | 3 | 3 | 106.37276 |
| P17097 | Zinc finger protein 7 OS=Homo sapiens C | ZNF7 | 77.887 | 1.5 | 1 | 1 | 60.031261 |
| P17098 | Zinc finger protein 8 OS=Homo sapiens C | ZNF8 | 64.971 | 4.2 | 2 | 2 | 88.922844 |
| P31277 | Homeobox protein Hox-D11 OS=Homo s | HOXD11 | 35.198 | 2.1 | 1 | 1 | 190.54832 |
| P33552 | Cyclin-dependent kinases regulatory subu | CKS2 | 9.861 | 12.7 | 1 | 1 | 176.03932 |
| P35542 | Serum amyloid A-4 protein OS=Homo sa | SAA4 | 14.747 | 6.2 | 1 | 1 | 115.3739 |
| P36915 | Guanine nucleotide-binding protein-like 1 | GNL1 | 68.661 | 12.2 | 6 | 6 | 96.874802 |
| P42696 | RNA-binding protein 34 OS=Homo sapie | RBM34 | 48.565 | 15.6 | 6 | 6 | 206.39699 |
| P49406 | Large ribosomal subunit protein bL19m C | MRPL19 | 33.535 | 33.6 | 10 | 10 | 2023.213 |
| P49750 | YLP motif-containing protein 1 OS=Hom | YLPM1 | 241.646 | 23.4 | 50 | 50 | 1888.9468 |
| P56385 | ATP synthase subunit e, mitochondrial OS | ATP5ME | 7.934 | 60.9 | 4 | 4 | 154.41205 |
| P57775 | F-box/WD repeat-containing protein 4 OS | FBXW4 | 46.338 | 5.1 | 2 | 2 | 74.125061 |
| P58317 | Zinc finger protein 121 OS=Homo sapiens | ZNF121 | 44.695 | 38.7 | 13 | 8 | 513.48529 |
| P58557 | Endoribonuclease YbeY OS=Homo sapie | YBEY | 19.299 | 15.6 | 3 | 3 | 147.79973 |
| P82663 | Small ribosomal subunit protein mS25 OS | MRPS25 | 20.117 | 35.3 | 5 | 5 | 728.10193 |
| P82673 | Small ribosomal subunit protein mS35 OS | MRPS35 | 36.845 | 25.1 | 7 | 7 | 462.92474 |
| P82675 | Small ribosomal subunit protein uS5m OS | MRPS5 | 48.007 | 21.6 | 9 | 9 | 758.66992 |
| P82912 | Small ribosomal subunit protein uS11m O | MRPS11 | 20.616 | 22.2 | 5 | 5 | 479.64728 |
| P82921 | Small ribosomal subunit protein bS21m O | MRPS21 | 10.689 | 33.3 | 3 | 3 | 173.98651 |
| P82933 | Small ribosomal subunit protein uS9m OS | MRPS9 | 45.835 | 20.7 | 9 | 9 | 972.77887 |
| Q02509 | Otoconin-90 OS=Homo sapiens OX=960 | OC90 | 51.728 | 1.5 | 1 | 1 | 148.39952 |
| Q03701 | CCAAT/enhancer-binding protein zeta OS | CEBPZ | 120.974 | 16.4 | 16 | 16 | 269.03375 |
| Q05048 | Cleavage stimulation factor subunit 1 OS= | CSTF1 | 48.358 | 14.4 | 5 | 5 | 222.44638 |
| Q0VDF9 | Heat shock 70 kDa protein 14 OS=Homo | HSPA14 | 54.795 | 19.6 | 8 | 8 | 291.99277 |
| Q12872 | Splicing factor, suppressor of white-aprico | SFSWAP | 104.822 | 10 | 9 | 9 | 140.62926 |
| Q12894 | Interferon-related developmental regulato | IFRD2 | 48.048 | 11.5 | 7 | 7 | 266.19894 |
| Q12926 | ELAV-like protein 2 OS=Homo sapiens C | ELAVL2 | 39.504 | 26.5 | 10 | 2 | 2771.1067 |
| Q13442 | 28 kDa heat- and acid-stable phosphoprot | PDAP1 | 20.631 | 7.2 | 1 | 1 | 103.15976 |
| Q15181 | Inorganic pyrophosphatase OS=Homo sap | PPA1 | 32.661 | 15.6 | 4 | 4 | 332.41333 |
| Q15390 | Mitochondrial fission regulator 1 OS=Ho | MTFR1 | 37.001 | 7.2 | 2 | 2 | 138.32336 |
| Q15404 | Ras suppressor protein 1 OS=Homo sapie | RSU1 | 31.541 | 2.5 | 1 | 1 | 53.434765 |
| Q15695 | Putative U2 small nuclear ribonucleoprote | ZRSR2P1 | 57.644 | 11.3 | 5 | 5 | 89.936676 |
| Q15773 | Myeloid leukemia factor 2 OS=Homo sap | MLF2 | 28.148 | 11.7 | 3 | 3 | 94.886551 |
| Q17RY6 | Lymphocyte antigen 6K OS=Homo sapien | LY6K | 18.673 | 24.8 | 4 | 4 | 180.71783 |
| Q2NKX9 | UPF0561 protein C2orf68 OS=Homo sap | C2orf68 | 18.751 | 18.1 | 3 | 3 | 235.33386 |

| | | | | | | | |
|---|---|---|---|---|---|---|---|
| Q2NL82 | Pre-rRNA-processing protein TSR1 homc | TSR1 | 91.811 | 34.2 | 25 | 25 | 3634.2097 |
| Q2T9L4 | Inhibitory synaptic factor 1 OS=Homo saj | INSYN1 | 31.929 | 2.4 | 1 | 1 | 918.21442 |
| Q3B726 | DNA-directed RNA polymerase I subunit | POLR1F | 37.433 | 3.6 | 1 | 1 | 229.6629 |
| Q3KQU3 | MAP7 domain-containing protein 1 OS=H | MAP7D1 | 92.82 | 18.1 | 14 | 13 | 395.58588 |
| Q3T8J9 | GON-4-like protein OS=Homo sapiens O | GON4L | 248.621 | 1.3 | 2 | 1 | 51.560593 |
| Q3ZCT1 | Zinc finger protein 260 OS=Homo sapien | ZNF260 | 47.223 | 4.6 | 2 | 2 | 30.255358 |
| Q4U2R6 | Large ribosomal subunit protein mL51 OS | MRPL51 | 15.095 | 8.6 | 1 | 1 | 654.01514 |
| Q53HC5 | Kelch-like protein 26 OS=Homo sapiens ( | KLHL26 | 68.14 | 5.7 | 3 | 3 | 83.71991 |
| Q567V2 | Mpv17-like protein 2 OS=Homo sapiens ( | MPV17L2 | 23.181 | 13.6 | 2 | 2 | 112.26791 |
| Q58FF6 | Putative heat shock protein HSP 90-beta 4 | HSP90AB4 | 58.265 | 2.2 | 1 | 1 | 209.31064 |
| Q58FF8 | Putative heat shock protein HSP 90-beta 2 | HSP90AB2 | 44.349 | 3.9 | 1 | 1 | 53.884033 |
| Q5H9U9 | Probable ATP-dependent RNA helicase D | DDX60L | 197.674 | 16.8 | 26 | 24 | 316.81668 |
| Q5JSZ5 | Protein PRRC2B OS=Homo sapiens OX= | PRRC2B | 242.967 | 32.3 | 63 | 60 | 2054.4646 |
| Q5T653 | Large ribosomal subunit protein uL2m OS | MRPL2 | 33.301 | 17.7 | 4 | 4 | 141.12941 |
| Q5T8P6 | RNA-binding protein 26 OS=Homo sapie | RBM26 | 113.598 | 23.3 | 22 | 21 | 533.50415 |
| Q5TEC6 | Histone H3-7 OS=Homo sapiens OX=96( | H3-7 | 15.431 | 30.9 | 4 | 1 | 5894.4272 |
| Q5VU97 | VWFA and cache domain-containing prot | CACHD1 | 142.29 | 0.5 | 1 | 1 | 396.91205 |
| Q5VV52 | Zinc finger protein 691 OS=Homo sapien | ZNF691 | 35.81 | 8.3 | 2 | 2 | 123.22382 |
| Q658Y4 | Protein FAM91A1 OS=Homo sapiens OX | FAM91A1 | 93.909 | 28.4 | 23 | 22 | 2005.0059 |
| Q69YN2 | CWF19-like protein 1 OS=Homo sapiens | CWF19L1 | 60.62 | 8.2 | 4 | 4 | 80.251488 |
| Q6IPR3 | tRNA wybutosine-synthesizing protein 3 l | TYW3 | 29.795 | 12.4 | 3 | 3 | 87.84745 |
| Q6NUK4 | Receptor expression-enhancing protein 3 ( | REEP3 | 29.264 | 42 | 10 | 10 | 393.12616 |
| Q6NVV7 | Cysteine-rich DPF motif domain-containi | CDPF1 | 13.878 | 11.4 | 1 | 1 | 61.923687 |
| Q6NW34 | Nucleolus and neural progenitor protein C | NEPRO | 64.552 | 12.5 | 6 | 6 | 102.10736 |
| Q6NXE6 | Armadillo repeat-containing protein 6 OS | ARMC6 | 54.142 | 3.8 | 2 | 2 | 66.686813 |
| Q6P1L8 | Large ribosomal subunit protein uL14m C | MRPL14 | 15.948 | 22.1 | 3 | 3 | 594.64258 |
| Q6P1R4 | tRNA-dihydrouridine(16/17) synthase [N | DUS1L | 53.231 | 6.8 | 3 | 3 | 55.140488 |
| Q6P582 | Mitotic-spindle organizing protein 2A OS | MZT2A | 16.221 | 10.1 | 1 | 1 | 19.785173 |
| Q6PJG2 | Mitotic deacetylase-associated SANT don | MIDEAS | 114.989 | 19.7 | 17 | 17 | 287.54239 |
| Q6UXM1 | Leucine-rich repeats and immunoglobulin | LRIG3 | 123.434 | 3 | 3 | 3 | 23.413998 |
| Q6ZNB6 | NF-X1-type zinc finger protein NFXL1 O | NFXL1 | 101.34 | 24.5 | 20 | 20 | 534.04211 |
| Q6ZWK6 | Transmembrane protease serine 11F OS=l | TMPRSS11 | 49.41 | 1.8 | 1 | 1 | 216.41034 |
| Q7L2K0 | Tubulin epsilon and delta complex protei | TEDC2 | 46.403 | 4.2 | 2 | 2 | 86.559708 |
| Q7L3T8 | Probable proline--tRNA ligase, mitochonc | PARS2 | 53.263 | 16.6 | 7 | 7 | 154.73756 |
| Q7Z2T5 | TRMT1-like protein OS=Homo sapiens C | TRMT1L | 81.748 | 28.4 | 21 | 21 | 826.59668 |

| | | | | | | | |
|---|---|---|---|---|---|---|---|
| Q7Z2W9 | Large ribosomal subunit protein bL21m C | MRPL21 | 22.815 | 22.9 | 5 | 5 | 193.83043 |
| Q7Z408 | CUB and sushi domain-containing protein | CSMD2 | 380.04 | 0.4 | 1 | 1 | 36.24794 |
| Q7Z4Q2 | HEAT repeat-containing protein 3 OS=Ho | HEATR3 | 74.583 | 15.1 | 9 | 9 | 142.52815 |
| Q7Z7A3 | Cytoplasmic tRNA 2-thiolation protein 1 ( | CTU1 | 36.45 | 16.7 | 5 | 5 | 150.15112 |
| Q7Z7F7 | Large ribosomal subunit protein mL55 OS | MRPL55 | 15.129 | 7 | 1 | 1 | 96.019264 |
| Q86SX3 | Tubulin epsilon and delta complex protein | TEDC1 | 54.232 | 18.2 | 8 | 8 | 186.60266 |
| Q86TN4 | tRNA 2'-phosphotransferase 1 OS=Homo | TRPT1 | 27.742 | 9.5 | 2 | 2 | 94.703598 |
| Q86U38 | Nucleolar protein 9 OS=Homo sapiens OX | NOP9 | 69.438 | 12.6 | 7 | 7 | 288.04996 |
| Q86XH1 | Dynein regulatory complex protein 11 OS | IQCA1 | 95.342 | 1 | 1 | 1 | 138.09633 |
| Q86Y79 | Peptidyl-tRNA hydrolase OS=Homo sapie | PTRH1 | 22.937 | 43.9 | 8 | 8 | 812.32379 |
| Q8IUR0 | Trafficking protein particle complex subu | TRAPPC5 | 20.784 | 14.9 | 3 | 3 | 40.729618 |
| Q8IWR0 | Zinc finger CCCH domain-containing pro | ZC3H7A | 110.538 | 27 | 25 | 25 | 1524.1835 |
| Q8IWY8 | Zinc finger and SCAN domain-containing | ZSCAN29 | 96.72 | 3.5 | 3 | 2 | 45.183594 |
| Q8IX18 | Probable ATP-dependent RNA helicase D | DHX40 | 88.561 | 18.5 | 13 | 13 | 183.24843 |
| Q8IY67 | Ribonucleoprotein PTB-binding 1 OS=Ho | RAVER1 | 63.878 | 24.8 | 11 | 11 | 718.00531 |
| Q8IYL2 | Probable tRNA (uracil-O(2)-)-methyltrans | TRMT44 | 84.629 | 2.8 | 2 | 2 | 35.162613 |
| Q8N0T1 | Ribosomal biogenesis factor OS=Homo sa | RBIS | 11.457 | 55 | 6 | 6 | 219.79419 |
| Q8N0Z8 | tRNA pseudouridine synthase-like 1 OS=I | PUSL1 | 33.233 | 9.9 | 2 | 2 | 23.097847 |
| Q8N143 | B-cell CLL/lymphoma 6 member B protei | BCL6B | 51.532 | 2.3 | 1 | 1 | 57.560783 |
| Q8N1G4 | Leucine-rich repeat-containing protein 47 | LRRC47 | 63.473 | 18.5 | 9 | 9 | 788.65802 |
| Q8N3J9 | Zinc finger protein 664 OS=Homo sapiens | ZNF664 | 30.284 | 5 | 1 | 1 | 33.493053 |
| Q8N3X1 | Formin-binding protein 4 OS=Homo sapie | FNBP4 | 110.266 | 12 | 12 | 12 | 258.85773 |
| Q8N442 | Translation factor GUF1, mitochondrial C | GUF1 | 74.328 | 6.7 | 4 | 4 | 82.682846 |
| Q8N4T4 | Rho guanine nucleotide exchange factor 3 | ARHGEF39 | 38.296 | 11.6 | 4 | 4 | 207.09247 |
| Q8N5L8 | Ribonuclease P protein subunit p25-like p | RPP25L | 17.632 | 29.4 | 6 | 6 | 459.23532 |
| Q8N5N7 | Large ribosomal subunit protein mL50 OS | MRPL50 | 18.325 | 46.8 | 7 | 7 | 1496.0156 |
| Q8N7A1 | Kelch domain-containing protein 1 OS=H | KLHDC1 | 46.72 | 1.7 | 1 | 1 | 249.73116 |
| Q8N8A2 | Serine/threonine-protein phosphatase 6 re | ANKRD44 | 107.604 | 7.5 | 6 | 6 | 60.043633 |
| Q8N8A6 | ATP-dependent RNA helicase DDX51 OS | DDX51 | 72.458 | 5.6 | 3 | 3 | 67.67569 |
| Q8N954 | G patch domain-containing protein 11 OS | GPATCH11 | 33.278 | 20.4 | 6 | 6 | 277.13199 |
| Q8NAF0 | Zinc finger protein 579 OS=Homo sapiens | ZNF579 | 60.51 | 2.7 | 1 | 1 | 42.363445 |
| Q8NB91 | Fanconi anemia group B protein OS=Hom | FANCB | 97.727 | 7.5 | 6 | 6 | 75.213417 |
| Q8NBA8 | tRNA-uridine aminocarboxypropyltransfe | DTWD2 | 33.416 | 7.7 | 2 | 2 | 179.34015 |
| Q8NC96 | Adaptin ear-binding coat-associated prote | NECAP1 | 29.738 | 9.1 | 2 | 2 | 210.40735 |
| Q8ND82 | Zinc finger protein 280C OS=Homo sapie | ZNF280C | 83.096 | 3.3 | 2 | 2 | 28.586994 |

| | | | | | | | |
|---|---|---|---|---|---|---|---|
| Q8NDX6 | Zinc finger protein 740 OS=Homo sapien: | ZNF740 | 21.857 | 33.2 | 5 | 5 | 71.72715 |
| Q8NEF9 | Serum response factor-binding protein 1 ( | SRFBP1 | 48.634 | 27.5 | 11 | 11 | 326.9267 |
| Q8NFI3 | Cytosolic endo-beta-N-acetylglucosamini | ENGASE | 83.987 | 11.7 | 8 | 8 | 191.59106 |
| Q8TBB5 | Kelch domain-containing protein 4 OS=H | KLHDC4 | 57.892 | 26.9 | 11 | 11 | 252.31482 |
| Q8TBF4 | Zinc finger CCHC-type and RNA-binding | ZCRB1 | 24.592 | 14.3 | 2 | 2 | 21.521833 |
| Q8TCC3 | Large ribosomal subunit protein uL30m C | MRPL30 | 18.546 | 6.8 | 1 | 1 | 198.98955 |
| Q8TDN6 | Ribosome biogenesis protein BRX1 homc | BRIX1 | 41.402 | 24.6 | 8 | 8 | 652.87012 |
| Q8TF39 | Zinc finger protein 483 OS=Homo sapien: | ZNF483 | 85.099 | 1.2 | 1 | 1 | 59.405617 |
| Q8TF68 | Zinc finger protein 384 OS=Homo sapien: | ZNF384 | 63.22 | 15.1 | 7 | 6 | 218.82506 |
| Q8WUA2 | Peptidyl-prolyl cis-trans isomerase-like 4 | PPIL4 | 57.226 | 19.5 | 9 | 9 | 541.34497 |
| Q8WUF8 | Cotranscriptional regulator FAM172A OS | FAM172A | 47.973 | 2.4 | 1 | 1 | 35.028233 |
| Q8WUH1 | Protein Churchill OS=Homo sapiens OX= | CHURC1 | 12.887 | 16.1 | 1 | 1 | 89.320572 |
| Q8WUZ0 | B-cell CLL/lymphoma 7 protein family m | BCL7C | 23.469 | 5.5 | 1 | 1 | 196.77083 |
| Q8WV60 | Pentatricopeptide repeat-containing protei | PTCD2 | 43.968 | 18 | 8 | 8 | 743.15417 |
| Q8WVI0 | Ubiquinol-cytochrome c reductase comple | UQCC5 | 8.697 | 15.7 | 1 | 1 | 498.0004 |
| Q8WYQ9 | Zinc finger CCHC domain-containing pro | ZCCHC14 | 100.043 | 2.4 | 2 | 2 | 163.07582 |
| Q92610 | Zinc finger protein 592 OS=Homo sapien: | ZNF592 | 137.529 | 1 | 1 | 1 | 90.247765 |
| Q93073 | Selenocysteine insertion sequence-binding | SECISBP2L | 121.776 | 16.8 | 16 | 15 | 1198.6118 |
| Q969Q0 | Ribosomal protein eL42-like OS=Homo s | RPL36AL | 12.469 | 9.4 | 2 | 2 | 981.11487 |
| Q969W8 | Zinc finger protein 566 OS=Homo sapien: | ZNF566 | 49.22 | 6.9 | 3 | 3 | 107.02306 |
| Q96A35 | Large ribosomal subunit protein uL24m C | MRPL24 | 24.915 | 45.4 | 10 | 10 | 1328.8048 |
| Q96A47 | Insulin gene enhancer protein ISL-2 OS=I | ISL2 | 39.769 | 35.9 | 11 | 8 | 594.79834 |
| Q96BU1 | S100P-binding protein OS=Homo sapiens | S100PBP | 45.582 | 8.1 | 3 | 3 | 112.93413 |
| Q96C19 | EF-hand domain-containing protein D2 O | EFHD2 | 26.698 | 11.2 | 2 | 2 | 120.89711 |
| Q96C28 | Zinc finger protein 707 OS=Homo sapien: | ZNF707 | 43.088 | 6.7 | 2 | 2 | 44.82299 |
| Q96CW6 | Probable RNA polymerase II nuclear loca | SLC7A6OS | 35.028 | 10 | 3 | 3 | 177.16074 |
| Q96DY7 | Mdm2-binding protein OS=Homo sapiens | MTBP | 102.194 | 2.8 | 2 | 2 | 66.75795 |
| Q96EK7 | Constitutive coactivator of peroxisome pr | FAM120B | 103.784 | 25.6 | 19 | 19 | 223.59433 |
| Q96GA3 | Protein LTV1 homolog OS=Homo sapien | LTV1 | 54.855 | 33.1 | 13 | 13 | 1595.6743 |
| Q96GC5 | Large ribosomal subunit protein mL48 OS | MRPL48 | 23.935 | 5.2 | 1 | 1 | 201.82947 |
| Q96GY0 | Zinc finger C2HC domain-containing prot | ZC2HC1A | 35.093 | 36 | 13 | 13 | 570.24139 |
| Q96HE9 | Proline-rich protein 11 OS=Homo sapiens | PRR11 | 40.086 | 16.7 | 5 | 5 | 284.90668 |
| Q96JN8 | Neuralized-like protein 4 OS=Homo sapie | NEURL4 | 166.907 | 3.8 | 5 | 5 | 73.25737 |
| Q96KR1 | Zinc finger RNA-binding protein OS=Hoi | ZFR | 117.013 | 32.5 | 28 | 28 | 987.36884 |
| Q96M89 | Coiled-coil domain-containing protein 13 | CCDC138 | 76.219 | 1.5 | 1 | 1 | 84.303452 |

| | | | | | | | |
|---|---|---|---|---|---|---|---|
| Q96MX6 | Dynein axonemal assembly factor 10 OS= | DNAAF10 | 39.741 | 2.8 | 1 | 1 | 40.25872 |
| Q96PE2 | Rho guanine nucleotide exchange factor 1 | ARHGEF17 | 221.673 | 1.3 | 2 | 2 | 63.733459 |
| Q96SL8 | Flt3-interacting zinc finger protein 1 OS= | FIZ1 | 51.996 | 11.7 | 5 | 5 | 182.02725 |
| Q99442 | Translocation protein SEC62 OS=Homo s | SEC62 | 45.862 | 5.3 | 2 | 2 | 92.122032 |
| Q99805 | Transmembrane 9 superfamily member 2 | TM9SF2 | 75.776 | 1.8 | 1 | 1 | 118.45199 |
| Q99811 | Paired mesoderm homeobox protein 2 OS | PRRX2 | 27.08 | 22.5 | 5 | 3 | 198.64903 |
| Q9BPW4 | Apolipoprotein L4 OS=Homo sapiens OX | APOL4 | 39.165 | 5.7 | 2 | 2 | 60.727394 |
| Q9BQC6 | Large ribosomal subunit protein mL63 OS | MRPL57 | 12.267 | 12.7 | 1 | 1 | 48.989429 |
| Q9BRJ2 | Large ribosomal subunit protein mL45 OS | MRPL45 | 35.351 | 26.5 | 9 | 9 | 688.74744 |
| Q9BRR8 | G patch domain-containing protein 1 OS= | GPATCH1 | 103.345 | 2.1 | 2 | 2 | 23.635105 |
| Q9BRT6 | Protein LLP homolog OS=Homo sapiens | LLPH | 15.226 | 15.5 | 2 | 2 | 316.78275 |
| Q9BS18 | Anaphase-promoting complex subunit 13 | ANAPC13 | 8.522 | 13.5 | 1 | 1 | 63.529984 |
| Q9BSF4 | Mitochondrial import inner membrane tra | TIMM29 | 29.234 | 9.6 | 2 | 2 | 117.96422 |
| Q9BSF8 | BTB/POZ domain-containing protein 10 ( | BTBD10 | 53.78 | 2.1 | 1 | 1 | 82.783112 |
| Q9BSJ5 | Uncharacterized protein C17orf80 OS=Ho | C17orf80 | 67.316 | 11.8 | 6 | 6 | 61.469776 |
| Q9BUT9 | MAPK regulated corepressor interacting p | MCRIP2 | 17.828 | 34.4 | 5 | 5 | 828.24701 |
| Q9BVC5 | Ashwin OS=Homo sapiens OX=9606 GN | C2orf49 | 25.859 | 5.6 | 1 | 1 | 145.98071 |
| Q9BWU0 | Kanadaptin OS=Homo sapiens OX=9606 | SLC4A1AP | 88.815 | 11.1 | 8 | 8 | 156.38795 |
| Q9BXK1 | Krueppel-like factor 16 OS=Homo sapien | KLF16 | 25.431 | 19.8 | 7 | 7 | 228.39365 |
| Q9BYC8 | Large ribosomal subunit protein bL32m C | MRPL32 | 21.405 | 17 | 3 | 3 | 268.5719 |
| Q9BYC9 | Large ribosomal subunit protein bL20m C | MRPL20 | 17.443 | 14.1 | 2 | 2 | 68.850044 |
| Q9BYD1 | Large ribosomal subunit protein uL13m C | MRPL13 | 20.692 | 38.8 | 7 | 7 | 94.904648 |
| Q9BYD2 | Large ribosomal subunit protein bL9m OS | MRPL9 | 30.244 | 18.4 | 5 | 5 | 764.23993 |
| Q9BYD3 | Large ribosomal subunit protein uL4m OS | MRPL4 | 34.92 | 31.2 | 8 | 8 | 984.23676 |
| Q9BYN8 | Small ribosomal subunit protein mS26 OS | MRPS26 | 24.212 | 32.2 | 8 | 8 | 1744.136 |
| Q9BZE1 | Large ribosomal subunit protein mL37 OS | MRPL37 | 48.118 | 9.7 | 3 | 3 | 92.095032 |
| Q9C075 | Keratin, type I cytoskeletal 23 OS=Homo | KRT23 | 48.132 | 2.6 | 1 | 1 | 177.39774 |
| Q9C0D7 | Probable ribonuclease ZC3H12C OS=Hon | ZC3H12C | 99.341 | 18.1 | 15 | 9 | 258.61505 |
| Q9C0H5 | Rho GTPase-activating protein 39 OS=Ho | ARHGAP39 | 121.286 | 8.1 | 7 | 7 | 78.971611 |
| Q9GZP4 | PITH domain-containing protein 1 OS=Ho | PITHD1 | 24.178 | 14.2 | 2 | 2 | 50.875816 |
| Q9GZR7 | ATP-dependent RNA helicase DDX24 OS | DDX24 | 96.332 | 17.1 | 13 | 13 | 416.34357 |
| Q9H061 | Transmembrane protein 126A OS=Homo | TMEM126A | 21.528 | 5.1 | 1 | 1 | 295.9201 |
| Q9H0U9 | Testis-specific Y-encoded-like protein 1 ( | TSPYL1 | 49.193 | 12.6 | 5 | 5 | 181.99211 |
| Q9H0X4 | Protein FAM234A OS=Homo sapiens OX | FAM234A | 59.66 | 1.8 | 1 | 1 | 36.459484 |
| Q9H1A3 | Protein-L-histidine N-pros-methyltransfer | METTL9 | 36.537 | 17.6 | 5 | 5 | 538.05859 |

| | | | | | | | |
|---|---|---|---|---|---|---|---|
| Q9H2W6 | Large ribosomal subunit protein mL46 OS | MRPL46 | 31.706 | 27.2 | 7 | 7 | 332.53842 |
| Q9H3C7 | Gametogenetin-binding protein 2 OS=Hor | GGNBP2 | 79.087 | 28.3 | 16 | 16 | 2301.0662 |
| Q9H501 | ESF1 homolog OS=Homo sapiens OX=9( | ESF1 | 98.797 | 17.5 | 13 | 13 | 233.55043 |
| Q9H5L6 | DNA transposase THAP9 OS=Homo sapi | THAP9 | 103.411 | 1 | 1 | 1 | 20.930315 |
| Q9H6F5 | Coiled-coil domain-containing protein 86 | CCDC86 | 40.236 | 23.1 | 9 | 9 | 525.92566 |
| Q9H8H0 | Nucleolar protein 11 OS=Homo sapiens C | NOL11 | 81.124 | 21.4 | 14 | 14 | 255.08604 |
| Q9H967 | WD repeat-containing protein 76 OS=Hor | WDR76 | 69.77 | 13.3 | 8 | 8 | 88.226707 |
| Q9H9Y2 | Ribosome production factor 1 OS=Homo | RPF1 | 40.111 | 10.9 | 3 | 3 | 100.1419 |
| Q9HA92 | Radical S-adenosyl methionine domain-co | RSAD1 | 48.714 | 5 | 2 | 2 | 76.044487 |
| Q9HCN8 | Stromal cell-derived factor 2-like protein | SDF2L1 | 23.599 | 5.9 | 1 | 1 | 186.7784 |
| Q9HD33 | Large ribosomal subunit protein uL29m C | MRPL47 | 29.451 | 38.8 | 10 | 10 | 696.50928 |
| Q9NP92 | Large ribosomal subunit protein mL65 OS | MRPS30 | 50.365 | 9.8 | 4 | 4 | 89.306824 |
| Q9NPA3 | Mid1-interacting protein 1 OS=Homo sap | MID1IP1 | 20.202 | 25.7 | 3 | 3 | 55.382278 |
| Q9NQS1 | Cell death regulator Aven OS=Homo sapi | AVEN | 38.507 | 22.9 | 6 | 6 | 281.67346 |
| Q9NRM2 | Zinc finger protein 277 OS=Homo sapien: | ZNF277 | 52.788 | 16.9 | 7 | 7 | 306.28366 |
| Q9NRX2 | Large ribosomal subunit protein bL17m C | MRPL17 | 20.051 | 32.6 | 7 | 7 | 782.14337 |
| Q9NSI2 | Ribosome biogenesis protein SLX9 homo | SLX9 | 25.457 | 29.6 | 7 | 7 | 1208.2122 |
| Q9NTZ6 | RNA-binding protein 12 OS=Homo sapie | RBM12 | 97.396 | 3.9 | 4 | 4 | 85.135574 |
| Q9NUQ6 | SPATS2-like protein OS=Homo sapiens ( | SPATS2L | 61.729 | 31.7 | 18 | 18 | 3059.4141 |
| Q9NUQ8 | ATP-binding cassette sub-family F memb | ABCF3 | 79.745 | 16.6 | 11 | 11 | 110.14736 |
| Q9NVE5 | Ubiquitin carboxyl-terminal hydrolase 40 | USP40 | 140.13 | 6.5 | 7 | 7 | 128.22188 |
| Q9NVR7 | TBCC domain-containing protein 1 OS=H | TBCCD1 | 63.531 | 4.1 | 2 | 2 | 72.322617 |
| Q9NVS2 | Large ribosomal subunit protein mL66 OS | MRPS18A | 22.184 | 9.2 | 2 | 2 | 79.865669 |
| Q9NVX7 | Kelch repeat and BTB domain-containing | KBTBD4 | 58.144 | 4.4 | 2 | 2 | 77.517303 |
| Q9NWA0 | Mediator of RNA polymerase II transcript | MED9 | 16.403 | 12.3 | 1 | 1 | 81.968719 |
| Q9NWB6 | Arginine and glutamate-rich protein 1 OS= | ARGLU1 | 33.217 | 16.8 | 6 | 6 | 590.29535 |
| Q9NX20 | Large ribosomal subunit protein uL16m C | MRPL16 | 28.45 | 23.5 | 5 | 5 | 428.55469 |
| Q9NXF1 | Testis-expressed protein 10 OS=Homo sa | TEX10 | 105.675 | 8 | 7 | 7 | 89.440346 |
| Q9NYL9 | Tropomodulin-3 OS=Homo sapiens OX=9 | TMOD3 | 39.595 | 28.4 | 10 | 9 | 397.03122 |
| Q9NZW5 | Protein PALS2 OS=Homo sapiens OX=9( | PALS2 | 61.117 | 4.1 | 2 | 1 | 37.640316 |
| Q9P015 | Large ribosomal subunit protein uL15m C | MRPL15 | 33.42 | 28.7 | 10 | 10 | 1197.3463 |
| Q9P031 | Thyroid transcription factor 1-associated p | CCDC59 | 28.67 | 20.7 | 5 | 5 | 313.23578 |
| Q9P0J6 | Large ribosomal subunit protein bL36m C | MRPL36 | 11.785 | 7.8 | 1 | 1 | 675.13409 |
| Q9P0M9 | Large ribosomal subunit protein bL27m C | MRPL27 | 16.073 | 29.7 | 4 | 4 | 154.08653 |
| Q9P1Y6 | PHD and RING finger domain-containing | PHRF1 | 178.667 | 2.2 | 3 | 3 | 89.178619 |

| | | | | | | | |
|---|---|---|---|---|---|---|---|
| Q9P206 | Uncharacterized protein KIAA1522 OS=I | KIAA1522 | 107.095 | 19.8 | 15 | 15 | 962.02881 |
| Q9P242 | Neuronal tyrosine-phosphorylated phosph | NYAP2 | 70.548 | 4.9 | 2 | 2 | 38.953056 |
| Q9P265 | Disco-interacting protein 2 homolog B OS | DIP2B | 171.492 | 15.9 | 20 | 19 | 208.40086 |
| Q9P2F8 | Signal-induced proliferation-associated 1- | SIPA1L2 | 190.438 | 3.5 | 5 | 5 | 70.870949 |
| Q9P2Z0 | THAP domain-containing protein 10 OS= | THAP10 | 28.351 | 11.3 | 2 | 2 | 25.788248 |
| Q9UHA3 | Probable ribosome biogenesis protein RLI | RSL24D1 | 19.622 | 11.7 | 2 | 2 | 251.0448 |
| Q9UHR6 | Zinc finger HIT domain-containing protei | ZNHIT2 | 42.884 | 6.2 | 2 | 2 | 64.610001 |
| Q9UHW5 | GPN-loop GTPase 3 OS=Homo sapiens ( | GPN3 | 32.762 | 9.2 | 2 | 2 | 104.55589 |
| Q9UJK0 | 18S rRNA aminocarboxypropyltransferas | TSR3 | 33.596 | 22.8 | 6 | 6 | 391.94797 |
| Q9UKD2 | mRNA turnover protein 4 homolog OS=H | MRTO4 | 27.561 | 41 | 12 | 12 | 2001.5552 |
| Q9UKJ3 | G patch domain-containing protein 8 OS= | GPATCH8 | 164.198 | 7 | 9 | 8 | 107.67581 |
| Q9UKZ1 | CCR4-NOT transcription complex subuni | CNOT11 | 55.216 | 12.2 | 4 | 4 | 94.765839 |
| Q9ULL5 | Proline-rich protein 12 OS=Homo sapiens | PRR12 | 211.045 | 2.5 | 5 | 5 | 98.80098 |
| Q9UN30 | Sex comb on midleg-like protein 1 OS=H | SCML1 | 37.448 | 2.1 | 1 | 1 | 98.736298 |
| Q9UNK9 | Protein angel homolog 1 OS=Homo sapie | ANGEL1 | 75.276 | 4.3 | 3 | 3 | 119.80428 |
| Q9Y2D9 | Zinc finger protein 652 OS=Homo sapien | ZNF652 | 69.745 | 7.4 | 4 | 1 | 83.428452 |
| Q9Y2L6 | FERM domain-containing protein 4B OS= | FRMD4B | 118.047 | 0.8 | 1 | 1 | 85.395363 |
| Q9Y2Q9 | Small ribosomal subunit protein bS1m OS | MRPS28 | 20.843 | 19.8 | 4 | 4 | 1600.4128 |
| Q9Y2R5 | Small ribosomal subunit protein uS17m O | MRPS17 | 14.503 | 22.3 | 2 | 2 | 1039.5516 |
| Q9Y2R9 | Small ribosomal subunit protein uS7m OS | MRPS7 | 28.135 | 52.1 | 11 | 11 | 2916.9165 |
| Q9Y305 | Acyl-coenzyme A thioesterase 9, mitocho | ACOT9 | 49.902 | 29.4 | 13 | 13 | 525.52142 |
| Q9Y3C1 | Nucleolar protein 16 OS=Homo sapiens ( | NOP16 | 21.189 | 38.8 | 8 | 8 | 1489.8859 |
| Q9Y3X0 | Coiled-coil domain-containing protein 9 ( | CCDC9 | 59.703 | 22 | 14 | 14 | 997.37885 |
| Q9Y421 | Protein FAM32A OS=Homo sapiens OX= | FAM32A | 13.179 | 35.7 | 4 | 4 | 186.84418 |
| Q9Y4F5 | Centrosomal protein of 170 kDa protein E | CEP170B | 171.689 | 10.6 | 14 | 14 | 102.79034 |
| Q9Y4P3 | Transducin beta-like protein 2 OS=Homo | TBL2 | 49.798 | 43.4 | 19 | 19 | 1549.7645 |
| Q9Y5J9 | Mitochondrial import inner membrane tra | TIMM8B | 9.344 | 26.5 | 2 | 2 | 60.9035 |
| Q9Y5M8 | Signal recognition particle receptor subun | SRPRB | 29.703 | 36.5 | 10 | 10 | 591.07819 |
| Q9Y6V7 | Probable ATP-dependent RNA helicase D | DDX49 | 54.227 | 33.5 | 17 | 17 | 10277.279 |
| A0A075B6I | Probable non-functional immunoglobulin | IGKV3-7 | 12.784 | 7.8 | 1 | 1 | 5571.3477 |
| A0A075B6( | Immunoglobulin heavy variable 3-64 OS= | IGHV3-64 | 12.891 | 9.3 | 1 | 1 | 47.864841 |
| A0A075B6I | Probable non-functional immunoglobulin | IGKV2D-24 | 13.079 | 18.3 | 2 | 1 | 1534.0225 |
| A0A075B6! | Immunoglobulin kappa variable 2D-29 OS | IGKV2D-29 | 13.144 | 20 | 2 | 1 | 17160.646 |
| A0A087WS | Immunoglobulin kappa variable 3D-15 OS | IGKV3D-15 | 12.535 | 7.8 | 1 | 1 | 237.53035 |
| A0A0A0MS | Immunoglobulin heavy variable 1-45 OS= | IGHV1-45 | 13.509 | 9.4 | 1 | 1 | 699.29346 |

| | | | | | | | |
|---|---|---|---|---|---|---|---|
| A0A0A0MS | Immunoglobulin heavy variable 3-49 OS= | IGHV3-49 | 13.056 | 16.8 | 2 | 1 | 151.27322 |
| A0A0B4J1X | Immunoglobulin heavy variable 3-74 OS= | IGHV3-74 | 12.84 | 24.8 | 3 | 0 | 130.90549 |
| A0A0B4J2I | Immunoglobulin heavy variable 1-69D OS | IGHV1-69D | 12.661 | 9.4 | 1 | 1 | 126.08485 |
| A0A0C4DH | Immunoglobulin kappa variable 3D-20 OS | IGKV3D-2( | 12.516 | 13.8 | 1 | 1 | 74.455872 |
| A0A0C4DH | Immunoglobulin heavy variable 1-18 OS= | IGHV1-18 | 12.821 | 9.4 | 1 | 1 | 29.451866 |
| A0A0J9YX | Immunoglobulin heavy variable 3-64D OS | IGHV3-64D | 12.823 | 9.4 | 1 | 1 | 1238.1682 |
| A8MWD9 | Putative small nuclear ribonucleoprotein ( | SNRPGP15 | 8.545 | 25 | 2 | 2 | 5084.6938 |
| B5ME19 | Eukaryotic translation initiation factor 3 s | EIF3CL | 105.473 | 24.6 | 24 | 24 | 3218.04 |
| B9A064 | Immunoglobulin lambda-like polypeptide | IGLL5 | 23.064 | 3.7 | 1 | 1 | 317.33917 |
| E7ETH6 | Zinc finger protein 587B OS=Homo sapie | ZNF587B | 45.542 | 4.5 | 2 | 2 | 32.200081 |
| L0R819 | ASNSD1 upstream open reading frame pr | ASDURF | 11.25 | 24 | 2 | 2 | 88.439545 |
| O15037 | Protein KHNYN OS=Homo sapiens OX= | KHNYN | 74.534 | 14.3 | 8 | 8 | 128.84662 |
| O15050 | TPR and ankyrin repeat-containing protei | TRANK1 | 336.222 | 5.2 | 13 | 13 | 90.619545 |
| O43824 | Putative GTP-binding protein 6 OS=Hom | GTPBP6 | 56.898 | 25.2 | 12 | 12 | 271.42413 |
| O60287 | Nucleolar pre-ribosomal-associated protei | URB1 | 254.39 | 2.2 | 4 | 4 | 73.365364 |
| O75794 | Cell division cycle protein 123 homolog ( | CDC123 | 39.135 | 25.9 | 7 | 7 | 238.27646 |
| O95478 | Ribosome biogenesis protein NSA2 homo | NSA2 | 30.066 | 28.8 | 7 | 7 | 463.29959 |
| P0CG12 | Decreased expression in renal and prostat | DERPC | 51.392 | 4.8 | 2 | 2 | 58.916264 |
| P0DPB5 | Protein POLR1D, isoform 2 OS=Homo sa | POLR1D | 14.333 | 16.4 | 2 | 1 | 75.042015 |
| P16415 | Zinc finger protein 823 OS=Homo sapien | ZNF823 | 70.271 | 1.5 | 1 | 1 | 105.39409 |
| P17017 | Zinc finger protein 14 OS=Homo sapiens | ZNF14 | 75.354 | 5.3 | 3 | 3 | 28.398222 |
| P52943 | Cysteine-rich protein 2 OS=Homo sapiens | CRIP2 | 22.493 | 20.2 | 2 | 2 | 60.034828 |
| P78312 | Protein FAM193A OS=Homo sapiens OX | FAM193A | 139.988 | 14.8 | 16 | 16 | 314.36987 |
| P82664 | Small ribosomal subunit protein uS10m O | MRPS10 | 23 | 25.4 | 4 | 4 | 423.99323 |
| P86790 | Vacuolar fusion protein CCZ1 homolog B | CCZ1B | 55.867 | 6.8 | 3 | 3 | 49.792488 |
| Q08AF3 | Schlafen family member 5 OS=Homo sap | SLFN5 | 101.055 | 3 | 3 | 3 | 49.017685 |
| Q15032 | R3H domain-containing protein 1 OS=Ho | R3HDM1 | 120.697 | 12.8 | 13 | 13 | 3224.5225 |
| Q15434 | RNA-binding motif, single-stranded-inter | RBMS2 | 43.959 | 12.5 | 4 | 4 | 2464.1929 |
| Q15527 | Surfeit locus protein 2 OS=Homo sapiens | SURF2 | 29.618 | 18.8 | 5 | 5 | 90.035408 |
| Q1ED39 | Lysine-rich nucleolar protein 1 OS=Homo | KNOP1 | 51.59 | 19.9 | 9 | 9 | 178.05962 |
| Q1KMD3 | Heterogeneous nuclear ribonucleoprotein | HNRNPUL2 | 85.105 | 22.8 | 16 | 16 | 753.29553 |
| Q2KHR3 | Glutamine and serine-rich protein 1 OS=H | QSER1 | 189.973 | 0.8 | 1 | 1 | 88.378395 |
| Q2M296 | Methenyltetrahydrofolate synthase domain | MTHFSD | 42.173 | 24.5 | 8 | 8 | 256.89124 |
| Q2TB10 | Zinc finger protein 800 OS=Homo sapien | ZNF800 | 75.236 | 4.4 | 3 | 3 | 56.634918 |
| Q499Z4 | Zinc finger protein 672 OS=Homo sapien | ZNF672 | 50.225 | 14.6 | 6 | 6 | 114.44088 |

| | | | | | | | |
|---|---|---|---|---|---|---|---|
| Q49AG3 | Zinc finger BED domain-containing prote | ZBED5 | 78.911 | 11.7 | 8 | 8 | 58.987225 |
| Q4VXU2 | Polyadenylate-binding protein 1-like OS= | PABPC1L | 68.393 | 1.6 | 1 | 1 | 39.50544 |
| Q56VL3 | OCIA domain-containing protein 2 OS=H | OCIAD2 | 16.954 | 6.5 | 1 | 1 | 12.699386 |
| Q5SWX8 | Protein odr-4 homolog OS=Homo sapiens | ODR4 | 51.104 | 5.9 | 2 | 2 | 34.181801 |
| Q5SYE7 | NHS-like protein 1 OS=Homo sapiens OX | NHSL1 | 170.669 | 0.9 | 1 | 1 | 169.12663 |
| Q5T1J5 | Putative coiled-coil-helix-coiled-coil-heli | CHCHD2P9 | 15.49 | 15.9 | 1 | 1 | 70.143463 |
| Q5T3I0 | G patch domain-containing protein 4 OS= | GPATCH4 | 50.382 | 25.6 | 10 | 10 | 1148.0273 |
| Q5VTE6 | Protein angel homolog 2 OS=Homo sapie | ANGEL2 | 62.339 | 3.9 | 2 | 2 | 44.93795 |
| Q5VTL8 | Pre-mRNA-splicing factor 38B OS=Hom | PRPF38B | 64.468 | 20.1 | 10 | 10 | 1977.8298 |
| Q5VWQ0 | Lysine-specific demethylase 9 OS=Homo | RSBN1 | 90.073 | 16.6 | 11 | 9 | 165.54767 |
| Q5VYV7 | Protein SLX4IP OS=Homo sapiens OX=9 | SLX4IP | 45.552 | 6.9 | 2 | 2 | 60.902199 |
| Q6AI08 | HEAT repeat-containing protein 6 OS=H | HEATR6 | 128.782 | 3.5 | 3 | 3 | 81.905548 |
| Q6DD87 | Zinc finger protein 787 OS=Homo sapien | ZNF787 | 40.429 | 35.3 | 10 | 10 | 1015.6679 |
| Q6DKI1 | Ribosomal protein uL30-like OS=Homo s | RPL7L1 | 29.67 | 29.4 | 8 | 8 | 1073.8909 |
| Q6ICC9 | Retrotransposon Gag-like protein 6 OS=H | RTL6 | 26.155 | 8.8 | 2 | 2 | 62.193554 |
| Q6ICG6 | Uncharacterized protein KIAA0930 OS=H | KIAA0930 | 45.795 | 18.3 | 7 | 7 | 90.728523 |
| Q6IQ21 | Zinc finger protein 770 OS=Homo sapien | ZNF770 | 80.008 | 2.7 | 2 | 2 | 67.793945 |
| Q6NW29 | RWD domain-containing protein 4 OS=H | RWDD4 | 21.251 | 10.6 | 2 | 2 | 127.99886 |
| Q6NX49 | Zinc finger protein 544 OS=Homo sapien | ZNF544 | 81.742 | 1.8 | 1 | 1 | 49.873787 |
| Q6P161 | Large ribosomal subunit protein mL54 OS | MRPL54 | 15.82 | 7.2 | 1 | 1 | 1300.0587 |
| Q6PCB5 | Lysine-specific demethylase RSBN1L OS | RSBN1L | 94.87 | 20.9 | 15 | 15 | 356.36847 |
| Q6PK04 | Coiled-coil domain-containing protein 13 | CCDC137 | 33.232 | 33.2 | 10 | 10 | 862.33044 |
| Q6Y2X3 | DnaJ homolog subfamily C member 14 O | DNAJC14 | 78.569 | 10.1 | 7 | 7 | 112.40831 |
| Q6ZMW3 | Echinoderm microtubule-associated prote | EML6 | 217.9 | 0.6 | 1 | 1 | 57.675968 |
| Q6ZN06 | Zinc finger protein 813 OS=Homo sapien | ZNF813 | 71.721 | 1.5 | 1 | 0 | 238.07899 |
| Q6ZUT6 | Coiled-coil domain-containing protein 9B | CCDC9B | 57.325 | 6.4 | 3 | 3 | 68.427551 |
| Q7Z309 | PABIR family member 2 OS=Homo sapie | PABIR2 | 26.929 | 4.5 | 1 | 1 | 35.20293 |
| Q7Z422 | SUZ domain-containing protein 1 OS=Ho | SZRD1 | 16.998 | 32.2 | 4 | 4 | 131.9003 |
| Q7Z736 | Pleckstrin homology domain-containing fa | PLEKHH3 | 85.318 | 12.2 | 8 | 8 | 90.32122 |
| Q7Z7K0 | COX assembly mitochondrial protein hom | CMC1 | 12.49 | 9.4 | 1 | 1 | 116.1627 |
| Q86VU5 | Catechol O-methyltransferase domain-con | COMTD1 | 28.809 | 5.7 | 1 | 1 | 114.93988 |
| Q86WR7 | Proline and serine-rich protein 2 OS=Hon | PROSER2 | 45.802 | 16.3 | 6 | 6 | 163.11267 |
| Q86XN6 | Zinc finger protein 761 OS=Homo sapien | ZNF761 | 87.717 | 7.5 | 5 | 1 | 226.51012 |
| Q86XZ4 | Spermatogenesis-associated serine-rich pr | SPATS2 | 59.545 | 32.3 | 15 | 15 | 3594.011 |
| Q86YQ8 | Copine-8 OS=Homo sapiens OX=9606 G | CPNE8 | 63.108 | 2.3 | 1 | 1 | 86.005714 |

| | | | | | | | |
|---|---|---|---|---|---|---|---|
| Q8IXT5 | RNA-binding protein 12B OS=Homo sap | RBM12B | 118.103 | 40.9 | 36 | 36 | 1295.3567 |
| Q8IXX5 | Transmembrane protein 183A OS=Homo | TMEM183A | 42.827 | 11.7 | 4 | 2 | 86.435654 |
| Q8IYL3 | UPF0688 protein C1orf174 OS=Homo sa | C1orf174 | 25.977 | 5.8 | 1 | 1 | 62.94471 |
| Q8IYN0 | Zinc finger protein 100 OS=Homo sapien | ZNF100 | 62.745 | 2.2 | 1 | 0 | 20.154261 |
| Q8N531 | F-box/LRR-repeat protein 6 OS=Homo sa | FBXL6 | 58.559 | 14.3 | 6 | 6 | 127.97021 |
| Q8N567 | Zinc finger CCHC domain-containing pro | ZCCHC9 | 30.477 | 24 | 6 | 6 | 441.87744 |
| Q8N5C6 | S1 RNA-binding domain-containing prote | SRBD1 | 111.776 | 33.9 | 30 | 30 | 618.69604 |
| Q8N5I9 | NOP protein chaperone 1 OS=Homo sapi | NOPCHAP1 | 20.124 | 21.6 | 4 | 4 | 139.91269 |
| Q8N6N3 | UPF0690 protein C1orf52 OS=Homo sap | C1orf52 | 20.599 | 19.2 | 3 | 3 | 43.095352 |
| Q8N8D1 | Programmed cell death protein 7 OS=Hor | PDCD7 | 54.7 | 2.9 | 1 | 1 | 48.277287 |
| Q8N9T8 | Protein KRI1 homolog OS=Homo sapiens | KRI1 | 82.599 | 13.9 | 8 | 8 | 188.51103 |
| Q8NB46 | Serine/threonine-protein phosphatase 6 re | ANKRD52 | 115.077 | 8.3 | 8 | 8 | 284.97961 |
| Q8NE00 | Transmembrane protein 104 OS=Homo sa | TMEM104 | 55.679 | 1.4 | 1 | 1 | 219.38 |
| Q8NEM2 | SHC SH2 domain-binding protein 1 OS=I | SHCBP1 | 75.691 | 29.6 | 17 | 17 | 1595.3481 |
| Q8TAV0 | Protein FAM76A OS=Homo sapiens OX= | FAM76A | 35.05 | 3.6 | 1 | 1 | 50.660046 |
| Q8TBR7 | TLC domain-containing protein 3A OS=H | TLCD3A | 29.383 | 10.9 | 2 | 2 | 62.084862 |
| Q8WUH6 | Transmembrane protein 263 OS=Homo sa | TMEM263 | 11.749 | 12.9 | 1 | 1 | 93.660889 |
| Q8WUY9 | DEP domain-containing protein 1B OS=H | DEPDC1B | 61.772 | 13.2 | 7 | 6 | 103.47694 |
| Q8WXE0 | Caskin-2 OS=Homo sapiens OX=9606 GI | CASKIN2 | 126.784 | 3.2 | 3 | 2 | 53.175423 |
| Q92545 | Transmembrane protein 131 OS=Homo sa | TMEM131 | 205.138 | 6.5 | 9 | 9 | 151.3945 |
| Q92576 | PHD finger protein 3 OS=Homo sapiens ( | PHF3 | 229.482 | 9.8 | 18 | 18 | 219.6871 |
| Q93075 | Putative deoxyribonuclease TATDN2 OS | TATDN2 | 85.024 | 10 | 7 | 7 | 82.907722 |
| Q969Y0 | NXPE family member 3 OS=Homo sapien | NXPE3 | 63.8 | 4.5 | 2 | 2 | 44.206478 |
| Q96C57 | Protein CUSTOS OS=Homo sapiens OX= | CUSTOS | 28.171 | 22.1 | 4 | 4 | 127.05176 |
| Q96E39 | RNA binding motif protein, X-linked-like | RBMXL1 | 42.142 | 7.2 | 3 | 3 | 94.368484 |
| Q96EI5 | Transcription elongation factor A protein- | TCEAL4 | 24.648 | 23.3 | 5 | 4 | 68.751816 |
| Q96EL2 | Small ribosomal subunit protein uS3m OS | MRPS24 | 19.016 | 28.7 | 4 | 4 | 494.74475 |
| Q96ET8 | Golgi apparatus membrane protein TVP23 | TVP23C | 31.105 | 6.5 | 2 | 2 | 61.47039 |
| Q96EV2 | RNA-binding protein 33 OS=Homo sapie | RBM33 | 129.986 | 12.6 | 12 | 12 | 504.08286 |
| Q96F63 | Coiled-coil domain-containing protein 97 | CCDC97 | 38.947 | 2.9 | 1 | 1 | 72.521042 |
| Q96H12 | Myb/SANT-like DNA-binding domain-co | MSANTD3 | 32.364 | 11.6 | 3 | 3 | 29.760391 |
| Q96HP4 | Oxidoreductase NAD-binding domain-coi | OXNAD1 | 34.855 | 3.5 | 1 | 1 | 124.21304 |
| Q96JF6 | Zinc finger protein 594 OS=Homo sapien | ZNF594 | 93.907 | 1.1 | 1 | 1 | 282.70392 |
| Q96ME7 | Zinc finger protein 512 OS=Homo sapien | ZNF512 | 64.683 | 10.8 | 5 | 5 | 121.83868 |
| Q96MN5 | Transcription elongation factor A N-termi | TCEANC2 | 24.151 | 14.9 | 3 | 3 | 60.751572 |

| | | | | | | | |
|---|---|---|---|---|---|---|---|
| Q96N46 | Tetratricopeptide repeat protein 14 OS=H | TTC14 | 88.319 | 7 | 4 | 4 | 45.002121 |
| Q96NA8 | t-SNARE domain-containing protein 1 OS | TSNARE1 | 55.949 | 6 | 2 | 2 | 41.460155 |
| Q96S19 | Methyltransferase-like 26 OS=Homo sapi | METTL26 | 22.579 | 6.9 | 1 | 1 | 40.136288 |
| Q99676 | Zinc finger protein 184 OS=Homo sapiens | ZNF184 | 86.175 | 1.6 | 1 | 1 | 47.541809 |
| Q9BQ48 | Large ribosomal subunit protein bL34m C | MRPL34 | 10.165 | 23.9 | 2 | 2 | 487.74405 |
| Q9BQ75 | Protein CMSS1 OS=Homo sapiens OX=9 | CMSS1 | 31.885 | 25.8 | 9 | 9 | 2830.4229 |
| Q9BQE6 | LBH domain-containing protein 1 OS=Ho | LBHD1 | 31.566 | 4.5 | 1 | 1 | 101.84477 |
| Q9BR77 | Coiled-coil domain-containing protein 77 | CCDC77 | 57.487 | 16.8 | 8 | 8 | 296.02982 |
| Q9BSC4 | Nucleolar protein 10 OS=Homo sapiens C | NOL10 | 80.302 | 11 | 8 | 8 | 146.53421 |
| Q9BTF0 | THUMP domain-containing protein 2 OS= | THUMPD2 | 56.434 | 4.4 | 2 | 2 | 15.033356 |
| Q9BTY7 | Protein HGH1 homolog OS=Homo sapien | HGH1 | 42.13 | 12.1 | 3 | 3 | 33.067509 |
| Q9BWG4 | Single-stranded DNA-binding protein 4 O | SSBP4 | 39.389 | 7 | 2 | 1 | 291.88068 |
| Q9BXW7 | Haloacid dehalogenase-like hydrolase dor | HDHD5 | 46.322 | 2.8 | 1 | 1 | 74.65815 |
| Q9BXY0 | Protein MAK16 homolog OS=Homo sapi | MAK16 | 35.369 | 19.7 | 5 | 5 | 171.57944 |
| Q9BYB4 | Guanine nucleotide-binding protein subun | GNB1L | 35.618 | 30.9 | 8 | 8 | 146.47302 |
| Q9BYD6 | Large ribosomal subunit protein uL1m OS | MRPL1 | 36.909 | 13.8 | 5 | 5 | 335.45154 |
| Q9C0B9 | Zinc finger CCHC domain-containing pro | ZCCHC2 | 125.936 | 9.5 | 7 | 7 | 379.09949 |
| Q9GZU3 | Transmembrane protein 39B OS=Homo s | TMEM39B | 56.274 | 3.5 | 1 | 1 | 98.15789 |
| Q9H568 | Actin-like protein 8 OS=Homo sapiens O | ACTL8 | 41.361 | 12.3 | 4 | 4 | 68.756866 |
| Q9H6Y2 | WD repeat-containing protein 55 OS=Hoi | WDR55 | 42.071 | 28.2 | 8 | 8 | 188.20892 |
| Q9H857 | 5'-nucleotidase domain-containing protein | NT5DC2 | 60.719 | 9.6 | 5 | 5 | 172.57437 |
| Q9H8G1 | Zinc finger protein 430 OS=Homo sapiens | ZNF430 | 66.319 | 8.1 | 4 | 0 | 134.42282 |
| Q9H8G2 | Caspase activity and apoptosis inhibitor 1 | CAAP1 | 38.368 | 10.2 | 3 | 3 | 43.969429 |
| Q9H9L3 | Interferon-stimulated 20 kDa exonuclease | ISG20L2 | 39.154 | 24.4 | 8 | 8 | 489.78201 |
| Q9HAH1 | Zinc finger protein 556 OS=Homo sapiens | ZNF556 | 51.582 | 5 | 2 | 2 | 39.201538 |
| Q9HC44 | Vasculin-like protein 1 OS=Homo sapiens | GPBP1L1 | 52.302 | 26.2 | 11 | 11 | 577.78809 |
| Q9HCJ3 | Ribonucleoprotein PTB-binding 2 OS=Ho | RAVER2 | 74.339 | 14 | 6 | 6 | 140.87807 |
| Q9HD45 | Transmembrane 9 superfamily member 3 | TM9SF3 | 67.889 | 1.9 | 1 | 1 | 26.04467 |
| Q9NPI7 | Lysine-rich coiled-coil protein 1 OS=Hon | KRCC1 | 30.98 | 17 | 4 | 4 | 95.307602 |
| Q9NQX6 | Zinc finger protein 331 OS=Homo sapiens | ZNF331 | 53.739 | 2.4 | 1 | 1 | 37.239258 |
| Q9NW61 | Pleckstrin homology domain-containing fa | PLEKHJ1 | 17.552 | 9.4 | 1 | 1 | 214.57098 |
| Q9NW82 | WD repeat-containing protein 70 OS=Hoi | WDR70 | 73.202 | 5.8 | 3 | 3 | 53.220654 |
| Q9NX05 | Constitutive coactivator of PPAR-gamma- | FAM120C | 120.577 | 24.6 | 23 | 23 | 687.20416 |
| Q9NXD2 | Myotubularin-related protein 10 OS=Hon | MTMR10 | 88.274 | 6.8 | 4 | 4 | 84.744514 |
| Q9NXH8 | Torsin-4A OS=Homo sapiens OX=9606 ( | TOR4A | 46.915 | 7.1 | 2 | 2 | 75.514442 |

| | | | | | | | |
|---|---|---|---|---|---|---|---|
| Q9NXT0 | Zinc finger protein 586 OS=Homo sapiens | ZNF586 | 46.414 | 5 | 1 | 1 | 60.97506 |
| Q9P016 | Thymocyte nuclear protein 1 OS=Homo s | THYN1 | 25.698 | 36 | 8 | 8 | 406.41257 |
| Q9P227 | Rho GTPase-activating protein 23 OS=Hc | ARHGAP23 | 162.192 | 5.7 | 7 | 7 | 72.962784 |
| Q9P2B7 | Cilia- and flagella-associated protein 97 C | CFAP97 | 59.476 | 15.4 | 8 | 8 | 205.39989 |
| Q9P2F5 | Storkhead-box protein 2 OS=Homo sapiei | STOX2 | 102.672 | 5 | 4 | 4 | 85.409637 |
| Q9P2N5 | RNA-binding protein 27 OS=Homo sapie | RBM27 | 118.718 | 19.9 | 20 | 20 | 944.94629 |
| Q9UBI9 | Headcase protein homolog OS=Homo sap | HECA | 58.837 | 22.1 | 11 | 11 | 251.10327 |
| Q9UG63 | ATP-binding cassette sub-family F memb | ABCF2 | 71.291 | 35.3 | 22 | 22 | 2767.1995 |
| Q9UJJ7 | RNA pseudouridylate synthase domain-cc | RPUSD1 | 34.756 | 2.9 | 1 | 1 | 95.231499 |
| Q9UJN7 | Zinc finger protein 391 OS=Homo sapiens | ZNF391 | 40.66 | 7.8 | 3 | 2 | 98.876228 |
| Q9UJT0 | Tubulin epsilon chain OS=Homo sapiens | TUBE1 | 52.932 | 6.9 | 3 | 3 | 40.881798 |
| Q9UL59 | Zinc finger protein 214 OS=Homo sapiens | ZNF214 | 70.992 | 1.5 | 1 | 1 | 87.82222 |
| Q9UNX3 | Ribosomal protein uL24-like OS=Homo s | RPL26L1 | 17.257 | 11.7 | 2 | 2 | 1406.9385 |
| Q9Y291 | Small ribosomal subunit protein mS33 OS | MRPS33 | 12.629 | 38.7 | 4 | 4 | 114.65418 |
| Q9Y3B9 | RRP15-like protein OS=Homo sapiens O | RRP15 | 31.485 | 32.3 | 9 | 9 | 479.71573 |
| Q9Y3D5 | Small ribosomal subunit protein bS18m O | MRPS18C | 15.85 | 14.1 | 2 | 2 | 436.02884 |
| Q9Y3T6 | R3H and coiled-coil domain-containing pı | R3HCC1 | 49.092 | 13 | 5 | 5 | 105.71871 |
| Q9Y426 | C2 domain-containing protein 2 OS=Hom | C2CD2 | 75.534 | 4.5 | 2 | 2 | 119.04793 |
| Q9Y485 | DmX-like protein 1 OS=Homo sapiens O | DMXL1 | 337.839 | 0.7 | 2 | 2 | 59.049641 |
| Q9Y4E6 | WD repeat-containing protein 7 OS=Hom | WDR7 | 163.811 | 1.1 | 1 | 1 | 29.274349 |
| A0A087WU | Neuroblastoma breakpoint family member | NBPF19 | 440.409 | 0.3 | 1 | 1 | 80.557526 |
| A1L429 | G antigen 12B/C/D/E OS=Homo sapiens | GAGE12B | 12.925 | 16.2 | 2 | 1 | 806.81891 |
| L0R6Q1 | SLC35A4 upstream open reading frame p | SLC35A4 | 11.133 | 9.7 | 1 | 1 | 88.833687 |
| P0DMU7 | Cancer/testis antigen family 45 member A | CT45A6 | 21.232 | 7.9 | 1 | 1 | 113.94914 |
| Q13670 | Putative postmeiotic segregation increased | PMS2P11 | 28.555 | 5.2 | 1 | 1 | 72.353851 |
| Q2TB18 | Protein asteroid homolog 1 OS=Homo sap | ASTE1 | 77.094 | 7.5 | 5 | 5 | 137.46451 |
| Q32NC0 | UPF0711 protein C18orf21 OS=Homo sa | C18orf21 | 24.827 | 6.4 | 1 | 1 | 20.989891 |
| Q49AM3 | Tetratricopeptide repeat protein 31 OS=H | TTC31 | 57.105 | 18.1 | 8 | 8 | 161.94214 |
| Q5BKY9 | Protein FAM133B OS=Homo sapiens OX | FAM133B | 28.385 | 7.3 | 2 | 2 | 149.49649 |
| Q5JUR7 | Testis-expressed protein 30 OS=Homo sa | TEX30 | 25.585 | 8.4 | 2 | 2 | 124.69713 |
| Q5R3I4 | Tetratricopeptide repeat protein 38 OS=H | TTC38 | 52.788 | 14.5 | 6 | 6 | 132.31094 |
| Q5RGS3 | Protein FAM74A1 OS=Homo sapiens OX | FAM74A1 | 14.779 | 6.3 | 1 | 1 | 175.85281 |
| Q5SVZ6 | Zinc finger MYM-type protein 1 OS=Hon | ZMYM1 | 128.717 | 4.6 | 5 | 5 | 46.565025 |
| Q5T6C5 | Ataxin-7-like protein 2 OS=Homo sapiens | ATXN7L2 | 77.181 | 5 | 3 | 3 | 139.43123 |
| Q5T8I3 | EEIG family member 2 OS=Homo sapien | EEIG2 | 39.309 | 17.8 | 6 | 6 | 93.281403 |

| | | | | | | | |
|---|---|---|---|---|---|---|---|
| Q69YL0 | Protein NCBP2AS2 OS=Homo sapiens O | NCBP2AS2 | 10.891 | 28.3 | 2 | 2 | 78.783875 |
| Q6DCA0 | AMMECR1-like protein OS=Homo sapie | AMMECR1 | 34.5 | 18.1 | 5 | 4 | 153.45181 |
| Q6P1M9 | Armadillo repeat-containing X-linked pro | ARMCX5 | 62.346 | 4.8 | 2 | 2 | 56.286068 |
| Q6P3X3 | Tetratricopeptide repeat protein 27 OS=H | TTC27 | 96.633 | 10.9 | 8 | 8 | 88.3927 |
| Q6P3X8 | PiggyBac transposable element-derived pr | PGBD2 | 68.012 | 1.2 | 1 | 1 | 19.251831 |
| Q6P5R6 | Ribosomal protein eL22-like OS=Homo s | RPL22L1 | 14.607 | 19.7 | 2 | 2 | 102.1027 |
| Q6ZSJ8 | Uncharacterized protein C1orf122 OS=Ho | C1orf122 | 11.471 | 33.6 | 3 | 3 | 324.8085 |
| Q6ZSR9 | Uncharacterized protein FLJ45252 OS=H | -- | 37.977 | 2.8 | 1 | 1 | 171.94524 |
| Q7Z5U6 | WD repeat-containing protein 53 OS=Hor | WDR53 | 38.99 | 3.4 | 1 | 1 | 63.561935 |
| Q7Z6I8 | UPF0461 protein C5orf24 OS=Homo sap | C5orf24 | 20.132 | 12.8 | 2 | 2 | 66.660339 |
| Q86VQ1 | Glucocorticoid-induced transcript 1 protei | GLCCI1 | 58.024 | 9.9 | 4 | 4 | 37.255508 |
| Q86X53 | Glutamate-rich protein 1 OS=Homo sapie | ERICH1 | 48.985 | 8.6 | 3 | 3 | 97.952927 |
| Q8IVB5 | LIX1-like protein OS=Homo sapiens OX= | LIX1L | 36.563 | 4.2 | 1 | 1 | 18.203917 |
| Q8IZQ5 | Selenoprotein H OS=Homo sapiens OX=9 | SELENOH | 13.454 | 9.8 | 1 | 1 | 38.639462 |
| Q8N2G6 | Zinc finger CCHC domain-containing pro | ZCCHC24 | 26.955 | 16.6 | 3 | 3 | 173.12302 |
| Q8NBP0 | Tetratricopeptide repeat protein 13 OS=H | TTC13 | 96.814 | 13.6 | 10 | 10 | 108.78894 |
| Q8NCJ5 | SPRY domain-containing protein 3 OS=H | SPRYD3 | 49.694 | 18.3 | 7 | 7 | 144.3726 |
| Q8ND24 | RING finger protein 214 OS=Homo sapie | RNF214 | 77.668 | 19.5 | 12 | 12 | 187.97273 |
| Q8WUY1 | Protein THEM6 OS=Homo sapiens OX=9 | THEM6 | 23.865 | 13 | 3 | 3 | 150.00024 |
| Q8WW59 | SPRY domain-containing protein 4 OS=H | SPRYD4 | 23.099 | 4.8 | 1 | 1 | 16.671032 |
| Q96A19 | Coiled-coil domain-containing protein 10 | CCDC102A | 62.596 | 14.5 | 7 | 7 | 211.28221 |
| Q96B54 | Zinc finger protein 428 OS=Homo sapiens | ZNF428 | 20.481 | 20.7 | 3 | 3 | 321.61679 |
| Q96B70 | Leukocyte receptor cluster member 9 OS= | LENG9 | 53.167 | 21 | 7 | 7 | 119.34654 |
| Q96CP2 | FLYWCH family member 2 OS=Homo sa | FLYWCH2 | 14.564 | 26.4 | 3 | 2 | 64.951195 |
| Q96D70 | R3H domain-containing protein 4 OS=Ho | R3HDM4 | 30.35 | 4.5 | 1 | 1 | 94.744278 |
| Q96H35 | Probable RNA-binding protein 18 OS=Ho | RBM18 | 21.649 | 20.5 | 3 | 3 | 88.344498 |
| Q9BRJ6 | Uncharacterized protein C7orf50 OS=Hor | C7orf50 | 22.084 | 27.3 | 4 | 4 | 2101.0579 |
| Q9BRP1 | Programmed cell death protein 2-like OS= | PDCD2L | 39.417 | 9.5 | 3 | 3 | 374.97134 |
| Q9BVG4 | Protein PBDC1 OS=Homo sapiens OX=9 | PBDC1 | 26.057 | 9 | 2 | 2 | 38.042034 |
| Q9H1X3 | DnaJ homolog subfamily C member 25 O | DNAJC25 | 42.404 | 8.6 | 3 | 3 | 78.154274 |
| Q9H4I3 | TraB domain-containing protein OS=Hom | TRABD | 42.322 | 2.1 | 1 | 1 | 37.429432 |
| Q9H8W3 | Protein FAM204A OS=Homo sapiens OX | FAM204A | 27.022 | 18.5 | 4 | 4 | 210.5063 |
| Q9NWW7 | Uncharacterized protein C2orf42 OS=Hor | C2orf42 | 64.077 | 8.7 | 4 | 4 | 43.306614 |
| Q9NX00 | Transmembrane protein 160 OS=Homo sa | TMEM160 | 19.658 | 11.2 | 2 | 2 | 162.696 |
| Q9NX31 | Oxidative stress-responsive serine-rich pro | OSER1 | 31.78 | 30.5 | 6 | 6 | 155.20323 |

| | | | | | | | |
|---|---|---|---|---|---|---|---|
| Q9UNZ5 | Leydig cell tumor 10 kDa protein homolo; | C19orf53 | 10.577 | 17.2 | 3 | 3 | 1077.9611 |
| Q9Y5V0 | Zinc finger protein 706 OS=Homo sapien: | ZNF706 | 8.498 | 30.3 | 2 | 2 | 1232.563 |
| A0A1W2P( | Myb/SANT-like DNA-binding domain-co | MSANTD7 | 39.761 | 4.7 | 1 | 1 | 25.592131 |
| E9PRG8 | Uncharacterized protein C11orf98 OS=H( | C11orf98 | 14.234 | 29.3 | 4 | 4 | 1821.967 |
| Q96BQ5 | Coiled-coil domain-containing protein 12' | CCDC127 | 30.835 | 11.9 | 3 | 3 | 49.998001 |
| Q9UF83 | Uncharacterized protein DKFZp434B061 | -- | 59.412 | 8 | 4 | 4 | 152.30038 |

| Sequence | Length | Protein accession | Protein description | Gene name | MW [kDa] | Charge | Score | Unique [yes/no] | PEP | m/z | Intensity PINK1 |
|---|---|---|---|---|---|---|---|---|---|---|---|
| LADFGFGNFYK | 11 | A0A0B4J2F2 | Putative serine/threonine-protein kinase S | SIK1B | 84.931 | 2 | 53.488495 | yes | 9.722E-08 | 639.81134 | 72.787872 |
| GFAFVEYESHR | 11 | A0AV96 | RNA-binding protein 47 OS=Homo sapie | RBM47 | 64.1 | 2 | 52.953346 | no | 1.028E-07 | 671.31476 | 3315.4282 |
| SFGQFNPGCVER | 12 | A0AV96 | RNA-binding protein 47 OS=Homo sapie | RBM47 | 64.1 | 2 | 52.953346 | yes | 9.722E-08 | 699.31696 | 3013.1924 |
| DYAFVHFTSR | 10 | A0AV96 | RNA-binding protein 47 OS=Homo sapie | RBM47 | 64.1 | 2 | 52.953346 | yes | 9.722E-08 | 621.79877 | 2682.3418 |
| PITPVYTVAPNVQ | 14 | A0AV96 | RNA-binding protein 47 OS=Homo sapie | RBM47 | 64.1 | 2 | 52.953346 | yes | 1.164E-07 | 777.93555 | 2166.9014 |
| LLGVCCSVDNCR | 12 | A0AV96 | RNA-binding protein 47 OS=Homo sapie | RBM47 | 64.1 | 2 | 52.953346 | yes | 1.073E-07 | 726.82324 | 1688.7045 |
| NLMIETTEDTIK | 12 | A0AV96 | RNA-binding protein 47 OS=Homo sapie | RBM47 | 64.1 | 2 | 52.953346 | yes | 1.135E-07 | 704.35541 | 1250.0859 |
| EEILEEIAK | 9 | A0AV96 | RNA-binding protein 47 OS=Homo sapie | RBM47 | 64.1 | 2 | 52.953346 | yes | 9.722E-08 | 537.28986 | 729.85437 |
| GYAFVMYCHK | 10 | A0AV96 | RNA-binding protein 47 OS=Homo sapie | RBM47 | 64.1 | 2 | 52.953346 | yes | 9.722E-08 | 638.28607 | 665.73657 |
| NLMIETTEDTIKK | 13 | A0AV96 | RNA-binding protein 47 OS=Homo sapie | RBM47 | 64.1 | 2 | 52.953346 | yes | 1.328E-07 | 768.40289 | 524.96893 |
| ELNNYEIR | 8 | A0AV96 | RNA-binding protein 47 OS=Homo sapie | RBM47 | 64.1 | 2 | 52.953346 | yes | 1.646E-07 | 525.76434 | 261.95929 |
| ELNNYEIRPGR | 11 | A0AV96 | RNA-binding protein 47 OS=Homo sapie | RBM47 | 64.1 | 2 | 52.953346 | yes | 9.81E-08 | 680.85205 | 58.966919 |
| ELNNYEIRPGR | 11 | A0AV96 | RNA-binding protein 47 OS=Homo sapie | RBM47 | 64.1 | 2 | 52.953346 | yes | 1.207E-07 | 454.23712 | 58.966919 |
| ELNNYEIRPGR | 11 | A0AV96 | RNA-binding protein 47 OS=Homo sapie | RBM47 | 64.1 | 3 | 52.953346 | yes | 9.81E-08 | 680.85205 | 58.966919 |
| ELNNYEIRPGR | 11 | A0AV96 | RNA-binding protein 47 OS=Homo sapie | RBM47 | 64.1 | 3 | 52.953346 | yes | 1.207E-07 | 454.23712 | 58.966919 |
| KREEILEEIAK | 11 | A0AV96 | RNA-binding protein 47 OS=Homo sapie | RBM47 | 64.1 | 2 | 52.953346 | yes | 1.183E-07 | 679.38794 | 85.670349 |
| REEILEEIAK | 10 | A0AV96 | RNA-binding protein 47 OS=Homo sapie | RBM47 | 64.1 | 2 | 52.953346 | yes | 2.118E-07 | 615.34045 | 74.53756 |
| TGYSMVQENGQR | 12 | A0AV96 | RNA-binding protein 47 OS=Homo sapie | RBM47 | 64.1 | 2 | 52.953346 | yes | 9.728E-08 | 685.31189 | 61.844128 |
| VPEGVAGAPNEA/ | 20 | A0AV96 | RNA-binding protein 47 OS=Homo sapie | RBM47 | 64.1 | 3 | 52.953346 | yes | 1.396E-07 | 670.02258 | 54.43853 |
| YGGPPPGWEGPHI | 15 | A0AV96 | RNA-binding protein 47 OS=Homo sapie | RBM47 | 64.1 | 2 | 52.953346 | yes | 2.277E-07 | 816.38934 | 45.411911 |
| SSSSLLASPGHISV | 15 | A0FGR8 | Extended synaptotagmin-2 OS=Homo sap | ESYT2 | 102.358 | 2 | 53.759075 | yes | 9.722E-08 | 735.40155 | 292.64297 |
| VLVALASEELAK | 12 | A0FGR8 | Extended synaptotagmin-2 OS=Homo sap | ESYT2 | 102.358 | 2 | 53.759075 | yes | 1.274E-07 | 621.86884 | 272.83389 |
| ITVPLVSEVQIAQL | 15 | A0FGR8 | Extended synaptotagmin-2 OS=Homo sap | ESYT2 | 102.358 | 2 | 53.759075 | yes | 1.233E-07 | 833.49854 | 222.86749 |
| IHFIEAQDLQGK | 12 | A0FGR8 | Extended synaptotagmin-2 OS=Homo sap | ESYT2 | 102.358 | 2 | 53.759075 | yes | 9.857E-08 | 699.87244 | 187.1181 |
| ALALLEDEER | 10 | A0FGR8 | Extended synaptotagmin-2 OS=Homo sap | ESYT2 | 102.358 | 2 | 53.759075 | yes | 1.02E-07 | 579.80371 | 184.07851 |
| VGNQIFQSR | 9 | A0FGR8 | Extended synaptotagmin-2 OS=Homo sap | ESYT2 | 102.358 | 2 | 53.759075 | yes | 3.478E-07 | 524.78033 | 167.09012 |
| SIQIHGTMR | 9 | A0FGR8 | Extended synaptotagmin-2 OS=Homo sap | ESYT2 | 102.358 | 2 | 53.759075 | yes | 8.885E-07 | 521.77673 | 137.24048 |
| LIVVVHACR | 9 | A0FGR8 | Extended synaptotagmin-2 OS=Homo sap | ESYT2 | 102.358 | 2 | 53.759075 | yes | 1.198E-07 | 533.81311 | 108.09221 |
| NLIAFSEDGSDPY\ | 15 | A0FGR8 | Extended synaptotagmin-2 OS=Homo sap | ESYT2 | 102.358 | 2 | 53.759075 | yes | 9.722E-08 | 841.90466 | 70.899727 |
| RQDLEVEVR | 9 | A0FGR8 | Extended synaptotagmin-2 OS=Homo sap | ESYT2 | 102.358 | 2 | 53.759075 | yes | 1.15E-07 | 572.30945 | 62.85656 |
| VDVGQQPLR | 9 | A0FGR8 | Extended synaptotagmin-2 OS=Homo sap | ESYT2 | 102.358 | 2 | 53.759075 | yes | 1.125E-07 | 506.28271 | 62.761063 |
| GANTHLSTFSFTK | 13 | A0FGR8 | Extended synaptotagmin-2 OS=Homo sap | ESYT2 | 102.358 | 2 | 53.759075 | yes | 9.722E-08 | 705.85425 | 57.506828 |
| LLDEWFTLDEVPK | 13 | A0FGR8 | Extended synaptotagmin-2 OS=Homo sap | ESYT2 | 102.358 | 2 | 53.759075 | yes | 3.425E-05 | 802.91394 | 38.097168 |
| ACDLPAWVHFPD | 15 | A0FGR8 | Extended synaptotagmin-2 OS=Homo sap | ESYT2 | 102.358 | 3 | 53.759075 | yes | 3.378E-07 | 605.28241 | 24.008066 |
| LMHIQPPK | 8 | A0JLT2 | Mediator of RNA polymerase II transcrip | MED19 | 26.274 | 2 | 42.998386 | yes | 2.776E-07 | 482.27585 | 63.606518 |
| SGAGCGPFYLMR | 12 | A0JLT2 | Mediator of RNA polymerase II transcrip | MED19 | 26.274 | 2 | 42.998386 | yes | 2.833E-07 | 658.29974 | 44.547989 |
| PSSSDTFFR | 9 | A0JNW5 | Bridge-like lipid transfer protein family m | BLTP3B | 164.199 | 2 | 18.483032 | yes | 5.313E-06 | 522.24329 | 448.94446 |

| | | | | | | | | | | | |
|---|---|---|---|---|---|---|---|---|---|---|---|
| LGDGEVFFLK | 10 | A1A4S6 | Rho GTPase-activating protein 10 OS=Ho | ARHGAP1( | 89.375 | 2 | 43.18329 | yes | 2.695E-07 | 562.80298 | 61.345829 |
| PNFLAPLPHEVTQI | 15 | A1X283 | SH3 and PX domain-containing protein 2 | SH3PXD2E | 101.58 | 3 | 46.06591 | yes | 5.238E-07 | 577.98956 | 308.1055 |
| SVPVPLQEAPQQR | 13 | A1X283 | SH3 and PX domain-containing protein 2 | SH3PXD2E | 101.58 | 2 | 46.06591 | yes | 1.663E-07 | 724.89642 | 51.92136 |
| VMDWYQTR | 8 | A2RRP1 | NBAS subunit of NRZ tethering complex | NBAS | 268.572 | 2 | 36.399487 | yes | 7.036E-07 | 557.75293 | 643.17908 |
| EISPATPNMQK | 11 | A2RUB6 | Coiled-coil domain-containing protein 66 | CCDC66 | 109.412 | 2 | 49.908825 | yes | 1.914E-07 | 608.30554 | 49.829796 |
| SVLNEHQETSK | 11 | A2RUB6 | Coiled-coil domain-containing protein 66 | CCDC66 | 109.412 | 2 | 49.908825 | yes | 9.989E-08 | 636.31494 | 38.979942 |
| SMTALLDPAQIEEI | 14 | A2RUB6 | Coiled-coil domain-containing protein 66 | CCDC66 | 109.412 | 2 | 49.908825 | yes | 4.271E-07 | 787.39813 | 24.744469 |
| NCLPVNTPQSR | 11 | A2VDJ0 | Transmembrane protein 131-like OS=Hor | TMEM131I | 179.34 | 2 | 42.267178 | yes | 6.023E-07 | 643.31952 | 52.128841 |
| YASGINVNLQK | 11 | A2VDJ0 | Transmembrane protein 131-like OS=Hor | TMEM131I | 179.34 | 2 | 42.267178 | yes | 3.114E-07 | 603.82751 | 46.944782 |
| GCYLESDDVLR | 11 | A2VDJ0 | Transmembrane protein 131-like OS=Hor | TMEM131I | 179.34 | 2 | 42.267178 | yes | 5.493E-06 | 663.80334 | 39.839867 |
| LVLLDGIHR | 9 | A3KMH1 | von Willebrand factor A domain-containi | VWA8 | 214.825 | 2 | 52.41734 | yes | 1.089E-07 | 518.31909 | 107.14443 |
| VIALGLPVPR | 10 | A3KMH1 | von Willebrand factor A domain-containi | VWA8 | 214.825 | 2 | 52.41734 | yes | 1.412E-07 | 517.83966 | 45.11084 |
| VNAGTLAVLQR | 11 | A3KMH1 | von Willebrand factor A domain-containi | VWA8 | 214.825 | 2 | 52.41734 | yes | 9.722E-08 | 571.33801 | 44.938984 |
| ILVAQGGLGGK | 11 | A4D1E9 | GTP-binding protein 10 OS=Homo sapiei | GTPBP10 | 42.933 | 2 | 53.209286 | yes | 4.321E-07 | 506.81113 | 486.07074 |
| TVEFQHIIPISAVTC | 22 | A4D1E9 | GTP-binding protein 10 OS=Homo sapiei | GTPBP10 | 42.933 | 3 | 53.209286 | yes | 1.082E-07 | 804.10034 | 291.30539 |
| SSLLSCVSHAK | 11 | A4D1E9 | GTP-binding protein 10 OS=Homo sapiei | GTPBP10 | 42.933 | 2 | 53.209286 | yes | 1.436E-07 | 594.80573 | 274.8703 |
| LIADVGLVGFPNA | 15 | A4D1E9 | GTP-binding protein 10 OS=Homo sapiei | GTPBP10 | 42.933 | 2 | 53.209286 | yes | 1.211E-07 | 735.91937 | 261.56955 |
| MDLPDAQDK | 9 | A4D1E9 | GTP-binding protein 10 OS=Homo sapiei | GTPBP10 | 42.933 | 2 | 53.209286 | yes | 9.722E-08 | 516.73694 | 239.49959 |
| GGDVWVVAQNR | 11 | A4D1E9 | GTP-binding protein 10 OS=Homo sapiei | GTPBP10 | 42.933 | 2 | 53.209286 | yes | 1.022E-07 | 600.80963 | 224.04515 |
| DFLHLFEK | 8 | A4D1E9 | GTP-binding protein 10 OS=Homo sapiei | GTPBP10 | 42.933 | 2 | 53.209286 | yes | 9.722E-08 | 524.77673 | 172.94026 |
| PAIADYAFTTLKPI | 17 | A4D1E9 | GTP-binding protein 10 OS=Homo sapiei | GTPBP10 | 42.933 | 3 | 53.209286 | yes | 1.287E-07 | 612.33649 | 108.22336 |
| FHELMSQLQNPK | 12 | A4D1E9 | GTP-binding protein 10 OS=Homo sapiei | GTPBP10 | 42.933 | 2 | 53.209286 | yes | 9.722E-08 | 736.37177 | 89.005577 |
| LLTNFLPLK | 9 | A4D1E9 | GTP-binding protein 10 OS=Homo sapiei | GTPBP10 | 42.933 | 2 | 53.209286 | yes | 1.335E-07 | 529.83405 | 63.897072 |
| QISVADLPGLIEGA | 18 | A4D1E9 | GTP-binding protein 10 OS=Homo sapiei | GTPBP10 | 42.933 | 3 | 53.209286 | yes | 4.964E-07 | 631.66815 | 48.58947 |
| EELQTKPALLAVN | 14 | A4D1E9 | GTP-binding protein 10 OS=Homo sapiei | GTPBP10 | 42.933 | 2 | 53.209286 | yes | 1.379E-07 | 777.44849 | 44.67588 |
| TAFETIILLTK | 11 | A4D1E9 | GTP-binding protein 10 OS=Homo sapiei | GTPBP10 | 42.933 | 2 | 53.209286 | yes | 3.724E-07 | 625.37372 | 25.104815 |
| ANLNALDR | 8 | A5A3E0 | POTE ankyrin domain family member F ( | POTEF | 121.445 | 2 | 48.42318 | no | 1.15E-07 | 443.74069 | 1578.1166 |
| NLEFQGPLEISR | 12 | A5D8V6 | Vacuolar protein sorting-associated protei | VPS37C | 38.659 | 2 | 25.35781 | yes | 3.295E-06 | 701.86987 | 58.857521 |
| IVEPPENIQEK | 11 | A5YKK6 | CCR4-NOT transcription complex subuni | CNOT1 | 266.939 | 2 | 55.33279 | yes | 1.667E-06 | 648.3457 | 870.50427 |
| NIIVQFGVR | 9 | A5YKK6 | CCR4-NOT transcription complex subuni | CNOT1 | 266.939 | 2 | 55.33279 | yes | 1.801E-07 | 523.31128 | 752.09796 |
| AVQELVHPVVDR | 12 | A5YKK6 | CCR4-NOT transcription complex subuni | CNOT1 | 266.939 | 2 | 55.33279 | yes | 9.722E-08 | 681.38043 | 744.72552 |
| LATEFELR | 8 | A5YKK6 | CCR4-NOT transcription complex subuni | CNOT1 | 266.939 | 2 | 55.33279 | yes | 4.385E-06 | 489.76639 | 587.4585 |
| IAMTTCEQIVR | 11 | A5YKK6 | CCR4-NOT transcription complex subuni | CNOT1 | 266.939 | 2 | 55.33279 | yes | 9.85E-08 | 661.3338 | 566.19849 |
| GNAPEGLPQLMEV | 15 | A5YKK6 | CCR4-NOT transcription complex subuni | CNOT1 | 266.939 | 2 | 55.33279 | yes | 2.18E-07 | 805.42194 | 555.14899 |
| SPVTFLSDLR | 10 | A5YKK6 | CCR4-NOT transcription complex subuni | CNOT1 | 266.939 | 2 | 55.33279 | yes | 9.789E-08 | 567.81134 | 554.40961 |
| NLILSAFPR | 9 | A5YKK6 | CCR4-NOT transcription complex subuni | CNOT1 | 266.939 | 2 | 55.33279 | yes | 2.328E-07 | 515.80585 | 504.50507 |
| LAEVGQYEQVK | 11 | A5YKK6 | CCR4-NOT transcription complex subuni | CNOT1 | 266.939 | 2 | 55.33279 | yes | 9.722E-08 | 632.33258 | 466.27225 |
| VDMLSEINIAPR | 12 | A5YKK6 | CCR4-NOT transcription complex subuni | CNOT1 | 266.939 | 2 | 55.33279 | yes | 9.722E-08 | 679.36084 | 450.75513 |
| ILLVDER | 7 | A5YKK6 | CCR4-NOT transcription complex subuni | CNOT1 | 266.939 | 2 | 55.33279 | yes | 1.794E-07 | 429.25818 | 399.12567 |

| | | | | | | | | | | | |
|---|---|---|---|---|---|---|---|---|---|---|---|
| GAFGVGQEQIDAF | 15 | A5YKK6 | CCR4-NOT transcription complex subuni | CNOT1 | 266.939 | 2 | 55.33279 | yes | 1.043E-07 | 790.40936 | 398.24976 |
| GLHNVVYGIQR | 11 | A5YKK6 | CCR4-NOT transcription complex subuni | CNOT1 | 266.939 | 2 | 55.33279 | yes | 1.421E-07 | 628.34894 | 355.02286 |
| NKPILHTDLDVK | 12 | A5YKK6 | CCR4-NOT transcription complex subuni | CNOT1 | 266.939 | 2 | 55.33279 | yes | 1.816E-07 | 696.89594 | 174.35165 |
| NKPILHTDLDVK | 12 | A5YKK6 | CCR4-NOT transcription complex subuni | CNOT1 | 266.939 | 2 | 55.33279 | yes | 9.722E-08 | 464.93304 | 174.35165 |
| NKPILHTDLDVK | 12 | A5YKK6 | CCR4-NOT transcription complex subuni | CNOT1 | 266.939 | 3 | 55.33279 | yes | 1.816E-07 | 696.89594 | 174.35165 |
| NKPILHTDLDVK | 12 | A5YKK6 | CCR4-NOT transcription complex subuni | CNOT1 | 266.939 | 3 | 55.33279 | yes | 9.722E-08 | 464.93304 | 174.35165 |
| ILTNFTGVMPPQFI | 14 | A5YKK6 | CCR4-NOT transcription complex subuni | CNOT1 | 266.939 | 2 | 55.33279 | yes | 4.863E-07 | 796.92889 | 340.31912 |
| MYYFGIAALDR | 11 | A5YKK6 | CCR4-NOT transcription complex subuni | CNOT1 | 266.939 | 2 | 55.33279 | yes | 9.722E-08 | 660.32629 | 312.90488 |
| CPSILGGLAPEK | 12 | A5YKK6 | CCR4-NOT transcription complex subuni | CNOT1 | 266.939 | 2 | 55.33279 | yes | 1.338E-07 | 621.33154 | 308.87759 |
| MQGSITTPGSIALA | 25 | A5YKK6 | CCR4-NOT transcription complex subuni | CNOT1 | 266.939 | 3 | 55.33279 | yes | 1.683E-07 | 823.10394 | 275.55292 |
| YVLEALR | 7 | A5YKK6 | CCR4-NOT transcription complex subuni | CNOT1 | 266.939 | 2 | 55.33279 | yes | 1.957E-07 | 432.2529 | 274.85379 |
| MVLNETYR | 8 | A5YKK6 | CCR4-NOT transcription complex subuni | CNOT1 | 266.939 | 2 | 55.33279 | yes | 1.6E-07 | 513.25787 | 258.01581 |
| QLAVYEEFAR | 10 | A5YKK6 | CCR4-NOT transcription complex subuni | CNOT1 | 266.939 | 2 | 55.33279 | yes | 9.819E-08 | 613.31421 | 247.92593 |
| MLAHTPQQK | 9 | A5YKK6 | CCR4-NOT transcription complex subuni | CNOT1 | 266.939 | 2 | 55.33279 | yes | 1.001E-07 | 527.27911 | 203.13092 |
| NLDEQLSAPK | 10 | A5YKK6 | CCR4-NOT transcription complex subuni | CNOT1 | 266.939 | 2 | 55.33279 | yes | 2.945E-07 | 557.79059 | 201.78743 |
| LFQSVAQCCMGQI | 13 | A5YKK6 | CCR4-NOT transcription complex subuni | CNOT1 | 266.939 | 2 | 55.33279 | yes | 1.203E-07 | 778.85455 | 199.98796 |
| ASQQEIQHIVNR | 12 | A5YKK6 | CCR4-NOT transcription complex subuni | CNOT1 | 266.939 | 2 | 55.33279 | yes | 1.041E-07 | 711.87604 | 191.30226 |
| DFALDSEESR | 10 | A5YKK6 | CCR4-NOT transcription complex subuni | CNOT1 | 266.939 | 2 | 55.33279 | yes | 1.007E-07 | 584.75946 | 184.12888 |
| LGTSGLNQPTFQQ | 15 | A5YKK6 | CCR4-NOT transcription complex subuni | CNOT1 | 266.939 | 2 | 55.33279 | yes | 1.022E-07 | 803.41522 | 181.62137 |
| EPLLMSISTNLK | 12 | A5YKK6 | CCR4-NOT transcription complex subuni | CNOT1 | 266.939 | 2 | 55.33279 | yes | 1.031E-07 | 673.37341 | 179.32971 |
| LPDPFTPNLK | 10 | A5YKK6 | CCR4-NOT transcription complex subuni | CNOT1 | 266.939 | 2 | 55.33279 | yes | 4.616E-07 | 571.31622 | 169.28294 |
| GQQELLYVVPFVA | 14 | A5YKK6 | CCR4-NOT transcription complex subuni | CNOT1 | 266.939 | 2 | 55.33279 | yes | 1.063E-07 | 795.94812 | 166.50517 |
| TVTVTRPTGVSFK | 13 | A5YKK6 | CCR4-NOT transcription complex subuni | CNOT1 | 266.939 | 2 | 55.33279 | yes | 4.251E-07 | 696.89594 | 82.360687 |
| TVTVTRPTGVSFK | 13 | A5YKK6 | CCR4-NOT transcription complex subuni | CNOT1 | 266.939 | 2 | 55.33279 | yes | 4.194E-06 | 464.93304 | 82.360687 |
| TVTVTRPTGVSFK | 13 | A5YKK6 | CCR4-NOT transcription complex subuni | CNOT1 | 266.939 | 3 | 55.33279 | yes | 4.251E-07 | 696.89594 | 82.360687 |
| TVTVTRPTGVSFK | 13 | A5YKK6 | CCR4-NOT transcription complex subuni | CNOT1 | 266.939 | 3 | 55.33279 | yes | 4.194E-06 | 464.93304 | 82.360687 |
| NLFEEYR | 7 | A5YKK6 | CCR4-NOT transcription complex subuni | CNOT1 | 266.939 | 2 | 55.33279 | yes | 1.454E-07 | 485.73508 | 154.442 |
| QAWATDDVAQIYI | 14 | A5YKK6 | CCR4-NOT transcription complex subuni | CNOT1 | 266.939 | 2 | 55.33279 | yes | 1.022E-07 | 812.38611 | 139.69493 |
| CYHNLDAFVR | 10 | A5YKK6 | CCR4-NOT transcription complex subuni | CNOT1 | 266.939 | 2 | 55.33279 | yes | 9.722E-08 | 647.80347 | 69.268356 |
| CYHNLDAFVR | 10 | A5YKK6 | CCR4-NOT transcription complex subuni | CNOT1 | 266.939 | 2 | 55.33279 | yes | 1.441E-07 | 432.20474 | 69.268356 |
| CYHNLDAFVR | 10 | A5YKK6 | CCR4-NOT transcription complex subuni | CNOT1 | 266.939 | 3 | 55.33279 | yes | 9.722E-08 | 647.80347 | 69.268356 |
| CYHNLDAFVR | 10 | A5YKK6 | CCR4-NOT transcription complex subuni | CNOT1 | 266.939 | 3 | 55.33279 | yes | 1.441E-07 | 432.20474 | 69.268356 |
| EVFNCMLR | 8 | A5YKK6 | CCR4-NOT transcription complex subuni | CNOT1 | 266.939 | 2 | 55.33279 | yes | 9.722E-08 | 534.75189 | 136.57361 |
| QKLPDLLR | 8 | A5YKK6 | CCR4-NOT transcription complex subuni | CNOT1 | 266.939 | 2 | 55.33279 | yes | 1.675E-07 | 491.80585 | 135.09853 |
| DAIAALGLLQK | 11 | A5YKK6 | CCR4-NOT transcription complex subuni | CNOT1 | 266.939 | 2 | 55.33279 | yes | 1.34E-07 | 556.83734 | 125.26849 |
| YRECHLLVLK | 10 | A5YKK6 | CCR4-NOT transcription complex subuni | CNOT1 | 266.939 | 3 | 55.33279 | yes | 3.604E-07 | 444.2482 | 116.48089 |
| SNYEAMIDR | 9 | A5YKK6 | CCR4-NOT transcription complex subuni | CNOT1 | 266.939 | 2 | 55.33279 | yes | 9.722E-08 | 549.74786 | 108.92279 |
| YCDPVVLTYQAEI | 13 | A5YKK6 | CCR4-NOT transcription complex subuni | CNOT1 | 266.939 | 2 | 55.33279 | yes | 9.722E-08 | 807.38507 | 107.57864 |
| KDFALDSEESR | 11 | A5YKK6 | CCR4-NOT transcription complex subuni | CNOT1 | 266.939 | 2 | 55.33279 | yes | 2.696E-07 | 648.80695 | 105.36219 |

| | | | | | | | | | | | |
|---|---|---|---|---|---|---|---|---|---|---|---|
| EIDDEANSYFQR | 12 | A5YKK6 | CCR4-NOT transcription complex subuni | CNOT1 | 266.939 | 2 | 55.33279 | yes | 9.722E-08 | 743.82587 | 105.31226 |
| AYGSPWCNK | 9 | A5YKK6 | CCR4-NOT transcription complex subuni | CNOT1 | 266.939 | 2 | 55.33279 | yes | 1.938E-06 | 541.73981 | 102.90836 |
| CPVVLAPLLYPEK | 13 | A5YKK6 | CCR4-NOT transcription complex subuni | CNOT1 | 266.939 | 2 | 55.33279 | yes | 1.041E-07 | 749.92053 | 94.646095 |
| YLAPFLR | 7 | A5YKK6 | CCR4-NOT transcription complex subuni | CNOT1 | 266.939 | 2 | 55.33279 | yes | 2.54E-07 | 440.258 | 93.520958 |
| GSTPSMSTITHSAF | 30 | A5YKK6 | CCR4-NOT transcription complex subuni | CNOT1 | 266.939 | 4 | 55.33279 | yes | 9.722E-08 | 808.63159 | 89.817017 |
| DVPPSINTTNIDTL | 23 | A5YKK6 | CCR4-NOT transcription complex subuni | CNOT1 | 266.939 | 3 | 55.33279 | yes | 6.994E-07 | 838.42914 | 88.689423 |
| NVELTKPMQILYK | 13 | A5YKK6 | CCR4-NOT transcription complex subuni | CNOT1 | 266.939 | 2 | 55.33279 | yes | 9.728E-08 | 788.94202 | 87.534653 |
| YNVEAVELLIR | 11 | A5YKK6 | CCR4-NOT transcription complex subuni | CNOT1 | 266.939 | 2 | 55.33279 | yes | 3.366E-07 | 659.87189 | 74.442192 |
| NLTAGMAMITCR | 12 | A5YKK6 | CCR4-NOT transcription complex subuni | CNOT1 | 266.939 | 2 | 55.33279 | yes | 1.483E-07 | 669.81995 | 67.248428 |
| SLLEVVVLSR | 10 | A5YKK6 | CCR4-NOT transcription complex subuni | CNOT1 | 266.939 | 2 | 55.33279 | yes | 1.724E-07 | 557.84515 | 66.930687 |
| VEELKETVK | 9 | A5YKK6 | CCR4-NOT transcription complex subuni | CNOT1 | 266.939 | 2 | 55.33279 | yes | 1.585E-07 | 537.80573 | 65.300926 |
| NLGHWLGMITLAF | 13 | A5YKK6 | CCR4-NOT transcription complex subuni | CNOT1 | 266.939 | 2 | 55.33279 | yes | 9.722E-08 | 727.40283 | 61.1297 |
| FEIEVLCK | 8 | A5YKK6 | CCR4-NOT transcription complex subuni | CNOT1 | 266.939 | 2 | 55.33279 | yes | 1.577E-07 | 519.27045 | 60.052887 |
| SLKPAPHLFAQLSI | 14 | A5YKK6 | CCR4-NOT transcription complex subuni | CNOT1 | 266.939 | 3 | 55.33279 | yes | 5.218E-07 | 512.96808 | 29.057278 |
| SLKPAPHLFAQLSI | 14 | A5YKK6 | CCR4-NOT transcription complex subuni | CNOT1 | 266.939 | 3 | 55.33279 | yes | 2.83E-07 | 768.94849 | 29.057278 |
| SLKPAPHLFAQLSI | 14 | A5YKK6 | CCR4-NOT transcription complex subuni | CNOT1 | 266.939 | 2 | 55.33279 | yes | 5.218E-07 | 512.96808 | 29.057278 |
| SLKPAPHLFAQLSI | 14 | A5YKK6 | CCR4-NOT transcription complex subuni | CNOT1 | 266.939 | 2 | 55.33279 | yes | 2.83E-07 | 768.94849 | 29.057278 |
| TDDLITR | 7 | A5YKK6 | CCR4-NOT transcription complex subuni | CNOT1 | 266.939 | 2 | 55.33279 | yes | 3.084E-07 | 417.2218 | 54.739212 |
| ECHLLVLK | 8 | A5YKK6 | CCR4-NOT transcription complex subuni | CNOT1 | 266.939 | 2 | 55.33279 | yes | 2.149E-07 | 506.28641 | 44.963921 |
| RYCDPVVLTYQAI | 14 | A5YKK6 | CCR4-NOT transcription complex subuni | CNOT1 | 266.939 | 3 | 55.33279 | yes | 2.93E-07 | 590.62616 | 40.183041 |
| CLFSHVDFSGDGK | 13 | A5YKK6 | CCR4-NOT transcription complex subuni | CNOT1 | 266.939 | 2 | 55.33279 | yes | 2.888E-07 | 734.8299 | 19.551893 |
| CLFSHVDFSGDGK | 13 | A5YKK6 | CCR4-NOT transcription complex subuni | CNOT1 | 266.939 | 2 | 55.33279 | yes | 2.554E-05 | 490.22235 | 19.551893 |
| CLFSHVDFSGDGK | 13 | A5YKK6 | CCR4-NOT transcription complex subuni | CNOT1 | 266.939 | 3 | 55.33279 | yes | 2.888E-07 | 734.8299 | 19.551893 |
| CLFSHVDFSGDGK | 13 | A5YKK6 | CCR4-NOT transcription complex subuni | CNOT1 | 266.939 | 3 | 55.33279 | yes | 2.554E-05 | 490.22235 | 19.551893 |
| PMQILYK | 7 | A5YKK6 | CCR4-NOT transcription complex subuni | CNOT1 | 266.939 | 2 | 55.33279 | yes | 2.831E-07 | 446.75168 | 18.662235 |
| LMILPDPEKPVR | 12 | A6NHR9 | Structural maintenance of chromosomes f | SMCHD1 | 226.374 | 2 | 55.723537 | yes | 1.128E-07 | 704.40503 | 104.51112 |
| LMILPDPEKPVR | 12 | A6NHR9 | Structural maintenance of chromosomes f | SMCHD1 | 226.374 | 2 | 55.723537 | yes | 1.218E-07 | 469.93912 | 104.51112 |
| LMILPDPEKPVR | 12 | A6NHR9 | Structural maintenance of chromosomes f | SMCHD1 | 226.374 | 3 | 55.723537 | yes | 1.128E-07 | 704.40503 | 104.51112 |
| LMILPDPEKPVR | 12 | A6NHR9 | Structural maintenance of chromosomes f | SMCHD1 | 226.374 | 3 | 55.723537 | yes | 1.218E-07 | 469.93912 | 104.51112 |
| GVITRPDLPSK | 11 | A6NHR9 | Structural maintenance of chromosomes f | SMCHD1 | 226.374 | 2 | 55.723537 | yes | 9.722E-08 | 591.8457 | 148.84895 |
| VQLISGPPAK | 10 | A6NHR9 | Structural maintenance of chromosomes f | SMCHD1 | 226.374 | 2 | 55.723537 | yes | 9.722E-08 | 505.30566 | 146.43388 |
| QQVLPLDSIYK | 11 | A6NHR9 | Structural maintenance of chromosomes f | SMCHD1 | 226.374 | 2 | 55.723537 | yes | 3.694E-07 | 652.36646 | 144.4971 |
| GSNEEDTDTPLFIC | 15 | A6NHR9 | Structural maintenance of chromosomes f | SMCHD1 | 226.374 | 2 | 55.723537 | yes | 9.722E-08 | 811.88086 | 123.66801 |
| LQIFSVEGQK | 10 | A6NHR9 | Structural maintenance of chromosomes f | SMCHD1 | 226.374 | 2 | 55.723537 | yes | 1.318E-07 | 574.81915 | 121.53666 |
| GLAPIECYNR | 10 | A6NHR9 | Structural maintenance of chromosomes f | SMCHD1 | 226.374 | 2 | 55.723537 | yes | 2.815E-07 | 596.7926 | 115.03602 |
| EINITSALAEK | 11 | A6NHR9 | Structural maintenance of chromosomes f | SMCHD1 | 226.374 | 2 | 55.723537 | yes | 1.007E-07 | 594.82715 | 104.17171 |
| IAHLAQIEDDR | 11 | A6NHR9 | Structural maintenance of chromosomes f | SMCHD1 | 226.374 | 2 | 55.723537 | yes | 1.004E-07 | 640.83331 | 102.81969 |
| SLNSDISYFGVGGI | 14 | A6NHR9 | Structural maintenance of chromosomes f | SMCHD1 | 226.374 | 2 | 55.723537 | yes | 9.722E-08 | 722.35938 | 98.647728 |
| DLNSQLEYLR | 10 | A6NHR9 | Structural maintenance of chromosomes f | SMCHD1 | 226.374 | 2 | 55.723537 | yes | 1.233E-07 | 625.82239 | 91.60701 |

| | | | | | | | | | | | |
|---|---|---|---|---|---|---|---|---|---|---|---|
| TTFQENTQSISVR | 13 | A6NHR9 | Structural maintenance of chromosomes f | SMCHD1 | 226.374 | 2 | 55.723537 | yes | 9.722E-08 | 755.87842 | 85.131714 |
| QAVFFVGQSAR | 11 | A6NHR9 | Structural maintenance of chromosomes f | SMCHD1 | 226.374 | 2 | 55.723537 | yes | 1.858E-07 | 605.32239 | 68.901321 |
| ACVCQTLGISPEEK | 14 | A6NHR9 | Structural maintenance of chromosomes f | SMCHD1 | 226.374 | 2 | 55.723537 | yes | 2.308E-07 | 796.37634 | 66.02137 |
| EHLLQGLLPDVQV | 18 | A6NHR9 | Structural maintenance of chromosomes f | SMCHD1 | 226.374 | 3 | 55.723537 | yes | 1.871E-07 | 658.37372 | 56.419327 |
| AHVEGDGVVEGII | 14 | A6NHR9 | Structural maintenance of chromosomes f | SMCHD1 | 226.374 | 2 | 55.723537 | yes | 9.722E-08 | 725.88605 | 53.770927 |
| KYVEDEMAR | 9 | A6NHR9 | Structural maintenance of chromosomes f | SMCHD1 | 226.374 | 2 | 55.723537 | yes | 7.751E-07 | 570.7713 | 47.587841 |
| TLPLFYGSIVR | 11 | A6NHR9 | Structural maintenance of chromosomes f | SMCHD1 | 226.374 | 2 | 55.723537 | yes | 2.093E-07 | 633.36627 | 44.661694 |
| NYFHLWTR | 8 | A6NHR9 | Structural maintenance of chromosomes f | SMCHD1 | 226.374 | 2 | 55.723537 | yes | 3.453E-06 | 568.78546 | 40.492752 |
| EKDQLSQSIVMYK | 13 | A6NHR9 | Structural maintenance of chromosomes f | SMCHD1 | 226.374 | 2 | 55.723537 | yes | 7.606E-07 | 784.90308 | 39.590031 |
| LSEQEELKK | 9 | A6NHR9 | Structural maintenance of chromosomes f | SMCHD1 | 226.374 | 2 | 55.723537 | yes | 4.122E-07 | 552.30078 | 35.210361 |
| FLHHLIIEEK | 10 | A6NHR9 | Structural maintenance of chromosomes f | SMCHD1 | 226.374 | 2 | 55.723537 | yes | 2.348E-07 | 639.86389 | 31.426092 |
| MISKPADSQDVHE | 18 | A6NHR9 | Structural maintenance of chromosomes f | SMCHD1 | 226.374 | 3 | 55.723537 | yes | 6.478E-07 | 666.35059 | 24.789227 |
| LLPSSHVAR | 9 | A6NHR9 | Structural maintenance of chromosomes f | SMCHD1 | 226.374 | 2 | 55.723537 | yes | 3.432E-07 | 490.28781 | 24.57202 |
| LYFKPIGDPVFAR | 13 | A6NHR9 | Structural maintenance of chromosomes f | SMCHD1 | 226.374 | 3 | 55.723537 | yes | 2.154E-06 | 508.2854 | 19.599478 |
| LQVVLNESNADTY | 21 | A6NHR9 | Structural maintenance of chromosomes f | SMCHD1 | 226.374 | 3 | 55.723537 | yes | 3.783E-06 | 758.06927 | 17.610516 |
| VYPPVPEQLQLR | 12 | A7E2V4 | Zinc finger SWIM domain-containing prc | ZSWIM8 | 197.298 | 2 | 53.580544 | yes | 5.141E-07 | 719.90625 | 142.25533 |
| LAVLDPALSPQR | 12 | A7E2V4 | Zinc finger SWIM domain-containing prc | ZSWIM8 | 197.298 | 2 | 53.580544 | yes | 6.023E-07 | 640.37207 | 136.94112 |
| APAFHQLVQR | 10 | A7E2V4 | Zinc finger SWIM domain-containing prc | ZSWIM8 | 197.298 | 2 | 53.580544 | yes | 2.093E-07 | 583.82507 | 68.375084 |
| APVSESLSR | 9 | A7E2V4 | Zinc finger SWIM domain-containing prc | ZSWIM8 | 197.298 | 2 | 53.580544 | yes | 2.997E-07 | 473.25363 | 66.522743 |
| VGMLALEMLGR | 11 | A7E2V4 | Zinc finger SWIM domain-containing prc | ZSWIM8 | 197.298 | 2 | 53.580544 | yes | 9.722E-08 | 595.3252 | 63.104305 |
| GLGEGVPSSQR | 11 | A7E2V4 | Zinc finger SWIM domain-containing prc | ZSWIM8 | 197.298 | 2 | 53.580544 | yes | 4.196E-07 | 543.78052 | 59.596222 |
| LAYQESEVAALLK | 13 | A7E2V4 | Zinc finger SWIM domain-containing prc | ZSWIM8 | 197.298 | 2 | 53.580544 | yes | 1.717E-07 | 717.89557 | 26.187778 |
| VFVVEAAVR | 9 | A8MT69 | Centromere protein X OS=Homo sapiens | CENPX | 8.96 | 2 | 50.52883 | yes | 9.789E-08 | 495.29257 | 394.31296 |
| VDVDQLEK | 8 | A8MT69 | Centromere protein X OS=Homo sapiens | CENPX | 8.96 | 2 | 50.52883 | yes | 1.019E-06 | 473.24802 | 199.2327 |
| IPAPQCDITK | 10 | A8MV65 | Transcription cofactor vestigial-like prote | VGLL3 | 36.009 | 2 | 48.49806 | yes | 1.615E-07 | 571.79736 | 403.19519 |
| ALGQAITLHPESAI | 16 | A8MV65 | Transcription cofactor vestigial-like prote | VGLL3 | 36.009 | 2 | 48.49806 | yes | 1.72E-07 | 818.45685 | 251.38765 |
| MQDSLEVTLPSK | 12 | A8MV65 | Transcription cofactor vestigial-like prote | VGLL3 | 36.009 | 2 | 48.49806 | yes | 1.139E-07 | 674.34485 | 199.04312 |
| MGLTPLWR | 8 | A8MV65 | Transcription cofactor vestigial-like prote | VGLL3 | 36.009 | 2 | 48.49806 | yes | 2.393E-07 | 487.26804 | 145.90674 |
| NITWTLSNLCR | 11 | A9QM74 | Importin subunit alpha-8 OS=Homo sapie | KPNA7 | 56.938 | 2 | 48.743385 | yes | 1.108E-07 | 689.35083 | 4740.5049 |
| FLEDKPQFISR | 11 | B2RTY4 | Unconventional myosin-IXa OS=Homo s | MYO9A | 292.706 | 2 | 48.25136 | yes | 3.087E-06 | 690.36951 | 190.24768 |
| GVYSVIDQLSR | 11 | B2RTY4 | Unconventional myosin-IXa OS=Homo s | MYO9A | 292.706 | 2 | 48.25136 | yes | 3.472E-07 | 618.83276 | 80.987778 |
| IALQEDTNR | 9 | B2RTY4 | Unconventional myosin-IXa OS=Homo s | MYO9A | 292.706 | 2 | 48.25136 | yes | 4.298E-06 | 530.27509 | 74.498421 |
| WSTELVPEGLQSP | 14 | B2RTY4 | Unconventional myosin-IXa OS=Homo s | MYO9A | 292.706 | 2 | 48.25136 | yes | 3.266E-07 | 799.91229 | 68.763809 |
| IILLQSTCR | 9 | B2RTY4 | Unconventional myosin-IXa OS=Homo s | MYO9A | 292.706 | 2 | 48.25136 | yes | 1.173E-07 | 552.31573 | 66.366913 |
| GNYPGPSSPVVVR | 13 | B2RTY4 | Unconventional myosin-IXa OS=Homo s | MYO9A | 292.706 | 2 | 48.25136 | yes | 5.318E-07 | 664.8515 | 62.883621 |
| IGECLVAPSSYQSK | 14 | B2RTY4 | Unconventional myosin-IXa OS=Homo s | MYO9A | 292.706 | 2 | 48.25136 | yes | 6.368E-07 | 769.87958 | 55.523098 |
| FSDVLVLR | 8 | B2RTY4 | Unconventional myosin-IXa OS=Homo s | MYO9A | 292.706 | 2 | 48.25136 | yes | 2.166E-06 | 474.7793 | 25.412333 |
| GELNVLGSLSLK | 12 | B2RTY4 | Unconventional myosin-IXa OS=Homo s | MYO9A | 292.706 | 2 | 48.25136 | yes | 2.669E-07 | 615.35864 | 20.755514 |
| GLVEEYVEK | 9 | C9JLW8 | Mapk-regulated corepressor-interacting p | MCRIP1 | 10.921 | 2 | 50.053547 | yes | 1.376E-07 | 533.27679 | 780.70538 |

| | | | | | | | | | | | |
|---|---|---|---|---|---|---|---|---|---|---|---|
| TFKPIDLSDLK | 11 | C9JLW8 | Mapk-regulated corepressor-interacting pı | MCRIP1 | 10.921 | 2 | 50.053547 | yes | 7.245E-07 | 638.86102 | 439.86691 |
| FIYEAWQGVER | 11 | C9JLW8 | Mapk-regulated corepressor-interacting pı | MCRIP1 | 10.921 | 2 | 50.053547 | yes | 9.921E-08 | 699.34607 | 259.82217 |
| EAVLSLYR | 8 | L0R8F8 | Mitochondrial ribosome and complex I as | MIEF1 | 8.445 | 2 | 45.760315 | yes | 1.749E-07 | 475.76892 | 487.30414 |
| ESISDVGFGMLK | 12 | O00116 | Alkyldihydroxyacetonephosphate synthas | AGPS | 72.912 | 2 | 52.530342 | yes | 1.46E-07 | 641.82104 | 77.207863 |
| TIISLDTSQMNR | 12 | O00116 | Alkyldihydroxyacetonephosphate synthas | AGPS | 72.912 | 2 | 52.530342 | yes | 1.098E-07 | 689.85339 | 69.386551 |
| EWIQNTLGVNVEI | 14 | O00116 | Alkyldihydroxyacetonephosphate synthas | AGPS | 72.912 | 2 | 52.530342 | yes | 9.722E-08 | 833.93103 | 24.682892 |
| ETNISYSQEADDR | 13 | O00116 | Alkyldihydroxyacetonephosphate synthas | AGPS | 72.912 | 2 | 52.530342 | yes | 1.673E-06 | 764.33154 | 23.89431 |
| IRPVPEYQK | 9 | O00116 | Alkyldihydroxyacetonephosphate synthas | AGPS | 72.912 | 2 | 52.530342 | yes | 3.867E-07 | 565.32184 | 23.215351 |
| PLVETIFER | 9 | O00139 | Kinesin-like protein KIF2A OS=Homo sa | KIF2A | 79.955 | 2 | 54.07929 | yes | 1.166E-07 | 552.30841 | 2384.5894 |
| SHAVFQIILR | 10 | O00139 | Kinesin-like protein KIF2A OS=Homo sa | KIF2A | 79.955 | 2 | 54.07929 | yes | 9.722E-08 | 592.35095 | 1303.1086 |
| LIDIGNSCR | 9 | O00139 | Kinesin-like protein KIF2A OS=Homo sa | KIF2A | 79.955 | 2 | 54.07929 | yes | 4.371E-07 | 524.26624 | 1105.9678 |
| AVFQESIR | 8 | O00139 | Kinesin-like protein KIF2A OS=Homo sa | KIF2A | 79.955 | 2 | 54.07929 | yes | 4.726E-07 | 475.25873 | 858.34863 |
| DVVMVHEPK | 9 | O00139 | Kinesin-like protein KIF2A OS=Homo sa | KIF2A | 79.955 | 2 | 54.07929 | yes | 1.677E-07 | 527.2735 | 709.11511 |
| FSLIDLAGNER | 11 | O00139 | Kinesin-like protein KIF2A OS=Homo sa | KIF2A | 79.955 | 2 | 54.07929 | yes | 1.157E-07 | 617.82495 | 697.17377 |
| LQQQELR | 7 | O00139 | Kinesin-like protein KIF2A OS=Homo sa | KIF2A | 79.955 | 2 | 54.07929 | yes | 1.048E-07 | 457.75635 | 660.4151 |
| AALQEEEQASK | 11 | O00139 | Kinesin-like protein KIF2A OS=Homo sa | KIF2A | 79.955 | 2 | 54.07929 | yes | 9.745E-08 | 602.2962 | 621.01984 |
| QQVQVVGLQER | 11 | O00139 | Kinesin-like protein KIF2A OS=Homo sa | KIF2A | 79.955 | 2 | 54.07929 | yes | 1.128E-07 | 642.35693 | 599.01849 |
| YLENQTFR | 8 | O00139 | Kinesin-like protein KIF2A OS=Homo sa | KIF2A | 79.955 | 2 | 54.07929 | yes | 1.514E-07 | 535.76691 | 409.60767 |
| DLDVITIPSK | 10 | O00139 | Kinesin-like protein KIF2A OS=Homo sa | KIF2A | 79.955 | 2 | 54.07929 | yes | 1.121E-07 | 550.81354 | 265.33243 |
| ELTVDPTAAGDVI | 13 | O00139 | Kinesin-like protein KIF2A OS=Homo sa | KIF2A | 79.955 | 2 | 54.07929 | yes | 1.695E-07 | 672.34369 | 222.13148 |
| DSFIGENSR | 9 | O00139 | Kinesin-like protein KIF2A OS=Homo sa | KIF2A | 79.955 | 2 | 54.07929 | no | 1.711E-07 | 512.73834 | 207.70511 |
| NKPHTPFR | 8 | O00139 | Kinesin-like protein KIF2A OS=Homo sa | KIF2A | 79.955 | 2 | 54.07929 | yes | 2.081E-07 | 498.77234 | 167.54254 |
| THTMGGDFSGK | 11 | O00139 | Kinesin-like protein KIF2A OS=Homo sa | KIF2A | 79.955 | 2 | 54.07929 | yes | 1.414E-07 | 569.2533 | 140.12656 |
| PLTTADPIDEHR | 12 | O00139 | Kinesin-like protein KIF2A OS=Homo sa | KIF2A | 79.955 | 2 | 54.07929 | yes | 9.722E-08 | 682.84387 | 114.90663 |
| VFDLLNR | 7 | O00139 | Kinesin-like protein KIF2A OS=Homo sa | KIF2A | 79.955 | 2 | 54.07929 | yes | 9.722E-08 | 438.75052 | 69.337234 |
| GIYALAAR | 8 | O00139 | Kinesin-like protein KIF2A OS=Homo sa | KIF2A | 79.955 | 2 | 54.07929 | yes | 1.385E-06 | 417.74524 | 50.983814 |
| KEFGPPSR | 8 | O00139 | Kinesin-like protein KIF2A OS=Homo sa | KIF2A | 79.955 | 2 | 54.07929 | yes | 2.953E-07 | 459.24561 | 50.83419 |
| PIMHHPPNQIDDLI | 25 | O00139 | Kinesin-like protein KIF2A OS=Homo sa | KIF2A | 79.955 | 4 | 54.07929 | yes | 1.056E-06 | 710.59497 | 46.312424 |
| GMATCFAYGQTG | 15 | O00139 | Kinesin-like protein KIF2A OS=Homo sa | KIF2A | 79.955 | 2 | 54.07929 | no | 1.059E-07 | 768.33447 | 41.248344 |
| SVICVEGNIASGK | 13 | O00142 | Thymidine kinase 2, mitochondrial OS=H | TK2 | 31.005 | 2 | 40.39803 | yes | 4.045E-07 | 667.34265 | 67.787186 |
| TNPETCYQR | 9 | O00142 | Thymidine kinase 2, mitochondrial OS=H | TK2 | 31.005 | 2 | 40.39803 | yes | 5.675E-07 | 584.75623 | 23.038349 |
| ILVATNLFGR | 10 | O00148 | ATP-dependent RNA helicase DDX39A ( | DDX39A | 49.13 | 2 | 53.15926 | no | 9.854E-08 | 552.33221 | 1133.7986 |
| MLEQLDMR | 8 | O00148 | ATP-dependent RNA helicase DDX39A ( | DDX39A | 49.13 | 2 | 53.15926 | no | 1.438E-07 | 518.25171 | 524.48596 |
| GLAITFVSDENDAI | 14 | O00148 | ATP-dependent RNA helicase DDX39A ( | DDX39A | 49.13 | 2 | 53.15926 | no | 1.359E-07 | 740.36993 | 519.76898 |
| GSYVSIHSSGFR | 12 | O00148 | ATP-dependent RNA helicase DDX39A ( | DDX39A | 49.13 | 2 | 53.15926 | no | 9.722E-08 | 648.82019 | 387.7077 |
| VSVFFGGLSIK | 11 | O00148 | ATP-dependent RNA helicase DDX39A ( | DDX39A | 49.13 | 2 | 53.15926 | yes | 9.722E-08 | 577.33441 | 204.91434 |
| HFVLDECDK | 9 | O00148 | ATP-dependent RNA helicase DDX39A ( | DDX39A | 49.13 | 2 | 53.15926 | yes | 3.514E-07 | 581.76349 | 131.18298 |
| LLSVEGSTLR | 10 | O00159 | Unconventional myosin-Ic OS=Homo sap | MYO1C | 121.682 | 2 | 54.194435 | yes | 1.876E-07 | 537.81134 | 620.24237 |
| TLFATEDALEVR | 12 | O00159 | Unconventional myosin-Ic OS=Homo sap | MYO1C | 121.682 | 2 | 54.194435 | yes | 9.722E-08 | 682.85645 | 612.14771 |

| | | | | | | | | | | | |
|---|---|---|---|---|---|---|---|---|---|---|---|
| NPIMSQCFDR | 10 | O00159 | Unconventional myosin-Ic OS=Homo sap | MYO1C | 121.682 | 2 | 54.194435 | yes | 1.144E-07 | 634.28156 | 395.53754 |
| SLCPETWPTWAGI | 13 | O00159 | Unconventional myosin-Ic OS=Homo sap | MYO1C | 121.682 | 2 | 54.194435 | yes | 1.473E-07 | 780.86682 | 362.2616 |
| YLGLLENLR | 9 | O00159 | Unconventional myosin-Ic OS=Homo sap | MYO1C | 121.682 | 2 | 54.194435 | yes | 9.722E-08 | 545.81641 | 357.6655 |
| PQDGVAVLVR | 10 | O00159 | Unconventional myosin-Ic OS=Homo sap | MYO1C | 121.682 | 2 | 54.194435 | yes | 1.338E-07 | 527.30621 | 285.85388 |
| DQAVMISGESGAC | 14 | O00159 | Unconventional myosin-Ic OS=Homo sap | MYO1C | 121.682 | 2 | 54.194435 | yes | 9.809E-08 | 675.3219 | 277.94919 |
| DNYPQSVPR | 9 | O00159 | Unconventional myosin-Ic OS=Homo sap | MYO1C | 121.682 | 2 | 54.194435 | yes | 1.724E-07 | 538.26196 | 272.16449 |
| SAICIQSWWR | 10 | O00159 | Unconventional myosin-Ic OS=Homo sap | MYO1C | 121.682 | 2 | 54.194435 | yes | 9.722E-08 | 653.82172 | 269.32007 |
| KYEAFLQR | 8 | O00159 | Unconventional myosin-Ic OS=Homo sap | MYO1C | 121.682 | 2 | 54.194435 | yes | 1.025E-07 | 527.78766 | 225.55063 |
| LGTDEISPR | 9 | O00159 | Unconventional myosin-Ic OS=Homo sap | MYO1C | 121.682 | 2 | 54.194435 | yes | 2.182E-07 | 494.25891 | 218.79504 |
| VTMESALTAR | 10 | O00159 | Unconventional myosin-Ic OS=Homo sap | MYO1C | 121.682 | 2 | 54.194435 | yes | 1.117E-06 | 539.78168 | 194.43616 |
| LLQSNPVLEAFGN | 15 | O00159 | Unconventional myosin-Ic OS=Homo sap | MYO1C | 121.682 | 2 | 54.194435 | yes | 1.133E-06 | 800.93829 | 188.11609 |
| IICDLVEEK | 9 | O00159 | Unconventional myosin-Ic OS=Homo sap | MYO1C | 121.682 | 2 | 54.194435 | yes | 1.024E-07 | 559.79175 | 167.25761 |
| NPQSYLYLVK | 10 | O00159 | Unconventional myosin-Ic OS=Homo sap | MYO1C | 121.682 | 2 | 54.194435 | yes | 3.367E-07 | 612.83478 | 160.23131 |
| TSFLLNLR | 8 | O00159 | Unconventional myosin-Ic OS=Homo sap | MYO1C | 121.682 | 2 | 54.194435 | yes | 1.233E-07 | 482.28473 | 146.12395 |
| NGHLAVVAPR | 10 | O00159 | Unconventional myosin-Ic OS=Homo sap | MYO1C | 121.682 | 2 | 54.194435 | yes | 5.345E-07 | 517.29871 | 144.96506 |
| LLQFYAETCPAPEI | 14 | O00159 | Unconventional myosin-Ic OS=Homo sap | MYO1C | 121.682 | 2 | 54.194435 | yes | 1.001E-07 | 847.914 | 132.61928 |
| YMDVQFDFK | 9 | O00159 | Unconventional myosin-Ic OS=Homo sap | MYO1C | 121.682 | 2 | 54.194435 | yes | 3.633E-07 | 596.77081 | 56.476387 |
| CPENAFFLDHVR | 12 | O00159 | Unconventional myosin-Ic OS=Homo sap | MYO1C | 121.682 | 2 | 54.194435 | yes | 5.054E-07 | 752.8537 | 48.781155 |
| HLGYKPEEYK | 10 | O00159 | Unconventional myosin-Ic OS=Homo sap | MYO1C | 121.682 | 2 | 54.194435 | yes | 1.572E-07 | 632.32202 | 48.344189 |
| ITKPDGIVEER | 11 | O00165 | HCLS1-associated protein X-1 OS=Homo | HAX1 | 31.621 | 2 | 50.171677 | yes | 9.869E-08 | 628.84589 | 133.05392 |
| IFGGVLESDAR | 11 | O00165 | HCLS1-associated protein X-1 OS=Homo | HAX1 | 31.621 | 2 | 50.171677 | yes | 3.235E-07 | 582.3064 | 114.23473 |
| VLELDPALAPVVS | 14 | O00170 | AH receptor-interacting protein OS=Hom | AIP | 37.665 | 2 | 49.183086 | yes | 1.061E-07 | 739.9325 | 93.981895 |
| TEQDLALGTDAEC | 15 | O00186 | Syntaxin-binding protein 3 OS=Homo sap | STXBP3 | 67.765 | 2 | 53.39742 | yes | 9.722E-08 | 788.37848 | 30.573503 |
| YVCEADGFWTSSK | 13 | O00187 | Mannan-binding lectin serine protease 2 ( | MASP2 | 75.703 | 2 | 46.248936 | yes | 1.611E-07 | 775.33502 | 34.54718 |
| TVIGSVLLR | 9 | O00203 | AP-3 complex subunit beta-1 OS=Homo : | AP3B1 | 121.321 | 2 | 53.897743 | yes | 2.071E-07 | 479.3082 | 158.10909 |
| LLDSITVPVAR | 11 | O00203 | AP-3 complex subunit beta-1 OS=Homo : | AP3B1 | 121.321 | 2 | 53.897743 | yes | 5.488E-07 | 592.3559 | 134.16078 |
| EASADLSPYVR | 11 | O00203 | AP-3 complex subunit beta-1 OS=Homo : | AP3B1 | 121.321 | 2 | 53.897743 | yes | 1.312E-07 | 604.30133 | 97.660789 |
| NTKPLLQSR | 9 | O00203 | AP-3 complex subunit beta-1 OS=Homo : | AP3B1 | 121.321 | 2 | 53.897743 | no | 8.706E-07 | 528.81165 | 95.270264 |
| QFAAATIQTIGR | 12 | O00203 | AP-3 complex subunit beta-1 OS=Homo : | AP3B1 | 121.321 | 2 | 53.897743 | yes | 9.722E-08 | 638.85406 | 75.851105 |
| SFTSEDDLVK | 10 | O00203 | AP-3 complex subunit beta-1 OS=Homo : | AP3B1 | 121.321 | 2 | 53.897743 | yes | 1.816E-06 | 570.7746 | 58.979321 |
| LYSLDPEQK | 9 | O00203 | AP-3 complex subunit beta-1 OS=Homo : | AP3B1 | 121.321 | 2 | 53.897743 | yes | 4.901E-07 | 546.78223 | 52.170509 |
| EGSTAQLIINTEK | 13 | O00203 | AP-3 complex subunit beta-1 OS=Homo : | AP3B1 | 121.321 | 2 | 53.897743 | yes | 1.133E-06 | 702.37244 | 50.57534 |
| TQFVSPWK | 8 | O00203 | AP-3 complex subunit beta-1 OS=Homo : | AP3B1 | 121.321 | 2 | 53.897743 | yes | 3.556E-06 | 496.76364 | 45.602631 |
| LLQMQPAQHGEIII | 14 | O00203 | AP-3 complex subunit beta-1 OS=Homo : | AP3B1 | 121.321 | 2 | 53.897743 | no | 1.346E-07 | 803.44269 | 42.407398 |
| ASILWLIGENCER | 13 | O00203 | AP-3 complex subunit beta-1 OS=Homo : | AP3B1 | 121.321 | 2 | 53.897743 | yes | 6.495E-07 | 780.89557 | 41.189167 |
| NASELFPAVVK | 11 | O00203 | AP-3 complex subunit beta-1 OS=Homo : | AP3B1 | 121.321 | 2 | 53.897743 | yes | 1.163E-07 | 587.82697 | 31.33522 |
| KLVYVYLVR | 9 | O00203 | AP-3 complex subunit beta-1 OS=Homo : | AP3B1 | 121.321 | 2 | 53.897743 | no | 2.355E-07 | 576.8606 | 29.256191 |
| TVHSGSLMLVTVE | 15 | O00203 | AP-3 complex subunit beta-1 OS=Homo : | AP3B1 | 121.321 | 3 | 53.897743 | yes | 2.555E-07 | 538.63586 | 23.079136 |
| EPATINYPFEK | 11 | O00217 | NADH dehydrogenase [ubiquinone] iron- | NDUFS8 | 23.706 | 2 | 49.54917 | yes | 2.725E-07 | 654.82715 | 208.68219 |

| | | | | | | | | | | |
|---|---|---|---|---|---|---|---|---|---|---|
| YVNMQDPEMDMI | 12 | O00217 | NADH dehydrogenase [ubiquinone] iron- | NDUFS8 | 23.706 | 2 | 49.54917 | yes | 1.022E-07 | 750.81201 | 67.388145 |
| TALHLAVETQER | 12 | O00221 | NF-kappa-B inhibitor epsilon OS=Homo | NFKBIE | 52.864 | 2 | 52.99933 | yes | 9.722E-08 | 684.36749 | 81.267899 |
| TGQAAELGGLLK | 12 | O00231 | 26S proteasome non-ATPase regulatory s | PSMD11 | 47.464 | 2 | 37.457783 | yes | 0.000227 | 579.3299 | 38.787022 |
| EASIDILHSIVK | 12 | O00231 | 26S proteasome non-ATPase regulatory s | PSMD11 | 47.464 | 2 | 37.457783 | yes | 1.346E-06 | 662.8772 | 35.227013 |
| DDPIISTHLAK | 11 | O00231 | 26S proteasome non-ATPase regulatory s | PSMD11 | 47.464 | 2 | 37.457783 | yes | 5.975E-07 | 605.32733 | 30.989517 |
| GLLGCNIIPLQR | 12 | O00233 | 26S proteasome non-ATPase regulatory s | PSMD9 | 24.682 | 2 | 44.58788 | yes | 2.139E-07 | 677.38721 | 45.532452 |
| PLVVQVPTFAR | 11 | O00257 | E3 SUMO-protein ligase CBX4 OS=Hon | CBX4 | 61.368 | 2 | 54.3758 | yes | 1.083E-07 | 613.86639 | 605.25641 |
| LLIAFQNR | 8 | O00257 | E3 SUMO-protein ligase CBX4 OS=Hon | CBX4 | 61.368 | 2 | 54.3758 | yes | 1.64E-07 | 487.79272 | 443.05371 |
| EAPSPTCPDLGAK | 13 | O00257 | E3 SUMO-protein ligase CBX4 OS=Hon | CBX4 | 61.368 | 2 | 54.3758 | yes | 1.204E-07 | 671.81903 | 395.45871 |
| SNVLTGLQDSSTD | 15 | O00257 | E3 SUMO-protein ligase CBX4 OS=Hon | CBX4 | 61.368 | 2 | 54.3758 | yes | 1.122E-07 | 803.88702 | 380.20917 |
| MYDLQYQGGHK | 11 | O00257 | E3 SUMO-protein ligase CBX4 OS=Hon | CBX4 | 61.368 | 2 | 54.3758 | yes | 9.722E-08 | 670.30859 | 224.83621 |
| SGEVAEGEAR | 10 | O00257 | E3 SUMO-protein ligase CBX4 OS=Hon | CBX4 | 61.368 | 2 | 54.3758 | yes | 1.054E-07 | 502.73581 | 178.35837 |
| GYLGAVKPLAGA | 18 | O00257 | E3 SUMO-protein ligase CBX4 OS=Hon | CBX4 | 61.368 | 2 | 54.3758 | yes | 1.059E-07 | 799.45667 | 73.194145 |
| GYLGAVKPLAGA | 18 | O00257 | E3 SUMO-protein ligase CBX4 OS=Hon | CBX4 | 61.368 | 2 | 54.3758 | yes | 2.165E-07 | 533.30688 | 73.194145 |
| GYLGAVKPLAGA | 18 | O00257 | E3 SUMO-protein ligase CBX4 OS=Hon | CBX4 | 61.368 | 3 | 54.3758 | yes | 1.059E-07 | 799.45667 | 73.194145 |
| GYLGAVKPLAGA | 18 | O00257 | E3 SUMO-protein ligase CBX4 OS=Hon | CBX4 | 61.368 | 3 | 54.3758 | yes | 2.165E-07 | 533.30688 | 73.194145 |
| QEQLMGYR | 8 | O00257 | E3 SUMO-protein ligase CBX4 OS=Hon | CBX4 | 61.368 | 2 | 54.3758 | yes | 3.05E-07 | 512.74768 | 97.456635 |
| PKPLVVQVPTFAR | 13 | O00257 | E3 SUMO-protein ligase CBX4 OS=Hon | CBX4 | 61.368 | 3 | 54.3758 | yes | 6.867E-07 | 484.62927 | 64.950806 |
| SISTPTCLGGSPAA | 16 | O00257 | E3 SUMO-protein ligase CBX4 OS=Hon | CBX4 | 61.368 | 2 | 54.3758 | yes | 3.361E-07 | 802.39087 | 61.419975 |
| EAFGEQPLQLTTK | 13 | O00257 | E3 SUMO-protein ligase CBX4 OS=Hon | CBX4 | 61.368 | 2 | 54.3758 | yes | 2.587E-07 | 731.38281 | 44.599613 |
| HHPYQPDPK | 9 | O00257 | E3 SUMO-protein ligase CBX4 OS=Hon | CBX4 | 61.368 | 2 | 54.3758 | yes | 1.633E-06 | 559.77252 | 37.846558 |
| LGYWNQQMVPIK | 12 | O00267 | Transcription elongation factor SPT5 OS= | SUPT5H | 121 | 2 | 56.451385 | yes | 1.313E-07 | 738.88702 | 221.26672 |
| ETFQVLNMYGK | 11 | O00267 | Transcription elongation factor SPT5 OS= | SUPT5H | 121 | 2 | 56.451385 | yes | 1.056E-07 | 665.32904 | 216.35857 |
| DMLEFPAQELR | 11 | O00267 | Transcription elongation factor SPT5 OS= | SUPT5H | 121 | 2 | 56.451385 | yes | 3.547E-07 | 674.83191 | 180.57599 |
| MDLDEQLK | 8 | O00267 | Transcription elongation factor SPT5 OS= | SUPT5H | 121 | 2 | 56.451385 | yes | 9.722E-08 | 496.24188 | 166.00754 |
| SVTGGMCSVYLK | 12 | O00267 | Transcription elongation factor SPT5 OS= | SUPT5H | 121 | 2 | 56.451385 | yes | 1.508E-07 | 651.31506 | 155.48428 |
| SVVAPEHVK | 9 | O00267 | Transcription elongation factor SPT5 OS= | SUPT5H | 121 | 2 | 56.451385 | yes | 2.911E-07 | 483.27435 | 145.10222 |
| FIAYQFTDTPLQIK | 14 | O00267 | Transcription elongation factor SPT5 OS= | SUPT5H | 121 | 2 | 56.451385 | yes | 9.722E-08 | 842.95087 | 114.99192 |
| FAVALDSEQNNIH | 15 | O00267 | Transcription elongation factor SPT5 OS= | SUPT5H | 121 | 2 | 56.451385 | yes | 9.722E-08 | 842.93628 | 109.8752 |
| SFAFLHCK | 8 | O00267 | Transcription elongation factor SPT5 OS= | SUPT5H | 121 | 2 | 56.451385 | yes | 5.868E-07 | 505.24985 | 82.639862 |
| TPMYGSQTPLQDC | 15 | O00267 | Transcription elongation factor SPT5 OS= | SUPT5H | 121 | 2 | 56.451385 | yes | 8.455E-07 | 819.38306 | 57.64035 |
| EEELGEYYMK | 10 | O00267 | Transcription elongation factor SPT5 OS= | SUPT5H | 121 | 2 | 56.451385 | yes | 1.896E-07 | 645.78156 | 34.60553 |
| TPHYGSQTPLHDG | 15 | O00267 | Transcription elongation factor SPT5 OS= | SUPT5H | 121 | 2 | 56.451385 | yes | 7.165E-07 | 826.8924 | 21.538822 |
| SFAMSAVITEGVK | 21 | O00267 | Transcription elongation factor SPT5 OS= | SUPT5H | 121 | 3 | 56.451385 | yes | 4.197E-07 | 746.396 | 21.114016 |
| QQQLLNQQVLEQI | 13 | O00287 | Regulatory factor X-associated protein OS | RFXAP | 28.233 | 2 | 42.457214 | yes | 3.02E-07 | 812.94189 | 59.585121 |
| LEESADNILSIVK | 13 | O00287 | Regulatory factor X-associated protein OS | RFXAP | 28.233 | 2 | 42.457214 | yes | 4.396E-06 | 715.8905 | 25.280886 |
| GVTFNVTTVDTK | 12 | O00299 | Chloride intracellular channel protein 1 O | CLIC1 | 26.923 | 2 | 46.00664 | yes | 1.679E-07 | 641.33789 | 76.704803 |
| VIGTLLGTVDK | 11 | O00303 | Eukaryotic translation initiation factor 3 s | EIF3F | 37.564 | 2 | 54.63323 | yes | 1.783E-07 | 558.33716 | 3066.7422 |
| VIGLSSDLQQVGG | 17 | O00303 | Eukaryotic translation initiation factor 3 s | EIF3F | 37.564 | 2 | 54.63323 | yes | 1.29E-07 | 829.44702 | 2762.6665 |

| | | | | | | | | | | |
|---|---|---|---|---|---|---|---|---|---|---|
| TMGVMFTPLTVK | 12 | O00303 | Eukaryotic translation initiation factor 3 s | EIF3F | 37.564 | 2 | 54.63323 | yes | 1.037E-07 | 662.85382 | 1725.3538 |
| YAYYDTER | 8 | O00303 | Eukaryotic translation initiation factor 3 s | EIF3F | 37.564 | 2 | 54.63323 | yes | 1.611E-07 | 540.73529 | 863.2373 |
| FLMSLVNQVPK | 11 | O00303 | Eukaryotic translation initiation factor 3 s | EIF3F | 37.564 | 2 | 54.63323 | yes | 1.222E-07 | 638.36011 | 829.14014 |
| AYVSTLMGVPGR | 12 | O00303 | Eukaryotic translation initiation factor 3 s | EIF3F | 37.564 | 2 | 54.63323 | yes | 3.297E-06 | 625.83173 | 300.88123 |
| HSVEVTNCFSVPH | 28 | O00303 | Eukaryotic translation initiation factor 3 s | EIF3F | 37.564 | 4 | 54.63323 | yes | 9.722E-08 | 802.35651 | 131.94896 |
| LHPVILASIVDSYE | 15 | O00303 | Eukaryotic translation initiation factor 3 s | EIF3F | 37.564 | 3 | 54.63323 | yes | 9.722E-08 | 571.31744 | 52.661968 |
| EAPNPIHLTVDTSI | 18 | O00303 | Eukaryotic translation initiation factor 3 s | EIF3F | 37.564 | 3 | 54.63323 | yes | 1.163E-07 | 654.67285 | 40.297886 |
| LLQFVTGTCR | 10 | O00308 | NEDD4-like E3 ubiquitin-protein ligase \ | WWP2 | 98.913 | 2 | 54.64761 | no | 9.722E-08 | 597.8186 | 250.51364 |
| TTPATGEQSPGAR | 13 | O00308 | NEDD4-like E3 ubiquitin-protein ligase \ | WWP2 | 98.913 | 2 | 54.64761 | yes | 1.248E-06 | 636.81256 | 125.96572 |
| FYYVDHNTR | 9 | O00308 | NEDD4-like E3 ubiquitin-protein ligase \ | WWP2 | 98.913 | 2 | 54.64761 | yes | 9.722E-08 | 607.78308 | 114.17902 |
| NELLGTASVNLSN | 16 | O00308 | NEDD4-like E3 ubiquitin-protein ligase \ | WWP2 | 98.913 | 2 | 54.64761 | yes | 2.727E-06 | 836.46741 | 57.899113 |
| NYEQWQSQR | 9 | O00308 | NEDD4-like E3 ubiquitin-protein ligase \ | WWP2 | 98.913 | 2 | 54.64761 | yes | 7.174E-07 | 619.78107 | 49.863266 |
| SHTCFNR | 7 | O00308 | NEDD4-like E3 ubiquitin-protein ligase \ | WWP2 | 98.913 | 2 | 54.64761 | no | 4.534E-07 | 461.20343 | 40.845509 |
| VTEENKEEYIMLL | 17 | O00308 | NEDD4-like E3 ubiquitin-protein ligase \ | WWP2 | 98.913 | 3 | 54.64761 | yes | 2.313E-06 | 723.68927 | 31.563663 |
| LYEAVPQLSNVFK | 13 | O00311 | Cell division cycle 7-related protein kinas | CDC7 | 63.889 | 2 | 52.728718 | yes | 3.946E-07 | 754.41138 | 399.93536 |
| EGSVGLSVQR | 10 | O00311 | Cell division cycle 7-related protein kinas | CDC7 | 63.889 | 2 | 52.728718 | yes | 1.962E-07 | 516.27765 | 365.44244 |
| LLDLNPASR | 9 | O00311 | Cell division cycle 7-related protein kinas | CDC7 | 63.889 | 2 | 52.728718 | yes | 1.493E-07 | 499.7851 | 287.39273 |
| IHQFGIVHR | 9 | O00311 | Cell division cycle 7-related protein kinas | CDC7 | 63.889 | 2 | 52.728718 | yes | 9.722E-08 | 553.81451 | 258.98755 |
| DVKPSNFLYNR | 11 | O00311 | Cell division cycle 7-related protein kinas | CDC7 | 63.889 | 2 | 52.728718 | yes | 1.618E-07 | 676.8515 | 251.04552 |
| VCSICLSR | 8 | O00311 | Cell division cycle 7-related protein kinas | CDC7 | 63.889 | 2 | 52.728718 | yes | 3.972E-07 | 497.74405 | 168.99068 |
| IPLSGPVPK | 9 | O00311 | Cell division cycle 7-related protein kinas | CDC7 | 63.889 | 2 | 52.728718 | yes | 3.166E-07 | 454.28421 | 71.089363 |
| KYALVDFGLAQG | 17 | O00311 | Cell division cycle 7-related protein kinas | CDC7 | 63.889 | 3 | 52.728718 | yes | 9.722E-08 | 621.99152 | 58.856358 |
| NFNIHSSISHESPA | 16 | O00311 | Cell division cycle 7-related protein kinas | CDC7 | 63.889 | 3 | 52.728718 | yes | 2.397E-06 | 589.6322 | 46.899731 |
| FVQSEAQQER | 10 | O00311 | Cell division cycle 7-related protein kinas | CDC7 | 63.889 | 2 | 52.728718 | yes | 2.247E-07 | 611.29657 | 41.989742 |
| ILNYSQNDLIGQSI | 21 | O00327 | Basic helix-loop-helix ARNT-like protein | BMAL1 | 68.762 | 3 | 41.110504 | yes | 9.54E-07 | 826.76031 | 144.35751 |
| FVFVDQR | 7 | O00327 | Basic helix-loop-helix ARNT-like protein | BMAL1 | 68.762 | 2 | 41.110504 | yes | 4.898E-07 | 455.74271 | 67.530014 |
| AADGFLFVVGCDI | 13 | O00327 | Basic helix-loop-helix ARNT-like protein | BMAL1 | 68.762 | 2 | 41.110504 | yes | 3.671E-07 | 713.84283 | 52.168034 |
| ILFVSESVFK | 10 | O00327 | Basic helix-loop-helix ARNT-like protein | BMAL1 | 68.762 | 2 | 41.110504 | yes | 3.661E-06 | 584.83423 | 33.599049 |
| LAGAPSEDPQFPK | 13 | O00391 | Sulfhydryl oxidase 1 OS=Homo sapiens ( | QSOX1 | 82.578 | 2 | 43.62168 | yes | 2.87E-06 | 678.84332 | 83.701416 |
| DTGAALLAESR | 11 | O00391 | Sulfhydryl oxidase 1 OS=Homo sapiens ( | QSOX1 | 82.578 | 2 | 43.62168 | yes | 2.514E-07 | 552.28821 | 64.258331 |
| KFVAVLAK | 8 | O00391 | Sulfhydryl oxidase 1 OS=Homo sapiens ( | QSOX1 | 82.578 | 2 | 43.62168 | yes | 4.668E-07 | 438.28928 | 25.151295 |
| FYFHDGVR | 8 | O00410 | Importin-5 OS=Homo sapiens OX=9606 | IPO5 | 123.63 | 2 | 54.743523 | yes | 9.823E-08 | 520.75104 | 220.302 |
| EGFVEYTEQVVK | 12 | O00410 | Importin-5 OS=Homo sapiens OX=9606 | IPO5 | 123.63 | 2 | 54.743523 | no | 1.137E-07 | 714.35626 | 204.32024 |
| VAAAESMPLLLEC | 15 | O00410 | Importin-5 OS=Homo sapiens OX=9606 | IPO5 | 123.63 | 2 | 54.743523 | yes | 2.023E-07 | 815.91821 | 186.46207 |
| FLFDSVSSQNVGL | 14 | O00410 | Importin-5 OS=Homo sapiens OX=9606 | IPO5 | 123.63 | 2 | 54.743523 | yes | 3.235E-07 | 784.90698 | 117.04507 |
| VCDIAAELAR | 10 | O00410 | Importin-5 OS=Homo sapiens OX=9606 | IPO5 | 123.63 | 2 | 54.743523 | yes | 9.722E-08 | 559.28717 | 107.90519 |
| TIECISLIGLAVGK | 14 | O00410 | Importin-5 OS=Homo sapiens OX=9606 | IPO5 | 123.63 | 2 | 54.743523 | yes | 0.0001616 | 737.4209 | 93.125473 |
| HLHSIMVLK | 9 | O00410 | Importin-5 OS=Homo sapiens OX=9606 | IPO5 | 123.63 | 2 | 54.743523 | yes | 1.767E-07 | 539.31549 | 84.531418 |
| LCGDTSLNNMQR | 12 | O00410 | Importin-5 OS=Homo sapiens OX=9606 | IPO5 | 123.63 | 2 | 54.743523 | yes | 1.112E-07 | 704.81903 | 76.096779 |

| | | | | | | | | | | | |
|---|---|---|---|---|---|---|---|---|---|---|---|
| ENVNATENCISAV | 15 | O00410 | Importin-5 OS=Homo sapiens OX=9606 | IPO5 | 123.63 | 2 | 54.743523 | yes | 9.722E-08 | 803.38049 | 73.254616 |
| YLRPHLEATLQLSI | 15 | O00410 | Importin-5 OS=Homo sapiens OX=9606 | IPO5 | 123.63 | 3 | 54.743523 | yes | 1.03E-07 | 594.6806 | 68.323547 |
| QAEETYENIPGQSI | 14 | O00410 | Importin-5 OS=Homo sapiens OX=9606 | IPO5 | 123.63 | 2 | 54.743523 | yes | 9.722E-08 | 797.37317 | 66.156265 |
| STACQMLVCYAK | 12 | O00410 | Importin-5 OS=Homo sapiens OX=9606 | IPO5 | 123.63 | 2 | 54.743523 | yes | 9.722E-08 | 716.32507 | 63.093929 |
| QMAAVLLR | 8 | O00410 | Importin-5 OS=Homo sapiens OX=9606 | IPO5 | 123.63 | 2 | 54.743523 | yes | 2.115E-07 | 451.26804 | 62.035583 |
| EHIMQMLQNPDW | 13 | O00410 | Importin-5 OS=Homo sapiens OX=9606 | IPO5 | 123.63 | 2 | 54.743523 | yes | 3.39E-07 | 835.3949 | 56.86668 |
| LQELIQK | 7 | O00410 | Importin-5 OS=Homo sapiens OX=9606 | IPO5 | 123.63 | 2 | 54.743523 | yes | 6.119E-07 | 436.26602 | 35.450798 |
| ITFLLQAIR | 9 | O00410 | Importin-5 OS=Homo sapiens OX=9606 | IPO5 | 123.63 | 2 | 54.743523 | yes | 3.648E-06 | 537.83716 | 29.988979 |
| VAQVLEGFITR | 11 | O00411 | DNA-directed RNA polymerase, mitocho | POLRMT | 138.621 | 2 | 53.14511 | yes | 5.548E-07 | 616.85352 | 1301.5168 |
| PGAFDLEQVK | 10 | O00411 | DNA-directed RNA polymerase, mitocho | POLRMT | 138.621 | 2 | 53.14511 | yes | 1.389E-07 | 552.29022 | 1107.6487 |
| AIQHWLTESAR | 11 | O00411 | DNA-directed RNA polymerase, mitocho | POLRMT | 138.621 | 2 | 53.14511 | yes | 9.722E-08 | 656.34387 | 1088.8689 |
| IHLVNLTGLK | 10 | O00411 | DNA-directed RNA polymerase, mitocho | POLRMT | 138.621 | 2 | 53.14511 | yes | 1.568E-07 | 554.34784 | 1014.4078 |
| LSLAQHLR | 8 | O00411 | DNA-directed RNA polymerase, mitocho | POLRMT | 138.621 | 2 | 53.14511 | yes | 1.062E-07 | 469.28253 | 980.45013 |
| ALLEFAQGR | 9 | O00411 | DNA-directed RNA polymerase, mitocho | POLRMT | 138.621 | 2 | 53.14511 | yes | 2.373E-07 | 502.77982 | 786.40161 |
| EQPWPLPVQMEL( | 14 | O00411 | DNA-directed RNA polymerase, mitocho | POLRMT | 138.621 | 2 | 53.14511 | yes | 2.613E-07 | 826.42926 | 665.06104 |
| VCVVSVEKPTLPS | 14 | O00411 | DNA-directed RNA polymerase, mitocho | POLRMT | 138.621 | 3 | 53.14511 | yes | 2.224E-07 | 514.95685 | 327.66266 |
| VCVVSVEKPTLPS | 14 | O00411 | DNA-directed RNA polymerase, mitocho | POLRMT | 138.621 | 3 | 53.14511 | yes | 9.892E-08 | 771.93164 | 327.66266 |
| VCVVSVEKPTLPS | 14 | O00411 | DNA-directed RNA polymerase, mitocho | POLRMT | 138.621 | 2 | 53.14511 | yes | 2.224E-07 | 514.95685 | 327.66266 |
| VCVVSVEKPTLPS | 14 | O00411 | DNA-directed RNA polymerase, mitocho | POLRMT | 138.621 | 2 | 53.14511 | yes | 9.892E-08 | 771.93164 | 327.66266 |
| LHSEPILQDLSR | 12 | O00411 | DNA-directed RNA polymerase, mitocho | POLRMT | 138.621 | 2 | 53.14511 | yes | 1.147E-07 | 704.38318 | 653.3075 |
| QAPESPWEEQLAR | 13 | O00411 | DNA-directed RNA polymerase, mitocho | POLRMT | 138.621 | 2 | 53.14511 | yes | 3.176E-07 | 770.87317 | 554.79016 |
| ILEASQLK | 8 | O00411 | DNA-directed RNA polymerase, mitocho | POLRMT | 138.621 | 2 | 53.14511 | yes | 9.741E-08 | 451.2713 | 540.4361 |
| QTVMTVVYGVTR | 12 | O00411 | DNA-directed RNA polymerase, mitocho | POLRMT | 138.621 | 2 | 53.14511 | yes | 9.722E-08 | 677.3634 | 529.56512 |
| QMAGCLEDCTR | 11 | O00411 | DNA-directed RNA polymerase, mitocho | POLRMT | 138.621 | 2 | 53.14511 | yes | 5.365E-07 | 670.77319 | 474.57993 |
| QLHMELASR | 9 | O00411 | DNA-directed RNA polymerase, mitocho | POLRMT | 138.621 | 2 | 53.14511 | yes | 2.102E-07 | 542.78204 | 463.63385 |
| PHPAYVQLLEK | 11 | O00411 | DNA-directed RNA polymerase, mitocho | POLRMT | 138.621 | 2 | 53.14511 | yes | 1.726E-07 | 647.86133 | 460.78409 |
| CLEQMSQEGLK | 11 | O00411 | DNA-directed RNA polymerase, mitocho | POLRMT | 138.621 | 2 | 53.14511 | yes | 9.722E-08 | 661.80762 | 458.84903 |
| PLGPHGLDWLK | 11 | O00411 | DNA-directed RNA polymerase, mitocho | POLRMT | 138.621 | 2 | 53.14511 | yes | 9.722E-08 | 616.84296 | 220.65768 |
| PLGPHGLDWLK | 11 | O00411 | DNA-directed RNA polymerase, mitocho | POLRMT | 138.621 | 2 | 53.14511 | yes | 2.404E-05 | 411.56439 | 220.65768 |
| PLGPHGLDWLK | 11 | O00411 | DNA-directed RNA polymerase, mitocho | POLRMT | 138.621 | 3 | 53.14511 | yes | 9.722E-08 | 616.84296 | 220.65768 |
| PLGPHGLDWLK | 11 | O00411 | DNA-directed RNA polymerase, mitocho | POLRMT | 138.621 | 3 | 53.14511 | yes | 2.404E-05 | 411.56439 | 220.65768 |
| VSGQVQALQNHY | 13 | O00411 | DNA-directed RNA polymerase, mitocho | POLRMT | 138.621 | 2 | 53.14511 | yes | 5.061E-07 | 750.38934 | 423.54697 |
| QLQAESVSEVVVN | 14 | O00411 | DNA-directed RNA polymerase, mitocho | POLRMT | 138.621 | 2 | 53.14511 | yes | 4.691E-07 | 779.41522 | 332.41916 |
| NVQQIGILK | 9 | O00411 | DNA-directed RNA polymerase, mitocho | POLRMT | 138.621 | 2 | 53.14511 | yes | 1.988E-07 | 506.81113 | 205.77515 |
| LQMPFQSGEFK | 11 | O00411 | DNA-directed RNA polymerase, mitocho | POLRMT | 138.621 | 2 | 53.14511 | yes | 4.264E-07 | 656.32373 | 182.97353 |
| TLQCLFEK | 8 | O00411 | DNA-directed RNA polymerase, mitocho | POLRMT | 138.621 | 2 | 53.14511 | yes | 2.681E-07 | 519.76807 | 154.12834 |
| VFWLPHNMDFR | 11 | O00411 | DNA-directed RNA polymerase, mitocho | POLRMT | 138.621 | 2 | 53.14511 | yes | 9.722E-08 | 731.35846 | 140.87373 |
| LVPVLYHVYSFR | 12 | O00411 | DNA-directed RNA polymerase, mitocho | POLRMT | 138.621 | 2 | 53.14511 | yes | 6.011E-07 | 746.91919 | 135.22556 |
| FCSEPQK | 7 | O00411 | DNA-directed RNA polymerase, mitocho | POLRMT | 138.621 | 2 | 53.14511 | yes | 1.427E-06 | 448.20255 | 122.91933 |

| | | | | | | | | | | | |
|---|---|---|---|---|---|---|---|---|---|---|---|
| QDQDAGTIER | 10 | O00411 | DNA-directed RNA polymerase, mitocho | POLRMT | 138.621 | 2 | 53.14511 | yes | 9.722E-08 | 566.76508 | 117.71548 |
| LPECGSGDGSLQPI | 15 | O00411 | DNA-directed RNA polymerase, mitocho | POLRMT | 138.621 | 2 | 53.14511 | yes | 2.447E-06 | 785.36993 | 83.359474 |
| CCLLTDQLPLAHH | 22 | O00411 | DNA-directed RNA polymerase, mitocho | POLRMT | 138.621 | 4 | 53.14511 | yes | 1.258E-06 | 655.09552 | 75.465897 |
| ETLQAVPK | 8 | O00411 | DNA-directed RNA polymerase, mitocho | POLRMT | 138.621 | 2 | 53.14511 | yes | 1.214E-06 | 443.25565 | 36.501823 |
| NVQQIGILKPHPAY | 20 | O00411 | DNA-directed RNA polymerase, mitocho | POLRMT | 138.621 | 3 | 53.14511 | yes | 5.541E-07 | 763.44238 | 36.0938 |
| KYLCLLASDAEVP | 19 | O00411 | DNA-directed RNA polymerase, mitocho | POLRMT | 138.621 | 3 | 53.14511 | yes | 4.193E-07 | 744.37817 | 29.661592 |
| KDWGHVELLEVLO | 15 | O00411 | DNA-directed RNA polymerase, mitocho | POLRMT | 138.621 | 3 | 53.14511 | yes | 2.923E-07 | 598.32831 | 26.599016 |
| GLTFVSVHDCYWT | 26 | O00411 | DNA-directed RNA polymerase, mitocho | POLRMT | 138.621 | 4 | 53.14511 | yes | 3.475E-06 | 763.85199 | 11.520315 |
| EAEMLFTGGYGLE | 14 | O00418 | Eukaryotic elongation factor 2 kinase OS= | EEF2K | 82.145 | 2 | 44.73534 | yes | 2.087E-07 | 772.86871 | 53.329788 |
| ICESMGLAPFDLSI | 15 | O00418 | Eukaryotic elongation factor 2 kinase OS= | EEF2K | 82.145 | 2 | 44.73534 | yes | 2.671E-06 | 846.90784 | 42.842484 |
| VTQEEIK | 7 | O00422 | Histone deacetylase complex subunit SAI | SAP18 | 17.562 | 2 | 50.257 | yes | 6.054E-07 | 423.73199 | 159.86371 |
| TCPLLLR | 7 | O00422 | Histone deacetylase complex subunit SAI | SAP18 | 17.562 | 2 | 50.257 | yes | 2.197E-07 | 436.75476 | 149.28059 |
| GTDDSMTLQSQK | 12 | O00422 | Histone deacetylase complex subunit SAI | SAP18 | 17.562 | 2 | 50.257 | yes | 9.825E-08 | 655.79828 | 84.632088 |
| KGTDDSMTLQSQI | 13 | O00422 | Histone deacetylase complex subunit SAI | SAP18 | 17.562 | 2 | 50.257 | yes | 4.774E-07 | 719.84576 | 32.811516 |
| GTHFNFAIVFTDVI | 14 | O00422 | Histone deacetylase complex subunit SAI | SAP18 | 17.562 | 2 | 50.257 | yes | 1.618E-06 | 798.41449 | 31.6136 |
| LISWSGNYQK | 10 | O00423 | Echinoderm microtubule-associated prote | EML1 | 89.861 | 2 | 41.462543 | yes | 3.497E-07 | 598.30896 | 54.390907 |
| MVIITGPPEAQFK | 13 | O00425 | Insulin-like growth factor 2 mRNA-bindi | IGF2BP3 | 63.706 | 2 | 51.539543 | no | 9.722E-08 | 715.88922 | 19552.877 |
| MVIITGPPEAQFK | 13 | O00425 | Insulin-like growth factor 2 mRNA-bindi | IGF2BP3 | 63.706 | 2 | 51.539543 | no | 2.381E-07 | 723.88672 | 19552.877 |
| SITILSTPEGTSAAC | 16 | O00425 | Insulin-like growth factor 2 mRNA-bindi | IGF2BP3 | 63.706 | 2 | 51.539543 | yes | 1.21E-07 | 818.41656 | 18552.854 |
| IPVSGPFLVK | 10 | O00425 | Insulin-like growth factor 2 mRNA-bindi | IGF2BP3 | 63.706 | 2 | 51.539543 | yes | 2.87E-07 | 528.82623 | 14461.611 |
| SILEIMHK | 8 | O00425 | Insulin-like growth factor 2 mRNA-bindi | IGF2BP3 | 63.706 | 2 | 51.539543 | yes | 9.861E-08 | 485.77313 | 6927.2656 |
| SILEIMHK | 8 | O00425 | Insulin-like growth factor 2 mRNA-bindi | IGF2BP3 | 63.706 | 2 | 51.539543 | yes | 2.701E-06 | 493.7706 | 6927.2656 |
| ILAHNNFVGR | 10 | O00425 | Insulin-like growth factor 2 mRNA-bindi | IGF2BP3 | 63.706 | 2 | 51.539543 | no | 1.795E-07 | 570.81726 | 13347.284 |
| IKEENFVSPK | 10 | O00425 | Insulin-like growth factor 2 mRNA-bindi | IGF2BP3 | 63.706 | 2 | 51.539543 | yes | 1.385E-07 | 595.8244 | 12117.851 |
| ALQSGPPQSR | 10 | O00425 | Insulin-like growth factor 2 mRNA-bindi | IGF2BP3 | 63.706 | 2 | 51.539543 | yes | 2.716E-06 | 520.77783 | 11863.576 |
| DQTPDENDQVVVI | 13 | O00425 | Insulin-like growth factor 2 mRNA-bindi | IGF2BP3 | 63.706 | 2 | 51.539543 | yes | 5.41E-07 | 743.85461 | 8296.5449 |
| IQEILTQVK | 9 | O00425 | Insulin-like growth factor 2 mRNA-bindi | IGF2BP3 | 63.706 | 2 | 51.539543 | yes | 1.76E-07 | 536.32404 | 7736.6826 |
| IAPAEAPDAK | 10 | O00425 | Insulin-like growth factor 2 mRNA-bindi | IGF2BP3 | 63.706 | 2 | 51.539543 | yes | 1.117E-07 | 491.76382 | 4642.5806 |
| KIQEILTQVK | 10 | O00425 | Insulin-like growth factor 2 mRNA-bindi | IGF2BP3 | 63.706 | 2 | 51.539543 | yes | 1.028E-07 | 600.37152 | 4310.7012 |
| QKPCDLPLR | 9 | O00425 | Insulin-like growth factor 2 mRNA-bindi | IGF2BP3 | 63.706 | 2 | 51.539543 | yes | 1.465E-07 | 563.80548 | 4125.7622 |
| IELHGKPIEVEHSV | 16 | O00425 | Insulin-like growth factor 2 mRNA-bindi | IGF2BP3 | 63.706 | 3 | 51.539543 | yes | 9.722E-08 | 604.672 | 1554.2429 |
| IELHGKPIEVEHSV | 16 | O00425 | Insulin-like growth factor 2 mRNA-bindi | IGF2BP3 | 63.706 | 3 | 51.539543 | yes | 9.759E-08 | 453.7558 | 1554.2429 |
| IELHGKPIEVEHSV | 16 | O00425 | Insulin-like growth factor 2 mRNA-bindi | IGF2BP3 | 63.706 | 4 | 51.539543 | yes | 9.722E-08 | 604.672 | 1554.2429 |
| IELHGKPIEVEHSV | 16 | O00425 | Insulin-like growth factor 2 mRNA-bindi | IGF2BP3 | 63.706 | 4 | 51.539543 | yes | 9.759E-08 | 453.7558 | 1554.2429 |
| FTEEIPLK | 8 | O00425 | Insulin-like growth factor 2 mRNA-bindi | IGF2BP3 | 63.706 | 2 | 51.539543 | yes | 2.691E-07 | 488.77112 | 2764.3679 |
| LNGFQLENFTLK | 12 | O00425 | Insulin-like growth factor 2 mRNA-bindi | IGF2BP3 | 63.706 | 2 | 51.539543 | yes | 1.105E-07 | 712.38263 | 2003.1594 |
| ITGHFYACQVAQR | 13 | O00425 | Insulin-like growth factor 2 mRNA-bindi | IGF2BP3 | 63.706 | 2 | 51.539543 | yes | 1.186E-07 | 775.88025 | 1992.6608 |
| PIEVEHSVPK | 10 | O00425 | Insulin-like growth factor 2 mRNA-bindi | IGF2BP3 | 63.706 | 2 | 51.539543 | yes | 1.465E-07 | 567.81134 | 1490.9366 |
| AEEEIMK | 7 | O00425 | Insulin-like growth factor 2 mRNA-bindi | IGF2BP3 | 63.706 | 2 | 51.539543 | yes | 3.662E-07 | 425.20477 | 636.58203 |

| | | | | | | | | | | | |
|---|---|---|---|---|---|---|---|---|---|---|---|
| TVNELQNLSSAEV | 17 | O00425 | Insulin-like growth factor 2 mRNA-bindi | IGF2BP3 | 63.706 | 3 | 51.539543 | yes | 1.006E-07 | 619.0022 | 390.76263 |
| EENFVSPK | 8 | O00425 | Insulin-like growth factor 2 mRNA-bindi | IGF2BP3 | 63.706 | 2 | 51.539543 | yes | 1.402E-07 | 475.23489 | 149.38971 |
| VAYIPDEMAAQQI | 19 | O00425 | Insulin-like growth factor 2 mRNA-bindi | IGF2BP3 | 63.706 | 3 | 51.539543 | yes | 1.108E-07 | 723.69678 | 109.5885 |
| EEVKLEAHIR | 10 | O00425 | Insulin-like growth factor 2 mRNA-bindi | IGF2BP3 | 63.706 | 2 | 51.539543 | no | 2.171E-07 | 612.34076 | 84.760406 |
| QLNPINLTER | 10 | O00442 | RNA 3'-terminal phosphate cyclase OS=F | RTCA | 39.337 | 2 | 52.40247 | yes | 1.84E-07 | 599.33295 | 623.95032 |
| VEVDGSIMEGGGQ | 16 | O00442 | RNA 3'-terminal phosphate cyclase OS=F | RTCA | 39.337 | 2 | 52.40247 | yes | 9.722E-08 | 830.42218 | 551.3587 |
| AFVAGVLPFK | 10 | O00442 | RNA 3'-terminal phosphate cyclase OS=F | RTCA | 39.337 | 2 | 52.40247 | yes | 2.317E-07 | 524.81311 | 356.50082 |
| DLYVNIQPVQEPK | 13 | O00442 | RNA 3'-terminal phosphate cyclase OS=F | RTCA | 39.337 | 2 | 52.40247 | yes | 1.253E-07 | 771.91174 | 293.08643 |
| FGFIFNCDIK | 10 | O00442 | RNA 3'-terminal phosphate cyclase OS=F | RTCA | 39.337 | 2 | 52.40247 | yes | 9.722E-08 | 630.80774 | 258.84131 |
| TGPVTLHTQTAIHI | 20 | O00442 | RNA 3'-terminal phosphate cyclase OS=F | RTCA | 39.337 | 3 | 52.40247 | yes | 1.06E-07 | 721.72339 | 210.71268 |
| GMVELVPASDTLF | 13 | O00443 | Phosphatidylinositol 4-phosphate 3-kinas | PIK3C2A | 190.68 | 2 | 50.214096 | yes | 9.849E-08 | 694.36615 | 77.31826 |
| LIESSLGSIATK | 12 | O00443 | Phosphatidylinositol 4-phosphate 3-kinas | PIK3C2A | 190.68 | 2 | 50.214096 | yes | 3.208E-07 | 609.85065 | 30.81596 |
| DPWDAVLLEER | 11 | O00443 | Phosphatidylinositol 4-phosphate 3-kinas | PIK3C2A | 190.68 | 2 | 50.214096 | yes | 1.841E-07 | 671.83551 | 22.643652 |
| DLVTEDGADPNPY | 15 | O00443 | Phosphatidylinositol 4-phosphate 3-kinas | PIK3C2A | 190.68 | 2 | 50.214096 | yes | 2.11E-06 | 816.89124 | 17.486479 |
| LAVQQVEEAQQLI | 13 | O00461 | Golgi integral membrane protein 4 OS=H | GOLIM4 | 81.881 | 2 | 38.18467 | yes | 5.4E-07 | 756.41248 | 38.257431 |
| SALAAAQTQVAEY | 14 | O00461 | Golgi integral membrane protein 4 OS=H | GOLIM4 | 81.881 | 2 | 38.18467 | yes | 1.327E-06 | 725.88043 | 20.67922 |
| VTLMLLDQSGK | 11 | O00463 | TNF receptor-associated factor 5 OS=Hoi | TRAF5 | 64.406 | 2 | 30.178556 | yes | 2.262E-06 | 602.83392 | 31.425241 |
| LELGIGPGAATR | 12 | O00468 | Agrin OS=Homo sapiens OX=9606 GN= | AGRN | 217.32 | 2 | 43.739616 | yes | 4.277E-07 | 577.83002 | 73.514633 |
| SAGDVDTLAFDGF | 13 | O00468 | Agrin OS=Homo sapiens OX=9606 GN= | AGRN | 217.32 | 2 | 43.739616 | yes | 2.466E-07 | 662.31238 | 47.298733 |
| LALFNPDVCWDR | 12 | O00483 | Cytochrome c oxidase subunit NDUFA4 ( | NDUFA4 | 9.37 | 2 | 51.783634 | yes | 1.317E-07 | 753.36389 | 1393.161 |
| FYSVNVDYSK | 10 | O00483 | Cytochrome c oxidase subunit NDUFA4 ( | NDUFA4 | 9.37 | 2 | 51.783634 | yes | 9.722E-08 | 611.29297 | 1065.4635 |
| LGPNDQYK | 8 | O00483 | Cytochrome c oxidase subunit NDUFA4 ( | NDUFA4 | 9.37 | 2 | 51.783634 | yes | 1.268E-06 | 467.73508 | 441.49698 |
| AVAVVVDPIQSVK | 13 | O00487 | 26S proteasome non-ATPase regulatory s | PSMD14 | 34.578 | 2 | 52.448162 | yes | 9.722E-08 | 662.89539 | 184.24059 |
| YFIDSTNLK | 9 | O00488 | Zinc finger protein 593 OS=Homo sapien | ZNF593 | 15.2 | 2 | 48.206978 | yes | 1.181E-07 | 550.78479 | 1207.5948 |
| QLSVEPYSQEEAE | 14 | O00488 | Zinc finger protein 593 OS=Homo sapien | ZNF593 | 15.2 | 2 | 48.206978 | yes | 4.806E-07 | 832.89172 | 373.2532 |
| EAAWAISNLTISGF | 14 | O00505 | Importin subunit alpha-4 OS=Homo sapie | KPNA3 | 57.811 | 2 | 54.381546 | no | 5.219E-07 | 744.89392 | 1232.4674 |
| NPPIDDLIK | 9 | O00505 | Importin subunit alpha-4 OS=Homo sapie | KPNA3 | 57.811 | 2 | 54.381546 | no | 1.251E-07 | 512.78729 | 263.02542 |
| AENPSLENHR | 10 | O00505 | Importin subunit alpha-4 OS=Homo sapie | KPNA3 | 57.811 | 2 | 54.381546 | yes | 1.443E-07 | 604.78638 | 246.99155 |
| DYVISLGVVK | 10 | O00505 | Importin subunit alpha-4 OS=Homo sapie | KPNA3 | 57.811 | 2 | 54.381546 | no | 9.722E-08 | 546.8186 | 147.91628 |
| DSQVVQVVLDGLI | 13 | O00505 | Importin subunit alpha-4 OS=Homo sapie | KPNA3 | 57.811 | 2 | 54.381546 | yes | 9.722E-08 | 700.39319 | 135.71732 |
| NEVTVELR | 8 | O00505 | Importin subunit alpha-4 OS=Homo sapie | KPNA3 | 57.811 | 2 | 54.381546 | yes | 1.506E-06 | 480.26144 | 126.00645 |
| RDEHLLK | 7 | O00505 | Importin subunit alpha-4 OS=Homo sapie | KPNA3 | 57.811 | 2 | 54.381546 | no | 4.326E-07 | 455.75888 | 89.694496 |
| NVTWVIVNLCR | 11 | O00505 | Importin subunit alpha-4 OS=Homo sapie | KPNA3 | 57.811 | 2 | 54.381546 | yes | 1.354E-07 | 687.37152 | 62.723331 |
| SGILPILVK | 9 | O00505 | Importin subunit alpha-4 OS=Homo sapie | KPNA3 | 57.811 | 2 | 54.381546 | yes | 3.88E-06 | 470.31549 | 7.1041393 |
| SACEGDSELLSR | 12 | O00522 | Krev interaction trapped protein 1 OS=H | KRIT1 | 84.349 | 2 | 51.236618 | yes | 9.722E-08 | 662.2959 | 57.516762 |
| EASLFIVPSVVK | 12 | O00522 | Krev interaction trapped protein 1 OS=H | KRIT1 | 84.349 | 2 | 51.236618 | yes | 2.21E-06 | 644.87921 | 27.293798 |
| LEGQAQAEAK | 10 | O00541 | Pescadillo homolog OS=Homo sapiens O | PES1 | 68.003 | 2 | 53.23156 | yes | 2.669E-07 | 522.76965 | 2297.7058 |
| LAALSASLAR | 10 | O00541 | Pescadillo homolog OS=Homo sapiens O | PES1 | 68.003 | 2 | 53.23156 | yes | 2.099E-07 | 486.79547 | 2015.7905 |
| LQLSLADFR | 9 | O00541 | Pescadillo homolog OS=Homo sapiens O | PES1 | 68.003 | 2 | 53.23156 | yes | 2.419E-07 | 531.80072 | 1632.0598 |

| | | | | | | | | | | | |
|---|---|---|---|---|---|---|---|---|---|---|---|
| SLCIGATYDVTDSI | 14 | O00541 | Pescadillo homolog OS=Homo sapiens O | PES1 | 68.003 | 2 | 53.23156 | yes | 1.073E-07 | 779.36432 | 1474.0562 |
| FLLHEPIVNK | 10 | O00541 | Pescadillo homolog OS=Homo sapiens O | PES1 | 68.003 | 2 | 53.23156 | yes | 1.855E-06 | 605.35315 | 1466.7085 |
| SEWNTVER | 8 | O00541 | Pescadillo homolog OS=Homo sapiens O | PES1 | 68.003 | 2 | 53.23156 | yes | 1.602E-07 | 510.74088 | 947.26776 |
| CHVQTIQLCR | 10 | O00541 | Pescadillo homolog OS=Homo sapiens O | PES1 | 68.003 | 2 | 53.23156 | yes | 1.121E-07 | 657.82391 | 852.27496 |
| YPTFIDALR | 9 | O00541 | Pescadillo homolog OS=Homo sapiens O | PES1 | 68.003 | 2 | 53.23156 | yes | 2.036E-07 | 548.29529 | 604.13068 |
| GIYPHEPK | 8 | O00541 | Pescadillo homolog OS=Homo sapiens O | PES1 | 68.003 | 2 | 53.23156 | yes | 1.615E-07 | 470.74799 | 381.90796 |
| LAALEEQR | 8 | O00541 | Pescadillo homolog OS=Homo sapiens O | PES1 | 68.003 | 2 | 53.23156 | yes | 1.165E-07 | 465.25616 | 353.8537 |
| KVFLSIK | 7 | O00541 | Pescadillo homolog OS=Homo sapiens O | PES1 | 68.003 | 2 | 53.23156 | yes | 5.211E-07 | 417.776 | 231.27914 |
| LTVEFMHYIIAAR | 13 | O00541 | Pescadillo homolog OS=Homo sapiens O | PES1 | 68.003 | 2 | 53.23156 | yes | 1.128E-07 | 782.42126 | 177.64529 |
| LAQEEESEAK | 10 | O00541 | Pescadillo homolog OS=Homo sapiens O | PES1 | 68.003 | 2 | 53.23156 | yes | 1.031E-07 | 567.26965 | 116.57019 |
| GSATNYITR | 9 | O00541 | Pescadillo homolog OS=Homo sapiens O | PES1 | 68.003 | 2 | 53.23156 | yes | 3.322E-07 | 491.75125 | 108.27864 |
| RLTVEFMHYIIAAI | 14 | O00541 | Pescadillo homolog OS=Homo sapiens O | PES1 | 68.003 | 3 | 53.23156 | yes | 1.013E-07 | 573.98364 | 74.683464 |
| LAQEEESEAKR | 11 | O00541 | Pescadillo homolog OS=Homo sapiens O | PES1 | 68.003 | 2 | 53.23156 | yes | 9.722E-08 | 645.32025 | 68.103638 |
| KLFEGLK | 7 | O00541 | Pescadillo homolog OS=Homo sapiens O | PES1 | 68.003 | 2 | 53.23156 | yes | 9.793E-08 | 417.75781 | 65.206482 |
| LKDNKPNYK | 9 | O00541 | Pescadillo homolog OS=Homo sapiens O | PES1 | 68.003 | 2 | 53.23156 | yes | 1.252E-06 | 560.31146 | 51.752003 |
| KLQLSLADFR | 10 | O00541 | Pescadillo homolog OS=Homo sapiens O | PES1 | 68.003 | 2 | 53.23156 | yes | 9.722E-08 | 595.84821 | 46.40234 |
| SLMDHTIPEV | 10 | O00560 | Syntenin-1 OS=Homo sapiens OX=9606 | SDCBP | 32.445 | 2 | 51.007175 | yes | 1.392E-07 | 571.28156 | 1476.0342 |
| DSTGHVGFIFK | 11 | O00560 | Syntenin-1 OS=Homo sapiens OX=9606 | SDCBP | 32.445 | 2 | 51.007175 | yes | 1.118E-07 | 604.30896 | 784.26349 |
| PSSINYMVAPVTG | 19 | O00560 | Syntenin-1 OS=Homo sapiens OX=9606 | SDCBP | 32.445 | 3 | 51.007175 | yes | 9.722E-08 | 664.00696 | 105.09644 |
| NGLLTEHNICEINC | 21 | O00560 | Syntenin-1 OS=Homo sapiens OX=9606 | SDCBP | 32.445 | 3 | 51.007175 | yes | 1.062E-07 | 779.40576 | 91.425331 |
| LSLAEIYEQEYIK | 13 | O00566 | U3 small nucleolar ribonucleoprotein pro | MPHOSPH | 78.864 | 2 | 48.03362 | yes | 2.935E-06 | 799.91925 | 174.40471 |
| LSEMEAYLENIEK | 13 | O00566 | U3 small nucleolar ribonucleoprotein pro | MPHOSPH | 78.864 | 2 | 48.03362 | yes | 1.464E-07 | 784.87927 | 168.53012 |
| IHGSPLQK | 8 | O00566 | U3 small nucleolar ribonucleoprotein pro | MPHOSPH | 78.864 | 2 | 48.03362 | yes | 1.863E-07 | 440.25598 | 160.81241 |
| MMDSLFLK | 8 | O00566 | U3 small nucleolar ribonucleoprotein pro | MPHOSPH | 78.864 | 2 | 48.03362 | yes | 1.231E-07 | 492.74829 | 117.6267 |
| LTLDHEK | 7 | O00566 | U3 small nucleolar ribonucleoprotein pro | MPHOSPH | 78.864 | 2 | 48.03362 | yes | 1.211E-07 | 428.23218 | 102.25391 |
| LDALSNFHFIPKPP | 19 | O00566 | U3 small nucleolar ribonucleoprotein pro | MPHOSPH | 78.864 | 3 | 48.03362 | yes | 8.828E-07 | 721.40515 | 50.51474 |
| TAEEENPEHVEIQI | 14 | O00566 | U3 small nucleolar ribonucleoprotein pro | MPHOSPH | 78.864 | 2 | 48.03362 | yes | 1.704E-07 | 826.89172 | 35.219639 |
| KPWQLQGEVTAQ | 13 | O00566 | U3 small nucleolar ribonucleoprotein pro | MPHOSPH | 78.864 | 2 | 48.03362 | yes | 1.941E-07 | 756.91211 | 21.883404 |
| RPENSLLEETLHFI | 18 | O00566 | U3 small nucleolar ribonucleoprotein pro | MPHOSPH | 78.864 | 3 | 48.03362 | yes | 2.675E-06 | 721.70288 | 17.006252 |
| LSFYETGEIPR | 11 | O00567 | Nucleolar protein 56 OS=Homo sapiens ( | NOP56 | 66.05 | 2 | 54.808876 | yes | 1.048E-07 | 656.33258 | 1112.3391 |
| EAMVQAEEAAAE | 15 | O00567 | Nucleolar protein 56 OS=Homo sapiens ( | NOP56 | 66.05 | 2 | 54.808876 | yes | 1.418E-07 | 809.89069 | 614.88135 |
| IINDNATYCR | 10 | O00567 | Nucleolar protein 56 OS=Homo sapiens ( | NOP56 | 66.05 | 2 | 54.808876 | yes | 9.893E-08 | 620.29297 | 389.09879 |
| YPASTVQILGAEK | 13 | O00567 | Nucleolar protein 56 OS=Homo sapiens ( | NOP56 | 66.05 | 2 | 54.808876 | yes | 9.977E-08 | 688.87463 | 336.92093 |
| LEELTMDGAK | 10 | O00567 | Nucleolar protein 56 OS=Homo sapiens ( | NOP56 | 66.05 | 2 | 54.808876 | yes | 2.144E-07 | 553.77356 | 311.31381 |
| VLLGVGDPK | 9 | O00567 | Nucleolar protein 56 OS=Homo sapiens ( | NOP56 | 66.05 | 2 | 54.808876 | yes | 1.611E-07 | 449.27383 | 269.72906 |
| LHFHNLVK | 8 | O00567 | Nucleolar protein 56 OS=Homo sapiens ( | NOP56 | 66.05 | 2 | 54.808876 | yes | 1.063E-07 | 504.29291 | 222.70825 |
| EWYGYHFPELVK | 12 | O00567 | Nucleolar protein 56 OS=Homo sapiens ( | NOP56 | 66.05 | 2 | 54.808876 | yes | 3.582E-07 | 784.38263 | 101.65082 |
| EWYGYHFPELVK | 12 | O00567 | Nucleolar protein 56 OS=Homo sapiens ( | NOP56 | 66.05 | 2 | 54.808876 | yes | 2.736E-07 | 523.25751 | 101.65082 |
| EWYGYHFPELVK | 12 | O00567 | Nucleolar protein 56 OS=Homo sapiens ( | NOP56 | 66.05 | 3 | 54.808876 | yes | 3.582E-07 | 784.38263 | 101.65082 |

| | | | | | | | | | | | |
|---|---|---|---|---|---|---|---|---|---|---|---|
| EWYGYHFPELVK | 12 | O00567 | Nucleolar protein 56 OS=Homo sapiens ( | NOP56 | 66.05 | 3 | 54.808876 | yes | 2.736E-07 | 523.25751 | 101.65082 |
| LIAHAGSLTNLAK | 13 | O00567 | Nucleolar protein 56 OS=Homo sapiens ( | NOP56 | 66.05 | 2 | 54.808876 | yes | 9.722E-08 | 654.88531 | 82.267159 |
| LIAHAGSLTNLAK | 13 | O00567 | Nucleolar protein 56 OS=Homo sapiens ( | NOP56 | 66.05 | 2 | 54.808876 | yes | 1.102E-06 | 436.92599 | 82.267159 |
| LIAHAGSLTNLAK | 13 | O00567 | Nucleolar protein 56 OS=Homo sapiens ( | NOP56 | 66.05 | 3 | 54.808876 | yes | 9.722E-08 | 654.88531 | 82.267159 |
| LIAHAGSLTNLAK | 13 | O00567 | Nucleolar protein 56 OS=Homo sapiens ( | NOP56 | 66.05 | 3 | 54.808876 | yes | 1.102E-06 | 436.92599 | 82.267159 |
| QSLHTYLR | 8 | O00567 | Nucleolar protein 56 OS=Homo sapiens ( | NOP56 | 66.05 | 2 | 54.808876 | yes | 2.07E-07 | 509.27744 | 160.17839 |
| LAQFIGNR | 8 | O00567 | Nucleolar protein 56 OS=Homo sapiens ( | NOP56 | 66.05 | 2 | 54.808876 | yes | 3.826E-07 | 459.76141 | 157.017 |
| ELNEDKLEK | 9 | O00567 | Nucleolar protein 56 OS=Homo sapiens ( | NOP56 | 66.05 | 2 | 54.808876 | yes | 9.722E-08 | 559.29041 | 152.31111 |
| YGLIFHSTFIGR | 12 | O00567 | Nucleolar protein 56 OS=Homo sapiens ( | NOP56 | 66.05 | 2 | 54.808876 | yes | 5.298E-07 | 705.88007 | 101.18268 |
| GLTDLSACK | 9 | O00567 | Nucleolar protein 56 OS=Homo sapiens ( | NOP56 | 66.05 | 2 | 54.808876 | yes | 9.997E-06 | 482.74203 | 15.190578 |
| ELAVQIYEEAR | 11 | O00571 | ATP-dependent RNA helicase DDX3X O | DDX3X | 73.244 | 2 | 55.443253 | no | 3.615E-07 | 660.84332 | 47211.82 |
| QYPISLVLAPTR | 12 | O00571 | ATP-dependent RNA helicase DDX3X O | DDX3X | 73.244 | 2 | 55.443253 | no | 1.47E-07 | 679.39557 | 37525.75 |
| VGNLGLATSFFNE | 14 | O00571 | ATP-dependent RNA helicase DDX3X O | DDX3X | 73.244 | 2 | 55.443253 | yes | 9.779E-08 | 762.89392 | 18325.154 |
| VGNLGLATSFFNE | 14 | O00571 | ATP-dependent RNA helicase DDX3X O | DDX3X | 73.244 | 2 | 55.443253 | yes | 1.11E-07 | 508.9317 | 18325.154 |
| VGNLGLATSFFNE | 14 | O00571 | ATP-dependent RNA helicase DDX3X O | DDX3X | 73.244 | 3 | 55.443253 | yes | 9.779E-08 | 762.89392 | 18325.154 |
| VGNLGLATSFFNE | 14 | O00571 | ATP-dependent RNA helicase DDX3X O | DDX3X | 73.244 | 3 | 55.443253 | yes | 1.11E-07 | 508.9317 | 18325.154 |
| YLVLDEADR | 9 | O00571 | ATP-dependent RNA helicase DDX3X O | DDX3X | 73.244 | 2 | 55.443253 | no | 1.568E-07 | 547.27985 | 35919.934 |
| SPILVATAVAAR | 12 | O00571 | ATP-dependent RNA helicase DDX3X O | DDX3X | 73.244 | 2 | 55.443253 | no | 6.538E-07 | 584.85608 | 34039.559 |
| VVWVEESDKR | 10 | O00571 | ATP-dependent RNA helicase DDX3X O | DDX3X | 73.244 | 2 | 55.443253 | yes | 6.431E-07 | 623.82495 | 32248.123 |
| GCHLLVATPGR | 11 | O00571 | ATP-dependent RNA helicase DDX3X O | DDX3X | 73.244 | 2 | 55.443253 | no | 1.173E-07 | 590.81641 | 30979.244 |
| DSLTLVFVETK | 11 | O00571 | ATP-dependent RNA helicase DDX3X O | DDX3X | 73.244 | 2 | 55.443253 | yes | 3.662E-07 | 626.34521 | 26852.359 |
| DLMACAQTGSGK | 12 | O00571 | ATP-dependent RNA helicase DDX3X O | DDX3X | 73.244 | 2 | 55.443253 | no | 1.21E-07 | 619.77881 | 9609.9502 |
| DLMACAQTGSGK | 12 | O00571 | ATP-dependent RNA helicase DDX3X O | DDX3X | 73.244 | 2 | 55.443253 | no | 1.503E-07 | 627.77631 | 9609.9502 |
| VRPCVVYGGADIC | 17 | O00571 | ATP-dependent RNA helicase DDX3X O | DDX3X | 73.244 | 3 | 55.443253 | no | 1.134E-07 | 630.00012 | 12860.354 |
| DKDAYSSFGSR | 11 | O00571 | ATP-dependent RNA helicase DDX3X O | DDX3X | 73.244 | 2 | 55.443253 | no | 3.475E-07 | 616.78076 | 11625.871 |
| SDYDGIGSR | 9 | O00571 | ATP-dependent RNA helicase DDX3X O | DDX3X | 73.244 | 2 | 55.443253 | yes | 1.174E-07 | 485.21725 | 9444.7539 |
| YTRPTPVQK | 9 | O00571 | ATP-dependent RNA helicase DDX3X O | DDX3X | 73.244 | 2 | 55.443253 | no | 1.474E-07 | 545.30621 | 7631.46 |
| QSSGASSSSFSSSR | 14 | O00571 | ATP-dependent RNA helicase DDX3X O | DDX3X | 73.244 | 2 | 55.443253 | yes | 9.889E-08 | 681.30005 | 7499.8906 |
| VGSTSENITQK | 11 | O00571 | ATP-dependent RNA helicase DDX3X O | DDX3X | 73.244 | 2 | 55.443253 | no | 9.722E-08 | 582.29877 | 7450.9048 |
| QEVPSWLENMAY | 17 | O00571 | ATP-dependent RNA helicase DDX3X O | DDX3X | 73.244 | 3 | 55.443253 | no | 1.401E-07 | 720.99841 | 6114.8994 |
| MLDMGFEPQIR | 11 | O00571 | ATP-dependent RNA helicase DDX3X O | DDX3X | 73.244 | 2 | 55.443253 | no | 9.723E-08 | 676.8205 | 1859.824 |
| MLDMGFEPQIR | 11 | O00571 | ATP-dependent RNA helicase DDX3X O | DDX3X | 73.244 | 2 | 55.443253 | no | 9.722E-08 | 676.8205 | 1859.824 |
| MLDMGFEPQIR | 11 | O00571 | ATP-dependent RNA helicase DDX3X O | DDX3X | 73.244 | 2 | 55.443253 | no | 9.722E-08 | 684.81793 | 1859.824 |
| MLDMGFEPQIR | 11 | O00571 | ATP-dependent RNA helicase DDX3X O | DDX3X | 73.244 | 2 | 55.443253 | no | 9.967E-08 | 668.82306 | 1859.824 |
| IVEQDTMPPK | 10 | O00571 | ATP-dependent RNA helicase DDX3X O | DDX3X | 73.244 | 2 | 55.443253 | no | 1.621E-06 | 579.29718 | 2817.031 |
| IVEQDTMPPK | 10 | O00571 | ATP-dependent RNA helicase DDX3X O | DDX3X | 73.244 | 2 | 55.443253 | no | 9.722E-08 | 587.29462 | 2817.031 |
| HTMMFSATFPK | 11 | O00571 | ATP-dependent RNA helicase DDX3X O | DDX3X | 73.244 | 2 | 55.443253 | no | 2.526E-07 | 649.30701 | 1858.4607 |
| HTMMFSATFPK | 11 | O00571 | ATP-dependent RNA helicase DDX3X O | DDX3X | 73.244 | 2 | 55.443253 | no | 9.867E-08 | 657.3045 | 1858.4607 |
| HTMMFSATFPK | 11 | O00571 | ATP-dependent RNA helicase DDX3X O | DDX3X | 73.244 | 2 | 55.443253 | no | 1.23E-07 | 657.3045 | 1858.4607 |

| | | | | | | | | | | | |
|---|---|---|---|---|---|---|---|---|---|---|---|
| LVDMMER | 7 | O00571 | ATP-dependent RNA helicase DDX3X O | DDX3X | 73.244 | 2 | 55.443253 | no | 1.041E-07 | 447.2146 | 2743.2625 |
| LVDMMER | 7 | O00571 | ATP-dependent RNA helicase DDX3X O | DDX3X | 73.244 | 2 | 55.443253 | no | 1.536E-07 | 455.21207 | 2743.2625 |
| YIPPHLR | 7 | O00571 | ATP-dependent RNA helicase DDX3X O | DDX3X | 73.244 | 2 | 55.443253 | no | 2.473E-07 | 448.26105 | 4508.1841 |
| EIQMLAR | 7 | O00571 | ATP-dependent RNA helicase DDX3X O | DDX3X | 73.244 | 2 | 55.443253 | no | 3.286E-07 | 430.73657 | 3073.9937 |
| EEALHQFR | 8 | O00571 | ATP-dependent RNA helicase DDX3X O | DDX3X | 73.244 | 2 | 55.443253 | no | 1.181E-07 | 515.25928 | 1956.4282 |
| DREEALHQFR | 10 | O00571 | ATP-dependent RNA helicase DDX3X O | DDX3X | 73.244 | 2 | 55.443253 | no | 1.033E-07 | 650.8233 | 1928.3481 |
| GLDISNVK | 8 | O00571 | ATP-dependent RNA helicase DDX3X O | DDX3X | 73.244 | 2 | 55.443253 | yes | 1.002E-07 | 423.23999 | 1793.1185 |
| VVWVEESDK | 9 | O00571 | ATP-dependent RNA helicase DDX3X O | DDX3X | 73.244 | 2 | 55.443253 | yes | 1.451E-07 | 545.77441 | 1533.6503 |
| KQYPISLVLAPTR | 13 | O00571 | ATP-dependent RNA helicase DDX3X O | DDX3X | 73.244 | 2 | 55.443253 | no | 9.722E-08 | 743.44305 | 689.69763 |
| KQYPISLVLAPTR | 13 | O00571 | ATP-dependent RNA helicase DDX3X O | DDX3X | 73.244 | 2 | 55.443253 | no | 2.128E-07 | 495.96445 | 689.69763 |
| KQYPISLVLAPTR | 13 | O00571 | ATP-dependent RNA helicase DDX3X O | DDX3X | 73.244 | 3 | 55.443253 | no | 9.722E-08 | 743.44305 | 689.69763 |
| KQYPISLVLAPTR | 13 | O00571 | ATP-dependent RNA helicase DDX3X O | DDX3X | 73.244 | 3 | 55.443253 | no | 2.128E-07 | 495.96445 | 689.69763 |
| RIVEQDTMPPK | 11 | O00571 | ATP-dependent RNA helicase DDX3X O | DDX3X | 73.244 | 2 | 55.443253 | no | 1.187E-07 | 657.34772 | 1150.5026 |
| DAYSSFGSR | 9 | O00571 | ATP-dependent RNA helicase DDX3X O | DDX3X | 73.244 | 2 | 55.443253 | no | 3.758E-07 | 495.21979 | 651.12927 |
| KGADSLEDFLYHE | 24 | O00571 | ATP-dependent RNA helicase DDX3X O | DDX3X | 73.244 | 4 | 55.443253 | no | 2.466E-07 | 686.06195 | 199.98758 |
| RDLMACAQTGSGI | 13 | O00571 | ATP-dependent RNA helicase DDX3X O | DDX3X | 73.244 | 2 | 55.443253 | no | 1.081E-07 | 697.82941 | 113.9236 |
| WCDKSDEDDWSK | 19 | O00571 | ATP-dependent RNA helicase DDX3X O | DDX3X | 73.244 | 4 | 55.443253 | yes | 3.877E-06 | 587.51282 | 106.83786 |
| LEQELFSGGNTGIN | 17 | O00571 | ATP-dependent RNA helicase DDX3X O | DDX3X | 73.244 | 3 | 55.443253 | no | 9.722E-08 | 628.31091 | 87.495712 |
| PCVVYGGADIGQQ | 15 | O00571 | ATP-dependent RNA helicase DDX3X O | DDX3X | 73.244 | 2 | 55.443253 | no | 1.131E-07 | 816.91174 | 69.348686 |
| ILYNPLQGQK | 10 | O00622 | CCN family member 1 OS=Homo sapiens | CCN1 | 42.027 | 2 | 51.755 | yes | 1.005E-07 | 587.33496 | 404.00745 |
| GLECNFGASSTALI | 14 | O00622 | CCN family member 1 OS=Homo sapiens | CCN1 | 42.027 | 2 | 51.755 | yes | 1.624E-07 | 727.85083 | 258.54736 |
| CAPGVGLVR | 9 | O00622 | CCN family member 1 OS=Homo sapiens | CCN1 | 42.027 | 2 | 51.755 | yes | 1.885E-07 | 464.75528 | 239.61823 |
| CCTPQLTR | 8 | O00622 | CCN family member 1 OS=Homo sapiens | CCN1 | 42.027 | 2 | 51.755 | yes | 5.553E-07 | 518.23914 | 234.50879 |
| ELGFDASEVELTR | 13 | O00622 | CCN family member 1 OS=Homo sapiens | CCN1 | 42.027 | 2 | 51.755 | yes | 1.611E-07 | 733.36212 | 225.71861 |
| FTYAGCLSVK | 10 | O00622 | CCN family member 1 OS=Homo sapiens | CCN1 | 42.027 | 2 | 51.755 | yes | 1.343E-06 | 573.28662 | 155.33282 |
| YCGSCVDGR | 9 | O00622 | CCN family member 1 OS=Homo sapiens | CCN1 | 42.027 | 2 | 51.755 | yes | 1.191E-07 | 537.21057 | 116.31826 |
| NVMMIQSCK | 9 | O00622 | CCN family member 1 OS=Homo sapiens | CCN1 | 42.027 | 2 | 51.755 | yes | 4.767E-07 | 555.75885 | 90.776093 |
| CEDGETFSK | 9 | O00622 | CCN family member 1 OS=Homo sapiens | CCN1 | 42.027 | 2 | 51.755 | yes | 5.483E-07 | 536.71625 | 78.198578 |
| CIVQTTSWSQCSK | 13 | O00622 | CCN family member 1 OS=Homo sapiens | CCN1 | 42.027 | 2 | 51.755 | yes | 9.722E-08 | 792.8609 | 50.167618 |
| MVEPQYQELK | 10 | O00625 | Pirin OS=Homo sapiens OX=9606 GN=P | PIR | 32.114 | 2 | 49.691334 | yes | 2.419E-07 | 632.81573 | 132.86133 |
| KVTLSVLSR | 9 | O00625 | Pirin OS=Homo sapiens OX=9606 GN=P | PIR | 32.114 | 2 | 49.691334 | yes | 1.01E-07 | 501.81894 | 91.128281 |
| SEEIPKPSK | 9 | O00625 | Pirin OS=Homo sapiens OX=9606 GN=P | PIR | 32.114 | 2 | 49.691334 | yes | 1.305E-07 | 507.77695 | 63.902569 |
| NEVVVELR | 8 | O00629 | Importin subunit alpha-3 OS=Homo sapie | KPNA4 | 57.887 | 2 | 50.3529 | yes | 7.02E-07 | 479.27182 | 352.57486 |
| NVTWVMVNLCR | 11 | O00629 | Importin subunit alpha-3 OS=Homo sapie | KPNA4 | 57.887 | 2 | 50.3529 | yes | 9.813E-08 | 696.34973 | 235.2202 |
| SGILPILVHCLER | 13 | O00629 | Importin subunit alpha-3 OS=Homo sapie | KPNA4 | 57.887 | 2 | 50.3529 | yes | 1.024E-07 | 753.9267 | 198.55338 |
| SGVFNVSELVR | 11 | O00712 | Nuclear factor 1 B-type OS=Homo sapien | NFIB | 47.442 | 2 | 50.230244 | yes | 3.54E-07 | 603.82751 | 1483.355 |
| DQDMSSPTTMK | 11 | O00712 | Nuclear factor 1 B-type OS=Homo sapien | NFIB | 47.442 | 2 | 50.230244 | yes | 1.305E-07 | 620.76282 | 876.73462 |
| NPPGYLEDSFVK | 12 | O00712 | Nuclear factor 1 B-type OS=Homo sapien | NFIB | 47.442 | 2 | 50.230244 | yes | 1.326E-07 | 683.33789 | 835.99664 |
| DELLSEKPEIK | 11 | O00712 | Nuclear factor 1 B-type OS=Homo sapien | NFIB | 47.442 | 2 | 50.230244 | yes | 1.104E-07 | 650.85339 | 651.85095 |

| | | | | | | | | | | | |
|---|---|---|---|---|---|---|---|---|---|---|---|
| AVKDELLSEKPEIK | 14 | O00712 | Nuclear factor 1 B-type OS=Homo sapien | NFIB | 47.442 | 3 | 50.230244 | yes | 9.91E-08 | 533.63818 | 212.79637 |
| AVKDELLSEKPEIK | 14 | O00712 | Nuclear factor 1 B-type OS=Homo sapien | NFIB | 47.442 | 3 | 50.230244 | yes | 9.848E-08 | 799.95361 | 212.79637 |
| AVKDELLSEKPEIK | 14 | O00712 | Nuclear factor 1 B-type OS=Homo sapien | NFIB | 47.442 | 2 | 50.230244 | yes | 9.91E-08 | 533.63818 | 212.79637 |
| AVKDELLSEKPEIK | 14 | O00712 | Nuclear factor 1 B-type OS=Homo sapien | NFIB | 47.442 | 2 | 50.230244 | yes | 9.848E-08 | 799.95361 | 212.79637 |
| AIAYTWFNLQAR | 12 | O00712 | Nuclear factor 1 B-type OS=Homo sapien | NFIB | 47.442 | 2 | 50.230244 | yes | 2.272E-07 | 727.383 | 285.56134 |
| LSTFPQHHHPGIPC | 23 | O00712 | Nuclear factor 1 B-type OS=Homo sapien | NFIB | 47.442 | 4 | 50.230244 | yes | 9.367E-07 | 619.58057 | 88.093567 |
| LSTFPQHHHPGIPC | 23 | O00712 | Nuclear factor 1 B-type OS=Homo sapien | NFIB | 47.442 | 4 | 50.230244 | yes | 2.676E-06 | 825.77167 | 88.093567 |
| LSTFPQHHHPGIPC | 23 | O00712 | Nuclear factor 1 B-type OS=Homo sapien | NFIB | 47.442 | 3 | 50.230244 | yes | 9.367E-07 | 619.58057 | 88.093567 |
| LSTFPQHHHPGIPC | 23 | O00712 | Nuclear factor 1 B-type OS=Homo sapien | NFIB | 47.442 | 3 | 50.230244 | yes | 2.676E-06 | 825.77167 | 88.093567 |
| KHPCCVLSNPDQK | 13 | O00712 | Nuclear factor 1 B-type OS=Homo sapien | NFIB | 47.442 | 2 | 50.230244 | yes | 1.368E-07 | 791.87689 | 106.35837 |
| PLFSSASPQDSSPR | 14 | O00712 | Nuclear factor 1 B-type OS=Homo sapien | NFIB | 47.442 | 2 | 50.230244 | yes | 1.128E-07 | 738.35986 | 67.47686 |
| SPHCTNPALCVQP | 21 | O00712 | Nuclear factor 1 B-type OS=Homo sapien | NFIB | 47.442 | 4 | 50.230244 | yes | 3.106E-06 | 596.3006 | 35.992485 |
| HPCCVLSNPDQK | 12 | O00712 | Nuclear factor 1 B-type OS=Homo sapien | NFIB | 47.442 | 2 | 50.230244 | yes | 6.707E-07 | 727.82941 | 31.424316 |
| KPEKPLFSSASPQD | 18 | O00712 | Nuclear factor 1 B-type OS=Homo sapien | NFIB | 47.442 | 3 | 50.230244 | yes | 2.985E-07 | 653.33746 | 16.826937 |
| LAYVTYQDIR | 10 | O00716 | Transcription factor E2F3 OS=Homo sapi | E2F3 | 49.162 | 2 | 49.93572 | no | 1.032E-07 | 621.3299 | 167.61981 |
| RLELGESGHQYLS | 17 | O00716 | Transcription factor E2F3 OS=Homo sapi | E2F3 | 49.162 | 3 | 49.93572 | yes | 5.014E-07 | 634.66205 | 111.87165 |
| EVTELSQEEK | 10 | O00716 | Transcription factor E2F3 OS=Homo sapi | E2F3 | 49.162 | 2 | 49.93572 | yes | 9.975E-08 | 596.29059 | 100.15572 |
| YDTSLGLLTK | 10 | O00716 | Transcription factor E2F3 OS=Homo sapi | E2F3 | 49.162 | 2 | 49.93572 | no | 6.597E-07 | 555.80573 | 72.284714 |
| AGLLQQPPALGR | 12 | O00716 | Transcription factor E2F3 OS=Homo sapi | E2F3 | 49.162 | 2 | 49.93572 | yes | 2.918E-07 | 610.85913 | 58.666077 |
| LLTEDSENQR | 10 | O00716 | Transcription factor E2F3 OS=Homo sapi | E2F3 | 49.162 | 2 | 49.93572 | yes | 3.015E-07 | 602.79382 | 29.52574 |
| LVTVWSAPNYCYI | 13 | O00743 | Serine/threonine-protein phosphatase 6 ca | PPP6C | 35.145 | 2 | 56.043232 | yes | 1.114E-07 | 814.89813 | 735.50531 |
| VTNEFVHINNLK | 12 | O00743 | Serine/threonine-protein phosphatase 6 ca | PPP6C | 35.145 | 2 | 56.043232 | yes | 9.722E-08 | 714.38568 | 250.20224 |
| VTNEFVHINNLK | 12 | O00743 | Serine/threonine-protein phosphatase 6 ca | PPP6C | 35.145 | 2 | 56.043232 | yes | 5.692E-07 | 476.5929 | 250.20224 |
| VTNEFVHINNLK | 12 | O00743 | Serine/threonine-protein phosphatase 6 ca | PPP6C | 35.145 | 3 | 56.043232 | yes | 9.722E-08 | 714.38568 | 250.20224 |
| VTNEFVHINNLK | 12 | O00743 | Serine/threonine-protein phosphatase 6 ca | PPP6C | 35.145 | 3 | 56.043232 | yes | 5.692E-07 | 476.5929 | 250.20224 |
| CGNIASIMVFK | 11 | O00743 | Serine/threonine-protein phosphatase 6 ca | PPP6C | 35.145 | 2 | 56.043232 | yes | 9.722E-08 | 620.31488 | 199.13733 |
| YGNANAWR | 8 | O00743 | Serine/threonine-protein phosphatase 6 ca | PPP6C | 35.145 | 2 | 56.043232 | yes | 1.873E-07 | 476.22522 | 169.36136 |
| AHQLVHEGYK | 10 | O00743 | Serine/threonine-protein phosphatase 6 ca | PPP6C | 35.145 | 2 | 56.043232 | yes | 1.779E-07 | 591.3067 | 104.09718 |
| YLPENDLK | 8 | O00743 | Serine/threonine-protein phosphatase 6 ca | PPP6C | 35.145 | 2 | 56.043232 | yes | 2.47E-05 | 496.25839 | 102.92706 |
| TFEEFQELHNK | 11 | O00750 | Phosphatidylinositol 4-phosphate 3-kinas | PIK3C2B | 184.768 | 2 | 52.038326 | yes | 4.375E-06 | 711.33844 | 415.36856 |
| LIESSLGSVATK | 12 | O00750 | Phosphatidylinositol 4-phosphate 3-kinas | PIK3C2B | 184.768 | 2 | 52.038326 | yes | 2.785E-07 | 602.84283 | 113.14851 |
| LSFQNVDPLGENII | 14 | O00750 | Phosphatidylinositol 4-phosphate 3-kinas | PIK3C2B | 184.768 | 2 | 52.038326 | yes | 1.621E-07 | 801.41772 | 89.599014 |
| GMVEMIPNAETLR | 13 | O00750 | Phosphatidylinositol 4-phosphate 3-kinas | PIK3C2B | 184.768 | 2 | 52.038326 | yes | 4.343E-07 | 730.86542 | 77.736969 |
| ISDVFLCR | 8 | O00750 | Phosphatidylinositol 4-phosphate 3-kinas | PIK3C2B | 184.768 | 2 | 52.038326 | yes | 3.176E-07 | 505.26041 | 73.776672 |
| YECYLDSPLVR | 11 | O00750 | Phosphatidylinositol 4-phosphate 3-kinas | PIK3C2B | 184.768 | 2 | 52.038326 | yes | 2.069E-07 | 707.83722 | 46.084991 |
| LPLSPSLLVK | 10 | O00750 | Phosphatidylinositol 4-phosphate 3-kinas | PIK3C2B | 184.768 | 2 | 52.038326 | yes | 1.623E-07 | 533.84717 | 42.469364 |
| ESLYWLTDADK | 11 | O00750 | Phosphatidylinositol 4-phosphate 3-kinas | PIK3C2B | 184.768 | 2 | 52.038326 | yes | 9.722E-08 | 670.82208 | 40.205761 |
| GSTLDLSDLEAEK | 13 | O00767 | Stearoyl-CoA desaturase OS=Homo sapie | SCD | 41.523 | 2 | 50.363136 | yes | 9.818E-08 | 689.34082 | 430.67252 |
| FSETHADPHNSR | 12 | O00767 | Stearoyl-CoA desaturase OS=Homo sapie | SCD | 41.523 | 2 | 50.363136 | yes | 9.81E-08 | 699.31329 | 44.108288 |

| | | | | | | | | | | | |
|---|---|---|---|---|---|---|---|---|---|---|---|
| ECLAETER | 8 | O14519 | Cyclin-dependent kinase 2-associated pro | CDK2AP1 | 12.365 | 2 | 49.144264 | yes | 1.098E-07 | 504.22675 | 110.1045 |
| EIRPTYAGSK | 10 | O14519 | Cyclin-dependent kinase 2-associated pro | CDK2AP1 | 12.365 | 2 | 49.144264 | yes | 1.064E-07 | 561.30109 | 57.619484 |
| AGIALNDNFVK | 11 | O14556 | Glyceraldehyde-3-phosphate dehydrogena | GAPDHS | 44.502 | 2 | 49.239986 | yes | 1.054E-07 | 581.31677 | 395.08673 |
| LSVNSHFMK | 9 | O14561 | Acyl carrier protein, mitochondrial OS=H | NDUFAB1 | 17.418 | 2 | 51.79194 | yes | 9.722E-08 | 531.77368 | 300.08771 |
| LMCPQEIVDYIAD | 14 | O14561 | Acyl carrier protein, mitochondrial OS=H | NDUFAB1 | 17.418 | 2 | 51.79194 | yes | 5.505E-07 | 847.91003 | 214.47713 |
| DIKPENILVDR | 11 | O14578 | Citron Rho-interacting kinase OS=Homo | CIT | 231.431 | 2 | 52.25101 | yes | 9.722E-08 | 656.367 | 174.42639 |
| AEILALQQALK | 11 | O14578 | Citron Rho-interacting kinase OS=Homo | CIT | 231.431 | 2 | 52.25101 | yes | 9.722E-08 | 599.36371 | 172.92834 |
| ASTEATELLQNIR | 13 | O14578 | Citron Rho-interacting kinase OS=Homo | CIT | 231.431 | 2 | 52.25101 | yes | 1.462E-07 | 723.38336 | 154.2841 |
| ESQLTALQAAR | 11 | O14578 | Citron Rho-interacting kinase OS=Homo | CIT | 231.431 | 2 | 52.25101 | yes | 1.857E-07 | 594.32257 | 132.77864 |
| ALQLLHDIR | 9 | O14578 | Citron Rho-interacting kinase OS=Homo | CIT | 231.431 | 2 | 52.25101 | yes | 2.779E-07 | 539.82202 | 132.59377 |
| LTQGLQEALDR | 11 | O14578 | Citron Rho-interacting kinase OS=Homo | CIT | 231.431 | 2 | 52.25101 | yes | 1.05E-06 | 622.33569 | 110.17906 |
| EQEYQAQVEEMR | 12 | O14578 | Citron Rho-interacting kinase OS=Homo | CIT | 231.431 | 2 | 52.25101 | yes | 1.053E-07 | 770.34082 | 103.04206 |
| LAANSSLFTQR | 11 | O14578 | Citron Rho-interacting kinase OS=Homo | CIT | 231.431 | 2 | 52.25101 | yes | 3.527E-07 | 604.32513 | 93.542709 |
| AYLDIPNPR | 9 | O14578 | Citron Rho-interacting kinase OS=Homo | CIT | 231.431 | 2 | 52.25101 | yes | 2.54E-07 | 529.7851 | 84.46978 |
| LMMNQLEEDLVS | 14 | O14578 | Citron Rho-interacting kinase OS=Homo | CIT | 231.431 | 2 | 52.25101 | yes | 1.002E-07 | 824.90527 | 80.820534 |
| CAELEEALQK | 10 | O14578 | Citron Rho-interacting kinase OS=Homo | CIT | 231.431 | 2 | 52.25101 | yes | 3.294E-05 | 595.78973 | 73.501358 |
| SPFAEGTSAR | 10 | O14578 | Citron Rho-interacting kinase OS=Homo | CIT | 231.431 | 2 | 52.25101 | yes | 1.248E-07 | 511.74872 | 71.286568 |
| QLTELQLSLQER | 12 | O14578 | Citron Rho-interacting kinase OS=Homo | CIT | 231.431 | 2 | 52.25101 | yes | 1.994E-06 | 729.40155 | 65.495491 |
| LQQQMDLQK | 9 | O14578 | Citron Rho-interacting kinase OS=Homo | CIT | 231.431 | 2 | 52.25101 | yes | 4.492E-05 | 566.29498 | 65.376221 |
| SESVVSGLDSPAK | 13 | O14578 | Citron Rho-interacting kinase OS=Homo | CIT | 231.431 | 2 | 52.25101 | yes | 2.182E-06 | 638.32501 | 64.9049 |
| AQEEMISELR | 10 | O14578 | Citron Rho-interacting kinase OS=Homo | CIT | 231.431 | 2 | 52.25101 | yes | 3.927E-07 | 603.29517 | 64.220879 |
| NSPPPFVPTLK | 11 | O14578 | Citron Rho-interacting kinase OS=Homo | CIT | 231.431 | 2 | 52.25101 | yes | 6.055E-07 | 598.83734 | 62.785843 |
| LIDFLQAK | 8 | O14578 | Citron Rho-interacting kinase OS=Homo | CIT | 231.431 | 2 | 52.25101 | yes | 3.822E-06 | 474.28165 | 60.079803 |
| TLYLLAPSFPDK | 12 | O14578 | Citron Rho-interacting kinase OS=Homo | CIT | 231.431 | 2 | 52.25101 | yes | 9.722E-08 | 682.87665 | 58.924522 |
| DQGKPEVGEYAK | 12 | O14578 | Citron Rho-interacting kinase OS=Homo | CIT | 231.431 | 2 | 52.25101 | yes | 4.609E-06 | 660.82513 | 34.026928 |
| MFADYLAHESR | 11 | O14579 | Coatomer subunit epsilon OS=Homo sapi | COPE | 34.483 | 2 | 53.35003 | yes | 9.722E-08 | 670.30859 | 189.31909 |
| MFADYLAHESR | 11 | O14579 | Coatomer subunit epsilon OS=Homo sapi | COPE | 34.483 | 2 | 53.35003 | yes | 9.82E-08 | 447.20816 | 189.31909 |
| MFADYLAHESR | 11 | O14579 | Coatomer subunit epsilon OS=Homo sapi | COPE | 34.483 | 3 | 53.35003 | yes | 9.722E-08 | 670.30859 | 189.31909 |
| MFADYLAHESR | 11 | O14579 | Coatomer subunit epsilon OS=Homo sapi | COPE | 34.483 | 3 | 53.35003 | yes | 9.82E-08 | 447.20816 | 189.31909 |
| WEAAEGLLQEALI | 14 | O14579 | Coatomer subunit epsilon OS=Homo sapi | COPE | 34.483 | 2 | 53.35003 | yes | 9.722E-08 | 786.89886 | 138.67499 |
| LFGALTPLEPR | 11 | O14617 | AP-3 complex subunit delta-1 OS=Homo | AP3D1 | 130.158 | 2 | 53.428 | yes | 9.722E-08 | 607.35059 | 281.284 |
| LYASILQQK | 9 | O14617 | AP-3 complex subunit delta-1 OS=Homo | AP3D1 | 130.158 | 2 | 53.428 | yes | 2.787E-06 | 532.31097 | 226.8925 |
| ILIEDSDQNLK | 11 | O14617 | AP-3 complex subunit delta-1 OS=Homo | AP3D1 | 130.158 | 2 | 53.428 | yes | 2.964E-07 | 644.3432 | 206.23448 |
| LLESGDLSMSSIK | 13 | O14617 | AP-3 complex subunit delta-1 OS=Homo | AP3D1 | 130.158 | 2 | 53.428 | yes | 9.722E-08 | 690.35797 | 181.44676 |
| GMELSVLDSLNAF | 13 | O14617 | AP-3 complex subunit delta-1 OS=Homo | AP3D1 | 130.158 | 2 | 53.428 | yes | 2.853E-07 | 702.86121 | 138.7662 |
| YISQCIDEIK | 10 | O14617 | AP-3 complex subunit delta-1 OS=Homo | AP3D1 | 130.158 | 2 | 53.428 | yes | 1.311E-06 | 634.81317 | 134.23518 |
| ALDLLYGMVSK | 11 | O14617 | AP-3 complex subunit delta-1 OS=Homo | AP3D1 | 130.158 | 2 | 53.428 | yes | 2.512E-07 | 605.33105 | 91.934906 |
| ALDIDLDKPLADSI | 15 | O14617 | AP-3 complex subunit delta-1 OS=Homo | AP3D1 | 130.158 | 2 | 53.428 | yes | 1.751E-07 | 821.93036 | 77.206131 |
| VPGLPMSDQYVK | 12 | O14617 | AP-3 complex subunit delta-1 OS=Homo | AP3D1 | 130.158 | 2 | 53.428 | yes | 2.712E-05 | 667.34467 | 62.615669 |

| | | | | | | | | | | | |
|---|---|---|---|---|---|---|---|---|---|---|---|
| ASCILQLVK | 9 | O14617 | AP-3 complex subunit delta-1 OS=Homo | AP3D1 | 130.158 | 2 | 53.428 | yes | 1.164E-07 | 516.2995 | 62.256153 |
| LMTSSTNNWVLIK | 13 | O14617 | AP-3 complex subunit delta-1 OS=Homo | AP3D1 | 130.158 | 2 | 53.428 | yes | 9.722E-08 | 753.90289 | 60.949837 |
| AVFHEEEQR | 9 | O14617 | AP-3 complex subunit delta-1 OS=Homo | AP3D1 | 130.158 | 2 | 53.428 | yes | 5.578E-06 | 572.77271 | 44.411972 |
| MSFQNLLAK | 9 | O14617 | AP-3 complex subunit delta-1 OS=Homo | AP3D1 | 130.158 | 2 | 53.428 | yes | 2.744E-07 | 526.28387 | 40.857262 |
| SIQGHHVCLLVK | 12 | O14617 | AP-3 complex subunit delta-1 OS=Homo | AP3D1 | 130.158 | 2 | 53.428 | yes | 1.495E-06 | 695.88483 | 32.967133 |
| LNSGLGQLILK | 11 | O14639 | Actin-binding LIM protein 1 OS=Homo s | ABLIM1 | 87.688 | 2 | 46.80465 | yes | 1.69E-07 | 578.35846 | 144.64929 |
| YDSPINSASHIPSSI | 15 | O14639 | Actin-binding LIM protein 1 OS=Homo s | ABLIM1 | 87.688 | 2 | 46.80465 | yes | 1.947E-06 | 801.89154 | 138.74568 |
| GVSMPNMLEPK | 11 | O14639 | Actin-binding LIM protein 1 OS=Homo s | ABLIM1 | 87.688 | 2 | 46.80465 | yes | 1.466E-07 | 601.79901 | 137.98572 |
| HSYTPTTSR | 9 | O14639 | Actin-binding LIM protein 1 OS=Homo s | ABLIM1 | 87.688 | 2 | 46.80465 | yes | 5.677E-07 | 525.25415 | 131.51746 |
| QSLGESPR | 8 | O14639 | Actin-binding LIM protein 1 OS=Homo s | ABLIM1 | 87.688 | 2 | 46.80465 | yes | 4.531E-07 | 437.22488 | 93.985107 |
| EIFGMSIQEFDR | 12 | O14639 | Actin-binding LIM protein 1 OS=Homo s | ABLIM1 | 87.688 | 2 | 46.80465 | yes | 1.499E-07 | 736.34796 | 90.193085 |
| GDGGIYIGSIMK | 12 | O14640 | Segment polarity protein dishevelled hom | DVL1 | 75.187 | 2 | 46.788273 | no | 1.47E-07 | 605.81049 | 391.60959 |
| VMQLPDSGLEIR | 12 | O14640 | Segment polarity protein dishevelled hom | DVL1 | 75.187 | 2 | 46.788273 | no | 2.419E-07 | 679.36084 | 149.86777 |
| SMDQDFGVVK | 10 | O14640 | Segment polarity protein dishevelled hom | DVL1 | 75.187 | 2 | 46.788273 | no | 5.829E-07 | 563.26587 | 139.22221 |
| YGTSPCSSAVTR | 12 | O14640 | Segment polarity protein dishevelled hom | DVL1 | 75.187 | 2 | 46.788273 | yes | 6.208E-07 | 643.29572 | 36.628414 |
| EIVSQTGPISLTVA | 15 | O14640 | Segment polarity protein dishevelled hom | DVL1 | 75.187 | 2 | 46.788273 | no | 4.493E-07 | 771.94049 | 14.536201 |
| NVLSNRPVHAYK | 12 | O14640 | Segment polarity protein dishevelled hom | DVL1 | 75.187 | 2 | 46.788273 | no | 2.388E-07 | 699.38605 | 10.376789 |
| AMAAPESGLEVR | 12 | O14641 | Segment polarity protein dishevelled hom | DVL2 | 78.948 | 2 | 39.005253 | yes | 4.849E-07 | 615.81097 | 186.4095 |
| DIVHKPGPIVLTVA | 15 | O14641 | Segment polarity protein dishevelled hom | DVL2 | 78.948 | 2 | 39.005253 | yes | 2.299E-06 | 793.98505 | 33.730698 |
| DIVHKPGPIVLTVA | 15 | O14641 | Segment polarity protein dishevelled hom | DVL2 | 78.948 | 2 | 39.005253 | yes | 2.088E-06 | 529.65912 | 33.730698 |
| DIVHKPGPIVLTVA | 15 | O14641 | Segment polarity protein dishevelled hom | DVL2 | 78.948 | 3 | 39.005253 | yes | 2.299E-06 | 793.98505 | 33.730698 |
| DIVHKPGPIVLTVA | 15 | O14641 | Segment polarity protein dishevelled hom | DVL2 | 78.948 | 3 | 39.005253 | yes | 2.088E-06 | 529.65912 | 33.730698 |
| GSVEEDILER | 10 | O14646 | Chromodomain-helicase-DNA-binding pr | CHD1 | 196.688 | 2 | 53.88361 | yes | 9.968E-08 | 573.78552 | 670.86969 |
| SSPSILAVQR | 10 | O14646 | Chromodomain-helicase-DNA-binding pr | CHD1 | 196.688 | 2 | 53.88361 | yes | 1.058E-07 | 529.30365 | 611.68854 |
| ELEEIYMLPR | 10 | O14646 | Chromodomain-helicase-DNA-binding pr | CHD1 | 196.688 | 2 | 53.88361 | no | 1.233E-07 | 646.83136 | 535.41278 |
| MDPDLSLTHK | 10 | O14646 | Chromodomain-helicase-DNA-binding pr | CHD1 | 196.688 | 2 | 53.88361 | yes | 1.936E-07 | 578.78699 | 520.11963 |
| NWEEIIPEDQR | 11 | O14646 | Chromodomain-helicase-DNA-binding pr | CHD1 | 196.688 | 2 | 53.88361 | yes | 9.722E-08 | 714.84131 | 423.55035 |
| AAHFDIDWGK | 10 | O14646 | Chromodomain-helicase-DNA-binding pr | CHD1 | 196.688 | 2 | 53.88361 | yes | 9.722E-08 | 580.28015 | 267.03354 |
| NKEPGEIQYLIK | 12 | O14646 | Chromodomain-helicase-DNA-binding pr | CHD1 | 196.688 | 2 | 53.88361 | yes | 3.365E-07 | 716.39575 | 260.47043 |
| STPEHTWSSR | 10 | O14646 | Chromodomain-helicase-DNA-binding pr | CHD1 | 196.688 | 2 | 53.88361 | yes | 9.962E-08 | 594.27563 | 233.07651 |
| IGDHITECLK | 10 | O14646 | Chromodomain-helicase-DNA-binding pr | CHD1 | 196.688 | 2 | 53.88361 | yes | 1.489E-07 | 593.30023 | 227.05182 |
| MVLDHLVIQR | 10 | O14646 | Chromodomain-helicase-DNA-binding pr | CHD1 | 196.688 | 2 | 53.88361 | no | 9.885E-08 | 612.3501 | 109.7997 |
| MVLDHLVIQR | 10 | O14646 | Chromodomain-helicase-DNA-binding pr | CHD1 | 196.688 | 2 | 53.88361 | no | 4.463E-07 | 408.56915 | 109.7997 |
| MVLDHLVIQR | 10 | O14646 | Chromodomain-helicase-DNA-binding pr | CHD1 | 196.688 | 3 | 53.88361 | no | 9.885E-08 | 612.3501 | 109.7997 |
| MVLDHLVIQR | 10 | O14646 | Chromodomain-helicase-DNA-binding pr | CHD1 | 196.688 | 3 | 53.88361 | no | 4.463E-07 | 408.56915 | 109.7997 |
| KQPSYIGGHEGLEI | 15 | O14646 | Chromodomain-helicase-DNA-binding pr | CHD1 | 196.688 | 3 | 53.88361 | yes | 1.166E-07 | 561.96527 | 214.62547 |
| LKNDDSLLYK | 10 | O14646 | Chromodomain-helicase-DNA-binding pr | CHD1 | 196.688 | 2 | 53.88361 | no | 3.4E-07 | 604.82971 | 213.78384 |
| FQACIDEYFSR | 11 | O14646 | Chromodomain-helicase-DNA-binding pr | CHD1 | 196.688 | 2 | 53.88361 | yes | 9.722E-08 | 718.31915 | 169.31841 |
| GNSCILADEMGLG | 14 | O14646 | Chromodomain-helicase-DNA-binding pr | CHD1 | 196.688 | 2 | 53.88361 | yes | 1.073E-07 | 732.84473 | 167.86639 |

| | | | | | | | | | | | |
|---|---|---|---|---|---|---|---|---|---|---|---|
| QEALQHLIR | 9 | O14646 | Chromodomain-helicase-DNA-binding pr | CHD1 | 196.688 | 2 | 53.88361 | yes | 1.043E-07 | 554.31708 | 153.22202 |
| EYGYASLHK | 9 | O14646 | Chromodomain-helicase-DNA-binding pr | CHD1 | 196.688 | 2 | 53.88361 | yes | 1.943E-07 | 534.26147 | 153.17593 |
| SAAGYPDYYCK | 11 | O14646 | Chromodomain-helicase-DNA-binding pr | CHD1 | 196.688 | 2 | 53.88361 | yes | 1.013E-07 | 647.77405 | 123.88472 |
| FTEFDAR | 7 | O14646 | Chromodomain-helicase-DNA-binding pr | CHD1 | 196.688 | 2 | 53.88361 | no | 9.722E-08 | 443.20868 | 118.06885 |
| KNLWIFVSK | 9 | O14646 | Chromodomain-helicase-DNA-binding pr | CHD1 | 196.688 | 2 | 53.88361 | yes | 1.346E-07 | 567.83716 | 111.38527 |
| ILPDDPDKKPQAK | 13 | O14646 | Chromodomain-helicase-DNA-binding pr | CHD1 | 196.688 | 3 | 53.88361 | yes | 6.94E-07 | 488.94006 | 103.04839 |
| LVISHEEELIPLHK | 14 | O14646 | Chromodomain-helicase-DNA-binding pr | CHD1 | 196.688 | 2 | 53.88361 | yes | 1.2E-07 | 828.9696 | 44.317318 |
| LVISHEEELIPLHK | 14 | O14646 | Chromodomain-helicase-DNA-binding pr | CHD1 | 196.688 | 2 | 53.88361 | yes | 5.333E-05 | 552.98218 | 44.317318 |
| LVISHEEELIPLHK | 14 | O14646 | Chromodomain-helicase-DNA-binding pr | CHD1 | 196.688 | 3 | 53.88361 | yes | 1.2E-07 | 828.9696 | 44.317318 |
| LVISHEEELIPLHK | 14 | O14646 | Chromodomain-helicase-DNA-binding pr | CHD1 | 196.688 | 3 | 53.88361 | yes | 5.333E-05 | 552.98218 | 44.317318 |
| FSSWEDFEEEHGK | 13 | O14646 | Chromodomain-helicase-DNA-binding pr | CHD1 | 196.688 | 2 | 53.88361 | yes | 9.722E-08 | 813.83899 | 80.435852 |
| LLITGTPLQNSLK | 13 | O14646 | Chromodomain-helicase-DNA-binding pr | CHD1 | 196.688 | 2 | 53.88361 | no | 1.024E-07 | 699.42175 | 74.590263 |
| QYTIPCHTK | 9 | O14646 | Chromodomain-helicase-DNA-binding pr | CHD1 | 196.688 | 2 | 53.88361 | yes | 9.903E-08 | 574.28186 | 70.147873 |
| QATVNVSYK | 9 | O14646 | Chromodomain-helicase-DNA-binding pr | CHD1 | 196.688 | 2 | 53.88361 | yes | 2.153E-07 | 505.26929 | 57.801449 |
| ELEPFLLR | 8 | O14646 | Chromodomain-helicase-DNA-binding pr | CHD1 | 196.688 | 2 | 53.88361 | yes | 2.149E-07 | 508.79239 | 57.448078 |
| NLWIFVSK | 8 | O14646 | Chromodomain-helicase-DNA-binding pr | CHD1 | 196.688 | 2 | 53.88361 | no | 1.067E-07 | 503.78964 | 49.936707 |
| YHSDWQMDHR | 10 | O14646 | Chromodomain-helicase-DNA-binding pr | CHD1 | 196.688 | 2 | 53.88361 | yes | 1.217E-06 | 687.78583 | 37.417049 |
| QPSYIGGHEGLELI | 14 | O14646 | Chromodomain-helicase-DNA-binding pr | CHD1 | 196.688 | 2 | 53.88361 | yes | 4.559E-07 | 778.39679 | 32.194019 |
| GTVEEEIIER | 10 | O14647 | Chromodomain-helicase-DNA-binding pr | CHD2 | 211.344 | 2 | 52.281593 | yes | 1.127E-07 | 587.80115 | 195.35172 |
| DLVEGFTDAEIR | 12 | O14647 | Chromodomain-helicase-DNA-binding pr | CHD2 | 211.344 | 2 | 52.281593 | yes | 9.722E-08 | 682.83826 | 165.08197 |
| AYSDQEHIK | 9 | O14647 | Chromodomain-helicase-DNA-binding pr | CHD2 | 211.344 | 2 | 52.281593 | yes | 2.425E-07 | 545.76184 | 135.55589 |
| DWDEIIPEEQR | 11 | O14647 | Chromodomain-helicase-DNA-binding pr | CHD2 | 211.344 | 2 | 52.281593 | yes | 9.722E-08 | 715.33331 | 110.91779 |
| FGLPLER | 7 | O14647 | Chromodomain-helicase-DNA-binding pr | CHD2 | 211.344 | 2 | 52.281593 | yes | 2.033E-07 | 416.23981 | 104.41439 |
| NNSVILADEMGLG | 14 | O14647 | Chromodomain-helicase-DNA-binding pr | CHD2 | 211.344 | 2 | 52.281593 | yes | 3.308E-07 | 730.87433 | 98.92215 |
| MWEEYPDVYGVR | 12 | O14647 | Chromodomain-helicase-DNA-binding pr | CHD2 | 211.344 | 2 | 52.281593 | yes | 1.295E-07 | 772.34796 | 88.533417 |
| ISGVQVNVK | 9 | O14647 | Chromodomain-helicase-DNA-binding pr | CHD2 | 211.344 | 2 | 52.281593 | yes | 3.609E-06 | 472.2822 | 84.655212 |
| GLNVQEQLEHTR | 12 | O14647 | Chromodomain-helicase-DNA-binding pr | CHD2 | 211.344 | 2 | 52.281593 | yes | 1.083E-07 | 712.36804 | 75.079437 |
| VLEPFLLR | 8 | O14647 | Chromodomain-helicase-DNA-binding pr | CHD2 | 211.344 | 2 | 52.281593 | yes | 6.801E-07 | 493.8053 | 37.055668 |
| TFLQDLAR | 8 | O14681 | Etoposide-induced protein 2.4 homolog C | EI24 | 38.965 | 2 | 46.43603 | yes | 1.664E-07 | 482.26654 | 120.91282 |
| DSIWGICTISK | 11 | O14681 | Etoposide-induced protein 2.4 homolog C | EI24 | 38.965 | 2 | 46.43603 | yes | 1.559E-07 | 640.32117 | 61.625156 |
| SSTVASLDTDSTK | 13 | O14713 | Integrin beta-1-binding protein 1 OS=Ho | ITGB1BP1 | 21.783 | 2 | 43.517426 | yes | 2.556E-07 | 656.31738 | 90.875504 |
| AGIPCPTVVLLK | 12 | O14730 | Serine/threonine-protein kinase RIO3 OS | RIOK3 | 59.094 | 2 | 50.63512 | yes | 9.733E-08 | 634.37579 | 239.30774 |
| VSISFENYR | 9 | O14730 | Serine/threonine-protein kinase RIO3 OS | RIOK3 | 59.094 | 2 | 50.63512 | yes | 1.71E-07 | 557.78003 | 227.6282 |
| LSNHVFNALK | 10 | O14730 | Serine/threonine-protein kinase RIO3 OS | RIOK3 | 59.094 | 2 | 50.63512 | yes | 2.898E-07 | 571.81946 | 22.905058 |
| PTLQLPLANDGGS | 14 | O14733 | Dual specificity mitogen-activated protei | MAP2K7 | 47.485 | 2 | 54.150898 | yes | 3.584E-07 | 719.88605 | 226.72906 |
| LLEHSFIK | 8 | O14733 | Dual specificity mitogen-activated protei | MAP2K7 | 47.485 | 2 | 54.150898 | yes | 1.021E-07 | 493.78711 | 207.77397 |
| LCDFGISGR | 9 | O14733 | Dual specificity mitogen-activated protei | MAP2K7 | 47.485 | 2 | 54.150898 | yes | 3.103E-07 | 512.74768 | 184.37859 |
| QTGYLTIGGQR | 11 | O14733 | Dual specificity mitogen-activated protei | MAP2K7 | 47.485 | 2 | 54.150898 | yes | 9.722E-08 | 597.31732 | 139.01918 |
| DVKPSNILLDER | 12 | O14733 | Dual specificity mitogen-activated protei | MAP2K7 | 47.485 | 2 | 54.150898 | yes | 9.722E-08 | 699.883 | 68.708908 |

| | | | | | | | | | | | |
|---|---|---|---|---|---|---|---|---|---|---|---|
| HMLGLPSTLFTPR | 13 | O14733 | Dual specificity mitogen-activated proteir | MAP2K7 | 47.485 | 2 | 54.150898 | yes | 1.742E-07 | 735.40033 | 55.009888 |
| ILMDLDVVLK | 10 | O14733 | Dual specificity mitogen-activated proteir | MAP2K7 | 47.485 | 2 | 54.150898 | yes | 1.672E-07 | 579.84375 | 50.6059 |
| SMESIEIDQK | 10 | O14733 | Dual specificity mitogen-activated proteir | MAP2K7 | 47.485 | 2 | 54.150898 | yes | 1.476E-06 | 590.28174 | 40.472294 |
| LPVLYQVER | 9 | O14734 | Acyl-coenzyme A thioesterase 8 OS=Hon | ACOT8 | 35.915 | 2 | 45.399925 | yes | 1.861E-07 | 558.82422 | 110.47009 |
| VSEQGLIEILK | 11 | O14737 | Programmed cell death protein 5 OS=Hor | PDCD5 | 14.286 | 2 | 35.87763 | yes | 7.695E-07 | 614.86102 | 10.914261 |
| LSPWIRPDSK | 10 | O14744 | Protein arginine N-methyltransferase 5 O: | PRMT5 | 72.684 | 2 | 53.757236 | yes | 1.678E-07 | 599.83258 | 1054.245 |
| VPLVAPEDLR | 10 | O14744 | Protein arginine N-methyltransferase 5 O: | PRMT5 | 72.684 | 2 | 53.757236 | yes | 1.187E-07 | 554.82166 | 906.89819 |
| GPLVNASLR | 9 | O14744 | Protein arginine N-methyltransferase 5 O: | PRMT5 | 72.684 | 2 | 53.757236 | yes | 3.423E-07 | 463.77454 | 516.95868 |
| YSQYQQAIYK | 10 | O14744 | Protein arginine N-methyltransferase 5 O: | PRMT5 | 72.684 | 2 | 53.757236 | yes | 9.722E-08 | 646.31952 | 311.61639 |
| DWNTLIVGK | 9 | O14744 | Protein arginine N-methyltransferase 5 O: | PRMT5 | 72.684 | 2 | 53.757236 | yes | 3.713E-07 | 523.28748 | 306.1705 |
| DPMIDNNR | 8 | O14744 | Protein arginine N-methyltransferase 5 O: | PRMT5 | 72.684 | 2 | 53.757236 | yes | 1.859E-07 | 487.72165 | 193.98399 |
| EFIQEPAK | 8 | O14744 | Protein arginine N-methyltransferase 5 O: | PRMT5 | 72.684 | 2 | 53.757236 | yes | 3.224E-07 | 481.25311 | 191.08597 |
| LHNFHQLSAPQPC | 21 | O14744 | Protein arginine N-methyltransferase 5 O: | PRMT5 | 72.684 | 4 | 53.757236 | yes | 1.123E-07 | 634.55792 | 160.13013 |
| KVWYEWAVTAPV | 22 | O14744 | Protein arginine N-methyltransferase 5 O: | PRMT5 | 72.684 | 4 | 53.757236 | yes | 1.179E-06 | 636.32196 | 49.819244 |
| KVWYEWAVTAPV | 22 | O14744 | Protein arginine N-methyltransferase 5 O: | PRMT5 | 72.684 | 4 | 53.757236 | yes | 1.088E-07 | 848.09351 | 49.819244 |
| KVWYEWAVTAPV | 22 | O14744 | Protein arginine N-methyltransferase 5 O: | PRMT5 | 72.684 | 3 | 53.757236 | yes | 1.179E-06 | 636.32196 | 49.819244 |
| KVWYEWAVTAPV | 22 | O14744 | Protein arginine N-methyltransferase 5 O: | PRMT5 | 72.684 | 3 | 53.757236 | yes | 1.088E-07 | 848.09351 | 49.819244 |
| VPEEEKDTNVQVI | 20 | O14744 | Protein arginine N-methyltransferase 5 O: | PRMT5 | 72.684 | 3 | 53.757236 | yes | 2.349E-07 | 728.71582 | 93.96801 |
| VWYEWAVTAPVC | 21 | O14744 | Protein arginine N-methyltransferase 5 O: | PRMT5 | 72.684 | 3 | 53.757236 | yes | 3.569E-07 | 805.3952 | 84.526199 |
| EGQTICVR | 8 | O14744 | Protein arginine N-methyltransferase 5 O: | PRMT5 | 72.684 | 2 | 53.757236 | yes | 1.002E-07 | 481.73984 | 67.446671 |
| IAVALEIGADLPSN | 19 | O14744 | Protein arginine N-methyltransferase 5 O: | PRMT5 | 72.684 | 3 | 53.757236 | yes | 7.226E-07 | 668.36884 | 15.595256 |
| IEPDIGMPEPDAQF | 14 | O14757 | Serine/threonine-protein kinase Chk1 OS: | CHEK1 | 54.434 | 2 | 52.343708 | yes | 8.967E-07 | 784.37469 | 158.07365 |
| ILVENPSAR | 9 | O14757 | Serine/threonine-protein kinase Chk1 OS: | CHEK1 | 54.434 | 2 | 52.343708 | yes | 1.191E-07 | 499.7851 | 129.19557 |
| VTSGGVSESPSGFS | 15 | O14757 | Serine/threonine-protein kinase Chk1 OS: | CHEK1 | 54.434 | 2 | 52.343708 | yes | 2.236E-07 | 713.34644 | 71.725876 |
| MCGTLPYVAPELL | 14 | O14757 | Serine/threonine-protein kinase Chk1 OS: | CHEK1 | 54.434 | 2 | 52.343708 | yes | 9.722E-08 | 796.41473 | 68.220123 |
| VNLLEMDDK | 9 | O14757 | Serine/threonine-protein kinase Chk1 OS: | CHEK1 | 54.434 | 2 | 52.343708 | yes | 1.522E-07 | 538.76825 | 52.956493 |
| ISDFGLATVFR | 11 | O14757 | Serine/threonine-protein kinase Chk1 OS: | CHEK1 | 54.434 | 2 | 52.343708 | yes | 1.874E-07 | 613.3324 | 46.724434 |
| VFFYNPTTR | 9 | O14776 | Transcription elongation regulator 1 OS= | TCERG1 | 123.902 | 2 | 53.842712 | yes | 9.722E-08 | 572.79291 | 260.45218 |
| WESGSLLER | 9 | O14776 | Transcription elongation regulator 1 OS= | TCERG1 | 123.902 | 2 | 53.842712 | yes | 2.508E-07 | 538.77222 | 143.80273 |
| YLVLDCVPEER | 11 | O14776 | Transcription elongation regulator 1 OS= | TCERG1 | 123.902 | 2 | 53.842712 | yes | 1.326E-07 | 696.84503 | 130.77138 |
| EALFNEFVAAAR | 12 | O14776 | Transcription elongation regulator 1 OS= | TCERG1 | 123.902 | 2 | 53.842712 | yes | 1.422E-07 | 669.34607 | 108.08291 |
| HPTPTMLSIQK | 11 | O14776 | Transcription elongation regulator 1 OS= | TCERG1 | 123.902 | 2 | 53.842712 | yes | 1.362E-07 | 626.83954 | 82.521156 |
| LFNEHIEALTK | 11 | O14776 | Transcription elongation regulator 1 OS= | TCERG1 | 123.902 | 2 | 53.842712 | yes | 9.722E-08 | 657.85626 | 70.716431 |
| TYYYNNR | 7 | O14776 | Transcription elongation regulator 1 OS= | TCERG1 | 123.902 | 2 | 53.842712 | yes | 1.249E-05 | 497.22488 | 57.378437 |
| GPPPPPTASEPTR | 13 | O14776 | Transcription elongation regulator 1 OS= | TCERG1 | 123.902 | 2 | 53.842712 | yes | 2.77E-07 | 652.33569 | 53.305759 |
| ESAWTKPDGVK | 11 | O14776 | Transcription elongation regulator 1 OS= | TCERG1 | 123.902 | 2 | 53.842712 | yes | 9.722E-08 | 609.31171 | 52.812645 |
| TLESTWEKPQELK | 13 | O14776 | Transcription elongation regulator 1 OS= | TCERG1 | 123.902 | 2 | 53.842712 | yes | 3.748E-06 | 794.91449 | 24.759109 |
| VELECETIK | 9 | O14777 | Kinetochore protein NDC80 homolog OS | NDC80 | 73.913 | 2 | 46.55204 | yes | 3.48E-06 | 560.78137 | 395.96332 |
| ELLNETEEEINK | 12 | O14777 | Kinetochore protein NDC80 homolog OS | NDC80 | 73.913 | 2 | 46.55204 | yes | 1.53E-07 | 730.85938 | 41.976875 |

| | | | | | | | | | | | |
|---|---|---|---|---|---|---|---|---|---|---|---|
| NSQLGIFSSSEK | 12 | O14777 | Kinetochore protein NDC80 homolog OS | NDC80 | 73.913 | 2 | 46.55204 | yes | 1.939E-06 | 648.82513 | 38.03862 |
| TIFGNLNPVWEEK | 13 | O14795 | Protein unc-13 homolog B OS=Homo sap | UNC13B | 180.679 | 2 | 53.572197 | yes | 9.722E-08 | 773.89862 | 131.13287 |
| VLMQYADILSK | 11 | O14795 | Protein unc-13 homolog B OS=Homo sap | UNC13B | 180.679 | 2 | 53.572197 | yes | 2.368E-07 | 640.84961 | 111.42257 |
| SPQQLLLQGSSR | 12 | O14795 | Protein unc-13 homolog B OS=Homo sap | UNC13B | 180.679 | 2 | 53.572197 | yes | 9.722E-08 | 657.36224 | 103.9057 |
| NKPEIFEVIR | 10 | O14795 | Protein unc-13 homolog B OS=Homo sap | UNC13B | 180.679 | 2 | 53.572197 | yes | 1.147E-07 | 622.85352 | 89.991936 |
| ESDDFLGQTIIEVR | 14 | O14795 | Protein unc-13 homolog B OS=Homo sap | UNC13B | 180.679 | 2 | 53.572197 | no | 3.411E-07 | 811.40704 | 77.361885 |
| CQDLLNADCLQR | 12 | O14795 | Protein unc-13 homolog B OS=Homo sap | UNC13B | 180.679 | 2 | 53.572197 | no | 3.939E-07 | 753.3454 | 65.960213 |
| SADYMNLHFK | 10 | O14795 | Protein unc-13 homolog B OS=Homo sap | UNC13B | 180.679 | 2 | 53.572197 | no | 9.964E-08 | 613.28717 | 63.558922 |
| VLGLAVMPLR | 10 | O14795 | Protein unc-13 homolog B OS=Homo sap | UNC13B | 180.679 | 2 | 53.572197 | yes | 1.094E-06 | 534.83356 | 56.390594 |
| TLSGEMDVWYNL | 14 | O14795 | Protein unc-13 homolog B OS=Homo sap | UNC13B | 180.679 | 2 | 53.572197 | no | 9.81E-08 | 842.89801 | 47.550835 |
| TSLKDEELK | 9 | O14795 | Protein unc-13 homolog B OS=Homo sap | UNC13B | 180.679 | 2 | 53.572197 | yes | 2.594E-06 | 531.78748 | 36.341496 |
| ITITVVCAQGLQAI | 14 | O14795 | Protein unc-13 homolog B OS=Homo sap | UNC13B | 180.679 | 2 | 53.572197 | yes | 6.789E-07 | 751.42395 | 32.971214 |
| LLVEGDNLR | 9 | O14802 | DNA-directed RNA polymerase III subun | POLR3A | 155.642 | 2 | 50.81428 | yes | 2.536E-07 | 514.79041 | 270.49509 |
| LAPVYLSNR | 9 | O14802 | DNA-directed RNA polymerase III subun | POLR3A | 155.642 | 2 | 50.81428 | yes | 2.551E-07 | 516.79547 | 250.44389 |
| NELILTTESIMK | 12 | O14802 | DNA-directed RNA polymerase III subun | POLR3A | 155.642 | 2 | 50.81428 | yes | 1.539E-07 | 696.37616 | 166.27646 |
| AVMATHGVK | 9 | O14802 | DNA-directed RNA polymerase III subun | POLR3A | 155.642 | 2 | 50.81428 | yes | 3.29E-07 | 457.24985 | 154.2516 |
| GFSIGIGDVTPGQC | 17 | O14802 | DNA-directed RNA polymerase III subun | POLR3A | 155.642 | 2 | 50.81428 | yes | 1.539E-07 | 829.95923 | 135.69916 |
| VVVQGIPEVSR | 11 | O14802 | DNA-directed RNA polymerase III subun | POLR3A | 155.642 | 2 | 50.81428 | yes | 1.001E-07 | 591.8457 | 124.32812 |
| AVFPCPSEPALSK | 13 | O14802 | DNA-directed RNA polymerase III subun | POLR3A | 155.642 | 2 | 50.81428 | yes | 9.307E-07 | 701.85541 | 110.7925 |
| TVISPDPNLR | 10 | O14802 | DNA-directed RNA polymerase III subun | POLR3A | 155.642 | 2 | 50.81428 | yes | 3.634E-07 | 556.30896 | 107.72556 |
| ESVLMLASFEK | 11 | O14802 | DNA-directed RNA polymerase III subun | POLR3A | 155.642 | 2 | 50.81428 | yes | 2.744E-07 | 627.32593 | 101.65779 |
| YELLNAGYK | 9 | O14802 | DNA-directed RNA polymerase III subun | POLR3A | 155.642 | 2 | 50.81428 | yes | 6.282E-07 | 535.77948 | 89.685097 |
| AEALVLMGTK | 10 | O14802 | DNA-directed RNA polymerase III subun | POLR3A | 155.642 | 2 | 50.81428 | yes | 3.556E-07 | 516.79156 | 87.293633 |
| AVIHIDEQSGK | 11 | O14802 | DNA-directed RNA polymerase III subun | POLR3A | 155.642 | 2 | 50.81428 | yes | 3.845E-07 | 598.81714 | 83.692757 |
| VPDGFENR | 8 | O14802 | DNA-directed RNA polymerase III subun | POLR3A | 155.642 | 2 | 50.81428 | yes | 4.861E-07 | 467.22488 | 64.977753 |
| SGTNEDDLTMK | 11 | O14802 | DNA-directed RNA polymerase III subun | POLR3A | 155.642 | 2 | 50.81428 | yes | 9.722E-08 | 605.76642 | 59.574612 |
| RPGLTYLQK | 9 | O14802 | DNA-directed RNA polymerase III subun | POLR3A | 155.642 | 2 | 50.81428 | yes | 1.746E-06 | 538.31659 | 59.571651 |
| SNSPLTMALCGSK | 13 | O14802 | DNA-directed RNA polymerase III subun | POLR3A | 155.642 | 2 | 50.81428 | yes | 3.164E-06 | 683.32867 | 58.090714 |
| ACQIIASILVGK | 12 | O14802 | DNA-directed RNA polymerase III subun | POLR3A | 155.642 | 2 | 50.81428 | yes | 3.148E-07 | 636.87085 | 42.270081 |
| LSIMAHLAR | 9 | O14802 | DNA-directed RNA polymerase III subun | POLR3A | 155.642 | 2 | 50.81428 | yes | 1.505E-06 | 506.29202 | 40.964397 |
| CDEYIEALNTGK | 12 | O14802 | DNA-directed RNA polymerase III subun | POLR3A | 155.642 | 2 | 50.81428 | yes | 2.608E-07 | 706.82178 | 28.138897 |
| TCCHIMLSQEEK | 12 | O14802 | DNA-directed RNA polymerase III subun | POLR3A | 155.642 | 2 | 50.81428 | yes | 1.071E-07 | 768.33618 | 22.12759 |
| LPPPTILKPVTLWT | 16 | O14802 | DNA-directed RNA polymerase III subun | POLR3A | 155.642 | 3 | 50.81428 | yes | 2.388E-06 | 587.69397 | 17.392021 |
| ILNPEEIEK | 9 | O14818 | Proteasome subunit alpha type-7 OS=Hor | PSMA7 | 27.887 | 2 | 48.588505 | yes | 2.613E-07 | 542.79785 | 488.93912 |
| ALLEVVQSGGK | 11 | O14818 | Proteasome subunit alpha type-7 OS=Hor | PSMA7 | 27.887 | 2 | 48.588505 | no | 3.97E-07 | 550.81915 | 223.036 |
| LTVEDPVTVEYITI | 14 | O14818 | Proteasome subunit alpha type-7 OS=Hor | PSMA7 | 27.887 | 2 | 48.588505 | no | 1.127E-07 | 817.93542 | 185.8114 |
| AITVFSPDGHLFQV | 22 | O14818 | Proteasome subunit alpha type-7 OS=Hor | PSMA7 | 27.887 | 3 | 48.588505 | no | 1.921E-06 | 817.08478 | 111.10124 |
| KICALDDNVCMAI | 21 | O14818 | Proteasome subunit alpha type-7 OS=Hor | PSMA7 | 27.887 | 3 | 48.588505 | yes | 4.2E-07 | 771.36591 | 38.88361 |
| TAAANAAAGAAEI | 16 | O14828 | Secretory carrier-associated membrane pr | SCAMP3 | 38.287 | 2 | 47.03996 | yes | 1.415E-07 | 738.8631 | 375.6362 |

| | | | | | | | | | | | |
|---|---|---|---|---|---|---|---|---|---|---|---|
| AQQEFAAGVFSNF | 16 | O14828 | Secretory carrier-associated membrane pr | SCAMP3 | 38.287 | 2 | 47.03996 | yes | 6.084E-07 | 846.42865 | 113.98425 |
| ELQHAALGGTATR | 13 | O14828 | Secretory carrier-associated membrane pr | SCAMP3 | 38.287 | 2 | 47.03996 | yes | 1.459E-07 | 662.85205 | 67.596703 |
| YLQQELPVR | 9 | O14874 | [3-methyl-2-oxobutanoate dehydrogenase | BCKDK | 46.361 | 2 | 41.80093 | yes | 3.495E-07 | 573.31934 | 106.2001 |
| DQADEAQYCQLV | 13 | O14874 | [3-methyl-2-oxobutanoate dehydrogenase | BCKDK | 46.361 | 2 | 41.80093 | yes | 3.334E-07 | 798.35956 | 34.678463 |
| SPCQTDVLR | 9 | O14879 | Interferon-induced protein with tetratrico | IFIT3 | 55.985 | 2 | 53.81647 | yes | 5.721E-07 | 538.26367 | 127.75562 |
| EAEGEQFVEEALE | 14 | O14879 | Interferon-induced protein with tetratrico | IFIT3 | 55.985 | 2 | 53.81647 | yes | 9.722E-08 | 804.37543 | 123.00331 |
| QAIELSPDNQYVK | 13 | O14879 | Interferon-induced protein with tetratrico | IFIT3 | 55.985 | 2 | 53.81647 | yes | 8.988E-07 | 752.88574 | 101.33514 |
| LSDAQIYVDK | 10 | O14879 | Interferon-induced protein with tetratrico | IFIT3 | 55.985 | 2 | 53.81647 | yes | 2.527E-07 | 576.30078 | 82.078468 |
| VCNQIEFLNTEFK | 13 | O14879 | Interferon-induced protein with tetratrico | IFIT3 | 55.985 | 2 | 53.81647 | yes | 9.722E-08 | 821.4007 | 48.854313 |
| ALEEKPNNPEFSSC | 28 | O14879 | Interferon-induced protein with tetratrico | IFIT3 | 55.985 | 4 | 53.81647 | yes | 8.656E-07 | 788.63428 | 36.994389 |
| VLYAYGYYTGEPS | 14 | O14880 | Glutathione S-transferase 3, mitochondria | MGST3 | 16.517 | 2 | 51.09447 | yes | 1.418E-07 | 805.8905 | 1298.115 |
| IASGLGLAWIVGR | 13 | O14880 | Glutathione S-transferase 3, mitochondria | MGST3 | 16.517 | 2 | 51.09447 | yes | 9.722E-08 | 656.89044 | 818.59637 |
| TPQEYLR | 7 | O14893 | Gem-associated protein 2 OS=Homo sapi | GEMIN2 | 31.586 | 2 | 49.871166 | yes | 1.966E-07 | 453.73761 | 864.26184 |
| SQQLDSNVTMPK | 12 | O14893 | Gem-associated protein 2 OS=Homo sapi | GEMIN2 | 31.586 | 2 | 49.871166 | yes | 1.001E-07 | 674.33228 | 469.30746 |
| RVQIEAAQCPDVV | 20 | O14893 | Gem-associated protein 2 OS=Homo sapi | GEMIN2 | 31.586 | 3 | 49.871166 | yes | 1.368E-07 | 746.06335 | 178.74141 |
| VQIEAAQCPDVVV | 19 | O14893 | Gem-associated protein 2 OS=Homo sapi | GEMIN2 | 31.586 | 3 | 49.871166 | yes | 4.845E-07 | 694.0296 | 39.944702 |
| KIPGWQAEVGK | 11 | O14901 | Krueppel-like factor 11 OS=Homo sapien | KLF11 | 55.14 | 2 | 42.219845 | yes | 1.096E-06 | 606.84039 | 124.52026 |
| ACTATDVLQSSAV | 16 | O14901 | Krueppel-like factor 11 OS=Homo sapien | KLF11 | 55.14 | 2 | 42.219845 | yes | 3.136E-07 | 824.9198 | 46.587086 |
| NYVCSFPGCR | 10 | O14901 | Krueppel-like factor 11 OS=Homo sapien | KLF11 | 55.14 | 2 | 42.219845 | yes | 2.847E-06 | 630.26843 | 45.351547 |
| NPDELAEALDER | 12 | O14908 | PDZ domain-containing protein GIPC1 O | GIPC1 | 36.05 | 2 | 49.68459 | yes | 7.616E-07 | 686.323 | 76.201218 |
| VDDLLESYMGIR | 12 | O14908 | PDZ domain-containing protein GIPC1 O | GIPC1 | 36.05 | 2 | 49.68459 | no | 1.011E-07 | 705.85028 | 65.537148 |
| GPATVEDLPSAFEI | 15 | O14908 | PDZ domain-containing protein GIPC1 O | GIPC1 | 36.05 | 2 | 49.68459 | yes | 1.727E-07 | 795.38831 | 60.91637 |
| ETQNMAWSKPR | 11 | O14925 | Mitochondrial import inner membrane tra | TIMM23 | 21.944 | 2 | 54.394524 | yes | 9.722E-08 | 674.32733 | 87.812599 |
| LLVTDMSDAEQYI | 13 | O14929 | Histone acetyltransferase type B catalytic | HAT1 | 49.541 | 2 | 47.930523 | yes | 2.128E-07 | 770.8692 | 236.06047 |
| LCQDLPCFSR | 10 | O14929 | Histone acetyltransferase type B catalytic | HAT1 | 49.541 | 2 | 47.930523 | yes | 1.23E-07 | 648.29718 | 200.80661 |
| RVYEILR | 7 | O14929 | Histone acetyltransferase type B catalytic | HAT1 | 49.541 | 2 | 47.930523 | yes | 2.409E-06 | 474.78491 | 55.506042 |
| CNTNTAIELK | 10 | O14929 | Histone acetyltransferase type B catalytic | HAT1 | 49.541 | 2 | 47.930523 | yes | 3.169E-07 | 582.28992 | 29.580648 |
| MLMLDPAER | 9 | O14936 | Peripheral plasma membrane protein CAS | CASK | 105.123 | 2 | 41.88319 | yes | 3.296E-07 | 538.2674 | 130.49898 |
| AYPHVFTK | 8 | O14949 | Cytochrome b-c1 complex subunit 8 OS= | UQCRQ | 9.907 | 2 | 49.930477 | yes | 9.975E-08 | 481.75836 | 450.20944 |
| KNPAAYENDK | 10 | O14949 | Cytochrome b-c1 complex subunit 8 OS= | UQCRQ | 9.907 | 2 | 49.930477 | yes | 7.345E-07 | 575.2804 | 372.24033 |
| ELLTTMGDR | 9 | O14950 | Myosin regulatory light chain 12B OS=H | MYL12B | 19.78 | 2 | 51.79153 | no | 1.022E-07 | 518.26062 | 4995.8174 |
| ELLTTMGDR | 9 | O14950 | Myosin regulatory light chain 12B OS=H | MYL12B | 19.78 | 2 | 51.79153 | no | 3.363E-07 | 526.25806 | 4995.8174 |
| EAFNMIDQNR | 10 | O14950 | Myosin regulatory light chain 12B OS=H | MYL12B | 19.78 | 2 | 51.79153 | no | 3.841E-07 | 619.28516 | 7069.5786 |
| FTDEEVDELYR | 11 | O14950 | Myosin regulatory light chain 12B OS=H | MYL12B | 19.78 | 2 | 51.79153 | yes | 1.466E-07 | 708.31989 | 6542.501 |
| GNFNYIEFTR | 10 | O14950 | Myosin regulatory light chain 12B OS=H | MYL12B | 19.78 | 2 | 51.79153 | yes | 1.128E-07 | 630.80402 | 6229.5508 |
| LNGTDPEDVIR | 11 | O14950 | Myosin regulatory light chain 12B OS=H | MYL12B | 19.78 | 2 | 51.79153 | no | 3.696E-07 | 614.81201 | 1875.4973 |
| EDLHDMLASLGK | 12 | O14950 | Myosin regulatory light chain 12B OS=H | MYL12B | 19.78 | 2 | 51.79153 | no | 9.722E-08 | 664.82941 | 193.73961 |
| ALQNAVTTFVNR | 12 | O14964 | Hepatocyte growth factor-regulated tyrosi | HGS | 86.193 | 2 | 53.80612 | yes | 9.722E-08 | 667.36475 | 31.619755 |
| AGVEHQLR | 8 | O14965 | Aurora kinase A OS=Homo sapiens OX= | AURKA | 45.824 | 2 | 51.714043 | yes | 1.744E-07 | 455.24869 | 3404.1643 |

| | | | | | | | | | | | |
|---|---|---|---|---|---|---|---|---|---|---|---|
| VEFTFPDFVTEGAI | 14 | O14965 | Aurora kinase A OS=Homo sapiens OX=' | AURKA | 45.824 | 2 | 51.714043 | yes | 1.866E-07 | 807.89355 | 3211.0313 |
| EVEIQSHLR | 9 | O14965 | Aurora kinase A OS=Homo sapiens OX=' | AURKA | 45.824 | 2 | 51.714043 | yes | 9.996E-08 | 555.79877 | 2905.8027 |
| VLCPSNSSQR | 10 | O14965 | Aurora kinase A OS=Homo sapiens OX=' | AURKA | 45.824 | 2 | 51.714043 | yes | 6.734E-07 | 574.27985 | 2529.4224 |
| QLQATSVPHPVSR | 13 | O14965 | Aurora kinase A OS=Homo sapiens OX=' | AURKA | 45.824 | 2 | 51.714043 | yes | 9.819E-08 | 710.38879 | 869.69397 |
| QLQATSVPHPVSR | 13 | O14965 | Aurora kinase A OS=Homo sapiens OX=' | AURKA | 45.824 | 2 | 51.714043 | yes | 2.336E-07 | 473.92828 | 869.69397 |
| QLQATSVPHPVSR | 13 | O14965 | Aurora kinase A OS=Homo sapiens OX=' | AURKA | 45.824 | 3 | 51.714043 | yes | 9.819E-08 | 710.38879 | 869.69397 |
| QLQATSVPHPVSR | 13 | O14965 | Aurora kinase A OS=Homo sapiens OX=' | AURKA | 45.824 | 3 | 51.714043 | yes | 2.336E-07 | 473.92828 | 869.69397 |
| FGNVYLAR | 8 | O14965 | Aurora kinase A OS=Homo sapiens OX=' | AURKA | 45.824 | 2 | 51.714043 | no | 1.045E-07 | 470.25598 | 1569.1591 |
| IADFGWSVHAPSS | 14 | O14965 | Aurora kinase A OS=Homo sapiens OX=' | AURKA | 45.824 | 2 | 51.714043 | yes | 1.074E-07 | 765.37842 | 1499.3125 |
| LYGYFHDATR | 10 | O14965 | Aurora kinase A OS=Homo sapiens OX=' | AURKA | 45.824 | 2 | 51.714043 | yes | 1.634E-07 | 621.79877 | 1249.8097 |
| DIKPENLLLGSAGI | 16 | O14965 | Aurora kinase A OS=Homo sapiens OX=' | AURKA | 45.824 | 2 | 51.714043 | yes | 2.862E-07 | 848.9776 | 421.91202 |
| DIKPENLLLGSAGI | 16 | O14965 | Aurora kinase A OS=Homo sapiens OX=' | AURKA | 45.824 | 2 | 51.714043 | yes | 1.48E-05 | 566.32086 | 421.91202 |
| DIKPENLLLGSAGI | 16 | O14965 | Aurora kinase A OS=Homo sapiens OX=' | AURKA | 45.824 | 3 | 51.714043 | yes | 2.862E-07 | 848.9776 | 421.91202 |
| DIKPENLLLGSAGI | 16 | O14965 | Aurora kinase A OS=Homo sapiens OX=' | AURKA | 45.824 | 3 | 51.714043 | yes | 1.48E-05 | 566.32086 | 421.91202 |
| QWALEDFEIGR | 11 | O14965 | Aurora kinase A OS=Homo sapiens OX=' | AURKA | 45.824 | 2 | 51.714043 | yes | 9.722E-08 | 682.33569 | 545.06903 |
| SKENCISGPVK | 11 | O14965 | Aurora kinase A OS=Homo sapiens OX=' | AURKA | 45.824 | 2 | 51.714043 | yes | 7.212E-07 | 609.81097 | 354.86319 |
| EVLEHPWITANSSI | 14 | O14965 | Aurora kinase A OS=Homo sapiens OX=' | AURKA | 45.824 | 2 | 51.714043 | yes | 1.26E-07 | 805.91229 | 229.08849 |
| ENCISGPVK | 9 | O14965 | Aurora kinase A OS=Homo sapiens OX=' | AURKA | 45.824 | 2 | 51.714043 | yes | 1.87E-07 | 502.2475 | 181.00528 |
| HNPSQRPMLR | 10 | O14965 | Aurora kinase A OS=Homo sapiens OX=' | AURKA | 45.824 | 3 | 51.714043 | yes | 6.161E-07 | 412.55237 | 76.220566 |
| TATYITELANALSY | 18 | O14965 | Aurora kinase A OS=Homo sapiens OX=' | AURKA | 45.824 | 3 | 51.714043 | yes | 2.021E-07 | 681.66656 | 24.699514 |
| LQLWDIAGQER | 11 | O14966 | Ras-related protein Rab-7L1 OS=Homo s | RAB29 | 23.156 | 2 | 53.33579 | no | 9.722E-08 | 664.8515 | 167.98097 |
| VLQWSDYEIVR | 11 | O14966 | Ras-related protein Rab-7L1 OS=Homo s | RAB29 | 23.156 | 2 | 53.33579 | yes | 3.942E-07 | 704.367 | 122.99889 |
| VLVVGDAAVGK | 11 | O14966 | Ras-related protein Rab-7L1 OS=Homo s | RAB29 | 23.156 | 2 | 53.33579 | yes | 2.329E-07 | 514.31097 | 79.804382 |
| CDLSPWAVSR | 10 | O14966 | Ras-related protein Rab-7L1 OS=Homo s | RAB29 | 23.156 | 2 | 53.33579 | yes | 1.065E-07 | 595.78479 | 70.930077 |
| LAYVAPTIPR | 10 | O14974 | Protein phosphatase 1 regulatory subunit | PPP1R12A | 115.281 | 2 | 53.904285 | yes | 1.067E-07 | 550.82678 | 606.10248 |
| STQGVTLTDLQEA | 15 | O14974 | Protein phosphatase 1 regulatory subunit | PPP1R12A | 115.281 | 2 | 53.904285 | yes | 9.79E-08 | 810.40979 | 240.92691 |
| LYEQILAENEK | 11 | O14974 | Protein phosphatase 1 regulatory subunit | PPP1R12A | 115.281 | 2 | 53.904285 | yes | 1.534E-07 | 675.35101 | 224.54979 |
| LWAEDSTEK | 9 | O14974 | Protein phosphatase 1 regulatory subunit | PPP1R12A | 115.281 | 2 | 53.904285 | yes | 4.201E-07 | 539.75623 | 189.58911 |
| TGSYGALAEITASI | 14 | O14974 | Protein phosphatase 1 regulatory subunit | PPP1R12A | 115.281 | 2 | 53.904285 | yes | 9.722E-08 | 684.85388 | 141.54968 |
| ILVDNLCDMEMVI | 14 | O14974 | Protein phosphatase 1 regulatory subunit | PPP1R12A | 115.281 | 2 | 53.904285 | yes | 1.489E-07 | 847.40155 | 126.29721 |
| SGGTALHVAAAK | 12 | O14974 | Protein phosphatase 1 regulatory subunit | PPP1R12A | 115.281 | 2 | 53.904285 | yes | 9.728E-08 | 541.80127 | 91.182465 |
| LASTSDIEEK | 10 | O14974 | Protein phosphatase 1 regulatory subunit | PPP1R12A | 115.281 | 2 | 53.904285 | yes | 3.122E-07 | 546.7746 | 90.213943 |
| TYDETYQR | 8 | O14974 | Protein phosphatase 1 regulatory subunit | PPP1R12A | 115.281 | 2 | 53.904285 | yes | 2.238E-07 | 538.23816 | 88.055557 |
| SLLSSTSTTTK | 11 | O14974 | Protein phosphatase 1 regulatory subunit | PPP1R12A | 115.281 | 2 | 53.904285 | yes | 2.935E-07 | 563.30353 | 70.258774 |
| YETSSTSAGDR | 11 | O14974 | Protein phosphatase 1 regulatory subunit | PPP1R12A | 115.281 | 2 | 53.904285 | yes | 2.555E-07 | 587.25458 | 66.137802 |
| SGSYSYLEER | 10 | O14974 | Protein phosphatase 1 regulatory subunit | PPP1R12A | 115.281 | 2 | 53.904285 | yes | 4.378E-07 | 595.76984 | 63.148643 |
| LKDENGALIR | 10 | O14974 | Protein phosphatase 1 regulatory subunit | PPP1R12A | 115.281 | 2 | 53.904285 | no | 1.577E-07 | 564.8222 | 55.432037 |
| ITTGSSSAGTQSST | 17 | O14974 | Protein phosphatase 1 regulatory subunit | PPP1R12A | 115.281 | 2 | 53.904285 | yes | 9.722E-08 | 821.38739 | 48.452599 |
| AQLHDTNMELTDI | 14 | O14974 | Protein phosphatase 1 regulatory subunit | PPP1R12A | 115.281 | 2 | 53.904285 | yes | 4.45E-07 | 814.90106 | 37.460033 |

| | | | | | | | | | | | |
|---|---|---|---|---|---|---|---|---|---|---|---|
| IITYDLIK | 8 | O14975 | Long-chain fatty acid transport protein 2 ( | SLC27A2 | 70.313 | 2 | 51.216 | yes | 6.776E-06 | 489.79715 | 84.91333 |
| MTLVEEGFNPAVI | 14 | O14975 | Long-chain fatty acid transport protein 2 ( | SLC27A2 | 70.313 | 2 | 51.216 | yes | 9.722E-08 | 774.41052 | 74.808853 |
| DETLTYAQVDR | 11 | O14975 | Long-chain fatty acid transport protein 2 ( | SLC27A2 | 70.313 | 2 | 51.216 | yes | 1.1E-06 | 655.81476 | 41.484917 |
| FGEVVDCTIK | 10 | O14979 | Heterogeneous nuclear ribonucleoprotein | HNRNPDL | 46.438 | 2 | 51.86694 | no | 9.722E-08 | 584.28937 | 46805.363 |
| MFIGGLSWDTSK | 12 | O14979 | Heterogeneous nuclear ribonucleoprotein | HNRNPDL | 46.438 | 2 | 51.86694 | yes | 2.088E-07 | 671.32904 | 15308.187 |
| MFIGGLSWDTSK | 12 | O14979 | Heterogeneous nuclear ribonucleoprotein | HNRNPDL | 46.438 | 2 | 51.86694 | yes | 9.728E-08 | 679.32648 | 15308.187 |
| KDLTEYLSR | 9 | O14979 | Heterogeneous nuclear ribonucleoprotein | HNRNPDL | 46.438 | 2 | 51.86694 | yes | 9.722E-08 | 562.80096 | 3641.0881 |
| DLTEYLSR | 8 | O14979 | Heterogeneous nuclear ribonucleoprotein | HNRNPDL | 46.438 | 2 | 51.86694 | yes | 1.069E-07 | 498.75348 | 3316.4204 |
| YHQIGSGK | 8 | O14979 | Heterogeneous nuclear ribonucleoprotein | HNRNPDL | 46.438 | 2 | 51.86694 | yes | 3.754E-07 | 445.22995 | 421.34943 |
| QLAPLLPSLAPSSA | 15 | O14979 | Heterogeneous nuclear ribonucleoprotein | HNRNPDL | 46.438 | 2 | 51.86694 | yes | 2.02E-07 | 760.94342 | 328.73694 |
| GFGFVLFK | 8 | O14979 | Heterogeneous nuclear ribonucleoprotein | HNRNPDL | 46.438 | 2 | 51.86694 | no | 2.573E-07 | 457.76038 | 283.73453 |
| RGFCFITYTDEEPV | 16 | O14979 | Heterogeneous nuclear ribonucleoprotein | HNRNPDL | 46.438 | 3 | 51.86694 | yes | 1.084E-07 | 663.99615 | 55.104607 |
| KVFVGGLSPDTSE | 17 | O14979 | Heterogeneous nuclear ribonucleoprotein | HNRNPDL | 46.438 | 3 | 51.86694 | yes | 1.441E-07 | 611.99127 | 18.282551 |
| MAQEVLTHLK | 10 | O14980 | Exportin-1 OS=Homo sapiens OX=9606 | XPO1 | 123.386 | 2 | 51.410103 | yes | 9.722E-08 | 585.32098 | 159.80804 |
| VDTILEFSQNMNT | 14 | O14980 | Exportin-1 OS=Homo sapiens OX=9606 | XPO1 | 123.386 | 2 | 51.410103 | yes | 5.802E-07 | 820.40344 | 91.69838 |
| IYLDMLNVYK | 10 | O14980 | Exportin-1 OS=Homo sapiens OX=9606 | XPO1 | 123.386 | 2 | 51.410103 | yes | 1.422E-07 | 636.33887 | 62.532829 |
| DLLGLCEQK | 9 | O14980 | Exportin-1 OS=Homo sapiens OX=9606 | XPO1 | 123.386 | 2 | 51.410103 | yes | 4.901E-07 | 538.27625 | 55.294216 |
| NVDILKDPETVK | 12 | O14980 | Exportin-1 OS=Homo sapiens OX=9606 | XPO1 | 123.386 | 2 | 51.410103 | yes | 2.306E-07 | 685.87994 | 50.509842 |
| TSESLCQNNMVIL | 14 | O14980 | Exportin-1 OS=Homo sapiens OX=9606 | XPO1 | 123.386 | 2 | 51.410103 | yes | 4.251E-07 | 818.90527 | 48.388382 |
| SDLSSSSGSLSLSH | 22 | O15013 | Rho guanine nucleotide exchange factor 1 | ARHGEF1( | 151.612 | 3 | 40.302776 | yes | 4.098E-07 | 739.68414 | 21.378105 |
| HPEAILTPMPEGLS | 20 | O15013 | Rho guanine nucleotide exchange factor 1 | ARHGEF1( | 151.612 | 3 | 40.302776 | yes | 3.034E-06 | 744.05981 | 14.339953 |
| FVTPVPGPQGK | 11 | O15014 | Zinc finger protein 609 OS=Homo sapien | ZNF609 | 151.191 | 2 | 52.740902 | yes | 1.571E-07 | 563.81641 | 954.46307 |
| TYVGTLLDCTR | 11 | O15014 | Zinc finger protein 609 OS=Homo sapien | ZNF609 | 151.191 | 2 | 52.740902 | yes | 1.185E-07 | 649.8241 | 507.61078 |
| LPGQAPEGLK | 10 | O15014 | Zinc finger protein 609 OS=Homo sapien | ZNF609 | 151.191 | 2 | 52.740902 | yes | 1.279E-07 | 505.28748 | 479.51727 |
| QAEMDPILWYR | 11 | O15014 | Zinc finger protein 609 OS=Homo sapien | ZNF609 | 151.191 | 2 | 52.740902 | yes | 1.086E-07 | 711.34772 | 465.99585 |
| FCDSPTSDLEMR | 12 | O15014 | Zinc finger protein 609 OS=Homo sapien | ZNF609 | 151.191 | 2 | 52.740902 | yes | 9.722E-08 | 729.30542 | 402.27518 |
| HDWAPPR | 7 | O15014 | Zinc finger protein 609 OS=Homo sapien | ZNF609 | 151.191 | 2 | 52.740902 | no | 2.411E-07 | 439.71701 | 364.04239 |
| LSDASHLSK | 9 | O15014 | Zinc finger protein 609 OS=Homo sapien | ZNF609 | 151.191 | 2 | 52.740902 | yes | 2.893E-07 | 479.25363 | 347.29562 |
| QSLEQQQR | 8 | O15014 | Zinc finger protein 609 OS=Homo sapien | ZNF609 | 151.191 | 2 | 52.740902 | yes | 9.722E-08 | 508.75961 | 324.82034 |
| TNSMGSATGPLPG | 15 | O15014 | Zinc finger protein 609 OS=Homo sapien | ZNF609 | 151.191 | 2 | 52.740902 | yes | 4.707E-07 | 709.85083 | 319.11462 |
| VEPTVLDR | 8 | O15014 | Zinc finger protein 609 OS=Homo sapien | ZNF609 | 151.191 | 2 | 52.740902 | yes | 1.263E-07 | 464.75854 | 246.26897 |
| ALDILQQHASHYK | 13 | O15014 | Zinc finger protein 609 OS=Homo sapien | ZNF609 | 151.191 | 2 | 52.740902 | yes | 9.722E-08 | 762.40192 | 232.2137 |
| SGDGANAGGLVA | 17 | O15014 | Zinc finger protein 609 OS=Homo sapien | ZNF609 | 151.191 | 2 | 52.740902 | yes | 9.722E-08 | 734.89136 | 195.19154 |
| DPGVLQPVPLGGR | 13 | O15014 | Zinc finger protein 609 OS=Homo sapien | ZNF609 | 151.191 | 2 | 52.740902 | yes | 3.116E-07 | 652.86969 | 177.61609 |
| MWTYVYPAK | 9 | O15014 | Zinc finger protein 609 OS=Homo sapien | ZNF609 | 151.191 | 2 | 52.740902 | yes | 9.722E-08 | 579.78625 | 154.82758 |
| VKNDICEEK | 9 | O15014 | Zinc finger protein 609 OS=Homo sapien | ZNF609 | 151.191 | 2 | 52.740902 | yes | 9.795E-08 | 567.77661 | 147.35902 |
| ELESPLTPGK | 10 | O15014 | Zinc finger protein 609 OS=Homo sapien | ZNF609 | 151.191 | 2 | 52.740902 | yes | 1.233E-06 | 535.79004 | 94.855949 |
| LVEPHSPSPSSK | 12 | O15014 | Zinc finger protein 609 OS=Homo sapien | ZNF609 | 151.191 | 2 | 52.740902 | yes | 3.203E-06 | 632.83026 | 79.652039 |
| TNSPAYSDISDAGI | 24 | O15014 | Zinc finger protein 609 OS=Homo sapien | ZNF609 | 151.191 | 3 | 52.740902 | yes | 8.962E-07 | 814.36987 | 0.30336 |

| | | | | | | | | | | | |
|---|---|---|---|---|---|---|---|---|---|---|---|
| NMGPLPVDMAR | 11 | O15021 | Microtubule-associated serine/threonine-p | MAST4 | 284.098 | 2 | 49.61298 | no | 1.018E-07 | 600.79681 | 160.98701 |
| ASMQEAPPLGSLL | 14 | O15021 | Microtubule-associated serine/threonine-p | MAST4 | 284.098 | 2 | 49.61298 | yes | 3.263E-07 | 721.38959 | 42.423061 |
| TMATMGDTLASR | 12 | O15027 | Protein transport protein Sec16A OS=Ho | SEC16A | 251.894 | 2 | 55.509995 | yes | 1.001E-07 | 627.79449 | 3499.8352 |
| FSVPHVCAR | 9 | O15027 | Protein transport protein Sec16A OS=Ho | SEC16A | 251.894 | 2 | 55.509995 | yes | 1.248E-07 | 536.77148 | 2773.635 |
| LFDPQLK | 7 | O15027 | Protein transport protein Sec16A OS=Ho | SEC16A | 251.894 | 2 | 55.509995 | yes | 3.276E-07 | 430.74747 | 2514.4377 |
| YVDVLNPSGTQR | 12 | O15027 | Protein transport protein Sec16A OS=Ho | SEC16A | 251.894 | 2 | 55.509995 | yes | 1.223E-07 | 674.84644 | 2441.3271 |
| FATNEAIQR | 9 | O15027 | Protein transport protein Sec16A OS=Ho | SEC16A | 251.894 | 2 | 55.509995 | yes | 1.142E-07 | 525.27234 | 2353.2793 |
| HLLQEAR | 7 | O15027 | Protein transport protein Sec16A OS=Ho | SEC16A | 251.894 | 2 | 55.509995 | yes | 1.172E-07 | 433.74579 | 1980.963 |
| VDVINFAQNK | 10 | O15027 | Protein transport protein Sec16A OS=Ho | SEC16A | 251.894 | 2 | 55.509995 | yes | 9.722E-08 | 574.30896 | 1827.6655 |
| IVNHWASPELR | 11 | O15027 | Protein transport protein Sec16A OS=Ho | SEC16A | 251.894 | 2 | 55.509995 | yes | 1.174E-07 | 661.35419 | 761.14893 |
| IVNHWASPELR | 11 | O15027 | Protein transport protein Sec16A OS=Ho | SEC16A | 251.894 | 2 | 55.509995 | yes | 1.264E-07 | 441.23856 | 761.14893 |
| IVNHWASPELR | 11 | O15027 | Protein transport protein Sec16A OS=Ho | SEC16A | 251.894 | 3 | 55.509995 | yes | 1.174E-07 | 661.35419 | 761.14893 |
| IVNHWASPELR | 11 | O15027 | Protein transport protein Sec16A OS=Ho | SEC16A | 251.894 | 3 | 55.509995 | yes | 1.264E-07 | 441.23856 | 761.14893 |
| ELLLYGR | 7 | O15027 | Protein transport protein Sec16A OS=Ho | SEC16A | 251.894 | 2 | 55.509995 | yes | 2.295E-07 | 432.2529 | 1449.4967 |
| NQWVNLNEPEEEI | 13 | O15027 | Protein transport protein Sec16A OS=Ho | SEC16A | 251.894 | 2 | 55.509995 | yes | 1.011E-07 | 814.88116 | 1439.1251 |
| QALQSTPLGSSSK | 13 | O15027 | Protein transport protein Sec16A OS=Ho | SEC16A | 251.894 | 2 | 55.509995 | yes | 1.07E-07 | 652.34625 | 1403.0217 |
| GLANPEPAPEPK | 12 | O15027 | Protein transport protein Sec16A OS=Ho | SEC16A | 251.894 | 2 | 55.509995 | yes | 1.165E-07 | 610.31952 | 1312.3591 |
| SVFWASSPYR | 10 | O15027 | Protein transport protein Sec16A OS=Ho | SEC16A | 251.894 | 2 | 55.509995 | yes | 9.722E-08 | 600.29584 | 1227.2808 |
| GGIGASENLENPPI | 14 | O15027 | Protein transport protein Sec16A OS=Ho | SEC16A | 251.894 | 2 | 55.509995 | yes | 1.002E-07 | 691.84918 | 1150.9426 |
| KGESWFFR | 8 | O15027 | Protein transport protein Sec16A OS=Ho | SEC16A | 251.894 | 2 | 55.509995 | yes | 2.258E-07 | 528.76672 | 1054.5991 |
| NPSSAAPVQSR | 11 | O15027 | Protein transport protein Sec16A OS=Ho | SEC16A | 251.894 | 2 | 55.509995 | yes | 9.722E-08 | 557.28601 | 972.68622 |
| MPAASTCCGDEK | 12 | O15027 | Protein transport protein Sec16A OS=Ho | SEC16A | 251.894 | 2 | 55.509995 | yes | 1.477E-07 | 663.75977 | 968.26556 |
| DAQGQPGLER | 10 | O15027 | Protein transport protein Sec16A OS=Ho | SEC16A | 251.894 | 2 | 55.509995 | yes | 2.755E-07 | 535.76489 | 852.17462 |
| CLQNENLIDK | 10 | O15027 | Protein transport protein Sec16A OS=Ho | SEC16A | 251.894 | 2 | 55.509995 | yes | 2.993E-07 | 623.80847 | 772.13483 |
| KAPPPPPTSMPK | 12 | O15027 | Protein transport protein Sec16A OS=Ho | SEC16A | 251.894 | 2 | 55.509995 | yes | 2.921E-07 | 624.34448 | 682.22791 |
| YRPYDGAASAYA( | 16 | O15027 | Protein transport protein Sec16A OS=Ho | SEC16A | 251.894 | 3 | 55.509995 | yes | 1.074E-07 | 622.62372 | 633.45416 |
| SHLVGVKPFEADR | 13 | O15027 | Protein transport protein Sec16A OS=Ho | SEC16A | 251.894 | 3 | 55.509995 | yes | 1.474E-07 | 485.59653 | 289.46661 |
| SHLVGVKPFEADR | 13 | O15027 | Protein transport protein Sec16A OS=Ho | SEC16A | 251.894 | 3 | 55.509995 | yes | 1.926E-07 | 727.89117 | 289.46661 |
| SHLVGVKPFEADR | 13 | O15027 | Protein transport protein Sec16A OS=Ho | SEC16A | 251.894 | 2 | 55.509995 | yes | 1.474E-07 | 485.59653 | 289.46661 |
| SHLVGVKPFEADR | 13 | O15027 | Protein transport protein Sec16A OS=Ho | SEC16A | 251.894 | 2 | 55.509995 | yes | 1.926E-07 | 727.89117 | 289.46661 |
| KTEAYLPDDK | 10 | O15027 | Protein transport protein Sec16A OS=Ho | SEC16A | 251.894 | 2 | 55.509995 | yes | 9.887E-08 | 590.29822 | 549.99023 |
| ANHSSHQEDTYG/ | 20 | O15027 | Protein transport protein Sec16A OS=Ho | SEC16A | 251.894 | 3 | 55.509995 | yes | 1.152E-07 | 750.34546 | 468.64282 |
| TLENPVNVYNPSH | 28 | O15027 | Protein transport protein Sec16A OS=Ho | SEC16A | 251.894 | 4 | 55.509995 | yes | 9.722E-08 | 759.37079 | 451.7821 |
| FANLTPSR | 8 | O15027 | Protein transport protein Sec16A OS=Ho | SEC16A | 251.894 | 2 | 55.509995 | yes | 1.301E-07 | 453.24561 | 388.8754 |
| TVPDSEAPPGWDI | 13 | O15027 | Protein transport protein Sec16A OS=Ho | SEC16A | 251.894 | 2 | 55.509995 | yes | 1.968E-07 | 713.8335 | 359.56326 |
| EKPEEESLAAPTW | 22 | O15027 | Protein transport protein Sec16A OS=Ho | SEC16A | 251.894 | 4 | 55.509995 | yes | 1.155E-07 | 648.08813 | 256.8656 |
| EHSAFGDRPEK | 11 | O15027 | Protein transport protein Sec16A OS=Ho | SEC16A | 251.894 | 2 | 55.509995 | yes | 3.619E-07 | 636.802 | 255.58035 |
| TEAYLPDDK | 9 | O15027 | Protein transport protein Sec16A OS=Ho | SEC16A | 251.894 | 2 | 55.509995 | yes | 1.824E-06 | 526.25073 | 248.63754 |
| SHNVAAGSYEAPI | 28 | O15027 | Protein transport protein Sec16A OS=Ho | SEC16A | 251.894 | 4 | 55.509995 | yes | 5.76E-07 | 742.84668 | 234.5038 |

| | | | | | | | | | | | |
|---|---|---|---|---|---|---|---|---|---|---|---|
| PASALVNPLAR | 11 | O15027 | Protein transport protein Sec16A OS=Hoı | SEC16A | 251.894 | 2 | 55.509995 | yes | 9.722E-08 | 554.82727 | 221.54292 |
| SSPPVLQGPAPAGI | 28 | O15027 | Protein transport protein Sec16A OS=Hoı | SEC16A | 251.894 | 4 | 55.509995 | yes | 3.07E-07 | 714.63202 | 211.16449 |
| KAGSALPGFANSP. | 35 | O15027 | Protein transport protein Sec16A OS=Hoı | SEC16A | 251.894 | 4 | 55.509995 | yes | 1.363E-07 | 829.19391 | 200.20552 |
| YWCDAEYDAYR | 11 | O15027 | Protein transport protein Sec16A OS=Hoı | SEC16A | 251.894 | 2 | 55.509995 | yes | 9.722E-08 | 756.29846 | 195.83496 |
| VLSSAASLPGSELI | 17 | O15027 | Protein transport protein Sec16A OS=Hoı | SEC16A | 251.894 | 2 | 55.509995 | yes | 9.722E-08 | 829.44141 | 174.44151 |
| SLHSAHSLASR | 11 | O15027 | Protein transport protein Sec16A OS=Hoı | SEC16A | 251.894 | 2 | 55.509995 | yes | 1.041E-07 | 583.30725 | 171.30002 |
| LVLIGSNHSLPFLK | 14 | O15027 | Protein transport protein Sec16A OS=Hoı | SEC16A | 251.894 | 2 | 55.509995 | yes | 9.809E-08 | 769.45868 | 82.44175 |
| LVLIGSNHSLPFLK | 14 | O15027 | Protein transport protein Sec16A OS=Hoı | SEC16A | 251.894 | 2 | 55.509995 | yes | 1.671E-06 | 513.30823 | 82.44175 |
| LVLIGSNHSLPFLK | 14 | O15027 | Protein transport protein Sec16A OS=Hoı | SEC16A | 251.894 | 3 | 55.509995 | yes | 9.809E-08 | 769.45868 | 82.44175 |
| LVLIGSNHSLPFLK | 14 | O15027 | Protein transport protein Sec16A OS=Hoı | SEC16A | 251.894 | 3 | 55.509995 | yes | 1.671E-06 | 513.30823 | 82.44175 |
| NEHRPASALVNPL | 15 | O15027 | Protein transport protein Sec16A OS=Hoı | SEC16A | 251.894 | 3 | 55.509995 | yes | 9.722E-08 | 548.96912 | 150.04166 |
| YGPLPGPAVPR | 11 | O15027 | Protein transport protein Sec16A OS=Hoı | SEC16A | 251.894 | 2 | 55.509995 | yes | 4.368E-07 | 562.31659 | 133.7804 |
| FYQQVTK | 7 | O15027 | Protein transport protein Sec16A OS=Hoı | SEC16A | 251.894 | 2 | 55.509995 | yes | 1.288E-06 | 457.24252 | 127.63178 |
| APPPPPTSMPK | 11 | O15027 | Protein transport protein Sec16A OS=Hoı | SEC16A | 251.894 | 2 | 55.509995 | yes | 2.212E-07 | 560.297 | 102.81557 |
| QGYPEGYYSSK | 11 | O15027 | Protein transport protein Sec16A OS=Hoı | SEC16A | 251.894 | 2 | 55.509995 | yes | 3.505E-06 | 639.78552 | 78.072281 |
| PQPPVVQPPEEAM | 19 | O15027 | Protein transport protein Sec16A OS=Hoı | SEC16A | 251.894 | 3 | 55.509995 | yes | 1.968E-06 | 688.00562 | 59.907982 |
| LAEMGLATQAFHY | 19 | O15027 | Protein transport protein Sec16A OS=Hoı | SEC16A | 251.894 | 3 | 55.509995 | yes | 6.001E-07 | 708.67993 | 55.621784 |
| DALESAMK | 8 | O15027 | Protein transport protein Sec16A OS=Hoı | SEC16A | 251.894 | 2 | 55.509995 | yes | 2.459E-07 | 432.71021 | 46.058304 |
| CSSMSSLSR | 9 | O15027 | Protein transport protein Sec16A OS=Hoı | SEC16A | 251.894 | 2 | 55.509995 | yes | 9.722E-08 | 507.72079 | 40.522285 |
| SSLSSHSHQSQIYR | 14 | O15027 | Protein transport protein Sec16A OS=Hoı | SEC16A | 251.894 | 2 | 55.509995 | yes | 1.043E-07 | 808.8924 | 23.714312 |
| NLLALIHR | 8 | O15042 | U2 snRNP-associated SURP motif-contai | U2SURP | 118.292 | 2 | 54.840206 | yes | 1.301E-07 | 475.30072 | 6540.2495 |
| LYSILQGDSPTK | 12 | O15042 | U2 snRNP-associated SURP motif-contai | U2SURP | 118.292 | 2 | 54.840206 | yes | 1.127E-07 | 661.35352 | 5484.9717 |
| LCQIFSDLNATYR | 13 | O15042 | U2 snRNP-associated SURP motif-contai | U2SURP | 118.292 | 2 | 54.840206 | yes | 9.965E-08 | 800.89307 | 4366.394 |
| DKLEEILR | 8 | O15042 | U2 snRNP-associated SURP motif-contai | U2SURP | 118.292 | 2 | 54.840206 | yes | 1.117E-07 | 508.29276 | 3451.9529 |
| LYLVSDVLYNSSA | 14 | O15042 | U2 snRNP-associated SURP motif-contai | U2SURP | 118.292 | 2 | 54.840206 | yes | 1.144E-07 | 786.41943 | 3407.1699 |
| FEPPQSDSDGQR | 12 | O15042 | U2 snRNP-associated SURP motif-contai | U2SURP | 118.292 | 2 | 54.840206 | yes | 3.194E-06 | 681.79968 | 2347.7542 |
| EINNPMFR | 8 | O15042 | U2 snRNP-associated SURP motif-contai | U2SURP | 118.292 | 2 | 54.840206 | yes | 2.093E-07 | 510.75021 | 1855.8809 |
| VVIPTER | 7 | O15042 | U2 snRNP-associated SURP motif-contai | U2SURP | 118.292 | 2 | 54.840206 | yes | 1.169E-07 | 407.24509 | 1751.2001 |
| NCGFVAFMNR | 10 | O15042 | U2 snRNP-associated SURP motif-contai | U2SURP | 118.292 | 2 | 54.840206 | yes | 1.12E-07 | 608.2735 | 1665.2841 |
| ESLCDSPHQNLSR | 13 | O15042 | U2 snRNP-associated SURP motif-contai | U2SURP | 118.292 | 3 | 54.840206 | yes | 2.539E-07 | 514.90369 | 614.4198 |
| ESLCDSPHQNLSR | 13 | O15042 | U2 snRNP-associated SURP motif-contai | U2SURP | 118.292 | 3 | 54.840206 | yes | 1.165E-07 | 771.85193 | 614.4198 |
| ESLCDSPHQNLSR | 13 | O15042 | U2 snRNP-associated SURP motif-contai | U2SURP | 118.292 | 2 | 54.840206 | yes | 2.539E-07 | 514.90369 | 614.4198 |
| ESLCDSPHQNLSR | 13 | O15042 | U2 snRNP-associated SURP motif-contai | U2SURP | 118.292 | 2 | 54.840206 | yes | 1.165E-07 | 771.85193 | 614.4198 |
| FQDELESGK | 9 | O15042 | U2 snRNP-associated SURP motif-contai | U2SURP | 118.292 | 2 | 54.840206 | yes | 1.163E-07 | 526.74835 | 967.37903 |
| TIQGHLQSENFK | 12 | O15042 | U2 snRNP-associated SURP motif-contai | U2SURP | 118.292 | 3 | 54.840206 | yes | 2.129E-07 | 467.90887 | 481.33887 |
| TIQGHLQSENFK | 12 | O15042 | U2 snRNP-associated SURP motif-contai | U2SURP | 118.292 | 3 | 54.840206 | yes | 6.375E-07 | 701.35968 | 481.33887 |
| TIQGHLQSENFK | 12 | O15042 | U2 snRNP-associated SURP motif-contai | U2SURP | 118.292 | 2 | 54.840206 | yes | 2.129E-07 | 467.90887 | 481.33887 |
| TIQGHLQSENFK | 12 | O15042 | U2 snRNP-associated SURP motif-contai | U2SURP | 118.292 | 2 | 54.840206 | yes | 6.375E-07 | 701.35968 | 481.33887 |
| MIEFVVR | 7 | O15042 | U2 snRNP-associated SURP motif-contai | U2SURP | 118.292 | 2 | 54.840206 | yes | 1.269E-07 | 447.2493 | 953.36469 |

| | | | | | | | | | | | |
|---|---|---|---|---|---|---|---|---|---|---|---|
| VANASYYR | 8 | O15042 | U2 snRNP-associated SURP motif-contai | U2SURP | 118.292 | 2 | 54.840206 | yes | 5.734E-07 | 472.23523 | 937.31586 |
| KPGQSFQEQVEHY | 14 | O15042 | U2 snRNP-associated SURP motif-contai | U2SURP | 118.292 | 3 | 54.840206 | yes | 9.926E-08 | 578.28491 | 819.48456 |
| YSEMSEEKR | 9 | O15042 | U2 snRNP-associated SURP motif-contai | U2SURP | 118.292 | 2 | 54.840206 | yes | 1.152E-07 | 579.75842 | 593.64227 |
| IYKPSSR | 7 | O15042 | U2 snRNP-associated SURP motif-contai | U2SURP | 118.292 | 2 | 54.840206 | yes | 1.172E-07 | 425.74271 | 488.33038 |
| LKNPNAPMLPPPK | 13 | O15042 | U2 snRNP-associated SURP motif-contai | U2SURP | 118.292 | 2 | 54.840206 | yes | 1.164E-07 | 708.90521 | 193.95773 |
| LKNPNAPMLPPPK | 13 | O15042 | U2 snRNP-associated SURP motif-contai | U2SURP | 118.292 | 2 | 54.840206 | yes | 3.072E-07 | 472.93924 | 193.95773 |
| LKNPNAPMLPPPK | 13 | O15042 | U2 snRNP-associated SURP motif-contai | U2SURP | 118.292 | 3 | 54.840206 | yes | 1.164E-07 | 708.90521 | 193.95773 |
| LKNPNAPMLPPPK | 13 | O15042 | U2 snRNP-associated SURP motif-contai | U2SURP | 118.292 | 3 | 54.840206 | yes | 3.072E-07 | 472.93924 | 193.95773 |
| YSEMSEEK | 8 | O15042 | U2 snRNP-associated SURP motif-contai | U2SURP | 118.292 | 2 | 54.840206 | yes | 2.51E-07 | 501.70786 | 362.09445 |
| KSNLELFK | 8 | O15042 | U2 snRNP-associated SURP motif-contai | U2SURP | 118.292 | 2 | 54.840206 | yes | 1.626E-07 | 489.78458 | 314.17084 |
| TLSQAIVK | 8 | O15042 | U2 snRNP-associated SURP motif-contai | U2SURP | 118.292 | 2 | 54.840206 | yes | 4.996E-07 | 430.26602 | 262.86853 |
| SEEHHLYSNPIK | 12 | O15042 | U2 snRNP-associated SURP motif-contai | U2SURP | 118.292 | 2 | 54.840206 | yes | 9.722E-08 | 727.35718 | 70.472939 |
| SEEHHLYSNPIK | 12 | O15042 | U2 snRNP-associated SURP motif-contai | U2SURP | 118.292 | 2 | 54.840206 | yes | 1.121E-07 | 485.24051 | 70.472939 |
| SEEHHLYSNPIK | 12 | O15042 | U2 snRNP-associated SURP motif-contai | U2SURP | 118.292 | 3 | 54.840206 | yes | 9.722E-08 | 727.35718 | 70.472939 |
| SEEHHLYSNPIK | 12 | O15042 | U2 snRNP-associated SURP motif-contai | U2SURP | 118.292 | 3 | 54.840206 | yes | 1.121E-07 | 485.24051 | 70.472939 |
| EGPMFEAMIMNR | 12 | O15042 | U2 snRNP-associated SURP motif-contai | U2SURP | 118.292 | 2 | 54.840206 | yes | 9.722E-08 | 713.31982 | 140.71082 |
| MIMSFEMK | 8 | O15042 | U2 snRNP-associated SURP motif-contai | U2SURP | 118.292 | 2 | 54.840206 | yes | 5.252E-07 | 508.73431 | 100.82971 |
| SLDDDLDGVPLDA | 19 | O15042 | U2 snRNP-associated SURP motif-contai | U2SURP | 118.292 | 3 | 54.840206 | yes | 1.42E-07 | 678.32343 | 18.991241 |
| AQAQAQDAELVR | 12 | O15049 | NEDD4-binding protein 3 OS=Homo sap | N4BP3 | 60.47 | 2 | 43.637836 | yes | 2.251E-05 | 650.33624 | 69.008713 |
| EIQGGYMDMYR | 11 | O15049 | NEDD4-binding protein 3 OS=Homo sap | N4BP3 | 60.47 | 2 | 43.637836 | yes | 2.508E-07 | 681.79449 | 44.60363 |
| EGFLGPPASR | 10 | O15054 | Lysine-specific demethylase 6B OS=Hom | KDM6B | 176.633 | 2 | 48.3462 | yes | 3.525E-07 | 515.76947 | 266.15375 |
| FGTNIDLSDAK | 11 | O15054 | Lysine-specific demethylase 6B OS=Hom | KDM6B | 176.633 | 2 | 48.3462 | yes | 1.162E-07 | 590.79584 | 199.88106 |
| LNLGLFSTK | 9 | O15054 | Lysine-specific demethylase 6B OS=Hom | KDM6B | 176.633 | 2 | 48.3462 | yes | 3.028E-06 | 496.79239 | 73.528 |
| VLGNLDLQSEEIQ | 15 | O15054 | Lysine-specific demethylase 6B OS=Hom | KDM6B | 176.633 | 2 | 48.3462 | yes | 3.002E-05 | 835.93903 | 46.917175 |
| SAGLQGVVVLEQ | 14 | O15054 | Lysine-specific demethylase 6B OS=Hom | KDM6B | 176.633 | 2 | 48.3462 | yes | 1.039E-06 | 759.91736 | 22.463305 |
| ENLLLEEELR | 10 | O15061 | Synemin OS=Homo sapiens OX=9606 G | SYNM | 172.868 | 2 | 50.844982 | yes | 1.328E-07 | 629.33789 | 112.77117 |
| ETVQLYEDEVR | 11 | O15061 | Synemin OS=Homo sapiens OX=9606 G | SYNM | 172.868 | 2 | 50.844982 | yes | 9.722E-08 | 690.83569 | 56.564781 |
| TFSPTYGLLR | 10 | O15061 | Synemin OS=Homo sapiens OX=9606 G | SYNM | 172.868 | 2 | 50.844982 | yes | 2.387E-06 | 577.81384 | 14.551657 |
| AFSITQGLLK | 10 | O15067 | Phosphoribosylformylglycinamidine synt | PFAS | 144.735 | 2 | 44.84655 | yes | 7.905E-07 | 539.31879 | 60.524254 |
| ELSDPAGAIIYTSR | 14 | O15067 | Phosphoribosylformylglycinamidine synt | PFAS | 144.735 | 2 | 44.84655 | yes | 2.048E-07 | 746.88574 | 48.599907 |
| CPACFVGTITGDR | 13 | O15067 | Phosphoribosylformylglycinamidine synt | PFAS | 144.735 | 2 | 44.84655 | yes | 1.831E-06 | 727.33179 | 43.200207 |
| SPVLHFYVR | 9 | O15067 | Phosphoribosylformylglycinamidine synt | PFAS | 144.735 | 2 | 44.84655 | yes | 3.972E-06 | 559.31128 | 33.59539 |
| VSQDLER | 7 | O15078 | Centrosomal protein of 290 kDa OS=Hon | CEP290 | 290.386 | 2 | 45.309242 | yes | 1.889E-07 | 423.71942 | 185.82582 |
| TMIDLTEFR | 9 | O15078 | Centrosomal protein of 290 kDa OS=Hon | CEP290 | 290.386 | 2 | 45.309242 | yes | 4.487E-06 | 563.28406 | 31.531128 |
| AVASCSEEAVQVI | 15 | O15084 | Serine/threonine-protein phosphatase 6 re | ANKRD28 | 112.966 | 2 | 52.42774 | yes | 1.594E-07 | 802.42163 | 801.05414 |
| GAVTGHEECVDAI | 19 | O15084 | Serine/threonine-protein phosphatase 6 re | ANKRD28 | 112.966 | 3 | 52.42774 | yes | 1.02E-07 | 664.65833 | 664.93335 |
| GHETSALLILEK | 12 | O15084 | Serine/threonine-protein phosphatase 6 re | ANKRD28 | 112.966 | 2 | 52.42774 | yes | 1.077E-07 | 655.86938 | 345.82724 |
| WLTPLHR | 7 | O15084 | Serine/threonine-protein phosphatase 6 re | ANKRD28 | 112.966 | 2 | 52.42774 | yes | 9.796E-08 | 461.76651 | 328.35526 |
| GHVECVDVLINQC | 19 | O15084 | Serine/threonine-protein phosphatase 6 re | ANKRD28 | 112.966 | 3 | 52.42774 | yes | 1.233E-07 | 684.37 | 308.71082 |

| | | | | | | | | | | | |
|---|---|---|---|---|---|---|---|---|---|---|---|
| NTALHLACSK | 10 | O15084 | Serine/threonine-protein phosphatase 6 re | ANKRD28 | 112.966 | 2 | 52.42774 | no | 2.274E-07 | 557.78729 | 306.20779 |
| TPLDLAAFK | 9 | O15084 | Serine/threonine-protein phosphatase 6 re | ANKRD28 | 112.966 | 2 | 52.42774 | yes | 9.722E-08 | 488.27911 | 279.0459 |
| EDVNFQDNEK | 10 | O15084 | Serine/threonine-protein phosphatase 6 re | ANKRD28 | 112.966 | 2 | 52.42774 | yes | 9.722E-08 | 619.2702 | 222.96581 |
| TVSFEALPIMR | 11 | O15084 | Serine/threonine-protein phosphatase 6 re | ANKRD28 | 112.966 | 2 | 52.42774 | yes | 1.76E-07 | 632.34192 | 204.61079 |
| NLINATNAALQTPI | 19 | O15084 | Serine/threonine-protein phosphatase 6 re | ANKRD28 | 112.966 | 3 | 52.42774 | yes | 1.064E-07 | 663.37274 | 200.83925 |
| SYTPLHAAASSGM | 18 | O15084 | Serine/threonine-protein phosphatase 6 re | ANKRD28 | 112.966 | 3 | 52.42774 | yes | 1.241E-07 | 606.98547 | 191.14879 |
| YGHELLINTLITSG | 19 | O15084 | Serine/threonine-protein phosphatase 6 re | ANKRD28 | 112.966 | 3 | 52.42774 | no | 1.284E-07 | 673.02484 | 149.3929 |
| TALHHAAFSGHGE | 16 | O15084 | Serine/threonine-protein phosphatase 6 re | ANKRD28 | 112.966 | 3 | 52.42774 | yes | 4.08E-07 | 564.94733 | 61.21059 |
| TALHHAAFSGHGE | 16 | O15084 | Serine/threonine-protein phosphatase 6 re | ANKRD28 | 112.966 | 3 | 52.42774 | yes | 1.602E-07 | 846.91736 | 61.21059 |
| TALHHAAFSGHGE | 16 | O15084 | Serine/threonine-protein phosphatase 6 re | ANKRD28 | 112.966 | 2 | 52.42774 | yes | 4.08E-07 | 564.94733 | 61.21059 |
| TALHHAAFSGHGE | 16 | O15084 | Serine/threonine-protein phosphatase 6 re | ANKRD28 | 112.966 | 2 | 52.42774 | yes | 1.602E-07 | 846.91736 | 61.21059 |
| NGNTPLHIAAR | 11 | O15084 | Serine/threonine-protein phosphatase 6 re | ANKRD28 | 112.966 | 2 | 52.42774 | yes | 7.169E-07 | 582.31763 | 48.846542 |
| TCLHAAAAGGNLI | 28 | O15084 | Serine/threonine-protein phosphatase 6 re | ANKRD28 | 112.966 | 4 | 52.42774 | yes | 1.196E-06 | 747.37732 | 48.744156 |
| GCTPLHYAATSDT | 16 | O15084 | Serine/threonine-protein phosphatase 6 re | ANKRD28 | 112.966 | 2 | 52.42774 | yes | 1.006E-07 | 847.37793 | 35.245605 |
| LLVSHGAEVTCK | 12 | O15084 | Serine/threonine-protein phosphatase 6 re | ANKRD28 | 112.966 | 3 | 52.42774 | yes | 4.394E-07 | 438.5676 | 29.693893 |
| AIHWAAYMGHIEV | 15 | O15084 | Serine/threonine-protein phosphatase 6 re | ANKRD28 | 112.966 | 3 | 52.42774 | yes | 1.348E-06 | 575.63623 | 20.21052 |
| GENVVLLGEIDLEI | 14 | O15116 | U6 snRNA-associated Sm-like protein LS | LSM1 | 15.18 | 2 | 48.883064 | yes | 1.091E-07 | 764.41687 | 679.38312 |
| TMEPEEEVEDDPK | 13 | O15119 | T-box transcription factor TBX3 OS=Hor | TBX3 | 79.389 | 2 | 52.081203 | yes | 9.722E-08 | 774.32452 | 73.865166 |
| LVSGLEAKPDR | 11 | O15119 | T-box transcription factor TBX3 OS=Hor | TBX3 | 79.389 | 2 | 52.081203 | yes | 1.605E-07 | 592.83533 | 46.067242 |
| EAATSELQSIQR | 12 | O15119 | T-box transcription factor TBX3 OS=Hor | TBX3 | 79.389 | 2 | 52.081203 | yes | 1.737E-07 | 666.84131 | 43.571152 |
| KEACWTVSNITAC | 15 | O15131 | Importin subunit alpha-6 OS=Homo sapie | KPNA5 | 60.667 | 3 | 38.663857 | yes | 5.067E-07 | 569.61414 | 103.18683 |
| TWKPTLVILR | 10 | O15143 | Actin-related protein 2/3 complex subunit | ARPC1B | 40.95 | 2 | 47.702503 | yes | 1.313E-07 | 613.88464 | 179.85165 |
| NSVSQISVLSGGK | 13 | O15143 | Actin-related protein 2/3 complex subunit | ARPC1B | 40.95 | 2 | 47.702503 | yes | 1.276E-07 | 638.34882 | 48.880421 |
| YFQFQEEGK | 9 | O15144 | Actin-related protein 2/3 complex subunit | ARPC2 | 34.334 | 2 | 54.5837 | yes | 9.722E-08 | 588.27203 | 128.57391 |
| MILLEVNNR | 9 | O15144 | Actin-related protein 2/3 complex subunit | ARPC2 | 34.334 | 2 | 54.5837 | yes | 9.722E-08 | 551.30786 | 96.103775 |
| AYLQQLR | 7 | O15145 | Actin-related protein 2/3 complex subunit | ARPC3 | 20.547 | 2 | 24.709694 | yes | 3.405E-06 | 446.25598 | 103.04248 |
| QLGHLCDLTIR | 11 | O15156 | Zinc finger and BTB domain-containing p | ZBTB7B | 58.027 | 2 | 43.293068 | yes | 2.651E-07 | 663.35333 | 38.608238 |
| QLGHLCDLTIR | 11 | O15156 | Zinc finger and BTB domain-containing p | ZBTB7B | 58.027 | 2 | 43.293068 | yes | 6.181E-07 | 442.57132 | 38.608238 |
| QLGHLCDLTIR | 11 | O15156 | Zinc finger and BTB domain-containing p | ZBTB7B | 58.027 | 3 | 43.293068 | yes | 2.651E-07 | 663.35333 | 38.608238 |
| QLGHLCDLTIR | 11 | O15156 | Zinc finger and BTB domain-containing p | ZBTB7B | 58.027 | 3 | 43.293068 | yes | 6.181E-07 | 442.57132 | 38.608238 |
| ANHLVPEVPTVPA | 26 | O15156 | Zinc finger and BTB domain-containing p | ZBTB7B | 58.027 | 4 | 43.293068 | yes | 9.864E-07 | 714.36469 | 53.284344 |
| FSPVATASYR | 10 | O15160 | DNA-directed RNA polymerases I and III | POLR1C | 39.25 | 2 | 53.195923 | yes | 5.148E-07 | 549.78253 | 418.74283 |
| AASQAVEEMR | 10 | O15160 | DNA-directed RNA polymerases I and III | POLR1C | 39.25 | 2 | 53.195923 | yes | 9.722E-08 | 567.26642 | 374.9512 |
| ILLAEVPTMAVEK | 13 | O15160 | DNA-directed RNA polymerases I and III | POLR1C | 39.25 | 2 | 53.195923 | yes | 3.749E-07 | 707.40472 | 346.61035 |
| CFSPGVIEVQEVQ | 15 | O15160 | DNA-directed RNA polymerases I and III | POLR1C | 39.25 | 2 | 53.195923 | yes | 9.722E-08 | 838.91925 | 212.49182 |
| RFLDELDAVQMD | 12 | O15160 | DNA-directed RNA polymerases I and III | POLR1C | 39.25 | 2 | 53.195923 | yes | 3.989E-07 | 726.3454 | 163.62878 |
| VVLGEFGVR | 9 | O15160 | DNA-directed RNA polymerases I and III | POLR1C | 39.25 | 2 | 53.195923 | yes | 9.727E-08 | 488.28473 | 163.02243 |
| DSSDPNELYVNHK | 13 | O15160 | DNA-directed RNA polymerases I and III | POLR1C | 39.25 | 2 | 53.195923 | yes | 9.722E-08 | 759.34698 | 62.740208 |
| LLLETAAPR | 9 | O15169 | Axin-1 OS=Homo sapiens OX=9606 GN= | AXIN1 | 95.635 | 2 | 52.638855 | yes | 2.48E-07 | 492.29785 | 188.15498 |

| | | | | | | | | | | | |
|---|---|---|---|---|---|---|---|---|---|---|---|
| FAEELIHR | 8 | O15169 | Axin-1 OS=Homo sapiens OX=9606 GN= | AXIN1 | 95.635 | 2 | 52.638855 | yes | 4.963E-07 | 507.772 | 168.33128 |
| IMQWIIEGEK | 10 | O15169 | Axin-1 OS=Homo sapiens OX=9606 GN= | AXIN1 | 95.635 | 2 | 52.638855 | yes | 9.819E-08 | 623.82867 | 109.49113 |
| VPLPHIPR | 8 | O15169 | Axin-1 OS=Homo sapiens OX=9606 GN= | AXIN1 | 95.635 | 2 | 52.638855 | yes | 2.627E-06 | 464.78998 | 105.52995 |
| PLSLEHPWAGPQL | 14 | O15169 | Axin-1 OS=Homo sapiens OX=9606 GN= | AXIN1 | 95.635 | 3 | 52.638855 | yes | 9.722E-08 | 534.29132 | 103.06964 |
| SDIYLEYTR | 9 | O15169 | Axin-1 OS=Homo sapiens OX=9606 GN= | AXIN1 | 95.635 | 2 | 52.638855 | yes | 3.759E-06 | 580.28516 | 71.11573 |
| EDEAVLPVFEEK | 12 | O15169 | Axin-1 OS=Homo sapiens OX=9606 GN= | AXIN1 | 95.635 | 2 | 52.638855 | yes | 5.998E-07 | 702.84827 | 69.968925 |
| KLEPCDSNEEK | 11 | O15169 | Axin-1 OS=Homo sapiens OX=9606 GN= | AXIN1 | 95.635 | 2 | 52.638855 | yes | 1.505E-07 | 674.80609 | 59.684349 |
| CVDMGCAGLR | 10 | O15169 | Axin-1 OS=Homo sapiens OX=9606 GN= | AXIN1 | 95.635 | 2 | 52.638855 | yes | 1.03E-07 | 569.74371 | 55.767609 |
| ELGENMSDEELR | 12 | O15182 | Centrin-3 OS=Homo sapiens OX=9606 G | CETN3 | 19.55 | 2 | 55.732807 | yes | 9.722E-08 | 711.31427 | 78.202286 |
| EELALLDGSNVVF | 14 | O15212 | Prefoldin subunit 6 OS=Homo sapiens O | PFDN6 | 14.583 | 2 | 50.089104 | yes | 3.992E-07 | 767.41156 | 116.36374 |
| ETLAQLQQEFQR | 12 | O15212 | Prefoldin subunit 6 OS=Homo sapiens O | PFDN6 | 14.583 | 2 | 50.089104 | yes | 9.904E-08 | 745.88354 | 96.52845 |
| LEAQLTENNIVK | 12 | O15212 | Prefoldin subunit 6 OS=Homo sapiens O | PFDN6 | 14.583 | 2 | 50.089104 | yes | 3.021E-06 | 686.37756 | 27.476784 |
| LMCEINVMEAVR | 12 | O15213 | WD repeat-containing protein 46 OS=Ho | WDR46 | 68.072 | 2 | 52.14307 | yes | 1.17E-07 | 732.854 | 225.55209 |
| VPAELICLDPR | 11 | O15213 | WD repeat-containing protein 46 OS=Ho | WDR46 | 68.072 | 2 | 52.14307 | yes | 1.825E-07 | 641.84485 | 176.77824 |
| ALAEVDVISLEQG | 14 | O15213 | WD repeat-containing protein 46 OS=Ho | WDR46 | 68.072 | 2 | 52.14307 | yes | 3.277E-07 | 736.40375 | 169.54201 |
| HFDLNLR | 7 | O15213 | WD repeat-containing protein 46 OS=Ho | WDR46 | 68.072 | 2 | 52.14307 | yes | 1.961E-07 | 457.74579 | 154.77792 |
| FLHSEALLAVAQN | 14 | O15213 | WD repeat-containing protein 46 OS=Ho | WDR46 | 68.072 | 3 | 52.14307 | yes | 9.722E-08 | 523.62299 | 60.307873 |
| FLHSEALLAVAQN | 14 | O15213 | WD repeat-containing protein 46 OS=Ho | WDR46 | 68.072 | 3 | 52.14307 | yes | 1.14E-07 | 784.93079 | 60.307873 |
| FLHSEALLAVAQN | 14 | O15213 | WD repeat-containing protein 46 OS=Ho | WDR46 | 68.072 | 2 | 52.14307 | yes | 9.722E-08 | 523.62299 | 60.307873 |
| FLHSEALLAVAQN | 14 | O15213 | WD repeat-containing protein 46 OS=Ho | WDR46 | 68.072 | 2 | 52.14307 | yes | 1.14E-07 | 784.93079 | 60.307873 |
| GHVAALDWVTK | 11 | O15213 | WD repeat-containing protein 46 OS=Ho | WDR46 | 68.072 | 2 | 52.14307 | yes | 2.442E-07 | 598.82477 | 107.5062 |
| LEVAEAEEEETSIK | 14 | O15213 | WD repeat-containing protein 46 OS=Ho | WDR46 | 68.072 | 2 | 52.14307 | yes | 9.722E-08 | 788.88306 | 95.19223 |
| METAPKPGK | 9 | O15213 | WD repeat-containing protein 46 OS=Ho | WDR46 | 68.072 | 2 | 52.14307 | yes | 1.291E-06 | 500.76022 | 41.415611 |
| LLTDGYACEVR | 11 | O15226 | NF-kappa-B-repressing factor OS=Homo | NKRF | 77.673 | 2 | 51.347385 | yes | 3.676E-07 | 648.81628 | 983.59924 |
| TNPEYIYAPLK | 11 | O15226 | NF-kappa-B-repressing factor OS=Homo | NKRF | 77.673 | 2 | 51.347385 | yes | 1.128E-07 | 654.84534 | 787.96893 |
| EGLGLDVER | 9 | O15226 | NF-kappa-B-repressing factor OS=Homo | NKRF | 77.673 | 2 | 51.347385 | yes | 1.602E-07 | 494.25891 | 745.35645 |
| INYTYMLTR | 9 | O15226 | NF-kappa-B-repressing factor OS=Homo | NKRF | 77.673 | 2 | 51.347385 | yes | 1.51E-07 | 587.79987 | 742.89551 |
| EDNIGNQLLR | 10 | O15226 | NF-kappa-B-repressing factor OS=Homo | NKRF | 77.673 | 2 | 51.347385 | yes | 9.722E-08 | 586.30695 | 729.76196 |
| TQPTYPSVK | 9 | O15226 | NF-kappa-B-repressing factor OS=Homo | NKRF | 77.673 | 2 | 51.347385 | yes | 8.238E-07 | 510.77167 | 527.21692 |
| TQPTVINNLK | 10 | O15226 | NF-kappa-B-repressing factor OS=Homo | NKRF | 77.673 | 2 | 51.347385 | yes | 1.564E-07 | 564.32458 | 517.17786 |
| CQNIYLTTGYAGS | 14 | O15226 | NF-kappa-B-repressing factor OS=Homo | NKRF | 77.673 | 2 | 51.347385 | yes | 1.028E-07 | 788.3772 | 473.93665 |
| HLSTCDGQNPPK | 12 | O15226 | NF-kappa-B-repressing factor OS=Homo | NKRF | 77.673 | 2 | 51.347385 | yes | 9.809E-08 | 677.31445 | 267.7728 |
| YEAAGEAVK | 9 | O15226 | NF-kappa-B-repressing factor OS=Homo | NKRF | 77.673 | 2 | 51.347385 | yes | 1.012E-07 | 469.23489 | 252.47911 |
| SESHTDLTFSR | 11 | O15226 | NF-kappa-B-repressing factor OS=Homo | NKRF | 77.673 | 2 | 51.347385 | yes | 1.088E-07 | 640.29932 | 247.90082 |
| MGWTGGGLGK | 10 | O15226 | NF-kappa-B-repressing factor OS=Homo | NKRF | 77.673 | 2 | 51.347385 | yes | 3.307E-07 | 482.23947 | 243.28891 |
| KEDLLDQLK | 9 | O15226 | NF-kappa-B-repressing factor OS=Homo | NKRF | 77.673 | 2 | 51.347385 | yes | 1.283E-07 | 551.31116 | 196.28375 |
| YEMMPNR | 7 | O15226 | NF-kappa-B-repressing factor OS=Homo | NKRF | 77.673 | 2 | 51.347385 | yes | 1.166E-07 | 470.70441 | 135.66434 |
| QEDPYGPQTK | 10 | O15226 | NF-kappa-B-repressing factor OS=Homo | NKRF | 77.673 | 2 | 51.347385 | yes | 9.722E-08 | 581.7724 | 123.13966 |
| DASGQPIFNASAK | 13 | O15226 | NF-kappa-B-repressing factor OS=Homo | NKRF | 77.673 | 2 | 51.347385 | yes | 2.643E-07 | 653.32532 | 95.14019 |

| | | | | | | | | | | | |
|---|---|---|---|---|---|---|---|---|---|---|---|
| EDLLDQLK | 8 | O15226 | NF-kappa-B-repressing factor OS=Homo | NKRF | 77.673 | 2 | 51.347385 | yes | 2.022E-06 | 487.26367 | 77.362923 |
| KTQPTVINNLK | 11 | O15226 | NF-kappa-B-repressing factor OS=Homo | NKRF | 77.673 | 2 | 51.347385 | yes | 1.314E-07 | 628.37207 | 62.293224 |
| EVNEQTHFASMPF | 13 | O15226 | NF-kappa-B-repressing factor OS=Homo | NKRF | 77.673 | 2 | 51.347385 | yes | 9.722E-08 | 773.35938 | 60.233757 |
| KGAVEDVISR | 10 | O15226 | NF-kappa-B-repressing factor OS=Homo | NKRF | 77.673 | 2 | 51.347385 | yes | 3.341E-07 | 537.30109 | 51.359207 |
| SGEGIREPISVK | 12 | O15226 | NF-kappa-B-repressing factor OS=Homo | NKRF | 77.673 | 2 | 51.347385 | yes | 5.277E-07 | 636.35132 | 40.352333 |
| VILESEVIAEAVGV | 15 | O15226 | NF-kappa-B-repressing factor OS=Homo | NKRF | 77.673 | 2 | 51.347385 | yes | 7.474E-07 | 778.45068 | 16.439863 |
| EEVSEILDEMSHK | 13 | O15228 | Dihydroxyacetone phosphate acyltransfera | GNPAT | 77.188 | 2 | 52.261436 | yes | 9.722E-08 | 773.3587 | 77.509979 |
| AIQEHPVVLLPSHI | 14 | O15228 | Dihydroxyacetone phosphate acyltransfera | GNPAT | 77.188 | 3 | 52.261436 | yes | 2.024E-07 | 532.63873 | 69.122452 |
| NQLLNIFVR | 9 | O15228 | Dihydroxyacetone phosphate acyltransfera | GNPAT | 77.188 | 2 | 52.261436 | yes | 3.311E-07 | 558.82983 | 37.452251 |
| GALDQLCGAGGLC | 14 | O15230 | Laminin subunit alpha-5 OS=Homo sapie | LAMA5 | 399.738 | 2 | 30.836817 | yes | 2.044E-06 | 724.34265 | 40.512444 |
| GLFFEHDLR | 9 | O15234 | Protein CASC3 OS=Homo sapiens OX=9 | CASC3 | 76.278 | 2 | 56.03233 | yes | 1.354E-07 | 567.29053 | 10737.906 |
| QELIALYGYDIR | 12 | O15234 | Protein CASC3 OS=Homo sapiens OX=9 | CASC3 | 76.278 | 2 | 56.03233 | yes | 2.931E-07 | 727.38794 | 6079.7617 |
| KNPAYIPR | 8 | O15234 | Protein CASC3 OS=Homo sapiens OX=9 | CASC3 | 76.278 | 2 | 56.03233 | yes | 3.136E-07 | 479.77707 | 5213.3105 |
| MEEMGVQGGR | 10 | O15234 | Protein CASC3 OS=Homo sapiens OX=9 | CASC3 | 76.278 | 2 | 56.03233 | yes | 1.15E-07 | 547.24188 | 2587.3154 |
| MEEMGVQGGR | 10 | O15234 | Protein CASC3 OS=Homo sapiens OX=9 | CASC3 | 76.278 | 2 | 56.03233 | yes | 2.03E-07 | 555.23932 | 2587.3154 |
| HLDDDEDR | 8 | O15234 | Protein CASC3 OS=Homo sapiens OX=9 | CASC3 | 76.278 | 2 | 56.03233 | yes | 4.701E-07 | 507.70978 | 1690.2048 |
| GQTQEEEVRPK | 11 | O15234 | Protein CASC3 OS=Homo sapiens OX=9 | CASC3 | 76.278 | 2 | 56.03233 | yes | 1.468E-07 | 650.82825 | 1459.6906 |
| AGFRPVEAGGQHC | 15 | O15234 | Protein CASC3 OS=Homo sapiens OX=9 | CASC3 | 76.278 | 3 | 56.03233 | yes | 4.718E-07 | 499.25528 | 1402.257 |
| KGLFFEHDLR | 10 | O15234 | Protein CASC3 OS=Homo sapiens OX=9 | CASC3 | 76.278 | 2 | 56.03233 | yes | 9.722E-08 | 631.33801 | 422.91605 |
| KGLFFEHDLR | 10 | O15234 | Protein CASC3 OS=Homo sapiens OX=9 | CASC3 | 76.278 | 2 | 56.03233 | yes | 3.637E-07 | 421.22778 | 422.91605 |
| KGLFFEHDLR | 10 | O15234 | Protein CASC3 OS=Homo sapiens OX=9 | CASC3 | 76.278 | 3 | 56.03233 | yes | 9.722E-08 | 631.33801 | 422.91605 |
| KGLFFEHDLR | 10 | O15234 | Protein CASC3 OS=Homo sapiens OX=9 | CASC3 | 76.278 | 3 | 56.03233 | yes | 3.637E-07 | 421.22778 | 422.91605 |
| HQGLGGTLPPR | 11 | O15234 | Protein CASC3 OS=Homo sapiens OX=9 | CASC3 | 76.278 | 2 | 56.03233 | yes | 3.286E-06 | 566.8147 | 820.01129 |
| GMPNHIHMGAGPI | 18 | O15234 | Protein CASC3 OS=Homo sapiens OX=9 | CASC3 | 76.278 | 4 | 56.03233 | yes | 8.714E-07 | 490.23727 | 584.5119 |
| LEQTSVR | 7 | O15234 | Protein CASC3 OS=Homo sapiens OX=9 | CASC3 | 76.278 | 2 | 56.03233 | yes | 2.215E-07 | 416.7298 | 543.21173 |
| SAHNPDDIKPR | 11 | O15234 | Protein CASC3 OS=Homo sapiens OX=9 | CASC3 | 76.278 | 2 | 56.03233 | yes | 1.822E-06 | 625.31781 | 504.28177 |
| NPAYIPR | 7 | O15234 | Protein CASC3 OS=Homo sapiens OX=9 | CASC3 | 76.278 | 2 | 56.03233 | yes | 5.412E-07 | 415.72958 | 291.29282 |
| RTPQPVTIKPPPPE | 18 | O15234 | Protein CASC3 OS=Homo sapiens OX=9 | CASC3 | 76.278 | 4 | 56.03233 | yes | 3.334E-07 | 500.29282 | 240.7388 |
| FREDEQAPK | 9 | O15234 | Protein CASC3 OS=Homo sapiens OX=9 | CASC3 | 76.278 | 2 | 56.03233 | yes | 5.453E-07 | 560.27509 | 187.5139 |
| QSGDGQESTEPVE | 15 | O15234 | Protein CASC3 OS=Homo sapiens OX=9 | CASC3 | 76.278 | 2 | 56.03233 | yes | 1.156E-07 | 802.85535 | 172.54475 |
| GQTQEEEVR | 9 | O15234 | Protein CASC3 OS=Homo sapiens OX=9 | CASC3 | 76.278 | 2 | 56.03233 | yes | 2.592E-06 | 538.25433 | 126.05449 |
| WEHDKFR | 7 | O15234 | Protein CASC3 OS=Homo sapiens OX=9 | CASC3 | 76.278 | 2 | 56.03233 | yes | 2.357E-06 | 509.24869 | 54.827682 |
| GGGSCSGSAGGGC | 21 | O15234 | Protein CASC3 OS=Homo sapiens OX=9 | CASC3 | 76.278 | 3 | 56.03233 | yes | 9.767E-08 | 593.92737 | 40.923103 |
| DPSPEADAPVLGSI | 16 | O15234 | Protein CASC3 OS=Homo sapiens OX=9 | CASC3 | 76.278 | 2 | 56.03233 | yes | 9.722E-08 | 804.89124 | 28.289429 |
| LGQIYQSWLDK | 11 | O15258 | Protein RER1 OS=Homo sapiens OX=96 | RER1 | 22.958 | 2 | 52.782753 | yes | 1.162E-07 | 675.85626 | 426.39435 |
| SEGDSVGESVHGK | 19 | O15258 | Protein RER1 OS=Homo sapiens OX=96 | RER1 | 22.958 | 3 | 52.782753 | yes | 9.722E-08 | 677.66412 | 55.871666 |
| SEGDSVGESVHGK | 13 | O15258 | Protein RER1 OS=Homo sapiens OX=96 | RER1 | 22.958 | 2 | 52.782753 | yes | 9.722E-08 | 665.2995 | 39.310974 |
| FDVLLAEHK | 9 | O15265 | Ataxin-7 OS=Homo sapiens OX=9606 GI | ATXN7 | 95.452 | 2 | 31.296106 | yes | 1.889E-06 | 536.29529 | 53.720921 |
| SFVIDHQR | 8 | O15269 | Serine palmitoyltransferase 1 OS=Homo s | SPTLC1 | 52.744 | 2 | 51.980556 | yes | 1.346E-07 | 501.26178 | 268.05792 |

| | | | | | | | | | | | |
|---|---|---|---|---|---|---|---|---|---|---|---|
| VVVTVEQTEEELE | 14 | O15269 | Serine palmitoyltransferase 1 OS=Homo s | SPTLC1 | 52.744 | 2 | 51.980556 | yes | 1.142E-07 | 830.42542 | 261.15897 |
| LLQEIVDQCMNR | 12 | O15269 | Serine palmitoyltransferase 1 OS=Homo s | SPTLC1 | 52.744 | 2 | 51.980556 | yes | 1.266E-07 | 759.87378 | 193.14935 |
| HNDMADLER | 9 | O15269 | Serine palmitoyltransferase 1 OS=Homo s | SPTLC1 | 52.744 | 2 | 51.980556 | yes | 9.722E-08 | 550.7431 | 114.90854 |
| RGDIVFVDR | 9 | O15269 | Serine palmitoyltransferase 1 OS=Homo s | SPTLC1 | 52.744 | 2 | 51.980556 | yes | 3.843E-07 | 538.79602 | 68.581467 |
| ALQGISGLK | 9 | O15269 | Serine palmitoyltransferase 1 OS=Homo s | SPTLC1 | 52.744 | 2 | 51.980556 | yes | 4.398E-06 | 443.77145 | 21.413725 |
| FCLSAAHTK | 9 | O15270 | Serine palmitoyltransferase 2 OS=Homo s | SPTLC2 | 62.925 | 2 | 52.18946 | yes | 2.801E-07 | 517.75806 | 96.111771 |
| VLEEYGAGVCSTR | 13 | O15270 | Serine palmitoyltransferase 2 OS=Homo s | SPTLC2 | 62.925 | 2 | 52.18946 | yes | 1.327E-07 | 720.84302 | 93.688927 |
| EIDEVGDLLQLK | 12 | O15270 | Serine palmitoyltransferase 2 OS=Homo s | SPTLC2 | 62.925 | 2 | 52.18946 | yes | 9.722E-08 | 686.37195 | 83.884827 |
| ECVQQLAENTR | 11 | O15270 | Serine palmitoyltransferase 2 OS=Homo s | SPTLC2 | 62.925 | 2 | 52.18946 | yes | 9.953E-08 | 674.3197 | 82.219215 |
| DSGNIPEAIASYR | 13 | O15294 | UDP-N-acetylglucosamine--peptide N-ac | OGT | 116.925 | 2 | 52.95161 | yes | 3.409E-07 | 696.84131 | 929.10291 |
| LVSIVADQLEK | 11 | O15294 | UDP-N-acetylglucosamine--peptide N-ac | OGT | 116.925 | 2 | 52.95161 | yes | 9.722E-08 | 607.85315 | 670.65002 |
| EQGNIEEAVR | 10 | O15294 | UDP-N-acetylglucosamine--peptide N-ac | OGT | 116.925 | 2 | 52.95161 | yes | 1.658E-07 | 572.78326 | 460.67093 |
| SAHFSTLAIK | 10 | O15294 | UDP-N-acetylglucosamine--peptide N-ac | OGT | 116.925 | 2 | 52.95161 | yes | 1.163E-06 | 537.80072 | 322.89435 |
| GQLQEAIEHYR | 11 | O15294 | UDP-N-acetylglucosamine--peptide N-ac | OGT | 116.925 | 2 | 52.95161 | yes | 1.16E-07 | 672.33875 | 290.5892 |
| LQEALMHYK | 9 | O15294 | UDP-N-acetylglucosamine--peptide N-ac | OGT | 116.925 | 2 | 52.95161 | yes | 1.01E-07 | 566.79462 | 259.43228 |
| HCMQLWR | 7 | O15294 | UDP-N-acetylglucosamine--peptide N-ac | OGT | 116.925 | 2 | 52.95161 | yes | 1.187E-07 | 515.73914 | 228.61513 |
| IIFSPVAPK | 9 | O15294 | UDP-N-acetylglucosamine--peptide N-ac | OGT | 116.925 | 2 | 52.95161 | yes | 4.166E-07 | 486.29987 | 202.75385 |
| EYQAGDFEAAER | 12 | O15294 | UDP-N-acetylglucosamine--peptide N-ac | OGT | 116.925 | 2 | 52.95161 | yes | 9.722E-08 | 693.30206 | 186.07724 |
| QYTMELER | 8 | O15294 | UDP-N-acetylglucosamine--peptide N-ac | OGT | 116.925 | 2 | 52.95161 | yes | 9.722E-08 | 535.25275 | 153.92474 |
| ASSVGNVADSTEP | 16 | O15294 | UDP-N-acetylglucosamine--peptide N-ac | OGT | 116.925 | 2 | 52.95161 | yes | 1.037E-07 | 830.91046 | 111.27998 |
| LGTDLEYLK | 9 | O15294 | UDP-N-acetylglucosamine--peptide N-ac | OGT | 116.925 | 2 | 52.95161 | yes | 1.155E-07 | 526.28711 | 107.87189 |
| INVLHKPPYEHPK | 13 | O15294 | UDP-N-acetylglucosamine--peptide N-ac | OGT | 116.925 | 3 | 52.95161 | yes | 1.348E-06 | 524.62793 | 49.667038 |
| INVLHKPPYEHPK | 13 | O15294 | UDP-N-acetylglucosamine--peptide N-ac | OGT | 116.925 | 3 | 52.95161 | yes | 2.902E-06 | 786.43829 | 49.667038 |
| INVLHKPPYEHPK | 13 | O15294 | UDP-N-acetylglucosamine--peptide N-ac | OGT | 116.925 | 2 | 52.95161 | yes | 1.348E-06 | 524.62793 | 49.667038 |
| INVLHKPPYEHPK | 13 | O15294 | UDP-N-acetylglucosamine--peptide N-ac | OGT | 116.925 | 2 | 52.95161 | yes | 2.902E-06 | 786.43829 | 49.667038 |
| ISSPLFNTK | 9 | O15294 | UDP-N-acetylglucosamine--peptide N-ac | OGT | 116.925 | 2 | 52.95161 | yes | 2.699E-07 | 503.78201 | 99.002541 |
| AFLDSLPDVK | 10 | O15294 | UDP-N-acetylglucosamine--peptide N-ac | OGT | 116.925 | 2 | 52.95161 | yes | 4.986E-07 | 552.80042 | 54.23806 |
| ASSVGNVADSTEP | 15 | O15294 | UDP-N-acetylglucosamine--peptide N-ac | OGT | 116.925 | 2 | 52.95161 | yes | 1.073E-06 | 752.85992 | 46.980743 |
| NRQEYEDIAVK | 11 | O15294 | UDP-N-acetylglucosamine--peptide N-ac | OGT | 116.925 | 2 | 52.95161 | yes | 2.237E-07 | 682.84387 | 39.155872 |
| GQMLIGPDGR | 10 | O15304 | Apoptosis regulatory protein Siva OS=Hc | SIVA1 | 18.695 | 2 | 52.02885 | yes | 9.722E-08 | 522.26874 | 359.52695 |
| SCPFADVAPLQLK | 13 | O15304 | Apoptosis regulatory protein Siva OS=Hc | SIVA1 | 18.695 | 2 | 52.02885 | yes | 9.85E-08 | 723.37646 | 170.08099 |
| ALCGQCVR | 8 | O15304 | Apoptosis regulatory protein Siva OS=Hc | SIVA1 | 18.695 | 2 | 52.02885 | yes | 9.853E-08 | 482.22858 | 101.55502 |
| GTPLTNTEDVLK | 12 | O15318 | DNA-directed RNA polymerase III subun | POLR3G | 25.915 | 2 | 46.751 | yes | 2.224E-07 | 644.3432 | 144.66966 |
| TGEGEEYMLALK | 12 | O15318 | DNA-directed RNA polymerase III subun | POLR3G | 25.915 | 2 | 46.751 | yes | 1.479E-07 | 670.82373 | 43.641167 |
| EIPIEPAPHLR | 11 | O15344 | E3 ubiquitin-protein ligase Midline-1 OS | MID1 | 75.251 | 2 | 49.55727 | yes | 1.117E-07 | 636.35895 | 98.627258 |
| NTELETLLAK | 10 | O15344 | E3 ubiquitin-protein ligase Midline-1 OS | MID1 | 75.251 | 2 | 49.55727 | yes | 1.021E-07 | 566.31641 | 70.436729 |
| AFDANTMTSAEK | 12 | O15344 | E3 ubiquitin-protein ligase Midline-1 OS | MID1 | 75.251 | 2 | 49.55727 | yes | 4.665E-07 | 643.2901 | 54.704067 |
| LAQQIANCK | 9 | O15344 | E3 ubiquitin-protein ligase Midline-1 OS | MID1 | 75.251 | 2 | 49.55727 | yes | 3.936E-06 | 523.27661 | 54.212143 |
| ASVSGPNSPSETR | 13 | O15344 | E3 ubiquitin-protein ligase Midline-1 OS | MID1 | 75.251 | 2 | 49.55727 | yes | 2.598E-06 | 644.81006 | 24.446236 |

| | | | | | | | | | | | |
|---|---|---|---|---|---|---|---|---|---|---|---|
| QNLESNLTNLIK | 12 | O15344 | E3 ubiquitin-protein ligase Midline-1 OS | MID1 | 75.251 | 2 | 49.55727 | yes | 2.537E-07 | 693.883 | 17.318529 |
| AYTGFSSNSER | 11 | O15355 | Protein phosphatase 1G OS=Homo sapier | PPM1G | 59.272 | 2 | 48.36894 | yes | 2.147E-07 | 609.77289 | 126.63875 |
| AIGDHFYK | 8 | O15355 | Protein phosphatase 1G OS=Homo sapier | PPM1G | 59.272 | 2 | 48.36894 | yes | 5.031E-06 | 475.74017 | 107.55044 |
| ALDMSYDHKPEDI | 18 | O15355 | Protein phosphatase 1G OS=Homo sapier | PPM1G | 59.272 | 3 | 48.36894 | yes | 4.112E-07 | 706.6651 | 101.87267 |
| ALEDAFLAIDAK | 12 | O15355 | Protein phosphatase 1G OS=Homo sapier | PPM1G | 59.272 | 2 | 48.36894 | yes | 1.157E-07 | 638.84283 | 57.459148 |
| ELAQIAGRPTEDEI | 18 | O15355 | Protein phosphatase 1G OS=Homo sapier | PPM1G | 59.272 | 3 | 48.36894 | yes | 3.176E-07 | 686.67102 | 36.45821 |
| ASGLGEAGMSAW | 14 | O15357 | Phosphatidylinositol 3,4,5-trisphosphate : | INPPL1 | 138.599 | 2 | 48.236053 | yes | 1.198E-07 | 703.34827 | 71.810844 |
| TLSEVDYAPAGPA | 14 | O15357 | Phosphatidylinositol 3,4,5-trisphosphate : | INPPL1 | 138.599 | 2 | 48.236053 | yes | 1.175E-07 | 723.86481 | 68.538879 |
| VLSGLEILSK | 10 | O15357 | Phosphatidylinositol 3,4,5-trisphosphate : | INPPL1 | 138.599 | 2 | 48.236053 | yes | 8.774E-07 | 529.82642 | 61.147713 |
| DTLCSCLSPGLIK | 13 | O15360 | Fanconi anemia group A protein OS=Hor | FANCA | 162.776 | 2 | 46.206673 | yes | 1.985E-06 | 732.3653 | 42.659576 |
| GLLNACLR | 8 | O15360 | Fanconi anemia group A protein OS=Hor | FANCA | 162.776 | 2 | 46.206673 | yes | 1.622E-07 | 458.75528 | 41.640938 |
| VAFIESLK | 8 | O15360 | Fanconi anemia group A protein OS=Hor | FANCA | 162.776 | 2 | 46.206673 | yes | 2.946E-06 | 453.76837 | 24.462482 |
| SVYSWDIVVQR | 11 | O15371 | Eukaryotic translation initiation factor 3 s | EIF3D | 63.973 | 2 | 53.41931 | yes | 9.927E-08 | 676.35388 | 3321.3796 |
| EEMDFPQLMK | 10 | O15371 | Eukaryotic translation initiation factor 3 s | EIF3D | 63.973 | 2 | 53.41931 | yes | 9.975E-08 | 634.28851 | 3128.7849 |
| WTCCALLAGSEYI | 14 | O15371 | Eukaryotic translation initiation factor 3 s | EIF3D | 63.973 | 2 | 53.41931 | yes | 1.108E-07 | 836.39709 | 1629.0684 |
| IFHTVTTTDDPVIR | 14 | O15371 | Eukaryotic translation initiation factor 3 s | EIF3D | 63.973 | 2 | 53.41931 | yes | 1.095E-07 | 807.92792 | 1426.0299 |
| TLNEWDSR | 8 | O15371 | Eukaryotic translation initiation factor 3 s | EIF3D | 63.973 | 2 | 53.41931 | yes | 1.42E-07 | 510.74088 | 1113.0347 |
| VADWTGATYQDK | 12 | O15371 | Eukaryotic translation initiation factor 3 s | EIF3D | 63.973 | 2 | 53.41931 | yes | 9.722E-08 | 677.81732 | 1034.6915 |
| CVIDICMK | 8 | O15371 | Eukaryotic translation initiation factor 3 s | EIF3D | 63.973 | 2 | 53.41931 | yes | 1.398E-07 | 519.74268 | 418.65936 |
| GAVIATELK | 9 | O15371 | Eukaryotic translation initiation factor 3 s | EIF3D | 63.973 | 2 | 53.41931 | yes | 1.223E-07 | 451.2713 | 264.78195 |
| HVILGTQQFK | 10 | O15371 | Eukaryotic translation initiation factor 3 s | EIF3D | 63.973 | 2 | 53.41931 | yes | 9.722E-08 | 585.83514 | 178.90829 |
| HCNGVDWR | 8 | O15371 | Eukaryotic translation initiation factor 3 s | EIF3D | 63.973 | 2 | 53.41931 | yes | 3.929E-06 | 522.22742 | 92.708435 |
| NMLQFNLQILPK | 12 | O15371 | Eukaryotic translation initiation factor 3 s | EIF3D | 63.973 | 2 | 53.41931 | yes | 1.847E-07 | 729.91052 | 70.153458 |
| NLAMEATYINHNI | 19 | O15371 | Eukaryotic translation initiation factor 3 s | EIF3D | 63.973 | 3 | 53.41931 | yes | 2.138E-07 | 770.69666 | 60.972149 |
| GEPPLPEEDLSK | 12 | O15372 | Eukaryotic translation initiation factor 3 s | EIF3H | 39.931 | 2 | 58.159416 | yes | 1.214E-07 | 655.82733 | 2701.5688 |
| HELLSLASSNHLGI | 14 | O15372 | Eukaryotic translation initiation factor 3 s | EIF3H | 39.931 | 3 | 58.159416 | yes | 1.415E-07 | 502.60721 | 871.32013 |
| HELLSLASSNHLGI | 14 | O15372 | Eukaryotic translation initiation factor 3 s | EIF3H | 39.931 | 3 | 58.159416 | yes | 9.722E-08 | 753.40717 | 871.32013 |
| HELLSLASSNHLGI | 14 | O15372 | Eukaryotic translation initiation factor 3 s | EIF3H | 39.931 | 2 | 58.159416 | yes | 1.415E-07 | 502.60721 | 871.32013 |
| HELLSLASSNHLGI | 14 | O15372 | Eukaryotic translation initiation factor 3 s | EIF3H | 39.931 | 2 | 58.159416 | yes | 9.722E-08 | 753.40717 | 871.32013 |
| EFTAQNLGK | 9 | O15372 | Eukaryotic translation initiation factor 3 s | EIF3H | 39.931 | 2 | 58.159416 | yes | 1.615E-07 | 504.26144 | 1350.3616 |
| QVQIDGLVVLK | 11 | O15372 | Eukaryotic translation initiation factor 3 s | EIF3H | 39.931 | 2 | 58.159416 | yes | 1.043E-07 | 606.37152 | 1037.9557 |
| SAVADKHELLSLA | 20 | O15372 | Eukaryotic translation initiation factor 3 s | EIF3H | 39.931 | 3 | 58.159416 | yes | 1.127E-07 | 693.03943 | 427.66342 |
| SAVADKHELLSLA | 20 | O15372 | Eukaryotic translation initiation factor 3 s | EIF3H | 39.931 | 3 | 58.159416 | yes | 1.344E-07 | 520.03137 | 427.66342 |
| SAVADKHELLSLA | 20 | O15372 | Eukaryotic translation initiation factor 3 s | EIF3H | 39.931 | 4 | 58.159416 | yes | 1.127E-07 | 693.03943 | 427.66342 |
| SAVADKHELLSLA | 20 | O15372 | Eukaryotic translation initiation factor 3 s | EIF3H | 39.931 | 4 | 58.159416 | yes | 1.344E-07 | 520.03137 | 427.66342 |
| VDEMSQDIVK | 10 | O15372 | Eukaryotic translation initiation factor 3 s | EIF3H | 39.931 | 2 | 58.159416 | yes | 4.616E-07 | 582.28424 | 733.75409 |
| LFKPPQPPAR | 10 | O15372 | Eukaryotic translation initiation factor 3 s | EIF3H | 39.931 | 2 | 58.159416 | yes | 3.977E-07 | 575.84021 | 583.948 |
| NLQLLMDR | 8 | O15372 | Eukaryotic translation initiation factor 3 s | EIF3H | 39.931 | 2 | 58.159416 | yes | 3.784E-07 | 501.77368 | 482.29608 |
| LFMAQALQEYNN | 12 | O15372 | Eukaryotic translation initiation factor 3 s | EIF3H | 39.931 | 2 | 58.159416 | yes | 9.722E-08 | 721.34265 | 391.42584 |

| | | | | | | | | | | | |
|---|---|---|---|---|---|---|---|---|---|---|---|
| EKDFSPEALK | 10 | O15372 | Eukaryotic translation initiation factor 3 s | EIF3H | 39.931 | 2 | 58.159416 | yes | 9.722E-08 | 582.30078 | 274.27509 |
| TAQGSLSLK | 9 | O15372 | Eukaryotic translation initiation factor 3 s | EIF3H | 39.931 | 2 | 58.159416 | yes | 2.067E-07 | 452.75854 | 173.69203 |
| DFSPEALKK | 9 | O15372 | Eukaryotic translation initiation factor 3 s | EIF3H | 39.931 | 2 | 58.159416 | yes | 1.421E-06 | 517.77948 | 171.8163 |
| HVNIDHLHVGWY | 23 | O15372 | Eukaryotic translation initiation factor 3 s | EIF3H | 39.931 | 4 | 58.159416 | yes | 3.091E-06 | 695.58984 | 66.531235 |
| EGTGSTATSSSSTA | 19 | O15372 | Eukaryotic translation initiation factor 3 s | EIF3H | 39.931 | 2 | 58.159416 | yes | 0.0001034 | 814.3739 | 1.6742973 |
| QYLDQIR | 7 | O15379 | Histone deacetylase 3 OS=Homo sapiens | HDAC3 | 48.848 | 2 | 38.113937 | yes | 4.023E-05 | 468.25089 | 66.207886 |
| LGCFNLSIR | 9 | O15379 | Histone deacetylase 3 OS=Homo sapiens | HDAC3 | 48.848 | 2 | 38.113937 | yes | 5.45E-07 | 540.28693 | 53.665363 |
| YVDIGVLASDLQR | 13 | O15381 | Nuclear valosin-containing protein-like O | NVL | 95.051 | 2 | 54.214695 | yes | 7.999E-07 | 724.89081 | 164.91306 |
| FEDVGGNDMTLK | 12 | O15381 | Nuclear valosin-containing protein-like O | NVL | 95.051 | 2 | 54.214695 | yes | 3.904E-07 | 663.30573 | 129.1282 |
| VVNQLLTEMDGLI | 15 | O15381 | Nuclear valosin-containing protein-like O | NVL | 95.051 | 2 | 54.214695 | yes | 9.722E-08 | 844.43781 | 126.41909 |
| NLTELEDEHLAK | 12 | O15381 | Nuclear valosin-containing protein-like O | NVL | 95.051 | 2 | 54.214695 | yes | 1.363E-07 | 706.35681 | 110.41573 |
| VFSIISSEK | 9 | O15381 | Nuclear valosin-containing protein-like O | NVL | 95.051 | 2 | 54.214695 | yes | 9.722E-08 | 505.28186 | 80.631821 |
| VAAPEIVSGVSGES | 17 | O15381 | Nuclear valosin-containing protein-like O | NVL | 95.051 | 2 | 54.214695 | yes | 2.325E-06 | 843.93085 | 73.95591 |
| AVANESGLNFISVI | 14 | O15381 | Nuclear valosin-containing protein-like O | NVL | 95.051 | 2 | 54.214695 | yes | 1.074E-07 | 724.89081 | 51.27737 |
| KPITEIQDSK | 10 | O15381 | Nuclear valosin-containing protein-like O | NVL | 95.051 | 2 | 54.214695 | yes | 4.179E-06 | 579.8219 | 51.262665 |
| EELTMAILAPVR | 12 | O15381 | Nuclear valosin-containing protein-like O | NVL | 95.051 | 2 | 54.214695 | yes | 1.187E-06 | 671.8736 | 35.541046 |
| KLPPGGLPGIDLSD | 16 | O15391 | Transcription factor YY2 OS=Homo sapi | YY2 | 41.348 | 2 | 31.861898 | yes | 1.701E-06 | 802.45636 | 258.4491 |
| HSSGCAFLSVK | 11 | O15392 | Baculoviral IAP repeat-containing protein | BIRC5 | 16.389 | 2 | 48.18798 | yes | 1.185E-07 | 596.7926 | 111.50111 |
| QFEELTLGEFLK | 12 | O15392 | Baculoviral IAP repeat-containing protein | BIRC5 | 16.389 | 2 | 48.18798 | yes | 1.796E-07 | 727.38232 | 57.337055 |
| KQFEELTLGEFLK | 13 | O15392 | Baculoviral IAP repeat-containing protein | BIRC5 | 16.389 | 2 | 48.18798 | yes | 3.768E-07 | 791.42981 | 25.378799 |
| IINFAPSLLR | 10 | O15397 | Importin-8 OS=Homo sapiens OX=9606 | IPO8 | 119.939 | 2 | 45.333046 | yes | 4.361E-07 | 572.34784 | 66.021019 |
| IAAENELNQSYK | 12 | O15397 | Importin-8 OS=Homo sapiens OX=9606 | IPO8 | 119.939 | 2 | 45.333046 | yes | 2.254E-07 | 690.34369 | 46.422493 |
| ETENDDVTNVIQK | 13 | O15397 | Importin-8 OS=Homo sapiens OX=9606 | IPO8 | 119.939 | 2 | 45.333046 | yes | 2.032E-06 | 752.85992 | 25.85568 |
| TYAVGIQQVLLK | 12 | O15397 | Importin-8 OS=Homo sapiens OX=9606 | IPO8 | 119.939 | 2 | 45.333046 | yes | 1.881E-07 | 666.89795 | 22.700335 |
| GGAVVDEGPTGVI | 13 | O15427 | Monocarboxylate transporter 4 OS=Homo | SLC16A3 | 49.47 | 2 | 50.24099 | yes | 1.21E-07 | 593.30914 | 851.54181 |
| PAAGFVAGLGK | 11 | O15427 | Monocarboxylate transporter 4 OS=Homo | SLC16A3 | 49.47 | 2 | 50.24099 | yes | 9.839E-08 | 494.28473 | 559.03113 |
| LHKPPADSGVDLR | 13 | O15427 | Monocarboxylate transporter 4 OS=Homo | SLC16A3 | 49.47 | 2 | 50.24099 | yes | 4.164E-07 | 702.88336 | 242.5625 |
| LHKPPADSGVDLR | 13 | O15427 | Monocarboxylate transporter 4 OS=Homo | SLC16A3 | 49.47 | 2 | 50.24099 | yes | 3.784E-07 | 468.92465 | 242.5625 |
| LHKPPADSGVDLR | 13 | O15427 | Monocarboxylate transporter 4 OS=Homo | SLC16A3 | 49.47 | 3 | 50.24099 | yes | 4.164E-07 | 702.88336 | 242.5625 |
| LHKPPADSGVDLR | 13 | O15427 | Monocarboxylate transporter 4 OS=Homo | SLC16A3 | 49.47 | 3 | 50.24099 | yes | 3.784E-07 | 468.92465 | 242.5625 |
| EVEHFLK | 7 | O15427 | Monocarboxylate transporter 4 OS=Homo | SLC16A3 | 49.47 | 2 | 50.24099 | yes | 1.747E-07 | 451.24252 | 282.97928 |
| EPQPEVAAAEEEK | 13 | O15427 | Monocarboxylate transporter 4 OS=Homo | SLC16A3 | 49.47 | 2 | 50.24099 | yes | 1.068E-07 | 713.83844 | 234.8266 |
| PLVVTAQPGSGPPI | 17 | O15427 | Monocarboxylate transporter 4 OS=Homo | SLC16A3 | 49.47 | 3 | 50.24099 | yes | 1.161E-07 | 572.65656 | 108.1145 |
| LLDLSVFR | 8 | O15427 | Monocarboxylate transporter 4 OS=Homo | SLC16A3 | 49.47 | 2 | 50.24099 | yes | 1.02E-07 | 481.78711 | 73.416557 |
| PLVVTAQPGSGPPI | 14 | O15427 | Monocarboxylate transporter 4 OS=Homo | SLC16A3 | 49.47 | 2 | 50.24099 | yes | 6.317E-07 | 688.38824 | 22.058258 |
| TLSLEAPAR | 9 | O15440 | ATP-binding cassette sub-family C memb | ABCC5 | 160.661 | 2 | 46.380676 | yes | 3.23E-07 | 479.27182 | 90.714462 |
| FFDTTPTGR | 9 | O15440 | ATP-binding cassette sub-family C memb | ABCC5 | 160.661 | 2 | 46.380676 | no | 1.575E-07 | 521.2536 | 82.174805 |
| ECIAQLPLK | 9 | O15440 | ATP-binding cassette sub-family C memb | ABCC5 | 160.661 | 2 | 46.380676 | yes | 8.539E-07 | 536.297 | 73.72654 |
| ISDIGLADLR | 10 | O15440 | ATP-binding cassette sub-family C memb | ABCC5 | 160.661 | 2 | 46.380676 | yes | 2.486E-06 | 536.80347 | 61.467354 |

| | | | | | | | | | | | |
|---|---|---|---|---|---|---|---|---|---|---|---|
| FCAFGGNPPVTGP | 14 | O15446 | DNA-directed RNA polymerase I subunit | POLR1G | 54.986 | 2 | 50.620384 | yes | 9.734E-08 | 738.85626 | 624.85559 |
| HVPLSGSQIVK | 11 | O15446 | DNA-directed RNA polymerase I subunit | POLR1G | 54.986 | 2 | 50.620384 | yes | 2.711E-07 | 582.84039 | 264.1326 |
| SALAPNLLTSGK | 12 | O15446 | DNA-directed RNA polymerase I subunit | POLR1G | 54.986 | 2 | 50.620384 | yes | 2.982E-06 | 586.33771 | 228.28018 |
| VEPLEEAIPLPPTK | 14 | O15446 | DNA-directed RNA polymerase I subunit | POLR1G | 54.986 | 2 | 50.620384 | yes | 7.648E-07 | 766.93213 | 205.83253 |
| SSWLEEDDDPVVA | 14 | O15460 | Prolyl 4-hydroxylase subunit alpha-2 OS= | P4HA2 | 60.902 | 2 | 51.717228 | yes | 1.294E-07 | 809.37317 | 56.329254 |
| LLSLDPSHER | 10 | O15460 | Prolyl 4-hydroxylase subunit alpha-2 OS= | P4HA2 | 60.902 | 2 | 51.717228 | yes | 9.722E-08 | 583.81183 | 49.945827 |
| YYDVMSDEEIER | 12 | O15460 | Prolyl 4-hydroxylase subunit alpha-2 OS= | P4HA2 | 60.902 | 2 | 51.717228 | yes | 4.797E-07 | 774.82977 | 37.278133 |
| SPSEDPLTR | 9 | O15479 | Melanoma-associated antigen B2 OS=Ho | MAGEB2 | 35.278 | 2 | 53.746128 | yes | 5.249E-07 | 501.24854 | 3287.374 |
| FQFLWGPR | 8 | O15479 | Melanoma-associated antigen B2 OS=Ho | MAGEB2 | 35.278 | 2 | 53.746128 | yes | 4.034E-06 | 525.7796 | 1798.6267 |
| QVPSSDPPR | 9 | O15479 | Melanoma-associated antigen B2 OS=Ho | MAGEB2 | 35.278 | 2 | 53.746128 | yes | 9.722E-08 | 491.75125 | 1154.5688 |
| APTTAAAAAAGVS | 16 | O15479 | Melanoma-associated antigen B2 OS=Ho | MAGEB2 | 35.278 | 2 | 53.746128 | yes | 9.722E-08 | 687.86481 | 588.86542 |
| SGSLVQFLLYK | 11 | O15479 | Melanoma-associated antigen B2 OS=Ho | MAGEB2 | 35.278 | 2 | 53.746128 | yes | 1.002E-07 | 627.85828 | 572.81055 |
| EHFPEILK | 8 | O15479 | Melanoma-associated antigen B2 OS=Ho | MAGEB2 | 35.278 | 2 | 53.746128 | yes | 8.266E-07 | 506.77673 | 522.59192 |
| ASEGLSVVFGLELI | 15 | O15479 | Melanoma-associated antigen B2 OS=Ho | MAGEB2 | 35.278 | 2 | 53.746128 | yes | 9.722E-08 | 781.92487 | 437.85974 |
| KSGSLVQFLLYK | 12 | O15479 | Melanoma-associated antigen B2 OS=Ho | MAGEB2 | 35.278 | 2 | 53.746128 | yes | 2.268E-07 | 691.90576 | 94.38382 |
| KASEGLSVVFGLE | 16 | O15479 | Melanoma-associated antigen B2 OS=Ho | MAGEB2 | 35.278 | 3 | 53.746128 | yes | 9.722E-08 | 564.31732 | 34.439091 |
| KASEGLSVVFGLE | 16 | O15479 | Melanoma-associated antigen B2 OS=Ho | MAGEB2 | 35.278 | 3 | 53.746128 | yes | 5.548E-07 | 845.97235 | 34.439091 |
| KASEGLSVVFGLE | 16 | O15479 | Melanoma-associated antigen B2 OS=Ho | MAGEB2 | 35.278 | 2 | 53.746128 | yes | 9.722E-08 | 564.31732 | 34.439091 |
| KASEGLSVVFGLE | 16 | O15479 | Melanoma-associated antigen B2 OS=Ho | MAGEB2 | 35.278 | 2 | 53.746128 | yes | 5.548E-07 | 845.97235 | 34.439091 |
| VNGTTPCAFPTHY | 22 | O15479 | Melanoma-associated antigen B2 OS=Ho | MAGEB2 | 35.278 | 4 | 53.746128 | yes | 4.299E-07 | 634.797 | 47.402943 |
| FREHFPEILK | 10 | O15479 | Melanoma-associated antigen B2 OS=Ho | MAGEB2 | 35.278 | 3 | 53.746128 | yes | 8.519E-07 | 439.24344 | 27.795317 |
| AICQFFLQGR | 10 | O15504 | Nucleoporin NUP42 OS=Homo sapiens ( | NUP42 | 44.872 | 2 | 51.159542 | yes | 1.675E-07 | 620.31879 | 546.82056 |
| YSNVIQPSSFSK | 12 | O15504 | Nucleoporin NUP42 OS=Homo sapiens ( | NUP42 | 44.872 | 2 | 51.159542 | yes | 9.722E-08 | 678.84332 | 176.8817 |
| KLLEGIVK | 8 | O15504 | Nucleoporin NUP42 OS=Homo sapiens ( | NUP42 | 44.872 | 2 | 51.159542 | yes | 6.44E-07 | 450.29987 | 103.89898 |
| DQEKPYFSSFDSG. | 18 | O15504 | Nucleoporin NUP42 OS=Homo sapiens ( | NUP42 | 44.872 | 3 | 51.159542 | yes | 1.055E-07 | 679.30457 | 84.114532 |
| ALAAGGVGSIVR | 12 | O15511 | Actin-related protein 2/3 complex subunit | ARPC5 | 16.321 | 2 | 46.101387 | yes | 6.366E-07 | 535.81946 | 126.21541 |
| QGNMTAALQAAL | 13 | O15511 | Actin-related protein 2/3 complex subunit | ARPC5 | 16.321 | 2 | 46.101387 | yes | 1.653E-07 | 658.85321 | 40.000057 |
| FSQGQQLVTK | 10 | O15516 | Circadian locomoter output cycles protein | CLOCK | 95.304 | 2 | 48.964573 | yes | 6.421E-07 | 568.30896 | 59.429905 |
| ELGIEESLPETAAD | 15 | O15516 | Circadian locomoter output cycles protein | CLOCK | 95.304 | 2 | 48.964573 | yes | 1.083E-07 | 801.39886 | 28.676334 |
| EITAQSDASEIR | 12 | O15516 | Circadian locomoter output cycles protein | CLOCK | 95.304 | 2 | 48.964573 | yes | 0.0003485 | 660.3255 | 27.811108 |
| GKPGYFSER | 9 | O15523 | ATP-dependent RNA helicase DDX3Y O | DDX3Y | 73.154 | 2 | 22.786482 | yes | 3.788E-06 | 520.7616 | 52.053806 |
| VSLMPATLAR | 10 | O15533 | Tapasin OS=Homo sapiens OX=9606 GN | TAPBP | 47.571 | 2 | 46.61124 | yes | 1.126E-06 | 529.80499 | 132.65894 |
| WASGLTPAQNCPF | 13 | O15533 | Tapasin OS=Homo sapiens OX=9606 GN | TAPBP | 47.571 | 2 | 46.61124 | yes | 1.519E-06 | 729.35132 | 52.823685 |
| RPGALLLR | 8 | O15533 | Tapasin OS=Homo sapiens OX=9606 GN | TAPBP | 47.571 | 2 | 46.61124 | yes | 1.515E-07 | 448.29544 | 44.336414 |
| AVDQVCTFLFK | 11 | O15541 | E3 ubiquitin-protein ligase RNF113A OS | RNF113A | 38.788 | 2 | 52.68445 | yes | 9.722E-08 | 664.33936 | 379.79092 |
| SAKPVGPEDMGAT | 22 | O15541 | E3 ubiquitin-protein ligase RNF113A OS | RNF113A | 38.788 | 3 | 52.68445 | yes | 2.577E-07 | 770.03864 | 344.19641 |
| HGWQIER | 7 | O15541 | E3 ubiquitin-protein ligase RNF113A OS | RNF113A | 38.788 | 2 | 52.68445 | yes | 2.641E-07 | 463.23557 | 179.92555 |
| WDYQPDICK | 9 | O15541 | E3 ubiquitin-protein ligase RNF113A OS | RNF113A | 38.788 | 2 | 52.68445 | no | 1.981E-07 | 612.7713 | 159.79648 |
| DTSMGNASSGMV | 13 | O15541 | E3 ubiquitin-protein ligase RNF113A OS | RNF113A | 38.788 | 2 | 52.68445 | yes | 1.778E-07 | 656.78467 | 147.155 |

| | | | | | | | | | | | |
|---|---|---|---|---|---|---|---|---|---|---|---|
| QSFQNPVVTK | 10 | O15541 | E3 ubiquitin-protein ligase RNF113A OS | RNF113A | 38.788 | 2 | 52.68445 | yes | 9.722E-08 | 574.30896 | 128.47984 |
| HYFCESCALQHFR | 13 | O15541 | E3 ubiquitin-protein ligase RNF113A OS | RNF113A | 38.788 | 3 | 52.68445 | yes | 3.329E-07 | 585.58905 | 108.99653 |
| VTHNPMIQK | 9 | O15541 | E3 ubiquitin-protein ligase RNF113A OS | RNF113A | 38.788 | 2 | 52.68445 | yes | 1.542E-07 | 534.28693 | 54.857685 |
| DAQAIFER | 8 | O15541 | E3 ubiquitin-protein ligase RNF113A OS | RNF113A | 38.788 | 2 | 52.68445 | yes | 9.809E-08 | 475.24051 | 25.604027 |
| VESLLVTAISK | 11 | O43143 | ATP-dependent RNA helicase DHX15 O: | DHX15 | 90.933 | 2 | 56.327045 | yes | 1.114E-07 | 580.35028 | 8539.4932 |
| EAMNDPLLER | 10 | O43143 | ATP-dependent RNA helicase DHX15 O: | DHX15 | 90.933 | 2 | 56.327045 | yes | 1.226E-07 | 594.28992 | 2969.9316 |
| EAMNDPLLER | 10 | O43143 | ATP-dependent RNA helicase DHX15 O: | DHX15 | 90.933 | 2 | 56.327045 | yes | 8.938E-07 | 602.28735 | 2969.9316 |
| SNLGSVVLQLK | 11 | O43143 | ATP-dependent RNA helicase DHX15 O: | DHX15 | 90.933 | 2 | 56.327045 | yes | 9.733E-08 | 579.34808 | 5216.7798 |
| EVDDLGPEVGDIK | 13 | O43143 | ATP-dependent RNA helicase DHX15 O: | DHX15 | 90.933 | 2 | 56.327045 | yes | 9.722E-08 | 693.34338 | 4867.7383 |
| LQLPVWEYK | 9 | O43143 | ATP-dependent RNA helicase DHX15 O: | DHX15 | 90.933 | 2 | 56.327045 | yes | 1.398E-07 | 588.3266 | 4802.5479 |
| VIVMSATLDAGK | 12 | O43143 | ATP-dependent RNA helicase DHX15 O: | DHX15 | 90.933 | 2 | 56.327045 | yes | 9.722E-08 | 602.83392 | 4377.9258 |
| TEMQDNTYPEILR | 13 | O43143 | ATP-dependent RNA helicase DHX15 O: | DHX15 | 90.933 | 2 | 56.327045 | yes | 9.823E-08 | 805.37994 | 3721.8936 |
| SLMSADNVR | 9 | O43143 | ATP-dependent RNA helicase DHX15 O: | DHX15 | 90.933 | 2 | 56.327045 | yes | 1.108E-07 | 496.74512 | 1596.9922 |
| SLMSADNVR | 9 | O43143 | ATP-dependent RNA helicase DHX15 O: | DHX15 | 90.933 | 2 | 56.327045 | yes | 3.452E-07 | 504.74258 | 1596.9922 |
| LDLGEDYPSGK | 11 | O43143 | ATP-dependent RNA helicase DHX15 O: | DHX15 | 90.933 | 2 | 56.327045 | yes | 1.003E-07 | 597.28784 | 2555.7798 |
| TCTDIKPEWLVK | 12 | O43143 | ATP-dependent RNA helicase DHX15 O: | DHX15 | 90.933 | 2 | 56.327045 | yes | 4.729E-07 | 745.38959 | 2551.4072 |
| FAHIDGDHLTLLN | 19 | O43143 | ATP-dependent RNA helicase DHX15 O: | DHX15 | 90.933 | 3 | 56.327045 | yes | 9.722E-08 | 737.71637 | 985.58063 |
| FAHIDGDHLTLLN | 19 | O43143 | ATP-dependent RNA helicase DHX15 O: | DHX15 | 90.933 | 3 | 56.327045 | yes | 1.05E-07 | 553.53912 | 985.58063 |
| FAHIDGDHLTLLN | 19 | O43143 | ATP-dependent RNA helicase DHX15 O: | DHX15 | 90.933 | 4 | 56.327045 | yes | 9.722E-08 | 737.71637 | 985.58063 |
| FAHIDGDHLTLLN | 19 | O43143 | ATP-dependent RNA helicase DHX15 O: | DHX15 | 90.933 | 4 | 56.327045 | yes | 1.05E-07 | 553.53912 | 985.58063 |
| HQSFVLVGETGSG | 14 | O43143 | ATP-dependent RNA helicase DHX15 O: | DHX15 | 90.933 | 2 | 56.327045 | yes | 1.209E-07 | 723.3728 | 1493.2231 |
| VAAMSVAQR | 9 | O43143 | ATP-dependent RNA helicase DHX15 O: | DHX15 | 90.933 | 2 | 56.327045 | yes | 1.287E-07 | 466.75275 | 1094.5222 |
| DYLEAAIR | 8 | O43143 | ATP-dependent RNA helicase DHX15 O: | DHX15 | 90.933 | 2 | 56.327045 | yes | 1.128E-07 | 475.75073 | 1048.9238 |
| IFEPPPPK | 8 | O43143 | ATP-dependent RNA helicase DHX15 O: | DHX15 | 90.933 | 2 | 56.327045 | yes | 1.382E-07 | 462.76309 | 926.08478 |
| TGHYLTVK | 8 | O43143 | ATP-dependent RNA helicase DHX15 O: | DHX15 | 90.933 | 2 | 56.327045 | yes | 1.009E-07 | 459.7558 | 882.89063 |
| DYYINIR | 7 | O43143 | ATP-dependent RNA helicase DHX15 O: | DHX15 | 90.933 | 2 | 56.327045 | yes | 1.347E-06 | 478.74545 | 471.55756 |
| ALVTGYFMQVAH | 15 | O43143 | ATP-dependent RNA helicase DHX15 O: | DHX15 | 90.933 | 3 | 56.327045 | yes | 2.112E-07 | 578.97137 | 347.92386 |
| FEDCSSAK | 8 | O43143 | ATP-dependent RNA helicase DHX15 O: | DHX15 | 90.933 | 2 | 56.327045 | yes | 9.722E-08 | 472.19492 | 296.61575 |
| ASTNAMLISAGLPI | 16 | O43143 | ATP-dependent RNA helicase DHX15 O: | DHX15 | 90.933 | 2 | 56.327045 | yes | 9.757E-08 | 792.44489 | 271.45654 |
| RLQLPVWEYK | 10 | O43143 | ATP-dependent RNA helicase DHX15 O: | DHX15 | 90.933 | 2 | 56.327045 | yes | 9.722E-08 | 666.37714 | 168.26025 |
| RGVACTQPR | 9 | O43143 | ATP-dependent RNA helicase DHX15 O: | DHX15 | 90.933 | 2 | 56.327045 | yes | 2.652E-06 | 522.77197 | 116.26637 |
| HRLDLGEDYPSGK | 13 | O43143 | ATP-dependent RNA helicase DHX15 O: | DHX15 | 90.933 | 2 | 56.327045 | yes | 9.722E-08 | 743.86786 | 102.86955 |
| FTDILVR | 7 | O43143 | ATP-dependent RNA helicase DHX15 O: | DHX15 | 90.933 | 2 | 56.327045 | yes | 1.148E-06 | 432.2529 | 93.519348 |
| RVAAMSVAQR | 10 | O43143 | ATP-dependent RNA helicase DHX15 O: | DHX15 | 90.933 | 2 | 56.327045 | yes | 2.214E-07 | 544.80328 | 50.754639 |
| IRVESLLVTAISK | 13 | O43143 | ATP-dependent RNA helicase DHX15 O: | DHX15 | 90.933 | 2 | 56.327045 | yes | 2.38E-06 | 714.94287 | 49.491673 |
| DRFTDILVR | 9 | O43143 | ATP-dependent RNA helicase DHX15 O: | DHX15 | 90.933 | 2 | 56.327045 | yes | 1.441E-07 | 567.81696 | 41.649647 |
| MQALEPYPANESS | 14 | O43148 | mRNA cap guanine-N7 methyltransferase | RNMT | 54.845 | 2 | 51.928482 | yes | 4.235E-07 | 782.8692 | 1502.4301 |
| LVCTDIADVSVK | 12 | O43148 | mRNA cap guanine-N7 methyltransferase | RNMT | 54.845 | 2 | 51.928482 | yes | 9.722E-08 | 660.34741 | 1122.3188 |
| KLDPEIVPEEK | 11 | O43148 | mRNA cap guanine-N7 methyltransferase | RNMT | 54.845 | 2 | 51.928482 | yes | 9.722E-08 | 648.8559 | 739.66174 |

| | | | | | | | | | | | |
|---|---|---|---|---|---|---|---|---|---|---|---|
| SVLIGEFLEK | 10 | O43148 | mRNA cap guanine-N7 methyltransferase | RNMT | 54.845 | 2 | 51.928482 | yes | 1.025E-07 | 567.82391 | 672.03467 |
| IALEDVPEK | 9 | O43148 | mRNA cap guanine-N7 methyltransferase | RNMT | 54.845 | 2 | 51.928482 | yes | 9.968E-08 | 507.27933 | 349.60947 |
| TFLEFYEEK | 9 | O43148 | mRNA cap guanine-N7 methyltransferase | RNMT | 54.845 | 2 | 51.928482 | yes | 9.935E-08 | 603.28986 | 296.94781 |
| KGDYPLFGCK | 10 | O43148 | mRNA cap guanine-N7 methyltransferase | RNMT | 54.845 | 2 | 51.928482 | yes | 9.728E-08 | 592.79205 | 266.83215 |
| VDDYEHAAK | 9 | O43148 | mRNA cap guanine-N7 methyltransferase | RNMT | 54.845 | 2 | 51.928482 | yes | 2.958E-07 | 524.24072 | 178.49196 |
| KIALEDVPEK | 10 | O43148 | mRNA cap guanine-N7 methyltransferase | RNMT | 54.845 | 2 | 51.928482 | yes | 1.425E-07 | 571.32678 | 171.37144 |
| DITVLDLGCGK | 11 | O43148 | mRNA cap guanine-N7 methyltransferase | RNMT | 54.845 | 2 | 51.928482 | yes | 9.729E-08 | 595.80792 | 76.481262 |
| EFEDDLVK | 8 | O43148 | mRNA cap guanine-N7 methyltransferase | RNMT | 54.845 | 2 | 51.928482 | yes | 1.433E-07 | 497.74002 | 66.673058 |
| KEFEDDLVK | 9 | O43148 | mRNA cap guanine-N7 methyltransferase | RNMT | 54.845 | 2 | 51.928482 | yes | 3.265E-06 | 561.78748 | 50.886276 |
| VLKPGGLLK | 9 | O43159 | Ribosomal RNA-processing protein 8 OS | RRP8 | 50.715 | 2 | 52.35274 | yes | 1.603E-07 | 462.81567 | 151.09842 |
| AQLSGLQLQPCLY | 14 | O43159 | Ribosomal RNA-processing protein 8 OS | RRP8 | 50.715 | 2 | 52.35274 | yes | 9.722E-08 | 809.93469 | 86.491837 |
| PAGGYQTITGR | 11 | O43164 | E3 ubiquitin-protein ligase Praja-2 OS=H | PJA2 | 78.215 | 2 | 54.432915 | yes | 2.543E-07 | 560.79089 | 177.84525 |
| AGDDYEVLELDD\ | 15 | O43164 | E3 ubiquitin-protein ligase Praja-2 OS=H | PJA2 | 78.215 | 2 | 54.432915 | yes | 9.722E-08 | 839.39636 | 155.69362 |
| HAYVSFKPCMTR | 12 | O43164 | E3 ubiquitin-protein ligase Praja-2 OS=H | PJA2 | 78.215 | 3 | 54.432915 | yes | 3.171E-07 | 499.57608 | 62.751072 |
| HAYVSFKPCMTR | 12 | O43164 | E3 ubiquitin-protein ligase Praja-2 OS=H | PJA2 | 78.215 | 3 | 54.432915 | yes | 3.386E-07 | 748.86047 | 62.751072 |
| HAYVSFKPCMTR | 12 | O43164 | E3 ubiquitin-protein ligase Praja-2 OS=H | PJA2 | 78.215 | 2 | 54.432915 | yes | 3.171E-07 | 499.57608 | 62.751072 |
| HAYVSFKPCMTR | 12 | O43164 | E3 ubiquitin-protein ligase Praja-2 OS=H | PJA2 | 78.215 | 2 | 54.432915 | yes | 3.386E-07 | 748.86047 | 62.751072 |
| DEFEELDSVPLVK | 13 | O43164 | E3 ubiquitin-protein ligase Praja-2 OS=H | PJA2 | 78.215 | 2 | 54.432915 | yes | 1.371E-07 | 760.37994 | 87.071487 |
| LISSSQVDQETGFN | 15 | O43164 | E3 ubiquitin-protein ligase Praja-2 OS=H | PJA2 | 78.215 | 2 | 54.432915 | yes | 1.78E-07 | 840.91302 | 84.814125 |
| NHGSSPEQVVRPK | 13 | O43164 | E3 ubiquitin-protein ligase Praja-2 OS=H | PJA2 | 78.215 | 3 | 54.432915 | yes | 1.51E-07 | 478.9198 | 61.70293 |
| EPAAMDQESGK | 11 | O43164 | E3 ubiquitin-protein ligase Praja-2 OS=H | PJA2 | 78.215 | 2 | 54.432915 | yes | 4.251E-05 | 581.75586 | 41.870358 |
| DDIATELPCHHFFI | 24 | O43164 | E3 ubiquitin-protein ligase Praja-2 OS=H | PJA2 | 78.215 | 4 | 54.432915 | yes | 6.159E-07 | 745.36646 | 31.031162 |
| LVGDQEDHGSAK | 12 | O43167 | Zinc finger and BTB domain-containing p | ZBTB24 | 78.283 | 2 | 47.655685 | yes | 1.284E-07 | 628.29932 | 241.01024 |
| SELAAEEEIQLR | 12 | O43167 | Zinc finger and BTB domain-containing p | ZBTB24 | 78.283 | 2 | 47.655685 | yes | 2.152E-07 | 694.35681 | 92.114334 |
| SSLQTHIR | 8 | O43167 | Zinc finger and BTB domain-containing p | ZBTB24 | 78.283 | 2 | 47.655685 | yes | 6.666E-07 | 471.26178 | 68.038857 |
| FMDVSQLK | 8 | O43167 | Zinc finger and BTB domain-containing p | ZBTB24 | 78.283 | 2 | 47.655685 | yes | 6.776E-07 | 484.24951 | 66.334717 |
| YNHFLAIHQR | 10 | O43167 | Zinc finger and BTB domain-containing p | ZBTB24 | 78.283 | 2 | 47.655685 | yes | 3.645E-07 | 649.84125 | 62.946804 |
| STEQILATAQFLK | 13 | O43167 | Zinc finger and BTB domain-containing p | ZBTB24 | 78.283 | 2 | 47.655685 | yes | 3.156E-06 | 725.401 | 48.969379 |
| NILSVVGTDALK | 12 | O43172 | U4/U6 small nuclear ribonucleoprotein P | PRPF4 | 58.45 | 2 | 54.564404 | yes | 9.722E-08 | 615.35864 | 1410.4877 |
| LWIANYSLPR | 10 | O43172 | U4/U6 small nuclear ribonucleoprotein P | PRPF4 | 58.45 | 2 | 54.564404 | yes | 1.03E-07 | 616.84296 | 702.80688 |
| QINVSTDDSEVK | 12 | O43172 | U4/U6 small nuclear ribonucleoprotein P | PRPF4 | 58.45 | 2 | 54.564404 | yes | 3.424E-07 | 667.82532 | 602.46027 |
| DVNLASCAADGS\ | 14 | O43172 | U4/U6 small nuclear ribonucleoprotein P | PRPF4 | 58.45 | 2 | 54.564404 | yes | 9.722E-08 | 703.83264 | 510.52399 |
| MLATACWSGLCK | 12 | O43172 | U4/U6 small nuclear ribonucleoprotein P | PRPF4 | 58.45 | 2 | 54.564404 | yes | 9.88E-08 | 699.32233 | 507.56931 |
| VMWHPSGR | 8 | O43172 | U4/U6 small nuclear ribonucleoprotein P | PRPF4 | 58.45 | 2 | 54.564404 | yes | 1.821E-07 | 485.23981 | 492.58441 |
| ALGEPITLFGEGPA | 16 | O43172 | U4/U6 small nuclear ribonucleoprotein P | PRPF4 | 58.45 | 2 | 54.564404 | yes | 3.242E-07 | 828.93323 | 480.87231 |
| QAEVLAEFER | 10 | O43172 | U4/U6 small nuclear ribonucleoprotein P | PRPF4 | 58.45 | 2 | 54.564404 | yes | 1.039E-07 | 596.30383 | 409.25192 |
| TSQMQELHK | 9 | O43172 | U4/U6 small nuclear ribonucleoprotein P | PRPF4 | 58.45 | 2 | 54.564404 | yes | 9.722E-08 | 551.27148 | 397.81564 |
| FLGTTCYDR | 9 | O43172 | U4/U6 small nuclear ribonucleoprotein P | PRPF4 | 58.45 | 2 | 54.564404 | yes | 9.757E-08 | 566.75824 | 373.2005 |
| GHNTNVGAIVFHP | 14 | O43172 | U4/U6 small nuclear ribonucleoprotein P | PRPF4 | 58.45 | 3 | 54.564404 | yes | 1.341E-07 | 497.60028 | 179.02365 |

| | | | | | | | | | | | |
|---|---|---|---|---|---|---|---|---|---|---|---|
| GHNTNVGAIVFHP | 14 | O43172 | U4/U6 small nuclear ribonucleoprotein P | PRPF4 | 58.45 | 3 | 54.564404 | yes | 2.355E-07 | 745.89679 | 179.02365 |
| GHNTNVGAIVFHP | 14 | O43172 | U4/U6 small nuclear ribonucleoprotein P | PRPF4 | 58.45 | 2 | 54.564404 | yes | 1.341E-07 | 497.60028 | 179.02365 |
| GHNTNVGAIVFHP | 14 | O43172 | U4/U6 small nuclear ribonucleoprotein P | PRPF4 | 58.45 | 2 | 54.564404 | yes | 2.355E-07 | 745.89679 | 179.02365 |
| IWTHPGWSPLK | 11 | O43172 | U4/U6 small nuclear ribonucleoprotein P | PRPF4 | 58.45 | 2 | 54.564404 | yes | 1.292E-07 | 661.3562 | 176.12982 |
| IWTHPGWSPLK | 11 | O43172 | U4/U6 small nuclear ribonucleoprotein P | PRPF4 | 58.45 | 2 | 54.564404 | yes | 9.728E-08 | 441.2399 | 176.12982 |
| IWTHPGWSPLK | 11 | O43172 | U4/U6 small nuclear ribonucleoprotein P | PRPF4 | 58.45 | 3 | 54.564404 | yes | 1.292E-07 | 661.3562 | 176.12982 |
| IWTHPGWSPLK | 11 | O43172 | U4/U6 small nuclear ribonucleoprotein P | PRPF4 | 58.45 | 3 | 54.564404 | yes | 9.728E-08 | 441.2399 | 176.12982 |
| LWSLDSDEPVADI | 19 | O43172 | U4/U6 small nuclear ribonucleoprotein P | PRPF4 | 58.45 | 3 | 54.564404 | yes | 1.56E-07 | 713.68329 | 344.02127 |
| TKAPDDLVAPVVI | 13 | O43172 | U4/U6 small nuclear ribonucleoprotein P | PRPF4 | 58.45 | 2 | 54.564404 | yes | 1.639E-07 | 676.89282 | 220.20651 |
| CVYTIPAHQNLVT | 16 | O43172 | U4/U6 small nuclear ribonucleoprotein P | PRPF4 | 58.45 | 3 | 54.564404 | yes | 1.409E-07 | 600.65393 | 183.74937 |
| LWSVPDCNLLHTI | 14 | O43172 | U4/U6 small nuclear ribonucleoprotein P | PRPF4 | 58.45 | 3 | 54.564404 | yes | 1.15E-07 | 575.3031 | 160.20076 |
| APDDLVAPVVK | 11 | O43172 | U4/U6 small nuclear ribonucleoprotein P | PRPF4 | 58.45 | 2 | 54.564404 | yes | 9.722E-08 | 562.32153 | 143.66006 |
| CIMFLEGHLK | 10 | O43172 | U4/U6 small nuclear ribonucleoprotein P | PRPF4 | 58.45 | 2 | 54.564404 | yes | 9.722E-08 | 624.31738 | 65.918938 |
| QIPQATASMK | 10 | O43175 | D-3-phosphoglycerate dehydrogenase OS | PHGDH | 56.651 | 2 | 52.61781 | yes | 2.934E-07 | 537.78424 | 245.22122 |
| AGTGVDNVDLEA | 15 | O43175 | D-3-phosphoglycerate dehydrogenase OS | PHGDH | 56.651 | 2 | 52.61781 | yes | 9.81E-08 | 744.8681 | 244.91367 |
| VLISDSLDPCCR | 12 | O43175 | D-3-phosphoglycerate dehydrogenase OS | PHGDH | 56.651 | 2 | 52.61781 | yes | 1.434E-07 | 717.83942 | 212.61119 |
| ILQDGGLQVVEK | 12 | O43175 | D-3-phosphoglycerate dehydrogenase OS | PHGDH | 56.651 | 2 | 52.61781 | yes | 9.722E-08 | 649.86938 | 183.9082 |
| GGIVDEGALLR | 11 | O43175 | D-3-phosphoglycerate dehydrogenase OS | PHGDH | 56.651 | 2 | 52.61781 | yes | 1.75E-07 | 550.30896 | 182.181 |
| GTIQVITQGTSLK | 13 | O43175 | D-3-phosphoglycerate dehydrogenase OS | PHGDH | 56.651 | 2 | 52.61781 | yes | 1.199E-07 | 673.38794 | 170.6003 |
| CGEEIAVQFVDMV | 14 | O43175 | D-3-phosphoglycerate dehydrogenase OS | PHGDH | 56.651 | 2 | 52.61781 | yes | 1.149E-07 | 812.8891 | 83.68232 |
| ALQSGQCAGAALI | 21 | O43175 | D-3-phosphoglycerate dehydrogenase OS | PHGDH | 56.651 | 3 | 52.61781 | yes | 1.623E-06 | 739.69165 | 23.506392 |
| FNQVLDQFGEK | 11 | O43242 | 26S proteasome non-ATPase regulatory s | PSMD3 | 60.978 | 2 | 55.325096 | yes | 2.841E-07 | 662.83026 | 586.37085 |
| DGVIEASINHEK | 12 | O43242 | 26S proteasome non-ATPase regulatory s | PSMD3 | 60.978 | 2 | 55.325096 | yes | 1.3E-07 | 656.33063 | 239.43788 |
| EMIDIYSTR | 9 | O43242 | 26S proteasome non-ATPase regulatory s | PSMD3 | 60.978 | 2 | 55.325096 | yes | 4.952E-07 | 564.27368 | 225.42751 |
| SVFPEQANNNEWA | 14 | O43242 | 26S proteasome non-ATPase regulatory s | PSMD3 | 60.978 | 2 | 55.325096 | yes | 1.046E-07 | 831.38696 | 169.5779 |
| ISLADIAQK | 9 | O43242 | 26S proteasome non-ATPase regulatory s | PSMD3 | 60.978 | 2 | 55.325096 | yes | 9.722E-08 | 479.78201 | 139.75003 |
| LQLDSPEDAEFIVA | 15 | O43242 | 26S proteasome non-ATPase regulatory s | PSMD3 | 60.978 | 2 | 55.325096 | yes | 9.722E-08 | 837.93286 | 133.89299 |
| EPQLAFHQR | 9 | O43242 | 26S proteasome non-ATPase regulatory s | PSMD3 | 60.978 | 2 | 55.325096 | yes | 3.473E-07 | 563.29364 | 129.03206 |
| AIQLEYSEAR | 10 | O43242 | 26S proteasome non-ATPase regulatory s | PSMD3 | 60.978 | 2 | 55.325096 | yes | 9.722E-08 | 590.30383 | 106.32089 |
| AVQGFFTSNNATR | 13 | O43242 | 26S proteasome non-ATPase regulatory s | PSMD3 | 60.978 | 2 | 55.325096 | yes | 6.887E-07 | 706.84949 | 93.13884 |
| KAPQHTAVGFK | 11 | O43242 | 26S proteasome non-ATPase regulatory s | PSMD3 | 60.978 | 2 | 55.325096 | yes | 3.669E-06 | 592.33276 | 90.243782 |
| AKPPPGGGEQEPP | 24 | O43242 | 26S proteasome non-ATPase regulatory s | PSMD3 | 60.978 | 3 | 55.325096 | yes | 3.163E-07 | 817.40552 | 52.781765 |
| ELDTVTLEDIK | 11 | O43242 | 26S proteasome non-ATPase regulatory s | PSMD3 | 60.978 | 2 | 55.325096 | yes | 2.891E-07 | 638.33759 | 48.685055 |
| GEPEAPPASGLASI | 14 | O43248 | Homeobox protein Hox-C11 OS=Homo s | HOXC11 | 33.748 | 2 | 50.111828 | yes | 9.9E-08 | 669.83606 | 330.88998 |
| NSVLPQAFDR | 10 | O43248 | Homeobox protein Hox-C11 OS=Homo s | HOXC11 | 33.748 | 2 | 50.111828 | yes | 9.938E-08 | 573.79877 | 297.24963 |
| MLNLTDR | 7 | O43248 | Homeobox protein Hox-C11 OS=Homo s | HOXC11 | 33.748 | 2 | 50.111828 | no | 2.036E-07 | 431.7262 | 231.04971 |
| EVSYGLEPSGK | 11 | O43248 | Homeobox protein Hox-C11 OS=Homo s | HOXC11 | 33.748 | 2 | 50.111828 | yes | 9.968E-08 | 583.29041 | 166.27446 |
| EFFFNVYINK | 10 | O43248 | Homeobox protein Hox-C11 OS=Homo s | HOXC11 | 33.748 | 2 | 50.111828 | no | 4.569E-07 | 660.83478 | 68.200607 |
| QISYPYSAQVPPVI | 14 | O43248 | Homeobox protein Hox-C11 OS=Homo s | HOXC11 | 33.748 | 2 | 50.111828 | yes | 3.567E-06 | 802.92517 | 24.88102 |

| | | | | | | | | | | | |
|---|---|---|---|---|---|---|---|---|---|---|---|
| GFGFVTFENSADA | 15 | O43251 | RNA binding protein fox-1 homolog 2 OS | RBFOX2 | 41.374 | 2 | 52.001644 | no | 1.073E-07 | 816.8681 | 2096.8062 |
| LHVSNIPFR | 9 | O43251 | RNA binding protein fox-1 homolog 2 OS | RBFOX2 | 41.374 | 2 | 52.001644 | no | 2.623E-07 | 541.8089 | 1927.1674 |
| LHGTVVEGR | 9 | O43251 | RNA binding protein fox-1 homolog 2 OS | RBFOX2 | 41.374 | 2 | 52.001644 | no | 4.671E-07 | 484.26962 | 1122.9297 |
| FRDPDLR | 7 | O43251 | RNA binding protein fox-1 homolog 2 OS | RBFOX2 | 41.374 | 2 | 52.001644 | no | 5.161E-06 | 459.74323 | 966.7951 |
| YAQPATATAATAA | 26 | O43251 | RNA binding protein fox-1 homolog 2 OS | RBFOX2 | 41.374 | 3 | 52.001644 | yes | 1.795E-07 | 811.38739 | 810.08405 |
| QMFGQFGK | 8 | O43251 | RNA binding protein fox-1 homolog 2 OS | RBFOX2 | 41.374 | 2 | 52.001644 | no | 2.488E-07 | 471.72873 | 589.1145 |
| KIEVNNATAR | 10 | O43251 | RNA binding protein fox-1 homolog 2 OS | RBFOX2 | 41.374 | 2 | 52.001644 | no | 9.722E-08 | 558.31201 | 448.97495 |
| ILDVEIIFNER | 11 | O43251 | RNA binding protein fox-1 homolog 2 OS | RBFOX2 | 41.374 | 2 | 52.001644 | no | 1.066E-07 | 680.8772 | 284.32062 |
| EKLHGTVVEGR | 11 | O43251 | RNA binding protein fox-1 homolog 2 OS | RBFOX2 | 41.374 | 2 | 52.001644 | no | 5.799E-07 | 612.83838 | 30.17054 |
| SLPSAVYCIEDK | 12 | O43290 | U4/U6.U5 tri-snRNP-associated protein 1 | SART1 | 90.255 | 2 | 50.240757 | yes | 1.164E-07 | 691.33704 | 546.32928 |
| TLGEIPTYGLAGNI | 14 | O43290 | U4/U6.U5 tri-snRNP-associated protein 1 | SART1 | 90.255 | 2 | 50.240757 | yes | 1.424E-07 | 731.38843 | 512.05457 |
| IEYVDETGR | 9 | O43290 | U4/U6.U5 tri-snRNP-associated protein 1 | SART1 | 90.255 | 2 | 50.240757 | yes | 1.081E-07 | 541.26166 | 248.07388 |
| QLQQLQQLR | 9 | O43290 | U4/U6.U5 tri-snRNP-associated protein 1 | SART1 | 90.255 | 2 | 50.240757 | yes | 1.022E-07 | 577.83563 | 235.69569 |
| GAIVFNATSEFCR | 13 | O43290 | U4/U6.U5 tri-snRNP-associated protein 1 | SART1 | 90.255 | 2 | 50.240757 | yes | 1.354E-07 | 736.35358 | 201.21553 |
| YDEELEGERPHSFI | 14 | O43290 | U4/U6.U5 tri-snRNP-associated protein 1 | SART1 | 90.255 | 3 | 50.240757 | yes | 2.284E-07 | 588.60431 | 185.87706 |
| TPYIVLSGSGK | 11 | O43290 | U4/U6.U5 tri-snRNP-associated protein 1 | SART1 | 90.255 | 2 | 50.240757 | yes | 2.135E-07 | 561.31366 | 120.60589 |
| GLLETTVQK | 9 | O43290 | U4/U6.U5 tri-snRNP-associated protein 1 | SART1 | 90.255 | 2 | 50.240757 | yes | 9.839E-08 | 494.78729 | 119.35047 |
| EEQEELMDFER | 11 | O43290 | U4/U6.U5 tri-snRNP-associated protein 1 | SART1 | 90.255 | 2 | 50.240757 | yes | 9.992E-08 | 727.80884 | 118.62453 |
| KLDEEALLK | 9 | O43290 | U4/U6.U5 tri-snRNP-associated protein 1 | SART1 | 90.255 | 2 | 50.240757 | yes | 2.058E-07 | 529.80823 | 112.68651 |
| TSSGDASSLSIEETI | 16 | O43290 | U4/U6.U5 tri-snRNP-associated protein 1 | SART1 | 90.255 | 2 | 50.240757 | yes | 3.212E-07 | 813.37866 | 106.25344 |
| LQAQSLSTVGPR | 12 | O43290 | U4/U6.U5 tri-snRNP-associated protein 1 | SART1 | 90.255 | 2 | 50.240757 | yes | 2.796E-07 | 628.8515 | 104.24567 |
| QQQDFSASSTTILD | 21 | O43290 | U4/U6.U5 tri-snRNP-associated protein 1 | SART1 | 90.255 | 3 | 50.240757 | yes | 1.617E-06 | 793.38721 | 70.676994 |
| EGETMILTLK | 10 | O43290 | U4/U6.U5 tri-snRNP-associated protein 1 | SART1 | 90.255 | 2 | 50.240757 | yes | 1.599E-07 | 567.80737 | 70.129631 |
| DGYKPDVK | 8 | O43290 | U4/U6.U5 tri-snRNP-associated protein 1 | SART1 | 90.255 | 2 | 50.240757 | yes | 5.65E-07 | 461.23746 | 60.475861 |
| LGLKPLEVNAIK | 12 | O43290 | U4/U6.U5 tri-snRNP-associated protein 1 | SART1 | 90.255 | 2 | 50.240757 | yes | 3.081E-06 | 647.90826 | 18.747698 |
| LGLKPLEVNAIK | 12 | O43290 | U4/U6.U5 tri-snRNP-associated protein 1 | SART1 | 90.255 | 2 | 50.240757 | yes | 2.296E-07 | 432.27463 | 18.747698 |
| LGLKPLEVNAIK | 12 | O43290 | U4/U6.U5 tri-snRNP-associated protein 1 | SART1 | 90.255 | 3 | 50.240757 | yes | 3.081E-06 | 647.90826 | 18.747698 |
| LGLKPLEVNAIK | 12 | O43290 | U4/U6.U5 tri-snRNP-associated protein 1 | SART1 | 90.255 | 3 | 50.240757 | yes | 2.296E-07 | 432.27463 | 18.747698 |
| GLAAALLLCQNK | 12 | O43290 | U4/U6.U5 tri-snRNP-associated protein 1 | SART1 | 90.255 | 2 | 50.240757 | yes | 1.193E-06 | 636.36066 | 27.85692 |
| EKEVVVR | 7 | O43290 | U4/U6.U5 tri-snRNP-associated protein 1 | SART1 | 90.255 | 2 | 50.240757 | yes | 0.0001602 | 429.7558 | 2.4417319 |
| LQGVECGLR | 9 | O43293 | Death-associated protein kinase 3 OS=Ho | DAPK3 | 52.536 | 2 | 53.60934 | yes | 1.301E-07 | 516.26874 | 212.37367 |
| VLEEAAAAEEGLR | 13 | O43293 | Death-associated protein kinase 3 OS=Ho | DAPK3 | 52.536 | 2 | 53.60934 | yes | 1.001E-07 | 679.3515 | 83.37719 |
| ESLTEDEATQFLK | 13 | O43293 | Death-associated protein kinase 3 OS=Ho | DAPK3 | 52.536 | 2 | 53.60934 | yes | 9.722E-08 | 755.86719 | 56.428131 |
| MTIAQSLEHSWIK | 13 | O43293 | Death-associated protein kinase 3 OS=Ho | DAPK3 | 52.536 | 2 | 53.60934 | yes | 9.85E-08 | 772.40051 | 15.338593 |
| SFVCTECGQVFR | 12 | O43296 | Zinc finger protein 264 OS=Homo sapien | ZNF264 | 70.587 | 2 | 46.989254 | yes | 1.425E-07 | 745.33179 | 46.365837 |
| PFTSGQTSVTLR | 12 | O43296 | Zinc finger protein 264 OS=Homo sapien | ZNF264 | 70.587 | 2 | 46.989254 | yes | 6.357E-07 | 647.34351 | 26.783909 |
| VEFPDFSSTILQK | 13 | O43298 | Zinc finger and BTB domain-containing p | ZBTB43 | 52.631 | 2 | 51.62491 | yes | 1.668E-07 | 755.89301 | 671.5282 |
| IVLPDVMNPR | 10 | O43298 | Zinc finger and BTB domain-containing p | ZBTB43 | 52.631 | 2 | 51.62491 | yes | 9.722E-08 | 577.32355 | 611.71973 |
| VFENILLSSYTGR | 13 | O43298 | Zinc finger and BTB domain-containing p | ZBTB43 | 52.631 | 2 | 51.62491 | yes | 1.629E-07 | 749.89862 | 360.91837 |

| | | | | | | | | | | | |
|---|---|---|---|---|---|---|---|---|---|---|---|
| WIHVKPER | 8 | O43298 | Zinc finger and BTB domain-containing p | ZBTB43 | 52.631 | 2 | 51.62491 | yes | 1.732E-07 | 532.80365 | 311.85043 |
| QQGQLCDVSIVVQ | 18 | O43298 | Zinc finger and BTB domain-containing p | ZBTB43 | 52.631 | 3 | 51.62491 | yes | 5.457E-07 | 695.36169 | 124.85664 |
| SDGNLIGHR | 9 | O43298 | Zinc finger and BTB domain-containing p | ZBTB43 | 52.631 | 2 | 51.62491 | yes | 4.134E-07 | 484.74905 | 110.18113 |
| VLVCPIYTCLR | 11 | O43310 | CBP80/20-dependent translation initiation | CTIF | 67.587 | 2 | 48.110977 | yes | 2.002E-07 | 697.37018 | 164.28021 |
| MLCPSESMLTR | 11 | O43310 | CBP80/20-dependent translation initiation | CTIF | 67.587 | 2 | 48.110977 | yes | 1.197E-07 | 662.80652 | 158.81036 |
| LDFTQFHR | 8 | O43310 | CBP80/20-dependent translation initiation | CTIF | 67.587 | 2 | 48.110977 | yes | 4.675E-07 | 532.26959 | 72.531746 |
| SPDTLAPVASER | 12 | O43310 | CBP80/20-dependent translation initiation | CTIF | 67.587 | 2 | 48.110977 | yes | 2.483E-07 | 621.81989 | 68.738075 |
| ELLQSQDVK | 9 | O43310 | CBP80/20-dependent translation initiation | CTIF | 67.587 | 2 | 48.110977 | yes | 1.215E-06 | 530.28766 | 56.481777 |
| ALADIVIPQEYGIT | 15 | O43314 | Inositol hexakisphosphate and diphospho | PPIP5K2 | 140.407 | 2 | 49.50046 | yes | 1.025E-07 | 815.95618 | 84.713562 |
| IPDIYDCIK | 9 | O43314 | Inositol hexakisphosphate and diphospho | PPIP5K2 | 140.407 | 2 | 49.50046 | yes | 9.04E-07 | 568.78644 | 23.909983 |
| FIVDLTVQEK | 10 | O43324 | Eukaryotic translation elongation factor 1 | EEF1E1 | 19.811 | 2 | 54.72056 | yes | 9.81E-08 | 596.33459 | 301.54904 |
| EYLLGSTAEEK | 11 | O43324 | Eukaryotic translation elongation factor 1 | EEF1E1 | 19.811 | 2 | 54.72056 | yes | 9.722E-08 | 620.30878 | 300.77435 |
| AIVQQWLEYR | 10 | O43324 | Eukaryotic translation elongation factor 1 | EEF1E1 | 19.811 | 2 | 54.72056 | yes | 1.396E-07 | 653.35114 | 219.9949 |
| DLNSYLEDK | 9 | O43324 | Eukaryotic translation elongation factor 1 | EEF1E1 | 19.811 | 2 | 54.72056 | yes | 2.904E-06 | 548.76147 | 31.952517 |
| SLPAPQDNDFLSR | 13 | O43353 | Receptor-interacting serine/threonine-pro | RIPK2 | 61.195 | 2 | 51.645576 | yes | 3.239E-07 | 730.36243 | 412.68472 |
| CLIELEPVLR | 10 | O43353 | Receptor-interacting serine/threonine-pro | RIPK2 | 61.195 | 2 | 51.645576 | yes | 9.722E-08 | 621.34973 | 314.62674 |
| LQPGIAQQWIQSK | 13 | O43353 | Receptor-interacting serine/threonine-pro | RIPK2 | 61.195 | 2 | 51.645576 | yes | 3.231E-07 | 748.91461 | 262.27789 |
| EDYELVSTKPTR | 12 | O43353 | Receptor-interacting serine/threonine-pro | RIPK2 | 61.195 | 2 | 51.645576 | yes | 9.884E-08 | 719.36462 | 158.26617 |
| SPSLNLLQNK | 10 | O43353 | Receptor-interacting serine/threonine-pro | RIPK2 | 61.195 | 2 | 51.645576 | yes | 9.752E-08 | 557.31677 | 122.21754 |
| MMSLSQSR | 8 | O43353 | Receptor-interacting serine/threonine-pro | RIPK2 | 61.195 | 2 | 51.645576 | yes | 1.888E-07 | 470.22296 | 116.7327 |
| LQSVSSAIHLCDK | 13 | O43353 | Receptor-interacting serine/threonine-pro | RIPK2 | 61.195 | 2 | 51.645576 | yes | 1.339E-07 | 729.37451 | 95.049194 |
| KTEYPDVAWPLR | 12 | O43353 | Receptor-interacting serine/threonine-pro | RIPK2 | 61.195 | 2 | 51.645576 | yes | 4.14E-07 | 737.88806 | 60.739712 |
| HLHIHTPLLDSER | 13 | O43353 | Receptor-interacting serine/threonine-pro | RIPK2 | 61.195 | 3 | 51.645576 | yes | 1.055E-07 | 523.28284 | 54.962524 |
| NQVAEMLGHK | 10 | O43379 | WD repeat-containing protein 62 OS=Ho | WDR62 | 165.954 | 2 | 45.921707 | yes | 1.149E-06 | 563.78729 | 82.165451 |
| DIQMISCGADK | 11 | O43379 | WD repeat-containing protein 62 OS=Ho | WDR62 | 165.954 | 2 | 45.921707 | yes | 3.68E-07 | 619.28119 | 58.8745 |
| LIHVLNVEK | 9 | O43379 | WD repeat-containing protein 62 OS=Ho | WDR62 | 165.954 | 2 | 45.921707 | yes | 5.894E-07 | 532.82678 | 42.127052 |
| LSISTQFLSSLQK | 13 | O43379 | WD repeat-containing protein 62 OS=Ho | WDR62 | 165.954 | 2 | 45.921707 | yes | 1.703E-07 | 726.40887 | 37.827923 |
| VLVSSGQVDTGQG | 16 | O43379 | WD repeat-containing protein 62 OS=Ho | WDR62 | 165.954 | 2 | 45.921707 | yes | 7.049E-07 | 836.93427 | 18.40765 |
| TGYTLDVTTGQR | 12 | O43390 | Heterogeneous nuclear ribonucleoprotein | HNRNPR | 70.944 | 2 | 54.54753 | no | 1.658E-07 | 656.33063 | 19146.17 |
| GFCFLEYEDHK | 11 | O43390 | Heterogeneous nuclear ribonucleoprotein | HNRNPR | 70.944 | 2 | 54.54753 | no | 1.164E-07 | 722.81372 | 7348.1826 |
| DYAFVHFEDR | 10 | O43390 | Heterogeneous nuclear ribonucleoprotein | HNRNPR | 70.944 | 2 | 54.54753 | yes | 1.011E-07 | 649.79364 | 2189.2529 |
| DYAFVHFEDR | 10 | O43390 | Heterogeneous nuclear ribonucleoprotein | HNRNPR | 70.944 | 2 | 54.54753 | yes | 1.206E-07 | 433.53152 | 2189.2529 |
| DYAFVHFEDR | 10 | O43390 | Heterogeneous nuclear ribonucleoprotein | HNRNPR | 70.944 | 3 | 54.54753 | yes | 1.011E-07 | 649.79364 | 2189.2529 |
| DYAFVHFEDR | 10 | O43390 | Heterogeneous nuclear ribonucleoprotein | HNRNPR | 70.944 | 3 | 54.54753 | yes | 1.206E-07 | 433.53152 | 2189.2529 |
| NLATTVTEEILEK | 13 | O43390 | Heterogeneous nuclear ribonucleoprotein | HNRNPR | 70.944 | 2 | 54.54753 | yes | 1.073E-07 | 730.89575 | 4190.1714 |
| AGPIWDLR | 8 | O43390 | Heterogeneous nuclear ribonucleoprotein | HNRNPR | 70.944 | 2 | 54.54753 | no | 2.739E-07 | 464.25598 | 3107.7524 |
| LCDSYEIRPGK | 11 | O43390 | Heterogeneous nuclear ribonucleoprotein | HNRNPR | 70.944 | 2 | 54.54753 | yes | 1.247E-07 | 669.32953 | 2926.1897 |
| SAFLCGVMK | 9 | O43390 | Heterogeneous nuclear ribonucleoprotein | HNRNPR | 70.944 | 2 | 54.54753 | no | 1.353E-07 | 506.75134 | 1458.6824 |
| SAFLCGVMK | 9 | O43390 | Heterogeneous nuclear ribonucleoprotein | HNRNPR | 70.944 | 2 | 54.54753 | no | 9.722E-08 | 514.74884 | 1458.6824 |

| | | | | | | | | | | | |
|---|---|---|---|---|---|---|---|---|---|---|---|
| GYAFITFCGK | 10 | O43390 | Heterogeneous nuclear ribonucleoprotein | HNRNPR | 70.944 | 2 | 54.54753 | yes | 9.893E-08 | 582.28131 | 2174.6565 |
| STAYEDYYYHPPP | 14 | O43390 | Heterogeneous nuclear ribonucleoprotein | HNRNPR | 70.944 | 3 | 54.54753 | yes | 1.511E-07 | 586.93011 | 2149.2075 |
| TLIEAGLPQK | 10 | O43390 | Heterogeneous nuclear ribonucleoprotein | HNRNPR | 70.944 | 2 | 54.54753 | yes | 9.722E-08 | 535.31622 | 1860.6603 |
| DLYEDELVPLFEK | 13 | O43390 | Heterogeneous nuclear ribonucleoprotein | HNRNPR | 70.944 | 2 | 54.54753 | yes | 9.722E-08 | 805.40344 | 1678.1152 |
| LMMDPLSGQNR | 11 | O43390 | Heterogeneous nuclear ribonucleoprotein | HNRNPR | 70.944 | 2 | 54.54753 | yes | 9.722E-08 | 631.30499 | 813.09308 |
| LMMDPLSGQNR | 11 | O43390 | Heterogeneous nuclear ribonucleoprotein | HNRNPR | 70.944 | 2 | 54.54753 | yes | 9.926E-08 | 639.30249 | 813.09308 |
| ENILEEFSK | 9 | O43390 | Heterogeneous nuclear ribonucleoprotein | HNRNPR | 70.944 | 2 | 54.54753 | yes | 1.577E-07 | 554.77966 | 1357.6956 |
| TKENILEEFSK | 11 | O43390 | Heterogeneous nuclear ribonucleoprotein | HNRNPR | 70.944 | 2 | 54.54753 | yes | 1.11E-07 | 669.35101 | 1256.5968 |
| LKDYAFVHFEDR | 12 | O43390 | Heterogeneous nuclear ribonucleoprotein | HNRNPR | 70.944 | 2 | 54.54753 | yes | 1.028E-07 | 770.38318 | 511.79861 |
| LKDYAFVHFEDR | 12 | O43390 | Heterogeneous nuclear ribonucleoprotein | HNRNPR | 70.944 | 2 | 54.54753 | yes | 2.221E-07 | 513.92456 | 511.79861 |
| LKDYAFVHFEDR | 12 | O43390 | Heterogeneous nuclear ribonucleoprotein | HNRNPR | 70.944 | 3 | 54.54753 | yes | 1.028E-07 | 770.38318 | 511.79861 |
| LKDYAFVHFEDR | 12 | O43390 | Heterogeneous nuclear ribonucleoprotein | HNRNPR | 70.944 | 3 | 54.54753 | yes | 2.221E-07 | 513.92456 | 511.79861 |
| LFVGSIPK | 8 | O43390 | Heterogeneous nuclear ribonucleoprotein | HNRNPR | 70.944 | 2 | 54.54753 | no | 3.911E-07 | 430.76566 | 882.5481 |
| LCDSYEIR | 8 | O43390 | Heterogeneous nuclear ribonucleoprotein | HNRNPR | 70.944 | 2 | 54.54753 | yes | 3.182E-07 | 528.24493 | 687.71588 |
| ESDLSHVQNK | 10 | O43390 | Heterogeneous nuclear ribonucleoprotein | HNRNPR | 70.944 | 2 | 54.54753 | yes | 9.935E-08 | 578.78326 | 660.52643 |
| EIEGEEIEIVLAKPI | 17 | O43390 | Heterogeneous nuclear ribonucleoprotein | HNRNPR | 70.944 | 3 | 54.54753 | yes | 9.722E-08 | 637.01031 | 297.69626 |
| EEEEPMDTSSVTH | 18 | O43390 | Heterogeneous nuclear ribonucleoprotein | HNRNPR | 70.944 | 3 | 54.54753 | yes | 4.271E-07 | 716.97241 | 264.62927 |
| KADGYNQPDSK | 11 | O43390 | Heterogeneous nuclear ribonucleoprotein | HNRNPR | 70.944 | 2 | 54.54753 | no | 4.647E-07 | 611.78857 | 234.42374 |
| YGGPPPDSVYSGV | 25 | O43390 | Heterogeneous nuclear ribonucleoprotein | HNRNPR | 70.944 | 3 | 54.54753 | yes | 1.166E-07 | 836.41992 | 59.312592 |
| VQESTKGPDEAK | 12 | O43390 | Heterogeneous nuclear ribonucleoprotein | HNRNPR | 70.944 | 2 | 54.54753 | yes | 1.78E-07 | 644.82263 | 21.82394 |
| ADGYNQPDSK | 10 | O43390 | Heterogeneous nuclear ribonucleoprotein | HNRNPR | 70.944 | 2 | 54.54753 | no | 5.28E-07 | 547.74109 | 21.164181 |
| LFEAVEEGR | 9 | O43395 | U4/U6 small nuclear ribonucleoprotein Pı | PRPF3 | 77.529 | 2 | 51.019325 | yes | 2.079E-07 | 525.26672 | 1101.6163 |
| VLGTEAVQDPTK | 12 | O43395 | U4/U6 small nuclear ribonucleoprotein Pı | PRPF3 | 77.529 | 2 | 51.019325 | yes | 1.341E-07 | 629.33789 | 813.96564 |
| LQAEISQAAR | 10 | O43395 | U4/U6 small nuclear ribonucleoprotein Pı | PRPF3 | 77.529 | 2 | 51.019325 | yes | 5.924E-07 | 543.79877 | 730.58344 |
| LGLMPPPEPK | 10 | O43395 | U4/U6 small nuclear ribonucleoprotein Pı | PRPF3 | 77.529 | 2 | 51.019325 | yes | 5.139E-06 | 539.80188 | 672.11304 |
| IEANAGQLYLTGV | 18 | O43395 | U4/U6 small nuclear ribonucleoprotein Pı | PRPF3 | 77.529 | 3 | 51.019325 | yes | 1.067E-07 | 642.3667 | 449.32111 |
| EIELTHR | 7 | O43395 | U4/U6 small nuclear ribonucleoprotein Pı | PRPF3 | 77.529 | 2 | 51.019325 | yes | 4.583E-07 | 449.24307 | 379.37015 |
| QLSFISPPTPQPK | 13 | O43395 | U4/U6 small nuclear ribonucleoprotein Pı | PRPF3 | 77.529 | 2 | 51.019325 | yes | 4.776E-07 | 720.39825 | 243.94508 |
| DVNVVVVEGGPK | 12 | O43395 | U4/U6 small nuclear ribonucleoprotein Pı | PRPF3 | 77.529 | 2 | 51.019325 | yes | 9.722E-08 | 606.33514 | 242.35948 |
| QCPTENMAR | 9 | O43395 | U4/U6 small nuclear ribonucleoprotein Pı | PRPF3 | 77.529 | 2 | 51.019325 | yes | 1.625E-07 | 553.7395 | 224.3443 |
| CVLVWEGTAK | 10 | O43395 | U4/U6 small nuclear ribonucleoprotein Pı | PRPF3 | 77.529 | 2 | 51.019325 | yes | 1.02E-07 | 581.79987 | 163.50578 |
| QMMEAATR | 8 | O43395 | U4/U6 small nuclear ribonucleoprotein Pı | PRPF3 | 77.529 | 2 | 51.019325 | yes | 5.847E-07 | 469.21515 | 141.90778 |
| WDEQTSNTK | 9 | O43395 | U4/U6 small nuclear ribonucleoprotein Pı | PRPF3 | 77.529 | 2 | 51.019325 | yes | 4.162E-07 | 554.7489 | 97.380264 |
| IKWDEQTSNTK | 11 | O43395 | U4/U6 small nuclear ribonucleoprotein Pı | PRPF3 | 77.529 | 2 | 51.019325 | yes | 2.204E-07 | 675.33844 | 77.072388 |
| LKEDISQGVHISVY | 15 | O43395 | U4/U6 small nuclear ribonucleoprotein Pı | PRPF3 | 77.529 | 3 | 51.019325 | yes | 3.504E-07 | 581.98444 | 75.679863 |
| RELDELKPWIEK | 12 | O43395 | U4/U6 small nuclear ribonucleoprotein Pı | PRPF3 | 77.529 | 3 | 51.019325 | yes | 3.421E-07 | 519.28748 | 51.825233 |
| DQTKPTPLILDEQ( | 15 | O43395 | U4/U6 small nuclear ribonucleoprotein Pı | PRPF3 | 77.529 | 3 | 51.019325 | yes | 1.575E-07 | 570.97223 | 47.907303 |
| EDISQGVHISVYR | 13 | O43395 | U4/U6 small nuclear ribonucleoprotein Pı | PRPF3 | 77.529 | 2 | 51.019325 | yes | 0.0002969 | 751.88354 | 18.384529 |
| ELDELKPWIEK | 11 | O43395 | U4/U6 small nuclear ribonucleoprotein Pı | PRPF3 | 77.529 | 2 | 51.019325 | yes | 4.397E-07 | 700.37701 | 14.489195 |

| | | | | | | | | | | | |
|---|---|---|---|---|---|---|---|---|---|---|---|
| DIFSEQHLK | 9 | O43422 | 52 kDa repressor of the inhibitor of the pr | THAP12 | 87.704 | 2 | 54.418327 | yes | 1.132E-07 | 558.78784 | 1549.21 |
| STQSDLAFFR | 10 | O43422 | 52 kDa repressor of the inhibitor of the pr | THAP12 | 87.704 | 2 | 54.418327 | yes | 1.102E-07 | 586.29077 | 1166.3281 |
| QMLEICESCIR | 11 | O43422 | 52 kDa repressor of the inhibitor of the pr | THAP12 | 87.704 | 2 | 54.418327 | yes | 9.919E-08 | 719.828 | 1057.2349 |
| FETTAVNTLFCSK | 13 | O43422 | 52 kDa repressor of the inhibitor of the pr | THAP12 | 87.704 | 2 | 54.418327 | yes | 9.722E-08 | 759.3689 | 993.66064 |
| FVDESHNLR | 9 | O43422 | 52 kDa repressor of the inhibitor of the pr | THAP12 | 87.704 | 2 | 54.418327 | yes | 9.935E-08 | 558.77527 | 940.01215 |
| GQAYIVSSGFSSK | 13 | O43422 | 52 kDa repressor of the inhibitor of the pr | THAP12 | 87.704 | 2 | 54.418327 | yes | 1.013E-07 | 665.83551 | 889.86853 |
| CLSLVPSVMGQLK | 13 | O43422 | 52 kDa repressor of the inhibitor of the pr | THAP12 | 87.704 | 2 | 54.418327 | yes | 1.584E-07 | 716.38855 | 801.23511 |
| HFETSMICR | 9 | O43422 | 52 kDa repressor of the inhibitor of the pr | THAP12 | 87.704 | 2 | 54.418327 | yes | 1.26E-07 | 590.7655 | 651.422 |
| FHTMITEK | 8 | O43422 | 52 kDa repressor of the inhibitor of the pr | THAP12 | 87.704 | 2 | 54.418327 | yes | 2.673E-06 | 503.75494 | 615.8833 |
| DNAIPTIFDLTSHL | 20 | O43422 | 52 kDa repressor of the inhibitor of the pr | THAP12 | 87.704 | 3 | 54.418327 | yes | 1.101E-06 | 754.71356 | 611.87799 |
| FNTSEEHHADMYI | 13 | O43422 | 52 kDa repressor of the inhibitor of the pr | THAP12 | 87.704 | 3 | 54.418327 | yes | 1.073E-07 | 546.23181 | 290.53333 |
| FNTSEEHHADMYI | 13 | O43422 | 52 kDa repressor of the inhibitor of the pr | THAP12 | 87.704 | 3 | 54.418327 | yes | 9.968E-08 | 818.84406 | 290.53333 |
| FNTSEEHHADMYI | 13 | O43422 | 52 kDa repressor of the inhibitor of the pr | THAP12 | 87.704 | 2 | 54.418327 | yes | 1.073E-07 | 546.23181 | 290.53333 |
| FNTSEEHHADMYI | 13 | O43422 | 52 kDa repressor of the inhibitor of the pr | THAP12 | 87.704 | 2 | 54.418327 | yes | 9.968E-08 | 818.84406 | 290.53333 |
| WNNYIAGR | 8 | O43422 | 52 kDa repressor of the inhibitor of the pr | THAP12 | 87.704 | 2 | 54.418327 | yes | 1.272E-07 | 497.24869 | 571.30432 |
| INSGEEVLR | 9 | O43422 | 52 kDa repressor of the inhibitor of the pr | THAP12 | 87.704 | 2 | 54.418327 | yes | 1.529E-07 | 508.77219 | 397.62619 |
| HDLDLMVDTYIK | 12 | O43422 | 52 kDa repressor of the inhibitor of the pr | THAP12 | 87.704 | 2 | 54.418327 | yes | 1.234E-07 | 731.86597 | 391.51901 |
| WGLNMEYCR | 9 | O43422 | 52 kDa repressor of the inhibitor of the pr | THAP12 | 87.704 | 2 | 54.418327 | yes | 1.905E-07 | 614.7655 | 367.05081 |
| VLCILPVMK | 9 | O43422 | 52 kDa repressor of the inhibitor of the pr | THAP12 | 87.704 | 2 | 54.418327 | yes | 9.766E-08 | 536.81653 | 317.83896 |
| SSNLALLNINFDIK | 14 | O43422 | 52 kDa repressor of the inhibitor of the pr | THAP12 | 87.704 | 2 | 54.418327 | yes | 9.722E-08 | 781.43286 | 287.78378 |
| EICHSQWTGR | 10 | O43422 | 52 kDa repressor of the inhibitor of the pr | THAP12 | 87.704 | 2 | 54.418327 | yes | 9.722E-08 | 637.29077 | 268.68158 |
| NTLTDQR | 7 | O43422 | 52 kDa repressor of the inhibitor of the pr | THAP12 | 87.704 | 2 | 54.418327 | yes | 2.207E-07 | 424.21704 | 171.74565 |
| SDLPNPDTLSAELI | 17 | O43422 | 52 kDa repressor of the inhibitor of the pr | THAP12 | 87.704 | 3 | 54.418327 | yes | 1.279E-07 | 670.98279 | 136.17944 |
| IDETSEQEQK | 10 | O43422 | 52 kDa repressor of the inhibitor of the pr | THAP12 | 87.704 | 2 | 54.418327 | yes | 1.199E-07 | 603.77789 | 129.57576 |
| IKELSEDEIR | 10 | O43422 | 52 kDa repressor of the inhibitor of the pr | THAP12 | 87.704 | 2 | 54.418327 | yes | 9.722E-08 | 616.33008 | 110.22572 |
| SELPTDNSETVENT | 14 | O43422 | 52 kDa repressor of the inhibitor of the pr | THAP12 | 87.704 | 2 | 54.418327 | yes | 9.722E-08 | 768.33899 | 99.697746 |
| GKDIELPSTIYEAL | 20 | O43422 | 52 kDa repressor of the inhibitor of the pr | THAP12 | 87.704 | 3 | 54.418327 | yes | 2.289E-07 | 751.41058 | 54.541412 |
| ADLEDKTPDQLNK | 13 | O43422 | 52 kDa repressor of the inhibitor of the pr | THAP12 | 87.704 | 2 | 54.418327 | yes | 2.948E-06 | 743.8728 | 32.269821 |
| ETLSVPTVEHIIQE | 16 | O43422 | 52 kDa repressor of the inhibitor of the pr | THAP12 | 87.704 | 3 | 54.418327 | yes | 2.01E-07 | 612.67529 | 17.517672 |
| IDLPNEEVK | 9 | O43426 | Synaptojanin-1 OS=Homo sapiens OX=9 | SYNJ1 | 173.104 | 2 | 31.48566 | yes | 1.825E-06 | 528.78223 | 56.859047 |
| MDQYFNQMEK | 10 | O43432 | Eukaryotic translation initiation factor 4 g | EIF4G3 | 176.653 | 2 | 54.382847 | no | 1.086E-07 | 667.28119 | 3219.7827 |
| MDQYFNQMEK | 10 | O43432 | Eukaryotic translation initiation factor 4 g | EIF4G3 | 176.653 | 2 | 54.382847 | no | 9.975E-08 | 675.27869 | 3219.7827 |
| MDQYFNQMEK | 10 | O43432 | Eukaryotic translation initiation factor 4 g | EIF4G3 | 176.653 | 2 | 54.382847 | no | 9.722E-08 | 675.27869 | 3219.7827 |
| MLTEAIMHDCVVI | 13 | O43432 | Eukaryotic translation initiation factor 4 g | EIF4G3 | 176.653 | 2 | 54.382847 | no | 1.391E-07 | 773.87494 | 2892.1311 |
| MLTEAIMHDCVVI | 13 | O43432 | Eukaryotic translation initiation factor 4 g | EIF4G3 | 176.653 | 2 | 54.382847 | no | 7.907E-07 | 781.87238 | 2892.1311 |
| MLTEAIMHDCVVI | 13 | O43432 | Eukaryotic translation initiation factor 4 g | EIF4G3 | 176.653 | 2 | 54.382847 | no | 1.122E-07 | 516.25238 | 2892.1311 |
| MLTEAIMHDCVVI | 13 | O43432 | Eukaryotic translation initiation factor 4 g | EIF4G3 | 176.653 | 2 | 54.382847 | no | 9.722E-08 | 781.87238 | 2892.1311 |
| MLTEAIMHDCVVI | 13 | O43432 | Eukaryotic translation initiation factor 4 g | EIF4G3 | 176.653 | 3 | 54.382847 | no | 1.391E-07 | 773.87494 | 2892.1311 |
| MLTEAIMHDCVVI | 13 | O43432 | Eukaryotic translation initiation factor 4 g | EIF4G3 | 176.653 | 3 | 54.382847 | no | 7.907E-07 | 781.87238 | 2892.1311 |

| | | | | | | | | | | | |
|---|---|---|---|---|---|---|---|---|---|---|---|
| MLTEAIMHDCVVI | 13 | O43432 | Eukaryotic translation initiation factor 4 g | EIF4G3 | 176.653 | 3 | 54.382847 | no | 1.122E-07 | 516.25238 | 2892.1311 |
| MLTEAIMHDCVVI | 13 | O43432 | Eukaryotic translation initiation factor 4 g | EIF4G3 | 176.653 | 3 | 54.382847 | no | 9.722E-08 | 781.87238 | 2892.1311 |
| GPDFTPAFADFGR | 13 | O43432 | Eukaryotic translation initiation factor 4 g | EIF4G3 | 176.653 | 2 | 54.382847 | yes | 1.165E-07 | 699.32788 | 1746.7913 |
| VLEEDESIR | 9 | O43432 | Eukaryotic translation initiation factor 4 g | EIF4G3 | 176.653 | 2 | 54.382847 | yes | 9.722E-08 | 545.27478 | 1466.635 |
| VGVESTLER | 9 | O43432 | Eukaryotic translation initiation factor 4 g | EIF4G3 | 176.653 | 2 | 54.382847 | yes | 1.182E-07 | 495.26672 | 1394.0557 |
| LSATESIVEIVK | 12 | O43432 | Eukaryotic translation initiation factor 4 g | EIF4G3 | 176.653 | 2 | 54.382847 | yes | 1.098E-07 | 644.87158 | 1331.0059 |
| LDQPANLLR | 9 | O43432 | Eukaryotic translation initiation factor 4 g | EIF4G3 | 176.653 | 2 | 54.382847 | yes | 1.51E-07 | 520.2984 | 1241.1127 |
| LTPQMFNQLMK | 11 | O43432 | Eukaryotic translation initiation factor 4 g | EIF4G3 | 176.653 | 2 | 54.382847 | yes | 1.388E-07 | 675.84906 | 1176.8883 |
| ELEAASAPEER | 11 | O43432 | Eukaryotic translation initiation factor 4 g | EIF4G3 | 176.653 | 2 | 54.382847 | yes | 1.434E-07 | 601.28839 | 1054.4965 |
| LCNWVSR | 7 | O43432 | Eukaryotic translation initiation factor 4 g | EIF4G3 | 176.653 | 2 | 54.382847 | yes | 1.736E-07 | 467.73181 | 903.18781 |
| AAIIADSSTFR | 11 | O43432 | Eukaryotic translation initiation factor 4 g | EIF4G3 | 176.653 | 2 | 54.382847 | yes | 1.086E-07 | 576.3064 | 871.5423 |
| VPMADKPGNTVN | 14 | O43432 | Eukaryotic translation initiation factor 4 g | EIF4G3 | 176.653 | 3 | 54.382847 | yes | 3.398E-07 | 515.93292 | 399.14276 |
| VPMADKPGNTVN | 14 | O43432 | Eukaryotic translation initiation factor 4 g | EIF4G3 | 176.653 | 3 | 54.382847 | yes | 1.206E-07 | 773.39575 | 399.14276 |
| VPMADKPGNTVN | 14 | O43432 | Eukaryotic translation initiation factor 4 g | EIF4G3 | 176.653 | 2 | 54.382847 | yes | 3.398E-07 | 515.93292 | 399.14276 |
| VPMADKPGNTVN | 14 | O43432 | Eukaryotic translation initiation factor 4 g | EIF4G3 | 176.653 | 2 | 54.382847 | yes | 1.206E-07 | 773.39575 | 399.14276 |
| ITKPTIDEK | 9 | O43432 | Eukaryotic translation initiation factor 4 g | EIF4G3 | 176.653 | 2 | 54.382847 | yes | 1.443E-07 | 522.80042 | 760.77405 |
| QLTGGVDVER | 10 | O43432 | Eukaryotic translation initiation factor 4 g | EIF4G3 | 176.653 | 2 | 54.382847 | yes | 9.722E-08 | 537.2829 | 716.84351 |
| AALSEEELER | 10 | O43432 | Eukaryotic translation initiation factor 4 g | EIF4G3 | 176.653 | 2 | 54.382847 | yes | 1.065E-07 | 573.78552 | 662.91791 |
| QVSGLTVDTEER | 12 | O43432 | Eukaryotic translation initiation factor 4 g | EIF4G3 | 176.653 | 2 | 54.382847 | yes | 9.722E-08 | 667.33331 | 647.82098 |
| DLLDNQSQEEQR | 12 | O43432 | Eukaryotic translation initiation factor 4 g | EIF4G3 | 176.653 | 2 | 54.382847 | yes | 9.838E-08 | 737.84204 | 527.38226 |
| TTEEMLEAELELK | 13 | O43432 | Eukaryotic translation initiation factor 4 g | EIF4G3 | 176.653 | 2 | 54.382847 | yes | 1.007E-07 | 768.37909 | 515.38074 |
| MSQGFHPER | 9 | O43432 | Eukaryotic translation initiation factor 4 g | EIF4G3 | 176.653 | 2 | 54.382847 | yes | 9.722E-08 | 544.75073 | 459.17194 |
| VDEGGWNTVQGA | 13 | O43432 | Eukaryotic translation initiation factor 4 g | EIF4G3 | 176.653 | 2 | 54.382847 | yes | 2.702E-07 | 680.82825 | 452.33786 |
| ELSAEELYK | 9 | O43432 | Eukaryotic translation initiation factor 4 g | EIF4G3 | 176.653 | 2 | 54.382847 | yes | 1.313E-07 | 541.27423 | 418.90915 |
| DKADDDVFEK | 10 | O43432 | Eukaryotic translation initiation factor 4 g | EIF4G3 | 176.653 | 2 | 54.382847 | yes | 1.138E-07 | 591.26965 | 411.20435 |
| IEEQEEQR | 8 | O43432 | Eukaryotic translation initiation factor 4 g | EIF4G3 | 176.653 | 2 | 54.382847 | yes | 1.022E-07 | 530.7489 | 397.19269 |
| LHDELEEAK | 9 | O43432 | Eukaryotic translation initiation factor 4 g | EIF4G3 | 176.653 | 2 | 54.382847 | yes | 1.004E-07 | 542.26947 | 308.46933 |
| KIITVSVK | 8 | O43432 | Eukaryotic translation initiation factor 4 g | EIF4G3 | 176.653 | 2 | 54.382847 | yes | 2.873E-07 | 444.29987 | 305.52853 |
| GVPLLNVGSR | 10 | O43432 | Eukaryotic translation initiation factor 4 g | EIF4G3 | 176.653 | 2 | 54.382847 | yes | 5.14E-07 | 506.3009 | 305.24847 |
| VLESEQDK | 8 | O43432 | Eukaryotic translation initiation factor 4 g | EIF4G3 | 176.653 | 2 | 54.382847 | yes | 4.947E-07 | 474.23764 | 281.75443 |
| EKNDKPLPSATAR | 13 | O43432 | Eukaryotic translation initiation factor 4 g | EIF4G3 | 176.653 | 3 | 54.382847 | yes | 3.723E-07 | 476.2598 | 132.55608 |
| EKNDKPLPSATAR | 13 | O43432 | Eukaryotic translation initiation factor 4 g | EIF4G3 | 176.653 | 3 | 54.382847 | yes | 9.651E-07 | 713.88605 | 132.55608 |
| EKNDKPLPSATAR | 13 | O43432 | Eukaryotic translation initiation factor 4 g | EIF4G3 | 176.653 | 2 | 54.382847 | yes | 3.723E-07 | 476.2598 | 132.55608 |
| EKNDKPLPSATAR | 13 | O43432 | Eukaryotic translation initiation factor 4 g | EIF4G3 | 176.653 | 2 | 54.382847 | yes | 9.651E-07 | 713.88605 | 132.55608 |
| NDKPLPSATAR | 11 | O43432 | Eukaryotic translation initiation factor 4 g | EIF4G3 | 176.653 | 2 | 54.382847 | yes | 2.907E-06 | 585.31732 | 213.49869 |
| GVIDLVFEK | 9 | O43432 | Eukaryotic translation initiation factor 4 g | EIF4G3 | 176.653 | 2 | 54.382847 | yes | 8.564E-07 | 510.29224 | 196.25159 |
| CELSSPR | 7 | O43432 | Eukaryotic translation initiation factor 4 g | EIF4G3 | 176.653 | 2 | 54.382847 | yes | 3.998E-06 | 424.70016 | 189.91319 |
| YLDSDTEK | 8 | O43432 | Eukaryotic translation initiation factor 4 g | EIF4G3 | 176.653 | 2 | 54.382847 | yes | 2.673E-06 | 485.72183 | 186.49484 |
| EAMQCVEELNAQ | 20 | O43432 | Eukaryotic translation initiation factor 4 g | EIF4G3 | 176.653 | 3 | 54.382847 | yes | 1.241E-07 | 781.7243 | 165.69475 |

| | | | | | | | | | | | |
|---|---|---|---|---|---|---|---|---|---|---|---|
| SPVPAQIAITVPK | 13 | O43432 | Eukaryotic translation initiation factor 4 g | EIF4G3 | 176.653 | 2 | 54.382847 | yes | 2.222E-07 | 660.89795 | 141.07008 |
| NPTPPIGR | 8 | O43432 | Eukaryotic translation initiation factor 4 g | EIF4G3 | 176.653 | 2 | 54.382847 | yes | 4.922E-07 | 426.24033 | 137.86278 |
| ELQALYALQASIV | 14 | O43432 | Eukaryotic translation initiation factor 4 g | EIF4G3 | 176.653 | 2 | 54.382847 | yes | 1.192E-07 | 773.94562 | 133.31015 |
| KVQQLMTK | 8 | O43432 | Eukaryotic translation initiation factor 4 g | EIF4G3 | 176.653 | 2 | 54.382847 | yes | 1.171E-07 | 488.28641 | 123.05671 |
| AENAWKPSQK | 10 | O43432 | Eukaryotic translation initiation factor 4 g | EIF4G3 | 176.653 | 2 | 54.382847 | yes | 1.147E-07 | 579.79877 | 106.80249 |
| QEEKPKPDPVLK | 12 | O43432 | Eukaryotic translation initiation factor 4 g | EIF4G3 | 176.653 | 2 | 54.382847 | yes | 9.244E-07 | 704.39575 | 48.890709 |
| QEEKPKPDPVLK | 12 | O43432 | Eukaryotic translation initiation factor 4 g | EIF4G3 | 176.653 | 2 | 54.382847 | yes | 5.28E-07 | 469.93292 | 48.890709 |
| QEEKPKPDPVLK | 12 | O43432 | Eukaryotic translation initiation factor 4 g | EIF4G3 | 176.653 | 3 | 54.382847 | yes | 9.244E-07 | 704.39575 | 48.890709 |
| QEEKPKPDPVLK | 12 | O43432 | Eukaryotic translation initiation factor 4 g | EIF4G3 | 176.653 | 3 | 54.382847 | yes | 5.28E-07 | 469.93292 | 48.890709 |
| KELSAEELYK | 10 | O43432 | Eukaryotic translation initiation factor 4 g | EIF4G3 | 176.653 | 2 | 54.382847 | yes | 5.123E-07 | 605.32172 | 95.666122 |
| KAENAWKPSQK | 11 | O43432 | Eukaryotic translation initiation factor 4 g | EIF4G3 | 176.653 | 2 | 54.382847 | yes | 1.555E-07 | 643.84619 | 91.714188 |
| TCLSEDAK | 8 | O43432 | Eukaryotic translation initiation factor 4 g | EIF4G3 | 176.653 | 2 | 54.382847 | yes | 1.379E-07 | 462.21057 | 84.920334 |
| LHDELEEAKDK | 11 | O43432 | Eukaryotic translation initiation factor 4 g | EIF4G3 | 176.653 | 2 | 54.382847 | yes | 5.918E-07 | 663.83044 | 82.918129 |
| NHDEESLECLCR | 12 | O43432 | Eukaryotic translation initiation factor 4 g | EIF4G3 | 176.653 | 2 | 54.382847 | no | 9.722E-08 | 781.32214 | 77.382744 |
| ELTIEFSK | 8 | O43432 | Eukaryotic translation initiation factor 4 g | EIF4G3 | 176.653 | 2 | 54.382847 | yes | 3.365E-07 | 483.76077 | 67.033691 |
| AEEELSIDK | 9 | O43432 | Eukaryotic translation initiation factor 4 g | EIF4G3 | 176.653 | 2 | 54.382847 | yes | 2.139E-07 | 517.25604 | 60.289398 |
| RSPVPAQIAITVPK | 14 | O43432 | Eukaryotic translation initiation factor 4 g | EIF4G3 | 176.653 | 2 | 54.382847 | yes | 1.847E-07 | 738.94849 | 54.778633 |
| KVGALWR | 7 | O43432 | Eukaryotic translation initiation factor 4 g | EIF4G3 | 176.653 | 2 | 54.382847 | yes | 4.051E-06 | 415.25577 | 54.203148 |
| ADDDVFEK | 8 | O43432 | Eukaryotic translation initiation factor 4 g | EIF4G3 | 176.653 | 2 | 54.382847 | yes | 1.423E-07 | 469.70871 | 51.266289 |
| ELTIEFSKPLLPVG | 15 | O43432 | Eukaryotic translation initiation factor 4 g | EIF4G3 | 176.653 | 3 | 54.382847 | yes | 1.585E-07 | 566.9978 | 40.651505 |
| ESAKPEISAMSAH | 15 | O43432 | Eukaryotic translation initiation factor 4 g | EIF4G3 | 176.653 | 2 | 54.382847 | yes | 2.536E-07 | 800.88544 | 33.021286 |
| FIGELFK | 7 | O43432 | Eukaryotic translation initiation factor 4 g | EIF4G3 | 176.653 | 2 | 54.382847 | no | 1.637E-07 | 427.24454 | 30.988998 |
| QRVPILLK | 8 | O43432 | Eukaryotic translation initiation factor 4 g | EIF4G3 | 176.653 | 2 | 54.382847 | yes | 1.58E-05 | 483.82657 | 13.311254 |
| DCDPEDWPR | 9 | O43435 | T-box transcription factor TBX1 OS=Hor | TBX1 | 43.133 | 2 | 38.81539 | yes | 4.97E-07 | 595.23254 | 52.068096 |
| FTAVTAYQNHR | 11 | O43435 | T-box transcription factor TBX1 OS=Hor | TBX1 | 43.133 | 2 | 38.81539 | yes | 5.321E-07 | 654.32819 | 45.496777 |
| ARMEQTIADVK | 11 | O43439 | Protein CBFA2T2 OS=Homo sapiens OX | CBFA2T2 | 67.133 | 2 | 37.089626 | yes | 6.31E-07 | 631.33209 | 113.06348 |
| IIDGLLVMR | 9 | O43447 | Peptidyl-prolyl cis-trans isomerase H OS= | PPIH | 19.209 | 2 | 52.48714 | yes | 2.202E-07 | 515.30988 | 798.3064 |
| IELFADVVPK | 10 | O43447 | Peptidyl-prolyl cis-trans isomerase H OS= | PPIH | 19.209 | 2 | 52.48714 | yes | 1.109E-07 | 565.82642 | 461.12921 |
| GPFADENFK | 9 | O43447 | Peptidyl-prolyl cis-trans isomerase H OS= | PPIH | 19.209 | 2 | 52.48714 | yes | 2.821E-07 | 512.74036 | 338.98859 |
| QFCTGEFR | 8 | O43447 | Peptidyl-prolyl cis-trans isomerase H OS= | PPIH | 19.209 | 2 | 52.48714 | yes | 9.722E-08 | 522.73199 | 159.40462 |
| KIENVPTGPNNKP | 14 | O43447 | Peptidyl-prolyl cis-trans isomerase H OS= | PPIH | 19.209 | 3 | 52.48714 | yes | 2.18E-07 | 512.62286 | 40.653629 |
| AQLTAVTPDTR | 11 | O43464 | Serine protease HTRA2, mitochondrial O | HTRA2 | 48.841 | 2 | 34.751595 | yes | 9.479E-07 | 586.81714 | 188.5739 |
| SQYNFIADVVEK | 12 | O43464 | Serine protease HTRA2, mitochondrial O | HTRA2 | 48.841 | 2 | 34.751595 | yes | 3.068E-06 | 706.85645 | 33.350311 |
| TQSQLAVQIR | 10 | O43464 | Serine protease HTRA2, mitochondrial O | HTRA2 | 48.841 | 2 | 34.751595 | yes | 3.902E-06 | 572.32764 | 10.555243 |
| TTPTLGLEEVLSSR | 14 | O43474 | Krueppel-like factor 4 OS=Homo sapiens | KLF4 | 54.671 | 2 | 48.626858 | yes | 1.122E-07 | 751.90668 | 932.8244 |
| TATHTCDYAGCGI | 13 | O43474 | Krueppel-like factor 4 OS=Homo sapiens | KLF4 | 54.671 | 2 | 48.626858 | yes | 5.77E-07 | 721.29535 | 20.820801 |
| APHLQLIEGK | 10 | O43491 | Band 4.1-like protein 2 OS=Homo sapien | EPB41L2 | 112.589 | 2 | 34.34331 | yes | 4.119E-05 | 553.32184 | 44.941082 |
| LVSPEQPPK | 9 | O43491 | Band 4.1-like protein 2 OS=Homo sapien | EPB41L2 | 112.589 | 2 | 34.34331 | yes | 1.028E-06 | 497.78201 | 17.485817 |
| NAWGNLSYADLIT | 14 | O43524 | Forkhead box protein O3 OS=Homo sapi | FOXO3 | 71.277 | 2 | 53.254234 | no | 9.722E-08 | 797.4046 | 172.67406 |

| | | | | | | | | | | | |
|---|---|---|---|---|---|---|---|---|---|---|---|
| SSDELDAWTDFR | 12 | O43524 | Forkhead box protein O3 OS=Homo sapi | FOXO3 | 71.277 | 2 | 53.254234 | yes | 9.722E-08 | 721.31512 | 136.17447 |
| SSWWIINPDGGK | 12 | O43524 | Forkhead box protein O3 OS=Homo sapi | FOXO3 | 71.277 | 2 | 53.254234 | yes | 1.588E-07 | 680.33826 | 122.5776 |
| LEIESNICR | 9 | O43572 | A-kinase anchor protein 10, mitochondria | AKAP10 | 73.819 | 2 | 34.763687 | yes | 9.456E-07 | 567.28461 | 156.70978 |
| LDADYPLR | 8 | O43583 | Density-regulated protein OS=Homo sapi | DENR | 22.093 | 2 | 48.156563 | yes | 6.668E-07 | 481.75073 | 158.53027 |
| VCGLATFEIDLK | 12 | O43583 | Density-regulated protein OS=Homo sapi | DENR | 22.093 | 2 | 48.156563 | yes | 1.19E-07 | 683.35779 | 148.70796 |
| LTVENSPK | 8 | O43583 | Density-regulated protein OS=Homo sapi | DENR | 22.093 | 2 | 48.156563 | yes | 2.588E-06 | 444.24527 | 115.11248 |
| AADISESSGADCK | 13 | O43583 | Density-regulated protein OS=Homo sapi | DENR | 22.093 | 2 | 48.156563 | yes | 3.967E-07 | 676.78534 | 98.162407 |
| NLLTPLMEK | 9 | O43592 | Exportin-T OS=Homo sapiens OX=9606 | XPOT | 109.965 | 2 | 50.75294 | yes | 9.723E-08 | 529.79938 | 85.810936 |
| DGPVGFADFVYK | 12 | O43592 | Exportin-T OS=Homo sapiens OX=9606 | XPOT | 109.965 | 2 | 50.75294 | yes | 4.51E-07 | 657.8219 | 74.907372 |
| YSELTTVQQQLIR | 13 | O43592 | Exportin-T OS=Homo sapiens OX=9606 | XPOT | 109.965 | 2 | 50.75294 | yes | 1.136E-07 | 789.92792 | 55.803692 |
| NAQEALQAIETK | 12 | O43592 | Exportin-T OS=Homo sapiens OX=9606 | XPOT | 109.965 | 2 | 50.75294 | yes | 6.085E-07 | 658.34625 | 17.975315 |
| YDESDNAFIR | 10 | O43615 | Mitochondrial import inner membrane tra | TIMM44 | 51.356 | 2 | 49.652546 | yes | 1.013E-07 | 615.27527 | 32.213516 |
| KTPEELLR | 8 | O43633 | Charged multivesicular body protein 2a C | CHMP2A | 25.104 | 2 | 46.545223 | yes | 3.584E-07 | 493.28748 | 335.88544 |
| TPEELLR | 7 | O43633 | Charged multivesicular body protein 2a C | CHMP2A | 25.104 | 2 | 46.545223 | yes | 1.532E-07 | 429.23999 | 190.28694 |
| ANIQAVSLK | 9 | O43633 | Charged multivesicular body protein 2a C | CHMP2A | 25.104 | 2 | 46.545223 | yes | 1.62E-07 | 472.2822 | 108.03176 |
| QAEIMDMK | 8 | O43633 | Charged multivesicular body protein 2a C | CHMP2A | 25.104 | 2 | 46.545223 | yes | 3.504E-06 | 483.22516 | 42.36895 |
| KAEAAASALADAI | 19 | O43633 | Charged multivesicular body protein 2a C | CHMP2A | 25.104 | 3 | 46.545223 | yes | 5.606E-06 | 639.64471 | 24.026302 |
| SPYLFSCGEDK | 11 | O43660 | Pleiotropic regulator 1 OS=Homo sapiens | PLRG1 | 57.194 | 2 | 51.915287 | yes | 1.303E-07 | 651.78717 | 247.69554 |
| HSVHTLVFR | 9 | O43660 | Pleiotropic regulator 1 OS=Homo sapiens | PLRG1 | 57.194 | 2 | 51.915287 | yes | 9.728E-08 | 548.30652 | 236.63086 |
| NEYGPVLHMPTSK | 13 | O43660 | Pleiotropic regulator 1 OS=Homo sapiens | PLRG1 | 57.194 | 2 | 51.915287 | yes | 4.093E-07 | 736.86377 | 159.93909 |
| GPQNATDSYVHK | 12 | O43660 | Pleiotropic regulator 1 OS=Homo sapiens | PLRG1 | 57.194 | 2 | 51.915287 | yes | 2.274E-07 | 658.81512 | 132.61475 |
| CQAAEPQIITGSHD | 18 | O43660 | Pleiotropic regulator 1 OS=Homo sapiens | PLRG1 | 57.194 | 3 | 51.915287 | yes | 3.67E-07 | 666.6618 | 123.55459 |
| LSLTGHISTVR | 11 | O43660 | Pleiotropic regulator 1 OS=Homo sapiens | PLRG1 | 57.194 | 2 | 51.915287 | yes | 9.722E-08 | 592.34332 | 116.3474 |
| HYTFASGSPDNIK | 13 | O43660 | Pleiotropic regulator 1 OS=Homo sapiens | PLRG1 | 57.194 | 2 | 51.915287 | yes | 3.279E-06 | 718.84387 | 77.850243 |
| VISGHLGWVR | 10 | O43660 | Pleiotropic regulator 1 OS=Homo sapiens | PLRG1 | 57.194 | 2 | 51.915287 | yes | 1.672E-07 | 562.3222 | 65.7883 |
| LRNEYGPVLHMPT | 15 | O43660 | Pleiotropic regulator 1 OS=Homo sapiens | PLRG1 | 57.194 | 3 | 51.915287 | yes | 1.736E-07 | 581.30664 | 50.096806 |
| PVPLDEESHK | 10 | O43660 | Pleiotropic regulator 1 OS=Homo sapiens | PLRG1 | 57.194 | 2 | 51.915287 | yes | 3.072E-07 | 575.79059 | 32.208126 |
| EDDTATEETHPVS | 20 | O43660 | Pleiotropic regulator 1 OS=Homo sapiens | PLRG1 | 57.194 | 3 | 51.915287 | yes | 2.958E-07 | 775.71301 | 31.057081 |
| LLTAEADK | 8 | O43660 | Pleiotropic regulator 1 OS=Homo sapiens | PLRG1 | 57.194 | 2 | 51.915287 | yes | 1.097E-06 | 430.73984 | 28.955561 |
| LQIPEEER | 8 | O43663 | Protein regulator of cytokinesis 1 OS=Ho | PRC1 | 71.607 | 2 | 55.604584 | yes | 1.602E-07 | 507.26672 | 3610.9944 |
| REEFVSIK | 8 | O43663 | Protein regulator of cytokinesis 1 OS=Ho | PRC1 | 71.607 | 2 | 55.604584 | yes | 4.849E-07 | 504.27966 | 2458.0459 |
| IELWEQEHSK | 10 | O43663 | Protein regulator of cytokinesis 1 OS=Ho | PRC1 | 71.607 | 2 | 55.604584 | yes | 9.889E-08 | 649.82239 | 2330.0029 |
| EIWELIGIPEDQR | 13 | O43663 | Protein regulator of cytokinesis 1 OS=Ho | PRC1 | 71.607 | 2 | 55.604584 | yes | 9.892E-08 | 799.41467 | 1948.8524 |
| LFLEFER | 7 | O43663 | Protein regulator of cytokinesis 1 OS=Ho | PRC1 | 71.607 | 2 | 55.604584 | yes | 1.248E-07 | 477.25818 | 1908.3037 |
| SEVLAEESIVCLQK | 14 | O43663 | Protein regulator of cytokinesis 1 OS=Ho | PRC1 | 71.607 | 2 | 55.604584 | yes | 1.7E-07 | 802.91364 | 1797.1903 |
| EAVATIMSGSK | 11 | O43663 | Protein regulator of cytokinesis 1 OS=Ho | PRC1 | 71.607 | 2 | 55.604584 | yes | 8.088E-07 | 547.28156 | 1712.97 |
| QTETEMLYGSAPR | 13 | O43663 | Protein regulator of cytokinesis 1 OS=Ho | PRC1 | 71.607 | 2 | 55.604584 | yes | 2.782E-07 | 741.84833 | 1652.5358 |
| SQNEAVCEGLR | 11 | O43663 | Protein regulator of cytokinesis 1 OS=Ho | PRC1 | 71.607 | 2 | 55.604584 | yes | 9.722E-08 | 631.79333 | 1313.6727 |
| TQVELMR | 7 | O43663 | Protein regulator of cytokinesis 1 OS=Ho | PRC1 | 71.607 | 2 | 55.604584 | yes | 4.946E-07 | 438.73401 | 993.10437 |

| | | | | | | | | | | | |
|---|---|---|---|---|---|---|---|---|---|---|---|
| ELLDMMIAEEESL | 14 | O43663 | Protein regulator of cytokinesis 1 OS=Ho | PRC1 | 71.607 | 2 | 55.604584 | yes | 9.809E-08 | 825.90186 | 960.96283 |
| ALQLEVDR | 8 | O43663 | Protein regulator of cytokinesis 1 OS=Ho | PRC1 | 71.607 | 2 | 55.604584 | yes | 9.975E-08 | 472.26401 | 805.53821 |
| ELFEGVQK | 8 | O43663 | Protein regulator of cytokinesis 1 OS=Ho | PRC1 | 71.607 | 2 | 55.604584 | yes | 5.139E-07 | 475.25311 | 715.86987 |
| PIFGGTVYHSPVSF | 14 | O43663 | Protein regulator of cytokinesis 1 OS=Ho | PRC1 | 71.607 | 2 | 55.604584 | yes | 1.637E-07 | 758.89899 | 283.60693 |
| PIFGGTVYHSPVSF | 14 | O43663 | Protein regulator of cytokinesis 1 OS=Ho | PRC1 | 71.607 | 2 | 55.604584 | yes | 5.721E-07 | 506.2684 | 283.60693 |
| PIFGGTVYHSPVSF | 14 | O43663 | Protein regulator of cytokinesis 1 OS=Ho | PRC1 | 71.607 | 3 | 55.604584 | yes | 1.637E-07 | 758.89899 | 283.60693 |
| PIFGGTVYHSPVSF | 14 | O43663 | Protein regulator of cytokinesis 1 OS=Ho | PRC1 | 71.607 | 3 | 55.604584 | yes | 5.721E-07 | 506.2684 | 283.60693 |
| KVIEAIR | 7 | O43663 | Protein regulator of cytokinesis 1 OS=Ho | PRC1 | 71.607 | 2 | 55.604584 | yes | 9.722E-08 | 414.76871 | 517.09192 |
| DPSLSDSSTVGLQI | 14 | O43663 | Protein regulator of cytokinesis 1 OS=Ho | PRC1 | 71.607 | 2 | 55.604584 | yes | 1.109E-07 | 731.36261 | 448.7352 |
| LNTTTMSNATANS | 16 | O43663 | Protein regulator of cytokinesis 1 OS=Ho | PRC1 | 71.607 | 2 | 55.604584 | yes | 1.152E-07 | 841.41235 | 409.28833 |
| QHVTTLR | 7 | O43663 | Protein regulator of cytokinesis 1 OS=Ho | PRC1 | 71.607 | 2 | 55.604584 | yes | 1.547E-06 | 427.74579 | 323.05267 |
| SDATSGILNSTNIQ | 15 | O43663 | Protein regulator of cytokinesis 1 OS=Ho | PRC1 | 71.607 | 2 | 55.604584 | yes | 1.026E-07 | 754.36536 | 141.4337 |
| RSEVLAEESIVCLQ | 15 | O43663 | Protein regulator of cytokinesis 1 OS=Ho | PRC1 | 71.607 | 3 | 55.604584 | yes | 9.722E-08 | 587.6452 | 139.0528 |
| ALQLEVDRLEELK | 13 | O43663 | Protein regulator of cytokinesis 1 OS=Ho | PRC1 | 71.607 | 2 | 55.604584 | yes | 1.173E-07 | 778.43811 | 95.366287 |
| RGLAPNTPGK | 10 | O43663 | Protein regulator of cytokinesis 1 OS=Ho | PRC1 | 71.607 | 2 | 55.604584 | yes | 1.054E-07 | 505.79071 | 90.447205 |
| NYYEVHK | 7 | O43663 | Protein regulator of cytokinesis 1 OS=Ho | PRC1 | 71.607 | 2 | 55.604584 | yes | 4.312E-06 | 476.7298 | 84.044167 |
| LPPSGSKPVAASTC | 17 | O43663 | Protein regulator of cytokinesis 1 OS=Ho | PRC1 | 71.607 | 2 | 55.604584 | yes | 1.535E-06 | 822.42468 | 53.670776 |
| KQTETEMLYGSAF | 14 | O43663 | Protein regulator of cytokinesis 1 OS=Ho | PRC1 | 71.607 | 2 | 55.604584 | yes | 9.825E-08 | 805.89575 | 51.703453 |
| ETIDAVPNAIPGR | 13 | O43670 | BUB3-interacting and GLEBS motif-cont | ZNF207 | 50.751 | 2 | 51.21952 | yes | 1.368E-07 | 676.86206 | 3190.2358 |
| LIHPDEDISLEER | 13 | O43670 | BUB3-interacting and GLEBS motif-cont | ZNF207 | 50.751 | 2 | 51.21952 | yes | 1.455E-07 | 783.39392 | 1754 |
| ILIQHQK | 7 | O43670 | BUB3-interacting and GLEBS motif-cont | ZNF207 | 50.751 | 2 | 51.21952 | yes | 1.223E-07 | 440.27417 | 1272.6553 |
| PATLTTTSATSK | 12 | O43670 | BUB3-interacting and GLEBS motif-cont | ZNF207 | 50.751 | 2 | 51.21952 | yes | 9.722E-08 | 589.81677 | 251.97902 |
| QLKPWCWYCNR | 11 | O43670 | BUB3-interacting and GLEBS motif-cont | ZNF207 | 50.751 | 2 | 51.21952 | yes | 1.287E-07 | 805.8714 | 112.51172 |
| PAASITSKPATLTT | 20 | O43670 | BUB3-interacting and GLEBS motif-cont | ZNF207 | 50.751 | 3 | 51.21952 | yes | 7.109E-07 | 645.35284 | 10.304014 |
| TMAVLQIEAEK | 11 | O43674 | NADH dehydrogenase [ubiquinone] 1 bet | NDUFB5 | 21.751 | 2 | 50.08436 | yes | 9.909E-08 | 616.83136 | 239.08881 |
| GDGPWYYYETIDK | 13 | O43674 | NADH dehydrogenase [ubiquinone] 1 bet | NDUFB5 | 21.751 | 2 | 50.08436 | yes | 1.042E-07 | 803.85663 | 137.23863 |
| ANPDLPILIR | 10 | O43678 | NADH dehydrogenase [ubiquinone] 1 alp | NDUFA2 | 10.922 | 2 | 26.976917 | yes | 3.035E-06 | 561.33752 | 28.364439 |
| LLNFPTIVER | 10 | O43681 | ATPase GET3 OS=Homo sapiens OX=96 | GET3 | 38.793 | 2 | 40.56462 | yes | 3.958E-07 | 601.35059 | 58.666294 |
| NQGSELSGVISSAC | 16 | O43683 | Mitotic checkpoint serine/threonine-prote | BUB1 | 122.376 | 2 | 36.053753 | yes | 7.46E-07 | 826.38324 | 22.328293 |
| VAVEYLDPSPEVQ | 14 | O43684 | Mitotic checkpoint protein BUB3 OS=Hc | BUB3 | 37.155 | 2 | 52.615936 | yes | 1.025E-07 | 787.40906 | 4901.3623 |
| LYDVPANSMR | 10 | O43684 | Mitotic checkpoint protein BUB3 OS=Hc | BUB3 | 37.155 | 2 | 52.615936 | yes | 9.722E-08 | 583.28717 | 2338.1855 |
| LNQPPEDGISSVK | 13 | O43684 | Mitotic checkpoint protein BUB3 OS=Hc | BUB3 | 37.155 | 2 | 52.615936 | yes | 1.355E-07 | 692.35938 | 2206.6956 |
| QGYVLSSIEGR | 11 | O43684 | Mitotic checkpoint protein BUB3 OS=Hc | BUB3 | 37.155 | 2 | 52.615936 | yes | 9.722E-08 | 604.81714 | 2141.0513 |
| VYTLSVSGDR | 10 | O43684 | Mitotic checkpoint protein BUB3 OS=Hc | BUB3 | 37.155 | 2 | 52.615936 | yes | 2.397E-07 | 548.78528 | 1925.7721 |
| VLVWDLR | 7 | O43684 | Mitotic checkpoint protein BUB3 OS=Hc | BUB3 | 37.155 | 2 | 52.615936 | yes | 2.462E-07 | 450.76871 | 896.37347 |
| NMGYVQQR | 8 | O43684 | Mitotic checkpoint protein BUB3 OS=Hc | BUB3 | 37.155 | 2 | 52.615936 | yes | 1.166E-07 | 498.23999 | 686.87891 |
| TPCNAGTFSQPEK | 13 | O43684 | Mitotic checkpoint protein BUB3 OS=Hc | BUB3 | 37.155 | 2 | 52.615936 | yes | 1.035E-07 | 718.82739 | 633.9953 |
| YQHTGAVLDCAFY | 28 | O43684 | Mitotic checkpoint protein BUB3 OS=Hc | BUB3 | 37.155 | 4 | 52.615936 | yes | 1.001E-07 | 797.62274 | 566.49469 |
| QVTDAETKPK | 10 | O43684 | Mitotic checkpoint protein BUB3 OS=Hc | BUB3 | 37.155 | 2 | 52.615936 | yes | 1.02E-07 | 558.7984 | 281.73514 |

| | | | | | | | | | | | |
|---|---|---|---|---|---|---|---|---|---|---|---|
| MHDLNTDQENLV | 20 | O43684 | Mitotic checkpoint protein BUB3 OS=Hc | BUB3 | 37.155 | 4 | 52.615936 | yes | 9.722E-08 | 569.77356 | 78.790169 |
| CVEYCPEVNVMV | 21 | O43684 | Mitotic checkpoint protein BUB3 OS=Hc | BUB3 | 37.155 | 3 | 52.615936 | yes | 2.608E-07 | 834.37634 | 43.49535 |
| GISQEQMQEFR | 11 | O43707 | Alpha-actinin-4 OS=Homo sapiens OX=9 | ACTN4 | 104.855 | 2 | 49.013344 | yes | 1.098E-07 | 676.81677 | 192.71025 |
| TIPWLEDR | 8 | O43707 | Alpha-actinin-4 OS=Homo sapiens OX=9 | ACTN4 | 104.855 | 2 | 49.013344 | yes | 4.257E-07 | 515.27185 | 162.01814 |
| AGTQIENIDEDFR | 13 | O43707 | Alpha-actinin-4 OS=Homo sapiens OX=9 | ACTN4 | 104.855 | 2 | 49.013344 | yes | 1.076E-07 | 754.3548 | 107.16431 |
| ACLISLGYDVEND | 14 | O43707 | Alpha-actinin-4 OS=Homo sapiens OX=9 | ACTN4 | 104.855 | 2 | 49.013344 | yes | 5.483E-07 | 812.88544 | 65.026505 |
| LVSIGAEEIVDGNA | 15 | O43707 | Alpha-actinin-4 OS=Homo sapiens OX=9 | ACTN4 | 104.855 | 2 | 49.013344 | yes | 0.0001197 | 757.90668 | 55.66576 |
| QLEAIDQLHLEYA | 14 | O43707 | Alpha-actinin-4 OS=Homo sapiens OX=9 | ACTN4 | 104.855 | 2 | 49.013344 | yes | 1.17E-07 | 835.94104 | 48.743774 |
| AGFSGGMVVDYP | 16 | O43709 | Probable 18S rRNA (guanine-N(7))-meth | BUD23 | 31.881 | 2 | 50.66373 | yes | 9.729E-08 | 800.3772 | 481.10034 |
| MIDIQTR | 7 | O43709 | Probable 18S rRNA (guanine-N(7))-meth | BUD23 | 31.881 | 2 | 50.66373 | yes | 2.061E-07 | 438.73401 | 310.96039 |
| AVLQLYPENSEQL | 22 | O43709 | Probable 18S rRNA (guanine-N(7))-meth | BUD23 | 31.881 | 3 | 50.66373 | yes | 1.151E-07 | 830.44263 | 191.02455 |
| EVRPDTQYTGR | 11 | O43709 | Probable 18S rRNA (guanine-N(7))-meth | BUD23 | 31.881 | 2 | 50.66373 | yes | 2.813E-07 | 661.32837 | 55.373207 |
| VQICELEK | 8 | O43711 | T-cell leukemia homeobox protein 3 OS= | TLX3 | 31.868 | 2 | 39.636967 | yes | 4.468E-07 | 509.7655 | 143.20013 |
| GWEAFLNAPEANI | 13 | O43719 | HIV Tat-specific factor 1 OS=Homo sapi | HTATSF1 | 85.853 | 2 | 48.02909 | yes | 1.211E-07 | 737.8573 | 90.744026 |
| LLDEDEIR | 8 | O43719 | HIV Tat-specific factor 1 OS=Homo sapi | HTATSF1 | 85.853 | 2 | 48.02909 | yes | 4.874E-07 | 501.75876 | 68.482491 |
| EEEYVAPPR | 9 | O43734 | E3 ubiquitin ligase TRAF3IP2 OS=Homo | TRAF3IP2 | 64.667 | 2 | 45.262695 | yes | 2.611E-07 | 545.26422 | 194.73016 |
| TSNLPEELR | 9 | O43734 | E3 ubiquitin ligase TRAF3IP2 OS=Homo | TRAF3IP2 | 64.667 | 2 | 45.262695 | yes | 1.962E-07 | 529.77747 | 125.99728 |
| MMQIEFIK | 8 | O43734 | E3 ubiquitin ligase TRAF3IP2 OS=Homo | TRAF3IP2 | 64.667 | 2 | 45.262695 | yes | 1.904E-07 | 520.26941 | 101.0441 |
| TSNLPEELRK | 10 | O43734 | E3 ubiquitin ligase TRAF3IP2 OS=Homo | TRAF3IP2 | 64.667 | 2 | 45.262695 | yes | 3.475E-07 | 593.82495 | 80.538651 |
| DPFYDMLATR | 10 | O43739 | Cytohesin-3 OS=Homo sapiens OX=960 | CYTH3 | 46.349 | 2 | 41.277714 | yes | 3.589E-07 | 614.78699 | 39.705078 |
| TVIGDYLGER | 10 | O43739 | Cytohesin-3 OS=Homo sapiens OX=960 | CYTH3 | 46.349 | 2 | 41.277714 | yes | 2.61E-06 | 561.79309 | 35.084469 |
| AALCAVHVIR | 10 | O43747 | AP-1 complex subunit gamma-1 OS=Hon | AP1G1 | 91.352 | 2 | 41.887257 | yes | 3.294E-07 | 555.31604 | 53.280899 |
| KLVPQLVR | 8 | O43747 | AP-1 complex subunit gamma-1 OS=Hon | AP1G1 | 91.352 | 2 | 41.887257 | yes | 5.147E-07 | 476.81876 | 39.7048 |
| ADCASGIFLAAEK | 13 | O43747 | AP-1 complex subunit gamma-1 OS=Hon | AP1G1 | 91.352 | 2 | 41.887257 | yes | 2.06E-06 | 676.82941 | 33.54501 |
| VLAINILGR | 9 | O43747 | AP-1 complex subunit gamma-1 OS=Hon | AP1G1 | 91.352 | 2 | 41.887257 | yes | 2.118E-05 | 484.81619 | 27.805515 |
| EPFPTIYVDSQK | 12 | O43776 | Asparagine--tRNA ligase, cytoplasmic OS | NARS1 | 62.943 | 2 | 50.834206 | yes | 9.766E-08 | 712.35883 | 369.4173 |
| LMTDTINEPILLCR | 14 | O43776 | Asparagine--tRNA ligase, cytoplasmic OS | NARS1 | 62.943 | 2 | 50.834206 | yes | 9.722E-08 | 844.93915 | 329.05353 |
| IGALEGYR | 8 | O43776 | Asparagine--tRNA ligase, cytoplasmic OS | NARS1 | 62.943 | 2 | 50.834206 | yes | 2.66E-07 | 439.74017 | 289.31439 |
| IFDSEEILAGYK | 12 | O43776 | Asparagine--tRNA ligase, cytoplasmic OS | NARS1 | 62.943 | 2 | 50.834206 | yes | 3.689E-07 | 692.85339 | 259.28787 |
| VFGWVHR | 7 | O43776 | Asparagine--tRNA ligase, cytoplasmic OS | NARS1 | 62.943 | 2 | 50.834206 | yes | 3.928E-07 | 450.74557 | 233.30882 |
| LEDLVCDVVDR | 11 | O43776 | Asparagine--tRNA ligase, cytoplasmic OS | NARS1 | 62.943 | 2 | 50.834206 | yes | 2.906E-07 | 666.82684 | 176.14519 |
| VLAELYVSDR | 10 | O43776 | Asparagine--tRNA ligase, cytoplasmic OS | NARS1 | 62.943 | 2 | 50.834206 | yes | 1.092E-07 | 582.81659 | 94.334511 |
| MNYSDAIVWLK | 11 | O43776 | Asparagine--tRNA ligase, cytoplasmic OS | NARS1 | 62.943 | 2 | 50.834206 | yes | 1.988E-07 | 670.33936 | 66.250046 |
| VEQLFQEFGNR | 11 | O43781 | Dual specificity tyrosine-phosphorylation | DYRK3 | 65.714 | 2 | 45.4353 | yes | 1.85E-07 | 683.84113 | 65.147659 |
| VIDFGSSCFEYQK | 13 | O43781 | Dual specificity tyrosine-phosphorylation | DYRK3 | 65.714 | 2 | 45.4353 | yes | 7.532E-07 | 790.35852 | 13.992859 |
| SMVVSHPHLVAEA | 15 | O43791 | Speckle-type POZ protein OS=Homo sap | SPOP | 42.133 | 3 | 54.906086 | yes | 1.008E-07 | 565.95935 | 310.97864 |
| SMVVSHPHLVAEA | 15 | O43791 | Speckle-type POZ protein OS=Homo sap | SPOP | 42.133 | 3 | 54.906086 | yes | 9.722E-08 | 848.43542 | 310.97864 |
| SMVVSHPHLVAEA | 15 | O43791 | Speckle-type POZ protein OS=Homo sap | SPOP | 42.133 | 2 | 54.906086 | yes | 1.008E-07 | 565.95935 | 310.97864 |
| SMVVSHPHLVAEA | 15 | O43791 | Speckle-type POZ protein OS=Homo sap | SPOP | 42.133 | 2 | 54.906086 | yes | 9.722E-08 | 848.43542 | 310.97864 |

| | | | | | | | | | | | |
|---|---|---|---|---|---|---|---|---|---|---|---|
| MADDLLAAADK | 11 | O43791 | Speckle-type POZ protein OS=Homo sap | SPOP | 42.133 | 2 | 54.906086 | yes | 1.883E-07 | 567.27899 | 303.96982 |
| VEINDVEPEVFK | 12 | O43791 | Speckle-type POZ protein OS=Homo sap | SPOP | 42.133 | 2 | 54.906086 | yes | 9.722E-08 | 709.36414 | 289.63361 |
| TQAVDFINYHASD | 21 | O43791 | Speckle-type POZ protein OS=Homo sap | SPOP | 42.133 | 3 | 54.906086 | yes | 5.277E-07 | 794.38513 | 131.34296 |
| SSTFSSGANDK | 11 | O43791 | Speckle-type POZ protein OS=Homo sap | SPOP | 42.133 | 2 | 54.906086 | yes | 9.787E-07 | 550.74634 | 83.586967 |
| DFLLDEANGLLPD | 15 | O43791 | Speckle-type POZ protein OS=Homo sap | SPOP | 42.133 | 2 | 54.906086 | no | 1.355E-07 | 837.91467 | 74.253738 |
| EMMCFIYTGK | 10 | O43791 | Speckle-type POZ protein OS=Homo sap | SPOP | 42.133 | 2 | 54.906086 | yes | 2.709E-07 | 640.2796 | 63.765701 |
| LADELGGLWENSI | 13 | O43791 | Speckle-type POZ protein OS=Homo sap | SPOP | 42.133 | 2 | 54.906086 | yes | 9.722E-08 | 730.36243 | 63.491306 |
| LEDLATLIQK | 10 | O43795 | Unconventional myosin-Ib OS=Homo sa | MYO1B | 131.986 | 2 | 53.541748 | yes | 9.722E-08 | 572.33459 | 383.72968 |
| YLGLLENVR | 9 | O43795 | Unconventional myosin-Ib OS=Homo sa | MYO1B | 131.986 | 2 | 53.541748 | no | 1.545E-07 | 538.80859 | 379.66351 |
| YNYLSLDSAK | 10 | O43795 | Unconventional myosin-Ib OS=Homo sa | MYO1B | 131.986 | 2 | 53.541748 | yes | 4.245E-07 | 587.29297 | 288.49881 |
| GDPLGGVISNYLL | 15 | O43795 | Unconventional myosin-Ib OS=Homo sa | MYO1B | 131.986 | 2 | 53.541748 | yes | 9.722E-08 | 787.92487 | 171.60925 |
| AAHIFNEALVCHQ | 15 | O43795 | Unconventional myosin-Ib OS=Homo sa | MYO1B | 131.986 | 3 | 53.541748 | yes | 9.722E-08 | 593.64215 | 151.03433 |
| FLNDTSLPHSCFR | 13 | O43795 | Unconventional myosin-Ib OS=Homo sa | MYO1B | 131.986 | 2 | 53.541748 | yes | 1.022E-07 | 797.37756 | 74.889175 |
| FLNDTSLPHSCFR | 13 | O43795 | Unconventional myosin-Ib OS=Homo sa | MYO1B | 131.986 | 2 | 53.541748 | yes | 4.622E-07 | 531.92078 | 74.889175 |
| FLNDTSLPHSCFR | 13 | O43795 | Unconventional myosin-Ib OS=Homo sa | MYO1B | 131.986 | 3 | 53.541748 | yes | 1.022E-07 | 797.37756 | 74.889175 |
| FLNDTSLPHSCFR | 13 | O43795 | Unconventional myosin-Ib OS=Homo sa | MYO1B | 131.986 | 3 | 53.541748 | yes | 4.622E-07 | 531.92078 | 74.889175 |
| VLYQVEGFVDK | 11 | O43795 | Unconventional myosin-Ib OS=Homo sa | MYO1B | 131.986 | 2 | 53.541748 | yes | 1.392E-07 | 648.84534 | 135.5591 |
| SSALVIQSYIR | 11 | O43795 | Unconventional myosin-Ib OS=Homo sa | MYO1B | 131.986 | 2 | 53.541748 | yes | 1.088E-07 | 618.85095 | 133.1857 |
| IIIAEVVNK | 9 | O43795 | Unconventional myosin-Ib OS=Homo sa | MYO1B | 131.986 | 2 | 53.541748 | yes | 2.329E-07 | 499.81586 | 123.57152 |
| QAYEPCLER | 9 | O43795 | Unconventional myosin-Ib OS=Homo sa | MYO1B | 131.986 | 2 | 53.541748 | yes | 9.885E-08 | 583.26898 | 107.83049 |
| LGNIEFKPESR | 11 | O43795 | Unconventional myosin-Ib OS=Homo sa | MYO1B | 131.986 | 2 | 53.541748 | yes | 9.722E-08 | 645.34607 | 107.42488 |
| IYEFTLQR | 8 | O43795 | Unconventional myosin-Ib OS=Homo sa | MYO1B | 131.986 | 2 | 53.541748 | yes | 9.722E-08 | 535.28748 | 105.19925 |
| THFLLMK | 7 | O43795 | Unconventional myosin-Ib OS=Homo sa | MYO1B | 131.986 | 2 | 53.541748 | yes | 2.793E-07 | 445.25186 | 85.394264 |
| LQQIFIELTLK | 11 | O43795 | Unconventional myosin-Ib OS=Homo sa | MYO1B | 131.986 | 2 | 53.541748 | no | 9.722E-08 | 673.40814 | 78.041969 |
| SQIVIAAWYR | 10 | O43795 | Unconventional myosin-Ib OS=Homo sa | MYO1B | 131.986 | 2 | 53.541748 | yes | 1.514E-06 | 603.83514 | 74.80101 |
| VSTTLNVAQAYY | 14 | O43795 | Unconventional myosin-Ib OS=Homo sa | MYO1B | 131.986 | 2 | 53.541748 | yes | 9.722E-08 | 778.90698 | 67.761169 |
| YMDIEFDFK | 9 | O43795 | Unconventional myosin-Ib OS=Homo sa | MYO1B | 131.986 | 2 | 53.541748 | no | 1.346E-07 | 604.27063 | 64.146492 |
| RPPTAGSQFK | 10 | O43795 | Unconventional myosin-Ib OS=Homo sa | MYO1B | 131.986 | 2 | 53.541748 | yes | 3.015E-07 | 544.79602 | 50.054111 |
| NILYLLHQR | 9 | O43808 | Peroxisomal membrane protein PMP34 O | SLC25A17 | 34.567 | 2 | 51.55386 | yes | 9.722E-08 | 585.34314 | 261.8873 |
| EEGLLAPYR | 9 | O43808 | Peroxisomal membrane protein PMP34 O | SLC25A17 | 34.567 | 2 | 51.55386 | yes | 2.179E-07 | 524.2771 | 223.28271 |
| TTHMVLLEIIK | 11 | O43808 | Peroxisomal membrane protein PMP34 O | SLC25A17 | 34.567 | 2 | 51.55386 | yes | 1.128E-07 | 649.38104 | 115.5255 |
| NEDIVPTNYK | 10 | O43808 | Peroxisomal membrane protein PMP34 O | SLC25A17 | 34.567 | 2 | 51.55386 | yes | 1.963E-07 | 596.79584 | 73.140823 |
| LTAATFTVMGLK | 12 | O43808 | Peroxisomal membrane protein PMP34 O | SLC25A17 | 34.567 | 2 | 51.55386 | yes | 1.231E-07 | 626.85211 | 48.310108 |
| GIIDAFHQIIR | 11 | O43808 | Peroxisomal membrane protein PMP34 O | SLC25A17 | 34.567 | 2 | 51.55386 | yes | 9.722E-08 | 641.86694 | 40.261471 |
| LFLVQLQEK | 9 | O43809 | Cleavage and polyadenylation specificity | NUDT21 | 26.228 | 2 | 50.15975 | yes | 1.361E-07 | 559.33441 | 6607.792 |
| KLFLVQLQEK | 10 | O43809 | Cleavage and polyadenylation specificity | NUDT21 | 26.228 | 2 | 50.15975 | yes | 1.504E-07 | 623.3819 | 1429.1125 |
| RTVEGVLIVHEHR | 13 | O43809 | Cleavage and polyadenylation specificity | NUDT21 | 26.228 | 3 | 50.15975 | yes | 2.119E-07 | 515.62671 | 587.02838 |
| LMTEILGR | 8 | O43809 | Cleavage and polyadenylation specificity | NUDT21 | 26.228 | 2 | 50.15975 | yes | 1.903E-07 | 466.76532 | 532.42902 |
| MREEFDK | 7 | O43809 | Cleavage and polyadenylation specificity | NUDT21 | 26.228 | 2 | 50.15975 | yes | 1.763E-07 | 477.7211 | 314.33676 |

| | | | | | | | | | | | |
|---|---|---|---|---|---|---|---|---|---|---|---|
| YIQQTKPLTLER | 12 | O43809 | Cleavage and polyadenylation specificity | NUDT21 | 26.228 | 2 | 50.15975 | yes | 9.881E-08 | 745.4223 | 222.64001 |
| PNFEPPQYPYIPAF | 19 | O43809 | Cleavage and polyadenylation specificity | NUDT21 | 26.228 | 3 | 50.15975 | yes | 1.645E-07 | 746.39661 | 99.863182 |
| PNFEPPQYPYIPAF | 19 | O43809 | Cleavage and polyadenylation specificity | NUDT21 | 26.228 | 3 | 50.15975 | yes | 5.532E-07 | 560.04932 | 99.863182 |
| PNFEPPQYPYIPAF | 19 | O43809 | Cleavage and polyadenylation specificity | NUDT21 | 26.228 | 4 | 50.15975 | yes | 1.645E-07 | 746.39661 | 99.863182 |
| PNFEPPQYPYIPAF | 19 | O43809 | Cleavage and polyadenylation specificity | NUDT21 | 26.228 | 4 | 50.15975 | yes | 5.532E-07 | 560.04932 | 99.863182 |
| PNFEPPQYPYIPAF | 17 | O43809 | Cleavage and polyadenylation specificity | NUDT21 | 26.228 | 3 | 50.15975 | yes | 2.826E-06 | 671.34741 | 23.752295 |
| SLAEGYFDAAGR | 12 | O43813 | Glutathione S-transferase LANCL1 OS=H | LANCL1 | 45.284 | 2 | 49.14024 | yes | 1.154E-07 | 628.79895 | 349.40054 |
| ELLQQMER | 8 | O43813 | Glutathione S-transferase LANCL1 OS=H | LANCL1 | 45.284 | 2 | 49.14024 | yes | 1.064E-07 | 523.76862 | 296.45343 |
| LTPEFSQR | 8 | O43813 | Glutathione S-transferase LANCL1 OS=H | LANCL1 | 45.284 | 2 | 49.14024 | yes | 1.879E-07 | 489.25616 | 285.03119 |
| AFPNPYADYNK | 11 | O43813 | Glutathione S-transferase LANCL1 OS=H | LANCL1 | 45.284 | 2 | 49.14024 | yes | 1.402E-07 | 650.30383 | 186.09851 |
| LKEDVLEQR | 9 | O43818 | U3 small nucleolar RNA-interacting prote | RRP9 | 51.841 | 2 | 48.011265 | yes | 3.641E-07 | 565.31421 | 451.91077 |
| AFEEDQVAGR | 10 | O43818 | U3 small nucleolar RNA-interacting prote | RRP9 | 51.841 | 2 | 48.011265 | yes | 3.001E-07 | 561.26471 | 243.98752 |
| LYLEQLR | 7 | O43818 | U3 small nucleolar RNA-interacting prote | RRP9 | 51.841 | 2 | 48.011265 | yes | 6.12E-07 | 467.77145 | 170.20261 |
| LWQCGEGFR | 9 | O43818 | U3 small nucleolar RNA-interacting prote | RRP9 | 51.841 | 2 | 48.011265 | yes | 2.091E-07 | 576.76636 | 114.63277 |
| GTHQLYSTSHDR | 12 | O43818 | U3 small nucleolar RNA-interacting prote | RRP9 | 51.841 | 2 | 48.011265 | yes | 2.814E-07 | 701.32892 | 49.722382 |
| KLHVIPR | 7 | O43818 | U3 small nucleolar RNA-interacting prote | RRP9 | 51.841 | 2 | 48.011265 | yes | 3.332E-07 | 431.7847 | 46.750076 |
| NSVCIIPLR | 9 | O43818 | U3 small nucleolar RNA-interacting prote | RRP9 | 51.841 | 2 | 48.011265 | yes | 1.214E-07 | 536.30261 | 21.134125 |
| LLGLTGSTK | 9 | O43819 | Protein SCO2 homolog, mitochondrial OS | SCO2 | 29.81 | 2 | 45.867386 | yes | 1.719E-07 | 445.2713 | 117.14239 |
| SAEQISDSVR | 10 | O43819 | Protein SCO2 homolog, mitochondrial OS | SCO2 | 29.81 | 2 | 45.867386 | yes | 5.293E-07 | 546.27002 | 80.291161 |
| TVEFLQEYIVNR | 12 | O43823 | A-kinase anchor protein 8 OS=Homo sap | AKAP8 | 76.109 | 2 | 52.39945 | yes | 1.159E-07 | 755.89862 | 5467.3516 |
| GIGQEHFFK | 9 | O43823 | A-kinase anchor protein 8 OS=Homo sap | AKAP8 | 76.109 | 2 | 52.39945 | yes | 4.132E-07 | 531.77197 | 3908.9604 |
| IQFACSVCK | 9 | O43823 | A-kinase anchor protein 8 OS=Homo sap | AKAP8 | 76.109 | 2 | 52.39945 | yes | 1.617E-07 | 556.76501 | 3813.1614 |
| RQELMEK | 7 | O43823 | A-kinase anchor protein 8 OS=Homo sap | AKAP8 | 76.109 | 2 | 52.39945 | yes | 1.247E-07 | 467.24475 | 2217.9993 |
| QFQLYEEPDTK | 11 | O43823 | A-kinase anchor protein 8 OS=Homo sap | AKAP8 | 76.109 | 2 | 52.39945 | yes | 9.722E-08 | 699.33282 | 1984.0116 |
| ETAKPKPDPFK | 11 | O43823 | A-kinase anchor protein 8 OS=Homo sap | AKAP8 | 76.109 | 2 | 52.39945 | yes | 1.061E-07 | 629.34552 | 676.08295 |
| ETAKPKPDPFK | 11 | O43823 | A-kinase anchor protein 8 OS=Homo sap | AKAP8 | 76.109 | 2 | 52.39945 | yes | 2.826E-07 | 419.89944 | 676.08295 |
| ETAKPKPDPFK | 11 | O43823 | A-kinase anchor protein 8 OS=Homo sap | AKAP8 | 76.109 | 3 | 52.39945 | yes | 1.061E-07 | 629.34552 | 676.08295 |
| ETAKPKPDPFK | 11 | O43823 | A-kinase anchor protein 8 OS=Homo sap | AKAP8 | 76.109 | 3 | 52.39945 | yes | 2.826E-07 | 419.89944 | 676.08295 |
| SFDDEEIQK | 9 | O43823 | A-kinase anchor protein 8 OS=Homo sap | AKAP8 | 76.109 | 2 | 52.39945 | yes | 2.635E-07 | 555.7511 | 852.5376 |
| FQPFESYDSR | 10 | O43823 | A-kinase anchor protein 8 OS=Homo sap | AKAP8 | 76.109 | 2 | 52.39945 | yes | 1.536E-06 | 638.28564 | 580.63269 |
| GGSGGGGEGIQDR | 13 | O43823 | A-kinase anchor protein 8 OS=Homo sap | AKAP8 | 76.109 | 2 | 52.39945 | yes | 9.722E-08 | 573.76031 | 570.45856 |
| SEAAEAGNGAETM | 23 | O43823 | A-kinase anchor protein 8 OS=Homo sap | AKAP8 | 76.109 | 3 | 52.39945 | yes | 1.555E-07 | 751.32922 | 423.66992 |
| VAPAPAAADAEVI | 20 | O43823 | A-kinase anchor protein 8 OS=Homo sap | AKAP8 | 76.109 | 3 | 52.39945 | yes | 1.165E-07 | 657.31647 | 205.89331 |
| KQFQLYEEPDTK | 12 | O43823 | A-kinase anchor protein 8 OS=Homo sap | AKAP8 | 76.109 | 2 | 52.39945 | yes | 3.081E-07 | 763.38031 | 47.15778 |
| GEDELCDSGR | 10 | O43823 | A-kinase anchor protein 8 OS=Homo sap | AKAP8 | 76.109 | 2 | 52.39945 | yes | 9.722E-08 | 569.22748 | 37.417286 |
| NGSFGGQYSECR | 12 | O43823 | A-kinase anchor protein 8 OS=Homo sap | AKAP8 | 76.109 | 2 | 52.39945 | yes | 1.464E-07 | 681.28058 | 26.820698 |
| GEKEDEDEDVK | 11 | O43823 | A-kinase anchor protein 8 OS=Homo sap | AKAP8 | 76.109 | 2 | 52.39945 | yes | 3.036E-06 | 646.77808 | 5.532361 |
| DMGGYSTTTDFIK | 13 | O43837 | Isocitrate dehydrogenase [NAD] subunit l | IDH3B | 42.184 | 2 | 47.8465 | yes | 1.247E-07 | 718.3241 | 62.664307 |
| HNNLDLVIIR | 10 | O43837 | Isocitrate dehydrogenase [NAD] subunit l | IDH3B | 42.184 | 2 | 47.8465 | yes | 1.252E-07 | 603.85132 | 28.310421 |

| | | | | | | | | | | |
|---|---|---|---|---|---|---|---|---|---|---|
| FHLISPLIQK | 10 | O43847 | Nardilysin OS=Homo sapiens OX=9606 ( | NRDC | 131.701 | 2 | 45.730453 | yes | 1.77E-07 | 598.36353 | 55.407429 |
| AFDCPETEYPVK | 12 | O43847 | Nardilysin OS=Homo sapiens OX=9606 ( | NRDC | 131.701 | 2 | 45.730453 | yes | 1.759E-07 | 728.32666 | 51.452152 |
| IVNTPQGCLWYK | 12 | O43847 | Nardilysin OS=Homo sapiens OX=9606 ( | NRDC | 131.701 | 2 | 45.730453 | yes | 2.193E-07 | 739.87665 | 42.810947 |
| SQLFVEGLVQGNV | 23 | O43847 | Nardilysin OS=Homo sapiens OX=9606 ( | NRDC | 131.701 | 3 | 45.730453 | yes | 3.429E-06 | 843.75623 | 13.823788 |
| TFDQLTPEESK | 11 | O43852 | Calumenin OS=Homo sapiens OX=9606 | CALU | 37.107 | 2 | 53.778797 | yes | 2.165E-07 | 647.81171 | 11766.352 |
| DIVVQETMEDIDK | 13 | O43852 | Calumenin OS=Homo sapiens OX=9606 | CALU | 37.107 | 2 | 53.778797 | yes | 2.722E-07 | 767.8689 | 4980.0332 |
| DIVVQETMEDIDK | 13 | O43852 | Calumenin OS=Homo sapiens OX=9606 | CALU | 37.107 | 2 | 53.778797 | yes | 6.039E-07 | 775.86633 | 4980.0332 |
| HLVYESDQNK | 10 | O43852 | Calumenin OS=Homo sapiens OX=9606 | CALU | 37.107 | 2 | 53.778797 | yes | 1.245E-07 | 616.79895 | 8995.9268 |
| VHHEPQLSDK | 10 | O43852 | Calumenin OS=Homo sapiens OX=9606 | CALU | 37.107 | 2 | 53.778797 | yes | 1.26E-07 | 595.30164 | 7996.1055 |
| WIYEDVER | 8 | O43852 | Calumenin OS=Homo sapiens OX=9606 | CALU | 37.107 | 2 | 53.778797 | yes | 1.151E-07 | 555.26672 | 7218.6064 |
| EEFTAFLHPEEYD | 16 | O43852 | Calumenin OS=Homo sapiens OX=9606 | CALU | 37.107 | 3 | 53.778797 | yes | 9.722E-08 | 683.63641 | 6169.186 |
| EQFVEFR | 7 | O43852 | Calumenin OS=Homo sapiens OX=9606 | CALU | 37.107 | 2 | 53.778797 | yes | 2.287E-07 | 477.73761 | 4486.1826 |
| MADKDGDLIATK | 12 | O43852 | Calumenin OS=Homo sapiens OX=9606 | CALU | 37.107 | 2 | 53.778797 | yes | 9.935E-08 | 639.32391 | 1369.4076 |
| MADKDGDLIATK | 12 | O43852 | Calumenin OS=Homo sapiens OX=9606 | CALU | 37.107 | 2 | 53.778797 | yes | 1.098E-07 | 647.32141 | 1369.4076 |
| MADKDGDLIATK | 12 | O43852 | Calumenin OS=Homo sapiens OX=9606 | CALU | 37.107 | 2 | 53.778797 | yes | 7.022E-07 | 426.5517 | 1369.4076 |
| MADKDGDLIATK | 12 | O43852 | Calumenin OS=Homo sapiens OX=9606 | CALU | 37.107 | 3 | 53.778797 | yes | 9.935E-08 | 639.32391 | 1369.4076 |
| MADKDGDLIATK | 12 | O43852 | Calumenin OS=Homo sapiens OX=9606 | CALU | 37.107 | 3 | 53.778797 | yes | 1.098E-07 | 647.32141 | 1369.4076 |
| MADKDGDLIATK | 12 | O43852 | Calumenin OS=Homo sapiens OX=9606 | CALU | 37.107 | 3 | 53.778797 | yes | 7.022E-07 | 426.5517 | 1369.4076 |
| IDGDKDGFVTVDE | 15 | O43852 | Calumenin OS=Homo sapiens OX=9606 | CALU | 37.107 | 2 | 53.778797 | yes | 1.926E-07 | 825.91467 | 2473.3713 |
| DGFVTVDELK | 10 | O43852 | Calumenin OS=Homo sapiens OX=9606 | CALU | 37.107 | 2 | 53.778797 | yes | 9.722E-08 | 561.78748 | 1198.8265 |
| GHDLNEDGLVSW | 16 | O43852 | Calumenin OS=Homo sapiens OX=9606 | CALU | 37.107 | 3 | 53.778797 | yes | 4.687E-07 | 630.95502 | 257.76044 |
| HLVYESDQNKDG | 13 | O43852 | Calumenin OS=Homo sapiens OX=9606 | CALU | 37.107 | 3 | 53.778797 | yes | 5.322E-07 | 511.58282 | 62.262783 |
| HLVYESDQNKDG | 13 | O43852 | Calumenin OS=Homo sapiens OX=9606 | CALU | 37.107 | 3 | 53.778797 | yes | 9.722E-08 | 766.87061 | 62.262783 |
| HLVYESDQNKDG | 13 | O43852 | Calumenin OS=Homo sapiens OX=9606 | CALU | 37.107 | 2 | 53.778797 | yes | 5.322E-07 | 511.58282 | 62.262783 |
| HLVYESDQNKDG | 13 | O43852 | Calumenin OS=Homo sapiens OX=9606 | CALU | 37.107 | 2 | 53.778797 | yes | 9.722E-08 | 766.87061 | 62.262783 |
| MDKEETKDWILPS | 23 | O43852 | Calumenin OS=Homo sapiens OX=9606 | CALU | 37.107 | 4 | 53.778797 | yes | 9.975E-08 | 688.06573 | 49.154762 |
| DRVHHEPQLSDK | 12 | O43852 | Calumenin OS=Homo sapiens OX=9606 | CALU | 37.107 | 2 | 53.778797 | yes | 2.743E-06 | 730.86566 | 20.197975 |
| GIVEESVTGVHR | 12 | O43865 | S-adenosylhomocysteine hydrolase-like p | AHCYL1 | 58.952 | 2 | 54.11108 | no | 9.722E-08 | 641.84113 | 151.10112 |
| LCVPAMNVNDSV | 14 | O43865 | S-adenosylhomocysteine hydrolase-like p | AHCYL1 | 58.952 | 2 | 54.11108 | no | 1.787E-07 | 774.38147 | 121.58688 |
| QVDVVITCTGNK | 12 | O43865 | S-adenosylhomocysteine hydrolase-like p | AHCYL1 | 58.952 | 2 | 54.11108 | yes | 9.722E-08 | 667.34265 | 78.212936 |
| FDNLYCCR | 8 | O43865 | S-adenosylhomocysteine hydrolase-like p | AHCYL1 | 58.952 | 2 | 54.11108 | no | 2.649E-06 | 574.23657 | 57.714912 |
| QIQFADDMQEFTK | 13 | O43865 | S-adenosylhomocysteine hydrolase-like p | AHCYL1 | 58.952 | 2 | 54.11108 | yes | 9.759E-08 | 800.8692 | 38.785164 |
| LMEEDPAFR | 9 | O43896 | Kinesin-like protein KIF1C OS=Homo sa | KIF1C | 122.947 | 2 | 55.391193 | yes | 1.085E-07 | 554.26062 | 1765.6177 |
| ELLEQQGIDIK | 11 | O43896 | Kinesin-like protein KIF1C OS=Homo sa | KIF1C | 122.947 | 2 | 55.391193 | yes | 9.722E-08 | 643.35352 | 1557.433 |
| YPPYTTPPR | 9 | O43896 | Kinesin-like protein KIF1C OS=Homo sa | KIF1C | 122.947 | 2 | 55.391193 | yes | 1.846E-07 | 546.27966 | 1333.0557 |
| EALLAEMGVAVR | 12 | O43896 | Kinesin-like protein KIF1C OS=Homo sa | KIF1C | 122.947 | 2 | 55.391193 | yes | 9.722E-08 | 629.84485 | 1300.729 |
| VGQVDMDIK | 9 | O43896 | Kinesin-like protein KIF1C OS=Homo sa | KIF1C | 122.947 | 2 | 55.391193 | yes | 3.194E-07 | 502.75769 | 1256.3657 |
| AEIEALAALK | 10 | O43896 | Kinesin-like protein KIF1C OS=Homo sa | KIF1C | 122.947 | 2 | 55.391193 | yes | 1.17E-07 | 514.80298 | 1251.5873 |
| FVPPHDCK | 8 | O43896 | Kinesin-like protein KIF1C OS=Homo sa | KIF1C | 122.947 | 2 | 55.391193 | yes | 1.418E-06 | 500.23947 | 1129.158 |

| | | | | | | | | | | | |
|---|---|---|---|---|---|---|---|---|---|---|---|
| VISALADMQSK | 11 | O43896 | Kinesin-like protein KIF1C OS=Homo sa | KIF1C | 122.947 | 2 | 55.391193 | yes | 1.691E-07 | 581.81049 | 1117.7362 |
| LQDLENQYR | 9 | O43896 | Kinesin-like protein KIF1C OS=Homo sa | KIF1C | 122.947 | 2 | 55.391193 | yes | 9.722E-08 | 589.79364 | 967.32159 |
| SHAVFTIVFTQR | 12 | O43896 | Kinesin-like protein KIF1C OS=Homo sa | KIF1C | 122.947 | 2 | 55.391193 | yes | 9.722E-08 | 703.383 | 953.73505 |
| EDGGTVGVFSPK | 12 | O43896 | Kinesin-like protein KIF1C OS=Homo sa | KIF1C | 122.947 | 2 | 55.391193 | yes | 9.722E-08 | 596.79584 | 817.42114 |
| EQHCLFR | 7 | O43896 | Kinesin-like protein KIF1C OS=Homo sa | KIF1C | 122.947 | 2 | 55.391193 | yes | 5.409E-07 | 495.23471 | 813.22302 |
| CNAIINEDPNAR | 12 | O43896 | Kinesin-like protein KIF1C OS=Homo sa | KIF1C | 122.947 | 2 | 55.391193 | yes | 9.839E-08 | 693.82513 | 739.22394 |
| LTGQFIR | 7 | O43896 | Kinesin-like protein KIF1C OS=Homo sa | KIF1C | 122.947 | 2 | 55.391193 | yes | 9.722E-08 | 417.74524 | 707.14496 |
| EHPILGPYVQDLSI | 14 | O43896 | Kinesin-like protein KIF1C OS=Homo sa | KIF1C | 122.947 | 2 | 55.391193 | yes | 3.744E-07 | 798.42505 | 554.20746 |
| LTGILQEVK | 9 | O43896 | Kinesin-like protein KIF1C OS=Homo sa | KIF1C | 122.947 | 2 | 55.391193 | yes | 2.123E-07 | 500.80551 | 502.10901 |
| GAGSAQPEPQHFQ | 15 | O43896 | Kinesin-like protein KIF1C OS=Homo sa | KIF1C | 122.947 | 2 | 55.391193 | yes | 1.002E-07 | 789.8866 | 239.62328 |
| GAGSAQPEPQHFQ | 15 | O43896 | Kinesin-like protein KIF1C OS=Homo sa | KIF1C | 122.947 | 2 | 55.391193 | yes | 3.554E-07 | 526.92682 | 239.62328 |
| GAGSAQPEPQHFQ | 15 | O43896 | Kinesin-like protein KIF1C OS=Homo sa | KIF1C | 122.947 | 3 | 55.391193 | yes | 1.002E-07 | 789.8866 | 239.62328 |
| GAGSAQPEPQHFQ | 15 | O43896 | Kinesin-like protein KIF1C OS=Homo sa | KIF1C | 122.947 | 3 | 55.391193 | yes | 3.554E-07 | 526.92682 | 239.62328 |
| TYGKPDGPGDAW | 13 | O43896 | Kinesin-like protein KIF1C OS=Homo sa | KIF1C | 122.947 | 2 | 55.391193 | yes | 1.121E-07 | 710.33624 | 206.55637 |
| TYGKPDGPGDAW | 13 | O43896 | Kinesin-like protein KIF1C OS=Homo sa | KIF1C | 122.947 | 2 | 55.391193 | yes | 9.722E-08 | 473.89325 | 206.55637 |
| TYGKPDGPGDAW | 13 | O43896 | Kinesin-like protein KIF1C OS=Homo sa | KIF1C | 122.947 | 3 | 55.391193 | yes | 1.121E-07 | 710.33624 | 206.55637 |
| TYGKPDGPGDAW | 13 | O43896 | Kinesin-like protein KIF1C OS=Homo sa | KIF1C | 122.947 | 3 | 55.391193 | yes | 9.722E-08 | 473.89325 | 206.55637 |
| LVTEPLVLK | 9 | O43896 | Kinesin-like protein KIF1C OS=Homo sa | KIF1C | 122.947 | 2 | 55.391193 | yes | 1.002E-07 | 506.32608 | 407.33228 |
| EQLPPTTVQTIVK | 13 | O43896 | Kinesin-like protein KIF1C OS=Homo sa | KIF1C | 122.947 | 2 | 55.391193 | yes | 5.706E-07 | 727.41669 | 357.95883 |
| EICYEVALADFR | 12 | O43896 | Kinesin-like protein KIF1C OS=Homo sa | KIF1C | 122.947 | 2 | 55.391193 | yes | 9.722E-08 | 743.35577 | 344.49384 |
| KSDFIPYR | 8 | O43896 | Kinesin-like protein KIF1C OS=Homo sa | KIF1C | 122.947 | 2 | 55.391193 | yes | 9.795E-08 | 513.27435 | 279.52301 |
| CHDQLTGLDSEK | 12 | O43896 | Kinesin-like protein KIF1C OS=Homo sa | KIF1C | 122.947 | 2 | 55.391193 | yes | 9.722E-08 | 701.81702 | 253.65508 |
| KHNSYPQPPQPYP | 21 | O43896 | Kinesin-like protein KIF1C OS=Homo sa | KIF1C | 122.947 | 4 | 55.391193 | yes | 3.31E-07 | 603.81348 | 241.19153 |
| ELQEEVAR | 8 | O43896 | Kinesin-like protein KIF1C OS=Homo sa | KIF1C | 122.947 | 2 | 55.391193 | yes | 1.155E-07 | 487.2511 | 201.33344 |
| VYQIPQR | 7 | O43896 | Kinesin-like protein KIF1C OS=Homo sa | KIF1C | 122.947 | 2 | 55.391193 | yes | 5.237E-07 | 452.25598 | 101.31285 |
| TPHLVNLNEDPLM | 22 | O43896 | Kinesin-like protein KIF1C OS=Homo sa | KIF1C | 122.947 | 4 | 55.391193 | yes | 9.722E-08 | 659.83575 | 76.373726 |
| HNSYPQPPQPYPA | 20 | O43896 | Kinesin-like protein KIF1C OS=Homo sa | KIF1C | 122.947 | 4 | 55.391193 | yes | 2.818E-07 | 571.78973 | 64.362701 |
| DRELQALR | 8 | O43896 | Kinesin-like protein KIF1C OS=Homo sa | KIF1C | 122.947 | 2 | 55.391193 | yes | 9.722E-08 | 500.78033 | 62.870773 |
| VRPFNAR | 7 | O43896 | Kinesin-like protein KIF1C OS=Homo sa | KIF1C | 122.947 | 2 | 55.391193 | yes | 1.238E-07 | 430.24847 | 42.986698 |
| TAMIAALSPADIN | 22 | O43896 | Kinesin-like protein KIF1C OS=Homo sa | KIF1C | 122.947 | 3 | 55.391193 | yes | 5.396E-07 | 794.07367 | 33.73917 |
| DVWDTVGEEEGG | 25 | O43896 | Kinesin-like protein KIF1C OS=Homo sa | KIF1C | 122.947 | 3 | 55.391193 | yes | 4.982E-07 | 788.99854 | 31.560274 |
| RVYQIPQR | 8 | O43896 | Kinesin-like protein KIF1C OS=Homo sa | KIF1C | 122.947 | 2 | 55.391193 | yes | 6.663E-07 | 530.30652 | 25.213257 |
| LYADSDSGDDSDK | 14 | O43896 | Kinesin-like protein KIF1C OS=Homo sa | KIF1C | 122.947 | 2 | 55.391193 | yes | 9.17E-07 | 772.32898 | 17.659513 |
| HSMPELGLR | 9 | O43900 | Prickle planar cell polarity protein 3 OS= | PRICKLE3 | 68.61 | 2 | 46.389626 | yes | 2.146E-06 | 520.2713 | 142.61029 |
| DPLVSEGGPR | 10 | O43900 | Prickle planar cell polarity protein 3 OS= | PRICKLE3 | 68.61 | 2 | 46.389626 | yes | 2.397E-07 | 513.76434 | 121.67477 |
| LISDFQR | 7 | O43900 | Prickle planar cell polarity protein 3 OS= | PRICKLE3 | 68.61 | 2 | 46.389626 | yes | 1.572E-07 | 439.74017 | 98.734985 |
| QIGGGDIAVFASR | 13 | O43900 | Prickle planar cell polarity protein 3 OS= | PRICKLE3 | 68.61 | 2 | 46.389626 | yes | 2.968E-07 | 645.84369 | 39.435051 |
| EEHAVHAVPVDLI | 14 | O43900 | Prickle planar cell polarity protein 3 OS= | PRICKLE3 | 68.61 | 2 | 46.389626 | yes | 1.013E-06 | 800.90753 | 13.937463 |
| VVVVWNSPK | 9 | O43909 | Exostosin-like 3 OS=Homo sapiens OX= | EXTL3 | 104.75 | 2 | 46.34253 | yes | 7.624E-07 | 514.30035 | 74.285591 |

| | | | | | | | | | | | |
|---|---|---|---|---|---|---|---|---|---|---|---|
| VTEVHFLLR | 9 | O43909 | Exostosin-like 3 OS=Homo sapiens OX=! | EXTL3 | 104.75 | 2 | 46.34253 | yes | 1.585E-07 | 557.3244 | 51.396446 |
| HDEIMFGFR | 9 | O43909 | Exostosin-like 3 OS=Homo sapiens OX=! | EXTL3 | 104.75 | 2 | 46.34253 | yes | 2.807E-06 | 576.26874 | 43.539921 |
| LLLEQILNK | 9 | O43913 | Origin recognition complex subunit 5 OS | ORC5 | 50.284 | 2 | 34.167725 | yes | 1.065E-06 | 542.34222 | 87.115799 |
| EWIECAHGIGYTR | 13 | O43920 | NADH dehydrogenase [ubiquinone] iron- | NDUFS5 | 12.518 | 3 | 48.32855 | yes | 1.164E-07 | 531.2489 | 45.71159 |
| PFLDIQK | 7 | O43920 | NADH dehydrogenase [ubiquinone] iron- | NDUFS5 | 12.518 | 2 | 48.32855 | yes | 1.406E-06 | 430.74747 | 40.323807 |
| EQLSLPAEFPDK | 12 | O43929 | Origin recognition complex subunit 4 OS | ORC4 | 50.378 | 2 | 46.80511 | yes | 5.581E-07 | 687.35101 | 127.11743 |
| WNENVQYLSEDR | 12 | O43929 | Origin recognition complex subunit 4 OS | ORC4 | 50.378 | 2 | 46.80511 | yes | 3.203E-07 | 776.85498 | 57.908348 |
| TALHGESNSVLIIG | 16 | O43929 | Origin recognition complex subunit 4 OS | ORC4 | 50.378 | 3 | 46.80511 | yes | 1.466E-07 | 555.30902 | 39.91761 |
| DQLFSQPPVLR | 11 | O43933 | Peroxisomal ATPase PEX1 OS=Homo sa | PEX1 | 142.868 | 2 | 51.77739 | yes | 9.722E-08 | 650.35645 | 837.13696 |
| TGILLYGPPGTGK | 13 | O43933 | Peroxisomal ATPase PEX1 OS=Homo sa | PEX1 | 142.868 | 2 | 51.77739 | yes | 1.276E-07 | 637.36115 | 582.01581 |
| LVLTTLDFQK | 10 | O43933 | Peroxisomal ATPase PEX1 OS=Homo sa | PEX1 | 142.868 | 2 | 51.77739 | yes | 1.418E-07 | 589.34497 | 458.97742 |
| EFSLSIVHSWEK | 12 | O43933 | Peroxisomal ATPase PEX1 OS=Homo sa | PEX1 | 142.868 | 2 | 51.77739 | yes | 1.025E-07 | 731.37225 | 398.63571 |
| TLLAGVIAR | 9 | O43933 | Peroxisomal ATPase PEX1 OS=Homo sa | PEX1 | 142.868 | 2 | 51.77739 | yes | 1.069E-07 | 457.2951 | 393.24438 |
| YIGASEQAVR | 10 | O43933 | Peroxisomal ATPase PEX1 OS=Homo sa | PEX1 | 142.868 | 2 | 51.77739 | yes | 1.6E-07 | 547.28546 | 382.66821 |
| IGGLHEVR | 8 | O43933 | Peroxisomal ATPase PEX1 OS=Homo sa | PEX1 | 142.868 | 2 | 51.77739 | yes | 1.722E-07 | 440.7536 | 374.10504 |
| NVEVLHLGK | 9 | O43933 | Peroxisomal ATPase PEX1 OS=Homo sa | PEX1 | 142.868 | 2 | 51.77739 | yes | 2.393E-07 | 504.79547 | 341.44821 |
| LAHALNDMIK | 10 | O43933 | Peroxisomal ATPase PEX1 OS=Homo sa | PEX1 | 142.868 | 2 | 51.77739 | yes | 4.019E-07 | 563.30786 | 293.0629 |
| QLMSLVAGLR | 10 | O43933 | Peroxisomal ATPase PEX1 OS=Homo sa | PEX1 | 142.868 | 2 | 51.77739 | yes | 1.049E-07 | 544.31824 | 283.21509 |
| TTIQVLLDPMVK | 12 | O43933 | Peroxisomal ATPase PEX1 OS=Homo sa | PEX1 | 142.868 | 2 | 51.77739 | yes | 2.159E-07 | 679.3916 | 250.15211 |
| NFAELYESFQNPK | 13 | O43933 | Peroxisomal ATPase PEX1 OS=Homo sa | PEX1 | 142.868 | 2 | 51.77739 | yes | 1.097E-07 | 793.87793 | 230.43054 |
| CEILCNVIK | 9 | O43933 | Peroxisomal ATPase PEX1 OS=Homo sa | PEX1 | 142.868 | 2 | 51.77739 | yes | 1.451E-07 | 574.79376 | 205.44833 |
| QILMDTIQLPAK | 12 | O43933 | Peroxisomal ATPase PEX1 OS=Homo sa | PEX1 | 142.868 | 2 | 51.77739 | yes | 1.02E-07 | 685.88922 | 198.5183 |
| LNIEMHAVVR | 10 | O43933 | Peroxisomal ATPase PEX1 OS=Homo sa | PEX1 | 142.868 | 2 | 51.77739 | yes | 5.359E-07 | 591.3266 | 102.72559 |
| VWIPDDLR | 8 | O43933 | Peroxisomal ATPase PEX1 OS=Homo sa | PEX1 | 142.868 | 2 | 51.77739 | yes | 3.641E-05 | 507.27435 | 93.122154 |
| LSSLGGVNSLGVS | 25 | O43933 | Peroxisomal ATPase PEX1 OS=Homo sa | PEX1 | 142.868 | 3 | 51.77739 | yes | 1.004E-07 | 845.12927 | 89.894142 |
| DFTVLVDR | 8 | O43933 | Peroxisomal ATPase PEX1 OS=Homo sa | PEX1 | 142.868 | 2 | 51.77739 | yes | 1.088E-07 | 482.75854 | 85.040596 |
| PSISEDDWK | 9 | O43933 | Peroxisomal ATPase PEX1 OS=Homo sa | PEX1 | 142.868 | 2 | 51.77739 | yes | 3.464E-07 | 538.74835 | 71.61705 |
| YPELFANLPIR | 11 | O43933 | Peroxisomal ATPase PEX1 OS=Homo sa | PEX1 | 142.868 | 2 | 51.77739 | yes | 1.488E-06 | 666.86914 | 68.361328 |
| LAISQSHLMTALG | 16 | O43933 | Peroxisomal ATPase PEX1 OS=Homo sa | PEX1 | 142.868 | 3 | 51.77739 | yes | 4.685E-07 | 579.3139 | 59.247742 |
| NQSGTMFRPGQK | 12 | O43933 | Peroxisomal ATPase PEX1 OS=Homo sa | PEX1 | 142.868 | 2 | 51.77739 | yes | 1.87E-07 | 675.83276 | 31.72401 |
| LQESVMEASR | 10 | O60216 | Double-strand-break repair protein rad21 | RAD21 | 71.69 | 2 | 49.2508 | yes | 5.737E-06 | 575.28204 | 146.95894 |
| YLLADCNEAFIK | 12 | O60216 | Double-strand-break repair protein rad21 | RAD21 | 71.69 | 2 | 49.2508 | yes | 2.266E-07 | 728.86066 | 112.79944 |
| IWLAAHWDK | 9 | O60216 | Double-strand-break repair protein rad21 | RAD21 | 71.69 | 2 | 49.2508 | yes | 1.281E-07 | 570.30347 | 102.72318 |
| CLTPLVPEDLR | 11 | O60216 | Double-strand-break repair protein rad21 | RAD21 | 71.69 | 2 | 49.2508 | yes | 1.053E-07 | 656.8501 | 69.800415 |
| DVIDEPIIEEPSR | 13 | O60216 | Double-strand-break repair protein rad21 | RAD21 | 71.69 | 2 | 49.2508 | yes | 1.168E-07 | 756.38306 | 59.491245 |
| PGVVDLPEENR | 11 | O60216 | Double-strand-break repair protein rad21 | RAD21 | 71.69 | 2 | 49.2508 | yes | 2.646E-07 | 612.81458 | 54.965851 |
| TGAESISLLELCR | 13 | O60216 | Double-strand-break repair protein rad21 | RAD21 | 71.69 | 2 | 49.2508 | yes | 1.085E-07 | 724.87433 | 47.673218 |
| TSGHLLLGVVR | 11 | O60216 | Double-strand-break repair protein rad21 | RAD21 | 71.69 | 2 | 49.2508 | no | 1.066E-07 | 576.34839 | 44.886147 |
| AHVFECNLESSVE | 19 | O60216 | Double-strand-break repair protein rad21 | RAD21 | 71.69 | 3 | 49.2508 | yes | 6.386E-07 | 716.021 | 25.28944 |

| | | | | | | | | | | | |
|---|---|---|---|---|---|---|---|---|---|---|---|
| YMTDGMLLR | 9 | O60231 | Pre-mRNA-splicing factor ATP-dependen | DHX16 | 119.264 | 2 | 53.76582 | no | 1.774E-07 | 550.2674 | 2995.0371 |
| YMTDGMLLR | 9 | O60231 | Pre-mRNA-splicing factor ATP-dependen | DHX16 | 119.264 | 2 | 53.76582 | no | 1.435E-07 | 558.26483 | 2995.0371 |
| DTDTLDLSGPAR | 12 | O60231 | Pre-mRNA-splicing factor ATP-dependen | DHX16 | 119.264 | 2 | 53.76582 | yes | 4.031E-07 | 630.80695 | 200.06146 |
| EQLEGLLER | 9 | O60231 | Pre-mRNA-splicing factor ATP-dependen | DHX16 | 119.264 | 2 | 53.76582 | yes | 3.74E-07 | 543.79309 | 166.84444 |
| LGNEVGYSIR | 10 | O60231 | Pre-mRNA-splicing factor ATP-dependen | DHX16 | 119.264 | 2 | 53.76582 | yes | 1.045E-07 | 554.29327 | 135.29634 |
| MAELPVDPMLSK | 12 | O60231 | Pre-mRNA-splicing factor ATP-dependen | DHX16 | 119.264 | 2 | 53.76582 | yes | 9.85E-08 | 665.84088 | 131.29659 |
| IFQPTPPGAR | 10 | O60231 | Pre-mRNA-splicing factor ATP-dependen | DHX16 | 119.264 | 2 | 53.76582 | yes | 2.021E-06 | 542.3009 | 117.77298 |
| LQDLEER | 7 | O60231 | Pre-mRNA-splicing factor ATP-dependen | DHX16 | 119.264 | 2 | 53.76582 | yes | 1.947E-07 | 451.73254 | 110.22742 |
| VLVASATMDTAR | 12 | O60231 | Pre-mRNA-splicing factor ATP-dependen | DHX16 | 119.264 | 2 | 53.76582 | yes | 9.722E-08 | 617.82666 | 96.718864 |
| FEDCTSER | 8 | O60231 | Pre-mRNA-splicing factor ATP-dependen | DHX16 | 119.264 | 2 | 53.76582 | yes | 8.236E-07 | 522.20856 | 88.313164 |
| CTSAEEFVQR | 10 | O60231 | Pre-mRNA-splicing factor ATP-dependen | DHX16 | 119.264 | 2 | 53.76582 | yes | 3.134E-07 | 613.77716 | 75.978859 |
| AITAGYFYHTAR | 12 | O60231 | Pre-mRNA-splicing factor ATP-dependen | DHX16 | 119.264 | 2 | 53.76582 | yes | 9.722E-08 | 685.84619 | 70.138435 |
| HVAQFLIGTAQR | 12 | O60231 | Pre-mRNA-splicing factor ATP-dependen | DHX16 | 119.264 | 2 | 53.76582 | yes | 1.076E-07 | 670.87531 | 64.601318 |
| FSTFFDDAPVFR | 12 | O60231 | Pre-mRNA-splicing factor ATP-dependen | DHX16 | 119.264 | 2 | 53.76582 | yes | 4.3E-07 | 724.84589 | 44.742077 |
| AVVEKPAR | 8 | O60231 | Pre-mRNA-splicing factor ATP-dependen | DHX16 | 119.264 | 2 | 53.76582 | yes | 8.558E-07 | 435.26379 | 33.570148 |
| TSLGNVVLLLK | 11 | O60231 | Pre-mRNA-splicing factor ATP-dependen | DHX16 | 119.264 | 2 | 53.76582 | yes | 5.261E-07 | 578.86865 | 26.297253 |
| LMGDYLLR | 8 | O60232 | Protein ZNRD2 OS=Homo sapiens OX=9 | ZNRD2 | 21.475 | 2 | 49.833546 | yes | 1.002E-07 | 490.76532 | 104.82904 |
| LPTVPMLGLTQR | 12 | O60244 | Mediator of RNA polymerase II transcrip | MED14 | 160.607 | 2 | 51.7216 | yes | 4.68E-07 | 663.38409 | 136.43468 |
| LYECVLEFAR | 10 | O60244 | Mediator of RNA polymerase II transcrip | MED14 | 160.607 | 2 | 51.7216 | yes | 1.169E-07 | 650.32373 | 101.56824 |
| TFLNMFVDSNQDA | 14 | O60244 | Mediator of RNA polymerase II transcrip | MED14 | 160.607 | 2 | 51.7216 | yes | 9.722E-08 | 829.38556 | 96.113869 |
| LVEGFYPAPGLK | 12 | O60244 | Mediator of RNA polymerase II transcrip | MED14 | 160.607 | 2 | 51.7216 | yes | 1.843E-07 | 645.85828 | 92.590736 |
| DGAYSLFDNSK | 11 | O60244 | Mediator of RNA polymerase II transcrip | MED14 | 160.607 | 2 | 51.7216 | yes | 3.252E-06 | 608.77765 | 88.384453 |
| IIPWIQQLK | 9 | O60244 | Mediator of RNA polymerase II transcrip | MED14 | 160.607 | 2 | 51.7216 | yes | 2.491E-07 | 569.85278 | 83.374374 |
| NMYCIDIYCR | 10 | O60244 | Mediator of RNA polymerase II transcrip | MED14 | 160.607 | 2 | 51.7216 | yes | 1.235E-07 | 704.29633 | 81.380188 |
| LELFPDQATQLK | 12 | O60244 | Mediator of RNA polymerase II transcrip | MED14 | 160.607 | 2 | 51.7216 | yes | 4.48E-07 | 701.88245 | 76.336792 |
| WGDLVQVER | 9 | O60244 | Mediator of RNA polymerase II transcrip | MED14 | 160.607 | 2 | 51.7216 | yes | 9.872E-08 | 551.28802 | 70.325966 |
| LLIDSVHAR | 9 | O60244 | Mediator of RNA polymerase II transcrip | MED14 | 160.607 | 2 | 51.7216 | yes | 1.144E-07 | 512.3009 | 66.651817 |
| ENIQDLVFR | 9 | O60244 | Mediator of RNA polymerase II transcrip | MED14 | 160.607 | 2 | 51.7216 | yes | 2.308E-06 | 567.30109 | 51.465229 |
| HVYLTYENLLSEP | 17 | O60244 | Mediator of RNA polymerase II transcrip | MED14 | 160.607 | 3 | 51.7216 | yes | 9.722E-08 | 649.67053 | 46.562855 |
| LEILVEDK | 8 | O60244 | Mediator of RNA polymerase II transcrip | MED14 | 160.607 | 2 | 51.7216 | yes | 3.45E-07 | 479.7764 | 38.88578 |
| LILCYGTTK | 9 | O60244 | Mediator of RNA polymerase II transcrip | MED14 | 160.607 | 2 | 51.7216 | yes | 6.663E-07 | 534.79156 | 31.550652 |
| NITCITCTDVR | 11 | O60260 | E3 ubiquitin-protein ligase parkin OS=Hc | PRKN | 51.641 | 2 | 51.618065 | yes | 9.85E-08 | 676.81848 | 1254.8154 |
| ILGEEQYNR | 9 | O60260 | E3 ubiquitin-protein ligase parkin OS=Hc | PRKN | 51.641 | 2 | 51.618065 | yes | 1.103E-07 | 561.2829 | 984.2782 |
| PGCGAGLLPEPDQ | 14 | O60260 | E3 ubiquitin-protein ligase parkin OS=Hc | PRKN | 51.641 | 2 | 51.618065 | yes | 1.483E-07 | 733.85645 | 586.46136 |
| CHVPVEK | 7 | O60260 | E3 ubiquitin-protein ligase parkin OS=Hc | PRKN | 51.641 | 2 | 51.618065 | yes | 3.002E-06 | 434.72092 | 565.03473 |
| HVICLDCFHLYCV | 15 | O60260 | E3 ubiquitin-protein ligase parkin OS=Hc | PRKN | 51.641 | 3 | 51.618065 | yes | 1.167E-07 | 664.97955 | 381.32831 |
| QGVPADQLR | 9 | O60260 | E3 ubiquitin-protein ligase parkin OS=Hc | PRKN | 51.641 | 2 | 51.618065 | yes | 3.307E-07 | 492.26706 | 355.37402 |
| SPVLVFQCNSR | 11 | O60260 | E3 ubiquitin-protein ligase parkin OS=Hc | PRKN | 51.641 | 2 | 51.618065 | yes | 1.3E-07 | 653.83228 | 328.16122 |
| VCMGDHWFDV | 10 | O60260 | E3 ubiquitin-protein ligase parkin OS=Hc | PRKN | 51.641 | 2 | 51.618065 | yes | 1.286E-07 | 633.25751 | 268.99985 |

| | | | | | | | | | | | |
|---|---|---|---|---|---|---|---|---|---|---|---|
| SIYNSFYVYCK | 11 | O60260 | E3 ubiquitin-protein ligase parkin OS=Hc | PRKN | 51.641 | 2 | 51.618065 | yes | 9.722E-08 | 722.33429 | 219.66829 |
| VDLSSSVLPGDSV | 24 | O60260 | E3 ubiquitin-protein ligase parkin OS=Hc | PRKN | 51.641 | 3 | 51.618065 | yes | 1.922E-07 | 813.1026 | 120.88696 |
| ETSVALHLIATNSF | 14 | O60260 | E3 ubiquitin-protein ligase parkin OS=Hc | PRKN | 51.641 | 2 | 51.618065 | yes | 2.737E-07 | 756.41248 | 100.77381 |
| YQQYGAEECVLQI | 20 | O60260 | E3 ubiquitin-protein ligase parkin OS=Hc | PRKN | 51.641 | 3 | 51.618065 | yes | 2.942E-07 | 786.69403 | 66.944809 |
| CGAHPTSDKETSV | 23 | O60260 | E3 ubiquitin-protein ligase parkin OS=Hc | PRKN | 51.641 | 3 | 51.618065 | yes | 1.161E-06 | 822.41156 | 36.265678 |
| LDSIVIQQGR | 10 | O60264 | SWI/SNF-related matrix-associated actin- | SMARCA5 | 121.906 | 2 | 53.060192 | no | 1.149E-07 | 564.8222 | 667.81042 |
| YLVIDEAHR | 9 | O60264 | SWI/SNF-related matrix-associated actin- | SMARCA5 | 121.906 | 2 | 53.060192 | no | 1.187E-07 | 558.29584 | 631.71973 |
| NIPGPHMVLVPK | 12 | O60264 | SWI/SNF-related matrix-associated actin- | SMARCA5 | 121.906 | 2 | 53.060192 | no | 9.727E-08 | 651.37354 | 285.84634 |
| VLIFSQMTR | 9 | O60264 | SWI/SNF-related matrix-associated actin- | SMARCA5 | 121.906 | 2 | 53.060192 | no | 1.064E-07 | 547.80499 | 267.8259 |
| ANYAVDAYFR | 10 | O60264 | SWI/SNF-related matrix-associated actin- | SMARCA5 | 121.906 | 2 | 53.060192 | no | 1.204E-07 | 595.28546 | 200.18474 |
| EIQEPDPTYEEK | 12 | O60264 | SWI/SNF-related matrix-associated actin- | SMARCA5 | 121.906 | 2 | 53.060192 | yes | 9.722E-08 | 739.33826 | 198.54025 |
| CNTLITLIER | 10 | O60264 | SWI/SNF-related matrix-associated actin- | SMARCA5 | 121.906 | 2 | 53.060192 | yes | 5.891E-07 | 616.83698 | 198.50223 |
| FITDNTVEER | 10 | O60264 | SWI/SNF-related matrix-associated actin- | SMARCA5 | 121.906 | 2 | 53.060192 | yes | 4.755E-07 | 612.29877 | 191.99359 |
| APFHQLR | 7 | O60264 | SWI/SNF-related matrix-associated actin- | SMARCA5 | 121.906 | 2 | 53.060192 | no | 1.083E-07 | 434.74304 | 180.79335 |
| LLTQGFTNWNK | 11 | O60264 | SWI/SNF-related matrix-associated actin- | SMARCA5 | 121.906 | 2 | 53.060192 | yes | 1.908E-07 | 661.34857 | 163.70494 |
| NPELPNAAQAQK | 12 | O60264 | SWI/SNF-related matrix-associated actin- | SMARCA5 | 121.906 | 2 | 53.060192 | yes | 9.667E-07 | 640.83331 | 158.98682 |
| LLNILMQLR | 9 | O60264 | SWI/SNF-related matrix-associated actin- | SMARCA5 | 121.906 | 2 | 53.060192 | no | 9.722E-08 | 557.3443 | 139.58794 |
| CNELQDIEK | 9 | O60264 | SWI/SNF-related matrix-associated actin- | SMARCA5 | 121.906 | 2 | 53.060192 | no | 6.558E-07 | 574.76624 | 124.32945 |
| STLHNWMSEFK | 11 | O60264 | SWI/SNF-related matrix-associated actin- | SMARCA5 | 121.906 | 2 | 53.060192 | yes | 1.282E-07 | 690.32428 | 101.70743 |
| QNLLSVGDYR | 10 | O60264 | SWI/SNF-related matrix-associated actin- | SMARCA5 | 121.906 | 2 | 53.060192 | yes | 9.722E-08 | 582.80402 | 82.243103 |
| HGATHVFASK | 10 | O60264 | SWI/SNF-related matrix-associated actin- | SMARCA5 | 121.906 | 2 | 53.060192 | no | 3.732E-07 | 527.77509 | 80.185913 |
| ENVYDELR | 8 | O60264 | SWI/SNF-related matrix-associated actin- | SMARCA5 | 121.906 | 2 | 53.060192 | yes | 2.093E-07 | 519.24854 | 73.526657 |
| FEDSPSYVK | 9 | O60264 | SWI/SNF-related matrix-associated actin- | SMARCA5 | 121.906 | 2 | 53.060192 | yes | 3.324E-07 | 536.2533 | 67.325829 |
| MVVLDKLLPK | 10 | O60264 | SWI/SNF-related matrix-associated actin- | SMARCA5 | 121.906 | 2 | 53.060192 | yes | 7.263E-05 | 578.36212 | 51.601601 |
| LDGQTPHDER | 10 | O60264 | SWI/SNF-related matrix-associated actin- | SMARCA5 | 121.906 | 2 | 53.060192 | yes | 9.722E-08 | 584.27307 | 50.92355 |
| QDSINAYNEPNSTI | 14 | O60264 | SWI/SNF-related matrix-associated actin- | SMARCA5 | 121.906 | 2 | 53.060192 | yes | 1.155E-07 | 790.86298 | 24.984343 |
| FLEMQSK | 7 | O60281 | Zinc finger protein 292 OS=Homo sapien | ZNF292 | 304.816 | 2 | 37.058487 | yes | 6.342E-07 | 449.72058 | 134.2377 |
| VVLQSEVASLR | 11 | O60292 | Signal-induced proliferation-associated 1- | SIPA1L3 | 194.61 | 2 | 47.718414 | yes | 1.272E-07 | 600.85095 | 121.58987 |
| STISASELSLADGR | 14 | O60292 | Signal-induced proliferation-associated 1- | SIPA1L3 | 194.61 | 2 | 47.718414 | yes | 3.774E-07 | 703.85974 | 60.434719 |
| LQEESQAASEQLR | 13 | O60292 | Signal-induced proliferation-associated 1- | SIPA1L3 | 194.61 | 2 | 47.718414 | yes | 3.09E-07 | 744.8681 | 33.198517 |
| FCSVMPANSGLAL | 14 | O60293 | Zinc finger C3H1 domain-containing prot | ZFC3H1 | 226.356 | 2 | 49.30542 | yes | 2.007E-07 | 761.87891 | 156.8219 |
| YNNLDLFR | 8 | O60293 | Zinc finger C3H1 domain-containing prot | ZFC3H1 | 226.356 | 2 | 49.30542 | yes | 2.935E-07 | 527.76947 | 95.624512 |
| DQLLFEASEGGK | 12 | O60293 | Zinc finger C3H1 domain-containing prot | ZFC3H1 | 226.356 | 2 | 49.30542 | yes | 4.431E-07 | 647.3197 | 91.833664 |
| LLALQSASK | 9 | O60293 | Zinc finger C3H1 domain-containing prot | ZFC3H1 | 226.356 | 2 | 49.30542 | yes | 3.323E-07 | 465.78458 | 75.26886 |
| KLQEQIHR | 8 | O60293 | Zinc finger C3H1 domain-containing prot | ZFC3H1 | 226.356 | 2 | 49.30542 | yes | 2.638E-07 | 526.30402 | 64.277328 |
| SNLSIVSINTVSQPI | 15 | O60293 | Zinc finger C3H1 domain-containing prot | ZFC3H1 | 226.356 | 2 | 49.30542 | yes | 3.413E-06 | 807.94409 | 61.900051 |
| MPSSSLSESSPR | 12 | O60293 | Zinc finger C3H1 domain-containing prot | ZFC3H1 | 226.356 | 2 | 49.30542 | yes | 1.083E-07 | 632.79553 | 61.160397 |
| SANDGIVAEYLK | 12 | O60293 | Zinc finger C3H1 domain-containing prot | ZFC3H1 | 226.356 | 2 | 49.30542 | yes | 1.047E-07 | 640.33008 | 58.958572 |
| STNSVFGGLESMIF | 14 | O60293 | Zinc finger C3H1 domain-containing prot | ZFC3H1 | 226.356 | 2 | 49.30542 | yes | 2.898E-07 | 735.3689 | 57.699043 |

| | | | | | | | | | | | |
|---|---|---|---|---|---|---|---|---|---|---|---|
| IAIAAEIVLK | 10 | O60293 | Zinc finger C3H1 domain-containing prot | ZFC3H1 | 226.356 | 2 | 49.30542 | yes | 2.858E-07 | 520.83936 | 55.060158 |
| LPLCASLWK | 9 | O60293 | Zinc finger C3H1 domain-containing prot | ZFC3H1 | 226.356 | 2 | 49.30542 | yes | 6.303E-07 | 544.30206 | 52.737907 |
| ACTDESLAVEER | 12 | O60293 | Zinc finger C3H1 domain-containing prot | ZFC3H1 | 226.356 | 2 | 49.30542 | yes | 1.331E-07 | 690.30902 | 50.777481 |
| ISTSDILSEK | 10 | O60293 | Zinc finger C3H1 domain-containing prot | ZFC3H1 | 226.356 | 2 | 49.30542 | yes | 1.958E-06 | 546.79279 | 47.85955 |
| KLGEDEEELSELQI | 15 | O60293 | Zinc finger C3H1 domain-containing prot | ZFC3H1 | 226.356 | 3 | 49.30542 | yes | 2.628E-07 | 596.63892 | 44.819649 |
| LGEDEEELSELQLI | 14 | O60293 | Zinc finger C3H1 domain-containing prot | ZFC3H1 | 226.356 | 2 | 49.30542 | yes | 7.73E-07 | 830.40723 | 38.743309 |
| MLEFLMGAAK | 10 | O60293 | Zinc finger C3H1 domain-containing prot | ZFC3H1 | 226.356 | 2 | 49.30542 | yes | 8.639E-07 | 555.78796 | 38.05917 |
| NPQNAEVFYHMC | 13 | O60293 | Zinc finger C3H1 domain-containing prot | ZFC3H1 | 226.356 | 2 | 49.30542 | yes | 7.212E-07 | 819.36359 | 36.388031 |
| FYDPSNDNPSR | 11 | O60293 | Zinc finger C3H1 domain-containing prot | ZFC3H1 | 226.356 | 2 | 49.30542 | yes | 2.317E-07 | 656.28363 | 34.417198 |
| NTVEKPELFLGLK | 13 | O60293 | Zinc finger C3H1 domain-containing prot | ZFC3H1 | 226.356 | 2 | 49.30542 | yes | 1.172E-07 | 744.42706 | 12.831323 |
| VGVPTVDLDAQGI | 13 | O60306 | RNA helicase aquarius OS=Homo sapien | AQR | 171.296 | 2 | 51.6011 | yes | 1.593E-07 | 663.85425 | 324.31567 |
| IFTQLEEFR | 9 | O60306 | RNA helicase aquarius OS=Homo sapien | AQR | 171.296 | 2 | 51.6011 | yes | 1.063E-07 | 591.81134 | 262.76691 |
| ASLCNLYNWR | 10 | O60306 | RNA helicase aquarius OS=Homo sapien | AQR | 171.296 | 2 | 51.6011 | yes | 1.477E-07 | 648.81134 | 153.54063 |
| IMALDIDER | 9 | O60306 | RNA helicase aquarius OS=Homo sapien | AQR | 171.296 | 2 | 51.6011 | yes | 2.613E-07 | 538.27625 | 134.73376 |
| GPYPYNQPK | 9 | O60306 | RNA helicase aquarius OS=Homo sapien | AQR | 171.296 | 2 | 51.6011 | yes | 1.136E-05 | 532.26398 | 126.19232 |
| RPFIEQVGLVYVR | 13 | O60306 | RNA helicase aquarius OS=Homo sapien | AQR | 171.296 | 3 | 51.6011 | yes | 2.591E-07 | 525.97168 | 56.369972 |
| RPFIEQVGLVYVR | 13 | O60306 | RNA helicase aquarius OS=Homo sapien | AQR | 171.296 | 3 | 51.6011 | yes | 3.175E-06 | 788.45392 | 56.369972 |
| RPFIEQVGLVYVR | 13 | O60306 | RNA helicase aquarius OS=Homo sapien | AQR | 171.296 | 2 | 51.6011 | yes | 2.591E-07 | 525.97168 | 56.369972 |
| RPFIEQVGLVYVR | 13 | O60306 | RNA helicase aquarius OS=Homo sapien | AQR | 171.296 | 2 | 51.6011 | yes | 3.175E-06 | 788.45392 | 56.369972 |
| AYLMSICCMVNEI | 13 | O60306 | RNA helicase aquarius OS=Homo sapien | AQR | 171.296 | 2 | 51.6011 | yes | 1.479E-07 | 809.85846 | 111.77962 |
| YSNMEQSLFTR | 11 | O60306 | RNA helicase aquarius OS=Homo sapien | AQR | 171.296 | 2 | 51.6011 | yes | 2.283E-07 | 688.31915 | 93.13382 |
| SVPLSEPVTMDK | 12 | O60306 | RNA helicase aquarius OS=Homo sapien | AQR | 171.296 | 2 | 51.6011 | yes | 3.066E-07 | 651.83411 | 92.295372 |
| VNYVLAR | 7 | O60306 | RNA helicase aquarius OS=Homo sapien | AQR | 171.296 | 2 | 51.6011 | yes | 3.131E-07 | 417.74524 | 73.436569 |
| GCEIQGMLDDK | 11 | O60306 | RNA helicase aquarius OS=Homo sapien | AQR | 171.296 | 2 | 51.6011 | yes | 1.156E-07 | 633.27869 | 66.991661 |
| ADVTINLNVR | 10 | O60306 | RNA helicase aquarius OS=Homo sapien | AQR | 171.296 | 2 | 51.6011 | yes | 6.817E-07 | 557.81439 | 63.037739 |
| KFWNLIK | 7 | O60306 | RNA helicase aquarius OS=Homo sapien | AQR | 171.296 | 2 | 51.6011 | yes | 9.722E-08 | 474.78693 | 61.392586 |
| LGHGEEELETEK | 12 | O60306 | RNA helicase aquarius OS=Homo sapien | AQR | 171.296 | 2 | 51.6011 | yes | 1.505E-07 | 685.82532 | 52.824711 |
| ENVPAWEIFK | 10 | O60306 | RNA helicase aquarius OS=Homo sapien | AQR | 171.296 | 2 | 51.6011 | yes | 0.0002562 | 616.81915 | 48.490952 |
| NTIQFTHTQIEAIR | 14 | O60306 | RNA helicase aquarius OS=Homo sapien | AQR | 171.296 | 2 | 51.6011 | yes | 9.722E-08 | 836.44427 | 41.182037 |
| LGLYIFAR | 8 | O60306 | RNA helicase aquarius OS=Homo sapien | AQR | 171.296 | 2 | 51.6011 | yes | 3.813E-07 | 476.78436 | 36.887573 |
| DADVEDEDTEEAI | 13 | O60306 | RNA helicase aquarius OS=Homo sapien | AQR | 171.296 | 2 | 51.6011 | yes | 1.556E-07 | 733.29425 | 27.566761 |
| CGNNPLIGRPNK | 12 | O60306 | RNA helicase aquarius OS=Homo sapien | AQR | 171.296 | 2 | 51.6011 | yes | 3.015E-07 | 670.34863 | 24.634722 |
| SLVVGTPSPTLSR | 13 | O60307 | Microtubule-associated serine/threonine-p | MAST3 | 143.138 | 2 | 50.696125 | yes | 9.728E-08 | 657.37482 | 128.56154 |
| DLKPDNLLITSLGI | 16 | O60307 | Microtubule-associated serine/threonine-p | MAST3 | 143.138 | 3 | 50.696125 | yes | 1.426E-07 | 593.01215 | 116.18577 |
| SPSPLPGHPPAPAI | 14 | O60307 | Microtubule-associated serine/threonine-p | MAST3 | 143.138 | 3 | 50.696125 | yes | 4.043E-06 | 460.9176 | 80.17765 |
| EGCTVSPETISLNV | 15 | O60313 | Dynamin-like 120 kDa protein, mitochon | OPA1 | 111.631 | 2 | 49.662064 | yes | 5.715E-07 | 817.40869 | 150.97581 |
| EVLEDFAEDGEK | 12 | O60313 | Dynamin-like 120 kDa protein, mitochon | OPA1 | 111.631 | 2 | 49.662064 | yes | 1.012E-07 | 690.81189 | 94.045525 |
| NLSLAVSDCFWK | 12 | O60313 | Dynamin-like 120 kDa protein, mitochon | OPA1 | 111.631 | 2 | 49.662064 | yes | 4.819E-07 | 720.35303 | 54.469086 |
| WNDFAEDSLR | 10 | O60313 | Dynamin-like 120 kDa protein, mitochon | OPA1 | 111.631 | 2 | 49.662064 | yes | 7.56E-07 | 626.78326 | 45.111656 |

| | | | | | | | | | | | |
|---|---|---|---|---|---|---|---|---|---|---|---|
| IQQIIEGK | 8 | O60313 | Dynamin-like 120 kDa protein, mitochon | OPA1 | 111.631 | 2 | 49.662064 | yes | 3.599E-07 | 464.77673 | 38.904915 |
| SIVTDLVSQMDPH | 15 | O60313 | Dynamin-like 120 kDa protein, mitochon | OPA1 | 111.631 | 2 | 49.662064 | yes | 6.407E-06 | 827.91449 | 25.331541 |
| TSVLEMIAQAR | 11 | O60313 | Dynamin-like 120 kDa protein, mitochon | OPA1 | 111.631 | 2 | 49.662064 | yes | 2.62E-06 | 609.82916 | 21.57119 |
| AFPAAPCCVDVSD | 14 | O60318 | Germinal-center associated nuclear protei | MCM3AP | 218.405 | 2 | 54.427135 | yes | 3.128E-06 | 782.84778 | 67.400124 |
| DMAIFWHR | 8 | O60318 | Germinal-center associated nuclear protei | MCM3AP | 218.405 | 2 | 54.427135 | yes | 1.357E-07 | 538.26074 | 57.193428 |
| GLLDLGHAGR | 10 | O60318 | Germinal-center associated nuclear protei | MCM3AP | 218.405 | 2 | 54.427135 | yes | 1.739E-06 | 504.7829 | 55.545326 |
| SAFLEPEGLSK | 11 | O60318 | Germinal-center associated nuclear protei | MCM3AP | 218.405 | 2 | 54.427135 | yes | 2.702E-06 | 589.30859 | 48.360622 |
| EVEDGLMLQDLV | 15 | O60318 | Germinal-center associated nuclear protei | MCM3AP | 218.405 | 2 | 54.427135 | yes | 3.619E-07 | 823.91895 | 36.855175 |
| TFVGTCLDMCPEK | 13 | O60318 | Germinal-center associated nuclear protei | MCM3AP | 218.405 | 2 | 54.427135 | yes | 9.722E-08 | 779.34094 | 26.088085 |
| ISLVDLAGSER | 11 | O60333 | Kinesin-like protein KIF1B OS=Homo sa | KIF1B | 204.477 | 2 | 52.113922 | no | 1.192E-07 | 580.31952 | 4466.25 |
| IIAELNETWEEK | 12 | O60333 | Kinesin-like protein KIF1B OS=Homo sa | KIF1B | 204.477 | 2 | 52.113922 | no | 9.722E-08 | 737.87482 | 2077.6904 |
| IMSTPGGEEAIER | 13 | O60333 | Kinesin-like protein KIF1B OS=Homo sa | KIF1B | 204.477 | 2 | 52.113922 | yes | 9.899E-08 | 695.33759 | 916.02289 |
| EEADLLLEQQR | 11 | O60333 | Kinesin-like protein KIF1B OS=Homo sa | KIF1B | 204.477 | 2 | 52.113922 | no | 9.733E-08 | 672.34369 | 666.19037 |
| EALLAEMGVAIR | 12 | O60333 | Kinesin-like protein KIF1B OS=Homo sa | KIF1B | 204.477 | 2 | 52.113922 | yes | 9.722E-08 | 636.85266 | 492.3215 |
| EDGGTLGVFSPK | 12 | O60333 | Kinesin-like protein KIF1B OS=Homo sa | KIF1B | 204.477 | 2 | 52.113922 | no | 9.729E-08 | 603.80371 | 433.47217 |
| EKEEADLLLEQQR | 13 | O60333 | Kinesin-like protein KIF1B OS=Homo sa | KIF1B | 204.477 | 2 | 52.113922 | no | 9.809E-08 | 800.91248 | 409.95715 |
| EEHCIFR | 7 | O60333 | Kinesin-like protein KIF1B OS=Homo sa | KIF1B | 204.477 | 2 | 52.113922 | yes | 2.649E-07 | 495.72672 | 352.4212 |
| VSQPVQLR | 8 | O60333 | Kinesin-like protein KIF1B OS=Homo sa | KIF1B | 204.477 | 2 | 52.113922 | yes | 2.284E-07 | 463.77454 | 296.03842 |
| RQDIVLSGAHIK | 12 | O60333 | Kinesin-like protein KIF1B OS=Homo sa | KIF1B | 204.477 | 3 | 52.113922 | yes | 7.638E-07 | 446.26138 | 294.95502 |
| EHPLLGPYVEDLS | 14 | O60333 | Kinesin-like protein KIF1B OS=Homo sa | KIF1B | 204.477 | 2 | 52.113922 | no | 9.728E-08 | 798.91705 | 281.86588 |
| LQEMEILYK | 9 | O60333 | Kinesin-like protein KIF1B OS=Homo sa | KIF1B | 204.477 | 2 | 52.113922 | yes | 1.594E-07 | 583.80994 | 266.71286 |
| ELKEEVTR | 8 | O60333 | Kinesin-like protein KIF1B OS=Homo sa | KIF1B | 204.477 | 2 | 52.113922 | yes | 8.081E-07 | 502.27457 | 198.41673 |
| SHAVFTIVFTQK | 12 | O60333 | Kinesin-like protein KIF1B OS=Homo sa | KIF1B | 204.477 | 2 | 52.113922 | yes | 1.131E-07 | 689.37988 | 194.73886 |
| KTDFIPYR | 8 | O60333 | Kinesin-like protein KIF1B OS=Homo sa | KIF1B | 204.477 | 2 | 52.113922 | no | 3.108E-06 | 520.28217 | 194.31567 |
| SNSGEVIVTLEPCE | 15 | O60333 | Kinesin-like protein KIF1B OS=Homo sa | KIF1B | 204.477 | 2 | 52.113922 | yes | 9.722E-08 | 845.40924 | 183.68886 |
| CNAVINEDPNAK | 12 | O60333 | Kinesin-like protein KIF1B OS=Homo sa | KIF1B | 204.477 | 2 | 52.113922 | yes | 9.722E-08 | 672.81427 | 98.58638 |
| QDIVLSGAHIK | 11 | O60333 | Kinesin-like protein KIF1B OS=Homo sa | KIF1B | 204.477 | 2 | 52.113922 | yes | 4.972E-07 | 590.83789 | 80.811066 |
| TVAATNMNETSSR | 13 | O60333 | Kinesin-like protein KIF1B OS=Homo sa | KIF1B | 204.477 | 2 | 52.113922 | no | 1.164E-07 | 699.31989 | 74.150909 |
| VYNDIGK | 7 | O60333 | Kinesin-like protein KIF1B OS=Homo sa | KIF1B | 204.477 | 2 | 52.113922 | yes | 1.284E-07 | 404.71362 | 46.593414 |
| RLQEMEILYK | 10 | O60333 | Kinesin-like protein KIF1B OS=Homo sa | KIF1B | 204.477 | 2 | 52.113922 | yes | 3.399E-07 | 661.86047 | 31.146164 |
| GIFGSSAVPQPK | 12 | O60341 | Lysine-specific histone demethylase 1A C | KDM1A | 92.903 | 2 | 45.253853 | yes | 5.467E-07 | 594.32458 | 99.996704 |
| VVLCFDR | 7 | O60341 | Lysine-specific histone demethylase 1A C | KDM1A | 92.903 | 2 | 45.253853 | yes | 1.908E-07 | 454.73657 | 44.978626 |
| EASEVKPPR | 9 | O60341 | Lysine-specific histone demethylase 1A C | KDM1A | 92.903 | 2 | 45.253853 | yes | 2.227E-06 | 506.77472 | 40.324913 |
| HGLINFGIYK | 10 | O60341 | Lysine-specific histone demethylase 1A C | KDM1A | 92.903 | 2 | 45.253853 | yes | 3.542E-07 | 581.3244 | 33.07354 |
| CDAVLCTLPLGVL | 14 | O60341 | Lysine-specific histone demethylase 1A C | KDM1A | 92.903 | 2 | 45.253853 | yes | 1.964E-06 | 779.92023 | 26.027275 |
| IITQVFEMDISK | 12 | O60343 | TBC1 domain family member 4 OS=Hom | TBC1D4 | 146.564 | 2 | 46.959747 | yes | 2.486E-07 | 712.37872 | 66.9617 |
| MKPVSDQEENELV | 18 | O60343 | TBC1 domain family member 4 OS=Hom | TBC1D4 | 146.564 | 3 | 46.959747 | yes | 2.302E-07 | 717.38068 | 59.608898 |
| LGSVDSFER | 9 | O60343 | TBC1 domain family member 4 OS=Hom | TBC1D4 | 146.564 | 2 | 46.959747 | yes | 2.225E-07 | 505.2511 | 58.37608 |
| SLTSSLENIFSR | 12 | O60343 | TBC1 domain family member 4 OS=Hom | TBC1D4 | 146.564 | 2 | 46.959747 | yes | 1.431E-07 | 677.35406 | 38.215504 |

| | | | | | | | | | | | |
|---|---|---|---|---|---|---|---|---|---|---|---|
| HASAPSHVQPSDS | 15 | O60343 | TBC1 domain family member 4 OS=Hom | TBC1D4 | 146.564 | 3 | 46.959747 | yes | 2.606E-06 | 526.24988 | 31.417061 |
| APSSLIDDCMEK | 12 | O60343 | TBC1 domain family member 4 OS=Hom | TBC1D4 | 146.564 | 2 | 46.959747 | yes | 2.953E-07 | 683.30487 | 22.639973 |
| NTLPDMNTSEMEI | 13 | O60343 | TBC1 domain family member 4 OS=Hom | TBC1D4 | 146.564 | 2 | 46.959747 | yes | 3.177E-06 | 755.3316 | 18.872492 |
| VAPLVQSLCAK | 11 | O60427 | Acyl-CoA (8-3)-desaturase OS=Homo saj | FADS1 | 51.965 | 2 | 45.847916 | yes | 6.999E-07 | 593.33667 | 133.63844 |
| YFTWDEVAQR | 10 | O60427 | Acyl-CoA (8-3)-desaturase OS=Homo saj | FADS1 | 51.965 | 2 | 45.847916 | yes | 1.725E-07 | 657.80933 | 69.729698 |
| AQYLALPER | 9 | O60479 | Homeobox protein DLX-3 OS=Homo sap | DLX3 | 31.739 | 2 | 51.46781 | yes | 2.318E-07 | 530.79291 | 215.51904 |
| EQPLPAQDPVSVK | 13 | O60479 | Homeobox protein DLX-3 OS=Homo sap | DLX3 | 31.739 | 2 | 51.46781 | yes | 9.722E-08 | 704.37756 | 204.49429 |
| AELAAQLGLTQTQ | 15 | O60479 | Homeobox protein DLX-3 OS=Homo sap | DLX3 | 31.739 | 2 | 51.46781 | yes | 2.171E-07 | 785.9436 | 40.824734 |
| TIYSSYQLAALQR | 13 | O60479 | Homeobox protein DLX-3 OS=Homo sap | DLX3 | 31.739 | 2 | 51.46781 | yes | 4.532E-06 | 757.40411 | 22.79495 |
| NTIAIFCETR | 10 | O60488 | Long-chain-fatty-acid--CoA ligase 4 OS= | ACSL4 | 79.189 | 2 | 42.65622 | yes | 2.935E-07 | 612.80573 | 112.81335 |
| WMNYLEVNR | 9 | O60488 | Long-chain-fatty-acid--CoA ligase 4 OS= | ACSL4 | 79.189 | 2 | 42.65622 | yes | 2.931E-07 | 612.79517 | 44.922131 |
| LTLLAQQK | 8 | O60488 | Long-chain-fatty-acid--CoA ligase 4 OS= | ACSL4 | 79.189 | 2 | 42.65622 | yes | 2.796E-06 | 457.78711 | 39.059929 |
| IGYSSPLTLSDQSS | 15 | O60488 | Long-chain-fatty-acid--CoA ligase 4 OS= | ACSL4 | 79.189 | 2 | 42.65622 | yes | 3.406E-07 | 791.90161 | 29.54377 |
| GDDGIFDDNFIEEI | 14 | O60493 | Sorting nexin-3 OS=Homo sapiens OX=9 | SNX3 | 18.763 | 2 | 49.19806 | yes | 1.06E-07 | 821.35498 | 83.756172 |
| YSDFEWLR | 8 | O60493 | Sorting nexin-3 OS=Homo sapiens OX=9 | SNX3 | 18.763 | 2 | 49.19806 | yes | 4.59E-07 | 558.26147 | 17.744717 |
| WCELLQESAAVGI | 13 | O60499 | Syntaxin-10 OS=Homo sapiens OX=960( | STX10 | 28.115 | 2 | 42.383865 | yes | 2.371E-06 | 759.87213 | 57.390419 |
| SPSDLLDASAVSA | 16 | O60499 | Syntaxin-10 OS=Homo sapiens OX=960( | STX10 | 28.115 | 2 | 42.383865 | yes | 3.056E-07 | 788.89429 | 45.882431 |
| DHMVSPTAVAFLI | 14 | O60499 | Syntaxin-10 OS=Homo sapiens OX=960( | STX10 | 28.115 | 3 | 42.383865 | yes | 2.893E-06 | 524.9328 | 29.811335 |
| FLCGVVEGFYGRP | 20 | O60502 | Protein O-GlcNAcase OS=Homo sapiens | OGA | 102.916 | 4 | 32.777035 | yes | 1.416E-06 | 615.30945 | 45.796143 |
| TGFSFPTQEPR | 11 | O60504 | Vinexin OS=Homo sapiens OX=9606 GN | SORBS3 | 75.342 | 2 | 50.27617 | yes | 9.82E-08 | 633.80933 | 254.09538 |
| HPSSPSALR | 9 | O60504 | Vinexin OS=Homo sapiens OX=9606 GN | SORBS3 | 75.342 | 2 | 50.27617 | yes | 2.802E-06 | 476.25397 | 177.86528 |
| PQTQNLGTPGPAL | 17 | O60504 | Vinexin OS=Homo sapiens OX=9606 GN | SORBS3 | 75.342 | 3 | 50.27617 | yes | 2.378E-07 | 587.63947 | 97.338295 |
| LMMDPLTGLNR | 11 | O60506 | Heterogeneous nuclear ribonucleoprotein | SYNCRIP | 69.603 | 2 | 51.577126 | yes | 9.938E-08 | 630.82556 | 8859.8574 |
| LMMDPLTGLNR | 11 | O60506 | Heterogeneous nuclear ribonucleoprotein | SYNCRIP | 69.603 | 2 | 51.577126 | yes | 1.191E-07 | 638.82306 | 8859.8574 |
| LMMDPLTGLNR | 11 | O60506 | Heterogeneous nuclear ribonucleoprotein | SYNCRIP | 69.603 | 2 | 51.577126 | yes | 9.722E-08 | 638.82306 | 8859.8574 |
| DYAFIHFDER | 10 | O60506 | Heterogeneous nuclear ribonucleoprotein | SYNCRIP | 69.603 | 2 | 51.577126 | yes | 1.141E-07 | 656.80145 | 11992.522 |
| LYNNHEIR | 8 | O60506 | Heterogeneous nuclear ribonucleoprotein | SYNCRIP | 69.603 | 2 | 51.577126 | yes | 1.536E-07 | 529.77252 | 8561.1396 |
| NLANTVTEEILEK | 13 | O60506 | Heterogeneous nuclear ribonucleoprotein | SYNCRIP | 69.603 | 2 | 51.577126 | yes | 9.819E-08 | 737.39337 | 8147.2627 |
| EQILEEFSK | 9 | O60506 | Heterogeneous nuclear ribonucleoprotein | SYNCRIP | 69.603 | 2 | 51.577126 | yes | 1.156E-07 | 561.78748 | 7946.8809 |
| HIGVCISVANNR | 12 | O60506 | Heterogeneous nuclear ribonucleoprotein | SYNCRIP | 69.603 | 2 | 51.577126 | yes | 2.262E-07 | 670.34863 | 5962.2612 |
| GYAFVTFCTK | 10 | O60506 | Heterogeneous nuclear ribonucleoprotein | SYNCRIP | 69.603 | 2 | 51.577126 | yes | 4.886E-07 | 597.28662 | 4341.6309 |
| DLEGENIEIVFAKP | 18 | O60506 | Heterogeneous nuclear ribonucleoprotein | SYNCRIP | 69.603 | 3 | 51.577126 | yes | 6.191E-07 | 681.35284 | 4152.7705 |
| DSDLSHVQNK | 10 | O60506 | Heterogeneous nuclear ribonucleoprotein | SYNCRIP | 69.603 | 2 | 51.577126 | yes | 1.439E-07 | 571.77545 | 3979.6069 |
| LKDYAFIHFDER | 12 | O60506 | Heterogeneous nuclear ribonucleoprotein | SYNCRIP | 69.603 | 3 | 51.577126 | yes | 3.203E-07 | 518.59644 | 1533.2295 |
| LKDYAFIHFDER | 12 | O60506 | Heterogeneous nuclear ribonucleoprotein | SYNCRIP | 69.603 | 3 | 51.577126 | yes | 1.48E-07 | 777.39099 | 1533.2295 |
| LKDYAFIHFDER | 12 | O60506 | Heterogeneous nuclear ribonucleoprotein | SYNCRIP | 69.603 | 2 | 51.577126 | yes | 3.203E-07 | 518.59644 | 1533.2295 |
| LKDYAFIHFDER | 12 | O60506 | Heterogeneous nuclear ribonucleoprotein | SYNCRIP | 69.603 | 2 | 51.577126 | yes | 1.48E-07 | 777.39099 | 1533.2295 |
| LDEIYVAGLVAHS | 18 | O60506 | Heterogeneous nuclear ribonucleoprotein | SYNCRIP | 69.603 | 3 | 51.577126 | yes | 1.002E-07 | 672.34088 | 2646.1287 |
| TKEQILEEFSK | 11 | O60506 | Heterogeneous nuclear ribonucleoprotein | SYNCRIP | 69.603 | 2 | 51.577126 | yes | 1.398E-07 | 676.35883 | 890.24286 |

| | | | | | | | | | | | |
|---|---|---|---|---|---|---|---|---|---|---|---|
| VAEKLDEIYVAGL | 22 | O60506 | Heterogeneous nuclear ribonucleoprotein | SYNCRIP | 69.603 | 3 | 51.577126 | yes | 1.129E-07 | 814.75519 | 214.55382 |
| EFNEDGALAVLQC | 15 | O60506 | Heterogeneous nuclear ribonucleoprotein | SYNCRIP | 69.603 | 3 | 51.577126 | yes | 9.722E-08 | 570.28955 | 36.711464 |
| LWDTETGQCISR | 12 | O60508 | Pre-mRNA-processing factor 17 OS=Hon | CDC40 | 65.522 | 2 | 51.68414 | yes | 1.441E-07 | 733.34064 | 312.75543 |
| VITCGWDGLIK | 11 | O60508 | Pre-mRNA-processing factor 17 OS=Hon | CDC40 | 65.522 | 2 | 51.68414 | yes | 6.678E-07 | 631.33411 | 209.12683 |
| NQGLTVFETGQK | 12 | O60508 | Pre-mRNA-processing factor 17 OS=Hon | CDC40 | 65.522 | 2 | 51.68414 | yes | 1.108E-07 | 661.34094 | 197.4747 |
| SYLHIPQDVGVNL | 14 | O60508 | Pre-mRNA-processing factor 17 OS=Hon | CDC40 | 65.522 | 2 | 51.68414 | yes | 9.722E-08 | 805.9361 | 197.19817 |
| LWEVYGER | 8 | O60508 | Pre-mRNA-processing factor 17 OS=Hon | CDC40 | 65.522 | 2 | 51.68414 | yes | 1.399E-06 | 526.26398 | 182.36804 |
| PSLAVAVDSAPEV | 16 | O60508 | Pre-mRNA-processing factor 17 OS=Hon | CDC40 | 65.522 | 2 | 51.68414 | yes | 1.013E-07 | 776.93268 | 170.61584 |
| SGEIVQEYDR | 10 | O60508 | Pre-mRNA-processing factor 17 OS=Hon | CDC40 | 65.522 | 2 | 51.68414 | yes | 9.689E-07 | 598.28314 | 162.6489 |
| EDLETGVHLDPAV | 14 | O60508 | Pre-mRNA-processing factor 17 OS=Hon | CDC40 | 65.522 | 2 | 51.68414 | yes | 6.898E-07 | 761.89099 | 63.913334 |
| EDLETGVHLDPAV | 14 | O60508 | Pre-mRNA-processing factor 17 OS=Hon | CDC40 | 65.522 | 2 | 51.68414 | yes | 5.576E-06 | 508.26309 | 63.913334 |
| EDLETGVHLDPAV | 14 | O60508 | Pre-mRNA-processing factor 17 OS=Hon | CDC40 | 65.522 | 3 | 51.68414 | yes | 6.898E-07 | 761.89099 | 63.913334 |
| EDLETGVHLDPAV | 14 | O60508 | Pre-mRNA-processing factor 17 OS=Hon | CDC40 | 65.522 | 3 | 51.68414 | yes | 5.576E-06 | 508.26309 | 63.913334 |
| YIGSVEEAEK | 10 | O60508 | Pre-mRNA-processing factor 17 OS=Hon | CDC40 | 65.522 | 2 | 51.68414 | yes | 6.177E-07 | 562.77716 | 119.52509 |
| HLGAVNTIVFVDE | 15 | O60508 | Pre-mRNA-processing factor 17 OS=Hon | CDC40 | 65.522 | 3 | 51.68414 | yes | 1.191E-07 | 561.96527 | 90.921326 |
| IVQWDIR | 7 | O60508 | Pre-mRNA-processing factor 17 OS=Hon | CDC40 | 65.522 | 2 | 51.68414 | yes | 1.449E-06 | 465.26379 | 67.747116 |
| VWEWDIPVDFK | 11 | O60508 | Pre-mRNA-processing factor 17 OS=Hon | CDC40 | 65.522 | 2 | 51.68414 | yes | 1.699E-07 | 717.35864 | 54.73159 |
| CPLPAADSLMHLT | 14 | O60508 | Pre-mRNA-processing factor 17 OS=Hon | CDC40 | 65.522 | 3 | 51.68414 | yes | 1.611E-07 | 518.59869 | 41.981079 |
| VCIGAVWHPHETS | 14 | O60508 | Pre-mRNA-processing factor 17 OS=Hon | CDC40 | 65.522 | 2 | 51.68414 | yes | 9.935E-08 | 810.90118 | 40.904926 |
| LFPLSGHLLLSCSM | 17 | O60508 | Pre-mRNA-processing factor 17 OS=Hon | CDC40 | 65.522 | 3 | 51.68414 | yes | 5.275E-07 | 659.99133 | 32.199612 |
| WLACQSMDNQILI | 19 | O60508 | Pre-mRNA-processing factor 17 OS=Hon | CDC40 | 65.522 | 3 | 51.68414 | yes | 9.763E-05 | 755.7016 | 7.5823665 |
| AAEPLLTLER | 10 | O60524 | Ribosome quality control complex subuni | NEMF | 122.955 | 2 | 55.345207 | yes | 1.192E-07 | 556.81915 | 706.32953 |
| LEALQQAQEIDK | 12 | O60524 | Ribosome quality control complex subuni | NEMF | 122.955 | 2 | 55.345207 | yes | 1.163E-07 | 693.36719 | 674.4231 |
| VSAPNLLNVK | 10 | O60524 | Ribosome quality control complex subuni | NEMF | 122.955 | 2 | 55.345207 | yes | 2.663E-07 | 527.81641 | 658.15558 |
| ATLLLESGIR | 10 | O60524 | Ribosome quality control complex subuni | NEMF | 122.955 | 2 | 55.345207 | yes | 2.916E-07 | 536.82166 | 621.45483 |
| EAQAQGDPVASAI | 14 | O60524 | Ribosome quality control complex subuni | NEMF | 122.955 | 2 | 55.345207 | yes | 9.722E-08 | 692.85699 | 432.67926 |
| LTEIVASAPK | 10 | O60524 | Ribosome quality control complex subuni | NEMF | 122.955 | 2 | 55.345207 | yes | 2.38E-07 | 514.80298 | 211.6207 |
| TALNSFMHSK | 10 | O60524 | Ribosome quality control complex subuni | NEMF | 122.955 | 2 | 55.345207 | yes | 9.722E-08 | 568.28186 | 181.75858 |
| GELIEMNLQIVDR | 13 | O60524 | Ribosome quality control complex subuni | NEMF | 122.955 | 2 | 55.345207 | yes | 1.489E-07 | 765.40326 | 180.84933 |
| DQQQNEIIVK | 10 | O60524 | Ribosome quality control complex subuni | NEMF | 122.955 | 2 | 55.345207 | yes | 9.809E-08 | 607.82239 | 165.24286 |
| KVYWFEK | 7 | O60524 | Ribosome quality control complex subuni | NEMF | 122.955 | 2 | 55.345207 | yes | 2.113E-07 | 500.26855 | 153.95508 |
| LQKPDFK | 7 | O60524 | Ribosome quality control complex subuni | NEMF | 122.955 | 2 | 55.345207 | yes | 3.331E-07 | 438.2529 | 147.18146 |
| KLPSDSGDLEALE( | 15 | O60524 | Ribosome quality control complex subuni | NEMF | 122.955 | 2 | 55.345207 | yes | 2.327E-07 | 779.90161 | 146.44946 |
| NMMPSSFAMK | 10 | O60524 | Ribosome quality control complex subuni | NEMF | 122.955 | 2 | 55.345207 | yes | 9.909E-08 | 572.25342 | 139.77342 |
| LQTNHVTMLLR | 11 | O60524 | Ribosome quality control complex subuni | NEMF | 122.955 | 3 | 55.345207 | yes | 1.423E-07 | 442.58347 | 65.090775 |
| LQTNHVTMLLR | 11 | O60524 | Ribosome quality control complex subuni | NEMF | 122.955 | 3 | 55.345207 | yes | 9.722E-08 | 663.37152 | 65.090775 |
| LQTNHVTMLLR | 11 | O60524 | Ribosome quality control complex subuni | NEMF | 122.955 | 2 | 55.345207 | yes | 1.423E-07 | 442.58347 | 65.090775 |
| LQTNHVTMLLR | 11 | O60524 | Ribosome quality control complex subuni | NEMF | 122.955 | 2 | 55.345207 | yes | 9.722E-08 | 663.37152 | 65.090775 |
| EVQTVTSIQK | 10 | O60524 | Ribosome quality control complex subuni | NEMF | 122.955 | 2 | 55.345207 | yes | 1.673E-07 | 566.81403 | 109.20464 |

| | | | | | | | | | | | |
|---|---|---|---|---|---|---|---|---|---|---|---|
| NPTGEPIPPR | 10 | O60524 | Ribosome quality control complex subuni | NEMF | 122.955 | 2 | 55.345207 | yes | 3.151E-06 | 539.28802 | 108.71438 |
| YPLDHAR | 7 | O60524 | Ribosome quality control complex subuni | NEMF | 122.955 | 2 | 55.345207 | yes | 6.485E-07 | 436.22467 | 107.29912 |
| AVLAELNASLLGM | 14 | O60524 | Ribosome quality control complex subuni | NEMF | 122.955 | 2 | 55.345207 | yes | 1.341E-07 | 729.41089 | 99.117393 |
| VNNVYDVDNK | 10 | O60524 | Ribosome quality control complex subuni | NEMF | 122.955 | 2 | 55.345207 | yes | 2.46E-06 | 590.28564 | 80.73616 |
| FRTDEADDVK | 10 | O60524 | Ribosome quality control complex subuni | NEMF | 122.955 | 2 | 55.345207 | yes | 3.204E-06 | 598.28314 | 67.846886 |
| GYEVHGIVR | 9 | O60547 | GDP-mannose 4,6 dehydratase OS=Homo | GMDS | 41.95 | 2 | 49.334404 | yes | 4.235E-07 | 515.27747 | 118.0239 |
| VAFDELVR | 8 | O60547 | GDP-mannose 4,6 dehydratase OS=Homo | GMDS | 41.95 | 2 | 49.334404 | yes | 1.043E-07 | 474.76108 | 111.67503 |
| IINEVKPTEIYNLG | 20 | O60547 | GDP-mannose 4,6 dehydratase OS=Homo | GMDS | 41.95 | 3 | 49.334404 | yes | 3.171E-07 | 751.74609 | 70.373589 |
| NPQAHIEGNMK | 11 | O60547 | GDP-mannose 4,6 dehydratase OS=Homo | GMDS | 41.95 | 2 | 49.334404 | yes | 2.535E-05 | 619.80096 | 16.575851 |
| MVEINFLCVHK | 11 | O60551 | Glycylpeptide N-tetradecanoyltransferase | NMT2 | 56.981 | 2 | 42.602173 | yes | 2.954E-07 | 695.35449 | 98.247017 |
| MPIEGSENPERPFL | 16 | O60563 | Cyclin-T1 OS=Homo sapiens OX=9606 ( | CCNT1 | 80.685 | 3 | 53.290997 | yes | 3.229E-06 | 624.97687 | 247.63351 |
| WSNWEIPVSTDGK | 13 | O60563 | Cyclin-T1 OS=Homo sapiens OX=9606 ( | CCNT1 | 80.685 | 2 | 53.290997 | no | 1.403E-07 | 759.86481 | 143.21162 |
| EQLENSPSR | 9 | O60563 | Cyclin-T1 OS=Homo sapiens OX=9606 ( | CCNT1 | 80.685 | 2 | 53.290997 | yes | 4.922E-07 | 530.2569 | 110.44629 |
| HNSVEDSVTK | 10 | O60563 | Cyclin-T1 OS=Homo sapiens OX=9606 ( | CCNT1 | 80.685 | 2 | 53.290997 | yes | 9.722E-08 | 558.27002 | 100.14267 |
| KLEHVIK | 7 | O60563 | Cyclin-T1 OS=Homo sapiens OX=9606 ( | CCNT1 | 80.685 | 2 | 53.290997 | no | 5.99E-07 | 433.77655 | 91.293068 |
| TYSLSSSFSSSSTF | 15 | O60563 | Cyclin-T1 OS=Homo sapiens OX=9606 ( | CCNT1 | 80.685 | 2 | 53.290997 | yes | 2.249E-07 | 792.36285 | 59.20005 |
| GPSEETGGAVFDH | 16 | O60563 | Cyclin-T1 OS=Homo sapiens OX=9606 ( | CCNT1 | 80.685 | 2 | 53.290997 | yes | 3.582E-07 | 799.87592 | 23.86401 |
| LVGPEEALSPGEAI | 14 | O60568 | Multifunctional procollagen lysine hydro | PLOD3 | 84.786 | 2 | 52.815052 | yes | 2.173E-07 | 712.87262 | 116.66785 |
| LYLDPGLR | 8 | O60568 | Multifunctional procollagen lysine hydro | PLOD3 | 84.786 | 2 | 52.815052 | yes | 2.436E-07 | 473.77145 | 114.81012 |
| YDCVISSPR | 9 | O60568 | Multifunctional procollagen lysine hydro | PLOD3 | 84.786 | 2 | 52.815052 | yes | 1.022E-07 | 548.75824 | 103.13066 |
| DMAMDLCR | 8 | O60568 | Multifunctional procollagen lysine hydro | PLOD3 | 84.786 | 2 | 52.815052 | yes | 3.423E-07 | 506.20645 | 69.685097 |
| GIFLHLSNQHEFGI | 14 | O60568 | Multifunctional procollagen lysine hydro | PLOD3 | 84.786 | 3 | 52.815052 | yes | 9.722E-08 | 552.28644 | 58.997288 |
| LSLNLDHK | 8 | O60568 | Multifunctional procollagen lysine hydro | PLOD3 | 84.786 | 2 | 52.815052 | yes | 4.591E-07 | 470.26654 | 46.428482 |
| FQEDIISIWNK | 11 | O60573 | Eukaryotic translation initiation factor 4E | EIF4E2 | 28.362 | 2 | 53.139183 | yes | 1.001E-07 | 696.86151 | 1949.2574 |
| VLNLPPNTIMEYK | 13 | O60573 | Eukaryotic translation initiation factor 4E | EIF4E2 | 28.362 | 2 | 53.139183 | yes | 1.414E-07 | 766.41309 | 1123.3636 |
| QIGTFASVEQFWR | 13 | O60573 | Eukaryotic translation initiation factor 4E | EIF4E2 | 28.362 | 2 | 53.139183 | yes | 1.353E-07 | 784.89642 | 508.34763 |
| FYSHMVR | 7 | O60573 | Eukaryotic translation initiation factor 4E | EIF4E2 | 28.362 | 2 | 53.139183 | yes | 1.051E-07 | 470.22891 | 433.37881 |
| LLFQNLWK | 8 | O60573 | Eukaryotic translation initiation factor 4E | EIF4E2 | 28.362 | 2 | 53.139183 | yes | 1.95E-07 | 531.31073 | 360.30469 |
| EGIKPMWEDDANI | 13 | O60573 | Eukaryotic translation initiation factor 4E | EIF4E2 | 28.362 | 2 | 53.139183 | yes | 5.416E-07 | 766.85614 | 124.31293 |
| EGIKPMWEDDANI | 13 | O60573 | Eukaryotic translation initiation factor 4E | EIF4E2 | 28.362 | 2 | 53.139183 | yes | 3.039E-06 | 511.57318 | 124.31293 |
| EGIKPMWEDDANI | 13 | O60573 | Eukaryotic translation initiation factor 4E | EIF4E2 | 28.362 | 3 | 53.139183 | yes | 5.416E-07 | 766.85614 | 124.31293 |
| EGIKPMWEDDANI | 13 | O60573 | Eukaryotic translation initiation factor 4E | EIF4E2 | 28.362 | 3 | 53.139183 | yes | 3.039E-06 | 511.57318 | 124.31293 |
| PTSSQSYEQNIK | 12 | O60573 | Eukaryotic translation initiation factor 4E | EIF4E2 | 28.362 | 2 | 53.139183 | yes | 9.722E-08 | 691.33331 | 123.2953 |
| PGDLTGHSDFHLF | 14 | O60573 | Eukaryotic translation initiation factor 4E | EIF4E2 | 28.362 | 2 | 53.139183 | yes | 3.532E-07 | 785.88605 | 66.331322 |
| DIVVPSLNVAK | 11 | O60645 | Exocyst complex component 3 OS=Homo | EXOC3 | 85.568 | 2 | 45.941578 | yes | 1.993E-07 | 577.84259 | 63.879257 |
| EAVATAVQR | 9 | O60645 | Exocyst complex component 3 OS=Homo | EXOC3 | 85.568 | 2 | 45.941578 | yes | 1.697E-07 | 472.76163 | 55.564964 |
| NIFSVPEIVR | 10 | O60645 | Exocyst complex component 3 OS=Homo | EXOC3 | 85.568 | 2 | 45.941578 | yes | 5.673E-06 | 587.33496 | 19.315294 |
| IQDYHILTR | 9 | O60674 | Tyrosine-protein kinase JAK2 OS=Homo | JAK2 | 130.674 | 2 | 29.593576 | yes | 2.443E-06 | 579.81696 | 21.490927 |
| ALSAEAALAR | 10 | O60678 | Protein arginine N-methyltransferase 3 OS | PRMT3 | 59.904 | 2 | 56.097218 | yes | 1.529E-06 | 486.77728 | 158.13646 |

| | | | | | | | | | | |
|---|---|---|---|---|---|---|---|---|---|---|
| VVFSTGPQSTK | 11 | O60678 | Protein arginine N-methyltransferase 3 O | PRMT3 | 59.904 | 2 | 56.097218 | yes | 9.722E-08 | 575.80878 | 102.30901 |
| QQTPCLFCNR | 10 | O60678 | Protein arginine N-methyltransferase 3 O | PRMT3 | 59.904 | 2 | 56.097218 | yes | 9.722E-08 | 662.30029 | 78.181023 |
| HGLEFYGYIK | 10 | O60678 | Protein arginine N-methyltransferase 3 O | PRMT3 | 59.904 | 2 | 56.097218 | yes | 1.611E-07 | 613.81384 | 46.046448 |
| LEDTITLIK | 9 | O60678 | Protein arginine N-methyltransferase 3 O | PRMT3 | 59.904 | 2 | 56.097218 | yes | 1.961E-07 | 523.31061 | 44.214348 |
| LFTSAEETFSHCK | 13 | O60678 | Protein arginine N-methyltransferase 3 O | PRMT3 | 59.904 | 2 | 56.097218 | yes | 1.598E-07 | 778.85614 | 27.952616 |
| IAFWDDVYGFK | 11 | O60678 | Protein arginine N-methyltransferase 3 O | PRMT3 | 59.904 | 2 | 56.097218 | yes | 2.497E-07 | 680.83228 | 15.008013 |
| NAVWALSNLCR | 11 | O60684 | Importin subunit alpha-7 OS=Homo sapi | KPNA6 | 60.03 | 2 | 54.920666 | no | 5.089E-07 | 652.3324 | 535.70557 |
| EPSPPIDEVINTPR | 14 | O60684 | Importin subunit alpha-7 OS=Homo sapi | KPNA6 | 60.03 | 2 | 54.920666 | yes | 9.722E-08 | 782.4043 | 294.9971 |
| IQAVIDSGVCR | 11 | O60684 | Importin subunit alpha-7 OS=Homo sapi | KPNA6 | 60.03 | 2 | 54.920666 | no | 1.893E-07 | 609.31897 | 200.9669 |
| EEEGIQLR | 8 | O60684 | Importin subunit alpha-7 OS=Homo sapi | KPNA6 | 60.03 | 2 | 54.920666 | no | 4.095E-07 | 487.2511 | 139.37643 |
| LVELLMHNDYK | 11 | O60684 | Importin subunit alpha-7 OS=Homo sapi | KPNA6 | 60.03 | 2 | 54.920666 | no | 9.727E-08 | 687.85791 | 95.54985 |
| EACWTISNITAGNI | 14 | O60684 | Importin subunit alpha-7 OS=Homo sapi | KPNA6 | 60.03 | 2 | 54.920666 | no | 1.017E-07 | 796.87793 | 94.575905 |
| NNALNPEEMR | 10 | O60684 | Importin subunit alpha-7 OS=Homo sapi | KPNA6 | 60.03 | 2 | 54.920666 | yes | 9.938E-08 | 594.27728 | 82.67717 |
| VTVVDVNESR | 10 | O60701 | UDP-glucose 6-dehydrogenase OS=Hom | UGDH | 55.025 | 2 | 44.675053 | yes | 2.228E-07 | 559.29602 | 100.88388 |
| ASVGFGGSCFQK | 12 | O60701 | UDP-glucose 6-dehydrogenase OS=Hom | UGDH | 55.025 | 2 | 44.675053 | yes | 5.621E-07 | 622.79004 | 92.834068 |
| EADLVFISVNTPTK | 14 | O60701 | UDP-glucose 6-dehydrogenase OS=Hom | UGDH | 55.025 | 2 | 44.675053 | yes | 7.69E-07 | 767.41156 | 91.673813 |
| IIDSLFNTVTDK | 12 | O60701 | UDP-glucose 6-dehydrogenase OS=Hom | UGDH | 55.025 | 2 | 44.675053 | yes | 3.399E-07 | 683.36664 | 90.468735 |
| ILTTNTWSSELSK | 13 | O60701 | UDP-glucose 6-dehydrogenase OS=Hom | UGDH | 55.025 | 2 | 44.675053 | yes | 2.109E-07 | 740.38812 | 55.527637 |
| NLFFSTNIDDAIK | 13 | O60701 | UDP-glucose 6-dehydrogenase OS=Hom | UGDH | 55.025 | 2 | 44.675053 | yes | 3.751E-07 | 749.38281 | 45.905441 |
| GYELLFQPEVVR | 12 | O60716 | Catenin delta-1 OS=Homo sapiens OX=9 | CTNND1 | 108.171 | 2 | 53.005108 | yes | 1.087E-07 | 725.39044 | 2005.936 |
| NCDGVPALVR | 10 | O60716 | Catenin delta-1 OS=Homo sapiens OX=9 | CTNND1 | 108.171 | 2 | 53.005108 | yes | 1.173E-07 | 550.77948 | 875.20435 |
| QDVYGPQPQVR | 11 | O60716 | Catenin delta-1 OS=Homo sapiens OX=9 | CTNND1 | 108.171 | 2 | 53.005108 | yes | 9.722E-08 | 643.828 | 685.50464 |
| LVENCVCLLR | 10 | O60716 | Catenin delta-1 OS=Homo sapiens OX=9 | CTNND1 | 108.171 | 2 | 53.005108 | yes | 1.622E-07 | 638.33105 | 515.06567 |
| NLSYQVHR | 8 | O60716 | Catenin delta-1 OS=Homo sapiens OX=9 | CTNND1 | 108.171 | 2 | 53.005108 | yes | 1.799E-07 | 508.76724 | 500.92053 |
| VGGSSVDLHR | 10 | O60716 | Catenin delta-1 OS=Homo sapiens OX=9 | CTNND1 | 108.171 | 2 | 53.005108 | yes | 1.765E-07 | 513.76996 | 492.84753 |
| GSLASLDSLR | 10 | O60716 | Catenin delta-1 OS=Homo sapiens OX=9 | CTNND1 | 108.171 | 2 | 53.005108 | yes | 1.073E-07 | 509.78 | 432.09747 |
| SGDLGDMEPLK | 11 | O60716 | Catenin delta-1 OS=Homo sapiens OX=9 | CTNND1 | 108.171 | 2 | 53.005108 | yes | 1.792E-07 | 581.27643 | 340.31805 |
| IYISLLK | 7 | O60716 | Catenin delta-1 OS=Homo sapiens OX=9 | CTNND1 | 108.171 | 2 | 53.005108 | yes | 1.503E-06 | 425.27585 | 312.98343 |
| EEIQMSNMGSNTK | 13 | O60716 | Catenin delta-1 OS=Homo sapiens OX=9 | CTNND1 | 108.171 | 2 | 53.005108 | yes | 9.722E-08 | 734.82397 | 286.56635 |
| EVHLGACGALK | 11 | O60716 | Catenin delta-1 OS=Homo sapiens OX=9 | CTNND1 | 108.171 | 2 | 53.005108 | yes | 1.082E-07 | 577.80298 | 267.49957 |
| FVGDADLER | 9 | O60716 | Catenin delta-1 OS=Homo sapiens OX=9 | CTNND1 | 108.171 | 2 | 53.005108 | yes | 3.788E-07 | 511.2511 | 235.4919 |
| FHPEPYGLEDDQR | 13 | O60716 | Catenin delta-1 OS=Homo sapiens OX=9 | CTNND1 | 108.171 | 2 | 53.005108 | yes | 1.437E-07 | 801.86279 | 209.15826 |
| SLDNNYSTPNER | 12 | O60716 | Catenin delta-1 OS=Homo sapiens OX=9 | CTNND1 | 108.171 | 2 | 53.005108 | yes | 9.722E-08 | 705.31824 | 192.46956 |
| ALSAIADLLTNEHI | 15 | O60716 | Catenin delta-1 OS=Homo sapiens OX=9 | CTNND1 | 108.171 | 2 | 53.005108 | yes | 1.615E-07 | 826.93378 | 176.33855 |
| KEVHLGACGALK | 12 | O60716 | Catenin delta-1 OS=Homo sapiens OX=9 | CTNND1 | 108.171 | 2 | 53.005108 | yes | 9.729E-08 | 641.85046 | 74.781052 |
| KEVHLGACGALK | 12 | O60716 | Catenin delta-1 OS=Homo sapiens OX=9 | CTNND1 | 108.171 | 2 | 53.005108 | yes | 1.088E-07 | 428.23605 | 74.781052 |
| KEVHLGACGALK | 12 | O60716 | Catenin delta-1 OS=Homo sapiens OX=9 | CTNND1 | 108.171 | 3 | 53.005108 | yes | 9.729E-08 | 641.85046 | 74.781052 |
| KEVHLGACGALK | 12 | O60716 | Catenin delta-1 OS=Homo sapiens OX=9 | CTNND1 | 108.171 | 3 | 53.005108 | yes | 1.088E-07 | 428.23605 | 74.781052 |
| AAALVLQTIWGYK | 13 | O60716 | Catenin delta-1 OS=Homo sapiens OX=9 | CTNND1 | 108.171 | 2 | 53.005108 | yes | 1.32E-07 | 717.41119 | 147.98128 |

| | | | | | | | | | | | |
|---|---|---|---|---|---|---|---|---|---|---|---|
| SNAAAYLQHLCYI | 13 | O60716 | Catenin delta-1 OS=Homo sapiens OX=9 | CTNND1 | 108.171 | 2 | 53.005108 | yes | 2.522E-07 | 783.87775 | 120.38483 |
| YRPSMEGYR | 9 | O60716 | Catenin delta-1 OS=Homo sapiens OX=9 | CTNND1 | 108.171 | 2 | 53.005108 | yes | 9.968E-08 | 579.77167 | 102.13176 |
| KPIEDPANDTVDFI | 15 | O60716 | Catenin delta-1 OS=Homo sapiens OX=9 | CTNND1 | 108.171 | 2 | 53.005108 | yes | 1.32E-07 | 843.42267 | 98.348846 |
| TPAILEASAGAIQN | 19 | O60716 | Catenin delta-1 OS=Homo sapiens OX=9 | CTNND1 | 108.171 | 3 | 53.005108 | yes | 8.64E-07 | 638.33514 | 48.458492 |
| ALSQLAEVEEK | 11 | O60749 | Sorting nexin-2 OS=Homo sapiens OX=9 | SNX2 | 58.472 | 2 | 44.22994 | yes | 2.272E-07 | 608.82465 | 54.88567 |
| EGNFDIVSGTR | 11 | O60762 | Dolichol-phosphate mannosyltransferase : | DPM1 | 29.635 | 2 | 50.303654 | yes | 9.819E-08 | 597.79114 | 1610.6907 |
| YSVLLPTYNER | 11 | O60762 | Dolichol-phosphate mannosyltransferase : | DPM1 | 29.635 | 2 | 50.303654 | yes | 2.04E-07 | 677.8537 | 1524.5076 |
| FIPEFIR | 7 | O60762 | Dolichol-phosphate mannosyltransferase : | DPM1 | 29.635 | 2 | 50.303654 | yes | 2.814E-07 | 461.26328 | 1444.5848 |
| LGGNEIVSFLK | 11 | O60762 | Dolichol-phosphate mannosyltransferase : | DPM1 | 29.635 | 2 | 50.303654 | yes | 1.138E-07 | 588.83478 | 1316.6595 |
| HATGNYIIIMDADI | 19 | O60762 | Dolichol-phosphate mannosyltransferase : | DPM1 | 29.635 | 3 | 50.303654 | yes | 9.893E-08 | 711.6897 | 980.35083 |
| GYVFQMEMIVR | 11 | O60762 | Dolichol-phosphate mannosyltransferase : | DPM1 | 29.635 | 2 | 50.303654 | yes | 1.457E-07 | 686.84125 | 207.41623 |
| GNGGVYGWDLK | 11 | O60762 | Dolichol-phosphate mannosyltransferase : | DPM1 | 29.635 | 2 | 50.303654 | yes | 1.108E-07 | 583.28546 | 111.9928 |
| KLGLGTAYIHGMK | 13 | O60762 | Dolichol-phosphate mannosyltransferase : | DPM1 | 29.635 | 2 | 50.303654 | yes | 1.355E-07 | 694.88959 | 21.316517 |
| MAYEYADER | 9 | O60783 | Small ribosomal subunit protein uS14m C | MRPS14 | 15.139 | 2 | 51.741074 | yes | 1.146E-07 | 574.23987 | 280.35004 |
| ILQDVADEEIAALI | 15 | O60783 | Small ribosomal subunit protein uS14m C | MRPS14 | 15.139 | 2 | 51.741074 | yes | 9.722E-08 | 826.94635 | 156.01195 |
| HLADHGQLSGIQR | 13 | O60783 | Small ribosomal subunit protein uS14m C | MRPS14 | 15.139 | 3 | 51.741074 | yes | 1.673E-07 | 477.91989 | 63.821041 |
| HLADHGQLSGIQR | 13 | O60783 | Small ribosomal subunit protein uS14m C | MRPS14 | 15.139 | 3 | 51.741074 | yes | 9.902E-08 | 716.37622 | 63.821041 |
| HLADHGQLSGIQR | 13 | O60783 | Small ribosomal subunit protein uS14m C | MRPS14 | 15.139 | 2 | 51.741074 | yes | 1.673E-07 | 477.91989 | 63.821041 |
| HLADHGQLSGIQR | 13 | O60783 | Small ribosomal subunit protein uS14m C | MRPS14 | 15.139 | 2 | 51.741074 | yes | 9.902E-08 | 716.37622 | 63.821041 |
| ESYSVYVYK | 9 | O60814 | Histone H2B type 1-K OS=Homo sapiens | H2BC12 | 13.891 | 2 | 48.328526 | yes | 1.164E-07 | 569.27679 | 3535.1079 |
| KESYSVYVYK | 10 | O60814 | Histone H2B type 1-K OS=Homo sapiens | H2BC12 | 13.891 | 2 | 48.328526 | yes | 1.46E-07 | 633.32428 | 2778.7649 |
| LEPVIMELER | 10 | O60825 | 6-phosphofructo-2-kinase/fructose-2,6-bi: | PFKFB2 | 58.477 | 2 | 56.909042 | no | 1.108E-07 | 614.83392 | 154.82655 |
| LNVEAVNTHR | 10 | O60825 | 6-phosphofructo-2-kinase/fructose-2,6-bi: | PFKFB2 | 58.477 | 2 | 56.909042 | yes | 9.722E-08 | 576.80963 | 150.07487 |
| NSFTPLSSSNTIR | 13 | O60825 | 6-phosphofructo-2-kinase/fructose-2,6-bi: | PFKFB2 | 58.477 | 2 | 56.909042 | yes | 2.493E-07 | 712.36243 | 97.581787 |
| YLNWIGVPTK | 10 | O60825 | 6-phosphofructo-2-kinase/fructose-2,6-bi: | PFKFB2 | 58.477 | 2 | 56.909042 | no | 1.021E-07 | 595.83203 | 93.23481 |
| PLKPLSPLR | 9 | O60825 | 6-phosphofructo-2-kinase/fructose-2,6-bi: | PFKFB2 | 58.477 | 2 | 56.909042 | yes | 9.935E-08 | 510.83185 | 87.827148 |
| FLEEQEITDLK | 11 | O60825 | 6-phosphofructo-2-kinase/fructose-2,6-bi: | PFKFB2 | 58.477 | 2 | 56.909042 | yes | 1.006E-06 | 682.85083 | 82.989479 |
| YPEEFALR | 8 | O60825 | 6-phosphofructo-2-kinase/fructose-2,6-bi: | PFKFB2 | 58.477 | 2 | 56.909042 | yes | 1.428E-07 | 512.75854 | 73.022484 |
| LTPVAYGCK | 9 | O60825 | 6-phosphofructo-2-kinase/fructose-2,6-bi: | PFKFB2 | 58.477 | 2 | 56.909042 | no | 1.05E-07 | 504.76279 | 67.545647 |
| IVYYLMNIHVQPR | 13 | O60825 | 6-phosphofructo-2-kinase/fructose-2,6-bi: | PFKFB2 | 58.477 | 2 | 56.909042 | no | 4.054E-07 | 823.44781 | 58.337631 |
| YPGGESYQDLVQI | 13 | O60825 | 6-phosphofructo-2-kinase/fructose-2,6-bi: | PFKFB2 | 58.477 | 2 | 56.909042 | yes | 1.791E-07 | 756.35986 | 57.396355 |
| QGNVLVISHQAVN | 14 | O60825 | 6-phosphofructo-2-kinase/fructose-2,6-bi: | PFKFB2 | 58.477 | 2 | 56.909042 | yes | 9.722E-08 | 776.42487 | 48.586674 |
| HGESEFNLLGK | 11 | O60825 | 6-phosphofructo-2-kinase/fructose-2,6-bi: | PFKFB2 | 58.477 | 2 | 56.909042 | yes | 3.654E-07 | 615.80933 | 45.170254 |
| ENVMEDFLK | 9 | O60825 | 6-phosphofructo-2-kinase/fructose-2,6-bi: | PFKFB2 | 58.477 | 2 | 56.909042 | yes | 8.042E-07 | 562.76825 | 44.952793 |
| DMILNFAEQNSFK | 13 | O60825 | 6-phosphofructo-2-kinase/fructose-2,6-bi: | PFKFB2 | 58.477 | 2 | 56.909042 | yes | 1.361E-05 | 778.87433 | 9.9701328 |
| PLPVALQTR | 9 | O60828 | Polyglutamine-binding protein 1 OS=Hor | PQBP1 | 30.473 | 2 | 50.646496 | yes | 2.22E-07 | 497.80585 | 429.96475 |
| PYPSPGAVLR | 10 | O60828 | Polyglutamine-binding protein 1 OS=Hor | PQBP1 | 30.473 | 2 | 50.646496 | yes | 1.211E-07 | 528.79547 | 185.61427 |
| TGADTTAAGPLFQ | 15 | O60828 | Polyglutamine-binding protein 1 OS=Hor | PQBP1 | 30.473 | 2 | 50.646496 | yes | 9.732E-08 | 767.38641 | 185.30841 |
| GKEDPWNSITSGA | 22 | O60830 | Mitochondrial import inner membrane tra | TIMM17B | 18.274 | 3 | 49.093296 | yes | 2.193E-07 | 738.72198 | 115.63132 |

| | | | | | | | | | | | |
|---|---|---|---|---|---|---|---|---|---|---|---|
| YTAQQFR | 7 | O60830 | Mitochondrial import inner membrane tra | TIMM17B | 18.274 | 2 | 49.093296 | yes | 2.961E-07 | 457.22995 | 108.48863 |
| DGTPAPGYPSYQC | 15 | O60830 | Mitochondrial import inner membrane tra | TIMM17B | 18.274 | 2 | 49.093296 | yes | 1.069E-07 | 840.8681 | 91.909874 |
| VTGCLIVCIER | 11 | O60832 | H/ACA ribonucleoprotein complex subun | DKC1 | 57.675 | 2 | 53.73567 | yes | 4.006E-07 | 660.34412 | 1337.9584 |
| RVVYPLEK | 8 | O60832 | H/ACA ribonucleoprotein complex subun | DKC1 | 57.675 | 2 | 53.73567 | yes | 5.178E-07 | 502.30038 | 1061.6787 |
| LDTSQWPLLLK | 11 | O60832 | H/ACA ribonucleoprotein complex subun | DKC1 | 57.675 | 2 | 53.73567 | yes | 1.073E-07 | 657.37683 | 930.90881 |
| DSAVNAICYGAK | 12 | O60832 | H/ACA ribonucleoprotein complex subun | DKC1 | 57.675 | 2 | 53.73567 | yes | 6.566E-07 | 634.8006 | 866.34137 |
| APQVVAEAAK | 10 | O60832 | H/ACA ribonucleoprotein complex subun | DKC1 | 57.675 | 2 | 53.73567 | yes | 1.001E-07 | 492.27966 | 765.84613 |
| TTHYTPLACGSNP | 15 | O60832 | H/ACA ribonucleoprotein complex subun | DKC1 | 57.675 | 2 | 53.73567 | yes | 2.959E-07 | 830.41162 | 708.33087 |
| LHNAIEGGTQLSR | 13 | O60832 | H/ACA ribonucleoprotein complex subun | DKC1 | 57.675 | 2 | 53.73567 | yes | 9.722E-08 | 698.37061 | 617.76007 |
| MIEYDPER | 8 | O60832 | H/ACA ribonucleoprotein complex subun | DKC1 | 57.675 | 2 | 53.73567 | yes | 1.442E-07 | 526.7395 | 406.77716 |
| QEYVDYSESAK | 11 | O60832 | H/ACA ribonucleoprotein complex subun | DKC1 | 57.675 | 2 | 53.73567 | yes | 9.81E-08 | 659.79352 | 289.92978 |
| DHMVTMHDVLDA | 20 | O60832 | H/ACA ribonucleoprotein complex subun | DKC1 | 57.675 | 3 | 53.73567 | yes | 1.055E-07 | 823.37555 | 243.66133 |
| EIGDYIR | 7 | O60832 | H/ACA ribonucleoprotein complex subun | DKC1 | 57.675 | 2 | 53.73567 | yes | 1.02E-07 | 433.22433 | 205.4043 |
| KEVVAEVVK | 9 | O60832 | H/ACA ribonucleoprotein complex subun | DKC1 | 57.675 | 2 | 53.73567 | yes | 1.994E-07 | 500.80551 | 204.67021 |
| EVVAEVVK | 8 | O60832 | H/ACA ribonucleoprotein complex subun | DKC1 | 57.675 | 2 | 53.73567 | yes | 9.722E-08 | 436.75803 | 188.11386 |
| VVYPLEK | 7 | O60832 | H/ACA ribonucleoprotein complex subun | DKC1 | 57.675 | 2 | 53.73567 | yes | 3.443E-07 | 424.24982 | 158.8871 |
| ALETLTGALFQRPI | 20 | O60832 | H/ACA ribonucleoprotein complex subun | DKC1 | 57.675 | 3 | 53.73567 | yes | 1.084E-07 | 703.7525 | 130.38257 |
| TTHYTPLACGSNP | 16 | O60832 | H/ACA ribonucleoprotein complex subun | DKC1 | 57.675 | 3 | 53.73567 | yes | 9.809E-08 | 605.97717 | 117.03008 |
| TGFINLDKPSNPSS | 22 | O60832 | H/ACA ribonucleoprotein complex subun | DKC1 | 57.675 | 4 | 53.73567 | yes | 7.492E-07 | 617.57367 | 94.962166 |
| HGKPTDSTPATWK | 13 | O60832 | H/ACA ribonucleoprotein complex subun | DKC1 | 57.675 | 2 | 53.73567 | yes | 3.333E-07 | 713.35968 | 85.295319 |
| MIEYDPERR | 9 | O60832 | H/ACA ribonucleoprotein complex subun | DKC1 | 57.675 | 2 | 53.73567 | yes | 9.722E-08 | 604.79004 | 84.28154 |
| IMLPGVLR | 8 | O60832 | H/ACA ribonucleoprotein complex subun | DKC1 | 57.675 | 2 | 53.73567 | yes | 2.623E-07 | 449.78076 | 66.975632 |
| ADAEVIILPK | 10 | O60832 | H/ACA ribonucleoprotein complex subun | DKC1 | 57.675 | 2 | 53.73567 | yes | 5.252E-06 | 555.82391 | 38.731491 |
| PPLIAAVK | 8 | O60832 | H/ACA ribonucleoprotein complex subun | DKC1 | 57.675 | 2 | 53.73567 | yes | 2.733E-07 | 404.76819 | 23.941738 |
| LLQAQGVEVPSK | 12 | O60841 | Eukaryotic translation initiation factor 5B | EIF5B | 138.827 | 2 | 52.75793 | yes | 1.151E-07 | 634.86407 | 2261.9546 |
| ILPQYIFNSR | 10 | O60841 | Eukaryotic translation initiation factor 5B | EIF5B | 138.827 | 2 | 52.75793 | yes | 1.451E-07 | 625.84821 | 1894.2246 |
| TSEVPYAGINIGPV | 16 | O60841 | Eukaryotic translation initiation factor 5B | EIF5B | 138.827 | 2 | 52.75793 | yes | 2.016E-07 | 841.44904 | 821.31097 |
| TSEVPYAGINIGPV | 16 | O60841 | Eukaryotic translation initiation factor 5B | EIF5B | 138.827 | 2 | 52.75793 | yes | 9.722E-08 | 561.30176 | 821.31097 |
| TSEVPYAGINIGPV | 16 | O60841 | Eukaryotic translation initiation factor 5B | EIF5B | 138.827 | 3 | 52.75793 | yes | 2.016E-07 | 841.44904 | 821.31097 |
| TSEVPYAGINIGPV | 16 | O60841 | Eukaryotic translation initiation factor 5B | EIF5B | 138.827 | 3 | 52.75793 | yes | 9.722E-08 | 561.30176 | 821.31097 |
| APIICVLGHVDTGI | 14 | O60841 | Eukaryotic translation initiation factor 5B | EIF5B | 138.827 | 2 | 52.75793 | yes | 5.904E-07 | 740.40302 | 1416.8973 |
| LAHCEELR | 8 | O60841 | Eukaryotic translation initiation factor 5B | EIF5B | 138.827 | 2 | 52.75793 | yes | 1.29E-07 | 514.25311 | 1334.4185 |
| DAQEMADSLGVR | 12 | O60841 | Eukaryotic translation initiation factor 5B | EIF5B | 138.827 | 2 | 52.75793 | yes | 5.464E-07 | 646.30096 | 1226.9042 |
| IEPIPGESPK | 10 | O60841 | Eukaryotic translation initiation factor 5B | EIF5B | 138.827 | 2 | 52.75793 | yes | 1.018E-07 | 533.7926 | 1190.5214 |
| HIAVFPCK | 8 | O60841 | Eukaryotic translation initiation factor 5B | EIF5B | 138.827 | 2 | 52.75793 | yes | 3.417E-07 | 486.26019 | 1186.5103 |
| VTVDTGVIPASEEI | 14 | O60841 | Eukaryotic translation initiation factor 5B | EIF5B | 138.827 | 2 | 52.75793 | yes | 1.229E-07 | 722.88013 | 1102.6772 |
| HFEATDILVSK | 11 | O60841 | Eukaryotic translation initiation factor 5B | EIF5B | 138.827 | 2 | 52.75793 | yes | 2.394E-07 | 630.33514 | 965.63049 |
| ELEELSLEAQGIK | 13 | O60841 | Eukaryotic translation initiation factor 5B | EIF5B | 138.827 | 2 | 52.75793 | yes | 1.287E-07 | 729.88794 | 897.40747 |
| SEDSTKDDIDLDAI | 25 | O60841 | Eukaryotic translation initiation factor 5B | EIF5B | 138.827 | 3 | 52.75793 | yes | 9.722E-08 | 835.73279 | 611.46564 |

| | | | | | | | | | | | |
|---|---|---|---|---|---|---|---|---|---|---|---|
| GLLLPPPMK | 9 | O60841 | Eukaryotic translation initiation factor 5B | EIF5B | 138.827 | 2 | 52.75793 | yes | 3.842E-07 | 483.29626 | 534.41711 |
| DELIHELK | 8 | O60841 | Eukaryotic translation initiation factor 5B | EIF5B | 138.827 | 2 | 52.75793 | yes | 1.304E-07 | 498.77167 | 527.60107 |
| KIPQQLESK | 9 | O60841 | Eukaryotic translation initiation factor 5B | EIF5B | 138.827 | 2 | 52.75793 | yes | 4.762E-07 | 535.81384 | 438.88046 |
| IPGMLIIDTPGHESI | 19 | O60841 | Eukaryotic translation initiation factor 5B | EIF5B | 138.827 | 3 | 52.75793 | yes | 9.722E-08 | 699.69812 | 404.82022 |
| KSPDSDVAATLK | 12 | O60841 | Eukaryotic translation initiation factor 5B | EIF5B | 138.827 | 2 | 52.75793 | yes | 9.733E-08 | 616.33008 | 398.81567 |
| HTHVQDGEAGGIT | 31 | O60841 | Eukaryotic translation initiation factor 5B | EIF5B | 138.827 | 4 | 52.75793 | yes | 1.11E-07 | 814.91028 | 357.52753 |
| KFEEETVK | 8 | O60841 | Eukaryotic translation initiation factor 5B | EIF5B | 138.827 | 2 | 52.75793 | yes | 1.841E-07 | 505.26367 | 326.34341 |
| QGTPMCVPSK | 10 | O60841 | Eukaryotic translation initiation factor 5B | EIF5B | 138.827 | 2 | 52.75793 | yes | 2.88E-07 | 552.76245 | 321.52402 |
| CPFIVALNK | 9 | O60841 | Eukaryotic translation initiation factor 5B | EIF5B | 138.827 | 2 | 52.75793 | yes | 9.722E-08 | 531.29425 | 287.51627 |
| EVEAAQGVK | 9 | O60841 | Eukaryotic translation initiation factor 5B | EIF5B | 138.827 | 2 | 52.75793 | yes | 3.329E-06 | 465.74817 | 245.12213 |
| KQDFDEDDILK | 11 | O60841 | Eukaryotic translation initiation factor 5B | EIF5B | 138.827 | 2 | 52.75793 | yes | 9.722E-08 | 683.33026 | 244.49658 |
| TLAGLPLLVAYK | 12 | O60841 | Eukaryotic translation initiation factor 5B | EIF5B | 138.827 | 2 | 52.75793 | yes | 6.875E-07 | 629.89209 | 232.2662 |
| DPIVMGVTVEAGC | 15 | O60841 | Eukaryotic translation initiation factor 5B | EIF5B | 138.827 | 2 | 52.75793 | yes | 2.54E-07 | 771.91345 | 203.70276 |
| NTKDEFEER | 9 | O60841 | Eukaryotic translation initiation factor 5B | EIF5B | 138.827 | 2 | 52.75793 | yes | 1.067E-07 | 584.26746 | 194.86847 |
| ASVMLEHDPQYA | 20 | O60841 | Eukaryotic translation initiation factor 5B | EIF5B | 138.827 | 3 | 52.75793 | yes | 3.661E-07 | 758.72443 | 180.90733 |
| AMQEALAK | 8 | O60841 | Eukaryotic translation initiation factor 5B | EIF5B | 138.827 | 2 | 52.75793 | yes | 2.916E-07 | 431.22858 | 177.78113 |
| QDFDEDDILK | 10 | O60841 | Eukaryotic translation initiation factor 5B | EIF5B | 138.827 | 2 | 52.75793 | yes | 9.722E-08 | 619.28278 | 103.15186 |
| SDWQLIVELK | 10 | O60841 | Eukaryotic translation initiation factor 5B | EIF5B | 138.827 | 2 | 52.75793 | yes | 9.722E-08 | 615.84009 | 99.734428 |
| AETPTAAEDDNEC | 16 | O60841 | Eukaryotic translation initiation factor 5B | EIF5B | 138.827 | 2 | 52.75793 | yes | 1.756E-07 | 845.87378 | 98.998375 |
| SPDSDVAATLK | 11 | O60841 | Eukaryotic translation initiation factor 5B | EIF5B | 138.827 | 2 | 52.75793 | yes | 1.176E-07 | 552.28259 | 81.57106 |
| EDEIPVLKDELIHE | 16 | O60841 | Eukaryotic translation initiation factor 5B | EIF5B | 138.827 | 3 | 52.75793 | yes | 1.261E-07 | 640.68231 | 20.692703 |
| LRAPIICVLGHVDT | 16 | O60841 | Eukaryotic translation initiation factor 5B | EIF5B | 138.827 | 3 | 52.75793 | yes | 2.862E-06 | 583.6662 | 18.763229 |
| TDYWLQPEIIVK | 12 | O60870 | DNA/RNA-binding protein KIN17 OS=H | KIN | 45.374 | 2 | 54.76121 | yes | 9.722E-08 | 752.90594 | 809.78357 |
| EQEVPTFTELSR | 12 | O60870 | DNA/RNA-binding protein KIN17 OS=H | KIN | 45.374 | 2 | 54.76121 | yes | 1.328E-07 | 718.35681 | 419.26736 |
| FIEEQVR | 7 | O60870 | DNA/RNA-binding protein KIN17 OS=H | KIN | 45.374 | 2 | 54.76121 | yes | 1.371E-07 | 460.74545 | 418.98209 |
| GNEGTLESINEK | 12 | O60870 | DNA/RNA-binding protein KIN17 OS=H | KIN | 45.374 | 2 | 54.76121 | yes | 9.722E-08 | 645.81226 | 377.76257 |
| TFSATIVIETGPLK | 14 | O60870 | DNA/RNA-binding protein KIN17 OS=H | KIN | 45.374 | 2 | 54.76121 | yes | 9.722E-08 | 738.91907 | 369.85468 |
| SALDEIMEIEEEK | 13 | O60870 | DNA/RNA-binding protein KIN17 OS=H | KIN | 45.374 | 2 | 54.76121 | yes | 1.232E-07 | 768.3609 | 288.24893 |
| LDQTHLETVIPAPC | 15 | O60870 | DNA/RNA-binding protein KIN17 OS=H | KIN | 45.374 | 2 | 54.76121 | yes | 1.05E-07 | 809.9436 | 121.38013 |
| LDQTHLETVIPAPC | 15 | O60870 | DNA/RNA-binding protein KIN17 OS=H | KIN | 45.374 | 2 | 54.76121 | yes | 9.722E-08 | 540.29816 | 121.38013 |
| LDQTHLETVIPAPC | 15 | O60870 | DNA/RNA-binding protein KIN17 OS=H | KIN | 45.374 | 3 | 54.76121 | yes | 1.05E-07 | 809.9436 | 121.38013 |
| LDQTHLETVIPAPC | 15 | O60870 | DNA/RNA-binding protein KIN17 OS=H | KIN | 45.374 | 3 | 54.76121 | yes | 9.722E-08 | 540.29816 | 121.38013 |
| RVEGIQYEDISK | 12 | O60870 | DNA/RNA-binding protein KIN17 OS=H | KIN | 45.374 | 2 | 54.76121 | yes | 4.243E-07 | 718.87262 | 130.7356 |
| WYCQMCQK | 8 | O60870 | DNA/RNA-binding protein KIN17 OS=H | KIN | 45.374 | 2 | 54.76121 | yes | 3.943E-07 | 602.24084 | 114.00005 |
| VEGIQYEDISK | 11 | O60870 | DNA/RNA-binding protein KIN17 OS=H | KIN | 45.374 | 2 | 54.76121 | yes | 1.966E-07 | 640.82208 | 95.951027 |
| NDFLELLR | 8 | O60870 | DNA/RNA-binding protein KIN17 OS=H | KIN | 45.374 | 2 | 54.76121 | yes | 9.722E-08 | 510.27966 | 59.178722 |
| ILVLNGGYR | 9 | O60870 | DNA/RNA-binding protein KIN17 OS=H | KIN | 45.374 | 2 | 54.76121 | yes | 1.288E-07 | 502.798 | 56.624146 |
| SSTLGPSALK | 10 | O60870 | DNA/RNA-binding protein KIN17 OS=H | KIN | 45.374 | 2 | 54.76121 | yes | 5.533E-07 | 480.77167 | 34.091156 |
| VIEPGCVR | 8 | O60884 | DnaJ homolog subfamily A member 2 OS | DNAJA2 | 45.746 | 2 | 54.5787 | yes | 3.12E-06 | 465.24728 | 1310.5219 |

| | | | | | | | | | | | |
|---|---|---|---|---|---|---|---|---|---|---|---|
| ILEVHVDK | 8 | O60884 | DnaJ homolog subfamily A member 2 OS | DNAJA2 | 45.746 | 2 | 54.5787 | yes | 2.479E-07 | 476.77673 | 877.95758 |
| NVLCSACSGQGGI | 13 | O60884 | DnaJ homolog subfamily A member 2 OS | DNAJA2 | 45.746 | 2 | 54.5787 | yes | 9.722E-08 | 669.30048 | 491.32379 |
| EISFAYEVLSNPEK | 14 | O60884 | DnaJ homolog subfamily A member 2 OS | DNAJA2 | 45.746 | 2 | 54.5787 | yes | 9.919E-08 | 813.40649 | 412.13873 |
| GEDMMHPLK | 9 | O60884 | DnaJ homolog subfamily A member 2 OS | DNAJA2 | 45.746 | 2 | 54.5787 | yes | 1.721E-07 | 529.2439 | 328.76895 |
| GEGMPQYR | 8 | O60884 | DnaJ homolog subfamily A member 2 OS | DNAJA2 | 45.746 | 2 | 54.5787 | yes | 9.722E-08 | 469.21344 | 317.37064 |
| DGNDLHMTYK | 10 | O60884 | DnaJ homolog subfamily A member 2 OS | DNAJA2 | 45.746 | 2 | 54.5787 | yes | 9.997E-08 | 597.26642 | 277.02625 |
| VSLEDLYNGK | 10 | O60884 | DnaJ homolog subfamily A member 2 OS | DNAJA2 | 45.746 | 2 | 54.5787 | yes | 9.722E-08 | 569.29297 | 207.8049 |
| TQSSLVPALTDFVI | 14 | O60888 | Protein CutA OS=Homo sapiens OX=960 | CUTA | 19.117 | 2 | 46.0755 | yes | 1.66E-07 | 767.41718 | 405.42188 |
| LGAIFLEGVTVK | 12 | O60942 | mRNA-capping enzyme OS=Homo sapie | RNGTT | 68.557 | 2 | 52.29881 | yes | 9.722E-08 | 623.8739 | 90.10643 |
| MGLLVDLTNTSR | 12 | O60942 | mRNA-capping enzyme OS=Homo sapie | RNGTT | 68.557 | 2 | 52.29881 | yes | 2.71E-06 | 660.35303 | 61.86631 |
| LLDLKPYK | 8 | O60942 | mRNA-capping enzyme OS=Homo sapie | RNGTT | 68.557 | 2 | 52.29881 | yes | 1.214E-07 | 495.30515 | 47.967373 |
| FNSQPVGDCDFNV | 14 | O60942 | mRNA-capping enzyme OS=Homo sapie | RNGTT | 68.557 | 2 | 52.29881 | yes | 2.073E-06 | 827.86755 | 28.052914 |
| EVSHEMDGLIFQP | 16 | O60942 | mRNA-capping enzyme OS=Homo sapie | RNGTT | 68.557 | 3 | 52.29881 | yes | 9.722E-08 | 596.62592 | 19.04006 |
| YHIQQAQR | 8 | O75030 | Microphthalmia-associated transcription f | MITF | 58.796 | 2 | 43.953133 | yes | 4.201E-06 | 522.27271 | 154.46672 |
| QYLSTTLANK | 10 | O75030 | Microphthalmia-associated transcription f | MITF | 58.796 | 2 | 43.953133 | yes | 4.334E-07 | 569.80878 | 105.52332 |
| LQAAQFMQQR | 10 | O75030 | Microphthalmia-associated transcription f | MITF | 58.796 | 2 | 43.953133 | yes | 2.381E-07 | 610.81384 | 78.870613 |
| VQTHLENPTK | 10 | O75030 | Microphthalmia-associated transcription f | MITF | 58.796 | 2 | 43.953133 | yes | 3.632E-07 | 583.81183 | 73.648766 |
| VSGSQVEVNDIK | 12 | O75044 | SLIT-ROBO Rho GTPase-activating prot | SRGAP2 | 120.871 | 2 | 55.82155 | no | 9.722E-08 | 637.83295 | 388.45404 |
| VLNELYSVMK | 10 | O75044 | SLIT-ROBO Rho GTPase-activating prot | SRGAP2 | 120.871 | 2 | 55.82155 | no | 2.249E-07 | 598.32318 | 252.51653 |
| EVGQQLQDDLMK | 12 | O75044 | SLIT-ROBO Rho GTPase-activating prot | SRGAP2 | 120.871 | 2 | 55.82155 | no | 9.722E-08 | 702.3454 | 204.57899 |
| TFLSAELNLEQSK | 13 | O75044 | SLIT-ROBO Rho GTPase-activating prot | SRGAP2 | 120.871 | 2 | 55.82155 | no | 9.91E-08 | 740.38812 | 182.53334 |
| GLEHPLFPK | 9 | O75044 | SLIT-ROBO Rho GTPase-activating prot | SRGAP2 | 120.871 | 2 | 55.82155 | yes | 2.388E-07 | 519.29254 | 171.59186 |
| ELEGPVYSR | 9 | O75044 | SLIT-ROBO Rho GTPase-activating prot | SRGAP2 | 120.871 | 2 | 55.82155 | yes | 5.308E-07 | 525.26672 | 164.30247 |
| FVQVSEDSGR | 10 | O75044 | SLIT-ROBO Rho GTPase-activating prot | SRGAP2 | 120.871 | 2 | 55.82155 | no | 2.303E-07 | 562.27252 | 159.69266 |
| SEIEVISEPPEEK | 13 | O75044 | SLIT-ROBO Rho GTPase-activating prot | SRGAP2 | 120.871 | 2 | 55.82155 | yes | 9.722E-08 | 743.36957 | 134.4725 |
| CLDQQCELR | 9 | O75044 | SLIT-ROBO Rho GTPase-activating prot | SRGAP2 | 120.871 | 2 | 55.82155 | no | 1.123E-06 | 611.27118 | 94.165352 |
| PSSQPIMSQSLPK | 13 | O75044 | SLIT-ROBO Rho GTPase-activating prot | SRGAP2 | 120.871 | 2 | 55.82155 | yes | 3.029E-07 | 700.36615 | 90.154839 |
| DKEIIAEYDTQVK | 13 | O75044 | SLIT-ROBO Rho GTPase-activating prot | SRGAP2 | 120.871 | 2 | 55.82155 | no | 9.722E-08 | 776.39868 | 84.838249 |
| SPDSTANVR | 9 | O75044 | SLIT-ROBO Rho GTPase-activating prot | SRGAP2 | 120.871 | 2 | 55.82155 | no | 4.398E-07 | 473.73306 | 72.606903 |
| GASLLLYQR | 9 | O75044 | SLIT-ROBO Rho GTPase-activating prot | SRGAP2 | 120.871 | 2 | 55.82155 | yes | 2.132E-07 | 510.79547 | 67.113014 |
| ANQQETEQFYFTK | 13 | O75044 | SLIT-ROBO Rho GTPase-activating prot | SRGAP2 | 120.871 | 2 | 55.82155 | yes | 1.4E-07 | 817.3783 | 56.295349 |
| KAEIEMDYSR | 10 | O75044 | SLIT-ROBO Rho GTPase-activating prot | SRGAP2 | 120.871 | 2 | 55.82155 | no | 3.345E-07 | 621.29517 | 53.363102 |
| KQDSSQAIPLVVE | 17 | O75044 | SLIT-ROBO Rho GTPase-activating prot | SRGAP2 | 120.871 | 3 | 55.82155 | yes | 1.762E-07 | 644.00696 | 48.532951 |
| KVLLVLPK | 8 | O75044 | SLIT-ROBO Rho GTPase-activating prot | SRGAP2 | 120.871 | 2 | 55.82155 | yes | 9.722E-08 | 455.3284 | 38.866848 |
| TIIIQHENIFPSPR | 14 | O75044 | SLIT-ROBO Rho GTPase-activating prot | SRGAP2 | 120.871 | 3 | 55.82155 | yes | 1.642E-06 | 555.64215 | 27.82951 |
| KGASLLLYQR | 10 | O75044 | SLIT-ROBO Rho GTPase-activating prot | SRGAP2 | 120.871 | 2 | 55.82155 | yes | 7.567E-05 | 574.84296 | 23.950382 |
| ALLILEGGQK | 10 | O75063 | Glycosaminoglycan xylosylkinase OS=H | FAM20B | 46.433 | 2 | 53.104984 | yes | 3.634E-07 | 521.31879 | 1013.753 |
| SILAPLYQCCIIR | 13 | O75063 | Glycosaminoglycan xylosylkinase OS=H | FAM20B | 46.433 | 2 | 53.104984 | yes | 9.81E-08 | 803.92584 | 768.23456 |
| SFGNPSLDER | 10 | O75063 | Glycosaminoglycan xylosylkinase OS=H | FAM20B | 46.433 | 2 | 53.104984 | yes | 5.239E-07 | 561.26471 | 404.57813 |

| | | | | | | | | | | | |
|---|---|---|---|---|---|---|---|---|---|---|---|
| WEYDESYCDAVK | 12 | O75063 | Glycosaminoglycan xylosylkinase OS=H | FAM20B | 46.433 | 2 | 53.104984 | yes | 9.722E-08 | 782.81665 | 270.28699 |
| HNAEVAAFHLDR | 12 | O75063 | Glycosaminoglycan xylosylkinase OS=H | FAM20B | 46.433 | 3 | 53.104984 | yes | 9.722E-08 | 460.56534 | 105.04124 |
| HNAEVAAFHLDR | 12 | O75063 | Glycosaminoglycan xylosylkinase OS=H | FAM20B | 46.433 | 3 | 53.104984 | yes | 3.415E-06 | 690.34436 | 105.04124 |
| HNAEVAAFHLDR | 12 | O75063 | Glycosaminoglycan xylosylkinase OS=H | FAM20B | 46.433 | 2 | 53.104984 | yes | 9.722E-08 | 460.56534 | 105.04124 |
| HNAEVAAFHLDR | 12 | O75063 | Glycosaminoglycan xylosylkinase OS=H | FAM20B | 46.433 | 2 | 53.104984 | yes | 3.415E-06 | 690.34436 | 105.04124 |
| LLSVLATVK | 9 | O75063 | Glycosaminoglycan xylosylkinase OS=H | FAM20B | 46.433 | 2 | 53.104984 | yes | 3.147E-07 | 472.31296 | 208.78502 |
| TSPYDSGPR | 9 | O75063 | Glycosaminoglycan xylosylkinase OS=H | FAM20B | 46.433 | 2 | 53.104984 | yes | 1.083E-06 | 490.2276 | 175.4388 |
| HHYESFQDDEGAS | 22 | O75063 | Glycosaminoglycan xylosylkinase OS=H | FAM20B | 46.433 | 3 | 53.104984 | yes | 9.722E-08 | 845.0603 | 153.65054 |
| KTSPYDSGPR | 10 | O75063 | Glycosaminoglycan xylosylkinase OS=H | FAM20B | 46.433 | 2 | 53.104984 | yes | 1.617E-07 | 554.27509 | 82.944305 |
| EVYPEETPELGAV | 20 | O75063 | Glycosaminoglycan xylosylkinase OS=H | FAM20B | 46.433 | 3 | 53.104984 | yes | 1.777E-07 | 735.01849 | 77.497498 |
| SAMAHDPISPVLSI | 23 | O75063 | Glycosaminoglycan xylosylkinase OS=H | FAM20B | 46.433 | 4 | 53.104984 | yes | 1.443E-07 | 618.55505 | 59.650169 |
| DHVVEGEPYAGYI | 14 | O75063 | Glycosaminoglycan xylosylkinase OS=H | FAM20B | 46.433 | 2 | 53.104984 | yes | 9.722E-08 | 803.86023 | 38.738686 |
| VFASLPQVER | 10 | O75083 | WD repeat-containing protein 1 OS=Hon | WDR1 | 66.194 | 2 | 46.81873 | yes | 5.879E-07 | 573.31934 | 148.36969 |
| NIDNPALADIYTEH | 22 | O75083 | WD repeat-containing protein 1 OS=Hon | WDR1 | 66.194 | 3 | 46.81873 | yes | 1.464E-07 | 806.75189 | 55.220955 |
| YAPSGFYIASGDV | 16 | O75083 | WD repeat-containing protein 1 OS=Hon | WDR1 | 66.194 | 2 | 46.81873 | yes | 1.197E-06 | 809.89099 | 46.737221 |
| AAALAHLDR | 9 | O75106 | Retina-specific copper amine oxidase OS | AOC2 | 83.674 | 2 | 45.381977 | yes | 1.866E-07 | 469.26434 | 200.16373 |
| ENLLLSDSPSCR | 12 | O75116 | Rho-associated protein kinase 2 OS=Hon | ROCK2 | 160.9 | 2 | 50.11236 | yes | 9.899E-08 | 695.83521 | 162.57025 |
| NSLCFPEDAEISK | 13 | O75116 | Rho-associated protein kinase 2 OS=Hon | ROCK2 | 160.9 | 2 | 50.11236 | yes | 2.203E-07 | 755.34814 | 110.14851 |
| DVKPDNMLLDK | 11 | O75116 | Rho-associated protein kinase 2 OS=Hon | ROCK2 | 160.9 | 2 | 50.11236 | no | 3.703E-07 | 644.33429 | 103.39326 |
| IFQILYANEGESK | 13 | O75116 | Rho-associated protein kinase 2 OS=Hon | ROCK2 | 160.9 | 2 | 50.11236 | yes | 1.043E-07 | 756.39069 | 99.952522 |
| EFINLQSALESER | 13 | O75116 | Rho-associated protein kinase 2 OS=Hon | ROCK2 | 160.9 | 2 | 50.11236 | yes | 2.072E-07 | 768.38867 | 77.202934 |
| LKDEEISAAAIK | 12 | O75116 | Rho-associated protein kinase 2 OS=Hon | ROCK2 | 160.9 | 2 | 50.11236 | yes | 9.926E-08 | 644.36139 | 69.272522 |
| SIAEEQYSDLEK | 12 | O75116 | Rho-associated protein kinase 2 OS=Hon | ROCK2 | 160.9 | 2 | 50.11236 | yes | 1.365E-07 | 706.33301 | 64.413963 |
| LEGWLSLPVR | 10 | O75116 | Rho-associated protein kinase 2 OS=Hon | ROCK2 | 160.9 | 2 | 50.11236 | yes | 1.722E-07 | 585.33752 | 39.614971 |
| NLICAFLTDR | 10 | O75116 | Rho-associated protein kinase 2 OS=Hon | ROCK2 | 160.9 | 2 | 50.11236 | no | 4.059E-07 | 611.8161 | 39.256451 |
| NIDNFLNR | 8 | O75116 | Rho-associated protein kinase 2 OS=Hon | ROCK2 | 160.9 | 2 | 50.11236 | yes | 4.976E-07 | 503.25925 | 38.915348 |
| IELQMTLDSK | 10 | O75116 | Rho-associated protein kinase 2 OS=Hon | ROCK2 | 160.9 | 2 | 50.11236 | yes | 3.244E-06 | 589.3103 | 33.828102 |
| GAFGEVQLVR | 10 | O75116 | Rho-associated protein kinase 2 OS=Hon | ROCK2 | 160.9 | 2 | 50.11236 | no | 1.988E-06 | 538.2984 | 27.769571 |
| VLTTTALSTVSSGV | 16 | O75122 | CLIP-associating protein 2 OS=Homo sa | CLASP2 | 141.064 | 2 | 54.38389 | yes | 1.26E-07 | 810.45178 | 1647.7477 |
| VLTTTALSTVSSGV | 16 | O75122 | CLIP-associating protein 2 OS=Homo sa | CLASP2 | 141.064 | 2 | 54.38389 | yes | 4.627E-07 | 540.63696 | 1647.7477 |
| VLTTTALSTVSSGV | 16 | O75122 | CLIP-associating protein 2 OS=Homo sa | CLASP2 | 141.064 | 3 | 54.38389 | yes | 1.26E-07 | 810.45178 | 1647.7477 |
| VLTTTALSTVSSGV | 16 | O75122 | CLIP-associating protein 2 OS=Homo sa | CLASP2 | 141.064 | 3 | 54.38389 | yes | 4.627E-07 | 540.63696 | 1647.7477 |
| FTVDQTQTPSLK | 12 | O75122 | CLIP-associating protein 2 OS=Homo sa | CLASP2 | 141.064 | 2 | 54.38389 | yes | 1.231E-07 | 682.85645 | 1430.1229 |
| VMATSGCAAIR | 11 | O75122 | CLIP-associating protein 2 OS=Homo sa | CLASP2 | 141.064 | 2 | 54.38389 | yes | 9.722E-08 | 568.78119 | 1161.2045 |
| LLHNHLR | 7 | O75122 | CLIP-associating protein 2 OS=Homo sa | CLASP2 | 141.064 | 2 | 54.38389 | yes | 1.747E-07 | 451.76959 | 1033.592 |
| SDIDVNAAAGAK | 12 | O75122 | CLIP-associating protein 2 OS=Homo sa | CLASP2 | 141.064 | 2 | 54.38389 | yes | 9.935E-08 | 566.28564 | 959.16071 |
| EGGAGAVDEDDFI | 14 | O75122 | CLIP-associating protein 2 OS=Homo sa | CLASP2 | 141.064 | 2 | 54.38389 | yes | 1.084E-07 | 711.82281 | 942.63568 |
| LSAPLAGMGNAK | 12 | O75122 | CLIP-associating protein 2 OS=Homo sa | CLASP2 | 141.064 | 2 | 54.38389 | yes | 9.722E-08 | 565.30536 | 910.82153 |
| ASSLPGSLQR | 10 | O75122 | CLIP-associating protein 2 OS=Homo sa | CLASP2 | 141.064 | 2 | 54.38389 | yes | 9.739E-08 | 508.28018 | 906.12134 |

| | | | | | | | | | | | |
|---|---|---|---|---|---|---|---|---|---|---|---|
| QTEDVAEVLNR | 11 | O75122 | CLIP-associating protein 2 OS=Homo sa | CLASP2 | 141.064 | 2 | 54.38389 | yes | 3.169E-07 | 637.32275 | 889.31421 |
| GVTEAIQNFSFR | 12 | O75122 | CLIP-associating protein 2 OS=Homo sa | CLASP2 | 141.064 | 2 | 54.38389 | yes | 9.722E-08 | 684.84894 | 863.40259 |
| VLVNSASAQK | 10 | O75122 | CLIP-associating protein 2 OS=Homo sa | CLASP2 | 141.064 | 2 | 54.38389 | yes | 3.212E-07 | 508.79037 | 668.37262 |
| GIHDADAEAR | 10 | O75122 | CLIP-associating protein 2 OS=Homo sa | CLASP2 | 141.064 | 2 | 54.38389 | yes | 9.722E-08 | 527.74927 | 620.88959 |
| HAAVLVETIK | 10 | O75122 | CLIP-associating protein 2 OS=Homo sa | CLASP2 | 141.064 | 2 | 54.38389 | yes | 2.105E-07 | 540.82422 | 532.43262 |
| IALYELMK | 8 | O75122 | CLIP-associating protein 2 OS=Homo sa | CLASP2 | 141.064 | 2 | 54.38389 | yes | 1.241E-07 | 490.77789 | 441.81555 |
| VITWTTEPK | 9 | O75122 | CLIP-associating protein 2 OS=Homo sa | CLASP2 | 141.064 | 2 | 54.38389 | yes | 1.149E-07 | 537.7951 | 420.48792 |
| AGGDATDSSQTAL | 16 | O75122 | CLIP-associating protein 2 OS=Homo sa | CLASP2 | 141.064 | 2 | 54.38389 | yes | 9.722E-08 | 775.8501 | 419.60626 |
| KTYMGLR | 7 | O75122 | CLIP-associating protein 2 OS=Homo sa | CLASP2 | 141.064 | 2 | 54.38389 | yes | 9.935E-08 | 434.73911 | 328.75269 |
| AFTDVPSIQIYSSR | 14 | O75122 | CLIP-associating protein 2 OS=Homo sa | CLASP2 | 141.064 | 2 | 54.38389 | yes | 1.857E-07 | 792.40686 | 302.33273 |
| WSTANPSTVAGR | 12 | O75122 | CLIP-associating protein 2 OS=Homo sa | CLASP2 | 141.064 | 2 | 54.38389 | yes | 1.083E-07 | 623.81238 | 251.54286 |
| QMDPGDFINSSETI | 14 | O75122 | CLIP-associating protein 2 OS=Homo sa | CLASP2 | 141.064 | 2 | 54.38389 | yes | 9.722E-08 | 798.85156 | 241.172 |
| SQEDMNEPLKR | 11 | O75122 | CLIP-associating protein 2 OS=Homo sa | CLASP2 | 141.064 | 2 | 54.38389 | yes | 9.722E-08 | 673.82208 | 230.23012 |
| SSGSVASLPQSDR | 13 | O75122 | CLIP-associating protein 2 OS=Homo sa | CLASP2 | 141.064 | 2 | 54.38389 | yes | 9.722E-08 | 645.81787 | 218.12206 |
| NYAELTVMK | 9 | O75122 | CLIP-associating protein 2 OS=Homo sa | CLASP2 | 141.064 | 2 | 54.38389 | yes | 3.132E-06 | 534.77332 | 182.81837 |
| MFADPHGK | 8 | O75122 | CLIP-associating protein 2 OS=Homo sa | CLASP2 | 141.064 | 2 | 54.38389 | yes | 1.568E-07 | 451.71307 | 151.12289 |
| YIETLAK | 7 | O75122 | CLIP-associating protein 2 OS=Homo sa | CLASP2 | 141.064 | 2 | 54.38389 | yes | 1.609E-07 | 419.23947 | 136.32784 |
| SQEDMNEPLK | 10 | O75122 | CLIP-associating protein 2 OS=Homo sa | CLASP2 | 141.064 | 2 | 54.38389 | yes | 9.722E-08 | 595.77155 | 135.06563 |
| ASLLHSMPTHSSPI | 14 | O75122 | CLIP-associating protein 2 OS=Homo sa | CLASP2 | 141.064 | 2 | 54.38389 | yes | 9.722E-08 | 760.88556 | 123.80211 |
| LIPLITSNCTSK | 12 | O75122 | CLIP-associating protein 2 OS=Homo sa | CLASP2 | 141.064 | 2 | 54.38389 | yes | 1.21E-07 | 673.87103 | 113.32404 |
| ELEETLNK | 8 | O75122 | CLIP-associating protein 2 OS=Homo sa | CLASP2 | 141.064 | 2 | 54.38389 | yes | 6.869E-06 | 488.2533 | 75.100464 |
| SLLVAGAAQYDCI | 18 | O75122 | CLIP-associating protein 2 OS=Homo sa | CLASP2 | 141.064 | 3 | 54.38389 | yes | 1.691E-07 | 699.35089 | 63.860573 |
| MVSQSQPGSR | 10 | O75122 | CLIP-associating protein 2 OS=Homo sa | CLASP2 | 141.064 | 2 | 54.38389 | yes | 1.229E-07 | 538.76129 | 61.028553 |
| SLQTYLK | 7 | O75122 | CLIP-associating protein 2 OS=Homo sa | CLASP2 | 141.064 | 2 | 54.38389 | yes | 9.701E-07 | 426.7449 | 45.798061 |
| EILSDDKHDWDQI | 13 | O75122 | CLIP-associating protein 2 OS=Homo sa | CLASP2 | 141.064 | 2 | 54.38389 | yes | 1.092E-07 | 828.88428 | 44.507523 |
| KIALYELMK | 9 | O75122 | CLIP-associating protein 2 OS=Homo sa | CLASP2 | 141.064 | 2 | 54.38389 | yes | 3.194E-07 | 554.82538 | 34.264286 |
| KGIHDADAEAR | 11 | O75122 | CLIP-associating protein 2 OS=Homo sa | CLASP2 | 141.064 | 2 | 54.38389 | yes | 9.722E-08 | 591.79675 | 31.07811 |
| RYESYGMHSDDD. | 24 | O75122 | CLIP-associating protein 2 OS=Homo sa | CLASP2 | 141.064 | 4 | 54.38389 | yes | 2.091E-07 | 678.26489 | 26.961924 |
| LTQEESFSVWDEH | 15 | O75122 | CLIP-associating protein 2 OS=Homo sa | CLASP2 | 141.064 | 3 | 54.38389 | yes | 1.345E-07 | 627.95996 | 26.149473 |
| EACITVAHLSTVL( | 16 | O75122 | CLIP-associating protein 2 OS=Homo sa | CLASP2 | 141.064 | 3 | 54.38389 | yes | 4.046E-06 | 571.63818 | 14.442513 |
| LMLSLGLQPSR | 11 | O75127 | Pentatricopeptide repeat-containing prote | PTCD1 | 78.856 | 2 | 51.033485 | yes | 9.991E-08 | 607.84991 | 2538.3992 |
| EEATVLQPPVSR | 12 | O75127 | Pentatricopeptide repeat-containing prote | PTCD1 | 78.856 | 2 | 51.033485 | yes | 1.23E-07 | 663.35663 | 2042.2316 |
| DGLQLLTDMK | 10 | O75127 | Pentatricopeptide repeat-containing prote | PTCD1 | 78.856 | 2 | 51.033485 | yes | 1.073E-07 | 567.29718 | 1184.3914 |
| SAAQFHNLR | 9 | O75127 | Pentatricopeptide repeat-containing prote | PTCD1 | 78.856 | 2 | 51.033485 | yes | 2.875E-07 | 522.27271 | 1044.7289 |
| VPVNEVVIR | 9 | O75127 | Pentatricopeptide repeat-containing prote | PTCD1 | 78.856 | 2 | 51.033485 | yes | 4.028E-07 | 512.8111 | 986.50378 |
| AFNLYNQMK | 9 | O75127 | Pentatricopeptide repeat-containing prote | PTCD1 | 78.856 | 2 | 51.033485 | yes | 9.722E-08 | 564.77893 | 897.52295 |
| LVEALDLFER | 10 | O75127 | Pentatricopeptide repeat-containing prote | PTCD1 | 78.856 | 2 | 51.033485 | yes | 1.136E-07 | 602.83228 | 842.86377 |
| NTPYWYFLQCK | 11 | O75127 | Pentatricopeptide repeat-containing prote | PTCD1 | 78.856 | 2 | 51.033485 | yes | 1.084E-07 | 760.35559 | 471.55078 |
| DCGLGDPQVASEI | 18 | O75127 | Pentatricopeptide repeat-containing prote | PTCD1 | 78.856 | 3 | 51.033485 | yes | 1.01E-07 | 656.67883 | 412.09198 |

| | | | | | | | | | | | |
|---|---|---|---|---|---|---|---|---|---|---|---|
| RDEQMEPEPK | 10 | O75127 | Pentatricopeptide repeat-containing prote | PTCD1 | 78.856 | 2 | 51.033485 | yes | 1.216E-07 | 629.79022 | 370.61169 |
| QWLTVMPAEETPI | 17 | O75127 | Pentatricopeptide repeat-containing prote | PTCD1 | 78.856 | 3 | 51.033485 | yes | 1.098E-07 | 693.34363 | 331.51624 |
| SQVTPNTHIYSALI | 19 | O75127 | Pentatricopeptide repeat-containing prote | PTCD1 | 78.856 | 3 | 51.033485 | yes | 1.002E-07 | 690.3761 | 290.42233 |
| DSYNLLLVAAR | 11 | O75127 | Pentatricopeptide repeat-containing prote | PTCD1 | 78.856 | 2 | 51.033485 | yes | 1.124E-07 | 617.84314 | 248.79251 |
| MCLDVFK | 7 | O75127 | Pentatricopeptide repeat-containing prote | PTCD1 | 78.856 | 2 | 51.033485 | yes | 2.545E-07 | 456.71951 | 205.72739 |
| QLEFAAQYPPTFD | 14 | O75127 | Pentatricopeptide repeat-containing prote | PTCD1 | 78.856 | 2 | 51.033485 | yes | 1.142E-07 | 841.91229 | 202.46173 |
| YALQVWR | 7 | O75127 | Pentatricopeptide repeat-containing prote | PTCD1 | 78.856 | 2 | 51.033485 | yes | 2.019E-07 | 468.25851 | 153.09276 |
| LGDLEGAK | 8 | O75127 | Pentatricopeptide repeat-containing prote | PTCD1 | 78.856 | 2 | 51.033485 | yes | 1.049E-06 | 401.7189 | 114.31258 |
| DSALQSALK | 9 | O75127 | Pentatricopeptide repeat-containing prote | PTCD1 | 78.856 | 2 | 51.033485 | yes | 9.722E-08 | 466.75601 | 97.231804 |
| KLNYTYLISILK | 12 | O75127 | Pentatricopeptide repeat-containing prote | PTCD1 | 78.856 | 2 | 51.033485 | yes | 1.077E-07 | 734.94232 | 94.765205 |
| NFELNLK | 7 | O75127 | Pentatricopeptide repeat-containing prote | PTCD1 | 78.856 | 2 | 51.033485 | yes | 1.426E-07 | 439.24252 | 62.788097 |
| GHVVTEETFSFLLI | 20 | O75127 | Pentatricopeptide repeat-containing prote | PTCD1 | 78.856 | 3 | 51.033485 | yes | 4.864E-06 | 771.04144 | 35.517181 |
| ASTMDVTVVLPSC | 16 | O75128 | Protein cordon-bleu OS=Homo sapiens O | COBL | 135.617 | 2 | 46.563183 | yes | 1.527E-07 | 823.93713 | 58.768784 |
| TSSQYVASAIAK | 12 | O75128 | Protein cordon-bleu OS=Homo sapiens O | COBL | 135.617 | 2 | 46.563183 | yes | 3.869E-06 | 613.32477 | 32.564781 |
| SDPYLEFHK | 9 | O75131 | Copine-3 OS=Homo sapiens OX=9606 G | CPNE3 | 60.131 | 2 | 49.043583 | yes | 3.91E-06 | 568.27454 | 88.587425 |
| FGVYDIDNK | 9 | O75131 | Copine-3 OS=Homo sapiens OX=9606 G | CPNE3 | 60.131 | 2 | 49.043583 | yes | 8.686E-07 | 535.76129 | 88.187286 |
| SSPVEFECINEK | 12 | O75131 | Copine-3 OS=Homo sapiens OX=9606 G | CPNE3 | 60.131 | 2 | 49.043583 | yes | 1.315E-06 | 719.82959 | 75.891846 |
| VVLFEMEAR | 9 | O75131 | Copine-3 OS=Homo sapiens OX=9606 G | CPNE3 | 60.131 | 2 | 49.043583 | yes | 6.259E-07 | 547.28918 | 73.366554 |
| SPLGEVAIR | 9 | O75131 | Copine-3 OS=Homo sapiens OX=9606 G | CPNE3 | 60.131 | 2 | 49.043583 | yes | 8.658E-07 | 471.27435 | 54.463867 |
| LYGPTNFSPIINHV | 16 | O75131 | Copine-3 OS=Homo sapiens OX=9606 G | CPNE3 | 60.131 | 3 | 49.043583 | yes | 1.074E-07 | 600.32483 | 37.152374 |
| VALNVSCANLLDK | 13 | O75131 | Copine-3 OS=Homo sapiens OX=9606 G | CPNE3 | 60.131 | 2 | 49.043583 | yes | 2.17E-07 | 708.87939 | 35.573044 |
| AEDGPGASLALR | 12 | O75147 | Obscurin-like protein 1 OS=Homo sapien | OBSL1 | 206.947 | 2 | 54.887028 | yes | 1.133E-07 | 578.80145 | 1062.3416 |
| VVLTCELSR | 9 | O75147 | Obscurin-like protein 1 OS=Homo sapien | OBSL1 | 206.947 | 2 | 54.887028 | yes | 2.115E-07 | 538.79205 | 1035.4226 |
| VVLTCELCR | 9 | O75147 | Obscurin-like protein 1 OS=Homo sapien | OBSL1 | 206.947 | 2 | 54.887028 | yes | 1.454E-07 | 575.29138 | 1008.0408 |
| AGGLPEPTLYWEK | 13 | O75147 | Obscurin-like protein 1 OS=Homo sapien | OBSL1 | 206.947 | 2 | 54.887028 | yes | 9.938E-08 | 730.87463 | 942.0423 |
| VSLTFTTSER | 10 | O75147 | Obscurin-like protein 1 OS=Homo sapien | OBSL1 | 206.947 | 2 | 54.887028 | yes | 9.722E-08 | 570.7984 | 854.62927 |
| ICTVSGHGR | 9 | O75147 | Obscurin-like protein 1 OS=Homo sapien | OBSL1 | 206.947 | 2 | 54.887028 | yes | 9.722E-08 | 493.74545 | 635.55884 |
| DGEEVVESPALLL | 15 | O75147 | Obscurin-like protein 1 OS=Homo sapien | OBSL1 | 206.947 | 2 | 54.887028 | yes | 4.804E-07 | 813.93286 | 538.01868 |
| ADDDGIYLCEMR | 12 | O75147 | Obscurin-like protein 1 OS=Homo sapien | OBSL1 | 206.947 | 2 | 54.887028 | yes | 1.413E-07 | 729.30542 | 517.44653 |
| VDFPATWYK | 9 | O75147 | Obscurin-like protein 1 OS=Homo sapien | OBSL1 | 206.947 | 2 | 54.887028 | yes | 3.749E-07 | 563.78204 | 513.75647 |
| VVSGAEAELK | 10 | O75147 | Obscurin-like protein 1 OS=Homo sapien | OBSL1 | 206.947 | 2 | 54.887028 | yes | 5.433E-07 | 501.77695 | 378.8425 |
| SPHVVFHGSAHLV | 17 | O75147 | Obscurin-like protein 1 OS=Homo sapien | OBSL1 | 206.947 | 3 | 54.887028 | yes | 9.722E-08 | 604.66028 | 323.07559 |
| HSPPGPPILAEMFK | 14 | O75147 | Obscurin-like protein 1 OS=Homo sapien | OBSL1 | 206.947 | 2 | 54.887028 | yes | 1.238E-07 | 760.90015 | 294.84344 |
| DRDGGFVLK | 9 | O75147 | Obscurin-like protein 1 OS=Homo sapien | OBSL1 | 206.947 | 2 | 54.887028 | yes | 2.94E-07 | 503.76944 | 263.13467 |
| VEESELLVVK | 10 | O75147 | Obscurin-like protein 1 OS=Homo sapien | OBSL1 | 206.947 | 2 | 54.887028 | yes | 1.592E-07 | 572.8266 | 239.44087 |
| GLYVCAAR | 8 | O75147 | Obscurin-like protein 1 OS=Homo sapien | OBSL1 | 206.947 | 2 | 54.887028 | yes | 1.011E-07 | 455.23419 | 226.5992 |
| EDAGEVTFSLGNS | 14 | O75147 | Obscurin-like protein 1 OS=Homo sapien | OBSL1 | 206.947 | 2 | 54.887028 | yes | 9.722E-08 | 741.34698 | 212.21306 |
| DGMALDEVWDSS | 20 | O75147 | Obscurin-like protein 1 OS=Homo sapien | OBSL1 | 206.947 | 3 | 54.887028 | yes | 1.084E-07 | 744.34436 | 192.46527 |
| KYEQIEEGTVR | 11 | O75147 | Obscurin-like protein 1 OS=Homo sapien | OBSL1 | 206.947 | 2 | 54.887028 | yes | 1.115E-07 | 676.34625 | 164.20195 |

| | | | | | | | | | | | |
|---|---|---|---|---|---|---|---|---|---|---|---|
| EHGIAVLECK | 10 | O75147 | Obscurin-like protein 1 OS=Homo sapien | OBSL1 | 206.947 | 2 | 54.887028 | yes | 1.934E-06 | 578.29498 | 125.42941 |
| SNEPEGQVEPGAL | 14 | O75147 | Obscurin-like protein 1 OS=Homo sapien | OBSL1 | 206.947 | 2 | 54.887028 | yes | 2.464E-07 | 741.86279 | 123.57722 |
| NSAGQTLSAVQLH | 15 | O75147 | Obscurin-like protein 1 OS=Homo sapien | OBSL1 | 206.947 | 2 | 54.887028 | yes | 1.499E-07 | 776.92572 | 91.751366 |
| GGQQLAASER | 10 | O75147 | Obscurin-like protein 1 OS=Homo sapien | OBSL1 | 206.947 | 2 | 54.887028 | yes | 4.037E-07 | 508.75961 | 78.649277 |
| FTRPLQDVEGR | 11 | O75147 | Obscurin-like protein 1 OS=Homo sapien | OBSL1 | 206.947 | 2 | 54.887028 | yes | 1.424E-07 | 659.34912 | 67.361488 |
| DGEELPVICQSSSC | 26 | O75147 | Obscurin-like protein 1 OS=Homo sapien | OBSL1 | 206.947 | 4 | 54.887028 | yes | 1.512E-07 | 698.09821 | 63.768791 |
| VLYCQAK | 7 | O75147 | Obscurin-like protein 1 OS=Homo sapien | OBSL1 | 206.947 | 2 | 54.887028 | yes | 1.046E-07 | 441.23111 | 61.469746 |
| YEQIEEGTVR | 10 | O75147 | Obscurin-like protein 1 OS=Homo sapien | OBSL1 | 206.947 | 2 | 54.887028 | yes | 9.722E-08 | 612.29877 | 53.069546 |
| VVLTCELCRPWAE | 15 | O75147 | Obscurin-like protein 1 OS=Homo sapien | OBSL1 | 206.947 | 3 | 54.887028 | yes | 2.806E-07 | 629.99042 | 29.640434 |
| DGQKVEESELLVV | 14 | O75147 | Obscurin-like protein 1 OS=Homo sapien | OBSL1 | 206.947 | 2 | 54.887028 | yes | 2.282E-07 | 786.92761 | 27.95554 |
| TEGVSAFFGVTVQ | 23 | O75147 | Obscurin-like protein 1 OS=Homo sapien | OBSL1 | 206.947 | 3 | 54.887028 | yes | 3.291E-06 | 823.09204 | 23.969917 |
| EMAPVPGTTTTTT | 16 | O75150 | E3 ubiquitin-protein ligase BRE1B OS=H | RNF40 | 113.679 | 2 | 48.741154 | yes | 3.337E-06 | 810.91107 | 65.175385 |
| TEVIQLEDTLAQVI | 14 | O75150 | E3 ubiquitin-protein ligase BRE1B OS=H | RNF40 | 113.679 | 2 | 48.741154 | no | 1.108E-07 | 807.93848 | 50.470875 |
| SQVDAQLLTVQK | 12 | O75150 | E3 ubiquitin-protein ligase BRE1B OS=H | RNF40 | 113.679 | 2 | 48.741154 | yes | 3.429E-06 | 665.37225 | 9.8211737 |
| VLNVTNLEFSDTR | 13 | O75151 | Lysine-specific demethylase PHF2 OS=H | PHF2 | 120.775 | 2 | 40.18739 | yes | 6.048E-07 | 754.39117 | 159.93906 |
| TSQPMAPGVFLTQ | 14 | O75151 | Lysine-specific demethylase PHF2 OS=H | PHF2 | 120.775 | 2 | 40.18739 | yes | 5.142E-07 | 766.89813 | 138.07248 |
| DEYEYVSDDGELK | 13 | O75151 | Lysine-specific demethylase PHF2 OS=H | PHF2 | 120.775 | 2 | 40.18739 | yes | 5.542E-07 | 781.33069 | 28.526264 |
| MSSFVEPPDIVK | 12 | O75151 | Lysine-specific demethylase PHF2 OS=H | PHF2 | 120.775 | 2 | 40.18739 | yes | 8.308E-07 | 674.84448 | 27.944021 |
| EFVDYYYSTNR | 11 | O75151 | Lysine-specific demethylase PHF2 OS=H | PHF2 | 120.775 | 2 | 40.18739 | yes | 4.163E-07 | 728.82263 | 24.185921 |
| TVVLPPIVASR | 11 | O75152 | Zinc finger CCCH domain-containing prc | ZC3H11A | 89.132 | 2 | 56.309666 | yes | 1.504E-07 | 576.36096 | 3131.4385 |
| SVTVPEAENPR | 11 | O75152 | Zinc finger CCCH domain-containing prc | ZC3H11A | 89.132 | 2 | 56.309666 | no | 1.498E-07 | 599.80676 | 2116.7888 |
| LSVQSNPSPQLR | 12 | O75152 | Zinc finger CCCH domain-containing prc | ZC3H11A | 89.132 | 2 | 56.309666 | no | 9.81E-08 | 663.36224 | 1867.4884 |
| TLEEILLER | 9 | O75152 | Zinc finger CCCH domain-containing prc | ZC3H11A | 89.132 | 2 | 56.309666 | yes | 9.781E-08 | 558.31897 | 1795.6056 |
| TVLPTVPESPEEEV | 15 | O75152 | Zinc finger CCCH domain-containing prc | ZC3H11A | 89.132 | 2 | 56.309666 | yes | 2.229E-07 | 827.43274 | 1490.2522 |
| SMQEVHIK | 8 | O75152 | Zinc finger CCCH domain-containing prc | ZC3H11A | 89.132 | 2 | 56.309666 | yes | 1.87E-07 | 486.25256 | 1066.4691 |
| YVDGLFLPPSK | 11 | O75152 | Zinc finger CCCH domain-containing prc | ZC3H11A | 89.132 | 2 | 56.309666 | no | 1.326E-07 | 618.33716 | 1024.3546 |
| QGEGSSGVSSLLLI | 23 | O75152 | Zinc finger CCCH domain-containing prc | ZC3H11A | 89.132 | 3 | 56.309666 | yes | 2.207E-07 | 767.72888 | 830.71619 |
| EASGETTGVDITK | 13 | O75152 | Zinc finger CCCH domain-containing prc | ZC3H11A | 89.132 | 2 | 56.309666 | yes | 1.108E-07 | 654.31989 | 777.89178 |
| KVEAPETNIDK | 11 | O75152 | Zinc finger CCCH domain-containing prc | ZC3H11A | 89.132 | 2 | 56.309666 | yes | 9.722E-08 | 622.33008 | 756.85028 |
| ASQLSVQQNK | 10 | O75152 | Zinc finger CCCH domain-containing prc | ZC3H11A | 89.132 | 2 | 56.309666 | no | 1.119E-07 | 551.7962 | 728.36554 |
| LNCAFHHNR | 9 | O75152 | Zinc finger CCCH domain-containing prc | ZC3H11A | 89.132 | 2 | 56.309666 | no | 1.34E-07 | 584.77509 | 693.6369 |
| VEVETSGIGDSLLN | 16 | O75152 | Zinc finger CCCH domain-containing prc | ZC3H11A | 89.132 | 2 | 56.309666 | yes | 3.296E-07 | 830.4436 | 599.15088 |
| QGECLNFGIK | 10 | O75152 | Zinc finger CCCH domain-containing prc | ZC3H11A | 89.132 | 2 | 56.309666 | yes | 1.105E-07 | 583.28717 | 528.38605 |
| FSAGGDSDPPLK | 12 | O75152 | Zinc finger CCCH domain-containing prc | ZC3H11A | 89.132 | 2 | 56.309666 | yes | 1.662E-07 | 595.78802 | 491.91098 |
| CAAQTLEK | 8 | O75152 | Zinc finger CCCH domain-containing prc | ZC3H11A | 89.132 | 2 | 56.309666 | yes | 3.906E-07 | 460.72894 | 424.25827 |
| QGEEPLVR | 8 | O75152 | Zinc finger CCCH domain-containing prc | ZC3H11A | 89.132 | 2 | 56.309666 | no | 1.08E-07 | 464.24835 | 408.26422 |
| TFSEVLAEK | 9 | O75152 | Zinc finger CCCH domain-containing prc | ZC3H11A | 89.132 | 2 | 56.309666 | yes | 1.664E-07 | 512.27148 | 365.69327 |
| TPTLQPTPEVHNGI | 15 | O75152 | Zinc finger CCCH domain-containing prc | ZC3H11A | 89.132 | 3 | 56.309666 | no | 1.001E-07 | 553.96527 | 169.0397 |
| TPTLQPTPEVHNGI | 15 | O75152 | Zinc finger CCCH domain-containing prc | ZC3H11A | 89.132 | 3 | 56.309666 | no | 1.35E-07 | 830.44427 | 169.0397 |

| | | | | | | | | | | |
|---|---|---|---|---|---|---|---|---|---|---|
| TPTLQPTPEVHNG | 15 | O75152 | Zinc finger CCCH domain-containing prc | ZC3H11A | 89.132 | 2 | 56.309666 | no | 1.001E-07 | 553.96527 | 169.0397 |
| TPTLQPTPEVHNG | 15 | O75152 | Zinc finger CCCH domain-containing prc | ZC3H11A | 89.132 | 2 | 56.309666 | no | 1.35E-07 | 830.44427 | 169.0397 |
| LGMSADPDNEDA | 15 | O75152 | Zinc finger CCCH domain-containing prc | ZC3H11A | 89.132 | 2 | 56.309666 | yes | 1E-07 | 789.83307 | 323.86469 |
| FSAGGDSDPPLKR | 13 | O75152 | Zinc finger CCCH domain-containing prc | ZC3H11A | 89.132 | 2 | 56.309666 | yes | 1.354E-07 | 673.83856 | 312.44223 |
| VEAPETNIDK | 10 | O75152 | Zinc finger CCCH domain-containing prc | ZC3H11A | 89.132 | 2 | 56.309666 | yes | 9.722E-08 | 558.28259 | 191.39552 |
| TLEEIKLEK | 9 | O75152 | Zinc finger CCCH domain-containing prc | ZC3H11A | 89.132 | 2 | 56.309666 | yes | 2.613E-07 | 551.82135 | 185.46632 |
| GDVASCNTQVAEI | 24 | O75152 | Zinc finger CCCH domain-containing prc | ZC3H11A | 89.132 | 3 | 56.309666 | yes | 2.463E-07 | 828.43561 | 154.70547 |
| AAVAVVPLVSEDI | 13 | O75152 | Zinc finger CCCH domain-containing prc | ZC3H11A | 89.132 | 2 | 56.309666 | yes | 9.722E-08 | 649.37177 | 139.47267 |
| PLSSSSVLQEPPAK | 14 | O75152 | Zinc finger CCCH domain-containing prc | ZC3H11A | 89.132 | 2 | 56.309666 | no | 9.722E-08 | 720.39069 | 97.981102 |
| PVLTAVPGITR | 11 | O75152 | Zinc finger CCCH domain-containing prc | ZC3H11A | 89.132 | 2 | 56.309666 | yes | 4.111E-07 | 562.34534 | 66.651596 |
| KQGEGSSGVSSLL | 24 | O75152 | Zinc finger CCCH domain-containing prc | ZC3H11A | 89.132 | 3 | 56.309666 | yes | 3.784E-07 | 810.42719 | 61.825691 |
| NLQEGNEVDSQSS | 15 | O75152 | Zinc finger CCCH domain-containing prc | ZC3H11A | 89.132 | 2 | 56.309666 | yes | 9.722E-08 | 838.39771 | 43.373619 |
| KFSAGGDSDPPLK | 13 | O75152 | Zinc finger CCCH domain-containing prc | ZC3H11A | 89.132 | 2 | 56.309666 | yes | 3.599E-07 | 659.83551 | 41.48798 |
| AVEMHAAVIAAV | 27 | O75152 | Zinc finger CCCH domain-containing prc | ZC3H11A | 89.132 | 4 | 56.309666 | yes | 5.072E-05 | 683.37695 | 3.1438038 |
| VSFFDVTYR | 9 | O75161 | Nephrocystin-4 OS=Homo sapiens OX=9 | NPHP4 | 157.599 | 2 | 39.884563 | yes | 4.328E-07 | 567.28491 | 22.80262 |
| VYNEVPTFQLEVP | 14 | O75165 | DnaJ homolog subfamily C member 13 O | DNAJC13 | 254.415 | 2 | 53.391956 | yes | 3.849E-07 | 831.94049 | 1642.0549 |
| VLALTETCLVER | 12 | O75165 | DnaJ homolog subfamily C member 13 O | DNAJC13 | 254.415 | 2 | 53.391956 | yes | 1.091E-07 | 702.38177 | 957.80933 |
| LHLFASEQR | 9 | O75165 | DnaJ homolog subfamily C member 13 O | DNAJC13 | 254.415 | 2 | 53.391956 | yes | 5.73E-07 | 550.79602 | 919.28101 |
| YIIGMLER | 8 | O75165 | DnaJ homolog subfamily C member 13 O | DNAJC13 | 254.415 | 2 | 53.391956 | yes | 1.499E-07 | 497.77313 | 873.13635 |
| ADTVGLACEAINR | 13 | O75165 | DnaJ homolog subfamily C member 13 O | DNAJC13 | 254.415 | 2 | 53.391956 | yes | 9.722E-08 | 695.3432 | 772.09741 |
| LAVASVTEILK | 11 | O75165 | DnaJ homolog subfamily C member 13 O | DNAJC13 | 254.415 | 2 | 53.391956 | yes | 1.054E-07 | 572.35284 | 712.29156 |
| SYEFFNELYHR | 11 | O75165 | DnaJ homolog subfamily C member 13 O | DNAJC13 | 254.415 | 2 | 53.391956 | yes | 1.018E-07 | 752.84644 | 651.0976 |
| GQGTEFNLTFR | 11 | O75165 | DnaJ homolog subfamily C member 13 O | DNAJC13 | 254.415 | 2 | 53.391956 | yes | 1.023E-07 | 635.31476 | 601.84869 |
| YECLAEEIK | 9 | O75165 | DnaJ homolog subfamily C member 13 O | DNAJC13 | 254.415 | 2 | 53.391956 | yes | 4.198E-07 | 577.77356 | 598.0213 |
| FLPSVFMDAMR | 11 | O75165 | DnaJ homolog subfamily C member 13 O | DNAJC13 | 254.415 | 2 | 53.391956 | yes | 1.246E-07 | 657.32269 | 591.09167 |
| YAGYPMLIR | 9 | O75165 | DnaJ homolog subfamily C member 13 O | DNAJC13 | 254.415 | 2 | 53.391956 | yes | 1.614E-07 | 542.28644 | 581.62378 |
| EMMLEHFK | 8 | O75165 | DnaJ homolog subfamily C member 13 O | DNAJC13 | 254.415 | 2 | 53.391956 | yes | 2.379E-07 | 532.74884 | 580.66132 |
| TELLTEALR | 9 | O75165 | DnaJ homolog subfamily C member 13 O | DNAJC13 | 254.415 | 2 | 53.391956 | yes | 1.727E-07 | 523.29803 | 565.25916 |
| YNPGSESECIGHFI | 14 | O75165 | DnaJ homolog subfamily C member 13 O | DNAJC13 | 254.415 | 2 | 53.391956 | yes | 1.792E-07 | 812.85663 | 556.81769 |
| TQSILFNR | 8 | O75165 | DnaJ homolog subfamily C member 13 O | DNAJC13 | 254.415 | 2 | 53.391956 | yes | 3.179E-07 | 489.772 | 552.3385 |
| TYGSEFVYSDHAK | 13 | O75165 | DnaJ homolog subfamily C member 13 O | DNAJC13 | 254.415 | 2 | 53.391956 | yes | 1.014E-07 | 752.34119 | 550.99371 |
| TITMETSDDLLFSI | 14 | O75165 | DnaJ homolog subfamily C member 13 O | DNAJC13 | 254.415 | 2 | 53.391956 | yes | 1.017E-07 | 800.8924 | 543.28119 |
| EQSELVAQALK | 11 | O75165 | DnaJ homolog subfamily C member 13 O | DNAJC13 | 254.415 | 2 | 53.391956 | yes | 1.065E-07 | 608.33258 | 532.49023 |
| IFIAQPAWVLR | 11 | O75165 | DnaJ homolog subfamily C member 13 O | DNAJC13 | 254.415 | 2 | 53.391956 | yes | 9.722E-08 | 657.39008 | 456.41238 |
| IEANWDLFYYR | 11 | O75165 | DnaJ homolog subfamily C member 13 O | DNAJC13 | 254.415 | 2 | 53.391956 | yes | 9.722E-08 | 745.35913 | 449.39746 |
| ITEMPSIIK | 9 | O75165 | DnaJ homolog subfamily C member 13 O | DNAJC13 | 254.415 | 2 | 53.391956 | yes | 9.722E-08 | 516.29388 | 431.29416 |
| CYSVAAQFEECR | 12 | O75165 | DnaJ homolog subfamily C member 13 O | DNAJC13 | 254.415 | 2 | 53.391956 | yes | 1.014E-07 | 760.31885 | 423.23608 |
| VIQAMNHR | 8 | O75165 | DnaJ homolog subfamily C member 13 O | DNAJC13 | 254.415 | 2 | 53.391956 | yes | 1.476E-07 | 484.75836 | 411.12296 |
| CVAVLTR | 7 | O75165 | DnaJ homolog subfamily C member 13 O | DNAJC13 | 254.415 | 2 | 53.391956 | yes | 2.537E-07 | 409.73129 | 409.28583 |

| | | | | | | | | | | | |
|---|---|---|---|---|---|---|---|---|---|---|---|
| VLCSYDYR | 8 | O75165 | DnaJ homolog subfamily C member 13 O | DNAJC13 | 254.415 | 2 | 53.391956 | yes | 9.722E-08 | 538.2475 | 384.46317 |
| AYEFLCTK | 8 | O75165 | DnaJ homolog subfamily C member 13 O | DNAJC13 | 254.415 | 2 | 53.391956 | yes | 2.063E-07 | 516.24695 | 348.97006 |
| VIHALSENELCVR | 13 | O75165 | DnaJ homolog subfamily C member 13 O | DNAJC13 | 254.415 | 2 | 53.391956 | yes | 1.075E-07 | 770.40106 | 348.95947 |
| ENINQKPVVLR | 11 | O75165 | DnaJ homolog subfamily C member 13 O | DNAJC13 | 254.415 | 2 | 53.391956 | yes | 9.722E-08 | 655.383 | 173.04982 |
| ENINQKPVVLR | 11 | O75165 | DnaJ homolog subfamily C member 13 O | DNAJC13 | 254.415 | 2 | 53.391956 | yes | 8.739E-07 | 437.25775 | 173.04982 |
| ENINQKPVVLR | 11 | O75165 | DnaJ homolog subfamily C member 13 O | DNAJC13 | 254.415 | 3 | 53.391956 | yes | 9.722E-08 | 655.383 | 173.04982 |
| ENINQKPVVLR | 11 | O75165 | DnaJ homolog subfamily C member 13 O | DNAJC13 | 254.415 | 3 | 53.391956 | yes | 8.739E-07 | 437.25775 | 173.04982 |
| EPLEFEQYLNLR | 12 | O75165 | DnaJ homolog subfamily C member 13 O | DNAJC13 | 254.415 | 2 | 53.391956 | yes | 1.163E-07 | 775.89612 | 339.23541 |
| IGDYYLR | 7 | O75165 | DnaJ homolog subfamily C member 13 O | DNAJC13 | 254.415 | 2 | 53.391956 | yes | 1.122E-07 | 450.23471 | 313.63593 |
| FPDWPIKDPVK | 11 | O75165 | DnaJ homolog subfamily C member 13 O | DNAJC13 | 254.415 | 2 | 53.391956 | yes | 4.249E-07 | 671.36371 | 312.47061 |
| LFQHPSMAIIK | 11 | O75165 | DnaJ homolog subfamily C member 13 O | DNAJC13 | 254.415 | 2 | 53.391956 | yes | 3.042E-07 | 642.86029 | 128.64151 |
| LFQHPSMAIIK | 11 | O75165 | DnaJ homolog subfamily C member 13 O | DNAJC13 | 254.415 | 2 | 53.391956 | yes | 1.137E-06 | 428.90927 | 128.64151 |
| LFQHPSMAIIK | 11 | O75165 | DnaJ homolog subfamily C member 13 O | DNAJC13 | 254.415 | 3 | 53.391956 | yes | 3.042E-07 | 642.86029 | 128.64151 |
| LFQHPSMAIIK | 11 | O75165 | DnaJ homolog subfamily C member 13 O | DNAJC13 | 254.415 | 3 | 53.391956 | yes | 1.137E-06 | 428.90927 | 128.64151 |
| CHEEIGPFTDTR | 12 | O75165 | DnaJ homolog subfamily C member 13 O | DNAJC13 | 254.415 | 2 | 53.391956 | yes | 9.722E-08 | 731.32501 | 235.36609 |
| DQDNAIIRPLPK | 12 | O75165 | DnaJ homolog subfamily C member 13 O | DNAJC13 | 254.415 | 2 | 53.391956 | yes | 1.079E-07 | 690.38568 | 232.71194 |
| SNLIWNFK | 8 | O75165 | DnaJ homolog subfamily C member 13 O | DNAJC13 | 254.415 | 2 | 53.391956 | yes | 2.601E-07 | 511.27692 | 228.69601 |
| DLACFYTTK | 9 | O75165 | DnaJ homolog subfamily C member 13 O | DNAJC13 | 254.415 | 2 | 53.391956 | yes | 9.722E-08 | 559.763 | 225.65799 |
| SEETNQQEVANSL | 15 | O75165 | DnaJ homolog subfamily C member 13 O | DNAJC13 | 254.415 | 2 | 53.391956 | yes | 9.722E-08 | 824.39465 | 225.2233 |
| KEPLEFEQYLNLR | 13 | O75165 | DnaJ homolog subfamily C member 13 O | DNAJC13 | 254.415 | 3 | 53.391956 | yes | 1.504E-07 | 560.29816 | 111.94205 |
| KEPLEFEQYLNLR | 13 | O75165 | DnaJ homolog subfamily C member 13 O | DNAJC13 | 254.415 | 3 | 53.391956 | yes | 1.224E-07 | 839.9436 | 111.94205 |
| KEPLEFEQYLNLR | 13 | O75165 | DnaJ homolog subfamily C member 13 O | DNAJC13 | 254.415 | 2 | 53.391956 | yes | 1.504E-07 | 560.29816 | 111.94205 |
| KEPLEFEQYLNLR | 13 | O75165 | DnaJ homolog subfamily C member 13 O | DNAJC13 | 254.415 | 2 | 53.391956 | yes | 1.224E-07 | 839.9436 | 111.94205 |
| ELIVGEIFVR | 10 | O75165 | DnaJ homolog subfamily C member 13 O | DNAJC13 | 254.415 | 2 | 53.391956 | yes | 1.05E-07 | 587.84515 | 208.60403 |
| KFLENLLEK | 9 | O75165 | DnaJ homolog subfamily C member 13 O | DNAJC13 | 254.415 | 2 | 53.391956 | yes | 1.175E-07 | 567.33191 | 204.52213 |
| SAIDHAGNYIGISL | 15 | O75165 | DnaJ homolog subfamily C member 13 O | DNAJC13 | 254.415 | 2 | 53.391956 | yes | 1.079E-07 | 793.91791 | 196.42032 |
| MQELALSEGALPR | 13 | O75165 | DnaJ homolog subfamily C member 13 O | DNAJC13 | 254.415 | 2 | 53.391956 | yes | 4.427E-07 | 707.87158 | 194.19237 |
| QLCDTLR | 7 | O75165 | DnaJ homolog subfamily C member 13 O | DNAJC13 | 254.415 | 2 | 53.391956 | yes | 9.722E-08 | 453.2291 | 174.46143 |
| HKEDLQPYK | 9 | O75165 | DnaJ homolog subfamily C member 13 O | DNAJC13 | 254.415 | 2 | 53.391956 | yes | 9.722E-08 | 579.30109 | 172.6721 |
| EWYFGNADK | 9 | O75165 | DnaJ homolog subfamily C member 13 O | DNAJC13 | 254.415 | 2 | 53.391956 | yes | 9.722E-08 | 565.2511 | 169.45389 |
| VEMALEALR | 9 | O75165 | DnaJ homolog subfamily C member 13 O | DNAJC13 | 254.415 | 2 | 53.391956 | yes | 8.161E-07 | 516.28131 | 79.703247 |
| VEMALEALR | 9 | O75165 | DnaJ homolog subfamily C member 13 O | DNAJC13 | 254.415 | 2 | 53.391956 | yes | 6.245E-06 | 524.27881 | 79.703247 |
| AGFLAFTQLPK | 11 | O75165 | DnaJ homolog subfamily C member 13 O | DNAJC13 | 254.415 | 2 | 53.391956 | yes | 1.491E-07 | 596.83984 | 144.33987 |
| ILVDLLTLAHLHV | 15 | O75165 | DnaJ homolog subfamily C member 13 O | DNAJC13 | 254.415 | 3 | 53.391956 | yes | 2.637E-07 | 567.34546 | 113.29221 |
| IAMDQYGK | 8 | O75165 | DnaJ homolog subfamily C member 13 O | DNAJC13 | 254.415 | 2 | 53.391956 | yes | 1.152E-06 | 463.22601 | 113.07616 |
| EELKDTLESEMR | 12 | O75165 | DnaJ homolog subfamily C member 13 O | DNAJC13 | 254.415 | 2 | 53.391956 | yes | 4.989E-07 | 740.35339 | 102.87454 |
| HLHTAMFTISSDQ | 14 | O75165 | DnaJ homolog subfamily C member 13 O | DNAJC13 | 254.415 | 2 | 53.391956 | yes | 9.722E-08 | 822.40155 | 49.895954 |
| HLHTAMFTISSDQ | 14 | O75165 | DnaJ homolog subfamily C member 13 O | DNAJC13 | 254.415 | 2 | 53.391956 | yes | 3.426E-06 | 548.60345 | 49.895954 |
| HLHTAMFTISSDQ | 14 | O75165 | DnaJ homolog subfamily C member 13 O | DNAJC13 | 254.415 | 3 | 53.391956 | yes | 9.722E-08 | 822.40155 | 49.895954 |

| | | | | | | | | | | | |
|---|---|---|---|---|---|---|---|---|---|---|---|
| HLHTAMFTISSDQI | 14 | O75165 | DnaJ homolog subfamily C member 13 O | DNAJC13 | 254.415 | 3 | 53.391956 | yes | 3.426E-06 | 548.60345 | 49.895954 |
| CLCLQALAIVYGR | 13 | O75165 | DnaJ homolog subfamily C member 13 O | DNAJC13 | 254.415 | 2 | 53.391956 | yes | 9.967E-08 | 768.90491 | 98.226135 |
| DTLESEMR | 8 | O75165 | DnaJ homolog subfamily C member 13 O | DNAJC13 | 254.415 | 2 | 53.391956 | yes | 3.942E-07 | 490.72131 | 95.226982 |
| NQQDNPEANWK | 11 | O75165 | DnaJ homolog subfamily C member 13 O | DNAJC13 | 254.415 | 2 | 53.391956 | yes | 9.722E-08 | 672.30237 | 90.76564 |
| ENGLEVLQEAFSR | 13 | O75165 | DnaJ homolog subfamily C member 13 O | DNAJC13 | 254.415 | 2 | 53.391956 | yes | 9.722E-08 | 746.37555 | 90.341187 |
| HLVGLWTADNAT | 18 | O75165 | DnaJ homolog subfamily C member 13 O | DNAJC13 | 254.415 | 3 | 53.391956 | yes | 4.824E-07 | 646.68628 | 66.529655 |
| FLENLLEK | 8 | O75165 | DnaJ homolog subfamily C member 13 O | DNAJC13 | 254.415 | 2 | 53.391956 | yes | 1.692E-07 | 503.28439 | 43.082077 |
| VDLVLMHWR | 9 | O75165 | DnaJ homolog subfamily C member 13 O | DNAJC13 | 254.415 | 2 | 53.391956 | yes | 3.06E-07 | 584.81842 | 41.333954 |
| PLQSIPQLK | 9 | O75165 | DnaJ homolog subfamily C member 13 O | DNAJC13 | 254.415 | 2 | 53.391956 | yes | 5.434E-07 | 512.31348 | 39.360531 |
| AMASLETIGPLMN | 16 | O75165 | DnaJ homolog subfamily C member 13 O | DNAJC13 | 254.415 | 2 | 53.391956 | yes | 2.07E-07 | 832.41443 | 37.437622 |
| IANAVVQNLER | 11 | O75170 | Serine/threonine-protein phosphatase 6 re | PPP6R2 | 104.942 | 2 | 51.87435 | yes | 9.722E-08 | 613.84619 | 81.710114 |
| LVELIHPSQDEDR | 13 | O75170 | Serine/threonine-protein phosphatase 6 re | PPP6R2 | 104.942 | 2 | 51.87435 | yes | 2.023E-07 | 775.8941 | 23.511173 |
| MLDNDSILVDAIR | 13 | O75175 | CCR4-NOT transcription complex subuni | CNOT3 | 81.872 | 2 | 52.97408 | yes | 9.722E-08 | 737.88214 | 270.59946 |
| VSEGVEQFEDIWQ | 14 | O75175 | CCR4-NOT transcription complex subuni | CNOT3 | 81.872 | 2 | 52.97408 | yes | 1.253E-07 | 847.40704 | 255.70909 |
| TWVASNEIK | 9 | O75175 | CCR4-NOT transcription complex subuni | CNOT3 | 81.872 | 2 | 52.97408 | yes | 5.5E-07 | 524.2771 | 232.69357 |
| KLIETQMER | 9 | O75175 | CCR4-NOT transcription complex subuni | CNOT3 | 81.872 | 2 | 52.97408 | yes | 9.722E-08 | 574.31061 | 177.76361 |
| APEPLSSLK | 9 | O75175 | CCR4-NOT transcription complex subuni | CNOT3 | 81.872 | 2 | 52.97408 | yes | 9.722E-08 | 471.26874 | 105.22163 |
| MLETILR | 7 | O75175 | CCR4-NOT transcription complex subuni | CNOT3 | 81.872 | 2 | 52.97408 | yes | 3.438E-07 | 438.25458 | 100.51425 |
| LIETQMER | 8 | O75175 | CCR4-NOT transcription complex subuni | CNOT3 | 81.872 | 2 | 52.97408 | yes | 5.911E-07 | 510.26315 | 98.951134 |
| STDSEVSQSPAK | 12 | O75175 | CCR4-NOT transcription complex subuni | CNOT3 | 81.872 | 2 | 52.97408 | yes | 1.164E-07 | 618.29114 | 86.886871 |
| EGFTFEYR | 8 | O75175 | CCR4-NOT transcription complex subuni | CNOT3 | 81.872 | 2 | 52.97408 | yes | 3.696E-07 | 524.74036 | 81.692825 |
| GKGEVTQQTIQK | 12 | O75177 | Calcium-responsive transactivator OS=H | SS18L1 | 42.991 | 2 | 52.236763 | yes | 1.017E-07 | 658.86206 | 160.78539 |
| APQTAPSAQQQR | 12 | O75177 | Calcium-responsive transactivator OS=H | SS18L1 | 42.991 | 2 | 52.236763 | yes | 9.722E-08 | 641.82855 | 117.03643 |
| TAECTQYQQILHR | 13 | O75177 | Calcium-responsive transactivator OS=H | SS18L1 | 42.991 | 2 | 52.236763 | yes | 9.766E-08 | 824.39905 | 90.842583 |
| GEVTQQTIQK | 10 | O75177 | Calcium-responsive transactivator OS=H | SS18L1 | 42.991 | 2 | 52.236763 | yes | 2.855E-07 | 566.30383 | 67.16674 |
| TEVVSLLLDR | 10 | O75179 | Ankyrin repeat domain-containing proteir | ANKRD17 | 274.258 | 2 | 54.921856 | yes | 9.792E-08 | 572.83228 | 754.95502 |
| KVPVPIGTER | 10 | O75179 | Ankyrin repeat domain-containing proteir | ANKRD17 | 274.258 | 2 | 54.921856 | yes | 1.705E-07 | 548.32965 | 656.9397 |
| KQPSVLVTFPK | 11 | O75179 | Ankyrin repeat domain-containing proteir | ANKRD17 | 274.258 | 2 | 54.921856 | yes | 2.523E-07 | 622.37408 | 482.28818 |
| VSVPSTVISR | 10 | O75179 | Ankyrin repeat domain-containing proteir | ANKRD17 | 274.258 | 2 | 54.921856 | yes | 5.573E-07 | 522.80603 | 391.72916 |
| FCELLIGR | 8 | O75179 | Ankyrin repeat domain-containing proteir | ANKRD17 | 274.258 | 2 | 54.921856 | yes | 1.123E-07 | 504.27078 | 356.92328 |
| QEVVELLLAR | 10 | O75179 | Ankyrin repeat domain-containing proteir | ANKRD17 | 274.258 | 2 | 54.921856 | yes | 9.809E-08 | 585.34808 | 347.11884 |
| KVSVPSTVISR | 11 | O75179 | Ankyrin repeat domain-containing proteir | ANKRD17 | 274.258 | 2 | 54.921856 | yes | 9.82E-08 | 586.85352 | 275.5578 |
| KLEEIEAK | 8 | O75179 | Ankyrin repeat domain-containing proteir | ANKRD17 | 274.258 | 2 | 54.921856 | yes | 9.722E-08 | 480.27402 | 253.5386 |
| LQQVDPVLLK | 10 | O75179 | Ankyrin repeat domain-containing proteir | ANKRD17 | 274.258 | 2 | 54.921856 | yes | 9.722E-08 | 576.85297 | 248.56073 |
| EFTGAHIDIDK | 11 | O75179 | Ankyrin repeat domain-containing proteir | ANKRD17 | 274.258 | 2 | 54.921856 | yes | 1.113E-07 | 623.30914 | 209.27748 |
| PMSDPGVFSQHQA | 16 | O75179 | Ankyrin repeat domain-containing proteir | ANKRD17 | 274.258 | 3 | 54.921856 | yes | 2.585E-07 | 606.27502 | 185.52831 |
| CHLCMESIVQAK | 12 | O75179 | Ankyrin repeat domain-containing proteir | ANKRD17 | 274.258 | 2 | 54.921856 | yes | 3.323E-07 | 738.34381 | 182.86362 |
| ENFELQAAQEK | 11 | O75179 | Ankyrin repeat domain-containing proteir | ANKRD17 | 274.258 | 2 | 54.921856 | yes | 9.722E-08 | 653.81732 | 162.82571 |
| VCDLLLK | 7 | O75179 | Ankyrin repeat domain-containing proteir | ANKRD17 | 274.258 | 2 | 54.921856 | yes | 5.353E-07 | 430.74915 | 143.69507 |

| | | | | | | | | | | |
|---|---|---|---|---|---|---|---|---|---|---|
| QATQLINALIK | 11 | O75179 | Ankyrin repeat domain-containing proteir | ANKRD17 | 274.258 | 2 | 54.921856 | yes | 1.986E-07 | 606.86914 | 134.4007 |
| EVNQFPSDSECMR | 13 | O75179 | Ankyrin repeat domain-containing proteir | ANKRD17 | 274.258 | 2 | 54.921856 | yes | 1.001E-07 | 799.83234 | 124.61307 |
| LTELEQR | 7 | O75179 | Ankyrin repeat domain-containing proteir | ANKRD17 | 274.258 | 2 | 54.921856 | yes | 8.315E-07 | 444.74289 | 124.26485 |
| NASILLEELDLEK | 13 | O75179 | Ankyrin repeat domain-containing proteir | ANKRD17 | 274.258 | 2 | 54.921856 | yes | 9.722E-08 | 743.90363 | 123.28577 |
| APSPAPSSVPLGSE | 15 | O75179 | Ankyrin repeat domain-containing proteir | ANKRD17 | 274.258 | 2 | 54.921856 | yes | 2.331E-07 | 712.375 | 78.743553 |
| NSASVQNSSVAVL | 19 | O75179 | Ankyrin repeat domain-containing proteir | ANKRD17 | 274.258 | 3 | 54.921856 | yes | 9.722E-08 | 652.0166 | 53.112137 |
| DPDKEIDELIPK | 12 | O75179 | Ankyrin repeat domain-containing proteir | ANKRD17 | 274.258 | 2 | 54.921856 | yes | 6.124E-07 | 706.36938 | 51.857765 |
| SLAEACSEGDVNA | 15 | O75179 | Ankyrin repeat domain-containing proteir | ANKRD17 | 274.258 | 2 | 54.921856 | yes | 9.722E-08 | 789.36487 | 45.867142 |
| NKENFELQAAQEK | 13 | O75179 | Ankyrin repeat domain-containing proteir | ANKRD17 | 274.258 | 2 | 54.921856 | yes | 3.835E-07 | 774.88629 | 38.655148 |
| APGFRPPLQR | 10 | O75179 | Ankyrin repeat domain-containing proteir | ANKRD17 | 274.258 | 2 | 54.921856 | yes | 1.164E-07 | 569.82764 | 36.408119 |
| VPVQALPMVVPP( | 29 | O75179 | Ankyrin repeat domain-containing proteir | ANKRD17 | 274.258 | 4 | 54.921856 | yes | 2.379E-07 | 769.43225 | 34.79636 |
| AGHVCTVQFLISK | 13 | O75179 | Ankyrin repeat domain-containing proteir | ANKRD17 | 274.258 | 2 | 54.921856 | yes | 2.309E-05 | 730.38995 | 24.006786 |
| VDWCNPGMGNPM | 15 | O75179 | Ankyrin repeat domain-containing proteir | ANKRD17 | 274.258 | 3 | 54.921856 | yes | 3.198E-06 | 595.26569 | 19.189003 |
| VDYYEVLGVQR | 11 | O75190 | DnaJ homolog subfamily B member 6 OS | DNAJB6 | 36.087 | 2 | 51.904755 | yes | 1.279E-07 | 670.84589 | 1105.3282 |
| VEVEEDGQLK | 10 | O75190 | DnaJ homolog subfamily B member 6 OS | DNAJB6 | 36.087 | 2 | 51.904755 | no | 1.063E-07 | 573.28784 | 753.3562 |
| QVAEAYEVLSDAI | 13 | O75190 | DnaJ homolog subfamily B member 6 OS | DNAJB6 | 36.087 | 2 | 51.904755 | no | 9.722E-08 | 711.85919 | 597.79639 |
| APGPWDPLASAA( | 15 | O75190 | DnaJ homolog subfamily B member 6 OS | DNAJB6 | 36.087 | 2 | 51.904755 | yes | 9.722E-08 | 725.88806 | 506.45901 |
| HAPHCLSEEEGEQ | 17 | O75190 | DnaJ homolog subfamily B member 6 OS | DNAJB6 | 36.087 | 4 | 51.904755 | yes | 2.134E-07 | 512.48163 | 67.984726 |
| HAPHCLSEEEGEQ | 17 | O75190 | DnaJ homolog subfamily B member 6 OS | DNAJB6 | 36.087 | 4 | 51.904755 | yes | 2.886E-07 | 682.97308 | 67.984726 |
| HAPHCLSEEEGEQ | 17 | O75190 | DnaJ homolog subfamily B member 6 OS | DNAJB6 | 36.087 | 3 | 51.904755 | yes | 2.134E-07 | 512.48163 | 67.984726 |
| HAPHCLSEEEGEQ | 17 | O75190 | DnaJ homolog subfamily B member 6 OS | DNAJB6 | 36.087 | 3 | 51.904755 | yes | 2.886E-07 | 682.97308 | 67.984726 |
| NPDDVFR | 7 | O75190 | DnaJ homolog subfamily B member 6 OS | DNAJB6 | 36.087 | 2 | 51.904755 | yes | 1.709E-07 | 431.70633 | 75.497139 |
| MVDYYEVLGVQR | 12 | O75190 | DnaJ homolog subfamily B member 6 OS | DNAJB6 | 36.087 | 2 | 51.904755 | yes | 2.58E-06 | 736.36615 | 17.290491 |
| LDDLVNWAR | 9 | O75251 | NADH dehydrogenase [ubiquinone] iron- | NDUFS7 | 23.564 | 2 | 51.656372 | yes | 9.722E-08 | 551.28802 | 443.20847 |
| QSDVMIVAGTLTN | 14 | O75251 | NADH dehydrogenase [ubiquinone] iron- | NDUFS7 | 23.564 | 2 | 51.656372 | yes | 1.031E-07 | 738.88995 | 265.95081 |
| KVYDQMPEPR | 10 | O75251 | NADH dehydrogenase [ubiquinone] iron- | NDUFS7 | 23.564 | 2 | 51.656372 | yes | 4.554E-07 | 631.81354 | 56.876709 |
| APGFAHLAGLDK | 12 | O75306 | NADH dehydrogenase [ubiquinone] iron- | NDUFS2 | 52.546 | 2 | 56.08079 | yes | 9.722E-08 | 598.82477 | 147.43077 |
| LDELEELLTNNR | 12 | O75306 | NADH dehydrogenase [ubiquinone] iron- | NDUFS2 | 52.546 | 2 | 56.08079 | yes | 1.795E-07 | 729.87537 | 88.569695 |
| VLFGEITR | 8 | O75306 | NADH dehydrogenase [ubiquinone] iron- | NDUFS2 | 52.546 | 2 | 56.08079 | yes | 8.449E-07 | 467.77145 | 85.162392 |
| GEFGVYLVSDGSS | 14 | O75306 | NADH dehydrogenase [ubiquinone] iron- | NDUFS2 | 52.546 | 2 | 56.08079 | yes | 1.174E-07 | 736.85443 | 82.5401 |
| LVMELSGEMVR | 11 | O75306 | NADH dehydrogenase [ubiquinone] iron- | NDUFS2 | 52.546 | 2 | 56.08079 | yes | 1.446E-07 | 632.32544 | 65.015701 |
| MFEFYER | 7 | O75306 | NADH dehydrogenase [ubiquinone] iron- | NDUFS2 | 52.546 | 2 | 56.08079 | yes | 2.68E-07 | 511.22601 | 56.762527 |
| VVFPLELR | 8 | O75317 | Ubiquitin carboxyl-terminal hydrolase 12 | USP12 | 42.858 | 2 | 35.72381 | yes | 7.907E-07 | 486.79749 | 61.808121 |
| QNYIEDLQNGPIR | 13 | O75319 | RNA/RNP complex-1-interacting phosph | DUSP11 | 43.711 | 2 | 43.189384 | yes | 2.693E-07 | 780.39429 | 63.371792 |
| TGYLICR | 7 | O75319 | RNA/RNP complex-1-interacting phosph | DUSP11 | 43.711 | 2 | 43.189384 | yes | 4.324E-07 | 441.72873 | 45.555279 |
| FILEQQER | 8 | O75330 | Hyaluronan mediated motility receptor O | HMMR | 84.101 | 2 | 52.432926 | yes | 9.881E-08 | 531.78253 | 2343.521 |
| LKDENSQLK | 9 | O75330 | Hyaluronan mediated motility receptor O | HMMR | 84.101 | 2 | 52.432926 | yes | 2.895E-07 | 537.79309 | 1822.3871 |
| ELQIDSLLQQEK | 12 | O75330 | Hyaluronan mediated motility receptor O | HMMR | 84.101 | 2 | 52.432926 | yes | 1.006E-07 | 722.38812 | 1404.2811 |
| LENSSLQEK | 9 | O75330 | Hyaluronan mediated motility receptor O | HMMR | 84.101 | 2 | 52.432926 | yes | 2.125E-07 | 524.26947 | 1334.7225 |

| | | | | | | | | | | | |
|---|---|---|---|---|---|---|---|---|---|---|---|
| MLLDLQTK | 8 | O75330 | Hyaluronan mediated motility receptor O! | HMMR | 84.101 | 2 | 52.432926 | yes | 1.45E-07 | 481.27298 | 1197.1913 |
| LDIAQLEENLK | 11 | O75330 | Hyaluronan mediated motility receptor O! | HMMR | 84.101 | 2 | 52.432926 | yes | 9.722E-08 | 643.35352 | 1022.0926 |
| ELSSSLHQK | 9 | O75330 | Hyaluronan mediated motility receptor O! | HMMR | 84.101 | 2 | 52.432926 | yes | 9.722E-08 | 514.77222 | 851.67413 |
| QSLEENIVILSK | 12 | O75330 | Hyaluronan mediated motility receptor O! | HMMR | 84.101 | 2 | 52.432926 | yes | 9.789E-08 | 686.88776 | 805.72058 |
| ITDLQNQLK | 9 | O75330 | Hyaluronan mediated motility receptor O! | HMMR | 84.101 | 2 | 52.432926 | yes | 1.86E-07 | 536.80347 | 724.65552 |
| QLIELTR | 7 | O75330 | Hyaluronan mediated motility receptor O! | HMMR | 84.101 | 2 | 52.432926 | yes | 3.824E-07 | 436.76364 | 720.8692 |
| IQDLETELEK | 10 | O75330 | Hyaluronan mediated motility receptor O! | HMMR | 84.101 | 2 | 52.432926 | yes | 1.365E-07 | 609.31665 | 665.37878 |
| LLYEELYNK | 9 | O75330 | Hyaluronan mediated motility receptor O! | HMMR | 84.101 | 2 | 52.432926 | yes | 1.71E-07 | 592.81354 | 664.11145 |
| ILSLELMK | 8 | O75330 | Hyaluronan mediated motility receptor O! | HMMR | 84.101 | 2 | 52.432926 | yes | 2.68E-07 | 473.78574 | 616.62177 |
| LLGHQNLK | 8 | O75330 | Hyaluronan mediated motility receptor O! | HMMR | 84.101 | 2 | 52.432926 | yes | 9.809E-08 | 461.77707 | 587.56775 |
| QVEDLNVK | 8 | O75330 | Hyaluronan mediated motility receptor O! | HMMR | 84.101 | 2 | 52.432926 | yes | 1.154E-07 | 472.75601 | 549.56647 |
| APMECQESWK | 10 | O75330 | Hyaluronan mediated motility receptor O! | HMMR | 84.101 | 2 | 52.432926 | yes | 9.849E-08 | 633.26813 | 544.83594 |
| YKLDIAQLEENLK | 13 | O75330 | Hyaluronan mediated motility receptor O! | HMMR | 84.101 | 2 | 52.432926 | yes | 3.694E-07 | 788.93268 | 247.74843 |
| YKLDIAQLEENLK | 13 | O75330 | Hyaluronan mediated motility receptor O! | HMMR | 84.101 | 2 | 52.432926 | yes | 6.168E-07 | 526.29089 | 247.74843 |
| YKLDIAQLEENLK | 13 | O75330 | Hyaluronan mediated motility receptor O! | HMMR | 84.101 | 3 | 52.432926 | yes | 3.694E-07 | 788.93268 | 247.74843 |
| YKLDIAQLEENLK | 13 | O75330 | Hyaluronan mediated motility receptor O! | HMMR | 84.101 | 3 | 52.432926 | yes | 6.168E-07 | 526.29089 | 247.74843 |
| EITVSFLQK | 9 | O75330 | Hyaluronan mediated motility receptor O! | HMMR | 84.101 | 2 | 52.432926 | yes | 3.715E-07 | 532.80298 | 484.30936 |
| EKNDEILSLK | 10 | O75330 | Hyaluronan mediated motility receptor O! | HMMR | 84.101 | 2 | 52.432926 | yes | 1.058E-07 | 594.82715 | 441.85782 |
| TSLSANNATLEK | 12 | O75330 | Hyaluronan mediated motility receptor O! | HMMR | 84.101 | 2 | 52.432926 | yes | 2.351E-07 | 624.82513 | 413.15646 |
| QTLDELDK | 8 | O75330 | Hyaluronan mediated motility receptor O! | HMMR | 84.101 | 2 | 52.432926 | yes | 1.427E-07 | 481.24548 | 291.49298 |
| LCSFQEEMVK | 10 | O75330 | Hyaluronan mediated motility receptor O! | HMMR | 84.101 | 2 | 52.432926 | yes | 9.729E-08 | 635.79395 | 265.16342 |
| ENTTAELTEEINK | 13 | O75330 | Hyaluronan mediated motility receptor O! | HMMR | 84.101 | 2 | 52.432926 | yes | 9.722E-08 | 746.3623 | 226.95313 |
| NAEDVQHQILATE | 21 | O75330 | Hyaluronan mediated motility receptor O! | HMMR | 84.101 | 3 | 52.432926 | yes | 1.065E-07 | 811.05432 | 226.20064 |
| ALTASEIEDLK | 11 | O75330 | Hyaluronan mediated motility receptor O! | HMMR | 84.101 | 2 | 52.432926 | yes | 1.014E-06 | 595.31915 | 203.35013 |
| QNLNVDKDTTLPA | 16 | O75330 | Hyaluronan mediated motility receptor O! | HMMR | 84.101 | 3 | 52.432926 | yes | 4.086E-07 | 581.63922 | 167.99313 |
| NLFEEELK | 8 | O75330 | Hyaluronan mediated motility receptor O! | HMMR | 84.101 | 2 | 52.432926 | yes | 2.578E-07 | 511.26367 | 135.01137 |
| QQEEDFRK | 8 | O75330 | Hyaluronan mediated motility receptor O! | HMMR | 84.101 | 2 | 52.432926 | yes | 3.086E-07 | 540.25946 | 132.55705 |
| RIQDLETELEK | 11 | O75330 | Hyaluronan mediated motility receptor O! | HMMR | 84.101 | 2 | 52.432926 | yes | 9.722E-08 | 687.36719 | 115.86874 |
| EKNLFEEELK | 10 | O75330 | Hyaluronan mediated motility receptor O! | HMMR | 84.101 | 2 | 52.432926 | yes | 2.226E-07 | 639.83246 | 110.15855 |
| SSAAHTQATLLLQ | 15 | O75330 | Hyaluronan mediated motility receptor O! | HMMR | 84.101 | 2 | 52.432926 | yes | 9.722E-08 | 799.43085 | 84.070839 |
| EHNENLNAEMQN | 14 | O75330 | Hyaluronan mediated motility receptor O! | HMMR | 84.101 | 2 | 52.432926 | yes | 9.722E-08 | 842.39142 | 63.534431 |
| QTLDELDKLQQK | 12 | O75330 | Hyaluronan mediated motility receptor O! | HMMR | 84.101 | 2 | 52.432926 | yes | 2.669E-07 | 729.89355 | 44.421761 |
| LSDQFHDILIR | 11 | O75340 | Programmed cell death protein 6 OS=Hor | PDCD6 | 21.869 | 2 | 50.94894 | yes | 9.722E-08 | 678.86713 | 351.23715 |
| AGVNFSEFTGVWI | 13 | O75340 | Programmed cell death protein 6 OS=Hor | PDCD6 | 21.869 | 2 | 50.94894 | yes | 2.077E-07 | 721.35913 | 248.51343 |
| QALSGFGYR | 9 | O75340 | Programmed cell death protein 6 OS=Hor | PDCD6 | 21.869 | 2 | 50.94894 | yes | 2.621E-07 | 499.75635 | 203.76643 |
| DNSGMIDKNELK | 12 | O75340 | Programmed cell death protein 6 OS=Hor | PDCD6 | 21.869 | 2 | 50.94894 | yes | 1.354E-07 | 682.32971 | 71.871307 |
| EEAQAEIEQYR | 11 | O75348 | V-type proton ATPase subunit G 1 OS=H | ATP6V1G1 | 13.758 | 2 | 41.705017 | yes | 3.379E-07 | 683.31769 | 74.500687 |
| SSVVALDVDQPGA | 16 | O75362 | Zinc finger protein 217 OS=Homo sapien | ZNF217 | 115.272 | 2 | 34.808582 | yes | 9.373E-07 | 845.9234 | 74.720299 |
| SAVETQANNLICR | 13 | O75362 | Zinc finger protein 217 OS=Homo sapien | ZNF217 | 115.272 | 2 | 34.808582 | yes | 2.583E-06 | 738.36719 | 25.186014 |

| | | | | | | | | | | | |
|---|---|---|---|---|---|---|---|---|---|---|---|
| EFVEAVLELR | 10 | O75367 | Core histone macro-H2A.1 OS=Homo sa | MACROH2 | 39.184 | 2 | 54.540104 | yes | 2.847E-07 | 602.83228 | 466.86478 |
| NCLALADDKK | 10 | O75367 | Core histone macro-H2A.1 OS=Homo sa | MACROH2 | 39.184 | 2 | 54.540104 | yes | 9.819E-08 | 574.29242 | 228.33813 |
| QTAAQLILK | 9 | O75367 | Core histone macro-H2A.1 OS=Homo sa | MACROH2 | 39.184 | 2 | 54.540104 | yes | 1.531E-07 | 493.30566 | 153.29611 |
| AISSYFVSTMSSSII | 15 | O75367 | Core histone macro-H2A.1 OS=Homo sa | MACROH2 | 39.184 | 2 | 54.540104 | yes | 9.722E-08 | 804.40289 | 116.31873 |
| GKLEAIITPPPAK | 13 | O75367 | Core histone macro-H2A.1 OS=Homo sa | MACROH2 | 39.184 | 2 | 54.540104 | yes | 9.722E-08 | 667.90576 | 66.769562 |
| GVTIASGGVLPNIH | 20 | O75367 | Core histone macro-H2A.1 OS=Homo sa | MACROH2 | 39.184 | 3 | 54.540104 | yes | 2.195E-07 | 662.71759 | 46.07188 |
| HILLAVANDEELN | 17 | O75367 | Core histone macro-H2A.1 OS=Homo sa | MACROH2 | 39.184 | 3 | 54.540104 | no | 9.722E-08 | 645.02997 | 42.341263 |
| APSVATVGSICDLI | 16 | O75369 | Filamin-B OS=Homo sapiens OX=9606 ( | FLNB | 278.165 | 2 | 46.52464 | yes | 1.537E-07 | 822.93488 | 68.706818 |
| VVASGPGLEHGK | 12 | O75369 | Filamin-B OS=Homo sapiens OX=9606 ( | FLNB | 278.165 | 2 | 46.52464 | yes | 1.693E-07 | 575.81439 | 36.017387 |
| FVPQEMGVHTVSV | 14 | O75369 | Filamin-B OS=Homo sapiens OX=9606 ( | FLNB | 278.165 | 2 | 46.52464 | yes | 5.43E-07 | 779.40833 | 34.835636 |
| SPFEVQVGPEAGM | 15 | O75369 | Filamin-B OS=Homo sapiens OX=9606 ( | FLNB | 278.165 | 2 | 46.52464 | yes | 3.897E-06 | 802.39288 | 26.755882 |
| SPFTVGVAAPLDL | 15 | O75369 | Filamin-B OS=Homo sapiens OX=9606 ( | FLNB | 278.165 | 2 | 46.52464 | yes | 4.7E-07 | 751.41669 | 25.219313 |
| VVPCLVTPVTGR | 12 | O75369 | Filamin-B OS=Homo sapiens OX=9606 ( | FLNB | 278.165 | 2 | 46.52464 | yes | 3.179E-06 | 649.36847 | 25.121225 |
| MDGTYACSYTPVI | 13 | O75369 | Filamin-B OS=Homo sapiens OX=9606 ( | FLNB | 278.165 | 2 | 46.52464 | yes | 4.707E-06 | 746.82599 | 23.626034 |
| VAVTEGCQPSR | 11 | O75369 | Filamin-B OS=Homo sapiens OX=9606 ( | FLNB | 278.165 | 2 | 46.52464 | yes | 7.854E-07 | 602.29297 | 17.851486 |
| GAGIGGLGITVEGI | 18 | O75369 | Filamin-B OS=Homo sapiens OX=9606 ( | FLNB | 278.165 | 2 | 46.52464 | yes | 4.483E-05 | 814.92816 | 16.944363 |
| LGSAPEVLLVR | 11 | O75382 | Tripartite motif-containing protein 3 OS= | TRIM3 | 80.83 | 2 | 53.312595 | yes | 1.009E-07 | 577.35059 | 816.09106 |
| VLQSQLDTLR | 10 | O75382 | Tripartite motif-containing protein 3 OS= | TRIM3 | 80.83 | 2 | 53.312595 | yes | 9.722E-08 | 586.83533 | 662.73236 |
| WVSIFSPEGK | 10 | O75382 | Tripartite motif-containing protein 3 OS= | TRIM3 | 80.83 | 2 | 53.312595 | yes | 1.003E-07 | 575.3006 | 311.08316 |
| QALVSDLETICGAI | 14 | O75382 | Tripartite motif-containing protein 3 OS= | TRIM3 | 80.83 | 2 | 53.312595 | yes | 1.142E-07 | 752.88745 | 268.45218 |
| VLPCLHTFCER | 11 | O75382 | Tripartite motif-containing protein 3 OS= | TRIM3 | 80.83 | 2 | 53.312595 | no | 8.638E-07 | 716.34723 | 262.285 |
| QFLVCSICLDR | 11 | O75382 | Tripartite motif-containing protein 3 OS= | TRIM3 | 80.83 | 2 | 53.312595 | yes | 1.414E-07 | 705.84705 | 181.08369 |
| DNPIEDELVFR | 11 | O75382 | Tripartite motif-containing protein 3 OS= | TRIM3 | 80.83 | 2 | 53.312595 | yes | 4.217E-07 | 673.83295 | 158.74571 |
| AEITGPDGTR | 10 | O75382 | Tripartite motif-containing protein 3 OS= | TRIM3 | 80.83 | 2 | 53.312595 | yes | 3.826E-07 | 508.754 | 132.50117 |
| PLSCPNHEGK | 10 | O75382 | Tripartite motif-containing protein 3 OS= | TRIM3 | 80.83 | 2 | 53.312595 | yes | 9.742E-08 | 569.7691 | 124.20448 |
| GEFTNLQGVSAAS | 16 | O75382 | Tripartite motif-containing protein 3 OS= | TRIM3 | 80.83 | 2 | 53.312595 | yes | 1.928E-07 | 790.88678 | 88.470749 |
| QALVGQPASLTVT | 15 | O75382 | Tripartite motif-containing protein 3 OS= | TRIM3 | 80.83 | 2 | 53.312595 | yes | 9.722E-08 | 757.43286 | 83.502487 |
| LPVPVVDHK | 9 | O75382 | Tripartite motif-containing protein 3 OS= | TRIM3 | 80.83 | 2 | 53.312595 | yes | 4.477E-07 | 502.30038 | 71.763191 |
| RPSSMYSTGGK | 11 | O75382 | Tripartite motif-containing protein 3 OS= | TRIM3 | 80.83 | 2 | 53.312595 | yes | 4.663E-07 | 585.78223 | 43.683239 |
| CLQNYIPAQSLTLS | 19 | O75382 | Tripartite motif-containing protein 3 OS= | TRIM3 | 80.83 | 3 | 53.312595 | yes | 2.908E-07 | 760.70264 | 39.30793 |
| NEIVVTDFHNHSV | 14 | O75382 | Tripartite motif-containing protein 3 OS= | TRIM3 | 80.83 | 2 | 53.312595 | yes | 9.903E-08 | 819.91534 | 38.426994 |
| NGTYELVYTAR | 11 | O75382 | Tripartite motif-containing protein 3 OS= | TRIM3 | 80.83 | 2 | 53.312595 | yes | 2.943E-07 | 643.82239 | 31.465443 |
| REDSPGPEVQPMD | 14 | O75382 | Tripartite motif-containing protein 3 OS= | TRIM3 | 80.83 | 2 | 53.312595 | yes | 3.208E-07 | 792.86975 | 16.500191 |
| TMENLVELHNK | 11 | O75386 | Tubby-related protein 3 OS=Homo sapier | TULP3 | 49.643 | 2 | 53.704006 | yes | 1.109E-07 | 664.33734 | 164.34099 |
| NDPDYIVMQFGR | 12 | O75386 | Tubby-related protein 3 OS=Homo sapier | TULP3 | 49.643 | 2 | 53.704006 | yes | 9.722E-08 | 727.84027 | 156.83025 |
| TANYLISIDPVDLS | 15 | O75386 | Tubby-related protein 3 OS=Homo sapier | TULP3 | 49.643 | 2 | 53.704006 | yes | 2.56E-07 | 838.94635 | 155.15057 |
| GLVGAAHTR | 9 | O75386 | Tubby-related protein 3 OS=Homo sapier | TULP3 | 49.643 | 2 | 53.704006 | yes | 1.414E-07 | 441.25122 | 69.165253 |
| GLFPTYYMYLEK | 12 | O75386 | Tubby-related protein 3 OS=Homo sapier | TULP3 | 49.643 | 2 | 53.704006 | yes | 9.969E-06 | 762.87579 | 20.984716 |
| IVPNVLLEQGK | 11 | O75390 | Citrate synthase, mitochondrial OS=Hom | CS | 51.713 | 2 | 39.995758 | yes | 4.265E-07 | 605.36371 | 86.279228 |

| | | | | | | | | | | | |
|---|---|---|---|---|---|---|---|---|---|---|---|
| IMVANIEEVLQR | 12 | O75396 | Vesicle-trafficking protein SEC22b OS=I | SEC22B | 24.741 | 2 | 51.901554 | yes | 9.722E-08 | 707.88977 | 235.6088 |
| NLGSINTELQDVQ | 14 | O75396 | Vesicle-trafficking protein SEC22b OS=I | SEC22B | 24.741 | 2 | 51.901554 | yes | 1.332E-07 | 793.91028 | 130.81572 |
| GEALSALDSK | 10 | O75396 | Vesicle-trafficking protein SEC22b OS=I | SEC22B | 24.741 | 2 | 51.901554 | yes | 1.46E-07 | 495.75876 | 86.595207 |
| VLLTMIAR | 8 | O75396 | Vesicle-trafficking protein SEC22b OS=I | SEC22B | 24.741 | 2 | 51.901554 | yes | 4.572E-06 | 458.78604 | 43.204517 |
| STTLDAGNIK | 10 | O75400 | Pre-mRNA-processing factor 40 homolog | PRPF40A | 108.805 | 2 | 51.824417 | yes | 1.108E-07 | 510.27203 | 236.62448 |
| LAFNSLLEK | 9 | O75400 | Pre-mRNA-processing factor 40 homolog | PRPF40A | 108.805 | 2 | 51.824417 | yes | 1.182E-07 | 517.79767 | 204.7056 |
| FLENHEK | 7 | O75400 | Pre-mRNA-processing factor 40 homolog | PRPF40A | 108.805 | 2 | 51.824417 | yes | 2.203E-07 | 458.7298 | 180.71091 |
| EPAFEDITLESER | 13 | O75400 | Pre-mRNA-processing factor 40 homolog | PRPF40A | 108.805 | 2 | 51.824417 | yes | 5.3E-07 | 768.36481 | 177.02142 |
| EDALICFEEHIR | 12 | O75400 | Pre-mRNA-processing factor 40 homolog | PRPF40A | 108.805 | 2 | 51.824417 | no | 9.722E-08 | 766.36414 | 111.54155 |
| QETVADFTPK | 10 | O75400 | Pre-mRNA-processing factor 40 homolog | PRPF40A | 108.805 | 2 | 51.824417 | yes | 7.178E-07 | 568.28516 | 98.830032 |
| FYVEDLK | 7 | O75400 | Pre-mRNA-processing factor 40 homolog | PRPF40A | 108.805 | 2 | 51.824417 | yes | 4.776E-07 | 457.23691 | 38.282578 |
| RVPSNASWEQAM | 13 | O75400 | Pre-mRNA-processing factor 40 homolog | PRPF40A | 108.805 | 2 | 51.824417 | yes | 3.103E-07 | 752.37225 | 8.7647114 |
| HSAAATALPLSHG | 16 | O75420 | GRB10-interacting GYF protein 1 OS=H | GIGYF1 | 114.602 | 3 | 52.66475 | yes | 1.165E-07 | 510.94269 | 233.49265 |
| HSAAATALPLSHG | 16 | O75420 | GRB10-interacting GYF protein 1 OS=H | GIGYF1 | 114.602 | 3 | 52.66475 | yes | 9.722E-08 | 765.9104 | 233.49265 |
| HSAAATALPLSHG | 16 | O75420 | GRB10-interacting GYF protein 1 OS=H | GIGYF1 | 114.602 | 2 | 52.66475 | yes | 1.165E-07 | 510.94269 | 233.49265 |
| HSAAATALPLSHG | 16 | O75420 | GRB10-interacting GYF protein 1 OS=H | GIGYF1 | 114.602 | 2 | 52.66475 | yes | 9.722E-08 | 765.9104 | 233.49265 |
| GGDQNLLPTMSR | 12 | O75420 | GRB10-interacting GYF protein 1 OS=H | GIGYF1 | 114.602 | 2 | 52.66475 | yes | 2.261E-07 | 644.81934 | 265.49231 |
| TLLELQLEGER | 11 | O75420 | GRB10-interacting GYF protein 1 OS=H | GIGYF1 | 114.602 | 2 | 52.66475 | yes | 1.088E-07 | 650.85901 | 258.24652 |
| SIEEGDGAFGR | 11 | O75420 | GRB10-interacting GYF protein 1 OS=H | GIGYF1 | 114.602 | 2 | 52.66475 | yes | 9.722E-08 | 569.26221 | 150.07889 |
| GCDEGFQPLGEVI | 14 | O75420 | GRB10-interacting GYF protein 1 OS=H | GIGYF1 | 114.602 | 2 | 52.66475 | yes | 1.695E-07 | 774.87177 | 143.64323 |
| ENKVPEELQDK | 11 | O75420 | GRB10-interacting GYF protein 1 OS=H | GIGYF1 | 114.602 | 2 | 52.66475 | yes | 1.883E-07 | 664.83826 | 141.54225 |
| SGGGSSGLGLWED | 16 | O75420 | GRB10-interacting GYF protein 1 OS=H | GIGYF1 | 114.602 | 2 | 52.66475 | yes | 2.675E-07 | 774.37042 | 137.16127 |
| EVESPYDVHDYIR | 13 | O75420 | GRB10-interacting GYF protein 1 OS=H | GIGYF1 | 114.602 | 2 | 52.66475 | yes | 1.805E-07 | 811.3783 | 121.10344 |
| EEMLALYVK | 9 | O75420 | GRB10-interacting GYF protein 1 OS=H | GIGYF1 | 114.602 | 2 | 52.66475 | yes | 9.728E-08 | 548.29138 | 116.0972 |
| SSPSLSDSYSHLSG | 15 | O75420 | GRB10-interacting GYF protein 1 OS=H | GIGYF1 | 114.602 | 2 | 52.66475 | yes | 4.106E-07 | 790.37097 | 102.94859 |
| ELTPLPPQEEK | 11 | O75420 | GRB10-interacting GYF protein 1 OS=H | GIGYF1 | 114.602 | 2 | 52.66475 | yes | 1.1E-06 | 640.84027 | 75.403702 |
| NFSLSVNSVAVLR | 13 | O75420 | GRB10-interacting GYF protein 1 OS=H | GIGYF1 | 114.602 | 2 | 52.66475 | yes | 1.347E-07 | 703.39355 | 69.689308 |
| CGFEEGGAGPR | 11 | O75420 | GRB10-interacting GYF protein 1 OS=H | GIGYF1 | 114.602 | 2 | 52.66475 | yes | 1.565E-07 | 568.7431 | 62.945023 |
| QQEIILWR | 8 | O75436 | Vacuolar protein sorting-associated prote | VPS26A | 38.17 | 2 | 50.014282 | yes | 9.934E-08 | 543.30872 | 62.005417 |
| VNLLQIVR | 8 | O75438 | NADH dehydrogenase [ubiquinone] 1 bet | NDUFB1 | 6.962 | 2 | 28.769236 | yes | 2.67E-06 | 477.80838 | 28.136671 |
| TILGPTENIK | 10 | O75439 | Mitochondrial-processing peptidase subu | PMPCB | 54.367 | 2 | 47.907234 | yes | 3.617E-07 | 543.31366 | 216.7614 |
| LCTSVTESEVAR | 12 | O75439 | Mitochondrial-processing peptidase subu | PMPCB | 54.367 | 2 | 47.907234 | yes | 1.375E-07 | 676.32971 | 182.63228 |
| IPIPELEAR | 9 | O75439 | Mitochondrial-processing peptidase subu | PMPCB | 54.367 | 2 | 47.907234 | yes | 1.882E-07 | 519.3031 | 126.13209 |
| DLVDYITTHYK | 11 | O75439 | Mitochondrial-processing peptidase subu | PMPCB | 54.367 | 2 | 47.907234 | yes | 1.234E-07 | 684.3457 | 124.35165 |
| IVLAAAGGVSHDE | 19 | O75439 | Mitochondrial-processing peptidase subu | PMPCB | 54.367 | 3 | 47.907234 | yes | 1.833E-07 | 631.35443 | 109.82944 |
| VTCLESGLR | 9 | O75439 | Mitochondrial-processing peptidase subu | PMPCB | 54.367 | 2 | 47.907234 | yes | 2.564E-07 | 517.76862 | 63.958408 |
| QMLCYNR | 7 | O75439 | Mitochondrial-processing peptidase subu | PMPCB | 54.367 | 2 | 47.907234 | yes | 4.688E-06 | 492.72314 | 57.560127 |
| RIPIPELEAR | 10 | O75439 | Mitochondrial-processing peptidase subu | PMPCB | 54.367 | 2 | 47.907234 | yes | 4.671E-06 | 597.3537 | 19.919865 |
| LPTRPGLNK | 9 | O75446 | Histone deacetylase complex subunit SAI | SAP30 | 23.307 | 2 | 47.043377 | yes | 1.414E-07 | 498.30347 | 26.148865 |

| | | | | | | | | | | | |
|---|---|---|---|---|---|---|---|---|---|---|---|
| IYIPLPSAK | 9 | O75449 | Katanin p60 ATPase-containing subunit A | KATNA1 | 55.965 | 2 | 49.1186 | yes | 5.923E-07 | 501.30515 | 212.39371 |
| TTFFNVSSSTLTSK | 14 | O75449 | Katanin p60 ATPase-containing subunit A | KATNA1 | 55.965 | 2 | 49.1186 | yes | 1.318E-07 | 760.38556 | 186.02678 |
| IEGLTPEEIR | 10 | O75449 | Katanin p60 ATPase-containing subunit A | KATNA1 | 55.965 | 2 | 49.1186 | yes | 2.439E-07 | 578.81403 | 164.36121 |
| WQQVWQEINVEA | 13 | O75449 | Katanin p60 ATPase-containing subunit A | KATNA1 | 55.965 | 2 | 49.1186 | yes | 1.473E-07 | 829.42029 | 114.6927 |
| WDDIADLVEAK | 11 | O75449 | Katanin p60 ATPase-containing subunit A | KATNA1 | 55.965 | 2 | 49.1186 | yes | 1.066E-07 | 637.81677 | 84.105331 |
| SPAAVTEPETNK | 12 | O75449 | Katanin p60 ATPase-containing subunit A | KATNA1 | 55.965 | 2 | 49.1186 | yes | 2.027E-07 | 622.31189 | 73.180618 |
| LLFEMAR | 7 | O75449 | Katanin p60 ATPase-containing subunit A | KATNA1 | 55.965 | 2 | 49.1186 | no | 3.781E-07 | 440.24149 | 36.140141 |
| VDEVPDGAVKPPT | 15 | O75475 | PC4 and SFRS1-interacting protein OS=H | PSIP1 | 60.104 | 2 | 51.914093 | yes | 1.083E-07 | 783.41211 | 1482.9574 |
| VDEVPDGAVKPPT | 15 | O75475 | PC4 and SFRS1-interacting protein OS=H | PSIP1 | 60.104 | 2 | 51.914093 | yes | 2.303E-07 | 522.61053 | 1482.9574 |
| VDEVPDGAVKPPT | 15 | O75475 | PC4 and SFRS1-interacting protein OS=H | PSIP1 | 60.104 | 3 | 51.914093 | yes | 1.083E-07 | 783.41211 | 1482.9574 |
| VDEVPDGAVKPPT | 15 | O75475 | PC4 and SFRS1-interacting protein OS=H | PSIP1 | 60.104 | 3 | 51.914093 | yes | 2.303E-07 | 522.61053 | 1482.9574 |
| DFKPGDLIFAK | 11 | O75475 | PC4 and SFRS1-interacting protein OS=H | PSIP1 | 60.104 | 2 | 51.914093 | yes | 1.673E-07 | 625.84259 | 2390.8982 |
| GFNEGLWEIDNNP | 14 | O75475 | PC4 and SFRS1-interacting protein OS=H | PSIP1 | 60.104 | 2 | 51.914093 | yes | 1.07E-07 | 816.88629 | 2174.1975 |
| GYPHWPAR | 8 | O75475 | PC4 and SFRS1-interacting protein OS=H | PSIP1 | 60.104 | 2 | 51.914093 | no | 3.73E-07 | 492.24594 | 1843.7046 |
| VSQVIMEK | 8 | O75475 | PC4 and SFRS1-interacting protein OS=H | PSIP1 | 60.104 | 2 | 51.914093 | yes | 2.044E-07 | 467.25732 | 579.82092 |
| IDNLDVNR | 8 | O75475 | PC4 and SFRS1-interacting protein OS=H | PSIP1 | 60.104 | 2 | 51.914093 | yes | 2.885E-07 | 479.75125 | 406.48712 |
| HTEMITTLK | 9 | O75475 | PC4 and SFRS1-interacting protein OS=H | PSIP1 | 60.104 | 2 | 51.914093 | yes | 2.273E-07 | 537.28662 | 402.0141 |
| QSNASSDVEVEEK | 13 | O75475 | PC4 and SFRS1-interacting protein OS=H | PSIP1 | 60.104 | 2 | 51.914093 | yes | 9.722E-08 | 711.32318 | 379.97195 |
| LPIFFFGTHETAFL( | 17 | O75475 | PC4 and SFRS1-interacting protein OS=H | PSIP1 | 60.104 | 3 | 51.914093 | yes | 1.087E-07 | 641.34521 | 264.38049 |
| DIFPYSENK | 9 | O75475 | PC4 and SFRS1-interacting protein OS=H | PSIP1 | 60.104 | 2 | 51.914093 | yes | 4.004E-07 | 556.7666 | 251.27451 |
| GRPAATEVK | 9 | O75475 | PC4 and SFRS1-interacting protein OS=H | PSIP1 | 60.104 | 2 | 51.914093 | yes | 8.065E-07 | 464.76416 | 211.92278 |
| AVDITTPK | 8 | O75475 | PC4 and SFRS1-interacting protein OS=H | PSIP1 | 60.104 | 2 | 51.914093 | yes | 1.357E-07 | 422.74237 | 69.720482 |
| QVETEEAGVVTTA | 20 | O75475 | PC4 and SFRS1-interacting protein OS=H | PSIP1 | 60.104 | 3 | 51.914093 | yes | 2.967E-07 | 683.02332 | 45.804653 |
| KGFNEGLWEIDNN | 15 | O75475 | PC4 and SFRS1-interacting protein OS=H | PSIP1 | 60.104 | 3 | 51.914093 | yes | 3.152E-06 | 587.62494 | 22.933475 |
| SVQTTLQTDEVK | 12 | O75477 | Erlin-1 OS=Homo sapiens OX=9606 GN= | ERLIN1 | 39.172 | 2 | 55.022003 | no | 9.722E-08 | 674.85138 | 93.19838 |
| ISEIEDAAFLAR | 12 | O75477 | Erlin-1 OS=Homo sapiens OX=9606 GN= | ERLIN1 | 39.172 | 2 | 55.022003 | no | 9.722E-08 | 667.8512 | 80.443436 |
| DLNLMAPGLTIQA | 15 | O75477 | Erlin-1 OS=Homo sapiens OX=9606 GN= | ERLIN1 | 39.172 | 2 | 55.022003 | yes | 1.003E-07 | 806.448 | 73.827721 |
| EALEPSGENVIQNI | 14 | O75477 | Erlin-1 OS=Homo sapiens OX=9606 GN= | ERLIN1 | 39.172 | 2 | 55.022003 | yes | 2.151E-06 | 764.38611 | 59.660324 |
| SLVDLTAVDVPTR | 13 | O75489 | NADH dehydrogenase [ubiquinone] iron- | NDUFS3 | 30.242 | 2 | 51.340527 | yes | 9.722E-08 | 693.38538 | 249.33385 |
| DFPLSGYVELR | 11 | O75489 | NADH dehydrogenase [ubiquinone] iron- | NDUFS3 | 30.242 | 2 | 51.340527 | yes | 1.022E-07 | 648.33514 | 103.9082 |
| DAEDALHNLDR | 11 | O75494 | Serine/arginine-rich splicing factor 10 OS | SRSF10 | 31.301 | 2 | 51.45538 | yes | 1.017E-07 | 634.79694 | 3161.5227 |
| GFAYVQFEDVR | 11 | O75494 | Serine/arginine-rich splicing factor 10 OS | SRSF10 | 31.301 | 2 | 51.45538 | no | 9.722E-08 | 665.82495 | 1136.9133 |
| QIEIQFAQGDR | 11 | O75494 | Serine/arginine-rich splicing factor 10 OS | SRSF10 | 31.301 | 2 | 51.45538 | no | 9.722E-08 | 652.83331 | 852.06006 |
| SFDYNYR | 7 | O75494 | Serine/arginine-rich splicing factor 10 OS | SRSF10 | 31.301 | 2 | 51.45538 | yes | 2.024E-07 | 482.71161 | 426.58298 |
| YLRPPNTSLFVR | 12 | O75494 | Serine/arginine-rich splicing factor 10 OS | SRSF10 | 31.301 | 2 | 51.45538 | yes | 1.497E-07 | 731.91187 | 324.95691 |
| MIQPSASGSLVGR | 13 | O75496 | Geminin OS=Homo sapiens OX=9606 Gl | GMNN | 23.566 | 2 | 44.59276 | yes | 2.138E-07 | 651.84534 | 160.71582 |
| NLGGVTQESFDLM | 15 | O75496 | Geminin OS=Homo sapiens OX=9606 Gl | GMNN | 23.566 | 2 | 44.59276 | yes | 4.357E-07 | 826.42163 | 14.032823 |
| ATEMLIFGK | 9 | O75521 | Enoyl-CoA delta isomerase 2 OS=Homo s | ECI2 | 43.586 | 2 | 27.296558 | yes | 2.979E-06 | 505.27298 | 14.014915 |
| WAQEDLLEEQK | 11 | O75528 | Transcriptional adapter 3 OS=Homo sapi( | TADA3 | 48.903 | 2 | 48.853306 | yes | 2.356E-06 | 694.83826 | 186.23265 |

| | | | | | | | | | | | |
|---|---|---|---|---|---|---|---|---|---|---|---|
| MADNEVMDAFR | 11 | O75528 | Transcriptional adapter 3 OS=Homo sapi | TADA3 | 48.903 | 2 | 48.853306 | yes | 1.095E-07 | 649.77881 | 111.64786 |
| GLMGPLTELDTK | 12 | O75528 | Transcriptional adapter 3 OS=Homo sapi | TADA3 | 48.903 | 2 | 48.853306 | yes | 2.026E-07 | 637.83667 | 78.92112 |
| ESGADGASTSPR | 12 | O75528 | Transcriptional adapter 3 OS=Homo sapi | TADA3 | 48.903 | 2 | 48.853306 | yes | 2.461E-06 | 567.7547 | 30.392931 |
| LLAAGFDNSCIK | 12 | O75529 | TAF5-like RNA polymerase II p300/CBP | TAF5L | 66.156 | 2 | 51.388634 | yes | 3.258E-07 | 654.83447 | 131.16183 |
| TDYQLYASGSSSR | 13 | O75529 | TAF5-like RNA polymerase II p300/CBP | TAF5L | 66.156 | 2 | 51.388634 | yes | 9.722E-08 | 717.82843 | 69.858368 |
| LQEDSYNYLIR | 11 | O75529 | TAF5-like RNA polymerase II p300/CBP | TAF5L | 66.156 | 2 | 51.388634 | yes | 1.34E-07 | 707.35406 | 52.47752 |
| GHCGPVYSTR | 10 | O75529 | TAF5-like RNA polymerase II p300/CBP | TAF5L | 66.156 | 2 | 51.388634 | yes | 2.376E-06 | 567.26147 | 46.040073 |
| YLASAGEDQR | 10 | O75529 | TAF5-like RNA polymerase II p300/CBP | TAF5L | 66.156 | 2 | 51.388634 | yes | 4.212E-06 | 555.26471 | 36.862877 |
| EGDPLVFATVGSN | 14 | O75530 | Polycomb protein EED OS=Homo sapien | EED | 50.198 | 2 | 53.28866 | yes | 1.117E-07 | 731.37024 | 2299.7429 |
| MLALGNQVGK | 10 | O75530 | Polycomb protein EED OS=Homo sapien | EED | 50.198 | 2 | 53.28866 | yes | 1.073E-07 | 515.78931 | 1062.0282 |
| IHFPDFSTR | 9 | O75530 | Polycomb protein EED OS=Homo sapien | EED | 50.198 | 2 | 53.28866 | yes | 3.647E-07 | 560.28271 | 889.37427 |
| DEVLSADYDLLGE | 14 | O75530 | Polycomb protein EED OS=Homo sapien | EED | 50.198 | 2 | 53.28866 | yes | 9.722E-08 | 783.88031 | 580.00189 |
| IINPITMQCIK | 11 | O75530 | Polycomb protein EED OS=Homo sapien | EED | 50.198 | 2 | 53.28866 | yes | 1.223E-07 | 665.86469 | 557.46741 |
| EVSTAPAGTDMPA | 15 | O75530 | Polycomb protein EED OS=Homo sapien | EED | 50.198 | 2 | 53.28866 | yes | 1.165E-07 | 723.35065 | 446.61371 |
| IKPSESNVTILGR | 13 | O75530 | Polycomb protein EED OS=Homo sapien | EED | 50.198 | 2 | 53.28866 | yes | 1.475E-07 | 707.40662 | 216.32874 |
| IKPSESNVTILGR | 13 | O75530 | Polycomb protein EED OS=Homo sapien | EED | 50.198 | 2 | 53.28866 | yes | 2.534E-06 | 471.94019 | 216.32874 |
| IKPSESNVTILGR | 13 | O75530 | Polycomb protein EED OS=Homo sapien | EED | 50.198 | 3 | 53.28866 | yes | 1.475E-07 | 707.40662 | 216.32874 |
| IKPSESNVTILGR | 13 | O75530 | Polycomb protein EED OS=Homo sapien | EED | 50.198 | 3 | 53.28866 | yes | 2.534E-06 | 471.94019 | 216.32874 |
| WLGDLILSK | 9 | O75530 | Polycomb protein EED OS=Homo sapien | EED | 50.198 | 2 | 53.28866 | yes | 3.132E-06 | 522.80804 | 331.73474 |
| LYVWDLEVEDPHI | 13 | O75530 | Polycomb protein EED OS=Homo sapien | EED | 50.198 | 2 | 53.28866 | yes | 1.064E-07 | 821.90924 | 303.05219 |
| HYVGHGNAINELK | 13 | O75530 | Polycomb protein EED OS=Homo sapien | EED | 50.198 | 2 | 53.28866 | yes | 9.722E-08 | 726.37311 | 82.637695 |
| HYVGHGNAINELK | 13 | O75530 | Polycomb protein EED OS=Homo sapien | EED | 50.198 | 2 | 53.28866 | yes | 1.959E-07 | 484.5845 | 82.637695 |
| HYVGHGNAINELK | 13 | O75530 | Polycomb protein EED OS=Homo sapien | EED | 50.198 | 3 | 53.28866 | yes | 9.722E-08 | 726.37311 | 82.637695 |
| HYVGHGNAINELK | 13 | O75530 | Polycomb protein EED OS=Homo sapien | EED | 50.198 | 3 | 53.28866 | yes | 1.959E-07 | 484.5845 | 82.637695 |
| FDYSQCDIWYMR | 12 | O75530 | Polycomb protein EED OS=Homo sapien | EED | 50.198 | 2 | 53.28866 | yes | 9.722E-08 | 842.35016 | 160.58844 |
| ESYDYNPNK | 9 | O75530 | Polycomb protein EED OS=Homo sapien | EED | 50.198 | 2 | 53.28866 | yes | 9.893E-08 | 565.24347 | 123.91424 |
| VTLYECHSQGEIR | 13 | O75530 | Polycomb protein EED OS=Homo sapien | EED | 50.198 | 2 | 53.28866 | yes | 9.808E-08 | 796.38031 | 120.71136 |
| FSMDFWQK | 8 | O75530 | Polycomb protein EED OS=Homo sapien | EED | 50.198 | 2 | 53.28866 | yes | 2.284E-07 | 544.74713 | 120.14795 |
| SCENAIVCWKPGK | 13 | O75530 | Polycomb protein EED OS=Homo sapien | EED | 50.198 | 2 | 53.28866 | yes | 2.492E-07 | 774.86853 | 112.10118 |
| TNRPFISQK | 9 | O75530 | Polycomb protein EED OS=Homo sapien | EED | 50.198 | 2 | 53.28866 | yes | 1.151E-07 | 545.80383 | 98.214348 |
| SCENAIVCWK | 10 | O75530 | Polycomb protein EED OS=Homo sapien | EED | 50.198 | 2 | 53.28866 | yes | 9.722E-08 | 633.78394 | 94.967781 |
| NYVDCVR | 7 | O75530 | Polycomb protein EED OS=Homo sapien | EED | 50.198 | 2 | 53.28866 | yes | 4.718E-07 | 463.21344 | 70.364098 |
| IMSCGMDHSLK | 11 | O75530 | Polycomb protein EED OS=Homo sapien | EED | 50.198 | 2 | 53.28866 | yes | 4.507E-07 | 639.78558 | 67.830589 |
| DPNLLLSVSK | 10 | O75530 | Polycomb protein EED OS=Homo sapien | EED | 50.198 | 2 | 53.28866 | yes | 5.912E-07 | 543.31366 | 33.789845 |
| KDEDLFR | 7 | O75531 | Barrier-to-autointegration factor OS=Hon | BANF1 | 10.059 | 2 | 54.38556 | yes | 2.221E-07 | 461.73508 | 1214.4001 |
| DCFGCLR | 7 | O75531 | Barrier-to-autointegration factor OS=Hon | BANF1 | 10.059 | 2 | 54.38556 | yes | 1.581E-07 | 464.19421 | 1107.6815 |
| DFVAEPMGEKPV( | 24 | O75531 | Barrier-to-autointegration factor OS=Hon | BANF1 | 10.059 | 3 | 54.38556 | yes | 2.006E-07 | 800.75415 | 412.52502 |
| HRDFVAEPMGEKI | 26 | O75531 | Barrier-to-autointegration factor OS=Hon | BANF1 | 10.059 | 4 | 54.38556 | yes | 1.157E-07 | 674.10742 | 265.73386 |
| PVGSLAGIGEVLGI | 14 | O75531 | Barrier-to-autointegration factor OS=Hon | BANF1 | 10.059 | 2 | 54.38556 | yes | 9.722E-08 | 648.8797 | 192.4894 |

| | | | | | | | | | | |
|---|---|---|---|---|---|---|---|---|---|---|
| EWLKDTCGANAK | 12 | O75531 | Barrier-to-autointegration factor OS=Hon | BANF1 | 10.059 | 2 | 54.38556 | yes | 2.748E-07 | 696.83246 | 44.110138 |
| AIVNVIGMHK | 10 | O75533 | Splicing factor 3B subunit 1 OS=Homo sa | SF3B1 | 145.831 | 2 | 56.280064 | yes | 1.003E-07 | 541.31299 | 507.72089 |
| AIVNVIGMHK | 10 | O75533 | Splicing factor 3B subunit 1 OS=Homo sa | SF3B1 | 145.831 | 2 | 56.280064 | yes | 9.882E-08 | 549.31042 | 507.72089 |
| VQENCIDLVGR | 11 | O75533 | Splicing factor 3B subunit 1 OS=Homo sa | SF3B1 | 145.831 | 2 | 56.280064 | yes | 9.927E-08 | 651.82715 | 970.14648 |
| GGDSIGETPTPGAS | 15 | O75533 | Splicing factor 3B subunit 1 OS=Homo sa | SF3B1 | 145.831 | 2 | 56.280064 | yes | 9.927E-08 | 687.33081 | 731.60834 |
| VLPPPAGYVPIR | 12 | O75533 | Splicing factor 3B subunit 1 OS=Homo sa | SF3B1 | 145.831 | 2 | 56.280064 | yes | 1.114E-06 | 639.88208 | 698.72797 |
| QQAADLISR | 9 | O75533 | Splicing factor 3B subunit 1 OS=Homo sa | SF3B1 | 145.831 | 2 | 56.280064 | yes | 2.565E-07 | 501.27237 | 671.96545 |
| QLVDTTVELANK | 12 | O75533 | Splicing factor 3B subunit 1 OS=Homo sa | SF3B1 | 145.831 | 2 | 56.280064 | yes | 1.423E-07 | 665.86426 | 531.45117 |
| DTPGHGSGWAETI | 14 | O75533 | Splicing factor 3B subunit 1 OS=Homo sa | SF3B1 | 145.831 | 2 | 56.280064 | yes | 4.715E-07 | 734.33423 | 457.60843 |
| THEDIEAQIR | 10 | O75533 | Splicing factor 3B subunit 1 OS=Homo sa | SF3B1 | 145.831 | 2 | 56.280064 | yes | 9.722E-08 | 606.30438 | 452.22504 |
| ATVNTFGYIAK | 11 | O75533 | Splicing factor 3B subunit 1 OS=Homo sa | SF3B1 | 145.831 | 2 | 56.280064 | yes | 9.722E-08 | 592.81915 | 403.61411 |
| IVDDLKDEAEQYR | 13 | O75533 | Splicing factor 3B subunit 1 OS=Homo sa | SF3B1 | 145.831 | 2 | 56.280064 | yes | 9.935E-08 | 797.39136 | 188.83405 |
| IVDDLKDEAEQYR | 13 | O75533 | Splicing factor 3B subunit 1 OS=Homo sa | SF3B1 | 145.831 | 2 | 56.280064 | yes | 2.72E-07 | 531.92999 | 188.83405 |
| IVDDLKDEAEQYR | 13 | O75533 | Splicing factor 3B subunit 1 OS=Homo sa | SF3B1 | 145.831 | 3 | 56.280064 | yes | 9.935E-08 | 797.39136 | 188.83405 |
| IVDDLKDEAEQYR | 13 | O75533 | Splicing factor 3B subunit 1 OS=Homo sa | SF3B1 | 145.831 | 3 | 56.280064 | yes | 2.72E-07 | 531.92999 | 188.83405 |
| IADREDEYK | 9 | O75533 | Splicing factor 3B subunit 1 OS=Homo sa | SF3B1 | 145.831 | 2 | 56.280064 | yes | 4.978E-07 | 569.7724 | 249.93193 |
| EFQSPDEEMKK | 11 | O75533 | Splicing factor 3B subunit 1 OS=Homo sa | SF3B1 | 145.831 | 2 | 56.280064 | yes | 9.722E-08 | 684.31104 | 230.29057 |
| IMGNLGAADIDHK | 13 | O75533 | Splicing factor 3B subunit 1 OS=Homo sa | SF3B1 | 145.831 | 2 | 56.280064 | yes | 1.318E-07 | 677.84283 | 95.835732 |
| IMGNLGAADIDHK | 13 | O75533 | Splicing factor 3B subunit 1 OS=Homo sa | SF3B1 | 145.831 | 2 | 56.280064 | yes | 3.363E-07 | 452.23096 | 95.835732 |
| IMGNLGAADIDHK | 13 | O75533 | Splicing factor 3B subunit 1 OS=Homo sa | SF3B1 | 145.831 | 3 | 56.280064 | yes | 1.318E-07 | 677.84283 | 95.835732 |
| IMGNLGAADIDHK | 13 | O75533 | Splicing factor 3B subunit 1 OS=Homo sa | SF3B1 | 145.831 | 3 | 56.280064 | yes | 3.363E-07 | 452.23096 | 95.835732 |
| IYNSIYIGSQDALIA | 19 | O75533 | Splicing factor 3B subunit 1 OS=Homo sa | SF3B1 | 145.831 | 3 | 56.280064 | yes | 9.722E-08 | 732.04791 | 191.32515 |
| TMIISPER | 8 | O75533 | Splicing factor 3B subunit 1 OS=Homo sa | SF3B1 | 145.831 | 2 | 56.280064 | yes | 1.475E-07 | 473.75494 | 184.04932 |
| KLTATPTPLGGMT | 22 | O75533 | Splicing factor 3B subunit 1 OS=Homo sa | SF3B1 | 145.831 | 3 | 56.280064 | yes | 2.003E-07 | 797.05945 | 85.284981 |
| KLTATPTPLGGMT | 22 | O75533 | Splicing factor 3B subunit 1 OS=Homo sa | SF3B1 | 145.831 | 3 | 56.280064 | yes | 3.599E-07 | 598.04645 | 85.284981 |
| KLTATPTPLGGMT | 22 | O75533 | Splicing factor 3B subunit 1 OS=Homo sa | SF3B1 | 145.831 | 4 | 56.280064 | yes | 2.003E-07 | 797.05945 | 85.284981 |
| KLTATPTPLGGMT | 22 | O75533 | Splicing factor 3B subunit 1 OS=Homo sa | SF3B1 | 145.831 | 4 | 56.280064 | yes | 3.599E-07 | 598.04645 | 85.284981 |
| PDIDNMDEYVR | 11 | O75533 | Splicing factor 3B subunit 1 OS=Homo sa | SF3B1 | 145.831 | 2 | 56.280064 | yes | 9.722E-08 | 683.80084 | 139.67537 |
| VPELNVQNGVLK | 12 | O75533 | Splicing factor 3B subunit 1 OS=Homo sa | SF3B1 | 145.831 | 2 | 56.280064 | yes | 9.722E-08 | 655.37738 | 132.1338 |
| TEILPPFFK | 9 | O75533 | Splicing factor 3B subunit 1 OS=Homo sa | SF3B1 | 145.831 | 2 | 56.280064 | yes | 1.515E-07 | 546.31042 | 120.95129 |
| LDPFADGGK | 9 | O75533 | Splicing factor 3B subunit 1 OS=Homo sa | SF3B1 | 145.831 | 2 | 56.280064 | yes | 3.654E-07 | 460.22961 | 105.12016 |
| TYMDVMR | 7 | O75533 | Splicing factor 3B subunit 1 OS=Homo sa | SF3B1 | 145.831 | 2 | 56.280064 | yes | 1.107E-07 | 458.20679 | 90.020546 |
| IYNDDKNTYIR | 11 | O75533 | Splicing factor 3B subunit 1 OS=Homo sa | SF3B1 | 145.831 | 2 | 56.280064 | yes | 2.522E-07 | 707.85168 | 71.906288 |
| QCCGTDGVEANYI | 14 | O75533 | Splicing factor 3B subunit 1 OS=Homo sa | SF3B1 | 145.831 | 2 | 56.280064 | yes | 9.722E-08 | 807.84796 | 71.792068 |
| WDETPGR | 7 | O75533 | Splicing factor 3B subunit 1 OS=Homo sa | SF3B1 | 145.831 | 2 | 56.280064 | yes | 1.318E-07 | 430.69849 | 49.691692 |
| WDQTADQTPGATI | 14 | O75533 | Splicing factor 3B subunit 1 OS=Homo sa | SF3B1 | 145.831 | 2 | 56.280064 | yes | 3.324E-07 | 758.35736 | 48.783676 |
| KLSSWDQAETPGH | 18 | O75533 | Splicing factor 3B subunit 1 OS=Homo sa | SF3B1 | 145.831 | 4 | 56.280064 | yes | 3.599E-07 | 503.25644 | 35.666775 |
| EVMLILIR | 8 | O75533 | Splicing factor 3B subunit 1 OS=Homo sa | SF3B1 | 145.831 | 2 | 56.280064 | yes | 9.968E-08 | 493.80698 | 26.174854 |
| MLQYCLQGLFHPA | 14 | O75533 | Splicing factor 3B subunit 1 OS=Homo sa | SF3B1 | 145.831 | 3 | 56.280064 | yes | 3.387E-05 | 578.62531 | 14.627187 |

| | | | | | | | | | | | |
|---|---|---|---|---|---|---|---|---|---|---|---|
| EFQSPDEEMK | 10 | O75533 | Splicing factor 3B subunit 1 OS=Homo sa | SF3B1 | 145.831 | 2 | 56.280064 | yes | 2.975E-06 | 620.26355 | 10.421307 |
| VTLLEGDHVR | 10 | O75534 | Cold shock domain-containing protein E1 | CSDE1 | 88.885 | 2 | 52.250637 | yes | 1.23E-07 | 569.81439 | 8093.3594 |
| LLTSYGFIQCSER | 13 | O75534 | Cold shock domain-containing protein E1 | CSDE1 | 88.885 | 2 | 52.250637 | yes | 1.075E-07 | 787.38757 | 7127.0947 |
| EMGVIAAMR | 9 | O75534 | Cold shock domain-containing protein E1 | CSDE1 | 88.885 | 2 | 52.250637 | yes | 1.367E-07 | 489.24899 | 2512.23 |
| EMGVIAAMR | 9 | O75534 | Cold shock domain-containing protein E1 | CSDE1 | 88.885 | 2 | 52.250637 | yes | 3.15E-07 | 497.24643 | 2512.23 |
| DNFGFIETANHDK | 13 | O75534 | Cold shock domain-containing protein E1 | CSDE1 | 88.885 | 2 | 52.250637 | yes | 1.078E-07 | 754.34424 | 4899.186 |
| VGDDVEFEVSSDR | 13 | O75534 | Cold shock domain-containing protein E1 | CSDE1 | 88.885 | 2 | 52.250637 | yes | 1.059E-07 | 727.32568 | 4616.3403 |
| GEVYPFGIVGMAN | 14 | O75534 | Cold shock domain-containing protein E1 | CSDE1 | 88.885 | 2 | 52.250637 | yes | 1.339E-07 | 741.37646 | 4454.4536 |
| IKQEILPEER | 10 | O75534 | Cold shock domain-containing protein E1 | CSDE1 | 88.885 | 2 | 52.250637 | yes | 9.722E-08 | 627.85626 | 4275.231 |
| DQFGFINYEVGDS | 14 | O75534 | Cold shock domain-containing protein E1 | CSDE1 | 88.885 | 2 | 52.250637 | yes | 1.535E-07 | 809.8728 | 3862.4221 |
| NITLDDASAPR | 11 | O75534 | Cold shock domain-containing protein E1 | CSDE1 | 88.885 | 2 | 52.250637 | yes | 9.722E-08 | 586.79895 | 3612.4141 |
| CQGVVCAMK | 9 | O75534 | Cold shock domain-containing protein E1 | CSDE1 | 88.885 | 2 | 52.250637 | yes | 1.154E-07 | 526.73792 | 2270.2332 |
| EIFFHYSEFK | 10 | O75534 | Cold shock domain-containing protein E1 | CSDE1 | 88.885 | 2 | 52.250637 | yes | 1.314E-07 | 673.8244 | 1096.9138 |
| EIFFHYSEFK | 10 | O75534 | Cold shock domain-containing protein E1 | CSDE1 | 88.885 | 2 | 52.250637 | yes | 6.687E-07 | 449.55203 | 1096.9138 |
| EIFFHYSEFK | 10 | O75534 | Cold shock domain-containing protein E1 | CSDE1 | 88.885 | 3 | 52.250637 | yes | 1.314E-07 | 673.8244 | 1096.9138 |
| EIFFHYSEFK | 10 | O75534 | Cold shock domain-containing protein E1 | CSDE1 | 88.885 | 3 | 52.250637 | yes | 6.687E-07 | 449.55203 | 1096.9138 |
| INFVIDNNK | 9 | O75534 | Cold shock domain-containing protein E1 | CSDE1 | 88.885 | 2 | 52.250637 | yes | 1.24E-07 | 538.79041 | 2064.3079 |
| ELPFGDKDTK | 10 | O75534 | Cold shock domain-containing protein E1 | CSDE1 | 88.885 | 2 | 52.250637 | yes | 1.311E-07 | 575.29297 | 1925.265 |
| AVAAPRPDR | 9 | O75534 | Cold shock domain-containing protein E1 | CSDE1 | 88.885 | 2 | 52.250637 | yes | 3.597E-07 | 476.76978 | 1860.5631 |
| EAFGFIER | 8 | O75534 | Cold shock domain-containing protein E1 | CSDE1 | 88.885 | 2 | 52.250637 | yes | 1.369E-07 | 484.74545 | 1756.7773 |
| IKVDFVIPK | 9 | O75534 | Cold shock domain-containing protein E1 | CSDE1 | 88.885 | 2 | 52.250637 | yes | 1.236E-07 | 529.83405 | 1640.1777 |
| LLGYVATLK | 9 | O75534 | Cold shock domain-containing protein E1 | CSDE1 | 88.885 | 2 | 52.250637 | yes | 1.408E-07 | 489.30515 | 1492.9082 |
| EAEDGIIAYDDCG | 15 | O75534 | Cold shock domain-containing protein E1 | CSDE1 | 88.885 | 2 | 52.250637 | yes | 1.318E-07 | 827.86688 | 1326.3064 |
| QEILPEER | 8 | O75534 | Cold shock domain-containing protein E1 | CSDE1 | 88.885 | 2 | 52.250637 | yes | 2.33E-07 | 507.26672 | 1256.9476 |
| KLFFHVK | 7 | O75534 | Cold shock domain-containing protein E1 | CSDE1 | 88.885 | 2 | 52.250637 | yes | 1.858E-07 | 459.78162 | 740.68579 |
| NQNDPLPGR | 9 | O75534 | Cold shock domain-containing protein E1 | CSDE1 | 88.885 | 2 | 52.250637 | yes | 1.488E-07 | 505.75433 | 621.55707 |
| FNISTDR | 7 | O75534 | Cold shock domain-containing protein E1 | CSDE1 | 88.885 | 2 | 52.250637 | yes | 1.144E-07 | 426.71414 | 441.39508 |
| LTIAFQAK | 8 | O75534 | Cold shock domain-containing protein E1 | CSDE1 | 88.885 | 2 | 52.250637 | yes | 9.774E-08 | 446.26855 | 401.11749 |
| GTVSFHSHSDHR | 12 | O75534 | Cold shock domain-containing protein E1 | CSDE1 | 88.885 | 2 | 52.250637 | yes | 2.521E-07 | 683.81598 | 392.85745 |
| DVEGSTSPQIGDK | 13 | O75534 | Cold shock domain-containing protein E1 | CSDE1 | 88.885 | 2 | 52.250637 | yes | 9.904E-08 | 666.8175 | 148.54723 |
| MNGQVVCAVPHN | 16 | O75534 | Cold shock domain-containing protein E1 | CSDE1 | 88.885 | 3 | 52.250637 | yes | 5.467E-07 | 594.95905 | 145.54778 |
| EATFSNPK | 8 | O75534 | Cold shock domain-containing protein E1 | CSDE1 | 88.885 | 2 | 52.250637 | yes | 2.327E-07 | 447.2218 | 131.60646 |
| VEFSISDK | 8 | O75534 | Cold shock domain-containing protein E1 | CSDE1 | 88.885 | 2 | 52.250637 | yes | 1.18E-07 | 462.73727 | 112.06628 |
| VDFVIPK | 7 | O75534 | Cold shock domain-containing protein E1 | CSDE1 | 88.885 | 2 | 52.250637 | yes | 2.98E-07 | 409.24454 | 81.410156 |
| RLLGYVATLK | 10 | O75534 | Cold shock domain-containing protein E1 | CSDE1 | 88.885 | 2 | 52.250637 | yes | 1.803E-07 | 567.35571 | 57.548901 |
| VGDDVEFEVSSDR | 14 | O75534 | Cold shock domain-containing protein E1 | CSDE1 | 88.885 | 2 | 52.250637 | yes | 1.724E-07 | 805.37628 | 45.761147 |
| CSACNVWR | 8 | O75534 | Cold shock domain-containing protein E1 | CSDE1 | 88.885 | 2 | 52.250637 | yes | 4.307E-07 | 526.72363 | 40.78606 |
| QRPGQQVATCVR | 12 | O75534 | Cold shock domain-containing protein E1 | CSDE1 | 88.885 | 2 | 52.250637 | yes | 2.422E-07 | 700.36481 | 27.16185 |
| AAEAAINILK | 10 | O75569 | Interferon-inducible double-stranded RNA | PRKRA | 34.405 | 2 | 49.96428 | yes | 3.133E-07 | 507.30313 | 2266.6411 |

| | | | | | | | | | | | |
|---|---|---|---|---|---|---|---|---|---|---|---|
| VTVGDITCTGEGT: | 15 | O75569 | Interferon-inducible double-stranded RN/ | PRKRA | 34.405 | 2 | 49.96428 | yes | 1E-07 | 762.86414 | 1650.3861 |
| SDVQIHVPTFTFR | 13 | O75569 | Interferon-inducible double-stranded RN/ | PRKRA | 34.405 | 2 | 49.96428 | yes | 1.003E-07 | 773.90424 | 909.26526 |
| EDSGTFSLGK | 10 | O75569 | Interferon-inducible double-stranded RN/ | PRKRA | 34.405 | 2 | 49.96428 | yes | 1.073E-07 | 520.74835 | 699.6062 |
| TPIQVLHEYGMK | 12 | O75569 | Interferon-inducible double-stranded RN/ | PRKRA | 34.405 | 2 | 49.96428 | yes | 9.967E-08 | 708.37122 | 259.99405 |
| TPIQVLHEYGMK | 12 | O75569 | Interferon-inducible double-stranded RN/ | PRKRA | 34.405 | 2 | 49.96428 | yes | 1.079E-07 | 472.58322 | 259.99405 |
| TPIQVLHEYGMK | 12 | O75569 | Interferon-inducible double-stranded RN/ | PRKRA | 34.405 | 3 | 49.96428 | yes | 9.967E-08 | 708.37122 | 259.99405 |
| TPIQVLHEYGMK | 12 | O75569 | Interferon-inducible double-stranded RN/ | PRKRA | 34.405 | 3 | 49.96428 | yes | 1.079E-07 | 472.58322 | 259.99405 |
| LPEYTLSQEGGPAI | 15 | O75569 | Interferon-inducible double-stranded RN/ | PRKRA | 34.405 | 2 | 49.96428 | yes | 1.316E-07 | 813.90973 | 209.1254 |
| LPEYTLSQEGGPAI | 15 | O75569 | Interferon-inducible double-stranded RN/ | PRKRA | 34.405 | 2 | 49.96428 | yes | 3.194E-07 | 542.94226 | 209.1254 |
| LPEYTLSQEGGPAI | 15 | O75569 | Interferon-inducible double-stranded RN/ | PRKRA | 34.405 | 3 | 49.96428 | yes | 1.316E-07 | 813.90973 | 209.1254 |
| LPEYTLSQEGGPAI | 15 | O75569 | Interferon-inducible double-stranded RN/ | PRKRA | 34.405 | 3 | 49.96428 | yes | 3.194E-07 | 542.94226 | 209.1254 |
| AEAPPLER | 8 | O75569 | Interferon-inducible double-stranded RN/ | PRKRA | 34.405 | 2 | 49.96428 | yes | 5.345E-07 | 441.73761 | 290.98337 |
| NQLNPIGSLQELAI | 19 | O75569 | Interferon-inducible double-stranded RN/ | PRKRA | 34.405 | 3 | 49.96428 | yes | 2.323E-07 | 728.38715 | 249.87717 |
| NIPVYECER | 9 | O75569 | Interferon-inducible double-stranded RN/ | PRKRA | 34.405 | 2 | 49.96428 | yes | 1.823E-07 | 590.27679 | 231.92119 |
| FSNISPENHISLTN\ | 28 | O75569 | Interferon-inducible double-stranded RN/ | PRKRA | 34.405 | 4 | 49.96428 | yes | 6.871E-07 | 791.39392 | 20.496475 |
| IEVSNLDGSLR | 11 | O75581 | Low-density lipoprotein receptor-related ן | LRP6 | 180.43 | 2 | 52.17074 | yes | 9.722E-08 | 601.82239 | 150.86069 |
| EASALDFDVTDNR | 13 | O75581 | Low-density lipoprotein receptor-related ן | LRP6 | 180.43 | 2 | 52.17074 | yes | 3.87E-07 | 726.83368 | 126.04134 |
| ELNLQEYR | 8 | O75581 | Low-density lipoprotein receptor-related ן | LRP6 | 180.43 | 2 | 52.17074 | yes | 2.7E-07 | 532.77222 | 103.64024 |
| ILISEDLEEPR | 11 | O75581 | Low-density lipoprotein receptor-related ן | LRP6 | 180.43 | 2 | 52.17074 | yes | 9.722E-08 | 657.35101 | 98.05648 |
| NLYWADTGTNR | 11 | O75581 | Low-density lipoprotein receptor-related ן | LRP6 | 180.43 | 2 | 52.17074 | no | 2.181E-07 | 655.80981 | 84.922623 |
| LYWTDSETNR | 10 | O75581 | Low-density lipoprotein receptor-related ן | LRP6 | 180.43 | 2 | 52.17074 | no | 1.333E-07 | 642.79639 | 61.105728 |
| NLYWTDTGTDR | 11 | O75581 | Low-density lipoprotein receptor-related ן | LRP6 | 180.43 | 2 | 52.17074 | no | 3.284E-07 | 671.30713 | 46.525032 |
| DGATELLLLAR | 11 | O75581 | Low-density lipoprotein receptor-related ן | LRP6 | 180.43 | 2 | 52.17074 | yes | 5.083E-06 | 586.33771 | 33.937096 |
| EIHSDIFSPMDIHA | 19 | O75581 | Low-density lipoprotein receptor-related ן | LRP6 | 180.43 | 3 | 52.17074 | yes | 2.35E-07 | 753.36017 | 25.28318 |
| VRQVKVLGWK | 10 | O75592 | E3 ubiquitin-protein ligase MYCBP2 OS | MYCBP2 | 513.637 | 3 | 54.83225 | yes | 5.522E-07 | 404.92404 | 458.6687 |
| FSHPSLCLR | 9 | O75592 | E3 ubiquitin-protein ligase MYCBP2 OS | MYCBP2 | 513.637 | 2 | 54.83225 | yes | 2.799E-07 | 558.78455 | 285.94788 |
| CSILSPELALPTGSI | 15 | O75592 | E3 ubiquitin-protein ligase MYCBP2 OS | MYCBP2 | 513.637 | 2 | 54.83225 | yes | 9.722E-08 | 800.92181 | 276.61279 |
| GSTPTQISEIIIK | 13 | O75592 | E3 ubiquitin-protein ligase MYCBP2 OS | MYCBP2 | 513.637 | 2 | 54.83225 | yes | 1.076E-07 | 693.89557 | 255.2571 |
| ATWIGASGDQTFL | 14 | O75592 | E3 ubiquitin-protein ligase MYCBP2 OS | MYCBP2 | 513.637 | 2 | 54.83225 | yes | 9.758E-08 | 761.88605 | 223.79332 |
| VTFPECVR | 8 | O75592 | E3 ubiquitin-protein ligase MYCBP2 OS | MYCBP2 | 513.637 | 2 | 54.83225 | yes | 4.102E-07 | 504.25256 | 223.43658 |
| HPSPELSR | 8 | O75592 | E3 ubiquitin-protein ligase MYCBP2 OS | MYCBP2 | 513.637 | 2 | 54.83225 | yes | 2.952E-07 | 461.74069 | 180.39258 |
| DAIYSDSSSLVTDI | 15 | O75592 | E3 ubiquitin-protein ligase MYCBP2 OS | MYCBP2 | 513.637 | 2 | 54.83225 | yes | 3.327E-07 | 807.39886 | 172.37416 |
| SDSYTLDPDTLR | 12 | O75592 | E3 ubiquitin-protein ligase MYCBP2 OS | MYCBP2 | 513.637 | 2 | 54.83225 | yes | 4.969E-07 | 691.82532 | 171.99857 |
| HCVLEWQK | 8 | O75592 | E3 ubiquitin-protein ligase MYCBP2 OS | MYCBP2 | 513.637 | 2 | 54.83225 | yes | 2.705E-07 | 550.2713 | 169.26768 |
| VYNATSDADLK | 11 | O75592 | E3 ubiquitin-protein ligase MYCBP2 OS | MYCBP2 | 513.637 | 2 | 54.83225 | yes | 5.242E-07 | 598.79333 | 168.86761 |
| VYTCEIYPVSATGI | 14 | O75592 | E3 ubiquitin-protein ligase MYCBP2 OS | MYCBP2 | 513.637 | 2 | 54.83225 | yes | 1.791E-07 | 794.38977 | 166.60121 |
| GTTITGTAGTTVGI | 14 | O75592 | E3 ubiquitin-protein ligase MYCBP2 OS | MYCBP2 | 513.637 | 2 | 54.83225 | yes | 1.474E-07 | 632.84082 | 165.53015 |
| IAGQISASVAQQR | 13 | O75592 | E3 ubiquitin-protein ligase MYCBP2 OS | MYCBP2 | 513.637 | 2 | 54.83225 | yes | 9.722E-08 | 664.86768 | 155.58076 |
| IYTAALNDR | 9 | O75592 | E3 ubiquitin-protein ligase MYCBP2 OS | MYCBP2 | 513.637 | 2 | 54.83225 | yes | 2.29E-07 | 518.77472 | 144.00497 |

| | | | | | | | | | | |
|---|---|---|---|---|---|---|---|---|---|---|
| ELANLGGVCAAI | 15 | O75592 | E3 ubiquitin-protein ligase MYCBP2 OS | MYCBP2 | 513.637 | 2 | 54.83225 | yes | 9.926E-08 | 759.39435 | 143.8833 |
| LSQSEENISGMTSF | 15 | O75592 | E3 ubiquitin-protein ligase MYCBP2 OS | MYCBP2 | 513.637 | 2 | 54.83225 | yes | 9.722E-08 | 843.39362 | 141.99809 |
| VNSGDTEVGSSLL | 14 | O75592 | E3 ubiquitin-protein ligase MYCBP2 OS | MYCBP2 | 513.637 | 2 | 54.83225 | yes | 1.021E-07 | 717.36517 | 140.30396 |
| SEFDGDLQSQLLSI | 14 | O75592 | E3 ubiquitin-protein ligase MYCBP2 OS | MYCBP2 | 513.637 | 2 | 54.83225 | yes | 9.722E-08 | 783.88593 | 138.21924 |
| VYENYSFEELR | 11 | O75592 | E3 ubiquitin-protein ligase MYCBP2 OS | MYCBP2 | 513.637 | 2 | 54.83225 | yes | 1.22E-07 | 724.83826 | 133.6492 |
| MPCLYLQTLAR | 11 | O75592 | E3 ubiquitin-protein ligase MYCBP2 OS | MYCBP2 | 513.637 | 2 | 54.83225 | yes | 1.658E-07 | 683.35449 | 131.1507 |
| ALSVVSTVVR | 10 | O75592 | E3 ubiquitin-protein ligase MYCBP2 OS | MYCBP2 | 513.637 | 2 | 54.83225 | yes | 9.722E-08 | 515.81641 | 128.80841 |
| EHMVGIIFSR | 10 | O75592 | E3 ubiquitin-protein ligase MYCBP2 OS | MYCBP2 | 513.637 | 2 | 54.83225 | yes | 9.722E-08 | 594.81335 | 126.81042 |
| KMPLTEPLR | 9 | O75592 | E3 ubiquitin-protein ligase MYCBP2 OS | MYCBP2 | 513.637 | 2 | 54.83225 | yes | 9.262E-07 | 542.81281 | 126.21114 |
| ELICGACSDVSR | 12 | O75592 | E3 ubiquitin-protein ligase MYCBP2 OS | MYCBP2 | 513.637 | 2 | 54.83225 | yes | 9.722E-08 | 683.80811 | 126.08076 |
| YVSVHVDNSR | 10 | O75592 | E3 ubiquitin-protein ligase MYCBP2 OS | MYCBP2 | 513.637 | 2 | 54.83225 | yes | 1.089E-07 | 588.29382 | 123.44241 |
| SYSVVASEYDK | 11 | O75592 | E3 ubiquitin-protein ligase MYCBP2 OS | MYCBP2 | 513.637 | 2 | 54.83225 | yes | 9.978E-08 | 624.29315 | 120.91317 |
| GTESLSASESLILK | 14 | O75592 | E3 ubiquitin-protein ligase MYCBP2 OS | MYCBP2 | 513.637 | 2 | 54.83225 | yes | 1.198E-07 | 717.88794 | 119.69546 |
| GHYQLLLSGR | 10 | O75592 | E3 ubiquitin-protein ligase MYCBP2 OS | MYCBP2 | 513.637 | 2 | 54.83225 | yes | 1.842E-07 | 572.31708 | 119.15885 |
| GQIPQILYYR | 10 | O75592 | E3 ubiquitin-protein ligase MYCBP2 OS | MYCBP2 | 513.637 | 2 | 54.83225 | yes | 4.574E-07 | 625.84821 | 117.62122 |
| ITFGFISCPICK | 12 | O75592 | E3 ubiquitin-protein ligase MYCBP2 OS | MYCBP2 | 513.637 | 2 | 54.83225 | yes | 9.722E-08 | 721.86218 | 107.75142 |
| MGVASTEEETQAV | 15 | O75592 | E3 ubiquitin-protein ligase MYCBP2 OS | MYCBP2 | 513.637 | 2 | 54.83225 | yes | 9.722E-08 | 805.87366 | 107.59984 |
| VDLHEGCGR | 9 | O75592 | E3 ubiquitin-protein ligase MYCBP2 OS | MYCBP2 | 513.637 | 2 | 54.83225 | yes | 1.471E-07 | 521.74036 | 106.95562 |
| MPLTEPLR | 8 | O75592 | E3 ubiquitin-protein ligase MYCBP2 OS | MYCBP2 | 513.637 | 2 | 54.83225 | yes | 6.307E-07 | 478.76532 | 98.160538 |
| GHFVTQVAMGK | 11 | O75592 | E3 ubiquitin-protein ligase MYCBP2 OS | MYCBP2 | 513.637 | 2 | 54.83225 | yes | 9.722E-08 | 587.80548 | 93.918358 |
| CGWPTTITVQTK | 12 | O75592 | E3 ubiquitin-protein ligase MYCBP2 OS | MYCBP2 | 513.637 | 2 | 54.83225 | yes | 3.738E-07 | 696.35303 | 92.812737 |
| ALLSSPEGEEK | 11 | O75592 | E3 ubiquitin-protein ligase MYCBP2 OS | MYCBP2 | 513.637 | 2 | 54.83225 | yes | 9.722E-08 | 580.29572 | 91.351959 |
| FYNDPAGYAMNR | 12 | O75592 | E3 ubiquitin-protein ligase MYCBP2 OS | MYCBP2 | 513.637 | 2 | 54.83225 | yes | 9.722E-08 | 709.81152 | 88.735619 |
| YQSLGIK | 7 | O75592 | E3 ubiquitin-protein ligase MYCBP2 OS | MYCBP2 | 513.637 | 2 | 54.83225 | yes | 1.511E-06 | 404.73181 | 84.820221 |
| MSLQQDQAK | 9 | O75592 | E3 ubiquitin-protein ligase MYCBP2 OS | MYCBP2 | 513.637 | 2 | 54.83225 | yes | 3.655E-07 | 524.75824 | 82.621346 |
| HIGWVTSELPGGD | 18 | O75592 | E3 ubiquitin-protein ligase MYCBP2 OS | MYCBP2 | 513.637 | 3 | 54.83225 | yes | 1.885E-07 | 658.34619 | 76.289642 |
| QHSILPAR | 8 | O75592 | E3 ubiquitin-protein ligase MYCBP2 OS | MYCBP2 | 513.637 | 2 | 54.83225 | yes | 2.676E-07 | 461.26688 | 75.060997 |
| INHIVLK | 7 | O75592 | E3 ubiquitin-protein ligase MYCBP2 OS | MYCBP2 | 513.637 | 2 | 54.83225 | yes | 4.001E-06 | 418.77127 | 72.259209 |
| LISAHSSLSK | 10 | O75592 | E3 ubiquitin-protein ligase MYCBP2 OS | MYCBP2 | 513.637 | 2 | 54.83225 | yes | 1.215E-05 | 521.79822 | 68.168785 |
| PILDVPYWNAK | 11 | O75592 | E3 ubiquitin-protein ligase MYCBP2 OS | MYCBP2 | 513.637 | 2 | 54.83225 | yes | 2.356E-07 | 658.3559 | 66.73748 |
| SENLENTVIIPDIK | 14 | O75592 | E3 ubiquitin-protein ligase MYCBP2 OS | MYCBP2 | 513.637 | 2 | 54.83225 | yes | 7.845E-06 | 792.92761 | 65.022842 |
| ELWCYNAVVADA | 13 | O75592 | E3 ubiquitin-protein ligase MYCBP2 OS | MYCBP2 | 513.637 | 2 | 54.83225 | yes | 2.776E-07 | 783.87213 | 62.986565 |
| DLHIISTGFDEESA | 20 | O75592 | E3 ubiquitin-protein ligase MYCBP2 OS | MYCBP2 | 513.637 | 3 | 54.83225 | yes | 3.208E-07 | 701.0235 | 57.444611 |
| HIIGLVPASISEPPP | 17 | O75592 | E3 ubiquitin-protein ligase MYCBP2 OS | MYCBP2 | 513.637 | 3 | 54.83225 | yes | 3.369E-06 | 601.34521 | 57.137032 |
| QQTSEPVHILK | 11 | O75592 | E3 ubiquitin-protein ligase MYCBP2 OS | MYCBP2 | 513.637 | 2 | 54.83225 | yes | 1.192E-07 | 640.35388 | 56.992073 |
| QAMSPSVAECAR | 12 | O75592 | E3 ubiquitin-protein ligase MYCBP2 OS | MYCBP2 | 513.637 | 2 | 54.83225 | yes | 9.803E-08 | 653.79755 | 54.037983 |
| LLAAVMSALCHTS | 15 | O75592 | E3 ubiquitin-protein ligase MYCBP2 OS | MYCBP2 | 513.637 | 2 | 54.83225 | yes | 9.722E-08 | 800.93109 | 53.799973 |
| IDEALINSHVLATS | 20 | O75592 | E3 ubiquitin-protein ligase MYCBP2 OS | MYCBP2 | 513.637 | 3 | 54.83225 | yes | 9.722E-08 | 720.05121 | 53.75959 |
| APLKDEQEMR | 10 | O75592 | E3 ubiquitin-protein ligase MYCBP2 OS | MYCBP2 | 513.637 | 2 | 54.83225 | yes | 1.334E-06 | 608.80316 | 53.436882 |

| | | | | | | | | | | | |
|---|---|---|---|---|---|---|---|---|---|---|---|
| GDGSKPLEPAK | 11 | O75592 | E3 ubiquitin-protein ligase MYCBP2 OS= | MYCBP2 | 513.637 | 2 | 54.83225 | yes | 1.351E-07 | 549.79309 | 50.998405 |
| WTSLELVK | 8 | O75592 | E3 ubiquitin-protein ligase MYCBP2 OS= | MYCBP2 | 513.637 | 2 | 54.83225 | yes | 2.106E-06 | 488.27911 | 48.96603 |
| SMMCPPGMHK | 10 | O75592 | E3 ubiquitin-protein ligase MYCBP2 OS= | MYCBP2 | 513.637 | 2 | 54.83225 | yes | 3.559E-07 | 588.24506 | 44.043892 |
| SHPSLQSEQIGIVK | 14 | O75592 | E3 ubiquitin-protein ligase MYCBP2 OS= | MYCBP2 | 513.637 | 2 | 54.83225 | yes | 3.604E-07 | 761.91486 | 43.021145 |
| ILSEGVDHCMVK | 12 | O75592 | E3 ubiquitin-protein ligase MYCBP2 OS= | MYCBP2 | 513.637 | 2 | 54.83225 | yes | 4.682E-07 | 694.33905 | 42.033051 |
| GFQFTATLLDLER | 13 | O75592 | E3 ubiquitin-protein ligase MYCBP2 OS= | MYCBP2 | 513.637 | 2 | 54.83225 | yes | 9.722E-08 | 755.89862 | 39.152176 |
| FPLLVAHMEK | 10 | O75592 | E3 ubiquitin-protein ligase MYCBP2 OS= | MYCBP2 | 513.637 | 2 | 54.83225 | yes | 1.431E-07 | 592.82843 | 36.078804 |
| SHAALHILGCLDTI | 21 | O75592 | E3 ubiquitin-protein ligase MYCBP2 OS= | MYCBP2 | 513.637 | 3 | 54.83225 | yes | 5.524E-07 | 760.06091 | 36.074444 |
| GIPIGMLVLGNK | 12 | O75592 | E3 ubiquitin-protein ligase MYCBP2 OS= | MYCBP2 | 513.637 | 2 | 54.83225 | yes | 9.838E-07 | 606.36267 | 27.531553 |
| DECNVVEVVAR | 11 | O75607 | Nucleoplasmin-3 OS=Homo sapiens OX= | NPM3 | 19.344 | 2 | 49.834553 | yes | 1.066E-07 | 645.31134 | 488.181 |
| VPAPVTMDSFFFG | 21 | O75607 | Nucleoplasmin-3 OS=Homo sapiens OX= | NPM3 | 19.344 | 3 | 49.834553 | yes | 1.004E-07 | 785.70142 | 75.951912 |
| NHDHQEIAVPVAN | 15 | O75607 | Nucleoplasmin-3 OS=Homo sapiens OX= | NPM3 | 19.344 | 2 | 49.834553 | yes | 1.002E-07 | 842.94189 | 55.853985 |
| YSQIPSVLVMNK | 12 | O75616 | GTPase Era, mitochondrial OS=Homo sa | ERAL1 | 48.351 | 2 | 53.474068 | yes | 1.43E-07 | 689.8736 | 1505.0989 |
| CQALGVITEK | 10 | O75616 | GTPase Era, mitochondrial OS=Homo sa | ERAL1 | 48.351 | 2 | 53.474068 | yes | 1.336E-07 | 559.79736 | 1200.0483 |
| STLSNQLLGR | 10 | O75616 | GTPase Era, mitochondrial OS=Homo sa | ERAL1 | 48.351 | 2 | 53.474068 | yes | 1.362E-07 | 544.80658 | 1016.4182 |
| VVLLGAPNAGK | 11 | O75616 | GTPase Era, mitochondrial OS=Homo sa | ERAL1 | 48.351 | 2 | 53.474068 | yes | 2.804E-07 | 519.81897 | 1014.7213 |
| NQLSPQLLR | 9 | O75616 | GTPase Era, mitochondrial OS=Homo sa | ERAL1 | 48.351 | 2 | 53.474068 | yes | 1.108E-07 | 534.81165 | 661.13904 |
| EIFMLSALSQEDVI | 14 | O75616 | GTPase Era, mitochondrial OS=Homo sa | ERAL1 | 48.351 | 2 | 53.474068 | yes | 9.722E-08 | 805.41071 | 363.55508 |
| KVFPVSR | 7 | O75616 | GTPase Era, mitochondrial OS=Homo sa | ERAL1 | 48.351 | 2 | 53.474068 | yes | 2.018E-07 | 416.75562 | 337.5527 |
| HHLELSLLEDPWK | 13 | O75616 | GTPase Era, mitochondrial OS=Homo sa | ERAL1 | 48.351 | 2 | 53.474068 | yes | 3.057E-07 | 808.92517 | 180.31265 |
| DPNTQSVGNPQR | 12 | O75616 | GTPase Era, mitochondrial OS=Homo sa | ERAL1 | 48.351 | 2 | 53.474068 | yes | 9.722E-08 | 656.81567 | 118.45994 |
| IGWPHFK | 7 | O75616 | GTPase Era, mitochondrial OS=Homo sa | ERAL1 | 48.351 | 2 | 53.474068 | yes | 1.878E-07 | 442.74249 | 104.00448 |
| QAFHSHPGTHCPS | 17 | O75616 | GTPase Era, mitochondrial OS=Homo sa | ERAL1 | 48.351 | 4 | 53.474068 | yes | 2.817E-06 | 465.2258 | 34.361507 |
| EIDLLLGQTDDTR | 13 | O75643 | U5 small nuclear ribonucleoprotein 200 k | SNRNP200 | 244.508 | 2 | 55.753304 | yes | 9.927E-08 | 744.88068 | 1099.8445 |
| FYDDAIVSQK | 10 | O75643 | U5 small nuclear ribonucleoprotein 200 k | SNRNP200 | 244.508 | 2 | 55.753304 | yes | 1.001E-07 | 593.29297 | 654.25909 |
| PVPLEQTYVGITEI | 14 | O75643 | U5 small nuclear ribonucleoprotein 200 k | SNRNP200 | 244.508 | 2 | 55.753304 | yes | 9.722E-08 | 787.42725 | 616.03687 |
| GNIIISTPEK | 10 | O75643 | U5 small nuclear ribonucleoprotein 200 k | SNRNP200 | 244.508 | 2 | 55.753304 | yes | 7.385E-07 | 536.30585 | 505.60223 |
| FQIMNEIVYEK | 11 | O75643 | U5 small nuclear ribonucleoprotein 200 k | SNRNP200 | 244.508 | 2 | 55.753304 | yes | 1.781E-07 | 707.35779 | 468.95551 |
| DEPTGEVLSLVGK | 13 | O75643 | U5 small nuclear ribonucleoprotein 200 k | SNRNP200 | 244.508 | 2 | 55.753304 | yes | 1.811E-07 | 672.35626 | 462.45163 |
| YAQAGFEGFK | 10 | O75643 | U5 small nuclear ribonucleoprotein 200 k | SNRNP200 | 244.508 | 2 | 55.753304 | yes | 7.822E-07 | 559.26929 | 424.66577 |
| YHVLVNLGK | 9 | O75643 | U5 small nuclear ribonucleoprotein 200 k | SNRNP200 | 244.508 | 2 | 55.753304 | yes | 1.017E-07 | 521.80585 | 406.60352 |
| KPVIVFVPSR | 10 | O75643 | U5 small nuclear ribonucleoprotein 200 k | SNRNP200 | 244.508 | 2 | 55.753304 | yes | 9.819E-08 | 571.35822 | 380.94962 |
| VVLLTGETSTDLK | 13 | O75643 | U5 small nuclear ribonucleoprotein 200 k | SNRNP200 | 244.508 | 2 | 55.753304 | yes | 9.722E-08 | 688.38757 | 348.065 |
| NQVLVFVHSR | 10 | O75643 | U5 small nuclear ribonucleoprotein 200 k | SNRNP200 | 244.508 | 2 | 55.753304 | yes | 9.722E-08 | 599.8382 | 343.66422 |
| ANSNLVLQADR | 11 | O75643 | U5 small nuclear ribonucleoprotein 200 k | SNRNP200 | 244.508 | 2 | 55.753304 | yes | 9.898E-08 | 600.82019 | 311.90366 |
| NIEMTQEDVR | 10 | O75643 | U5 small nuclear ribonucleoprotein 200 k | SNRNP200 | 244.508 | 2 | 55.753304 | yes | 9.722E-08 | 617.79022 | 308.93393 |
| AIFEIVLNR | 9 | O75643 | U5 small nuclear ribonucleoprotein 200 k | SNRNP200 | 244.508 | 2 | 55.753304 | yes | 3.582E-07 | 537.81897 | 297.19739 |
| SPTLYGISHDDLK | 13 | O75643 | U5 small nuclear ribonucleoprotein 200 k | SNRNP200 | 244.508 | 2 | 55.753304 | yes | 1.619E-07 | 723.36719 | 296.26315 |
| NSAFESLYQDK | 11 | O75643 | U5 small nuclear ribonucleoprotein 200 k | SNRNP200 | 244.508 | 2 | 55.753304 | yes | 9.809E-08 | 651.30408 | 296.11661 |

| | | | | | | | | | | | |
|---|---|---|---|---|---|---|---|---|---|---|---|
| WLSCETQLPVSFR | 13 | O75643 | U5 small nuclear ribonucleoprotein 200 k | SNRNP200 | 244.508 | 2 | 55.753304 | yes | 1.328E-07 | 811.90338 | 293.98657 |
| GPVLEALVAR | 10 | O75643 | U5 small nuclear ribonucleoprotein 200 k | SNRNP200 | 244.508 | 2 | 55.753304 | yes | 1.37E-07 | 512.8111 | 289.45221 |
| EGSASTEVLR | 10 | O75643 | U5 small nuclear ribonucleoprotein 200 k | SNRNP200 | 244.508 | 2 | 55.753304 | yes | 1.823E-07 | 524.76709 | 284.14856 |
| MLLQSSEGR | 9 | O75643 | U5 small nuclear ribonucleoprotein 200 k | SNRNP200 | 244.508 | 2 | 55.753304 | yes | 1.911E-07 | 510.76077 | 263.25406 |
| KLPEEVVK | 8 | O75643 | U5 small nuclear ribonucleoprotein 200 k | SNRNP200 | 244.508 | 2 | 55.753304 | yes | 4.284E-07 | 471.28696 | 245.33388 |
| FLHCTEK | 7 | O75643 | U5 small nuclear ribonucleoprotein 200 k | SNRNP200 | 244.508 | 2 | 55.753304 | yes | 3.409E-07 | 467.7262 | 238.40504 |
| TNLLLQAHLSR | 11 | O75643 | U5 small nuclear ribonucleoprotein 200 k | SNRNP200 | 244.508 | 2 | 55.753304 | yes | 9.977E-08 | 633.36987 | 226.56294 |
| LELSVHLQPITR | 12 | O75643 | U5 small nuclear ribonucleoprotein 200 k | SNRNP200 | 244.508 | 2 | 55.753304 | yes | 4.078E-07 | 703.41174 | 111.89684 |
| LELSVHLQPITR | 12 | O75643 | U5 small nuclear ribonucleoprotein 200 k | SNRNP200 | 244.508 | 2 | 55.753304 | yes | 3.248E-07 | 469.27692 | 111.89684 |
| LELSVHLQPITR | 12 | O75643 | U5 small nuclear ribonucleoprotein 200 k | SNRNP200 | 244.508 | 3 | 55.753304 | yes | 4.078E-07 | 703.41174 | 111.89684 |
| LELSVHLQPITR | 12 | O75643 | U5 small nuclear ribonucleoprotein 200 k | SNRNP200 | 244.508 | 3 | 55.753304 | yes | 3.248E-07 | 469.27692 | 111.89684 |
| ESIEEPSAK | 9 | O75643 | U5 small nuclear ribonucleoprotein 200 k | SNRNP200 | 244.508 | 2 | 55.753304 | yes | 1.915E-07 | 495.24292 | 219.45164 |
| DIDAFWLQR | 9 | O75643 | U5 small nuclear ribonucleoprotein 200 k | SNRNP200 | 244.508 | 2 | 55.753304 | yes | 1.134E-07 | 582.29584 | 217.46129 |
| SLVQEMVGSFGK | 12 | O75643 | U5 small nuclear ribonucleoprotein 200 k | SNRNP200 | 244.508 | 2 | 55.753304 | yes | 9.722E-08 | 641.32904 | 213.91399 |
| MWQSMCPLR | 9 | O75643 | U5 small nuclear ribonucleoprotein 200 k | SNRNP200 | 244.508 | 2 | 55.753304 | yes | 9.722E-08 | 604.77228 | 210.73743 |
| KADEVLEILK | 10 | O75643 | U5 small nuclear ribonucleoprotein 200 k | SNRNP200 | 244.508 | 2 | 55.753304 | yes | 4.221E-07 | 579.34247 | 209.70885 |
| LIILDEIHLLHDDR | 14 | O75643 | U5 small nuclear ribonucleoprotein 200 k | SNRNP200 | 244.508 | 3 | 55.753304 | yes | 1.652E-07 | 572.32104 | 205.53967 |
| EEEKLELQK | 9 | O75643 | U5 small nuclear ribonucleoprotein 200 k | SNRNP200 | 244.508 | 2 | 55.753304 | yes | 1.078E-06 | 573.30603 | 178.88368 |
| VFSLSSEFK | 9 | O75643 | U5 small nuclear ribonucleoprotein 200 k | SNRNP200 | 244.508 | 2 | 55.753304 | yes | 1.185E-07 | 522.27405 | 178.72815 |
| TLVEDLFADK | 10 | O75643 | U5 small nuclear ribonucleoprotein 200 k | SNRNP200 | 244.508 | 2 | 55.753304 | yes | 1.282E-07 | 575.80316 | 175.94307 |
| SGGPVVVLVQLEF | 13 | O75643 | U5 small nuclear ribonucleoprotein 200 k | SNRNP200 | 244.508 | 2 | 55.753304 | yes | 9.722E-08 | 676.89844 | 162.84621 |
| PVPLELHIQGFNIS | 19 | O75643 | U5 small nuclear ribonucleoprotein 200 k | SNRNP200 | 244.508 | 3 | 55.753304 | yes | 2.354E-07 | 729.72717 | 75.288956 |
| PVPLELHIQGFNIS | 19 | O75643 | U5 small nuclear ribonucleoprotein 200 k | SNRNP200 | 244.508 | 3 | 55.753304 | yes | 2.685E-07 | 547.54718 | 75.288956 |
| PVPLELHIQGFNIS | 19 | O75643 | U5 small nuclear ribonucleoprotein 200 k | SNRNP200 | 244.508 | 4 | 55.753304 | yes | 2.354E-07 | 729.72717 | 75.288956 |
| PVPLELHIQGFNIS | 19 | O75643 | U5 small nuclear ribonucleoprotein 200 k | SNRNP200 | 244.508 | 4 | 55.753304 | yes | 2.685E-07 | 547.54718 | 75.288956 |
| TICAEFAILR | 10 | O75643 | U5 small nuclear ribonucleoprotein 200 k | SNRNP200 | 244.508 | 2 | 55.753304 | yes | 1.26E-07 | 597.32098 | 139.36481 |
| FLYQLHETEK | 10 | O75643 | U5 small nuclear ribonucleoprotein 200 k | SNRNP200 | 244.508 | 2 | 55.753304 | yes | 9.722E-08 | 654.33514 | 138.18614 |
| IIYIAPMR | 8 | O75643 | U5 small nuclear ribonucleoprotein 200 k | SNRNP200 | 244.508 | 2 | 55.753304 | yes | 1.46E-07 | 488.78604 | 134.56342 |
| TNVALMCMLR | 10 | O75643 | U5 small nuclear ribonucleoprotein 200 k | SNRNP200 | 244.508 | 2 | 55.753304 | yes | 9.722E-08 | 604.80103 | 127.44137 |
| GLFYFDNSFR | 10 | O75643 | U5 small nuclear ribonucleoprotein 200 k | SNRNP200 | 244.508 | 2 | 55.753304 | yes | 9.722E-08 | 633.30109 | 123.17651 |
| GDPLLDQR | 8 | O75643 | U5 small nuclear ribonucleoprotein 200 k | SNRNP200 | 244.508 | 2 | 55.753304 | yes | 3.015E-06 | 457.24051 | 117.5319 |
| FNDPHVK | 7 | O75643 | U5 small nuclear ribonucleoprotein 200 k | SNRNP200 | 244.508 | 2 | 55.753304 | yes | 8.234E-07 | 428.71924 | 117.28625 |
| YPNIELSYEVVDK | 13 | O75643 | U5 small nuclear ribonucleoprotein 200 k | SNRNP200 | 244.508 | 2 | 55.753304 | yes | 9.722E-08 | 784.89575 | 98.416229 |
| YVHLFPK | 7 | O75643 | U5 small nuclear ribonucleoprotein 200 k | SNRNP200 | 244.508 | 2 | 55.753304 | yes | 3.636E-06 | 452.258 | 68.716393 |
| PFGSEEQLLPVEK | 13 | O75643 | U5 small nuclear ribonucleoprotein 200 k | SNRNP200 | 244.508 | 2 | 55.753304 | yes | 1.13E-06 | 736.88519 | 63.855186 |
| DTLGLFLR | 8 | O75643 | U5 small nuclear ribonucleoprotein 200 k | SNRNP200 | 244.508 | 2 | 55.753304 | yes | 1.524E-07 | 467.77145 | 60.431973 |
| HINMDGTINVDDF | 14 | O75643 | U5 small nuclear ribonucleoprotein 200 k | SNRNP200 | 244.508 | 2 | 55.753304 | yes | 1.743E-07 | 809.88013 | 47.528496 |
| RLDLVHTAALMLI | 14 | O75643 | U5 small nuclear ribonucleoprotein 200 k | SNRNP200 | 244.508 | 3 | 55.753304 | yes | 1.468E-07 | 532.63611 | 41.723152 |
| KVVLLTGETSTDL | 14 | O75643 | U5 small nuclear ribonucleoprotein 200 k | SNRNP200 | 244.508 | 2 | 55.753304 | yes | 4.992E-07 | 752.43506 | 41.466862 |

| | | | | | | | | | | | |
|---|---|---|---|---|---|---|---|---|---|---|---|
| PVYHAITK | 8 | O75643 | U5 small nuclear ribonucleoprotein 200 k | SNRNP200 | 244.508 | 2 | 55.753304 | yes | 2.64E-07 | 464.76617 | 36.341644 |
| KGYEEVHVPALKI | 14 | O75643 | U5 small nuclear ribonucleoprotein 200 k | SNRNP200 | 244.508 | 3 | 55.753304 | yes | 9.722E-08 | 532.30322 | 29.380083 |
| EEGWWVVIGDAK | 12 | O75643 | U5 small nuclear ribonucleoprotein 200 k | SNRNP200 | 244.508 | 2 | 55.753304 | yes | 1.332E-07 | 694.84589 | 29.12685 |
| LPDMLNAEIVLGN | 18 | O75643 | U5 small nuclear ribonucleoprotein 200 k | SNRNP200 | 244.508 | 3 | 55.753304 | yes | 6.239E-05 | 647.01544 | 17.423418 |
| ADEVLEILK | 9 | O75643 | U5 small nuclear ribonucleoprotein 200 k | SNRNP200 | 244.508 | 2 | 55.753304 | yes | 1.637E-05 | 515.29498 | 15.928264 |
| VACAEEWQESR | 11 | O75663 | TIP41-like protein OS=Homo sapiens OX | TIPRL | 31.444 | 2 | 46.590103 | yes | 1.521E-07 | 682.79858 | 49.531811 |
| VVSEQPQVGTLEE | 14 | O75665 | Centriole and centriolar satellite protein C | OFD1 | 116.672 | 2 | 44.665554 | yes | 2.112E-07 | 785.90723 | 59.876427 |
| IGPVAQDLLQR | 11 | O75676 | Ribosomal protein S6 kinase alpha-4 OS= | RPS6KA4 | 85.606 | 2 | 53.654568 | yes | 9.722E-08 | 605.35114 | 263.93039 |
| SELDVGNFAEEFTI | 14 | O75676 | Ribosomal protein S6 kinase alpha-4 OS= | RPS6KA4 | 85.606 | 2 | 53.654568 | yes | 9.892E-08 | 807.37573 | 149.19373 |
| GGELLEHIR | 9 | O75676 | Ribosomal protein S6 kinase alpha-4 OS= | RPS6KA4 | 85.606 | 2 | 53.654568 | yes | 1.156E-07 | 512.28271 | 136.22131 |
| LGAGPQGAQEVR | 12 | O75676 | Ribosomal protein S6 kinase alpha-4 OS= | RPS6KA4 | 85.606 | 2 | 53.654568 | yes | 9.722E-08 | 591.81494 | 124.06781 |
| SLVSAVSFMHEEA | 18 | O75676 | Ribosomal protein S6 kinase alpha-4 OS= | RPS6KA4 | 85.606 | 3 | 53.654568 | yes | 4.192E-07 | 652.33167 | 111.37437 |
| TPDVLESSGPAVR | 13 | O75676 | Ribosomal protein S6 kinase alpha-4 OS= | RPS6KA4 | 85.606 | 2 | 53.654568 | yes | 5.05E-07 | 664.34625 | 83.176468 |
| IIDFGFAR | 8 | O75676 | Ribosomal protein S6 kinase alpha-4 OS= | RPS6KA4 | 85.606 | 2 | 53.654568 | no | 1.37E-07 | 469.75836 | 78.401604 |
| EPALGQGSFSVCR | 13 | O75676 | Ribosomal protein S6 kinase alpha-4 OS= | RPS6KA4 | 85.606 | 2 | 53.654568 | yes | 2.145E-07 | 704.33789 | 56.001999 |
| KIPAPFRPQIR | 11 | O75676 | Ribosomal protein S6 kinase alpha-4 OS= | RPS6KA4 | 85.606 | 3 | 53.654568 | yes | 1.823E-07 | 441.60669 | 26.289343 |
| FLASDLDQK | 9 | O75691 | Small subunit processome component 20 | UTP20 | 318.386 | 2 | 55.286522 | yes | 3.494E-06 | 518.7691 | 435.39694 |
| LGTVLPVIEK | 10 | O75691 | Small subunit processome component 20 | UTP20 | 318.386 | 2 | 55.286522 | yes | 1.171E-07 | 534.83679 | 360.10956 |
| YILDYPLGDK | 10 | O75691 | Small subunit processome component 20 | UTP20 | 318.386 | 2 | 55.286522 | yes | 1.167E-07 | 598.81354 | 319.51392 |
| HLPEPVAIK | 9 | O75691 | Small subunit processome component 20 | UTP20 | 318.386 | 2 | 55.286522 | yes | 6.194E-07 | 502.30038 | 281.03992 |
| NIQGTITGDILPR | 13 | O75691 | Small subunit processome component 20 | UTP20 | 318.386 | 2 | 55.286522 | yes | 9.722E-08 | 699.39099 | 265.5994 |
| QAELVPATVNDYI | 13 | O75691 | Small subunit processome component 20 | UTP20 | 318.386 | 2 | 55.286522 | yes | 9.722E-08 | 738.37805 | 254.48357 |
| LEAAYSPR | 8 | O75691 | Small subunit processome component 20 | UTP20 | 318.386 | 2 | 55.286522 | yes | 6.522E-07 | 453.73761 | 242.53661 |
| LGNVNIDIIHR | 11 | O75691 | Small subunit processome component 20 | UTP20 | 318.386 | 2 | 55.286522 | yes | 9.722E-08 | 632.36206 | 232.67151 |
| LAVSAQSEPAR | 11 | O75691 | Small subunit processome component 20 | UTP20 | 318.386 | 2 | 55.286522 | yes | 9.722E-08 | 564.80402 | 220.53461 |
| LLVPEIDEVMR | 11 | O75691 | Small subunit processome component 20 | UTP20 | 318.386 | 2 | 55.286522 | yes | 1.116E-07 | 657.36029 | 214.68549 |
| ELNSTYSEQDPLLI | 14 | O75691 | Small subunit processome component 20 | UTP20 | 318.386 | 2 | 55.286522 | yes | 9.722E-08 | 818.90686 | 195.74918 |
| LAALNVTEK | 9 | O75691 | Small subunit processome component 20 | UTP20 | 318.386 | 2 | 55.286522 | yes | 1.361E-07 | 479.78201 | 179.45848 |
| FLDQSLGEQVVK | 12 | O75691 | Small subunit processome component 20 | UTP20 | 318.386 | 2 | 55.286522 | yes | 2.553E-07 | 681.86682 | 177.84924 |
| ELNLTEHFGK | 10 | O75691 | Small subunit processome component 20 | UTP20 | 318.386 | 2 | 55.286522 | yes | 1.209E-07 | 594.3064 | 174.4465 |
| VVNDEEVVR | 9 | O75691 | Small subunit processome component 20 | UTP20 | 318.386 | 2 | 55.286522 | yes | 1.807E-07 | 529.77747 | 150.35513 |
| LQANISTGVSK | 11 | O75691 | Small subunit processome component 20 | UTP20 | 318.386 | 2 | 55.286522 | yes | 1.535E-07 | 559.31421 | 145.29514 |
| LGSESQYSPTPLLK | 14 | O75691 | Small subunit processome component 20 | UTP20 | 318.386 | 2 | 55.286522 | yes | 2.085E-07 | 760.40375 | 97.564049 |
| LPPQSCLLLPPTPV | 15 | O75691 | Small subunit processome component 20 | UTP20 | 318.386 | 2 | 55.286522 | yes | 1.223E-06 | 844.48181 | 92.544746 |
| ADLFPILMR | 9 | O75691 | Small subunit processome component 20 | UTP20 | 318.386 | 2 | 55.286522 | yes | 2.035E-07 | 538.30206 | 90.1063 |
| FLGAVAMDLGIDK | 13 | O75691 | Small subunit processome component 20 | UTP20 | 318.386 | 2 | 55.286522 | yes | 2.273E-07 | 675.36029 | 89.674896 |
| ALEFVTNPDIAAK | 13 | O75691 | Small subunit processome component 20 | UTP20 | 318.386 | 2 | 55.286522 | yes | 2.74E-07 | 694.87463 | 89.511642 |
| ELQTTLVR | 8 | O75691 | Small subunit processome component 20 | UTP20 | 318.386 | 2 | 55.286522 | yes | 5.104E-07 | 480.27966 | 87.490295 |
| ITLDCIMTYK | 10 | O75691 | Small subunit processome component 20 | UTP20 | 318.386 | 2 | 55.286522 | yes | 9.722E-08 | 629.31451 | 76.917213 |

| | | | | | | | | | | | |
|---|---|---|---|---|---|---|---|---|---|---|---|
| FETFQTITSYIK | 12 | O75691 | Small subunit processome component 20 | UTP20 | 318.386 | 2 | 55.286522 | yes | 1.045E-07 | 739.38232 | 69.881119 |
| HPHVLPYR | 8 | O75691 | Small subunit processome component 20 | UTP20 | 318.386 | 2 | 55.286522 | yes | 4.735E-07 | 509.78268 | 67.61763 |
| ELSPLFLR | 8 | O75691 | Small subunit processome component 20 | UTP20 | 318.386 | 2 | 55.286522 | yes | 6.457E-07 | 487.78711 | 66.94001 |
| ELQNDMTDEK | 10 | O75691 | Small subunit processome component 20 | UTP20 | 318.386 | 2 | 55.286522 | yes | 9.722E-08 | 611.76642 | 59.148312 |
| TNCCESSEQIK | 11 | O75691 | Small subunit processome component 20 | UTP20 | 318.386 | 2 | 55.286522 | yes | 1.684E-07 | 678.28192 | 57.53006 |
| HLDDINFDVR | 10 | O75691 | Small subunit processome component 20 | UTP20 | 318.386 | 2 | 55.286522 | yes | 9.174E-06 | 622.30695 | 51.488194 |
| LRPNLEFMLAQLN | 20 | O75691 | Small subunit processome component 20 | UTP20 | 318.386 | 3 | 55.286522 | yes | 8.737E-07 | 811.07776 | 43.347607 |
| GQNFHLVVNCFK | 12 | O75691 | Small subunit processome component 20 | UTP20 | 318.386 | 2 | 55.286522 | yes | 5.025E-05 | 731.86664 | 35.283272 |
| RLGTVLPVIEK | 11 | O75691 | Small subunit processome component 20 | UTP20 | 318.386 | 2 | 55.286522 | yes | 1.821E-07 | 612.88733 | 20.306755 |
| ANELLQR | 7 | O75694 | Nuclear pore complex protein Nup155 OS | NUP155 | 155.2 | 2 | 59.25273 | yes | 1.912E-07 | 422.23779 | 336.52231 |
| LLQVASPFLEPHL | 15 | O75694 | Nuclear pore complex protein Nup155 OS | NUP155 | 155.2 | 3 | 59.25273 | yes | 9.781E-08 | 573.66992 | 271.41254 |
| IQLQIQETLQR | 11 | O75694 | Nuclear pore complex protein Nup155 OS | NUP155 | 155.2 | 2 | 59.25273 | yes | 1.355E-07 | 685.39355 | 234.10181 |
| IMGNIWDASLVVE | 14 | O75694 | Nuclear pore complex protein Nup155 OS | NUP155 | 155.2 | 2 | 59.25273 | yes | 9.722E-08 | 801.91907 | 230.18552 |
| ISLQAIQQLVR | 11 | O75694 | Nuclear pore complex protein Nup155 OS | NUP155 | 155.2 | 2 | 59.25273 | yes | 1.172E-07 | 634.88788 | 224.62311 |
| ELSDSVTLSSSDR | 13 | O75694 | Nuclear pore complex protein Nup155 OS | NUP155 | 155.2 | 2 | 59.25273 | yes | 9.722E-08 | 698.33356 | 201.06421 |
| LYFSTCPFR | 9 | O75694 | Nuclear pore complex protein Nup155 OS | NUP155 | 155.2 | 2 | 59.25273 | yes | 9.81E-08 | 595.7868 | 196.8551 |
| YVENPSQVLNCER | 13 | O75694 | Nuclear pore complex protein Nup155 OS | NUP155 | 155.2 | 2 | 59.25273 | yes | 1.043E-07 | 804.37775 | 194.16104 |
| LYGEFADPFK | 10 | O75694 | Nuclear pore complex protein Nup155 OS | NUP155 | 155.2 | 2 | 59.25273 | yes | 2.351E-06 | 593.7926 | 189.90761 |
| LLCEHQFTIIVAEL | 16 | O75694 | Nuclear pore complex protein Nup155 OS | NUP155 | 155.2 | 3 | 59.25273 | yes | 5.837E-07 | 648.02039 | 137.83954 |
| LLEVYDQLFK | 10 | O75694 | Nuclear pore complex protein Nup155 OS | NUP155 | 155.2 | 2 | 59.25273 | yes | 9.722E-08 | 634.35028 | 117.14365 |
| NSQFAGGPLGNPN | 17 | O75694 | Nuclear pore complex protein Nup155 OS | NUP155 | 155.2 | 2 | 59.25273 | yes | 1.073E-07 | 837.41571 | 103.87353 |
| HGEPEEDIVGLQA | 17 | O75694 | Nuclear pore complex protein Nup155 OS | NUP155 | 155.2 | 3 | 59.25273 | yes | 2.041E-07 | 651.98157 | 78.182091 |
| KPGPPVLSSDPNM | 28 | O75694 | Nuclear pore complex protein Nup155 OS | NUP155 | 155.2 | 4 | 59.25273 | yes | 3.265E-06 | 773.12592 | 77.39785 |
| CITDTLQELVNQSI | 14 | O75694 | Nuclear pore complex protein Nup155 OS | NUP155 | 155.2 | 2 | 59.25273 | yes | 9.722E-08 | 824.91418 | 57.7299 |
| DPQGLGLHFYK | 11 | O75694 | Nuclear pore complex protein Nup155 OS | NUP155 | 155.2 | 2 | 59.25273 | yes | 3.422E-07 | 637.83002 | 56.438671 |
| SRDPFWNR | 8 | O75694 | Nuclear pore complex protein Nup155 OS | NUP155 | 155.2 | 2 | 59.25273 | yes | 2.273E-06 | 539.26489 | 46.078812 |
| LADMHSTEISLQQ | 14 | O75694 | Nuclear pore complex protein Nup155 OS | NUP155 | 155.2 | 2 | 59.25273 | yes | 1.557E-07 | 814.90668 | 37.190281 |
| VASVSQNAIVSAA | 18 | O75694 | Nuclear pore complex protein Nup155 OS | NUP155 | 155.2 | 3 | 59.25273 | yes | 4.504E-07 | 576.65149 | 35.988098 |
| PAVAILSFK | 9 | O75717 | WD repeat and HMG-box DNA-binding p | WDHD1 | 125.968 | 2 | 36.544743 | yes | 6.87E-07 | 473.29202 | 14.154386 |
| AFYECLAACEGSR | 13 | O75718 | Cartilage-associated protein OS=Homo sa | CRTAP | 46.562 | 2 | 49.556614 | yes | 1.021E-07 | 767.32666 | 73.166069 |
| DSEAFCHR | 8 | O75718 | Cartilage-associated protein OS=Homo sa | CRTAP | 46.562 | 2 | 49.556614 | yes | 1.757E-07 | 511.21143 | 52.760006 |
| AIAAAHTFLLK | 11 | O75718 | Cartilage-associated protein OS=Homo sa | CRTAP | 46.562 | 2 | 49.556614 | yes | 2.244E-07 | 578.34784 | 40.66069 |
| LQVAGEITTGPR | 12 | O75746 | Electrogenic aspartate/glutamate antiporte | SLC25A12 | 74.762 | 2 | 53.512764 | no | 9.727E-08 | 621.34607 | 1216.5347 |
| LTVNDFVR | 8 | O75746 | Electrogenic aspartate/glutamate antiporte | SLC25A12 | 74.762 | 2 | 53.512764 | no | 9.894E-08 | 482.26654 | 668.87006 |
| AGQTTYSGVIDCF | 14 | O75746 | Electrogenic aspartate/glutamate antiporte | SLC25A12 | 74.762 | 2 | 53.512764 | no | 9.722E-08 | 787.867 | 324.63043 |
| IYSTLAGTR | 9 | O75746 | Electrogenic aspartate/glutamate antiporte | SLC25A12 | 74.762 | 2 | 53.512764 | no | 1.742E-07 | 491.27182 | 235.16722 |
| YEGFFGLYR | 9 | O75746 | Electrogenic aspartate/glutamate antiporte | SLC25A12 | 74.762 | 2 | 53.512764 | no | 9.722E-08 | 576.27966 | 203.8949 |
| YLGLYNDPNSNPK | 13 | O75746 | Electrogenic aspartate/glutamate antiporte | SLC25A12 | 74.762 | 2 | 53.512764 | yes | 9.722E-08 | 747.86481 | 145.62152 |
| LATATFAGIENK | 12 | O75746 | Electrogenic aspartate/glutamate antiporte | SLC25A12 | 74.762 | 2 | 53.512764 | yes | 9.961E-08 | 618.33514 | 133.73822 |

| | | | | | | | | | | | |
|---|---|---|---|---|---|---|---|---|---|---|---|
| GSGSVVGELMYK | 12 | O75746 | Electrogenic aspartate/glutamate antiporte | SLC25A12 | 74.762 | 2 | 53.512764 | yes | 1.047E-07 | 613.80792 | 106.92671 |
| KIYSTLAGTR | 10 | O75746 | Electrogenic aspartate/glutamate antiporte | SLC25A12 | 74.762 | 2 | 53.512764 | no | 1.571E-07 | 555.31934 | 73.520607 |
| LTLADIER | 8 | O75746 | Electrogenic aspartate/glutamate antiporte | SLC25A12 | 74.762 | 2 | 53.512764 | yes | 9.722E-08 | 465.76639 | 65.936974 |
| SPSVAVVQPK | 10 | O75746 | Electrogenic aspartate/glutamate antiporte | SLC25A12 | 74.762 | 2 | 53.512764 | yes | 2.628E-07 | 506.29529 | 62.937592 |
| ASLEFALFQEGSR | 13 | O75808 | Calpain-15 OS=Homo sapiens OX=9606 | CAPN15 | 117.315 | 2 | 42.299763 | yes | 4.389E-07 | 727.86737 | 98.182785 |
| AAQAFLSDWTASK | 13 | O75808 | Calpain-15 OS=Homo sapiens OX=9606 | CAPN15 | 117.315 | 2 | 42.299763 | yes | 3.097E-07 | 698.34882 | 47.276165 |
| EAGFLMGASCGGQ | 16 | O75808 | Calpain-15 OS=Homo sapiens OX=9606 | CAPN15 | 117.315 | 2 | 42.299763 | yes | 2.564E-06 | 793.84161 | 22.036831 |
| LSLTMGGVQAR | 11 | O75815 | Breast cancer anti-estrogen resistance pro | BCAR3 | 92.566 | 2 | 50.43439 | yes | 6.079E-07 | 566.81079 | 97.80864 |
| TGSEPALSPAVVR | 13 | O75815 | Breast cancer anti-estrogen resistance pro | BCAR3 | 92.566 | 2 | 50.43439 | yes | 9.809E-08 | 642.35132 | 37.385933 |
| FNQILTALSR | 10 | O75815 | Breast cancer anti-estrogen resistance pro | BCAR3 | 92.566 | 2 | 50.43439 | yes | 0.0001501 | 581.83258 | 11.334805 |
| GAVEAELDPVEYT | 15 | O75817 | Ribonuclease P protein subunit p20 OS=H | POP7 | 15.651 | 2 | 50.094162 | yes | 9.903E-08 | 831.42267 | 3047.7944 |
| NNSAIHIR | 8 | O75817 | Ribonuclease P protein subunit p20 OS=H | POP7 | 15.651 | 2 | 50.094162 | yes | 1.731E-07 | 462.75412 | 774.05127 |
| RPNDIYVNMK | 10 | O75817 | Ribonuclease P protein subunit p20 OS=H | POP7 | 15.651 | 2 | 50.094162 | yes | 1.373E-07 | 625.32153 | 241.3333 |
| YQIQEHQPK | 9 | O75818 | Ribonuclease P protein subunit p40 OS=H | RPP40 | 41.834 | 2 | 56.35921 | yes | 1.522E-07 | 585.79877 | 179.55379 |
| FDFLLAWHK | 9 | O75818 | Ribonuclease P protein subunit p40 OS=H | RPP40 | 41.834 | 2 | 56.35921 | yes | 2.824E-06 | 588.81366 | 130.67757 |
| NLVMNTGPYYFVI | 13 | O75818 | Ribonuclease P protein subunit p40 OS=H | RPP40 | 41.834 | 2 | 56.35921 | yes | 9.722E-08 | 773.39215 | 122.9281 |
| HLVQTHYYNYR | 11 | O75818 | Ribonuclease P protein subunit p40 OS=H | RPP40 | 41.834 | 3 | 56.35921 | yes | 2.078E-07 | 498.58099 | 50.893982 |
| HLVQTHYYNYR | 11 | O75818 | Ribonuclease P protein subunit p40 OS=H | RPP40 | 41.834 | 3 | 56.35921 | yes | 9.722E-08 | 747.36786 | 50.893982 |
| HLVQTHYYNYR | 11 | O75818 | Ribonuclease P protein subunit p40 OS=H | RPP40 | 41.834 | 2 | 56.35921 | yes | 2.078E-07 | 498.58099 | 50.893982 |
| HLVQTHYYNYR | 11 | O75818 | Ribonuclease P protein subunit p40 OS=H | RPP40 | 41.834 | 2 | 56.35921 | yes | 9.722E-08 | 747.36786 | 50.893982 |
| LILSLDKDTYEETC | 25 | O75818 | Ribonuclease P protein subunit p40 OS=H | RPP40 | 41.834 | 4 | 56.35921 | yes | 1.592E-06 | 698.60291 | 72.989075 |
| TGSEESTMMSYFS | 14 | O75818 | Ribonuclease P protein subunit p40 OS=H | RPP40 | 41.834 | 2 | 56.35921 | yes | 4.086E-07 | 792.83148 | 34.310314 |
| ELAEQLGLSTGEK | 13 | O75821 | Eukaryotic translation initiation factor 3 s | EIF3G | 35.612 | 2 | 49.959564 | yes | 1.984E-07 | 687.85919 | 1439.3811 |
| ETDLQELFR | 9 | O75821 | Eukaryotic translation initiation factor 3 s | EIF3G | 35.612 | 2 | 49.959564 | yes | 5.827E-07 | 575.79059 | 1387.9594 |
| GFAFISFHR | 9 | O75821 | Eukaryotic translation initiation factor 3 s | EIF3G | 35.612 | 2 | 49.959564 | yes | 1.876E-07 | 541.28253 | 980.83746 |
| VTNLSEDTR | 9 | O75821 | Eukaryotic translation initiation factor 3 s | EIF3G | 35.612 | 2 | 49.959564 | yes | 4.45E-07 | 517.75928 | 769.75256 |
| LPGELEPVQATQN | 14 | O75821 | Eukaryotic translation initiation factor 3 s | EIF3G | 35.612 | 2 | 49.959564 | yes | 9.968E-08 | 762.40686 | 508.96515 |
| GIPLATGDTSPEPE | 25 | O75821 | Eukaryotic translation initiation factor 3 s | EIF3G | 35.612 | 3 | 49.959564 | yes | 2.633E-07 | 822.11597 | 299.67505 |
| AIAGVSGFGYDHL | 21 | O75821 | Eukaryotic translation initiation factor 3 s | EIF3G | 35.612 | 3 | 49.959564 | yes | 8.746E-07 | 754.06354 | 92.84227 |
| RADDNATIR | 9 | O75821 | Eukaryotic translation initiation factor 3 s | EIF3G | 35.612 | 2 | 49.959564 | yes | 3.135E-07 | 516.26508 | 71.18399 |
| VLTPEEQLADK | 11 | O75822 | Eukaryotic translation initiation factor 3 s | EIF3J | 29.063 | 2 | 55.194336 | yes | 2.597E-07 | 621.83246 | 367.57489 |
| ITNSLTVLCSEK | 12 | O75822 | Eukaryotic translation initiation factor 3 s | EIF3J | 29.063 | 2 | 55.194336 | yes | 9.722E-08 | 682.85815 | 110.65662 |
| EEVMVLER | 8 | O75909 | Cyclin-K OS=Homo sapiens OX=9606 G | CCNK | 64.24 | 2 | 48.42288 | yes | 1.57E-07 | 502.75769 | 220.70409 |
| LGLHYDTLATGIIY | 17 | O75909 | Cyclin-K OS=Homo sapiens OX=9606 G | CCNK | 64.24 | 3 | 48.42288 | yes | 2.112E-07 | 664.02301 | 121.24601 |
| YVTGACCLFLAGK | 13 | O75909 | Cyclin-K OS=Homo sapiens OX=9606 G | CCNK | 64.24 | 2 | 48.42288 | yes | 1.15E-07 | 730.35724 | 82.506042 |
| AWDDFFPGSDR | 11 | O75915 | PRA1 family protein 3 OS=Homo sapiens | ARL6IP5 | 21.615 | 2 | 37.93526 | yes | 5.581E-07 | 656.78326 | 35.847301 |
| LTADPDSEIATTSL | 15 | O75925 | E3 SUMO-protein ligase PIAS1 OS=Hon | PIAS1 | 71.836 | 2 | 52.63667 | no | 9.722E-08 | 795.40448 | 224.38376 |
| VSLLCPLGK | 9 | O75925 | E3 SUMO-protein ligase PIAS1 OS=Hon | PIAS1 | 71.836 | 2 | 52.63667 | yes | 1.404E-07 | 493.78879 | 178.97519 |
| QLSSTVLLQR | 10 | O75925 | E3 SUMO-protein ligase PIAS1 OS=Hon | PIAS1 | 71.836 | 2 | 52.63667 | yes | 1.456E-07 | 572.83789 | 100.24281 |

| | | | | | | | | | | | |
|---|---|---|---|---|---|---|---|---|---|---|---|
| CDFTVQVQLR | 10 | O75925 | E3 SUMO-protein ligase PIAS1 OS=Hon | PIAS1 | 71.836 | 2 | 52.63667 | yes | 1.979E-06 | 633.31897 | 68.67466 |
| YCTDCDEIQFK | 11 | O75925 | E3 SUMO-protein ligase PIAS1 OS=Hon | PIAS1 | 71.836 | 2 | 52.63667 | yes | 1.418E-07 | 739.79999 | 65.712654 |
| RPSRPINITSLVR | 13 | O75925 | E3 SUMO-protein ligase PIAS1 OS=Hon | PIAS1 | 71.836 | 3 | 52.63667 | yes | 8.676E-06 | 503.63885 | 24.508892 |
| VSELQVLLGYAGR | 13 | O75925 | E3 SUMO-protein ligase PIAS1 OS=Hon | PIAS1 | 71.836 | 2 | 52.63667 | yes | 9.722E-08 | 702.89594 | 24.256966 |
| EMESNWVSLVSK | 12 | O75934 | Pre-mRNA-splicing factor SPF27 OS=Hc | BCAS2 | 26.132 | 2 | 55.151943 | yes | 9.968E-08 | 704.84247 | 1431.3835 |
| TIVQLENEIYQIK | 13 | O75934 | Pre-mRNA-splicing factor SPF27 OS=Hc | BCAS2 | 26.132 | 2 | 55.151943 | yes | 9.754E-08 | 795.94049 | 1135.0844 |
| EAAAALVEEETR | 12 | O75934 | Pre-mRNA-splicing factor SPF27 OS=Hc | BCAS2 | 26.132 | 2 | 55.151943 | yes | 9.809E-08 | 644.82263 | 1025.563 |
| HIQDLNWQR | 9 | O75934 | Pre-mRNA-splicing factor SPF27 OS=Hc | BCAS2 | 26.132 | 2 | 55.151943 | yes | 1E-07 | 605.30981 | 603.83575 |
| QPIELLSMK | 9 | O75934 | Pre-mRNA-splicing factor SPF27 OS=Hc | BCAS2 | 26.132 | 2 | 55.151943 | yes | 9.722E-08 | 529.79938 | 483.96725 |
| KHIQDLNWQR | 10 | O75934 | Pre-mRNA-splicing factor SPF27 OS=Hc | BCAS2 | 26.132 | 2 | 55.151943 | yes | 9.722E-08 | 669.3573 | 89.849693 |
| RYELPAPSSGQK | 12 | O75934 | Pre-mRNA-splicing factor SPF27 OS=Hc | BCAS2 | 26.132 | 2 | 55.151943 | yes | 2.029E-07 | 666.84894 | 82.674591 |
| EAAAALVEEETRR | 13 | O75934 | Pre-mRNA-splicing factor SPF27 OS=Hc | BCAS2 | 26.132 | 2 | 55.151943 | yes | 4.548E-07 | 722.87317 | 45.968758 |
| SIFASPESVTGK | 12 | O75940 | Survival of motor neuron-related-splicing | SMNDC1 | 26.712 | 2 | 48.750732 | yes | 1.127E-07 | 611.81934 | 691.19336 |
| DLQEVIELTK | 10 | O75940 | Survival of motor neuron-related-splicing | SMNDC1 | 26.712 | 2 | 48.750732 | yes | 1.108E-07 | 594.32953 | 614.30231 |
| WQQFNNR | 7 | O75940 | Survival of motor neuron-related-splicing | SMNDC1 | 26.712 | 2 | 48.750732 | yes | 1.341E-07 | 496.73849 | 457.91385 |
| EMIAQQR | 7 | O75940 | Survival of motor neuron-related-splicing | SMNDC1 | 26.712 | 2 | 48.750732 | yes | 2.107E-07 | 438.22382 | 350.60202 |
| VGVGTCGIADK | 11 | O75940 | Survival of motor neuron-related-splicing | SMNDC1 | 26.712 | 2 | 48.750732 | yes | 3.4E-07 | 538.77386 | 132.76134 |
| IKELEQER | 8 | O75940 | Survival of motor neuron-related-splicing | SMNDC1 | 26.712 | 2 | 48.750732 | yes | 2.132E-07 | 522.78784 | 128.35191 |
| KDLQEVIELTK | 11 | O75940 | Survival of motor neuron-related-splicing | SMNDC1 | 26.712 | 2 | 48.750732 | yes | 2.338E-07 | 658.37701 | 38.977573 |
| PMTQYQDTSK | 10 | O75940 | Survival of motor neuron-related-splicing | SMNDC1 | 26.712 | 2 | 48.750732 | yes | 1.544E-07 | 599.77405 | 37.088802 |
| VQVQVVER | 8 | O75955 | Flotillin-1 OS=Homo sapiens OX=9606 ( | FLOT1 | 47.356 | 2 | 51.940598 | yes | 1.301E-07 | 478.77982 | 288.40421 |
| VTGEVLDILTR | 11 | O75955 | Flotillin-1 OS=Homo sapiens OX=9606 ( | FLOT1 | 47.356 | 2 | 51.940598 | yes | 9.722E-08 | 608.35083 | 138.6981 |
| AQQVAVQEQEIAR | 13 | O75955 | Flotillin-1 OS=Homo sapiens OX=9606 ( | FLOT1 | 47.356 | 2 | 51.940598 | yes | 1.001E-07 | 735.38898 | 116.83665 |
| ITLVSSGSGTMGA | 15 | O75955 | Flotillin-1 OS=Homo sapiens OX=9606 ( | FLOT1 | 47.356 | 2 | 51.940598 | yes | 2.399E-07 | 690.36359 | 109.37846 |
| AQADLAYQLQVA | 13 | O75955 | Flotillin-1 OS=Homo sapiens OX=9606 ( | FLOT1 | 47.356 | 2 | 51.940598 | yes | 4.084E-06 | 709.88556 | 98.928467 |
| HGVPISVTGIAQVI | 14 | O75955 | Flotillin-1 OS=Homo sapiens OX=9606 ( | FLOT1 | 47.356 | 2 | 51.940598 | yes | 2.165E-07 | 703.41174 | 92.455032 |
| SQLIMQAEAEAAS | 15 | O75955 | Flotillin-1 OS=Homo sapiens OX=9606 ( | FLOT1 | 47.356 | 2 | 51.940598 | yes | 9.722E-08 | 802.40906 | 84.748375 |
| TEAEIAHIALETLE | 18 | O75955 | Flotillin-1 OS=Homo sapiens OX=9606 ( | FLOT1 | 47.356 | 3 | 51.940598 | yes | 1.23E-07 | 673.34821 | 73.543518 |
| VFVLPCIQQIQR | 12 | O75955 | Flotillin-1 OS=Homo sapiens OX=9606 ( | FLOT1 | 47.356 | 2 | 51.940598 | yes | 1.31E-07 | 750.92139 | 42.783424 |
| ISLNTLTLNVK | 11 | O75955 | Flotillin-1 OS=Homo sapiens OX=9606 ( | FLOT1 | 47.356 | 2 | 51.940598 | yes | 1.735E-07 | 608.36902 | 38.99789 |
| LLIQLADGFCEK | 12 | O75962 | Triple functional domain protein OS=Hor | TRIO | 346.901 | 2 | 48.466164 | yes | 1.144E-07 | 703.87103 | 63.637508 |
| APIEDLDLEGQK | 12 | O75962 | Triple functional domain protein OS=Hor | TRIO | 346.901 | 2 | 48.466164 | yes | 6.633E-07 | 664.34064 | 61.230125 |
| VQDLLEFLHEK | 11 | O75962 | Triple functional domain protein OS=Hor | TRIO | 346.901 | 2 | 48.466164 | yes | 3.564E-07 | 685.86938 | 60.750393 |
| YMSNVDSWCK | 10 | O75962 | Triple functional domain protein OS=Hor | TRIO | 346.901 | 2 | 48.466164 | yes | 5.331E-07 | 645.26813 | 36.86731 |
| MFDWITHNK | 9 | O75962 | Triple functional domain protein OS=Hor | TRIO | 346.901 | 2 | 48.466164 | yes | 6.745E-06 | 596.28442 | 33.486839 |
| DFFLGELEK | 9 | O75962 | Triple functional domain protein OS=Hor | TRIO | 346.901 | 2 | 48.466164 | yes | 3.666E-06 | 549.2793 | 24.690674 |
| THQSALQVQQK | 11 | O75962 | Triple functional domain protein OS=Hor | TRIO | 346.901 | 2 | 48.466164 | no | 1.687E-06 | 634.34131 | 10.638392 |
| QVLGWIR | 7 | O75962 | Triple functional domain protein OS=Hor | TRIO | 346.901 | 2 | 48.466164 | no | 3.29E-06 | 436.26105 | 9.3695793 |
| GELVGGLDIVK | 11 | O76003 | Glutaredoxin-3 OS=Homo sapiens OX=9 | GLRX3 | 37.433 | 2 | 29.651995 | yes | 2.425E-06 | 550.32153 | 67.645546 |

| | | | | | | | | | | | |
|---|---|---|---|---|---|---|---|---|---|---|---|
| ENGELLPILR | 10 | O76003 | Glutaredoxin-3 OS=Homo sapiens OX=9 | GLRX3 | 37.433 | 2 | 29.651995 | yes | 5.191E-06 | 577.3324 | 10.914615 |
| EINDCIGGTVLNIS | 15 | O76021 | Ribosomal L1 domain-containing protein | RSL1D1 | 54.973 | 2 | 54.352448 | yes | 1.402E-07 | 816.91669 | 1820.9209 |
| TVSQIISLQTLK | 12 | O76021 | Ribosomal L1 domain-containing protein | RSL1D1 | 54.973 | 2 | 54.352448 | yes | 1.072E-07 | 665.90063 | 1367.847 |
| KVPVSVNLLSK | 11 | O76021 | Ribosomal L1 domain-containing protein | RSL1D1 | 54.973 | 2 | 54.352448 | yes | 2.466E-07 | 592.37408 | 1266.3723 |
| AVDALLTHCK | 10 | O76021 | Ribosomal L1 domain-containing protein | RSL1D1 | 54.973 | 2 | 54.352448 | yes | 9.722E-08 | 564.29755 | 965.02325 |
| SDSEDICLFTK | 11 | O76021 | Ribosomal L1 domain-containing protein | RSL1D1 | 54.973 | 2 | 54.352448 | yes | 1.11E-07 | 657.79773 | 804.69037 |
| LLPSLIGR | 8 | O76021 | Ribosomal L1 domain-containing protein | RSL1D1 | 54.973 | 2 | 54.352448 | yes | 1.101E-07 | 434.78436 | 400.32486 |
| VPVSVNLLSK | 10 | O76021 | Ribosomal L1 domain-containing protein | RSL1D1 | 54.973 | 2 | 54.352448 | yes | 1.218E-07 | 528.3266 | 337.73151 |
| DDVAPESGDTTVK | 13 | O76021 | Ribosomal L1 domain-containing protein | RSL1D1 | 54.973 | 2 | 54.352448 | yes | 1.127E-07 | 667.30951 | 190.98143 |
| LTLPHSIR | 8 | O76021 | Ribosomal L1 domain-containing protein | RSL1D1 | 54.973 | 2 | 54.352448 | yes | 9.722E-08 | 468.78491 | 188.52798 |
| ALPASETPK | 9 | O76021 | Ribosomal L1 domain-containing protein | RSL1D1 | 54.973 | 2 | 54.352448 | yes | 1.66E-06 | 457.25311 | 83.41066 |
| KALPASETPK | 10 | O76021 | Ribosomal L1 domain-containing protein | RSL1D1 | 54.973 | 2 | 54.352448 | yes | 2.553E-07 | 521.3006 | 76.958672 |
| KAVDALLTHCK | 11 | O76021 | Ribosomal L1 domain-containing protein | RSL1D1 | 54.973 | 2 | 54.352448 | yes | 9.722E-08 | 628.34497 | 71.652283 |
| IYNNIPANLR | 10 | O76031 | ATP-dependent Clp protease ATP-bindin | CLPX | 69.224 | 2 | 53.47943 | yes | 3.238E-06 | 594.3302 | 122.22569 |
| AQQGIVFLDEVDK | 13 | O76031 | ATP-dependent Clp protease ATP-bindin | CLPX | 69.224 | 2 | 53.47943 | yes | 1.979E-07 | 731.38281 | 119.35899 |
| IGSVPGIHQLR | 11 | O76031 | ATP-dependent Clp protease ATP-bindin | CLPX | 69.224 | 2 | 53.47943 | yes | 1.987E-07 | 588.84601 | 105.1751 |
| TLVQILTEPR | 10 | O76031 | ATP-dependent Clp protease ATP-bindin | CLPX | 69.224 | 2 | 53.47943 | yes | 3.689E-07 | 585.34808 | 66.596092 |
| SNILLLGPTGSGK | 13 | O76031 | ATP-dependent Clp protease ATP-bindin | CLPX | 69.224 | 2 | 53.47943 | yes | 9.722E-08 | 628.86407 | 66.561684 |
| LPVVVPLHSLDEK | 13 | O76031 | ATP-dependent Clp protease ATP-bindin | CLPX | 69.224 | 2 | 53.47943 | yes | 1.317E-07 | 723.42175 | 65.273987 |
| DVGGEGVQQGLLI | 13 | O76031 | ATP-dependent Clp protease ATP-bindin | CLPX | 69.224 | 2 | 53.47943 | yes | 9.722E-08 | 650.34882 | 60.560337 |
| LLQDANYNVEK | 11 | O76031 | ATP-dependent Clp protease ATP-bindin | CLPX | 69.224 | 2 | 53.47943 | yes | 9.722E-08 | 653.83551 | 53.50407 |
| GGEVLDSSHDDIK | 13 | O76031 | ATP-dependent Clp protease ATP-bindin | CLPX | 69.224 | 2 | 53.47943 | yes | 2.282E-07 | 686.323 | 43.473671 |
| VLGPLPSEQMK | 11 | O76039 | Cyclin-dependent kinase-like 5 OS=Hom | CDKL5 | 107.52 | 2 | 54.177586 | yes | 1.68E-07 | 599.82867 | 228.42601 |
| HLYNDPVPR | 9 | O76039 | Cyclin-dependent kinase-like 5 OS=Hom | CDKL5 | 107.52 | 2 | 54.177586 | yes | 1.004E-07 | 555.78815 | 150.29953 |
| FPAVNHPQSLER | 12 | O76039 | Cyclin-dependent kinase-like 5 OS=Hom | CDKL5 | 107.52 | 3 | 54.177586 | yes | 2.97E-07 | 465.57736 | 70.390388 |
| FPAVNHPQSLER | 12 | O76039 | Cyclin-dependent kinase-like 5 OS=Hom | CDKL5 | 107.52 | 3 | 54.177586 | yes | 9.722E-08 | 697.86243 | 70.390388 |
| FPAVNHPQSLER | 12 | O76039 | Cyclin-dependent kinase-like 5 OS=Hom | CDKL5 | 107.52 | 2 | 54.177586 | yes | 2.97E-07 | 465.57736 | 70.390388 |
| FPAVNHPQSLER | 12 | O76039 | Cyclin-dependent kinase-like 5 OS=Hom | CDKL5 | 107.52 | 2 | 54.177586 | yes | 9.722E-08 | 697.86243 | 70.390388 |
| GLDGSLSIGQGMA | 15 | O76039 | Cyclin-dependent kinase-like 5 OS=Hom | CDKL5 | 107.52 | 2 | 54.177586 | yes | 1.79E-07 | 716.86426 | 126.08934 |
| DIQNLSVGLPR | 11 | O76039 | Cyclin-dependent kinase-like 5 OS=Hom | CDKL5 | 107.52 | 2 | 54.177586 | yes | 1.811E-07 | 606.34076 | 117.01841 |
| SATEGPSYSEQLG | 15 | O76039 | Cyclin-dependent kinase-like 5 OS=Hom | CDKL5 | 107.52 | 2 | 54.177586 | yes | 9.722E-08 | 762.86249 | 97.499435 |
| ISDLQTQSQPLK | 12 | O76039 | Cyclin-dependent kinase-like 5 OS=Hom | CDKL5 | 107.52 | 2 | 54.177586 | yes | 2.205E-07 | 679.36975 | 95.296021 |
| SPELLLGAPYGK | 12 | O76039 | Cyclin-dependent kinase-like 5 OS=Hom | CDKL5 | 107.52 | 2 | 54.177586 | yes | 1.084E-07 | 622.8479 | 93.377953 |
| IHPLSQASGGSSNI | 15 | O76039 | Cyclin-dependent kinase-like 5 OS=Hom | CDKL5 | 107.52 | 2 | 54.177586 | yes | 1.37E-07 | 762.3999 | 25.75515 |
| DIKPENLLISHNDV | 16 | O76039 | Cyclin-dependent kinase-like 5 OS=Hom | CDKL5 | 107.52 | 3 | 54.177586 | yes | 4.335E-07 | 616.67902 | 25.375942 |
| SVLSEGHQR | 9 | O76071 | Probable cytosolic iron-sulfur protein ass | CIAO1 | 37.841 | 2 | 50.876728 | yes | 9.722E-08 | 506.76215 | 637.30719 |
| SVAWAPSGNLLAT | 16 | O76071 | Probable cytosolic iron-sulfur protein ass | CIAO1 | 37.841 | 2 | 50.876728 | yes | 2.119E-07 | 845.42249 | 322.06812 |
| IWGTEGDSWICK | 12 | O76071 | Probable cytosolic iron-sulfur protein ass | CIAO1 | 37.841 | 2 | 50.876728 | yes | 1E-07 | 726.33484 | 218.13345 |
| HVVWHPSQELLAS | 21 | O76071 | Probable cytosolic iron-sulfur protein ass | CIAO1 | 37.841 | 3 | 50.876728 | yes | 9.85E-08 | 794.729 | 177.95631 |

| | | | | | | | | | | | |
|---|---|---|---|---|---|---|---|---|---|---|---|
| MKDSLVLLGR | 10 | O76071 | Probable cytosolic iron-sulfur protein ass | CIAO1 | 37.841 | 2 | 50.876728 | yes | 1.847E-07 | 566.33136 | 159.63864 |
| YQRPEGL | 7 | O76071 | Probable cytosolic iron-sulfur protein ass | CIAO1 | 37.841 | 2 | 50.876728 | yes | 1.585E-07 | 431.72452 | 137.18652 |
| DSLVLLGR | 8 | O76071 | Probable cytosolic iron-sulfur protein ass | CIAO1 | 37.841 | 2 | 50.876728 | yes | 5.379E-07 | 436.76364 | 77.015846 |
| ELYGQVLYR | 9 | O76094 | Signal recognition particle subunit SRP72 | SRP72 | 74.607 | 2 | 53.326366 | yes | 9.722E-08 | 570.80603 | 542.14282 |
| YDECLAVYR | 9 | O76094 | Signal recognition particle subunit SRP72 | SRP72 | 74.607 | 2 | 53.326366 | yes | 9.722E-08 | 594.7713 | 445.82626 |
| HLPSSDSMSLK | 11 | O76094 | Signal recognition particle subunit SRP72 | SRP72 | 74.607 | 2 | 53.326366 | yes | 4.011E-07 | 601.29773 | 419.4303 |
| EAISDLQQLWK | 11 | O76094 | Signal recognition particle subunit SRP72 | SRP72 | 74.607 | 2 | 53.326366 | yes | 9.722E-08 | 665.8537 | 393.77686 |
| SPAHLSLIR | 9 | O76094 | Signal recognition particle subunit SRP72 | SRP72 | 74.607 | 2 | 53.326366 | yes | 3.27E-07 | 497.29562 | 349.60455 |
| EALNVINTHTK | 11 | O76094 | Signal recognition particle subunit SRP72 | SRP72 | 74.607 | 2 | 53.326366 | yes | 1.046E-07 | 620.33826 | 332.91586 |
| TEEALQLYNQIIK | 13 | O76094 | Signal recognition particle subunit SRP72 | SRP72 | 74.607 | 2 | 53.326366 | yes | 9.722E-08 | 781.92487 | 258.565 |
| VLANNSLSFEK | 11 | O76094 | Signal recognition particle subunit SRP72 | SRP72 | 74.607 | 2 | 53.326366 | yes | 1.186E-07 | 611.32733 | 243.40575 |
| DDVTALHCK | 9 | O76094 | Signal recognition particle subunit SRP72 | SRP72 | 74.607 | 2 | 53.326366 | yes | 9.742E-08 | 529.75043 | 235.64639 |
| LTNAEGVEFK | 10 | O76094 | Signal recognition particle subunit SRP72 | SRP72 | 74.607 | 2 | 53.326366 | yes | 1.532E-07 | 554.28766 | 232.27097 |
| QLQAIEFNK | 9 | O76094 | Signal recognition particle subunit SRP72 | SRP72 | 74.607 | 2 | 53.326366 | yes | 9.722E-08 | 545.79822 | 197.28348 |
| TNLSAVVAAQSNV | 15 | O76094 | Signal recognition particle subunit SRP72 | SRP72 | 74.607 | 2 | 53.326366 | yes | 9.847E-08 | 809.41522 | 159.29221 |
| NSQDDYDEER | 10 | O76094 | Signal recognition particle subunit SRP72 | SRP72 | 74.607 | 2 | 53.326366 | yes | 9.722E-08 | 635.74457 | 130.84293 |
| GTQGATAGASSEL | 17 | O76094 | Signal recognition particle subunit SRP72 | SRP72 | 74.607 | 2 | 53.326366 | yes | 1.232E-07 | 775.86829 | 116.99595 |
| ALLAMYTNQAEQ | 14 | O76094 | Signal recognition particle subunit SRP72 | SRP72 | 74.607 | 2 | 53.326366 | yes | 9.722E-08 | 834.89526 | 110.65726 |
| VVCLIQNGSFK | 11 | O76094 | Signal recognition particle subunit SRP72 | SRP72 | 74.607 | 2 | 53.326366 | yes | 2.033E-07 | 632.83954 | 84.934044 |
| SSESLPAAAEEAPI | 16 | O94761 | ATP-dependent DNA helicase Q4 OS=H | RECQL4 | 133.068 | 2 | 54.270714 | yes | 3.787E-07 | 820.89172 | 414.50336 |
| LGSLDPGWLQR | 11 | O94761 | ATP-dependent DNA helicase Q4 OS=H | RECQL4 | 133.068 | 2 | 54.270714 | yes | 9.722E-08 | 621.33551 | 291.97034 |
| TTGQAGGGLR | 10 | O94761 | ATP-dependent DNA helicase Q4 OS=H | RECQL4 | 133.068 | 2 | 54.270714 | yes | 2.845E-07 | 459.24359 | 235.7332 |
| VSDEPPQLPEPQPI | 14 | O94761 | ATP-dependent DNA helicase Q4 OS=H | RECQL4 | 133.068 | 2 | 54.270714 | yes | 2.206E-07 | 794.90192 | 185.03075 |
| LQDWEDQVR | 9 | O94761 | ATP-dependent DNA helicase Q4 OS=H | RECQL4 | 133.068 | 2 | 54.270714 | yes | 6.223E-07 | 594.78583 | 119.83734 |
| FQNLDSIIIYCNR | 13 | O94761 | ATP-dependent DNA helicase Q4 OS=H | RECQL4 | 133.068 | 2 | 54.270714 | yes | 2.909E-07 | 828.41412 | 111.58119 |
| LVDSMGWELASV | 13 | O94761 | ATP-dependent DNA helicase Q4 OS=H | RECQL4 | 133.068 | 2 | 54.270714 | yes | 9.722E-08 | 731.87158 | 88.871391 |
| YHGLSPSSQAR | 11 | O94761 | ATP-dependent DNA helicase Q4 OS=H | RECQL4 | 133.068 | 2 | 54.270714 | yes | 7.641E-07 | 601.79926 | 70.37999 |
| CHSEVPDFLGAPK | 13 | O94761 | ATP-dependent DNA helicase Q4 OS=H | RECQL4 | 133.068 | 2 | 54.270714 | yes | 5.965E-06 | 728.84808 | 44.281769 |
| SLCYQLPALLYSR | 13 | O94761 | ATP-dependent DNA helicase Q4 OS=H | RECQL4 | 133.068 | 2 | 54.270714 | yes | 1.926E-06 | 792.41614 | 33.060539 |
| DTDQALLTLLQGK | 13 | O94761 | ATP-dependent DNA helicase Q4 OS=H | RECQL4 | 133.068 | 2 | 54.270714 | yes | 5.112E-07 | 708.39069 | 29.480312 |
| TCLHAAWVPGSG | 14 | O94761 | ATP-dependent DNA helicase Q4 OS=H | RECQL4 | 133.068 | 3 | 54.270714 | yes | 3.242E-06 | 490.24197 | 10.811985 |
| LLEEALSSNR | 10 | O94762 | ATP-dependent DNA helicase Q5 OS=H | RECQL5 | 108.859 | 2 | 56.591457 | yes | 1.183E-07 | 566.30383 | 288.67877 |
| GLSGCGIVYCR | 11 | O94762 | ATP-dependent DNA helicase Q5 OS=H | RECQL5 | 108.859 | 2 | 56.591457 | yes | 9.722E-08 | 621.29193 | 204.29407 |
| ANLFYDVQFK | 10 | O94762 | ATP-dependent DNA helicase Q5 OS=H | RECQL5 | 108.859 | 2 | 56.591457 | yes | 1.243E-06 | 622.81915 | 187.0448 |
| DVFVCMPTGAGK | 12 | O94762 | ATP-dependent DNA helicase Q5 OS=H | RECQL5 | 108.859 | 2 | 56.591457 | yes | 9.722E-08 | 641.30194 | 119.54115 |
| SLCYQLPALLAK | 12 | O94762 | ATP-dependent DNA helicase Q5 OS=H | RECQL5 | 108.859 | 2 | 56.591457 | yes | 1.104E-07 | 688.88397 | 115.57861 |
| DQGTPEVQPTPAK | 13 | O94762 | ATP-dependent DNA helicase Q5 OS=H | RECQL5 | 108.859 | 2 | 56.591457 | yes | 1.795E-07 | 684.34369 | 98.633484 |
| AVELEHETFR | 10 | O94762 | ATP-dependent DNA helicase Q5 OS=H | RECQL5 | 108.859 | 2 | 56.591457 | yes | 9.722E-08 | 615.80933 | 89.771881 |
| EACEQLAIELSCR | 13 | O94762 | ATP-dependent DNA helicase Q5 OS=H | RECQL5 | 108.859 | 2 | 56.591457 | yes | 3.019E-07 | 789.86615 | 83.105507 |

| | | | | | | | | | | | |
|---|---|---|---|---|---|---|---|---|---|---|---|
| SMAGYYQESGR | 11 | O94762 | ATP-dependent DNA helicase Q5 OS=H( | RECQL5 | 108.859 | 2 | 56.591457 | yes | 9.722E-08 | 624.76935 | 79.164207 |
| IEEFVPPDENCPLK | 14 | O94762 | ATP-dependent DNA helicase Q5 OS=H( | RECQL5 | 108.859 | 2 | 56.591457 | yes | 7.481E-07 | 843.90582 | 77.324142 |
| TPLQESATMAVVK | 13 | O94762 | ATP-dependent DNA helicase Q5 OS=H( | RECQL5 | 108.859 | 2 | 56.591457 | yes | 9.722E-08 | 687.86853 | 69.750946 |
| DQVSFLIR | 8 | O94762 | ATP-dependent DNA helicase Q5 OS=H( | RECQL5 | 108.859 | 2 | 56.591457 | yes | 1.444E-06 | 489.27435 | 61.458958 |
| EWNLFYQK | 8 | O94762 | ATP-dependent DNA helicase Q5 OS=H( | RECQL5 | 108.859 | 2 | 56.591457 | yes | 4.301E-07 | 564.27966 | 61.19146 |
| TLVQNDWMEEK | 11 | O94762 | ATP-dependent DNA helicase Q5 OS=H( | RECQL5 | 108.859 | 2 | 56.591457 | yes | 1.93E-07 | 696.82684 | 57.39085 |
| HLSHLLTQK | 9 | O94762 | ATP-dependent DNA helicase Q5 OS=H( | RECQL5 | 108.859 | 2 | 56.591457 | yes | 3.212E-07 | 538.81421 | 54.548447 |
| VFGFDSFK | 8 | O94762 | ATP-dependent DNA helicase Q5 OS=H( | RECQL5 | 108.859 | 2 | 56.591457 | yes | 3.942E-07 | 473.73709 | 31.033779 |
| TSDIFEADIANDVK | 14 | O94763 | Unconventional prefoldin RPB5 interacto | URI1 | 59.833 | 2 | 50.868378 | yes | 9.932E-08 | 769.37268 | 113.70171 |
| MSDAAGDIVDIR | 12 | O94763 | Unconventional prefoldin RPB5 interacto | URI1 | 59.833 | 2 | 50.868378 | yes | 1.104E-07 | 631.80591 | 95.734657 |
| VEFTEDLQK | 9 | O94763 | Unconventional prefoldin RPB5 interacto | URI1 | 59.833 | 2 | 50.868378 | yes | 9.722E-08 | 554.77966 | 76.84552 |
| EFVSPSLTPPPAIAI | 24 | O94763 | Unconventional prefoldin RPB5 interacto | URI1 | 59.833 | 3 | 50.868378 | yes | 1.475E-06 | 846.46362 | 20.955494 |
| NSTGSGHSAQELP' | 16 | O94763 | Unconventional prefoldin RPB5 interacto | URI1 | 59.833 | 3 | 50.868378 | yes | 8.654E-05 | 552.27631 | 14.389824 |
| ISQLQAELR | 9 | O94766 | Galactosylgalactosylxylosylprotein 3-beta | B3GAT3 | 37.122 | 2 | 41.42979 | yes | 3.514E-07 | 529.30365 | 241.57745 |
| GVSVWPVGLVGG | 14 | O94766 | Galactosylgalactosylxylosylprotein 3-beta | B3GAT3 | 37.122 | 2 | 41.42979 | yes | 8.122E-07 | 698.409 | 59.31982 |
| TLLADQGEIR | 10 | O94776 | Metastasis-associated protein MTA2 OS= | MTA2 | 75.024 | 2 | 51.38954 | yes | 9.722E-08 | 558.3064 | 520.93115 |
| QVYIPTYTK | 9 | O94776 | Metastasis-associated protein MTA2 OS= | MTA2 | 75.024 | 2 | 51.38954 | yes | 4.9E-06 | 556.80298 | 324.12808 |
| AMSTLVPQGGPVI | 15 | O94776 | Metastasis-associated protein MTA2 OS= | MTA2 | 75.024 | 2 | 51.38954 | yes | 1.041E-07 | 793.41309 | 264.95444 |
| LNPADAPNPVVFV | 16 | O94776 | Metastasis-associated protein MTA2 OS= | MTA2 | 75.024 | 2 | 51.38954 | yes | 1.684E-07 | 826.95398 | 249.17303 |
| QIDQFLVVAR | 10 | O94776 | Metastasis-associated protein MTA2 OS= | MTA2 | 75.024 | 2 | 51.38954 | no | 9.915E-08 | 594.84039 | 213.56656 |
| VWDPDNPLTDR | 11 | O94776 | Metastasis-associated protein MTA2 OS= | MTA2 | 75.024 | 2 | 51.38954 | yes | 9.809E-08 | 664.3175 | 178.06897 |
| RIEELNK | 7 | O94776 | Metastasis-associated protein MTA2 OS= | MTA2 | 75.024 | 2 | 51.38954 | no | 2.63E-07 | 451.25873 | 155.81372 |
| DISSSLNSLADSNA | 15 | O94776 | Metastasis-associated protein MTA2 OS= | MTA2 | 75.024 | 2 | 51.38954 | yes | 1.77E-07 | 775.37628 | 149.70319 |
| ALTHLEMR | 8 | O94776 | Metastasis-associated protein MTA2 OS= | MTA2 | 75.024 | 2 | 51.38954 | yes | 2.501E-07 | 485.76056 | 121.0696 |
| QTFLLQTTK | 9 | O94776 | Metastasis-associated protein MTA2 OS= | MTA2 | 75.024 | 2 | 51.38954 | yes | 1.33E-07 | 540.30841 | 94.964058 |
| RPNLPLK | 7 | O94776 | Metastasis-associated protein MTA2 OS= | MTA2 | 75.024 | 2 | 51.38954 | yes | 1.929E-06 | 419.26889 | 73.524437 |
| DITLFHAMDTLQR | 13 | O94776 | Metastasis-associated protein MTA2 OS= | MTA2 | 75.024 | 2 | 51.38954 | yes | 3.22E-07 | 780.89557 | 15.515372 |
| GVVENYNDEEVSI | 14 | O94782 | Ubiquitin carboxyl-terminal hydrolase 1 ( | USP1 | 88.208 | 2 | 44.305077 | yes | 2.245E-07 | 811.88647 | 36.354057 |
| MPEVITIHLK | 10 | O94782 | Ubiquitin carboxyl-terminal hydrolase 1 ( | USP1 | 88.208 | 2 | 44.305077 | yes | 2.964E-07 | 590.84155 | 24.777678 |
| FFDTELENLER | 11 | O94804 | Serine/threonine-protein kinase 10 OS=H | STK10 | 112.136 | 2 | 48.533024 | yes | 1.135E-07 | 706.83826 | 30.510921 |
| CHLLVEHETQK | 11 | O94804 | Serine/threonine-protein kinase 10 OS=H | STK10 | 112.136 | 2 | 48.533024 | no | 1.845E-06 | 697.34827 | 10.771919 |
| LSNVSLPGPGLSVI | 15 | O94806 | Serine/threonine-protein kinase D3 OS=H | PRKD3 | 100.471 | 2 | 52.410637 | yes | 1.368E-07 | 746.92773 | 503.22153 |
| LCDFGFAR | 8 | O94806 | Serine/threonine-protein kinase D3 OS=H | PRKD3 | 100.471 | 2 | 52.410637 | no | 1.457E-07 | 493.23166 | 313.64963 |
| SVVGTPAYLAPEV | 15 | O94806 | Serine/threonine-protein kinase D3 OS=H | PRKD3 | 100.471 | 2 | 52.410637 | no | 8.245E-07 | 786.44324 | 222.69061 |
| YITHESDDAR | 10 | O94806 | Serine/threonine-protein kinase D3 OS=H | PRKD3 | 100.471 | 2 | 52.410637 | no | 1.941E-07 | 603.77289 | 147.24303 |
| CLTLFQNESGSK | 12 | O94806 | Serine/threonine-protein kinase D3 OS=H | PRKD3 | 100.471 | 2 | 52.410637 | yes | 9.722E-08 | 692.33228 | 123.20616 |
| EGWMVHYTSR | 10 | O94806 | Serine/threonine-protein kinase D3 OS=H | PRKD3 | 100.471 | 2 | 52.410637 | yes | 1.018E-07 | 633.29022 | 100.29027 |
| MFFLDPSDLDVER | 19 | O94806 | Serine/threonine-protein kinase D3 OS=H | PRKD3 | 100.471 | 3 | 52.410637 | yes | 1.517E-07 | 752.35608 | 72.957764 |
| EIPLSEILR | 9 | O94806 | Serine/threonine-protein kinase D3 OS=H | PRKD3 | 100.471 | 2 | 52.410637 | yes | 3.31E-06 | 535.31622 | 46.415771 |

| | | | | | | | | | | | |
|---|---|---|---|---|---|---|---|---|---|---|---|
| GLDDTEEPSPPEDI | 14 | O94806 | Serine/threonine-protein kinase D3 OS=H | PRKD3 | 100.471 | 2 | 52.410637 | yes | 2.93E-06 | 764.83612 | 43.060867 |
| FPECGFFGMYDK | 12 | O94806 | Serine/threonine-protein kinase D3 OS=H | PRKD3 | 100.471 | 2 | 52.410637 | yes | 3.207E-06 | 749.3125 | 12.419304 |
| HSVPQIR | 7 | O94822 | E3 ubiquitin-protein ligase listerin OS=H | LTN1 | 200.553 | 2 | 51.253426 | yes | 3.466E-07 | 418.74051 | 55.109184 |
| LADISINYVNER | 12 | O94822 | E3 ubiquitin-protein ligase listerin OS=H | LTN1 | 200.553 | 2 | 51.253426 | yes | 9.722E-08 | 703.86737 | 45.59787 |
| NDYLIGDVYVER | 12 | O94822 | E3 ubiquitin-protein ligase listerin OS=H | LTN1 | 200.553 | 2 | 51.253426 | yes | 2.247E-06 | 728.35938 | 43.760712 |
| GLELISK | 7 | O94826 | Mitochondrial import receptor subunit TC | TOMM70 | 67.455 | 1 | 24.120823 | yes | 3.512E-06 | 759.46106 | 25.951998 |
| IVASMWDSLGEEQ | 14 | O94842 | TOX high mobility group box family men | TOX4 | 66.195 | 2 | 49.566868 | no | 1.197E-07 | 796.88489 | 1413.3738 |
| INLQQQPPPLQIK | 13 | O94842 | TOX high mobility group box family men | TOX4 | 66.195 | 2 | 49.566868 | yes | 1.182E-07 | 758.94592 | 1216.7921 |
| GQNPNATFGEVSK | 13 | O94842 | TOX high mobility group box family men | TOX4 | 66.195 | 2 | 49.566868 | no | 7.251E-07 | 674.82825 | 953.03577 |
| SGCENPPIVSK | 11 | O94842 | TOX high mobility group box family men | TOX4 | 66.195 | 2 | 49.566868 | yes | 1.021E-07 | 594.28992 | 570.04883 |
| NSNTVVFVK | 9 | O94842 | TOX high mobility group box family men | TOX4 | 66.195 | 2 | 49.566868 | yes | 1.274E-07 | 504.27966 | 76.655243 |
| DVFLAWVASR | 10 | O94842 | TOX high mobility group box family men | TOX4 | 66.195 | 2 | 49.566868 | yes | 1.392E-07 | 582.31403 | 73.196617 |
| SVPLPTLK | 8 | O94842 | TOX high mobility group box family men | TOX4 | 66.195 | 2 | 49.566868 | yes | 1.581E-07 | 427.77094 | 70.4394 |
| LLPQTTPENINK | 12 | O94851 | [F-actin]-monooxygenase MICAL2 OS=H | MICAL2 | 219.062 | 2 | 53.428032 | yes | 4.108E-07 | 684.38007 | 355.84061 |
| CLIVGGGPCGLR | 12 | O94851 | [F-actin]-monooxygenase MICAL2 OS=H | MICAL2 | 219.062 | 2 | 53.428032 | yes | 3.925E-07 | 629.82336 | 215.45012 |
| SGLALCAIIHR | 11 | O94851 | [F-actin]-monooxygenase MICAL2 OS=H | MICAL2 | 219.062 | 2 | 53.428032 | no | 9.722E-08 | 605.8399 | 137.29945 |
| ELEHYPLER | 9 | O94851 | [F-actin]-monooxygenase MICAL2 OS=H | MICAL2 | 219.062 | 2 | 53.428032 | yes | 4.559E-07 | 593.29858 | 122.62814 |
| TAIELAYLGAK | 11 | O94851 | [F-actin]-monooxygenase MICAL2 OS=H | MICAL2 | 219.062 | 2 | 53.428032 | yes | 1.083E-07 | 575.32935 | 115.01394 |
| CSICATTLR | 9 | O94851 | [F-actin]-monooxygenase MICAL2 OS=H | MICAL2 | 219.062 | 2 | 53.428032 | yes | 4.492E-07 | 541.26007 | 102.14754 |
| NFEQYTLDPGTR | 12 | O94851 | [F-actin]-monooxygenase MICAL2 OS=H | MICAL2 | 219.062 | 2 | 53.428032 | yes | 6.435E-07 | 720.84131 | 54.411747 |
| EETGIDLENIVYYK | 14 | O94851 | [F-actin]-monooxygenase MICAL2 OS=H | MICAL2 | 219.062 | 2 | 53.428032 | yes | 1.202E-07 | 843.41705 | 53.990501 |
| GFTNLDEPSNFSSF | 14 | O94851 | [F-actin]-monooxygenase MICAL2 OS=H | MICAL2 | 219.062 | 2 | 53.428032 | yes | 2.229E-07 | 785.86023 | 53.376724 |
| EEEATWQEQEAPF | 13 | O94851 | [F-actin]-monooxygenase MICAL2 OS=H | MICAL2 | 219.062 | 2 | 53.428032 | yes | 9.722E-08 | 801.85516 | 28.87509 |
| SSISNNYLNLTFPR | 14 | O94851 | [F-actin]-monooxygenase MICAL2 OS=H | MICAL2 | 219.062 | 2 | 53.428032 | yes | 1.693E-06 | 813.41772 | 27.838596 |
| DCTHYFVMTAK | 11 | O94851 | [F-actin]-monooxygenase MICAL2 OS=H | MICAL2 | 219.062 | 2 | 53.428032 | yes | 2.093E-07 | 686.80487 | 14.37718 |
| VEEISGVAFIFNQK | 14 | O94851 | [F-actin]-monooxygenase MICAL2 OS=H | MICAL2 | 219.062 | 2 | 53.428032 | no | 2.455E-07 | 790.91956 | 11.099684 |
| YWGEIPISSSQTNR | 14 | O94864 | STAGA complex 65 subunit gamma OS= | SUPT7L | 46.193 | 2 | 49.05429 | yes | 1.66E-07 | 819.39954 | 189.93045 |
| HSDPESDFYR | 10 | O94864 | STAGA complex 65 subunit gamma OS= | SUPT7L | 46.193 | 2 | 49.05429 | yes | 1.073E-07 | 626.76508 | 88.860138 |
| FFADDTQAALTK | 12 | O94874 | E3 UFM1-protein ligase 1 OS=Homo sap | UFL1 | 89.596 | 2 | 50.34761 | yes | 9.818E-08 | 664.33008 | 84.499641 |
| LQESGQVTISELCK | 14 | O94874 | E3 UFM1-protein ligase 1 OS=Homo sap | UFL1 | 89.596 | 2 | 50.34761 | yes | 1.084E-07 | 796.40344 | 32.365322 |
| SVFMSSTTSASGT( | 15 | O94874 | E3 UFM1-protein ligase 1 OS=Homo sap | UFL1 | 89.596 | 2 | 50.34761 | yes | 1.498E-06 | 738.34338 | 28.349318 |
| VDLPGSSTTLTK | 12 | O94875 | Sorbin and SH3 domain-containing protei | SORBS2 | 124.108 | 2 | 52.70148 | yes | 2.486E-07 | 609.83246 | 395.49408 |
| SYSSTLTDMGR | 11 | O94875 | Sorbin and SH3 domain-containing protei | SORBS2 | 124.108 | 2 | 52.70148 | yes | 1.525E-07 | 609.27698 | 304.52109 |
| PASLYQSSIDR | 11 | O94875 | Sorbin and SH3 domain-containing protei | SORBS2 | 124.108 | 2 | 52.70148 | yes | 9.722E-08 | 618.81458 | 175.11224 |
| PMSSASMASDFR | 12 | O94875 | Sorbin and SH3 domain-containing protei | SORBS2 | 124.108 | 2 | 52.70148 | yes | 9.722E-08 | 643.77881 | 139.10754 |
| IHSLSSNKPQR | 11 | O94875 | Sorbin and SH3 domain-containing protei | SORBS2 | 124.108 | 2 | 52.70148 | yes | 2.225E-07 | 633.8493 | 105.13097 |
| TQTYRPLSK | 9 | O94875 | Sorbin and SH3 domain-containing protei | SORBS2 | 124.108 | 2 | 52.70148 | yes | 9.861E-08 | 547.30365 | 100.31177 |
| QGIFPVSYVEVVK | 13 | O94875 | Sorbin and SH3 domain-containing protei | SORBS2 | 124.108 | 2 | 52.70148 | yes | 9.722E-08 | 732.90851 | 98.470261 |
| KGDTVYILR | 9 | O94875 | Sorbin and SH3 domain-containing protei | SORBS2 | 124.108 | 2 | 52.70148 | yes | 1.527E-07 | 532.80859 | 87.983482 |

| | | | | | | | | | | | |
|---|---|---|---|---|---|---|---|---|---|---|---|
| ESDVIDVMEK | 10 | O94875 | Sorbin and SH3 domain-containing protei | SORBS2 | 124.108 | 2 | 52.70148 | yes | 2.286E-07 | 582.77631 | 69.464081 |
| VGIFPISYVEK | 11 | O94875 | Sorbin and SH3 domain-containing protei | SORBS2 | 124.108 | 2 | 52.70148 | yes | 9.968E-08 | 626.35284 | 52.751236 |
| RVDQNWYEGK | 10 | O94875 | Sorbin and SH3 domain-containing protei | SORBS2 | 124.108 | 2 | 52.70148 | yes | 1.737E-07 | 647.81238 | 46.684391 |
| SNEDSLILEK | 10 | O94880 | PHD finger protein 14 OS=Homo sapiens | PHF14 | 106.987 | 2 | 53.786118 | yes | 9.722E-08 | 574.29572 | 72.082214 |
| MIQIQENMAEQK | 12 | O94880 | PHD finger protein 14 OS=Homo sapiens | PHF14 | 106.987 | 2 | 53.786118 | yes | 1.159E-07 | 731.85504 | 49.200615 |
| SEGQGIWALLGR | 12 | O94880 | PHD finger protein 14 OS=Homo sapiens | PHF14 | 106.987 | 2 | 53.786118 | yes | 9.722E-08 | 643.84619 | 46.908813 |
| TGVCISCDAGMCR | 13 | O94880 | PHD finger protein 14 OS=Homo sapiens | PHF14 | 106.987 | 2 | 53.786118 | yes | 4.51E-07 | 743.79889 | 46.283276 |
| CDGQVLLTQVWK | 12 | O94887 | FERM, ARHGEF and pleckstrin domain- | FARP2 | 119.889 | 2 | 50.723103 | yes | 1.606E-07 | 723.87415 | 1237.0499 |
| KLEMYGIR | 8 | O94887 | FERM, ARHGEF and pleckstrin domain- | FARP2 | 119.889 | 2 | 50.723103 | yes | 1.839E-07 | 505.27859 | 1049.8239 |
| ILEFHQK | 7 | O94887 | FERM, ARHGEF and pleckstrin domain- | FARP2 | 119.889 | 2 | 50.723103 | yes | 1.436E-07 | 457.75836 | 787.42957 |
| DSSSSLTDPQVSY | 15 | O94887 | FERM, ARHGEF and pleckstrin domain- | FARP2 | 119.889 | 2 | 50.723103 | yes | 1.239E-07 | 806.88867 | 764.42908 |
| HVGQTPAESDFQV | 18 | O94887 | FERM, ARHGEF and pleckstrin domain- | FARP2 | 119.889 | 3 | 50.723103 | yes | 4.507E-07 | 666.34106 | 722.03522 |
| ICVEYHTFFR | 10 | O94887 | FERM, ARHGEF and pleckstrin domain- | FARP2 | 119.889 | 2 | 50.723103 | yes | 1.211E-07 | 686.32935 | 666.67596 |
| QSISFPEGLR | 10 | O94887 | FERM, ARHGEF and pleckstrin domain- | FARP2 | 119.889 | 2 | 50.723103 | yes | 9.926E-08 | 567.30109 | 537.38873 |
| LLDNTMEIFDIEPK | 14 | O94887 | FERM, ARHGEF and pleckstrin domain- | FARP2 | 119.889 | 2 | 50.723103 | yes | 9.809E-08 | 839.42383 | 495.49188 |
| YLFALQLK | 8 | O94887 | FERM, ARHGEF and pleckstrin domain- | FARP2 | 119.889 | 2 | 50.723103 | yes | 3.206E-07 | 498.29987 | 348.85291 |
| LGAQTPVGVSTLE | 21 | O94887 | FERM, ARHGEF and pleckstrin domain- | FARP2 | 119.889 | 3 | 50.723103 | yes | 1.777E-07 | 712.06708 | 199.49982 |
| LLDQPKPK | 8 | O94887 | FERM, ARHGEF and pleckstrin domain- | FARP2 | 119.889 | 2 | 50.723103 | yes | 1.074E-07 | 469.78711 | 171.05246 |
| IQLAVSHMGVLVF | 18 | O94887 | FERM, ARHGEF and pleckstrin domain- | FARP2 | 119.889 | 3 | 50.723103 | yes | 9.727E-08 | 643.69238 | 105.9681 |
| WMEVIQGASSSAC | 14 | O94887 | FERM, ARHGEF and pleckstrin domain- | FARP2 | 119.889 | 2 | 50.723103 | yes | 2.083E-07 | 739.85645 | 103.85089 |
| GVAGTSHFR | 9 | O94887 | FERM, ARHGEF and pleckstrin domain- | FARP2 | 119.889 | 2 | 50.723103 | yes | 1.783E-07 | 466.24084 | 87.945541 |
| QLVDYFK | 7 | O94887 | FERM, ARHGEF and pleckstrin domain- | FARP2 | 119.889 | 2 | 50.723103 | yes | 1.473E-07 | 456.7449 | 76.728889 |
| LHPEVHGPYQDTL | 20 | O94887 | FERM, ARHGEF and pleckstrin domain- | FARP2 | 119.889 | 3 | 50.723103 | yes | 2.645E-07 | 770.06219 | 35.944427 |
| LHPEVHGPYQDTL | 20 | O94887 | FERM, ARHGEF and pleckstrin domain- | FARP2 | 119.889 | 3 | 50.723103 | yes | 2.93E-07 | 577.79846 | 35.944427 |
| LHPEVHGPYQDTL | 20 | O94887 | FERM, ARHGEF and pleckstrin domain- | FARP2 | 119.889 | 4 | 50.723103 | yes | 2.645E-07 | 770.06219 | 35.944427 |
| LHPEVHGPYQDTL | 20 | O94887 | FERM, ARHGEF and pleckstrin domain- | FARP2 | 119.889 | 4 | 50.723103 | yes | 2.93E-07 | 577.79846 | 35.944427 |
| GEIEGTYR | 8 | O94887 | FERM, ARHGEF and pleckstrin domain- | FARP2 | 119.889 | 2 | 50.723103 | yes | 5.819E-06 | 483.72998 | 70.310188 |
| LSFGLEDEPLETAT | 15 | O94905 | Erlin-2 OS=Homo sapiens OX=9606 GN | ERLIN2 | 37.84 | 2 | 42.751896 | yes | 2.89E-07 | 825.41705 | 42.951744 |
| VAQVAEITYGQK | 12 | O94905 | Erlin-2 OS=Homo sapiens OX=9606 GN | ERLIN2 | 37.84 | 2 | 42.751896 | yes | 3.903E-07 | 653.8537 | 24.190571 |
| ESLEALLQR | 9 | O94906 | Pre-mRNA-processing factor 6 OS=Hom | PRPF6 | 106.925 | 2 | 53.67132 | yes | 4.066E-07 | 529.79565 | 291.49246 |
| LEWVQDNIR | 9 | O94906 | Pre-mRNA-processing factor 6 OS=Hom | PRPF6 | 106.925 | 2 | 53.67132 | yes | 1.113E-07 | 586.80658 | 272.51398 |
| NPGLWLESVR | 10 | O94906 | Pre-mRNA-processing factor 6 OS=Hom | PRPF6 | 106.925 | 2 | 53.67132 | yes | 2.782E-07 | 585.81696 | 251.91058 |
| AAVELEEPEDAR | 12 | O94906 | Pre-mRNA-processing factor 6 OS=Hom | PRPF6 | 106.925 | 2 | 53.67132 | yes | 1.902E-07 | 664.82007 | 230.34711 |
| WLAGDVPAAR | 10 | O94906 | Pre-mRNA-processing factor 6 OS=Hom | PRPF6 | 106.925 | 2 | 53.67132 | yes | 2.201E-07 | 528.28528 | 211.69653 |
| AGSVATCQAVMR | 12 | O94906 | Pre-mRNA-processing factor 6 OS=Hom | PRPF6 | 106.925 | 2 | 53.67132 | yes | 2.474E-07 | 625.80261 | 207.58641 |
| EQWIQDAEECDR | 12 | O94906 | Pre-mRNA-processing factor 6 OS=Hom | PRPF6 | 106.925 | 2 | 53.67132 | yes | 1.001E-07 | 789.82806 | 152.26886 |
| SEDVWLEAAR | 10 | O94906 | Pre-mRNA-processing factor 6 OS=Hom | PRPF6 | 106.925 | 2 | 53.67132 | yes | 9.722E-08 | 588.28821 | 120.46484 |
| HIWITAAK | 8 | O94906 | Pre-mRNA-processing factor 6 OS=Hom | PRPF6 | 106.925 | 2 | 53.67132 | yes | 9.722E-08 | 470.27417 | 118.02743 |
| LETYENAR | 8 | O94906 | Pre-mRNA-processing factor 6 OS=Hom | PRPF6 | 106.925 | 2 | 53.67132 | yes | 7.409E-07 | 498.24326 | 112.10304 |

| | | | | | | | | | | | |
|---|---|---|---|---|---|---|---|---|---|---|---|
| ALEHVPNSVR | 10 | O94906 | Pre-mRNA-processing factor 6 OS=Hom | PRPF6 | 106.925 | 2 | 53.67132 | yes | 1.872E-07 | 561.3067 | 101.68877 |
| LESENDEYER | 10 | O94906 | Pre-mRNA-processing factor 6 OS=Hom | PRPF6 | 106.925 | 2 | 53.67132 | yes | 2.09E-07 | 642.27295 | 54.38308 |
| HYEDFPK | 7 | O94906 | Pre-mRNA-processing factor 6 OS=Hom | PRPF6 | 106.925 | 2 | 53.67132 | yes | 2.522E-06 | 468.21652 | 52.579414 |
| AEVLWLMGAK | 10 | O94906 | Pre-mRNA-processing factor 6 OS=Hom | PRPF6 | 106.925 | 2 | 53.67132 | yes | 2.683E-07 | 559.30737 | 36.157318 |
| NLTQEQLIR | 9 | O94913 | Pre-mRNA cleavage complex 2 protein P | PCF11 | 173.051 | 2 | 56.063362 | yes | 9.722E-08 | 557.81439 | 402.44641 |
| LSVDANLQIPK | 11 | O94913 | Pre-mRNA cleavage complex 2 protein P | PCF11 | 173.051 | 2 | 56.063362 | yes | 1.02E-07 | 599.34552 | 330.24414 |
| EIVSLIEAQTAK | 12 | O94913 | Pre-mRNA cleavage complex 2 protein P | PCF11 | 173.051 | 2 | 56.063362 | yes | 9.722E-08 | 651.3692 | 317.3306 |
| FDIPLGLQGTR | 11 | O94913 | Pre-mRNA cleavage complex 2 protein P | PCF11 | 173.051 | 2 | 56.063362 | yes | 8.765E-07 | 608.83789 | 242.24953 |
| QLLELQQK | 8 | O94913 | Pre-mRNA cleavage complex 2 protein P | PCF11 | 173.051 | 2 | 56.063362 | yes | 1.85E-07 | 500.29529 | 167.02922 |
| FAGLDTNQR | 9 | O94913 | Pre-mRNA cleavage complex 2 protein P | PCF11 | 173.051 | 2 | 56.063362 | yes | 3.135E-07 | 511.25671 | 146.48705 |
| DYQSSLEDLTFNSI | 14 | O94913 | Pre-mRNA cleavage complex 2 protein P | PCF11 | 173.051 | 2 | 56.063362 | yes | 9.722E-08 | 823.88086 | 142.25485 |
| NELQEPCDSPK | 11 | O94913 | Pre-mRNA cleavage complex 2 protein P | PCF11 | 173.051 | 2 | 56.063362 | yes | 1.737E-07 | 658.79297 | 136.68622 |
| LASGEITQDDFLV | 19 | O94913 | Pre-mRNA cleavage complex 2 protein P | PCF11 | 173.051 | 3 | 56.063362 | yes | 1.021E-07 | 714.05139 | 134.88258 |
| LYTGIQCYSCGMR | 13 | O94913 | Pre-mRNA cleavage complex 2 protein P | PCF11 | 173.051 | 2 | 56.063362 | yes | 9.722E-08 | 804.85199 | 123.55695 |
| KLELELEQAK | 10 | O94913 | Pre-mRNA cleavage complex 2 protein P | PCF11 | 173.051 | 2 | 56.063362 | yes | 9.809E-08 | 600.84534 | 113.37753 |
| QLFQYQEGVR | 10 | O94913 | Pre-mRNA cleavage complex 2 protein P | PCF11 | 173.051 | 2 | 56.063362 | yes | 9.794E-08 | 634.32513 | 97.634056 |
| ENVENWQSSK | 10 | O94913 | Pre-mRNA cleavage complex 2 protein P | PCF11 | 173.051 | 2 | 56.063362 | yes | 9.722E-08 | 610.78076 | 70.134468 |
| STWDEIFPLKK | 11 | O94913 | Pre-mRNA cleavage complex 2 protein P | PCF11 | 173.051 | 2 | 56.063362 | yes | 1.355E-07 | 682.36646 | 51.818996 |
| VHEEVVLK | 8 | O94913 | Pre-mRNA cleavage complex 2 protein P | PCF11 | 173.051 | 2 | 56.063362 | yes | 1.679E-07 | 476.77673 | 47.626007 |
| LHVSQIPPMAVK | 12 | O94913 | Pre-mRNA cleavage complex 2 protein P | PCF11 | 173.051 | 2 | 56.063362 | yes | 4.064E-07 | 660.37885 | 42.814632 |
| STWDEIFPLK | 10 | O94913 | Pre-mRNA cleavage complex 2 protein P | PCF11 | 173.051 | 2 | 56.063362 | yes | 5.081E-07 | 618.31897 | 21.659302 |
| GEEEVFLIGK | 10 | O94916 | Nuclear factor of activated T-cells 5 OS=I | NFAT5 | 165.764 | 2 | 51.533318 | yes | 1.88E-06 | 560.79785 | 68.806702 |
| AMQVESCSSAVG | 16 | O94916 | Nuclear factor of activated T-cells 5 OS=I | NFAT5 | 165.764 | 2 | 51.533318 | yes | 9.722E-08 | 841.38525 | 62.281006 |
| VIFQENVSDENSW | 14 | O94916 | Nuclear factor of activated T-cells 5 OS=I | NFAT5 | 165.764 | 2 | 51.533318 | yes | 2.68E-07 | 847.90466 | 57.962795 |
| SEDVTPMEVTAEK | 13 | O94916 | Nuclear factor of activated T-cells 5 OS=I | NFAT5 | 165.764 | 2 | 51.533318 | yes | 3.885E-05 | 718.33472 | 12.236326 |
| QLTSNTVQQHPST | 15 | O94916 | Nuclear factor of activated T-cells 5 OS=I | NFAT5 | 165.764 | 3 | 51.533318 | yes | 4.023E-06 | 555.95667 | 11.479334 |
| APPVVGGFNSNV | 15 | O94923 | D-glucuronyl C5-epimerase OS=Homo sa | GLCE | 70.102 | 2 | 31.320671 | yes | 1.88E-06 | 715.37537 | 63.961792 |
| GCFMANVADK | 10 | O94923 | D-glucuronyl C5-epimerase OS=Homo sa | GLCE | 70.102 | 2 | 31.320671 | yes | 2.589E-06 | 556.74683 | 40.781418 |
| DGPGETDAFGNSE | 15 | O94925 | Glutaminase kidney isoform, mitochondri | GLS | 73.462 | 2 | 48.867107 | yes | 1.911E-07 | 740.81299 | 21.074322 |
| CVQSNIVLLTQAF | 14 | O94925 | Glutaminase kidney isoform, mitochondri | GLS | 73.462 | 2 | 48.867107 | yes | 1.093E-07 | 824.94562 | 18.322941 |
| GSSFIALSHK | 10 | O94927 | HAUS augmin-like complex subunit 5 OS | HAUS5 | 71.683 | 2 | 48.32057 | yes | 1.982E-07 | 523.7851 | 218.02672 |
| VVPTFEAVAPQSR | 13 | O94927 | HAUS augmin-like complex subunit 5 OS | HAUS5 | 71.683 | 2 | 48.32057 | yes | 4.106E-07 | 700.88025 | 72.960426 |
| SPGEVLALVQR | 11 | O94927 | HAUS augmin-like complex subunit 5 OS | HAUS5 | 71.683 | 2 | 48.32057 | yes | 2.652E-07 | 584.83789 | 65.312813 |
| KLELEAAVTR | 10 | O94927 | HAUS augmin-like complex subunit 5 OS | HAUS5 | 71.683 | 2 | 48.32057 | yes | 1.164E-07 | 565.3324 | 61.402336 |
| ASELLLPAAASLR | 13 | O94927 | HAUS augmin-like complex subunit 5 OS | HAUS5 | 71.683 | 2 | 48.32057 | yes | 1.141E-06 | 656.38519 | 49.880867 |
| QLVEETQEQVR | 11 | O94927 | HAUS augmin-like complex subunit 5 OS | HAUS5 | 71.683 | 2 | 48.32057 | yes | 6.797E-07 | 679.84918 | 24.3876 |
| SVLIPVYGVSK | 11 | O94955 | Rho-related BTB domain-containing prot | RHOBTB3 | 69.414 | 2 | 49.379913 | yes | 3.668E-07 | 581.34753 | 98.594704 |
| SPTDIQDSSIIR | 12 | O94955 | Rho-related BTB domain-containing prot | RHOBTB3 | 69.414 | 2 | 49.379913 | yes | 0.0001134 | 666.34369 | 83.724792 |
| GSCVSTTEGIQLAI | 14 | O94955 | Rho-related BTB domain-containing prot | RHOBTB3 | 69.414 | 2 | 49.379913 | yes | 1.039E-07 | 725.86395 | 36.034885 |

| | | | | | | | | | | | |
|---|---|---|---|---|---|---|---|---|---|---|---|
| EFELQSLSLQR | 11 | O94964 | Protein SOGA1 OS=Homo sapiens OX=9 | SOGA1 | 159.76 | 2 | 53.227787 | yes | 9.722E-08 | 675.35663 | 521.44232 |
| AAQTGQVDGELIR | 13 | O94964 | Protein SOGA1 OS=Homo sapiens OX=9 | SOGA1 | 159.76 | 2 | 53.227787 | yes | 1.818E-07 | 679.35718 | 477.31598 |
| LVEEEANLLSR | 11 | O94964 | Protein SOGA1 OS=Homo sapiens OX=9 | SOGA1 | 159.76 | 2 | 53.227787 | yes | 9.722E-08 | 636.84332 | 384.8367 |
| NMLVQESQQFK | 11 | O94964 | Protein SOGA1 OS=Homo sapiens OX=9 | SOGA1 | 159.76 | 2 | 53.227787 | yes | 1.192E-07 | 676.33734 | 348.87903 |
| LCLDNECADFR | 11 | O94964 | Protein SOGA1 OS=Homo sapiens OX=9 | SOGA1 | 159.76 | 2 | 53.227787 | yes | 1.222E-07 | 706.80029 | 348.68863 |
| VLLSNLQR | 8 | O94964 | Protein SOGA1 OS=Homo sapiens OX=9 | SOGA1 | 159.76 | 2 | 53.227787 | yes | 1.118E-06 | 471.79019 | 331.54047 |
| YGIVQEFFR | 9 | O94964 | Protein SOGA1 OS=Homo sapiens OX=9 | SOGA1 | 159.76 | 2 | 53.227787 | yes | 4.247E-07 | 579.80072 | 286.22726 |
| GLQEQLSQER | 10 | O94964 | Protein SOGA1 OS=Homo sapiens OX=9 | SOGA1 | 159.76 | 2 | 53.227787 | yes | 1.076E-07 | 594.30438 | 251.62975 |
| LIHAILVR | 8 | O94964 | Protein SOGA1 OS=Homo sapiens OX=9 | SOGA1 | 159.76 | 2 | 53.227787 | yes | 2.072E-07 | 467.81348 | 227.18636 |
| TVSVGLQTDPPR | 12 | O94964 | Protein SOGA1 OS=Homo sapiens OX=9 | SOGA1 | 159.76 | 2 | 53.227787 | yes | 9.722E-08 | 635.34351 | 214.5892 |
| SLWNLHQGK | 9 | O94964 | Protein SOGA1 OS=Homo sapiens OX=9 | SOGA1 | 159.76 | 2 | 53.227787 | yes | 3.233E-07 | 541.79071 | 194.78221 |
| TLVQEDSADLK | 11 | O94964 | Protein SOGA1 OS=Homo sapiens OX=9 | SOGA1 | 159.76 | 2 | 53.227787 | yes | 1.356E-07 | 609.81427 | 187.84332 |
| LQIGELSGK | 9 | O94964 | Protein SOGA1 OS=Homo sapiens OX=9 | SOGA1 | 159.76 | 2 | 53.227787 | yes | 5.027E-07 | 472.7742 | 183.44417 |
| CDLASCQSTR | 10 | O94964 | Protein SOGA1 OS=Homo sapiens OX=9 | SOGA1 | 159.76 | 2 | 53.227787 | yes | 1.073E-07 | 599.25299 | 178.72536 |
| LEEENEELR | 9 | O94964 | Protein SOGA1 OS=Homo sapiens OX=9 | SOGA1 | 159.76 | 2 | 53.227787 | yes | 1.001E-07 | 580.77515 | 178.39238 |
| LSVLQQELNAFTR | 13 | O94964 | Protein SOGA1 OS=Homo sapiens OX=9 | SOGA1 | 159.76 | 2 | 53.227787 | yes | 1.477E-07 | 759.91736 | 145.74896 |
| SSAELEDQNK | 10 | O94964 | Protein SOGA1 OS=Homo sapiens OX=9 | SOGA1 | 159.76 | 2 | 53.227787 | yes | 1.622E-07 | 560.75946 | 141.19128 |
| DLPEWEPR | 8 | O94964 | Protein SOGA1 OS=Homo sapiens OX=9 | SOGA1 | 159.76 | 2 | 53.227787 | yes | 6.165E-07 | 521.2536 | 132.93005 |
| LTMLPQVGK | 9 | O94964 | Protein SOGA1 OS=Homo sapiens OX=9 | SOGA1 | 159.76 | 2 | 53.227787 | yes | 1.248E-07 | 493.78879 | 132.63791 |
| EAEVLPGLR | 9 | O94964 | Protein SOGA1 OS=Homo sapiens OX=9 | SOGA1 | 159.76 | 2 | 53.227787 | yes | 1.896E-07 | 492.27966 | 131.68489 |
| GHTSQMELK | 9 | O94964 | Protein SOGA1 OS=Homo sapiens OX=9 | SOGA1 | 159.76 | 2 | 53.227787 | yes | 4.408E-07 | 515.75293 | 126.98267 |
| WLCNFSR | 7 | O94964 | Protein SOGA1 OS=Homo sapiens OX=9 | SOGA1 | 159.76 | 2 | 53.227787 | yes | 2.408E-07 | 491.73181 | 126.41301 |
| AEDNDSYASEIK | 12 | O94964 | Protein SOGA1 OS=Homo sapiens OX=9 | SOGA1 | 159.76 | 2 | 53.227787 | yes | 2.892E-07 | 671.29388 | 121.99403 |
| KADAVLGCSVK | 11 | O94964 | Protein SOGA1 OS=Homo sapiens OX=9 | SOGA1 | 159.76 | 2 | 53.227787 | yes | 9.722E-08 | 574.31061 | 121.92387 |
| LHEAPDNSEGPR | 12 | O94964 | Protein SOGA1 OS=Homo sapiens OX=9 | SOGA1 | 159.76 | 2 | 53.227787 | yes | 9.722E-08 | 661.31018 | 112.36259 |
| ESAEADGLAESSY | 15 | O94964 | Protein SOGA1 OS=Homo sapiens OX=9 | SOGA1 | 159.76 | 2 | 53.227787 | yes | 9.734E-08 | 771.33936 | 108.6548 |
| LLLNELAK | 8 | O94964 | Protein SOGA1 OS=Homo sapiens OX=9 | SOGA1 | 159.76 | 2 | 53.227787 | yes | 2E-07 | 457.28949 | 90.018166 |
| SLYGDLDSALSAE | 22 | O94964 | Protein SOGA1 OS=Homo sapiens OX=9 | SOGA1 | 159.76 | 3 | 53.227787 | yes | 3.945E-07 | 773.03625 | 86.091606 |
| TGDLDSKPDPSR | 12 | O94964 | Protein SOGA1 OS=Homo sapiens OX=9 | SOGA1 | 159.76 | 2 | 53.227787 | yes | 9.722E-08 | 644.31238 | 84.274422 |
| IERPCCSPK | 9 | O94964 | Protein SOGA1 OS=Homo sapiens OX=9 | SOGA1 | 159.76 | 2 | 53.227787 | yes | 1.21E-06 | 573.77338 | 70.672066 |
| GLTELQQQFAK | 11 | O94964 | Protein SOGA1 OS=Homo sapiens OX=9 | SOGA1 | 159.76 | 2 | 53.227787 | yes | 1.002E-07 | 631.84058 | 65.077469 |
| APSPTSSAGEEGTK | 14 | O94964 | Protein SOGA1 OS=Homo sapiens OX=9 | SOGA1 | 159.76 | 2 | 53.227787 | yes | 4.249E-07 | 659.80969 | 59.109261 |
| ELYLLMEEEEINA( | 19 | O94964 | Protein SOGA1 OS=Homo sapiens OX=9 | SOGA1 | 159.76 | 3 | 53.227787 | yes | 2.952E-07 | 769.02661 | 52.545681 |
| RLGVPVVHDK | 10 | O94964 | Protein SOGA1 OS=Homo sapiens OX=9 | SOGA1 | 159.76 | 2 | 53.227787 | yes | 4.769E-07 | 560.33527 | 26.30381 |
| LTHDASNIHTSTPF | 14 | O94967 | WD repeat-containing protein 47 OS=Ho | WDR47 | 101.95 | 2 | 39.330853 | yes | 4.649E-07 | 775.38953 | 10.491266 |
| TSPDLVPMGDWTS | 14 | O94973 | AP-2 complex subunit alpha-2 OS=Homo | AP2A2 | 103.961 | 2 | 55.042244 | yes | 4.595E-07 | 781.36938 | 932.32312 |
| TTQIGCLLR | 9 | O94973 | AP-2 complex subunit alpha-2 OS=Homo | AP2A2 | 103.961 | 2 | 55.042244 | yes | 9.723E-08 | 531.29224 | 517.86615 |
| FHLCSVPTR | 9 | O94973 | AP-2 complex subunit alpha-2 OS=Homo | AP2A2 | 103.961 | 2 | 55.042244 | yes | 2.736E-07 | 558.78455 | 356.9165 |
| THIETVINALK | 11 | O94973 | AP-2 complex subunit alpha-2 OS=Homo | AP2A2 | 103.961 | 2 | 55.042244 | yes | 9.787E-08 | 619.85876 | 282.44141 |

| | | | | | | | | | | | |
|---|---|---|---|---|---|---|---|---|---|---|---|
| EMAEAFAGEIPK | 12 | O94973 | AP-2 complex subunit alpha-2 OS=Homc | AP2A2 | 103.961 | 2 | 55.042244 | yes | 9.722E-08 | 646.81317 | 244.8651 |
| LEPNLQAQMYR | 11 | O94973 | AP-2 complex subunit alpha-2 OS=Homc | AP2A2 | 103.961 | 2 | 55.042244 | yes | 9.819E-08 | 681.84534 | 236.69334 |
| AVDLLYAMCDR | 11 | O94973 | AP-2 complex subunit alpha-2 OS=Homc | AP2A2 | 103.961 | 2 | 55.042244 | yes | 4.583E-07 | 663.81268 | 230.49887 |
| LLQCYPPPDPAVR | 13 | O94973 | AP-2 complex subunit alpha-2 OS=Homc | AP2A2 | 103.961 | 2 | 55.042244 | yes | 9.85E-08 | 763.3952 | 220.87386 |
| MFIFYGNK | 8 | O94973 | AP-2 complex subunit alpha-2 OS=Homc | AP2A2 | 103.961 | 2 | 55.042244 | yes | 2.76E-07 | 510.25458 | 173.42937 |
| TSVSLAVSR | 9 | O94973 | AP-2 complex subunit alpha-2 OS=Homc | AP2A2 | 103.961 | 2 | 55.042244 | yes | 3.886E-07 | 460.26401 | 93.45401 |
| QLSNPQQEVQNIFI | 14 | O94973 | AP-2 complex subunit alpha-2 OS=Homc | AP2A2 | 103.961 | 2 | 55.042244 | yes | 9.796E-08 | 836.93628 | 85.464195 |
| LTECLETILNK | 11 | O94973 | AP-2 complex subunit alpha-2 OS=Homc | AP2A2 | 103.961 | 2 | 55.042244 | yes | 9.722E-08 | 667.35522 | 83.212555 |
| SSPLIQFHLLHSK | 13 | O94973 | AP-2 complex subunit alpha-2 OS=Homc | AP2A2 | 103.961 | 2 | 55.042244 | yes | 9.722E-08 | 753.92499 | 76.794662 |
| VIQIVINR | 8 | O94973 | AP-2 complex subunit alpha-2 OS=Homc | AP2A2 | 103.961 | 2 | 55.042244 | yes | 1.627E-07 | 477.80838 | 45.886387 |
| NQLWPESSTSDVV | 14 | O94986 | Centrosomal protein of 152 kDa OS=Hon | CEP152 | 195.627 | 2 | 53.90249 | yes | 1.177E-07 | 809.39697 | 1334.8386 |
| LEELQTLCK | 9 | O94986 | Centrosomal protein of 152 kDa OS=Hon | CEP152 | 195.627 | 2 | 53.90249 | yes | 1.063E-07 | 567.29718 | 548.24207 |
| ETELQTCLDQSR | 12 | O94986 | Centrosomal protein of 152 kDa OS=Hon | CEP152 | 195.627 | 2 | 53.90249 | yes | 9.722E-08 | 740.34082 | 497.42773 |
| VVEELIEENNDMK | 13 | O94986 | Centrosomal protein of 152 kDa OS=Hon | CEP152 | 195.627 | 2 | 53.90249 | yes | 9.722E-08 | 781.37433 | 436.79785 |
| LLETPISSK | 9 | O94986 | Centrosomal protein of 152 kDa OS=Hon | CEP152 | 195.627 | 2 | 53.90249 | yes | 9.819E-08 | 494.28967 | 432.93954 |
| CTIQQNLEQEK | 11 | O94986 | Centrosomal protein of 152 kDa OS=Hon | CEP152 | 195.627 | 2 | 53.90249 | yes | 9.722E-08 | 695.83521 | 412.48056 |
| ALETQIQALK | 10 | O94986 | Centrosomal protein of 152 kDa OS=Hon | CEP152 | 195.627 | 2 | 53.90249 | yes | 9.849E-08 | 557.82697 | 363.72754 |
| EMAIMIEEQK | 10 | O94986 | Centrosomal protein of 152 kDa OS=Hon | CEP152 | 195.627 | 2 | 53.90249 | yes | 6.097E-06 | 611.29633 | 321.30719 |
| EQEDICSR | 8 | O94986 | Centrosomal protein of 152 kDa OS=Hon | CEP152 | 195.627 | 2 | 53.90249 | yes | 1.93E-07 | 518.72186 | 294.26944 |
| LSSTLPSSVCQQPS | 15 | O94986 | Centrosomal protein of 152 kDa OS=Hon | CEP152 | 195.627 | 2 | 53.90249 | yes | 2.375E-07 | 823.91199 | 288.10605 |
| NLDATVTALK | 10 | O94986 | Centrosomal protein of 152 kDa OS=Hon | CEP152 | 195.627 | 2 | 53.90249 | yes | 1.55E-07 | 523.29803 | 249.57597 |
| VNEEQMIK | 8 | O94986 | Centrosomal protein of 152 kDa OS=Hon | CEP152 | 195.627 | 2 | 53.90249 | yes | 2.04E-07 | 495.74988 | 241.80293 |
| SQGLDVQEPPVK | 12 | O94986 | Centrosomal protein of 152 kDa OS=Hon | CEP152 | 195.627 | 2 | 53.90249 | yes | 9.722E-08 | 648.84332 | 239.97485 |
| QQLFEAYER | 9 | O94986 | Centrosomal protein of 152 kDa OS=Hon | CEP152 | 195.627 | 2 | 53.90249 | yes | 9.722E-08 | 592.29077 | 236.83105 |
| QEFNNQATNVIK | 12 | O94986 | Centrosomal protein of 152 kDa OS=Hon | CEP152 | 195.627 | 2 | 53.90249 | yes | 1.089E-07 | 703.35718 | 223.5916 |
| EKDNLELTLR | 10 | O94986 | Centrosomal protein of 152 kDa OS=Hon | CEP152 | 195.627 | 2 | 53.90249 | yes | 9.722E-08 | 615.83807 | 221.31731 |
| QLENLIEK | 8 | O94986 | Centrosomal protein of 152 kDa OS=Hon | CEP152 | 195.627 | 2 | 53.90249 | yes | 1.071E-07 | 493.77948 | 178.20978 |
| ISFAVSEAK | 9 | O94986 | Centrosomal protein of 152 kDa OS=Hon | CEP152 | 195.627 | 2 | 53.90249 | yes | 1.161E-07 | 476.26093 | 173.1299 |
| PQGTLEIPSESVK | 13 | O94986 | Centrosomal protein of 152 kDa OS=Hon | CEP152 | 195.627 | 2 | 53.90249 | yes | 1.761E-07 | 692.86957 | 167.32463 |
| WMSVQYFEK | 9 | O94986 | Centrosomal protein of 152 kDa OS=Hon | CEP152 | 195.627 | 2 | 53.90249 | yes | 9.722E-08 | 609.28662 | 158.56699 |
| NEEVPVVIR | 9 | O94986 | Centrosomal protein of 152 kDa OS=Hon | CEP152 | 195.627 | 2 | 53.90249 | yes | 2.754E-07 | 527.79822 | 156.94438 |
| EQHESIVMGLTK | 12 | O94986 | Centrosomal protein of 152 kDa OS=Hon | CEP152 | 195.627 | 2 | 53.90249 | yes | 1.022E-07 | 686.35046 | 147.23073 |
| IEDLHQVK | 8 | O94986 | Centrosomal protein of 152 kDa OS=Hon | CEP152 | 195.627 | 2 | 53.90249 | yes | 1.204E-07 | 491.27182 | 130.55637 |
| CNPLMESENAASE | 14 | O94986 | Centrosomal protein of 152 kDa OS=Hon | CEP152 | 195.627 | 2 | 53.90249 | yes | 1.662E-07 | 790.33997 | 127.30238 |
| IQEQNEQDYR | 10 | O94986 | Centrosomal protein of 152 kDa OS=Hon | CEP152 | 195.627 | 2 | 53.90249 | yes | 1.359E-07 | 661.80219 | 125.75404 |
| ALQEELTELKDEIS | 21 | O94986 | Centrosomal protein of 152 kDa OS=Hon | CEP152 | 195.627 | 3 | 53.90249 | yes | 9.728E-08 | 794.07587 | 118.91502 |
| KIEDLHQVK | 9 | O94986 | Centrosomal protein of 152 kDa OS=Hon | CEP152 | 195.627 | 2 | 53.90249 | yes | 9.722E-08 | 555.31934 | 100.37093 |
| HLVSQLQNDLK | 11 | O94986 | Centrosomal protein of 152 kDa OS=Hon | CEP152 | 195.627 | 2 | 53.90249 | yes | 9.722E-08 | 647.85931 | 98.548622 |
| ISVNCGHPSR | 10 | O94986 | Centrosomal protein of 152 kDa OS=Hon | CEP152 | 195.627 | 2 | 53.90249 | yes | 1.498E-07 | 563.77472 | 89.078094 |

| | | | | | | | | | | | |
|---|---|---|---|---|---|---|---|---|---|---|---|
| EIQLEAQIK | 9 | O94986 | Centrosomal protein of 152 kDa OS=Hon | CEP152 | 195.627 | 2 | 53.90249 | yes | 6.389E-06 | 536.30585 | 88.049149 |
| CGNVWEENR | 9 | O94986 | Centrosomal protein of 152 kDa OS=Hon | CEP152 | 195.627 | 2 | 53.90249 | yes | 1.188E-07 | 582.24854 | 70.080498 |
| QLLEIMSTCSSK | 12 | O94986 | Centrosomal protein of 152 kDa OS=Hon | CEP152 | 195.627 | 2 | 53.90249 | yes | 9.766E-08 | 698.84418 | 63.302452 |
| SELENMR | 7 | O94986 | Centrosomal protein of 152 kDa OS=Hon | CEP152 | 195.627 | 2 | 53.90249 | yes | 2.629E-07 | 439.70547 | 60.031006 |
| INEVLAAAK | 9 | O94986 | Centrosomal protein of 152 kDa OS=Hon | CEP152 | 195.627 | 2 | 53.90249 | yes | 3.668E-07 | 464.77673 | 45.349869 |
| WEEQHEVSVNK | 11 | O94986 | Centrosomal protein of 152 kDa OS=Hon | CEP152 | 195.627 | 2 | 53.90249 | yes | 3.859E-07 | 692.82825 | 36.130207 |
| ETHLEFQFGDGSC | 14 | O94986 | Centrosomal protein of 152 kDa OS=Hon | CEP152 | 195.627 | 2 | 53.90249 | yes | 9.722E-08 | 827.86194 | 35.356483 |
| EWTMQEAK | 8 | O94986 | Centrosomal protein of 152 kDa OS=Hon | CEP152 | 195.627 | 2 | 53.90249 | yes | 3.161E-07 | 511.73422 | 33.666874 |
| KWEEQHEVSVNK | 12 | O94986 | Centrosomal protein of 152 kDa OS=Hon | CEP152 | 195.627 | 2 | 53.90249 | yes | 5.218E-07 | 756.87573 | 21.972551 |
| KYEEQVLSLQK | 11 | O94986 | Centrosomal protein of 152 kDa OS=Hon | CEP152 | 195.627 | 2 | 53.90249 | yes | 5.501E-06 | 682.87463 | 21.811844 |
| QKTELLLQK | 9 | O94986 | Centrosomal protein of 152 kDa OS=Hon | CEP152 | 195.627 | 2 | 53.90249 | yes | 7.302E-05 | 550.83734 | 16.752117 |
| AENLQLLTENELH | 14 | O94992 | Protein HEXIM1 OS=Homo sapiens OX= | HEXIM1 | 40.624 | 2 | 53.572224 | yes | 9.722E-08 | 840.43921 | 354.11768 |
| AENLQLLTENELH | 14 | O94992 | Protein HEXIM1 OS=Homo sapiens OX= | HEXIM1 | 40.624 | 2 | 53.572224 | yes | 8.008E-07 | 560.62854 | 354.11768 |
| AENLQLLTENELH | 14 | O94992 | Protein HEXIM1 OS=Homo sapiens OX= | HEXIM1 | 40.624 | 3 | 53.572224 | yes | 9.722E-08 | 840.43921 | 354.11768 |
| AENLQLLTENELH | 14 | O94992 | Protein HEXIM1 OS=Homo sapiens OX= | HEXIM1 | 40.624 | 3 | 53.572224 | yes | 8.008E-07 | 560.62854 | 354.11768 |
| ELELELDR | 8 | O94992 | Protein HEXIM1 OS=Homo sapiens OX= | HEXIM1 | 40.624 | 2 | 53.572224 | yes | 9.722E-08 | 508.76657 | 219.93198 |
| YHTESLQNMSK | 11 | O94992 | Protein HEXIM1 OS=Homo sapiens OX= | HEXIM1 | 40.624 | 2 | 53.572224 | yes | 9.722E-08 | 669.31134 | 154.55219 |
| EYLELEK | 7 | O94992 | Protein HEXIM1 OS=Homo sapiens OX= | HEXIM1 | 40.624 | 2 | 53.572224 | yes | 5.436E-07 | 462.23965 | 77.208939 |
| HITDSGLAAR | 10 | O95049 | Tight junction protein ZO-3 OS=Homo sa | TJP3 | 101.397 | 2 | 50.167458 | yes | 3.945E-07 | 520.77783 | 184.41212 |
| HALLDVTPSAIER | 13 | O95049 | Tight junction protein ZO-3 OS=Homo sa | TJP3 | 101.397 | 2 | 50.167458 | yes | 1.236E-07 | 711.39099 | 59.083138 |
| HALLDVTPSAIER | 13 | O95049 | Tight junction protein ZO-3 OS=Homo sa | TJP3 | 101.397 | 2 | 50.167458 | yes | 1.002E-07 | 474.59641 | 59.083138 |
| HALLDVTPSAIER | 13 | O95049 | Tight junction protein ZO-3 OS=Homo sa | TJP3 | 101.397 | 3 | 50.167458 | yes | 1.236E-07 | 711.39099 | 59.083138 |
| HALLDVTPSAIER | 13 | O95049 | Tight junction protein ZO-3 OS=Homo sa | TJP3 | 101.397 | 3 | 50.167458 | yes | 1.002E-07 | 474.59641 | 59.083138 |
| AEQLASLEAAQR | 12 | O95049 | Tight junction protein ZO-3 OS=Homo sa | TJP3 | 101.397 | 2 | 50.167458 | yes | 2.656E-07 | 643.83856 | 114.60386 |
| VGDSFYIR | 8 | O95049 | Tight junction protein ZO-3 OS=Homo sa | TJP3 | 101.397 | 2 | 50.167458 | yes | 3.822E-07 | 478.74545 | 105.6438 |
| ISAHQGAQVDSR | 12 | O95049 | Tight junction protein ZO-3 OS=Homo sa | TJP3 | 101.397 | 2 | 50.167458 | yes | 1.414E-07 | 634.82074 | 99.949867 |
| GFGIAISGGR | 10 | O95049 | Tight junction protein ZO-3 OS=Homo sa | TJP3 | 101.397 | 2 | 50.167458 | yes | 2.061E-06 | 467.75888 | 88.345642 |
| AVGVGPGSSAGSN | 15 | O95049 | Tight junction protein ZO-3 OS=Homo sa | TJP3 | 101.397 | 2 | 50.167458 | yes | 1.133E-07 | 643.82599 | 87.795517 |
| LGSQIFIK | 8 | O95049 | Tight junction protein ZO-3 OS=Homo sa | TJP3 | 101.397 | 2 | 50.167458 | yes | 9.873E-08 | 453.27637 | 84.362015 |
| ESSYDIYR | 8 | O95049 | Tight junction protein ZO-3 OS=Homo sa | TJP3 | 101.397 | 2 | 50.167458 | yes | 1.122E-05 | 516.73529 | 69.883682 |
| VPSSQSMEDR | 10 | O95049 | Tight junction protein ZO-3 OS=Homo sa | TJP3 | 101.397 | 2 | 50.167458 | yes | 4.501E-07 | 568.25604 | 56.005299 |
| PAPAATYER | 9 | O95059 | Ribonuclease P protein subunit p14 OS=I | RPP14 | 13.694 | 2 | 49.856846 | yes | 1.001E-07 | 488.24835 | 344.03418 |
| NPSEYHYMK | 9 | O95059 | Ribonuclease P protein subunit p14 OS=I | RPP14 | 13.694 | 2 | 49.856846 | yes | 2.078E-07 | 584.75824 | 93.916039 |
| LWSSLTLLGSYK | 12 | O95059 | Ribonuclease P protein subunit p14 OS=I | RPP14 | 13.694 | 2 | 49.856846 | yes | 3.179E-06 | 684.38208 | 52.167721 |
| TAALGPDSMGGPV | 15 | O95070 | Protein YIF1A OS=Homo sapiens OX=9 | YIF1A | 32.012 | 2 | 48.22437 | yes | 1.215E-07 | 713.36139 | 147.63622 |
| YFFAVDTAYVAK | 12 | O95070 | Protein YIF1A OS=Homo sapiens OX=9 | YIF1A | 32.012 | 2 | 48.22437 | yes | 1.178E-07 | 697.85315 | 37.617962 |
| VATWVDETLSSVA | 15 | O95071 | E3 ubiquitin-protein ligase UBR5 OS=Hc | UBR5 | 309.353 | 2 | 53.983456 | yes | 1.495E-07 | 796.91193 | 81.920502 |
| WLDGASFDNER | 11 | O95071 | E3 ubiquitin-protein ligase UBR5 OS=Hc | UBR5 | 309.353 | 2 | 53.983456 | yes | 9.722E-08 | 655.29401 | 76.005646 |
| IVLLSANSIR | 10 | O95071 | E3 ubiquitin-protein ligase UBR5 OS=Hc | UBR5 | 309.353 | 2 | 53.983456 | yes | 3.906E-07 | 543.33752 | 74.379951 |

| | | | | | | | | | | | |
|---|---|---|---|---|---|---|---|---|---|---|---|
| LLTATNLVTLPNSI | 14 | O95071 | E3 ubiquitin-protein ligase UBR5 OS=Hc | UBR5 | 309.353 | 2 | 53.983456 | yes | 1.356E-07 | 756.94086 | 52.7141 |
| TEILAVNVDSK | 11 | O95071 | E3 ubiquitin-protein ligase UBR5 OS=Hc | UBR5 | 309.353 | 2 | 53.983456 | yes | 1.553E-06 | 594.82715 | 51.578499 |
| YCIFDLATGK | 10 | O95071 | E3 ubiquitin-protein ligase UBR5 OS=Hc | UBR5 | 309.353 | 2 | 53.983456 | yes | 4.336E-07 | 594.29187 | 38.973656 |
| EVVFVEDVK | 9 | O95071 | E3 ubiquitin-protein ligase UBR5 OS=Hc | UBR5 | 309.353 | 2 | 53.983456 | yes | 1.405E-07 | 532.28711 | 38.775871 |
| PSQGLYSSSASSGK | 14 | O95071 | E3 ubiquitin-protein ligase UBR5 OS=Hc | UBR5 | 309.353 | 2 | 53.983456 | yes | 2.495E-06 | 678.3255 | 10.739939 |
| LSLELFGR | 8 | O95071 | E3 ubiquitin-protein ligase UBR5 OS=Hc | UBR5 | 309.353 | 2 | 53.983456 | yes | 4.382E-05 | 467.77145 | 5.6043849 |
| MVHGVYK | 7 | O95139 | NADH dehydrogenase [ubiquinone] 1 bet | NDUFB6 | 15.49 | 2 | 45.668995 | yes | 1.779E-07 | 417.22055 | 56.371101 |
| SATLCIAYLMK | 11 | O95147 | Dual specificity protein phosphatase 14 O | DUSP14 | 22.255 | 2 | 20.498846 | yes | 4.444E-06 | 635.83032 | 31.735893 |
| VTNLFCFEHR | 10 | O95159 | Zinc finger protein-like 1 OS=Homo sapi | ZFPL1 | 34.114 | 2 | 51.706367 | yes | 9.722E-08 | 661.81915 | 168.67065 |
| LATVNWAR | 8 | O95159 | Zinc finger protein-like 1 OS=Homo sapi | ZFPL1 | 34.114 | 2 | 51.706367 | yes | 1.378E-07 | 465.76141 | 152.21057 |
| LCNIPLASR | 9 | O95159 | Zinc finger protein-like 1 OS=Homo sapi | ZFPL1 | 34.114 | 2 | 51.706367 | yes | 2.516E-07 | 522.28693 | 136.48689 |
| SVVLQLADGQIFK | 13 | O95163 | Elongator complex protein 1 OS=Homo s | ELP1 | 150.255 | 2 | 54.358864 | yes | 9.722E-08 | 709.40613 | 630.63074 |
| TMYPAPVTSSVYL | 15 | O95163 | Elongator complex protein 1 OS=Homo s | ELP1 | 150.255 | 2 | 54.358864 | yes | 1.05E-07 | 836.42419 | 534.0658 |
| IVTVVPQDTK | 10 | O95163 | Elongator complex protein 1 OS=Homo s | ELP1 | 150.255 | 2 | 54.358864 | yes | 9.722E-08 | 550.32153 | 378.03259 |
| IDLVCDAMR | 9 | O95163 | Elongator complex protein 1 OS=Homo s | ELP1 | 150.255 | 2 | 54.358864 | yes | 1.798E-07 | 546.76245 | 346.58096 |
| TTPELEIVLQK | 11 | O95163 | Elongator complex protein 1 OS=Homo s | ELP1 | 150.255 | 2 | 54.358864 | yes | 1.365E-07 | 635.86627 | 330.68454 |
| ALSAFLTCGNWK | 12 | O95163 | Elongator complex protein 1 OS=Homo s | ELP1 | 150.255 | 2 | 54.358864 | yes | 1.453E-07 | 684.34247 | 312.85828 |
| AVMESINPHK | 10 | O95163 | Elongator complex protein 1 OS=Homo s | ELP1 | 150.255 | 2 | 54.358864 | yes | 1.065E-07 | 563.28967 | 276.24603 |
| AFEDTLQLMER | 11 | O95163 | Elongator complex protein 1 OS=Homo s | ELP1 | 150.255 | 2 | 54.358864 | yes | 9.722E-08 | 676.82941 | 250.34366 |
| VFLGNVETFIK | 11 | O95163 | Elongator complex protein 1 OS=Homo s | ELP1 | 150.255 | 2 | 54.358864 | yes | 1.277E-07 | 633.85828 | 172.12462 |
| GNLEVVHHR | 9 | O95163 | Elongator complex protein 1 OS=Homo s | ELP1 | 150.255 | 2 | 54.358864 | yes | 2.372E-06 | 530.78595 | 151.6106 |
| IVSLMWDPVTPYR | 13 | O95163 | Elongator complex protein 1 OS=Homo s | ELP1 | 150.255 | 2 | 54.358864 | yes | 9.722E-08 | 788.91327 | 144.82948 |
| TEQGTVLIGSEHGI | 22 | O95163 | Elongator complex protein 1 OS=Homo s | ELP1 | 150.255 | 3 | 54.358864 | yes | 1.823E-07 | 779.4079 | 105.37375 |
| DIQGPGNPQCFSLI | 14 | O95163 | Elongator complex protein 1 OS=Homo s | ELP1 | 150.255 | 2 | 54.358864 | yes | 1.062E-07 | 794.88049 | 98.68087 |
| ETQFHGSEGR | 10 | O95163 | Elongator complex protein 1 OS=Homo s | ELP1 | 150.255 | 2 | 54.358864 | yes | 3.695E-07 | 574.25995 | 97.073334 |
| DEVYHILK | 8 | O95163 | Elongator complex protein 1 OS=Homo s | ELP1 | 150.255 | 2 | 54.358864 | yes | 2.354E-07 | 508.7742 | 96.196808 |
| CGDCPSADPTVK | 12 | O95163 | Elongator complex protein 1 OS=Homo s | ELP1 | 150.255 | 2 | 54.358864 | yes | 2.457E-07 | 653.77374 | 88.123192 |
| QSLSFSTCGK | 10 | O95163 | Elongator complex protein 1 OS=Homo s | ELP1 | 150.255 | 2 | 54.358864 | yes | 1.39E-07 | 557.76349 | 85.634087 |
| LGAVGGSGFK | 10 | O95163 | Elongator complex protein 1 OS=Homo s | ELP1 | 150.255 | 2 | 54.358864 | yes | 5.431E-07 | 446.74799 | 74.62513 |
| QALCVAAQLNFTK | 13 | O95163 | Elongator complex protein 1 OS=Homo s | ELP1 | 150.255 | 2 | 54.358864 | yes | 5.395E-07 | 732.38739 | 60.539749 |
| TSVPVLDAELFIPP | 15 | O95163 | Elongator complex protein 1 OS=Homo s | ELP1 | 150.255 | 2 | 54.358864 | yes | 4.335E-07 | 813.46106 | 54.677616 |
| DKNLYNEALK | 10 | O95163 | Elongator complex protein 1 OS=Homo s | ELP1 | 150.255 | 2 | 54.358864 | yes | 3.432E-07 | 604.31952 | 50.306606 |
| KTTPELEIVLQK | 12 | O95163 | Elongator complex protein 1 OS=Homo s | ELP1 | 150.255 | 2 | 54.358864 | yes | 1.839E-07 | 699.91376 | 33.62228 |
| INLNLIYDHNPK | 12 | O95163 | Elongator complex protein 1 OS=Homo s | ELP1 | 150.255 | 2 | 54.358864 | yes | 1.021E-07 | 727.39355 | 26.307022 |
| GLIENPALLR | 10 | O95168 | NADH dehydrogenase [ubiquinone] 1 bet | NDUFB4 | 15.209 | 2 | 48.55 | yes | 2.5E-07 | 548.32965 | 686.62988 |
| EYLLQYNDPNR | 11 | O95168 | NADH dehydrogenase [ubiquinone] 1 bet | NDUFB4 | 15.209 | 2 | 48.55 | yes | 1.132E-07 | 712.84387 | 322.25787 |
| TINVYPNFR | 9 | O95168 | NADH dehydrogenase [ubiquinone] 1 bet | NDUFB4 | 15.209 | 2 | 48.55 | yes | 3.277E-06 | 562.2984 | 44.966473 |
| AMYLPDTLSPADQ | 15 | O95202 | Mitochondrial proton/calcium exchanger | LETM1 | 83.355 | 2 | 41.994545 | yes | 3.244E-07 | 831.92401 | 38.389893 |
| TGYGGGFNER | 10 | O95218 | Zinc finger Ran-binding domain-containi | ZRANB2 | 37.405 | 2 | 53.507324 | yes | 1.002E-07 | 529.23853 | 711.61707 |

| | | | | | | | | | | | |
|---|---|---|---|---|---|---|---|---|---|---|---|
| TCSNVNWAR | 9 | O95218 | Zinc finger Ran-binding domain-containi | ZRANB2 | 37.405 | 2 | 53.507324 | yes | 9.723E-08 | 554.25366 | 566.4231 |
| GLFSANDWQCK | 11 | O95218 | Zinc finger Ran-binding domain-containi | ZRANB2 | 37.405 | 2 | 53.507324 | yes | 2.991E-07 | 663.30078 | 548.97009 |
| CGNVNFAR | 8 | O95218 | Zinc finger Ran-binding domain-containi | ZRANB2 | 37.405 | 2 | 53.507324 | yes | 2.233E-07 | 469.21906 | 456.3988 |
| VSDGDWICPDKK | 12 | O95218 | Zinc finger Ran-binding domain-containi | ZRANB2 | 37.405 | 2 | 53.507324 | yes | 9.722E-08 | 710.33228 | 380.45398 |
| ENVEYIER | 8 | O95218 | Zinc finger Ran-binding domain-containi | ZRANB2 | 37.405 | 2 | 53.507324 | yes | 1.29E-07 | 526.25635 | 300.38644 |
| SECNMCNTPK | 10 | O95218 | Zinc finger Ran-binding domain-containi | ZRANB2 | 37.405 | 2 | 53.507324 | yes | 1.075E-07 | 620.74139 | 178.87892 |
| LDEDEDEDDADLS | 14 | O95218 | Zinc finger Ran-binding domain-containi | ZRANB2 | 37.405 | 2 | 53.507324 | yes | 9.722E-08 | 804.82336 | 101.14137 |
| RSECNMCNTPK | 11 | O95218 | Zinc finger Ran-binding domain-containi | ZRANB2 | 37.405 | 2 | 53.507324 | yes | 1.18E-07 | 698.79193 | 72.987877 |
| YNLDASEEEDSNK | 13 | O95218 | Zinc finger Ran-binding domain-containi | ZRANB2 | 37.405 | 2 | 53.507324 | yes | 9.722E-08 | 757.31805 | 70.512558 |
| EESDGEYDEFGR | 12 | O95218 | Zinc finger Ran-binding domain-containi | ZRANB2 | 37.405 | 2 | 53.507324 | yes | 9.722E-08 | 716.77856 | 34.32608 |
| GLDPLASEDTSR | 12 | O95229 | ZW10 interactor OS=Homo sapiens OX= | ZWINT | 31.294 | 2 | 47.522438 | yes | 1.312E-07 | 630.80695 | 74.724808 |
| YLQSLLAEVER | 11 | O95232 | Luc7-like protein 3 OS=Homo sapiens O | LUC7L3 | 51.467 | 2 | 49.450554 | yes | 1.064E-07 | 660.86151 | 2210.6221 |
| STTSTIESFAAQEK | 14 | O95232 | Luc7-like protein 3 OS=Homo sapiens O | LUC7L3 | 51.467 | 2 | 49.450554 | yes | 1.393E-07 | 750.36481 | 1310.2087 |
| VDDHLMGK | 8 | O95232 | Luc7-like protein 3 OS=Homo sapiens O | LUC7L3 | 51.467 | 2 | 49.450554 | yes | 3.037E-07 | 457.72366 | 546.18542 |
| VEEAQGMMK | 9 | O95232 | Luc7-like protein 3 OS=Homo sapiens O | LUC7L3 | 51.467 | 2 | 49.450554 | yes | 3.556E-07 | 511.7359 | 209.748 |
| QMEVCEVCGAFLI | 20 | O95232 | Luc7-like protein 3 OS=Homo sapiens O | LUC7L3 | 51.467 | 3 | 49.450554 | yes | 3.604E-07 | 757.01849 | 162.69618 |
| WDHESVCK | 8 | O95232 | Luc7-like protein 3 OS=Homo sapiens O | LUC7L3 | 51.467 | 2 | 49.450554 | yes | 1.423E-07 | 530.72949 | 82.849716 |
| LVEQLKEER | 9 | O95232 | Luc7-like protein 3 OS=Homo sapiens O | LUC7L3 | 51.467 | 2 | 49.450554 | yes | 1.029E-07 | 572.32202 | 27.4419 |
| ISELSLCDLAGSER | 14 | O95235 | Kinesin-like protein KIF20A OS=Homo s | KIF20A | 100.278 | 2 | 54.506832 | yes | 9.722E-08 | 775.37994 | 153.9794 |
| VFQGFFTGR | 9 | O95235 | Kinesin-like protein KIF20A OS=Homo s | KIF20A | 100.278 | 2 | 54.506832 | yes | 9.722E-08 | 529.77454 | 107.06023 |
| LGESLQSAER | 10 | O95235 | Kinesin-like protein KIF20A OS=Homo s | KIF20A | 100.278 | 2 | 54.506832 | yes | 2.322E-07 | 545.2804 | 89.994476 |
| EAGNINTSLHTLGI | 14 | O95235 | Kinesin-like protein KIF20A OS=Homo s | KIF20A | 100.278 | 2 | 54.506832 | yes | 1.733E-07 | 741.8866 | 80.515533 |
| THTIQGTIK | 9 | O95235 | Kinesin-like protein KIF20A OS=Homo s | KIF20A | 100.278 | 2 | 54.506832 | yes | 1.034E-06 | 499.7851 | 79.644981 |
| AACIAEQYHTVLK | 13 | O95235 | Kinesin-like protein KIF20A OS=Homo s | KIF20A | 100.278 | 2 | 54.506832 | yes | 1.153E-07 | 752.38483 | 74.134308 |
| LAASASTQQLQEV | 14 | O95235 | Kinesin-like protein KIF20A OS=Homo s | KIF20A | 100.278 | 2 | 54.506832 | yes | 1.272E-07 | 737.39899 | 68.09462 |
| QALTTCDDILIK | 12 | O95235 | Kinesin-like protein KIF20A OS=Homo s | KIF20A | 100.278 | 2 | 54.506832 | yes | 5.366E-07 | 695.86597 | 64.751633 |
| ELLEEMYEEK | 10 | O95235 | Kinesin-like protein KIF20A OS=Homo s | KIF20A | 100.278 | 2 | 54.506832 | yes | 3.148E-07 | 656.80249 | 63.642895 |
| IEELEALLQEAR | 12 | O95235 | Kinesin-like protein KIF20A OS=Homo s | KIF20A | 100.278 | 2 | 54.506832 | yes | 4.857E-07 | 707.38281 | 47.41761 |
| QNLVPFR | 7 | O95235 | Kinesin-like protein KIF20A OS=Homo s | KIF20A | 100.278 | 2 | 54.506832 | yes | 4.971E-07 | 437.2507 | 39.77227 |
| EELLQVVEAMK | 11 | O95235 | Kinesin-like protein KIF20A OS=Homo s | KIF20A | 100.278 | 2 | 54.506832 | yes | 1.748E-07 | 644.84448 | 35.749386 |
| ILHLQGEGDIVPK | 13 | O95235 | Kinesin-like protein KIF20A OS=Homo s | KIF20A | 100.278 | 2 | 54.506832 | yes | 1.192E-07 | 709.90375 | 30.614578 |
| QQVPSEDSMEK | 11 | O95235 | Kinesin-like protein KIF20A OS=Homo s | KIF20A | 100.278 | 2 | 54.506832 | yes | 2.812E-06 | 639.28754 | 19.956104 |
| IENVETLVLQAPK | 13 | O95235 | Kinesin-like protein KIF20A OS=Homo s | KIF20A | 100.278 | 2 | 54.506832 | yes | 3.723E-05 | 727.41669 | 19.128063 |
| EQWCSEHLDTQK | 12 | O95235 | Kinesin-like protein KIF20A OS=Homo s | KIF20A | 100.278 | 2 | 54.506832 | yes | 9.722E-08 | 780.841 | 13.056612 |
| LTLLQVASR | 9 | O95239 | Chromosome-associated kinesin KIF4A ( | KIF4A | 139.882 | 2 | 52.836567 | yes | 1.35E-07 | 500.81113 | 701.83972 |
| ILAQDVAQLK | 10 | O95239 | Chromosome-associated kinesin KIF4A ( | KIF4A | 139.882 | 2 | 52.836567 | yes | 1.101E-07 | 549.82953 | 564.94629 |
| SAQIADLQQK | 10 | O95239 | Chromosome-associated kinesin KIF4A ( | KIF4A | 139.882 | 2 | 52.836567 | no | 2.779E-06 | 551.29858 | 373.14575 |
| GLSEAAGQTAQMI | 15 | O95239 | Chromosome-associated kinesin KIF4A ( | KIF4A | 139.882 | 2 | 52.836567 | yes | 9.819E-08 | 781.38556 | 361.81635 |
| HLNDLLEDR | 9 | O95239 | Chromosome-associated kinesin KIF4A ( | KIF4A | 139.882 | 2 | 52.836567 | no | 3.832E-06 | 562.78839 | 311.59351 |

| | | | | | | | | | | | |
|---|---|---|---|---|---|---|---|---|---|---|---|
| YLIGELVSSK | 10 | O95239 | Chromosome-associated kinesin KIF4A ( | KIF4A | 139.882 | 2 | 52.836567 | no | 1.28E-07 | 554.81604 | 298.5545 |
| NKPIVNIDPQTAEI | 18 | O95239 | Chromosome-associated kinesin KIF4A ( | KIF4A | 139.882 | 3 | 52.836567 | yes | 1.061E-07 | 682.04437 | 296.33374 |
| EMCDVEQVLSK | 11 | O95239 | Chromosome-associated kinesin KIF4A ( | KIF4A | 139.882 | 2 | 52.836567 | yes | 1.106E-07 | 669.30743 | 280.36325 |
| LVETLEDQELK | 11 | O95239 | Chromosome-associated kinesin KIF4A ( | KIF4A | 139.882 | 2 | 52.836567 | no | 1.066E-07 | 658.85083 | 230.78955 |
| LQELEGQIADLK | 12 | O95239 | Chromosome-associated kinesin KIF4A ( | KIF4A | 139.882 | 2 | 52.836567 | yes | 9.85E-08 | 678.87207 | 230.19484 |
| QIESLETEMEFR | 12 | O95239 | Chromosome-associated kinesin KIF4A ( | KIF4A | 139.882 | 2 | 52.836567 | yes | 9.734E-08 | 756.35596 | 211.47601 |
| WENIATILEAK | 11 | O95239 | Chromosome-associated kinesin KIF4A ( | KIF4A | 139.882 | 2 | 52.836567 | yes | 9.882E-08 | 644.35083 | 193.3416 |
| DKEVIQLK | 8 | O95239 | Chromosome-associated kinesin KIF4A ( | KIF4A | 139.882 | 2 | 52.836567 | no | 5.123E-07 | 486.78986 | 183.18045 |
| SSDAFTTQHALR | 12 | O95239 | Chromosome-associated kinesin KIF4A ( | KIF4A | 139.882 | 2 | 52.836567 | yes | 1.033E-07 | 667.32837 | 180.04065 |
| FLEQSMDIEDLK | 12 | O95239 | Chromosome-associated kinesin KIF4A ( | KIF4A | 139.882 | 2 | 52.836567 | no | 5.298E-07 | 734.35541 | 174.44829 |
| GQVSESEDSITK | 12 | O95239 | Chromosome-associated kinesin KIF4A ( | KIF4A | 139.882 | 2 | 52.836567 | yes | 9.722E-08 | 640.30426 | 161.82353 |
| NQSLVEENEK | 10 | O95239 | Chromosome-associated kinesin KIF4A ( | KIF4A | 139.882 | 2 | 52.836567 | no | 8.719E-07 | 595.28839 | 144.36655 |
| TVASTAMNSQSSR | 13 | O95239 | Chromosome-associated kinesin KIF4A ( | KIF4A | 139.882 | 2 | 52.836567 | no | 9.968E-08 | 670.31714 | 121.845 |
| CQDEELEK | 8 | O95239 | Chromosome-associated kinesin KIF4A ( | KIF4A | 139.882 | 2 | 52.836567 | yes | 9.819E-08 | 525.72406 | 98.984856 |
| EKEELVLELQTAK | 13 | O95239 | Chromosome-associated kinesin KIF4A ( | KIF4A | 139.882 | 2 | 52.836567 | yes | 1.354E-07 | 765.42468 | 94.09697 |
| EQQLLSTLK | 9 | O95239 | Chromosome-associated kinesin KIF4A ( | KIF4A | 139.882 | 2 | 52.836567 | yes | 5.677E-07 | 530.30585 | 59.583691 |
| ELELEVINLQK | 11 | O95239 | Chromosome-associated kinesin KIF4A ( | KIF4A | 139.882 | 2 | 52.836567 | yes | 1.046E-07 | 664.37701 | 53.813198 |
| EVCEQNQQLLR | 11 | O95239 | Chromosome-associated kinesin KIF4A ( | KIF4A | 139.882 | 2 | 52.836567 | yes | 9.722E-08 | 708.84863 | 49.586819 |
| GYNATVLAYGQT( | 16 | O95239 | Chromosome-associated kinesin KIF4A ( | KIF4A | 139.882 | 2 | 52.836567 | no | 8.471E-07 | 793.8941 | 26.64547 |
| PLVPKEISEGCQM( | 30 | O95239 | Chromosome-associated kinesin KIF4A ( | KIF4A | 139.882 | 4 | 52.836567 | yes | 0.000143 | 825.91327 | 10.726283 |
| ADAESEPVAQK | 11 | O95243 | Methyl-CpG-binding domain protein 4 O: | MBD4 | 66.051 | 2 | 54.779037 | yes | 1.299E-06 | 572.77765 | 347.51367 |
| LVPDPPNDLR | 10 | O95243 | Methyl-CpG-binding domain protein 4 O: | MBD4 | 66.051 | 2 | 54.779037 | yes | 9.722E-08 | 568.30896 | 328.44223 |
| EDVAMELER | 9 | O95243 | Methyl-CpG-binding domain protein 4 O: | MBD4 | 66.051 | 2 | 54.779037 | yes | 1.091E-07 | 546.25549 | 287.22321 |
| FDVYFISPQGLK | 12 | O95243 | Methyl-CpG-binding domain protein 4 O: | MBD4 | 66.051 | 2 | 54.779037 | yes | 3.382E-07 | 707.37427 | 284.1647 |
| EDEGVDDVNFR | 11 | O95243 | Methyl-CpG-binding domain protein 4 O: | MBD4 | 66.051 | 2 | 54.779037 | yes | 9.722E-08 | 647.78094 | 234.06271 |
| YEDTFLESEEIGTK | 14 | O95243 | Methyl-CpG-binding domain protein 4 O: | MBD4 | 66.051 | 2 | 54.779037 | yes | 1.036E-07 | 830.88306 | 205.18518 |
| VGEDEEQMMIK | 11 | O95243 | Methyl-CpG-binding domain protein 4 O: | MBD4 | 66.051 | 2 | 54.779037 | yes | 9.722E-08 | 654.79413 | 193.30667 |
| IFQEDTIPR | 9 | O95243 | Methyl-CpG-binding domain protein 4 O: | MBD4 | 66.051 | 2 | 54.779037 | yes | 1.281E-06 | 559.79565 | 168.06693 |
| SSLANYLHK | 9 | O95243 | Methyl-CpG-binding domain protein 4 O: | MBD4 | 66.051 | 2 | 54.779037 | yes | 3.027E-06 | 516.77728 | 167.92339 |
| YPSAEVAR | 8 | O95243 | Methyl-CpG-binding domain protein 4 O: | MBD4 | 66.051 | 2 | 54.779037 | yes | 4.749E-07 | 446.7298 | 152.84648 |
| YPIELHGIGK | 10 | O95243 | Methyl-CpG-binding domain protein 4 O: | MBD4 | 66.051 | 2 | 54.779037 | yes | 1.072E-07 | 563.81641 | 139.38548 |
| GAAPTVTSSER | 11 | O95243 | Methyl-CpG-binding domain protein 4 O: | MBD4 | 66.051 | 2 | 54.779037 | yes | 3.942E-07 | 538.27252 | 134.53003 |
| KEDVAMELER | 10 | O95243 | Methyl-CpG-binding domain protein 4 O: | MBD4 | 66.051 | 2 | 54.779037 | yes | 4.754E-06 | 610.30298 | 93.400703 |
| SVPCGWER | 8 | O95243 | Methyl-CpG-binding domain protein 4 O: | MBD4 | 66.051 | 2 | 54.779037 | yes | 2.388E-06 | 495.72672 | 90.177177 |
| GTTGLESLSLGDR | 13 | O95243 | Methyl-CpG-binding domain protein 4 O: | MBD4 | 66.051 | 2 | 54.779037 | yes | 1.91E-06 | 674.34119 | 72.128593 |
| SLSSGSNFCSEQK | 13 | O95243 | Methyl-CpG-binding domain protein 4 O: | MBD4 | 66.051 | 2 | 54.779037 | yes | 9.722E-08 | 715.81445 | 53.315281 |
| EHLHTDILK | 9 | O95243 | Methyl-CpG-binding domain protein 4 O: | MBD4 | 66.051 | 2 | 54.779037 | yes | 2.404E-06 | 553.30365 | 51.547215 |
| GLSNFTSTHLLLK | 13 | O95243 | Methyl-CpG-binding domain protein 4 O: | MBD4 | 66.051 | 2 | 54.779037 | yes | 5.118E-07 | 715.90375 | 21.823864 |
| LGLGTLSSSLSR | 12 | O95248 | Myotubularin-related protein 5 OS=Hom( | SBF1 | 208.444 | 2 | 53.80546 | yes | 1.225E-07 | 595.84058 | 175.77861 |

| | | | | | | | | | | | |
|---|---|---|---|---|---|---|---|---|---|---|---|
| SALDVASEQR | 10 | O95248 | Myotubularin-related protein 5 OS=Hom | SBF1 | 208.444 | 2 | 53.80546 | yes | 2.184E-07 | 538.27252 | 168.51344 |
| TFLLDSDYER | 10 | O95248 | Myotubularin-related protein 5 OS=Hom | SBF1 | 208.444 | 2 | 53.80546 | no | 9.848E-08 | 629.80115 | 158.01292 |
| EWLSFGHR | 8 | O95248 | Myotubularin-related protein 5 OS=Hom | SBF1 | 208.444 | 2 | 53.80546 | no | 1.525E-07 | 516.25653 | 157.25761 |
| GSGEGQGQILQR | 12 | O95248 | Myotubularin-related protein 5 OS=Hom | SBF1 | 208.444 | 2 | 53.80546 | yes | 3.506E-07 | 615.31531 | 121.5726 |
| MSYLLLPLDSSK | 12 | O95248 | Myotubularin-related protein 5 OS=Hom | SBF1 | 208.444 | 2 | 53.80546 | yes | 1.612E-07 | 683.86798 | 108.69276 |
| QVIQTPLADSLPVS | 15 | O95248 | Myotubularin-related protein 5 OS=Hom | SBF1 | 208.444 | 2 | 53.80546 | yes | 9.722E-08 | 812.45685 | 103.79512 |
| VYLLPDGR | 8 | O95248 | Myotubularin-related protein 5 OS=Hom | SBF1 | 208.444 | 2 | 53.80546 | yes | 1.247E-07 | 466.76364 | 101.2682 |
| CSVALLFR | 8 | O95248 | Myotubularin-related protein 5 OS=Hom | SBF1 | 208.444 | 2 | 53.80546 | yes | 9.722E-08 | 483.26547 | 90.901894 |
| TTVPSGPPMTAILE | 15 | O95248 | Myotubularin-related protein 5 OS=Hom | SBF1 | 208.444 | 2 | 53.80546 | yes | 3.624E-06 | 785.41888 | 88.873131 |
| LFAQLLQGYR | 10 | O95248 | Myotubularin-related protein 5 OS=Hom | SBF1 | 208.444 | 2 | 53.80546 | yes | 1.949E-07 | 604.84296 | 84.278099 |
| VVWPCYDSCPR | 11 | O95248 | Myotubularin-related protein 5 OS=Hom | SBF1 | 208.444 | 2 | 53.80546 | yes | 4.48E-07 | 719.81573 | 52.561028 |
| VCTESGVTSDHLK | 13 | O95248 | Myotubularin-related protein 5 OS=Hom | SBF1 | 208.444 | 2 | 53.80546 | yes | 1.04E-07 | 716.84045 | 49.561218 |
| GLVEDDFLMK | 10 | O95248 | Myotubularin-related protein 5 OS=Hom | SBF1 | 208.444 | 2 | 53.80546 | yes | 3.617E-06 | 583.79175 | 44.400795 |
| AVLDHQQFDFVVI | 13 | O95248 | Myotubularin-related protein 5 OS=Hom | SBF1 | 208.444 | 2 | 53.80546 | yes | 9.722E-08 | 787.40973 | 42.360153 |
| WCLHVVR | 7 | O95248 | Myotubularin-related protein 5 OS=Hom | SBF1 | 208.444 | 2 | 53.80546 | yes | 3.211E-06 | 485.258 | 42.236145 |
| IELGLLYEEK | 10 | O95248 | Myotubularin-related protein 5 OS=Hom | SBF1 | 208.444 | 2 | 53.80546 | yes | 1.326E-06 | 603.83447 | 38.800449 |
| HVQELAEQLYK | 11 | O95248 | Myotubularin-related protein 5 OS=Hom | SBF1 | 208.444 | 2 | 53.80546 | yes | 9.722E-08 | 679.35913 | 38.156979 |
| LETELGQPAER | 11 | O95248 | Myotubularin-related protein 5 OS=Hom | SBF1 | 208.444 | 2 | 53.80546 | yes | 1.239E-07 | 621.81989 | 37.930721 |
| KLLPAVLR | 8 | O95248 | Myotubularin-related protein 5 OS=Hom | SBF1 | 208.444 | 2 | 53.80546 | yes | 2.944E-07 | 455.31583 | 37.523121 |
| MTMSSLVER | 9 | O95248 | Myotubularin-related protein 5 OS=Hom | SBF1 | 208.444 | 2 | 53.80546 | yes | 4.346E-06 | 527.25702 | 23.602526 |
| LQGQITEGSNMIK | 13 | O95251 | Histone acetyltransferase KAT7 OS=Hon | KAT7 | 70.643 | 2 | 50.782448 | yes | 9.819E-08 | 709.86902 | 450.02219 |
| SQQQPTPVTPK | 11 | O95251 | Histone acetyltransferase KAT7 OS=Hon | KAT7 | 70.643 | 2 | 50.782448 | yes | 1.012E-07 | 605.82495 | 320.24478 |
| IYCQNLCLLAK | 11 | O95251 | Histone acetyltransferase KAT7 OS=Hon | KAT7 | 70.643 | 2 | 50.782448 | no | 1.002E-07 | 698.3598 | 288.83072 |
| LYMCEFCLK | 9 | O95251 | Histone acetyltransferase KAT7 OS=Hon | KAT7 | 70.643 | 2 | 50.782448 | yes | 1.398E-07 | 632.28217 | 257.09616 |
| HPPGDEIYR | 9 | O95251 | Histone acetyltransferase KAT7 OS=Hon | KAT7 | 70.643 | 2 | 50.782448 | yes | 8.387E-07 | 542.26453 | 223.54028 |
| FHESYNFNMK | 10 | O95251 | Histone acetyltransferase KAT7 OS=Hon | KAT7 | 70.643 | 2 | 50.782448 | yes | 9.722E-08 | 658.79004 | 187.09196 |
| TMDPSCLK | 8 | O95251 | Histone acetyltransferase KAT7 OS=Hon | KAT7 | 70.643 | 2 | 50.782448 | yes | 9.608E-07 | 476.21735 | 175.41861 |
| PLSDLGLISYR | 11 | O95251 | Histone acetyltransferase KAT7 OS=Hon | KAT7 | 70.643 | 2 | 50.782448 | yes | 4.455E-06 | 617.34552 | 161.62257 |
| RQDLIDEWIAK | 11 | O95251 | Histone acetyltransferase KAT7 OS=Hon | KAT7 | 70.643 | 2 | 50.782448 | yes | 1.23E-07 | 693.87244 | 134.07394 |
| QDLIDEWIAK | 10 | O95251 | Histone acetyltransferase KAT7 OS=Hon | KAT7 | 70.643 | 2 | 50.782448 | yes | 1.232E-06 | 615.8219 | 79.53109 |
| YLHNFQGK | 8 | O95251 | Histone acetyltransferase KAT7 OS=Hon | KAT7 | 70.643 | 2 | 50.782448 | yes | 1.073E-07 | 503.75888 | 65.161774 |
| VGSPERPLSDLGLI | 17 | O95251 | Histone acetyltransferase KAT7 OS=Hon | KAT7 | 70.643 | 3 | 50.782448 | yes | 5.534E-07 | 620.33887 | 48.484612 |
| KGSISVFEVDGK | 12 | O95251 | Histone acetyltransferase KAT7 OS=Hon | KAT7 | 70.643 | 2 | 50.782448 | yes | 3.078E-07 | 633.34045 | 40.43 |
| CPTPGCNSLGHLT | 15 | O95251 | Histone acetyltransferase KAT7 OS=Hon | KAT7 | 70.643 | 2 | 50.782448 | yes | 1.001E-06 | 799.87433 | 31.970268 |
| DSGSDLSHRPK | 11 | O95251 | Histone acetyltransferase KAT7 OS=Hon | KAT7 | 70.643 | 2 | 50.782448 | yes | 9.977E-08 | 599.79419 | 27.033939 |
| GQELGALR | 8 | O95255 | ATP-binding cassette sub-family C memb | ABCC6 | 164.907 | 2 | 35.30181 | yes | 8.536E-07 | 422.23779 | 733.05389 |
| TTELYSPQLGTK | 12 | O95263 | High affinity cAMP-specific and IBMX-i | PDE8B | 98.979 | 2 | 40.8388 | yes | 3.813E-07 | 669.35101 | 42.556061 |
| FTPLHEAAAK | 10 | O95271 | Poly [ADP-ribose] polymerase tankyrase- | TNKS | 142.04 | 2 | 54.705795 | no | 9.728E-08 | 542.79291 | 412.1655 |
| SAVDMAPTPELR | 12 | O95271 | Poly [ADP-ribose] polymerase tankyrase- | TNKS | 142.04 | 2 | 54.705795 | yes | 9.722E-08 | 643.8241 | 361.77841 |

| | | | | | | | | | | | |
|---|---|---|---|---|---|---|---|---|---|---|---|
| EGDTDIQDLLR | 11 | O95271 | Poly [ADP-ribose] polymerase tankyrase- | TNKS | 142.04 | 2 | 54.705795 | yes | 1.009E-07 | 637.81476 | 281.3634 |
| DGGNAGGIFNR | 11 | O95271 | Poly [ADP-ribose] polymerase tankyrase- | TNKS | 142.04 | 2 | 54.705795 | yes | 1.084E-07 | 539.25726 | 274.68021 |
| SLGLEHLR | 8 | O95271 | Poly [ADP-ribose] polymerase tankyrase- | TNKS | 142.04 | 2 | 54.705795 | yes | 3.561E-06 | 462.76672 | 269.55057 |
| EIGINAYGHR | 10 | O95271 | Poly [ADP-ribose] polymerase tankyrase- | TNKS | 142.04 | 2 | 54.705795 | no | 1.881E-07 | 565.29108 | 245.63863 |
| LVDAANVNAK | 10 | O95271 | Poly [ADP-ribose] polymerase tankyrase- | TNKS | 142.04 | 2 | 54.705795 | yes | 9.819E-08 | 507.78256 | 244.41171 |
| AALAGHLQTCR | 11 | O95271 | Poly [ADP-ribose] polymerase tankyrase- | TNKS | 142.04 | 2 | 54.705795 | yes | 9.733E-08 | 599.31146 | 236.1586 |
| GHSLLQAAR | 9 | O95271 | Poly [ADP-ribose] polymerase tankyrase- | TNKS | 142.04 | 2 | 54.705795 | no | 1.107E-07 | 476.76978 | 209.45532 |
| LCTPENINCR | 10 | O95271 | Poly [ADP-ribose] polymerase tankyrase- | TNKS | 142.04 | 2 | 54.705795 | yes | 9.722E-08 | 638.7923 | 208.70081 |
| DNWNYTPLHEAA | 14 | O95271 | Poly [ADP-ribose] polymerase tankyrase- | TNKS | 142.04 | 2 | 54.705795 | no | 9.722E-08 | 836.40991 | 194.61549 |
| TQLCALLLAHGAD | 17 | O95271 | Poly [ADP-ribose] polymerase tankyrase- | TNKS | 142.04 | 3 | 54.705795 | yes | 1.675E-07 | 613.98669 | 185.13612 |
| HGLALPEGDGSR | 12 | O95271 | Poly [ADP-ribose] polymerase tankyrase- | TNKS | 142.04 | 2 | 54.705795 | yes | 9.722E-08 | 604.80457 | 162.79491 |
| QVTELLLR | 8 | O95271 | Poly [ADP-ribose] polymerase tankyrase- | TNKS | 142.04 | 2 | 54.705795 | yes | 4.271E-07 | 486.29785 | 158.30898 |
| TLALEIINFK | 10 | O95271 | Poly [ADP-ribose] polymerase tankyrase- | TNKS | 142.04 | 2 | 54.705795 | yes | 9.722E-08 | 581.34753 | 152.52069 |
| MAHAPPGHHSVIC | 14 | O95271 | Poly [ADP-ribose] polymerase tankyrase- | TNKS | 142.04 | 3 | 54.705795 | yes | 3.34E-07 | 489.58597 | 125.6769 |
| STPLHLAAGYNR | 12 | O95271 | Poly [ADP-ribose] polymerase tankyrase- | TNKS | 142.04 | 2 | 54.705795 | no | 6.907E-07 | 650.34387 | 116.77924 |
| HSTPLHFAAGYNR | 13 | O95271 | Poly [ADP-ribose] polymerase tankyrase- | TNKS | 142.04 | 3 | 54.705795 | yes | 9.964E-08 | 490.91275 | 101.78628 |
| YNTCVNATDK | 10 | O95271 | Poly [ADP-ribose] polymerase tankyrase- | TNKS | 142.04 | 2 | 54.705795 | yes | 9.722E-08 | 593.26385 | 82.263466 |
| SSPLHFAAGFGR | 12 | O95271 | Poly [ADP-ribose] polymerase tankyrase- | TNKS | 142.04 | 2 | 54.705795 | yes | 4.198E-07 | 623.82001 | 81.399078 |
| HGASVNVADLWK | 12 | O95271 | Poly [ADP-ribose] polymerase tankyrase- | TNKS | 142.04 | 2 | 54.705795 | yes | 9.722E-08 | 648.83838 | 65.544571 |
| GKYEICK | 7 | O95271 | Poly [ADP-ribose] polymerase tankyrase- | TNKS | 142.04 | 2 | 54.705795 | no | 1.073E-07 | 449.22858 | 64.949745 |
| QLCSSQNVNCR | 11 | O95271 | Poly [ADP-ribose] polymerase tankyrase- | TNKS | 142.04 | 2 | 54.705795 | yes | 9.823E-08 | 683.30353 | 58.3881 |
| ELLEACR | 7 | O95271 | Poly [ADP-ribose] polymerase tankyrase- | TNKS | 142.04 | 2 | 54.705795 | yes | 3.867E-07 | 445.72366 | 56.942265 |
| IVQLLLQHGADVH | 15 | O95271 | Poly [ADP-ribose] polymerase tankyrase- | TNKS | 142.04 | 2 | 54.705795 | no | 2.256E-07 | 821.4754 | 23.555676 |
| SFLQFSTMK | 9 | O95271 | Poly [ADP-ribose] polymerase tankyrase- | TNKS | 142.04 | 2 | 54.705795 | yes | 2.216E-07 | 544.77588 | 22.513931 |
| GGLIPLHNAASYG | 23 | O95271 | Poly [ADP-ribose] polymerase tankyrase- | TNKS | 142.04 | 3 | 54.705795 | yes | 1.041E-07 | 781.77765 | 15.761884 |
| KNYYTQK | 7 | O95279 | Potassium channel subfamily K member 5 | KCNK5 | 55.13 | 2 | 19.475641 | yes | 4.837E-06 | 472.74545 | 124.07233 |
| VEQVLSLEPQHEL | 14 | O95292 | Vesicle-associated membrane protein-ass | VAPB | 27.229 | 2 | 54.66766 | yes | 9.722E-08 | 824.94885 | 280.60095 |
| TVQSNSPISALAPT | 16 | O95292 | Vesicle-associated membrane protein-ass | VAPB | 27.229 | 2 | 54.66766 | yes | 3.948E-07 | 785.92542 | 79.024498 |
| TETPIVSK | 8 | O95292 | Vesicle-associated membrane protein-ass | VAPB | 27.229 | 2 | 54.66766 | yes | 2.837E-06 | 437.74765 | 32.430035 |
| SLSSSLDDTEVKK | 13 | O95292 | Vesicle-associated membrane protein-ass | VAPB | 27.229 | 2 | 54.66766 | yes | 9.722E-08 | 704.86194 | 31.187107 |
| EQIDNLATELCR | 12 | O95295 | SNARE-associated protein Snapin OS=H | SNAPIN | 14.874 | 2 | 47.959522 | yes | 2.492E-07 | 731.35376 | 56.741127 |
| VVLVNNILQNAQE | 14 | O95295 | SNARE-associated protein Snapin OS=H | SNAPIN | 14.874 | 2 | 47.959522 | yes | 1.224E-07 | 805.45465 | 50.088444 |
| VALDLDPYVK | 10 | O95295 | SNARE-associated protein Snapin OS=H | SNAPIN | 14.874 | 2 | 47.959522 | yes | 0.0001383 | 566.81604 | 12.032446 |
| LQSWLYSSR | 9 | O95299 | NADH dehydrogenase [ubiquinone] 1 alp | NDUFA10 | 40.751 | 2 | 49.95487 | yes | 1.267E-07 | 570.29584 | 911.94891 |
| YSPGYNTEVGDK | 12 | O95299 | NADH dehydrogenase [ubiquinone] 1 alp | NDUFA10 | 40.751 | 2 | 49.95487 | yes | 1.082E-07 | 665.30151 | 498.72946 |
| VITVDGNICTGK | 12 | O95299 | NADH dehydrogenase [ubiquinone] 1 alp | NDUFA10 | 40.751 | 2 | 49.95487 | yes | 1.284E-07 | 638.83191 | 480.14584 |
| TFLPEMSEK | 9 | O95299 | NADH dehydrogenase [ubiquinone] 1 alp | NDUFA10 | 40.751 | 2 | 49.95487 | yes | 1.456E-07 | 541.26532 | 413.96518 |
| CEVLQYSAR | 9 | O95299 | NADH dehydrogenase [ubiquinone] 1 alp | NDUFA10 | 40.751 | 2 | 49.95487 | yes | 9.968E-08 | 563.27148 | 294.48825 |
| ITSAYLQDIENAYK | 14 | O95299 | NADH dehydrogenase [ubiquinone] 1 alp | NDUFA10 | 40.751 | 2 | 49.95487 | yes | 1.21E-07 | 814.91193 | 205.90761 |

| | | | | | | | | | | | |
|---|---|---|---|---|---|---|---|---|---|---|---|
| VVEDIEYLK | 9 | O95299 | NADH dehydrogenase [ubiquinone] 1 alp | NDUFA10 | 40.751 | 2 | 49.95487 | yes | 1.199E-07 | 554.30023 | 192.29263 |
| YGMWHFLLGDK | 11 | O95299 | NADH dehydrogenase [ubiquinone] 1 alp | NDUFA10 | 40.751 | 2 | 49.95487 | yes | 1.557E-07 | 683.83429 | 86.254707 |
| KVVEDIEYLK | 10 | O95299 | NADH dehydrogenase [ubiquinone] 1 alp | NDUFA10 | 40.751 | 2 | 49.95487 | yes | 2.129E-07 | 618.34772 | 52.457703 |
| FDKGPWLK | 8 | O95299 | NADH dehydrogenase [ubiquinone] 1 alp | NDUFA10 | 40.751 | 2 | 49.95487 | yes | 1.332E-06 | 495.77402 | 46.176071 |
| LLQYSDALEHLLT | 22 | O95299 | NADH dehydrogenase [ubiquinone] 1 alp | NDUFA10 | 40.751 | 3 | 49.95487 | yes | 2.268E-07 | 819.11121 | 42.677528 |
| LYNVVVDTEVTGI | 13 | O95347 | Structural maintenance of chromosomes p | SMC2 | 135.657 | 2 | 57.004215 | yes | 9.722E-08 | 718.88519 | 397.6633 |
| ENLTELSGGQR | 11 | O95347 | Structural maintenance of chromosomes p | SMC2 | 135.657 | 2 | 57.004215 | yes | 9.722E-08 | 602.30182 | 340.56433 |
| EVITAQDTVIK | 11 | O95347 | Structural maintenance of chromosomes p | SMC2 | 135.657 | 2 | 57.004215 | yes | 2.578E-07 | 608.84283 | 317.99249 |
| ALEEELAGLK | 10 | O95347 | Structural maintenance of chromosomes p | SMC2 | 135.657 | 2 | 57.004215 | yes | 9.722E-08 | 536.79785 | 246.05392 |
| AMNVLTEAEER | 11 | O95347 | Structural maintenance of chromosomes p | SMC2 | 135.657 | 2 | 57.004215 | yes | 9.722E-08 | 631.80591 | 231.67296 |
| FVDGVSTVAR | 10 | O95347 | Structural maintenance of chromosomes p | SMC2 | 135.657 | 2 | 57.004215 | yes | 2.947E-07 | 525.78253 | 222.05316 |
| SLEDALAEAQR | 11 | O95347 | Structural maintenance of chromosomes p | SMC2 | 135.657 | 2 | 57.004215 | yes | 9.722E-08 | 601.8042 | 186.90552 |
| TEEADLLQTK | 10 | O95347 | Structural maintenance of chromosomes p | SMC2 | 135.657 | 2 | 57.004215 | yes | 2.71E-07 | 574.29572 | 186.01451 |
| TIEESEETLK | 10 | O95347 | Structural maintenance of chromosomes p | SMC2 | 135.657 | 2 | 57.004215 | yes | 1.541E-07 | 589.79297 | 144.91121 |
| NQALNIAWQK | 10 | O95347 | Structural maintenance of chromosomes p | SMC2 | 135.657 | 2 | 57.004215 | yes | 9.722E-08 | 593.32239 | 130.41246 |
| QQLEAVNEAIK | 11 | O95347 | Structural maintenance of chromosomes p | SMC2 | 135.657 | 2 | 57.004215 | yes | 9.722E-08 | 621.83807 | 125.69707 |
| EGMFNNANVLFK | 12 | O95347 | Structural maintenance of chromosomes p | SMC2 | 135.657 | 2 | 57.004215 | yes | 9.722E-08 | 692.3399 | 121.92353 |
| DKETGGILR | 9 | O95347 | Structural maintenance of chromosomes p | SMC2 | 135.657 | 2 | 57.004215 | yes | 9.632E-07 | 494.77472 | 117.91021 |
| TILEEEITPTIQK | 13 | O95347 | Structural maintenance of chromosomes p | SMC2 | 135.657 | 2 | 57.004215 | yes | 9.722E-08 | 757.91925 | 113.71394 |
| ASNLQDLVYK | 10 | O95347 | Structural maintenance of chromosomes p | SMC2 | 135.657 | 2 | 57.004215 | yes | 1.894E-07 | 575.80878 | 111.67537 |
| ILTTIEDLDQK | 11 | O95347 | Structural maintenance of chromosomes p | SMC2 | 135.657 | 2 | 57.004215 | yes | 2.031E-07 | 644.85339 | 106.96281 |
| LKETYEALLAR | 11 | O95347 | Structural maintenance of chromosomes p | SMC2 | 135.657 | 2 | 57.004215 | yes | 1.341E-07 | 653.87189 | 84.619339 |
| FAYKDPEK | 8 | O95347 | Structural maintenance of chromosomes p | SMC2 | 135.657 | 2 | 57.004215 | yes | 1.315E-05 | 499.25311 | 77.292267 |
| LQEELSENDKK | 11 | O95347 | Structural maintenance of chromosomes p | SMC2 | 135.657 | 2 | 57.004215 | yes | 9.722E-08 | 666.83569 | 62.186806 |
| ITDGLHALQEASN | 14 | O95347 | Structural maintenance of chromosomes p | SMC2 | 135.657 | 2 | 57.004215 | yes | 4.587E-06 | 748.88879 | 56.825657 |
| QQEVEAITLELEEI | 16 | O95347 | Structural maintenance of chromosomes p | SMC2 | 135.657 | 3 | 57.004215 | yes | 2.526E-07 | 643.34937 | 37.674309 |
| YTIIPLNK | 8 | O95347 | Structural maintenance of chromosomes p | SMC2 | 135.657 | 2 | 57.004215 | yes | 2.346E-06 | 481.28949 | 35.249409 |
| VAQNLVGPDNVH | 26 | O95347 | Structural maintenance of chromosomes p | SMC2 | 135.657 | 4 | 57.004215 | yes | 2.173E-06 | 715.89465 | 19.721344 |
| DGESLQLFEQSSR | 13 | O95363 | Phenylalanine--tRNA ligase, mitochondri | FARS2 | 52.357 | 2 | 54.818733 | yes | 1.101E-07 | 748.3548 | 916.28186 |
| HIHQALQEAAVQL | 19 | O95363 | Phenylalanine--tRNA ligase, mitochondri | FARS2 | 52.357 | 3 | 54.818733 | yes | 1.117E-07 | 690.37988 | 808.66217 |
| APGSVVELLGK | 11 | O95363 | Phenylalanine--tRNA ligase, mitochondri | FARS2 | 52.357 | 2 | 54.818733 | yes | 1.519E-07 | 535.31622 | 659.41241 |
| LFWCEDER | 8 | O95363 | Phenylalanine--tRNA ligase, mitochondri | FARS2 | 52.357 | 2 | 54.818733 | yes | 9.935E-08 | 577.75043 | 486.42874 |
| HELFAGIK | 8 | O95363 | Phenylalanine--tRNA ligase, mitochondri | FARS2 | 52.357 | 2 | 54.818733 | yes | 1.078E-07 | 457.75836 | 457.08243 |
| LMAHLFGDELEIR | 13 | O95363 | Phenylalanine--tRNA ligase, mitochondri | FARS2 | 52.357 | 2 | 54.818733 | yes | 9.722E-08 | 772.40051 | 451.32831 |
| LAMILYDIPDIR | 12 | O95363 | Phenylalanine--tRNA ligase, mitochondri | FARS2 | 52.357 | 2 | 54.818733 | yes | 2.307E-06 | 716.89703 | 358.42963 |
| IGWAFGLGLER | 11 | O95363 | Phenylalanine--tRNA ligase, mitochondri | FARS2 | 52.357 | 2 | 54.818733 | yes | 1.456E-07 | 609.83514 | 291.51749 |
| QFCVSNINQK | 10 | O95363 | Phenylalanine--tRNA ligase, mitochondri | FARS2 | 52.357 | 2 | 54.818733 | yes | 9.809E-08 | 619.30334 | 283.62326 |
| SYPQDDHSNLTR | 12 | O95363 | Phenylalanine--tRNA ligase, mitochondri | FARS2 | 52.357 | 2 | 54.818733 | yes | 2.586E-06 | 716.82623 | 263.29211 |
| QETHTMEAVK | 10 | O95363 | Phenylalanine--tRNA ligase, mitochondri | FARS2 | 52.357 | 2 | 54.818733 | yes | 9.722E-08 | 587.28204 | 220.86108 |

| | | | | | | | | | | | |
|---|---|---|---|---|---|---|---|---|---|---|---|
| LVEFDLK | 7 | O95363 | Phenylalanine--tRNA ligase, mitochondri | FARS2 | 52.357 | 2 | 54.818733 | yes | 9.722E-08 | 432.24728 | 182.70151 |
| KGDNYYLNR | 9 | O95363 | Phenylalanine--tRNA ligase, mitochondri | FARS2 | 52.357 | 2 | 54.818733 | yes | 2.352E-07 | 571.78308 | 139.07567 |
| SVLAACSQYFK | 11 | O95365 | Zinc finger and BTB domain-containing p | ZBTB7A | 61.44 | 2 | 50.89581 | yes | 9.722E-08 | 637.31592 | 271.28595 |
| HTGEKPYLCQQCC | 23 | O95365 | Zinc finger and BTB domain-containing p | ZBTB7A | 61.44 | 4 | 50.89581 | yes | 1.125E-07 | 677.81409 | 31.194714 |
| AIFQTIQNR | 9 | O95373 | Importin-7 OS=Homo sapiens OX=9606 | IPO7 | 119.517 | 2 | 55.684402 | yes | 2.807E-07 | 545.80383 | 501.86005 |
| LLEVIILQCK | 10 | O95373 | Importin-7 OS=Homo sapiens OX=9606 | IPO7 | 119.517 | 2 | 55.684402 | no | 1.116E-07 | 614.8703 | 368.39514 |
| SDQNLQTALELTR | 13 | O95373 | Importin-7 OS=Homo sapiens OX=9606 | IPO7 | 119.517 | 2 | 55.684402 | yes | 1.345E-07 | 744.88629 | 332.34488 |
| ETENDDLTNVIQK | 13 | O95373 | Importin-7 OS=Homo sapiens OX=9606 | IPO7 | 119.517 | 2 | 55.684402 | yes | 2.612E-07 | 759.86774 | 195.55692 |
| NMITQYWPDR | 10 | O95373 | Importin-7 OS=Homo sapiens OX=9606 | IPO7 | 119.517 | 2 | 55.684402 | yes | 9.722E-08 | 662.31116 | 144.21094 |
| QLQDIATLADQR | 12 | O95373 | Importin-7 OS=Homo sapiens OX=9606 | IPO7 | 119.517 | 2 | 55.684402 | yes | 2.373E-07 | 686.36499 | 130.97926 |
| YLEMIYSMCK | 10 | O95373 | Importin-7 OS=Homo sapiens OX=9606 | IPO7 | 119.517 | 2 | 55.684402 | yes | 9.722E-08 | 669.30054 | 114.57751 |
| WALHILAR | 8 | O95373 | Importin-7 OS=Homo sapiens OX=9606 | IPO7 | 119.517 | 2 | 55.684402 | yes | 1.514E-07 | 490.29544 | 94.112244 |
| VQLTTCIHHIIK | 12 | O95373 | Importin-7 OS=Homo sapiens OX=9606 | IPO7 | 119.517 | 2 | 55.684402 | yes | 1.903E-07 | 731.91357 | 28.892918 |
| KWALHILAR | 9 | O95373 | Importin-7 OS=Homo sapiens OX=9606 | IPO7 | 119.517 | 2 | 55.684402 | yes | 4.662E-07 | 554.3429 | 27.144529 |
| EYNEFAEVFLK | 11 | O95373 | Importin-7 OS=Homo sapiens OX=9606 | IPO7 | 119.517 | 2 | 55.684402 | yes | 9.722E-08 | 694.84027 | 24.729004 |
| VLTGVAGEDAECI | 16 | O95373 | Importin-7 OS=Homo sapiens OX=9606 | IPO7 | 119.517 | 2 | 55.684402 | yes | 1.022E-06 | 814.39087 | 23.579187 |
| EPTEEEMEAYR | 11 | O95391 | Pre-mRNA-splicing factor SLU7 OS=Ho | SLU7 | 68.387 | 2 | 52.832832 | yes | 9.722E-08 | 692.29028 | 507.33188 |
| GSEVHLQADPTK | 12 | O95391 | Pre-mRNA-splicing factor SLU7 OS=Ho | SLU7 | 68.387 | 2 | 52.832832 | yes | 1.026E-07 | 641.32532 | 502.63239 |
| LVEQANSPK | 9 | O95391 | Pre-mRNA-splicing factor SLU7 OS=Ho | SLU7 | 68.387 | 2 | 52.832832 | yes | 1.724E-07 | 493.26929 | 421.5965 |
| QRPDDPMASFLGC | 13 | O95391 | Pre-mRNA-splicing factor SLU7 OS=Ho | SLU7 | 68.387 | 2 | 52.832832 | yes | 9.722E-08 | 731.3432 | 395.87854 |
| IREDIAK | 7 | O95391 | Pre-mRNA-splicing factor SLU7 OS=Ho | SLU7 | 68.387 | 2 | 52.832832 | yes | 2.601E-06 | 422.74799 | 331.09363 |
| NPDEVSYAGDNFV | 14 | O95391 | Pre-mRNA-splicing factor SLU7 OS=Ho | SLU7 | 68.387 | 2 | 52.832832 | yes | 3.187E-07 | 791.86023 | 308.40799 |
| SATVVDAVNAAPI | 17 | O95391 | Pre-mRNA-splicing factor SLU7 OS=Ho | SLU7 | 68.387 | 2 | 52.832832 | yes | 4.017E-07 | 814.92816 | 273.49557 |
| QFSSSGEWYK | 10 | O95391 | Pre-mRNA-splicing factor SLU7 OS=Ho | SLU7 | 68.387 | 2 | 52.832832 | yes | 9.734E-08 | 609.7749 | 208.73795 |
| ENPYANAGK | 9 | O95391 | Pre-mRNA-splicing factor SLU7 OS=Ho | SLU7 | 68.387 | 2 | 52.832832 | yes | 1.095E-06 | 482.23016 | 166.39166 |
| EMSLEEPKK | 9 | O95391 | Pre-mRNA-splicing factor SLU7 OS=Ho | SLU7 | 68.387 | 2 | 52.832832 | yes | 1.735E-07 | 545.77606 | 147.05075 |
| LQEELASGK | 9 | O95391 | Pre-mRNA-splicing factor SLU7 OS=Ho | SLU7 | 68.387 | 2 | 52.832832 | yes | 4.194E-07 | 487.76129 | 146.0808 |
| YSYCTGEAGK | 10 | O95391 | Pre-mRNA-splicing factor SLU7 OS=Ho | SLU7 | 68.387 | 2 | 52.832832 | yes | 9.722E-08 | 568.23987 | 140.01956 |
| CCHSFFK | 7 | O95391 | Pre-mRNA-splicing factor SLU7 OS=Ho | SLU7 | 68.387 | 2 | 52.832832 | yes | 1.996E-07 | 493.20456 | 130.51428 |
| ALNAEEAR | 8 | O95391 | Pre-mRNA-splicing factor SLU7 OS=Ho | SLU7 | 68.387 | 2 | 52.832832 | yes | 1.735E-07 | 437.22488 | 124.72897 |
| IVEEYAK | 7 | O95391 | Pre-mRNA-splicing factor SLU7 OS=Ho | SLU7 | 68.387 | 2 | 52.832832 | yes | 6.141E-07 | 426.2291 | 107.54018 |
| NLDPNSAYYDPK | 12 | O95391 | Pre-mRNA-splicing factor SLU7 OS=Ho | SLU7 | 68.387 | 2 | 52.832832 | yes | 9.722E-08 | 698.82263 | 79.82431 |
| WNGYNPEEHMK | 11 | O95391 | Pre-mRNA-splicing factor SLU7 OS=Ho | SLU7 | 68.387 | 2 | 52.832832 | yes | 9.722E-08 | 702.80371 | 59.719902 |
| KPQTLMELHQEK | 12 | O95391 | Pre-mRNA-splicing factor SLU7 OS=Ho | SLU7 | 68.387 | 2 | 52.832832 | yes | 6.134E-07 | 741.3927 | 55.050507 |
| KELEEQR | 7 | O95391 | Pre-mRNA-splicing factor SLU7 OS=Ho | SLU7 | 68.387 | 2 | 52.832832 | yes | 3.836E-05 | 466.24582 | 24.066448 |
| LVDPVAGSGGPGS | 14 | O95400 | CD2 antigen cytoplasmic tail-binding pro | CD2BP2 | 37.647 | 2 | 50.95548 | yes | 9.722E-08 | 634.83331 | 148.80792 |
| GDGLVDVMWEYK | 12 | O95400 | CD2 antigen cytoplasmic tail-binding pro | CD2BP2 | 37.647 | 2 | 50.95548 | yes | 2.195E-07 | 706.33173 | 105.83167 |
| LSGLADQMVAR | 11 | O95400 | CD2 antigen cytoplasmic tail-binding pro | CD2BP2 | 37.647 | 2 | 50.95548 | yes | 5.078E-07 | 580.80823 | 104.40083 |
| GNLGVYQETR | 10 | O95400 | CD2 antigen cytoplasmic tail-binding pro | CD2BP2 | 37.647 | 2 | 50.95548 | yes | 1.791E-07 | 568.78839 | 82.818558 |

| | | | | | | | | | | | |
|---|---|---|---|---|---|---|---|---|---|---|---|
| RIDFDLYT | 8 | O95400 | CD2 antigen cytoplasmic tail-binding pro | CD2BP2 | 37.647 | 2 | 50.95548 | yes | 6.485E-07 | 521.76385 | 75.568733 |
| CSDCLVPNEVR | 11 | O95405 | Zinc finger FYVE domain-containing pro | ZFYVE9 | 156.403 | 2 | 45.3673 | yes | 4.997E-06 | 674.80286 | 61.126072 |
| VFCASCCSLK | 10 | O95405 | Zinc finger FYVE domain-containing pro | ZFYVE9 | 156.403 | 2 | 45.3673 | yes | 2.816E-06 | 616.26703 | 55.774323 |
| SECYSNIYEQR | 11 | O95405 | Zinc finger FYVE domain-containing pro | ZFYVE9 | 156.403 | 2 | 45.3673 | yes | 1.871E-07 | 724.80914 | 50.027176 |
| LNEMNDSQVNEEI | 13 | O95405 | Zinc finger FYVE domain-containing pro | ZFYVE9 | 156.403 | 2 | 45.3673 | yes | 1.922E-07 | 775.34357 | 13.08955 |
| LGLPGEVFGR | 10 | O95409 | Zinc finger protein ZIC 2 OS=Homo sapi | ZIC2 | 55.006 | 2 | 50.403362 | yes | 4.392E-07 | 522.79547 | 580.70697 |
| WIDPEQLSNPK | 11 | O95409 | Zinc finger protein ZIC 2 OS=Homo sapi | ZIC2 | 55.006 | 2 | 50.403362 | yes | 9.809E-08 | 663.83807 | 362.51385 |
| SYTHPSSLR | 9 | O95409 | Zinc finger protein ZIC 2 OS=Homo sapi | ZIC2 | 55.006 | 2 | 50.403362 | no | 2.898E-07 | 524.26453 | 231.40831 |
| DASSLYPGTETMG | 15 | O95425 | Supervillin OS=Homo sapiens OX=9606 | SVIL | 247.747 | 2 | 37.288868 | yes | 6.125E-07 | 799.37994 | 59.689953 |
| GTSALMTVELDEE | 14 | O95425 | Supervillin OS=Homo sapiens OX=9606 | SVIL | 247.747 | 2 | 37.288868 | yes | 1.971E-06 | 775.87195 | 38.4394 |
| SNEEEETSDSSLEK | 14 | O95425 | Supervillin OS=Homo sapiens OX=9606 | SVIL | 247.747 | 2 | 37.288868 | yes | 7.203E-07 | 792.33136 | 12.046308 |
| SGYGPSDGPSYGR | 13 | O95429 | BAG family molecular chaperone regulat | BAG4 | 49.595 | 2 | 51.880955 | yes | 9.722E-08 | 650.28363 | 586.94836 |
| SSGNSPTPVSR | 11 | O95429 | BAG family molecular chaperone regulat | BAG4 | 49.595 | 2 | 51.880955 | yes | 1.26E-07 | 544.7702 | 459.15228 |
| IIHVLEK | 7 | O95429 | BAG family molecular chaperone regulat | BAG4 | 49.595 | 2 | 51.880955 | yes | 2.292E-07 | 426.27109 | 356.80127 |
| VQYLEQEVEEFVC | 14 | O95429 | BAG family molecular chaperone regulat | BAG4 | 49.595 | 2 | 51.880955 | yes | 1.372E-07 | 848.92505 | 234.83537 |
| IQAILEK | 7 | O95429 | BAG family molecular chaperone regulat | BAG4 | 49.595 | 2 | 51.880955 | yes | 2.989E-07 | 407.75528 | 213.16273 |
| SVPQSGPTVR | 10 | O95429 | BAG family molecular chaperone regulat | BAG4 | 49.595 | 2 | 51.880955 | yes | 2.709E-06 | 514.28015 | 103.98735 |
| KIIHVLEK | 8 | O95429 | BAG family molecular chaperone regulat | BAG4 | 49.595 | 2 | 51.880955 | yes | 2.585E-07 | 490.31857 | 68.926208 |
| EAMGIYISTLK | 11 | O95433 | Activator of 90 kDa heat shock protein A | AHSA1 | 38.275 | 2 | 48.523067 | yes | 1.136E-07 | 613.32849 | 70.105728 |
| DEPDTNLVALMK | 12 | O95433 | Activator of 90 kDa heat shock protein A | AHSA1 | 38.275 | 2 | 48.523067 | yes | 2.722E-07 | 673.33704 | 68.59333 |
| ETFLTSPEELYR | 12 | O95433 | Activator of 90 kDa heat shock protein A | AHSA1 | 38.275 | 2 | 48.523067 | yes | 2.332E-07 | 742.867 | 67.388054 |
| IQTYAEYMGR | 10 | O95453 | Poly(A)-specific ribonuclease PARN OS= | PARN | 73.451 | 2 | 52.872498 | yes | 9.722E-08 | 616.29242 | 433.35696 |
| IEDLLQSEENK | 11 | O95453 | Poly(A)-specific ribonuclease PARN OS= | PARN | 73.451 | 2 | 52.872498 | yes | 9.722E-08 | 659.33026 | 416.92542 |
| EMTTCVFPR | 9 | O95453 | Poly(A)-specific ribonuclease PARN OS= | PARN | 73.451 | 2 | 52.872498 | yes | 1.047E-07 | 570.76245 | 375.61621 |
| EQEELNDAVGFSR | 13 | O95453 | Poly(A)-specific ribonuclease PARN OS= | PARN | 73.451 | 2 | 52.872498 | yes | 9.722E-08 | 747.34698 | 318.48969 |
| NNSFTAPSTVGK | 12 | O95453 | Poly(A)-specific ribonuclease PARN OS= | PARN | 73.451 | 2 | 52.872498 | yes | 9.722E-08 | 611.80676 | 277.99042 |
| LIEPFFNK | 8 | O95453 | Poly(A)-specific ribonuclease PARN OS= | PARN | 73.451 | 2 | 52.872498 | yes | 2.565E-07 | 504.28165 | 251.43196 |
| NLDLEPCTGFQR | 12 | O95453 | Poly(A)-specific ribonuclease PARN OS= | PARN | 73.451 | 2 | 52.872498 | yes | 4.969E-07 | 725.3432 | 241.66838 |
| DIINNTSLAELEK | 13 | O95453 | Poly(A)-specific ribonuclease PARN OS= | PARN | 73.451 | 2 | 52.872498 | yes | 9.9E-08 | 730.38556 | 227.75073 |
| GIHVETLETEK | 11 | O95453 | Poly(A)-specific ribonuclease PARN OS= | PARN | 73.451 | 2 | 52.872498 | yes | 9.722E-08 | 628.33008 | 179.74811 |
| VIHAIANSGK | 10 | O95453 | Poly(A)-specific ribonuclease PARN OS= | PARN | 73.451 | 2 | 52.872498 | yes | 2.103E-07 | 505.29309 | 106.37582 |
| KLIYQTLSWK | 10 | O95453 | Poly(A)-specific ribonuclease PARN OS= | PARN | 73.451 | 2 | 52.872498 | yes | 2.838E-07 | 640.37408 | 88.593475 |
| LIYQTLSWK | 9 | O95453 | Poly(A)-specific ribonuclease PARN OS= | PARN | 73.451 | 2 | 52.872498 | yes | 5.189E-07 | 576.3266 | 61.097546 |
| LMASTQPFK | 9 | O95453 | Poly(A)-specific ribonuclease PARN OS= | PARN | 73.451 | 2 | 52.872498 | yes | 1.009E-07 | 511.7706 | 58.188416 |
| RDHVLHVTFPK | 11 | O95453 | Poly(A)-specific ribonuclease PARN OS= | PARN | 73.451 | 3 | 52.872498 | yes | 5.611E-07 | 450.25433 | 35.915306 |
| KIEAEIVR | 8 | O95470 | Sphingosine-1-phosphate lyase 1 OS=Ho | SGPL1 | 63.524 | 2 | 51.743683 | yes | 4.189E-07 | 479.29001 | 462.33347 |
| ALPSQGLSSSAVLI | 15 | O95470 | Sphingosine-1-phosphate lyase 1 OS=Ho | SGPL1 | 63.524 | 2 | 51.743683 | yes | 1.512E-07 | 743.90924 | 293.70996 |
| GVTSISADTHK | 11 | O95470 | Sphingosine-1-phosphate lyase 1 OS=Ho | SGPL1 | 63.524 | 2 | 51.743683 | yes | 1.356E-07 | 558.28821 | 256.285 |
| EYSSMDAFWQEGI | 13 | O95470 | Sphingosine-1-phosphate lyase 1 OS=Ho | SGPL1 | 63.524 | 2 | 51.743683 | yes | 9.722E-08 | 803.33557 | 232.89383 |

| | | | | | | | | | | | |
|---|---|---|---|---|---|---|---|---|---|---|---|
| GSSLVLYSDK | 10 | O95470 | Sphingosine-1-phosphate lyase 1 OS=Hoi | SGPL1 | 63.524 | 2 | 51.743683 | yes | 1.622E-06 | 534.78223 | 182.74388 |
| AGYPLEHPFDFR | 12 | O95470 | Sphingosine-1-phosphate lyase 1 OS=Hoi | SGPL1 | 63.524 | 2 | 51.743683 | yes | 4.054E-07 | 724.8515 | 111.39371 |
| ESVTQIMK | 8 | O95470 | Sphingosine-1-phosphate lyase 1 OS=Hoi | SGPL1 | 63.524 | 2 | 51.743683 | yes | 6.092E-07 | 468.24695 | 85.568367 |
| ASGTVYSGEEK | 11 | O95470 | Sphingosine-1-phosphate lyase 1 OS=Hoi | SGPL1 | 63.524 | 2 | 51.743683 | yes | 1.512E-07 | 564.2644 | 60.710743 |
| PSTDLLMLK | 9 | O95470 | Sphingosine-1-phosphate lyase 1 OS=Hoi | SGPL1 | 63.524 | 2 | 51.743683 | yes | 6.393E-07 | 509.28607 | 51.913326 |
| VDNLSDEGALNISI | 15 | O95486 | Protein transport protein Sec24A OS=Hoi | SEC24A | 119.75 | 2 | 35.127335 | yes | 9.393E-07 | 809.38934 | 42.776958 |
| DIHMTPSTDFYK | 12 | O95486 | Protein transport protein Sec24A OS=Hoi | SEC24A | 119.75 | 2 | 35.127335 | yes | 8.818E-07 | 727.83466 | 23.645494 |
| AEFFQHLIEDR | 11 | O95487 | Protein transport protein Sec24B OS=Hoi | SEC24B | 137.418 | 2 | 52.56922 | yes | 3.301E-07 | 702.84894 | 87.118523 |
| MIHPNLYR | 8 | O95487 | Protein transport protein Sec24B OS=Hoi | SEC24B | 137.418 | 2 | 52.56922 | yes | 9.722E-08 | 522.27637 | 86.477478 |
| SQPLVHLMK | 9 | O95487 | Protein transport protein Sec24B OS=Hoi | SEC24B | 137.418 | 2 | 52.56922 | yes | 1.554E-07 | 526.79968 | 66.994316 |
| LTDEGAVHVNDR | 12 | O95487 | Protein transport protein Sec24B OS=Hoi | SEC24B | 137.418 | 2 | 52.56922 | yes | 9.722E-08 | 663.32587 | 65.4189 |
| DLTQLPVITSNTIV | 15 | O95487 | Protein transport protein Sec24B OS=Hoi | SEC24B | 137.418 | 2 | 52.56922 | yes | 1.275E-07 | 835.47778 | 58.756149 |
| SAPAGSSHPAASA | 14 | O95487 | Protein transport protein Sec24B OS=Hoi | SEC24B | 137.418 | 2 | 52.56922 | yes | 2.208E-06 | 654.81818 | 24.039574 |
| DLCCHLSYK | 9 | O95497 | Pantetheinase OS=Homo sapiens OX=96( | VNN1 | 57.012 | 2 | 41.345356 | yes | 3.556E-07 | 598.26538 | 266.98541 |
| LLPETVSPSAPSWI | 14 | O95503 | Chromobox protein homolog 6 OS=Homo | CBX6 | 43.899 | 2 | 47.005516 | yes | 1.422E-07 | 770.41193 | 107.99857 |
| MELSAVGER | 9 | O95503 | Chromobox protein homolog 6 OS=Homo | CBX6 | 43.899 | 2 | 47.005516 | no | 1.738E-07 | 517.25275 | 95.369339 |
| EFCNPEDFEK | 10 | O95503 | Chromobox protein homolog 6 OS=Homo | CBX6 | 43.899 | 2 | 47.005516 | yes | 2.238E-07 | 657.76898 | 93.652718 |
| ELSAVGER | 8 | O95503 | Chromobox protein homolog 6 OS=Homo | CBX6 | 43.899 | 2 | 47.005516 | no | 3.325E-06 | 430.72726 | 57.889378 |
| GAGGGGAGQGAG | 16 | O95503 | Chromobox protein homolog 6 OS=Homo | CBX6 | 43.899 | 2 | 47.005516 | yes | 1.544E-07 | 614.31305 | 25.301762 |
| IMLSPEAR | 8 | O95544 | NAD kinase OS=Homo sapiens OX=960( | NADK | 49.229 | 2 | 49.75953 | yes | 4.976E-07 | 458.74966 | 180.19617 |
| SLSASPALGSTK | 12 | O95544 | NAD kinase OS=Homo sapiens OX=960( | NADK | 49.229 | 2 | 49.75953 | yes | 1.095E-07 | 559.80621 | 176.21571 |
| NTAWVSFDGR | 10 | O95544 | NAD kinase OS=Homo sapiens OX=960( | NADK | 49.229 | 2 | 49.75953 | yes | 2.396E-07 | 576.77527 | 142.83685 |
| PIVVPAGVELK | 11 | O95544 | NAD kinase OS=Homo sapiens OX=960( | NADK | 49.229 | 2 | 49.75953 | yes | 1.004E-07 | 561.3501 | 95.897415 |
| SLHGPCPVTTFGPI | 14 | O95544 | NAD kinase OS=Homo sapiens OX=960( | NADK | 49.229 | 3 | 49.75953 | yes | 1.475E-06 | 499.92215 | 69.411087 |
| ELCTHLMEENMIV | 17 | O95544 | NAD kinase OS=Homo sapiens OX=960( | NADK | 49.229 | 3 | 49.75953 | yes | 4.447E-06 | 713.3371 | 48.948818 |
| PETISEPK | 8 | O95551 | Tyrosyl-DNA phosphodiesterase 2 OS=H | TDP2 | 40.93 | 2 | 35.512608 | yes | 9.076E-07 | 450.73727 | 143.34508 |
| SQEIIPFPSTK | 11 | O95551 | Tyrosyl-DNA phosphodiesterase 2 OS=H | TDP2 | 40.93 | 2 | 35.512608 | yes | 1.96E-06 | 623.83752 | 80.355721 |
| SLDLLGLEK | 9 | O95551 | Tyrosyl-DNA phosphodiesterase 2 OS=H | TDP2 | 40.93 | 2 | 35.512608 | yes | 8.211E-07 | 494.28967 | 20.78299 |
| LRPLYNHPAGPR | 12 | O95563 | Mitochondrial pyruvate carrier 2 OS=Hor | MPC2 | 14.279 | 3 | 46.059483 | yes | 1.664E-07 | 464.2616 | 51.730103 |
| EAVLIDPVLETAPI | 14 | O95571 | Persulfide dioxygenase ETHE1, mitochor | ETHE1 | 27.873 | 2 | 51.566845 | yes | 9.81E-08 | 761.92743 | 262.11676 |
| LTLSCEEFVK | 10 | O95571 | Persulfide dioxygenase ETHE1, mitochor | ETHE1 | 27.873 | 2 | 51.566845 | yes | 1.937E-07 | 613.3103 | 118.14698 |
| SLLPGCQSVISR | 12 | O95571 | Persulfide dioxygenase ETHE1, mitochor | ETHE1 | 27.873 | 2 | 51.566845 | yes | 9.722E-08 | 658.85321 | 103.50906 |
| IMGNLNLPK | 9 | O95571 | Persulfide dioxygenase ETHE1, mitochor | ETHE1 | 27.873 | 2 | 51.566845 | yes | 1.4E-07 | 500.28641 | 64.365028 |
| NTPLCDSFVFR | 11 | O95573 | Fatty acid CoA ligase Acsl3 OS=Homo sa | ACSL3 | 80.421 | 2 | 52.73989 | yes | 1.809E-07 | 678.32428 | 426.93555 |
| IGYSSPQTLADQSS | 15 | O95573 | Fatty acid CoA ligase Acsl3 OS=Homo sa | ACSL3 | 80.421 | 2 | 52.73989 | yes | 9.722E-08 | 791.39136 | 213.61137 |
| LQAGEYVSLGK | 11 | O95573 | Fatty acid CoA ligase Acsl3 OS=Homo sa | ACSL3 | 80.421 | 2 | 52.73989 | no | 1.215E-07 | 582.81659 | 203.10522 |
| VLSEAAISASLEK | 13 | O95573 | Fatty acid CoA ligase Acsl3 OS=Homo sa | ACSL3 | 80.421 | 2 | 52.73989 | yes | 9.722E-08 | 659.36664 | 197.2695 |
| LKDIVSLVPR | 10 | O95573 | Fatty acid CoA ligase Acsl3 OS=Homo sa | ACSL3 | 80.421 | 2 | 52.73989 | yes | 9.722E-08 | 570.36096 | 173.9825 |
| THYQADIER | 9 | O95573 | Fatty acid CoA ligase Acsl3 OS=Homo sa | ACSL3 | 80.421 | 2 | 52.73989 | yes | 1.636E-07 | 566.77271 | 171.55629 |

| | | | | | | | | | | | |
|---|---|---|---|---|---|---|---|---|---|---|---|
| VGAPLVCCEIK | 11 | O95573 | Fatty acid CoA ligase Acsl3 OS=Homo sa | ACSL3 | 80.421 | 2 | 52.73989 | yes | 5.713E-06 | 623.32013 | 154.35265 |
| LLLCGGAPLSATT( | 15 | O95573 | Fatty acid CoA ligase Acsl3 OS=Homo sa | ACSL3 | 80.421 | 2 | 52.73989 | yes | 9.722E-08 | 779.4245 | 129.10248 |
| SLLGGNIR | 8 | O95573 | Fatty acid CoA ligase Acsl3 OS=Homo sa | ACSL3 | 80.421 | 2 | 52.73989 | yes | 3.519E-07 | 415.24814 | 110.34782 |
| GVMISHSNIIAGIT( | 19 | O95573 | Fatty acid CoA ligase Acsl3 OS=Homo sa | ACSL3 | 80.421 | 3 | 52.73989 | yes | 2.249E-07 | 653.00464 | 87.822678 |
| GIIVHTMAAVEAL | 16 | O95573 | Fatty acid CoA ligase Acsl3 OS=Homo sa | ACSL3 | 80.421 | 2 | 52.73989 | yes | 9.722E-08 | 790.94507 | 79.649498 |
| ADFFEDENGQR | 11 | O95573 | Fatty acid CoA ligase Acsl3 OS=Homo sa | ACSL3 | 80.421 | 2 | 52.73989 | yes | 1.854E-07 | 664.28113 | 73.486923 |
| HIITVDGKPPTWSI | 17 | O95573 | Fatty acid CoA ligase Acsl3 OS=Homo sa | ACSL3 | 80.421 | 3 | 52.73989 | yes | 9.722E-08 | 651.34741 | 46.479706 |
| AALNLPEAASYDE | 15 | O95602 | DNA-directed RNA polymerase I subunit | POLR1A | 194.812 | 2 | 50.191776 | yes | 2.036E-07 | 809.90723 | 175.35896 |
| TALSAVDMTQR | 11 | O95602 | DNA-directed RNA polymerase I subunit | POLR1A | 194.812 | 2 | 50.191776 | yes | 9.851E-08 | 596.80316 | 110.51212 |
| DSEPLGIEEAQIGK | 14 | O95602 | DNA-directed RNA polymerase I subunit | POLR1A | 194.812 | 2 | 50.191776 | yes | 2.735E-07 | 743.37518 | 87.804886 |
| SLPCFEPYEFTPR | 13 | O95602 | DNA-directed RNA polymerase I subunit | POLR1A | 194.812 | 2 | 50.191776 | yes | 4.814E-07 | 821.88214 | 86.463692 |
| TPMMSVPVLNTK | 12 | O95602 | DNA-directed RNA polymerase I subunit | POLR1A | 194.812 | 2 | 50.191776 | yes | 2.512E-07 | 659.34888 | 53.227562 |
| VDDYSQEWAAQT | 14 | O95602 | DNA-directed RNA polymerase I subunit | POLR1A | 194.812 | 2 | 50.191776 | yes | 3.209E-07 | 835.37067 | 19.928871 |
| LLTEQLSQR | 9 | O95613 | Pericentrin OS=Homo sapiens OX=9606 | PCNT | 378.038 | 2 | 53.577557 | yes | 2.982E-07 | 544.30896 | 168.39552 |
| AAGSDADHLR | 10 | O95613 | Pericentrin OS=Homo sapiens OX=9606 | PCNT | 378.038 | 2 | 53.577557 | yes | 2.364E-06 | 506.74396 | 152.60788 |
| LEEMNINIR | 9 | O95613 | Pericentrin OS=Homo sapiens OX=9606 | PCNT | 378.038 | 2 | 53.577557 | yes | 3.748E-06 | 566.29498 | 115.65523 |
| AQELALLQSR | 10 | O95613 | Pericentrin OS=Homo sapiens OX=9606 | PCNT | 378.038 | 2 | 53.577557 | yes | 1.658E-07 | 564.8222 | 112.47026 |
| LDMQNSQTAVSLI | 13 | O95613 | Pericentrin OS=Homo sapiens OX=9606 | PCNT | 378.038 | 2 | 53.577557 | yes | 9.722E-08 | 731.86957 | 107.44204 |
| AELALELHK | 9 | O95613 | Pericentrin OS=Homo sapiens OX=9606 | PCNT | 378.038 | 2 | 53.577557 | yes | 7.362E-07 | 512.29529 | 104.57436 |
| ATIENLQENQK | 11 | O95613 | Pericentrin OS=Homo sapiens OX=9606 | PCNT | 378.038 | 2 | 53.577557 | yes | 1.533E-07 | 644.33063 | 104.07582 |
| VQLSLLQTELK | 11 | O95613 | Pericentrin OS=Homo sapiens OX=9606 | PCNT | 378.038 | 2 | 53.577557 | yes | 1.065E-07 | 636.38208 | 100.57885 |
| EKEEVVLR | 8 | O95613 | Pericentrin OS=Homo sapiens OX=9606 | PCNT | 378.038 | 2 | 53.577557 | yes | 2.296E-07 | 501.28494 | 99.879463 |
| LLAAEQTVVR | 10 | O95613 | Pericentrin OS=Homo sapiens OX=9606 | PCNT | 378.038 | 2 | 53.577557 | yes | 3.452E-07 | 550.32715 | 98.897041 |
| SLCEVQQEVLQLR | 13 | O95613 | Pericentrin OS=Homo sapiens OX=9606 | PCNT | 378.038 | 2 | 53.577557 | yes | 3.639E-07 | 801.41943 | 93.239021 |
| ESFLMSPESVR | 11 | O95613 | Pericentrin OS=Homo sapiens OX=9606 | PCNT | 378.038 | 2 | 53.577557 | yes | 1.115E-07 | 641.31079 | 89.220848 |
| ALQSQLEEEQLR | 12 | O95613 | Pericentrin OS=Homo sapiens OX=9606 | PCNT | 378.038 | 2 | 53.577557 | yes | 1.464E-07 | 722.37555 | 87.382187 |
| NEEILHLNLK | 10 | O95613 | Pericentrin OS=Homo sapiens OX=9606 | PCNT | 378.038 | 2 | 53.577557 | yes | 9.722E-08 | 611.84314 | 82.971428 |
| EGANLLSMLK | 10 | O95613 | Pericentrin OS=Homo sapiens OX=9606 | PCNT | 378.038 | 2 | 53.577557 | yes | 6.44E-07 | 538.29443 | 77.743515 |
| NLEIDALNQR | 10 | O95613 | Pericentrin OS=Homo sapiens OX=9606 | PCNT | 378.038 | 2 | 53.577557 | yes | 9.722E-08 | 593.31476 | 75.313034 |
| ELEEENTSLK | 10 | O95613 | Pericentrin OS=Homo sapiens OX=9606 | PCNT | 378.038 | 2 | 53.577557 | yes | 6.515E-07 | 596.29059 | 72.677361 |
| SSLLSEIQALR | 11 | O95613 | Pericentrin OS=Homo sapiens OX=9606 | PCNT | 378.038 | 2 | 53.577557 | yes | 2.169E-06 | 608.84845 | 70.253792 |
| LGGVLPDSTSK | 11 | O95613 | Pericentrin OS=Homo sapiens OX=9606 | PCNT | 378.038 | 2 | 53.577557 | yes | 3.784E-06 | 537.29547 | 60.782162 |
| CEALLAQER | 9 | O95613 | Pericentrin OS=Homo sapiens OX=9606 | PCNT | 378.038 | 2 | 53.577557 | yes | 2.571E-05 | 545.27148 | 59.110626 |
| LVLELESLR | 9 | O95613 | Pericentrin OS=Homo sapiens OX=9606 | PCNT | 378.038 | 2 | 53.577557 | yes | 1.755E-07 | 536.32404 | 58.670208 |
| IQEFEAALK | 9 | O95613 | Pericentrin OS=Homo sapiens OX=9606 | PCNT | 378.038 | 2 | 53.577557 | yes | 3.73E-06 | 524.78729 | 58.342941 |
| QELSELHEQLLAR | 13 | O95613 | Pericentrin OS=Homo sapiens OX=9606 | PCNT | 378.038 | 3 | 53.577557 | yes | 3.194E-07 | 522.61426 | 53.499695 |
| EQEMQGVELQPR | 12 | O95613 | Pericentrin OS=Homo sapiens OX=9606 | PCNT | 378.038 | 2 | 53.577557 | yes | 1.418E-07 | 722.34845 | 53.276085 |
| TQHESELEQLR | 11 | O95613 | Pericentrin OS=Homo sapiens OX=9606 | PCNT | 378.038 | 2 | 53.577557 | yes | 9.722E-08 | 685.33893 | 52.729828 |
| QLQQAAQEQAALI | 13 | O95613 | Pericentrin OS=Homo sapiens OX=9606 | PCNT | 378.038 | 2 | 53.577557 | yes | 9.722E-08 | 727.88916 | 52.539883 |

| | | | | | | | | | | | |
|---|---|---|---|---|---|---|---|---|---|---|---|
| TVNDWTSSNEK | 11 | O95613 | Pericentrin OS=Homo sapiens OX=9606 | PCNT | 378.038 | 2 | 53.577557 | yes | 7.154E-07 | 640.79132 | 52.165241 |
| LQSEMEKNAQIVK | 13 | O95613 | Pericentrin OS=Homo sapiens OX=9606 | PCNT | 378.038 | 2 | 53.577557 | yes | 3.328E-07 | 767.4007 | 52.123695 |
| EDSALCGGGDICK | 13 | O95613 | Pericentrin OS=Homo sapiens OX=9606 | PCNT | 378.038 | 2 | 53.577557 | yes | 1.82E-07 | 691.28973 | 50.437183 |
| NLESHHQAAIEK | 12 | O95613 | Pericentrin OS=Homo sapiens OX=9606 | PCNT | 378.038 | 3 | 53.577557 | yes | 8.707E-07 | 459.56876 | 44.364182 |
| EDCALQLMLAR | 11 | O95613 | Pericentrin OS=Homo sapiens OX=9606 | PCNT | 378.038 | 2 | 53.577557 | yes | 9.722E-08 | 660.32593 | 42.401733 |
| LSGSDLGGHSSLLI | 15 | O95613 | Pericentrin OS=Homo sapiens OX=9606 | PCNT | 378.038 | 2 | 53.577557 | yes | 5.183E-06 | 764.39172 | 42.323318 |
| ADVNLSHSER | 10 | O95613 | Pericentrin OS=Homo sapiens OX=9606 | PCNT | 378.038 | 2 | 53.577557 | yes | 3.593E-06 | 564.27563 | 38.72551 |
| EEFQQEIQRLEGQI | 15 | O95613 | Pericentrin OS=Homo sapiens OX=9606 | PCNT | 378.038 | 3 | 53.577557 | yes | 3.183E-05 | 634.99377 | 38.426853 |
| EKETALTELR | 10 | O95613 | Pericentrin OS=Homo sapiens OX=9606 | PCNT | 378.038 | 2 | 53.577557 | yes | 4.109E-07 | 595.32477 | 36.685387 |
| LSLSNMHTAQLEL | 20 | O95613 | Pericentrin OS=Homo sapiens OX=9606 | PCNT | 378.038 | 3 | 53.577557 | yes | 6.635E-07 | 747.39496 | 33.912048 |
| VLEDCQLPK | 9 | O95613 | Pericentrin OS=Homo sapiens OX=9606 | PCNT | 378.038 | 2 | 53.577557 | yes | 1.144E-07 | 551.28406 | 32.794128 |
| QVLMQEEEIK | 10 | O95613 | Pericentrin OS=Homo sapiens OX=9606 | PCNT | 378.038 | 2 | 53.577557 | yes | 3.395E-07 | 623.82104 | 26.898756 |
| VDLVAQVK | 8 | O95613 | Pericentrin OS=Homo sapiens OX=9606 | PCNT | 378.038 | 2 | 53.577557 | yes | 3.172E-07 | 436.26602 | 26.71652 |
| SLTASQDPEHSLTI | 25 | O95613 | Pericentrin OS=Homo sapiens OX=9606 | PCNT | 378.038 | 4 | 53.577557 | yes | 5.991E-07 | 733.62036 | 26.409151 |
| EKEVSATLK | 9 | O95613 | Pericentrin OS=Homo sapiens OX=9606 | PCNT | 378.038 | 2 | 53.577557 | yes | 4.338E-07 | 502.78476 | 21.405201 |
| FSAEQDAFLQEAQ | 18 | O95613 | Pericentrin OS=Homo sapiens OX=9606 | PCNT | 378.038 | 3 | 53.577557 | yes | 2.261E-06 | 702.32764 | 18.339113 |
| GSAVDASVQEESP | 16 | O95613 | Pericentrin OS=Homo sapiens OX=9606 | PCNT | 378.038 | 2 | 53.577557 | yes | 3.939E-07 | 802.3941 | 17.629271 |
| LHESYGNPLR | 10 | O95619 | YEATS domain-containing protein 4 OS= | YEATS4 | 26.5 | 2 | 49.658016 | yes | 2.422E-07 | 593.3042 | 267.577 |
| LFQSDTNAMLGK | 12 | O95619 | YEATS domain-containing protein 4 OS= | YEATS4 | 26.5 | 2 | 49.658016 | yes | 1.013E-07 | 662.83191 | 158.6333 |
| QLTLGAYK | 8 | O95619 | YEATS domain-containing protein 4 OS= | YEATS4 | 26.5 | 2 | 49.658016 | yes | 3.462E-06 | 447.25818 | 102.78128 |
| GVTIVKPIVYGNV | 15 | O95619 | YEATS domain-containing protein 4 OS= | YEATS4 | 26.5 | 2 | 49.658016 | yes | 1.022E-07 | 793.47485 | 90.080757 |
| HETEFAELEVK | 11 | O95619 | YEATS domain-containing protein 4 OS= | YEATS4 | 26.5 | 2 | 49.658016 | yes | 2.754E-07 | 666.32751 | 68.062851 |
| NEFTITHVLIPK | 12 | O95630 | STAM-binding protein OS=Homo sapien | STAMBP | 48.077 | 2 | 37.5038 | yes | 5.934E-07 | 706.40082 | 37.786404 |
| GHPDLSVMYR | 10 | O95633 | Follistatin-related protein 3 OS=Homo sa | FSTL3 | 27.664 | 2 | 52.16846 | yes | 2.748E-07 | 587.78729 | 219.99681 |
| LQVCGSDGATYR | 12 | O95633 | Follistatin-related protein 3 OS=Homo sa | FSTL3 | 27.664 | 2 | 52.16846 | yes | 9.722E-08 | 663.80896 | 143.55391 |
| SCEHVVCPR | 9 | O95633 | Follistatin-related protein 3 OS=Homo sa | FSTL3 | 27.664 | 2 | 52.16846 | yes | 1.14E-07 | 572.25531 | 40.35202 |
| CECAPDCSGLPAR | 13 | O95633 | Follistatin-related protein 3 OS=Homo sa | FSTL3 | 27.664 | 2 | 52.16846 | yes | 9.722E-08 | 746.80249 | 36.720737 |
| PQSCVVDQTGSAI | 18 | O95633 | Follistatin-related protein 3 OS=Homo sa | FSTL3 | 27.664 | 3 | 52.16846 | yes | 3.375E-07 | 687.30841 | 29.293402 |
| PLEQVTCYK | 9 | O95639 | Cleavage and polyadenylation specificity | CPSF4 | 30.255 | 2 | 51.16785 | yes | 9.722E-08 | 569.28406 | 129.24205 |
| SGAAVCEFFLK | 11 | O95639 | Cleavage and polyadenylation specificity | CPSF4 | 30.255 | 2 | 51.16785 | yes | 3.105E-07 | 614.80518 | 128.339 |
| MPECYFYSK | 9 | O95639 | Cleavage and polyadenylation specificity | CPSF4 | 30.255 | 2 | 51.16785 | no | 1.69E-07 | 612.75684 | 111.1555 |
| KGDQCEFLHEYDI | 15 | O95639 | Cleavage and polyadenylation specificity | CPSF4 | 30.255 | 3 | 51.16785 | yes | 2.599E-07 | 634.27832 | 81.958817 |
| GLGACTLLGSPR | 12 | O95644 | Nuclear factor of activated T-cells, cytopl | NFATC1 | 101.243 | 2 | 44.35521 | yes | 2.225E-07 | 601.32153 | 94.431648 |
| ASVTEESWLGAR | 12 | O95644 | Nuclear factor of activated T-cells, cytopl | NFATC1 | 101.243 | 2 | 44.35521 | yes | 4.295E-06 | 653.32532 | 85.818108 |
| TTLEQPPSVALK | 12 | O95644 | Nuclear factor of activated T-cells, cytopl | NFATC1 | 101.243 | 2 | 44.35521 | yes | 1.825E-06 | 642.36389 | 50.071808 |
| SAQELPLVEK | 10 | O95644 | Nuclear factor of activated T-cells, cytopl | NFATC1 | 101.243 | 2 | 44.35521 | yes | 2.343E-06 | 557.31116 | 39.912045 |
| VLGSVSQFEEFGR | 13 | O95672 | Endothelin-converting enzyme-like 1 OS= | ECEL1 | 87.792 | 2 | 50.378517 | yes | 3.98E-07 | 727.86737 | 156.424 |
| LEELDWMDAETR | 12 | O95672 | Endothelin-converting enzyme-like 1 OS= | ECEL1 | 87.792 | 2 | 50.378517 | yes | 9.81E-08 | 754.34033 | 141.8443 |
| VLSLLGADAVEQK | 13 | O95672 | Endothelin-converting enzyme-like 1 OS= | ECEL1 | 87.792 | 2 | 50.378517 | yes | 6.689E-07 | 671.88245 | 128.16728 |

| | | | | | | | | | | | |
|---|---|---|---|---|---|---|---|---|---|---|---|
| IDQDGLTLPER | 11 | O95672 | Endothelin-converting enzyme-like 1 OS= | ECEL1 | 87.792 | 2 | 50.378517 | yes | 1.108E-07 | 628.8277 | 92.47995 |
| TLYLAQDEDSEK | 12 | O95672 | Endothelin-converting enzyme-like 1 OS= | ECEL1 | 87.792 | 2 | 50.378517 | yes | 6.266E-07 | 706.33301 | 63.772377 |
| VVVVLSEHLSPPFI | 14 | O95672 | Endothelin-converting enzyme-like 1 OS= | ECEL1 | 87.792 | 2 | 50.378517 | yes | 1.418E-07 | 789.95374 | 44.374821 |
| FSIQLSVK | 8 | O95672 | Endothelin-converting enzyme-like 1 OS= | ECEL1 | 87.792 | 2 | 50.378517 | yes | 3.419E-07 | 461.27383 | 25.424519 |
| EHGPEHPLPR | 10 | O95672 | Endothelin-converting enzyme-like 1 OS= | ECEL1 | 87.792 | 2 | 50.378517 | yes | 1.895E-06 | 584.79651 | 23.160065 |
| SQSIYLQVLTDK | 12 | O95672 | Endothelin-converting enzyme-like 1 OS= | ECEL1 | 87.792 | 2 | 50.378517 | yes | 5.08E-06 | 697.87994 | 23.067518 |
| NNVGGLLGPAK | 11 | O95677 | Eyes absent homolog 4 OS=Homo sapien | EYA4 | 69.506 | 2 | 49.742382 | no | 2.702E-06 | 520.2984 | 54.117931 |
| MEDSQDLNEQSVI | 13 | O95677 | Eyes absent homolog 4 OS=Homo sapien | EYA4 | 69.506 | 2 | 49.742382 | yes | 1.006E-07 | 782.84338 | 19.11068 |
| IGSFLSNR | 8 | O95684 | Centrosomal protein 43 OS=Homo sapier | CEP43 | 43.065 | 2 | 52.592503 | yes | 1.125E-07 | 447.24561 | 153.72292 |
| TPLVNESLK | 9 | O95684 | Centrosomal protein 43 OS=Homo sapier | CEP43 | 43.065 | 2 | 52.592503 | yes | 1.467E-07 | 500.78729 | 124.28567 |
| AATAAAVVAEED' | 16 | O95684 | Centrosomal protein 43 OS=Homo sapier | CEP43 | 43.065 | 2 | 52.592503 | yes | 1.874E-07 | 829.91522 | 116.52726 |
| SGLSSLAGAPSLK | 13 | O95684 | Centrosomal protein 43 OS=Homo sapier | CEP43 | 43.065 | 2 | 52.592503 | yes | 1.011E-07 | 594.33514 | 114.57818 |
| AAVFLALEEQEK | 12 | O95684 | Centrosomal protein 43 OS=Homo sapier | CEP43 | 43.065 | 2 | 52.592503 | yes | 2.312E-07 | 674.36139 | 114.26348 |
| DLLVQTLENSGVL | 15 | O95684 | Centrosomal protein 43 OS=Homo sapier | CEP43 | 43.065 | 2 | 52.592503 | yes | 9.722E-08 | 835.95721 | 51.4412 |
| AVVLEYFTR | 9 | O95707 | Ribonuclease P protein subunit p29 OS=I | POP4 | 25.425 | 2 | 50.81947 | yes | 1.113E-07 | 549.3031 | 739.19025 |
| LFDIKPEQQR | 10 | O95707 | Ribonuclease P protein subunit p29 OS=I | POP4 | 25.425 | 2 | 50.81947 | yes | 2.871E-07 | 637.34857 | 541.29358 |
| SVIYHALSQK | 10 | O95707 | Ribonuclease P protein subunit p29 OS=I | POP4 | 25.425 | 2 | 50.81947 | yes | 9.722E-08 | 573.31934 | 324.5723 |
| ADLHGAIISVTK | 12 | O95707 | Ribonuclease P protein subunit p29 OS=I | POP4 | 25.425 | 2 | 50.81947 | yes | 9.809E-08 | 612.85095 | 231.2934 |
| ILLSLDGNLAK | 11 | O95714 | E3 ubiquitin-protein ligase HERC2 OS=I | HERC2 | 527.228 | 2 | 55.270603 | yes | 1.002E-07 | 578.85046 | 147.74242 |
| FTVYPIMPAAGPK | 13 | O95714 | E3 ubiquitin-protein ligase HERC2 OS=I | HERC2 | 527.228 | 2 | 55.270603 | yes | 3.67E-07 | 696.37323 | 105.91637 |
| SAQGTSAPLLPLL( | 15 | O95714 | E3 ubiquitin-protein ligase HERC2 OS=I | HERC2 | 527.228 | 2 | 55.270603 | yes | 9.901E-08 | 776.44629 | 100.41616 |
| NNEEVTLIR | 9 | O95714 | E3 ubiquitin-protein ligase HERC2 OS=I | HERC2 | 527.228 | 2 | 55.270603 | yes | 2.982E-07 | 544.29077 | 95.087593 |
| IHGLILLGR | 9 | O95714 | E3 ubiquitin-protein ligase HERC2 OS=I | HERC2 | 527.228 | 2 | 55.270603 | yes | 1.001E-07 | 496.32419 | 88.334427 |
| QFEYEDPIVR | 10 | O95714 | E3 ubiquitin-protein ligase HERC2 OS=I | HERC2 | 527.228 | 2 | 55.270603 | yes | 2.247E-07 | 648.31696 | 85.314201 |
| VFSWGEGDDGK | 11 | O95714 | E3 ubiquitin-protein ligase HERC2 OS=I | HERC2 | 527.228 | 2 | 55.270603 | yes | 1.11E-07 | 598.76459 | 82.977005 |
| VIGELGEDGWIR | 12 | O95714 | E3 ubiquitin-protein ligase HERC2 OS=I | HERC2 | 527.228 | 2 | 55.270603 | no | 1.04E-06 | 672.35132 | 75.365013 |
| INEPGQSAVFCGR | 13 | O95714 | E3 ubiquitin-protein ligase HERC2 OS=I | HERC2 | 527.228 | 2 | 55.270603 | yes | 5.546E-06 | 717.84332 | 72.894623 |
| LLTTEFGQSININR | 14 | O95714 | E3 ubiquitin-protein ligase HERC2 OS=I | HERC2 | 527.228 | 2 | 55.270603 | yes | 9.722E-08 | 803.43341 | 70.331955 |
| LEGIDTILK | 9 | O95714 | E3 ubiquitin-protein ligase HERC2 OS=I | HERC2 | 527.228 | 2 | 55.270603 | no | 2.748E-06 | 501.29752 | 69.131706 |
| AYEEVCEGDVGK | 12 | O95714 | E3 ubiquitin-protein ligase HERC2 OS=I | HERC2 | 527.228 | 2 | 55.270603 | yes | 9.722E-08 | 678.29285 | 64.185692 |
| SVVTGDVHGTPAT | 14 | O95714 | E3 ubiquitin-protein ligase HERC2 OS=I | HERC2 | 527.228 | 2 | 55.270603 | yes | 1.486E-07 | 684.8595 | 63.879917 |
| LQDLDVVK | 8 | O95714 | E3 ubiquitin-protein ligase HERC2 OS=I | HERC2 | 527.228 | 2 | 55.270603 | yes | 1.298E-06 | 465.26874 | 63.229389 |
| LLLCCLLK | 8 | O95714 | E3 ubiquitin-protein ligase HERC2 OS=I | HERC2 | 527.228 | 2 | 55.270603 | no | 1.146E-07 | 516.80084 | 60.770023 |
| DVDLIPPFNR | 10 | O95714 | E3 ubiquitin-protein ligase HERC2 OS=I | HERC2 | 527.228 | 2 | 55.270603 | no | 7.806E-07 | 593.31677 | 60.510292 |
| ADFLSNDDYAVY' | 14 | O95714 | E3 ubiquitin-protein ligase HERC2 OS=I | HERC2 | 527.228 | 2 | 55.270603 | yes | 5.8E-07 | 824.38611 | 46.985424 |
| TGSPLSLNLAEPV' | 15 | O95714 | E3 ubiquitin-protein ligase HERC2 OS=I | HERC2 | 527.228 | 2 | 55.270603 | yes | 9.722E-08 | 806.44067 | 42.381683 |
| QAFAGQVDLDLLI | 13 | O95714 | E3 ubiquitin-protein ligase HERC2 OS=I | HERC2 | 527.228 | 2 | 55.270603 | no | 9.722E-08 | 723.39099 | 41.205208 |
| MLVESGTTDK | 10 | O95714 | E3 ubiquitin-protein ligase HERC2 OS=I | HERC2 | 527.228 | 2 | 55.270603 | yes | 1.982E-07 | 540.76569 | 40.434978 |
| IDSLTGLGVVK | 11 | O95714 | E3 ubiquitin-protein ligase HERC2 OS=I | HERC2 | 527.228 | 2 | 55.270603 | yes | 3.091E-07 | 551.32935 | 39.29842 |

| | | | | | | | | | | | |
|---|---|---|---|---|---|---|---|---|---|---|---|
| APVHSSMFR | 9 | O95714 | E3 ubiquitin-protein ligase HERC2 OS=I | HERC2 | 527.228 | 2 | 55.270603 | yes | 9.22E-07 | 516.25818 | 38.40752 |
| VECGSQFSVALTK | 13 | O95714 | E3 ubiquitin-protein ligase HERC2 OS=I | HERC2 | 527.228 | 2 | 55.270603 | yes | 6.273E-07 | 713.35577 | 37.660477 |
| DGGFWTVIDGK | 11 | O95714 | E3 ubiquitin-protein ligase HERC2 OS=I | HERC2 | 527.228 | 2 | 55.270603 | yes | 2.705E-07 | 597.79309 | 18.525253 |
| VACGSSHSVAWT | 31 | O95714 | E3 ubiquitin-protein ligase HERC2 OS=I | HERC2 | 527.228 | 4 | 55.270603 | yes | 2.946E-06 | 828.16345 | 18.524199 |
| DFVIQVLDK | 9 | O95714 | E3 ubiquitin-protein ligase HERC2 OS=I | HERC2 | 527.228 | 2 | 55.270603 | yes | 3.56E-06 | 538.80298 | 17.621813 |
| VQWDTGSTNSYR | 12 | O95714 | E3 ubiquitin-protein ligase HERC2 OS=I | HERC2 | 527.228 | 2 | 55.270603 | no | 1.131E-06 | 707.3233 | 13.975663 |
| ALSIENSQLMEQLI | 14 | O95751 | Protein LDOC1 OS=Homo sapiens OX=9 | LDOC1 | 16.969 | 2 | 50.960495 | yes | 9.722E-08 | 816.42468 | 120.60863 |
| EGQEDQGLTK | 10 | O95758 | Polypyrimidine tract-binding protein 3 OS | PTBP3 | 59.69 | 2 | 52.392937 | no | 9.722E-08 | 552.76202 | 14266.339 |
| DFSNSPLHR | 9 | O95758 | Polypyrimidine tract-binding protein 3 OS | PTBP3 | 59.69 | 2 | 52.392937 | yes | 2.064E-07 | 536.76215 | 2892.7607 |
| HQAVQLPR | 8 | O95758 | Polypyrimidine tract-binding protein 3 OS | PTBP3 | 59.69 | 2 | 52.392937 | yes | 9.92E-07 | 474.77234 | 2551.5747 |
| NLFIEAGCSVK | 11 | O95758 | Polypyrimidine tract-binding protein 3 OS | PTBP3 | 59.69 | 2 | 52.392937 | yes | 2.294E-07 | 619.31592 | 1259.8137 |
| SQPVYIQYSNHR | 12 | O95758 | Polypyrimidine tract-binding protein 3 OS | PTBP3 | 59.69 | 2 | 52.392937 | yes | 2.93E-07 | 746.37061 | 830.32666 |
| TDNLPNQAR | 9 | O95758 | Polypyrimidine tract-binding protein 3 OS | PTBP3 | 59.69 | 2 | 52.392937 | yes | 9.722E-08 | 514.75958 | 309.37924 |
| VTNLLMLK | 8 | O95758 | Polypyrimidine tract-binding protein 3 OS | PTBP3 | 59.69 | 2 | 52.392937 | no | 1.794E-07 | 474.28336 | 276.82983 |
| KIPCDVTEAEIISL( | 20 | O95758 | Polypyrimidine tract-binding protein 3 OS | PTBP3 | 59.69 | 3 | 52.392937 | yes | 9.722E-08 | 729.72919 | 152.79176 |
| ASPDLVPMGEWT/ | 14 | O95782 | AP-2 complex subunit alpha-1 OS=Homo | AP2A1 | 107.546 | 2 | 52.254997 | yes | 1.021E-07 | 765.37451 | 1613.4246 |
| ALQVGCLLR | 9 | O95782 | AP-2 complex subunit alpha-1 OS=Homo | AP2A1 | 107.546 | 2 | 52.254997 | yes | 1.039E-07 | 515.2973 | 1314.2128 |
| LLQCYPPPEDAAV | 14 | O95782 | AP-2 complex subunit alpha-1 OS=Homo | AP2A1 | 107.546 | 2 | 52.254997 | yes | 2.25E-07 | 800.90564 | 1020.5544 |
| IAGDYVSEEVWYI | 13 | O95782 | AP-2 complex subunit alpha-1 OS=Homo | AP2A1 | 107.546 | 2 | 52.254997 | no | 9.722E-08 | 793.87793 | 980.90845 |
| EMGEAFAADIPR | 12 | O95782 | AP-2 complex subunit alpha-1 OS=Homo | AP2A1 | 107.546 | 2 | 52.254997 | yes | 9.809E-08 | 653.80847 | 925.63885 |
| AADLLYAMCDR | 11 | O95782 | AP-2 complex subunit alpha-1 OS=Homo | AP2A1 | 107.546 | 2 | 52.254997 | yes | 9.722E-08 | 649.797 | 837.97491 |
| LVECLETVLNK | 11 | O95782 | AP-2 complex subunit alpha-1 OS=Homo | AP2A1 | 107.546 | 2 | 52.254997 | yes | 9.838E-08 | 659.35779 | 775.0296 |
| YLETADYAIR | 10 | O95782 | AP-2 complex subunit alpha-1 OS=Homo | AP2A1 | 107.546 | 2 | 52.254997 | yes | 9.722E-08 | 607.80621 | 724.05225 |
| YGGAPQALTLK | 11 | O95782 | AP-2 complex subunit alpha-1 OS=Homo | AP2A1 | 107.546 | 2 | 52.254997 | yes | 1.436E-07 | 559.81384 | 707.71136 |
| FHLCSVATR | 9 | O95782 | AP-2 complex subunit alpha-1 OS=Homo | AP2A1 | 107.546 | 2 | 52.254997 | yes | 9.722E-08 | 545.77673 | 690.79279 |
| GLAVFISDIR | 10 | O95782 | AP-2 complex subunit alpha-1 OS=Homo | AP2A1 | 107.546 | 2 | 52.254997 | no | 3.169E-07 | 545.81641 | 684.06354 |
| THIDTVINALK | 11 | O95782 | AP-2 complex subunit alpha-1 OS=Homo | AP2A1 | 107.546 | 2 | 52.254997 | yes | 3.926E-07 | 612.85095 | 614.77997 |
| QSAALCLLR | 9 | O95782 | AP-2 complex subunit alpha-1 OS=Homo | AP2A1 | 107.546 | 2 | 52.254997 | no | 5.566E-07 | 516.28693 | 595.48004 |
| VLQIVTNR | 8 | O95782 | AP-2 complex subunit alpha-1 OS=Homo | AP2A1 | 107.546 | 2 | 52.254997 | yes | 1.225E-07 | 471.79019 | 587.41846 |
| LEPNAQAQMYR | 11 | O95782 | AP-2 complex subunit alpha-1 OS=Homo | AP2A1 | 107.546 | 2 | 52.254997 | yes | 1.142E-07 | 660.8219 | 490.12579 |
| MYLFYGNK | 8 | O95782 | AP-2 complex subunit alpha-1 OS=Homo | AP2A1 | 107.546 | 2 | 52.254997 | yes | 2.045E-07 | 518.25208 | 407.34238 |
| HLCELLAQQF | 10 | O95782 | AP-2 complex subunit alpha-1 OS=Homo | AP2A1 | 107.546 | 2 | 52.254997 | yes | 2.164E-07 | 629.8161 | 404.16681 |
| TCVSLAVSR | 9 | O95782 | AP-2 complex subunit alpha-1 OS=Homo | AP2A1 | 107.546 | 2 | 52.254997 | yes | 3.454E-07 | 496.76331 | 402.40125 |
| QLSLPQQEAQK | 11 | O95782 | AP-2 complex subunit alpha-1 OS=Homo | AP2A1 | 107.546 | 2 | 52.254997 | yes | 1.021E-07 | 635.34351 | 392.56296 |
| YLALESMCTLASS | 21 | O95782 | AP-2 complex subunit alpha-1 OS=Homo | AP2A1 | 107.546 | 3 | 52.254997 | no | 1.041E-07 | 791.70819 | 346.43155 |
| FINLFPETK | 9 | O95782 | AP-2 complex subunit alpha-1 OS=Homo | AP2A1 | 107.546 | 2 | 52.254997 | yes | 8.546E-07 | 554.80548 | 334.31073 |
| ACNQLGQFLQHR | 12 | O95782 | AP-2 complex subunit alpha-1 OS=Homo | AP2A1 | 107.546 | 2 | 52.254997 | no | 1.065E-07 | 736.36481 | 319.52917 |
| QIVSEMLR | 8 | O95782 | AP-2 complex subunit alpha-1 OS=Homo | AP2A1 | 107.546 | 2 | 52.254997 | yes | 8.388E-07 | 488.26822 | 308.08917 |
| ANHPMDAEVTK | 11 | O95782 | AP-2 complex subunit alpha-1 OS=Homo | AP2A1 | 107.546 | 2 | 52.254997 | yes | 1.062E-07 | 606.78754 | 208.3927 |

| | | | | | | | | | | | |
|---|---|---|---|---|---|---|---|---|---|---|---|
| KGPGAGSALDDGI | 13 | O95782 | AP-2 complex subunit alpha-1 OS=Homc | AP2A1 | 107.546 | 2 | 52.254997 | yes | 1.557E-06 | 600.802 | 170.96132 |
| GPGAGSALDDGR | 12 | O95782 | AP-2 complex subunit alpha-1 OS=Homc | AP2A1 | 107.546 | 2 | 52.254997 | yes | 1.888E-07 | 536.75452 | 136.47701 |
| ALLLSTYIK | 9 | O95782 | AP-2 complex subunit alpha-1 OS=Homc | AP2A1 | 107.546 | 2 | 52.254997 | no | 2.321E-06 | 511.31824 | 70.456993 |
| DDVQGYAAK | 9 | O95782 | AP-2 complex subunit alpha-1 OS=Homc | AP2A1 | 107.546 | 2 | 52.254997 | no | 1.308E-06 | 483.72998 | 67.640701 |
| VAAQVDGGAQVQ | 21 | O95782 | AP-2 complex subunit alpha-1 OS=Homc | AP2A1 | 107.546 | 3 | 52.254997 | yes | 3.296E-07 | 756.73035 | 53.931538 |
| NPTFMCLALHCIA | 18 | O95782 | AP-2 complex subunit alpha-1 OS=Homc | AP2A1 | 107.546 | 3 | 52.254997 | yes | 5.53E-07 | 687.66577 | 18.938499 |
| TYIQTELPFK | 10 | O95785 | Protein Wiz OS=Homo sapiens OX=960€ | WIZ | 178.675 | 2 | 53.217556 | yes | 2.23E-06 | 620.33459 | 171.98489 |
| CEFCGEFFENR | 11 | O95785 | Protein Wiz OS=Homo sapiens OX=960€ | WIZ | 178.675 | 2 | 53.217556 | yes | 9.722E-08 | 747.79248 | 168.67415 |
| ELSLTPITGAK | 11 | O95785 | Protein Wiz OS=Homo sapiens OX=960€ | WIZ | 178.675 | 2 | 53.217556 | yes | 9.753E-07 | 565.32678 | 162.84941 |
| MMGGAGPGSSLE. | 14 | O95785 | Protein Wiz OS=Homo sapiens OX=960€ | WIZ | 178.675 | 2 | 53.217556 | yes | 2.72E-07 | 660.80536 | 130.29369 |
| PLDAPAVNK | 9 | O95785 | Protein Wiz OS=Homo sapiens OX=960€ | WIZ | 178.675 | 2 | 53.217556 | yes | 3.011E-06 | 462.76108 | 85.688263 |
| PSATGYLGSVAAK | 13 | O95785 | Protein Wiz OS=Homo sapiens OX=960€ | WIZ | 178.675 | 2 | 53.217556 | yes | 1.595E-07 | 611.32733 | 67.775703 |
| SPSDLHISPLAK | 12 | O95785 | Protein Wiz OS=Homo sapiens OX=960€ | WIZ | 178.675 | 2 | 53.217556 | yes | 9.722E-08 | 632.84845 | 60.232555 |
| PGGPPNPPGPSPK | 13 | O95785 | Protein Wiz OS=Homo sapiens OX=960€ | WIZ | 178.675 | 2 | 53.217556 | yes | 5.856E-07 | 599.81439 | 44.680809 |
| KTPLALAGSPTPK | 13 | O95785 | Protein Wiz OS=Homo sapiens OX=960€ | WIZ | 178.675 | 2 | 53.217556 | yes | 1.471E-06 | 640.88226 | 18.771517 |
| VSVGEFVGEGEGK | 13 | O95793 | Double-stranded RNA-binding protein St | STAU1 | 63.183 | 2 | 54.35891 | yes | 9.722E-08 | 647.3197 | 2201.4165 |
| NLPVNFEVAR | 10 | O95793 | Double-stranded RNA-binding protein St | STAU1 | 63.183 | 2 | 54.35891 | yes | 1.165E-07 | 579.81696 | 2043.2913 |
| NAAIAVLEELKK | 12 | O95793 | Double-stranded RNA-binding protein St | STAU1 | 63.183 | 2 | 54.35891 | yes | 2.529E-07 | 649.88757 | 1731.2775 |
| VQGFQVEYK | 9 | O95793 | Double-stranded RNA-binding protein St | STAU1 | 63.183 | 2 | 54.35891 | yes | 1.443E-07 | 549.28491 | 1531.5995 |
| ILQNEPLPER | 10 | O95793 | Double-stranded RNA-binding protein St | STAU1 | 63.183 | 2 | 54.35891 | yes | 9.722E-08 | 604.83533 | 1480.2444 |
| EPEYTLLTER | 10 | O95793 | Double-stranded RNA-binding protein St | STAU1 | 63.183 | 2 | 54.35891 | yes | 1.726E-07 | 625.81677 | 1380.1874 |
| LLSELDQQSTEMP | 14 | O95793 | Double-stranded RNA-binding protein St | STAU1 | 63.183 | 2 | 54.35891 | yes | 1.073E-07 | 823.90637 | 1330.8728 |
| PSEQLDYLSR | 10 | O95793 | Double-stranded RNA-binding protein St | STAU1 | 63.183 | 2 | 54.35891 | yes | 9.722E-08 | 604.30133 | 1228.8927 |
| KLPPLPAVER | 10 | O95793 | Double-stranded RNA-binding protein St | STAU1 | 63.183 | 2 | 54.35891 | yes | 3.106E-06 | 560.34784 | 941.46844 |
| VPQAQPTKPALK | 12 | O95793 | Double-stranded RNA-binding protein St | STAU1 | 63.183 | 2 | 54.35891 | yes | 1.043E-07 | 639.38245 | 365.43253 |
| VPQAQPTKPALK | 12 | O95793 | Double-stranded RNA-binding protein St | STAU1 | 63.183 | 2 | 54.35891 | yes | 2.114E-07 | 426.59073 | 365.43253 |
| VPQAQPTKPALK | 12 | O95793 | Double-stranded RNA-binding protein St | STAU1 | 63.183 | 3 | 54.35891 | yes | 1.043E-07 | 639.38245 | 365.43253 |
| VPQAQPTKPALK | 12 | O95793 | Double-stranded RNA-binding protein St | STAU1 | 63.183 | 3 | 54.35891 | yes | 2.114E-07 | 426.59073 | 365.43253 |
| ESEEENLNK | 9 | O95793 | Double-stranded RNA-binding protein St | STAU1 | 63.183 | 2 | 54.35891 | yes | 1.508E-07 | 546.24622 | 642.83533 |
| EKEPEYTLLTER | 12 | O95793 | Double-stranded RNA-binding protein St | STAU1 | 63.183 | 2 | 54.35891 | yes | 9.758E-08 | 754.38556 | 614.82117 |
| NAAIAVLEELK | 11 | O95793 | Double-stranded RNA-binding protein St | STAU1 | 63.183 | 2 | 54.35891 | yes | 1.575E-07 | 585.84009 | 547.63092 |
| SEISQVFEIALK | 12 | O95793 | Double-stranded RNA-binding protein St | STAU1 | 63.183 | 2 | 54.35891 | yes | 3.451E-07 | 682.37701 | 427.14481 |
| NNISSGHVPHGPL | 15 | O95793 | Double-stranded RNA-binding protein St | STAU1 | 63.183 | 3 | 54.35891 | yes | 3.247E-07 | 529.27795 | 198.35115 |
| NNISSGHVPHGPL | 15 | O95793 | Double-stranded RNA-binding protein St | STAU1 | 63.183 | 3 | 54.35891 | yes | 9.722E-08 | 793.41333 | 198.35115 |
| NNISSGHVPHGPL | 15 | O95793 | Double-stranded RNA-binding protein St | STAU1 | 63.183 | 2 | 54.35891 | yes | 3.247E-07 | 529.27795 | 198.35115 |
| NNISSGHVPHGPL | 15 | O95793 | Double-stranded RNA-binding protein St | STAU1 | 63.183 | 2 | 54.35891 | yes | 9.722E-08 | 793.41333 | 198.35115 |
| NEFVSLINCSSQPP | 22 | O95793 | Double-stranded RNA-binding protein St | STAU1 | 63.183 | 3 | 54.35891 | yes | 2.737E-07 | 799.74591 | 362.86014 |
| ELLYGGTSPTAETI | 16 | O95793 | Double-stranded RNA-binding protein St | STAU1 | 63.183 | 2 | 54.35891 | yes | 1.173E-07 | 846.95636 | 342.08459 |
| MQSTYNYNMR | 10 | O95793 | Double-stranded RNA-binding protein St | STAU1 | 63.183 | 2 | 54.35891 | yes | 1.504E-07 | 654.27899 | 254.72667 |

| | | | | | | | | | | | |
|---|---|---|---|---|---|---|---|---|---|---|---|
| TGNGPMSVCGR | 11 | O95793 | Double-stranded RNA-binding protein St | STAU1 | 63.183 | 2 | 54.35891 | yes | 5.116E-07 | 568.25281 | 216.92572 |
| TKPIVKPQTSPEY( | 22 | O95793 | Double-stranded RNA-binding protein St | STAU1 | 63.183 | 4 | 54.35891 | yes | 9.903E-08 | 603.3327 | 181.0251 |
| MPYLSHQQLPAGI | 33 | O95793 | Double-stranded RNA-binding protein St | STAU1 | 63.183 | 5 | 54.35891 | yes | 9.862E-08 | 704.96967 | 146.12828 |
| KNAAIAVLEELK | 12 | O95793 | Double-stranded RNA-binding protein St | STAU1 | 63.183 | 2 | 54.35891 | yes | 9.722E-08 | 649.88757 | 105.06212 |
| NNISSGHVPHGPL' | 25 | O95793 | Double-stranded RNA-binding protein St | STAU1 | 63.183 | 4 | 54.35891 | yes | 1.93E-07 | 694.35468 | 76.922417 |
| SQVQVQVQNPSA/ | 22 | O95793 | Double-stranded RNA-binding protein St | STAU1 | 63.183 | 3 | 54.35891 | yes | 1.523E-07 | 780.08356 | 68.601845 |
| LPPLPAVER | 9 | O95793 | Double-stranded RNA-binding protein St | STAU1 | 63.183 | 2 | 54.35891 | yes | 2.975E-06 | 496.30038 | 7.4017744 |
| GALCHLELK | 9 | O95801 | Tetratricopeptide repeat protein 4 OS=Ho | TTC4 | 44.679 | 2 | 51.968307 | yes | 1.508E-07 | 520.78149 | 205.53076 |
| STLLQVLQHQR | 11 | O95801 | Tetratricopeptide repeat protein 4 OS=Ho | TTC4 | 44.679 | 2 | 51.968307 | yes | 9.788E-08 | 661.88062 | 174.12489 |
| CADPDLNAVLYTN | 14 | O95801 | Tetratricopeptide repeat protein 4 OS=Ho | TTC4 | 44.679 | 2 | 51.968307 | yes | 9.722E-08 | 811.38556 | 109.58943 |
| AAAQYYLGNFR | 11 | O95801 | Tetratricopeptide repeat protein 4 OS=Ho | TTC4 | 44.679 | 2 | 51.968307 | yes | 2.081E-07 | 637.31982 | 68.585968 |
| GGFHEDQWEK | 10 | O95801 | Tetratricopeptide repeat protein 4 OS=Ho | TTC4 | 44.679 | 2 | 51.968307 | yes | 7.797E-07 | 616.7702 | 42.617188 |
| NQNEALLQAIK | 11 | O95801 | Tetratricopeptide repeat protein 4 OS=Ho | TTC4 | 44.679 | 2 | 51.968307 | yes | 1.549E-06 | 621.34607 | 30.696381 |
| TLQQNAESR | 9 | O95816 | BAG family molecular chaperone regulat | BAG2 | 23.772 | 2 | 52.42223 | yes | 1.269E-07 | 523.76489 | 3473.1042 |
| FQSIVIGCALEDQK | 14 | O95816 | BAG family molecular chaperone regulat | BAG2 | 23.772 | 2 | 52.42223 | yes | 1.125E-07 | 804.40851 | 2623.9912 |
| FLDDLGNAK | 9 | O95816 | BAG family molecular chaperone regulat | BAG2 | 23.772 | 2 | 52.42223 | yes | 1.678E-07 | 496.75601 | 1309.9875 |
| EAATAVEQEK | 10 | O95816 | BAG family molecular chaperone regulat | BAG2 | 23.772 | 2 | 52.42223 | yes | 1.046E-07 | 538.26691 | 1191.64 |
| NPQQQESLK | 9 | O95816 | BAG family molecular chaperone regulat | BAG2 | 23.772 | 2 | 52.42223 | yes | 1.03E-07 | 536.27509 | 1158.676 |
| LLESLDQLELR | 11 | O95816 | BAG family molecular chaperone regulat | BAG2 | 23.772 | 2 | 52.42223 | yes | 1.133E-07 | 664.87463 | 1113.193 |
| RLETLLR | 7 | O95816 | BAG family molecular chaperone regulat | BAG2 | 23.772 | 2 | 52.42223 | yes | 2.113E-07 | 450.78491 | 989.16583 |
| TLTVEVSVETIR | 12 | O95816 | BAG family molecular chaperone regulat | BAG2 | 23.772 | 2 | 52.42223 | yes | 1.033E-06 | 673.87994 | 184.94641 |
| EELNLTANR | 9 | O95816 | BAG family molecular chaperone regulat | BAG2 | 23.772 | 2 | 52.42223 | yes | 9.722E-08 | 530.27509 | 111.8539 |
| SHLMSLYSACSSE' | 22 | O95816 | BAG family molecular chaperone regulat | BAG2 | 23.772 | 3 | 52.42223 | yes | 3.903E-07 | 810.37897 | 76.365547 |
| ELLALDSVDPEGR | 13 | O95817 | BAG family molecular chaperone regulat | BAG3 | 61.595 | 2 | 51.94595 | yes | 3.86E-07 | 707.36462 | 1046.1637 |
| VQGLEQAVDNFE( | 14 | O95817 | BAG family molecular chaperone regulat | BAG3 | 61.595 | 2 | 51.94595 | yes | 9.722E-08 | 767.3808 | 496.62228 |
| EGHPVYPQLRPGY | 26 | O95817 | BAG family molecular chaperone regulat | BAG3 | 61.595 | 4 | 51.94595 | yes | 1.088E-07 | 735.38849 | 287.07986 |
| SSLGSHQLPR | 10 | O95817 | BAG family molecular chaperone regulat | BAG3 | 61.595 | 2 | 51.94595 | yes | 1.081E-07 | 541.29108 | 205.30731 |
| EGHPVYPQLR | 10 | O95817 | BAG family molecular chaperone regulat | BAG3 | 61.595 | 2 | 51.94595 | yes | 1.65E-07 | 598.31458 | 194.18372 |
| EVDSKPVSQKPPP | 17 | O95817 | BAG family molecular chaperone regulat | BAG3 | 61.595 | 3 | 51.94595 | yes | 9.977E-08 | 616.99493 | 190.04553 |
| KVQTILEK | 8 | O95817 | BAG family molecular chaperone regulat | BAG3 | 61.595 | 2 | 51.94595 | yes | 9.977E-08 | 479.80023 | 180.80208 |
| YLMIEEYLTK | 10 | O95817 | BAG family molecular chaperone regulat | BAG3 | 61.595 | 2 | 51.94595 | yes | 1.151E-07 | 651.83612 | 168.89984 |
| PGYIPIPVLHEGAE | 16 | O95817 | BAG family molecular chaperone regulat | BAG3 | 61.595 | 3 | 51.94595 | yes | 2.051E-07 | 587.98096 | 154.96532 |
| PAAQPSFHQAQK | 12 | O95817 | BAG family molecular chaperone regulat | BAG3 | 61.595 | 2 | 51.94595 | yes | 1.03E-07 | 655.336 | 130.79778 |
| SSTPLHSPSPIR | 12 | O95817 | BAG family molecular chaperone regulat | BAG3 | 61.595 | 2 | 51.94595 | yes | 9.722E-08 | 639.84369 | 108.08833 |
| QCGQVAAAAAAQ | 21 | O95817 | BAG family molecular chaperone regulat | BAG3 | 61.595 | 3 | 51.94595 | yes | 9.366E-07 | 692.00098 | 103.27916 |
| GMPETTQPDK | 10 | O95817 | BAG family molecular chaperone regulat | BAG3 | 61.595 | 2 | 51.94595 | yes | 1.503E-07 | 552.25549 | 89.646446 |
| GYISIPVIHEQNVT | 15 | O95817 | BAG family molecular chaperone regulat | BAG3 | 61.595 | 3 | 51.94595 | yes | 2.212E-06 | 575.98096 | 14.541855 |
| LVDFGVSAQLDR | 12 | O95819 | Mitogen-activated protein kinase kinase k | MAP4K4 | 142.101 | 2 | 51.30719 | no | 1.197E-07 | 660.35132 | 576.71393 |
| GQNVLLTENAEVK | 13 | O95819 | Mitogen-activated protein kinase kinase k | MAP4K4 | 142.101 | 2 | 51.30719 | no | 1.231E-07 | 707.88043 | 357.74768 |

| | | | | | | | | | | |
|---|---|---|---|---|---|---|---|---|---|---|
| RLEELER | 7 | O95819 | Mitogen-activated protein kinase kinase k | MAP4K4 | 142.101 | 2 | 51.30719 | no | 1.01E-07 | 472.76163 | 330.15344 |
| SLVDIDLSSLR | 11 | O95819 | Mitogen-activated protein kinase kinase k | MAP4K4 | 142.101 | 2 | 51.30719 | yes | 9.722E-08 | 609.34045 | 259.76035 |
| QQLLQEQQLR | 10 | O95819 | Mitogen-activated protein kinase kinase k | MAP4K4 | 142.101 | 2 | 51.30719 | yes | 1.074E-07 | 642.35693 | 182.33783 |
| PSTEQLLK | 8 | O95819 | Mitogen-activated protein kinase kinase k | MAP4K4 | 142.101 | 2 | 51.30719 | yes | 2.24E-06 | 458.26093 | 163.94046 |
| SNQTMGWGEK | 10 | O95819 | Mitogen-activated protein kinase kinase k | MAP4K4 | 142.101 | 2 | 51.30719 | yes | 1.678E-07 | 569.2533 | 114.85017 |
| ILHNDPEVEK | 10 | O95819 | Mitogen-activated protein kinase kinase k | MAP4K4 | 142.101 | 2 | 51.30719 | no | 9.722E-08 | 597.31171 | 114.20807 |
| AVEDVRPPHK | 10 | O95819 | Mitogen-activated protein kinase kinase k | MAP4K4 | 142.101 | 2 | 51.30719 | yes | 9.722E-08 | 574.31458 | 110.59428 |
| QTQSASSTLQK | 11 | O95819 | Mitogen-activated protein kinase kinase k | MAP4K4 | 142.101 | 2 | 51.30719 | yes | 2.232E-07 | 589.8042 | 106.37315 |
| EDWIAYISR | 9 | O95819 | Mitogen-activated protein kinase kinase k | MAP4K4 | 142.101 | 2 | 51.30719 | yes | 1.417E-07 | 576.78784 | 94.787933 |
| SGGSSQVYFMTLC | 14 | O95819 | Mitogen-activated protein kinase kinase k | MAP4K4 | 142.101 | 2 | 51.30719 | yes | 1.877E-07 | 745.35883 | 75.314941 |
| FFSFIEGCLVK | 11 | O95819 | Mitogen-activated protein kinase kinase k | MAP4K4 | 142.101 | 2 | 51.30719 | yes | 2.411E-07 | 673.8443 | 60.625809 |
| AASSLNLSNGETES | 16 | O95819 | Mitogen-activated protein kinase kinase k | MAP4K4 | 142.101 | 2 | 51.30719 | yes | 3.037E-07 | 803.8996 | 57.096352 |
| PLLVDLTVEEGQR | 13 | O95819 | Mitogen-activated protein kinase kinase k | MAP4K4 | 142.101 | 2 | 51.30719 | no | 2.061E-07 | 734.90393 | 39.125359 |
| KGSVVNVNPTNTI | 22 | O95819 | Mitogen-activated protein kinase kinase k | MAP4K4 | 142.101 | 4 | 51.30719 | yes | 1.752E-07 | 603.0675 | 36.297657 |
| VMDVTEDEEEEIK | 20 | O95819 | Mitogen-activated protein kinase kinase k | MAP4K4 | 142.101 | 3 | 51.30719 | yes | 3.315E-06 | 803.05884 | 31.341507 |
| GSVVNVNPTNTR | 12 | O95819 | Mitogen-activated protein kinase kinase k | MAP4K4 | 142.101 | 2 | 51.30719 | no | 1.893E-07 | 629.33093 | 28.995403 |
| ELWFSDDPNVTK | 12 | O95831 | Apoptosis-inducing factor 1, mitochondri | AIFM1 | 66.901 | 2 | 52.443317 | yes | 9.995E-08 | 725.84607 | 409.51831 |
| TGGLEIDSDFGGFI | 14 | O95831 | Apoptosis-inducing factor 1, mitochondri | AIFM1 | 66.901 | 2 | 52.443317 | yes | 1.053E-07 | 735.84662 | 402.95581 |
| ISGLGLTPEQK | 11 | O95831 | Apoptosis-inducing factor 1, mitochondri | AIFM1 | 66.901 | 2 | 52.443317 | yes | 9.722E-08 | 571.8244 | 398.22165 |
| AALSASEGEEVPQ | 15 | O95831 | Apoptosis-inducing factor 1, mitochondri | AIFM1 | 66.901 | 2 | 52.443317 | yes | 9.918E-08 | 765.86774 | 286.66766 |
| ILPEYLSNWTMEK | 13 | O95831 | Apoptosis-inducing factor 1, mitochondri | AIFM1 | 66.901 | 2 | 52.443317 | yes | 1.352E-07 | 812.40802 | 244.49681 |
| LNDGSQITYEK | 11 | O95831 | Apoptosis-inducing factor 1, mitochondri | AIFM1 | 66.901 | 2 | 52.443317 | yes | 1.53E-07 | 634.31189 | 188.77852 |
| VMPNAIVQSVGVS | 16 | O95831 | Apoptosis-inducing factor 1, mitochondri | AIFM1 | 66.901 | 2 | 52.443317 | yes | 2.079E-07 | 786.92432 | 137.45247 |
| LFNIHED | 7 | O95831 | Apoptosis-inducing factor 1, mitochondri | AIFM1 | 66.901 | 2 | 52.443317 | yes | 4.3E-07 | 444.21652 | 89.701164 |
| ALGTEVIQLFPEK | 13 | O95831 | Apoptosis-inducing factor 1, mitochondri | AIFM1 | 66.901 | 2 | 52.443317 | yes | 1.248E-07 | 722.90594 | 74.458321 |
| APSHVPFLLIGGGT | 21 | O95831 | Apoptosis-inducing factor 1, mitochondri | AIFM1 | 66.901 | 3 | 52.443317 | yes | 1.203E-07 | 675.37402 | 39.056023 |
| MGIFPESLAR | 10 | O95835 | Serine/threonine-protein kinase LATS1 O | LATS1 | 126.87 | 2 | 45.19325 | yes | 5.157E-07 | 560.79462 | 40.000595 |
| LSPEASDLIIK | 11 | O95835 | Serine/threonine-protein kinase LATS1 O | LATS1 | 126.87 | 2 | 45.19325 | yes | 1.93E-07 | 593.3399 | 39.686535 |
| VAELTSLSDEDSGI | 14 | O95863 | Zinc finger protein SNAI1 OS=Homo sap | SNAI1 | 29.084 | 2 | 48.502914 | yes | 1.138E-07 | 725.849 | 177.76973 |
| EYLSLGALK | 9 | O95863 | Zinc finger protein SNAI1 OS=Homo sap | SNAI1 | 29.084 | 2 | 48.502914 | yes | 4.455E-07 | 497.28439 | 42.437454 |
| LLSCVLGPR | 9 | O95870 | Phosphatidylserine lipase ABHD16A OS= | ABHD16A | 63.244 | 2 | 34.658398 | yes | 9.651E-07 | 507.79187 | 283.70187 |
| MLYPGSVYLLQK | 12 | O95870 | Phosphatidylserine lipase ABHD16A OS= | ABHD16A | 63.244 | 2 | 34.658398 | yes | 1.526E-05 | 706.38635 | 38.215248 |
| AATPQVAAELEK | 12 | O95900 | Pseudouridylate synthase TRUB2, mitoch | TRUB2 | 36.695 | 2 | 52.52968 | yes | 1.723E-07 | 614.33258 | 2278.821 |
| TQWDLTNIQDAIR | 13 | O95900 | Pseudouridylate synthase TRUB2, mitoch | TRUB2 | 36.695 | 2 | 52.52968 | yes | 9.722E-08 | 787.4021 | 1790.8606 |
| TTYDHVTR | 8 | O95900 | Pseudouridylate synthase TRUB2, mitoch | TRUB2 | 36.695 | 2 | 52.52968 | yes | 2.153E-07 | 496.74344 | 1140.1263 |
| ALVMYSNLDLK | 11 | O95900 | Pseudouridylate synthase TRUB2, mitoch | TRUB2 | 36.695 | 2 | 52.52968 | yes | 9.819E-08 | 633.84174 | 993.41125 |
| TTAVCTQVR | 9 | O95900 | Pseudouridylate synthase TRUB2, mitoch | TRUB2 | 36.695 | 2 | 52.52968 | yes | 9.843E-08 | 518.26624 | 838.30219 |
| FLLGPMEGSEEK | 12 | O95900 | Pseudouridylate synthase TRUB2, mitoch | TRUB2 | 36.695 | 2 | 52.52968 | yes | 1.73E-07 | 668.82629 | 712.3479 |
| DTVELQLLK | 9 | O95900 | Pseudouridylate synthase TRUB2, mitoch | TRUB2 | 36.695 | 2 | 52.52968 | yes | 2.374E-06 | 529.80823 | 631.33936 |

| | | | | | | | | | | | |
|---|---|---|---|---|---|---|---|---|---|---|---|
| SPMLITGIR | 9 | O95900 | Pseudouridylate synthase TRUB2, mitoch | TRUB2 | 36.695 | 2 | 52.52968 | yes | 1.738E-07 | 494.28641 | 626.83856 |
| LLTDMYNAHLTK | 12 | O95900 | Pseudouridylate synthase TRUB2, mitoch | TRUB2 | 36.695 | 2 | 52.52968 | yes | 1.25E-07 | 710.36865 | 290.28482 |
| LLTDMYNAHLTK | 12 | O95900 | Pseudouridylate synthase TRUB2, mitoch | TRUB2 | 36.695 | 2 | 52.52968 | yes | 5.728E-06 | 473.91489 | 290.28482 |
| LLTDMYNAHLTK | 12 | O95900 | Pseudouridylate synthase TRUB2, mitoch | TRUB2 | 36.695 | 3 | 52.52968 | yes | 1.25E-07 | 710.36865 | 290.28482 |
| LLTDMYNAHLTK | 12 | O95900 | Pseudouridylate synthase TRUB2, mitoch | TRUB2 | 36.695 | 3 | 52.52968 | yes | 5.728E-06 | 473.91489 | 290.28482 |
| TQEAYEMAVR | 10 | O95900 | Pseudouridylate synthase TRUB2, mitoch | TRUB2 | 36.695 | 2 | 52.52968 | yes | 9.722E-08 | 599.28204 | 579.46313 |
| LHGLFAVYKPPGL | 14 | O95900 | Pseudouridylate synthase TRUB2, mitoch | TRUB2 | 36.695 | 3 | 52.52968 | yes | 9.862E-08 | 513.97302 | 180.6738 |
| LHGLFAVYKPPGL | 14 | O95900 | Pseudouridylate synthase TRUB2, mitoch | TRUB2 | 36.695 | 3 | 52.52968 | yes | 1.945E-06 | 770.45593 | 180.6738 |
| LHGLFAVYKPPGL | 14 | O95900 | Pseudouridylate synthase TRUB2, mitoch | TRUB2 | 36.695 | 2 | 52.52968 | yes | 9.862E-08 | 513.97302 | 180.6738 |
| LHGLFAVYKPPGL | 14 | O95900 | Pseudouridylate synthase TRUB2, mitoch | TRUB2 | 36.695 | 2 | 52.52968 | yes | 1.945E-06 | 770.45593 | 180.6738 |
| SLSPGLDTK | 9 | O95900 | Pseudouridylate synthase TRUB2, mitoch | TRUB2 | 36.695 | 2 | 52.52968 | yes | 9.722E-08 | 459.25055 | 328.9115 |
| KLVHEIGLELK | 11 | O95900 | Pseudouridylate synthase TRUB2, mitoch | TRUB2 | 36.695 | 2 | 52.52968 | yes | 1.076E-07 | 639.89264 | 149.09792 |
| KLVHEIGLELK | 11 | O95900 | Pseudouridylate synthase TRUB2, mitoch | TRUB2 | 36.695 | 2 | 52.52968 | yes | 9.777E-08 | 426.93085 | 149.09792 |
| KLVHEIGLELK | 11 | O95900 | Pseudouridylate synthase TRUB2, mitoch | TRUB2 | 36.695 | 3 | 52.52968 | yes | 1.076E-07 | 639.89264 | 149.09792 |
| KLVHEIGLELK | 11 | O95900 | Pseudouridylate synthase TRUB2, mitoch | TRUB2 | 36.695 | 3 | 52.52968 | yes | 9.777E-08 | 426.93085 | 149.09792 |
| LVHEIGLELK | 10 | O95900 | Pseudouridylate synthase TRUB2, mitoch | TRUB2 | 36.695 | 2 | 52.52968 | yes | 1.098E-07 | 575.84515 | 266.92111 |
| ILAVIQGSHQK | 11 | O95900 | Pseudouridylate synthase TRUB2, mitoch | TRUB2 | 36.695 | 2 | 52.52968 | yes | 8.901E-05 | 597.3537 | 16.264853 |
| GGVPAHMFGVTK | 12 | O95905 | Protein ecdysoneless homolog OS=Homo | ECD | 72.759 | 2 | 51.952465 | yes | 9.722E-08 | 600.81335 | 151.76282 |
| LPPPSDPQYR | 10 | O95905 | Protein ecdysoneless homolog OS=Homo | ECD | 72.759 | 2 | 51.952465 | yes | 3.51E-07 | 585.30109 | 95.581635 |
| LLEISAEDAER | 11 | O95926 | Pre-mRNA-splicing factor SYF2 OS=Hor | SYF2 | 28.723 | 2 | 48.149506 | yes | 1.192E-07 | 623.3197 | 104.35349 |
| GKPDLNTALPVR | 12 | O95983 | Methyl-CpG-binding domain protein 3 O | MBD3 | 32.845 | 2 | 50.75512 | yes | 9.723E-08 | 640.86969 | 53.214115 |
| KQEELVQQVR | 10 | O95983 | Methyl-CpG-binding domain protein 3 O | MBD3 | 32.845 | 2 | 50.75512 | yes | 2.049E-07 | 628.8515 | 34.657871 |
| YLGGSMDLSTFDF | 14 | O95983 | Methyl-CpG-binding domain protein 3 O | MBD3 | 32.845 | 2 | 50.75512 | yes | 9.728E-08 | 804.87177 | 17.271679 |
| IDAELVLPTIR | 11 | O95985 | DNA topoisomerase 3-beta-1 OS=Homo | TOP3B | 96.662 | 2 | 53.2078 | yes | 4.438E-07 | 620.36902 | 4444.1392 |
| SATEAELGGDAWI | 13 | O95985 | DNA topoisomerase 3-beta-1 OS=Homo | TOP3B | 96.662 | 2 | 53.2078 | yes | 9.722E-08 | 681.81787 | 3654.6458 |
| VDPAELFSQAPTEI | 14 | O95985 | DNA topoisomerase 3-beta-1 OS=Homo | TOP3B | 96.662 | 2 | 53.2078 | yes | 4.393E-07 | 766.38556 | 3198.4255 |
| LKPTNLGIVLVHG | 16 | O95985 | DNA topoisomerase 3-beta-1 OS=Homo | TOP3B | 96.662 | 3 | 53.2078 | yes | 2.107E-07 | 605.68909 | 1188.3098 |
| LKPTNLGIVLVHG | 16 | O95985 | DNA topoisomerase 3-beta-1 OS=Homo | TOP3B | 96.662 | 3 | 53.2078 | yes | 9.722E-08 | 454.51865 | 1188.3098 |
| LKPTNLGIVLVHG | 16 | O95985 | DNA topoisomerase 3-beta-1 OS=Homo | TOP3B | 96.662 | 4 | 53.2078 | yes | 2.107E-07 | 605.68909 | 1188.3098 |
| LKPTNLGIVLVHG | 16 | O95985 | DNA topoisomerase 3-beta-1 OS=Homo | TOP3B | 96.662 | 4 | 53.2078 | yes | 9.722E-08 | 454.51865 | 1188.3098 |
| EIAQMFLNMTK | 11 | O95985 | DNA topoisomerase 3-beta-1 OS=Homo | TOP3B | 96.662 | 2 | 53.2078 | yes | 1.362E-07 | 663.33325 | 2127.4575 |
| YLQSTISFR | 9 | O95985 | DNA topoisomerase 3-beta-1 OS=Homo | TOP3B | 96.662 | 2 | 53.2078 | yes | 1.02E-07 | 557.79822 | 1740.8142 |
| QLNLIAQGK | 9 | O95985 | DNA topoisomerase 3-beta-1 OS=Homo | TOP3B | 96.662 | 2 | 53.2078 | yes | 9.236E-07 | 492.79547 | 1294.3398 |
| GDAFPVGEVK | 10 | O95985 | DNA topoisomerase 3-beta-1 OS=Homo | TOP3B | 96.662 | 2 | 53.2078 | yes | 2.278E-07 | 509.76382 | 1285.218 |
| EAQVEATSR | 9 | O95985 | DNA topoisomerase 3-beta-1 OS=Homo | TOP3B | 96.662 | 2 | 53.2078 | yes | 2.188E-07 | 495.74619 | 1273.4858 |
| SLLLDWDR | 8 | O95985 | DNA topoisomerase 3-beta-1 OS=Homo | TOP3B | 96.662 | 2 | 53.2078 | yes | 2.346E-07 | 509.27182 | 1143.8152 |
| TETTHYPENFDLK | 13 | O95985 | DNA topoisomerase 3-beta-1 OS=Homo | TOP3B | 96.662 | 2 | 53.2078 | yes | 9.722E-08 | 797.8728 | 954.32635 |
| LYEYITR | 7 | O95985 | DNA topoisomerase 3-beta-1 OS=Homo | TOP3B | 96.662 | 2 | 53.2078 | yes | 5.89E-07 | 479.25565 | 931.66986 |
| NYVTVESGR | 9 | O95985 | DNA topoisomerase 3-beta-1 OS=Homo | TOP3B | 96.662 | 2 | 53.2078 | yes | 1.532E-07 | 512.75653 | 830.57678 |

| | | | | | | | | | | | |
|---|---|---|---|---|---|---|---|---|---|---|---|
| IQSFKPETYWVLQ | 15 | O95985 | DNA topoisomerase 3-beta-1 OS=Homo : | TOP3B | 96.662 | 3 | 53.2078 | yes | 3.918E-06 | 613.33307 | 650.92584 |
| HFIATVSHDCK | 11 | O95985 | DNA topoisomerase 3-beta-1 OS=Homo : | TOP3B | 96.662 | 2 | 53.2078 | yes | 5.072E-07 | 657.81659 | 524.51941 |
| QQANHPYWADTV | 13 | O95985 | DNA topoisomerase 3-beta-1 OS=Homo : | TOP3B | 96.662 | 3 | 53.2078 | yes | 5.452E-07 | 519.91968 | 177.96881 |
| QQANHPYWADTV | 13 | O95985 | DNA topoisomerase 3-beta-1 OS=Homo : | TOP3B | 96.662 | 3 | 53.2078 | yes | 9.809E-08 | 779.37585 | 177.96881 |
| QQANHPYWADTV | 13 | O95985 | DNA topoisomerase 3-beta-1 OS=Homo : | TOP3B | 96.662 | 2 | 53.2078 | yes | 5.452E-07 | 519.91968 | 177.96881 |
| QQANHPYWADTV | 13 | O95985 | DNA topoisomerase 3-beta-1 OS=Homo : | TOP3B | 96.662 | 2 | 53.2078 | yes | 9.809E-08 | 779.37585 | 177.96881 |
| TVLMVAEKPSLAQ | 17 | O95985 | DNA topoisomerase 3-beta-1 OS=Homo : | TOP3B | 96.662 | 3 | 53.2078 | yes | 1.128E-07 | 596.00958 | 335.1499 |
| GLNGACSVHEYTC | 21 | O95985 | DNA topoisomerase 3-beta-1 OS=Homo : | TOP3B | 96.662 | 3 | 53.2078 | yes | 1.154E-07 | 741.02002 | 310.39294 |
| WDKVDPAELFSQ/ | 17 | O95985 | DNA topoisomerase 3-beta-1 OS=Homo : | TOP3B | 96.662 | 3 | 53.2078 | yes | 1.002E-07 | 654.32654 | 281.06799 |
| IGPELFTCSGK | 11 | O95985 | DNA topoisomerase 3-beta-1 OS=Homo : | TOP3B | 96.662 | 2 | 53.2078 | yes | 9.729E-08 | 604.80261 | 242.06296 |
| GCDYIVLWLDCDI | 13 | O95985 | DNA topoisomerase 3-beta-1 OS=Homo : | TOP3B | 96.662 | 2 | 53.2078 | yes | 1.006E-07 | 828.87347 | 221.48877 |
| PSLAQSIAK | 9 | O95985 | DNA topoisomerase 3-beta-1 OS=Homo : | TOP3B | 96.662 | 2 | 53.2078 | yes | 1.976E-07 | 457.76892 | 202.62213 |
| PLALNTVEMLR | 11 | O95985 | DNA topoisomerase 3-beta-1 OS=Homo : | TOP3B | 96.662 | 2 | 53.2078 | yes | 9.141E-07 | 628.85522 | 105.85249 |
| QVLGHTLDVFK | 11 | O95985 | DNA topoisomerase 3-beta-1 OS=Homo : | TOP3B | 96.662 | 3 | 53.2078 | yes | 1.06E-07 | 419.57144 | 87.863274 |
| LLAEGINRPR | 10 | O95985 | DNA topoisomerase 3-beta-1 OS=Homo : | TOP3B | 96.662 | 2 | 53.2078 | yes | 4.277E-07 | 569.8382 | 60.816856 |
| GHDAGDHPPITPM | 14 | O95985 | DNA topoisomerase 3-beta-1 OS=Homo : | TOP3B | 96.662 | 3 | 53.2078 | yes | 1.343E-07 | 491.56989 | 28.501497 |
| GHDAGDHPPITPM | 14 | O95985 | DNA topoisomerase 3-beta-1 OS=Homo : | TOP3B | 96.662 | 3 | 53.2078 | yes | 5.11E-07 | 736.8512 | 28.501497 |
| GHDAGDHPPITPM | 14 | O95985 | DNA topoisomerase 3-beta-1 OS=Homo : | TOP3B | 96.662 | 2 | 53.2078 | yes | 1.343E-07 | 491.56989 | 28.501497 |
| GHDAGDHPPITPM | 14 | O95985 | DNA topoisomerase 3-beta-1 OS=Homo : | TOP3B | 96.662 | 2 | 53.2078 | yes | 5.11E-07 | 736.8512 | 28.501497 |
| CNVVAHCFENAHI | 13 | O95985 | DNA topoisomerase 3-beta-1 OS=Homo : | TOP3B | 96.662 | 2 | 53.2078 | yes | 9.722E-08 | 807.35663 | 35.187065 |
| KGHDAGDHPPITP | 15 | O95985 | DNA topoisomerase 3-beta-1 OS=Homo : | TOP3B | 96.662 | 2 | 53.2078 | yes | 1.747E-07 | 800.89868 | 34.34631 |
| GTPIVDGLAPEDM | 15 | O95995 | Dynein regulatory complex subunit 4 OS= | GAS8 | 56.356 | 2 | 53.559223 | yes | 9.722E-08 | 765.37946 | 146.77211 |
| DLQYELAQVCK | 11 | O95995 | Dynein regulatory complex subunit 4 OS= | GAS8 | 56.356 | 2 | 53.559223 | yes | 3.619E-07 | 683.83722 | 62.343506 |
| KFTAAIQEVQQK | 12 | O95995 | Dynein regulatory complex subunit 4 OS= | GAS8 | 56.356 | 2 | 53.559223 | yes | 6.064E-07 | 695.88806 | 30.681219 |
| DLQWEHEVLEQR | 12 | O95995 | Dynein regulatory complex subunit 4 OS= | GAS8 | 56.356 | 2 | 53.559223 | yes | 6.295E-07 | 791.38641 | 23.029737 |
| SSVPASDDAYPEII | 15 | O95997 | Securin OS=Homo sapiens OX=9606 GN | PTTG1 | 22.025 | 2 | 46.423862 | no | 1.562E-07 | 804.37543 | 129.80176 |
| LFQLGPPSPVK | 11 | O95997 | Securin OS=Homo sapiens OX=9606 GN | PTTG1 | 22.025 | 2 | 46.423862 | no | 1.062E-06 | 591.84772 | 81.03643 |
| TFDAPPALPK | 10 | O95997 | Securin OS=Homo sapiens OX=9606 GN | PTTG1 | 22.025 | 2 | 46.423862 | yes | 4.621E-05 | 528.78986 | 48.466129 |
| TENTIFSSTTLPR | 13 | O95999 | B-cell lymphoma/leukemia 10 OS=Homo | BCL10 | 26.252 | 2 | 50.650425 | yes | 9.732E-08 | 733.87793 | 159.33429 |
| LLDYLQENPK | 10 | O95999 | B-cell lymphoma/leukemia 10 OS=Homo | BCL10 | 26.252 | 2 | 50.650425 | yes | 2.776E-07 | 616.82971 | 74.029465 |
| GLDTLVESIR | 10 | O95999 | B-cell lymphoma/leukemia 10 OS=Homo | BCL10 | 26.252 | 2 | 50.650425 | yes | 1.385E-07 | 551.80878 | 49.511803 |
| VDQEIINIMQDR | 12 | O96000 | NADH dehydrogenase [ubiquinone] 1 bet | NDUFB10 | 20.777 | 2 | 48.465626 | yes | 1.144E-07 | 737.37195 | 323.5394 |
| EVEQFTQVAK | 10 | O96000 | NADH dehydrogenase [ubiquinone] 1 bet | NDUFB10 | 20.777 | 2 | 48.465626 | yes | 2.202E-06 | 589.80621 | 62.231209 |
| EGQNYQQNCIK | 11 | O96000 | NADH dehydrogenase [ubiquinone] 1 bet | NDUFB10 | 20.777 | 2 | 48.465626 | yes | 1.823E-07 | 691.31189 | 19.895164 |
| GSVDSNWIVGATI | 15 | O96008 | Mitochondrial import receptor subunit TC | TOMM40 | 37.894 | 2 | 54.970596 | yes | 9.722E-08 | 788.4043 | 129.45618 |
| ELFPIQMEGVK | 11 | O96008 | Mitochondrial import receptor subunit TC | TOMM40 | 37.894 | 2 | 54.970596 | yes | 2.422E-06 | 645.84174 | 79.89888 |
| FVNWQVDGEYR | 11 | O96008 | Mitochondrial import receptor subunit TC | TOMM40 | 37.894 | 2 | 54.970596 | yes | 2.16E-07 | 706.83331 | 38.008656 |
| GHPPLLLDVVR | 11 | O96011 | Peroxisomal membrane protein 11B OS=l | PEX11B | 28.432 | 2 | 41.53554 | yes | 5.974E-07 | 608.36407 | 161.80919 |
| AVHLSDVVLR | 10 | O96011 | Peroxisomal membrane protein 11B OS=l | PEX11B | 28.432 | 2 | 41.53554 | yes | 3.485E-06 | 554.82727 | 57.806751 |

| | | | | | | | | | | | |
|---|---|---|---|---|---|---|---|---|---|---|---|
| LQVLLLAR | 8 | O96011 | Peroxisomal membrane protein 11B OS=1 | PEX11B | 28.432 | 2 | 41.53554 | yes | 5.4E-07 | 463.31329 | 48.083305 |
| ALYFACDNVLWA | 14 | O96011 | Peroxisomal membrane protein 11B OS=1 | PEX11B | 28.432 | 2 | 41.53554 | yes | 3.46E-07 | 814.40051 | 36.323814 |
| IGEGSTGIVCIATV | 15 | O96013 | Serine/threonine-protein kinase PAK 4 O | PAK4 | 64.073 | 2 | 50.106075 | yes | 9.902E-08 | 766.90869 | 65.690964 |
| VDFPTAIGMVVER | 13 | O96019 | Actin-like protein 6A OS=Homo sapiens | ACTL6A | 47.461 | 2 | 51.30797 | yes | 2.527E-07 | 717.37646 | 1167.8394 |
| LIANNTTVER | 10 | O96019 | Actin-like protein 6A OS=Homo sapiens | ACTL6A | 47.461 | 2 | 51.30797 | yes | 1.127E-07 | 565.81183 | 1104.025 |
| DDGSTLMEIDGDK | 13 | O96019 | Actin-like protein 6A OS=Homo sapiens | ACTL6A | 47.461 | 2 | 51.30797 | yes | 9.723E-08 | 698.30084 | 887.80725 |
| SPLAGDFITMQCR | 13 | O96019 | Actin-like protein 6A OS=Homo sapiens | ACTL6A | 47.461 | 2 | 51.30797 | yes | 9.819E-08 | 748.35522 | 702.8808 |
| IPEGLFDPSNVK | 12 | O96019 | Actin-like protein 6A OS=Homo sapiens | ACTL6A | 47.461 | 2 | 51.30797 | no | 9.722E-08 | 658.34827 | 544.49554 |
| QGGPTYYIDTNAL | 14 | O96019 | Actin-like protein 6A OS=Homo sapiens | ACTL6A | 47.461 | 2 | 51.30797 | yes | 2.48E-07 | 784.88879 | 375.12146 |
| STGLILDSGATHTT | 30 | O96019 | Actin-like protein 6A OS=Homo sapiens | ACTL6A | 47.461 | 4 | 51.30797 | yes | 1.235E-07 | 773.66602 | 295.50534 |
| AGYAGEDCPK | 10 | O96019 | Actin-like protein 6A OS=Homo sapiens | ACTL6A | 47.461 | 2 | 51.30797 | no | 2.287E-06 | 534.22675 | 244.42038 |
| ENMEAISPLK | 10 | O96019 | Actin-like protein 6A OS=Homo sapiens | ACTL6A | 47.461 | 2 | 51.30797 | yes | 3.771E-07 | 566.28937 | 200.64909 |
| NGMVEDWDSFQA | 19 | O96019 | Actin-like protein 6A OS=Homo sapiens | ACTL6A | 47.461 | 3 | 51.30797 | yes | 3.801E-07 | 757.01251 | 66.187241 |
| DDGSTLMEIDGDK | 15 | O96019 | Actin-like protein 6A OS=Homo sapiens | ACTL6A | 47.461 | 2 | 51.30797 | yes | 2.393E-07 | 790.85907 | 41.404037 |
| ETFYLAQDFFDR | 12 | O96020 | G1/S-specific cyclin-E2 OS=Homo sapie | CCNE2 | 46.758 | 2 | 41.25651 | yes | 3.6E-07 | 776.35938 | 61.69743 |
| SLVAFEGEGQFEK | 13 | O96028 | Histone-lysine N-methyltransferase NSD2 | NSD2 | 152.259 | 2 | 57.30725 | yes | 9.722E-08 | 720.85388 | 202.26173 |
| TPSCEVNR | 8 | O96028 | Histone-lysine N-methyltransferase NSD2 | NSD2 | 152.259 | 2 | 57.30725 | yes | 1.362E-07 | 481.72165 | 155.80127 |
| SQWSLLSEK | 9 | O96028 | Histone-lysine N-methyltransferase NSD2 | NSD2 | 152.259 | 2 | 57.30725 | yes | 9.531E-07 | 539.28241 | 126.30727 |
| ASTAASSALGFSK | 13 | O96028 | Histone-lysine N-methyltransferase NSD2 | NSD2 | 152.259 | 2 | 57.30725 | yes | 2.542E-07 | 599.30914 | 119.77412 |
| AQLSSSLQEGVMQ | 14 | O96028 | Histone-lysine N-methyltransferase NSD2 | NSD2 | 152.259 | 2 | 57.30725 | yes | 9.722E-08 | 753.38507 | 105.09753 |
| FTFLYVGDQLHLN | 18 | O96028 | Histone-lysine N-methyltransferase NSD2 | NSD2 | 152.259 | 3 | 57.30725 | yes | 1.19E-07 | 697.37384 | 85.76683 |
| LCSSAETLESHPDI | 16 | O96028 | Histone-lysine N-methyltransferase NSD2 | NSD2 | 152.259 | 3 | 57.30725 | yes | 1.216E-07 | 581.94543 | 56.857719 |
| NLSDACKPLK | 10 | O96028 | Histone-lysine N-methyltransferase NSD2 | NSD2 | 152.259 | 2 | 57.30725 | yes | 1.006E-06 | 573.3028 | 51.809517 |
| VIGSGCNLDSAR | 12 | P00338 | L-lactate dehydrogenase A chain OS=Hor | LDHA | 36.689 | 2 | 49.8646 | no | 3.417E-07 | 624.80371 | 192.48895 |
| VTLTSEEEAR | 10 | P00338 | L-lactate dehydrogenase A chain OS=Hor | LDHA | 36.689 | 2 | 49.8646 | yes | 1.001E-07 | 567.78552 | 112.4026 |
| SADTLWGIQK | 10 | P00338 | L-lactate dehydrogenase A chain OS=Hor | LDHA | 36.689 | 2 | 49.8646 | yes | 2.505E-07 | 559.79565 | 108.38259 |
| DQLIYNLLK | 9 | P00338 | L-lactate dehydrogenase A chain OS=Hor | LDHA | 36.689 | 2 | 49.8646 | yes | 5.149E-07 | 560.32404 | 48.760632 |
| TLHPDLGTDKDK | 12 | P00338 | L-lactate dehydrogenase A chain OS=Hor | LDHA | 36.689 | 2 | 49.8646 | yes | 1.017E-07 | 670.34625 | 29.469618 |
| QVVESAYEVIK | 11 | P00338 | L-lactate dehydrogenase A chain OS=Hor | LDHA | 36.689 | 2 | 49.8646 | yes | 2.894E-06 | 632.84283 | 9.9454288 |
| YNLGLDLR | 8 | P00367 | Glutamate dehydrogenase 1, mitochondria | GLUD1 | 61.398 | 2 | 50.329308 | yes | 6.367E-06 | 482.26654 | 152.67665 |
| ALASLMTYK | 9 | P00367 | Glutamate dehydrogenase 1, mitochondria | GLUD1 | 61.398 | 2 | 50.329308 | yes | 9.819E-08 | 499.27298 | 91.224121 |
| YSTDVSVDEVK | 11 | P00367 | Glutamate dehydrogenase 1, mitochondria | GLUD1 | 61.398 | 2 | 50.329308 | yes | 1.39E-07 | 621.29846 | 88.814552 |
| DDGSWEVIEGYR | 12 | P00367 | Glutamate dehydrogenase 1, mitochondria | GLUD1 | 61.398 | 2 | 50.329308 | yes | 9.927E-08 | 713.31769 | 64.274841 |
| MVEGFFDR | 8 | P00367 | Glutamate dehydrogenase 1, mitochondria | GLUD1 | 61.398 | 2 | 50.329308 | yes | 7.089E-06 | 500.73148 | 24.368996 |
| ALLTPVAIAAGR | 12 | P00390 | Glutathione reductase, mitochondrial OS= | GSR | 56.258 | 2 | 41.660854 | yes | 3.401E-07 | 576.85858 | 108.74495 |
| VVLPIEAPIR | 10 | P00403 | Cytochrome c oxidase subunit 2 OS=Hon | MT-CO2 | 25.566 | 2 | 47.25096 | yes | 1.369E-07 | 553.85022 | 385.00342 |
| DIFTGLIGPMK | 11 | P00450 | Ceruloplasmin OS=Homo sapiens OX=96 | CP | 122.206 | 2 | 51.31302 | yes | 1.076E-07 | 596.32574 | 715.83533 |
| DIFTGLIGPMK | 11 | P00450 | Ceruloplasmin OS=Homo sapiens OX=96 | CP | 122.206 | 2 | 51.31302 | yes | 2.307E-07 | 604.32318 | 715.83533 |
| IYHSHIDAPK | 10 | P00450 | Ceruloplasmin OS=Homo sapiens OX=96 | CP | 122.206 | 2 | 51.31302 | yes | 1.969E-07 | 590.80908 | 927.64606 |

| | | | | | | | | | | | |
|---|---|---|---|---|---|---|---|---|---|---|---|
| MYYSAVDPTK | 10 | P00450 | Ceruloplasmin OS=Homo sapiens OX=9( | CP | 122.206 | 2 | 51.31302 | yes | 1.104E-07 | 587.77606 | 874.37134 |
| DNEDFQESNR | 10 | P00450 | Ceruloplasmin OS=Homo sapiens OX=9( | CP | 122.206 | 2 | 51.31302 | yes | 9.722E-08 | 627.25507 | 463.81958 |
| TYCSEPEK | 8 | P00450 | Ceruloplasmin OS=Homo sapiens OX=9( | CP | 122.206 | 2 | 51.31302 | yes | 9.722E-08 | 507.21585 | 320.97269 |
| MFTTAPDQVDK | 11 | P00450 | Ceruloplasmin OS=Homo sapiens OX=9( | CP | 122.206 | 2 | 51.31302 | yes | 6.699E-07 | 634.79498 | 95.0616 |
| MFTTAPDQVDK | 11 | P00450 | Ceruloplasmin OS=Homo sapiens OX=9( | CP | 122.206 | 2 | 51.31302 | yes | 2.187E-07 | 626.79755 | 95.0616 |
| GTYIPVPIVSELQS( | 16 | P00488 | Coagulation factor XIII A chain OS=Hon | F13A1 | 83.269 | 2 | 45.872528 | yes | 1.717E-07 | 844.46692 | 149.52461 |
| VIGGDDLSTLTGK | 13 | P00492 | Hypoxanthine-guanine phosphoribosyltra | HPRT1 | 24.58 | 2 | 50.973934 | yes | 3.072E-06 | 638.3432 | 443.89951 |
| NVLIVEDIIDTGK | 13 | P00492 | Hypoxanthine-guanine phosphoribosyltra | HPRT1 | 24.58 | 2 | 50.973934 | yes | 9.722E-08 | 714.90088 | 81.900665 |
| EMGGHHIVALCVI | 14 | P00492 | Hypoxanthine-guanine phosphoribosyltra | HPRT1 | 24.58 | 3 | 50.973934 | yes | 1.721E-07 | 521.94263 | 35.94952 |
| NYVVTDHGSCVR | 12 | P00533 | Epidermal growth factor receptor OS=Ho | EGFR | 134.278 | 2 | 50.294453 | yes | 9.819E-08 | 703.8277 | 13.374226 |
| IQLINNMLDK | 10 | P00558 | Phosphoglycerate kinase 1 OS=Homo sap | PGK1 | 44.615 | 2 | 52.884624 | yes | 9.722E-08 | 601.33411 | 75.74086 |
| SGIECQLWR | 9 | P00734 | Prothrombin OS=Homo sapiens OX=960 | F2 | 70.037 | 2 | 50.968105 | yes | 9.722E-08 | 574.77948 | 903.76764 |
| SPQELLCGASLISD | 15 | P00734 | Prothrombin OS=Homo sapiens OX=960 | F2 | 70.037 | 2 | 50.968105 | yes | 1.215E-07 | 823.41431 | 106.57318 |
| YGFYTHVFR | 9 | P00734 | Prothrombin OS=Homo sapiens OX=960 | F2 | 70.037 | 2 | 50.968105 | yes | 1.585E-06 | 595.29309 | 79.894524 |
| GDACEGDSGGPFV | 15 | P00734 | Prothrombin OS=Homo sapiens OX=960 | F2 | 70.037 | 2 | 50.968105 | yes | 1.51E-06 | 763.81616 | 46.946682 |
| VGYVSGWGR | 9 | P00738 | Haptoglobin OS=Homo sapiens OX=960( | HP | 45.206 | 2 | 48.77938 | yes | 2.197E-07 | 490.75107 | 1653.92 |
| LPECEAVCGK | 10 | P00738 | Haptoglobin OS=Homo sapiens OX=960( | HP | 45.206 | 2 | 48.77938 | no | 1.105E-07 | 581.7652 | 467.65997 |
| LPECEAVCGKPK | 12 | P00738 | Haptoglobin OS=Homo sapiens OX=960( | HP | 45.206 | 2 | 48.77938 | no | 1.461E-07 | 694.33905 | 435.70267 |
| SCAVAEYGVYVK | 12 | P00738 | Haptoglobin OS=Homo sapiens OX=960( | HP | 45.206 | 2 | 48.77938 | no | 1.335E-07 | 673.32648 | 105.08411 |
| EAQLPVIENK | 10 | P00747 | Plasminogen OS=Homo sapiens OX=960 | PLG | 90.569 | 2 | 50.990627 | yes | 1.544E-07 | 570.81659 | 2844.887 |
| PNKPGVYVR | 9 | P00747 | Plasminogen OS=Homo sapiens OX=960 | PLG | 90.569 | 2 | 50.990627 | yes | 2.663E-07 | 515.29565 | 1407.2385 |
| YILQGVTSWGLGC | 15 | P00747 | Plasminogen OS=Homo sapiens OX=960 | PLG | 90.569 | 2 | 50.990627 | no | 1.102E-07 | 840.92993 | 810.59363 |
| WEYCNLK | 7 | P00747 | Plasminogen OS=Homo sapiens OX=960 | PLG | 90.569 | 2 | 50.990627 | yes | 2.637E-07 | 506.73148 | 459.90903 |
| TPENFPCK | 8 | P00747 | Plasminogen OS=Homo sapiens OX=960 | PLG | 90.569 | 2 | 50.990627 | yes | 3.841E-07 | 496.72894 | 294.24011 |
| DKYILQGVTSWGI | 17 | P00747 | Plasminogen OS=Homo sapiens OX=960 | PLG | 90.569 | 3 | 50.990627 | no | 9.722E-08 | 641.99634 | 27.090343 |
| LITTQQWLIK | 10 | P00846 | ATP synthase subunit a OS=Homo sapier | MT-ATP6 | 24.818 | 2 | 53.35717 | yes | 9.722E-08 | 622.37408 | 134.57587 |
| YLLGTSLAR | 9 | P00966 | Argininosuccinate synthase OS=Homo sa | ASS1 | 46.531 | 2 | 51.08464 | yes | 1.122E-07 | 497.29001 | 139.84627 |
| VFIEDVSR | 8 | P00966 | Argininosuccinate synthase OS=Homo sa | ASS1 | 46.531 | 2 | 51.08464 | yes | 9.722E-08 | 482.75854 | 118.51721 |
| NQAPPGLYTK | 10 | P00966 | Argininosuccinate synthase OS=Homo sa | ASS1 | 46.531 | 2 | 51.08464 | yes | 2.676E-06 | 544.79041 | 91.539978 |
| FELSCYSLAPQIK | 13 | P00966 | Argininosuccinate synthase OS=Homo sa | ASS1 | 46.531 | 2 | 51.08464 | yes | 4.416E-07 | 778.3949 | 88.674156 |
| KVFIEDVSR | 9 | P00966 | Argininosuccinate synthase OS=Homo sa | ASS1 | 46.531 | 2 | 51.08464 | yes | 2.69E-07 | 546.80603 | 88.571976 |
| IDIVENR | 7 | P00966 | Argininosuccinate synthase OS=Homo sa | ASS1 | 46.531 | 2 | 51.08464 | yes | 8.046E-07 | 429.73761 | 88.01709 |
| MPEFYNR | 7 | P00966 | Argininosuccinate synthase OS=Homo sa | ASS1 | 46.531 | 2 | 51.08464 | yes | 9.625E-07 | 478.71838 | 67.75544 |
| APNTPDILEIEFK | 13 | P00966 | Argininosuccinate synthase OS=Homo sa | ASS1 | 46.531 | 2 | 51.08464 | yes | 5.425E-07 | 743.89301 | 60.968353 |
| QHGIPIPVTPK | 11 | P00966 | Argininosuccinate synthase OS=Homo sa | ASS1 | 46.531 | 2 | 51.08464 | yes | 5.787E-07 | 593.85077 | 40.433704 |
| EGEFSTCFTELQR | 13 | P00973 | 2'-5'-oligoadenylate synthase 1 OS=Homc | OAS1 | 46.029 | 2 | 48.885685 | yes | 1.09E-07 | 802.35651 | 61.820007 |
| LPGIVAEGR | 9 | P01008 | Antithrombin-III OS=Homo sapiens OX= | SERPINC1 | 52.603 | 2 | 50.05581 | yes | 2.357E-07 | 456.26907 | 1079.0801 |
| TSDQIHFFFAK | 11 | P01008 | Antithrombin-III OS=Homo sapiens OX= | SERPINC1 | 52.603 | 2 | 50.05581 | yes | 9.919E-08 | 670.83533 | 365.89343 |
| RVWELSK | 7 | P01008 | Antithrombin-III OS=Homo sapiens OX= | SERPINC1 | 52.603 | 2 | 50.05581 | yes | 1.577E-07 | 459.26379 | 262.28381 |

| | | | | | | | | | | | |
|---|---|---|---|---|---|---|---|---|---|---|---|
| DEELSCTVVELK | 12 | P01011 | Alpha-1-antichymotrypsin OS=Homo sap | SERPINA3 | 47.651 | 2 | 44.67303 | yes | 2.11E-07 | 711.34503 | 50.523445 |
| YGAATFTR | 8 | P01023 | Alpha-2-macroglobulin OS=Homo sapien | A2M | 163.291 | 2 | 46.485046 | no | 1.546E-07 | 443.72452 | 396.51437 |
| LPPNVVEESAR | 11 | P01023 | Alpha-2-macroglobulin OS=Homo sapien | A2M | 163.291 | 2 | 46.485046 | yes | 4.16E-07 | 605.82495 | 124.06527 |
| CCEDGMR | 7 | P01024 | Complement C3 OS=Homo sapiens OX=' | C3 | 187.149 | 2 | 53.579216 | yes | 1.165E-07 | 464.15952 | 695.37494 |
| TIYTPGSTVLYR | 12 | P01024 | Complement C3 OS=Homo sapiens OX=' | C3 | 187.149 | 2 | 53.579216 | yes | 9.722E-08 | 685.86938 | 445.7309 |
| ADIGCTPGSGK | 11 | P01024 | Complement C3 OS=Homo sapiens OX=' | C3 | 187.149 | 2 | 53.579216 | yes | 1.102E-07 | 531.74786 | 133.48729 |
| LVAYYTLIGASGQ | 14 | P01024 | Complement C3 OS=Homo sapiens OX=' | C3 | 187.149 | 2 | 53.579216 | yes | 5.03E-07 | 756.41449 | 112.23015 |
| ACEPGVDYVYK | 11 | P01024 | Complement C3 OS=Homo sapiens OX=' | C3 | 187.149 | 2 | 53.579216 | yes | 5.578E-07 | 650.79755 | 98.574112 |
| EVVADSVWVDVK | 12 | P01024 | Complement C3 OS=Homo sapiens OX=' | C3 | 187.149 | 2 | 53.579216 | yes | 9.722E-08 | 673.35352 | 95.531868 |
| LPYSVVR | 7 | P01024 | Complement C3 OS=Homo sapiens OX=' | C3 | 187.149 | 2 | 53.579216 | yes | 1.223E-07 | 417.24762 | 71.05265 |
| RPPGFSPFR | 9 | P01042 | Kininogen-1 OS=Homo sapiens OX=960 | KNG1 | 71.958 | 2 | 46.59948 | yes | 1.518E-07 | 530.78796 | 99.45343 |
| TEPFDDFLFPASSR | 14 | P01100 | Protein c-Fos OS=Homo sapiens OX=96( | FOS | 40.696 | 2 | 40.715393 | yes | 3.88E-07 | 814.88318 | 51.03791 |
| LISEEDLLR | 9 | P01106 | Myc proto-oncogene protein OS=Homo s | MYC | 50.566 | 2 | 50.301617 | yes | 2.538E-07 | 544.30334 | 1118.5863 |
| DQIPELENNEK | 11 | P01106 | Myc proto-oncogene protein OS=Homo s | MYC | 50.566 | 2 | 50.301617 | yes | 9.819E-08 | 664.82007 | 583.73773 |
| THNVLER | 7 | P01106 | Myc proto-oncogene protein OS=Homo s | MYC | 50.566 | 2 | 50.301617 | yes | 1.032E-07 | 434.73541 | 242.59595 |
| ATAYILSVQAEEQ | 14 | P01106 | Myc proto-oncogene protein OS=Homo s | MYC | 50.566 | 2 | 50.301617 | yes | 2.253E-07 | 775.90668 | 163.29823 |
| KFELLPTPPLSPSR | 14 | P01106 | Myc proto-oncogene protein OS=Homo s | MYC | 50.566 | 2 | 50.301617 | no | 1.358E-07 | 791.45361 | 115.73034 |
| LASYQAAR | 8 | P01106 | Myc proto-oncogene protein OS=Homo s | MYC | 50.566 | 2 | 50.301617 | yes | 2.049E-07 | 440.23779 | 101.503 |
| LVVVGAGGVGK | 11 | P01111 | GTPase NRas OS=Homo sapiens OX=96 | NRAS | 21.23 | 2 | 48.636784 | no | 1.818E-07 | 478.30038 | 150.7728 |
| SYGIPFIETSAK | 12 | P01111 | GTPase NRas OS=Homo sapiens OX=96 | NRAS | 21.23 | 2 | 48.636784 | yes | 1.121E-07 | 656.84283 | 126.08788 |
| IYWSDLSQR | 9 | P01130 | Low-density lipoprotein receptor OS=Ho | LDLR | 95.377 | 2 | 53.23331 | yes | 4.25E-07 | 584.29327 | 137.23254 |
| SEYTSLIPNLR | 11 | P01130 | Low-density lipoprotein receptor OS=Ho | LDLR | 95.377 | 2 | 53.23331 | yes | 9.722E-08 | 646.84589 | 112.49105 |
| FTCACPDGMLLAR | 13 | P01130 | Low-density lipoprotein receptor OS=Ho | LDLR | 95.377 | 2 | 53.23331 | yes | 2.47E-06 | 756.34381 | 94.252754 |
| AHGVSSYDTVISR | 13 | P01130 | Low-density lipoprotein receptor OS=Ho | LDLR | 95.377 | 2 | 53.23331 | yes | 3.188E-07 | 696.3493 | 56.649204 |
| GIVEECCFR | 9 | P01344 | Insulin-like growth factor II OS=Homo sa | IGF2 | 20.141 | 2 | 45.706856 | yes | 1.767E-07 | 585.25751 | 1309.2108 |
| FVYHLSDLCK | 10 | P01591 | Immunoglobulin J chain OS=Homo sapie | JCHAIN | 18.099 | 2 | 54.02805 | yes | 9.722E-08 | 641.31842 | 192.97881 |
| EEQFNSTFR | 9 | P01859 | Immunoglobulin heavy constant gamma 2 | IGHG2 | 43.806 | 2 | 49.32374 | yes | 1.044E-07 | 579.26471 | 2575.3381 |
| WAAVVVPSGEEQ | 13 | P01889 | HLA class I histocompatibility antigen, B | HLA-B | 40.461 | 2 | 50.57223 | no | 9.759E-08 | 714.36749 | 156.02086 |
| FISVGYVDDTQFV | 14 | P01889 | HLA class I histocompatibility antigen, B | HLA-B | 40.461 | 2 | 50.57223 | no | 1.42E-07 | 823.41467 | 96.089188 |
| SWTAADTAAQITQ | 14 | P01889 | HLA class I histocompatibility antigen, B | HLA-B | 40.461 | 2 | 50.57223 | no | 2.81E-07 | 760.3786 | 72.820442 |
| YTCHVQHEGLPK | 12 | P01889 | HLA class I histocompatibility antigen, B | HLA-B | 40.461 | 2 | 50.57223 | no | 7.081E-07 | 734.8537 | 9.3565588 |
| LHVDPENFR | 9 | P02042 | Hemoglobin subunit delta OS=Homo sapi | HBD | 16.056 | 2 | 42.066154 | yes | 3.209E-07 | 563.78564 | 852.44843 |
| LLVVYPWTQR | 10 | P02100 | Hemoglobin subunit epsilon OS=Homo sa | HBE1 | 16.203 | 2 | 49.09762 | no | 4.235E-07 | 637.86639 | 2501.0552 |
| LHVDPENFK | 9 | P02100 | Hemoglobin subunit epsilon OS=Homo sa | HBE1 | 16.203 | 2 | 49.09762 | yes | 1.068E-07 | 549.78253 | 352.39197 |
| SLSQQIENIR | 10 | P02452 | Collagen alpha-1(I) chain OS=Homo sapi | COL1A1 | 138.912 | 2 | 54.229694 | yes | 9.722E-08 | 594.32257 | 1795.6404 |
| GQAGVMGFPGPK | 12 | P02452 | Collagen alpha-1(I) chain OS=Homo sapi | COL1A1 | 138.912 | 2 | 54.229694 | yes | 3.345E-07 | 581.28967 | 1445.4012 |
| NSVAYMDQQTGN | 14 | P02452 | Collagen alpha-1(I) chain OS=Homo sapi | COL1A1 | 138.912 | 2 | 54.229694 | yes | 4.357E-07 | 784.87231 | 385.2077 |
| NSVAYMDQQTGN | 14 | P02452 | Collagen alpha-1(I) chain OS=Homo sapi | COL1A1 | 138.912 | 2 | 54.229694 | yes | 1.763E-07 | 792.86975 | 385.2077 |
| ALLLQGSNEIEIR | 13 | P02452 | Collagen alpha-1(I) chain OS=Homo sapi | COL1A1 | 138.912 | 2 | 54.229694 | yes | 4.031E-06 | 728.41193 | 400.57993 |

| | | | | | | | | | | | |
|---|---|---|---|---|---|---|---|---|---|---|---|
| GPPGPMGPPGLAG | 20 | P02452 | Collagen alpha-1(I) chain OS=Homo sapi | COL1A1 | 138.912 | 3 | 54.229694 | yes | 5.406E-07 | 600.96143 | 376.23447 |
| ADDANVVR | 8 | P02452 | Collagen alpha-1(I) chain OS=Homo sapi | COL1A1 | 138.912 | 2 | 54.229694 | yes | 1.506E-07 | 430.21704 | 332.43402 |
| DLEVDTTLK | 9 | P02452 | Collagen alpha-1(I) chain OS=Homo sapi | COL1A1 | 138.912 | 2 | 54.229694 | yes | 2.174E-07 | 517.27423 | 205.86891 |
| DRDLEVDTTLK | 11 | P02452 | Collagen alpha-1(I) chain OS=Homo sapi | COL1A1 | 138.912 | 2 | 54.229694 | yes | 9.722E-08 | 652.83826 | 65.176804 |
| MCHSDWK | 7 | P02452 | Collagen alpha-1(I) chain OS=Homo sapi | COL1A1 | 138.912 | 2 | 54.229694 | yes | 1.05E-07 | 482.19421 | 36.394547 |
| FTYTVLEDGCTK | 12 | P02461 | Collagen alpha-1(III) chain OS=Homo sa | COL3A1 | 138.565 | 2 | 53.30145 | yes | 4.988E-07 | 717.33447 | 2906.8342 |
| LMGSNEGEFK | 10 | P02461 | Collagen alpha-1(III) chain OS=Homo sa | COL3A1 | 138.565 | 2 | 53.30145 | yes | 1.457E-07 | 556.25806 | 2222.1106 |
| SGEYWVDPNQGC | 13 | P02461 | Collagen alpha-1(III) chain OS=Homo sa | COL3A1 | 138.565 | 2 | 53.30145 | yes | 1.018E-07 | 770.33026 | 2212.0405 |
| NSIAYMDQASGNV | 14 | P02461 | Collagen alpha-1(III) chain OS=Homo sa | COL3A1 | 138.565 | 2 | 53.30145 | yes | 9.722E-08 | 749.35376 | 1396.9639 |
| FCHPELK | 7 | P02461 | Collagen alpha-1(III) chain OS=Homo sa | COL3A1 | 138.565 | 2 | 53.30145 | yes | 2.673E-07 | 465.72873 | 894.82373 |
| SVNGQIESLISPDG | 16 | P02461 | Collagen alpha-1(III) chain OS=Homo sa | COL3A1 | 138.565 | 2 | 53.30145 | yes | 1.622E-07 | 829.92084 | 620.06439 |
| ILYHGYSLLYVQG | 16 | P02462 | Collagen alpha-1(IV) chain OS=Homo sa | COL4A1 | 160.611 | 3 | 50.240543 | yes | 9.839E-08 | 642.33539 | 225.49792 |
| AHGQDLGTAGSCI | 14 | P02462 | Collagen alpha-1(IV) chain OS=Homo sa | COL4A1 | 160.611 | 2 | 50.240543 | no | 2.167E-07 | 721.84387 | 64.329842 |
| VTMQNLNDR | 9 | P02533 | Keratin, type I cytoskeletal 14 OS=Homo | KRT14 | 51.562 | 2 | 50.946518 | no | 1.613E-07 | 545.7691 | 955.24548 |
| VTMQNLNDR | 9 | P02533 | Keratin, type I cytoskeletal 14 OS=Homo | KRT14 | 51.562 | 2 | 50.946518 | no | 1.321E-07 | 553.7666 | 955.24548 |
| LAADDFR | 7 | P02533 | Keratin, type I cytoskeletal 14 OS=Homo | KRT14 | 51.562 | 2 | 50.946518 | no | 2.47E-07 | 404.2034 | 1104.7639 |
| VLDELTLAR | 9 | P02533 | Keratin, type I cytoskeletal 14 OS=Homo | KRT14 | 51.562 | 2 | 50.946518 | no | 3.299E-07 | 515.3006 | 439.51904 |
| LEQEIATYR | 9 | P02533 | Keratin, type I cytoskeletal 14 OS=Homo | KRT14 | 51.562 | 2 | 50.946518 | no | 9.722E-08 | 561.79309 | 181.50882 |
| ALEEANADLEVK | 12 | P02533 | Keratin, type I cytoskeletal 14 OS=Homo | KRT14 | 51.562 | 2 | 50.946518 | no | 1.797E-07 | 651.33282 | 114.19952 |
| TKYETELNLR | 10 | P02533 | Keratin, type I cytoskeletal 14 OS=Homo | KRT14 | 51.562 | 2 | 50.946518 | yes | 1.05E-07 | 633.83807 | 74.139534 |
| APSTYGGGLSVSS | 15 | P02533 | Keratin, type I cytoskeletal 14 OS=Homo | KRT14 | 51.562 | 2 | 50.946518 | yes | 3.023E-07 | 713.35205 | 67.705635 |
| ASLENSLEETK | 11 | P02533 | Keratin, type I cytoskeletal 14 OS=Homo | KRT14 | 51.562 | 2 | 50.946518 | no | 2.004E-07 | 610.80389 | 57.609837 |
| EVATNSELVQSGK | 13 | P02533 | Keratin, type I cytoskeletal 14 OS=Homo | KRT14 | 51.562 | 2 | 50.946518 | no | 0.0001185 | 681.349 | 4.9632764 |
| EYQELMNVK | 9 | P02538 | Keratin, type II cytoskeletal 6A OS=Hom | KRT6A | 60.045 | 2 | 55.147278 | no | 1.043E-07 | 577.28156 | 3182.2451 |
| LALDVEIATYR | 11 | P02538 | Keratin, type II cytoskeletal 6A OS=Hom | KRT6A | 60.045 | 2 | 55.147278 | no | 9.722E-08 | 632.35083 | 951.84113 |
| AQYEEIAQR | 9 | P02538 | Keratin, type II cytoskeletal 6A OS=Hom | KRT6A | 60.045 | 2 | 55.147278 | no | 1.011E-07 | 554.27509 | 569.5257 |
| WTLLQEQGTK | 10 | P02538 | Keratin, type II cytoskeletal 6A OS=Hom | KRT6A | 60.045 | 2 | 55.147278 | no | 1.356E-07 | 602.32202 | 354.94977 |
| LLEGEECR | 8 | P02538 | Keratin, type II cytoskeletal 6A OS=Hom | KRT6A | 60.045 | 2 | 55.147278 | no | 1.536E-07 | 503.23712 | 332.8479 |
| NKYEDEINKR | 10 | P02538 | Keratin, type II cytoskeletal 6A OS=Hom | KRT6A | 60.045 | 2 | 55.147278 | no | 1.157E-07 | 654.83075 | 289.97195 |
| ADTLTDEINFLR | 12 | P02538 | Keratin, type II cytoskeletal 6A OS=Hom | KRT6A | 60.045 | 2 | 55.147278 | no | 9.722E-08 | 704.35938 | 157.09344 |
| QLDSIVGER | 9 | P02538 | Keratin, type II cytoskeletal 6A OS=Hom | KRT6A | 60.045 | 2 | 55.147278 | no | 1.065E-07 | 508.77219 | 153.63899 |
| TAAENEFVTLK | 11 | P02538 | Keratin, type II cytoskeletal 6A OS=Hom | KRT6A | 60.045 | 2 | 55.147278 | no | 9.722E-08 | 611.81934 | 115.23479 |
| NKLEGLEDALQK | 12 | P02538 | Keratin, type II cytoskeletal 6A OS=Hom | KRT6A | 60.045 | 2 | 55.147278 | no | 6.458E-07 | 679.36975 | 101.02728 |
| AIGGGLSSVGGGS | 17 | P02538 | Keratin, type II cytoskeletal 6A OS=Hom | KRT6A | 60.045 | 2 | 55.147278 | no | 4.24E-07 | 724.39117 | 101.02475 |
| FASFIDKVR | 9 | P02538 | Keratin, type II cytoskeletal 6A OS=Hom | KRT6A | 60.045 | 2 | 55.147278 | no | 1.324E-07 | 541.80328 | 86.62973 |
| AEAESWYQTK | 10 | P02538 | Keratin, type II cytoskeletal 6A OS=Hom | KRT6A | 60.045 | 2 | 55.147278 | no | 1.194E-07 | 606.78021 | 84.121162 |
| NKYEDEINK | 9 | P02538 | Keratin, type II cytoskeletal 6A OS=Hom | KRT6A | 60.045 | 2 | 55.147278 | no | 1.257E-07 | 576.78021 | 73.390739 |
| QCANLQAAIADAE | 15 | P02538 | Keratin, type II cytoskeletal 6A OS=Hom | KRT6A | 60.045 | 2 | 55.147278 | no | 1.844E-06 | 829.89941 | 47.931831 |
| KLLEGEECR | 9 | P02538 | Keratin, type II cytoskeletal 6A OS=Hom | KRT6A | 60.045 | 2 | 55.147278 | no | 1.483E-07 | 567.28461 | 33.155212 |

| | | | | | | | | | | | |
|---|---|---|---|---|---|---|---|---|---|---|---|
| NLDLDSIIAEVK | 12 | P02538 | Keratin, type II cytoskeletal 6A OS=Hom | KRT6A | 60.045 | 2 | 55.147278 | no | 1.426E-07 | 665.36664 | 25.68791 |
| ISIGGGSCAISGGY( | 17 | P02538 | Keratin, type II cytoskeletal 6A OS=Hom | KRT6A | 60.045 | 2 | 55.147278 | no | 2.506E-05 | 799.88318 | 13.874951 |
| EAALSTALSEK | 11 | P02545 | Prelamin-A/C OS=Homo sapiens OX=96 | LMNA | 74.14 | 2 | 55.27941 | yes | 9.722E-08 | 560.29822 | 1031.3363 |
| TALINSTGEEVAM | 14 | P02545 | Prelamin-A/C OS=Homo sapiens OX=96 | LMNA | 74.14 | 2 | 55.27941 | yes | 1.09E-07 | 746.3772 | 931.29974 |
| LADALQELR | 9 | P02545 | Prelamin-A/C OS=Homo sapiens OX=96 | LMNA | 74.14 | 2 | 55.27941 | yes | 1.346E-07 | 514.79041 | 854.53064 |
| AAYEAELGDAR | 11 | P02545 | Prelamin-A/C OS=Homo sapiens OX=96 | LMNA | 74.14 | 2 | 55.27941 | yes | 9.722E-08 | 583.27783 | 758.37555 |
| ITESEEVVSR | 10 | P02545 | Prelamin-A/C OS=Homo sapiens OX=96 | LMNA | 74.14 | 2 | 55.27941 | yes | 1.574E-07 | 574.79333 | 564.60364 |
| EGDLIAAQAR | 10 | P02545 | Prelamin-A/C OS=Homo sapiens OX=96 | LMNA | 74.14 | 2 | 55.27941 | yes | 3.845E-07 | 522.27765 | 498.1445 |
| TLEGELHDLR | 10 | P02545 | Prelamin-A/C OS=Homo sapiens OX=96 | LMNA | 74.14 | 2 | 55.27941 | yes | 1.265E-06 | 591.80933 | 485.65802 |
| IDSLSAQLSQLQK | 13 | P02545 | Prelamin-A/C OS=Homo sapiens OX=96 | LMNA | 74.14 | 2 | 55.27941 | yes | 1.022E-07 | 715.89612 | 472.7627 |
| SNEDQSMGNWQII | 13 | P02545 | Prelamin-A/C OS=Homo sapiens OX=96 | LMNA | 74.14 | 2 | 55.27941 | yes | 1.206E-07 | 768.841 | 464.40796 |
| EDLQELNDR | 9 | P02545 | Prelamin-A/C OS=Homo sapiens OX=96 | LMNA | 74.14 | 2 | 55.27941 | yes | 9.722E-08 | 566.26746 | 443.49667 |
| LLEGEEER | 8 | P02545 | Prelamin-A/C OS=Homo sapiens OX=96 | LMNA | 74.14 | 2 | 55.27941 | no | 1.342E-07 | 487.7431 | 394.94745 |
| SGAQASSTPLSPTF | 14 | P02545 | Prelamin-A/C OS=Homo sapiens OX=96 | LMNA | 74.14 | 2 | 55.27941 | yes | 1.595E-07 | 680.3468 | 366.71722 |
| QLQDEMLR | 8 | P02545 | Prelamin-A/C OS=Homo sapiens OX=96 | LMNA | 74.14 | 2 | 55.27941 | yes | 9.722E-08 | 516.76074 | 354.58044 |
| SVGGSGGGSFGDN | 17 | P02545 | Prelamin-A/C OS=Homo sapiens OX=96 | LMNA | 74.14 | 2 | 55.27941 | yes | 3.886E-07 | 783.87897 | 284.97562 |
| AQNTWGCGNSLR | 12 | P02545 | Prelamin-A/C OS=Homo sapiens OX=96 | LMNA | 74.14 | 2 | 55.27941 | yes | 9.722E-08 | 682.31219 | 243.88713 |
| TVLCGTCGQPADK | 13 | P02545 | Prelamin-A/C OS=Homo sapiens OX=96 | LMNA | 74.14 | 2 | 55.27941 | yes | 9.722E-08 | 703.82379 | 243.1331 |
| LAVYIDR | 7 | P02545 | Prelamin-A/C OS=Homo sapiens OX=96 | LMNA | 74.14 | 2 | 55.27941 | yes | 2.526E-07 | 425.24509 | 238.2135 |
| LEAALGEAK | 9 | P02545 | Prelamin-A/C OS=Homo sapiens OX=96 | LMNA | 74.14 | 2 | 55.27941 | yes | 3.247E-07 | 451.25311 | 190.74368 |
| EELDFQK | 7 | P02545 | Prelamin-A/C OS=Homo sapiens OX=96 | LMNA | 74.14 | 2 | 55.27941 | yes | 1.846E-07 | 454.72162 | 151.2876 |
| LKDLEALLNSK | 11 | P02545 | Prelamin-A/C OS=Homo sapiens OX=96 | LMNA | 74.14 | 2 | 55.27941 | yes | 9.722E-08 | 622.36646 | 147.61932 |
| KLLEGEEER | 9 | P02545 | Prelamin-A/C OS=Homo sapiens OX=96 | LMNA | 74.14 | 2 | 55.27941 | no | 9.722E-08 | 551.79059 | 134.39578 |
| LQLELSK | 7 | P02545 | Prelamin-A/C OS=Homo sapiens OX=96 | LMNA | 74.14 | 2 | 55.27941 | yes | 1.182E-06 | 415.75275 | 111.96918 |
| KQLQDEMLR | 9 | P02545 | Prelamin-A/C OS=Homo sapiens OX=96 | LMNA | 74.14 | 2 | 55.27941 | yes | 9.722E-08 | 580.80823 | 103.91596 |
| KEGDLIAAQAR | 11 | P02545 | Prelamin-A/C OS=Homo sapiens OX=96 | LMNA | 74.14 | 2 | 55.27941 | yes | 1.387E-07 | 586.32513 | 102.4147 |
| QNGDDPLLTYR | 11 | P02545 | Prelamin-A/C OS=Homo sapiens OX=96 | LMNA | 74.14 | 2 | 55.27941 | yes | 9.926E-08 | 646.3175 | 92.365211 |
| SLETENAGLR | 10 | P02545 | Prelamin-A/C OS=Homo sapiens OX=96 | LMNA | 74.14 | 2 | 55.27941 | yes | 2.829E-07 | 545.2804 | 89.461311 |
| DLEDSLAR | 8 | P02545 | Prelamin-A/C OS=Homo sapiens OX=96 | LMNA | 74.14 | 2 | 55.27941 | yes | 1.897E-07 | 459.72998 | 77.612343 |
| DLEALLNSK | 9 | P02545 | Prelamin-A/C OS=Homo sapiens OX=96 | LMNA | 74.14 | 2 | 55.27941 | yes | 1.058E-07 | 501.77695 | 61.346493 |
| AQHEDQVEQYK | 11 | P02545 | Prelamin-A/C OS=Homo sapiens OX=96 | LMNA | 74.14 | 2 | 55.27941 | yes | 3.344E-07 | 687.81787 | 37.976894 |
| VAVEEVDEEGK | 11 | P02545 | Prelamin-A/C OS=Homo sapiens OX=96 | LMNA | 74.14 | 2 | 55.27941 | yes | 1.955E-07 | 602.29059 | 24.690619 |
| DALSSVQESQVAQ | 16 | P02656 | Apolipoprotein C-III OS=Homo sapiens ( | APOC3 | 10.853 | 3 | 45.597393 | yes | 2.199E-07 | 572.9552 | 61.846821 |
| TAKDALSSVQESQ | 19 | P02656 | Apolipoprotein C-III OS=Homo sapiens ( | APOC3 | 10.853 | 3 | 45.597393 | yes | 1.801E-07 | 673.01514 | 33.485924 |
| RLEVDIDIK | 9 | P02671 | Fibrinogen alpha chain OS=Homo sapien | FGA | 94.974 | 2 | 52.414295 | yes | 7.701E-07 | 550.81915 | 3867.5815 |
| LEVDIDIK | 8 | P02671 | Fibrinogen alpha chain OS=Homo sapien | FGA | 94.974 | 2 | 52.414295 | yes | 1.106E-07 | 472.76859 | 219.5809 |
| RLEVDIDIKIR | 11 | P02671 | Fibrinogen alpha chain OS=Homo sapien | FGA | 94.974 | 3 | 52.414295 | yes | 9.722E-08 | 457.27692 | 145.40353 |
| IRPFFPQQ | 8 | P02675 | Fibrinogen beta chain OS=Homo sapiens | FGB | 55.929 | 2 | 50.981125 | yes | 1.562E-07 | 516.78491 | 6693.8413 |
| ECEEIIR | 7 | P02675 | Fibrinogen beta chain OS=Homo sapiens | FGB | 55.929 | 2 | 50.981125 | yes | 3.591E-07 | 474.72638 | 6577.585 |

| | | | | | | | | | | | |
|---|---|---|---|---|---|---|---|---|---|---|---|
| TPCTVSCNIPVVSC | 15 | P02675 | Fibrinogen beta chain OS=Homo sapiens | FGB | 55.929 | 2 | 50.981125 | yes | 9.722E-08 | 809.90002 | 6342.979 |
| DNDGWLTSDPR | 11 | P02675 | Fibrinogen beta chain OS=Homo sapiens | FGB | 55.929 | 2 | 50.981125 | yes | 2.839E-06 | 638.28363 | 5942.209 |
| QDGSVDFGR | 9 | P02675 | Fibrinogen beta chain OS=Homo sapiens | FGB | 55.929 | 2 | 50.981125 | yes | 1.353E-07 | 490.72525 | 3422.4636 |
| EDGGGWWYNR | 10 | P02675 | Fibrinogen beta chain OS=Homo sapiens | FGB | 55.929 | 2 | 50.981125 | yes | 1.048E-07 | 620.26251 | 2204.1182 |
| HGTDDGVVWMNV | 13 | P02675 | Fibrinogen beta chain OS=Homo sapiens | FGB | 55.929 | 2 | 50.981125 | yes | 1.128E-07 | 772.8512 | 719.3573 |
| HGTDDGVVWMNV | 13 | P02675 | Fibrinogen beta chain OS=Homo sapiens | FGB | 55.929 | 2 | 50.981125 | yes | 5.28E-06 | 515.56989 | 719.3573 |
| HGTDDGVVWMNV | 13 | P02675 | Fibrinogen beta chain OS=Homo sapiens | FGB | 55.929 | 2 | 50.981125 | yes | 2.645E-07 | 780.84863 | 719.3573 |
| HGTDDGVVWMNV | 13 | P02675 | Fibrinogen beta chain OS=Homo sapiens | FGB | 55.929 | 3 | 50.981125 | yes | 1.128E-07 | 772.8512 | 719.3573 |
| HGTDDGVVWMNV | 13 | P02675 | Fibrinogen beta chain OS=Homo sapiens | FGB | 55.929 | 3 | 50.981125 | yes | 5.28E-06 | 515.56989 | 719.3573 |
| HGTDDGVVWMNV | 13 | P02675 | Fibrinogen beta chain OS=Homo sapiens | FGB | 55.929 | 3 | 50.981125 | yes | 2.645E-07 | 780.84863 | 719.3573 |
| SILENLR | 7 | P02675 | Fibrinogen beta chain OS=Homo sapiens | FGB | 55.929 | 2 | 50.981125 | yes | 4.352E-07 | 422.74799 | 106.39928 |
| ECEEIIRK | 8 | P02675 | Fibrinogen beta chain OS=Homo sapiens | FGB | 55.929 | 2 | 50.981125 | yes | 3.018E-07 | 538.77386 | 36.120811 |
| GSWYSMR | 7 | P02675 | Fibrinogen beta chain OS=Homo sapiens | FGB | 55.929 | 2 | 50.981125 | yes | 5.324E-07 | 451.69489 | 33.921276 |
| YLQEIYNSNNQK | 12 | P02679 | Fibrinogen gamma chain OS=Homo sapie | FGG | 51.512 | 2 | 52.16527 | yes | 9.722E-08 | 757.36774 | 5173.0093 |
| VAQLEAQCQEPCK | 13 | P02679 | Fibrinogen gamma chain OS=Homo sapie | FGG | 51.512 | 2 | 52.16527 | yes | 1.894E-07 | 780.8609 | 2263.4368 |
| DNCCILDER | 9 | P02679 | Fibrinogen gamma chain OS=Homo sapie | FGG | 51.512 | 2 | 52.16527 | yes | 1.048E-07 | 597.74756 | 955.96881 |
| KNWIQYK | 7 | P02679 | Fibrinogen gamma chain OS=Homo sapie | FGG | 51.512 | 2 | 52.16527 | yes | 9.819E-08 | 490.27164 | 304.9101 |
| WYSMKKTTMK | 10 | P02679 | Fibrinogen gamma chain OS=Homo sapie | FGG | 51.512 | 2 | 52.16527 | yes | 1.007E-05 | 660.32794 | 183.16891 |
| EHSSLAFWK | 9 | P02749 | Beta-2-glycoprotein 1 OS=Homo sapiens | APOH | 38.299 | 2 | 45.194485 | yes | 1.929E-07 | 552.77728 | 471.22534 |
| YEKPGSPPR | 9 | P02751 | Fibronectin OS=Homo sapiens OX=9606 | FN1 | 272.321 | 2 | 52.674675 | yes | 9.733E-08 | 515.76947 | 1962.7953 |
| TFYSCTTEGR | 10 | P02751 | Fibronectin OS=Homo sapiens OX=9606 | FN1 | 272.321 | 2 | 52.674675 | yes | 1.202E-07 | 611.26385 | 1263.3995 |
| FTNIGPDTMR | 10 | P02751 | Fibronectin OS=Homo sapiens OX=9606 | FN1 | 272.321 | 2 | 52.674675 | yes | 9.926E-08 | 576.2793 | 1052.3357 |
| TYHVGEQWQK | 10 | P02751 | Fibronectin OS=Homo sapiens OX=9606 | FN1 | 272.321 | 2 | 52.674675 | yes | 9.819E-08 | 638.30945 | 928.6651 |
| PQAPITGYR | 9 | P02751 | Fibronectin OS=Homo sapiens OX=9606 | FN1 | 272.321 | 2 | 52.674675 | yes | 1.112E-07 | 501.772 | 736.68793 |
| DSMIWDCTCIGAG | 14 | P02751 | Fibronectin OS=Homo sapiens OX=9606 | FN1 | 272.321 | 2 | 52.674675 | yes | 1.113E-07 | 821.34454 | 321.29285 |
| DSMIWDCTCIGAG | 14 | P02751 | Fibronectin OS=Homo sapiens OX=9606 | FN1 | 272.321 | 2 | 52.674675 | yes | 9.723E-08 | 829.34198 | 321.29285 |
| SYTITGLQPGTDYI | 14 | P02751 | Fibronectin OS=Homo sapiens OX=9606 | FN1 | 272.321 | 2 | 52.674675 | yes | 9.722E-08 | 772.38556 | 577.66669 |
| HYQINQQWER | 10 | P02751 | Fibronectin OS=Homo sapiens OX=9606 | FN1 | 272.321 | 2 | 52.674675 | yes | 9.722E-08 | 701.33655 | 525.25293 |
| YQCYCYGR | 8 | P02751 | Fibronectin OS=Homo sapiens OX=9606 | FN1 | 272.321 | 2 | 52.674675 | yes | 1.197E-07 | 585.22876 | 500.35718 |
| YEVSVYALK | 9 | P02751 | Fibronectin OS=Homo sapiens OX=9606 | FN1 | 272.321 | 2 | 52.674675 | yes | 2.489E-07 | 536.28967 | 481.50159 |
| VTDATETTITISWR | 14 | P02751 | Fibronectin OS=Homo sapiens OX=9606 | FN1 | 272.321 | 2 | 52.674675 | yes | 2.689E-07 | 797.40961 | 375.78018 |
| EATIPGHLNSYTIK | 14 | P02751 | Fibronectin OS=Homo sapiens OX=9606 | FN1 | 272.321 | 2 | 52.674675 | yes | 2.034E-07 | 772.40936 | 184.35547 |
| EATIPGHLNSYTIK | 14 | P02751 | Fibronectin OS=Homo sapiens OX=9606 | FN1 | 272.321 | 2 | 52.674675 | yes | 2.898E-07 | 515.27533 | 184.35547 |
| EATIPGHLNSYTIK | 14 | P02751 | Fibronectin OS=Homo sapiens OX=9606 | FN1 | 272.321 | 3 | 52.674675 | yes | 2.034E-07 | 772.40936 | 184.35547 |
| EATIPGHLNSYTIK | 14 | P02751 | Fibronectin OS=Homo sapiens OX=9606 | FN1 | 272.321 | 3 | 52.674675 | yes | 2.898E-07 | 515.27533 | 184.35547 |
| GATYNVIVEALK | 12 | P02751 | Fibronectin OS=Homo sapiens OX=9606 | FN1 | 272.321 | 2 | 52.674675 | yes | 2.841E-07 | 639.35864 | 321.65906 |
| GEWTCKPIAEK | 11 | P02751 | Fibronectin OS=Homo sapiens OX=9606 | FN1 | 272.321 | 2 | 52.674675 | yes | 1.048E-07 | 659.82666 | 305.79688 |
| VGDTYERPK | 9 | P02751 | Fibronectin OS=Homo sapiens OX=9606 | FN1 | 272.321 | 2 | 52.674675 | yes | 4.651E-07 | 532.77222 | 272.10394 |
| VDVIPVNLPGEHG | 15 | P02751 | Fibronectin OS=Homo sapiens OX=9606 | FN1 | 272.321 | 2 | 52.674675 | yes | 1.632E-06 | 815.43903 | 126.93192 |

| | | | | | | | | | | | |
|---|---|---|---|---|---|---|---|---|---|---|---|
| VDVIPVNLPGEHG | 15 | P02751 | Fibronectin OS=Homo sapiens OX=9606 | FN1 | 272.321 | 2 | 52.674675 | yes | 1.882E-07 | 543.96173 | 126.93192 |
| VDVIPVNLPGEHG | 15 | P02751 | Fibronectin OS=Homo sapiens OX=9606 | FN1 | 272.321 | 3 | 52.674675 | yes | 1.632E-06 | 815.43903 | 126.93192 |
| VDVIPVNLPGEHG | 15 | P02751 | Fibronectin OS=Homo sapiens OX=9606 | FN1 | 272.321 | 3 | 52.674675 | yes | 1.882E-07 | 543.96173 | 126.93192 |
| PAQGVVTTLENVS | 16 | P02751 | Fibronectin OS=Homo sapiens OX=9606 | FN1 | 272.321 | 2 | 52.674675 | yes | 9.722E-08 | 832.95197 | 224.99722 |
| STTPDITGYR | 10 | P02751 | Fibronectin OS=Homo sapiens OX=9606 | FN1 | 272.321 | 2 | 52.674675 | yes | 9.722E-08 | 555.7749 | 202.73743 |
| WSRPQAPITGYR | 12 | P02751 | Fibronectin OS=Homo sapiens OX=9606 | FN1 | 272.321 | 3 | 52.674675 | yes | 3.928E-07 | 477.92123 | 156.9653 |
| GLAFTDVDVDSIK | 13 | P02751 | Fibronectin OS=Homo sapiens OX=9606 | FN1 | 272.321 | 2 | 52.674675 | yes | 9.722E-08 | 690.35626 | 137.81023 |
| DLQFVEVTDVK | 11 | P02751 | Fibronectin OS=Homo sapiens OX=9606 | FN1 | 272.321 | 2 | 52.674675 | yes | 1.031E-07 | 646.84027 | 136.03204 |
| LGVRPSQGGEAPR | 13 | P02751 | Fibronectin OS=Homo sapiens OX=9606 | FN1 | 272.321 | 2 | 52.674675 | yes | 9.809E-08 | 662.36005 | 115.58226 |
| GNLLQCICTGNGR | 13 | P02751 | Fibronectin OS=Homo sapiens OX=9606 | FN1 | 272.321 | 2 | 52.674675 | yes | 1.9E-06 | 731.84808 | 104.61863 |
| WCGTTQNYDADQ | 13 | P02751 | Fibronectin OS=Homo sapiens OX=9606 | FN1 | 272.321 | 2 | 52.674675 | yes | 9.722E-08 | 793.83063 | 103.05232 |
| STATISGLKPGVDY | 24 | P02751 | Fibronectin OS=Homo sapiens OX=9606 | FN1 | 272.321 | 3 | 52.674675 | yes | 1.499E-07 | 824.11108 | 96.575699 |
| IAWESPQGQVSR | 12 | P02751 | Fibronectin OS=Homo sapiens OX=9606 | FN1 | 272.321 | 2 | 52.674675 | yes | 1.114E-07 | 679.34656 | 95.782516 |
| SSPVVIDASTAIDA | 19 | P02751 | Fibronectin OS=Homo sapiens OX=9606 | FN1 | 272.321 | 3 | 52.674675 | yes | 1.673E-06 | 638.33734 | 66.394997 |
| WCHDNGVNYK | 10 | P02751 | Fibronectin OS=Homo sapiens OX=9606 | FN1 | 272.321 | 2 | 52.674675 | yes | 1.622E-07 | 646.77747 | 57.763714 |
| ECLQTCR | 7 | P02760 | Protein AMBP OS=Homo sapiens OX=9( | AMBP | 39 | 2 | 47.649994 | yes | 1.285E-07 | 483.71021 | 1747.7454 |
| YLYEIAR | 7 | P02768 | Albumin OS=Homo sapiens OX=9606 G | ALB | 69.367 | 2 | 53.101555 | yes | 2.009E-07 | 464.25037 | 76108.344 |
| KVPQVSTPTLVEV | 15 | P02768 | Albumin OS=Homo sapiens OX=9606 G | ALB | 69.367 | 2 | 53.101555 | yes | 4.145E-07 | 820.47253 | 26394.34 |
| KVPQVSTPTLVEV | 15 | P02768 | Albumin OS=Homo sapiens OX=9606 G | ALB | 69.367 | 2 | 53.101555 | yes | 1.863E-07 | 547.31744 | 26394.34 |
| KVPQVSTPTLVEV | 15 | P02768 | Albumin OS=Homo sapiens OX=9606 G | ALB | 69.367 | 3 | 53.101555 | yes | 4.145E-07 | 820.47253 | 26394.34 |
| KVPQVSTPTLVEV | 15 | P02768 | Albumin OS=Homo sapiens OX=9606 G | ALB | 69.367 | 3 | 53.101555 | yes | 1.863E-07 | 547.31744 | 26394.34 |
| YICENQDSISSK | 12 | P02768 | Albumin OS=Homo sapiens OX=9606 G | ALB | 69.367 | 2 | 53.101555 | yes | 4.066E-07 | 722.32465 | 42296.629 |
| CCTESLVNR | 9 | P02768 | Albumin OS=Homo sapiens OX=9606 G | ALB | 69.367 | 2 | 53.101555 | yes | 1.845E-07 | 569.75262 | 34352.809 |
| VPQVSTPTLVEVS | 14 | P02768 | Albumin OS=Homo sapiens OX=9606 G | ALB | 69.367 | 2 | 53.101555 | yes | 1.189E-06 | 756.42505 | 4586.7925 |
| CASLQKFGER | 10 | P02768 | Albumin OS=Homo sapiens OX=9606 G | ALB | 69.367 | 2 | 53.101555 | yes | 9.722E-08 | 598.29803 | 3967.5815 |
| AFKAWAVAR | 9 | P02768 | Albumin OS=Homo sapiens OX=9606 G | ALB | 69.367 | 2 | 53.101555 | yes | 1.26E-07 | 510.29291 | 3633.7417 |
| SLHTLFGDK | 9 | P02768 | Albumin OS=Homo sapiens OX=9606 G | ALB | 69.367 | 2 | 53.101555 | yes | 1.897E-07 | 509.27182 | 2522.4045 |
| FQNALLVR | 8 | P02768 | Albumin OS=Homo sapiens OX=9606 G | ALB | 69.367 | 2 | 53.101555 | yes | 1.441E-07 | 480.78491 | 2267.9651 |
| NECFLQHK | 8 | P02768 | Albumin OS=Homo sapiens OX=9606 G | ALB | 69.367 | 2 | 53.101555 | yes | 2.097E-07 | 538.25311 | 724.1806 |
| YLYEIARR | 8 | P02768 | Albumin OS=Homo sapiens OX=9606 G | ALB | 69.367 | 2 | 53.101555 | yes | 1.188E-07 | 542.3009 | 310.29172 |
| VTKCCTESLVNR | 12 | P02768 | Albumin OS=Homo sapiens OX=9606 G | ALB | 69.367 | 2 | 53.101555 | yes | 1.133E-07 | 733.85815 | 283.3895 |
| YICENQDSISSKLK | 14 | P02768 | Albumin OS=Homo sapiens OX=9606 G | ALB | 69.367 | 2 | 53.101555 | yes | 1.256E-07 | 842.91418 | 260.94556 |
| SSGLPNIPVQTISR | 14 | P02786 | Transferrin receptor protein 1 OS=Homo | TFRC | 84.872 | 2 | 54.903893 | yes | 5.56E-07 | 734.90955 | 148.37355 |
| LAQMFSDMVLK | 11 | P02786 | Transferrin receptor protein 1 OS=Homo | TFRC | 84.872 | 2 | 54.903893 | yes | 9.722E-08 | 641.83032 | 133.40591 |
| LDSTDFTGTIK | 11 | P02786 | Transferrin receptor protein 1 OS=Homo | TFRC | 84.872 | 2 | 54.903893 | yes | 9.975E-08 | 599.30353 | 71.275986 |
| AAAEVAGQFVIK | 12 | P02786 | Transferrin receptor protein 1 OS=Homo | TFRC | 84.872 | 2 | 54.903893 | yes | 9.722E-08 | 602.34021 | 68.989876 |
| SGVGTALLLK | 10 | P02786 | Transferrin receptor protein 1 OS=Homo | TFRC | 84.872 | 2 | 54.903893 | yes | 4.081E-07 | 479.80023 | 62.039177 |
| VSASPLLYTLIEK | 13 | P02786 | Transferrin receptor protein 1 OS=Homo | TFRC | 84.872 | 2 | 54.903893 | yes | 9.722E-08 | 717.41614 | 46.255135 |
| DSAQNSVIIVDK | 12 | P02786 | Transferrin receptor protein 1 OS=Homo | TFRC | 84.872 | 2 | 54.903893 | yes | 9.722E-08 | 644.84082 | 38.995205 |

| | | | | | | | | | | | |
|---|---|---|---|---|---|---|---|---|---|---|---|
| ILNIFGVIK | 9 | P02786 | Transferrin receptor protein 1 OS=Homo | TFRC | 84.872 | 2 | 54.903893 | yes | 1.952E-06 | 508.82877 | 26.958366 |
| LFGNMEGDCPSDV | 14 | P02786 | Transferrin receptor protein 1 OS=Homo | TFRC | 84.872 | 2 | 54.903893 | yes | 9.79E-08 | 828.34509 | 26.710976 |
| YYGYTGAFR | 9 | P02788 | Lactotransferrin OS=Homo sapiens OX=9 | LTF | 78.182 | 2 | 48.167458 | yes | 1.188E-07 | 549.25616 | 3947.6409 |
| CGLVPVLAENYK | 12 | P02788 | Lactotransferrin OS=Homo sapiens OX=9 | LTF | 78.182 | 2 | 48.167458 | yes | 1.3E-07 | 681.85791 | 3561.7639 |
| RLWWLDLK | 8 | P02790 | Hemopexin OS=Homo sapiens OX=9606 | HPX | 51.677 | 2 | 33.127293 | yes | 1.317E-06 | 565.32947 | 215.61758 |
| KLNQALLDLHALC | 16 | P02792 | Ferritin light chain OS=Homo sapiens OX | FTL | 20.02 | 3 | 42.499874 | yes | 3E-07 | 574.00031 | 17.392036 |
| MGAPESGLAEYLF | 15 | P02794 | Ferritin heavy chain OS=Homo sapiens O | FTH1 | 21.226 | 2 | 46.908722 | yes | 1.443E-07 | 814.38727 | 108.98978 |
| ELGDHVTNLR | 10 | P02794 | Ferritin heavy chain OS=Homo sapiens O | FTH1 | 21.226 | 2 | 46.908722 | yes | 1.493E-07 | 577.30164 | 97.568771 |
| CAQGCICK | 8 | P02795 | Metallothionein-2 OS=Homo sapiens OX | MT2A | 6.043 | 2 | 34.312706 | yes | 1.034E-06 | 498.70663 | 245.603 |
| NYNKPWEPK | 9 | P03928 | ATP synthase protein 8 OS=Homo sapien | MT-ATP8 | 7.992 | 2 | 49.426723 | yes | 1.032E-07 | 588.29584 | 692.00671 |
| ICSLHSLPPQS | 11 | P03928 | ATP synthase protein 8 OS=Homo sapien | MT-ATP8 | 7.992 | 2 | 49.426723 | yes | 7.664E-07 | 619.81354 | 420.24081 |
| MLNTNYHLPPSPK | 13 | P03928 | ATP synthase protein 8 OS=Homo sapien | MT-ATP8 | 7.992 | 3 | 49.426723 | yes | 2.266E-06 | 504.59402 | 76.207726 |
| MLNTNYHLPPSPK | 13 | P03928 | ATP synthase protein 8 OS=Homo sapien | MT-ATP8 | 7.992 | 3 | 49.426723 | yes | 2.873E-06 | 756.38739 | 76.207726 |
| MLNTNYHLPPSPK | 13 | P03928 | ATP synthase protein 8 OS=Homo sapien | MT-ATP8 | 7.992 | 2 | 49.426723 | yes | 2.266E-06 | 504.59402 | 76.207726 |
| MLNTNYHLPPSPK | 13 | P03928 | ATP synthase protein 8 OS=Homo sapien | MT-ATP8 | 7.992 | 2 | 49.426723 | yes | 2.873E-06 | 756.38739 | 76.207726 |
| EDVYVVGTVLR | 11 | P04003 | C4b-binding protein alpha chain OS=Hor | C4BPA | 67.034 | 2 | 32.353996 | yes | 1.543E-06 | 625.34296 | 32.760994 |
| AVRPGYPK | 8 | P04004 | Vitronectin OS=Homo sapiens OX=9606 | VTN | 54.306 | 2 | 50.43947 | yes | 1.144E-07 | 444.25851 | 270.71359 |
| DVWGIEGPIDAAF | 15 | P04004 | Vitronectin OS=Homo sapiens OX=9606 | VTN | 54.306 | 2 | 50.43947 | yes | 9.809E-08 | 823.91229 | 251.98067 |
| PFDAFTDLK | 9 | P04004 | Vitronectin OS=Homo sapiens OX=9606 | VTN | 54.306 | 2 | 50.43947 | yes | 4.359E-06 | 527.26624 | 36.798782 |
| FSTVAGESGSADT | 15 | P04040 | Catalase OS=Homo sapiens OX=9606 GN | CAT | 59.757 | 2 | 35.58378 | yes | 8.105E-07 | 742.3548 | 116.56654 |
| LAFCDICQK | 9 | P04049 | RAF proto-oncogene serine/threonine-prc | RAF1 | 73.052 | 2 | 46.70714 | yes | 1.495E-07 | 577.77026 | 142.84364 |
| VPTMCVDWSNIR | 12 | P04049 | RAF proto-oncogene serine/threonine-prc | RAF1 | 73.052 | 2 | 46.70714 | yes | 1.49E-07 | 739.34998 | 142.32991 |
| VVDPTPEQFQAFR | 13 | P04049 | RAF proto-oncogene serine/threonine-prc | RAF1 | 73.052 | 2 | 46.70714 | yes | 6.471E-07 | 767.38843 | 116.66845 |
| IGSGSFGTVYK | 11 | P04049 | RAF proto-oncogene serine/threonine-prc | RAF1 | 73.052 | 2 | 46.70714 | no | 2.795E-07 | 558.29022 | 100.73536 |
| GILAADESTGSIAK | 14 | P04075 | Fructose-bisphosphate aldolase A OS=Hc | ALDOA | 39.421 | 2 | 49.922962 | yes | 1.375E-07 | 666.85388 | 1404.8478 |
| PYQYPALTPEQK | 12 | P04075 | Fructose-bisphosphate aldolase A OS=Hc | ALDOA | 39.421 | 2 | 49.922962 | yes | 3.234E-07 | 717.86682 | 134.35204 |
| ELSDIAHR | 8 | P04075 | Fructose-bisphosphate aldolase A OS=Hc | ALDOA | 39.421 | 2 | 49.922962 | yes | 9.976E-08 | 470.74597 | 87.070724 |
| ADDGRPFPQVIK | 12 | P04075 | Fructose-bisphosphate aldolase A OS=Hc | ALDOA | 39.421 | 2 | 49.922962 | yes | 1.028E-07 | 671.85931 | 67.158775 |
| LQSIGTENTEENR | 13 | P04075 | Fructose-bisphosphate aldolase A OS=Hc | ALDOA | 39.421 | 2 | 49.922962 | yes | 2.045E-07 | 745.85773 | 55.118309 |
| SQVVAGTNYFIK | 12 | P04080 | Cystatin-B OS=Homo sapiens OX=9606 | CSTB | 11.14 | 2 | 52.035816 | yes | 9.81E-08 | 663.85626 | 520.16638 |
| VHVGDEDFVHLR | 12 | P04080 | Cystatin-B OS=Homo sapiens OX=9606 | CSTB | 11.14 | 2 | 52.035816 | yes | 9.722E-08 | 711.85986 | 100.54613 |
| AKHDELTYF | 9 | P04080 | Cystatin-B OS=Homo sapiens OX=9606 | CSTB | 11.14 | 2 | 52.035816 | yes | 5.809E-05 | 562.27454 | 38.362442 |
| TEVIPPLIENR | 11 | P04114 | Apolipoprotein B-100 OS=Homo sapiens | APOB | 515.605 | 2 | 42.80753 | yes | 2.865E-07 | 640.86407 | 179.00606 |
| LDNIYSSDK | 9 | P04114 | Apolipoprotein B-100 OS=Homo sapiens | APOB | 515.605 | 2 | 42.80753 | yes | 2.058E-06 | 527.75623 | 122.76432 |
| VSASSPSLAVASQ! | 17 | P04150 | Glucocorticoid receptor OS=Homo sapier | NR3C1 | 85.66 | 2 | 45.607693 | yes | 2.88E-07 | 810.90741 | 75.605049 |
| SSASTAVSAAPTEI | 14 | P04150 | Glucocorticoid receptor OS=Homo sapier | NR3C1 | 85.66 | 2 | 45.607693 | yes | 1.798E-07 | 653.82788 | 44.063194 |
| SQELFDEIR | 9 | P04150 | Glucocorticoid receptor OS=Homo sapier | NR3C1 | 85.66 | 2 | 45.607693 | yes | 2.559E-06 | 568.78278 | 40.39135 |
| VVEQMCITQYER | 12 | P04156 | Major prion protein OS=Homo sapiens O | PRNP | 27.662 | 2 | 52.384003 | yes | 9.722E-08 | 778.36578 | 73.744514 |
| YPGQGSPGGNR | 11 | P04156 | Major prion protein OS=Homo sapiens O | PRNP | 27.662 | 2 | 52.384003 | yes | 1.157E-07 | 545.25726 | 63.308636 |

| | | | | | | | | | |
|---|---|---|---|---|---|---|---|---|---|
| LTAVCMECFR | 10 | P04183 | Thymidine kinase, cytosolic OS=Homo sa | TK1 | 25.469 | 2 | 56.283703 | yes | 9.893E-08 | 643.78815 | 482.10776 |
| GQIQVILGPMFSGI | 14 | P04183 | Thymidine kinase, cytosolic OS=Homo sa | TK1 | 25.469 | 2 | 56.283703 | yes | 9.722E-08 | 737.90796 | 186.2587 |
| NTMEALPACLLR | 12 | P04183 | Thymidine kinase, cytosolic OS=Homo sa | TK1 | 25.469 | 2 | 56.283703 | yes | 3.321E-07 | 694.85486 | 165.58679 |
| SCINLPTVLPGSPS | 15 | P04183 | Thymidine kinase, cytosolic OS=Homo sa | TK1 | 25.469 | 2 | 56.283703 | yes | 9.722E-08 | 806.42419 | 124.21649 |
| RFQIAQYK | 8 | P04183 | Thymidine kinase, cytosolic OS=Homo sa | TK1 | 25.469 | 2 | 56.283703 | yes | 9.733E-08 | 527.29565 | 108.15507 |
| TVIVAALDGTFQR | 13 | P04183 | Thymidine kinase, cytosolic OS=Homo sa | TK1 | 25.469 | 2 | 56.283703 | yes | 2.805E-07 | 695.88806 | 106.98698 |
| LFAPQQILQCSPAN | 14 | P04183 | Thymidine kinase, cytosolic OS=Homo sa | TK1 | 25.469 | 2 | 56.283703 | yes | 3.431E-06 | 793.90338 | 99.457268 |
| YSSSFCTHDR | 10 | P04183 | Thymidine kinase, cytosolic OS=Homo sa | TK1 | 25.469 | 2 | 56.283703 | yes | 9.894E-08 | 630.25909 | 97.953346 |
| ENCPVPGKPGEAV | 16 | P04183 | Thymidine kinase, cytosolic OS=Homo sa | TK1 | 25.469 | 2 | 56.283703 | yes | 1.751E-07 | 826.41467 | 15.720692 |
| FLEQQNQVLQTK | 12 | P04264 | Keratin, type II cytoskeletal 1 OS=Homo | KRT1 | 66.039 | 2 | 54.46101 | no | 1.061E-07 | 738.3963 | 2558.9978 |
| IEISELNR | 8 | P04264 | Keratin, type II cytoskeletal 1 OS=Homo | KRT1 | 66.039 | 2 | 54.46101 | no | 1.512E-07 | 487.26929 | 2494.9238 |
| LNDLEDALQQAK | 12 | P04264 | Keratin, type II cytoskeletal 1 OS=Homo | KRT1 | 66.039 | 2 | 54.46101 | yes | 1.004E-07 | 679.3515 | 1904.441 |
| WELLQQVDTSTR | 12 | P04264 | Keratin, type II cytoskeletal 1 OS=Homo | KRT1 | 66.039 | 2 | 54.46101 | yes | 9.722E-08 | 738.37805 | 1596.4247 |
| TNAENEFVTIK | 11 | P04264 | Keratin, type II cytoskeletal 1 OS=Homo | KRT1 | 66.039 | 2 | 54.46101 | yes | 1.142E-07 | 633.32227 | 1488.7196 |
| TLLEGEESR | 9 | P04264 | Keratin, type II cytoskeletal 1 OS=Homo | KRT1 | 66.039 | 2 | 54.46101 | yes | 2.94E-07 | 517.26166 | 939.30536 |
| SLNNQFASFIDK | 12 | P04264 | Keratin, type II cytoskeletal 1 OS=Homo | KRT1 | 66.039 | 2 | 54.46101 | yes | 2.885E-07 | 692.34882 | 902.77802 |
| AEAESLYQSK | 10 | P04264 | Keratin, type II cytoskeletal 1 OS=Homo | KRT1 | 66.039 | 2 | 54.46101 | yes | 1.041E-07 | 563.27478 | 811.60791 |
| DYQELMNTK | 9 | P04264 | Keratin, type II cytoskeletal 1 OS=Homo | KRT1 | 66.039 | 2 | 54.46101 | yes | 1.063E-07 | 571.26337 | 265.81799 |
| DYQELMNTK | 9 | P04264 | Keratin, type II cytoskeletal 1 OS=Homo | KRT1 | 66.039 | 2 | 54.46101 | yes | 5.641E-07 | 579.2608 | 265.81799 |
| NMQDMVEDYR | 10 | P04264 | Keratin, type II cytoskeletal 1 OS=Homo | KRT1 | 66.039 | 2 | 54.46101 | yes | 3.057E-07 | 650.76843 | 265.50204 |
| NMQDMVEDYR | 10 | P04264 | Keratin, type II cytoskeletal 1 OS=Homo | KRT1 | 66.039 | 2 | 54.46101 | yes | 2.333E-06 | 658.76593 | 265.50204 |
| DVDGAYMTK | 9 | P04264 | Keratin, type II cytoskeletal 1 OS=Homo | KRT1 | 66.039 | 2 | 54.46101 | yes | 5.071E-07 | 500.22623 | 455.89429 |
| SKAEAESLYQSK | 12 | P04264 | Keratin, type II cytoskeletal 1 OS=Homo | KRT1 | 66.039 | 2 | 54.46101 | yes | 9.741E-08 | 670.83826 | 396.94928 |
| YEELQITAGR | 10 | P04264 | Keratin, type II cytoskeletal 1 OS=Homo | KRT1 | 66.039 | 2 | 54.46101 | no | 9.722E-08 | 590.30383 | 228.61789 |
| SGGGFSSGSAGIIN | 17 | P04264 | Keratin, type II cytoskeletal 1 OS=Homo | KRT1 | 66.039 | 2 | 54.46101 | yes | 9.722E-08 | 829.40009 | 196.06497 |
| SLNNQFASFIDKVI | 14 | P04264 | Keratin, type II cytoskeletal 1 OS=Homo | KRT1 | 66.039 | 2 | 54.46101 | yes | 9.722E-08 | 819.93353 | 65.748978 |
| THNLEPYFESFINN | 16 | P04264 | Keratin, type II cytoskeletal 1 OS=Homo | KRT1 | 66.039 | 3 | 54.46101 | yes | 9.722E-08 | 665.33038 | 61.819077 |
| LRSEIDNVK | 9 | P04264 | Keratin, type II cytoskeletal 1 OS=Homo | KRT1 | 66.039 | 2 | 54.46101 | yes | 1.141E-07 | 537.30109 | 60.630177 |
| NSKIEISELNR | 11 | P04264 | Keratin, type II cytoskeletal 1 OS=Homo | KRT1 | 66.039 | 2 | 54.46101 | yes | 1.931E-07 | 651.85425 | 32.563244 |
| TNAENEFVTIKK | 12 | P04264 | Keratin, type II cytoskeletal 1 OS=Homo | KRT1 | 66.039 | 2 | 54.46101 | yes | 1.293E-07 | 697.36975 | 23.511417 |
| GALQNIIPASTGAA | 15 | P04406 | Glyceraldehyde-3-phosphate dehydrogena | GAPDH | 36.054 | 2 | 52.947353 | yes | 2.439E-07 | 706.3988 | 6543.9678 |
| VIISAPSADAPMFV | 21 | P04406 | Glyceraldehyde-3-phosphate dehydrogena | GAPDH | 36.054 | 3 | 52.947353 | yes | 1.058E-07 | 738.37463 | 989.84595 |
| VIISAPSADAPMFV | 21 | P04406 | Glyceraldehyde-3-phosphate dehydrogena | GAPDH | 36.054 | 3 | 52.947353 | yes | 1.532E-06 | 743.70624 | 989.84595 |
| VIISAPSADAPMFV | 21 | P04406 | Glyceraldehyde-3-phosphate dehydrogena | GAPDH | 36.054 | 3 | 52.947353 | yes | 1.107E-06 | 743.70624 | 989.84595 |
| VPTANVSVVDLTC | 14 | P04406 | Glyceraldehyde-3-phosphate dehydrogena | GAPDH | 36.054 | 2 | 52.947353 | yes | 9.997E-08 | 765.90088 | 2838.0571 |
| LVINGNPITIFQER | 14 | P04406 | Glyceraldehyde-3-phosphate dehydrogena | GAPDH | 36.054 | 2 | 52.947353 | yes | 1.456E-07 | 807.4541 | 804.41455 |
| VVDLMAHMASK | 11 | P04406 | Glyceraldehyde-3-phosphate dehydrogena | GAPDH | 36.054 | 2 | 52.947353 | yes | 9.722E-08 | 601.30701 | 710.09595 |
| RVIISAPSADAPMI | 22 | P04406 | Glyceraldehyde-3-phosphate dehydrogena | GAPDH | 36.054 | 3 | 52.947353 | yes | 1.32E-07 | 790.40833 | 213.29086 |
| WGDAGAEYVVES | 21 | P04406 | Glyceraldehyde-3-phosphate dehydrogena | GAPDH | 36.054 | 3 | 52.947353 | yes | 1.121E-07 | 759.6842 | 87.385979 |

| | | | | | | | | | | | |
|---|---|---|---|---|---|---|---|---|---|---|---|
| VGVNGFGR | 8 | P04406 | Glyceraldehyde-3-phosphate dehydrogena | GAPDH | 36.054 | 2 | 52.947353 | yes | 2.18E-06 | 403.21939 | 58.547638 |
| FIAVGYVDDTQFV | 14 | P04439 | HLA class I histocompatibility antigen, A | HLA-A | 40.841 | 2 | 44.553608 | yes | 2.152E-07 | 815.41718 | 739.61365 |
| SMVAVMDSDTTG | 13 | P04632 | Calpain small subunit 1 OS=Homo sapier | CAPNS1 | 28.316 | 2 | 44.60369 | yes | 2.134E-07 | 671.30487 | 152.15987 |
| LFDQAFGLPR | 10 | P04792 | Heat shock protein beta-1 OS=Homo sapi | HSPB1 | 22.783 | 2 | 50.926804 | yes | 1.446E-07 | 582.31403 | 3812.7498 |
| QLSSGVSEIR | 10 | P04792 | Heat shock protein beta-1 OS=Homo sapi | HSPB1 | 22.783 | 2 | 50.926804 | yes | 1.349E-07 | 538.29077 | 1769.693 |
| DWYPHSR | 7 | P04792 | Heat shock protein beta-1 OS=Homo sapi | HSPB1 | 22.783 | 2 | 50.926804 | yes | 3.32E-07 | 480.71976 | 687.63116 |
| QDEHGYISR | 9 | P04792 | Heat shock protein beta-1 OS=Homo sapi | HSPB1 | 22.783 | 2 | 50.926804 | yes | 1.776E-07 | 552.75708 | 534.06799 |
| TKDGVVEITGK | 11 | P04792 | Heat shock protein beta-1 OS=Homo sapi | HSPB1 | 22.783 | 2 | 50.926804 | yes | 9.722E-08 | 573.8219 | 491.10565 |
| AQLGGPEAAK | 10 | P04792 | Heat shock protein beta-1 OS=Homo sapi | HSPB1 | 22.783 | 2 | 50.926804 | yes | 2.721E-07 | 471.25616 | 424.94424 |
| PLPPAAIESPAVAA | 19 | P04792 | Heat shock protein beta-1 OS=Homo sapi | HSPB1 | 22.783 | 3 | 50.926804 | yes | 2.712E-06 | 626.67554 | 56.973518 |
| GPSWDPFR | 8 | P04792 | Heat shock protein beta-1 OS=Homo sapi | HSPB1 | 22.783 | 2 | 50.926804 | yes | 2.406E-07 | 481.22995 | 44.249641 |
| SEDLLDYGPFR | 11 | P04843 | Dolichyl-diphosphooligosaccharide--prot | RPN1 | 68.57 | 2 | 53.006268 | yes | 9.722E-08 | 656.31439 | 890.77411 |
| TILPAAAQDVYYR | 13 | P04843 | Dolichyl-diphosphooligosaccharide--prot | RPN1 | 68.57 | 2 | 53.006268 | yes | 2.249E-07 | 740.89337 | 866.48883 |
| TVDLSSHLAK | 10 | P04843 | Dolichyl-diphosphooligosaccharide--prot | RPN1 | 68.57 | 2 | 53.006268 | yes | 9.722E-08 | 535.79565 | 677.31311 |
| ALTSEIALLQSR | 12 | P04843 | Dolichyl-diphosphooligosaccharide--prot | RPN1 | 68.57 | 2 | 53.006268 | yes | 1.452E-07 | 651.37482 | 628.66144 |
| LPVALDPGAK | 10 | P04843 | Dolichyl-diphosphooligosaccharide--prot | RPN1 | 68.57 | 2 | 53.006268 | yes | 1.9E-07 | 490.79239 | 608.05469 |
| DVPAYSQDTFK | 11 | P04843 | Dolichyl-diphosphooligosaccharide--prot | RPN1 | 68.57 | 2 | 53.006268 | yes | 1.585E-07 | 635.80115 | 428.91626 |
| NIEIDSPYEISR | 12 | P04843 | Dolichyl-diphosphooligosaccharide--prot | RPN1 | 68.57 | 2 | 53.006268 | yes | 9.722E-08 | 718.35681 | 415.63089 |
| VACITEQVLTLVN | 14 | P04843 | Dolichyl-diphosphooligosaccharide--prot | RPN1 | 68.57 | 2 | 53.006268 | yes | 9.722E-08 | 794.44238 | 377.28842 |
| TEGSDLCDR | 9 | P04843 | Dolichyl-diphosphooligosaccharide--prot | RPN1 | 68.57 | 2 | 53.006268 | yes | 1.32E-07 | 526.7193 | 314.21613 |
| QPDSGISSIR | 10 | P04843 | Dolichyl-diphosphooligosaccharide--prot | RPN1 | 68.57 | 2 | 53.006268 | yes | 9.722E-08 | 530.27509 | 310.08893 |
| IDHILDAL | 8 | P04843 | Dolichyl-diphosphooligosaccharide--prot | RPN1 | 68.57 | 2 | 53.006268 | yes | 2.512E-07 | 455.25565 | 286.49216 |
| LAHLGVQVK | 9 | P04843 | Dolichyl-diphosphooligosaccharide--prot | RPN1 | 68.57 | 2 | 53.006268 | yes | 1.602E-07 | 482.80054 | 245.64827 |
| HFDETVNR | 8 | P04843 | Dolichyl-diphosphooligosaccharide--prot | RPN1 | 68.57 | 2 | 53.006268 | yes | 1.987E-07 | 509.24106 | 241.52632 |
| VTAEVVLAHLGG( | 17 | P04843 | Dolichyl-diphosphooligosaccharide--prot | RPN1 | 68.57 | 2 | 53.006268 | yes | 1.717E-07 | 827.44958 | 100.06482 |
| VTAEVVLAHLGG( | 17 | P04843 | Dolichyl-diphosphooligosaccharide--prot | RPN1 | 68.57 | 2 | 53.006268 | yes | 1.464E-06 | 551.96881 | 100.06482 |
| VTAEVVLAHLGG( | 17 | P04843 | Dolichyl-diphosphooligosaccharide--prot | RPN1 | 68.57 | 3 | 53.006268 | yes | 1.717E-07 | 827.44958 | 100.06482 |
| VTAEVVLAHLGG( | 17 | P04843 | Dolichyl-diphosphooligosaccharide--prot | RPN1 | 68.57 | 3 | 53.006268 | yes | 1.464E-06 | 551.96881 | 100.06482 |
| GEDEEENNLEVR | 12 | P04843 | Dolichyl-diphosphooligosaccharide--prot | RPN1 | 68.57 | 2 | 53.006268 | yes | 9.722E-08 | 716.81299 | 162.39639 |
| IILPEGAK | 8 | P04843 | Dolichyl-diphosphooligosaccharide--prot | RPN1 | 68.57 | 2 | 53.006268 | yes | 8.368E-07 | 420.76309 | 146.20692 |
| VIEVSHWGNIAVE | 19 | P04843 | Dolichyl-diphosphooligosaccharide--prot | RPN1 | 68.57 | 3 | 53.006268 | yes | 2.097E-07 | 717.70752 | 124.92497 |
| LKTEGSDLCDR | 11 | P04843 | Dolichyl-diphosphooligosaccharide--prot | RPN1 | 68.57 | 2 | 53.006268 | yes | 1.247E-07 | 647.30884 | 72.558899 |
| FPLFGGWK | 8 | P04843 | Dolichyl-diphosphooligosaccharide--prot | RPN1 | 68.57 | 2 | 53.006268 | yes | 3.371E-06 | 476.258 | 67.3405 |
| NLVEQHIQDIVVH | 18 | P04843 | Dolichyl-diphosphooligosaccharide--prot | RPN1 | 68.57 | 3 | 53.006268 | yes | 1.796E-07 | 733.05151 | 17.890076 |
| TGQEVVFVAEPDN | 14 | P04844 | Dolichyl-diphosphooligosaccharide--prot | RPN2 | 69.284 | 2 | 58.109734 | yes | 9.722E-08 | 766.88318 | 135.57713 |
| YHVPVVVVPEGS/ | 23 | P04844 | Dolichyl-diphosphooligosaccharide--prot | RPN2 | 69.284 | 4 | 58.109734 | yes | 2.159E-07 | 626.57892 | 60.074627 |
| YHVPVVVVPEGS/ | 23 | P04844 | Dolichyl-diphosphooligosaccharide--prot | RPN2 | 69.284 | 4 | 58.109734 | yes | 4.007E-07 | 835.10284 | 60.074627 |
| YHVPVVVVPEGS/ | 23 | P04844 | Dolichyl-diphosphooligosaccharide--prot | RPN2 | 69.284 | 3 | 58.109734 | yes | 2.159E-07 | 626.57892 | 60.074627 |
| YHVPVVVVPEGS/ | 23 | P04844 | Dolichyl-diphosphooligosaccharide--prot | RPN2 | 69.284 | 3 | 58.109734 | yes | 4.007E-07 | 835.10284 | 60.074627 |

| | | | | | | | | | | | |
|---|---|---|---|---|---|---|---|---|---|---|---|
| QEIQHLFR | 8 | P04844 | Dolichyl-diphosphooligosaccharide--prot | RPN2 | 69.284 | 2 | 58.109734 | yes | 1.23E-07 | 535.79071 | 68.758095 |
| LLLLGAGESGK | 11 | P04899 | Guanine nucleotide-binding protein G(i) s | GNAI2 | 40.451 | 2 | 53.5467 | no | 9.722E-08 | 529.31622 | 355.13266 |
| TTGIVETHFTFK | 12 | P04899 | Guanine nucleotide-binding protein G(i) s | GNAI2 | 40.451 | 2 | 53.5467 | no | 9.722E-08 | 690.86151 | 160.77696 |
| LFDSICNNK | 9 | P04899 | Guanine nucleotide-binding protein G(i) s | GNAI2 | 40.451 | 2 | 53.5467 | yes | 5.078E-07 | 555.76605 | 157.06229 |
| EIYTHFTCATDTK | 13 | P04899 | Guanine nucleotide-binding protein G(i) s | GNAI2 | 40.451 | 2 | 53.5467 | yes | 1.673E-07 | 793.86139 | 44.652958 |
| APQTGIVDECCFR | 13 | P05019 | Insulin-like growth factor I OS=Homo saj | IGF1 | 21.842 | 2 | 52.403915 | yes | 9.722E-08 | 776.84778 | 373.5932 |
| RLEMYCAPLKPAK | 13 | P05019 | Insulin-like growth factor I OS=Homo saj | IGF1 | 21.842 | 3 | 52.403915 | yes | 1.331E-07 | 526.28278 | 108.97166 |
| RLEMYCAPLKPAK | 13 | P05019 | Insulin-like growth factor I OS=Homo saj | IGF1 | 21.842 | 3 | 52.403915 | yes | 2.807E-07 | 531.61444 | 108.97166 |
| RLEMYCAPLKPAK | 13 | P05019 | Insulin-like growth factor I OS=Homo saj | IGF1 | 21.842 | 3 | 52.403915 | yes | 6.503E-07 | 788.92053 | 108.97166 |
| RLEMYCAPLKPAK | 13 | P05019 | Insulin-like growth factor I OS=Homo saj | IGF1 | 21.842 | 2 | 52.403915 | yes | 1.331E-07 | 526.28278 | 108.97166 |
| RLEMYCAPLKPAK | 13 | P05019 | Insulin-like growth factor I OS=Homo saj | IGF1 | 21.842 | 2 | 52.403915 | yes | 2.807E-07 | 531.61444 | 108.97166 |
| RLEMYCAPLKPAK | 13 | P05019 | Insulin-like growth factor I OS=Homo saj | IGF1 | 21.842 | 2 | 52.403915 | yes | 6.503E-07 | 788.92053 | 108.97166 |
| GFYFNKPTGYGSS | 15 | P05019 | Insulin-like growth factor I OS=Homo saj | IGF1 | 21.842 | 2 | 52.403915 | yes | 3.486E-07 | 834.39429 | 54.034534 |
| RAPQTGIVDECCFI | 14 | P05019 | Insulin-like growth factor I OS=Homo saj | IGF1 | 21.842 | 3 | 52.403915 | yes | 9.722E-08 | 570.26801 | 23.470772 |
| LIIVEGCQR | 9 | P05023 | Sodium/potassium-transporting ATPase s | ATP1A1 | 112.897 | 2 | 51.447655 | no | 1.1E-07 | 544.30005 | 486.60986 |
| LNIPVSQVNPR | 11 | P05023 | Sodium/potassium-transporting ATPase s | ATP1A1 | 112.897 | 2 | 51.447655 | no | 1.623E-07 | 618.85657 | 316.63718 |
| AVAGDASESALLK | 13 | P05023 | Sodium/potassium-transporting ATPase s | ATP1A1 | 112.897 | 2 | 51.447655 | yes | 4.637E-07 | 616.33008 | 194.32939 |
| YHTEIVFAR | 9 | P05023 | Sodium/potassium-transporting ATPase s | ATP1A1 | 112.897 | 2 | 51.447655 | yes | 1.479E-07 | 568.2984 | 193.65848 |
| LSLDELHR | 8 | P05023 | Sodium/potassium-transporting ATPase s | ATP1A1 | 112.897 | 2 | 51.447655 | yes | 1.164E-07 | 491.76944 | 173.12386 |
| NMVPQQALVIR | 11 | P05023 | Sodium/potassium-transporting ATPase s | ATP1A1 | 112.897 | 2 | 51.447655 | no | 1.755E-06 | 634.86084 | 171.9857 |
| IVEIPFNSTNK | 11 | P05023 | Sodium/potassium-transporting ATPase s | ATP1A1 | 112.897 | 2 | 51.447655 | yes | 6.198E-07 | 631.34296 | 130.40445 |
| VIMVTGDHPITAK | 13 | P05023 | Sodium/potassium-transporting ATPase s | ATP1A1 | 112.897 | 2 | 51.447655 | no | 9.79E-08 | 691.37903 | 91.611404 |
| AVFQANQENLPILI | 14 | P05023 | Sodium/potassium-transporting ATPase s | ATP1A1 | 112.897 | 2 | 51.447655 | yes | 1.618E-07 | 792.94086 | 88.213142 |
| NIAFFSTNCVEGTA | 15 | P05023 | Sodium/potassium-transporting ATPase s | ATP1A1 | 112.897 | 2 | 51.447655 | yes | 2.179E-06 | 843.89886 | 77.802963 |
| DMTSEQLDDILK | 12 | P05023 | Sodium/potassium-transporting ATPase s | ATP1A1 | 112.897 | 2 | 51.447655 | yes | 9.722E-08 | 704.33722 | 75.394104 |
| NPNTSEPQHLLVM | 14 | P05023 | Sodium/potassium-transporting ATPase s | ATP1A1 | 112.897 | 2 | 51.447655 | yes | 1.161E-07 | 804.41412 | 73.88105 |
| ADIGVAMGIAGSD | 16 | P05023 | Sodium/potassium-transporting ATPase s | ATP1A1 | 112.897 | 2 | 51.447655 | no | 9.076E-07 | 745.87958 | 60.697212 |
| CSSILLHGK | 9 | P05023 | Sodium/potassium-transporting ATPase s | ATP1A1 | 112.897 | 2 | 51.447655 | yes | 5.45E-07 | 507.77368 | 38.991112 |
| VDNSSLTGESEPQ | 15 | P05023 | Sodium/potassium-transporting ATPase s | ATP1A1 | 112.897 | 2 | 51.447655 | no | 3.4E-05 | 810.37903 | 2.2033169 |
| YNPNVLPVQCTGK | 13 | P05026 | Sodium/potassium-transporting ATPase s | ATP1B1 | 35.062 | 2 | 45.61655 | yes | 1.795E-07 | 745.37701 | 77.949692 |
| VEAMLNDR | 8 | P05067 | Amyloid-beta precursor protein OS=Hom | APP | 86.944 | 2 | 47.93719 | no | 1.229E-07 | 474.23438 | 181.34642 |
| THPHFVIPYR | 10 | P05067 | Amyloid-beta precursor protein OS=Hom | APP | 86.944 | 3 | 47.93719 | yes | 6.103E-07 | 422.89624 | 48.070999 |
| THPHFVIPYR | 10 | P05067 | Amyloid-beta precursor protein OS=Hom | APP | 86.944 | 3 | 47.93719 | yes | 5.843E-07 | 633.84076 | 48.070999 |
| THPHFVIPYR | 10 | P05067 | Amyloid-beta precursor protein OS=Hom | APP | 86.944 | 2 | 47.93719 | yes | 6.103E-07 | 422.89624 | 48.070999 |
| THPHFVIPYR | 10 | P05067 | Amyloid-beta precursor protein OS=Hom | APP | 86.944 | 2 | 47.93719 | yes | 5.843E-07 | 633.84076 | 48.070999 |
| LALENYITALQAV | 18 | P05067 | Amyloid-beta precursor protein OS=Hom | APP | 86.944 | 3 | 47.93719 | yes | 5.942E-07 | 674.72131 | 58.296837 |
| TEESPASDEAGEK | 13 | P05114 | Non-histone chromosomal protein HMG- | HMGN1 | 10.659 | 2 | 40.785503 | yes | 3.842E-07 | 675.28882 | 44.161098 |
| EQGVLSFWR | 9 | P05141 | ADP/ATP translocase 2 OS=Homo sapier | SLC25A5 | 32.853 | 2 | 53.02103 | no | 1.637E-07 | 561.29053 | 15562.646 |
| GTDIMYTGTLDCW | 14 | P05141 | ADP/ATP translocase 2 OS=Homo sapier | SLC25A5 | 32.853 | 2 | 53.02103 | yes | 1.274E-07 | 844.87396 | 11537.225 |

| | | | | | | | | | | | |
|---|---|---|---|---|---|---|---|---|---|---|---|
| YFPTQALNFAFK | 12 | P05141 | ADP/ATP translocase 2 OS=Homo sapie | SLC25A5 | 32.853 | 2 | 53.02103 | no | 1.048E-07 | 723.87445 | 10780.385 |
| QIFLGGVDK | 9 | P05141 | ADP/ATP translocase 2 OS=Homo sapie | SLC25A5 | 32.853 | 2 | 53.02103 | no | 1.364E-07 | 488.77673 | 9603.0762 |
| TAVAPIER | 8 | P05141 | ADP/ATP translocase 2 OS=Homo sapie | SLC25A5 | 32.853 | 2 | 53.02103 | no | 1.543E-07 | 428.74799 | 6099.9297 |
| GAWSNVLR | 8 | P05141 | ADP/ATP translocase 2 OS=Homo sapie | SLC25A5 | 32.853 | 2 | 53.02103 | no | 1.466E-07 | 451.74579 | 1967.3885 |
| GLGDCLVK | 8 | P05141 | ADP/ATP translocase 2 OS=Homo sapie | SLC25A5 | 32.853 | 2 | 53.02103 | no | 4.248E-07 | 431.22858 | 1789.0338 |
| QIFLGGVDKR | 10 | P05141 | ADP/ATP translocase 2 OS=Homo sapie | SLC25A5 | 32.853 | 2 | 53.02103 | yes | 1.028E-07 | 566.82727 | 1520.9286 |
| TDAAVSFAK | 9 | P05141 | ADP/ATP translocase 2 OS=Homo sapie | SLC25A5 | 32.853 | 2 | 53.02103 | yes | 9.893E-08 | 476.24274 | 1428.9901 |
| GNLANVIR | 8 | P05141 | ADP/ATP translocase 2 OS=Homo sapie | SLC25A5 | 32.853 | 2 | 53.02103 | no | 1.655E-07 | 428.7536 | 761.97168 |
| DFLAGGVAAAISK | 13 | P05141 | ADP/ATP translocase 2 OS=Homo sapie | SLC25A5 | 32.853 | 2 | 53.02103 | yes | 5.296E-06 | 610.33771 | 424.96609 |
| AAYFGIYDTAK | 11 | P05141 | ADP/ATP translocase 2 OS=Homo sapie | SLC25A5 | 32.853 | 2 | 53.02103 | yes | 1.5E-07 | 610.30334 | 409.99173 |
| KGTDIMYTGTLDC | 15 | P05141 | ADP/ATP translocase 2 OS=Homo sapie | SLC25A5 | 32.853 | 3 | 53.02103 | yes | 9.722E-08 | 606.28339 | 188.10562 |
| MMMQSGR | 7 | P05141 | ADP/ATP translocase 2 OS=Homo sapie | SLC25A5 | 32.853 | 2 | 53.02103 | no | 1.006E-06 | 420.67987 | 26.280632 |
| GLYQGFNVSVQGI | 17 | P05141 | ADP/ATP translocase 2 OS=Homo sapie | SLC25A5 | 32.853 | 3 | 53.02103 | no | 2.766E-07 | 643.01941 | 21.83675 |
| LPYQCPK | 7 | P05156 | Complement factor I OS=Homo sapiens ( | CFI | 65.751 | 2 | 48.973362 | yes | 1.082E-07 | 453.23111 | 3107.1772 |
| GLETSLAECTFTK | 13 | P05156 | Complement factor I OS=Homo sapiens ( | CFI | 65.751 | 2 | 48.973362 | yes | 1.697E-07 | 728.85303 | 1504.5007 |
| NWYSDADVPASA | 13 | P05187 | Alkaline phosphatase, placental type OS= | ALPP | 57.954 | 2 | 50.05576 | no | 1.001E-07 | 726.33112 | 117.72939 |
| LGPEIPLAMDR | 11 | P05187 | Alkaline phosphatase, placental type OS= | ALPP | 57.954 | 2 | 50.05576 | yes | 6.069E-07 | 606.32629 | 98.254524 |
| NLVQEWLAK | 9 | P05187 | Alkaline phosphatase, placental type OS= | ALPP | 57.954 | 2 | 50.05576 | no | 2.53E-07 | 550.80859 | 90.577934 |
| GNFQTIGLSAAAR | 13 | P05187 | Alkaline phosphatase, placental type OS= | ALPP | 57.954 | 2 | 50.05576 | no | 1.873E-07 | 653.34912 | 84.66729 |
| GNEVISVMNR | 10 | P05187 | Alkaline phosphatase, placental type OS= | ALPP | 57.954 | 2 | 50.05576 | no | 1.352E-07 | 559.78479 | 82.996674 |
| GSSIFGLAPGK | 11 | P05187 | Alkaline phosphatase, placental type OS= | ALPP | 57.954 | 2 | 50.05576 | no | 1.124E-06 | 517.28748 | 61.820091 |
| ALTETIMFDDAIEF | 14 | P05187 | Alkaline phosphatase, placental type OS= | ALPP | 57.954 | 2 | 50.05576 | no | 1.507E-07 | 812.89801 | 39.379868 |
| GFFLFVEGGR | 10 | P05187 | Alkaline phosphatase, placental type OS= | ALPP | 57.954 | 2 | 50.05576 | no | 9.919E-08 | 564.79547 | 35.935139 |
| DEQLESLFQR | 10 | P05198 | Eukaryotic translation initiation factor 2 s | EIF2S1 | 36.113 | 2 | 51.828033 | yes | 1.446E-07 | 632.81201 | 4585.9648 |
| YVMTTTTLER | 10 | P05198 | Eukaryotic translation initiation factor 2 s | EIF2S1 | 36.113 | 2 | 51.828033 | yes | 9.722E-08 | 607.80792 | 3936.4036 |
| VVTDTDETELAR | 12 | P05198 | Eukaryotic translation initiation factor 2 s | EIF2S1 | 36.113 | 2 | 51.828033 | yes | 1.021E-07 | 674.83319 | 3481.2976 |
| EVLINNINR | 9 | P05198 | Eukaryotic translation initiation factor 2 s | EIF2S1 | 36.113 | 2 | 51.828033 | yes | 1.183E-07 | 542.80908 | 3384.5254 |
| AGLNCSTENMPIK | 13 | P05198 | Eukaryotic translation initiation factor 2 s | EIF2S1 | 36.113 | 2 | 51.828033 | yes | 1.122E-07 | 717.83942 | 1373.4453 |
| AGLNCSTENMPIK | 13 | P05198 | Eukaryotic translation initiation factor 2 s | EIF2S1 | 36.113 | 2 | 51.828033 | yes | 3.491E-07 | 725.83685 | 1373.4453 |
| FPEVEDVVMVNV | 13 | P05198 | Eukaryotic translation initiation factor 2 s | EIF2S1 | 36.113 | 2 | 51.828033 | yes | 2.811E-07 | 766.89252 | 2411.7749 |
| HVAEVLEYTK | 10 | P05198 | Eukaryotic translation initiation factor 2 s | EIF2S1 | 36.113 | 2 | 51.828033 | yes | 1.022E-07 | 594.81659 | 1439.1533 |
| NECVVVIR | 8 | P05198 | Eukaryotic translation initiation factor 2 s | EIF2S1 | 36.113 | 2 | 51.828033 | yes | 1.574E-07 | 494.76584 | 1300.1016 |
| VSPEEAIK | 8 | P05198 | Eukaryotic translation initiation factor 2 s | EIF2S1 | 36.113 | 2 | 51.828033 | yes | 1.763E-07 | 436.73984 | 1276.8433 |
| TEGLSVLSQAMAV | 15 | P05198 | Eukaryotic translation initiation factor 2 s | EIF2S1 | 36.113 | 2 | 51.828033 | yes | 9.722E-08 | 773.92908 | 837.45264 |
| GVFNVQMEPK | 10 | P05198 | Eukaryotic translation initiation factor 2 s | EIF2S1 | 36.113 | 2 | 51.828033 | yes | 1.528E-07 | 574.79205 | 713.87299 |
| RPGYGAYDAFK | 11 | P05198 | Eukaryotic translation initiation factor 2 s | EIF2S1 | 36.113 | 2 | 51.828033 | yes | 1.473E-07 | 622.80658 | 407.91617 |
| TVYSILR | 7 | P05198 | Eukaryotic translation initiation factor 2 s | EIF2S1 | 36.113 | 2 | 51.828033 | yes | 2.502E-07 | 426.2529 | 211.02782 |
| RLTPQAVK | 8 | P05198 | Eukaryotic translation initiation factor 2 s | EIF2S1 | 36.113 | 2 | 51.828033 | yes | 1.022E-07 | 456.78491 | 153.90526 |
| HAVSDPSILDSLDI | 19 | P05198 | Eukaryotic translation initiation factor 2 s | EIF2S1 | 36.113 | 3 | 51.828033 | yes | 1.157E-07 | 709.00629 | 149.8183 |

| | | | | | | | | | | | |
|---|---|---|---|---|---|---|---|---|---|---|---|
| RGVFNVQMEPK | 11 | P05198 | Eukaryotic translation initiation factor 2 s | EIF2S1 | 36.113 | 2 | 51.828033 | yes | 1.931E-07 | 652.84265 | 147.59621 |
| HAVSDPSILDSLDI | 28 | P05198 | Eukaryotic translation initiation factor 2 s | EIF2S1 | 36.113 | 4 | 51.828033 | yes | 1.001E-07 | 798.40485 | 141.4684 |
| RVSPEEAIK | 9 | P05198 | Eukaryotic translation initiation factor 2 s | EIF2S1 | 36.113 | 2 | 51.828033 | yes | 1.034E-07 | 514.79041 | 122.67815 |
| INLIAPPR | 8 | P05198 | Eukaryotic translation initiation factor 2 s | EIF2S1 | 36.113 | 2 | 51.828033 | yes | 1.108E-07 | 447.28198 | 95.892883 |
| PGYGAYDAFK | 10 | P05198 | Eukaryotic translation initiation factor 2 s | EIF2S1 | 36.113 | 2 | 51.828033 | yes | 2.5E-07 | 544.75598 | 53.170055 |
| HVAEVLEYTKDEC | 20 | P05198 | Eukaryotic translation initiation factor 2 s | EIF2S1 | 36.113 | 3 | 51.828033 | yes | 2.064E-07 | 812.07983 | 43.023308 |
| VSPEEAIKCEDK | 12 | P05198 | Eukaryotic translation initiation factor 2 s | EIF2S1 | 36.113 | 2 | 51.828033 | yes | 1.362E-07 | 702.8374 | 36.266384 |
| EGNNPAENGDAK | 12 | P05204 | Non-histone chromosomal protein HMG- | HMGN2 | 9.393 | 2 | 28.841269 | yes | 2.651E-06 | 608.26544 | 21.88023 |
| DGTFPLPIGESVTV | 16 | P05362 | Intercellular adhesion molecule 1 OS=Ho | ICAM1 | 57.826 | 2 | 52.74916 | yes | 2.967E-07 | 844.94635 | 263.75385 |
| VELAPLPSWQPVC | 14 | P05362 | Intercellular adhesion molecule 1 OS=Ho | ICAM1 | 57.826 | 2 | 52.74916 | yes | 9.722E-08 | 760.92719 | 85.226723 |
| ASVSVTAEDEGTQ | 14 | P05362 | Intercellular adhesion molecule 1 OS=Ho | ICAM1 | 57.826 | 2 | 52.74916 | yes | 2.684E-07 | 725.34442 | 35.130051 |
| GGSVLVTCSTSCD | 16 | P05362 | Intercellular adhesion molecule 1 OS=Ho | ICAM1 | 57.826 | 2 | 52.74916 | yes | 4.769E-07 | 848.38745 | 30.638464 |
| KVTVNVLSPR | 10 | P05362 | Intercellular adhesion molecule 1 OS=Ho | ICAM1 | 57.826 | 2 | 52.74916 | yes | 5.255E-07 | 556.84296 | 27.639565 |
| ILDSVGIEADDDR | 13 | P05387 | Large ribosomal subunit protein P2 OS=H | RPLP2 | 11.665 | 2 | 53.931633 | yes | 1.674E-07 | 709.34393 | 34045.594 |
| NIEDVIAQGIGK | 12 | P05387 | Large ribosomal subunit protein P2 OS=H | RPLP2 | 11.665 | 2 | 53.931633 | yes | 9.967E-08 | 628.84589 | 24710.656 |
| KILDSVGIEADDDI | 14 | P05387 | Large ribosomal subunit protein P2 OS=H | RPLP2 | 11.665 | 2 | 53.931633 | yes | 1.014E-07 | 773.39136 | 929.38641 |
| KILDSVGIEADDDI | 17 | P05387 | Large ribosomal subunit protein P2 OS=H | RPLP2 | 11.665 | 3 | 53.931633 | yes | 1.065E-07 | 634.33734 | 447.54779 |
| MRYVASYLLAAL( | 21 | P05387 | Large ribosomal subunit protein P2 OS=H | RPLP2 | 11.665 | 3 | 53.931633 | yes | 1.133E-07 | 719.37714 | 94.920059 |
| YVASYLLAALGGN | 19 | P05387 | Large ribosomal subunit protein P2 OS=H | RPLP2 | 11.665 | 3 | 53.931633 | yes | 1.029E-07 | 623.66327 | 93.374565 |
| ILDSVGIEADDDRI | 16 | P05387 | Large ribosomal subunit protein P2 OS=H | RPLP2 | 11.665 | 3 | 53.931633 | yes | 7.006E-07 | 591.63904 | 65.657883 |
| VISELNGKNIEDVI | 20 | P05387 | Large ribosomal subunit protein P2 OS=H | RPLP2 | 11.665 | 3 | 53.931633 | yes | 9.722E-08 | 699.72321 | 24.207575 |
| CFIVGADNVGSK | 12 | P05388 | Large ribosomal subunit protein uL10 OS | RPLP0 | 34.274 | 2 | 50.366493 | no | 1.043E-07 | 633.81097 | 27125.512 |
| IIQLLDDYPK | 10 | P05388 | Large ribosomal subunit protein uL10 OS | RPLP0 | 34.274 | 2 | 50.366493 | no | 9.81E-08 | 609.34247 | 23191.457 |
| TSFFQALGITTK | 12 | P05388 | Large ribosomal subunit protein uL10 OS | RPLP0 | 34.274 | 2 | 50.366493 | no | 9.968E-08 | 657.35864 | 17039.135 |
| GTIEILSDVQLIK | 13 | P05388 | Large ribosomal subunit protein uL10 OS | RPLP0 | 34.274 | 2 | 50.366493 | no | 1.159E-07 | 714.91907 | 9797.1084 |
| GNVGFVFTK | 9 | P05388 | Large ribosomal subunit protein uL10 OS | RPLP0 | 34.274 | 2 | 50.366493 | yes | 9.214E-07 | 484.76364 | 8099.7573 |
| EDLTEIR | 7 | P05388 | Large ribosomal subunit protein uL10 OS | RPLP0 | 34.274 | 2 | 50.366493 | no | 7.725E-07 | 438.22708 | 6295.4556 |
| GHLENNPALEK | 11 | P05388 | Large ribosomal subunit protein uL10 OS | RPLP0 | 34.274 | 2 | 50.366493 | no | 2.176E-07 | 611.31476 | 5551.5894 |
| AGAIAPCEVTVPA( | 22 | P05388 | Large ribosomal subunit protein uL10 OS | RPLP0 | 34.274 | 3 | 50.366493 | no | 1.083E-07 | 727.37207 | 1394.127 |
| NVASVCLQIGYPT | 27 | P05388 | Large ribosomal subunit protein uL10 OS | RPLP0 | 34.274 | 4 | 50.366493 | yes | 5.331E-07 | 736.64319 | 55.151138 |
| TGEMTVLGHVK | 11 | P05423 | DNA-directed RNA polymerase III subun | POLR3D | 44.396 | 2 | 51.836445 | yes | 1.647E-07 | 586.31061 | 120.5955 |
| LVCSPDFESLLDHI | 14 | P05423 | DNA-directed RNA polymerase III subun | POLR3D | 44.396 | 2 | 51.836445 | yes | 9.722E-08 | 830.40601 | 25.957895 |
| LVCSPDFESLLDHI | 14 | P05423 | DNA-directed RNA polymerase III subun | POLR3D | 44.396 | 2 | 51.836445 | yes | 4.367E-07 | 553.93976 | 25.957895 |
| LVCSPDFESLLDHI | 14 | P05423 | DNA-directed RNA polymerase III subun | POLR3D | 44.396 | 3 | 51.836445 | yes | 9.722E-08 | 830.40601 | 25.957895 |
| LVCSPDFESLLDHI | 14 | P05423 | DNA-directed RNA polymerase III subun | POLR3D | 44.396 | 3 | 51.836445 | yes | 4.367E-07 | 553.93976 | 25.957895 |
| EVTWEVLEGEVEI | 13 | P05455 | Lupus La protein OS=Homo sapiens OX= | SSB | 46.838 | 2 | 52.912483 | yes | 1.055E-07 | 773.88538 | 3380.1284 |
| GQVLNIQMR | 9 | P05455 | Lupus La protein OS=Homo sapiens OX= | SSB | 46.838 | 2 | 52.912483 | yes | 2.523E-07 | 529.79242 | 2634.3416 |
| FSGDLDDQTCR | 11 | P05455 | Lupus La protein OS=Homo sapiens OX= | SSB | 46.838 | 2 | 52.912483 | yes | 2.654E-07 | 657.27496 | 1446.4788 |
| IIEDQQESLNK | 11 | P05455 | Lupus La protein OS=Homo sapiens OX= | SSB | 46.838 | 2 | 52.912483 | yes | 1.396E-07 | 658.83826 | 1116.8512 |

| | | | | | | | | | | | |
|---|---|---|---|---|---|---|---|---|---|---|---|
| GFPTDATLDDIK | 12 | P05455 | Lupus La protein OS=Homo sapiens OX= | SSB | 46.838 | 2 | 52.912483 | yes | 2.035E-07 | 646.82208 | 1088.3583 |
| EDLHILFSNHGEIK | 14 | P05455 | Lupus La protein OS=Homo sapiens OX= | SSB | 46.838 | 2 | 52.912483 | yes | 1.002E-07 | 826.42554 | 939.62408 |
| AELMEISEDK | 10 | P05455 | Lupus La protein OS=Homo sapiens OX= | SSB | 46.838 | 2 | 52.912483 | yes | 1.013E-07 | 582.77631 | 709.08698 |
| SKAELMEISEDK | 12 | P05455 | Lupus La protein OS=Homo sapiens OX= | SSB | 46.838 | 2 | 52.912483 | yes | 9.722E-08 | 690.33978 | 404.79858 |
| KFVETPGQK | 9 | P05455 | Lupus La protein OS=Homo sapiens OX= | SSB | 46.838 | 2 | 52.912483 | yes | 9.81E-08 | 517.28748 | 279.50244 |
| PLPEVTDEYK | 10 | P05455 | Lupus La protein OS=Homo sapiens OX= | SSB | 46.838 | 2 | 52.912483 | yes | 1.816E-07 | 595.8006 | 232.17918 |
| RSPSKPLPEVTDEY | 15 | P05455 | Lupus La protein OS=Homo sapiens OX= | SSB | 46.838 | 3 | 52.912483 | yes | 1.175E-07 | 582.64044 | 217.4716 |
| ICHQIEYYFGDFNI | 16 | P05455 | Lupus La protein OS=Homo sapiens OX= | SSB | 46.838 | 3 | 52.912483 | yes | 9.722E-08 | 691.328 | 216.89133 |
| SPSKPLPEVTDEYI | 14 | P05455 | Lupus La protein OS=Homo sapiens OX= | SSB | 46.838 | 2 | 52.912483 | yes | 1.022E-07 | 795.40649 | 206.64665 |
| QKLEEDAEMK | 10 | P05455 | Lupus La protein OS=Homo sapiens OX= | SSB | 46.838 | 2 | 52.912483 | yes | 9.722E-08 | 610.79498 | 205.5593 |
| YKETDLLILFK | 11 | P05455 | Lupus La protein OS=Homo sapiens OX= | SSB | 46.838 | 2 | 52.912483 | yes | 1.371E-07 | 691.90015 | 120.093 |
| KIIEDQQESLNK | 12 | P05455 | Lupus La protein OS=Homo sapiens OX= | SSB | 46.838 | 2 | 52.912483 | yes | 2.781E-07 | 722.88574 | 73.684631 |
| DANNGNLQLR | 10 | P05455 | Lupus La protein OS=Homo sapiens OX= | SSB | 46.838 | 2 | 52.912483 | yes | 2.329E-07 | 557.78363 | 57.009396 |
| SPSKPLPEVTDEYI | 18 | P05455 | Lupus La protein OS=Homo sapiens OX= | SSB | 46.838 | 3 | 52.912483 | yes | 3.328E-07 | 682.68451 | 28.950794 |
| GSIFVVFDSIESAK | 14 | P05455 | Lupus La protein OS=Homo sapiens OX= | SSB | 46.838 | 2 | 52.912483 | yes | 1.977E-05 | 749.89301 | 2.2137306 |
| SNSNAVSAIPINK | 13 | P05549 | Transcription factor AP-2-alpha OS=Hon | TFAP2A | 48.063 | 2 | 52.81594 | yes | 1.061E-07 | 657.85425 | 3165.2654 |
| EFTDLLAQDR | 10 | P05549 | Transcription factor AP-2-alpha OS=Hon | TFAP2A | 48.063 | 2 | 52.81594 | no | 1.026E-07 | 604.30133 | 2950.9033 |
| VTVAEVQR | 8 | P05549 | Transcription factor AP-2-alpha OS=Hon | TFAP2A | 48.063 | 2 | 52.81594 | no | 2.123E-07 | 451.25873 | 2731.9302 |
| DFGYVCETEFPAK | 13 | P05549 | Transcription factor AP-2-alpha OS=Hon | TFAP2A | 48.063 | 2 | 52.81594 | no | 1.163E-06 | 781.84521 | 1166.3794 |
| GLPHQLSGLDPR | 12 | P05549 | Transcription factor AP-2-alpha OS=Hon | TFAP2A | 48.063 | 2 | 52.81594 | yes | 1.819E-06 | 645.35168 | 416.86694 |
| GLPHQLSGLDPR | 12 | P05549 | Transcription factor AP-2-alpha OS=Hon | TFAP2A | 48.063 | 2 | 52.81594 | yes | 4.433E-07 | 430.57019 | 416.86694 |
| GLPHQLSGLDPR | 12 | P05549 | Transcription factor AP-2-alpha OS=Hon | TFAP2A | 48.063 | 3 | 52.81594 | yes | 1.819E-06 | 645.35168 | 416.86694 |
| GLPHQLSGLDPR | 12 | P05549 | Transcription factor AP-2-alpha OS=Hon | TFAP2A | 48.063 | 3 | 52.81594 | yes | 4.433E-07 | 430.57019 | 416.86694 |
| IGLNLPAGR | 9 | P05549 | Transcription factor AP-2-alpha OS=Hon | TFAP2A | 48.063 | 2 | 52.81594 | no | 3.888E-07 | 455.77707 | 570.23206 |
| AVAEFLNR | 8 | P05549 | Transcription factor AP-2-alpha OS=Hon | TFAP2A | 48.063 | 2 | 52.81594 | yes | 2.581E-07 | 460.25345 | 363.87341 |
| QSQESGLLHTHR | 12 | P05549 | Transcription factor AP-2-alpha OS=Hon | TFAP2A | 48.063 | 2 | 52.81594 | yes | 9.722E-08 | 696.85254 | 349.98434 |
| AANVTLLTSLVEG | 20 | P05549 | Transcription factor AP-2-alpha OS=Hon | TFAP2A | 48.063 | 3 | 52.81594 | no | 9.722E-08 | 688.71979 | 331.61243 |
| LSLLSSTSK | 9 | P05549 | Transcription factor AP-2-alpha OS=Hon | TFAP2A | 48.063 | 2 | 52.81594 | no | 1.569E-07 | 468.27402 | 310.4494 |
| KNMLLATK | 8 | P05549 | Transcription factor AP-2-alpha OS=Hon | TFAP2A | 48.063 | 2 | 52.81594 | no | 1.672E-07 | 459.7757 | 182.71925 |
| YEDCEDR | 7 | P05549 | Transcription factor AP-2-alpha OS=Hon | TFAP2A | 48.063 | 2 | 52.81594 | yes | 1.108E-07 | 493.67963 | 101.74875 |
| YKVTVAEVQR | 10 | P05549 | Transcription factor AP-2-alpha OS=Hon | TFAP2A | 48.063 | 2 | 52.81594 | no | 7.814E-07 | 596.83789 | 25.331951 |
| TVQSLEIDLDSMR | 13 | P05783 | Keratin, type I cytoskeletal 18 OS=Homo | KRT18 | 48.058 | 2 | 54.98605 | yes | 5.58E-07 | 753.87708 | 2732.7996 |
| TVQSLEIDLDSMR | 13 | P05783 | Keratin, type I cytoskeletal 18 OS=Homo | KRT18 | 48.058 | 2 | 54.98605 | yes | 9.722E-08 | 761.87451 | 2732.7996 |
| AQYDELAR | 8 | P05783 | Keratin, type I cytoskeletal 18 OS=Homo | KRT18 | 48.058 | 2 | 54.98605 | yes | 1.009E-07 | 483.23798 | 3621.6755 |
| LEAEIATYR | 9 | P05783 | Keratin, type I cytoskeletal 18 OS=Homo | KRT18 | 48.058 | 2 | 54.98605 | yes | 1.446E-07 | 533.28241 | 3371.2275 |
| SLGSVQAPSYGAR | 13 | P05783 | Keratin, type I cytoskeletal 18 OS=Homo | KRT18 | 48.058 | 2 | 54.98605 | yes | 9.722E-08 | 646.83331 | 2989.0725 |
| ASLENSLR | 8 | P05783 | Keratin, type I cytoskeletal 18 OS=Homo | KRT18 | 48.058 | 2 | 54.98605 | yes | 9.722E-08 | 445.24051 | 2689.0596 |
| QAQEYEALLNIK | 12 | P05783 | Keratin, type I cytoskeletal 18 OS=Homo | KRT18 | 48.058 | 2 | 54.98605 | yes | 2.101E-07 | 710.37756 | 2411.1238 |
| STFSTNYR | 8 | P05783 | Keratin, type I cytoskeletal 18 OS=Homo | KRT18 | 48.058 | 2 | 54.98605 | yes | 2.975E-07 | 488.23016 | 1422.1887 |

| | | | | | | | | | | | |
|---|---|---|---|---|---|---|---|---|---|---|---|
| VKLEAEIATYR | 11 | P05783 | Keratin, type I cytoskeletal 18 OS=Homo | KRT18 | 48.058 | 2 | 54.98605 | yes | 1.087E-07 | 646.86407 | 1294.3269 |
| IVLQIDNAR | 9 | P05783 | Keratin, type I cytoskeletal 18 OS=Homo | KRT18 | 48.058 | 2 | 54.98605 | no | 9.722E-08 | 521.30621 | 1042.0983 |
| PVSSAASVYAGAC | 18 | P05783 | Keratin, type I cytoskeletal 18 OS=Homo | KRT18 | 48.058 | 2 | 54.98605 | yes | 1.004E-07 | 790.88678 | 427.17804 |
| PVSSAASVYAGAC | 18 | P05783 | Keratin, type I cytoskeletal 18 OS=Homo | KRT18 | 48.058 | 2 | 54.98605 | yes | 3.069E-07 | 527.59363 | 427.17804 |
| PVSSAASVYAGAC | 18 | P05783 | Keratin, type I cytoskeletal 18 OS=Homo | KRT18 | 48.058 | 3 | 54.98605 | yes | 1.004E-07 | 790.88678 | 427.17804 |
| PVSSAASVYAGAC | 18 | P05783 | Keratin, type I cytoskeletal 18 OS=Homo | KRT18 | 48.058 | 3 | 54.98605 | yes | 3.069E-07 | 527.59363 | 427.17804 |
| GGMGSGGLATGIA | 26 | P05783 | Keratin, type I cytoskeletal 18 OS=Homo | KRT18 | 48.058 | 3 | 54.98605 | yes | 1.641E-07 | 754.37195 | 658.70197 |
| VIDDTNITR | 9 | P05783 | Keratin, type I cytoskeletal 18 OS=Homo | KRT18 | 48.058 | 2 | 54.98605 | yes | 3.619E-07 | 523.77747 | 575.72247 |
| AQIFANTVDNAR | 12 | P05783 | Keratin, type I cytoskeletal 18 OS=Homo | KRT18 | 48.058 | 2 | 54.98605 | yes | 1.847E-07 | 660.33875 | 531.9884 |
| VKYETELAMR | 10 | P05783 | Keratin, type I cytoskeletal 18 OS=Homo | KRT18 | 48.058 | 2 | 54.98605 | yes | 9.819E-08 | 620.32373 | 458.44498 |
| LASYLDR | 7 | P05783 | Keratin, type I cytoskeletal 18 OS=Homo | KRT18 | 48.058 | 2 | 54.98605 | yes | 3.039E-07 | 419.2269 | 163.54176 |
| ETMQSLNDR | 9 | P05783 | Keratin, type I cytoskeletal 18 OS=Homo | KRT18 | 48.058 | 2 | 54.98605 | no | 1.112E-07 | 547.25073 | 158.23222 |
| QSVENDIHGLR | 11 | P05783 | Keratin, type I cytoskeletal 18 OS=Homo | KRT18 | 48.058 | 2 | 54.98605 | yes | 2.403E-07 | 634.32312 | 149.86188 |
| VVSETNDTK | 9 | P05783 | Keratin, type I cytoskeletal 18 OS=Homo | KRT18 | 48.058 | 2 | 54.98605 | yes | 5.251E-07 | 496.74838 | 141.27469 |
| YETELAMR | 8 | P05783 | Keratin, type I cytoskeletal 18 OS=Homo | KRT18 | 48.058 | 2 | 54.98605 | yes | 5.672E-07 | 506.74203 | 116.09385 |
| KVIDDTNITR | 10 | P05783 | Keratin, type I cytoskeletal 18 OS=Homo | KRT18 | 48.058 | 2 | 54.98605 | yes | 9.722E-08 | 587.82495 | 108.51888 |
| DWSHYFK | 7 | P05783 | Keratin, type I cytoskeletal 18 OS=Homo | KRT18 | 48.058 | 2 | 54.98605 | yes | 2.324E-07 | 491.72452 | 104.547 |
| LQLETEIEALK | 11 | P05783 | Keratin, type I cytoskeletal 18 OS=Homo | KRT18 | 48.058 | 2 | 54.98605 | yes | 5.497E-07 | 643.86377 | 37.863037 |
| LSELEAALQR | 10 | P05787 | Keratin, type II cytoskeletal 8 OS=Homo | KRT8 | 53.705 | 2 | 53.211 | yes | 9.9E-08 | 565.31421 | 5375.4536 |
| SLDMDSIIAEVK | 12 | P05787 | Keratin, type II cytoskeletal 8 OS=Homo | KRT8 | 53.705 | 2 | 53.211 | yes | 9.722E-08 | 660.83942 | 3178.9067 |
| YEELQSLAGK | 10 | P05787 | Keratin, type II cytoskeletal 8 OS=Homo | KRT8 | 53.705 | 2 | 53.211 | yes | 9.241E-07 | 569.29297 | 3150.6829 |
| LVSESSDVLPK | 11 | P05787 | Keratin, type II cytoskeletal 8 OS=Homo | KRT8 | 53.705 | 2 | 53.211 | yes | 6.119E-07 | 587.32172 | 2901.4233 |
| LQAEIEGLK | 9 | P05787 | Keratin, type II cytoskeletal 8 OS=Homo | KRT8 | 53.705 | 2 | 53.211 | yes | 2.389E-07 | 500.78729 | 2630.679 |
| ASLEAAIADAEQR | 13 | P05787 | Keratin, type II cytoskeletal 8 OS=Homo | KRT8 | 53.705 | 2 | 53.211 | yes | 1.303E-07 | 672.84131 | 2563.635 |
| AQYEDIANR | 9 | P05787 | Keratin, type II cytoskeletal 8 OS=Homo | KRT8 | 53.705 | 2 | 53.211 | no | 1.955E-07 | 540.25946 | 2342.7424 |
| AEAESMYQIK | 10 | P05787 | Keratin, type II cytoskeletal 8 OS=Homo | KRT8 | 53.705 | 2 | 53.211 | yes | 9.85E-08 | 585.27899 | 2163.502 |
| LESGMQNMSIHTK | 13 | P05787 | Keratin, type II cytoskeletal 8 OS=Homo | KRT8 | 53.705 | 2 | 53.211 | yes | 1.001E-07 | 738.35272 | 445.70731 |
| LESGMQNMSIHTK | 13 | P05787 | Keratin, type II cytoskeletal 8 OS=Homo | KRT8 | 53.705 | 2 | 53.211 | yes | 1.997E-07 | 492.57089 | 445.70731 |
| LESGMQNMSIHTK | 13 | P05787 | Keratin, type II cytoskeletal 8 OS=Homo | KRT8 | 53.705 | 3 | 53.211 | yes | 1.001E-07 | 738.35272 | 445.70731 |
| LESGMQNMSIHTK | 13 | P05787 | Keratin, type II cytoskeletal 8 OS=Homo | KRT8 | 53.705 | 3 | 53.211 | yes | 1.997E-07 | 492.57089 | 445.70731 |
| WSLLQQQK | 8 | P05787 | Keratin, type II cytoskeletal 8 OS=Homo | KRT8 | 53.705 | 2 | 53.211 | yes | 2.503E-07 | 515.78766 | 795.90259 |
| LEGLTDEINFLR | 12 | P05787 | Keratin, type II cytoskeletal 8 OS=Homo | KRT8 | 53.705 | 2 | 53.211 | yes | 9.722E-08 | 710.37756 | 595.77014 |
| TKTEISEMNR | 10 | P05787 | Keratin, type II cytoskeletal 8 OS=Homo | KRT8 | 53.705 | 2 | 53.211 | yes | 3.455E-07 | 604.8006 | 485.99234 |
| TEMENEFVLIK | 11 | P05787 | Keratin, type II cytoskeletal 8 OS=Homo | KRT8 | 53.705 | 2 | 53.211 | yes | 1.941E-07 | 676.84198 | 477.7287 |
| TEISEMNR | 8 | P05787 | Keratin, type II cytoskeletal 8 OS=Homo | KRT8 | 53.705 | 2 | 53.211 | yes | 4.718E-07 | 490.22931 | 167.54541 |
| DVDEAYMNK | 9 | P05787 | Keratin, type II cytoskeletal 8 OS=Homo | KRT8 | 53.705 | 2 | 53.211 | yes | 4.457E-06 | 542.73444 | 22.580671 |
| EGNGTVMGAELR | 12 | P05976 | Myosin light chain 1/3, skeletal muscle is | MYL1 | 21.146 | 2 | 48.98989 | yes | 1.08E-07 | 617.29822 | 638.09369 |
| LLIYWASTR | 9 | P06312 | Immunoglobulin kappa variable 4-1 OS=l | IGKV4-1 | 13.381 | 2 | 48.246056 | yes | 1.174E-07 | 561.81897 | 6217.5098 |
| NYLAWYQQK | 9 | P06312 | Immunoglobulin kappa variable 4-1 OS=l | IGKV4-1 | 13.381 | 2 | 48.246056 | yes | 2.287E-07 | 607.30365 | 2171.7036 |

| | | | | | | | | | | | |
|---|---|---|---|---|---|---|---|---|---|---|---|
| HVVPNEVVVQR | 11 | P06396 | Gelsolin OS=Homo sapiens OX=9606 GN | GSN | 85.698 | 2 | 48.91432 | yes | 1.087E-07 | 638.36206 | 87.0448 |
| VPFDAATLHTSTA | 28 | P06396 | Gelsolin OS=Homo sapiens OX=9606 GN | GSN | 85.698 | 4 | 48.91432 | yes | 2.759E-06 | 723.07489 | 23.035137 |
| LESEEEGVPSTAIR | 14 | P06493 | Cyclin-dependent kinase 1 OS=Homo sap | CDK1 | 34.096 | 2 | 51.938675 | yes | 9.722E-08 | 758.87811 | 664.16217 |
| SPEVLLGSAR | 10 | P06493 | Cyclin-dependent kinase 1 OS=Homo sap | CDK1 | 34.096 | 2 | 51.938675 | yes | 2.011E-07 | 514.79041 | 426.77057 |
| DLKPQNLLIDDK | 12 | P06493 | Cyclin-dependent kinase 1 OS=Homo sap | CDK1 | 34.096 | 2 | 51.938675 | yes | 3.521E-07 | 706.39319 | 215.74127 |
| NLDENGLDLLSK | 12 | P06493 | Cyclin-dependent kinase 1 OS=Homo sap | CDK1 | 34.096 | 2 | 51.938675 | yes | 8.17E-07 | 665.84607 | 132.03876 |
| HPNIVSLQDVLMQ | 16 | P06493 | Cyclin-dependent kinase 1 OS=Homo sap | CDK1 | 34.096 | 3 | 51.938675 | yes | 2.323E-07 | 617.98425 | 44.511574 |
| MLIYDPAK | 8 | P06493 | Cyclin-dependent kinase 1 OS=Homo sap | CDK1 | 34.096 | 2 | 51.938675 | yes | 9.327E-05 | 475.75443 | 42.613377 |
| IRLESEEEGVPSTA | 16 | P06493 | Cyclin-dependent kinase 1 OS=Homo sap | CDK1 | 34.096 | 3 | 51.938675 | yes | 2.504E-06 | 595.98291 | 39.432053 |
| WKPGSLASHVK | 11 | P06493 | Cyclin-dependent kinase 1 OS=Homo sap | CDK1 | 34.096 | 3 | 51.938675 | yes | 1.021E-07 | 403.89615 | 35.383656 |
| IGLFGGAGVGK | 11 | P06576 | ATP synthase subunit beta, mitochondrial | ATP5F1B | 56.56 | 2 | 53.258327 | yes | 1.691E-07 | 488.28473 | 325.29025 |
| IMNVIGEPIDER | 12 | P06576 | ATP synthase subunit beta, mitochondrial | ATP5F1B | 56.56 | 2 | 53.258327 | yes | 2.104E-06 | 693.35828 | 254.45438 |
| FTQAGSEVSALLG | 14 | P06576 | ATP synthase subunit beta, mitochondrial | ATP5F1B | 56.56 | 2 | 53.258327 | yes | 1.236E-07 | 718.38062 | 232.67996 |
| VVDLLAPYAK | 10 | P06576 | ATP synthase subunit beta, mitochondrial | ATP5F1B | 56.56 | 2 | 53.258327 | yes | 1.411E-07 | 544.82117 | 160.45667 |
| VALVYGQMNEPP | 15 | P06576 | ATP synthase subunit beta, mitochondrial | ATP5F1B | 56.56 | 2 | 53.258327 | yes | 6.79E-07 | 801.40887 | 159.36275 |
| IPVGPETLGR | 10 | P06576 | ATP synthase subunit beta, mitochondrial | ATP5F1B | 56.56 | 2 | 53.258327 | yes | 5.167E-07 | 519.80072 | 134.82497 |
| IMDPNIVGSEHYD | 16 | P06576 | ATP synthase subunit beta, mitochondrial | ATP5F1B | 56.56 | 3 | 53.258327 | yes | 1.861E-07 | 605.9613 | 97.112289 |
| FLSQPFQVAEVFT | 18 | P06576 | ATP synthase subunit beta, mitochondrial | ATP5F1B | 56.56 | 3 | 53.258327 | yes | 9.722E-08 | 675.00836 | 55.935654 |
| VALTGLTVAEYFR | 13 | P06576 | ATP synthase subunit beta, mitochondrial | ATP5F1B | 56.56 | 2 | 53.258327 | yes | 4.702E-06 | 720.39825 | 54.952728 |
| IPSAVGYQPTLATI | 21 | P06576 | ATP synthase subunit beta, mitochondrial | ATP5F1B | 56.56 | 3 | 53.258327 | yes | 1.78E-06 | 756.0329 | 25.020428 |
| LQDAEIAR | 8 | P06703 | Protein S100-A6 OS=Homo sapiens OX= | S100A6 | 10.18 | 2 | 46.520203 | yes | 2.404E-07 | 458.24835 | 1013.4776 |
| LMEDLDR | 7 | P06703 | Protein S100-A6 OS=Homo sapiens OX= | S100A6 | 10.18 | 2 | 46.520203 | yes | 1.537E-07 | 446.21567 | 159.18854 |
| IAIWTTECENR | 11 | P06730 | Eukaryotic translation initiation factor 4E | EIF4E | 25.098 | 2 | 48.905178 | yes | 1.654E-07 | 696.83246 | 10265.67 |
| DGIEPMWEDEK | 11 | P06730 | Eukaryotic translation initiation factor 4E | EIF4E | 25.098 | 2 | 48.905178 | yes | 1.21E-07 | 674.78992 | 4169.5425 |
| DGIEPMWEDEK | 11 | P06730 | Eukaryotic translation initiation factor 4E | EIF4E | 25.098 | 2 | 48.905178 | yes | 2.579E-06 | 682.78735 | 4169.5425 |
| IVIGYQSHADTATI | 14 | P06730 | Eukaryotic translation initiation factor 4E | EIF4E | 25.098 | 2 | 48.905178 | yes | 1.272E-07 | 752.39374 | 2269.8433 |
| IVIGYQSHADTATI | 14 | P06730 | Eukaryotic translation initiation factor 4E | EIF4E | 25.098 | 2 | 48.905178 | yes | 4.338E-07 | 501.93158 | 2269.8433 |
| IVIGYQSHADTATI | 14 | P06730 | Eukaryotic translation initiation factor 4E | EIF4E | 25.098 | 3 | 48.905178 | yes | 1.272E-07 | 752.39374 | 2269.8433 |
| IVIGYQSHADTATI | 14 | P06730 | Eukaryotic translation initiation factor 4E | EIF4E | 25.098 | 3 | 48.905178 | yes | 4.338E-07 | 501.93158 | 2269.8433 |
| TWQANLR | 7 | P06730 | Eukaryotic translation initiation factor 4E | EIF4E | 25.098 | 2 | 48.905178 | yes | 1.382E-07 | 444.73795 | 3411.1277 |
| EAVTHIGR | 8 | P06730 | Eukaryotic translation initiation factor 4E | EIF4E | 25.098 | 2 | 48.905178 | yes | 1.165E-07 | 441.74323 | 2377.2886 |
| TESNQEVANPEHY | 15 | P06730 | Eukaryotic translation initiation factor 4E | EIF4E | 25.098 | 3 | 48.905178 | yes | 1.088E-07 | 586.94794 | 639.13196 |
| DGIEPMWEDEKNI | 13 | P06730 | Eukaryotic translation initiation factor 4E | EIF4E | 25.098 | 2 | 48.905178 | yes | 1.135E-07 | 795.85889 | 454.27179 |
| VNQIGSVTESLQA | 15 | P06733 | Alpha-enolase OS=Homo sapiens OX=96 | ENO1 | 47.169 | 2 | 50.841423 | yes | 1.776E-07 | 817.41431 | 558.59912 |
| GNPTVEVDLFTSK | 13 | P06733 | Alpha-enolase OS=Homo sapiens OX=96 | ENO1 | 47.169 | 2 | 50.841423 | yes | 1.444E-07 | 703.86176 | 472.04205 |
| IGAEVYHNLK | 10 | P06733 | Alpha-enolase OS=Homo sapiens OX=96 | ENO1 | 47.169 | 2 | 50.841423 | yes | 1.219E-07 | 572.31146 | 452.82175 |
| YISPDQLADLYK | 12 | P06733 | Alpha-enolase OS=Homo sapiens OX=96 | ENO1 | 47.169 | 2 | 50.841423 | yes | 9.722E-08 | 713.36664 | 362.49292 |
| LMIEMDGTENK | 11 | P06733 | Alpha-enolase OS=Homo sapiens OX=96 | ENO1 | 47.169 | 2 | 50.841423 | yes | 1.104E-07 | 640.79669 | 356.37326 |
| SCNCLLLK | 8 | P06733 | Alpha-enolase OS=Homo sapiens OX=96 | ENO1 | 47.169 | 2 | 50.841423 | yes | 5.595E-07 | 504.25427 | 219.52707 |

| | | | | | | | | | | | |
|---|---|---|---|---|---|---|---|---|---|---|---|
| KLNVTEQEK | 9 | P06733 | Alpha-enolase OS=Homo sapiens OX=96 | ENO1 | 47.169 | 2 | 50.841423 | yes | 9.727E-08 | 544.80096 | 130.21082 |
| TIAPALVSK | 9 | P06733 | Alpha-enolase OS=Homo sapiens OX=96 | ENO1 | 47.169 | 2 | 50.841423 | yes | 2.954E-07 | 450.28165 | 81.411041 |
| VVIGMDVAASEFF | 14 | P06733 | Alpha-enolase OS=Homo sapiens OX=96 | ENO1 | 47.169 | 2 | 50.841423 | yes | 2.087E-07 | 770.89508 | 71.283371 |
| TFTTQETITNAETA | 15 | P06744 | Glucose-6-phosphate isomerase OS=Hom | GPI | 63.148 | 2 | 45.30222 | yes | 1.892E-07 | 828.40979 | 29.661901 |
| VHFITDTLSK | 10 | P06746 | DNA polymerase beta OS=Homo sapiens | POLB | 38.178 | 2 | 55.31288 | yes | 9.722E-08 | 580.81915 | 173.46286 |
| FMGVCQLPSK | 10 | P06746 | DNA polymerase beta OS=Homo sapiens | POLB | 38.178 | 2 | 55.31288 | yes | 2.645E-07 | 583.78845 | 139.40271 |
| LLHQVVEQLQK | 11 | P06746 | DNA polymerase beta OS=Homo sapiens | POLB | 38.178 | 2 | 55.31288 | yes | 9.722E-08 | 667.89319 | 113.07574 |
| QDDTSSSINFLTR | 13 | P06746 | DNA polymerase beta OS=Homo sapiens | POLB | 38.178 | 2 | 55.31288 | yes | 9.722E-08 | 742.3548 | 73.027428 |
| KFVDEGIK | 8 | P06746 | DNA polymerase beta OS=Homo sapiens | POLB | 38.178 | 2 | 55.31288 | yes | 4.223E-06 | 468.26346 | 45.814922 |
| DIFDYIQWK | 9 | P06746 | DNA polymerase beta OS=Homo sapiens | POLB | 38.178 | 2 | 55.31288 | yes | 3.155E-06 | 614.30585 | 13.662835 |
| VDNDENEHQLSLF | 13 | P06748 | Nucleophosmin OS=Homo sapiens OX=9 | NPM1 | 32.576 | 3 | 53.12892 | yes | 3.138E-07 | 523.58148 | 2437.2793 |
| VDNDENEHQLSLF | 13 | P06748 | Nucleophosmin OS=Homo sapiens OX=9 | NPM1 | 32.576 | 3 | 53.12892 | yes | 2.916E-07 | 784.86859 | 2437.2793 |
| VDNDENEHQLSLF | 13 | P06748 | Nucleophosmin OS=Homo sapiens OX=9 | NPM1 | 32.576 | 2 | 53.12892 | yes | 3.138E-07 | 523.58148 | 2437.2793 |
| VDNDENEHQLSLF | 13 | P06748 | Nucleophosmin OS=Homo sapiens OX=9 | NPM1 | 32.576 | 2 | 53.12892 | yes | 2.916E-07 | 784.86859 | 2437.2793 |
| GPSSVEDIK | 9 | P06748 | Nucleophosmin OS=Homo sapiens OX=9 | NPM1 | 32.576 | 2 | 53.12892 | yes | 1.604E-07 | 466.2402 | 3448.5022 |
| DELHIVEAEAMNY | 19 | P06748 | Nucleophosmin OS=Homo sapiens OX=9 | NPM1 | 32.576 | 3 | 53.12892 | yes | 4.225E-07 | 715.67712 | 3301.5474 |
| ADKDYHFK | 8 | P06748 | Nucleophosmin OS=Homo sapiens OX=9 | NPM1 | 32.576 | 2 | 53.12892 | yes | 6.464E-07 | 512.24835 | 2198.4421 |
| MQASIEK | 7 | P06748 | Nucleophosmin OS=Homo sapiens OX=9 | NPM1 | 32.576 | 2 | 53.12892 | yes | 2.041E-07 | 403.70746 | 1517.4777 |
| TVSLGAGAKDELF | 28 | P06748 | Nucleophosmin OS=Homo sapiens OX=9 | NPM1 | 32.576 | 4 | 53.12892 | yes | 9.722E-08 | 733.12073 | 696.78113 |
| PQNYLFGCELK | 11 | P06748 | Nucleophosmin OS=Homo sapiens OX=9 | NPM1 | 32.576 | 2 | 53.12892 | yes | 9.722E-08 | 684.83447 | 95.88382 |
| AADAEAEVASLNF | 13 | P06753 | Tropomyosin alpha-3 chain OS=Homo sa | TPM3 | 32.95 | 2 | 49.84652 | yes | 1.001E-07 | 658.82568 | 1917.4946 |
| LVIIEGDLER | 10 | P06753 | Tropomyosin alpha-3 chain OS=Homo sa | TPM3 | 32.95 | 2 | 49.84652 | yes | 1.317E-07 | 578.83228 | 1029.8441 |
| KLVIIEGDLER | 11 | P06753 | Tropomyosin alpha-3 chain OS=Homo sa | TPM3 | 32.95 | 2 | 49.84652 | yes | 1.293E-07 | 642.8797 | 504.86453 |
| LLLPGELAK | 9 | P06899 | Histone H2B type 1-J OS=Homo sapiens | H2BC11 | 13.905 | 2 | 48.87259 | no | 1.054E-06 | 477.30515 | 41985.824 |
| QVHPDTGISSK | 11 | P06899 | Histone H2B type 1-J OS=Homo sapiens | H2BC11 | 13.905 | 2 | 48.87259 | no | 1.379E-07 | 584.80145 | 15262.956 |
| KESYSIYVYK | 10 | P06899 | Histone H2B type 1-J OS=Homo sapiens | H2BC11 | 13.905 | 2 | 48.87259 | no | 1.092E-07 | 640.33209 | 871.42859 |
| ESYSIYVYK | 9 | P06899 | Histone H2B type 1-J OS=Homo sapiens | H2BC11 | 13.905 | 2 | 48.87259 | no | 1.854E-07 | 576.28461 | 759.9444 |
| LIAPVAEEEATVPI | 16 | P07195 | L-lactate dehydrogenase B chain OS=Hor | LDHB | 36.639 | 2 | 51.199825 | yes | 1.119E-07 | 847.9516 | 108.06194 |
| IVVVTAGVR | 9 | P07195 | L-lactate dehydrogenase B chain OS=Hor | LDHB | 36.639 | 2 | 51.199825 | yes | 9.722E-08 | 457.2951 | 101.98087 |
| GLTSVINQK | 9 | P07195 | L-lactate dehydrogenase B chain OS=Hor | LDHB | 36.639 | 2 | 51.199825 | yes | 7.691E-07 | 480.27966 | 53.28318 |
| MDSTANEVEAVK | 12 | P07237 | Protein disulfide-isomerase OS=Homo sa | P4HB | 57.117 | 2 | 52.898464 | yes | 9.809E-08 | 647.30322 | 235.50952 |
| YKPESEELTAER | 12 | P07237 | Protein disulfide-isomerase OS=Homo sa | P4HB | 57.117 | 2 | 52.898464 | yes | 2.023E-07 | 726.35425 | 188.43832 |
| NFEDVAFDEK | 10 | P07237 | Protein disulfide-isomerase OS=Homo sa | P4HB | 57.117 | 2 | 52.898464 | yes | 9.722E-08 | 607.27222 | 186.82898 |
| SNFAEALAAHK | 11 | P07237 | Protein disulfide-isomerase OS=Homo sa | P4HB | 57.117 | 2 | 52.898464 | yes | 4.767E-07 | 579.79877 | 171.47301 |
| HNQLPLVIEFTEQT | 17 | P07237 | Protein disulfide-isomerase OS=Homo sa | P4HB | 57.117 | 3 | 52.898464 | yes | 3.427E-07 | 655.68616 | 147.93901 |
| ITEFCHR | 7 | P07237 | Protein disulfide-isomerase OS=Homo sa | P4HB | 57.117 | 2 | 52.898464 | yes | 1.004E-07 | 481.72928 | 112.91847 |
| EADDIVNWLK | 10 | P07237 | Protein disulfide-isomerase OS=Homo sa | P4HB | 57.117 | 2 | 52.898464 | yes | 1.35E-06 | 601.80621 | 98.805573 |
| THILLFLPK | 9 | P07237 | Protein disulfide-isomerase OS=Homo sa | P4HB | 57.117 | 2 | 52.898464 | yes | 1.267E-07 | 541.34204 | 84.105835 |
| TVIDYNGER | 9 | P07237 | Protein disulfide-isomerase OS=Homo sa | P4HB | 57.117 | 2 | 52.898464 | yes | 1.075E-06 | 533.76184 | 67.36734 |

| | | | | | | | | | | |
|---|---|---|---|---|---|---|---|---|---|---|
| VHSFPTLK | 8 | P07237 | Protein disulfide-isomerase OS=Homo sa | P4HB | 57.117 | 2 | 52.898464 | yes | 2.663E-07 | 464.76617 | 55.845249 |
| LITLEEEMTK | 10 | P07237 | Protein disulfide-isomerase OS=Homo sa | P4HB | 57.117 | 2 | 52.898464 | yes | 7.901E-07 | 603.81793 | 45.238251 |
| VGENADSQIK | 10 | P07305 | Histone H1.0 OS=Homo sapiens OX=96( | H1-0 | 20.863 | 2 | 47.05175 | yes | 1.413E-07 | 530.76709 | 390.52219 |
| YSDMIVAAIQAEK | 13 | P07305 | Histone H1.0 OS=Homo sapiens OX=96( | H1-0 | 20.863 | 2 | 47.05175 | yes | 3.664E-07 | 719.86597 | 67.180763 |
| SLYYYIQQDTK | 11 | P07355 | Annexin A2 OS=Homo sapiens OX=960( | ANXA2 | 38.605 | 2 | 52.558792 | no | 1.051E-07 | 711.35101 | 2272.0786 |
| STVHEILCK | 9 | P07355 | Annexin A2 OS=Homo sapiens OX=960( | ANXA2 | 38.605 | 2 | 52.558792 | no | 9.728E-08 | 564.78955 | 2036.2533 |
| WISIMTER | 8 | P07355 | Annexin A2 OS=Homo sapiens OX=960( | ANXA2 | 38.605 | 2 | 52.558792 | no | 1.238E-07 | 518.26825 | 2000.9185 |
| SYSPYDMLESIR | 12 | P07355 | Annexin A2 OS=Homo sapiens OX=960( | ANXA2 | 38.605 | 2 | 52.558792 | no | 1.451E-07 | 730.83997 | 1915.4958 |
| QDIAFAYQR | 9 | P07355 | Annexin A2 OS=Homo sapiens OX=960( | ANXA2 | 38.605 | 2 | 52.558792 | yes | 9.849E-08 | 556.28015 | 1771.2683 |
| TPAQYDASELK | 11 | P07355 | Annexin A2 OS=Homo sapiens OX=960( | ANXA2 | 38.605 | 2 | 52.558792 | no | 1.128E-07 | 611.80115 | 1423.4229 |
| TNQELQEINR | 10 | P07355 | Annexin A2 OS=Homo sapiens OX=960( | ANXA2 | 38.605 | 2 | 52.558792 | no | 9.722E-08 | 622.81512 | 838.03314 |
| AYTNFDAER | 9 | P07355 | Annexin A2 OS=Homo sapiens OX=960( | ANXA2 | 38.605 | 2 | 52.558792 | no | 1.152E-07 | 543.74615 | 768.07867 |
| SVPHLQK | 7 | P07355 | Annexin A2 OS=Homo sapiens OX=960( | ANXA2 | 38.605 | 2 | 52.558792 | no | 9.722E-08 | 404.73743 | 579.07068 |
| GVDEVTIVNILTNI | 14 | P07355 | Annexin A2 OS=Homo sapiens OX=960( | ANXA2 | 38.605 | 2 | 52.558792 | yes | 9.722E-08 | 771.92792 | 402.47702 |
| KLMVALAK | 8 | P07355 | Annexin A2 OS=Homo sapiens OX=960( | ANXA2 | 38.605 | 2 | 52.558792 | no | 9.722E-08 | 437.28314 | 155.17496 |
| DALNIETAIK | 10 | P07355 | Annexin A2 OS=Homo sapiens OX=960( | ANXA2 | 38.605 | 2 | 52.558792 | no | 1.022E-07 | 544.30334 | 117.24324 |
| RAEDGSVIDYELII | 18 | P07355 | Annexin A2 OS=Homo sapiens OX=960( | ANXA2 | 38.605 | 3 | 52.558792 | yes | 5.987E-05 | 688.99927 | 58.650887 |
| CEGFVCAQTGR | 11 | P07358 | Complement component C8 beta chain O | C8B | 66.948 | 2 | 35.07069 | yes | 8.916E-07 | 642.77661 | 34.16618 |
| ALTVPELTQQVFD | 15 | P07437 | Tubulin beta chain OS=Homo sapiens O> | TUBB | 49.671 | 2 | 49.96019 | yes | 1.417E-06 | 830.45123 | 24984.41 |
| ISVYYNEATGGK | 12 | P07437 | Tubulin beta chain OS=Homo sapiens O> | TUBB | 49.671 | 2 | 49.96019 | yes | 9.967E-08 | 651.32227 | 8984.4766 |
| FWEVISDEHGIDPT | 27 | P07437 | Tubulin beta chain OS=Homo sapiens O> | TUBB | 49.671 | 4 | 49.96019 | yes | 2.197E-07 | 776.35736 | 4953.5518 |
| NKPGVYTK | 8 | P07477 | Serine protease 1 OS=Homo sapiens OX= | PRSS1 | 26.559 | 2 | 43.384525 | yes | 2.614E-07 | 453.7558 | 10679.297 |
| NHCGIASAASYPT | 14 | P07711 | Procathepsin L OS=Homo sapiens OX=9 | CTSL | 37.565 | 2 | 52.9541 | yes | 9.722E-08 | 724.33539 | 266.99121 |
| NSWGEEWGMGGY | 14 | P07711 | Procathepsin L OS=Homo sapiens OX=9 | CTSL | 37.565 | 2 | 52.9541 | yes | 9.722E-08 | 800.34845 | 77.388641 |
| TFVNITPAEVGVL | 16 | P07737 | Profilin-1 OS=Homo sapiens OX=9606 C | PFN1 | 15.055 | 2 | 53.11663 | yes | 2.834E-07 | 822.47198 | 176.6427 |
| DSLLQDGEFSMDL | 14 | P07737 | Profilin-1 OS=Homo sapiens OX=9606 C | PFN1 | 15.055 | 2 | 53.11663 | yes | 1.305E-07 | 813.37744 | 148.01581 |
| CYEMASHLR | 9 | P07737 | Profilin-1 OS=Homo sapiens OX=9606 C | PFN1 | 15.055 | 2 | 53.11663 | yes | 9.722E-08 | 583.75769 | 116.13374 |
| STGGAPTFNVTVT | 14 | P07737 | Profilin-1 OS=Homo sapiens OX=9606 C | PFN1 | 15.055 | 2 | 53.11663 | yes | 3.227E-07 | 690.36188 | 102.49551 |
| TLVLLMGK | 8 | P07737 | Profilin-1 OS=Homo sapiens OX=9606 C | PFN1 | 15.055 | 2 | 53.11663 | yes | 2.535E-07 | 437.77515 | 56.806534 |
| SSFYVNGLTLGGQ | 14 | P07737 | Profilin-1 OS=Homo sapiens OX=9606 C | PFN1 | 15.055 | 2 | 53.11663 | yes | 2.215E-06 | 735.883 | 36.125637 |
| IDYIAGLDSR | 10 | P07741 | Adenine phosphoribosyltransferase OS=H | APRT | 19.608 | 2 | 47.09949 | yes | 2.639E-07 | 561.79309 | 154.87044 |
| SFPDFPTPGVVFR | 13 | P07741 | Adenine phosphoribosyltransferase OS=H | APRT | 19.608 | 2 | 47.09949 | yes | 1.401E-07 | 733.37738 | 101.79001 |
| SLYDEVAAQGEVV | 14 | P07814 | Bifunctional glutamate/proline--tRNA lig | EPRS1 | 170.591 | 2 | 55.68973 | yes | 2.231E-07 | 768.38867 | 361.63809 |
| LNQWCNVVR | 9 | P07814 | Bifunctional glutamate/proline--tRNA lig | EPRS1 | 170.591 | 2 | 55.68973 | yes | 2.628E-07 | 594.80078 | 352.3255 |
| VAVQGDVVR | 9 | P07814 | Bifunctional glutamate/proline--tRNA lig | EPRS1 | 170.591 | 2 | 55.68973 | yes | 9.722E-08 | 471.772 | 319.93906 |
| LTVAENEAETK | 11 | P07814 | Bifunctional glutamate/proline--tRNA lig | EPRS1 | 170.591 | 2 | 55.68973 | yes | 9.722E-08 | 602.8064 | 312.21594 |
| GSQFGQSCCLR | 11 | P07814 | Bifunctional glutamate/proline--tRNA lig | EPRS1 | 170.591 | 2 | 55.68973 | yes | 9.722E-08 | 650.28204 | 283.15298 |
| DQDLEPGAPSMGA | 14 | P07814 | Bifunctional glutamate/proline--tRNA lig | EPRS1 | 170.591 | 2 | 55.68973 | yes | 1.105E-07 | 708.32721 | 279.82953 |
| AYVDDTPAEQMK | 12 | P07814 | Bifunctional glutamate/proline--tRNA lig | EPRS1 | 170.591 | 2 | 55.68973 | yes | 9.722E-08 | 684.31104 | 272.19351 |

| | | | | | | | | | | | |
|---|---|---|---|---|---|---|---|---|---|---|---|
| SCQFVAVR | 8 | P07814 | Bifunctional glutamate/proline--tRNA lig | EPRS1 | 170.591 | 2 | 55.68973 | yes | 1.206E-07 | 483.74493 | 264.05951 |
| MFEIVFEDPK | 10 | P07814 | Bifunctional glutamate/proline--tRNA lig | EPRS1 | 170.591 | 2 | 55.68973 | yes | 9.722E-08 | 627.80737 | 261.91739 |
| KGDIIQLQR | 9 | P07814 | Bifunctional glutamate/proline--tRNA lig | EPRS1 | 170.591 | 2 | 55.68973 | yes | 2.033E-07 | 535.81946 | 254.99435 |
| GNAAWQEQLK | 10 | P07814 | Bifunctional glutamate/proline--tRNA lig | EPRS1 | 170.591 | 2 | 55.68973 | yes | 9.722E-08 | 572.79089 | 242.70679 |
| PLCELQPGAK | 10 | P07814 | Bifunctional glutamate/proline--tRNA lig | EPRS1 | 170.591 | 2 | 55.68973 | yes | 1.002E-07 | 556.79205 | 217.91893 |
| LNLNNTVLSK | 10 | P07814 | Bifunctional glutamate/proline--tRNA lig | EPRS1 | 170.591 | 2 | 55.68973 | yes | 3.619E-07 | 558.32458 | 192.49863 |
| TELAEPIAIR | 10 | P07814 | Bifunctional glutamate/proline--tRNA lig | EPRS1 | 170.591 | 2 | 55.68973 | yes | 9.722E-08 | 556.81915 | 187.74834 |
| KPYIWEYSR | 9 | P07814 | Bifunctional glutamate/proline--tRNA lig | EPRS1 | 170.591 | 2 | 55.68973 | yes | 9.722E-08 | 621.31934 | 127.25804 |
| KVIDPVAPR | 9 | P07814 | Bifunctional glutamate/proline--tRNA lig | EPRS1 | 170.591 | 2 | 55.68973 | yes | 2.234E-07 | 497.80585 | 120.67201 |
| NSEPAGLETPEAK | 13 | P07814 | Bifunctional glutamate/proline--tRNA lig | EPRS1 | 170.591 | 2 | 55.68973 | yes | 5.702E-07 | 671.82788 | 118.93959 |
| FVELPGAEMGK | 11 | P07814 | Bifunctional glutamate/proline--tRNA lig | EPRS1 | 170.591 | 2 | 55.68973 | yes | 1.218E-06 | 589.29974 | 104.38788 |
| SLIGVEYKPVSAT( | 20 | P07814 | Bifunctional glutamate/proline--tRNA lig | EPRS1 | 170.591 | 3 | 55.68973 | yes | 1.135E-07 | 703.03546 | 100.27676 |
| NLQMWEEMK | 9 | P07814 | Bifunctional glutamate/proline--tRNA lig | EPRS1 | 170.591 | 2 | 55.68973 | yes | 3.958E-07 | 604.77557 | 92.886406 |
| INEAVECLLSLK | 12 | P07814 | Bifunctional glutamate/proline--tRNA lig | EPRS1 | 170.591 | 2 | 55.68973 | yes | 9.722E-08 | 694.87634 | 86.564072 |
| YYTLFGR | 7 | P07814 | Bifunctional glutamate/proline--tRNA lig | EPRS1 | 170.591 | 2 | 55.68973 | yes | 1.772E-07 | 460.23724 | 85.598618 |
| HEELMLGDPCLK | 12 | P07814 | Bifunctional glutamate/proline--tRNA lig | EPRS1 | 170.591 | 2 | 55.68973 | yes | 9.722E-08 | 721.34436 | 85.508804 |
| QFAYQNSWGLTTI | 13 | P07814 | Bifunctional glutamate/proline--tRNA lig | EPRS1 | 170.591 | 2 | 55.68973 | yes | 2.161E-07 | 786.38367 | 74.234863 |
| TIGVMTMVHGDN | 20 | P07814 | Bifunctional glutamate/proline--tRNA lig | EPRS1 | 170.591 | 3 | 55.68973 | yes | 4.76E-05 | 713.36884 | 58.836273 |
| THVADFAPEVAW | 15 | P07814 | Bifunctional glutamate/proline--tRNA lig | EPRS1 | 170.591 | 3 | 55.68973 | yes | 9.722E-08 | 566.95813 | 57.773865 |
| THMVVANTMEDF | 14 | P07814 | Bifunctional glutamate/proline--tRNA lig | EPRS1 | 170.591 | 2 | 55.68973 | yes | 9.722E-08 | 825.88434 | 47.183914 |
| AALLNQHYQVNFI | 13 | P07814 | Bifunctional glutamate/proline--tRNA lig | EPRS1 | 170.591 | 2 | 55.68973 | yes | 5.969E-07 | 773.41223 | 36.011108 |
| AIQGGTSHHLGQN | 16 | P07814 | Bifunctional glutamate/proline--tRNA lig | EPRS1 | 170.591 | 2 | 55.68973 | yes | 1.441E-06 | 841.42389 | 5.7813234 |
| TLTIVDTGIGMTK | 13 | P07900 | Heat shock protein HSP 90-alpha OS=Ho | HSP90AA1 | 84.66 | 2 | 56.433414 | no | 1.815E-07 | 675.37085 | 1507.7026 |
| ELHINLIPNK | 10 | P07900 | Heat shock protein HSP 90-alpha OS=Ho | HSP90AA1 | 84.66 | 2 | 56.433414 | no | 3.17E-07 | 595.84821 | 565.09686 |
| YYTSASGDEMVSI | 14 | P07900 | Heat shock protein HSP 90-alpha OS=Ho | HSP90AA1 | 84.66 | 2 | 56.433414 | yes | 9.722E-08 | 775.85577 | 414.90582 |
| HIYYITGETK | 10 | P07900 | Heat shock protein HSP 90-alpha OS=Ho | HSP90AA1 | 84.66 | 2 | 56.433414 | no | 9.722E-08 | 612.81659 | 360.95923 |
| ELISNSSDALDK | 12 | P07900 | Heat shock protein HSP 90-alpha OS=Ho | HSP90AA1 | 84.66 | 2 | 56.433414 | no | 2.703E-07 | 646.32245 | 299.95547 |
| LGIHEDSQNR | 10 | P07900 | Heat shock protein HSP 90-alpha OS=Ho | HSP90AA1 | 84.66 | 2 | 56.433414 | yes | 9.899E-08 | 584.78888 | 229.58286 |
| DQVANSAFVER | 11 | P07900 | Heat shock protein HSP 90-alpha OS=Ho | HSP90AA1 | 84.66 | 2 | 56.433414 | yes | 2.006E-07 | 618.30438 | 226.30496 |
| KLSELLR | 7 | P07900 | Heat shock protein HSP 90-alpha OS=Ho | HSP90AA1 | 84.66 | 2 | 56.433414 | yes | 9.786E-08 | 429.77402 | 205.90248 |
| RAPFDLFENR | 10 | P07900 | Heat shock protein HSP 90-alpha OS=Ho | HSP90AA1 | 84.66 | 2 | 56.433414 | yes | 2.838E-07 | 632.82526 | 131.84987 |
| CLELFTELAEDK | 12 | P07900 | Heat shock protein HSP 90-alpha OS=Ho | HSP90AA1 | 84.66 | 2 | 56.433414 | yes | 9.722E-08 | 734.35541 | 120.54177 |
| HLEINPDHSIIETLF | 15 | P07900 | Heat shock protein HSP 90-alpha OS=Ho | HSP90AA1 | 84.66 | 3 | 56.433414 | yes | 2.363E-07 | 596.3197 | 116.795 |
| DNSTMGYMAAK | 11 | P07900 | Heat shock protein HSP 90-alpha OS=Ho | HSP90AA1 | 84.66 | 2 | 56.433414 | yes | 9.722E-08 | 594.75482 | 101.42139 |
| FYEQFSK | 7 | P07900 | Heat shock protein HSP 90-alpha OS=Ho | HSP90AA1 | 84.66 | 2 | 56.433414 | yes | 1.789E-07 | 474.72672 | 100.43167 |
| APFDLFENR | 9 | P07900 | Heat shock protein HSP 90-alpha OS=Ho | HSP90AA1 | 84.66 | 2 | 56.433414 | yes | 1.106E-07 | 554.77472 | 59.290432 |
| TDTGEPMGR | 9 | P07900 | Heat shock protein HSP 90-alpha OS=Ho | HSP90AA1 | 84.66 | 2 | 56.433414 | yes | 9.007E-07 | 482.21365 | 42.129028 |
| ALLFVPR | 7 | P07900 | Heat shock protein HSP 90-alpha OS=Ho | HSP90AA1 | 84.66 | 2 | 56.433414 | yes | 1.298E-06 | 408.26053 | 34.468182 |
| HSQFIGYPITLFVE | 15 | P07900 | Heat shock protein HSP 90-alpha OS=Ho | HSP90AA1 | 84.66 | 3 | 56.433414 | no | 4.228E-07 | 593.65405 | 17.693306 |

| | | | | | | | | | | | |
|---|---|---|---|---|---|---|---|---|---|---|---|
| GFAFVQYVNER | 11 | P07910 | Heterogeneous nuclear ribonucleoprotein | HNRNPC | 33.67 | 2 | 49.51851 | yes | 1.028E-07 | 665.33295 | 9879.4688 |
| LKGDDLQAIK | 10 | P07910 | Heterogeneous nuclear ribonucleoprotein | HNRNPC | 33.67 | 2 | 49.51851 | no | 1.837E-07 | 550.81915 | 8836.7529 |
| VFIGNLNTLVVK | 12 | P07910 | Heterogeneous nuclear ribonucleoprotein | HNRNPC | 33.67 | 2 | 49.51851 | no | 1.94E-07 | 658.90045 | 8693.5586 |
| VPPPPPIAR | 9 | P07910 | Heterogeneous nuclear ribonucleoprotein | HNRNPC | 33.67 | 2 | 49.51851 | yes | 6.72E-07 | 472.28983 | 6099.3711 |
| MIAGQVLDINLAA | 16 | P07910 | Heterogeneous nuclear ribonucleoprotein | HNRNPC | 33.67 | 2 | 49.51851 | yes | 1.022E-07 | 841.96094 | 1870.5458 |
| MIAGQVLDINLAA | 16 | P07910 | Heterogeneous nuclear ribonucleoprotein | HNRNPC | 33.67 | 2 | 49.51851 | yes | 2.912E-06 | 561.64307 | 1870.5458 |
| MIAGQVLDINLAA | 16 | P07910 | Heterogeneous nuclear ribonucleoprotein | HNRNPC | 33.67 | 3 | 49.51851 | yes | 1.022E-07 | 841.96094 | 1870.5458 |
| MIAGQVLDINLAA | 16 | P07910 | Heterogeneous nuclear ribonucleoprotein | HNRNPC | 33.67 | 3 | 49.51851 | yes | 2.912E-06 | 561.64307 | 1870.5458 |
| KSDVEAIFSK | 10 | P07910 | Heterogeneous nuclear ribonucleoprotein | HNRNPC | 33.67 | 2 | 49.51851 | no | 1.211E-07 | 562.30334 | 3459.302 |
| VDSLLENLEK | 10 | P07910 | Heterogeneous nuclear ribonucleoprotein | HNRNPC | 33.67 | 2 | 49.51851 | no | 1.94E-07 | 580.3139 | 3021.468 |
| QKVDSLLENLEK | 12 | P07910 | Heterogeneous nuclear ribonucleoprotein | HNRNPC | 33.67 | 2 | 49.51851 | no | 1.536E-07 | 708.39069 | 1908.9185 |
| MYSYPAR | 7 | P07910 | Heterogeneous nuclear ribonucleoprotein | HNRNPC | 33.67 | 2 | 49.51851 | yes | 2.748E-07 | 444.20764 | 1632.5553 |
| GDDQLELIKDDEK | 13 | P07910 | Heterogeneous nuclear ribonucleoprotein | HNRNPC | 33.67 | 2 | 49.51851 | yes | 1.162E-07 | 759.37012 | 990.42358 |
| IVGCSVHK | 8 | P07910 | Heterogeneous nuclear ribonucleoprotein | HNRNPC | 33.67 | 2 | 49.51851 | no | 2.656E-07 | 450.242 | 901.66864 |
| AAVAGEDGR | 9 | P07910 | Heterogeneous nuclear ribonucleoprotein | HNRNPC | 33.67 | 2 | 49.51851 | no | 1.28E-07 | 423.20923 | 170.45578 |
| GDDQLELIK | 9 | P07910 | Heterogeneous nuclear ribonucleoprotein | HNRNPC | 33.67 | 2 | 49.51851 | yes | 1.229E-06 | 515.77441 | 154.45636 |
| SDVEAIFSK | 9 | P07910 | Heterogeneous nuclear ribonucleoprotein | HNRNPC | 33.67 | 2 | 49.51851 | no | 1.916E-07 | 498.25583 | 21.004093 |
| EQCEQLEK | 8 | P07919 | Cytochrome b-c1 complex subunit 6, mito | UQCRH | 10.739 | 2 | 48.163765 | no | 1.277E-07 | 532.23987 | 1046.6738 |
| LELCDER | 7 | P07919 | Cytochrome b-c1 complex subunit 6, mito | UQCRH | 10.739 | 2 | 48.163765 | yes | 1.189E-07 | 467.71857 | 534.91681 |
| ERLELCDER | 9 | P07919 | Cytochrome b-c1 complex subunit 6, mito | UQCRH | 10.739 | 2 | 48.163765 | yes | 8.661E-07 | 610.29041 | 329.65997 |
| SHTEEDCTEELFDI | 18 | P07919 | Cytochrome b-c1 complex subunit 6, mito | UQCRH | 10.739 | 3 | 48.163765 | yes | 7.255E-07 | 745.99188 | 60.609081 |
| ISGVIGPYR | 9 | P07942 | Laminin subunit beta-1 OS=Homo sapien | LAMB1 | 198.039 | 2 | 48.03071 | yes | 4.999E-07 | 481.27692 | 159.5452 |
| DILAQSPAAEPLK | 13 | P07942 | Laminin subunit beta-1 OS=Homo sapien | LAMB1 | 198.039 | 2 | 48.03071 | yes | 1.211E-07 | 676.87463 | 126.06345 |
| KVDDIICDSR | 10 | P07942 | Laminin subunit beta-1 OS=Homo sapien | LAMB1 | 198.039 | 2 | 48.03071 | yes | 3.577E-07 | 610.8006 | 43.788811 |
| YYYAVYDMVVR | 11 | P07942 | Laminin subunit beta-1 OS=Homo sapien | LAMB1 | 198.039 | 2 | 48.03071 | yes | 4.117E-06 | 721.34467 | 19.744999 |
| LIEDNEYTAR | 10 | P07947 | Tyrosine-protein kinase Yes OS=Homo sa | YES1 | 60.802 | 2 | 50.72903 | no | 1.223E-07 | 612.29877 | 252.92032 |
| VPYPGMVNR | 9 | P07947 | Tyrosine-protein kinase Yes OS=Homo sa | YES1 | 60.802 | 2 | 50.72903 | no | 1.267E-07 | 516.76837 | 199.23323 |
| WTAPEAALYGR | 11 | P07947 | Tyrosine-protein kinase Yes OS=Homo sa | YES1 | 60.802 | 2 | 50.72903 | no | 3.347E-07 | 617.81439 | 192.02737 |
| EVLEQVER | 8 | P07947 | Tyrosine-protein kinase Yes OS=Homo sa | YES1 | 60.802 | 2 | 50.72903 | no | 1.479E-07 | 501.26672 | 125.9883 |
| AANILVGENLVCK | 13 | P07947 | Tyrosine-protein kinase Yes OS=Homo sa | YES1 | 60.802 | 2 | 50.72903 | no | 9.727E-08 | 700.88196 | 105.7492 |
| LLLNPGNQR | 9 | P07947 | Tyrosine-protein kinase Yes OS=Homo sa | YES1 | 60.802 | 2 | 50.72903 | yes | 1.494E-07 | 512.79852 | 79.027199 |
| AANVLVSESLMCK | 13 | P07948 | Tyrosine-protein kinase Lyn OS=Homo sa | LYN | 58.575 | 2 | 49.302277 | yes | 4.525E-07 | 711.35999 | 253.34471 |
| EGFIPSNYVAK | 11 | P07948 | Tyrosine-protein kinase Lyn OS=Homo sa | LYN | 58.575 | 2 | 49.302277 | yes | 1.047E-07 | 612.81659 | 189.10489 |
| DFDPVHGDVIK | 11 | P07948 | Tyrosine-protein kinase Lyn OS=Homo sa | LYN | 58.575 | 2 | 49.302277 | yes | 2.442E-07 | 621.31171 | 122.78713 |
| VIEDNEYTAR | 10 | P07948 | Tyrosine-protein kinase Lyn OS=Homo sa | LYN | 58.575 | 2 | 49.302277 | no | 2.845E-07 | 605.29095 | 100.77815 |
| VLEEHGEWWK | 10 | P07948 | Tyrosine-protein kinase Lyn OS=Homo sa | LYN | 58.575 | 2 | 49.302277 | yes | 1.382E-07 | 656.81964 | 87.487694 |
| VENCPDELYDIMK | 13 | P07948 | Tyrosine-protein kinase Lyn OS=Homo sa | LYN | 58.575 | 2 | 49.302277 | yes | 1.833E-07 | 813.36292 | 85.904266 |
| TNADVMTALSQG | 14 | P07948 | Tyrosine-protein kinase Lyn OS=Homo sa | LYN | 58.575 | 2 | 49.302277 | yes | 3.059E-06 | 763.867 | 80.629333 |
| PVPESQLLPGQR | 12 | P07948 | Tyrosine-protein kinase Lyn OS=Homo sa | LYN | 58.575 | 2 | 49.302277 | yes | 2.705E-07 | 660.86713 | 70.671791 |

| | | | | | | | | | | | |
|---|---|---|---|---|---|---|---|---|---|---|---|
| ITFPCISDMIK | 11 | P07948 | Tyrosine-protein kinase Lyn OS=Homo sa | LYN | 58.575 | 2 | 49.302277 | yes | 9.236E-07 | 662.83563 | 39.890003 |
| LNTLETEEWFFK | 12 | P07948 | Tyrosine-protein kinase Lyn OS=Homo sa | LYN | 58.575 | 2 | 49.302277 | yes | 1.251E-07 | 778.88519 | 37.793964 |
| QLLAPGNSAGAFL | 15 | P07948 | Tyrosine-protein kinase Lyn OS=Homo sa | LYN | 58.575 | 2 | 49.302277 | yes | 1.105E-07 | 764.43573 | 16.787294 |
| IQLVEEELDR | 10 | P07951 | Tropomyosin beta chain OS=Homo sapie | TPM2 | 32.851 | 2 | 53.110035 | no | 1.362E-07 | 622.33008 | 5589.3457 |
| LVILEGELER | 10 | P07951 | Tropomyosin beta chain OS=Homo sapie | TPM2 | 32.851 | 2 | 53.110035 | no | 1.071E-07 | 585.84009 | 1257.4159 |
| EDKYEEEIK | 9 | P07951 | Tropomyosin beta chain OS=Homo sapie | TPM2 | 32.851 | 2 | 53.110035 | no | 9.722E-08 | 591.77985 | 1086.4823 |
| KLVILEGELER | 11 | P07951 | Tropomyosin beta chain OS=Homo sapie | TPM2 | 32.851 | 2 | 53.110035 | no | 9.722E-08 | 649.88757 | 547.96417 |
| ATDAEADVASLNI | 13 | P07951 | Tropomyosin beta chain OS=Homo sapie | TPM2 | 32.851 | 2 | 53.110035 | no | 9.722E-08 | 666.82312 | 382.8974 |
| KYEEVAR | 7 | P07951 | Tropomyosin beta chain OS=Homo sapie | TPM2 | 32.851 | 2 | 53.110035 | no | 2.87E-07 | 447.73761 | 319.17526 |
| QLEEEQQALQK | 11 | P07951 | Tropomyosin beta chain OS=Homo sapie | TPM2 | 32.851 | 2 | 53.110035 | yes | 1.001E-07 | 672.34369 | 291.1908 |
| CGDLEEELK | 9 | P07951 | Tropomyosin beta chain OS=Homo sapie | TPM2 | 32.851 | 2 | 53.110035 | no | 1.187E-07 | 546.7475 | 274.49225 |
| LDKENAIDR | 9 | P07951 | Tropomyosin beta chain OS=Homo sapie | TPM2 | 32.851 | 2 | 53.110035 | yes | 1.386E-06 | 537.2829 | 229.5743 |
| MELQEMQLK | 9 | P07951 | Tropomyosin beta chain OS=Homo sapie | TPM2 | 32.851 | 2 | 53.110035 | yes | 1.435E-07 | 575.28577 | 211.99348 |
| TIDDLEDEVYAQK | 13 | P07951 | Tropomyosin beta chain OS=Homo sapie | TPM2 | 32.851 | 2 | 53.110035 | yes | 3.34E-07 | 769.86469 | 57.37698 |
| KATDAEADVASLI | 14 | P07951 | Tropomyosin beta chain OS=Homo sapie | TPM2 | 32.851 | 2 | 53.110035 | no | 1.125E-07 | 730.87061 | 35.835285 |
| RIQLVEEELDR | 11 | P07951 | Tropomyosin beta chain OS=Homo sapie | TPM2 | 32.851 | 2 | 53.110035 | no | 5.433E-07 | 700.38062 | 30.97525 |
| VLLVQVDVK | 9 | P07992 | DNA excision repair protein ERCC-1 OS | ERCC1 | 32.563 | 2 | 44.92723 | yes | 3.508E-07 | 506.8237 | 49.994415 |
| LFDVLHEPFLK | 11 | P07992 | DNA excision repair protein ERCC-1 OS | ERCC1 | 32.563 | 2 | 44.92723 | yes | 2.022E-07 | 679.37933 | 49.001621 |
| AYEQKPADLLMEI | 13 | P07992 | DNA excision repair protein ERCC-1 OS | ERCC1 | 32.563 | 2 | 44.92723 | yes | 1.072E-06 | 768.39233 | 42.895878 |
| FQDLVDAVR | 9 | P07996 | Thrombospondin-1 OS=Homo sapiens O | THBS1 | 129.383 | 2 | 50.199272 | yes | 1.275E-07 | 531.78253 | 314.21954 |
| FVFGTTPEDILR | 12 | P07996 | Thrombospondin-1 OS=Homo sapiens O | THBS1 | 129.383 | 2 | 50.199272 | yes | 1.15E-07 | 697.86938 | 212.62485 |
| TIVTTLQDSIR | 11 | P07996 | Thrombospondin-1 OS=Homo sapiens O | THBS1 | 129.383 | 2 | 50.199272 | yes | 3.873E-07 | 623.8537 | 189.10985 |
| FQMIPLDPK | 9 | P07996 | Thrombospondin-1 OS=Homo sapiens O | THBS1 | 129.383 | 2 | 50.199272 | yes | 4.729E-07 | 544.79407 | 181.35677 |
| IMADSGPIYDK | 11 | P07996 | Thrombospondin-1 OS=Homo sapiens O | THBS1 | 129.383 | 2 | 50.199272 | yes | 3.467E-07 | 605.29462 | 172.06482 |
| AQLYIDCEK | 9 | P07996 | Thrombospondin-1 OS=Homo sapiens O | THBS1 | 129.383 | 2 | 50.199272 | yes | 1.197E-07 | 570.27368 | 165.42352 |
| GGVNDNFQGVLQ | 15 | P07996 | Thrombospondin-1 OS=Homo sapiens O | THBS1 | 129.383 | 2 | 50.199272 | yes | 1.005E-07 | 808.91058 | 163.99637 |
| DNCQYVYNVDQR | 12 | P07996 | Thrombospondin-1 OS=Homo sapiens O | THBS1 | 129.383 | 2 | 50.199272 | yes | 9.85E-08 | 787.33862 | 136.47844 |
| SITLFVQEDR | 10 | P07996 | Thrombospondin-1 OS=Homo sapiens O | THBS1 | 129.383 | 2 | 50.199272 | yes | 2.702E-07 | 604.31952 | 134.80084 |
| GTSQNDPNWVVR | 12 | P07996 | Thrombospondin-1 OS=Homo sapiens O | THBS1 | 129.383 | 2 | 50.199272 | yes | 1.647E-07 | 686.83386 | 119.36616 |
| CNYLGHYSDPMY | 13 | P07996 | Thrombospondin-1 OS=Homo sapiens O | THBS1 | 129.383 | 2 | 50.199272 | yes | 1.112E-07 | 838.35321 | 23.409258 |
| FACPECPK | 8 | P08047 | Transcription factor Sp1 OS=Homo sapie | SP1 | 80.693 | 2 | 50.70846 | yes | 1.05E-07 | 504.7175 | 176.32101 |
| PFMCTWSYCGK | 11 | P08047 | Transcription factor Sp1 OS=Homo sapie | SP1 | 80.693 | 2 | 50.70846 | yes | 9.728E-08 | 718.7934 | 57.75462 |
| KFACPECPK | 9 | P08047 | Transcription factor Sp1 OS=Homo sapie | SP1 | 80.693 | 2 | 50.70846 | yes | 1.107E-06 | 568.76501 | 50.466732 |
| EACTCPYCK | 9 | P08047 | Transcription factor Sp1 OS=Homo sapie | SP1 | 80.693 | 2 | 50.70846 | yes | 1.063E-07 | 594.72772 | 49.058109 |
| AVILQGSNDVELV | 19 | P08123 | Collagen alpha-2(I) chain OS=Homo sapi | COL1A2 | 129.314 | 3 | 52.5528 | yes | 9.722E-08 | 657.67621 | 243.10533 |
| LVIVGDGACGK | 11 | P08134 | Rho-related GTP-binding protein RhoC C | RHOC | 22.007 | 2 | 49.3114 | no | 1.046E-07 | 544.79205 | 313.80093 |
| ISAFGYLECSAK | 12 | P08134 | Rho-related GTP-binding protein RhoC C | RHOC | 22.007 | 2 | 49.3114 | yes | 3.805E-07 | 673.32648 | 149.34938 |
| HFCPNVPIILVGNK | 14 | P08134 | Rho-related GTP-binding protein RhoC C | RHOC | 22.007 | 2 | 49.3114 | no | 1.813E-07 | 804.43994 | 54.474144 |
| HFCPNVPIILVGNK | 14 | P08134 | Rho-related GTP-binding protein RhoC C | RHOC | 22.007 | 2 | 49.3114 | no | 2.291E-06 | 536.62909 | 54.474144 |

| | | | | | | | | | | | |
|---|---|---|---|---|---|---|---|---|---|---|---|
| HFCPNVPIILVGNK | 14 | P08134 | Rho-related GTP-binding protein RhoC C | RHOC | 22.007 | 3 | 49.3114 | no | 1.813E-07 | 804.43994 | 54.474144 |
| HFCPNVPIILVGNK | 14 | P08134 | Rho-related GTP-binding protein RhoC C | RHOC | 22.007 | 3 | 49.3114 | no | 2.291E-06 | 536.62909 | 54.474144 |
| EVFEMATR | 8 | P08134 | Rho-related GTP-binding protein RhoC C | RHOC | 22.007 | 2 | 49.3114 | no | 2.703E-07 | 491.73676 | 54.715332 |
| TCLLIVFSK | 9 | P08134 | Rho-related GTP-binding protein RhoC C | RHOC | 22.007 | 2 | 49.3114 | no | 2.063E-07 | 540.80975 | 50.655224 |
| LTCLQNLK | 8 | P08174 | Complement decay-accelerating factor OS | CD55 | 41.401 | 2 | 41.672295 | yes | 3.397E-07 | 495.27606 | 91.762695 |
| GLVLGPIHK | 9 | P08195 | Amino acid transporter heavy chain SLC3 | SLC3A2 | 67.994 | 2 | 51.92491 | yes | 3.127E-07 | 467.29764 | 149.91457 |
| ADLLLSTQPGR | 11 | P08195 | Amino acid transporter heavy chain SLC3 | SLC3A2 | 67.994 | 2 | 51.92491 | yes | 9.722E-08 | 585.82751 | 134.77739 |
| VAEDEAEAAAAI | 13 | P08195 | Amino acid transporter heavy chain SLC3 | SLC3A2 | 67.994 | 2 | 51.92491 | yes | 9.722E-08 | 623.30151 | 126.4634 |
| EVELNELEPEK | 11 | P08195 | Amino acid transporter heavy chain SLC3 | SLC3A2 | 67.994 | 2 | 51.92491 | yes | 1.095E-07 | 664.83264 | 80.011391 |
| EDFDSLLQSAK | 11 | P08195 | Amino acid transporter heavy chain SLC3 | SLC3A2 | 67.994 | 2 | 51.92491 | yes | 4.84E-07 | 626.8064 | 76.35305 |
| TFVLEVMGR | 9 | P08237 | ATP-dependent 6-phosphofructokinase, n | PFKM | 85.183 | 2 | 52.425312 | no | 9.722E-08 | 526.28387 | 501.47504 |
| TFVLEVMGR | 9 | P08237 | ATP-dependent 6-phosphofructokinase, n | PFKM | 85.183 | 2 | 52.425312 | no | 2.652E-07 | 534.28131 | 501.47504 |
| VTVLGHVQR | 9 | P08237 | ATP-dependent 6-phosphofructokinase, n | PFKM | 85.183 | 2 | 52.425312 | no | 2.432E-07 | 504.80109 | 718.62549 |
| VFFVHEGYQGLVI | 19 | P08237 | ATP-dependent 6-phosphofructokinase, n | PFKM | 85.183 | 3 | 52.425312 | yes | 1.058E-07 | 706.35321 | 650.36407 |
| ALVFQPVAELK | 11 | P08237 | ATP-dependent 6-phosphofructokinase, n | PFKM | 85.183 | 2 | 52.425312 | yes | 1.127E-07 | 607.86078 | 559.7865 |
| IGLIQGNR | 8 | P08237 | ATP-dependent 6-phosphofructokinase, n | PFKM | 85.183 | 2 | 52.425312 | yes | 1.821E-07 | 435.76141 | 483.38223 |
| LPLMECVQVTK | 11 | P08237 | ATP-dependent 6-phosphofructokinase, n | PFKM | 85.183 | 2 | 52.425312 | yes | 1.136E-07 | 659.34888 | 466.60825 |
| SFMNNWEVYK | 10 | P08237 | ATP-dependent 6-phosphofructokinase, n | PFKM | 85.183 | 2 | 52.425312 | yes | 1.169E-07 | 659.30023 | 251.41765 |
| DQTDFEHR | 8 | P08237 | ATP-dependent 6-phosphofructokinase, n | PFKM | 85.183 | 2 | 52.425312 | yes | 2.37E-07 | 524.22815 | 229.5705 |
| VGIFTGAR | 8 | P08237 | ATP-dependent 6-phosphofructokinase, n | PFKM | 85.183 | 2 | 52.425312 | yes | 9.975E-08 | 410.73743 | 213.75902 |
| IFANTPDSGCVLGI | 15 | P08237 | ATP-dependent 6-phosphofructokinase, n | PFKM | 85.183 | 2 | 52.425312 | yes | 1.047E-07 | 819.39233 | 184.9357 |
| KFDEALK | 7 | P08237 | ATP-dependent 6-phosphofructokinase, n | PFKM | 85.183 | 2 | 52.425312 | yes | 2.082E-07 | 425.73709 | 161.01814 |
| SEWSDLLSDLQK | 12 | P08237 | ATP-dependent 6-phosphofructokinase, n | PFKM | 85.183 | 2 | 52.425312 | yes | 9.987E-08 | 710.85138 | 113.37173 |
| LNIIIVAEGAIDK | 13 | P08237 | ATP-dependent 6-phosphofructokinase, n | PFKM | 85.183 | 2 | 52.425312 | yes | 2.12E-07 | 684.90851 | 90.829399 |
| YEIDLDTSDHAHL | 18 | P08237 | ATP-dependent 6-phosphofructokinase, n | PFKM | 85.183 | 3 | 52.425312 | yes | 1.813E-07 | 722.3468 | 63.915436 |
| VLVVHDGFEGLAI | 13 | P08237 | ATP-dependent 6-phosphofructokinase, n | PFKM | 85.183 | 2 | 52.425312 | yes | 9.722E-08 | 692.38519 | 63.676411 |
| SFEQISANITK | 11 | P08237 | ATP-dependent 6-phosphofructokinase, n | PFKM | 85.183 | 2 | 52.425312 | yes | 1.048E-07 | 619.32477 | 47.531456 |
| IMEIVDAITTTAQS | 17 | P08237 | ATP-dependent 6-phosphofructokinase, n | PFKM | 85.183 | 3 | 52.425312 | yes | 4.381E-07 | 638.66315 | 47.359638 |
| KSFEQISANITK | 12 | P08237 | ATP-dependent 6-phosphofructokinase, n | PFKM | 85.183 | 2 | 52.425312 | yes | 2.491E-07 | 683.37225 | 27.78722 |
| LLAHVRPPVSK | 11 | P08237 | ATP-dependent 6-phosphofructokinase, n | PFKM | 85.183 | 3 | 52.425312 | yes | 1.089E-06 | 406.25568 | 24.072392 |
| IDIIPNPQER | 10 | P08238 | Heat shock protein HSP 90-beta OS=Hon | HSP90AB1 | 83.265 | 2 | 55.02816 | no | 1.594E-07 | 597.82751 | 1917.3494 |
| ADLINNLGTIAK | 12 | P08238 | Heat shock protein HSP 90-beta OS=Hon | HSP90AB1 | 83.265 | 2 | 55.02816 | no | 9.809E-08 | 621.85626 | 1871.2103 |
| EDQTEYLEER | 10 | P08238 | Heat shock protein HSP 90-beta OS=Hon | HSP90AB1 | 83.265 | 2 | 55.02816 | no | 9.722E-08 | 656.28857 | 1679.1351 |
| EGLELPEDEEEK | 12 | P08238 | Heat shock protein HSP 90-beta OS=Hon | HSP90AB1 | 83.265 | 2 | 55.02816 | no | 1.57E-07 | 708.82245 | 1411.9319 |
| HFSVEGQLEFR | 11 | P08238 | Heat shock protein HSP 90-beta OS=Hon | HSP90AB1 | 83.265 | 2 | 55.02816 | no | 1.082E-07 | 674.83588 | 1404.9816 |
| SIYYITGESK | 10 | P08238 | Heat shock protein HSP 90-beta OS=Hon | HSP90AB1 | 83.265 | 2 | 55.02816 | no | 1.456E-07 | 580.79535 | 1349.241 |
| HLEINPDHPIVETL | 15 | P08238 | Heat shock protein HSP 90-beta OS=Hon | HSP90AB1 | 83.265 | 3 | 55.02816 | yes | 1.08E-07 | 594.9881 | 1290.9692 |
| ELISNASDALDK | 12 | P08238 | Heat shock protein HSP 90-beta OS=Hon | HSP90AB1 | 83.265 | 2 | 55.02816 | no | 9.722E-08 | 638.32501 | 1252.5005 |
| YIDQEELNK | 9 | P08238 | Heat shock protein HSP 90-beta OS=Hon | HSP90AB1 | 83.265 | 2 | 55.02816 | no | 2.88E-07 | 576.28259 | 1202.0326 |

| | | | | | | | | | | | |
|---|---|---|---|---|---|---|---|---|---|---|---|
| SLTNDWEDHLAVI | 13 | P08238 | Heat shock protein HSP 90-beta OS=Hon | HSP90AB1 | 83.265 | 2 | 55.02816 | no | 1.252E-07 | 764.37555 | 1041.6041 |
| ADHGEPIGR | 9 | P08238 | Heat shock protein HSP 90-beta OS=Hon | HSP90AB1 | 83.265 | 2 | 55.02816 | yes | 1.154E-07 | 476.23578 | 976.11407 |
| EQVANSAFVER | 11 | P08238 | Heat shock protein HSP 90-beta OS=Hon | HSP90AB1 | 83.265 | 2 | 55.02816 | no | 9.722E-08 | 625.31219 | 829.44983 |
| RAPFDLFENK | 10 | P08238 | Heat shock protein HSP 90-beta OS=Hon | HSP90AB1 | 83.265 | 2 | 55.02816 | no | 4.993E-07 | 618.8222 | 803.48175 |
| YESLTDPSK | 9 | P08238 | Heat shock protein HSP 90-beta OS=Hon | HSP90AB1 | 83.265 | 2 | 55.02816 | no | 4.15E-07 | 520.25073 | 585.45135 |
| HNDDEQYAWESS. | 20 | P08238 | Heat shock protein HSP 90-beta OS=Hon | HSP90AB1 | 83.265 | 3 | 55.02816 | no | 9.722E-08 | 752.65778 | 460.11731 |
| LGIHEDSTNR | 10 | P08238 | Heat shock protein HSP 90-beta OS=Hon | HSP90AB1 | 83.265 | 2 | 55.02816 | yes | 9.722E-08 | 571.28345 | 220.55424 |
| CLELFSELAEDK | 12 | P08238 | Heat shock protein HSP 90-beta OS=Hon | HSP90AB1 | 83.265 | 2 | 55.02816 | yes | 9.727E-08 | 727.3476 | 175.36584 |
| APFDLFENK | 9 | P08238 | Heat shock protein HSP 90-beta OS=Hon | HSP90AB1 | 83.265 | 2 | 55.02816 | no | 7.718E-07 | 540.77167 | 143.3362 |
| CLELFSELAEDKEI | 16 | P08238 | Heat shock protein HSP 90-beta OS=Hon | HSP90AB1 | 83.265 | 3 | 55.02816 | yes | 1.165E-07 | 663.31543 | 105.73214 |
| DNSTMGYMMAK | 11 | P08238 | Heat shock protein HSP 90-beta OS=Hon | HSP90AB1 | 83.265 | 2 | 55.02816 | no | 9.722E-08 | 624.75653 | 95.291962 |
| AKFENLCK | 8 | P08238 | Heat shock protein HSP 90-beta OS=Hon | HSP90AB1 | 83.265 | 2 | 55.02816 | no | 2.138E-07 | 505.26041 | 86.177704 |
| TLTLVDTGIGMTK | 13 | P08238 | Heat shock protein HSP 90-beta OS=Hon | HSP90AB1 | 83.265 | 2 | 55.02816 | yes | 6.103E-07 | 683.36835 | 64.157021 |
| RLSELLR | 7 | P08238 | Heat shock protein HSP 90-beta OS=Hon | HSP90AB1 | 83.265 | 2 | 55.02816 | yes | 6.5E-07 | 443.77707 | 56.850342 |
| FYEAFSK | 7 | P08238 | Heat shock protein HSP 90-beta OS=Hon | HSP90AB1 | 83.265 | 2 | 55.02816 | yes | 1.77E-06 | 446.21597 | 42.594116 |
| KHLEINPDHPIVET | 16 | P08238 | Heat shock protein HSP 90-beta OS=Hon | HSP90AB1 | 83.265 | 4 | 55.02816 | yes | 1.517E-07 | 478.51663 | 37.190147 |
| ALLFIPR | 7 | P08238 | Heat shock protein HSP 90-beta OS=Hon | HSP90AB1 | 83.265 | 2 | 55.02816 | yes | 7.458E-07 | 415.26834 | 29.12348 |
| NQGFDVVLVDTA( | 14 | P08240 | Signal recognition particle receptor subun | SRPRA | 69.812 | 2 | 53.523254 | yes | 2.894E-07 | 745.88354 | 476.93268 |
| ILTLTYVDK | 9 | P08240 | Signal recognition particle receptor subun | SRPRA | 69.812 | 2 | 53.523254 | yes | 2.94E-07 | 533.31317 | 457.31665 |
| LIDGIVLTK | 9 | P08240 | Signal recognition particle receptor subun | SRPRA | 69.812 | 2 | 53.523254 | yes | 1.376E-07 | 486.31042 | 450.65112 |
| LIDDVHR | 7 | P08240 | Signal recognition particle receptor subun | SRPRA | 69.812 | 2 | 53.523254 | yes | 4.608E-07 | 434.23779 | 404.39618 |
| VWELGGCANK | 10 | P08240 | Signal recognition particle receptor subun | SRPRA | 69.812 | 2 | 53.523254 | yes | 2.25E-07 | 567.27405 | 364.38983 |
| MQDNAPLMTALA | 13 | P08240 | Signal recognition particle receptor subun | SRPRA | 69.812 | 2 | 53.523254 | yes | 1.532E-07 | 702.35468 | 341.4856 |
| VMGTFSTVTSTVK | 13 | P08240 | Signal recognition particle receptor subun | SRPRA | 69.812 | 2 | 53.523254 | yes | 9.722E-08 | 679.35522 | 312.91455 |
| VGAAISMTYITSK | 13 | P08240 | Signal recognition particle receptor subun | SRPRA | 69.812 | 2 | 53.523254 | yes | 2.964E-07 | 671.35779 | 214.62949 |
| LSALHPPEK | 9 | P08240 | Signal recognition particle receptor subun | SRPRA | 69.812 | 2 | 53.523254 | yes | 9.722E-08 | 496.2822 | 213.07077 |
| SVLLQER | 7 | P08240 | Signal recognition particle receptor subun | SRPRA | 69.812 | 2 | 53.523254 | yes | 5.127E-07 | 422.74799 | 168.03943 |
| SGLPVGPENGVEL | 15 | P08240 | Signal recognition particle receptor subun | SRPRA | 69.812 | 2 | 53.523254 | yes | 2.826E-07 | 741.89355 | 163.86804 |
| GTLGGMFGMLK | 11 | P08240 | Signal recognition particle receptor subun | SRPRA | 69.812 | 2 | 53.523254 | yes | 1.524E-07 | 556.28558 | 140.41113 |
| ALADHSMAQTPR | 12 | P08240 | Signal recognition particle receptor subun | SRPRA | 69.812 | 2 | 53.523254 | yes | 1.3E-07 | 649.31952 | 95.52459 |
| GGNNSFTHEALTL | 14 | P08240 | Signal recognition particle receptor subun | SRPRA | 69.812 | 2 | 53.523254 | yes | 3.202E-06 | 744.87573 | 47.281364 |
| GGNNSFTHEALTL | 14 | P08240 | Signal recognition particle receptor subun | SRPRA | 69.812 | 2 | 53.523254 | yes | 2.457E-06 | 496.91956 | 47.281364 |
| GGNNSFTHEALTL | 14 | P08240 | Signal recognition particle receptor subun | SRPRA | 69.812 | 3 | 53.523254 | yes | 3.202E-06 | 744.87573 | 47.281364 |
| GGNNSFTHEALTL | 14 | P08240 | Signal recognition particle receptor subun | SRPRA | 69.812 | 3 | 53.523254 | yes | 2.457E-06 | 496.91956 | 47.281364 |
| TTAAHGLELR | 10 | P08243 | Asparagine synthetase [glutamine-hydroly | ASNS | 64.37 | 2 | 52.330677 | yes | 9.132E-07 | 534.79346 | 195.06548 |
| LFPGFEIETVK | 11 | P08243 | Asparagine synthetase [glutamine-hydroly | ASNS | 64.37 | 2 | 52.330677 | yes | 9.722E-08 | 640.35028 | 163.87152 |
| ILFNNAVK | 8 | P08243 | Asparagine synthetase [glutamine-hydroly | ASNS | 64.37 | 2 | 52.330677 | yes | 1.93E-07 | 459.77402 | 143.74812 |
| LAVVDPLFGMQPI | 14 | P08243 | Asparagine synthetase [glutamine-hydroly | ASNS | 64.37 | 2 | 52.330677 | yes | 2.628E-06 | 778.43689 | 130.40904 |
| ILQEYVEHQVDDA | 21 | P08243 | Asparagine synthetase [glutamine-hydroly | ASNS | 64.37 | 3 | 52.330677 | yes | 1.16E-07 | 802.04724 | 124.96674 |

| | | | | | | | | | | | |
|---|---|---|---|---|---|---|---|---|---|---|---|
| AMTEDGFLAVCSE | 15 | P08243 | Asparagine synthetase [glutamine-hydroly | ASNS | 64.37 | 2 | 52.330677 | yes | 9.823E-08 | 814.86835 | 90.72126 |
| EAFSDGITSVK | 11 | P08243 | Asparagine synthetase [glutamine-hydroly | ASNS | 64.37 | 2 | 52.330677 | yes | 2.143E-06 | 577.29041 | 88.65686 |
| ETFEDSNLIPK | 11 | P08243 | Asparagine synthetase [glutamine-hydroly | ASNS | 64.37 | 2 | 52.330677 | yes | 6.852E-07 | 646.82208 | 47.278332 |
| VDGEIILHLYDK | 12 | P08243 | Asparagine synthetase [glutamine-hydroly | ASNS | 64.37 | 2 | 52.330677 | yes | 7.999E-07 | 707.88245 | 35.414429 |
| VEPFLPGHYEVLD | 19 | P08243 | Asparagine synthetase [glutamine-hydroly | ASNS | 64.37 | 3 | 52.330677 | yes | 4.848E-06 | 718.05273 | 31.932552 |
| EILAELTGR | 9 | P08559 | Pyruvate dehydrogenase E1 component su | PDHA1 | 43.296 | 2 | 56.151543 | yes | 2.265E-07 | 501.28494 | 249.6636 |
| LEEGPPVTTVLTR | 13 | P08559 | Pyruvate dehydrogenase E1 component su | PDHA1 | 43.296 | 2 | 56.151543 | yes | 9.722E-08 | 706.39319 | 213.9799 |
| YGMGTSVER | 9 | P08559 | Pyruvate dehydrogenase E1 component su | PDHA1 | 43.296 | 2 | 56.151543 | yes | 1.526E-07 | 500.23184 | 182.73955 |
| GPILMELQTYR | 11 | P08559 | Pyruvate dehydrogenase E1 component su | PDHA1 | 43.296 | 2 | 56.151543 | no | 9.968E-08 | 660.85266 | 121.04055 |
| LPCIFICENNR | 11 | P08559 | Pyruvate dehydrogenase E1 component su | PDHA1 | 43.296 | 2 | 56.151543 | yes | 9.722E-08 | 718.34467 | 91.526932 |
| MVNSNLASVEELK | 13 | P08559 | Pyruvate dehydrogenase E1 component su | PDHA1 | 43.296 | 2 | 56.151543 | yes | 1.331E-07 | 717.3689 | 70.901474 |
| VDGMDILCVR | 10 | P08559 | Pyruvate dehydrogenase E1 component su | PDHA1 | 43.296 | 2 | 56.151543 | yes | 6.251E-07 | 589.28882 | 70.264153 |
| AHNQDLGLAGSCI | 15 | P08572 | Collagen alpha-2(IV) chain OS=Homo sa | COL4A2 | 167.554 | 3 | 44.168343 | yes | 2.295E-07 | 528.26331 | 79.205864 |
| AHNQDLGLAGSCI | 15 | P08572 | Collagen alpha-2(IV) chain OS=Homo sa | COL4A2 | 167.554 | 3 | 44.168343 | yes | 3.971E-07 | 791.89136 | 79.205864 |
| AHNQDLGLAGSCI | 15 | P08572 | Collagen alpha-2(IV) chain OS=Homo sa | COL4A2 | 167.554 | 2 | 44.168343 | yes | 2.295E-07 | 528.26331 | 79.205864 |
| AHNQDLGLAGSCI | 15 | P08572 | Collagen alpha-2(IV) chain OS=Homo sa | COL4A2 | 167.554 | 2 | 44.168343 | yes | 3.971E-07 | 791.89136 | 79.205864 |
| SVSIGYLLVK | 10 | P08572 | Collagen alpha-2(IV) chain OS=Homo sa | COL4A2 | 167.554 | 2 | 44.168343 | yes | 1.33E-06 | 539.82898 | 90.185394 |
| PGSPGLPGMPGR | 12 | P08572 | Collagen alpha-2(IV) chain OS=Homo sa | COL4A2 | 167.554 | 2 | 44.168343 | yes | 5.455E-07 | 569.78729 | 72.591888 |
| PYPNSEAAR | 9 | P08574 | Cytochrome c1, heme protein, mitochond | CYC1 | 35.422 | 2 | 48.410046 | yes | 7.589E-07 | 502.74344 | 1020.1095 |
| LAYRPPK | 7 | P08574 | Cytochrome c1, heme protein, mitochond | CYC1 | 35.422 | 2 | 48.410046 | yes | 1.364E-07 | 422.75562 | 682.48547 |
| HLVGVCYTEDEAK | 13 | P08574 | Cytochrome c1, heme protein, mitochond | CYC1 | 35.422 | 2 | 48.410046 | yes | 3.854E-07 | 760.85614 | 414.3811 |
| WASEPEHDHR | 10 | P08574 | Cytochrome c1, heme protein, mitochond | CYC1 | 35.422 | 2 | 48.410046 | yes | 1.276E-07 | 632.27869 | 254.60672 |
| AANNGALPPDLSY | 16 | P08574 | Cytochrome c1, heme protein, mitochond | CYC1 | 35.422 | 2 | 48.410046 | yes | 2.691E-07 | 835.94666 | 180.63516 |
| DVCTFLR | 7 | P08574 | Cytochrome c1, heme protein, mitochond | CYC1 | 35.422 | 2 | 48.410046 | yes | 1.152E-07 | 455.7262 | 175.28981 |
| LFDYFPK | 7 | P08574 | Cytochrome c1, heme protein, mitochond | CYC1 | 35.422 | 2 | 48.410046 | yes | 1.858E-07 | 465.242 | 67.564308 |
| ELGSSTNALR | 10 | P08579 | U2 small nuclear ribonucleoprotein B'' OS | SNRPB2 | 25.487 | 2 | 45.591427 | yes | 8.599E-07 | 524.27509 | 818.92029 |
| TVEQTATTTNK | 11 | P08579 | U2 small nuclear ribonucleoprotein B'' OS | SNRPB2 | 25.487 | 2 | 45.591427 | yes | 1.803E-07 | 597.30408 | 677.95837 |
| QLQGFPFYGK | 10 | P08579 | U2 small nuclear ribonucleoprotein B'' OS | SNRPB2 | 25.487 | 2 | 45.591427 | yes | 7.328E-07 | 592.80859 | 622.15186 |
| GQAFVIFK | 8 | P08579 | U2 small nuclear ribonucleoprotein B'' OS | SNRPB2 | 25.487 | 2 | 45.591427 | no | 2.258E-07 | 455.26328 | 182.37592 |
| PNHTIYINNMNDK | 13 | P08579 | U2 small nuclear ribonucleoprotein B'' OS | SNRPB2 | 25.487 | 2 | 45.591427 | yes | 2.913E-07 | 787.375 | 58.715912 |
| GYAFIEYEHER | 11 | P08621 | U1 small nuclear ribonucleoprotein 70 kD | SNRNP70 | 51.557 | 2 | 51.770782 | yes | 9.892E-08 | 707.32532 | 6845.7759 |
| DPIPYLPPLEK | 11 | P08621 | U1 small nuclear ribonucleoprotein 70 kD | SNRNP70 | 51.557 | 2 | 51.770782 | yes | 2.72E-07 | 641.35809 | 6042.2642 |
| QQEVETELK | 9 | P08621 | U1 small nuclear ribonucleoprotein 70 kD | SNRNP70 | 51.557 | 2 | 51.770782 | yes | 9.722E-08 | 552.28259 | 2265.3691 |
| VNYDTTESK | 9 | P08621 | U1 small nuclear ribonucleoprotein 70 kD | SNRNP70 | 51.557 | 2 | 51.770782 | yes | 1.086E-07 | 528.74585 | 2160.7893 |
| EFEVYGPIK | 9 | P08621 | U1 small nuclear ribonucleoprotein 70 kD | SNRNP70 | 51.557 | 2 | 51.770782 | yes | 1.641E-07 | 541.28186 | 1385.1108 |
| RGGADVNIR | 9 | P08621 | U1 small nuclear ribonucleoprotein 70 kD | SNRNP70 | 51.557 | 2 | 51.770782 | yes | 1.146E-07 | 479.26486 | 412.36142 |
| REFEVYGPIK | 10 | P08621 | U1 small nuclear ribonucleoprotein 70 kD | SNRNP70 | 51.557 | 2 | 51.770782 | yes | 2.154E-07 | 619.3324 | 394.86459 |
| VLVDVER | 7 | P08621 | U1 small nuclear ribonucleoprotein 70 kD | SNRNP70 | 51.557 | 2 | 51.770782 | yes | 2.49E-07 | 415.24252 | 392.7674 |
| IHMVYSK | 7 | P08621 | U1 small nuclear ribonucleoprotein 70 kD | SNRNP70 | 51.557 | 2 | 51.770782 | yes | 1.611E-07 | 439.23364 | 379.63846 |

| | | | | | | | | | | | |
|---|---|---|---|---|---|---|---|---|---|---|---|
| RQQEVETELK | 10 | P08621 | U1 small nuclear ribonucleoprotein 70 kD | SNRNP70 | 51.557 | 2 | 51.770782 | yes | 1.578E-07 | 630.33313 | 330.6814 |
| YDERPGPSPLPHR | 13 | P08621 | U1 small nuclear ribonucleoprotein 70 kD | SNRNP70 | 51.557 | 2 | 51.770782 | yes | 1.526E-07 | 760.88385 | 298.82669 |
| RVLVDVER | 8 | P08621 | U1 small nuclear ribonucleoprotein 70 kD | SNRNP70 | 51.557 | 2 | 51.770782 | yes | 9.722E-08 | 493.29309 | 265.56323 |
| DMHSAYK | 7 | P08621 | U1 small nuclear ribonucleoprotein 70 kD | SNRNP70 | 51.557 | 2 | 51.770782 | yes | 1.086E-07 | 426.18945 | 219.9928 |
| HHNQPYCGIAPYII | 14 | P08621 | U1 small nuclear ribonucleoprotein 70 kD | SNRNP70 | 51.557 | 3 | 51.770782 | yes | 9.894E-08 | 575.94745 | 155.4059 |
| GGGGGQDNGLEGI | 18 | P08621 | U1 small nuclear ribonucleoprotein 70 kD | SNRNP70 | 51.557 | 2 | 51.770782 | yes | 4.251E-07 | 830.36951 | 83.963219 |
| DELLGEKPEVK | 11 | P08651 | Nuclear factor 1 C-type OS=Homo sapien | NFIC | 55.675 | 2 | 53.91512 | yes | 1.004E-07 | 628.84027 | 2021.5947 |
| EDFVLSITGK | 10 | P08651 | Nuclear factor 1 C-type OS=Homo sapien | NFIC | 55.675 | 2 | 53.91512 | yes | 1.12E-07 | 554.79785 | 1966.4083 |
| AVKDELLGEKPEV | 14 | P08651 | Nuclear factor 1 C-type OS=Homo sapien | NFIC | 55.675 | 3 | 53.91512 | yes | 2.718E-07 | 518.96277 | 936.07593 |
| AVKDELLGEKPEV | 14 | P08651 | Nuclear factor 1 C-type OS=Homo sapien | NFIC | 55.675 | 3 | 53.91512 | yes | 9.722E-08 | 777.94049 | 936.07593 |
| AVKDELLGEKPEV | 14 | P08651 | Nuclear factor 1 C-type OS=Homo sapien | NFIC | 55.675 | 2 | 53.91512 | yes | 2.718E-07 | 518.96277 | 936.07593 |
| AVKDELLGEKPEV | 14 | P08651 | Nuclear factor 1 C-type OS=Homo sapien | NFIC | 55.675 | 2 | 53.91512 | yes | 9.722E-08 | 777.94049 | 936.07593 |
| SPFNSPSPQDSPR | 13 | P08651 | Nuclear factor 1 C-type OS=Homo sapien | NFIC | 55.675 | 2 | 53.91512 | yes | 1.364E-07 | 708.33112 | 1531.7152 |
| DIRPECR | 7 | P08651 | Nuclear factor 1 C-type OS=Homo sapien | NFIC | 55.675 | 2 | 53.91512 | yes | 8.863E-07 | 473.23218 | 1390.8878 |
| NWTEDMEGGISSP | 15 | P08651 | Nuclear factor 1 C-type OS=Homo sapien | NFIC | 55.675 | 2 | 53.91512 | yes | 1.065E-07 | 825.37744 | 663.59229 |
| NWTEDMEGGISSP | 15 | P08651 | Nuclear factor 1 C-type OS=Homo sapien | NFIC | 55.675 | 2 | 53.91512 | yes | 1.662E-07 | 833.37488 | 663.59229 |
| PVIAVHSGIAR | 11 | P08651 | Nuclear factor 1 C-type OS=Homo sapien | NFIC | 55.675 | 2 | 53.91512 | yes | 1.164E-07 | 560.33527 | 1193.3269 |
| KAPGCVLSNPDQK | 13 | P08651 | Nuclear factor 1 C-type OS=Homo sapien | NFIC | 55.675 | 2 | 53.91512 | yes | 4.589E-07 | 707.36139 | 949.25909 |
| APGCVLSNPDQK | 12 | P08651 | Nuclear factor 1 C-type OS=Homo sapien | NFIC | 55.675 | 2 | 53.91512 | yes | 9.722E-08 | 643.3139 | 748.78937 |
| SPHPSSALHFPTTS | 30 | P08651 | Nuclear factor 1 C-type OS=Homo sapien | NFIC | 55.675 | 5 | 53.91512 | yes | 1.112E-07 | 653.33868 | 255.24924 |
| SPHPSSALHFPTTS | 30 | P08651 | Nuclear factor 1 C-type OS=Homo sapien | NFIC | 55.675 | 5 | 53.91512 | yes | 1.479E-07 | 816.42157 | 255.24924 |
| SPHPSSALHFPTTS | 30 | P08651 | Nuclear factor 1 C-type OS=Homo sapien | NFIC | 55.675 | 4 | 53.91512 | yes | 1.112E-07 | 653.33868 | 255.24924 |
| SPHPSSALHFPTTS | 30 | P08651 | Nuclear factor 1 C-type OS=Homo sapien | NFIC | 55.675 | 4 | 53.91512 | yes | 1.479E-07 | 816.42157 | 255.24924 |
| TLPSTSSSGSK | 11 | P08651 | Nuclear factor 1 C-type OS=Homo sapien | NFIC | 55.675 | 2 | 53.91512 | yes | 4.656E-07 | 526.26691 | 508.2829 |
| LSSFTQHHR | 9 | P08651 | Nuclear factor 1 C-type OS=Homo sapien | NFIC | 55.675 | 2 | 53.91512 | yes | 2.651E-06 | 556.78345 | 401.71213 |
| YPPHLNPQDPLK | 12 | P08651 | Nuclear factor 1 C-type OS=Homo sapien | NFIC | 55.675 | 2 | 53.91512 | yes | 2.478E-06 | 709.875 | 240.0442 |
| AAQCGHPVLCVQI | 21 | P08651 | Nuclear factor 1 C-type OS=Homo sapien | NFIC | 55.675 | 3 | 53.91512 | yes | 1.002E-07 | 760.0614 | 55.445183 |
| AAQCGHPVLCVQI | 21 | P08651 | Nuclear factor 1 C-type OS=Homo sapien | NFIC | 55.675 | 3 | 53.91512 | yes | 3.88E-07 | 570.29785 | 55.445183 |
| AAQCGHPVLCVQI | 21 | P08651 | Nuclear factor 1 C-type OS=Homo sapien | NFIC | 55.675 | 4 | 53.91512 | yes | 1.002E-07 | 760.0614 | 55.445183 |
| AAQCGHPVLCVQI | 21 | P08651 | Nuclear factor 1 C-type OS=Homo sapien | NFIC | 55.675 | 4 | 53.91512 | yes | 3.88E-07 | 570.29785 | 55.445183 |
| MPSHCLSAQMLAI | 20 | P08651 | Nuclear factor 1 C-type OS=Homo sapien | NFIC | 55.675 | 3 | 53.91512 | yes | 1.163E-07 | 719.69702 | 67.548683 |
| LGDLYEEEMR | 10 | P08670 | Vimentin OS=Homo sapiens OX=9606 G | VIM | 53.652 | 2 | 52.919807 | yes | 1.369E-07 | 627.78717 | 5037.4102 |
| SLYASSPGGVYAT | 14 | P08670 | Vimentin OS=Homo sapiens OX=9606 G | VIM | 53.652 | 2 | 52.919807 | yes | 1.174E-07 | 714.8595 | 4778.5127 |
| LQEEMLQR | 8 | P08670 | Vimentin OS=Homo sapiens OX=9606 G | VIM | 53.652 | 2 | 52.919807 | yes | 1.902E-07 | 523.76862 | 4582.7192 |
| EEAENTLQSFR | 11 | P08670 | Vimentin OS=Homo sapiens OX=9606 G | VIM | 53.652 | 2 | 52.919807 | yes | 9.927E-08 | 662.31238 | 4054.2212 |
| MALDIEIATYR | 11 | P08670 | Vimentin OS=Homo sapiens OX=9606 G | VIM | 53.652 | 2 | 52.919807 | no | 1.128E-07 | 648.33685 | 1934.5857 |
| MALDIEIATYR | 11 | P08670 | Vimentin OS=Homo sapiens OX=9606 G | VIM | 53.652 | 2 | 52.919807 | no | 5.089E-07 | 656.33429 | 1934.5857 |
| VELQELNDR | 9 | P08670 | Vimentin OS=Homo sapiens OX=9606 G | VIM | 53.652 | 2 | 52.919807 | no | 1.108E-07 | 558.28821 | 3738.0393 |
| LQDEIQNMK | 9 | P08670 | Vimentin OS=Homo sapiens OX=9606 G | VIM | 53.652 | 2 | 52.919807 | yes | 1.003E-07 | 559.77917 | 1613.3195 |

| | | | | | | | | | | | |
|---|---|---|---|---|---|---|---|---|---|---|---|
| LQDEIQNMK | 9 | P08670 | Vimentin OS=Homo sapiens OX=9606 G | VIM | 53.652 | 2 | 52.919807 | yes | 2.23E-05 | 567.77661 | 1613.3195 |
| MFGGPGTASR | 10 | P08670 | Vimentin OS=Homo sapiens OX=9606 G | VIM | 53.652 | 2 | 52.919807 | yes | 1.446E-07 | 490.73456 | 2373.335 |
| QVQSLTCEVDALK | 13 | P08670 | Vimentin OS=Homo sapiens OX=9606 G | VIM | 53.652 | 2 | 52.919807 | yes | 2.058E-07 | 745.87958 | 2049.3369 |
| FADLSEAANR | 10 | P08670 | Vimentin OS=Homo sapiens OX=9606 G | VIM | 53.652 | 2 | 52.919807 | yes | 1.19E-07 | 547.26727 | 1901.6151 |
| NLQEAEEWYK | 10 | P08670 | Vimentin OS=Homo sapiens OX=9606 G | VIM | 53.652 | 2 | 52.919807 | no | 1.036E-07 | 655.30658 | 1881.3446 |
| EYQDLLNVK | 9 | P08670 | Vimentin OS=Homo sapiens OX=9606 G | VIM | 53.652 | 2 | 52.919807 | no | 1.067E-07 | 561.29547 | 1812.3169 |
| ETNLDSLPLVDTH | 15 | P08670 | Vimentin OS=Homo sapiens OX=9606 G | VIM | 53.652 | 2 | 52.919807 | yes | 1.211E-07 | 834.9256 | 726.90619 |
| ETNLDSLPLVDTH | 15 | P08670 | Vimentin OS=Homo sapiens OX=9606 G | VIM | 53.652 | 2 | 52.919807 | yes | 1.682E-07 | 556.95282 | 726.90619 |
| ETNLDSLPLVDTH | 15 | P08670 | Vimentin OS=Homo sapiens OX=9606 G | VIM | 53.652 | 3 | 52.919807 | yes | 1.211E-07 | 834.9256 | 726.90619 |
| ETNLDSLPLVDTH | 15 | P08670 | Vimentin OS=Homo sapiens OX=9606 G | VIM | 53.652 | 3 | 52.919807 | yes | 1.682E-07 | 556.95282 | 726.90619 |
| ILLAELEQLK | 10 | P08670 | Vimentin OS=Homo sapiens OX=9606 G | VIM | 53.652 | 2 | 52.919807 | yes | 1.364E-07 | 585.36066 | 1228.9341 |
| KVESLQEEIAFLK | 13 | P08670 | Vimentin OS=Homo sapiens OX=9606 G | VIM | 53.652 | 2 | 52.919807 | yes | 9.224E-07 | 767.42981 | 1026.8422 |
| DNLAEDIMR | 9 | P08670 | Vimentin OS=Homo sapiens OX=9606 G | VIM | 53.652 | 2 | 52.919807 | yes | 2.22E-07 | 538.75568 | 491.84381 |
| DNLAEDIMR | 9 | P08670 | Vimentin OS=Homo sapiens OX=9606 G | VIM | 53.652 | 2 | 52.919807 | yes | 1.081E-05 | 546.75311 | 491.84381 |
| TYSLGSALR | 9 | P08670 | Vimentin OS=Homo sapiens OX=9606 G | VIM | 53.652 | 2 | 52.919807 | yes | 2.411E-07 | 484.26401 | 673.53735 |
| QVDQLTNDK | 9 | P08670 | Vimentin OS=Homo sapiens OX=9606 G | VIM | 53.652 | 2 | 52.919807 | yes | 9.967E-08 | 530.76709 | 652.89331 |
| VESLQEEIAFLK | 12 | P08670 | Vimentin OS=Homo sapiens OX=9606 G | VIM | 53.652 | 2 | 52.919807 | yes | 1.25E-07 | 703.38232 | 508.24564 |
| EKLQEEMLQR | 10 | P08670 | Vimentin OS=Homo sapiens OX=9606 G | VIM | 53.652 | 2 | 52.919807 | yes | 1.084E-07 | 652.33734 | 431.31998 |
| QQYESVAAK | 9 | P08670 | Vimentin OS=Homo sapiens OX=9606 G | VIM | 53.652 | 2 | 52.919807 | yes | 1.002E-07 | 512.25891 | 396.82584 |
| TYSLGSALRPSTSR | 14 | P08670 | Vimentin OS=Homo sapiens OX=9606 G | VIM | 53.652 | 3 | 52.919807 | yes | 9.722E-08 | 499.26697 | 87.219254 |
| TYSLGSALRPSTSR | 14 | P08670 | Vimentin OS=Homo sapiens OX=9606 G | VIM | 53.652 | 3 | 52.919807 | yes | 1.6E-07 | 748.39679 | 87.219254 |
| TYSLGSALRPSTSR | 14 | P08670 | Vimentin OS=Homo sapiens OX=9606 G | VIM | 53.652 | 2 | 52.919807 | yes | 9.722E-08 | 499.26697 | 87.219254 |
| TYSLGSALRPSTSR | 14 | P08670 | Vimentin OS=Homo sapiens OX=9606 G | VIM | 53.652 | 2 | 52.919807 | yes | 1.6E-07 | 748.39679 | 87.219254 |
| QDVDNASLAR | 10 | P08670 | Vimentin OS=Homo sapiens OX=9606 G | VIM | 53.652 | 2 | 52.919807 | yes | 1.247E-07 | 544.7702 | 164.15271 |
| RQVDQLTNDK | 10 | P08670 | Vimentin OS=Homo sapiens OX=9606 G | VIM | 53.652 | 2 | 52.919807 | yes | 9.722E-08 | 608.81769 | 105.83701 |
| KLHEEEIQELQAQI | 35 | P08670 | Vimentin OS=Homo sapiens OX=9606 G | VIM | 53.652 | 5 | 52.919807 | yes | 9.722E-08 | 811.02771 | 45.08448 |
| VCEEIAIIPSK | 11 | P08708 | Small ribosomal subunit protein eS17 OS | RPS17 | 15.551 | 2 | 52.959507 | yes | 9.722E-08 | 629.83923 | 19439.154 |
| IAGYVTHLMK | 10 | P08708 | Small ribosomal subunit protein eS17 OS | RPS17 | 15.551 | 2 | 52.959507 | yes | 1.018E-07 | 566.81281 | 9668.0332 |
| IAGYVTHLMK | 10 | P08708 | Small ribosomal subunit protein eS17 OS | RPS17 | 15.551 | 2 | 52.959507 | yes | 1.772E-07 | 574.81024 | 9668.0332 |
| LGNDFHTNK | 9 | P08708 | Small ribosomal subunit protein eS17 OS | RPS17 | 15.551 | 2 | 52.959507 | yes | 2.894E-06 | 523.25671 | 11777.051 |
| RVCEEIAIIPSK | 12 | P08708 | Small ribosomal subunit protein eS17 OS | RPS17 | 15.551 | 2 | 52.959507 | yes | 4.32E-07 | 707.88977 | 7086.4663 |
| LLDFGSLSNLQVTQ | 22 | P08708 | Small ribosomal subunit protein eS17 OS | RPS17 | 15.551 | 3 | 52.959507 | yes | 9.974E-08 | 803.75427 | 941.13129 |
| LLDFGSLSNLQVTQ | 22 | P08708 | Small ribosomal subunit protein eS17 OS | RPS17 | 15.551 | 3 | 52.959507 | yes | 2.221E-07 | 809.08594 | 941.13129 |
| VCEEIAIIPSKK | 12 | P08708 | Small ribosomal subunit protein eS17 OS | RPS17 | 15.551 | 2 | 52.959507 | yes | 9.722E-08 | 693.88672 | 494.32318 |
| LGNDFHTNKR | 10 | P08708 | Small ribosomal subunit protein eS17 OS | RPS17 | 15.551 | 2 | 52.959507 | yes | 9.722E-08 | 601.30725 | 294.14453 |
| NKIAGYVTHLMK | 12 | P08708 | Small ribosomal subunit protein eS17 OS | RPS17 | 15.551 | 3 | 52.959507 | yes | 0.0001032 | 458.92358 | 194.10272 |
| RVCEEIAIIPSKK | 13 | P08708 | Small ribosomal subunit protein eS17 OS | RPS17 | 15.551 | 3 | 52.959507 | yes | 1.047E-07 | 514.96057 | 96.897446 |
| DNYVPEVSALDQE | 22 | P08708 | Small ribosomal subunit protein eS17 OS | RPS17 | 15.551 | 3 | 52.959507 | yes | 1.811E-07 | 830.40253 | 22.214752 |
| LALDIEIATYR | 11 | P08729 | Keratin, type II cytoskeletal 7 OS=Homo | KRT7 | 51.386 | 2 | 55.910484 | no | 9.819E-08 | 639.35864 | 3213.9436 |

| | | | | | | | | | | | |
|---|---|---|---|---|---|---|---|---|---|---|---|
| KLLEGEESR | 9 | P08729 | Keratin, type II cytoskeletal 7 OS=Homo | KRT7 | 51.386 | 2 | 55.910484 | no | 9.935E-08 | 530.78528 | 1763.7592 |
| LLEGEESR | 8 | P08729 | Keratin, type II cytoskeletal 7 OS=Homo | KRT7 | 51.386 | 2 | 55.910484 | no | 5.051E-07 | 466.73782 | 1070.8192 |
| SIHFSSPVFTSR | 12 | P08729 | Keratin, type II cytoskeletal 7 OS=Homo | KRT7 | 51.386 | 2 | 55.910484 | yes | 9.81E-08 | 703.85681 | 924.00867 |
| EVTINQSLLAPLR | 13 | P08729 | Keratin, type II cytoskeletal 7 OS=Homo | KRT7 | 51.386 | 2 | 55.910484 | yes | 9.882E-08 | 727.4223 | 756.32983 |
| FETLQAQAGK | 10 | P08729 | Keratin, type II cytoskeletal 7 OS=Homo | KRT7 | 51.386 | 2 | 55.910484 | yes | 1.375E-07 | 546.78784 | 624.05963 |
| LQAEIDNIK | 9 | P08729 | Keratin, type II cytoskeletal 7 OS=Homo | KRT7 | 51.386 | 2 | 55.910484 | yes | 1.12E-07 | 522.29022 | 559.49225 |
| LDADPSLQR | 9 | P08729 | Keratin, type II cytoskeletal 7 OS=Homo | KRT7 | 51.386 | 2 | 55.910484 | yes | 1.951E-07 | 507.76437 | 539.0755 |
| WTLLQEQK | 8 | P08729 | Keratin, type II cytoskeletal 7 OS=Homo | KRT7 | 51.386 | 2 | 55.910484 | yes | 2.828E-07 | 523.28748 | 458.78128 |
| AEAEAWYQTK | 10 | P08729 | Keratin, type II cytoskeletal 7 OS=Homo | KRT7 | 51.386 | 2 | 55.910484 | no | 1.202E-07 | 598.78278 | 328.21698 |
| GQLEALQVDGGR | 12 | P08729 | Keratin, type II cytoskeletal 7 OS=Homo | KRT7 | 51.386 | 2 | 55.910484 | yes | 5.067E-07 | 621.8255 | 251.85609 |
| PGGLGSSSLYGLG. | 16 | P08729 | Keratin, type II cytoskeletal 7 OS=Homo | KRT7 | 51.386 | 2 | 55.910484 | yes | 9.722E-08 | 739.88354 | 219.51306 |
| VDALNDEINFLR | 12 | P08729 | Keratin, type II cytoskeletal 7 OS=Homo | KRT7 | 51.386 | 2 | 55.910484 | yes | 1.088E-07 | 709.86737 | 208.67628 |
| QEELEAALQR | 10 | P08729 | Keratin, type II cytoskeletal 7 OS=Homo | KRT7 | 51.386 | 2 | 55.910484 | yes | 2.277E-07 | 593.80676 | 205.55447 |
| AKQEELEAALQR | 12 | P08729 | Keratin, type II cytoskeletal 7 OS=Homo | KRT7 | 51.386 | 2 | 55.910484 | yes | 1.028E-07 | 693.3728 | 193.79837 |
| SMQDVVEDFK | 10 | P08729 | Keratin, type II cytoskeletal 7 OS=Homo | KRT7 | 51.386 | 2 | 55.910484 | yes | 9.722E-08 | 599.27643 | 180.67899 |
| EYQELMSVK | 9 | P08729 | Keratin, type II cytoskeletal 7 OS=Homo | KRT7 | 51.386 | 2 | 55.910484 | no | 1.823E-07 | 563.77606 | 136.72023 |
| SLDLDGIIAEVK | 12 | P08729 | Keratin, type II cytoskeletal 7 OS=Homo | KRT7 | 51.386 | 2 | 55.910484 | yes | 1.342E-07 | 636.8559 | 120.10255 |
| SAYGGPVGAGIR | 12 | P08729 | Keratin, type II cytoskeletal 7 OS=Homo | KRT7 | 51.386 | 2 | 55.910484 | yes | 4.425E-06 | 552.79346 | 108.88866 |
| LPDIFEAQIAGLR | 13 | P08729 | Keratin, type II cytoskeletal 7 OS=Homo | KRT7 | 51.386 | 2 | 55.910484 | yes | 1.229E-07 | 721.90375 | 77.983131 |
| LEAAIAEAEER | 11 | P08729 | Keratin, type II cytoskeletal 7 OS=Homo | KRT7 | 51.386 | 2 | 55.910484 | yes | 2.092E-07 | 601.30658 | 74.828278 |
| KDVDAAYMSK | 10 | P08729 | Keratin, type II cytoskeletal 7 OS=Homo | KRT7 | 51.386 | 2 | 55.910484 | yes | 0.0001035 | 564.27368 | 8.1300497 |
| APSTYGGGLSVSSI | 14 | P08779 | Keratin, type I cytoskeletal 16 OS=Homo | KRT16 | 51.268 | 2 | 56.175457 | yes | 9.722E-08 | 669.83606 | 51.359066 |
| EVASNSELVQSSR | 13 | P08779 | Keratin, type I cytoskeletal 16 OS=Homo | KRT16 | 51.268 | 2 | 56.175457 | yes | 9.733E-08 | 703.34955 | 42.113598 |
| EVSSATNALR | 10 | P09012 | U1 small nuclear ribonucleoprotein A OS | SNRPA | 31.28 | 2 | 52.01242 | yes | 1.541E-07 | 524.27509 | 2662.3718 |
| SMQGFPFYDKPMI | 13 | P09012 | U1 small nuclear ribonucleoprotein A OS | SNRPA | 31.28 | 2 | 52.01242 | yes | 1.843E-07 | 802.37341 | 1508.8947 |
| ITQNNAMK | 8 | P09012 | U1 small nuclear ribonucleoprotein A OS | SNRPA | 31.28 | 2 | 52.01242 | yes | 1.002E-07 | 460.23694 | 1039.239 |
| SMQGFPFYDK | 10 | P09012 | U1 small nuclear ribonucleoprotein A OS | SNRPA | 31.28 | 2 | 52.01242 | yes | 9.722E-08 | 610.27625 | 632.05176 |
| PNHTIYINNLNEK | 13 | P09012 | U1 small nuclear ribonucleoprotein A OS | SNRPA | 31.28 | 2 | 52.01242 | yes | 1.08E-07 | 785.4046 | 554.51489 |
| HDIAFVEFDNEVQ | 18 | P09012 | U1 small nuclear ribonucleoprotein A OS | SNRPA | 31.28 | 3 | 52.01242 | yes | 9.848E-08 | 663.65356 | 533.21228 |
| SQETPATK | 8 | P09012 | U1 small nuclear ribonucleoprotein A OS | SNRPA | 31.28 | 2 | 52.01242 | yes | 3.473E-06 | 431.21927 | 32.642956 |
| LQLQAEER | 8 | P09038 | Fibroblast growth factor 2 OS=Homo sap | FGF2 | 30.77 | 2 | 51.834538 | yes | 2.224E-07 | 493.76691 | 531.94226 |
| CVTDECFFFER | 11 | P09038 | Fibroblast growth factor 2 OS=Homo sap | FGF2 | 30.77 | 2 | 51.834538 | yes | 9.722E-08 | 755.31049 | 62.318649 |
| LESNNYNTYR | 10 | P09038 | Fibroblast growth factor 2 OS=Homo sap | FGF2 | 30.77 | 2 | 51.834538 | yes | 1.004E-07 | 637.29401 | 45.052959 |
| KYTSWYVALK | 10 | P09038 | Fibroblast growth factor 2 OS=Homo sap | FGF2 | 30.77 | 2 | 51.834538 | yes | 1.548E-06 | 629.84515 | 39.516853 |
| QEDGSLCLVQFPS. | 14 | P09132 | Signal recognition particle 19 kDa proteir | SRP19 | 16.156 | 2 | 54.14435 | yes | 9.722E-08 | 818.39343 | 169.72745 |
| FICIYPAYLNNK | 12 | P09132 | Signal recognition particle 19 kDa proteir | SRP19 | 16.156 | 2 | 54.14435 | yes | 9.809E-08 | 758.38684 | 124.82735 |
| WMEEQAQSLIDK | 12 | P09234 | U1 small nuclear ribonucleoprotein C OS | SNRPC | 17.394 | 2 | 50.125343 | yes | 9.894E-08 | 739.35321 | 1063.0861 |
| WMEEQAQSLIDK | 12 | P09234 | U1 small nuclear ribonucleoprotein C OS | SNRPC | 17.394 | 2 | 50.125343 | yes | 1.232E-06 | 747.35065 | 1063.0861 |
| VRGEVAPDAK | 10 | P09382 | Galectin-1 OS=Homo sapiens OX=9606 ( | LGALS1 | 14.716 | 2 | 48.935806 | yes | 3.166E-07 | 521.28802 | 1767.5403 |

| | | | | | | | | | | | |
|---|---|---|---|---|---|---|---|---|---|---|---|
| DSNNLCLHFNPR | 12 | P09382 | Galectin-1 OS=Homo sapiens OX=9606 ( | LGALS1 | 14.716 | 2 | 48.935806 | yes | 1.264E-07 | 743.84644 | 1468.2906 |
| LPDGYEFK | 8 | P09382 | Galectin-1 OS=Homo sapiens OX=9606 ( | LGALS1 | 14.716 | 2 | 48.935806 | yes | 1.139E-07 | 484.73984 | 1394.6182 |
| DGGAWGTEQR | 10 | P09382 | Galectin-1 OS=Homo sapiens OX=9606 ( | LGALS1 | 14.716 | 2 | 48.935806 | yes | 1.084E-07 | 538.74139 | 1049.1639 |
| FNAHGDANTIVCN | 15 | P09382 | Galectin-1 OS=Homo sapiens OX=9606 ( | LGALS1 | 14.716 | 2 | 48.935806 | yes | 1.519E-07 | 824.3808 | 525.52441 |
| SFVLNLGK | 8 | P09382 | Galectin-1 OS=Homo sapiens OX=9606 ( | LGALS1 | 14.716 | 2 | 48.935806 | yes | 1.483E-07 | 439.26074 | 484.29318 |
| FKDPNAPK | 8 | P09429 | High mobility group protein B1 OS=Hom | HMGB1 | 24.894 | 2 | 54.50457 | no | 2.362E-07 | 458.74799 | 329.36032 |
| GEHPGLSIGDVAK | 13 | P09429 | High mobility group protein B1 OS=Hom | HMGB1 | 24.894 | 2 | 54.50457 | no | 1.074E-07 | 640.33569 | 147.5408 |
| MSSYAFFVQTCR | 12 | P09429 | High mobility group protein B1 OS=Hom | HMGB1 | 24.894 | 2 | 54.50457 | no | 9.722E-08 | 748.83667 | 143.97453 |
| RPPSAFFLFCSEYR | 16 | P09429 | High mobility group protein B1 OS=Hom | HMGB1 | 24.894 | 3 | 54.50457 | yes | 9.722E-08 | 668.00494 | 131.23476 |
| KHPDASVNFSEFSI | 14 | P09429 | High mobility group protein B1 OS=Hom | HMGB1 | 24.894 | 2 | 54.50457 | yes | 1.073E-07 | 796.88879 | 39.820045 |
| LGEMWNNTAADD | 13 | P09429 | High mobility group protein B1 OS=Hom | HMGB1 | 24.894 | 2 | 54.50457 | no | 5.35E-06 | 732.82483 | 28.794762 |
| IKGEHPGLSIGDVA | 15 | P09429 | High mobility group protein B1 OS=Hom | HMGB1 | 24.894 | 2 | 54.50457 | no | 0.0001146 | 760.92517 | 25.905458 |
| MEIQEIQLK | 9 | P09493 | Tropomyosin alpha-1 chain OS=Homo sa | TPM1 | 32.709 | 2 | 53.30372 | yes | 1.473E-07 | 566.30756 | 1709.0211 |
| LVIIESDLER | 10 | P09493 | Tropomyosin alpha-1 chain OS=Homo sa | TPM1 | 32.709 | 2 | 53.30372 | yes | 9.722E-08 | 593.83752 | 213.72597 |
| EDRYEEEIK | 9 | P09493 | Tropomyosin alpha-1 chain OS=Homo sa | TPM1 | 32.709 | 2 | 53.30372 | yes | 3.316E-07 | 605.78296 | 36.758736 |
| LQSEPESIR | 9 | P09496 | Clathrin light chain A OS=Homo sapiens | CLTA | 27.077 | 2 | 49.12297 | yes | 3.49E-07 | 529.77747 | 421.33237 |
| ELEEWYAR | 8 | P09496 | Clathrin light chain A OS=Homo sapiens | CLTA | 27.077 | 2 | 49.12297 | yes | 1.066E-07 | 548.25891 | 243.44214 |
| LCDFNPK | 7 | P09496 | Clathrin light chain A OS=Homo sapiens | CLTA | 27.077 | 2 | 49.12297 | yes | 4.441E-07 | 447.21292 | 202.55203 |
| LTQEPESIR | 9 | P09497 | Clathrin light chain B OS=Homo sapiens | CLTB | 25.191 | 2 | 47.386307 | yes | 1.563E-07 | 536.78528 | 312.72314 |
| VAQLCDFNPK | 10 | P09497 | Clathrin light chain B OS=Homo sapiens | CLTB | 25.191 | 2 | 47.386307 | yes | 1.33E-06 | 596.29498 | 194.8391 |
| DLEEWNQR | 8 | P09497 | Clathrin light chain B OS=Homo sapiens | CLTB | 25.191 | 2 | 47.386307 | yes | 1.341E-07 | 545.25165 | 189.3855 |
| LDEDLAAYCR | 10 | P09543 | 2',3'-cyclic-nucleotide 3'-phosphodiestera: | CNP | 47.579 | 2 | 53.25533 | yes | 9.722E-08 | 613.27954 | 537.24042 |
| AGQVFLEELGNHK | 13 | P09543 | 2',3'-cyclic-nucleotide 3'-phosphodiestera: | CNP | 47.579 | 2 | 53.25533 | yes | 1.55E-07 | 721.37537 | 355.52859 |
| ILVLDDTNHER | 11 | P09543 | 2',3'-cyclic-nucleotide 3'-phosphodiestera: | CNP | 47.579 | 2 | 53.25533 | yes | 2.819E-07 | 662.84644 | 328.81091 |
| APGAEEYAQQDVI | 14 | P09543 | 2',3'-cyclic-nucleotide 3'-phosphodiestera: | CNP | 47.579 | 2 | 53.25533 | yes | 1.119E-07 | 759.87537 | 247.00558 |
| NQWQLSADDLK | 11 | P09543 | 2',3'-cyclic-nucleotide 3'-phosphodiestera: | CNP | 47.579 | 2 | 53.25533 | yes | 1.061E-07 | 659.32532 | 237.82494 |
| MVSADAYK | 8 | P09543 | 2',3'-cyclic-nucleotide 3'-phosphodiestera: | CNP | 47.579 | 2 | 53.25533 | yes | 2.262E-06 | 442.71277 | 211.4848 |
| RPPGVLHCTTK | 11 | P09543 | 2',3'-cyclic-nucleotide 3'-phosphodiestera: | CNP | 47.579 | 3 | 53.25533 | yes | 1.839E-07 | 422.56427 | 103.26806 |
| RPPGVLHCTTK | 11 | P09543 | 2',3'-cyclic-nucleotide 3'-phosphodiestera: | CNP | 47.579 | 3 | 53.25533 | yes | 2.225E-07 | 633.34277 | 103.26806 |
| RPPGVLHCTTK | 11 | P09543 | 2',3'-cyclic-nucleotide 3'-phosphodiestera: | CNP | 47.579 | 2 | 53.25533 | yes | 1.839E-07 | 422.56427 | 103.26806 |
| RPPGVLHCTTK | 11 | P09543 | 2',3'-cyclic-nucleotide 3'-phosphodiestera: | CNP | 47.579 | 2 | 53.25533 | yes | 2.225E-07 | 633.34277 | 103.26806 |
| LSPTDNLPR | 9 | P09543 | 2',3'-cyclic-nucleotide 3'-phosphodiestera: | CNP | 47.579 | 2 | 53.25533 | yes | 1.984E-07 | 506.77472 | 183.22473 |
| GEEVGELSR | 9 | P09543 | 2',3'-cyclic-nucleotide 3'-phosphodiestera: | CNP | 47.579 | 2 | 53.25533 | yes | 1.096E-07 | 488.24072 | 134.82312 |
| MDLVTYFGK | 9 | P09543 | 2',3'-cyclic-nucleotide 3'-phosphodiestera: | CNP | 47.579 | 2 | 53.25533 | yes | 2.97E-07 | 537.27045 | 75.987679 |
| GAFSEEYKR | 9 | P09543 | 2',3'-cyclic-nucleotide 3'-phosphodiestera: | CNP | 47.579 | 2 | 53.25533 | yes | 3.854E-07 | 543.76434 | 53.144306 |
| AIFTGYYGK | 9 | P09543 | 2',3'-cyclic-nucleotide 3'-phosphodiestera: | CNP | 47.579 | 2 | 53.25533 | yes | 8.746E-07 | 510.26346 | 44.401463 |
| WQEVIPYTPAMQI | 13 | P09601 | Heme oxygenase 1 OS=Homo sapiens O> | HMOX1 | 32.819 | 2 | 40.860752 | yes | 3.801E-07 | 809.90594 | 103.14761 |
| VQDSAPVETPR | 11 | P09601 | Heme oxygenase 1 OS=Homo sapiens O> | HMOX1 | 32.819 | 2 | 40.860752 | yes | 7.243E-07 | 599.80676 | 59.83897 |
| TAGPGTTGQDR | 11 | P09629 | Homeobox protein Hox-B7 OS=Homo sa | HOXB7 | 24.015 | 2 | 46.008198 | yes | 4.156E-07 | 530.75452 | 139.34503 |

| | | | | | | | | | | | |
|---|---|---|---|---|---|---|---|---|---|---|---|
| DSDLAAESNFR | 11 | P09629 | Homeobox protein Hox-B7 OS=Homo sa | HOXB7 | 24.015 | 2 | 46.008198 | yes | 1.679E-07 | 612.7782 | 103.86549 |
| IEVIEIMTDR | 10 | P09651 | Heterogeneous nuclear ribonucleoprotein | HNRNPA1 | 38.747 | 2 | 51.898205 | no | 1.096E-07 | 609.82355 | 42029.512 |
| IEVIEIMTDR | 10 | P09651 | Heterogeneous nuclear ribonucleoprotein | HNRNPA1 | 38.747 | 2 | 51.898205 | no | 2.394E-07 | 617.82104 | 42029.512 |
| SSGPYGGGGQYFA | 14 | P09651 | Heterogeneous nuclear ribonucleoprotein | HNRNPA1 | 38.747 | 2 | 51.898205 | no | 1.482E-07 | 688.3175 | 43219.539 |
| KIFVGGIK | 8 | P09651 | Heterogeneous nuclear ribonucleoprotein | HNRNPA1 | 38.747 | 2 | 51.898205 | no | 1.092E-07 | 431.28146 | 15576.933 |
| DYFEQYGK | 8 | P09651 | Heterogeneous nuclear ribonucleoprotein | HNRNPA1 | 38.747 | 2 | 51.898205 | no | 1.849E-07 | 525.23236 | 10550.141 |
| SHFEQWGTLTDCV | 16 | P09651 | Heterogeneous nuclear ribonucleoprotein | HNRNPA1 | 38.747 | 3 | 51.898205 | no | 9.723E-08 | 655.96973 | 4493.7041 |
| SHFEQWGTLTDCV | 16 | P09651 | Heterogeneous nuclear ribonucleoprotein | HNRNPA1 | 38.747 | 3 | 51.898205 | no | 7.697E-07 | 661.30133 | 4493.7041 |
| GGGFGGNDNFGR | 12 | P09651 | Heterogeneous nuclear ribonucleoprotein | HNRNPA1 | 38.747 | 2 | 51.898205 | no | 1.082E-07 | 577.75232 | 6584.6455 |
| EDSQRPGAHLTVK | 13 | P09651 | Heterogeneous nuclear ribonucleoprotein | HNRNPA1 | 38.747 | 2 | 51.898205 | no | 1.022E-07 | 719.37585 | 5697.1934 |
| SESPKEPEQLR | 11 | P09651 | Heterogeneous nuclear ribonucleoprotein | HNRNPA1 | 38.747 | 3 | 51.898205 | yes | 1.577E-07 | 433.8895 | 830.07532 |
| SESPKEPEQLR | 11 | P09651 | Heterogeneous nuclear ribonucleoprotein | HNRNPA1 | 38.747 | 3 | 51.898205 | yes | 9.91E-08 | 650.33063 | 830.07532 |
| SESPKEPEQLR | 11 | P09651 | Heterogeneous nuclear ribonucleoprotein | HNRNPA1 | 38.747 | 2 | 51.898205 | yes | 1.577E-07 | 433.8895 | 830.07532 |
| SESPKEPEQLR | 11 | P09651 | Heterogeneous nuclear ribonucleoprotein | HNRNPA1 | 38.747 | 2 | 51.898205 | yes | 9.91E-08 | 650.33063 | 830.07532 |
| RGFAFVTFDDHDS | 16 | P09651 | Heterogeneous nuclear ribonucleoprotein | HNRNPA1 | 38.747 | 3 | 51.898205 | no | 6.104E-07 | 619.29181 | 1229.173 |
| NQGGYGGSSSSSS | 18 | P09651 | Heterogeneous nuclear ribonucleoprotein | HNRNPA1 | 38.747 | 2 | 51.898205 | no | 4.668E-07 | 847.8537 | 808.40186 |
| EDTEEHHLR | 9 | P09651 | Heterogeneous nuclear ribonucleoprotein | HNRNPA1 | 38.747 | 2 | 51.898205 | no | 4.677E-07 | 583.26526 | 616.80518 |
| QEMASASSSQR | 11 | P09651 | Heterogeneous nuclear ribonucleoprotein | HNRNPA1 | 38.747 | 2 | 51.898205 | no | 9.722E-08 | 591.2644 | 569.41736 |
| PGAHLTVK | 8 | P09651 | Heterogeneous nuclear ribonucleoprotein | HNRNPA1 | 38.747 | 2 | 51.898205 | no | 9.722E-08 | 411.74524 | 442.72113 |
| SSGPYGGGGQYFA | 16 | P09651 | Heterogeneous nuclear ribonucleoprotein | HNRNPA1 | 38.747 | 3 | 51.898205 | yes | 1.031E-07 | 543.59869 | 95.211456 |
| SSGPYGGGGQYFA | 16 | P09651 | Heterogeneous nuclear ribonucleoprotein | HNRNPA1 | 38.747 | 3 | 51.898205 | yes | 3.249E-07 | 814.89441 | 95.211456 |
| SSGPYGGGGQYFA | 16 | P09651 | Heterogeneous nuclear ribonucleoprotein | HNRNPA1 | 38.747 | 2 | 51.898205 | yes | 1.031E-07 | 543.59869 | 95.211456 |
| SSGPYGGGGQYFA | 16 | P09651 | Heterogeneous nuclear ribonucleoprotein | HNRNPA1 | 38.747 | 2 | 51.898205 | yes | 3.249E-07 | 814.89441 | 95.211456 |
| YHTVNGHNCEVR | 12 | P09651 | Heterogeneous nuclear ribonucleoprotein | HNRNPA1 | 38.747 | 3 | 51.898205 | no | 8.123E-07 | 495.89325 | 69.888222 |
| YHTVNGHNCEVR | 12 | P09651 | Heterogeneous nuclear ribonucleoprotein | HNRNPA1 | 38.747 | 3 | 51.898205 | no | 2.2E-06 | 743.33624 | 69.888222 |
| YHTVNGHNCEVR | 12 | P09651 | Heterogeneous nuclear ribonucleoprotein | HNRNPA1 | 38.747 | 2 | 51.898205 | no | 8.123E-07 | 495.89325 | 69.888222 |
| YHTVNGHNCEVR | 12 | P09651 | Heterogeneous nuclear ribonucleoprotein | HNRNPA1 | 38.747 | 2 | 51.898205 | no | 2.2E-06 | 743.33624 | 69.888222 |
| KLFIGGLSFETTDE | 17 | P09651 | Heterogeneous nuclear ribonucleoprotein | HNRNPA1 | 38.747 | 3 | 51.898205 | no | 9.722E-08 | 638.33868 | 95.689217 |
| SHFEQWGTLTDCV | 21 | P09651 | Heterogeneous nuclear ribonucleoprotein | HNRNPA1 | 38.747 | 3 | 51.898205 | no | 2.749E-07 | 841.05811 | 69.356888 |
| GFGFVTYATVEEV | 20 | P09651 | Heterogeneous nuclear ribonucleoprotein | HNRNPA1 | 38.747 | 3 | 51.898205 | no | 1.074E-07 | 722.00537 | 52.779892 |
| GLLQSGQIPGR | 11 | P09661 | U2 small nuclear ribonucleoprotein A' OS | SNRPA1 | 28.416 | 2 | 51.89585 | yes | 9.722E-08 | 563.32239 | 2216.719 |
| TFNPGAGLPTDK | 12 | P09661 | U2 small nuclear ribonucleoprotein A' OS | SNRPA1 | 28.416 | 2 | 51.89585 | yes | 9.722E-08 | 609.31171 | 1606.8981 |
| KLDGFPLLR | 9 | P09661 | U2 small nuclear ribonucleoprotein A' OS | SNRPA1 | 28.416 | 2 | 51.89585 | yes | 3.898E-07 | 529.82147 | 1306.7023 |
| TLLVNNNR | 8 | P09661 | U2 small nuclear ribonucleoprotein A' OS | SNRPA1 | 28.416 | 2 | 51.89585 | yes | 2.197E-07 | 472.26962 | 944.87915 |
| NAIANASTLAEVE | 14 | P09661 | U2 small nuclear ribonucleoprotein A' OS | SNRPA1 | 28.416 | 2 | 51.89585 | yes | 1.066E-07 | 729.88098 | 732.41034 |
| GGPSPGDVEAIK | 12 | P09661 | U2 small nuclear ribonucleoprotein A' OS | SNRPA1 | 28.416 | 2 | 51.89585 | yes | 7.148E-07 | 563.79059 | 574.13574 |
| SLTYLSILR | 9 | P09661 | U2 small nuclear ribonucleoprotein A' OS | SNRPA1 | 28.416 | 2 | 51.89585 | yes | 1.305E-07 | 533.31879 | 157.7114 |
| AYADFYR | 7 | P09669 | Cytochrome c oxidase subunit 6C OS=Hc | COX6C | 8.782 | 2 | 47.662777 | yes | 1.775E-07 | 453.21124 | 168.62823 |
| AGIFQSVK | 8 | P09669 | Cytochrome c oxidase subunit 6C OS=Hc | COX6C | 8.782 | 2 | 47.662777 | yes | 1.283E-07 | 425.24509 | 94.157547 |

| | | | | | | | | | | | |
|---|---|---|---|---|---|---|---|---|---|---|---|
| KAGIFQSVK | 9 | P09669 | Cytochrome c oxidase subunit 6C OS=Hc | COX6C | 8.782 | 2 | 47.662777 | yes | 4.229E-07 | 489.29257 | 74.977188 |
| APEVLPKPR | 9 | P09669 | Cytochrome c oxidase subunit 6C OS=Hc | COX6C | 8.782 | 2 | 47.662777 | yes | 1.53E-07 | 503.80585 | 70.819611 |
| IAPPEAPVTGYMF( | 15 | P09874 | Poly [ADP-ribose] polymerase 1 OS=Hor | PARP1 | 113.084 | 2 | 51.05493 | yes | 1.187E-07 | 789.40527 | 2923.866 |
| FYTLIPHDFGMK | 12 | P09874 | Poly [ADP-ribose] polymerase 1 OS=Hor | PARP1 | 113.084 | 2 | 51.05493 | yes | 1.299E-07 | 734.86829 | 1396.161 |
| FYTLIPHDFGMK | 12 | P09874 | Poly [ADP-ribose] polymerase 1 OS=Hor | PARP1 | 113.084 | 2 | 51.05493 | yes | 3.307E-07 | 490.24796 | 1396.161 |
| FYTLIPHDFGMK | 12 | P09874 | Poly [ADP-ribose] polymerase 1 OS=Hor | PARP1 | 113.084 | 3 | 51.05493 | yes | 1.299E-07 | 734.86829 | 1396.161 |
| FYTLIPHDFGMK | 12 | P09874 | Poly [ADP-ribose] polymerase 1 OS=Hor | PARP1 | 113.084 | 3 | 51.05493 | yes | 3.307E-07 | 490.24796 | 1396.161 |
| VVSEDFLQDVSAS | 15 | P09874 | Poly [ADP-ribose] polymerase 1 OS=Hor | PARP1 | 113.084 | 2 | 51.05493 | yes | 1.043E-07 | 812.90686 | 2512.2446 |
| TTNFAGILSQGLR | 13 | P09874 | Poly [ADP-ribose] polymerase 1 OS=Hor | PARP1 | 113.084 | 2 | 51.05493 | yes | 9.79E-08 | 689.37787 | 1469.5189 |
| TLGDFAAEYAK | 11 | P09874 | Poly [ADP-ribose] polymerase 1 OS=Hor | PARP1 | 113.084 | 2 | 51.05493 | yes | 1.288E-07 | 593.29297 | 1162.135 |
| GIYFADMVSK | 10 | P09874 | Poly [ADP-ribose] polymerase 1 OS=Hor | PARP1 | 113.084 | 2 | 51.05493 | yes | 1.262E-07 | 565.78119 | 1124.8781 |
| QQVPSGESAILDR | 13 | P09874 | Poly [ADP-ribose] polymerase 1 OS=Hor | PARP1 | 113.084 | 2 | 51.05493 | yes | 1.443E-07 | 700.36243 | 1036.6206 |
| LLWHGSR | 7 | P09874 | Poly [ADP-ribose] polymerase 1 OS=Hor | PARP1 | 113.084 | 2 | 51.05493 | yes | 1.139E-07 | 434.74304 | 995.35028 |
| AMVEYEIDLQK | 11 | P09874 | Poly [ADP-ribose] polymerase 1 OS=Hor | PARP1 | 113.084 | 2 | 51.05493 | yes | 9.722E-08 | 669.83411 | 932.90833 |
| MIFDVESMK | 9 | P09874 | Poly [ADP-ribose] polymerase 1 OS=Hor | PARP1 | 113.084 | 2 | 51.05493 | yes | 1.774E-07 | 550.26178 | 878.48865 |
| MVDPEKPQLGMII | 14 | P09874 | Poly [ADP-ribose] polymerase 1 OS=Hor | PARP1 | 113.084 | 2 | 51.05493 | yes | 1.002E-07 | 814.9104 | 798.58862 |
| MAIMVQSPMFDGI | 13 | P09874 | Poly [ADP-ribose] polymerase 1 OS=Hor | PARP1 | 113.084 | 2 | 51.05493 | yes | 1.017E-07 | 727.84564 | 722.63055 |
| AEPVEVVAPR | 10 | P09874 | Poly [ADP-ribose] polymerase 1 OS=Hor | PARP1 | 113.084 | 2 | 51.05493 | yes | 3.343E-07 | 533.79822 | 717.88348 |
| KLTVNPGTK | 9 | P09874 | Poly [ADP-ribose] polymerase 1 OS=Hor | PARP1 | 113.084 | 2 | 51.05493 | yes | 2.27E-07 | 479.29001 | 691.10333 |
| LQLLEDDKENR | 11 | P09874 | Poly [ADP-ribose] polymerase 1 OS=Hor | PARP1 | 113.084 | 2 | 51.05493 | yes | 9.935E-08 | 686.85699 | 562.0191 |
| KPPLLNNADSVQA | 14 | P09874 | Poly [ADP-ribose] polymerase 1 OS=Hor | PARP1 | 113.084 | 2 | 51.05493 | yes | 2.43E-07 | 747.91736 | 413.0527 |
| WYHPGCFVK | 9 | P09874 | Poly [ADP-ribose] polymerase 1 OS=Hor | PARP1 | 113.084 | 2 | 51.05493 | yes | 2.475E-07 | 597.28168 | 397.67078 |
| HPDVEVDGFSELR | 13 | P09874 | Poly [ADP-ribose] polymerase 1 OS=Hor | PARP1 | 113.084 | 2 | 51.05493 | yes | 1.022E-07 | 750.35986 | 388.41495 |
| PVQDLIK | 7 | P09874 | Poly [ADP-ribose] polymerase 1 OS=Hor | PARP1 | 113.084 | 2 | 51.05493 | yes | 2.32E-07 | 406.74747 | 331.25668 |
| EDAIEHFMK | 9 | P09874 | Poly [ADP-ribose] polymerase 1 OS=Hor | PARP1 | 113.084 | 2 | 51.05493 | yes | 1.691E-07 | 560.26062 | 286.2431 |
| GFSLLATEDK | 10 | P09874 | Poly [ADP-ribose] polymerase 1 OS=Hor | PARP1 | 113.084 | 2 | 51.05493 | yes | 1.866E-07 | 540.78223 | 234.38947 |
| LEQMPSKEDAIEH | 16 | P09874 | Poly [ADP-ribose] polymerase 1 OS=Hor | PARP1 | 113.084 | 3 | 51.05493 | yes | 1.115E-07 | 644.97797 | 214.19029 |
| ELLIFNK | 7 | P09874 | Poly [ADP-ribose] polymerase 1 OS=Hor | PARP1 | 113.084 | 2 | 51.05493 | yes | 1.922E-07 | 438.76309 | 150.14986 |
| DSEEAEIIR | 9 | P09874 | Poly [ADP-ribose] polymerase 1 OS=Hor | PARP1 | 113.084 | 2 | 51.05493 | yes | 1.429E-07 | 531.25909 | 144.58479 |
| KGDEVDGVDEVA | 13 | P09874 | Poly [ADP-ribose] polymerase 1 OS=Hor | PARP1 | 113.084 | 2 | 51.05493 | yes | 9.935E-08 | 680.83319 | 103.52905 |
| VVDRDSEEAEIIR | 13 | P09874 | Poly [ADP-ribose] polymerase 1 OS=Hor | PARP1 | 113.084 | 3 | 51.05493 | yes | 1.287E-07 | 510.93011 | 29.504356 |
| VVDRDSEEAEIIR | 13 | P09874 | Poly [ADP-ribose] polymerase 1 OS=Hor | PARP1 | 113.084 | 3 | 51.05493 | yes | 1.037E-07 | 765.89154 | 29.504356 |
| VVDRDSEEAEIIR | 13 | P09874 | Poly [ADP-ribose] polymerase 1 OS=Hor | PARP1 | 113.084 | 2 | 51.05493 | yes | 1.287E-07 | 510.93011 | 29.504356 |
| VVDRDSEEAEIIR | 13 | P09874 | Poly [ADP-ribose] polymerase 1 OS=Hor | PARP1 | 113.084 | 2 | 51.05493 | yes | 1.037E-07 | 765.89154 | 29.504356 |
| GGAAVDPDSGLEI | 20 | P09874 | Poly [ADP-ribose] polymerase 1 OS=Hor | PARP1 | 113.084 | 3 | 51.05493 | yes | 5.035E-07 | 663.66058 | 53.936943 |
| DPIDVNYEK | 9 | P09874 | Poly [ADP-ribose] polymerase 1 OS=Hor | PARP1 | 113.084 | 2 | 51.05493 | yes | 2.66E-07 | 546.76404 | 52.436722 |
| AQNDLIWNIKDEL | 14 | P09874 | Poly [ADP-ribose] polymerase 1 OS=Hor | PARP1 | 113.084 | 3 | 51.05493 | yes | 1.301E-07 | 567.3053 | 45.343269 |
| NTHATTHNAYDLI | 19 | P09874 | Poly [ADP-ribose] polymerase 1 OS=Hor | PARP1 | 113.084 | 3 | 51.05493 | yes | 6.217E-06 | 734.69904 | 43.846119 |
| VGTVIGSNK | 9 | P09874 | Poly [ADP-ribose] polymerase 1 OS=Hor | PARP1 | 113.084 | 2 | 51.05493 | yes | 5.518E-07 | 437.75327 | 41.975754 |

| | | | | | | | | | | | |
|---|---|---|---|---|---|---|---|---|---|---|---|
| ASLCISTK | 8 | P09874 | Poly [ADP-ribose] polymerase 1 OS=Hoı | PARP1 | 113.084 | 2 | 51.05493 | yes | 1.043E-07 | 440.23386 | 34.328049 |
| VFSATLGLVDIVK | 13 | P09874 | Poly [ADP-ribose] polymerase 1 OS=Hoı | PARP1 | 113.084 | 2 | 51.05493 | yes | 3.367E-07 | 681.40558 | 32.152145 |
| NTAEQFLSR | 9 | P09884 | DNA polymerase alpha catalytic subunit ( | POLA1 | 165.913 | 2 | 45.105724 | yes | 4.304E-06 | 533.26984 | 385.60443 |
| YSAEMPQLPQDLK | 13 | P09884 | DNA polymerase alpha catalytic subunit ( | POLA1 | 165.913 | 2 | 45.105724 | yes | 1.961E-07 | 760.37671 | 147.12305 |
| GPCWLEVK | 8 | P09884 | DNA polymerase alpha catalytic subunit ( | POLA1 | 165.913 | 2 | 45.105724 | yes | 3.019E-07 | 494.74966 | 52.296387 |
| ASPLTFTVQLSNK | 13 | P09884 | DNA polymerase alpha catalytic subunit ( | POLA1 | 165.913 | 2 | 45.105724 | yes | 7.144E-07 | 703.38794 | 49.990814 |
| LGDEDEEIDGDTN | 14 | P09884 | DNA polymerase alpha catalytic subunit ( | POLA1 | 165.913 | 2 | 45.105724 | yes | 1.245E-06 | 775.32867 | 24.278357 |
| PLAALVTYK | 9 | P09884 | DNA polymerase alpha catalytic subunit ( | POLA1 | 165.913 | 2 | 45.105724 | yes | 5.051E-07 | 488.2973 | 13.371191 |
| APGVTDVLR | 9 | P09913 | Interferon-induced protein with tetratricoj | IFIT2 | 54.633 | 2 | 43.25154 | yes | 3.012E-07 | 464.26654 | 49.848541 |
| LSDVQIYVDK | 10 | P09913 | Interferon-induced protein with tetratricoj | IFIT2 | 54.633 | 2 | 43.25154 | yes | 2.667E-07 | 590.31641 | 12.081763 |
| VLDQIEFLDTK | 11 | P09914 | Interferon-induced protein with tetratricoj | IFIT1 | 55.361 | 2 | 49.393036 | yes | 1.037E-07 | 660.8559 | 196.15416 |
| SAIFHFESAVEK | 12 | P09914 | Interferon-induced protein with tetratricoj | IFIT1 | 55.361 | 2 | 49.393036 | yes | 5.129E-07 | 682.84589 | 49.499355 |
| KPTFEVAHLDLAR | 13 | P09914 | Interferon-induced protein with tetratricoj | IFIT1 | 55.361 | 3 | 49.393036 | yes | 3.49E-07 | 499.61218 | 35.521072 |
| HLQLAIR | 7 | P0C0S5 | Histone H2A.Z OS=Homo sapiens OX=9 | H2AZ1 | 13.553 | 2 | 51.4326 | no | 1.477E-06 | 425.76651 | 63415.645 |
| GDEELDSLIK | 10 | P0C0S5 | Histone H2A.Z OS=Homo sapiens OX=9 | H2AZ1 | 13.553 | 2 | 51.4326 | yes | 9.722E-08 | 559.78241 | 657.17358 |
| AGNALTPELAPVQ | 15 | P0C1Z6 | TCF3 fusion partner OS=Homo sapiens C | TFPT | 28.279 | 2 | 44.497208 | yes | 2.172E-07 | 761.43542 | 85.29229 |
| VILDMEDK | 8 | P0C2W1 | F-box/SPRY domain-containing protein 1 | FBXO45 | 30.633 | 2 | 48.800926 | yes | 3.399E-07 | 481.7468 | 106.54505 |
| NPIAQSTDGAR | 11 | P0C2W1 | F-box/SPRY domain-containing protein 1 | FBXO45 | 30.633 | 2 | 48.800926 | yes | 1.103E-07 | 565.28345 | 62.74173 |
| TDILCNLPSYK | 11 | P0C2W1 | F-box/SPRY domain-containing protein 1 | FBXO45 | 30.633 | 2 | 48.800926 | yes | 2.074E-07 | 662.33429 | 46.062981 |
| LCPGSQGACAR | 11 | P0CW18 | Serine protease 56 OS=Homo sapiens OX | PRSS56 | 64.598 | 2 | 52.197342 | yes | 2.645E-07 | 588.76605 | 560.92957 |
| LAHQQCLQR | 9 | P0CW18 | Serine protease 56 OS=Homo sapiens OX | PRSS56 | 64.598 | 2 | 52.197342 | yes | 2.428E-07 | 577.2984 | 506.70877 |
| VPLLSTDTCR | 10 | P0CW18 | Serine protease 56 OS=Homo sapiens OX | PRSS56 | 64.598 | 2 | 52.197342 | yes | 1.695E-07 | 581.30023 | 474.09335 |
| LAPALALPAPALR | 13 | P0CW18 | Serine protease 56 OS=Homo sapiens OX | PRSS56 | 64.598 | 2 | 52.197342 | yes | 9.722E-08 | 637.4032 | 408.42566 |
| LLVQALQAFR | 10 | P0CW18 | Serine protease 56 OS=Homo sapiens OX | PRSS56 | 64.598 | 2 | 52.197342 | yes | 9.722E-08 | 579.85333 | 379.00015 |
| VAMEIQHR | 8 | P0CW18 | Serine protease 56 OS=Homo sapiens OX | PRSS56 | 64.598 | 2 | 52.197342 | yes | 2.892E-07 | 492.25818 | 207.24422 |
| GEQAEEVPVNR | 11 | P0CW18 | Serine protease 56 OS=Homo sapiens OX | PRSS56 | 64.598 | 2 | 52.197342 | yes | 9.722E-08 | 614.30182 | 203.27068 |
| LCAFYAR | 7 | P0CW18 | Serine protease 56 OS=Homo sapiens OX | PRSS56 | 64.598 | 2 | 52.197342 | yes | 3.25E-06 | 450.72345 | 132.83232 |
| HVAFSGLVGLEPA | 17 | P0CW18 | Serine protease 56 OS=Homo sapiens OX | PRSS56 | 64.598 | 3 | 52.197342 | yes | 9.722E-08 | 579.99304 | 128.49596 |
| ESPLHPAR | 8 | P0CW18 | Serine protease 56 OS=Homo sapiens OX | PRSS56 | 64.598 | 2 | 52.197342 | yes | 3.343E-07 | 453.74323 | 89.780014 |
| DWLQEQMSASSSI | 13 | P0CW18 | Serine protease 56 OS=Homo sapiens OX | PRSS56 | 64.598 | 2 | 52.197342 | yes | 2.877E-07 | 762.841 | 55.837631 |
| NAQELLGPR | 9 | P0CW18 | Serine protease 56 OS=Homo sapiens OX | PRSS56 | 64.598 | 2 | 52.197342 | yes | 2.688E-07 | 499.2749 | 55.800789 |
| SLAHTLLGLLR | 11 | P0CW18 | Serine protease 56 OS=Homo sapiens OX | PRSS56 | 64.598 | 2 | 52.197342 | yes | 3.971E-06 | 597.37189 | 23.663406 |
| VIVSSPWPAVVVA | 16 | P0DI83 | Ras-related protein Rab-34, isoform NAR | RAB34 | 21.119 | 2 | 52.458706 | yes | 1.312E-07 | 832.48016 | 140.05823 |
| VIVGTIRPR | 9 | P0DI83 | Ras-related protein Rab-34, isoform NAR | RAB34 | 21.119 | 2 | 52.458706 | yes | 9.722E-08 | 505.82709 | 45.376057 |
| VEIIANDQGNR | 11 | P0DMV8 | Heat shock 70 kDa protein 1A OS=Homo | HSPA1A | 70.053 | 2 | 51.94918 | no | 9.722E-08 | 614.81769 | 15544.507 |
| NQVALNPQNTVFI | 15 | P0DMV8 | Heat shock 70 kDa protein 1A OS=Homo | HSPA1A | 70.053 | 2 | 51.94918 | yes | 1.428E-07 | 829.92847 | 10404.866 |
| TTPSYVAFTDTER | 13 | P0DMV8 | Heat shock 70 kDa protein 1A OS=Homo | HSPA1A | 70.053 | 2 | 51.94918 | no | 9.809E-08 | 744.35425 | 10213.417 |
| FEELCSDLFR | 10 | P0DMV8 | Heat shock 70 kDa protein 1A OS=Homo | HSPA1A | 70.053 | 2 | 51.94918 | no | 9.85E-08 | 658.30298 | 10003.889 |
| AFYPEEISSMVLTK | 14 | P0DMV8 | Heat shock 70 kDa protein 1A OS=Homo | HSPA1A | 70.053 | 2 | 51.94918 | no | 5.464E-07 | 807.90784 | 2789.1836 |

| | | | | | | | | | | | |
|---|---|---|---|---|---|---|---|---|---|---|---|
| AFYPEEISSMVLTK | 14 | P0DMV8 | Heat shock 70 kDa protein 1A OS=Homo | HSPA1A | 70.053 | 2 | 51.94918 | no | 2.616E-07 | 815.90527 | 2789.1836 |
| IINEPTAAAIAYGL | 16 | P0DMV8 | Heat shock 70 kDa protein 1A OS=Homo | HSPA1A | 70.053 | 2 | 51.94918 | no | 1.473E-07 | 844.45435 | 5378.6538 |
| AQIHDLVLVGGST | 14 | P0DMV8 | Heat shock 70 kDa protein 1A OS=Homo | HSPA1A | 70.053 | 2 | 51.94918 | yes | 9.809E-08 | 733.40973 | 3444.5652 |
| DAGVIAGLNVLR | 12 | P0DMV8 | Heat shock 70 kDa protein 1A OS=Homo | HSPA1A | 70.053 | 2 | 51.94918 | no | 1.331E-07 | 599.35114 | 3337.4436 |
| FGDPVVQSDMK | 11 | P0DMV8 | Heat shock 70 kDa protein 1A OS=Homo | HSPA1A | 70.053 | 2 | 51.94918 | yes | 2.139E-07 | 611.79224 | 3229.8523 |
| LVNHFVEEFK | 10 | P0DMV8 | Heat shock 70 kDa protein 1A OS=Homo | HSPA1A | 70.053 | 2 | 51.94918 | yes | 1.532E-07 | 631.3324 | 1109.1589 |
| LVNHFVEEFK | 10 | P0DMV8 | Heat shock 70 kDa protein 1A OS=Homo | HSPA1A | 70.053 | 2 | 51.94918 | yes | 1.74E-07 | 421.22403 | 1109.1589 |
| LVNHFVEEFK | 10 | P0DMV8 | Heat shock 70 kDa protein 1A OS=Homo | HSPA1A | 70.053 | 3 | 51.94918 | yes | 1.532E-07 | 631.3324 | 1109.1589 |
| LVNHFVEEFK | 10 | P0DMV8 | Heat shock 70 kDa protein 1A OS=Homo | HSPA1A | 70.053 | 3 | 51.94918 | yes | 1.74E-07 | 421.22403 | 1109.1589 |
| KFGDPVVQSDMK | 12 | P0DMV8 | Heat shock 70 kDa protein 1A OS=Homo | HSPA1A | 70.053 | 2 | 51.94918 | yes | 9.968E-08 | 675.83972 | 1958.9122 |
| LLQDFFNGR | 9 | P0DMV8 | Heat shock 70 kDa protein 1A OS=Homo | HSPA1A | 70.053 | 2 | 51.94918 | yes | 5.837E-07 | 555.29053 | 689.24188 |
| SAVEDEGLK | 9 | P0DMV8 | Heat shock 70 kDa protein 1A OS=Homo | HSPA1A | 70.053 | 2 | 51.94918 | yes | 1.337E-07 | 474.23764 | 671.09015 |
| NALESYAFNMK | 11 | P0DMV8 | Heat shock 70 kDa protein 1A OS=Homo | HSPA1A | 70.053 | 2 | 51.94918 | no | 9.722E-08 | 644.30554 | 553.49329 |
| AAAIGIDLGTTYSC | 22 | P0DMV8 | Heat shock 70 kDa protein 1A OS=Homo | HSPA1A | 70.053 | 3 | 51.94918 | yes | 1.543E-07 | 755.71594 | 540.51874 |
| ATAGDTHLGGEDF | 16 | P0DMV8 | Heat shock 70 kDa protein 1A OS=Homo | HSPA1A | 70.053 | 2 | 51.94918 | no | 1.072E-07 | 838.36896 | 475.53345 |
| MVQEAEK | 7 | P0DMV8 | Heat shock 70 kDa protein 1A OS=Homo | HSPA1A | 70.053 | 2 | 51.94918 | no | 2.281E-07 | 417.70493 | 470.70392 |
| HWPFQVINDGDKI | 14 | P0DMV8 | Heat shock 70 kDa protein 1A OS=Homo | HSPA1A | 70.053 | 3 | 51.94918 | yes | 2.136E-07 | 560.95459 | 98.332375 |
| HWPFQVINDGDKI | 14 | P0DMV8 | Heat shock 70 kDa protein 1A OS=Homo | HSPA1A | 70.053 | 3 | 51.94918 | yes | 2.314E-07 | 840.92828 | 98.332375 |
| HWPFQVINDGDKI | 14 | P0DMV8 | Heat shock 70 kDa protein 1A OS=Homo | HSPA1A | 70.053 | 2 | 51.94918 | yes | 2.136E-07 | 560.95459 | 98.332375 |
| HWPFQVINDGDKI | 14 | P0DMV8 | Heat shock 70 kDa protein 1A OS=Homo | HSPA1A | 70.053 | 2 | 51.94918 | yes | 2.314E-07 | 840.92828 | 98.332375 |
| STLEPVEK | 8 | P0DMV8 | Heat shock 70 kDa protein 1A OS=Homo | HSPA1A | 70.053 | 2 | 51.94918 | no | 2.097E-07 | 451.74512 | 177.08472 |
| LVNHFVEEFKR | 11 | P0DMV8 | Heat shock 70 kDa protein 1A OS=Homo | HSPA1A | 70.053 | 2 | 51.94918 | yes | 9.723E-08 | 709.383 | 154.14099 |
| CQEVISWLDANTL | 16 | P0DMV8 | Heat shock 70 kDa protein 1A OS=Homo | HSPA1A | 70.053 | 3 | 51.94918 | yes | 4.59E-07 | 626.30847 | 27.556383 |
| ADQLTEEQIAEFK | 13 | P0DP23 | Calmodulin-1 OS=Homo sapiens OX=96 | CALM1 | 16.838 | 2 | 53.56798 | yes | 1.213E-07 | 782.38049 | 2825.6091 |
| MKDTDSEEEIR | 11 | P0DP23 | Calmodulin-1 OS=Homo sapiens OX=96 | CALM1 | 16.838 | 2 | 53.56798 | yes | 9.722E-08 | 676.80353 | 323.20377 |
| MKDTDSEEEIR | 11 | P0DP23 | Calmodulin-1 OS=Homo sapiens OX=96 | CALM1 | 16.838 | 2 | 53.56798 | yes | 2.266E-06 | 684.80103 | 323.20377 |
| DGNGYISAAELR | 12 | P0DP23 | Calmodulin-1 OS=Homo sapiens OX=96 | CALM1 | 16.838 | 2 | 53.56798 | yes | 9.722E-08 | 633.30969 | 514.58875 |
| DTDSEEEIR | 9 | P0DP23 | Calmodulin-1 OS=Homo sapiens OX=96 | CALM1 | 16.838 | 2 | 53.56798 | yes | 1.429E-07 | 547.23584 | 332.26437 |
| IIEQLSPQAATSR | 13 | P0DP91 | Chimeric ERCC6-PGBD3 protein OS=H | ERCC6 | 119.487 | 2 | 51.39102 | no | 1.014E-07 | 707.38843 | 863.37225 |
| VTAPPNDFFTVMR | 13 | P0DP91 | Chimeric ERCC6-PGBD3 protein OS=H | ERCC6 | 119.487 | 2 | 51.39102 | no | 1.664E-07 | 747.87415 | 531.7204 |
| IMLNEASGFEK | 11 | P0DP91 | Chimeric ERCC6-PGBD3 protein OS=H | ERCC6 | 119.487 | 2 | 51.39102 | no | 9.903E-08 | 619.80792 | 446.79425 |
| TGQMTPFGTQIPQ | 14 | P0DP91 | Chimeric ERCC6-PGBD3 protein OS=H | ERCC6 | 119.487 | 2 | 51.39102 | no | 3.196E-07 | 767.39014 | 339.80154 |
| VPLESDVALK | 10 | P0DP91 | Chimeric ERCC6-PGBD3 protein OS=H | ERCC6 | 119.487 | 2 | 51.39102 | no | 4.831E-07 | 535.80823 | 292.98178 |
| ADLTVQPVAGR | 11 | P0DP91 | Chimeric ERCC6-PGBD3 protein OS=H | ERCC6 | 119.487 | 2 | 51.39102 | no | 1.014E-07 | 563.81439 | 291.01129 |
| SVLDDLTSCTTSLI | 14 | P0DP91 | Chimeric ERCC6-PGBD3 protein OS=H | ERCC6 | 119.487 | 2 | 51.39102 | no | 1.192E-07 | 784.38525 | 234.82986 |
| TYDEKPVDFLEFR | 13 | P0DP91 | Chimeric ERCC6-PGBD3 protein OS=H | ERCC6 | 119.487 | 2 | 51.39102 | no | 9.722E-08 | 829.90668 | 195.19997 |
| HLQAILGGAEVK | 12 | P0DP91 | Chimeric ERCC6-PGBD3 protein OS=H | ERCC6 | 119.487 | 2 | 51.39102 | no | 1.53E-07 | 618.35895 | 156.36839 |
| TDVHNVLVSAAM | 13 | P0DP91 | Chimeric ERCC6-PGBD3 protein OS=H | ERCC6 | 119.487 | 2 | 51.39102 | no | 1.452E-07 | 706.86938 | 147.33682 |
| ASQLVDVEK | 9 | P0DP91 | Chimeric ERCC6-PGBD3 protein OS=H | ERCC6 | 119.487 | 2 | 51.39102 | no | 3.665E-07 | 494.7691 | 134.42389 |

| | | | | | | | | | | | |
|---|---|---|---|---|---|---|---|---|---|---|---|
| VYNQFMGGVDR | 11 | P0DP91 | Chimeric ERCC6-PGBD3 protein OS=Hc | ERCC6 | 119.487 | 2 | 51.39102 | no | 3.448E-07 | 643.30334 | 117.8605 |
| GVHLGLTSSEFK | 12 | P0DP91 | Chimeric ERCC6-PGBD3 protein OS=Hc | ERCC6 | 119.487 | 2 | 51.39102 | no | 1.736E-07 | 637.84058 | 116.0911 |
| KADLTVQPVAGR | 12 | P0DP91 | Chimeric ERCC6-PGBD3 protein OS=Hc | ERCC6 | 119.487 | 2 | 51.39102 | no | 9.758E-08 | 627.86188 | 109.59459 |
| YDGINHVIVK | 10 | P0DP91 | Chimeric ERCC6-PGBD3 protein OS=Hc | ERCC6 | 119.487 | 2 | 51.39102 | no | 9.917E-08 | 579.31934 | 79.401077 |
| GPALLHIDR | 9 | P0DP91 | Chimeric ERCC6-PGBD3 protein OS=Hc | ERCC6 | 119.487 | 2 | 51.39102 | no | 7.95E-07 | 496.28781 | 78.192436 |
| KAPAPVTPPAPVQ | 15 | P0DP91 | Chimeric ERCC6-PGBD3 protein OS=Hc | ERCC6 | 119.487 | 2 | 51.39102 | no | 4.147E-07 | 757.93811 | 52.414749 |
| YSNLYACSK | 9 | P0DP91 | Chimeric ERCC6-PGBD3 protein OS=Hc | ERCC6 | 119.487 | 2 | 51.39102 | no | 1.247E-07 | 553.25275 | 45.152111 |
| LSSMGHQATGTVI | 13 | P0DP91 | Chimeric ERCC6-PGBD3 protein OS=Hc | ERCC6 | 119.487 | 2 | 51.39102 | no | 1.266E-07 | 672.83807 | 36.512062 |
| YRDDGDEDYYK | 11 | P0DP91 | Chimeric ERCC6-PGBD3 protein OS=Hc | ERCC6 | 119.487 | 2 | 51.39102 | no | 1.381E-07 | 719.7915 | 27.659254 |
| TALEMVQAAGTDI | 13 | P0DPB6 | DNA-directed RNA polymerases I and III | POLR1D | 15.238 | 2 | 43.334934 | yes | 2.656E-07 | 681.83771 | 222.52625 |
| GTLPAVEPFQR | 11 | P0DPB6 | DNA-directed RNA polymerases I and III | POLR1D | 15.238 | 2 | 43.334934 | yes | 2.635E-07 | 607.83002 | 126.28233 |
| KPCAAEPALSAGS | 22 | P10074 | Telomere zinc finger-associated protein C | ZBTB48 | 77.054 | 3 | 51.133087 | yes | 9.722E-08 | 737.36719 | 59.313774 |
| TELQLHEAFK | 10 | P10074 | Telomere zinc finger-associated protein C | ZBTB48 | 77.054 | 2 | 51.133087 | yes | 2.194E-06 | 608.32202 | 51.417519 |
| VVVLGSGGVGK | 11 | P10114 | Ras-related protein Rap-2a OS=Homo sap | RAP2A | 20.616 | 2 | 48.03734 | yes | 1.21E-07 | 486.29785 | 89.313522 |
| YDPTIEDFYR | 10 | P10114 | Ras-related protein Rap-2a OS=Homo sap | RAP2A | 20.616 | 2 | 48.03734 | yes | 3.482E-06 | 659.80115 | 7.0777349 |
| GPMFELLPGESNK | 13 | P10124 | Serglycin OS=Homo sapiens OX=9606 G | SRGN | 17.652 | 2 | 39.63745 | yes | 4.467E-07 | 709.85284 | 82.467056 |
| LGLENAEALIR | 11 | P10155 | RNA-binding protein RO60 OS=Homo sa | RO60 | 60.671 | 2 | 54.870148 | yes | 1.12E-07 | 599.84314 | 398.44894 |
| GCEVIQEIK | 9 | P10155 | RNA-binding protein RO60 OS=Homo sa | RO60 | 60.671 | 2 | 54.870148 | yes | 1.492E-07 | 538.27625 | 176.74843 |
| FLCFGSEGGTYYIK | 14 | P10155 | RNA-binding protein RO60 OS=Homo sa | RO60 | 60.671 | 2 | 54.870148 | yes | 1.733E-07 | 821.38452 | 138.67334 |
| GGMALALAVTK | 11 | P10155 | RNA-binding protein RO60 OS=Homo sa | RO60 | 60.671 | 2 | 54.870148 | yes | 4.221E-07 | 516.2995 | 118.40627 |
| WRPDEEILK | 9 | P10155 | RNA-binding protein RO60 OS=Homo sa | RO60 | 60.671 | 2 | 54.870148 | yes | 3.037E-07 | 593.31677 | 104.57686 |
| LSHLKPSSEGLAIV | 16 | P10155 | RNA-binding protein RO60 OS=Homo sa | RO60 | 60.671 | 3 | 54.870148 | yes | 9.722E-08 | 560.66119 | 35.269196 |
| LSHLKPSSEGLAIV | 16 | P10155 | RNA-binding protein RO60 OS=Homo sa | RO60 | 60.671 | 3 | 54.870148 | yes | 2.93E-07 | 840.48816 | 35.269196 |
| LSHLKPSSEGLAIV | 16 | P10155 | RNA-binding protein RO60 OS=Homo sa | RO60 | 60.671 | 2 | 54.870148 | yes | 9.722E-08 | 560.66119 | 35.269196 |
| LSHLKPSSEGLAIV | 16 | P10155 | RNA-binding protein RO60 OS=Homo sa | RO60 | 60.671 | 2 | 54.870148 | yes | 2.93E-07 | 840.48816 | 35.269196 |
| IPTHLFTFIQFK | 12 | P10155 | RNA-binding protein RO60 OS=Homo sa | RO60 | 60.671 | 2 | 54.870148 | yes | 9.722E-08 | 746.42157 | 38.846664 |
| AYLEGTCVEWLR | 12 | P10321 | HLA class I histocompatibility antigen, C | HLA-C | 40.649 | 2 | 51.63984 | yes | 9.722E-08 | 748.86377 | 221.47856 |
| IGDFGLATVK | 10 | P10398 | Serine/threonine-protein kinase A-Raf OS | ARAF | 67.586 | 2 | 54.766422 | no | 5.98E-07 | 510.78986 | 303.27832 |
| VSQPTAEQAQAFK | 13 | P10398 | Serine/threonine-protein kinase A-Raf OS | ARAF | 67.586 | 2 | 54.766422 | yes | 2.124E-07 | 702.8595 | 163.70995 |
| TQADELPACLLSA | 15 | P10398 | Serine/threonine-protein kinase A-Raf OS | ARAF | 67.586 | 2 | 54.766422 | yes | 3.504E-06 | 808.40906 | 161.1026 |
| SNNIFLHEGLTVK | 13 | P10398 | Serine/threonine-protein kinase A-Raf OS | ARAF | 67.586 | 2 | 54.766422 | no | 1.175E-07 | 736.3988 | 114.53947 |
| IGTGSFGTVFR | 11 | P10398 | Serine/threonine-protein kinase A-Raf OS | ARAF | 67.586 | 2 | 54.766422 | yes | 1.443E-07 | 571.30365 | 95.742874 |
| DGMSVYDSLDK | 11 | P10398 | Serine/threonine-protein kinase A-Raf OS | ARAF | 67.586 | 2 | 54.766422 | yes | 1.187E-07 | 615.27136 | 92.59407 |
| VPTVCVDMSTNR | 12 | P10398 | Serine/threonine-protein kinase A-Raf OS | ARAF | 67.586 | 2 | 54.766422 | yes | 9.722E-08 | 689.82629 | 79.869293 |
| QTAQGMDYLHAK | 12 | P10398 | Serine/threonine-protein kinase A-Raf OS | ARAF | 67.586 | 2 | 54.766422 | no | 9.722E-08 | 681.82715 | 68.711372 |
| GLNQDCCVVYR | 11 | P10398 | Serine/threonine-protein kinase A-Raf OS | ARAF | 67.586 | 2 | 54.766422 | yes | 4.018E-07 | 692.31085 | 66.028679 |
| GYLSPDLSK | 9 | P10398 | Serine/threonine-protein kinase A-Raf OS | ARAF | 67.586 | 2 | 54.766422 | no | 1.69E-07 | 490.25839 | 60.979198 |
| WHGDVAVK | 8 | P10398 | Serine/threonine-protein kinase A-Raf OS | ARAF | 67.586 | 2 | 54.766422 | no | 1.972E-06 | 456.24033 | 53.01334 |
| FDMVQLIDVAR | 11 | P10398 | Serine/threonine-protein kinase A-Raf OS | ARAF | 67.586 | 2 | 54.766422 | yes | 9.722E-08 | 653.84485 | 44.770752 |

| | | | | | | | | | | | |
|---|---|---|---|---|---|---|---|---|---|---|---|
| DQIIFMVGR | 9 | P10398 | Serine/threonine-protein kinase A-Raf OS | ARAF | 67.586 | 2 | 54.766422 | no | 1.509E-06 | 539.78931 | 20.828878 |
| ALAAAGYDVEK | 11 | P10412 | Histone H1.4 OS=Homo sapiens OX=96( | H1-4 | 21.866 | 2 | 51.82133 | no | 3.76E-07 | 554.28766 | 22276.186 |
| SETAPAAPAAPAP. | 16 | P10412 | Histone H1.4 OS=Homo sapiens OX=96( | H1-4 | 21.866 | 2 | 51.82133 | yes | 3.926E-07 | 760.88318 | 19650.871 |
| ASGPPVSELITK | 12 | P10412 | Histone H1.4 OS=Homo sapiens OX=96( | H1-4 | 21.866 | 2 | 51.82133 | no | 1.002E-07 | 599.83752 | 15510.292 |
| KASGPPVSELITK | 13 | P10412 | Histone H1.4 OS=Homo sapiens OX=96( | H1-4 | 21.866 | 2 | 51.82133 | no | 1.649E-07 | 663.88501 | 4064.4648 |
| SGVSLAALK | 9 | P10412 | Histone H1.4 OS=Homo sapiens OX=96( | H1-4 | 21.866 | 2 | 51.82133 | no | 2.734E-07 | 423.25818 | 957.70996 |
| KALAAAGYDVEK | 12 | P10412 | Histone H1.4 OS=Homo sapiens OX=96( | H1-4 | 21.866 | 2 | 51.82133 | no | 9.722E-08 | 618.33514 | 525.68164 |
| SGVSLAALKK | 10 | P10412 | Histone H1.4 OS=Homo sapiens OX=96( | H1-4 | 21.866 | 2 | 51.82133 | no | 7.765E-07 | 487.30566 | 33.747826 |
| VPEANSSWMDTV. | 14 | P10515 | Dihydrolipoyllysine-residue acetyltransfe | DLAT | 68.997 | 2 | 49.28839 | yes | 6.737E-07 | 802.8905 | 103.16652 |
| DVPLGTPLCIIVEK | 14 | P10515 | Dihydrolipoyllysine-residue acetyltransfe | DLAT | 68.997 | 2 | 49.28839 | yes | 3.32E-07 | 777.43402 | 77.716003 |
| ILVAEGTR | 8 | P10515 | Dihydrolipoyllysine-residue acetyltransfe | DLAT | 68.997 | 2 | 49.28839 | yes | 1.048E-07 | 429.7558 | 74.211769 |
| LSEGDLLAEIETDK | 14 | P10515 | Dihydrolipoyllysine-residue acetyltransfe | DLAT | 68.997 | 2 | 49.28839 | yes | 2.966E-07 | 766.88812 | 56.515385 |
| GVETIANDVVSLA | 15 | P10515 | Dihydrolipoyllysine-residue acetyltransfe | DLAT | 68.997 | 2 | 49.28839 | yes | 2.745E-07 | 758.91449 | 54.322002 |
| ILVPEGTR | 8 | P10515 | Dihydrolipoyllysine-residue acetyltransfe | DLAT | 68.997 | 2 | 49.28839 | yes | 7.955E-07 | 442.76364 | 46.778057 |
| VFVSPLAK | 8 | P10515 | Dihydrolipoyllysine-residue acetyltransfe | DLAT | 68.997 | 2 | 49.28839 | yes | 1.247E-06 | 430.76566 | 20.267387 |
| SQCALEEYVR | 10 | P10589 | COUP transcription factor 1 OS=Homo sa | NR2F1 | 46.156 | 2 | 54.878754 | no | 9.722E-08 | 627.79279 | 176.72409 |
| TPIETLIR | 8 | P10589 | COUP transcription factor 1 OS=Homo sa | NR2F1 | 46.156 | 2 | 54.878754 | no | 9.722E-08 | 471.78458 | 119.30296 |
| LLFSAVEWAR | 10 | P10589 | COUP transcription factor 1 OS=Homo sa | NR2F1 | 46.156 | 2 | 54.878754 | yes | 1.157E-07 | 596.32965 | 33.058079 |
| CMPTFQFFK | 9 | P10599 | Thioredoxin OS=Homo sapiens OX=960( | TXN | 11.738 | 2 | 50.857746 | yes | 2.797E-07 | 603.27771 | 1729.9459 |
| TAFQEALDAAGDK | 13 | P10599 | Thioredoxin OS=Homo sapiens OX=960( | TXN | 11.738 | 2 | 50.857746 | yes | 1.022E-07 | 668.82263 | 1728.9373 |
| EKLEATINELV | 11 | P10599 | Thioredoxin OS=Homo sapiens OX=960( | TXN | 11.738 | 2 | 50.857746 | yes | 1.459E-07 | 629.84808 | 1446.4506 |
| MIKPFFHSLSEK | 12 | P10599 | Thioredoxin OS=Homo sapiens OX=960( | TXN | 11.738 | 2 | 50.857746 | yes | 1.375E-07 | 732.3894 | 470.43533 |
| PFFHSLSEK | 9 | P10599 | Thioredoxin OS=Homo sapiens OX=960( | TXN | 11.738 | 2 | 50.857746 | yes | 9.722E-08 | 546.27966 | 250.75415 |
| VFANPEDCVAFGK | 13 | P10620 | Microsomal glutathione S-transferase 1 O | MGST1 | 17.599 | 2 | 51.74148 | yes | 9.722E-08 | 727.34265 | 316.97531 |
| MMLMSTATAFYR | 12 | P10620 | Microsomal glutathione S-transferase 1 O | MGST1 | 17.599 | 2 | 51.74148 | yes | 9.722E-08 | 711.83258 | 55.942165 |
| EIGNIISDAMK | 11 | P10809 | 60 kDa heat shock protein, mitochondrial | HSPD1 | 61.055 | 2 | 52.786106 | yes | 1.678E-07 | 595.80792 | 1121.194 |
| TLNDELEIIEGMK | 13 | P10809 | 60 kDa heat shock protein, mitochondrial | HSPD1 | 61.055 | 2 | 52.786106 | yes | 1.153E-07 | 752.88184 | 993.47534 |
| ISSIQSIVPALEIAN. | 18 | P10809 | 60 kDa heat shock protein, mitochondrial | HSPD1 | 61.055 | 3 | 52.786106 | yes | 1.483E-07 | 640.36182 | 726.29364 |
| NAGVEGSLIVEK | 12 | P10809 | 60 kDa heat shock protein, mitochondrial | HSPD1 | 61.055 | 2 | 52.786106 | yes | 9.819E-08 | 608.33258 | 687.91534 |
| VTDALNATR | 9 | P10809 | 60 kDa heat shock protein, mitochondrial | HSPD1 | 61.055 | 2 | 52.786106 | yes | 9.722E-08 | 480.75909 | 670.0766 |
| CEFQDAYVLLSEK | 13 | P10809 | 60 kDa heat shock protein, mitochondrial | HSPD1 | 61.055 | 2 | 52.786106 | yes | 9.766E-08 | 801.37946 | 576.1543 |
| TVIIEQSWGSPK | 12 | P10809 | 60 kDa heat shock protein, mitochondrial | HSPD1 | 61.055 | 2 | 52.786106 | yes | 1.498E-07 | 672.86151 | 515.17603 |
| AAVEEGIVLGGGC | 17 | P10809 | 60 kDa heat shock protein, mitochondrial | HSPD1 | 61.055 | 2 | 52.786106 | yes | 1.084E-07 | 842.95618 | 212.67267 |
| AAVEEGIVLGGGC | 17 | P10809 | 60 kDa heat shock protein, mitochondrial | HSPD1 | 61.055 | 2 | 52.786106 | yes | 3.968E-07 | 562.30652 | 212.67267 |
| AAVEEGIVLGGGC | 17 | P10809 | 60 kDa heat shock protein, mitochondrial | HSPD1 | 61.055 | 3 | 52.786106 | yes | 1.084E-07 | 842.95618 | 212.67267 |
| AAVEEGIVLGGGC | 17 | P10809 | 60 kDa heat shock protein, mitochondrial | HSPD1 | 61.055 | 3 | 52.786106 | yes | 3.968E-07 | 562.30652 | 212.67267 |
| VGGTSDVEVNEK | 12 | P10809 | 60 kDa heat shock protein, mitochondrial | HSPD1 | 61.055 | 2 | 52.786106 | yes | 1.279E-07 | 617.30151 | 415.31635 |
| GANPVEIR | 8 | P10809 | 60 kDa heat shock protein, mitochondrial | HSPD1 | 61.055 | 2 | 52.786106 | yes | 1.065E-05 | 428.23779 | 283.65921 |
| IPAMTIAK | 8 | P10809 | 60 kDa heat shock protein, mitochondrial | HSPD1 | 61.055 | 2 | 52.786106 | yes | 1.163E-07 | 422.75168 | 258.54599 |

| | | | | | | | | | | | |
|---|---|---|---|---|---|---|---|---|---|---|---|
| LSDGVAVLK | 9 | P10809 | 60 kDa heat shock protein, mitochondrial | HSPD1 | 61.055 | 2 | 52.786106 | yes | 1.241E-07 | 451.2713 | 146.12898 |
| APGFGDNR | 8 | P10809 | 60 kDa heat shock protein, mitochondrial | HSPD1 | 61.055 | 2 | 52.786106 | yes | 1.177E-07 | 417.19867 | 141.52081 |
| GYISPYFINTSK | 12 | P10809 | 60 kDa heat shock protein, mitochondrial | HSPD1 | 61.055 | 2 | 52.786106 | yes | 9.722E-08 | 695.35608 | 94.814209 |
| VGLQVVAVK | 9 | P10809 | 60 kDa heat shock protein, mitochondrial | HSPD1 | 61.055 | 2 | 52.786106 | yes | 2.656E-07 | 456.79749 | 56.650787 |
| KPLVIIAEDVDGE/ | 22 | P10809 | 60 kDa heat shock protein, mitochondrial | HSPD1 | 61.055 | 3 | 52.786106 | yes | 9.766E-08 | 789.11609 | 56.097481 |
| QSKPVTTPEEIAQ\ | 23 | P10809 | 60 kDa heat shock protein, mitochondrial | HSPD1 | 61.055 | 3 | 52.786106 | yes | 7.598E-07 | 795.41498 | 35.081711 |
| VTTVASHTSDSDV | 23 | P10909 | Clusterin OS=Homo sapiens OX=9606 G | CLU | 52.495 | 3 | 35.44971 | yes | 8.307E-07 | 772.06396 | 13.552837 |
| ITPSYVAFTPEGER | 14 | P11021 | Endoplasmic reticulum chaperone BiP OS | HSPA5 | 72.333 | 2 | 50.764507 | yes | 1.159E-07 | 783.89355 | 3661.4199 |
| NQLTSNPENTVFD | 15 | P11021 | Endoplasmic reticulum chaperone BiP OS | HSPA5 | 72.333 | 2 | 50.764507 | yes | 1.198E-07 | 839.40759 | 3416.7117 |
| TFAPEEISAMVLTI | 14 | P11021 | Endoplasmic reticulum chaperone BiP OS | HSPA5 | 72.333 | 2 | 50.764507 | yes | 1.371E-07 | 768.90253 | 2655.6194 |
| DAGTIAGLNVMR | 12 | P11021 | Endoplasmic reticulum chaperone BiP OS | HSPA5 | 72.333 | 2 | 50.764507 | yes | 1.088E-07 | 609.31897 | 2234.521 |
| ELEEIVQPIISK | 12 | P11021 | Endoplasmic reticulum chaperone BiP OS | HSPA5 | 72.333 | 2 | 50.764507 | yes | 9.728E-08 | 699.39795 | 2110.925 |
| TWNDPSVQQDIK | 12 | P11021 | Endoplasmic reticulum chaperone BiP OS | HSPA5 | 72.333 | 2 | 50.764507 | yes | 1.422E-07 | 715.84918 | 1960.16 |
| VMEHFIK | 7 | P11021 | Endoplasmic reticulum chaperone BiP OS | HSPA5 | 72.333 | 2 | 50.764507 | yes | 1.201E-07 | 452.24149 | 1572.9137 |
| TKPYIQVDIGGGQ | 15 | P11021 | Endoplasmic reticulum chaperone BiP OS | HSPA5 | 72.333 | 2 | 50.764507 | yes | 1.165E-07 | 802.93579 | 772.82568 |
| TKPYIQVDIGGGQ | 15 | P11021 | Endoplasmic reticulum chaperone BiP OS | HSPA5 | 72.333 | 2 | 50.764507 | yes | 4.824E-07 | 535.62628 | 772.82568 |
| TKPYIQVDIGGGQ | 15 | P11021 | Endoplasmic reticulum chaperone BiP OS | HSPA5 | 72.333 | 3 | 50.764507 | yes | 1.165E-07 | 802.93579 | 772.82568 |
| TKPYIQVDIGGGQ | 15 | P11021 | Endoplasmic reticulum chaperone BiP OS | HSPA5 | 72.333 | 3 | 50.764507 | yes | 4.824E-07 | 535.62628 | 772.82568 |
| FEELNMDLFR | 10 | P11021 | Endoplasmic reticulum chaperone BiP OS | HSPA5 | 72.333 | 2 | 50.764507 | yes | 9.819E-08 | 657.31335 | 1494.6526 |
| NELESYAYSLK | 11 | P11021 | Endoplasmic reticulum chaperone BiP OS | HSPA5 | 72.333 | 2 | 50.764507 | yes | 9.843E-08 | 658.82208 | 1409.1309 |
| LTPEEIER | 8 | P11021 | Endoplasmic reticulum chaperone BiP OS | HSPA5 | 72.333 | 2 | 50.764507 | yes | 2.228E-07 | 493.76129 | 1016.7796 |
| VYEGERPLTK | 10 | P11021 | Endoplasmic reticulum chaperone BiP OS | HSPA5 | 72.333 | 2 | 50.764507 | yes | 1.061E-07 | 596.32202 | 998.58618 |
| AKFEELNMDLFR | 12 | P11021 | Endoplasmic reticulum chaperone BiP OS | HSPA5 | 72.333 | 2 | 50.764507 | yes | 1.572E-07 | 756.87939 | 917.65417 |
| VLEDSDLKK | 9 | P11021 | Endoplasmic reticulum chaperone BiP OS | HSPA5 | 72.333 | 2 | 50.764507 | yes | 1.388E-07 | 523.79004 | 518.04651 |
| ITITNDQNR | 9 | P11021 | Endoplasmic reticulum chaperone BiP OS | HSPA5 | 72.333 | 2 | 50.764507 | yes | 1.847E-07 | 537.78052 | 466.47263 |
| MKETAEAYLGK | 11 | P11021 | Endoplasmic reticulum chaperone BiP OS | HSPA5 | 72.333 | 2 | 50.764507 | yes | 1.157E-07 | 620.81573 | 370.32504 |
| ETAEAYLGK | 9 | P11021 | Endoplasmic reticulum chaperone BiP OS | HSPA5 | 72.333 | 2 | 50.764507 | yes | 8.116E-07 | 491.24802 | 323.40128 |
| KSDIDEIVLVGGST | 15 | P11021 | Endoplasmic reticulum chaperone BiP OS | HSPA5 | 72.333 | 3 | 50.764507 | yes | 9.722E-08 | 530.28955 | 152.97784 |
| KSDIDEIVLVGGST | 15 | P11021 | Endoplasmic reticulum chaperone BiP OS | HSPA5 | 72.333 | 3 | 50.764507 | yes | 9.819E-08 | 794.93066 | 152.97784 |
| KSDIDEIVLVGGST | 15 | P11021 | Endoplasmic reticulum chaperone BiP OS | HSPA5 | 72.333 | 2 | 50.764507 | yes | 9.722E-08 | 530.28955 | 152.97784 |
| KSDIDEIVLVGGST | 15 | P11021 | Endoplasmic reticulum chaperone BiP OS | HSPA5 | 72.333 | 2 | 50.764507 | yes | 9.819E-08 | 794.93066 | 152.97784 |
| SDIDEIVLVGGSTR | 14 | P11021 | Endoplasmic reticulum chaperone BiP OS | HSPA5 | 72.333 | 2 | 50.764507 | yes | 3.339E-07 | 730.88318 | 180.24333 |
| DNHLLGTFDLTGII | 18 | P11021 | Endoplasmic reticulum chaperone BiP OS | HSPA5 | 72.333 | 3 | 50.764507 | yes | 1.924E-07 | 645.34253 | 178.55998 |
| LSSEDKETMEK | 11 | P11021 | Endoplasmic reticulum chaperone BiP OS | HSPA5 | 72.333 | 2 | 50.764507 | yes | 1.162E-07 | 648.80304 | 134.87482 |
| VTHAVVTVPAYFI | 17 | P11021 | Endoplasmic reticulum chaperone BiP OS | HSPA5 | 72.333 | 3 | 50.764507 | yes | 1.339E-07 | 629.99524 | 130.56265 |
| IINEPTAAAIAYGL | 17 | P11021 | Endoplasmic reticulum chaperone BiP OS | HSPA5 | 72.333 | 3 | 50.764507 | yes | 4.554E-07 | 606.0036 | 50.880646 |
| SQECYFDPELYR | 12 | P11047 | Laminin subunit gamma-1 OS=Homo sap | LAMC1 | 177.603 | 2 | 52.695038 | yes | 9.722E-08 | 803.84576 | 160.55481 |
| QDIAVISDSYFPR | 13 | P11047 | Laminin subunit gamma-1 OS=Homo sap | LAMC1 | 177.603 | 2 | 52.695038 | yes | 1.897E-07 | 755.88043 | 150.52631 |
| LNTFGDEVFNDPK | 13 | P11047 | Laminin subunit gamma-1 OS=Homo sap | LAMC1 | 177.603 | 2 | 52.695038 | yes | 1.31E-07 | 748.35681 | 90.475647 |

| | | | | | | | | | | | |
|---|---|---|---|---|---|---|---|---|---|---|---|
| LKDYEDLR | 8 | P11047 | Laminin subunit gamma-1 OS=Homo sap | LAMC1 | 177.603 | 2 | 52.695038 | yes | 1.841E-07 | 526.27454 | 77.899948 |
| LSAEDLVLEGAGL | 14 | P11047 | Laminin subunit gamma-1 OS=Homo sap | LAMC1 | 177.603 | 2 | 52.695038 | yes | 1.035E-07 | 721.89612 | 54.23103 |
| CSCKPGVMGDKC | 14 | P11047 | Laminin subunit gamma-1 OS=Homo sap | LAMC1 | 177.603 | 3 | 52.695038 | yes | 1.661E-07 | 557.23529 | 43.180595 |
| DAGTIAGLNVLR | 12 | P11142 | Heat shock cognate 71 kDa protein OS=H | HSPA8 | 70.899 | 2 | 53.044407 | yes | 3.141E-07 | 600.34076 | 17867.193 |
| FEELNADLFR | 10 | P11142 | Heat shock cognate 71 kDa protein OS=H | HSPA8 | 70.899 | 2 | 53.044407 | no | 2.106E-07 | 627.31171 | 16722.023 |
| SFYPEEVSSMVLT | 14 | P11142 | Heat shock cognate 71 kDa protein OS=H | HSPA8 | 70.899 | 2 | 53.044407 | yes | 6.247E-07 | 808.89746 | 8289.8594 |
| SFYPEEVSSMVLT | 14 | P11142 | Heat shock cognate 71 kDa protein OS=H | HSPA8 | 70.899 | 2 | 53.044407 | yes | 4.824E-07 | 816.8949 | 8289.8594 |
| NQVAMNPTNTVF | 15 | P11142 | Heat shock cognate 71 kDa protein OS=H | HSPA8 | 70.899 | 2 | 53.044407 | yes | 6.445E-07 | 825.40125 | 7987.2759 |
| NQVAMNPTNTVF | 15 | P11142 | Heat shock cognate 71 kDa protein OS=H | HSPA8 | 70.899 | 2 | 53.044407 | yes | 1.101E-07 | 833.39868 | 7987.2759 |
| IINEPTAAAIAYGL | 16 | P11142 | Heat shock cognate 71 kDa protein OS=H | HSPA8 | 70.899 | 2 | 53.044407 | no | 1.142E-07 | 830.45123 | 7225.9341 |
| CNEIINWLDK | 10 | P11142 | Heat shock cognate 71 kDa protein OS=H | HSPA8 | 70.899 | 2 | 53.044407 | yes | 6.395E-07 | 652.81879 | 4726.4521 |
| FDDAVVQSDMK | 11 | P11142 | Heat shock cognate 71 kDa protein OS=H | HSPA8 | 70.899 | 2 | 53.044407 | yes | 1.039E-07 | 627.78717 | 2193.291 |
| FDDAVVQSDMK | 11 | P11142 | Heat shock cognate 71 kDa protein OS=H | HSPA8 | 70.899 | 2 | 53.044407 | yes | 2.572E-07 | 635.78461 | 2193.291 |
| MVNHFIAEFK | 10 | P11142 | Heat shock cognate 71 kDa protein OS=H | HSPA8 | 70.899 | 2 | 53.044407 | yes | 9.819E-08 | 618.31573 | 1400.3151 |
| MVNHFIAEFK | 10 | P11142 | Heat shock cognate 71 kDa protein OS=H | HSPA8 | 70.899 | 2 | 53.044407 | yes | 1.422E-07 | 412.54623 | 1400.3151 |
| MVNHFIAEFK | 10 | P11142 | Heat shock cognate 71 kDa protein OS=H | HSPA8 | 70.899 | 2 | 53.044407 | yes | 9.722E-08 | 626.31317 | 1400.3151 |
| MVNHFIAEFK | 10 | P11142 | Heat shock cognate 71 kDa protein OS=H | HSPA8 | 70.899 | 3 | 53.044407 | yes | 9.819E-08 | 618.31573 | 1400.3151 |
| MVNHFIAEFK | 10 | P11142 | Heat shock cognate 71 kDa protein OS=H | HSPA8 | 70.899 | 3 | 53.044407 | yes | 1.422E-07 | 412.54623 | 1400.3151 |
| MVNHFIAEFK | 10 | P11142 | Heat shock cognate 71 kDa protein OS=H | HSPA8 | 70.899 | 3 | 53.044407 | yes | 9.722E-08 | 626.31317 | 1400.3151 |
| SQIHDIVLVGGSTF | 14 | P11142 | Heat shock cognate 71 kDa protein OS=H | HSPA8 | 70.899 | 2 | 53.044407 | yes | 4.508E-07 | 741.40717 | 3546.9321 |
| RFDDAVVQSDMK | 12 | P11142 | Heat shock cognate 71 kDa protein OS=H | HSPA8 | 70.899 | 2 | 53.044407 | yes | 2.004E-07 | 705.83771 | 3163.4734 |
| NQTAEKEEFEHQC | 14 | P11142 | Heat shock cognate 71 kDa protein OS=H | HSPA8 | 70.899 | 3 | 53.044407 | yes | 1.108E-07 | 582.60785 | 3069.2483 |
| GPAVGIDLGTTYS | 22 | P11142 | Heat shock cognate 71 kDa protein OS=H | HSPA8 | 70.899 | 3 | 53.044407 | yes | 2.613E-07 | 755.04407 | 2413.6821 |
| HWPFMVVNDAGF | 12 | P11142 | Heat shock cognate 71 kDa protein OS=H | HSPA8 | 70.899 | 2 | 53.044407 | yes | 1.338E-07 | 714.8457 | 1080.317 |
| HWPFMVVNDAGF | 12 | P11142 | Heat shock cognate 71 kDa protein OS=H | HSPA8 | 70.899 | 2 | 53.044407 | yes | 2.445E-07 | 476.89957 | 1080.317 |
| HWPFMVVNDAGF | 12 | P11142 | Heat shock cognate 71 kDa protein OS=H | HSPA8 | 70.899 | 3 | 53.044407 | yes | 1.338E-07 | 714.8457 | 1080.317 |
| HWPFMVVNDAGF | 12 | P11142 | Heat shock cognate 71 kDa protein OS=H | HSPA8 | 70.899 | 3 | 53.044407 | yes | 2.445E-07 | 476.89957 | 1080.317 |
| EIAEAYLGK | 9 | P11142 | Heat shock cognate 71 kDa protein OS=H | HSPA8 | 70.899 | 2 | 53.044407 | yes | 1.003E-07 | 497.2662 | 1854.8076 |
| NSLESYAFNMK | 11 | P11142 | Heat shock cognate 71 kDa protein OS=H | HSPA8 | 70.899 | 2 | 53.044407 | yes | 1.232E-07 | 652.30298 | 1753.6232 |
| HWPFMVVNDAGF | 14 | P11142 | Heat shock cognate 71 kDa protein OS=H | HSPA8 | 70.899 | 3 | 53.044407 | yes | 5.341E-07 | 551.94879 | 1724.7905 |
| STAGDTHLGGEDF | 16 | P11142 | Heat shock cognate 71 kDa protein OS=H | HSPA8 | 70.899 | 2 | 53.044407 | no | 1.063E-07 | 846.36646 | 1265.614 |
| TVTNAVVTVPAYI | 18 | P11142 | Heat shock cognate 71 kDa protein OS=H | HSPA8 | 70.899 | 3 | 53.044407 | yes | 1.381E-07 | 661.33746 | 999.55225 |
| FELTGIPPAPR | 11 | P11142 | Heat shock cognate 71 kDa protein OS=H | HSPA8 | 70.899 | 2 | 53.044407 | yes | 3.39E-07 | 599.33496 | 709.32782 |
| SINPDEAVAYGAA | 23 | P11142 | Heat shock cognate 71 kDa protein OS=H | HSPA8 | 70.899 | 3 | 53.044407 | yes | 1.233E-07 | 754.05334 | 623.2652 |
| RNTTIPTK | 8 | P11142 | Heat shock cognate 71 kDa protein OS=H | HSPA8 | 70.899 | 2 | 53.044407 | no | 3.282E-07 | 465.772 | 541.5343 |
| LLQDFFNGK | 9 | P11142 | Heat shock cognate 71 kDa protein OS=H | HSPA8 | 70.899 | 2 | 53.044407 | no | 2.805E-07 | 541.28748 | 239.09927 |
| KVTILELFR | 9 | P11166 | Solute carrier family 2, facilitated glucose | SLC2A1 | 54.084 | 2 | 49.383163 | yes | 1.038E-07 | 559.85022 | 32.902245 |
| YYLCLQLR | 8 | P11171 | Protein 4.1 OS=Homo sapiens OX=9606 | EPB41 | 97.017 | 2 | 49.298576 | no | 1.047E-07 | 564.79718 | 67.88961 |
| IVITGDADIDHDQ | 20 | P11171 | Protein 4.1 OS=Homo sapiens OX=9606 | EPB41 | 97.017 | 3 | 49.298576 | yes | 4.763E-07 | 721.72668 | 32.067238 |

| | | | | | | | | | | | |
|---|---|---|---|---|---|---|---|---|---|---|---|
| TQTVTISDNANAV | 14 | P11171 | Protein 4.1 OS=Homo sapiens OX=9606 | EPB41 | 97.017 | 2 | 49.298576 | yes | 7.97E-07 | 731.3808 | 18.790792 |
| VTDAIVLLDR | 10 | P11172 | Uridine 5'-monophosphate synthase OS=I | UMPS | 52.222 | 2 | 36.950523 | yes | 6.446E-07 | 557.82697 | 103.15926 |
| LEAAEMYR | 8 | P11172 | Uridine 5'-monophosphate synthase OS=I | UMPS | 52.222 | 2 | 36.950523 | yes | 4.747E-06 | 491.73676 | 69.741852 |
| FADIGNTVK | 9 | P11172 | Uridine 5'-monophosphate synthase OS=I | UMPS | 52.222 | 2 | 36.950523 | yes | 2.412E-06 | 482.75854 | 28.453432 |
| IASWADLVNAHV | 20 | P11172 | Uridine 5'-monophosphate synthase OS=I | UMPS | 52.222 | 3 | 36.950523 | yes | 8.462E-06 | 673.70557 | 23.367767 |
| EGVECEVINMR | 11 | P11177 | Pyruvate dehydrogenase E1 component su | PDHB | 39.234 | 2 | 36.411385 | yes | 9.649E-07 | 668.30524 | 107.29465 |
| DAINQGMDEELER | 13 | P11177 | Pyruvate dehydrogenase E1 component su | PDHB | 39.234 | 2 | 36.411385 | yes | 7.022E-07 | 760.33832 | 49.949001 |
| LSDIGEGIR | 9 | P11182 | Lipoamide acyltransferase component of l | DBT | 53.488 | 2 | 54.14636 | yes | 4.627E-07 | 480.26144 | 175.71838 |
| VIDGATMSR | 9 | P11182 | Lipoamide acyltransferase component of l | DBT | 53.488 | 2 | 54.14636 | yes | 1E-07 | 475.24222 | 81.088409 |
| LAMENNIK | 8 | P11182 | Lipoamide acyltransferase component of l | DBT | 53.488 | 2 | 54.14636 | yes | 6.887E-07 | 466.74713 | 54.361565 |
| AQIMNVSWSADH | 13 | P11182 | Lipoamide acyltransferase component of l | DBT | 53.488 | 2 | 54.14636 | yes | 9.722E-08 | 757.86206 | 35.543034 |
| VEIMPPPPKPK | 11 | P11182 | Lipoamide acyltransferase component of l | DBT | 53.488 | 3 | 54.14636 | yes | 0.0001224 | 411.57391 | 24.124144 |
| AEQWNVNYVETS | 14 | P11233 | Ras-related protein Ral-A OS=Homo sapi | RALA | 23.567 | 2 | 48.807774 | yes | 1.102E-07 | 819.89154 | 69.971207 |
| VFFDLMR | 7 | P11233 | Ras-related protein Ral-A OS=Homo sapi | RALA | 23.567 | 2 | 48.807774 | no | 6.866E-06 | 464.24149 | 23.965485 |
| STLVLHDLLK | 10 | P11274 | Breakpoint cluster region protein OS=Ho | BCR | 142.82 | 2 | 50.87492 | no | 9.722E-08 | 569.84515 | 149.87776 |
| LASQLGVYR | 9 | P11274 | Breakpoint cluster region protein OS=Ho | BCR | 142.82 | 2 | 50.87492 | yes | 6.124E-07 | 503.78763 | 146.41 |
| VSGVATDIQALK | 12 | P11274 | Breakpoint cluster region protein OS=Ho | BCR | 142.82 | 2 | 50.87492 | yes | 3.372E-07 | 601.34296 | 136.85657 |
| AFVDNYGVAMEM | 15 | P11274 | Breakpoint cluster region protein OS=Ho | BCR | 142.82 | 2 | 50.87492 | yes | 9.728E-08 | 837.87872 | 108.67641 |
| SVGDIEQELER | 11 | P11274 | Breakpoint cluster region protein OS=Ho | BCR | 142.82 | 2 | 50.87492 | yes | 9.789E-08 | 637.81476 | 88.567307 |
| ISSLGSQAMQMER | 13 | P11274 | Breakpoint cluster region protein OS=Ho | BCR | 142.82 | 2 | 50.87492 | yes | 1.082E-06 | 719.34485 | 67.818275 |
| HCPVVVSEATIVG | 15 | P11274 | Breakpoint cluster region protein OS=Ho | BCR | 142.82 | 3 | 50.87492 | yes | 3.451E-07 | 541.62762 | 39.431606 |
| MIYLQTLLAK | 10 | P11274 | Breakpoint cluster region protein OS=Ho | BCR | 142.82 | 2 | 50.87492 | yes | 1.173E-05 | 597.35175 | 11.483852 |
| VIVPADPTLWSTD | 16 | P11308 | Transcriptional regulator ERG OS=Homo | ERG | 53.838 | 3 | 46.32728 | yes | 1.589E-07 | 602.65631 | 85.30851 |
| LTPSYNADILLSHL | 18 | P11308 | Transcriptional regulator ERG OS=Homo | ERG | 53.838 | 3 | 46.32728 | yes | 4.39E-06 | 709.38464 | 50.4888 |
| MTEEPLMCAYCV | 25 | P11310 | Medium-chain specific acyl-CoA dehydro | ACADM | 46.589 | 4 | 45.04447 | yes | 2.222E-07 | 672.30212 | 176.46783 |
| TGEYPVPLIR | 10 | P11310 | Medium-chain specific acyl-CoA dehydro | ACADM | 46.589 | 2 | 45.04447 | yes | 3.184E-06 | 572.82166 | 113.18498 |
| IYQIYEGTSQIQR | 13 | P11310 | Medium-chain specific acyl-CoA dehydro | ACADM | 46.589 | 2 | 45.04447 | yes | 1.982E-07 | 799.91229 | 102.73425 |
| IMPEDIIINCSK | 12 | P11387 | DNA topoisomerase 1 OS=Homo sapiens | TOP1 | 90.726 | 2 | 52.224533 | yes | 2.326E-06 | 716.86237 | 3201.1057 |
| AVAILCNHQR | 10 | P11387 | DNA topoisomerase 1 OS=Homo sapiens | TOP1 | 90.726 | 2 | 52.224533 | yes | 9.772E-08 | 591.31403 | 1913.9554 |
| YIMLNPSSR | 9 | P11387 | DNA topoisomerase 1 OS=Homo sapiens | TOP1 | 90.726 | 2 | 52.224533 | yes | 1.164E-07 | 540.77893 | 1420.4619 |
| ELTAPDENIPAK | 12 | P11387 | DNA topoisomerase 1 OS=Homo sapiens | TOP1 | 90.726 | 2 | 52.224533 | yes | 9.809E-08 | 649.33533 | 1192.3976 |
| LNYLDPR | 7 | P11387 | DNA topoisomerase 1 OS=Homo sapiens | TOP1 | 90.726 | 2 | 52.224533 | no | 3.021E-06 | 445.74017 | 1124.828 |
| NLQLFMENK | 9 | P11387 | DNA topoisomerase 1 OS=Homo sapiens | TOP1 | 90.726 | 2 | 52.224533 | no | 9.901E-08 | 568.79205 | 1076.814 |
| EYGFCIMDNHK | 11 | P11387 | DNA topoisomerase 1 OS=Homo sapiens | TOP1 | 90.726 | 2 | 52.224533 | yes | 1.001E-07 | 707.29993 | 993.84857 |
| CDFTQMSQYFK | 11 | P11387 | DNA topoisomerase 1 OS=Homo sapiens | TOP1 | 90.726 | 2 | 52.224533 | yes | 1.309E-07 | 727.80762 | 970.48724 |
| LEVQATDR | 8 | P11387 | DNA topoisomerase 1 OS=Homo sapiens | TOP1 | 90.726 | 2 | 52.224533 | yes | 1.844E-07 | 466.24582 | 829.05579 |
| VPSPPPGHK | 9 | P11387 | DNA topoisomerase 1 OS=Homo sapiens | TOP1 | 90.726 | 2 | 52.224533 | yes | 2.952E-07 | 458.25598 | 788.50122 |
| KWGVPIEK | 8 | P11387 | DNA topoisomerase 1 OS=Homo sapiens | TOP1 | 90.726 | 2 | 52.224533 | yes | 2.841E-07 | 478.78183 | 768.76855 |
| AEEVATFFAK | 10 | P11387 | DNA topoisomerase 1 OS=Homo sapiens | TOP1 | 90.726 | 2 | 52.224533 | yes | 1.05E-07 | 556.78479 | 718.7616 |

| | | | | | | | | | | |
|---|---|---|---|---|---|---|---|---|---|---|
| AVALYFIDK | 9 | P11387 | DNA topoisomerase 1 OS=Homo sapiens TOP1 | 90.726 | 2 | 52.224533 | no | 2.41E-07 | 520.29474 | 633.00458 |
| SMMNLQTK | 8 | P11387 | DNA topoisomerase 1 OS=Homo sapiens TOP1 | 90.726 | 2 | 52.224533 | yes | 1.796E-07 | 476.73315 | 632.06317 |
| TYNASITLQQQLK | 13 | P11387 | DNA topoisomerase 1 OS=Homo sapiens TOP1 | 90.726 | 2 | 52.224533 | yes | 1.001E-07 | 754.40936 | 627.72064 |
| DEDDADYKPK | 10 | P11387 | DNA topoisomerase 1 OS=Homo sapiens TOP1 | 90.726 | 2 | 52.224533 | yes | 9.722E-08 | 598.25928 | 609.56982 |
| QPEDDLFDR | 9 | P11387 | DNA topoisomerase 1 OS=Homo sapiens TOP1 | 90.726 | 2 | 52.224533 | yes | 9.722E-08 | 567.75671 | 587.03497 |
| MLDHEYTTK | 9 | P11387 | DNA topoisomerase 1 OS=Homo sapiens TOP1 | 90.726 | 2 | 52.224533 | no | 9.722E-08 | 569.26587 | 551.24762 |
| IEPPGLFR | 8 | P11387 | DNA topoisomerase 1 OS=Homo sapiens TOP1 | 90.726 | 2 | 52.224533 | no | 1.626E-07 | 464.76617 | 471.29767 |
| LEEQLMK | 7 | P11387 | DNA topoisomerase 1 OS=Homo sapiens TOP1 | 90.726 | 2 | 52.224533 | yes | 2.776E-07 | 445.73624 | 471.10672 |
| HLQDLMEGLTAK | 12 | P11387 | DNA topoisomerase 1 OS=Homo sapiens TOP1 | 90.726 | 2 | 52.224533 | yes | 9.722E-08 | 678.35303 | 433.69287 |
| NIITNLSK | 8 | P11387 | DNA topoisomerase 1 OS=Homo sapiens TOP1 | 90.726 | 2 | 52.224533 | yes | 2.069E-07 | 451.76892 | 396.29291 |
| EDIKPLK | 7 | P11387 | DNA topoisomerase 1 OS=Homo sapiens TOP1 | 90.726 | 2 | 52.224533 | yes | 2.507E-07 | 421.75275 | 305.11139 |
| KLEEEEDGK | 9 | P11387 | DNA topoisomerase 1 OS=Homo sapiens TOP1 | 90.726 | 2 | 52.224533 | yes | 9.722E-08 | 538.75897 | 265.39395 |
| LEVQATDREENK | 12 | P11387 | DNA topoisomerase 1 OS=Homo sapiens TOP1 | 90.726 | 2 | 52.224533 | yes | 1.084E-07 | 716.35736 | 228.72765 |
| KLEEEEDGKLK | 11 | P11387 | DNA topoisomerase 1 OS=Homo sapiens TOP1 | 90.726 | 2 | 52.224533 | yes | 9.925E-08 | 659.34845 | 180.80692 |
| DEPEDDGYFVPPK | 13 | P11387 | DNA topoisomerase 1 OS=Homo sapiens TOP1 | 90.726 | 2 | 52.224533 | yes | 2.874E-07 | 754.33301 | 178.87419 |
| VEHINLHPELDGQ | 24 | P11387 | DNA topoisomerase 1 OS=Homo sapiens TOP1 | 90.726 | 4 | 52.224533 | yes | 1.046E-07 | 707.60388 | 131.44992 |
| WGVPIEK | 7 | P11387 | DNA topoisomerase 1 OS=Homo sapiens TOP1 | 90.726 | 2 | 52.224533 | yes | 3.01E-07 | 414.73434 | 129.71822 |
| WWEEERYPEGIK | 12 | P11387 | DNA topoisomerase 1 OS=Homo sapiens TOP1 | 90.726 | 2 | 52.224533 | yes | 2.992E-07 | 811.38593 | 45.434669 |
| EDIKPLKR | 8 | P11387 | DNA topoisomerase 1 OS=Homo sapiens TOP1 | 90.726 | 2 | 52.224533 | yes | 4.448E-07 | 499.80328 | 16.492041 |
| TLAVSGLGVVGR | 12 | P11388 | DNA topoisomerase 2-alpha OS=Homo s TOP2A | 174.386 | 2 | 53.68906 | yes | 9.722E-08 | 564.84039 | 2039.6019 |
| SIPSMVDGLKPGQ | 14 | P11388 | DNA topoisomerase 2-alpha OS=Homo s TOP2A | 174.386 | 2 | 53.68906 | yes | 1.156E-06 | 742.89813 | 1791.6803 |
| TPPLITDYR | 9 | P11388 | DNA topoisomerase 2-alpha OS=Homo s TOP2A | 174.386 | 2 | 53.68906 | yes | 1.23E-07 | 538.29279 | 1626.797 |
| SFGSTCQLSEK | 11 | P11388 | DNA topoisomerase 2-alpha OS=Homo s TOP2A | 174.386 | 2 | 53.68906 | yes | 2.697E-07 | 622.28479 | 1317.2426 |
| FLEEFITPIVK | 11 | P11388 | DNA topoisomerase 2-alpha OS=Homo s TOP2A | 174.386 | 2 | 53.68906 | yes | 1.489E-07 | 668.38159 | 1024.238 |
| KYDTVLDILR | 10 | P11388 | DNA topoisomerase 2-alpha OS=Homo s TOP2A | 174.386 | 2 | 53.68906 | yes | 1.306E-07 | 618.35333 | 840.49298 |
| YSGPEDDAAISLAI | 16 | P11388 | DNA topoisomerase 2-alpha OS=Homo s TOP2A | 174.386 | 2 | 53.68906 | yes | 9.965E-08 | 835.89905 | 780.27948 |
| HVDYVADQIVTK | 12 | P11388 | DNA topoisomerase 2-alpha OS=Homo s TOP2A | 174.386 | 2 | 53.68906 | yes | 1.138E-07 | 694.36444 | 638.0625 |
| QTWMDNMGR | 9 | P11388 | DNA topoisomerase 2-alpha OS=Homo s TOP2A | 174.386 | 2 | 53.68906 | yes | 1.146E-07 | 569.74207 | 591.53925 |
| GFQQISFVNSIATS | 15 | P11388 | DNA topoisomerase 2-alpha OS=Homo s TOP2A | 174.386 | 2 | 53.68906 | yes | 1.503E-07 | 813.92792 | 587.5166 |
| IIIENKPK | 8 | P11388 | DNA topoisomerase 2-alpha OS=Homo s TOP2A | 174.386 | 2 | 53.68906 | yes | 3.261E-07 | 477.80276 | 555.02344 |
| WEVCLTMSEK | 10 | P11388 | DNA topoisomerase 2-alpha OS=Homo s TOP2A | 174.386 | 2 | 53.68906 | yes | 9.809E-08 | 641.79395 | 261.5394 |
| WEVCLTMSEK | 10 | P11388 | DNA topoisomerase 2-alpha OS=Homo s TOP2A | 174.386 | 2 | 53.68906 | yes | 7.506E-07 | 649.79138 | 261.5394 |
| IPNFDVR | 7 | P11388 | DNA topoisomerase 2-alpha OS=Homo s TOP2A | 174.386 | 2 | 53.68906 | yes | 3.278E-07 | 430.73486 | 513.12048 |
| TQMAEVLPSPR | 11 | P11388 | DNA topoisomerase 2-alpha OS=Homo s TOP2A | 174.386 | 2 | 53.68906 | yes | 1.215E-07 | 614.82135 | 503.60779 |
| SVVSDLEADDVK | 12 | P11388 | DNA topoisomerase 2-alpha OS=Homo s TOP2A | 174.386 | 2 | 53.68906 | yes | 3.184E-07 | 638.81702 | 457.0936 |
| DKLDETGNSLK | 11 | P11388 | DNA topoisomerase 2-alpha OS=Homo s TOP2A | 174.386 | 2 | 53.68906 | yes | 3.063E-07 | 610.31189 | 435.14798 |
| RDPALNSGVSQKP | 17 | P11388 | DNA topoisomerase 2-alpha OS=Homo s TOP2A | 174.386 | 3 | 53.68906 | yes | 1.066E-07 | 593.98309 | 403.18814 |
| EWLLGMLGAESAI | 13 | P11388 | DNA topoisomerase 2-alpha OS=Homo s TOP2A | 174.386 | 2 | 53.68906 | yes | 9.722E-08 | 702.86322 | 397.66251 |
| LQTSLTCNSMVLF | 20 | P11388 | DNA topoisomerase 2-alpha OS=Homo s TOP2A | 174.386 | 3 | 53.68906 | yes | 1.283E-07 | 775.04626 | 315.98224 |

| | | | | | | | | | | | |
|---|---|---|---|---|---|---|---|---|---|---|---|
| QTTLAFKPIK | 10 | P11388 | DNA topoisomerase 2-alpha OS=Homo s | TOP2A | 174.386 | 2 | 53.68906 | yes | 9.914E-08 | 573.84772 | 311.8829 |
| YDTVLDILR | 9 | P11388 | DNA topoisomerase 2-alpha OS=Homo s | TOP2A | 174.386 | 2 | 53.68906 | yes | 2.958E-07 | 554.30585 | 280.81619 |
| QDEQVGLPGK | 10 | P11388 | DNA topoisomerase 2-alpha OS=Homo s | TOP2A | 174.386 | 2 | 53.68906 | yes | 1.65E-07 | 535.77747 | 234.08084 |
| KEWLLGMLGAES | 14 | P11388 | DNA topoisomerase 2-alpha OS=Homo s | TOP2A | 174.386 | 2 | 53.68906 | yes | 1.021E-07 | 766.91071 | 226.70224 |
| GLLINFIHHNWPSI | 16 | P11388 | DNA topoisomerase 2-alpha OS=Homo s | TOP2A | 174.386 | 3 | 53.68906 | yes | 9.722E-08 | 644.03186 | 223.25171 |
| EQELDTLKR | 9 | P11388 | DNA topoisomerase 2-alpha OS=Homo s | TOP2A | 174.386 | 2 | 53.68906 | yes | 9.722E-08 | 566.30383 | 213.50922 |
| FTVETASR | 8 | P11388 | DNA topoisomerase 2-alpha OS=Homo s | TOP2A | 174.386 | 2 | 53.68906 | yes | 9.893E-08 | 455.73508 | 204.93559 |
| EKQDEQVGLPGK | 12 | P11388 | DNA topoisomerase 2-alpha OS=Homo s | TOP2A | 174.386 | 2 | 53.68906 | yes | 9.722E-08 | 664.34625 | 185.78223 |
| SYVDMYLK | 8 | P11388 | DNA topoisomerase 2-alpha OS=Homo s | TOP2A | 174.386 | 2 | 53.68906 | yes | 9.722E-08 | 509.74933 | 161.15601 |
| DPALNSGVSQKPD | 16 | P11388 | DNA topoisomerase 2-alpha OS=Homo s | TOP2A | 174.386 | 2 | 53.68906 | yes | 5.704E-07 | 812.42047 | 129.95787 |
| DKYGVFPLR | 9 | P11388 | DNA topoisomerase 2-alpha OS=Homo s | TOP2A | 174.386 | 2 | 53.68906 | yes | 1.295E-06 | 547.80328 | 119.88587 |
| KQTTLAFKPIK | 11 | P11388 | DNA topoisomerase 2-alpha OS=Homo s | TOP2A | 174.386 | 3 | 53.68906 | yes | 9.722E-08 | 425.59921 | 54.865673 |
| KQTTLAFKPIK | 11 | P11388 | DNA topoisomerase 2-alpha OS=Homo s | TOP2A | 174.386 | 3 | 53.68906 | yes | 1.051E-07 | 637.8952 | 54.865673 |
| KQTTLAFKPIK | 11 | P11388 | DNA topoisomerase 2-alpha OS=Homo s | TOP2A | 174.386 | 2 | 53.68906 | yes | 9.722E-08 | 425.59921 | 54.865673 |
| KQTTLAFKPIK | 11 | P11388 | DNA topoisomerase 2-alpha OS=Homo s | TOP2A | 174.386 | 2 | 53.68906 | yes | 1.051E-07 | 637.8952 | 54.865673 |
| LDETGNSLK | 9 | P11388 | DNA topoisomerase 2-alpha OS=Homo s | TOP2A | 174.386 | 2 | 53.68906 | yes | 1.456E-07 | 488.75092 | 92.835335 |
| YIFTMLSSLAR | 11 | P11388 | DNA topoisomerase 2-alpha OS=Homo s | TOP2A | 174.386 | 2 | 53.68906 | yes | 2.476E-06 | 651.34973 | 92.019798 |
| TDDEDFVPSDASP | 15 | P11388 | DNA topoisomerase 2-alpha OS=Homo s | TOP2A | 174.386 | 2 | 53.68906 | yes | 1.737E-06 | 810.35718 | 87.952721 |
| EYFADMK | 7 | P11388 | DNA topoisomerase 2-alpha OS=Homo s | TOP2A | 174.386 | 2 | 53.68906 | yes | 3.813E-07 | 452.19949 | 57.327377 |
| KNYEDEDSLK | 10 | P11388 | DNA topoisomerase 2-alpha OS=Homo s | TOP2A | 174.386 | 2 | 53.68906 | yes | 1.021E-07 | 620.78821 | 46.187149 |
| MQSLDKDIVALM | 14 | P11388 | DNA topoisomerase 2-alpha OS=Homo s | TOP2A | 174.386 | 2 | 53.68906 | yes | 2.249E-07 | 809.9364 | 36.952625 |
| GYLDDPTVPR | 10 | P11413 | Glucose-6-phosphate 1-dehydrogenase O | G6PD | 59.257 | 2 | 55.9609 | yes | 2.058E-07 | 566.78528 | 203.64996 |
| IIVEKPFGR | 9 | P11413 | Glucose-6-phosphate 1-dehydrogenase O | G6PD | 59.257 | 2 | 55.9609 | yes | 2.188E-07 | 529.82147 | 149.8161 |
| GGYFDEFGIIR | 11 | P11413 | Glucose-6-phosphate 1-dehydrogenase O | G6PD | 59.257 | 2 | 55.9609 | yes | 9.722E-08 | 637.31421 | 130.86482 |
| GPTEADELMK | 10 | P11413 | Glucose-6-phosphate 1-dehydrogenase O | G6PD | 59.257 | 2 | 55.9609 | yes | 6.143E-07 | 545.75787 | 98.342064 |
| EMVQNLMVLR | 10 | P11413 | Glucose-6-phosphate 1-dehydrogenase O | G6PD | 59.257 | 2 | 55.9609 | yes | 9.722E-08 | 616.82813 | 74.961189 |
| VGFQYEGTYK | 10 | P11413 | Glucose-6-phosphate 1-dehydrogenase O | G6PD | 59.257 | 2 | 55.9609 | yes | 9.722E-08 | 596.28766 | 63.573246 |
| LKLEDFFAR | 9 | P11413 | Glucose-6-phosphate 1-dehydrogenase O | G6PD | 59.257 | 2 | 55.9609 | yes | 6.923E-07 | 569.81641 | 62.341 |
| DNIACVILTFK | 11 | P11413 | Glucose-6-phosphate 1-dehydrogenase O | G6PD | 59.257 | 2 | 55.9609 | yes | 1.717E-06 | 647.34723 | 27.662777 |
| LTLDDIER | 8 | P11441 | Ubiquitin-like protein 4A OS=Homo sapi | UBL4A | 17.777 | 2 | 48.12299 | yes | 1.197E-07 | 487.76129 | 89.227135 |
| VLLEEGEAQR | 10 | P11441 | Ubiquitin-like protein 4A OS=Homo sapi | UBL4A | 17.777 | 2 | 48.12299 | yes | 1.47E-06 | 572.30383 | 50.989998 |
| ILADLEEENR | 10 | P11532 | Dystrophin OS=Homo sapiens OX=9606 | DMD | 426.75 | 2 | 50.783947 | yes | 3.793E-07 | 601.30658 | 854.66144 |
| ECPIIGFR | 8 | P11532 | Dystrophin OS=Homo sapiens OX=9606 | DMD | 426.75 | 2 | 50.783947 | yes | 1.287E-07 | 496.25513 | 575.36127 |
| MQILEDHNK | 9 | P11532 | Dystrophin OS=Homo sapiens OX=9606 | DMD | 426.75 | 2 | 50.783947 | no | 2.052E-07 | 564.2793 | 521.64795 |
| SDSSQPMLLR | 10 | P11532 | Dystrophin OS=Homo sapiens OX=9606 | DMD | 426.75 | 2 | 50.783947 | yes | 1.999E-07 | 567.28461 | 456.02383 |
| SPAQILISLESEER | 14 | P11532 | Dystrophin OS=Homo sapiens OX=9606 | DMD | 426.75 | 2 | 50.783947 | yes | 9.722E-08 | 786.41742 | 340.65253 |
| QLLEQPQAEAK | 11 | P11532 | Dystrophin OS=Homo sapiens OX=9606 | DMD | 426.75 | 2 | 50.783947 | yes | 1.561E-07 | 627.83807 | 164.82529 |
| TGIISLCK | 8 | P11532 | Dystrophin OS=Homo sapiens OX=9606 | DMD | 426.75 | 2 | 50.783947 | yes | 1.022E-07 | 446.25204 | 138.353 |
| NLQAEYDR | 8 | P11532 | Dystrophin OS=Homo sapiens OX=9606 | DMD | 426.75 | 2 | 50.783947 | yes | 4.841E-07 | 504.74088 | 109.22994 |

| | | | | | | | | | | | |
|---|---|---|---|---|---|---|---|---|---|---|---|
| QLESQLHR | 8 | P11532 | Dystrophin OS=Homo sapiens OX=9606 | DMD | 426.75 | 2 | 50.783947 | no | 1.8E-07 | 505.77252 | 100.04044 |
| DAELIAEAK | 9 | P11532 | Dystrophin OS=Homo sapiens OX=9606 | DMD | 426.75 | 2 | 50.783947 | yes | 3.512E-07 | 480.25583 | 87.181114 |
| QVASSTGFCDQR | 12 | P11532 | Dystrophin OS=Homo sapiens OX=9606 | DMD | 426.75 | 2 | 50.783947 | yes | 9.81E-08 | 678.30408 | 82.096321 |
| ALCLDLLSLSAACI | 22 | P11532 | Dystrophin OS=Homo sapiens OX=9606 | DMD | 426.75 | 3 | 50.783947 | yes | 1.891E-06 | 814.41028 | 25.615515 |
| GALALAQAVQR | 11 | P11586 | C-1-tetrahydrofolate synthase, cytoplasmi | MTHFD1 | 101.532 | 2 | 49.548557 | yes | 1.073E-07 | 549.32495 | 125.56617 |
| GDLNDCFIPCTPK | 13 | P11586 | C-1-tetrahydrofolate synthase, cytoplasmi | MTHFD1 | 101.532 | 2 | 49.548557 | yes | 1.398E-07 | 768.84473 | 101.68974 |
| YVVVTGITPTPLGI | 16 | P11586 | C-1-tetrahydrofolate synthase, cytoplasmi | MTHFD1 | 101.532 | 2 | 49.548557 | yes | 1.022E-07 | 815.95618 | 59.600285 |
| FSDIQIR | 7 | P11586 | C-1-tetrahydrofolate synthase, cytoplasmi | MTHFD1 | 101.532 | 2 | 49.548557 | yes | 2.594E-06 | 439.74017 | 38.399906 |
| TDPTTLTDEEINR | 13 | P11586 | C-1-tetrahydrofolate synthase, cytoplasmi | MTHFD1 | 101.532 | 2 | 49.548557 | yes | 5.491E-07 | 752.85992 | 38.207207 |
| TDTESELDLISR | 12 | P11586 | C-1-tetrahydrofolate synthase, cytoplasmi | MTHFD1 | 101.532 | 2 | 49.548557 | yes | 7.686E-06 | 689.83844 | 31.761276 |
| KITIGQAPTEK | 11 | P11586 | C-1-tetrahydrofolate synthase, cytoplasmi | MTHFD1 | 101.532 | 2 | 49.548557 | yes | 1.537E-06 | 593.34552 | 30.399151 |
| YVDGDLCPDGIR | 12 | P11717 | Cation-independent mannose-6-phosphat | IGF2R | 274.375 | 2 | 50.20913 | yes | 9.85E-08 | 690.31665 | 113.65907 |
| TVEACPVVR | 9 | P11717 | Cation-independent mannose-6-phosphat | IGF2R | 274.375 | 2 | 50.20913 | yes | 9.185E-07 | 515.77112 | 96.8564 |
| TASSVIELTCTK | 12 | P11717 | Cation-independent mannose-6-phosphat | IGF2R | 274.375 | 2 | 50.20913 | yes | 5.47E-07 | 655.33704 | 96.8013 |
| STAIFFYCDR | 10 | P11717 | Cation-independent mannose-6-phosphat | IGF2R | 274.375 | 2 | 50.20913 | yes | 2.625E-07 | 640.29242 | 68.686455 |
| YVDQVLQLVYK | 11 | P11717 | Cation-independent mannose-6-phosphat | IGF2R | 274.375 | 2 | 50.20913 | yes | 2.034E-07 | 684.38208 | 67.317909 |
| GCPEDAAVCAVDI | 13 | P11717 | Cation-independent mannose-6-phosphat | IGF2R | 274.375 | 2 | 50.20913 | yes | 1.769E-07 | 696.30011 | 65.431847 |
| GTQRPVFLK | 9 | P11717 | Cation-independent mannose-6-phosphat | IGF2R | 274.375 | 2 | 50.20913 | yes | 1.157E-07 | 523.31128 | 60.015724 |
| ETSDCSYLFEWR | 12 | P11717 | Cation-independent mannose-6-phosphat | IGF2R | 274.375 | 2 | 50.20913 | yes | 1.362E-07 | 796.83795 | 50.235638 |
| LSYYDGMIQLNYF | 13 | P11717 | Cation-independent mannose-6-phosphat | IGF2R | 274.375 | 2 | 50.20913 | yes | 5.193E-07 | 818.39539 | 45.602283 |
| PLNPVPGCNR | 10 | P11717 | Cation-independent mannose-6-phosphat | IGF2R | 274.375 | 2 | 50.20913 | yes | 1.404E-06 | 562.28748 | 45.356579 |
| FVCNDDVYSGPLK | 13 | P11717 | Cation-independent mannose-6-phosphat | IGF2R | 274.375 | 2 | 50.20913 | yes | 5.927E-07 | 757.35321 | 45.317104 |
| VPIDGPPIDIGR | 12 | P11717 | Cation-independent mannose-6-phosphat | IGF2R | 274.375 | 2 | 50.20913 | yes | 2.216E-07 | 624.85095 | 43.710289 |
| KHDLNPLIK | 9 | P11717 | Cation-independent mannose-6-phosphat | IGF2R | 274.375 | 2 | 50.20913 | yes | 9.082E-07 | 539.3244 | 35.563095 |
| DGGVCLLSGTK | 11 | P11717 | Cation-independent mannose-6-phosphat | IGF2R | 274.375 | 2 | 50.20913 | yes | 4.285E-07 | 553.77917 | 32.935921 |
| EVPCYVFDEELR | 12 | P11717 | Cation-independent mannose-6-phosphat | IGF2R | 274.375 | 2 | 50.20913 | yes | 1.547E-07 | 778.35852 | 16.283701 |
| GTADAFIVR | 9 | P11717 | Cation-independent mannose-6-phosphat | IGF2R | 274.375 | 2 | 50.20913 | yes | 6.542E-07 | 475.25873 | 13.653204 |
| PASGCEAETQTEEI | 15 | P11717 | Cation-independent mannose-6-phosphat | IGF2R | 274.375 | 2 | 50.20913 | yes | 2.182E-07 | 825.36981 | 9.9682999 |
| RLEAFEHPNVVR | 12 | P11802 | Cyclin-dependent kinase 4 OS=Homo sap | CDK4 | 33.73 | 3 | 49.827175 | yes | 2.016E-07 | 489.60028 | 173.89714 |
| LMDVCATSR | 9 | P11802 | Cyclin-dependent kinase 4 OS=Homo sap | CDK4 | 33.73 | 2 | 49.827175 | yes | 1.002E-07 | 526.74683 | 125.82674 |
| APPPGLPAETIK | 12 | P11802 | Cyclin-dependent kinase 4 OS=Homo sap | CDK4 | 33.73 | 2 | 49.827175 | yes | 1.362E-06 | 595.84259 | 80.013519 |
| DLKPENILVTSGGT | 16 | P11802 | Cyclin-dependent kinase 4 OS=Homo sap | CDK4 | 33.73 | 2 | 49.827175 | yes | 3.694E-07 | 835.96979 | 44.582024 |
| VTLVFEHVDQDLF | 13 | P11802 | Cyclin-dependent kinase 4 OS=Homo sap | CDK4 | 33.73 | 2 | 49.827175 | yes | 1.337E-07 | 785.91486 | 35.178169 |
| LNVEFALIHK | 10 | P11908 | Ribose-phosphate pyrophosphokinase 2 C | PRPS2 | 34.77 | 2 | 54.77418 | yes | 9.722E-08 | 592.34534 | 75.667351 |
| FSNQETSVEIGESV | 15 | P11908 | Ribose-phosphate pyrophosphokinase 2 C | PRPS2 | 34.77 | 2 | 54.77418 | yes | 9.722E-08 | 841.40503 | 30.964417 |
| PLYVALAQR | 9 | P11940 | Polyadenylate-binding protein 1 OS=Hon | PABPC1 | 70.671 | 2 | 51.73703 | no | 2.311E-07 | 515.80585 | 129901.58 |
| FSPAGPILSIR | 11 | P11940 | Polyadenylate-binding protein 1 OS=Hon | PABPC1 | 70.671 | 2 | 51.73703 | no | 1.739E-07 | 579.33752 | 113685.24 |
| ALDTMNFDVIK | 11 | P11940 | Polyadenylate-binding protein 1 OS=Hon | PABPC1 | 70.671 | 2 | 51.73703 | no | 1.18E-07 | 633.82355 | 35937.902 |
| ALDTMNFDVIK | 11 | P11940 | Polyadenylate-binding protein 1 OS=Hon | PABPC1 | 70.671 | 2 | 51.73703 | no | 1.674E-07 | 641.82104 | 35937.902 |

| | | | | | | | | | | | |
|---|---|---|---|---|---|---|---|---|---|---|---|
| EFSPFGTITSAK | 12 | P11940 | Polyadenylate-binding protein 1 OS=Hon | PABPC1 | 70.671 | 2 | 51.73703 | yes | 1.325E-07 | 642.82715 | 64608.754 |
| NLDDGIDDER | 10 | P11940 | Polyadenylate-binding protein 1 OS=Hon | PABPC1 | 70.671 | 2 | 51.73703 | no | 9.732E-08 | 581.25458 | 55701.332 |
| GFGFVCFSSPEEAT | 15 | P11940 | Polyadenylate-binding protein 1 OS=Hon | PABPC1 | 70.671 | 2 | 51.73703 | no | 1.102E-07 | 831.87708 | 40891.789 |
| NFGEDMDDER | 10 | P11940 | Polyadenylate-binding protein 1 OS=Hon | PABPC1 | 70.671 | 2 | 51.73703 | no | 1.298E-07 | 614.23279 | 16962.109 |
| NFGEDMDDER | 10 | P11940 | Polyadenylate-binding protein 1 OS=Hon | PABPC1 | 70.671 | 2 | 51.73703 | no | 9.722E-08 | 622.23022 | 16962.109 |
| SGVGNIFIK | 9 | P11940 | Polyadenylate-binding protein 1 OS=Hon | PABPC1 | 70.671 | 2 | 51.73703 | no | 1.577E-07 | 467.77145 | 32037.008 |
| MNGMLLNDR | 9 | P11940 | Polyadenylate-binding protein 1 OS=Hon | PABPC1 | 70.671 | 2 | 51.73703 | no | 1.197E-07 | 532.25482 | 9767.6875 |
| MNGMLLNDR | 9 | P11940 | Polyadenylate-binding protein 1 OS=Hon | PABPC1 | 70.671 | 2 | 51.73703 | no | 2.346E-07 | 540.25226 | 9767.6875 |
| MNGMLLNDR | 9 | P11940 | Polyadenylate-binding protein 1 OS=Hon | PABPC1 | 70.671 | 2 | 51.73703 | no | 2.494E-07 | 540.25226 | 9767.6875 |
| SKVDEAVAVLQAI | 16 | P11940 | Polyadenylate-binding protein 1 OS=Hon | PABPC1 | 70.671 | 3 | 51.73703 | no | 1.213E-07 | 565.31256 | 14366.51 |
| SKVDEAVAVLQAI | 16 | P11940 | Polyadenylate-binding protein 1 OS=Hon | PABPC1 | 70.671 | 3 | 51.73703 | no | 2.249E-07 | 847.46521 | 14366.51 |
| SKVDEAVAVLQAI | 16 | P11940 | Polyadenylate-binding protein 1 OS=Hon | PABPC1 | 70.671 | 2 | 51.73703 | no | 1.213E-07 | 565.31256 | 14366.51 |
| SKVDEAVAVLQAI | 16 | P11940 | Polyadenylate-binding protein 1 OS=Hon | PABPC1 | 70.671 | 2 | 51.73703 | no | 2.249E-07 | 847.46521 | 14366.51 |
| IVATKPLYVALAQ | 14 | P11940 | Polyadenylate-binding protein 1 OS=Hon | PABPC1 | 70.671 | 3 | 51.73703 | no | 1.728E-07 | 514.9837 | 13364.758 |
| IVATKPLYVALAQ | 14 | P11940 | Polyadenylate-binding protein 1 OS=Hon | PABPC1 | 70.671 | 3 | 51.73703 | no | 3.688E-07 | 771.97192 | 13364.758 |
| IVATKPLYVALAQ | 14 | P11940 | Polyadenylate-binding protein 1 OS=Hon | PABPC1 | 70.671 | 2 | 51.73703 | no | 1.728E-07 | 514.9837 | 13364.758 |
| IVATKPLYVALAQ | 14 | P11940 | Polyadenylate-binding protein 1 OS=Hon | PABPC1 | 70.671 | 2 | 51.73703 | no | 3.688E-07 | 771.97192 | 13364.758 |
| YQGVNLYVK | 9 | P11940 | Polyadenylate-binding protein 1 OS=Hon | PABPC1 | 70.671 | 2 | 51.73703 | no | 1.234E-07 | 542.29529 | 22570.613 |
| KEFSPFGTITSAK | 13 | P11940 | Polyadenylate-binding protein 1 OS=Hon | PABPC1 | 70.671 | 2 | 51.73703 | yes | 7.69E-07 | 706.87463 | 18620.846 |
| KSGVGNIFIK | 10 | P11940 | Polyadenylate-binding protein 1 OS=Hon | PABPC1 | 70.671 | 2 | 51.73703 | no | 1.554E-07 | 531.81897 | 14198.934 |
| PAAAAAAATPAVI | 13 | P11940 | Polyadenylate-binding protein 1 OS=Hon | PABPC1 | 70.671 | 2 | 51.73703 | yes | 1.106E-07 | 569.32239 | 13290.395 |
| EFTNVYIK | 8 | P11940 | Polyadenylate-binding protein 1 OS=Hon | PABPC1 | 70.671 | 2 | 51.73703 | no | 2.366E-07 | 507.26874 | 12700.673 |
| QAHLTNQYMQR | 11 | P11940 | Polyadenylate-binding protein 1 OS=Hon | PABPC1 | 70.671 | 2 | 51.73703 | yes | 1.007E-07 | 695.33826 | 5926.8706 |
| QAHLTNQYMQR | 11 | P11940 | Polyadenylate-binding protein 1 OS=Hon | PABPC1 | 70.671 | 2 | 51.73703 | yes | 9.75E-08 | 703.33569 | 5926.8706 |
| LFPLIQAMHPTLA( | 15 | P11940 | Polyadenylate-binding protein 1 OS=Hon | PABPC1 | 70.671 | 2 | 51.73703 | no | 1.683E-07 | 818.96582 | 3039.614 |
| LFPLIQAMHPTLA( | 15 | P11940 | Polyadenylate-binding protein 1 OS=Hon | PABPC1 | 70.671 | 2 | 51.73703 | no | 2.497E-07 | 546.31299 | 3039.614 |
| LFPLIQAMHPTLA( | 15 | P11940 | Polyadenylate-binding protein 1 OS=Hon | PABPC1 | 70.671 | 2 | 51.73703 | no | 1.307E-06 | 551.64459 | 3039.614 |
| LFPLIQAMHPTLA( | 15 | P11940 | Polyadenylate-binding protein 1 OS=Hon | PABPC1 | 70.671 | 2 | 51.73703 | no | 2.062E-07 | 826.96326 | 3039.614 |
| LFPLIQAMHPTLA( | 15 | P11940 | Polyadenylate-binding protein 1 OS=Hon | PABPC1 | 70.671 | 3 | 51.73703 | no | 1.683E-07 | 818.96582 | 3039.614 |
| LFPLIQAMHPTLA( | 15 | P11940 | Polyadenylate-binding protein 1 OS=Hon | PABPC1 | 70.671 | 3 | 51.73703 | no | 2.497E-07 | 546.31299 | 3039.614 |
| LFPLIQAMHPTLA( | 15 | P11940 | Polyadenylate-binding protein 1 OS=Hon | PABPC1 | 70.671 | 3 | 51.73703 | no | 1.307E-06 | 551.64459 | 3039.614 |
| LFPLIQAMHPTLA( | 15 | P11940 | Polyadenylate-binding protein 1 OS=Hon | PABPC1 | 70.671 | 3 | 51.73703 | no | 2.062E-07 | 826.96326 | 3039.614 |
| PASSQVPR | 8 | P11940 | Polyadenylate-binding protein 1 OS=Hon | PABPC1 | 70.671 | 2 | 51.73703 | no | 1.058E-07 | 421.22995 | 4258.5181 |
| LKDLFGK | 7 | P11940 | Polyadenylate-binding protein 1 OS=Hon | PABPC1 | 70.671 | 2 | 51.73703 | no | 1.93E-07 | 410.75 | 3605.2219 |
| VANTSTQTMGPR | 12 | P11940 | Polyadenylate-binding protein 1 OS=Hon | PABPC1 | 70.671 | 2 | 51.73703 | yes | 2.958E-07 | 631.81152 | 1521.3939 |
| VANTSTQTMGPR | 12 | P11940 | Polyadenylate-binding protein 1 OS=Hon | PABPC1 | 70.671 | 2 | 51.73703 | yes | 5.406E-07 | 639.80896 | 1521.3939 |
| VMTDESGK | 8 | P11940 | Polyadenylate-binding protein 1 OS=Hon | PABPC1 | 70.671 | 2 | 51.73703 | no | 1.993E-07 | 433.69983 | 999.97528 |
| VMTDESGK | 8 | P11940 | Polyadenylate-binding protein 1 OS=Hon | PABPC1 | 70.671 | 2 | 51.73703 | no | 2.617E-06 | 441.6973 | 999.97528 |
| VDEAVAVLQAHQ | 14 | P11940 | Polyadenylate-binding protein 1 OS=Hon | PABPC1 | 70.671 | 3 | 51.73703 | no | 9.968E-08 | 493.60358 | 568.06287 |

| | | | | | | | | | | | |
|---|---|---|---|---|---|---|---|---|---|---|---|
| VDEAVAVLQAHQ | 14 | P11940 | Polyadenylate-binding protein 1 OS=Hon | PABPC1 | 70.671 | 3 | 51.73703 | no | 1.256E-07 | 739.90173 | 568.06287 |
| VDEAVAVLQAHQ | 14 | P11940 | Polyadenylate-binding protein 1 OS=Hon | PABPC1 | 70.671 | 2 | 51.73703 | no | 9.968E-08 | 493.60358 | 568.06287 |
| VDEAVAVLQAHQ | 14 | P11940 | Polyadenylate-binding protein 1 OS=Hon | PABPC1 | 70.671 | 2 | 51.73703 | no | 1.256E-07 | 739.90173 | 568.06287 |
| AAYYPPSQIAQLR | 13 | P11940 | Polyadenylate-binding protein 1 OS=Hon | PABPC1 | 70.671 | 2 | 51.73703 | yes | 6.447E-07 | 739.39355 | 1056.7306 |
| AKEFTNVYIK | 10 | P11940 | Polyadenylate-binding protein 1 OS=Hon | PABPC1 | 70.671 | 2 | 51.73703 | no | 2.996E-07 | 606.83478 | 635.94025 |
| GFGFVSFER | 9 | P11940 | Polyadenylate-binding protein 1 OS=Hon | PABPC1 | 70.671 | 2 | 51.73703 | no | 4.319E-07 | 523.25873 | 274.82187 |
| AVTEMNGR | 8 | P11940 | Polyadenylate-binding protein 1 OS=Hon | PABPC1 | 70.671 | 2 | 51.73703 | no | 9.722E-08 | 439.21344 | 265.73776 |
| PHPFQNMPGAIR | 12 | P11940 | Polyadenylate-binding protein 1 OS=Hon | PABPC1 | 70.671 | 2 | 51.73703 | yes | 4.246E-07 | 682.84827 | 128.49057 |
| AAYYPPSQIAQLR | 17 | P11940 | Polyadenylate-binding protein 1 OS=Hon | PABPC1 | 70.671 | 3 | 51.73703 | yes | 3.34E-07 | 639.01099 | 93.632805 |
| PAAPRPPFSTMR | 12 | P11940 | Polyadenylate-binding protein 1 OS=Hon | PABPC1 | 70.671 | 3 | 51.73703 | yes | 6.028E-07 | 443.23618 | 46.443481 |
| SEGFDTYR | 8 | P12004 | Proliferating cell nuclear antigen OS=Hor | PCNA | 28.769 | 2 | 50.828472 | yes | 3.717E-07 | 487.71432 | 236.01927 |
| NLAMGVNLTSMS | 13 | P12004 | Proliferating cell nuclear antigen OS=Hor | PCNA | 28.769 | 2 | 50.828472 | yes | 2.417E-07 | 683.34686 | 210.44127 |
| IADMGHLK | 8 | P12004 | Proliferating cell nuclear antigen OS=Hor | PCNA | 28.769 | 2 | 50.828472 | yes | 4.871E-07 | 442.73657 | 191.78647 |
| CAGNEDIITLR | 11 | P12004 | Proliferating cell nuclear antigen OS=Hor | PCNA | 28.769 | 2 | 50.828472 | yes | 9.722E-08 | 631.3139 | 140.20773 |
| DLSHIGDAVVISC | 15 | P12004 | Proliferating cell nuclear antigen OS=Hor | PCNA | 28.769 | 2 | 50.828472 | yes | 2.543E-07 | 792.90613 | 102.37207 |
| YLNFFTK | 7 | P12004 | Proliferating cell nuclear antigen OS=Hor | PCNA | 28.769 | 2 | 50.828472 | yes | 3.651E-07 | 466.74747 | 33.739162 |
| GADIMYTGTVDC | 14 | P12235 | ADP/ATP translocase 1 OS=Homo sapie | SLC25A4 | 33.065 | 2 | 50.886562 | no | 1.248E-07 | 822.8609 | 1659.4335 |
| GADIMYTGTVDC | 14 | P12235 | ADP/ATP translocase 1 OS=Homo sapie | SLC25A4 | 33.065 | 2 | 50.886562 | no | 9.809E-08 | 830.85834 | 1659.4335 |
| AAYFGVYDTAK | 11 | P12235 | ADP/ATP translocase 1 OS=Homo sapie | SLC25A4 | 33.065 | 2 | 50.886562 | no | 1.006E-07 | 603.29547 | 541.12036 |
| QLFLGGVDR | 9 | P12235 | ADP/ATP translocase 1 OS=Homo sapie | SLC25A4 | 33.065 | 2 | 50.886562 | yes | 5.464E-07 | 502.77982 | 481.94189 |
| GDHAWSFLK | 9 | P12235 | ADP/ATP translocase 1 OS=Homo sapie | SLC25A4 | 33.065 | 2 | 50.886562 | yes | 9.938E-08 | 551.76947 | 316.48013 |
| EQGFLSFWR | 9 | P12235 | ADP/ATP translocase 1 OS=Homo sapie | SLC25A4 | 33.065 | 2 | 50.886562 | yes | 1.007E-07 | 585.29053 | 265.32489 |
| EFHGLGDCIIK | 11 | P12235 | ADP/ATP translocase 1 OS=Homo sapie | SLC25A4 | 33.065 | 2 | 50.886562 | yes | 9.722E-08 | 644.82135 | 144.17978 |
| TEQAISFAK | 9 | P12236 | ADP/ATP translocase 3 OS=Homo sapie | SLC25A6 | 32.867 | 2 | 48.111668 | yes | 1.197E-07 | 518.7691 | 375.84158 |
| GIVDCIVR | 8 | P12236 | ADP/ATP translocase 3 OS=Homo sapie | SLC25A6 | 32.867 | 2 | 48.111668 | yes | 5.596E-07 | 466.25513 | 127.57404 |
| MTEQAISFAK | 10 | P12236 | ADP/ATP translocase 3 OS=Homo sapie | SLC25A6 | 32.867 | 2 | 48.111668 | yes | 1.247E-07 | 584.28937 | 77.20343 |
| DFLAGGIAAAISK | 13 | P12236 | ADP/ATP translocase 3 OS=Homo sapie | SLC25A6 | 32.867 | 2 | 48.111668 | yes | 3.792E-06 | 617.34552 | 21.045111 |
| NLIDAGVDALR | 11 | P12268 | Inosine-5'-monophosphate dehydrogenase | IMPDH2 | 55.805 | 2 | 51.579716 | yes | 1.142E-07 | 578.81964 | 884.77411 |
| EANEILQR | 8 | P12268 | Inosine-5'-monophosphate dehydrogenase | IMPDH2 | 55.805 | 2 | 51.579716 | no | 2.436E-07 | 486.75909 | 638.37604 |
| VAQGVSGAVQDK | 12 | P12268 | Inosine-5'-monophosphate dehydrogenase | IMPDH2 | 55.805 | 2 | 51.579716 | yes | 1.736E-07 | 579.80933 | 474.24896 |
| EDLVVAPAGITLK | 13 | P12268 | Inosine-5'-monophosphate dehydrogenase | IMPDH2 | 55.805 | 2 | 51.579716 | yes | 1.248E-07 | 663.38739 | 391.54999 |
| DKYPNLQVIGGN | 20 | P12268 | Inosine-5'-monophosphate dehydrogenase | IMPDH2 | 55.805 | 3 | 51.579716 | yes | 1.02E-07 | 696.04791 | 337.2699 |
| QLLCGAAIGTHED | 17 | P12268 | Inosine-5'-monophosphate dehydrogenase | IMPDH2 | 55.805 | 3 | 51.579716 | yes | 1.028E-07 | 649.65112 | 267.56686 |
| TSSAQVEGGVHSL | 18 | P12268 | Inosine-5'-monophosphate dehydrogenase | IMPDH2 | 55.805 | 3 | 51.579716 | yes | 2.454E-07 | 639.30969 | 204.22482 |
| AMMYSGELK | 9 | P12268 | Inosine-5'-monophosphate dehydrogenase | IMPDH2 | 55.805 | 2 | 51.579716 | yes | 4.603E-07 | 515.24084 | 185.49866 |
| LVGIISSR | 8 | P12268 | Inosine-5'-monophosphate dehydrogenase | IMPDH2 | 55.805 | 2 | 51.579716 | yes | 2.915E-07 | 422.76617 | 168.56573 |
| YEQGFITDPVVLS | 15 | P12268 | Inosine-5'-monophosphate dehydrogenase | IMPDH2 | 55.805 | 2 | 51.579716 | yes | 2.07E-07 | 846.9458 | 96.320694 |
| HGFCGIPITDTGR | 13 | P12268 | Inosine-5'-monophosphate dehydrogenase | IMPDH2 | 55.805 | 2 | 51.579716 | yes | 6.618E-07 | 715.84589 | 77.858551 |
| REDLVVAPAGITL | 14 | P12268 | Inosine-5'-monophosphate dehydrogenase | IMPDH2 | 55.805 | 2 | 51.579716 | yes | 9.722E-08 | 741.43793 | 75.051361 |

| | | | | | | | | | | | |
|---|---|---|---|---|---|---|---|---|---|---|---|
| GMGSLDAMDK | 10 | P12268 | Inosine-5'-monophosphate dehydrogenase | IMPDH2 | 55.805 | 2 | 51.579716 | yes | 1.819E-07 | 512.72552 | 67.620132 |
| PQATAVYK | 8 | P12268 | Inosine-5'-monophosphate dehydrogenase | IMPDH2 | 55.805 | 2 | 51.579716 | yes | 1.491E-07 | 439.24252 | 57.918552 |
| FGVPVIADGGIQN | 19 | P12268 | Inosine-5'-monophosphate dehydrogenase | IMPDH2 | 55.805 | 3 | 51.579716 | yes | 4.74E-07 | 631.35114 | 52.074577 |
| IQQLTEEIGR | 10 | P12270 | Nucleoprotein TPR OS=Homo sapiens O | TPR | 267.293 | 2 | 49.79112 | yes | 1.015E-07 | 593.82495 | 111.49379 |
| SQEQILEILR | 10 | P12270 | Nucleoprotein TPR OS=Homo sapiens O | TPR | 267.293 | 2 | 49.79112 | yes | 3.737E-07 | 614.84845 | 89.051956 |
| LDELQASDVSVK | 12 | P12270 | Nucleoprotein TPR OS=Homo sapiens O | TPR | 267.293 | 2 | 49.79112 | yes | 1.951E-07 | 652.34064 | 82.643791 |
| ELQELQDSLNAER | 13 | P12270 | Nucleoprotein TPR OS=Homo sapiens O | TPR | 267.293 | 2 | 49.79112 | yes | 2.212E-07 | 772.88116 | 77.917274 |
| LSQELEYLTEDVK | 13 | P12270 | Nucleoprotein TPR OS=Homo sapiens O | TPR | 267.293 | 2 | 49.79112 | yes | 2.465E-07 | 783.8985 | 58.113708 |
| ASTALSNEQQAR | 12 | P12270 | Nucleoprotein TPR OS=Homo sapiens O | TPR | 267.293 | 2 | 49.79112 | yes | 3.837E-07 | 638.31799 | 56.13591 |
| ESAEFQTQLEK | 11 | P12270 | Nucleoprotein TPR OS=Homo sapiens O | TPR | 267.293 | 2 | 49.79112 | yes | 4.747E-07 | 655.31714 | 54.003216 |
| FLADQQSEIDGLK | 13 | P12270 | Nucleoprotein TPR OS=Homo sapiens O | TPR | 267.293 | 2 | 49.79112 | yes | 3.582E-06 | 732.37244 | 49.546379 |
| LESALTELEQLR | 12 | P12270 | Nucleoprotein TPR OS=Homo sapiens O | TPR | 267.293 | 2 | 49.79112 | yes | 1.003E-07 | 701.38281 | 43.890522 |
| TMAQHEELMK | 10 | P12270 | Nucleoprotein TPR OS=Homo sapiens O | TPR | 267.293 | 2 | 49.79112 | yes | 2.749E-07 | 609.28632 | 42.199352 |
| TDELLALGR | 9 | P12270 | Nucleoprotein TPR OS=Homo sapiens O | TPR | 267.293 | 2 | 49.79112 | yes | 2.861E-07 | 494.2771 | 41.252705 |
| VLLMELEEAR | 10 | P12270 | Nucleoprotein TPR OS=Homo sapiens O | TPR | 267.293 | 2 | 49.79112 | yes | 3.632E-07 | 601.82611 | 30.865547 |
| LEQGQAIDDLMPA | 15 | P12277 | Creatine kinase B-type OS=Homo sapiens | CKB | 42.645 | 2 | 50.931496 | yes | 7.972E-07 | 828.91669 | 196.23796 |
| VLTPELYAELR | 11 | P12277 | Creatine kinase B-type OS=Homo sapiens | CKB | 42.645 | 2 | 50.931496 | yes | 9.722E-08 | 652.36646 | 114.48339 |
| DLFDPIIEDR | 10 | P12277 | Creatine kinase B-type OS=Homo sapiens | CKB | 42.645 | 2 | 50.931496 | yes | 1.57E-07 | 616.81152 | 28.593685 |
| LYPPSAEYPDLR | 12 | P12532 | Creatine kinase U-type, mitochondrial OS | CKMT1A | 47.037 | 2 | 52.271065 | yes | 3.647E-07 | 710.85901 | 814.46851 |
| SFLIWVNEEDHTR | 13 | P12532 | Creatine kinase U-type, mitochondrial OS | CKMT1A | 47.037 | 2 | 52.271065 | yes | 9.722E-08 | 823.4021 | 262.55402 |
| GWEFMWNER | 9 | P12532 | Creatine kinase U-type, mitochondrial OS | CKMT1A | 47.037 | 2 | 52.271065 | no | 9.722E-08 | 627.77167 | 153.89238 |
| IPTPVIHTK | 9 | P12532 | Creatine kinase U-type, mitochondrial OS | CKMT1A | 47.037 | 2 | 52.271065 | yes | 1.216E-06 | 503.3082 | 138.63237 |
| VVVDALSGLK | 10 | P12532 | Creatine kinase U-type, mitochondrial OS | CKMT1A | 47.037 | 2 | 52.271065 | yes | 1.347E-07 | 500.80551 | 106.38045 |
| EGLLLWCQR | 9 | P12814 | Alpha-actinin-1 OS=Homo sapiens OX=9 | ACTN1 | 103.058 | 2 | 57.069122 | no | 9.722E-08 | 587.80548 | 393.74683 |
| GYEEWLLNEIR | 11 | P12814 | Alpha-actinin-1 OS=Homo sapiens OX=9 | ACTN1 | 103.058 | 2 | 57.069122 | no | 9.899E-08 | 711.35663 | 228.33151 |
| FAIQDISVEETSAK | 14 | P12814 | Alpha-actinin-1 OS=Homo sapiens OX=9 | ACTN1 | 103.058 | 2 | 57.069122 | no | 9.722E-08 | 769.39087 | 184.15881 |
| DLLLDPAWEK | 10 | P12814 | Alpha-actinin-1 OS=Homo sapiens OX=9 | ACTN1 | 103.058 | 2 | 57.069122 | no | 9.722E-08 | 600.31897 | 150.451 |
| CQLEINFNTLQTK | 13 | P12814 | Alpha-actinin-1 OS=Homo sapiens OX=9 | ACTN1 | 103.058 | 2 | 57.069122 | no | 9.722E-08 | 804.90613 | 145.61435 |
| TINEVENQILTR | 12 | P12814 | Alpha-actinin-1 OS=Homo sapiens OX=9 | ACTN1 | 103.058 | 2 | 57.069122 | no | 1.165E-07 | 715.38593 | 122.53725 |
| ASIHEAWTDGK | 11 | P12814 | Alpha-actinin-1 OS=Homo sapiens OX=9 | ACTN1 | 103.058 | 2 | 57.069122 | no | 1.087E-07 | 607.79364 | 100.14562 |
| DGLGFCALIHR | 11 | P12814 | Alpha-actinin-1 OS=Homo sapiens OX=9 | ACTN1 | 103.058 | 2 | 57.069122 | yes | 1.736E-07 | 629.82172 | 96.264198 |
| GISQEQMNEFR | 11 | P12814 | Alpha-actinin-1 OS=Homo sapiens OX=9 | ACTN1 | 103.058 | 2 | 57.069122 | yes | 5.362E-07 | 669.80896 | 95.845482 |
| TFTAWCNSHLR | 11 | P12814 | Alpha-actinin-1 OS=Homo sapiens OX=9 | ACTN1 | 103.058 | 2 | 57.069122 | no | 1.004E-07 | 696.82751 | 72.044716 |
| ATLPDADKER | 10 | P12814 | Alpha-actinin-1 OS=Homo sapiens OX=9 | ACTN1 | 103.058 | 2 | 57.069122 | yes | 1.178E-07 | 558.28821 | 48.908237 |
| AGTQIENIEEDFR | 13 | P12814 | Alpha-actinin-1 OS=Homo sapiens OX=9 | ACTN1 | 103.058 | 2 | 57.069122 | no | 1.272E-07 | 761.36261 | 28.292696 |
| LASDLLEWIR | 10 | P12814 | Alpha-actinin-1 OS=Homo sapiens OX=9 | ACTN1 | 103.058 | 2 | 57.069122 | no | 2.02E-07 | 608.34021 | 25.494793 |
| HRPELIDYGK | 10 | P12814 | Alpha-actinin-1 OS=Homo sapiens OX=9 | ACTN1 | 103.058 | 3 | 57.069122 | yes | 2.539E-06 | 409.88754 | 24.386066 |
| ICDQWDNLGALTQ | 14 | P12814 | Alpha-actinin-1 OS=Homo sapiens OX=9 | ACTN1 | 103.058 | 2 | 57.069122 | yes | 2.241E-06 | 831.40125 | 18.153027 |
| ALEDQLSEIK | 10 | P12882 | Myosin-1 OS=Homo sapiens OX=9606 C | MYH1 | 223.146 | 2 | 55.190514 | yes | 1.128E-07 | 573.30603 | 188.65869 |

| | | | | | | | | | | | |
|---|---|---|---|---|---|---|---|---|---|---|---|
| HDCDLLR | 7 | P12882 | Myosin-1 OS=Homo sapiens OX=9606 C | MYH1 | 223.146 | 2 | 55.190514 | no | 3.031E-07 | 464.7189 | 171.78854 |
| NLTEEMAGLDETL | 15 | P12882 | Myosin-1 OS=Homo sapiens OX=9606 C | MYH1 | 223.146 | 2 | 55.190514 | no | 9.722E-08 | 817.90076 | 101.2715 |
| DLEEATLQHEATA | 18 | P12882 | Myosin-1 OS=Homo sapiens OX=9606 C | MYH1 | 223.146 | 3 | 55.190514 | no | 1.184E-06 | 666.34814 | 30.389803 |
| TFNTSTGGLLLPSD | 16 | P12956 | X-ray repair cross-complementing protein | XRCC6 | 69.844 | 2 | 53.34053 | yes | 2.04E-07 | 826.43048 | 15080.245 |
| NIYVLQELDNPGA | 14 | P12956 | X-ray repair cross-complementing protein | XRCC6 | 69.844 | 2 | 53.34053 | yes | 1.144E-07 | 787.41467 | 13661.902 |
| ILELDQFK | 8 | P12956 | X-ray repair cross-complementing protein | XRCC6 | 69.844 | 2 | 53.34053 | yes | 1.554E-07 | 503.28439 | 12496.276 |
| DIISIAEDEDLR | 12 | P12956 | X-ray repair cross-complementing protein | XRCC6 | 69.844 | 2 | 53.34053 | yes | 1.001E-07 | 694.84882 | 10238.64 |
| EETEELKR | 8 | P12956 | X-ray repair cross-complementing protein | XRCC6 | 69.844 | 2 | 53.34053 | yes | 2.491E-07 | 517.26166 | 6791.5444 |
| ELVYPPDYNPEGK | 13 | P12956 | X-ray repair cross-complementing protein | XRCC6 | 69.844 | 2 | 53.34053 | yes | 9.241E-07 | 760.867 | 5841.9253 |
| LGSLVDEFK | 9 | P12956 | X-ray repair cross-complementing protein | XRCC6 | 69.844 | 2 | 53.34053 | yes | 1.925E-07 | 504.27402 | 5271.9321 |
| VHFEESSK | 8 | P12956 | X-ray repair cross-complementing protein | XRCC6 | 69.844 | 2 | 53.34053 | yes | 2.083E-07 | 481.73254 | 3946.0229 |
| KQELLEALTK | 10 | P12956 | X-ray repair cross-complementing protein | XRCC6 | 69.844 | 2 | 53.34053 | yes | 1.376E-07 | 586.8479 | 3808.8472 |
| VEYSEEELK | 9 | P12956 | X-ray repair cross-complementing protein | XRCC6 | 69.844 | 2 | 53.34053 | yes | 9.819E-08 | 563.26917 | 3419.9875 |
| KPGGFDISLFYR | 12 | P12956 | X-ray repair cross-complementing protein | XRCC6 | 69.844 | 2 | 53.34053 | yes | 1.542E-07 | 700.37207 | 2232.4746 |
| FDDPGLMLMGFK | 12 | P12956 | X-ray repair cross-complementing protein | XRCC6 | 69.844 | 2 | 53.34053 | yes | 9.722E-08 | 685.82782 | 741.03442 |
| FDDPGLMLMGFK | 12 | P12956 | X-ray repair cross-complementing protein | XRCC6 | 69.844 | 2 | 53.34053 | yes | 1.31E-07 | 693.82526 | 741.03442 |
| FDDPGLMLMGFK | 12 | P12956 | X-ray repair cross-complementing protein | XRCC6 | 69.844 | 2 | 53.34053 | yes | 3.344E-06 | 693.82526 | 741.03442 |
| DSLIFLVDASK | 11 | P12956 | X-ray repair cross-complementing protein | XRCC6 | 69.844 | 2 | 53.34053 | yes | 9.976E-08 | 604.33209 | 2093.7085 |
| IMLFTNEDNPHGN | 17 | P12956 | X-ray repair cross-complementing protein | XRCC6 | 69.844 | 3 | 53.34053 | yes | 5.54E-07 | 634.95984 | 1761.479 |
| QELLEALTK | 9 | P12956 | X-ray repair cross-complementing protein | XRCC6 | 69.844 | 2 | 53.34053 | yes | 1.796E-07 | 522.80042 | 802.15369 |
| DKNSVNFK | 8 | P12956 | X-ray repair cross-complementing protein | XRCC6 | 69.844 | 2 | 53.34053 | yes | 1.837E-07 | 476.24835 | 683.95825 |
| IMATPEQVGK | 10 | P12956 | X-ray repair cross-complementing protein | XRCC6 | 69.844 | 2 | 53.34053 | yes | 1.637E-07 | 537.28662 | 621.81348 |
| KMPFTEK | 7 | P12956 | X-ray repair cross-complementing protein | XRCC6 | 69.844 | 2 | 53.34053 | yes | 2.695E-07 | 440.73349 | 457.71606 |
| SQIYGSR | 7 | P12956 | X-ray repair cross-complementing protein | XRCC6 | 69.844 | 2 | 53.34053 | yes | 2.505E-07 | 405.70886 | 338.88412 |
| NIPPYFVALVPQEE | 20 | P12956 | X-ray repair cross-complementing protein | XRCC6 | 69.844 | 3 | 53.34053 | yes | 1.235E-07 | 782.06177 | 84.722321 |
| DIVISVGIYNLVQK | 14 | P12956 | X-ray repair cross-complementing protein | XRCC6 | 69.844 | 2 | 53.34053 | yes | 9.722E-08 | 780.95343 | 65.829643 |
| IISSDRDLLAVVFY | 18 | P12956 | X-ray repair cross-complementing protein | XRCC6 | 69.844 | 3 | 53.34053 | yes | 9.722E-08 | 676.03339 | 49.583565 |
| FTVPMLK | 7 | P12956 | X-ray repair cross-complementing protein | XRCC6 | 69.844 | 2 | 53.34053 | yes | 3.016E-07 | 418.24094 | 20.709076 |
| FTVPMLK | 7 | P12956 | X-ray repair cross-complementing protein | XRCC6 | 69.844 | 2 | 53.34053 | yes | 4.632E-07 | 426.2384 | 20.709076 |
| RFDDPGLMLMGFI | 13 | P12956 | X-ray repair cross-complementing protein | XRCC6 | 69.844 | 2 | 53.34053 | yes | 1.61E-06 | 763.87836 | 26.398655 |
| PGGFDISLFYR | 11 | P12956 | X-ray repair cross-complementing protein | XRCC6 | 69.844 | 2 | 53.34053 | yes | 1.373E-06 | 636.32458 | 22.227869 |
| QIILEKEETEELK | 13 | P12956 | X-ray repair cross-complementing protein | XRCC6 | 69.844 | 2 | 53.34053 | yes | 9.722E-08 | 801.43524 | 19.848017 |
| FFMGNQVLK | 9 | P13010 | X-ray repair cross-complementing protein | XRCC5 | 82.705 | 2 | 51.832966 | yes | 2.864E-07 | 542.28644 | 11814.289 |
| TDTLEDLFPTTK | 12 | P13010 | X-ray repair cross-complementing protein | XRCC5 | 82.705 | 2 | 51.832966 | yes | 1.26E-07 | 690.84827 | 11021.649 |
| LFQCLLHR | 8 | P13010 | X-ray repair cross-complementing protein | XRCC5 | 82.705 | 2 | 51.832966 | yes | 1.112E-07 | 543.79749 | 9797.8203 |
| LTIGSNLSIR | 10 | P13010 | X-ray repair cross-complementing protein | XRCC5 | 82.705 | 2 | 51.832966 | yes | 2.447E-07 | 537.31934 | 9059.8477 |
| EDIIQGFR | 8 | P13010 | X-ray repair cross-complementing protein | XRCC5 | 82.705 | 2 | 51.832966 | yes | 1.572E-07 | 489.25616 | 7845.2334 |
| CFSVLGFCK | 9 | P13010 | X-ray repair cross-complementing protein | XRCC5 | 82.705 | 2 | 51.832966 | yes | 1.376E-07 | 559.26208 | 5805.2881 |
| HIEIFTDLSSR | 11 | P13010 | X-ray repair cross-complementing protein | XRCC5 | 82.705 | 2 | 51.832966 | yes | 1.108E-07 | 659.34351 | 4966.6133 |

| | | | | | | | | | | | |
|---|---|---|---|---|---|---|---|---|---|---|---|
| LGGHGPSFPLK | 11 | P13010 | X-ray repair cross-complementing protein | XRCC5 | 82.705 | 2 | 51.832966 | yes | 9.819E-08 | 555.30872 | 4504.1123 |
| TLFPLIEAK | 9 | P13010 | X-ray repair cross-complementing protein | XRCC5 | 82.705 | 2 | 51.832966 | yes | 1.459E-07 | 516.31042 | 3807.9929 |
| VITMFVQR | 8 | P13010 | X-ray repair cross-complementing protein | XRCC5 | 82.705 | 2 | 51.832966 | yes | 1.993E-07 | 497.28113 | 1517.0045 |
| VITMFVQR | 8 | P13010 | X-ray repair cross-complementing protein | XRCC5 | 82.705 | 2 | 51.832966 | yes | 2.344E-07 | 505.27859 | 1517.0045 |
| SQLDIIIHSLK | 11 | P13010 | X-ray repair cross-complementing protein | XRCC5 | 82.705 | 2 | 51.832966 | yes | 2.677E-07 | 633.87445 | 2797.1912 |
| QYMFSSLK | 8 | P13010 | X-ray repair cross-complementing protein | XRCC5 | 82.705 | 2 | 51.832966 | yes | 1.365E-07 | 502.24951 | 1393.8832 |
| QYMFSSLK | 8 | P13010 | X-ray repair cross-complementing protein | XRCC5 | 82.705 | 2 | 51.832966 | yes | 2.85E-07 | 510.24695 | 1393.8832 |
| KVITMFVQR | 9 | P13010 | X-ray repair cross-complementing protein | XRCC5 | 82.705 | 2 | 51.832966 | yes | 1.106E-07 | 561.32861 | 2037.2126 |
| KTWTVVDAK | 9 | P13010 | X-ray repair cross-complementing protein | XRCC5 | 82.705 | 2 | 51.832966 | yes | 1.062E-07 | 524.29529 | 1331.0171 |
| HSIHWPCR | 8 | P13010 | X-ray repair cross-complementing protein | XRCC5 | 82.705 | 2 | 51.832966 | yes | 2.587E-07 | 546.76141 | 1098.7699 |
| VDEEQMK | 7 | P13010 | X-ray repair cross-complementing protein | XRCC5 | 82.705 | 2 | 51.832966 | yes | 9.722E-08 | 439.69983 | 1074.9819 |
| TWTVVDAK | 8 | P13010 | X-ray repair cross-complementing protein | XRCC5 | 82.705 | 2 | 51.832966 | yes | 2.312E-07 | 460.2478 | 671.70447 |
| RFFMGNQVLK | 10 | P13010 | X-ray repair cross-complementing protein | XRCC5 | 82.705 | 2 | 51.832966 | yes | 2.617E-07 | 620.33698 | 532.54779 |
| EEASGSSVTAEEA | 14 | P13010 | X-ray repair cross-complementing protein | XRCC5 | 82.705 | 2 | 51.832966 | yes | 1.329E-07 | 697.81769 | 394.83426 |
| VNPIQGLASK | 10 | P13073 | Cytochrome c oxidase subunit 4 isoform | COX4I1 | 19.577 | 2 | 51.945454 | yes | 1.922E-07 | 513.80072 | 197.9225 |
| SEDFSLPAYMDR | 12 | P13073 | Cytochrome c oxidase subunit 4 isoform | COX4I1 | 19.577 | 2 | 51.945454 | yes | 4.95E-07 | 715.81647 | 73.534836 |
| ASWSSLSMDEK | 11 | P13073 | Cytochrome c oxidase subunit 4 isoform | COX4I1 | 19.577 | 2 | 51.945454 | yes | 9.722E-08 | 620.77936 | 49.821396 |
| TDGGDPSGLLK | 11 | P13196 | 5-aminolevulinate synthase, non-specific, | ALAS1 | 70.581 | 2 | 43.611023 | yes | 1.569E-06 | 530.26947 | 85.414001 |
| SVSTFQYDR | 9 | P13196 | 5-aminolevulinate synthase, non-specific, | ALAS1 | 70.581 | 2 | 43.611023 | yes | 2.518E-07 | 551.76184 | 73.115318 |
| EVAETSAGPSVVS | 15 | P13196 | 5-aminolevulinate synthase, non-specific, | ALAS1 | 70.581 | 2 | 43.611023 | yes | 5.219E-07 | 730.38556 | 25.794065 |
| NTEVCDELMSR | 11 | P13196 | 5-aminolevulinate synthase, non-specific, | ALAS1 | 70.581 | 2 | 43.611023 | yes | 6.368E-07 | 677.2923 | 22.111668 |
| EFLFNPYLTR | 10 | P13378 | Homeobox protein Hox-D8 OS=Homo sa | HOXD8 | 31.911 | 2 | 54.79863 | no | 9.722E-08 | 650.34021 | 125.03669 |
| LDDCGLTEAR | 10 | P13489 | Ribonuclease inhibitor OS=Homo sapiens | RNH1 | 49.974 | 2 | 50.355385 | yes | 1.963E-07 | 575.26385 | 145.45462 |
| VNPALAELNLR | 11 | P13489 | Ribonuclease inhibitor OS=Homo sapiens | RNH1 | 49.974 | 2 | 50.355385 | yes | 3.273E-07 | 605.35114 | 134.61821 |
| ELSLAGNELGDEG | 15 | P13489 | Ribonuclease inhibitor OS=Homo sapiens | RNH1 | 49.974 | 2 | 50.355385 | yes | 9.81E-08 | 765.87335 | 131.72829 |
| ELCQGLGQPGSVL | 14 | P13489 | Ribonuclease inhibitor OS=Homo sapiens | RNH1 | 49.974 | 2 | 50.355385 | yes | 1.202E-07 | 757.39319 | 111.58654 |
| DSPCQLEALK | 10 | P13489 | Ribonuclease inhibitor OS=Homo sapiens | RNH1 | 49.974 | 2 | 50.355385 | yes | 1.925E-06 | 580.78442 | 65.368355 |
| ELTVSNNDINEAG | 15 | P13489 | Ribonuclease inhibitor OS=Homo sapiens | RNH1 | 49.974 | 2 | 50.355385 | yes | 1.381E-07 | 815.90521 | 54.057613 |
| FLLELQISNNR | 11 | P13489 | Ribonuclease inhibitor OS=Homo sapiens | RNH1 | 49.974 | 2 | 50.355385 | yes | 2.306E-07 | 673.875 | 35.30468 |
| HTCVTFLER | 9 | P13497 | Bone morphogenetic protein 1 OS=Homo | BMP1 | 111.249 | 2 | 43.407345 | yes | 2.605E-07 | 581.78729 | 44.575745 |
| CEAACGGFLTK | 11 | P13497 | Bone morphogenetic protein 1 OS=Homo | BMP1 | 111.249 | 2 | 43.407345 | yes | 3.724E-07 | 607.27063 | 40.99157 |
| CSCDPGYELAPDK | 13 | P13497 | Bone morphogenetic protein 1 OS=Homo | BMP1 | 111.249 | 2 | 43.407345 | yes | 2.699E-06 | 756.31067 | 30.282707 |
| FYSDNSVQR | 9 | P13497 | Bone morphogenetic protein 1 OS=Homo | BMP1 | 111.249 | 2 | 43.407345 | yes | 9.533E-07 | 558.25946 | 11.484206 |
| ERPQIGGTIK | 10 | P13498 | Cytochrome b-245 light chain OS=Homo | CYBA | 21.013 | 2 | 31.508673 | yes | 1.818E-06 | 549.81696 | 106.38649 |
| VNFTVDQIR | 9 | P13639 | Elongation factor 2 OS=Homo sapiens O | EEF2 | 95.339 | 2 | 52.4218 | yes | 1.392E-07 | 546.29584 | 1277.4854 |
| YEWDVAEAR | 9 | P13639 | Elongation factor 2 OS=Homo sapiens O | EEF2 | 95.339 | 2 | 52.4218 | yes | 1.082E-07 | 569.76184 | 1161.9586 |
| ETVSEESNVLCLS | 14 | P13639 | Elongation factor 2 OS=Homo sapiens O | EEF2 | 95.339 | 2 | 52.4218 | yes | 1.006E-07 | 797.88507 | 1053.0836 |
| EDLYLKPIQR | 10 | P13639 | Elongation factor 2 OS=Homo sapiens O | EEF2 | 95.339 | 2 | 52.4218 | yes | 9.819E-08 | 637.85876 | 956.2381 |
| IMGPNYTPGK | 10 | P13639 | Elongation factor 2 OS=Homo sapiens O | EEF2 | 95.339 | 2 | 52.4218 | yes | 9.926E-08 | 539.2735 | 857.99695 |

| | | | | | | | | | | |
|---|---|---|---|---|---|---|---|---|---|---|
| EGALCEENMR | 10 | P13639 | Elongation factor 2 OS=Homo sapiens O | EEF2 | 95.339 | 2 | 52.4218 | yes | 2.97E-07 | 604.75537 | 851.60059 |
| STLTDSLVCK | 10 | P13639 | Elongation factor 2 OS=Homo sapiens O | EEF2 | 95.339 | 2 | 52.4218 | yes | 1.655E-07 | 562.2868 | 831.33313 |
| IKPVLMMNK | 9 | P13639 | Elongation factor 2 OS=Homo sapiens O | EEF2 | 95.339 | 2 | 52.4218 | yes | 3.438E-07 | 537.31415 | 753.78717 |
| VFDAIMNFK | 9 | P13639 | Elongation factor 2 OS=Homo sapiens O | EEF2 | 95.339 | 2 | 52.4218 | yes | 1.148E-07 | 542.77844 | 714.88751 |
| GVQYLNEIK | 9 | P13639 | Elongation factor 2 OS=Homo sapiens O | EEF2 | 95.339 | 2 | 52.4218 | yes | 9.911E-08 | 532.29279 | 537.90796 |
| VFSGLVSTGLK | 11 | P13639 | Elongation factor 2 OS=Homo sapiens O | EEF2 | 95.339 | 2 | 52.4218 | yes | 1.46E-07 | 554.32404 | 507.26788 |
| DSVVAGFQWATK | 12 | P13639 | Elongation factor 2 OS=Homo sapiens O | EEF2 | 95.339 | 2 | 52.4218 | yes | 9.722E-08 | 654.83276 | 503.24271 |
| DLEEDHACIPIK | 12 | P13639 | Elongation factor 2 OS=Homo sapiens O | EEF2 | 95.339 | 2 | 52.4218 | yes | 1.073E-07 | 720.3454 | 202.29068 |
| DLEEDHACIPIK | 12 | P13639 | Elongation factor 2 OS=Homo sapiens O | EEF2 | 95.339 | 2 | 52.4218 | yes | 1.222E-07 | 480.56601 | 202.29068 |
| DLEEDHACIPIK | 12 | P13639 | Elongation factor 2 OS=Homo sapiens O | EEF2 | 95.339 | 3 | 52.4218 | yes | 1.073E-07 | 720.3454 | 202.29068 |
| DLEEDHACIPIK | 12 | P13639 | Elongation factor 2 OS=Homo sapiens O | EEF2 | 95.339 | 3 | 52.4218 | yes | 1.222E-07 | 480.56601 | 202.29068 |
| PSQVVAETR | 9 | P13639 | Elongation factor 2 OS=Homo sapiens O | EEF2 | 95.339 | 2 | 52.4218 | yes | 1.381E-07 | 493.76691 | 251.5685 |
| NMSVIAHVDHGK | 12 | P13639 | Elongation factor 2 OS=Homo sapiens O | EEF2 | 95.339 | 2 | 52.4218 | yes | 9.723E-08 | 654.3299 | 243.96356 |
| KVEDMMK | 7 | P13639 | Elongation factor 2 OS=Homo sapiens O | EEF2 | 95.339 | 2 | 52.4218 | yes | 9.722E-08 | 440.71698 | 234.43852 |
| QFAEMYVAK | 9 | P13639 | Elongation factor 2 OS=Homo sapiens O | EEF2 | 95.339 | 2 | 52.4218 | yes | 1.102E-07 | 543.76807 | 196.87471 |
| SDPVVSYR | 8 | P13639 | Elongation factor 2 OS=Homo sapiens O | EEF2 | 95.339 | 2 | 52.4218 | yes | 9.722E-08 | 461.73508 | 169.29054 |
| ARPFPDGLAEDIDI | 20 | P13639 | Elongation factor 2 OS=Homo sapiens O | EEF2 | 95.339 | 3 | 52.4218 | yes | 1.14E-06 | 715.03076 | 146.44742 |
| CLYASVLTAQPR | 12 | P13639 | Elongation factor 2 OS=Homo sapiens O | EEF2 | 95.339 | 2 | 52.4218 | yes | 1.578E-07 | 689.86102 | 139.24405 |
| EGIPALDNFLDK | 12 | P13639 | Elongation factor 2 OS=Homo sapiens O | EEF2 | 95.339 | 2 | 52.4218 | yes | 1.183E-07 | 666.3457 | 138.53998 |
| TILMMGR | 7 | P13639 | Elongation factor 2 OS=Homo sapiens O | EEF2 | 95.339 | 2 | 52.4218 | yes | 1.978E-07 | 411.22223 | 121.91387 |
| FSVSPVVR | 8 | P13639 | Elongation factor 2 OS=Homo sapiens O | EEF2 | 95.339 | 2 | 52.4218 | yes | 1.85E-07 | 445.75836 | 106.4157 |
| KSDPVVSYR | 9 | P13639 | Elongation factor 2 OS=Homo sapiens O | EEF2 | 95.339 | 2 | 52.4218 | yes | 8.199E-06 | 525.78253 | 87.546661 |
| GPLMMYISK | 9 | P13639 | Elongation factor 2 OS=Homo sapiens O | EEF2 | 95.339 | 2 | 52.4218 | yes | 1.966E-07 | 520.26941 | 71.246178 |
| GHVFEESQVAGTP | 18 | P13639 | Elongation factor 2 OS=Homo sapiens O | EEF2 | 95.339 | 3 | 52.4218 | yes | 1.218E-07 | 654.66449 | 45.879162 |
| DLEEDHACIPIKK | 13 | P13639 | Elongation factor 2 OS=Homo sapiens O | EEF2 | 95.339 | 2 | 52.4218 | yes | 1.965E-07 | 784.39288 | 27.248547 |
| LENEIQTYR | 9 | P13645 | Keratin, type I cytoskeletal 10 OS=Homo | KRT10 | 58.828 | 2 | 52.451595 | yes | 9.722E-08 | 583.29602 | 1651.5471 |
| SQYEQLAEQNR | 11 | P13645 | Keratin, type I cytoskeletal 10 OS=Homo | KRT10 | 58.828 | 2 | 52.451595 | yes | 9.722E-08 | 683.3233 | 1416.3424 |
| QSLEASLAETEGR | 13 | P13645 | Keratin, type I cytoskeletal 10 OS=Homo | KRT10 | 58.828 | 2 | 52.451595 | yes | 1.298E-07 | 695.84406 | 1271.3673 |
| ALEESNYELEGK | 12 | P13645 | Keratin, type I cytoskeletal 10 OS=Homo | KRT10 | 58.828 | 2 | 52.451595 | yes | 9.935E-08 | 691.3277 | 1117.0212 |
| VLDELTLTK | 9 | P13645 | Keratin, type I cytoskeletal 10 OS=Homo | KRT10 | 58.828 | 2 | 52.451595 | yes | 1.699E-07 | 516.3028 | 983.01416 |
| LKYENEVALR | 10 | P13645 | Keratin, type I cytoskeletal 10 OS=Homo | KRT10 | 58.828 | 2 | 52.451595 | yes | 1.402E-07 | 617.84314 | 856.54388 |
| QSVEADINGLR | 11 | P13645 | Keratin, type I cytoskeletal 10 OS=Homo | KRT10 | 58.828 | 2 | 52.451595 | no | 1.976E-07 | 601.31219 | 610.6781 |
| SLLEGEGSSGGGG | 14 | P13645 | Keratin, type I cytoskeletal 10 OS=Homo | KRT10 | 58.828 | 2 | 52.451595 | yes | 1.597E-07 | 631.80219 | 431.05576 |
| DAEAWFNEK | 9 | P13645 | Keratin, type I cytoskeletal 10 OS=Homo | KRT10 | 58.828 | 2 | 52.451595 | no | 9.823E-08 | 555.24854 | 375.1994 |
| YENEVALR | 8 | P13645 | Keratin, type I cytoskeletal 10 OS=Homo | KRT10 | 58.828 | 2 | 52.451595 | yes | 1.692E-06 | 497.25363 | 324.9584 |
| IKEWYEK | 7 | P13645 | Keratin, type I cytoskeletal 10 OS=Homo | KRT10 | 58.828 | 2 | 52.451595 | yes | 3.192E-07 | 498.26346 | 198.79463 |
| SQYEQLAEQNRK | 12 | P13645 | Keratin, type I cytoskeletal 10 OS=Homo | KRT10 | 58.828 | 2 | 52.451595 | yes | 2.601E-07 | 747.37079 | 77.290695 |
| NQILNLTTDNANII | 21 | P13645 | Keratin, type I cytoskeletal 10 OS=Homo | KRT10 | 58.828 | 3 | 52.451595 | yes | 9.722E-08 | 789.75909 | 67.897537 |
| RVLDELTLTK | 10 | P13645 | Keratin, type I cytoskeletal 10 OS=Homo | KRT10 | 58.828 | 2 | 52.451595 | yes | 3.793E-07 | 594.35333 | 35.509422 |

| | | | | | | | | | | | |
|---|---|---|---|---|---|---|---|---|---|---|---|
| AQYEEIANR | 9 | P13647 | Keratin, type II cytoskeletal 5 OS=Homo | KRT5 | 62.379 | 2 | 51.62419 | yes | 6.268E-07 | 547.26727 | 127.37054 |
| VSLAGACGVGGY( | 15 | P13647 | Keratin, type II cytoskeletal 5 OS=Homo | KRT5 | 62.379 | 2 | 51.62419 | yes | 2.437E-07 | 705.84332 | 125.44559 |
| LAELEEALQK | 10 | P13647 | Keratin, type II cytoskeletal 5 OS=Homo | KRT5 | 62.379 | 2 | 51.62419 | yes | 9.722E-08 | 572.31641 | 117.00761 |
| ISISTSGGSFR | 11 | P13647 | Keratin, type II cytoskeletal 5 OS=Homo | KRT5 | 62.379 | 2 | 51.62419 | yes | 9.722E-08 | 556.29077 | 77.017731 |
| TTAENEFVMLK | 11 | P13647 | Keratin, type II cytoskeletal 5 OS=Homo | KRT5 | 62.379 | 2 | 51.62419 | yes | 9.722E-08 | 649.81848 | 72.432625 |
| SFSTASAITPSVSR | 14 | P13647 | Keratin, type II cytoskeletal 5 OS=Homo | KRT5 | 62.379 | 2 | 51.62419 | yes | 3.212E-06 | 705.86481 | 60.551579 |
| VDALMDEINFMK | 12 | P13647 | Keratin, type II cytoskeletal 5 OS=Homo | KRT5 | 62.379 | 2 | 51.62419 | yes | 1.71E-07 | 713.34125 | 23.494196 |
| SAWLSGYENPVVS | 14 | P13674 | Prolyl 4-hydroxylase subunit alpha-1 OS= | P4HA1 | 61.05 | 2 | 53.383812 | yes | 9.809E-08 | 782.89136 | 278.59009 |
| SFLTAEDCFELGK | 13 | P13674 | Prolyl 4-hydroxylase subunit alpha-1 OS= | P4HA1 | 61.05 | 2 | 53.383812 | yes | 9.722E-08 | 758.85303 | 133.89388 |
| LLELDPEHQR | 10 | P13674 | Prolyl 4-hydroxylase subunit alpha-1 OS= | P4HA1 | 61.05 | 2 | 53.383812 | yes | 1.987E-07 | 625.33038 | 125.91161 |
| DPEGFVGHPVNAF | 14 | P13674 | Prolyl 4-hydroxylase subunit alpha-1 OS= | P4HA1 | 61.05 | 2 | 53.383812 | yes | 9.722E-08 | 757.37537 | 106.23521 |
| YFEYIMAK | 8 | P13674 | Prolyl 4-hydroxylase subunit alpha-1 OS= | P4HA1 | 61.05 | 2 | 53.383812 | yes | 3.413E-07 | 532.7597 | 104.1642 |
| LQDTYNLDTDTISI | 14 | P13674 | Prolyl 4-hydroxylase subunit alpha-1 OS= | P4HA1 | 61.05 | 2 | 53.383812 | yes | 9.736E-08 | 813.89648 | 74.351852 |
| VAYTEADYYHTEI | 20 | P13674 | Prolyl 4-hydroxylase subunit alpha-1 OS= | P4HA1 | 61.05 | 3 | 53.383812 | yes | 2.316E-07 | 830.38623 | 66.285484 |
| AEEDKLEQIK | 10 | P13674 | Prolyl 4-hydroxylase subunit alpha-1 OS= | P4HA1 | 61.05 | 2 | 53.383812 | yes | 1.653E-07 | 601.81677 | 57.310822 |
| HAACPVLVGNK | 11 | P13674 | Prolyl 4-hydroxylase subunit alpha-1 OS= | P4HA1 | 61.05 | 2 | 53.383812 | yes | 7.999E-07 | 583.31097 | 18.581724 |
| EAMALAPWGPVK | 12 | P13682 | Zinc finger protein 35 OS=Homo sapiens | ZNF35 | 59.09 | 2 | 37.825195 | yes | 5.67E-07 | 635.33667 | 43.42664 |
| NLQLLVPK | 8 | P13682 | Zinc finger protein 35 OS=Homo sapiens | ZNF35 | 59.09 | 2 | 37.825195 | yes | 4.61E-05 | 462.79749 | 11.186185 |
| CVLIFGVPSR | 10 | P13716 | Delta-aminolevulinic acid dehydratase OS | ALAD | 36.295 | 2 | 42.575012 | yes | 2.965E-07 | 574.31824 | 88.217407 |
| LAEVALAYAK | 10 | P13716 | Delta-aminolevulinic acid dehydratase OS | ALAD | 36.295 | 2 | 42.575012 | yes | 2.582E-06 | 524.80548 | 56.058308 |
| FASCFYGPFR | 10 | P13716 | Delta-aminolevulinic acid dehydratase OS | ALAD | 36.295 | 2 | 42.575012 | yes | 8.327E-07 | 626.28442 | 24.533056 |
| ISFDEFIK | 8 | P13796 | Plastin-2 OS=Homo sapiens OX=9606 G | LCP1 | 70.289 | 2 | 31.367062 | yes | 1.865E-06 | 499.76331 | 17.623938 |
| MINLSVPDTIDER | 13 | P13797 | Plastin-3 OS=Homo sapiens OX=9606 G | PLS3 | 70.812 | 2 | 54.917004 | no | 9.722E-08 | 751.87958 | 157.61855 |
| LSPEELLLR | 9 | P13797 | Plastin-3 OS=Homo sapiens OX=9606 G | PLS3 | 70.812 | 2 | 54.917004 | no | 1.916E-07 | 535.31622 | 66.366356 |
| NEALAALLR | 9 | P13797 | Plastin-3 OS=Homo sapiens OX=9606 G | PLS3 | 70.812 | 2 | 54.917004 | yes | 1.091E-07 | 485.78763 | 38.174969 |
| ANDDIIVNWVNR | 12 | P13797 | Plastin-3 OS=Homo sapiens OX=9606 G | PLS3 | 70.812 | 2 | 54.917004 | yes | 8.031E-07 | 714.86511 | 35.520786 |
| IGLFADIELSR | 11 | P13797 | Plastin-3 OS=Homo sapiens OX=9606 G | PLS3 | 70.812 | 2 | 54.917004 | no | 3.378E-06 | 617.34552 | 29.88418 |
| TQVELLEAPTPALI | 14 | P13807 | Glycogen [starch] synthase, muscle OS=F | GYS1 | 83.786 | 2 | 56.287796 | yes | 1.646E-07 | 755.42981 | 1794.7996 |
| IGLFNSSADR | 10 | P13807 | Glycogen [starch] synthase, muscle OS=F | GYS1 | 83.786 | 2 | 56.287796 | yes | 1.607E-07 | 540.27765 | 613.87683 |
| LYESLLVGSLPDM | 15 | P13807 | Glycogen [starch] synthase, muscle OS=F | GYS1 | 83.786 | 2 | 56.287796 | yes | 9.722E-08 | 839.93964 | 387.76242 |
| RLPVATIFTTHATL | 17 | P13807 | Glycogen [starch] synthase, muscle OS=F | GYS1 | 83.786 | 3 | 56.287796 | yes | 9.722E-08 | 623.03528 | 332.62064 |
| QLWDTANTVK | 10 | P13807 | Glycogen [starch] synthase, muscle OS=F | GYS1 | 83.786 | 2 | 56.287796 | yes | 9.722E-08 | 588.3064 | 310.72278 |
| TNNFNVETLK | 10 | P13807 | Glycogen [starch] synthase, muscle OS=F | GYS1 | 83.786 | 2 | 56.287796 | no | 1.022E-07 | 590.30383 | 214.20886 |
| GADVFLEALAR | 11 | P13807 | Glycogen [starch] synthase, muscle OS=F | GYS1 | 83.786 | 2 | 56.287796 | yes | 1.219E-07 | 581.31677 | 179.46213 |
| TLYFFIAGR | 9 | P13807 | Glycogen [starch] synthase, muscle OS=F | GYS1 | 83.786 | 2 | 56.287796 | yes | 9.722E-08 | 544.30035 | 177.6826 |
| VGGIYTVLQTK | 11 | P13807 | Glycogen [starch] synthase, muscle OS=F | GYS1 | 83.786 | 2 | 56.287796 | yes | 3.211E-07 | 589.84259 | 168.41527 |
| KLYESLLVGSLPDI | 16 | P13807 | Glycogen [starch] synthase, muscle OS=F | GYS1 | 83.786 | 3 | 56.287796 | yes | 1.304E-07 | 602.99384 | 123.3153 |
| GHFYGHLDFNLDK | 13 | P13807 | Glycogen [starch] synthase, muscle OS=F | GYS1 | 83.786 | 2 | 56.287796 | yes | 9.919E-08 | 781.87299 | 77.881042 |
| MLDKEDFTMMK | 11 | P13807 | Glycogen [starch] synthase, muscle OS=F | GYS1 | 83.786 | 2 | 56.287796 | yes | 9.722E-08 | 694.81659 | 54.549519 |

| | | | | | | | | | | | |
|---|---|---|---|---|---|---|---|---|---|---|---|
| APASVLPAATPR | 12 | P13861 | cAMP-dependent protein kinase type II-a | PRKAR2A | 45.519 | 2 | 55.69249 | yes | 9.85E-08 | 575.83258 | 1264.8158 |
| CLVMDVQAFER | 11 | P13861 | cAMP-dependent protein kinase type II-a | PRKAR2A | 45.519 | 2 | 55.69249 | yes | 9.79E-08 | 684.32593 | 953.02594 |
| LLGPCMDIMK | 10 | P13861 | cAMP-dependent protein kinase type II-a | PRKAR2A | 45.519 | 2 | 55.69249 | yes | 1.446E-07 | 589.29254 | 720.55322 |
| DGGNQEVEIAR | 11 | P13861 | cAMP-dependent protein kinase type II-a | PRKAR2A | 45.519 | 2 | 55.69249 | yes | 1.084E-07 | 594.28619 | 422.23001 |
| AASAYAVGDVK | 11 | P13861 | cAMP-dependent protein kinase type II-a | PRKAR2A | 45.519 | 2 | 55.69249 | yes | 1.022E-07 | 526.27454 | 393.70706 |
| GSFGELALMYNTP | 14 | P13861 | cAMP-dependent protein kinase type II-a | PRKAR2A | 45.519 | 2 | 55.69249 | no | 4.45E-07 | 778.38232 | 212.05446 |
| GTYDILVTK | 9 | P13861 | cAMP-dependent protein kinase type II-a | PRKAR2A | 45.519 | 2 | 55.69249 | yes | 1.474E-07 | 505.28186 | 180.36697 |
| GQYFGELALVTNK | 15 | P13861 | cAMP-dependent protein kinase type II-a | PRKAR2A | 45.519 | 3 | 55.69249 | no | 9.722E-08 | 564.97375 | 75.934479 |
| GQYFGELALVTNK | 15 | P13861 | cAMP-dependent protein kinase type II-a | PRKAR2A | 45.519 | 3 | 55.69249 | no | 1.069E-07 | 846.95703 | 75.934479 |
| GQYFGELALVTNK | 15 | P13861 | cAMP-dependent protein kinase type II-a | PRKAR2A | 45.519 | 2 | 55.69249 | no | 9.722E-08 | 564.97375 | 75.934479 |
| GQYFGELALVTNK | 15 | P13861 | cAMP-dependent protein kinase type II-a | PRKAR2A | 45.519 | 2 | 55.69249 | no | 1.069E-07 | 846.95703 | 75.934479 |
| QSLGHPPPEPGPDI | 14 | P13861 | cAMP-dependent protein kinase type II-a | PRKAR2A | 45.519 | 2 | 55.69249 | yes | 1.21E-07 | 742.36804 | 124.05229 |
| IVDVIGEK | 8 | P13861 | cAMP-dependent protein kinase type II-a | PRKAR2A | 45.519 | 2 | 55.69249 | yes | 2.151E-07 | 436.75803 | 51.2934 |
| NISHYEEQLVK | 11 | P13861 | cAMP-dependent protein kinase type II-a | PRKAR2A | 45.519 | 2 | 55.69249 | yes | 4.688E-07 | 680.34882 | 22.418774 |
| LQIEESSKPVR | 11 | P13984 | General transcription factor IIF subunit 2 | GTF2F2 | 28.381 | 2 | 47.731968 | yes | 1.27E-07 | 643.35913 | 23.622341 |
| ENELTYYCCK | 10 | P13987 | CD59 glycoprotein OS=Homo sapiens OX | CD59 | 14.178 | 2 | 43.424976 | yes | 2.598E-07 | 690.28394 | 26.828384 |
| TALATSPITVGDLC | 17 | P13994 | Probable splicing factor YJU2B OS=Hom | YJU2B | 44.803 | 2 | 46.475227 | yes | 1.549E-07 | 842.4856 | 82.474716 |
| KLSQGILIIR | 10 | P13994 | Probable splicing factor YJU2B OS=Hom | YJU2B | 44.803 | 2 | 46.475227 | yes | 2.136E-05 | 570.87677 | 50.291843 |
| LSQGILIIR | 9 | P13994 | Probable splicing factor YJU2B OS=Hom | YJU2B | 44.803 | 2 | 46.475227 | yes | 1.706E-07 | 506.82932 | 48.520386 |
| DVPESPQHAADTP | 14 | P13994 | Probable splicing factor YJU2B OS=Hom | YJU2B | 44.803 | 2 | 46.475227 | yes | 2.961E-07 | 746.35736 | 41.564007 |
| DDFALNSMLR | 10 | P13994 | Probable splicing factor YJU2B OS=Hom | YJU2B | 44.803 | 2 | 46.475227 | yes | 4.45E-07 | 591.28461 | 38.678905 |
| EGAAVIDVGINR | 12 | P13995 | Bifunctional methylenetetrahydrofolate de | MTHFD2 | 37.896 | 2 | 50.300568 | yes | 9.819E-08 | 607.33038 | 257.84314 |
| PASISEEELLNLINI | 15 | P13995 | Bifunctional methylenetetrahydrofolate de | MTHFD2 | 37.896 | 2 | 50.300568 | yes | 1.651E-07 | 835.45398 | 13.802101 |
| DLQENYCR | 8 | P14210 | Hepatocyte growth factor OS=Homo sapi | HGF | 83.134 | 2 | 32.790398 | yes | 1.412E-06 | 549.23767 | 70.239464 |
| GLTVVVAFR | 9 | P14324 | Farnesyl pyrophosphate synthase OS=Ho | FDPS | 48.276 | 2 | 49.858074 | yes | 4.7E-07 | 481.2951 | 61.637482 |
| ATPEQYQILK | 10 | P14324 | Farnesyl pyrophosphate synthase OS=Ho | FDPS | 48.276 | 2 | 49.858074 | yes | 2.548E-07 | 595.8244 | 47.559528 |
| CSWLVVQCLQR | 11 | P14324 | Farnesyl pyrophosphate synthase OS=Ho | FDPS | 48.276 | 2 | 49.858074 | yes | 1.001E-07 | 724.86047 | 46.331215 |
| ELLQDIGDTLSR | 12 | P14373 | Zinc finger protein RFP OS=Homo sapier | TRIM27 | 58.49 | 2 | 50.48253 | yes | 1.997E-07 | 680.35938 | 82.372566 |
| WTIGVCEDSVCR | 12 | P14373 | Zinc finger protein RFP OS=Homo sapier | TRIM27 | 58.49 | 2 | 50.48253 | yes | 9.804E-08 | 741.32922 | 75.106674 |
| AELLSLTQMER | 11 | P14373 | Zinc finger protein RFP OS=Homo sapier | TRIM27 | 58.49 | 2 | 50.48253 | yes | 5.016E-07 | 645.83972 | 73.136314 |
| HYWEVEVGDK | 10 | P14373 | Zinc finger protein RFP OS=Homo sapier | TRIM27 | 58.49 | 2 | 50.48253 | no | 4.904E-07 | 631.29602 | 35.195515 |
| LFQEDDEIPLYLK | 13 | P14406 | Cytochrome c oxidase subunit 7A2, mitoc | COX7A2 | 9.396 | 2 | 50.92091 | yes | 9.722E-08 | 811.91925 | 225.09439 |
| GNLYWTDWNR | 10 | P14543 | Nidogen-1 OS=Homo sapiens OX=9606 | NID1 | 136.378 | 2 | 49.161972 | no | 1.063E-07 | 662.80707 | 230.14412 |
| VIIGLAFDCVDK | 12 | P14543 | Nidogen-1 OS=Homo sapiens OX=9606 | NID1 | 136.378 | 2 | 49.161972 | yes | 1.441E-07 | 675.36029 | 130.66867 |
| ASLHGGEPTTIIR | 13 | P14543 | Nidogen-1 OS=Homo sapiens OX=9606 | NID1 | 136.378 | 2 | 49.161972 | yes | 5.336E-07 | 676.37006 | 82.554688 |
| APIIAVTR | 8 | P14618 | Pyruvate kinase PKM OS=Homo sapiens | PKM | 57.937 | 2 | 53.62051 | yes | 3.238E-07 | 420.76871 | 568.18677 |
| GDYPLEAVR | 9 | P14618 | Pyruvate kinase PKM OS=Homo sapiens | PKM | 57.937 | 2 | 53.62051 | yes | 1.306E-07 | 510.26144 | 488.17874 |
| NTGIICTIGPASR | 13 | P14618 | Pyruvate kinase PKM OS=Homo sapiens | PKM | 57.937 | 2 | 53.62051 | yes | 9.722E-08 | 680.35608 | 373.96954 |
| MQHLIAR | 7 | P14618 | Pyruvate kinase PKM OS=Homo sapiens | PKM | 57.937 | 2 | 53.62051 | yes | 1.34E-07 | 434.74472 | 306.93964 |

| | | | | | | | | | | | |
|---|---|---|---|---|---|---|---|---|---|---|---|
| ITLDNAYMEK | 10 | P14618 | Pyruvate kinase PKM OS=Homo sapiens | PKM | 57.937 | 2 | 53.62051 | yes | 2.226E-07 | 599.29462 | 286.49026 |
| GVNLPGAAVDLP/ | 17 | P14618 | Pyruvate kinase PKM OS=Homo sapiens | PKM | 57.937 | 2 | 53.62051 | yes | 2.027E-07 | 818.94885 | 250.58983 |
| GSGTAEVELK | 10 | P14618 | Pyruvate kinase PKM OS=Homo sapiens | PKM | 57.937 | 2 | 53.62051 | yes | 4.038E-07 | 495.75876 | 222.91461 |
| LDIDSPPITAR | 11 | P14618 | Pyruvate kinase PKM OS=Homo sapiens | PKM | 57.937 | 2 | 53.62051 | yes | 1.061E-07 | 599.32733 | 205.80626 |
| CCSGAIIVLTK | 11 | P14618 | Pyruvate kinase PKM OS=Homo sapiens | PKM | 57.937 | 2 | 53.62051 | yes | 9.722E-08 | 611.32013 | 189.59569 |
| PGSGFTNTMR | 10 | P14618 | Pyruvate kinase PKM OS=Homo sapiens | PKM | 57.937 | 2 | 53.62051 | yes | 7.769E-07 | 534.25055 | 171.73215 |
| IYVDDGLISLQVK | 13 | P14618 | Pyruvate kinase PKM OS=Homo sapiens | PKM | 57.937 | 2 | 53.62051 | yes | 1.289E-07 | 731.91119 | 168.75246 |
| PVAVALDTK | 9 | P14618 | Pyruvate kinase PKM OS=Homo sapiens | PKM | 57.937 | 2 | 53.62051 | yes | 1.627E-07 | 457.2713 | 161.95277 |
| KGDVVIVLTGWR | 12 | P14618 | Pyruvate kinase PKM OS=Homo sapiens | PKM | 57.937 | 2 | 53.62051 | yes | 9.722E-08 | 671.89569 | 106.41993 |
| TATESFASDPILYR | 14 | P14618 | Pyruvate kinase PKM OS=Homo sapiens | PKM | 57.937 | 2 | 53.62051 | yes | 9.722E-08 | 785.89099 | 105.61732 |
| AEGSDVANAVLD( | 25 | P14618 | Pyruvate kinase PKM OS=Homo sapiens | PKM | 57.937 | 3 | 53.62051 | yes | 9.722E-08 | 832.05273 | 64.639671 |
| RFDEILEASDGIM\ | 16 | P14618 | Pyruvate kinase PKM OS=Homo sapiens | PKM | 57.937 | 3 | 53.62051 | yes | 9.81E-08 | 607.97699 | 62.709362 |
| DPVQEAWAEDVD | 14 | P14618 | Pyruvate kinase PKM OS=Homo sapiens | PKM | 57.937 | 2 | 53.62051 | yes | 3.971E-07 | 821.88904 | 48.298065 |
| LNFSHGTHEYHAE | 16 | P14618 | Pyruvate kinase PKM OS=Homo sapiens | PKM | 57.937 | 3 | 53.62051 | yes | 3.142E-07 | 628.63934 | 32.269905 |
| GVVDSDDLPLNVS | 14 | P14625 | Endoplasmin OS=Homo sapiens OX=960 | HSP90B1 | 92.469 | 2 | 51.174812 | yes | 9.722E-08 | 743.3808 | 168.63516 |
| FAFQAEVNR | 9 | P14625 | Endoplasmin OS=Homo sapiens OX=960 | HSP90B1 | 92.469 | 2 | 51.174812 | yes | 4.241E-07 | 541.2749 | 160.3578 |
| SILFVPTSAPR | 11 | P14625 | Endoplasmin OS=Homo sapiens OX=960 | HSP90B1 | 92.469 | 2 | 51.174812 | yes | 3.149E-07 | 594.34277 | 111.74265 |
| EEASDYLELDTIK | 13 | P14625 | Endoplasmin OS=Homo sapiens OX=960 | HSP90B1 | 92.469 | 2 | 51.174812 | yes | 4.037E-07 | 763.36707 | 97.648636 |
| IYFMAGSSR | 9 | P14625 | Endoplasmin OS=Homo sapiens OX=960 | HSP90B1 | 92.469 | 2 | 51.174812 | yes | 5.051E-07 | 516.25256 | 83.481499 |
| LIINSLYK | 8 | P14625 | Endoplasmin OS=Homo sapiens OX=960 | HSP90B1 | 92.469 | 2 | 51.174812 | yes | 3.984E-07 | 482.2973 | 80.398499 |
| EAESSPFVER | 10 | P14625 | Endoplasmin OS=Homo sapiens OX=960 | HSP90B1 | 92.469 | 2 | 51.174812 | yes | 3.556E-07 | 575.7724 | 51.07481 |
| NLLHVTDTGVGM' | 14 | P14625 | Endoplasmin OS=Homo sapiens OX=960 | HSP90B1 | 92.469 | 2 | 51.174812 | yes | 1.382E-07 | 757.39319 | 25.00915 |
| PLPLHFLR | 8 | P14635 | G2/mitotic-specific cyclin-B1 OS=Homo | CCNB1 | 48.338 | 2 | 50.50433 | no | 2.816E-07 | 496.80563 | 343.33463 |
| NVVMVNQGLTK | 11 | P14635 | G2/mitotic-specific cyclin-B1 OS=Homo | CCNB1 | 48.338 | 2 | 50.50433 | yes | 1.318E-07 | 601.83173 | 198.98453 |
| RVPTAPAATSK | 11 | P14635 | G2/mitotic-specific cyclin-B1 OS=Homo | CCNB1 | 48.338 | 2 | 50.50433 | yes | 1.087E-07 | 549.81696 | 118.00576 |
| QLEEEQAVRPK | 11 | P14635 | G2/mitotic-specific cyclin-B1 OS=Homo | CCNB1 | 48.338 | 2 | 50.50433 | yes | 9.791E-08 | 663.85425 | 71.511215 |
| LPKPLEK | 7 | P14635 | G2/mitotic-specific cyclin-B1 OS=Homo | CCNB1 | 48.338 | 2 | 50.50433 | yes | 1.565E-07 | 412.76566 | 56.322689 |
| IGEVDVEQHTLAK | 13 | P14635 | G2/mitotic-specific cyclin-B1 OS=Homo | CCNB1 | 48.338 | 2 | 50.50433 | yes | 4.927E-06 | 719.88043 | 55.075996 |
| ALGQNPTNAEVLK | 13 | P14649 | Myosin light chain 6B OS=Homo sapiens | MYL6B | 22.764 | 2 | 51.17878 | no | 8.304E-07 | 677.86987 | 11701.797 |
| ILYSQCGDVMR | 11 | P14649 | Myosin light chain 6B OS=Homo sapiens | MYL6B | 22.764 | 2 | 51.17878 | no | 1.048E-07 | 671.31812 | 4769.1636 |
| ILYSQCGDVMR | 11 | P14649 | Myosin light chain 6B OS=Homo sapiens | MYL6B | 22.764 | 2 | 51.17878 | no | 9.722E-08 | 679.31555 | 4769.1636 |
| HVLTTLGEK | 9 | P14649 | Myosin light chain 6B OS=Homo sapiens | MYL6B | 22.764 | 2 | 51.17878 | yes | 9.722E-08 | 499.28748 | 170.97725 |
| TQEPPVDLSK | 10 | P14649 | Myosin light chain 6B OS=Homo sapiens | MYL6B | 22.764 | 2 | 51.17878 | yes | 1.696E-07 | 557.29297 | 100.33583 |
| AEPAVPQAPQK | 11 | P14649 | Myosin light chain 6B OS=Homo sapiens | MYL6B | 22.764 | 2 | 51.17878 | yes | 1.284E-07 | 568.30896 | 92.962158 |
| GQGTYEDYLEGFF | 13 | P14649 | Myosin light chain 6B OS=Homo sapiens | MYL6B | 22.764 | 2 | 51.17878 | yes | 1.272E-07 | 767.84406 | 51.567799 |
| TKAEPAVPQAPQK | 13 | P14649 | Myosin light chain 6B OS=Homo sapiens | MYL6B | 22.764 | 2 | 51.17878 | yes | 6.708E-07 | 682.88025 | 45.198132 |
| MLQHIDYR | 8 | P14678 | Small nuclear ribonucleoprotein-associate | SNRPB | 24.611 | 2 | 49.94659 | no | 3.921E-07 | 538.2713 | 2542.1289 |
| MLQHIDYR | 8 | P14678 | Small nuclear ribonucleoprotein-associate | SNRPB | 24.611 | 2 | 49.94659 | no | 1.002E-07 | 546.26874 | 2542.1289 |
| GENLVSMTVEGPF | 15 | P14678 | Small nuclear ribonucleoprotein-associate | SNRPB | 24.611 | 2 | 49.94659 | no | 9.968E-08 | 777.89526 | 1537.8914 |

| | | | | | | | | | | | |
|---|---|---|---|---|---|---|---|---|---|---|---|
| RVLGLVLLR | 9 | P14678 | Small nuclear ribonucleoprotein-associate | SNRPB | 24.611 | 2 | 49.94659 | no | 1.84E-07 | 519.86096 | 369.45282 |
| VLGLVLLR | 8 | P14678 | Small nuclear ribonucleoprotein-associate | SNRPB | 24.611 | 2 | 49.94659 | no | 2.599E-07 | 441.81039 | 240.40564 |
| VPLAGAAGGPGIG | 14 | P14678 | Small nuclear ribonucleoprotein-associate | SNRPB | 24.611 | 2 | 49.94659 | yes | 5.383E-07 | 596.84351 | 54.0075 |
| HMNLILCDCDEFR | 13 | P14678 | Small nuclear ribonucleoprotein-associate | SNRPB | 24.611 | 3 | 49.94659 | no | 4.958E-07 | 574.91809 | 9.229888 |
| VLLISDPTTDK | 11 | P14735 | Insulin-degrading enzyme OS=Homo sap | IDE | 117.969 | 2 | 50.811813 | yes | 9.722E-08 | 601.33734 | 202.84627 |
| GWVNTLVGGQK | 11 | P14735 | Insulin-degrading enzyme OS=Homo sap | IDE | 117.969 | 2 | 50.811813 | yes | 1.229E-07 | 579.81696 | 89.632629 |
| EMLAVDAPR | 9 | P14735 | Insulin-degrading enzyme OS=Homo sap | IDE | 117.969 | 2 | 50.811813 | yes | 2.779E-07 | 501.25787 | 75.776642 |
| PLLPSQLVR | 9 | P14735 | Insulin-degrading enzyme OS=Homo sap | IDE | 117.969 | 2 | 50.811813 | yes | 9.722E-08 | 511.82147 | 61.920727 |
| GGDISVCEWYQR | 12 | P14854 | Cytochrome c oxidase subunit 6B1 OS=H | COX6B1 | 10.193 | 2 | 45.193504 | yes | 1.93E-07 | 735.32751 | 99.343224 |
| TPASPVVHIR | 10 | P14866 | Heterogeneous nuclear ribonucleoprotein | HNRNPL | 64.133 | 2 | 50.66089 | yes | 1.541E-07 | 538.81421 | 15895.453 |
| MGPPVGGHR | 9 | P14866 | Heterogeneous nuclear ribonucleoprotein | HNRNPL | 64.133 | 2 | 50.66089 | yes | 2.756E-07 | 454.23196 | 8589.7119 |
| IEYAKPTR | 8 | P14866 | Heterogeneous nuclear ribonucleoprotein | HNRNPL | 64.133 | 2 | 50.66089 | yes | 4.616E-07 | 489.27435 | 8075.7373 |
| LCFSTAQHAS | 10 | P14866 | Heterogeneous nuclear ribonucleoprotein | HNRNPL | 64.133 | 2 | 50.66089 | yes | 1.026E-07 | 561.25586 | 6905.1606 |
| SSSGLLEWESK | 11 | P14866 | Heterogeneous nuclear ribonucleoprotein | HNRNPL | 64.133 | 2 | 50.66089 | yes | 3.411E-07 | 611.80115 | 6757.083 |
| AITHLNNNFMFGQ | 14 | P14866 | Heterogeneous nuclear ribonucleoprotein | HNRNPL | 64.133 | 2 | 50.66089 | yes | 1.487E-07 | 817.909 | 3341.0757 |
| NPNGPYPYTLK | 11 | P14866 | Heterogeneous nuclear ribonucleoprotein | HNRNPL | 64.133 | 2 | 50.66089 | yes | 2.102E-07 | 632.32202 | 2589.7993 |
| VFNVFCLYGNVEK | 13 | P14866 | Heterogeneous nuclear ribonucleoprotein | HNRNPL | 64.133 | 2 | 50.66089 | yes | 1.68E-07 | 794.89508 | 2056.959 |
| YYGGGSEGGR | 10 | P14866 | Heterogeneous nuclear ribonucleoprotein | HNRNPL | 64.133 | 2 | 50.66089 | yes | 2.563E-07 | 501.71741 | 1959.0313 |
| FSTPEQAAK | 9 | P14866 | Heterogeneous nuclear ribonucleoprotein | HNRNPL | 64.133 | 2 | 50.66089 | yes | 1.171E-07 | 489.74817 | 1277.7181 |
| ISRPGDSDDSR | 11 | P14866 | Heterogeneous nuclear ribonucleoprotein | HNRNPL | 64.133 | 2 | 50.66089 | yes | 1.046E-07 | 602.78125 | 1261.3293 |
| LKTDNAGDQHGG | 37 | P14866 | Heterogeneous nuclear ribonucleoprotein | HNRNPL | 64.133 | 5 | 50.66089 | yes | 1.417E-07 | 645.2821 | 204.17336 |
| LKTDNAGDQHGG | 37 | P14866 | Heterogeneous nuclear ribonucleoprotein | HNRNPL | 64.133 | 5 | 50.66089 | yes | 1.797E-06 | 806.35083 | 204.17336 |
| LKTDNAGDQHGG | 37 | P14866 | Heterogeneous nuclear ribonucleoprotein | HNRNPL | 64.133 | 4 | 50.66089 | yes | 1.417E-07 | 645.2821 | 204.17336 |
| LKTDNAGDQHGG | 37 | P14866 | Heterogeneous nuclear ribonucleoprotein | HNRNPL | 64.133 | 4 | 50.66089 | yes | 1.797E-06 | 806.35083 | 204.17336 |
| QPPLLGDHPAEYG | 38 | P14866 | Heterogeneous nuclear ribonucleoprotein | HNRNPL | 64.133 | 6 | 50.66089 | yes | 8.429E-07 | 696.64948 | 260.98911 |
| PGAAMVEMADGY | 16 | P14866 | Heterogeneous nuclear ribonucleoprotein | HNRNPL | 64.133 | 2 | 50.66089 | yes | 9.729E-08 | 826.87396 | 111.96313 |
| QPAIMPGQSYGLE | 20 | P14866 | Heterogeneous nuclear ribonucleoprotein | HNRNPL | 64.133 | 3 | 50.66089 | yes | 2.475E-07 | 729.99438 | 53.636765 |
| TDNAGDQHGGGG | 45 | P14866 | Heterogeneous nuclear ribonucleoprotein | HNRNPL | 64.133 | 5 | 50.66089 | yes | 1.123E-07 | 808.56696 | 40.846832 |
| SDALETLGFLNHY | 16 | P14866 | Heterogeneous nuclear ribonucleoprotein | HNRNPL | 64.133 | 3 | 50.66089 | yes | 7.193E-07 | 622.97333 | 16.324991 |
| TDNAGDQHGGGG | 35 | P14866 | Heterogeneous nuclear ribonucleoprotein | HNRNPL | 64.133 | 4 | 50.66089 | yes | 3.878E-07 | 746.05609 | 13.683558 |
| IHDPQLLTER | 10 | P14868 | Aspartate--tRNA ligase, cytoplasmic OS= | DARS1 | 57.137 | 2 | 52.92275 | yes | 9.722E-08 | 611.33295 | 272.52164 |
| ESIVDVEGVVR | 11 | P14868 | Aspartate--tRNA ligase, cytoplasmic OS= | DARS1 | 57.137 | 2 | 52.92275 | yes | 9.819E-08 | 601.32477 | 224.1563 |
| EIMDAAEDYAK | 11 | P14868 | Aspartate--tRNA ligase, cytoplasmic OS= | DARS1 | 57.137 | 2 | 52.92275 | yes | 9.722E-08 | 628.27917 | 219.16815 |
| LEYCEALAMLR | 11 | P14868 | Aspartate--tRNA ligase, cytoplasmic OS= | DARS1 | 57.137 | 2 | 52.92275 | yes | 9.722E-08 | 684.83612 | 215.77313 |
| QCFLVLR | 7 | P14868 | Aspartate--tRNA ligase, cytoplasmic OS= | DARS1 | 57.137 | 2 | 52.92275 | yes | 1.893E-07 | 468.26019 | 179.0629 |
| NNAYLAQSPQLYK | 13 | P14868 | Aspartate--tRNA ligase, cytoplasmic OS= | DARS1 | 57.137 | 2 | 52.92275 | yes | 2.552E-06 | 755.38843 | 147.47231 |
| VFSIGPVFR | 9 | P14868 | Aspartate--tRNA ligase, cytoplasmic OS= | DARS1 | 57.137 | 2 | 52.92275 | yes | 9.809E-08 | 511.2951 | 132.29497 |
| QQQFNVQALVAV | 18 | P14868 | Aspartate--tRNA ligase, cytoplasmic OS= | DARS1 | 57.137 | 3 | 52.92275 | yes | 9.722E-08 | 647.33765 | 130.71034 |
| GFVEIQTPK | 9 | P14868 | Aspartate--tRNA ligase, cytoplasmic OS= | DARS1 | 57.137 | 2 | 52.92275 | yes | 1.586E-07 | 509.78201 | 126.65082 |

| | | | | | | | | | | | |
|---|---|---|---|---|---|---|---|---|---|---|---|
| GEEILSGAQR | 10 | P14868 | Aspartate--tRNA ligase, cytoplasmic OS= | DARS1 | 57.137 | 2 | 52.92275 | yes | 2.705E-07 | 530.27509 | 108.84195 |
| ADEVVWVR | 8 | P14868 | Aspartate--tRNA ligase, cytoplasmic OS= | DARS1 | 57.137 | 2 | 52.92275 | yes | 2.047E-07 | 487.25873 | 107.48251 |
| FQTEIQTVNK | 10 | P14868 | Aspartate--tRNA ligase, cytoplasmic OS= | DARS1 | 57.137 | 2 | 52.92275 | yes | 4.368E-07 | 604.31952 | 100.99783 |
| FLEPTLR | 7 | P14868 | Aspartate--tRNA ligase, cytoplasmic OS= | DARS1 | 57.137 | 2 | 52.92275 | yes | 1.055E-07 | 438.2529 | 99.65715 |
| ALHHGIDLEK | 10 | P14868 | Aspartate--tRNA ligase, cytoplasmic OS= | DARS1 | 57.137 | 2 | 52.92275 | yes | 1.095E-07 | 566.80908 | 93.217896 |
| QTSMFPR | 7 | P14868 | Aspartate--tRNA ligase, cytoplasmic OS= | DARS1 | 57.137 | 2 | 52.92275 | yes | 6.385E-06 | 433.71307 | 81.214531 |
| QMCICADFEK | 10 | P14868 | Aspartate--tRNA ligase, cytoplasmic OS= | DARS1 | 57.137 | 2 | 52.92275 | yes | 2.886E-07 | 651.26978 | 45.987591 |
| QSNSYDMFMR | 10 | P14868 | Aspartate--tRNA ligase, cytoplasmic OS= | DARS1 | 57.137 | 2 | 52.92275 | yes | 1.006E-07 | 639.76575 | 36.069962 |
| LSDPDEVAR | 9 | P14921 | Protein C-ets-1 OS=Homo sapiens OX=9 | ETS1 | 50.408 | 2 | 49.220234 | yes | 3.939E-07 | 501.24854 | 114.83168 |
| YENDYPSVILR | 11 | P14921 | Protein C-ets-1 OS=Homo sapiens OX=9 | ETS1 | 50.408 | 2 | 49.220234 | yes | 1.058E-07 | 684.84332 | 101.68263 |
| LLNQPNQWPLVK | 12 | P14923 | Junction plakoglobin OS=Homo sapiens ( | JUP | 81.745 | 2 | 51.130486 | yes | 9.722E-08 | 725.41425 | 86.187859 |
| LLNDEDPVVVTK | 12 | P14923 | Junction plakoglobin OS=Homo sapiens ( | JUP | 81.745 | 2 | 51.130486 | yes | 2.867E-07 | 671.36664 | 82.973419 |
| NLALCPANHAPLQ | 20 | P14923 | Junction plakoglobin OS=Homo sapiens ( | JUP | 81.745 | 3 | 51.130486 | yes | 1.362E-07 | 719.05286 | 73.281929 |
| LNYGIPAIVK | 10 | P14923 | Junction plakoglobin OS=Homo sapiens ( | JUP | 81.745 | 2 | 51.130486 | yes | 1.075E-07 | 544.32916 | 58.868599 |
| VLSVCPSNKPAIVI | 23 | P14923 | Junction plakoglobin OS=Homo sapiens ( | JUP | 81.745 | 3 | 51.130486 | yes | 3.607E-07 | 776.08008 | 43.651337 |
| TMQNTSDLDTAR | 12 | P14923 | Junction plakoglobin OS=Homo sapiens ( | JUP | 81.745 | 2 | 51.130486 | yes | 7.815E-07 | 676.80914 | 38.059837 |
| MPILGLGTWK | 10 | P15121 | Aldo-keto reductase family 1 member B1 | AKR1B1 | 35.854 | 2 | 33.49517 | yes | 1.221E-06 | 558.31775 | 25.259211 |
| MDDPTVNWSNER | 12 | P15170 | Eukaryotic peptide chain release factor G | GSPT1 | 55.756 | 2 | 52.761395 | yes | 1.072E-07 | 732.31464 | 748.54974 |
| GQQLVMMPNK | 10 | P15170 | Eukaryotic peptide chain release factor G | GSPT1 | 55.756 | 2 | 52.761395 | no | 9.722E-08 | 573.29395 | 705.78394 |
| HLIVLINK | 8 | P15170 | Eukaryotic peptide chain release factor G | GSPT1 | 55.756 | 2 | 52.761395 | no | 1.393E-06 | 475.31329 | 580.48419 |
| TAGTICLETFK | 11 | P15170 | Eukaryotic peptide chain release factor G | GSPT1 | 55.756 | 2 | 52.761395 | no | 9.823E-08 | 620.81573 | 566.1452 |
| QDQVCIAR | 8 | P15170 | Eukaryotic peptide chain release factor G | GSPT1 | 55.756 | 2 | 52.761395 | no | 2.334E-07 | 495.24527 | 447.28955 |
| HFTILDAPGHK | 11 | P15170 | Eukaryotic peptide chain release factor G | GSPT1 | 55.756 | 2 | 52.761395 | no | 9.722E-08 | 618.3302 | 430.72205 |
| KGEFETGFEK | 10 | P15170 | Eukaryotic peptide chain release factor G | GSPT1 | 55.756 | 2 | 52.761395 | no | 3.121E-07 | 586.28516 | 287.54553 |
| DMGTVVLGK | 9 | P15170 | Eukaryotic peptide chain release factor G | GSPT1 | 55.756 | 2 | 52.761395 | no | 2.228E-07 | 460.24951 | 77.60891 |
| LESGSICK | 8 | P15170 | Eukaryotic peptide chain release factor G | GSPT1 | 55.756 | 2 | 52.761395 | yes | 3.614E-07 | 447.22348 | 45.703255 |
| EHVNVVFIGHVDA | 15 | P15170 | Eukaryotic peptide chain release factor G | GSPT1 | 55.756 | 2 | 52.761395 | no | 2.6E-06 | 810.92828 | 32.670589 |
| LLNVGFQEALK | 11 | P15291 | Beta-1,4-galactosyltransferase 1 OS=Hon | B4GALT1 | 43.921 | 2 | 49.06508 | yes | 1.073E-07 | 616.3559 | 45.943382 |
| QLLTLSSELSQAR | 13 | P15311 | Ezrin OS=Homo sapiens OX=9606 GN=I | EZR | 69.413 | 2 | 50.121124 | yes | 2.23E-07 | 723.40155 | 101.39027 |
| ELSEQIQR | 8 | P15311 | Ezrin OS=Homo sapiens OX=9606 GN=I | EZR | 69.413 | 2 | 50.121124 | yes | 2.262E-07 | 501.76437 | 82.859077 |
| DNAMLEYLK | 9 | P15311 | Ezrin OS=Homo sapiens OX=9606 GN=I | EZR | 69.413 | 2 | 50.121124 | yes | 1.43E-07 | 548.77081 | 78.513435 |
| SQEQLAAELAEYT | 15 | P15311 | Ezrin OS=Homo sapiens OX=9606 GN=I | EZR | 69.413 | 2 | 50.121124 | yes | 8.241E-07 | 826.41229 | 36.644592 |
| EVWYFGLHYVDN | 13 | P15311 | Ezrin OS=Homo sapiens OX=9606 GN=I | EZR | 69.413 | 2 | 50.121124 | yes | 9.896E-08 | 835.40411 | 33.676308 |
| LQAETEELEEEK | 12 | P15408 | Fos-related antigen 2 OS=Homo sapiens ( | FOSL2 | 35.194 | 2 | 47.885044 | yes | 1.238E-07 | 724.34357 | 114.26325 |
| SGGGSVGAVVVK | 12 | P15408 | Fos-related antigen 2 OS=Homo sapiens ( | FOSL2 | 35.194 | 2 | 47.885044 | yes | 3.304E-07 | 508.79037 | 51.960243 |
| FMQASEDLLK | 10 | P15531 | Nucleoside diphosphate kinase A OS=Ho | NME1 | 17.149 | 2 | 50.06575 | yes | 9.918E-08 | 591.29718 | 122.70119 |
| NIIHGSDSVESAEK | 14 | P15531 | Nucleoside diphosphate kinase A OS=Ho | NME1 | 17.149 | 2 | 50.06575 | yes | 1.823E-07 | 743.36261 | 46.166336 |
| EGHLSPDIVAEQK | 13 | P15559 | NAD(P)H dehydrogenase [quinone] 1 OS | NQO1 | 30.868 | 2 | 42.343147 | yes | 3.076E-07 | 711.86481 | 256.41034 |
| ATLIVHLR | 8 | P15622 | Zinc finger protein 250 OS=Homo sapien | ZNF250 | 63.468 | 2 | 51.75824 | yes | 9.722E-08 | 461.79526 | 134.24373 |

| | | | | | | | | | | | |
|---|---|---|---|---|---|---|---|---|---|---|---|
| AFNHSTVLR | 9 | P15622 | Zinc finger protein 250 OS=Homo sapien | ZNF250 | 63.468 | 2 | 51.75824 | yes | 4.24E-07 | 522.7829 | 68.385941 |
| TLLQHQR | 7 | P15622 | Zinc finger protein 250 OS=Homo sapien | ZNF250 | 63.468 | 2 | 51.75824 | yes | 2.156E-06 | 448.25906 | 57.139954 |
| GEDPWVLDR | 9 | P15622 | Zinc finger protein 250 OS=Homo sapien | ZNF250 | 63.468 | 2 | 51.75824 | yes | 1.67E-07 | 543.76434 | 48.903111 |
| AFSQYSVLIQHQR | 13 | P15622 | Zinc finger protein 250 OS=Homo sapien | ZNF250 | 63.468 | 2 | 51.75824 | yes | 2.875E-07 | 788.91516 | 41.183472 |
| AFSQSSVLSK | 10 | P15622 | Zinc finger protein 250 OS=Homo sapien | ZNF250 | 63.468 | 2 | 51.75824 | yes | 2.755E-06 | 527.28241 | 37.873104 |
| GTGIVSAPVPK | 11 | P15880 | Small ribosomal subunit protein uS5 OS= | RPS2 | 31.325 | 2 | 52.070168 | yes | 8.4E-07 | 513.3031 | 40261.086 |
| GCTATLGNFAK | 11 | P15880 | Small ribosomal subunit protein uS5 OS= | RPS2 | 31.325 | 2 | 52.070168 | yes | 4.511E-07 | 570.2793 | 39979.73 |
| IGKPHTVPCK | 10 | P15880 | Small ribosomal subunit protein uS5 OS= | RPS2 | 31.325 | 2 | 52.070168 | yes | 1.141E-07 | 568.81586 | 16023.551 |
| SPYQEFTDHLVK | 12 | P15880 | Small ribosomal subunit protein uS5 OS= | RPS2 | 31.325 | 2 | 52.070168 | yes | 9.722E-08 | 732.36188 | 15079.391 |
| AEDKEWMPVTK | 11 | P15880 | Small ribosomal subunit protein uS5 OS= | RPS2 | 31.325 | 2 | 52.070168 | yes | 1.213E-07 | 667.32648 | 5493.0249 |
| AEDKEWMPVTK | 11 | P15880 | Small ribosomal subunit protein uS5 OS= | RPS2 | 31.325 | 2 | 52.070168 | yes | 1.001E-07 | 675.32391 | 5493.0249 |
| EWMPVTK | 7 | P15880 | Small ribosomal subunit protein uS5 OS= | RPS2 | 31.325 | 2 | 52.070168 | yes | 1.458E-07 | 445.72568 | 4074.5881 |
| EWMPVTK | 7 | P15880 | Small ribosomal subunit protein uS5 OS= | RPS2 | 31.325 | 2 | 52.070168 | yes | 1.632E-07 | 453.72311 | 4074.5881 |
| AFVAIGDYNGHV( | 17 | P15880 | Small ribosomal subunit protein uS5 OS= | RPS2 | 31.325 | 3 | 52.070168 | yes | 2.845E-07 | 572.97375 | 167.10283 |
| TYSYLTPDLWK | 11 | P15880 | Small ribosomal subunit protein uS5 OS= | RPS2 | 31.325 | 2 | 52.070168 | yes | 1.402E-07 | 693.85065 | 123.05933 |
| LSIVPVRR | 8 | P15880 | Small ribosomal subunit protein uS5 OS= | RPS2 | 31.325 | 2 | 52.070168 | yes | 1.723E-07 | 470.30853 | 66.963272 |
| CSKEVATAIR | 10 | P15880 | Small ribosomal subunit protein uS5 OS= | RPS2 | 31.325 | 2 | 52.070168 | yes | 9.723E-08 | 567.80042 | 39.246616 |
| AITGFDDPFSGK | 12 | P15924 | Desmoplakin OS=Homo sapiens OX=96( | DSP | 331.774 | 2 | 53.746677 | yes | 9.823E-08 | 627.80371 | 2526.2278 |
| LLQLQEQMR | 9 | P15924 | Desmoplakin OS=Homo sapiens OX=96( | DSP | 331.774 | 2 | 53.746677 | yes | 1.602E-07 | 579.8186 | 2522.5847 |
| ISITEGIER | 9 | P15924 | Desmoplakin OS=Homo sapiens OX=96( | DSP | 331.774 | 2 | 53.746677 | yes | 2.173E-07 | 509.28238 | 2308.9797 |
| VLLQEEGTR | 9 | P15924 | Desmoplakin OS=Homo sapiens OX=96( | DSP | 331.774 | 2 | 53.746677 | yes | 1.041E-07 | 522.78784 | 2159.7034 |
| LASLEELKR | 9 | P15924 | Desmoplakin OS=Homo sapiens OX=96( | DSP | 331.774 | 2 | 53.746677 | yes | 1.031E-07 | 529.81384 | 2003.5897 |
| LLEAASVSSK | 10 | P15924 | Desmoplakin OS=Homo sapiens OX=96( | DSP | 331.774 | 2 | 53.746677 | yes | 1.422E-07 | 502.78476 | 1904.6682 |
| YGDGIQLTR | 9 | P15924 | Desmoplakin OS=Homo sapiens OX=96( | DSP | 331.774 | 2 | 53.746677 | yes | 1.01E-06 | 511.76691 | 1715.433 |
| LTVDSAIAR | 9 | P15924 | Desmoplakin OS=Homo sapiens OX=96( | DSP | 331.774 | 2 | 53.746677 | yes | 1.429E-07 | 473.27182 | 1687.3687 |
| SMVEDITGLR | 10 | P15924 | Desmoplakin OS=Homo sapiens OX=96( | DSP | 331.774 | 2 | 53.746677 | yes | 1.002E-07 | 560.78699 | 1630.3201 |
| SNKPIILR | 8 | P15924 | Desmoplakin OS=Homo sapiens OX=96( | DSP | 331.774 | 2 | 53.746677 | yes | 4.014E-07 | 470.80054 | 1255.816 |
| LPVDIAYK | 8 | P15924 | Desmoplakin OS=Homo sapiens OX=96( | DSP | 331.774 | 2 | 53.746677 | yes | 1.941E-07 | 459.76837 | 1232.5159 |
| GAGSIAGASASPK | 13 | P15924 | Desmoplakin OS=Homo sapiens OX=96( | DSP | 331.774 | 2 | 53.746677 | yes | 2.685E-07 | 537.2829 | 1195.7423 |
| TGSQYDIQDAIDK | 13 | P15924 | Desmoplakin OS=Homo sapiens OX=96( | DSP | 331.774 | 2 | 53.746677 | yes | 1.022E-07 | 727.34393 | 1179.134 |
| FLDQNLQK | 8 | P15924 | Desmoplakin OS=Homo sapiens OX=96( | DSP | 331.774 | 2 | 53.746677 | yes | 3.419E-07 | 503.27182 | 1137.9761 |
| IVQLKPR | 7 | P15924 | Desmoplakin OS=Homo sapiens OX=96( | DSP | 331.774 | 2 | 53.746677 | yes | 1.441E-07 | 427.28455 | 1055.1786 |
| EKYSLVEAK | 9 | P15924 | Desmoplakin OS=Homo sapiens OX=96( | DSP | 331.774 | 2 | 53.746677 | yes | 7.317E-07 | 533.7926 | 1031.6617 |
| DLIDFDDR | 8 | P15924 | Desmoplakin OS=Homo sapiens OX=96( | DSP | 331.774 | 2 | 53.746677 | yes | 3.003E-07 | 504.73526 | 1028.0713 |
| NLHSEISGK | 9 | P15924 | Desmoplakin OS=Homo sapiens OX=96( | DSP | 331.774 | 2 | 53.746677 | yes | 1.736E-07 | 492.75909 | 980.78107 |
| NMPLQHLLEQIK | 12 | P15924 | Desmoplakin OS=Homo sapiens OX=96( | DSP | 331.774 | 2 | 53.746677 | yes | 9.181E-07 | 732.40558 | 939.24438 |
| LQSLTENLTK | 10 | P15924 | Desmoplakin OS=Homo sapiens OX=96( | DSP | 331.774 | 2 | 53.746677 | yes | 1.53E-07 | 573.8219 | 912.42621 |
| IEPHTGLLLLSVQK | 14 | P15924 | Desmoplakin OS=Homo sapiens OX=96( | DSP | 331.774 | 2 | 53.746677 | yes | 2.234E-07 | 774.46143 | 878.17719 |
| LLEAQACTGGIIHF | 19 | P15924 | Desmoplakin OS=Homo sapiens OX=96( | DSP | 331.774 | 3 | 53.746677 | yes | 9.935E-08 | 665.68243 | 877.34857 |

| | | | | | | | | | | | |
|---|---|---|---|---|---|---|---|---|---|---|---|
| IEVLEEELR | 9 | P15924 | Desmoplakin OS=Homo sapiens OX=96( | DSP | 331.774 | 2 | 53.746677 | yes | 9.85E-08 | 565.30859 | 855.31226 |
| TMIQSPSGVILQEA | 20 | P15924 | Desmoplakin OS=Homo sapiens OX=96( | DSP | 331.774 | 3 | 53.746677 | yes | 1.455E-07 | 708.36865 | 853.92212 |
| ITNLTQQLEQASIV | 15 | P15924 | Desmoplakin OS=Homo sapiens OX=96( | DSP | 331.774 | 2 | 53.746677 | yes | 1.083E-07 | 843.47528 | 824.67444 |
| LNDSILQATEQR | 12 | P15924 | Desmoplakin OS=Homo sapiens OX=96( | DSP | 331.774 | 2 | 53.746677 | yes | 9.927E-08 | 694.36243 | 793.65106 |
| LTYEIEDEKR | 10 | P15924 | Desmoplakin OS=Homo sapiens OX=96( | DSP | 331.774 | 2 | 53.746677 | yes | 2.071E-07 | 648.32751 | 775.38678 |
| QLQNIIQATSR | 11 | P15924 | Desmoplakin OS=Homo sapiens OX=96( | DSP | 331.774 | 2 | 53.746677 | yes | 1.358E-07 | 636.35693 | 747.40216 |
| LPVEEAYK | 8 | P15924 | Desmoplakin OS=Homo sapiens OX=96( | DSP | 331.774 | 2 | 53.746677 | yes | 1.111E-07 | 474.75546 | 730.67273 |
| NQCTQVVQER | 10 | P15924 | Desmoplakin OS=Homo sapiens OX=96( | DSP | 331.774 | 2 | 53.746677 | yes | 9.903E-08 | 631.30133 | 707.46191 |
| IAELCANSIK | 10 | P15924 | Desmoplakin OS=Homo sapiens OX=96( | DSP | 331.774 | 2 | 53.746677 | yes | 1.546E-07 | 559.79736 | 698.78992 |
| IQSQFTDAQK | 10 | P15924 | Desmoplakin OS=Homo sapiens OX=96( | DSP | 331.774 | 2 | 53.746677 | yes | 4.712E-07 | 583.29602 | 679.69781 |
| ETQSQLETER | 10 | P15924 | Desmoplakin OS=Homo sapiens OX=96( | DSP | 331.774 | 2 | 53.746677 | yes | 1.451E-07 | 610.79132 | 670.6778 |
| DVLDGHLR | 8 | P15924 | Desmoplakin OS=Homo sapiens OX=96( | DSP | 331.774 | 2 | 53.746677 | yes | 1.074E-07 | 462.7485 | 666.71906 |
| GLPSPYNMSSAPG | 15 | P15924 | Desmoplakin OS=Homo sapiens OX=96( | DSP | 331.774 | 2 | 53.746677 | yes | 1.74E-07 | 760.86176 | 661.90363 |
| GIHNSIGDYR | 10 | P15924 | Desmoplakin OS=Homo sapiens OX=96( | DSP | 331.774 | 2 | 53.746677 | yes | 1.157E-07 | 566.2807 | 640.87732 |
| VVIVDPETNK | 10 | P15924 | Desmoplakin OS=Homo sapiens OX=96( | DSP | 331.774 | 2 | 53.746677 | yes | 9.722E-08 | 557.31116 | 615.63538 |
| LGIYEAMK | 8 | P15924 | Desmoplakin OS=Homo sapiens OX=96( | DSP | 331.774 | 2 | 53.746677 | yes | 1.465E-07 | 462.74658 | 610.05457 |
| EHLMLEEELR | 10 | P15924 | Desmoplakin OS=Homo sapiens OX=96( | DSP | 331.774 | 2 | 53.746677 | yes | 1.697E-07 | 649.8241 | 560.88318 |
| SQLQISNNR | 9 | P15924 | Desmoplakin OS=Homo sapiens OX=96( | DSP | 331.774 | 2 | 53.746677 | yes | 9.722E-08 | 530.2807 | 559.64923 |
| VQEQELTR | 8 | P15924 | Desmoplakin OS=Homo sapiens OX=96( | DSP | 331.774 | 2 | 53.746677 | yes | 9.733E-08 | 501.76437 | 558.68182 |
| DLKDEIVR | 8 | P15924 | Desmoplakin OS=Homo sapiens OX=96( | DSP | 331.774 | 2 | 53.746677 | yes | 1.317E-07 | 494.2771 | 533.71216 |
| AFIGFEGVK | 9 | P15924 | Desmoplakin OS=Homo sapiens OX=96( | DSP | 331.774 | 2 | 53.746677 | yes | 3.316E-07 | 484.26602 | 505.10248 |
| EMSVQEAYK | 9 | P15924 | Desmoplakin OS=Homo sapiens OX=96( | DSP | 331.774 | 2 | 53.746677 | yes | 2.605E-07 | 542.75262 | 490.12769 |
| ISTEEAIR | 8 | P15924 | Desmoplakin OS=Homo sapiens OX=96( | DSP | 331.774 | 2 | 53.746677 | yes | 1.075E-07 | 459.74817 | 448.73679 |
| KQQLEVELR | 9 | P15924 | Desmoplakin OS=Homo sapiens OX=96( | DSP | 331.774 | 2 | 53.746677 | yes | 9.81E-08 | 571.83002 | 446.60254 |
| ISTISSVR | 8 | P15924 | Desmoplakin OS=Homo sapiens OX=96( | DSP | 331.774 | 2 | 53.746677 | yes | 1.356E-07 | 431.75327 | 441.91925 |
| IKNDLNLK | 8 | P15924 | Desmoplakin OS=Homo sapiens OX=96( | DSP | 331.774 | 2 | 53.746677 | yes | 2.444E-06 | 479.29001 | 434.58981 |
| LQDTSSYAK | 9 | P15924 | Desmoplakin OS=Homo sapiens OX=96( | DSP | 331.774 | 2 | 53.746677 | yes | 9.722E-08 | 506.75092 | 431.51465 |
| CIDVHLK | 7 | P15924 | Desmoplakin OS=Homo sapiens OX=96( | DSP | 331.774 | 2 | 53.746677 | yes | 1.113E-07 | 442.73657 | 419.33075 |
| QLLQEQESVK | 10 | P15924 | Desmoplakin OS=Homo sapiens OX=96( | DSP | 331.774 | 2 | 53.746677 | yes | 1.507E-07 | 601.32477 | 404.88266 |
| YQAECSQFK | 9 | P15924 | Desmoplakin OS=Homo sapiens OX=96( | DSP | 331.774 | 2 | 53.746677 | yes | 9.722E-08 | 580.75568 | 390.14423 |
| HVTSECLGWMR | 11 | P15924 | Desmoplakin OS=Homo sapiens OX=96( | DSP | 331.774 | 2 | 53.746677 | yes | 1.084E-07 | 688.31592 | 387.63358 |
| WEYENELSK | 9 | P15924 | Desmoplakin OS=Homo sapiens OX=96( | DSP | 331.774 | 2 | 53.746677 | yes | 9.783E-08 | 599.27478 | 379.61133 |
| QMGQPCDAYQK | 11 | P15924 | Desmoplakin OS=Homo sapiens OX=96( | DSP | 331.774 | 2 | 53.746677 | yes | 1.048E-07 | 663.2843 | 370.40994 |
| QQIQNDLNQWK | 11 | P15924 | Desmoplakin OS=Homo sapiens OX=96( | DSP | 331.774 | 2 | 53.746677 | yes | 1.152E-07 | 707.8573 | 352.28485 |
| NTLTQTTENLR | 11 | P15924 | Desmoplakin OS=Homo sapiens OX=96( | DSP | 331.774 | 2 | 53.746677 | yes | 1.199E-07 | 645.83606 | 347.74011 |
| GLIDYETFK | 9 | P15924 | Desmoplakin OS=Homo sapiens OX=96( | DSP | 331.774 | 2 | 53.746677 | yes | 9.722E-08 | 543.2793 | 344.7002 |
| ANSSATETINK | 11 | P15924 | Desmoplakin OS=Homo sapiens OX=96( | DSP | 331.774 | 2 | 53.746677 | yes | 2.854E-07 | 568.28314 | 341.16272 |
| LTEEETVCLDLDK | 13 | P15924 | Desmoplakin OS=Homo sapiens OX=96( | DSP | 331.774 | 2 | 53.746677 | yes | 2.674E-07 | 782.87421 | 332.15811 |
| NQFETEINITK | 11 | P15924 | Desmoplakin OS=Homo sapiens OX=96( | DSP | 331.774 | 2 | 53.746677 | yes | 5.072E-07 | 668.84082 | 327.79935 |

| | | | | | | | | | | | |
|---|---|---|---|---|---|---|---|---|---|---|---|
| AELIVQPELK | 10 | P15924 | Desmoplakin OS=Homo sapiens OX=96( | DSP | 331.774 | 2 | 53.746677 | yes | 3.432E-07 | 570.33716 | 318.15479 |
| NKYETEINITK | 11 | P15924 | Desmoplakin OS=Homo sapiens OX=96( | DSP | 331.774 | 2 | 53.746677 | yes | 1.003E-07 | 676.85645 | 315.71091 |
| QQIYAAEK | 8 | P15924 | Desmoplakin OS=Homo sapiens OX=96( | DSP | 331.774 | 2 | 53.746677 | yes | 1.434E-07 | 475.75073 | 304.96161 |
| LSSEVEALR | 9 | P15924 | Desmoplakin OS=Homo sapiens OX=96( | DSP | 331.774 | 2 | 53.746677 | yes | 3.203E-07 | 502.27457 | 296.57828 |
| RVVIVDPETNK | 11 | P15924 | Desmoplakin OS=Homo sapiens OX=96( | DSP | 331.774 | 2 | 53.746677 | yes | 1.795E-07 | 635.36169 | 291.79788 |
| VSYVQLK | 7 | P15924 | Desmoplakin OS=Homo sapiens OX=96( | DSP | 331.774 | 2 | 53.746677 | yes | 1.978E-07 | 418.74747 | 279.49243 |
| TIADLELHYQEFIR | 14 | P15924 | Desmoplakin OS=Homo sapiens OX=96( | DSP | 331.774 | 3 | 53.746677 | yes | 1.125E-07 | 583.3053 | 272.85461 |
| LLEAQIATGGIIDPI | 15 | P15924 | Desmoplakin OS=Homo sapiens OX=96( | DSP | 331.774 | 2 | 53.746677 | yes | 9.81E-08 | 769.94305 | 266.48563 |
| QKVEEELNR | 9 | P15924 | Desmoplakin OS=Homo sapiens OX=96( | DSP | 331.774 | 2 | 53.746677 | yes | 2.217E-07 | 572.80145 | 264.86295 |
| FLEFQYLTGGLVD | 19 | P15924 | Desmoplakin OS=Homo sapiens OX=96( | DSP | 331.774 | 3 | 53.746677 | yes | 1.316E-07 | 726.37238 | 261.854 |
| NNYDEEIISLK | 11 | P15924 | Desmoplakin OS=Homo sapiens OX=96( | DSP | 331.774 | 2 | 53.746677 | yes | 9.722E-08 | 669.33282 | 260.56702 |
| YIELLTR | 7 | P15924 | Desmoplakin OS=Homo sapiens OX=96( | DSP | 331.774 | 2 | 53.746677 | yes | 1.448E-07 | 454.26602 | 260.34088 |
| NSQGSEMFGDDDI | 13 | P15924 | Desmoplakin OS=Homo sapiens OX=96( | DSP | 331.774 | 2 | 53.746677 | yes | 1.037E-07 | 715.28046 | 259.38885 |
| QETWMLMELQK | 11 | P15924 | Desmoplakin OS=Homo sapiens OX=96( | DSP | 331.774 | 2 | 53.746677 | yes | 9.722E-08 | 718.84924 | 245.86035 |
| QVQNLVNK | 8 | P15924 | Desmoplakin OS=Homo sapiens OX=96( | DSP | 331.774 | 2 | 53.746677 | yes | 1.911E-07 | 471.772 | 236.20282 |
| KLEEELEGMR | 10 | P15924 | Desmoplakin OS=Homo sapiens OX=96( | DSP | 331.774 | 2 | 53.746677 | yes | 1.69E-07 | 617.31079 | 234.49232 |
| QDSLESMK | 8 | P15924 | Desmoplakin OS=Homo sapiens OX=96( | DSP | 331.774 | 2 | 53.746677 | yes | 1.522E-07 | 469.21841 | 228.72205 |
| NHYNEEMSNLR | 11 | P15924 | Desmoplakin OS=Homo sapiens OX=96( | DSP | 331.774 | 2 | 53.746677 | yes | 9.976E-08 | 703.80951 | 226.56403 |
| YSLVEAK | 7 | P15924 | Desmoplakin OS=Homo sapiens OX=96( | DSP | 331.774 | 2 | 53.746677 | yes | 2.244E-07 | 405.22382 | 216.51428 |
| KVSYVQLK | 8 | P15924 | Desmoplakin OS=Homo sapiens OX=96( | DSP | 331.774 | 2 | 53.746677 | yes | 1.489E-07 | 482.79492 | 209.03442 |
| TLELQGLINDLQR | 13 | P15924 | Desmoplakin OS=Homo sapiens OX=96( | DSP | 331.774 | 2 | 53.746677 | yes | 9.902E-08 | 756.92267 | 205.59189 |
| AEENALQQK | 9 | P15924 | Desmoplakin OS=Homo sapiens OX=96( | DSP | 331.774 | 2 | 53.746677 | yes | 3.715E-06 | 515.76184 | 177.33075 |
| VTQLTDR | 7 | P15924 | Desmoplakin OS=Homo sapiens OX=96( | DSP | 331.774 | 2 | 53.746677 | yes | 1.793E-07 | 416.7298 | 174.14709 |
| LKQESDQLVLNQH | 18 | P15924 | Desmoplakin OS=Homo sapiens OX=96( | DSP | 331.774 | 3 | 53.746677 | yes | 4.794E-07 | 684.02362 | 163.37119 |
| TTIHQLTMQK | 10 | P15924 | Desmoplakin OS=Homo sapiens OX=96( | DSP | 331.774 | 2 | 53.746677 | yes | 9.722E-08 | 600.82391 | 162.15073 |
| GIVDSITGQR | 10 | P15924 | Desmoplakin OS=Homo sapiens OX=96( | DSP | 331.774 | 2 | 53.746677 | yes | 9.722E-08 | 523.28546 | 150.66385 |
| KIVQLKPR | 8 | P15924 | Desmoplakin OS=Homo sapiens OX=96( | DSP | 331.774 | 2 | 53.746677 | yes | 1.588E-07 | 491.33203 | 145.07416 |
| ACGSEIMQK | 9 | P15924 | Desmoplakin OS=Homo sapiens OX=96( | DSP | 331.774 | 2 | 53.746677 | yes | 2.325E-07 | 512.23352 | 141.02919 |
| MSQLEVK | 7 | P15924 | Desmoplakin OS=Homo sapiens OX=96( | DSP | 331.774 | 2 | 53.746677 | yes | 2.267E-07 | 417.72311 | 135.80577 |
| FGDSNTVMR | 9 | P15924 | Desmoplakin OS=Homo sapiens OX=96( | DSP | 331.774 | 2 | 53.746677 | yes | 2.245E-07 | 513.7373 | 135.15796 |
| SLSEEIKR | 8 | P15924 | Desmoplakin OS=Homo sapiens OX=96( | DSP | 331.774 | 2 | 53.746677 | yes | 8.521E-07 | 481.26929 | 130.43927 |
| TVSVSEAIK | 9 | P15924 | Desmoplakin OS=Homo sapiens OX=96( | DSP | 331.774 | 2 | 53.746677 | yes | 2.376E-07 | 467.2662 | 124.80218 |
| KCLEDENAR | 9 | P15924 | Desmoplakin OS=Homo sapiens OX=96( | DSP | 331.774 | 2 | 53.746677 | yes | 9.095E-07 | 567.76404 | 111.90698 |
| CIKDEETGLCLLPI | 15 | P15924 | Desmoplakin OS=Homo sapiens OX=96( | DSP | 331.774 | 3 | 53.746677 | yes | 1.041E-07 | 596.97931 | 105.8417 |
| RVEEDIQQQK | 10 | P15924 | Desmoplakin OS=Homo sapiens OX=96( | DSP | 331.774 | 2 | 53.746677 | yes | 9.938E-08 | 636.83075 | 101.22166 |
| LTYEIEDEK | 9 | P15924 | Desmoplakin OS=Homo sapiens OX=96( | DSP | 331.774 | 2 | 53.746677 | yes | 9.722E-08 | 570.27698 | 98.99559 |
| LEEELEGMR | 9 | P15924 | Desmoplakin OS=Homo sapiens OX=96( | DSP | 331.774 | 2 | 53.746677 | yes | 9.934E-08 | 553.26337 | 98.13446 |
| LSLQDAVSQGVID | 19 | P15924 | Desmoplakin OS=Homo sapiens OX=96( | DSP | 331.774 | 3 | 53.746677 | yes | 1.196E-06 | 683.00897 | 88.790924 |
| RQDSLESMK | 9 | P15924 | Desmoplakin OS=Homo sapiens OX=96( | DSP | 331.774 | 2 | 53.746677 | yes | 9.722E-08 | 547.26898 | 82.367844 |

| | | | | | | | | | | | |
|---|---|---|---|---|---|---|---|---|---|---|---|
| NATILELR | 8 | P15924 | Desmoplakin OS=Homo sapiens OX=96( | DSP | 331.774 | 2 | 53.746677 | yes | 1.247E-06 | 465.27435 | 79.300858 |
| KFFDQYR | 7 | P15924 | Desmoplakin OS=Homo sapiens OX=96( | DSP | 331.774 | 2 | 53.746677 | yes | 1.633E-07 | 502.25345 | 74.044739 |
| DEETGLCLLPLK | 12 | P15924 | Desmoplakin OS=Homo sapiens OX=96( | DSP | 331.774 | 2 | 53.746677 | yes | 9.722E-08 | 694.36053 | 68.684875 |
| LEDELNR | 7 | P15924 | Desmoplakin OS=Homo sapiens OX=96( | DSP | 331.774 | 2 | 53.746677 | yes | 1.737E-07 | 444.7247 | 67.408325 |
| NFVDPVTK | 8 | P15924 | Desmoplakin OS=Homo sapiens OX=96( | DSP | 331.774 | 2 | 53.746677 | yes | 1.037E-07 | 460.2478 | 60.018982 |
| FLSEMLK | 7 | P15924 | Desmoplakin OS=Homo sapiens OX=96( | DSP | 331.774 | 2 | 53.746677 | yes | 1.481E-06 | 434.23587 | 55.847115 |
| SVQNDSQAIAEVL | 17 | P15924 | Desmoplakin OS=Homo sapiens OX=96( | DSP | 331.774 | 3 | 53.746677 | yes | 3.27E-07 | 619.66193 | 44.999466 |
| KIQSQFTDAQK | 11 | P15924 | Desmoplakin OS=Homo sapiens OX=96( | DSP | 331.774 | 2 | 53.746677 | yes | 2.205E-07 | 647.34351 | 37.793789 |
| WLPYEAGQR | 9 | P15924 | Desmoplakin OS=Homo sapiens OX=96( | DSP | 331.774 | 2 | 53.746677 | yes | 5.094E-07 | 560.28271 | 36.503063 |
| STLEAETR | 8 | P15924 | Desmoplakin OS=Homo sapiens OX=96( | DSP | 331.774 | 2 | 53.746677 | yes | 1.975E-05 | 453.72998 | 28.707117 |
| KTGSQYDIQDAIDI | 14 | P15924 | Desmoplakin OS=Homo sapiens OX=96( | DSP | 331.774 | 2 | 53.746677 | yes | 1.178E-06 | 791.39136 | 18.32312 |
| NSQGSEMFGDDDI | 14 | P15924 | Desmoplakin OS=Homo sapiens OX=96( | DSP | 331.774 | 2 | 53.746677 | yes | 2.049E-07 | 793.33099 | 15.155088 |
| IDDMTAAPMDVR | 12 | P15927 | Replication protein A 32 kDa subunit OS | RPA2 | 29.247 | 2 | 52.980335 | yes | 9.722E-08 | 667.80762 | 531.77307 |
| APTNIVYK | 8 | P15927 | Replication protein A 32 kDa subunit OS | RPA2 | 29.247 | 2 | 52.980335 | yes | 1.389E-07 | 453.25818 | 271.85028 |
| HMSVSSIK | 8 | P15927 | Replication protein A 32 kDa subunit OS | RPA2 | 29.247 | 2 | 52.980335 | yes | 2.873E-06 | 444.73401 | 110.72042 |
| EVDSGNDIYGNPII | 14 | P16035 | Metalloproteinase inhibitor 2 OS=Homo s | TIMP2 | 24.4 | 2 | 47.465374 | yes | 1.324E-07 | 760.86499 | 64.217613 |
| YGFIEGHVVIPR | 12 | P16070 | CD44 antigen OS=Homo sapiens OX=96 | CD44 | 81.538 | 3 | 48.078125 | yes | 1.203E-07 | 462.92245 | 530.20093 |
| ALSIGFETCR | 10 | P16070 | CD44 antigen OS=Homo sapiens OX=96 | CD44 | 81.538 | 2 | 48.078125 | yes | 3.269E-05 | 577.28717 | 104.20389 |
| TEAADLCK | 8 | P16070 | CD44 antigen OS=Homo sapiens OX=96 | CD44 | 81.538 | 2 | 48.078125 | yes | 2.145E-06 | 454.2131 | 81.432304 |
| SQEMVHLVNK | 10 | P16070 | CD44 antigen OS=Homo sapiens OX=96 | CD44 | 81.538 | 2 | 48.078125 | yes | 1.063E-06 | 592.80823 | 32.710793 |
| VAVLENQNK | 9 | P16220 | Cyclic AMP-responsive element-binding | CREB1 | 35.137 | 2 | 47.384983 | no | 1.415E-07 | 507.78256 | 261.452 |
| ILNDLSSDAPGVPI | 14 | P16220 | Cyclic AMP-responsive element-binding | CREB1 | 35.137 | 2 | 47.384983 | yes | 1.341E-07 | 727.38593 | 126.80122 |
| LTLLQETHR | 9 | P16383 | Intron Large complex component GCFC2 | GCFC2 | 89.385 | 2 | 43.94502 | yes | 2.384E-07 | 555.81696 | 88.317253 |
| IYVENLIDCLNEK | 13 | P16383 | Intron Large complex component GCFC2 | GCFC2 | 89.385 | 2 | 43.94502 | yes | 3.123E-06 | 811.90839 | 27.770132 |
| VQLIDWNPLK | 10 | P16383 | Intron Large complex component GCFC2 | GCFC2 | 89.385 | 2 | 43.94502 | yes | 6.666E-07 | 613.35059 | 21.470531 |
| IPDAAFIQAAR | 11 | P16383 | Intron Large complex component GCFC2 | GCFC2 | 89.385 | 2 | 43.94502 | yes | 3.995E-06 | 586.82477 | 20.32901 |
| LLNPLIR | 7 | P16383 | Intron Large complex component GCFC2 | GCFC2 | 89.385 | 2 | 43.94502 | yes | 5.387E-06 | 419.77908 | 15.53547 |
| ATGPPVSELITK | 12 | P16401 | Histone H1.5 OS=Homo sapiens OX=96( | H1-5 | 22.581 | 2 | 50.735462 | yes | 1.426E-07 | 606.84534 | 10983.721 |
| ALAAGGYDVEK | 11 | P16401 | Histone H1.5 OS=Homo sapiens OX=96( | H1-5 | 22.581 | 2 | 50.735462 | yes | 1.219E-07 | 547.27985 | 8589.8115 |
| SETAPAETATPAP\ | 16 | P16401 | Histone H1.5 OS=Homo sapiens OX=96( | H1-5 | 22.581 | 2 | 50.735462 | yes | 1.354E-07 | 820.9043 | 1226.7638 |
| SETAPAETATPAP\ | 16 | P16401 | Histone H1.5 OS=Homo sapiens OX=96( | H1-5 | 22.581 | 2 | 50.735462 | yes | 1.014E-07 | 799.89905 | 1226.7638 |
| KATGPPVSELITK | 13 | P16401 | Histone H1.5 OS=Homo sapiens OX=96( | H1-5 | 22.581 | 2 | 50.735462 | yes | 4.565E-07 | 670.89282 | 1397.2987 |
| NGLSLAALK | 9 | P16401 | Histone H1.5 OS=Homo sapiens OX=96( | H1-5 | 22.581 | 2 | 50.735462 | yes | 9.723E-08 | 443.77145 | 107.12981 |
| KALAAGGYDVEK | 12 | P16401 | Histone H1.5 OS=Homo sapiens OX=96( | H1-5 | 22.581 | 2 | 50.735462 | yes | 1.159E-07 | 611.32733 | 100.87691 |
| SETAPAAPAAAPP. | 16 | P16403 | Histone H1.2 OS=Homo sapiens OX=96( | H1-2 | 21.365 | 2 | 51.38997 | yes | 1.341E-07 | 760.88318 | 4429.8003 |
| SETAPAAPAAAPP. | 16 | P16403 | Histone H1.2 OS=Homo sapiens OX=96( | H1-2 | 21.365 | 2 | 51.38997 | yes | 9.722E-08 | 739.87793 | 4429.8003 |
| KPAAATVTK | 9 | P16403 | Histone H1.2 OS=Homo sapiens OX=96( | H1-2 | 21.365 | 2 | 51.38997 | yes | 2.436E-06 | 443.77145 | 586.6806 |
| EVGETLLYYGCR | 12 | P16435 | NADPH--cytochrome P450 reductase OS | POR | 76.69 | 2 | 42.224743 | yes | 5.498E-07 | 730.34796 | 50.10186 |
| NPFLAAVTTNR | 11 | P16435 | NADPH--cytochrome P450 reductase OS | POR | 76.69 | 2 | 42.224743 | yes | 3.134E-07 | 602.32764 | 47.959091 |

| | | | | | | | | | | | |
|---|---|---|---|---|---|---|---|---|---|---|---|
| IGIFGQDEDVTSK | 13 | P16615 | Sarcoplasmic/endoplasmic reticulum calc | ATP2A2 | 114.757 | 2 | 51.591972 | yes | 1.802E-07 | 704.85138 | 261.07745 |
| VDQSILTGESVSVI | 15 | P16615 | Sarcoplasmic/endoplasmic reticulum calc | ATP2A2 | 114.757 | 2 | 51.591972 | no | 9.722E-08 | 787.93542 | 260.53552 |
| VGEATETALTCLV | 15 | P16615 | Sarcoplasmic/endoplasmic reticulum calc | ATP2A2 | 114.757 | 2 | 51.591972 | no | 2.26E-07 | 810.91107 | 222.13457 |
| EFDELNPSAQR | 11 | P16615 | Sarcoplasmic/endoplasmic reticulum calc | ATP2A2 | 114.757 | 2 | 51.591972 | yes | 1.072E-07 | 653.30713 | 121.86656 |
| MNVFDTELK | 9 | P16615 | Sarcoplasmic/endoplasmic reticulum calc | ATP2A2 | 114.757 | 2 | 51.591972 | yes | 1.065E-07 | 548.77081 | 115.58829 |
| NMLFSGTNIAAGK | 13 | P16615 | Sarcoplasmic/endoplasmic reticulum calc | ATP2A2 | 114.757 | 2 | 51.591972 | no | 2.756E-07 | 662.3399 | 73.898148 |
| EFTLEFSR | 8 | P16615 | Sarcoplasmic/endoplasmic reticulum calc | ATP2A2 | 114.757 | 2 | 51.591972 | no | 5.756E-07 | 514.75598 | 68.259857 |
| TVEEVLGHFGVNE | 23 | P16615 | Sarcoplasmic/endoplasmic reticulum calc | ATP2A2 | 114.757 | 3 | 51.591972 | yes | 9.968E-08 | 824.75873 | 63.445438 |
| TGTLTTNQMSVCR | 13 | P16615 | Sarcoplasmic/endoplasmic reticulum calc | ATP2A2 | 114.757 | 2 | 51.591972 | no | 1.519E-07 | 734.84778 | 55.081013 |
| IEVASSVK | 8 | P16615 | Sarcoplasmic/endoplasmic reticulum calc | ATP2A2 | 114.757 | 2 | 51.591972 | yes | 1.053E-07 | 416.74237 | 53.570045 |
| DIVPGDIVEIAVGD | 15 | P16615 | Sarcoplasmic/endoplasmic reticulum calc | ATP2A2 | 114.757 | 2 | 51.591972 | yes | 3.183E-06 | 770.41687 | 53.190075 |
| AIYNNMK | 7 | P16615 | Sarcoplasmic/endoplasmic reticulum calc | ATP2A2 | 114.757 | 2 | 51.591972 | no | 5.729E-07 | 427.21545 | 45.556595 |
| EEMHLEDSANFIK | 13 | P16615 | Sarcoplasmic/endoplasmic reticulum calc | ATP2A2 | 114.757 | 2 | 51.591972 | yes | 2.4E-07 | 781.86139 | 43.557346 |
| AMGVVVATGVNT | 16 | P16615 | Sarcoplasmic/endoplasmic reticulum calc | ATP2A2 | 114.757 | 2 | 51.591972 | yes | 1.014E-06 | 773.41888 | 33.60482 |
| VSFYQLSHFLQCK | 13 | P16615 | Sarcoplasmic/endoplasmic reticulum calc | ATP2A2 | 114.757 | 2 | 51.591972 | yes | 2.646E-07 | 828.91376 | 12.044636 |
| EDVFVHQTAIK | 11 | P16989 | Y-box-binding protein 3 OS=Homo sapie | YBX3 | 40.09 | 2 | 48.576538 | no | 1.209E-07 | 643.84058 | 15871.82 |
| EDVFVHQTAIK | 11 | P16989 | Y-box-binding protein 3 OS=Homo sapie | YBX3 | 40.09 | 2 | 48.576538 | no | 1.49E-07 | 429.56284 | 15871.82 |
| EDVFVHQTAIK | 11 | P16989 | Y-box-binding protein 3 OS=Homo sapie | YBX3 | 40.09 | 3 | 48.576538 | no | 1.209E-07 | 643.84058 | 15871.82 |
| EDVFVHQTAIK | 11 | P16989 | Y-box-binding protein 3 OS=Homo sapie | YBX3 | 40.09 | 3 | 48.576538 | no | 1.49E-07 | 429.56284 | 15871.82 |
| VLPHPNR | 7 | P16989 | Y-box-binding protein 3 OS=Homo sapie | YBX3 | 40.09 | 2 | 48.576538 | yes | 1.235E-07 | 416.74304 | 16672.682 |
| IQAGEIGEMK | 10 | P16989 | Y-box-binding protein 3 OS=Homo sapie | YBX3 | 40.09 | 2 | 48.576538 | yes | 2.587E-07 | 538.27625 | 5068.5488 |
| IQAGEIGEMK | 10 | P16989 | Y-box-binding protein 3 OS=Homo sapie | YBX3 | 40.09 | 2 | 48.576538 | yes | 2.442E-07 | 546.27368 | 5068.5488 |
| NGYGFINR | 8 | P16989 | Y-box-binding protein 3 OS=Homo sapie | YBX3 | 40.09 | 2 | 48.576538 | no | 1.941E-07 | 470.73541 | 8093.4932 |
| NPTYRPR | 7 | P16989 | Y-box-binding protein 3 OS=Homo sapie | YBX3 | 40.09 | 2 | 48.576538 | yes | 2.027E-07 | 452.24341 | 3684.8584 |
| RPRPPNAPSQDGK | 13 | P16989 | Y-box-binding protein 3 OS=Homo sapie | YBX3 | 40.09 | 3 | 48.576538 | yes | 1.445E-07 | 473.91989 | 2652.3875 |
| FPPYHVGQTFDR | 12 | P16989 | Y-box-binding protein 3 OS=Homo sapie | YBX3 | 40.09 | 2 | 48.576538 | yes | 2.279E-06 | 732.35693 | 611.20709 |
| RFPPYHVGQTFDR | 13 | P16989 | Y-box-binding protein 3 OS=Homo sapie | YBX3 | 40.09 | 3 | 48.576538 | yes | 3.991E-07 | 540.60742 | 607.6322 |
| NDTKEDVFVHQTA | 15 | P16989 | Y-box-binding protein 3 OS=Homo sapie | YBX3 | 40.09 | 3 | 48.576538 | no | 1.128E-07 | 582.30035 | 235.45662 |
| PPNAPSQDGK | 10 | P16989 | Y-box-binding protein 3 OS=Homo sapie | YBX3 | 40.09 | 2 | 48.576538 | yes | 6.159E-07 | 505.74872 | 112.44695 |
| PRPPNAPSQDGK | 12 | P16989 | Y-box-binding protein 3 OS=Homo sapie | YBX3 | 40.09 | 3 | 48.576538 | yes | 1.295E-07 | 421.8862 | 50.382103 |
| PAPAVGEAEDK | 11 | P16989 | Y-box-binding protein 3 OS=Homo sapie | YBX3 | 40.09 | 2 | 48.576538 | yes | 3.292E-06 | 542.26947 | 31.985439 |
| AFSYSSVLR | 9 | P17020 | Zinc finger protein 16 OS=Homo sapiens | ZNF16 | 76.472 | 2 | 54.152725 | yes | 2.725E-07 | 515.27185 | 347.68356 |
| SVLIQHQR | 8 | P17020 | Zinc finger protein 16 OS=Homo sapiens | ZNF16 | 76.472 | 2 | 54.152725 | no | 1.052E-07 | 490.78543 | 272.16196 |
| TFQGNPDLIQR | 11 | P17020 | Zinc finger protein 16 OS=Homo sapiens | ZNF16 | 76.472 | 2 | 54.152725 | yes | 9.722E-08 | 644.83588 | 230.88541 |
| SSNLIQHQR | 9 | P17020 | Zinc finger protein 16 OS=Homo sapiens | ZNF16 | 76.472 | 2 | 54.152725 | yes | 1.827E-07 | 541.7887 | 192.38185 |
| AFSQSAHLR | 9 | P17020 | Zinc finger protein 16 OS=Homo sapiens | ZNF16 | 76.472 | 2 | 54.152725 | yes | 5.999E-07 | 508.76724 | 113.70735 |
| TFSQSSTLIQHQR | 13 | P17020 | Zinc finger protein 16 OS=Homo sapiens | ZNF16 | 76.472 | 2 | 54.152725 | yes | 1.119E-07 | 766.89441 | 113.29948 |
| AFSQSSSLIQHR | 12 | P17020 | Zinc finger protein 16 OS=Homo sapiens | ZNF16 | 76.472 | 2 | 54.152725 | yes | 1.214E-07 | 680.85205 | 105.16781 |
| DLDCNGFDSR | 10 | P17020 | Zinc finger protein 16 OS=Homo sapiens | ZNF16 | 76.472 | 2 | 54.152725 | yes | 1.217E-06 | 599.74329 | 68.055855 |

| | | | | | | | | | | | |
|---|---|---|---|---|---|---|---|---|---|---|---|
| TEDKEFLHK | 9 | P17020 | Zinc finger protein 16 OS=Homo sapiens | ZNF16 | 76.472 | 2 | 54.152725 | yes | 9.11E-06 | 573.79309 | 53.703197 |
| GFSQSSHLIQHQIII | 19 | P17020 | Zinc finger protein 16 OS=Homo sapiens | ZNF16 | 76.472 | 3 | 54.152725 | yes | 1.732E-06 | 725.70667 | 36.934998 |
| PYVCSECGK | 9 | P17020 | Zinc finger protein 16 OS=Homo sapiens | ZNF16 | 76.472 | 2 | 54.152725 | no | 3.726E-07 | 550.23096 | 35.500946 |
| QFGYQDSPGPR | 11 | P17028 | Zinc finger protein 24 OS=Homo sapiens | ZNF24 | 42.156 | 2 | 52.04522 | yes | 9.722E-08 | 626.29126 | 470.8782 |
| SSILVQHQR | 9 | P17028 | Zinc finger protein 24 OS=Homo sapiens | ZNF24 | 42.156 | 2 | 52.04522 | yes | 9.903E-08 | 534.30145 | 439.75604 |
| SYSQSSNLFR | 10 | P17028 | Zinc finger protein 24 OS=Homo sapiens | ZNF24 | 42.156 | 2 | 52.04522 | yes | 9.722E-08 | 594.78583 | 415.38901 |
| HFSQGSALILHQR | 13 | P17028 | Zinc finger protein 24 OS=Homo sapiens | ZNF24 | 42.156 | 3 | 52.04522 | yes | 1.065E-07 | 498.60391 | 379.96051 |
| YGETCFPK | 8 | P17028 | Zinc finger protein 24 OS=Homo sapiens | ZNF24 | 42.156 | 2 | 52.04522 | yes | 1.26E-06 | 501.22348 | 111.09032 |
| AFSQNSGLINHQR | 13 | P17028 | Zinc finger protein 24 OS=Homo sapiens | ZNF24 | 42.156 | 2 | 52.04522 | yes | 9.722E-08 | 736.37366 | 107.98607 |
| QHICDECGK | 9 | P17028 | Zinc finger protein 24 OS=Homo sapiens | ZNF24 | 42.156 | 2 | 52.04522 | yes | 2.805E-07 | 573.73694 | 55.6894 |
| WASWELHSLR | 10 | P17028 | Zinc finger protein 24 OS=Homo sapiens | ZNF24 | 42.156 | 2 | 52.04522 | yes | 9.722E-08 | 642.82782 | 29.053617 |
| LWLRPETHTK | 10 | P17028 | Zinc finger protein 24 OS=Homo sapiens | ZNF24 | 42.156 | 2 | 52.04522 | yes | 4.115E-07 | 640.85913 | 20.477352 |
| SHGVLLGR | 8 | P17036 | Zinc finger protein 3 OS=Homo sapiens ( | ZNF3 | 50.916 | 2 | 41.729744 | yes | 3.367E-07 | 419.74832 | 301.89029 |
| LPVGDRPHK | 9 | P17036 | Zinc finger protein 3 OS=Homo sapiens ( | ZNF3 | 50.916 | 2 | 41.729744 | yes | 3.424E-07 | 509.79324 | 196.21632 |
| TSDLIQHQR | 9 | P17036 | Zinc finger protein 3 OS=Homo sapiens ( | ZNF3 | 50.916 | 2 | 41.729744 | yes | 3.766E-07 | 549.28851 | 83.686516 |
| PICVEQGQK | 9 | P17081 | Rho-related GTP-binding protein RhoQ ( | RHOQ | 22.66 | 2 | 19.623302 | yes | 4.775E-06 | 529.76862 | 41.656784 |
| KQPPVSPGTALVG | 16 | P17096 | High mobility group protein HMG-I/HM( | HMGA1 | 11.677 | 2 | 49.763115 | yes | 1.966E-07 | 797.4516 | 3217.9397 |
| KQPPVSPGTALVG | 16 | P17096 | High mobility group protein HMG-I/HM( | HMGA1 | 11.677 | 2 | 49.763115 | yes | 1.004E-07 | 531.97015 | 3217.9397 |
| KQPPVSPGTALVG | 16 | P17096 | High mobility group protein HMG-I/HM( | HMGA1 | 11.677 | 3 | 49.763115 | yes | 1.966E-07 | 797.4516 | 3217.9397 |
| KQPPVSPGTALVG | 16 | P17096 | High mobility group protein HMG-I/HM( | HMGA1 | 11.677 | 3 | 49.763115 | yes | 1.004E-07 | 531.97015 | 3217.9397 |
| EPSEVPTPK | 9 | P17096 | High mobility group protein HMG-I/HM( | HMGA1 | 11.677 | 2 | 49.763115 | yes | 1.124E-07 | 492.25583 | 2027.4385 |
| QPPVSPGTALVGS( | 15 | P17096 | High mobility group protein HMG-I/HM( | HMGA1 | 11.677 | 2 | 49.763115 | yes | 4.514E-07 | 733.40411 | 330.2825 |
| LPLHTLTSSTPVVI | 16 | P17152 | Transmembrane protein 11, mitochondria | TMEM11 | 21.541 | 3 | 49.77356 | yes | 1.004E-07 | 578.34882 | 103.5674 |
| APGGLSLHDYK | 11 | P17275 | Transcription factor JunB OS=Homo sapi | JUNB | 35.88 | 2 | 51.933395 | yes | 9.722E-08 | 579.30109 | 468.07971 |
| AENAGLSSTAGLL | 14 | P17275 | Transcription factor JunB OS=Homo sapi | JUNB | 35.88 | 2 | 51.933395 | yes | 9.722E-08 | 680.36499 | 388.62576 |
| LASSELER | 8 | P17275 | Transcription factor JunB OS=Homo sapi | JUNB | 35.88 | 2 | 51.933395 | yes | 9.914E-08 | 452.74036 | 319.71686 |
| LLKPSLAVNLADP | 15 | P17275 | Transcription factor JunB OS=Homo sapi | JUNB | 35.88 | 2 | 51.933395 | yes | 9.81E-08 | 835.48541 | 126.34023 |
| LLKPSLAVNLADP | 15 | P17275 | Transcription factor JunB OS=Homo sapi | JUNB | 35.88 | 2 | 51.933395 | yes | 1.686E-07 | 557.32605 | 126.34023 |
| LLKPSLAVNLADP | 15 | P17275 | Transcription factor JunB OS=Homo sapi | JUNB | 35.88 | 3 | 51.933395 | yes | 9.81E-08 | 835.48541 | 126.34023 |
| LLKPSLAVNLADP | 15 | P17275 | Transcription factor JunB OS=Homo sapi | JUNB | 35.88 | 3 | 51.933395 | yes | 1.686E-07 | 557.32605 | 126.34023 |
| EEPQTVPEAR | 10 | P17275 | Transcription factor JunB OS=Homo sapi | JUNB | 35.88 | 2 | 51.933395 | yes | 9.789E-08 | 578.28564 | 132.27937 |
| FLESLPEEEQQR | 12 | P17480 | Nucleolar transcription factor 1 OS=Hom | UBTF | 89.406 | 2 | 53.20042 | yes | 4.507E-07 | 752.86755 | 1495.7194 |
| WVEISNEVR | 9 | P17480 | Nucleolar transcription factor 1 OS=Hom | UBTF | 89.406 | 2 | 53.20042 | yes | 1.087E-07 | 566.29327 | 1096.8295 |
| LHPEMSNLDLTK | 12 | P17480 | Nucleolar transcription factor 1 OS=Hom | UBTF | 89.406 | 2 | 53.20042 | yes | 1.838E-07 | 699.35828 | 618.15808 |
| FREDHPDLIQNAK | 13 | P17480 | Nucleolar transcription factor 1 OS=Hom | UBTF | 89.406 | 3 | 53.20042 | yes | 1.804E-07 | 528.27057 | 295.20477 |
| FREDHPDLIQNAK | 13 | P17480 | Nucleolar transcription factor 1 OS=Hom | UBTF | 89.406 | 3 | 53.20042 | yes | 9.787E-08 | 791.90228 | 295.20477 |
| FREDHPDLIQNAK | 13 | P17480 | Nucleolar transcription factor 1 OS=Hom | UBTF | 89.406 | 2 | 53.20042 | yes | 1.804E-07 | 528.27057 | 295.20477 |
| FREDHPDLIQNAK | 13 | P17480 | Nucleolar transcription factor 1 OS=Hom | UBTF | 89.406 | 2 | 53.20042 | yes | 9.787E-08 | 791.90228 | 295.20477 |
| MVLCSQQWK | 9 | P17480 | Nucleolar transcription factor 1 OS=Hom | UBTF | 89.406 | 2 | 53.20042 | yes | 1.004E-07 | 590.28607 | 587.5827 |

| | | | | | | | | | | | |
|---|---|---|---|---|---|---|---|---|---|---|---|
| NNLPSNDSSK | 10 | P17480 | Nucleolar transcription factor 1 OS=Hom | UBTF | 89.406 | 2 | 53.20042 | yes | 1.053E-07 | 538.25433 | 463.88361 |
| HPELNISEEGITK | 13 | P17480 | Nucleolar transcription factor 1 OS=Hom | UBTF | 89.406 | 2 | 53.20042 | yes | 5.983E-07 | 733.87793 | 424.63364 |
| YIQDFQR | 7 | P17480 | Nucleolar transcription factor 1 OS=Hom | UBTF | 89.406 | 2 | 53.20042 | yes | 1.187E-07 | 485.24307 | 356.79871 |
| MWNDLSEK | 8 | P17480 | Nucleolar transcription factor 1 OS=Hom | UBTF | 89.406 | 2 | 53.20042 | yes | 1.022E-07 | 511.73422 | 353.92957 |
| TPQQLWYTHEK | 11 | P17480 | Nucleolar transcription factor 1 OS=Hom | UBTF | 89.406 | 2 | 53.20042 | yes | 9.722E-08 | 715.85681 | 298.28699 |
| KHPDFPK | 7 | P17480 | Nucleolar transcription factor 1 OS=Hom | UBTF | 89.406 | 2 | 53.20042 | yes | 2.338E-07 | 434.73743 | 267.38101 |
| KSDIPEKPK | 9 | P17480 | Nucleolar transcription factor 1 OS=Hom | UBTF | 89.406 | 2 | 53.20042 | yes | 9.733E-08 | 521.3006 | 178.51187 |
| AMEMTWNNMEK | 11 | P17480 | Nucleolar transcription factor 1 OS=Hom | UBTF | 89.406 | 2 | 53.20042 | yes | 9.81E-08 | 692.78833 | 174.30469 |
| KEYEEIMR | 8 | P17480 | Nucleolar transcription factor 1 OS=Hom | UBTF | 89.406 | 2 | 53.20042 | yes | 9.789E-08 | 549.26843 | 172.61928 |
| GKLPESPK | 8 | P17480 | Nucleolar transcription factor 1 OS=Hom | UBTF | 89.406 | 2 | 53.20042 | yes | 4.058E-06 | 428.25037 | 170.4435 |
| DYEVELLR | 8 | P17480 | Nucleolar transcription factor 1 OS=Hom | UBTF | 89.406 | 2 | 53.20042 | yes | 3.397E-07 | 518.7691 | 156.48946 |
| TTESHMDWEK | 10 | P17480 | Nucleolar transcription factor 1 OS=Hom | UBTF | 89.406 | 2 | 53.20042 | yes | 9.733E-08 | 632.26917 | 149.52971 |
| RPVSAMFIFSEEK | 13 | P17480 | Nucleolar transcription factor 1 OS=Hom | UBTF | 89.406 | 2 | 53.20042 | yes | 9.809E-08 | 770.89508 | 95.915108 |
| EDHPDLIQNAK | 11 | P17480 | Nucleolar transcription factor 1 OS=Hom | UBTF | 89.406 | 2 | 53.20042 | yes | 1.12E-07 | 640.3175 | 46.474987 |
| RAEEIWQQSVIGD | 17 | P17480 | Nucleolar transcription factor 1 OS=Hom | UBTF | 89.406 | 3 | 53.20042 | yes | 7.68E-07 | 678.68494 | 40.030598 |
| AAPCDYGPAPAFY | 14 | P17509 | Homeobox protein Hox-B6 OS=Homo sa | HOXB6 | 25.433 | 2 | 45.559155 | yes | 1.812E-07 | 778.35358 | 69.011528 |
| TDSVIIADQTPTPT | 15 | P17544 | Cyclic AMP-dependent transcription fact | ATF7 | 51.757 | 2 | 42.312508 | yes | 3.091E-07 | 807.92029 | 92.820267 |
| PVLISTPTPTIVR | 13 | P17544 | Cyclic AMP-dependent transcription fact | ATF7 | 51.757 | 2 | 42.312508 | yes | 3.59E-07 | 697.42432 | 52.909199 |
| NLLQVDLTK | 9 | P17612 | cAMP-dependent protein kinase catalytic | PRKACA | 40.59 | 2 | 53.198307 | yes | 1.584E-07 | 522.30841 | 462.89767 |
| TLGTGSFGR | 9 | P17612 | cAMP-dependent protein kinase catalytic | PRKACA | 40.59 | 2 | 53.198307 | yes | 1.615E-07 | 448.23523 | 429.2002 |
| FSEPHAR | 7 | P17612 | cAMP-dependent protein kinase catalytic | PRKACA | 40.59 | 2 | 53.198307 | yes | 2.014E-07 | 422.20901 | 351.47928 |
| KVEAPFIPK | 9 | P17612 | cAMP-dependent protein kinase catalytic | PRKACA | 40.59 | 2 | 53.198307 | no | 2.959E-07 | 514.81061 | 288.83936 |
| FPSHFSSDLK | 10 | P17612 | cAMP-dependent protein kinase catalytic | PRKACA | 40.59 | 2 | 53.198307 | yes | 2.89E-06 | 582.78784 | 156.74281 |
| WFATTDWIAIYQR | 13 | P17612 | cAMP-dependent protein kinase catalytic | PRKACA | 40.59 | 2 | 53.198307 | yes | 9.722E-08 | 835.91992 | 116.08478 |
| QIEHTLNEK | 9 | P17612 | cAMP-dependent protein kinase catalytic | PRKACA | 40.59 | 2 | 53.198307 | yes | 9.722E-08 | 556.29077 | 108.04351 |
| AKEDFLK | 7 | P17612 | cAMP-dependent protein kinase catalytic | PRKACA | 40.59 | 2 | 53.198307 | yes | 2.388E-07 | 425.73709 | 102.22948 |
| YLNQDYEALR | 10 | P17655 | Calpain-2 catalytic subunit OS=Homo sa | CAPN2 | 79.996 | 2 | 52.87485 | yes | 9.722E-08 | 642.81458 | 138.85394 |
| SDGFSIETCK | 10 | P17655 | Calpain-2 catalytic subunit OS=Homo sa | CAPN2 | 79.996 | 2 | 52.87485 | yes | 6.668E-07 | 572.25299 | 104.9901 |
| SDTFINLR | 8 | P17655 | Calpain-2 catalytic subunit OS=Homo sa | CAPN2 | 79.996 | 2 | 52.87485 | yes | 1.435E-06 | 483.25616 | 84.569588 |
| IMVDMLDSDGSG | 13 | P17655 | Calpain-2 catalytic subunit OS=Homo sa | CAPN2 | 79.996 | 2 | 52.87485 | yes | 1.108E-07 | 684.31268 | 77.835464 |
| MPCQLHQVIVAR | 12 | P17655 | Calpain-2 catalytic subunit OS=Homo sa | CAPN2 | 79.996 | 2 | 52.87485 | yes | 1.516E-07 | 726.38416 | 35.11359 |
| NPWGEVEWTGR | 11 | P17655 | Calpain-2 catalytic subunit OS=Homo sa | CAPN2 | 79.996 | 2 | 52.87485 | yes | 7.404E-05 | 665.81238 | 12.085909 |
| VLELTAENER | 10 | P17676 | CCAAT/enhancer-binding protein beta O | CEBPB | 36.106 | 2 | 52.9845 | yes | 1.367E-07 | 587.30914 | 636.64276 |
| QLPEPLLASSGHC | 13 | P17676 | CCAAT/enhancer-binding protein beta O | CEBPB | 36.106 | 2 | 52.9845 | yes | 9.722E-08 | 704.84808 | 297.7258 |
| KPAEYGYVSLGR | 12 | P17676 | CCAAT/enhancer-binding protein beta O | CEBPB | 36.106 | 2 | 52.9845 | yes | 1.832E-07 | 670.35388 | 231.81441 |
| KVEQLSR | 7 | P17676 | CCAAT/enhancer-binding protein beta O | CEBPB | 36.106 | 2 | 52.9845 | yes | 3.81E-07 | 430.25345 | 164.57646 |
| AEPGFEPADCK | 11 | P17676 | CCAAT/enhancer-binding protein beta O | CEBPB | 36.106 | 2 | 52.9845 | yes | 1.169E-07 | 610.76624 | 58.314438 |
| GALHPGCFAPLHP | 25 | P17676 | CCAAT/enhancer-binding protein beta O | CEBPB | 36.106 | 4 | 52.9845 | yes | 2.997E-06 | 648.59521 | 16.115313 |
| MGLIQTPDQLR | 11 | P17706 | Tyrosine-protein phosphatase non-recept | PTPN2 | 48.474 | 2 | 48.948612 | yes | 1.084E-07 | 636.34247 | 247.39488 |

| | | | | | | | | | | | |
|---|---|---|---|---|---|---|---|---|---|---|---|
| FSYMAIIEGAK | 11 | P17706 | Tyrosine-protein phosphatase non-recepto | PTPN2 | 48.474 | 2 | 48.948612 | yes | 1.212E-06 | 615.31537 | 74.669975 |
| MQDTMEENSESAI | 14 | P17706 | Tyrosine-protein phosphatase non-recepto | PTPN2 | 48.474 | 2 | 48.948612 | yes | 1.755E-06 | 820.84814 | 19.866381 |
| GLGLSPDLVVCR | 12 | P17812 | CTP synthase 1 OS=Homo sapiens OX=9 | CTPS1 | 66.691 | 2 | 51.880295 | yes | 1.128E-07 | 643.35028 | 518.87323 |
| FVGQDVEGER | 10 | P17812 | CTP synthase 1 OS=Homo sapiens OX=9 | CTPS1 | 66.691 | 2 | 51.880295 | yes | 4.286E-07 | 568.27252 | 297.58594 |
| GIIASSVGTILK | 12 | P17812 | CTP synthase 1 OS=Homo sapiens OX=9 | CTPS1 | 66.691 | 2 | 51.880295 | yes | 9.782E-08 | 579.85828 | 262.57867 |
| IQAIAWAR | 8 | P17812 | CTP synthase 1 OS=Homo sapiens OX=9 | CTPS1 | 66.691 | 2 | 51.880295 | yes | 1.767E-07 | 464.77179 | 216.67509 |
| YILVTGGVISGIGK | 14 | P17812 | CTP synthase 1 OS=Homo sapiens OX=9 | CTPS1 | 66.691 | 2 | 51.880295 | no | 2.225E-07 | 688.91101 | 149.32747 |
| LYGDADYLEER | 11 | P17812 | CTP synthase 1 OS=Homo sapiens OX=9 | CTPS1 | 66.691 | 2 | 51.880295 | yes | 9.722E-08 | 672.30933 | 144.36407 |
| LLETCSIALVGK | 12 | P17812 | CTP synthase 1 OS=Homo sapiens OX=9 | CTPS1 | 66.691 | 2 | 51.880295 | yes | 4.427E-07 | 652.36816 | 131.28513 |
| CSNPLDTSVK | 10 | P17812 | CTP synthase 1 OS=Homo sapiens OX=9 | CTPS1 | 66.691 | 2 | 51.880295 | yes | 1.456E-07 | 560.7688 | 97.403687 |
| SCGLHVTSIK | 10 | P17812 | CTP synthase 1 OS=Homo sapiens OX=9 | CTPS1 | 66.691 | 2 | 51.880295 | yes | 1.238E-07 | 551.28967 | 95.760696 |
| FSDSYASVIK | 10 | P17812 | CTP synthase 1 OS=Homo sapiens OX=9 | CTPS1 | 66.691 | 2 | 51.880295 | yes | 1.062E-07 | 558.78223 | 95.186157 |
| IYQYVINK | 8 | P17812 | CTP synthase 1 OS=Homo sapiens OX=9 | CTPS1 | 66.691 | 2 | 51.880295 | yes | 3.709E-06 | 520.79242 | 52.58112 |
| KLYGDADYLEER | 12 | P17812 | CTP synthase 1 OS=Homo sapiens OX=9 | CTPS1 | 66.691 | 2 | 51.880295 | yes | 8.678E-07 | 736.35681 | 43.979816 |
| RLDLPIER | 8 | P17812 | CTP synthase 1 OS=Homo sapiens OX=9 | CTPS1 | 66.691 | 2 | 51.880295 | yes | 3.205E-06 | 506.3009 | 39.701611 |
| APILIATDVASR | 12 | P17844 | Probable ATP-dependent RNA helicase D | DDX5 | 69.149 | 2 | 53.686756 | no | 1.09E-07 | 613.85876 | 69408.625 |
| GDGPICLVLAPTR | 13 | P17844 | Probable ATP-dependent RNA helicase D | DDX5 | 69.149 | 2 | 53.686756 | no | 8.526E-07 | 684.86884 | 44901.996 |
| TTYLVLDEADR | 11 | P17844 | Probable ATP-dependent RNA helicase D | DDX5 | 69.149 | 2 | 53.686756 | yes | 1.425E-07 | 648.32751 | 35041.113 |
| DGWPAMGIHGDK | 12 | P17844 | Probable ATP-dependent RNA helicase D | DDX5 | 69.149 | 2 | 53.686756 | yes | 2.061E-07 | 642.29553 | 16060.414 |
| DGWPAMGIHGDK | 12 | P17844 | Probable ATP-dependent RNA helicase D | DDX5 | 69.149 | 2 | 53.686756 | yes | 1.379E-07 | 650.29297 | 16060.414 |
| GVEICIATPGR | 11 | P17844 | Probable ATP-dependent RNA helicase D | DDX5 | 69.149 | 2 | 53.686756 | no | 9.722E-08 | 586.80823 | 31655.357 |
| QTLMWSATWPK | 11 | P17844 | Probable ATP-dependent RNA helicase D | DDX5 | 69.149 | 2 | 53.686756 | no | 1.108E-07 | 674.83954 | 13722.091 |
| QTLMWSATWPK | 11 | P17844 | Probable ATP-dependent RNA helicase D | DDX5 | 69.149 | 2 | 53.686756 | no | 9.722E-08 | 682.83698 | 13722.091 |
| TAQEVETYR | 9 | P17844 | Probable ATP-dependent RNA helicase D | DDX5 | 69.149 | 2 | 53.686756 | yes | 9.882E-08 | 548.76709 | 24604.506 |
| LLQLVEDR | 8 | P17844 | Probable ATP-dependent RNA helicase D | DDX5 | 69.149 | 2 | 53.686756 | yes | 4.786E-07 | 493.28748 | 24020.615 |
| NFYQEHPDLAR | 11 | P17844 | Probable ATP-dependent RNA helicase D | DDX5 | 69.149 | 2 | 53.686756 | yes | 9.722E-08 | 695.33093 | 11659.195 |
| NFYQEHPDLAR | 11 | P17844 | Probable ATP-dependent RNA helicase D | DDX5 | 69.149 | 2 | 53.686756 | yes | 1.496E-07 | 463.88971 | 11659.195 |
| NFYQEHPDLAR | 11 | P17844 | Probable ATP-dependent RNA helicase D | DDX5 | 69.149 | 3 | 53.686756 | yes | 9.722E-08 | 695.33093 | 11659.195 |
| NFYQEHPDLAR | 11 | P17844 | Probable ATP-dependent RNA helicase D | DDX5 | 69.149 | 3 | 53.686756 | yes | 1.496E-07 | 463.88971 | 11659.195 |
| LIDFLECGK | 9 | P17844 | Probable ATP-dependent RNA helicase D | DDX5 | 69.149 | 2 | 53.686756 | yes | 1.692E-07 | 547.78119 | 21918.561 |
| LMEEIMSEK | 9 | P17844 | Probable ATP-dependent RNA helicase D | DDX5 | 69.149 | 2 | 53.686756 | yes | 1.219E-07 | 555.26447 | 5948.1865 |
| LMEEIMSEK | 9 | P17844 | Probable ATP-dependent RNA helicase D | DDX5 | 69.149 | 2 | 53.686756 | yes | 9.819E-08 | 563.26196 | 5948.1865 |
| LMEEIMSEK | 9 | P17844 | Probable ATP-dependent RNA helicase D | DDX5 | 69.149 | 2 | 53.686756 | yes | 1.098E-07 | 563.26196 | 5948.1865 |
| WNLDELPK | 8 | P17844 | Probable ATP-dependent RNA helicase D | DDX5 | 69.149 | 2 | 53.686756 | yes | 2.182E-07 | 507.76639 | 15436.403 |
| EANQAINPK | 9 | P17844 | Probable ATP-dependent RNA helicase D | DDX5 | 69.149 | 2 | 53.686756 | yes | 2.142E-06 | 492.75909 | 14842.343 |
| DWVLNEFK | 8 | P17844 | Probable ATP-dependent RNA helicase D | DDX5 | 69.149 | 2 | 53.686756 | yes | 1.388E-07 | 525.76636 | 11925.01 |
| STCIYGGAPK | 10 | P17844 | Probable ATP-dependent RNA helicase D | DDX5 | 69.149 | 2 | 53.686756 | no | 6.58E-07 | 527.25531 | 10406.561 |
| QVSDLISVLR | 10 | P17844 | Probable ATP-dependent RNA helicase D | DDX5 | 69.149 | 2 | 53.686756 | yes | 9.722E-08 | 565.3324 | 9262.3369 |
| TIVFVETK | 8 | P17844 | Probable ATP-dependent RNA helicase D | DDX5 | 69.149 | 2 | 53.686756 | yes | 1.409E-07 | 468.77368 | 8488.9922 |

| | | | | | | | | | | | |
|---|---|---|---|---|---|---|---|---|---|---|---|
| TGTAYTFFTPNNIK | 14 | P17844 | Probable ATP-dependent RNA helicase D | DDX5 | 69.149 | 2 | 53.686756 | yes | 9.722E-08 | 787.89612 | 7023.3696 |
| IVDQIRPDR | 9 | P17844 | Probable ATP-dependent RNA helicase D | DDX5 | 69.149 | 2 | 53.686756 | no | 2.049E-07 | 556.31458 | 6814.7349 |
| QLAEDFLK | 8 | P17844 | Probable ATP-dependent RNA helicase D | DDX5 | 69.149 | 2 | 53.686756 | yes | 1.072E-07 | 482.26093 | 6607.7881 |
| KIVDQIRPDR | 10 | P17844 | Probable ATP-dependent RNA helicase D | DDX5 | 69.149 | 3 | 53.686756 | no | 3.117E-07 | 413.91046 | 2885.1648 |
| KIVDQIRPDR | 10 | P17844 | Probable ATP-dependent RNA helicase D | DDX5 | 69.149 | 3 | 53.686756 | no | 1.101E-07 | 620.36206 | 2885.1648 |
| KIVDQIRPDR | 10 | P17844 | Probable ATP-dependent RNA helicase D | DDX5 | 69.149 | 2 | 53.686756 | no | 3.117E-07 | 413.91046 | 2885.1648 |
| KIVDQIRPDR | 10 | P17844 | Probable ATP-dependent RNA helicase D | DDX5 | 69.149 | 2 | 53.686756 | no | 1.101E-07 | 620.36206 | 2885.1648 |
| RCDELTR | 7 | P17844 | Probable ATP-dependent RNA helicase D | DDX5 | 69.149 | 2 | 53.686756 | yes | 9.722E-08 | 475.22964 | 4158.6709 |
| GLDVEDVK | 8 | P17844 | Probable ATP-dependent RNA helicase D | DDX5 | 69.149 | 2 | 53.686756 | no | 1.842E-07 | 437.72946 | 3958.6682 |
| TLSYLLPAIVHINH | 20 | P17844 | Probable ATP-dependent RNA helicase D | DDX5 | 69.149 | 3 | 53.686756 | yes | 1.448E-07 | 787.77411 | 3802.4221 |
| RDGWPAMGIHGD | 13 | P17844 | Probable ATP-dependent RNA helicase D | DDX5 | 69.149 | 3 | 53.686756 | yes | 1.895E-07 | 480.56647 | 835.62048 |
| RDGWPAMGIHGD | 13 | P17844 | Probable ATP-dependent RNA helicase D | DDX5 | 69.149 | 3 | 53.686756 | yes | 1.837E-07 | 720.34607 | 835.62048 |
| RDGWPAMGIHGD | 13 | P17844 | Probable ATP-dependent RNA helicase D | DDX5 | 69.149 | 2 | 53.686756 | yes | 1.895E-07 | 480.56647 | 835.62048 |
| RDGWPAMGIHGD | 13 | P17844 | Probable ATP-dependent RNA helicase D | DDX5 | 69.149 | 2 | 53.686756 | yes | 1.837E-07 | 720.34607 | 835.62048 |
| KFGNPGEK | 8 | P17844 | Probable ATP-dependent RNA helicase D | DDX5 | 69.149 | 2 | 53.686756 | yes | 2.285E-07 | 438.73233 | 1570.9006 |
| RTAQEVETYR | 10 | P17844 | Probable ATP-dependent RNA helicase D | DDX5 | 69.149 | 2 | 53.686756 | yes | 1.093E-07 | 626.81769 | 720.84174 |
| ELAQQVQQVAAE | 15 | P17844 | Probable ATP-dependent RNA helicase D | DDX5 | 69.149 | 3 | 53.686756 | yes | 1.061E-07 | 598.29309 | 673.31824 |
| RTTYLVLDEADR | 12 | P17844 | Probable ATP-dependent RNA helicase D | DDX5 | 69.149 | 2 | 53.686756 | yes | 1.197E-07 | 726.37805 | 445.91214 |
| RGGFNTFR | 8 | P17844 | Probable ATP-dependent RNA helicase D | DDX5 | 69.149 | 2 | 53.686756 | yes | 1.521E-07 | 477.74884 | 401.74359 |
| KWNLDELPK | 9 | P17844 | Probable ATP-dependent RNA helicase D | DDX5 | 69.149 | 2 | 53.686756 | yes | 9.722E-08 | 571.81384 | 332.40753 |
| SGYSSDRDR | 9 | P17844 | Probable ATP-dependent RNA helicase D | DDX5 | 69.149 | 2 | 53.686756 | yes | 1.118E-07 | 542.73633 | 170.267 |
| DRGFGAPR | 8 | P17844 | Probable ATP-dependent RNA helicase D | DDX5 | 69.149 | 2 | 53.686756 | yes | 2.339E-07 | 438.22775 | 168.3886 |
| FVINYDYPNSSED | 17 | P17844 | Probable ATP-dependent RNA helicase D | DDX5 | 69.149 | 3 | 53.686756 | yes | 1.857E-07 | 711.32886 | 74.468628 |
| KIVDQIR | 7 | P17844 | Probable ATP-dependent RNA helicase D | DDX5 | 69.149 | 2 | 53.686756 | no | 6.726E-07 | 436.27164 | 68.536552 |
| SKEITVR | 7 | P17844 | Probable ATP-dependent RNA helicase D | DDX5 | 69.149 | 2 | 53.686756 | yes | 3.663E-05 | 416.74799 | 34.057682 |
| GYSSLLKR | 8 | P17844 | Probable ATP-dependent RNA helicase D | DDX5 | 69.149 | 2 | 53.686756 | yes | 3.637E-07 | 462.26907 | 26.946701 |
| TIVFVETKR | 9 | P17844 | Probable ATP-dependent RNA helicase D | DDX5 | 69.149 | 2 | 53.686756 | yes | 4.79E-07 | 546.82422 | 24.113899 |
| VFANAPDSACVIG | 15 | P17858 | ATP-dependent 6-phosphofructokinase, li | PFKL | 85.019 | 2 | 52.646847 | yes | 2.01E-07 | 781.40576 | 487.65417 |
| GQLESIVENIR | 11 | P17858 | ATP-dependent 6-phosphofructokinase, li | PFKL | 85.019 | 2 | 52.646847 | yes | 9.938E-08 | 629.34351 | 407.21381 |
| LPLMECVQMTK | 11 | P17858 | ATP-dependent 6-phosphofructokinase, li | PFKL | 85.019 | 2 | 52.646847 | yes | 9.722E-08 | 675.3349 | 344.94611 |
| FDEATQLR | 8 | P17858 | ATP-dependent 6-phosphofructokinase, li | PFKL | 85.019 | 2 | 52.646847 | yes | 9.975E-08 | 490.24582 | 147.66718 |
| AVAFSPVTELK | 11 | P17858 | ATP-dependent 6-phosphofructokinase, li | PFKL | 85.019 | 2 | 52.646847 | yes | 1.305E-07 | 581.32935 | 119.62 |
| AMLWLSEK | 8 | P17858 | ATP-dependent 6-phosphofructokinase, li | PFKL | 85.019 | 2 | 52.646847 | yes | 9.722E-08 | 489.25986 | 109.90767 |
| VNVEHMTEK | 9 | P17858 | ATP-dependent 6-phosphofructokinase, li | PFKL | 85.019 | 2 | 52.646847 | yes | 1.212E-07 | 543.76605 | 104.80723 |
| TGISHGHTVYVVH | 21 | P17858 | ATP-dependent 6-phosphofructokinase, li | PFKL | 85.019 | 3 | 52.646847 | yes | 4.081E-07 | 742.04303 | 51.880127 |
| TGISHGHTVYVVH | 21 | P17858 | ATP-dependent 6-phosphofructokinase, li | PFKL | 85.019 | 3 | 52.646847 | yes | 1.55E-07 | 556.78412 | 51.880127 |
| TGISHGHTVYVVH | 21 | P17858 | ATP-dependent 6-phosphofructokinase, li | PFKL | 85.019 | 4 | 52.646847 | yes | 4.081E-07 | 742.04303 | 51.880127 |
| TGISHGHTVYVVH | 21 | P17858 | ATP-dependent 6-phosphofructokinase, li | PFKL | 85.019 | 4 | 52.646847 | yes | 1.55E-07 | 556.78412 | 51.880127 |
| GGSFENNWNIYK | 12 | P17858 | ATP-dependent 6-phosphofructokinase, li | PFKL | 85.019 | 2 | 52.646847 | yes | 9.722E-08 | 714.83075 | 89.961357 |

| | | | | | | | | | | | |
|---|---|---|---|---|---|---|---|---|---|---|---|
| EQWWLSLR | 8 | P17858 | ATP-dependent 6-phosphofructokinase, li | PFKL | 85.019 | 2 | 52.646847 | yes | 1.047E-07 | 559.29309 | 65.790222 |
| LLAHQKPPK | 9 | P17858 | ATP-dependent 6-phosphofructokinase, li | PFKL | 85.019 | 2 | 52.646847 | yes | 2.616E-07 | 516.32166 | 63.063374 |
| IQVLVEPDHFK | 11 | P17931 | Galectin-3 OS=Homo sapiens OX=9606 ( | LGALS3 | 26.153 | 2 | 51.60235 | yes | 9.722E-08 | 662.86664 | 1313.7213 |
| GNDVAFHFNPR | 11 | P17931 | Galectin-3 OS=Homo sapiens OX=9606 ( | LGALS3 | 26.153 | 2 | 51.60235 | yes | 1.31E-07 | 637.30725 | 949.57831 |
| QSVFPFESGKPFK | 13 | P17931 | Galectin-3 OS=Homo sapiens OX=9606 ( | LGALS3 | 26.153 | 2 | 51.60235 | yes | 1.428E-07 | 749.39044 | 203.4248 |
| QSVFPFESGKPFK | 13 | P17931 | Galectin-3 OS=Homo sapiens OX=9606 ( | LGALS3 | 26.153 | 2 | 51.60235 | yes | 8.796E-07 | 499.92941 | 203.4248 |
| QSVFPFESGKPFK | 13 | P17931 | Galectin-3 OS=Homo sapiens OX=9606 ( | LGALS3 | 26.153 | 3 | 51.60235 | yes | 1.428E-07 | 749.39044 | 203.4248 |
| QSVFPFESGKPFK | 13 | P17931 | Galectin-3 OS=Homo sapiens OX=9606 ( | LGALS3 | 26.153 | 3 | 51.60235 | yes | 8.796E-07 | 499.92941 | 203.4248 |
| MLITILGTVKPNAN | 15 | P17931 | Galectin-3 OS=Homo sapiens OX=9606 ( | LGALS3 | 26.153 | 3 | 51.60235 | yes | 1.197E-07 | 547.6554 | 69.736671 |
| MLITILGTVKPNAN | 15 | P17931 | Galectin-3 OS=Homo sapiens OX=9606 ( | LGALS3 | 26.153 | 3 | 51.60235 | yes | 1.046E-07 | 820.97943 | 69.736671 |
| MLITILGTVKPNAN | 15 | P17931 | Galectin-3 OS=Homo sapiens OX=9606 ( | LGALS3 | 26.153 | 2 | 51.60235 | yes | 1.197E-07 | 547.6554 | 69.736671 |
| MLITILGTVKPNAN | 15 | P17931 | Galectin-3 OS=Homo sapiens OX=9606 ( | LGALS3 | 26.153 | 2 | 51.60235 | yes | 1.046E-07 | 820.97943 | 69.736671 |
| KLNEISK | 7 | P17931 | Galectin-3 OS=Homo sapiens OX=9606 ( | LGALS3 | 26.153 | 2 | 51.60235 | yes | 3.171E-07 | 416.25037 | 62.413284 |
| CEPCDAR | 7 | P17936 | Insulin-like growth factor-binding protein | IGFBP3 | 31.675 | 2 | 52.74288 | yes | 1.745E-07 | 454.17346 | 935.96252 |
| EMEDTLNHLK | 10 | P17936 | Insulin-like growth factor-binding protein | IGFBP3 | 31.675 | 2 | 52.74288 | yes | 1.382E-07 | 615.29517 | 245.18239 |
| EMEDTLNHLK | 10 | P17936 | Insulin-like growth factor-binding protein | IGFBP3 | 31.675 | 2 | 52.74288 | yes | 9.722E-08 | 623.2926 | 245.18239 |
| PLQALLDGR | 9 | P17936 | Insulin-like growth factor-binding protein | IGFBP3 | 31.675 | 2 | 52.74288 | yes | 2.206E-07 | 491.78763 | 357.15408 |
| LKPGDLVGVNK | 11 | P17980 | 26S proteasome regulatory subunit 6A OS | PSMC3 | 49.204 | 2 | 48.247673 | yes | 5.597E-07 | 570.34277 | 130.16518 |
| MSTEEIIQR | 9 | P17980 | 26S proteasome regulatory subunit 6A OS | PSMC3 | 49.204 | 2 | 48.247673 | yes | 1.117E-06 | 553.77917 | 99.70816 |
| VDILDPALLR | 10 | P17980 | 26S proteasome regulatory subunit 6A OS | PSMC3 | 49.204 | 2 | 48.247673 | yes | 6.573E-07 | 562.83734 | 93.144165 |
| GVLMYGPPGTGK | 12 | P17980 | 26S proteasome regulatory subunit 6A OS | PSMC3 | 49.204 | 2 | 48.247673 | yes | 4.94E-07 | 588.80774 | 77.728195 |
| GATELTHEDYMEC | 20 | P17980 | 26S proteasome regulatory subunit 6A OS | PSMC3 | 49.204 | 3 | 48.247673 | yes | 1.174E-07 | 745.35968 | 63.626656 |
| VTHELQAMK | 9 | P17980 | 26S proteasome regulatory subunit 6A OS | PSMC3 | 49.204 | 2 | 48.247673 | yes | 4.357E-07 | 528.77893 | 49.929165 |
| LLEVEHPAAK | 10 | P17987 | T-complex protein 1 subunit alpha OS=H | TCP1 | 60.344 | 2 | 54.691174 | yes | 1.033E-07 | 553.81384 | 375.84332 |
| VLCELADLQDK | 11 | P17987 | T-complex protein 1 subunit alpha OS=H | TCP1 | 60.344 | 2 | 54.691174 | yes | 9.722E-08 | 652.33173 | 360.38205 |
| IHPTSVISGYR | 11 | P17987 | T-complex protein 1 subunit alpha OS=H | TCP1 | 60.344 | 2 | 54.691174 | yes | 6.767E-07 | 615.33551 | 325.49567 |
| IACLDFSLQK | 10 | P17987 | T-complex protein 1 subunit alpha OS=H | TCP1 | 60.344 | 2 | 54.691174 | yes | 9.722E-08 | 597.81299 | 287.59088 |
| GANDFMCDEMER | 12 | P17987 | T-complex protein 1 subunit alpha OS=H | TCP1 | 60.344 | 2 | 54.691174 | yes | 9.935E-08 | 737.77344 | 284.20209 |
| YFVEAGAMAVR | 11 | P17987 | T-complex protein 1 subunit alpha OS=H | TCP1 | 60.344 | 2 | 54.691174 | yes | 1.676E-07 | 607.30536 | 268.25424 |
| ICDDELILIK | 10 | P17987 | T-complex protein 1 subunit alpha OS=H | TCP1 | 60.344 | 2 | 54.691174 | yes | 2.05E-07 | 616.33374 | 253.1902 |
| QAGVFEPTIVK | 11 | P17987 | T-complex protein 1 subunit alpha OS=H | TCP1 | 60.344 | 2 | 54.691174 | yes | 2.664E-07 | 594.83478 | 220.21118 |
| SLHDALCVVK | 10 | P17987 | T-complex protein 1 subunit alpha OS=H | TCP1 | 60.344 | 2 | 54.691174 | yes | 9.722E-08 | 571.30536 | 199.87852 |
| LGVQVVITDPEK | 12 | P17987 | T-complex protein 1 subunit alpha OS=H | TCP1 | 60.344 | 2 | 54.691174 | yes | 9.722E-08 | 649.37177 | 185.50203 |
| EQLAIAEFAR | 10 | P17987 | T-complex protein 1 subunit alpha OS=H | TCP1 | 60.344 | 2 | 54.691174 | yes | 1.026E-07 | 574.30896 | 149.41331 |
| DCLINAAK | 8 | P17987 | T-complex protein 1 subunit alpha OS=H | TCP1 | 60.344 | 2 | 54.691174 | yes | 8.085E-07 | 452.73148 | 127.9968 |
| YPVNSVNILK | 10 | P17987 | T-complex protein 1 subunit alpha OS=H | TCP1 | 60.344 | 2 | 54.691174 | yes | 3.148E-07 | 573.82953 | 126.12569 |
| SQNVMAAASIANI | 15 | P17987 | T-complex protein 1 subunit alpha OS=H | TCP1 | 60.344 | 2 | 54.691174 | yes | 9.809E-08 | 758.91125 | 83.928017 |
| AFHNEAQVNPER | 12 | P17987 | T-complex protein 1 subunit alpha OS=H | TCP1 | 60.344 | 2 | 54.691174 | yes | 9.819E-08 | 706.33929 | 75.325378 |
| FATEAAITILR | 11 | P17987 | T-complex protein 1 subunit alpha OS=H | TCP1 | 60.344 | 2 | 54.691174 | yes | 3.34E-06 | 603.34808 | 70.706635 |

| | | | | | | | | | | | |
|---|---|---|---|---|---|---|---|---|---|---|---|
| HGSYEDAVHSGAI | 15 | P17987 | T-complex protein 1 subunit alpha OS=H | TCP1 | 60.344 | 2 | 54.691174 | yes | 5.368E-07 | 786.33966 | 56.168324 |
| TSASIILR | 8 | P17987 | T-complex protein 1 subunit alpha OS=H | TCP1 | 60.344 | 2 | 54.691174 | yes | 7.926E-07 | 430.76364 | 42.674496 |
| SSLGPVGLDK | 10 | P17987 | T-complex protein 1 subunit alpha OS=H | TCP1 | 60.344 | 2 | 54.691174 | yes | 3.9E-06 | 486.77167 | 10.154914 |
| SGSWAAIYQDIR | 12 | P18031 | Tyrosine-protein phosphatase non-recepto | PTPN1 | 49.967 | 2 | 55.947327 | yes | 9.722E-08 | 683.84113 | 308.98651 |
| DVSPFDHSR | 9 | P18031 | Tyrosine-protein phosphatase non-recepto | PTPN1 | 49.967 | 2 | 55.947327 | yes | 2.286E-07 | 530.24634 | 177.22607 |
| EMIFEDTNLK | 10 | P18031 | Tyrosine-protein phosphatase non-recepto | PTPN1 | 49.967 | 2 | 55.947327 | yes | 1.102E-07 | 620.29993 | 175.99844 |
| MGLIQTADQLR | 11 | P18031 | Tyrosine-protein phosphatase non-recepto | PTPN1 | 49.967 | 2 | 55.947327 | yes | 9.722E-08 | 623.33466 | 127.53556 |
| FSYLAVIEGAK | 11 | P18031 | Tyrosine-protein phosphatase non-recepto | PTPN1 | 49.967 | 2 | 55.947327 | yes | 9.722E-08 | 599.32935 | 94.889679 |
| FIMGDSSVQDQWI | 13 | P18031 | Tyrosine-protein phosphatase non-recepto | PTPN1 | 49.967 | 2 | 55.947327 | yes | 9.722E-08 | 770.85864 | 87.851105 |
| LTLISEDIK | 9 | P18031 | Tyrosine-protein phosphatase non-recepto | PTPN1 | 49.967 | 2 | 55.947327 | yes | 1.097E-07 | 516.3028 | 73.574425 |
| ESGSLSPEHGPVV | 22 | P18031 | Tyrosine-protein phosphatase non-recepto | PTPN1 | 49.967 | 3 | 55.947327 | yes | 1.065E-07 | 744.69904 | 51.400146 |
| QLELENLTTQETR | 13 | P18031 | Tyrosine-protein phosphatase non-recepto | PTPN1 | 49.967 | 2 | 55.947327 | yes | 1.201E-07 | 787.90466 | 45.100883 |
| EVPLPAGIYNLDD | 15 | P18074 | General transcription and DNA repair fac | ERCC2 | 86.91 | 2 | 51.062817 | yes | 1.31E-07 | 828.9458 | 214.5761 |
| ENDFLTFDAMR | 11 | P18074 | General transcription and DNA repair fac | ERCC2 | 86.91 | 2 | 51.062817 | yes | 9.722E-08 | 679.80591 | 120.92462 |
| TAEHFLGFLR | 10 | P18074 | General transcription and DNA repair fac | ERCC2 | 86.91 | 2 | 51.062817 | yes | 1.063E-07 | 595.81946 | 110.69698 |
| NLCIHPEVTPLR | 12 | P18074 | General transcription and DNA repair fac | ERCC2 | 86.91 | 2 | 51.062817 | yes | 9.771E-08 | 724.88757 | 97.501549 |
| GAILLSVAR | 9 | P18074 | General transcription and DNA repair fac | ERCC2 | 86.91 | 2 | 51.062817 | yes | 4.001E-07 | 450.28726 | 68.779266 |
| TDYGLMVFADK | 11 | P18074 | General transcription and DNA repair fac | ERCC2 | 86.91 | 2 | 51.062817 | yes | 1.001E-07 | 630.30249 | 62.420418 |
| GHGVLEMPSGTGI | 13 | P18074 | General transcription and DNA repair fac | ERCC2 | 86.91 | 2 | 51.062817 | yes | 4.45E-07 | 635.31647 | 56.20327 |
| LPFLGLALSSR | 11 | P18074 | General transcription and DNA repair fac | ERCC2 | 86.91 | 2 | 51.062817 | yes | 1.93E-07 | 587.35315 | 40.989235 |
| VIEELRK | 7 | P18074 | General transcription and DNA repair fac | ERCC2 | 86.91 | 2 | 51.062817 | yes | 7.395E-07 | 443.77145 | 40.110584 |
| CHSLTASYVR | 10 | P18074 | General transcription and DNA repair fac | ERCC2 | 86.91 | 2 | 51.062817 | yes | 3.046E-06 | 597.29022 | 35.193134 |
| GFTIIIEPFDDR | 12 | P18074 | General transcription and DNA repair fac | ERCC2 | 86.91 | 2 | 51.062817 | yes | 1.735E-06 | 711.86682 | 14.52536 |
| VMLYPSR | 7 | P18077 | Large ribosomal subunit protein eL33 OS | RPL35A | 12.538 | 2 | 50.894516 | yes | 2.003E-07 | 433.23364 | 4672.2217 |
| VMLYPSR | 7 | P18077 | Large ribosomal subunit protein eL33 OS | RPL35A | 12.538 | 2 | 50.894516 | yes | 1.137E-07 | 441.23111 | 4672.2217 |
| IEGVYAR | 7 | P18077 | Large ribosomal subunit protein eL33 OS | RPL35A | 12.538 | 2 | 50.894516 | yes | 1.023E-07 | 404.22159 | 7006.3555 |
| EHTALLK | 7 | P18077 | Large ribosomal subunit protein eL33 OS | RPL35A | 12.538 | 2 | 50.894516 | yes | 2.9E-06 | 406.23724 | 5175.1602 |
| NNTVTPGGKPNK | 12 | P18077 | Large ribosomal subunit protein eL33 OS | RPL35A | 12.538 | 2 | 50.894516 | yes | 4.601E-07 | 613.828 | 177.79735 |
| NNTVTPGGKPNK | 12 | P18077 | Large ribosomal subunit protein eL33 OS | RPL35A | 12.538 | 2 | 50.894516 | yes | 4.607E-06 | 409.55444 | 177.79735 |
| NNTVTPGGKPNK | 12 | P18077 | Large ribosomal subunit protein eL33 OS | RPL35A | 12.538 | 3 | 50.894516 | yes | 4.601E-07 | 613.828 | 177.79735 |
| NNTVTPGGKPNK | 12 | P18077 | Large ribosomal subunit protein eL33 OS | RPL35A | 12.538 | 3 | 50.894516 | yes | 4.607E-06 | 409.55444 | 177.79735 |
| RCAYVYK | 7 | P18077 | Large ribosomal subunit protein eL33 OS | RPL35A | 12.538 | 2 | 50.894516 | yes | 9.722E-08 | 480.242 | 235.10638 |
| MLLVDELR | 8 | P18085 | ADP-ribosylation factor 4 OS=Homo sap | ARF4 | 20.511 | 2 | 54.282097 | yes | 9.722E-08 | 494.77844 | 407.54498 |
| NICFTVWDVGGQI | 14 | P18085 | ADP-ribosylation factor 4 OS=Homo sap | ARF4 | 20.511 | 2 | 54.282097 | yes | 9.722E-08 | 833.88574 | 195.96173 |
| QDLPNAMAISEMT | 15 | P18085 | ADP-ribosylation factor 4 OS=Homo sap | ARF4 | 20.511 | 2 | 54.282097 | yes | 9.722E-08 | 832.3869 | 174.4133 |
| IQEVADELQK | 10 | P18085 | ADP-ribosylation factor 4 OS=Homo sap | ARF4 | 20.511 | 2 | 54.282097 | yes | 1.917E-07 | 586.81152 | 89.87706 |
| DAVLLLFANK | 10 | P18085 | ADP-ribosylation factor 4 OS=Homo sap | ARF4 | 20.511 | 2 | 54.282097 | yes | 1.479E-06 | 552.3266 | 32.564781 |
| EVPAVPETLK | 10 | P18124 | Large ribosomal subunit protein uL30 OS | RPL7 | 29.226 | 2 | 53.44788 | yes | 2.991E-07 | 541.80823 | 29934.15 |
| AGNFYVPAEPK | 11 | P18124 | Large ribosomal subunit protein uL30 OS | RPL7 | 29.226 | 2 | 53.44788 | yes | 3.921E-07 | 596.80347 | 10652.976 |

| | | | | | | | | | | | |
|---|---|---|---|---|---|---|---|---|---|---|---|
| KVLQLLR | 7 | P18124 | Large ribosomal subunit protein uL30 OS | RPL7 | 29.226 | 2 | 53.44788 | yes | 1.926E-07 | 435.3002 | 5739.3774 |
| IVEPYIAWGYPNLI | 14 | P18124 | Large ribosomal subunit protein uL30 OS | RPL7 | 29.226 | 2 | 53.44788 | yes | 9.722E-08 | 831.94812 | 4616.3774 |
| SVNELIYK | 8 | P18124 | Large ribosomal subunit protein uL30 OS | RPL7 | 29.226 | 2 | 53.44788 | yes | 1.213E-07 | 483.26874 | 3952.2197 |
| IALTDNALIAR | 11 | P18124 | Large ribosomal subunit protein uL30 OS | RPL7 | 29.226 | 2 | 53.44788 | yes | 9.927E-08 | 585.8457 | 3659.5762 |
| KAGNFYVPAEPK | 12 | P18124 | Large ribosomal subunit protein uL30 OS | RPL7 | 29.226 | 2 | 53.44788 | yes | 1.025E-07 | 660.85095 | 1743.5437 |
| KAGNFYVPAEPK | 12 | P18124 | Large ribosomal subunit protein uL30 OS | RPL7 | 29.226 | 2 | 53.44788 | yes | 9.722E-08 | 440.90308 | 1743.5437 |
| KAGNFYVPAEPK | 12 | P18124 | Large ribosomal subunit protein uL30 OS | RPL7 | 29.226 | 3 | 53.44788 | yes | 1.025E-07 | 660.85095 | 1743.5437 |
| KAGNFYVPAEPK | 12 | P18124 | Large ribosomal subunit protein uL30 OS | RPL7 | 29.226 | 3 | 53.44788 | yes | 9.722E-08 | 440.90308 | 1743.5437 |
| RNFAELK | 7 | P18124 | Large ribosomal subunit protein uL30 OS | RPL7 | 29.226 | 2 | 53.44788 | yes | 3.263E-07 | 439.24814 | 2368.4456 |
| TTHFVEGGDAGNI | 13 | P18124 | Large ribosomal subunit protein uL30 OS | RPL7 | 29.226 | 2 | 53.44788 | yes | 1.469E-07 | 680.81567 | 2180.1157 |
| QIFNGTFVK | 9 | P18124 | Large ribosomal subunit protein uL30 OS | RPL7 | 29.226 | 2 | 53.44788 | yes | 2.431E-07 | 527.29004 | 2076.0815 |
| EVPAVPETLKK | 11 | P18124 | Large ribosomal subunit protein uL30 OS | RPL7 | 29.226 | 2 | 53.44788 | yes | 1.319E-07 | 605.85571 | 1655.1498 |
| ASINMLR | 7 | P18124 | Large ribosomal subunit protein uL30 OS | RPL7 | 29.226 | 2 | 53.44788 | yes | 1.043E-07 | 402.72345 | 1456.7972 |
| KTTHFVEGGDAGN | 14 | P18124 | Large ribosomal subunit protein uL30 OS | RPL7 | 29.226 | 2 | 53.44788 | yes | 9.722E-08 | 744.8631 | 192.37921 |
| KEVPAVPETLK | 11 | P18124 | Large ribosomal subunit protein uL30 OS | RPL7 | 29.226 | 2 | 53.44788 | yes | 9.722E-08 | 605.85571 | 155.06909 |
| TTHFVEGGDAGNI | 19 | P18124 | Large ribosomal subunit protein uL30 OS | RPL7 | 29.226 | 3 | 53.44788 | yes | 1.177E-07 | 705.99829 | 31.565186 |
| YGIICMEDLIHEIY | 18 | P18124 | Large ribosomal subunit protein uL30 OS | RPL7 | 29.226 | 3 | 53.44788 | yes | 5.661E-07 | 724.0235 | 24.136768 |
| LGHDPYR | 7 | P18583 | Protein SON OS=Homo sapiens OX=960 | SON | 263.83 | 2 | 54.27649 | yes | 3.231E-07 | 429.21686 | 329.2421 |
| FDSEPSAVALELPT | 15 | P18583 | Protein SON OS=Homo sapiens OX=960 | SON | 263.83 | 2 | 54.27649 | yes | 1.075E-07 | 816.41742 | 297.77954 |
| EADSVYGEWVPVI | 14 | P18583 | Protein SON OS=Homo sapiens OX=960 | SON | 263.83 | 2 | 54.27649 | yes | 9.722E-08 | 804.38306 | 242.55733 |
| IEEVLSGVLDTELF | 14 | P18583 | Protein SON OS=Homo sapiens OX=960 | SON | 263.83 | 2 | 54.27649 | yes | 1.438E-07 | 786.92761 | 193.9516 |
| AGIEGPLLASDVGI | 14 | P18583 | Protein SON OS=Homo sapiens OX=960 | SON | 263.83 | 2 | 54.27649 | yes | 1.164E-07 | 677.86987 | 166.90427 |
| SMMSPMAER | 9 | P18583 | Protein SON OS=Homo sapiens OX=960 | SON | 263.83 | 2 | 54.27649 | yes | 2.752E-07 | 520.22211 | 101.03041 |
| CVSVQTDPTDEIPT | 15 | P18583 | Protein SON OS=Homo sapiens OX=960 | SON | 263.83 | 2 | 54.27649 | yes | 9.779E-08 | 845.40363 | 100.67071 |
| SMMSSYSAADR | 11 | P18583 | Protein SON OS=Homo sapiens OX=960 | SON | 263.83 | 2 | 54.27649 | yes | 1.536E-07 | 603.24988 | 76.394905 |
| AQLLEIAK | 8 | P18583 | Protein SON OS=Homo sapiens OX=960 | SON | 263.83 | 2 | 54.27649 | yes | 2.877E-07 | 443.27383 | 63.833717 |
| EIQQELSSGR | 10 | P18583 | Protein SON OS=Homo sapiens OX=960 | SON | 263.83 | 2 | 54.27649 | yes | 9.938E-08 | 573.79114 | 61.439911 |
| SVESTSPEPSK | 11 | P18583 | Protein SON OS=Homo sapiens OX=960 | SON | 263.83 | 2 | 54.27649 | yes | 2.112E-07 | 574.27753 | 59.827099 |
| SMMSSYTADR | 10 | P18583 | Protein SON OS=Homo sapiens OX=960 | SON | 263.83 | 2 | 54.27649 | yes | 3.839E-07 | 574.7392 | 50.884964 |
| SAASPVVSSMPER | 13 | P18583 | Protein SON OS=Homo sapiens OX=960 | SON | 263.83 | 2 | 54.27649 | yes | 2.83E-07 | 659.32703 | 45.284962 |
| EFPVSSGSQHR | 11 | P18583 | Protein SON OS=Homo sapiens OX=960 | SON | 263.83 | 2 | 54.27649 | yes | 4.319E-07 | 615.79675 | 38.701439 |
| ESDQTLAALLSPK | 13 | P18583 | Protein SON OS=Homo sapiens OX=960 | SON | 263.83 | 2 | 54.27649 | yes | 2.066E-07 | 686.86957 | 38.442425 |
| GIEFTTASTLSLVN | 15 | P18583 | Protein SON OS=Homo sapiens OX=960 | SON | 263.83 | 2 | 54.27649 | yes | 2.19E-07 | 790.93018 | 26.135803 |
| DTEEPLPVK | 9 | P18583 | Protein SON OS=Homo sapiens OX=960 | SON | 263.83 | 2 | 54.27649 | yes | 1.154E-05 | 514.26892 | 25.04818 |
| TQIVYSDDVYK | 11 | P18615 | Negative elongation factor E OS=Homo s | NELFE | 43.24 | 2 | 54.82922 | yes | 9.921E-08 | 665.8299 | 1359.6349 |
| SLYESFVSSSDR | 12 | P18615 | Negative elongation factor E OS=Homo s | NELFE | 43.24 | 2 | 54.82922 | yes | 9.722E-08 | 688.82007 | 1246.8857 |
| SFDWGYEER | 9 | P18615 | Negative elongation factor E OS=Homo s | NELFE | 43.24 | 2 | 54.82922 | yes | 1.121E-07 | 594.75146 | 786.89349 |
| SISADDDLQESSR | 13 | P18615 | Negative elongation factor E OS=Homo s | NELFE | 43.24 | 2 | 54.82922 | yes | 9.722E-08 | 711.8208 | 738.48175 |
| KQPMLDAATGK | 11 | P18615 | Negative elongation factor E OS=Homo s | NELFE | 43.24 | 2 | 54.82922 | yes | 1.381E-07 | 580.31061 | 395.45624 |

| | | | | | | | | | | |
|---|---|---|---|---|---|---|---|---|---|---|
| NCAFVTYEK | 9 | P18615 | Negative elongation factor E OS=Homo s | NELFE | 43.24 | 2 | 54.82922 | yes | 1.277E-07 | 566.26062 | 368.80435 |
| KGNTLYVYGEDM | 18 | P18615 | Negative elongation factor E OS=Homo s | NELFE | 43.24 | 3 | 54.82922 | yes | 5.32E-07 | 691.02246 | 74.679092 |
| YSLDPENPTK | 10 | P18621 | Large ribosomal subunit protein uL22 OS | RPL17 | 21.398 | 2 | 49.739304 | yes | 1.046E-07 | 582.28259 | 33595.109 |
| EQIVPKPEEEVAQI | 14 | P18621 | Large ribosomal subunit protein uL22 OS | RPL17 | 21.398 | 3 | 49.739304 | yes | 6.198E-07 | 541.95776 | 14626.521 |
| EQIVPKPEEEVAQI | 14 | P18621 | Large ribosomal subunit protein uL22 OS | RPL17 | 21.398 | 3 | 49.739304 | yes | 5.306E-07 | 812.43304 | 14626.521 |
| EQIVPKPEEEVAQI | 14 | P18621 | Large ribosomal subunit protein uL22 OS | RPL17 | 21.398 | 2 | 49.739304 | yes | 6.198E-07 | 541.95776 | 14626.521 |
| EQIVPKPEEEVAQI | 14 | P18621 | Large ribosomal subunit protein uL22 OS | RPL17 | 21.398 | 2 | 49.739304 | yes | 5.306E-07 | 812.43304 | 14626.521 |
| QWGWTQGR | 8 | P18621 | Large ribosomal subunit protein uL22 OS | RPL17 | 21.398 | 2 | 49.739304 | yes | 1.797E-07 | 509.74631 | 23065.766 |
| SAEFLLHMLK | 10 | P18621 | Large ribosomal subunit protein uL22 OS | RPL17 | 21.398 | 2 | 49.739304 | yes | 1.006E-07 | 594.82593 | 10828.776 |
| SAEFLLHMLK | 10 | P18621 | Large ribosomal subunit protein uL22 OS | RPL17 | 21.398 | 2 | 49.739304 | yes | 1.317E-07 | 602.82336 | 10828.776 |
| NAESNAELK | 9 | P18621 | Large ribosomal subunit protein uL22 OS | RPL17 | 21.398 | 2 | 49.739304 | yes | 1.105E-07 | 488.24072 | 7239.5439 |
| SMVPVQVQLDVPY | 15 | P18754 | Regulator of chromosome condensation C | RCC1 | 44.97 | 2 | 51.07188 | yes | 9.722E-08 | 819.46838 | 196.56108 |
| LGLGEGAEEK | 10 | P18754 | Regulator of chromosome condensation C | RCC1 | 44.97 | 2 | 51.07188 | yes | 3.636E-07 | 501.75876 | 109.12414 |
| KKPALVSIPEDVV( | 27 | P18754 | Regulator of chromosome condensation C | RCC1 | 44.97 | 4 | 51.07188 | yes | 3.032E-06 | 716.88068 | 48.482998 |
| DTSVEGSEMVPGK | 13 | P18754 | Regulator of chromosome condensation C | RCC1 | 44.97 | 2 | 51.07188 | yes | 7.527E-07 | 668.30847 | 47.464706 |
| VVLSVSSGGQHTV | 17 | P18754 | Regulator of chromosome condensation C | RCC1 | 44.97 | 3 | 51.07188 | yes | 3.387E-06 | 575.00861 | 26.812435 |
| NVGSDIAVLR | 10 | P18850 | Cyclic AMP-dependent transcription facto | ATF6 | 74.585 | 2 | 53.27881 | yes | 3.819E-07 | 522.29584 | 225.05513 |
| PLLLPAAPK | 9 | P18850 | Cyclic AMP-dependent transcription facto | ATF6 | 74.585 | 2 | 53.27881 | yes | 9.935E-08 | 460.3024 | 130.72699 |
| QQPIISLQPAPTK | 13 | P18850 | Cyclic AMP-dependent transcription facto | ATF6 | 74.585 | 2 | 53.27881 | yes | 1.074E-07 | 710.91156 | 116.49393 |
| QLDEVVSENQR | 11 | P18850 | Cyclic AMP-dependent transcription facto | ATF6 | 74.585 | 2 | 53.27881 | yes | 9.722E-08 | 658.82568 | 90.894264 |
| MNPSVSPANQR | 11 | P18850 | Cyclic AMP-dependent transcription facto | ATF6 | 74.585 | 2 | 53.27881 | yes | 3.534E-06 | 600.79315 | 52.717342 |
| EAQDTSDGIIQK | 12 | P18850 | Cyclic AMP-dependent transcription facto | ATF6 | 74.585 | 2 | 53.27881 | yes | 5.35E-06 | 652.82007 | 19.533459 |
| TISVVLQLEK | 10 | P18887 | DNA repair protein XRCC1 OS=Homo sa | XRCC1 | 69.498 | 2 | 52.0416 | yes | 1.559E-07 | 565.34497 | 1189.6909 |
| TPATAPVPAR | 10 | P18887 | DNA repair protein XRCC1 OS=Homo sa | XRCC1 | 69.498 | 2 | 52.0416 | yes | 2.249E-07 | 490.77982 | 1169.4127 |
| LLPHQLYGVVPQA | 13 | P18887 | DNA repair protein XRCC1 OS=Homo sa | XRCC1 | 69.498 | 2 | 52.0416 | yes | 1.958E-07 | 717.90881 | 992.21753 |
| DSPFGLSFVR | 10 | P18887 | DNA repair protein XRCC1 OS=Homo sa | XRCC1 | 69.498 | 2 | 52.0416 | yes | 9.722E-08 | 562.79041 | 869.95007 |
| EWVLDCHR | 8 | P18887 | DNA repair protein XRCC1 OS=Homo sa | XRCC1 | 69.498 | 2 | 52.0416 | yes | 9.722E-08 | 557.75854 | 588.50214 |
| HVVSCSSQDSTHC | 19 | P18887 | DNA repair protein XRCC1 OS=Homo sa | XRCC1 | 69.498 | 3 | 52.0416 | yes | 1.194E-07 | 724.6651 | 489.85001 |
| DSTHLICAFANTPK | 14 | P18887 | DNA repair protein XRCC1 OS=Homo sa | XRCC1 | 69.498 | 2 | 52.0416 | yes | 3.862E-07 | 787.88519 | 239.51573 |
| DSTHLICAFANTPK | 14 | P18887 | DNA repair protein XRCC1 OS=Homo sa | XRCC1 | 69.498 | 2 | 52.0416 | yes | 7.059E-07 | 525.59259 | 239.51573 |
| DSTHLICAFANTPK | 14 | P18887 | DNA repair protein XRCC1 OS=Homo sa | XRCC1 | 69.498 | 3 | 52.0416 | yes | 3.862E-07 | 787.88519 | 239.51573 |
| DSTHLICAFANTPK | 14 | P18887 | DNA repair protein XRCC1 OS=Homo sa | XRCC1 | 69.498 | 3 | 52.0416 | yes | 7.059E-07 | 525.59259 | 239.51573 |
| WIYSCNEK | 8 | P18887 | DNA repair protein XRCC1 OS=Homo sa | XRCC1 | 69.498 | 2 | 52.0416 | yes | 2.925E-07 | 550.2475 | 446.44751 |
| YRPDWTR | 7 | P18887 | DNA repair protein XRCC1 OS=Homo sa | XRCC1 | 69.498 | 2 | 52.0416 | yes | 9.723E-08 | 497.24869 | 352.12677 |
| HFFLYGEFPGDER | 13 | P18887 | DNA repair protein XRCC1 OS=Homo sa | XRCC1 | 69.498 | 2 | 52.0416 | yes | 9.722E-08 | 807.3728 | 138.37306 |
| HFFLYGEFPGDER | 13 | P18887 | DNA repair protein XRCC1 OS=Homo sa | XRCC1 | 69.498 | 2 | 52.0416 | yes | 4.605E-07 | 538.58429 | 138.37306 |
| HFFLYGEFPGDER | 13 | P18887 | DNA repair protein XRCC1 OS=Homo sa | XRCC1 | 69.498 | 3 | 52.0416 | yes | 9.722E-08 | 807.3728 | 138.37306 |
| HFFLYGEFPGDER | 13 | P18887 | DNA repair protein XRCC1 OS=Homo sa | XRCC1 | 69.498 | 3 | 52.0416 | yes | 4.605E-07 | 538.58429 | 138.37306 |
| KTPSKPPAQLSPSV | 16 | P18887 | DNA repair protein XRCC1 OS=Homo sa | XRCC1 | 69.498 | 3 | 52.0416 | yes | 1.298E-07 | 554.65765 | 75.510147 |

| | | | | | | | | | | | |
|---|---|---|---|---|---|---|---|---|---|---|---|
| KTPSKPPAQLSPS\ | 16 | P18887 | DNA repair protein XRCC1 OS=Homo sa | XRCC1 | 69.498 | 3 | 52.0416 | yes | 2.249E-06 | 831.48285 | 75.510147 |
| KTPSKPPAQLSPS\ | 16 | P18887 | DNA repair protein XRCC1 OS=Homo sa | XRCC1 | 69.498 | 2 | 52.0416 | yes | 1.298E-07 | 554.65765 | 75.510147 |
| KTPSKPPAQLSPS\ | 16 | P18887 | DNA repair protein XRCC1 OS=Homo sa | XRCC1 | 69.498 | 2 | 52.0416 | yes | 2.249E-06 | 831.48285 | 75.510147 |
| KEWVLDCHR | 9 | P18887 | DNA repair protein XRCC1 OS=Homo sa | XRCC1 | 69.498 | 2 | 52.0416 | yes | 1.15E-07 | 621.80603 | 96.149399 |
| IVCSQPYSK | 9 | P18887 | DNA repair protein XRCC1 OS=Homo sa | XRCC1 | 69.498 | 2 | 52.0416 | yes | 2.02E-07 | 541.271 | 22.752361 |
| GFGFVDFNSEEDA | 14 | P19338 | Nucleolin OS=Homo sapiens OX=9606 ( | NCL | 76.615 | 2 | 51.604534 | yes | 9.722E-08 | 781.34393 | 29295.164 |
| GLSEDTTEETLK | 12 | P19338 | Nucleolin OS=Homo sapiens OX=9606 ( | NCL | 76.615 | 2 | 51.604534 | yes | 9.722E-08 | 661.8197 | 17127.432 |
| NDLAVVDVR | 9 | P19338 | Nucleolin OS=Homo sapiens OX=9606 ( | NCL | 76.615 | 2 | 51.604534 | yes | 1.894E-07 | 500.77472 | 14511.825 |
| FGYVDFESAEDLE | 14 | P19338 | Nucleolin OS=Homo sapiens OX=9606 ( | NCL | 76.615 | 2 | 51.604534 | yes | 1.897E-07 | 824.8725 | 14100.283 |
| SISLYYTGEK | 10 | P19338 | Nucleolin OS=Homo sapiens OX=9606 ( | NCL | 76.615 | 2 | 51.604534 | yes | 1.138E-07 | 580.79535 | 11229.516 |
| GYAFIEFASFEDAI | 14 | P19338 | Nucleolin OS=Homo sapiens OX=9606 ( | NCL | 76.615 | 2 | 51.604534 | yes | 9.722E-08 | 797.87482 | 10767.807 |
| TGISDVFAK | 9 | P19338 | Nucleolin OS=Homo sapiens OX=9606 ( | NCL | 76.615 | 2 | 51.604534 | yes | 1.257E-07 | 469.25311 | 10619.068 |
| LELQGPR | 7 | P19338 | Nucleolin OS=Homo sapiens OX=9606 ( | NCL | 76.615 | 2 | 51.604534 | yes | 1.182E-07 | 406.73486 | 7604.6563 |
| EVFEDAAEIR | 10 | P19338 | Nucleolin OS=Homo sapiens OX=9606 ( | NCL | 76.615 | 2 | 51.604534 | yes | 1.997E-07 | 589.78802 | 7029.4458 |
| VTLDWAKPK | 9 | P19338 | Nucleolin OS=Homo sapiens OX=9606 ( | NCL | 76.615 | 2 | 51.604534 | yes | 1.075E-07 | 529.30566 | 5791.4883 |
| GIAYIEFK | 8 | P19338 | Nucleolin OS=Homo sapiens OX=9606 ( | NCL | 76.615 | 2 | 51.604534 | yes | 3.022E-07 | 470.76056 | 5294.5898 |
| EAMEDGEIDGNK | 12 | P19338 | Nucleolin OS=Homo sapiens OX=9606 ( | NCL | 76.615 | 2 | 51.604534 | yes | 2.712E-06 | 654.27466 | 2514.0298 |
| EAMEDGEIDGNK | 12 | P19338 | Nucleolin OS=Homo sapiens OX=9606 ( | NCL | 76.615 | 2 | 51.604534 | yes | 9.722E-08 | 662.27209 | 2514.0298 |
| ALELTGLK | 8 | P19338 | Nucleolin OS=Homo sapiens OX=9606 ( | NCL | 76.615 | 2 | 51.604534 | yes | 1.382E-07 | 422.76056 | 3130.0002 |
| VTQDELK | 7 | P19338 | Nucleolin OS=Homo sapiens OX=9606 ( | NCL | 76.615 | 2 | 51.604534 | yes | 1.596E-07 | 416.72418 | 3108.4956 |
| KVAVATPAK | 9 | P19338 | Nucleolin OS=Homo sapiens OX=9606 ( | NCL | 76.615 | 2 | 51.604534 | yes | 1.119E-07 | 442.78183 | 1008.4614 |
| KVVVSPTK | 8 | P19338 | Nucleolin OS=Homo sapiens OX=9606 ( | NCL | 76.615 | 2 | 51.604534 | yes | 1.071E-07 | 429.27637 | 859.71875 |
| ESFDGSVR | 8 | P19338 | Nucleolin OS=Homo sapiens OX=9606 ( | NCL | 76.615 | 2 | 51.604534 | yes | 2.338E-07 | 448.70905 | 502.13721 |
| EAMEDGEIDGNK\ | 19 | P19338 | Nucleolin OS=Homo sapiens OX=9606 ( | NCL | 76.615 | 3 | 51.604534 | yes | 2.258E-07 | 707.66504 | 329.36075 |
| QGTEIDGR | 8 | P19338 | Nucleolin OS=Homo sapiens OX=9606 ( | NCL | 76.615 | 2 | 51.604534 | yes | 1.197E-07 | 438.21451 | 227.23552 |
| EALNSCNK | 8 | P19338 | Nucleolin OS=Homo sapiens OX=9606 ( | NCL | 76.615 | 2 | 51.604534 | yes | 1.898E-07 | 468.21619 | 171.71095 |
| KFGYVDFESAEDL | 15 | P19338 | Nucleolin OS=Homo sapiens OX=9606 ( | NCL | 76.615 | 3 | 51.604534 | yes | 1.422E-07 | 592.9491 | 138.0778 |
| EAMEDGEIDGNK\ | 21 | P19338 | Nucleolin OS=Homo sapiens OX=9606 ( | NCL | 76.615 | 3 | 51.604534 | yes | 1.881E-07 | 782.71423 | 105.28895 |
| ALVATPGKK | 9 | P19338 | Nucleolin OS=Homo sapiens OX=9606 ( | NCL | 76.615 | 2 | 51.604534 | yes | 4.464E-07 | 442.78183 | 65.496864 |
| GQNQDYR | 7 | P19338 | Nucleolin OS=Homo sapiens OX=9606 ( | NCL | 76.615 | 2 | 51.604534 | yes | 1.565E-07 | 440.69904 | 26.637142 |
| MISGMYLGEIVR | 12 | P19367 | Hexokinase-1 OS=Homo sapiens OX=96( | HK1 | 102.486 | 2 | 45.39144 | yes | 1.864E-07 | 684.85437 | 56.894119 |
| PYANQPTVR | 9 | P19387 | DNA-directed RNA polymerase II subuni | POLR2C | 31.442 | 2 | 43.552364 | yes | 8.048E-07 | 523.2749 | 143.10066 |
| ITELTDENVK | 10 | P19387 | DNA-directed RNA polymerase II subuni | POLR2C | 31.442 | 2 | 43.552364 | yes | 2.542E-07 | 581.30353 | 60.901836 |
| FIIENTDLAVANSII | 15 | P19387 | DNA-directed RNA polymerase II subuni | POLR2C | 31.442 | 2 | 43.552364 | yes | 1.024E-06 | 838.45435 | 50.821239 |
| ALIVVQQGMTPSA | 14 | P19388 | DNA-directed RNA polymerases I, II, and | POLR2E | 24.552 | 2 | 51.757557 | yes | 9.722E-08 | 721.9054 | 278.69821 |
| TIMQLCHDR | 9 | P19388 | DNA-directed RNA polymerases I, II, and | POLR2E | 24.552 | 2 | 51.757557 | yes | 9.722E-08 | 587.27881 | 225.61705 |
| EEVTELLAR | 9 | P19388 | DNA-directed RNA polymerases I, II, and | POLR2E | 24.552 | 2 | 51.757557 | yes | 1.31E-07 | 530.28766 | 147.72641 |
| MQEENITR | 8 | P19388 | DNA-directed RNA polymerases I, II, and | POLR2E | 24.552 | 2 | 51.757557 | yes | 6.278E-07 | 510.74258 | 133.97507 |
| QSLVDMAPK | 9 | P19388 | DNA-directed RNA polymerases I, II, and | POLR2E | 24.552 | 2 | 51.757557 | yes | 6.209E-07 | 494.76022 | 124.25614 |

| | | | | | | | | | | | |
|---|---|---|---|---|---|---|---|---|---|---|---|
| IIRPSETAGR | 10 | P19388 | DNA-directed RNA polymerases I, II, and | POLR2E | 24.552 | 2 | 51.757557 | yes | 5.465E-07 | 550.31458 | 53.029705 |
| AAAVLPVLDLAQI | 13 | P19404 | NADH dehydrogenase [ubiquinone] flavo | NDUFV2 | 27.392 | 2 | 32.397774 | yes | 1.53E-06 | 668.901 | 54.317524 |
| LGLTATLVR | 9 | P19447 | General transcription and DNA repair fac | ERCC3 | 89.278 | 2 | 51.764156 | yes | 1.983E-07 | 472.30038 | 567.71863 |
| SLVGVTAACTVR | 12 | P19447 | General transcription and DNA repair fac | ERCC3 | 89.278 | 2 | 51.764156 | yes | 2.716E-07 | 617.33466 | 310.84225 |
| MQILQNFK | 8 | P19447 | General transcription and DNA repair fac | ERCC3 | 89.278 | 2 | 51.764156 | yes | 4.696E-07 | 511.27859 | 307.89969 |
| FLVDQGYSFK | 10 | P19447 | General transcription and DNA repair fac | ERCC3 | 89.278 | 2 | 51.764156 | yes | 1.43E-06 | 602.30585 | 195.02771 |
| VLAATDLDAEEEV | 21 | P19447 | General transcription and DNA repair fac | ERCC3 | 89.278 | 3 | 51.764156 | yes | 1.293E-07 | 726.69019 | 187.62889 |
| MWSTIDDSQICR | 12 | P19447 | General transcription and DNA repair fac | ERCC3 | 89.278 | 2 | 51.764156 | yes | 1.35E-07 | 756.33447 | 158.52905 |
| SGVIVLPCGAGK | 12 | P19447 | General transcription and DNA repair fac | ERCC3 | 89.278 | 2 | 51.764156 | yes | 2.258E-07 | 579.32098 | 137.46365 |
| LAGMEEEDLAFST | 14 | P19447 | General transcription and DNA repair fac | ERCC3 | 89.278 | 2 | 51.764156 | yes | 2.703E-07 | 770.86359 | 133.1438 |
| IVDLNFLIGPK | 11 | P19447 | General transcription and DNA repair fac | ERCC3 | 89.278 | 2 | 51.764156 | yes | 1.524E-07 | 614.86865 | 121.82111 |
| EEQQQLLQK | 9 | P19447 | General transcription and DNA repair fac | ERCC3 | 89.278 | 2 | 51.764156 | yes | 3.058E-06 | 572.30383 | 113.17909 |
| LCTVSYGK | 8 | P19447 | General transcription and DNA repair fac | ERCC3 | 89.278 | 2 | 51.764156 | yes | 2.265E-07 | 464.23386 | 89.215164 |
| VLTIVQAHCK | 10 | P19447 | General transcription and DNA repair fac | ERCC3 | 89.278 | 2 | 51.764156 | yes | 1.375E-07 | 584.82898 | 74.922974 |
| ILLYTMNPNK | 10 | P19447 | General transcription and DNA repair fac | ERCC3 | 89.278 | 2 | 51.764156 | yes | 1.225E-07 | 603.83118 | 67.586823 |
| IIVFADNVFALK | 12 | P19447 | General transcription and DNA repair fac | ERCC3 | 89.278 | 2 | 51.764156 | yes | 3.086E-06 | 675.39502 | 61.8013 |
| ACQFLIK | 7 | P19447 | General transcription and DNA repair fac | ERCC3 | 89.278 | 2 | 51.764156 | yes | 2.782E-07 | 440.24149 | 56.053043 |
| CLVLGNSAVSVEQ | 15 | P19447 | General transcription and DNA repair fac | ERCC3 | 89.278 | 2 | 51.764156 | yes | 9.722E-08 | 845.43506 | 50.631832 |
| VGDTSFDLPEANV | 24 | P19447 | General transcription and DNA repair fac | ERCC3 | 89.278 | 3 | 51.764156 | yes | 7.737E-07 | 833.75397 | 44.291496 |
| DKPIGCSVAISTYS | 21 | P19447 | General transcription and DNA repair fac | ERCC3 | 89.278 | 3 | 51.764156 | yes | 3.344E-07 | 756.04504 | 42.524521 |
| CIHLEYPLLAEYDI | 15 | P19447 | General transcription and DNA repair fac | ERCC3 | 89.278 | 3 | 51.764156 | yes | 5.345E-07 | 646.98547 | 23.211657 |
| YFVESCHPDVIQH | 21 | P19447 | General transcription and DNA repair fac | ERCC3 | 89.278 | 4 | 51.764156 | yes | 2.682E-06 | 642.07843 | 18.057617 |
| LQVALGELR | 9 | P19474 | E3 ubiquitin-protein ligase TRIM21 OS= | TRIM21 | 54.17 | 2 | 51.56556 | yes | 1.623E-07 | 499.80328 | 21847.832 |
| NFLVEEEQR | 9 | P19474 | E3 ubiquitin-protein ligase TRIM21 OS= | TRIM21 | 54.17 | 2 | 51.56556 | yes | 9.85E-08 | 582.28821 | 15353.715 |
| LHLFCEK | 7 | P19474 | E3 ubiquitin-protein ligase TRIM21 OS= | TRIM21 | 54.17 | 2 | 51.56556 | yes | 1.346E-07 | 473.74438 | 8188.0737 |
| ALCWVCAQSR | 10 | P19474 | E3 ubiquitin-protein ligase TRIM21 OS= | TRIM21 | 54.17 | 2 | 51.56556 | yes | 9.722E-08 | 625.79205 | 7583.0127 |
| IHAEFVQQK | 9 | P19474 | E3 ubiquitin-protein ligase TRIM21 OS= | TRIM21 | 54.17 | 2 | 51.56556 | yes | 2.466E-07 | 550.2984 | 7377.522 |
| SVCHVPGLK | 9 | P19474 | E3 ubiquitin-protein ligase TRIM21 OS= | TRIM21 | 54.17 | 2 | 51.56556 | yes | 3.118E-07 | 498.7684 | 6249.7612 |
| EAWDLGVCR | 9 | P19474 | E3 ubiquitin-protein ligase TRIM21 OS= | TRIM21 | 54.17 | 2 | 51.56556 | yes | 1.353E-07 | 553.25836 | 4941.2368 |
| FDSYPMVLGAQHI | 17 | P19474 | E3 ubiquitin-protein ligase TRIM21 OS= | TRIM21 | 54.17 | 3 | 51.56556 | yes | 2.287E-07 | 640.97357 | 4751.2344 |
| HYWEVDVTGK | 10 | P19474 | E3 ubiquitin-protein ligase TRIM21 OS= | TRIM21 | 54.17 | 2 | 51.56556 | yes | 1.711E-07 | 617.29858 | 4328.3091 |
| QLANMVNNLK | 10 | P19474 | E3 ubiquitin-protein ligase TRIM21 OS= | TRIM21 | 54.17 | 2 | 51.56556 | yes | 1.029E-07 | 572.81079 | 2882.7473 |
| GHFLLSSK | 8 | P19474 | E3 ubiquitin-protein ligase TRIM21 OS= | TRIM21 | 54.17 | 2 | 51.56556 | yes | 1.467E-07 | 444.75052 | 2782.5164 |
| DLDITSPELR | 10 | P19474 | E3 ubiquitin-protein ligase TRIM21 OS= | TRIM21 | 54.17 | 2 | 51.56556 | yes | 1.002E-07 | 579.80371 | 2385.0166 |
| GGGSVCPVCR | 10 | P19474 | E3 ubiquitin-protein ligase TRIM21 OS= | TRIM21 | 54.17 | 2 | 51.56556 | yes | 4.097E-07 | 524.73676 | 2273.5007 |
| QLQELEKDER | 10 | P19474 | E3 ubiquitin-protein ligase TRIM21 OS= | TRIM21 | 54.17 | 2 | 51.56556 | yes | 1.023E-07 | 644.33063 | 876.70288 |
| QLQELEKDER | 10 | P19474 | E3 ubiquitin-protein ligase TRIM21 OS= | TRIM21 | 54.17 | 2 | 51.56556 | yes | 1.362E-07 | 429.8895 | 876.70288 |
| QLQELEKDER | 10 | P19474 | E3 ubiquitin-protein ligase TRIM21 OS= | TRIM21 | 54.17 | 3 | 51.56556 | yes | 1.023E-07 | 644.33063 | 876.70288 |
| QLQELEKDER | 10 | P19474 | E3 ubiquitin-protein ligase TRIM21 OS= | TRIM21 | 54.17 | 3 | 51.56556 | yes | 1.362E-07 | 429.8895 | 876.70288 |

| | | | | | | | | | | | |
|---|---|---|---|---|---|---|---|---|---|---|---|
| KGHFLLSSK | 9 | P19474 | E3 ubiquitin-protein ligase TRIM21 OS= | TRIM21 | 54.17 | 2 | 51.56556 | yes | 2.193E-07 | 508.798 | 1036.3424 |
| DHAMVPLEEAAQ | 17 | P19474 | E3 ubiquitin-protein ligase TRIM21 OS= | TRIM21 | 54.17 | 3 | 51.56556 | yes | 4.093E-07 | 663.30707 | 1002.8914 |
| LEVEIAIK | 8 | P19474 | E3 ubiquitin-protein ligase TRIM21 OS= | TRIM21 | 54.17 | 2 | 51.56556 | yes | 1.693E-07 | 457.78149 | 943.047 |
| EISQEAR | 7 | P19474 | E3 ubiquitin-protein ligase TRIM21 OS= | TRIM21 | 54.17 | 2 | 51.56556 | yes | 9.935E-08 | 416.71161 | 931.11072 |
| PFFSPGFNDGGK | 12 | P19474 | E3 ubiquitin-protein ligase TRIM21 OS= | TRIM21 | 54.17 | 2 | 51.56556 | yes | 1.624E-07 | 635.29858 | 864.41705 |
| LGDTQQSIPGNEEI | 14 | P19474 | E3 ubiquitin-protein ligase TRIM21 OS= | TRIM21 | 54.17 | 2 | 51.56556 | yes | 5.176E-07 | 772.37097 | 640.11621 |
| QLQELEK | 7 | P19474 | E3 ubiquitin-protein ligase TRIM21 OS= | TRIM21 | 54.17 | 2 | 51.56556 | yes | 1.025E-07 | 444.24527 | 569.11981 |
| TCAVHITLDPDTAI | 24 | P19474 | E3 ubiquitin-protein ligase TRIM21 OS= | TRIM21 | 54.17 | 4 | 51.56556 | yes | 1.561E-07 | 699.10516 | 333.70834 |
| LAQQSQALQELISI | 18 | P19474 | E3 ubiquitin-protein ligase TRIM21 OS= | TRIM21 | 54.17 | 3 | 51.56556 | yes | 1.08E-07 | 700.04657 | 233.16496 |
| FAAHISPAQGSPKI | 23 | P19484 | Transcription factor EB OS=Homo sapier | TFEB | 52.865 | 4 | 45.063095 | yes | 4.448E-07 | 560.80383 | 64.774071 |
| ELGMLIPK | 8 | P19484 | Transcription factor EB OS=Homo sapier | TFEB | 52.865 | 2 | 45.063095 | yes | 1.976E-07 | 450.7648 | 23.142159 |
| IGDFGLVTSLK | 11 | P19525 | Interferon-induced, double-stranded RNA | EIF2AK2 | 62.095 | 2 | 53.066578 | yes | 1.503E-07 | 575.32935 | 6028.002 |
| LAYLQILSEETSVK | 14 | P19525 | Interferon-induced, double-stranded RNA | EIF2AK2 | 62.095 | 2 | 53.066578 | yes | 9.722E-08 | 797.44037 | 4952.1846 |
| YMSPEQISSQDYG | 14 | P19525 | Interferon-induced, double-stranded RNA | EIF2AK2 | 62.095 | 2 | 53.066578 | yes | 2.715E-07 | 816.86414 | 2189.8955 |
| YMSPEQISSQDYG | 14 | P19525 | Interferon-induced, double-stranded RNA | EIF2AK2 | 62.095 | 2 | 53.066578 | yes | 1.405E-07 | 824.86157 | 2189.8955 |
| GVDYIHSK | 8 | P19525 | Interferon-induced, double-stranded RNA | EIF2AK2 | 62.095 | 2 | 53.066578 | yes | 1.703E-07 | 459.73761 | 3272.9043 |
| CLFIQMEFCDK | 11 | P19525 | Interferon-induced, double-stranded RNA | EIF2AK2 | 62.095 | 2 | 53.066578 | yes | 1.076E-07 | 745.82745 | 1454.3348 |
| CLFIQMEFCDK | 11 | P19525 | Interferon-induced, double-stranded RNA | EIF2AK2 | 62.095 | 2 | 53.066578 | yes | 4.707E-07 | 753.82495 | 1454.3348 |
| DLKPSNIFLVDTK | 13 | P19525 | Interferon-induced, double-stranded RNA | EIF2AK2 | 62.095 | 2 | 53.066578 | yes | 1.945E-07 | 745.41669 | 2673.877 |
| GTLEQWIEK | 9 | P19525 | Interferon-induced, double-stranded RNA | EIF2AK2 | 62.095 | 2 | 53.066578 | yes | 9.934E-08 | 552.29022 | 2336.2434 |
| FTFQVIIDGR | 10 | P19525 | Interferon-induced, double-stranded RNA | EIF2AK2 | 62.095 | 2 | 53.066578 | yes | 4.582E-07 | 598.32715 | 1821.0262 |
| EIELIGSGGFGQVF | 15 | P19525 | Interferon-induced, double-stranded RNA | EIF2AK2 | 62.095 | 2 | 53.066578 | yes | 9.722E-08 | 790.91956 | 1626.5789 |
| EYSIGTGSTK | 10 | P19525 | Interferon-induced, double-stranded RNA | EIF2AK2 | 62.095 | 2 | 53.066578 | yes | 2.301E-07 | 521.75623 | 1410.301 |
| DGIISDIFDKK | 11 | P19525 | Interferon-induced, double-stranded RNA | EIF2AK2 | 62.095 | 2 | 53.066578 | yes | 2.04E-07 | 625.83496 | 1116.1309 |
| KPEDRPNTSEILR | 13 | P19525 | Interferon-induced, double-stranded RNA | EIF2AK2 | 62.095 | 3 | 53.066578 | yes | 3.404E-07 | 518.94598 | 300.18665 |
| KPEDRPNTSEILR | 13 | P19525 | Interferon-induced, double-stranded RNA | EIF2AK2 | 62.095 | 3 | 53.066578 | yes | 2.084E-07 | 777.91534 | 300.18665 |
| KPEDRPNTSEILR | 13 | P19525 | Interferon-induced, double-stranded RNA | EIF2AK2 | 62.095 | 2 | 53.066578 | yes | 3.404E-07 | 518.94598 | 300.18665 |
| KPEDRPNTSEILR | 13 | P19525 | Interferon-induced, double-stranded RNA | EIF2AK2 | 62.095 | 2 | 53.066578 | yes | 2.084E-07 | 777.91534 | 300.18665 |
| YQELPNSGPPHDR | 13 | P19525 | Interferon-induced, double-stranded RNA | EIF2AK2 | 62.095 | 2 | 53.066578 | yes | 3.623E-07 | 755.35767 | 214.65927 |
| YQELPNSGPPHDR | 13 | P19525 | Interferon-induced, double-stranded RNA | EIF2AK2 | 62.095 | 2 | 53.066578 | yes | 9.722E-08 | 503.90753 | 214.65927 |
| YQELPNSGPPHDR | 13 | P19525 | Interferon-induced, double-stranded RNA | EIF2AK2 | 62.095 | 3 | 53.066578 | yes | 3.623E-07 | 755.35767 | 214.65927 |
| YQELPNSGPPHDR | 13 | P19525 | Interferon-induced, double-stranded RNA | EIF2AK2 | 62.095 | 3 | 53.066578 | yes | 9.722E-08 | 503.90753 | 214.65927 |
| DGIISDIFDK | 10 | P19525 | Interferon-induced, double-stranded RNA | EIF2AK2 | 62.095 | 2 | 53.066578 | yes | 1.464E-07 | 561.78748 | 359.48669 |
| FDLPDMK | 7 | P19525 | Interferon-induced, double-stranded RNA | EIF2AK2 | 62.095 | 2 | 53.066578 | yes | 9.46E-07 | 433.20984 | 303.50961 |
| PNTSEILR | 8 | P19525 | Interferon-induced, double-stranded RNA | EIF2AK2 | 62.095 | 2 | 53.066578 | yes | 1.976E-07 | 465.25616 | 225.29974 |
| RFTFQVIIDGR | 11 | P19525 | Interferon-induced, double-stranded RNA | EIF2AK2 | 62.095 | 2 | 53.066578 | yes | 1.048E-07 | 676.37769 | 113.04662 |
| PSNIFLVDTK | 10 | P19525 | Interferon-induced, double-stranded RNA | EIF2AK2 | 62.095 | 2 | 53.066578 | yes | 3.254E-06 | 567.31366 | 41.661583 |
| EFPEGEGR | 8 | P19525 | Interferon-induced, double-stranded RNA | EIF2AK2 | 62.095 | 2 | 53.066578 | yes | 6.955E-07 | 460.70905 | 37.412407 |
| YHLQQAR | 7 | P19532 | Transcription factor E3 OS=Homo sapien | TFE3 | 61.522 | 2 | 49.620174 | yes | 1.037E-07 | 458.24341 | 393.64279 |

| | | | | | | | | | | | |
|---|---|---|---|---|---|---|---|---|---|---|---|
| QYLSTTLGPK | 10 | P19532 | Transcription factor E3 OS=Homo sapien | TFE3 | 61.522 | 2 | 49.620174 | yes | 1.022E-07 | 554.30585 | 344.99765 |
| DNHNLIER | 8 | P19532 | Transcription factor E3 OS=Homo sapien | TFE3 | 61.522 | 2 | 49.620174 | no | 1.017E-07 | 505.75433 | 177.54614 |
| AASDPLLSSVSPAV | 16 | P19532 | Transcription factor E3 OS=Homo sapien | TFE3 | 61.522 | 2 | 49.620174 | yes | 1.245E-07 | 764.91449 | 166.08369 |
| VQTHLENPTR | 10 | P19532 | Transcription factor E3 OS=Homo sapien | TFE3 | 61.522 | 2 | 49.620174 | yes | 2.498E-07 | 597.81494 | 91.029167 |
| DGVEASAEGPR | 11 | P19532 | Transcription factor E3 OS=Homo sapien | TFE3 | 61.522 | 2 | 49.620174 | yes | 4.487E-07 | 544.25433 | 86.56311 |
| AVFVLLEER | 9 | P19532 | Transcription factor E3 OS=Homo sapien | TFE3 | 61.522 | 2 | 49.620174 | yes | 8.823E-06 | 538.31097 | 59.768734 |
| ELGTLIPK | 8 | P19532 | Transcription factor E3 OS=Homo sapien | TFE3 | 61.522 | 2 | 49.620174 | no | 2.442E-07 | 435.76837 | 22.765179 |
| VLIIGGGDGGVLR | 13 | P19623 | Spermidine synthase OS=Homo sapiens ( | SRM | 33.825 | 2 | 40.373085 | yes | 4.059E-07 | 613.36676 | 136.10327 |
| YQDILVFR | 8 | P19623 | Spermidine synthase OS=Homo sapiens ( | SRM | 33.825 | 2 | 40.373085 | yes | 5.951E-07 | 527.29004 | 27.419222 |
| LIDTVKDPVSK | 11 | P19784 | Casein kinase II subunit alpha' OS=Homo | CSNK2A2 | 41.214 | 2 | 54.02791 | yes | 8.911E-07 | 607.85315 | 473.10681 |
| EAMEHPYFYPVVI | 13 | P19784 | Casein kinase II subunit alpha' OS=Homo | CSNK2A2 | 41.214 | 2 | 54.02791 | yes | 1.687E-07 | 805.38959 | 92.692924 |
| ALDYCHSK | 8 | P19784 | Casein kinase II subunit alpha' OS=Homo | CSNK2A2 | 41.214 | 2 | 54.02791 | yes | 7.042E-07 | 497.22656 | 74.575302 |
| HLVSPEALDLLDK | 13 | P19784 | Casein kinase II subunit alpha' OS=Homo | CSNK2A2 | 41.214 | 2 | 54.02791 | yes | 9.722E-08 | 725.401 | 61.156876 |
| WENFIHSENR | 10 | P19784 | Casein kinase II subunit alpha' OS=Homo | CSNK2A2 | 41.214 | 2 | 54.02791 | yes | 4.542E-07 | 666.31 | 57.308632 |
| VLGTEELYGYLK | 12 | P19784 | Casein kinase II subunit alpha' OS=Homo | CSNK2A2 | 41.214 | 2 | 54.02791 | yes | 1.804E-07 | 692.87158 | 51.957642 |
| FYNQVSTPLLR | 11 | P19823 | Inter-alpha-trypsin inhibitor heavy chain I | ITIH2 | 106.464 | 2 | 36.223057 | yes | 7.247E-07 | 669.36426 | 57.00037 |
| MVTGDNINTAR | 11 | P20020 | Plasma membrane calcium-transporting A | ATP2B1 | 134.685 | 2 | 55.40208 | no | 1.506E-06 | 596.29297 | 56.514 |
| IDESSLTGESDHVI | 14 | P20020 | Plasma membrane calcium-transporting A | ATP2B1 | 134.685 | 2 | 55.40208 | yes | 9.722E-08 | 758.85992 | 42.547886 |
| IFDIDEAEEGVK | 12 | P20042 | Eukaryotic translation initiation factor 2 s | EIF2S2 | 38.389 | 2 | 52.13827 | yes | 9.938E-08 | 682.83264 | 4133.3467 |
| DYTYEELLNR | 10 | P20042 | Eukaryotic translation initiation factor 2 s | EIF2S2 | 38.389 | 2 | 52.13827 | yes | 3.926E-07 | 658.31189 | 3426.5703 |
| TSFVNFTDICK | 11 | P20042 | Eukaryotic translation initiation factor 2 s | EIF2S2 | 38.389 | 2 | 52.13827 | yes | 1.317E-07 | 666.31866 | 2327.5593 |
| EYVTCHTCR | 9 | P20042 | Eukaryotic translation initiation factor 2 s | EIF2S2 | 38.389 | 2 | 52.13827 | yes | 1.396E-07 | 613.25806 | 1463.413 |
| KFVMKPPQVVR | 11 | P20042 | Eukaryotic translation initiation factor 2 s | EIF2S2 | 38.389 | 3 | 52.13827 | yes | 9.722E-08 | 443.60056 | 542.0542 |
| KFVMKPPQVVR | 11 | P20042 | Eukaryotic translation initiation factor 2 s | EIF2S2 | 38.389 | 3 | 52.13827 | yes | 9.722E-08 | 664.89722 | 542.0542 |
| KFVMKPPQVVR | 11 | P20042 | Eukaryotic translation initiation factor 2 s | EIF2S2 | 38.389 | 2 | 52.13827 | yes | 9.722E-08 | 443.60056 | 542.0542 |
| KFVMKPPQVVR | 11 | P20042 | Eukaryotic translation initiation factor 2 s | EIF2S2 | 38.389 | 2 | 52.13827 | yes | 9.722E-08 | 664.89722 | 542.0542 |
| SGDEMIFDPTMSK | 13 | P20042 | Eukaryotic translation initiation factor 2 s | EIF2S2 | 38.389 | 2 | 52.13827 | yes | 4.034E-07 | 750.32324 | 951.2475 |
| QIENVLR | 7 | P20042 | Eukaryotic translation initiation factor 2 s | EIF2S2 | 38.389 | 2 | 52.13827 | yes | 4.327E-07 | 436.25345 | 480.26996 |
| LYFLQCETCHSR | 12 | P20042 | Eukaryotic translation initiation factor 2 s | EIF2S2 | 38.389 | 2 | 52.13827 | yes | 2.314E-07 | 807.36359 | 421.19992 |
| FPDEDEILEK | 10 | P20042 | Eukaryotic translation initiation factor 2 s | EIF2S2 | 38.389 | 2 | 52.13827 | yes | 7.533E-07 | 617.79553 | 337.01263 |
| KIFDIDEAEEGVK | 13 | P20042 | Eukaryotic translation initiation factor 2 s | EIF2S2 | 38.389 | 2 | 52.13827 | yes | 1.679E-07 | 746.88013 | 218.24368 |
| SPDTILQK | 8 | P20042 | Eukaryotic translation initiation factor 2 s | EIF2S2 | 38.389 | 2 | 52.13827 | yes | 2.658E-07 | 451.25311 | 205.13895 |
| FPDEDEILEKDEAI | 21 | P20042 | Eukaryotic translation initiation factor 2 s | EIF2S2 | 38.389 | 3 | 52.13827 | yes | 1.394E-07 | 841.05365 | 126.68344 |
| EKNPDMVAGEK | 11 | P20042 | Eukaryotic translation initiation factor 2 s | EIF2S2 | 38.389 | 2 | 52.13827 | yes | 1.072E-07 | 609.29517 | 95.486763 |
| DEALEDEDNKK | 11 | P20042 | Eukaryotic translation initiation factor 2 s | EIF2S2 | 38.389 | 2 | 52.13827 | yes | 1.459E-07 | 653.29388 | 68.944901 |
| KTSFVNFTDICK | 12 | P20042 | Eukaryotic translation initiation factor 2 s | EIF2S2 | 38.389 | 2 | 52.13827 | yes | 1.282E-07 | 730.36615 | 55.169411 |
| FVMKPPQVVR | 10 | P20042 | Eukaryotic translation initiation factor 2 s | EIF2S2 | 38.389 | 2 | 52.13827 | yes | 3.871E-06 | 600.84973 | 46.999962 |
| DLEADEEDTR | 10 | P20042 | Eukaryotic translation initiation factor 2 s | EIF2S2 | 38.389 | 2 | 52.13827 | yes | 1.271E-06 | 596.75183 | 31.339756 |
| MSGDEMIFDPTMS | 14 | P20042 | Eukaryotic translation initiation factor 2 s | EIF2S2 | 38.389 | 2 | 52.13827 | yes | 2.296E-07 | 815.84351 | 27.350323 |

| | | | | | | | | | | | |
|---|---|---|---|---|---|---|---|---|---|---|---|
| DEALEDEDNK | 10 | P20042 | Eukaryotic translation initiation factor 2 s | EIF2S2 | 38.389 | 2 | 52.13827 | yes | 5.585E-05 | 589.2464 | 21.330811 |
| PAANFDAIR | 9 | P20073 | Annexin A7 OS=Homo sapiens OX=960( | ANXA7 | 52.74 | 2 | 49.600372 | yes | 1.561E-07 | 487.75635 | 149.25638 |
| VLIEILCTR | 9 | P20073 | Annexin A7 OS=Homo sapiens OX=960( | ANXA7 | 52.74 | 2 | 49.600372 | yes | 1.734E-07 | 558.82593 | 135.09204 |
| GAGTDDSTLVR | 11 | P20073 | Annexin A7 OS=Homo sapiens OX=960( | ANXA7 | 52.74 | 2 | 49.600372 | yes | 3.372E-07 | 546.27002 | 110.1534 |
| TLGTMIAGDTSGD | 15 | P20073 | Annexin A7 OS=Homo sapiens OX=960( | ANXA7 | 52.74 | 2 | 49.600372 | yes | 1.467E-07 | 779.36432 | 98.046478 |
| SDTSGHFER | 9 | P20073 | Annexin A7 OS=Homo sapiens OX=960( | ANXA7 | 52.74 | 2 | 49.600372 | yes | 2.546E-07 | 518.22815 | 82.68013 |
| LLVSMCQGNR | 10 | P20073 | Annexin A7 OS=Homo sapiens OX=960( | ANXA7 | 52.74 | 2 | 49.600372 | yes | 1.392E-07 | 589.29443 | 74.398552 |
| SEIDLVQIK | 9 | P20073 | Annexin A7 OS=Homo sapiens OX=960( | ANXA7 | 52.74 | 2 | 49.600372 | yes | 1.487E-06 | 522.80042 | 72.187263 |
| GFGTDEQAIVDVV | 16 | P20073 | Annexin A7 OS=Homo sapiens OX=960( | ANXA7 | 52.74 | 2 | 49.600372 | yes | 1.019E-07 | 845.9234 | 47.454189 |
| IQNMVGSCDVK | 11 | P20226 | TATA-box-binding protein OS=Homo sa | TBP | 37.699 | 2 | 44.715332 | yes | 2.093E-07 | 625.797 | 92.214691 |
| EDALAFNSAISLPC | 16 | P20248 | Cyclin-A2 OS=Homo sapiens OX=9606 ( | CCNA2 | 48.551 | 2 | 42.62288 | yes | 4.093E-07 | 829.43085 | 93.503403 |
| MEHLVLK | 7 | P20248 | Cyclin-A2 OS=Homo sapiens OX=9606 ( | CCNA2 | 48.551 | 2 | 42.62288 | yes | 2.945E-07 | 435.2493 | 37.497051 |
| TGYTLESLKPCLM | 21 | P20248 | Cyclin-A2 OS=Homo sapiens OX=9606 ( | CCNA2 | 48.551 | 3 | 42.62288 | yes | 6.246E-05 | 843.09058 | 17.472263 |
| GGCPGGEATLSQP | 16 | P20290 | Transcription factor BTF3 OS=Homo sap | BTF3 | 22.168 | 2 | 49.326557 | yes | 6.132E-07 | 790.87793 | 339.25082 |
| ETIMNQEK | 8 | P20290 | Transcription factor BTF3 OS=Homo sap | BTF3 | 22.168 | 2 | 49.326557 | yes | 6.281E-07 | 496.7395 | 125.27433 |
| VQASLAANTFTIT( | 19 | P20290 | Transcription factor BTF3 OS=Homo sap | BTF3 | 22.168 | 3 | 49.326557 | yes | 1.043E-07 | 654.00934 | 49.688423 |
| TGAPAQADSR | 10 | P20290 | Transcription factor BTF3 OS=Homo sap | BTF3 | 22.168 | 2 | 49.326557 | yes | 1.24E-06 | 487.23853 | 48.504013 |
| KLGVNNISGIEEVN | 29 | P20290 | Transcription factor BTF3 OS=Homo sap | BTF3 | 22.168 | 4 | 49.326557 | yes | 8.04E-06 | 804.41327 | 23.841341 |
| FEIWDTAGQER | 11 | P20339 | Ras-related protein Rab-5A OS=Homo sa | RAB5A | 23.659 | 2 | 43.069065 | no | 5.937E-07 | 676.3175 | 94.401001 |
| QASPNIVIALSGNK | 14 | P20339 | Ras-related protein Rab-5A OS=Homo sa | RAB5A | 23.659 | 2 | 43.069065 | yes | 9.87E-07 | 706.3988 | 51.854324 |
| YHSLAPMYYR | 10 | P20339 | Ras-related protein Rab-5A OS=Homo sa | RAB5A | 23.659 | 2 | 43.069065 | no | 2.745E-07 | 650.81079 | 39.107079 |
| VAAALPGMESTQI | 14 | P20340 | Ras-related protein Rab-6A OS=Homo sa | RAB6A | 23.593 | 2 | 47.59877 | yes | 3.148E-07 | 723.35626 | 52.355629 |
| GSDVIIMLVGNK | 12 | P20340 | Ras-related protein Rab-6A OS=Homo sa | RAB6A | 23.593 | 2 | 47.59877 | no | 1.296E-07 | 623.34723 | 23.609121 |
| STLIGEDVNPLIK | 13 | P20585 | DNA mismatch repair protein Msh3 OS=] | MSH3 | 127.412 | 2 | 53.508392 | yes | 9.722E-08 | 699.89557 | 417.811 |
| FFGEDAEIAAR | 11 | P20585 | DNA mismatch repair protein Msh3 OS=] | MSH3 | 127.412 | 2 | 53.508392 | yes | 1.155E-07 | 613.29602 | 367.66638 |
| NSAVSCIPTDWVK | 13 | P20585 | DNA mismatch repair protein Msh3 OS=] | MSH3 | 127.412 | 2 | 53.508392 | yes | 1.202E-07 | 738.86121 | 268.22501 |
| NLEILQNQTDMK | 12 | P20585 | DNA mismatch repair protein Msh3 OS=] | MSH3 | 127.412 | 2 | 53.508392 | yes | 1.681E-07 | 723.86652 | 219.802 |
| EFCCDSALPQSR | 12 | P20585 | DNA mismatch repair protein Msh3 OS=] | MSH3 | 127.412 | 2 | 53.508392 | yes | 1.1E-07 | 735.31104 | 175.28894 |
| ATSQSLVILDELGF | 14 | P20585 | DNA mismatch repair protein Msh3 OS=] | MSH3 | 127.412 | 2 | 53.508392 | yes | 9.722E-08 | 751.41467 | 147.13347 |
| GLCSIYHK | 8 | P20585 | DNA mismatch repair protein Msh3 OS=] | MSH3 | 127.412 | 2 | 53.508392 | yes | 1.701E-07 | 489.24728 | 143.89476 |
| VMIITGPNMGGK | 12 | P20585 | DNA mismatch repair protein Msh3 OS=] | MSH3 | 127.412 | 2 | 53.508392 | yes | 7.125E-07 | 609.32269 | 127.96535 |
| VQTESLQER | 9 | P20585 | DNA mismatch repair protein Msh3 OS=] | MSH3 | 127.412 | 2 | 53.508392 | yes | 6.184E-07 | 545.2804 | 124.80025 |
| ATSVSVQDDR | 10 | P20585 | DNA mismatch repair protein Msh3 OS=] | MSH3 | 127.412 | 2 | 53.508392 | yes | 1.031E-07 | 539.26221 | 111.66433 |
| DAVLCVECGYK | 11 | P20585 | DNA mismatch repair protein Msh3 OS=] | MSH3 | 127.412 | 2 | 53.508392 | yes | 1.506E-07 | 657.29688 | 95.227669 |
| WVTQPLLK | 8 | P20585 | DNA mismatch repair protein Msh3 OS=] | MSH3 | 127.412 | 2 | 53.508392 | yes | 2.606E-06 | 492.79749 | 79.991829 |
| EGGSDLGMSGNSE | 15 | P20585 | DNA mismatch repair protein Msh3 OS=] | MSH3 | 127.412 | 2 | 53.508392 | yes | 9.723E-08 | 732.8172 | 65.194832 |
| MLSKPENFK | 9 | P20585 | DNA mismatch repair protein Msh3 OS=] | MSH3 | 127.412 | 2 | 53.508392 | yes | 6.041E-07 | 547.28918 | 58.457256 |
| DLSDFPLIK | 9 | P20585 | DNA mismatch repair protein Msh3 OS=] | MSH3 | 127.412 | 2 | 53.508392 | yes | 6.208E-07 | 524.28967 | 47.880001 |
| KDEIQGVIDEIR | 12 | P20585 | DNA mismatch repair protein Msh3 OS=] | MSH3 | 127.412 | 2 | 53.508392 | yes | 1.139E-05 | 707.88043 | 38.197395 |

| | | | | | | | | | | | |
|---|---|---|---|---|---|---|---|---|---|---|---|
| DTTLFDLSQFGSSN | 22 | P20585 | DNA mismatch repair protein Msh3 OS=I | MSH3 | 127.412 | 3 | 53.508392 | yes | 2.367E-07 | 823.72247 | 26.749969 |
| KWVTQPLLK | 9 | P20585 | DNA mismatch repair protein Msh3 OS=I | MSH3 | 127.412 | 2 | 53.508392 | yes | 1.916E-07 | 556.84497 | 20.502014 |
| GAVYSFDPVGSYQ | 14 | P20618 | Proteasome subunit beta type-1 OS=Hom | PSMB1 | 26.49 | 2 | 48.907635 | yes | 2.227E-07 | 773.37024 | 163.32072 |
| LSEGFSIHTR | 10 | P20618 | Proteasome subunit beta type-1 OS=Hom | PSMB1 | 26.49 | 2 | 48.907635 | yes | 1.088E-07 | 573.79877 | 80.021324 |
| LNDFASTVR | 9 | P20674 | Cytochrome c oxidase subunit 5A, mitoch | COX5A | 16.763 | 2 | 45.579357 | yes | 1.807E-07 | 511.76691 | 123.26364 |
| ALYETELADAR | 11 | P20700 | Lamin-B1 OS=Homo sapiens OX=9606 ( | LMNB1 | 66.409 | 2 | 53.87309 | yes | 9.722E-08 | 626.31439 | 348.72366 |
| LALDMEISAYR | 11 | P20700 | Lamin-B1 OS=Homo sapiens OX=9606 ( | LMNB1 | 66.409 | 2 | 53.87309 | yes | 1.204E-07 | 641.32904 | 310.70798 |
| IESLSSQLSNLQK | 13 | P20700 | Lamin-B1 OS=Homo sapiens OX=9606 ( | LMNB1 | 66.409 | 2 | 53.87309 | yes | 1.279E-07 | 723.89355 | 177.85127 |
| CQSLTEDLEFR | 11 | P20700 | Lamin-B1 OS=Homo sapiens OX=9606 ( | LMNB1 | 66.409 | 2 | 53.87309 | yes | 1.114E-07 | 699.3219 | 147.83713 |
| NSQGEEVAQR | 10 | P20700 | Lamin-B1 OS=Homo sapiens OX=9606 ( | LMNB1 | 66.409 | 2 | 53.87309 | yes | 1.757E-07 | 559.26526 | 111.77661 |
| LSSEMNTSTVNSA | 14 | P20700 | Lamin-B1 OS=Homo sapiens OX=9606 ( | LMNB1 | 66.409 | 2 | 53.87309 | yes | 9.968E-08 | 748.85413 | 104.15307 |
| SMYEEEINETR | 11 | P20700 | Lamin-B1 OS=Homo sapiens OX=9606 ( | LMNB1 | 66.409 | 2 | 53.87309 | yes | 9.722E-08 | 700.80353 | 96.779579 |
| LAQALHEMR | 9 | P20700 | Lamin-B1 OS=Homo sapiens OX=9606 ( | LMNB1 | 66.409 | 2 | 53.87309 | yes | 9.722E-08 | 534.78455 | 95.607864 |
| DAALATALGDK | 11 | P20700 | Lamin-B1 OS=Homo sapiens OX=9606 ( | LMNB1 | 66.409 | 2 | 53.87309 | yes | 3.128E-06 | 523.27985 | 82.415833 |
| QLADETLLK | 9 | P20700 | Lamin-B1 OS=Homo sapiens OX=9606 ( | LMNB1 | 66.409 | 2 | 53.87309 | yes | 7.081E-07 | 515.7926 | 65.49437 |
| AGGPTTPLSPTR | 12 | P20700 | Lamin-B1 OS=Homo sapiens OX=9606 ( | LMNB1 | 66.409 | 2 | 53.87309 | yes | 1.375E-07 | 577.81183 | 62.56459 |
| EELEQTYHAK | 10 | P20700 | Lamin-B1 OS=Homo sapiens OX=9606 ( | LMNB1 | 66.409 | 2 | 53.87309 | yes | 2.678E-06 | 624.29877 | 38.848125 |
| LYKEELEQTYHAK | 13 | P20700 | Lamin-B1 OS=Homo sapiens OX=9606 ( | LMNB1 | 66.409 | 2 | 53.87309 | yes | 1.772E-06 | 826.41992 | 14.527311 |
| YQTLELEK | 8 | P20719 | Homeobox protein Hox-A5 OS=Homo sa | HOXA5 | 29.346 | 2 | 55.727352 | no | 1.588E-07 | 512.27148 | 392.63608 |
| SSYFVNSFCGR | 11 | P20719 | Homeobox protein Hox-A5 OS=Homo sa | HOXA5 | 29.346 | 2 | 55.727352 | yes | 9.722E-08 | 683.29822 | 223.3604 |
| SYAASASAAPAEP | 14 | P20719 | Homeobox protein Hox-A5 OS=Homo sa | HOXA5 | 29.346 | 2 | 55.727352 | yes | 2.523E-07 | 674.82825 | 155.09521 |
| RIEIAHALCLSER | 13 | P20719 | Homeobox protein Hox-A5 OS=Homo sa | HOXA5 | 29.346 | 3 | 55.727352 | no | 3.425E-07 | 523.28394 | 79.018852 |
| LHISHDNIGGPEGK | 14 | P20719 | Homeobox protein Hox-A5 OS=Homo sa | HOXA5 | 29.346 | 2 | 55.727352 | yes | 9.722E-08 | 737.37585 | 77.514076 |
| YGYGYNGMDLSV | 14 | P20719 | Homeobox protein Hox-A5 OS=Homo sa | HOXA5 | 29.346 | 2 | 55.727352 | yes | 2.398E-07 | 776.34845 | 57.09568 |
| NSLSNSSGASADA | 21 | P20719 | Homeobox protein Hox-A5 OS=Homo sa | HOXA5 | 29.346 | 3 | 55.727352 | yes | 3.472E-06 | 669.31049 | 26.516798 |
| PASNMVIVDVK | 11 | P20742 | Pregnancy zone protein OS=Homo sapien | PZP | 163.863 | 2 | 37.70853 | yes | 5.765E-07 | 594.8183 | 114.8447 |
| LVNLFQQGGR | 10 | P20749 | B-cell lymphoma 3 protein OS=Homo saj | BCL3 | 47.584 | 2 | 53.042473 | yes | 2.776E-07 | 566.31708 | 102.59543 |
| LLVTAGASPMALD | 14 | P20749 | B-cell lymphoma 3 protein OS=Homo saj | BCL3 | 47.584 | 2 | 53.042473 | yes | 9.722E-08 | 707.88977 | 71.941727 |
| AAGLPGAALPLR | 12 | P20749 | B-cell lymphoma 3 protein OS=Homo saj | BCL3 | 47.584 | 2 | 53.042473 | yes | 1.46E-06 | 553.83765 | 71.221161 |
| ELDIYNNLR | 9 | P20749 | B-cell lymphoma 3 protein OS=Homo saj | BCL3 | 47.584 | 2 | 53.042473 | yes | 1.88E-06 | 575.29858 | 71.072327 |
| GAAGLVVPLDPLR | 13 | P20749 | B-cell lymphoma 3 protein OS=Homo saj | BCL3 | 47.584 | 2 | 53.042473 | yes | 9.722E-08 | 639.38245 | 57.003166 |
| QLLCGAAVGTR | 11 | P20839 | Inosine-5'-monophosphate dehydrogenase | IMPDH1 | 55.406 | 2 | 47.4775 | yes | 4.377E-07 | 573.30841 | 179.94955 |
| IAQGVSGSIQDK | 12 | P20839 | Inosine-5'-monophosphate dehydrogenase | IMPDH1 | 55.406 | 2 | 47.4775 | yes | 1.32E-07 | 601.82239 | 150.49678 |
| NLIDAGVDGLR | 11 | P20839 | Inosine-5'-monophosphate dehydrogenase | IMPDH1 | 55.406 | 2 | 47.4775 | yes | 2.384E-07 | 571.81183 | 146.91376 |
| TMSAQIEGGVHGL | 18 | P20839 | Inosine-5'-monophosphate dehydrogenase | IMPDH1 | 55.406 | 3 | 47.4775 | yes | 2.948E-07 | 648.64752 | 73.870461 |
| IELVVAPAGVTLK | 13 | P20839 | Inosine-5'-monophosphate dehydrogenase | IMPDH1 | 55.406 | 2 | 47.4775 | yes | 1.996E-06 | 655.40814 | 17.594999 |
| ENPGSWFSEFK | 11 | P20908 | Collagen alpha-1(V) chain OS=Homo saj | COL5A1 | 183.56 | 2 | 49.343735 | yes | 1.043E-07 | 664.30133 | 269.88434 |
| GPAGPMGLTGR | 11 | P20908 | Collagen alpha-1(V) chain OS=Homo saj | COL5A1 | 183.56 | 2 | 49.343735 | yes | 1.878E-07 | 507.26349 | 200.34256 |
| ALVDGCATK | 9 | P20908 | Collagen alpha-1(V) chain OS=Homo saj | COL5A1 | 183.56 | 2 | 49.343735 | yes | 4.093E-07 | 467.73676 | 143.20976 |

| | | | | | | | | | | | |
|---|---|---|---|---|---|---|---|---|---|---|---|
| RPLGTQQNPAR | 11 | P20908 | Collagen alpha-1(V) chain OS=Homo sap | COL5A1 | 183.56 | 2 | 49.343735 | yes | 9.129E-07 | 619.34161 | 21.474928 |
| IEEGTYGVVYR | 11 | P21127 | Cyclin-dependent kinase 11B OS=Homo | CDK11B | 92.62 | 2 | 52.310764 | yes | 9.722E-08 | 643.32477 | 149.40103 |
| EGFPITSLR | 9 | P21127 | Cyclin-dependent kinase 11B OS=Homo | CDK11B | 92.62 | 2 | 52.310764 | yes | 1.002E-07 | 510.27966 | 145.15222 |
| YLPALQGCR | 9 | P21127 | Cyclin-dependent kinase 11B OS=Homo | CDK11B | 92.62 | 2 | 52.310764 | yes | 1.261E-07 | 539.27911 | 135.29245 |
| TLMIQLLR | 8 | P21127 | Cyclin-dependent kinase 11B OS=Homo | CDK11B | 92.62 | 2 | 52.310764 | yes | 9.722E-08 | 494.30463 | 105.72617 |
| SVEEFQCLNR | 10 | P21127 | Cyclin-dependent kinase 11B OS=Homo | CDK11B | 92.62 | 2 | 52.310764 | yes | 9.722E-08 | 641.29822 | 90.652245 |
| FGALLSDQGFDLM | 15 | P21127 | Cyclin-dependent kinase 11B OS=Homo | CDK11B | 92.62 | 2 | 52.310764 | yes | 1.303E-06 | 828.40851 | 85.674339 |
| EIVVGSNMDK | 10 | P21127 | Cyclin-dependent kinase 11B OS=Homo | CDK11B | 92.62 | 2 | 52.310764 | yes | 1.288E-07 | 546.27368 | 76.70874 |
| MTFSEHPYNNLR | 12 | P21127 | Cyclin-dependent kinase 11B OS=Homo | CDK11B | 92.62 | 2 | 52.310764 | yes | 1.728E-07 | 754.8512 | 74.397324 |
| EKEGFPITSLR | 11 | P21127 | Cyclin-dependent kinase 11B OS=Homo | CDK11B | 92.62 | 2 | 52.310764 | yes | 1.038E-06 | 638.84845 | 53.066692 |
| IYIVMNYVEHDLK | 13 | P21127 | Cyclin-dependent kinase 11B OS=Homo | CDK11B | 92.62 | 2 | 52.310764 | yes | 1.874E-07 | 818.92383 | 47.445339 |
| DLLSDLQDISDSEF | 14 | P21127 | Cyclin-dependent kinase 11B OS=Homo | CDK11B | 92.62 | 2 | 52.310764 | yes | 4.071E-05 | 803.38373 | 14.383075 |
| VDIIENQVMDFR | 12 | P21266 | Glutathione S-transferase Mu 3 OS=Hom | GSTM3 | 26.56 | 2 | 48.660942 | yes | 4.349E-06 | 739.86902 | 88.869102 |
| LLLEFTDTSYEEK | 13 | P21266 | Glutathione S-transferase Mu 3 OS=Hom | GSTM3 | 26.56 | 2 | 48.660942 | yes | 1.118E-07 | 794.39307 | 65.400345 |
| GFGFGQGAGALVI | 15 | P21291 | Cysteine and glycine-rich protein 1 OS=H | CSRP1 | 20.568 | 2 | 53.5489 | yes | 6.819E-07 | 717.34406 | 1968.1941 |
| GYGYGQGAGTLST | 15 | P21291 | Cysteine and glycine-rich protein 1 OS=H | CSRP1 | 20.568 | 2 | 53.5489 | yes | 9.722E-08 | 737.84406 | 738.90741 |
| CSQAVYAAEK | 10 | P21291 | Cysteine and glycine-rich protein 1 OS=H | CSRP1 | 20.568 | 2 | 53.5489 | yes | 9.722E-08 | 563.76349 | 376.82886 |
| GLESTTLADK | 10 | P21291 | Cysteine and glycine-rich protein 1 OS=H | CSRP1 | 20.568 | 2 | 53.5489 | yes | 3.398E-07 | 517.77185 | 151.12491 |
| KNLDSTTVAVHGI | 18 | P21291 | Cysteine and glycine-rich protein 1 OS=H | CSRP1 | 20.568 | 3 | 53.5489 | yes | 3.677E-07 | 688.67371 | 66.345421 |
| NLDSTTVAVHGEE | 17 | P21291 | Cysteine and glycine-rich protein 1 OS=H | CSRP1 | 20.568 | 3 | 53.5489 | yes | 1.58E-07 | 645.9754 | 36.769676 |
| AGNNMLLVGVHG | 14 | P21333 | Filamin-A OS=Homo sapiens OX=9606 ( | FLNA | 280.739 | 2 | 52.846935 | yes | 0.0001869 | 717.88538 | 837.61255 |
| AGNNMLLVGVHG | 14 | P21333 | Filamin-A OS=Homo sapiens OX=9606 ( | FLNA | 280.739 | 2 | 52.846935 | yes | 2.824E-06 | 478.92599 | 837.61255 |
| AGNNMLLVGVHG | 14 | P21333 | Filamin-A OS=Homo sapiens OX=9606 ( | FLNA | 280.739 | 3 | 52.846935 | yes | 0.0001869 | 717.88538 | 837.61255 |
| AGNNMLLVGVHG | 14 | P21333 | Filamin-A OS=Homo sapiens OX=9606 ( | FLNA | 280.739 | 3 | 52.846935 | yes | 2.824E-06 | 478.92599 | 837.61255 |
| SPFSVAVSPSLDLS | 15 | P21333 | Filamin-A OS=Homo sapiens OX=9606 ( | FLNA | 280.739 | 2 | 52.846935 | yes | 1.485E-07 | 767.41156 | 327.98691 |
| AWGPGLEGGVVG | 13 | P21333 | Filamin-A OS=Homo sapiens OX=9606 ( | FLNA | 280.739 | 2 | 52.846935 | yes | 2.018E-07 | 613.83002 | 322.19589 |
| VEPGLGADNSVVI | 13 | P21333 | Filamin-A OS=Homo sapiens OX=9606 ( | FLNA | 280.739 | 2 | 52.846935 | yes | 4.517E-07 | 656.84644 | 294.61325 |
| GTVEPQLEAR | 10 | P21333 | Filamin-A OS=Homo sapiens OX=9606 ( | FLNA | 280.739 | 2 | 52.846935 | yes | 1.752E-07 | 550.29077 | 255.21304 |
| AYGPGIEPTGNMV | 14 | P21333 | Filamin-A OS=Homo sapiens OX=9606 ( | FLNA | 280.739 | 2 | 52.846935 | yes | 1.301E-07 | 717.35828 | 249.84074 |
| YAPSEAGLHEMDI | 14 | P21333 | Filamin-A OS=Homo sapiens OX=9606 ( | FLNA | 280.739 | 2 | 52.846935 | yes | 9.722E-08 | 794.87482 | 230.00487 |
| VPVHDVTDASK | 11 | P21333 | Filamin-A OS=Homo sapiens OX=9606 ( | FLNA | 280.739 | 2 | 52.846935 | yes | 9.722E-08 | 584.30383 | 183.07684 |
| IANLQTDLSDGLR | 13 | P21333 | Filamin-A OS=Homo sapiens OX=9606 ( | FLNA | 280.739 | 2 | 52.846935 | yes | 1.526E-07 | 708.37805 | 173.15175 |
| CSGPGLSPGMVR | 12 | P21333 | Filamin-A OS=Homo sapiens OX=9606 ( | FLNA | 280.739 | 2 | 52.846935 | yes | 9.722E-08 | 609.29193 | 137.99802 |
| AFGPGLQGGSAGS | 16 | P21333 | Filamin-A OS=Homo sapiens OX=9606 ( | FLNA | 280.739 | 2 | 52.846935 | yes | 2.139E-07 | 715.36279 | 130.1591 |
| KGEITGEVR | 9 | P21333 | Filamin-A OS=Homo sapiens OX=9606 ( | FLNA | 280.739 | 2 | 52.846935 | yes | 3.381E-07 | 494.77472 | 128.69312 |
| ANLPQSFQVDTSK | 13 | P21333 | Filamin-A OS=Homo sapiens OX=9606 ( | FLNA | 280.739 | 2 | 52.846935 | yes | 1.257E-06 | 717.86481 | 96.768295 |
| ALTQTGGPHVK | 11 | P21333 | Filamin-A OS=Homo sapiens OX=9606 ( | FLNA | 280.739 | 2 | 52.846935 | yes | 1.531E-07 | 554.80908 | 94.411858 |
| VTAQGPGLEPSGN | 17 | P21333 | Filamin-A OS=Homo sapiens OX=9606 ( | FLNA | 280.739 | 2 | 52.846935 | yes | 1.212E-07 | 826.93378 | 93.75547 |
| LDVQFSGLTK | 10 | P21333 | Filamin-A OS=Homo sapiens OX=9606 ( | FLNA | 280.739 | 2 | 52.846935 | yes | 9.819E-08 | 554.30585 | 92.058556 |

| | | | | | | | | | | | |
|---|---|---|---|---|---|---|---|---|---|---|---|
| LTVSSLQESGLK | 12 | P21333 | Filamin-A OS=Homo sapiens OX=9606 ( | FLNA | 280.739 | 2 | 52.846935 | yes | 9.819E-08 | 631.35352 | 85.131332 |
| TPCEEILVK | 9 | P21333 | Filamin-A OS=Homo sapiens OX=9606 ( | FLNA | 280.739 | 2 | 52.846935 | yes | 2.705E-07 | 544.78644 | 78.604706 |
| GAGTGGLGLAVE( | 18 | P21333 | Filamin-A OS=Homo sapiens OX=9606 ( | FLNA | 280.739 | 2 | 52.846935 | yes | 2.006E-07 | 785.90723 | 77.746407 |
| EAGAGGLAIAVEG | 16 | P21333 | Filamin-A OS=Homo sapiens OX=9606 ( | FLNA | 280.739 | 2 | 52.846935 | yes | 1.047E-07 | 713.88043 | 68.05719 |
| LPQLPITNFSR | 11 | P21333 | Filamin-A OS=Homo sapiens OX=9606 ( | FLNA | 280.739 | 2 | 52.846935 | yes | 4.287E-07 | 643.36676 | 59.60957 |
| VHGPGIQSGTTNK | 16 | P21333 | Filamin-A OS=Homo sapiens OX=9606 ( | FLNA | 280.739 | 3 | 52.846935 | yes | 4.493E-07 | 545.62512 | 29.614613 |
| VHGPGIQSGTTNK | 16 | P21333 | Filamin-A OS=Homo sapiens OX=9606 ( | FLNA | 280.739 | 3 | 52.846935 | yes | 2.51E-06 | 817.93408 | 29.614613 |
| VHGPGIQSGTTNK | 16 | P21333 | Filamin-A OS=Homo sapiens OX=9606 ( | FLNA | 280.739 | 2 | 52.846935 | yes | 4.493E-07 | 545.62512 | 29.614613 |
| VHGPGIQSGTTNK | 16 | P21333 | Filamin-A OS=Homo sapiens OX=9606 ( | FLNA | 280.739 | 2 | 52.846935 | yes | 2.51E-06 | 817.93408 | 29.614613 |
| LKPGAPLRPK | 10 | P21333 | Filamin-A OS=Homo sapiens OX=9606 ( | FLNA | 280.739 | 2 | 52.846935 | yes | 1.365E-07 | 538.85059 | 57.034966 |
| GKLDVQFSGLTK | 12 | P21333 | Filamin-A OS=Homo sapiens OX=9606 ( | FLNA | 280.739 | 2 | 52.846935 | yes | 1.051E-07 | 646.86407 | 42.038738 |
| ETGEHLVHVK | 10 | P21333 | Filamin-A OS=Homo sapiens OX=9606 ( | FLNA | 280.739 | 2 | 52.846935 | yes | 7.468E-07 | 574.80658 | 39.614708 |
| AHVVPCFDASK | 11 | P21333 | Filamin-A OS=Homo sapiens OX=9606 ( | FLNA | 280.739 | 2 | 52.846935 | yes | 8.889E-07 | 615.80042 | 38.227711 |
| AEAGVPAEFSIWT | 14 | P21333 | Filamin-A OS=Homo sapiens OX=9606 ( | FLNA | 280.739 | 2 | 52.846935 | yes | 5.086E-07 | 767.38843 | 34.547066 |
| DAPQDFHPDR | 10 | P21333 | Filamin-A OS=Homo sapiens OX=9606 ( | FLNA | 280.739 | 2 | 52.846935 | yes | 4.569E-07 | 599.26782 | 26.399118 |
| TGVAVNKPAEFTV | 16 | P21333 | Filamin-A OS=Homo sapiens OX=9606 ( | FLNA | 280.739 | 2 | 52.846935 | yes | 3.898E-07 | 823.94104 | 17.908958 |
| VDINTEDLEDGTC | 14 | P21333 | Filamin-A OS=Homo sapiens OX=9606 ( | FLNA | 280.739 | 2 | 52.846935 | yes | 4.633E-07 | 818.85962 | 17.121912 |
| TLSIFYQAGTSK | 12 | P21359 | Neurofibromin OS=Homo sapiens OX=9( | NF1 | 319.373 | 2 | 50.003273 | yes | 9.935E-08 | 658.34827 | 45.338188 |
| PVEWVQAVVSR | 11 | P21359 | Neurofibromin OS=Homo sapiens OX=9( | NF1 | 319.373 | 2 | 50.003273 | yes | 2.299E-07 | 635.35114 | 35.774464 |
| LFDLVDGFAESTK | 13 | P21359 | Neurofibromin OS=Homo sapiens OX=9( | NF1 | 319.373 | 2 | 50.003273 | yes | 1.165E-07 | 721.36414 | 31.529377 |
| LLLCNPR | 7 | P21359 | Neurofibromin OS=Homo sapiens OX=9( | NF1 | 319.373 | 2 | 50.003273 | yes | 5.741E-07 | 443.25238 | 27.070608 |
| NIEVPFKPAR | 10 | P21399 | Cytoplasmic aconitate hydratase OS=Hon | ACO1 | 98.399 | 2 | 51.270386 | yes | 1.435E-07 | 585.83514 | 232.28831 |
| YLQAVGMFR | 9 | P21399 | Cytoplasmic aconitate hydratase OS=Hon | ACO1 | 98.399 | 2 | 51.270386 | yes | 5.728E-07 | 542.78406 | 101.6032 |
| QYVIPGMFK | 9 | P21399 | Cytoplasmic aconitate hydratase OS=Hon | ACO1 | 98.399 | 2 | 51.270386 | yes | 1.57E-07 | 541.78882 | 89.026604 |
| VLLEAAIR | 8 | P21399 | Cytoplasmic aconitate hydratase OS=Hon | ACO1 | 98.399 | 2 | 51.270386 | yes | 1.165E-07 | 442.78183 | 78.530685 |
| YTIIIPENLKPQMK | 14 | P21399 | Cytoplasmic aconitate hydratase OS=Hon | ACO1 | 98.399 | 2 | 51.270386 | yes | 1.068E-07 | 844.4762 | 34.854618 |
| YTIIIPENLKPQMK | 14 | P21399 | Cytoplasmic aconitate hydratase OS=Hon | ACO1 | 98.399 | 2 | 51.270386 | yes | 1.169E-07 | 563.31989 | 34.854618 |
| YTIIIPENLKPQMK | 14 | P21399 | Cytoplasmic aconitate hydratase OS=Hon | ACO1 | 98.399 | 3 | 51.270386 | yes | 1.068E-07 | 844.4762 | 34.854618 |
| YTIIIPENLKPQMK | 14 | P21399 | Cytoplasmic aconitate hydratase OS=Hon | ACO1 | 98.399 | 3 | 51.270386 | yes | 1.169E-07 | 563.31989 | 34.854618 |
| YQQAGLPLIVLAG | 14 | P21399 | Cytoplasmic aconitate hydratase OS=Hon | ACO1 | 98.399 | 2 | 51.270386 | yes | 9.722E-08 | 735.93756 | 62.900768 |
| TVVPCCSGPK | 10 | P21399 | Cytoplasmic aconitate hydratase OS=Hon | ACO1 | 98.399 | 2 | 51.270386 | yes | 1.553E-06 | 552.76245 | 41.249077 |
| MASTDTPMAR | 10 | P21580 | Tumor necrosis factor alpha-induced prot | TNFAIP3 | 89.614 | 2 | 51.369095 | yes | 6.538E-07 | 540.74426 | 256.3992 |
| FQNTIPCLGR | 10 | P21580 | Tumor necrosis factor alpha-induced prot | TNFAIP3 | 89.614 | 2 | 51.369095 | yes | 2.45E-07 | 603.30841 | 245.5916 |
| CQACLQDVTR | 10 | P21580 | Tumor necrosis factor alpha-induced prot | TNFAIP3 | 89.614 | 2 | 51.369095 | yes | 1.541E-07 | 625.78442 | 243.07594 |
| SLESGSNFAPLK | 12 | P21580 | Tumor necrosis factor alpha-induced prot | TNFAIP3 | 89.614 | 2 | 51.369095 | yes | 9.722E-08 | 625.32477 | 224.87384 |
| RPIIVISDK | 9 | P21580 | Tumor necrosis factor alpha-induced prot | TNFAIP3 | 89.614 | 2 | 51.369095 | yes | 1.045E-07 | 520.82678 | 181.16675 |
| AGCVYFGTPENK | 12 | P21580 | Tumor necrosis factor alpha-induced prot | TNFAIP3 | 89.614 | 2 | 51.369095 | yes | 1.318E-07 | 671.80847 | 167.65456 |
| YPIVLGYDSHHFV | 19 | P21580 | Tumor necrosis factor alpha-induced prot | TNFAIP3 | 89.614 | 3 | 51.369095 | yes | 3.177E-07 | 733.40515 | 151.27101 |
| TCQFCPQFR | 9 | P21580 | Tumor necrosis factor alpha-induced prot | TNFAIP3 | 89.614 | 2 | 51.369095 | yes | 9.722E-08 | 622.271 | 149.13531 |

| | | | | | | | | | | | |
|---|---|---|---|---|---|---|---|---|---|---|---|
| PGPEGLPGMALGA | 15 | P21580 | Tumor necrosis factor alpha-induced prot | TNFAIP3 | 89.614 | 2 | 51.369095 | yes | 2.537E-07 | 705.36389 | 142.30559 |
| GFCTLCFIEYR | 11 | P21580 | Tumor necrosis factor alpha-induced prot | TNFAIP3 | 89.614 | 2 | 51.369095 | yes | 1.164E-07 | 733.3338 | 127.86505 |
| NIQATLESQK | 10 | P21580 | Tumor necrosis factor alpha-induced prot | TNFAIP3 | 89.614 | 2 | 51.369095 | yes | 2.348E-07 | 566.30383 | 126.95854 |
| VHFLTDPENEMK | 12 | P21580 | Tumor necrosis factor alpha-induced prot | TNFAIP3 | 89.614 | 2 | 51.369095 | yes | 8.818E-07 | 730.34796 | 92.986023 |
| WQLESLK | 7 | P21580 | Tumor necrosis factor alpha-induced prot | TNFAIP3 | 89.614 | 2 | 51.369095 | yes | 1.637E-07 | 452.25037 | 92.402771 |
| TFNGICSTCFK | 11 | P21580 | Tumor necrosis factor alpha-induced prot | TNFAIP3 | 89.614 | 2 | 51.369095 | yes | 4.539E-06 | 667.797 | 84.706795 |
| CFIEAQNQR | 9 | P21580 | Tumor necrosis factor alpha-induced prot | TNFAIP3 | 89.614 | 2 | 51.369095 | yes | 4.82E-07 | 583.2746 | 83.681976 |
| AGNDAPAGCLSQA | 15 | P21580 | Tumor necrosis factor alpha-induced prot | TNFAIP3 | 89.614 | 2 | 51.369095 | yes | 3.143E-07 | 729.84131 | 63.212467 |
| NWNDEWDNLIK | 11 | P21580 | Tumor necrosis factor alpha-induced prot | TNFAIP3 | 89.614 | 2 | 51.369095 | yes | 3.331E-07 | 723.83606 | 59.965187 |
| YTLEMFR | 7 | P21580 | Tumor necrosis factor alpha-induced prot | TNFAIP3 | 89.614 | 2 | 51.369095 | yes | 1.108E-07 | 480.23639 | 58.206852 |
| APACDHFGNAK | 11 | P21580 | Tumor necrosis factor alpha-induced prot | TNFAIP3 | 89.614 | 2 | 51.369095 | yes | 1.105E-07 | 594.26672 | 51.83778 |
| WQENSEQGR | 9 | P21580 | Tumor necrosis factor alpha-induced prot | TNFAIP3 | 89.614 | 2 | 51.369095 | yes | 4.307E-07 | 567.25214 | 48.865463 |
| VIVNPMDLETIR | 12 | P21675 | Transcription initiation factor TFIID subu | TAF1 | 214.714 | 2 | 35.19353 | yes | 8.707E-07 | 700.38434 | 17.75909 |
| YQIDPDACFSAK | 12 | P21796 | Voltage-dependent anion-selective chann | VDAC1 | 30.773 | 2 | 48.641502 | yes | 2.773E-07 | 707.81903 | 102.26602 |
| LTFDSSFSPNTGK | 13 | P21796 | Voltage-dependent anion-selective chann | VDAC1 | 30.773 | 2 | 48.641502 | yes | 2.883E-07 | 700.83826 | 95.157585 |
| LTLSALLDGK | 10 | P21796 | Voltage-dependent anion-selective chann | VDAC1 | 30.773 | 2 | 48.641502 | yes | 1.12E-07 | 515.81079 | 82.666016 |
| WTEYGLTFTEK | 11 | P21796 | Voltage-dependent anion-selective chann | VDAC1 | 30.773 | 2 | 48.641502 | yes | 9.086E-07 | 687.83246 | 29.444746 |
| LQDPFSLYR | 9 | P21912 | Succinate dehydrogenase [ubiquinone] irc | SDHB | 31.63 | 2 | 47.71997 | yes | 3.212E-07 | 569.79822 | 185.70093 |
| QQYLQSIEER | 10 | P21912 | Succinate dehydrogenase [ubiquinone] irc | SDHB | 31.63 | 2 | 47.71997 | yes | 1.642E-07 | 647.32532 | 170.11702 |
| CGPMVLDALIK | 11 | P21912 | Succinate dehydrogenase [ubiquinone] irc | SDHB | 31.63 | 2 | 47.71997 | yes | 1.272E-07 | 608.82507 | 93.560562 |
| DLVPDLSNFYAQY | 14 | P21912 | Succinate dehydrogenase [ubiquinone] irc | SDHB | 31.63 | 2 | 47.71997 | yes | 4.594E-07 | 836.91449 | 61.500443 |
| KYDVDTLDMVFLI | 16 | P21964 | Catechol O-methyltransferase OS=Homo | COMT | 30.038 | 3 | 53.53492 | yes | 9.722E-08 | 675.66437 | 68.755692 |
| SVPLCILYEK | 10 | P21980 | Protein-glutamine gamma-glutamyltransfe | TGM2 | 77.329 | 2 | 51.48398 | yes | 2.665E-07 | 611.33105 | 418.7561 |
| SEGTYCCGPVPVR | 13 | P21980 | Protein-glutamine gamma-glutamyltransfe | TGM2 | 77.329 | 2 | 51.48398 | yes | 1.139E-07 | 741.32922 | 312.01443 |
| TVEIPDPVEAGEEV | 15 | P21980 | Protein-glutamine gamma-glutamyltransfe | TGM2 | 77.329 | 2 | 51.48398 | yes | 1.058E-07 | 806.40924 | 263.82629 |
| LVVNFESDK | 9 | P21980 | Protein-glutamine gamma-glutamyltransfe | TGM2 | 77.329 | 2 | 51.48398 | yes | 2.723E-07 | 525.77692 | 235.90498 |
| MDLLPLHMGLHK | 12 | P21980 | Protein-glutamine gamma-glutamyltransfe | TGM2 | 77.329 | 2 | 51.48398 | yes | 9.977E-08 | 702.87817 | 193.90874 |
| TVSYNGILGPECGT | 15 | P21980 | Protein-glutamine gamma-glutamyltransfe | TGM2 | 77.329 | 2 | 51.48398 | yes | 3.751E-07 | 798.39032 | 153.85611 |
| CLGIPTR | 7 | P21980 | Protein-glutamine gamma-glutamyltransfe | TGM2 | 77.329 | 2 | 51.48398 | no | 1.734E-07 | 408.72345 | 120.78877 |
| YPEGSSEER | 9 | P21980 | Protein-glutamine gamma-glutamyltransfe | TGM2 | 77.329 | 2 | 51.48398 | no | 9.722E-08 | 527.22778 | 117.66048 |
| NEFGEIQGDK | 10 | P21980 | Protein-glutamine gamma-glutamyltransfe | TGM2 | 77.329 | 2 | 51.48398 | yes | 2.479E-06 | 568.76459 | 96.360512 |
| GQPFWLTLHFEGR | 13 | P21980 | Protein-glutamine gamma-glutamyltransfe | TGM2 | 77.329 | 2 | 51.48398 | yes | 1.577E-07 | 794.40698 | 63.275528 |
| VQLVVGDGR | 9 | P22061 | Protein-L-isoaspartate(D-aspartate) O-me | PCMT1 | 24.637 | 2 | 52.359665 | yes | 2.115E-07 | 471.772 | 2825.406 |
| VFEVMLATDR | 10 | P22061 | Protein-L-isoaspartate(D-aspartate) O-me | PCMT1 | 24.637 | 2 | 52.359665 | yes | 1.15E-07 | 590.80518 | 2625.6057 |
| VIGIDHIK | 8 | P22061 | Protein-L-isoaspartate(D-aspartate) O-me | PCMT1 | 24.637 | 2 | 52.359665 | yes | 3.881E-07 | 447.77402 | 1907.3809 |
| KDDPTLLSSGR | 11 | P22061 | Protein-L-isoaspartate(D-aspartate) O-me | PCMT1 | 24.637 | 2 | 52.359665 | yes | 1.133E-07 | 594.81458 | 1080.4258 |
| ALDVGSGSGILTA( | 17 | P22061 | Protein-L-isoaspartate(D-aspartate) O-me | PCMT1 | 24.637 | 2 | 52.359665 | yes | 1.021E-07 | 847.93018 | 472.8653 |
| SGGASHSELIHNLF | 14 | P22061 | Protein-L-isoaspartate(D-aspartate) O-me | PCMT1 | 24.637 | 2 | 52.359665 | yes | 9.722E-08 | 739.37897 | 291.64966 |
| TDKVFEVMLATDI | 13 | P22061 | Protein-L-isoaspartate(D-aspartate) O-me | PCMT1 | 24.637 | 2 | 52.359665 | yes | 1.084E-07 | 762.88995 | 234.41617 |

| | | | | | | | | | | | |
|---|---|---|---|---|---|---|---|---|---|---|---|
| DDPTLLSSGR | 10 | P22061 | Protein-L-isoaspartate(D-aspartate) O-me | PCMT1 | 24.637 | 2 | 52.359665 | yes | 2.946E-06 | 530.76709 | 42.370041 |
| MKPLMGVIYVPLT | 17 | P22061 | Protein-L-isoaspartate(D-aspartate) O-me | PCMT1 | 24.637 | 3 | 52.359665 | yes | 2.658E-07 | 654.69824 | 35.761986 |
| PLMGVIYVPLTDK | 13 | P22061 | Protein-L-isoaspartate(D-aspartate) O-me | PCMT1 | 24.637 | 2 | 52.359665 | yes | 7.912E-06 | 723.40729 | 35.084873 |
| TNIIPVIEDAR | 11 | P22087 | rRNA 2'-O-methyltransferase fibrillarin C | FBL | 33.785 | 2 | 51.686462 | yes | 1.102E-07 | 620.84845 | 4713.5073 |
| HEGVFICR | 8 | P22087 | rRNA 2'-O-methyltransferase fibrillarin C | FBL | 33.785 | 2 | 51.686462 | yes | 1.69E-07 | 509.25037 | 1718.9287 |
| NVMVEPHR | 8 | P22087 | rRNA 2'-O-methyltransferase fibrillarin C | FBL | 33.785 | 2 | 51.686462 | yes | 2.165E-07 | 491.25037 | 609.21381 |
| NVMVEPHR | 8 | P22087 | rRNA 2'-O-methyltransferase fibrillarin C | FBL | 33.785 | 2 | 51.686462 | yes | 6.819E-07 | 499.24783 | 609.21381 |
| NLVPGESVYGEK | 12 | P22087 | rRNA 2'-O-methyltransferase fibrillarin C | FBL | 33.785 | 2 | 51.686462 | yes | 2.511E-07 | 646.33008 | 1090.3149 |
| VSISEGDDKIEYR | 13 | P22087 | rRNA 2'-O-methyltransferase fibrillarin C | FBL | 33.785 | 2 | 51.686462 | yes | 9.722E-08 | 755.8728 | 601.34082 |
| DHAVVVGVYRPPI | 14 | P22087 | rRNA 2'-O-methyltransferase fibrillarin C | FBL | 33.785 | 2 | 51.686462 | yes | 9.722E-08 | 767.43042 | 256.22705 |
| DHAVVVGVYRPPI | 14 | P22087 | rRNA 2'-O-methyltransferase fibrillarin C | FBL | 33.785 | 2 | 51.686462 | yes | 3.326E-07 | 511.95605 | 256.22705 |
| DHAVVVGVYRPPI | 14 | P22087 | rRNA 2'-O-methyltransferase fibrillarin C | FBL | 33.785 | 3 | 51.686462 | yes | 9.722E-08 | 767.43042 | 256.22705 |
| DHAVVVGVYRPPI | 14 | P22087 | rRNA 2'-O-methyltransferase fibrillarin C | FBL | 33.785 | 3 | 51.686462 | yes | 3.326E-07 | 511.95605 | 256.22705 |
| LAAAILGGVDQIHI | 19 | P22087 | rRNA 2'-O-methyltransferase fibrillarin C | FBL | 33.785 | 3 | 51.686462 | yes | 1.655E-07 | 624.70703 | 257.24509 |
| NGGHFVISIK | 10 | P22087 | rRNA 2'-O-methyltransferase fibrillarin C | FBL | 33.785 | 2 | 51.686462 | yes | 1.973E-07 | 536.3009 | 197.81078 |
| ANCIDSTASAEAV | 19 | P22087 | rRNA 2'-O-methyltransferase fibrillarin C | FBL | 33.785 | 3 | 51.686462 | yes | 6.377E-07 | 657.31055 | 60.181686 |
| LAAAILGGVDQIHI | 15 | P22087 | rRNA 2'-O-methyltransferase fibrillarin C | FBL | 33.785 | 3 | 51.686462 | no | 2.029E-07 | 506.97159 | 44.938145 |
| IVALNAHTFLR | 11 | P22087 | rRNA 2'-O-methyltransferase fibrillarin C | FBL | 33.785 | 3 | 51.686462 | no | 6.83E-07 | 418.91544 | 27.115021 |
| ILSGPFVQK | 9 | P22102 | Trifunctional purine biosynthetic protein a | GART | 107.768 | 2 | 52.148193 | yes | 2.231E-07 | 494.79492 | 192.93201 |
| EEAWVIGSVVAR | 12 | P22102 | Trifunctional purine biosynthetic protein a | GART | 107.768 | 2 | 52.148193 | yes | 2.023E-07 | 658.35388 | 102.36124 |
| ENLISALEEAK | 11 | P22102 | Trifunctional purine biosynthetic protein a | GART | 107.768 | 2 | 52.148193 | yes | 9.722E-08 | 608.82465 | 95.666878 |
| QVLVAPGNAGTA( | 16 | P22102 | Trifunctional purine biosynthetic protein a | GART | 107.768 | 2 | 52.148193 | yes | 1.02E-07 | 801.40125 | 62.127335 |
| HGIPTAQWK | 9 | P22102 | Trifunctional purine biosynthetic protein a | GART | 107.768 | 2 | 52.148193 | yes | 4.014E-06 | 519.27997 | 57.513493 |
| LGVDLDAQTWR | 11 | P22102 | Trifunctional purine biosynthetic protein a | GART | 107.768 | 2 | 52.148193 | yes | 9.848E-08 | 637.33038 | 49.317661 |
| AIAFLQQPR | 9 | P22102 | Trifunctional purine biosynthetic protein a | GART | 107.768 | 2 | 52.148193 | yes | 8.51E-06 | 522.30347 | 48.572788 |
| TFNCGVGAVLVVS | 14 | P22102 | Trifunctional purine biosynthetic protein a | GART | 107.768 | 2 | 52.148193 | yes | 5.123E-07 | 725.88977 | 40.312576 |
| EVTPEGLQMVK | 11 | P22234 | Bifunctional phosphoribosylaminoimidaz | PAICS | 47.08 | 2 | 49.528732 | yes | 1.022E-07 | 615.82355 | 108.11523 |
| EIVLADVIDNDSW | 14 | P22234 | Bifunctional phosphoribosylaminoimidaz | PAICS | 47.08 | 2 | 49.528732 | yes | 5.536E-07 | 822.91504 | 59.43903 |
| LAGTQPLEVLEAV | 15 | P22314 | Ubiquitin-like modifier-activating enzyme | UBA1 | 117.849 | 2 | 50.695286 | yes | 9.728E-08 | 812.45685 | 83.958199 |
| AENYDIPSADR | 11 | P22314 | Ubiquitin-like modifier-activating enzyme | UBA1 | 117.849 | 2 | 50.695286 | yes | 6.109E-07 | 625.78601 | 66.764664 |
| YDGQVAVFGSDL( | 15 | P22314 | Ubiquitin-like modifier-activating enzyme | UBA1 | 117.849 | 2 | 50.695286 | yes | 3.895E-07 | 828.39923 | 62.462517 |
| LQTSSVLVSGLR | 12 | P22314 | Ubiquitin-like modifier-activating enzyme | UBA1 | 117.849 | 2 | 50.695286 | yes | 4.885E-07 | 630.36951 | 44.166443 |
| KPLLESGTLGTK | 12 | P22314 | Ubiquitin-like modifier-activating enzyme | UBA1 | 117.849 | 2 | 50.695286 | yes | 4.042E-07 | 622.36646 | 32.886215 |
| TFIAIKPDGVQR | 12 | P22392 | Nucleoside diphosphate kinase B OS=Ho | NME2 | 17.299 | 2 | 50.398666 | no | 9.809E-08 | 672.88531 | 227.04684 |
| NIIHGSDSVK | 10 | P22392 | Nucleoside diphosphate kinase B OS=Ho | NME2 | 17.299 | 2 | 50.398666 | no | 2.186E-07 | 535.28546 | 153.35674 |
| VMLGETNPADSKI | 17 | P22392 | Nucleoside diphosphate kinase B OS=Ho | NME2 | 17.299 | 3 | 50.398666 | no | 1.026E-07 | 595.97699 | 121.81012 |
| SCAHDWVYE | 9 | P22392 | Nucleoside diphosphate kinase B OS=Ho | NME2 | 17.299 | 2 | 50.398666 | no | 1.295E-07 | 583.73218 | 106.06152 |
| DRPFFPGLVK | 10 | P22392 | Nucleoside diphosphate kinase B OS=Ho | NME2 | 17.299 | 2 | 50.398666 | no | 9.819E-08 | 588.33221 | 95.341019 |
| INNWIVQLSK | 10 | P22415 | Upstream stimulatory factor 1 OS=Homo | USF1 | 33.539 | 2 | 46.654697 | yes | 1.503E-07 | 607.84821 | 44.685394 |

| | | | | | | | | | | | |
|---|---|---|---|---|---|---|---|---|---|---|---|
| IDTIEIITDR | 10 | P22626 | Heterogeneous nuclear ribonucleoprotein: | HNRNPA2 | 37.43 | 2 | 52.344246 | yes | 1.301E-07 | 594.82715 | 64748.84 |
| LTDCVVMR | 8 | P22626 | Heterogeneous nuclear ribonucleoprotein: | HNRNPA2 | 37.43 | 2 | 52.344246 | yes | 1.415E-07 | 497.24643 | 30389.547 |
| LTDCVVMR | 8 | P22626 | Heterogeneous nuclear ribonucleoprotein: | HNRNPA2 | 37.43 | 2 | 52.344246 | yes | 1.278E-07 | 505.2439 | 30389.547 |
| GGGGNFGPGPGSN | 15 | P22626 | Heterogeneous nuclear ribonucleoprotein: | HNRNPA2 | 37.43 | 2 | 52.344246 | yes | 9.759E-08 | 689.31836 | 39827.875 |
| TLETVPLER | 9 | P22626 | Heterogeneous nuclear ribonucleoprotein: | HNRNPA2 | 37.43 | 2 | 52.344246 | yes | 1.571E-07 | 529.29803 | 31180.65 |
| GGNFGFGDSR | 10 | P22626 | Heterogeneous nuclear ribonucleoprotein: | HNRNPA2 | 37.43 | 2 | 52.344246 | yes | 1.821E-07 | 507.2254 | 24683.027 |
| QEMQEVQSSR | 10 | P22626 | Heterogeneous nuclear ribonucleoprotein: | HNRNPA2 | 37.43 | 2 | 52.344246 | yes | 9.722E-08 | 611.28003 | 7454.0713 |
| QEMQEVQSSR | 10 | P22626 | Heterogeneous nuclear ribonucleoprotein: | HNRNPA2 | 37.43 | 2 | 52.344246 | yes | 9.722E-08 | 619.27753 | 7454.0713 |
| EESGKPGAHVTVK | 13 | P22626 | Heterogeneous nuclear ribonucleoprotein: | HNRNPA2 | 37.43 | 2 | 52.344246 | yes | 9.877E-08 | 669.85425 | 6602.6675 |
| EESGKPGAHVTVK | 13 | P22626 | Heterogeneous nuclear ribonucleoprotein: | HNRNPA2 | 37.43 | 2 | 52.344246 | yes | 2.263E-07 | 446.90524 | 6602.6675 |
| EESGKPGAHVTVK | 13 | P22626 | Heterogeneous nuclear ribonucleoprotein: | HNRNPA2 | 37.43 | 3 | 52.344246 | yes | 9.877E-08 | 669.85425 | 6602.6675 |
| EESGKPGAHVTVK | 13 | P22626 | Heterogeneous nuclear ribonucleoprotein: | HNRNPA2 | 37.43 | 3 | 52.344246 | yes | 2.263E-07 | 446.90524 | 6602.6675 |
| DYFEEYGK | 8 | P22626 | Heterogeneous nuclear ribonucleoprotein: | HNRNPA2 | 37.43 | 2 | 52.344246 | yes | 2.035E-07 | 525.72437 | 9438.0586 |
| NYYEQWGK | 8 | P22626 | Heterogeneous nuclear ribonucleoprotein: | HNRNPA2 | 37.43 | 2 | 52.344246 | yes | 1.632E-07 | 544.24579 | 5383.0781 |
| KLFVGGIK | 8 | P22626 | Heterogeneous nuclear ribonucleoprotein: | HNRNPA2 | 37.43 | 2 | 52.344246 | yes | 2.189E-07 | 431.28146 | 3701.6982 |
| KLFIGGLSFETTEE | 17 | P22626 | Heterogeneous nuclear ribonucleoprotein: | HNRNPA2 | 37.43 | 3 | 52.344246 | yes | 1.087E-07 | 643.01056 | 1165.9248 |
| YHTINGHNAEVR | 12 | P22626 | Heterogeneous nuclear ribonucleoprotein: | HNRNPA2 | 37.43 | 2 | 52.344246 | yes | 9.851E-08 | 705.84729 | 298.00702 |
| YHTINGHNAEVR | 12 | P22626 | Heterogeneous nuclear ribonucleoprotein: | HNRNPA2 | 37.43 | 2 | 52.344246 | yes | 9.863E-07 | 470.9006 | 298.00702 |
| YHTINGHNAEVR | 12 | P22626 | Heterogeneous nuclear ribonucleoprotein: | HNRNPA2 | 37.43 | 3 | 52.344246 | yes | 9.851E-08 | 705.84729 | 298.00702 |
| YHTINGHNAEVR | 12 | P22626 | Heterogeneous nuclear ribonucleoprotein: | HNRNPA2 | 37.43 | 3 | 52.344246 | yes | 9.863E-07 | 470.9006 | 298.00702 |
| GGSDGYGSGR | 10 | P22626 | Heterogeneous nuclear ribonucleoprotein: | HNRNPA2 | 37.43 | 2 | 52.344246 | yes | 5.206E-07 | 456.69394 | 150.13956 |
| SRGFGFVTFSSMA | 22 | P22626 | Heterogeneous nuclear ribonucleoprotein: | HNRNPA2 | 37.43 | 3 | 52.344246 | yes | 3.807E-07 | 770.03314 | 114.9198 |
| LTDCVVMRDPASI | 13 | P22626 | Heterogeneous nuclear ribonucleoprotein: | HNRNPA2 | 37.43 | 2 | 52.344246 | yes | 3.591E-07 | 746.36835 | 25.33626 |
| LTDCVVMRDPASI | 13 | P22626 | Heterogeneous nuclear ribonucleoprotein: | HNRNPA2 | 37.43 | 2 | 52.344246 | yes | 3.14E-07 | 754.36578 | 25.33626 |
| NMGGPYGGGNYG | 25 | P22626 | Heterogeneous nuclear ribonucleoprotein: | HNRNPA2 | 37.43 | 3 | 52.344246 | yes | 3.517E-07 | 730.63995 | 46.330036 |
| PGAHVTVK | 8 | P22626 | Heterogeneous nuclear ribonucleoprotein: | HNRNPA2 | 37.43 | 2 | 52.344246 | yes | 1.179E-07 | 404.73743 | 44.266106 |
| AVAFQNPQTHVIE | 21 | P22695 | Cytochrome b-c1 complex subunit 2, mitc | UQCRC2 | 48.443 | 3 | 54.620495 | yes | 1.367E-07 | 784.07324 | 196.33191 |
| NALANPLYCPDYF | 13 | P22695 | Cytochrome b-c1 complex subunit 2, mitc | UQCRC2 | 48.443 | 2 | 54.620495 | yes | 1.438E-07 | 783.87213 | 149.50693 |
| TIAQGNLSNTDVQ | 16 | P22695 | Cytochrome b-c1 complex subunit 2, mitc | UQCRC2 | 48.443 | 2 | 54.620495 | yes | 9.722E-08 | 815.9234 | 145.81892 |
| ENMAYTVECLR | 11 | P22695 | Cytochrome b-c1 complex subunit 2, mitc | UQCRC2 | 48.443 | 2 | 54.620495 | yes | 9.722E-08 | 693.31305 | 139.92322 |
| YEDFSNLGTTHLLI | 14 | P22695 | Cytochrome b-c1 complex subunit 2, mitc | UQCRC2 | 48.443 | 2 | 54.620495 | yes | 1.11E-07 | 833.41522 | 134.07693 |
| QVAEQFLNMR | 10 | P22695 | Cytochrome b-c1 complex subunit 2, mitc | UQCRC2 | 48.443 | 2 | 54.620495 | yes | 7.847E-07 | 618.31372 | 53.798405 |
| GDPYPQEVSATVQ | 14 | P22830 | Ferrochelatase, mitochondrial OS=Homo | FECH | 47.863 | 2 | 45.861385 | yes | 1.721E-07 | 759.87537 | 92.935455 |
| DLMTLPIQNK | 10 | P22830 | Ferrochelatase, mitochondrial OS=Homo | FECH | 47.863 | 2 | 45.861385 | yes | 6.584E-07 | 586.8208 | 67.73896 |
| SLEVWGSQEALEE | 15 | P23025 | DNA repair protein complementing XP-A | XPA | 31.368 | 2 | 45.84526 | yes | 1.726E-07 | 838.41229 | 169.59409 |
| LGESCINTVGSFR | 13 | P23142 | Fibulin-1 OS=Homo sapiens OX=9606 G | FBLN1 | 77.215 | 3 | 53.79689 | yes | 7.662E-07 | 480.56976 | 324.05292 |
| GYHLNEEGTR | 10 | P23142 | Fibulin-1 OS=Homo sapiens OX=9606 G | FBLN1 | 77.215 | 2 | 53.79689 | yes | 1.458E-07 | 588.27563 | 343.23654 |
| EFTRPEEIIFLR | 12 | P23142 | Fibulin-1 OS=Homo sapiens OX=9606 G | FBLN1 | 77.215 | 3 | 53.79689 | yes | 9.722E-08 | 517.28394 | 157.23758 |
| YMDGMTVGVVR | 11 | P23142 | Fibulin-1 OS=Homo sapiens OX=9606 G | FBLN1 | 77.215 | 2 | 53.79689 | yes | 1.272E-07 | 614.29663 | 63.449936 |

| | | | | | | | | | | | |
|---|---|---|---|---|---|---|---|---|---|---|---|
| CCHCCLLGR | 9 | P23142 | Fibulin-1 OS=Homo sapiens OX=9606 G | FBLN1 | 77.215 | 2 | 53.79689 | yes | 1.158E-07 | 618.24866 | 39.47554 |
| CLAFECPENYR | 11 | P23142 | Fibulin-1 OS=Homo sapiens OX=9606 G | FBLN1 | 77.215 | 2 | 53.79689 | yes | 9.728E-08 | 729.81067 | 37.066101 |
| MVQEQCCHSQLEI | 28 | P23142 | Fibulin-1 OS=Homo sapiens OX=9606 G | FBLN1 | 77.215 | 4 | 53.79689 | yes | 3.216E-07 | 836.61945 | 32.306705 |
| MTAEEMASDELK | 12 | P23193 | Transcription elongation factor A protein | TCEA1 | 33.97 | 2 | 50.454243 | yes | 9.809E-08 | 677.79688 | 125.55293 |
| DTYVSSFPR | 9 | P23193 | Transcription elongation factor A protein | TCEA1 | 33.97 | 2 | 50.454243 | yes | 3.403E-07 | 536.25891 | 117.52461 |
| TGGTQTDLFTCGK | 13 | P23193 | Transcription elongation factor A protein | TCEA1 | 33.97 | 2 | 50.454243 | no | 1.282E-07 | 693.3219 | 56.513264 |
| NCTYTQVQTR | 10 | P23193 | Transcription elongation factor A protein | TCEA1 | 33.97 | 2 | 50.454243 | no | 1.022E-07 | 635.7959 | 54.131847 |
| QSTDEEVTSLAK | 12 | P23193 | Transcription elongation factor A protein | TCEA1 | 33.97 | 2 | 50.454243 | yes | 2.303E-07 | 654.31989 | 43.263538 |
| CSEGVFLLTTTPR | 13 | P23246 | Splicing factor, proline- and glutamine-ric | SFPQ | 76.15 | 2 | 52.789658 | yes | 1.778E-07 | 740.87683 | 60532.734 |
| SPPPGMGLNQNR | 12 | P23246 | Splicing factor, proline- and glutamine-ric | SFPQ | 76.15 | 2 | 52.789658 | yes | 2.3E-07 | 634.31421 | 27383.412 |
| SPPPGMGLNQNR | 12 | P23246 | Splicing factor, proline- and glutamine-ric | SFPQ | 76.15 | 2 | 52.789658 | yes | 1.507E-07 | 642.31171 | 27383.412 |
| AELDDTPMR | 9 | P23246 | Splicing factor, proline- and glutamine-ric | SFPQ | 76.15 | 2 | 52.789658 | yes | 1.145E-07 | 524.24243 | 26883.584 |
| AELDDTPMR | 9 | P23246 | Splicing factor, proline- and glutamine-ric | SFPQ | 76.15 | 2 | 52.789658 | yes | 1.323E-07 | 532.23987 | 26883.584 |
| YGEPGEVFINK | 11 | P23246 | Splicing factor, proline- and glutamine-ric | SFPQ | 76.15 | 2 | 52.789658 | yes | 1.603E-07 | 626.81403 | 46681.711 |
| FATHAAALSVR | 11 | P23246 | Splicing factor, proline- and glutamine-ric | SFPQ | 76.15 | 2 | 52.789658 | yes | 1.219E-07 | 572.31708 | 37549.512 |
| GMGPGTPAGYGR | 12 | P23246 | Splicing factor, proline- and glutamine-ric | SFPQ | 76.15 | 2 | 52.789658 | yes | 2.284E-07 | 560.76385 | 15211.068 |
| GMGPGTPAGYGR | 12 | P23246 | Splicing factor, proline- and glutamine-ric | SFPQ | 76.15 | 2 | 52.789658 | yes | 2.184E-07 | 568.76129 | 15211.068 |
| MPGGPKPGGGPGI | 25 | P23246 | Splicing factor, proline- and glutamine-ric | SFPQ | 76.15 | 4 | 52.789658 | yes | 1.233E-07 | 593.31165 | 9178.1289 |
| MPGGPKPGGGPGI | 25 | P23246 | Splicing factor, proline- and glutamine-ric | SFPQ | 76.15 | 4 | 52.789658 | yes | 3.824E-07 | 597.31036 | 9178.1289 |
| MPGGPKPGGGPGI | 25 | P23246 | Splicing factor, proline- and glutamine-ric | SFPQ | 76.15 | 4 | 52.789658 | yes | 2.63E-07 | 474.85077 | 9178.1289 |
| MPGGPKPGGGPGI | 25 | P23246 | Splicing factor, proline- and glutamine-ric | SFPQ | 76.15 | 4 | 52.789658 | yes | 1.008E-07 | 790.74646 | 9178.1289 |
| MPGGPKPGGGPGI | 25 | P23246 | Splicing factor, proline- and glutamine-ric | SFPQ | 76.15 | 4 | 52.789658 | yes | 2.926E-06 | 796.07806 | 9178.1289 |
| MPGGPKPGGGPGI | 25 | P23246 | Splicing factor, proline- and glutamine-ric | SFPQ | 76.15 | 5 | 52.789658 | yes | 1.233E-07 | 593.31165 | 9178.1289 |
| MPGGPKPGGGPGI | 25 | P23246 | Splicing factor, proline- and glutamine-ric | SFPQ | 76.15 | 5 | 52.789658 | yes | 3.824E-07 | 597.31036 | 9178.1289 |
| MPGGPKPGGGPGI | 25 | P23246 | Splicing factor, proline- and glutamine-ric | SFPQ | 76.15 | 5 | 52.789658 | yes | 2.63E-07 | 474.85077 | 9178.1289 |
| MPGGPKPGGGPGI | 25 | P23246 | Splicing factor, proline- and glutamine-ric | SFPQ | 76.15 | 5 | 52.789658 | yes | 1.008E-07 | 790.74646 | 9178.1289 |
| MPGGPKPGGGPGI | 25 | P23246 | Splicing factor, proline- and glutamine-ric | SFPQ | 76.15 | 5 | 52.789658 | yes | 2.926E-06 | 796.07806 | 9178.1289 |
| MPGGPKPGGGPGI | 25 | P23246 | Splicing factor, proline- and glutamine-ric | SFPQ | 76.15 | 3 | 52.789658 | yes | 1.233E-07 | 593.31165 | 9178.1289 |
| MPGGPKPGGGPGI | 25 | P23246 | Splicing factor, proline- and glutamine-ric | SFPQ | 76.15 | 3 | 52.789658 | yes | 3.824E-07 | 597.31036 | 9178.1289 |
| MPGGPKPGGGPGI | 25 | P23246 | Splicing factor, proline- and glutamine-ric | SFPQ | 76.15 | 3 | 52.789658 | yes | 2.63E-07 | 474.85077 | 9178.1289 |
| MPGGPKPGGGPGI | 25 | P23246 | Splicing factor, proline- and glutamine-ric | SFPQ | 76.15 | 3 | 52.789658 | yes | 1.008E-07 | 790.74646 | 9178.1289 |
| MPGGPKPGGGPGI | 25 | P23246 | Splicing factor, proline- and glutamine-ric | SFPQ | 76.15 | 3 | 52.789658 | yes | 2.926E-06 | 796.07806 | 9178.1289 |
| FGQGGAGPVGGQ( | 15 | P23246 | Splicing factor, proline- and glutamine-ric | SFPQ | 76.15 | 2 | 52.789658 | yes | 1.163E-07 | 671.33655 | 26294.705 |
| MGYMDPR | 7 | P23246 | Splicing factor, proline- and glutamine-ric | SFPQ | 76.15 | 2 | 52.789658 | yes | 5.462E-07 | 435.18585 | 5995.7407 |
| MGYMDPR | 7 | P23246 | Splicing factor, proline- and glutamine-ric | SFPQ | 76.15 | 2 | 52.789658 | yes | 1.296E-06 | 443.18332 | 5995.7407 |
| MGYMDPR | 7 | P23246 | Splicing factor, proline- and glutamine-ric | SFPQ | 76.15 | 2 | 52.789658 | yes | 3.187E-06 | 443.18332 | 5995.7407 |
| PGGGPGLSTPGGH | 19 | P23246 | Splicing factor, proline- and glutamine-ric | SFPQ | 76.15 | 4 | 52.789658 | yes | 2.405E-06 | 451.49066 | 8533.3281 |
| PGGGPGLSTPGGH | 19 | P23246 | Splicing factor, proline- and glutamine-ric | SFPQ | 76.15 | 4 | 52.789658 | yes | 1.271E-06 | 601.65179 | 8533.3281 |
| PGGGPGLSTPGGH | 19 | P23246 | Splicing factor, proline- and glutamine-ric | SFPQ | 76.15 | 3 | 52.789658 | yes | 2.405E-06 | 451.49066 | 8533.3281 |

| | | | | | | | | | | | |
|---|---|---|---|---|---|---|---|---|---|---|---|
| PGGGPGLSTPGGH | 19 | P23246 | Splicing factor, proline- and glutamine-ric | SFPQ | 76.15 | 3 | 52.789658 | yes | 1.271E-06 | 601.65179 | 8533.3281 |
| EMEEQMR | 7 | P23246 | Splicing factor, proline- and glutamine-ric | SFPQ | 76.15 | 2 | 52.789658 | yes | 1.71E-07 | 476.69678 | 6537.6807 |
| EMEEQMR | 7 | P23246 | Splicing factor, proline- and glutamine-ric | SFPQ | 76.15 | 2 | 52.789658 | yes | 1.766E-07 | 484.69424 | 6537.6807 |
| DKLESEMEDAYHI | 20 | P23246 | Splicing factor, proline- and glutamine-ric | SFPQ | 76.15 | 4 | 52.789658 | yes | 9.722E-08 | 607.7854 | 3762.9089 |
| DKLESEMEDAYHI | 20 | P23246 | Splicing factor, proline- and glutamine-ric | SFPQ | 76.15 | 4 | 52.789658 | yes | 4.72E-07 | 611.78412 | 3762.9089 |
| GREEYEGPNK | 10 | P23246 | Splicing factor, proline- and glutamine-ric | SFPQ | 76.15 | 2 | 52.789658 | yes | 1.837E-07 | 589.77545 | 5709.2285 |
| EEEMMIR | 7 | P23246 | Splicing factor, proline- and glutamine-ric | SFPQ | 76.15 | 2 | 52.789658 | yes | 2.159E-07 | 469.20953 | 2256.0791 |
| EEEMMIR | 7 | P23246 | Splicing factor, proline- and glutamine-ric | SFPQ | 76.15 | 2 | 52.789658 | yes | 8.869E-05 | 477.20697 | 2256.0791 |
| RMEELHNQEMQK | 12 | P23246 | Splicing factor, proline- and glutamine-ric | SFPQ | 76.15 | 3 | 52.789658 | yes | 1.468E-07 | 524.91345 | 1459.9216 |
| RMEELHNQEMQK | 12 | P23246 | Splicing factor, proline- and glutamine-ric | SFPQ | 76.15 | 3 | 52.789658 | yes | 1.076E-07 | 786.86652 | 1459.9216 |
| RMEELHNQEMQK | 12 | P23246 | Splicing factor, proline- and glutamine-ric | SFPQ | 76.15 | 3 | 52.789658 | yes | 1.425E-07 | 530.24506 | 1459.9216 |
| RMEELHNQEMQK | 12 | P23246 | Splicing factor, proline- and glutamine-ric | SFPQ | 76.15 | 3 | 52.789658 | yes | 4.239E-06 | 794.86395 | 1459.9216 |
| RMEELHNQEMQK | 12 | P23246 | Splicing factor, proline- and glutamine-ric | SFPQ | 76.15 | 2 | 52.789658 | yes | 1.468E-07 | 524.91345 | 1459.9216 |
| RMEELHNQEMQK | 12 | P23246 | Splicing factor, proline- and glutamine-ric | SFPQ | 76.15 | 2 | 52.789658 | yes | 1.076E-07 | 786.86652 | 1459.9216 |
| RMEELHNQEMQK | 12 | P23246 | Splicing factor, proline- and glutamine-ric | SFPQ | 76.15 | 2 | 52.789658 | yes | 1.425E-07 | 530.24506 | 1459.9216 |
| RMEELHNQEMQK | 12 | P23246 | Splicing factor, proline- and glutamine-ric | SFPQ | 76.15 | 2 | 52.789658 | yes | 4.239E-06 | 794.86395 | 1459.9216 |
| MEELHNQEMQK | 11 | P23246 | Splicing factor, proline- and glutamine-ric | SFPQ | 76.15 | 2 | 52.789658 | yes | 1.558E-07 | 708.81592 | 1250.0444 |
| MEELHNQEMQK | 11 | P23246 | Splicing factor, proline- and glutamine-ric | SFPQ | 76.15 | 2 | 52.789658 | yes | 1.061E-07 | 472.87973 | 1250.0444 |
| MEELHNQEMQK | 11 | P23246 | Splicing factor, proline- and glutamine-ric | SFPQ | 76.15 | 2 | 52.789658 | yes | 9.722E-08 | 716.81342 | 1250.0444 |
| MEELHNQEMQK | 11 | P23246 | Splicing factor, proline- and glutamine-ric | SFPQ | 76.15 | 2 | 52.789658 | yes | 2.42E-06 | 716.81342 | 1250.0444 |
| MEELHNQEMQK | 11 | P23246 | Splicing factor, proline- and glutamine-ric | SFPQ | 76.15 | 3 | 52.789658 | yes | 1.558E-07 | 708.81592 | 1250.0444 |
| MEELHNQEMQK | 11 | P23246 | Splicing factor, proline- and glutamine-ric | SFPQ | 76.15 | 3 | 52.789658 | yes | 1.061E-07 | 472.87973 | 1250.0444 |
| MEELHNQEMQK | 11 | P23246 | Splicing factor, proline- and glutamine-ric | SFPQ | 76.15 | 3 | 52.789658 | yes | 9.722E-08 | 716.81342 | 1250.0444 |
| MEELHNQEMQK | 11 | P23246 | Splicing factor, proline- and glutamine-ric | SFPQ | 76.15 | 3 | 52.789658 | yes | 2.42E-06 | 716.81342 | 1250.0444 |
| REEEMMIR | 8 | P23246 | Splicing factor, proline- and glutamine-ric | SFPQ | 76.15 | 2 | 52.789658 | yes | 1.231E-07 | 547.26007 | 2877.1499 |
| LFVGNLPADITEDI | 17 | P23246 | Splicing factor, proline- and glutamine-ric | SFPQ | 76.15 | 3 | 52.789658 | yes | 1.611E-07 | 655.34229 | 2271.1292 |
| ISDSEGFK | 8 | P23246 | Splicing factor, proline- and glutamine-ric | SFPQ | 76.15 | 2 | 52.789658 | yes | 1.404E-07 | 441.71381 | 2103.949 |
| PVIVEPLEQLDDEI | 19 | P23246 | Splicing factor, proline- and glutamine-ric | SFPQ | 76.15 | 3 | 52.789658 | yes | 2.473E-07 | 712.36334 | 1078.8353 |
| QHHPPYHQQHHQ | 22 | P23246 | Splicing factor, proline- and glutamine-ric | SFPQ | 76.15 | 4 | 52.789658 | yes | 1.15E-07 | 601.53925 | 420.9375 |
| QHHPPYHQQHHQ | 22 | P23246 | Splicing factor, proline- and glutamine-ric | SFPQ | 76.15 | 4 | 52.789658 | yes | 2.114E-07 | 481.43283 | 420.9375 |
| QHHPPYHQQHHQ | 22 | P23246 | Splicing factor, proline- and glutamine-ric | SFPQ | 76.15 | 5 | 52.789658 | yes | 1.15E-07 | 601.53925 | 420.9375 |
| QHHPPYHQQHHQ | 22 | P23246 | Splicing factor, proline- and glutamine-ric | SFPQ | 76.15 | 5 | 52.789658 | yes | 2.114E-07 | 481.43283 | 420.9375 |
| RREEEMMIR | 9 | P23246 | Splicing factor, proline- and glutamine-ric | SFPQ | 76.15 | 2 | 52.789658 | yes | 9.722E-08 | 625.31061 | 499.97723 |
| SLDEMEK | 7 | P23246 | Splicing factor, proline- and glutamine-ric | SFPQ | 76.15 | 2 | 52.789658 | yes | 1.257E-07 | 426.1944 | 339.46869 |
| EEYEGPNK | 8 | P23246 | Splicing factor, proline- and glutamine-ric | SFPQ | 76.15 | 2 | 52.789658 | yes | 1.863E-07 | 483.21417 | 280.27802 |
| LESEMEDAYHEH | 18 | P23246 | Splicing factor, proline- and glutamine-ric | SFPQ | 76.15 | 4 | 52.789658 | yes | 5.526E-07 | 547.00494 | 278.50464 |
| GPMGPGPGQSGPK | 13 | P23246 | Splicing factor, proline- and glutamine-ric | SFPQ | 76.15 | 2 | 52.789658 | yes | 6.676E-07 | 583.78479 | 252.44778 |
| GGLHDFR | 7 | P23246 | Splicing factor, proline- and glutamine-ric | SFPQ | 76.15 | 2 | 52.789658 | yes | 1.025E-07 | 401.20374 | 244.75766 |
| FPPLGGGGGIGYEA | 33 | P23246 | Splicing factor, proline- and glutamine-ric | SFPQ | 76.15 | 4 | 52.789658 | yes | 1.144E-07 | 799.61462 | 111.75299 |

| | | | | | | | | | | |
|---|---|---|---|---|---|---|---|---|---|---|
| FPPLGGGGGIGYEA | 33 | P23246 | Splicing factor, proline- and glutamine-ric | SFPQ | 76.15 | 4 | 52.789658 | yes | 3.444E-06 | 803.61334 | 111.75299 |
| MPGGPKPGGGPGI | 21 | P23246 | Splicing factor, proline- and glutamine-ric | SFPQ | 76.15 | 3 | 52.789658 | yes | 1.422E-07 | 628.32458 | 188.36577 |
| MEELHNQEMQKR | 12 | P23246 | Splicing factor, proline- and glutamine-ric | SFPQ | 76.15 | 2 | 52.789658 | yes | 2.696E-06 | 786.86652 | 13.549305 |
| AVLLDLEPR | 9 | P23258 | Tubulin gamma-1 chain OS=Homo sapier | TUBG1 | 51.17 | 2 | 52.501022 | no | 2.54E-07 | 513.3031 | 2231.156 |
| IHEDIFDIIDR | 11 | P23258 | Tubulin gamma-1 chain OS=Homo sapier | TUBG1 | 51.17 | 2 | 52.501022 | no | 1.313E-07 | 693.35663 | 1380.5345 |
| EAFLEQFR | 8 | P23258 | Tubulin gamma-1 chain OS=Homo sapier | TUBG1 | 51.17 | 2 | 52.501022 | yes | 2.333E-07 | 520.26398 | 1379.2859 |
| KTTVLDVMR | 9 | P23258 | Tubulin gamma-1 chain OS=Homo sapier | TUBG1 | 51.17 | 2 | 52.501022 | no | 1.606E-07 | 531.80243 | 1344.213 |
| SPYLPSAHR | 9 | P23258 | Tubulin gamma-1 chain OS=Homo sapier | TUBG1 | 51.17 | 2 | 52.501022 | no | 1.465E-07 | 514.26959 | 804.46332 |
| KSPYLPSAHR | 10 | P23258 | Tubulin gamma-1 chain OS=Homo sapier | TUBG1 | 51.17 | 2 | 52.501022 | no | 9.722E-08 | 578.31708 | 584.65125 |
| TTVLDVMR | 8 | P23258 | Tubulin gamma-1 chain OS=Homo sapier | TUBG1 | 51.17 | 2 | 52.501022 | no | 2.969E-07 | 467.75494 | 384.30121 |
| LTQNADCVVVLDI | 18 | P23258 | Tubulin gamma-1 chain OS=Homo sapier | TUBG1 | 51.17 | 3 | 52.501022 | no | 1.248E-07 | 672.67749 | 357.97372 |
| VSGLMMANHTSIS | 18 | P23258 | Tubulin gamma-1 chain OS=Homo sapier | TUBG1 | 51.17 | 3 | 52.501022 | yes | 1.629E-07 | 660.66077 | 188.49585 |
| DNFDEMDTSR | 10 | P23258 | Tubulin gamma-1 chain OS=Homo sapier | TUBG1 | 51.17 | 2 | 52.501022 | yes | 1.464E-07 | 615.2406 | 128.89783 |
| DVFFYQADDEHYI | 15 | P23258 | Tubulin gamma-1 chain OS=Homo sapier | TUBG1 | 51.17 | 3 | 52.501022 | no | 1.602E-06 | 638.96008 | 106.23158 |
| KDVFFYQADDEHY | 16 | P23258 | Tubulin gamma-1 chain OS=Homo sapier | TUBG1 | 51.17 | 3 | 52.501022 | no | 6.129E-07 | 681.65839 | 24.618988 |
| TVDNFVALATGEK | 13 | P23284 | Peptidyl-prolyl cis-trans isomerase B OS= | PPIB | 23.743 | 2 | 50.330154 | yes | 9.819E-08 | 682.85645 | 851.02832 |
| VLEGMEVVR | 9 | P23284 | Peptidyl-prolyl cis-trans isomerase B OS= | PPIB | 23.743 | 2 | 50.330154 | yes | 1.959E-07 | 516.28131 | 578.32739 |
| DFMIQGGDFTR | 11 | P23284 | Peptidyl-prolyl cis-trans isomerase B OS= | PPIB | 23.743 | 2 | 50.330154 | yes | 1.168E-07 | 643.79535 | 494.84695 |
| IGDEDVGR | 8 | P23284 | Peptidyl-prolyl cis-trans isomerase B OS= | PPIB | 23.743 | 2 | 50.330154 | yes | 2.607E-07 | 430.70905 | 392.65601 |
| HYGPGWVSMANA | 14 | P23284 | Peptidyl-prolyl cis-trans isomerase B OS= | PPIB | 23.743 | 2 | 50.330154 | yes | 4.841E-07 | 737.84845 | 58.366432 |
| DTNGSQFFITTVK | 13 | P23284 | Peptidyl-prolyl cis-trans isomerase B OS= | PPIB | 23.743 | 2 | 50.330154 | yes | 1.384E-06 | 729.36719 | 46.5896 |
| ALTSQLTDEELAQ | 15 | P23368 | NAD-dependent malic enzyme, mitochon | ME2 | 65.444 | 2 | 44.010265 | yes | 2.357E-07 | 816.41541 | 105.99948 |
| GKPLMLNPR | 9 | P23368 | NAD-dependent malic enzyme, mitochon | ME2 | 65.444 | 2 | 44.010265 | yes | 4.863E-07 | 513.29987 | 83.092201 |
| TQQYDDLIDEFMK | 13 | P23368 | NAD-dependent malic enzyme, mitochon | ME2 | 65.444 | 2 | 44.010265 | yes | 5.057E-07 | 823.37433 | 32.648472 |
| DIIACGFDINK | 11 | P23381 | Tryptophan--tRNA ligase, cytoplasmic OS | WARS1 | 53.166 | 2 | 39.09368 | yes | 4.793E-07 | 633.31335 | 18.005459 |
| TEIIILATR | 9 | P23396 | Small ribosomal subunit protein uS3 OS= | RPS3 | 26.689 | 2 | 52.50722 | yes | 1.948E-07 | 515.31879 | 62638.613 |
| FGFPEGSVELYAEI | 14 | P23396 | Small ribosomal subunit protein uS3 OS= | RPS3 | 26.689 | 2 | 52.50722 | yes | 2.809E-07 | 786.88269 | 37168.031 |
| AELNEFLTR | 9 | P23396 | Small ribosomal subunit protein uS3 OS= | RPS3 | 26.689 | 2 | 52.50722 | yes | 2.131E-07 | 546.78784 | 35549.594 |
| GLCAIAQAESLR | 12 | P23396 | Small ribosomal subunit protein uS3 OS= | RPS3 | 26.689 | 2 | 52.50722 | yes | 1.906E-07 | 644.83752 | 34292.719 |
| DEILPTTPISEQK | 13 | P23396 | Small ribosomal subunit protein uS3 OS= | RPS3 | 26.689 | 2 | 52.50722 | yes | 2.12E-07 | 735.88794 | 26967.529 |
| ELAEDGYSGVEVF | 13 | P23396 | Small ribosomal subunit protein uS3 OS= | RPS3 | 26.689 | 2 | 52.50722 | yes | 9.722E-08 | 712.33862 | 25274.857 |
| GGKPEPPAMPQPV | 16 | P23396 | Small ribosomal subunit protein uS3 OS= | RPS3 | 26.689 | 2 | 52.50722 | yes | 1.293E-07 | 787.40576 | 9382.9834 |
| GGKPEPPAMPQPV | 16 | P23396 | Small ribosomal subunit protein uS3 OS= | RPS3 | 26.689 | 2 | 52.50722 | yes | 7.714E-07 | 795.40326 | 9382.9834 |
| IMLPWDPTGK | 10 | P23396 | Small ribosomal subunit protein uS3 OS= | RPS3 | 26.689 | 2 | 52.50722 | yes | 1.841E-07 | 579.30481 | 5911.7656 |
| IMLPWDPTGK | 10 | P23396 | Small ribosomal subunit protein uS3 OS= | RPS3 | 26.689 | 2 | 52.50722 | yes | 1.238E-06 | 587.30225 | 5911.7656 |
| KFVADGIFK | 9 | P23396 | Small ribosomal subunit protein uS3 OS= | RPS3 | 26.689 | 2 | 52.50722 | yes | 1.499E-07 | 512.79492 | 10771.694 |
| KPLPDHVSIVEPK | 13 | P23396 | Small ribosomal subunit protein uS3 OS= | RPS3 | 26.689 | 3 | 52.50722 | yes | 3.004E-07 | 486.94867 | 5258.3608 |
| KPLPDHVSIVEPK | 13 | P23396 | Small ribosomal subunit protein uS3 OS= | RPS3 | 26.689 | 3 | 52.50722 | yes | 2.602E-07 | 729.91937 | 5258.3608 |
| KPLPDHVSIVEPK | 13 | P23396 | Small ribosomal subunit protein uS3 OS= | RPS3 | 26.689 | 2 | 52.50722 | yes | 3.004E-07 | 486.94867 | 5258.3608 |

| | | | | | | | | | | |
|---|---|---|---|---|---|---|---|---|---|---|
| KPLPDHVSIVEPK | 13 | P23396 | Small ribosomal subunit protein uS3 OS= | RPS3 | 26.689 | 2 | 52.50722 | yes | 2.602E-07 | 729.91937 | 5258.3608 |
| FIMESGAK | 8 | P23396 | Small ribosomal subunit protein uS3 OS= | RPS3 | 26.689 | 2 | 52.50722 | yes | 2.875E-07 | 441.72311 | 8535.1533 |
| ACYGVLR | 7 | P23396 | Small ribosomal subunit protein uS3 OS= | RPS3 | 26.689 | 2 | 52.50722 | yes | 1.952E-07 | 419.71564 | 5659.3618 |
| TQNVLGEK | 8 | P23396 | Small ribosomal subunit protein uS3 OS= | RPS3 | 26.689 | 2 | 52.50722 | yes | 1.685E-07 | 444.74289 | 5235.2524 |
| ELTAVVQK | 8 | P23396 | Small ribosomal subunit protein uS3 OS= | RPS3 | 26.689 | 2 | 52.50722 | yes | 1.175E-07 | 444.26346 | 2331.3313 |
| RFGFPEGSVELYA | 15 | P23396 | Small ribosomal subunit protein uS3 OS= | RPS3 | 26.689 | 3 | 52.50722 | yes | 1.536E-07 | 576.95789 | 745.79871 |
| GCEVVVSGK | 9 | P23396 | Small ribosomal subunit protein uS3 OS= | RPS3 | 26.689 | 2 | 52.50722 | yes | 1.737E-07 | 467.73676 | 431.4693 |
| RACYGVLR | 8 | P23396 | Small ribosomal subunit protein uS3 OS= | RPS3 | 26.689 | 2 | 52.50722 | yes | 9.722E-08 | 497.76617 | 192.34171 |
| YKLLGGLAVR | 10 | P23396 | Small ribosomal subunit protein uS3 OS= | RPS3 | 26.689 | 2 | 52.50722 | yes | 3.384E-07 | 545.34259 | 66.01947 |
| FVDGLMIHSGDPV | 22 | P23396 | Small ribosomal subunit protein uS3 OS= | RPS3 | 26.689 | 3 | 52.50722 | yes | 2.992E-06 | 823.40204 | 6.2445903 |
| IPWIAPECVEDSK | 13 | P23458 | Tyrosine-protein kinase JAK1 OS=Homo | JAK1 | 133.278 | 2 | 52.94874 | yes | 1.082E-07 | 772.37671 | 377.60748 |
| LSDPGIPITVLSR | 13 | P23458 | Tyrosine-protein kinase JAK1 OS=Homo | JAK1 | 133.278 | 2 | 52.94874 | yes | 3.243E-07 | 684.39825 | 344.13611 |
| DLGEGHFGK | 9 | P23458 | Tyrosine-protein kinase JAK1 OS=Homo | JAK1 | 133.278 | 2 | 52.94874 | no | 1.707E-07 | 480.2327 | 326.65955 |
| NVLVESEHQVK | 11 | P23458 | Tyrosine-protein kinase JAK1 OS=Homo | JAK1 | 133.278 | 2 | 52.94874 | yes | 2.26E-07 | 641.34351 | 288.90332 |
| CLAPIRDPK | 9 | P23458 | Tyrosine-protein kinase JAK1 OS=Homo | JAK1 | 133.278 | 2 | 52.94874 | yes | 1.641E-07 | 535.2948 | 265.6333 |
| LIMEFLPSGSLK | 12 | P23458 | Tyrosine-protein kinase JAK1 OS=Homo | JAK1 | 133.278 | 2 | 52.94874 | yes | 9.722E-08 | 667.87305 | 248.46884 |
| YLATLETLTK | 10 | P23458 | Tyrosine-protein kinase JAK1 OS=Homo | JAK1 | 133.278 | 2 | 52.94874 | yes | 1.223E-07 | 576.82916 | 243.67993 |
| LEEQNPDIVSEK | 12 | P23458 | Tyrosine-protein kinase JAK1 OS=Homo | JAK1 | 133.278 | 2 | 52.94874 | yes | 1.363E-07 | 700.84882 | 196.59419 |
| ELADLMTR | 8 | P23458 | Tyrosine-protein kinase JAK1 OS=Homo | JAK1 | 133.278 | 2 | 52.94874 | yes | 9.722E-08 | 474.7446 | 194.36723 |
| DLVQGEHLGR | 10 | P23458 | Tyrosine-protein kinase JAK1 OS=Homo | JAK1 | 133.278 | 2 | 52.94874 | yes | 1.995E-07 | 562.29639 | 184.36748 |
| HIVYLYGVCVR | 11 | P23458 | Tyrosine-protein kinase JAK1 OS=Homo | JAK1 | 133.278 | 2 | 52.94874 | yes | 9.722E-08 | 689.86865 | 169.3035 |
| TICDSSVSTHDLK | 13 | P23458 | Tyrosine-protein kinase JAK1 OS=Homo | JAK1 | 133.278 | 2 | 52.94874 | yes | 2.191E-07 | 731.84576 | 153.01393 |
| DLVHGNVCTK | 10 | P23458 | Tyrosine-protein kinase JAK1 OS=Homo | JAK1 | 133.278 | 2 | 52.94874 | yes | 1.001E-07 | 571.78479 | 145.16455 |
| EGIDSECGPFIK | 12 | P23458 | Tyrosine-protein kinase JAK1 OS=Homo | JAK1 | 133.278 | 2 | 52.94874 | yes | 9.722E-08 | 676.31354 | 142.60654 |
| LGSGEYTAEELCIF | 14 | P23458 | Tyrosine-protein kinase JAK1 OS=Homo | JAK1 | 133.278 | 2 | 52.94874 | yes | 1.014E-07 | 799.37994 | 127.10226 |
| NFQIEVQK | 8 | P23458 | Tyrosine-protein kinase JAK1 OS=Homo | JAK1 | 133.278 | 2 | 52.94874 | yes | 4.032E-07 | 503.27182 | 127.07112 |
| YDPEGDNTGEQV | 15 | P23458 | Tyrosine-protein kinase JAK1 OS=Homo | JAK1 | 133.278 | 2 | 52.94874 | yes | 1.381E-07 | 811.37067 | 118.25077 |
| QLASALSYLEDK | 12 | P23458 | Tyrosine-protein kinase JAK1 OS=Homo | JAK1 | 133.278 | 2 | 52.94874 | yes | 9.819E-08 | 669.35101 | 106.63557 |
| YSLHGSDR | 8 | P23458 | Tyrosine-protein kinase JAK1 OS=Homo | JAK1 | 133.278 | 2 | 52.94874 | yes | 1.675E-07 | 467.7225 | 102.66228 |
| KSDVLTTPWK | 10 | P23458 | Tyrosine-protein kinase JAK1 OS=Homo | JAK1 | 133.278 | 2 | 52.94874 | yes | 4.115E-07 | 587.82697 | 97.773216 |
| NLYHENIVK | 9 | P23458 | Tyrosine-protein kinase JAK1 OS=Homo | JAK1 | 133.278 | 2 | 52.94874 | yes | 2.115E-07 | 565.30365 | 80.313446 |
| QEGSEEGMYVLR | 12 | P23458 | Tyrosine-protein kinase JAK1 OS=Homo | JAK1 | 133.278 | 2 | 52.94874 | yes | 9.722E-08 | 699.3219 | 79.176796 |
| CRPVTPSCK | 9 | P23458 | Tyrosine-protein kinase JAK1 OS=Homo | JAK1 | 133.278 | 2 | 52.94874 | yes | 1.28E-06 | 552.76807 | 76.900627 |
| KPATEVDPTHFEK | 13 | P23458 | Tyrosine-protein kinase JAK1 OS=Homo | JAK1 | 133.278 | 2 | 52.94874 | yes | 1.698E-06 | 749.88043 | 60.296921 |
| MIGPTHGQMTVTI | 13 | P23458 | Tyrosine-protein kinase JAK1 OS=Homo | JAK1 | 133.278 | 3 | 52.94874 | yes | 8.363E-07 | 476.90771 | 56.984749 |
| THIYSGTLMDYK | 12 | P23458 | Tyrosine-protein kinase JAK1 OS=Homo | JAK1 | 133.278 | 2 | 52.94874 | yes | 5.477E-07 | 714.84503 | 53.110847 |
| MQLPELPK | 8 | P23458 | Tyrosine-protein kinase JAK1 OS=Homo | JAK1 | 133.278 | 2 | 52.94874 | yes | 5.114E-07 | 478.2677 | 46.817581 |
| TDNISFMLK | 9 | P23458 | Tyrosine-protein kinase JAK1 OS=Homo | JAK1 | 133.278 | 2 | 52.94874 | yes | 2.738E-07 | 534.77332 | 33.467812 |
| SLKPESGGNHIADI | 15 | P23458 | Tyrosine-protein kinase JAK1 OS=Homo | JAK1 | 133.278 | 2 | 52.94874 | yes | 1.618E-07 | 783.41772 | 33.262589 |

| | | | | | | | | | | | |
|---|---|---|---|---|---|---|---|---|---|---|---|
| KCWEFQPSNR | 10 | P23458 | Tyrosine-protein kinase JAK1 OS=Homo | JAK1 | 133.278 | 2 | 52.94874 | yes | 2.684E-06 | 676.31421 | 26.614481 |
| VVYNVLSELEK | 11 | P23497 | Nuclear autoantigen Sp-100 OS=Homo sa | SP100 | 100.417 | 2 | 51.631054 | no | 1.21E-07 | 646.85846 | 205.60532 |
| MFTEDQGVDDR | 11 | P23497 | Nuclear autoantigen Sp-100 OS=Homo sa | SP100 | 100.417 | 2 | 51.631054 | yes | 9.722E-08 | 656.77734 | 122.39952 |
| GFENVIHDK | 9 | P23497 | Nuclear autoantigen Sp-100 OS=Homo sa | SP100 | 100.417 | 2 | 51.631054 | yes | 3.035E-07 | 529.76691 | 82.473366 |
| MFEDSQDSCR | 10 | P23497 | Nuclear autoantigen Sp-100 OS=Homo sa | SP100 | 100.417 | 2 | 51.631054 | yes | 9.722E-08 | 637.74243 | 60.078976 |
| ALDIAENEMPGLM | 14 | P23526 | Adenosylhomocysteinase OS=Homo sapi | AHCY | 47.717 | 2 | 45.30569 | yes | 5.572E-07 | 780.38147 | 102.32147 |
| VPAINVNDSVTK | 12 | P23526 | Adenosylhomocysteinase OS=Homo sapi | AHCY | 47.717 | 2 | 45.30569 | yes | 1.891E-07 | 628.84589 | 74.081116 |
| FDNLYGCR | 8 | P23526 | Adenosylhomocysteinase OS=Homo sapi | AHCY | 47.717 | 2 | 45.30569 | yes | 6.734E-07 | 522.73199 | 41.059319 |
| YPQLLPGIR | 9 | P23526 | Adenosylhomocysteinase OS=Homo sapi | AHCY | 47.717 | 2 | 45.30569 | yes | 2.512E-06 | 528.81366 | 30.097523 |
| YALYDATYETK | 11 | P23528 | Cofilin-1 OS=Homo sapiens OX=9606 G | CFL1 | 18.503 | 2 | 49.99421 | no | 9.937E-08 | 669.31665 | 1649.0251 |
| AVLFCLSEDKK | 11 | P23528 | Cofilin-1 OS=Homo sapiens OX=9606 G | CFL1 | 18.503 | 2 | 49.99421 | yes | 1.672E-07 | 655.34467 | 1169.5106 |
| LGGSAVISLEGKPI | 14 | P23528 | Cofilin-1 OS=Homo sapiens OX=9606 G | CFL1 | 18.503 | 2 | 49.99421 | yes | 5.057E-07 | 670.89282 | 837.23151 |
| MLPDKDCR | 8 | P23528 | Cofilin-1 OS=Homo sapiens OX=9606 G | CFL1 | 18.503 | 2 | 49.99421 | yes | 2.242E-06 | 517.74152 | 497.72928 |
| NIILEEGK | 8 | P23528 | Cofilin-1 OS=Homo sapiens OX=9606 G | CFL1 | 18.503 | 2 | 49.99421 | yes | 2.563E-07 | 458.26093 | 228.2067 |
| AVLFCLSEDK | 10 | P23528 | Cofilin-1 OS=Homo sapiens OX=9606 G | CFL1 | 18.503 | 2 | 49.99421 | yes | 1.889E-07 | 591.29718 | 113.85654 |
| HELQANCYEEVK | 12 | P23528 | Cofilin-1 OS=Homo sapiens OX=9606 G | CFL1 | 18.503 | 2 | 49.99421 | yes | 3.789E-07 | 760.34595 | 113.8544 |
| KSSTPEEVK | 9 | P23528 | Cofilin-1 OS=Homo sapiens OX=9606 G | CFL1 | 18.503 | 2 | 49.99421 | yes | 2.757E-06 | 502.76657 | 34.349957 |
| VDVADQAQDK | 10 | P23588 | Eukaryotic translation initiation factor 4B | EIF4B | 69.152 | 2 | 55.653385 | yes | 1.21E-07 | 544.76459 | 1503.7079 |
| VDVADQAQDKDR | 12 | P23588 | Eukaryotic translation initiation factor 4B | EIF4B | 69.152 | 2 | 55.653385 | yes | 1.367E-07 | 680.32861 | 746.58728 |
| AASIFGGAKPVDT. | 16 | P23588 | Eukaryotic translation initiation factor 4B | EIF4B | 69.152 | 3 | 55.653385 | yes | 9.722E-08 | 511.27908 | 341.88309 |
| AASIFGGAKPVDT. | 16 | P23588 | Eukaryotic translation initiation factor 4B | EIF4B | 69.152 | 3 | 55.653385 | yes | 9.85E-08 | 766.41498 | 341.88309 |
| AASIFGGAKPVDT. | 16 | P23588 | Eukaryotic translation initiation factor 4B | EIF4B | 69.152 | 2 | 55.653385 | yes | 9.722E-08 | 511.27908 | 341.88309 |
| AASIFGGAKPVDT. | 16 | P23588 | Eukaryotic translation initiation factor 4B | EIF4B | 69.152 | 2 | 55.653385 | yes | 9.85E-08 | 766.41498 | 341.88309 |
| DSDKTDTDWR | 10 | P23588 | Eukaryotic translation initiation factor 4B | EIF4B | 69.152 | 2 | 55.653385 | yes | 9.722E-08 | 619.76782 | 653.88263 |
| SILPTAPR | 8 | P23588 | Eukaryotic translation initiation factor 4B | EIF4B | 69.152 | 2 | 55.653385 | yes | 9.722E-08 | 427.75836 | 359.06165 |
| SLENETLNK | 9 | P23588 | Eukaryotic translation initiation factor 4B | EIF4B | 69.152 | 2 | 55.653385 | yes | 4.147E-07 | 524.26947 | 334.08716 |
| RGDDSFGDK | 9 | P23588 | Eukaryotic translation initiation factor 4B | EIF4B | 69.152 | 2 | 55.653385 | yes | 1.688E-07 | 498.72269 | 210.89415 |
| YEDRYDR | 7 | P23588 | Eukaryotic translation initiation factor 4B | EIF4B | 69.152 | 2 | 55.653385 | yes | 3.067E-07 | 508.72525 | 202.40404 |
| VAPAQPSEEGPGR | 13 | P23588 | Eukaryotic translation initiation factor 4B | EIF4B | 69.152 | 2 | 55.653385 | yes | 2.748E-07 | 647.82294 | 175.47543 |
| GLNISAVR | 8 | P23588 | Eukaryotic translation initiation factor 4B | EIF4B | 69.152 | 2 | 55.653385 | yes | 5.087E-07 | 415.24814 | 114.09962 |
| ARPATDSFDDYPP. | 14 | P23588 | Eukaryotic translation initiation factor 4B | EIF4B | 69.152 | 2 | 55.653385 | yes | 3.506E-06 | 804.3761 | 110.21378 |
| IRVDVADQAQDK | 12 | P23588 | Eukaryotic translation initiation factor 4B | EIF4B | 69.152 | 2 | 55.653385 | yes | 9.722E-08 | 679.35718 | 55.770573 |
| DENKVDGMNAPK | 12 | P23588 | Eukaryotic translation initiation factor 4B | EIF4B | 69.152 | 2 | 55.653385 | yes | 9.722E-08 | 659.30884 | 54.35463 |
| TISLTDFLAEDGGT | 23 | P23588 | Eukaryotic translation initiation factor 4B | EIF4B | 69.152 | 3 | 55.653385 | yes | 1.197E-07 | 759.36914 | 38.175049 |
| VDVADQAQDKDR | 15 | P23588 | Eukaryotic translation initiation factor 4B | EIF4B | 69.152 | 3 | 55.653385 | yes | 2.156E-06 | 582.60651 | 25.558928 |
| VSYSPAHAR | 9 | P23769 | Endothelial transcription factor GATA-2 ( | GATA2 | 50.5 | 2 | 53.262913 | yes | 1.733E-07 | 494.25397 | 310.24414 |
| ECVNCGATATPLW | 14 | P23769 | Endothelial transcription factor GATA-2 ( | GATA2 | 50.5 | 2 | 53.262913 | no | 9.722E-08 | 817.87433 | 278.55087 |
| GAECFEELSK | 10 | P23769 | Endothelial transcription factor GATA-2 ( | GATA2 | 50.5 | 2 | 53.262913 | yes | 2.633E-07 | 585.2608 | 63.883827 |
| GALIVLEGVDR | 11 | P23919 | Thymidylate kinase OS=Homo sapiens O. | DTYMK | 23.82 | 2 | 48.95493 | yes | 1.083E-07 | 571.3324 | 120.58646 |

| | | | | | | | | | | | |
|---|---|---|---|---|---|---|---|---|---|---|---|
| LVEALCAAGHR | 11 | P23919 | Thymidylate kinase OS=Homo sapiens O | DTYMK | 23.82 | 2 | 48.95493 | yes | 3.387E-06 | 598.81384 | 45.961739 |
| LNSAIIYDR | 9 | P23921 | Ribonucleoside-diphosphate reductase lar | RRM1 | 90.071 | 2 | 53.757793 | yes | 1.106E-07 | 532.79041 | 270.02036 |
| SAGGIGVAVSCIR | 13 | P23921 | Ribonucleoside-diphosphate reductase lar | RRM1 | 90.071 | 2 | 53.757793 | yes | 4.496E-07 | 623.83228 | 222.99385 |
| NTAAMVCSLENR | 12 | P23921 | Ribonucleoside-diphosphate reductase lar | RRM1 | 90.071 | 2 | 53.757793 | yes | 4.265E-07 | 683.3161 | 164.17462 |
| HPDYAILAAR | 10 | P23921 | Ribonucleoside-diphosphate reductase lar | RRM1 | 90.071 | 2 | 53.757793 | yes | 5.454E-07 | 563.80383 | 141.33685 |
| IAVSNLHK | 8 | P23921 | Ribonucleoside-diphosphate reductase lar | RRM1 | 90.071 | 2 | 53.757793 | yes | 9.722E-08 | 441.26379 | 108.77444 |
| DFSYNYFGFK | 10 | P23921 | Ribonucleoside-diphosphate reductase lar | RRM1 | 90.071 | 2 | 53.757793 | yes | 9.722E-08 | 644.28766 | 89.360489 |
| LTSMHFYGWK | 10 | P23921 | Ribonucleoside-diphosphate reductase lar | RRM1 | 90.071 | 2 | 53.757793 | yes | 1.006E-07 | 635.30786 | 84.343613 |
| DDSIEGIYDTLK | 12 | P23921 | Ribonucleoside-diphosphate reductase lar | RRM1 | 90.071 | 2 | 53.757793 | yes | 9.722E-08 | 684.83008 | 77.207748 |
| GLWHEEMK | 8 | P23921 | Ribonucleoside-diphosphate reductase lar | RRM1 | 90.071 | 2 | 53.757793 | yes | 5.243E-06 | 515.24475 | 59.330898 |
| STLDIVLANK | 10 | P23921 | Ribonucleoside-diphosphate reductase lar | RRM1 | 90.071 | 2 | 53.757793 | yes | 2.997E-07 | 537.31366 | 41.592278 |
| SNQQNLGTIK | 10 | P23921 | Ribonucleoside-diphosphate reductase lar | RRM1 | 90.071 | 2 | 53.757793 | yes | 3.249E-07 | 551.7962 | 40.372261 |
| VLSGEFQIVNPHLI | 15 | P23921 | Ribonucleoside-diphosphate reductase lar | RRM1 | 90.071 | 2 | 53.757793 | yes | 8.894E-05 | 847.48541 | 13.468156 |
| ATYDGNHDTFR | 11 | P24390 | ER lumen protein-retaining receptor 1 OS | KDELR1 | 24.543 | 2 | 50.977993 | yes | 9.722E-08 | 648.78381 | 31.808434 |
| LVPVGYGIK | 9 | P24534 | Elongation factor 1-beta OS=Homo sapie | EEF1B2 | 24.764 | 2 | 51.64396 | yes | 2.758E-07 | 473.29202 | 22333.936 |
| SPAGLQVLNDYLA | 15 | P24534 | Elongation factor 1-beta OS=Homo sapie | EEF1B2 | 24.764 | 2 | 51.64396 | yes | 1.113E-07 | 802.41992 | 14954.224 |
| SIQADGLVWGSSK | 13 | P24534 | Elongation factor 1-beta OS=Homo sapie | EEF1B2 | 24.764 | 2 | 51.64396 | yes | 1.026E-07 | 674.34882 | 11906.94 |
| LAQYESK | 7 | P24534 | Elongation factor 1-beta OS=Homo sapie | EEF1B2 | 24.764 | 2 | 51.64396 | yes | 1.165E-07 | 419.7189 | 3777.835 |
| PWDDETDMAK | 10 | P24534 | Elongation factor 1-beta OS=Homo sapie | EEF1B2 | 24.764 | 2 | 51.64396 | yes | 9.722E-08 | 604.25043 | 25.16827 |
| HYLFDVQR | 8 | P24539 | ATP synthase F(0) complex subunit B1, r | ATP5PB | 28.909 | 2 | 52.718124 | yes | 9.722E-08 | 539.27747 | 681.42603 |
| YGPFVADFADK | 11 | P24539 | ATP synthase F(0) complex subunit B1, r | ATP5PB | 28.909 | 2 | 52.718124 | yes | 1.043E-07 | 615.29547 | 558.68518 |
| NNIAMALEVTYR | 12 | P24539 | ATP synthase F(0) complex subunit B1, r | ATP5PB | 28.909 | 2 | 52.718124 | yes | 1.114E-07 | 697.85846 | 156.01643 |
| LAQLEEAK | 8 | P24539 | ATP synthase F(0) complex subunit B1, r | ATP5PB | 28.909 | 2 | 52.718124 | yes | 4.231E-07 | 451.25311 | 151.86296 |
| LDYHISVQNMMR | 12 | P24539 | ATP synthase F(0) complex subunit B1, r | ATP5PB | 28.909 | 2 | 52.718124 | yes | 1.345E-07 | 753.86322 | 139.01553 |
| HVVQSISTQQEK | 12 | P24539 | ATP synthase F(0) complex subunit B1, r | ATP5PB | 28.909 | 2 | 52.718124 | yes | 1.113E-07 | 692.36499 | 111.37807 |
| EQEHMINWVEK | 11 | P24539 | ATP synthase F(0) complex subunit B1, r | ATP5PB | 28.909 | 2 | 52.718124 | yes | 1.113E-07 | 721.84027 | 87.291473 |
| QASIQHIQNAIDTE | 15 | P24539 | ATP synthase F(0) complex subunit B1, r | ATP5PB | 28.909 | 3 | 52.718124 | yes | 6.615E-07 | 565.96021 | 42.354652 |
| QASIQHIQNAIDTE | 15 | P24539 | ATP synthase F(0) complex subunit B1, r | ATP5PB | 28.909 | 3 | 52.718124 | yes | 1.604E-07 | 848.43665 | 42.354652 |
| QASIQHIQNAIDTE | 15 | P24539 | ATP synthase F(0) complex subunit B1, r | ATP5PB | 28.909 | 2 | 52.718124 | yes | 6.615E-07 | 565.96021 | 42.354652 |
| QASIQHIQNAIDTE | 15 | P24539 | ATP synthase F(0) complex subunit B1, r | ATP5PB | 28.909 | 2 | 52.718124 | yes | 1.604E-07 | 848.43665 | 42.354652 |
| HMEASLQELK | 10 | P24593 | Insulin-like growth factor-binding protein | IGFBP5 | 30.571 | 2 | 45.727234 | yes | 1.76E-07 | 593.30023 | 104.49757 |
| SVLFVCLGNICR | 12 | P24666 | Low molecular weight phosphotyrosine p | ACP1 | 18.043 | 2 | 51.122765 | yes | 9.722E-08 | 719.37067 | 908.09698 |
| LVTDQNISENWR | 12 | P24666 | Low molecular weight phosphotyrosine p | ACP1 | 18.043 | 2 | 51.122765 | yes | 9.809E-08 | 737.86786 | 725.37048 |
| SPIAEAVFR | 9 | P24666 | Low molecular weight phosphotyrosine p | ACP1 | 18.043 | 2 | 51.122765 | yes | 1.183E-07 | 495.27435 | 650.26611 |
| IELLGSYDPQK | 11 | P24666 | Low molecular weight phosphotyrosine p | ACP1 | 18.043 | 2 | 51.122765 | yes | 1.614E-07 | 631.83496 | 586.89557 |
| HGIPMSHVAR | 10 | P24666 | Low molecular weight phosphotyrosine p | ACP1 | 18.043 | 2 | 51.122765 | yes | 9.795E-08 | 552.79022 | 280.95908 |
| FGNEVIPVTVTVK | 13 | P24752 | Acetyl-CoA acetyltransferase, mitochondı | ACAT1 | 45.2 | 2 | 46.635536 | yes | 1.508E-07 | 701.90063 | 82.661804 |
| FTDEEVDEMYR | 11 | P24844 | Myosin regulatory light polypeptide 9 OS | MYL9 | 19.828 | 2 | 32.008625 | yes | 1.653E-06 | 717.2981 | 38.418186 |
| VEEFGVVSNTR | 11 | P24863 | Cyclin-C OS=Homo sapiens OX=9606 G | CCNC | 33.243 | 2 | 49.99367 | yes | 9.937E-08 | 618.81458 | 148.44788 |

| | | | | | | | | | | |
|---|---|---|---|---|---|---|---|---|---|---|
| LIAAATSVLK | 10 | P24863 | Cyclin-C OS=Homo sapiens OX=9606 G | CCNC | 33.243 | 2 | 49.99367 | yes | 4.002E-06 | 493.81586 | 53.608185 |
| VVLPCNLLR | 9 | P24928 | DNA-directed RNA polymerase II subuni | POLR2A | 217.177 | 2 | 54.885693 | yes | 1.151E-07 | 542.3208 | 383.74673 |
| FGVEQPEGDEDLT | 14 | P24928 | DNA-directed RNA polymerase II subuni | POLR2A | 217.177 | 2 | 54.885693 | yes | 1.44E-07 | 782.3623 | 331.00891 |
| YSPTSPTYSPTSPK | 14 | P24928 | DNA-directed RNA polymerase II subuni | POLR2A | 217.177 | 2 | 54.885693 | yes | 2.289E-07 | 756.86444 | 321.34384 |
| VIFPTGDSK | 9 | P24928 | DNA-directed RNA polymerase II subuni | POLR2A | 217.177 | 2 | 54.885693 | yes | 1.503E-07 | 482.26093 | 304.67416 |
| LLVDSNNPK | 9 | P24928 | DNA-directed RNA polymerase II subuni | POLR2A | 217.177 | 2 | 54.885693 | yes | 2.949E-07 | 500.2771 | 272.10175 |
| NSINQVVQLR | 10 | P24928 | DNA-directed RNA polymerase II subuni | POLR2A | 217.177 | 2 | 54.885693 | yes | 1.878E-07 | 585.83313 | 236.0061 |
| ISDEECFVLGMEPI | 14 | P24928 | DNA-directed RNA polymerase II subuni | POLR2A | 217.177 | 2 | 54.885693 | yes | 9.722E-08 | 841.38165 | 216.13126 |
| AEIQELAMVPR | 11 | P24928 | DNA-directed RNA polymerase II subuni | POLR2A | 217.177 | 2 | 54.885693 | yes | 9.722E-08 | 628.83698 | 182.34218 |
| IFHINPR | 7 | P24928 | DNA-directed RNA polymerase II subuni | POLR2A | 217.177 | 2 | 54.885693 | yes | 5.693E-07 | 448.7587 | 178.04597 |
| SLGTSAGSLVHISY | 23 | P24928 | DNA-directed RNA polymerase II subuni | POLR2A | 217.177 | 3 | 54.885693 | yes | 3.243E-07 | 815.41278 | 172.99823 |
| EGLIDTAVK | 9 | P24928 | DNA-directed RNA polymerase II subuni | POLR2A | 217.177 | 2 | 54.885693 | yes | 2.626E-07 | 473.2662 | 168.74905 |
| VQFGVLSPDELK | 12 | P24928 | DNA-directed RNA polymerase II subuni | POLR2A | 217.177 | 2 | 54.885693 | yes | 1.13E-07 | 666.36389 | 162.90692 |
| LEHTTLR | 7 | P24928 | DNA-directed RNA polymerase II subuni | POLR2A | 217.177 | 2 | 54.885693 | yes | 9.728E-08 | 435.24561 | 147.36201 |
| TYQDIQNTIK | 10 | P24928 | DNA-directed RNA polymerase II subuni | POLR2A | 217.177 | 2 | 54.885693 | yes | 3.198E-07 | 612.31696 | 136.18123 |
| LTHVYDLCK | 9 | P24928 | DNA-directed RNA polymerase II subuni | POLR2A | 217.177 | 2 | 54.885693 | yes | 1.557E-07 | 574.79205 | 116.14647 |
| LPSDLHPIK | 9 | P24928 | DNA-directed RNA polymerase II subuni | POLR2A | 217.177 | 2 | 54.885693 | yes | 2.372E-07 | 510.29785 | 108.21023 |
| PSDLHLQTGYK | 11 | P24928 | DNA-directed RNA polymerase II subuni | POLR2A | 217.177 | 2 | 54.885693 | yes | 1.084E-07 | 629.82495 | 103.84007 |
| ILPWSTFR | 8 | P24928 | DNA-directed RNA polymerase II subuni | POLR2A | 217.177 | 2 | 54.885693 | yes | 3.37E-06 | 510.28726 | 102.1488 |
| PVFHVGFLVK | 10 | P24928 | DNA-directed RNA polymerase II subuni | POLR2A | 217.177 | 2 | 54.885693 | yes | 9.722E-08 | 571.83966 | 101.67414 |
| IIITEDGEFK | 10 | P24928 | DNA-directed RNA polymerase II subuni | POLR2A | 217.177 | 2 | 54.885693 | yes | 9.722E-08 | 582.81097 | 99.327065 |
| KLTMEQIAEK | 10 | P24928 | DNA-directed RNA polymerase II subuni | POLR2A | 217.177 | 2 | 54.885693 | yes | 9.722E-08 | 595.82611 | 95.835159 |
| ISPWLLR | 7 | P24928 | DNA-directed RNA polymerase II subuni | POLR2A | 217.177 | 2 | 54.885693 | yes | 1.839E-07 | 442.77127 | 93.882431 |
| YSPTSPTYSPTTPK | 14 | P24928 | DNA-directed RNA polymerase II subuni | POLR2A | 217.177 | 2 | 54.885693 | yes | 2.423E-07 | 763.87231 | 93.366226 |
| VPQPAILKPR | 10 | P24928 | DNA-directed RNA polymerase II subuni | POLR2A | 217.177 | 2 | 54.885693 | yes | 3.46E-07 | 559.85583 | 90.509811 |
| HLALLCDTMTCR | 12 | P24928 | DNA-directed RNA polymerase II subuni | POLR2A | 217.177 | 2 | 54.885693 | yes | 7.022E-07 | 745.84924 | 90.006752 |
| MQEEEEVVDK | 10 | P24928 | DNA-directed RNA polymerase II subuni | POLR2A | 217.177 | 2 | 54.885693 | yes | 3.119E-07 | 618.27667 | 84.962143 |
| SGLELYAEWK | 10 | P24928 | DNA-directed RNA polymerase II subuni | POLR2A | 217.177 | 2 | 54.885693 | yes | 5.548E-07 | 598.30334 | 84.121391 |
| MDDDVFLR | 8 | P24928 | DNA-directed RNA polymerase II subuni | POLR2A | 217.177 | 2 | 54.885693 | yes | 2.162E-07 | 505.73422 | 70.853638 |
| RVDFSAR | 7 | P24928 | DNA-directed RNA polymerase II subuni | POLR2A | 217.177 | 2 | 54.885693 | yes | 6.79E-07 | 425.73013 | 63.245663 |
| HMCDGDIVIFNR | 12 | P24928 | DNA-directed RNA polymerase II subuni | POLR2A | 217.177 | 2 | 54.885693 | yes | 2.393E-07 | 738.83978 | 48.633678 |
| AHNNELEPTPGNT | 15 | P24928 | DNA-directed RNA polymerase II subuni | POLR2A | 217.177 | 2 | 54.885693 | yes | 9.722E-08 | 831.91333 | 35.840256 |
| DVLSNAHIQNELE | 14 | P24928 | DNA-directed RNA polymerase II subuni | POLR2A | 217.177 | 2 | 54.885693 | yes | 9.722E-08 | 819.41571 | 24.216747 |
| LDTETEGVPSTAIR | 14 | P24941 | Cyclin-dependent kinase 2 OS=Homo sap | CDK2 | 33.93 | 2 | 52.710926 | yes | 9.722E-08 | 744.88068 | 444.28787 |
| IGEGTYGVVYK | 11 | P24941 | Cyclin-dependent kinase 2 OS=Homo sap | CDK2 | 33.93 | 2 | 52.710926 | no | 9.722E-08 | 593.31116 | 390.79535 |
| LLDVIHTENK | 10 | P24941 | Cyclin-dependent kinase 2 OS=Homo sap | CDK2 | 33.93 | 2 | 52.710926 | yes | 9.722E-08 | 591.3299 | 283.02451 |
| VVPPLDEDGR | 10 | P24941 | Cyclin-dependent kinase 2 OS=Homo sap | CDK2 | 33.93 | 2 | 52.710926 | yes | 3.992E-07 | 548.78528 | 107.72895 |
| ALFPGDSEIDQLFR | 14 | P24941 | Cyclin-dependent kinase 2 OS=Homo sap | CDK2 | 33.93 | 2 | 52.710926 | no | 1.001E-07 | 804.40686 | 80.183388 |
| ELNHPNIVK | 9 | P24941 | Cyclin-dependent kinase 2 OS=Homo sap | CDK2 | 33.93 | 2 | 52.710926 | yes | 8.32E-07 | 532.2984 | 73.234856 |

| | | | | | | | | | | | |
|---|---|---|---|---|---|---|---|---|---|---|---|
| APEILLGCK | 9 | P24941 | Cyclin-dependent kinase 2 OS=Homo sap | CDK2 | 33.93 | 2 | 52.710926 | yes | 1.235E-07 | 500.77844 | 64.508934 |
| MIAMGSAAALR | 11 | P25054 | Adenomatous polyposis coli protein OS=I | APC | 311.646 | 2 | 57.930008 | no | 3.813E-07 | 546.28864 | 236.44955 |
| TLIYQMAPAVSK | 12 | P25054 | Adenomatous polyposis coli protein OS=I | APC | 311.646 | 2 | 57.930008 | yes | 4.937E-07 | 661.36285 | 204.29333 |
| IEDCPINNPR | 10 | P25054 | Adenomatous polyposis coli protein OS=I | APC | 311.646 | 2 | 57.930008 | yes | 1.064E-07 | 614.29297 | 173.77144 |
| NVSSLIATNEDHR | 13 | P25054 | Adenomatous polyposis coli protein OS=I | APC | 311.646 | 2 | 57.930008 | yes | 9.802E-08 | 728.36298 | 155.14053 |
| VLHLLEQIR | 9 | P25054 | Adenomatous polyposis coli protein OS=I | APC | 311.646 | 2 | 57.930008 | yes | 1.25E-07 | 560.84552 | 147.21165 |
| ESTGYLEELEK | 11 | P25054 | Adenomatous polyposis coli protein OS=I | APC | 311.646 | 2 | 57.930008 | yes | 9.722E-08 | 649.31152 | 146.76015 |
| LNSFIQVDAPDQK | 13 | P25054 | Adenomatous polyposis coli protein OS=I | APC | 311.646 | 2 | 57.930008 | yes | 1.229E-07 | 737.88043 | 139.7872 |
| PSQIPTPVNNNTK | 13 | P25054 | Adenomatous polyposis coli protein OS=I | APC | 311.646 | 2 | 57.930008 | yes | 4.95E-07 | 705.3728 | 128.37122 |
| ALMECALEVK | 10 | P25054 | Adenomatous polyposis coli protein OS=I | APC | 311.646 | 2 | 57.930008 | yes | 1.266E-07 | 582.29358 | 114.87296 |
| TGSSSSILSASSESS | 17 | P25054 | Adenomatous polyposis coli protein OS=I | APC | 311.646 | 2 | 57.930008 | yes | 1.454E-07 | 822.38397 | 107.77973 |
| LSFDEEHR | 8 | P25054 | Adenomatous polyposis coli protein OS=I | APC | 311.646 | 2 | 57.930008 | yes | 1.419E-07 | 516.74091 | 105.32327 |
| SGECSPVPMGSFPI | 14 | P25054 | Adenomatous polyposis coli protein OS=I | APC | 311.646 | 2 | 57.930008 | yes | 1.259E-07 | 754.33704 | 95.027557 |
| QLLQSQATEAER | 12 | P25054 | Adenomatous polyposis coli protein OS=I | APC | 311.646 | 2 | 57.930008 | yes | 1.371E-07 | 687.35461 | 93.547272 |
| SLLLADLDK | 9 | P25054 | Adenomatous polyposis coli protein OS=I | APC | 311.646 | 2 | 57.930008 | yes | 3.056E-07 | 494.28967 | 90.560593 |
| HVSFTPGDDMPR | 12 | P25054 | Adenomatous polyposis coli protein OS=I | APC | 311.646 | 2 | 57.930008 | yes | 1.458E-06 | 679.81152 | 83.146973 |
| KPSQLPVYK | 9 | P25054 | Adenomatous polyposis coli protein OS=I | APC | 311.646 | 2 | 57.930008 | yes | 6.147E-07 | 530.31348 | 77.066727 |
| HVDQPIDYSLK | 11 | P25054 | Adenomatous polyposis coli protein OS=I | APC | 311.646 | 2 | 57.930008 | yes | 9.722E-08 | 657.83807 | 72.851006 |
| QPLQANMPSISR | 12 | P25054 | Adenomatous polyposis coli protein OS=I | APC | 311.646 | 2 | 57.930008 | yes | 1.525E-07 | 671.35083 | 67.538528 |
| ELNLDSSNFPGVK | 13 | P25054 | Adenomatous polyposis coli protein OS=I | APC | 311.646 | 2 | 57.930008 | yes | 7.628E-07 | 710.35938 | 66.802917 |
| RLTSHLGTK | 9 | P25054 | Adenomatous polyposis coli protein OS=I | APC | 311.646 | 2 | 57.930008 | yes | 9.165E-06 | 506.79855 | 58.609383 |
| LQGSSLSSESAR | 12 | P25054 | Adenomatous polyposis coli protein OS=I | APC | 311.646 | 2 | 57.930008 | yes | 2.803E-07 | 611.30713 | 44.105389 |
| ENNCLQTLLQHLK | 13 | P25054 | Adenomatous polyposis coli protein OS=I | APC | 311.646 | 2 | 57.930008 | yes | 1.137E-07 | 805.91962 | 43.570621 |
| SSGSTTELHCVTDI | 15 | P25054 | Adenomatous polyposis coli protein OS=I | APC | 311.646 | 2 | 57.930008 | yes | 7.992E-07 | 839.87048 | 29.59177 |
| SFHVEDTPVCFSR | 13 | P25054 | Adenomatous polyposis coli protein OS=I | APC | 311.646 | 2 | 57.930008 | yes | 9.722E-08 | 790.86176 | 26.35887 |
| SPSEGQTATTSPR | 13 | P25054 | Adenomatous polyposis coli protein OS=I | APC | 311.646 | 2 | 57.930008 | yes | 0.0001929 | 659.81531 | 11.7455 |
| QELEDNSNHLTK | 12 | P25054 | Adenomatous polyposis coli protein OS=I | APC | 311.646 | 2 | 57.930008 | yes | 9.722E-08 | 714.34167 | 10.219709 |
| SVDVILDDDLVDK | 13 | P25205 | DNA replication licensing factor MCM3 ( | MCM3 | 90.981 | 2 | 53.06903 | yes | 1.188E-07 | 723.37213 | 990.77881 |
| CSVLAAANPVYGI | 13 | P25205 | DNA replication licensing factor MCM3 ( | MCM3 | 90.981 | 2 | 53.06903 | yes | 9.743E-08 | 689.35083 | 562.67902 |
| VQVVGTYR | 8 | P25205 | DNA replication licensing factor MCM3 ( | MCM3 | 90.981 | 2 | 53.06903 | yes | 9.722E-08 | 461.26126 | 425.27716 |
| SQDSMSSDTAR | 11 | P25205 | DNA replication licensing factor MCM3 ( | MCM3 | 90.981 | 2 | 53.06903 | yes | 9.819E-08 | 592.74603 | 394.23495 |
| LIVNVNDLR | 9 | P25205 | DNA replication licensing factor MCM3 ( | MCM3 | 90.981 | 2 | 53.06903 | yes | 2.207E-07 | 528.31403 | 375.82285 |
| TPMENIGLQDSLLS | 15 | P25205 | DNA replication licensing factor MCM3 ( | MCM3 | 90.981 | 2 | 53.06903 | yes | 9.722E-08 | 837.42999 | 324.48886 |
| DAQPSFSAEDIAK | 13 | P25205 | DNA replication licensing factor MCM3 ( | MCM3 | 90.981 | 2 | 53.06903 | yes | 1.132E-07 | 689.82788 | 318.892 |
| GDINILLIGDPSVA | 15 | P25205 | DNA replication licensing factor MCM3 ( | MCM3 | 90.981 | 2 | 53.06903 | yes | 1.232E-07 | 762.93524 | 315.96936 |
| SVHYCPATK | 9 | P25205 | DNA replication licensing factor MCM3 ( | MCM3 | 90.981 | 2 | 53.06903 | yes | 2.782E-07 | 531.75549 | 308.0484 |
| GVVCIDEFDK | 10 | P25205 | DNA replication licensing factor MCM3 ( | MCM3 | 90.981 | 2 | 53.06903 | yes | 1.902E-07 | 591.27899 | 285.28552 |
| TAIHEVMEQGR | 11 | P25205 | DNA replication licensing factor MCM3 ( | MCM3 | 90.981 | 2 | 53.06903 | yes | 9.733E-08 | 635.81409 | 275.59216 |
| SLAPSIHGHDYVK | 13 | P25205 | DNA replication licensing factor MCM3 ( | MCM3 | 90.981 | 2 | 53.06903 | yes | 2.131E-07 | 712.37006 | 252.16577 |

| | | | | | | | | | | | |
|---|---|---|---|---|---|---|---|---|---|---|---|
| EISDHVLR | 8 | P25205 | DNA replication licensing factor MCM3 ( | MCM3 | 90.981 | 2 | 53.06903 | yes | 2.06E-07 | 484.76163 | 237.55728 |
| TVLIACNVK | 9 | P25205 | DNA replication licensing factor MCM3 ( | MCM3 | 90.981 | 2 | 53.06903 | yes | 4.65E-07 | 509.29172 | 213.34769 |
| EAHAQSIGMNR | 11 | P25205 | DNA replication licensing factor MCM3 ( | MCM3 | 90.981 | 2 | 53.06903 | yes | 1.102E-07 | 607.29077 | 200.78702 |
| YVLCTAPR | 8 | P25205 | DNA replication licensing factor MCM3 ( | MCM3 | 90.981 | 2 | 53.06903 | yes | 3.202E-07 | 490.25513 | 180.51389 |
| ELISDNQYR | 9 | P25205 | DNA replication licensing factor MCM3 ( | MCM3 | 90.981 | 2 | 53.06903 | yes | 3.992E-07 | 569.2804 | 166.37978 |
| CSLVRPK | 7 | P25205 | DNA replication licensing factor MCM3 ( | MCM3 | 90.981 | 2 | 53.06903 | yes | 3.316E-07 | 430.24457 | 164.32611 |
| APAGQLPR | 8 | P25205 | DNA replication licensing factor MCM3 ( | MCM3 | 90.981 | 2 | 53.06903 | yes | 2.231E-07 | 405.23505 | 86.447205 |
| HDNLLHGTK | 9 | P25205 | DNA replication licensing factor MCM3 ( | MCM3 | 90.981 | 2 | 53.06903 | yes | 1.874E-07 | 517.77252 | 80.024872 |
| SKDIFDQLAK | 10 | P25205 | DNA replication licensing factor MCM3 ( | MCM3 | 90.981 | 2 | 53.06903 | yes | 1.85E-07 | 582.81659 | 77.724861 |
| DFVASIDATYAK | 12 | P25205 | DNA replication licensing factor MCM3 ( | MCM3 | 90.981 | 2 | 53.06903 | yes | 4.296E-06 | 650.82465 | 75.831841 |
| DLENGSHIR | 9 | P25205 | DNA replication licensing factor MCM3 ( | MCM3 | 90.981 | 2 | 53.06903 | yes | 4.044E-07 | 520.75958 | 72.157677 |
| LLNNAFEELVAFQ | 14 | P25205 | DNA replication licensing factor MCM3 ( | MCM3 | 90.981 | 2 | 53.06903 | yes | 1.286E-07 | 832.44373 | 71.291481 |
| VALLDVFR | 8 | P25205 | DNA replication licensing factor MCM3 ( | MCM3 | 90.981 | 2 | 53.06903 | yes | 2.785E-07 | 466.78183 | 57.558907 |
| RLEAGAMVLADR | 12 | P25205 | DNA replication licensing factor MCM3 ( | MCM3 | 90.981 | 2 | 53.06903 | yes | 9.722E-08 | 651.35333 | 53.787312 |
| LGVGAATHVVIYD | 24 | P25325 | 3-mercaptopyruvate sulfurtransferase OS= | MPST | 33.179 | 3 | 49.946312 | yes | 9.968E-08 | 820.75543 | 104.01199 |
| TPVTLGSDQPLLR | 13 | P25391 | Laminin subunit alpha-1 OS=Homo sapie | LAMA1 | 337.084 | 2 | 54.09321 | yes | 1.706E-07 | 698.89337 | 339.66864 |
| ELATSASQSAVSTI | 15 | P25391 | Laminin subunit alpha-1 OS=Homo sapie | LAMA1 | 337.084 | 2 | 54.09321 | yes | 9.722E-08 | 760.89935 | 305.90717 |
| ISDISMEVGR | 10 | P25391 | Laminin subunit alpha-1 OS=Homo sapie | LAMA1 | 337.084 | 2 | 54.09321 | yes | 2.979E-07 | 553.77917 | 204.19778 |
| LEFPDFPIDDNR | 12 | P25391 | Laminin subunit alpha-1 OS=Homo sapie | LAMA1 | 337.084 | 2 | 54.09321 | yes | 9.722E-08 | 739.3515 | 203.28415 |
| LQMSITEIMEK | 11 | P25391 | Laminin subunit alpha-1 OS=Homo sapie | LAMA1 | 337.084 | 2 | 54.09321 | yes | 1.734E-07 | 661.83838 | 181.31261 |
| YYWAAPEAYLGN | 13 | P25391 | Laminin subunit alpha-1 OS=Homo sapie | LAMA1 | 337.084 | 2 | 54.09321 | yes | 1.026E-07 | 773.37225 | 150.85715 |
| VDAIGLELVDGK | 12 | P25391 | Laminin subunit alpha-1 OS=Homo sapie | LAMA1 | 337.084 | 2 | 54.09321 | yes | 5.452E-05 | 614.84283 | 150.06001 |
| SIGLWNYIER | 10 | P25391 | Laminin subunit alpha-1 OS=Homo sapie | LAMA1 | 337.084 | 2 | 54.09321 | yes | 2.258E-07 | 625.83002 | 149.08069 |
| ASYGQGLQQSR | 11 | P25391 | Laminin subunit alpha-1 OS=Homo sapie | LAMA1 | 337.084 | 2 | 54.09321 | yes | 9.722E-08 | 597.79675 | 140.53195 |
| VAFLWDLGSGSTF | 13 | P25391 | Laminin subunit alpha-1 OS=Homo sapie | LAMA1 | 337.084 | 2 | 54.09321 | yes | 9.783E-08 | 704.86462 | 132.2957 |
| LHFMFDLGK | 9 | P25391 | Laminin subunit alpha-1 OS=Homo sapie | LAMA1 | 337.084 | 2 | 54.09321 | yes | 1.67E-06 | 554.28644 | 119.31398 |
| LDKDPIYVGGLPR | 13 | P25391 | Laminin subunit alpha-1 OS=Homo sapie | LAMA1 | 337.084 | 2 | 54.09321 | yes | 2.117E-07 | 721.90375 | 118.03934 |
| KIPSQQDALGGR | 12 | P25391 | Laminin subunit alpha-1 OS=Homo sapie | LAMA1 | 337.084 | 2 | 54.09321 | yes | 1.026E-07 | 635.34912 | 117.99804 |
| CDACLLGYEGK | 11 | P25391 | Laminin subunit alpha-1 OS=Homo sapie | LAMA1 | 337.084 | 2 | 54.09321 | yes | 9.823E-08 | 643.28119 | 112.2931 |
| SPQVQSFDFSR | 11 | P25391 | Laminin subunit alpha-1 OS=Homo sapie | LAMA1 | 337.084 | 2 | 54.09321 | yes | 9.881E-08 | 649.31219 | 110.97151 |
| YLQESLLK | 8 | P25391 | Laminin subunit alpha-1 OS=Homo sapie | LAMA1 | 337.084 | 2 | 54.09321 | yes | 1.6E-07 | 497.28439 | 108.52892 |
| DFPDCVPCDCDLR | 13 | P25391 | Laminin subunit alpha-1 OS=Homo sapie | LAMA1 | 337.084 | 2 | 54.09321 | yes | 9.722E-08 | 834.82617 | 101.00109 |
| FGNIGSLSVK | 10 | P25391 | Laminin subunit alpha-1 OS=Homo sapie | LAMA1 | 337.084 | 2 | 54.09321 | yes | 1.108E-07 | 511.28748 | 100.58475 |
| LADVLYSGLENIR | 13 | P25391 | Laminin subunit alpha-1 OS=Homo sapie | LAMA1 | 337.084 | 2 | 54.09321 | yes | 9.722E-08 | 731.89862 | 94.712616 |
| LEGVAEETDNLQK | 13 | P25391 | Laminin subunit alpha-1 OS=Homo sapie | LAMA1 | 337.084 | 2 | 54.09321 | yes | 9.722E-08 | 723.35956 | 91.810753 |
| NAVDLVYR | 8 | P25391 | Laminin subunit alpha-1 OS=Homo sapie | LAMA1 | 337.084 | 2 | 54.09321 | yes | 4.081E-07 | 475.25873 | 88.575096 |
| VMTDLGSGPITLLT | 16 | P25391 | Laminin subunit alpha-1 OS=Homo sapie | LAMA1 | 337.084 | 2 | 54.09321 | yes | 2.112E-07 | 844.948 | 86.746628 |
| WHSIHVAR | 8 | P25391 | Laminin subunit alpha-1 OS=Homo sapie | LAMA1 | 337.084 | 2 | 54.09321 | yes | 7.833E-07 | 503.27249 | 82.138268 |
| QTNESLLILR | 10 | P25391 | Laminin subunit alpha-1 OS=Homo sapie | LAMA1 | 337.084 | 2 | 54.09321 | yes | 1.4E-07 | 593.84314 | 73.789963 |

| | | | | | | | | | | | |
|---|---|---|---|---|---|---|---|---|---|---|---|
| LSVELSIR | 8 | P25391 | Laminin subunit alpha-1 OS=Homo sapie | LAMA1 | 337.084 | 2 | 54.09321 | yes | 4.416E-07 | 458.77673 | 72.592712 |
| ACDQCSLGYR | 10 | P25391 | Laminin subunit alpha-1 OS=Homo sapie | LAMA1 | 337.084 | 2 | 54.09321 | yes | 2.969E-07 | 615.25549 | 64.199806 |
| KLPGIALELSELR | 13 | P25391 | Laminin subunit alpha-1 OS=Homo sapie | LAMA1 | 337.084 | 2 | 54.09321 | yes | 1.02E-07 | 719.93506 | 58.159924 |
| LHVQEEQNLTSEL | 18 | P25391 | Laminin subunit alpha-1 OS=Homo sapie | LAMA1 | 337.084 | 3 | 54.09321 | yes | 2.601E-07 | 698.37103 | 37.417133 |
| QVIYMDAPAPENC | 15 | P25391 | Laminin subunit alpha-1 OS=Homo sapie | LAMA1 | 337.084 | 2 | 54.09321 | yes | 2.789E-06 | 830.41162 | 34.808666 |
| TSSQNGVLLGISTA | 15 | P25391 | Laminin subunit alpha-1 OS=Homo sapie | LAMA1 | 337.084 | 2 | 54.09321 | yes | 2.393E-06 | 738.40686 | 30.5049 |
| AEDHAAEFQR | 10 | P25391 | Laminin subunit alpha-1 OS=Homo sapie | LAMA1 | 337.084 | 2 | 54.09321 | yes | 1.487E-07 | 587.26782 | 28.107981 |
| DFTQLHQNATLEL | 14 | P25391 | Laminin subunit alpha-1 OS=Homo sapie | LAMA1 | 337.084 | 2 | 54.09321 | yes | 1.704E-07 | 829.43085 | 16.505075 |
| TALIHDGLAR | 10 | P25398 | Small ribosomal subunit protein eS12 OS | RPS12 | 14.515 | 2 | 52.035583 | yes | 1.593E-07 | 533.80383 | 29479.988 |
| VVGCSCVVVK | 10 | P25398 | Small ribosomal subunit protein eS12 OS | RPS12 | 14.515 | 2 | 52.035583 | yes | 1.364E-07 | 553.78845 | 15597.053 |
| DVIEEYFK | 8 | P25398 | Small ribosomal subunit protein eS12 OS | RPS12 | 14.515 | 2 | 52.035583 | yes | 1.418E-07 | 521.75824 | 8827.9951 |
| KVVGCSCVVVK | 11 | P25398 | Small ribosomal subunit protein eS12 OS | RPS12 | 14.515 | 2 | 52.035583 | yes | 9.722E-08 | 617.83594 | 4944.0264 |
| LGEWVGLCK | 9 | P25398 | Small ribosomal subunit protein eS12 OS | RPS12 | 14.515 | 2 | 52.035583 | yes | 1.644E-07 | 531.27606 | 1948.5914 |
| KLGEWVGLCK | 10 | P25398 | Small ribosomal subunit protein eS12 OS | RPS12 | 14.515 | 2 | 52.035583 | yes | 1.305E-07 | 595.32355 | 1321.1552 |
| QAHLCVLASNCDI | 18 | P25398 | Small ribosomal subunit protein eS12 OS | RPS12 | 14.515 | 3 | 52.035583 | yes | 9.722E-08 | 712.3288 | 346.58276 |
| LVEALCAEHQINL | 15 | P25398 | Small ribosomal subunit protein eS12 OS | RPS12 | 14.515 | 3 | 52.035583 | yes | 1.083E-07 | 584.3222 | 164.78619 |
| DITSDSENSNFR | 12 | P25445 | Tumor necrosis factor receptor superfamil | FAS | 37.733 | 2 | 53.2753 | yes | 9.81E-08 | 692.80243 | 243.63264 |
| ECTLTSNTK | 9 | P25445 | Tumor necrosis factor receptor superfamil | FAS | 37.733 | 2 | 53.2753 | yes | 1.341E-07 | 527.24768 | 138.54224 |
| NWHQLHGK | 8 | P25445 | Tumor necrosis factor receptor superfamil | FAS | 37.733 | 2 | 53.2753 | yes | 1.972E-07 | 510.26212 | 115.82405 |
| ANLCTLAEK | 9 | P25445 | Tumor necrosis factor receptor superfamil | FAS | 37.733 | 2 | 53.2753 | yes | 9.722E-08 | 510.26315 | 81.174942 |
| IQTIILK | 7 | P25445 | Tumor necrosis factor receptor superfamil | FAS | 37.733 | 2 | 53.2753 | yes | 2.195E-06 | 414.78131 | 31.358498 |
| FSLDFNLR | 8 | P25490 | Transcriptional repressor protein YY1 OS | YY1 | 44.713 | 2 | 51.737293 | no | 4.507E-07 | 506.26654 | 1180.3899 |
| EEVVGGDDSDGLI | 13 | P25490 | Transcriptional repressor protein YY1 OS | YY1 | 44.713 | 2 | 51.737293 | yes | 1.429E-07 | 674.30475 | 975.96155 |
| HQLVHTGEKPFQC | 20 | P25490 | Transcriptional repressor protein YY1 OS | YY1 | 44.713 | 4 | 51.737293 | no | 2.13E-07 | 590.7782 | 349.48389 |
| QLAEFAR | 7 | P25490 | Transcriptional repressor protein YY1 OS | YY1 | 44.713 | 2 | 51.737293 | no | 9.722E-08 | 417.72705 | 258.91476 |
| FAQSTNLK | 8 | P25490 | Transcriptional repressor protein YY1 OS | YY1 | 44.713 | 2 | 51.737293 | no | 9.722E-08 | 454.74545 | 181.66124 |
| VHVCAECGK | 9 | P25490 | Transcriptional repressor protein YY1 OS | YY1 | 44.713 | 2 | 51.737293 | no | 1.946E-07 | 530.23914 | 178.55716 |
| SYLSGGAGAAGG( | 20 | P25490 | Transcriptional repressor protein YY1 OS | YY1 | 44.713 | 2 | 51.737293 | yes | 9.728E-08 | 832.38715 | 116.61623 |
| PFQCTFEGCGK | 11 | P25490 | Transcriptional repressor protein YY1 OS | YY1 | 44.713 | 2 | 51.737293 | no | 2.061E-07 | 665.78137 | 100.90831 |
| KFAQSTNLK | 9 | P25490 | Transcriptional repressor protein YY1 OS | YY1 | 44.713 | 2 | 51.737293 | no | 1.456E-07 | 518.79291 | 66.466911 |
| LPPGGIPGIDLSDPI | 15 | P25490 | Transcriptional repressor protein YY1 OS | YY1 | 44.713 | 2 | 51.737293 | no | 1.68E-06 | 738.40887 | 62.52528 |
| PYVCPFDGCNK | 11 | P25490 | Transcriptional repressor protein YY1 OS | YY1 | 44.713 | 2 | 51.737293 | yes | 8.889E-07 | 678.78918 | 42.299141 |
| KSYLSGGAGAAG( | 21 | P25490 | Transcriptional repressor protein YY1 OS | YY1 | 44.713 | 3 | 51.737293 | yes | 4.658E-07 | 597.95886 | 35.461201 |
| RFSLDFNLR | 9 | P25490 | Transcriptional repressor protein YY1 OS | YY1 | 44.713 | 2 | 51.737293 | no | 7.703E-07 | 584.31708 | 32.780056 |
| VSLEEIYSGCTK | 12 | P25685 | DnaJ homolog subfamily B member 1 OS | DNAJB1 | 38.045 | 2 | 54.282146 | no | 9.722E-08 | 693.33447 | 328.99396 |
| EIAEAYDVLSDPR | 13 | P25685 | DnaJ homolog subfamily B member 1 OS | DNAJB1 | 38.045 | 2 | 54.282146 | yes | 9.722E-08 | 739.36212 | 198.32184 |
| KVPGEGLPLPK | 11 | P25685 | DnaJ homolog subfamily B member 1 OS | DNAJB1 | 38.045 | 2 | 54.282146 | yes | 3.757E-07 | 567.84772 | 184.14081 |
| NPFDTFFGQR | 10 | P25685 | DnaJ homolog subfamily B member 1 OS | DNAJB1 | 38.045 | 2 | 54.282146 | yes | 9.722E-08 | 614.79089 | 134.00955 |
| GKDYYQTLGLAR | 12 | P25685 | DnaJ homolog subfamily B member 1 OS | DNAJB1 | 38.045 | 2 | 54.282146 | yes | 2.5E-07 | 692.86462 | 45.043221 |

| | | | | | | | | | | | |
|---|---|---|---|---|---|---|---|---|---|---|---|
| EAYPGDVFYLHSR | 13 | P25705 | ATP synthase subunit alpha, mitochondri | ATP5F1A | 59.751 | 2 | 52.077778 | yes | 9.885E-08 | 777.3728 | 711.04974 |
| TGTAEMSSILEER | 13 | P25705 | ATP synthase subunit alpha, mitochondri | ATP5F1A | 59.751 | 2 | 52.077778 | yes | 9.849E-08 | 712.34033 | 682.05969 |
| TGAIVDVPVGEEL | 16 | P25705 | ATP synthase subunit alpha, mitochondri | ATP5F1A | 59.751 | 2 | 52.077778 | yes | 1.656E-07 | 812.94885 | 634.35242 |
| VLSIGDGIAR | 10 | P25705 | ATP synthase subunit alpha, mitochondri | ATP5F1A | 59.751 | 2 | 52.077778 | yes | 3.683E-07 | 500.79294 | 618.29077 |
| LELAQYR | 7 | P25705 | ATP synthase subunit alpha, mitochondri | ATP5F1A | 59.751 | 2 | 52.077778 | yes | 6.789E-07 | 446.74799 | 611.9361 |
| VVDALGNAIDGK | 12 | P25705 | ATP synthase subunit alpha, mitochondri | ATP5F1A | 59.751 | 2 | 52.077778 | yes | 1.718E-07 | 586.31952 | 488.87338 |
| ILGADTSVDLEET( | 15 | P25705 | ATP synthase subunit alpha, mitochondri | ATP5F1A | 59.751 | 2 | 52.077778 | yes | 9.722E-08 | 788.39667 | 485.10367 |
| HALIIYDDLSK | 11 | P25705 | ATP synthase subunit alpha, mitochondri | ATP5F1A | 59.751 | 2 | 52.077778 | yes | 9.834E-08 | 644.35083 | 439.94775 |
| PAINVGLSVSR | 11 | P25705 | ATP synthase subunit alpha, mitochondri | ATP5F1A | 59.751 | 2 | 52.077778 | yes | 9.722E-08 | 556.82477 | 339.80661 |
| TSIAIDTIINQK | 12 | P25705 | ATP synthase subunit alpha, mitochondri | ATP5F1A | 59.751 | 2 | 52.077778 | yes | 1.721E-07 | 658.87463 | 307.16187 |
| AVDSLVPIGR | 10 | P25705 | ATP synthase subunit alpha, mitochondri | ATP5F1A | 59.751 | 2 | 52.077778 | yes | 2.377E-07 | 513.80072 | 248.50217 |
| LYCIYVAIGQK | 11 | P25705 | ATP synthase subunit alpha, mitochondri | ATP5F1A | 59.751 | 2 | 52.077778 | yes | 1.121E-07 | 664.35754 | 91.556854 |
| EPMQTGIK | 8 | P25705 | ATP synthase subunit alpha, mitochondri | ATP5F1A | 59.751 | 2 | 52.077778 | yes | 1.111E-06 | 452.23386 | 82.493431 |
| RLTDADAMK | 9 | P25705 | ATP synthase subunit alpha, mitochondri | ATP5F1A | 59.751 | 2 | 52.077778 | yes | 2.975E-06 | 510.76077 | 78.873077 |
| QMSLLLR | 7 | P25705 | ATP synthase subunit alpha, mitochondri | ATP5F1A | 59.751 | 2 | 52.077778 | yes | 4.574E-07 | 430.75476 | 59.096802 |
| NVQAEEMVEFSS( | 15 | P25705 | ATP synthase subunit alpha, mitochondri | ATP5F1A | 59.751 | 2 | 52.077778 | yes | 1.425E-06 | 834.40088 | 33.104958 |
| GIRPAINVGLSVSR | 14 | P25705 | ATP synthase subunit alpha, mitochondri | ATP5F1A | 59.751 | 3 | 52.077778 | yes | 2.492E-06 | 480.28781 | 28.381809 |
| ETLPAEQDLTTK | 12 | P25786 | Proteasome subunit alpha type-1 OS=Hor | PSMA1 | 29.556 | 2 | 51.48884 | yes | 9.722E-08 | 673.34589 | 84.568779 |
| ILHVDNHIGISIAGl | 20 | P25786 | Proteasome subunit alpha type-1 OS=Hor | PSMA1 | 29.556 | 4 | 51.48884 | yes | 2.708E-05 | 522.29053 | 5.9548607 |
| LLDEVFFSEK | 10 | P25789 | Proteasome subunit alpha type-4 OS=Hor | PSMA4 | 29.484 | 2 | 50.52979 | yes | 9.788E-08 | 613.81879 | 280.29904 |
| VEIATLTR | 8 | P25789 | Proteasome subunit alpha type-4 OS=Hor | PSMA4 | 29.484 | 2 | 50.52979 | yes | 1E-07 | 451.76892 | 116.49095 |
| ATCIGNNSAAAVS | 16 | P25789 | Proteasome subunit alpha type-4 OS=Hor | PSMA4 | 29.484 | 2 | 50.52979 | yes | 1.806E-07 | 804.39764 | 88.541443 |
| TTIFSPEGR | 9 | P25789 | Proteasome subunit alpha type-4 OS=Hor | PSMA4 | 29.484 | 2 | 50.52979 | yes | 4.464E-07 | 504.26144 | 49.755772 |
| ALTMEVIR | 8 | P25963 | NF-kappa-B inhibitor alpha OS=Homo sa | NFKBIA | 35.609 | 2 | 47.431156 | yes | 1.331E-07 | 466.76532 | 118.77834 |
| AQMVQEDLEK | 10 | P26038 | Moesin OS=Homo sapiens OX=9606 GN | MSN | 67.821 | 2 | 54.152756 | yes | 9.722E-08 | 595.78973 | 225.41302 |
| ALELEQER | 8 | P26038 | Moesin OS=Homo sapiens OX=9606 GN | MSN | 67.821 | 2 | 54.152756 | yes | 3.03E-07 | 494.25891 | 224.79918 |
| EDAVLEYLK | 9 | P26038 | Moesin OS=Homo sapiens OX=9606 GN | MSN | 67.821 | 2 | 54.152756 | yes | 1.12E-07 | 540.28461 | 156.23674 |
| KESPLLFK | 8 | P26038 | Moesin OS=Homo sapiens OX=9606 GN | MSN | 67.821 | 2 | 54.152756 | yes | 3.015E-06 | 481.28949 | 117.13446 |
| IQVWHEEHR | 9 | P26038 | Moesin OS=Homo sapiens OX=9606 GN | MSN | 67.821 | 2 | 54.152756 | yes | 3.6E-07 | 617.30981 | 105.05701 |
| EVWFFGLQYQDTI | 13 | P26038 | Moesin OS=Homo sapiens OX=9606 GN | MSN | 67.821 | 2 | 54.152756 | yes | 9.722E-08 | 830.90393 | 100.0246 |
| AQQELEEQTR | 10 | P26038 | Moesin OS=Homo sapiens OX=9606 GN | MSN | 67.821 | 2 | 54.152756 | yes | 1.511E-07 | 616.29932 | 67.957558 |
| EALLQASR | 8 | P26038 | Moesin OS=Homo sapiens OX=9606 GN | MSN | 67.821 | 2 | 54.152756 | yes | 1.43E-07 | 444.25089 | 63.428261 |
| TANDMIHAENMR | 12 | P26038 | Moesin OS=Homo sapiens OX=9606 GN | MSN | 67.821 | 2 | 54.152756 | yes | 1.221E-07 | 701.81372 | 46.659763 |
| INPESPADTCIPK | 13 | P26045 | Tyrosine-protein phosphatase non-recepto | PTPN3 | 103.99 | 2 | 47.19968 | yes | 5.305E-07 | 721.35321 | 118.14722 |
| GSLAPQDSDSEVS( | 16 | P26045 | Tyrosine-protein phosphatase non-recepto | PTPN3 | 103.99 | 2 | 47.19968 | yes | 3.888E-07 | 845.38739 | 112.88644 |
| YKDVLPYDTTR | 11 | P26045 | Tyrosine-protein phosphatase non-recepto | PTPN3 | 103.99 | 2 | 47.19968 | yes | 2.849E-06 | 685.8512 | 80.721802 |
| AMMVQTSSQYK | 11 | P26045 | Tyrosine-protein phosphatase non-recepto | PTPN3 | 103.99 | 2 | 47.19968 | yes | 1.554E-07 | 637.29938 | 74.443649 |
| QSEAESCYINIAR | 13 | P26045 | Tyrosine-protein phosphatase non-recepto | PTPN3 | 103.99 | 2 | 47.19968 | yes | 4.403E-07 | 770.85663 | 51.478352 |
| VIGGMVWNPAMR | 12 | P26045 | Tyrosine-protein phosphatase non-recepto | PTPN3 | 103.99 | 2 | 47.19968 | yes | 1.83E-07 | 665.84155 | 47.485733 |

| | | | | | | | | | | | |
|---|---|---|---|---|---|---|---|---|---|---|---|
| SLSVEHLETK | 10 | P26045 | Tyrosine-protein phosphatase non-recepto | PTPN3 | 103.99 | 2 | 47.19968 | yes | 5.22E-07 | 571.80621 | 47.089497 |
| FVCEAILR | 8 | P26045 | Tyrosine-protein phosphatase non-recepto | PTPN3 | 103.99 | 2 | 47.19968 | yes | 2.581E-07 | 504.27078 | 23.577642 |
| HLYFLQLK | 8 | P26045 | Tyrosine-protein phosphatase non-recepto | PTPN3 | 103.99 | 2 | 47.19968 | yes | 1.707E-07 | 531.31073 | 20.512983 |
| MDICEGR | 7 | P26045 | Tyrosine-protein phosphatase non-recepto | PTPN3 | 103.99 | 2 | 47.19968 | yes | 1.379E-07 | 440.68622 | 18.830221 |
| MPLVVSR | 7 | P26045 | Tyrosine-protein phosphatase non-recepto | PTPN3 | 103.99 | 2 | 47.19968 | yes | 3.559E-06 | 401.23621 | 17.309498 |
| NLVCTDLFTR | 10 | P26196 | Probable ATP-dependent RNA helicase D | DDX6 | 54.417 | 2 | 52.113106 | yes | 1.426E-07 | 619.81354 | 24657.506 |
| LAETYLHR | 8 | P26196 | Probable ATP-dependent RNA helicase D | DDX6 | 54.417 | 2 | 52.113106 | yes | 1.285E-07 | 501.772 | 17897.645 |
| VMATTGGTNLR | 11 | P26196 | Probable ATP-dependent RNA helicase D | DDX6 | 54.417 | 2 | 52.113106 | yes | 1.074E-07 | 560.7926 | 7730.9717 |
| VMATTGGTNLR | 11 | P26196 | Probable ATP-dependent RNA helicase D | DDX6 | 54.417 | 2 | 52.113106 | yes | 1.624E-07 | 568.79004 | 7730.9717 |
| SGAYLIPLLER | 11 | P26196 | Probable ATP-dependent RNA helicase D | DDX6 | 54.417 | 2 | 52.113106 | yes | 3.65E-07 | 616.3559 | 11625.184 |
| GVTQYYAYVTER | 12 | P26196 | Probable ATP-dependent RNA helicase D | DDX6 | 54.417 | 2 | 52.113106 | yes | 3.583E-07 | 725.35406 | 8691.4971 |
| GNEFEDYCLK | 10 | P26196 | Probable ATP-dependent RNA helicase D | DDX6 | 54.417 | 2 | 52.113106 | yes | 2.95E-07 | 637.77155 | 7688.4082 |
| LDDTVHVVIATPG | 14 | P26196 | Probable ATP-dependent RNA helicase D | DDX6 | 54.417 | 2 | 52.113106 | yes | 9.722E-08 | 746.90955 | 2997.9026 |
| LDDTVHVVIATPG | 14 | P26196 | Probable ATP-dependent RNA helicase D | DDX6 | 54.417 | 2 | 52.113106 | yes | 1.117E-07 | 498.27545 | 2997.9026 |
| LDDTVHVVIATPG | 14 | P26196 | Probable ATP-dependent RNA helicase D | DDX6 | 54.417 | 3 | 52.113106 | yes | 9.722E-08 | 746.90955 | 2997.9026 |
| LDDTVHVVIATPG | 14 | P26196 | Probable ATP-dependent RNA helicase D | DDX6 | 54.417 | 3 | 52.113106 | yes | 1.117E-07 | 498.27545 | 2997.9026 |
| DNIQAMVIVPTR | 12 | P26196 | Probable ATP-dependent RNA helicase D | DDX6 | 54.417 | 2 | 52.113106 | yes | 1.083E-07 | 678.86884 | 2741.843 |
| DNIQAMVIVPTR | 12 | P26196 | Probable ATP-dependent RNA helicase D | DDX6 | 54.417 | 2 | 52.113106 | yes | 2.315E-07 | 686.86627 | 2741.843 |
| VHCLNTLFSR | 10 | P26196 | Probable ATP-dependent RNA helicase D | DDX6 | 54.417 | 2 | 52.113106 | yes | 5.519E-07 | 623.82172 | 4747.915 |
| PSPIQEESIPIALSG | 16 | P26196 | Probable ATP-dependent RNA helicase D | DDX6 | 54.417 | 2 | 52.113106 | yes | 1.222E-07 | 847.45959 | 2922.6086 |
| VDHVQMIVLDEAI | 14 | P26196 | Probable ATP-dependent RNA helicase D | DDX6 | 54.417 | 2 | 52.113106 | yes | 1.296E-07 | 806.40601 | 2631.7048 |
| KDNIQAMVIVPTR | 13 | P26196 | Probable ATP-dependent RNA helicase D | DDX6 | 54.417 | 2 | 52.113106 | yes | 1.483E-07 | 742.91632 | 352.60059 |
| KDNIQAMVIVPTR | 13 | P26196 | Probable ATP-dependent RNA helicase D | DDX6 | 54.417 | 2 | 52.113106 | yes | 3.945E-07 | 495.61331 | 352.60059 |
| KDNIQAMVIVPTR | 13 | P26196 | Probable ATP-dependent RNA helicase D | DDX6 | 54.417 | 2 | 52.113106 | yes | 3.9E-07 | 500.94495 | 352.60059 |
| KDNIQAMVIVPTR | 13 | P26196 | Probable ATP-dependent RNA helicase D | DDX6 | 54.417 | 3 | 52.113106 | yes | 1.483E-07 | 742.91632 | 352.60059 |
| KDNIQAMVIVPTR | 13 | P26196 | Probable ATP-dependent RNA helicase D | DDX6 | 54.417 | 3 | 52.113106 | yes | 3.945E-07 | 495.61331 | 352.60059 |
| KDNIQAMVIVPTR | 13 | P26196 | Probable ATP-dependent RNA helicase D | DDX6 | 54.417 | 3 | 52.113106 | yes | 3.9E-07 | 500.94495 | 352.60059 |
| KISQLGYSCFYIHA | 15 | P26196 | Probable ATP-dependent RNA helicase D | DDX6 | 54.417 | 3 | 52.113106 | yes | 9.819E-08 | 605.64679 | 856.36798 |
| FMNSHLQK | 8 | P26196 | Probable ATP-dependent RNA helicase D | DDX6 | 54.417 | 2 | 52.113106 | yes | 2.094E-07 | 502.75275 | 599.82117 |
| ISQLGYSCFYIHAK | 14 | P26196 | Probable ATP-dependent RNA helicase D | DDX6 | 54.417 | 2 | 52.113106 | yes | 1.597E-07 | 843.91907 | 371.8891 |
| PYEINLMEELTLK | 13 | P26196 | Probable ATP-dependent RNA helicase D | DDX6 | 54.417 | 2 | 52.113106 | yes | 9.722E-08 | 796.91565 | 261.50821 |
| GNEFEDYCLKR | 11 | P26196 | Probable ATP-dependent RNA helicase D | DDX6 | 54.417 | 2 | 52.113106 | yes | 9.722E-08 | 715.82208 | 253.0318 |
| SIEEQLGTEIK | 11 | P26196 | Probable ATP-dependent RNA helicase D | DDX6 | 54.417 | 2 | 52.113106 | yes | 1.923E-07 | 623.8299 | 178.84111 |
| PLPAEEPER | 9 | P26358 | DNA (cytosine-5)-methyltransferase 1 OS | DNMT1 | 183.166 | 2 | 53.531235 | yes | 1.018E-07 | 519.26672 | 340.26224 |
| ISWVGEAVK | 9 | P26358 | DNA (cytosine-5)-methyltransferase 1 OS | DNMT1 | 183.166 | 2 | 53.531235 | yes | 1.499E-07 | 494.77673 | 337.21332 |
| FTEDSLLR | 8 | P26358 | DNA (cytosine-5)-methyltransferase 1 OS | DNMT1 | 183.166 | 2 | 53.531235 | yes | 1.841E-07 | 490.75601 | 318.42249 |
| LVMAGETTNSR | 11 | P26358 | DNA (cytosine-5)-methyltransferase 1 OS | DNMT1 | 183.166 | 2 | 53.531235 | yes | 1.369E-07 | 589.79535 | 312.92899 |
| GSNLDAPEPYR | 11 | P26358 | DNA (cytosine-5)-methyltransferase 1 OS | DNMT1 | 183.166 | 2 | 53.531235 | yes | 2.097E-07 | 609.79114 | 252.8159 |
| VLEQLEDLDSR | 11 | P26358 | DNA (cytosine-5)-methyltransferase 1 OS | DNMT1 | 183.166 | 2 | 53.531235 | yes | 1.615E-07 | 658.83826 | 238.23242 |

| | | | | | | | | | | | |
|---|---|---|---|---|---|---|---|---|---|---|---|
| SQGFPDTYR | 9 | P26358 | DNA (cytosine-5)-methyltransferase 1 OS | DNMT1 | 183.166 | 2 | 53.531235 | yes | 2.638E-07 | 535.74872 | 222.35448 |
| GAQYQPILR | 9 | P26358 | DNA (cytosine-5)-methyltransferase 1 OS | DNMT1 | 183.166 | 2 | 53.531235 | yes | 1.87E-07 | 523.29309 | 212.70726 |
| LAGVTLGQR | 9 | P26358 | DNA (cytosine-5)-methyltransferase 1 OS | DNMT1 | 183.166 | 2 | 53.531235 | yes | 1.3E-07 | 457.77454 | 199.17094 |
| VPTLAVPAISLPDI | 16 | P26358 | DNA (cytosine-5)-methyltransferase 1 OS | DNMT1 | 183.166 | 2 | 53.531235 | yes | 6.67E-07 | 831.97491 | 192.36589 |
| FFLLENVR | 8 | P26358 | DNA (cytosine-5)-methyltransferase 1 OS | DNMT1 | 183.166 | 2 | 53.531235 | yes | 9.722E-08 | 519.29254 | 185.59354 |
| NQLCDLETK | 9 | P26358 | DNA (cytosine-5)-methyltransferase 1 OS | DNMT1 | 183.166 | 2 | 53.531235 | yes | 1.084E-07 | 560.7688 | 175.3481 |
| AIILAAAPGEK | 11 | P26358 | DNA (cytosine-5)-methyltransferase 1 OS | DNMT1 | 183.166 | 2 | 53.531235 | yes | 1.908E-07 | 527.31879 | 168.04062 |
| SQACEPSEPEIEIK | 14 | P26358 | DNA (cytosine-5)-methyltransferase 1 OS | DNMT1 | 183.166 | 2 | 53.531235 | yes | 1.041E-07 | 808.87726 | 154.21631 |
| DTMSDLPEVR | 10 | P26358 | DNA (cytosine-5)-methyltransferase 1 OS | DNMT1 | 183.166 | 2 | 53.531235 | yes | 9.722E-08 | 581.77405 | 154.07635 |
| QTTITSHFAK | 10 | P26358 | DNA (cytosine-5)-methyltransferase 1 OS | DNMT1 | 183.166 | 2 | 53.531235 | yes | 1.452E-07 | 567.30109 | 134.03806 |
| LTCFSVYCK | 9 | P26358 | DNA (cytosine-5)-methyltransferase 1 OS | DNMT1 | 183.166 | 2 | 53.531235 | yes | 3.389E-07 | 589.27264 | 123.75716 |
| ACQLSVVVDDK | 11 | P26358 | DNA (cytosine-5)-methyltransferase 1 OS | DNMT1 | 183.166 | 2 | 53.531235 | yes | 9.85E-08 | 617.31079 | 102.04635 |
| HGHLCPIDTGLIEK | 14 | P26358 | DNA (cytosine-5)-methyltransferase 1 OS | DNMT1 | 183.166 | 3 | 53.531235 | yes | 1.006E-07 | 530.60834 | 100.01183 |
| DMSALVAAR | 9 | P26358 | DNA (cytosine-5)-methyltransferase 1 OS | DNMT1 | 183.166 | 2 | 53.531235 | yes | 2.601E-07 | 467.24475 | 98.905327 |
| QVGNAVPPPLAK | 12 | P26358 | DNA (cytosine-5)-methyltransferase 1 OS | DNMT1 | 183.166 | 2 | 53.531235 | yes | 1.066E-06 | 595.84821 | 91.783829 |
| EELSEEGYLAK | 11 | P26358 | DNA (cytosine-5)-methyltransferase 1 OS | DNMT1 | 183.166 | 2 | 53.531235 | yes | 4.254E-07 | 634.30627 | 91.527039 |
| SFEDPPNHAR | 10 | P26358 | DNA (cytosine-5)-methyltransferase 1 OS | DNMT1 | 183.166 | 2 | 53.531235 | yes | 1.381E-07 | 585.27032 | 82.086823 |
| GVCSCVEAGK | 10 | P26358 | DNA (cytosine-5)-methyltransferase 1 OS | DNMT1 | 183.166 | 2 | 53.531235 | yes | 5.107E-07 | 533.73645 | 77.630859 |
| VLYYSATK | 8 | P26358 | DNA (cytosine-5)-methyltransferase 1 OS | DNMT1 | 183.166 | 2 | 53.531235 | yes | 1.69E-07 | 472.75803 | 73.972183 |
| FYFLEAYNAK | 10 | P26358 | DNA (cytosine-5)-methyltransferase 1 OS | DNMT1 | 183.166 | 2 | 53.531235 | yes | 9.722E-08 | 633.31366 | 69.748703 |
| LSIFDANESGFESY | 21 | P26358 | DNA (cytosine-5)-methyltransferase 1 OS | DNMT1 | 183.166 | 3 | 53.531235 | yes | 3.844E-07 | 794.71313 | 64.593346 |
| LFGNILDK | 8 | P26358 | DNA (cytosine-5)-methyltransferase 1 OS | DNMT1 | 183.166 | 2 | 53.531235 | yes | 7.327E-07 | 460.26602 | 60.661194 |
| HNHWAGLYGR | 10 | P26358 | DNA (cytosine-5)-methyltransferase 1 OS | DNMT1 | 183.166 | 3 | 53.531235 | yes | 1.527E-07 | 404.20035 | 42.629436 |
| KVCIDAETLEVGD | 30 | P26358 | DNA (cytosine-5)-methyltransferase 1 OS | DNMT1 | 183.166 | 4 | 53.531235 | yes | 1.711E-06 | 838.17181 | 40.751835 |
| PIYDDDPSLEGGV | 16 | P26358 | DNA (cytosine-5)-methyltransferase 1 OS | DNMT1 | 183.166 | 2 | 53.531235 | yes | 1.022E-07 | 838.3941 | 40.28957 |
| KEELSEEGYLAK | 12 | P26358 | DNA (cytosine-5)-methyltransferase 1 OS | DNMT1 | 183.166 | 2 | 53.531235 | yes | 2.288E-07 | 698.35376 | 32.416477 |
| LKEEPDR | 7 | P26358 | DNA (cytosine-5)-methyltransferase 1 OS | DNMT1 | 183.166 | 2 | 53.531235 | yes | 3.709E-06 | 443.73508 | 30.177731 |
| TSFTQEQIEALEK | 13 | P26367 | Paired box protein Pax-6 OS=Homo sapi | PAX6 | 46.684 | 2 | 45.443134 | yes | 1.847E-07 | 762.38306 | 39.689548 |
| ELLTSFGPLK | 10 | P26368 | Splicing factor U2AF 65 kDa subunit OS | U2AF2 | 53.501 | 2 | 51.9563 | yes | 1.192E-07 | 552.8186 | 8934.2197 |
| LGGLTQAPGNPVL | 21 | P26368 | Splicing factor U2AF 65 kDa subunit OS | U2AF2 | 53.501 | 3 | 51.9563 | yes | 3.796E-07 | 711.72693 | 3553.4551 |
| RPHDYQPLPGMSE | 33 | P26368 | Splicing factor U2AF 65 kDa subunit OS | U2AF2 | 53.501 | 5 | 51.9563 | yes | 5.135E-07 | 712.75861 | 3204.8916 |
| IFVEFTSVFDCQK | 13 | P26368 | Splicing factor U2AF 65 kDa subunit OS | U2AF2 | 53.501 | 2 | 51.9563 | yes | 9.741E-08 | 810.39233 | 2948.1338 |
| PVDGVEVPGCGK | 12 | P26368 | Splicing factor U2AF 65 kDa subunit OS | U2AF2 | 53.501 | 2 | 51.9563 | yes | 9.722E-08 | 607.29773 | 2137.6492 |
| EEHGGLIR | 8 | P26368 | Splicing factor U2AF 65 kDa subunit OS | U2AF2 | 53.501 | 2 | 51.9563 | yes | 1.241E-07 | 455.74069 | 1657.0848 |
| NFAFLEFR | 8 | P26368 | Splicing factor U2AF 65 kDa subunit OS | U2AF2 | 53.501 | 2 | 51.9563 | yes | 2.077E-06 | 522.2691 | 1649.3815 |
| SVDETTQAMAFD | 22 | P26368 | Splicing factor U2AF 65 kDa subunit OS | U2AF2 | 53.501 | 3 | 51.9563 | yes | 9.928E-08 | 796.05804 | 692.38385 |
| YWDVPPPGFEHIT | 18 | P26368 | Splicing factor U2AF 65 kDa subunit OS | U2AF2 | 53.501 | 3 | 51.9563 | yes | 1.346E-05 | 735.68732 | 76.625748 |
| SIEIPRPVDGVEVP | 18 | P26368 | Splicing factor U2AF 65 kDa subunit OS | U2AF2 | 53.501 | 3 | 51.9563 | yes | 1.01E-06 | 636.99976 | 11.198334 |
| LATQLTGPVMPVF | 13 | P26373 | Large ribosomal subunit protein eL13 OS | RPL13 | 24.262 | 2 | 52.531616 | yes | 1.551E-07 | 691.89484 | 22629.756 |

| | | | | | | | | | | | |
|---|---|---|---|---|---|---|---|---|---|---|---|
| LATQLTGPVMPVF | 13 | P26373 | Large ribosomal subunit protein eL13 OS | RPL13 | 24.262 | 2 | 52.531616 | yes | 1E-07 | 699.89233 | 22629.756 |
| STESLQANVQR | 11 | P26373 | Large ribosomal subunit protein eL13 OS | RPL13 | 24.262 | 2 | 52.531616 | yes | 9.926E-08 | 616.81512 | 18177.666 |
| VATWFNQPAR | 10 | P26373 | Large ribosomal subunit protein eL13 OS | RPL13 | 24.262 | 2 | 52.531616 | yes | 1.163E-07 | 595.30927 | 16690.018 |
| GFSLEELR | 8 | P26373 | Large ribosomal subunit protein eL13 OS | RPL13 | 24.262 | 2 | 52.531616 | yes | 1.532E-07 | 475.75073 | 15074.337 |
| EAAEQDVEK | 9 | P26373 | Large ribosomal subunit protein eL13 OS | RPL13 | 24.262 | 2 | 52.531616 | yes | 1.427E-07 | 509.73801 | 13353.38 |
| TIGISVDPR | 9 | P26373 | Large ribosomal subunit protein eL13 OS | RPL13 | 24.262 | 2 | 52.531616 | yes | 1.785E-05 | 479.27182 | 10761.524 |
| VITEEEK | 7 | P26373 | Large ribosomal subunit protein eL13 OS | RPL13 | 24.262 | 2 | 52.531616 | yes | 1.216E-07 | 424.224 | 3042.0671 |
| PASGPIRPIVR | 11 | P26373 | Large ribosomal subunit protein eL13 OS | RPL13 | 24.262 | 2 | 52.531616 | yes | 2.343E-07 | 581.85638 | 1378.4193 |
| NGMVLKPHFHK | 11 | P26373 | Large ribosomal subunit protein eL13 OS | RPL13 | 24.262 | 2 | 52.531616 | yes | 1.012E-07 | 654.35571 | 622.90308 |
| NGMVLKPHFHK | 11 | P26373 | Large ribosomal subunit protein eL13 OS | RPL13 | 24.262 | 2 | 52.531616 | yes | 1.137E-07 | 436.57291 | 622.90308 |
| NGMVLKPHFHK | 11 | P26373 | Large ribosomal subunit protein eL13 OS | RPL13 | 24.262 | 3 | 52.531616 | yes | 1.012E-07 | 654.35571 | 622.90308 |
| NGMVLKPHFHK | 11 | P26373 | Large ribosomal subunit protein eL13 OS | RPL13 | 24.262 | 3 | 52.531616 | yes | 1.137E-07 | 436.57291 | 622.90308 |
| KGDSSAEELK | 10 | P26373 | Large ribosomal subunit protein eL13 OS | RPL13 | 24.262 | 2 | 52.531616 | yes | 1.758E-06 | 532.26691 | 284.77274 |
| AKEAAEQDVEK | 11 | P26373 | Large ribosomal subunit protein eL13 OS | RPL13 | 24.262 | 2 | 52.531616 | yes | 9.722E-08 | 609.30408 | 139.19783 |
| RVATWFNQPAR | 11 | P26373 | Large ribosomal subunit protein eL13 OS | RPL13 | 24.262 | 2 | 52.531616 | yes | 6.579E-06 | 673.3598 | 27.723423 |
| LMSNLDSNR | 9 | P26447 | Protein S100-A4 OS=Homo sapiens OX= | S100A4 | 11.729 | 2 | 51.30009 | yes | 3.162E-07 | 525.25586 | 693.01404 |
| ALDVMVSTFHK | 11 | P26447 | Protein S100-A4 OS=Homo sapiens OX= | S100A4 | 11.729 | 2 | 51.30009 | yes | 9.722E-08 | 624.32629 | 287.147 |
| RTDEAAFQK | 9 | P26447 | Protein S100-A4 OS=Homo sapiens OX= | S100A4 | 11.729 | 2 | 51.30009 | yes | 3.429E-06 | 533.26984 | 136.45108 |
| VYGAPQLQVEK | 11 | P26572 | Alpha-1,3-mannosyl-glycoprotein 2-beta- | MGAT1 | 50.879 | 2 | 46.364468 | yes | 3.857E-07 | 616.33771 | 146.54936 |
| GIVTFQFR | 8 | P26572 | Alpha-1,3-mannosyl-glycoprotein 2-beta- | MGAT1 | 50.879 | 2 | 46.364468 | yes | 1.579E-07 | 484.27164 | 45.478302 |
| LNQQFVHFTQLDL | 18 | P26572 | Alpha-1,3-mannosyl-glycoprotein 2-beta- | MGAT1 | 50.879 | 3 | 46.364468 | yes | 1.308E-05 | 750.72705 | 13.672573 |
| SEHPGLSIGDTAK | 13 | P26583 | High mobility group protein B2 OS=Hom | HMGB2 | 24.034 | 2 | 39.343613 | yes | 4.641E-07 | 656.33063 | 84.105812 |
| HQNVQLPR | 8 | P26599 | Polypyrimidine tract-binding protein 1 O | PTBP1 | 59.634 | 2 | 49.46054 | yes | 1.907E-06 | 496.27524 | 22798.184 |
| DYGNSPLHR | 9 | P26599 | Polypyrimidine tract-binding protein 1 O | PTBP1 | 59.634 | 2 | 49.46054 | yes | 1.356E-07 | 529.75433 | 13243.618 |
| GQPIYIQFSNHK | 12 | P26599 | Polypyrimidine tract-binding protein 1 O | PTBP1 | 59.634 | 2 | 49.46054 | yes | 2.39E-07 | 716.37262 | 8494.4434 |
| VLFSSNGGVVK | 11 | P26599 | Polypyrimidine tract-binding protein 1 O | PTBP1 | 59.634 | 2 | 49.46054 | yes | 1.84E-07 | 553.81384 | 6305.0518 |
| NNQFQALLQYADI | 20 | P26599 | Polypyrimidine tract-binding protein 1 O | PTBP1 | 59.634 | 3 | 49.46054 | yes | 3.019E-07 | 748.3783 | 4585.8262 |
| IAIPGLAGAGNSVI | 23 | P26599 | Polypyrimidine tract-binding protein 1 O | PTBP1 | 59.634 | 3 | 49.46054 | yes | 7.394E-07 | 759.09711 | 4388.8276 |
| KLPIDVTEGEVISL | 20 | P26599 | Polypyrimidine tract-binding protein 1 O | PTBP1 | 59.634 | 3 | 49.46054 | yes | 1.473E-07 | 704.73657 | 749.85419 |
| MDGIVPDIAVGTK | 13 | P26599 | Polypyrimidine tract-binding protein 1 O | PTBP1 | 59.634 | 2 | 49.46054 | yes | 1.028E-07 | 687.35266 | 71.512123 |
| MDGIVPDIAVGTK | 13 | P26599 | Polypyrimidine tract-binding protein 1 O | PTBP1 | 59.634 | 2 | 49.46054 | yes | 1.758E-07 | 679.35522 | 71.512123 |
| AEHDSILAEK | 10 | P26639 | Threonine--tRNA ligase 1, cytoplasmic O | TARS1 | 83.436 | 2 | 52.504055 | yes | 8.315E-06 | 556.78278 | 872.68427 |
| AELNPWPEYIYTR | 13 | P26639 | Threonine--tRNA ligase 1, cytoplasmic O | TARS1 | 83.436 | 2 | 52.504055 | yes | 2.73E-07 | 826.40936 | 125.56548 |
| FLGDIEVWDQAEK | 13 | P26639 | Threonine--tRNA ligase 1, cytoplasmic O | TARS1 | 83.436 | 2 | 52.504055 | yes | 1.073E-07 | 775.38031 | 120.45857 |
| CGPLIDLCR | 9 | P26639 | Threonine--tRNA ligase 1, cytoplasmic O | TARS1 | 83.436 | 2 | 52.504055 | yes | 1.269E-07 | 552.27045 | 113.34029 |
| MIAILTENYGGK | 12 | P26639 | Threonine--tRNA ligase 1, cytoplasmic O | TARS1 | 83.436 | 2 | 52.504055 | yes | 9.722E-08 | 655.34467 | 105.9187 |
| LADFGVLHR | 9 | P26639 | Threonine--tRNA ligase 1, cytoplasmic O | TARS1 | 83.436 | 2 | 52.504055 | yes | 3.988E-07 | 514.28778 | 102.28556 |
| IYGISFPDPK | 10 | P26639 | Threonine--tRNA ligase 1, cytoplasmic O | TARS1 | 83.436 | 2 | 52.504055 | yes | 1.11E-07 | 568.80298 | 68.674812 |
| WELNSGDGAFYGI | 14 | P26639 | Threonine--tRNA ligase 1, cytoplasmic O | TARS1 | 83.436 | 2 | 52.504055 | yes | 7.926E-07 | 770.85699 | 49.845181 |

| | | | | | | | | | | | |
|---|---|---|---|---|---|---|---|---|---|---|---|
| GCLDFLR | 7 | P26639 | Threonine--tRNA ligase 1, cytoplasmic O | TARS1 | 83.436 | 2 | 52.504055 | yes | 6.198E-07 | 440.72092 | 31.092024 |
| FNLTYVSHDGDDI | 13 | P26639 | Threonine--tRNA ligase 1, cytoplasmic O | TARS1 | 83.436 | 2 | 52.504055 | yes | 3.299E-07 | 755.84406 | 27.87455 |
| GIEDNPMVVPLCN | 14 | P26640 | Valine--tRNA ligase OS=Homo sapiens ( | VARS1 | 140.477 | 2 | 51.694195 | yes | 5.703E-07 | 807.39233 | 2673.7444 |
| SSAQDPQAVLGAL | 15 | P26640 | Valine--tRNA ligase OS=Homo sapiens ( | VARS1 | 140.477 | 2 | 51.694195 | yes | 1.11E-07 | 735.38898 | 2263.2629 |
| EVYLHAIVR | 9 | P26640 | Valine--tRNA ligase OS=Homo sapiens ( | VARS1 | 140.477 | 2 | 51.694195 | yes | 1.014E-07 | 550.31659 | 2174.1968 |
| LHEEGIIYR | 9 | P26640 | Valine--tRNA ligase OS=Homo sapiens ( | VARS1 | 140.477 | 2 | 51.694195 | yes | 1.02E-07 | 565.30365 | 2078.9851 |
| LSAAVTEAFVR | 11 | P26640 | Valine--tRNA ligase OS=Homo sapiens ( | VARS1 | 140.477 | 2 | 51.694195 | yes | 1.061E-07 | 582.32458 | 1959.4121 |
| ADFPAGIPECGTD | 16 | P26640 | Valine--tRNA ligase OS=Homo sapiens ( | VARS1 | 140.477 | 2 | 51.694195 | yes | 2.43E-07 | 845.39868 | 1943.1233 |
| VPLEVQEADEAK | 12 | P26640 | Valine--tRNA ligase OS=Homo sapiens ( | VARS1 | 140.477 | 2 | 51.694195 | yes | 9.722E-08 | 664.34064 | 1564.1133 |
| ILPEAHQR | 8 | P26640 | Valine--tRNA ligase OS=Homo sapiens ( | VARS1 | 140.477 | 2 | 51.694195 | yes | 1.022E-07 | 482.27216 | 1541.137 |
| LQQTEAELR | 9 | P26640 | Valine--tRNA ligase OS=Homo sapiens ( | VARS1 | 140.477 | 2 | 51.694195 | yes | 2.092E-07 | 544.29077 | 1417.9832 |
| ICLQPPPTSR | 10 | P26640 | Valine--tRNA ligase OS=Homo sapiens ( | VARS1 | 140.477 | 2 | 51.694195 | yes | 1.365E-07 | 584.81079 | 1352.0353 |
| CGEMAQAASAAV | 14 | P26640 | Valine--tRNA ligase OS=Homo sapiens ( | VARS1 | 140.477 | 2 | 51.694195 | yes | 1.022E-07 | 711.82684 | 1290.106 |
| AVLGEVVLYSGAI | 13 | P26640 | Valine--tRNA ligase OS=Homo sapiens ( | VARS1 | 140.477 | 2 | 51.694195 | yes | 9.723E-08 | 667.37738 | 1257.6824 |
| YVLDPPAR | 8 | P26640 | Valine--tRNA ligase OS=Homo sapiens ( | VARS1 | 140.477 | 2 | 51.694195 | yes | 2.414E-07 | 465.7558 | 1238.9774 |
| ALSPLEEWLR | 10 | P26640 | Valine--tRNA ligase OS=Homo sapiens ( | VARS1 | 140.477 | 2 | 51.694195 | yes | 1.25E-07 | 607.3324 | 1153.2665 |
| DVSGPMPDSYSPR | 13 | P26640 | Valine--tRNA ligase OS=Homo sapiens ( | VARS1 | 140.477 | 2 | 51.694195 | yes | 1.21E-07 | 704.31409 | 1089.6744 |
| EAFLQEVWK | 9 | P26640 | Valine--tRNA ligase OS=Homo sapiens ( | VARS1 | 140.477 | 2 | 51.694195 | yes | 1.165E-07 | 575.3006 | 961.42657 |
| QTLYTCLDVGLR | 12 | P26640 | Valine--tRNA ligase OS=Homo sapiens ( | VARS1 | 140.477 | 2 | 51.694195 | yes | 1.256E-07 | 719.87158 | 913.55591 |
| PLSHQPGPEAPALI | 15 | P26640 | Valine--tRNA ligase OS=Homo sapiens ( | VARS1 | 140.477 | 3 | 51.694195 | yes | 2.448E-07 | 513.61572 | 435.87225 |
| PLSHQPGPEAPALI | 15 | P26640 | Valine--tRNA ligase OS=Homo sapiens ( | VARS1 | 140.477 | 3 | 51.694195 | yes | 1.421E-07 | 769.91992 | 435.87225 |
| PLSHQPGPEAPALI | 15 | P26640 | Valine--tRNA ligase OS=Homo sapiens ( | VARS1 | 140.477 | 2 | 51.694195 | yes | 2.448E-07 | 513.61572 | 435.87225 |
| PLSHQPGPEAPALI | 15 | P26640 | Valine--tRNA ligase OS=Homo sapiens ( | VARS1 | 140.477 | 2 | 51.694195 | yes | 1.421E-07 | 769.91992 | 435.87225 |
| WFVTCVR | 7 | P26640 | Valine--tRNA ligase OS=Homo sapiens ( | VARS1 | 140.477 | 2 | 51.694195 | yes | 2.717E-06 | 484.24457 | 870.51593 |
| VQGSDSDEEVVV/ | 16 | P26640 | Valine--tRNA ligase OS=Homo sapiens ( | VARS1 | 140.477 | 2 | 51.694195 | yes | 9.722E-08 | 846.40778 | 821.20184 |
| HGLEAISIMDSR | 12 | P26640 | Valine--tRNA ligase OS=Homo sapiens ( | VARS1 | 140.477 | 2 | 51.694195 | yes | 1.341E-07 | 664.83502 | 748.8653 |
| IQQQQPPPGEK | 11 | P26640 | Valine--tRNA ligase OS=Homo sapiens ( | VARS1 | 140.477 | 2 | 51.694195 | yes | 2.832E-07 | 625.33038 | 658.64758 |
| TWHAWMDNIR | 10 | P26640 | Valine--tRNA ligase OS=Homo sapiens ( | VARS1 | 140.477 | 2 | 51.694195 | yes | 1.073E-07 | 665.31146 | 638.47681 |
| PNVSAANPR | 9 | P26640 | Valine--tRNA ligase OS=Homo sapiens ( | VARS1 | 140.477 | 2 | 51.694195 | yes | 1.306E-07 | 463.24615 | 627.60895 |
| LGSSLDWDR | 9 | P26640 | Valine--tRNA ligase OS=Homo sapiens ( | VARS1 | 140.477 | 2 | 51.694195 | yes | 2.049E-07 | 524.75653 | 590.53174 |
| IWNNVTR | 7 | P26640 | Valine--tRNA ligase OS=Homo sapiens ( | VARS1 | 140.477 | 2 | 51.694195 | yes | 1.025E-07 | 451.74579 | 568.35986 |
| KVDEAIALFQK | 11 | P26640 | Valine--tRNA ligase OS=Homo sapiens ( | VARS1 | 140.477 | 2 | 51.694195 | yes | 1.252E-07 | 631.36115 | 566.77747 |
| NVIHPFLSR | 9 | P26640 | Valine--tRNA ligase OS=Homo sapiens ( | VARS1 | 140.477 | 2 | 51.694195 | yes | 2.974E-07 | 541.8089 | 493.47467 |
| TPFPPPR | 7 | P26640 | Valine--tRNA ligase OS=Homo sapiens ( | VARS1 | 140.477 | 2 | 51.694195 | yes | 1.276E-07 | 406.22668 | 388.09283 |
| DINLDVNR | 8 | P26640 | Valine--tRNA ligase OS=Homo sapiens ( | VARS1 | 140.477 | 2 | 51.694195 | yes | 1.611E-07 | 479.75125 | 321.53494 |
| KAVLVALK | 8 | P26640 | Valine--tRNA ligase OS=Homo sapiens ( | VARS1 | 140.477 | 2 | 51.694195 | yes | 1.082E-07 | 421.29712 | 281.45059 |
| KLGSSLDWDR | 10 | P26640 | Valine--tRNA ligase OS=Homo sapiens ( | VARS1 | 140.477 | 2 | 51.694195 | yes | 1.226E-07 | 588.80402 | 264.76926 |
| SKDVVEPLLR | 10 | P26640 | Valine--tRNA ligase OS=Homo sapiens ( | VARS1 | 140.477 | 2 | 51.694195 | yes | 9.728E-08 | 578.34021 | 262.02219 |
| IETMLGDVAVAVI | 15 | P26640 | Valine--tRNA ligase OS=Homo sapiens ( | VARS1 | 140.477 | 3 | 51.694195 | yes | 2.165E-07 | 527.28857 | 257.25323 |

| | | | | | | | | | | | |
|---|---|---|---|---|---|---|---|---|---|---|---|
| TLLSVPGYK | 9 | P26640 | Valine--tRNA ligase OS=Homo sapiens ( | VARS1 | 140.477 | 2 | 51.694195 | yes | 5.159E-07 | 489.28696 | 161.56607 |
| ADYNLTR | 7 | P26640 | Valine--tRNA ligase OS=Homo sapiens ( | VARS1 | 140.477 | 2 | 51.694195 | yes | 9.809E-08 | 426.71414 | 139.74657 |
| KDVSGPMPDSYSF | 14 | P26640 | Valine--tRNA ligase OS=Homo sapiens ( | VARS1 | 140.477 | 2 | 51.694195 | yes | 5.253E-07 | 768.36157 | 57.651108 |
| KDVSGPMPDSYSF | 14 | P26640 | Valine--tRNA ligase OS=Homo sapiens ( | VARS1 | 140.477 | 2 | 51.694195 | yes | 9.722E-08 | 512.57678 | 57.651108 |
| KDVSGPMPDSYSF | 14 | P26640 | Valine--tRNA ligase OS=Homo sapiens ( | VARS1 | 140.477 | 3 | 51.694195 | yes | 5.253E-07 | 768.36157 | 57.651108 |
| KDVSGPMPDSYSF | 14 | P26640 | Valine--tRNA ligase OS=Homo sapiens ( | VARS1 | 140.477 | 3 | 51.694195 | yes | 9.722E-08 | 512.57678 | 57.651108 |
| SKDVVEPLLRPQW | 16 | P26640 | Valine--tRNA ligase OS=Homo sapiens ( | VARS1 | 140.477 | 3 | 51.694195 | yes | 1.047E-07 | 662.37042 | 113.53909 |
| STLYVSPHPDAFPS | 16 | P26640 | Valine--tRNA ligase OS=Homo sapiens ( | VARS1 | 140.477 | 3 | 51.694195 | yes | 4.65E-07 | 610.31244 | 105.66609 |
| YGEAGEGPGWGG | 16 | P26640 | Valine--tRNA ligase OS=Homo sapiens ( | VARS1 | 140.477 | 2 | 51.694195 | yes | 1.398E-07 | 799.36078 | 100.23859 |
| QLWWGHR | 7 | P26640 | Valine--tRNA ligase OS=Homo sapiens ( | VARS1 | 140.477 | 2 | 51.694195 | yes | 1.44E-07 | 491.75394 | 80.945206 |
| LVNWSCTLNSAISI | 20 | P26640 | Valine--tRNA ligase OS=Homo sapiens ( | VARS1 | 140.477 | 3 | 51.694195 | yes | 1.335E-07 | 764.72284 | 61.690571 |
| VEFGVLVSFAYK | 12 | P26640 | Valine--tRNA ligase OS=Homo sapiens ( | VARS1 | 140.477 | 2 | 51.694195 | yes | 2.685E-07 | 679.8714 | 46.725819 |
| FGLCAYMSQGR | 11 | P26640 | Valine--tRNA ligase OS=Homo sapiens ( | VARS1 | 140.477 | 2 | 51.694195 | yes | 3.305E-07 | 645.29193 | 34.096573 |
| DVVEPLLR | 8 | P26640 | Valine--tRNA ligase OS=Homo sapiens ( | VARS1 | 140.477 | 2 | 51.694195 | yes | 7.022E-07 | 470.77673 | 23.888321 |
| QVLEPSFR | 8 | P26641 | Elongation factor 1-gamma OS=Homo sa | EEF1G | 50.119 | 2 | 51.568462 | yes | 1.692E-07 | 488.26654 | 18951.508 |
| STFVLDEFKR | 10 | P26641 | Elongation factor 1-gamma OS=Homo sa | EEF1G | 50.119 | 2 | 51.568462 | yes | 9.722E-08 | 621.3299 | 16873.342 |
| ALIAAQYSGAQVR | 13 | P26641 | Elongation factor 1-gamma OS=Homo sa | EEF1G | 50.119 | 2 | 51.568462 | yes | 1.392E-07 | 674.37262 | 15787.18 |
| AAGTLYTYPENWI | 13 | P26641 | Elongation factor 1-gamma OS=Homo sa | EEF1G | 50.119 | 2 | 51.568462 | yes | 1.002E-07 | 792.37805 | 6928.5781 |
| AAGTLYTYPENWI | 13 | P26641 | Elongation factor 1-gamma OS=Homo sa | EEF1G | 50.119 | 2 | 51.568462 | yes | 2.299E-07 | 771.3728 | 6928.5781 |
| WFLTCINQPQFR | 12 | P26641 | Elongation factor 1-gamma OS=Homo sa | EEF1G | 50.119 | 2 | 51.568462 | yes | 3.076E-07 | 805.40082 | 12879.865 |
| EYFSWEGAFQHV( | 14 | P26641 | Elongation factor 1-gamma OS=Homo sa | EEF1G | 50.119 | 2 | 51.568462 | yes | 1.777E-07 | 842.89136 | 5084.8691 |
| EYFSWEGAFQHV( | 14 | P26641 | Elongation factor 1-gamma OS=Homo sa | EEF1G | 50.119 | 2 | 51.568462 | yes | 4.145E-07 | 562.26331 | 5084.8691 |
| EYFSWEGAFQHV( | 14 | P26641 | Elongation factor 1-gamma OS=Homo sa | EEF1G | 50.119 | 3 | 51.568462 | yes | 1.777E-07 | 842.89136 | 5084.8691 |
| EYFSWEGAFQHV( | 14 | P26641 | Elongation factor 1-gamma OS=Homo sa | EEF1G | 50.119 | 3 | 51.568462 | yes | 4.145E-07 | 562.26331 | 5084.8691 |
| AKDPFAHLPK | 10 | P26641 | Elongation factor 1-gamma OS=Homo sa | EEF1G | 50.119 | 2 | 51.568462 | yes | 9.722E-08 | 562.31659 | 7928.6343 |
| LDPGSEETQTLVR | 13 | P26641 | Elongation factor 1-gamma OS=Homo sa | EEF1G | 50.119 | 2 | 51.568462 | yes | 1.543E-07 | 722.86755 | 6181.9731 |
| KLDPGSEETQTLV | 14 | P26641 | Elongation factor 1-gamma OS=Homo sa | EEF1G | 50.119 | 2 | 51.568462 | yes | 1.163E-07 | 786.91504 | 2809.9678 |
| KLDPGSEETQTLV | 14 | P26641 | Elongation factor 1-gamma OS=Homo sa | EEF1G | 50.119 | 2 | 51.568462 | yes | 1.183E-07 | 524.9458 | 2809.9678 |
| KLDPGSEETQTLV | 14 | P26641 | Elongation factor 1-gamma OS=Homo sa | EEF1G | 50.119 | 3 | 51.568462 | yes | 1.163E-07 | 786.91504 | 2809.9678 |
| KLDPGSEETQTLV | 14 | P26641 | Elongation factor 1-gamma OS=Homo sa | EEF1G | 50.119 | 3 | 51.568462 | yes | 1.183E-07 | 524.9458 | 2809.9678 |
| KFAETQPK | 8 | P26641 | Elongation factor 1-gamma OS=Homo sa | EEF1G | 50.119 | 2 | 51.568462 | yes | 2.064E-07 | 474.76108 | 4645.4429 |
| STFVLDEFK | 9 | P26641 | Elongation factor 1-gamma OS=Homo sa | EEF1G | 50.119 | 2 | 51.568462 | yes | 1.849E-07 | 543.2793 | 4639.1274 |
| FAETQPK | 7 | P26641 | Elongation factor 1-gamma OS=Homo sa | EEF1G | 50.119 | 2 | 51.568462 | yes | 1.562E-07 | 410.71362 | 2570.8098 |
| KAAAPAPEEEMDI | 23 | P26641 | Elongation factor 1-gamma OS=Homo sa | EEF1G | 50.119 | 3 | 51.568462 | yes | 4.241E-07 | 829.04553 | 2459.6775 |
| DGWSLWYSEYR | 11 | P26641 | Elongation factor 1-gamma OS=Homo sa | EEF1G | 50.119 | 2 | 51.568462 | yes | 1.331E-07 | 731.32532 | 2113.2649 |
| ILGLLDAYLK | 10 | P26641 | Elongation factor 1-gamma OS=Homo sa | EEF1G | 50.119 | 2 | 51.568462 | yes | 1.014E-06 | 559.8446 | 2032.4503 |
| DPFAHLPK | 8 | P26641 | Elongation factor 1-gamma OS=Homo sa | EEF1G | 50.119 | 2 | 51.568462 | yes | 2.449E-07 | 462.75052 | 1535.6826 |
| VLSAPPHFHFGQT | 15 | P26641 | Elongation factor 1-gamma OS=Homo sa | EEF1G | 50.119 | 4 | 51.568462 | yes | 2.942E-07 | 427.7233 | 932.22546 |
| RILGLLDAYLK | 11 | P26641 | Elongation factor 1-gamma OS=Homo sa | EEF1G | 50.119 | 2 | 51.568462 | yes | 1.488E-07 | 637.8952 | 188.77036 |

| | | | | | | | | | | | |
|---|---|---|---|---|---|---|---|---|---|---|---|
| QAFPNTNR | 8 | P26641 | Elongation factor 1-gamma OS=Homo sa | EEF1G | 50.119 | 2 | 51.568462 | yes | 9.722E-08 | 474.23831 | 151.90182 |
| TFLVGER | 7 | P26641 | Elongation factor 1-gamma OS=Homo sa | EEF1G | 50.119 | 2 | 51.568462 | yes | 1.145E-07 | 411.22943 | 71.814133 |
| KYSNEDTLSVALP | 21 | P26641 | Elongation factor 1-gamma OS=Homo sa | EEF1G | 50.119 | 4 | 51.568462 | yes | 3.807E-05 | 648.06287 | 6.0359859 |
| LLAQTTLR | 8 | P27105 | Stomatin OS=Homo sapiens OX=9606 G | STOM | 31.731 | 2 | 51.629128 | yes | 9.722E-08 | 458.28473 | 92.151756 |
| NLSQILSDR | 9 | P27105 | Stomatin OS=Homo sapiens OX=9606 G | STOM | 31.731 | 2 | 51.629128 | yes | 1.818E-06 | 523.28546 | 84.302841 |
| YLQTLTTIAAEK | 12 | P27105 | Stomatin OS=Homo sapiens OX=9606 G | STOM | 31.731 | 2 | 51.629128 | yes | 2.676E-07 | 676.37701 | 35.942238 |
| YVYIAELLAHK | 11 | P27169 | Serum paraoxonase/arylesterase 1 OS=Hc | PON1 | 39.732 | 2 | 46.44927 | yes | 1.556E-07 | 660.37152 | 226.63841 |
| YLIANATNPESK | 12 | P27348 | 14-3-3 protein theta OS=Homo sapiens O | YWHAQ | 27.765 | 2 | 50.430782 | yes | 9.809E-08 | 660.84332 | 2477.2627 |
| EKVESELR | 8 | P27348 | 14-3-3 protein theta OS=Homo sapiens O | YWHAQ | 27.765 | 2 | 50.430782 | yes | 1.72E-07 | 495.26672 | 1442.6244 |
| YDDMATCMK | 9 | P27348 | 14-3-3 protein theta OS=Homo sapiens O | YWHAQ | 27.765 | 2 | 50.430782 | yes | 9.819E-08 | 567.71686 | 1049.2494 |
| YLAEVACGDDR | 11 | P27348 | 14-3-3 protein theta OS=Homo sapiens O | YWHAQ | 27.765 | 2 | 50.430782 | yes | 1.061E-07 | 634.78241 | 333.11053 |
| YLAEVACGDDRK | 12 | P27348 | 14-3-3 protein theta OS=Homo sapiens O | YWHAQ | 27.765 | 2 | 50.430782 | yes | 1.173E-07 | 698.8299 | 294.96915 |
| LFEVIETEK | 9 | P27448 | MAP/microtubule affinity-regulating kina | MARK3 | 84.429 | 2 | 51.9562 | no | 1.425E-07 | 554.30023 | 1901.4154 |
| TPLPTVNER | 9 | P27448 | MAP/microtubule affinity-regulating kina | MARK3 | 84.429 | 2 | 51.9562 | no | 2.093E-07 | 513.78253 | 1207.1619 |
| TQLNPTSLQK | 10 | P27448 | MAP/microtubule affinity-regulating kina | MARK3 | 84.429 | 2 | 51.9562 | no | 9.722E-08 | 565.31421 | 1008.0387 |
| QIVSAVQYCHQK | 12 | P27448 | MAP/microtubule affinity-regulating kina | MARK3 | 84.429 | 2 | 51.9562 | no | 9.722E-08 | 730.86938 | 748.17883 |
| AENLLLDADMNIK | 13 | P27448 | MAP/microtubule affinity-regulating kina | MARK3 | 84.429 | 2 | 51.9562 | no | 1.166E-07 | 730.37671 | 658.66553 |
| ILNHPNIVK | 9 | P27448 | MAP/microtubule affinity-regulating kina | MARK3 | 84.429 | 2 | 51.9562 | no | 2.388E-07 | 524.31909 | 496.80811 |
| SSTVPSSNTASGGI | 16 | P27448 | MAP/microtubule affinity-regulating kina | MARK3 | 84.429 | 2 | 51.9562 | yes | 1.661E-07 | 770.35699 | 386.69943 |
| GIAPASPMLGNAS | 17 | P27448 | MAP/microtubule affinity-regulating kina | MARK3 | 84.429 | 2 | 51.9562 | yes | 1.008E-07 | 819.91705 | 306.36984 |
| IADFGFSNEFTVGC | 15 | P27448 | MAP/microtubule affinity-regulating kina | MARK3 | 84.429 | 2 | 51.9562 | yes | 1.071E-07 | 794.88574 | 291.31274 |
| FLVLNPIK | 8 | P27448 | MAP/microtubule affinity-regulating kina | MARK3 | 84.429 | 2 | 51.9562 | yes | 2.111E-07 | 472.3024 | 194.29803 |
| ENSTIPDQR | 9 | P27448 | MAP/microtubule affinity-regulating kina | MARK3 | 84.429 | 2 | 51.9562 | yes | 3.762E-06 | 530.2569 | 144.19092 |
| VLDANNCDYEQR | 12 | P27448 | MAP/microtubule affinity-regulating kina | MARK3 | 84.429 | 2 | 51.9562 | yes | 9.722E-08 | 748.82538 | 120.34339 |
| YDEITATYLLLGR | 13 | P27448 | MAP/microtubule affinity-regulating kina | MARK3 | 84.429 | 2 | 51.9562 | yes | 9.722E-08 | 764.40631 | 112.95345 |
| SQTSTADSDLK | 11 | P27448 | MAP/microtubule affinity-regulating kina | MARK3 | 84.429 | 2 | 51.9562 | yes | 1.973E-07 | 576.77258 | 59.389248 |
| RGTLEQIMK | 9 | P27448 | MAP/microtubule affinity-regulating kina | MARK3 | 84.429 | 2 | 51.9562 | no | 1.901E-06 | 538.30005 | 56.835911 |
| TTSSMDPGDMMR | 12 | P27448 | MAP/microtubule affinity-regulating kina | MARK3 | 84.429 | 2 | 51.9562 | yes | 3.123E-06 | 664.76758 | 46.370186 |
| YSDHAGPAIPSVV. | 17 | P27448 | MAP/microtubule affinity-regulating kina | MARK3 | 84.429 | 3 | 51.9562 | yes | 4.306E-07 | 591.3053 | 43.547489 |
| WINAGHEEDELKP | 26 | P27448 | MAP/microtubule affinity-regulating kina | MARK3 | 84.429 | 4 | 51.9562 | yes | 2.212E-05 | 774.38336 | 33.915211 |
| WINAGHEEDELKP | 25 | P27448 | MAP/microtubule affinity-regulating kina | MARK3 | 84.429 | 4 | 51.9562 | yes | 2.522E-06 | 735.35809 | 21.270279 |
| SGFSAQQVATQAT | 15 | P27540 | Aryl hydrocarbon receptor nuclear translo | ARNT | 86.637 | 2 | 51.778557 | yes | 9.722E-08 | 747.88098 | 132.04347 |
| CGSSSVDPVSVNR | 13 | P27540 | Aryl hydrocarbon receptor nuclear translo | ARNT | 86.637 | 2 | 51.778557 | yes | 1E-07 | 682.31714 | 86.15815 |
| QQQQQTELDMVPC | 14 | P27540 | Aryl hydrocarbon receptor nuclear translo | ARNT | 86.637 | 2 | 51.778557 | yes | 1.73E-07 | 829.40173 | 70.886711 |
| FCLVAIGR | 8 | P27540 | Aryl hydrocarbon receptor nuclear translo | ARNT | 86.637 | 2 | 51.778557 | yes | 1.05E-07 | 468.26019 | 66.851639 |
| EQLSTSENALTGR | 13 | P27540 | Aryl hydrocarbon receptor nuclear translo | ARNT | 86.637 | 2 | 51.778557 | yes | 1.609E-07 | 703.34955 | 63.996471 |
| CVATVGYQPQELI | 15 | P27540 | Aryl hydrocarbon receptor nuclear translo | ARNT | 86.637 | 2 | 51.778557 | yes | 1.087E-07 | 831.92963 | 54.92498 |
| DGLASYNHSQVVC | 28 | P27540 | Aryl hydrocarbon receptor nuclear translo | ARNT | 86.637 | 4 | 51.778557 | yes | 4.392E-07 | 755.63342 | 34.707317 |
| EHVIEALR | 8 | P27635 | Large ribosomal subunit protein uL16 OS | RPL10 | 24.577 | 2 | 51.299465 | yes | 1.559E-07 | 483.772 | 36766.406 |

| | | | | | | | | | | | |
|---|---|---|---|---|---|---|---|---|---|---|---|
| VHIGQVIMSIR | 11 | P27635 | Large ribosomal subunit protein uL16 OS | RPL10 | 24.577 | 2 | 51.299465 | no | 1.048E-07 | 626.86334 | 10324.849 |
| VHIGQVIMSIR | 11 | P27635 | Large ribosomal subunit protein uL16 OS | RPL10 | 24.577 | 2 | 51.299465 | no | 9.722E-08 | 634.86084 | 10324.849 |
| LIPDGCGVK | 9 | P27635 | Large ribosomal subunit protein uL16 OS | RPL10 | 24.577 | 2 | 51.299465 | yes | 3.406E-07 | 479.75494 | 20174.484 |
| MLSCAGADR | 9 | P27635 | Large ribosomal subunit protein uL16 OS | RPL10 | 24.577 | 2 | 51.299465 | no | 1.376E-07 | 490.71805 | 10000.039 |
| MLSCAGADR | 9 | P27635 | Large ribosomal subunit protein uL16 OS | RPL10 | 24.577 | 2 | 51.299465 | no | 6.167E-07 | 498.71552 | 10000.039 |
| GAFGKPQGTVAR | 12 | P27635 | Large ribosomal subunit protein uL16 OS | RPL10 | 24.577 | 2 | 51.299465 | no | 1.85E-07 | 594.82782 | 15866.968 |
| LHPFHVIR | 8 | P27635 | Large ribosomal subunit protein uL16 OS | RPL10 | 24.577 | 2 | 51.299465 | no | 2.015E-07 | 509.80087 | 10971.003 |
| FNADEFEDMVAEI | 13 | P27635 | Large ribosomal subunit protein uL16 OS | RPL10 | 24.577 | 2 | 51.299465 | yes | 3.403E-07 | 780.82977 | 496.849 |
| FNADEFEDMVAEI | 13 | P27635 | Large ribosomal subunit protein uL16 OS | RPL10 | 24.577 | 2 | 51.299465 | yes | 9.722E-08 | 772.83234 | 496.849 |
| RLIPDGCGVK | 10 | P27635 | Large ribosomal subunit protein uL16 OS | RPL10 | 24.577 | 2 | 51.299465 | yes | 1.674E-07 | 557.80548 | 179.91833 |
| LFSLELVDESGEIR | 14 | P27694 | Replication protein A 70 kDa DNA-bindi | RPA1 | 68.139 | 2 | 53.58033 | yes | 1.36E-07 | 803.9198 | 445.50388 |
| FFPLIEVNK | 9 | P27694 | Replication protein A 70 kDa DNA-bindi | RPA1 | 68.139 | 2 | 53.58033 | yes | 1.495E-07 | 553.81586 | 285.23062 |
| DSLVDIIGICK | 11 | P27694 | Replication protein A 70 kDa DNA-bindi | RPA1 | 68.139 | 2 | 53.58033 | yes | 1.012E-07 | 616.83136 | 182.94543 |
| NIYLMDTSGK | 10 | P27694 | Replication protein A 70 kDa DNA-bindi | RPA1 | 68.139 | 2 | 53.58033 | yes | 9.722E-08 | 571.28156 | 179.02081 |
| VGQLSEGAIAAIM | 15 | P27694 | Replication protein A 70 kDa DNA-bindi | RPA1 | 68.139 | 2 | 53.58033 | yes | 9.722E-08 | 758.41364 | 166.22426 |
| VVTATLWGEDAD | 13 | P27694 | Replication protein A 70 kDa DNA-bindi | RPA1 | 68.139 | 2 | 53.58033 | yes | 9.722E-08 | 702.85388 | 150.97934 |
| GDTNIKPILQVINI | 15 | P27694 | Replication protein A 70 kDa DNA-bindi | RPA1 | 68.139 | 2 | 53.58033 | yes | 1.778E-07 | 847.50159 | 72.468803 |
| GDTNIKPILQVINI | 15 | P27694 | Replication protein A 70 kDa DNA-bindi | RPA1 | 68.139 | 2 | 53.58033 | yes | 1.743E-07 | 565.33685 | 72.468803 |
| GDTNIKPILQVINI | 15 | P27694 | Replication protein A 70 kDa DNA-bindi | RPA1 | 68.139 | 3 | 53.58033 | yes | 1.778E-07 | 847.50159 | 72.468803 |
| GDTNIKPILQVINI | 15 | P27694 | Replication protein A 70 kDa DNA-bindi | RPA1 | 68.139 | 3 | 53.58033 | yes | 1.743E-07 | 565.33685 | 72.468803 |
| VKVETYNDESR | 11 | P27694 | Replication protein A 70 kDa DNA-bindi | RPA1 | 68.139 | 2 | 53.58033 | yes | 9.722E-08 | 670.32806 | 95.931419 |
| VVPIASLTPYQSK | 13 | P27694 | Replication protein A 70 kDa DNA-bindi | RPA1 | 68.139 | 2 | 53.58033 | yes | 1.374E-07 | 701.90063 | 77.396317 |
| FIVNTLK | 7 | P27694 | Replication protein A 70 kDa DNA-bindi | RPA1 | 68.139 | 2 | 53.58033 | yes | 9.819E-08 | 417.75781 | 71.542442 |
| VETYNDESR | 9 | P27694 | Replication protein A 70 kDa DNA-bindi | RPA1 | 68.139 | 2 | 53.58033 | yes | 4.375E-07 | 556.74634 | 64.259926 |
| PILQVINIR | 9 | P27694 | Replication protein A 70 kDa DNA-bindi | RPA1 | 68.139 | 2 | 53.58033 | yes | 3.142E-07 | 533.34259 | 37.476822 |
| ATVMDVKPVDYR | 12 | P27694 | Replication protein A 70 kDa DNA-bindi | RPA1 | 68.139 | 2 | 53.58033 | yes | 0.0001181 | 697.36084 | 36.862541 |
| CEKCDTEFPNFK | 12 | P27694 | Replication protein A 70 kDa DNA-bindi | RPA1 | 68.139 | 2 | 53.58033 | yes | 6.204E-07 | 787.83435 | 30.16143 |
| MVGQLSEGAIAAI | 16 | P27694 | Replication protein A 70 kDa DNA-bindi | RPA1 | 68.139 | 2 | 53.58033 | yes | 1.3E-07 | 823.93384 | 29.085949 |
| VIDQQNGLYR | 10 | P27694 | Replication protein A 70 kDa DNA-bindi | RPA1 | 68.139 | 2 | 53.58033 | yes | 1.981E-07 | 603.31732 | 23.958933 |
| EEAPDILCLQETK | 13 | P27695 | DNA-(apurinic or apyrimidinic site) endo | APEX1 | 35.555 | 2 | 46.928795 | yes | 2.618E-07 | 773.37689 | 148.22559 |
| ICSWNVDGLR | 10 | P27695 | DNA-(apurinic or apyrimidinic site) endo | APEX1 | 35.555 | 2 | 46.928795 | yes | 1.439E-07 | 610.29803 | 88.094528 |
| LLDTIGISQPQWR | 13 | P27708 | CAD protein OS=Homo sapiens OX=960 | CAD | 242.984 | 2 | 56.685856 | yes | 1.877E-07 | 763.91992 | 280.4823 |
| MVVMHPMPR | 9 | P27708 | CAD protein OS=Homo sapiens OX=960 | CAD | 242.984 | 2 | 56.685856 | yes | 1.296E-07 | 549.27448 | 249.10516 |
| LYLNETFSELR | 11 | P27708 | CAD protein OS=Homo sapiens OX=960 | CAD | 242.984 | 2 | 56.685856 | yes | 9.722E-08 | 692.85901 | 246.37482 |
| ILALDCGLK | 9 | P27708 | CAD protein OS=Homo sapiens OX=960 | CAD | 242.984 | 2 | 56.685856 | yes | 1.166E-07 | 501.78625 | 218.48886 |
| LGGAVLSFSEATS | 17 | P27708 | CAD protein OS=Homo sapiens OX=960 | CAD | 242.984 | 2 | 56.685856 | yes | 1.267E-07 | 840.94379 | 212.05843 |
| VLGTSPEAIDSAEN | 15 | P27708 | CAD protein OS=Homo sapiens OX=960 | CAD | 242.984 | 2 | 56.685856 | yes | 9.722E-08 | 779.88904 | 157.32019 |
| VMASMFYEVSTR | 12 | P27708 | CAD protein OS=Homo sapiens OX=960 | CAD | 242.984 | 2 | 56.685856 | yes | 9.728E-08 | 710.83362 | 146.11241 |
| ELSDLESAR | 9 | P27708 | CAD protein OS=Homo sapiens OX=960 | CAD | 242.984 | 2 | 56.685856 | yes | 1.088E-07 | 510.25381 | 142.61012 |

| | | | | | | | | | | | |
|---|---|---|---|---|---|---|---|---|---|---|---|
| MALLATVLGR | 10 | P27708 | CAD protein OS=Homo sapiens OX=960 | CAD | 242.984 | 2 | 56.685856 | yes | 3.591E-07 | 522.81537 | 129.67955 |
| PLVPEVSIK | 9 | P27708 | CAD protein OS=Homo sapiens OX=960 | CAD | 242.984 | 2 | 56.685856 | yes | 2.76E-07 | 491.30258 | 119.50008 |
| LSLDDLLQR | 9 | P27708 | CAD protein OS=Homo sapiens OX=960 | CAD | 242.984 | 2 | 56.685856 | yes | 2.231E-07 | 536.80347 | 118.18063 |
| VVLRGEVAYIDGQ | 25 | P27708 | CAD protein OS=Homo sapiens OX=960 | CAD | 242.984 | 4 | 56.685856 | yes | 4.493E-07 | 675.61835 | 117.65602 |
| QELGICPAVK | 10 | P27708 | CAD protein OS=Homo sapiens OX=960 | CAD | 242.984 | 2 | 56.685856 | yes | 1.87E-07 | 557.79987 | 112.17087 |
| VHVDCMTSQK | 10 | P27708 | CAD protein OS=Homo sapiens OX=960 | CAD | 242.984 | 2 | 56.685856 | yes | 9.722E-08 | 602.77606 | 111.38393 |
| QFCQTVGYPCVVI | 13 | P27708 | CAD protein OS=Homo sapiens OX=960 | CAD | 242.984 | 2 | 56.685856 | yes | 9.722E-08 | 807.38177 | 110.3829 |
| SILEQLAEK | 9 | P27708 | CAD protein OS=Homo sapiens OX=960 | CAD | 242.984 | 2 | 56.685856 | yes | 2.577E-07 | 515.7926 | 100.14729 |
| LALGIPLPELR | 11 | P27708 | CAD protein OS=Homo sapiens OX=960 | CAD | 242.984 | 2 | 56.685856 | yes | 5.494E-07 | 596.37665 | 99.631813 |
| VLGTPVETIELTED | 15 | P27708 | CAD protein OS=Homo sapiens OX=960 | CAD | 242.984 | 2 | 56.685856 | yes | 5.479E-07 | 836.4436 | 97.973801 |
| TSSSFAAAMAR | 11 | P27708 | CAD protein OS=Homo sapiens OX=960 | CAD | 242.984 | 2 | 56.685856 | yes | 5.524E-07 | 550.26367 | 87.537399 |
| ASDPGLPAEEPK | 12 | P27708 | CAD protein OS=Homo sapiens OX=960 | CAD | 242.984 | 2 | 56.685856 | yes | 7.807E-07 | 605.80115 | 85.071854 |
| AHWTPFEGQK | 10 | P27708 | CAD protein OS=Homo sapiens OX=960 | CAD | 242.984 | 2 | 56.685856 | yes | 1.049E-07 | 600.79346 | 83.578079 |
| VLGTPVETIELTED | 16 | P27708 | CAD protein OS=Homo sapiens OX=960 | CAD | 242.984 | 3 | 56.685856 | yes | 6.063E-07 | 609.99854 | 77.589996 |
| KEEILLIK | 8 | P27708 | CAD protein OS=Homo sapiens OX=960 | CAD | 242.984 | 2 | 56.685856 | yes | 2.097E-07 | 493.31824 | 68.33757 |
| LFVEALGQIGPAPI | 16 | P27708 | CAD protein OS=Homo sapiens OX=960 | CAD | 242.984 | 2 | 56.685856 | yes | 2.777E-06 | 825.48486 | 63.54459 |
| HPQPGAVELAAK | 12 | P27708 | CAD protein OS=Homo sapiens OX=960 | CAD | 242.984 | 2 | 56.685856 | yes | 3.984E-07 | 609.33551 | 61.741116 |
| SELLPTVR | 8 | P27708 | CAD protein OS=Homo sapiens OX=960 | CAD | 242.984 | 2 | 56.685856 | yes | 8.169E-07 | 457.76892 | 54.243305 |
| SVHICHVAR | 9 | P27708 | CAD protein OS=Homo sapiens OX=960 | CAD | 242.984 | 2 | 56.685856 | yes | 5.675E-07 | 539.78235 | 49.583412 |
| TLGVDLVALATR | 12 | P27708 | CAD protein OS=Homo sapiens OX=960 | CAD | 242.984 | 2 | 56.685856 | yes | 1.251E-07 | 614.86664 | 46.272148 |
| VNEISVEVDSDPR | 13 | P27708 | CAD protein OS=Homo sapiens OX=960 | CAD | 242.984 | 2 | 56.685856 | yes | 0.0001667 | 729.85718 | 40.472389 |
| VFNTGGAPR | 9 | P27708 | CAD protein OS=Homo sapiens OX=960 | CAD | 242.984 | 2 | 56.685856 | yes | 3.145E-06 | 459.74323 | 38.163651 |
| KNILLTIGSYK | 11 | P27708 | CAD protein OS=Homo sapiens OX=960 | CAD | 242.984 | 2 | 56.685856 | yes | 9.722E-08 | 625.37933 | 33.125702 |
| VTAVDWHFEEAV | 20 | P27708 | CAD protein OS=Homo sapiens OX=960 | CAD | 242.984 | 3 | 56.685856 | yes | 4.251E-06 | 781.35504 | 21.263426 |
| EQFLDGDGWTSR | 12 | P27797 | Calreticulin OS=Homo sapiens OX=9606 | CALR | 48.142 | 2 | 46.183723 | yes | 1.627E-07 | 705.81787 | 84.669342 |
| FYALSASFEPFSNK | 14 | P27797 | Calreticulin OS=Homo sapiens OX=9606 | CALR | 48.142 | 2 | 46.183723 | yes | 2.204E-06 | 804.39069 | 79.213882 |
| VHVIFNYK | 8 | P27797 | Calreticulin OS=Homo sapiens OX=9606 | CALR | 48.142 | 2 | 46.183723 | yes | 4.397E-07 | 510.28726 | 54.76545 |
| GQTLVVQFTVK | 11 | P27797 | Calreticulin OS=Homo sapiens OX=9606 | CALR | 48.142 | 2 | 46.183723 | yes | 6.87E-06 | 610.3559 | 40.803978 |
| GMEISPMCDK | 10 | P27815 | cAMP-specific 3',5'-cyclic phosphodiester | PDE4A | 98.144 | 2 | 41.19762 | yes | 3.628E-07 | 584.24579 | 15.370816 |
| DVTPPPETEVVLIK | 14 | P27816 | Microtubule-associated protein 4 OS=Ho | MAP4 | 121.005 | 2 | 54.509968 | yes | 1.427E-07 | 768.92963 | 2714.4478 |
| TDYIPLLDVDEK | 12 | P27816 | Microtubule-associated protein 4 OS=Ho | MAP4 | 121.005 | 2 | 54.509968 | yes | 1.809E-07 | 710.86395 | 2425.6365 |
| NVVLPTETEVAPA | 14 | P27816 | Microtubule-associated protein 4 OS=Ho | MAP4 | 121.005 | 2 | 54.509968 | yes | 1.727E-07 | 734.40631 | 2352.9839 |
| TTTLSGTAPAAGV | 17 | P27816 | Microtubule-associated protein 4 OS=Ho | MAP4 | 121.005 | 2 | 54.509968 | yes | 3.794E-07 | 793.43085 | 2217.6431 |
| DMAQLPETEIAPA | 14 | P27816 | Microtubule-associated protein 4 OS=Ho | MAP4 | 121.005 | 2 | 54.509968 | yes | 1.34E-07 | 757.38196 | 1047.0662 |
| DMAQLPETEIAPA | 14 | P27816 | Microtubule-associated protein 4 OS=Ho | MAP4 | 121.005 | 2 | 54.509968 | yes | 3.101E-06 | 765.37946 | 1047.0662 |
| DVPPLSETEATPVI | 16 | P27816 | Microtubule-associated protein 4 OS=Ho | MAP4 | 121.005 | 2 | 54.509968 | yes | 4.671E-07 | 846.95636 | 1843.5865 |
| DMSPLSETEMALC | 14 | P27816 | Microtubule-associated protein 4 OS=Ho | MAP4 | 121.005 | 2 | 54.509968 | yes | 1.409E-07 | 754.85217 | 1513.2158 |
| LATNTSAPDLK | 11 | P27816 | Microtubule-associated protein 4 OS=Ho | MAP4 | 121.005 | 2 | 54.509968 | yes | 1.639E-07 | 565.80621 | 1428.7002 |
| TKPLATTQPAK | 11 | P27816 | Microtubule-associated protein 4 OS=Ho | MAP4 | 121.005 | 2 | 54.509968 | yes | 1.84E-07 | 578.34021 | 1412.1193 |

| | | | | | | | | | | | |
|---|---|---|---|---|---|---|---|---|---|---|---|
| CSLPAEEDSVLEK | 13 | P27816 | Microtubule-associated protein 4 OS=Hoı | MAP4 | 121.005 | 2 | 54.509968 | yes | 1.365E-07 | 738.84796 | 1050.1216 |
| KPMSLASGLVPAA | 16 | P27816 | Microtubule-associated protein 4 OS=Hoı | MAP4 | 121.005 | 3 | 54.509968 | yes | 9.722E-08 | 521.96912 | 423.00861 |
| KPMSLASGLVPAA | 16 | P27816 | Microtubule-associated protein 4 OS=Hoı | MAP4 | 121.005 | 3 | 54.509968 | yes | 1.379E-07 | 782.45001 | 423.00861 |
| KPMSLASGLVPAA | 16 | P27816 | Microtubule-associated protein 4 OS=Hoı | MAP4 | 121.005 | 2 | 54.509968 | yes | 9.722E-08 | 521.96912 | 423.00861 |
| KPMSLASGLVPAA | 16 | P27816 | Microtubule-associated protein 4 OS=Hoı | MAP4 | 121.005 | 2 | 54.509968 | yes | 1.379E-07 | 782.45001 | 423.00861 |
| DMELPTEK | 8 | P27816 | Microtubule-associated protein 4 OS=Hoı | MAP4 | 121.005 | 2 | 54.509968 | yes | 9.722E-08 | 481.72861 | 833.62933 |
| DMEIAQTQK | 9 | P27816 | Microtubule-associated protein 4 OS=Hoı | MAP4 | 121.005 | 2 | 54.509968 | yes | 9.722E-08 | 532.25806 | 817.29742 |
| DMTLPPETNVILTI | 14 | P27816 | Microtubule-associated protein 4 OS=Hoı | MAP4 | 121.005 | 2 | 54.509968 | yes | 5.251E-07 | 786.42108 | 770.29956 |
| QPAPTTIGGLNK | 12 | P27816 | Microtubule-associated protein 4 OS=Hoı | MAP4 | 121.005 | 2 | 54.509968 | yes | 2.001E-07 | 598.83533 | 730.07294 |
| PSTTPFIDK | 9 | P27816 | Microtubule-associated protein 4 OS=Hoı | MAP4 | 121.005 | 2 | 54.509968 | yes | 2.213E-07 | 503.2662 | 713.39691 |
| VALSSETEVALAR | 13 | P27816 | Microtubule-associated protein 4 OS=Hoı | MAP4 | 121.005 | 2 | 54.509968 | yes | 9.722E-08 | 673.36975 | 658.51575 |
| HVPGGGNVQIQNI | 13 | P27816 | Microtubule-associated protein 4 OS=Hoı | MAP4 | 121.005 | 2 | 54.509968 | yes | 1.506E-07 | 674.36005 | 625.43176 |
| ALPLEAEVAPVK | 12 | P27816 | Microtubule-associated protein 4 OS=Hoı | MAP4 | 121.005 | 2 | 54.509968 | yes | 1.061E-07 | 618.86353 | 477.39893 |
| VGSLDNVGHLPA( | 17 | P27816 | Microtubule-associated protein 4 OS=Hoı | MAP4 | 121.005 | 2 | 54.509968 | yes | 9.727E-08 | 795.93353 | 442.11319 |
| DGVLTLANNVTP/ | 14 | P27816 | Microtubule-associated protein 4 OS=Hoı | MAP4 | 121.005 | 2 | 54.509968 | yes | 1.006E-07 | 706.89081 | 402.68408 |
| WPTETDVSSAK | 11 | P27816 | Microtubule-associated protein 4 OS=Hoı | MAP4 | 121.005 | 2 | 54.509968 | yes | 5.86E-07 | 610.79333 | 300.99835 |
| VSYSHIQSK | 9 | P27816 | Microtubule-associated protein 4 OS=Hoı | MAP4 | 121.005 | 2 | 54.509968 | yes | 3.011E-07 | 524.77472 | 252.50504 |
| DMQPSMESDMAL | 14 | P27816 | Microtubule-associated protein 4 OS=Hoı | MAP4 | 121.005 | 2 | 54.509968 | yes | 1.92E-07 | 791.3515 | 243.00044 |
| TQPTSLPK | 8 | P27816 | Microtubule-associated protein 4 OS=Hoı | MAP4 | 121.005 | 2 | 54.509968 | yes | 2.162E-07 | 436.2478 | 242.18248 |
| KVSYSHIQSK | 10 | P27816 | Microtubule-associated protein 4 OS=Hoı | MAP4 | 121.005 | 2 | 54.509968 | yes | 1.065E-07 | 588.8222 | 182.82256 |
| TTTAAAVASTGPS | 15 | P27816 | Microtubule-associated protein 4 OS=Hoı | MAP4 | 121.005 | 2 | 54.509968 | yes | 9.722E-08 | 689.35205 | 157.37801 |
| KCSLPAEEDSVLEI | 14 | P27816 | Microtubule-associated protein 4 OS=Hoı | MAP4 | 121.005 | 2 | 54.509968 | yes | 1.106E-07 | 802.89545 | 141.27956 |
| PLATTQPAK | 9 | P27816 | Microtubule-associated protein 4 OS=Hoı | MAP4 | 121.005 | 2 | 54.509968 | yes | 4.538E-07 | 463.76892 | 105.36139 |
| SPSTLLPK | 8 | P27816 | Microtubule-associated protein 4 OS=Hoı | MAP4 | 121.005 | 2 | 54.509968 | yes | 3.891E-06 | 421.75275 | 90.496811 |
| NVCLPPEMEVALT | 21 | P27816 | Microtubule-associated protein 4 OS=Hoı | MAP4 | 121.005 | 3 | 54.509968 | yes | 1.501E-07 | 785.06409 | 82.13031 |
| SVPADLSRPK | 10 | P27816 | Microtubule-associated protein 4 OS=Hoı | MAP4 | 121.005 | 2 | 54.509968 | yes | 1.084E-06 | 535.30365 | 43.32016 |
| DVKPKPIADAK | 11 | P27816 | Microtubule-associated protein 4 OS=Hoı | MAP4 | 121.005 | 2 | 54.509968 | yes | 6.346E-07 | 591.34808 | 37.983738 |
| GTLSGWILSK | 10 | P27824 | Calnexin OS=Homo sapiens OX=9606 G | CANX | 67.569 | 2 | 56.766937 | yes | 5.071E-07 | 531.3031 | 146.56258 |
| PVIDNPNYK | 9 | P27824 | Calnexin OS=Homo sapiens OX=9606 G | CANX | 67.569 | 2 | 56.766937 | yes | 3.214E-07 | 530.2771 | 114.50783 |
| AEEDEILNR | 9 | P27824 | Calnexin OS=Homo sapiens OX=9606 G | CANX | 67.569 | 2 | 56.766937 | yes | 5.397E-07 | 544.76459 | 93.517601 |
| IVDDWANDGWGL | 13 | P27824 | Calnexin OS=Homo sapiens OX=9606 G | CANX | 67.569 | 2 | 56.766937 | yes | 1.697E-07 | 744.8595 | 69.676323 |
| CESAPGCGVWQR | 12 | P27824 | Calnexin OS=Homo sapiens OX=9606 G | CANX | 67.569 | 2 | 56.766937 | yes | 9.722E-08 | 703.8006 | 69.355309 |
| LFQVEYAIEAIK | 12 | P28066 | Proteasome subunit alpha type-5 OS=Hor | PSMA5 | 26.412 | 2 | 51.891216 | yes | 9.722E-08 | 712.3952 | 179.99643 |
| ITSPLMEPSSIEK | 13 | P28066 | Proteasome subunit alpha type-5 OS=Hor | PSMA5 | 26.412 | 2 | 51.891216 | yes | 3.668E-06 | 716.3736 | 63.683342 |
| IVEIDAHIGCAMS( | 20 | P28066 | Proteasome subunit alpha type-5 OS=Hor | PSMA5 | 26.412 | 3 | 51.891216 | yes | 0.0001009 | 695.35535 | 14.817804 |
| SGLWAGGPAPGQI | 15 | P28070 | Proteasome subunit beta type-4 OS=Hom | PSMB4 | 29.205 | 2 | 43.42544 | yes | 2.598E-07 | 782.38879 | 315.85181 |
| FQIATVTEK | 9 | P28070 | Proteasome subunit beta type-4 OS=Hom | PSMB4 | 29.205 | 2 | 43.42544 | yes | 3.659E-06 | 518.78729 | 122.92552 |
| QVLGQMVIDEELL | 22 | P28070 | Proteasome subunit beta type-4 OS=Hom | PSMB4 | 29.205 | 3 | 43.42544 | yes | 6.441E-05 | 815.06891 | 16.992369 |
| LLANMVYQYK | 10 | P28074 | Proteasome subunit beta type-5 OS=Hom | PSMB5 | 28.481 | 2 | 36.61255 | yes | 6.798E-07 | 621.83118 | 52.546444 |

| | | | | | | | | | | | |
|---|---|---|---|---|---|---|---|---|---|---|---|
| ITELMQVLK | 9 | P28288 | ATP-binding cassette sub-family D memt | ABCD3 | 75.476 | 2 | 54.68571 | yes | 1.517E-07 | 537.815 | 997.29846 |
| DQVIYPDGR | 9 | P28288 | ATP-binding cassette sub-family D memt | ABCD3 | 75.476 | 2 | 54.68571 | yes | 1.136E-07 | 531.76434 | 861.68335 |
| LVEHLHNFILFR | 12 | P28288 | ATP-binding cassette sub-family D memt | ABCD3 | 75.476 | 3 | 54.68571 | yes | 9.722E-08 | 513.29279 | 204.71143 |
| LVEHLHNFILFR | 12 | P28288 | ATP-binding cassette sub-family D memt | ABCD3 | 75.476 | 3 | 54.68571 | yes | 9.722E-08 | 769.43555 | 204.71143 |
| LVEHLHNFILFR | 12 | P28288 | ATP-binding cassette sub-family D memt | ABCD3 | 75.476 | 2 | 54.68571 | yes | 9.722E-08 | 513.29279 | 204.71143 |
| LVEHLHNFILFR | 12 | P28288 | ATP-binding cassette sub-family D memt | ABCD3 | 75.476 | 2 | 54.68571 | yes | 9.722E-08 | 769.43555 | 204.71143 |
| PFLDLSHPR | 9 | P28288 | ATP-binding cassette sub-family D memt | ABCD3 | 75.476 | 2 | 54.68571 | yes | 1.846E-07 | 541.29309 | 385.89246 |
| PYMTLGTLR | 9 | P28288 | ATP-binding cassette sub-family D memt | ABCD3 | 75.476 | 2 | 54.68571 | yes | 5.096E-07 | 526.28387 | 307.99945 |
| QTVHSVFR | 8 | P28288 | ATP-binding cassette sub-family D memt | ABCD3 | 75.476 | 2 | 54.68571 | yes | 2.003E-07 | 487.26434 | 301.63992 |
| KGISDLVLK | 9 | P28288 | ATP-binding cassette sub-family D memt | ABCD3 | 75.476 | 2 | 54.68571 | yes | 9.722E-08 | 486.80804 | 214.66756 |
| DLNFEVR | 7 | P28288 | ATP-binding cassette sub-family D memt | ABCD3 | 75.476 | 2 | 54.68571 | yes | 1.466E-06 | 446.7298 | 183.34828 |
| TMVSQQEK | 8 | P28288 | ATP-binding cassette sub-family D memt | ABCD3 | 75.476 | 2 | 54.68571 | yes | 1.034E-05 | 475.73422 | 131.24942 |
| LFYVPQR | 7 | P28288 | ATP-binding cassette sub-family D memt | ABCD3 | 75.476 | 2 | 54.68571 | yes | 1.076E-07 | 461.76089 | 127.48851 |
| IANPDQLLTQDVE | 14 | P28288 | ATP-binding cassette sub-family D memt | ABCD3 | 75.476 | 2 | 54.68571 | yes | 9.722E-08 | 792.41742 | 113.79743 |
| SGANVLICGPNGC | 15 | P28288 | ATP-binding cassette sub-family D memt | ABCD3 | 75.476 | 2 | 54.68571 | yes | 1.693E-07 | 752.35577 | 40.632923 |
| FSMGFIDSIIAK | 12 | P28288 | ATP-binding cassette sub-family D memt | ABCD3 | 75.476 | 2 | 54.68571 | yes | 2.439E-06 | 672.84705 | 21.251787 |
| EYLDNVQLGHILE | 14 | P28288 | ATP-binding cassette sub-family D memt | ABCD3 | 75.476 | 3 | 54.68571 | yes | 5.034E-05 | 566.96515 | 15.3106 |
| LEEVNLNNIR | 10 | P28289 | Tropomodulin-1 OS=Homo sapiens OX= | TMOD1 | 40.57 | 2 | 49.753155 | yes | 3.198E-07 | 607.33038 | 86.066299 |
| LADLTGPIIPK | 11 | P28289 | Tropomodulin-1 OS=Homo sapiens OX= | TMOD1 | 40.57 | 2 | 49.753155 | yes | 1.426E-07 | 569.34753 | 49.707253 |
| VEMEIVSMLEK | 11 | P28289 | Tropomodulin-1 OS=Homo sapiens OX= | TMOD1 | 40.57 | 2 | 49.753155 | yes | 1.005E-07 | 654.33289 | 48.750137 |
| LEEVSPNLVR | 10 | P28331 | NADH-ubiquinone oxidoreductase 75 kD | NDUFS1 | 79.468 | 2 | 50.886463 | yes | 1.173E-07 | 578.32202 | 161.56142 |
| VLFLLGADGGCITI | 14 | P28331 | NADH-ubiquinone oxidoreductase 75 kD | NDUFS1 | 79.468 | 2 | 50.886463 | yes | 9.722E-08 | 746.40302 | 142.95006 |
| GNDMQVGTYIEK | 12 | P28331 | NADH-ubiquinone oxidoreductase 75 kD | NDUFS1 | 79.468 | 2 | 50.886463 | yes | 3.262E-06 | 677.81903 | 92.441048 |
| SATYVNTEGR | 10 | P28331 | NADH-ubiquinone oxidoreductase 75 kD | NDUFS1 | 79.468 | 2 | 50.886463 | yes | 1.035E-06 | 549.26471 | 76.762901 |
| FASEIAGVDDLGT | 16 | P28331 | NADH-ubiquinone oxidoreductase 75 kD | NDUFS1 | 79.468 | 2 | 50.886463 | yes | 1.15E-07 | 804.89685 | 58.142822 |
| MCLVEIEK | 8 | P28331 | NADH-ubiquinone oxidoreductase 75 kD | NDUFS1 | 79.468 | 2 | 50.886463 | yes | 4.163E-07 | 511.25647 | 53.769897 |
| GLLTYTSWEDALS | 14 | P28331 | NADH-ubiquinone oxidoreductase 75 kD | NDUFS1 | 79.468 | 2 | 50.886463 | yes | 1.125E-07 | 806.4043 | 26.189432 |
| IFEPILGEGR | 10 | P28340 | DNA polymerase delta catalytic subunit C | POLD1 | 123.632 | 2 | 52.96563 | yes | 4.477E-07 | 565.81384 | 423.12314 |
| FGVSSVAEAMAL( | 14 | P28340 | DNA polymerase delta catalytic subunit C | POLD1 | 123.632 | 2 | 52.96563 | yes | 9.722E-08 | 697.85846 | 308.44275 |
| VLSFDIECAGR | 11 | P28340 | DNA polymerase delta catalytic subunit C | POLD1 | 123.632 | 2 | 52.96563 | yes | 9.722E-08 | 633.81097 | 279.54813 |
| LALTLRPCAPILGA | 15 | P28340 | DNA polymerase delta catalytic subunit C | POLD1 | 123.632 | 3 | 52.96563 | yes | 2.843E-07 | 531.98846 | 123.67722 |
| LALTLRPCAPILGA | 15 | P28340 | DNA polymerase delta catalytic subunit C | POLD1 | 123.632 | 3 | 52.96563 | yes | 3.353E-07 | 797.47906 | 123.67722 |
| LALTLRPCAPILGA | 15 | P28340 | DNA polymerase delta catalytic subunit C | POLD1 | 123.632 | 2 | 52.96563 | yes | 2.843E-07 | 531.98846 | 123.67722 |
| LALTLRPCAPILGA | 15 | P28340 | DNA polymerase delta catalytic subunit C | POLD1 | 123.632 | 2 | 52.96563 | yes | 3.353E-07 | 797.47906 | 123.67722 |
| IDISQLVITK | 10 | P28340 | DNA polymerase delta catalytic subunit C | POLD1 | 123.632 | 2 | 52.96563 | yes | 2.049E-07 | 565.34497 | 228.20277 |
| LGLTEDQFIR | 10 | P28340 | DNA polymerase delta catalytic subunit C | POLD1 | 123.632 | 2 | 52.96563 | yes | 4.47E-07 | 596.32202 | 137.72449 |
| LWTQCQR | 7 | P28340 | DNA polymerase delta catalytic subunit C | POLD1 | 123.632 | 2 | 52.96563 | yes | 3.332E-07 | 496.24255 | 134.38728 |
| LMVLVNAVEMAR | 12 | P28340 | DNA polymerase delta catalytic subunit C | POLD1 | 123.632 | 2 | 52.96563 | yes | 3.379E-07 | 673.37018 | 127.36821 |
| ELTGPAVLAVELC | 15 | P28340 | DNA polymerase delta catalytic subunit C | POLD1 | 123.632 | 2 | 52.96563 | yes | 9.722E-08 | 807.92963 | 101.50223 |

| | | | | | | | | | | | |
|---|---|---|---|---|---|---|---|---|---|---|---|
| DLEDQEQLLR | 10 | P28340 | DNA polymerase delta catalytic subunit C | POLD1 | 123.632 | 2 | 52.96563 | yes | 9.722E-08 | 629.81732 | 101.34789 |
| VSANSVYGFTGAC | 16 | P28340 | DNA polymerase delta catalytic subunit C | POLD1 | 123.632 | 2 | 52.96563 | yes | 4.04E-07 | 792.90448 | 99.863541 |
| QAHVELAER | 9 | P28340 | DNA polymerase delta catalytic subunit C | POLD1 | 123.632 | 2 | 52.96563 | yes | 1.882E-07 | 526.77783 | 91.770088 |
| EVSHLNALEER | 11 | P28340 | DNA polymerase delta catalytic subunit C | POLD1 | 123.632 | 2 | 52.96563 | yes | 1.705E-07 | 648.83075 | 81.126045 |
| VVYGDTDSVMCR | 12 | P28340 | DNA polymerase delta catalytic subunit C | POLD1 | 123.632 | 2 | 52.96563 | yes | 1.204E-07 | 701.31049 | 64.549614 |
| VQTFPFLGR | 9 | P28340 | DNA polymerase delta catalytic subunit C | POLD1 | 123.632 | 2 | 52.96563 | yes | 2.672E-07 | 532.79803 | 63.123241 |
| DCPIFYMR | 8 | P28340 | DNA polymerase delta catalytic subunit C | POLD1 | 123.632 | 2 | 52.96563 | yes | 4.018E-07 | 551.24646 | 62.534626 |
| YAGLLFSSR | 9 | P28340 | DNA polymerase delta catalytic subunit C | POLD1 | 123.632 | 2 | 52.96563 | yes | 1.254E-07 | 507.27435 | 60.673641 |
| LPCLEISQSVTGFG | 15 | P28340 | DNA polymerase delta catalytic subunit C | POLD1 | 123.632 | 2 | 52.96563 | yes | 1.189E-07 | 832.42725 | 60.384167 |
| VAGLCSNIR | 9 | P28340 | DNA polymerase delta catalytic subunit C | POLD1 | 123.632 | 2 | 52.96563 | yes | 2.079E-07 | 495.26349 | 57.262573 |
| VTGVPLSYLLSR | 12 | P28340 | DNA polymerase delta catalytic subunit C | POLD1 | 123.632 | 2 | 52.96563 | yes | 1.781E-06 | 652.88226 | 47.92799 |
| DNCPLVANLVTAS | 15 | P28340 | DNA polymerase delta catalytic subunit C | POLD1 | 123.632 | 2 | 52.96563 | yes | 1.031E-07 | 821.93268 | 33.72118 |
| PSQFEEDLALMEE | 19 | P28340 | DNA polymerase delta catalytic subunit C | POLD1 | 123.632 | 3 | 52.96563 | yes | 2.035E-07 | 764.33667 | 23.617188 |
| CQGSLHEDVICTSI | 14 | P28340 | DNA polymerase delta catalytic subunit C | POLD1 | 123.632 | 2 | 52.96563 | yes | 1.96E-07 | 831.37213 | 8.3837051 |
| LVEFSAFLEQQR | 12 | P28347 | Transcriptional enhancer factor TEF-1 OS | TEAD1 | 47.946 | 2 | 51.256702 | yes | 9.722E-08 | 733.88556 | 98.444405 |
| GPQNAFFLVK | 10 | P28347 | Transcriptional enhancer factor TEF-1 OS | TEAD1 | 47.946 | 2 | 51.256702 | yes | 5.706E-07 | 560.8111 | 73.635979 |
| LLTQGFTNWTK | 11 | P28370 | Probable global transcription activator SN | SMARCA1 | 122.605 | 2 | 55.199486 | yes | 9.722E-08 | 654.85095 | 212.40898 |
| LITDNTVEER | 10 | P28370 | Probable global transcription activator SN | SMARCA1 | 122.605 | 2 | 55.199486 | yes | 9.809E-08 | 595.30658 | 172.91704 |
| FEVSPSYVK | 9 | P28370 | Probable global transcription activator SN | SMARCA1 | 122.605 | 2 | 55.199486 | yes | 2.087E-07 | 528.27405 | 170.99823 |
| LDGQTPHEER | 10 | P28370 | Probable global transcription activator SN | SMARCA1 | 122.605 | 2 | 55.199486 | yes | 3.521E-07 | 591.28088 | 137.87057 |
| ENMEIEER | 8 | P28370 | Probable global transcription activator SN | SMARCA1 | 122.605 | 2 | 55.199486 | yes | 3.285E-07 | 525.23206 | 118.88333 |
| NPDIPNPALAQR | 12 | P28370 | Probable global transcription activator SN | SMARCA1 | 122.605 | 2 | 55.199486 | yes | 1.555E-07 | 653.34912 | 109.35664 |
| QSLISAGDYR | 10 | P28370 | Probable global transcription activator SN | SMARCA1 | 122.605 | 2 | 55.199486 | yes | 2.301E-07 | 555.2829 | 100.35155 |
| CNTLISLIEK | 10 | P28370 | Probable global transcription activator SN | SMARCA1 | 122.605 | 2 | 55.199486 | yes | 1.611E-07 | 595.82611 | 93.133186 |
| DIDVLNSSGK | 10 | P28370 | Probable global transcription activator SN | SMARCA1 | 122.605 | 2 | 55.199486 | yes | 2.086E-07 | 524.26947 | 78.132759 |
| RFEFLLK | 7 | P28370 | Probable global transcription activator SN | SMARCA1 | 122.605 | 2 | 55.199486 | yes | 3.427E-07 | 476.78436 | 62.829842 |
| TEQEEDEELLSESF | 14 | P28370 | Probable global transcription activator SN | SMARCA1 | 122.605 | 2 | 55.199486 | yes | 1.002E-07 | 847.3736 | 50.957272 |
| STLHNWMNEFK | 11 | P28370 | Probable global transcription activator SN | SMARCA1 | 122.605 | 2 | 55.199486 | yes | 1.306E-07 | 703.82971 | 36.141708 |
| ICDFGLAR | 8 | P28482 | Mitogen-activated protein kinase 1 OS=H | MAPK1 | 41.39 | 2 | 53.18169 | no | 9.722E-08 | 476.23947 | 560.56995 |
| NYLLSLPHK | 9 | P28482 | Mitogen-activated protein kinase 1 OS=H | MAPK1 | 41.39 | 2 | 53.18169 | yes | 1.388E-07 | 542.8111 | 384.69736 |
| FDMELDDLPK | 10 | P28482 | Mitogen-activated protein kinase 1 OS=H | MAPK1 | 41.39 | 2 | 53.18169 | yes | 9.722E-08 | 611.78662 | 209.65326 |
| HENIIGINDIIR | 12 | P28482 | Mitogen-activated protein kinase 1 OS=H | MAPK1 | 41.39 | 2 | 53.18169 | yes | 9.722E-08 | 703.89117 | 189.65961 |
| MLTFNPHK | 8 | P28482 | Mitogen-activated protein kinase 1 OS=H | MAPK1 | 41.39 | 2 | 53.18169 | yes | 4.406E-07 | 494.25766 | 160.50987 |
| APTIEQMK | 8 | P28482 | Mitogen-activated protein kinase 1 OS=H | MAPK1 | 41.39 | 2 | 53.18169 | yes | 1.572E-07 | 459.24167 | 95.434265 |
| ISPFEHQTYCQR | 12 | P28482 | Mitogen-activated protein kinase 1 OS=H | MAPK1 | 41.39 | 2 | 53.18169 | no | 1.121E-07 | 783.36188 | 87.636848 |
| APEIMLNSK | 9 | P28482 | Mitogen-activated protein kinase 1 OS=H | MAPK1 | 41.39 | 2 | 53.18169 | no | 4.954E-07 | 501.76807 | 73.59465 |
| YIHSANVLHR | 10 | P28482 | Mitogen-activated protein kinase 1 OS=H | MAPK1 | 41.39 | 2 | 53.18169 | no | 9.722E-08 | 605.328 | 72.190277 |
| GLSNPSEVEVLR | 12 | P28702 | Retinoic acid receptor RXR-beta OS=Hor | RXRB | 56.922 | 2 | 48.116154 | yes | 1.197E-07 | 650.34882 | 108.67935 |
| AIILFNPDAK | 10 | P28702 | Retinoic acid receptor RXR-beta OS=Hor | RXRB | 56.922 | 2 | 48.116154 | yes | 2.253E-07 | 551.31879 | 77.743423 |

| | | | | | | | | | | | |
|---|---|---|---|---|---|---|---|---|---|---|---|
| LCAICGDR | 8 | P28702 | Retinoic acid receptor RXR-beta OS=Hoı | RXRB | 56.922 | 2 | 48.116154 | yes | 1.468E-06 | 482.72058 | 58.606491 |
| EGEAPAESESLLR | 13 | P28715 | DNA excision repair protein ERCC-5 OS | ERCC5 | 133.109 | 2 | 55.721054 | yes | 1.316E-07 | 694.33862 | 432.927 |
| GITNTLEESSSLK | 13 | P28715 | DNA excision repair protein ERCC-5 OS | ERCC5 | 133.109 | 2 | 55.721054 | yes | 1.867E-07 | 689.85663 | 266.76324 |
| NAPAAVDEGSISPI | 14 | P28715 | DNA excision repair protein ERCC-5 OS | ERCC5 | 133.109 | 2 | 55.721054 | yes | 1.429E-07 | 692.3468 | 265.68518 |
| VVSEDTSHYILIK | 13 | P28715 | DNA excision repair protein ERCC-5 OS | ERCC5 | 133.109 | 2 | 55.721054 | yes | 9.873E-08 | 752.40631 | 238.7193 |
| ELTPASPTCTNSVS | 15 | P28715 | DNA excision repair protein ERCC-5 OS | ERCC5 | 133.109 | 2 | 55.721054 | yes | 9.722E-08 | 796.38525 | 192.07259 |
| GSFLWGKPDLDK | 12 | P28715 | DNA excision repair protein ERCC-5 OS | ERCC5 | 133.109 | 2 | 55.721054 | yes | 9.722E-08 | 681.85626 | 190.01642 |
| MHGMSFDVK | 9 | P28715 | DNA excision repair protein ERCC-5 OS | ERCC5 | 133.109 | 2 | 55.721054 | yes | 8.053E-07 | 526.23859 | 188.46638 |
| LPLESAVVR | 9 | P28715 | DNA excision repair protein ERCC-5 OS | ERCC5 | 133.109 | 2 | 55.721054 | yes | 1.003E-07 | 492.29785 | 181.41621 |
| EAAASEIEAVSVAI | 16 | P28715 | DNA excision repair protein ERCC-5 OS | ERCC5 | 133.109 | 2 | 55.721054 | yes | 9.722E-08 | 817.90076 | 164.66316 |
| FSEWWHEAQK | 10 | P28715 | DNA excision repair protein ERCC-5 OS | ERCC5 | 133.109 | 2 | 55.721054 | yes | 2.253E-07 | 674.30945 | 146.06401 |
| HEILTDMK | 8 | P28715 | DNA excision repair protein ERCC-5 OS | ERCC5 | 133.109 | 2 | 55.721054 | yes | 1.217E-06 | 493.75241 | 134.11148 |
| QLDAQQTQLR | 10 | P28715 | DNA excision repair protein ERCC-5 OS | ERCC5 | 133.109 | 2 | 55.721054 | yes | 9.722E-08 | 600.82019 | 126.27644 |
| TDESLFPVLK | 10 | P28715 | DNA excision repair protein ERCC-5 OS | ERCC5 | 133.109 | 2 | 55.721054 | yes | 1.557E-07 | 574.81354 | 116.68546 |
| TVAEVDSESLPSSS | 15 | P28715 | DNA excision repair protein ERCC-5 OS | ERCC5 | 133.109 | 2 | 55.721054 | yes | 1.007E-07 | 768.37543 | 104.43056 |
| FDSSLLSSDDETK | 13 | P28715 | DNA excision repair protein ERCC-5 OS | ERCC5 | 133.109 | 2 | 55.721054 | yes | 9.722E-08 | 722.32794 | 85.938324 |
| LLECSGR | 7 | P28715 | DNA excision repair protein ERCC-5 OS | ERCC5 | 133.109 | 2 | 55.721054 | yes | 1.32E-07 | 417.71054 | 68.65181 |
| DRLPLESAVVR | 11 | P28715 | DNA excision repair protein ERCC-5 OS | ERCC5 | 133.109 | 2 | 55.721054 | yes | 1.041E-06 | 627.86188 | 62.318527 |
| NYLNQHIEHVQK | 12 | P28715 | DNA excision repair protein ERCC-5 OS | ERCC5 | 133.109 | 2 | 55.721054 | yes | 9.722E-08 | 761.89166 | 15.141222 |
| HCCPAGFR | 8 | P28799 | Progranulin OS=Homo sapiens OX=9606 | GRN | 63.545 | 2 | 52.310875 | yes | 1.116E-06 | 502.7131 | 402.88196 |
| QGWACCPYR | 9 | P28799 | Progranulin OS=Homo sapiens OX=9606 | GRN | 63.545 | 2 | 52.310875 | yes | 9.722E-08 | 599.25006 | 161.33434 |
| AVALSSSVMCPDA | 14 | P28799 | Progranulin OS=Homo sapiens OX=9606 | GRN | 63.545 | 2 | 52.310875 | yes | 2.63E-06 | 732.35272 | 21.335367 |
| EGGIDMDAFQER | 12 | P29083 | General transcription factor IIE subunit 1 | GTF2E1 | 49.453 | 2 | 54.642895 | yes | 9.722E-08 | 684.29846 | 188.54898 |
| ESTVQGAYGSEDM | 14 | P29083 | General transcription factor IIE subunit 1 | GTF2E1 | 49.453 | 2 | 54.642895 | yes | 2.741E-07 | 751.32739 | 37.762165 |
| FNEQIEPIYALLR | 13 | P29083 | General transcription factor IIE subunit 1 | GTF2E1 | 49.453 | 2 | 54.642895 | yes | 5.026E-05 | 803.43542 | 8.5009098 |
| SCQFSVDEEFQK | 12 | P29084 | Transcription initiation factor IIE subunit | GTF2E2 | 33.044 | 2 | 52.9233 | yes | 9.722E-08 | 752.32465 | 203.15662 |
| QWLMTEALVNNP | 13 | P29084 | Transcription initiation factor IIE subunit | GTF2E2 | 33.044 | 2 | 52.9233 | yes | 9.722E-08 | 772.40051 | 199.33324 |
| ALGDQILFVNRPDI | 14 | P29084 | Transcription initiation factor IIE subunit | GTF2E2 | 33.044 | 2 | 52.9233 | yes | 1.422E-07 | 793.43848 | 163.98485 |
| LLDQHDQR | 8 | P29084 | Transcription initiation factor IIE subunit | GTF2E2 | 33.044 | 2 | 52.9233 | yes | 1.991E-06 | 512.76215 | 102.58964 |
| AMIEVINHK | 9 | P29144 | Tripeptidyl-peptidase 2 OS=Homo sapien | TPP2 | 138.35 | 2 | 43.67131 | yes | 2.493E-07 | 527.78931 | 50.73177 |
| ACIDSNEDGDLSK | 13 | P29144 | Tripeptidyl-peptidase 2 OS=Homo sapien | TPP2 | 138.35 | 2 | 43.67131 | yes | 1.144E-06 | 712.30389 | 24.227413 |
| AYDYLVQNTSFAN | 14 | P29144 | Tripeptidyl-peptidase 2 OS=Homo sapien | TPP2 | 138.35 | 2 | 43.67131 | yes | 7.756E-06 | 817.39648 | 12.186722 |
| FCGVCTDGR | 9 | P29279 | CCN family member 2 OS=Homo sapiens | CCN2 | 38.092 | 2 | 38.151035 | yes | 5.423E-07 | 536.22095 | 33.579464 |
| VMGPGVSYLVR | 11 | P29353 | SHC-transforming protein 1 OS=Homo sa | SHC1 | 62.823 | 2 | 48.888897 | yes | 1.09E-07 | 589.32355 | 134.7429 |
| VEGGQLGGEEWTI | 13 | P29353 | SHC-transforming protein 1 OS=Homo sa | SHC1 | 62.823 | 2 | 48.888897 | yes | 7.669E-07 | 709.33893 | 97.080185 |
| EAISLVCEAVPGAI | 14 | P29353 | SHC-transforming protein 1 OS=Homo sa | SHC1 | 62.823 | 2 | 48.888897 | yes | 2.243E-05 | 722.37921 | 31.7533 |
| ALEPLEGLETMR | 12 | P29372 | DNA-3-methyladenine glycosylase OS=H | MPG | 32.869 | 2 | 53.74638 | yes | 9.926E-08 | 679.85284 | 871.40839 |
| GPLEPSEPAVVAA | 15 | P29372 | DNA-3-methyladenine glycosylase OS=H | MPG | 32.869 | 2 | 53.74638 | yes | 2.307E-07 | 732.3963 | 598.47327 |
| AFLGQVLVR | 9 | P29372 | DNA-3-methyladenine glycosylase OS=H | MPG | 32.869 | 2 | 53.74638 | yes | 1.483E-07 | 501.80838 | 383.79111 |

| | | | | | | | | | | | |
|---|---|---|---|---|---|---|---|---|---|---|---|
| VGVGHAGEWAR | 11 | P29372 | DNA-3-methyladenine glycosylase OS=H | MPG | 32.869 | 2 | 53.74638 | yes | 9.722E-08 | 569.79126 | 313.68466 |
| LGLEFFDQPAVPL/ | 15 | P29372 | DNA-3-methyladenine glycosylase OS=H | MPG | 32.869 | 2 | 53.74638 | yes | 9.722E-08 | 836.95648 | 278.63843 |
| GSPWVSVVDR | 10 | P29372 | DNA-3-methyladenine glycosylase OS=H | MPG | 32.869 | 2 | 53.74638 | yes | 6.058E-07 | 551.28802 | 225.50162 |
| LCQALAINK | 9 | P29372 | DNA-3-methyladenine glycosylase OS=H | MPG | 32.869 | 2 | 53.74638 | yes | 4.974E-07 | 515.78931 | 188.27634 |
| CLGPPTTPGPYR | 12 | P29372 | DNA-3-methyladenine glycosylase OS=H | MPG | 32.869 | 2 | 53.74638 | yes | 9.722E-08 | 658.32678 | 183.35854 |
| SIYFSSPK | 8 | P29372 | DNA-3-methyladenine glycosylase OS=H | MPG | 32.869 | 2 | 53.74638 | yes | 4.251E-07 | 464.74237 | 129.12277 |
| DLAQDEAVWLER | 12 | P29372 | DNA-3-methyladenine glycosylase OS=H | MPG | 32.869 | 2 | 53.74638 | yes | 7.397E-07 | 722.85699 | 89.469933 |
| IVETEAYLGPEDE/ | 18 | P29372 | DNA-3-methyladenine glycosylase OS=H | MPG | 32.869 | 3 | 53.74638 | yes | 1.223E-07 | 662.98499 | 65.336937 |
| AGQPHSSSDAAQ/ | 31 | P29372 | DNA-3-methyladenine glycosylase OS=H | MPG | 32.869 | 4 | 53.74638 | yes | 8.021E-07 | 779.09857 | 15.510251 |
| LTLSDPQQVTLDV | 15 | P29375 | Lysine-specific demethylase 5A OS=Hom | KDM5A | 192.096 | 2 | 40.940388 | yes | 5.961E-07 | 844.45038 | 41.63887 |
| LEALPQVESQVAA | 15 | P29375 | Lysine-specific demethylase 5A OS=Hom | KDM5A | 192.096 | 2 | 40.940388 | yes | 3.76E-07 | 791.43341 | 38.20816 |
| EAETCASVAQLLL | 15 | P29375 | Lysine-specific demethylase 5A OS=Hom | KDM5A | 192.096 | 2 | 40.940388 | yes | 1.567E-06 | 810.41907 | 22.117283 |
| NSTFSEIFK | 9 | P29401 | Transketolase OS=Homo sapiens OX=96 | TKT | 67.878 | 2 | 43.66208 | yes | 2.497E-07 | 536.7691 | 41.690086 |
| TPPGVSAPTEPLLC | 15 | P29558 | RNA-binding motif, single-stranded-inter | RBMS1 | 44.506 | 2 | 51.599903 | yes | 1.025E-07 | 783.91345 | 6936.8647 |
| GYGFVDFDSPAA/ | 15 | P29558 | RNA-binding motif, single-stranded-inter | RBMS1 | 44.506 | 2 | 51.599903 | no | 9.722E-08 | 786.87012 | 4796.5742 |
| GLPPHTTDQDLVK | 13 | P29558 | RNA-binding motif, single-stranded-inter | RBMS1 | 44.506 | 2 | 51.599903 | yes | 9.778E-08 | 710.87518 | 2011.2478 |
| ASGVQAQMAK | 10 | P29558 | RNA-binding motif, single-stranded-inter | RBMS1 | 44.506 | 2 | 51.599903 | no | 3.668E-07 | 495.75549 | 1983.6614 |
| LCQPYGK | 7 | P29558 | RNA-binding motif, single-stranded-inter | RBMS1 | 44.506 | 2 | 51.599903 | no | 1.556E-07 | 433.21545 | 1977.8522 |
| CEAVIGHFNGK | 11 | P29558 | RNA-binding motif, single-stranded-inter | RBMS1 | 44.506 | 2 | 51.599903 | yes | 9.722E-08 | 616.29803 | 708.52075 |
| PFGQVISTR | 9 | P29558 | RNA-binding motif, single-stranded-inter | RBMS1 | 44.506 | 2 | 51.599903 | no | 1.596E-07 | 502.77982 | 454.70135 |
| QQMYPQYATYYY | 19 | P29558 | RNA-binding motif, single-stranded-inter | RBMS1 | 44.506 | 3 | 51.599903 | yes | 1.916E-07 | 816.38409 | 143.36771 |
| LTELLER | 7 | P29597 | Non-receptor tyrosine-protein kinase TYI | TYK2 | 133.65 | 2 | 53.913025 | yes | 2.45E-07 | 437.25565 | 160.75679 |
| ELGAALQGCLLR | 12 | P29597 | Non-receptor tyrosine-protein kinase TYI | TYK2 | 133.65 | 2 | 53.913025 | yes | 9.722E-08 | 650.85571 | 145.64059 |
| HGIPLEEVAK | 10 | P29597 | Non-receptor tyrosine-protein kinase TYI | TYK2 | 133.65 | 2 | 53.913025 | yes | 9.722E-08 | 546.80603 | 134.61842 |
| LSDPGVGLGALSR | 13 | P29597 | Non-receptor tyrosine-protein kinase TYI | TYK2 | 133.65 | 2 | 53.913025 | yes | 1.334E-07 | 621.34607 | 94.641647 |
| SQAPDGMQSLR | 11 | P29597 | Non-receptor tyrosine-protein kinase TYI | TYK2 | 133.65 | 2 | 53.913025 | yes | 1.517E-07 | 595.28516 | 87.742836 |
| VPVCHLR | 7 | P29597 | Non-receptor tyrosine-protein kinase TYI | TYK2 | 133.65 | 2 | 53.913025 | yes | 4.388E-07 | 440.74472 | 84.702232 |
| EITQLSHLGQGTR | 13 | P29597 | Non-receptor tyrosine-protein kinase TYI | TYK2 | 133.65 | 2 | 53.913025 | yes | 4.209E-07 | 720.38367 | 62.106754 |
| LGLAEGTSPFIK | 12 | P29597 | Non-receptor tyrosine-protein kinase TYI | TYK2 | 133.65 | 2 | 53.913025 | yes | 1.438E-07 | 616.8479 | 59.271042 |
| DITHVVLK | 8 | P29597 | Non-receptor tyrosine-protein kinase TYI | TYK2 | 133.65 | 2 | 53.913025 | yes | 3.262E-06 | 462.7793 | 33.892014 |
| TLYHEHIIK | 9 | P29597 | Non-receptor tyrosine-protein kinase TYI | TYK2 | 133.65 | 2 | 53.913025 | yes | 2.928E-06 | 577.32184 | 20.852621 |
| IASLEVENQSLR | 12 | P29692 | Elongation factor 1-delta OS=Homo sapi | EEF1D | 31.122 | 2 | 51.86479 | yes | 1.067E-07 | 679.86737 | 11430.96 |
| LVPVGYGIR | 9 | P29692 | Elongation factor 1-delta OS=Homo sapi | EEF1D | 31.122 | 2 | 51.86479 | yes | 2.674E-07 | 487.2951 | 11264.916 |
| ATNFLAHEK | 9 | P29692 | Elongation factor 1-delta OS=Homo sapi | EEF1D | 31.122 | 2 | 51.86479 | yes | 1.328E-07 | 536.77472 | 10230.637 |
| GVVQELQQAISK | 12 | P29692 | Elongation factor 1-delta OS=Homo sapi | EEF1D | 31.122 | 2 | 51.86479 | yes | 9.722E-08 | 650.367 | 5492.9229 |
| SIQLDGLVWGASK | 13 | P29692 | Elongation factor 1-delta OS=Homo sapi | EEF1D | 31.122 | 2 | 51.86479 | yes | 1.027E-07 | 687.37482 | 5427.1694 |
| MATNFLAHEK | 10 | P29692 | Elongation factor 1-delta OS=Homo sapi | EEF1D | 31.122 | 2 | 51.86479 | yes | 9.919E-08 | 581.28967 | 1373.7227 |
| MATNFLAHEK | 10 | P29692 | Elongation factor 1-delta OS=Homo sapi | EEF1D | 31.122 | 2 | 51.86479 | yes | 2.764E-07 | 589.28717 | 1373.7227 |
| FYEQMNGPVAGA | 14 | P29692 | Elongation factor 1-delta OS=Homo sapi | EEF1D | 31.122 | 2 | 51.86479 | yes | 3.376E-07 | 763.85645 | 1280.6351 |

| | | | | | | | | | | | |
|---|---|---|---|---|---|---|---|---|---|---|---|
| FYEQMNGPVAGA | 14 | P29692 | Elongation factor 1-delta OS=Homo sapic | EEF1D | 31.122 | 2 | 51.86479 | yes | 1.138E-07 | 771.85394 | 1280.6351 |
| LQIQCVVEDDK | 11 | P29692 | Elongation factor 1-delta OS=Homo sapic | EEF1D | 31.122 | 2 | 51.86479 | no | 9.722E-08 | 673.83466 | 2455.5784 |
| ATAPQTQHVSPMI | 13 | P29692 | Elongation factor 1-delta OS=Homo sapic | EEF1D | 31.122 | 3 | 51.86479 | yes | 1.665E-07 | 475.24188 | 748.13007 |
| ATAPQTQHVSPMI | 13 | P29692 | Elongation factor 1-delta OS=Homo sapic | EEF1D | 31.122 | 3 | 51.86479 | yes | 1.038E-07 | 712.35919 | 748.13007 |
| ATAPQTQHVSPMI | 13 | P29692 | Elongation factor 1-delta OS=Homo sapic | EEF1D | 31.122 | 3 | 51.86479 | yes | 1.201E-07 | 720.35663 | 748.13007 |
| ATAPQTQHVSPMI | 13 | P29692 | Elongation factor 1-delta OS=Homo sapic | EEF1D | 31.122 | 2 | 51.86479 | yes | 1.665E-07 | 475.24188 | 748.13007 |
| ATAPQTQHVSPMI | 13 | P29692 | Elongation factor 1-delta OS=Homo sapic | EEF1D | 31.122 | 2 | 51.86479 | yes | 1.038E-07 | 712.35919 | 748.13007 |
| ATAPQTQHVSPMI | 13 | P29692 | Elongation factor 1-delta OS=Homo sapic | EEF1D | 31.122 | 2 | 51.86479 | yes | 1.201E-07 | 720.35663 | 748.13007 |
| SLAGSSGPGASSG | 24 | P29692 | Elongation factor 1-delta OS=Homo sapic | EEF1D | 31.122 | 3 | 51.86479 | yes | 1.388E-07 | 729.02087 | 1021.9304 |
| VGTDLLEEEITK | 12 | P29692 | Elongation factor 1-delta OS=Homo sapic | EEF1D | 31.122 | 2 | 51.86479 | yes | 1.753E-07 | 673.85614 | 623.27893 |
| QENGASVILR | 10 | P29692 | Elongation factor 1-delta OS=Homo sapic | EEF1D | 31.122 | 2 | 51.86479 | yes | 1.832E-07 | 543.79877 | 613.5459 |
| KLQIQCVVEDDK | 12 | P29692 | Elongation factor 1-delta OS=Homo sapic | EEF1D | 31.122 | 2 | 51.86479 | no | 9.934E-08 | 737.88214 | 400.64502 |
| FKYDDAER | 8 | P29692 | Elongation factor 1-delta OS=Homo sapic | EEF1D | 31.122 | 2 | 51.86479 | yes | 2.099E-07 | 522.24329 | 277.75314 |
| GEPAAAAAPEAGA | 18 | P29966 | Myristoylated alanine-rich C-kinase subst | MARCKS | 31.555 | 2 | 34.771633 | yes | 9.442E-07 | 811.90466 | 50.301857 |
| LPFPIIDDR | 9 | P30041 | Peroxiredoxin-6 OS=Homo sapiens OX= | PRDX6 | 25.035 | 2 | 49.523808 | yes | 6.65E-07 | 543.3031 | 415.20459 |
| VVISLQLTAEK | 11 | P30041 | Peroxiredoxin-6 OS=Homo sapiens OX= | PRDX6 | 25.035 | 2 | 49.523808 | yes | 1.339E-07 | 600.86353 | 320.31186 |
| VVFVFGPDKK | 10 | P30041 | Peroxiredoxin-6 OS=Homo sapiens OX= | PRDX6 | 25.035 | 2 | 49.523808 | yes | 3.304E-07 | 568.32916 | 93.691849 |
| DINAYNCEEPTEK | 13 | P30041 | Peroxiredoxin-6 OS=Homo sapiens OX= | PRDX6 | 25.035 | 2 | 49.523808 | yes | 3.716E-07 | 791.83813 | 76.588982 |
| FHDFLGDSWGILF | 17 | P30041 | Peroxiredoxin-6 OS=Homo sapiens OX= | PRDX6 | 25.035 | 3 | 49.523808 | yes | 1.022E-07 | 677.66724 | 71.672073 |
| VVFVFGPDK | 9 | P30041 | Peroxiredoxin-6 OS=Homo sapiens OX= | PRDX6 | 25.035 | 2 | 49.523808 | yes | 1.632E-07 | 504.28165 | 70.631363 |
| NFDEILR | 7 | P30041 | Peroxiredoxin-6 OS=Homo sapiens OX= | PRDX6 | 25.035 | 2 | 49.523808 | yes | 5.005E-07 | 453.73761 | 21.756193 |
| FSMVVQDGIVK | 11 | P30044 | Peroxiredoxin-5, mitochondrial OS=Hom | PRDX5 | 22.087 | 2 | 45.177948 | yes | 1.934E-07 | 611.82867 | 107.51529 |
| DYGVLLEGSGLAL | 14 | P30048 | Thioredoxin-dependent peroxide reductas | PRDX3 | 27.693 | 2 | 52.42316 | yes | 9.722E-08 | 731.89862 | 338.04495 |
| HLSVNDLPVGR | 11 | P30048 | Thioredoxin-dependent peroxide reductas | PRDX3 | 27.693 | 2 | 52.42316 | yes | 1.376E-07 | 603.83313 | 198.1057 |
| GTAVVNGEFK | 10 | P30048 | Thioredoxin-dependent peroxide reductas | PRDX3 | 27.693 | 2 | 52.42316 | yes | 1.503E-07 | 511.26929 | 103.61997 |
| AFQYVETHGEVCI | 31 | P30048 | Thioredoxin-dependent peroxide reductas | PRDX3 | 27.693 | 4 | 52.42316 | yes | 1.4E-07 | 847.15942 | 97.441818 |
| GLFIIDPNGVIK | 12 | P30048 | Thioredoxin-dependent peroxide reductas | PRDX3 | 27.693 | 2 | 52.42316 | yes | 4.689E-07 | 643.37933 | 52.305012 |
| IEANEALVK | 9 | P30049 | ATP synthase subunit delta, mitochondria | ATP5F1D | 17.49 | 2 | 54.79944 | yes | 1.757E-07 | 493.77948 | 235.35316 |
| AQAELVGTADEA | 14 | P30049 | ATP synthase subunit delta, mitochondria | ATP5F1D | 17.49 | 2 | 54.79944 | yes | 1.866E-07 | 716.35736 | 53.036446 |
| PGLVVVHAEDGT | 15 | P30049 | ATP synthase subunit delta, mitochondria | ATP5F1D | 17.49 | 2 | 54.79944 | yes | 9.722E-08 | 755.39899 | 50.687244 |
| IGPLGLSPK | 9 | P30050 | Large ribosomal subunit protein uL11 OS | RPL12 | 17.819 | 2 | 50.314846 | yes | 3.271E-07 | 441.27637 | 40669.855 |
| QAQIEVVPSASAL | 16 | P30050 | Large ribosomal subunit protein uL11 OS | RPL12 | 17.819 | 2 | 50.314846 | yes | 4.168E-07 | 833.99054 | 8151.793 |
| QAQIEVVPSASAL | 16 | P30050 | Large ribosomal subunit protein uL11 OS | RPL12 | 17.819 | 2 | 50.314846 | yes | 1.499E-07 | 556.32947 | 8151.793 |
| QAQIEVVPSASAL | 16 | P30050 | Large ribosomal subunit protein uL11 OS | RPL12 | 17.819 | 3 | 50.314846 | yes | 4.168E-07 | 833.99054 | 8151.793 |
| QAQIEVVPSASAL | 16 | P30050 | Large ribosomal subunit protein uL11 OS | RPL12 | 17.819 | 3 | 50.314846 | yes | 1.499E-07 | 556.32947 | 8151.793 |
| HSGNITFDEIVNIA | 15 | P30050 | Large ribosomal subunit protein uL11 OS | RPL12 | 17.819 | 3 | 50.314846 | yes | 9.819E-08 | 562.62506 | 7325.0942 |
| HSGNITFDEIVNIA | 15 | P30050 | Large ribosomal subunit protein uL11 OS | RPL12 | 17.819 | 3 | 50.314846 | yes | 4.527E-07 | 843.4339 | 7325.0942 |
| HSGNITFDEIVNIA | 15 | P30050 | Large ribosomal subunit protein uL11 OS | RPL12 | 17.819 | 2 | 50.314846 | yes | 9.819E-08 | 562.62506 | 7325.0942 |
| HSGNITFDEIVNIA | 15 | P30050 | Large ribosomal subunit protein uL11 OS | RPL12 | 17.819 | 2 | 50.314846 | yes | 4.527E-07 | 843.4339 | 7325.0942 |

| | | | | | | | | | | | |
|---|---|---|---|---|---|---|---|---|---|---|---|
| KVGDDIAK | 8 | P30050 | Large ribosomal subunit protein uL11 OS | RPL12 | 17.819 | 2 | 50.314846 | yes | 1.578E-07 | 423.23999 | 14433.519 |
| EILGTAQSVGCNV | 16 | P30050 | Large ribosomal subunit protein uL11 OS | RPL12 | 17.819 | 2 | 50.314846 | yes | 4.192E-07 | 838.40704 | 9237.8652 |
| CTGGEVGATSALA | 15 | P30050 | Large ribosomal subunit protein uL11 OS | RPL12 | 17.819 | 2 | 50.314846 | yes | 3.035E-07 | 709.85083 | 1057.365 |
| FDPNEIK | 7 | P30050 | Large ribosomal subunit protein uL11 OS | RPL12 | 17.819 | 2 | 50.314846 | yes | 3.857E-07 | 431.7189 | 166.39009 |
| FVMQEEFSR | 9 | P30101 | Protein disulfide-isomerase A3 OS=Homo | PDIA3 | 56.783 | 2 | 47.304703 | yes | 2.279E-07 | 586.77386 | 128.22211 |
| FLQDYFDGNLK | 11 | P30101 | Protein disulfide-isomerase A3 OS=Homo | PDIA3 | 56.783 | 2 | 47.304703 | yes | 3.671E-07 | 680.33258 | 92.056618 |
| SEPIPESNDGPVK | 13 | P30101 | Protein disulfide-isomerase A3 OS=Homo | PDIA3 | 56.783 | 2 | 47.304703 | yes | 1.358E-07 | 684.83569 | 88.331718 |
| LNFAVASR | 8 | P30101 | Protein disulfide-isomerase A3 OS=Homo | PDIA3 | 56.783 | 2 | 47.304703 | yes | 4.84E-07 | 439.24814 | 54.881462 |
| YGVSGYPTLK | 10 | P30101 | Protein disulfide-isomerase A3 OS=Homo | PDIA3 | 56.783 | 2 | 47.304703 | yes | 2.664E-07 | 542.78729 | 52.705063 |
| ELSDFISYLQR | 11 | P30101 | Protein disulfide-isomerase A3 OS=Homo | PDIA3 | 56.783 | 2 | 47.304703 | yes | 2.69E-07 | 685.8512 | 42.473476 |
| TADGIVSHLK | 10 | P30101 | Protein disulfide-isomerase A3 OS=Homo | PDIA3 | 56.783 | 2 | 47.304703 | yes | 3.453E-07 | 520.79041 | 41.784176 |
| DLLIAYYDVDYEK | 13 | P30101 | Protein disulfide-isomerase A3 OS=Homo | PDIA3 | 56.783 | 2 | 47.304703 | yes | 4.411E-06 | 810.39563 | 38.467449 |
| MDATANDVPSPYI | 15 | P30101 | Protein disulfide-isomerase A3 OS=Homo | PDIA3 | 56.783 | 2 | 47.304703 | yes | 6.547E-07 | 832.88287 | 26.375374 |
| TSACGLFSVCYPR | 13 | P30153 | Serine/threonine-protein phosphatase 2A | PPP2R1A | 65.309 | 2 | 53.81717 | no | 9.968E-08 | 759.34741 | 1271.0134 |
| IGPILDNSTLQSEV | 20 | P30153 | Serine/threonine-protein phosphatase 2A | PPP2R1A | 65.309 | 3 | 53.81717 | yes | 1.946E-07 | 732.08252 | 1117.1372 |
| EWAHATIIPK | 10 | P30153 | Serine/threonine-protein phosphatase 2A | PPP2R1A | 65.309 | 2 | 53.81717 | yes | 2.093E-07 | 583.32184 | 1094.0392 |
| ELVSDANQHVK | 11 | P30153 | Serine/threonine-protein phosphatase 2A | PPP2R1A | 65.309 | 2 | 53.81717 | yes | 9.722E-08 | 620.32001 | 633.75269 |
| MAGDPVANVR | 10 | P30153 | Serine/threonine-protein phosphatase 2A | PPP2R1A | 65.309 | 2 | 53.81717 | yes | 1.372E-07 | 515.26093 | 556.81506 |
| TDLVPAFQNLMK | 12 | P30153 | Serine/threonine-protein phosphatase 2A | PPP2R1A | 65.309 | 2 | 53.81717 | yes | 1.388E-07 | 688.86578 | 547.09924 |
| ENVIMSQILPCIK | 13 | P30153 | Serine/threonine-protein phosphatase 2A | PPP2R1A | 65.309 | 2 | 53.81717 | yes | 1.091E-07 | 772.91241 | 522.93439 |
| VLELDNVK | 8 | P30153 | Serine/threonine-protein phosphatase 2A | PPP2R1A | 65.309 | 2 | 53.81717 | yes | 2.51E-07 | 465.26874 | 507.54416 |
| NLCSDDTPMVR | 11 | P30153 | Serine/threonine-protein phosphatase 2A | PPP2R1A | 65.309 | 2 | 53.81717 | yes | 1.223E-07 | 654.28955 | 312.1004 |
| EFCENLSADCR | 11 | P30153 | Serine/threonine-protein phosphatase 2A | PPP2R1A | 65.309 | 2 | 53.81717 | yes | 1.025E-07 | 700.7821 | 291.25485 |
| NEDVQLR | 7 | P30153 | Serine/threonine-protein phosphatase 2A | PPP2R1A | 65.309 | 2 | 53.81717 | no | 1.877E-07 | 437.22488 | 157.06139 |
| VLAMSGDPNYLHI | 13 | P30153 | Serine/threonine-protein phosphatase 2A | PPP2R1A | 65.309 | 2 | 53.81717 | yes | 3.138E-07 | 736.86938 | 149.07881 |
| AISHEHSPSDLEAH | 20 | P30153 | Serine/threonine-protein phosphatase 2A | PPP2R1A | 65.309 | 3 | 53.81717 | yes | 2.247E-07 | 738.38318 | 66.898659 |
| AISHEHSPSDLEAH | 20 | P30153 | Serine/threonine-protein phosphatase 2A | PPP2R1A | 65.309 | 3 | 53.81717 | yes | 2.167E-07 | 554.03918 | 66.898659 |
| AISHEHSPSDLEAH | 20 | P30153 | Serine/threonine-protein phosphatase 2A | PPP2R1A | 65.309 | 4 | 53.81717 | yes | 2.247E-07 | 738.38318 | 66.898659 |
| AISHEHSPSDLEAH | 20 | P30153 | Serine/threonine-protein phosphatase 2A | PPP2R1A | 65.309 | 4 | 53.81717 | yes | 2.167E-07 | 554.03918 | 66.898659 |
| KLSTIALALGVER | 13 | P30153 | Serine/threonine-protein phosphatase 2A | PPP2R1A | 65.309 | 2 | 53.81717 | no | 9.722E-08 | 685.92194 | 52.069054 |
| KLSTIALALGVER | 13 | P30153 | Serine/threonine-protein phosphatase 2A | PPP2R1A | 65.309 | 2 | 53.81717 | no | 1.296E-07 | 457.61703 | 52.069054 |
| KLSTIALALGVER | 13 | P30153 | Serine/threonine-protein phosphatase 2A | PPP2R1A | 65.309 | 3 | 53.81717 | no | 9.722E-08 | 685.92194 | 52.069054 |
| KLSTIALALGVER | 13 | P30153 | Serine/threonine-protein phosphatase 2A | PPP2R1A | 65.309 | 3 | 53.81717 | no | 1.296E-07 | 457.61703 | 52.069054 |
| DKAVESLR | 8 | P30153 | Serine/threonine-protein phosphatase 2A | PPP2R1A | 65.309 | 2 | 53.81717 | no | 2.76E-07 | 459.25616 | 98.513 |
| LSTIALALGVER | 12 | P30153 | Serine/threonine-protein phosphatase 2A | PPP2R1A | 65.309 | 2 | 53.81717 | no | 3.089E-06 | 621.87445 | 43.528362 |
| LTQDQDVDVK | 10 | P30153 | Serine/threonine-protein phosphatase 2A | PPP2R1A | 65.309 | 2 | 53.81717 | yes | 9.722E-08 | 580.79333 | 21.794529 |
| VKEFCENLSADCR | 13 | P30153 | Serine/threonine-protein phosphatase 2A | PPP2R1A | 65.309 | 2 | 53.81717 | yes | 3.132E-06 | 814.36377 | 10.234884 |
| ELGENLPIEDR | 11 | P30154 | Serine/threonine-protein phosphatase 2A | PPP2R1B | 66.214 | 2 | 46.053757 | yes | 9.526E-07 | 642.82513 | 95.59185 |
| IGPILDTNALQGEV | 20 | P30154 | Serine/threonine-protein phosphatase 2A | PPP2R1B | 66.214 | 3 | 46.053757 | yes | 1.666E-07 | 711.74744 | 60.938248 |

| | | | | | | | | | | | |
|---|---|---|---|---|---|---|---|---|---|---|---|
| LGQDEDMDVK | 10 | P30154 | Serine/threonine-protein phosphatase 2A | PPP2R1B | 66.214 | 2 | 46.053757 | yes | 1.998E-07 | 575.25824 | 57.145428 |
| LAEGEQILSGGVFI | 15 | P30260 | Cell division cycle protein 27 homolog O | CDC27 | 91.868 | 2 | 54.101112 | yes | 1.008E-07 | 781.41467 | 325.03171 |
| SALQELEELK | 10 | P30260 | Cell division cycle protein 27 homolog O | CDC27 | 91.868 | 2 | 54.101112 | yes | 1.361E-07 | 580.3139 | 263.33002 |
| GYLALCSYNCK | 11 | P30260 | Cell division cycle protein 27 homolog O | CDC27 | 91.868 | 2 | 54.101112 | yes | 1.308E-07 | 674.80487 | 173.73486 |
| LDSSIISEGK | 10 | P30260 | Cell division cycle protein 27 homolog O | CDC27 | 91.868 | 2 | 54.101112 | yes | 4.024E-07 | 524.77966 | 166.16046 |
| AYFELSEYMQAEI | 13 | P30260 | Cell division cycle protein 27 homolog O | CDC27 | 91.868 | 2 | 54.101112 | yes | 9.722E-08 | 818.8692 | 141.12498 |
| CCVDLSK | 7 | P30260 | Cell division cycle protein 27 homolog O | CDC27 | 91.868 | 2 | 54.101112 | yes | 1.764E-06 | 441.19641 | 111.56519 |
| ASVLFANEK | 9 | P30260 | Cell division cycle protein 27 homolog O | CDC27 | 91.868 | 2 | 54.101112 | yes | 2.98E-07 | 489.76639 | 104.12268 |
| LNLESSNSK | 9 | P30260 | Cell division cycle protein 27 homolog O | CDC27 | 91.868 | 2 | 54.101112 | yes | 3.387E-06 | 496.25638 | 77.605232 |
| AAAEGLMSLLR | 11 | P30260 | Cell division cycle protein 27 homolog O | CDC27 | 91.868 | 2 | 54.101112 | yes | 2.364E-07 | 566.31317 | 60.131523 |
| FSLAEMHFQK | 10 | P30260 | Cell division cycle protein 27 homolog O | CDC27 | 91.868 | 2 | 54.101112 | yes | 1.196E-07 | 619.30536 | 59.471474 |
| VMIHPDPER | 9 | P30291 | Wee1-like protein kinase OS=Homo sapi | WEE1 | 71.598 | 2 | 53.279484 | yes | 3.12E-07 | 547.27661 | 498.82358 |
| IGSGEFGSVFK | 11 | P30291 | Wee1-like protein kinase OS=Homo sapi | WEE1 | 71.598 | 2 | 53.279484 | yes | 9.722E-08 | 564.29022 | 329.14691 |
| HSVLLSASR | 9 | P30291 | Wee1-like protein kinase OS=Homo sapi | WEE1 | 71.598 | 2 | 53.279484 | yes | 1.087E-07 | 485.27744 | 265.94717 |
| GSSLFMDTEK | 10 | P30291 | Wee1-like protein kinase OS=Homo sapi | WEE1 | 71.598 | 2 | 53.279484 | yes | 1.114E-07 | 557.75787 | 256.38831 |
| YTTEFHELEK | 10 | P30291 | Wee1-like protein kinase OS=Homo sapi | WEE1 | 71.598 | 2 | 53.279484 | yes | 9.722E-08 | 648.80896 | 241.47435 |
| IGDLGHVTR | 9 | P30291 | Wee1-like protein kinase OS=Homo sapi | WEE1 | 71.598 | 2 | 53.279484 | yes | 1.209E-07 | 484.26962 | 240.13379 |
| ITITESNMK | 9 | P30291 | Wee1-like protein kinase OS=Homo sapi | WEE1 | 71.598 | 2 | 53.279484 | yes | 2.03E-07 | 518.77081 | 149.22212 |
| ISSPQVEEGDSR | 12 | P30291 | Wee1-like protein kinase OS=Homo sapi | WEE1 | 71.598 | 2 | 53.279484 | yes | 6.187E-07 | 652.30988 | 82.478401 |
| SPAAPYFLGSSFSP | 16 | P30291 | Wee1-like protein kinase OS=Homo sapi | WEE1 | 71.598 | 2 | 53.279484 | yes | 4.68E-07 | 841.93048 | 64.574768 |
| DLLLQVGR | 8 | P30291 | Wee1-like protein kinase OS=Homo sapi | WEE1 | 71.598 | 2 | 53.279484 | yes | 2.112E-07 | 457.27692 | 40.67551 |
| LFDTPHTPK | 9 | P30291 | Wee1-like protein kinase OS=Homo sapi | WEE1 | 71.598 | 2 | 53.279484 | yes | 9.722E-08 | 528.27966 | 28.133312 |
| LDGCIYAIK | 9 | P30291 | Wee1-like protein kinase OS=Homo sapi | WEE1 | 71.598 | 2 | 53.279484 | yes | 2.45E-06 | 526.77588 | 26.416893 |
| LDTPDINIVLK | 11 | P30414 | NK-tumor recognition protein OS=Homo | NKTR | 165.677 | 2 | 47.53534 | yes | 1.31E-07 | 620.86102 | 141.19376 |
| DMPVVTAEPEPK | 12 | P30414 | NK-tumor recognition protein OS=Homo | NKTR | 165.677 | 2 | 47.53534 | yes | 2.167E-07 | 656.82629 | 125.30826 |
| VLTTVPEMKPQGI | 14 | P30414 | NK-tumor recognition protein OS=Homo | NKTR | 165.677 | 2 | 47.53534 | yes | 3.072E-07 | 784.94507 | 95.798141 |
| QDMATEHPQAEV | 14 | P30414 | NK-tumor recognition protein OS=Homo | NKTR | 165.677 | 2 | 47.53534 | yes | 2.953E-07 | 791.88013 | 25.814056 |
| TATITVLPQQPR | 12 | P30530 | Tyrosine-protein kinase receptor UFO OS | AXL | 98.337 | 2 | 45.369946 | yes | 2.149E-07 | 662.88281 | 124.8848 |
| YGEVFEPTVER | 11 | P30530 | Tyrosine-protein kinase receptor UFO OS | AXL | 98.337 | 2 | 45.369946 | yes | 2.225E-07 | 663.32227 | 102.16421 |
| FMADIASGMEYLS | 15 | P30530 | Tyrosine-protein kinase receptor UFO OS | AXL | 98.337 | 2 | 45.369946 | yes | 1.871E-07 | 832.38892 | 51.70137 |
| APLQGTLLGYR | 11 | P30530 | Tyrosine-protein kinase receptor UFO OS | AXL | 98.337 | 2 | 45.369946 | yes | 4.106E-05 | 594.84039 | 42.691605 |
| SELEDFLSEAVCM | 14 | P30530 | Tyrosine-protein kinase receptor UFO OS | AXL | 98.337 | 2 | 45.369946 | yes | 2.285E-07 | 829.37604 | 30.907707 |
| WIAIESLADR | 10 | P30530 | Tyrosine-protein kinase receptor UFO OS | AXL | 98.337 | 2 | 45.369946 | no | 2.365E-07 | 587.31677 | 25.739998 |
| FVHGEGLR | 8 | P30536 | Translocator protein OS=Homo sapiens C | TSPO | 18.828 | 2 | 35.937057 | yes | 7.618E-07 | 457.74579 | 389.51556 |
| GEWCGVELDEPLC | 14 | P30622 | CAP-Gly domain-containing linker protei | CLIP1 | 162.247 | 2 | 47.572884 | yes | 1.301E-07 | 794.8692 | 54.183208 |
| ITFDDYIACCVK | 12 | P30626 | Sorcin OS=Homo sapiens OX=9606 GN= | SRI | 21.677 | 2 | 44.00199 | yes | 2.361E-07 | 752.84418 | 67.191849 |
| DMSGTMGFNEFK | 12 | P30626 | Sorcin OS=Homo sapiens OX=9606 GN= | SRI | 21.677 | 2 | 44.00199 | yes | 4.406E-07 | 682.2865 | 42.99641 |
| LMVSMLDR | 8 | P30626 | Sorcin OS=Homo sapiens OX=9606 GN= | SRI | 21.677 | 2 | 44.00199 | yes | 3.205E-06 | 482.75134 | 28.548328 |
| YSLATGNWGDQK | 12 | P30876 | DNA-directed RNA polymerase II subuni | POLR2B | 133.897 | 2 | 52.172474 | yes | 1.028E-07 | 670.3175 | 154.36827 |

| | | | | | | | | | | | |
|---|---|---|---|---|---|---|---|---|---|---|---|
| LDDDGLIAPGVR | 12 | P30876 | DNA-directed RNA polymerase II subuni | POLR2B | 133.897 | 2 | 52.172474 | yes | 1.002E-07 | 620.83026 | 106.33171 |
| GPIQILNR | 8 | P30876 | DNA-directed RNA polymerase II subuni | POLR2B | 133.897 | 2 | 52.172474 | yes | 9.722E-08 | 455.77707 | 95.908623 |
| NLTYSAPLYVDITI | 14 | P30876 | DNA-directed RNA polymerase II subuni | POLR2B | 133.897 | 2 | 52.172474 | yes | 1.108E-07 | 799.42725 | 88.859474 |
| IVATLPYIK | 9 | P30876 | DNA-directed RNA polymerase II subuni | POLR2B | 133.897 | 2 | 52.172474 | yes | 9.182E-07 | 509.3208 | 78.254715 |
| VSGDDVIIGK | 10 | P30876 | DNA-directed RNA polymerase II subuni | POLR2B | 133.897 | 2 | 52.172474 | yes | 5.253E-07 | 501.77695 | 58.982857 |
| PTSTIWVSMLAR | 12 | P30876 | DNA-directed RNA polymerase II subuni | POLR2B | 133.897 | 2 | 52.172474 | yes | 2.656E-07 | 681.36591 | 58.666801 |
| GNEVLYNGFTGR | 12 | P30876 | DNA-directed RNA polymerase II subuni | POLR2B | 133.897 | 2 | 52.172474 | yes | 1.932E-07 | 663.8255 | 49.954994 |
| DCQIAHGAAQFLR | 13 | P30876 | DNA-directed RNA polymerase II subuni | POLR2B | 133.897 | 2 | 52.172474 | yes | 1.655E-07 | 743.86462 | 37.84959 |
| ICRPLLIVEK | 10 | P30876 | DNA-directed RNA polymerase II subuni | POLR2B | 133.897 | 2 | 52.172474 | yes | 3.494E-07 | 620.87592 | 31.769665 |
| PLLIVEK | 7 | P30876 | DNA-directed RNA polymerase II subuni | POLR2B | 133.897 | 2 | 52.172474 | yes | 2.819E-06 | 406.26804 | 25.266577 |
| QMDIIVSEVSMIR | 13 | P30876 | DNA-directed RNA polymerase II subuni | POLR2B | 133.897 | 2 | 52.172474 | yes | 2.773E-05 | 760.89417 | 18.301571 |
| GVIALCIEDGSIHR | 14 | P31040 | Succinate dehydrogenase [ubiquinone] fla | SDHA | 72.692 | 2 | 53.719486 | yes | 1.388E-07 | 770.40106 | 313.55481 |
| VGSVLQEGCGK | 11 | P31040 | Succinate dehydrogenase [ubiquinone] fla | SDHA | 72.692 | 2 | 53.719486 | yes | 1.684E-07 | 567.28461 | 215.87996 |
| TGHSLLHTLYGR | 12 | P31040 | Succinate dehydrogenase [ubiquinone] fla | SDHA | 72.692 | 3 | 53.719486 | yes | 1.678E-07 | 452.24573 | 81.18425 |
| TGHSLLHTLYGR | 12 | P31040 | Succinate dehydrogenase [ubiquinone] fla | SDHA | 72.692 | 3 | 53.719486 | yes | 1.104E-07 | 677.86493 | 81.18425 |
| TGHSLLHTLYGR | 12 | P31040 | Succinate dehydrogenase [ubiquinone] fla | SDHA | 72.692 | 2 | 53.719486 | yes | 1.678E-07 | 452.24573 | 81.18425 |
| TGHSLLHTLYGR | 12 | P31040 | Succinate dehydrogenase [ubiquinone] fla | SDHA | 72.692 | 2 | 53.719486 | yes | 1.104E-07 | 677.86493 | 81.18425 |
| NTVVATGGYGR | 11 | P31040 | Succinate dehydrogenase [ubiquinone] fla | SDHA | 72.692 | 2 | 53.719486 | yes | 3.84E-06 | 547.78308 | 85.076126 |
| TLNEADCATVPPA | 15 | P31040 | Succinate dehydrogenase [ubiquinone] fla | SDHA | 72.692 | 2 | 53.719486 | yes | 4.89E-07 | 814.40906 | 74.65757 |
| GEGGILINSQGER | 13 | P31040 | Succinate dehydrogenase [ubiquinone] fla | SDHA | 72.692 | 2 | 53.719486 | yes | 1.026E-07 | 665.34149 | 54.335934 |
| EPIPVLPTVHYNM | 21 | P31040 | Succinate dehydrogenase [ubiquinone] fla | SDHA | 72.692 | 3 | 53.719486 | yes | 3.928E-07 | 780.74005 | 37.245895 |
| WHFYDTVK | 8 | P31040 | Succinate dehydrogenase [ubiquinone] fla | SDHA | 72.692 | 2 | 53.719486 | yes | 1.282E-07 | 548.26654 | 34.722786 |
| HTLSYVDVGTGK | 12 | P31040 | Succinate dehydrogenase [ubiquinone] fla | SDHA | 72.692 | 2 | 53.719486 | yes | 1.923E-07 | 638.83026 | 33.049316 |
| KHTLSYVDVGTGI | 13 | P31040 | Succinate dehydrogenase [ubiquinone] fla | SDHA | 72.692 | 2 | 53.719486 | yes | 9.722E-08 | 702.87769 | 28.193157 |
| ACALSIEESCRPGI | 15 | P31040 | Succinate dehydrogenase [ubiquinone] fla | SDHA | 72.692 | 2 | 53.719486 | yes | 1.113E-07 | 846.88763 | 26.601751 |
| TGMILLAGEITSR | 13 | P31153 | S-adenosylmethionine synthase isoform ty | MAT2A | 43.661 | 2 | 54.2022 | yes | 9.722E-08 | 681.37646 | 572.97833 |
| FVIGGPQGDAGLT | 15 | P31153 | S-adenosylmethionine synthase isoform ty | MAT2A | 43.661 | 2 | 54.2022 | yes | 3.299E-07 | 722.8808 | 265.02185 |
| TAAYGHFGR | 9 | P31153 | S-adenosylmethionine synthase isoform ty | MAT2A | 43.661 | 2 | 54.2022 | yes | 1.015E-07 | 490.24084 | 204.89734 |
| NFDLRPGVIVR | 11 | P31153 | S-adenosylmethionine synthase isoform ty | MAT2A | 43.661 | 2 | 54.2022 | no | 9.903E-08 | 643.37238 | 102.40897 |
| NFDLRPGVIVR | 11 | P31153 | S-adenosylmethionine synthase isoform ty | MAT2A | 43.661 | 2 | 54.2022 | no | 9.722E-08 | 429.2507 | 102.40897 |
| NFDLRPGVIVR | 11 | P31153 | S-adenosylmethionine synthase isoform ty | MAT2A | 43.661 | 3 | 54.2022 | no | 9.903E-08 | 643.37238 | 102.40897 |
| NFDLRPGVIVR | 11 | P31153 | S-adenosylmethionine synthase isoform ty | MAT2A | 43.661 | 3 | 54.2022 | no | 9.722E-08 | 429.2507 | 102.40897 |
| VACETVAK | 8 | P31153 | S-adenosylmethionine synthase isoform ty | MAT2A | 43.661 | 2 | 54.2022 | yes | 1.937E-07 | 439.22601 | 134.4128 |
| TQVTVQYMQDR | 11 | P31153 | S-adenosylmethionine synthase isoform ty | MAT2A | 43.661 | 2 | 54.2022 | yes | 9.722E-08 | 684.83246 | 127.47906 |
| YLDEDTIYHLQPS( | 15 | P31153 | S-adenosylmethionine synthase isoform ty | MAT2A | 43.661 | 3 | 54.2022 | yes | 1.573E-07 | 602.96008 | 121.97952 |
| DSFPWEVPK | 9 | P31153 | S-adenosylmethionine synthase isoform ty | MAT2A | 43.661 | 2 | 54.2022 | yes | 2.425E-07 | 552.77167 | 81.442291 |
| IIVDTYGGWGAHG | 20 | P31153 | S-adenosylmethionine synthase isoform ty | MAT2A | 43.661 | 3 | 54.2022 | no | 2.782E-07 | 650.65497 | 44.4104 |
| SYPDYMYGSPGEI | 14 | P31274 | Homeobox protein Hox-C9 OS=Homo sa | HOXC9 | 29.248 | 2 | 47.080936 | yes | 1.747E-07 | 817.86139 | 235.67877 |
| APQTLPSPEADAL | 17 | P31274 | Homeobox protein Hox-C9 OS=Homo sa | HOXC9 | 29.248 | 2 | 47.080936 | yes | 1.405E-07 | 826.92816 | 193.93115 |

| | | | | | | | | | | | |
|---|---|---|---|---|---|---|---|---|---|---|---|
| FPATGAHPAAAR | 12 | P31274 | Homeobox protein Hox-C9 OS=Homo sa | HOXC9 | 29.248 | 2 | 47.080936 | yes | 5.795E-07 | 583.80688 | 74.614258 |
| VLGTSVESIMATEI | 15 | P31327 | Carbamoyl-phosphate synthase [ammonia | CPS1 | 164.94 | 2 | 54.611835 | yes | 9.79E-08 | 804.40088 | 690.87329 |
| GQNQPVLNITNK | 12 | P31327 | Carbamoyl-phosphate synthase [ammonia | CPS1 | 164.94 | 2 | 54.611835 | yes | 9.722E-08 | 663.36224 | 576.35339 |
| SLGQWLQEEK | 10 | P31327 | Carbamoyl-phosphate synthase [ammonia | CPS1 | 164.94 | 2 | 54.611835 | yes | 1.086E-07 | 609.31171 | 487.71927 |
| IMGTSPLQIDR | 11 | P31327 | Carbamoyl-phosphate synthase [ammonia | CPS1 | 164.94 | 2 | 54.611835 | yes | 1.105E-07 | 615.82916 | 448.6344 |
| AVNTLNEALEFAK | 13 | P31327 | Carbamoyl-phosphate synthase [ammonia | CPS1 | 164.94 | 2 | 54.611835 | yes | 9.722E-08 | 710.37756 | 415.52893 |
| AIDDNMSLDEIEK | 13 | P31327 | Carbamoyl-phosphate synthase [ammonia | CPS1 | 164.94 | 2 | 54.611835 | yes | 9.722E-08 | 746.8454 | 408.37692 |
| AADTIGYPVMIR | 12 | P31327 | Carbamoyl-phosphate synthase [ammonia | CPS1 | 164.94 | 2 | 54.611835 | yes | 1.186E-07 | 653.84485 | 387.80936 |
| VVAVDCGIK | 9 | P31327 | Carbamoyl-phosphate synthase [ammonia | CPS1 | 164.94 | 2 | 54.611835 | yes | 1.36E-07 | 480.76279 | 357.92001 |
| IAPSFAVESIEDAL | 15 | P31327 | Carbamoyl-phosphate synthase [ammonia | CPS1 | 164.94 | 2 | 54.611835 | yes | 2.035E-07 | 795.42468 | 354.28802 |
| CLGLTEAQTR | 10 | P31327 | Carbamoyl-phosphate synthase [ammonia | CPS1 | 164.94 | 2 | 54.611835 | yes | 1.356E-07 | 574.79004 | 352.16461 |
| VPAIYGVDTR | 10 | P31327 | Carbamoyl-phosphate synthase [ammonia | CPS1 | 164.94 | 2 | 54.611835 | yes | 4.172E-07 | 545.79822 | 341.69043 |
| TLGVDFIDVATK | 12 | P31327 | Carbamoyl-phosphate synthase [ammonia | CPS1 | 164.94 | 2 | 54.611835 | yes | 1.234E-06 | 639.85065 | 312.69971 |
| MCHPSIEGFTPR | 12 | P31327 | Carbamoyl-phosphate synthase [ammonia | CPS1 | 164.94 | 2 | 54.611835 | yes | 1.003E-07 | 716.32904 | 147.54514 |
| MCHPSIEGFTPR | 12 | P31327 | Carbamoyl-phosphate synthase [ammonia | CPS1 | 164.94 | 2 | 54.611835 | yes | 3.321E-07 | 477.88843 | 147.54514 |
| MCHPSIEGFTPR | 12 | P31327 | Carbamoyl-phosphate synthase [ammonia | CPS1 | 164.94 | 3 | 54.611835 | yes | 1.003E-07 | 716.32904 | 147.54514 |
| MCHPSIEGFTPR | 12 | P31327 | Carbamoyl-phosphate synthase [ammonia | CPS1 | 164.94 | 3 | 54.611835 | yes | 3.321E-07 | 477.88843 | 147.54514 |
| VMIGENVDEK | 10 | P31327 | Carbamoyl-phosphate synthase [ammonia | CPS1 | 164.94 | 2 | 54.611835 | yes | 3.683E-07 | 567.27899 | 294.94821 |
| FLGVAEQLHNEGF | 14 | P31327 | Carbamoyl-phosphate synthase [ammonia | CPS1 | 164.94 | 2 | 54.611835 | yes | 9.722E-08 | 794.90955 | 276.53003 |
| QNLIAEVSTK | 10 | P31327 | Carbamoyl-phosphate synthase [ammonia | CPS1 | 164.94 | 2 | 54.611835 | yes | 3.681E-07 | 551.80878 | 266.71597 |
| GNDVLVIECNLR | 12 | P31327 | Carbamoyl-phosphate synthase [ammonia | CPS1 | 164.94 | 2 | 54.611835 | yes | 6.961E-07 | 701.36139 | 258.47064 |
| SAYALGGLGSGICI | 16 | P31327 | Carbamoyl-phosphate synthase [ammonia | CPS1 | 164.94 | 2 | 54.611835 | yes | 9.722E-08 | 796.89612 | 253.97142 |
| VSGLLVLDYSK | 11 | P31327 | Carbamoyl-phosphate synthase [ammonia | CPS1 | 164.94 | 2 | 54.611835 | yes | 4.526E-07 | 597.34247 | 250.59796 |
| EIEYEVVR | 8 | P31327 | Carbamoyl-phosphate synthase [ammonia | CPS1 | 164.94 | 2 | 54.611835 | no | 2.244E-07 | 518.7691 | 250.05423 |
| VSQEHPVVLTK | 11 | P31327 | Carbamoyl-phosphate synthase [ammonia | CPS1 | 164.94 | 2 | 54.611835 | yes | 9.722E-08 | 618.85095 | 122.96913 |
| VSQEHPVVLTK | 11 | P31327 | Carbamoyl-phosphate synthase [ammonia | CPS1 | 164.94 | 2 | 54.611835 | yes | 3.69E-06 | 412.90308 | 122.96913 |
| VSQEHPVVLTK | 11 | P31327 | Carbamoyl-phosphate synthase [ammonia | CPS1 | 164.94 | 3 | 54.611835 | yes | 9.722E-08 | 618.85095 | 122.96913 |
| VSQEHPVVLTK | 11 | P31327 | Carbamoyl-phosphate synthase [ammonia | CPS1 | 164.94 | 3 | 54.611835 | yes | 3.69E-06 | 412.90308 | 122.96913 |
| AFAMTNQILVEK | 12 | P31327 | Carbamoyl-phosphate synthase [ammonia | CPS1 | 164.94 | 2 | 54.611835 | yes | 9.722E-08 | 682.86578 | 217.75931 |
| CEMASTGEVACFC | 22 | P31327 | Carbamoyl-phosphate synthase [ammonia | CPS1 | 164.94 | 3 | 54.611835 | yes | 1.823E-07 | 805.69745 | 189.56847 |
| GILIGIQQSFR | 11 | P31327 | Carbamoyl-phosphate synthase [ammonia | CPS1 | 164.94 | 2 | 54.611835 | yes | 1.353E-07 | 616.36151 | 183.01347 |
| GLNSESMTEETLK | 13 | P31327 | Carbamoyl-phosphate synthase [ammonia | CPS1 | 164.94 | 2 | 54.611835 | yes | 9.984E-08 | 719.84015 | 177.0246 |
| SVDYPCLLR | 9 | P31327 | Carbamoyl-phosphate synthase [ammonia | CPS1 | 164.94 | 2 | 54.611835 | yes | 1.066E-07 | 561.78424 | 172.3616 |
| IALGIPLPEIK | 11 | P31327 | Carbamoyl-phosphate synthase [ammonia | CPS1 | 164.94 | 2 | 54.611835 | yes | 9.759E-08 | 582.37354 | 132.54228 |
| DYNHWLATK | 9 | P31327 | Carbamoyl-phosphate synthase [ammonia | CPS1 | 164.94 | 2 | 54.611835 | yes | 1.594E-07 | 574.28015 | 128.75513 |
| FVHDNYVIR | 9 | P31327 | Carbamoyl-phosphate synthase [ammonia | CPS1 | 164.94 | 2 | 54.611835 | yes | 9.722E-08 | 581.80383 | 114.77093 |
| SVGEVMAIGR | 10 | P31327 | Carbamoyl-phosphate synthase [ammonia | CPS1 | 164.94 | 2 | 54.611835 | yes | 9.722E-08 | 509.77115 | 86.125954 |
| AQTAHIVLEDGTK | 13 | P31327 | Carbamoyl-phosphate synthase [ammonia | CPS1 | 164.94 | 2 | 54.611835 | yes | 9.722E-08 | 691.86737 | 83.474724 |
| GLNSESMTEETLK | 14 | P31327 | Carbamoyl-phosphate synthase [ammonia | CPS1 | 164.94 | 2 | 54.611835 | yes | 1.21E-07 | 797.89069 | 61.52058 |

| | | | | | | | | | | | |
|---|---|---|---|---|---|---|---|---|---|---|---|
| TFEESFQK | 8 | P31327 | Carbamoyl-phosphate synthase [ammonia | CPS1 | 164.94 | 2 | 54.611835 | yes | 2.912E-05 | 508.2402 | 28.699877 |
| VVAVDCGIKNNVI | 14 | P31327 | Carbamoyl-phosphate synthase [ammonia | CPS1 | 164.94 | 3 | 54.611835 | yes | 1.734E-06 | 519.62408 | 28.210094 |
| IFQEPTEPK | 9 | P31350 | Ribonucleoside-diphosphate reductase sul | RRM2 | 44.878 | 2 | 51.113605 | yes | 1.264E-06 | 544.78479 | 114.19129 |
| IEQEFLTEALPVK | 13 | P31350 | Ribonucleoside-diphosphate reductase sul | RRM2 | 44.878 | 2 | 51.113605 | yes | 9.722E-08 | 758.9165 | 82.967178 |
| ETEVIDPQDLLEGI | 14 | P31431 | Syndecan-4 OS=Homo sapiens OX=9606 | SDC4 | 21.642 | 2 | 52.80228 | yes | 9.722E-08 | 807.40448 | 732.80573 |
| ISPVEESEDVSNK | 13 | P31431 | Syndecan-4 OS=Homo sapiens OX=9606 | SDC4 | 21.642 | 2 | 52.80228 | yes | 9.722E-08 | 716.84375 | 467.67239 |
| KLEENEVIPK | 10 | P31431 | Syndecan-4 OS=Homo sapiens OX=9606 | SDC4 | 21.642 | 2 | 52.80228 | yes | 1.864E-07 | 599.83752 | 236.94157 |
| AGSGSQVPTEPK | 12 | P31431 | Syndecan-4 OS=Homo sapiens OX=9606 | SDC4 | 21.642 | 2 | 52.80228 | yes | 9.722E-08 | 579.29346 | 150.59084 |
| VNWATTPSSQK | 11 | P31483 | Cytotoxic granule associated RNA bindin | TIA1 | 42.964 | 2 | 50.42957 | no | 9.809E-08 | 609.80933 | 2290.3672 |
| TLYVGNLSR | 9 | P31483 | Cytotoxic granule associated RNA bindin | TIA1 | 42.964 | 2 | 50.42957 | no | 4.14E-07 | 511.7851 | 1775.4136 |
| QTFSPFGQIMEIR | 13 | P31483 | Cytotoxic granule associated RNA bindin | TIA1 | 42.964 | 2 | 50.42957 | no | 1.063E-07 | 777.3927 | 305.61429 |
| GYGFVSFFNK | 10 | P31483 | Cytotoxic granule associated RNA bindin | TIA1 | 42.964 | 2 | 50.42957 | yes | 1.088E-07 | 583.28748 | 133.86551 |
| DTSSSTVVSTQR | 12 | P31483 | Cytotoxic granule associated RNA bindin | TIA1 | 42.964 | 2 | 50.42957 | yes | 9.976E-08 | 634.30988 | 131.74327 |
| TIVITSHPGQIVK | 13 | P31689 | DnaJ homolog subfamily A member 1 OS | DNAJA1 | 44.869 | 2 | 53.117554 | yes | 1.157E-07 | 696.91412 | 2019.9154 |
| TIVITSHPGQIVK | 13 | P31689 | DnaJ homolog subfamily A member 1 OS | DNAJA1 | 44.869 | 2 | 53.117554 | yes | 2.156E-07 | 464.94516 | 2019.9154 |
| TIVITSHPGQIVK | 13 | P31689 | DnaJ homolog subfamily A member 1 OS | DNAJA1 | 44.869 | 3 | 53.117554 | yes | 1.157E-07 | 696.91412 | 2019.9154 |
| TIVITSHPGQIVK | 13 | P31689 | DnaJ homolog subfamily A member 1 OS | DNAJA1 | 44.869 | 3 | 53.117554 | yes | 2.156E-07 | 464.94516 | 2019.9154 |
| CVLNEGMPIYR | 11 | P31689 | DnaJ homolog subfamily A member 1 OS | DNAJA1 | 44.869 | 2 | 53.117554 | yes | 9.722E-08 | 676.32849 | 3269.592 |
| QISQAYEVLSDAK | 13 | P31689 | DnaJ homolog subfamily A member 1 OS | DNAJA1 | 44.869 | 2 | 53.117554 | yes | 3.34E-07 | 726.37244 | 2819.4487 |
| DHAVFTR | 7 | P31689 | DnaJ homolog subfamily A member 1 OS | DNAJA1 | 44.869 | 2 | 53.117554 | yes | 1.87E-07 | 423.21686 | 2020.7294 |
| ILEVHIDK | 8 | P31689 | DnaJ homolog subfamily A member 1 OS | DNAJA1 | 44.869 | 2 | 53.117554 | yes | 1.141E-07 | 483.78458 | 1850.1221 |
| NVICDKCEGR | 10 | P31689 | DnaJ homolog subfamily A member 1 OS | DNAJA1 | 44.869 | 2 | 53.117554 | yes | 9.722E-08 | 625.78442 | 1554.3718 |
| VNFPENGFLSPDK | 13 | P31689 | DnaJ homolog subfamily A member 1 OS | DNAJA1 | 44.869 | 2 | 53.117554 | yes | 1.178E-07 | 732.36188 | 1104.0969 |
| KILEVHIDK | 9 | P31689 | DnaJ homolog subfamily A member 1 OS | DNAJA1 | 44.869 | 2 | 53.117554 | yes | 1.581E-07 | 547.83203 | 91.849571 |
| NVVHQLSVTLEDI | 19 | P31689 | DnaJ homolog subfamily A member 1 OS | DNAJA1 | 44.869 | 3 | 53.117554 | yes | 9.722E-08 | 710.37103 | 67.315277 |
| ETTYYDVLGVK | 11 | P31689 | DnaJ homolog subfamily A member 1 OS | DNAJA1 | 44.869 | 2 | 53.117554 | yes | 4.555E-07 | 644.32697 | 63.793213 |
| HYNGEAYEDDEHI | 15 | P31689 | DnaJ homolog subfamily A member 1 OS | DNAJA1 | 44.869 | 3 | 53.117554 | yes | 1.314E-07 | 623.59094 | 37.152905 |
| ETTYYDVLGVKPN | 21 | P31689 | DnaJ homolog subfamily A member 1 OS | DNAJA1 | 44.869 | 4 | 53.117554 | yes | 3.25E-06 | 607.31671 | 18.468922 |
| ITDFGLCK | 8 | P31749 | RAC-alpha serine/threonine-protein kinas | AKT1 | 55.687 | 2 | 42.635246 | yes | 2.941E-07 | 477.24167 | 140.45848 |
| MVLAAAGGVEHQ | 20 | P31930 | Cytochrome b-c1 complex subunit 1, mitc | UQCRC1 | 52.646 | 3 | 51.185856 | yes | 1.473E-07 | 698.04547 | 76.555145 |
| LCTSATESEVAR | 12 | P31930 | Cytochrome b-c1 complex subunit 1, mitc | UQCRC1 | 52.646 | 2 | 51.185856 | yes | 1.121E-07 | 662.31409 | 72.631989 |
| ADLTEYLSTHYK | 12 | P31930 | Cytochrome b-c1 complex subunit 1, mitc | UQCRC1 | 52.646 | 2 | 51.185856 | yes | 9.722E-08 | 720.85388 | 42.925034 |
| EMQENDASMR | 10 | P31930 | Cytochrome b-c1 complex subunit 1, mitc | UQCRC1 | 52.646 | 3 | 51.185856 | yes | 2.889E-06 | 404.16574 | 3.6585667 |
| VHIDIGADGR | 10 | P31942 | Heterogeneous nuclear ribonucleoprotein | HNRNPH3 | 36.927 | 2 | 49.849194 | yes | 3.353E-07 | 526.77783 | 17461.373 |
| STGEAFVQFASK | 12 | P31942 | Heterogeneous nuclear ribonucleoprotein | HNRNPH3 | 36.927 | 2 | 49.849194 | yes | 3.988E-07 | 636.31696 | 13163.097 |
| DGMDNQGGYGSV | 14 | P31942 | Heterogeneous nuclear ribonucleoprotein | HNRNPH3 | 36.927 | 2 | 49.849194 | yes | 6.12E-07 | 706.79657 | 4773.7861 |
| GMGGHGYGGAGI | 26 | P31942 | Heterogeneous nuclear ribonucleoprotein | HNRNPH3 | 36.927 | 4 | 49.849194 | yes | 1.225E-07 | 636.52643 | 1195.1174 |
| GMGGHGYGGAGI | 26 | P31942 | Heterogeneous nuclear ribonucleoprotein | HNRNPH3 | 36.927 | 4 | 49.849194 | yes | 1.043E-07 | 509.42261 | 1195.1174 |
| GMGGHGYGGAGI | 26 | P31942 | Heterogeneous nuclear ribonucleoprotein | HNRNPH3 | 36.927 | 4 | 49.849194 | yes | 1.001E-07 | 640.52521 | 1195.1174 |

| | | | | | | | | | | | |
|---|---|---|---|---|---|---|---|---|---|---|---|
| GMGGHGYGGAGI | 26 | P31942 | Heterogeneous nuclear ribonucleoprotein | HNRNPH3 | 36.927 | 4 | 49.849194 | yes | 4.16E-07 | 512.62158 | 1195.1174 |
| GMGGHGYGGAGI | 26 | P31942 | Heterogeneous nuclear ribonucleoprotein | HNRNPH3 | 36.927 | 4 | 49.849194 | yes | 1.046E-07 | 640.52521 | 1195.1174 |
| GMGGHGYGGAGI | 26 | P31942 | Heterogeneous nuclear ribonucleoprotein | HNRNPH3 | 36.927 | 5 | 49.849194 | yes | 1.225E-07 | 636.52643 | 1195.1174 |
| GMGGHGYGGAGI | 26 | P31942 | Heterogeneous nuclear ribonucleoprotein | HNRNPH3 | 36.927 | 5 | 49.849194 | yes | 1.043E-07 | 509.42261 | 1195.1174 |
| GMGGHGYGGAGI | 26 | P31942 | Heterogeneous nuclear ribonucleoprotein | HNRNPH3 | 36.927 | 5 | 49.849194 | yes | 1.001E-07 | 640.52521 | 1195.1174 |
| GMGGHGYGGAGI | 26 | P31942 | Heterogeneous nuclear ribonucleoprotein | HNRNPH3 | 36.927 | 5 | 49.849194 | yes | 4.16E-07 | 512.62158 | 1195.1174 |
| GMGGHGYGGAGI | 26 | P31942 | Heterogeneous nuclear ribonucleoprotein | HNRNPH3 | 36.927 | 5 | 49.849194 | yes | 1.046E-07 | 640.52521 | 1195.1174 |
| ATGEADVEFVTHE | 21 | P31942 | Heterogeneous nuclear ribonucleoprotein | HNRNPH3 | 36.927 | 3 | 49.849194 | yes | 1.197E-07 | 726.67218 | 1415.4731 |
| ATGEADVEFVTHE | 21 | P31942 | Heterogeneous nuclear ribonucleoprotein | HNRNPH3 | 36.927 | 3 | 49.849194 | yes | 2.661E-07 | 732.00378 | 1415.4731 |
| GFYDPPRR | 8 | P31942 | Heterogeneous nuclear ribonucleoprotein | HNRNPH3 | 36.927 | 2 | 49.849194 | yes | 2.014E-07 | 504.2565 | 412.57623 |
| YIELFLNSTPGGGS | 26 | P31942 | Heterogeneous nuclear ribonucleoprotein | HNRNPH3 | 36.927 | 3 | 49.849194 | yes | 2.658E-07 | 841.38617 | 396.7623 |
| GFYDPPR | 7 | P31942 | Heterogeneous nuclear ribonucleoprotein | HNRNPH3 | 36.927 | 2 | 49.849194 | yes | 1.375E-07 | 426.20596 | 377.54031 |
| PGPYDRPIGGR | 11 | P31942 | Heterogeneous nuclear ribonucleoprotein | HNRNPH3 | 36.927 | 2 | 49.849194 | yes | 1.933E-07 | 592.81219 | 158.77156 |
| EIAENALGK | 9 | P31942 | Heterogeneous nuclear ribonucleoprotein | HNRNPH3 | 36.927 | 2 | 49.849194 | yes | 5.826E-07 | 472.75601 | 96.926651 |
| VHIEIGPDGR | 10 | P31943 | Heterogeneous nuclear ribonucleoprotein | HNRNPH1 | 49.23 | 2 | 54.823204 | no | 1.511E-07 | 546.79346 | 78949.859 |
| GLPWSCSADEVQF | 13 | P31943 | Heterogeneous nuclear ribonucleoprotein | HNRNPH1 | 49.23 | 2 | 54.823204 | yes | 4.506E-07 | 752.84607 | 48514.082 |
| SNNVEMDWVLK | 11 | P31943 | Heterogeneous nuclear ribonucleoprotein | HNRNPH1 | 49.23 | 2 | 54.823204 | yes | 2.035E-07 | 667.8241 | 18504.191 |
| SNNVEMDWVLK | 11 | P31943 | Heterogeneous nuclear ribonucleoprotein | HNRNPH1 | 49.23 | 2 | 54.823204 | yes | 1.141E-07 | 675.82153 | 18504.191 |
| DLNYCFSGMSDHI | 13 | P31943 | Heterogeneous nuclear ribonucleoprotein | HNRNPH1 | 49.23 | 2 | 54.823204 | no | 1.169E-07 | 801.32721 | 14388.183 |
| DLNYCFSGMSDHI | 13 | P31943 | Heterogeneous nuclear ribonucleoprotein | HNRNPH1 | 49.23 | 2 | 54.823204 | no | 9.734E-08 | 809.32465 | 14388.183 |
| MLGTEGGEGFVVI | 13 | P31943 | Heterogeneous nuclear ribonucleoprotein | HNRNPH1 | 49.23 | 2 | 54.823204 | yes | 1.56E-07 | 683.3396 | 4799.2993 |
| MLGTEGGEGFVVI | 13 | P31943 | Heterogeneous nuclear ribonucleoprotein | HNRNPH1 | 49.23 | 2 | 54.823204 | yes | 1.122E-07 | 662.33429 | 4799.2993 |
| MLGTEGGEGFVVI | 13 | P31943 | Heterogeneous nuclear ribonucleoprotein | HNRNPH1 | 49.23 | 2 | 54.823204 | yes | 9.722E-08 | 691.33704 | 4799.2993 |
| MLGTEGGEGFVVI | 13 | P31943 | Heterogeneous nuclear ribonucleoprotein | HNRNPH1 | 49.23 | 2 | 54.823204 | yes | 1.003E-07 | 670.33173 | 4799.2993 |
| THYDPPR | 7 | P31943 | Heterogeneous nuclear ribonucleoprotein | HNRNPH1 | 49.23 | 2 | 54.823204 | no | 1.456E-07 | 443.21429 | 13205.573 |
| YVELFLNSTAGAS | 20 | P31943 | Heterogeneous nuclear ribonucleoprotein | HNRNPH1 | 49.23 | 3 | 54.823204 | yes | 6.762E-07 | 714.67987 | 9820.4971 |
| VTGEADVEFATHE | 21 | P31943 | Heterogeneous nuclear ribonucleoprotein | HNRNPH1 | 49.23 | 3 | 54.823204 | yes | 1.464E-07 | 726.67218 | 3349.4685 |
| VTGEADVEFATHE | 21 | P31943 | Heterogeneous nuclear ribonucleoprotein | HNRNPH1 | 49.23 | 3 | 54.823204 | yes | 1.28E-07 | 732.00378 | 3349.4685 |
| MMLGTEGGEGFV | 14 | P31943 | Heterogeneous nuclear ribonucleoprotein | HNRNPH1 | 49.23 | 2 | 54.823204 | yes | 1.272E-07 | 748.8598 | 822.30072 |
| MMLGTEGGEGFV | 14 | P31943 | Heterogeneous nuclear ribonucleoprotein | HNRNPH1 | 49.23 | 2 | 54.823204 | yes | 4.561E-07 | 756.8573 | 822.30072 |
| MMLGTEGGEGFV | 14 | P31943 | Heterogeneous nuclear ribonucleoprotein | HNRNPH1 | 49.23 | 2 | 54.823204 | yes | 3.982E-07 | 756.8573 | 822.30072 |
| MMLGTEGGEGFV | 14 | P31943 | Heterogeneous nuclear ribonucleoprotein | HNRNPH1 | 49.23 | 2 | 54.823204 | yes | 4.239E-07 | 764.85474 | 822.30072 |
| HTGPNSPDTANDC | 16 | P31943 | Heterogeneous nuclear ribonucleoprotein | HNRNPH1 | 49.23 | 2 | 54.823204 | no | 1.022E-07 | 842.88733 | 1231.6731 |
| HTGPNSPDTANDC | 16 | P31943 | Heterogeneous nuclear ribonucleoprotein | HNRNPH1 | 49.23 | 2 | 54.823204 | no | 9.722E-08 | 562.26062 | 1231.6731 |
| HTGPNSPDTANDC | 16 | P31943 | Heterogeneous nuclear ribonucleoprotein | HNRNPH1 | 49.23 | 3 | 54.823204 | no | 1.022E-07 | 842.88733 | 1231.6731 |
| HTGPNSPDTANDC | 16 | P31943 | Heterogeneous nuclear ribonucleoprotein | HNRNPH1 | 49.23 | 3 | 54.823204 | no | 9.722E-08 | 562.26062 | 1231.6731 |
| EGRPSGEAFVELES | 19 | P31943 | Heterogeneous nuclear ribonucleoprotein | HNRNPH1 | 49.23 | 3 | 54.823204 | yes | 9.722E-08 | 702.99902 | 1006.9521 |
| GLPFGCSK | 8 | P31943 | Heterogeneous nuclear ribonucleoprotein | HNRNPH1 | 49.23 | 2 | 54.823204 | no | 1.108E-07 | 433.21545 | 507.87396 |
| KLMAMQR | 7 | P31943 | Heterogeneous nuclear ribonucleoprotein | HNRNPH1 | 49.23 | 2 | 54.823204 | no | 1.761E-07 | 439.24097 | 256.27826 |

| | | | | | | | | | | | |
|---|---|---|---|---|---|---|---|---|---|---|---|
| THYDPPRK | 8 | P31943 | Heterogeneous nuclear ribonucleoprotein | HNRNPH1 | 49.23 | 2 | 54.823204 | no | 1.28E-07 | 507.26178 | 191.93257 |
| PSGEAFVELESEDI | 16 | P31943 | Heterogeneous nuclear ribonucleoprotein | HNRNPH1 | 49.23 | 3 | 54.823204 | yes | 2.203E-07 | 588.94397 | 85.615738 |
| EMQPTHPIR | 9 | P31946 | 14-3-3 protein beta/alpha OS=Homo sapi | YWHAB | 28.083 | 2 | 55.459553 | no | 1.041E-07 | 554.78204 | 654.63562 |
| EMQPTHPIR | 9 | P31946 | 14-3-3 protein beta/alpha OS=Homo sapi | YWHAB | 28.083 | 2 | 55.459553 | no | 9.722E-08 | 562.77948 | 654.63562 |
| VISSIEQK | 8 | P31946 | 14-3-3 protein beta/alpha OS=Homo sapi | YWHAB | 28.083 | 2 | 55.459553 | no | 5.835E-07 | 452.26093 | 425.63867 |
| YLIPNATQPESK | 12 | P31946 | 14-3-3 protein beta/alpha OS=Homo sapi | YWHAB | 28.083 | 2 | 55.459553 | yes | 3.521E-07 | 680.85901 | 263.19498 |
| MKGDYFR | 7 | P31946 | 14-3-3 protein beta/alpha OS=Homo sapi | YWHAB | 28.083 | 2 | 55.459553 | no | 1.434E-07 | 458.72092 | 179.21466 |
| NLLSVAYK | 8 | P31946 | 14-3-3 protein beta/alpha OS=Homo sapi | YWHAB | 28.083 | 2 | 55.459553 | no | 1.643E-07 | 454.26602 | 104.17703 |
| YDDMAAAMK | 9 | P31946 | 14-3-3 protein beta/alpha OS=Homo sapi | YWHAB | 28.083 | 2 | 55.459553 | no | 9.729E-08 | 508.21481 | 75.938095 |
| AVTEQGHELSNEE | 14 | P31946 | 14-3-3 protein beta/alpha OS=Homo sapi | YWHAB | 28.083 | 2 | 55.459553 | yes | 4.922E-07 | 799.8739 | 17.889158 |
| YLAEVATGDDK | 11 | P31947 | 14-3-3 protein sigma OS=Homo sapiens ( | SFN | 27.775 | 2 | 50.52324 | yes | 1.056E-07 | 591.28784 | 130.7811 |
| SAYQEAMDISK | 11 | P31947 | 14-3-3 protein sigma OS=Homo sapiens ( | SFN | 27.775 | 2 | 50.52324 | yes | 9.789E-08 | 621.78717 | 74.882469 |
| TLLSDPTYR | 9 | P31948 | Stress-induced-phosphoprotein 1 OS=Ho | STIP1 | 62.64 | 2 | 48.02493 | yes | 4.373E-07 | 533.28241 | 87.857887 |
| LMDVGLIAIR | 10 | P31948 | Stress-induced-phosphoprotein 1 OS=Ho | STIP1 | 62.64 | 2 | 48.02493 | yes | 1.212E-07 | 550.82843 | 42.575798 |
| DPQALSEHLK | 10 | P31948 | Stress-induced-phosphoprotein 1 OS=Ho | STIP1 | 62.64 | 2 | 48.02493 | yes | 5.761E-07 | 569.29858 | 39.203037 |
| VFVTLTCAFR | 10 | P32121 | Beta-arrestin-2 OS=Homo sapiens OX=9 | ARRB2 | 46.106 | 2 | 50.053726 | yes | 1.2E-07 | 607.32355 | 131.02679 |
| EVLGILVSYR | 10 | P32121 | Beta-arrestin-2 OS=Homo sapiens OX=9 | ARRB2 | 46.106 | 2 | 50.053726 | yes | 1.075E-07 | 574.83734 | 124.04813 |
| ACGVDFEIR | 9 | P32121 | Beta-arrestin-2 OS=Homo sapiens OX=9 | ARRB2 | 46.106 | 2 | 50.053726 | yes | 9.921E-08 | 533.75293 | 79.292641 |
| SLHLEASLDK | 10 | P32121 | Beta-arrestin-2 OS=Homo sapiens OX=9 | ARRB2 | 46.106 | 2 | 50.053726 | yes | 4.537E-07 | 556.80096 | 69.590431 |
| EDLDVLGLSFR | 11 | P32121 | Beta-arrestin-2 OS=Homo sapiens OX=9 | ARRB2 | 46.106 | 2 | 50.053726 | yes | 5.591E-07 | 632.33258 | 40.188164 |
| EPGAAAASSAWK | 12 | P32242 | Homeobox protein OTX1 OS=Homo sapi | OTX1 | 37.328 | 2 | 52.00364 | yes | 9.722E-08 | 573.2829 | 661.74585 |
| LNFNSPDCLDYK | 12 | P32242 | Homeobox protein OTX1 OS=Homo sapi | OTX1 | 37.328 | 2 | 52.00364 | yes | 2.018E-07 | 743.33759 | 533.69452 |
| TRYPDIFMR | 9 | P32242 | Homeobox protein OTX1 OS=Homo sapi | OTX1 | 37.328 | 2 | 52.00364 | no | 9.795E-08 | 599.80548 | 143.26317 |
| INLPESR | 7 | P32242 | Homeobox protein OTX1 OS=Homo sapi | OTX1 | 37.328 | 2 | 52.00364 | no | 1.077E-06 | 414.73233 | 40.947929 |
| RPACTLKPECVQQ | 22 | P32320 | Cytidine deaminase OS=Homo sapiens O | CDA | 16.185 | 4 | 48.71478 | yes | 1.112E-07 | 654.58301 | 177.34071 |
| SLLINAVEASCIR | 13 | P32322 | Pyrroline-5-carboxylate reductase 1, mito | PYCR1 | 33.361 | 2 | 55.46571 | no | 9.722E-08 | 723.3927 | 136.77281 |
| CMTNTPVVVR | 10 | P32322 | Pyrroline-5-carboxylate reductase 1, mito | PYCR1 | 33.361 | 2 | 55.46571 | yes | 1.33E-07 | 588.79681 | 121.7252 |
| LGAQALLGAAK | 11 | P32322 | Pyrroline-5-carboxylate reductase 1, mito | PYCR1 | 33.361 | 2 | 55.46571 | yes | 9.608E-07 | 506.81113 | 36.005814 |
| MGFEPLAYR | 9 | P32754 | 4-hydroxyphenylpyruvate dioxygenase O | HPD | 44.935 | 2 | 51.335205 | yes | 2.144E-07 | 542.26825 | 82.76532 |
| ILVDYDEK | 8 | P32754 | 4-hydroxyphenylpyruvate dioxygenase O | HPD | 44.935 | 2 | 51.335205 | yes | 9.722E-08 | 497.75821 | 77.757248 |
| SIVVANYEESIK | 12 | P32754 | 4-hydroxyphenylpyruvate dioxygenase O | HPD | 44.935 | 2 | 51.335205 | yes | 1.12E-07 | 676.35883 | 73.158936 |
| TEDIITAIR | 9 | P32754 | 4-hydroxyphenylpyruvate dioxygenase O | HPD | 44.935 | 2 | 51.335205 | yes | 7.24E-07 | 516.29022 | 46.483425 |
| ENIDALEELK | 10 | P32754 | 4-hydroxyphenylpyruvate dioxygenase O | HPD | 44.935 | 2 | 51.335205 | yes | 1.177E-07 | 587.30353 | 45.958553 |
| GLEFLSVPSTYYK | 13 | P32754 | 4-hydroxyphenylpyruvate dioxygenase O | HPD | 44.935 | 2 | 51.335205 | yes | 6.893E-07 | 752.39014 | 40.317757 |
| LQESIEYEDLGK | 12 | P32780 | General transcription factor IIH subunit 1 | GTF2H1 | 62.032 | 2 | 53.79589 | yes | 2.436E-07 | 712.3512 | 419.22095 |
| HLYVAVGELLR | 11 | P32780 | General transcription factor IIH subunit 1 | GTF2H1 | 62.032 | 2 | 53.79589 | yes | 9.722E-08 | 635.36932 | 354.92203 |
| LHTWQSR | 7 | P32780 | General transcription factor IIH subunit 1 | GTF2H1 | 62.032 | 2 | 53.79589 | yes | 2.086E-07 | 464.24341 | 151.63539 |
| MLQEDPVLFQLYK | 13 | P32780 | General transcription factor IIH subunit 1 | GTF2H1 | 62.032 | 2 | 53.79589 | yes | 3.179E-07 | 812.42621 | 127.80428 |
| QDGALYLMAER | 11 | P32780 | General transcription factor IIH subunit 1 | GTF2H1 | 62.032 | 2 | 53.79589 | yes | 3.645E-07 | 633.81097 | 105.95933 |

| | | | | | | | | | | | |
|---|---|---|---|---|---|---|---|---|---|---|---|
| YAENVPHNMTEK | 12 | P32780 | General transcription factor IIH subunit 1 | GTF2H1 | 62.032 | 2 | 53.79589 | yes | 9.728E-08 | 716.8299 | 103.16904 |
| FNHHSAMVLAAG | 14 | P32780 | General transcription factor IIH subunit 1 | GTF2H1 | 62.032 | 3 | 53.79589 | yes | 2.041E-07 | 508.6015 | 67.444878 |
| LNVNATDSSSTSN | 15 | P32780 | General transcription factor IIH subunit 1 | GTF2H1 | 62.032 | 2 | 53.79589 | yes | 9.722E-08 | 787.8739 | 46.446926 |
| KQDGALYLMAER | 12 | P32780 | General transcription factor IIH subunit 1 | GTF2H1 | 62.032 | 2 | 53.79589 | yes | 1.161E-06 | 697.85846 | 41.444603 |
| DLLQQLLPK | 9 | P32780 | General transcription factor IIH subunit 1 | GTF2H1 | 62.032 | 2 | 53.79589 | yes | 2.154E-06 | 534.3266 | 31.52784 |
| FLDGIYVSEK | 10 | P32969 | Large ribosomal subunit protein uL6 OS= | RPL9 | 21.864 | 2 | 51.84279 | yes | 8.099E-07 | 585.80573 | 28280.902 |
| DFNHINVELSLLGI | 14 | P32969 | Large ribosomal subunit protein uL6 OS= | RPL9 | 21.864 | 2 | 51.84279 | yes | 1.808E-07 | 799.93048 | 5622.4829 |
| KFLDGIYVSEK | 11 | P32969 | Large ribosomal subunit protein uL6 OS= | RPL9 | 21.864 | 2 | 51.84279 | yes | 1.065E-07 | 649.85315 | 4997.4302 |
| TICSHVQNMIK | 11 | P32969 | Large ribosomal subunit protein uL6 OS= | RPL9 | 21.864 | 2 | 51.84279 | yes | 1.597E-07 | 665.83392 | 1530.5771 |
| TICSHVQNMIK | 11 | P32969 | Large ribosomal subunit protein uL6 OS= | RPL9 | 21.864 | 2 | 51.84279 | yes | 1.194E-07 | 444.22507 | 1530.5771 |
| TICSHVQNMIK | 11 | P32969 | Large ribosomal subunit protein uL6 OS= | RPL9 | 21.864 | 2 | 51.84279 | yes | 1.131E-07 | 673.83142 | 1530.5771 |
| TICSHVQNMIK | 11 | P32969 | Large ribosomal subunit protein uL6 OS= | RPL9 | 21.864 | 3 | 51.84279 | yes | 1.597E-07 | 665.83392 | 1530.5771 |
| TICSHVQNMIK | 11 | P32969 | Large ribosomal subunit protein uL6 OS= | RPL9 | 21.864 | 3 | 51.84279 | yes | 1.194E-07 | 444.22507 | 1530.5771 |
| TICSHVQNMIK | 11 | P32969 | Large ribosomal subunit protein uL6 OS= | RPL9 | 21.864 | 3 | 51.84279 | yes | 1.131E-07 | 673.83142 | 1530.5771 |
| MRPGVACSVSQA( | 14 | P32969 | Large ribosomal subunit protein uL6 OS= | RPL9 | 21.864 | 2 | 51.84279 | yes | 1.32E-07 | 759.87939 | 705.48346 |
| MRPGVACSVSQA( | 14 | P32969 | Large ribosomal subunit protein uL6 OS= | RPL9 | 21.864 | 2 | 51.84279 | yes | 9.819E-08 | 506.92203 | 705.48346 |
| MRPGVACSVSQA( | 14 | P32969 | Large ribosomal subunit protein uL6 OS= | RPL9 | 21.864 | 2 | 51.84279 | yes | 9.722E-08 | 512.25366 | 705.48346 |
| MRPGVACSVSQA( | 14 | P32969 | Large ribosomal subunit protein uL6 OS= | RPL9 | 21.864 | 2 | 51.84279 | yes | 3.539E-07 | 767.87689 | 705.48346 |
| MRPGVACSVSQA( | 14 | P32969 | Large ribosomal subunit protein uL6 OS= | RPL9 | 21.864 | 3 | 51.84279 | yes | 1.32E-07 | 759.87939 | 705.48346 |
| MRPGVACSVSQA( | 14 | P32969 | Large ribosomal subunit protein uL6 OS= | RPL9 | 21.864 | 3 | 51.84279 | yes | 9.819E-08 | 506.92203 | 705.48346 |
| MRPGVACSVSQA( | 14 | P32969 | Large ribosomal subunit protein uL6 OS= | RPL9 | 21.864 | 3 | 51.84279 | yes | 9.722E-08 | 512.25366 | 705.48346 |
| MRPGVACSVSQA( | 14 | P32969 | Large ribosomal subunit protein uL6 OS= | RPL9 | 21.864 | 3 | 51.84279 | yes | 3.539E-07 | 767.87689 | 705.48346 |
| SVYAHFPINVVIQI | 22 | P32969 | Large ribosomal subunit protein uL6 OS= | RPL9 | 21.864 | 3 | 51.84279 | yes | 1.144E-07 | 828.77972 | 808.19556 |
| KELATVR | 7 | P32969 | Large ribosomal subunit protein uL6 OS= | RPL9 | 21.864 | 2 | 51.84279 | yes | 9.722E-08 | 408.75052 | 745.87244 |
| PGVACSVSQAQK | 12 | P32969 | Large ribosomal subunit protein uL6 OS= | RPL9 | 21.864 | 2 | 51.84279 | yes | 1.088E-07 | 616.30859 | 639.21674 |
| TILSNQTVDIPENV | 19 | P32969 | Large ribosomal subunit protein uL6 OS= | RPL9 | 21.864 | 3 | 51.84279 | yes | 1.077E-07 | 705.05109 | 148.60674 |
| PEEGSGCSVR | 10 | P32970 | CD70 antigen OS=Homo sapiens OX=96 | CD70 | 21.119 | 2 | 50.62049 | yes | 9.734E-08 | 539.23511 | 133.53156 |
| SFLHGPELDK | 10 | P32970 | CD70 antigen OS=Homo sapiens OX=96 | CD70 | 21.119 | 2 | 50.62049 | yes | 4.958E-07 | 571.79565 | 124.89149 |
| NTDETFFGVQWVI | 14 | P32970 | CD70 antigen OS=Homo sapiens OX=96 | CD70 | 21.119 | 2 | 50.62049 | yes | 1.644E-07 | 848.40991 | 51.49765 |
| VLQPTVFPVVPR | 12 | P33121 | Long-chain-fatty-acid--CoA ligase 1 OS= | ACSL1 | 77.944 | 2 | 43.659355 | yes | 3.153E-07 | 676.40845 | 73.253159 |
| AILEDMVR | 8 | P33121 | Long-chain-fatty-acid--CoA ligase 1 OS= | ACSL1 | 77.944 | 2 | 43.659355 | no | 2.498E-07 | 473.75494 | 35.576206 |
| EYELLSDELNQK | 12 | P33176 | Kinesin-1 heavy chain OS=Homo sapiens | KIF5B | 109.685 | 2 | 55.776104 | yes | 9.722E-08 | 740.86194 | 434.59567 |
| LITDLQDQNQK | 11 | P33176 | Kinesin-1 heavy chain OS=Homo sapiens | KIF5B | 109.685 | 2 | 55.776104 | yes | 9.722E-08 | 658.34625 | 428.55389 |
| SLTEYLQNVEQK | 12 | P33176 | Kinesin-1 heavy chain OS=Homo sapiens | KIF5B | 109.685 | 2 | 55.776104 | yes | 9.722E-08 | 726.37244 | 343.38306 |
| TGAEGAVLDEAK | 12 | P33176 | Kinesin-1 heavy chain OS=Homo sapiens | KIF5B | 109.685 | 2 | 55.776104 | no | 9.722E-08 | 580.79333 | 296.7804 |
| TNLSVHEDK | 9 | P33176 | Kinesin-1 heavy chain OS=Homo sapiens | KIF5B | 109.685 | 2 | 55.776104 | no | 1.594E-07 | 521.76184 | 260.32913 |
| LHDPEGMGIIPR | 12 | P33176 | Kinesin-1 heavy chain OS=Homo sapiens | KIF5B | 109.685 | 2 | 55.776104 | yes | 1.109E-07 | 667.8479 | 250.07367 |
| TQMLDQEELLAST | 14 | P33176 | Kinesin-1 heavy chain OS=Homo sapiens | KIF5B | 109.685 | 2 | 55.776104 | yes | 9.722E-08 | 817.90637 | 248.80698 |
| MMLEQER | 7 | P33176 | Kinesin-1 heavy chain OS=Homo sapiens | KIF5B | 109.685 | 2 | 55.776104 | yes | 1.642E-07 | 468.7175 | 195.26305 |

| | | | | | | | | | | |
|---|---|---|---|---|---|---|---|---|---|---|
| STLLFGQR | 8 | P33176 | Kinesin-1 heavy chain OS=Homo sapiens | KIF5B | 109.685 | 2 | 55.776104 | yes | 1.586E-07 | 461.26126 | 188.61136 |
| AAEMMASLLK | 10 | P33176 | Kinesin-1 heavy chain OS=Homo sapiens | KIF5B | 109.685 | 2 | 55.776104 | yes | 2.127E-07 | 532.77759 | 186.67731 |
| ISFLENNLEQLTK | 13 | P33176 | Kinesin-1 heavy chain OS=Homo sapiens | KIF5B | 109.685 | 2 | 55.776104 | no | 1.083E-07 | 774.91705 | 168.34003 |
| GGGAFVQNSQPV/ | 15 | P33176 | Kinesin-1 heavy chain OS=Homo sapiens | KIF5B | 109.685 | 2 | 55.776104 | yes | 4.97E-07 | 743.89166 | 137.32503 |
| NTIQWLENELNR | 12 | P33176 | Kinesin-1 heavy chain OS=Homo sapiens | KIF5B | 109.685 | 2 | 55.776104 | yes | 9.722E-08 | 765.38898 | 135.45184 |
| FQGEDTVVIASK | 12 | P33176 | Kinesin-1 heavy chain OS=Homo sapiens | KIF5B | 109.685 | 2 | 55.776104 | yes | 1.095E-07 | 647.33789 | 131.14529 |
| KLFVQDLATR | 10 | P33176 | Kinesin-1 heavy chain OS=Homo sapiens | KIF5B | 109.685 | 2 | 55.776104 | yes | 1.998E-07 | 595.84821 | 123.33948 |
| ILQDSLGGNCR | 11 | P33176 | Kinesin-1 heavy chain OS=Homo sapiens | KIF5B | 109.685 | 2 | 55.776104 | no | 1.211E-07 | 616.80621 | 118.67645 |
| ELQTLHNLR | 9 | P33176 | Kinesin-1 heavy chain OS=Homo sapiens | KIF5B | 109.685 | 2 | 55.776104 | no | 9.722E-08 | 562.31458 | 91.178322 |
| DLAEIGIAVGNND' | 15 | P33176 | Kinesin-1 heavy chain OS=Homo sapiens | KIF5B | 109.685 | 2 | 55.776104 | yes | 1.261E-07 | 764.4043 | 90.658028 |
| LYLVDLAGSEK | 11 | P33176 | Kinesin-1 heavy chain OS=Homo sapiens | KIF5B | 109.685 | 2 | 55.776104 | no | 9.722E-08 | 604.33209 | 87.392799 |
| NGETVPIDEQFDK | 13 | P33176 | Kinesin-1 heavy chain OS=Homo sapiens | KIF5B | 109.685 | 2 | 55.776104 | yes | 1.4E-07 | 746.35175 | 82.719887 |
| DITLTNDKPATAIC | 24 | P33176 | Kinesin-1 heavy chain OS=Homo sapiens | KIF5B | 109.685 | 3 | 55.776104 | yes | 9.722E-08 | 844.76953 | 72.47966 |
| VSYFEIYLDK | 10 | P33176 | Kinesin-1 heavy chain OS=Homo sapiens | KIF5B | 109.685 | 2 | 55.776104 | no | 1.061E-07 | 638.8266 | 71.197578 |
| QAVEQQIQSHR | 11 | P33176 | Kinesin-1 heavy chain OS=Homo sapiens | KIF5B | 109.685 | 2 | 55.776104 | yes | 9.722E-08 | 662.34186 | 70.937836 |
| SHSIFLINVK | 10 | P33176 | Kinesin-1 heavy chain OS=Homo sapiens | KIF5B | 109.685 | 2 | 55.776104 | yes | 1.707E-06 | 579.33752 | 54.903641 |
| GGTLLSVTGEVEP | 14 | P33240 | Cleavage stimulation factor subunit 2 OS= | CSTF2 | 60.96 | 2 | 51.412815 | no | 9.722E-08 | 707.88043 | 137.8231 |
| PLMAEPR | 7 | P33240 | Cleavage stimulation factor subunit 2 OS= | CSTF2 | 60.96 | 2 | 51.412815 | yes | 4.094E-06 | 407.21802 | 62.481827 |
| TDIQIALPSGCYGR | 14 | P33316 | Deoxyuridine 5'-triphosphate nucleotidoh | DUT | 26.564 | 2 | 49.736835 | yes | 1.165E-07 | 775.88519 | 242.66762 |
| GNVGVVLFNFGK | 12 | P33316 | Deoxyuridine 5'-triphosphate nucleotidoh | DUT | 26.564 | 2 | 49.736835 | yes | 1.297E-07 | 625.84821 | 151.16565 |
| LSEHATAPTR | 10 | P33316 | Deoxyuridine 5'-triphosphate nucleotidoh | DUT | 26.564 | 2 | 49.736835 | yes | 4.68E-07 | 541.78308 | 90.13839 |
| IAQLICER | 8 | P33316 | Deoxyuridine 5'-triphosphate nucleotidoh | DUT | 26.564 | 2 | 49.736835 | yes | 3.458E-07 | 501.77368 | 87.478035 |
| HFIDVGAGVIDED' | 15 | P33316 | Deoxyuridine 5'-triphosphate nucleotidoh | DUT | 26.564 | 3 | 49.736835 | yes | 1.006E-07 | 569.27753 | 72.765137 |
| NLLTAAADAIER | 12 | P33897 | ATP-binding cassette sub-family D memt | ABCD1 | 82.937 | 2 | 50.82589 | yes | 9.722E-08 | 629.34351 | 113.9114 |
| DAGIALLSITHR | 12 | P33897 | ATP-binding cassette sub-family D memt | ABCD1 | 82.937 | 2 | 50.82589 | yes | 1.164E-07 | 633.86188 | 109.40656 |
| DQVIYPDSVEDMC | 14 | P33897 | ATP-binding cassette sub-family D memt | ABCD1 | 82.937 | 2 | 50.82589 | yes | 2.033E-07 | 847.88812 | 80.045479 |
| ELEDAQAGSGTIGI | 14 | P33897 | ATP-binding cassette sub-family D memt | ABCD1 | 82.937 | 2 | 50.82589 | yes | 1.946E-07 | 702.34167 | 65.221848 |
| EVTELAGYTAR | 11 | P33897 | ATP-binding cassette sub-family D memt | ABCD1 | 82.937 | 2 | 50.82589 | no | 1.565E-07 | 605.30914 | 64.003052 |
| LEQQLAGIPK | 10 | P33897 | ATP-binding cassette sub-family D memt | ABCD1 | 82.937 | 2 | 50.82589 | yes | 1.32E-07 | 548.82166 | 60.730274 |
| TFLSVYVAR | 9 | P33897 | ATP-binding cassette sub-family D memt | ABCD1 | 82.937 | 2 | 50.82589 | yes | 7.192E-07 | 528.29785 | 60.00066 |
| FGELVAEEAR | 10 | P33897 | ATP-binding cassette sub-family D memt | ABCD1 | 82.937 | 2 | 50.82589 | yes | 2.443E-07 | 560.78528 | 59.808968 |
| SSELIITDSITLK | 13 | P33981 | Dual specificity protein kinase TTK OS=l | TTK | 97.073 | 2 | 52.022446 | yes | 9.722E-08 | 710.4007 | 84.642937 |
| TDDSVVPCFMK | 11 | P33981 | Dual specificity protein kinase TTK OS=l | TTK | 97.073 | 2 | 52.022446 | yes | 3.311E-07 | 649.79138 | 74.895157 |
| VFQVLNEK | 8 | P33981 | Dual specificity protein kinase TTK OS=l | TTK | 97.073 | 2 | 52.022446 | yes | 1.022E-07 | 488.77673 | 56.112633 |
| ELTIDSIMNK | 10 | P33981 | Dual specificity protein kinase TTK OS=l | TTK | 97.073 | 2 | 52.022446 | yes | 3.283E-07 | 582.30249 | 54.150719 |
| EYQEPEVPESNQK | 13 | P33981 | Dual specificity protein kinase TTK OS=l | TTK | 97.073 | 2 | 52.022446 | yes | 1.246E-07 | 788.85992 | 50.573856 |
| NLSASTVLTAQESI | 24 | P33981 | Dual specificity protein kinase TTK OS=l | TTK | 97.073 | 3 | 52.022446 | yes | 1.352E-06 | 839.76123 | 47.28904 |
| NEDLTDELSLNK | 12 | P33981 | Dual specificity protein kinase TTK OS=l | TTK | 97.073 | 2 | 52.022446 | yes | 3.159E-07 | 695.83844 | 38.686092 |
| VPVNLLNSPDCDV | 14 | P33981 | Dual specificity protein kinase TTK OS=l | TTK | 97.073 | 2 | 52.022446 | yes | 2.678E-06 | 785.4007 | 14.416773 |

| | | | | | | | | | | | |
|---|---|---|---|---|---|---|---|---|---|---|---|
| ALADDDFLTVTGK | 13 | P33991 | DNA replication licensing factor MCM4 ( | MCM4 | 96.558 | 2 | 55.784466 | yes | 9.722E-08 | 683.34845 | 160.91602 |
| VEAIDVEEAK | 10 | P33991 | DNA replication licensing factor MCM4 ( | MCM4 | 96.558 | 2 | 55.784466 | yes | 8.17E-07 | 551.78497 | 106.46289 |
| THIDVIHYR | 9 | P33991 | DNA replication licensing factor MCM4 ( | MCM4 | 96.558 | 2 | 55.784466 | yes | 7.992E-07 | 577.30927 | 105.78284 |
| AEINILLCGDPGTS | 15 | P33991 | DNA replication licensing factor MCM4 ( | MCM4 | 96.558 | 2 | 55.784466 | yes | 2.916E-07 | 794.40601 | 104.83607 |
| LVIWGTDVNVAA( | 14 | P33991 | DNA replication licensing factor MCM4 ( | MCM4 | 96.558 | 2 | 55.784466 | yes | 1.915E-06 | 773.40833 | 62.528103 |
| KPDIYER | 7 | P33991 | DNA replication licensing factor MCM4 ( | MCM4 | 96.558 | 2 | 55.784466 | yes | 8.183E-07 | 460.74545 | 61.432953 |
| LGEINVIGEPFLNV | 20 | P33991 | DNA replication licensing factor MCM4 ( | MCM4 | 96.558 | 3 | 55.784466 | yes | 1.871E-07 | 765.73157 | 39.546238 |
| GILLQLFGGTR | 11 | P33991 | DNA replication licensing factor MCM4 ( | MCM4 | 96.558 | 2 | 55.784466 | yes | 1.081E-06 | 587.85077 | 28.713537 |
| LQELPDAVPHGEM | 15 | P33992 | DNA replication licensing factor MCM5 ( | MCM5 | 82.286 | 3 | 55.31454 | yes | 2.878E-07 | 563.61902 | 565.05548 |
| LQELPDAVPHGEM | 15 | P33992 | DNA replication licensing factor MCM5 ( | MCM5 | 82.286 | 3 | 55.31454 | yes | 1.317E-07 | 844.92487 | 565.05548 |
| LQELPDAVPHGEM | 15 | P33992 | DNA replication licensing factor MCM5 ( | MCM5 | 82.286 | 2 | 55.31454 | yes | 2.878E-07 | 563.61902 | 565.05548 |
| LQELPDAVPHGEM | 15 | P33992 | DNA replication licensing factor MCM5 ( | MCM5 | 82.286 | 2 | 55.31454 | yes | 1.317E-07 | 844.92487 | 565.05548 |
| LQPFATEADVEEA | 15 | P33992 | DNA replication licensing factor MCM5 ( | MCM5 | 82.286 | 2 | 55.31454 | yes | 9.81E-08 | 844.92816 | 986.73236 |
| IPGIIIAASAVR | 12 | P33992 | DNA replication licensing factor MCM5 ( | MCM5 | 82.286 | 2 | 55.31454 | yes | 1.315E-07 | 590.87427 | 944.4577 |
| SIAPSIFGGTDMK | 13 | P33992 | DNA replication licensing factor MCM5 ( | MCM5 | 82.286 | 2 | 55.31454 | yes | 1.001E-07 | 662.33429 | 886.59662 |
| CSVLAAANSVFGR | 13 | P33992 | DNA replication licensing factor MCM5 ( | MCM5 | 82.286 | 2 | 55.31454 | yes | 9.722E-08 | 676.34296 | 773.25757 |
| PGLEGYALPR | 10 | P33992 | DNA replication licensing factor MCM5 ( | MCM5 | 82.286 | 2 | 55.31454 | yes | 1.034E-06 | 536.79291 | 688.02338 |
| GEDNIDFMPTILSR | 14 | P33992 | DNA replication licensing factor MCM5 ( | MCM5 | 82.286 | 2 | 55.31454 | yes | 9.722E-08 | 804.39032 | 443.67255 |
| VTIMGIYSIK | 10 | P33992 | DNA replication licensing factor MCM5 ( | MCM5 | 82.286 | 2 | 55.31454 | yes | 1.863E-07 | 562.82281 | 439.93503 |
| NTLTNIAMR | 9 | P33992 | DNA replication licensing factor MCM5 ( | MCM5 | 82.286 | 2 | 55.31454 | yes | 9.722E-08 | 517.27661 | 412.77078 |
| VLGIQVDTDGSGR | 13 | P33992 | DNA replication licensing factor MCM5 ( | MCM5 | 82.286 | 2 | 55.31454 | yes | 9.79E-08 | 658.84387 | 398.91595 |
| AGITTTLNSR | 10 | P33992 | DNA replication licensing factor MCM5 ( | MCM5 | 82.286 | 2 | 55.31454 | yes | 5.989E-07 | 517.28546 | 372.03168 |
| SDMMSHLVK | 9 | P33992 | DNA replication licensing factor MCM5 ( | MCM5 | 82.286 | 2 | 55.31454 | yes | 2.521E-07 | 524.25171 | 369.01422 |
| LAALPNVYEVISK | 13 | P33992 | DNA replication licensing factor MCM5 ( | MCM5 | 82.286 | 2 | 55.31454 | yes | 2.687E-07 | 708.90851 | 318.56604 |
| AIACLLFGGSR | 11 | P33992 | DNA replication licensing factor MCM5 ( | MCM5 | 82.286 | 2 | 55.31454 | yes | 2.177E-07 | 582.81335 | 281.90845 |
| FAIGSQVSEHSIIK | 14 | P33992 | DNA replication licensing factor MCM5 ( | MCM5 | 82.286 | 2 | 55.31454 | yes | 9.758E-08 | 758.41193 | 266.19006 |
| CSPIGVYTSGK | 11 | P33992 | DNA replication licensing factor MCM5 ( | MCM5 | 82.286 | 2 | 55.31454 | yes | 3.172E-06 | 584.78699 | 250.59332 |
| RGDINLLMLGDPG | 16 | P33992 | DNA replication licensing factor MCM5 ( | MCM5 | 82.286 | 2 | 55.31454 | yes | 3.086E-07 | 835.94836 | 120.03987 |
| RGDINLLMLGDPG | 16 | P33992 | DNA replication licensing factor MCM5 ( | MCM5 | 82.286 | 2 | 55.31454 | yes | 1.675E-07 | 557.63464 | 120.03987 |
| RGDINLLMLGDPG | 16 | P33992 | DNA replication licensing factor MCM5 ( | MCM5 | 82.286 | 3 | 55.31454 | yes | 3.086E-07 | 835.94836 | 120.03987 |
| RGDINLLMLGDPG | 16 | P33992 | DNA replication licensing factor MCM5 ( | MCM5 | 82.286 | 3 | 55.31454 | yes | 1.675E-07 | 557.63464 | 120.03987 |
| VLQLMLR | 7 | P33992 | DNA replication licensing factor MCM5 ( | MCM5 | 82.286 | 2 | 55.31454 | yes | 3.445E-07 | 436.77295 | 232.9473 |
| CPLDPYFIMPDK | 12 | P33992 | DNA replication licensing factor MCM5 ( | MCM5 | 82.286 | 2 | 55.31454 | yes | 9.722E-08 | 748.35162 | 210.37842 |
| SFAGAVSPQEEEEI | 15 | P33992 | DNA replication licensing factor MCM5 ( | MCM5 | 82.286 | 2 | 55.31454 | yes | 9.722E-08 | 841.88647 | 171.10722 |
| QPAEHLQLLEEAA | 14 | P33992 | DNA replication licensing factor MCM5 ( | MCM5 | 82.286 | 3 | 55.31454 | yes | 3.475E-07 | 526.28247 | 57.593258 |
| QPAEHLQLLEEAA | 14 | P33992 | DNA replication licensing factor MCM5 ( | MCM5 | 82.286 | 3 | 55.31454 | yes | 9.722E-08 | 788.9201 | 57.593258 |
| QPAEHLQLLEEAA | 14 | P33992 | DNA replication licensing factor MCM5 ( | MCM5 | 82.286 | 2 | 55.31454 | yes | 3.475E-07 | 526.28247 | 57.593258 |
| QPAEHLQLLEEAA | 14 | P33992 | DNA replication licensing factor MCM5 ( | MCM5 | 82.286 | 2 | 55.31454 | yes | 9.722E-08 | 788.9201 | 57.593258 |
| CVDFQTLK | 8 | P33992 | DNA replication licensing factor MCM5 ( | MCM5 | 82.286 | 2 | 55.31454 | yes | 3.647E-07 | 505.75241 | 93.598999 |

| | | | | | | | | | | | |
|---|---|---|---|---|---|---|---|---|---|---|---|
| RLPDGLTR | 8 | P33992 | DNA replication licensing factor MCM5 ( | MCM5 | 82.286 | 2 | 55.31454 | yes | 1.012E-07 | 464.27216 | 90.030693 |
| GDINLLMLGDPGT | 15 | P33992 | DNA replication licensing factor MCM5 ( | MCM5 | 82.286 | 2 | 55.31454 | yes | 9.722E-08 | 757.89777 | 84.317688 |
| QLEAIVR | 7 | P33992 | DNA replication licensing factor MCM5 ( | MCM5 | 82.286 | 2 | 55.31454 | yes | 8.426E-07 | 414.75052 | 79.955795 |
| HMQLYCDR | 8 | P33992 | DNA replication licensing factor MCM5 ( | MCM5 | 82.286 | 2 | 55.31454 | yes | 1.017E-07 | 561.74457 | 69.524902 |
| EVADEVTRPR | 10 | P33992 | DNA replication licensing factor MCM5 ( | MCM5 | 82.286 | 2 | 55.31454 | yes | 9.819E-08 | 586.30695 | 60.808891 |
| NTLTNIAMRPGLE( | 19 | P33992 | DNA replication licensing factor MCM5 ( | MCM5 | 82.286 | 3 | 55.31454 | yes | 1.391E-07 | 696.37372 | 27.726469 |
| SRDPGMVR | 8 | P33993 | DNA replication licensing factor MCM7 ( | MCM7 | 81.308 | 2 | 51.452545 | yes | 4.431E-07 | 459.23471 | 256.40125 |
| GNINICLMGDPGV. | 15 | P33993 | DNA replication licensing factor MCM7 ( | MCM7 | 81.308 | 2 | 51.452545 | yes | 4.414E-07 | 779.88947 | 177.13943 |
| AGILTTLNAR | 10 | P33993 | DNA replication licensing factor MCM7 ( | MCM7 | 81.308 | 2 | 51.452545 | yes | 3.251E-07 | 515.30621 | 170.24249 |
| SQLLSYIDR | 9 | P33993 | DNA replication licensing factor MCM7 ( | MCM7 | 81.308 | 2 | 51.452545 | yes | 2.4E-07 | 547.79565 | 168.21886 |
| SPQNQYPAELMR | 12 | P33993 | DNA replication licensing factor MCM7 ( | MCM7 | 81.308 | 2 | 51.452545 | yes | 2.119E-06 | 717.3457 | 131.79788 |
| CSILAAANPAYGR | 13 | P33993 | DNA replication licensing factor MCM7 ( | MCM7 | 81.308 | 2 | 51.452545 | yes | 9.722E-08 | 682.34296 | 125.72558 |
| SLEQNIQLPAALLS | 15 | P33993 | DNA replication licensing factor MCM7 ( | MCM7 | 81.308 | 2 | 51.452545 | yes | 9.722E-08 | 826.97015 | 106.9201 |
| SITVLVEGENTR | 12 | P33993 | DNA replication licensing factor MCM7 ( | MCM7 | 81.308 | 2 | 51.452545 | yes | 4.513E-07 | 659.35406 | 84.627686 |
| LFADAVQELLPQY | 14 | P33993 | DNA replication licensing factor MCM7 ( | MCM7 | 81.308 | 2 | 51.452545 | yes | 5.118E-07 | 817.94305 | 83.900612 |
| FLQEFYQDDELGK | 13 | P33993 | DNA replication licensing factor MCM7 ( | MCM7 | 81.308 | 2 | 51.452545 | yes | 1.846E-07 | 816.38306 | 64.694923 |
| LAASIAPEIYGHED | 17 | P33993 | DNA replication licensing factor MCM7 ( | MCM7 | 81.308 | 3 | 51.452545 | yes | 3.895E-07 | 614.3316 | 62.690731 |
| LAASIAPEIYGHED | 16 | P33993 | DNA replication licensing factor MCM7 ( | MCM7 | 81.308 | 3 | 51.452545 | yes | 0.0001243 | 571.6333 | 53.119183 |
| QIAEEDFYEK | 10 | P33993 | DNA replication licensing factor MCM7 ( | MCM7 | 81.308 | 2 | 51.452545 | yes | 1.248E-07 | 636.29315 | 50.517391 |
| YGNQLVR | 7 | P33993 | DNA replication licensing factor MCM7 ( | MCM7 | 81.308 | 2 | 51.452545 | yes | 5.212E-07 | 425.23251 | 49.916035 |
| QVVQGLLSETYLE | 16 | P33993 | DNA replication licensing factor MCM7 ( | MCM7 | 81.308 | 3 | 51.452545 | yes | 5.548E-07 | 614.99512 | 36.082272 |
| SICSTTNIQCK | 11 | P34096 | Ribonuclease 4 OS=Homo sapiens OX=9 | RNASE4 | 16.841 | 2 | 51.561146 | yes | 9.722E-08 | 656.30524 | 82.426262 |
| VHLSDSER | 8 | P34741 | Syndecan-2 OS=Homo sapiens OX=9606 | SDC2 | 22.16 | 2 | 43.75333 | yes | 2.46E-07 | 471.7356 | 356.08871 |
| MDPAEEDTNVYTI | 14 | P34741 | Syndecan-2 OS=Homo sapiens OX=9606 | SDC2 | 22.16 | 2 | 43.75333 | yes | 3.772E-07 | 821.35107 | 29.830755 |
| VGLELIASENFASF | 14 | P34896 | Serine hydroxymethyltransferase, cytosoli | SHMT1 | 53.083 | 2 | 42.73521 | yes | 2.896E-07 | 753.40155 | 55.571369 |
| GLELIASENFCSR | 13 | P34897 | Serine hydroxymethyltransferase, mitocho | SHMT2 | 55.993 | 2 | 52.515347 | yes | 9.722E-08 | 748.36414 | 130.96205 |
| AMADALLER | 9 | P34897 | Serine hydroxymethyltransferase, mitocho | SHMT2 | 55.993 | 2 | 52.515347 | yes | 4.781E-07 | 495.25787 | 107.8258 |
| GYSLVSGGTDNHL | 21 | P34897 | Serine hydroxymethyltransferase, mitocho | SHMT2 | 55.993 | 3 | 52.515347 | yes | 9.722E-08 | 747.40594 | 100.90723 |
| TGLIDYNQLALTA | 14 | P34897 | Serine hydroxymethyltransferase, mitocho | SHMT2 | 55.993 | 2 | 52.515347 | yes | 9.722E-08 | 774.92267 | 84.831276 |
| AALEALGSCLNNK | 13 | P34897 | Serine hydroxymethyltransferase, mitocho | SHMT2 | 55.993 | 2 | 52.515347 | yes | 4.361E-07 | 680.84808 | 52.943642 |
| VLELVSITANK | 11 | P34897 | Serine hydroxymethyltransferase, mitocho | SHMT2 | 55.993 | 2 | 52.515347 | yes | 2.148E-07 | 593.85571 | 42.006222 |
| AFPMPGFDEH | 10 | P34897 | Serine hydroxymethyltransferase, mitocho | SHMT2 | 55.993 | 2 | 52.515347 | yes | 4.556E-06 | 574.2475 | 28.801125 |
| SVMDATQIAGLNC | 15 | P34932 | Heat shock 70 kDa protein 4 OS=Homo s | HSPA4 | 94.331 | 2 | 54.17608 | yes | 1.204E-07 | 824.91089 | 196.24438 |
| EDIYAVEIVGGATI | 14 | P34932 | Heat shock 70 kDa protein 4 OS=Homo s | HSPA4 | 94.331 | 2 | 54.17608 | yes | 5.167E-07 | 746.88574 | 131.65948 |
| VLATAFDTTLGGR | 13 | P34932 | Heat shock 70 kDa protein 4 OS=Homo s | HSPA4 | 94.331 | 2 | 54.17608 | yes | 3.566E-07 | 661.35913 | 67.789925 |
| AGGIETIANEYSDF | 14 | P34932 | Heat shock 70 kDa protein 4 OS=Homo s | HSPA4 | 94.331 | 2 | 54.17608 | yes | 9.722E-08 | 748.3548 | 59.404442 |
| AFSDPFVEAEK | 11 | P34932 | Heat shock 70 kDa protein 4 OS=Homo s | HSPA4 | 94.331 | 2 | 54.17608 | yes | 2.267E-07 | 620.29822 | 57.384296 |
| ELTSTCSPIISK | 12 | P34932 | Heat shock 70 kDa protein 4 OS=Homo s | HSPA4 | 94.331 | 2 | 54.17608 | yes | 3.082E-07 | 668.34485 | 51.443516 |
| GGFGEVCACQVR | 12 | P34947 | G protein-coupled receptor kinase 5 OS=I | GRK5 | 67.787 | 2 | 51.43217 | no | 9.722E-08 | 670.29773 | 70.743416 |

| | | | | | | | | | | | |
|---|---|---|---|---|---|---|---|---|---|---|---|
| DVLDIEQFSTVK | 12 | P34947 | G protein-coupled receptor kinase 5 OS=l | GRK5 | 67.787 | 2 | 51.43217 | no | 6.435E-07 | 697.36414 | 42.851681 |
| DLKPENILLDDYG | 16 | P34947 | G protein-coupled receptor kinase 5 OS=l | GRK5 | 67.787 | 3 | 51.43217 | yes | 3.306E-07 | 637.67053 | 42.370213 |
| VLETEEVYSHK | 11 | P34947 | G protein-coupled receptor kinase 5 OS=l | GRK5 | 67.787 | 2 | 51.43217 | yes | 1.175E-07 | 667.33533 | 40.763657 |
| VLEGNEQFINAAK | 13 | P35030 | Trypsin-3 OS=Homo sapiens OX=9606 C | PRSS3 | 32.529 | 2 | 47.89612 | yes | 1.236E-07 | 716.87518 | 6086.2832 |
| VAQVPLGPECSR | 12 | P35052 | Glypican-1 OS=Homo sapiens OX=9606 | GPC1 | 61.681 | 2 | 49.57897 | yes | 3.594E-07 | 656.83752 | 111.77221 |
| SFVQGLGVASDV\ | 14 | P35052 | Glypican-1 OS=Homo sapiens OX=9606 | GPC1 | 61.681 | 2 | 49.57897 | yes | 1.02E-07 | 717.39099 | 69.349487 |
| MALSTASDDR | 10 | P35052 | Glypican-1 OS=Homo sapiens OX=9606 | GPC1 | 61.681 | 2 | 49.57897 | yes | 4.559E-06 | 533.7453 | 58.91227 |
| GCLANQADLDAE\ | 14 | P35052 | Glypican-1 OS=Homo sapiens OX=9606 | GPC1 | 61.681 | 2 | 49.57897 | yes | 1.823E-07 | 809.86755 | 33.838371 |
| MAGWQSYVDNLM | 29 | P35080 | Profilin-2 OS=Homo sapiens OX=9606 C | PFN2 | 15.047 | 4 | 50.09258 | yes | 3.364E-06 | 847.84308 | 100.95511 |
| MAGWQSYVDNLM | 29 | P35080 | Profilin-2 OS=Homo sapiens OX=9606 C | PFN2 | 15.047 | 4 | 50.09258 | yes | 2.405E-06 | 843.84436 | 100.95511 |
| DSLYVDGDCTMD | 14 | P35080 | Profilin-2 OS=Homo sapiens OX=9606 C | PFN2 | 15.047 | 2 | 50.09258 | yes | 9.903E-08 | 830.35309 | 105.61487 |
| VLVFVMGK | 8 | P35080 | Profilin-2 OS=Homo sapiens OX=9606 C | PFN2 | 15.047 | 2 | 50.09258 | yes | 7.022E-07 | 446.76987 | 16.660913 |
| SQGMASLNLPAVS | 15 | P35221 | Catenin alpha-1 OS=Homo sapiens OX=9 | CTNNA1 | 100.072 | 2 | 52.194473 | yes | 2.287E-06 | 794.91125 | 247.14842 |
| TAVHAGNINFK | 11 | P35221 | Catenin alpha-1 OS=Homo sapiens OX=9 | CTNNA1 | 100.072 | 2 | 52.194473 | yes | 9.722E-08 | 607.31982 | 190.16797 |
| LLSNTVMPR | 9 | P35221 | Catenin alpha-1 OS=Homo sapiens OX=9 | CTNNA1 | 100.072 | 2 | 52.194473 | yes | 1.198E-06 | 515.78931 | 167.76614 |
| EELVAAVEDVR | 11 | P35221 | Catenin alpha-1 OS=Homo sapiens OX=9 | CTNNA1 | 100.072 | 2 | 52.194473 | no | 5.526E-07 | 615.32227 | 139.21199 |
| QIIVDPLSFSEER | 13 | P35221 | Catenin alpha-1 OS=Homo sapiens OX=9 | CTNNA1 | 100.072 | 2 | 52.194473 | yes | 5.142E-07 | 766.90137 | 127.57834 |
| CVIALQEK | 8 | P35221 | Catenin alpha-1 OS=Homo sapiens OX=9 | CTNNA1 | 100.072 | 2 | 52.194473 | yes | 3.843E-06 | 480.76279 | 118.89769 |
| AAAGEFADDPCSS | 15 | P35221 | Catenin alpha-1 OS=Homo sapiens OX=9 | CTNNA1 | 100.072 | 2 | 52.194473 | yes | 9.722E-08 | 762.83539 | 87.823006 |
| QDLLAYLQR | 9 | P35221 | Catenin alpha-1 OS=Homo sapiens OX=9 | CTNNA1 | 100.072 | 2 | 52.194473 | no | 2.086E-07 | 560.31146 | 62.514374 |
| LAQENMDLFK | 10 | P35221 | Catenin alpha-1 OS=Homo sapiens OX=9 | CTNNA1 | 100.072 | 2 | 52.194473 | yes | 9.722E-08 | 604.80261 | 59.849838 |
| IAEQVASFQEEK | 12 | P35221 | Catenin alpha-1 OS=Homo sapiens OX=9 | CTNNA1 | 100.072 | 2 | 52.194473 | yes | 1.146E-07 | 689.84607 | 49.800232 |
| LSVELTSSLFR | 11 | P35222 | Catenin beta-1 OS=Homo sapiens OX=9( | CTNNB1 | 85.497 | 2 | 49.501133 | yes | 3.525E-07 | 626.35083 | 137.76227 |
| LVQNCLWTLR | 10 | P35222 | Catenin beta-1 OS=Homo sapiens OX=9( | CTNNB1 | 85.497 | 2 | 49.501133 | no | 1.025E-07 | 651.85297 | 92.101021 |
| LLNDEDQVVVNK | 12 | P35222 | Catenin beta-1 OS=Homo sapiens OX=9( | CTNNB1 | 85.497 | 2 | 49.501133 | yes | 8.391E-07 | 693.36719 | 63.933609 |
| SPQMVSAIVR | 10 | P35222 | Catenin beta-1 OS=Homo sapiens OX=9( | CTNNB1 | 85.497 | 2 | 49.501133 | yes | 2.99E-05 | 544.30005 | 62.141598 |
| LHYGLPVVVK | 10 | P35222 | Catenin beta-1 OS=Homo sapiens OX=9( | CTNNB1 | 85.497 | 2 | 49.501133 | yes | 5.004E-06 | 562.84497 | 48.015259 |
| TSMGGTQQQFVE( | 15 | P35222 | Catenin beta-1 OS=Homo sapiens OX=9( | CTNNB1 | 85.497 | 2 | 49.501133 | yes | 1.161E-07 | 812.89105 | 47.389202 |
| VAAGVLCELAQDI | 13 | P35222 | Catenin beta-1 OS=Homo sapiens OX=9( | CTNNB1 | 85.497 | 2 | 49.501133 | no | 2.869E-07 | 687.35828 | 43.989677 |
| AAVMVHQLSK | 10 | P35222 | Catenin beta-1 OS=Homo sapiens OX=9( | CTNNB1 | 85.497 | 2 | 49.501133 | yes | 4.39E-07 | 542.30261 | 43.895306 |
| EAAEAIEAEGATA | 22 | P35222 | Catenin beta-1 OS=Homo sapiens OX=9( | CTNNB1 | 85.497 | 3 | 49.501133 | yes | 3.708E-07 | 760.38867 | 10.798688 |
| TRPLLNIR | 8 | P35226 | Polycomb complex protein BMI-1 OS=H | BMI1 | 36.949 | 2 | 54.13409 | yes | 3.467E-07 | 491.81146 | 48.686203 |
| DGLTNAGELESDS | 17 | P35226 | Polycomb complex protein BMI-1 OS=H | BMI1 | 36.949 | 2 | 54.13409 | yes | 9.722E-08 | 847.87122 | 30.034098 |
| VLPSITTEILK | 11 | P35232 | Prohibitin 1 OS=Homo sapiens OX=960( | PHB1 | 29.805 | 2 | 39.56684 | yes | 4.508E-07 | 607.37372 | 38.379505 |
| SCDFLSSFR | 9 | P35237 | Serpin B6 OS=Homo sapiens OX=9606 ( | SERPINB6 | 42.622 | 2 | 32.74133 | yes | 1.426E-06 | 559.75043 | 41.914886 |
| ITAWYAEHR | 9 | P35240 | Merlin OS=Homo sapiens OX=9606 GN= | NF2 | 69.691 | 2 | 49.567745 | yes | 3.002E-07 | 573.78815 | 162.39963 |
| EEATMANEALMR | 12 | P35240 | Merlin OS=Homo sapiens OX=9606 GN= | NF2 | 69.691 | 2 | 49.567745 | yes | 2.26E-07 | 683.31049 | 123.31573 |
| VLEAEVLALK | 10 | P35240 | Merlin OS=Homo sapiens OX=9606 GN= | NF2 | 69.691 | 2 | 49.567745 | yes | 1.557E-07 | 542.83423 | 120.17975 |
| ISFPWNEIR | 9 | P35240 | Merlin OS=Homo sapiens OX=9606 GN= | NF2 | 69.691 | 2 | 49.567745 | yes | 2.923E-07 | 581.30621 | 97.400658 |

| | | | | | | | | | | |
|---|---|---|---|---|---|---|---|---|---|---|
| HLQEQLNELK | 10 | P35240 | Merlin OS=Homo sapiens OX=9606 GN= | NF2 | 69.691 | 2 | 49.567745 | yes | 1.021E-07 | 626.33826 | 58.975136 |
| GFLAQEELLPK | 11 | P35240 | Merlin OS=Homo sapiens OX=9606 GN= | NF2 | 69.691 | 2 | 49.567745 | yes | 4.77E-07 | 622.8479 | 54.794231 |
| ARDEAEMEYLK | 11 | P35240 | Merlin OS=Homo sapiens OX=9606 GN= | NF2 | 69.691 | 2 | 49.567745 | yes | 4.239E-07 | 677.81903 | 36.483543 |
| AAEAEQEMQR | 10 | P35240 | Merlin OS=Homo sapiens OX=9606 GN= | NF2 | 69.691 | 2 | 49.567745 | yes | 6.541E-06 | 581.76147 | 33.943993 |
| IGFPWSEIR | 9 | P35241 | Radixin OS=Homo sapiens OX=9606 GN | RDX | 68.564 | 2 | 53.50245 | no | 9.722E-08 | 552.79547 | 903.55518 |
| FVIKPIDK | 8 | P35241 | Radixin OS=Homo sapiens OX=9606 GN | RDX | 68.564 | 2 | 53.50245 | no | 9.9E-08 | 480.29987 | 669.92511 |
| APDFVFYAPR | 10 | P35241 | Radixin OS=Homo sapiens OX=9606 GN | RDX | 68.564 | 2 | 53.50245 | no | 9.819E-08 | 591.80072 | 407.39673 |
| KAPDFVFYAPR | 11 | P35241 | Radixin OS=Homo sapiens OX=9606 GN | RDX | 68.564 | 2 | 53.50245 | no | 1.151E-07 | 655.84821 | 268.3331 |
| ALELDQER | 8 | P35241 | Radixin OS=Homo sapiens OX=9606 GN | RDX | 68.564 | 2 | 53.50245 | yes | 9.722E-08 | 487.2511 | 174.65099 |
| AFAAQEDLEK | 10 | P35241 | Radixin OS=Homo sapiens OX=9606 GN | RDX | 68.564 | 2 | 53.50245 | yes | 7.862E-07 | 561.27728 | 148.10262 |
| LFFLQVK | 7 | P35241 | Radixin OS=Homo sapiens OX=9606 GN | RDX | 68.564 | 2 | 53.50245 | no | 1.509E-07 | 447.776 | 143.65338 |
| EKEELMER | 8 | P35241 | Radixin OS=Homo sapiens OX=9606 GN | RDX | 68.564 | 2 | 53.50245 | no | 1.765E-07 | 532.25806 | 138.72621 |
| NQEQLAAELAEFT | 15 | P35241 | Radixin OS=Homo sapiens OX=9606 GN | RDX | 68.564 | 2 | 53.50245 | yes | 1.245E-07 | 831.92029 | 105.1815 |
| IALLEEAK | 8 | P35241 | Radixin OS=Homo sapiens OX=9606 GN | RDX | 68.564 | 2 | 53.50245 | yes | 4.099E-07 | 443.76584 | 38.297447 |
| EDSMMEYLK | 9 | P35241 | Radixin OS=Homo sapiens OX=9606 GN | RDX | 68.564 | 2 | 53.50245 | yes | 3.071E-05 | 573.24628 | 33.837116 |
| RKPDTIEVQQMK | 12 | P35241 | Radixin OS=Homo sapiens OX=9606 GN | RDX | 68.564 | 2 | 53.50245 | no | 1.026E-07 | 736.89813 | 23.201164 |
| MFILSDGEGK | 10 | P35244 | Replication protein A 14 kDa subunit OS= | RPA3 | 13.569 | 2 | 49.763615 | yes | 3.766E-07 | 548.77081 | 229.35643 |
| ATILCTSYVQFK | 12 | P35244 | Replication protein A 14 kDa subunit OS= | RPA3 | 13.569 | 2 | 49.763615 | yes | 1.004E-07 | 715.87103 | 133.27708 |
| IIEPLTSR | 8 | P35249 | Replication factor C subunit 4 OS=Homo | RFC4 | 39.682 | 2 | 53.658535 | yes | 2.084E-07 | 464.77673 | 1516.5094 |
| VITDIAGVIPAEK | 13 | P35249 | Replication factor C subunit 4 OS=Homo | RFC4 | 39.682 | 2 | 53.658535 | yes | 9.938E-08 | 663.38739 | 1213.9518 |
| VLELNASDER | 10 | P35249 | Replication factor C subunit 4 OS=Homo | RFC4 | 39.682 | 2 | 53.658535 | yes | 9.722E-08 | 573.29346 | 698.2171 |
| ISDEGIAYLVK | 11 | P35249 | Replication factor C subunit 4 OS=Homo | RFC4 | 39.682 | 2 | 53.658535 | yes | 1.678E-07 | 604.33209 | 626.48822 |
| ELFGPELFR | 9 | P35249 | Replication factor C subunit 4 OS=Homo | RFC4 | 39.682 | 2 | 53.658535 | yes | 1.22E-07 | 554.29529 | 618.39563 |
| NFAQLTVSGSR | 11 | P35249 | Replication factor C subunit 4 OS=Homo | RFC4 | 39.682 | 2 | 53.658535 | yes | 9.722E-08 | 590.30945 | 586.40094 |
| AKPVPWVEK | 9 | P35249 | Replication factor C subunit 4 OS=Homo | RFC4 | 39.682 | 2 | 53.658535 | yes | 1.108E-07 | 527.30823 | 524.14471 |
| GTSISTKPPLTK | 12 | P35249 | Replication factor C subunit 4 OS=Homo | RFC4 | 39.682 | 2 | 53.658535 | yes | 9.722E-08 | 615.35864 | 451.09003 |
| AITFLQSATR | 10 | P35249 | Replication factor C subunit 4 OS=Homo | RFC4 | 39.682 | 2 | 53.658535 | yes | 9.819E-08 | 554.31146 | 274.71268 |
| IDGVFAACQSGSFl | 15 | P35249 | Replication factor C subunit 4 OS=Homo | RFC4 | 39.682 | 2 | 53.658535 | yes | 1.341E-07 | 801.36688 | 272.66824 |
| GVAASAGSSGENK | 13 | P35249 | Replication factor C subunit 4 OS=Homo | RFC4 | 39.682 | 2 | 53.658535 | yes | 1.165E-07 | 567.77289 | 130.68945 |
| IDGVFAACQSGSFl | 21 | P35249 | Replication factor C subunit 4 OS=Homo | RFC4 | 39.682 | 3 | 53.658535 | yes | 7.848E-07 | 747.71216 | 82.184586 |
| FCLICNYVSR | 10 | P35249 | Replication factor C subunit 4 OS=Homo | RFC4 | 39.682 | 2 | 53.658535 | yes | 5.385E-07 | 666.31537 | 77.102005 |
| LTDAQILTR | 9 | P35250 | Replication factor C subunit 2 OS=Homo | RFC2 | 39.158 | 2 | 51.4284 | yes | 9.937E-08 | 515.79822 | 1578.556 |
| EGNVPNIIIAGPPG | 17 | P35250 | Replication factor C subunit 2 OS=Homo | RFC2 | 39.158 | 2 | 51.4284 | yes | 2.149E-07 | 817.44904 | 1386.5424 |
| LNEIVGNEDTVSR | 13 | P35250 | Replication factor C subunit 2 OS=Homo | RFC2 | 39.158 | 2 | 51.4284 | yes | 9.722E-08 | 723.36517 | 1087.7386 |
| TTSILCLAR | 9 | P35250 | Replication factor C subunit 2 OS=Homo | RFC2 | 39.158 | 2 | 51.4284 | yes | 1.455E-07 | 517.7868 | 924.52869 |
| DAMLELNASNDR | 12 | P35250 | Replication factor C subunit 2 OS=Homo | RFC2 | 39.158 | 2 | 51.4284 | yes | 1.102E-07 | 674.81171 | 766.24341 |
| FALACNASDK | 10 | P35250 | Replication factor C subunit 2 OS=Homo | RFC2 | 39.158 | 2 | 51.4284 | yes | 1.152E-07 | 548.75824 | 718.70923 |
| VCDEPHPLLVK | 11 | P35250 | Replication factor C subunit 2 OS=Homo | RFC2 | 39.158 | 2 | 51.4284 | yes | 2.739E-07 | 653.84485 | 264.91946 |
| VCDEPHPLLVK | 11 | P35250 | Replication factor C subunit 2 OS=Homo | RFC2 | 39.158 | 2 | 51.4284 | yes | 3.587E-06 | 436.2323 | 264.91946 |

| | | | | | | | | | | | |
|---|---|---|---|---|---|---|---|---|---|---|---|
| VCDEPHPLLVK | 11 | P35250 | Replication factor C subunit 2 OS=Homo | RFC2 | 39.158 | 3 | 51.4284 | yes | 2.739E-07 | 653.84485 | 264.91946 |
| VCDEPHPLLVK | 11 | P35250 | Replication factor C subunit 2 OS=Homo | RFC2 | 39.158 | 3 | 51.4284 | yes | 3.587E-06 | 436.2323 | 264.91946 |
| IIEPIQSR | 8 | P35250 | Replication factor C subunit 2 OS=Homo | RFC2 | 39.158 | 2 | 51.4284 | yes | 1.043E-07 | 478.2822 | 491.68027 |
| EIGYTHMK | 8 | P35250 | Replication factor C subunit 2 OS=Homo | RFC2 | 39.158 | 2 | 51.4284 | yes | 1.062E-07 | 489.73929 | 361.66486 |
| TFQMAEYLK | 9 | P35250 | Replication factor C subunit 2 OS=Homo | RFC2 | 39.158 | 2 | 51.4284 | yes | 1.022E-07 | 565.78119 | 344.763 |
| APGSAGHYELPWV | 15 | P35250 | Replication factor C subunit 2 OS=Homo | RFC2 | 39.158 | 2 | 51.4284 | yes | 1.789E-07 | 820.90698 | 102.47947 |
| LMNVIEK | 7 | P35250 | Replication factor C subunit 2 OS=Homo | RFC2 | 39.158 | 2 | 51.4284 | yes | 2.648E-07 | 423.7413 | 44.255161 |
| AIVAESLNNTSIK | 13 | P35251 | Replication factor C subunit 1 OS=Homo | RFC1 | 128.255 | 2 | 55.81233 | yes | 9.722E-08 | 680.37756 | 738.18323 |
| MGPFDVAR | 8 | P35251 | Replication factor C subunit 1 OS=Homo | RFC1 | 128.255 | 2 | 55.81233 | yes | 2.369E-07 | 446.72092 | 724.85706 |
| IPPPAMNEIILGAN | 18 | P35251 | Replication factor C subunit 1 OS=Homo | RFC1 | 128.255 | 3 | 55.81233 | yes | 2.139E-07 | 654.68744 | 606.04999 |
| AALLSGPPGVGK | 12 | P35251 | Replication factor C subunit 1 OS=Homo | RFC1 | 128.255 | 2 | 55.81233 | yes | 3.348E-07 | 533.81641 | 584.45264 |
| IPIICMCNDR | 10 | P35251 | Replication factor C subunit 1 OS=Homo | RFC1 | 128.255 | 2 | 55.81233 | yes | 2.606E-07 | 646.30139 | 536.93744 |
| LPVSSKPGK | 9 | P35251 | Replication factor C subunit 1 OS=Homo | RFC1 | 128.255 | 2 | 55.81233 | yes | 3.2E-07 | 456.7793 | 459.49698 |
| TLAMLDEEPK | 10 | P35251 | Replication factor C subunit 1 OS=Homo | RFC1 | 128.255 | 2 | 55.81233 | yes | 1.014E-07 | 573.78918 | 433.86569 |
| TIIGQQGDQSCANI | 14 | P35251 | Replication factor C subunit 1 OS=Homo | RFC1 | 128.255 | 2 | 55.81233 | yes | 9.722E-08 | 760.36212 | 390.54059 |
| IIDEDGLLNLIR | 12 | P35251 | Replication factor C subunit 1 OS=Homo | RFC1 | 128.255 | 2 | 55.81233 | yes | 9.766E-08 | 692.39575 | 333.80521 |
| SLVHYCFDLR | 10 | P35251 | Replication factor C subunit 1 OS=Homo | RFC1 | 128.255 | 2 | 55.81233 | yes | 1.022E-07 | 655.32153 | 305.66974 |
| IVQDLALHMSLR | 12 | P35251 | Replication factor C subunit 1 OS=Homo | RFC1 | 128.255 | 2 | 55.81233 | yes | 1.09E-07 | 698.39246 | 267.46652 |
| KFFGVIPSGK | 10 | P35251 | Replication factor C subunit 1 OS=Homo | RFC1 | 128.255 | 2 | 55.81233 | yes | 1.168E-07 | 540.31604 | 214.00455 |
| TVNMDYLSLLR | 11 | P35251 | Replication factor C subunit 1 OS=Homo | RFC1 | 128.255 | 2 | 55.81233 | yes | 9.722E-08 | 662.8501 | 207.94702 |
| QLQLDEDAELER | 12 | P35251 | Replication factor C subunit 1 OS=Homo | RFC1 | 128.255 | 2 | 55.81233 | yes | 9.722E-08 | 729.85718 | 186.08864 |
| KTNYLVMGR | 9 | P35251 | Replication factor C subunit 1 OS=Homo | RFC1 | 128.255 | 2 | 55.81233 | yes | 9.722E-08 | 541.2948 | 172.68187 |
| TNYQAYR | 7 | P35251 | Replication factor C subunit 1 OS=Homo | RFC1 | 128.255 | 2 | 55.81233 | yes | 1.311E-07 | 458.2196 | 169.25684 |
| HALIMDEVDGMA | 17 | P35251 | Replication factor C subunit 1 OS=Homo | RFC1 | 128.255 | 3 | 55.81233 | yes | 9.926E-08 | 624.94531 | 158.38153 |
| EQVAEETSGDSK | 12 | P35251 | Replication factor C subunit 1 OS=Homo | RFC1 | 128.255 | 2 | 55.81233 | yes | 9.722E-08 | 640.28607 | 155.22948 |
| YKPTSLK | 7 | P35251 | Replication factor C subunit 1 OS=Homo | RFC1 | 128.255 | 2 | 55.81233 | yes | 1.848E-07 | 418.74747 | 154.66418 |
| HLMLLSR | 7 | P35251 | Replication factor C subunit 1 OS=Homo | RFC1 | 128.255 | 2 | 55.81233 | yes | 1.795E-07 | 435.25491 | 152.58797 |
| FFGVIPSGK | 9 | P35251 | Replication factor C subunit 1 OS=Homo | RFC1 | 128.255 | 2 | 55.81233 | yes | 2.073E-07 | 476.26855 | 142.79233 |
| KLVSETVK | 8 | P35251 | Replication factor C subunit 1 OS=Homo | RFC1 | 128.255 | 2 | 55.81233 | yes | 2.014E-07 | 452.27911 | 125.74948 |
| QLHEDEEFAR | 10 | P35251 | Replication factor C subunit 1 OS=Homo | RFC1 | 128.255 | 2 | 55.81233 | yes | 1.15E-07 | 637.29401 | 123.11689 |
| TAQVSDER | 8 | P35251 | Replication factor C subunit 1 OS=Homo | RFC1 | 128.255 | 2 | 55.81233 | yes | 4.677E-07 | 453.21979 | 117.47472 |
| ESVSPEDSEK | 10 | P35251 | Replication factor C subunit 1 OS=Homo | RFC1 | 128.255 | 2 | 55.81233 | yes | 2.578E-07 | 553.74603 | 105.9286 |
| SKYEIAVETEMK | 12 | P35251 | Replication factor C subunit 1 OS=Homo | RFC1 | 128.255 | 2 | 55.81233 | yes | 2.019E-07 | 714.35797 | 94.396355 |
| NKPLSPIK | 8 | P35251 | Replication factor C subunit 1 OS=Homo | RFC1 | 128.255 | 2 | 55.81233 | yes | 2.193E-07 | 448.78183 | 81.439651 |
| TNYLVMGR | 8 | P35251 | Replication factor C subunit 1 OS=Homo | RFC1 | 128.255 | 2 | 55.81233 | yes | 9.627E-07 | 477.24728 | 72.51284 |
| VFAAGEETAHMSI | 16 | P35251 | Replication factor C subunit 1 OS=Homo | RFC1 | 128.255 | 3 | 55.81233 | yes | 1.756E-07 | 568.9469 | 72.04026 |
| QVLHNLSMWCAR | 12 | P35251 | Replication factor C subunit 1 OS=Homo | RFC1 | 128.255 | 3 | 55.81233 | yes | 1.554E-07 | 505.58334 | 32.418594 |
| QVLHNLSMWCAR | 12 | P35251 | Replication factor C subunit 1 OS=Homo | RFC1 | 128.255 | 3 | 55.81233 | yes | 1.477E-07 | 757.8714 | 32.418594 |
| QVLHNLSMWCAR | 12 | P35251 | Replication factor C subunit 1 OS=Homo | RFC1 | 128.255 | 2 | 55.81233 | yes | 1.554E-07 | 505.58334 | 32.418594 |

| | | | | | | | | | | | |
|---|---|---|---|---|---|---|---|---|---|---|---|
| QVLHNLSMWCAR | 12 | P35251 | Replication factor C subunit 1 OS=Homo | RFC1 | 128.255 | 2 | 55.81233 | yes | 1.477E-07 | 757.8714 | 32.418594 |
| GGIQELIGLIK | 11 | P35251 | Replication factor C subunit 1 OS=Homo | RFC1 | 128.255 | 2 | 55.81233 | yes | 5.036E-07 | 570.85297 | 47.08844 |
| KVFAAGEETAHM: | 17 | P35251 | Replication factor C subunit 1 OS=Homo | RFC1 | 128.255 | 3 | 55.81233 | yes | 2.983E-06 | 611.6452 | 45.42112 |
| ITVTSEVPFSK | 11 | P35268 | Large ribosomal subunit protein eL22 OS | RPL22 | 14.787 | 2 | 49.133495 | yes | 1.286E-07 | 604.33209 | 39058.371 |
| AGNLGGGVVTIER | 13 | P35268 | Large ribosomal subunit protein eL22 OS | RPL22 | 14.787 | 2 | 49.133495 | yes | 1.065E-07 | 621.84369 | 36332.805 |
| LDTGPQSLSGK | 11 | P35269 | General transcription factor IIF subunit 1 | GTF2F1 | 58.241 | 2 | 39.984554 | yes | 3.073E-06 | 551.79059 | 87.170792 |
| TLTAEEAEEEWER | 13 | P35269 | General transcription factor IIF subunit 1 | GTF2F1 | 58.241 | 2 | 39.984554 | yes | 4.271E-07 | 796.85736 | 24.045692 |
| TTPNSGDVQVTED | 16 | P35269 | General transcription factor IIF subunit 1 | GTF2F1 | 58.241 | 2 | 39.984554 | yes | 2.085E-06 | 844.90796 | 3.7955754 |
| CVVILPDSVR | 10 | P35520 | Cystathionine beta-synthase OS=Homo sa | CBS | 60.587 | 2 | 49.62882 | yes | 1.017E-07 | 579.32098 | 141.41365 |
| EPLWIRPDAPSR | 12 | P35520 | Cystathionine beta-synthase OS=Homo sa | CBS | 60.587 | 3 | 49.62882 | yes | 3.963E-06 | 479.59302 | 44.722775 |
| TLLDIDNTR | 9 | P35527 | Keratin, type I cytoskeletal 9 OS=Homo s | KRT9 | 62.065 | 2 | 52.731136 | yes | 2.139E-07 | 530.78528 | 1057.1594 |
| IKFEMEQNLR | 10 | P35527 | Keratin, type I cytoskeletal 9 OS=Homo s | KRT9 | 62.065 | 2 | 52.731136 | yes | 9.984E-08 | 654.34247 | 221.21559 |
| IKFEMEQNLR | 10 | P35527 | Keratin, type I cytoskeletal 9 OS=Homo s | KRT9 | 62.065 | 2 | 52.731136 | yes | 9.722E-08 | 662.3399 | 221.21559 |
| QGVDADINGLR | 11 | P35527 | Keratin, type I cytoskeletal 9 OS=Homo s | KRT9 | 62.065 | 2 | 52.731136 | yes | 1.035E-07 | 579.29907 | 437.27786 |
| VQALEEANNDLEN | 14 | P35527 | Keratin, type I cytoskeletal 9 OS=Homo s | KRT9 | 62.065 | 2 | 52.731136 | yes | 9.722E-08 | 793.88647 | 407.60614 |
| SGGGGGGGLGSGC | 16 | P35527 | Keratin, type I cytoskeletal 9 OS=Homo s | KRT9 | 62.065 | 2 | 52.731136 | yes | 9.722E-08 | 616.80255 | 389.17343 |
| HGVQELEIELQSQI | 16 | P35527 | Keratin, type I cytoskeletal 9 OS=Homo s | KRT9 | 62.065 | 3 | 52.731136 | yes | 9.722E-08 | 613.32666 | 286.00995 |
| DQIVDLTVGNNK | 12 | P35527 | Keratin, type I cytoskeletal 9 OS=Homo s | KRT9 | 62.065 | 2 | 52.731136 | yes | 9.722E-08 | 658.34625 | 237.9821 |
| FEMEQNLR | 8 | P35527 | Keratin, type I cytoskeletal 9 OS=Homo s | KRT9 | 62.065 | 2 | 52.731136 | yes | 1.304E-06 | 533.75293 | 155.98959 |
| FSSSSGYGGGSSR | 13 | P35527 | Keratin, type I cytoskeletal 9 OS=Homo s | KRT9 | 62.065 | 2 | 52.731136 | yes | 3.418E-07 | 618.26801 | 135.62851 |
| MTLDDFR | 7 | P35527 | Keratin, type I cytoskeletal 9 OS=Homo s | KRT9 | 62.065 | 2 | 52.731136 | yes | 1.367E-07 | 449.21039 | 132.651 |
| QEYEQLIAK | 9 | P35527 | Keratin, type I cytoskeletal 9 OS=Homo s | KRT9 | 62.065 | 2 | 52.731136 | yes | 2.01E-07 | 561.29547 | 129.41739 |
| QVLDNLTMEK | 10 | P35527 | Keratin, type I cytoskeletal 9 OS=Homo s | KRT9 | 62.065 | 2 | 52.731136 | yes | 9.722E-08 | 603.80536 | 115.15917 |
| HGVQELEIELQSQI | 17 | P35527 | Keratin, type I cytoskeletal 9 OS=Homo s | KRT9 | 62.065 | 3 | 52.731136 | yes | 1.952E-06 | 656.02496 | 73.160126 |
| CECEIGYELDR | 11 | P35555 | Fibrillin-1 OS=Homo sapiens OX=9606 ( | FBN1 | 312.298 | 2 | 50.316162 | yes | 9.819E-08 | 722.29761 | 71.522003 |
| LETCFLR | 7 | P35555 | Fibrillin-1 OS=Homo sapiens OX=9606 ( | FBN1 | 312.298 | 2 | 50.316162 | yes | 2.203E-07 | 469.74185 | 65.916725 |
| CSCSPGWIGDGIK | 13 | P35555 | Fibrillin-1 OS=Homo sapiens OX=9606 ( | FBN1 | 312.298 | 2 | 50.316162 | yes | 1.639E-07 | 718.81848 | 50.994602 |
| CECDMGFVPSADC | 14 | P35555 | Fibrillin-1 OS=Homo sapiens OX=9606 ( | FBN1 | 312.298 | 2 | 50.316162 | yes | 1.036E-07 | 786.81 | 36.023823 |
| VKPLLQVSR | 9 | P35579 | Myosin-9 OS=Homo sapiens OX=9606 C | MYH9 | 226.533 | 2 | 53.599823 | yes | 1.521E-07 | 520.33478 | 12140.817 |
| IMGIPEEEQMGLLI | 14 | P35579 | Myosin-9 OS=Homo sapiens OX=9606 C | MYH9 | 226.533 | 2 | 53.599823 | yes | 1.175E-07 | 808.41278 | 3643.3215 |
| IMGIPEEEQMGLLI | 14 | P35579 | Myosin-9 OS=Homo sapiens OX=9606 C | MYH9 | 226.533 | 2 | 53.599823 | yes | 4.796E-07 | 816.41022 | 3643.3215 |
| IMGIPEEEQMGLLI | 14 | P35579 | Myosin-9 OS=Homo sapiens OX=9606 C | MYH9 | 226.533 | 2 | 53.599823 | yes | 1.847E-06 | 816.41022 | 3643.3215 |
| ALELDSNLYR | 10 | P35579 | Myosin-9 OS=Homo sapiens OX=9606 C | MYH9 | 226.533 | 2 | 53.599823 | yes | 1.165E-07 | 597.31171 | 7869.9004 |
| LDPHLVLDQLR | 11 | P35579 | Myosin-9 OS=Homo sapiens OX=9606 C | MYH9 | 226.533 | 2 | 53.599823 | no | 1.556E-07 | 659.8775 | 6998.5259 |
| AGVLAHLEEER | 11 | P35579 | Myosin-9 OS=Homo sapiens OX=9606 C | MYH9 | 226.533 | 2 | 53.599823 | no | 9.722E-08 | 612.32257 | 6878.8789 |
| EQADFAIEALAK | 12 | P35579 | Myosin-9 OS=Homo sapiens OX=9606 C | MYH9 | 226.533 | 2 | 53.599823 | yes | 1.008E-07 | 653.33789 | 6714.3672 |
| NFINNPLAQADWA | 15 | P35579 | Myosin-9 OS=Homo sapiens OX=9606 C | MYH9 | 226.533 | 2 | 53.599823 | yes | 1.026E-07 | 836.92572 | 6457.3779 |
| LEVNLQAMK | 9 | P35579 | Myosin-9 OS=Homo sapiens OX=9606 C | MYH9 | 226.533 | 2 | 53.599823 | yes | 2.613E-07 | 523.28918 | 3050.8755 |
| LEVNLQAMK | 9 | P35579 | Myosin-9 OS=Homo sapiens OX=9606 C | MYH9 | 226.533 | 2 | 53.599823 | yes | 1.787E-07 | 531.28662 | 3050.8755 |

| | | | | | | | | | | | |
|---|---|---|---|---|---|---|---|---|---|---|---|
| HSQAVEELAEQLE | 16 | P35579 | Myosin-9 OS=Homo sapiens OX=9606 C | MYH9 | 226.533 | 3 | 53.599823 | yes | 1.146E-07 | 613.9743 | 5507.8486 |
| IAQLEEQLDNETK | 13 | P35579 | Myosin-9 OS=Homo sapiens OX=9606 C | MYH9 | 226.533 | 2 | 53.599823 | yes | 2.834E-07 | 765.88593 | 5288.0229 |
| EDQSILCTGESGA( | 15 | P35579 | Myosin-9 OS=Homo sapiens OX=9606 C | MYH9 | 226.533 | 2 | 53.599823 | no | 1.11E-07 | 776.35138 | 5121.0859 |
| KLVWVPSDK | 9 | P35579 | Myosin-9 OS=Homo sapiens OX=9606 C | MYH9 | 226.533 | 2 | 53.599823 | yes | 1.365E-07 | 536.31348 | 4883.0195 |
| DELADEIANSSGK | 13 | P35579 | Myosin-9 OS=Homo sapiens OX=9606 C | MYH9 | 226.533 | 2 | 53.599823 | yes | 9.722E-08 | 674.815 | 4755.981 |
| ALEQQVEEMK | 10 | P35579 | Myosin-9 OS=Homo sapiens OX=9606 C | MYH9 | 226.533 | 2 | 53.599823 | yes | 9.722E-08 | 602.79755 | 2105.1746 |
| ALEQQVEEMK | 10 | P35579 | Myosin-9 OS=Homo sapiens OX=9606 C | MYH9 | 226.533 | 2 | 53.599823 | yes | 2.734E-06 | 610.79498 | 2105.1746 |
| KEEELQAALAR | 11 | P35579 | Myosin-9 OS=Homo sapiens OX=9606 C | MYH9 | 226.533 | 2 | 53.599823 | no | 4.289E-07 | 629.34351 | 4180.207 |
| VSHLLGINVTDFTI | 14 | P35579 | Myosin-9 OS=Homo sapiens OX=9606 C | MYH9 | 226.533 | 2 | 53.599823 | yes | 1.21E-07 | 786.43066 | 4077.012 |
| DLEGLSQR | 8 | P35579 | Myosin-9 OS=Homo sapiens OX=9606 C | MYH9 | 226.533 | 2 | 53.599823 | yes | 1.674E-07 | 459.23798 | 4070.6689 |
| EMEAELEDER | 10 | P35579 | Myosin-9 OS=Homo sapiens OX=9606 C | MYH9 | 226.533 | 2 | 53.599823 | yes | 1.064E-07 | 625.76392 | 4015.5769 |
| TDLLLEPYNK | 10 | P35579 | Myosin-9 OS=Homo sapiens OX=9606 C | MYH9 | 226.533 | 2 | 53.599823 | yes | 9.965E-08 | 603.32428 | 3639.9399 |
| IAEFTTNLTEEEEK | 14 | P35579 | Myosin-9 OS=Homo sapiens OX=9606 C | MYH9 | 226.533 | 2 | 53.599823 | yes | 1.157E-07 | 827.39636 | 3600.5977 |
| EQLEEEEEAK | 10 | P35579 | Myosin-9 OS=Homo sapiens OX=9606 C | MYH9 | 226.533 | 2 | 53.599823 | yes | 1.197E-07 | 617.27771 | 3558.9941 |
| CIIPNHEK | 8 | P35579 | Myosin-9 OS=Homo sapiens OX=9606 C | MYH9 | 226.533 | 2 | 53.599823 | no | 1.743E-07 | 505.75803 | 3197.9609 |
| RGDLPFVVPR | 10 | P35579 | Myosin-9 OS=Homo sapiens OX=9606 C | MYH9 | 226.533 | 2 | 53.599823 | yes | 1.361E-07 | 578.33527 | 3016.418 |
| LQEMEGTVK | 9 | P35579 | Myosin-9 OS=Homo sapiens OX=9606 C | MYH9 | 226.533 | 2 | 53.599823 | yes | 1.499E-07 | 517.763 | 1504.6517 |
| LQEMEGTVK | 9 | P35579 | Myosin-9 OS=Homo sapiens OX=9606 C | MYH9 | 226.533 | 2 | 53.599823 | yes | 3.124E-07 | 525.76044 | 1504.6517 |
| ALEEAMEQK | 9 | P35579 | Myosin-9 OS=Homo sapiens OX=9606 C | MYH9 | 226.533 | 2 | 53.599823 | yes | 1.812E-07 | 524.75262 | 1479.2836 |
| ALEEAMEQK | 9 | P35579 | Myosin-9 OS=Homo sapiens OX=9606 C | MYH9 | 226.533 | 2 | 53.599823 | yes | 1.478E-06 | 532.75006 | 1479.2836 |
| NTNPNFVR | 8 | P35579 | Myosin-9 OS=Homo sapiens OX=9606 C | MYH9 | 226.533 | 2 | 53.599823 | no | 1.66E-07 | 481.24615 | 2825.3057 |
| DLEAHIDSANK | 11 | P35579 | Myosin-9 OS=Homo sapiens OX=9606 C | MYH9 | 226.533 | 2 | 53.599823 | yes | 9.903E-08 | 606.79639 | 2735.8354 |
| QIATLHAQVADMI | 13 | P35579 | Myosin-9 OS=Homo sapiens OX=9606 C | MYH9 | 226.533 | 2 | 53.599823 | yes | 4.932E-07 | 713.37958 | 1320.2932 |
| QIATLHAQVADMI | 13 | P35579 | Myosin-9 OS=Homo sapiens OX=9606 C | MYH9 | 226.533 | 2 | 53.599823 | yes | 4.708E-07 | 475.92215 | 1320.2932 |
| QIATLHAQVADMI | 13 | P35579 | Myosin-9 OS=Homo sapiens OX=9606 C | MYH9 | 226.533 | 3 | 53.599823 | yes | 4.932E-07 | 713.37958 | 1320.2932 |
| QIATLHAQVADMI | 13 | P35579 | Myosin-9 OS=Homo sapiens OX=9606 C | MYH9 | 226.533 | 3 | 53.599823 | yes | 4.708E-07 | 475.92215 | 1320.2932 |
| TEMEDLMSSK | 10 | P35579 | Myosin-9 OS=Homo sapiens OX=9606 C | MYH9 | 226.533 | 2 | 53.599823 | yes | 1.426E-07 | 585.75452 | 1301.7244 |
| TEMEDLMSSK | 10 | P35579 | Myosin-9 OS=Homo sapiens OX=9606 C | MYH9 | 226.533 | 2 | 53.599823 | yes | 6.307E-07 | 593.75195 | 1301.7244 |
| GDLPFVVPR | 9 | P35579 | Myosin-9 OS=Homo sapiens OX=9606 C | MYH9 | 226.533 | 2 | 53.599823 | yes | 1.587E-07 | 500.28473 | 2563.7957 |
| DVLLQVDDER | 10 | P35579 | Myosin-9 OS=Homo sapiens OX=9606 C | MYH9 | 226.533 | 2 | 53.599823 | yes | 1.69E-07 | 601.30658 | 2557.9514 |
| VVFQEFR | 7 | P35579 | Myosin-9 OS=Homo sapiens OX=9606 C | MYH9 | 226.533 | 2 | 53.599823 | yes | 1.323E-07 | 462.75052 | 2440.6011 |
| HEAMITDLEER | 11 | P35579 | Myosin-9 OS=Homo sapiens OX=9606 C | MYH9 | 226.533 | 2 | 53.599823 | yes | 9.722E-08 | 672.31665 | 2383.5947 |
| QTLENER | 7 | P35579 | Myosin-9 OS=Homo sapiens OX=9606 C | MYH9 | 226.533 | 2 | 53.599823 | yes | 1.146E-07 | 445.22232 | 2333.4001 |
| ADFCIIHYAGK | 11 | P35579 | Myosin-9 OS=Homo sapiens OX=9606 C | MYH9 | 226.533 | 2 | 53.599823 | no | 5.218E-07 | 647.8161 | 2330.4319 |
| SGFEPASLK | 9 | P35579 | Myosin-9 OS=Homo sapiens OX=9606 C | MYH9 | 226.533 | 2 | 53.599823 | yes | 9.796E-08 | 468.24527 | 2276.5276 |
| ELEDATETADAMI | 14 | P35579 | Myosin-9 OS=Homo sapiens OX=9606 C | MYH9 | 226.533 | 2 | 53.599823 | yes | 9.85E-08 | 783.34106 | 1131.6505 |
| ELEDATETADAMI | 14 | P35579 | Myosin-9 OS=Homo sapiens OX=9606 C | MYH9 | 226.533 | 2 | 53.599823 | yes | 1.345E-07 | 791.3385 | 1131.6505 |
| KFDQLLAEEK | 10 | P35579 | Myosin-9 OS=Homo sapiens OX=9606 C | MYH9 | 226.533 | 2 | 53.599823 | no | 9.722E-08 | 610.82971 | 2219.0498 |
| KLEMDLK | 7 | P35579 | Myosin-9 OS=Homo sapiens OX=9606 C | MYH9 | 226.533 | 2 | 53.599823 | yes | 6.621E-07 | 438.74658 | 2164.1016 |

| | | | | | | | | | | | |
|---|---|---|---|---|---|---|---|---|---|---|---|
| VEAQLQELQVK | 11 | P35579 | Myosin-9 OS=Homo sapiens OX=9606 C | MYH9 | 226.533 | 2 | 53.599823 | yes | 1.397E-07 | 642.86151 | 2074.1255 |
| QLEEAEEEAQR | 11 | P35579 | Myosin-9 OS=Homo sapiens OX=9606 C | MYH9 | 226.533 | 2 | 53.599823 | yes | 9.729E-08 | 666.30731 | 2063.043 |
| EQEVNILK | 8 | P35579 | Myosin-9 OS=Homo sapiens OX=9606 C | MYH9 | 226.533 | 2 | 53.599823 | yes | 6.736E-07 | 486.77167 | 2024.0392 |
| EMEAELEDERK | 11 | P35579 | Myosin-9 OS=Homo sapiens OX=9606 C | MYH9 | 226.533 | 2 | 53.599823 | yes | 9.722E-08 | 689.8114 | 1970.9053 |
| DMFQETMEAMR | 11 | P35579 | Myosin-9 OS=Homo sapiens OX=9606 C | MYH9 | 226.533 | 2 | 53.599823 | yes | 9.728E-08 | 694.78583 | 962.91565 |
| DMFQETMEAMR | 11 | P35579 | Myosin-9 OS=Homo sapiens OX=9606 C | MYH9 | 226.533 | 2 | 53.599823 | yes | 1.136E-07 | 702.78326 | 962.91565 |
| QACVLMIK | 8 | P35579 | Myosin-9 OS=Homo sapiens OX=9606 C | MYH9 | 226.533 | 2 | 53.599823 | yes | 1.222E-07 | 481.76172 | 1793.5928 |
| EEILAQAK | 8 | P35579 | Myosin-9 OS=Homo sapiens OX=9606 C | MYH9 | 226.533 | 2 | 53.599823 | yes | 1.808E-07 | 451.25311 | 1778.2244 |
| ITDVIIGFQACCR | 13 | P35579 | Myosin-9 OS=Homo sapiens OX=9606 C | MYH9 | 226.533 | 2 | 53.599823 | yes | 2.823E-07 | 776.88416 | 1736.7183 |
| DLGEELEALK | 10 | P35579 | Myosin-9 OS=Homo sapiens OX=9606 C | MYH9 | 226.533 | 2 | 53.599823 | no | 1.339E-07 | 558.79279 | 1560.9166 |
| QQQLTAMK | 8 | P35579 | Myosin-9 OS=Homo sapiens OX=9606 C | MYH9 | 226.533 | 2 | 53.599823 | no | 3.176E-07 | 474.25256 | 1553.1565 |
| CNGVLEGIR | 9 | P35579 | Myosin-9 OS=Homo sapiens OX=9606 C | MYH9 | 226.533 | 2 | 53.599823 | no | 1.846E-07 | 509.26093 | 1496.4047 |
| ASITALEAK | 9 | P35579 | Myosin-9 OS=Homo sapiens OX=9606 C | MYH9 | 226.533 | 2 | 53.599823 | yes | 1.388E-07 | 452.26093 | 1352.748 |
| FDQLLAEEK | 9 | P35579 | Myosin-9 OS=Homo sapiens OX=9606 C | MYH9 | 226.533 | 2 | 53.599823 | no | 1.528E-07 | 546.78223 | 1350.6389 |
| MEDSVGCLETAEE | 15 | P35579 | Myosin-9 OS=Homo sapiens OX=9606 C | MYH9 | 226.533 | 2 | 53.599823 | yes | 1.361E-07 | 848.87384 | 1297.2104 |
| TFHIFYYLLSGAGE | 17 | P35579 | Myosin-9 OS=Homo sapiens OX=9606 C | MYH9 | 226.533 | 3 | 53.599823 | yes | 1.33E-07 | 666.01575 | 1217.8392 |
| KTLEEEAK | 8 | P35579 | Myosin-9 OS=Homo sapiens OX=9606 C | MYH9 | 226.533 | 2 | 53.599823 | yes | 1.582E-07 | 474.25583 | 1101.0009 |
| LVWVPSDK | 8 | P35579 | Myosin-9 OS=Homo sapiens OX=9606 C | MYH9 | 226.533 | 2 | 53.599823 | yes | 1.365E-07 | 472.26602 | 1058.2896 |
| LKDVLLQVDDER | 12 | P35579 | Myosin-9 OS=Homo sapiens OX=9606 C | MYH9 | 226.533 | 2 | 53.599823 | yes | 1.036E-07 | 721.89612 | 1020.7967 |
| KVEAQLQELQVK | 12 | P35579 | Myosin-9 OS=Homo sapiens OX=9606 C | MYH9 | 226.533 | 2 | 53.599823 | yes | 1.02E-07 | 706.909 | 999.64172 |
| GALALEEK | 8 | P35579 | Myosin-9 OS=Homo sapiens OX=9606 C | MYH9 | 226.533 | 2 | 53.599823 | yes | 1.377E-07 | 415.73456 | 760.46948 |
| QSACNLEK | 8 | P35579 | Myosin-9 OS=Homo sapiens OX=9606 C | MYH9 | 226.533 | 2 | 53.599823 | yes | 1.95E-07 | 475.22403 | 735.0675 |
| KLEEEQIILEDQNC | 15 | P35579 | Myosin-9 OS=Homo sapiens OX=9606 C | MYH9 | 226.533 | 3 | 53.599823 | yes | 9.722E-08 | 630.31555 | 700.99908 |
| DKADFCIIHYAGK | 13 | P35579 | Myosin-9 OS=Homo sapiens OX=9606 C | MYH9 | 226.533 | 2 | 53.599823 | no | 9.893E-08 | 769.37701 | 591.37738 |
| REQEVNILK | 9 | P35579 | Myosin-9 OS=Homo sapiens OX=9606 C | MYH9 | 226.533 | 2 | 53.599823 | yes | 1.317E-07 | 564.8222 | 582.81226 |
| TLEEEAK | 7 | P35579 | Myosin-9 OS=Homo sapiens OX=9606 C | MYH9 | 226.533 | 2 | 53.599823 | yes | 9.722E-08 | 410.20834 | 512.92371 |
| VEDMAELTCLNEA | 20 | P35579 | Myosin-9 OS=Homo sapiens OX=9606 C | MYH9 | 226.533 | 3 | 53.599823 | no | 1.805E-07 | 762.70831 | 252.81285 |
| VEDMAELTCLNEA | 20 | P35579 | Myosin-9 OS=Homo sapiens OX=9606 C | MYH9 | 226.533 | 3 | 53.599823 | no | 1.969E-07 | 768.03998 | 252.81285 |
| SVHELEK | 7 | P35579 | Myosin-9 OS=Homo sapiens OX=9606 C | MYH9 | 226.533 | 2 | 53.599823 | yes | 4.206E-07 | 421.22433 | 488.35144 |
| LQQELDDLLVDLD | 16 | P35579 | Myosin-9 OS=Homo sapiens OX=9606 C | MYH9 | 226.533 | 3 | 53.599823 | yes | 1.166E-07 | 650.66907 | 484.47473 |
| KMEDSVGCLETAE | 17 | P35579 | Myosin-9 OS=Homo sapiens OX=9606 C | MYH9 | 226.533 | 3 | 53.599823 | yes | 1.414E-07 | 660.9837 | 425.96832 |
| MEDSVGCLETAEE | 16 | P35579 | Myosin-9 OS=Homo sapiens OX=9606 C | MYH9 | 226.533 | 3 | 53.599823 | yes | 9.722E-08 | 618.28534 | 373.55246 |
| EEELQAALAR | 10 | P35579 | Myosin-9 OS=Homo sapiens OX=9606 C | MYH9 | 226.533 | 2 | 53.599823 | no | 1.287E-07 | 565.29602 | 364.31519 |
| NKHEAMITDLEER | 13 | P35579 | Myosin-9 OS=Homo sapiens OX=9606 C | MYH9 | 226.533 | 2 | 53.599823 | yes | 1.075E-07 | 793.38556 | 360.66418 |
| RQQQLTAMK | 9 | P35579 | Myosin-9 OS=Homo sapiens OX=9606 C | MYH9 | 226.533 | 2 | 53.599823 | no | 1.061E-07 | 552.30316 | 352.64032 |
| GELANEVK | 8 | P35579 | Myosin-9 OS=Homo sapiens OX=9606 C | MYH9 | 226.533 | 2 | 53.599823 | yes | 1.8E-07 | 430.22961 | 328.34338 |
| QEEEMMAK | 8 | P35579 | Myosin-9 OS=Homo sapiens OX=9606 C | MYH9 | 226.533 | 2 | 53.599823 | yes | 1.136E-07 | 498.21225 | 267.02997 |
| LQLQEQLQAETEL | 21 | P35579 | Myosin-9 OS=Homo sapiens OX=9606 C | MYH9 | 226.533 | 3 | 53.599823 | yes | 9.722E-08 | 834.41113 | 232.16682 |
| FLSNGHVTIPGQQI | 15 | P35579 | Myosin-9 OS=Homo sapiens OX=9606 C | MYH9 | 226.533 | 2 | 53.599823 | yes | 1.468E-07 | 820.92316 | 225.34879 |

| | | | | | | | | | | |
|---|---|---|---|---|---|---|---|---|---|---|
| DFSALESQLQDTQ | 21 | P35579 | Myosin-9 OS=Homo sapiens OX=9606 C | MYH9 | 226.533 | 3 | 53.599823 | yes | 9.722E-08 | 831.72949 | 223.58803 |
| DVLLQVDDERR | 11 | P35579 | Myosin-9 OS=Homo sapiens OX=9606 C | MYH9 | 226.533 | 2 | 53.599823 | yes | 1.02E-07 | 679.35718 | 212.02042 |
| CQHLQAEK | 8 | P35579 | Myosin-9 OS=Homo sapiens OX=9606 C | MYH9 | 226.533 | 2 | 53.599823 | yes | 5.883E-07 | 507.24527 | 196.86304 |
| NCAAYLK | 7 | P35579 | Myosin-9 OS=Homo sapiens OX=9606 C | MYH9 | 226.533 | 2 | 53.599823 | no | 9.722E-08 | 420.20764 | 176.34076 |
| YEILTPNSIPK | 11 | P35579 | Myosin-9 OS=Homo sapiens OX=9606 C | MYH9 | 226.533 | 2 | 53.599823 | yes | 2.739E-07 | 637.85315 | 165.71225 |
| KMEDSVGCLETAI | 16 | P35579 | Myosin-9 OS=Homo sapiens OX=9606 C | MYH9 | 226.533 | 3 | 53.599823 | yes | 9.819E-08 | 608.95001 | 155.56985 |
| QELEEICHDLEAR | 13 | P35579 | Myosin-9 OS=Homo sapiens OX=9606 C | MYH9 | 226.533 | 2 | 53.599823 | yes | 1.002E-07 | 821.38049 | 135.15775 |
| RDLGEELEALK | 11 | P35579 | Myosin-9 OS=Homo sapiens OX=9606 C | MYH9 | 226.533 | 2 | 53.599823 | no | 9.722E-08 | 636.84332 | 83.448891 |
| MQQNIQELEEQLE | 19 | P35579 | Myosin-9 OS=Homo sapiens OX=9606 C | MYH9 | 226.533 | 3 | 53.599823 | yes | 2.177E-07 | 778.35693 | 66.084015 |
| NAEQYKDQADK | 11 | P35579 | Myosin-9 OS=Homo sapiens OX=9606 C | MYH9 | 226.533 | 2 | 53.599823 | yes | 1.426E-07 | 655.30457 | 65.209343 |
| KKVEAQLQELQVI | 13 | P35579 | Myosin-9 OS=Homo sapiens OX=9606 C | MYH9 | 226.533 | 2 | 53.599823 | yes | 1.467E-07 | 770.95648 | 64.796089 |
| NTDQASMPDNTA | 15 | P35579 | Myosin-9 OS=Homo sapiens OX=9606 C | MYH9 | 226.533 | 2 | 53.599823 | no | 9.722E-08 | 796.35449 | 57.826859 |
| VAAYDKLEK | 9 | P35579 | Myosin-9 OS=Homo sapiens OX=9606 C | MYH9 | 226.533 | 2 | 53.599823 | yes | 9.722E-08 | 518.78729 | 56.580669 |
| RQLEEAEEEAQR | 12 | P35579 | Myosin-9 OS=Homo sapiens OX=9606 C | MYH9 | 226.533 | 2 | 53.599823 | yes | 4.894E-07 | 744.35785 | 27.487431 |
| IIGLDQVAGMSETA | 20 | P35579 | Myosin-9 OS=Homo sapiens OX=9606 C | MYH9 | 226.533 | 3 | 53.599823 | yes | 3.154E-06 | 673.35907 | 24.661861 |
| SDLLLEGFNNYR | 12 | P35580 | Myosin-10 OS=Homo sapiens OX=9606 | MYH10 | 229 | 2 | 54.66976 | yes | 9.933E-08 | 720.8595 | 1218.8208 |
| VKPLLQVTR | 9 | P35580 | Myosin-10 OS=Homo sapiens OX=9606 | MYH10 | 229 | 2 | 54.66976 | no | 2.598E-07 | 527.34259 | 1066.7878 |
| ALELDPNLYR | 10 | P35580 | Myosin-10 OS=Homo sapiens OX=9606 | MYH10 | 229 | 2 | 54.66976 | no | 3.784E-07 | 602.32202 | 1000.7902 |
| EIFMQVEDER | 10 | P35580 | Myosin-10 OS=Homo sapiens OX=9606 | MYH10 | 229 | 2 | 54.66976 | yes | 9.722E-08 | 648.30048 | 798.1264 |
| LVWIPSER | 8 | P35580 | Myosin-10 OS=Homo sapiens OX=9606 | MYH10 | 229 | 2 | 54.66976 | yes | 2.178E-07 | 500.28473 | 757.09186 |
| AVIYNPATQADWT | 15 | P35580 | Myosin-10 OS=Homo sapiens OX=9606 | MYH10 | 229 | 2 | 54.66976 | yes | 1.833E-07 | 824.9201 | 691.22083 |
| ELDDATEANEGLS | 14 | P35580 | Myosin-10 OS=Homo sapiens OX=9606 | MYH10 | 229 | 2 | 54.66976 | yes | 3.982E-07 | 760.34717 | 651.56616 |
| ALEQQVEEMR | 10 | P35580 | Myosin-10 OS=Homo sapiens OX=9606 | MYH10 | 229 | 2 | 54.66976 | yes | 1.076E-07 | 616.8006 | 647.13293 |
| DLEAQIEAANK | 11 | P35580 | Myosin-10 OS=Homo sapiens OX=9606 | MYH10 | 229 | 2 | 54.66976 | yes | 1.417E-07 | 601.30658 | 585.05249 |
| EQADFAVEALAK | 12 | P35580 | Myosin-10 OS=Homo sapiens OX=9606 | MYH10 | 229 | 2 | 54.66976 | no | 1.207E-07 | 646.33008 | 527.2204 |
| NMDPLNDNVATLI | 19 | P35580 | Myosin-10 OS=Homo sapiens OX=9606 | MYH10 | 229 | 3 | 54.66976 | yes | 1.205E-07 | 714.00775 | 452.19724 |
| LEVNMQAMK | 9 | P35580 | Myosin-10 OS=Homo sapiens OX=9606 | MYH10 | 229 | 2 | 54.66976 | yes | 1.259E-07 | 532.2674 | 430.98386 |
| VEGELEEMER | 10 | P35580 | Myosin-10 OS=Homo sapiens OX=9606 | MYH10 | 229 | 2 | 54.66976 | yes | 1.272E-07 | 610.77679 | 384.38712 |
| LQELEGAVK | 9 | P35580 | Myosin-10 OS=Homo sapiens OX=9606 | MYH10 | 229 | 2 | 54.66976 | yes | 2.377E-07 | 493.77948 | 374.78754 |
| MEEEILLLEDQNSI | 14 | P35580 | Myosin-10 OS=Homo sapiens OX=9606 | MYH10 | 229 | 2 | 54.66976 | yes | 1.955E-07 | 845.91382 | 337.78876 |
| VDDDLGTIESLEEA | 15 | P35580 | Myosin-10 OS=Homo sapiens OX=9606 | MYH10 | 229 | 2 | 54.66976 | yes | 9.722E-08 | 817.3938 | 332.41925 |
| IGQLEEQLEQEAK | 13 | P35580 | Myosin-10 OS=Homo sapiens OX=9606 | MYH10 | 229 | 2 | 54.66976 | yes | 1.063E-07 | 757.88849 | 312.90161 |
| HATALEELSEQLEQ | 16 | P35580 | Myosin-10 OS=Homo sapiens OX=9606 | MYH10 | 229 | 3 | 54.66976 | yes | 9.722E-08 | 599.63904 | 311.42889 |
| KLVWIPSER | 9 | P35580 | Myosin-10 OS=Homo sapiens OX=9606 | MYH10 | 229 | 2 | 54.66976 | yes | 1.427E-07 | 564.33221 | 294.24362 |
| ELEAELEDER | 10 | P35580 | Myosin-10 OS=Homo sapiens OX=9606 | MYH10 | 229 | 2 | 54.66976 | yes | 1.2E-07 | 616.78571 | 286.07013 |
| TFHIFYQLLSGAGI | 17 | P35580 | Myosin-10 OS=Homo sapiens OX=9606 | MYH10 | 229 | 3 | 54.66976 | yes | 9.722E-08 | 654.34753 | 265.73105 |
| IAECSSQLAEEEEK | 14 | P35580 | Myosin-10 OS=Homo sapiens OX=9606 | MYH10 | 229 | 2 | 54.66976 | yes | 1.032E-07 | 811.86432 | 219.46248 |
| DLSEELEALK | 10 | P35580 | Myosin-10 OS=Homo sapiens OX=9606 | MYH10 | 229 | 2 | 54.66976 | yes | 2.635E-07 | 573.7981 | 207.26385 |
| TGLEDPER | 8 | P35580 | Myosin-10 OS=Homo sapiens OX=9606 | MYH10 | 229 | 2 | 54.66976 | yes | 1.233E-07 | 458.72217 | 203.2054 |

| | | | | | | | | | | | |
|---|---|---|---|---|---|---|---|---|---|---|---|
| VVSSVLQFGNISFI | 14 | P35580 | Myosin-10 OS=Homo sapiens OX=9606 | MYH10 | 229 | 2 | 54.66976 | yes | 1.202E-07 | 762.92468 | 196.8455 |
| NHEAQIQDMR | 10 | P35580 | Myosin-10 OS=Homo sapiens OX=9606 | MYH10 | 229 | 2 | 54.66976 | yes | 9.722E-08 | 621.28821 | 175.63841 |
| KHQQLLEEK | 9 | P35580 | Myosin-10 OS=Homo sapiens OX=9606 | MYH10 | 229 | 2 | 54.66976 | yes | 2.881E-06 | 576.8222 | 173.75105 |
| KEEELQGALAR | 11 | P35580 | Myosin-10 OS=Homo sapiens OX=9606 | MYH10 | 229 | 2 | 54.66976 | yes | 5.197E-07 | 622.33569 | 163.65797 |
| DELADEITNSASGI | 14 | P35580 | Myosin-10 OS=Homo sapiens OX=9606 | MYH10 | 229 | 2 | 54.66976 | yes | 9.722E-08 | 725.33881 | 159.14786 |
| FLSNGYIPIPGQQD | 15 | P35580 | Myosin-10 OS=Homo sapiens OX=9606 | MYH10 | 229 | 2 | 54.66976 | yes | 1.541E-07 | 838.93579 | 157.47565 |
| DHNIPGELER | 10 | P35580 | Myosin-10 OS=Homo sapiens OX=9606 | MYH10 | 229 | 2 | 54.66976 | yes | 1.737E-07 | 590.29126 | 146.8587 |
| ADMEDLMSSK | 10 | P35580 | Myosin-10 OS=Homo sapiens OX=9606 | MYH10 | 229 | 2 | 54.66976 | yes | 9.722E-08 | 563.74139 | 145.62463 |
| QQQLSALK | 8 | P35580 | Myosin-10 OS=Homo sapiens OX=9606 | MYH10 | 229 | 2 | 54.66976 | yes | 2.417E-07 | 458.26654 | 127.83379 |
| QEEELQAK | 8 | P35580 | Myosin-10 OS=Homo sapiens OX=9606 | MYH10 | 229 | 2 | 54.66976 | yes | 1.183E-07 | 487.7431 | 122.53714 |
| NQILQNEK | 8 | P35580 | Myosin-10 OS=Homo sapiens OX=9606 | MYH10 | 229 | 2 | 54.66976 | yes | 1.599E-07 | 493.76691 | 110.2761 |
| ELEAELEDERK | 11 | P35580 | Myosin-10 OS=Homo sapiens OX=9606 | MYH10 | 229 | 2 | 54.66976 | yes | 9.722E-08 | 680.83319 | 103.23695 |
| KQQQLSALK | 9 | P35580 | Myosin-10 OS=Homo sapiens OX=9606 | MYH10 | 229 | 2 | 54.66976 | yes | 3.144E-07 | 522.31403 | 97.711899 |
| LRVELAEK | 8 | P35580 | Myosin-10 OS=Homo sapiens OX=9606 | MYH10 | 229 | 2 | 54.66976 | yes | 1.074E-07 | 479.29001 | 96.087921 |
| DAASLESQLQDTQ | 21 | P35580 | Myosin-10 OS=Homo sapiens OX=9606 | MYH10 | 229 | 3 | 54.66976 | yes | 1.088E-07 | 802.05402 | 94.361206 |
| ATISALEAK | 9 | P35580 | Myosin-10 OS=Homo sapiens OX=9606 | MYH10 | 229 | 2 | 54.66976 | yes | 1.44E-07 | 452.26093 | 87.491814 |
| KDHNIPGELER | 11 | P35580 | Myosin-10 OS=Homo sapiens OX=9606 | MYH10 | 229 | 2 | 54.66976 | yes | 3.266E-07 | 654.33875 | 65.879715 |
| QVASNLEK | 8 | P35580 | Myosin-10 OS=Homo sapiens OX=9606 | MYH10 | 229 | 2 | 54.66976 | yes | 1.002E-06 | 444.74289 | 58.848331 |
| HQQLLEEK | 8 | P35580 | Myosin-10 OS=Homo sapiens OX=9606 | MYH10 | 229 | 2 | 54.66976 | yes | 2.218E-06 | 512.77472 | 58.225761 |
| KLDAQVQELHAK | 12 | P35580 | Myosin-10 OS=Homo sapiens OX=9606 | MYH10 | 229 | 2 | 54.66976 | yes | 1.191E-07 | 690.38568 | 58.148663 |
| DAEALSQR | 8 | P35580 | Myosin-10 OS=Homo sapiens OX=9606 | MYH10 | 229 | 2 | 54.66976 | yes | 1.721E-07 | 445.22232 | 53.508854 |
| VIQYLAHVASSHK | 13 | P35580 | Myosin-10 OS=Homo sapiens OX=9606 | MYH10 | 229 | 2 | 54.66976 | yes | 1.26E-07 | 726.90155 | 51.197186 |
| IVFQEFR | 7 | P35580 | Myosin-10 OS=Homo sapiens OX=9606 | MYH10 | 229 | 2 | 54.66976 | no | 2.587E-07 | 469.75836 | 44.182423 |
| LQNELDNVSTLLE | 18 | P35580 | Myosin-10 OS=Homo sapiens OX=9606 | MYH10 | 229 | 3 | 54.66976 | yes | 1.327E-06 | 691.69519 | 37.355583 |
| KLQAQMKDYQR | 11 | P35580 | Myosin-10 OS=Homo sapiens OX=9606 | MYH10 | 229 | 2 | 54.66976 | yes | 2.428E-05 | 712.86938 | 36.739502 |
| QVLALQSQLADTI | 13 | P35580 | Myosin-10 OS=Homo sapiens OX=9606 | MYH10 | 229 | 2 | 54.66976 | yes | 3.419E-07 | 707.89862 | 34.916355 |
| NKQEVMISDLEER | 13 | P35580 | Myosin-10 OS=Homo sapiens OX=9606 | MYH10 | 229 | 2 | 54.66976 | yes | 2.983E-06 | 795.89325 | 31.290751 |
| EQEVAELKK | 9 | P35580 | Myosin-10 OS=Homo sapiens OX=9606 | MYH10 | 229 | 2 | 54.66976 | yes | 1.889E-07 | 537.29547 | 27.527937 |
| NTDQASMPENTV | 15 | P35580 | Myosin-10 OS=Homo sapiens OX=9606 | MYH10 | 229 | 2 | 54.66976 | yes | 3.6E-07 | 817.37793 | 15.540915 |
| VFNYNTLER | 9 | P35606 | Coatomer subunit beta' OS=Homo sapien | COPB2 | 102.488 | 2 | 54.95824 | yes | 9.928E-08 | 578.29327 | 218.51128 |
| LPEAAFLAR | 9 | P35606 | Coatomer subunit beta' OS=Homo sapien | COPB2 | 102.488 | 2 | 54.95824 | yes | 3.729E-07 | 494.28473 | 174.9493 |
| QYPLVTPNEER | 11 | P35606 | Coatomer subunit beta' OS=Homo sapien | COPB2 | 102.488 | 2 | 54.95824 | yes | 1.836E-07 | 673.34094 | 140.85811 |
| IAYQLAVEAESEQ | 14 | P35606 | Coatomer subunit beta' OS=Homo sapien | COPB2 | 102.488 | 2 | 54.95824 | yes | 1.532E-07 | 789.90411 | 138.56937 |
| EAFVVEEWVK | 10 | P35606 | Coatomer subunit beta' OS=Homo sapien | COPB2 | 102.488 | 2 | 54.95824 | yes | 9.722E-08 | 618.31897 | 120.6291 |
| LESTLNYGMER | 11 | P35606 | Coatomer subunit beta' OS=Homo sapien | COPB2 | 102.488 | 2 | 54.95824 | yes | 2.69E-07 | 656.81372 | 105.70772 |
| RIEIQPK | 7 | P35606 | Coatomer subunit beta' OS=Homo sapien | COPB2 | 102.488 | 2 | 54.95824 | yes | 3.289E-07 | 442.27164 | 102.13907 |
| NWVVTGADDMQI | 13 | P35606 | Coatomer subunit beta' OS=Homo sapien | COPB2 | 102.488 | 2 | 54.95824 | yes | 2.558E-07 | 752.86426 | 100.91827 |
| IWHSSTYR | 8 | P35606 | Coatomer subunit beta' OS=Homo sapien | COPB2 | 102.488 | 2 | 54.95824 | yes | 7.556E-07 | 525.26178 | 97.771889 |
| VWCVASLR | 8 | P35606 | Coatomer subunit beta' OS=Homo sapien | COPB2 | 102.488 | 2 | 54.95824 | yes | 2.93E-07 | 495.76312 | 94.757896 |

| | | | | | | | | | | | |
|---|---|---|---|---|---|---|---|---|---|---|---|
| ETHADLWPAK | 10 | P35606 | Coatomer subunit beta' OS=Homo sapien | COPB2 | 102.488 | 2 | 54.95824 | yes | 2.199E-06 | 584.29327 | 72.884232 |
| EAFVVEEWVKETI | 20 | P35606 | Coatomer subunit beta' OS=Homo sapien | COPB2 | 102.488 | 3 | 54.95824 | yes | 1.016E-06 | 795.40222 | 63.496941 |
| QQALTVSTDPEHR | 13 | P35606 | Coatomer subunit beta' OS=Homo sapien | COPB2 | 102.488 | 2 | 54.95824 | yes | 5.333E-07 | 741.37079 | 42.391296 |
| TFEVCDLPVR | 10 | P35606 | Coatomer subunit beta' OS=Homo sapien | COPB2 | 102.488 | 2 | 54.95824 | yes | 1.234E-07 | 618.30811 | 35.079395 |
| VHMFEAHSDYIR | 12 | P35606 | Coatomer subunit beta' OS=Homo sapien | COPB2 | 102.488 | 3 | 54.95824 | yes | 3.662E-06 | 502.23825 | 25.86858 |
| VDENNPEYLR | 10 | P35611 | Alpha-adducin OS=Homo sapiens OX=9 | ADD1 | 80.956 | 2 | 53.25517 | yes | 2.337E-06 | 624.79639 | 335.93793 |
| INLQGDIVDR | 10 | P35611 | Alpha-adducin OS=Homo sapiens OX=9 | ADD1 | 80.956 | 2 | 53.25517 | yes | 1.507E-07 | 571.81183 | 132.01459 |
| GSEENLDEAR | 10 | P35611 | Alpha-adducin OS=Homo sapiens OX=9 | ADD1 | 80.956 | 2 | 53.25517 | yes | 1.959E-07 | 560.24927 | 71.228409 |
| WQIGEQEFEALMF | 13 | P35611 | Alpha-adducin OS=Homo sapiens OX=9 | ADD1 | 80.956 | 2 | 53.25517 | yes | 1.108E-07 | 818.89307 | 59.874054 |
| SPGSPVGEGTGSPI | 15 | P35611 | Alpha-adducin OS=Homo sapiens OX=9 | ADD1 | 80.956 | 2 | 53.25517 | yes | 9.722E-08 | 677.33588 | 28.971094 |
| SESVPPVTDWAW | 14 | P35613 | Basigin OS=Homo sapiens OX=9606 GN | BSG | 42.201 | 2 | 51.586594 | yes | 9.722E-08 | 832.90137 | 423.68741 |
| GSDQAIITLR | 10 | P35613 | Basigin OS=Homo sapiens OX=9606 GN | BSG | 42.201 | 2 | 51.586594 | yes | 1.013E-07 | 537.30109 | 269.00845 |
| FFVSSSQGR | 9 | P35613 | Basigin OS=Homo sapiens OX=9606 GN | BSG | 42.201 | 2 | 51.586594 | yes | 1.102E-07 | 507.75378 | 226.02217 |
| WDQLTLSQR | 9 | P35625 | Metalloproteinase inhibitor 3 OS=Homo | TIMP3 | 24.145 | 2 | 52.018764 | yes | 2.035E-07 | 573.79877 | 277.83231 |
| SCYYLPCFVTSK | 12 | P35625 | Metalloproteinase inhibitor 3 OS=Homo | TIMP3 | 24.145 | 2 | 52.018764 | yes | 9.722E-08 | 762.84674 | 263.22028 |
| MYTGLCNFVER | 11 | P35625 | Metalloproteinase inhibitor 3 OS=Homo | TIMP3 | 24.145 | 2 | 52.018764 | yes | 1.009E-07 | 695.31812 | 127.18945 |
| YHLGCNCK | 8 | P35625 | Metalloproteinase inhibitor 3 OS=Homo | TIMP3 | 24.145 | 2 | 52.018764 | yes | 7.083E-07 | 526.22601 | 86.040688 |
| LKGEATVSFDDPP | 16 | P35637 | RNA-binding protein FUS OS=Homo sap | FUS | 53.426 | 3 | 51.636475 | yes | 1.364E-07 | 554.61755 | 33251.133 |
| LKGEATVSFDDPP | 16 | P35637 | RNA-binding protein FUS OS=Homo sap | FUS | 53.426 | 3 | 51.636475 | yes | 1.187E-07 | 831.42267 | 33251.133 |
| LKGEATVSFDDPP | 16 | P35637 | RNA-binding protein FUS OS=Homo sap | FUS | 53.426 | 2 | 51.636475 | yes | 1.364E-07 | 554.61755 | 33251.133 |
| LKGEATVSFDDPP | 16 | P35637 | RNA-binding protein FUS OS=Homo sap | FUS | 53.426 | 2 | 51.636475 | yes | 1.187E-07 | 831.42267 | 33251.133 |
| TGQPMINLYTDR | 12 | P35637 | RNA-binding protein FUS OS=Homo sap | FUS | 53.426 | 2 | 51.636475 | yes | 1.166E-07 | 704.84808 | 31299.68 |
| TGQPMINLYTDR | 12 | P35637 | RNA-binding protein FUS OS=Homo sap | FUS | 53.426 | 2 | 51.636475 | yes | 1.074E-07 | 712.84558 | 31299.68 |
| GEATVSFDDPPSA | 14 | P35637 | RNA-binding protein FUS OS=Homo sap | FUS | 53.426 | 2 | 51.636475 | no | 9.722E-08 | 710.83319 | 48121.668 |
| AAIDWFDGK | 9 | P35637 | RNA-binding protein FUS OS=Homo sap | FUS | 53.426 | 2 | 51.636475 | no | 1.141E-07 | 511.75073 | 34197.316 |
| APKPDGPGGGPGC | 24 | P35637 | RNA-binding protein FUS OS=Homo sap | FUS | 53.426 | 4 | 51.636475 | yes | 1.017E-07 | 563.9989 | 1851.1307 |
| APKPDGPGGGPGC | 24 | P35637 | RNA-binding protein FUS OS=Homo sap | FUS | 53.426 | 4 | 51.636475 | yes | 1.121E-07 | 567.99762 | 1851.1307 |
| KTGQPMINLYTDR | 13 | P35637 | RNA-binding protein FUS OS=Homo sap | FUS | 53.426 | 2 | 51.636475 | yes | 1.388E-07 | 768.89557 | 1006.4547 |
| KTGQPMINLYTDR | 13 | P35637 | RNA-binding protein FUS OS=Homo sap | FUS | 53.426 | 2 | 51.636475 | yes | 2.233E-07 | 776.89307 | 1006.4547 |
| KTGQPMINLYTDR | 13 | P35637 | RNA-binding protein FUS OS=Homo sap | FUS | 53.426 | 2 | 51.636475 | yes | 3.082E-07 | 512.9328 | 1006.4547 |
| KTGQPMINLYTDR | 13 | P35637 | RNA-binding protein FUS OS=Homo sap | FUS | 53.426 | 3 | 51.636475 | yes | 1.388E-07 | 768.89557 | 1006.4547 |
| KTGQPMINLYTDR | 13 | P35637 | RNA-binding protein FUS OS=Homo sap | FUS | 53.426 | 3 | 51.636475 | yes | 2.233E-07 | 776.89307 | 1006.4547 |
| KTGQPMINLYTDR | 13 | P35637 | RNA-binding protein FUS OS=Homo sap | FUS | 53.426 | 3 | 51.636475 | yes | 3.082E-07 | 512.9328 | 1006.4547 |
| EFSGNPIK | 8 | P35637 | RNA-binding protein FUS OS=Homo sap | FUS | 53.426 | 2 | 51.636475 | yes | 2.949E-07 | 446.23218 | 599.60736 |
| APKPDGPGGGPGC | 25 | P35637 | RNA-binding protein FUS OS=Homo sap | FUS | 53.426 | 4 | 51.636475 | yes | 9.722E-08 | 603.02417 | 322.58063 |
| GGFPSGGGGGGGC | 15 | P35637 | RNA-binding protein FUS OS=Homo sap | FUS | 53.426 | 2 | 51.636475 | yes | 1.12E-07 | 638.29486 | 68.228394 |
| ETTESLHGDISSLK | 14 | P35658 | Nuclear pore complex protein Nup214 OS | NUP214 | 213.62 | 2 | 47.160694 | yes | 1.388E-07 | 758.87811 | 38.711159 |
| ASSTSLTSTQPTK | 13 | P35658 | Nuclear pore complex protein Nup214 OS | NUP214 | 213.62 | 2 | 47.160694 | yes | 1.529E-07 | 654.83569 | 34.551544 |
| SAQGSSSPVPSMV | 15 | P35658 | Nuclear pore complex protein Nup214 OS | NUP214 | 213.62 | 2 | 47.160694 | yes | 5.689E-07 | 745.36938 | 25.253075 |

| | | | | | | | | | | | |
|---|---|---|---|---|---|---|---|---|---|---|---|
| SICEVLDLER | 10 | P35659 | Protein DEK OS=Homo sapiens OX=960 | DEK | 42.675 | 2 | 53.555565 | yes | 9.81E-08 | 617.31079 | 1083.1799 |
| LTMQVSSLQR | 10 | P35659 | Protein DEK OS=Homo sapiens OX=960 | DEK | 42.675 | 2 | 53.555565 | yes | 1.066E-07 | 581.8161 | 1048.0607 |
| ILNFLMHPK | 9 | P35659 | Protein DEK OS=Homo sapiens OX=960 | DEK | 42.675 | 2 | 53.555565 | yes | 1.022E-07 | 556.81787 | 1015.2912 |
| LLASANLEEVTMK | 13 | P35659 | Protein DEK OS=Homo sapiens OX=960 | DEK | 42.675 | 2 | 53.555565 | yes | 1.004E-07 | 709.88159 | 723.48407 |
| NVGQFSGFPFEK | 12 | P35659 | Protein DEK OS=Homo sapiens OX=960 | DEK | 42.675 | 2 | 53.555565 | yes | 9.722E-08 | 678.83276 | 620.15588 |
| EPFTIAQGK | 9 | P35659 | Protein DEK OS=Homo sapiens OX=960 | DEK | 42.675 | 2 | 53.555565 | yes | 1.469E-07 | 495.76639 | 429.92834 |
| VYENYPTYDLTER | 13 | P35659 | Protein DEK OS=Homo sapiens OX=960 | DEK | 42.675 | 2 | 53.555565 | yes | 9.722E-08 | 831.88593 | 384.28702 |
| LLYNRPGTVSSLK | 13 | P35659 | Protein DEK OS=Homo sapiens OX=960 | DEK | 42.675 | 2 | 53.555565 | yes | 9.722E-08 | 724.41699 | 179.91666 |
| LLYNRPGTVSSLK | 13 | P35659 | Protein DEK OS=Homo sapiens OX=960 | DEK | 42.675 | 2 | 53.555565 | yes | 2.947E-07 | 483.28043 | 179.91666 |
| LLYNRPGTVSSLK | 13 | P35659 | Protein DEK OS=Homo sapiens OX=960 | DEK | 42.675 | 3 | 53.555565 | yes | 9.722E-08 | 724.41699 | 179.91666 |
| LLYNRPGTVSSLK | 13 | P35659 | Protein DEK OS=Homo sapiens OX=960 | DEK | 42.675 | 3 | 53.555565 | yes | 2.947E-07 | 483.28043 | 179.91666 |
| IHFFLSK | 7 | P35659 | Protein DEK OS=Homo sapiens OX=960 | DEK | 42.675 | 2 | 53.555565 | yes | 2.384E-07 | 446.258 | 312.33945 |
| KPPTDEELK | 9 | P35659 | Protein DEK OS=Homo sapiens OX=960 | DEK | 42.675 | 2 | 53.555565 | yes | 4.394E-07 | 528.78223 | 256.39703 |
| APYAPTELSR | 10 | P35713 | Transcription factor SOX-18 OS=Homo s | SOX18 | 40.892 | 2 | 48.377094 | yes | 1.316E-07 | 552.78784 | 187.88823 |
| DPGGCYGAPLAEA | 15 | P35713 | Transcription factor SOX-18 OS=Homo s | SOX18 | 40.892 | 2 | 48.377094 | yes | 1.156E-07 | 773.86957 | 104.43483 |
| TRPDAPGLPYHVA | 16 | P35713 | Transcription factor SOX-18 OS=Homo s | SOX18 | 40.892 | 3 | 48.377094 | yes | 3.019E-07 | 569.31763 | 84.34243 |
| IVLGYTEAELCTR | 13 | P35869 | Aryl hydrocarbon receptor OS=Homo sap | AHR | 96.148 | 2 | 50.98682 | yes | 1.212E-07 | 762.88995 | 196.15315 |
| GSGYQFIHAADMI | 21 | P35869 | Aryl hydrocarbon receptor OS=Homo sap | AHR | 96.148 | 3 | 50.98682 | yes | 9.722E-08 | 809.37207 | 121.60539 |
| LASLLPFPQDVINK | 14 | P35869 | Aryl hydrocarbon receptor OS=Homo sap | AHR | 96.148 | 2 | 50.98682 | yes | 1.897E-07 | 777.94812 | 107.02003 |
| TGESGMIVFR | 10 | P35869 | Aryl hydrocarbon receptor OS=Homo sap | AHR | 96.148 | 2 | 50.98682 | yes | 5.795E-07 | 548.77643 | 99.201981 |
| QVVVEPQQQLCQI | 13 | P35869 | Aryl hydrocarbon receptor OS=Homo sap | AHR | 96.148 | 2 | 50.98682 | yes | 1.108E-07 | 792.41412 | 88.794312 |
| WTWVQSNAR | 9 | P35869 | Aryl hydrocarbon receptor OS=Homo sap | AHR | 96.148 | 2 | 50.98682 | yes | 1.215E-05 | 574.28577 | 85.63372 |
| NDFSGEVDFR | 10 | P35869 | Aryl hydrocarbon receptor OS=Homo sap | AHR | 96.148 | 2 | 50.98682 | yes | 1.097E-07 | 593.26221 | 59.640965 |
| LDFTPIGCDAK | 11 | P35869 | Aryl hydrocarbon receptor OS=Homo sap | AHR | 96.148 | 2 | 50.98682 | yes | 2.702E-05 | 618.80011 | 17.940689 |
| LNDLEEALQQAK | 12 | P35908 | Keratin, type II cytoskeletal 2 epidermal ( | KRT2 | 65.433 | 2 | 53.632557 | yes | 1.019E-07 | 686.35938 | 608.25262 |
| YLDGLTAER | 9 | P35908 | Keratin, type II cytoskeletal 2 epidermal ( | KRT2 | 65.433 | 2 | 53.632557 | yes | 2.329E-07 | 519.26672 | 548.27838 |
| VDLLNQEIEFLK | 12 | P35908 | Keratin, type II cytoskeletal 2 epidermal ( | KRT2 | 65.433 | 2 | 53.632557 | yes | 9.722E-08 | 730.90338 | 410.38416 |
| GFSSGSAVVSGGS | 14 | P35908 | Keratin, type II cytoskeletal 2 epidermal ( | KRT2 | 65.433 | 2 | 53.632557 | yes | 2.136E-07 | 627.80731 | 388.6691 |
| TAAENDFVTLK | 11 | P35908 | Keratin, type II cytoskeletal 2 epidermal ( | KRT2 | 65.433 | 2 | 53.632557 | yes | 7.382E-07 | 604.81152 | 343.28851 |
| NVQDAIADAEQR | 12 | P35908 | Keratin, type II cytoskeletal 2 epidermal ( | KRT2 | 65.433 | 2 | 53.632557 | yes | 1.637E-07 | 665.3233 | 234.29523 |
| STSSFSCLSR | 10 | P35908 | Keratin, type II cytoskeletal 2 epidermal ( | KRT2 | 65.433 | 2 | 53.632557 | yes | 2.663E-07 | 566.25861 | 216.40474 |
| LQGEIAHVK | 9 | P35908 | Keratin, type II cytoskeletal 2 epidermal ( | KRT2 | 65.433 | 2 | 53.632557 | yes | 1.414E-07 | 497.78763 | 200.75005 |
| YEELQVTVGR | 10 | P35908 | Keratin, type II cytoskeletal 2 epidermal ( | KRT2 | 65.433 | 2 | 53.632557 | yes | 1.921E-07 | 597.31171 | 156.92302 |
| DYQELMNVK | 9 | P35908 | Keratin, type II cytoskeletal 2 epidermal ( | KRT2 | 65.433 | 2 | 53.632557 | no | 9.997E-08 | 570.27368 | 77.450569 |
| DYQELMNVK | 9 | P35908 | Keratin, type II cytoskeletal 2 epidermal ( | KRT2 | 65.433 | 2 | 53.632557 | no | 2.131E-07 | 578.27118 | 77.450569 |
| GGSISGGGYGSGG | 15 | P35908 | Keratin, type II cytoskeletal 2 epidermal ( | KRT2 | 65.433 | 2 | 53.632557 | yes | 3.071E-07 | 599.27838 | 154.5983 |
| DVDNAYMIK | 9 | P35908 | Keratin, type II cytoskeletal 2 epidermal ( | KRT2 | 65.433 | 2 | 53.632557 | yes | 2.733E-06 | 542.75262 | 59.5443 |
| HGGGGGGFGGGG | 16 | P35908 | Keratin, type II cytoskeletal 2 epidermal ( | KRT2 | 65.433 | 2 | 53.632557 | yes | 1.793E-07 | 660.79504 | 54.32251 |
| GVLLFGPPGTGK | 12 | P35998 | 26S proteasome regulatory subunit 7 OS= | PSMC2 | 48.634 | 2 | 44.203316 | yes | 2.283E-07 | 571.83203 | 177.4006 |

| | | | | | | | | | | |
|---|---|---|---|---|---|---|---|---|---|---|
| QTLQSEQPLQVAR | 13 | P35998 | 26S proteasome regulatory subunit 7 OS= | PSMC2 | 48.634 | 2 | 44.203316 | yes | 2.324E-07 | 749.4046 | 58.787781 |
| SVCTEAGMFAIR | 12 | P35998 | 26S proteasome regulatory subunit 7 OS= | PSMC2 | 48.634 | 2 | 44.203316 | yes | 5.799E-07 | 671.31812 | 56.859028 |
| ALDEGDIALLK | 11 | P35998 | 26S proteasome regulatory subunit 7 OS= | PSMC2 | 48.634 | 2 | 44.203316 | yes | 2.995E-06 | 579.32428 | 30.824469 |
| EVVETPLLHPER | 12 | P35998 | 26S proteasome regulatory subunit 7 OS= | PSMC2 | 48.634 | 3 | 44.203316 | yes | 3.549E-06 | 473.59277 | 26.886173 |
| ESAAINQILGR | 11 | P36402 | Transcription factor 7 OS=Homo sapiens | TCF7 | 41.642 | 2 | 45.280804 | yes | 1.899E-07 | 586.32513 | 63.42453 |
| HLLIGVSSDR | 10 | P36542 | ATP synthase subunit gamma, mitochond | ATP5F1C | 32.997 | 2 | 54.10322 | yes | 1.32E-07 | 548.80908 | 2151.2507 |
| THSDQFLVAFK | 11 | P36542 | ATP synthase subunit gamma, mitochond | ATP5F1C | 32.997 | 2 | 54.10322 | yes | 1.157E-07 | 646.83533 | 1952.0024 |
| SEVATLTAAGK | 11 | P36542 | ATP synthase subunit gamma, mitochond | ATP5F1C | 32.997 | 2 | 54.10322 | yes | 3.347E-07 | 524.28766 | 1775.0571 |
| EVMLVGIGDK | 10 | P36542 | ATP synthase subunit gamma, mitochond | ATP5F1C | 32.997 | 2 | 54.10322 | yes | 1.375E-07 | 530.789 | 849.84326 |
| EVMLVGIGDK | 10 | P36542 | ATP synthase subunit gamma, mitochond | ATP5F1C | 32.997 | 2 | 54.10322 | yes | 9.722E-08 | 538.78644 | 849.84326 |
| LTLTFNR | 7 | P36542 | ATP synthase subunit gamma, mitochond | ATP5F1C | 32.997 | 2 | 54.10322 | yes | 1.093E-07 | 432.75052 | 837.02881 |
| GLCGAIHSSIAK | 12 | P36542 | ATP synthase subunit gamma, mitochond | ATP5F1C | 32.997 | 2 | 54.10322 | yes | 6.026E-07 | 607.32153 | 245.8149 |
| IYGLGSLALYEK | 12 | P36542 | ATP synthase subunit gamma, mitochond | ATP5F1C | 32.997 | 2 | 54.10322 | yes | 9.722E-08 | 663.86884 | 223.72113 |
| KHLLIGVSSDR | 11 | P36542 | ATP synthase subunit gamma, mitochond | ATP5F1C | 32.997 | 2 | 54.10322 | yes | 9.722E-08 | 612.85657 | 105.89308 |
| IEEVPELPLVVEDK | 14 | P36578 | Large ribosomal subunit protein uL4 OS= | RPL4 | 47.698 | 2 | 54.021023 | yes | 2.035E-07 | 804.94019 | 49806.473 |
| PLISVYSEK | 9 | P36578 | Large ribosomal subunit protein uL4 OS= | RPL4 | 47.698 | 2 | 54.021023 | yes | 1.993E-07 | 518.28967 | 36915.875 |
| FCIWTESAFR | 10 | P36578 | Large ribosomal subunit protein uL4 OS= | RPL4 | 47.698 | 2 | 54.021023 | yes | 1.284E-07 | 658.80823 | 22604.07 |
| APIRPDIVNFVHTN | 16 | P36578 | Large ribosomal subunit protein uL4 OS= | RPL4 | 47.698 | 3 | 54.021023 | yes | 2.272E-07 | 621.35138 | 10614.853 |
| APIRPDIVNFVHTN | 16 | P36578 | Large ribosomal subunit protein uL4 OS= | RPL4 | 47.698 | 3 | 54.021023 | yes | 3.78E-07 | 466.26535 | 10614.853 |
| APIRPDIVNFVHTN | 16 | P36578 | Large ribosomal subunit protein uL4 OS= | RPL4 | 47.698 | 4 | 54.021023 | yes | 2.272E-07 | 621.35138 | 10614.853 |
| APIRPDIVNFVHTN | 16 | P36578 | Large ribosomal subunit protein uL4 OS= | RPL4 | 47.698 | 4 | 54.021023 | yes | 3.78E-07 | 466.26535 | 10614.853 |
| NIPGITLLNVSK | 12 | P36578 | Large ribosomal subunit protein uL4 OS= | RPL4 | 47.698 | 2 | 54.021023 | yes | 1.485E-07 | 634.88226 | 17919.824 |
| SGQGAFGNMCR | 11 | P36578 | Large ribosomal subunit protein uL4 OS= | RPL4 | 47.698 | 2 | 54.021023 | yes | 9.722E-08 | 592.75043 | 7562.8931 |
| SGQGAFGNMCR | 11 | P36578 | Large ribosomal subunit protein uL4 OS= | RPL4 | 47.698 | 2 | 54.021023 | yes | 9.722E-08 | 600.74786 | 7562.8931 |
| MINTDLSR | 8 | P36578 | Large ribosomal subunit protein uL4 OS= | RPL4 | 47.698 | 2 | 54.021023 | yes | 1.24E-07 | 475.24222 | 15059.126 |
| KLDELYGTWR | 10 | P36578 | Large ribosomal subunit protein uL4 OS= | RPL4 | 47.698 | 2 | 54.021023 | yes | 1.212E-07 | 640.83533 | 12511.165 |
| AAAAAAALQAK | 11 | P36578 | Large ribosomal subunit protein uL4 OS= | RPL4 | 47.698 | 2 | 54.021023 | yes | 1.1E-07 | 478.77982 | 11368.41 |
| GPCIIYNEDNGIIK | 14 | P36578 | Large ribosomal subunit protein uL4 OS= | RPL4 | 47.698 | 2 | 54.021023 | yes | 2.181E-07 | 803.4007 | 9714.2578 |
| SNYNLPMHK | 9 | P36578 | Large ribosomal subunit protein uL4 OS= | RPL4 | 47.698 | 2 | 54.021023 | yes | 3.702E-07 | 552.26874 | 4150.4937 |
| SNYNLPMHK | 9 | P36578 | Large ribosomal subunit protein uL4 OS= | RPL4 | 47.698 | 2 | 54.021023 | yes | 2.284E-07 | 560.26624 | 4150.4937 |
| LAPGGHVGR | 9 | P36578 | Large ribosomal subunit protein uL4 OS= | RPL4 | 47.698 | 2 | 54.021023 | yes | 1.284E-07 | 432.24594 | 7974.2734 |
| LDELYGTWR | 9 | P36578 | Large ribosomal subunit protein uL4 OS= | RPL4 | 47.698 | 2 | 54.021023 | yes | 1.02E-07 | 576.78784 | 7853.79 |
| NVTLPAVFK | 9 | P36578 | Large ribosomal subunit protein uL4 OS= | RPL4 | 47.698 | 2 | 54.021023 | yes | 2.141E-06 | 494.79492 | 328.12778 |
| IEEVPELPLVVEDK | 19 | P36578 | Large ribosomal subunit protein uL4 OS= | RPL4 | 47.698 | 3 | 54.021023 | yes | 1.465E-06 | 729.05945 | 113.12852 |
| ACARPLISVYSEK | 13 | P36578 | Large ribosomal subunit protein uL4 OS= | RPL4 | 47.698 | 2 | 54.021023 | yes | 4.327E-07 | 768.39795 | 112.25894 |
| QPYAVSELAGHQT | 22 | P36578 | Large ribosomal subunit protein uL4 OS= | RPL4 | 47.698 | 3 | 54.021023 | yes | 3.515E-07 | 778.03662 | 81.316193 |
| ILCFYGPPGVGK | 12 | P36776 | Lon protease homolog, mitochondrial OS | LONP1 | 106.49 | 2 | 45.04012 | yes | 7.514E-07 | 654.34448 | 35.634975 |
| TENPLILIDEVDK | 13 | P36776 | Lon protease homolog, mitochondrial OS | LONP1 | 106.49 | 2 | 45.04012 | yes | 2.747E-06 | 749.90363 | 25.098001 |
| KYLLQEQLK | 9 | P36776 | Lon protease homolog, mitochondrial OS | LONP1 | 106.49 | 2 | 45.04012 | yes | 1.195E-06 | 581.84515 | 18.023338 |

| | | | | | | | | | | | |
|---|---|---|---|---|---|---|---|---|---|---|---|
| QLEVEPEEPEAENI | 14 | P36776 | Lon protease homolog, mitochondrial OS | LONP1 | 106.49 | 2 | 45.04012 | yes | 1.984E-07 | 820.88611 | 15.327987 |
| HDLDLICR | 8 | P36873 | Serine/threonine-protein phosphatase PP1 | PPP1CC | 36.984 | 2 | 54.36372 | no | 1.952E-07 | 521.26093 | 2976.5793 |
| YPENFFLLR | 9 | P36873 | Serine/threonine-protein phosphatase PP1 | PPP1CC | 36.984 | 2 | 54.36372 | no | 1.279E-07 | 599.81641 | 2046.1187 |
| LNIDSIIQR | 9 | P36873 | Serine/threonine-protein phosphatase PP1 | PPP1CC | 36.984 | 2 | 54.36372 | yes | 2.369E-07 | 536.31146 | 2010.8719 |
| AHQVVEDGYEFFA | 14 | P36873 | Serine/threonine-protein phosphatase PP1 | PPP1CC | 36.984 | 2 | 54.36372 | no | 1.364E-07 | 820.39117 | 741.12708 |
| AHQVVEDGYEFFA | 14 | P36873 | Serine/threonine-protein phosphatase PP1 | PPP1CC | 36.984 | 2 | 54.36372 | no | 1.056E-07 | 547.26324 | 741.12708 |
| AHQVVEDGYEFFA | 14 | P36873 | Serine/threonine-protein phosphatase PP1 | PPP1CC | 36.984 | 3 | 54.36372 | no | 1.364E-07 | 820.39117 | 741.12708 |
| AHQVVEDGYEFFA | 14 | P36873 | Serine/threonine-protein phosphatase PP1 | PPP1CC | 36.984 | 3 | 54.36372 | no | 1.056E-07 | 547.26324 | 741.12708 |
| IYGFYDECK | 9 | P36873 | Serine/threonine-protein phosphatase PP1 | PPP1CC | 36.984 | 2 | 54.36372 | no | 2.918E-07 | 597.76044 | 977.08405 |
| ICGDIHGQYYDLLI | 14 | P36873 | Serine/threonine-protein phosphatase PP1 | PPP1CC | 36.984 | 3 | 54.36372 | no | 1.003E-07 | 574.94714 | 618.37256 |
| IKYPENFFLLR | 11 | P36873 | Serine/threonine-protein phosphatase PP1 | PPP1CC | 36.984 | 2 | 54.36372 | no | 1.406E-07 | 720.40588 | 576.52826 |
| NVQLQENEIR | 10 | P36873 | Serine/threonine-protein phosphatase PP1 | PPP1CC | 36.984 | 2 | 54.36372 | yes | 9.975E-08 | 621.8255 | 279.38074 |
| GNHECASINR | 10 | P36873 | Serine/threonine-protein phosphatase PP1 | PPP1CC | 36.984 | 2 | 54.36372 | no | 1.133E-07 | 579.25946 | 257.25363 |
| QSLETICLLLAYK | 13 | P36873 | Serine/threonine-protein phosphatase PP1 | PPP1CC | 36.984 | 2 | 54.36372 | no | 9.722E-08 | 776.42621 | 209.29697 |
| GVSFTFGAEVVAK | 13 | P36873 | Serine/threonine-protein phosphatase PP1 | PPP1CC | 36.984 | 2 | 54.36372 | no | 9.722E-08 | 656.35083 | 77.544106 |
| DVLGWGENDR | 10 | P36873 | Serine/threonine-protein phosphatase PP1 | PPP1CC | 36.984 | 2 | 54.36372 | yes | 1.646E-07 | 580.7702 | 41.539116 |
| EAEIYQTVMLR | 11 | P36897 | TGF-beta receptor type-1 OS=Homo sapi | TGFBR1 | 55.96 | 2 | 38.546364 | no | 5.146E-07 | 676.8476 | 71.424431 |
| IELPTTVK | 8 | P36897 | TGF-beta receptor type-1 OS=Homo sapi | TGFBR1 | 55.96 | 2 | 38.546364 | yes | 2.997E-06 | 450.77368 | 40.199829 |
| ILLYACR | 7 | P36954 | DNA-directed RNA polymerase II subuni | POLR2I | 14.524 | 2 | 51.48668 | yes | 1.81E-06 | 454.75476 | 131.51619 |
| FCQECNNMLYPK | 12 | P36954 | DNA-directed RNA polymerase II subuni | POLR2I | 14.524 | 2 | 51.48668 | yes | 9.722E-08 | 802.33875 | 83.63784 |
| LYYVCTAPHCGHI | 13 | P36954 | DNA-directed RNA polymerase II subuni | POLR2I | 14.524 | 2 | 51.48668 | yes | 1.382E-07 | 817.37177 | 58.250507 |
| ALHFLTR | 7 | P36956 | Sterol regulatory element-binding protein | SREBF1 | 121.676 | 2 | 41.204884 | yes | 3.638E-07 | 429.25323 | 56.33173 |
| AESGPASLTICEK | 13 | P36956 | Sterol regulatory element-binding protein | SREBF1 | 121.676 | 2 | 41.204884 | yes | 3.624E-07 | 681.83209 | 55.680916 |
| QQQPPAPAPAAQC | 17 | P36956 | Sterol regulatory element-binding protein | SREBF1 | 121.676 | 2 | 41.204884 | yes | 2.608E-06 | 846.42664 | 11.481316 |
| NVEAMNFADIER | 12 | P36957 | Dihydrolipoyllysine-residue succinyltrans | DLST | 48.756 | 2 | 55.859463 | yes | 9.722E-08 | 704.8299 | 67.826706 |
| GLVVPVIR | 8 | P36957 | Dihydrolipoyllysine-residue succinyltrans | DLST | 48.756 | 2 | 55.859463 | yes | 8.926E-07 | 426.78693 | 46.125038 |
| VLLESEQFLTELTF | 14 | P37108 | Signal recognition particle 14 kDa protein | SRP14 | 14.57 | 2 | 50.516193 | yes | 9.79E-08 | 839.45654 | 4419.9683 |
| FQMAYSNLLR | 10 | P37108 | Signal recognition particle 14 kDa protein | SRP14 | 14.57 | 2 | 50.516193 | yes | 1.526E-07 | 621.8186 | 1257.4135 |
| FQMAYSNLLR | 10 | P37108 | Signal recognition particle 14 kDa protein | SRP14 | 14.57 | 2 | 50.516193 | yes | 4.622E-07 | 629.8161 | 1257.4135 |
| KGTVEGFEPADNK | 13 | P37108 | Signal recognition particle 14 kDa protein | SRP14 | 14.57 | 2 | 50.516193 | yes | 1.027E-07 | 696.34369 | 2186.0542 |
| TSGSVYITLK | 10 | P37108 | Signal recognition particle 14 kDa protein | SRP14 | 14.57 | 2 | 50.516193 | yes | 1.281E-07 | 534.80042 | 1805.6251 |
| KISTVVSSK | 9 | P37108 | Signal recognition particle 14 kDa protein | SRP14 | 14.57 | 2 | 50.516193 | yes | 1.301E-07 | 474.78986 | 850.68085 |
| GTVEGFEPADNK | 12 | P37108 | Signal recognition particle 14 kDa protein | SRP14 | 14.57 | 2 | 50.516193 | yes | 2.965E-07 | 632.2962 | 668.39514 |
| AAAAAAAAAPAA | 29 | P37108 | Signal recognition particle 14 kDa protein | SRP14 | 14.57 | 3 | 50.516193 | yes | 1.069E-07 | 790.07495 | 281.13425 |
| ANMDGLKK | 8 | P37108 | Signal recognition particle 14 kDa protein | SRP14 | 14.57 | 2 | 50.516193 | yes | 1.13E-07 | 438.73401 | 162.92754 |
| LAENIDAQLK | 10 | P37198 | Nuclear pore glycoprotein p62 OS=Homo | NUP62 | 53.255 | 2 | 51.291348 | yes | 9.722E-08 | 557.80878 | 67.505455 |
| WSLELEDQER | 10 | P37198 | Nuclear pore glycoprotein p62 OS=Homo | NUP62 | 53.255 | 2 | 51.291348 | yes | 1.286E-07 | 652.80951 | 36.577877 |
| LSLEEFIR | 8 | P37235 | Hippocalcin-like protein 1 OS=Homo sap | HPCAL1 | 22.314 | 2 | 41.76186 | no | 7.401E-07 | 503.78201 | 50.216785 |
| SEMLEIVQAIYK | 12 | P37235 | Hippocalcin-like protein 1 OS=Homo sap | HPCAL1 | 22.314 | 2 | 41.76186 | yes | 3.352E-07 | 712.37872 | 39.74799 |

| | | | | | | | | | | |
|---|---|---|---|---|---|---|---|---|---|---|
| TQNLPNCQLISR | 12 | P37268 | Squalene synthase OS=Homo sapiens OX | FDFT1 | 48.116 | 2 | 48.937763 | yes | 4.534E-07 | 722.37225 | 52.688995 |
| YQTVIADICR | 10 | P37268 | Squalene synthase OS=Homo sapiens OX | FDFT1 | 48.116 | 2 | 48.937763 | yes | 1.084E-07 | 619.81354 | 45.204487 |
| MGIGMAEFLDK | 11 | P37268 | Squalene synthase OS=Homo sapiens OX | FDFT1 | 48.116 | 2 | 48.937763 | yes | 1.552E-05 | 606.29358 | 26.518541 |
| NVIGLQMGTNR | 11 | P37802 | Transgelin-2 OS=Homo sapiens OX=960 | TAGLN2 | 22.392 | 2 | 44.09163 | yes | 8.028E-07 | 601.81915 | 93.771904 |
| TLMNLGGLAVAR | 12 | P37802 | Transgelin-2 OS=Homo sapiens OX=960 | TAGLN2 | 22.392 | 2 | 44.09163 | yes | 3.14E-06 | 608.34753 | 68.485229 |
| QMEQISQFLQAAE | 14 | P37802 | Transgelin-2 OS=Homo sapiens OX=960 | TAGLN2 | 22.392 | 2 | 44.09163 | yes | 2.324E-07 | 839.91449 | 32.651543 |
| LSVISVEDPPQR | 12 | P38117 | Electron transfer flavoprotein subunit beta | ETFB | 27.844 | 2 | 50.70293 | yes | 9.728E-08 | 670.36444 | 108.03108 |
| LFIGGLNTETNEK | 13 | P38159 | RNA-binding motif protein, X chromosor | RBMX | 42.332 | 2 | 53.202946 | no | 1.21E-07 | 718.375 | 15611.541 |
| GFAFVTFESPADA | 14 | P38159 | RNA-binding motif protein, X chromosor | RBMX | 42.332 | 2 | 53.202946 | no | 1.417E-07 | 743.86426 | 7513.8433 |
| IVEVLLMK | 8 | P38159 | RNA-binding motif protein, X chromosor | RBMX | 42.332 | 2 | 53.202946 | no | 1.017E-07 | 472.79608 | 4020.8257 |
| GPPPSYGGSSR | 11 | P38159 | RNA-binding motif protein, X chromosor | RBMX | 42.332 | 2 | 53.202946 | no | 3.011E-07 | 531.25415 | 3749.0591 |
| REPLPSR | 7 | P38159 | RNA-binding motif protein, X chromosor | RBMX | 42.332 | 2 | 53.202946 | no | 1.776E-07 | 427.74579 | 3108.1772 |
| YDDYSSSR | 8 | P38159 | RNA-binding motif protein, X chromosor | RBMX | 42.332 | 2 | 53.202946 | no | 1.867E-07 | 496.70142 | 2385.4834 |
| DRDYSDHPSGGSY | 14 | P38159 | RNA-binding motif protein, X chromosor | RBMX | 42.332 | 3 | 53.202946 | no | 1.727E-07 | 537.89758 | 1547.2137 |
| DYSDHPSGGSYR | 12 | P38159 | RNA-binding motif protein, X chromosor | RBMX | 42.332 | 3 | 53.202946 | no | 2.193E-07 | 447.52158 | 460.68277 |
| DYSDHPSGGSYR | 12 | P38159 | RNA-binding motif protein, X chromosor | RBMX | 42.332 | 3 | 53.202946 | no | 1.022E-07 | 670.77875 | 460.68277 |
| DYSDHPSGGSYR | 12 | P38159 | RNA-binding motif protein, X chromosor | RBMX | 42.332 | 2 | 53.202946 | no | 2.193E-07 | 447.52158 | 460.68277 |
| DYSDHPSGGSYR | 12 | P38159 | RNA-binding motif protein, X chromosor | RBMX | 42.332 | 2 | 53.202946 | no | 1.022E-07 | 670.77875 | 460.68277 |
| DSYESYGNSR | 10 | P38159 | RNA-binding motif protein, X chromosor | RBMX | 42.332 | 2 | 53.202946 | no | 9.722E-08 | 589.24146 | 729.03088 |
| VEQATKPSFESGR | 13 | P38159 | RNA-binding motif protein, X chromosor | RBMX | 42.332 | 2 | 53.202946 | yes | 9.722E-08 | 718.36243 | 596.4895 |
| DVYLSPR | 7 | P38159 | RNA-binding motif protein, X chromosor | RBMX | 42.332 | 2 | 53.202946 | no | 2.945E-07 | 425.2269 | 423.17361 |
| VEADRPGK | 8 | P38159 | RNA-binding motif protein, X chromosor | RBMX | 42.332 | 2 | 53.202946 | no | 1.242E-07 | 457.24051 | 394.57242 |
| RGPPPPPR | 8 | P38159 | RNA-binding motif protein, X chromosor | RBMX | 42.332 | 2 | 53.202946 | yes | 1.106E-07 | 437.25632 | 382.00766 |
| SDLYSSGR | 8 | P38159 | RNA-binding motif protein, X chromosor | RBMX | 42.332 | 2 | 53.202946 | yes | 1.076E-07 | 442.70905 | 322.24823 |
| RDVYLSPR | 8 | P38159 | RNA-binding motif protein, X chromosor | RBMX | 42.332 | 2 | 53.202946 | no | 3.642E-06 | 503.27744 | 91.778824 |
| GNETGLITPNK | 11 | P38398 | Breast cancer type 1 susceptibility protein | BRCA1 | 207.721 | 2 | 54.46176 | yes | 2.705E-06 | 572.30383 | 1089.8243 |
| VGNIPSSTSALK | 12 | P38398 | Breast cancer type 1 susceptibility protein | BRCA1 | 207.721 | 2 | 54.46176 | yes | 3.185E-07 | 587.32733 | 448.72839 |
| LGVLQPEVYK | 10 | P38398 | Breast cancer type 1 susceptibility protein | BRCA1 | 207.721 | 2 | 54.46176 | yes | 2.156E-07 | 573.33191 | 330.82938 |
| TDAEFVCER | 9 | P38398 | Breast cancer type 1 susceptibility protein | BRCA1 | 207.721 | 2 | 54.46176 | yes | 1.034E-07 | 563.7453 | 276.98621 |
| FCLSSQFR | 8 | P38398 | Breast cancer type 1 susceptibility protein | BRCA1 | 207.721 | 2 | 54.46176 | yes | 1.998E-07 | 522.75018 | 268.38559 |
| CVSQCAAFENPK | 12 | P38398 | Breast cancer type 1 susceptibility protein | BRCA1 | 207.721 | 2 | 54.46176 | yes | 9.742E-08 | 705.81067 | 267.22879 |
| VEEVQNVINAMQI | 13 | P38398 | Breast cancer type 1 susceptibility protein | BRCA1 | 207.721 | 2 | 54.46176 | yes | 9.722E-08 | 751.38757 | 253.36104 |
| NPNPIESLEK | 10 | P38398 | Breast cancer type 1 susceptibility protein | BRCA1 | 207.721 | 2 | 54.46176 | yes | 2.093E-07 | 570.7984 | 234.88432 |
| HGLLQNPYR | 9 | P38398 | Breast cancer type 1 susceptibility protein | BRCA1 | 207.721 | 2 | 54.46176 | yes | 9.722E-08 | 549.29614 | 222.96188 |
| EFVNPSLPR | 9 | P38398 | Breast cancer type 1 susceptibility protein | BRCA1 | 207.721 | 2 | 54.46176 | yes | 5.146E-07 | 529.7851 | 213.17696 |
| MLNEHDFEVR | 10 | P38398 | Breast cancer type 1 susceptibility protein | BRCA1 | 207.721 | 2 | 54.46176 | yes | 2.156E-07 | 645.30078 | 209.10648 |
| RPTSGLHPEDFIK | 13 | P38398 | Breast cancer type 1 susceptibility protein | BRCA1 | 207.721 | 3 | 54.46176 | yes | 1.974E-07 | 499.60007 | 192.11093 |
| FSQLVEELLK | 10 | P38398 | Breast cancer type 1 susceptibility protein | BRCA1 | 207.721 | 2 | 54.46176 | yes | 9.722E-08 | 603.34247 | 190.29814 |
| VNNIPSQSTR | 10 | P38398 | Breast cancer type 1 susceptibility protein | BRCA1 | 207.721 | 2 | 54.46176 | yes | 1.584E-07 | 558.29382 | 179.92172 |

| | | | | | | | | | | | |
|---|---|---|---|---|---|---|---|---|---|---|---|
| HIHALELVVSR | 11 | P38398 | Breast cancer type 1 susceptibility protein | BRCA1 | 207.721 | 2 | 54.46176 | yes | 1.425E-06 | 637.37238 | 163.68652 |
| VDLNADPLCER | 11 | P38398 | Breast cancer type 1 susceptibility protein | BRCA1 | 207.721 | 2 | 54.46176 | yes | 2.319E-07 | 651.31134 | 151.81229 |
| EKPELTASTER | 11 | P38398 | Breast cancer type 1 susceptibility protein | BRCA1 | 207.721 | 2 | 54.46176 | yes | 3.635E-07 | 630.82513 | 150.35185 |
| DTMQHNLIK | 9 | P38398 | Breast cancer type 1 susceptibility protein | BRCA1 | 207.721 | 2 | 54.46176 | yes | 8.318E-07 | 550.28186 | 143.5614 |
| KVDLNADPLCER | 12 | P38398 | Breast cancer type 1 susceptibility protein | BRCA1 | 207.721 | 2 | 54.46176 | yes | 5.944E-07 | 715.35883 | 123.73903 |
| NSLNDCSNQVILA | 14 | P38398 | Breast cancer type 1 susceptibility protein | BRCA1 | 207.721 | 2 | 54.46176 | yes | 1.092E-07 | 788.39343 | 113.16946 |
| KADLAVQK | 8 | P38398 | Breast cancer type 1 susceptibility protein | BRCA1 | 207.721 | 2 | 54.46176 | yes | 1.234E-05 | 436.76364 | 111.64975 |
| ILECPICLELIK | 12 | P38398 | Breast cancer type 1 susceptibility protein | BRCA1 | 207.721 | 2 | 54.46176 | yes | 9.722E-08 | 750.91187 | 107.52255 |
| NTEENLLSLK | 10 | P38398 | Breast cancer type 1 susceptibility protein | BRCA1 | 207.721 | 2 | 54.46176 | yes | 9.722E-08 | 580.81152 | 100.82184 |
| DEISLDSAK | 9 | P38398 | Breast cancer type 1 susceptibility protein | BRCA1 | 207.721 | 2 | 54.46176 | yes | 1.493E-07 | 489.24292 | 97.58387 |
| NYPSQEELIK | 10 | P38398 | Breast cancer type 1 susceptibility protein | BRCA1 | 207.721 | 2 | 54.46176 | yes | 4.621E-07 | 610.81152 | 77.028778 |
| RHDSDTFPELK | 11 | P38398 | Breast cancer type 1 susceptibility protein | BRCA1 | 207.721 | 2 | 54.46176 | yes | 9.722E-08 | 672.83075 | 76.463341 |
| DEVSIIQSMGYR | 12 | P38398 | Breast cancer type 1 susceptibility protein | BRCA1 | 207.721 | 2 | 54.46176 | yes | 1.778E-07 | 699.34009 | 74.69915 |
| AEPISSSISNMELEI | 21 | P38398 | Breast cancer type 1 susceptibility protein | BRCA1 | 207.721 | 3 | 54.46176 | yes | 6.424E-06 | 771.7179 | 69.077553 |
| NLQLMEGK | 8 | P38398 | Breast cancer type 1 susceptibility protein | BRCA1 | 207.721 | 2 | 54.46176 | yes | 2.765E-07 | 466.74713 | 62.015007 |
| QSLPGSNCK | 9 | P38398 | Breast cancer type 1 susceptibility protein | BRCA1 | 207.721 | 2 | 54.46176 | yes | 3.086E-06 | 495.7373 | 57.968788 |
| KGPSQCPLCK | 10 | P38398 | Breast cancer type 1 susceptibility protein | BRCA1 | 207.721 | 2 | 54.46176 | yes | 2.88E-06 | 587.789 | 47.641739 |
| CDHIFCK | 7 | P38398 | Breast cancer type 1 susceptibility protein | BRCA1 | 207.721 | 2 | 54.46176 | yes | 3.059E-07 | 490.20987 | 46.205177 |
| HQSESQGVGLSDK | 13 | P38398 | Breast cancer type 1 susceptibility protein | BRCA1 | 207.721 | 2 | 54.46176 | yes | 9.722E-08 | 686.32861 | 43.343559 |
| YFLGIAGGK | 9 | P38398 | Breast cancer type 1 susceptibility protein | BRCA1 | 207.721 | 2 | 54.46176 | yes | 5.431E-07 | 463.26074 | 40.304691 |
| KYNQMPVR | 8 | P38398 | Breast cancer type 1 susceptibility protein | BRCA1 | 207.721 | 2 | 54.46176 | yes | 1.555E-06 | 518.27386 | 32.206226 |
| HSTVATECLSK | 11 | P38398 | Breast cancer type 1 susceptibility protein | BRCA1 | 207.721 | 2 | 54.46176 | yes | 3.455E-06 | 616.8006 | 28.687468 |
| HHITLTNLITEETTI | 19 | P38398 | Breast cancer type 1 susceptibility protein | BRCA1 | 207.721 | 3 | 54.46176 | yes | 1.832E-07 | 739.72803 | 21.961031 |
| NESNIKPVQTVNIT | 23 | P38398 | Breast cancer type 1 susceptibility protein | BRCA1 | 207.721 | 3 | 54.46176 | yes | 1.096E-05 | 814.11133 | 10.090931 |
| NSSTIIQNPVETPK | 14 | P38432 | Coilin OS=Homo sapiens OX=9606 GN= | COIL | 62.609 | 2 | 50.36804 | yes | 1.329E-07 | 764.4043 | 144.38585 |
| VVTDLISLIR | 10 | P38432 | Coilin OS=Homo sapiens OX=9606 GN= | COIL | 62.609 | 2 | 50.36804 | yes | 2.664E-07 | 564.85297 | 122.27652 |
| ATCGTVGDDNEEA | 14 | P38432 | Coilin OS=Homo sapiens OX=9606 GN= | COIL | 62.609 | 2 | 50.36804 | yes | 9.81E-08 | 733.80682 | 58.809013 |
| LQQLLPLK | 8 | P38435 | Vitamin K-dependent gamma-carboxylase | GGCX | 87.561 | 2 | 21.078754 | yes | 4.25E-06 | 476.81314 | 24.7803 |
| DFPELTMEVDGK | 12 | P38606 | V-type proton ATPase catalytic subunit A | ATP6V1A | 68.305 | 2 | 39.458866 | yes | 2.411E-06 | 690.82123 | 65.469612 |
| TALVANTSNMPVA | 15 | P38606 | V-type proton ATPase catalytic subunit A | ATP6V1A | 68.305 | 2 | 39.458866 | yes | 2.266E-06 | 758.40106 | 57.77784 |
| FTMVQVWPVR | 10 | P38606 | V-type proton ATPase catalytic subunit A | ATP6V1A | 68.305 | 2 | 39.458866 | yes | 4.573E-07 | 631.83936 | 27.38493 |
| DAGQISGLNVLR | 12 | P38646 | Stress-70 protein, mitochondrial OS=Hon | HSPA9 | 73.681 | 2 | 52.239925 | yes | 1.23E-07 | 621.84369 | 5268.0552 |
| NAVITVPAYFNDS | 15 | P38646 | Stress-70 protein, mitochondrial OS=Hon | HSPA9 | 73.681 | 2 | 52.239925 | yes | 1.429E-07 | 847.92847 | 4921.6514 |
| VINEPTAAALAYG | 16 | P38646 | Stress-70 protein, mitochondrial OS=Hon | HSPA9 | 73.681 | 2 | 52.239925 | yes | 1.172E-07 | 823.44342 | 4853.3975 |
| EQQIVIQSSGGLSK | 14 | P38646 | Stress-70 protein, mitochondrial OS=Hon | HSPA9 | 73.681 | 2 | 52.239925 | yes | 9.723E-08 | 737.39899 | 3814.5752 |
| AQFEGIVTDLIR | 12 | P38646 | Stress-70 protein, mitochondrial OS=Hon | HSPA9 | 73.681 | 2 | 52.239925 | yes | 9.722E-08 | 681.37482 | 3081.3757 |
| RYDDPEVQK | 9 | P38646 | Stress-70 protein, mitochondrial OS=Hon | HSPA9 | 73.681 | 2 | 52.239925 | yes | 1.084E-07 | 575.2804 | 2363.9331 |
| VLENAEGAR | 9 | P38646 | Stress-70 protein, mitochondrial OS=Hon | HSPA9 | 73.681 | 2 | 52.239925 | yes | 1.306E-07 | 479.75125 | 2038.6418 |
| VQQTVQDLFGR | 11 | P38646 | Stress-70 protein, mitochondrial OS=Hon | HSPA9 | 73.681 | 2 | 52.239925 | yes | 1.108E-07 | 645.84369 | 1829.5853 |

| | | | | | | | | | | | |
|---|---|---|---|---|---|---|---|---|---|---|---|
| ETAENYLGHTAK | 12 | P38646 | Stress-70 protein, mitochondrial OS=Hon | HSPA9 | 73.681 | 2 | 52.239925 | yes | 9.722E-08 | 667.32275 | 1645.9724 |
| SDIGEVILVGGMTI | 14 | P38646 | Stress-70 protein, mitochondrial OS=Hon | HSPA9 | 73.681 | 2 | 52.239925 | yes | 9.722E-08 | 723.8847 | 1426.1348 |
| DQLPADECNK | 10 | P38646 | Stress-70 protein, mitochondrial OS=Hon | HSPA9 | 73.681 | 2 | 52.239925 | yes | 1.703E-07 | 595.26135 | 1424.8192 |
| TTPSVVAFTADGE | 14 | P38646 | Stress-70 protein, mitochondrial OS=Hon | HSPA9 | 73.681 | 2 | 52.239925 | yes | 1.117E-07 | 725.86224 | 1396.3345 |
| QAVTNPNNTFYAT | 14 | P38646 | Stress-70 protein, mitochondrial OS=Hon | HSPA9 | 73.681 | 2 | 52.239925 | yes | 1.06E-07 | 784.88879 | 1080.3599 |
| QAASSLQQASLK | 12 | P38646 | Stress-70 protein, mitochondrial OS=Hon | HSPA9 | 73.681 | 2 | 52.239925 | yes | 1.594E-07 | 616.33569 | 1029.2458 |
| LLGQFTLIGIPPAPI | 15 | P38646 | Stress-70 protein, mitochondrial OS=Hon | HSPA9 | 73.681 | 2 | 52.239925 | yes | 3.376E-07 | 796.9798 | 943.85846 |
| AMQDAEVSK | 9 | P38646 | Stress-70 protein, mitochondrial OS=Hon | HSPA9 | 73.681 | 2 | 52.239925 | yes | 1.839E-07 | 489.73169 | 844.69263 |
| LYSPSQIGAFVLMI | 14 | P38646 | Stress-70 protein, mitochondrial OS=Hon | HSPA9 | 73.681 | 2 | 52.239925 | yes | 9.722E-08 | 777.42346 | 813.28882 |
| KDSETGENIR | 10 | P38646 | Stress-70 protein, mitochondrial OS=Hon | HSPA9 | 73.681 | 2 | 52.239925 | yes | 1.152E-07 | 574.78076 | 563.08093 |
| ERVEAVNMAEGII | 19 | P38646 | Stress-70 protein, mitochondrial OS=Hon | HSPA9 | 73.681 | 3 | 52.239925 | yes | 1.481E-07 | 714.68805 | 502.40497 |
| LKEEISK | 7 | P38646 | Stress-70 protein, mitochondrial OS=Hon | HSPA9 | 73.681 | 2 | 52.239925 | yes | 1.123E-07 | 423.75018 | 273.19583 |
| YDDPEVQK | 8 | P38646 | Stress-70 protein, mitochondrial OS=Hon | HSPA9 | 73.681 | 2 | 52.239925 | yes | 1.58E-07 | 497.22983 | 198.29646 |
| ETGVDLTK | 8 | P38646 | Stress-70 protein, mitochondrial OS=Hon | HSPA9 | 73.681 | 2 | 52.239925 | yes | 2.054E-07 | 431.72946 | 185.76601 |
| MKETAENYLGHTA | 14 | P38646 | Stress-70 protein, mitochondrial OS=Hon | HSPA9 | 73.681 | 3 | 52.239925 | yes | 1.276E-07 | 531.59607 | 90.418579 |
| MKETAENYLGHTA | 14 | P38646 | Stress-70 protein, mitochondrial OS=Hon | HSPA9 | 73.681 | 3 | 52.239925 | yes | 1.09E-07 | 796.8905 | 90.418579 |
| MKETAENYLGHTA | 14 | P38646 | Stress-70 protein, mitochondrial OS=Hon | HSPA9 | 73.681 | 2 | 52.239925 | yes | 1.276E-07 | 531.59607 | 90.418579 |
| MKETAENYLGHTA | 14 | P38646 | Stress-70 protein, mitochondrial OS=Hon | HSPA9 | 73.681 | 2 | 52.239925 | yes | 1.09E-07 | 796.8905 | 90.418579 |
| DSETGENIR | 9 | P38646 | Stress-70 protein, mitochondrial OS=Hon | HSPA9 | 73.681 | 2 | 52.239925 | yes | 1.847E-07 | 510.73325 | 171.56839 |
| ASNGDAWVEAHG | 13 | P38646 | Stress-70 protein, mitochondrial OS=Hon | HSPA9 | 73.681 | 2 | 52.239925 | yes | 1.138E-07 | 671.31274 | 169.40956 |
| DDIENMVK | 8 | P38646 | Stress-70 protein, mitochondrial OS=Hon | HSPA9 | 73.681 | 2 | 52.239925 | yes | 1.084E-07 | 482.22623 | 93.785599 |
| RETGVDLTK | 9 | P38646 | Stress-70 protein, mitochondrial OS=Hon | HSPA9 | 73.681 | 2 | 52.239925 | yes | 6.61E-07 | 509.78 | 74.637589 |
| MLVLDEADEMLN | 13 | P38919 | Eukaryotic initiation factor 4A-III OS=Hc | EIF4A3 | 46.872 | 2 | 54.21554 | yes | 1.496E-07 | 760.87036 | 7580.9268 |
| MLVLDEADEMLN | 13 | P38919 | Eukaryotic initiation factor 4A-III OS=Hc | EIF4A3 | 46.872 | 2 | 54.21554 | yes | 3.451E-07 | 768.86786 | 7580.9268 |
| VLISTDVWAR | 10 | P38919 | Eukaryotic initiation factor 4A-III OS=Hc | EIF4A3 | 46.872 | 2 | 54.21554 | yes | 9.935E-08 | 580.32715 | 13265.051 |
| ETQALILAPTR | 11 | P38919 | Eukaryotic initiation factor 4A-III OS=Hc | EIF4A3 | 46.872 | 2 | 54.21554 | yes | 1.276E-07 | 606.85095 | 10869.739 |
| EQIYDVYR | 8 | P38919 | Eukaryotic initiation factor 4A-III OS=Hc | EIF4A3 | 46.872 | 2 | 54.21554 | yes | 1.557E-07 | 543.26672 | 5846.707 |
| RDELTLEGIK | 10 | P38919 | Eukaryotic initiation factor 4A-III OS=Hc | EIF4A3 | 46.872 | 2 | 54.21554 | yes | 1.312E-07 | 587.32733 | 3363.0911 |
| KGVAINFVK | 9 | P38919 | Eukaryotic initiation factor 4A-III OS=Hc | EIF4A3 | 46.872 | 2 | 54.21554 | yes | 1.081E-07 | 488.30292 | 3137.4277 |
| KLDYGQHVVAGT | 15 | P38919 | Eukaryotic initiation factor 4A-III OS=Hc | EIF4A3 | 46.872 | 3 | 54.21554 | yes | 1.88E-07 | 533.28638 | 1448.0872 |
| KLDYGQHVVAGT | 15 | P38919 | Eukaryotic initiation factor 4A-III OS=Hc | EIF4A3 | 46.872 | 3 | 54.21554 | yes | 9.819E-08 | 799.4259 | 1448.0872 |
| KLDYGQHVVAGT | 15 | P38919 | Eukaryotic initiation factor 4A-III OS=Hc | EIF4A3 | 46.872 | 2 | 54.21554 | yes | 1.88E-07 | 533.28638 | 1448.0872 |
| KLDYGQHVVAGT | 15 | P38919 | Eukaryotic initiation factor 4A-III OS=Hc | EIF4A3 | 46.872 | 2 | 54.21554 | yes | 9.819E-08 | 799.4259 | 1448.0872 |
| QFFVAVER | 8 | P38919 | Eukaryotic initiation factor 4A-III OS=Hc | EIF4A3 | 46.872 | 2 | 54.21554 | yes | 2.419E-07 | 498.26907 | 2829.1479 |
| DVIAQSQSGTGK | 12 | P38919 | Eukaryotic initiation factor 4A-III OS=Hc | EIF4A3 | 46.872 | 2 | 54.21554 | yes | 9.722E-08 | 595.8042 | 2407.7546 |
| ELAVQIQK | 8 | P38919 | Eukaryotic initiation factor 4A-III OS=Hc | EIF4A3 | 46.872 | 2 | 54.21554 | yes | 1.765E-07 | 464.77673 | 1950.219 |
| LDYGQHVVAGTP( | 14 | P38919 | Eukaryotic initiation factor 4A-III OS=Hc | EIF4A3 | 46.872 | 2 | 54.21554 | yes | 9.927E-08 | 735.37842 | 862.71375 |
| EANFTVSSMHGDI | 16 | P38919 | Eukaryotic initiation factor 4A-III OS=Hc | EIF4A3 | 46.872 | 3 | 54.21554 | yes | 9.722E-08 | 593.59937 | 623.59583 |
| ATTATMATSGSAR | 13 | P38919 | Eukaryotic initiation factor 4A-III OS=Hc | EIF4A3 | 46.872 | 2 | 54.21554 | yes | 1.022E-07 | 634.30096 | 305.78815 |

| | | | | | | | | | | | |
|---|---|---|---|---|---|---|---|---|---|---|---|
| ATTATMATSGSAR | 13 | P38919 | Eukaryotic initiation factor 4A-III OS=Hc | EIF4A3 | 46.872 | 2 | 54.21554 | yes | 3.656E-07 | 642.29846 | 305.78815 |
| FMTDPIR | 7 | P38919 | Eukaryotic initiation factor 4A-III OS=Hc | EIF4A3 | 46.872 | 2 | 54.21554 | yes | 1.338E-07 | 440.2233 | 300.14282 |
| FMTDPIR | 7 | P38919 | Eukaryotic initiation factor 4A-III OS=Hc | EIF4A3 | 46.872 | 2 | 54.21554 | yes | 1.032E-07 | 448.22073 | 300.14282 |
| LLKEEDMTK | 9 | P38919 | Eukaryotic initiation factor 4A-III OS=Hc | EIF4A3 | 46.872 | 2 | 54.21554 | yes | 1.446E-07 | 553.79175 | 385.25494 |
| GVAINFVK | 8 | P38919 | Eukaryotic initiation factor 4A-III OS=Hc | EIF4A3 | 46.872 | 2 | 54.21554 | yes | 1.012E-07 | 424.25543 | 229.93326 |
| GRDVIAQSQSGTG | 14 | P38919 | Eukaryotic initiation factor 4A-III OS=Hc | EIF4A3 | 46.872 | 2 | 54.21554 | yes | 9.722E-08 | 702.36548 | 126.6829 |
| GLDVPQVSLIINYI | 19 | P38919 | Eukaryotic initiation factor 4A-III OS=Hc | EIF4A3 | 46.872 | 3 | 54.21554 | yes | 1.225E-07 | 714.05139 | 50.031944 |
| VEFETSEEVDVTP' | 21 | P38919 | Eukaryotic initiation factor 4A-III OS=Hc | EIF4A3 | 46.872 | 3 | 54.21554 | yes | 1.692E-07 | 801.37378 | 42.660023 |
| AMIVEFMASKK | 11 | P38935 | DNA-binding protein SMUBP-2 OS=Hoı | IGHMBP2 | 109.15 | 2 | 52.302925 | yes | 9.777E-08 | 643.82782 | 895.32068 |
| LLVTFEPR | 8 | P38935 | DNA-binding protein SMUBP-2 OS=Hoı | IGHMBP2 | 109.15 | 2 | 52.302925 | yes | 2.909E-07 | 487.78711 | 411.56992 |
| EGVLYAGSGTK | 11 | P38935 | DNA-binding protein SMUBP-2 OS=Hoı | IGHMBP2 | 109.15 | 2 | 52.302925 | yes | 2.233E-07 | 541.27985 | 303.66122 |
| ELAIIHGPPGTGK | 13 | P38935 | DNA-binding protein SMUBP-2 OS=Hoı | IGHMBP2 | 109.15 | 2 | 52.302925 | yes | 9.722E-08 | 645.36426 | 275.76227 |
| LAEEYGAR | 8 | P38935 | DNA-binding protein SMUBP-2 OS=Hoı | IGHMBP2 | 109.15 | 2 | 52.302925 | yes | 1.622E-07 | 454.72726 | 252.29498 |
| AMIVEFMASK | 10 | P38935 | DNA-binding protein SMUBP-2 OS=Hoı | IGHMBP2 | 109.15 | 2 | 52.302925 | yes | 1.031E-07 | 563.7854 | 246.92238 |
| HVAVICDSR | 9 | P38935 | DNA-binding protein SMUBP-2 OS=Hoı | IGHMBP2 | 109.15 | 2 | 52.302925 | yes | 3.497E-07 | 528.76636 | 237.5721 |
| QLDLLELER | 9 | P38935 | DNA-binding protein SMUBP-2 OS=Hoı | IGHMBP2 | 109.15 | 2 | 52.302925 | yes | 9.722E-08 | 564.81659 | 232.86795 |
| SWQENISLK | 9 | P38935 | DNA-binding protein SMUBP-2 OS=Hoı | IGHMBP2 | 109.15 | 2 | 52.302925 | yes | 9.722E-08 | 552.78784 | 227.63429 |
| MHQAIMR | 7 | P38935 | DNA-binding protein SMUBP-2 OS=Hoı | IGHMBP2 | 109.15 | 2 | 52.302925 | yes | 1.206E-07 | 443.72293 | 198.55023 |
| LVSLHIQALVDAG | 17 | P38935 | DNA-binding protein SMUBP-2 OS=Hoı | IGHMBP2 | 109.15 | 3 | 52.302925 | yes | 9.779E-08 | 587.01233 | 186.25362 |
| TLVEYFTQHGEVR | 13 | P38935 | DNA-binding protein SMUBP-2 OS=Hoı | IGHMBP2 | 109.15 | 2 | 52.302925 | yes | 1.056E-07 | 789.89917 | 183.51707 |
| LANDVTYR | 8 | P38935 | DNA-binding protein SMUBP-2 OS=Hoı | IGHMBP2 | 109.15 | 2 | 52.302925 | yes | 9.977E-08 | 476.24835 | 145.5235 |
| TLTVQYR | 7 | P38935 | DNA-binding protein SMUBP-2 OS=Hoı | IGHMBP2 | 109.15 | 2 | 52.302925 | yes | 9.851E-08 | 440.74799 | 143.02673 |
| VHQIAEEHGLR | 11 | P38935 | DNA-binding protein SMUBP-2 OS=Hoı | IGHMBP2 | 109.15 | 2 | 52.302925 | yes | 3.168E-07 | 644.84149 | 112.34665 |
| KGEVGFLAEDR | 11 | P38935 | DNA-binding protein SMUBP-2 OS=Hoı | IGHMBP2 | 109.15 | 2 | 52.302925 | yes | 3.824E-07 | 610.81714 | 65.781647 |
| HPELEIK | 7 | P38935 | DNA-binding protein SMUBP-2 OS=Hoı | IGHMBP2 | 109.15 | 2 | 52.302925 | yes | 2.804E-07 | 433.24252 | 65.065292 |
| ASAAVESFVTK | 11 | P38935 | DNA-binding protein SMUBP-2 OS=Hoı | IGHMBP2 | 109.15 | 2 | 52.302925 | yes | 9.131E-05 | 576.30078 | 48.381409 |
| KDIDQVFVK | 9 | P38935 | DNA-binding protein SMUBP-2 OS=Hoı | IGHMBP2 | 109.15 | 2 | 52.302925 | yes | 3.996E-07 | 546.30841 | 45.996483 |
| EKEAVILSFVR | 11 | P38935 | DNA-binding protein SMUBP-2 OS=Hoı | IGHMBP2 | 109.15 | 2 | 52.302925 | yes | 1.162E-07 | 645.87445 | 45.773796 |
| YCLSHHLPEIHGC( | 16 | P38935 | DNA-binding protein SMUBP-2 OS=Hoı | IGHMBP2 | 109.15 | 4 | 52.302925 | yes | 9.15E-07 | 491.97681 | 28.547283 |
| GHPATDLPTEEDFI | 22 | P38935 | DNA-binding protein SMUBP-2 OS=Hoı | IGHMBP2 | 109.15 | 3 | 52.302925 | yes | 2.083E-06 | 766.38141 | 26.887205 |
| DVNQQEFVR | 9 | P39019 | Small ribosomal subunit protein eS19 OS | RPS19 | 16.061 | 2 | 50.241646 | yes | 1.345E-07 | 567.78052 | 29966.182 |
| VLQALEGLK | 9 | P39019 | Small ribosomal subunit protein eS19 OS | RPS19 | 16.061 | 2 | 50.241646 | yes | 1.4E-07 | 485.80023 | 25577.381 |
| VPEWVDTVK | 9 | P39019 | Small ribosomal subunit protein eS19 OS | RPS19 | 16.061 | 2 | 50.241646 | yes | 1.222E-07 | 536.78729 | 12200.309 |
| LKVPEWVDTVK | 11 | P39019 | Small ribosomal subunit protein eS19 OS | RPS19 | 16.061 | 2 | 50.241646 | yes | 1.951E-07 | 657.37683 | 5608.4868 |
| LKVPEWVDTVK | 11 | P39019 | Small ribosomal subunit protein eS19 OS | RPS19 | 16.061 | 2 | 50.241646 | yes | 1.001E-07 | 438.58698 | 5608.4868 |
| LKVPEWVDTVK | 11 | P39019 | Small ribosomal subunit protein eS19 OS | RPS19 | 16.061 | 3 | 50.241646 | yes | 1.951E-07 | 657.37683 | 5608.4868 |
| LKVPEWVDTVK | 11 | P39019 | Small ribosomal subunit protein eS19 OS | RPS19 | 16.061 | 3 | 50.241646 | yes | 1.001E-07 | 438.58698 | 5608.4868 |
| NGVMPSHFSR | 10 | P39019 | Small ribosomal subunit protein eS19 OS | RPS19 | 16.061 | 2 | 50.241646 | yes | 2.244E-07 | 566.27185 | 4526.1538 |
| NGVMPSHFSR | 10 | P39019 | Small ribosomal subunit protein eS19 OS | RPS19 | 16.061 | 2 | 50.241646 | yes | 1.686E-07 | 574.26929 | 4526.1538 |

| | | | | | | | | | | |
|---|---|---|---|---|---|---|---|---|---|---|
| IAGQVAAANK | 10 | P39019 | Small ribosomal subunit protein eS19 OS | RPS19 | 16.061 | 2 | 50.241646 | yes | 2.548E-07 | 471.772 | 5008.5518 |
| RVLQALEGLK | 10 | P39019 | Small ribosomal subunit protein eS19 OS | RPS19 | 16.061 | 2 | 50.241646 | yes | 1.582E-07 | 563.85077 | 1942.0071 |
| KLTPQGQR | 8 | P39019 | Small ribosomal subunit protein eS19 OS | RPS19 | 16.061 | 2 | 50.241646 | yes | 9.839E-08 | 464.27216 | 891.87451 |
| IAGQVAAANKK | 11 | P39019 | Small ribosomal subunit protein eS19 OS | RPS19 | 16.061 | 2 | 50.241646 | yes | 1.695E-07 | 535.81946 | 377.3511 |
| VACIGAWHPAR | 11 | P39023 | Large ribosomal subunit protein uL3 OS= | RPL3 | 46.109 | 2 | 50.056194 | no | 1.361E-07 | 619.31659 | 31772.471 |
| HGSLGFLPR | 9 | P39023 | Large ribosomal subunit protein uL3 OS= | RPL3 | 46.109 | 2 | 50.056194 | yes | 2.457E-06 | 492.27469 | 27338.035 |
| VIAHTQMR | 8 | P39023 | Large ribosomal subunit protein uL3 OS= | RPL3 | 46.109 | 2 | 50.056194 | yes | 1.251E-07 | 478.26074 | 18271.045 |
| AGMTHIVR | 8 | P39023 | Large ribosomal subunit protein uL3 OS= | RPL3 | 46.109 | 2 | 50.056194 | yes | 1.828E-07 | 442.74219 | 10281.465 |
| TVFAEHISDECK | 12 | P39023 | Large ribosomal subunit protein uL3 OS= | RPL3 | 46.109 | 2 | 50.056194 | yes | 1.046E-07 | 718.32971 | 7010.3457 |
| IGQGYLIK | 8 | P39023 | Large ribosomal subunit protein uL3 OS= | RPL3 | 46.109 | 2 | 50.056194 | yes | 1.197E-07 | 446.26855 | 6692.7593 |
| FQTMEEK | 7 | P39023 | Large ribosomal subunit protein uL3 OS= | RPL3 | 46.109 | 2 | 50.056194 | yes | 2.75E-07 | 456.71021 | 4763.0703 |
| NNASTDYDLSDK | 12 | P39023 | Large ribosomal subunit protein uL3 OS= | RPL3 | 46.109 | 2 | 50.056194 | yes | 9.919E-08 | 671.7915 | 2771.5159 |
| KYCQVIR | 7 | P39023 | Large ribosomal subunit protein uL3 OS= | RPL3 | 46.109 | 2 | 50.056194 | yes | 3.707E-07 | 483.76312 | 2326.8376 |
| AHLMEIQVNGGTV | 16 | P39023 | Large ribosomal subunit protein uL3 OS= | RPL3 | 46.109 | 3 | 50.056194 | yes | 1.104E-07 | 566.29443 | 653.2085 |
| AHLMEIQVNGGTV | 16 | P39023 | Large ribosomal subunit protein uL3 OS= | RPL3 | 46.109 | 3 | 50.056194 | yes | 1.84E-07 | 848.93799 | 653.2085 |
| AHLMEIQVNGGTV | 16 | P39023 | Large ribosomal subunit protein uL3 OS= | RPL3 | 46.109 | 2 | 50.056194 | yes | 1.104E-07 | 566.29443 | 653.2085 |
| AHLMEIQVNGGTV | 16 | P39023 | Large ribosomal subunit protein uL3 OS= | RPL3 | 46.109 | 2 | 50.056194 | yes | 1.84E-07 | 848.93799 | 653.2085 |
| KAFMGPLK | 8 | P39023 | Large ribosomal subunit protein uL3 OS= | RPL3 | 46.109 | 2 | 50.056194 | yes | 1.144E-07 | 446.25967 | 959.35596 |
| KVACIGAWHPAR | 12 | P39023 | Large ribosomal subunit protein uL3 OS= | RPL3 | 46.109 | 2 | 50.056194 | no | 1.147E-07 | 683.36407 | 918.01074 |
| DDPSKPVHLTAFL | 16 | P39023 | Large ribosomal subunit protein uL3 OS= | RPL3 | 46.109 | 3 | 50.056194 | yes | 2.274E-07 | 596.64905 | 491.29565 |
| AFMGPLKK | 8 | P39023 | Large ribosomal subunit protein uL3 OS= | RPL3 | 46.109 | 2 | 50.056194 | yes | 1.742E-07 | 446.25967 | 379.21677 |
| PVHLTAFLGYK | 11 | P39023 | Large ribosomal subunit protein uL3 OS= | RPL3 | 46.109 | 2 | 50.056194 | yes | 1.742E-07 | 623.35315 | 169.54517 |
| PVHLTAFLGYK | 11 | P39023 | Large ribosomal subunit protein uL3 OS= | RPL3 | 46.109 | 2 | 50.056194 | yes | 1.332E-07 | 415.90451 | 169.54517 |
| PVHLTAFLGYK | 11 | P39023 | Large ribosomal subunit protein uL3 OS= | RPL3 | 46.109 | 3 | 50.056194 | yes | 1.742E-07 | 623.35315 | 169.54517 |
| PVHLTAFLGYK | 11 | P39023 | Large ribosomal subunit protein uL3 OS= | RPL3 | 46.109 | 3 | 50.056194 | yes | 1.332E-07 | 415.90451 | 169.54517 |
| KWQDEDGK | 8 | P39023 | Large ribosomal subunit protein uL3 OS= | RPL3 | 46.109 | 2 | 50.056194 | yes | 3.851E-07 | 503.23544 | 193.55571 |
| KSLLVQTK | 8 | P39023 | Large ribosomal subunit protein uL3 OS= | RPL3 | 46.109 | 2 | 50.056194 | yes | 1.004E-07 | 458.79492 | 87.904022 |
| SINPLGGFVHYGE | 22 | P39023 | Large ribosomal subunit protein uL3 OS= | RPL3 | 46.109 | 3 | 50.056194 | yes | 1.2E-07 | 813.07886 | 69.682587 |
| TVFAEHISDECKR | 13 | P39023 | Large ribosomal subunit protein uL3 OS= | RPL3 | 46.109 | 2 | 50.056194 | yes | 4.786E-07 | 796.38031 | 21.701012 |
| LQDLYSIVR | 9 | P39060 | Collagen alpha-1(XVIII) chain OS=Hom | COL18A1 | 178.188 | 2 | 28.688858 | yes | 2.689E-06 | 553.81384 | 66.626709 |
| TADDPSLSLIK | 11 | P39656 | Dolichyl-diphosphooligosaccharide--prot | DDOST | 50.801 | 2 | 52.082024 | yes | 9.722E-08 | 580.3139 | 242.94974 |
| TLVLLDNLNVR | 11 | P39656 | Dolichyl-diphosphooligosaccharide--prot | DDOST | 50.801 | 2 | 52.082024 | yes | 2.919E-07 | 635.37988 | 236.47998 |
| LPDVYGVFQFK | 11 | P39656 | Dolichyl-diphosphooligosaccharide--prot | DDOST | 50.801 | 2 | 52.082024 | yes | 1.922E-07 | 656.85046 | 156.28313 |
| NTLLIAGLQAR | 11 | P39656 | Dolichyl-diphosphooligosaccharide--prot | DDOST | 50.801 | 2 | 52.082024 | yes | 1.058E-07 | 585.3537 | 155.56259 |
| ELGSECGIEFDEEK | 14 | P39656 | Dolichyl-diphosphooligosaccharide--prot | DDOST | 50.801 | 2 | 52.082024 | yes | 9.722E-08 | 821.35107 | 95.67305 |
| ETHSLFFR | 8 | P39656 | Dolichyl-diphosphooligosaccharide--prot | DDOST | 50.801 | 2 | 52.082024 | yes | 1.938E-07 | 518.76416 | 95.034584 |
| SSLNPILFR | 9 | P39656 | Dolichyl-diphosphooligosaccharide--prot | DDOST | 50.801 | 2 | 52.082024 | yes | 8.174E-07 | 523.80328 | 50.276794 |
| YSQTGNYELAVAI | 15 | P39656 | Dolichyl-diphosphooligosaccharide--prot | DDOST | 50.801 | 2 | 52.082024 | yes | 2.411E-07 | 836.42047 | 30.896631 |
| IKDLSTIEPLK | 11 | P39687 | Acidic leucine-rich nuclear phosphoprote | ANP32A | 28.586 | 2 | 38.797176 | no | 4.98E-07 | 628.87665 | 20.973162 |

| | | | | | | | | | | |
|---|---|---|---|---|---|---|---|---|---|---|
| WSEPNEEELIK | 11 | P39748 | Flap endonuclease 1 OS=Homo sapiens C | FEN1 | 42.593 | 2 | 48.870552 | yes | 1.092E-07 | 687.33282 | 91.778282 |
| VLGLSQGSVSDMI | 15 | P39880 | Homeobox protein cut-like 1 OS=Homo s | CUX1 | 164.188 | 2 | 54.43856 | no | 9.722E-08 | 774.90613 | 139.9082 |
| VQSLQTALEK | 10 | P39880 | Homeobox protein cut-like 1 OS=Homo s | CUX1 | 164.188 | 2 | 54.43856 | no | 1.464E-07 | 558.81659 | 138.11717 |
| MQLWLNDPNNVE | 13 | P39880 | Homeobox protein cut-like 1 OS=Homo s | CUX1 | 164.188 | 2 | 54.43856 | yes | 9.722E-08 | 800.89307 | 67.838577 |
| SQLQGPSSSEYWK | 13 | P39880 | Homeobox protein cut-like 1 OS=Homo s | CUX1 | 164.188 | 2 | 54.43856 | yes | 9.722E-08 | 748.85443 | 62.111553 |
| ELDATATVLANR | 12 | P39880 | Homeobox protein cut-like 1 OS=Homo s | CUX1 | 164.188 | 2 | 54.43856 | no | 2.681E-07 | 637.34094 | 61.239079 |
| EIAQLVEDVQR | 11 | P39880 | Homeobox protein cut-like 1 OS=Homo s | CUX1 | 164.188 | 2 | 54.43856 | no | 5.368E-07 | 650.34882 | 61.045708 |
| EAEAAFLNVYK | 11 | P39880 | Homeobox protein cut-like 1 OS=Homo s | CUX1 | 164.188 | 2 | 54.43856 | no | 3.976E-07 | 627.8219 | 59.041412 |
| TSTVINWFHNYR | 12 | P39880 | Homeobox protein cut-like 1 OS=Homo s | CUX1 | 164.188 | 2 | 54.43856 | yes | 1.374E-07 | 769.38098 | 54.726593 |
| PLEVLLLEK | 9 | P39880 | Homeobox protein cut-like 1 OS=Homo s | CUX1 | 164.188 | 2 | 54.43856 | no | 8.754E-07 | 527.33136 | 51.717804 |
| TIEDLATQLNLK | 12 | P39880 | Homeobox protein cut-like 1 OS=Homo s | CUX1 | 164.188 | 2 | 54.43856 | yes | 3.468E-07 | 679.87994 | 43.066189 |
| QFLSDEQNILALR | 13 | P39880 | Homeobox protein cut-like 1 OS=Homo s | CUX1 | 164.188 | 2 | 54.43856 | yes | 2.559E-07 | 773.91486 | 41.466133 |
| SMEFAPSEGAGTQ | 17 | P39880 | Homeobox protein cut-like 1 OS=Homo s | CUX1 | 164.188 | 2 | 54.43856 | no | 2.775E-06 | 848.87781 | 40.626053 |
| LQNDFAEK | 8 | P39880 | Homeobox protein cut-like 1 OS=Homo s | CUX1 | 164.188 | 2 | 54.43856 | no | 6.697E-07 | 482.74036 | 38.624165 |
| LIEQYELR | 8 | P40222 | Alpha-taxilin OS=Homo sapiens OX=96( | TXLNA | 61.892 | 2 | 54.579647 | yes | 1.142E-07 | 532.29279 | 446.98929 |
| YAELLEEHR | 9 | P40222 | Alpha-taxilin OS=Homo sapiens OX=96( | TXLNA | 61.892 | 2 | 54.579647 | yes | 1.83E-07 | 580.29077 | 437.07715 |
| FEEFQNTLSK | 10 | P40222 | Alpha-taxilin OS=Homo sapiens OX=96( | TXLNA | 61.892 | 2 | 54.579647 | yes | 2.487E-07 | 621.80371 | 405.15588 |
| LQQAQEMLK | 9 | P40222 | Alpha-taxilin OS=Homo sapiens OX=96( | TXLNA | 61.892 | 2 | 54.579647 | yes | 1.28E-07 | 544.79205 | 369.91827 |
| QENMELAER | 9 | P40222 | Alpha-taxilin OS=Homo sapiens OX=96( | TXLNA | 61.892 | 2 | 54.579647 | yes | 1.371E-07 | 560.25861 | 240.91203 |
| SKLESLCR | 8 | P40222 | Alpha-taxilin OS=Homo sapiens OX=96( | TXLNA | 61.892 | 2 | 54.579647 | no | 1.874E-07 | 496.76331 | 234.14275 |
| DKELEGLQVK | 10 | P40222 | Alpha-taxilin OS=Homo sapiens OX=96( | TXLNA | 61.892 | 2 | 54.579647 | yes | 9.722E-08 | 579.8219 | 206.58426 |
| QQLALYTEK | 9 | P40222 | Alpha-taxilin OS=Homo sapiens OX=96( | TXLNA | 61.892 | 2 | 54.579647 | yes | 1.734E-07 | 547.29803 | 108.12196 |
| KLIEQYELR | 9 | P40222 | Alpha-taxilin OS=Homo sapiens OX=96( | TXLNA | 61.892 | 2 | 54.579647 | yes | 9.484E-07 | 596.34021 | 78.609566 |
| ALLEMAEEK | 9 | P40222 | Alpha-taxilin OS=Homo sapiens OX=96( | TXLNA | 61.892 | 2 | 54.579647 | yes | 2.311E-07 | 517.26532 | 65.772736 |
| QSQLVQEK | 8 | P40222 | Alpha-taxilin OS=Homo sapiens OX=96( | TXLNA | 61.892 | 2 | 54.579647 | yes | 1.24E-06 | 480.26144 | 47.44157 |
| EVTSHFQVTLNDI( | 23 | P40222 | Alpha-taxilin OS=Homo sapiens OX=96( | TXLNA | 61.892 | 4 | 54.579647 | yes | 5.576E-07 | 699.83972 | 5.2513518 |
| TEVNSGFFYK | 10 | P40227 | T-complex protein 1 subunit zeta OS=Ho | CCT6A | 58.025 | 2 | 48.097248 | no | 2.667E-07 | 596.28766 | 183.37509 |
| ETLIDVAR | 8 | P40227 | T-complex protein 1 subunit zeta OS=Ho | CCT6A | 58.025 | 2 | 48.097248 | yes | 2.687E-07 | 458.75854 | 138.42786 |
| ALQFLEEVK | 9 | P40227 | T-complex protein 1 subunit zeta OS=Ho | CCT6A | 58.025 | 2 | 48.097248 | yes | 2.927E-06 | 538.80298 | 113.60281 |
| AQAALAVNISAAR | 13 | P40227 | T-complex protein 1 subunit zeta OS=Ho | CCT6A | 58.025 | 2 | 48.097248 | yes | 2.136E-07 | 628.3595 | 105.381 |
| QADLYISEGLHPR | 13 | P40227 | T-complex protein 1 subunit zeta OS=Ho | CCT6A | 58.025 | 2 | 48.097248 | no | 1.2E-07 | 749.88605 | 72.843491 |
| GFVVINQK | 8 | P40227 | T-complex protein 1 subunit zeta OS=Ho | CCT6A | 58.025 | 2 | 48.097248 | no | 2.652E-07 | 452.76617 | 65.095909 |
| HTLTQIK | 7 | P40227 | T-complex protein 1 subunit zeta OS=Ho | CCT6A | 58.025 | 2 | 48.097248 | yes | 1.885E-07 | 420.75052 | 40.646488 |
| MVSIIHR | 7 | P40424 | Pre-B-cell leukemia transcription factor 1 | PBX1 | 46.626 | 2 | 49.35886 | no | 1.041E-07 | 428.2471 | 64.738411 |
| GAQEEEPTDPQLM | 14 | P40424 | Pre-B-cell leukemia transcription factor 1 | PBX1 | 46.626 | 2 | 49.35886 | yes | 1.564E-07 | 800.86725 | 57.592976 |
| LDNMLLAEGVAGI | 15 | P40424 | Pre-B-cell leukemia transcription factor 1 | PBX1 | 46.626 | 2 | 49.35886 | no | 6.932E-07 | 778.90308 | 42.325661 |
| YQAVTATLEEK | 11 | P40429 | Large ribosomal subunit protein uL13 OS | RPL13A | 23.578 | 2 | 52.00163 | no | 3.045E-07 | 626.82465 | 35558.176 |
| VFDGIPPPYDK | 11 | P40429 | Large ribosomal subunit protein uL13 OS | RPL13A | 23.578 | 2 | 52.00163 | no | 3.856E-07 | 624.31897 | 13223.331 |
| CEGINISGNFYR | 12 | P40429 | Large ribosomal subunit protein uL13 OS | RPL13A | 23.578 | 2 | 52.00163 | yes | 9.722E-08 | 715.33008 | 12381.938 |

| | | | | | | | | | | | |
|---|---|---|---|---|---|---|---|---|---|---|---|
| LAHEVGWK | 8 | P40429 | Large ribosomal subunit protein uL13 OS | RPL13A | 23.578 | 2 | 52.00163 | yes | 1.248E-07 | 470.25598 | 7331.7461 |
| KFAYLGR | 7 | P40429 | Large ribosomal subunit protein uL13 OS | RPL13A | 23.578 | 2 | 52.00163 | yes | 2.068E-07 | 427.7478 | 6270.7915 |
| AEVQVLVLDGR | 11 | P40429 | Large ribosomal subunit protein uL13 OS | RPL13A | 23.578 | 2 | 52.00163 | yes | 9.722E-08 | 599.84314 | 1559.0936 |
| AEVQVLVLDGR | 11 | P40429 | Large ribosomal subunit protein uL13 OS | RPL13A | 23.578 | 2 | 52.00163 | yes | 1.156E-07 | 620.84845 | 1559.0936 |
| MVVPAALK | 8 | P40429 | Large ribosomal subunit protein uL13 OS | RPL13A | 23.578 | 2 | 52.00163 | no | 2.471E-07 | 414.75421 | 973.45819 |
| IDKYTEVLK | 9 | P40429 | Large ribosomal subunit protein uL13 OS | RPL13A | 23.578 | 2 | 52.00163 | yes | 1.043E-07 | 554.81604 | 842.30756 |
| ILILGLDGAGK | 11 | P40616 | ADP-ribosylation factor-like protein 1 OS | ARL1 | 20.418 | 2 | 51.571793 | yes | 1.956E-07 | 535.33441 | 239.22745 |
| FQVWDLGGQTSIR | 13 | P40616 | ADP-ribosylation factor-like protein 1 OS | ARL1 | 20.418 | 2 | 51.571793 | yes | 9.722E-08 | 753.88861 | 147.08191 |
| AILVVFANK | 9 | P40616 | ADP-ribosylation factor-like protein 1 OS | ARL1 | 20.418 | 2 | 51.571793 | yes | 1.001E-07 | 487.8053 | 85.056862 |
| SELVAMLEEEELR | 14 | P40616 | ADP-ribosylation factor-like protein 1 OS | ARL1 | 20.418 | 2 | 51.571793 | yes | 4.41E-07 | 838.43219 | 37.648155 |
| TLPNASTVDNIR | 12 | P40692 | DNA mismatch repair protein Mlh1 OS=I | MLH1 | 84.601 | 2 | 53.74515 | yes | 1.61E-07 | 650.84644 | 353.48608 |
| IAAGEVIQR | 9 | P40692 | DNA mismatch repair protein Mlh1 OS=I | MLH1 | 84.601 | 2 | 53.74515 | yes | 9.968E-08 | 478.77982 | 217.25244 |
| ELIEIGCEDK | 10 | P40692 | DNA mismatch repair protein Mlh1 OS=I | MLH1 | 84.601 | 2 | 53.74515 | yes | 9.722E-08 | 603.28955 | 161.19357 |
| EMIENCLDAK | 10 | P40692 | DNA mismatch repair protein Mlh1 OS=I | MLH1 | 84.601 | 2 | 53.74515 | yes | 9.554E-07 | 611.77576 | 127.56591 |
| SIFGNAVSR | 9 | P40692 | DNA mismatch repair protein Mlh1 OS=I | MLH1 | 84.601 | 2 | 53.74515 | yes | 7.415E-06 | 475.75635 | 123.38016 |
| LDAFLQPLSK | 10 | P40692 | DNA mismatch repair protein Mlh1 OS=I | MLH1 | 84.601 | 2 | 53.74515 | yes | 9.722E-08 | 566.32404 | 116.64871 |
| EMTAACTPR | 9 | P40692 | DNA mismatch repair protein Mlh1 OS=I | MLH1 | 84.601 | 2 | 53.74515 | yes | 1.047E-07 | 518.73114 | 116.13601 |
| NQSLEGDTTK | 10 | P40692 | DNA mismatch repair protein Mlh1 OS=I | MLH1 | 84.601 | 2 | 53.74515 | yes | 2.01E-07 | 546.76202 | 113.52128 |
| LYLLNTTK | 8 | P40692 | DNA mismatch repair protein Mlh1 OS=I | MLH1 | 84.601 | 2 | 53.74515 | yes | 1.697E-07 | 483.28696 | 78.719658 |
| PLSSQPQAIVTEDK | 14 | P40692 | DNA mismatch repair protein Mlh1 OS=I | MLH1 | 84.601 | 2 | 53.74515 | yes | 1.717E-07 | 756.89886 | 70.131279 |
| STSIQVIVK | 9 | P40692 | DNA mismatch repair protein Mlh1 OS=I | MLH1 | 84.601 | 2 | 53.74515 | yes | 3.115E-07 | 487.79767 | 61.725708 |
| VYAHQMVR | 8 | P40692 | DNA mismatch repair protein Mlh1 OS=I | MLH1 | 84.601 | 2 | 53.74515 | yes | 2.401E-07 | 502.26074 | 61.072666 |
| WTVEHIVYK | 9 | P40692 | DNA mismatch repair protein Mlh1 OS=I | MLH1 | 84.601 | 2 | 53.74515 | yes | 1.823E-07 | 587.81641 | 44.308334 |
| HFTEDGNILQLAN | 19 | P40692 | DNA mismatch repair protein Mlh1 OS=I | MLH1 | 84.601 | 3 | 53.74515 | yes | 2.985E-07 | 734.37939 | 35.994881 |
| KEDLDIVCER | 10 | P40692 | DNA mismatch repair protein Mlh1 OS=I | MLH1 | 84.601 | 2 | 53.74515 | yes | 9.722E-08 | 638.81372 | 34.709377 |
| STTSLTSSSTSGSSI | 16 | P40692 | DNA mismatch repair protein Mlh1 OS=I | MLH1 | 84.601 | 2 | 53.74515 | yes | 3.762E-07 | 766.84973 | 21.009329 |
| KFNILGTNTK | 10 | P40763 | Signal transducer and activator of transcri | STAT3 | 88.068 | 2 | 45.698647 | yes | 1.769E-07 | 568.32715 | 112.88949 |
| TQIQSVEPYTK | 11 | P40763 | Signal transducer and activator of transcri | STAT3 | 88.068 | 2 | 45.698647 | yes | 3.234E-07 | 647.33789 | 53.620087 |
| TLTDEELADWK | 11 | P40763 | Signal transducer and activator of transcri | STAT3 | 88.068 | 2 | 45.698647 | yes | 3.519E-06 | 660.81952 | 50.550602 |
| GLSIEQLTTLAEK | 13 | P40763 | Signal transducer and activator of transcri | STAT3 | 88.068 | 2 | 45.698647 | yes | 1.854E-07 | 701.89301 | 49.524986 |
| FPELNYQLK | 9 | P40763 | Signal transducer and activator of transcri | STAT3 | 88.068 | 2 | 45.698647 | yes | 4.059E-06 | 576.30841 | 25.715141 |
| NLNPVFGGSGPAL | 17 | P40818 | Ubiquitin carboxyl-terminal hydrolase 8 ( | USP8 | 127.524 | 2 | 41.216232 | yes | 3.619E-07 | 835.45465 | 47.979736 |
| PQIPAER | 7 | P40818 | Ubiquitin carboxyl-terminal hydrolase 8 ( | USP8 | 127.524 | 2 | 41.216232 | yes | 7.444E-07 | 405.72705 | 32.16045 |
| GCDVVVIPAGVPR | 13 | P40926 | Malate dehydrogenase, mitochondrial OS | MDH2 | 35.504 | 2 | 50.39586 | yes | 9.81E-08 | 669.86353 | 66.269539 |
| IQEAGTEVVK | 10 | P40926 | Malate dehydrogenase, mitochondrial OS | MDH2 | 35.504 | 2 | 50.39586 | yes | 3.886E-07 | 537.29547 | 54.259556 |
| FGPLTPELMVPR | 12 | P40937 | Replication factor C subunit 5 OS=Homo | RFC5 | 38.497 | 2 | 52.816498 | yes | 1.054E-07 | 678.87085 | 1153.2444 |
| GPILSFASTR | 10 | P40937 | Replication factor C subunit 5 OS=Homo | RFC5 | 38.497 | 2 | 52.816498 | yes | 2.647E-07 | 524.79291 | 1048.0183 |
| ALVTLSSGDMR | 11 | P40937 | Replication factor C subunit 5 OS=Homo | RFC5 | 38.497 | 2 | 52.816498 | yes | 1.249E-07 | 575.30023 | 1026.4542 |
| IIPALQSR | 8 | P40937 | Replication factor C subunit 5 OS=Homo | RFC5 | 38.497 | 2 | 52.816498 | yes | 1.612E-07 | 449.27945 | 811.97925 |

| | | | | | | | | | | | |
|---|---|---|---|---|---|---|---|---|---|---|---|
| LPHLLLYGPPGTGI | 14 | P40937 | Replication factor C subunit 5 OS=Homo | RFC5 | 38.497 | 2 | 52.816498 | yes | 9.722E-08 | 731.9245 | 778.62115 |
| VDFPSSVR | 8 | P40937 | Replication factor C subunit 5 OS=Homo | RFC5 | 38.497 | 2 | 52.816498 | yes | 2.286E-07 | 453.73761 | 434.39725 |
| MADIEYR | 7 | P40937 | Replication factor C subunit 5 OS=Homo | RFC5 | 38.497 | 2 | 52.816498 | yes | 9.901E-08 | 449.21039 | 397.58878 |
| ALNILQSTNMAFG | 14 | P40937 | Replication factor C subunit 5 OS=Homo | RFC5 | 38.497 | 2 | 52.816498 | yes | 2.498E-07 | 754.40051 | 287.1593 |
| TSTILACAK | 9 | P40937 | Replication factor C subunit 5 OS=Homo | RFC5 | 38.497 | 2 | 52.816498 | yes | 4.135E-07 | 482.76022 | 165.96246 |
| LEHVVEEEK | 9 | P40937 | Replication factor C subunit 5 OS=Homo | RFC5 | 38.497 | 2 | 52.816498 | yes | 2.745E-07 | 556.28516 | 120.4845 |
| LEHVVEEEKVDISI | 18 | P40937 | Replication factor C subunit 5 OS=Homo | RFC5 | 38.497 | 3 | 52.816498 | yes | 9.722E-08 | 696.00513 | 73.361488 |
| FCLICNYLSK | 10 | P40937 | Replication factor C subunit 5 OS=Homo | RFC5 | 38.497 | 2 | 52.816498 | yes | 4.272E-07 | 659.32013 | 64.580101 |
| EFGSMVLELNASD | 15 | P40937 | Replication factor C subunit 5 OS=Homo | RFC5 | 38.497 | 2 | 52.816498 | yes | 1.886E-06 | 841.88812 | 15.655544 |
| EGLNLPSQLAHR | 12 | P40938 | Replication factor C subunit 3 OS=Homo | RFC3 | 40.557 | 2 | 51.723465 | yes | 1.24E-07 | 667.86243 | 1468.7854 |
| ALLMCEACR | 9 | P40938 | Replication factor C subunit 3 OS=Homo | RFC3 | 40.557 | 2 | 51.723465 | yes | 1.02E-07 | 562.25647 | 918.3595 |
| ETANAIVSQQTPQI | 14 | P40938 | Replication factor C subunit 3 OS=Homo | RFC3 | 40.557 | 2 | 51.723465 | yes | 9.819E-08 | 771.89716 | 861.29468 |
| GEVAQMAAYYEH | 13 | P40938 | Replication factor C subunit 3 OS=Homo | RFC3 | 40.557 | 2 | 51.723465 | yes | 1E-07 | 762.84863 | 665.92523 |
| VVLLTEVDK | 9 | P40938 | Replication factor C subunit 3 OS=Homo | RFC3 | 40.557 | 2 | 51.723465 | yes | 4.628E-07 | 508.30533 | 610.91071 |
| LILCCNSTSK | 10 | P40938 | Replication factor C subunit 3 OS=Homo | RFC3 | 40.557 | 2 | 51.723465 | yes | 1.17E-07 | 598.29413 | 539.33093 |
| IEHQTITTPSK | 11 | P40938 | Replication factor C subunit 3 OS=Homo | RFC3 | 40.557 | 2 | 51.723465 | yes | 1.405E-07 | 627.83807 | 416.1145 |
| VVIQEMLK | 8 | P40938 | Replication factor C subunit 3 OS=Homo | RFC3 | 40.557 | 2 | 51.723465 | yes | 2.207E-07 | 480.28336 | 356.91748 |
| ELYGVGVEK | 9 | P40938 | Replication factor C subunit 3 OS=Homo | RFC3 | 40.557 | 2 | 51.723465 | yes | 1.122E-07 | 497.2662 | 241.00159 |
| AIYHLEAFVAK | 11 | P40938 | Replication factor C subunit 3 OS=Homo | RFC3 | 40.557 | 2 | 51.723465 | yes | 1.15E-07 | 631.35059 | 229.17648 |
| TVAQSQQLETNSQ | 14 | P40938 | Replication factor C subunit 3 OS=Homo | RFC3 | 40.557 | 2 | 51.723465 | yes | 9.722E-08 | 795.39752 | 156.33017 |
| KFMEDGLEGMMF | 12 | P40938 | Replication factor C subunit 3 OS=Homo | RFC3 | 40.557 | 2 | 51.723465 | yes | 5.293E-07 | 717.80872 | 87.644043 |
| YRPCSLGR | 8 | P40938 | Replication factor C subunit 3 OS=Homo | RFC3 | 40.557 | 2 | 51.723465 | yes | 4.849E-07 | 504.75583 | 51.389225 |
| DAQHALR | 7 | P40938 | Replication factor C subunit 3 OS=Homo | RFC3 | 40.557 | 2 | 51.723465 | yes | 3.716E-07 | 405.71448 | 49.631248 |
| FGGGNPELLTQMV | 15 | P40939 | Trifunctional enzyme subunit alpha, mito | HADHA | 83 | 2 | 54.732292 | yes | 2.199E-07 | 789.40326 | 567.90436 |
| GFYIYQEGVK | 10 | P40939 | Trifunctional enzyme subunit alpha, mito | HADHA | 83 | 2 | 54.732292 | yes | 9.722E-08 | 602.30585 | 520.17188 |
| FVDLYGAQK | 9 | P40939 | Trifunctional enzyme subunit alpha, mito | HADHA | 83 | 2 | 54.732292 | yes | 2.386E-07 | 520.77417 | 475.16385 |
| TLQEVTQLSQEAQ | 14 | P40939 | Trifunctional enzyme subunit alpha, mito | HADHA | 83 | 2 | 54.732292 | yes | 9.722E-08 | 815.9234 | 362.41864 |
| DATLTALDR | 9 | P40939 | Trifunctional enzyme subunit alpha, mito | HADHA | 83 | 2 | 54.732292 | yes | 2.102E-07 | 488.25891 | 287.93076 |
| DTSASAVAVGLK | 12 | P40939 | Trifunctional enzyme subunit alpha, mito | HADHA | 83 | 2 | 54.732292 | yes | 1.888E-07 | 559.80621 | 285.16458 |
| VIGMHYFSPVDK | 12 | P40939 | Trifunctional enzyme subunit alpha, mito | HADHA | 83 | 2 | 54.732292 | yes | 9.722E-08 | 696.85266 | 247.81813 |
| ILQEGVDPK | 9 | P40939 | Trifunctional enzyme subunit alpha, mito | HADHA | 83 | 2 | 54.732292 | yes | 2.267E-07 | 499.77948 | 224.14044 |
| MQLLEIITTEK | 11 | P40939 | Trifunctional enzyme subunit alpha, mito | HADHA | 83 | 2 | 54.732292 | yes | 9.722E-08 | 667.86542 | 92.538536 |
| MQLLEIITTEK | 11 | P40939 | Trifunctional enzyme subunit alpha, mito | HADHA | 83 | 2 | 54.732292 | yes | 9.722E-08 | 659.86798 | 92.538536 |
| THINYGVK | 8 | P40939 | Trifunctional enzyme subunit alpha, mito | HADHA | 83 | 2 | 54.732292 | yes | 9.987E-08 | 466.25345 | 182.28659 |
| SEVSSDEDIQFR | 12 | P40939 | Trifunctional enzyme subunit alpha, mito | HADHA | 83 | 2 | 54.732292 | yes | 9.722E-08 | 706.32043 | 165.62318 |
| HVAEDLGK | 8 | P40939 | Trifunctional enzyme subunit alpha, mito | HADHA | 83 | 2 | 54.732292 | yes | 4.045E-07 | 434.7298 | 150.70273 |
| DGPGFYTTR | 9 | P40939 | Trifunctional enzyme subunit alpha, mito | HADHA | 83 | 2 | 54.732292 | yes | 1.971E-06 | 507.23798 | 116.01173 |
| GLYPAPLK | 8 | P40939 | Trifunctional enzyme subunit alpha, mito | HADHA | 83 | 2 | 54.732292 | yes | 9.722E-08 | 429.75781 | 57.757591 |
| ALMGLYHGQVLC | 13 | P40939 | Trifunctional enzyme subunit alpha, mito | HADHA | 83 | 2 | 54.732292 | yes | 1.318E-07 | 745.38635 | 54.066418 |

| | | | | | | | | | | | |
|---|---|---|---|---|---|---|---|---|---|---|---|
| CLAPMMSEVIR | 11 | P40939 | Trifunctional enzyme subunit alpha, mito | HADHA | 83 | 2 | 54.732292 | yes | 1.071E-06 | 653.81946 | 47.475121 |
| IVLTNPVCTEVGEI | 14 | P41091 | Eukaryotic translation initiation factor 2 s | EIF2S3 | 51.11 | 2 | 54.354008 | no | 1.4E-07 | 779.91089 | 6245.0044 |
| NEVLMVNIGSLST | 16 | P41091 | Eukaryotic translation initiation factor 2 s | EIF2S3 | 51.11 | 2 | 54.354008 | no | 3.484E-07 | 823.93018 | 2325.332 |
| NEVLMVNIGSLST | 16 | P41091 | Eukaryotic translation initiation factor 2 s | EIF2S3 | 51.11 | 2 | 54.354008 | no | 9.722E-08 | 831.92761 | 2325.332 |
| HILILQNK | 8 | P41091 | Eukaryotic translation initiation factor 2 s | EIF2S3 | 51.11 | 2 | 54.354008 | no | 4.782E-07 | 489.80838 | 3801.6504 |
| LIGWGQIR | 8 | P41091 | Eukaryotic translation initiation factor 2 s | EIF2S3 | 51.11 | 2 | 54.354008 | no | 4.162E-07 | 471.77963 | 3760.5552 |
| AGGEAGVTLGQPI | 16 | P41091 | Eukaryotic translation initiation factor 2 s | EIF2S3 | 51.11 | 2 | 54.354008 | no | 1.204E-07 | 796.41296 | 1214.686 |
| AGGEAGVTLGQPI | 16 | P41091 | Eukaryotic translation initiation factor 2 s | EIF2S3 | 51.11 | 2 | 54.354008 | no | 9.818E-08 | 775.40771 | 1214.686 |
| AGGEAGVTLGQPI | 16 | P41091 | Eukaryotic translation initiation factor 2 s | EIF2S3 | 51.11 | 2 | 54.354008 | no | 5.497E-07 | 531.27777 | 1214.686 |
| AGGEAGVTLGQPI | 16 | P41091 | Eukaryotic translation initiation factor 2 s | EIF2S3 | 51.11 | 2 | 54.354008 | no | 6.761E-07 | 517.27423 | 1214.686 |
| AGGEAGVTLGQPI | 16 | P41091 | Eukaryotic translation initiation factor 2 s | EIF2S3 | 51.11 | 3 | 54.354008 | no | 1.204E-07 | 796.41296 | 1214.686 |
| AGGEAGVTLGQPI | 16 | P41091 | Eukaryotic translation initiation factor 2 s | EIF2S3 | 51.11 | 3 | 54.354008 | no | 9.818E-08 | 775.40771 | 1214.686 |
| AGGEAGVTLGQPI | 16 | P41091 | Eukaryotic translation initiation factor 2 s | EIF2S3 | 51.11 | 3 | 54.354008 | no | 5.497E-07 | 531.27777 | 1214.686 |
| AGGEAGVTLGQPI | 16 | P41091 | Eukaryotic translation initiation factor 2 s | EIF2S3 | 51.11 | 3 | 54.354008 | no | 6.761E-07 | 517.27423 | 1214.686 |
| AISGVHTVR | 9 | P41091 | Eukaryotic translation initiation factor 2 s | EIF2S3 | 51.11 | 2 | 54.354008 | no | 1.108E-07 | 470.27216 | 2674.9924 |
| QDLTTLDVTK | 10 | P41091 | Eukaryotic translation initiation factor 2 s | EIF2S3 | 51.11 | 2 | 54.354008 | no | 1.087E-07 | 567.30603 | 2513.3647 |
| FKNELER | 7 | P41091 | Eukaryotic translation initiation factor 2 s | EIF2S3 | 51.11 | 2 | 54.354008 | no | 1.17E-07 | 468.25089 | 2388.2761 |
| VGQEIEVR | 8 | P41091 | Eukaryotic translation initiation factor 2 s | EIF2S3 | 51.11 | 2 | 54.354008 | yes | 1.196E-07 | 465.25616 | 2365.0747 |
| LTPLSHEVISR | 11 | P41091 | Eukaryotic translation initiation factor 2 s | EIF2S3 | 51.11 | 2 | 54.354008 | no | 5.826E-07 | 626.35645 | 1047.074 |
| LTPLSHEVISR | 11 | P41091 | Eukaryotic translation initiation factor 2 s | EIF2S3 | 51.11 | 2 | 54.354008 | no | 1.227E-06 | 417.90671 | 1047.074 |
| LTPLSHEVISR | 11 | P41091 | Eukaryotic translation initiation factor 2 s | EIF2S3 | 51.11 | 3 | 54.354008 | no | 5.826E-07 | 626.35645 | 1047.074 |
| LTPLSHEVISR | 11 | P41091 | Eukaryotic translation initiation factor 2 s | EIF2S3 | 51.11 | 3 | 54.354008 | no | 1.227E-06 | 417.90671 | 1047.074 |
| YNIEVVCEYIVK | 12 | P41091 | Eukaryotic translation initiation factor 2 s | EIF2S3 | 51.11 | 2 | 54.354008 | no | 1.303E-07 | 764.8894 | 1513.1617 |
| QATINIGTIGHVAI | 16 | P41091 | Eukaryotic translation initiation factor 2 s | EIF2S3 | 51.11 | 3 | 54.354008 | no | 1.165E-07 | 539.63373 | 694.80054 |
| QATINIGTIGHVAI | 16 | P41091 | Eukaryotic translation initiation factor 2 s | EIF2S3 | 51.11 | 3 | 54.354008 | no | 9.722E-08 | 808.94702 | 694.80054 |
| QATINIGTIGHVAI | 16 | P41091 | Eukaryotic translation initiation factor 2 s | EIF2S3 | 51.11 | 2 | 54.354008 | no | 1.165E-07 | 539.63373 | 694.80054 |
| QATINIGTIGHVAI | 16 | P41091 | Eukaryotic translation initiation factor 2 s | EIF2S3 | 51.11 | 2 | 54.354008 | no | 9.722E-08 | 808.94702 | 694.80054 |
| LMCKPIFSK | 9 | P41091 | Eukaryotic translation initiation factor 2 s | EIF2S3 | 51.11 | 2 | 54.354008 | yes | 1.073E-07 | 562.30377 | 1073.1537 |
| VGQEIEVRPGIVSK | 14 | P41091 | Eukaryotic translation initiation factor 2 s | EIF2S3 | 51.11 | 3 | 54.354008 | yes | 1.575E-07 | 504.29111 | 448.76111 |
| VGQEIEVRPGIVSK | 14 | P41091 | Eukaryotic translation initiation factor 2 s | EIF2S3 | 51.11 | 3 | 54.354008 | yes | 9.722E-08 | 755.93304 | 448.76111 |
| VGQEIEVRPGIVSK | 14 | P41091 | Eukaryotic translation initiation factor 2 s | EIF2S3 | 51.11 | 2 | 54.354008 | yes | 1.575E-07 | 504.29111 | 448.76111 |
| VGQEIEVRPGIVSK | 14 | P41091 | Eukaryotic translation initiation factor 2 s | EIF2S3 | 51.11 | 2 | 54.354008 | yes | 9.722E-08 | 755.93304 | 448.76111 |
| LDDPSCPRPECYR | 13 | P41091 | Eukaryotic translation initiation factor 2 s | EIF2S3 | 51.11 | 2 | 54.354008 | yes | 1.041E-07 | 832.86139 | 814.51984 |
| RGVTIKPTVDDD | 12 | P41091 | Eukaryotic translation initiation factor 2 s | EIF2S3 | 51.11 | 2 | 54.354008 | no | 5.34E-07 | 658.34625 | 601.99207 |
| GGVAGGSILK | 10 | P41091 | Eukaryotic translation initiation factor 2 s | EIF2S3 | 51.11 | 2 | 54.354008 | no | 5.427E-07 | 429.7558 | 321.96335 |
| VASGSTATAAAGP | 18 | P41134 | DNA-binding protein inhibitor ID-1 OS= | ID1 | 16.133 | 2 | 41.112316 | yes | 3.67E-07 | 810.40649 | 35.396069 |
| LIEPEEVAR | 9 | P41161 | ETS translocation variant 5 OS=Homo sa | ETV5 | 57.838 | 2 | 48.356422 | no | 1.74E-07 | 528.29022 | 506.4545 |
| LYFDDTCVVPER | 12 | P41161 | ETS translocation variant 5 OS=Homo sa | ETV5 | 57.838 | 2 | 48.356422 | yes | 1.16E-07 | 757.35321 | 103.22366 |
| GRPVIDR | 7 | P41161 | ETS translocation variant 5 OS=Homo sa | ETV5 | 57.838 | 2 | 48.356422 | yes | 3.681E-07 | 406.74051 | 58.220798 |

| | | | | | | | | | | | |
|---|---|---|---|---|---|---|---|---|---|---|---|
| MLMPQDIMAYR | 11 | P41182 | B-cell lymphoma 6 protein OS=Homo sap | BCL6 | 78.847 | 2 | 46.799423 | yes | 6.266E-07 | 684.82727 | 191.16925 |
| FVQVAHLR | 8 | P41182 | B-cell lymphoma 6 protein OS=Homo sap | BCL6 | 78.847 | 2 | 46.799423 | no | 1.467E-07 | 485.28506 | 128.45406 |
| LSASGEDSTIPQAS | 15 | P41182 | B-cell lymphoma 6 protein OS=Homo sap | BCL6 | 78.847 | 2 | 46.799423 | yes | 3.332E-07 | 759.87335 | 123.93774 |
| MPVANPFPK | 9 | P41182 | B-cell lymphoma 6 protein OS=Homo sap | BCL6 | 78.847 | 2 | 46.799423 | yes | 5.534E-07 | 500.76785 | 113.40897 |
| SDMHYSVAEGLKI | 19 | P41182 | B-cell lymphoma 6 protein OS=Homo sap | BCL6 | 78.847 | 3 | 46.799423 | yes | 4.806E-07 | 662.99493 | 101.97964 |
| HASDVLLNLNR | 11 | P41182 | B-cell lymphoma 6 protein OS=Homo sap | BCL6 | 78.847 | 2 | 46.799423 | yes | 2.003E-07 | 626.34387 | 98.79026 |
| EVVENNLPLR | 10 | P41182 | B-cell lymphoma 6 protein OS=Homo sap | BCL6 | 78.847 | 2 | 46.799423 | yes | 3.225E-07 | 591.82751 | 23.638145 |
| AHVLIHTGEKPYP | 20 | P41182 | B-cell lymphoma 6 protein OS=Homo sap | BCL6 | 78.847 | 4 | 46.799423 | yes | 4.79E-07 | 585.2912 | 15.398672 |
| SDCQPNSPTESCSS | 15 | P41182 | B-cell lymphoma 6 protein OS=Homo sap | BCL6 | 78.847 | 2 | 46.799423 | yes | 4.951E-05 | 842.33289 | 11.378807 |
| LFDDDETGK | 9 | P41208 | Centrin-2 OS=Homo sapiens OX=9606 G | CETN2 | 19.739 | 2 | 44.264 | yes | 2.259E-07 | 520.23254 | 44.792374 |
| ICPIDITLAQR | 11 | P41214 | Eukaryotic translation initiation factor 2D | EIF2D | 64.707 | 2 | 48.930817 | yes | 1.085E-07 | 650.35809 | 317.65625 |
| LDPILCDCILEK | 12 | P41214 | Eukaryotic translation initiation factor 2D | EIF2D | 64.707 | 2 | 48.930817 | yes | 3.867E-07 | 744.87549 | 217.75073 |
| LPWDSLLTR | 9 | P41214 | Eukaryotic translation initiation factor 2D | EIF2D | 64.707 | 2 | 48.930817 | yes | 1.509E-07 | 550.80859 | 132.19057 |
| GVESIVAVDWK | 11 | P41214 | Eukaryotic translation initiation factor 2D | EIF2D | 64.707 | 2 | 48.930817 | yes | 2.857E-07 | 601.8244 | 72.198067 |
| GDLCAISLVGNR | 12 | P41214 | Eukaryotic translation initiation factor 2D | EIF2D | 64.707 | 2 | 48.930817 | yes | 1.783E-07 | 637.82971 | 70.098763 |
| GSVQMDSTLQGDI | 14 | P41214 | Eukaryotic translation initiation factor 2D | EIF2D | 64.707 | 2 | 48.930817 | yes | 9.338E-07 | 762.84271 | 66.431656 |
| CQASTTVNPAPGA | 14 | P41214 | Eukaryotic translation initiation factor 2D | EIF2D | 64.707 | 2 | 48.930817 | yes | 3.634E-07 | 701.3432 | 45.102234 |
| IIECTHCGCR | 10 | P41223 | Protein BUD31 homolog OS=Homo sapi | BUD31 | 17 | 2 | 50.797626 | yes | 9.722E-08 | 653.2785 | 108.41352 |
| QGYENLCCLR | 10 | P41223 | Protein BUD31 homolog OS=Homo sapi | BUD31 | 17 | 2 | 50.797626 | yes | 1.076E-07 | 656.7923 | 105.54913 |
| KVESLWPIFR | 10 | P41223 | Protein BUD31 homolog OS=Homo sapi | BUD31 | 17 | 2 | 50.797626 | yes | 6.441E-07 | 637.86639 | 35.92794 |
| AMIENFNAK | 9 | P41227 | N-alpha-acetyltransferase 10 OS=Homo s | NAA10 | 26.459 | 2 | 43.905697 | no | 1.449E-06 | 519.25787 | 119.10339 |
| GNSPPSSGEACR | 12 | P41227 | N-alpha-acetyltransferase 10 OS=Homo s | NAA10 | 26.459 | 2 | 43.905697 | yes | 3.94E-07 | 609.76202 | 76.557892 |
| YYADGEDAYAMK | 12 | P41227 | N-alpha-acetyltransferase 10 OS=Homo s | NAA10 | 26.459 | 2 | 43.905697 | no | 3.888E-07 | 698.78992 | 64.554756 |
| DLSEVSETTESTDV | 15 | P41227 | N-alpha-acetyltransferase 10 OS=Homo s | NAA10 | 26.459 | 2 | 43.905697 | yes | 2.398E-07 | 820.38086 | 54.604141 |
| HVVLGAIENK | 10 | P41227 | N-alpha-acetyltransferase 10 OS=Homo s | NAA10 | 26.459 | 2 | 43.905697 | yes | 3.398E-07 | 540.31403 | 51.274811 |
| YPLFEGQETGK | 11 | P41250 | Glycine--tRNA ligase OS=Homo sapiens | GARS1 | 83.166 | 2 | 52.03979 | yes | 3.495E-07 | 634.81152 | 162.15953 |
| TVNVVQFEPSK | 11 | P41250 | Glycine--tRNA ligase OS=Homo sapiens | GARS1 | 83.166 | 2 | 52.03979 | yes | 9.722E-08 | 624.33514 | 102.87278 |
| NNIIQTWR | 8 | P41250 | Glycine--tRNA ligase OS=Homo sapiens | GARS1 | 83.166 | 2 | 52.03979 | yes | 2.487E-06 | 522.7829 | 79.355759 |
| LPFAAAQIGNSFR | 13 | P41250 | Glycine--tRNA ligase OS=Homo sapiens | GARS1 | 83.166 | 2 | 52.03979 | yes | 1.417E-07 | 696.37512 | 67.065689 |
| VPLVAEKPLK | 10 | P41250 | Glycine--tRNA ligase OS=Homo sapiens | GARS1 | 83.166 | 2 | 52.03979 | yes | 5.073E-07 | 547.3526 | 60.212311 |
| SCYDLSCHAR | 10 | P41250 | Glycine--tRNA ligase OS=Homo sapiens | GARS1 | 83.166 | 2 | 52.03979 | yes | 9.722E-08 | 634.76099 | 49.813019 |
| GEFTIETEGK | 10 | P41250 | Glycine--tRNA ligase OS=Homo sapiens | GARS1 | 83.166 | 2 | 52.03979 | yes | 2.19E-07 | 555.76929 | 47.792377 |
| VLIDPVSVQDK | 11 | P41252 | Isoleucine--tRNA ligase, cytoplasmic OS= | IARS1 | 144.499 | 2 | 52.777588 | yes | 1.202E-07 | 606.84534 | 245.96104 |
| LYLINSPVVR | 10 | P41252 | Isoleucine--tRNA ligase, cytoplasmic OS= | IARS1 | 144.499 | 2 | 52.777588 | yes | 1.088E-07 | 587.35315 | 200.94682 |
| AVPSWFVR | 8 | P41252 | Isoleucine--tRNA ligase, cytoplasmic OS= | IARS1 | 144.499 | 2 | 52.777588 | yes | 1.278E-07 | 481.26636 | 119.72021 |
| AEPDHMVLGK | 10 | P41252 | Isoleucine--tRNA ligase, cytoplasmic OS= | IARS1 | 144.499 | 2 | 52.777588 | yes | 1.004E-07 | 548.77643 | 119.28319 |
| MGITEYNNQCR | 11 | P41252 | Isoleucine--tRNA ligase, cytoplasmic OS= | IARS1 | 144.499 | 2 | 52.777588 | yes | 1.26E-07 | 693.30048 | 108.94464 |
| KVTLSTDK | 8 | P41252 | Isoleucine--tRNA ligase, cytoplasmic OS= | IARS1 | 144.499 | 2 | 52.777588 | yes | 1.146E-07 | 446.26093 | 91.137726 |
| QLSSEELEQFQK | 12 | P41252 | Isoleucine--tRNA ligase, cytoplasmic OS= | IARS1 | 144.499 | 2 | 52.777588 | yes | 9.722E-08 | 733.36212 | 83.418785 |

| | | | | | | | | | | |
|---|---|---|---|---|---|---|---|---|---|---|
| VCMDFNIIR | 9 | P41252 | Isoleucine--tRNA ligase, cytoplasmic OS= | IARS1 | 144.499 | 2 | 52.777588 | yes | 9.722E-08 | 584.28607 | 77.478371 |
| LYTVVPR | 7 | P41252 | Isoleucine--tRNA ligase, cytoplasmic OS= | IARS1 | 144.499 | 2 | 52.777588 | yes | 1.355E-07 | 424.25543 | 75.636414 |
| WIDFDNDYK | 9 | P41252 | Isoleucine--tRNA ligase, cytoplasmic OS= | IARS1 | 144.499 | 2 | 52.777588 | yes | 1.121E-07 | 608.26947 | 73.376472 |
| NYPDPVSIIQK | 11 | P41252 | Isoleucine--tRNA ligase, cytoplasmic OS= | IARS1 | 144.499 | 2 | 52.777588 | yes | 3.059E-06 | 637.34296 | 72.946938 |
| VREELIDK | 8 | P41252 | Isoleucine--tRNA ligase, cytoplasmic OS= | IARS1 | 144.499 | 2 | 52.777588 | yes | 3.434E-07 | 501.28494 | 67.03891 |
| LESDYEILER | 10 | P41252 | Isoleucine--tRNA ligase, cytoplasmic OS= | IARS1 | 144.499 | 2 | 52.777588 | yes | 9.722E-08 | 633.81427 | 62.784794 |
| NNDLCYWVPELV | 13 | P41252 | Isoleucine--tRNA ligase, cytoplasmic OS= | IARS1 | 144.499 | 2 | 52.777588 | yes | 2.475E-06 | 839.40631 | 59.523029 |
| ESVDHLTIPSR | 11 | P41252 | Isoleucine--tRNA ligase, cytoplasmic OS= | IARS1 | 144.499 | 2 | 52.777588 | yes | 6.23E-07 | 627.32788 | 58.198536 |
| GSELEITLTR | 10 | P41252 | Isoleucine--tRNA ligase, cytoplasmic OS= | IARS1 | 144.499 | 2 | 52.777588 | yes | 3.262E-07 | 559.80621 | 53.515919 |
| EIVVIHQDPEALK | 13 | P41252 | Isoleucine--tRNA ligase, cytoplasmic OS= | IARS1 | 144.499 | 2 | 52.777588 | yes | 1.466E-07 | 745.91431 | 52.800632 |
| SVVTSIFGVK | 10 | P41252 | Isoleucine--tRNA ligase, cytoplasmic OS= | IARS1 | 144.499 | 2 | 52.777588 | yes | 2.954E-07 | 518.80548 | 43.288254 |
| YAHQSGFHVDR | 11 | P41252 | Isoleucine--tRNA ligase, cytoplasmic OS= | IARS1 | 144.499 | 2 | 52.777588 | yes | 1.959E-06 | 658.81018 | 21.358562 |
| ASSSLGLQDFDLL | 14 | P41743 | Protein kinase C iota type OS=Homo sapi | PRKCI | 68.263 | 2 | 51.503494 | yes | 1.12E-07 | 761.39899 | 673.60779 |
| LVTIECGR | 8 | P41743 | Protein kinase C iota type OS=Homo sapi | PRKCI | 68.263 | 2 | 51.503494 | yes | 1.137E-07 | 474.25256 | 673.2597 |
| LDNVLLDSEGHIK | 13 | P41743 | Protein kinase C iota type OS=Homo sapi | PRKCI | 68.263 | 2 | 51.503494 | yes | 9.722E-08 | 726.88831 | 634.59131 |
| LTDYGMCK | 8 | P41743 | Protein kinase C iota type OS=Homo sapi | PRKCI | 68.263 | 2 | 51.503494 | no | 1.697E-07 | 494.21735 | 325.9834 |
| LGCHPQTGFADIQ | 19 | P41743 | Protein kinase C iota type OS=Homo sapi | PRKCI | 68.263 | 4 | 51.503494 | yes | 6.298E-07 | 547.01538 | 311.3468 |
| AHCAICTDR | 9 | P41743 | Protein kinase C iota type OS=Homo sapi | PRKCI | 68.263 | 2 | 51.503494 | yes | 4.053E-07 | 552.23969 | 282.30603 |
| NVDWDMMEQK | 10 | P41743 | Protein kinase C iota type OS=Homo sapi | PRKCI | 68.263 | 2 | 51.503494 | yes | 9.722E-08 | 648.27338 | 272.43649 |
| HVFEQASNHPFLV | 24 | P41743 | Protein kinase C iota type OS=Homo sapi | PRKCI | 68.263 | 4 | 51.503494 | yes | 2.906E-07 | 707.58966 | 56.028931 |
| SFLNKDPK | 8 | P41743 | Protein kinase C iota type OS=Homo sapi | PRKCI | 68.263 | 2 | 51.503494 | yes | 1.65E-06 | 474.76108 | 52.999817 |
| DSSTMSHTVAGGC | 22 | P41743 | Protein kinase C iota type OS=Homo sapi | PRKCI | 68.263 | 4 | 51.503494 | yes | 1.342E-06 | 553.24646 | 38.218346 |
| VMAGALEGDLFIG | 15 | P42025 | Beta-centractin OS=Homo sapiens OX=9 | ACTR1B | 42.294 | 2 | 42.129272 | yes | 3.179E-07 | 759.40527 | 318.29428 |
| VLVEPDAGAGVA | 15 | P42126 | Enoyl-CoA delta isomerase 1, mitochondr | ECI1 | 32.816 | 2 | 48.89645 | yes | 2.496E-07 | 728.3974 | 285.89581 |
| DADVQNFVSFISK | 13 | P42126 | Enoyl-CoA delta isomerase 1, mitochondr | ECI1 | 32.816 | 2 | 48.89645 | yes | 1.089E-07 | 735.36719 | 31.419207 |
| SSTPLPTISSSAENT | 16 | P42166 | Lamina-associated polypeptide 2, isoform | TMPO | 75.493 | 2 | 51.963207 | no | 4.97E-07 | 824.41284 | 2885.8716 |
| YGVNPGPIVGTTR | 13 | P42166 | Lamina-associated polypeptide 2, isoform | TMPO | 75.493 | 2 | 51.963207 | no | 1.684E-07 | 665.85931 | 2790.3518 |
| GGTLFGGEVCK | 11 | P42166 | Lamina-associated polypeptide 2, isoform | TMPO | 75.493 | 2 | 51.963207 | yes | 9.722E-08 | 562.77386 | 1447.8207 |
| EPLVATNLPGR | 11 | P42166 | Lamina-associated polypeptide 2, isoform | TMPO | 75.493 | 2 | 51.963207 | yes | 1.401E-07 | 583.83002 | 1339.5183 |
| PEFLEDPSVLTK | 12 | P42166 | Lamina-associated polypeptide 2, isoform | TMPO | 75.493 | 2 | 51.963207 | no | 1.423E-07 | 687.86121 | 1092.4905 |
| SGIQPLCPER | 10 | P42166 | Lamina-associated polypeptide 2, isoform | TMPO | 75.493 | 2 | 51.963207 | yes | 3.551E-07 | 578.7926 | 926.65619 |
| FQETEFLSPPR | 11 | P42166 | Lamina-associated polypeptide 2, isoform | TMPO | 75.493 | 2 | 51.963207 | yes | 2.248E-06 | 675.83807 | 580.97949 |
| SHISDQSPLSSK | 12 | P42166 | Lamina-associated polypeptide 2, isoform | TMPO | 75.493 | 2 | 51.963207 | yes | 9.85E-08 | 643.32275 | 574.67609 |
| THQALGILSK | 10 | P42166 | Lamina-associated polypeptide 2, isoform | TMPO | 75.493 | 2 | 51.963207 | yes | 1.104E-07 | 534.31403 | 526.07391 |
| VIEEEWQQVDR | 11 | P42166 | Lamina-associated polypeptide 2, isoform | TMPO | 75.493 | 2 | 51.963207 | yes | 1.135E-07 | 715.84918 | 381.53082 |
| QLPSLACK | 8 | P42166 | Lamina-associated polypeptide 2, isoform | TMPO | 75.493 | 2 | 51.963207 | yes | 1.427E-07 | 458.74966 | 339.20599 |
| NRPPLPAGTNSK | 12 | P42166 | Lamina-associated polypeptide 2, isoform | TMPO | 75.493 | 2 | 51.963207 | no | 1.467E-07 | 626.34387 | 277.43295 |
| AYEAAASALQIAT | 20 | P42166 | Lamina-associated polypeptide 2, isoform | TMPO | 75.493 | 3 | 51.963207 | yes | 1.521E-07 | 678.69336 | 267.04602 |
| EPTPVLGSGAAAA | 15 | P42166 | Lamina-associated polypeptide 2, isoform | TMPO | 75.493 | 2 | 51.963207 | no | 1.073E-07 | 677.35968 | 252.25063 |

| | | | | | | | | | | | |
|---|---|---|---|---|---|---|---|---|---|---|---|
| DIVENICGR | 9 | P42166 | Lamina-associated polypeptide 2, isoform | TMPO | 75.493 | 2 | 51.963207 | yes | 2.763E-07 | 538.26367 | 142.1951 |
| MAAHTMGNATVC | 13 | P42166 | Lamina-associated polypeptide 2, isoform | TMPO | 75.493 | 2 | 51.963207 | yes | 2.776E-07 | 658.81354 | 68.320877 |
| MAAHTMGNATVC | 13 | P42166 | Lamina-associated polypeptide 2, isoform | TMPO | 75.493 | 2 | 51.963207 | yes | 1.273E-07 | 439.5448 | 68.320877 |
| MAAHTMGNATVC | 13 | P42166 | Lamina-associated polypeptide 2, isoform | TMPO | 75.493 | 3 | 51.963207 | yes | 2.776E-07 | 658.81354 | 68.320877 |
| MAAHTMGNATVC | 13 | P42166 | Lamina-associated polypeptide 2, isoform | TMPO | 75.493 | 3 | 51.963207 | yes | 1.273E-07 | 439.5448 | 68.320877 |
| EATQILSVPK | 10 | P42166 | Lamina-associated polypeptide 2, isoform | TMPO | 75.493 | 2 | 51.963207 | yes | 1.133E-07 | 543.31366 | 124.05019 |
| NLFISCK | 7 | P42166 | Lamina-associated polypeptide 2, isoform | TMPO | 75.493 | 2 | 51.963207 | yes | 1.074E-07 | 441.23111 | 100.3283 |
| GPPDFSSDEER | 11 | P42166 | Lamina-associated polypeptide 2, isoform | TMPO | 75.493 | 2 | 51.963207 | no | 6.868E-07 | 618.26239 | 56.236897 |
| PLELSDFR | 8 | P42167 | Lamina-associated polypeptide 2, isoform | TMPO | 50.671 | 2 | 49.296597 | yes | 1.047E-07 | 488.75854 | 1176.9603 |
| VETSEHFR | 8 | P42167 | Lamina-associated polypeptide 2, isoform | TMPO | 50.671 | 2 | 49.296597 | yes | 1.388E-07 | 502.74344 | 739.44415 |
| HASPILPITEFSDIP | 16 | P42167 | Lamina-associated polypeptide 2, isoform | TMPO | 50.671 | 3 | 49.296597 | yes | 6.265E-07 | 598.32458 | 684.06982 |
| YVPLADVK | 8 | P42167 | Lamina-associated polypeptide 2, isoform | TMPO | 50.671 | 2 | 49.296597 | yes | 2.28E-07 | 452.76056 | 540.34265 |
| RVEHNQSYSQAGI | 24 | P42167 | Lamina-associated polypeptide 2, isoform | TMPO | 50.671 | 4 | 49.296597 | yes | 1.439E-07 | 671.31519 | 154.88293 |
| YTYEHDPITK | 10 | P42224 | Signal transducer and activator of transcri | STAT1 | 87.335 | 2 | 34.611507 | yes | 9.743E-07 | 633.80371 | 34.233913 |
| VGEIHLLR | 8 | P42226 | Signal transducer and activator of transcri | STAT6 | 94.136 | 2 | 49.03914 | yes | 1.074E-07 | 468.78491 | 57.343933 |
| TLVTSCFLVEK | 11 | P42226 | Signal transducer and activator of transcri | STAT6 | 94.136 | 2 | 49.03914 | yes | 1.03E-05 | 648.84705 | 52.289955 |
| TVLQMDELK | 9 | P42285 | Exosome RNA helicase MTR4 OS=Homo | MTREX | 117.805 | 2 | 53.995205 | yes | 2.883E-07 | 538.78644 | 841.22607 |
| LTEQLAGPLR | 10 | P42285 | Exosome RNA helicase MTR4 OS=Homo | MTREX | 117.805 | 2 | 53.995205 | yes | 9.81E-08 | 549.31934 | 698.39313 |
| LGFATSSDVIEMK | 13 | P42285 | Exosome RNA helicase MTR4 OS=Homo | MTREX | 117.805 | 2 | 53.995205 | yes | 1.032E-07 | 699.35266 | 519.32111 |
| VEEINPEYMLEK | 12 | P42285 | Exosome RNA helicase MTR4 OS=Homo | MTREX | 117.805 | 2 | 53.995205 | yes | 9.918E-08 | 747.36322 | 493.2673 |
| EDNFNTAMQVLR | 12 | P42285 | Exosome RNA helicase MTR4 OS=Homo | MTREX | 117.805 | 2 | 53.995205 | yes | 1.169E-07 | 719.3432 | 471.95349 |
| DVDFEGTDEPIFGI | 14 | P42285 | Exosome RNA helicase MTR4 OS=Homo | MTREX | 117.805 | 2 | 53.995205 | yes | 9.99E-08 | 784.85938 | 440.17111 |
| IVIPNEESVVIYYK | 14 | P42285 | Exosome RNA helicase MTR4 OS=Homo | MTREX | 117.805 | 2 | 53.995205 | yes | 1.255E-07 | 833.45856 | 406.66653 |
| TVLFTNAR | 8 | P42285 | Exosome RNA helicase MTR4 OS=Homo | MTREX | 117.805 | 2 | 53.995205 | yes | 9.967E-08 | 461.26126 | 365.68396 |
| YCLPFLQPGR | 10 | P42285 | Exosome RNA helicase MTR4 OS=Homo | MTREX | 117.805 | 2 | 53.995205 | yes | 1.172E-07 | 625.82117 | 283.20102 |
| VEAFEHR | 7 | P42285 | Exosome RNA helicase MTR4 OS=Homo | MTREX | 117.805 | 2 | 53.995205 | yes | 1.422E-07 | 444.22214 | 248.17082 |
| DLRPVDNR | 8 | P42285 | Exosome RNA helicase MTR4 OS=Homo | MTREX | 117.805 | 2 | 53.995205 | yes | 5.216E-06 | 492.76471 | 217.26109 |
| RLEELLR | 7 | P42285 | Exosome RNA helicase MTR4 OS=Homo | MTREX | 117.805 | 2 | 53.995205 | yes | 1.084E-07 | 464.78235 | 201.51453 |
| EIEEYIHKPK | 10 | P42285 | Exosome RNA helicase MTR4 OS=Homo | MTREX | 117.805 | 2 | 53.995205 | yes | 1.263E-07 | 643.34296 | 162.58029 |
| SFYQFQHYR | 9 | P42285 | Exosome RNA helicase MTR4 OS=Homo | MTREX | 117.805 | 2 | 53.995205 | yes | 9.991E-08 | 638.29889 | 159.48825 |
| LDFNTDEEK | 9 | P42285 | Exosome RNA helicase MTR4 OS=Homo | MTREX | 117.805 | 2 | 53.995205 | yes | 9.809E-08 | 555.7511 | 158.5663 |
| GIVILMVDEK | 10 | P42285 | Exosome RNA helicase MTR4 OS=Homo | MTREX | 117.805 | 2 | 53.995205 | yes | 9.722E-08 | 558.82031 | 153.53882 |
| MTDVFEGSIIR | 11 | P42285 | Exosome RNA helicase MTR4 OS=Homo | MTREX | 117.805 | 2 | 53.995205 | yes | 1.379E-07 | 634.32117 | 142.23654 |
| DCEAYALQMTK | 11 | P42285 | Exosome RNA helicase MTR4 OS=Homo | MTREX | 117.805 | 2 | 53.995205 | yes | 5.022E-07 | 665.29431 | 129.54404 |
| EAIQCVDNNQSVL | 22 | P42285 | Exosome RNA helicase MTR4 OS=Homo | MTREX | 117.805 | 3 | 53.995205 | yes | 1.805E-07 | 776.71313 | 120.98293 |
| KLPQVEHVLPLLK | 13 | P42285 | Exosome RNA helicase MTR4 OS=Homo | MTREX | 117.805 | 3 | 53.995205 | yes | 4.231E-07 | 505.32034 | 108.92232 |
| EYPFILDAFQR | 11 | P42285 | Exosome RNA helicase MTR4 OS=Homo | MTREX | 117.805 | 2 | 53.995205 | yes | 1.76E-07 | 699.85626 | 87.48069 |
| LPQVEHVLPLLK | 12 | P42285 | Exosome RNA helicase MTR4 OS=Homo | MTREX | 117.805 | 2 | 53.995205 | yes | 9.722E-08 | 693.42938 | 66.917557 |
| WISSGEYIQMSGR | 13 | P42285 | Exosome RNA helicase MTR4 OS=Homo | MTREX | 117.805 | 2 | 53.995205 | yes | 1.261E-07 | 757.35883 | 64.114594 |

| | | | | | | | | | | | |
|---|---|---|---|---|---|---|---|---|---|---|---|
| GIGIHHGGLLPILK | 14 | P42285 | Exosome RNA helicase MTR4 OS=Hom | MTREX | 117.805 | 2 | 53.995205 | yes | 1.026E-07 | 712.94043 | 28.939539 |
| GIGIHHGGLLPILK | 14 | P42285 | Exosome RNA helicase MTR4 OS=Hom | MTREX | 117.805 | 2 | 53.995205 | yes | 1.784E-06 | 475.62939 | 28.939539 |
| GIGIHHGGLLPILK | 14 | P42285 | Exosome RNA helicase MTR4 OS=Hom | MTREX | 117.805 | 3 | 53.995205 | yes | 1.026E-07 | 712.94043 | 28.939539 |
| GIGIHHGGLLPILK | 14 | P42285 | Exosome RNA helicase MTR4 OS=Hom | MTREX | 117.805 | 3 | 53.995205 | yes | 1.784E-06 | 475.62939 | 28.939539 |
| VQSVETVEGCTHE | 28 | P42285 | Exosome RNA helicase MTR4 OS=Hom | MTREX | 117.805 | 4 | 53.995205 | yes | 5.688E-07 | 792.39856 | 53.525227 |
| VIFTSPIK | 8 | P42285 | Exosome RNA helicase MTR4 OS=Hom | MTREX | 117.805 | 2 | 53.995205 | yes | 2.891E-06 | 452.77875 | 50.567493 |
| NSATEAAKPAKPD | 15 | P42285 | Exosome RNA helicase MTR4 OS=Hom | MTREX | 117.805 | 3 | 53.995205 | yes | 1.142E-07 | 519.60199 | 29.358456 |
| KDCEAYALQMTK | 12 | P42285 | Exosome RNA helicase MTR4 OS=Hom | MTREX | 117.805 | 2 | 53.995205 | yes | 1.756E-07 | 729.3418 | 18.282967 |
| ELDTLNNEIVDLQ | 14 | P42566 | Epidermal growth factor receptor substrat | EPS15 | 98.656 | 2 | 52.69476 | yes | 9.722E-08 | 836.43103 | 120.16505 |
| NIIGSSPVADFSAIK | 15 | P42566 | Epidermal growth factor receptor substrat | EPS15 | 98.656 | 2 | 52.69476 | yes | 2.56E-07 | 759.91174 | 103.32389 |
| LPVDILGR | 8 | P42566 | Epidermal growth factor receptor substrat | EPS15 | 98.656 | 2 | 52.69476 | yes | 2.324E-07 | 441.77402 | 60.712383 |
| SATSSSVSNVVITK | 14 | P42566 | Epidermal growth factor receptor substrat | EPS15 | 98.656 | 2 | 52.69476 | yes | 1.78E-07 | 690.37244 | 55.844471 |
| GSDPFASDCFFR | 12 | P42566 | Epidermal growth factor receptor substrat | EPS15 | 98.656 | 2 | 52.69476 | yes | 9.809E-08 | 703.29572 | 53.708378 |
| TSEVQDLQDEVQF | 13 | P42566 | Epidermal growth factor receptor substrat | EPS15 | 98.656 | 2 | 52.69476 | yes | 1.199E-07 | 773.87079 | 42.536583 |
| QEFFVALR | 8 | P42566 | Epidermal growth factor receptor substrat | EPS15 | 98.656 | 2 | 52.69476 | yes | 1.922E-07 | 505.27692 | 37.712662 |
| SFSSAPPLILTCSAI | 16 | P42568 | Protein AF-9 OS=Homo sapiens OX=960 | MLLT3 | 63.352 | 2 | 43.805397 | yes | 2.439E-07 | 847.42694 | 108.90061 |
| DLLHPSPEEEK | 11 | P42677 | Small ribosomal subunit protein eS27 OS | RPS27 | 9.462 | 2 | 38.86218 | yes | 4.939E-07 | 647.3197 | 20696.057 |
| DLLHPSPEEEKR | 12 | P42677 | Small ribosomal subunit protein eS27 OS | RPS27 | 9.462 | 2 | 38.86218 | yes | 1.298E-06 | 725.37024 | 1023.0005 |
| GNLVNFLR | 8 | P42679 | Megakaryocyte-associated tyrosine-protei | MATK | 56.469 | 2 | 52.481617 | yes | 5.356E-07 | 466.76926 | 131.59746 |
| LSLMPWFHGK | 10 | P42679 | Megakaryocyte-associated tyrosine-protei | MATK | 56.469 | 2 | 52.481617 | no | 9.722E-08 | 608.3208 | 125.77985 |
| MQHENLVR | 8 | P42679 | Megakaryocyte-associated tyrosine-protei | MATK | 56.469 | 2 | 52.481617 | yes | 2.928E-06 | 513.76111 | 88.591942 |
| HHTSGQEGLLAAC | 16 | P42679 | Megakaryocyte-associated tyrosine-protei | MATK | 56.469 | 3 | 52.481617 | yes | 1.594E-07 | 539.95337 | 67.368645 |
| AWHPPPVSAR | 10 | P42679 | Megakaryocyte-associated tyrosine-protei | MATK | 56.469 | 2 | 52.481617 | yes | 2.679E-06 | 559.29871 | 48.677391 |
| ELGSGLFGVVR | 11 | P42680 | Tyrosine-protein kinase Tec OS=Homo sa | TEC | 73.582 | 2 | 48.456024 | yes | 1.286E-07 | 567.31934 | 105.86026 |
| YNLFESSIR | 9 | P42680 | Tyrosine-protein kinase Tec OS=Homo sa | TEC | 73.582 | 2 | 48.456024 | yes | 1.146E-07 | 564.78784 | 89.065987 |
| YVLDDQYTSSSGA | 14 | P42680 | Tyrosine-protein kinase Tec OS=Homo sa | TEC | 73.582 | 2 | 48.456024 | yes | 8.027E-07 | 767.35699 | 44.744022 |
| GCLLNFLR | 8 | P42680 | Tyrosine-protein kinase Tec OS=Homo sa | TEC | 73.582 | 2 | 48.456024 | yes | 2.428E-07 | 496.77094 | 40.40126 |
| PSFEDLLR | 8 | P42680 | Tyrosine-protein kinase Tec OS=Homo sa | TEC | 73.582 | 2 | 48.456024 | yes | 5.874E-07 | 488.75854 | 32.835655 |
| TYVLNPLAPEFIPR | 14 | P42694 | Probable helicase with zinc finger domain | HELZ | 218.971 | 2 | 53.675266 | yes | 9.722E-08 | 815.45361 | 249.20468 |
| VLSECIEGVK | 10 | P42694 | Probable helicase with zinc finger domain | HELZ | 218.971 | 2 | 53.675266 | yes | 1.865E-07 | 567.29718 | 196.06915 |
| LTETLSEDTLAGR | 13 | P42694 | Probable helicase with zinc finger domain | HELZ | 218.971 | 2 | 53.675266 | yes | 3.4E-07 | 703.36212 | 190.78362 |
| PSVVYPSTK | 9 | P42694 | Probable helicase with zinc finger domain | HELZ | 218.971 | 2 | 53.675266 | yes | 1.339E-07 | 489.26874 | 181.88638 |
| IHGSVALETLR | 11 | P42694 | Probable helicase with zinc finger domain | HELZ | 218.971 | 2 | 53.675266 | yes | 1.188E-06 | 598.34332 | 150.54834 |
| FLEQGICR | 8 | P42694 | Probable helicase with zinc finger domain | HELZ | 218.971 | 2 | 53.675266 | yes | 4.882E-07 | 511.75803 | 142.23367 |
| LSDVNVER | 8 | P42694 | Probable helicase with zinc finger domain | HELZ | 218.971 | 2 | 53.675266 | yes | 2.165E-06 | 466.24582 | 137.17209 |
| VLFLSTVR | 8 | P42694 | Probable helicase with zinc finger domain | HELZ | 218.971 | 2 | 53.675266 | yes | 1.017E-07 | 467.78964 | 123.633 |
| LLNTAITR | 8 | P42694 | Probable helicase with zinc finger domain | HELZ | 218.971 | 2 | 53.675266 | yes | 2.167E-07 | 451.27692 | 116.09167 |
| LQPVSTILAK | 10 | P42694 | Probable helicase with zinc finger domain | HELZ | 218.971 | 2 | 53.675266 | yes | 1.186E-07 | 535.33441 | 112.75107 |
| ILLCENYR | 8 | P42694 | Probable helicase with zinc finger domain | HELZ | 218.971 | 2 | 53.675266 | yes | 3.104E-07 | 540.77893 | 102.62232 |

| | | | | | | | | | | | |
|---|---|---|---|---|---|---|---|---|---|---|---|
| APPKPRPPPEQAK | 13 | P42694 | Probable helicase with zinc finger domair | HELZ | 218.971 | 3 | 53.675266 | yes | 9.722E-08 | 471.60513 | 78.777298 |
| AVLCCMQLK | 9 | P42694 | Probable helicase with zinc finger domair | HELZ | 218.971 | 2 | 53.675266 | yes | 2.389E-07 | 561.77704 | 77.048042 |
| SCEQACESLK | 10 | P42694 | Probable helicase with zinc finger domair | HELZ | 218.971 | 2 | 53.675266 | yes | 1.117E-07 | 606.25519 | 74.743248 |
| WMNSLSPEK | 9 | P42694 | Probable helicase with zinc finger domair | HELZ | 218.971 | 2 | 53.675266 | yes | 1.427E-06 | 546.26312 | 69.187103 |
| QLSGSYMETLIEK | 13 | P42694 | Probable helicase with zinc finger domair | HELZ | 218.971 | 2 | 53.675266 | yes | 9.722E-08 | 749.87653 | 68.419937 |
| TVKPPEDQLK | 10 | P42694 | Probable helicase with zinc finger domair | HELZ | 218.971 | 2 | 53.675266 | yes | 1.754E-06 | 577.8244 | 64.419243 |
| VNAVYLLPVPK | 11 | P42694 | Probable helicase with zinc finger domair | HELZ | 218.971 | 2 | 53.675266 | yes | 2.13E-07 | 606.87115 | 57.617588 |
| DYLHPYVEAGNP( | 15 | P42694 | Probable helicase with zinc finger domair | HELZ | 218.971 | 3 | 53.675266 | yes | 6.8E-07 | 577.28125 | 48.032295 |
| VEHNPDLSVTVST | 14 | P42694 | Probable helicase with zinc finger domair | HELZ | 218.971 | 2 | 53.675266 | yes | 3.971E-07 | 763.39648 | 47.08313 |
| HVLGEGLAK | 9 | P42694 | Probable helicase with zinc finger domair | HELZ | 218.971 | 2 | 53.675266 | yes | 4.534E-07 | 462.26907 | 45.199577 |
| LHDLLYIEEIAQYK | 14 | P42694 | Probable helicase with zinc finger domair | HELZ | 218.971 | 3 | 53.675266 | yes | 1.117E-07 | 583.31366 | 39.959789 |
| LPLCEMHYALDR | 12 | P42694 | Probable helicase with zinc finger domair | HELZ | 218.971 | 2 | 53.675266 | yes | 6.972E-05 | 759.3656 | 6.5917883 |
| APLAVTSQVINCR | 13 | P42695 | Condensin-2 complex subunit D3 OS=Hc | NCAPD3 | 168.891 | 2 | 53.4428 | yes | 9.722E-08 | 714.88501 | 189.98233 |
| ILLQHICAK | 9 | P42695 | Condensin-2 complex subunit D3 OS=Hc | NCAPD3 | 168.891 | 2 | 53.4428 | yes | 1.881E-07 | 548.3208 | 164.48387 |
| ELEVCEDVAVR | 11 | P42695 | Condensin-2 complex subunit D3 OS=Hc | NCAPD3 | 168.891 | 2 | 53.4428 | yes | 9.755E-07 | 659.81903 | 116.71596 |
| ELMHYLR | 7 | P42695 | Condensin-2 complex subunit D3 OS=Hc | NCAPD3 | 168.891 | 2 | 53.4428 | yes | 8.072E-07 | 481.24985 | 102.29994 |
| EDLWILQDQCR | 11 | P42695 | Condensin-2 complex subunit D3 OS=Hc | NCAPD3 | 168.891 | 2 | 53.4428 | yes | 9.722E-08 | 738.35101 | 94.056419 |
| YQEQLVQEQELAK | 13 | P42695 | Condensin-2 complex subunit D3 OS=Hc | NCAPD3 | 168.891 | 2 | 53.4428 | yes | 3.858E-07 | 803.40961 | 86.167877 |
| FLLEHFTDEQR | 11 | P42695 | Condensin-2 complex subunit D3 OS=Hc | NCAPD3 | 168.891 | 2 | 53.4428 | yes | 9.938E-08 | 717.85425 | 79.336884 |
| QLASELEYDMK | 11 | P42695 | Condensin-2 complex subunit D3 OS=Hc | NCAPD3 | 168.891 | 2 | 53.4428 | yes | 9.722E-08 | 663.81592 | 72.079262 |
| AHAIITLGK | 9 | P42695 | Condensin-2 complex subunit D3 OS=Hc | NCAPD3 | 168.891 | 2 | 53.4428 | yes | 1.671E-07 | 462.28726 | 59.971291 |
| NAIFHLLK | 8 | P42695 | Condensin-2 complex subunit D3 OS=Hc | NCAPD3 | 168.891 | 2 | 53.4428 | yes | 1.344E-06 | 478.28983 | 57.56617 |
| SNTLPISLQSIR | 12 | P42704 | Leucine-rich PPR motif-containing protei | LRPPRC | 157.906 | 2 | 55.360527 | yes | 4.799E-07 | 664.88025 | 3243.0166 |
| AGYPQYVSEILEK | 13 | P42704 | Leucine-rich PPR motif-containing protei | LRPPRC | 157.906 | 2 | 55.360527 | yes | 9.314E-07 | 748.88519 | 2301.9744 |
| MEEANIQPNR | 10 | P42704 | Leucine-rich PPR motif-containing protei | LRPPRC | 157.906 | 2 | 55.360527 | yes | 9.722E-08 | 601.28516 | 2000.3164 |
| SELHLMDR | 8 | P42704 | Leucine-rich PPR motif-containing protei | LRPPRC | 157.906 | 2 | 55.360527 | yes | 2.792E-07 | 500.74765 | 1952.6987 |
| AGDMENAENILTV | 15 | P42704 | Leucine-rich PPR motif-containing protei | LRPPRC | 157.906 | 2 | 55.360527 | yes | 1.046E-07 | 832.39252 | 972.93463 |
| AGDMENAENILTV | 15 | P42704 | Leucine-rich PPR motif-containing protei | LRPPRC | 157.906 | 2 | 55.360527 | yes | 1.172E-07 | 840.39001 | 972.93463 |
| IPENIYR | 7 | P42704 | Leucine-rich PPR motif-containing protei | LRPPRC | 157.906 | 2 | 55.360527 | yes | 1.815E-06 | 452.74799 | 1918.5698 |
| LQDAINILK | 9 | P42704 | Leucine-rich PPR motif-containing protei | LRPPRC | 157.906 | 2 | 55.360527 | yes | 1.14E-07 | 514.31097 | 1686.0239 |
| ALYEHLTAK | 9 | P42704 | Leucine-rich PPR motif-containing protei | LRPPRC | 157.906 | 2 | 55.360527 | yes | 4.292E-07 | 523.28748 | 1678.7731 |
| NVQGIIEILK | 10 | P42704 | Leucine-rich PPR motif-containing protei | LRPPRC | 157.906 | 2 | 55.360527 | yes | 1.427E-07 | 563.84515 | 1578.8461 |
| IQEENVIPR | 9 | P42704 | Leucine-rich PPR motif-containing protei | LRPPRC | 157.906 | 2 | 55.360527 | yes | 9.722E-08 | 549.30109 | 1567.1899 |
| AFAETHIK | 8 | P42704 | Leucine-rich PPR motif-containing protei | LRPPRC | 157.906 | 2 | 55.360527 | yes | 9.722E-08 | 458.74799 | 1546.3499 |
| SCGSLLPELK | 10 | P42704 | Leucine-rich PPR motif-containing protei | LRPPRC | 157.906 | 2 | 55.360527 | yes | 1.883E-07 | 552.29187 | 1480.5286 |
| LIITQVR | 7 | P42704 | Leucine-rich PPR motif-containing protei | LRPPRC | 157.906 | 2 | 55.360527 | yes | 1.146E-07 | 421.77655 | 1475.8079 |
| TVLDQQQTPSR | 11 | P42704 | Leucine-rich PPR motif-containing protei | LRPPRC | 157.906 | 2 | 55.360527 | yes | 1.01E-07 | 636.83075 | 1245.5389 |
| GFTLNDAANSR | 11 | P42704 | Leucine-rich PPR motif-containing protei | LRPPRC | 157.906 | 2 | 55.360527 | yes | 9.976E-08 | 583.28345 | 1237.3619 |
| VYLQNEYK | 8 | P42704 | Leucine-rich PPR motif-containing protei | LRPPRC | 157.906 | 2 | 55.360527 | yes | 1.579E-07 | 528.77167 | 1116.2512 |

| | | | | | | | | | | | |
|---|---|---|---|---|---|---|---|---|---|---|---|
| SVLELIPELNEK | 12 | P42704 | Leucine-rich PPR motif-containing protei | LRPPRC | 157.906 | 2 | 55.360527 | yes | 1.446E-07 | 692.39014 | 1095.8098 |
| KIIETPGIR | 9 | P42704 | Leucine-rich PPR motif-containing protei | LRPPRC | 157.906 | 2 | 55.360527 | yes | 1.697E-07 | 513.81897 | 1013.58 |
| ISNQFDWALMR | 11 | P42704 | Leucine-rich PPR motif-containing protei | LRPPRC | 157.906 | 2 | 55.360527 | yes | 9.935E-08 | 690.84009 | 495.11526 |
| ISNQFDWALMR | 11 | P42704 | Leucine-rich PPR motif-containing protei | LRPPRC | 157.906 | 2 | 55.360527 | yes | 3.465E-07 | 698.83752 | 495.11526 |
| CVANNQVETLEK | 12 | P42704 | Leucine-rich PPR motif-containing protei | LRPPRC | 157.906 | 2 | 55.360527 | yes | 2.477E-07 | 702.84302 | 972.45313 |
| DAHLLVESK | 9 | P42704 | Leucine-rich PPR motif-containing protei | LRPPRC | 157.906 | 2 | 55.360527 | yes | 9.789E-08 | 506.2771 | 970.31067 |
| LDDLFLK | 7 | P42704 | Leucine-rich PPR motif-containing protei | LRPPRC | 157.906 | 2 | 55.360527 | yes | 1.107E-07 | 432.24728 | 903.80914 |
| GAYDIFLNAK | 10 | P42704 | Leucine-rich PPR motif-containing protei | LRPPRC | 157.906 | 2 | 55.360527 | yes | 1.029E-07 | 556.29279 | 844.0838 |
| LIASYCNVGDIEG/ | 16 | P42704 | Leucine-rich PPR motif-containing protei | LRPPRC | 157.906 | 2 | 55.360527 | yes | 9.722E-08 | 848.91418 | 802.00769 |
| MVFINNIALAQIK | 13 | P42704 | Leucine-rich PPR motif-containing protei | LRPPRC | 157.906 | 2 | 55.360527 | yes | 1.027E-07 | 737.92615 | 790.51007 |
| QYFHQLEK | 8 | P42704 | Leucine-rich PPR motif-containing protei | LRPPRC | 157.906 | 2 | 55.360527 | yes | 9.937E-08 | 546.77728 | 762.71204 |
| DIQEESTFSSR | 11 | P42704 | Leucine-rich PPR motif-containing protei | LRPPRC | 157.906 | 2 | 55.360527 | yes | 9.722E-08 | 649.79657 | 736.86377 |
| EEAYNSLMK | 9 | P42704 | Leucine-rich PPR motif-containing protei | LRPPRC | 157.906 | 2 | 55.360527 | yes | 1.318E-07 | 542.75262 | 669.60138 |
| IWDTLQK | 7 | P42704 | Leucine-rich PPR motif-containing protei | LRPPRC | 157.906 | 2 | 55.360527 | yes | 3.498E-06 | 452.25037 | 668.46735 |
| LDSSAVLDTGK | 11 | P42704 | Leucine-rich PPR motif-containing protei | LRPPRC | 157.906 | 2 | 55.360527 | yes | 9.728E-08 | 553.29041 | 647.09595 |
| VEDALNLK | 8 | P42704 | Leucine-rich PPR motif-containing protei | LRPPRC | 157.906 | 2 | 55.360527 | yes | 1.272E-07 | 451.25311 | 551.32617 |
| IIETPGIR | 8 | P42704 | Leucine-rich PPR motif-containing protei | LRPPRC | 157.906 | 2 | 55.360527 | yes | 9.819E-08 | 449.77145 | 480.86966 |
| VIEEQLEPAVEK | 12 | P42704 | Leucine-rich PPR motif-containing protei | LRPPRC | 157.906 | 2 | 55.360527 | yes | 1.401E-07 | 692.37195 | 430.15198 |
| LLAEILR | 7 | P42704 | Leucine-rich PPR motif-containing protei | LRPPRC | 157.906 | 2 | 55.360527 | yes | 1.789E-07 | 414.27109 | 415.30518 |
| GDVENIEVVQK | 11 | P42704 | Leucine-rich PPR motif-containing protei | LRPPRC | 157.906 | 2 | 55.360527 | yes | 9.722E-08 | 615.32227 | 373.61179 |
| NLLESYHVPELIK | 13 | P42704 | Leucine-rich PPR motif-containing protei | LRPPRC | 157.906 | 2 | 55.360527 | yes | 9.722E-08 | 777.92993 | 184.13205 |
| NLLESYHVPELIK | 13 | P42704 | Leucine-rich PPR motif-containing protei | LRPPRC | 157.906 | 2 | 55.360527 | yes | 1.004E-07 | 518.95575 | 184.13205 |
| NLLESYHVPELIK | 13 | P42704 | Leucine-rich PPR motif-containing protei | LRPPRC | 157.906 | 3 | 55.360527 | yes | 9.722E-08 | 777.92993 | 184.13205 |
| NLLESYHVPELIK | 13 | P42704 | Leucine-rich PPR motif-containing protei | LRPPRC | 157.906 | 3 | 55.360527 | yes | 1.004E-07 | 518.95575 | 184.13205 |
| FSPTDFLAK | 9 | P42704 | Leucine-rich PPR motif-containing protei | LRPPRC | 157.906 | 2 | 55.360527 | yes | 5.245E-07 | 513.26874 | 349.30316 |
| KGAYDIFLNAK | 11 | P42704 | Leucine-rich PPR motif-containing protei | LRPPRC | 157.906 | 2 | 55.360527 | yes | 1.346E-07 | 620.34021 | 317.01089 |
| IHDVLCK | 7 | P42704 | Leucine-rich PPR motif-containing protei | LRPPRC | 157.906 | 2 | 55.360527 | yes | 3.095E-07 | 442.73657 | 306.51791 |
| LANQFAIYK | 9 | P42704 | Leucine-rich PPR motif-containing protei | LRPPRC | 157.906 | 2 | 55.360527 | yes | 1.21E-07 | 534.29785 | 305.65341 |
| MLNGLEDSIGLSK | 13 | P42704 | Leucine-rich PPR motif-containing protei | LRPPRC | 157.906 | 2 | 55.360527 | yes | 9.733E-08 | 688.85815 | 290.80283 |
| HSLNSSSASTTEPL | 17 | P42704 | Leucine-rich PPR motif-containing protei | LRPPRC | 157.906 | 3 | 55.360527 | yes | 1.017E-07 | 612.61841 | 216.4581 |
| DQMYYNLLK | 9 | P42704 | Leucine-rich PPR motif-containing protei | LRPPRC | 157.906 | 2 | 55.360527 | yes | 1.037E-07 | 594.29187 | 204.68312 |
| EKDIQEESTFSSR | 13 | P42704 | Leucine-rich PPR motif-containing protei | LRPPRC | 157.906 | 2 | 55.360527 | yes | 1.02E-07 | 778.36536 | 198.74579 |
| SSLLLGFR | 8 | P42704 | Leucine-rich PPR motif-containing protei | LRPPRC | 157.906 | 2 | 55.360527 | yes | 1.006E-06 | 446.76617 | 197.45399 |
| LQWFCDR | 7 | P42704 | Leucine-rich PPR motif-containing protei | LRPPRC | 157.906 | 2 | 55.360527 | yes | 4.562E-07 | 512.73712 | 181.47275 |
| KVIEEQLEPAVEK | 13 | P42704 | Leucine-rich PPR motif-containing protei | LRPPRC | 157.906 | 2 | 55.360527 | yes | 1.777E-07 | 756.41943 | 76.384529 |
| KVIEEQLEPAVEK | 13 | P42704 | Leucine-rich PPR motif-containing protei | LRPPRC | 157.906 | 2 | 55.360527 | yes | 1.362E-06 | 504.61536 | 76.384529 |
| KVIEEQLEPAVEK | 13 | P42704 | Leucine-rich PPR motif-containing protei | LRPPRC | 157.906 | 3 | 55.360527 | yes | 1.777E-07 | 756.41943 | 76.384529 |
| KVIEEQLEPAVEK | 13 | P42704 | Leucine-rich PPR motif-containing protei | LRPPRC | 157.906 | 3 | 55.360527 | yes | 1.362E-06 | 504.61536 | 76.384529 |
| EKDVLIK | 7 | P42704 | Leucine-rich PPR motif-containing protei | LRPPRC | 157.906 | 2 | 55.360527 | yes | 1.949E-07 | 422.76056 | 149.87094 |

| | | | | | | | | | | | |
|---|---|---|---|---|---|---|---|---|---|---|---|
| VIQALAMK | 8 | P42704 | Leucine-rich PPR motif-containing protei | LRPPRC | 157.906 | 2 | 55.360527 | yes | 3.341E-07 | 437.26495 | 131.35735 |
| RYASLLK | 7 | P42704 | Leucine-rich PPR motif-containing protei | LRPPRC | 157.906 | 2 | 55.360527 | yes | 3.58E-07 | 425.76089 | 108.69721 |
| QLHEAIVTLGLAEI | 29 | P42704 | Leucine-rich PPR motif-containing protei | LRPPRC | 157.906 | 4 | 55.360527 | yes | 1.046E-07 | 789.93805 | 104.19736 |
| SVLELIPELNEKEE | 21 | P42704 | Leucine-rich PPR motif-containing protei | LRPPRC | 157.906 | 3 | 55.360527 | yes | 1.088E-07 | 817.08923 | 96.325493 |
| HDKVEDALNLK | 11 | P42704 | Leucine-rich PPR motif-containing protei | LRPPRC | 157.906 | 2 | 55.360527 | yes | 9.722E-08 | 641.34351 | 83.445129 |
| AKYESDMVTGGY | 21 | P42704 | Leucine-rich PPR motif-containing protei | LRPPRC | 157.906 | 3 | 55.360527 | yes | 2.231E-07 | 798.04132 | 73.109505 |
| EEGFPIRPHYFWPI | 18 | P42704 | Leucine-rich PPR motif-containing protei | LRPPRC | 157.906 | 4 | 55.360527 | yes | 1.724E-07 | 554.04681 | 57.491074 |
| EKNVQGIIEILK | 12 | P42704 | Leucine-rich PPR motif-containing protei | LRPPRC | 157.906 | 2 | 55.360527 | yes | 2.742E-07 | 692.41394 | 35.142902 |
| VEDALNLKEEFDR | 13 | P42704 | Leucine-rich PPR motif-containing protei | LRPPRC | 157.906 | 2 | 55.360527 | yes | 2.726E-07 | 789.39392 | 34.712204 |
| YKPLDLRPK | 9 | P42766 | Large ribosomal subunit protein uL29 OS | RPL35 | 14.552 | 2 | 50.75205 | yes | 1.537E-07 | 565.34003 | 27198.83 |
| VELSQLR | 7 | P42766 | Large ribosomal subunit protein uL29 OS | RPL35 | 14.552 | 2 | 50.75205 | yes | 1.986E-07 | 422.74799 | 25830.346 |
| VLTVINQTQK | 10 | P42766 | Large ribosomal subunit protein uL29 OS | RPL35 | 14.552 | 2 | 50.75205 | yes | 9.723E-08 | 572.34021 | 5524.1572 |
| PLDLRPK | 7 | P42766 | Large ribosomal subunit protein uL29 OS | RPL35 | 14.552 | 2 | 50.75205 | yes | 1.305E-07 | 419.76089 | 4814.6201 |
| QLDDLKVELSQLR | 13 | P42766 | Large ribosomal subunit protein uL29 OS | RPL35 | 14.552 | 2 | 50.75205 | yes | 1.088E-07 | 778.93579 | 825.35211 |
| KKEELLK | 7 | P42766 | Large ribosomal subunit protein uL29 OS | RPL35 | 14.552 | 2 | 50.75205 | yes | 2.042E-07 | 444.28165 | 778.80219 |
| KYKPLDLR | 8 | P42766 | Large ribosomal subunit protein uL29 OS | RPL35 | 14.552 | 2 | 50.75205 | yes | 5.119E-07 | 516.81366 | 171.54897 |
| LPVDLAEELGHR | 12 | P42771 | Cyclin-dependent kinase inhibitor 2A OS | CDKN2A | 16.533 | 2 | 50.987885 | yes | 2.143E-07 | 674.86462 | 330.18488 |
| EGFLDTLVVLHR | 12 | P42771 | Cyclin-dependent kinase inhibitor 2A OS | CDKN2A | 16.533 | 2 | 50.987885 | no | 9.722E-08 | 699.89063 | 100.90391 |
| TAPYVVTGSVDQT | 15 | P43034 | Platelet-activating factor acetylhydrolase | PAFAH1B | 46.638 | 2 | 44.8149 | yes | 2.059E-07 | 782.91449 | 70.59343 |
| SQAFIEMETR | 10 | P43243 | Matrin-3 OS=Homo sapiens OX=9606 G | MATR3 | 94.624 | 2 | 52.519684 | yes | 1.182E-07 | 606.28992 | 10669.566 |
| GPLPLSSQHR | 10 | P43243 | Matrin-3 OS=Homo sapiens OX=9606 G | MATR3 | 94.624 | 2 | 52.519684 | yes | 3.326E-07 | 546.30145 | 8842.7412 |
| EDAMAMVDHCLK | 12 | P43243 | Matrin-3 OS=Homo sapiens OX=9606 G | MATR3 | 94.624 | 2 | 52.519684 | yes | 1.583E-07 | 710.30688 | 3458.0186 |
| EDAMAMVDHCLK | 12 | P43243 | Matrin-3 OS=Homo sapiens OX=9606 G | MATR3 | 94.624 | 2 | 52.519684 | yes | 2.011E-07 | 718.30438 | 3458.0186 |
| TEEGPTLSYGR | 11 | P43243 | Matrin-3 OS=Homo sapiens OX=9606 G | MATR3 | 94.624 | 2 | 52.519684 | yes | 1.197E-07 | 605.29095 | 5617.8618 |
| SFQQSSLSR | 9 | P43243 | Matrin-3 OS=Homo sapiens OX=9606 G | MATR3 | 94.624 | 2 | 52.519684 | yes | 2.255E-07 | 520.26196 | 4568.7744 |
| LCSLFYTNEEVAK | 13 | P43243 | Matrin-3 OS=Homo sapiens OX=9606 G | MATR3 | 94.624 | 2 | 52.519684 | yes | 2.951E-07 | 787.38196 | 4007.7708 |
| VVHIMDFQR | 9 | P43243 | Matrin-3 OS=Homo sapiens OX=9606 G | MATR3 | 94.624 | 2 | 52.519684 | yes | 4.307E-07 | 572.80023 | 1928.1475 |
| VVHIMDFQR | 9 | P43243 | Matrin-3 OS=Homo sapiens OX=9606 G | MATR3 | 94.624 | 2 | 52.519684 | yes | 1.274E-07 | 580.79767 | 1928.1475 |
| GPGPLQER | 8 | P43243 | Matrin-3 OS=Homo sapiens OX=9606 G | MATR3 | 94.624 | 2 | 52.519684 | yes | 5.84E-07 | 427.22995 | 3833.9236 |
| SQESGYYDR | 9 | P43243 | Matrin-3 OS=Homo sapiens OX=9606 G | MATR3 | 94.624 | 2 | 52.519684 | yes | 1.108E-07 | 552.73328 | 2504.8699 |
| GPSLNPVLDYDHC | 15 | P43243 | Matrin-3 OS=Homo sapiens OX=9606 G | MATR3 | 94.624 | 2 | 52.519684 | yes | 2.212E-07 | 813.89716 | 2256.53 |
| NTHCSSLPHYQK | 12 | P43243 | Matrin-3 OS=Homo sapiens OX=9606 G | MATR3 | 94.624 | 2 | 52.519684 | yes | 9.722E-08 | 736.341 | 1020.9702 |
| MDYEDDRLR | 9 | P43243 | Matrin-3 OS=Homo sapiens OX=9606 G | MATR3 | 94.624 | 2 | 52.519684 | yes | 2.082E-07 | 606.76935 | 687.61737 |
| DDYTIPDEYR | 10 | P43243 | Matrin-3 OS=Homo sapiens OX=9606 G | MATR3 | 94.624 | 2 | 52.519684 | yes | 9.722E-08 | 643.7804 | 681.38025 |
| CRDDSFFGETSHN | 16 | P43243 | Matrin-3 OS=Homo sapiens OX=9606 G | MATR3 | 94.624 | 3 | 52.519684 | yes | 1.004E-07 | 667.28326 | 383.63556 |
| GYPHLCSICDLPVI | 17 | P43243 | Matrin-3 OS=Homo sapiens OX=9606 G | MATR3 | 94.624 | 3 | 52.519684 | yes | 1.321E-07 | 666.31714 | 378.94882 |
| VIHLSNLPHSGYSI | 19 | P43243 | Matrin-3 OS=Homo sapiens OX=9606 G | MATR3 | 94.624 | 4 | 52.519684 | yes | 2.076E-07 | 510.02454 | 148.01201 |
| VIHLSNLPHSGYSI | 19 | P43243 | Matrin-3 OS=Homo sapiens OX=9606 G | MATR3 | 94.624 | 4 | 52.519684 | yes | 2.83E-07 | 679.69696 | 148.01201 |
| VIHLSNLPHSGYSI | 19 | P43243 | Matrin-3 OS=Homo sapiens OX=9606 G | MATR3 | 94.624 | 3 | 52.519684 | yes | 2.076E-07 | 510.02454 | 148.01201 |

| | | | | | | | | | | | |
|---|---|---|---|---|---|---|---|---|---|---|---|
| VIHLSNLPHSGYSI | 19 | P43243 | Matrin-3 OS=Homo sapiens OX=9606 G | MATR3 | 94.624 | 3 | 52.519684 | yes | 2.83E-07 | 679.69696 | 148.01201 |
| FDSEYER | 7 | P43243 | Matrin-3 OS=Homo sapiens OX=9606 G | MATR3 | 94.624 | 2 | 52.519684 | yes | 2.161E-07 | 473.20105 | 234.56332 |
| DDSFFGETSHNYH | 14 | P43243 | Matrin-3 OS=Homo sapiens OX=9606 G | MATR3 | 94.624 | 2 | 52.519684 | yes | 9.757E-08 | 842.35535 | 197.42397 |
| RTEEGPTLSYGR | 12 | P43243 | Matrin-3 OS=Homo sapiens OX=9606 G | MATR3 | 94.624 | 2 | 52.519684 | yes | 9.722E-08 | 683.34149 | 88.551506 |
| RTEEGPTLSYGR | 12 | P43243 | Matrin-3 OS=Homo sapiens OX=9606 G | MATR3 | 94.624 | 2 | 52.519684 | yes | 5.227E-07 | 455.89676 | 88.551506 |
| RTEEGPTLSYGR | 12 | P43243 | Matrin-3 OS=Homo sapiens OX=9606 G | MATR3 | 94.624 | 3 | 52.519684 | yes | 9.722E-08 | 683.34149 | 88.551506 |
| RTEEGPTLSYGR | 12 | P43243 | Matrin-3 OS=Homo sapiens OX=9606 G | MATR3 | 94.624 | 3 | 52.519684 | yes | 5.227E-07 | 455.89676 | 88.551506 |
| EWSQHINGASHSR | 13 | P43243 | Matrin-3 OS=Homo sapiens OX=9606 G | MATR3 | 94.624 | 2 | 52.519684 | yes | 1.022E-07 | 754.85309 | 130.28931 |
| RGAPPSSNIEDFHC | 18 | P43243 | Matrin-3 OS=Homo sapiens OX=9606 G | MATR3 | 94.624 | 3 | 52.519684 | yes | 3.214E-07 | 645.67426 | 86.712494 |
| IGPYQPNVPVGIDY | 18 | P43243 | Matrin-3 OS=Homo sapiens OX=9606 G | MATR3 | 94.624 | 3 | 52.519684 | yes | 1.198E-07 | 657.03131 | 60.516331 |
| GNLGAGNGNLQGI | 14 | P43243 | Matrin-3 OS=Homo sapiens OX=9606 G | MATR3 | 94.624 | 2 | 52.519684 | yes | 9.722E-08 | 662.83948 | 49.761547 |
| VDKIEELDQENEA | 20 | P43243 | Matrin-3 OS=Homo sapiens OX=9606 G | MATR3 | 94.624 | 3 | 52.519684 | yes | 9.722E-08 | 753.04468 | 23.207327 |
| ECVLPGGETAGDM | 15 | P43246 | DNA mismatch repair protein Msh2 OS=I | MSH2 | 104.744 | 2 | 54.864136 | yes | 1.846E-07 | 760.8396 | 485.5015 |
| MQSTLISAAR | 10 | P43246 | DNA mismatch repair protein Msh2 OS=I | MSH2 | 104.744 | 2 | 54.864136 | yes | 1.985E-07 | 539.28967 | 355.75446 |
| NNSFVNEIISR | 11 | P43246 | DNA mismatch repair protein Msh2 OS=I | MSH2 | 104.744 | 2 | 54.864136 | yes | 9.722E-08 | 646.83331 | 273.49548 |
| LDSSAQFGYYFR | 12 | P43246 | DNA mismatch repair protein Msh2 OS=I | MSH2 | 104.744 | 2 | 54.864136 | yes | 2.867E-07 | 727.34094 | 253.8519 |
| LTSLNEEYTK | 10 | P43246 | DNA mismatch repair protein Msh2 OS=I | MSH2 | 104.744 | 2 | 54.864136 | yes | 1.012E-07 | 599.30353 | 252.13147 |
| QVGVGYVDSIQR | 12 | P43246 | DNA mismatch repair protein Msh2 OS=I | MSH2 | 104.744 | 2 | 54.864136 | yes | 9.722E-08 | 660.84894 | 213.95102 |
| QTLQEDLLR | 9 | P43246 | DNA mismatch repair protein Msh2 OS=I | MSH2 | 104.744 | 2 | 54.864136 | yes | 2.32E-07 | 558.3064 | 172.20212 |
| ETLQLESAAEVGF | 15 | P43246 | DNA mismatch repair protein Msh2 OS=I | MSH2 | 104.744 | 2 | 54.864136 | yes | 9.722E-08 | 824.93066 | 122.55885 |
| GGILITER | 8 | P43246 | DNA mismatch repair protein Msh2 OS=I | MSH2 | 104.744 | 2 | 54.864136 | yes | 2.667E-06 | 429.7558 | 116.77924 |
| IIQEFLSK | 8 | P43246 | DNA mismatch repair protein Msh2 OS=I | MSH2 | 104.744 | 2 | 54.864136 | yes | 9.722E-08 | 489.28696 | 109.26518 |
| DSLIIIDELGR | 11 | P43246 | DNA mismatch repair protein Msh2 OS=I | MSH2 | 104.744 | 2 | 54.864136 | yes | 9.722E-08 | 622.34827 | 90.627388 |
| ENDWYLAYK | 9 | P43246 | DNA mismatch repair protein Msh2 OS=I | MSH2 | 104.744 | 2 | 54.864136 | yes | 3.792E-07 | 601.27985 | 85.762901 |
| TEYEEAQDAIVK | 12 | P43246 | DNA mismatch repair protein Msh2 OS=I | MSH2 | 104.744 | 2 | 54.864136 | yes | 1.108E-07 | 698.33557 | 80.445145 |
| DIYQDLNR | 8 | P43246 | DNA mismatch repair protein Msh2 OS=I | MSH2 | 104.744 | 2 | 54.864136 | yes | 4.809E-07 | 518.75653 | 77.702354 |
| QAANLQDCYR | 10 | P43246 | DNA mismatch repair protein Msh2 OS=I | MSH2 | 104.744 | 2 | 54.864136 | yes | 2.141E-07 | 619.78278 | 73.483337 |
| MNFESFVK | 8 | P43246 | DNA mismatch repair protein Msh2 OS=I | MSH2 | 104.744 | 2 | 54.864136 | yes | 2.565E-07 | 501.24167 | 56.064472 |
| NKTEYEEAQDAIV | 14 | P43246 | DNA mismatch repair protein Msh2 OS=I | MSH2 | 104.744 | 2 | 54.864136 | yes | 2.96E-06 | 819.40448 | 9.5239258 |
| AADCISEPVNR | 11 | P43304 | Glycerol-3-phosphate dehydrogenase, mit | GPD2 | 80.853 | 2 | 29.291235 | yes | 2.53E-06 | 616.29041 | 40.974022 |
| GTEDFIVESLDASF | 15 | P43307 | Translocon-associated protein subunit alp | SSR1 | 32.236 | 2 | 50.559086 | yes | 9.809E-08 | 843.40448 | 467.88806 |
| GEDFPANNIVK | 11 | P43307 | Translocon-associated protein subunit alp | SSR1 | 32.236 | 2 | 50.559086 | yes | 9.975E-08 | 602.30383 | 300.40088 |
| FLVGFTNK | 8 | P43307 | Translocon-associated protein subunit alp | SSR1 | 32.236 | 2 | 50.559086 | yes | 1.895E-07 | 463.26074 | 174.5278 |
| PFGLVINLNYK | 11 | P43307 | Translocon-associated protein subunit alp | SSR1 | 32.236 | 2 | 50.559086 | yes | 9.766E-08 | 639.36627 | 112.62376 |
| QATFEYSFIPAEPM | 17 | P43307 | Translocon-associated protein subunit alp | SSR1 | 32.236 | 3 | 50.559086 | yes | 1.673E-07 | 634.30145 | 22.685583 |
| DPLGDTLR | 8 | P43353 | Aldehyde dehydrogenase family 3 membe | ALDH3B1 | 51.84 | 2 | 47.213314 | yes | 4.136E-07 | 464.74036 | 547.27136 |
| HLTPVTLELGGK | 12 | P43353 | Aldehyde dehydrogenase family 3 membe | ALDH3B1 | 51.84 | 2 | 47.213314 | no | 1.376E-07 | 632.86664 | 430.29672 |
| LEDLVPPPAIIDK | 13 | P43354 | Nuclear receptor subfamily 4 group A me | NR4A2 | 66.591 | 2 | 50.958977 | yes | 9.722E-08 | 710.40833 | 169.37091 |
| TLCTQGLQR | 9 | P43354 | Nuclear receptor subfamily 4 group A me | NR4A2 | 66.591 | 2 | 50.958977 | no | 1.528E-06 | 538.77948 | 92.016106 |

| | | | | | | | | | | | |
|---|---|---|---|---|---|---|---|---|---|---|---|
| KPASMGFPGLQIG | 29 | P43354 | Nuclear receptor subfamily 4 group A me | NR4A2 | 66.591 | 4 | 50.958977 | yes | 2.116E-07 | 754.89728 | 84.319511 |
| KLLTQDLVQEK | 11 | P43355 | Melanoma-associated antigen 1 OS=Hom | MAGEA1 | 34.343 | 2 | 53.924397 | yes | 2.93E-07 | 657.88501 | 128.03246 |
| ASESLQLVFGIDVI | 14 | P43355 | Melanoma-associated antigen 1 OS=Hom | MAGEA1 | 34.343 | 2 | 53.924397 | yes | 9.722E-08 | 753.41412 | 88.83194 |
| HCFPEIFGK | 9 | P43355 | Melanoma-associated antigen 1 OS=Hom | MAGEA1 | 34.343 | 2 | 53.924397 | yes | 4.416E-07 | 567.77368 | 65.653778 |
| YEFLWGPR | 8 | P43361 | Melanoma-associated antigen 8 OS=Hom | MAGEA8 | 35.215 | 2 | 48.126263 | no | 1.196E-07 | 534.2691 | 309.10721 |
| AEMLESVIK | 9 | P43361 | Melanoma-associated antigen 8 OS=Hom | MAGEA8 | 35.215 | 2 | 48.126263 | no | 2.316E-07 | 510.27573 | 67.949104 |
| AEEGLQAQGEAPC | 26 | P43361 | Melanoma-associated antigen 8 OS=Hom | MAGEA8 | 35.215 | 4 | 48.126263 | yes | 8.795E-06 | 685.58398 | 45.81633 |
| VLVIVMTTR | 9 | P43378 | Tyrosine-protein phosphatase non-recepto | PTPN9 | 68.021 | 2 | 34.141697 | yes | 1.07E-06 | 516.31775 | 19.749256 |
| FASENDLPEWK | 11 | P43487 | Ran-specific GTPase-activating protein O | RANBP1 | 23.311 | 2 | 50.12587 | yes | 1.13E-07 | 668.31439 | 125.82823 |
| TLEEDEEELFK | 11 | P43487 | Ran-specific GTPase-activating protein O | RANBP1 | 23.311 | 2 | 50.12587 | yes | 9.894E-08 | 691.32208 | 26.681761 |
| VYSYFECR | 8 | P43490 | Nicotinamide phosphoribosyltransferase ( | NAMPT | 55.522 | 2 | 41.049667 | yes | 3.703E-07 | 562.2475 | 37.995071 |
| TPAGNFVTLEEGK | 13 | P43490 | Nicotinamide phosphoribosyltransferase ( | NAMPT | 55.522 | 2 | 41.049667 | yes | 4.153E-07 | 681.84863 | 32.598839 |
| GVLMYGPPGCGK | 12 | P43686 | 26S proteasome regulatory subunit 6B OS | PSMC4 | 47.367 | 2 | 38.236637 | yes | 3.103E-06 | 618.29919 | 42.702049 |
| ELLKPNASVALHK | 13 | P43686 | 26S proteasome regulatory subunit 6B OS | PSMC4 | 47.367 | 3 | 38.236637 | yes | 5.362E-07 | 473.94879 | 36.171387 |
| YGALVICETSEQK | 13 | P43897 | Elongation factor Ts, mitochondrial OS=I | TSFM | 35.391 | 2 | 48.556442 | yes | 1.13E-07 | 749.36633 | 271.76993 |
| LGENMILK | 8 | P43897 | Elongation factor Ts, mitochondrial OS=I | TSFM | 35.391 | 2 | 48.556442 | yes | 1.69E-07 | 459.25986 | 236.43196 |
| ALETCGGDLK | 10 | P43897 | Elongation factor Ts, mitochondrial OS=I | TSFM | 35.391 | 2 | 48.556442 | yes | 2.97E-06 | 532.25806 | 98.38028 |
| QAEIWLHK | 8 | P43897 | Elongation factor Ts, mitochondrial OS=I | TSFM | 35.391 | 2 | 48.556442 | yes | 1.591E-07 | 512.78235 | 75.547874 |
| DQLALAIGK | 9 | P43897 | Elongation factor Ts, mitochondrial OS=I | TSFM | 35.391 | 2 | 48.556442 | yes | 6.596E-07 | 464.77673 | 30.509365 |
| FQLLVQQVALGTM | 20 | P43897 | Elongation factor Ts, mitochondrial OS=I | TSFM | 35.391 | 3 | 48.556442 | yes | 3.686E-07 | 782.74255 | 29.361094 |
| YQLDPTASISAK | 12 | P45880 | Voltage-dependent anion-selective chann | VDAC2 | 31.567 | 2 | 53.568787 | yes | 4.357E-07 | 647.33789 | 445.70596 |
| LTFDTTFSPNTGK | 13 | P45880 | Voltage-dependent anion-selective chann | VDAC2 | 31.567 | 2 | 53.568787 | yes | 2.066E-07 | 714.85388 | 256.39957 |
| LTLSALVDGK | 10 | P45880 | Voltage-dependent anion-selective chann | VDAC2 | 31.567 | 2 | 53.568787 | yes | 3.096E-07 | 508.80295 | 230.2327 |
| WCEYGLTFTEK | 11 | P45880 | Voltage-dependent anion-selective chann | VDAC2 | 31.567 | 2 | 53.568787 | yes | 9.722E-08 | 717.32391 | 77.084015 |
| CPQIVIAFYEER | 12 | P45973 | Chromobox protein homolog 5 OS=Hom | CBX5 | 22.225 | 2 | 52.323673 | no | 9.766E-08 | 762.87939 | 643.0188 |
| SNFSNSADDIK | 11 | P45973 | Chromobox protein homolog 5 OS=Hom | CBX5 | 22.225 | 2 | 52.323673 | yes | 9.722E-08 | 599.27277 | 267.61804 |
| LTWHAYPEDAENI | 13 | P45973 | Chromobox protein homolog 5 OS=Hom | CBX5 | 22.225 | 2 | 52.323673 | yes | 9.722E-08 | 787.36774 | 187.92151 |
| GFSEEHNTWEPEK | 13 | P45973 | Chromobox protein homolog 5 OS=Hom | CBX5 | 22.225 | 2 | 52.323673 | yes | 9.789E-08 | 795.34698 | 134.05331 |
| WKDTDEADLVLA | 13 | P45973 | Chromobox protein homolog 5 OS=Hom | CBX5 | 22.225 | 2 | 52.323673 | yes | 1.095E-07 | 752.38812 | 133.55525 |
| NLDCPELISEFMK | 13 | P45973 | Chromobox protein homolog 5 OS=Hom | CBX5 | 22.225 | 2 | 52.323673 | yes | 1.166E-07 | 798.37585 | 98.640533 |
| VIEQLGTPSAEFMI | 14 | P45984 | Mitogen-activated protein kinase 9 OS=H | MAPK9 | 48.14 | 2 | 43.35043 | yes | 2.628E-07 | 775.40015 | 129.34427 |
| APEVILGMGYK | 11 | P45984 | Mitogen-activated protein kinase 9 OS=H | MAPK9 | 48.14 | 2 | 43.35043 | no | 8.426E-07 | 589.31793 | 99.895409 |
| LDLLGNLPGSK | 11 | P46013 | Proliferation marker protein Ki-67 OS=H | MKI67 | 358.694 | 2 | 58.091614 | yes | 1.476E-07 | 563.82697 | 902.24957 |
| AQALEDLAGFK | 11 | P46013 | Proliferation marker protein Ki-67 OS=H | MKI67 | 358.694 | 2 | 58.091614 | yes | 1.455E-06 | 581.80878 | 664.96875 |
| LDLAGTLPGSK | 11 | P46013 | Proliferation marker protein Ki-67 OS=H | MKI67 | 358.694 | 2 | 58.091614 | yes | 1.656E-07 | 536.30585 | 620.78802 |
| LDLPGNLPGSK | 11 | P46013 | Proliferation marker protein Ki-67 OS=H | MKI67 | 358.694 | 2 | 58.091614 | yes | 1.488E-07 | 555.81134 | 437.61945 |
| AQSLVISPPAPSPR | 14 | P46013 | Proliferation marker protein Ki-67 OS=H | MKI67 | 358.694 | 2 | 58.091614 | yes | 1.05E-07 | 710.40137 | 424.78833 |
| EEAQSLEDLAGFK | 13 | P46013 | Proliferation marker protein Ki-67 OS=H | MKI67 | 358.694 | 2 | 58.091614 | yes | 9.722E-08 | 718.84882 | 417.77634 |
| LDLTENLTGSK | 11 | P46013 | Proliferation marker protein Ki-67 OS=H | MKI67 | 358.694 | 2 | 58.091614 | yes | 1.326E-07 | 595.81677 | 344.35931 |

| | | | | | | | | | | | |
|---|---|---|---|---|---|---|---|---|---|---|---|
| SGASEANLIVAK | 12 | P46013 | Proliferation marker protein Ki-67 OS=H | MKI67 | 358.694 | 2 | 58.091614 | yes | 1.175E-07 | 580.31952 | 311.10602 |
| CAPMSDLTDLK | 11 | P46013 | Proliferation marker protein Ki-67 OS=H | MKI67 | 358.694 | 2 | 58.091614 | yes | 2.978E-07 | 625.79138 | 289.49518 |
| GQNLLQTQDHAK | 12 | P46013 | Proliferation marker protein Ki-67 OS=H | MKI67 | 358.694 | 2 | 58.091614 | yes | 9.722E-08 | 676.84949 | 287.19644 |
| HGDVITIIDR | 10 | P46013 | Proliferation marker protein Ki-67 OS=H | MKI67 | 358.694 | 2 | 58.091614 | yes | 9.722E-08 | 569.81439 | 284.18231 |
| LDQPGNLPGSNR | 12 | P46013 | Proliferation marker protein Ki-67 OS=H | MKI67 | 358.694 | 2 | 58.091614 | yes | 1.203E-07 | 634.32312 | 282.15454 |
| AQPLEDLAGLK | 11 | P46013 | Proliferation marker protein Ki-67 OS=H | MKI67 | 358.694 | 2 | 58.091614 | yes | 9.722E-08 | 577.8244 | 277.20563 |
| SEETNTEIVECILK | 14 | P46013 | Proliferation marker protein Ki-67 OS=H | MKI67 | 358.694 | 2 | 58.091614 | yes | 9.722E-08 | 832.90601 | 258.3367 |
| AFMGTPVQK | 9 | P46013 | Proliferation marker protein Ki-67 OS=H | MKI67 | 358.694 | 2 | 58.091614 | yes | 2.624E-06 | 489.75751 | 252.72641 |
| DINTFVGTPVEK | 12 | P46013 | Proliferation marker protein Ki-67 OS=H | MKI67 | 358.694 | 2 | 58.091614 | yes | 1.493E-07 | 660.3457 | 248.61563 |
| VEPVGDVVSTR | 11 | P46013 | Proliferation marker protein Ki-67 OS=H | MKI67 | 358.694 | 2 | 58.091614 | yes | 1.992E-07 | 579.31171 | 233.49435 |
| LTQTSGETTHTHTI | 20 | P46013 | Proliferation marker protein Ki-67 OS=H | MKI67 | 358.694 | 3 | 58.091614 | yes | 1.819E-07 | 699.99432 | 230.71609 |
| NINTFVETPVQK | 12 | P46013 | Proliferation marker protein Ki-67 OS=H | MKI67 | 358.694 | 2 | 58.091614 | yes | 9.722E-08 | 695.37225 | 223.69156 |
| QILDPAASVTGSR | 13 | P46013 | Proliferation marker protein Ki-67 OS=H | MKI67 | 358.694 | 2 | 58.091614 | yes | 4.759E-07 | 657.85425 | 220.49115 |
| ADVEGELLACR | 11 | P46013 | Proliferation marker protein Ki-67 OS=H | MKI67 | 358.694 | 2 | 58.091614 | yes | 4.013E-07 | 616.8006 | 174.36922 |
| QTPAPAASVTGSR | 13 | P46013 | Proliferation marker protein Ki-67 OS=H | MKI67 | 358.694 | 2 | 58.091614 | yes | 9.81E-08 | 621.8255 | 173.06784 |
| EDLSGIAEMFK | 11 | P46013 | Proliferation marker protein Ki-67 OS=H | MKI67 | 358.694 | 2 | 58.091614 | yes | 9.722E-08 | 620.29993 | 169.89005 |
| DIIIFVGTPVQK | 12 | P46013 | Proliferation marker protein Ki-67 OS=H | MKI67 | 358.694 | 2 | 58.091614 | yes | 1.161E-07 | 665.39246 | 163.53476 |
| SVPTTQCLDNSK | 12 | P46013 | Proliferation marker protein Ki-67 OS=H | MKI67 | 358.694 | 2 | 58.091614 | yes | 9.722E-08 | 675.3219 | 147.82999 |
| KADVEEEFLALR | 12 | P46013 | Proliferation marker protein Ki-67 OS=H | MKI67 | 358.694 | 2 | 58.091614 | yes | 1.139E-07 | 710.37756 | 123.80169 |
| SPQSDPADTPTNTI | 14 | P46013 | Proliferation marker protein Ki-67 OS=H | MKI67 | 358.694 | 2 | 58.091614 | yes | 1.225E-07 | 729.83899 | 119.27241 |
| DINTFLGTPVQK | 12 | P46013 | Proliferation marker protein Ki-67 OS=H | MKI67 | 358.694 | 2 | 58.091614 | yes | 9.728E-08 | 666.86151 | 112.70989 |
| QILDSAASLTGSK | 13 | P46013 | Proliferation marker protein Ki-67 OS=H | MKI67 | 358.694 | 2 | 58.091614 | yes | 9.722E-08 | 645.84863 | 111.26418 |
| VLNNFISNQK | 10 | P46013 | Proliferation marker protein Ki-67 OS=H | MKI67 | 358.694 | 2 | 58.091614 | yes | 2.396E-06 | 588.8222 | 99.813156 |
| SSPELEDTATSSK | 13 | P46013 | Proliferation marker protein Ki-67 OS=H | MKI67 | 358.694 | 2 | 58.091614 | yes | 1.714E-07 | 676.31482 | 98.5625 |
| SLVMHTPPVLK | 11 | P46013 | Proliferation marker protein Ki-67 OS=H | MKI67 | 358.694 | 2 | 58.091614 | yes | 2.252E-07 | 611.35486 | 98.28138 |
| SPQPDPVDTPASTI | 14 | P46013 | Proliferation marker protein Ki-67 OS=H | MKI67 | 358.694 | 2 | 58.091614 | yes | 1.584E-07 | 720.35425 | 93.271011 |
| EKAEALEDLVGFK | 13 | P46013 | Proliferation marker protein Ki-67 OS=H | MKI67 | 358.694 | 2 | 58.091614 | yes | 1.624E-07 | 724.88519 | 92.035759 |
| SPPPESVDTPTSTK | 14 | P46013 | Proliferation marker protein Ki-67 OS=H | MKI67 | 358.694 | 2 | 58.091614 | yes | 8.938E-07 | 721.85413 | 82.475807 |
| LTQTSGETTHTDK | 20 | P46013 | Proliferation marker protein Ki-67 OS=H | MKI67 | 358.694 | 3 | 58.091614 | yes | 1.723E-07 | 705.67822 | 78.668938 |
| QESGSEIHVEVK | 12 | P46013 | Proliferation marker protein Ki-67 OS=H | MKI67 | 358.694 | 2 | 58.091614 | yes | 1.573E-07 | 671.33588 | 77.599564 |
| IEPAEELNSNDMK | 13 | P46013 | Proliferation marker protein Ki-67 OS=H | MKI67 | 358.694 | 2 | 58.091614 | yes | 3.183E-06 | 745.34558 | 70.882401 |
| EELLAVGK | 8 | P46013 | Proliferation marker protein Ki-67 OS=H | MKI67 | 358.694 | 2 | 58.091614 | yes | 2.232E-07 | 429.75018 | 70.042786 |
| SQHDILQMICSK | 12 | P46013 | Proliferation marker protein Ki-67 OS=H | MKI67 | 358.694 | 2 | 58.091614 | yes | 1.398E-07 | 730.35522 | 64.267029 |
| EEVLPVGK | 8 | P46013 | Proliferation marker protein Ki-67 OS=H | MKI67 | 358.694 | 2 | 58.091614 | yes | 2.922E-06 | 435.75018 | 64.255844 |
| KVDVEEEFFALR | 12 | P46013 | Proliferation marker protein Ki-67 OS=H | MKI67 | 358.694 | 2 | 58.091614 | yes | 1.083E-07 | 741.38538 | 63.556107 |
| ADVEEEFLAFR | 11 | P46013 | Proliferation marker protein Ki-67 OS=H | MKI67 | 358.694 | 2 | 58.091614 | yes | 2.322E-07 | 663.32227 | 60.848991 |
| TSPEMDIQNPDDG | 15 | P46013 | Proliferation marker protein Ki-67 OS=H | MKI67 | 358.694 | 2 | 58.091614 | yes | 3.907E-07 | 823.35974 | 60.421505 |
| VSGNPQVHIK | 10 | P46013 | Proliferation marker protein Ki-67 OS=H | MKI67 | 358.694 | 2 | 58.091614 | yes | 1.401E-07 | 539.80383 | 59.985302 |
| DIVEELSALK | 10 | P46013 | Proliferation marker protein Ki-67 OS=H | MKI67 | 358.694 | 2 | 58.091614 | yes | 9.722E-08 | 558.81097 | 54.798351 |

| | | | | | | | | | | | |
|---|---|---|---|---|---|---|---|---|---|---|---|
| SPPPESMDTPTSTF | 14 | P46013 | Proliferation marker protein Ki-67 OS=H | MKI67 | 358.694 | 2 | 58.091614 | yes | 2.137E-07 | 751.8432 | 54.774765 |
| KADTEEEFLAFR | 12 | P46013 | Proliferation marker protein Ki-67 OS=H | MKI67 | 358.694 | 2 | 58.091614 | yes | 9.722E-08 | 728.35938 | 54.341228 |
| KPVGEVHSQFSTG | 25 | P46013 | Proliferation marker protein Ki-67 OS=H | MKI67 | 358.694 | 4 | 58.091614 | yes | 3.344E-07 | 666.84454 | 54.095379 |
| GEAGNSDSMCLR | 12 | P46013 | Proliferation marker protein Ki-67 OS=H | MKI67 | 358.694 | 2 | 58.091614 | yes | 9.722E-08 | 648.76898 | 53.037762 |
| SSEPVVIMK | 9 | P46013 | Proliferation marker protein Ki-67 OS=H | MKI67 | 358.694 | 2 | 58.091614 | yes | 1.312E-05 | 495.27045 | 52.473141 |
| ADVEEEFLALR | 11 | P46013 | Proliferation marker protein Ki-67 OS=H | MKI67 | 358.694 | 2 | 58.091614 | yes | 9.722E-08 | 646.33008 | 50.653385 |
| KADVEEESLALR | 12 | P46013 | Proliferation marker protein Ki-67 OS=H | MKI67 | 358.694 | 2 | 58.091614 | yes | 2.126E-07 | 680.35938 | 41.475716 |
| EIERPFETYK | 10 | P46013 | Proliferation marker protein Ki-67 OS=H | MKI67 | 358.694 | 2 | 58.091614 | yes | 9.722E-08 | 656.33258 | 40.915878 |
| AMDTPKPAGGDEI | 13 | P46013 | Proliferation marker protein Ki-67 OS=H | MKI67 | 358.694 | 2 | 58.091614 | yes | 1.95E-06 | 658.81116 | 32.530865 |
| SPPPELTDTATSTK | 14 | P46013 | Proliferation marker protein Ki-67 OS=H | MKI67 | 358.694 | 2 | 58.091614 | yes | 6.2E-05 | 722.86194 | 32.244041 |
| ASQPDLVDTPTSSI | 18 | P46013 | Proliferation marker protein Ki-67 OS=H | MKI67 | 358.694 | 3 | 58.091614 | yes | 2.903E-06 | 632.6618 | 26.354933 |
| QMLDPANYGTGM | 14 | P46013 | Proliferation marker protein Ki-67 OS=H | MKI67 | 358.694 | 2 | 58.091614 | yes | 3.012E-06 | 791.85303 | 23.394979 |
| EALEGILIR | 9 | P46019 | Phosphorylase b kinase regulatory subuni | PHKA2 | 138.409 | 2 | 47.741123 | yes | 1.268E-07 | 507.30313 | 55.471851 |
| LEGNTVGVEAAR | 12 | P46060 | Ran GTPase-activating protein 1 OS=Hor | RANGAP1 | 63.542 | 2 | 52.86459 | yes | 8.528E-07 | 608.32001 | 265.54056 |
| TQVAGGQLSFK | 11 | P46060 | Ran GTPase-activating protein 1 OS=Hor | RANGAP1 | 63.542 | 2 | 52.86459 | yes | 3.582E-07 | 568.30896 | 140.39809 |
| GAVAIADAIR | 10 | P46060 | Ran GTPase-activating protein 1 OS=Hor | RANGAP1 | 63.542 | 2 | 52.86459 | yes | 1.028E-07 | 478.77982 | 131.40779 |
| ILAAALTECHR | 11 | P46060 | Ran GTPase-activating protein 1 OS=Hor | RANGAP1 | 63.542 | 2 | 52.86459 | yes | 1.039E-07 | 627.83478 | 126.14585 |
| QVEVINFGDCLVR | 13 | P46060 | Ran GTPase-activating protein 1 OS=Hor | RANGAP1 | 63.542 | 2 | 52.86459 | yes | 2.82E-07 | 774.89557 | 123.59238 |
| VINLNDNTFTEK | 12 | P46060 | Ran GTPase-activating protein 1 OS=Hor | RANGAP1 | 63.542 | 2 | 52.86459 | yes | 9.722E-08 | 704.35938 | 120.75417 |
| ELNLSFCEIK | 10 | P46060 | Ran GTPase-activating protein 1 OS=Hor | RANGAP1 | 63.542 | 2 | 52.86459 | yes | 1.726E-07 | 626.81573 | 116.81449 |
| EIEDFDSLEALR | 12 | P46060 | Ran GTPase-activating protein 1 OS=Hor | RANGAP1 | 63.542 | 2 | 52.86459 | yes | 1.106E-07 | 718.84882 | 108.26741 |
| LENDGATALAEAF | 14 | P46060 | Ran GTPase-activating protein 1 OS=Hor | RANGAP1 | 63.542 | 2 | 52.86459 | yes | 9.728E-08 | 739.36774 | 106.64185 |
| MAVQDAVDALM( | 13 | P46060 | Ran GTPase-activating protein 1 OS=Hor | RANGAP1 | 63.542 | 2 | 52.86459 | yes | 9.722E-08 | 710.35217 | 98.46933 |
| HSLLQTLYK | 9 | P46060 | Ran GTPase-activating protein 1 OS=Hor | RANGAP1 | 63.542 | 2 | 52.86459 | yes | 9.81E-08 | 551.81641 | 38.298531 |
| LLVHMGLLK | 9 | P46060 | Ran GTPase-activating protein 1 OS=Hor | RANGAP1 | 63.542 | 2 | 52.86459 | yes | 4.248E-06 | 512.32281 | 18.927559 |
| VAGVVAPTLPR | 11 | P46063 | ATP-dependent DNA helicase Q1 OS=H | RECQL | 73.458 | 2 | 54.929905 | yes | 2.123E-07 | 540.33221 | 2516.3372 |
| ISSMVVMENVGQ( | 14 | P46063 | ATP-dependent DNA helicase Q1 OS=H | RECQL | 73.458 | 2 | 54.929905 | yes | 9.826E-08 | 775.38928 | 1262.4025 |
| LIYVTPEK | 8 | P46063 | ATP-dependent DNA helicase Q1 OS=H | RECQL | 73.458 | 2 | 54.929905 | yes | 1.828E-07 | 481.78149 | 1144.929 |
| SMENYYQESGR | 11 | P46063 | ATP-dependent DNA helicase Q1 OS=H | RECQL | 73.458 | 2 | 54.929905 | yes | 1.009E-07 | 682.28278 | 901.02246 |
| GQSGIIYCFSQK | 12 | P46063 | ATP-dependent DNA helicase Q1 OS=H | RECQL | 73.458 | 2 | 54.929905 | yes | 9.722E-08 | 694.33734 | 878.65576 |
| WVHAEMVNK | 9 | P46063 | ATP-dependent DNA helicase Q1 OS=H | RECQL | 73.458 | 2 | 54.929905 | yes | 1.063E-07 | 557.27911 | 871.16107 |
| LYEMVSYCQNISK | 13 | P46063 | ATP-dependent DNA helicase Q1 OS=H | RECQL | 73.458 | 2 | 54.929905 | yes | 1.106E-07 | 817.88129 | 621.92102 |
| QLGISATMLNASS | 15 | P46063 | ATP-dependent DNA helicase Q1 OS=H | RECQL | 73.458 | 2 | 54.929905 | yes | 9.809E-08 | 754.39288 | 570.19531 |
| EVFLVMPTGGGK | 12 | P46063 | ATP-dependent DNA helicase Q1 OS=H | RECQL | 73.458 | 2 | 54.929905 | yes | 5.352E-07 | 617.82867 | 438.52371 |
| VKDILQNVFK | 10 | P46063 | ATP-dependent DNA helicase Q1 OS=H | RECQL | 73.458 | 2 | 54.929905 | yes | 9.741E-08 | 602.35846 | 334.55814 |
| FVIHHSMSK | 9 | P46063 | ATP-dependent DNA helicase Q1 OS=H | RECQL | 73.458 | 2 | 54.929905 | yes | 1.608E-07 | 543.28168 | 241.17505 |
| QQELIQK | 7 | P46063 | ATP-dependent DNA helicase Q1 OS=H | RECQL | 73.458 | 2 | 54.929905 | yes | 3.217E-06 | 443.75327 | 188.36824 |
| AANMLQQSGSK | 11 | P46063 | ATP-dependent DNA helicase Q1 OS=H | RECQL | 73.458 | 2 | 54.929905 | yes | 1.732E-07 | 567.78223 | 166.06299 |
| FRPLQLETINVTM. | 16 | P46063 | ATP-dependent DNA helicase Q1 OS=H | RECQL | 73.458 | 3 | 54.929905 | yes | 9.722E-08 | 606.66962 | 127.79373 |

| | | | | | | | | | | | |
|---|---|---|---|---|---|---|---|---|---|---|---|
| QFPNASLIGLTATA | 24 | P46063 | ATP-dependent DNA helicase Q1 OS=Ho | RECQL | 73.458 | 3 | 54.929905 | yes | 9.79E-08 | 837.77826 | 87.269196 |
| PNLYYEVR | 8 | P46063 | ATP-dependent DNA helicase Q1 OS=Ho | RECQL | 73.458 | 2 | 54.929905 | yes | 3.051E-06 | 527.27185 | 81.685593 |
| KNITEYCR | 8 | P46063 | ATP-dependent DNA helicase Q1 OS=Ho | RECQL | 73.458 | 2 | 54.929905 | yes | 3.917E-07 | 542.26624 | 63.634216 |
| IIAHFLIQQYLK | 12 | P46063 | ATP-dependent DNA helicase Q1 OS=Ho | RECQL | 73.458 | 2 | 54.929905 | yes | 1.537E-07 | 743.94269 | 56.31493 |
| LVPTGLDFGQEGF | 15 | P46087 | Probable 28S rRNA (cytosine(4447)-C(5) | NOP2 | 89.302 | 2 | 52.263023 | yes | 1.525E-07 | 818.9201 | 1515.2473 |
| ELLLSAIDSVNATS | 15 | P46087 | Probable 28S rRNA (cytosine(4447)-C(5) | NOP2 | 89.302 | 2 | 52.263023 | yes | 1.006E-07 | 780.92761 | 1213.45 |
| VLLDAPCSGTGVIS | 15 | P46087 | Probable 28S rRNA (cytosine(4447)-C(5) | NOP2 | 89.302 | 2 | 52.263023 | yes | 1.103E-07 | 758.90564 | 990.11945 |
| GADSELSTVPSVTI | 14 | P46087 | Probable 28S rRNA (cytosine(4447)-C(5) | NOP2 | 89.302 | 2 | 52.263023 | yes | 9.819E-08 | 695.85663 | 879.00757 |
| IQDIVGILR | 9 | P46087 | Probable 28S rRNA (cytosine(4447)-C(5) | NOP2 | 89.302 | 2 | 52.263023 | yes | 9.722E-08 | 513.81897 | 849.67285 |
| DLAQALINR | 9 | P46087 | Probable 28S rRNA (cytosine(4447)-C(5) | NOP2 | 89.302 | 2 | 52.263023 | yes | 1.464E-07 | 507.29056 | 843.98273 |
| KGPQSLFNAPR | 11 | P46087 | Probable 28S rRNA (cytosine(4447)-C(5) | NOP2 | 89.302 | 2 | 52.263023 | yes | 6.983E-07 | 607.83563 | 831.13672 |
| FLPAVSDENSK | 11 | P46087 | Probable 28S rRNA (cytosine(4447)-C(5) | NOP2 | 89.302 | 2 | 52.263023 | yes | 9.722E-08 | 603.80371 | 568.63568 |
| QQLPEQPFEK | 10 | P46087 | Probable 28S rRNA (cytosine(4447)-C(5) | NOP2 | 89.302 | 2 | 52.263023 | yes | 9.722E-08 | 622.31952 | 479.67795 |
| LGVTNTIISHYDGF | 14 | P46087 | Probable 28S rRNA (cytosine(4447)-C(5) | NOP2 | 89.302 | 3 | 52.263023 | yes | 2.38E-07 | 515.93884 | 220.92064 |
| LGVTNTIISHYDGF | 14 | P46087 | Probable 28S rRNA (cytosine(4447)-C(5) | NOP2 | 89.302 | 3 | 52.263023 | yes | 9.728E-08 | 773.4046 | 220.92064 |
| LGVTNTIISHYDGF | 14 | P46087 | Probable 28S rRNA (cytosine(4447)-C(5) | NOP2 | 89.302 | 2 | 52.263023 | yes | 2.38E-07 | 515.93884 | 220.92064 |
| LGVTNTIISHYDGF | 14 | P46087 | Probable 28S rRNA (cytosine(4447)-C(5) | NOP2 | 89.302 | 2 | 52.263023 | yes | 9.728E-08 | 773.4046 | 220.92064 |
| GISAGAVQTAGK | 12 | P46087 | Probable 28S rRNA (cytosine(4447)-C(5) | NOP2 | 89.302 | 2 | 52.263023 | yes | 4.855E-07 | 530.29327 | 417.1683 |
| SVVGNLHR | 8 | P46087 | Probable 28S rRNA (cytosine(4447)-C(5) | NOP2 | 89.302 | 2 | 52.263023 | yes | 9.722E-08 | 441.25122 | 405.87048 |
| NTGVILANDANAE | 14 | P46087 | Probable 28S rRNA (cytosine(4447)-C(5) | NOP2 | 89.302 | 2 | 52.263023 | yes | 3.647E-07 | 729.37079 | 403.22363 |
| ILDMCCAPGGK | 11 | P46087 | Probable 28S rRNA (cytosine(4447)-C(5) | NOP2 | 89.302 | 2 | 52.263023 | yes | 9.882E-08 | 611.27484 | 377.37137 |
| GPQPPTVSPIR | 11 | P46087 | Probable 28S rRNA (cytosine(4447)-C(5) | NOP2 | 89.302 | 2 | 52.263023 | yes | 2.097E-07 | 574.82477 | 229.6671 |
| GTDTQTPAVLSPSI | 14 | P46087 | Probable 28S rRNA (cytosine(4447)-C(5) | NOP2 | 89.302 | 2 | 52.263023 | yes | 2.185E-07 | 701.36462 | 174.88091 |
| GPQSLFNAPR | 10 | P46087 | Probable 28S rRNA (cytosine(4447)-C(5) | NOP2 | 89.302 | 2 | 52.263023 | yes | 3.28E-07 | 543.78815 | 123.51529 |
| TSYMAQLMK | 9 | P46087 | Probable 28S rRNA (cytosine(4447)-C(5) | NOP2 | 89.302 | 2 | 52.263023 | yes | 1.709E-07 | 536.7619 | 115.25151 |
| GVNLDPLGK | 9 | P46087 | Probable 28S rRNA (cytosine(4447)-C(5) | NOP2 | 89.302 | 2 | 52.263023 | yes | 8.774E-07 | 456.76108 | 108.53377 |
| RLGSVEAPK | 9 | P46087 | Probable 28S rRNA (cytosine(4447)-C(5) | NOP2 | 89.302 | 2 | 52.263023 | yes | 1.695E-07 | 478.77982 | 73.396759 |
| TALALEVGDIVK | 12 | P46109 | Crk-like protein OS=Homo sapiens OX=9 | CRKL | 33.777 | 2 | 52.15059 | yes | 9.722E-08 | 614.86102 | 197.53952 |
| LVEDLQEELSSR | 12 | P46199 | Translation initiation factor IF-2, mitocho | MTIF2 | 81.318 | 2 | 53.968254 | yes | 9.722E-08 | 709.36212 | 64.542175 |
| GLVTTAIIQR | 10 | P46199 | Translation initiation factor IF-2, mitocho | MTIF2 | 81.318 | 2 | 53.968254 | yes | 1.585E-06 | 536.32965 | 10.579438 |
| LINLVGESLR | 10 | P46379 | Large proline-rich protein BAG6 OS=Hor | BAG6 | 119.409 | 2 | 46.808475 | yes | 1.465E-07 | 557.33496 | 129.29494 |
| SQLSCVVVDDIER | 13 | P46459 | Vesicle-fusing ATPase OS=Homo sapiens | NSF | 82.595 | 2 | 53.15439 | yes | 9.722E-08 | 760.37469 | 36.333908 |
| DVKPSNVLINK | 11 | P46734 | Dual specificity mitogen-activated protein | MAP2K3 | 39.319 | 2 | 50.602467 | yes | 9.748E-08 | 613.85876 | 269.15634 |
| QVVEEPSPQLPAD | 14 | P46734 | Dual specificity mitogen-activated protein | MAP2K3 | 39.319 | 2 | 50.602467 | yes | 1.138E-07 | 782.90192 | 241.41548 |
| HAQSGTIMAVK | 11 | P46734 | Dual specificity mitogen-activated protein | MAP2K3 | 39.319 | 2 | 50.602467 | yes | 1.272E-07 | 571.80298 | 223.13539 |
| TFITIGDR | 8 | P46734 | Dual specificity mitogen-activated protein | MAP2K3 | 39.319 | 2 | 50.602467 | yes | 6.511E-07 | 461.75327 | 185.28152 |
| KTDIAAFVK | 9 | P46734 | Dual specificity mitogen-activated protein | MAP2K3 | 39.319 | 2 | 50.602467 | yes | 2.162E-07 | 496.79239 | 112.16368 |
| ISCMSKPPAPNPTF | 16 | P46734 | Dual specificity mitogen-activated protein | MAP2K3 | 39.319 | 3 | 50.602467 | yes | 1.34E-07 | 583.96399 | 111.61534 |
| LLMDLDINMR | 10 | P46734 | Dual specificity mitogen-activated protein | MAP2K3 | 39.319 | 2 | 50.602467 | yes | 4.515E-07 | 617.32013 | 100.18607 |

| | | | | | | | | | | | |
|---|---|---|---|---|---|---|---|---|---|---|---|
| VCLESAVELPK | 11 | P46736 | Lys-63-specific deubiquitinase BRCC36 ( | BRCC3 | 36.072 | 2 | 53.87677 | yes | 9.748E-08 | 622.83136 | 666.5564 |
| ILCQEEQDAYR | 11 | P46736 | Lys-63-specific deubiquitinase BRCC36 ( | BRCC3 | 36.072 | 2 | 53.87677 | yes | 9.722E-08 | 712.82739 | 360.32446 |
| FAYTGTEMR | 9 | P46736 | Lys-63-specific deubiquitinase BRCC36 ( | BRCC3 | 36.072 | 2 | 53.87677 | yes | 1.028E-07 | 538.2475 | 275.53708 |
| VLYTCFQSIQAQK | 13 | P46736 | Lys-63-specific deubiquitinase BRCC36 ( | BRCC3 | 36.072 | 2 | 53.87677 | yes | 9.722E-08 | 793.40576 | 251.42868 |
| IVHIHSVIILR | 11 | P46736 | Lys-63-specific deubiquitinase BRCC36 ( | BRCC3 | 36.072 | 2 | 53.87677 | yes | 9.722E-08 | 650.41663 | 240.28693 |
| IHSLTHLDSVTK | 12 | P46736 | Lys-63-specific deubiquitinase BRCC36 ( | BRCC3 | 36.072 | 2 | 53.87677 | yes | 9.722E-08 | 675.87244 | 106.33249 |
| DRVEISPEQLSAAS | 19 | P46736 | Lys-63-specific deubiquitinase BRCC36 ( | BRCC3 | 36.072 | 3 | 53.87677 | yes | 1.434E-07 | 696.67828 | 77.753914 |
| IEIPIHIVPHVTIGK | 15 | P46736 | Lys-63-specific deubiquitinase BRCC36 ( | BRCC3 | 36.072 | 2 | 53.87677 | yes | 2.493E-07 | 833.50616 | 33.820713 |
| VVGWYHSHPHIT\ | 21 | P46736 | Lys-63-specific deubiquitinase BRCC36 ( | BRCC3 | 36.072 | 4 | 53.87677 | yes | 3.182E-07 | 627.82257 | 22.035135 |
| LWTLVSEQTR | 10 | P46776 | Large ribosomal subunit protein uL15 OS | RPL27A | 16.562 | 2 | 50.30582 | yes | 1.04E-07 | 616.83533 | 61072.133 |
| TGAAPIIDVVR | 11 | P46776 | Large ribosomal subunit protein uL15 OS | RPL27A | 16.562 | 2 | 50.30582 | yes | 1.238E-07 | 556.32715 | 41318.098 |
| NQSFCPTVNLDK | 12 | P46776 | Large ribosomal subunit protein uL15 OS | RPL27A | 16.562 | 2 | 50.30582 | yes | 1.501E-07 | 711.83771 | 28838.156 |
| YHPGYFGK | 8 | P46776 | Large ribosomal subunit protein uL15 OS | RPL27A | 16.562 | 2 | 50.30582 | yes | 1.403E-07 | 484.73486 | 15098.282 |
| RNQSFCPTVNLDK | 13 | P46776 | Large ribosomal subunit protein uL15 OS | RPL27A | 16.562 | 2 | 50.30582 | yes | 9.819E-08 | 789.88831 | 339.91061 |
| NSVTPDMMEEMY | 13 | P46777 | Large ribosomal subunit protein uL18 OS | RPL5 | 34.363 | 2 | 53.316948 | yes | 2.667E-07 | 787.83038 | 6185.6646 |
| NSVTPDMMEEMY | 13 | P46777 | Large ribosomal subunit protein uL18 OS | RPL5 | 34.363 | 2 | 53.316948 | yes | 2.091E-07 | 795.82788 | 6185.6646 |
| EFNAEVHR | 8 | P46777 | Large ribosomal subunit protein uL18 OS | RPL5 | 34.363 | 2 | 53.316948 | yes | 1.388E-07 | 501.24359 | 9364.543 |
| HIMGQNVADYMR | 12 | P46777 | Large ribosomal subunit protein uL18 OS | RPL5 | 34.363 | 2 | 53.316948 | yes | 3.991E-07 | 717.83447 | 3971.8953 |
| HIMGQNVADYMR | 12 | P46777 | Large ribosomal subunit protein uL18 OS | RPL5 | 34.363 | 2 | 53.316948 | yes | 9.722E-08 | 725.83191 | 3971.8953 |
| YLMEEDEDAYK | 11 | P46777 | Large ribosomal subunit protein uL18 OS | RPL5 | 34.363 | 2 | 53.316948 | yes | 1.395E-07 | 703.29504 | 2779.9431 |
| YLMEEDEDAYK | 11 | P46777 | Large ribosomal subunit protein uL18 OS | RPL5 | 34.363 | 2 | 53.316948 | yes | 9.722E-08 | 711.29248 | 2779.9431 |
| YLMEEDEDAYKK | 12 | P46777 | Large ribosomal subunit protein uL18 OS | RPL5 | 34.363 | 2 | 53.316948 | yes | 1.006E-07 | 767.34253 | 2435.9617 |
| YLMEEDEDAYKK | 12 | P46777 | Large ribosomal subunit protein uL18 OS | RPL5 | 34.363 | 2 | 53.316948 | yes | 2.329E-07 | 775.33997 | 2435.9617 |
| IEGDMIVCAAYAH | 17 | P46777 | Large ribosomal subunit protein uL18 OS | RPL5 | 34.363 | 3 | 53.316948 | yes | 1.082E-07 | 639.64636 | 1340.2233 |
| IEGDMIVCAAYAH | 17 | P46777 | Large ribosomal subunit protein uL18 OS | RPL5 | 34.363 | 3 | 53.316948 | yes | 2.253E-06 | 644.97797 | 1340.2233 |
| RFPGYDSESK | 10 | P46777 | Large ribosomal subunit protein uL18 OS | RPL5 | 34.363 | 2 | 53.316948 | yes | 1.778E-07 | 593.2804 | 2502.9761 |
| QFSQYIK | 7 | P46777 | Large ribosomal subunit protein uL18 OS | RPL5 | 34.363 | 2 | 53.316948 | yes | 2.85E-07 | 457.24252 | 1841.5342 |
| DIICQIAYAR | 10 | P46777 | Large ribosomal subunit protein uL18 OS | RPL5 | 34.363 | 2 | 53.316948 | yes | 1.318E-07 | 611.8161 | 1760.5345 |
| ENPVYEK | 7 | P46777 | Large ribosomal subunit protein uL18 OS | RPL5 | 34.363 | 2 | 53.316948 | yes | 2.685E-06 | 439.71634 | 1447.6931 |
| GAVDGGLSIPHSTI | 14 | P46777 | Large ribosomal subunit protein uL18 OS | RPL5 | 34.363 | 2 | 53.316948 | yes | 1.414E-07 | 669.85425 | 1238.5841 |
| FPGYDSESK | 9 | P46777 | Large ribosomal subunit protein uL18 OS | RPL5 | 34.363 | 2 | 53.316948 | yes | 6.519E-07 | 515.2298 | 574.2348 |
| VGLTNYAAAYCT( | 18 | P46777 | Large ribosomal subunit protein uL18 OS | RPL5 | 34.363 | 3 | 53.316948 | yes | 9.722E-08 | 643.00836 | 526.79132 |
| RLVIQDK | 7 | P46777 | Large ribosomal subunit protein uL18 OS | RPL5 | 34.363 | 2 | 53.316948 | yes | 1.937E-07 | 436.27164 | 416.0748 |
| KQFSQYIK | 8 | P46777 | Large ribosomal subunit protein uL18 OS | RPL5 | 34.363 | 2 | 53.316948 | yes | 1.585E-07 | 521.29004 | 144.70189 |
| KHIMGQNVADYM | 13 | P46777 | Large ribosomal subunit protein uL18 OS | RPL5 | 34.363 | 2 | 53.316948 | yes | 2.418E-06 | 781.88196 | 34.133987 |
| HGVVPLATYMR | 11 | P46778 | Large ribosomal subunit protein eL21 OS | RPL21 | 18.565 | 2 | 49.361473 | yes | 1.291E-07 | 622.33441 | 7784.792 |
| HGVVPLATYMR | 11 | P46778 | Large ribosomal subunit protein eL21 OS | RPL21 | 18.565 | 2 | 49.361473 | yes | 1.14E-07 | 630.33191 | 7784.792 |
| KGDIVDIK | 8 | P46778 | Large ribosomal subunit protein eL21 OS | RPL21 | 18.565 | 2 | 49.361473 | yes | 1.175E-07 | 444.26346 | 8918.6387 |
| VYNVTQHAVGIV\ | 15 | P46778 | Large ribosomal subunit protein eL21 OS | RPL21 | 18.565 | 2 | 49.361473 | yes | 1.219E-07 | 820.95959 | 2865.8989 |

| | | | | | | | | | | |
|---|---|---|---|---|---|---|---|---|---|---|
| VYNVTQHAVGIV\ | 15 | P46778 | Large ribosomal subunit protein eL21 OS RPL21 | 18.565 | 2 | 49.361473 | yes | 2.679E-07 | 547.64215 | 2865.8989 |
| VYNVTQHAVGIV\ | 15 | P46778 | Large ribosomal subunit protein eL21 OS RPL21 | 18.565 | 3 | 49.361473 | yes | 1.219E-07 | 820.95959 | 2865.8989 |
| VYNVTQHAVGIV\ | 15 | P46778 | Large ribosomal subunit protein eL21 OS RPL21 | 18.565 | 3 | 49.361473 | yes | 2.679E-07 | 547.64215 | 2865.8989 |
| EKGTWVQLK | 9 | P46778 | Large ribosomal subunit protein eL21 OS RPL21 | 18.565 | 2 | 49.361473 | yes | 1.104E-07 | 544.80859 | 1775.6224 |
| KHGVVPLATYMR | 12 | P46778 | Large ribosomal subunit protein eL21 OS RPL21 | 18.565 | 2 | 49.361473 | yes | 2.624E-07 | 686.3819 | 374.52295 |
| KHGVVPLATYMR | 12 | P46778 | Large ribosomal subunit protein eL21 OS RPL21 | 18.565 | 2 | 49.361473 | yes | 2.294E-07 | 457.92371 | 374.52295 |
| KHGVVPLATYMR | 12 | P46778 | Large ribosomal subunit protein eL21 OS RPL21 | 18.565 | 3 | 49.361473 | yes | 2.624E-07 | 686.3819 | 374.52295 |
| KHGVVPLATYMR | 12 | P46778 | Large ribosomal subunit protein eL21 OS RPL21 | 18.565 | 3 | 49.361473 | yes | 2.294E-07 | 457.92371 | 374.52295 |
| GTWVQLK | 7 | P46778 | Large ribosomal subunit protein eL21 OS RPL21 | 18.565 | 2 | 49.361473 | yes | 1.041E-07 | 416.23981 | 531.62048 |
| SAHLQWMVVR | 10 | P46779 | Large ribosomal subunit protein eL28 OS RPL28 | 15.748 | 2 | 52.785774 | yes | 1.004E-07 | 634.83203 | 7164.8569 |
| SAHLQWMVVR | 10 | P46779 | Large ribosomal subunit protein eL28 OS RPL28 | 15.748 | 2 | 52.785774 | yes | 9.722E-08 | 642.82953 | 7164.8569 |
| SAHLQWMVVR | 10 | P46779 | Large ribosomal subunit protein eL28 OS RPL28 | 15.748 | 2 | 52.785774 | yes | 1.562E-07 | 613.82678 | 7164.8569 |
| KPATSYVR | 8 | P46779 | Large ribosomal subunit protein eL28 OS RPL28 | 15.748 | 2 | 52.785774 | yes | 1.001E-07 | 461.26126 | 20752.373 |
| QTYSTEPNNLK | 11 | P46779 | Large ribosomal subunit protein eL28 OS RPL28 | 15.748 | 2 | 52.785774 | yes | 1.108E-07 | 647.81732 | 9611.457 |
| TVGVEPAADGK | 11 | P46779 | Large ribosomal subunit protein eL28 OS RPL28 | 15.748 | 2 | 52.785774 | yes | 1.637E-07 | 522.27203 | 8956.1377 |
| NCSSFLIK | 8 | P46779 | Large ribosomal subunit protein eL28 OS RPL28 | 15.748 | 2 | 52.785774 | yes | 3.557E-07 | 484.74713 | 6467.6855 |
| KTVGVEPAADGK | 12 | P46779 | Large ribosomal subunit protein eL28 OS RPL28 | 15.748 | 2 | 52.785774 | yes | 1.017E-07 | 586.31952 | 4737.1367 |
| YNGLIHR | 7 | P46779 | Large ribosomal subunit protein eL28 OS RPL28 | 15.748 | 2 | 52.785774 | yes | 1.274E-07 | 436.74051 | 2347.4082 |
| SQKPVMVK | 8 | P46779 | Large ribosomal subunit protein eL28 OS RPL28 | 15.748 | 2 | 52.785774 | yes | 1.572E-07 | 458.76785 | 325.41254 |
| SQKPVMVK | 8 | P46779 | Large ribosomal subunit protein eL28 OS RPL28 | 15.748 | 2 | 52.785774 | yes | 2.03E-07 | 466.76532 | 325.41254 |
| NCSSFLIKR | 9 | P46779 | Large ribosomal subunit protein eL28 OS RPL28 | 15.748 | 2 | 52.785774 | yes | 9.722E-08 | 562.79767 | 85.794769 |
| QVVNIPSFIVR | 11 | P46781 | Small ribosomal subunit protein uS4 OS= RPS9 | 22.592 | 2 | 50.542587 | yes | 1.166E-07 | 636.37714 | 56527.531 |
| LIGEYGLR | 8 | P46781 | Small ribosomal subunit protein uS4 OS= RPS9 | 22.592 | 2 | 50.542587 | yes | 2.748E-07 | 460.76364 | 50840.375 |
| HIDFSLR | 7 | P46781 | Small ribosomal subunit protein uS4 OS= RPS9 | 22.592 | 2 | 50.542587 | yes | 6.926E-07 | 444.24033 | 37576.285 |
| LFEGNALLR | 9 | P46781 | Small ribosomal subunit protein uS4 OS= RPS9 | 22.592 | 2 | 50.542587 | yes | 6.718E-05 | 516.79547 | 34420.145 |
| IEDFLER | 7 | P46781 | Small ribosomal subunit protein uS4 OS= RPS9 | 22.592 | 2 | 50.542587 | yes | 1.582E-07 | 461.23746 | 26107.693 |
| IGVLDEGK | 8 | P46781 | Small ribosomal subunit protein uS4 OS= RPS9 | 22.592 | 2 | 50.542587 | yes | 1.478E-07 | 415.73456 | 23514.693 |
| ELLTLDEK | 8 | P46781 | Small ribosomal subunit protein uS4 OS= RPS9 | 22.592 | 2 | 50.542587 | yes | 1.468E-07 | 480.76605 | 21014.086 |
| LDYILGLK | 8 | P46781 | Small ribosomal subunit protein uS4 OS= RPS9 | 22.592 | 2 | 50.542587 | yes | 1.263E-07 | 467.78403 | 20773.041 |
| LQTQVFK | 7 | P46781 | Small ribosomal subunit protein uS4 OS= RPS9 | 22.592 | 2 | 50.542587 | yes | 1.628E-07 | 432.2529 | 15221.721 |
| KQVVNIPSFIVR | 12 | P46781 | Small ribosomal subunit protein uS4 OS= RPS9 | 22.592 | 2 | 50.542587 | yes | 1.448E-07 | 700.42462 | 4402.707 |
| KQVVNIPSFIVR | 12 | P46781 | Small ribosomal subunit protein uS4 OS= RPS9 | 22.592 | 2 | 50.542587 | yes | 1.565E-07 | 467.28552 | 4402.707 |
| KQVVNIPSFIVR | 12 | P46781 | Small ribosomal subunit protein uS4 OS= RPS9 | 22.592 | 3 | 50.542587 | yes | 1.448E-07 | 700.42462 | 4402.707 |
| KQVVNIPSFIVR | 12 | P46781 | Small ribosomal subunit protein uS4 OS= RPS9 | 22.592 | 3 | 50.542587 | yes | 1.565E-07 | 467.28552 | 4402.707 |
| ELLTLDEKDPR | 11 | P46781 | Small ribosomal subunit protein uS4 OS= RPS9 | 22.592 | 2 | 50.542587 | yes | 2.785E-07 | 664.85645 | 4108.563 |
| ELLTLDEKDPR | 11 | P46781 | Small ribosomal subunit protein uS4 OS= RPS9 | 22.592 | 2 | 50.542587 | yes | 1.458E-07 | 443.57339 | 4108.563 |
| ELLTLDEKDPR | 11 | P46781 | Small ribosomal subunit protein uS4 OS= RPS9 | 22.592 | 3 | 50.542587 | yes | 2.785E-07 | 664.85645 | 4108.563 |
| ELLTLDEKDPR | 11 | P46781 | Small ribosomal subunit protein uS4 OS= RPS9 | 22.592 | 3 | 50.542587 | yes | 1.458E-07 | 443.57339 | 4108.563 |
| RLQTQVFK | 8 | P46781 | Small ribosomal subunit protein uS4 OS= RPS9 | 22.592 | 2 | 50.542587 | yes | 7.035E-07 | 510.30347 | 7083.4932 |

| | | | | | | | | | | | |
|---|---|---|---|---|---|---|---|---|---|---|---|
| SRLDQELK | 8 | P46781 | Small ribosomal subunit protein uS4 OS= | RPS9 | 22.592 | 2 | 50.542587 | yes | 2.178E-07 | 494.77472 | 5076.1328 |
| KTYVTPR | 7 | P46781 | Small ribosomal subunit protein uS4 OS= | RPS9 | 22.592 | 2 | 50.542587 | yes | 1.93E-07 | 432.75052 | 4443.4458 |
| MKLDYILGLK | 10 | P46781 | Small ribosomal subunit protein uS4 OS= | RPS9 | 22.592 | 2 | 50.542587 | yes | 1.076E-07 | 597.35175 | 1575.8716 |
| MKLDYILGLK | 10 | P46781 | Small ribosomal subunit protein uS4 OS= | RPS9 | 22.592 | 2 | 50.542587 | yes | 9.779E-08 | 605.34924 | 1575.8716 |
| SPYGGGRPGR | 10 | P46781 | Small ribosomal subunit protein uS4 OS= | RPS9 | 22.592 | 2 | 50.542587 | yes | 2.662E-07 | 502.25705 | 802.1571 |
| VNQAIWLLCTGAF | 13 | P46782 | Small ribosomal subunit protein uS7 OS= | RPS5 | 22.877 | 2 | 52.855354 | yes | 9.722E-08 | 751.40082 | 20762.176 |
| YLPHSAGR | 8 | P46782 | Small ribosomal subunit protein uS7 OS= | RPS5 | 22.877 | 2 | 52.855354 | yes | 2.784E-06 | 450.73795 | 20172.84 |
| QAVDVSPLR | 9 | P46782 | Small ribosomal subunit protein uS7 OS= | RPS5 | 22.877 | 2 | 52.855354 | yes | 1.335E-07 | 492.77728 | 15928.376 |
| AQCPIVER | 8 | P46782 | Small ribosomal subunit protein uS7 OS= | RPS5 | 22.877 | 2 | 52.855354 | yes | 8.411E-07 | 486.75021 | 15063.356 |
| TIAECLADELINAA | 15 | P46782 | Small ribosomal subunit protein uS7 OS= | RPS5 | 22.877 | 2 | 52.855354 | yes | 1.176E-07 | 816.41907 | 6202.4229 |
| LTNSMMMHGR | 10 | P46782 | Small ribosomal subunit protein uS7 OS= | RPS5 | 22.877 | 2 | 52.855354 | yes | 9.722E-08 | 589.2674 | 1530.4965 |
| LTNSMMMHGR | 10 | P46782 | Small ribosomal subunit protein uS7 OS= | RPS5 | 22.877 | 2 | 52.855354 | yes | 9.722E-08 | 597.26483 | 1530.4965 |
| LTNSMMMHGR | 10 | P46782 | Small ribosomal subunit protein uS7 OS= | RPS5 | 22.877 | 2 | 52.855354 | yes | 8.93E-07 | 597.26483 | 1530.4965 |
| KAQCPIVER | 9 | P46782 | Small ribosomal subunit protein uS7 OS= | RPS5 | 22.877 | 2 | 52.855354 | yes | 1.644E-07 | 550.79767 | 4071.7495 |
| RQAVDVSPLR | 10 | P46782 | Small ribosomal subunit protein uS7 OS= | RPS5 | 22.877 | 2 | 52.855354 | yes | 5.517E-05 | 570.82782 | 1171.1749 |
| GSSNSYAIK | 9 | P46782 | Small ribosomal subunit protein uS7 OS= | RPS5 | 22.877 | 2 | 52.855354 | yes | 1.325E-07 | 463.73254 | 37.0103 |
| NVPNLHVMK | 9 | P46783 | Small ribosomal subunit protein eS10 OS | RPS10 | 18.898 | 2 | 50.25909 | no | 3.216E-07 | 526.28949 | 13214.573 |
| NVPNLHVMK | 9 | P46783 | Small ribosomal subunit protein eS10 OS | RPS10 | 18.898 | 2 | 50.25909 | no | 4.987E-07 | 534.28693 | 13214.573 |
| AEAGAGSATEFQF | 14 | P46783 | Small ribosomal subunit protein eS10 OS | RPS10 | 18.898 | 2 | 50.25909 | no | 1.136E-07 | 721.33893 | 12077.473 |
| HPELADK | 7 | P46783 | Small ribosomal subunit protein eS10 OS | RPS10 | 18.898 | 2 | 50.25909 | no | 9.825E-08 | 405.21124 | 5523.085 |
| KAEAGAGSATEFQ | 15 | P46783 | Small ribosomal subunit protein eS10 OS | RPS10 | 18.898 | 3 | 50.25909 | no | 1.898E-07 | 523.9267 | 2279.333 |
| KAEAGAGSATEFQ | 15 | P46783 | Small ribosomal subunit protein eS10 OS | RPS10 | 18.898 | 3 | 50.25909 | no | 1.077E-07 | 785.38641 | 2279.333 |
| KAEAGAGSATEFQ | 15 | P46783 | Small ribosomal subunit protein eS10 OS | RPS10 | 18.898 | 2 | 50.25909 | no | 1.898E-07 | 523.9267 | 2279.333 |
| KAEAGAGSATEFQ | 15 | P46783 | Small ribosomal subunit protein eS10 OS | RPS10 | 18.898 | 2 | 50.25909 | no | 1.077E-07 | 785.38641 | 2279.333 |
| GLEGERPAR | 9 | P46783 | Small ribosomal subunit protein eS10 OS | RPS10 | 18.898 | 2 | 50.25909 | yes | 6.078E-07 | 492.76471 | 3446.2534 |
| DYLHLPPEIVPATL | 15 | P46783 | Small ribosomal subunit protein eS10 OS | RPS10 | 18.898 | 3 | 50.25909 | yes | 1.211E-07 | 578.65765 | 2276.9697 |
| HFYWYLTNEGIQY | 15 | P46783 | Small ribosomal subunit protein eS10 OS | RPS10 | 18.898 | 3 | 50.25909 | yes | 1.127E-07 | 668.33185 | 2070.7207 |
| GEADRDTYR | 9 | P46783 | Small ribosomal subunit protein eS10 OS | RPS10 | 18.898 | 2 | 50.25909 | yes | 1.709E-07 | 541.7467 | 410.46167 |
| KDVHMPK | 7 | P46783 | Small ribosomal subunit protein eS10 OS | RPS10 | 18.898 | 2 | 50.25909 | no | 1.23E-07 | 427.73129 | 192.55357 |
| LQLLLLHTR | 9 | P46939 | Utrophin OS=Homo sapiens OX=9606 G | UTRN | 394.467 | 2 | 34.740776 | yes | 9.499E-07 | 553.85583 | 29.030697 |
| IFYPETTDIYDR | 12 | P46940 | Ras GTPase-activating-like protein IQGA | IQGAP1 | 189.252 | 2 | 54.70373 | yes | 1.348E-07 | 766.867 | 798.57458 |
| ALQSPALGLR | 10 | P46940 | Ras GTPase-activating-like protein IQGA | IQGAP1 | 189.252 | 2 | 54.70373 | yes | 1.443E-07 | 513.30872 | 564.41534 |
| FDVPGDENAEMD | 14 | P46940 | Ras GTPase-activating-like protein IQGA | IQGAP1 | 189.252 | 2 | 54.70373 | yes | 2.553E-07 | 783.33044 | 560.85785 |
| LGLAPQIQDLYGK | 13 | P46940 | Ras GTPase-activating-like protein IQGA | IQGAP1 | 189.252 | 2 | 54.70373 | yes | 1.446E-07 | 708.39825 | 377.40222 |
| FLSAIVSSVDK | 11 | P46940 | Ras GTPase-activating-like protein IQGA | IQGAP1 | 189.252 | 2 | 54.70373 | yes | 1.183E-06 | 583.32678 | 341.70346 |
| TCLDNLASK | 9 | P46940 | Ras GTPase-activating-like protein IQGA | IQGAP1 | 189.252 | 2 | 54.70373 | yes | 1.429E-07 | 511.25278 | 289.87729 |
| ATFYGEQVDYYK | 12 | P46940 | Ras GTPase-activating-like protein IQGA | IQGAP1 | 189.252 | 2 | 54.70373 | yes | 9.722E-08 | 742.34064 | 266.81525 |
| TLQALQIPAAK | 11 | P46940 | Ras GTPase-activating-like protein IQGA | IQGAP1 | 189.252 | 2 | 54.70373 | yes | 9.81E-08 | 577.35059 | 259.69177 |
| ATGLHFR | 7 | P46940 | Ras GTPase-activating-like protein IQGA | IQGAP1 | 189.252 | 2 | 54.70373 | yes | 1.157E-07 | 401.22192 | 248.48997 |

| | | | | | | | | | | | |
|---|---|---|---|---|---|---|---|---|---|---|---|
| EQLWLANEGLITR | 13 | P46940 | Ras GTPase-activating-like protein IQGA | IQGAP1 | 189.252 | 2 | 54.70373 | yes | 9.938E-08 | 771.91736 | 244.4328 |
| VDFTEEEINNMK | 12 | P46940 | Ras GTPase-activating-like protein IQGA | IQGAP1 | 189.252 | 2 | 54.70373 | yes | 9.728E-08 | 734.83484 | 243.57635 |
| LTAEEMDER | 9 | P46940 | Ras GTPase-activating-like protein IQGA | IQGAP1 | 189.252 | 2 | 54.70373 | yes | 3.792E-07 | 547.24512 | 242.21355 |
| GLQQQNSDWYLK | 12 | P46940 | Ras GTPase-activating-like protein IQGA | IQGAP1 | 189.252 | 2 | 54.70373 | yes | 1.136E-07 | 740.36499 | 217.36969 |
| TEVSLTLTNK | 10 | P46940 | Ras GTPase-activating-like protein IQGA | IQGAP1 | 189.252 | 2 | 54.70373 | yes | 2.812E-07 | 553.30859 | 203.52518 |
| KLTELGTVDPK | 11 | P46940 | Ras GTPase-activating-like protein IQGA | IQGAP1 | 189.252 | 2 | 54.70373 | yes | 9.722E-08 | 600.84534 | 166.89262 |
| LTELGTVDPK | 10 | P46940 | Ras GTPase-activating-like protein IQGA | IQGAP1 | 189.252 | 2 | 54.70373 | yes | 3.368E-07 | 536.79785 | 161.77313 |
| TALQEEIK | 8 | P46940 | Ras GTPase-activating-like protein IQGA | IQGAP1 | 189.252 | 2 | 54.70373 | no | 3.675E-07 | 466.25839 | 159.27635 |
| IPADTFAALK | 10 | P46940 | Ras GTPase-activating-like protein IQGA | IQGAP1 | 189.252 | 2 | 54.70373 | yes | 3.962E-07 | 523.79767 | 153.89407 |
| DHINDIIK | 8 | P46940 | Ras GTPase-activating-like protein IQGA | IQGAP1 | 189.252 | 2 | 54.70373 | yes | 3.766E-07 | 484.26401 | 138.78577 |
| QIPAITCIQSQWR | 13 | P46940 | Ras GTPase-activating-like protein IQGA | IQGAP1 | 189.252 | 2 | 54.70373 | yes | 1.197E-07 | 800.91687 | 138.67949 |
| EQLSDMMMINK | 11 | P46940 | Ras GTPase-activating-like protein IQGA | IQGAP1 | 189.252 | 2 | 54.70373 | yes | 5.796E-07 | 670.3064 | 137.50427 |
| ALESGDVNTVWK | 12 | P46940 | Ras GTPase-activating-like protein IQGA | IQGAP1 | 189.252 | 2 | 54.70373 | yes | 1.003E-07 | 659.83551 | 135.8649 |
| PHYGSVLDNER | 11 | P46940 | Ras GTPase-activating-like protein IQGA | IQGAP1 | 189.252 | 2 | 54.70373 | yes | 9.722E-08 | 643.80981 | 131.22977 |
| IIGNLLYYR | 9 | P46940 | Ras GTPase-activating-like protein IQGA | IQGAP1 | 189.252 | 2 | 54.70373 | yes | 1.166E-07 | 562.82678 | 121.77051 |
| RIPADTFAALK | 11 | P46940 | Ras GTPase-activating-like protein IQGA | IQGAP1 | 189.252 | 2 | 54.70373 | yes | 3.105E-07 | 601.84821 | 111.85713 |
| TLINAEDPPMVVV | 14 | P46940 | Ras GTPase-activating-like protein IQGA | IQGAP1 | 189.252 | 2 | 54.70373 | yes | 6.204E-07 | 777.42145 | 106.22549 |
| FNVDEYSDLVTLT | 14 | P46940 | Ras GTPase-activating-like protein IQGA | IQGAP1 | 189.252 | 2 | 54.70373 | yes | 9.722E-08 | 822.4118 | 104.98269 |
| FPDAGEDELLK | 11 | P46940 | Ras GTPase-activating-like protein IQGA | IQGAP1 | 189.252 | 2 | 54.70373 | yes | 4.723E-07 | 617.30353 | 98.987617 |
| LQQTYAALNSK | 11 | P46940 | Ras GTPase-activating-like protein IQGA | IQGAP1 | 189.252 | 2 | 54.70373 | yes | 9.722E-08 | 618.83276 | 94.151199 |
| LPYDVTPEQALAH | 17 | P46940 | Ras GTPase-activating-like protein IQGA | IQGAP1 | 189.252 | 3 | 54.70373 | yes | 1.357E-06 | 646.99841 | 88.49408 |
| TDPVDIYK | 8 | P46940 | Ras GTPase-activating-like protein IQGA | IQGAP1 | 189.252 | 2 | 54.70373 | yes | 2.746E-06 | 475.74512 | 87.665848 |
| QSGQTDPLQK | 10 | P46940 | Ras GTPase-activating-like protein IQGA | IQGAP1 | 189.252 | 2 | 54.70373 | yes | 3.492E-07 | 551.2804 | 43.487167 |
| SWVNQMESQTGE | 15 | P46940 | Ras GTPase-activating-like protein IQGA | IQGAP1 | 189.252 | 2 | 54.70373 | yes | 1.456E-07 | 841.37793 | 41.16016 |
| LGNFFSPK | 8 | P46940 | Ras GTPase-activating-like protein IQGA | IQGAP1 | 189.252 | 2 | 54.70373 | yes | 4.993E-07 | 455.24509 | 36.863106 |
| LIFQMPQNK | 9 | P46940 | Ras GTPase-activating-like protein IQGA | IQGAP1 | 189.252 | 2 | 54.70373 | no | 1.663E-07 | 559.80499 | 33.604565 |
| MREEVITLIR | 10 | P46940 | Ras GTPase-activating-like protein IQGA | IQGAP1 | 189.252 | 2 | 54.70373 | yes | 6.189E-07 | 630.36066 | 27.082687 |
| YGIQMPAFSK | 10 | P46940 | Ras GTPase-activating-like protein IQGA | IQGAP1 | 189.252 | 2 | 54.70373 | yes | 3.142E-07 | 571.28918 | 22.518499 |
| LVVLATPQVSDSM | 14 | P46976 | Glycogenin-1 OS=Homo sapiens OX=96( | GYG1 | 39.384 | 2 | 48.532257 | yes | 4.874E-07 | 758.41364 | 1251.3634 |
| VKPWNYTYDPK | 11 | P46976 | Glycogenin-1 OS=Homo sapiens OX=96( | GYG1 | 39.384 | 2 | 48.532257 | yes | 1.135E-07 | 705.85626 | 272.02771 |
| RPELGVTLTK | 10 | P46976 | Glycogenin-1 OS=Homo sapiens OX=96( | GYG1 | 39.384 | 2 | 48.532257 | yes | 2.313E-07 | 557.33496 | 261.63684 |
| GALVLGSSLK | 10 | P46976 | Glycogenin-1 OS=Homo sapiens OX=96( | GYG1 | 39.384 | 2 | 48.532257 | yes | 1.548E-07 | 472.79239 | 258.36511 |
| VVHFLGR | 7 | P46976 | Glycogenin-1 OS=Homo sapiens OX=96( | GYG1 | 39.384 | 2 | 48.532257 | yes | 5.402E-07 | 414.24796 | 126.36747 |
| FLAEEGFYK | 9 | P46977 | Dolichyl-diphosphooligosaccharide--prot | STT3A | 80.53 | 2 | 46.95019 | yes | 9.334E-07 | 552.27405 | 87.016006 |
| VGQAMASTEEK | 11 | P46977 | Dolichyl-diphosphooligosaccharide--prot | STT3A | 80.53 | 2 | 46.95019 | yes | 1.434E-07 | 575.77405 | 72.64769 |
| IIFDDFR | 7 | P46977 | Dolichyl-diphosphooligosaccharide--prot | STT3A | 80.53 | 2 | 46.95019 | yes | 3.194E-07 | 463.24252 | 52.301224 |
| ENDYYTPTGEFR | 12 | P46977 | Dolichyl-diphosphooligosaccharide--prot | STT3A | 80.53 | 2 | 46.95019 | yes | 7.919E-07 | 746.323 | 39.467194 |
| EIMFHVSTK | 9 | P47736 | Rap1 GTPase-activating protein 1 OS=Hc | RAP1GAP | 73.362 | 2 | 41.914722 | yes | 3.281E-07 | 546.28131 | 68.840874 |
| LLLNNDNLLR | 10 | P47755 | F-actin-capping protein subunit alpha-2 C | CAPZA2 | 32.95 | 2 | 54.334038 | no | 9.85E-08 | 599.35114 | 4148.9702 |

| | | | | | | | | | | | |
|---|---|---|---|---|---|---|---|---|---|---|---|
| FTITPSTTQVVGILI | 15 | P47755 | F-actin-capping protein subunit alpha-2 C | CAPZA2 | 32.95 | 2 | 54.334038 | yes | 2.666E-07 | 802.96655 | 962.59607 |
| ADLEEQLSDEEK | 12 | P47755 | F-actin-capping protein subunit alpha-2 C | CAPZA2 | 32.95 | 2 | 54.334038 | yes | 1.856E-07 | 724.32538 | 126.93831 |
| IDGQQTIIACIESH( | 19 | P47755 | F-actin-capping protein subunit alpha-2 C | CAPZA2 | 32.95 | 3 | 54.334038 | yes | 9.722E-08 | 729.70026 | 108.12588 |
| TSVETALR | 8 | P47755 | F-actin-capping protein subunit alpha-2 C | CAPZA2 | 32.95 | 2 | 54.334038 | yes | 9.722E-08 | 438.74289 | 97.027443 |
| EHYPNGVCTVYGI | 13 | P47755 | F-actin-capping protein subunit alpha-2 C | CAPZA2 | 32.95 | 2 | 54.334038 | yes | 1.396E-07 | 762.35101 | 88.075394 |
| KIDGQQTIIACIESI | 20 | P47755 | F-actin-capping protein subunit alpha-2 C | CAPZA2 | 32.95 | 3 | 54.334038 | yes | 9.722E-08 | 772.39862 | 54.3475 |
| IQVHYYEDGNVQI | 17 | P47755 | F-actin-capping protein subunit alpha-2 C | CAPZA2 | 32.95 | 3 | 54.334038 | no | 1.594E-07 | 677.00946 | 35.206387 |
| YDPPLEDGAMPSA | 14 | P47756 | F-actin-capping protein subunit beta OS= | CAPZB | 30.629 | 2 | 50.72378 | yes | 1.27E-07 | 759.84833 | 5957.1685 |
| SGSGTMNLGGSLT | 14 | P47756 | F-actin-capping protein subunit beta OS= | CAPZB | 30.629 | 2 | 50.72378 | yes | 9.728E-08 | 669.32751 | 1860.7996 |
| SGSGTMNLGGSLT | 14 | P47756 | F-actin-capping protein subunit beta OS= | CAPZB | 30.629 | 2 | 50.72378 | yes | 1.092E-07 | 677.32501 | 1860.7996 |
| DYLLCDYNR | 9 | P47756 | F-actin-capping protein subunit beta OS= | CAPZB | 30.629 | 2 | 50.72378 | yes | 9.728E-08 | 616.27423 | 3307.438 |
| GCWDSIHVVEVQI | 14 | P47756 | F-actin-capping protein subunit beta OS= | CAPZB | 30.629 | 2 | 50.72378 | yes | 9.818E-08 | 843.40125 | 784.13159 |
| GCWDSIHVVEVQI | 14 | P47756 | F-actin-capping protein subunit beta OS= | CAPZB | 30.629 | 2 | 50.72378 | yes | 1.41E-07 | 562.60327 | 784.13159 |
| GCWDSIHVVEVQI | 14 | P47756 | F-actin-capping protein subunit beta OS= | CAPZB | 30.629 | 3 | 50.72378 | yes | 9.818E-08 | 843.40125 | 784.13159 |
| GCWDSIHVVEVQI | 14 | P47756 | F-actin-capping protein subunit beta OS= | CAPZB | 30.629 | 3 | 50.72378 | yes | 1.41E-07 | 562.60327 | 784.13159 |
| STLNEIYFGK | 10 | P47756 | F-actin-capping protein subunit beta OS= | CAPZB | 30.629 | 2 | 50.72378 | yes | 1.025E-07 | 586.30334 | 848.69476 |
| SDQQLDCALDLMI | 13 | P47756 | F-actin-capping protein subunit beta OS= | CAPZB | 30.629 | 2 | 50.72378 | yes | 9.727E-08 | 803.86365 | 815.96136 |
| LTSTVMLWLQTNI | 13 | P47756 | F-actin-capping protein subunit beta OS= | CAPZB | 30.629 | 2 | 50.72378 | yes | 1.007E-07 | 767.91852 | 791.34149 |
| NDLVEALKR | 9 | P47756 | F-actin-capping protein subunit beta OS= | CAPZB | 30.629 | 2 | 50.72378 | yes | 9.789E-08 | 529.30365 | 412.20038 |
| NDLVEALK | 8 | P47756 | F-actin-capping protein subunit beta OS= | CAPZB | 30.629 | 2 | 50.72378 | yes | 4.752E-07 | 451.25311 | 291.26428 |
| LEVEANNAFDQYI | 13 | P47756 | F-actin-capping protein subunit beta OS= | CAPZB | 30.629 | 2 | 50.72378 | yes | 2.398E-07 | 784.87061 | 195.98705 |
| QMEKDETVSDCSI | 20 | P47756 | F-actin-capping protein subunit beta OS= | CAPZB | 30.629 | 4 | 50.72378 | yes | 6.673E-07 | 572.51648 | 47.704605 |
| QMEKDETVSDCSI | 20 | P47756 | F-actin-capping protein subunit beta OS= | CAPZB | 30.629 | 4 | 50.72378 | yes | 1.5E-07 | 763.01959 | 47.704605 |
| QMEKDETVSDCSI | 20 | P47756 | F-actin-capping protein subunit beta OS= | CAPZB | 30.629 | 3 | 50.72378 | yes | 6.673E-07 | 572.51648 | 47.704605 |
| QMEKDETVSDCSI | 20 | P47756 | F-actin-capping protein subunit beta OS= | CAPZB | 30.629 | 3 | 50.72378 | yes | 1.5E-07 | 763.01959 | 47.704605 |
| RLPPQQIEK | 9 | P47756 | F-actin-capping protein subunit beta OS= | CAPZB | 30.629 | 2 | 50.72378 | yes | 1.662E-05 | 554.82727 | 84.602333 |
| IKGCWDSIHVVEV | 16 | P47756 | F-actin-capping protein subunit beta OS= | CAPZB | 30.629 | 3 | 50.72378 | yes | 2.441E-06 | 642.99628 | 18.851097 |
| EEIFGPVQPILK | 12 | P47895 | Retinaldehyde dehydrogenase 3 OS=Hom | ALDH1A3 | 56.109 | 2 | 38.870155 | yes | 4.935E-07 | 685.38995 | 29.497816 |
| VFVEEQVYSEFVR | 13 | P47895 | Retinaldehyde dehydrogenase 3 OS=Hom | ALDH1A3 | 56.109 | 2 | 38.870155 | yes | 1.348E-05 | 815.90924 | 13.80782 |
| GPSGCVESLEVTC | 14 | P47897 | Glutamine--tRNA ligase OS=Homo sapie | QARS1 | 87.799 | 2 | 51.66968 | yes | 2.207E-07 | 775.85052 | 720.24878 |
| NEVDMQVLHLLG | 14 | P47897 | Glutamine--tRNA ligase OS=Homo sapie | QARS1 | 87.799 | 2 | 51.66968 | yes | 1.334E-07 | 796.92688 | 512.41455 |
| LAWGQPVGLR | 10 | P47897 | Glutamine--tRNA ligase OS=Homo sapie | QARS1 | 87.799 | 2 | 51.66968 | yes | 1.229E-07 | 548.81671 | 422.79263 |
| EAATQAQQTLGST | 16 | P47897 | Glutamine--tRNA ligase OS=Homo sapie | QARS1 | 87.799 | 2 | 51.66968 | yes | 1.176E-07 | 831.42065 | 325.89847 |
| SLDIQVPNFPADET | 15 | P47897 | Glutamine--tRNA ligase OS=Homo sapie | QARS1 | 87.799 | 2 | 51.66968 | yes | 9.938E-08 | 837.42267 | 280.82126 |
| ATGILLYGLASR | 12 | P47897 | Glutamine--tRNA ligase OS=Homo sapie | QARS1 | 87.799 | 2 | 51.66968 | yes | 1.684E-07 | 617.86133 | 172.48235 |
| ILQLVATGAVR | 11 | P47897 | Glutamine--tRNA ligase OS=Homo sapie | QARS1 | 87.799 | 2 | 51.66968 | yes | 1.074E-07 | 570.85858 | 140.99876 |
| AINFNFGYAK | 10 | P47897 | Glutamine--tRNA ligase OS=Homo sapie | QARS1 | 87.799 | 2 | 51.66968 | yes | 9.722E-08 | 572.79291 | 123.69573 |
| LNLHYAVVSK | 10 | P47897 | Glutamine--tRNA ligase OS=Homo sapie | QARS1 | 87.799 | 2 | 51.66968 | yes | 1.404E-07 | 572.32965 | 112.68369 |
| LGYFSVDPDSHQC | 14 | P47897 | Glutamine--tRNA ligase OS=Homo sapie | QARS1 | 87.799 | 2 | 51.66968 | yes | 2.84E-07 | 775.36774 | 103.22506 |

| | | | | | | | | | | | |
|---|---|---|---|---|---|---|---|---|---|---|---|
| TDFKEEPEPGFK | 12 | P47897 | Glutamine--tRNA ligase OS=Homo sapie | QARS1 | 87.799 | 2 | 51.66968 | yes | 3.513E-07 | 712.34064 | 98.518463 |
| GHNTLPSPWR | 10 | P47897 | Glutamine--tRNA ligase OS=Homo sapie | QARS1 | 87.799 | 2 | 51.66968 | yes | 2.436E-07 | 582.79907 | 92.218895 |
| AMAVLESLR | 9 | P47897 | Glutamine--tRNA ligase OS=Homo sapie | QARS1 | 87.799 | 2 | 51.66968 | yes | 4.241E-07 | 495.27606 | 88.410019 |
| GLAYVCHQR | 9 | P47897 | Glutamine--tRNA ligase OS=Homo sapie | QARS1 | 87.799 | 2 | 51.66968 | yes | 9.34E-07 | 552.27435 | 77.608749 |
| FSEGEATLR | 9 | P47897 | Glutamine--tRNA ligase OS=Homo sapie | QARS1 | 87.799 | 2 | 51.66968 | yes | 5.795E-07 | 505.2511 | 74.598694 |
| VIITNFPAAK | 10 | P47897 | Glutamine--tRNA ligase OS=Homo sapie | QARS1 | 87.799 | 2 | 51.66968 | yes | 6.861E-07 | 537.32129 | 74.184448 |
| VGVTVAQTTMEPI | 20 | P47897 | Glutamine--tRNA ligase OS=Homo sapie | QARS1 | 87.799 | 3 | 51.66968 | yes | 1.958E-06 | 737.7135 | 67.876343 |
| DRPMEESLLLFEA | 15 | P47897 | Glutamine--tRNA ligase OS=Homo sapie | QARS1 | 87.799 | 3 | 51.66968 | yes | 2.584E-07 | 612.97095 | 37.737637 |
| HTGYVIELQHVVK | 13 | P47897 | Glutamine--tRNA ligase OS=Homo sapie | QARS1 | 87.799 | 3 | 51.66968 | yes | 2.379E-07 | 508.28406 | 14.943827 |
| HTGYVIELQHVVK | 13 | P47897 | Glutamine--tRNA ligase OS=Homo sapie | QARS1 | 87.799 | 3 | 51.66968 | yes | 3.862E-07 | 761.92249 | 14.943827 |
| HTGYVIELQHVVK | 13 | P47897 | Glutamine--tRNA ligase OS=Homo sapie | QARS1 | 87.799 | 2 | 51.66968 | yes | 2.379E-07 | 508.28406 | 14.943827 |
| HTGYVIELQHVVK | 13 | P47897 | Glutamine--tRNA ligase OS=Homo sapie | QARS1 | 87.799 | 2 | 51.66968 | yes | 3.862E-07 | 761.92249 | 14.943827 |
| AQAAAPASVPAQ | 15 | P47914 | Large ribosomal subunit protein eL29 OS | RPL29 | 17.753 | 2 | 48.0294 | yes | 1.211E-07 | 689.37787 | 29269.055 |
| LAYIAHPK | 8 | P47914 | Large ribosomal subunit protein eL29 OS | RPL29 | 17.753 | 2 | 48.0294 | yes | 1.249E-07 | 456.76871 | 12042.688 |
| DALHLGFPR | 9 | P47974 | mRNA decay activator protein ZFP36L2 | ZFP36L2 | 51.063 | 2 | 54.081955 | yes | 1.136E-07 | 513.27997 | 1426.0103 |
| EPSGGGGTALLNK | 13 | P47974 | mRNA decay activator protein ZFP36L2 | ZFP36L2 | 51.063 | 2 | 54.081955 | yes | 4.921E-07 | 600.81458 | 538.62183 |
| SLANLNLNNMLDI | 13 | P47974 | mRNA decay activator protein ZFP36L2 | ZFP36L2 | 51.063 | 2 | 54.081955 | yes | 9.722E-08 | 730.38232 | 399.09543 |
| CHFIHNADER | 10 | P47974 | mRNA decay activator protein ZFP36L2 | ZFP36L2 | 51.063 | 2 | 54.081955 | yes | 1.157E-07 | 649.78839 | 272.38974 |
| SQHLLHLQQQQK | 12 | P47974 | mRNA decay activator protein ZFP36L2 | ZFP36L2 | 51.063 | 2 | 54.081955 | yes | 9.81E-08 | 744.40753 | 154.13011 |
| HSASNLHALAHPA | 22 | P47974 | mRNA decay activator protein ZFP36L2 | ZFP36L2 | 51.063 | 3 | 54.081955 | yes | 1.335E-07 | 741.69562 | 16.574812 |
| VPDFSEYR | 8 | P47985 | Cytochrome b-c1 complex subunit Rieske | UQCRFS1 | 29.668 | 2 | 46.200882 | no | 1.623E-07 | 506.74036 | 99.647659 |
| EIEQEAAVELSQLI | 14 | P47985 | Cytochrome b-c1 complex subunit Rieske | UQCRFS1 | 29.668 | 2 | 46.200882 | yes | 1.719E-07 | 807.92029 | 54.295376 |
| DPQHDLDR | 8 | P47985 | Cytochrome b-c1 complex subunit Rieske | UQCRFS1 | 29.668 | 2 | 46.200882 | no | 3.079E-06 | 498.23068 | 39.738663 |
| SFLSQGQVLK | 10 | P48047 | ATP synthase subunit O, mitochondrial C | ATP5PO | 23.278 | 2 | 51.51765 | yes | 1.565E-07 | 553.81384 | 780.48767 |
| TDPSILGGMIVR | 12 | P48047 | ATP synthase subunit O, mitochondrial C | ATP5PO | 23.278 | 2 | 51.51765 | yes | 9.722E-08 | 629.84485 | 515.51196 |
| VAASVLNPYVK | 11 | P48047 | ATP synthase subunit O, mitochondrial C | ATP5PO | 23.278 | 2 | 51.51765 | yes | 9.722E-08 | 580.83734 | 487.21912 |
| YATALYSAASK | 11 | P48047 | ATP synthase subunit O, mitochondrial C | ATP5PO | 23.278 | 2 | 51.51765 | yes | 1.006E-07 | 573.29547 | 384.12491 |
| LVRPPVQVYGIEG | 14 | P48047 | ATP synthase subunit O, mitochondrial C | ATP5PO | 23.278 | 3 | 51.51765 | yes | 1.5E-07 | 528.30701 | 182.29758 |
| LVRPPVQVYGIEG | 14 | P48047 | ATP synthase subunit O, mitochondrial C | ATP5PO | 23.278 | 3 | 51.51765 | yes | 1.7E-07 | 791.95685 | 182.29758 |
| LVRPPVQVYGIEG | 14 | P48047 | ATP synthase subunit O, mitochondrial C | ATP5PO | 23.278 | 2 | 51.51765 | yes | 1.5E-07 | 528.30701 | 182.29758 |
| LVRPPVQVYGIEG | 14 | P48047 | ATP synthase subunit O, mitochondrial C | ATP5PO | 23.278 | 2 | 51.51765 | yes | 1.7E-07 | 791.95685 | 182.29758 |
| SLNDITAK | 8 | P48047 | ATP synthase subunit O, mitochondrial C | ATP5PO | 23.278 | 2 | 51.51765 | yes | 1.284E-07 | 431.23746 | 249.61514 |
| YVDMSVK | 7 | P48047 | ATP synthase subunit O, mitochondrial C | ATP5PO | 23.278 | 2 | 51.51765 | yes | 1.038E-07 | 421.20984 | 26.16046 |
| VIEGDVVSALNK | 12 | P48059 | LIM and senescent cell antigen-like-conta | LIMS1 | 37.252 | 2 | 49.28859 | yes | 1.048E-07 | 622.34827 | 211.06538 |
| FVEFDMKPVCK | 11 | P48059 | LIM and senescent cell antigen-like-conta | LIMS1 | 37.252 | 2 | 49.28859 | no | 1.721E-07 | 700.34106 | 96.691994 |
| MGVPICGACR | 10 | P48059 | LIM and senescent cell antigen-like-conta | LIMS1 | 37.252 | 2 | 49.28859 | no | 1.155E-05 | 560.75665 | 40.810204 |
| AMNNSWHPECFR | 12 | P48059 | LIM and senescent cell antigen-like-conta | LIMS1 | 37.252 | 3 | 49.28859 | yes | 2.848E-06 | 516.88721 | 6.8417826 |
| QSNVEVPFFPAR | 12 | P48200 | Iron-responsive element-binding protein 2 | IREB2 | 105.059 | 2 | 56.43033 | yes | 1.102E-07 | 695.85931 | 292.8692 |
| AVLAESYEK | 9 | P48200 | Iron-responsive element-binding protein 2 | IREB2 | 105.059 | 2 | 56.43033 | yes | 2.582E-07 | 505.26367 | 91.738174 |

| | | | | | | | | | | | |
|---|---|---|---|---|---|---|---|---|---|---|---|
| SDFQACLNEK | 10 | P48200 | Iron-responsive element-binding protein 2 | IREB2 | 105.059 | 2 | 56.43033 | yes | 9.722E-08 | 606.27167 | 79.142639 |
| TIHFPSGQTLDVFE | 21 | P48200 | Iron-responsive element-binding protein 2 | IREB2 | 105.059 | 3 | 56.43033 | yes | 1.66E-06 | 798.73737 | 74.145576 |
| DIVSIHYEGSEYK | 13 | P48200 | Iron-responsive element-binding protein 2 | IREB2 | 105.059 | 2 | 56.43033 | yes | 3.746E-07 | 770.36993 | 69.509888 |
| VEEEHVILSMFK | 12 | P48200 | Iron-responsive element-binding protein 2 | IREB2 | 105.059 | 2 | 56.43033 | yes | 1.066E-07 | 730.87634 | 47.854847 |
| LESMETYLK | 9 | P48200 | Iron-responsive element-binding protein 2 | IREB2 | 105.059 | 2 | 56.43033 | yes | 6.148E-06 | 557.27844 | 43.783516 |
| EGIPLIILAGK | 11 | P48200 | Iron-responsive element-binding protein 2 | IREB2 | 105.059 | 2 | 56.43033 | yes | 2.398E-07 | 562.35791 | 41.318203 |
| KEDVMNILDWK | 11 | P48200 | Iron-responsive element-binding protein 2 | IREB2 | 105.059 | 2 | 56.43033 | yes | 2.54E-07 | 695.85541 | 27.732605 |
| QGDLVTCGILSGN | 14 | P48200 | Iron-responsive element-binding protein 2 | IREB2 | 105.059 | 2 | 56.43033 | yes | 2.971E-07 | 731.37195 | 25.519791 |
| HGGLLNFVAR | 10 | P48200 | Iron-responsive element-binding protein 2 | IREB2 | 105.059 | 2 | 56.43033 | yes | 5.278E-05 | 542.30652 | 17.984489 |
| GAVLAQGQGDGT | 15 | P48382 | DNA-binding protein RFX5 OS=Homo sa | RFX5 | 65.323 | 2 | 38.92032 | yes | 4.902E-07 | 694.36243 | 17.611843 |
| ENVNLAQIR | 9 | P48444 | Coatomer subunit delta OS=Homo sapien | ARCN1 | 57.211 | 2 | 54.717022 | yes | 1.232E-07 | 528.79346 | 232.40765 |
| YVYQPMEK | 8 | P48444 | Coatomer subunit delta OS=Homo sapien | ARCN1 | 57.211 | 2 | 54.717022 | yes | 4.718E-07 | 529.25476 | 167.867 |
| VLLAAAVCTK | 10 | P48444 | Coatomer subunit delta OS=Homo sapien | ARCN1 | 57.211 | 2 | 54.717022 | yes | 2.567E-07 | 523.30737 | 133.90257 |
| TFTEMDSHEEK | 11 | P48444 | Coatomer subunit delta OS=Homo sapien | ARCN1 | 57.211 | 2 | 54.717022 | yes | 9.722E-08 | 677.28497 | 103.48813 |
| NTLEWCLPVIDAK | 13 | P48444 | Coatomer subunit delta OS=Homo sapien | ARCN1 | 57.211 | 2 | 54.717022 | yes | 1.316E-07 | 779.90033 | 99.850441 |
| LYMVLITTK | 9 | P48444 | Coatomer subunit delta OS=Homo sapien | ARCN1 | 57.211 | 2 | 54.717022 | yes | 9.722E-08 | 541.31995 | 91.722984 |
| SEGETIMSSSMGK | 13 | P48444 | Coatomer subunit delta OS=Homo sapien | ARCN1 | 57.211 | 2 | 54.717022 | yes | 1.063E-07 | 672.29449 | 83.946892 |
| NSNILEDLETLR | 12 | P48444 | Coatomer subunit delta OS=Homo sapien | ARCN1 | 57.211 | 2 | 54.717022 | yes | 9.722E-08 | 708.87012 | 73.03775 |
| LFTAESLIGLK | 11 | P48444 | Coatomer subunit delta OS=Homo sapien | ARCN1 | 57.211 | 2 | 54.717022 | yes | 9.722E-08 | 596.35284 | 58.940208 |
| IEGLLAAFPK | 10 | P48444 | Coatomer subunit delta OS=Homo sapien | ARCN1 | 57.211 | 2 | 54.717022 | yes | 4.673E-06 | 529.81586 | 57.779533 |
| FSTETTFLVDK | 11 | P48444 | Coatomer subunit delta OS=Homo sapien | ARCN1 | 57.211 | 2 | 54.717022 | yes | 1.128E-07 | 644.32697 | 49.140034 |
| QFVEMTR | 7 | P48444 | Coatomer subunit delta OS=Homo sapien | ARCN1 | 57.211 | 2 | 54.717022 | yes | 4.48E-07 | 455.7262 | 45.834888 |
| VTQVDGNSPVR | 11 | P48444 | Coatomer subunit delta OS=Homo sapien | ARCN1 | 57.211 | 2 | 54.717022 | yes | 1.718E-07 | 586.30695 | 43.493942 |
| QHTFVETESVR | 11 | P48444 | Coatomer subunit delta OS=Homo sapien | ARCN1 | 57.211 | 2 | 54.717022 | yes | 1.397E-06 | 666.83075 | 41.503323 |
| IVAIGTSDLDK | 11 | P48507 | Glutamate--cysteine ligase regulatory sub | GCLM | 30.728 | 2 | 46.71475 | yes | 1.96E-07 | 566.31641 | 271.3385 |
| LFIVESNSSSSTR | 13 | P48507 | Glutamate--cysteine ligase regulatory sub | GCLM | 30.728 | 2 | 46.71475 | yes | 1.489E-07 | 713.86224 | 150.19196 |
| EEFPTLQAAGDQD | 14 | P48634 | Protein PRRC2A OS=Homo sapiens OX= | PRRC2A | 228.864 | 2 | 54.516624 | yes | 1.325E-07 | 774.86249 | 7025.106 |
| LLQGRPPLDFYPP( | 23 | P48634 | Protein PRRC2A OS=Homo sapiens OX= | PRRC2A | 228.864 | 4 | 54.516624 | yes | 1.701E-07 | 628.85101 | 2735.9006 |
| LLQGRPPLDFYPP( | 23 | P48634 | Protein PRRC2A OS=Homo sapiens OX= | PRRC2A | 228.864 | 4 | 54.516624 | yes | 1.113E-07 | 838.13226 | 2735.9006 |
| LLQGRPPLDFYPP( | 23 | P48634 | Protein PRRC2A OS=Homo sapiens OX= | PRRC2A | 228.864 | 3 | 54.516624 | yes | 1.701E-07 | 628.85101 | 2735.9006 |
| LLQGRPPLDFYPP( | 23 | P48634 | Protein PRRC2A OS=Homo sapiens OX= | PRRC2A | 228.864 | 3 | 54.516624 | yes | 1.113E-07 | 838.13226 | 2735.9006 |
| LIPGPLSPVAR | 11 | P48634 | Protein PRRC2A OS=Homo sapiens OX= | PRRC2A | 228.864 | 2 | 54.516624 | yes | 6.822E-07 | 560.34784 | 5088.5527 |
| SLEIQKPAVAPR | 12 | P48634 | Protein PRRC2A OS=Homo sapiens OX= | PRRC2A | 228.864 | 2 | 54.516624 | yes | 1.253E-07 | 654.88531 | 2282.8486 |
| SLEIQKPAVAPR | 12 | P48634 | Protein PRRC2A OS=Homo sapiens OX= | PRRC2A | 228.864 | 2 | 54.516624 | yes | 1.087E-07 | 436.92599 | 2282.8486 |
| SLEIQKPAVAPR | 12 | P48634 | Protein PRRC2A OS=Homo sapiens OX= | PRRC2A | 228.864 | 3 | 54.516624 | yes | 1.253E-07 | 654.88531 | 2282.8486 |
| SLEIQKPAVAPR | 12 | P48634 | Protein PRRC2A OS=Homo sapiens OX= | PRRC2A | 228.864 | 3 | 54.516624 | yes | 1.087E-07 | 436.92599 | 2282.8486 |
| VDLYQQASPPDAI | 14 | P48634 | Protein PRRC2A OS=Homo sapiens OX= | PRRC2A | 228.864 | 2 | 54.516624 | yes | 9.722E-08 | 786.90448 | 4216.4189 |
| GNDPNVSLVPK | 11 | P48634 | Protein PRRC2A OS=Homo sapiens OX= | PRRC2A | 228.864 | 2 | 54.516624 | yes | 1E-07 | 570.3064 | 3555.8743 |
| YSSLNLFDTYK | 11 | P48634 | Protein PRRC2A OS=Homo sapiens OX= | PRRC2A | 228.864 | 2 | 54.516624 | yes | 3.912E-07 | 675.83246 | 3198.3752 |

| | | | | | | | | | | | |
|---|---|---|---|---|---|---|---|---|---|---|---|
| LKAEPAAPPAAPS' | 23 | P48634 | Protein PRRC2A OS=Homo sapiens OX= | PRRC2A | 228.864 | 3 | 54.516624 | yes | 1.587E-07 | 725.7439 | 3126.2783 |
| ATPSTYSGVFR | 11 | P48634 | Protein PRRC2A OS=Homo sapiens OX= | PRRC2A | 228.864 | 2 | 54.516624 | yes | 9.915E-08 | 593.29858 | 3099.0347 |
| RPPAAPENTPLVPS | 17 | P48634 | Protein PRRC2A OS=Homo sapiens OX= | PRRC2A | 228.864 | 3 | 54.516624 | yes | 1.193E-07 | 577.32471 | 2980.6162 |
| WMMIPPYVDPR | 11 | P48634 | Protein PRRC2A OS=Homo sapiens OX= | PRRC2A | 228.864 | 2 | 54.516624 | yes | 2.165E-07 | 702.84375 | 2478.1338 |
| PLPWPPGSDEVAK | 13 | P48634 | Protein PRRC2A OS=Homo sapiens OX= | PRRC2A | 228.864 | 2 | 54.516624 | yes | 5.915E-07 | 696.86151 | 2395.8125 |
| WIPKPWER | 8 | P48634 | Protein PRRC2A OS=Homo sapiens OX= | PRRC2A | 228.864 | 2 | 54.516624 | yes | 1.763E-07 | 556.30603 | 2238.2788 |
| LSSNLGGPGSSR | 12 | P48634 | Protein PRRC2A OS=Homo sapiens OX= | PRRC2A | 228.864 | 2 | 54.516624 | yes | 1.187E-07 | 566.29126 | 2172.4373 |
| FPLEEPGPR | 9 | P48634 | Protein PRRC2A OS=Homo sapiens OX= | PRRC2A | 228.864 | 2 | 54.516624 | yes | 1.583E-06 | 521.27185 | 1852.0005 |
| GPDELEGPDSK | 11 | P48634 | Protein PRRC2A OS=Homo sapiens OX= | PRRC2A | 228.864 | 2 | 54.516624 | yes | 1.076E-07 | 572.26184 | 1834.1637 |
| QGSVTAPGGHPR | 12 | P48634 | Protein PRRC2A OS=Homo sapiens OX= | PRRC2A | 228.864 | 2 | 54.516624 | yes | 9.722E-08 | 582.29944 | 1827.5321 |
| QSSSEISLAVER | 12 | P48634 | Protein PRRC2A OS=Homo sapiens OX= | PRRC2A | 228.864 | 2 | 54.516624 | yes | 1.128E-07 | 653.33588 | 1693.269 |
| GVGSGGQGPPPPR | 13 | P48634 | Protein PRRC2A OS=Homo sapiens OX= | PRRC2A | 228.864 | 2 | 54.516624 | yes | 2.914E-06 | 581.80182 | 1548.8055 |
| EQPLPPGPIGTER | 13 | P48634 | Protein PRRC2A OS=Homo sapiens OX= | PRRC2A | 228.864 | 2 | 54.516624 | yes | 1.12E-07 | 695.86987 | 1547.7303 |
| ELTASVTEAIPVSR | 14 | P48634 | Protein PRRC2A OS=Homo sapiens OX= | PRRC2A | 228.864 | 2 | 54.516624 | yes | 1.173E-07 | 736.90137 | 1375.8683 |
| PPPAPATR | 8 | P48634 | Protein PRRC2A OS=Homo sapiens OX= | PRRC2A | 228.864 | 2 | 54.516624 | yes | 4.095E-07 | 403.72958 | 1330.5714 |
| EGTLTQVPLAPPPI | 24 | P48634 | Protein PRRC2A OS=Homo sapiens OX= | PRRC2A | 228.864 | 3 | 54.516624 | yes | 2.74E-07 | 773.42163 | 1148.3934 |
| GDKEPGLPPPR | 11 | P48634 | Protein PRRC2A OS=Homo sapiens OX= | PRRC2A | 228.864 | 2 | 54.516624 | yes | 9.722E-08 | 581.81439 | 1106.5024 |
| AELHPVELKPFQD | 16 | P48634 | Protein PRRC2A OS=Homo sapiens OX= | PRRC2A | 228.864 | 3 | 54.516624 | yes | 1.02E-07 | 648.00714 | 1063.7335 |
| EAPPPVLLTPK | 11 | P48634 | Protein PRRC2A OS=Homo sapiens OX= | PRRC2A | 228.864 | 2 | 54.516624 | yes | 1.372E-07 | 581.34753 | 895.30511 |
| QPPSQGLGYPK | 11 | P48634 | Protein PRRC2A OS=Homo sapiens OX= | PRRC2A | 228.864 | 2 | 54.516624 | yes | 2.252E-07 | 586.30896 | 642.69739 |
| GGGTGGPNHPPAP | 14 | P48634 | Protein PRRC2A OS=Homo sapiens OX= | PRRC2A | 228.864 | 2 | 54.516624 | yes | 4.376E-07 | 636.31561 | 620.95367 |
| PFPASLGR | 8 | P48634 | Protein PRRC2A OS=Homo sapiens OX= | PRRC2A | 228.864 | 2 | 54.516624 | yes | 1.157E-07 | 422.73743 | 563.00116 |
| GIPPEEPGAPPR | 12 | P48634 | Protein PRRC2A OS=Homo sapiens OX= | PRRC2A | 228.864 | 2 | 54.516624 | yes | 2.255E-07 | 608.81964 | 528.99927 |
| PPPMPPMNFDPR | 12 | P48634 | Protein PRRC2A OS=Homo sapiens OX= | PRRC2A | 228.864 | 2 | 54.516624 | yes | 2.086E-07 | 698.33105 | 484.56503 |
| GPAGNWGPPGDY | 15 | P48634 | Protein PRRC2A OS=Homo sapiens OX= | PRRC2A | 228.864 | 2 | 54.516624 | yes | 1.443E-07 | 778.34985 | 476.25092 |
| KPPTGPLPPSK | 11 | P48634 | Protein PRRC2A OS=Homo sapiens OX= | PRRC2A | 228.864 | 2 | 54.516624 | yes | 9.722E-08 | 559.83203 | 468.44592 |
| KGNSPNSEPPTPK | 13 | P48634 | Protein PRRC2A OS=Homo sapiens OX= | PRRC2A | 228.864 | 2 | 54.516624 | yes | 9.722E-08 | 676.84387 | 430.79846 |
| ALYPGALGR | 9 | P48634 | Protein PRRC2A OS=Homo sapiens OX= | PRRC2A | 228.864 | 2 | 54.516624 | yes | 9.847E-07 | 459.26379 | 430.1376 |
| QAADEDDKGMR | 11 | P48634 | Protein PRRC2A OS=Homo sapiens OX= | PRRC2A | 228.864 | 2 | 54.516624 | yes | 5.195E-07 | 618.26965 | 372.36307 |
| TAWAETSRPPETE | 28 | P48634 | Protein PRRC2A OS=Homo sapiens OX= | PRRC2A | 228.864 | 5 | 54.516624 | yes | 5.103E-07 | 602.9165 | 183.89566 |
| TAWAETSRPPETE | 28 | P48634 | Protein PRRC2A OS=Homo sapiens OX= | PRRC2A | 228.864 | 5 | 54.516624 | yes | 3.249E-07 | 753.3938 | 183.89566 |
| TAWAETSRPPETE | 28 | P48634 | Protein PRRC2A OS=Homo sapiens OX= | PRRC2A | 228.864 | 4 | 54.516624 | yes | 5.103E-07 | 602.9165 | 183.89566 |
| TAWAETSRPPETE | 28 | P48634 | Protein PRRC2A OS=Homo sapiens OX= | PRRC2A | 228.864 | 4 | 54.516624 | yes | 3.249E-07 | 753.3938 | 183.89566 |
| SWAQASVTHGAH | 17 | P48634 | Protein PRRC2A OS=Homo sapiens OX= | PRRC2A | 228.864 | 2 | 54.516624 | yes | 9.809E-08 | 847.39313 | 299.61676 |
| LVVGDSLK | 8 | P48634 | Protein PRRC2A OS=Homo sapiens OX= | PRRC2A | 228.864 | 2 | 54.516624 | yes | 1.733E-07 | 415.75275 | 284.2988 |
| KYSSLNLFDTYK | 12 | P48634 | Protein PRRC2A OS=Homo sapiens OX= | PRRC2A | 228.864 | 2 | 54.516624 | yes | 1.049E-07 | 739.87994 | 219.05884 |
| SEGSEYEEIPK | 11 | P48634 | Protein PRRC2A OS=Homo sapiens OX= | PRRC2A | 228.864 | 2 | 54.516624 | yes | 9.722E-08 | 634.28809 | 212.76355 |
| KPPTGPLPPSKEPL | 15 | P48634 | Protein PRRC2A OS=Homo sapiens OX= | PRRC2A | 228.864 | 3 | 54.516624 | yes | 6.147E-07 | 529.31525 | 205.35338 |
| KQSSSEISLAVER | 13 | P48634 | Protein PRRC2A OS=Homo sapiens OX= | PRRC2A | 228.864 | 3 | 54.516624 | yes | 9.882E-08 | 478.59134 | 102.43511 |

| | | | | | | | | | | | |
|---|---|---|---|---|---|---|---|---|---|---|---|
| KQSSSEISLAVER | 13 | P48634 | Protein PRRC2A OS=Homo sapiens OX= | PRRC2A | 228.864 | 3 | 54.516624 | yes | 9.722E-08 | 717.38336 | 102.43511 |
| KQSSSEISLAVER | 13 | P48634 | Protein PRRC2A OS=Homo sapiens OX= | PRRC2A | 228.864 | 2 | 54.516624 | yes | 9.882E-08 | 478.59134 | 102.43511 |
| KQSSSEISLAVER | 13 | P48634 | Protein PRRC2A OS=Homo sapiens OX= | PRRC2A | 228.864 | 2 | 54.516624 | yes | 9.722E-08 | 717.38336 | 102.43511 |
| GNSPNSEPPTPK | 12 | P48634 | Protein PRRC2A OS=Homo sapiens OX= | PRRC2A | 228.864 | 2 | 54.516624 | yes | 1.717E-07 | 612.79639 | 180.18994 |
| PGPPVQFGTSDK | 12 | P48634 | Protein PRRC2A OS=Homo sapiens OX= | PRRC2A | 228.864 | 2 | 54.516624 | yes | 7.196E-06 | 615.31171 | 165.17506 |
| PPLDFYPPGVHPS( | 18 | P48634 | Protein PRRC2A OS=Homo sapiens OX= | PRRC2A | 228.864 | 3 | 54.516624 | yes | 3.488E-07 | 649.01587 | 144.19725 |
| SFSSQRPGMER | 11 | P48634 | Protein PRRC2A OS=Homo sapiens OX= | PRRC2A | 228.864 | 2 | 54.516624 | yes | 2.619E-07 | 641.30383 | 141.37917 |
| KEPPKEETAQLTG | 18 | P48634 | Protein PRRC2A OS=Homo sapiens OX= | PRRC2A | 228.864 | 4 | 54.516624 | yes | 1.776E-07 | 485.25363 | 128.66875 |
| SEGSEYEEIPKR | 12 | P48634 | Protein PRRC2A OS=Homo sapiens OX= | PRRC2A | 228.864 | 2 | 54.516624 | yes | 2.856E-07 | 712.33862 | 99.100853 |
| PGTPSLHPYR | 10 | P48634 | Protein PRRC2A OS=Homo sapiens OX= | PRRC2A | 228.864 | 2 | 54.516624 | yes | 2.424E-07 | 562.79602 | 97.925858 |
| SFSGLNSR | 8 | P48634 | Protein PRRC2A OS=Homo sapiens OX= | PRRC2A | 228.864 | 2 | 54.516624 | yes | 1.135E-06 | 434.2196 | 97.552025 |
| DSAGVSPFPPK | 11 | P48634 | Protein PRRC2A OS=Homo sapiens OX= | PRRC2A | 228.864 | 2 | 54.516624 | yes | 6.338E-07 | 551.28241 | 95.535522 |
| KPELLQEESLPPPI | 33 | P48634 | Protein PRRC2A OS=Homo sapiens OX= | PRRC2A | 228.864 | 5 | 54.516624 | yes | 1.975E-07 | 697.35681 | 88.1464 |
| SDSGGSSSEPFDR | 13 | P48634 | Protein PRRC2A OS=Homo sapiens OX= | PRRC2A | 228.864 | 2 | 54.516624 | yes | 9.722E-08 | 664.2735 | 74.683693 |
| GGLQPSGPPQFPPY | 15 | P48634 | Protein PRRC2A OS=Homo sapiens OX= | PRRC2A | 228.864 | 2 | 54.516624 | yes | 8.424E-07 | 799.40973 | 54.645565 |
| RPPPAPHDGDR | 11 | P48634 | Protein PRRC2A OS=Homo sapiens OX= | PRRC2A | 228.864 | 3 | 54.516624 | yes | 3.549E-06 | 405.539 | 51.596287 |
| VNSGLSSDPHFEEI | 19 | P48634 | Protein PRRC2A OS=Homo sapiens OX= | PRRC2A | 228.864 | 3 | 54.516624 | yes | 1.1E-07 | 685.6582 | 35.415184 |
| LVEPVGRPSILK | 12 | P48634 | Protein PRRC2A OS=Homo sapiens OX= | PRRC2A | 228.864 | 2 | 54.516624 | yes | 1.718E-06 | 654.40588 | 20.821285 |
| ESAEQSSGPGPSLI | 14 | P48634 | Protein PRRC2A OS=Homo sapiens OX= | PRRC2A | 228.864 | 2 | 54.516624 | yes | 3.527E-06 | 701.33386 | 4.0013041 |
| SLHDALCVIR | 10 | P48643 | T-complex protein 1 subunit epsilon OS= | CCT5 | 59.672 | 2 | 54.205784 | yes | 9.722E-08 | 592.31622 | 290.71646 |
| VAIEHLDK | 8 | P48643 | T-complex protein 1 subunit epsilon OS= | CCT5 | 59.672 | 2 | 54.205784 | yes | 1.2E-07 | 462.76108 | 243.3222 |
| MLVIEQCK | 8 | P48643 | T-complex protein 1 subunit epsilon OS= | CCT5 | 59.672 | 2 | 54.205784 | yes | 2.658E-07 | 510.76447 | 172.97133 |
| EKFEEMIQQIK | 11 | P48643 | T-complex protein 1 subunit epsilon OS= | CCT5 | 59.672 | 2 | 54.205784 | yes | 9.722E-08 | 711.86853 | 148.71425 |
| ISDSVLVDIK | 10 | P48643 | T-complex protein 1 subunit epsilon OS= | CCT5 | 59.672 | 2 | 54.205784 | yes | 9.722E-08 | 544.81354 | 125.56215 |
| LMGLEALK | 8 | P48643 | T-complex protein 1 subunit epsilon OS= | CCT5 | 59.672 | 2 | 54.205784 | yes | 7.059E-07 | 437.75696 | 121.12376 |
| EMNPALGIDCLHK | 13 | P48643 | T-complex protein 1 subunit epsilon OS= | CCT5 | 59.672 | 2 | 54.205784 | yes | 7.95E-07 | 749.36304 | 118.59084 |
| DFSHPQMPK | 9 | P48643 | T-complex protein 1 subunit epsilon OS= | CCT5 | 59.672 | 2 | 54.205784 | yes | 2.053E-06 | 543.75549 | 107.5028 |
| QQISLATQMVR | 11 | P48643 | T-complex protein 1 subunit epsilon OS= | CCT5 | 59.672 | 2 | 54.205784 | yes | 9.722E-08 | 637.8479 | 76.777214 |
| QQHVIETLIGK | 11 | P48643 | T-complex protein 1 subunit epsilon OS= | CCT5 | 59.672 | 2 | 54.205784 | yes | 9.722E-08 | 633.36426 | 65.316238 |
| DTEPLIQTAK | 10 | P48643 | T-complex protein 1 subunit epsilon OS= | CCT5 | 59.672 | 2 | 54.205784 | yes | 0.0002044 | 558.30078 | 25.081272 |
| ASMGTLAFDEYGI | 13 | P48643 | T-complex protein 1 subunit epsilon OS= | CCT5 | 59.672 | 2 | 54.205784 | yes | 5.59E-05 | 730.32971 | 16.248653 |
| AVLQFTPASWTY\ | 14 | P48651 | Phosphatidylserine synthase 1 OS=Homo | PTDSS1 | 55.528 | 2 | 53.572964 | yes | 1.018E-07 | 819.93555 | 586.8493 |
| QYYAYLTDTQCK | 12 | P48651 | Phosphatidylserine synthase 1 OS=Homo | PTDSS1 | 55.528 | 2 | 53.572964 | yes | 9.722E-08 | 777.35065 | 479.94287 |
| ILFIGGITAPTVR | 13 | P48651 | Phosphatidylserine synthase 1 OS=Homo | PTDSS1 | 55.528 | 2 | 53.572964 | yes | 1.048E-07 | 679.41376 | 441.44379 |
| TYHWASFK | 8 | P48651 | Phosphatidylserine synthase 1 OS=Homo | PTDSS1 | 55.528 | 2 | 53.572964 | yes | 1.462E-07 | 520.25342 | 210.16628 |
| SLMYWLDPNLR | 11 | P48651 | Phosphatidylserine synthase 1 OS=Homo | PTDSS1 | 55.528 | 2 | 53.572964 | yes | 9.722E-08 | 704.35809 | 151.15985 |
| FGQDLFSK | 8 | P48651 | Phosphatidylserine synthase 1 OS=Homo | PTDSS1 | 55.528 | 2 | 53.572964 | yes | 3.347E-07 | 471.23999 | 49.516392 |
| LELQQLQAER | 10 | P48681 | Nestin OS=Homo sapiens OX=9606 GN= | NES | 177.439 | 2 | 50.928104 | yes | 4.549E-07 | 614.33826 | 120.0453 |
| GPPAPAPEVEELAI | 14 | P48681 | Nestin OS=Homo sapiens OX=9606 GN= | NES | 177.439 | 2 | 50.928104 | yes | 2.687E-06 | 716.87518 | 108.85152 |

| | | | | | | | | | | | |
|---|---|---|---|---|---|---|---|---|---|---|---|
| GEGEGQIWGLVEK | 13 | P48681 | Nestin OS=Homo sapiens OX=9606 GN= | NES | 177.439 | 2 | 50.928104 | yes | 3.718E-07 | 701.35406 | 73.713684 |
| SLEEQDQETLR | 11 | P48681 | Nestin OS=Homo sapiens OX=9606 GN= | NES | 177.439 | 2 | 50.928104 | yes | 9.722E-08 | 674.323 | 60.449215 |
| DVEVVRPLEK | 10 | P48681 | Nestin OS=Homo sapiens OX=9606 GN= | NES | 177.439 | 2 | 50.928104 | yes | 8.452E-07 | 592.33771 | 59.387604 |
| EAVGQLKPTGK | 11 | P48681 | Nestin OS=Homo sapiens OX=9606 GN= | NES | 177.439 | 2 | 50.928104 | yes | 1.744E-07 | 564.32458 | 38.494164 |
| NQEFLQAR | 8 | P48681 | Nestin OS=Homo sapiens OX=9606 GN= | NES | 177.439 | 2 | 50.928104 | yes | 4.765E-07 | 503.25925 | 38.393002 |
| EDTQTLQSLQK | 11 | P48681 | Nestin OS=Homo sapiens OX=9606 GN= | NES | 177.439 | 2 | 50.928104 | yes | 3.859E-07 | 645.83044 | 36.006096 |
| EIQDSQVPLEK | 11 | P48681 | Nestin OS=Homo sapiens OX=9606 GN= | NES | 177.439 | 2 | 50.928104 | yes | 2.396E-06 | 643.33533 | 26.496 |
| FEEAPDYMYLR | 11 | P48729 | Casein kinase I isoform alpha OS=Homo | CSNK1A1 | 38.915 | 2 | 51.232506 | yes | 1.363E-07 | 717.32391 | 1143.0123 |
| TVLMLADQMISR | 12 | P48729 | Casein kinase I isoform alpha OS=Homo | CSNK1A1 | 38.915 | 2 | 51.232506 | no | 9.722E-08 | 689.36505 | 1108.9525 |
| DIKPDNFLMGIGR | 13 | P48729 | Casein kinase I isoform alpha OS=Homo | CSNK1A1 | 38.915 | 2 | 51.232506 | yes | 1.482E-07 | 738.38739 | 619.42743 |
| HPQLLYESK | 9 | P48729 | Casein kinase I isoform alpha OS=Homo | CSNK1A1 | 38.915 | 2 | 51.232506 | no | 9.722E-08 | 557.79822 | 556.95343 |
| IEYVHTK | 7 | P48729 | Casein kinase I isoform alpha OS=Homo | CSNK1A1 | 38.915 | 2 | 51.232506 | no | 9.722E-08 | 445.24252 | 547.00214 |
| MSTPVEVLCK | 10 | P48729 | Casein kinase I isoform alpha OS=Homo | CSNK1A1 | 38.915 | 2 | 51.232506 | no | 9.727E-08 | 582.29358 | 313.32675 |
| TSLPWQGLK | 9 | P48729 | Casein kinase I isoform alpha OS=Homo | CSNK1A1 | 38.915 | 2 | 51.232506 | yes | 4.748E-07 | 515.29004 | 258.46976 |
| YASINAHLGIEQSR | 14 | P48729 | Casein kinase I isoform alpha OS=Homo | CSNK1A1 | 38.915 | 2 | 51.232506 | no | 6.368E-07 | 779.90228 | 120.46853 |
| GAPVNISSSDLTGR | 14 | P48730 | Casein kinase I isoform delta OS=Homo s | CSNK1D | 47.331 | 2 | 49.838768 | yes | 1.304E-07 | 687.35461 | 1335.3657 |
| MMQGGVGIPTIR | 12 | P48730 | Casein kinase I isoform delta OS=Homo s | CSNK1D | 47.331 | 2 | 49.838768 | yes | 1.002E-07 | 630.33356 | 1099.6959 |
| VETEDQVLATFCG | 14 | P48740 | Mannan-binding lectin serine protease 1 ( | MASP1 | 79.247 | 2 | 51.669094 | yes | 9.722E-08 | 812.88544 | 109.47717 |
| FSEINPDMK | 9 | P48960 | Adhesion G protein-coupled receptor E5 ( | ADGRE5 | 91.87 | 2 | 32.701973 | yes | 1.438E-06 | 540.75513 | 37.785538 |
| FLGTSGQNVSDIFF | 14 | P49005 | DNA polymerase delta subunit 2 OS=Hor | POLD2 | 51.29 | 2 | 52.078934 | yes | 9.722E-08 | 770.89136 | 56.634098 |
| AMPLQPSILR | 10 | P49005 | DNA polymerase delta subunit 2 OS=Hor | POLD2 | 51.29 | 2 | 52.078934 | yes | 1.145E-06 | 563.32605 | 24.242792 |
| VPVATYTNSSQPF | 14 | P49005 | DNA polymerase delta subunit 2 OS=Hor | POLD2 | 51.29 | 2 | 52.078934 | yes | 6.834E-07 | 783.89917 | 22.523703 |
| DVKPENLLYTSK | 12 | P49137 | MAP kinase-activated protein kinase 2 OS | MAPKAPK | 45.568 | 2 | 35.73907 | yes | 7.885E-07 | 703.87994 | 27.992748 |
| GIKPVTLELGGK | 12 | P49189 | 4-trimethylaminobutyraldehyde dehydrog | ALDH9A1 | 53.802 | 2 | 35.28011 | yes | 8.569E-07 | 606.37152 | 28.978949 |
| AFLIEEQK | 8 | P49207 | Large ribosomal subunit protein eL34 OS | RPL34 | 13.293 | 2 | 51.27728 | yes | 3.034E-07 | 489.26874 | 25502.303 |
| IVYLYTK | 7 | P49207 | Large ribosomal subunit protein eL34 OS | RPL34 | 13.293 | 2 | 51.27728 | yes | 1.984E-07 | 450.26547 | 6810.082 |
| LSYNTASNK | 9 | P49207 | Large ribosomal subunit protein eL34 OS | RPL34 | 13.293 | 2 | 51.27728 | yes | 9.722E-08 | 499.2511 | 5662.1768 |
| RLSYNTASNK | 10 | P49207 | Large ribosomal subunit protein eL34 OS | RPL34 | 13.293 | 2 | 51.27728 | yes | 1.009E-07 | 577.30164 | 1551.7026 |
| AYGGSMCAK | 9 | P49207 | Large ribosomal subunit protein eL34 OS | RPL34 | 13.293 | 2 | 51.27728 | yes | 3.016E-07 | 472.70187 | 1414.5437 |
| SACGVCPGR | 9 | P49207 | Large ribosomal subunit protein eL34 OS | RPL34 | 13.293 | 2 | 51.27728 | yes | 3.558E-06 | 482.21039 | 553.39795 |
| MVVPGLDGAQIPR | 13 | P49327 | Fatty acid synthase OS=Homo sapiens OX | FASN | 273.427 | 2 | 54.59156 | yes | 1.317E-07 | 676.8714 | 379.1268 |
| HGLYLPTR | 8 | P49327 | Fatty acid synthase OS=Homo sapiens OX | FASN | 273.427 | 2 | 54.59156 | yes | 1.005E-07 | 478.76926 | 355.45978 |
| LSIPTYGLQCTR | 12 | P49327 | Fatty acid synthase OS=Homo sapiens OX | FASN | 273.427 | 2 | 54.59156 | yes | 2.291E-07 | 704.86627 | 263.63153 |
| ACLDTAVENMPSI | 14 | P49327 | Fatty acid synthase OS=Homo sapiens OX | FASN | 273.427 | 2 | 54.59156 | yes | 9.722E-08 | 774.87347 | 229.40172 |
| TGTVSLEVR | 9 | P49327 | Fatty acid synthase OS=Homo sapiens OX | FASN | 273.427 | 2 | 54.59156 | yes | 2.48E-06 | 481.26929 | 221.79485 |
| GLVQALQTK | 9 | P49327 | Fatty acid synthase OS=Homo sapiens OX | FASN | 273.427 | 2 | 54.59156 | yes | 1.086E-07 | 479.29001 | 214.89569 |
| LGMLSPEGTCK | 11 | P49327 | Fatty acid synthase OS=Homo sapiens OX | FASN | 273.427 | 2 | 54.59156 | yes | 4.611E-07 | 596.7887 | 200.60709 |
| SEGVVAVLLTK | 11 | P49327 | Fatty acid synthase OS=Homo sapiens OX | FASN | 273.427 | 2 | 54.59156 | yes | 1.564E-07 | 558.33716 | 199.22745 |
| DPSQQELPR | 9 | P49327 | Fatty acid synthase OS=Homo sapiens OX | FASN | 273.427 | 2 | 54.59156 | yes | 1.009E-07 | 535.26727 | 198.85971 |

| | | | | | | | | | | | |
|---|---|---|---|---|---|---|---|---|---|---|---|
| FDASFFGVHPK | 11 | P49327 | Fatty acid synthase OS=Homo sapiens OX | FASN | 273.427 | 2 | 54.59156 | yes | 9.862E-08 | 626.31146 | 184.01645 |
| SLLVNPEGPTLMR | 13 | P49327 | Fatty acid synthase OS=Homo sapiens OX | FASN | 273.427 | 2 | 54.59156 | yes | 1.333E-06 | 713.88977 | 179.29669 |
| ELNLVLSVR | 9 | P49327 | Fatty acid synthase OS=Homo sapiens OX | FASN | 273.427 | 2 | 54.59156 | yes | 1.247E-07 | 521.81641 | 153.03751 |
| VLQGDLVMNVYR | 12 | P49327 | Fatty acid synthase OS=Homo sapiens OX | FASN | 273.427 | 2 | 54.59156 | yes | 9.81E-08 | 703.87665 | 148.33098 |
| GTPLISPLIK | 10 | P49327 | Fatty acid synthase OS=Homo sapiens OX | FASN | 273.427 | 2 | 54.59156 | yes | 1.562E-07 | 519.83154 | 139.73732 |
| YHGNVMLLR | 9 | P49327 | Fatty acid synthase OS=Homo sapiens OX | FASN | 273.427 | 2 | 54.59156 | yes | 4.939E-07 | 551.79492 | 139.67668 |
| LQVVDQPLPVR | 11 | P49327 | Fatty acid synthase OS=Homo sapiens OX | FASN | 273.427 | 2 | 54.59156 | yes | 2.438E-07 | 632.37463 | 138.48222 |
| EVWALVQAGIR | 11 | P49327 | Fatty acid synthase OS=Homo sapiens OX | FASN | 273.427 | 2 | 54.59156 | yes | 9.722E-08 | 621.3537 | 130.40727 |
| SLYQSAGVAPESF | 24 | P49327 | Fatty acid synthase OS=Homo sapiens OX | FASN | 273.427 | 3 | 54.59156 | yes | 1.456E-07 | 848.07471 | 125.88086 |
| AALQEELQLCK | 11 | P49327 | Fatty acid synthase OS=Homo sapiens OX | FASN | 273.427 | 2 | 54.59156 | yes | 2.212E-07 | 651.83972 | 121.75175 |
| VVEVLAGHGHLYS | 14 | P49327 | Fatty acid synthase OS=Homo sapiens OX | FASN | 273.427 | 3 | 54.59156 | yes | 9.722E-08 | 512.94757 | 57.645546 |
| VVEVLAGHGHLYS | 14 | P49327 | Fatty acid synthase OS=Homo sapiens OX | FASN | 273.427 | 3 | 54.59156 | yes | 6.558E-07 | 768.91772 | 57.645546 |
| VVEVLAGHGHLYS | 14 | P49327 | Fatty acid synthase OS=Homo sapiens OX | FASN | 273.427 | 2 | 54.59156 | yes | 9.722E-08 | 512.94757 | 57.645546 |
| VVEVLAGHGHLYS | 14 | P49327 | Fatty acid synthase OS=Homo sapiens OX | FASN | 273.427 | 2 | 54.59156 | yes | 6.558E-07 | 768.91772 | 57.645546 |
| QAHTMDPQLR | 10 | P49327 | Fatty acid synthase OS=Homo sapiens OX | FASN | 273.427 | 2 | 54.59156 | yes | 5.148E-07 | 598.79565 | 114.84114 |
| GVDLVLNSLAEEK | 13 | P49327 | Fatty acid synthase OS=Homo sapiens OX | FASN | 273.427 | 2 | 54.59156 | yes | 1.667E-07 | 693.87738 | 105.85928 |
| VAAAVDLIIK | 10 | P49327 | Fatty acid synthase OS=Homo sapiens OX | FASN | 273.427 | 2 | 54.59156 | yes | 2.274E-07 | 506.8237 | 93.426376 |
| ADEASELACPTPK | 13 | P49327 | Fatty acid synthase OS=Homo sapiens OX | FASN | 273.427 | 2 | 54.59156 | yes | 2.334E-07 | 694.82178 | 89.078094 |
| VSVHVIEGDHR | 11 | P49327 | Fatty acid synthase OS=Homo sapiens OX | FASN | 273.427 | 2 | 54.59156 | yes | 9.722E-08 | 624.32819 | 78.534508 |
| VGDPQELNGITR | 12 | P49327 | Fatty acid synthase OS=Homo sapiens OX | FASN | 273.427 | 2 | 54.59156 | yes | 1.13E-06 | 649.83856 | 67.3675 |
| SFYGSTLFLCR | 11 | P49327 | Fatty acid synthase OS=Homo sapiens OX | FASN | 273.427 | 2 | 54.59156 | yes | 9.722E-08 | 675.82916 | 64.516769 |
| EDGLAQQQTQLNI | 14 | P49327 | Fatty acid synthase OS=Homo sapiens OX | FASN | 273.427 | 2 | 54.59156 | yes | 1.237E-06 | 807.41571 | 63.110207 |
| QEPLLIGSTK | 10 | P49327 | Fatty acid synthase OS=Homo sapiens OX | FASN | 273.427 | 2 | 54.59156 | yes | 2.114E-07 | 543.31366 | 61.440022 |
| VVVQVLAEEPEAV | 15 | P49327 | Fatty acid synthase OS=Homo sapiens OX | FASN | 273.427 | 2 | 54.59156 | yes | 1.154E-07 | 811.9718 | 55.329895 |
| FPQLDSTSFANSR | 13 | P49327 | Fatty acid synthase OS=Homo sapiens OX | FASN | 273.427 | 2 | 54.59156 | yes | 1.304E-05 | 735.35461 | 47.15881 |
| CTVFHGAQVEDAI | 14 | P49327 | Fatty acid synthase OS=Homo sapiens OX | FASN | 273.427 | 2 | 54.59156 | yes | 1.791E-06 | 818.88049 | 44.852741 |
| DGLLENQTPEFFQI | 19 | P49327 | Fatty acid synthase OS=Homo sapiens OX | FASN | 273.427 | 3 | 54.59156 | yes | 1.632E-07 | 755.70007 | 44.111179 |
| VLEALLPLK | 9 | P49327 | Fatty acid synthase OS=Homo sapiens OX | FASN | 273.427 | 2 | 54.59156 | yes | 1.952E-07 | 498.32861 | 40.810177 |
| VTVAGGVHISGLH | 19 | P49327 | Fatty acid synthase OS=Homo sapiens OX | FASN | 273.427 | 3 | 54.59156 | yes | 2.509E-07 | 630.00604 | 37.667549 |
| VYATILNAGTNTD | 16 | P49327 | Fatty acid synthase OS=Homo sapiens OX | FASN | 273.427 | 2 | 54.59156 | yes | 7.694E-07 | 842.93066 | 34.728218 |
| AINCATSGVVGLV | 17 | P49327 | Fatty acid synthase OS=Homo sapiens OX | FASN | 273.427 | 3 | 54.59156 | yes | 5.841E-07 | 601.97833 | 31.795319 |
| GMGLSLMR | 8 | P49327 | Fatty acid synthase OS=Homo sapiens OX | FASN | 273.427 | 2 | 54.59156 | yes | 3.068E-07 | 432.72513 | 29.699482 |
| VTAIHIDPATHR | 12 | P49327 | Fatty acid synthase OS=Homo sapiens OX | FASN | 273.427 | 2 | 54.59156 | yes | 9.722E-08 | 665.86493 | 20.926208 |
| PLFLVHPIEGSTTV | 21 | P49327 | Fatty acid synthase OS=Homo sapiens OX | FASN | 273.427 | 4 | 54.59156 | yes | 3.214E-06 | 577.81665 | 14.185351 |
| GILADEDSSRPVW | 15 | P49327 | Fatty acid synthase OS=Homo sapiens OX | FASN | 273.427 | 2 | 54.59156 | yes | 5.28E-07 | 843.44647 | 11.146296 |
| QIEGTGISMSACR | 13 | P49336 | Cyclin-dependent kinase 8 OS=Homo sap | CDK8 | 53.284 | 2 | 52.601322 | no | 3.933E-05 | 705.32922 | 136.84726 |
| VFLSHADR | 8 | P49336 | Cyclin-dependent kinase 8 OS=Homo sap | CDK8 | 53.284 | 2 | 52.601322 | yes | 5.282E-07 | 472.75107 | 104.26577 |
| APELLLGAR | 9 | P49336 | Cyclin-dependent kinase 8 OS=Homo sap | CDK8 | 53.284 | 2 | 52.601322 | no | 9.722E-08 | 470.28473 | 60.783543 |
| IPGGIIEDSCVLR | 13 | P49368 | T-complex protein 1 subunit gamma OS= | CCT3 | 60.534 | 2 | 52.07971 | yes | 9.722E-08 | 714.87939 | 357.33685 |

| | | | | | | | | | | | |
|---|---|---|---|---|---|---|---|---|---|---|---|
| TAVETAVLLLR | 11 | P49368 | T-complex protein 1 subunit gamma OS= | CCT3 | 60.534 | 2 | 52.07971 | yes | 1.02E-07 | 593.36371 | 333.95883 |
| IVLLDSSLEYK | 11 | P49368 | T-complex protein 1 subunit gamma OS= | CCT3 | 60.534 | 2 | 52.07971 | yes | 1.542E-07 | 640.36084 | 279.84732 |
| WSSLACNIALDAV | 14 | P49368 | T-complex protein 1 subunit gamma OS= | CCT3 | 60.534 | 2 | 52.07971 | yes | 1.012E-07 | 774.39795 | 261.5213 |
| TLIQNCGASTIR | 12 | P49368 | T-complex protein 1 subunit gamma OS= | CCT3 | 60.534 | 2 | 52.07971 | yes | 9.722E-08 | 667.34827 | 245.06911 |
| ALDDMISTLK | 10 | P49368 | T-complex protein 1 subunit gamma OS= | CCT3 | 60.534 | 2 | 52.07971 | yes | 2.212E-07 | 553.79175 | 225.99174 |
| AVAQALEVIPR | 11 | P49368 | T-complex protein 1 subunit gamma OS= | CCT3 | 60.534 | 2 | 52.07971 | yes | 1.054E-07 | 583.84821 | 224.66515 |
| ELGIWEPLAVK | 11 | P49368 | T-complex protein 1 subunit gamma OS= | CCT3 | 60.534 | 2 | 52.07971 | yes | 6.66E-07 | 627.85828 | 204.63991 |
| IDDIVSGHK | 9 | P49368 | T-complex protein 1 subunit gamma OS= | CCT3 | 60.534 | 2 | 52.07971 | yes | 3.394E-07 | 492.26144 | 157.58194 |
| VQSGNINAAK | 10 | P49368 | T-complex protein 1 subunit gamma OS= | CCT3 | 60.534 | 2 | 52.07971 | yes | 5.074E-06 | 501.27237 | 149.0761 |
| EIQVQHPAAK | 10 | P49368 | T-complex protein 1 subunit gamma OS= | CCT3 | 60.534 | 2 | 52.07971 | yes | 1.476E-07 | 560.80908 | 135.62883 |
| NLQDAMQVCR | 10 | P49368 | T-complex protein 1 subunit gamma OS= | CCT3 | 60.534 | 2 | 52.07971 | yes | 2.035E-07 | 617.78699 | 123.47263 |
| IGDEYFTFITDCK | 13 | P49368 | T-complex protein 1 subunit gamma OS= | CCT3 | 60.534 | 2 | 52.07971 | yes | 3.548E-07 | 804.86615 | 103.03219 |
| AMTGVEQWPYR | 11 | P49368 | T-complex protein 1 subunit gamma OS= | CCT3 | 60.534 | 2 | 52.07971 | yes | 2.951E-07 | 669.31897 | 83.872696 |
| PVLVLSQNTK | 10 | P49368 | T-complex protein 1 subunit gamma OS= | CCT3 | 60.534 | 2 | 52.07971 | yes | 1.541E-07 | 549.82953 | 71.465332 |
| ACTILLR | 7 | P49368 | T-complex protein 1 subunit gamma OS= | CCT3 | 60.534 | 2 | 52.07971 | yes | 4.26E-07 | 423.74692 | 70.026985 |
| IVSRPEELR | 9 | P49368 | T-complex protein 1 subunit gamma OS= | CCT3 | 60.534 | 2 | 52.07971 | yes | 3.334E-07 | 549.81696 | 66.755714 |
| KGESQTDIEITR | 12 | P49368 | T-complex protein 1 subunit gamma OS= | CCT3 | 60.534 | 2 | 52.07971 | yes | 9.809E-08 | 688.85443 | 59.51712 |
| EDDVGTGAGLLEI | 14 | P49368 | T-complex protein 1 subunit gamma OS= | CCT3 | 60.534 | 2 | 52.07971 | yes | 5.703E-07 | 708.86444 | 27.825397 |
| EILGIIVSYK | 10 | P49407 | Beta-arrestin-1 OS=Homo sapiens OX=9 | ARRB1 | 47.066 | 2 | 50.35449 | yes | 9.81E-08 | 567.8421 | 107.77237 |
| ACGVDYEVK | 9 | P49407 | Beta-arrestin-1 OS=Homo sapiens OX=9 | ARRB1 | 47.066 | 2 | 50.35449 | yes | 2.416E-06 | 520.7395 | 85.356895 |
| AFCAENLEEK | 10 | P49407 | Beta-arrestin-1 OS=Homo sapiens OX=9 | ARRB1 | 47.066 | 2 | 50.35449 | yes | 9.995E-08 | 605.77405 | 50.059944 |
| EDLDVLGLTFR | 11 | P49407 | Beta-arrestin-1 OS=Homo sapiens OX=9 | ARRB1 | 47.066 | 2 | 50.35449 | yes | 1.902E-07 | 639.34045 | 31.869902 |
| GTVVTGTLER | 10 | P49411 | Elongation factor Tu, mitochondrial OS= | TUFM | 49.875 | 2 | 52.68331 | yes | 1.637E-07 | 516.78784 | 3372.3235 |
| AEAGDNLGALVR | 12 | P49411 | Elongation factor Tu, mitochondrial OS= | TUFM | 49.875 | 2 | 52.68331 | yes | 2.868E-07 | 593.31476 | 3027.5322 |
| ELAMPGEDLK | 10 | P49411 | Elongation factor Tu, mitochondrial OS= | TUFM | 49.875 | 2 | 52.68331 | yes | 1.642E-07 | 551.77606 | 2214.3057 |
| LLDAVDTYIPVPA | 14 | P49411 | Elongation factor Tu, mitochondrial OS= | TUFM | 49.875 | 2 | 52.68331 | yes | 1.095E-07 | 771.92993 | 1745.2379 |
| TVVTGIEMFHK | 11 | P49411 | Elongation factor Tu, mitochondrial OS= | TUFM | 49.875 | 2 | 52.68331 | yes | 1.003E-07 | 631.33411 | 716.11792 |
| TVVTGIEMFHK | 11 | P49411 | Elongation factor Tu, mitochondrial OS= | TUFM | 49.875 | 2 | 52.68331 | yes | 1.814E-06 | 421.22516 | 716.11792 |
| TVVTGIEMFHK | 11 | P49411 | Elongation factor Tu, mitochondrial OS= | TUFM | 49.875 | 3 | 52.68331 | yes | 1.003E-07 | 631.33411 | 716.11792 |
| TVVTGIEMFHK | 11 | P49411 | Elongation factor Tu, mitochondrial OS= | TUFM | 49.875 | 3 | 52.68331 | yes | 1.814E-06 | 421.22516 | 716.11792 |
| VEAQVYILSK | 10 | P49411 | Elongation factor Tu, mitochondrial OS= | TUFM | 49.875 | 2 | 52.68331 | yes | 9.825E-08 | 575.32935 | 1407.0442 |
| TTLTAAITK | 9 | P49411 | Elongation factor Tu, mitochondrial OS= | TUFM | 49.875 | 2 | 52.68331 | yes | 1.202E-07 | 460.27658 | 1230.4554 |
| ELLTEFGYK | 9 | P49411 | Elongation factor Tu, mitochondrial OS= | TUFM | 49.875 | 2 | 52.68331 | yes | 1.255E-07 | 550.28711 | 1127.0533 |
| YEEIDNAPEER | 11 | P49411 | Elongation factor Tu, mitochondrial OS= | TUFM | 49.875 | 2 | 52.68331 | yes | 1.118E-07 | 682.80188 | 876.67328 |
| QIGVEHVVVYVNI | 13 | P49411 | Elongation factor Tu, mitochondrial OS= | TUFM | 49.875 | 2 | 52.68331 | yes | 9.968E-08 | 742.41699 | 278.46014 |
| QIGVEHVVVYVNI | 13 | P49411 | Elongation factor Tu, mitochondrial OS= | TUFM | 49.875 | 2 | 52.68331 | yes | 9.809E-08 | 495.28043 | 278.46014 |
| QIGVEHVVVYVNI | 13 | P49411 | Elongation factor Tu, mitochondrial OS= | TUFM | 49.875 | 3 | 52.68331 | yes | 9.968E-08 | 742.41699 | 278.46014 |
| QIGVEHVVVYVNI | 13 | P49411 | Elongation factor Tu, mitochondrial OS= | TUFM | 49.875 | 3 | 52.68331 | yes | 9.809E-08 | 495.28043 | 278.46014 |
| GITINAAHVEYSTA | 16 | P49411 | Elongation factor Tu, mitochondrial OS= | TUFM | 49.875 | 2 | 52.68331 | yes | 9.722E-08 | 837.4339 | 191.97789 |

| | | | | | | | | | | | |
|---|---|---|---|---|---|---|---|---|---|---|---|
| GITINAAHVEYSTA | 16 | P49411 | Elongation factor Tu, mitochondrial OS=I | TUFM | 49.875 | 2 | 52.68331 | yes | 4.348E-07 | 558.62506 | 191.97789 |
| GITINAAHVEYSTA | 16 | P49411 | Elongation factor Tu, mitochondrial OS=I | TUFM | 49.875 | 3 | 52.68331 | yes | 9.722E-08 | 837.4339 | 191.97789 |
| GITINAAHVEYSTA | 16 | P49411 | Elongation factor Tu, mitochondrial OS=I | TUFM | 49.875 | 3 | 52.68331 | yes | 4.348E-07 | 558.62506 | 191.97789 |
| GDECELLGHSK | 11 | P49411 | Elongation factor Tu, mitochondrial OS=I | TUFM | 49.875 | 2 | 52.68331 | yes | 9.722E-08 | 622.78241 | 322.85388 |
| GLVMVKPGSIKPH | 15 | P49411 | Elongation factor Tu, mitochondrial OS=I | TUFM | 49.875 | 3 | 52.68331 | yes | 3.001E-07 | 540.32025 | 104.49762 |
| GLVMVKPGSIKPH | 15 | P49411 | Elongation factor Tu, mitochondrial OS=I | TUFM | 49.875 | 3 | 52.68331 | yes | 1.402E-06 | 809.97668 | 104.49762 |
| GLVMVKPGSIKPH | 15 | P49411 | Elongation factor Tu, mitochondrial OS=I | TUFM | 49.875 | 2 | 52.68331 | yes | 3.001E-07 | 540.32025 | 104.49762 |
| GLVMVKPGSIKPH | 15 | P49411 | Elongation factor Tu, mitochondrial OS=I | TUFM | 49.875 | 2 | 52.68331 | yes | 1.402E-06 | 809.97668 | 104.49762 |
| EHLLLAR | 7 | P49411 | Elongation factor Tu, mitochondrial OS=I | TUFM | 49.875 | 2 | 52.68331 | yes | 2.543E-07 | 426.25851 | 202.43541 |
| HYAHTDCPGHADY | 15 | P49411 | Elongation factor Tu, mitochondrial OS=I | TUFM | 49.875 | 4 | 52.68331 | yes | 1.692E-07 | 443.44678 | 190.983 |
| DLEKPFLLPVEAV | 19 | P49411 | Elongation factor Tu, mitochondrial OS=I | TUFM | 49.875 | 3 | 52.68331 | yes | 1.365E-07 | 710.39288 | 160.24133 |
| KYEEIDNAPEER | 12 | P49411 | Elongation factor Tu, mitochondrial OS=I | TUFM | 49.875 | 2 | 52.68331 | yes | 9.758E-08 | 746.84937 | 126.43615 |
| IILPPEK | 7 | P49411 | Elongation factor Tu, mitochondrial OS=I | TUFM | 49.875 | 2 | 52.68331 | yes | 1.662E-07 | 405.26019 | 106.35028 |
| DKPHVNVGTIGHV | 17 | P49411 | Elongation factor Tu, mitochondrial OS=I | TUFM | 49.875 | 4 | 52.68331 | yes | 3.103E-07 | 453.23932 | 97.749969 |
| GLVMVKPGSIK | 11 | P49411 | Elongation factor Tu, mitochondrial OS=I | TUFM | 49.875 | 2 | 52.68331 | yes | 9.722E-08 | 564.84412 | 67.904884 |
| KGDECELLGHSK | 12 | P49411 | Elongation factor Tu, mitochondrial OS=I | TUFM | 49.875 | 2 | 52.68331 | yes | 9.722E-08 | 686.8299 | 62.315533 |
| VPTTLAEYCVK | 11 | P49427 | Ubiquitin-conjugating enzyme E2 R1 OS= | CDC34 | 26.737 | 2 | 51.178444 | yes | 9.722E-08 | 640.83136 | 125.48709 |
| GLQEEPVEGFR | 11 | P49427 | Ubiquitin-conjugating enzyme E2 R1 OS= | CDC34 | 26.737 | 2 | 51.178444 | yes | 2.079E-07 | 630.81458 | 107.78888 |
| PLVPSSQK | 8 | P49427 | Ubiquitin-conjugating enzyme E2 R1 OS= | CDC34 | 26.737 | 2 | 51.178444 | yes | 1.673E-07 | 428.25037 | 36.68755 |
| STHLLIR | 7 | P49450 | Histone H3-like centromeric protein A OS | CENPA | 15.991 | 2 | 52.37685 | yes | 9.722E-08 | 420.25851 | 117.1261 |
| EIQELEQLLSSER | 13 | P49454 | Centromere protein F OS=Homo sapiens | CENPF | 357.528 | 2 | 53.483513 | yes | 2.18E-07 | 787.40704 | 149.96327 |
| AENLVLSTNLR | 11 | P49454 | Centromere protein F OS=Homo sapiens | CENPF | 357.528 | 2 | 53.483513 | yes | 4.799E-07 | 615.34607 | 79.541245 |
| LALSPLSLGK | 10 | P49454 | Centromere protein F OS=Homo sapiens | CENPF | 357.528 | 2 | 53.483513 | yes | 3.931E-07 | 499.81586 | 74.371941 |
| ELSQLQAAQEK | 11 | P49454 | Centromere protein F OS=Homo sapiens | CENPF | 357.528 | 2 | 53.483513 | yes | 1.732E-07 | 622.8277 | 62.576546 |
| SLLGIDTEDAIQGR | 14 | P49454 | Centromere protein F OS=Homo sapiens | CENPF | 357.528 | 2 | 53.483513 | yes | 1.685E-07 | 744.38867 | 53.081959 |
| LMEICESLEK | 10 | P49454 | Centromere protein F OS=Homo sapiens | CENPF | 357.528 | 2 | 53.483513 | yes | 3.98E-07 | 626.30164 | 50.454319 |
| LLTQMESEK | 9 | P49454 | Centromere protein F OS=Homo sapiens | CENPF | 357.528 | 2 | 53.483513 | yes | 1.711E-06 | 539.77606 | 49.432453 |
| GVEELESLCEVYR | 13 | P49454 | Centromere protein F OS=Homo sapiens | CENPF | 357.528 | 2 | 53.483513 | yes | 1.25E-07 | 791.87451 | 48.281849 |
| QFQLDSLEAALQK | 13 | P49454 | Centromere protein F OS=Homo sapiens | CENPF | 357.528 | 2 | 53.483513 | yes | 9.722E-08 | 745.89612 | 47.31287 |
| IQELEGQLDK | 10 | P49454 | Centromere protein F OS=Homo sapiens | CENPF | 357.528 | 2 | 53.483513 | yes | 4.935E-06 | 586.81152 | 36.2939 |
| SFQTLDQECIQMK | 13 | P49454 | Centromere protein F OS=Homo sapiens | CENPF | 357.528 | 2 | 53.483513 | yes | 2.352E-06 | 814.37634 | 26.166727 |
| NQIQDQEQLVSK | 12 | P49454 | Centromere protein F OS=Homo sapiens | CENPF | 357.528 | 2 | 53.483513 | yes | 3.197E-06 | 715.36774 | 19.659016 |
| FHSQLMR | 7 | P49458 | Signal recognition particle 9 kDa protein | SRP9 | 10.112 | 2 | 48.88817 | yes | 1.585E-07 | 459.73434 | 10944.132 |
| VTDDLVCLVYK | 11 | P49458 | Signal recognition particle 9 kDa protein | SRP9 | 10.112 | 2 | 48.88817 | yes | 1.248E-07 | 662.84448 | 7073.5278 |
| PQYQTWEEFSR | 11 | P49458 | Signal recognition particle 9 kDa protein | SRP9 | 10.112 | 2 | 48.88817 | yes | 1.165E-07 | 735.83606 | 5071.6152 |
| LYLADPMK | 8 | P49458 | Signal recognition particle 9 kDa protein | SRP9 | 10.112 | 2 | 48.88817 | yes | 3.405E-07 | 475.75443 | 2267.9785 |
| LYLADPMK | 8 | P49458 | Signal recognition particle 9 kDa protein | SRP9 | 10.112 | 2 | 48.88817 | yes | 1.454E-06 | 483.75189 | 2267.9785 |
| HSDGNLCVK | 9 | P49458 | Signal recognition particle 9 kDa protein | SRP9 | 10.112 | 2 | 48.88817 | yes | 1.09E-07 | 515.24274 | 3728.0745 |
| ITCLCQVPQNAAN | 14 | P49588 | Alanine--tRNA ligase, cytoplasmic OS=H | AARS1 | 106.811 | 2 | 51.886856 | yes | 9.819E-08 | 822.90088 | 363.04224 |

| | | | | | | | | | | | |
|---|---|---|---|---|---|---|---|---|---|---|---|
| AVFDETYPDPVR | 12 | P49588 | Alanine--tRNA ligase, cytoplasmic OS=H | AARS1 | 106.811 | 2 | 51.886856 | yes | 2.174E-07 | 704.84082 | 340.02319 |
| GLVVDMDGFEEEI | 13 | P49588 | Alanine--tRNA ligase, cytoplasmic OS=H | AARS1 | 106.811 | 2 | 51.886856 | yes | 9.809E-08 | 748.34033 | 306.35751 |
| VGAEDADGIDMA' | 14 | P49588 | Alanine--tRNA ligase, cytoplasmic OS=H | AARS1 | 106.811 | 2 | 51.886856 | yes | 1.464E-07 | 741.83008 | 279.36203 |
| VGDQVWLFIDEPR | 13 | P49588 | Alanine--tRNA ligase, cytoplasmic OS=H | AARS1 | 106.811 | 2 | 51.886856 | yes | 9.722E-08 | 787.40411 | 256.80328 |
| SIDTGMGLER | 10 | P49588 | Alanine--tRNA ligase, cytoplasmic OS=H | AARS1 | 106.811 | 2 | 51.886856 | yes | 9.722E-08 | 539.76349 | 213.68311 |
| DIINEEEVQFLK | 12 | P49588 | Alanine--tRNA ligase, cytoplasmic OS=H | AARS1 | 106.811 | 2 | 51.886856 | yes | 1.009E-07 | 738.88269 | 147.46054 |
| GAMSTQQIK | 9 | P49588 | Alanine--tRNA ligase, cytoplasmic OS=H | AARS1 | 106.811 | 2 | 51.886856 | yes | 2.851E-07 | 482.25003 | 132.23595 |
| ASEWVQQVSGLM | 15 | P49588 | Alanine--tRNA ligase, cytoplasmic OS=H | AARS1 | 106.811 | 2 | 51.886856 | yes | 9.722E-08 | 817.89575 | 127.04497 |
| AVYTQDCPLAAAI | 13 | P49588 | Alanine--tRNA ligase, cytoplasmic OS=H | AARS1 | 106.811 | 2 | 51.886856 | yes | 1.127E-07 | 704.35046 | 120.54421 |
| KVMDDLDR | 8 | P49588 | Alanine--tRNA ligase, cytoplasmic OS=H | AARS1 | 106.811 | 2 | 51.886856 | yes | 5.686E-07 | 496.2475 | 116.69801 |
| NSSHAGAFVIVTEI | 18 | P49588 | Alanine--tRNA ligase, cytoplasmic OS=H | AARS1 | 106.811 | 3 | 51.886856 | yes | 1.376E-07 | 615.32312 | 113.82604 |
| LVSVLQNK | 8 | P49588 | Alanine--tRNA ligase, cytoplasmic OS=H | AARS1 | 106.811 | 2 | 51.886856 | yes | 2.691E-07 | 450.7793 | 110.69962 |
| SVLGEADQK | 9 | P49588 | Alanine--tRNA ligase, cytoplasmic OS=H | AARS1 | 106.811 | 2 | 51.886856 | yes | 2.023E-06 | 473.74564 | 94.237907 |
| PIFLNTIDPSHPMA | 15 | P49588 | Alanine--tRNA ligase, cytoplasmic OS=H | AARS1 | 106.811 | 3 | 51.886856 | yes | 3.496E-07 | 560.96411 | 84.250595 |
| EADGILKPLPK | 11 | P49588 | Alanine--tRNA ligase, cytoplasmic OS=H | AARS1 | 106.811 | 2 | 51.886856 | yes | 2.947E-07 | 590.85046 | 77.274231 |
| QIWQNLGLDDTK | 12 | P49588 | Alanine--tRNA ligase, cytoplasmic OS=H | AARS1 | 106.811 | 2 | 51.886856 | yes | 1.004E-07 | 715.86737 | 74.204231 |
| KAEEIANEMIEAAI | 14 | P49588 | Alanine--tRNA ligase, cytoplasmic OS=H | AARS1 | 106.811 | 2 | 51.886856 | yes | 3.852E-07 | 773.8927 | 25.702021 |
| VDDSSEDKTEFTV | 14 | P49588 | Alanine--tRNA ligase, cytoplasmic OS=H | AARS1 | 106.811 | 2 | 51.886856 | yes | 5.578E-07 | 800.37286 | 19.90423 |
| DNILPELGVR | 10 | P49589 | Cysteine--tRNA ligase, cytoplasmic OS=I | CARS1 | 85.474 | 2 | 55.129692 | yes | 1.11E-07 | 563.31677 | 738.02258 |
| APVDITGQFEK | 11 | P49589 | Cysteine--tRNA ligase, cytoplasmic OS=I | CARS1 | 85.474 | 2 | 55.129692 | yes | 4.086E-07 | 602.81403 | 725.48993 |
| VSEYVPEIVNFVQ | 14 | P49589 | Cysteine--tRNA ligase, cytoplasmic OS=I | CARS1 | 85.474 | 2 | 55.129692 | yes | 2.859E-07 | 825.94049 | 626.35474 |
| LVPEAVGDQK | 10 | P49589 | Cysteine--tRNA ligase, cytoplasmic OS=I | CARS1 | 85.474 | 2 | 55.129692 | yes | 1.02E-07 | 528.29022 | 502.33356 |
| FWEGDFHR | 8 | P49589 | Cysteine--tRNA ligase, cytoplasmic OS=I | CARS1 | 85.474 | 2 | 55.129692 | yes | 9.722E-08 | 547.24615 | 374.6012 |
| LHLYNSLTR | 9 | P49589 | Cysteine--tRNA ligase, cytoplasmic OS=I | CARS1 | 85.474 | 2 | 55.129692 | yes | 9.897E-08 | 558.81165 | 351.85986 |
| IQHAVQLATEPLEI | 14 | P49589 | Cysteine--tRNA ligase, cytoplasmic OS=I | CARS1 | 85.474 | 2 | 55.129692 | yes | 1.103E-07 | 788.93829 | 151.53195 |
| IQHAVQLATEPLEI | 14 | P49589 | Cysteine--tRNA ligase, cytoplasmic OS=I | CARS1 | 85.474 | 2 | 55.129692 | yes | 1.8E-07 | 526.29462 | 151.53195 |
| IQHAVQLATEPLEI | 14 | P49589 | Cysteine--tRNA ligase, cytoplasmic OS=I | CARS1 | 85.474 | 3 | 55.129692 | yes | 1.103E-07 | 788.93829 | 151.53195 |
| IQHAVQLATEPLEI | 14 | P49589 | Cysteine--tRNA ligase, cytoplasmic OS=I | CARS1 | 85.474 | 3 | 55.129692 | yes | 1.8E-07 | 526.29462 | 151.53195 |
| EVFIPQDGK | 9 | P49589 | Cysteine--tRNA ligase, cytoplasmic OS=I | CARS1 | 85.474 | 2 | 55.129692 | yes | 1.3E-06 | 516.77167 | 280.55243 |
| IPPSEMFLSETDK | 13 | P49589 | Cysteine--tRNA ligase, cytoplasmic OS=I | CARS1 | 85.474 | 2 | 55.129692 | yes | 7.519E-07 | 747.36322 | 200.03307 |
| FLNEFFLNVK | 10 | P49589 | Cysteine--tRNA ligase, cytoplasmic OS=I | CARS1 | 85.474 | 2 | 55.129692 | yes | 2.325E-07 | 635.84515 | 162.48065 |
| ALVSQCNLYMAA' | 13 | P49589 | Cysteine--tRNA ligase, cytoplasmic OS=I | CARS1 | 85.474 | 2 | 55.129692 | yes | 9.722E-08 | 748.87103 | 151.8197 |
| VPEILQLSDALR | 12 | P49589 | Cysteine--tRNA ligase, cytoplasmic OS=I | CARS1 | 85.474 | 2 | 55.129692 | yes | 9.722E-08 | 677.39044 | 142.66531 |
| WGEEEAELNK | 10 | P49589 | Cysteine--tRNA ligase, cytoplasmic OS=I | CARS1 | 85.474 | 2 | 55.129692 | yes | 2.276E-07 | 602.77765 | 132.77376 |
| DMEALNVLPPDVI | 15 | P49589 | Cysteine--tRNA ligase, cytoplasmic OS=I | CARS1 | 85.474 | 2 | 55.129692 | yes | 9.722E-08 | 841.94275 | 127.71844 |
| FEDHEGLPTVVK | 12 | P49589 | Cysteine--tRNA ligase, cytoplasmic OS=I | CARS1 | 85.474 | 2 | 55.129692 | yes | 9.722E-08 | 685.8512 | 118.06968 |
| ALQEGEGDLSISAI | 15 | P49589 | Cysteine--tRNA ligase, cytoplasmic OS=I | CARS1 | 85.474 | 2 | 55.129692 | yes | 9.722E-08 | 780.87866 | 97.456833 |
| LAFLMHSWK | 9 | P49589 | Cysteine--tRNA ligase, cytoplasmic OS=I | CARS1 | 85.474 | 2 | 55.129692 | yes | 7.52E-07 | 566.80225 | 72.349754 |
| TVMEEMR | 7 | P49589 | Cysteine--tRNA ligase, cytoplasmic OS=I | CARS1 | 85.474 | 2 | 55.129692 | yes | 1.802E-06 | 448.20422 | 70.480644 |

| | | | | | | | | | | | |
|---|---|---|---|---|---|---|---|---|---|---|---|
| RSPNDFALWK | 10 | P49589 | Cysteine--tRNA ligase, cytoplasmic OS=I | CARS1 | 85.474 | 2 | 55.129692 | yes | 4.135E-07 | 617.32239 | 30.397825 |
| IFYIVEQR | 8 | P49590 | Histidine--tRNA ligase, mitochondrial OS | HARS2 | 56.889 | 2 | 53.776875 | yes | 5.364E-07 | 534.29785 | 147.17065 |
| TTETQVFVATPQK | 13 | P49590 | Histidine--tRNA ligase, mitochondrial OS | HARS2 | 56.889 | 2 | 53.776875 | yes | 2.093E-07 | 725.38281 | 146.85164 |
| LIAELWDSGIK | 11 | P49590 | Histidine--tRNA ligase, mitochondrial OS | HARS2 | 56.889 | 2 | 53.776875 | yes | 2.542E-07 | 622.8479 | 139.76733 |
| YDGLVGMFDPK | 11 | P49590 | Histidine--tRNA ligase, mitochondrial OS | HARS2 | 56.889 | 2 | 53.776875 | no | 1.038E-07 | 621.29718 | 130.30827 |
| IVDGMFAVCGVPE | 15 | P49590 | Histidine--tRNA ligase, mitochondrial OS | HARS2 | 56.889 | 2 | 53.776875 | yes | 9.722E-08 | 804.89166 | 120.07244 |
| YDLTVPFAR | 9 | P49590 | Histidine--tRNA ligase, mitochondrial OS | HARS2 | 56.889 | 2 | 53.776875 | no | 1.422E-07 | 541.28748 | 118.03529 |
| YGEDSGLMYDLK | 12 | P49590 | Histidine--tRNA ligase, mitochondrial OS | HARS2 | 56.889 | 2 | 53.776875 | yes | 9.898E-08 | 695.81342 | 111.89999 |
| ISFDLSLAR | 9 | P49590 | Histidine--tRNA ligase, mitochondrial OS | HARS2 | 56.889 | 2 | 53.776875 | no | 1.617E-07 | 511.28748 | 97.267914 |
| ENFVAEIQK | 9 | P49590 | Histidine--tRNA ligase, mitochondrial OS | HARS2 | 56.889 | 2 | 53.776875 | yes | 6.658E-07 | 539.28241 | 88.580444 |
| DQGGELLSLR | 10 | P49590 | Histidine--tRNA ligase, mitochondrial OS | HARS2 | 56.889 | 2 | 53.776875 | no | 4.471E-07 | 544.29077 | 80.568573 |
| AHQEKPNFIIK | 11 | P49590 | Histidine--tRNA ligase, mitochondrial OS | HARS2 | 56.889 | 2 | 53.776875 | yes | 3.842E-07 | 662.87225 | 34.143547 |
| SLPAVQSDER | 10 | P49643 | DNA primase large subunit OS=Homo sa | PRIM2 | 58.806 | 2 | 48.082287 | yes | 1.441E-07 | 551.2804 | 115.44883 |
| EQEIVASSPSLSGL | 15 | P49643 | DNA primase large subunit OS=Homo sa | PRIM2 | 58.806 | 2 | 48.082287 | yes | 1.202E-07 | 772.91193 | 104.49046 |
| ISLDQIDLLSTK | 12 | P49643 | DNA primase large subunit OS=Homo sa | PRIM2 | 58.806 | 2 | 48.082287 | yes | 3.35E-07 | 673.38232 | 55.966476 |
| VYLEDGFAYVPLK | 13 | P49643 | DNA primase large subunit OS=Homo sa | PRIM2 | 58.806 | 2 | 48.082287 | yes | 4.055E-07 | 757.40051 | 49.717461 |
| SVENLGVSYVK | 11 | P49643 | DNA primase large subunit OS=Homo sa | PRIM2 | 58.806 | 2 | 48.082287 | yes | 8.593E-07 | 597.8219 | 32.42342 |
| DVKPDNFLMGLGI | 13 | P49674 | Casein kinase I isoform epsilon OS=Hom | CSNK1E | 47.316 | 2 | 53.728302 | no | 2.626E-07 | 717.37646 | 951.19482 |
| FDDKPDYSYLR | 11 | P49674 | Casein kinase I isoform epsilon OS=Hom | CSNK1E | 47.316 | 2 | 53.728302 | no | 2.457E-07 | 709.83295 | 448.09464 |
| FDDKPDYSYLR | 11 | P49674 | Casein kinase I isoform epsilon OS=Hom | CSNK1E | 47.316 | 2 | 53.728302 | no | 2.014E-07 | 473.55774 | 448.09464 |
| FDDKPDYSYLR | 11 | P49674 | Casein kinase I isoform epsilon OS=Hom | CSNK1E | 47.316 | 3 | 53.728302 | no | 2.457E-07 | 709.83295 | 448.09464 |
| FDDKPDYSYLR | 11 | P49674 | Casein kinase I isoform epsilon OS=Hom | CSNK1E | 47.316 | 3 | 53.728302 | no | 2.014E-07 | 473.55774 | 448.09464 |
| IEYIHSK | 7 | P49674 | Casein kinase I isoform epsilon OS=Hom | CSNK1E | 47.316 | 2 | 53.728302 | no | 4.264E-07 | 445.24252 | 820.14703 |
| TVLLLADQMISR | 12 | P49674 | Casein kinase I isoform epsilon OS=Hom | CSNK1E | 47.316 | 2 | 53.728302 | no | 9.722E-08 | 680.38684 | 721.96387 |
| MSTPIEVLCK | 10 | P49674 | Casein kinase I isoform epsilon OS=Hom | CSNK1E | 47.316 | 2 | 53.728302 | no | 1.302E-06 | 589.30139 | 579.27075 |
| GAPANVSSSDLTG | 14 | P49674 | Casein kinase I isoform epsilon OS=Hom | CSNK1E | 47.316 | 2 | 53.728302 | yes | 3.54E-07 | 666.33112 | 421.82855 |
| SAAEPVASTPASR | 13 | P49674 | Casein kinase I isoform epsilon OS=Hom | CSNK1E | 47.316 | 2 | 53.728302 | yes | 1.324E-07 | 622.3175 | 308.58005 |
| THQHIPYR | 8 | P49674 | Casein kinase I isoform epsilon OS=Hom | CSNK1E | 47.316 | 2 | 53.728302 | no | 1.431E-07 | 526.27521 | 288.39975 |
| ALPPGPPTGATAN | 14 | P49674 | Casein kinase I isoform epsilon OS=Hom | CSNK1E | 47.316 | 2 | 53.728302 | yes | 1.844E-06 | 660.35693 | 247.37225 |
| MMQGGVGIPSIK | 12 | P49674 | Casein kinase I isoform epsilon OS=Hom | CSNK1E | 47.316 | 2 | 53.728302 | yes | 9.722E-08 | 609.32269 | 167.19153 |
| HPQLHIESK | 9 | P49674 | Casein kinase I isoform epsilon OS=Hom | CSNK1E | 47.316 | 2 | 53.728302 | no | 1.276E-07 | 544.79602 | 156.30334 |
| IPASQTSVPFDHLC | 15 | P49674 | Casein kinase I isoform epsilon OS=Hom | CSNK1E | 47.316 | 2 | 53.728302 | yes | 1.457E-06 | 798.92267 | 95.81945 |
| TKHPQLHIESK | 11 | P49674 | Casein kinase I isoform epsilon OS=Hom | CSNK1E | 47.316 | 2 | 53.728302 | no | 9.722E-08 | 659.36731 | 88.75296 |
| ASDNQGVPVLVLA | 15 | P49703 | ADP-ribosylation factor-like protein 4D ( | ARL4D | 22.156 | 2 | 49.033604 | yes | 1.075E-07 | 762.92267 | 48.174187 |
| SDLGVHLR | 8 | P49711 | Transcriptional repressor CTCF OS=Hom | CTCF | 82.786 | 2 | 49.732197 | yes | 2.227E-07 | 448.75107 | 132.85388 |
| YALIQHQK | 8 | P49711 | Transcriptional repressor CTCF OS=Hom | CTCF | 82.786 | 2 | 49.732197 | no | 1.612E-07 | 500.78235 | 129.08711 |
| YCDAVFHER | 9 | P49711 | Transcriptional repressor CTCF OS=Hom | CTCF | 82.786 | 2 | 49.732197 | yes | 2.96E-07 | 598.76129 | 124.79426 |
| FHCPHCDTVIAR | 12 | P49711 | Transcriptional repressor CTCF OS=Hom | CTCF | 82.786 | 3 | 49.732197 | yes | 2.812E-06 | 504.89935 | 110.34689 |
| CSMCDYASVEVSI | 13 | P49711 | Transcriptional repressor CTCF OS=Hom | CTCF | 82.786 | 2 | 49.732197 | yes | 3.574E-07 | 768.31238 | 61.326328 |

| | | | | | | | | | | | |
|---|---|---|---|---|---|---|---|---|---|---|---|
| KSDLGVHLR | 9 | P49711 | Transcriptional repressor CTCF OS=Hom | CTCF | 82.786 | 2 | 49.732197 | yes | 3.824E-07 | 512.79852 | 54.118263 |
| PFQCSLCSYASR | 12 | P49711 | Transcriptional repressor CTCF OS=Hom | CTCF | 82.786 | 2 | 49.732197 | yes | 1.006E-07 | 738.32391 | 23.753839 |
| LVELSAENEK | 10 | P49716 | CCAAT/enhancer-binding protein delta C | CEBPD | 28.468 | 2 | 46.21531 | yes | 7.125E-07 | 566.29822 | 90.49865 |
| AGGAGPLELLPGG | 16 | P49716 | CCAAT/enhancer-binding protein delta C | CEBPD | 28.468 | 2 | 46.21531 | yes | 1.619E-07 | 716.89899 | 46.280815 |
| VAASNIVQMK | 10 | P49721 | Proteasome subunit beta type-2 OS=Hom | PSMB2 | 22.837 | 2 | 43.999855 | yes | 2.362E-07 | 530.79462 | 90.728859 |
| DTDLDGFPDEK | 11 | P49747 | Cartilage oligomeric matrix protein OS=H | COMP | 82.861 | 2 | 36.966747 | yes | 6.431E-07 | 626.2724 | 41.998047 |
| VPSENVLGEVGSG | 15 | P49748 | Very long-chain specific acyl-CoA dehyd | ACADVL | 70.391 | 2 | 48.023968 | yes | 1.212E-07 | 759.89355 | 30.817316 |
| LKPLEVELR | 9 | P49755 | Transmembrane emp24 domain-containin | TMED10 | 24.976 | 2 | 43.90896 | yes | 3.778E-07 | 548.83984 | 231.27773 |
| ITDSAGHILYSK | 12 | P49755 | Transmembrane emp24 domain-containin | TMED10 | 24.976 | 2 | 43.90896 | yes | 1.31E-06 | 652.84589 | 61.715454 |
| IPDQLVILDMK | 11 | P49755 | Transmembrane emp24 domain-containin | TMED10 | 24.976 | 2 | 43.90896 | yes | 2.397E-07 | 642.86523 | 52.887695 |
| LLHDLQIGEK | 10 | P49756 | RNA-binding protein 25 OS=Homo sapie | RBM25 | 100.186 | 2 | 52.548294 | yes | 9.722E-08 | 583.3324 | 236.11456 |
| EDINAIEMEEDKR | 13 | P49756 | RNA-binding protein 25 OS=Homo sapie | RBM25 | 100.186 | 2 | 52.548294 | yes | 9.722E-08 | 796.36707 | 176.67096 |
| LLAEGHPDPDAEL | 15 | P49756 | RNA-binding protein 25 OS=Homo sapie | RBM25 | 100.186 | 2 | 52.548294 | yes | 1.007E-06 | 830.91809 | 164.29295 |
| LPVDSVFNK | 9 | P49756 | RNA-binding protein 25 OS=Homo sapie | RBM25 | 100.186 | 2 | 52.548294 | yes | 5.646E-07 | 509.78201 | 158.46669 |
| LVPLDYGEDDKN/ | 15 | P49756 | RNA-binding protein 25 OS=Homo sapie | RBM25 | 100.186 | 2 | 52.548294 | yes | 1.881E-07 | 839.42017 | 135.84166 |
| IRPWINK | 7 | P49756 | RNA-binding protein 25 OS=Homo sapie | RBM25 | 100.186 | 2 | 52.548294 | yes | 7.654E-07 | 463.78217 | 93.426552 |
| FEDEDSDDVPR | 11 | P49756 | RNA-binding protein 25 OS=Homo sapie | RBM25 | 100.186 | 2 | 52.548294 | yes | 3.74E-07 | 662.27039 | 61.257462 |
| LQAFGFCEYK | 10 | P49756 | RNA-binding protein 25 OS=Homo sapie | RBM25 | 100.186 | 2 | 52.548294 | yes | 9.722E-08 | 631.79736 | 55.494278 |
| LGASNSPGQPNSV | 14 | P49756 | RNA-binding protein 25 OS=Homo sapie | RBM25 | 100.186 | 2 | 52.548294 | yes | 2.958E-07 | 678.3493 | 44.842464 |
| LVPLDYGEDDK | 11 | P49756 | RNA-binding protein 25 OS=Homo sapie | RBM25 | 100.186 | 2 | 52.548294 | yes | 1.054E-06 | 632.30878 | 13.130093 |
| HAPIEQLAR | 9 | P49757 | Protein numb homolog OS=Homo sapien | NUMB | 70.804 | 2 | 53.285904 | yes | 4.566E-07 | 517.79071 | 530.42242 |
| DLIVDQTIEK | 10 | P49757 | Protein numb homolog OS=Homo sapien | NUMB | 70.804 | 2 | 53.285904 | yes | 1.998E-07 | 587.32172 | 264.09729 |
| AVLWVSADGLR | 11 | P49757 | Protein numb homolog OS=Homo sapien | NUMB | 70.804 | 2 | 53.285904 | yes | 2.812E-07 | 593.83258 | 256.21133 |
| YLGHVEVDESR | 11 | P49757 | Protein numb homolog OS=Homo sapien | NUMB | 70.804 | 2 | 53.285904 | yes | 5.437E-07 | 652.3175 | 250.26781 |
| INELPSTMQR | 10 | P49757 | Protein numb homolog OS=Homo sapien | NUMB | 70.804 | 2 | 53.285904 | yes | 9.908E-08 | 594.80573 | 186.5378 |
| TNPSPTNPFSSDLQ | 15 | P49757 | Protein numb homolog OS=Homo sapien | NUMB | 70.804 | 2 | 53.285904 | yes | 1.13E-07 | 816.89685 | 124.39703 |
| AFSYICR | 7 | P49757 | Protein numb homolog OS=Homo sapien | NUMB | 70.804 | 2 | 53.285904 | no | 1.02E-07 | 458.72092 | 105.57512 |
| VTTATEQAER | 10 | P49757 | Protein numb homolog OS=Homo sapien | NUMB | 70.804 | 2 | 53.285904 | yes | 1.246E-07 | 553.27783 | 104.52266 |
| LSHAVGCAFAACL | 15 | P49757 | Protein numb homolog OS=Homo sapien | NUMB | 70.804 | 2 | 53.285904 | no | 9.722E-08 | 831.39795 | 65.87178 |
| VSFCAPDR | 8 | P49757 | Protein numb homolog OS=Homo sapien | NUMB | 70.804 | 2 | 53.285904 | no | 1.185E-07 | 476.22128 | 51.890732 |
| GMHICEDAVK | 10 | P49757 | Protein numb homolog OS=Homo sapien | NUMB | 70.804 | 2 | 53.285904 | yes | 9.892E-08 | 580.26538 | 44.097321 |
| ETNPWAHAPDAAI | 14 | P49757 | Protein numb homolog OS=Homo sapien | NUMB | 70.804 | 2 | 53.285904 | yes | 3.453E-06 | 761.35767 | 23.099756 |
| WICHCFMAVK | 10 | P49757 | Protein numb homolog OS=Homo sapien | NUMB | 70.804 | 2 | 53.285904 | yes | 2.097E-06 | 676.30902 | 12.947255 |
| YEIVDTLGEGAFGI | 14 | P49759 | Dual specificity protein kinase CLK1 OS= | CLK1 | 57.291 | 2 | 52.211346 | no | 9.722E-08 | 749.87482 | 258.24121 |
| VVECIDHK | 8 | P49759 | Dual specificity protein kinase CLK1 OS= | CLK1 | 57.291 | 2 | 52.211346 | yes | 1.368E-05 | 500.25003 | 95.973907 |
| LDWDEHSSAGR | 11 | P49759 | Dual specificity protein kinase CLK1 OS= | CLK1 | 57.291 | 2 | 52.211346 | yes | 1.02E-07 | 636.78381 | 93.60685 |
| TLINPDIK | 8 | P49759 | Dual specificity protein kinase CLK1 OS= | CLK1 | 57.291 | 2 | 52.211346 | yes | 8.949E-07 | 457.2713 | 92.075546 |
| LFDLIQK | 7 | P49759 | Dual specificity protein kinase CLK1 OS= | CLK1 | 57.291 | 2 | 52.211346 | yes | 5.971E-07 | 438.76309 | 85.180756 |
| EFMLSQDVEHER | 12 | P49759 | Dual specificity protein kinase CLK1 OS= | CLK1 | 57.291 | 2 | 52.211346 | yes | 1.292E-07 | 760.34595 | 72.773338 |

| | | | | | | | | | | | |
|---|---|---|---|---|---|---|---|---|---|---|---|
| NDYTQGCEPGHR | 12 | P49759 | Dual specificity protein kinase CLK1 OS= | CLK1 | 57.291 | 2 | 52.211346 | yes | 1.641E-07 | 717.29675 | 40.659645 |
| MCDSHYLESR | 10 | P49759 | Dual specificity protein kinase CLK1 OS= | CLK1 | 57.291 | 2 | 52.211346 | yes | 3.021E-07 | 649.26862 | 32.42421 |
| ILGPIPSR | 8 | P49760 | Dual specificity protein kinase CLK2 OS= | CLK2 | 60.091 | 2 | 50.93933 | yes | 2.436E-07 | 426.76871 | 272.8779 |
| DNNYLPYPIHQVR | 13 | P49760 | Dual specificity protein kinase CLK2 OS= | CLK2 | 60.091 | 2 | 50.93933 | yes | 1.968E-07 | 814.9126 | 156.9697 |
| HMAFQLCQAVK | 11 | P49760 | Dual specificity protein kinase CLK2 OS= | CLK2 | 60.091 | 2 | 50.93933 | yes | 1.283E-07 | 666.83118 | 141.67856 |
| LEINVLEK | 8 | P49760 | Dual specificity protein kinase CLK2 OS= | CLK2 | 60.091 | 2 | 50.93933 | yes | 8.043E-07 | 479.28439 | 112.56936 |
| YEIVSTLGEGTFGI | 14 | P49760 | Dual specificity protein kinase CLK2 OS= | CLK2 | 60.091 | 2 | 50.93933 | yes | 9.722E-08 | 764.88574 | 98.859261 |
| VVDFGSATFDHEH | 21 | P49760 | Dual specificity protein kinase CLK2 OS= | CLK2 | 60.091 | 4 | 50.93933 | yes | 4.824E-07 | 586.28448 | 31.538584 |
| YEIVGNLGEGTFG | 14 | P49761 | Dual specificity protein kinase CLK3 OS= | CLK3 | 73.516 | 2 | 51.088825 | yes | 1.409E-07 | 742.375 | 874.60095 |
| MLEFDPAQR | 9 | P49761 | Dual specificity protein kinase CLK3 OS= | CLK3 | 73.516 | 2 | 51.088825 | yes | 9.927E-08 | 553.76862 | 874.29041 |
| IGDWLQER | 8 | P49761 | Dual specificity protein kinase CLK3 OS= | CLK3 | 73.516 | 2 | 51.088825 | yes | 3.297E-07 | 508.76163 | 525.33929 |
| VVECLDHAR | 9 | P49761 | Dual specificity protein kinase CLK3 OS= | CLK3 | 73.516 | 2 | 51.088825 | yes | 1.605E-07 | 549.77167 | 443.09369 |
| GGLVWDENSSDGI | 13 | P49761 | Dual specificity protein kinase CLK3 OS= | CLK3 | 73.516 | 2 | 51.088825 | yes | 9.722E-08 | 696.31293 | 220.49742 |
| VADFGSATFDHEH | 21 | P49761 | Dual specificity protein kinase CLK3 OS= | CLK3 | 73.516 | 3 | 51.088825 | yes | 2.287E-07 | 771.37335 | 109.12475 |
| VADFGSATFDHEH | 21 | P49761 | Dual specificity protein kinase CLK3 OS= | CLK3 | 73.516 | 3 | 51.088825 | yes | 2.685E-06 | 578.7818 | 109.12475 |
| VADFGSATFDHEH | 21 | P49761 | Dual specificity protein kinase CLK3 OS= | CLK3 | 73.516 | 4 | 51.088825 | yes | 2.287E-07 | 771.37335 | 109.12475 |
| VADFGSATFDHEH | 21 | P49761 | Dual specificity protein kinase CLK3 OS= | CLK3 | 73.516 | 4 | 51.088825 | yes | 2.685E-06 | 578.7818 | 109.12475 |
| EHLVMMEK | 8 | P49761 | Dual specificity protein kinase CLK3 OS= | CLK3 | 73.516 | 2 | 51.088825 | yes | 1.107E-06 | 508.74881 | 116.5071 |
| FLHENQLTHTDLK | 32 | P49761 | Dual specificity protein kinase CLK3 OS= | CLK3 | 73.516 | 5 | 51.088825 | yes | 1.429E-07 | 780.79059 | 100.11173 |
| HMAYQLCHALR | 11 | P49761 | Dual specificity protein kinase CLK3 OS= | CLK3 | 73.516 | 3 | 51.088825 | yes | 3.859E-07 | 467.22891 | 65.038895 |
| ILGPIPSHMIHR | 12 | P49761 | Dual specificity protein kinase CLK3 OS= | CLK3 | 73.516 | 3 | 51.088825 | yes | 4.228E-07 | 457.5957 | 35.611179 |
| LMSELYHPDDHVI | 13 | P49770 | Translation initiation factor eIF-2B subun | EIF2B2 | 38.99 | 2 | 50.422462 | yes | 9.935E-08 | 784.87433 | 120.33366 |
| MTAAQPSETTVGN | 16 | P49770 | Translation initiation factor eIF-2B subun | EIF2B2 | 38.99 | 2 | 50.422462 | yes | 3.306E-06 | 846.90582 | 87.989838 |
| WSNAGELMELIR | 12 | P49770 | Translation initiation factor eIF-2B subun | EIF2B2 | 38.99 | 2 | 50.422462 | yes | 6.432E-07 | 709.85846 | 56.064838 |
| SDESDQQESLHK | 12 | P49770 | Translation initiation factor eIF-2B subun | EIF2B2 | 38.99 | 2 | 50.422462 | yes | 9.809E-08 | 701.80768 | 28.395027 |
| PSLTPSGEFR | 10 | P49790 | Nuclear pore complex protein Nup153 OS | NUP153 | 153.939 | 2 | 55.98675 | yes | 1.756E-06 | 545.78003 | 244.20255 |
| TTYGGAAAAVR | 11 | P49790 | Nuclear pore complex protein Nup153 OS | NUP153 | 153.939 | 2 | 55.98675 | yes | 9.819E-08 | 519.27234 | 205.07205 |
| SSFNLGTIETK | 11 | P49790 | Nuclear pore complex protein Nup153 OS | NUP153 | 153.939 | 2 | 55.98675 | yes | 9.722E-08 | 598.81152 | 167.98438 |
| TSQLGDSPFYPGK | 13 | P49790 | Nuclear pore complex protein Nup153 OS | NUP153 | 153.939 | 2 | 55.98675 | yes | 9.722E-08 | 698.84082 | 167.7747 |
| TTLSASGTGFGDK | 13 | P49790 | Nuclear pore complex protein Nup153 OS | NUP153 | 153.939 | 2 | 55.98675 | yes | 1.092E-07 | 621.30408 | 130.94878 |
| SGIDITDFQAK | 11 | P49790 | Nuclear pore complex protein Nup153 OS | NUP153 | 153.939 | 2 | 55.98675 | yes | 2.613E-07 | 597.80371 | 120.34807 |
| FVASKPLEEEEME | 19 | P49790 | Nuclear pore complex protein Nup153 OS | NUP153 | 153.939 | 3 | 55.98675 | yes | 1.002E-07 | 724.38165 | 98.478165 |
| NTPYQAPVR | 9 | P49790 | Nuclear pore complex protein Nup153 OS | NUP153 | 153.939 | 2 | 55.98675 | yes | 1.421E-07 | 523.2749 | 96.675697 |
| CQPVFSFGNSEQTI | 14 | P49790 | Nuclear pore complex protein Nup153 OS | NUP153 | 153.939 | 2 | 55.98675 | yes | 5.118E-07 | 814.87231 | 88.876022 |
| IPSIVSSPLNSPLDI | 15 | P49790 | Nuclear pore complex protein Nup153 OS | NUP153 | 153.939 | 2 | 55.98675 | yes | 9.722E-08 | 797.9436 | 77.260376 |
| NVFSSSGTSFSGR | 13 | P49790 | Nuclear pore complex protein Nup153 OS | NUP153 | 153.939 | 2 | 55.98675 | yes | 4.211E-07 | 666.81256 | 69.96035 |
| QQHQGILSR | 9 | P49790 | Nuclear pore complex protein Nup153 OS | NUP153 | 153.939 | 2 | 55.98675 | yes | 3.305E-07 | 533.79126 | 55.772499 |
| ASDKDITVSK | 10 | P49790 | Nuclear pore complex protein Nup153 OS | NUP153 | 153.939 | 2 | 55.98675 | yes | 1.391E-07 | 532.28516 | 48.690197 |
| QQEPVTSTSLVFGI | 14 | P49790 | Nuclear pore complex protein Nup153 OS | NUP153 | 153.939 | 2 | 55.98675 | yes | 5.373E-07 | 760.90137 | 48.408775 |

| | | | | | | | | | | | |
|---|---|---|---|---|---|---|---|---|---|---|---|
| QLSAQSYGVTSST. | 15 | P49790 | Nuclear pore complex protein Nup153 OS | NUP153 | 153.939 | 2 | 55.98675 | yes | 9.722E-08 | 778.38916 | 44.366474 |
| PLEEEEMEVPVLP. | 14 | P49790 | Nuclear pore complex protein Nup153 OS | NUP153 | 153.939 | 2 | 55.98675 | yes | 3.152E-06 | 819.9184 | 35.510849 |
| NIVPGWLQR | 9 | P49790 | Nuclear pore complex protein Nup153 OS | NUP153 | 153.939 | 2 | 55.98675 | yes | 7.285E-07 | 541.8089 | 32.819458 |
| CVACETPKPGTCV | 14 | P49790 | Nuclear pore complex protein Nup153 OS | NUP153 | 153.939 | 2 | 55.98675 | yes | 3.679E-07 | 803.87292 | 25.426199 |
| VQMTSPSSTGSPM | 15 | P49790 | Nuclear pore complex protein Nup153 OS | NUP153 | 153.939 | 2 | 55.98675 | yes | 3.574E-07 | 792.87347 | 19.6099 |
| TAENFRALCTGEK | 13 | P49792 | E3 SUMO-protein ligase RanBP2 OS=Hc | RANBP2 | 358.2 | 2 | 55.77997 | no | 1.248E-07 | 748.86176 | 223.07637 |
| PIPGGQTIGPR | 11 | P49792 | E3 SUMO-protein ligase RanBP2 OS=Hc | RANBP2 | 358.2 | 2 | 55.77997 | yes | 2.778E-06 | 546.81165 | 160.70107 |
| EGQWDCSVCLVR | 12 | P49792 | E3 SUMO-protein ligase RanBP2 OS=Hc | RANBP2 | 358.2 | 2 | 55.77997 | yes | 9.722E-08 | 754.83466 | 158.95512 |
| SGFEGMFIR | 9 | P49792 | E3 SUMO-protein ligase RanBP2 OS=Hc | RANBP2 | 358.2 | 2 | 55.77997 | yes | 2.985E-07 | 522.25256 | 118.05438 |
| KLPVPLESVK | 10 | P49792 | E3 SUMO-protein ligase RanBP2 OS=Hc | RANBP2 | 358.2 | 2 | 55.77997 | no | 7.208E-07 | 555.3501 | 105.40337 |
| ELLQSFDSALQSVI | 14 | P49792 | E3 SUMO-protein ligase RanBP2 OS=Hc | RANBP2 | 358.2 | 2 | 55.77997 | yes | 1.317E-07 | 782.91449 | 103.23804 |
| SVFGTPTLETANK | 13 | P49792 | E3 SUMO-protein ligase RanBP2 OS=Hc | RANBP2 | 358.2 | 2 | 55.77997 | yes | 4.165E-07 | 682.85645 | 94.181976 |
| LLVQHEINTLR | 11 | P49792 | E3 SUMO-protein ligase RanBP2 OS=Hc | RANBP2 | 358.2 | 2 | 55.77997 | no | 9.722E-08 | 668.39081 | 92.493378 |
| YIASVQGSTPSPR | 13 | P49792 | E3 SUMO-protein ligase RanBP2 OS=Hc | RANBP2 | 358.2 | 2 | 55.77997 | yes | 1.054E-07 | 681.85425 | 87.078453 |
| FGISEPGNQEK | 11 | P49792 | E3 SUMO-protein ligase RanBP2 OS=Hc | RANBP2 | 358.2 | 2 | 55.77997 | no | 4.325E-07 | 603.29346 | 78.581009 |
| VIPDFVCQGGDITI | 14 | P49792 | E3 SUMO-protein ligase RanBP2 OS=Hc | RANBP2 | 358.2 | 2 | 55.77997 | yes | 0.0001341 | 774.88995 | 77.649826 |
| KVEHLAVR | 8 | P49792 | E3 SUMO-protein ligase RanBP2 OS=Hc | RANBP2 | 358.2 | 2 | 55.77997 | no | 3.509E-07 | 476.29034 | 73.5494 |
| TGEEDEEEFFCNR | 13 | P49792 | E3 SUMO-protein ligase RanBP2 OS=Hc | RANBP2 | 358.2 | 2 | 55.77997 | yes | 9.722E-08 | 831.32281 | 69.673538 |
| ELVGPPLAETVFTI | 15 | P49792 | E3 SUMO-protein ligase RanBP2 OS=Hc | RANBP2 | 358.2 | 2 | 55.77997 | yes | 1.39E-07 | 799.44543 | 68.240791 |
| HVVFGFVK | 8 | P49792 | E3 SUMO-protein ligase RanBP2 OS=Hc | RANBP2 | 358.2 | 2 | 55.77997 | yes | 2.93E-07 | 466.77127 | 60.30994 |
| LQDVADSFK | 9 | P49792 | E3 SUMO-protein ligase RanBP2 OS=Hc | RANBP2 | 358.2 | 2 | 55.77997 | no | 1.35E-06 | 511.76129 | 58.894035 |
| SGSSFVHQASFK | 12 | P49792 | E3 SUMO-protein ligase RanBP2 OS=Hc | RANBP2 | 358.2 | 2 | 55.77997 | yes | 1.077E-07 | 641.31476 | 57.940517 |
| EIDTDSTSQGESK | 13 | P49792 | E3 SUMO-protein ligase RanBP2 OS=Hc | RANBP2 | 358.2 | 2 | 55.77997 | yes | 9.722E-08 | 698.80737 | 55.354359 |
| VELVTGEEDEK | 11 | P49792 | E3 SUMO-protein ligase RanBP2 OS=Hc | RANBP2 | 358.2 | 2 | 55.77997 | yes | 4.894E-07 | 624.30371 | 54.886402 |
| TGSGLNSFYDQR | 12 | P49792 | E3 SUMO-protein ligase RanBP2 OS=Hc | RANBP2 | 358.2 | 2 | 55.77997 | yes | 1.645E-06 | 672.81256 | 44.284077 |
| IAVAVLEETTR | 11 | P49792 | E3 SUMO-protein ligase RanBP2 OS=Hc | RANBP2 | 358.2 | 2 | 55.77997 | no | 1.235E-07 | 601.34296 | 39.433903 |
| GEAGQNLLEMMA | 15 | P49792 | E3 SUMO-protein ligase RanBP2 OS=Hc | RANBP2 | 358.2 | 2 | 55.77997 | no | 9.722E-08 | 847.86841 | 39.074245 |
| STSGEGFQFGK | 11 | P49792 | E3 SUMO-protein ligase RanBP2 OS=Hc | RANBP2 | 358.2 | 2 | 55.77997 | no | 1.677E-06 | 572.76709 | 30.832787 |
| MGQHSSNVQWR | 11 | P49792 | E3 SUMO-protein ligase RanBP2 OS=Hc | RANBP2 | 358.2 | 2 | 55.77997 | yes | 5.513E-06 | 665.30945 | 17.236795 |
| DGFGFTICCDSPVI | 14 | P49796 | Regulator of G-protein signaling 3 OS=H | RGS3 | 132.336 | 2 | 49.803627 | yes | 1.09E-07 | 815.85303 | 63.427032 |
| CVELAHEIR | 9 | P49796 | Regulator of G-protein signaling 3 OS=H | RGS3 | 132.336 | 2 | 49.803627 | yes | 1.865E-06 | 563.78729 | 62.125446 |
| VLLLHIIEGK | 10 | P49796 | Regulator of G-protein signaling 3 OS=H | RGS3 | 132.336 | 2 | 49.803627 | yes | 1.003E-07 | 567.86591 | 50.381649 |
| FCVLYLAEK | 9 | P49796 | Regulator of G-protein signaling 3 OS=H | RGS3 | 132.336 | 2 | 49.803627 | yes | 3.378E-06 | 571.79938 | 33.043861 |
| QSGLIGCMSFGVK | 13 | P49796 | Regulator of G-protein signaling 3 OS=H | RGS3 | 132.336 | 2 | 49.803627 | yes | 1.59E-06 | 692.34161 | 25.035913 |
| LLEGCLVGGR | 10 | P49821 | NADH dehydrogenase [ubiquinone] flavc | NDUFV1 | 50.818 | 2 | 43.388035 | yes | 2.613E-07 | 537.29224 | 86.089584 |
| GPDWILGEIK | 10 | P49821 | NADH dehydrogenase [ubiquinone] flavc | NDUFV1 | 50.818 | 2 | 43.388035 | yes | 4.336E-07 | 564.30841 | 65.283249 |
| LCDSGELVAIK | 11 | P49841 | Glycogen synthase kinase-3 beta OS=Hor | GSK3B | 46.745 | 2 | 46.619038 | yes | 1.838E-06 | 602.81573 | 395.79047 |
| LLEYTPTAR | 9 | P49841 | Glycogen synthase kinase-3 beta OS=Hor | GSK3B | 46.745 | 2 | 46.619038 | yes | 1.792E-07 | 532.29279 | 292.73999 |
| YFFYSSGEK | 9 | P49841 | Glycogen synthase kinase-3 beta OS=Hor | GSK3B | 46.745 | 2 | 46.619038 | no | 1.897E-07 | 564.25586 | 184.09666 |

| | | | | | | | | | | | |
|---|---|---|---|---|---|---|---|---|---|---|---|
| LYMYQLFR | 8 | P49841 | Glycogen synthase kinase-3 beta OS=Hor | GSK3B | 46.745 | 2 | 46.619038 | yes | 2.735E-07 | 567.29425 | 115.61066 |
| LCDFGSAK | 8 | P49841 | Glycogen synthase kinase-3 beta OS=Hor | GSK3B | 46.745 | 2 | 46.619038 | no | 3.542E-07 | 449.21039 | 100.53795 |
| EMNPNYTEFK | 10 | P49841 | Glycogen synthase kinase-3 beta OS=Hor | GSK3B | 46.745 | 2 | 46.619038 | no | 2.726E-06 | 636.78192 | 85.789383 |
| KLDHCNIVR | 9 | P49841 | Glycogen synthase kinase-3 beta OS=Hor | GSK3B | 46.745 | 2 | 46.619038 | no | 1.512E-07 | 577.80859 | 38.829208 |
| VVYIFGPPVK | 10 | P49915 | GMP synthase [glutamine-hydrolyzing] C | GMPS | 76.716 | 2 | 50.387524 | yes | 2.949E-07 | 559.83405 | 172.44641 |
| ELGLPEELVSR | 11 | P49915 | GMP synthase [glutamine-hydrolyzing] C | GMPS | 76.716 | 2 | 50.387524 | yes | 2.393E-07 | 621.34045 | 114.55521 |
| HPFPGPGLAIR | 11 | P49915 | GMP synthase [glutamine-hydrolyzing] C | GMPS | 76.716 | 2 | 50.387524 | yes | 4.074E-07 | 581.33002 | 91.328957 |
| TLNMTTSPEEK | 11 | P49915 | GMP synthase [glutamine-hydrolyzing] C | GMPS | 76.716 | 2 | 50.387524 | yes | 9.81E-08 | 625.80029 | 65.891106 |
| SYSYVCGISSK | 11 | P49915 | GMP synthase [glutamine-hydrolyzing] C | GMPS | 76.716 | 2 | 50.387524 | yes | 2.36E-07 | 625.78973 | 60.967751 |
| EEVVLLTHGDSVD | 14 | P49915 | GMP synthase [glutamine-hydrolyzing] C | GMPS | 76.716 | 2 | 50.387524 | yes | 1.135E-07 | 770.8963 | 46.035294 |
| VICAEEPYICK | 11 | P49915 | GMP synthase [glutamine-hydrolyzing] C | GMPS | 76.716 | 2 | 50.387524 | yes | 6.501E-07 | 691.32819 | 23.8169 |
| TQIIQDFLR | 9 | P49916 | DNA ligase 3 OS=Homo sapiens OX=96( | LIG3 | 112.907 | 2 | 53.9544 | yes | 1.071E-07 | 567.31934 | 2061.8657 |
| VIQEGLEGLVLK | 12 | P49916 | DNA ligase 3 OS=Homo sapiens OX=96( | LIG3 | 112.907 | 2 | 53.9544 | yes | 1.055E-07 | 649.38995 | 2008.4075 |
| EDEQQQALQDIAS | 14 | P49916 | DNA ligase 3 OS=Homo sapiens OX=96( | LIG3 | 112.907 | 2 | 53.9544 | yes | 9.722E-08 | 815.88702 | 1891.9724 |
| ALDLADMITR | 10 | P49916 | DNA ligase 3 OS=Homo sapiens OX=96( | LIG3 | 112.907 | 2 | 53.9544 | yes | 1.145E-07 | 559.79736 | 1720.9166 |
| AAVWEITGAEFSK | 13 | P49916 | DNA ligase 3 OS=Homo sapiens OX=96( | LIG3 | 112.907 | 2 | 53.9544 | yes | 1.46E-07 | 704.85901 | 1388.4718 |
| IYYPDFIVPDPK | 12 | P49916 | DNA ligase 3 OS=Homo sapiens OX=96( | LIG3 | 112.907 | 2 | 53.9544 | yes | 9.903E-08 | 733.88196 | 1230.2111 |
| LQNELDMVK | 9 | P49916 | DNA ligase 3 OS=Homo sapiens OX=96( | LIG3 | 112.907 | 2 | 53.9544 | yes | 1.199E-07 | 545.28406 | 1216.9414 |
| DLEQGDVSETIR | 12 | P49916 | DNA ligase 3 OS=Homo sapiens OX=96( | LIG3 | 112.907 | 2 | 53.9544 | yes | 9.722E-08 | 681.33081 | 1077.9615 |
| VVPNPFSESGGDM | 14 | P49916 | DNA ligase 3 OS=Homo sapiens OX=96( | LIG3 | 112.907 | 2 | 53.9544 | yes | 1.902E-07 | 732.3454 | 860.31305 |
| SLKPVLPHK | 9 | P49916 | DNA ligase 3 OS=Homo sapiens OX=96( | LIG3 | 112.907 | 2 | 53.9544 | yes | 3.115E-07 | 509.82404 | 828.76703 |
| LTTTGQVTSPVK | 12 | P49916 | DNA ligase 3 OS=Homo sapiens OX=96( | LIG3 | 112.907 | 2 | 53.9544 | yes | 4.646E-07 | 616.34827 | 741.84912 |
| LYLPPSTPDFSR | 12 | P49916 | DNA ligase 3 OS=Homo sapiens OX=96( | LIG3 | 112.907 | 2 | 53.9544 | yes | 3.172E-06 | 696.86151 | 691.745 |
| NLQDVVER | 8 | P49916 | DNA ligase 3 OS=Homo sapiens OX=96( | LIG3 | 112.907 | 2 | 53.9544 | yes | 9.809E-08 | 486.75909 | 670.52728 |
| SEAHTADGISIR | 12 | P49916 | DNA ligase 3 OS=Homo sapiens OX=96( | LIG3 | 112.907 | 2 | 53.9544 | yes | 1.206E-07 | 628.81512 | 596.06018 |
| GASFVTSTNPR | 11 | P49916 | DNA ligase 3 OS=Homo sapiens OX=96( | LIG3 | 112.907 | 2 | 53.9544 | yes | 1.303E-07 | 568.78839 | 585.18408 |
| VLLDIFTGVR | 10 | P49916 | DNA ligase 3 OS=Homo sapiens OX=96( | LIG3 | 112.907 | 2 | 53.9544 | yes | 1.104E-07 | 566.83984 | 568.22107 |
| TGKPLPFGTLGVH | 14 | P49916 | DNA ligase 3 OS=Homo sapiens OX=96( | LIG3 | 112.907 | 3 | 53.9544 | yes | 1.382E-07 | 484.61716 | 255.49084 |
| TGKPLPFGTLGVH | 14 | P49916 | DNA ligase 3 OS=Homo sapiens OX=96( | LIG3 | 112.907 | 3 | 53.9544 | yes | 9.79E-08 | 726.42212 | 255.49084 |
| TGKPLPFGTLGVH | 14 | P49916 | DNA ligase 3 OS=Homo sapiens OX=96( | LIG3 | 112.907 | 2 | 53.9544 | yes | 1.382E-07 | 484.61716 | 255.49084 |
| TGKPLPFGTLGVH | 14 | P49916 | DNA ligase 3 OS=Homo sapiens OX=96( | LIG3 | 112.907 | 2 | 53.9544 | yes | 9.79E-08 | 726.42212 | 255.49084 |
| VLHNAQEVEK | 10 | P49916 | DNA ligase 3 OS=Homo sapiens OX=96( | LIG3 | 112.907 | 2 | 53.9544 | yes | 3.468E-07 | 583.81183 | 473.54599 |
| FLHDNMVEIPNR | 12 | P49916 | DNA ligase 3 OS=Homo sapiens OX=96( | LIG3 | 112.907 | 2 | 53.9544 | yes | 2.95E-07 | 742.86938 | 169.30325 |
| FLHDNMVEIPNR | 12 | P49916 | DNA ligase 3 OS=Homo sapiens OX=96( | LIG3 | 112.907 | 2 | 53.9544 | yes | 1.974E-07 | 495.582 | 169.30325 |
| FLHDNMVEIPNR | 12 | P49916 | DNA ligase 3 OS=Homo sapiens OX=96( | LIG3 | 112.907 | 3 | 53.9544 | yes | 2.95E-07 | 742.86938 | 169.30325 |
| FLHDNMVEIPNR | 12 | P49916 | DNA ligase 3 OS=Homo sapiens OX=96( | LIG3 | 112.907 | 3 | 53.9544 | yes | 1.974E-07 | 495.582 | 169.30325 |
| LCAMVADNPSYN | 14 | P49916 | DNA ligase 3 OS=Homo sapiens OX=96( | LIG3 | 112.907 | 2 | 53.9544 | yes | 9.722E-08 | 792.36328 | 328.33978 |
| FCVDYAK | 7 | P49916 | DNA ligase 3 OS=Homo sapiens OX=96( | LIG3 | 112.907 | 2 | 53.9544 | yes | 1.004E-07 | 451.70746 | 290.08459 |
| LLLPGVIK | 8 | P49916 | DNA ligase 3 OS=Homo sapiens OX=96( | LIG3 | 112.907 | 2 | 53.9544 | yes | 1.874E-07 | 426.7995 | 278.37787 |

| | | | | | | | | | | | |
|---|---|---|---|---|---|---|---|---|---|---|---|
| AADETLCQTK | 10 | P49916 | DNA ligase 3 OS=Homo sapiens OX=96( | LIG3 | 112.907 | 2 | 53.9544 | yes | 9.9E-08 | 568.76624 | 260.27811 |
| CAGGHDDATLAR | 12 | P49916 | DNA ligase 3 OS=Homo sapiens OX=96( | LIG3 | 112.907 | 2 | 53.9544 | yes | 1.113E-07 | 622.27783 | 121.62611 |
| CAGGHDDATLAR | 12 | P49916 | DNA ligase 3 OS=Homo sapiens OX=96( | LIG3 | 112.907 | 2 | 53.9544 | yes | 1.145E-07 | 415.18765 | 121.62611 |
| CAGGHDDATLAR | 12 | P49916 | DNA ligase 3 OS=Homo sapiens OX=96( | LIG3 | 112.907 | 3 | 53.9544 | yes | 1.113E-07 | 622.27783 | 121.62611 |
| CAGGHDDATLAR | 12 | P49916 | DNA ligase 3 OS=Homo sapiens OX=96( | LIG3 | 112.907 | 3 | 53.9544 | yes | 1.145E-07 | 415.18765 | 121.62611 |
| SATNLPQLK | 9 | P49916 | DNA ligase 3 OS=Homo sapiens OX=96( | LIG3 | 112.907 | 2 | 53.9544 | yes | 5.863E-07 | 486.27966 | 185.11592 |
| KIEDLTELEGWEE] | 19 | P49916 | DNA ligase 3 OS=Homo sapiens OX=96( | LIG3 | 112.907 | 3 | 53.9544 | yes | 1.128E-07 | 773.70068 | 169.8936 |
| IFNCNPDDMAR | 11 | P49916 | DNA ligase 3 OS=Homo sapiens OX=96( | LIG3 | 112.907 | 2 | 53.9544 | yes | 9.722E-08 | 676.78973 | 168.37437 |
| VLHNAQEVEKEP( | 15 | P49916 | DNA ligase 3 OS=Homo sapiens OX=96( | LIG3 | 112.907 | 3 | 53.9544 | yes | 9.81E-08 | 578.6358 | 132.8297 |
| EQITQHIADLSSK | 13 | P49916 | DNA ligase 3 OS=Homo sapiens OX=96( | LIG3 | 112.907 | 2 | 53.9544 | yes | 2.805E-07 | 735.38336 | 131.64587 |
| IMFSEMK | 7 | P49916 | DNA ligase 3 OS=Homo sapiens OX=96( | LIG3 | 112.907 | 2 | 53.9544 | yes | 2.079E-07 | 443.21408 | 130.23578 |
| HVLDALDPNAYE/ | 15 | P49916 | DNA ligase 3 OS=Homo sapiens OX=96( | LIG3 | 112.907 | 3 | 53.9544 | yes | 1.648E-07 | 568.28601 | 110.36714 |
| VFFEQSK | 7 | P49916 | DNA ligase 3 OS=Homo sapiens OX=96( | LIG3 | 112.907 | 2 | 53.9544 | yes | 1.457E-07 | 442.72925 | 87.940651 |
| IRDDKDWK | 8 | P49916 | DNA ligase 3 OS=Homo sapiens OX=96( | LIG3 | 112.907 | 2 | 53.9544 | yes | 1.672E-07 | 538.28015 | 87.381424 |
| NGDHFSYFSR | 10 | P49916 | DNA ligase 3 OS=Homo sapiens OX=96( | LIG3 | 112.907 | 2 | 53.9544 | yes | 2.243E-07 | 615.27032 | 86.406296 |
| GSAGDGFHGDVY] | 16 | P49916 | DNA ligase 3 OS=Homo sapiens OX=96( | LIG3 | 112.907 | 2 | 53.9544 | yes | 9.81E-08 | 811.8941 | 58.216522 |
| KLCAMVADNPSY | 15 | P49916 | DNA ligase 3 OS=Homo sapiens OX=96( | LIG3 | 112.907 | 3 | 53.9544 | yes | 1.967E-07 | 571.27625 | 25.896334 |
| KAADETLCQTK | 11 | P49916 | DNA ligase 3 OS=Homo sapiens OX=96( | LIG3 | 112.907 | 2 | 53.9544 | yes | 6.75E-06 | 632.81372 | 20.013798 |
| IDISPVLLQK | 10 | P49959 | Double-strand break repair protein MRE1 | MRE11 | 80.594 | 2 | 51.381256 | yes | 1.043E-07 | 563.34753 | 264.82932 |
| IPLHTVR | 7 | P49959 | Double-strand break repair protein MRE1 | MRE11 | 80.594 | 2 | 51.381256 | yes | 9.893E-08 | 418.26105 | 184.53255 |
| IALYGLGSIPDER | 13 | P49959 | Double-strand break repair protein MRE1 | MRE11 | 80.594 | 2 | 51.381256 | yes | 9.722E-08 | 702.38007 | 115.59167 |
| NVQLSLLTER | 10 | P49959 | Double-strand break repair protein MRE1 | MRE11 | 80.594 | 2 | 51.381256 | yes | 1.093E-07 | 586.83533 | 115.58035 |
| VTQAIQSFCLEK | 12 | P49959 | Double-strand break repair protein MRE1 | MRE11 | 80.594 | 2 | 51.381256 | yes | 3.401E-07 | 712.36615 | 104.98942 |
| DIIHFFR | 7 | P49959 | Double-strand break repair protein MRE1 | MRE11 | 80.594 | 2 | 51.381256 | yes | 9.976E-08 | 474.25851 | 96.143875 |
| TLHTCLELLR | 10 | P49959 | Double-strand break repair protein MRE1 | MRE11 | 80.594 | 2 | 51.381256 | yes | 9.722E-08 | 628.34497 | 88.48571 |
| IEEMLENAER | 10 | P49959 | Double-strand break repair protein MRE1 | MRE11 | 80.594 | 2 | 51.381256 | yes | 1.544E-06 | 617.2926 | 72.851913 |
| NTNEEDDEVR | 10 | P49959 | Double-strand break repair protein MRE1 | MRE11 | 80.594 | 2 | 51.381256 | yes | 6.368E-07 | 610.75494 | 66.532623 |
| GNDTFVTLDEILR | 13 | P49959 | Double-strand break repair protein MRE1 | MRE11 | 80.594 | 2 | 51.381256 | yes | 2.458E-06 | 746.88574 | 66.209015 |
| LGNSHQPEKPLVR | 13 | P49959 | Double-strand break repair protein MRE1 | MRE11 | 80.594 | 3 | 51.381256 | yes | 1.311E-07 | 492.27567 | 28.222445 |
| IAEFAFEYAR | 10 | P50213 | Isocitrate dehydrogenase [NAD] subunit a | IDH3A | 39.592 | 2 | 39.365 | yes | 8.583E-07 | 608.80347 | 88.178635 |
| TPYTDVNIVTIR | 12 | P50213 | Isocitrate dehydrogenase [NAD] subunit a | IDH3A | 39.592 | 2 | 39.365 | yes | 4.628E-07 | 696.38007 | 54.714226 |
| MDPAASSCMR | 10 | P50221 | Homeobox protein MOX-1 OS=Homo sa | MEOX1 | 27.997 | 2 | 28.14901 | yes | 2.813E-06 | 584.23322 | 37.751915 |
| TLTSGGHAEHEGK | 28 | P50238 | Cysteine-rich protein 1 OS=Homo sapien | CRIP1 | 8.533 | 4 | 51.751194 | yes | 2.442E-07 | 780.09955 | 541.6521 |
| TLTSGGHAEHEGK | 28 | P50238 | Cysteine-rich protein 1 OS=Homo sapien | CRIP1 | 8.533 | 4 | 51.751194 | yes | 3.844E-06 | 624.28113 | 541.6521 |
| TLTSGGHAEHEGK | 28 | P50238 | Cysteine-rich protein 1 OS=Homo sapien | CRIP1 | 8.533 | 4 | 51.751194 | yes | 1.575E-07 | 784.09827 | 541.6521 |
| TLTSGGHAEHEGK | 28 | P50238 | Cysteine-rich protein 1 OS=Homo sapien | CRIP1 | 8.533 | 5 | 51.751194 | yes | 2.442E-07 | 780.09955 | 541.6521 |
| TLTSGGHAEHEGK | 28 | P50238 | Cysteine-rich protein 1 OS=Homo sapien | CRIP1 | 8.533 | 5 | 51.751194 | yes | 3.844E-06 | 624.28113 | 541.6521 |
| TLTSGGHAEHEGK | 28 | P50238 | Cysteine-rich protein 1 OS=Homo sapien | CRIP1 | 8.533 | 5 | 51.751194 | yes | 1.575E-07 | 784.09827 | 541.6521 |
| DWHRPCLK | 8 | P50238 | Cysteine-rich protein 1 OS=Homo sapien | CRIP1 | 8.533 | 2 | 51.751194 | yes | 8.226E-07 | 556.27692 | 1058.5648 |

| | | | | | | | | | | | |
|---|---|---|---|---|---|---|---|---|---|---|---|
| EVYFAER | 7 | P50238 | Cysteine-rich protein 1 OS=Homo sapien: | CRIP1 | 8.533 | 2 | 51.751194 | yes | 9.722E-08 | 457.22433 | 371.77969 |
| PYCNHPCYAAMF( | 15 | P50238 | Cysteine-rich protein 1 OS=Homo sapien: | CRIP1 | 8.533 | 3 | 51.751194 | yes | 9.722E-08 | 604.9267 | 236.88646 |
| GGAESHTFK | 9 | P50238 | Cysteine-rich protein 1 OS=Homo sapien: | CRIP1 | 8.533 | 2 | 51.751194 | yes | 1.668E-07 | 467.22488 | 188.0276 |
| CNKEVYFAER | 10 | P50238 | Cysteine-rich protein 1 OS=Homo sapien: | CRIP1 | 8.533 | 2 | 51.751194 | yes | 9.722E-08 | 658.30859 | 130.69545 |
| YNIPHGPVVGSTR | 13 | P50402 | Emerin OS=Homo sapiens OX=9606 GN | EMD | 28.994 | 2 | 53.404797 | yes | 1.167E-07 | 698.87024 | 1512.6252 |
| YNIPHGPVVGSTR | 13 | P50402 | Emerin OS=Homo sapiens OX=9606 GN | EMD | 28.994 | 2 | 53.404797 | yes | 1.131E-07 | 466.24924 | 1512.6252 |
| YNIPHGPVVGSTR | 13 | P50402 | Emerin OS=Homo sapiens OX=9606 GN | EMD | 28.994 | 3 | 53.404797 | yes | 1.167E-07 | 698.87024 | 1512.6252 |
| YNIPHGPVVGSTR | 13 | P50402 | Emerin OS=Homo sapiens OX=9606 GN | EMD | 28.994 | 3 | 53.404797 | yes | 1.131E-07 | 466.24924 | 1512.6252 |
| IFEYETQR | 8 | P50402 | Emerin OS=Homo sapiens OX=9606 GN | EMD | 28.994 | 2 | 53.404797 | yes | 1.466E-07 | 543.26672 | 1908.2126 |
| GDADMYDLPK | 10 | P50402 | Emerin OS=Homo sapiens OX=9606 GN | EMD | 28.994 | 2 | 53.404797 | yes | 2.894E-07 | 562.75006 | 1575.1198 |
| KEDALLYQSK | 10 | P50402 | Emerin OS=Homo sapiens OX=9606 GN | EMD | 28.994 | 2 | 53.404797 | yes | 9.722E-08 | 597.8219 | 1164.4663 |
| DSAYQSITHYR | 11 | P50402 | Emerin OS=Homo sapiens OX=9606 GN | EMD | 28.994 | 2 | 53.404797 | yes | 2.18E-07 | 670.81512 | 1090.2161 |
| RYNIPHGPVVGST | 14 | P50402 | Emerin OS=Homo sapiens OX=9606 GN | EMD | 28.994 | 3 | 53.404797 | yes | 1.012E-07 | 518.28296 | 765.2395 |
| ERPMYGR | 7 | P50402 | Emerin OS=Homo sapiens OX=9606 GN | EMD | 28.994 | 2 | 53.404797 | yes | 8.706E-07 | 454.724 | 521.2887 |
| KIFEYETQR | 9 | P50402 | Emerin OS=Homo sapiens OX=9606 GN | EMD | 28.994 | 2 | 53.404797 | yes | 9.991E-08 | 607.31421 | 427.94614 |
| DSAYQSITHYRPV: | 17 | P50402 | Emerin OS=Homo sapiens OX=9606 GN | EMD | 28.994 | 3 | 53.404797 | yes | 1.611E-07 | 646.65363 | 411.26407 |
| LSPPSSSAASSYSF: | 21 | P50402 | Emerin OS=Homo sapiens OX=9606 GN | EMD | 28.994 | 3 | 53.404797 | yes | 5.331E-07 | 721.00629 | 75.71991 |
| MTALAQDFAVGL( | 15 | P50416 | Carnitine O-palmitoyltransferase 1, liver i | CPT1A | 88.368 | 2 | 53.90078 | yes | 1.203E-07 | 773.90594 | 793.63867 |
| LAALTAGDR | 9 | P50416 | Carnitine O-palmitoyltransferase 1, liver i | CPT1A | 88.368 | 2 | 53.90078 | yes | 9.722E-08 | 444.25089 | 763.24854 |
| YLAVESPFLK | 10 | P50416 | Carnitine O-palmitoyltransferase 1, liver i | CPT1A | 88.368 | 2 | 53.90078 | yes | 1.758E-07 | 583.82642 | 720.24194 |
| LAMTGSGIDR | 10 | P50416 | Carnitine O-palmitoyltransferase 1, liver i | CPT1A | 88.368 | 2 | 53.90078 | yes | 1.152E-07 | 510.76077 | 664.05627 |
| GDINPNIPYPTR | 12 | P50416 | Carnitine O-palmitoyltransferase 1, liver i | CPT1A | 88.368 | 2 | 53.90078 | yes | 9.862E-08 | 678.84894 | 642.14099 |
| IDPSLGIIAK | 10 | P50416 | Carnitine O-palmitoyltransferase 1, liver i | CPT1A | 88.368 | 2 | 53.90078 | yes | 4.208E-07 | 513.81335 | 638.45538 |
| AMVDPAQTVEQR | 12 | P50416 | Carnitine O-palmitoyltransferase 1, liver i | CPT1A | 88.368 | 2 | 53.90078 | yes | 1.004E-07 | 672.83246 | 613.8396 |
| EVLSEPWR | 8 | P50416 | Carnitine O-palmitoyltransferase 1, liver i | CPT1A | 88.368 | 2 | 53.90078 | yes | 7.164E-07 | 508.26401 | 575.39441 |
| ILDNTSEPQPGEAF | 14 | P50416 | Carnitine O-palmitoyltransferase 1, liver i | CPT1A | 88.368 | 2 | 53.90078 | yes | 1.664E-07 | 763.87592 | 567.97357 |
| VWLYHDGR | 8 | P50416 | Carnitine O-palmitoyltransferase 1, liver i | CPT1A | 88.368 | 2 | 53.90078 | yes | 1.443E-07 | 523.26434 | 554.38239 |
| LSHEALR | 7 | P50416 | Carnitine O-palmitoyltransferase 1, liver i | CPT1A | 88.368 | 2 | 53.90078 | yes | 2.165E-07 | 413.23251 | 450.55386 |
| SCTTESCDFVR | 11 | P50416 | Carnitine O-palmitoyltransferase 1, liver i | CPT1A | 88.368 | 2 | 53.90078 | yes | 9.722E-08 | 681.27667 | 443.04233 |
| SEDPDTSMDSYAK | 13 | P50416 | Carnitine O-palmitoyltransferase 1, liver i | CPT1A | 88.368 | 2 | 53.90078 | yes | 4.334E-07 | 723.29047 | 403.12704 |
| EMEQQMQR | 8 | P50416 | Carnitine O-palmitoyltransferase 1, liver i | CPT1A | 88.368 | 2 | 53.90078 | yes | 9.722E-08 | 540.23407 | 380.81775 |
| FSCPETDSHR | 10 | P50416 | Carnitine O-palmitoyltransferase 1, liver i | CPT1A | 88.368 | 2 | 53.90078 | yes | 6.121E-07 | 618.25909 | 335.89514 |
| FCLTYEASMTR | 11 | P50416 | Carnitine O-palmitoyltransferase 1, liver i | CPT1A | 88.368 | 2 | 53.90078 | no | 9.722E-08 | 689.81012 | 269.26318 |
| TLETANCMSSQTK | 13 | P50416 | Carnitine O-palmitoyltransferase 1, liver i | CPT1A | 88.368 | 2 | 53.90078 | yes | 9.722E-08 | 735.83179 | 222.17728 |
| HLFCLYVVSK | 10 | P50416 | Carnitine O-palmitoyltransferase 1, liver i | CPT1A | 88.368 | 2 | 53.90078 | yes | 8.129E-07 | 633.33917 | 176.20828 |
| IPGEETDTIQHMR | 13 | P50416 | Carnitine O-palmitoyltransferase 1, liver i | CPT1A | 88.368 | 2 | 53.90078 | yes | 1.452E-07 | 763.867 | 174.54134 |
| IWMGMVK | 7 | P50416 | Carnitine O-palmitoyltransferase 1, liver i | CPT1A | 88.368 | 2 | 53.90078 | yes | 8.924E-07 | 432.72714 | 33.825287 |
| GVVEVTHDLQK | 11 | P50454 | Serpin H1 OS=Homo sapiens OX=9606 ( | SERPINH1 | 46.441 | 2 | 48.128864 | yes | 1.551E-07 | 612.83276 | 237.92139 |
| LYGPSSVSFADDF | 15 | P50454 | Serpin H1 OS=Homo sapiens OX=9606 ( | SERPINH1 | 46.441 | 2 | 48.128864 | yes | 1.418E-07 | 830.4043 | 189.75797 |

| | | | | | | | | | | | |
|---|---|---|---|---|---|---|---|---|---|---|---|
| LQIVEMPLAHK | 11 | P50454 | Serpin H1 OS=Homo sapiens OX=9606 ( | SERPINH1 | 46.441 | 2 | 48.128864 | yes | 1.306E-07 | 639.86554 | 188.89642 |
| AVLSAEQLR | 9 | P50454 | Serpin H1 OS=Homo sapiens OX=9606 ( | SERPINH1 | 46.441 | 2 | 48.128864 | yes | 5.199E-07 | 493.7851 | 177.26013 |
| HLAGLGLTEAIDK | 13 | P50454 | Serpin H1 OS=Homo sapiens OX=9606 ( | SERPINH1 | 46.441 | 2 | 48.128864 | yes | 1.195E-07 | 669.37482 | 119.41959 |
| LSSLIILMPHHVEP | 17 | P50454 | Serpin H1 OS=Homo sapiens OX=9606 ( | SERPINH1 | 46.441 | 3 | 48.128864 | yes | 2.813E-06 | 662.03979 | 94.069359 |
| GEGGAMSENIWST | 15 | P50542 | Peroxisomal targeting signal 1 receptor O | PEX5 | 70.865 | 2 | 54.716564 | yes | 1.053E-07 | 804.37775 | 144.34552 |
| ALELQPGYIR | 10 | P50542 | Peroxisomal targeting signal 1 receptor O | PEX5 | 70.865 | 2 | 54.716564 | yes | 1.953E-07 | 580.32715 | 143.96049 |
| LQAELEEMAK | 10 | P50542 | Peroxisomal targeting signal 1 receptor O | PEX5 | 70.865 | 2 | 54.716564 | yes | 3.973E-07 | 581.29462 | 112.36421 |
| LWLGEPEGTATDR | 13 | P50542 | Peroxisomal targeting signal 1 receptor O | PEX5 | 70.865 | 2 | 54.716564 | yes | 4.937E-07 | 722.85699 | 107.77785 |
| GYQFEEENPLR | 11 | P50542 | Peroxisomal targeting signal 1 receptor O | PEX5 | 70.865 | 2 | 54.716564 | yes | 9.722E-08 | 691.32275 | 92.160469 |
| WAEEYLEQSEEK | 12 | P50542 | Peroxisomal targeting signal 1 receptor O | PEX5 | 70.865 | 2 | 54.716564 | yes | 4.213E-06 | 770.84375 | 73.164627 |
| PVNTSALDMEFER | 13 | P50542 | Peroxisomal targeting signal 1 receptor O | PEX5 | 70.865 | 2 | 54.716564 | yes | 9.722E-08 | 754.85614 | 69.933426 |
| ELVEAECGGANPL | 15 | P50542 | Peroxisomal targeting signal 1 receptor O | PEX5 | 70.865 | 2 | 54.716564 | yes | 1.365E-07 | 809.38422 | 59.352768 |
| SAIESDVDFWDK | 12 | P50542 | Peroxisomal targeting signal 1 receptor O | PEX5 | 70.865 | 2 | 54.716564 | yes | 9.722E-08 | 706.32245 | 56.841541 |
| ELFLAAVR | 8 | P50542 | Peroxisomal targeting signal 1 receptor O | PEX5 | 70.865 | 2 | 54.716564 | yes | 3.008E-06 | 459.77402 | 52.114941 |
| QIGEGQVSLESGA( | 17 | P50542 | Peroxisomal targeting signal 1 receptor O | PEX5 | 70.865 | 2 | 54.716564 | yes | 3.136E-07 | 816.40283 | 7.6237535 |
| VSSDLQHATAQLS | 17 | P50548 | ETS domain-containing transcription fact | ERF | 58.703 | 3 | 49.4262 | yes | 1.032E-07 | 631.32556 | 178.56226 |
| GEGPGEAGGPLTP | 14 | P50548 | ETS domain-containing transcription fact | ERF | 58.703 | 2 | 49.4262 | yes | 3.58E-07 | 647.82294 | 98.131134 |
| ARPPGPPDLGAFR | 13 | P50548 | ETS domain-containing transcription fact | ERF | 58.703 | 2 | 49.4262 | yes | 3.09E-06 | 675.86749 | 71.427261 |
| NFDVGHVPIR | 10 | P50579 | Methionine aminopeptidase 2 OS=Homo | METAP2 | 52.892 | 2 | 50.924297 | yes | 1.557E-07 | 577.30927 | 886.75586 |
| ALDQASEEIWNDF | 14 | P50579 | Methionine aminopeptidase 2 OS=Homo | METAP2 | 52.892 | 2 | 50.924297 | yes | 9.722E-08 | 847.39447 | 686.74463 |
| TYQVKPIR | 8 | P50579 | Methionine aminopeptidase 2 OS=Homo | METAP2 | 52.892 | 2 | 50.924297 | yes | 9.849E-08 | 502.798 | 507.88977 |
| IDFGTHISGR | 10 | P50579 | Methionine aminopeptidase 2 OS=Homo | METAP2 | 52.892 | 2 | 50.924297 | yes | 2.48E-07 | 551.78564 | 482.99896 |
| IIDCAFTVTFNPK | 13 | P50579 | Methionine aminopeptidase 2 OS=Homo | METAP2 | 52.892 | 2 | 50.924297 | yes | 1.2E-07 | 763.38959 | 274.04126 |
| PGMTMIEICEK | 11 | P50579 | Methionine aminopeptidase 2 OS=Homo | METAP2 | 52.892 | 2 | 50.924297 | yes | 1.946E-07 | 654.80347 | 105.11724 |
| KYVMSWIK | 8 | P50579 | Methionine aminopeptidase 2 OS=Homo | METAP2 | 52.892 | 2 | 50.924297 | yes | 4.831E-07 | 527.79132 | 39.553715 |
| APELLFGAR | 9 | P50613 | Cyclin-dependent kinase 7 OS=Homo sap | CDK7 | 39.039 | 2 | 52.70547 | yes | 1.876E-07 | 487.27692 | 286.12918 |
| DNSLVLTPSHIK | 12 | P50613 | Cyclin-dependent kinase 7 OS=Homo sap | CDK7 | 39.039 | 2 | 52.70547 | yes | 1.197E-07 | 662.367 | 242.19951 |
| VPFLPGDSDLDQL' | 15 | P50613 | Cyclin-dependent kinase 7 OS=Homo sap | CDK7 | 39.039 | 2 | 52.70547 | yes | 1.16E-07 | 836.93066 | 151.16493 |
| LDFLGEGQFATVY | 14 | P50613 | Cyclin-dependent kinase 7 OS=Homo sap | CDK7 | 39.039 | 2 | 52.70547 | yes | 9.722E-08 | 794.40631 | 100.57195 |
| TEALEQGGLPK | 11 | P50613 | Cyclin-dependent kinase 7 OS=Homo sap | CDK7 | 39.039 | 2 | 52.70547 | yes | 9.975E-08 | 571.80621 | 73.400917 |
| AYTHQVVTR | 9 | P50613 | Cyclin-dependent kinase 7 OS=Homo sap | CDK7 | 39.039 | 2 | 52.70547 | yes | 1.188E-06 | 537.78815 | 44.30661 |
| YFSNRPGPTPGCQ | 25 | P50613 | Cyclin-dependent kinase 7 OS=Homo sap | CDK7 | 39.039 | 4 | 52.70547 | yes | 7.662E-07 | 722.11011 | 42.029778 |
| STQPLTFTTATFAA | 16 | P50616 | Protein Tob1 OS=Homo sapiens OX=960 | TOB1 | 38.155 | 2 | 42.762867 | yes | 2.885E-07 | 843.44086 | 22.317057 |
| WLCPSTKPGEKPS | 25 | P50748 | Kinetochore-associated protein 1 OS=Ho | KNTC1 | 250.749 | 4 | 31.890842 | yes | 1.691E-06 | 740.6181 | 72.624908 |
| LLVLDPAQR | 9 | P50750 | Cyclin-dependent kinase 9 OS=Homo sap | CDK9 | 42.778 | 2 | 50.49054 | yes | 1.151E-06 | 512.8111 | 323.35339 |
| NPATTNQTEFER | 12 | P50750 | Cyclin-dependent kinase 9 OS=Homo sap | CDK9 | 42.778 | 2 | 50.49054 | yes | 9.796E-08 | 704.32861 | 97.977943 |
| GSQITQQSTNQSR | 13 | P50750 | Cyclin-dependent kinase 9 OS=Homo sap | CDK9 | 42.778 | 2 | 50.49054 | yes | 1.035E-07 | 717.85022 | 60.871212 |
| EGFPITALR | 9 | P50750 | Cyclin-dependent kinase 9 OS=Homo sap | CDK9 | 42.778 | 2 | 50.49054 | yes | 1.463E-06 | 502.2822 | 43.970863 |
| GMLSTHLTSMFEY | 18 | P50750 | Cyclin-dependent kinase 9 OS=Homo sap | CDK9 | 42.778 | 3 | 50.49054 | yes | 2.626E-07 | 684.34113 | 38.717228 |

| | | | | | | | | | | | |
|---|---|---|---|---|---|---|---|---|---|---|---|
| VAYVSFGPHAGK | 12 | P50914 | Large ribosomal subunit protein eL14 OS | RPL14 | 23.432 | 2 | 56.39677 | yes | 1.059E-07 | 616.82477 | 26338.467 |
| CMQLTDFILK | 10 | P50914 | Large ribosomal subunit protein eL14 OS | RPL14 | 23.432 | 2 | 56.39677 | yes | 9.819E-08 | 634.82251 | 4143.2256 |
| CMQLTDFILK | 10 | P50914 | Large ribosomal subunit protein eL14 OS | RPL14 | 23.432 | 2 | 56.39677 | yes | 2.272E-07 | 642.81995 | 4143.2256 |
| ALVDGPCTQVR | 11 | P50914 | Large ribosomal subunit protein eL14 OS | RPL14 | 23.432 | 2 | 56.39677 | yes | 1.838E-06 | 608.31116 | 6976.1284 |
| RFVEVGR | 7 | P50914 | Large ribosomal subunit protein eL14 OS | RPL14 | 23.432 | 2 | 56.39677 | yes | 2.153E-07 | 431.74832 | 1889.58 |
| LVAIVDVIDQNR | 12 | P50914 | Large ribosomal subunit protein eL14 OS | RPL14 | 23.432 | 2 | 56.39677 | yes | 9.722E-08 | 677.88806 | 124.01537 |
| GSTDNLMDDIER | 12 | P50990 | T-complex protein 1 subunit theta OS=Ho | CCT8 | 59.621 | 2 | 53.552917 | yes | 9.728E-08 | 683.30121 | 241.46646 |
| NVGLDIEAEVPAV | 14 | P50990 | T-complex protein 1 subunit theta OS=Ho | CCT8 | 59.621 | 2 | 53.552917 | yes | 2.458E-07 | 727.3985 | 233.52209 |
| LATNAAVTVLR | 11 | P50990 | T-complex protein 1 subunit theta OS=Ho | CCT8 | 59.621 | 2 | 53.552917 | yes | 4.381E-07 | 564.84039 | 166.41298 |
| TVGATALPR | 9 | P50990 | T-complex protein 1 subunit theta OS=Ho | CCT8 | 59.621 | 2 | 53.552917 | yes | 2.33E-06 | 443.26126 | 143.87003 |
| LFVTNDAATILR | 12 | P50990 | T-complex protein 1 subunit theta OS=Ho | CCT8 | 59.621 | 2 | 53.552917 | yes | 9.722E-08 | 667.37738 | 140.96222 |
| FAEAFEAIPR | 10 | P50990 | T-complex protein 1 subunit theta OS=Ho | CCT8 | 59.621 | 2 | 53.552917 | yes | 2.188E-07 | 575.79822 | 97.306015 |
| VDQIIMAK | 8 | P50990 | T-complex protein 1 subunit theta OS=Ho | CCT8 | 59.621 | 2 | 53.552917 | yes | 2.868E-06 | 459.25986 | 90.906342 |
| MVINHLEK | 8 | P50990 | T-complex protein 1 subunit theta OS=Ho | CCT8 | 59.621 | 2 | 53.552917 | yes | 1.426E-07 | 492.27078 | 89.617172 |
| AIADTGANVVVTC | 15 | P50990 | T-complex protein 1 subunit theta OS=Ho | CCT8 | 59.621 | 2 | 53.552917 | yes | 1.153E-06 | 686.87518 | 89.087662 |
| LYAVHQEGNK | 10 | P50990 | T-complex protein 1 subunit theta OS=Ho | CCT8 | 59.621 | 2 | 53.552917 | yes | 3.196E-07 | 579.79877 | 79.111496 |
| GEENLMDAQVK | 11 | P50990 | T-complex protein 1 subunit theta OS=Ho | CCT8 | 59.621 | 2 | 53.552917 | yes | 9.722E-08 | 617.2926 | 78.600639 |
| YNIMLVR | 7 | P50990 | T-complex protein 1 subunit theta OS=Ho | CCT8 | 59.621 | 2 | 53.552917 | yes | 2.556E-06 | 454.75476 | 61.10162 |
| ELEVQHPAAK | 10 | P50990 | T-complex protein 1 subunit theta OS=Ho | CCT8 | 59.621 | 2 | 53.552917 | yes | 9.722E-08 | 561.30109 | 54.868088 |
| APGFAQMLK | 9 | P50990 | T-complex protein 1 subunit theta OS=Ho | CCT8 | 59.621 | 2 | 53.552917 | yes | 5.83E-07 | 481.76004 | 49.464314 |
| DMLEAGILDTYLG | 14 | P50990 | T-complex protein 1 subunit theta OS=Ho | CCT8 | 59.621 | 2 | 53.552917 | yes | 1.007E-07 | 769.89215 | 42.229408 |
| VADMALHYANK | 11 | P50990 | T-complex protein 1 subunit theta OS=Ho | CCT8 | 59.621 | 2 | 53.552917 | yes | 9.936E-07 | 616.80823 | 23.725403 |
| EDGAISTIVLR | 11 | P50990 | T-complex protein 1 subunit theta OS=Ho | CCT8 | 59.621 | 2 | 53.552917 | yes | 1.516E-07 | 587.32733 | 15.637742 |
| VIDPATATSVDLR | 13 | P50991 | T-complex protein 1 subunit delta OS=Ho | CCT4 | 57.925 | 2 | 53.356117 | yes | 1.151E-07 | 679.36975 | 648.85565 |
| SIHDALCVIR | 10 | P50991 | T-complex protein 1 subunit delta OS=Ho | CCT4 | 57.925 | 2 | 53.356117 | yes | 2.344E-07 | 592.31622 | 213.44962 |
| TLSGMESYCVR | 11 | P50991 | T-complex protein 1 subunit delta OS=Ho | CCT4 | 57.925 | 2 | 53.356117 | yes | 2.77E-07 | 651.79449 | 195.07101 |
| EDIEFICK | 8 | P50991 | T-complex protein 1 subunit delta OS=Ho | CCT4 | 57.925 | 2 | 53.356117 | yes | 9.722E-08 | 527.24969 | 151.50987 |
| TGCNVLLIQK | 10 | P50991 | T-complex protein 1 subunit delta OS=Ho | CCT4 | 57.925 | 2 | 53.356117 | yes | 1.041E-06 | 573.32098 | 134.07501 |
| GDVTITNDGATILK | 14 | P50991 | T-complex protein 1 subunit delta OS=Ho | CCT4 | 57.925 | 2 | 53.356117 | yes | 9.809E-08 | 709.38031 | 128.93002 |
| GIHPTIISESFQK | 13 | P50991 | T-complex protein 1 subunit delta OS=Ho | CCT4 | 57.925 | 2 | 53.356117 | yes | 2.562E-07 | 728.89337 | 125.89452 |
| ETLLNSATTSLNSK | 14 | P50991 | T-complex protein 1 subunit delta OS=Ho | CCT4 | 57.925 | 2 | 53.356117 | yes | 9.722E-08 | 739.88849 | 115.3011 |
| ALIAGGGAPEIELA | 16 | P50991 | T-complex protein 1 subunit delta OS=Ho | CCT4 | 57.925 | 2 | 53.356117 | yes | 9.919E-08 | 775.94867 | 104.47264 |
| GIEILTDMSR | 10 | P50991 | T-complex protein 1 subunit delta OS=Ho | CCT4 | 57.925 | 2 | 53.356117 | yes | 1.737E-07 | 567.7948 | 96.270813 |
| LVIEEAER | 8 | P50991 | T-complex protein 1 subunit delta OS=Ho | CCT4 | 57.925 | 2 | 53.356117 | yes | 9.722E-08 | 479.76382 | 88.521286 |
| QMQVLHPAAR | 10 | P50991 | T-complex protein 1 subunit delta OS=Ho | CCT4 | 57.925 | 2 | 53.356117 | yes | 2.838E-07 | 575.8111 | 88.079659 |
| LGGTIDDCELVEG | 19 | P50991 | T-complex protein 1 subunit delta OS=Ho | CCT4 | 57.925 | 3 | 53.356117 | yes | 3.528E-06 | 687.35754 | 3.1513827 |
| CTVDVPEDLR | 10 | P51114 | RNA-binding protein FXR1 OS=Homo sa | FXR1 | 69.721 | 2 | 52.797337 | yes | 1.374E-07 | 602.28735 | 44192.582 |
| ANDQEPCGWWLA | 13 | P51114 | RNA-binding protein FXR1 OS=Homo sa | FXR1 | 69.721 | 2 | 52.797337 | yes | 1.204E-07 | 787.85645 | 25364.357 |
| IFYHPETTQLMILS | 21 | P51114 | RNA-binding protein FXR1 OS=Homo sa | FXR1 | 69.721 | 3 | 52.797337 | yes | 9.732E-08 | 793.74701 | 10773.271 |

| | | | | | | | | | | |
|---|---|---|---|---|---|---|---|---|---|---|
| IFYHPETTQLMILS | 21 | P51114 | RNA-binding protein FXR1 OS=Homo sa | FXR1 | 69.721 | 3 | 52.797337 | yes | 9.804E-08 | 799.07861 | 10773.271 |
| QVPFNEVR | 8 | P51114 | RNA-binding protein FXR1 OS=Homo sa | FXR1 | 69.721 | 2 | 52.797337 | yes | 1.247E-06 | 494.76416 | 14502.334 |
| LPPPPDIK | 8 | P51114 | RNA-binding protein FXR1 OS=Homo sa | FXR1 | 69.721 | 2 | 52.797337 | yes | 1.422E-07 | 438.76309 | 13991.664 |
| IYGESADAVK | 10 | P51114 | RNA-binding protein FXR1 OS=Homo sa | FXR1 | 69.721 | 2 | 52.797337 | yes | 2.035E-07 | 526.7666 | 13337.671 |
| QLAAAFHEEFVVR | 13 | P51114 | RNA-binding protein FXR1 OS=Homo sa | FXR1 | 69.721 | 2 | 52.797337 | yes | 1.477E-07 | 758.89899 | 5738.1851 |
| QLAAAFHEEFVVR | 13 | P51114 | RNA-binding protein FXR1 OS=Homo sa | FXR1 | 69.721 | 2 | 52.797337 | yes | 1.34E-07 | 506.2684 | 5738.1851 |
| QLAAAFHEEFVVR | 13 | P51114 | RNA-binding protein FXR1 OS=Homo sa | FXR1 | 69.721 | 3 | 52.797337 | yes | 1.477E-07 | 758.89899 | 5738.1851 |
| QLAAAFHEEFVVR | 13 | P51114 | RNA-binding protein FXR1 OS=Homo sa | FXR1 | 69.721 | 3 | 52.797337 | yes | 1.34E-07 | 506.2684 | 5738.1851 |
| VNILSDMHLR | 10 | P51114 | RNA-binding protein FXR1 OS=Homo sa | FXR1 | 69.721 | 2 | 52.797337 | yes | 1.106E-06 | 599.32404 | 3660.5649 |
| VNILSDMHLR | 10 | P51114 | RNA-binding protein FXR1 OS=Homo sa | FXR1 | 69.721 | 2 | 52.797337 | yes | 9.722E-08 | 607.32153 | 3660.5649 |
| EDLMGLAIGTHGS | 19 | P51114 | RNA-binding protein FXR1 OS=Homo sa | FXR1 | 69.721 | 3 | 52.797337 | yes | 1.837E-07 | 671.00568 | 6136.793 |
| EDGMVPFVFVGTI | 13 | P51114 | RNA-binding protein FXR1 OS=Homo sa | FXR1 | 69.721 | 2 | 52.797337 | yes | 4.572E-07 | 713.35779 | 2297.4924 |
| EDGMVPFVFVGTI | 13 | P51114 | RNA-binding protein FXR1 OS=Homo sa | FXR1 | 69.721 | 2 | 52.797337 | yes | 2.575E-07 | 721.35522 | 2297.4924 |
| KDELSDWSLAGEI | 18 | P51114 | RNA-binding protein FXR1 OS=Homo sa | FXR1 | 69.721 | 3 | 52.797337 | yes | 2.406E-07 | 698.65051 | 4242.4287 |
| AINGPTSASGDDIS | 15 | P51114 | RNA-binding protein FXR1 OS=Homo sa | FXR1 | 69.721 | 2 | 52.797337 | yes | 2.075E-07 | 716.84937 | 3167.1797 |
| EISEGDEVEVYSR | 13 | P51114 | RNA-binding protein FXR1 OS=Homo sa | FXR1 | 69.721 | 2 | 52.797337 | yes | 9.729E-08 | 756.34662 | 2848.8809 |
| AELTVEVR | 8 | P51114 | RNA-binding protein FXR1 OS=Homo sa | FXR1 | 69.721 | 2 | 52.797337 | yes | 1.16E-07 | 479.76382 | 1190.1726 |
| DVIEEHGPSEK | 11 | P51114 | RNA-binding protein FXR1 OS=Homo sa | FXR1 | 69.721 | 2 | 52.797337 | yes | 2.438E-07 | 620.2962 | 1177.2804 |
| KEISEGDEVEVYSI | 14 | P51114 | RNA-binding protein FXR1 OS=Homo sa | FXR1 | 69.721 | 3 | 52.797337 | yes | 9.803E-08 | 547.2652 | 405.58234 |
| KEISEGDEVEVYSI | 14 | P51114 | RNA-binding protein FXR1 OS=Homo sa | FXR1 | 69.721 | 3 | 52.797337 | yes | 1.031E-07 | 820.3941 | 405.58234 |
| KEISEGDEVEVYSI | 14 | P51114 | RNA-binding protein FXR1 OS=Homo sa | FXR1 | 69.721 | 2 | 52.797337 | yes | 9.803E-08 | 547.2652 | 405.58234 |
| KEISEGDEVEVYSI | 14 | P51114 | RNA-binding protein FXR1 OS=Homo sa | FXR1 | 69.721 | 2 | 52.797337 | yes | 1.031E-07 | 820.3941 | 405.58234 |
| ESIGNVQVLLEYH | 18 | P51114 | RNA-binding protein FXR1 OS=Homo sa | FXR1 | 69.721 | 3 | 52.797337 | yes | 1.304E-07 | 697.04913 | 522.03638 |
| DELSDWSLAGEDI | 14 | P51114 | RNA-binding protein FXR1 OS=Homo sa | FXR1 | 69.721 | 2 | 52.797337 | yes | 9.809E-08 | 804.3446 | 453.71167 |
| RVNILSDMHLR | 11 | P51114 | RNA-binding protein FXR1 OS=Homo sa | FXR1 | 69.721 | 2 | 52.797337 | yes | 1.075E-07 | 677.37463 | 310.87225 |
| KVPGVTAIELDED | 18 | P51114 | RNA-binding protein FXR1 OS=Homo sa | FXR1 | 69.721 | 3 | 52.797337 | yes | 9.722E-08 | 650.00555 | 309.70813 |
| IEGDNENKLPR | 11 | P51114 | RNA-binding protein FXR1 OS=Homo sa | FXR1 | 69.721 | 2 | 52.797337 | yes | 9.81E-08 | 642.83075 | 306.56281 |
| LRPVNQNK | 8 | P51114 | RNA-binding protein FXR1 OS=Homo sa | FXR1 | 69.721 | 2 | 52.797337 | yes | 1.191E-06 | 484.78543 | 286.84647 |
| EENTQEAAVLNG | 14 | P51114 | RNA-binding protein FXR1 OS=Homo sa | FXR1 | 69.721 | 2 | 52.797337 | yes | 4.653E-07 | 730.8468 | 171.92708 |
| DVHEDSLTVVFEN | 19 | P51114 | RNA-binding protein FXR1 OS=Homo sa | FXR1 | 69.721 | 3 | 52.797337 | yes | 2.521E-07 | 772.02936 | 71.654907 |
| LQIDEQLR | 8 | P51116 | RNA-binding protein FXR2 OS=Homo sa | FXR2 | 74.224 | 2 | 53.14861 | no | 1.215E-07 | 507.78256 | 61581.949 |
| ASLLGDMHFR | 10 | P51116 | RNA-binding protein FXR2 OS=Homo sa | FXR2 | 74.224 | 2 | 53.14861 | yes | 1.58E-07 | 573.78986 | 6597.9731 |
| ASLLGDMHFR | 10 | P51116 | RNA-binding protein FXR2 OS=Homo sa | FXR2 | 74.224 | 2 | 53.14861 | yes | 1.026E-06 | 581.78729 | 6597.9731 |
| IYGETPEACR | 10 | P51116 | RNA-binding protein FXR2 OS=Homo sa | FXR2 | 74.224 | 2 | 53.14861 | yes | 3.229E-07 | 598.27423 | 12395.499 |
| SYLEFSEDSVQVPI | 14 | P51116 | RNA-binding protein FXR2 OS=Homo sa | FXR2 | 74.224 | 2 | 53.14861 | yes | 1.602E-07 | 828.39923 | 9876.2129 |
| QLAAAFQEEFTVR | 13 | P51116 | RNA-binding protein FXR2 OS=Homo sa | FXR2 | 74.224 | 2 | 53.14861 | yes | 1.241E-07 | 755.38843 | 8714.7646 |
| YNSSSISSVLK | 11 | P51116 | RNA-binding protein FXR2 OS=Homo sa | FXR2 | 74.224 | 2 | 53.14861 | yes | 9.722E-08 | 592.81152 | 8052.7651 |
| EDLMGLAIGTHGA | 19 | P51116 | RNA-binding protein FXR2 OS=Homo sa | FXR2 | 74.224 | 3 | 53.14861 | no | 2.432E-07 | 665.67407 | 3827.9485 |
| EDLMGLAIGTHGA | 19 | P51116 | RNA-binding protein FXR2 OS=Homo sa | FXR2 | 74.224 | 3 | 53.14861 | no | 2.356E-05 | 671.00568 | 3827.9485 |

| | | | | | | | | | | | |
|---|---|---|---|---|---|---|---|---|---|---|---|
| ANEQEPCGWWLA | 13 | P51116 | RNA-binding protein FXR2 OS=Homo sa | FXR2 | 74.224 | 2 | 53.14861 | yes | 2.07E-07 | 808.86737 | 7451.4814 |
| EITEGDEVEVYSR | 13 | P51116 | RNA-binding protein FXR2 OS=Homo sa | FXR2 | 74.224 | 2 | 53.14861 | yes | 9.933E-08 | 763.35449 | 6749.5337 |
| VTMAVPEDLR | 10 | P51116 | RNA-binding protein FXR2 OS=Homo sa | FXR2 | 74.224 | 2 | 53.14861 | yes | 1.618E-07 | 565.79736 | 6209.2124 |
| QIPFGDVR | 8 | P51116 | RNA-binding protein FXR2 OS=Homo sa | FXR2 | 74.224 | 2 | 53.14861 | yes | 1.814E-06 | 466.25345 | 4596.856 |
| LRPVNPNPLATK | 12 | P51116 | RNA-binding protein FXR2 OS=Homo sa | FXR2 | 74.224 | 2 | 53.14861 | yes | 5.203E-07 | 660.39331 | 2102.7354 |
| LRPVNPNPLATK | 12 | P51116 | RNA-binding protein FXR2 OS=Homo sa | FXR2 | 74.224 | 2 | 53.14861 | yes | 1.732E-07 | 440.59799 | 2102.7354 |
| LRPVNPNPLATK | 12 | P51116 | RNA-binding protein FXR2 OS=Homo sa | FXR2 | 74.224 | 3 | 53.14861 | yes | 5.203E-07 | 660.39331 | 2102.7354 |
| LRPVNPNPLATK | 12 | P51116 | RNA-binding protein FXR2 OS=Homo sa | FXR2 | 74.224 | 3 | 53.14861 | yes | 1.732E-07 | 440.59799 | 2102.7354 |
| TKPSEDSLSGQK | 12 | P51116 | RNA-binding protein FXR2 OS=Homo sa | FXR2 | 74.224 | 2 | 53.14861 | yes | 1.445E-07 | 638.82263 | 4119.4883 |
| LPPPADYNK | 9 | P51116 | RNA-binding protein FXR2 OS=Homo sa | FXR2 | 74.224 | 2 | 53.14861 | yes | 1.736E-06 | 507.76639 | 3806.5911 |
| QRPPLER | 7 | P51116 | RNA-binding protein FXR2 OS=Homo sa | FXR2 | 74.224 | 2 | 53.14861 | yes | 8.595E-07 | 448.25906 | 3250.0474 |
| QIGLGFRPPGSGR | 13 | P51116 | RNA-binding protein FXR2 OS=Homo sa | FXR2 | 74.224 | 3 | 53.14861 | yes | 6.824E-07 | 447.91772 | 692.73871 |
| QIGLGFRPPGSGR | 13 | P51116 | RNA-binding protein FXR2 OS=Homo sa | FXR2 | 74.224 | 3 | 53.14861 | yes | 3.657E-07 | 671.37292 | 692.73871 |
| QIGLGFRPPGSGR | 13 | P51116 | RNA-binding protein FXR2 OS=Homo sa | FXR2 | 74.224 | 2 | 53.14861 | yes | 6.824E-07 | 447.91772 | 692.73871 |
| QIGLGFRPPGSGR | 13 | P51116 | RNA-binding protein FXR2 OS=Homo sa | FXR2 | 74.224 | 2 | 53.14861 | yes | 3.657E-07 | 671.37292 | 692.73871 |
| AGPGDRDPPTR | 11 | P51116 | RNA-binding protein FXR2 OS=Homo sa | FXR2 | 74.224 | 2 | 53.14861 | yes | 5.202E-07 | 569.78363 | 1022.2296 |
| TDGSISGDR | 9 | P51116 | RNA-binding protein FXR2 OS=Homo sa | FXR2 | 74.224 | 2 | 53.14861 | yes | 1.383E-07 | 454.20941 | 836.73792 |
| QPVTVADYISR | 11 | P51116 | RNA-binding protein FXR2 OS=Homo sa | FXR2 | 74.224 | 2 | 53.14861 | yes | 1.054E-07 | 624.83276 | 542.53711 |
| VRVEGDNDK | 9 | P51116 | RNA-binding protein FXR2 OS=Homo sa | FXR2 | 74.224 | 2 | 53.14861 | yes | 1.269E-07 | 516.25946 | 449.16705 |
| KVPGVTAIELGEE | 18 | P51116 | RNA-binding protein FXR2 OS=Homo sa | FXR2 | 74.224 | 3 | 53.14861 | yes | 1.741E-07 | 669.67871 | 389.56348 |
| TGGPAYGPSSDVS | 23 | P51116 | RNA-binding protein FXR2 OS=Homo sa | FXR2 | 74.224 | 3 | 53.14861 | yes | 2.061E-07 | 771.68744 | 125.06587 |
| AGYSTDESSSSSLI | 17 | P51116 | RNA-binding protein FXR2 OS=Homo sa | FXR2 | 74.224 | 3 | 53.14861 | yes | 2.045E-05 | 585.93085 | 11.609873 |
| QASPNIVIALAGNI | 14 | P51148 | Ras-related protein Rab-5C OS=Homo sa | RAB5C | 23.483 | 2 | 52.515816 | yes | 9.722E-08 | 698.40137 | 56.153751 |
| GVDLQENNPASR | 12 | P51148 | Ras-related protein Rab-5C OS=Homo sa | RAB5C | 23.483 | 2 | 52.515816 | yes | 5.838E-07 | 650.31799 | 40.915905 |
| LVTMQIWDTAGQI | 14 | P51149 | Ras-related protein Rab-7a OS=Homo sa | RAB7A | 23.49 | 2 | 48.13722 | yes | 1.194E-07 | 824.41162 | 114.00082 |
| DATNVAAAFEEA | 14 | P51151 | Ras-related protein Rab-9A OS=Homo sa | RAB9A | 22.838 | 2 | 53.96519 | yes | 9.722E-08 | 732.35986 | 245.2278 |
| EPESFPFVILGNK | 13 | P51151 | Ras-related protein Rab-9A OS=Homo sa | RAB9A | 22.838 | 2 | 53.96519 | yes | 2.266E-06 | 738.89032 | 96.801567 |
| SDHLIQTDTVNLH | 14 | P51151 | Ras-related protein Rab-9A OS=Homo sa | RAB9A | 22.838 | 2 | 53.96519 | yes | 2.838E-07 | 824.92371 | 31.53908 |
| QVSTEEAQAWCR | 12 | P51151 | Ras-related protein Rab-9A OS=Homo sa | RAB9A | 22.838 | 2 | 53.96519 | yes | 9.728E-08 | 732.83044 | 23.813084 |
| LLLIGDSGVGK | 11 | P51153 | Ras-related protein Rab-13 OS=Homo sa | RAB13 | 22.775 | 2 | 50.708828 | no | 9.728E-08 | 536.32404 | 1028.5134 |
| SSMNVDEAFSSLA | 14 | P51153 | Ras-related protein Rab-13 OS=Homo sa | RAB13 | 22.775 | 2 | 50.708828 | yes | 1.199E-07 | 757.3512 | 454.11002 |
| SFENIQNWMK | 10 | P51153 | Ras-related protein Rab-13 OS=Homo sa | RAB13 | 22.775 | 2 | 50.708828 | yes | 1.063E-07 | 648.80573 | 346.50055 |
| AYDHLFK | 7 | P51153 | Ras-related protein Rab-13 OS=Homo sa | RAB13 | 22.775 | 2 | 50.708828 | yes | 2.396E-07 | 447.22943 | 183.69632 |
| LQVWDTAGQER | 11 | P51153 | Ras-related protein Rab-13 OS=Homo sa | RAB13 | 22.775 | 2 | 50.708828 | yes | 2.168E-07 | 651.8255 | 130.19026 |
| GAMGIILVYDITDI | 15 | P51153 | Ras-related protein Rab-13 OS=Homo sa | RAB13 | 22.775 | 2 | 50.708828 | yes | 1.64E-06 | 819.42639 | 51.391216 |
| GSPLGEVVEQGITI | 14 | P51398 | Small ribosomal subunit protein mS29 O | DAP3 | 45.567 | 2 | 55.299023 | yes | 8.881E-07 | 721.38593 | 2830.9333 |
| FDQPLEASTWLK | 12 | P51398 | Small ribosomal subunit protein mS29 O | DAP3 | 45.567 | 2 | 55.299023 | yes | 1.197E-07 | 717.86682 | 2439.9197 |
| AYLPQELLGK | 10 | P51398 | Small ribosomal subunit protein mS29 O | DAP3 | 45.567 | 2 | 55.299023 | yes | 2.045E-07 | 566.32404 | 1545.3618 |
| TFSEACLMVR | 10 | P51398 | Small ribosomal subunit protein mS29 O | DAP3 | 45.567 | 2 | 55.299023 | yes | 9.809E-08 | 607.28882 | 586.73981 |

| | | | | | | | | | | | |
|---|---|---|---|---|---|---|---|---|---|---|---|
| TFSEACLMVR | 10 | P51398 | Small ribosomal subunit protein mS29 OS | DAP3 | 45.567 | 2 | 55.299023 | yes | 5.341E-07 | 615.28632 | 586.73981 |
| NDWHGGAIVSALS | 22 | P51398 | Small ribosomal subunit protein mS29 OS | DAP3 | 45.567 | 3 | 55.299023 | yes | 1.318E-07 | 781.07312 | 246.2674 |
| NDWHGGAIVSALS | 22 | P51398 | Small ribosomal subunit protein mS29 OS | DAP3 | 45.567 | 3 | 55.299023 | yes | 6.58E-07 | 586.05664 | 246.2674 |
| NDWHGGAIVSALS | 22 | P51398 | Small ribosomal subunit protein mS29 OS | DAP3 | 45.567 | 4 | 55.299023 | yes | 1.318E-07 | 781.07312 | 246.2674 |
| NDWHGGAIVSALS | 22 | P51398 | Small ribosomal subunit protein mS29 OS | DAP3 | 45.567 | 4 | 55.299023 | yes | 6.58E-07 | 586.05664 | 246.2674 |
| NTSFAYPAIR | 10 | P51398 | Small ribosomal subunit protein mS29 OS | DAP3 | 45.567 | 2 | 55.299023 | yes | 4.878E-07 | 570.29584 | 441.89926 |
| KAYLPQELLGK | 11 | P51398 | Small ribosomal subunit protein mS29 OS | DAP3 | 45.567 | 2 | 55.299023 | yes | 9.728E-08 | 630.37152 | 324.79758 |
| TLSLCHVIHFCAK | 13 | P51398 | Small ribosomal subunit protein mS29 OS | DAP3 | 45.567 | 3 | 55.299023 | yes | 1.075E-07 | 529.27075 | 137.45874 |
| TLSLCHVIHFCAK | 13 | P51398 | Small ribosomal subunit protein mS29 OS | DAP3 | 45.567 | 3 | 55.299023 | yes | 9.722E-08 | 793.40253 | 137.45874 |
| TLSLCHVIHFCAK | 13 | P51398 | Small ribosomal subunit protein mS29 OS | DAP3 | 45.567 | 2 | 55.299023 | yes | 1.075E-07 | 529.27075 | 137.45874 |
| TLSLCHVIHFCAK | 13 | P51398 | Small ribosomal subunit protein mS29 OS | DAP3 | 45.567 | 2 | 55.299023 | yes | 9.722E-08 | 793.40253 | 137.45874 |
| SPIAPEELALVHNL | 15 | P51398 | Small ribosomal subunit protein mS29 OS | DAP3 | 45.567 | 3 | 55.299023 | yes | 3.322E-07 | 553.64569 | 238.70795 |
| YLLYGEK | 7 | P51398 | Small ribosomal subunit protein mS29 OS | DAP3 | 45.567 | 2 | 55.299023 | yes | 1.025E-07 | 443.23947 | 231.50197 |
| DLLQSSYNK | 9 | P51398 | Small ribosomal subunit protein mS29 OS | DAP3 | 45.567 | 2 | 55.299023 | yes | 2.663E-07 | 534.27203 | 195.75421 |
| EDKSPIAPEELALV | 18 | P51398 | Small ribosomal subunit protein mS29 OS | DAP3 | 45.567 | 3 | 55.299023 | yes | 3.612E-07 | 677.7005 | 112.14227 |
| NDWHGGAIVSALS | 20 | P51398 | Small ribosomal subunit protein mS29 OS | DAP3 | 45.567 | 3 | 55.299023 | yes | 9.722E-08 | 696.68848 | 110.11067 |
| NATDAVGIVLK | 11 | P51398 | Small ribosomal subunit protein mS29 OS | DAP3 | 45.567 | 2 | 55.299023 | yes | 1.676E-07 | 550.81915 | 94.205093 |
| RLNVAITR | 8 | P51530 | DNA replication ATP-dependent helicase | DNA2 | 120.415 | 2 | 49.223095 | yes | 2.613E-07 | 471.79581 | 1885.7058 |
| ALGMSESLFK | 10 | P51530 | DNA replication ATP-dependent helicase | DNA2 | 120.415 | 2 | 49.223095 | yes | 2.471E-06 | 541.78119 | 87.793823 |
| DYTLIVGMPGTGK | 13 | P51530 | DNA replication ATP-dependent helicase | DNA2 | 120.415 | 2 | 49.223095 | yes | 1.262E-06 | 676.34998 | 67.841888 |
| IINDLLAR | 8 | P51530 | DNA replication ATP-dependent helicase | DNA2 | 120.415 | 2 | 49.223095 | yes | 2.263E-07 | 464.28473 | 40.721832 |
| LQELAFQTIQEIR | 13 | P51530 | DNA replication ATP-dependent helicase | DNA2 | 120.415 | 2 | 49.223095 | yes | 1.057E-07 | 794.93829 | 40.142876 |
| YCSQIGNCALYSR | 13 | P51530 | DNA replication ATP-dependent helicase | DNA2 | 120.415 | 2 | 49.223095 | yes | 4.958E-07 | 796.35321 | 21.5576 |
| GQPLPETLQLAVQ | 15 | P51531 | Probable global transcription activator SN | SMARCA2 | 181.28 | 2 | 56.144314 | yes | 1.317E-07 | 789.94611 | 226.07211 |
| DVDYSDALTEK | 11 | P51531 | Probable global transcription activator SN | SMARCA2 | 181.28 | 2 | 56.144314 | yes | 9.722E-08 | 628.28809 | 149.19785 |
| QMNAIIDTVINYK | 13 | P51531 | Probable global transcription activator SN | SMARCA2 | 181.28 | 2 | 56.144314 | yes | 9.722E-08 | 761.90033 | 120.96422 |
| ILLDPNSEEVSEK | 13 | P51531 | Probable global transcription activator SN | SMARCA2 | 181.28 | 2 | 56.144314 | yes | 1.665E-07 | 736.87756 | 119.91643 |
| DTTLETALNSK | 11 | P51531 | Probable global transcription activator SN | SMARCA2 | 181.28 | 2 | 56.144314 | yes | 2.775E-07 | 596.8064 | 101.85387 |
| GILLTDGSEK | 10 | P51531 | Probable global transcription activator SN | SMARCA2 | 181.28 | 2 | 56.144314 | yes | 1.194E-07 | 516.78223 | 87.893738 |
| GIVEDIHCGSMK | 12 | P51531 | Probable global transcription activator SN | SMARCA2 | 181.28 | 2 | 56.144314 | yes | 1.108E-07 | 673.31555 | 60.268749 |
| GSQSYYTVAHAIS | 15 | P51531 | Probable global transcription activator SN | SMARCA2 | 181.28 | 2 | 56.144314 | yes | 9.723E-08 | 834.90247 | 16.738281 |
| GLDPVEILQER | 11 | P51532 | Transcription activator BRG1 OS=Homo | SMARCA4 | 184.646 | 2 | 55.26119 | yes | 5.102E-07 | 634.84589 | 1933.3547 |
| YMIVDEGHR | 9 | P51532 | Transcription activator BRG1 OS=Homo | SMARCA4 | 184.646 | 2 | 55.26119 | no | 1.342E-07 | 560.26624 | 1149.1777 |
| ELPEYYELIR | 10 | P51532 | Transcription activator BRG1 OS=Homo | SMARCA4 | 184.646 | 2 | 55.26119 | no | 9.722E-08 | 662.84283 | 1070.7893 |
| VIQAGMFDQK | 10 | P51532 | Transcription activator BRG1 OS=Homo | SMARCA4 | 184.646 | 2 | 55.26119 | no | 1.08E-07 | 568.79205 | 865.55762 |
| LTQVLNTHYVAPF | 13 | P51532 | Transcription activator BRG1 OS=Homo | SMARCA4 | 184.646 | 2 | 55.26119 | no | 1.065E-07 | 756.4201 | 405.63483 |
| LTQVLNTHYVAPF | 13 | P51532 | Transcription activator BRG1 OS=Homo | SMARCA4 | 184.646 | 2 | 55.26119 | no | 1.747E-07 | 504.61581 | 405.63483 |
| LTQVLNTHYVAPF | 13 | P51532 | Transcription activator BRG1 OS=Homo | SMARCA4 | 184.646 | 3 | 55.26119 | no | 1.065E-07 | 756.4201 | 405.63483 |
| LTQVLNTHYVAPF | 13 | P51532 | Transcription activator BRG1 OS=Homo | SMARCA4 | 184.646 | 3 | 55.26119 | no | 1.747E-07 | 504.61581 | 405.63483 |

| | | | | | | | | | | | |
|---|---|---|---|---|---|---|---|---|---|---|---|
| EVDYSDSLTEK | 11 | P51532 | Transcription activator BRG1 OS=Homo | SMARCA4 | 184.646 | 2 | 55.26119 | yes | 1.161E-07 | 643.29333 | 657.8255 |
| ITPIQKPR | 8 | P51532 | Transcription activator BRG1 OS=Homo | SMARCA4 | 184.646 | 2 | 55.26119 | yes | 2.207E-07 | 476.80054 | 628.71429 |
| TLMNTIMQLR | 10 | P51532 | Transcription activator BRG1 OS=Homo | SMARCA4 | 184.646 | 2 | 55.26119 | no | 1.388E-07 | 610.82813 | 620.60089 |
| DTALETALNAK | 11 | P51532 | Transcription activator BRG1 OS=Homo | SMARCA4 | 184.646 | 2 | 55.26119 | yes | 7.847E-07 | 573.80371 | 614.86188 |
| VDLNEEETILIIR | 13 | P51532 | Transcription activator BRG1 OS=Homo | SMARCA4 | 184.646 | 2 | 55.26119 | no | 1.427E-07 | 778.93018 | 465.9357 |
| LCTVNSVEEK | 10 | P51532 | Transcription activator BRG1 OS=Homo | SMARCA4 | 184.646 | 2 | 55.26119 | no | 1.873E-07 | 589.78973 | 461.71588 |
| LLLTGTPLQNK | 11 | P51532 | Transcription activator BRG1 OS=Homo | SMARCA4 | 184.646 | 2 | 55.26119 | yes | 2.609E-07 | 599.36371 | 453.2598 |
| LTCEEEEEK | 9 | P51532 | Transcription activator BRG1 OS=Homo | SMARCA4 | 184.646 | 2 | 55.26119 | no | 9.722E-08 | 583.74774 | 445.51294 |
| DSDAGSSTPTTSTI | 14 | P51532 | Transcription activator BRG1 OS=Homo | SMARCA4 | 184.646 | 2 | 55.26119 | yes | 3.34E-07 | 691.80511 | 434.93201 |
| LMEEDELPSWIIK | 13 | P51532 | Transcription activator BRG1 OS=Homo | SMARCA4 | 184.646 | 2 | 55.26119 | no | 9.722E-08 | 801.90784 | 418.85184 |
| CDMSALQR | 8 | P51532 | Transcription activator BRG1 OS=Homo | SMARCA4 | 184.646 | 2 | 55.26119 | yes | 9.934E-08 | 490.71805 | 324.20074 |
| QDVDDEYGVSQA | 15 | P51532 | Transcription activator BRG1 OS=Homo | SMARCA4 | 184.646 | 2 | 55.26119 | yes | 2.02E-07 | 833.38934 | 313.25308 |
| HEEEFDLFMR | 10 | P51532 | Transcription activator BRG1 OS=Homo | SMARCA4 | 184.646 | 2 | 55.26119 | yes | 1.79E-07 | 676.8006 | 284.41959 |
| GVLLTDGSEK | 10 | P51532 | Transcription activator BRG1 OS=Homo | SMARCA4 | 184.646 | 2 | 55.26119 | yes | 1.822E-07 | 509.77438 | 281.52997 |
| GQPLPDHLQMAV( | 15 | P51532 | Transcription activator BRG1 OS=Homo | SMARCA4 | 184.646 | 2 | 55.26119 | yes | 1.627E-07 | 809.92212 | 123.59776 |
| GQPLPDHLQMAV( | 15 | P51532 | Transcription activator BRG1 OS=Homo | SMARCA4 | 184.646 | 2 | 55.26119 | yes | 7.137E-05 | 540.28387 | 123.59776 |
| GQPLPDHLQMAV( | 15 | P51532 | Transcription activator BRG1 OS=Homo | SMARCA4 | 184.646 | 3 | 55.26119 | yes | 1.627E-07 | 809.92212 | 123.59776 |
| GQPLPDHLQMAV( | 15 | P51532 | Transcription activator BRG1 OS=Homo | SMARCA4 | 184.646 | 3 | 55.26119 | yes | 7.137E-05 | 540.28387 | 123.59776 |
| EVEAQLPEK | 9 | P51532 | Transcription activator BRG1 OS=Homo | SMARCA4 | 184.646 | 2 | 55.26119 | yes | 2.72E-07 | 521.77441 | 228.19595 |
| LIPPQPTGR | 9 | P51532 | Transcription activator BRG1 OS=Homo | SMARCA4 | 184.646 | 2 | 55.26119 | yes | 7.663E-07 | 489.79019 | 197.75288 |
| VIHVESGK | 8 | P51532 | Transcription activator BRG1 OS=Homo | SMARCA4 | 184.646 | 2 | 55.26119 | yes | 1.814E-07 | 434.74799 | 189.97507 |
| LMAEDEEGYRK | 11 | P51532 | Transcription activator BRG1 OS=Homo | SMARCA4 | 184.646 | 2 | 55.26119 | no | 9.722E-08 | 670.81116 | 181.36397 |
| STPDPPLGGTPR | 12 | P51532 | Transcription activator BRG1 OS=Homo | SMARCA4 | 184.646 | 2 | 55.26119 | yes | 4.355E-07 | 618.81458 | 179.88525 |
| QLSEVFIQLPSR | 12 | P51532 | Transcription activator BRG1 OS=Homo | SMARCA4 | 184.646 | 2 | 55.26119 | no | 1.043E-07 | 708.89594 | 179.62054 |
| QEVVVCMR | 8 | P51532 | Transcription activator BRG1 OS=Homo | SMARCA4 | 184.646 | 2 | 55.26119 | yes | 5.073E-07 | 510.75189 | 170.94543 |
| ILTGTDAPK | 9 | P51532 | Transcription activator BRG1 OS=Homo | SMARCA4 | 184.646 | 2 | 55.26119 | yes | 2.275E-07 | 458.26093 | 150.39439 |
| HQEYLNSILQHAK | 13 | P51532 | Transcription activator BRG1 OS=Homo | SMARCA4 | 184.646 | 2 | 55.26119 | no | 9.722E-08 | 790.9126 | 82.841843 |
| AFVPQLR | 7 | P51532 | Transcription activator BRG1 OS=Homo | SMARCA4 | 184.646 | 2 | 55.26119 | yes | 1.315E-07 | 415.7478 | 72.236 |
| LMAEDEEGYR | 10 | P51532 | Transcription activator BRG1 OS=Homo | SMARCA4 | 184.646 | 2 | 55.26119 | no | 9.722E-08 | 606.76373 | 62.244549 |
| GPTPFNQNQLHQL | 14 | P51532 | Transcription activator BRG1 OS=Homo | SMARCA4 | 184.646 | 2 | 55.26119 | yes | 2.94E-07 | 825.42896 | 53.807068 |
| QSALMVNGVLK | 11 | P51532 | Transcription activator BRG1 OS=Homo | SMARCA4 | 184.646 | 2 | 55.26119 | yes | 5.598E-07 | 580.3288 | 19.878794 |
| IPDPDSDDVSEVD. | 15 | P51532 | Transcription activator BRG1 OS=Homo | SMARCA4 | 184.646 | 2 | 55.26119 | yes | 1.118E-06 | 815.36554 | 8.6820602 |
| HTVTMIPGDGIGPI | 20 | P51553 | Isocitrate dehydrogenase [NAD] subunit g | IDH3G | 42.795 | 3 | 45.443634 | yes | 1.847E-07 | 715.71136 | 25.763578 |
| EEFGAEPELAVSAI | 16 | P51570 | Galactokinase OS=Homo sapiens OX=96 | GALK1 | 42.273 | 2 | 45.52183 | yes | 6.483E-07 | 829.90466 | 65.51207 |
| QPQVAELLAEAR | 12 | P51570 | Galactokinase OS=Homo sapiens OX=96 | GALK1 | 42.273 | 2 | 45.52183 | yes | 5.303E-07 | 662.86462 | 58.528622 |
| HSLASSEYPVR | 11 | P51570 | Galactokinase OS=Homo sapiens OX=96 | GALK1 | 42.273 | 2 | 45.52183 | yes | 1.823E-07 | 623.31476 | 50.632019 |
| TAQAAAALR | 9 | P51570 | Galactokinase OS=Homo sapiens OX=96 | GALK1 | 42.273 | 2 | 45.52183 | yes | 5.161E-07 | 436.75107 | 21.725437 |
| VQNMALYADVGC | 13 | P51571 | Translocon-associated protein subunit del | SSR4 | 18.999 | 2 | 53.993607 | yes | 9.722E-08 | 683.34521 | 456.42316 |
| FFDEESYSLLR | 11 | P51571 | Translocon-associated protein subunit del | SSR4 | 18.999 | 2 | 53.993607 | yes | 9.722E-08 | 703.33533 | 415.29407 |

| | | | | | | | | | | | |
|---|---|---|---|---|---|---|---|---|---|---|---|
| YQVSWSLDHK | 10 | P51571 | Translocon-associated protein subunit del | SSR4 | 18.999 | 2 | 53.993607 | yes | 4.25E-07 | 631.81183 | 147.37573 |
| LQAAVDGPMDK | 11 | P51572 | B-cell receptor-associated protein 31 OS= | BCAP31 | 27.992 | 2 | 48.116215 | yes | 1.197E-07 | 572.78699 | 147.44261 |
| LDVGNAEVK | 9 | P51572 | B-cell receptor-associated protein 31 OS= | BCAP31 | 27.992 | 2 | 48.116215 | yes | 8.99E-07 | 472.75601 | 101.65972 |
| TQPAVATAATAAI | 14 | P51608 | Methyl-CpG-binding protein 2 OS=Homo | MECP2 | 52.441 | 2 | 50.99473 | yes | 2.633E-07 | 665.35406 | 216.86043 |
| AATSEGVQVK | 10 | P51608 | Methyl-CpG-binding protein 2 OS=Homo | MECP2 | 52.441 | 2 | 50.99473 | yes | 1.046E-06 | 495.26672 | 65.978836 |
| SEDQDLQGLK | 10 | P51608 | Methyl-CpG-binding protein 2 OS=Homo | MECP2 | 52.441 | 2 | 50.99473 | yes | 4.877E-07 | 566.77765 | 59.80447 |
| AEGGGATTSTQVM | 16 | P51608 | Methyl-CpG-binding protein 2 OS=Homo | MECP2 | 52.441 | 2 | 50.99473 | yes | 9.722E-08 | 775.39813 | 50.959763 |
| SVQETVLPIK | 10 | P51608 | Methyl-CpG-binding protein 2 OS=Homo | MECP2 | 52.441 | 2 | 50.99473 | yes | 9.847E-08 | 557.32935 | 43.977837 |
| SKVELIAYFEK | 11 | P51608 | Methyl-CpG-binding protein 2 OS=Homo | MECP2 | 52.441 | 2 | 50.99473 | yes | 2.967E-06 | 663.86884 | 27.177761 |
| EVVKPLLVSTLGE | 14 | P51608 | Methyl-CpG-binding protein 2 OS=Homo | MECP2 | 52.441 | 2 | 50.99473 | yes | 1.966E-06 | 756.45581 | 26.259785 |
| KPGSVVAAAAAEA | 14 | P51608 | Methyl-CpG-binding protein 2 OS=Homo | MECP2 | 52.441 | 2 | 50.99473 | yes | 9.98E-07 | 635.36169 | 17.806633 |
| SPISVPGGSALISNI | 17 | P51610 | Host cell factor 1 OS=Homo sapiens OX= | HCFC1 | 208.733 | 2 | 48.499634 | yes | 2.311E-07 | 798.95142 | 160.53403 |
| VVGWSGPVPR | 10 | P51610 | Host cell factor 1 OS=Homo sapiens OX= | HCFC1 | 208.733 | 2 | 48.499634 | yes | 1.225E-07 | 527.29565 | 141.53777 |
| LVIYGGMSGCR | 11 | P51610 | Host cell factor 1 OS=Homo sapiens OX= | HCFC1 | 208.733 | 2 | 48.499634 | yes | 1.173E-07 | 606.79681 | 91.547188 |
| VASSPVMVSNPAT | 14 | P51610 | Host cell factor 1 OS=Homo sapiens OX= | HCFC1 | 208.733 | 2 | 48.499634 | yes | 1.139E-07 | 708.3692 | 72.785583 |
| GAPGQPGTILR | 11 | P51610 | Host cell factor 1 OS=Homo sapiens OX= | HCFC1 | 208.733 | 2 | 48.499634 | yes | 4.415E-07 | 533.80383 | 70.558449 |
| CYLFGGLANDSED | 15 | P51610 | Host cell factor 1 OS=Homo sapiens OX= | HCFC1 | 208.733 | 2 | 48.499634 | yes | 1.727E-07 | 843.37744 | 58.343853 |
| SPAFVQLAPLSSK | 13 | P51610 | Host cell factor 1 OS=Homo sapiens OX= | HCFC1 | 208.733 | 2 | 48.499634 | yes | 1.209E-07 | 672.8797 | 52.912399 |
| YSNDLYELQASR | 12 | P51610 | Host cell factor 1 OS=Homo sapiens OX= | HCFC1 | 208.733 | 2 | 48.499634 | yes | 6.166E-07 | 729.84662 | 49.169273 |
| LGHSFSLVGNK | 11 | P51610 | Host cell factor 1 OS=Homo sapiens OX= | HCFC1 | 208.733 | 2 | 48.499634 | yes | 5.21E-07 | 579.81696 | 47.05183 |
| LYIWSGR | 7 | P51610 | Host cell factor 1 OS=Homo sapiens OX= | HCFC1 | 208.733 | 2 | 48.499634 | yes | 3.524E-06 | 447.74524 | 37.311756 |
| IATGHGQQGVTQV | 16 | P51610 | Host cell factor 1 OS=Homo sapiens OX= | HCFC1 | 208.733 | 2 | 48.499634 | yes | 1.098E-06 | 818.46246 | 33.088036 |
| ESHTAVVYTEK | 11 | P51610 | Host cell factor 1 OS=Homo sapiens OX= | HCFC1 | 208.733 | 2 | 48.499634 | yes | 2.605E-06 | 632.31439 | 24.044224 |
| GTLAYLPEEYIK | 12 | P51617 | Interleukin-1 receptor-associated kinase 1 | IRAK1 | 76.537 | 2 | 55.17361 | yes | 9.722E-08 | 698.87158 | 91.299477 |
| LDILLGTAR | 9 | P51617 | Interleukin-1 receptor-associated kinase 1 | IRAK1 | 76.537 | 2 | 55.17361 | yes | 2.619E-07 | 486.29785 | 90.785652 |
| TASVLWPWINR | 11 | P51617 | Interleukin-1 receptor-associated kinase 1 | IRAK1 | 76.537 | 2 | 55.17361 | yes | 1.388E-07 | 671.86694 | 69.160187 |
| QSFLTEVEQLSR | 12 | P51617 | Interleukin-1 receptor-associated kinase 1 | IRAK1 | 76.537 | 2 | 55.17361 | yes | 4.45E-07 | 718.87262 | 62.825428 |
| SSNVLLDER | 9 | P51617 | Interleukin-1 receptor-associated kinase 1 | IRAK1 | 76.537 | 2 | 55.17361 | yes | 9.623E-07 | 516.76965 | 51.223228 |
| GTHNFSEELK | 10 | P51617 | Interleukin-1 receptor-associated kinase 1 | IRAK1 | 76.537 | 2 | 55.17361 | yes | 4.815E-06 | 581.2804 | 23.197079 |
| AIQFLHQDSPSLIH | 18 | P51617 | Interleukin-1 receptor-associated kinase 1 | IRAK1 | 76.537 | 3 | 55.17361 | yes | 4.778E-05 | 673.69348 | 0.76231 |
| LQQLEALR | 8 | P51648 | Aldehyde dehydrogenase family 3 membe | ALDH3A2 | 54.848 | 2 | 51.26532 | yes | 1.075E-07 | 485.78763 | 422.94446 |
| IAFGGETDEATR | 12 | P51648 | Aldehyde dehydrogenase family 3 membe | ALDH3A2 | 54.848 | 2 | 51.26532 | yes | 9.728E-08 | 633.8017 | 415.20688 |
| NVDEAINFINER | 12 | P51648 | Aldehyde dehydrogenase family 3 membe | ALDH3A2 | 54.848 | 2 | 51.26532 | yes | 9.722E-08 | 717.35461 | 379.6506 |
| YIAPTVLTDVDPK | 13 | P51648 | Aldehyde dehydrogenase family 3 membe | ALDH3A2 | 54.848 | 2 | 51.26532 | yes | 1.232E-07 | 716.39014 | 350.31107 |
| FDHIFYTGNTAVG | 14 | P51648 | Aldehyde dehydrogenase family 3 membe | ALDH3A2 | 54.848 | 2 | 51.26532 | yes | 9.91E-08 | 785.38843 | 119.24607 |
| EKDILTAIAADLCK | 14 | P51648 | Aldehyde dehydrogenase family 3 membe | ALDH3A2 | 54.848 | 2 | 51.26532 | yes | 1.002E-07 | 780.9187 | 69.067139 |
| EKPLALYVFSHNH | 14 | P51648 | Aldehyde dehydrogenase family 3 membe | ALDH3A2 | 54.848 | 3 | 51.26532 | yes | 2.634E-07 | 561.63861 | 58.215279 |
| ILSLLEGQK | 9 | P51648 | Aldehyde dehydrogenase family 3 membe | ALDH3A2 | 54.848 | 2 | 51.26532 | yes | 2.696E-06 | 500.80551 | 41.560551 |
| IDVVVNNAGILR | 12 | P51659 | Peroxisomal multifunctional enzyme type | HSD17B4 | 79.687 | 2 | 45.208145 | yes | 1.924E-07 | 641.8775 | 25.5301 |

| | | | | | | | | | | | |
|---|---|---|---|---|---|---|---|---|---|---|---|
| EVMSTPVTCLR | 11 | P51798 | H(+)/Cl(-) exchange transporter 7 OS=Hc | CLCN7 | 88.68 | 2 | 36.098423 | yes | 7.402E-07 | 646.8205 | 69.730347 |
| HCDEVGFNAEEAI | 17 | P51808 | Dynein light chain Tctex-type 3 OS=Hom | DYNLT3 | 13.062 | 3 | 34.301964 | yes | 1.036E-06 | 656.96716 | 40.919525 |
| DLKPENILLDEEGI | 16 | P51812 | Ribosomal protein S6 kinase alpha-3 OS= | RPS6KA3 | 83.736 | 3 | 55.531567 | no | 1.551E-07 | 621.66681 | 451.05249 |
| ADPSQFELLK | 10 | P51812 | Ribosomal protein S6 kinase alpha-3 OS= | RPS6KA3 | 83.736 | 2 | 55.531567 | no | 9.722E-08 | 574.30334 | 170.90733 |
| GAMAATYSALNR | 12 | P51812 | Ribosomal protein S6 kinase alpha-3 OS= | RPS6KA3 | 83.736 | 2 | 55.531567 | yes | 9.722E-08 | 613.30334 | 166.72009 |
| EDIGVGSYSVCK | 12 | P51812 | Ribosomal protein S6 kinase alpha-3 OS= | RPS6KA3 | 83.736 | 2 | 55.531567 | no | 9.722E-08 | 657.30573 | 162.62456 |
| DILVEVNHPFIVK | 13 | P51812 | Ribosomal protein S6 kinase alpha-3 OS= | RPS6KA3 | 83.736 | 2 | 55.531567 | no | 1.076E-07 | 761.93506 | 162.48106 |
| ICDFGFAK | 8 | P51812 | Ribosomal protein S6 kinase alpha-3 OS= | RPS6KA3 | 83.736 | 2 | 55.531567 | no | 9.729E-08 | 479.22858 | 151.29724 |
| YVYVVTELMK | 10 | P51812 | Ribosomal protein S6 kinase alpha-3 OS= | RPS6KA3 | 83.736 | 2 | 55.531567 | yes | 9.722E-08 | 622.83337 | 141.5674 |
| NQSPVLEPVGR | 11 | P51812 | Ribosomal protein S6 kinase alpha-3 OS= | RPS6KA3 | 83.736 | 2 | 55.531567 | yes | 9.754E-08 | 598.32513 | 138.33997 |
| MLHVDPHQR | 9 | P51812 | Ribosomal protein S6 kinase alpha-3 OS= | RPS6KA3 | 83.736 | 2 | 55.531567 | no | 1.15E-07 | 566.78766 | 99.032631 |
| EIAITHHVK | 9 | P51812 | Ribosomal protein S6 kinase alpha-3 OS= | RPS6KA3 | 83.736 | 2 | 55.531567 | yes | 2.844E-07 | 524.3009 | 77.146301 |
| DLKPSNILYVDES( | 20 | P51812 | Ribosomal protein S6 kinase alpha-3 OS= | RPS6KA3 | 83.736 | 3 | 55.531567 | yes | 9.722E-08 | 749.38147 | 76.503845 |
| LGAGPDGVEEIKR | 13 | P51812 | Ribosomal protein S6 kinase alpha-3 OS= | RPS6KA3 | 83.736 | 2 | 55.531567 | yes | 9.722E-08 | 670.86206 | 73.660576 |
| NSIQFTDGYEVK | 12 | P51812 | Ribosomal protein S6 kinase alpha-3 OS= | RPS6KA3 | 83.736 | 2 | 55.531567 | yes | 5.489E-07 | 700.83826 | 73.111732 |
| PLAQLADPWQK | 11 | P51812 | Ribosomal protein S6 kinase alpha-3 OS= | RPS6KA3 | 83.736 | 2 | 55.531567 | yes | 1.775E-06 | 633.8457 | 58.381241 |
| LTAALVLR | 8 | P51812 | Ribosomal protein S6 kinase alpha-3 OS= | RPS6KA3 | 83.736 | 2 | 55.531567 | yes | 6.807E-07 | 428.78436 | 57.016853 |
| EASAVLFTITK | 11 | P51812 | Ribosomal protein S6 kinase alpha-3 OS= | RPS6KA3 | 83.736 | 2 | 55.531567 | yes | 3.541E-07 | 590.33459 | 43.235592 |
| LTDFGLSK | 8 | P51812 | Ribosomal protein S6 kinase alpha-3 OS= | RPS6KA3 | 83.736 | 2 | 55.531567 | no | 7.125E-07 | 440.74237 | 37.844074 |
| EIHPPFKPATGRPE | 26 | P51812 | Ribosomal protein S6 kinase alpha-3 OS= | RPS6KA3 | 83.736 | 4 | 55.531567 | yes | 3.71E-06 | 760.12372 | 13.191756 |
| EPGLLPYGSR | 10 | P51825 | AF4/FMR2 family member 1 OS=Homo : | AFF1 | 131.422 | 2 | 53.374023 | yes | 3.452E-06 | 544.79041 | 108.49826 |
| SAYSVYSETVDLII | 14 | P51825 | AF4/FMR2 family member 1 OS=Homo : | AFF1 | 131.422 | 2 | 53.374023 | yes | 9.722E-08 | 787.90106 | 87.357544 |
| SFSDATAPTQEK | 12 | P51825 | AF4/FMR2 family member 1 OS=Homo : | AFF1 | 131.422 | 2 | 53.374023 | yes | 9.722E-08 | 641.30151 | 68.635422 |
| ITLDLLSR | 8 | P51825 | AF4/FMR2 family member 1 OS=Homo : | AFF1 | 131.422 | 2 | 53.374023 | yes | 3.169E-06 | 465.78458 | 64.977798 |
| IFAVLCMR | 8 | P51825 | AF4/FMR2 family member 1 OS=Homo : | AFF1 | 131.422 | 2 | 53.374023 | yes | 1.475E-07 | 505.26971 | 62.880733 |
| APPEAPHPGK | 10 | P51825 | AF4/FMR2 family member 1 OS=Homo : | AFF1 | 131.422 | 2 | 53.374023 | yes | 1.557E-07 | 500.76416 | 56.203758 |
| WQLDNWLTK | 9 | P51825 | AF4/FMR2 family member 1 OS=Homo : | AFF1 | 131.422 | 2 | 53.374023 | yes | 1.847E-06 | 602.31146 | 42.855762 |
| DRLPLPLR | 8 | P51825 | AF4/FMR2 family member 1 OS=Homo : | AFF1 | 131.422 | 2 | 53.374023 | yes | 7.375E-07 | 490.306 | 35.343918 |
| YPILENPEILR | 11 | P51946 | Cyclin-H OS=Homo sapiens OX=9606 G | CCNH | 37.644 | 2 | 49.15634 | yes | 1.376E-07 | 678.8797 | 124.74426 |
| KTADDFLNR | 9 | P51946 | Cyclin-H OS=Homo sapiens OX=9606 G | CCNH | 37.644 | 2 | 49.15634 | yes | 3.861E-07 | 540.27765 | 119.12682 |
| LLEFCSVFK | 9 | P51946 | Cyclin-H OS=Homo sapiens OX=9606 G | CCNH | 37.644 | 2 | 49.15634 | yes | 3.031E-07 | 571.79938 | 66.679207 |
| TADDFLNR | 8 | P51946 | Cyclin-H OS=Homo sapiens OX=9606 G | CCNH | 37.644 | 2 | 49.15634 | yes | 6.776E-07 | 476.23016 | 56.5947 |
| TRYPILENPEILR | 13 | P51946 | Cyclin-H OS=Homo sapiens OX=9606 G | CCNH | 37.644 | 3 | 49.15634 | yes | 4.934E-06 | 538.63849 | 54.845917 |
| HWTFSSEEQLAR | 12 | P51946 | Cyclin-H OS=Homo sapiens OX=9606 G | CCNH | 37.644 | 2 | 49.15634 | yes | 1.063E-07 | 745.8548 | 51.814957 |
| LGYLNHVR | 8 | P51948 | CDK-activating kinase assembly factor M | MNAT1 | 35.823 | 2 | 46.093117 | yes | 2.359E-07 | 486.27469 | 95.91803 |
| EQEELEEALEVER | 13 | P51948 | CDK-activating kinase assembly factor M | MNAT1 | 35.823 | 2 | 46.093117 | yes | 1.655E-07 | 801.8783 | 61.672127 |
| QYLDEEFVLR | 10 | P51955 | Serine/threonine-protein kinase Nek2 OS: | NEK2 | 51.764 | 2 | 53.98266 | yes | 1.084E-07 | 656.33258 | 165.69641 |
| QMLVSEVNLLR | 11 | P51955 | Serine/threonine-protein kinase Nek2 OS: | NEK2 | 51.764 | 2 | 53.98266 | yes | 1.129E-07 | 651.36591 | 139.74673 |
| DLKPANVFLDGK | 12 | P51955 | Serine/threonine-protein kinase Nek2 OS: | NEK2 | 51.764 | 2 | 53.98266 | yes | 3.716E-07 | 658.86407 | 134.08278 |

| | | | | | | | | | | | |
|---|---|---|---|---|---|---|---|---|---|---|---|
| ELDYGSMTEAEK | 12 | P51955 | Serine/threonine-protein kinase Nek2 OS= | NEK2 | 51.764 | 2 | 53.98266 | yes | 1.625E-07 | 686.80048 | 123.36774 |
| YSDELNEIITR | 11 | P51955 | Serine/threonine-protein kinase Nek2 OS= | NEK2 | 51.764 | 2 | 53.98266 | yes | 9.722E-08 | 676.83826 | 104.13158 |
| VMTQLTLALK | 10 | P51955 | Serine/threonine-protein kinase Nek2 OS= | NEK2 | 51.764 | 2 | 53.98266 | yes | 9.722E-08 | 559.33612 | 57.541859 |
| ILNHDTSFAK | 10 | P51955 | Serine/threonine-protein kinase Nek2 OS= | NEK2 | 51.764 | 2 | 53.98266 | yes | 2.756E-07 | 573.30109 | 53.309254 |
| SQDSSPVLSELK | 12 | P51955 | Serine/threonine-protein kinase Nek2 OS= | NEK2 | 51.764 | 2 | 53.98266 | yes | 6.926E-07 | 645.33282 | 48.515293 |
| SDGGHTVLHR | 10 | P51955 | Serine/threonine-protein kinase Nek2 OS= | NEK2 | 51.764 | 2 | 53.98266 | yes | 2.131E-07 | 539.77307 | 41.528858 |
| AAEQEAQLLSQLK | 13 | P51957 | Serine/threonine-protein kinase Nek4 OS= | NEK4 | 94.598 | 2 | 39.757812 | yes | 4.398E-07 | 714.88831 | 46.800781 |
| DYSPELAELIR | 11 | P51957 | Serine/threonine-protein kinase Nek4 OS= | NEK4 | 94.598 | 2 | 39.757812 | yes | 3.457E-06 | 653.33789 | 41.854813 |
| NVPLATDLIR | 10 | P51959 | Cyclin-G1 OS=Homo sapiens OX=9606 ( | CCNG1 | 34.074 | 2 | 55.706585 | yes | 9.722E-08 | 556.32715 | 88.211739 |
| CVELTEGIECLQK | 13 | P51959 | Cyclin-G1 OS=Homo sapiens OX=9606 ( | CCNG1 | 34.074 | 2 | 55.706585 | yes | 1.077E-06 | 789.87872 | 73.730644 |
| LLHQLNALLEQES | 14 | P51959 | Cyclin-G1 OS=Homo sapiens OX=9606 ( | CCNG1 | 34.074 | 2 | 55.706585 | yes | 1.564E-07 | 832.45996 | 53.574955 |
| ITHLPTIPEMVP | 12 | P51959 | Cyclin-G1 OS=Homo sapiens OX=9606 ( | CCNG1 | 34.074 | 2 | 55.706585 | yes | 5.415E-06 | 674.37067 | 36.79287 |
| HLGCVGLSCFYLA | 15 | P51959 | Cyclin-G1 OS=Homo sapiens OX=9606 ( | CCNG1 | 34.074 | 3 | 55.706585 | yes | 3.588E-06 | 575.29346 | 30.02445 |
| DNWSPALTISK | 11 | P51965 | Ubiquitin-conjugating enzyme E2 E1 OS= | UBE2E1 | 21.405 | 2 | 41.540188 | yes | 3.458E-07 | 616.31952 | 54.196674 |
| CALDFFR | 7 | P51970 | NADH dehydrogenase [ubiquinone] 1 alp | NDUFA8 | 20.106 | 2 | 47.186806 | yes | 1.381E-07 | 464.72092 | 54.552246 |
| PGIVELPTLEELK | 13 | P51970 | NADH dehydrogenase [ubiquinone] 1 alp | NDUFA8 | 20.106 | 2 | 47.186806 | yes | 3.507E-06 | 719.41357 | 36.695847 |
| PRPDPSPEIEGDLQ | 21 | P51970 | NADH dehydrogenase [ubiquinone] 1 alp | NDUFA8 | 20.106 | 3 | 47.186806 | yes | 3.512E-06 | 752.70496 | 18.656017 |
| LGWVRPDLGELSK | 13 | P51970 | NADH dehydrogenase [ubiquinone] 1 alp | NDUFA8 | 20.106 | 2 | 47.186806 | yes | 3.179E-06 | 735.40918 | 3.8863034 |
| WGTLTDCVVMR | 11 | P51991 | Heterogeneous nuclear ribonucleoprotein | HNRNPA3 | 39.595 | 2 | 52.220684 | yes | 1.197E-07 | 669.32068 | 13236.661 |
| WGTLTDCVVMR | 11 | P51991 | Heterogeneous nuclear ribonucleoprotein | HNRNPA3 | 39.595 | 2 | 52.220684 | yes | 1.002E-07 | 677.31812 | 13236.661 |
| IETIEVMEDR | 10 | P51991 | Heterogeneous nuclear ribonucleoprotein | HNRNPA3 | 39.595 | 2 | 52.220684 | yes | 1.21E-07 | 617.80286 | 12612.345 |
| IETIEVMEDR | 10 | P51991 | Heterogeneous nuclear ribonucleoprotein | HNRNPA3 | 39.595 | 2 | 52.220684 | yes | 1.503E-07 | 625.80029 | 12612.345 |
| EDTEEYNLR | 9 | P51991 | Heterogeneous nuclear ribonucleoprotein | HNRNPA3 | 39.595 | 2 | 52.220684 | yes | 1.083E-07 | 584.75946 | 18659.031 |
| MEVKPPPGRPQPD | 16 | P51991 | Heterogeneous nuclear ribonucleoprotein | HNRNPA3 | 39.595 | 3 | 52.220684 | yes | 2.381E-06 | 597.3053 | 13754.614 |
| EDSVKPGAHLTVK | 13 | P51991 | Heterogeneous nuclear ribonucleoprotein | HNRNPA3 | 39.595 | 2 | 52.220684 | yes | 1.693E-07 | 690.87769 | 2433.9663 |
| EDSVKPGAHLTVK | 13 | P51991 | Heterogeneous nuclear ribonucleoprotein | HNRNPA3 | 39.595 | 2 | 52.220684 | yes | 1.074E-07 | 460.9209 | 2433.9663 |
| EDSVKPGAHLTVK | 13 | P51991 | Heterogeneous nuclear ribonucleoprotein | HNRNPA3 | 39.595 | 3 | 52.220684 | yes | 1.693E-07 | 690.87769 | 2433.9663 |
| EDSVKPGAHLTVK | 13 | P51991 | Heterogeneous nuclear ribonucleoprotein | HNRNPA3 | 39.595 | 3 | 52.220684 | yes | 1.074E-07 | 460.9209 | 2433.9663 |
| MEVKPPPGR | 9 | P51991 | Heterogeneous nuclear ribonucleoprotein | HNRNPA3 | 39.595 | 2 | 52.220684 | yes | 1.542E-07 | 526.78149 | 4570.668 |
| GFGFVTYSCVEEV | 20 | P51991 | Heterogeneous nuclear ribonucleoprotein | HNRNPA3 | 39.595 | 3 | 52.220684 | yes | 1.011E-07 | 756.99561 | 1334.9977 |
| QEMQSAGSQR | 10 | P51991 | Heterogeneous nuclear ribonucleoprotein | HNRNPA3 | 39.595 | 2 | 52.220684 | yes | 9.722E-08 | 561.25385 | 1304.5898 |
| YHTINGHNCEVK | 12 | P51991 | Heterogeneous nuclear ribonucleoprotein | HNRNPA3 | 39.595 | 2 | 52.220684 | yes | 9.757E-08 | 736.341 | 583.88196 |
| YHTINGHNCEVK | 12 | P51991 | Heterogeneous nuclear ribonucleoprotein | HNRNPA3 | 39.595 | 2 | 52.220684 | yes | 2.114E-07 | 491.22974 | 583.88196 |
| YHTINGHNCEVK | 12 | P51991 | Heterogeneous nuclear ribonucleoprotein | HNRNPA3 | 39.595 | 3 | 52.220684 | yes | 9.757E-08 | 736.341 | 583.88196 |
| YHTINGHNCEVK | 12 | P51991 | Heterogeneous nuclear ribonucleoprotein | HNRNPA3 | 39.595 | 3 | 52.220684 | yes | 2.114E-07 | 491.22974 | 583.88196 |
| RGEEGHDPKEPEQ | 15 | P51991 | Heterogeneous nuclear ribonucleoprotein | HNRNPA3 | 39.595 | 4 | 52.220684 | yes | 2.495E-07 | 444.97104 | 1065.3729 |
| RGFAFVTFDDHDT | 16 | P51991 | Heterogeneous nuclear ribonucleoprotein | HNRNPA3 | 39.595 | 3 | 52.220684 | yes | 1.003E-07 | 623.96375 | 975.41083 |
| GEEGHDPKEPEQL | 14 | P51991 | Heterogeneous nuclear ribonucleoprotein | HNRNPA3 | 39.595 | 3 | 52.220684 | yes | 2.296E-07 | 540.92529 | 366.33017 |
| GEEGHDPKEPEQL | 14 | P51991 | Heterogeneous nuclear ribonucleoprotein | HNRNPA3 | 39.595 | 3 | 52.220684 | yes | 3.011E-07 | 810.88428 | 366.33017 |

| | | | | | | | | | | |
|---|---|---|---|---|---|---|---|---|---|---|
| GEEGHDPKEPEQL | 14 | P51991 | Heterogeneous nuclear ribonucleoprotein | HNRNPA3 | 39.595 | 2 | 52.220684 | yes | 2.296E-07 | 540.92529 | 366.33017 |
| GEEGHDPKEPEQL | 14 | P51991 | Heterogeneous nuclear ribonucleoprotein | HNRNPA3 | 39.595 | 2 | 52.220684 | yes | 3.011E-07 | 810.88428 | 366.33017 |
| GGNFGGGGGNFGI | 13 | P51991 | Heterogeneous nuclear ribonucleoprotein | HNRNPA3 | 39.595 | 2 | 52.220684 | yes | 1.181E-07 | 577.26031 | 207.41685 |
| KLFIGGLSFETTDD | 17 | P51991 | Heterogeneous nuclear ribonucleoprotein | HNRNPA3 | 39.595 | 3 | 52.220684 | yes | 1.208E-06 | 633.66681 | 27.492321 |
| MVPAGMGAGLER | 12 | P52272 | Heterogeneous nuclear ribonucleoprotein | HNRNPM | 77.516 | 2 | 52.973534 | yes | 1.262E-07 | 594.79681 | 9634.3359 |
| MVPAGMGAGLER | 12 | P52272 | Heterogeneous nuclear ribonucleoprotein | HNRNPM | 77.516 | 2 | 52.973534 | yes | 1.921E-07 | 602.79431 | 9634.3359 |
| MGAGMGFGLER | 11 | P52272 | Heterogeneous nuclear ribonucleoprotein | HNRNPM | 77.516 | 2 | 52.973534 | yes | 1.065E-07 | 563.26263 | 6394.6875 |
| MGAGMGFGLER | 11 | P52272 | Heterogeneous nuclear ribonucleoprotein | HNRNPM | 77.516 | 2 | 52.973534 | yes | 2.921E-07 | 571.26007 | 6394.6875 |
| MGAGMGFGLER | 11 | P52272 | Heterogeneous nuclear ribonucleoprotein | HNRNPM | 77.516 | 2 | 52.973534 | yes | 1.001E-07 | 571.26007 | 6394.6875 |
| AFITNIPFDVK | 11 | P52272 | Heterogeneous nuclear ribonucleoprotein | HNRNPM | 77.516 | 2 | 52.973534 | yes | 9.741E-08 | 632.85046 | 19066.92 |
| MGPAMGPALGAG | 15 | P52272 | Heterogeneous nuclear ribonucleoprotein | HNRNPM | 77.516 | 2 | 52.973534 | yes | 4.963E-07 | 714.36035 | 5918.875 |
| MGPAMGPALGAG | 15 | P52272 | Heterogeneous nuclear ribonucleoprotein | HNRNPM | 77.516 | 2 | 52.973534 | yes | 9.975E-08 | 722.35779 | 5918.875 |
| MGPAMGPALGAG | 15 | P52272 | Heterogeneous nuclear ribonucleoprotein | HNRNPM | 77.516 | 2 | 52.973534 | yes | 2.297E-07 | 722.35779 | 5918.875 |
| MGLAMGGGGGAS | 15 | P52272 | Heterogeneous nuclear ribonucleoprotein | HNRNPM | 77.516 | 2 | 52.973534 | yes | 2.213E-06 | 692.31085 | 3536.4141 |
| MGLAMGGGGGAS | 15 | P52272 | Heterogeneous nuclear ribonucleoprotein | HNRNPM | 77.516 | 2 | 52.973534 | yes | 2.245E-07 | 700.30829 | 3536.4141 |
| MGLAMGGGGGAS | 15 | P52272 | Heterogeneous nuclear ribonucleoprotein | HNRNPM | 77.516 | 2 | 52.973534 | yes | 1.019E-07 | 700.30829 | 3536.4141 |
| FESPEVAER | 9 | P52272 | Heterogeneous nuclear ribonucleoprotein | HNRNPM | 77.516 | 2 | 52.973534 | yes | 1.621E-07 | 532.25635 | 9613.0234 |
| MGANNLER | 8 | P52272 | Heterogeneous nuclear ribonucleoprotein | HNRNPM | 77.516 | 2 | 52.973534 | yes | 1.206E-07 | 452.7189 | 4757.4292 |
| MGANNLER | 8 | P52272 | Heterogeneous nuclear ribonucleoprotein | HNRNPM | 77.516 | 2 | 52.973534 | yes | 1.466E-07 | 460.71637 | 4757.4292 |
| MGAGLGHGMDR | 11 | P52272 | Heterogeneous nuclear ribonucleoprotein | HNRNPM | 77.516 | 2 | 52.973534 | yes | 1.267E-07 | 551.25006 | 2747.9827 |
| MGAGLGHGMDR | 11 | P52272 | Heterogeneous nuclear ribonucleoprotein | HNRNPM | 77.516 | 2 | 52.973534 | yes | 9.722E-08 | 559.2475 | 2747.9827 |
| MGAGLGHGMDR | 11 | P52272 | Heterogeneous nuclear ribonucleoprotein | HNRNPM | 77.516 | 2 | 52.973534 | yes | 1.086E-07 | 559.2475 | 2747.9827 |
| EVFSMAGVVVR | 11 | P52272 | Heterogeneous nuclear ribonucleoprotein | HNRNPM | 77.516 | 2 | 52.973534 | yes | 1.141E-07 | 597.32098 | 3117.1443 |
| EVFSMAGVVVR | 11 | P52272 | Heterogeneous nuclear ribonucleoprotein | HNRNPM | 77.516 | 2 | 52.973534 | yes | 1.177E-07 | 605.31842 | 3117.1443 |
| KACQIFVR | 8 | P52272 | Heterogeneous nuclear ribonucleoprotein | HNRNPM | 77.516 | 2 | 52.973534 | yes | 2.615E-07 | 511.28421 | 6212.5103 |
| ADILEDKDGK | 10 | P52272 | Heterogeneous nuclear ribonucleoprotein | HNRNPM | 77.516 | 2 | 52.973534 | yes | 1.128E-07 | 552.28259 | 5918.3188 |
| ACQIFVR | 7 | P52272 | Heterogeneous nuclear ribonucleoprotein | HNRNPM | 77.516 | 2 | 52.973534 | yes | 2.776E-07 | 447.23672 | 5733.936 |
| FEPYANPTK | 9 | P52272 | Heterogeneous nuclear ribonucleoprotein | HNRNPM | 77.516 | 2 | 52.973534 | yes | 3.435E-07 | 533.76385 | 4319.686 |
| LGSTVFVANLDYK | 13 | P52272 | Heterogeneous nuclear ribonucleoprotein | HNRNPM | 77.516 | 2 | 52.973534 | yes | 1.133E-07 | 713.88245 | 4100.2808 |
| INEILSNALK | 10 | P52272 | Heterogeneous nuclear ribonucleoprotein | HNRNPM | 77.516 | 2 | 52.973534 | yes | 1.038E-07 | 557.82697 | 3673.8218 |
| MGPLGLDHMASSI | 15 | P52272 | Heterogeneous nuclear ribonucleoprotein | HNRNPM | 77.516 | 2 | 52.973534 | yes | 2.745E-07 | 807.39233 | 1003.6302 |
| MGPLGLDHMASSI | 15 | P52272 | Heterogeneous nuclear ribonucleoprotein | HNRNPM | 77.516 | 2 | 52.973534 | yes | 3.49E-07 | 538.59735 | 1003.6302 |
| MGPLGLDHMASSI | 15 | P52272 | Heterogeneous nuclear ribonucleoprotein | HNRNPM | 77.516 | 2 | 52.973534 | yes | 1.863E-07 | 815.38983 | 1003.6302 |
| MGPLGLDHMASSI | 15 | P52272 | Heterogeneous nuclear ribonucleoprotein | HNRNPM | 77.516 | 3 | 52.973534 | yes | 2.745E-07 | 807.39233 | 1003.6302 |
| MGPLGLDHMASSI | 15 | P52272 | Heterogeneous nuclear ribonucleoprotein | HNRNPM | 77.516 | 3 | 52.973534 | yes | 3.49E-07 | 538.59735 | 1003.6302 |
| MGPLGLDHMASSI | 15 | P52272 | Heterogeneous nuclear ribonucleoprotein | HNRNPM | 77.516 | 3 | 52.973534 | yes | 1.863E-07 | 815.38983 | 1003.6302 |
| QGGGGGGGSVPGI | 15 | P52272 | Heterogeneous nuclear ribonucleoprotein | HNRNPM | 77.516 | 2 | 52.973534 | yes | 4.873E-07 | 642.81818 | 2005.4661 |
| DKFNECGHVLYAI | 15 | P52272 | Heterogeneous nuclear ribonucleoprotein | HNRNPM | 77.516 | 3 | 52.973534 | yes | 1.075E-07 | 603.62604 | 1593.3972 |
| NLPFDFTWK | 9 | P52272 | Heterogeneous nuclear ribonucleoprotein | HNRNPM | 77.516 | 2 | 52.973534 | yes | 1.606E-07 | 584.29529 | 1591.3677 |

| | | | | | | | | | | | |
|---|---|---|---|---|---|---|---|---|---|---|---|
| MGLVMDR | 7 | P52272 | Heterogeneous nuclear ribonucleoprotein | HNRNPM | 77.516 | 2 | 52.973534 | yes | 1.17E-07 | 411.20404 | 765.94458 |
| MGLVMDR | 7 | P52272 | Heterogeneous nuclear ribonucleoprotein | HNRNPM | 77.516 | 2 | 52.973534 | yes | 1.561E-07 | 419.20151 | 765.94458 |
| MGANSLER | 8 | P52272 | Heterogeneous nuclear ribonucleoprotein | HNRNPM | 77.516 | 2 | 52.973534 | yes | 1.192E-07 | 439.21344 | 1254.8624 |
| GNFGGSFAGSFGG | 23 | P52272 | Heterogeneous nuclear ribonucleoprotein | HNRNPM | 77.516 | 3 | 52.973534 | yes | 1.662E-07 | 678.98914 | 1133.6462 |
| MGQTMER | 7 | P52272 | Heterogeneous nuclear ribonucleoprotein | HNRNPM | 77.516 | 2 | 52.973534 | yes | 2.474E-07 | 426.68875 | 435.86972 |
| MGQTMER | 7 | P52272 | Heterogeneous nuclear ribonucleoprotein | HNRNPM | 77.516 | 2 | 52.973534 | yes | 2.673E-06 | 434.68622 | 435.86972 |
| FNECGHVLYADIK | 13 | P52272 | Heterogeneous nuclear ribonucleoprotein | HNRNPM | 77.516 | 2 | 52.973534 | yes | 1.069E-07 | 783.37451 | 415.89075 |
| FNECGHVLYADIK | 13 | P52272 | Heterogeneous nuclear ribonucleoprotein | HNRNPM | 77.516 | 2 | 52.973534 | yes | 1.481E-07 | 522.58539 | 415.89075 |
| FNECGHVLYADIK | 13 | P52272 | Heterogeneous nuclear ribonucleoprotein | HNRNPM | 77.516 | 3 | 52.973534 | yes | 1.069E-07 | 783.37451 | 415.89075 |
| FNECGHVLYADIK | 13 | P52272 | Heterogeneous nuclear ribonucleoprotein | HNRNPM | 77.516 | 3 | 52.973534 | yes | 1.481E-07 | 522.58539 | 415.89075 |
| MGLSMER | 7 | P52272 | Heterogeneous nuclear ribonucleoprotein | HNRNPM | 77.516 | 2 | 52.973534 | yes | 2.1E-07 | 412.19366 | 719.42487 |
| GDFFPPERPQQLPI | 37 | P52272 | Heterogeneous nuclear ribonucleoprotein | HNRNPM | 77.516 | 5 | 52.973534 | yes | 1.277E-07 | 764.58948 | 286.29858 |
| GIGMGNIGPAGMC | 23 | P52272 | Heterogeneous nuclear ribonucleoprotein | HNRNPM | 77.516 | 3 | 52.973534 | yes | 1.473E-07 | 726.68835 | 269.90689 |
| AAGVEAAAEVAA | 16 | P52272 | Heterogeneous nuclear ribonucleoprotein | HNRNPM | 77.516 | 2 | 52.973534 | yes | 1.942E-07 | 771.90411 | 243.34482 |
| GCAVVEFK | 8 | P52272 | Heterogeneous nuclear ribonucleoprotein | HNRNPM | 77.516 | 2 | 52.973534 | yes | 4.759E-07 | 455.22858 | 216.9407 |
| HSLSGRPLK | 9 | P52272 | Heterogeneous nuclear ribonucleoprotein | HNRNPM | 77.516 | 2 | 52.973534 | yes | 1.097E-07 | 497.79324 | 155.12584 |
| GGNRFEPYANPTK | 13 | P52272 | Heterogeneous nuclear ribonucleoprotein | HNRNPM | 77.516 | 2 | 52.973534 | yes | 5.067E-07 | 725.8573 | 68.779243 |
| GGNRFEPYANPTK | 13 | P52272 | Heterogeneous nuclear ribonucleoprotein | HNRNPM | 77.516 | 2 | 52.973534 | yes | 1.182E-06 | 484.24063 | 68.779243 |
| GGNRFEPYANPTK | 13 | P52272 | Heterogeneous nuclear ribonucleoprotein | HNRNPM | 77.516 | 3 | 52.973534 | yes | 5.067E-07 | 725.8573 | 68.779243 |
| GGNRFEPYANPTK | 13 | P52272 | Heterogeneous nuclear ribonucleoprotein | HNRNPM | 77.516 | 3 | 52.973534 | yes | 1.182E-06 | 484.24063 | 68.779243 |
| PQQLPHGLGGIGM | 29 | P52272 | Heterogeneous nuclear ribonucleoprotein | HNRNPM | 77.516 | 4 | 52.973534 | yes | 1.866E-07 | 719.1264 | 77.336349 |
| FGSGMNMGR | 9 | P52272 | Heterogeneous nuclear ribonucleoprotein | HNRNPM | 77.516 | 2 | 52.973534 | yes | 1.477E-06 | 478.70749 | 32.358917 |
| LLGASELPIVTPAL | 15 | P52292 | Importin subunit alpha-1 OS=Homo sapie | KPNA2 | 57.862 | 2 | 53.375225 | yes | 1.716E-07 | 775.46924 | 5723.8789 |
| NNQGTVNWSVDD | 15 | P52292 | Importin subunit alpha-1 OS=Homo sapie | KPNA2 | 57.862 | 2 | 53.375225 | yes | 1.177E-07 | 844.91559 | 3004.9985 |
| EKQPPIDNIIR | 11 | P52292 | Importin subunit alpha-1 OS=Homo sapie | KPNA2 | 57.862 | 2 | 53.375225 | yes | 1.362E-07 | 661.875 | 2966.6365 |
| EATWTMSNITAGR | 13 | P52292 | Importin subunit alpha-1 OS=Homo sapie | KPNA2 | 57.862 | 2 | 53.375225 | yes | 9.81E-08 | 719.3432 | 1421.6433 |
| EATWTMSNITAGR | 13 | P52292 | Importin subunit alpha-1 OS=Homo sapie | KPNA2 | 57.862 | 2 | 53.375225 | yes | 1.733E-07 | 727.34064 | 1421.6433 |
| LSIMIEECGGLDK | 13 | P52292 | Importin subunit alpha-1 OS=Homo sapie | KPNA2 | 57.862 | 2 | 53.375225 | yes | 9.722E-08 | 732.8573 | 658.53003 |
| LSIMIEECGGLDK | 13 | P52292 | Importin subunit alpha-1 OS=Homo sapie | KPNA2 | 57.862 | 2 | 53.375225 | yes | 9.722E-08 | 740.85474 | 658.53003 |
| IEVNVELR | 8 | P52292 | Importin subunit alpha-1 OS=Homo sapie | KPNA2 | 57.862 | 2 | 53.375225 | yes | 5.14E-07 | 486.27966 | 1213.4708 |
| STNENANTPAAR | 12 | P52292 | Importin subunit alpha-1 OS=Homo sapie | KPNA2 | 57.862 | 2 | 53.375225 | yes | 9.722E-08 | 644.2998 | 1185.558 |
| QPPIDNIIR | 9 | P52292 | Importin subunit alpha-1 OS=Homo sapie | KPNA2 | 57.862 | 2 | 53.375225 | yes | 1.001E-07 | 533.30621 | 875.44885 |
| KDDQMLK | 7 | P52292 | Importin subunit alpha-1 OS=Homo sapie | KPNA2 | 57.862 | 2 | 53.375225 | yes | 5.994E-07 | 439.22601 | 332.83774 |
| RIEVNVELR | 9 | P52292 | Importin subunit alpha-1 OS=Homo sapie | KPNA2 | 57.862 | 2 | 53.375225 | yes | 6.315E-07 | 564.3302 | 287.86484 |
| IEALQNHENESVY | 14 | P52292 | Importin subunit alpha-1 OS=Homo sapie | KPNA2 | 57.862 | 2 | 53.375225 | yes | 7.87E-07 | 837.4101 | 224.35962 |
| TGVVPQLVK | 9 | P52292 | Importin subunit alpha-1 OS=Homo sapie | KPNA2 | 57.862 | 2 | 53.375225 | yes | 1.621E-07 | 470.79492 | 213.60828 |
| FVSFLGR | 7 | P52292 | Importin subunit alpha-1 OS=Homo sapie | KPNA2 | 57.862 | 2 | 53.375225 | yes | 2.861E-07 | 413.23453 | 191.59637 |
| ASLSLIEK | 8 | P52292 | Importin subunit alpha-1 OS=Homo sapie | KPNA2 | 57.862 | 2 | 53.375225 | yes | 9.728E-08 | 430.75803 | 164.19624 |
| LSIMIEECGGLDKI | 27 | P52292 | Importin subunit alpha-1 OS=Homo sapie | KPNA2 | 57.862 | 4 | 53.375225 | yes | 1.745E-07 | 780.63104 | 38.030918 |

| | | | | | | | | | | | |
|---|---|---|---|---|---|---|---|---|---|---|---|
| IQAVIDAGVCR | 11 | P52294 | Importin subunit alpha-5 OS=Homo sapie | KPNA1 | 60.222 | 2 | 53.97813 | yes | 1.683E-07 | 601.32153 | 495.93744 |
| SPEQQLSATQK | 11 | P52294 | Importin subunit alpha-5 OS=Homo sapie | KPNA1 | 60.222 | 2 | 53.97813 | yes | 9.722E-08 | 608.81201 | 361.14594 |
| SDSYVELSQYR | 11 | P52298 | Nuclear cap-binding protein subunit 2 OS | NCBP2 | 18.002 | 2 | 50.724003 | yes | 9.727E-08 | 673.81476 | 8364.3506 |
| TACGFCFVEYYSR | 13 | P52298 | Nuclear cap-binding protein subunit 2 OS | NCBP2 | 18.002 | 2 | 50.724003 | yes | 1.865E-06 | 830.35016 | 5314.1812 |
| KIIMGLDK | 8 | P52298 | Nuclear cap-binding protein subunit 2 OS | NCBP2 | 18.002 | 2 | 50.724003 | yes | 1.445E-07 | 459.27808 | 1240.7545 |
| ADAENAMR | 8 | P52298 | Nuclear cap-binding protein subunit 2 OS | NCBP2 | 18.002 | 2 | 50.724003 | no | 1.727E-07 | 439.19525 | 1040.2667 |
| LSAYVSYGGLLMF | 13 | P52434 | DNA-directed RNA polymerases I, II, and | POLR2H | 17.144 | 2 | 52.645325 | yes | 9.722E-08 | 715.37903 | 125.0703 |
| ADQFEYVMYGK | 11 | P52434 | DNA-directed RNA polymerases I, II, and | POLR2H | 17.144 | 2 | 52.645325 | yes | 9.722E-08 | 675.80536 | 117.60985 |
| LHCESESFK | 9 | P52434 | DNA-directed RNA polymerases I, II, and | POLR2H | 17.144 | 2 | 52.645325 | yes | 3.153E-06 | 568.75568 | 115.2914 |
| INPELNQK | 8 | P52564 | Dual specificity mitogen-activated proteir | MAP2K6 | 37.493 | 2 | 49.936245 | no | 6.122E-07 | 478.26401 | 163.25117 |
| ADDLEPIMELGR | 12 | P52564 | Dual specificity mitogen-activated proteir | MAP2K6 | 37.493 | 2 | 49.936245 | yes | 9.975E-08 | 679.83466 | 68.908714 |
| ACISIGNQNFEVK | 13 | P52564 | Dual specificity mitogen-activated proteir | MAP2K6 | 37.493 | 2 | 49.936245 | yes | 9.915E-07 | 740.36664 | 54.139713 |
| TDYMVGSYGPR | 11 | P52565 | Rho GDP-dissociation inhibitor 1 OS=Hc | ARHGDIA | 23.208 | 2 | 45.36389 | yes | 1.873E-07 | 623.28204 | 61.651733 |
| TEMDWVLK | 8 | P52597 | Heterogeneous nuclear ribonucleoprotein | HNRNPF | 45.672 | 2 | 50.64138 | yes | 1.426E-07 | 511.25479 | 17991.727 |
| TEMDWVLK | 8 | P52597 | Heterogeneous nuclear ribonucleoprotein | HNRNPF | 45.672 | 2 | 50.64138 | yes | 2.081E-07 | 519.25226 | 17991.727 |
| SYSDPPLK | 8 | P52597 | Heterogeneous nuclear ribonucleoprotein | HNRNPF | 45.672 | 2 | 50.64138 | yes | 1.776E-07 | 453.73199 | 24579.07 |
| MLGPEGGEGFVVF | 13 | P52597 | Heterogeneous nuclear ribonucleoprotein | HNRNPF | 45.672 | 2 | 50.64138 | yes | 1.515E-07 | 681.3421 | 6201.9717 |
| MLGPEGGEGFVVF | 13 | P52597 | Heterogeneous nuclear ribonucleoprotein | HNRNPF | 45.672 | 2 | 50.64138 | yes | 1.102E-07 | 689.3396 | 6201.9717 |
| MLGPEGGEGFVVF | 13 | P52597 | Heterogeneous nuclear ribonucleoprotein | HNRNPF | 45.672 | 2 | 50.64138 | yes | 2.716E-07 | 660.33685 | 6201.9717 |
| VTGEADVEFATHE | 21 | P52597 | Heterogeneous nuclear ribonucleoprotein | HNRNPF | 45.672 | 3 | 50.64138 | yes | 2.45E-07 | 731.34406 | 16791.826 |
| RYIGIVK | 7 | P52597 | Heterogeneous nuclear ribonucleoprotein | HNRNPF | 45.672 | 2 | 50.64138 | yes | 2.868E-07 | 424.77127 | 14086.897 |
| DLSYCLSGMYDHI | 13 | P52597 | Heterogeneous nuclear ribonucleoprotein | HNRNPF | 45.672 | 2 | 50.64138 | yes | 9.919E-08 | 808.84521 | 6810.9907 |
| DLSYCLSGMYDHI | 13 | P52597 | Heterogeneous nuclear ribonucleoprotein | HNRNPF | 45.672 | 2 | 50.64138 | yes | 9.733E-08 | 816.84271 | 6810.9907 |
| GLPFGCTK | 8 | P52597 | Heterogeneous nuclear ribonucleoprotein | HNRNPF | 45.672 | 2 | 50.64138 | yes | 2.722E-07 | 440.2233 | 8496.3438 |
| HSGPNSADSANDC | 16 | P52597 | Heterogeneous nuclear ribonucleoprotein | HNRNPF | 45.672 | 3 | 50.64138 | yes | 1.348E-07 | 544.245 | 3365.3982 |
| HSGPNSADSANDC | 16 | P52597 | Heterogeneous nuclear ribonucleoprotein | HNRNPF | 45.672 | 3 | 50.64138 | yes | 1.025E-07 | 815.86383 | 3365.3982 |
| HSGPNSADSANDC | 16 | P52597 | Heterogeneous nuclear ribonucleoprotein | HNRNPF | 45.672 | 2 | 50.64138 | yes | 1.348E-07 | 544.245 | 3365.3982 |
| HSGPNSADSANDC | 16 | P52597 | Heterogeneous nuclear ribonucleoprotein | HNRNPF | 45.672 | 2 | 50.64138 | yes | 1.025E-07 | 815.86383 | 3365.3982 |
| PGPYDRPGTAR | 11 | P52597 | Heterogeneous nuclear ribonucleoprotein | HNRNPF | 45.672 | 2 | 50.64138 | yes | 3.047E-07 | 593.80182 | 1598.3308 |
| SSQEEVR | 7 | P52597 | Heterogeneous nuclear ribonucleoprotein | HNRNPF | 45.672 | 2 | 50.64138 | yes | 1.211E-07 | 417.70123 | 246.51276 |
| FMSVQRPGPYDRF | 17 | P52597 | Heterogeneous nuclear ribonucleoprotein | HNRNPF | 45.672 | 4 | 50.64138 | yes | 2.537E-07 | 484.4968 | 126.27059 |
| LANLPEEVIQK | 11 | P52701 | DNA mismatch repair protein Msh6 OS=] | MSH6 | 152.786 | 2 | 56.638954 | yes | 2.258E-07 | 627.35864 | 497.89008 |
| AYCVLVTGPNMG | 14 | P52701 | DNA mismatch repair protein Msh6 OS=] | MSH6 | 152.786 | 2 | 56.638954 | yes | 2.101E-07 | 733.86017 | 318.99832 |
| FFIGQFSDDR | 10 | P52701 | DNA mismatch repair protein Msh6 OS=] | MSH6 | 152.786 | 2 | 56.638954 | yes | 9.722E-08 | 616.29077 | 285.86649 |
| STVDAEAVHK | 10 | P52701 | DNA mismatch repair protein Msh6 OS=] | MSH6 | 152.786 | 2 | 56.638954 | yes | 3.092E-07 | 528.76965 | 200.39026 |
| FPDLTVELNR | 10 | P52701 | DNA mismatch repair protein Msh6 OS=] | MSH6 | 152.786 | 2 | 56.638954 | yes | 2.533E-07 | 602.32202 | 198.97583 |
| IIGIMEEVADGFK | 13 | P52701 | DNA mismatch repair protein Msh6 OS=] | MSH6 | 152.786 | 2 | 56.638954 | yes | 1.265E-07 | 711.37085 | 179.4093 |
| VEQTETPEMMEAI | 13 | P52701 | DNA mismatch repair protein Msh6 OS=] | MSH6 | 152.786 | 2 | 56.638954 | yes | 1.547E-07 | 775.84491 | 174.76324 |
| GMTSESDSIGLTPC | 16 | P52701 | DNA mismatch repair protein Msh6 OS=] | MSH6 | 152.786 | 2 | 56.638954 | yes | 1.197E-07 | 804.87469 | 174.29437 |

| | | | | | | | | | | | |
|---|---|---|---|---|---|---|---|---|---|---|---|
| TIVYWGIGR | 9 | P52701 | DNA mismatch repair protein Msh6 OS=] | MSH6 | 152.786 | 2 | 56.638954 | yes | 4.339E-07 | 532.79803 | 169.55476 |
| WDTAFDHEK | 9 | P52701 | DNA mismatch repair protein Msh6 OS=] | MSH6 | 152.786 | 2 | 56.638954 | yes | 1.311E-07 | 574.75397 | 166.89745 |
| YQLEIPENFTTR | 12 | P52701 | DNA mismatch repair protein Msh6 OS=] | MSH6 | 152.786 | 2 | 56.638954 | yes | 9.722E-08 | 755.88043 | 153.77357 |
| SQNFDLVICYK | 11 | P52701 | DNA mismatch repair protein Msh6 OS=] | MSH6 | 152.786 | 2 | 56.638954 | yes | 1.166E-07 | 693.83972 | 135.40523 |
| GGHFYSAKPEILR | 13 | P52701 | DNA mismatch repair protein Msh6 OS=] | MSH6 | 152.786 | 2 | 56.638954 | yes | 6.598E-07 | 737.89368 | 66.103569 |
| GGHFYSAKPEILR | 13 | P52701 | DNA mismatch repair protein Msh6 OS=] | MSH6 | 152.786 | 2 | 56.638954 | yes | 9.722E-08 | 492.26489 | 66.103569 |
| GGHFYSAKPEILR | 13 | P52701 | DNA mismatch repair protein Msh6 OS=] | MSH6 | 152.786 | 3 | 56.638954 | yes | 6.598E-07 | 737.89368 | 66.103569 |
| GGHFYSAKPEILR | 13 | P52701 | DNA mismatch repair protein Msh6 OS=] | MSH6 | 152.786 | 3 | 56.638954 | yes | 9.722E-08 | 492.26489 | 66.103569 |
| AGFDSDYDQALAI | 15 | P52701 | DNA mismatch repair protein Msh6 OS=] | MSH6 | 152.786 | 2 | 56.638954 | yes | 1.047E-07 | 828.87866 | 128.96022 |
| AYGVCFVDTSLGK | 13 | P52701 | DNA mismatch repair protein Msh6 OS=] | MSH6 | 152.786 | 2 | 56.638954 | yes | 1.138E-07 | 708.84503 | 119.15858 |
| VHVQFFDDSPTR | 12 | P52701 | DNA mismatch repair protein Msh6 OS=] | MSH6 | 152.786 | 2 | 56.638954 | yes | 1.281E-07 | 724.35187 | 117.63552 |
| LSDGIGVMLPQVL | 14 | P52701 | DNA mismatch repair protein Msh6 OS=] | MSH6 | 152.786 | 2 | 56.638954 | yes | 1.496E-07 | 735.42346 | 110.06689 |
| TLLEEEYFR | 9 | P52701 | DNA mismatch repair protein Msh6 OS=] | MSH6 | 152.786 | 2 | 56.638954 | yes | 1.025E-07 | 600.30078 | 105.59808 |
| ENEQSLLEYLEK | 12 | P52701 | DNA mismatch repair protein Msh6 OS=] | MSH6 | 152.786 | 2 | 56.638954 | yes | 9.722E-08 | 747.86975 | 105.57834 |
| ISEVVELLK | 9 | P52701 | DNA mismatch repair protein Msh6 OS=] | MSH6 | 152.786 | 2 | 56.638954 | yes | 2.192E-07 | 515.31317 | 103.66821 |
| LDAIEDLMVVPDK | 13 | P52701 | DNA mismatch repair protein Msh6 OS=] | MSH6 | 152.786 | 2 | 56.638954 | yes | 1.166E-07 | 729.38147 | 87.882484 |
| GTATFDGTAIANA | 16 | P52701 | DNA mismatch repair protein Msh6 OS=] | MSH6 | 152.786 | 2 | 56.638954 | yes | 6.978E-07 | 768.40686 | 87.835037 |
| QSTLYSFFPK | 10 | P52701 | DNA mismatch repair protein Msh6 OS=] | MSH6 | 152.786 | 2 | 56.638954 | yes | 1.915E-07 | 609.31366 | 82.09436 |
| AIMYEETTYSK | 11 | P52701 | DNA mismatch repair protein Msh6 OS=] | MSH6 | 152.786 | 2 | 56.638954 | yes | 3.607E-06 | 668.31049 | 77.698921 |
| KTGLITPK | 8 | P52701 | DNA mismatch repair protein Msh6 OS=] | MSH6 | 152.786 | 2 | 56.638954 | yes | 2.92E-07 | 429.27637 | 54.181747 |
| VDTCHTPFGK | 10 | P52701 | DNA mismatch repair protein Msh6 OS=] | MSH6 | 152.786 | 2 | 56.638954 | yes | 8.349E-07 | 581.27148 | 24.215504 |
| ISQETEQR | 8 | P52732 | Kinesin-like protein KIF11 OS=Homo sa | KIF11 | 119.159 | 2 | 48.937492 | yes | 3.404E-07 | 495.74619 | 75.14473 |
| EEYITSALESTEEK | 14 | P52732 | Kinesin-like protein KIF11 OS=Homo sa | KIF11 | 119.159 | 2 | 48.937492 | yes | 1.084E-07 | 814.88049 | 53.43681 |
| SYLYPSTLVR | 10 | P52732 | Kinesin-like protein KIF11 OS=Homo sa | KIF11 | 119.159 | 2 | 48.937492 | yes | 2.734E-06 | 599.82697 | 46.531132 |
| EAGNINQSLLTLGI | 14 | P52732 | Kinesin-like protein KIF11 OS=Homo sa | KIF11 | 119.159 | 2 | 48.937492 | yes | 5.254E-07 | 743.4046 | 43.339485 |
| GLPITITESDIR | 12 | P52756 | RNA-binding protein 5 OS=Homo sapien | RBM5 | 92.154 | 2 | 52.991856 | yes | 1.217E-07 | 657.86682 | 1437.4591 |
| NSFQPVNSLR | 10 | P52756 | RNA-binding protein 5 OS=Homo sapien | RBM5 | 92.154 | 2 | 52.991856 | yes | 2.272E-07 | 581.3042 | 1106.5754 |
| LSEQELEALELR | 12 | P52756 | RNA-binding protein 5 OS=Homo sapien | RBM5 | 92.154 | 2 | 52.991856 | yes | 1.387E-07 | 715.38031 | 851.92926 |
| QQLIPELVR | 9 | P52756 | RNA-binding protein 5 OS=Homo sapien | RBM5 | 92.154 | 2 | 52.991856 | yes | 1.437E-07 | 548.32965 | 561.54175 |
| HIAMHYSNPR | 10 | P52756 | RNA-binding protein 5 OS=Homo sapien | RBM5 | 92.154 | 3 | 52.991856 | yes | 1.536E-07 | 409.20135 | 243.30013 |
| HIAMHYSNPR | 10 | P52756 | RNA-binding protein 5 OS=Homo sapien | RBM5 | 92.154 | 3 | 52.991856 | yes | 9.722E-08 | 613.2984 | 243.30013 |
| HIAMHYSNPR | 10 | P52756 | RNA-binding protein 5 OS=Homo sapien | RBM5 | 92.154 | 2 | 52.991856 | yes | 1.536E-07 | 409.20135 | 243.30013 |
| HIAMHYSNPR | 10 | P52756 | RNA-binding protein 5 OS=Homo sapien | RBM5 | 92.154 | 2 | 52.991856 | yes | 9.722E-08 | 613.2984 | 243.30013 |
| DLVLSDGNR | 9 | P52756 | RNA-binding protein 5 OS=Homo sapien | RBM5 | 92.154 | 2 | 52.991856 | yes | 1.188E-06 | 494.75653 | 484.89923 |
| GSAYGLSGADSYK | 13 | P52756 | RNA-binding protein 5 OS=Homo sapien | RBM5 | 92.154 | 2 | 52.991856 | yes | 9.722E-08 | 638.2962 | 449.01776 |
| CQGITAPIEAQVR | 13 | P52756 | RNA-binding protein 5 OS=Homo sapien | RBM5 | 92.154 | 2 | 52.991856 | yes | 1.091E-07 | 721.87463 | 424.8866 |
| CCLNNFR | 7 | P52756 | RNA-binding protein 5 OS=Homo sapien | RBM5 | 92.154 | 2 | 52.991856 | yes | 1.736E-07 | 492.21292 | 371.09177 |
| FEDWLCNK | 8 | P52756 | RNA-binding protein 5 OS=Homo sapien | RBM5 | 92.154 | 2 | 52.991856 | yes | 7.783E-07 | 556.2475 | 348.10071 |
| MLQAMGWR | 8 | P52756 | RNA-binding protein 5 OS=Homo sapien | RBM5 | 92.154 | 2 | 52.991856 | yes | 9.722E-08 | 496.74387 | 338.47363 |

| | | | | | | | | | | | |
|---|---|---|---|---|---|---|---|---|---|---|---|
| KDLVLSDGNR | 10 | P52756 | RNA-binding protein 5 OS=Homo sapien | RBM5 | 92.154 | 2 | 52.991856 | yes | 1.539E-07 | 558.80402 | 323.28726 |
| ESAAADAGFALFE | 14 | P52756 | RNA-binding protein 5 OS=Homo sapien | RBM5 | 92.154 | 2 | 52.991856 | yes | 1.006E-07 | 713.84607 | 319.38318 |
| KCQGITAPIEAQVI | 14 | P52756 | RNA-binding protein 5 OS=Homo sapien | RBM5 | 92.154 | 3 | 52.991856 | yes | 9.722E-08 | 524.28387 | 140.57098 |
| KCQGITAPIEAQVI | 14 | P52756 | RNA-binding protein 5 OS=Homo sapien | RBM5 | 92.154 | 3 | 52.991856 | yes | 2.888E-07 | 785.92212 | 140.57098 |
| KCQGITAPIEAQVI | 14 | P52756 | RNA-binding protein 5 OS=Homo sapien | RBM5 | 92.154 | 2 | 52.991856 | yes | 9.722E-08 | 524.28387 | 140.57098 |
| KCQGITAPIEAQVI | 14 | P52756 | RNA-binding protein 5 OS=Homo sapien | RBM5 | 92.154 | 2 | 52.991856 | yes | 2.888E-07 | 785.92212 | 140.57098 |
| HDISDER | 7 | P52756 | RNA-binding protein 5 OS=Homo sapien | RBM5 | 92.154 | 2 | 52.991856 | yes | 2.2E-07 | 436.19885 | 229.19966 |
| DGIDHSNIGNK | 11 | P52756 | RNA-binding protein 5 OS=Homo sapien | RBM5 | 92.154 | 2 | 52.991856 | yes | 1.128E-07 | 585.28088 | 173.61502 |
| QFDAGTVNYEQPT | 14 | P52756 | RNA-binding protein 5 OS=Homo sapien | RBM5 | 92.154 | 2 | 52.991856 | yes | 9.722E-08 | 799.3783 | 158.80589 |
| MACLLCR | 7 | P52756 | RNA-binding protein 5 OS=Homo sapien | RBM5 | 92.154 | 2 | 52.991856 | yes | 2.591E-07 | 462.21661 | 150.75673 |
| TIGVDFAK | 8 | P52756 | RNA-binding protein 5 OS=Homo sapien | RBM5 | 92.154 | 2 | 52.991856 | yes | 2.231E-07 | 425.73709 | 108.52441 |
| YGSIIDR | 7 | P52756 | RNA-binding protein 5 OS=Homo sapien | RBM5 | 92.154 | 2 | 52.991856 | yes | 2.816E-07 | 412.21906 | 82.260162 |
| LTEAKPVDK | 9 | P52815 | Large ribosomal subunit protein bL12m C | MRPL12 | 21.349 | 2 | 49.529453 | yes | 2.851E-07 | 500.78729 | 3055.1672 |
| KLVESLPQEIK | 11 | P52815 | Large ribosomal subunit protein bL12m C | MRPL12 | 21.349 | 2 | 49.529453 | yes | 1.154E-07 | 642.38208 | 1036.8149 |
| LVESLPQEIK | 10 | P52815 | Large ribosomal subunit protein bL12m C | MRPL12 | 21.349 | 2 | 49.529453 | yes | 1.051E-07 | 578.33459 | 911.06763 |
| NYIQGINLVQAK | 12 | P52815 | Large ribosomal subunit protein bL12m C | MRPL12 | 21.349 | 2 | 49.529453 | yes | 1.061E-07 | 680.88281 | 728.87091 |
| CEALAGAPLDNAF | 14 | P52815 | Large ribosomal subunit protein bL12m C | MRPL12 | 21.349 | 2 | 49.529453 | yes | 1.022E-07 | 713.85339 | 212.13586 |
| FTITPPTAQVVGVI | 15 | P52907 | F-actin-capping protein subunit alpha-1 C | CAPZA1 | 32.923 | 2 | 52.71526 | yes | 9.536E-07 | 785.96381 | 2034.7715 |
| EASDPQPEEADGG | 15 | P52907 | F-actin-capping protein subunit alpha-1 C | CAPZA1 | 32.923 | 2 | 52.71526 | yes | 1.674E-07 | 771.84955 | 1201.9513 |
| TIDGQQTIIACIESH | 20 | P52907 | F-actin-capping protein subunit alpha-1 C | CAPZA1 | 32.923 | 3 | 52.71526 | yes | 1.606E-07 | 772.05475 | 771.16669 |
| IEGYEDQVLITEHC | 20 | P52907 | F-actin-capping protein subunit alpha-1 C | CAPZA1 | 32.923 | 3 | 52.71526 | yes | 1.705E-07 | 749.02924 | 639.36023 |
| ESCDSALR | 8 | P52907 | F-actin-capping protein subunit alpha-1 C | CAPZA1 | 32.923 | 2 | 52.71526 | yes | 3.236E-07 | 469.20581 | 191.97311 |
| DHYSNGFCTVYAI | 13 | P52907 | F-actin-capping protein subunit alpha-1 C | CAPZA1 | 32.923 | 2 | 52.71526 | yes | 9.722E-08 | 781.34064 | 61.840118 |
| KEASDPQPEEADG | 16 | P52907 | F-actin-capping protein subunit alpha-1 C | CAPZA1 | 32.923 | 2 | 52.71526 | yes | 4.938E-07 | 835.89703 | 21.551605 |
| GECIVSDFTIGR | 12 | P52948 | Nuclear pore complex protein Nup98-Nu | NUP98 | 197.58 | 2 | 52.14499 | yes | 1.075E-07 | 677.32703 | 269.38974 |
| EASGDLPEAQIVK | 13 | P52948 | Nuclear pore complex protein Nup98-Nu | NUP98 | 197.58 | 2 | 52.14499 | yes | 1.412E-07 | 678.85388 | 176.46059 |
| PIPQTPESAGNK | 12 | P52948 | Nuclear pore complex protein Nup98-Nu | NUP98 | 197.58 | 2 | 52.14499 | yes | 1.279E-07 | 619.82239 | 163.24153 |
| GFLEDLAPPER | 11 | P52948 | Nuclear pore complex protein Nup98-Nu | NUP98 | 197.58 | 2 | 52.14499 | yes | 9.722E-08 | 622.31952 | 162.68765 |
| LADINYEGR | 9 | P52948 | Nuclear pore complex protein Nup98-Nu | NUP98 | 197.58 | 2 | 52.14499 | yes | 3.745E-07 | 525.76434 | 95.337288 |
| PVWQLSEK | 8 | P52948 | Nuclear pore complex protein Nup98-Nu | NUP98 | 197.58 | 2 | 52.14499 | yes | 1.696E-07 | 493.76892 | 93.725075 |
| DVCFHLLK | 8 | P52948 | Nuclear pore complex protein Nup98-Nu | NUP98 | 197.58 | 2 | 52.14499 | yes | 4.261E-07 | 516.27075 | 88.799278 |
| LPMPEDYAMDELI | 13 | P52948 | Nuclear pore complex protein Nup98-Nu | NUP98 | 197.58 | 2 | 52.14499 | yes | 9.722E-08 | 790.36017 | 83.162224 |
| LYQTPLELK | 9 | P52948 | Nuclear pore complex protein Nup98-Nu | NUP98 | 197.58 | 2 | 52.14499 | yes | 8.36E-07 | 552.8186 | 48.003563 |
| LEDYQANR | 8 | P52948 | Nuclear pore complex protein Nup98-Nu | NUP98 | 197.58 | 2 | 52.14499 | yes | 6.34E-07 | 504.74088 | 32.896538 |
| ADTSQEICSPR | 11 | P52948 | Nuclear pore complex protein Nup98-Nu | NUP98 | 197.58 | 2 | 52.14499 | yes | 3.571E-05 | 632.28534 | 32.028942 |
| HQCITAMK | 8 | P52948 | Nuclear pore complex protein Nup98-Nu | NUP98 | 197.58 | 2 | 52.14499 | yes | 7.286E-05 | 494.73877 | 30.189409 |
| NLNNSNLFSPVNR | 13 | P52948 | Nuclear pore complex protein Nup98-Nu | NUP98 | 197.58 | 2 | 52.14499 | yes | 9.722E-08 | 744.88135 | 28.600681 |
| DILNLEQQQR | 10 | P52952 | Homeobox protein Nkx-2.5 OS=Homo sa | NKX2-5 | 34.918 | 2 | 51.313324 | yes | 9.722E-08 | 628.83331 | 482.26382 |
| IAVPVLVR | 8 | P52952 | Homeobox protein Nkx-2.5 OS=Homo sa | NKX2-5 | 34.918 | 2 | 51.313324 | yes | 9.81E-08 | 433.79474 | 392.56573 |

| | | | | | | | | | | | |
|---|---|---|---|---|---|---|---|---|---|---|---|
| VLFSQAQVYELER | 13 | P52952 | Homeobox protein Nkx-2.5 OS=Homo sa | NKX2-5 | 34.918 | 2 | 51.313324 | no | 1.407E-07 | 791.41718 | 264.97922 |
| AYSDPDPAKDPR | 12 | P52952 | Homeobox protein Nkx-2.5 OS=Homo sa | NKX2-5 | 34.918 | 2 | 51.313324 | yes | 1.021E-07 | 666.31494 | 245.20581 |
| CASAFPAAPAFYPI | 14 | P52952 | Homeobox protein Nkx-2.5 OS=Homo sa | NKX2-5 | 34.918 | 2 | 51.313324 | yes | 2.483E-07 | 763.36646 | 229.80228 |
| TQLQLDER | 8 | P53007 | Tricarboxylate transport protein, mitochoı | SLC25A1 | 34.013 | 2 | 52.16528 | yes | 9.722E-08 | 501.76437 | 1800.7271 |
| GTYQGLTATVLK | 12 | P53007 | Tricarboxylate transport protein, mitochoı | SLC25A1 | 34.013 | 2 | 52.16528 | yes | 1.361E-07 | 626.35083 | 1780.1156 |
| SHGVLGLYR | 9 | P53007 | Tricarboxylate transport protein, mitochoı | SLC25A1 | 34.013 | 2 | 52.16528 | yes | 1.186E-07 | 501.27997 | 1704.0792 |
| GFFHGVR | 7 | P53007 | Tricarboxylate transport protein, mitochoı | SLC25A1 | 34.013 | 2 | 52.16528 | yes | 2.721E-07 | 410.21664 | 1359.7538 |
| FIHDQTSPNPK | 11 | P53007 | Tricarboxylate transport protein, mitochoı | SLC25A1 | 34.013 | 2 | 52.16528 | yes | 1.001E-07 | 642.32257 | 548.81073 |
| FIHDQTSPNPK | 11 | P53007 | Tricarboxylate transport protein, mitochoı | SLC25A1 | 34.013 | 2 | 52.16528 | yes | 3.925E-07 | 428.55081 | 548.81073 |
| FIHDQTSPNPK | 11 | P53007 | Tricarboxylate transport protein, mitochoı | SLC25A1 | 34.013 | 3 | 52.16528 | yes | 1.001E-07 | 642.32257 | 548.81073 |
| FIHDQTSPNPK | 11 | P53007 | Tricarboxylate transport protein, mitochoı | SLC25A1 | 34.013 | 3 | 52.16528 | yes | 3.925E-07 | 428.55081 | 548.81073 |
| NTWDCGLQILK | 11 | P53007 | Tricarboxylate transport protein, mitochoı | SLC25A1 | 34.013 | 2 | 52.16528 | yes | 9.727E-08 | 674.3399 | 661.58521 |
| FFVMTSLR | 8 | P53007 | Tricarboxylate transport protein, mitochoı | SLC25A1 | 34.013 | 2 | 52.16528 | yes | 2.294E-07 | 500.76785 | 634.63794 |
| FGMFEFLSNHMR | 12 | P53007 | Tricarboxylate transport protein, mitochoı | SLC25A1 | 34.013 | 2 | 52.16528 | yes | 9.722E-08 | 758.34723 | 429.74841 |
| TECYGYALGDATF | 13 | P53041 | Serine/threonine-protein phosphatase 5 O | PPP5C | 56.879 | 2 | 48.52253 | yes | 2.927E-07 | 738.82483 | 60.891556 |
| LSTLVETTLK | 10 | P53041 | Serine/threonine-protein phosphatase 5 O | PPP5C | 56.879 | 2 | 48.52253 | yes | 1.136E-07 | 552.82916 | 38.855495 |
| FSIAPSSLDPSNR | 13 | P53350 | Serine/threonine-protein kinase PLK1 OS | PLK1 | 68.255 | 2 | 53.505043 | yes | 9.862E-08 | 695.85168 | 1642.7209 |
| MSMEISIHR | 9 | P53350 | Serine/threonine-protein kinase PLK1 OS | PLK1 | 68.255 | 2 | 53.505043 | yes | 1.27E-07 | 552.27045 | 1000.9409 |
| LGNLFLNEDLEVK | 13 | P53350 | Serine/threonine-protein kinase PLK1 OS | PLK1 | 68.255 | 2 | 53.505043 | yes | 1.039E-07 | 752.40631 | 714.30701 |
| LSLLEEYGCCK | 11 | P53350 | Serine/threonine-protein kinase PLK1 OS | PLK1 | 68.255 | 2 | 53.505043 | yes | 9.722E-08 | 686.31781 | 688.31378 |
| MLQTDPTAR | 9 | P53350 | Serine/threonine-protein kinase PLK1 OS | PLK1 | 68.255 | 2 | 53.505043 | yes | 2.343E-07 | 516.76074 | 620.52332 |
| NYMSEHLLK | 9 | P53350 | Serine/threonine-protein kinase PLK1 OS | PLK1 | 68.255 | 2 | 53.505043 | yes | 1.602E-07 | 567.78424 | 435.53571 |
| QIVLGCQYLHR | 11 | P53350 | Serine/threonine-protein kinase PLK1 OS | PLK1 | 68.255 | 2 | 53.505043 | yes | 1.003E-07 | 693.86914 | 334.20959 |
| IGDFGLATK | 9 | P53350 | Serine/threonine-protein kinase PLK1 OS | PLK1 | 68.255 | 2 | 53.505043 | yes | 1.105E-07 | 461.25565 | 327.17426 |
| EKEEPVVR | 8 | P53350 | Serine/threonine-protein kinase PLK1 OS | PLK1 | 68.255 | 2 | 53.505043 | yes | 2.991E-06 | 493.26929 | 324.70572 |
| VEYDGER | 7 | P53350 | Serine/threonine-protein kinase PLK1 OS | PLK1 | 68.255 | 2 | 53.505043 | yes | 9.722E-08 | 434.19577 | 314.60138 |
| CFEISDADTK | 10 | P53350 | Serine/threonine-protein kinase PLK1 OS | PLK1 | 68.255 | 2 | 53.505043 | yes | 1.076E-07 | 593.25824 | 261.3826 |
| HINPVAASLIQK | 12 | P53350 | Serine/threonine-protein kinase PLK1 OS | PLK1 | 68.255 | 2 | 53.505043 | yes | 9.809E-08 | 645.88007 | 250.12355 |
| KPLTVLNK | 8 | P53350 | Serine/threonine-protein kinase PLK1 OS | PLK1 | 68.255 | 2 | 53.505043 | yes | 1.923E-07 | 456.79749 | 220.50531 |
| GLENPLPERPR | 11 | P53350 | Serine/threonine-protein kinase PLK1 OS | PLK1 | 68.255 | 2 | 53.505043 | yes | 9.722E-08 | 639.35168 | 82.743858 |
| GLENPLPERPR | 11 | P53350 | Serine/threonine-protein kinase PLK1 OS | PLK1 | 68.255 | 2 | 53.505043 | yes | 4.631E-07 | 426.57019 | 82.743858 |
| GLENPLPERPR | 11 | P53350 | Serine/threonine-protein kinase PLK1 OS | PLK1 | 68.255 | 3 | 53.505043 | yes | 9.722E-08 | 639.35168 | 82.743858 |
| GLENPLPERPR | 11 | P53350 | Serine/threonine-protein kinase PLK1 OS | PLK1 | 68.255 | 3 | 53.505043 | yes | 4.631E-07 | 426.57019 | 82.743858 |
| SLLELHK | 7 | P53350 | Serine/threonine-protein kinase PLK1 OS | PLK1 | 68.255 | 2 | 53.505043 | yes | 1.939E-07 | 420.2529 | 133.70349 |
| LPITCLTIPPR | 11 | P53350 | Serine/threonine-protein kinase PLK1 OS | PLK1 | 68.255 | 2 | 53.505043 | yes | 2.982E-07 | 640.87341 | 130.20584 |
| EIPEVLVDPR | 10 | P53350 | Serine/threonine-protein kinase PLK1 OS | PLK1 | 68.255 | 2 | 53.505043 | yes | 1.633E-07 | 583.8244 | 88.045105 |
| DGTESYLTVSSHPI | 18 | P53350 | Serine/threonine-protein kinase PLK1 OS | PLK1 | 68.255 | 3 | 53.505043 | yes | 1.116E-07 | 655.97894 | 51.257092 |
| IIDFGLAHK | 9 | P53355 | Death-associated protein kinase 1 OS=Hc | DAPK1 | 160.047 | 2 | 39.367817 | yes | 4.627E-07 | 507.29257 | 110.41029 |
| LIMGIGHR | 8 | P53396 | ATP-citrate synthase OS=Homo sapiens ( | ACLY | 120.84 | 2 | 51.936665 | yes | 9.722E-08 | 448.76038 | 135.0816 |

| | | | | | | | | | | | |
|---|---|---|---|---|---|---|---|---|---|---|---|
| SGGMSNELNNIISF | 14 | P53396 | ATP-citrate synthase OS=Homo sapiens ( | ACLY | 120.84 | 2 | 51.936665 | yes | 2.98E-07 | 746.36462 | 93.68573 |
| AFDSGIIPMEFVNK | 14 | P53396 | ATP-citrate synthase OS=Homo sapiens ( | ACLY | 120.84 | 2 | 51.936665 | yes | 1.242E-07 | 784.3949 | 80.506363 |
| EHEVTIFVR | 9 | P53396 | ATP-citrate synthase OS=Homo sapiens ( | ACLY | 120.84 | 2 | 51.936665 | yes | 1.841E-07 | 565.30365 | 71.56279 |
| VDATADYICK | 10 | P53396 | ATP-citrate synthase OS=Homo sapiens ( | ACLY | 120.84 | 2 | 51.936665 | yes | 4.308E-07 | 578.27118 | 57.362751 |
| AVQGMLDFDYVC | 14 | P53396 | ATP-citrate synthase OS=Homo sapiens ( | ACLY | 120.84 | 2 | 51.936665 | yes | 1.759E-07 | 830.87653 | 51.85051 |
| MIVVLGEIGGTEEY | 15 | P53396 | ATP-citrate synthase OS=Homo sapiens ( | ACLY | 120.84 | 2 | 51.936665 | yes | 5.111E-06 | 819.42639 | 42.537262 |
| SMGFIGHYLDQK | 12 | P53396 | ATP-citrate synthase OS=Homo sapiens ( | ACLY | 120.84 | 2 | 51.936665 | yes | 1.496E-07 | 698.3399 | 18.639191 |
| EETLTPEEEEK | 11 | P53539 | Protein FosB OS=Homo sapiens OX=960 | FOSB | 35.928 | 2 | 53.732475 | yes | 9.722E-08 | 667.30389 | 225.92052 |
| MFQAFPGDYDSGS | 14 | P53539 | Protein FosB OS=Homo sapiens OX=960 | FOSB | 35.928 | 2 | 53.732475 | yes | 1.273E-06 | 810.34338 | 25.186447 |
| LLSSEDIEGMR | 11 | P53582 | Methionine aminopeptidase 1 OS=Homo | METAP1 | 43.216 | 2 | 50.36654 | yes | 1.269E-07 | 625.30829 | 155.5526 |
| LFHTAPNVPHYAK | 13 | P53582 | Methionine aminopeptidase 1 OS=Homo | METAP1 | 43.216 | 3 | 50.36654 | yes | 1.144E-07 | 498.93326 | 64.182144 |
| LFHTAPNVPHYAK | 13 | P53582 | Methionine aminopeptidase 1 OS=Homo | METAP1 | 43.216 | 3 | 50.36654 | yes | 9.81E-08 | 747.89624 | 64.182144 |
| LFHTAPNVPHYAK | 13 | P53582 | Methionine aminopeptidase 1 OS=Homo | METAP1 | 43.216 | 2 | 50.36654 | yes | 1.144E-07 | 498.93326 | 64.182144 |
| LFHTAPNVPHYAK | 13 | P53582 | Methionine aminopeptidase 1 OS=Homo | METAP1 | 43.216 | 2 | 50.36654 | yes | 9.81E-08 | 747.89624 | 64.182144 |
| LRPHYPLMPTR | 11 | P53582 | Methionine aminopeptidase 1 OS=Homo | METAP1 | 43.216 | 3 | 50.36654 | yes | 8.137E-07 | 460.92383 | 60.683826 |
| LVQTTYECLMQAI | 21 | P53582 | Methionine aminopeptidase 1 OS=Homo | METAP1 | 43.216 | 3 | 50.36654 | yes | 2.734E-07 | 798.08362 | 39.498421 |
| EVLDVAAGMIKPC | 30 | P53582 | Methionine aminopeptidase 1 OS=Homo | METAP1 | 43.216 | 4 | 50.36654 | yes | 2.435E-05 | 804.66925 | 12.100859 |
| SDAPDTLLLEK | 11 | P53611 | Geranylgeranyl transferase type-2 subunit | RABGGTB | 36.925 | 2 | 43.468708 | yes | 2.578E-07 | 601.31915 | 89.291153 |
| SLGEIPIVESEIK | 13 | P53618 | Coatomer subunit beta OS=Homo sapiens | COPB1 | 107.143 | 2 | 56.02267 | yes | 2.238E-07 | 707.39539 | 325.07153 |
| EDIQSVMTEIR | 11 | P53618 | Coatomer subunit beta OS=Homo sapiens | COPB1 | 107.143 | 2 | 56.02267 | yes | 9.723E-08 | 660.82684 | 306.19901 |
| SQGMALSLGDK | 11 | P53618 | Coatomer subunit beta OS=Homo sapiens | COPB1 | 107.143 | 2 | 56.02267 | yes | 2.892E-07 | 553.77917 | 225.05061 |
| CIYNLLQSSSPAVK | 14 | P53618 | Coatomer subunit beta OS=Homo sapiens | COPB1 | 107.143 | 2 | 56.02267 | yes | 2.258E-07 | 790.41107 | 210.0376 |
| AAAQCYIDLIIK | 12 | P53618 | Coatomer subunit beta OS=Homo sapiens | COPB1 | 107.143 | 2 | 56.02267 | yes | 9.722E-08 | 689.8736 | 209.71165 |
| GALWILGEYCSTK | 13 | P53618 | Coatomer subunit beta OS=Homo sapiens | COPB1 | 107.143 | 2 | 56.02267 | yes | 9.722E-08 | 749.37396 | 138.3651 |
| MLEVFHAIK | 9 | P53618 | Coatomer subunit beta OS=Homo sapiens | COPB1 | 107.143 | 2 | 56.02267 | yes | 1.49E-07 | 544.30206 | 117.02756 |
| EAELLEPLMPAIR | 13 | P53618 | Coatomer subunit beta OS=Homo sapiens | COPB1 | 107.143 | 2 | 56.02267 | yes | 1.525E-06 | 741.40527 | 116.51646 |
| DLQHPNEFIR | 10 | P53618 | Coatomer subunit beta OS=Homo sapiens | COPB1 | 107.143 | 2 | 56.02267 | yes | 9.722E-08 | 634.82275 | 106.10072 |
| KPITDDDVDR | 10 | P53618 | Coatomer subunit beta OS=Homo sapiens | COPB1 | 107.143 | 2 | 56.02267 | yes | 1.12E-07 | 587.29095 | 103.87396 |
| LVEKPSPLTLAPHI | 19 | P53618 | Coatomer subunit beta OS=Homo sapiens | COPB1 | 107.143 | 3 | 56.02267 | yes | 1.115E-07 | 697.39301 | 100.88748 |
| TLQLALDLVSSR | 12 | P53618 | Coatomer subunit beta OS=Homo sapiens | COPB1 | 107.143 | 2 | 56.02267 | yes | 9.722E-08 | 658.38263 | 70.142952 |
| QMWAEFEWENK | 11 | P53618 | Coatomer subunit beta OS=Homo sapiens | COPB1 | 107.143 | 2 | 56.02267 | yes | 2.032E-07 | 749.32703 | 63.782013 |
| NVEELVIVLK | 10 | P53618 | Coatomer subunit beta OS=Homo sapiens | COPB1 | 107.143 | 2 | 56.02267 | yes | 1.034E-07 | 578.35284 | 62.411388 |
| QSLSHMLSAK | 10 | P53618 | Coatomer subunit beta OS=Homo sapiens | COPB1 | 107.143 | 2 | 56.02267 | yes | 6.326E-07 | 551.28967 | 42.839916 |
| LVTEMGTYATQSA | 21 | P53618 | Coatomer subunit beta OS=Homo sapiens | COPB1 | 107.143 | 3 | 56.02267 | yes | 1.784E-07 | 743.37909 | 34.454903 |
| DADSITLFDVQQK | 13 | P53621 | Coatomer subunit alpha OS=Homo sapier | COPA | 138.346 | 2 | 54.994915 | yes | 1.673E-07 | 740.36993 | 579.19977 |
| VWDISGLR | 8 | P53621 | Coatomer subunit alpha OS=Homo sapier | COPA | 138.346 | 2 | 54.994915 | yes | 7.262E-07 | 473.26126 | 447.73703 |
| TLDLPIYVTR | 10 | P53621 | Coatomer subunit alpha OS=Homo sapier | COPA | 138.346 | 2 | 54.994915 | yes | 1.263E-07 | 595.84259 | 418.48563 |
| VLTIDPTEFK | 10 | P53621 | Coatomer subunit alpha OS=Homo sapier | COPA | 138.346 | 2 | 54.994915 | yes | 2.069E-07 | 581.82135 | 384.9805 |
| QQPLFVSGGDDYK | 13 | P53621 | Coatomer subunit alpha OS=Homo sapier | COPA | 138.346 | 2 | 54.994915 | yes | 3.802E-07 | 727.3515 | 272.27725 |

| | | | | | | | | | | | |
|---|---|---|---|---|---|---|---|---|---|---|---|
| YAVTTGDHGIIR | 12 | P53621 | Coatomer subunit alpha OS=Homo sapier | COPA | 138.346 | 2 | 54.994915 | yes | 9.893E-08 | 651.84369 | 265.25842 |
| DVAVMQLR | 8 | P53621 | Coatomer subunit alpha OS=Homo sapier | COPA | 138.346 | 2 | 54.994915 | yes | 2.077E-06 | 466.25513 | 262.14279 |
| AWEVDTCR | 8 | P53621 | Coatomer subunit alpha OS=Homo sapier | COPA | 138.346 | 2 | 54.994915 | yes | 4.242E-07 | 518.72949 | 256.81525 |
| LQLCYQLTTVGK | 12 | P53621 | Coatomer subunit alpha OS=Homo sapier | COPA | 138.346 | 2 | 54.994915 | yes | 1.145E-07 | 712.38434 | 235.14725 |
| GITGVDLFGTTDA | 16 | P53621 | Coatomer subunit alpha OS=Homo sapier | COPA | 138.346 | 2 | 54.994915 | yes | 9.722E-08 | 796.93018 | 225.03419 |
| GFFEGTIASK | 10 | P53621 | Coatomer subunit alpha OS=Homo sapier | COPA | 138.346 | 2 | 54.994915 | yes | 4.509E-07 | 528.77167 | 202.54184 |
| EYIVGLSVETER | 12 | P53621 | Coatomer subunit alpha OS=Homo sapier | COPA | 138.346 | 2 | 54.994915 | yes | 1.326E-07 | 697.86176 | 195.2159 |
| QEIAEAQQLITICR | 14 | P53621 | Coatomer subunit alpha OS=Homo sapier | COPA | 138.346 | 2 | 54.994915 | yes | 3.603E-07 | 836.93799 | 193.45886 |
| SSGLTAVWVAR | 11 | P53621 | Coatomer subunit alpha OS=Homo sapier | COPA | 138.346 | 2 | 54.994915 | yes | 9.722E-08 | 573.81696 | 192.72871 |
| CPLSGACYSPEFK | 13 | P53621 | Coatomer subunit alpha OS=Homo sapier | COPA | 138.346 | 2 | 54.994915 | yes | 1.616E-07 | 758.33398 | 172.46954 |
| QELILSNSEDK | 11 | P53621 | Coatomer subunit alpha OS=Homo sapier | COPA | 138.346 | 2 | 54.994915 | yes | 1.942E-07 | 638.32501 | 170.34169 |
| GNNVYCLDR | 9 | P53621 | Coatomer subunit alpha OS=Homo sapier | COPA | 138.346 | 2 | 54.994915 | yes | 9.722E-08 | 555.75348 | 145.55562 |
| KYDEVLHMVR | 10 | P53621 | Coatomer subunit alpha OS=Homo sapier | COPA | 138.346 | 3 | 54.994915 | yes | 2.9E-07 | 430.56055 | 143.31146 |
| DADSQNPDAPEGK | 13 | P53621 | Coatomer subunit alpha OS=Homo sapier | COPA | 138.346 | 2 | 54.994915 | yes | 9.722E-08 | 672.28912 | 143.10104 |
| KLDALCNIHENIR | 13 | P53621 | Coatomer subunit alpha OS=Homo sapier | COPA | 138.346 | 3 | 54.994915 | yes | 1.313E-07 | 532.6156 | 135.65909 |
| LGEVALLQGNHQI | 20 | P53621 | Coatomer subunit alpha OS=Homo sapier | COPA | 138.346 | 3 | 54.994915 | yes | 1.284E-07 | 786.72797 | 122.00323 |
| TALNLFFK | 8 | P53621 | Coatomer subunit alpha OS=Homo sapier | COPA | 138.346 | 2 | 54.994915 | yes | 2.124E-07 | 477.27637 | 75.942741 |
| LNDLIQR | 7 | P53621 | Coatomer subunit alpha OS=Homo sapier | COPA | 138.346 | 2 | 54.994915 | yes | 9.462E-06 | 436.25345 | 66.574333 |
| GVNWAAFHPTMP | 21 | P53621 | Coatomer subunit alpha OS=Homo sapier | COPA | 138.346 | 3 | 54.994915 | yes | 2.328E-07 | 752.04065 | 64.175117 |
| YDEVLHMVR | 9 | P53621 | Coatomer subunit alpha OS=Homo sapier | COPA | 138.346 | 2 | 54.994915 | yes | 1.341E-07 | 581.28967 | 48.575996 |
| MHSLLIK | 7 | P53621 | Coatomer subunit alpha OS=Homo sapier | COPA | 138.346 | 2 | 54.994915 | yes | 8.216E-06 | 421.25186 | 43.531967 |
| LLELGPKPEVAQQ | 15 | P53621 | Coatomer subunit alpha OS=Homo sapier | COPA | 138.346 | 2 | 54.994915 | yes | 1.483E-07 | 839.97797 | 40.803593 |
| YVIWSADMSHVA | 16 | P53621 | Coatomer subunit alpha OS=Homo sapier | COPA | 138.346 | 3 | 54.994915 | yes | 3.617E-07 | 601.98694 | 40.643215 |
| VQVPNCDEIFYAG | 20 | P53621 | Coatomer subunit alpha OS=Homo sapier | COPA | 138.346 | 3 | 54.994915 | yes | 8.262E-07 | 760.38782 | 39.392231 |
| HAIVICNR | 8 | P53621 | Coatomer subunit alpha OS=Homo sapier | COPA | 138.346 | 2 | 54.994915 | yes | 3.4E-06 | 491.76617 | 38.048241 |
| LVGQSIIAYLQK | 12 | P53621 | Coatomer subunit alpha OS=Homo sapier | COPA | 138.346 | 2 | 54.994915 | yes | 6.666E-07 | 666.89795 | 36.280632 |
| SGAWDESGVFIYT | 19 | P53621 | Coatomer subunit alpha OS=Homo sapier | COPA | 138.346 | 3 | 54.994915 | yes | 1.825E-07 | 704.67261 | 35.629753 |
| DMSGHYQNALYL | 18 | P53621 | Coatomer subunit alpha OS=Homo sapier | COPA | 138.346 | 3 | 54.994915 | yes | 1.366E-06 | 685.64606 | 33.003357 |
| TTFFHHEYPWILS | 21 | P53621 | Coatomer subunit alpha OS=Homo sapier | COPA | 138.346 | 4 | 54.994915 | yes | 2.162E-07 | 641.81055 | 32.393116 |
| QLFLQTYAR | 9 | P53621 | Coatomer subunit alpha OS=Homo sapier | COPA | 138.346 | 2 | 54.994915 | yes | 1.209E-06 | 570.31403 | 31.642857 |
| ALDDKNCWEK | 10 | P53621 | Coatomer subunit alpha OS=Homo sapier | COPA | 138.346 | 2 | 54.994915 | yes | 1.297E-07 | 639.79279 | 31.029682 |
| CLFTLLGHLDYIR | 13 | P53621 | Coatomer subunit alpha OS=Homo sapier | COPA | 138.346 | 2 | 54.994915 | yes | 1.902E-07 | 810.93195 | 25.637163 |
| GHYNNVSCAVFHI | 14 | P53621 | Coatomer subunit alpha OS=Homo sapier | COPA | 138.346 | 2 | 54.994915 | yes | 1.835E-07 | 829.38623 | 10.74925 |
| NVHGINFVSPVR | 12 | P53634 | Dipeptidyl peptidase 1 OS=Homo sapiens | CTSC | 51.854 | 3 | 39.503586 | yes | 4.547E-07 | 446.91409 | 43.862865 |
| GQFSTDELVAEVE | 14 | P53675 | Clathrin heavy chain 2 OS=Homo sapiens | CLTCL1 | 187.03 | 2 | 50.703396 | no | 1.493E-07 | 776.38049 | 866.22089 |
| IVLDNSVFSEHR | 12 | P53675 | Clathrin heavy chain 2 OS=Homo sapiens | CLTCL1 | 187.03 | 2 | 50.703396 | no | 9.766E-08 | 708.36749 | 542.49103 |
| TLQIFNIEMK | 10 | P53675 | Clathrin heavy chain 2 OS=Homo sapiens | CLTCL1 | 187.03 | 2 | 50.703396 | no | 9.783E-08 | 618.83649 | 492.16757 |
| YIEIYVQK | 8 | P53675 | Clathrin heavy chain 2 OS=Homo sapiens | CLTCL1 | 187.03 | 2 | 50.703396 | no | 2.779E-05 | 528.29224 | 416.31812 |
| DPHLACVAYER | 11 | P53675 | Clathrin heavy chain 2 OS=Homo sapiens | CLTCL1 | 187.03 | 2 | 50.703396 | no | 1.131E-07 | 665.81409 | 303.48996 |

| | | | | | | | | | | | |
|---|---|---|---|---|---|---|---|---|---|---|---|
| VVGAMQLYSVDR | 12 | P53675 | Clathrin heavy chain 2 OS=Homo sapiens | CLTCL1 | 187.03 | 2 | 50.703396 | no | 9.728E-08 | 669.34772 | 206.20081 |
| AHIAQLCEK | 9 | P53675 | Clathrin heavy chain 2 OS=Homo sapiens | CLTCL1 | 187.03 | 2 | 50.703396 | no | 2.144E-07 | 535.27661 | 162.3799 |
| LTDQLPLIIVCDR | 13 | P53675 | Clathrin heavy chain 2 OS=Homo sapiens | CLTCL1 | 187.03 | 2 | 50.703396 | no | 1.219E-07 | 778.42926 | 132.1312 |
| QNLQLCVQVASK | 12 | P53675 | Clathrin heavy chain 2 OS=Homo sapiens | CLTCL1 | 187.03 | 2 | 50.703396 | yes | 1.88E-07 | 694.37177 | 95.797508 |
| LECSEELGDLVK | 12 | P53675 | Clathrin heavy chain 2 OS=Homo sapiens | CLTCL1 | 187.03 | 2 | 50.703396 | no | 1.796E-07 | 696.33978 | 92.397514 |
| GNNWWAQSVELC | 13 | P53675 | Clathrin heavy chain 2 OS=Homo sapiens | CLTCL1 | 187.03 | 2 | 50.703396 | yes | 4.678E-06 | 796.36975 | 49.853973 |
| AHMGMFTELAILY | 15 | P53675 | Clathrin heavy chain 2 OS=Homo sapiens | CLTCL1 | 187.03 | 3 | 50.703396 | no | 3.096E-07 | 571.28973 | 16.838539 |
| FILIQNR | 7 | P53680 | AP-2 complex subunit sigma OS=Homo s | AP2S1 | 17.018 | 2 | 53.8677 | yes | 1.348E-07 | 452.27417 | 2521.0723 |
| HTNFVEFR | 8 | P53680 | AP-2 complex subunit sigma OS=Homo s | AP2S1 | 17.018 | 2 | 53.8677 | yes | 5.078E-07 | 525.26178 | 1297.3054 |
| LIEEVHAVVTVR | 12 | P53680 | AP-2 complex subunit sigma OS=Homo s | AP2S1 | 17.018 | 2 | 53.8677 | yes | 1.139E-07 | 682.89844 | 445.63156 |
| WYMQFDDDEK | 10 | P53680 | AP-2 complex subunit sigma OS=Homo s | AP2S1 | 17.018 | 2 | 53.8677 | yes | 9.722E-08 | 688.77679 | 138.83266 |
| WYMQFDDDEK | 10 | P53680 | AP-2 complex subunit sigma OS=Homo s | AP2S1 | 17.018 | 2 | 53.8677 | yes | 9.108E-07 | 696.77429 | 138.83266 |
| YVIDYYDGGEVNI | 13 | P53701 | Holocytochrome c-type synthase OS=Hor | HCCS | 30.602 | 2 | 46.021973 | yes | 1.675E-07 | 767.85663 | 61.837669 |
| AYEYVECPIR | 10 | P53701 | Holocytochrome c-type synthase OS=Hor | HCCS | 30.602 | 2 | 46.021973 | yes | 1.678E-07 | 650.30554 | 54.496559 |
| LVVFDAR | 7 | P53803 | DNA-directed RNA polymerases I, II, and | POLR2K | 7.005 | 2 | 54.862534 | yes | 9.722E-08 | 410.23981 | 120.92618 |
| SVPVVPSFVANDR | 13 | P53804 | E3 ubiquitin-protein ligase TTC3 OS=Ho | TTC3 | 229.869 | 2 | 46.432114 | yes | 3.191E-06 | 693.87244 | 92.073746 |
| LQLNPAAR | 8 | P53804 | E3 ubiquitin-protein ligase TTC3 OS=Ho | TTC3 | 229.869 | 2 | 46.432114 | yes | 1.591E-06 | 441.76141 | 64.061935 |
| FSSSSPLTLPADLK | 14 | P53804 | E3 ubiquitin-protein ligase TTC3 OS=Ho | TTC3 | 229.869 | 2 | 46.432114 | yes | 5.101E-07 | 731.89301 | 62.901318 |
| FVVIDNCIALK | 11 | P53804 | E3 ubiquitin-protein ligase TTC3 OS=Ho | TTC3 | 229.869 | 2 | 46.432114 | yes | 1.56E-07 | 646.35754 | 46.001549 |
| VLSTEDYTTCFSSF | 14 | P53804 | E3 ubiquitin-protein ligase TTC3 OS=Ho | TTC3 | 229.869 | 2 | 46.432114 | yes | 4.792E-07 | 833.37488 | 29.900219 |
| SSQGSPSVVVAPSI | 15 | P53804 | E3 ubiquitin-protein ligase TTC3 OS=Ho | TTC3 | 229.869 | 2 | 46.432114 | yes | 8.12E-07 | 713.88043 | 10.911736 |
| GVAQPLAQLR | 10 | P53814 | Smoothelin OS=Homo sapiens OX=9606 | SMTN | 99.059 | 2 | 44.836918 | yes | 7.503E-07 | 526.81421 | 62.843277 |
| ESTPLASGPSSFQR | 14 | P53814 | Smoothelin OS=Homo sapiens OX=9606 | SMTN | 99.059 | 2 | 44.836918 | yes | 3.703E-07 | 732.35986 | 60.836712 |
| LLEVTADLAER | 11 | P53814 | Smoothelin OS=Homo sapiens OX=9606 | SMTN | 99.059 | 2 | 44.836918 | yes | 2.051E-07 | 615.34045 | 52.820621 |
| AQEIEAATLAGR | 12 | P53814 | Smoothelin OS=Homo sapiens OX=9606 | SMTN | 99.059 | 2 | 44.836918 | yes | 4.873E-07 | 615.32788 | 46.157478 |
| ETPAAQSPTR | 10 | P53814 | Smoothelin OS=Homo sapiens OX=9606 | SMTN | 99.059 | 2 | 44.836918 | yes | 9.658E-07 | 529.26727 | 30.542753 |
| DLHDANTDLIGR | 12 | P53985 | Monocarboxylate transporter 1 OS=Homo | SLC16A1 | 53.945 | 2 | 54.251705 | yes | 4.793E-07 | 670.33368 | 733.36877 |
| PSMGLVANTK | 10 | P53985 | Monocarboxylate transporter 1 OS=Homo | SLC16A1 | 53.945 | 2 | 54.251705 | yes | 1.806E-06 | 509.2735 | 131.25726 |
| ESKEEETSIDVAGI | 20 | P53985 | Monocarboxylate transporter 1 OS=Homo | SLC16A1 | 53.945 | 3 | 54.251705 | yes | 9.722E-08 | 730.69727 | 102.07219 |
| LNDMYGDYK | 9 | P53985 | Monocarboxylate transporter 1 OS=Homo | SLC16A1 | 53.945 | 2 | 54.251705 | yes | 5.453E-07 | 559.74481 | 43.042877 |
| TNQIGTVNDR | 10 | P53990 | IST1 homolog OS=Homo sapiens OX=96 | IST1 | 39.751 | 2 | 47.60001 | yes | 1.296E-07 | 559.28345 | 72.551086 |
| FGLIQSMK | 8 | P53990 | IST1 homolog OS=Homo sapiens OX=96 | IST1 | 39.751 | 2 | 47.60001 | yes | 3.859E-07 | 462.25458 | 61.71582 |
| SDVLQPGAEVTTD | 15 | P53992 | Protein transport protein Sec24C OS=Hor | SEC24C | 118.325 | 2 | 55.025925 | yes | 9.938E-08 | 801.88391 | 379.19489 |
| GTEPFVTGVR | 10 | P53992 | Protein transport protein Sec24C OS=Hor | SEC24C | 118.325 | 2 | 55.025925 | yes | 1.405E-07 | 531.78253 | 303.19632 |
| VVGFDAVMR | 9 | P53992 | Protein transport protein Sec24C OS=Hor | SEC24C | 118.325 | 2 | 55.025925 | yes | 4.32E-07 | 497.26294 | 228.36035 |
| TLFQPQTGAYQTL | 15 | P53992 | Protein transport protein Sec24C OS=Hor | SEC24C | 118.325 | 2 | 55.025925 | yes | 1.318E-07 | 833.9436 | 167.717 |
| YASFQVENDQER | 12 | P53992 | Protein transport protein Sec24C OS=Hor | SEC24C | 118.325 | 2 | 55.025925 | yes | 1.001E-07 | 743.33386 | 111.1297 |
| CTSYNIPCTSDMA | 14 | P53992 | Protein transport protein Sec24C OS=Hor | SEC24C | 118.325 | 2 | 55.025925 | yes | 1.381E-07 | 824.34424 | 88.793633 |
| NCETDTLINYMAK | 13 | P53992 | Protein transport protein Sec24C OS=Hor | SEC24C | 118.325 | 2 | 55.025925 | yes | 9.722E-08 | 786.85529 | 87.298752 |

| | | | | | | | | | | | |
|---|---|---|---|---|---|---|---|---|---|---|---|
| GQVPPLVTTNFLV | 14 | P53992 | Protein transport protein Sec24C OS=Hoı | SEC24C | 118.325 | 2 | 55.025925 | yes | 1.456E-07 | 756.94287 | 86.971085 |
| LLPVYLNCVLK | 11 | P53992 | Protein transport protein Sec24C OS=Hoı | SEC24C | 118.325 | 2 | 55.025925 | yes | 9.722E-08 | 666.39142 | 85.992981 |
| SPVESTTEPPAVR | 13 | P53992 | Protein transport protein Sec24C OS=Hoı | SEC24C | 118.325 | 2 | 55.025925 | yes | 9.722E-08 | 685.3515 | 68.280731 |
| LNEESGALLQCAL | 22 | P53992 | Protein transport protein Sec24C OS=Hoı | SEC24C | 118.325 | 3 | 55.025925 | yes | 1.087E-07 | 818.72992 | 51.309101 |
| QAQVPLAAVIKPL | 15 | P53992 | Protein transport protein Sec24C OS=Hoı | SEC24C | 118.325 | 3 | 55.025925 | yes | 5.081E-07 | 525.6629 | 13.995234 |
| EQISDIDDAVR | 11 | P53999 | Activated RNA polymerase II transcriptio | SUB1 | 14.396 | 2 | 51.54738 | yes | 9.722E-08 | 630.80695 | 408.94287 |
| GISLNPEQWSQLK | 13 | P53999 | Activated RNA polymerase II transcriptio | SUB1 | 14.396 | 2 | 51.54738 | yes | 9.722E-08 | 750.3963 | 253.51132 |
| EYWMDPEGEMKP | 14 | P53999 | Activated RNA polymerase II transcriptio | SUB1 | 14.396 | 3 | 51.54738 | yes | 1.106E-07 | 575.58508 | 43.791637 |
| YLAEAQGTYEELQ | 14 | P54098 | DNA polymerase subunit gamma-1 OS=H | POLG | 139.563 | 2 | 42.40342 | yes | 1.3E-06 | 835.90466 | 71.944008 |
| ISSQMVVWLPR | 11 | P54098 | DNA polymerase subunit gamma-1 OS=H | POLG | 139.563 | 2 | 42.40342 | yes | 3.046E-07 | 658.36322 | 43.678982 |
| GLGTGTLYIAESR | 13 | P54105 | Methylosome subunit pICln OS=Homo sa | CLNS1A | 26.216 | 2 | 53.76711 | yes | 9.722E-08 | 669.35663 | 246.61449 |
| SFPPPGPAEGLLR | 13 | P54105 | Methylosome subunit pICln OS=Homo sa | CLNS1A | 26.216 | 2 | 53.76711 | yes | 9.345E-07 | 669.36426 | 154.59384 |
| LISTLENLYER | 11 | P54132 | RecQ-like DNA helicase BLM OS=Homo | BLM | 159.001 | 2 | 55.402374 | yes | 9.934E-08 | 675.86682 | 309.14789 |
| PLFNTHLQK | 9 | P54132 | RecQ-like DNA helicase BLM OS=Homo | BLM | 159.001 | 2 | 55.402374 | yes | 9.722E-08 | 549.30872 | 298.7775 |
| EVVCTTQNTPTVK | 13 | P54132 | RecQ-like DNA helicase BLM OS=Homo | BLM | 159.001 | 2 | 55.402374 | yes | 1.838E-07 | 738.87177 | 253.32626 |
| TSSDNNVSVTNVS | 16 | P54132 | RecQ-like DNA helicase BLM OS=Homo | BLM | 159.001 | 2 | 55.402374 | yes | 1.152E-07 | 811.40503 | 213.41992 |
| LLTEVDFNK | 9 | P54132 | RecQ-like DNA helicase BLM OS=Homo | BLM | 159.001 | 2 | 55.402374 | yes | 1.681E-07 | 539.7926 | 208.27777 |
| FQSLSFPHTK | 10 | P54132 | RecQ-like DNA helicase BLM OS=Homo | BLM | 159.001 | 2 | 55.402374 | yes | 6.985E-07 | 596.31146 | 207.50079 |
| SVEGYYQESGR | 11 | P54132 | RecQ-like DNA helicase BLM OS=Homo | BLM | 159.001 | 2 | 55.402374 | yes | 9.722E-08 | 637.78601 | 199.79721 |
| DLLSKPEK | 8 | P54132 | RecQ-like DNA helicase BLM OS=Homo | BLM | 159.001 | 2 | 55.402374 | yes | 1.938E-07 | 465.26874 | 186.16635 |
| SFVTPPQSHFVR | 12 | P54132 | RecQ-like DNA helicase BLM OS=Homo | BLM | 159.001 | 2 | 55.402374 | yes | 3.194E-06 | 701.36731 | 159.92535 |
| LTSLDIPATYLTGD | 15 | P54132 | RecQ-like DNA helicase BLM OS=Homo | BLM | 159.001 | 2 | 55.402374 | yes | 9.728E-08 | 804.42999 | 156.69232 |
| HTASINDLER | 10 | P54132 | RecQ-like DNA helicase BLM OS=Homo | BLM | 159.001 | 2 | 55.402374 | yes | 3.357E-07 | 578.29126 | 142.0887 |
| VAFDCLEWIR | 10 | P54132 | RecQ-like DNA helicase BLM OS=Homo | BLM | 159.001 | 2 | 55.402374 | yes | 9.722E-08 | 654.82391 | 141.49689 |
| LIDTIPDDK | 9 | P54132 | RecQ-like DNA helicase BLM OS=Homo | BLM | 159.001 | 2 | 55.402374 | yes | 3.471E-06 | 515.27679 | 133.04056 |
| SLIVDQVQK | 9 | P54132 | RecQ-like DNA helicase BLM OS=Homo | BLM | 159.001 | 2 | 55.402374 | yes | 7.838E-07 | 515.3006 | 126.96152 |
| ECDTMADTLQR | 11 | P54132 | RecQ-like DNA helicase BLM OS=Homo | BLM | 159.001 | 2 | 55.402374 | yes | 9.722E-08 | 670.28448 | 117.38219 |
| YGAEVISVLQK | 11 | P54132 | RecQ-like DNA helicase BLM OS=Homo | BLM | 159.001 | 2 | 55.402374 | yes | 9.722E-08 | 603.84009 | 107.27621 |
| VDFMETENSSSVK | 13 | P54132 | RecQ-like DNA helicase BLM OS=Homo | BLM | 159.001 | 2 | 55.402374 | yes | 1.777E-07 | 736.83234 | 100.19379 |
| TDSEATNIYLQLSK | 14 | P54132 | RecQ-like DNA helicase BLM OS=Homo | BLM | 159.001 | 2 | 55.402374 | yes | 9.834E-08 | 791.90161 | 97.228722 |
| FVIHASLPK | 9 | P54132 | RecQ-like DNA helicase BLM OS=Homo | BLM | 159.001 | 2 | 55.402374 | yes | 4.409E-07 | 506.30292 | 94.197357 |
| NLEEAELHSTEK | 12 | P54132 | RecQ-like DNA helicase BLM OS=Homo | BLM | 159.001 | 2 | 55.402374 | yes | 9.722E-08 | 700.33862 | 83.462585 |
| SSSIIGSSSASHTSQ | 23 | P54132 | RecQ-like DNA helicase BLM OS=Homo | BLM | 159.001 | 3 | 55.402374 | yes | 2.303E-07 | 718.00861 | 58.991817 |
| LLDCGNELLQQR | 12 | P54132 | RecQ-like DNA helicase BLM OS=Homo | BLM | 159.001 | 2 | 55.402374 | yes | 2.405E-07 | 729.87213 | 57.48925 |
| FPSVPVMALTATA | 16 | P54132 | RecQ-like DNA helicase BLM OS=Homo | BLM | 159.001 | 2 | 55.402374 | yes | 2.831E-06 | 836.448 | 35.565266 |
| FVQEHSSSQGMR | 12 | P54132 | RecQ-like DNA helicase BLM OS=Homo | BLM | 159.001 | 2 | 55.402374 | yes | 1.745E-07 | 696.81989 | 26.464474 |
| STIIGESISR | 10 | P54136 | Arginine--tRNA ligase, cytoplasmic OS=I | RARS1 | 75.379 | 2 | 55.594933 | yes | 3.314E-07 | 531.79309 | 510.20132 |
| LMDLLGEGLK | 10 | P54136 | Arginine--tRNA ligase, cytoplasmic OS=I | RARS1 | 75.379 | 2 | 55.594933 | yes | 1.001E-07 | 544.80463 | 411.72839 |
| VEIAGPGFINVHLF | 14 | P54136 | Arginine--tRNA ligase, cytoplasmic OS=I | RARS1 | 75.379 | 2 | 55.594933 | yes | 1.673E-07 | 761.43042 | 201.96606 |

| | | | | | | | | | | | |
|---|---|---|---|---|---|---|---|---|---|---|---|
| VEIAGPGFINVHLF | 14 | P54136 | Arginine--tRNA ligase, cytoplasmic OS=l | RARS1 | 75.379 | 2 | 55.594933 | yes | 4.611E-07 | 507.95605 | 201.96606 |
| VEIAGPGFINVHLF | 14 | P54136 | Arginine--tRNA ligase, cytoplasmic OS=l | RARS1 | 75.379 | 3 | 55.594933 | yes | 1.673E-07 | 761.43042 | 201.96606 |
| VEIAGPGFINVHLF | 14 | P54136 | Arginine--tRNA ligase, cytoplasmic OS=l | RARS1 | 75.379 | 3 | 55.594933 | yes | 4.611E-07 | 507.95605 | 201.96606 |
| IVFVPGCSIPLTIVK | 15 | P54136 | Arginine--tRNA ligase, cytoplasmic OS=l | RARS1 | 75.379 | 2 | 55.594933 | yes | 1.055E-07 | 821.98364 | 262.2655 |
| AYQCVVLLQGK | 11 | P54136 | Arginine--tRNA ligase, cytoplasmic OS=l | RARS1 | 75.379 | 2 | 55.594933 | yes | 3.23E-07 | 639.84735 | 249.49146 |
| NMINIISR | 8 | P54136 | Arginine--tRNA ligase, cytoplasmic OS=l | RARS1 | 75.379 | 2 | 55.594933 | yes | 2.554E-07 | 480.7684 | 232.71011 |
| SDGGYTYDTSDLA | 16 | P54136 | Arginine--tRNA ligase, cytoplasmic OS=l | RARS1 | 75.379 | 2 | 55.594933 | yes | 1.122E-07 | 838.88611 | 223.97272 |
| LFEFAGYDVLR | 11 | P54136 | Arginine--tRNA ligase, cytoplasmic OS=l | RARS1 | 75.379 | 2 | 55.594933 | yes | 9.722E-08 | 665.34552 | 203.92108 |
| EMHVGHLR | 8 | P54136 | Arginine--tRNA ligase, cytoplasmic OS=l | RARS1 | 75.379 | 2 | 55.594933 | yes | 1.434E-07 | 489.75055 | 196.70119 |
| LANIDEEMLQK | 11 | P54136 | Arginine--tRNA ligase, cytoplasmic OS=l | RARS1 | 75.379 | 2 | 55.594933 | yes | 1.116E-07 | 652.33173 | 190.78844 |
| ILLDHEK | 7 | P54136 | Arginine--tRNA ligase, cytoplasmic OS=l | RARS1 | 75.379 | 2 | 55.594933 | yes | 9.722E-08 | 434.25037 | 188.69681 |
| LLQQEEEIK | 9 | P54136 | Arginine--tRNA ligase, cytoplasmic OS=l | RARS1 | 75.379 | 2 | 55.594933 | yes | 5.322E-07 | 565.30859 | 188.57074 |
| IYDALDVSLIER | 12 | P54136 | Arginine--tRNA ligase, cytoplasmic OS=l | RARS1 | 75.379 | 2 | 55.594933 | yes | 9.722E-08 | 703.87994 | 173.17973 |
| YADLSHNR | 8 | P54136 | Arginine--tRNA ligase, cytoplasmic OS=l | RARS1 | 75.379 | 2 | 55.594933 | yes | 2.443E-06 | 488.23578 | 151.63293 |
| GFDILGIKPVQR | 12 | P54136 | Arginine--tRNA ligase, cytoplasmic OS=l | RARS1 | 75.379 | 2 | 55.594933 | yes | 9.722E-08 | 671.89569 | 116.8736 |
| MDVLVSECSAR | 11 | P54136 | Arginine--tRNA ligase, cytoplasmic OS=l | RARS1 | 75.379 | 2 | 55.594933 | yes | 9.824E-08 | 654.79974 | 116.86015 |
| GESFYQDR | 8 | P54136 | Arginine--tRNA ligase, cytoplasmic OS=l | RARS1 | 75.379 | 2 | 55.594933 | yes | 9.634E-07 | 501.21979 | 80.284164 |
| LQEVFGHAIK | 10 | P54136 | Arginine--tRNA ligase, cytoplasmic OS=l | RARS1 | 75.379 | 2 | 55.594933 | yes | 9.722E-08 | 571.32184 | 78.92054 |
| LNDYIFSFDK | 10 | P54136 | Arginine--tRNA ligase, cytoplasmic OS=l | RARS1 | 75.379 | 2 | 55.594933 | yes | 3.269E-07 | 631.30859 | 37.222103 |
| MLLCEAVAAVMA | 13 | P54136 | Arginine--tRNA ligase, cytoplasmic OS=l | RARS1 | 75.379 | 2 | 55.594933 | yes | 6.079E-06 | 703.86383 | 34.072845 |
| SDLYFVPLEGSK | 12 | P54259 | Atrophin-1 OS=Homo sapiens OX=9606 | ATN1 | 125.415 | 2 | 42.986412 | yes | 2.782E-07 | 677.84808 | 41.195972 |
| TLDAQFENDER | 11 | P54274 | Telomeric repeat-binding factor 1 OS=Hc | TERF1 | 50.246 | 2 | 54.070305 | yes | 9.722E-08 | 669.30206 | 309.96777 |
| LLMIISQK | 8 | P54274 | Telomeric repeat-binding factor 1 OS=Hc | TERF1 | 50.246 | 2 | 54.070305 | yes | 1.048E-07 | 473.2937 | 239.80592 |
| TIYICQFLTR | 10 | P54274 | Telomeric repeat-binding factor 1 OS=Hc | TERF1 | 50.246 | 2 | 54.070305 | yes | 9.722E-08 | 657.84735 | 160.27611 |
| IFGDPNSHMPFK | 12 | P54274 | Telomeric repeat-binding factor 1 OS=Hc | TERF1 | 50.246 | 2 | 54.070305 | yes | 9.722E-08 | 695.33466 | 131.05386 |
| LHEEIQNLIK | 10 | P54274 | Telomeric repeat-binding factor 1 OS=Hc | TERF1 | 50.246 | 2 | 54.070305 | yes | 1.427E-07 | 618.85095 | 129.06856 |
| SYVNYVLSEK | 10 | P54274 | Telomeric repeat-binding factor 1 OS=Hc | TERF1 | 50.246 | 2 | 54.070305 | yes | 1.446E-07 | 601.30859 | 124.42244 |
| IQAIAVCMENGNF | 14 | P54274 | Telomeric repeat-binding factor 1 OS=Hc | TERF1 | 50.246 | 2 | 54.070305 | yes | 2.45E-07 | 797.88947 | 56.075493 |
| QAWLWEEDK | 9 | P54274 | Telomeric repeat-binding factor 1 OS=Hc | TERF1 | 50.246 | 2 | 54.070305 | yes | 2.84E-06 | 602.78528 | 55.234989 |
| NSAEAIIHGLSSLT. | 19 | P54274 | Telomeric repeat-binding factor 1 OS=Hc | TERF1 | 50.246 | 3 | 54.070305 | yes | 1.697E-07 | 681.02136 | 49.50938 |
| LDELLQSQIEK | 11 | P54277 | PMS1 protein homolog 1 OS=Homo sapi | PMS1 | 105.831 | 2 | 44.923344 | yes | 2.023E-07 | 658.35883 | 53.852562 |
| TEVMLLNPYR | 10 | P54277 | PMS1 protein homolog 1 OS=Homo sapi | PMS1 | 105.831 | 2 | 44.923344 | yes | 2.144E-06 | 618.32629 | 27.499052 |
| TSLIGMFDSDVNK | 13 | P54278 | Mismatch repair endonuclease PMS2 OS= | PMS2 | 95.797 | 2 | 48.625694 | yes | 2.923E-07 | 713.84778 | 139.17441 |
| VVPLDFSMSSLAK | 13 | P54278 | Mismatch repair endonuclease PMS2 OS= | PMS2 | 95.797 | 2 | 48.625694 | yes | 5.078E-07 | 697.37341 | 106.18518 |
| ENIGSVFGQK | 10 | P54278 | Mismatch repair endonuclease PMS2 OS= | PMS2 | 95.797 | 2 | 48.625694 | yes | 7.961E-06 | 539.78003 | 79.04158 |
| VLPQPTNLATPNT | 14 | P54278 | Mismatch repair endonuclease PMS2 OS= | PMS2 | 95.797 | 2 | 48.625694 | yes | 2.812E-06 | 747.41974 | 66.235794 |
| GMLSSSTSGAISDK | 14 | P54278 | Mismatch repair endonuclease PMS2 OS= | PMS2 | 95.797 | 2 | 48.625694 | yes | 2.69E-07 | 670.82178 | 59.991844 |
| SVHQICSGQVVLS | 19 | P54278 | Mismatch repair endonuclease PMS2 OS= | PMS2 | 95.797 | 3 | 48.625694 | yes | 1.122E-07 | 671.69812 | 50.592133 |
| QFFFINR | 7 | P54278 | Mismatch repair endonuclease PMS2 OS= | PMS2 | 95.797 | 2 | 48.625694 | yes | 1.069E-06 | 486.25851 | 41.824921 |

| | | | | | | | | | | | |
|---|---|---|---|---|---|---|---|---|---|---|---|
| EAVSSSHGPSDPTI | 15 | P54278 | Mismatch repair endonuclease PMS2 OS= | PMS2 | 95.797 | 2 | 48.625694 | yes | 2.068E-06 | 771.34497 | 16.071444 |
| SSIAEVQSEIER | 12 | P54284 | Voltage-dependent L-type calcium chann | CACNB3 | 54.532 | 2 | 49.646954 | yes | 4.606E-07 | 674.34119 | 89.71946 |
| PVVLVGPSLK | 10 | P54284 | Voltage-dependent L-type calcium chann | CACNB3 | 54.532 | 2 | 49.646954 | no | 1.014E-07 | 504.82623 | 67.261261 |
| PLYSVTVK | 8 | P54578 | Ubiquitin carboxyl-terminal hydrolase 14 | USP14 | 56.07 | 2 | 38.532917 | yes | 5.154E-07 | 453.76837 | 19.858801 |
| FDVINLAAEK | 10 | P54619 | 5'-AMP-activated protein kinase subunit g | PRKAG1 | 37.58 | 2 | 54.35056 | no | 1.786E-07 | 560.30585 | 121.94462 |
| LVVVDENDVVK | 11 | P54619 | 5'-AMP-activated protein kinase subunit g | PRKAG1 | 37.58 | 2 | 54.35056 | yes | 1.672E-07 | 614.84283 | 95.454758 |
| LVEAEVHR | 8 | P54619 | 5'-AMP-activated protein kinase subunit g | PRKAG1 | 37.58 | 2 | 54.35056 | yes | 3.454E-06 | 476.76416 | 82.842766 |
| VSALPVVDEK | 10 | P54619 | 5'-AMP-activated protein kinase subunit g | PRKAG1 | 37.58 | 2 | 54.35056 | yes | 2.291E-05 | 528.80042 | 63.238232 |
| SALVQIYELEEHK | 13 | P54619 | 5'-AMP-activated protein kinase subunit g | PRKAG1 | 37.58 | 2 | 54.35056 | yes | 9.722E-08 | 779.90924 | 43.471657 |
| DLKPENVLLDAHM | 16 | P54646 | 5'-AMP-activated protein kinase catalytic | PRKAA2 | 62.32 | 3 | 49.53203 | no | 1.022E-07 | 603.3172 | 91.218048 |
| GGVFYIPEYLNR | 12 | P54646 | 5'-AMP-activated protein kinase catalytic | PRKAA2 | 62.32 | 2 | 49.53203 | yes | 1.166E-07 | 714.36951 | 48.197601 |
| PVTALEYTFSR | 11 | P54709 | Sodium/potassium-transporting ATPase s | ATP1B3 | 31.513 | 2 | 45.598236 | yes | 1.801E-07 | 642.33514 | 49.561691 |
| IDIDPEETVK | 10 | P54727 | UV excision repair protein RAD23 homo | RAD23B | 43.172 | 2 | 44.922153 | yes | 2.023E-07 | 579.7981 | 153.72838 |
| EQVIAALR | 8 | P54727 | UV excision repair protein RAD23 homo | RAD23B | 43.172 | 2 | 44.922153 | yes | 7.317E-07 | 450.26907 | 81.422768 |
| AVEYLLMGIPGDR | 13 | P54727 | UV excision repair protein RAD23 homo | RAD23B | 43.172 | 2 | 44.922153 | yes | 3.524E-07 | 717.37646 | 40.396603 |
| MLATLEPEQR | 10 | P54886 | Delta-1-pyrroline-5-carboxylate synthase | ALDH18A | 87.303 | 2 | 52.352444 | yes | 9.722E-08 | 594.30811 | 158.03751 |
| EAVQLVNTR | 9 | P54886 | Delta-1-pyrroline-5-carboxylate synthase | ALDH18A | 87.303 | 2 | 52.352444 | yes | 5.293E-07 | 515.28802 | 96.961075 |
| FASYLTFSPSEVK | 13 | P54886 | Delta-1-pyrroline-5-carboxylate synthase | ALDH18A | 87.303 | 2 | 52.352444 | yes | 1.823E-07 | 738.37445 | 95.976227 |
| HEILLSQSVR | 10 | P54886 | Delta-1-pyrroline-5-carboxylate synthase | ALDH18A | 87.303 | 2 | 52.352444 | yes | 1.001E-07 | 591.33551 | 76.49028 |
| GDECGLALGR | 10 | P54886 | Delta-1-pyrroline-5-carboxylate synthase | ALDH18A | 87.303 | 2 | 52.352444 | yes | 4.003E-07 | 524.24805 | 51.327339 |
| GPVGLEGLLTTK | 12 | P54886 | Delta-1-pyrroline-5-carboxylate synthase | ALDH18A | 87.303 | 2 | 52.352444 | yes | 4.627E-05 | 592.8479 | 11.186328 |
| VLTLSDDLER | 10 | P55010 | Eukaryotic translation initiation factor 5 ( | EIF5 | 49.223 | 2 | 38.953526 | yes | 4.881E-07 | 580.81152 | 143.72462 |
| EQCVLSEEQIR | 11 | P55055 | Oxysterols receptor LXR-beta OS=Homo | NR1H2 | 50.974 | 2 | 32.658085 | yes | 1.451E-06 | 695.83521 | 105.42067 |
| VIVPNMEFR | 9 | P55060 | Exportin-2 OS=Homo sapiens OX=9606 | CSE1L | 110.417 | 2 | 51.148705 | yes | 4.164E-07 | 552.79718 | 286.2211 |
| IIIPEIQK | 8 | P55060 | Exportin-2 OS=Homo sapiens OX=9606 | CSE1L | 110.417 | 2 | 51.148705 | yes | 3.107E-07 | 477.30515 | 258.06412 |
| LLQAFLER | 8 | P55060 | Exportin-2 OS=Homo sapiens OX=9606 | CSE1L | 110.417 | 2 | 51.148705 | yes | 2.649E-07 | 495.29257 | 237.23483 |
| IHLAQSLHK | 9 | P55060 | Exportin-2 OS=Homo sapiens OX=9606 | CSE1L | 110.417 | 2 | 51.148705 | yes | 1.063E-07 | 523.8089 | 152.60458 |
| QLSDAISIIGR | 11 | P55060 | Exportin-2 OS=Homo sapiens OX=9606 | CSE1L | 110.417 | 2 | 51.148705 | yes | 9.722E-08 | 586.83533 | 150.47115 |
| IPGLLGVFQK | 10 | P55060 | Exportin-2 OS=Homo sapiens OX=9606 | CSE1L | 110.417 | 2 | 51.148705 | yes | 2.89E-07 | 536.33167 | 102.22861 |
| GSNTIASAAADK | 12 | P55060 | Exportin-2 OS=Homo sapiens OX=9606 | CSE1L | 110.417 | 2 | 51.148705 | yes | 3.772E-06 | 553.27783 | 100.58859 |
| SQICDNAALYAQK | 13 | P55060 | Exportin-2 OS=Homo sapiens OX=9606 | CSE1L | 110.417 | 2 | 51.148705 | yes | 1.257E-07 | 741.35626 | 91.033203 |
| TLDPDPAIR | 9 | P55060 | Exportin-2 OS=Homo sapiens OX=9606 | CSE1L | 110.417 | 2 | 51.148705 | yes | 4.439E-07 | 499.26929 | 88.433647 |
| SANVNEFPVLK | 11 | P55060 | Exportin-2 OS=Homo sapiens OX=9606 | CSE1L | 110.417 | 2 | 51.148705 | yes | 3.602E-07 | 609.3299 | 76.439697 |
| YDEEFQR | 7 | P55060 | Exportin-2 OS=Homo sapiens OX=9606 | CSE1L | 110.417 | 2 | 51.148705 | yes | 3.556E-07 | 493.71432 | 31.581039 |
| MFGMVLEK | 8 | P55060 | Exportin-2 OS=Homo sapiens OX=9606 | CSE1L | 110.417 | 2 | 51.148705 | yes | 2.8E-07 | 477.74301 | 30.609499 |
| FQSGDFHVINGVL | 14 | P55060 | Exportin-2 OS=Homo sapiens OX=9606 | CSE1L | 110.417 | 2 | 51.148705 | yes | 3.34E-06 | 794.91516 | 11.405482 |
| EMVELPLR | 8 | P55072 | Transitional endoplasmic reticulum ATPa | VCP | 89.322 | 2 | 53.548664 | yes | 3.137E-07 | 493.7706 | 203.54185 |
| EVDIGIPDATGR | 12 | P55072 | Transitional endoplasmic reticulum ATPa | VCP | 89.322 | 2 | 53.548664 | yes | 2.726E-07 | 621.81989 | 190.78413 |
| VINQILTEMDGMS | 15 | P55072 | Transitional endoplasmic reticulum ATPa | VCP | 89.322 | 2 | 53.548664 | yes | 1.653E-07 | 840.42078 | 109.26474 |

| | | | | | | | | | | | |
|---|---|---|---|---|---|---|---|---|---|---|---|
| GVLFYGPPGCGK | 12 | P55072 | Transitional endoplasmic reticulum ATPa | VCP | 89.322 | 2 | 53.548664 | yes | 9.722E-08 | 626.31317 | 95.729347 |
| WALSQSNPSALR | 12 | P55072 | Transitional endoplasmic reticulum ATPa | VCP | 89.322 | 2 | 53.548664 | yes | 1.003E-07 | 665.34912 | 89.091202 |
| MDELQLFR | 8 | P55072 | Transitional endoplasmic reticulum ATPa | VCP | 89.322 | 2 | 53.548664 | yes | 1.174E-07 | 526.26569 | 86.543167 |
| KGDIFLVR | 8 | P55072 | Transitional endoplasmic reticulum ATPa | VCP | 89.322 | 2 | 53.548664 | yes | 1.072E-07 | 474.28726 | 77.606255 |
| LAGESESNLR | 10 | P55072 | Transitional endoplasmic reticulum ATPa | VCP | 89.322 | 2 | 53.548664 | yes | 5.373E-07 | 538.27252 | 68.37162 |
| AIGVKPPR | 8 | P55072 | Transitional endoplasmic reticulum ATPa | VCP | 89.322 | 2 | 53.548664 | yes | 2.346E-07 | 419.26889 | 61.205952 |
| LEILQIHTK | 9 | P55072 | Transitional endoplasmic reticulum ATPa | VCP | 89.322 | 2 | 53.548664 | yes | 1.376E-07 | 547.83203 | 59.190956 |
| GGNIGDGGGAADI | 13 | P55072 | Transitional endoplasmic reticulum ATPa | VCP | 89.322 | 2 | 53.548664 | yes | 8.08E-07 | 558.75507 | 44.542564 |
| LGDVISIQPCPDVK | 14 | P55072 | Transitional endoplasmic reticulum ATPa | VCP | 89.322 | 2 | 53.548664 | yes | 3.171E-07 | 770.90564 | 43.874325 |
| AIANECQANFISIK | 14 | P55072 | Transitional endoplasmic reticulum ATPa | VCP | 89.322 | 2 | 53.548664 | yes | 2.064E-07 | 789.90088 | 36.986008 |
| ETVVEVPQVTWEI | 22 | P55072 | Transitional endoplasmic reticulum ATPa | VCP | 89.322 | 3 | 53.548664 | yes | 4.737E-05 | 833.4306 | 27.673201 |
| LKPVFIR | 7 | P55081 | Microfibrillar-associated protein 1 OS=H | MFAP1 | 51.959 | 2 | 51.55038 | yes | 9.722E-08 | 436.78946 | 392.4758 |
| QPPIQSTAGAVPVI | 14 | P55081 | Microfibrillar-associated protein 1 OS=H | MFAP1 | 51.959 | 2 | 51.55038 | yes | 9.722E-08 | 710.89899 | 172.4286 |
| GAFFMDEDEEVYI | 13 | P55081 | Microfibrillar-associated protein 1 OS=H | MFAP1 | 51.959 | 2 | 51.55038 | yes | 9.722E-08 | 790.33472 | 113.58974 |
| DVFERPSAK | 9 | P55081 | Microfibrillar-associated protein 1 OS=H | MFAP1 | 51.959 | 2 | 51.55038 | yes | 6.509E-07 | 524.77472 | 79.692162 |
| ISEDVEER | 8 | P55081 | Microfibrillar-associated protein 1 OS=H | MFAP1 | 51.959 | 2 | 51.55038 | yes | 1.9E-06 | 488.73273 | 79.430176 |
| AALTGLLHR | 9 | P55084 | Trifunctional enzyme subunit beta, mitocl | HADHB | 51.295 | 2 | 52.812073 | yes | 1.122E-07 | 476.29034 | 451.22684 |
| DLMPHDLAR | 9 | P55084 | Trifunctional enzyme subunit beta, mitocl | HADHB | 51.295 | 2 | 52.812073 | yes | 6.636E-07 | 534.26874 | 433.53989 |
| LAAAFAVSR | 9 | P55084 | Trifunctional enzyme subunit beta, mitocl | HADHB | 51.295 | 2 | 52.812073 | yes | 1.573E-07 | 453.26379 | 343.08466 |
| TPFLLSGTSYK | 11 | P55084 | Trifunctional enzyme subunit beta, mitocl | HADHB | 51.295 | 2 | 52.812073 | yes | 9.722E-08 | 607.32678 | 286.14435 |
| AMDSDWFAENYM | 14 | P55084 | Trifunctional enzyme subunit beta, mitocl | HADHB | 51.295 | 2 | 52.812073 | yes | 9.722E-08 | 846.84271 | 204.29018 |
| DQLLLGPTYATPK | 13 | P55084 | Trifunctional enzyme subunit beta, mitocl | HADHB | 51.295 | 2 | 52.812073 | yes | 9.722E-08 | 708.89032 | 192.39458 |
| LEQDEYALR | 9 | P55084 | Trifunctional enzyme subunit beta, mitocl | HADHB | 51.295 | 2 | 52.812073 | yes | 1.128E-07 | 568.78278 | 136.99942 |
| AQDEGLLSDVVPF | 14 | P55084 | Trifunctional enzyme subunit beta, mitocl | HADHB | 51.295 | 2 | 52.812073 | yes | 9.722E-08 | 759.39594 | 124.62068 |
| EAALGAGFSDK | 11 | P55084 | Trifunctional enzyme subunit beta, mitocl | HADHB | 51.295 | 2 | 52.812073 | yes | 1.874E-06 | 533.26422 | 102.28062 |
| NVVVVDGVR | 9 | P55084 | Trifunctional enzyme subunit beta, mitocl | HADHB | 51.295 | 2 | 52.812073 | yes | 2.685E-07 | 478.77982 | 75.811607 |
| ALAMGYKPK | 9 | P55084 | Trifunctional enzyme subunit beta, mitocl | HADHB | 51.295 | 2 | 52.812073 | yes | 9.733E-08 | 489.7757 | 45.763512 |
| KAQDEGLLSDVVI | 15 | P55084 | Trifunctional enzyme subunit beta, mitocl | HADHB | 51.295 | 2 | 52.812073 | yes | 2.526E-07 | 823.44342 | 37.396305 |
| SIVTNMDK | 8 | P55160 | Nck-associated protein 1-like OS=Homo : | NCKAP1L | 128.154 | 2 | 22.247904 | yes | 0.0001141 | 462.22876 | 70.729431 |
| VTEDLFSSLK | 10 | P55160 | Nck-associated protein 1-like OS=Homo : | NCKAP1L | 128.154 | 2 | 22.247904 | yes | 3.919E-06 | 569.80316 | 15.651498 |
| LAAGDQLLSVDGI | 13 | P55196 | Afadin OS=Homo sapiens OX=9606 GN= | AFDN | 206.805 | 2 | 54.694424 | yes | 9.722E-08 | 657.85425 | 104.97223 |
| TPAAIPATPVAVS( | 17 | P55196 | Afadin OS=Homo sapiens OX=9606 GN= | AFDN | 206.805 | 2 | 54.694424 | yes | 1.399E-07 | 844.98834 | 103.32897 |
| KEPEIITVTLK | 11 | P55196 | Afadin OS=Homo sapiens OX=9606 GN= | AFDN | 206.805 | 2 | 54.694424 | yes | 2.25E-06 | 635.88446 | 37.689682 |
| LIAEYDQR | 8 | P55199 | RNA polymerase II elongation factor ELI | ELL | 68.265 | 2 | 42.358746 | yes | 2.003E-06 | 504.26144 | 81.201218 |
| KAELLLR | 7 | P55199 | RNA polymerase II elongation factor ELI | ELL | 68.265 | 2 | 42.358746 | yes | 1.813E-06 | 421.77655 | 32.987061 |
| SGASAVSGGSGVS | 15 | P55199 | RNA polymerase II elongation factor ELI | ELL | 68.265 | 2 | 42.358746 | yes | 4.625E-07 | 653.82092 | 31.368227 |
| NDFNAEYSEYR | 11 | P55199 | RNA polymerase II elongation factor ELI | ELL | 68.265 | 2 | 42.358746 | yes | 3.069E-07 | 704.29419 | 19.946278 |
| ITVCATDDSYQK | 12 | P55199 | RNA polymerase II elongation factor ELI | ELL | 68.265 | 2 | 42.358746 | yes | 4.695E-07 | 700.82178 | 11.40613 |
| RCLQSPSR | 8 | P55201 | Peregrin OS=Homo sapiens OX=9606 GI | BRPF1 | 137.499 | 2 | 47.257824 | yes | 2.736E-06 | 502.25873 | 173.01353 |

| | | | | | | | | | | | |
|---|---|---|---|---|---|---|---|---|---|---|---|
| MVEVDLHGR | 9 | P55201 | Peregrin OS=Homo sapiens OX=9606 GI | BRPF1 | 137.499 | 2 | 47.257824 | yes | 1.001E-06 | 528.26874 | 107.8851 |
| LGEQMTQEAR | 10 | P55201 | Peregrin OS=Homo sapiens OX=9606 GI | BRPF1 | 137.499 | 2 | 47.257824 | yes | 2.149E-07 | 581.77966 | 94.823479 |
| LQTHLQSQR | 9 | P55201 | Peregrin OS=Homo sapiens OX=9606 GI | BRPF1 | 137.499 | 2 | 47.257824 | yes | 1.367E-07 | 555.80438 | 88.291199 |
| LPEVVYR | 7 | P55201 | Peregrin OS=Homo sapiens OX=9606 GI | BRPF1 | 137.499 | 2 | 47.257824 | yes | 2.647E-06 | 438.2529 | 46.211147 |
| YAVLYQPLFDK | 11 | P55209 | Nucleosome assembly protein 1-like 1 OS | NAP1L1 | 45.375 | 2 | 48.418945 | yes | 1.151E-07 | 678.86353 | 1599.7546 |
| FYEEVHDLER | 10 | P55209 | Nucleosome assembly protein 1-like 1 OS | NAP1L1 | 45.375 | 2 | 48.418945 | no | 1.218E-07 | 668.81201 | 1081.5481 |
| KYAVLYQPLFDK | 12 | P55209 | Nucleosome assembly protein 1-like 1 OS | NAP1L1 | 45.375 | 2 | 48.418945 | yes | 1.105E-06 | 742.91101 | 344.15656 |
| MGFTEVTPVTGAS | 15 | P55265 | Double-stranded RNA-specific adenosine | ADAR | 136.067 | 2 | 53.272297 | yes | 1.48E-07 | 783.40326 | 2129.6482 |
| MGFTEVTPVTGAS | 15 | P55265 | Double-stranded RNA-specific adenosine | ADAR | 136.067 | 2 | 53.272297 | yes | 6.924E-07 | 791.4007 | 2129.6482 |
| SPVTTLLECMHK | 12 | P55265 | Double-stranded RNA-specific adenosine | ADAR | 136.067 | 2 | 53.272297 | yes | 9.161E-07 | 708.35468 | 1815.1125 |
| YLNTNPVGGLLEY | 15 | P55265 | Double-stranded RNA-specific adenosine | ADAR | 136.067 | 2 | 53.272297 | yes | 2.662E-07 | 840.44122 | 1666.1677 |
| VLIGENEK | 8 | P55265 | Double-stranded RNA-specific adenosine | ADAR | 136.067 | 2 | 53.272297 | yes | 1.169E-07 | 451.25311 | 1403.287 |
| HYPVFENPK | 9 | P55265 | Double-stranded RNA-specific adenosine | ADAR | 136.067 | 2 | 53.272297 | yes | 1.139E-07 | 565.7851 | 1316.7804 |
| LVDQSGPPHEPK | 12 | P55265 | Double-stranded RNA-specific adenosine | ADAR | 136.067 | 2 | 53.272297 | yes | 1.443E-07 | 652.33569 | 1232.7126 |
| ILAAIIMK | 8 | P55265 | Double-stranded RNA-specific adenosine | ADAR | 136.067 | 2 | 53.272297 | yes | 9.851E-08 | 436.78552 | 1087.2814 |
| FLYSELMK | 8 | P55265 | Double-stranded RNA-specific adenosine | ADAR | 136.067 | 2 | 53.272297 | yes | 2.471E-07 | 515.76752 | 1050.5829 |
| QGTTPPIWHLTDK | 13 | P55265 | Double-stranded RNA-specific adenosine | ADAR | 136.067 | 2 | 53.272297 | yes | 1.066E-07 | 747.39099 | 1045.5973 |
| AMTILLEEAK | 10 | P55265 | Double-stranded RNA-specific adenosine | ADAR | 136.067 | 2 | 53.272297 | yes | 9.722E-08 | 559.80994 | 962.63104 |
| GVDCLSSHFQELS | 21 | P55265 | Double-stranded RNA-specific adenosine | ADAR | 136.067 | 3 | 53.272297 | yes | 8.57E-07 | 847.05542 | 647.53772 |
| HPFIVNHPK | 9 | P55265 | Double-stranded RNA-specific adenosine | ADAR | 136.067 | 2 | 53.272297 | yes | 1.214E-07 | 544.80365 | 637.73859 |
| KIGELVR | 7 | P55265 | Double-stranded RNA-specific adenosine | ADAR | 136.067 | 2 | 53.272297 | yes | 2.483E-07 | 407.76089 | 586.00464 |
| GQLPEAPVIGK | 11 | P55265 | Double-stranded RNA-specific adenosine | ADAR | 136.067 | 2 | 53.272297 | yes | 1.523E-07 | 554.82166 | 527.2384 |
| LKPPVHYNGPSK | 12 | P55265 | Double-stranded RNA-specific adenosine | ADAR | 136.067 | 2 | 53.272297 | yes | 3.505E-07 | 668.87225 | 259.5293 |
| LKPPVHYNGPSK | 12 | P55265 | Double-stranded RNA-specific adenosine | ADAR | 136.067 | 2 | 53.272297 | yes | 9.722E-08 | 446.25058 | 259.5293 |
| LKPPVHYNGPSK | 12 | P55265 | Double-stranded RNA-specific adenosine | ADAR | 136.067 | 3 | 53.272297 | yes | 3.505E-07 | 668.87225 | 259.5293 |
| LKPPVHYNGPSK | 12 | P55265 | Double-stranded RNA-specific adenosine | ADAR | 136.067 | 3 | 53.272297 | yes | 9.722E-08 | 446.25058 | 259.5293 |
| DINAVLIDMER | 11 | P55265 | Double-stranded RNA-specific adenosine | ADAR | 136.067 | 2 | 53.272297 | yes | 9.893E-08 | 644.83191 | 517.31494 |
| GETVNDCHAEIISF | 14 | P55265 | Double-stranded RNA-specific adenosine | ADAR | 136.067 | 2 | 53.272297 | yes | 9.722E-08 | 800.87286 | 254.61331 |
| GETVNDCHAEIISF | 14 | P55265 | Double-stranded RNA-specific adenosine | ADAR | 136.067 | 2 | 53.272297 | yes | 1.065E-07 | 534.25098 | 254.61331 |
| GETVNDCHAEIISF | 14 | P55265 | Double-stranded RNA-specific adenosine | ADAR | 136.067 | 3 | 53.272297 | yes | 9.722E-08 | 800.87286 | 254.61331 |
| GETVNDCHAEIISF | 14 | P55265 | Double-stranded RNA-specific adenosine | ADAR | 136.067 | 3 | 53.272297 | yes | 1.065E-07 | 534.25098 | 254.61331 |
| QEAADAALR | 9 | P55265 | Double-stranded RNA-specific adenosine | ADAR | 136.067 | 2 | 53.272297 | yes | 1.501E-07 | 472.74344 | 472.12302 |
| KLTECQLK | 8 | P55265 | Double-stranded RNA-specific adenosine | ADAR | 136.067 | 2 | 53.272297 | yes | 2.628E-07 | 510.28134 | 443.08035 |
| DSEDMGVVVSLG | 16 | P55265 | Double-stranded RNA-specific adenosine | ADAR | 136.067 | 2 | 53.272297 | yes | 1.007E-07 | 818.3858 | 381.28381 |
| FLEELGEGK | 9 | P55265 | Double-stranded RNA-specific adenosine | ADAR | 136.067 | 2 | 53.272297 | yes | 1.047E-07 | 511.26367 | 257.3186 |
| YQQPGPGSSPSSFI | 16 | P55265 | Double-stranded RNA-specific adenosine | ADAR | 136.067 | 2 | 53.272297 | yes | 6.765E-07 | 846.93323 | 255.02745 |
| VENGQEPVIK | 10 | P55265 | Double-stranded RNA-specific adenosine | ADAR | 136.067 | 2 | 53.272297 | yes | 9.722E-08 | 556.80096 | 226.04831 |
| TMLLLSR | 7 | P55265 | Double-stranded RNA-specific adenosine | ADAR | 136.067 | 2 | 53.272297 | yes | 1.259E-07 | 417.2493 | 219.00206 |
| KNIFLLFK | 8 | P55265 | Double-stranded RNA-specific adenosine | ADAR | 136.067 | 2 | 53.272297 | yes | 3.246E-07 | 511.82349 | 164.52757 |

| | | | | | | | | | | | |
|---|---|---|---|---|---|---|---|---|---|---|---|
| AIMEMPSFYSHGL | 15 | P55265 | Double-stranded RNA-specific adenosine | ADAR | 136.067 | 3 | 53.272297 | yes | 1.402E-07 | 579.28131 | 154.69919 |
| FQVVINGR | 8 | P55265 | Double-stranded RNA-specific adenosine | ADAR | 136.067 | 2 | 53.272297 | yes | 4.372E-06 | 466.76926 | 131.63774 |
| SVTLGYLFSQGHL | 15 | P55265 | Double-stranded RNA-specific adenosine | ADAR | 136.067 | 3 | 53.272297 | yes | 9.722E-08 | 560.30188 | 65.240921 |
| SVTLGYLFSQGHL | 15 | P55265 | Double-stranded RNA-specific adenosine | ADAR | 136.067 | 3 | 53.272297 | yes | 4.866E-07 | 839.94922 | 65.240921 |
| SVTLGYLFSQGHL | 15 | P55265 | Double-stranded RNA-specific adenosine | ADAR | 136.067 | 2 | 53.272297 | yes | 9.722E-08 | 560.30188 | 65.240921 |
| SVTLGYLFSQGHL | 15 | P55265 | Double-stranded RNA-specific adenosine | ADAR | 136.067 | 2 | 53.272297 | yes | 4.866E-07 | 839.94922 | 65.240921 |
| QMAAEEAMK | 9 | P55265 | Double-stranded RNA-specific adenosine | ADAR | 136.067 | 2 | 53.272297 | yes | 3.84E-06 | 504.72809 | 111.29106 |
| FPVLLASSTR | 10 | P55265 | Double-stranded RNA-specific adenosine | ADAR | 136.067 | 2 | 53.272297 | yes | 2.529E-07 | 545.81641 | 102.19118 |
| KILAAIIMK | 9 | P55265 | Double-stranded RNA-specific adenosine | ADAR | 136.067 | 2 | 53.272297 | yes | 5.185E-07 | 500.83301 | 101.50697 |
| TLPLTGSTFHDQIA | 19 | P55265 | Double-stranded RNA-specific adenosine | ADAR | 136.067 | 3 | 53.272297 | yes | 6.731E-07 | 709.03351 | 82.690575 |
| DMGYGNWISK | 10 | P55265 | Double-stranded RNA-specific adenosine | ADAR | 136.067 | 2 | 53.272297 | yes | 9.722E-08 | 585.76605 | 80.573196 |
| KDSEDMGVVVSL | 17 | P55265 | Double-stranded RNA-specific adenosine | ADAR | 136.067 | 3 | 53.272297 | yes | 1.309E-07 | 588.62457 | 72.634651 |
| FQYCVAVGAQTFI | 20 | P55265 | Double-stranded RNA-specific adenosine | ADAR | 136.067 | 3 | 53.272297 | yes | 3.249E-07 | 715.35425 | 70.079178 |
| EAGTPPLWK | 9 | P55265 | Double-stranded RNA-specific adenosine | ADAR | 136.067 | 2 | 53.272297 | yes | 5.817E-07 | 499.76892 | 48.899475 |
| IQNVVTSFAPQR | 12 | P55268 | Laminin subunit beta-2 OS=Homo sapien | LAMB2 | 195.982 | 2 | 45.671947 | yes | 1.778E-07 | 680.37262 | 65.260361 |
| LGIVQGIVGAR | 11 | P55268 | Laminin subunit beta-2 OS=Homo sapien | LAMB2 | 195.982 | 2 | 45.671947 | yes | 8.253E-07 | 541.83765 | 35.202675 |
| HSLSFNDCFVK | 11 | P55317 | Hepatocyte nuclear factor 3-alpha OS=Hc | FOXA1 | 49.148 | 2 | 48.7974 | yes | 1.103E-07 | 677.31647 | 100.70558 |
| FASGGCDNLIK | 11 | P55735 | Protein SEC13 homolog OS=Homo sapie | SEC13 | 35.541 | 2 | 53.111008 | yes | 1.108E-07 | 591.28461 | 1769.938 |
| LEAHSDWVR | 9 | P55735 | Protein SEC13 homolog OS=Homo sapie | SEC13 | 35.541 | 2 | 53.111008 | yes | 1.302E-07 | 556.77783 | 1586.5786 |
| NGGQILIADLR | 11 | P55735 | Protein SEC13 homolog OS=Homo sapie | SEC13 | 35.541 | 2 | 53.111008 | yes | 9.722E-08 | 585.33551 | 1126.8569 |
| LATCSSDR | 8 | P55735 | Protein SEC13 homolog OS=Homo sapie | SEC13 | 35.541 | 2 | 53.111008 | yes | 1.305E-07 | 455.20837 | 146.47661 |
| DVAWAPSIGLPTS | 23 | P55735 | Protein SEC13 homolog OS=Homo sapie | SEC13 | 35.541 | 3 | 53.111008 | yes | 3.145E-06 | 797.05328 | 113.34267 |
| GHEGPVWQVAWA | 27 | P55735 | Protein SEC13 homolog OS=Homo sapie | SEC13 | 35.541 | 4 | 53.111008 | yes | 9.722E-08 | 776.10907 | 105.22211 |
| EEQKLEAHSDWVI | 13 | P55735 | Protein SEC13 homolog OS=Homo sapie | SEC13 | 35.541 | 3 | 53.111008 | yes | 1.301E-07 | 542.9339 | 92.637787 |
| EENGTWEK | 8 | P55735 | Protein SEC13 homolog OS=Homo sapie | SEC13 | 35.541 | 2 | 53.111008 | yes | 3.846E-07 | 496.71964 | 65.154182 |
| LWKEEEDGQWK | 11 | P55735 | Protein SEC13 homolog OS=Homo sapie | SEC13 | 35.541 | 2 | 53.111008 | yes | 8.026E-07 | 724.34625 | 31.864199 |
| LLDLVQQSCNYK | 12 | P55769 | NHP2-like protein 1 OS=Homo sapiens C | SNU13 | 14.174 | 2 | 48.800583 | yes | 5.513E-07 | 740.87683 | 1299.6177 |
| PVIACSVTIK | 10 | P55769 | NHP2-like protein 1 OS=Homo sapiens C | SNU13 | 14.174 | 2 | 48.800583 | yes | 1.103E-07 | 544.31262 | 1243.2849 |
| AYPLADAHLTK | 11 | P55769 | NHP2-like protein 1 OS=Homo sapiens C | SNU13 | 14.174 | 2 | 48.800583 | yes | 7.817E-07 | 600.32458 | 968.95068 |
| QQIQSIQQSIER | 12 | P55769 | NHP2-like protein 1 OS=Homo sapiens C | SNU13 | 14.174 | 2 | 48.800583 | yes | 1.493E-07 | 729.38898 | 604.59277 |
| NVPYVFVR | 8 | P55769 | NHP2-like protein 1 OS=Homo sapiens C | SNU13 | 14.174 | 2 | 48.800583 | yes | 4.522E-07 | 497.27945 | 47.665829 |
| SPVYLTVLK | 9 | P55786 | Puromycin-sensitive aminopeptidase OS= | NPEPPS | 103.277 | 2 | 32.310406 | yes | 1.556E-06 | 510.31042 | 22.307293 |
| GLPWSCSADEVMI | 13 | P55795 | Heterogeneous nuclear ribonucleoprotein | HNRNPH2 | 49.264 | 2 | 50.91975 | yes | 2.167E-07 | 754.33704 | 8953.2686 |
| GLPWSCSADEVMI | 13 | P55795 | Heterogeneous nuclear ribonucleoprotein | HNRNPH2 | 49.264 | 2 | 50.91975 | yes | 1.405E-07 | 762.33447 | 8953.2686 |
| SNSVEMDWVLK | 11 | P55795 | Heterogeneous nuclear ribonucleoprotein | HNRNPH2 | 49.264 | 2 | 50.91975 | yes | 1.262E-07 | 654.31866 | 6006.3184 |
| SNSVEMDWVLK | 11 | P55795 | Heterogeneous nuclear ribonucleoprotein | HNRNPH2 | 49.264 | 2 | 50.91975 | yes | 9.722E-08 | 662.3161 | 6006.3184 |
| VTGEADVEFATHE | 21 | P55795 | Heterogeneous nuclear ribonucleoprotein | HNRNPH2 | 49.264 | 3 | 50.91975 | yes | 2.26E-07 | 721.34052 | 1606.8313 |
| VTGEADVEFATHE | 21 | P55795 | Heterogeneous nuclear ribonucleoprotein | HNRNPH2 | 49.264 | 3 | 50.91975 | yes | 1.267E-07 | 726.67218 | 1606.8313 |
| EGRPSGEAFVELE | 19 | P55795 | Heterogeneous nuclear ribonucleoprotein | HNRNPH2 | 49.264 | 3 | 50.91975 | yes | 4.599E-07 | 707.67096 | 1784.72 |

| | | | | | | | | | | | |
|---|---|---|---|---|---|---|---|---|---|---|---|
| YGDGGSSFQSTTG | 19 | P55795 | Heterogeneous nuclear ribonucleoprotein | HNRNPH2 | 49.264 | 4 | 50.91975 | yes | 3.468E-06 | 521.72321 | 4.4303365 |
| VNLFTDFDK | 9 | P55884 | Eukaryotic translation initiation factor 3 s | EIF3B | 92.482 | 2 | 53.821922 | yes | 1.248E-07 | 549.77692 | 2216.5134 |
| FAVLHGEAPR | 10 | P55884 | Eukaryotic translation initiation factor 3 s | EIF3B | 92.482 | 2 | 53.821922 | yes | 1.075E-07 | 548.79852 | 1943.9896 |
| YWLEEAECR | 9 | P55884 | Eukaryotic translation initiation factor 3 s | EIF3B | 92.482 | 2 | 53.821922 | yes | 3.571E-07 | 628.27423 | 1884.5265 |
| ITNDFYPEEDGK | 12 | P55884 | Eukaryotic translation initiation factor 3 s | EIF3B | 92.482 | 2 | 53.821922 | yes | 9.819E-08 | 714.31989 | 1833.3228 |
| GTQGVVTNFEIFR | 13 | P55884 | Eukaryotic translation initiation factor 3 s | EIF3B | 92.482 | 2 | 53.821922 | yes | 1.183E-06 | 734.38318 | 1790.7789 |
| MAQELYMEQK | 10 | P55884 | Eukaryotic translation initiation factor 3 s | EIF3B | 92.482 | 2 | 53.821922 | yes | 1.065E-07 | 635.79395 | 1410.5797 |
| YVVTSVSWWSHK | 12 | P55884 | Eukaryotic translation initiation factor 3 s | EIF3B | 92.482 | 2 | 53.821922 | yes | 1.025E-07 | 739.875 | 1264.1638 |
| VTLMQLPTR | 9 | P55884 | Eukaryotic translation initiation factor 3 s | EIF3B | 92.482 | 2 | 53.821922 | yes | 1.47E-07 | 529.80499 | 1205.0173 |
| GTYLATFHQR | 10 | P55884 | Eukaryotic translation initiation factor 3 s | EIF3B | 92.482 | 2 | 53.821922 | yes | 9.722E-08 | 597.3067 | 927.85681 |
| ISVSFYHVK | 9 | P55884 | Eukaryotic translation initiation factor 3 s | EIF3B | 92.482 | 2 | 53.821922 | yes | 1.121E-07 | 540.29785 | 750.48047 |
| YMTISDEWDIPEK | 13 | P55884 | Eukaryotic translation initiation factor 3 s | EIF3B | 92.482 | 2 | 53.821922 | yes | 2.472E-06 | 813.87146 | 748.59857 |
| DQYSVIFESGDR | 12 | P55884 | Eukaryotic translation initiation factor 3 s | EIF3B | 92.482 | 2 | 53.821922 | yes | 9.722E-08 | 708.3255 | 689.99628 |
| QVPVDVVEMK | 10 | P55884 | Eukaryotic translation initiation factor 3 s | EIF3B | 92.482 | 2 | 53.821922 | yes | 9.975E-08 | 572.30756 | 639.1745 |
| NLFNVVDCK | 9 | P55884 | Eukaryotic translation initiation factor 3 s | EIF3B | 92.482 | 2 | 53.821922 | yes | 1.121E-07 | 554.77643 | 602.67627 |
| VDNAYWLWTFQC | 13 | P55884 | Eukaryotic translation initiation factor 3 s | EIF3B | 92.482 | 2 | 53.821922 | yes | 9.722E-08 | 828.40192 | 385.63022 |
| DPVSIEER | 8 | P55884 | Eukaryotic translation initiation factor 3 s | EIF3B | 92.482 | 2 | 53.821922 | yes | 3.494E-07 | 472.73782 | 339.39331 |
| TSIFWNDVK | 9 | P55884 | Eukaryotic translation initiation factor 3 s | EIF3B | 92.482 | 2 | 53.821922 | yes | 2.312E-07 | 555.28491 | 324.45505 |
| WTETYVR | 7 | P55884 | Eukaryotic translation initiation factor 3 s | EIF3B | 92.482 | 2 | 53.821922 | yes | 1.862E-07 | 477.73761 | 226.93283 |
| GFHCESSAHWPIFI | 14 | P55884 | Eukaryotic translation initiation factor 3 s | EIF3B | 92.482 | 3 | 53.821922 | yes | 1.107E-07 | 568.26465 | 218.49707 |
| TMMEDFR | 7 | P55884 | Eukaryotic translation initiation factor 3 s | EIF3B | 92.482 | 2 | 53.821922 | yes | 9.722E-08 | 465.19641 | 195.40099 |
| GYIFLEYASPAHA | 18 | P55884 | Eukaryotic translation initiation factor 3 s | EIF3B | 92.482 | 3 | 53.821922 | yes | 9.722E-08 | 651.00354 | 162.70386 |
| ETIIAFAWEPNGSK | 14 | P55884 | Eukaryotic translation initiation factor 3 s | EIF3B | 92.482 | 2 | 53.821922 | yes | 1.002E-07 | 781.89612 | 159.12952 |
| PRPPTLLSQEQIK | 13 | P55884 | Eukaryotic translation initiation factor 3 s | EIF3B | 92.482 | 2 | 53.821922 | yes | 1.31E-07 | 753.93555 | 73.885345 |
| PRPPTLLSQEQIK | 13 | P55884 | Eukaryotic translation initiation factor 3 s | EIF3B | 92.482 | 2 | 53.821922 | yes | 3.368E-07 | 502.95947 | 73.885345 |
| PRPPTLLSQEQIK | 13 | P55884 | Eukaryotic translation initiation factor 3 s | EIF3B | 92.482 | 3 | 53.821922 | yes | 1.31E-07 | 753.93555 | 73.885345 |
| PRPPTLLSQEQIK | 13 | P55884 | Eukaryotic translation initiation factor 3 s | EIF3B | 92.482 | 3 | 53.821922 | yes | 3.368E-07 | 502.95947 | 73.885345 |
| GIALWGGEK | 9 | P55884 | Eukaryotic translation initiation factor 3 s | EIF3B | 92.482 | 2 | 53.821922 | yes | 8.015E-07 | 465.7558 | 116.9356 |
| TMMEDFRK | 8 | P55884 | Eukaryotic translation initiation factor 3 s | EIF3B | 92.482 | 2 | 53.821922 | yes | 6.808E-07 | 529.2439 | 91.073105 |
| DRFCQLLWR | 9 | P55884 | Eukaryotic translation initiation factor 3 s | EIF3B | 92.482 | 2 | 53.821922 | yes | 6.872E-07 | 647.32971 | 84.482407 |
| DFSPSGIFGAFQR | 13 | P56134 | ATP synthase subunit f, mitochondrial OS | ATP5MF | 10.918 | 2 | 51.15601 | yes | 9.722E-08 | 714.84894 | 644.70264 |
| ASVGECPAPVPVK | 13 | P56134 | ATP synthase subunit f, mitochondrial OS | ATP5MF | 10.918 | 2 | 51.15601 | yes | 1.682E-07 | 676.8476 | 99.897697 |
| LEDPGADSEK | 10 | P56177 | Homeobox protein DLX-1 OS=Homo sap | DLX1 | 27.32 | 2 | 54.732193 | yes | 2.45E-07 | 530.74329 | 305.64865 |
| TIYSSLQLQALNR | 13 | P56177 | Homeobox protein DLX-1 OS=Homo sap | DLX1 | 27.32 | 2 | 54.732193 | yes | 9.722E-08 | 753.91736 | 149.74539 |
| FQQTQYLALPER | 12 | P56177 | Homeobox protein DLX-1 OS=Homo sap | DLX1 | 27.32 | 2 | 54.732193 | no | 1.401E-07 | 747.39099 | 114.85693 |
| STVVEGGEVR | 10 | P56177 | Homeobox protein DLX-1 OS=Homo sap | DLX1 | 27.32 | 2 | 54.732193 | yes | 9.722E-08 | 516.76965 | 93.641342 |
| AELAASLGLTQTQ | 15 | P56177 | Homeobox protein DLX-1 OS=Homo sap | DLX1 | 27.32 | 2 | 54.732193 | no | 1.121E-07 | 765.4303 | 88.80529 |
| TMTTMPESLNSPV | 16 | P56177 | Homeobox protein DLX-1 OS=Homo sap | DLX1 | 27.32 | 2 | 54.732193 | yes | 2.914E-07 | 840.40259 | 30.576263 |
| TQYLALPER | 9 | P56178 | Homeobox protein DLX-5 OS=Homo sap | DLX5 | 31.54 | 2 | 49.98112 | yes | 2.943E-07 | 545.79822 | 449.31244 |

| | | | | | | | | | | | |
|---|---|---|---|---|---|---|---|---|---|---|---|
| TIYSSFQLAALQR | 13 | P56178 | Homeobox protein DLX-5 OS=Homo sap | DLX5 | 31.54 | 2 | 49.98112 | yes | 9.939E-08 | 749.40662 | 183.55626 |
| DSLVLNNITR | 10 | P56182 | Ribosomal RNA processing protein 1 hor | RRP1 | 52.84 | 2 | 55.595093 | yes | 7.073E-07 | 572.81964 | 726.27234 |
| SEKPPAGSICR | 11 | P56182 | Ribosomal RNA processing protein 1 hor | RRP1 | 52.84 | 2 | 55.595093 | yes | 3.064E-06 | 601.30334 | 605.74219 |
| AAGGFTHDELLK | 12 | P56182 | Ribosomal RNA processing protein 1 hor | RRP1 | 52.84 | 2 | 55.595093 | yes | 9.722E-08 | 629.82495 | 438.39569 |
| MVLNESLK | 8 | P56182 | Ribosomal RNA processing protein 1 hor | RRP1 | 52.84 | 2 | 55.595093 | yes | 1.142E-07 | 467.25732 | 413.42856 |
| VQLPPEIQLAQR | 12 | P56182 | Ribosomal RNA processing protein 1 hor | RRP1 | 52.84 | 2 | 55.595093 | yes | 2.145E-07 | 696.40393 | 396.18839 |
| AEAGEQPGTAER | 12 | P56182 | Ribosomal RNA processing protein 1 hor | RRP1 | 52.84 | 2 | 55.595093 | yes | 9.722E-08 | 608.28363 | 180.43568 |
| RGVGADPEAR | 10 | P56182 | Ribosomal RNA processing protein 1 hor | RRP1 | 52.84 | 2 | 55.595093 | yes | 5.037E-07 | 514.26758 | 152.17097 |
| LAGNEQVTR | 9 | P56182 | Ribosomal RNA processing protein 1 hor | RRP1 | 52.84 | 2 | 55.595093 | yes | 1.382E-07 | 494.26453 | 151.00298 |
| FIDPFCR | 7 | P56182 | Ribosomal RNA processing protein 1 hor | RRP1 | 52.84 | 2 | 55.595093 | yes | 5.158E-07 | 477.72873 | 114.11726 |
| MQGWEER | 7 | P56182 | Ribosomal RNA processing protein 1 hor | RRP1 | 52.84 | 2 | 55.595093 | yes | 1.316E-05 | 468.20563 | 63.26553 |
| AEPEAGEEQAGDI | 14 | P56182 | Ribosomal RNA processing protein 1 hor | RRP1 | 52.84 | 2 | 55.595093 | yes | 3.204E-07 | 737.30804 | 47.951206 |
| TKDSLVLNNITR | 12 | P56182 | Ribosomal RNA processing protein 1 hor | RRP1 | 52.84 | 2 | 55.595093 | yes | 2.288E-07 | 687.39099 | 42.82996 |
| LFEMASR | 7 | P56182 | Ribosomal RNA processing protein 1 hor | RRP1 | 52.84 | 2 | 55.595093 | yes | 1.719E-07 | 427.21545 | 27.13673 |
| GFVLQDTVEQLR | 12 | P56192 | Methionine--tRNA ligase, cytoplasmic O! | MARS1 | 101.116 | 2 | 55.544018 | yes | 1.329E-07 | 702.87769 | 248.60365 |
| LENDQIESLR | 10 | P56192 | Methionine--tRNA ligase, cytoplasmic O! | MARS1 | 101.116 | 2 | 55.544018 | yes | 3.404E-07 | 608.81201 | 202.56288 |
| KGEDVLGSVR | 10 | P56192 | Methionine--tRNA ligase, cytoplasmic O! | MARS1 | 101.116 | 2 | 55.544018 | yes | 4.3E-07 | 530.29327 | 100.81409 |
| ALEEGLTPQEICDI | 14 | P56192 | Methionine--tRNA ligase, cytoplasmic O! | MARS1 | 101.116 | 2 | 55.544018 | yes | 9.722E-08 | 801.88763 | 96.15258 |
| QLAVAEGKPPEAP | 14 | P56192 | Methionine--tRNA ligase, cytoplasmic O! | MARS1 | 101.116 | 2 | 55.544018 | yes | 9.722E-08 | 717.90118 | 83.763565 |
| ADKNEVAAEVAK | 12 | P56192 | Methionine--tRNA ligase, cytoplasmic O! | MARS1 | 101.116 | 2 | 55.544018 | yes | 9.722E-08 | 622.8277 | 53.724091 |
| NPEQVDLYQFMAI | 13 | P56192 | Methionine--tRNA ligase, cytoplasmic O! | MARS1 | 101.116 | 2 | 55.544018 | yes | 9.722E-08 | 791.88214 | 48.821075 |
| RLEEWLGR | 8 | P56192 | Methionine--tRNA ligase, cytoplasmic O! | MARS1 | 101.116 | 2 | 55.544018 | yes | 0.00017 | 529.79071 | 38.916103 |
| PAVVETVTTAK | 11 | P56192 | Methionine--tRNA ligase, cytoplasmic O! | MARS1 | 101.116 | 2 | 55.544018 | yes | 1.456E-07 | 558.31897 | 28.382315 |
| YHIIHADIYR | 10 | P56192 | Methionine--tRNA ligase, cytoplasmic O! | MARS1 | 101.116 | 3 | 55.544018 | yes | 1.442E-06 | 434.23141 | 28.276678 |
| AEVLISTVGPEDC\ | 22 | P56192 | Methionine--tRNA ligase, cytoplasmic O! | MARS1 | 101.116 | 3 | 55.544018 | yes | 3.47E-06 | 809.7699 | 20.296089 |
| ELAIQISK | 8 | P56199 | Integrin alpha-1 OS=Homo sapiens OX=! | ITGA1 | 130.848 | 2 | 22.970036 | yes | 3.747E-06 | 451.2713 | 67.751099 |
| GPYICALCAK | 10 | P56270 | Myc-associated zinc finger protein OS=H | MAZ | 48.608 | 2 | 54.179516 | yes | 9.723E-08 | 576.78064 | 220.34134 |
| VPCHVCGK | 8 | P56270 | Myc-associated zinc finger protein OS=H | MAZ | 48.608 | 2 | 54.179516 | yes | 2.287E-07 | 478.72568 | 174.62654 |
| QVHSTERPFK | 10 | P56270 | Myc-associated zinc finger protein OS=H | MAZ | 48.608 | 2 | 54.179516 | yes | 1.28E-07 | 614.82526 | 169.06256 |
| CEKCEAAFATK | 11 | P56270 | Myc-associated zinc finger protein OS=H | MAZ | 48.608 | 2 | 54.179516 | yes | 9.722E-08 | 657.79449 | 121.18005 |
| VHSQGPHHVCELC | 15 | P56270 | Myc-associated zinc finger protein OS=H | MAZ | 48.608 | 4 | 54.179516 | yes | 5.176E-07 | 451.21097 | 78.257278 |
| MLSSAYISDHMK | 12 | P56270 | Myc-associated zinc finger protein OS=H | MAZ | 48.608 | 2 | 54.179516 | yes | 3.662E-06 | 691.82581 | 44.929165 |
| SFSRPDHLNSHVR | 13 | P56270 | Myc-associated zinc finger protein OS=H | MAZ | 48.608 | 2 | 54.179516 | yes | 1.549E-06 | 776.3924 | 42.064243 |
| QAGLSYIR | 8 | P56381 | ATP synthase subunit epsilon, mitochond | ATP5F1E | 5.78 | 2 | 28.157084 | yes | 2.811E-06 | 454.25345 | 179.26093 |
| LTLPALQQR | 9 | P56524 | Histone deacetylase 4 OS=Homo sapiens | HDAC4 | 119.04 | 2 | 32.812233 | yes | 1.406E-06 | 520.31659 | 33.411053 |
| LNEAQPSTIATSMI | 14 | P56537 | Eukaryotic translation initiation factor 6 ( | EIF6 | 26.6 | 2 | 51.62206 | yes | 1.815E-07 | 759.88269 | 592.59662 |
| LNEAQPSTIATSMI | 14 | P56537 | Eukaryotic translation initiation factor 6 ( | EIF6 | 26.6 | 2 | 51.62206 | yes | 4.435E-07 | 767.88013 | 592.59662 |
| NSLPDTVQIR | 10 | P56537 | Eukaryotic translation initiation factor 6 ( | EIF6 | 26.6 | 2 | 51.62206 | yes | 2.72E-07 | 571.81183 | 957.29993 |
| ETEEILADVLK | 11 | P56537 | Eukaryotic translation initiation factor 6 ( | EIF6 | 26.6 | 2 | 51.62206 | yes | 9.722E-08 | 630.34009 | 514.75476 |

| | | | | | | | | | | | |
|---|---|---|---|---|---|---|---|---|---|---|---|
| HGLLVPNNTTDQE | 18 | P56537 | Eukaryotic translation initiation factor 6 ( | EIF6 | 26.6 | 3 | 51.62206 | yes | 1.012E-07 | 695.6994 | 394.21112 |
| ASFENNCEIGCFAI | 14 | P56537 | Eukaryotic translation initiation factor 6 ( | EIF6 | 26.6 | 2 | 51.62206 | yes | 9.722E-08 | 823.85052 | 357.71289 |
| LSALGNVTTCNDY | 23 | P56537 | Eukaryotic translation initiation factor 6 ( | EIF6 | 26.6 | 3 | 51.62206 | yes | 3.351E-07 | 848.42346 | 260.12698 |
| GETLGLIGFGR | 11 | P56545 | C-terminal-binding protein 2 OS=Homo s | CTBP2 | 48.945 | 2 | 56.62551 | yes | 9.722E-08 | 560.31146 | 280.86292 |
| PLVALLDGR | 9 | P56545 | C-terminal-binding protein 2 OS=Homo s | CTBP2 | 48.945 | 2 | 56.62551 | no | 3.147E-07 | 477.29257 | 259.70529 |
| QGAFLVNAAR | 10 | P56545 | C-terminal-binding protein 2 OS=Homo s | CTBP2 | 48.945 | 2 | 56.62551 | yes | 2.25E-07 | 523.79071 | 194.34541 |
| VQSVEQIR | 8 | P56545 | C-terminal-binding protein 2 OS=Homo s | CTBP2 | 48.945 | 2 | 56.62551 | no | 1.832E-07 | 479.76944 | 184.56242 |
| IGSGYDNVDIK | 11 | P56545 | C-terminal-binding protein 2 OS=Homo s | CTBP2 | 48.945 | 2 | 56.62551 | yes | 9.722E-08 | 590.79584 | 106.71864 |
| DCTVEMPILK | 10 | P56545 | C-terminal-binding protein 2 OS=Homo s | CTBP2 | 48.945 | 2 | 56.62551 | no | 9.722E-08 | 603.29889 | 100.43471 |
| NTWLYQALR | 9 | P56545 | C-terminal-binding protein 2 OS=Homo s | CTBP2 | 48.945 | 2 | 56.62551 | yes | 1.781E-07 | 582.81165 | 69.326729 |
| YATDLLSVALNR | 12 | P56937 | 3-keto-steroid reductase/17-beta-hydroxy: | HSD17B7 | 38.207 | 2 | 36.91993 | yes | 6.477E-07 | 668.367 | 44.32972 |
| LVFIGDTLSR | 10 | P56945 | Breast cancer anti-estrogen resistance pro | BCAR1 | 93.372 | 2 | 46.6863 | yes | 3.765E-07 | 560.82166 | 108.32941 |
| SAVGNAAHTSDR | 12 | P56945 | Breast cancer anti-estrogen resistance pro | BCAR1 | 93.372 | 2 | 46.6863 | yes | 1.496E-07 | 593.284 | 32.280312 |
| ALLANALTSALR | 12 | P57088 | Transmembrane protein 33 OS=Homo sa | TMEM33 | 27.979 | 2 | 52.23263 | yes | 9.934E-08 | 607.36676 | 1171.0941 |
| LDTAMWLSR | 9 | P57088 | Transmembrane protein 33 OS=Homo sa | TMEM33 | 27.979 | 2 | 52.23263 | yes | 1.058E-07 | 546.77893 | 631.2793 |
| PACPLFVR | 8 | P57088 | Transmembrane protein 33 OS=Homo sa | TMEM33 | 27.979 | 2 | 52.23263 | yes | 2.441E-07 | 480.26019 | 617.79596 |
| LPHFQLSR | 8 | P57088 | Transmembrane protein 33 OS=Homo sa | TMEM33 | 27.979 | 2 | 52.23263 | yes | 1.079E-07 | 499.28253 | 434.61957 |
| IVVEHIIMK | 9 | P57088 | Transmembrane protein 33 OS=Homo sa | TMEM33 | 27.979 | 2 | 52.23263 | yes | 1.276E-07 | 541.32556 | 384.38513 |
| LSANQQNILK | 10 | P57088 | Transmembrane protein 33 OS=Homo sa | TMEM33 | 27.979 | 2 | 52.23263 | yes | 9.722E-08 | 564.8222 | 379.16168 |
| LCLQSIAFISR | 11 | P57088 | Transmembrane protein 33 OS=Homo sa | TMEM33 | 27.979 | 2 | 52.23263 | yes | 3.244E-07 | 654.36066 | 374.30222 |
| TLFNELR | 7 | P57088 | Transmembrane protein 33 OS=Homo sa | TMEM33 | 27.979 | 2 | 52.23263 | yes | 9.722E-08 | 446.74799 | 275.81619 |
| GSNSLPLLR | 9 | P57088 | Transmembrane protein 33 OS=Homo sa | TMEM33 | 27.979 | 2 | 52.23263 | yes | 1.883E-07 | 478.77982 | 149.00424 |
| IVVEHIIMKPACPL | 17 | P57088 | Transmembrane protein 33 OS=Homo sa | TMEM33 | 27.979 | 3 | 52.23263 | yes | 1.232E-07 | 674.7179 | 55.931316 |
| IVVEHIIMKPACPL | 17 | P57088 | Transmembrane protein 33 OS=Homo sa | TMEM33 | 27.979 | 3 | 52.23263 | yes | 1.982E-07 | 680.0495 | 55.931316 |
| LLETVIDVSTADR | 13 | P57678 | Gem-associated protein 4 OS=Homo sapi | GEMIN4 | 120.038 | 2 | 53.446934 | yes | 1.496E-07 | 716.38812 | 6481.9458 |
| YLPALDEFPHPPK | 13 | P57678 | Gem-associated protein 4 OS=Homo sapi | GEMIN4 | 120.038 | 2 | 53.446934 | yes | 2.455E-06 | 762.39825 | 2829.4346 |
| SFFEGHFK | 8 | P57678 | Gem-associated protein 4 OS=Homo sapi | GEMIN4 | 120.038 | 2 | 53.446934 | yes | 3.154E-07 | 499.74017 | 2731.5688 |
| SIAEGIGPEER | 11 | P57678 | Gem-associated protein 4 OS=Homo sapi | GEMIN4 | 120.038 | 2 | 53.446934 | yes | 9.722E-08 | 579.29346 | 2616.5671 |
| EWGEELQAVLR | 11 | P57678 | Gem-associated protein 4 OS=Homo sapi | GEMIN4 | 120.038 | 2 | 53.446934 | yes | 1.078E-07 | 665.34351 | 2613.376 |
| CEVPATLFEICK | 12 | P57678 | Gem-associated protein 4 OS=Homo sapi | GEMIN4 | 120.038 | 2 | 53.446934 | yes | 4.762E-07 | 733.85455 | 2556.6067 |
| TNPSVSSLLQR | 11 | P57678 | Gem-associated protein 4 OS=Homo sapi | GEMIN4 | 120.038 | 2 | 53.446934 | yes | 1.204E-07 | 601.33038 | 2275.9993 |
| LDVEEVDLSLR | 11 | P57678 | Gem-associated protein 4 OS=Homo sapi | GEMIN4 | 120.038 | 2 | 53.446934 | yes | 3.458E-07 | 644.3432 | 1623.6399 |
| MCSLAVVNLGTHI | 13 | P57678 | Gem-associated protein 4 OS=Homo sapi | GEMIN4 | 120.038 | 2 | 53.446934 | yes | 9.809E-08 | 715.36816 | 1614.1399 |
| QVVSELAECVR | 11 | P57678 | Gem-associated protein 4 OS=Homo sapi | GEMIN4 | 120.038 | 2 | 53.446934 | yes | 1.066E-07 | 645.32953 | 1520.7245 |
| LLCGSLTR | 8 | P57678 | Gem-associated protein 4 OS=Homo sapi | GEMIN4 | 120.038 | 2 | 53.446934 | yes | 2.362E-07 | 460.25513 | 1280.2217 |
| IFIQTLEANACR | 12 | P57678 | Gem-associated protein 4 OS=Homo sapi | GEMIN4 | 120.038 | 2 | 53.446934 | yes | 2.646E-07 | 718.37177 | 1093.9475 |
| GLTQIQSR | 8 | P57678 | Gem-associated protein 4 OS=Homo sapi | GEMIN4 | 120.038 | 2 | 53.446934 | yes | 4.752E-07 | 451.75635 | 858.64642 |
| GHPQDPLLSQFSAI | 17 | P57678 | Gem-associated protein 4 OS=Homo sapi | GEMIN4 | 120.038 | 3 | 53.446934 | yes | 2.436E-06 | 621.97717 | 426.05609 |
| GHPQDPLLSQFSAI | 17 | P57678 | Gem-associated protein 4 OS=Homo sapi | GEMIN4 | 120.038 | 3 | 53.446934 | yes | 2.907E-07 | 627.30884 | 426.05609 |

| | | | | | | | | | | | |
|---|---|---|---|---|---|---|---|---|---|---|---|
| KLEQLDWTVGLR | 12 | P57678 | Gem-associated protein 4 OS=Homo sapi | GEMIN4 | 120.038 | 2 | 53.446934 | yes | 1.125E-07 | 729.40918 | 716.23383 |
| LVIVHPEVTVK | 11 | P57678 | Gem-associated protein 4 OS=Homo sapi | GEMIN4 | 120.038 | 2 | 53.446934 | yes | 1.825E-07 | 617.3819 | 319.65894 |
| LVIVHPEVTVK | 11 | P57678 | Gem-associated protein 4 OS=Homo sapi | GEMIN4 | 120.038 | 2 | 53.446934 | yes | 2.492E-07 | 411.92371 | 319.65894 |
| LVIVHPEVTVK | 11 | P57678 | Gem-associated protein 4 OS=Homo sapi | GEMIN4 | 120.038 | 3 | 53.446934 | yes | 1.825E-07 | 617.3819 | 319.65894 |
| LVIVHPEVTVK | 11 | P57678 | Gem-associated protein 4 OS=Homo sapi | GEMIN4 | 120.038 | 3 | 53.446934 | yes | 2.492E-07 | 411.92371 | 319.65894 |
| TFQFLCSHSCR | 11 | P57678 | Gem-associated protein 4 OS=Homo sapi | GEMIN4 | 120.038 | 2 | 53.446934 | yes | 9.722E-08 | 721.81885 | 637.92072 |
| VLQPHPVTPSDTE | 15 | P57678 | Gem-associated protein 4 OS=Homo sapi | GEMIN4 | 120.038 | 2 | 53.446934 | yes | 8.337E-07 | 838.93378 | 605.03052 |
| HMEVCYIFASEK | 12 | P57678 | Gem-associated protein 4 OS=Homo sapi | GEMIN4 | 120.038 | 2 | 53.446934 | yes | 2.935E-07 | 757.34436 | 478.05167 |
| EISSAAAHSQPFAV | 15 | P57678 | Gem-associated protein 4 OS=Homo sapi | GEMIN4 | 120.038 | 2 | 53.446934 | yes | 1.046E-07 | 815.4046 | 229.61938 |
| EISSAAAHSQPFAV | 15 | P57678 | Gem-associated protein 4 OS=Homo sapi | GEMIN4 | 120.038 | 2 | 53.446934 | yes | 2.717E-07 | 543.93884 | 229.61938 |
| EISSAAAHSQPFAV | 15 | P57678 | Gem-associated protein 4 OS=Homo sapi | GEMIN4 | 120.038 | 3 | 53.446934 | yes | 1.046E-07 | 815.4046 | 229.61938 |
| EISSAAAHSQPFAV | 15 | P57678 | Gem-associated protein 4 OS=Homo sapi | GEMIN4 | 120.038 | 3 | 53.446934 | yes | 2.717E-07 | 543.93884 | 229.61938 |
| LEQLDWTVGLR | 11 | P57678 | Gem-associated protein 4 OS=Homo sapi | GEMIN4 | 120.038 | 2 | 53.446934 | yes | 1.141E-07 | 665.36169 | 370.9086 |
| SSQGTSYDSYR | 11 | P57678 | Gem-associated protein 4 OS=Homo sapi | GEMIN4 | 120.038 | 2 | 53.446934 | yes | 9.722E-08 | 625.76782 | 327.50381 |
| DVSLTSLAK | 9 | P57678 | Gem-associated protein 4 OS=Homo sapi | GEMIN4 | 120.038 | 2 | 53.446934 | yes | 9.722E-08 | 467.2662 | 286.8674 |
| LPSETIFVGCEFLH | 18 | P57678 | Gem-associated protein 4 OS=Homo sapi | GEMIN4 | 120.038 | 3 | 53.446934 | yes | 1.004E-07 | 723.3822 | 171.12943 |
| AIPESQIR | 8 | P57678 | Gem-associated protein 4 OS=Homo sapi | GEMIN4 | 120.038 | 2 | 53.446934 | yes | 1.325E-07 | 457.25873 | 133.06781 |
| VGRPIVEALR | 10 | P57678 | Gem-associated protein 4 OS=Homo sapi | GEMIN4 | 120.038 | 2 | 53.446934 | yes | 1.389E-07 | 555.34314 | 64.456696 |
| KMCSLAVVNLGTI | 14 | P57678 | Gem-associated protein 4 OS=Homo sapi | GEMIN4 | 120.038 | 2 | 53.446934 | yes | 1.079E-07 | 779.41565 | 54.594055 |
| ALIIIWAK | 8 | P57678 | Gem-associated protein 4 OS=Homo sapi | GEMIN4 | 120.038 | 2 | 53.446934 | yes | 1.009E-07 | 464.30493 | 28.716507 |
| KALIIIWAK | 9 | P57678 | Gem-associated protein 4 OS=Homo sapi | GEMIN4 | 120.038 | 2 | 53.446934 | yes | 1.385E-06 | 528.35242 | 27.492867 |
| YGVIYSTPLPEK | 12 | P57682 | Krueppel-like factor 3 OS=Homo sapiens | KLF3 | 38.829 | 2 | 48.615196 | yes | 1.122E-07 | 683.86627 | 54.583584 |
| QICVVMLESPPK | 12 | P57721 | Poly(rC)-binding protein 3 OS=Homo sap | PCBP3 | 39.466 | 2 | 41.26371 | yes | 3.596E-07 | 700.86743 | 4037.5339 |
| TVVEALFQR | 9 | P57740 | Nuclear pore complex protein Nup107 OS | NUP107 | 106.375 | 2 | 51.118023 | yes | 9.722E-08 | 531.80072 | 174.57915 |
| THQMVLLR | 8 | P57740 | Nuclear pore complex protein Nup107 OS | NUP107 | 106.375 | 2 | 51.118023 | yes | 1.043E-07 | 499.28421 | 165.37061 |
| SGFGEISSPVIR | 12 | P57740 | Nuclear pore complex protein Nup107 OS | NUP107 | 106.375 | 2 | 51.118023 | yes | 9.819E-08 | 624.83276 | 141.85425 |
| MAEDELFNR | 9 | P57740 | Nuclear pore complex protein Nup107 OS | NUP107 | 106.375 | 2 | 51.118023 | yes | 1.101E-07 | 562.75568 | 84.189461 |
| GHLDALTADVK | 11 | P57740 | Nuclear pore complex protein Nup107 OS | NUP107 | 106.375 | 2 | 51.118023 | yes | 2.555E-07 | 570.3064 | 54.019207 |
| AIYAALSGNLK | 11 | P57740 | Nuclear pore complex protein Nup107 OS | NUP107 | 106.375 | 2 | 51.118023 | yes | 8.431E-06 | 560.82166 | 32.683178 |
| KYEMDFGIWK | 10 | P57740 | Nuclear pore complex protein Nup107 OS | NUP107 | 106.375 | 2 | 51.118023 | yes | 4.159E-07 | 658.8208 | 25.886379 |
| AMDDYSVIGR | 10 | P57772 | Selenocysteine-specific elongation factor | EEFSEC | 65.305 | 2 | 50.53275 | yes | 9.787E-08 | 563.76349 | 236.2606 |
| DLTPAVTDNDEAI | 15 | P57772 | Selenocysteine-specific elongation factor | EEFSEC | 65.305 | 2 | 50.53275 | yes | 4.3E-07 | 816.3916 | 186.55919 |
| IHIPGGLSPESK | 12 | P57772 | Selenocysteine-specific elongation factor | EEFSEC | 65.305 | 2 | 50.53275 | yes | 1.624E-07 | 617.84314 | 140.69296 |
| LGICVTQFDPK | 11 | P57772 | Selenocysteine-specific elongation factor | EEFSEC | 65.305 | 2 | 50.53275 | yes | 4.563E-07 | 639.33154 | 105.19908 |
| YVFDTHK | 7 | P57772 | Selenocysteine-specific elongation factor | EEFSEC | 65.305 | 2 | 50.53275 | yes | 7.805E-07 | 455.2269 | 102.4199 |
| ETNIQLFVGLK | 11 | P57772 | Selenocysteine-specific elongation factor | EEFSEC | 65.305 | 2 | 50.53275 | yes | 1.197E-07 | 631.36115 | 80.987778 |
| LAFHGILLHGLEDI | 14 | P57772 | Selenocysteine-specific elongation factor | EEFSEC | 65.305 | 3 | 50.53275 | yes | 9.978E-08 | 530.9632 | 80.23436 |
| FHITVGHETVMGR | 13 | P57772 | Selenocysteine-specific elongation factor | EEFSEC | 65.305 | 3 | 50.53275 | yes | 1.188E-07 | 495.254 | 74.83152 |
| GAPIIPVAAK | 10 | P57772 | Selenocysteine-specific elongation factor | EEFSEC | 65.305 | 2 | 50.53275 | yes | 1.401E-07 | 468.79749 | 42.401062 |

| | | | | | | | | | | | |
|---|---|---|---|---|---|---|---|---|---|---|---|
| QQWALVEFEKPV1 | 16 | P57772 | Selenocysteine-specific elongation factor | EEFSEC | 65.305 | 3 | 50.53275 | yes | 5.016E-06 | 663.34015 | 30.983791 |
| RPFRPPPAHFQPGI | 14 | P58012 | Forkhead box protein L2 OS=Homo sapie | FOXL2 | 38.773 | 3 | 47.21188 | yes | 1.376E-07 | 544.63544 | 120.30508 |
| RPFRPPPAHFQPGI | 14 | P58012 | Forkhead box protein L2 OS=Homo sapie | FOXL2 | 38.773 | 3 | 47.21188 | yes | 1.444E-07 | 408.72839 | 120.30508 |
| RPFRPPPAHFQPGI | 14 | P58012 | Forkhead box protein L2 OS=Homo sapie | FOXL2 | 38.773 | 4 | 47.21188 | yes | 1.376E-07 | 544.63544 | 120.30508 |
| RPFRPPPAHFQPGI | 14 | P58012 | Forkhead box protein L2 OS=Homo sapie | FOXL2 | 38.773 | 4 | 47.21188 | yes | 1.444E-07 | 408.72839 | 120.30508 |
| GLAGPAASYGPYT | 14 | P58012 | Forkhead box protein L2 OS=Homo sapie | FOXL2 | 38.773 | 2 | 47.21188 | yes | 6.139E-07 | 690.84894 | 77.54689 |
| GLFGAGGAAGGC( | 29 | P58012 | Forkhead box protein L2 OS=Homo sapie | FOXL2 | 38.773 | 3 | 47.21188 | yes | 1.612E-07 | 847.07263 | 52.742115 |
| TLQEVTEMDSVK | 12 | P58107 | Epiplakin OS=Homo sapiens OX=9606 C | EPPK1 | 555.659 | 2 | 34.53997 | yes | 4.219E-06 | 690.33978 | 41.134636 |
| GFFDPNTHENLTY | 18 | P58107 | Epiplakin OS=Homo sapiens OX=9606 C | EPPK1 | 555.659 | 3 | 34.53997 | yes | 9.884E-07 | 722.03223 | 19.475811 |
| ELLLQPVTISR | 11 | P59998 | Actin-related protein 2/3 complex subunit | ARPC4 | 19.668 | 2 | 43.810303 | yes | 5.055E-07 | 634.88226 | 90.063416 |
| IVAEEFLK | 8 | P59998 | Actin-related protein 2/3 complex subunit | ARPC4 | 19.668 | 2 | 43.810303 | yes | 1.527E-06 | 474.77368 | 74.135483 |
| AENFFILR | 8 | P59998 | Actin-related protein 2/3 complex subunit | ARPC4 | 19.668 | 2 | 43.810303 | yes | 2.438E-07 | 505.27692 | 35.91378 |
| LIHIPINNIIVGG | 13 | P60059 | Protein transport protein Sec61 subunit ga | SEC61G | 7.742 | 2 | 47.84983 | yes | 1.247E-07 | 686.91919 | 74.33139 |
| TATPQQAQEVHEK | 13 | P60174 | Triosephosphate isomerase OS=Homo sa | TPI1 | 26.67 | 2 | 32.354652 | yes | 1.543E-06 | 733.86536 | 40.052074 |
| NLYSDDIPHALR | 12 | P60228 | Eukaryotic translation initiation factor 3 s | EIF3E | 52.221 | 2 | 50.949917 | yes | 1.006E-07 | 707.35968 | 1785.7843 |
| NLYSDDIPHALR | 12 | P60228 | Eukaryotic translation initiation factor 3 s | EIF3E | 52.221 | 2 | 50.949917 | yes | 2.107E-07 | 471.90887 | 1785.7843 |
| NLYSDDIPHALR | 12 | P60228 | Eukaryotic translation initiation factor 3 s | EIF3E | 52.221 | 3 | 50.949917 | yes | 1.006E-07 | 707.35968 | 1785.7843 |
| NLYSDDIPHALR | 12 | P60228 | Eukaryotic translation initiation factor 3 s | EIF3E | 52.221 | 3 | 50.949917 | yes | 2.107E-07 | 471.90887 | 1785.7843 |
| MFEDPETTR | 9 | P60228 | Eukaryotic translation initiation factor 3 s | EIF3E | 52.221 | 2 | 50.949917 | yes | 1.747E-07 | 563.24768 | 3281.5769 |
| YLTTAVITNK | 10 | P60228 | Eukaryotic translation initiation factor 3 s | EIF3E | 52.221 | 2 | 50.949917 | yes | 9.926E-08 | 562.32153 | 2133.8684 |
| QLQAETEPIVK | 11 | P60228 | Eukaryotic translation initiation factor 3 s | EIF3E | 52.221 | 2 | 50.949917 | yes | 9.761E-08 | 628.34827 | 1990.0906 |
| MLFDYLADK | 9 | P60228 | Eukaryotic translation initiation factor 3 s | EIF3E | 52.221 | 2 | 50.949917 | yes | 2.058E-07 | 558.2757 | 1653.8245 |
| LNMTPEEAER | 10 | P60228 | Eukaryotic translation initiation factor 3 s | EIF3E | 52.221 | 2 | 50.949917 | yes | 9.722E-08 | 595.27954 | 974.34686 |
| IAHFLDR | 7 | P60228 | Eukaryotic translation initiation factor 3 s | EIF3E | 52.221 | 2 | 50.949917 | yes | 5.252E-07 | 436.24286 | 920.30109 |
| LFIFETFCR | 9 | P60228 | Eukaryotic translation initiation factor 3 s | EIF3E | 52.221 | 2 | 50.949917 | yes | 9.781E-08 | 616.81024 | 891.1217 |
| QEYLDTLYR | 9 | P60228 | Eukaryotic translation initiation factor 3 s | EIF3E | 52.221 | 2 | 50.949917 | yes | 9.722E-08 | 600.7984 | 851.52814 |
| SQMLAMNIEK | 10 | P60228 | Eukaryotic translation initiation factor 3 s | EIF3E | 52.221 | 2 | 50.949917 | yes | 2.02E-07 | 582.7912 | 661.57642 |
| VLVPATDR | 8 | P60228 | Eukaryotic translation initiation factor 3 s | EIF3E | 52.221 | 2 | 50.949917 | yes | 1.551E-07 | 435.7558 | 649.68799 |
| VIQQESYTYK | 10 | P60228 | Eukaryotic translation initiation factor 3 s | EIF3E | 52.221 | 2 | 50.949917 | yes | 9.823E-08 | 629.81934 | 541.9519 |
| WIVNLIR | 7 | P60228 | Eukaryotic translation initiation factor 3 s | EIF3E | 52.221 | 2 | 50.949917 | yes | 2.117E-07 | 457.28455 | 330.71118 |
| RTTVVAQLK | 9 | P60228 | Eukaryotic translation initiation factor 3 s | EIF3E | 52.221 | 2 | 50.949917 | yes | 1.416E-07 | 508.31656 | 212.04955 |
| TTSAGTGGMWR | 11 | P60468 | Protein transport protein Sec61 subunit be | SEC61B | 9.975 | 2 | 51.3959 | yes | 9.722E-08 | 562.76129 | 54.549507 |
| DFIIFEAAPQETR | 13 | P60510 | Serine/threonine-protein phosphatase 4 ca | PPP4C | 35.08 | 2 | 45.4653 | yes | 4.355E-07 | 768.88831 | 81.451126 |
| EILVEESNVQR | 11 | P60510 | Serine/threonine-protein phosphatase 4 ca | PPP4C | 35.08 | 2 | 45.4653 | yes | 1.841E-07 | 658.34625 | 60.355751 |
| HVLVTLGEK | 9 | P60660 | Myosin light polypeptide 6 OS=Homo sa | MYL6 | 16.931 | 2 | 51.63636 | yes | 1.455E-07 | 498.29785 | 6862.8325 |
| CDFTEDQTAEFK | 12 | P60660 | Myosin light polypeptide 6 OS=Homo sa | MYL6 | 16.931 | 2 | 51.63636 | yes | 9.722E-08 | 766.81415 | 4650.3833 |
| DQGTYEDYVEGLI | 13 | P60660 | Myosin light polypeptide 6 OS=Homo sa | MYL6 | 16.931 | 2 | 51.63636 | yes | 9.722E-08 | 772.8468 | 1757.0011 |
| AGFAGDDAPR | 10 | P60709 | Actin, cytoplasmic 1 OS=Homo sapiens ( | ACTB | 41.737 | 2 | 51.744602 | no | 1.423E-07 | 488.72778 | 34801.453 |
| GYSFTTTAER | 10 | P60709 | Actin, cytoplasmic 1 OS=Homo sapiens ( | ACTB | 41.737 | 2 | 51.744602 | yes | 1.074E-07 | 566.76709 | 30149.836 |

| | | | | | | | | | | | |
|---|---|---|---|---|---|---|---|---|---|---|---|
| EITALAPSTMK | 11 | P60709 | Actin, cytoplasmic 1 OS=Homo sapiens ( | ACTB | 41.737 | 2 | 51.744602 | no | 1.078E-07 | 581.31281 | 7321.7861 |
| EITALAPSTMK | 11 | P60709 | Actin, cytoplasmic 1 OS=Homo sapiens ( | ACTB | 41.737 | 2 | 51.744602 | no | 6.771E-07 | 589.3103 | 7321.7861 |
| DSYVGDEAQSK | 11 | P60709 | Actin, cytoplasmic 1 OS=Homo sapiens ( | ACTB | 41.737 | 2 | 51.744602 | no | 9.722E-08 | 599.76477 | 12465.204 |
| HQGVMVGMGQK | 11 | P60709 | Actin, cytoplasmic 1 OS=Homo sapiens ( | ACTB | 41.737 | 2 | 51.744602 | no | 1.022E-07 | 586.28918 | 3274.3838 |
| HQGVMVGMGQK | 11 | P60709 | Actin, cytoplasmic 1 OS=Homo sapiens ( | ACTB | 41.737 | 2 | 51.744602 | no | 2.942E-07 | 594.28662 | 3274.3838 |
| HQGVMVGMGQK | 11 | P60709 | Actin, cytoplasmic 1 OS=Homo sapiens ( | ACTB | 41.737 | 2 | 51.744602 | no | 9.722E-08 | 594.28662 | 3274.3838 |
| HQGVMVGMGQK | 11 | P60709 | Actin, cytoplasmic 1 OS=Homo sapiens ( | ACTB | 41.737 | 2 | 51.744602 | no | 5.056E-07 | 602.28406 | 3274.3838 |
| IWHHTFYNELR | 11 | P60709 | Actin, cytoplasmic 1 OS=Homo sapiens ( | ACTB | 41.737 | 3 | 51.744602 | no | 2.361E-07 | 505.92123 | 3382.3943 |
| IWHHTFYNELR | 11 | P60709 | Actin, cytoplasmic 1 OS=Homo sapiens ( | ACTB | 41.737 | 3 | 51.744602 | no | 9.722E-08 | 758.37823 | 3382.3943 |
| IWHHTFYNELR | 11 | P60709 | Actin, cytoplasmic 1 OS=Homo sapiens ( | ACTB | 41.737 | 2 | 51.744602 | no | 2.361E-07 | 505.92123 | 3382.3943 |
| IWHHTFYNELR | 11 | P60709 | Actin, cytoplasmic 1 OS=Homo sapiens ( | ACTB | 41.737 | 2 | 51.744602 | no | 9.722E-08 | 758.37823 | 3382.3943 |
| QEYDESGPSIVHR | 13 | P60709 | Actin, cytoplasmic 1 OS=Homo sapiens ( | ACTB | 41.737 | 2 | 51.744602 | no | 3.846E-07 | 758.85498 | 2235.697 |
| QEYDESGPSIVHR | 13 | P60709 | Actin, cytoplasmic 1 OS=Homo sapiens ( | ACTB | 41.737 | 2 | 51.744602 | no | 2.135E-06 | 506.23907 | 2235.697 |
| QEYDESGPSIVHR | 13 | P60709 | Actin, cytoplasmic 1 OS=Homo sapiens ( | ACTB | 41.737 | 3 | 51.744602 | no | 3.846E-07 | 758.85498 | 2235.697 |
| QEYDESGPSIVHR | 13 | P60709 | Actin, cytoplasmic 1 OS=Homo sapiens ( | ACTB | 41.737 | 3 | 51.744602 | no | 2.135E-06 | 506.23907 | 2235.697 |
| VAPEEHPVLLTEA | 18 | P60709 | Actin, cytoplasmic 1 OS=Homo sapiens ( | ACTB | 41.737 | 3 | 51.744602 | yes | 2.685E-06 | 652.02631 | 3856.9841 |
| DLTDYLMK | 8 | P60709 | Actin, cytoplasmic 1 OS=Homo sapiens ( | ACTB | 41.737 | 2 | 51.744602 | no | 2.212E-07 | 499.7468 | 981.37476 |
| DLTDYLMK | 8 | P60709 | Actin, cytoplasmic 1 OS=Homo sapiens ( | ACTB | 41.737 | 2 | 51.744602 | no | 5.35E-07 | 507.74426 | 981.37476 |
| TTGIVMDSGDGVT | 30 | P60709 | Actin, cytoplasmic 1 OS=Homo sapiens ( | ACTB | 41.737 | 4 | 51.744602 | yes | 1.39E-07 | 796.65906 | 742.08325 |
| TTGIVMDSGDGVT | 30 | P60709 | Actin, cytoplasmic 1 OS=Homo sapiens ( | ACTB | 41.737 | 4 | 51.744602 | yes | 1.726E-07 | 800.65778 | 742.08325 |
| AVFPSIVGR | 9 | P60709 | Actin, cytoplasmic 1 OS=Homo sapiens ( | ACTB | 41.737 | 2 | 51.744602 | no | 3.405E-07 | 473.27945 | 575.80261 |
| KDLYANTVLSGGT | 22 | P60709 | Actin, cytoplasmic 1 OS=Homo sapiens ( | ACTB | 41.737 | 3 | 51.744602 | yes | 2.621E-07 | 781.7265 | 349.06174 |
| AVFPSIVGRPR | 11 | P60709 | Actin, cytoplasmic 1 OS=Homo sapiens ( | ACTB | 41.737 | 2 | 51.744602 | no | 1.963E-07 | 599.85638 | 35.161751 |
| DLYANTVLSGGTT | 21 | P60709 | Actin, cytoplasmic 1 OS=Homo sapiens ( | ACTB | 41.737 | 3 | 51.744602 | yes | 1.628E-06 | 739.0282 | 22.851788 |
| LQMEAPHIIVGTP( | 15 | P60842 | Eukaryotic initiation factor 4A-I OS=Hon | EIF4A1 | 46.154 | 3 | 51.687397 | yes | 1.284E-07 | 540.29596 | 1079.7195 |
| LQMEAPHIIVGTP( | 15 | P60842 | Eukaryotic initiation factor 4A-I OS=Hon | EIF4A1 | 46.154 | 3 | 51.687397 | yes | 1.483E-07 | 809.94031 | 1079.7195 |
| LQMEAPHIIVGTP( | 15 | P60842 | Eukaryotic initiation factor 4A-I OS=Hon | EIF4A1 | 46.154 | 3 | 51.687397 | yes | 3.285E-07 | 545.62762 | 1079.7195 |
| LQMEAPHIIVGTP( | 15 | P60842 | Eukaryotic initiation factor 4A-I OS=Hon | EIF4A1 | 46.154 | 3 | 51.687397 | yes | 3.088E-06 | 817.93781 | 1079.7195 |
| LQMEAPHIIVGTP( | 15 | P60842 | Eukaryotic initiation factor 4A-I OS=Hon | EIF4A1 | 46.154 | 2 | 51.687397 | yes | 1.284E-07 | 540.29596 | 1079.7195 |
| LQMEAPHIIVGTP( | 15 | P60842 | Eukaryotic initiation factor 4A-I OS=Hon | EIF4A1 | 46.154 | 2 | 51.687397 | yes | 1.483E-07 | 809.94031 | 1079.7195 |
| LQMEAPHIIVGTP( | 15 | P60842 | Eukaryotic initiation factor 4A-I OS=Hon | EIF4A1 | 46.154 | 2 | 51.687397 | yes | 3.285E-07 | 545.62762 | 1079.7195 |
| LQMEAPHIIVGTP( | 15 | P60842 | Eukaryotic initiation factor 4A-I OS=Hon | EIF4A1 | 46.154 | 2 | 51.687397 | yes | 3.088E-06 | 817.93781 | 1079.7195 |
| DQIYDIFQK | 9 | P60842 | Eukaryotic initiation factor 4A-I OS=Hon | EIF4A1 | 46.154 | 2 | 51.687397 | yes | 1.403E-07 | 585.29547 | 1854.8772 |
| KEELTLEGIR | 10 | P60842 | Eukaryotic initiation factor 4A-I OS=Hon | EIF4A1 | 46.154 | 2 | 51.687397 | yes | 4.987E-07 | 594.33514 | 1444.412 |
| GVAINMVTEEDKI | 13 | P60842 | Eukaryotic initiation factor 4A-I OS=Hon | EIF4A1 | 46.154 | 2 | 51.687397 | yes | 9.757E-08 | 731.37195 | 1007.7407 |
| ATQALVLAPTR | 11 | P60842 | Eukaryotic initiation factor 4A-I OS=Hon | EIF4A1 | 46.154 | 2 | 51.687397 | yes | 9.722E-08 | 570.84039 | 744.88141 |
| EELTLEGIR | 9 | P60842 | Eukaryotic initiation factor 4A-I OS=Hon | EIF4A1 | 46.154 | 2 | 51.687397 | yes | 1.019E-07 | 530.28766 | 727.79077 |
| VVMALGDYMGAS | 23 | P60842 | Eukaryotic initiation factor 4A-I OS=Hon | EIF4A1 | 46.154 | 3 | 51.687397 | yes | 9.758E-08 | 813.70605 | 200.19362 |
| VVMALGDYMGAS | 23 | P60842 | Eukaryotic initiation factor 4A-I OS=Hon | EIF4A1 | 46.154 | 3 | 51.687397 | yes | 2.81E-07 | 819.03772 | 200.19362 |

| | | | | | | | | | | | |
|---|---|---|---|---|---|---|---|---|---|---|---|
| VVMALGDYMGAS | 23 | P60842 | Eukaryotic initiation factor 4A-I OS=Hon | EIF4A1 | 46.154 | 3 | 51.687397 | yes | 2.947E-07 | 819.03772 | 200.19362 |
| DFTVSAMHGDMD | 14 | P60842 | Eukaryotic initiation factor 4A-I OS=Hon | EIF4A1 | 46.154 | 2 | 51.687397 | yes | 1.245E-07 | 791.33722 | 500.92337 |
| KGVAINMVTEEDI | 14 | P60842 | Eukaryotic initiation factor 4A-I OS=Hon | EIF4A1 | 46.154 | 3 | 51.687397 | yes | 1.845E-07 | 530.61536 | 232.11246 |
| KGVAINMVTEEDI | 14 | P60842 | Eukaryotic initiation factor 4A-I OS=Hon | EIF4A1 | 46.154 | 3 | 51.687397 | yes | 9.789E-08 | 795.41943 | 232.11246 |
| KGVAINMVTEEDI | 14 | P60842 | Eukaryotic initiation factor 4A-I OS=Hon | EIF4A1 | 46.154 | 2 | 51.687397 | yes | 1.845E-07 | 530.61536 | 232.11246 |
| KGVAINMVTEEDI | 14 | P60842 | Eukaryotic initiation factor 4A-I OS=Hon | EIF4A1 | 46.154 | 2 | 51.687397 | yes | 9.789E-08 | 795.41943 | 232.11246 |
| TPVEPEVAIHR | 11 | P60866 | Small ribosomal subunit protein uS10 OS | RPS20 | 13.373 | 2 | 50.301743 | yes | 2.08E-07 | 624.34076 | 45838.211 |
| VCADLIR | 7 | P60866 | Small ribosomal subunit protein uS10 OS | RPS20 | 13.373 | 2 | 50.301743 | yes | 1.292E-07 | 423.72873 | 26018.607 |
| LIDLHSPSEIVK | 12 | P60866 | Small ribosomal subunit protein uS10 OS | RPS20 | 13.373 | 2 | 50.301743 | yes | 1.697E-07 | 675.88501 | 1087.7454 |
| LIDLHSPSEIVK | 12 | P60866 | Small ribosomal subunit protein uS10 OS | RPS20 | 13.373 | 2 | 50.301743 | yes | 9.819E-08 | 450.92575 | 1087.7454 |
| LIDLHSPSEIVK | 12 | P60866 | Small ribosomal subunit protein uS10 OS | RPS20 | 13.373 | 3 | 50.301743 | yes | 1.697E-07 | 675.88501 | 1087.7454 |
| LIDLHSPSEIVK | 12 | P60866 | Small ribosomal subunit protein uS10 OS | RPS20 | 13.373 | 3 | 50.301743 | yes | 9.819E-08 | 450.92575 | 1087.7454 |
| RLIDLHSPSEIVK | 13 | P60866 | Small ribosomal subunit protein uS10 OS | RPS20 | 13.373 | 3 | 50.301743 | yes | 9.819E-08 | 502.95947 | 279.78766 |
| RLIDLHSPSEIVK | 13 | P60866 | Small ribosomal subunit protein uS10 OS | RPS20 | 13.373 | 3 | 50.301743 | yes | 1.306E-07 | 753.93555 | 279.78766 |
| RLIDLHSPSEIVK | 13 | P60866 | Small ribosomal subunit protein uS10 OS | RPS20 | 13.373 | 2 | 50.301743 | yes | 9.819E-08 | 502.95947 | 279.78766 |
| RLIDLHSPSEIVK | 13 | P60866 | Small ribosomal subunit protein uS10 OS | RPS20 | 13.373 | 2 | 50.301743 | yes | 1.306E-07 | 753.93555 | 279.78766 |
| VTAVIPCFPYAR | 12 | P60891 | Ribose-phosphate pyrophosphokinase 1 C | PRPS1 | 34.835 | 2 | 52.882977 | no | 1.607E-07 | 697.36847 | 2395.3977 |
| LNVDFALIHK | 10 | P60891 | Ribose-phosphate pyrophosphokinase 1 C | PRPS1 | 34.835 | 2 | 52.882977 | yes | 1.193E-07 | 585.33752 | 1159.9175 |
| NCTIVSPDAGGAK | 13 | P60891 | Ribose-phosphate pyrophosphokinase 1 C | PRPS1 | 34.835 | 2 | 52.882977 | yes | 2.556E-07 | 645.31134 | 302.63745 |
| IFSGSSHQDLSQK | 13 | P60891 | Ribose-phosphate pyrophosphokinase 1 C | PRPS1 | 34.835 | 2 | 52.882977 | no | 9.722E-08 | 717.35461 | 242.02542 |
| VYAILTHGIFSGPA | 17 | P60891 | Ribose-phosphate pyrophosphokinase 1 C | PRPS1 | 34.835 | 3 | 52.882977 | no | 3.127E-07 | 601.33685 | 236.11951 |
| INNACFEAVVVTN | 20 | P60891 | Ribose-phosphate pyrophosphokinase 1 C | PRPS1 | 34.835 | 3 | 52.882977 | yes | 9.722E-08 | 754.70721 | 128.84132 |
| MVLVGDVK | 8 | P60891 | Ribose-phosphate pyrophosphokinase 1 C | PRPS1 | 34.835 | 2 | 52.882977 | no | 1.284E-07 | 430.74915 | 73.656815 |
| AINQGGLTSVAVR | 13 | P60900 | Proteasome subunit alpha type-6 OS=Hor | PSMA6 | 27.4 | 2 | 40.925613 | yes | 3.768E-07 | 643.36475 | 62.873459 |
| EFPGFLENQK | 10 | P60903 | Protein S100-A10 OS=Homo sapiens OX | S100A10 | 11.204 | 2 | 51.982662 | yes | 9.722E-08 | 604.80096 | 497.04718 |
| PSQMEHAMETMM | 17 | P60903 | Protein S100-A10 OS=Homo sapiens OX | S100A10 | 11.204 | 3 | 51.982662 | yes | 1.759E-07 | 694.96832 | 39.746582 |
| YVECSALTQK | 10 | P60953 | Cell division control protein 42 homolog | CDC42 | 21.259 | 2 | 45.186207 | yes | 5.553E-07 | 599.79224 | 181.75655 |
| QKPITPETAEK | 11 | P60953 | Cell division control protein 42 homolog | CDC42 | 21.259 | 2 | 45.186207 | yes | 4.188E-07 | 621.34045 | 157.97035 |
| TPFLLVGTQIDLR | 13 | P60953 | Cell division control protein 42 homolog | CDC42 | 21.259 | 2 | 45.186207 | yes | 1.931E-07 | 736.92719 | 58.903858 |
| WVPEITHHCPK | 11 | P60953 | Cell division control protein 42 homolog | CDC42 | 21.259 | 2 | 45.186207 | yes | 3.979E-07 | 702.34808 | 33.012131 |
| YALYDASFETK | 11 | P60981 | Destrin OS=Homo sapiens OX=9606 GN | DSTN | 18.506 | 2 | 50.70428 | yes | 9.728E-08 | 654.31134 | 605.84814 |
| HFVGMLPEK | 9 | P60981 | Destrin OS=Homo sapiens OX=9606 GN | DSTN | 18.506 | 2 | 50.70428 | yes | 1.752E-07 | 529.27856 | 532.47058 |
| AVIFCLSADKK | 11 | P60981 | Destrin OS=Homo sapiens OX=9606 GN | DSTN | 18.506 | 2 | 50.70428 | yes | 1.256E-07 | 626.34192 | 107.96886 |
| KCSTPEEIK | 9 | P60981 | Destrin OS=Homo sapiens OX=9606 GN | DSTN | 18.506 | 2 | 50.70428 | yes | 3.313E-07 | 546.27368 | 86.822563 |
| MNTVLSR | 7 | P61009 | Signal peptidase complex subunit 3 OS=I | SPCS3 | 20.314 | 2 | 37.373116 | yes | 6.05E-07 | 418.71838 | 65.015831 |
| SAIDLEEMASGLN | 14 | P61011 | Signal recognition particle subunit SRP5 | SRP54 | 55.705 | 2 | 43.741398 | yes | 2.465E-07 | 739.36377 | 56.554848 |
| DVQELLTQYTK | 11 | P61011 | Signal recognition particle subunit SRP5 | SRP54 | 55.705 | 2 | 43.741398 | yes | 6.191E-07 | 669.35101 | 20.811115 |
| TASNVEEAFINTAI | 14 | P61019 | Ras-related protein Rab-2A OS=Homo sa | RAB2A | 23.546 | 2 | 50.94709 | yes | 1.256E-07 | 747.87537 | 50.20895 |
| LQIWDTAGQESFR | 13 | P61019 | Ras-related protein Rab-2A OS=Homo sa | RAB2A | 23.546 | 2 | 50.94709 | no | 9.722E-08 | 775.88354 | 44.252472 |

| | | | | | | | | | | | |
|---|---|---|---|---|---|---|---|---|---|---|---|
| THLMSESEWR | 10 | P61024 | Cyclin-dependent kinases regulatory subu | CKS1B | 9.661 | 2 | 39.42855 | yes | 4.591E-07 | 638.29297 | 25.387562 |
| TSAQHALTSVSR | 12 | P61073 | C-X-C chemokine receptor type 4 OS=Hc | CXCR4 | 39.746 | 2 | 51.77084 | yes | 9.722E-08 | 629.33093 | 141.42879 |
| SQWSPALTISK | 11 | P61077 | Ubiquitin-conjugating enzyme E2 D3 OS | UBE2D3 | 16.688 | 2 | 48.75836 | yes | 1.108E-07 | 609.3299 | 149.37094 |
| TNEAQAIETAR | 11 | P61088 | Ubiquitin-conjugating enzyme E2 N OS= | UBE2N | 17.138 | 2 | 21.97414 | yes | 3.99E-06 | 602.30182 | 51.825077 |
| WSPALQIR | 8 | P61088 | Ubiquitin-conjugating enzyme E2 N OS= | UBE2N | 17.138 | 2 | 21.97414 | yes | 5.611E-06 | 485.77707 | 27.746443 |
| TGENVEDAFLEAA | 14 | P61106 | Ras-related protein Rab-14 OS=Homo saj | RAB14 | 23.897 | 2 | 55.510956 | yes | 9.722E-08 | 747.35956 | 103.48236 |
| NLTNPNTVIILIGNI | 15 | P61106 | Ras-related protein Rab-14 OS=Homo saj | RAB14 | 23.897 | 2 | 55.510956 | yes | 9.722E-08 | 812.47504 | 47.780518 |
| SCLLHQFTEK | 10 | P61106 | Ras-related protein Rab-14 OS=Homo saj | RAB14 | 23.897 | 2 | 55.510956 | yes | 9.809E-08 | 631.81354 | 26.189648 |
| EFSIDVGYER | 10 | P61158 | Actin-related protein 3 OS=Homo sapiens | ACTR3 | 47.372 | 2 | 54.31653 | yes | 1.809E-07 | 607.78802 | 88.827461 |
| NIVLSGGSTMFR | 12 | P61158 | Actin-related protein 3 OS=Homo sapiens | ACTR3 | 47.372 | 2 | 54.31653 | yes | 9.722E-08 | 641.33466 | 76.985458 |
| YSYVCPDLVK | 10 | P61158 | Actin-related protein 3 OS=Homo sapiens | ACTR3 | 47.372 | 2 | 54.31653 | yes | 3.272E-07 | 622.30499 | 55.029861 |
| LPACVVDCGTGYT | 14 | P61158 | Actin-related protein 3 OS=Homo sapiens | ACTR3 | 47.372 | 2 | 54.31653 | yes | 6.768E-07 | 770.86035 | 54.807663 |
| HGIVEDWDLMER | 12 | P61158 | Actin-related protein 3 OS=Homo sapiens | ACTR3 | 47.372 | 2 | 54.31653 | yes | 2.59E-07 | 750.35101 | 26.02396 |
| DLMVGDEASELR | 12 | P61160 | Actin-related protein 2 OS=Homo sapiens | ACTR2 | 44.761 | 2 | 52.143753 | yes | 9.722E-08 | 667.81647 | 316.5582 |
| LCYVGYNIEQEQK | 13 | P61160 | Actin-related protein 2 OS=Homo sapiens | ACTR2 | 44.761 | 2 | 52.143753 | yes | 1.467E-07 | 822.39032 | 104.85886 |
| HIVLSGGSTMYPG | 17 | P61160 | Actin-related protein 2 OS=Homo sapiens | ACTR2 | 44.761 | 3 | 52.143753 | yes | 6.321E-07 | 591.31018 | 57.686363 |
| DKDNFWMTR | 9 | P61160 | Actin-related protein 2 OS=Homo sapiens | ACTR2 | 44.761 | 2 | 52.143753 | yes | 1.327E-07 | 606.77698 | 36.802711 |
| ILMVGLDAAGK | 11 | P61204 | ADP-ribosylation factor 3 OS=Homo sap | ARF3 | 20.601 | 2 | 49.1203 | no | 4.872E-07 | 544.31262 | 2025.6521 |
| HYFQNTQGLIFVV | 18 | P61204 | ADP-ribosylation factor 3 OS=Homo sap | ARF3 | 20.601 | 3 | 49.1203 | no | 1.198E-07 | 718.35187 | 604.72406 |
| MLAEDELR | 8 | P61204 | ADP-ribosylation factor 3 OS=Homo sap | ARF3 | 20.601 | 2 | 49.1203 | yes | 1.066E-07 | 488.74203 | 557.66187 |
| NISFTVWDVGGQD | 14 | P61204 | ADP-ribosylation factor 3 OS=Homo sap | ARF3 | 20.601 | 2 | 49.1203 | yes | 2.965E-07 | 783.38336 | 474.30106 |
| DAVLLVFANK | 10 | P61204 | ADP-ribosylation factor 3 OS=Homo sap | ARF3 | 20.601 | 2 | 49.1203 | no | 3.793E-07 | 545.31879 | 342.09109 |
| QDLPNAMNAAEIT | 15 | P61204 | ADP-ribosylation factor 3 OS=Homo sap | ARF3 | 20.601 | 2 | 49.1203 | yes | 1.073E-07 | 815.89069 | 270.57651 |
| GSELQNYFTK | 10 | P61221 | ATP-binding cassette sub-family E memb | ABCE1 | 67.315 | 2 | 51.572453 | yes | 1.025E-07 | 593.79059 | 1791.9612 |
| FLSQLEITFR | 10 | P61221 | ATP-binding cassette sub-family E memb | ABCE1 | 67.315 | 2 | 51.572453 | yes | 1.75E-07 | 627.34808 | 1655.4812 |
| LKPDEGGEVPVLN | 20 | P61221 | ATP-binding cassette sub-family E memb | ABCE1 | 67.315 | 3 | 51.572453 | yes | 1.436E-07 | 733.06696 | 812.4469 |
| LKPDEGGEVPVLN | 20 | P61221 | ATP-binding cassette sub-family E memb | ABCE1 | 67.315 | 3 | 51.572453 | yes | 5.773E-07 | 550.05206 | 812.4469 |
| LKPDEGGEVPVLN | 20 | P61221 | ATP-binding cassette sub-family E memb | ABCE1 | 67.315 | 4 | 51.572453 | yes | 1.436E-07 | 733.06696 | 812.4469 |
| LKPDEGGEVPVLN | 20 | P61221 | ATP-binding cassette sub-family E memb | ABCE1 | 67.315 | 4 | 51.572453 | yes | 5.773E-07 | 550.05206 | 812.4469 |
| GTVGSILDR | 9 | P61221 | ATP-binding cassette sub-family E memb | ABCE1 | 67.315 | 2 | 51.572453 | yes | 2.93E-07 | 459.25616 | 1401.3812 |
| FACAVVCIQK | 10 | P61221 | ATP-binding cassette sub-family E memb | ABCE1 | 67.315 | 2 | 51.572453 | yes | 9.722E-08 | 598.30176 | 1344.007 |
| NVEDLSGGELQR | 12 | P61221 | ATP-binding cassette sub-family E memb | ABCE1 | 67.315 | 2 | 51.572453 | yes | 9.838E-08 | 658.82568 | 1238.1614 |
| LCIEVTPQSK | 10 | P61221 | ATP-binding cassette sub-family E memb | ABCE1 | 67.315 | 2 | 51.572453 | yes | 1.168E-07 | 587.81049 | 1160.5055 |
| VIVFDGVPSK | 10 | P61221 | ATP-binding cassette sub-family E memb | ABCE1 | 67.315 | 2 | 51.572453 | yes | 6.046E-07 | 530.80548 | 626.84814 |
| VAETANEEEVKK | 12 | P61221 | ATP-binding cassette sub-family E memb | ABCE1 | 67.315 | 2 | 51.572453 | yes | 9.976E-08 | 673.84357 | 601.05267 |
| IAIVNHDK | 8 | P61221 | ATP-binding cassette sub-family E memb | ABCE1 | 67.315 | 2 | 51.572453 | yes | 9.788E-08 | 455.26126 | 577.9259 |
| YCANAFK | 7 | P61221 | ATP-binding cassette sub-family E memb | ABCE1 | 67.315 | 2 | 51.572453 | yes | 3.171E-07 | 437.1998 | 281.68936 |
| AIIKPQYVDQIPK | 13 | P61221 | ATP-binding cassette sub-family E memb | ABCE1 | 67.315 | 2 | 51.572453 | yes | 9.728E-08 | 756.94287 | 111.31606 |
| AIIKPQYVDQIPK | 13 | P61221 | ATP-binding cassette sub-family E memb | ABCE1 | 67.315 | 2 | 51.572453 | yes | 7.091E-07 | 504.96432 | 111.31606 |

| | | | | | | | | | | | |
|---|---|---|---|---|---|---|---|---|---|---|---|
| AIIKPQYVDQIPK | 13 | P61221 | ATP-binding cassette sub-family E memb | ABCE1 | 67.315 | 3 | 51.572453 | yes | 9.728E-08 | 756.94287 | 111.31606 |
| AIIKPQYVDQIPK | 13 | P61221 | ATP-binding cassette sub-family E memb | ABCE1 | 67.315 | 3 | 51.572453 | yes | 7.091E-07 | 504.96432 | 111.31606 |
| VALALCLGK | 9 | P61221 | ATP-binding cassette sub-family E memb | ABCE1 | 67.315 | 2 | 51.572453 | yes | 9.823E-08 | 472.78351 | 165.40169 |
| NTVANSPQTLLAG | 16 | P61221 | ATP-binding cassette sub-family E memb | ABCE1 | 67.315 | 2 | 51.572453 | yes | 1.05E-07 | 829.93018 | 71.386162 |
| NTVANSPQTLLAG | 16 | P61221 | ATP-binding cassette sub-family E memb | ABCE1 | 67.315 | 2 | 51.572453 | yes | 3.328E-06 | 837.92761 | 71.386162 |
| PGEVLGLVGTNGI | 15 | P61221 | ATP-binding cassette sub-family E memb | ABCE1 | 67.315 | 2 | 51.572453 | yes | 1.53E-07 | 705.90118 | 116.72195 |
| DAYTHPQFVTDVN | 14 | P61221 | ATP-binding cassette sub-family E memb | ABCE1 | 67.315 | 2 | 51.572453 | yes | 9.722E-08 | 826.39288 | 85.751686 |
| VAETANEEEVK | 11 | P61221 | ATP-binding cassette sub-family E memb | ABCE1 | 67.315 | 2 | 51.572453 | yes | 9.935E-08 | 609.79608 | 52.287376 |
| SLINPDR | 7 | P61221 | ATP-binding cassette sub-family E memb | ABCE1 | 67.315 | 2 | 51.572453 | yes | 5.438E-07 | 407.72452 | 39.95805 |
| QNALLEQQVR | 10 | P61244 | Protein max OS=Homo sapiens OX=960( | MAX | 18.275 | 2 | 48.73022 | yes | 1.11E-07 | 599.83057 | 554.29199 |
| DSFHSLR | 7 | P61244 | Protein max OS=Homo sapiens OX=960( | MAX | 18.275 | 2 | 48.73022 | yes | 3.478E-07 | 431.21429 | 469.41498 |
| ATEYIQYMR | 9 | P61244 | Protein max OS=Homo sapiens OX=960( | MAX | 18.275 | 2 | 48.73022 | yes | 1.357E-07 | 587.78168 | 417.43811 |
| NHTHQQDIDDLKF | 13 | P61244 | Protein max OS=Homo sapiens OX=960( | MAX | 18.275 | 2 | 48.73022 | yes | 3.253E-06 | 810.39789 | 27.316002 |
| APAMFNIR | 8 | P61247 | Small ribosomal subunit protein eS1 OS= | RPS3A | 29.945 | 2 | 52.055485 | yes | 1.483E-07 | 460.24457 | 46607.418 |
| TTDGYLLR | 8 | P61247 | Small ribosomal subunit protein eS1 OS= | RPS3A | 29.945 | 2 | 52.055485 | yes | 2.766E-06 | 469.75073 | 46563.094 |
| LMELHGEGSSSGK | 13 | P61247 | Small ribosomal subunit protein eS1 OS= | RPS3A | 29.945 | 2 | 52.055485 | yes | 9.722E-08 | 666.31665 | 4967.7754 |
| LMELHGEGSSSGK | 13 | P61247 | Small ribosomal subunit protein eS1 OS= | RPS3A | 29.945 | 2 | 52.055485 | yes | 2.295E-07 | 444.54684 | 4967.7754 |
| LMELHGEGSSSGK | 13 | P61247 | Small ribosomal subunit protein eS1 OS= | RPS3A | 29.945 | 2 | 52.055485 | yes | 9.722E-08 | 674.31409 | 4967.7754 |
| LMELHGEGSSSGK | 13 | P61247 | Small ribosomal subunit protein eS1 OS= | RPS3A | 29.945 | 2 | 52.055485 | yes | 2.112E-07 | 449.87848 | 4967.7754 |
| LMELHGEGSSSGK | 13 | P61247 | Small ribosomal subunit protein eS1 OS= | RPS3A | 29.945 | 3 | 52.055485 | yes | 9.722E-08 | 666.31665 | 4967.7754 |
| LMELHGEGSSSGK | 13 | P61247 | Small ribosomal subunit protein eS1 OS= | RPS3A | 29.945 | 3 | 52.055485 | yes | 2.295E-07 | 444.54684 | 4967.7754 |
| LMELHGEGSSSGK | 13 | P61247 | Small ribosomal subunit protein eS1 OS= | RPS3A | 29.945 | 3 | 52.055485 | yes | 9.722E-08 | 674.31409 | 4967.7754 |
| LMELHGEGSSSGK | 13 | P61247 | Small ribosomal subunit protein eS1 OS= | RPS3A | 29.945 | 3 | 52.055485 | yes | 2.112E-07 | 449.87848 | 4967.7754 |
| MMEIMTR | 7 | P61247 | Small ribosomal subunit protein eS1 OS= | RPS3A | 29.945 | 2 | 52.055485 | yes | 1.606E-07 | 456.211 | 4108.6797 |
| MMEIMTR | 7 | P61247 | Small ribosomal subunit protein eS1 OS= | RPS3A | 29.945 | 2 | 52.055485 | yes | 1.165E-07 | 464.20847 | 4108.6797 |
| MMEIMTR | 7 | P61247 | Small ribosomal subunit protein eS1 OS= | RPS3A | 29.945 | 2 | 52.055485 | yes | 1.192E-07 | 464.20847 | 4108.6797 |
| MMEIMTR | 7 | P61247 | Small ribosomal subunit protein eS1 OS= | RPS3A | 29.945 | 2 | 52.055485 | yes | 1.049E-07 | 464.20847 | 4108.6797 |
| LFCVGFTK | 8 | P61247 | Small ribosomal subunit protein eS1 OS= | RPS3A | 29.945 | 2 | 52.055485 | yes | 1.272E-07 | 486.25458 | 9678.5498 |
| NCLTNFHGMDLTF | 13 | P61247 | Small ribosomal subunit protein eS1 OS= | RPS3A | 29.945 | 2 | 52.055485 | yes | 9.849E-08 | 789.86121 | 3952.916 |
| NCLTNFHGMDLTF | 13 | P61247 | Small ribosomal subunit protein eS1 OS= | RPS3A | 29.945 | 2 | 52.055485 | yes | 9.722E-08 | 797.8587 | 3952.916 |
| WQTMIEAHVDVK | 12 | P61247 | Small ribosomal subunit protein eS1 OS= | RPS3A | 29.945 | 2 | 52.055485 | yes | 3.175E-06 | 728.86627 | 3759.8455 |
| WQTMIEAHVDVK | 12 | P61247 | Small ribosomal subunit protein eS1 OS= | RPS3A | 29.945 | 2 | 52.055485 | yes | 3.184E-06 | 486.24661 | 3759.8455 |
| WQTMIEAHVDVK | 12 | P61247 | Small ribosomal subunit protein eS1 OS= | RPS3A | 29.945 | 3 | 52.055485 | yes | 3.175E-06 | 728.86627 | 3759.8455 |
| WQTMIEAHVDVK | 12 | P61247 | Small ribosomal subunit protein eS1 OS= | RPS3A | 29.945 | 3 | 52.055485 | yes | 3.184E-06 | 486.24661 | 3759.8455 |
| EVQTNDLK | 8 | P61247 | Small ribosomal subunit protein eS1 OS= | RPS3A | 29.945 | 2 | 52.055485 | yes | 6.387E-07 | 473.74564 | 5774.0171 |
| KDWYDVK | 7 | P61247 | Small ribosomal subunit protein eS1 OS= | RPS3A | 29.945 | 2 | 52.055485 | yes | 7.039E-07 | 477.23999 | 4044.3289 |
| DKMCSMVK | 8 | P61247 | Small ribosomal subunit protein eS1 OS= | RPS3A | 29.945 | 2 | 52.055485 | yes | 1.123E-07 | 499.72702 | 3845.0786 |
| LITEDVQGK | 9 | P61247 | Small ribosomal subunit protein eS1 OS= | RPS3A | 29.945 | 2 | 52.055485 | yes | 2.956E-07 | 501.77695 | 1827.2754 |
| KWQTMIEAHVDV | 13 | P61247 | Small ribosomal subunit protein eS1 OS= | RPS3A | 29.945 | 2 | 52.055485 | yes | 2.081E-06 | 792.91376 | 527.16486 |

| | | | | | | | | | | | |
|---|---|---|---|---|---|---|---|---|---|---|---|
| KWQTMIEAHVDV | 13 | P61247 | Small ribosomal subunit protein eS1 OS= | RPS3A | 29.945 | 2 | 52.055485 | yes | 2.789E-06 | 800.91125 | 527.16486 |
| KVVDPFSK | 8 | P61247 | Small ribosomal subunit protein eS1 OS= | RPS3A | 29.945 | 2 | 52.055485 | yes | 1.657E-07 | 460.26602 | 633.37701 |
| LIPDSIGK | 8 | P61247 | Small ribosomal subunit protein eS1 OS= | RPS3A | 29.945 | 2 | 52.055485 | yes | 8.177E-07 | 421.75275 | 425.77475 |
| ADGYEPPVQESV | 12 | P61247 | Small ribosomal subunit protein eS1 OS= | RPS3A | 29.945 | 2 | 52.055485 | yes | 1.596E-07 | 645.79608 | 408.71851 |
| ACQSIYPLHDVFV | 14 | P61247 | Small ribosomal subunit protein eS1 OS= | RPS3A | 29.945 | 3 | 52.055485 | yes | 1.443E-07 | 568.95575 | 67.095444 |
| LFCVGFTKK | 9 | P61247 | Small ribosomal subunit protein eS1 OS= | RPS3A | 29.945 | 2 | 52.055485 | yes | 5.732E-05 | 550.30206 | 22.595453 |
| FNPFVTSDR | 9 | P61254 | Large ribosomal subunit protein uL24 OS | RPL26 | 17.259 | 2 | 50.465675 | no | 1.353E-07 | 541.76691 | 54875.098 |
| KYVIYIER | 8 | P61254 | Large ribosomal subunit protein uL24 OS | RPL26 | 17.259 | 2 | 50.465675 | no | 1.281E-07 | 542.31348 | 21956.66 |
| YKEETIEK | 8 | P61254 | Large ribosomal subunit protein uL24 OS | RPL26 | 17.259 | 2 | 50.465675 | yes | 1.011E-06 | 520.26892 | 18472.816 |
| YVIYIER | 7 | P61254 | Large ribosomal subunit protein uL24 OS | RPL26 | 17.259 | 2 | 50.465675 | no | 2.825E-07 | 478.26602 | 18239.559 |
| HFNAPSHIR | 9 | P61254 | Large ribosomal subunit protein uL24 OS | RPL26 | 17.259 | 2 | 50.465675 | yes | 2.254E-07 | 539.7807 | 17870.674 |
| KDDEVQVVR | 9 | P61254 | Large ribosomal subunit protein uL24 OS | RPL26 | 17.259 | 2 | 50.465675 | no | 1.981E-07 | 544.29077 | 12723.171 |
| IMSSPLSK | 8 | P61254 | Large ribosomal subunit protein uL24 OS | RPL26 | 17.259 | 2 | 50.465675 | no | 1.878E-07 | 431.73877 | 2662.8892 |
| IMSSPLSK | 8 | P61254 | Large ribosomal subunit protein uL24 OS | RPL26 | 17.259 | 2 | 50.465675 | no | 2.42E-07 | 439.73624 | 2662.8892 |
| KIMSSPLSK | 9 | P61254 | Large ribosomal subunit protein uL24 OS | RPL26 | 17.259 | 2 | 50.465675 | no | 3.04E-06 | 495.78625 | 3337.0422 |
| DDEVQVVR | 8 | P61254 | Large ribosomal subunit protein uL24 OS | RPL26 | 17.259 | 2 | 50.465675 | no | 1.146E-07 | 480.24326 | 1009.3 |
| ANGTTVHVGIHPS | 14 | P61254 | Large ribosomal subunit protein uL24 OS | RPL26 | 17.259 | 2 | 50.465675 | no | 9.809E-08 | 709.38098 | 678.6756 |
| TVESEAASYLDQIS | 15 | P61289 | Proteasome activator complex subunit 3 ( | PSME3 | 29.507 | 2 | 52.352882 | yes | 1.107E-06 | 834.90741 | 161.79697 |
| YPHVEDYR | 8 | P61289 | Proteasome activator complex subunit 3 ( | PSME3 | 29.507 | 2 | 52.352882 | yes | 2.716E-07 | 539.75128 | 93.654243 |
| LDECEEAFQGTK | 12 | P61289 | Proteasome activator complex subunit 3 ( | PSME3 | 29.507 | 2 | 52.352882 | yes | 9.722E-08 | 713.8114 | 49.615269 |
| LLELDSFLK | 9 | P61289 | Proteasome activator complex subunit 3 ( | PSME3 | 29.507 | 2 | 52.352882 | yes | 2.602E-07 | 539.31317 | 48.170055 |
| VFVMPNGMLK | 10 | P61289 | Proteasome activator complex subunit 3 ( | PSME3 | 29.507 | 2 | 52.352882 | yes | 4.876E-06 | 568.30377 | 27.796963 |
| VLNSYWVGEDST | 14 | P61313 | Large ribosomal subunit protein eL15 OS | RPL15 | 24.147 | 2 | 50.33394 | yes | 1.114E-07 | 830.8963 | 21725.406 |
| FHHTIGGSR | 9 | P61313 | Large ribosomal subunit protein eL15 OS | RPL15 | 24.147 | 2 | 50.33394 | yes | 1.244E-07 | 506.25958 | 14239.177 |
| SLQSVAEER | 9 | P61313 | Large ribosomal subunit protein eL15 OS | RPL15 | 24.147 | 2 | 50.33394 | yes | 1.068E-07 | 509.76181 | 11365.813 |
| QGYVIYR | 7 | P61313 | Large ribosomal subunit protein eL15 OS | RPL15 | 24.147 | 2 | 50.33394 | yes | 1.212E-07 | 449.74271 | 6059.5015 |
| RNPDTQWITK | 10 | P61313 | Large ribosomal subunit protein eL15 OS | RPL15 | 24.147 | 2 | 50.33394 | yes | 2.97E-07 | 629.83057 | 4062.4883 |
| NPDTQWITK | 9 | P61313 | Large ribosomal subunit protein eL15 OS | RPL15 | 24.147 | 2 | 50.33394 | yes | 1.147E-07 | 551.78003 | 2474.4419 |
| QLSALHR | 7 | P61313 | Large ribosomal subunit protein eL15 OS | RPL15 | 24.147 | 2 | 50.33394 | yes | 1.938E-07 | 412.74051 | 1175.7437 |
| GATYGKPVHHGV | 16 | P61313 | Large ribosomal subunit protein eL15 OS | RPL15 | 24.147 | 3 | 50.33394 | yes | 1.949E-07 | 569.30927 | 1097.7031 |
| NPDTQWITKPVHK | 13 | P61313 | Large ribosomal subunit protein eL15 OS | RPL15 | 24.147 | 2 | 50.33394 | yes | 3.089E-07 | 782.41754 | 837.55157 |
| YIQELWR | 7 | P61313 | Large ribosomal subunit protein eL15 OS | RPL15 | 24.147 | 2 | 50.33394 | yes | 1.931E-07 | 504.26907 | 444.5401 |
| RNTLQLHR | 8 | P61313 | Large ribosomal subunit protein eL15 OS | RPL15 | 24.147 | 2 | 50.33394 | yes | 1.591E-07 | 519.30176 | 355.51935 |
| NTLQLHR | 7 | P61313 | Large ribosomal subunit protein eL15 OS | RPL15 | 24.147 | 2 | 50.33394 | yes | 9.819E-08 | 441.25122 | 147.33627 |
| RNPDTQWITKPVH | 14 | P61313 | Large ribosomal subunit protein eL15 OS | RPL15 | 24.147 | 3 | 50.33394 | yes | 2.981E-06 | 573.98114 | 82.289444 |
| PVHHGVNQLK | 10 | P61313 | Large ribosomal subunit protein eL15 OS | RPL15 | 24.147 | 2 | 50.33394 | yes | 2.472E-05 | 564.81726 | 57.136768 |
| SVMEELKR | 8 | P61326 | Protein mago nashi homolog OS=Homo s | MAGOH | 17.164 | 2 | 55.507977 | yes | 2.127E-07 | 496.26569 | 8955.7168 |
| IGSLIDVNQSK | 11 | P61326 | Protein mago nashi homolog OS=Homo s | MAGOH | 17.164 | 2 | 55.507977 | yes | 1.204E-07 | 587.32733 | 8852.8447 |
| YANNSNYK | 8 | P61326 | Protein mago nashi homolog OS=Homo s | MAGOH | 17.164 | 2 | 55.507977 | yes | 3.6E-07 | 487.22232 | 1326.7952 |

| | | | | | | | | | | | |
|---|---|---|---|---|---|---|---|---|---|---|---|
| QELEIVIGDEHISF1 | 18 | P61326 | Protein mago nashi homolog OS=Homo s | MAGOH | 17.164 | 3 | 55.507977 | yes | 1.567E-07 | 682.68451 | 928.06561 |
| IIDDSEITK | 9 | P61326 | Protein mago nashi homolog OS=Homo s | MAGOH | 17.164 | 2 | 55.507977 | yes | 9.885E-08 | 517.27423 | 643.86316 |
| EDDALWPPPDR | 11 | P61326 | Protein mago nashi homolog OS=Homo s | MAGOH | 17.164 | 2 | 55.507977 | yes | 9.722E-08 | 655.8042 | 592.64294 |
| VFYYLVQDLK | 10 | P61326 | Protein mago nashi homolog OS=Homo s | MAGOH | 17.164 | 2 | 55.507977 | yes | 3.344E-07 | 644.35284 | 42.733673 |
| YSVDIPLDK | 9 | P61353 | Large ribosomal subunit protein eL27 OS | RPL27 | 15.798 | 2 | 54.104626 | yes | 7.351E-07 | 525.2793 | 16339.997 |
| VYNYNHLMPTR | 11 | P61353 | Large ribosomal subunit protein eL27 OS | RPL27 | 15.798 | 2 | 54.104626 | yes | 9.758E-08 | 704.34552 | 5024.397 |
| VYNYNHLMPTR | 11 | P61353 | Large ribosomal subunit protein eL27 OS | RPL27 | 15.798 | 2 | 54.104626 | yes | 2.795E-07 | 469.89944 | 5024.397 |
| VYNYNHLMPTR | 11 | P61353 | Large ribosomal subunit protein eL27 OS | RPL27 | 15.798 | 2 | 54.104626 | yes | 2.581E-07 | 712.34296 | 5024.397 |
| VYNYNHLMPTR | 11 | P61353 | Large ribosomal subunit protein eL27 OS | RPL27 | 15.798 | 2 | 54.104626 | yes | 1.682E-07 | 475.23108 | 5024.397 |
| VYNYNHLMPTR | 11 | P61353 | Large ribosomal subunit protein eL27 OS | RPL27 | 15.798 | 3 | 54.104626 | yes | 9.758E-08 | 704.34552 | 5024.397 |
| VYNYNHLMPTR | 11 | P61353 | Large ribosomal subunit protein eL27 OS | RPL27 | 15.798 | 3 | 54.104626 | yes | 2.795E-07 | 469.89944 | 5024.397 |
| VYNYNHLMPTR | 11 | P61353 | Large ribosomal subunit protein eL27 OS | RPL27 | 15.798 | 3 | 54.104626 | yes | 2.581E-07 | 712.34296 | 5024.397 |
| VYNYNHLMPTR | 11 | P61353 | Large ribosomal subunit protein eL27 OS | RPL27 | 15.798 | 3 | 54.104626 | yes | 1.682E-07 | 475.23108 | 5024.397 |
| DVFRDPALK | 9 | P61353 | Large ribosomal subunit protein eL27 OS | RPL27 | 15.798 | 2 | 54.104626 | yes | 4.177E-07 | 530.79291 | 7315.9663 |
| NIDDGTSDRPYSH. | 21 | P61353 | Large ribosomal subunit protein eL27 OS | RPL27 | 15.798 | 4 | 54.104626 | yes | 1.301E-07 | 568.7793 | 3050.1709 |
| VVLVLAGR | 8 | P61353 | Large ribosomal subunit protein eL27 OS | RPL27 | 15.798 | 2 | 54.104626 | yes | 1.209E-07 | 413.77908 | 1696.9032 |
| PYSHALVAGIDR | 12 | P61353 | Large ribosomal subunit protein eL27 OS | RPL27 | 15.798 | 2 | 54.104626 | yes | 3.662E-07 | 649.84619 | 325.17896 |
| PYSHALVAGIDR | 12 | P61353 | Large ribosomal subunit protein eL27 OS | RPL27 | 15.798 | 2 | 54.104626 | yes | 9.722E-08 | 433.56656 | 325.17896 |
| PYSHALVAGIDR | 12 | P61353 | Large ribosomal subunit protein eL27 OS | RPL27 | 15.798 | 3 | 54.104626 | yes | 3.662E-07 | 649.84619 | 325.17896 |
| PYSHALVAGIDR | 12 | P61353 | Large ribosomal subunit protein eL27 OS | RPL27 | 15.798 | 3 | 54.104626 | yes | 9.722E-08 | 433.56656 | 325.17896 |
| KVTAAMGK | 8 | P61353 | Large ribosomal subunit protein eL27 OS | RPL27 | 15.798 | 2 | 54.104626 | yes | 9.722E-08 | 403.23364 | 76.581573 |
| DYIRLYGIK | 9 | P61371 | Insulin gene enhancer protein ISL-1 OS=I | ISL1 | 39.036 | 2 | 27.175863 | yes | 3.001E-06 | 570.82422 | 38.848701 |
| AVGIWHCGSCMK | 12 | P61513 | Large ribosomal subunit protein eL43 OS | RPL37A | 10.276 | 2 | 53.187225 | yes | 9.722E-08 | 703.31232 | 3645.8872 |
| AVGIWHCGSCMK | 12 | P61513 | Large ribosomal subunit protein eL43 OS | RPL37A | 10.276 | 2 | 53.187225 | yes | 9.722E-08 | 711.30975 | 3645.8872 |
| AVGIWHCGSCMK | 12 | P61513 | Large ribosomal subunit protein eL43 OS | RPL37A | 10.276 | 2 | 53.187225 | yes | 1.043E-07 | 469.21063 | 3645.8872 |
| AVGIWHCGSCMK | 12 | P61513 | Large ribosomal subunit protein eL43 OS | RPL37A | 10.276 | 3 | 53.187225 | yes | 9.722E-08 | 703.31232 | 3645.8872 |
| AVGIWHCGSCMK | 12 | P61513 | Large ribosomal subunit protein eL43 OS | RPL37A | 10.276 | 3 | 53.187225 | yes | 9.722E-08 | 711.30975 | 3645.8872 |
| AVGIWHCGSCMK | 12 | P61513 | Large ribosomal subunit protein eL43 OS | RPL37A | 10.276 | 3 | 53.187225 | yes | 1.043E-07 | 469.21063 | 3645.8872 |
| YTCSFCGK | 8 | P61513 | Large ribosomal subunit protein eL43 OS | RPL37A | 10.276 | 2 | 53.187225 | no | 1.512E-07 | 511.70715 | 7117.2124 |
| IEISQHAK | 8 | P61513 | Large ribosomal subunit protein eL43 OS | RPL37A | 10.276 | 2 | 53.187225 | yes | 1.12E-07 | 463.25873 | 3918.0034 |
| KIEISQHAK | 9 | P61513 | Large ribosomal subunit protein eL43 OS | RPL37A | 10.276 | 2 | 53.187225 | yes | 2.022E-07 | 527.30621 | 3237.5889 |
| IGAFGYMECSAK | 12 | P61586 | Transforming protein RhoA OS=Homo sa | RHOA | 21.769 | 2 | 46.911205 | yes | 1.443E-07 | 667.29938 | 9.4949217 |
| SIMDPNQNVK | 10 | P61587 | Rho-related GTP-binding protein RhoE C | RND3 | 27.369 | 2 | 42.034027 | yes | 3.084E-06 | 573.28461 | 56.985603 |
| TDVSTLVELSNHR | 13 | P61587 | Rho-related GTP-binding protein RhoE C | RND3 | 27.369 | 2 | 42.034027 | yes | 3.225E-07 | 735.88098 | 38.966125 |
| TALLHVFAK | 9 | P61587 | Rho-related GTP-binding protein RhoE C | RND3 | 27.369 | 2 | 42.034027 | yes | 2.588E-06 | 500.30292 | 32.712906 |
| IIEVGDTPK | 9 | P61619 | Protein transport protein Sec61 subunit al | SEC61A1 | 52.265 | 2 | 50.839516 | no | 1.211E-06 | 486.27402 | 283.10291 |
| GQYNTYPIK | 9 | P61619 | Protein transport protein Sec61 subunit al | SEC61A1 | 52.265 | 2 | 50.839516 | yes | 1.014E-07 | 542.2771 | 244.84653 |
| AFSPTTVNTGR | 11 | P61619 | Protein transport protein Sec61 subunit al | SEC61A1 | 52.265 | 2 | 50.839516 | yes | 2.251E-07 | 575.7962 | 200.41193 |
| EQQMVMR | 7 | P61619 | Protein transport protein Sec61 subunit al | SEC61A1 | 52.265 | 2 | 50.839516 | no | 9.722E-08 | 461.21768 | 116.65884 |

| | | | | | | | | | | | |
|---|---|---|---|---|---|---|---|---|---|---|---|
| KLDEQYQK | 8 | P61758 | Prefoldin subunit 3 OS=Homo sapiens O: | VBP1 | 22.626 | 2 | 41.26765 | yes | 1.652E-06 | 526.27454 | 76.801811 |
| DQFTTTEVNMAR | 12 | P61758 | Prefoldin subunit 3 OS=Homo sapiens O: | VBP1 | 22.626 | 2 | 41.26765 | yes | 3.594E-07 | 706.82739 | 65.886574 |
| NLDSLEEDLDFLR | 13 | P61758 | Prefoldin subunit 3 OS=Homo sapiens O: | VBP1 | 22.626 | 2 | 41.26765 | yes | 3.6E-05 | 789.88593 | 7.6547532 |
| VNHVTLSQPK | 10 | P61769 | Beta-2-microglobulin OS=Homo sapiens | B2M | 13.715 | 2 | 45.93358 | yes | 1.699E-07 | 561.81696 | 58.320831 |
| FLEEYLSSTPQR | 12 | P61803 | Dolichyl-diphosphooligosaccharide--prot | DAD1 | 12.497 | 2 | 31.36451 | yes | 1.866E-06 | 735.36719 | 64.877243 |
| KYNWSAK | 7 | P61927 | Large ribosomal subunit protein eL37 OS | RPL37 | 11.078 | 2 | 42.606686 | yes | 2.952E-07 | 448.73486 | 2931.5134 |
| VLSVPESTPFTAVI | 15 | P61960 | Ubiquitin-fold modifier 1 OS=Homo sapi | UFM1 | 9.118 | 2 | 47.145763 | yes | 1.39E-07 | 794.45325 | 249.71428 |
| VNLVSGHVK | 9 | P61962 | DDB1- and CUL4-associated factor 7 OS | DCAF7 | 38.927 | 2 | 53.230476 | yes | 1.001E-07 | 476.78235 | 652.28815 |
| HLEHSTIIYEDPQH | 19 | P61962 | DDB1- and CUL4-associated factor 7 OS | DCAF7 | 38.927 | 4 | 53.230476 | yes | 2.042E-07 | 584.55402 | 171.44019 |
| HLEHSTIIYEDPQH | 19 | P61962 | DDB1- and CUL4-associated factor 7 OS | DCAF7 | 38.927 | 4 | 53.230476 | yes | 9.722E-08 | 467.84467 | 171.44019 |
| HLEHSTIIYEDPQH | 19 | P61962 | DDB1- and CUL4-associated factor 7 OS | DCAF7 | 38.927 | 4 | 53.230476 | yes | 1.345E-07 | 779.06958 | 171.44019 |
| HLEHSTIIYEDPQH | 19 | P61962 | DDB1- and CUL4-associated factor 7 OS | DCAF7 | 38.927 | 5 | 53.230476 | yes | 2.042E-07 | 584.55402 | 171.44019 |
| HLEHSTIIYEDPQH | 19 | P61962 | DDB1- and CUL4-associated factor 7 OS | DCAF7 | 38.927 | 5 | 53.230476 | yes | 9.722E-08 | 467.84467 | 171.44019 |
| HLEHSTIIYEDPQH | 19 | P61962 | DDB1- and CUL4-associated factor 7 OS | DCAF7 | 38.927 | 5 | 53.230476 | yes | 1.345E-07 | 779.06958 | 171.44019 |
| HLEHSTIIYEDPQH | 19 | P61962 | DDB1- and CUL4-associated factor 7 OS | DCAF7 | 38.927 | 3 | 53.230476 | yes | 2.042E-07 | 584.55402 | 171.44019 |
| HLEHSTIIYEDPQH | 19 | P61962 | DDB1- and CUL4-associated factor 7 OS | DCAF7 | 38.927 | 3 | 53.230476 | yes | 9.722E-08 | 467.84467 | 171.44019 |
| HLEHSTIIYEDPQH | 19 | P61962 | DDB1- and CUL4-associated factor 7 OS | DCAF7 | 38.927 | 3 | 53.230476 | yes | 1.345E-07 | 779.06958 | 171.44019 |
| LECLLNNNK | 9 | P61962 | DDB1- and CUL4-associated factor 7 OS | DCAF7 | 38.927 | 2 | 53.230476 | yes | 1.647E-07 | 559.28717 | 458.20496 |
| LALGSFVEEYNNK | 13 | P61962 | DDB1- and CUL4-associated factor 7 OS | DCAF7 | 38.927 | 2 | 53.230476 | yes | 1.345E-07 | 742.375 | 434.65295 |
| GVYPDLLATSGDY | 15 | P61962 | DDB1- and CUL4-associated factor 7 OS | DCAF7 | 38.927 | 2 | 53.230476 | yes | 2.146E-06 | 820.41992 | 373.99246 |
| DMFASVGADGSVI | 13 | P61962 | DDB1- and CUL4-associated factor 7 OS | DCAF7 | 38.927 | 2 | 53.230476 | yes | 1.402E-07 | 656.30353 | 363.66632 |
| TQLIAHDK | 8 | P61962 | DDB1- and CUL4-associated factor 7 OS | DCAF7 | 38.927 | 2 | 53.230476 | yes | 6.329E-07 | 463.25873 | 333.4711 |
| VPCTPVAR | 8 | P61962 | DDB1- and CUL4-associated factor 7 OS | DCAF7 | 38.927 | 2 | 53.230476 | yes | 2.323E-07 | 450.242 | 213.46091 |
| NTFDHPYPTTK | 11 | P61962 | DDB1- and CUL4-associated factor 7 OS | DCAF7 | 38.927 | 2 | 53.230476 | yes | 9.722E-08 | 660.81458 | 197.86925 |
| EVYDIAFSR | 9 | P61962 | DDB1- and CUL4-associated factor 7 OS | DCAF7 | 38.927 | 2 | 53.230476 | yes | 1.912E-07 | 550.27454 | 181.09628 |
| LMWIPDTK | 8 | P61962 | DDB1- and CUL4-associated factor 7 OS | DCAF7 | 38.927 | 2 | 53.230476 | yes | 1.293E-07 | 502.2677 | 91.518898 |
| DGSLIVSSSYDGL( | 15 | P61964 | WD repeat-containing protein 5 OS=Hon | WDR5 | 36.589 | 2 | 53.847324 | yes | 2.97E-07 | 814.88287 | 397.90372 |
| FSPNGEWLASSSA | 15 | P61964 | WD repeat-containing protein 5 OS=Hon | WDR5 | 36.589 | 2 | 53.847324 | yes | 2.969E-07 | 798.37042 | 230.58032 |
| IWDTASGQCLK | 11 | P61964 | WD repeat-containing protein 5 OS=Hon | WDR5 | 36.589 | 2 | 53.847324 | yes | 1.375E-07 | 639.81097 | 174.82259 |
| TLIDDDNPPVSFVI | 14 | P61964 | WD repeat-containing protein 5 OS=Hon | WDR5 | 36.589 | 2 | 53.847324 | yes | 1.537E-07 | 780.40125 | 174.63245 |
| YCIFANFSVTGGK | 13 | P61964 | WD repeat-containing protein 5 OS=Hon | WDR5 | 36.589 | 2 | 53.847324 | no | 9.722E-08 | 732.35303 | 88.656853 |
| IWDVSSGK | 8 | P61964 | WD repeat-containing protein 5 OS=Hon | WDR5 | 36.589 | 2 | 53.847324 | yes | 2.866E-07 | 446.23218 | 84.398705 |
| TLPAHSDPVSAVH | 16 | P61964 | WD repeat-containing protein 5 OS=Hon | WDR5 | 36.589 | 3 | 53.847324 | yes | 1.205E-07 | 583.30066 | 53.521629 |
| AQPTPSSSATQSKI | 24 | P61964 | WD repeat-containing protein 5 OS=Hon | WDR5 | 36.589 | 3 | 53.847324 | yes | 1.364E-07 | 833.44647 | 43.529354 |
| AQPTPSSSATQSK | 13 | P61964 | WD repeat-containing protein 5 OS=Hon | WDR5 | 36.589 | 2 | 53.847324 | yes | 5.942E-07 | 645.32025 | 13.027825 |
| AQGNVYHLK | 9 | P61968 | LIM domain transcription factor LMO4 C | LMO4 | 17.994 | 2 | 52.37479 | yes | 6.835E-07 | 515.27747 | 96.195473 |
| CFTCSTCR | 8 | P61968 | LIM domain transcription factor LMO4 C | LMO4 | 17.994 | 2 | 52.37479 | yes | 9.722E-08 | 546.20697 | 46.542931 |
| IDEPLEGSEDR | 11 | P61978 | Heterogeneous nuclear ribonucleoprotein | HNRNPK | 50.977 | 2 | 52.66764 | yes | 1.358E-06 | 630.29114 | 42668.77 |
| GSDFDCELR | 9 | P61978 | Heterogeneous nuclear ribonucleoprotein | HNRNPK | 50.977 | 2 | 52.66764 | yes | 1.592E-07 | 549.72968 | 29896.127 |

| | | | | | | | | | | | |
|---|---|---|---|---|---|---|---|---|---|---|---|
| VVLIGGKPDR | 10 | P61978 | Heterogeneous nuclear ribonucleoprotein | HNRNPK | 50.977 | 2 | 52.66764 | yes | 1.737E-07 | 527.3244 | 27999.145 |
| NTDEMVELR | 9 | P61978 | Heterogeneous nuclear ribonucleoprotein | HNRNPK | 50.977 | 2 | 52.66764 | yes | 1.113E-07 | 553.76099 | 11268.834 |
| NTDEMVELR | 9 | P61978 | Heterogeneous nuclear ribonucleoprotein | HNRNPK | 50.977 | 2 | 52.66764 | yes | 2.125E-07 | 561.75842 | 11268.834 |
| NLPLPPPPPPR | 11 | P61978 | Heterogeneous nuclear ribonucleoprotein | HNRNPK | 50.977 | 2 | 52.66764 | yes | 2.623E-06 | 597.85333 | 16106.785 |
| LFQECCPHSTDR | 12 | P61978 | Heterogeneous nuclear ribonucleoprotein | HNRNPK | 50.977 | 2 | 52.66764 | yes | 9.722E-08 | 775.32977 | 5390.4463 |
| LFQECCPHSTDR | 12 | P61978 | Heterogeneous nuclear ribonucleoprotein | HNRNPK | 50.977 | 2 | 52.66764 | yes | 9.934E-08 | 517.22223 | 5390.4463 |
| LFQECCPHSTDR | 12 | P61978 | Heterogeneous nuclear ribonucleoprotein | HNRNPK | 50.977 | 3 | 52.66764 | yes | 9.722E-08 | 775.32977 | 5390.4463 |
| LFQECCPHSTDR | 12 | P61978 | Heterogeneous nuclear ribonucleoprotein | HNRNPK | 50.977 | 3 | 52.66764 | yes | 9.934E-08 | 517.22223 | 5390.4463 |
| GGDLMAYDR | 9 | P61978 | Heterogeneous nuclear ribonucleoprotein | HNRNPK | 50.977 | 2 | 52.66764 | yes | 1.123E-07 | 499.22403 | 5237.5029 |
| GGDLMAYDR | 9 | P61978 | Heterogeneous nuclear ribonucleoprotein | HNRNPK | 50.977 | 2 | 52.66764 | yes | 1.422E-07 | 507.22147 | 5237.5029 |
| LLIHQSLAGGIIGV | 15 | P61978 | Heterogeneous nuclear ribonucleoprotein | HNRNPK | 50.977 | 2 | 52.66764 | yes | 1.443E-07 | 759.97192 | 4616.335 |
| LLIHQSLAGGIIGV | 15 | P61978 | Heterogeneous nuclear ribonucleoprotein | HNRNPK | 50.977 | 2 | 52.66764 | yes | 1.216E-07 | 506.98373 | 4616.335 |
| LLIHQSLAGGIIGV | 15 | P61978 | Heterogeneous nuclear ribonucleoprotein | HNRNPK | 50.977 | 3 | 52.66764 | yes | 1.443E-07 | 759.97192 | 4616.335 |
| LLIHQSLAGGIIGV | 15 | P61978 | Heterogeneous nuclear ribonucleoprotein | HNRNPK | 50.977 | 3 | 52.66764 | yes | 1.216E-07 | 506.98373 | 4616.335 |
| IILDLISESPIK | 12 | P61978 | Heterogeneous nuclear ribonucleoprotein | HNRNPK | 50.977 | 2 | 52.66764 | yes | 1.003E-07 | 670.9054 | 7172.6582 |
| DYDDMSPR | 8 | P61978 | Heterogeneous nuclear ribonucleoprotein | HNRNPK | 50.977 | 2 | 52.66764 | yes | 1.2E-07 | 499.69785 | 5254.7822 |
| RPAEDMEEEQAFK | 13 | P61978 | Heterogeneous nuclear ribonucleoprotein | HNRNPK | 50.977 | 2 | 52.66764 | yes | 1.084E-07 | 790.35651 | 1151.8589 |
| RPAEDMEEEQAFK | 13 | P61978 | Heterogeneous nuclear ribonucleoprotein | HNRNPK | 50.977 | 2 | 52.66764 | yes | 2.373E-07 | 798.35394 | 1151.8589 |
| ENTQTTIK | 8 | P61978 | Heterogeneous nuclear ribonucleoprotein | HNRNPK | 50.977 | 2 | 52.66764 | yes | 3.656E-07 | 467.74564 | 1626.6204 |
| RDYDDMSPR | 9 | P61978 | Heterogeneous nuclear ribonucleoprotein | HNRNPK | 50.977 | 2 | 52.66764 | yes | 9.99E-08 | 577.74841 | 1074.6969 |
| DLAGSIIGK | 9 | P61978 | Heterogeneous nuclear ribonucleoprotein | HNRNPK | 50.977 | 2 | 52.66764 | yes | 1.276E-07 | 437.25565 | 820.38141 |
| PAEDMEEEQAFK | 12 | P61978 | Heterogeneous nuclear ribonucleoprotein | HNRNPK | 50.977 | 2 | 52.66764 | yes | 9.722E-08 | 712.30591 | 455.39572 |
| GRPVGFPMR | 9 | P61978 | Heterogeneous nuclear ribonucleoprotein | HNRNPK | 50.977 | 2 | 52.66764 | yes | 3.244E-07 | 508.77655 | 194.07625 |
| RPAEDMEEEQAFK | 14 | P61978 | Heterogeneous nuclear ribonucleoprotein | HNRNPK | 50.977 | 3 | 52.66764 | yes | 3.034E-06 | 579.2738 | 22.990421 |
| METEQPEETFPNT | 21 | P61978 | Heterogeneous nuclear ribonucleoprotein | HNRNPK | 50.977 | 3 | 52.66764 | yes | 2.873E-05 | 819.68481 | 16.127859 |
| EHMQPTHPIR | 10 | P61981 | 14-3-3 protein gamma OS=Homo sapiens | YWHAG | 28.303 | 3 | 53.233208 | yes | 1.227E-07 | 415.87674 | 264.5231 |
| EHMQPTHPIR | 10 | P61981 | 14-3-3 protein gamma OS=Homo sapiens | YWHAG | 28.303 | 3 | 53.233208 | yes | 9.937E-08 | 623.31146 | 264.5231 |
| EHMQPTHPIR | 10 | P61981 | 14-3-3 protein gamma OS=Homo sapiens | YWHAG | 28.303 | 2 | 53.233208 | yes | 1.227E-07 | 415.87674 | 264.5231 |
| EHMQPTHPIR | 10 | P61981 | 14-3-3 protein gamma OS=Homo sapiens | YWHAG | 28.303 | 2 | 53.233208 | yes | 9.937E-08 | 623.31146 | 264.5231 |
| NVTELNEPLSNEE | 14 | P61981 | 14-3-3 protein gamma OS=Homo sapiens | YWHAG | 28.303 | 2 | 53.233208 | yes | 9.722E-08 | 822.39722 | 272.19333 |
| ATVVESSEK | 9 | P61981 | 14-3-3 protein gamma OS=Homo sapiens | YWHAG | 28.303 | 2 | 53.233208 | yes | 2.532E-07 | 475.24548 | 228.20789 |
| AYSEAHEISK | 10 | P61981 | 14-3-3 protein gamma OS=Homo sapiens | YWHAG | 28.303 | 2 | 53.233208 | yes | 1.117E-07 | 567.7749 | 222.37692 |
| MKGDYYR | 7 | P61981 | 14-3-3 protein gamma OS=Homo sapiens | YWHAG | 28.303 | 2 | 53.233208 | no | 2.223E-07 | 466.71838 | 196.10158 |
| YLAEVATGEK | 10 | P61981 | 14-3-3 protein gamma OS=Homo sapiens | YWHAG | 28.303 | 2 | 53.233208 | yes | 1.002E-07 | 540.78223 | 150.50609 |
| NCSETQYESK | 10 | P61981 | 14-3-3 protein gamma OS=Homo sapiens | YWHAG | 28.303 | 2 | 53.233208 | yes | 9.722E-08 | 623.25623 | 136.83679 |
| YLDIHER | 7 | P62072 | Mitochondrial import inner membrane tra | TIMM10 | 10.333 | 2 | 48.643566 | yes | 1.985E-07 | 473.24307 | 131.94481 |
| LTELSMQDEELMK | 13 | P62072 | Mitochondrial import inner membrane tra | TIMM10 | 10.333 | 2 | 48.643566 | yes | 1.12E-07 | 783.87311 | 52.83667 |
| HVVFIAQR | 8 | P62081 | Small ribosomal subunit protein eS7 OS= | RPS7 | 22.127 | 2 | 48.78619 | yes | 1.15E-07 | 485.28506 | 33097.547 |
| VETFSGVYK | 9 | P62081 | Small ribosomal subunit protein eS7 OS= | RPS7 | 22.127 | 2 | 48.78619 | yes | 1.418E-07 | 515.26624 | 22814.707 |

| | | | | | | | | | | | |
|---|---|---|---|---|---|---|---|---|---|---|---|
| IVKPNGEKPDEFES | 30 | P62081 | Small ribosomal subunit protein eS7 OS= | RPS7 | 22.127 | 4 | 48.78619 | yes | 2.508E-07 | 833.4248 | 736.25732 |
| KAIIIFVPVPQLK | 13 | P62081 | Small ribosomal subunit protein eS7 OS= | RPS7 | 22.127 | 2 | 48.78619 | yes | 1.104E-07 | 733.47888 | 341.24722 |
| LNLDSIIGR | 9 | P62136 | Serine/threonine-protein phosphatase PP1 | PPP1CA | 37.513 | 2 | 47.0928 | yes | 6.666E-07 | 500.79294 | 1443.0468 |
| YGQFSGLNPGGR | 12 | P62136 | Serine/threonine-protein phosphatase PP1 | PPP1CA | 37.513 | 2 | 47.0928 | yes | 1.787E-07 | 626.80707 | 1319.5026 |
| LLEVQGSRPGK | 11 | P62136 | Serine/threonine-protein phosphatase PP1 | PPP1CA | 37.513 | 2 | 47.0928 | yes | 2.189E-07 | 592.34332 | 534.64111 |
| NVQLTENEIR | 10 | P62136 | Serine/threonine-protein phosphatase PP1 | PPP1CA | 37.513 | 2 | 47.0928 | yes | 1.402E-07 | 608.32001 | 320.35144 |
| YQYGGLNSGR | 10 | P62140 | Serine/threonine-protein phosphatase PP1 | PPP1CB | 37.187 | 2 | 52.591743 | yes | 1.074E-07 | 557.76746 | 890.38263 |
| IVQMTEAEVR | 10 | P62140 | Serine/threonine-protein phosphatase PP1 | PPP1CB | 37.187 | 2 | 52.591743 | yes | 1.194E-07 | 588.30811 | 628.21533 |
| ICGDIHGQYTDLLI | 14 | P62140 | Serine/threonine-protein phosphatase PP1 | PPP1CB | 37.187 | 2 | 52.591743 | yes | 1.183E-07 | 830.90924 | 284.8299 |
| ICGDIHGQYTDLLI | 14 | P62140 | Serine/threonine-protein phosphatase PP1 | PPP1CB | 37.187 | 2 | 52.591743 | yes | 1.063E-07 | 554.27527 | 284.8299 |
| ICGDIHGQYTDLLI | 14 | P62140 | Serine/threonine-protein phosphatase PP1 | PPP1CB | 37.187 | 3 | 52.591743 | yes | 1.183E-07 | 830.90924 | 284.8299 |
| ICGDIHGQYTDLLI | 14 | P62140 | Serine/threonine-protein phosphatase PP1 | PPP1CB | 37.187 | 3 | 52.591743 | yes | 1.063E-07 | 554.27527 | 284.8299 |
| GVSFTFGADVVSK | 13 | P62140 | Serine/threonine-protein phosphatase PP1 | PPP1CB | 37.187 | 2 | 52.591743 | yes | 1.394E-06 | 657.34045 | 386.48163 |
| ADGELNVDSLITR | 13 | P62140 | Serine/threonine-protein phosphatase PP1 | PPP1CB | 37.187 | 2 | 52.591743 | yes | 9.722E-08 | 722.86755 | 309.70432 |
| LKPEVVEELTR | 11 | P62166 | Neuronal calcium sensor 1 OS=Homo sap | NCS1 | 21.879 | 2 | 51.153503 | yes | 9.722E-08 | 656.8772 | 97.201035 |
| GVILYGPPGTGK | 12 | P62191 | 26S proteasome regulatory subunit 4 OS= | PSMC1 | 49.185 | 2 | 51.061565 | yes | 9.722E-08 | 579.82953 | 124.78631 |
| KYEPPVPTR | 9 | P62191 | 26S proteasome regulatory subunit 4 OS= | PSMC1 | 49.185 | 2 | 51.061565 | yes | 2.273E-06 | 543.80072 | 39.046112 |
| IEELQLIVNDK | 11 | P62195 | 26S proteasome regulatory subunit 8 OS= | PSMC5 | 45.626 | 2 | 47.986225 | yes | 1.219E-07 | 657.3692 | 159.93628 |
| GVCTEAGMYALR | 12 | P62195 | 26S proteasome regulatory subunit 8 OS= | PSMC5 | 45.626 | 2 | 47.986225 | yes | 3.03E-07 | 664.3103 | 80.51368 |
| IAELMPGASGAEV | 14 | P62195 | 26S proteasome regulatory subunit 8 OS= | PSMC5 | 45.626 | 2 | 47.986225 | yes | 3.667E-07 | 686.86066 | 73.951172 |
| LTPEEEEILNK | 11 | P62241 | Small ribosomal subunit protein eS8 OS= | RPS8 | 24.206 | 2 | 50.73727 | yes | 1.52E-07 | 657.84302 | 43125.102 |
| NCIVLIDSTPYR | 12 | P62241 | Small ribosomal subunit protein eS8 OS= | RPS8 | 24.206 | 2 | 50.73727 | yes | 2.898E-07 | 725.87158 | 38792.609 |
| LLACIASR | 8 | P62241 | Small ribosomal subunit protein eS8 OS= | RPS8 | 24.206 | 2 | 50.73727 | yes | 1.559E-07 | 452.25766 | 30553.48 |
| ISSLLEEQFQQGK | 13 | P62241 | Small ribosomal subunit protein eS8 OS= | RPS8 | 24.206 | 2 | 50.73727 | yes | 9.723E-08 | 753.89355 | 29888.373 |
| ELEFYLR | 7 | P62241 | Small ribosomal subunit protein eS8 OS= | RPS8 | 24.206 | 2 | 50.73727 | yes | 2.971E-07 | 485.25565 | 11861.3 |
| ADGYVLEGK | 9 | P62241 | Small ribosomal subunit protein eS8 OS= | RPS8 | 24.206 | 2 | 50.73727 | yes | 1.422E-07 | 476.24274 | 8580.4395 |
| QWYESHYALPLGI | 13 | P62241 | Small ribosomal subunit protein eS8 OS= | RPS8 | 24.206 | 2 | 50.73727 | yes | 1.557E-07 | 810.40192 | 3572.6475 |
| QWYESHYALPLGI | 13 | P62241 | Small ribosomal subunit protein eS8 OS= | RPS8 | 24.206 | 2 | 50.73727 | yes | 1.128E-07 | 540.6037 | 3572.6475 |
| QWYESHYALPLGI | 13 | P62241 | Small ribosomal subunit protein eS8 OS= | RPS8 | 24.206 | 3 | 50.73727 | yes | 1.557E-07 | 810.40192 | 3572.6475 |
| QWYESHYALPLGI | 13 | P62241 | Small ribosomal subunit protein eS8 OS= | RPS8 | 24.206 | 3 | 50.73727 | yes | 1.128E-07 | 540.6037 | 3572.6475 |
| LTPEEEEILNKK | 12 | P62241 | Small ribosomal subunit protein eS8 OS= | RPS8 | 24.206 | 2 | 50.73727 | yes | 2.636E-07 | 721.8905 | 1093.8573 |
| LLACIASRPGQCGI | 14 | P62241 | Small ribosomal subunit protein eS8 OS= | RPS8 | 24.206 | 2 | 50.73727 | yes | 1.695E-07 | 779.9007 | 1007.3788 |
| IIDVVYNASNNEL | 15 | P62241 | Small ribosomal subunit protein eS8 OS= | RPS8 | 24.206 | 3 | 50.73727 | yes | 1.686E-07 | 573.64056 | 347.85446 |
| YELGRPAANTK | 11 | P62241 | Small ribosomal subunit protein eS8 OS= | RPS8 | 24.206 | 2 | 50.73727 | yes | 9.758E-08 | 610.32513 | 73.588364 |
| WQNNLLPSR | 9 | P62244 | Small ribosomal subunit protein uS8 OS= | RPS15A | 14.84 | 2 | 50.738586 | yes | 3.569E-07 | 564.30145 | 49470.809 |
| HGYIGEFEIIDDHR | 14 | P62244 | Small ribosomal subunit protein uS8 OS= | RPS15A | 14.84 | 3 | 50.738586 | yes | 1.088E-07 | 567.60577 | 46680.895 |
| IVVNLTGR | 8 | P62244 | Small ribosomal subunit protein uS8 OS= | RPS15A | 14.84 | 2 | 50.738586 | yes | 4.109E-07 | 436.27164 | 32289.779 |
| MNVLADALK | 9 | P62244 | Small ribosomal subunit protein uS8 OS= | RPS15A | 14.84 | 2 | 50.738586 | yes | 1.775E-07 | 487.7706 | 11997.588 |
| MNVLADALK | 9 | P62244 | Small ribosomal subunit protein uS8 OS= | RPS15A | 14.84 | 2 | 50.738586 | yes | 2.333E-07 | 495.76807 | 11997.588 |

| | | | | | | | | | | |
|---|---|---|---|---|---|---|---|---|---|---|
| QVLIRPCSK | 9 | P62244 | Small ribosomal subunit protein uS8 OS= RPS15A | 14.84 | 2 | 50.738586 | yes | 1.106E-07 | 550.81586 | 12244.101 |
| QFGFIVLTTSAGIM | 20 | P62244 | Small ribosomal subunit protein uS8 OS= RPS15A | 14.84 | 3 | 50.738586 | yes | 9.723E-08 | 741.36853 | 2688.8972 |
| QFGFIVLTTSAGIM | 20 | P62244 | Small ribosomal subunit protein uS8 OS= RPS15A | 14.84 | 3 | 50.738586 | yes | 3.356E-07 | 746.7002 | 2688.8972 |
| AGKIVVNLTGR | 11 | P62244 | Small ribosomal subunit protein uS8 OS= RPS15A | 14.84 | 2 | 50.738586 | yes | 5.099E-07 | 564.34839 | 56.271297 |
| FLTVMMK | 7 | P62244 | Small ribosomal subunit protein uS8 OS= RPS15A | 14.84 | 2 | 50.738586 | yes | 2.102E-06 | 443.23227 | 43.737679 |
| PLEMIEPR | 8 | P62249 | Small ribosomal subunit protein uS9 OS= RPS16 | 16.446 | 2 | 49.6987 | yes | 1.804E-07 | 492.76279 | 24035.822 |
| PLEMIEPR | 8 | P62249 | Small ribosomal subunit protein uS9 OS= RPS16 | 16.446 | 2 | 49.6987 | yes | 6.46E-06 | 500.76022 | 24035.822 |
| GPLQSVQVFGR | 11 | P62249 | Small ribosomal subunit protein uS9 OS= RPS16 | 16.446 | 2 | 49.6987 | yes | 1.009E-07 | 594.3302 | 41864.043 |
| GGGHVAQIYAIR | 12 | P62249 | Small ribosomal subunit protein uS9 OS= RPS16 | 16.446 | 2 | 49.6987 | yes | 1.094E-07 | 621.34113 | 30803.145 |
| DILIQYDR | 8 | P62249 | Small ribosomal subunit protein uS9 OS= RPS16 | 16.446 | 2 | 49.6987 | yes | 1.362E-07 | 518.2771 | 22357.811 |
| TLLVADPR | 8 | P62249 | Small ribosomal subunit protein uS9 OS= RPS16 | 16.446 | 2 | 49.6987 | yes | 5.358E-07 | 442.76364 | 16486.596 |
| LLEPVLLLGK | 10 | P62249 | Small ribosomal subunit protein uS9 OS= RPS16 | 16.446 | 2 | 49.6987 | yes | 4.825E-07 | 547.86279 | 14396.674 |
| ALVAYYQK | 8 | P62249 | Small ribosomal subunit protein uS9 OS= RPS16 | 16.446 | 2 | 49.6987 | yes | 1.87E-07 | 478.26602 | 5566.8379 |
| VNGRPLEMIEPR | 12 | P62249 | Small ribosomal subunit protein uS9 OS= RPS16 | 16.446 | 2 | 49.6987 | yes | 1.735E-07 | 705.87976 | 367.172 |
| EIKDILIQYDR | 11 | P62249 | Small ribosomal subunit protein uS9 OS= RPS16 | 16.446 | 2 | 49.6987 | yes | 2.662E-07 | 703.38794 | 119.0767 |
| EIKDILIQYDR | 11 | P62249 | Small ribosomal subunit protein uS9 OS= RPS16 | 16.446 | 2 | 49.6987 | yes | 6.41E-07 | 469.26105 | 119.0767 |
| EIKDILIQYDR | 11 | P62249 | Small ribosomal subunit protein uS9 OS= RPS16 | 16.446 | 3 | 49.6987 | yes | 2.662E-07 | 703.38794 | 119.0767 |
| EIKDILIQYDR | 11 | P62249 | Small ribosomal subunit protein uS9 OS= RPS16 | 16.446 | 3 | 49.6987 | yes | 6.41E-07 | 469.26105 | 119.0767 |
| VKGGGHVAQIYAI | 14 | P62249 | Small ribosomal subunit protein uS9 OS= RPS16 | 16.446 | 3 | 49.6987 | yes | 1.462E-07 | 490.2843 | 82.956924 |
| YLAEFATGNDR | 11 | P62258 | 14-3-3 protein epsilon OS=Homo sapiens YWHAE | 29.174 | 2 | 53.09024 | yes | 9.722E-08 | 628.79895 | 754.18518 |
| VAGMDVELTVEEI | 13 | P62258 | 14-3-3 protein epsilon OS=Homo sapiens YWHAE | 29.174 | 2 | 53.09024 | yes | 1.355E-07 | 724.35852 | 545.47375 |
| IISSIEQK | 8 | P62258 | 14-3-3 protein epsilon OS=Homo sapiens YWHAE | 29.174 | 2 | 53.09024 | yes | 2.438E-06 | 459.26874 | 412.72827 |
| EAAENSLVAYK | 11 | P62258 | 14-3-3 protein epsilon OS=Homo sapiens YWHAE | 29.174 | 2 | 53.09024 | yes | 1.075E-07 | 597.80371 | 365.01602 |
| HLIPAANTGESK | 12 | P62258 | 14-3-3 protein epsilon OS=Homo sapiens YWHAE | 29.174 | 2 | 53.09024 | yes | 2.038E-05 | 619.33038 | 238.12819 |
| YDEMVESMK | 9 | P62258 | 14-3-3 protein epsilon OS=Homo sapiens YWHAE | 29.174 | 2 | 53.09024 | yes | 2.546E-07 | 566.23846 | 179.74965 |
| MDDREDLVYQAK | 12 | P62258 | 14-3-3 protein epsilon OS=Homo sapiens YWHAE | 29.174 | 2 | 53.09024 | yes | 9.722E-08 | 762.85358 | 132.20305 |
| QMVETELK | 8 | P62258 | 14-3-3 protein epsilon OS=Homo sapiens YWHAE | 29.174 | 2 | 53.09024 | yes | 1.453E-07 | 489.25223 | 107.88268 |
| LICCDILDVLDK | 12 | P62258 | 14-3-3 protein epsilon OS=Homo sapiens YWHAE | 29.174 | 2 | 53.09024 | yes | 3.576E-07 | 738.87549 | 75.980728 |
| IEDVTPIPSDSTR | 13 | P62263 | Small ribosomal subunit protein uS11 OS RPS14 | 16.273 | 2 | 55.680573 | yes | 9.722E-08 | 715.36212 | 82643.289 |
| TPGPGAQSALR | 11 | P62263 | Small ribosomal subunit protein uS11 OS RPS14 | 16.273 | 2 | 55.680573 | yes | 1.888E-07 | 527.78564 | 40101.371 |
| ADRDESSPYAAMI | 21 | P62263 | Small ribosomal subunit protein uS11 OS RPS14 | 16.273 | 3 | 55.680573 | yes | 1.156E-07 | 755.69031 | 17743.68 |
| ADRDESSPYAAMI | 21 | P62263 | Small ribosomal subunit protein uS11 OS RPS14 | 16.273 | 3 | 55.680573 | yes | 6.436E-07 | 761.02197 | 17743.68 |
| ELGITALHIK | 10 | P62263 | Small ribosomal subunit protein uS11 OS RPS14 | 16.273 | 2 | 55.680573 | yes | 1.108E-07 | 547.83203 | 754.60437 |
| CKELGITALHIK | 12 | P62263 | Small ribosomal subunit protein uS11 OS RPS14 | 16.273 | 3 | 55.680573 | yes | 9.722E-08 | 461.599 | 347.99612 |
| CKELGITALHIK | 12 | P62263 | Small ribosomal subunit protein uS11 OS RPS14 | 16.273 | 3 | 55.680573 | yes | 9.722E-08 | 691.89484 | 347.99612 |
| CKELGITALHIK | 12 | P62263 | Small ribosomal subunit protein uS11 OS RPS14 | 16.273 | 2 | 55.680573 | yes | 9.722E-08 | 461.599 | 347.99612 |
| CKELGITALHIK | 12 | P62263 | Small ribosomal subunit protein uS11 OS RPS14 | 16.273 | 2 | 55.680573 | yes | 9.722E-08 | 691.89484 | 347.99612 |
| TKTPGPGAQSALR | 13 | P62263 | Small ribosomal subunit protein uS11 OS RPS14 | 16.273 | 2 | 55.680573 | yes | 1.405E-07 | 642.35693 | 256.69083 |
| DESSPYAAMLAAC | 18 | P62263 | Small ribosomal subunit protein uS11 OS RPS14 | 16.273 | 3 | 55.680573 | yes | 2.92E-07 | 641.63525 | 33.754799 |

| | | | | | | | | | | | |
|---|---|---|---|---|---|---|---|---|---|---|---|
| ANPFGGASHAK | 11 | P62266 | Small ribosomal subunit protein uS12 OS | RPS23 | 15.808 | 2 | 47.78988 | yes | 1.259E-07 | 528.76471 | 37226.688 |
| GHAVGDIPGVR | 11 | P62266 | Small ribosomal subunit protein uS12 OS | RPS23 | 15.808 | 2 | 47.78988 | yes | 1.284E-07 | 539.29364 | 27429.379 |
| AHLGTALK | 8 | P62266 | Small ribosomal subunit protein uS12 OS | RPS23 | 15.808 | 2 | 47.78988 | yes | 2.298E-07 | 405.74524 | 14272.312 |
| VANVSLLALYK | 11 | P62266 | Small ribosomal subunit protein uS12 OS | RPS23 | 15.808 | 2 | 47.78988 | yes | 1.313E-07 | 595.86078 | 11649.539 |
| KGHAVGDIPGVR | 12 | P62266 | Small ribosomal subunit protein uS12 OS | RPS23 | 15.808 | 2 | 47.78988 | yes | 2.063E-07 | 603.34113 | 3425.616 |
| KGHAVGDIPGVR | 12 | P62266 | Small ribosomal subunit protein uS12 OS | RPS23 | 15.808 | 2 | 47.78988 | yes | 2.767E-07 | 402.56317 | 3425.616 |
| KGHAVGDIPGVR | 12 | P62266 | Small ribosomal subunit protein uS12 OS | RPS23 | 15.808 | 3 | 47.78988 | yes | 2.063E-07 | 603.34113 | 3425.616 |
| KGHAVGDIPGVR | 12 | P62266 | Small ribosomal subunit protein uS12 OS | RPS23 | 15.808 | 3 | 47.78988 | yes | 2.767E-07 | 402.56317 | 3425.616 |
| KAHLGTALK | 9 | P62266 | Small ribosomal subunit protein uS12 OS | RPS23 | 15.808 | 2 | 47.78988 | yes | 2.482E-07 | 469.79272 | 3970.3071 |
| IPDWFLNR | 8 | P62269 | Small ribosomal subunit protein uS13 OS | RPS18 | 17.719 | 2 | 51.402775 | yes | 2.105E-07 | 530.78235 | 59952.559 |
| VITIMQNPR | 9 | P62269 | Small ribosomal subunit protein uS13 OS | RPS18 | 17.719 | 2 | 51.402775 | yes | 1.526E-07 | 536.30261 | 27066.35 |
| VITIMQNPR | 9 | P62269 | Small ribosomal subunit protein uS13 OS | RPS18 | 17.719 | 2 | 51.402775 | yes | 3.533E-07 | 544.30005 | 27066.35 |
| YAHVVLR | 7 | P62269 | Small ribosomal subunit protein uS13 OS | RPS18 | 17.719 | 2 | 51.402775 | yes | 3.112E-07 | 429.25323 | 34248.074 |
| AGELTEDEVER | 11 | P62269 | Small ribosomal subunit protein uS13 OS | RPS18 | 17.719 | 2 | 51.402775 | yes | 9.722E-08 | 624.29114 | 33820.746 |
| LREDLER | 7 | P62269 | Small ribosomal subunit protein uS13 OS | RPS18 | 17.719 | 2 | 51.402775 | yes | 9.926E-08 | 465.75378 | 12264.375 |
| KADIDLTK | 8 | P62269 | Small ribosomal subunit protein uS13 OS | RPS18 | 17.719 | 2 | 51.402775 | yes | 2E-07 | 452.26093 | 11722.364 |
| YSQVLANGLDNK | 12 | P62269 | Small ribosomal subunit protein uS13 OS | RPS18 | 17.719 | 2 | 51.402775 | yes | 1.022E-07 | 661.34094 | 11366.484 |
| IAFAITAIK | 9 | P62269 | Small ribosomal subunit protein uS13 OS | RPS18 | 17.719 | 2 | 51.402775 | yes | 1.992E-07 | 474.29987 | 6088.0596 |
| RAGELTEDEVER | 12 | P62269 | Small ribosomal subunit protein uS13 OS | RPS18 | 17.719 | 2 | 51.402775 | yes | 1.028E-07 | 702.34167 | 1047.6113 |
| RYAHVVLR | 8 | P62269 | Small ribosomal subunit protein uS13 OS | RPS18 | 17.719 | 2 | 51.402775 | yes | 1.108E-07 | 507.3038 | 904.99652 |
| VLNTNIDGR | 9 | P62269 | Small ribosomal subunit protein uS13 OS | RPS18 | 17.719 | 2 | 51.402775 | yes | 2.717E-06 | 501.27237 | 886.52545 |
| KIAFAITAIK | 10 | P62269 | Small ribosomal subunit protein uS13 OS | RPS18 | 17.719 | 2 | 51.402775 | yes | 9.722E-08 | 538.34735 | 175.50316 |
| KYGLNMCR | 8 | P62273 | Small ribosomal subunit protein uS14 OS | RPS29 | 6.677 | 2 | 51.531677 | yes | 1.629E-07 | 521.25208 | 5540.8535 |
| KYGLNMCR | 8 | P62273 | Small ribosomal subunit protein uS14 OS | RPS29 | 6.677 | 2 | 51.531677 | yes | 1.75E-06 | 529.24951 | 5540.8535 |
| GHQQLYWSHPR | 11 | P62273 | Small ribosomal subunit protein uS14 OS | RPS29 | 6.677 | 2 | 51.531677 | yes | 9.722E-08 | 704.84711 | 1612.0565 |
| GHQQLYWSHPR | 11 | P62273 | Small ribosomal subunit protein uS14 OS | RPS29 | 6.677 | 2 | 51.531677 | yes | 3.344E-06 | 725.85236 | 1612.0565 |
| YGLNMCR | 7 | P62273 | Small ribosomal subunit protein uS14 OS | RPS29 | 6.677 | 2 | 51.531677 | yes | 3.774E-06 | 457.20456 | 27.320969 |
| GLTPSQIGVILR | 12 | P62277 | Small ribosomal subunit protein uS15 OS | RPS13 | 17.223 | 2 | 52.346756 | yes | 7.589E-07 | 627.38245 | 52262.281 |
| GLSQSALPYR | 10 | P62277 | Small ribosomal subunit protein uS15 OS | RPS13 | 17.223 | 2 | 52.346756 | yes | 1.176E-07 | 546.29584 | 47551.195 |
| DSHGVAQVR | 9 | P62277 | Small ribosomal subunit protein uS15 OS | RPS13 | 17.223 | 2 | 52.346756 | yes | 1.682E-07 | 484.74905 | 36077.859 |
| LILIESR | 7 | P62277 | Small ribosomal subunit protein uS15 OS | RPS13 | 17.223 | 2 | 52.346756 | yes | 3.074E-06 | 422.26855 | 20653.186 |
| KGLTPSQIGVILR | 13 | P62277 | Small ribosomal subunit protein uS15 OS | RPS13 | 17.223 | 2 | 52.346756 | yes | 1.083E-07 | 691.42993 | 2148.3328 |
| KGLTPSQIGVILR | 13 | P62277 | Small ribosomal subunit protein uS15 OS | RPS13 | 17.223 | 2 | 52.346756 | yes | 2.712E-07 | 461.28903 | 2148.3328 |
| KGLTPSQIGVILR | 13 | P62277 | Small ribosomal subunit protein uS15 OS | RPS13 | 17.223 | 3 | 52.346756 | yes | 1.083E-07 | 691.42993 | 2148.3328 |
| KGLTPSQIGVILR | 13 | P62277 | Small ribosomal subunit protein uS15 OS | RPS13 | 17.223 | 3 | 52.346756 | yes | 2.712E-07 | 461.28903 | 2148.3328 |
| VLPPNWK | 7 | P62277 | Small ribosomal subunit protein uS15 OS | RPS13 | 17.223 | 2 | 52.346756 | yes | 2.75E-06 | 427.25015 | 2144.9124 |
| GLAPDLPEDLYHL | 15 | P62277 | Small ribosomal subunit protein uS15 OS | RPS13 | 17.223 | 2 | 52.346756 | yes | 1.221E-07 | 847.46161 | 998.26086 |
| GLAPDLPEDLYHL | 15 | P62277 | Small ribosomal subunit protein uS15 OS | RPS13 | 17.223 | 2 | 52.346756 | yes | 1.467E-07 | 565.31018 | 998.26086 |
| GLAPDLPEDLYHL | 15 | P62277 | Small ribosomal subunit protein uS15 OS | RPS13 | 17.223 | 3 | 52.346756 | yes | 1.221E-07 | 847.46161 | 998.26086 |

| | | | | | | | | | | | |
|---|---|---|---|---|---|---|---|---|---|---|---|
| GLAPDLPEDLYHL | 15 | P62277 | Small ribosomal subunit protein uS15 OS | RPS13 | 17.223 | 3 | 52.346756 | yes | 1.467E-07 | 565.31018 | 998.26086 |
| SVPTWLK | 7 | P62277 | Small ribosomal subunit protein uS15 OS | RPS13 | 17.223 | 1 | 52.346756 | yes | 1.169E-07 | 830.47705 | 362.24161 |
| SVPTWLK | 7 | P62277 | Small ribosomal subunit protein uS15 OS | RPS13 | 17.223 | 1 | 52.346756 | yes | 9.722E-08 | 415.74216 | 362.24161 |
| SVPTWLK | 7 | P62277 | Small ribosomal subunit protein uS15 OS | RPS13 | 17.223 | 2 | 52.346756 | yes | 1.169E-07 | 830.47705 | 362.24161 |
| SVPTWLK | 7 | P62277 | Small ribosomal subunit protein uS15 OS | RPS13 | 17.223 | 2 | 52.346756 | yes | 9.722E-08 | 415.74216 | 362.24161 |
| RVLPPNWK | 8 | P62277 | Small ribosomal subunit protein uS15 OS | RPS13 | 17.223 | 2 | 52.346756 | yes | 1.294E-06 | 505.30072 | 340.76474 |
| CPFTGNVSIR | 10 | P62280 | Small ribosomal subunit protein uS17 OS | RPS11 | 18.431 | 2 | 50.692535 | yes | 1.298E-07 | 575.78729 | 36719.223 |
| DYLHYIR | 7 | P62280 | Small ribosomal subunit protein uS17 OS | RPS11 | 18.431 | 2 | 50.692535 | yes | 2.421E-06 | 490.25345 | 31563.26 |
| NMSVHLSPCFR | 11 | P62280 | Small ribosomal subunit protein uS17 OS | RPS11 | 18.431 | 2 | 50.692535 | yes | 3.208E-07 | 674.31848 | 7702.0151 |
| NMSVHLSPCFR | 11 | P62280 | Small ribosomal subunit protein uS17 OS | RPS11 | 18.431 | 2 | 50.692535 | yes | 1.657E-07 | 449.88141 | 7702.0151 |
| NMSVHLSPCFR | 11 | P62280 | Small ribosomal subunit protein uS17 OS | RPS11 | 18.431 | 2 | 50.692535 | yes | 5.736E-07 | 682.31592 | 7702.0151 |
| NMSVHLSPCFR | 11 | P62280 | Small ribosomal subunit protein uS17 OS | RPS11 | 18.431 | 3 | 50.692535 | yes | 3.208E-07 | 674.31848 | 7702.0151 |
| NMSVHLSPCFR | 11 | P62280 | Small ribosomal subunit protein uS17 OS | RPS11 | 18.431 | 3 | 50.692535 | yes | 1.657E-07 | 449.88141 | 7702.0151 |
| NMSVHLSPCFR | 11 | P62280 | Small ribosomal subunit protein uS17 OS | RPS11 | 18.431 | 3 | 50.692535 | yes | 5.736E-07 | 682.31592 | 7702.0151 |
| VLLGETGK | 8 | P62280 | Small ribosomal subunit protein uS17 OS | RPS11 | 18.431 | 2 | 50.692535 | yes | 1.122E-07 | 408.7449 | 12438.148 |
| EAIEGTYIDK | 10 | P62280 | Small ribosomal subunit protein uS17 OS | RPS11 | 18.431 | 2 | 50.692535 | yes | 2.199E-07 | 569.78497 | 11502.168 |
| QPTIFQNK | 8 | P62280 | Small ribosomal subunit protein uS17 OS | RPS11 | 18.431 | 2 | 50.692535 | yes | 7.626E-07 | 488.26654 | 11139.867 |
| EAIEGTYIDKK | 11 | P62280 | Small ribosomal subunit protein uS17 OS | RPS11 | 18.431 | 2 | 50.692535 | yes | 9.728E-08 | 633.83246 | 10732.221 |
| DVQIGDIVTVGEC | 14 | P62280 | Small ribosomal subunit protein uS17 OS | RPS11 | 18.431 | 2 | 50.692535 | yes | 1.002E-07 | 780.88794 | 6305.9717 |
| KCPFTGNVSIR | 11 | P62280 | Small ribosomal subunit protein uS17 OS | RPS11 | 18.431 | 2 | 50.692535 | yes | 1.297E-07 | 639.83478 | 1743.4572 |
| RVLLGETGK | 9 | P62280 | Small ribosomal subunit protein uS17 OS | RPS11 | 18.431 | 2 | 50.692535 | yes | 1.087E-07 | 486.79547 | 1040.6521 |
| RDYLHYIR | 8 | P62280 | Small ribosomal subunit protein uS17 OS | RPS11 | 18.431 | 2 | 50.692535 | yes | 1.33E-07 | 568.30402 | 505.93848 |
| ILSGVVTK | 8 | P62280 | Small ribosomal subunit protein uS17 OS | RPS11 | 18.431 | 2 | 50.692535 | yes | 5.075E-07 | 408.76309 | 163.06563 |
| DVQIGDIVTVGEC | 18 | P62280 | Small ribosomal subunit protein uS17 OS | RPS11 | 18.431 | 3 | 50.692535 | yes | 1.335E-07 | 662.68231 | 87.57634 |
| NIGLGFK | 7 | P62280 | Small ribosomal subunit protein uS17 OS | RPS11 | 18.431 | 1 | 50.692535 | yes | 2.762E-06 | 748.43518 | 59.55024 |
| IQVWLYEQVNMR | 12 | P62304 | Small nuclear ribonucleoprotein E OS=H | SNRPE | 10.804 | 2 | 51.06214 | yes | 9.722E-08 | 789.90851 | 2347.4929 |
| VMVQPINLIFR | 11 | P62304 | Small nuclear ribonucleoprotein E OS=H | SNRPE | 10.804 | 2 | 51.06214 | yes | 1.31E-06 | 665.38922 | 248.17862 |
| VMVQPINLIFR | 11 | P62304 | Small nuclear ribonucleoprotein E OS=H | SNRPE | 10.804 | 2 | 51.06214 | yes | 6.316E-07 | 673.38666 | 248.17862 |
| GDNITLLQSVSN | 12 | P62304 | Small nuclear ribonucleoprotein E OS=H | SNRPE | 10.804 | 2 | 51.06214 | yes | 2.45E-07 | 630.82513 | 363.9772 |
| CNNVLYIR | 8 | P62306 | Small nuclear ribonucleoprotein F OS=H | SNRPF | 9.726 | 2 | 53.250412 | yes | 1.776E-07 | 526.2713 | 2296.1807 |
| PFLNGLTGK | 9 | P62306 | Small nuclear ribonucleoprotein F OS=H | SNRPF | 9.726 | 2 | 53.250412 | yes | 1.3E-07 | 473.77145 | 311.5224 |
| LKWGMEYK | 8 | P62306 | Small nuclear ribonucleoprotein F OS=H | SNRPF | 9.726 | 2 | 53.250412 | yes | 5.627E-07 | 527.77313 | 198.86104 |
| GVEEEEEDGEMR | 12 | P62306 | Small nuclear ribonucleoprotein F OS=H | SNRPF | 9.726 | 2 | 53.250412 | yes | 9.722E-08 | 704.78027 | 91.846382 |
| PFLNGLTGKPVMV | 14 | P62306 | Small nuclear ribonucleoprotein F OS=H | SNRPF | 9.726 | 3 | 53.250412 | yes | 2.54E-07 | 500.9584 | 26.860519 |
| GVEEEEEDGEMRI | 13 | P62306 | Small nuclear ribonucleoprotein F OS=H | SNRPF | 9.726 | 2 | 53.250412 | yes | 1.623E-07 | 769.30157 | 21.610765 |
| GDGVVLVAPPLR | 12 | P62310 | U6 snRNA-associated Sm-like protein LS | LSM3 | 11.846 | 2 | 44.288124 | yes | 2.25E-07 | 596.85608 | 1702.0432 |
| GNNVLYISTQK | 11 | P62312 | U6 snRNA-associated Sm-like protein LS | LSM6 | 9.128 | 2 | 52.10183 | yes | 2.49E-07 | 618.83276 | 1050.0359 |
| QTPSDFLK | 8 | P62312 | U6 snRNA-associated Sm-like protein LS | LSM6 | 9.128 | 2 | 52.10183 | yes | 9.722E-08 | 468.24527 | 411.85059 |
| KQTPSDFLK | 9 | P62312 | U6 snRNA-associated Sm-like protein LS | LSM6 | 9.128 | 2 | 52.10183 | yes | 9.722E-08 | 532.29279 | 90.046371 |

| | | | | | | | | | | | |
|---|---|---|---|---|---|---|---|---|---|---|---|
| QIIGRPVVVK | 10 | P62312 | U6 snRNA-associated Sm-like protein LS | LSM6 | 9.128 | 2 | 52.10183 | yes | 3.524E-06 | 554.86371 | 3.2913523 |
| EPVQLETLSIR | 11 | P62314 | Small nuclear ribonucleoprotein Sm D1 C | SNRPD1 | 13.282 | 2 | 52.245045 | yes | 4.119E-07 | 642.86151 | 1394.2994 |
| LSHETVTIELK | 11 | P62314 | Small nuclear ribonucleoprotein Sm D1 C | SNRPD1 | 13.282 | 2 | 52.245045 | yes | 9.722E-08 | 635.35608 | 998.41443 |
| NREPVQLETLSIR | 13 | P62314 | Small nuclear ribonucleoprotein Sm D1 C | SNRPD1 | 13.282 | 3 | 52.245045 | yes | 1.998E-07 | 518.95813 | 357.76556 |
| NREPVQLETLSIR | 13 | P62314 | Small nuclear ribonucleoprotein Sm D1 C | SNRPD1 | 13.282 | 3 | 52.245045 | yes | 1.28E-07 | 777.93353 | 357.76556 |
| NREPVQLETLSIR | 13 | P62314 | Small nuclear ribonucleoprotein Sm D1 C | SNRPD1 | 13.282 | 2 | 52.245045 | yes | 1.998E-07 | 518.95813 | 357.76556 |
| NREPVQLETLSIR | 13 | P62314 | Small nuclear ribonucleoprotein Sm D1 C | SNRPD1 | 13.282 | 2 | 52.245045 | yes | 1.28E-07 | 777.93353 | 357.76556 |
| NGTQVHGTITGVD | 21 | P62314 | Small nuclear ribonucleoprotein Sm D1 C | SNRPD1 | 13.282 | 3 | 52.245045 | yes | 9.722E-08 | 737.03918 | 377.93854 |
| SEMTPEELQK | 10 | P62316 | Small nuclear ribonucleoprotein Sm D2 C | SNRPD2 | 13.527 | 2 | 49.481632 | yes | 1.928E-07 | 596.28174 | 3163.2173 |
| NNTQVLINCR | 10 | P62316 | Small nuclear ribonucleoprotein Sm D2 C | SNRPD2 | 13.527 | 2 | 49.481632 | yes | 1.16E-07 | 616.31421 | 3095.6084 |
| GDSVIVVLR | 9 | P62316 | Small nuclear ribonucleoprotein Sm D2 C | SNRPD2 | 13.527 | 2 | 49.481632 | yes | 1.183E-07 | 479.29001 | 2538.4541 |
| SLLNKPK | 7 | P62316 | Small nuclear ribonucleoprotein Sm D2 C | SNRPD2 | 13.527 | 2 | 49.481632 | yes | 2.071E-07 | 421.26074 | 2025.6727 |
| HCNMVLENVK | 10 | P62316 | Small nuclear ribonucleoprotein Sm D2 C | SNRPD2 | 13.527 | 2 | 49.481632 | yes | 2.199E-07 | 622.29974 | 1648.0066 |
| EMWTEVPK | 8 | P62316 | Small nuclear ribonucleoprotein Sm D2 C | SNRPD2 | 13.527 | 2 | 49.481632 | yes | 1.026E-07 | 510.24695 | 166.6517 |
| VAQLEQVYIR | 10 | P62318 | Small nuclear ribonucleoprotein Sm D3 C | SNRPD3 | 13.917 | 2 | 48.993614 | yes | 1.079E-07 | 609.8457 | 8119.9829 |
| FLILPDMLK | 9 | P62318 | Small nuclear ribonucleoprotein Sm D3 C | SNRPD3 | 13.917 | 2 | 48.993614 | yes | 2.058E-07 | 553.31995 | 156.30458 |
| VLHEAEGHIVTCE | 21 | P62318 | Small nuclear ribonucleoprotein Sm D3 C | SNRPD3 | 13.917 | 4 | 48.993614 | yes | 1.465E-07 | 604.29059 | 74.338104 |
| ILMLGLDAAGK | 11 | P62330 | ADP-ribosylation factor 6 OS=Homo sap | ARF6 | 20.083 | 2 | 43.722027 | yes | 2.473E-07 | 551.32043 | 296.39615 |
| QDLPDAMKPHEIQ | 15 | P62330 | ADP-ribosylation factor 6 OS=Homo sap | ARF6 | 20.083 | 3 | 43.722027 | yes | 2.475E-07 | 593.62958 | 112.1265 |
| VALDMTTLTIMR | 12 | P62333 | 26S proteasome regulatory subunit 10B C | PSMC6 | 44.173 | 2 | 41.846992 | yes | 5.747E-07 | 682.86743 | 52.124741 |
| GCLLYGPPGTGK | 12 | P62333 | 26S proteasome regulatory subunit 10B C | PSMC6 | 44.173 | 2 | 41.846992 | yes | 3.313E-07 | 610.31061 | 47.928017 |
| AGVNTVTTLVENK | 13 | P62424 | Large ribosomal subunit protein eL8 OS= | RPL7A | 29.996 | 2 | 55.034126 | yes | 4.178E-07 | 673.36975 | 32355.951 |
| KVVNPLFEK | 9 | P62424 | Large ribosomal subunit protein eL8 OS= | RPL7A | 29.996 | 2 | 55.034126 | yes | 1.296E-07 | 537.32129 | 22077.311 |
| HWGGNVLGPK | 10 | P62424 | Large ribosomal subunit protein eL8 OS= | RPL7A | 29.996 | 2 | 55.034126 | yes | 2.058E-07 | 532.78546 | 20437.895 |
| NFGIGQDIQPK | 11 | P62424 | Large ribosomal subunit protein eL8 OS= | RPL7A | 29.996 | 2 | 55.034126 | yes | 1.187E-07 | 608.81964 | 12104.143 |
| VPPAINQFTQALDI | 14 | P62424 | Large ribosomal subunit protein eL8 OS= | RPL7A | 29.996 | 2 | 55.034126 | yes | 3.02E-06 | 785.42285 | 7946.7959 |
| QTATQLLK | 8 | P62424 | Large ribosomal subunit protein eL8 OS= | RPL7A | 29.996 | 2 | 55.034126 | yes | 1.248E-07 | 451.76892 | 6346.5693 |
| KVAPAPAVVK | 10 | P62424 | Large ribosomal subunit protein eL8 OS= | RPL7A | 29.996 | 2 | 55.034126 | yes | 1.767E-07 | 490.31857 | 5673.8496 |
| VVNPLFEK | 8 | P62424 | Large ribosomal subunit protein eL8 OS= | RPL7A | 29.996 | 2 | 55.034126 | yes | 1.681E-07 | 473.27383 | 3186.7468 |
| VAPAPAVVK | 9 | P62424 | Large ribosomal subunit protein eL8 OS= | RPL7A | 29.996 | 2 | 55.034126 | yes | 3.584E-07 | 426.27109 | 2790.5166 |
| MGVPYCIIK | 9 | P62424 | Large ribosomal subunit protein eL8 OS= | RPL7A | 29.996 | 2 | 55.034126 | yes | 4.24E-07 | 540.78265 | 601.49152 |
| KMGVPYCIIK | 10 | P62424 | Large ribosomal subunit protein eL8 OS= | RPL7A | 29.996 | 2 | 55.034126 | yes | 9.809E-08 | 604.83014 | 427.81277 |
| KTCTTVAFTQVNS | 16 | P62424 | Large ribosomal subunit protein eL8 OS= | RPL7A | 29.996 | 3 | 55.034126 | yes | 9.793E-08 | 610.29639 | 294.54208 |
| RHWGGNVLGPK | 11 | P62424 | Large ribosomal subunit protein eL8 OS= | RPL7A | 29.996 | 2 | 55.034126 | yes | 9.918E-08 | 610.836 | 89.811264 |
| RHWGGNVLGPK | 11 | P62424 | Large ribosomal subunit protein eL8 OS= | RPL7A | 29.996 | 2 | 55.034126 | yes | 1.777E-07 | 407.55975 | 89.811264 |
| RHWGGNVLGPK | 11 | P62424 | Large ribosomal subunit protein eL8 OS= | RPL7A | 29.996 | 3 | 55.034126 | yes | 9.918E-08 | 610.836 | 89.811264 |
| RHWGGNVLGPK | 11 | P62424 | Large ribosomal subunit protein eL8 OS= | RPL7A | 29.996 | 3 | 55.034126 | yes | 1.777E-07 | 407.55975 | 89.811264 |
| KTCTTVAFTQVNS | 21 | P62424 | Large ribosomal subunit protein eL8 OS= | RPL7A | 29.996 | 3 | 55.034126 | yes | 2.564E-07 | 757.05463 | 63.57613 |
| AGVNTVTTLVENK | 14 | P62424 | Large ribosomal subunit protein eL8 OS= | RPL7A | 29.996 | 2 | 55.034126 | yes | 9.722E-08 | 737.41718 | 55.453529 |

| | | | | | | | | | | | |
|---|---|---|---|---|---|---|---|---|---|---|---|
| TCTTVAFTQVNSE | 15 | P62424 | Large ribosomal subunit protein eL8 OS= | RPL7A | 29.996 | 3 | 55.034126 | yes | 2.961E-07 | 567.59802 | 28.674883 |
| AQIWDTAGQER | 11 | P62491 | Ras-related protein Rab-11A OS=Homo s | RAB11A | 24.394 | 2 | 50.75047 | yes | 1.2E-07 | 637.80981 | 60.115963 |
| VVLIGDSGVGK | 11 | P62491 | Ras-related protein Rab-11A OS=Homo s | RAB11A | 24.394 | 2 | 50.75047 | yes | 9.723E-08 | 522.30841 | 54.208141 |
| LSVLGAITSVQQR | 13 | P62495 | Eukaryotic peptide chain release factor su | ETF1 | 49.031 | 2 | 52.39877 | yes | 9.81E-08 | 686.40137 | 416.32434 |
| GFGGIGGILR | 10 | P62495 | Eukaryotic peptide chain release factor su | ETF1 | 49.031 | 2 | 52.39877 | yes | 9.927E-08 | 473.77707 | 260.88803 |
| YCFGVEDTLK | 10 | P62495 | Eukaryotic peptide chain release factor su | ETF1 | 49.031 | 2 | 52.39877 | yes | 1.12E-07 | 616.2868 | 112.47882 |
| FGATLEIVTDK | 11 | P62495 | Eukaryotic peptide chain release factor su | ETF1 | 49.031 | 2 | 52.39877 | yes | 2.74E-07 | 597.32428 | 106.05777 |
| YFDEISQDTGK | 11 | P62495 | Eukaryotic peptide chain release factor su | ETF1 | 49.031 | 2 | 52.39877 | yes | 1.18E-07 | 651.79608 | 97.604309 |
| TELSQSDMFDQR | 12 | P62495 | Eukaryotic peptide chain release factor su | ETF1 | 49.031 | 2 | 52.39877 | yes | 9.722E-08 | 728.82227 | 76.932343 |
| TSEVNCYR | 8 | P62633 | CCHC-type zinc finger nucleic acid bindi | CNBP | 19.463 | 2 | 51.22902 | yes | 1.027E-07 | 514.72693 | 20966.893 |
| CYSCGEFGHIQK | 12 | P62633 | CCHC-type zinc finger nucleic acid bindi | CNBP | 19.463 | 2 | 51.22902 | yes | 1.384E-07 | 743.3161 | 4707.4995 |
| CYSCGEFGHIQK | 12 | P62633 | CCHC-type zinc finger nucleic acid bindi | CNBP | 19.463 | 2 | 51.22902 | yes | 9.722E-08 | 495.87982 | 4707.4995 |
| CYSCGEFGHIQK | 12 | P62633 | CCHC-type zinc finger nucleic acid bindi | CNBP | 19.463 | 3 | 51.22902 | yes | 1.384E-07 | 743.3161 | 4707.4995 |
| CYSCGEFGHIQK | 12 | P62633 | CCHC-type zinc finger nucleic acid bindi | CNBP | 19.463 | 3 | 51.22902 | yes | 9.722E-08 | 495.87982 | 4707.4995 |
| CGETGHVAINCSK | 13 | P62633 | CCHC-type zinc finger nucleic acid bindi | CNBP | 19.463 | 2 | 51.22902 | yes | 1.283E-07 | 716.81903 | 3890.5981 |
| CGETGHVAINCSK | 13 | P62633 | CCHC-type zinc finger nucleic acid bindi | CNBP | 19.463 | 2 | 51.22902 | yes | 2.874E-07 | 478.21512 | 3890.5981 |
| CGETGHVAINCSK | 13 | P62633 | CCHC-type zinc finger nucleic acid bindi | CNBP | 19.463 | 3 | 51.22902 | yes | 1.283E-07 | 716.81903 | 3890.5981 |
| CGETGHVAINCSK | 13 | P62633 | CCHC-type zinc finger nucleic acid bindi | CNBP | 19.463 | 3 | 51.22902 | yes | 2.874E-07 | 478.21512 | 3890.5981 |
| CGESGHLAR | 9 | P62633 | CCHC-type zinc finger nucleic acid bindi | CNBP | 19.463 | 2 | 51.22902 | yes | 6.687E-07 | 493.72726 | 2112.8198 |
| EQCCYNCGKPGHI | 15 | P62633 | CCHC-type zinc finger nucleic acid bindi | CNBP | 19.463 | 4 | 51.22902 | yes | 1.742E-07 | 463.20273 | 632.14313 |
| SSNECFKCGR | 10 | P62633 | CCHC-type zinc finger nucleic acid bindi | CNBP | 19.463 | 2 | 51.22902 | yes | 1.196E-07 | 643.76624 | 176.77399 |
| EQCCYNCGK | 9 | P62633 | CCHC-type zinc finger nucleic acid bindi | CNBP | 19.463 | 2 | 51.22902 | yes | 1.12E-07 | 609.72046 | 150.01553 |
| CGESGHLAK | 9 | P62633 | CCHC-type zinc finger nucleic acid bindi | CNBP | 19.463 | 2 | 51.22902 | yes | 4.048E-07 | 479.72418 | 106.15583 |
| CYRCGETGHVAIN | 16 | P62633 | CCHC-type zinc finger nucleic acid bindi | CNBP | 19.463 | 3 | 51.22902 | yes | 1.136E-07 | 637.94678 | 50.114944 |
| GNKPWISLPR | 10 | P62701 | Small ribosomal subunit protein eS4, X is | RPS4X | 29.598 | 2 | 52.283 | yes | 1.644E-07 | 584.33527 | 61316.012 |
| LTGVFAPR | 8 | P62701 | Small ribosomal subunit protein eS4, X is | RPS4X | 29.598 | 2 | 52.283 | no | 1.367E-06 | 430.75308 | 37392.301 |
| GIPHLVTHDAR | 11 | P62701 | Small ribosomal subunit protein eS4, X is | RPS4X | 29.598 | 2 | 52.283 | no | 1.785E-07 | 608.33325 | 13154.414 |
| GIPHLVTHDAR | 11 | P62701 | Small ribosomal subunit protein eS4, X is | RPS4X | 29.598 | 2 | 52.283 | no | 9.722E-08 | 405.89127 | 13154.414 |
| GIPHLVTHDAR | 11 | P62701 | Small ribosomal subunit protein eS4, X is | RPS4X | 29.598 | 3 | 52.283 | no | 1.785E-07 | 608.33325 | 13154.414 |
| GIPHLVTHDAR | 11 | P62701 | Small ribosomal subunit protein eS4, X is | RPS4X | 29.598 | 3 | 52.283 | no | 9.722E-08 | 405.89127 | 13154.414 |
| VNDTIQIDLETGK | 13 | P62701 | Small ribosomal subunit protein eS4, X is | RPS4X | 29.598 | 2 | 52.283 | yes | 1.275E-07 | 723.37775 | 23796.865 |
| YALTGDEVK | 9 | P62701 | Small ribosomal subunit protein eS4, X is | RPS4X | 29.598 | 2 | 52.283 | no | 9.722E-08 | 498.25583 | 16593.543 |
| LSNIFVIGK | 9 | P62701 | Small ribosomal subunit protein eS4, X is | RPS4X | 29.598 | 2 | 52.283 | yes | 1.699E-07 | 495.80276 | 15675.235 |
| LTIAEER | 7 | P62701 | Small ribosomal subunit protein eS4, X is | RPS4X | 29.598 | 2 | 52.283 | no | 1.001E-07 | 416.23218 | 10704.88 |
| HPGSFDVVHVK | 11 | P62701 | Small ribosomal subunit protein eS4, X is | RPS4X | 29.598 | 2 | 52.283 | no | 1.198E-07 | 611.32239 | 5074.6533 |
| HPGSFDVVHVK | 11 | P62701 | Small ribosomal subunit protein eS4, X is | RPS4X | 29.598 | 2 | 52.283 | no | 6.49E-07 | 407.884 | 5074.6533 |
| HPGSFDVVHVK | 11 | P62701 | Small ribosomal subunit protein eS4, X is | RPS4X | 29.598 | 3 | 52.283 | no | 1.198E-07 | 611.32239 | 5074.6533 |
| HPGSFDVVHVK | 11 | P62701 | Small ribosomal subunit protein eS4, X is | RPS4X | 29.598 | 3 | 52.283 | no | 6.49E-07 | 407.884 | 5074.6533 |
| YPDPLIK | 7 | P62701 | Small ribosomal subunit protein eS4, X is | RPS4X | 29.598 | 2 | 52.283 | no | 1.375E-07 | 423.242 | 3055.5437 |

| | | | | | | | | | | | |
|---|---|---|---|---|---|---|---|---|---|---|---|
| YPDPLIK | 7 | P62701 | Small ribosomal subunit protein eS4, X is | RPS4X | 29.598 | 2 | 52.283 | no | 9.722E-08 | 845.47675 | 3055.5437 |
| YPDPLIK | 7 | P62701 | Small ribosomal subunit protein eS4, X is | RPS4X | 29.598 | 1 | 52.283 | no | 1.375E-07 | 423.242 | 3055.5437 |
| YPDPLIK | 7 | P62701 | Small ribosomal subunit protein eS4, X is | RPS4X | 29.598 | 1 | 52.283 | no | 9.722E-08 | 845.47675 | 3055.5437 |
| ERHPGSFDVVHVK | 13 | P62701 | Small ribosomal subunit protein eS4, X is | RPS4X | 29.598 | 3 | 52.283 | no | 2.069E-07 | 502.93192 | 1453.6583 |
| ERHPGSFDVVHVK | 13 | P62701 | Small ribosomal subunit protein eS4, X is | RPS4X | 29.598 | 3 | 52.283 | no | 1.352E-07 | 753.89423 | 1453.6583 |
| ERHPGSFDVVHVK | 13 | P62701 | Small ribosomal subunit protein eS4, X is | RPS4X | 29.598 | 2 | 52.283 | no | 2.069E-07 | 502.93192 | 1453.6583 |
| ERHPGSFDVVHVK | 13 | P62701 | Small ribosomal subunit protein eS4, X is | RPS4X | 29.598 | 2 | 52.283 | no | 1.352E-07 | 753.89423 | 1453.6583 |
| DANGNSFATR | 10 | P62701 | Small ribosomal subunit protein eS4, X is | RPS4X | 29.598 | 2 | 52.283 | no | 1.771E-07 | 526.74139 | 275.35208 |
| FDTGNLCMVTGG/ | 17 | P62701 | Small ribosomal subunit protein eS4, X is | RPS4X | 29.598 | 3 | 52.283 | yes | 1.93E-07 | 594.9469 | 210.23846 |
| YALTGDEVKK | 10 | P62701 | Small ribosomal subunit protein eS4, X is | RPS4X | 29.598 | 2 | 52.283 | no | 9.722E-08 | 562.30334 | 208.49005 |
| CGNQAAIMELDDT | 15 | P62714 | Serine/threonine-protein phosphatase 2A | PPP2CB | 35.576 | 2 | 51.01616 | no | 9.722E-08 | 839.8924 | 642.27631 |
| YSFLQFDPAPR | 11 | P62714 | Serine/threonine-protein phosphatase 2A | PPP2CB | 35.576 | 2 | 51.01616 | no | 1.127E-07 | 670.83533 | 638.62457 |
| ELDQWVEQLNECI | 13 | P62714 | Serine/threonine-protein phosphatase 2A | PPP2CB | 35.576 | 2 | 51.01616 | yes | 1.167E-07 | 845.89069 | 164.2852 |
| KYGNANVWK | 9 | P62714 | Serine/threonine-protein phosphatase 2A | PPP2CB | 35.576 | 2 | 51.01616 | no | 4.647E-07 | 540.28528 | 73.089279 |
| QEYDEAGPSIVHR | 13 | P62736 | Actin, aortic smooth muscle OS=Homo sa | ACTA2 | 42.009 | 3 | 37.15023 | no | 6.252E-07 | 500.90744 | 1553.5259 |
| LAPDYDALDVANI | 13 | P62750 | Large ribosomal subunit protein uL23 OS | RPL23A | 17.696 | 2 | 51.674854 | yes | 2.34E-07 | 702.85388 | 42847.313 |
| FPLTTESAMK | 10 | P62750 | Large ribosomal subunit protein uL23 OS | RPL23A | 17.696 | 2 | 51.674854 | yes | 2.614E-06 | 562.78644 | 14490.998 |
| FPLTTESAMK | 10 | P62750 | Large ribosomal subunit protein uL23 OS | RPL23A | 17.696 | 2 | 51.674854 | yes | 5.408E-07 | 570.78394 | 14490.998 |
| LYDIDVAK | 8 | P62750 | Large ribosomal subunit protein uL23 OS | RPL23A | 17.696 | 2 | 51.674854 | yes | 8.287E-07 | 468.75546 | 14528.363 |
| KLYDIDVAK | 9 | P62750 | Large ribosomal subunit protein uL23 OS | RPL23A | 17.696 | 2 | 51.674854 | yes | 1.8E-07 | 532.80298 | 13538.339 |
| VNTLIRPDGEK | 11 | P62750 | Large ribosomal subunit protein uL23 OS | RPL23A | 17.696 | 2 | 51.674854 | yes | 9.722E-08 | 621.34607 | 6537.6411 |
| VNTLIRPDGEK | 11 | P62750 | Large ribosomal subunit protein uL23 OS | RPL23A | 17.696 | 2 | 51.674854 | yes | 4.672E-07 | 414.56647 | 6537.6411 |
| VNTLIRPDGEK | 11 | P62750 | Large ribosomal subunit protein uL23 OS | RPL23A | 17.696 | 3 | 51.674854 | yes | 9.722E-08 | 621.34607 | 6537.6411 |
| VNTLIRPDGEK | 11 | P62750 | Large ribosomal subunit protein uL23 OS | RPL23A | 17.696 | 3 | 51.674854 | yes | 4.672E-07 | 414.56647 | 6537.6411 |
| IEDNNTLVFIVDVI | 14 | P62750 | Large ribosomal subunit protein uL23 OS | RPL23A | 17.696 | 2 | 51.674854 | yes | 1.108E-07 | 809.93799 | 3010.988 |
| NKLDHYAIIK | 10 | P62750 | Large ribosomal subunit protein uL23 OS | RPL23A | 17.696 | 2 | 51.674854 | yes | 9.722E-08 | 607.84821 | 1390.5966 |
| NKLDHYAIIK | 10 | P62750 | Large ribosomal subunit protein uL23 OS | RPL23A | 17.696 | 2 | 51.674854 | yes | 1.003E-07 | 405.5679 | 1390.5966 |
| NKLDHYAIIK | 10 | P62750 | Large ribosomal subunit protein uL23 OS | RPL23A | 17.696 | 3 | 51.674854 | yes | 9.722E-08 | 607.84821 | 1390.5966 |
| NKLDHYAIIK | 10 | P62750 | Large ribosomal subunit protein uL23 OS | RPL23A | 17.696 | 3 | 51.674854 | yes | 1.003E-07 | 405.5679 | 1390.5966 |
| LDHYAIIK | 8 | P62750 | Large ribosomal subunit protein uL23 OS | RPL23A | 17.696 | 2 | 51.674854 | yes | 2.018E-07 | 486.7793 | 2512.0322 |
| VNTLIRPDGEKK | 12 | P62750 | Large ribosomal subunit protein uL23 OS | RPL23A | 17.696 | 2 | 51.674854 | yes | 1.004E-07 | 685.39355 | 75.528488 |
| KIEDNNTLVFIVD\ | 15 | P62750 | Large ribosomal subunit protein uL23 OS | RPL23A | 17.696 | 3 | 51.674854 | yes | 1.022E-07 | 582.99274 | 48.773853 |
| LNISFPATGCQK | 12 | P62753 | Small ribosomal subunit protein eS6 OS= | RPS6 | 28.681 | 2 | 51.836098 | yes | 1.108E-07 | 668.3399 | 49754.574 |
| MATEVAADALGE | 15 | P62753 | Small ribosomal subunit protein eS6 OS= | RPS6 | 28.681 | 2 | 51.836098 | yes | 1.537E-07 | 810.88232 | 14488.473 |
| MATEVAADALGE | 15 | P62753 | Small ribosomal subunit protein eS6 OS= | RPS6 | 28.681 | 2 | 51.836098 | yes | 1.07E-07 | 818.87982 | 14488.473 |
| MATEVAADALGE | 15 | P62753 | Small ribosomal subunit protein eS6 OS= | RPS6 | 28.681 | 2 | 51.836098 | yes | 2.102E-07 | 540.92401 | 14488.473 |
| MATEVAADALGE | 15 | P62753 | Small ribosomal subunit protein eS6 OS= | RPS6 | 28.681 | 3 | 51.836098 | yes | 1.537E-07 | 810.88232 | 14488.473 |
| MATEVAADALGE | 15 | P62753 | Small ribosomal subunit protein eS6 OS= | RPS6 | 28.681 | 3 | 51.836098 | yes | 1.07E-07 | 818.87982 | 14488.473 |
| MATEVAADALGE | 15 | P62753 | Small ribosomal subunit protein eS6 OS= | RPS6 | 28.681 | 3 | 51.836098 | yes | 2.102E-07 | 540.92401 | 14488.473 |

| | | | | | | | | | | | |
|---|---|---|---|---|---|---|---|---|---|---|---|
| DIPGLTDTTVPR | 12 | P62753 | Small ribosomal subunit protein eS6 OS= | RPS6 | 28.681 | 2 | 51.836098 | yes | 1.539E-07 | 642.84332 | 37051.902 |
| LIEVDDER | 8 | P62753 | Small ribosomal subunit protein eS6 OS= | RPS6 | 28.681 | 2 | 51.836098 | yes | 1.692E-07 | 494.75092 | 31376.467 |
| QGVLTHGR | 8 | P62753 | Small ribosomal subunit protein eS6 OS= | RPS6 | 28.681 | 2 | 51.836098 | yes | 1.81E-07 | 434.24341 | 14403.449 |
| LIEVDDERK | 9 | P62753 | Small ribosomal subunit protein eS6 OS= | RPS6 | 28.681 | 2 | 51.836098 | yes | 9.974E-08 | 558.7984 | 5430.8374 |
| NKEEAAEYAK | 10 | P62753 | Small ribosomal subunit protein eS6 OS= | RPS6 | 28.681 | 2 | 51.836098 | yes | 9.722E-08 | 576.78021 | 3032.9766 |
| KLFNLSK | 7 | P62753 | Small ribosomal subunit protein eS6 OS= | RPS6 | 28.681 | 2 | 51.836098 | yes | 2.664E-07 | 425.26328 | 351.03448 |
| VFLENVIR | 8 | P62805 | Histone H4 OS=Homo sapiens OX=9606 | H4C1 | 11.368 | 2 | 52.815487 | yes | 1.542E-07 | 495.29257 | 46596.652 |
| ISGLIYEETR | 10 | P62805 | Histone H4 OS=Homo sapiens OX=9606 | H4C1 | 11.368 | 2 | 52.815487 | yes | 9.722E-08 | 590.81403 | 27839.943 |
| DAVTYTEHAK | 10 | P62805 | Histone H4 OS=Homo sapiens OX=9606 | H4C1 | 11.368 | 2 | 52.815487 | yes | 9.722E-08 | 567.7749 | 20223.336 |
| TVTAMDVVYALK | 12 | P62805 | Histone H4 OS=Homo sapiens OX=9606 | H4C1 | 11.368 | 2 | 52.815487 | yes | 1.114E-07 | 655.85486 | 7206.4604 |
| TVTAMDVVYALK | 12 | P62805 | Histone H4 OS=Homo sapiens OX=9606 | H4C1 | 11.368 | 2 | 52.815487 | yes | 1.146E-07 | 663.85229 | 7206.4604 |
| DNIQGITKPAIR | 12 | P62805 | Histone H4 OS=Homo sapiens OX=9606 | H4C1 | 11.368 | 2 | 52.815487 | yes | 1.078E-07 | 663.38043 | 3599.2964 |
| DNIQGITKPAIR | 12 | P62805 | Histone H4 OS=Homo sapiens OX=9606 | H4C1 | 11.368 | 2 | 52.815487 | yes | 1.728E-07 | 442.58939 | 3599.2964 |
| DNIQGITKPAIR | 12 | P62805 | Histone H4 OS=Homo sapiens OX=9606 | H4C1 | 11.368 | 3 | 52.815487 | yes | 1.078E-07 | 663.38043 | 3599.2964 |
| DNIQGITKPAIR | 12 | P62805 | Histone H4 OS=Homo sapiens OX=9606 | H4C1 | 11.368 | 3 | 52.815487 | yes | 1.728E-07 | 442.58939 | 3599.2964 |
| KTVTAMDVVYAL | 13 | P62805 | Histone H4 OS=Homo sapiens OX=9606 | H4C1 | 11.368 | 2 | 52.815487 | yes | 1.821E-07 | 719.90234 | 959.92102 |
| LVLVGDGGTGK | 11 | P62826 | GTP-binding nuclear protein Ran OS=Ho | RAN | 24.424 | 2 | 50.506798 | yes | 1.837E-07 | 508.29276 | 3894.6743 |
| FNVWDTAGQEK | 11 | P62826 | GTP-binding nuclear protein Ran OS=Ho | RAN | 24.424 | 2 | 50.506798 | yes | 9.81E-08 | 647.80676 | 1715.5519 |
| VCENIPIVLCGNK | 13 | P62826 | GTP-binding nuclear protein Ran OS=Ho | RAN | 24.424 | 2 | 50.506798 | yes | 2.776E-07 | 758.38654 | 1476.5585 |
| NLQYYDISAK | 10 | P62826 | GTP-binding nuclear protein Ran OS=Ho | RAN | 24.424 | 2 | 50.506798 | yes | 3.653E-07 | 607.80621 | 1421.9943 |
| YVATLGVEVHPL\ | 18 | P62826 | GTP-binding nuclear protein Ran OS=Ho | RAN | 24.424 | 3 | 50.506798 | yes | 1.418E-07 | 684.70569 | 949.28094 |
| HLTGEFEK | 8 | P62826 | GTP-binding nuclear protein Ran OS=Ho | RAN | 24.424 | 2 | 50.506798 | yes | 3.319E-07 | 480.74289 | 780.00916 |
| AAQGEPQVQFK | 11 | P62826 | GTP-binding nuclear protein Ran OS=Ho | RAN | 24.424 | 2 | 50.506798 | yes | 9.79E-08 | 622.81714 | 200.85191 |
| NVPNWHR | 7 | P62826 | GTP-binding nuclear protein Ran OS=Ho | RAN | 24.424 | 2 | 50.506798 | yes | 8.138E-06 | 461.73575 | 72.357452 |
| PFLWLARK | 8 | P62826 | GTP-binding nuclear protein Ran OS=Ho | RAN | 24.424 | 2 | 50.506798 | yes | 1.719E-05 | 515.81348 | 56.346741 |
| PFLWLAR | 7 | P62826 | GTP-binding nuclear protein Ran OS=Ho | RAN | 24.424 | 2 | 50.506798 | yes | 4.931E-07 | 451.76599 | 41.706749 |
| RHLTGEFEK | 9 | P62826 | GTP-binding nuclear protein Ran OS=Ho | RAN | 24.424 | 2 | 50.506798 | yes | 2.042E-06 | 558.79346 | 40.676228 |
| KYVATLGVEVHPI | 19 | P62826 | GTP-binding nuclear protein Ran OS=Ho | RAN | 24.424 | 3 | 50.506798 | yes | 5.544E-06 | 727.40399 | 17.940332 |
| KNLQYYDISAK | 11 | P62826 | GTP-binding nuclear protein Ran OS=Ho | RAN | 24.424 | 2 | 50.506798 | yes | 4.778E-06 | 671.8537 | 12.368112 |
| VHPAVVIR | 8 | P62829 | Large ribosomal subunit protein uL14 OS | RPL23 | 14.866 | 2 | 47.831985 | yes | 2.579E-06 | 445.78217 | 23126.949 |
| LPAAGVGDMVMA | 15 | P62829 | Large ribosomal subunit protein uL14 OS | RPL23 | 14.866 | 2 | 47.831985 | yes | 1.421E-07 | 730.38599 | 5411.9741 |
| LPAAGVGDMVMA | 15 | P62829 | Large ribosomal subunit protein uL14 OS | RPL23 | 14.866 | 2 | 47.831985 | yes | 2.471E-07 | 738.38348 | 5411.9741 |
| LPAAGVGDMVMA | 15 | P62829 | Large ribosomal subunit protein uL14 OS | RPL23 | 14.866 | 2 | 47.831985 | yes | 4.325E-07 | 738.38348 | 5411.9741 |
| LPAAGVGDMVMA | 15 | P62829 | Large ribosomal subunit protein uL14 OS | RPL23 | 14.866 | 2 | 47.831985 | yes | 2.336E-07 | 746.38092 | 5411.9741 |
| ECADLWPR | 8 | P62829 | Large ribosomal subunit protein uL14 OS | RPL23 | 14.866 | 2 | 47.831985 | yes | 3.563E-07 | 523.73981 | 14235.165 |
| GSAITGPVAK | 10 | P62829 | Large ribosomal subunit protein uL14 OS | RPL23 | 14.866 | 2 | 47.831985 | yes | 1.666E-07 | 450.76108 | 7399.5229 |
| KVHPAVVIR | 9 | P62829 | Large ribosomal subunit protein uL14 OS | RPL23 | 14.866 | 2 | 47.831985 | yes | 1.249E-07 | 509.82965 | 7338.8042 |
| ISLGLPVGAVINCA | 20 | P62829 | Large ribosomal subunit protein uL14 OS | RPL23 | 14.866 | 3 | 47.831985 | yes | 1.417E-07 | 657.35071 | 74.85482 |
| KDGVFLYFEDNAC | 19 | P62829 | Large ribosomal subunit protein uL14 OS | RPL23 | 14.866 | 3 | 47.831985 | yes | 1.673E-06 | 714.70038 | 52.469158 |

| | | | | | | | | | | |
|---|---|---|---|---|---|---|---|---|---|---|
| YDPTIEDSYR | 10 | P62834 | Ras-related protein Rap-1A OS=Homo sa | RAP1A | 20.988 | 2 | 47.332066 | no | 4.195E-07 | 629.78296 | 214.54335 |
| LVVLGSGGVGK | 11 | P62834 | Ras-related protein Rap-1A OS=Homo sa | RAP1A | 20.988 | 2 | 47.332066 | no | 1.353E-07 | 493.30566 | 176.6568 |
| SALTVQFVQGIFVI | 15 | P62834 | Ras-related protein Rap-1A OS=Homo sa | RAP1A | 20.988 | 2 | 47.332066 | no | 1.725E-07 | 833.46417 | 154.8123 |
| QWCNCAFLESSAK | 13 | P62834 | Ras-related protein Rap-1A OS=Homo sa | RAP1A | 20.988 | 2 | 47.332066 | yes | 1.602E-07 | 800.84778 | 57.334072 |
| EAPPMEKPEVVK | 12 | P62841 | Small ribosomal subunit protein uS19 OS | RPS15 | 17.041 | 2 | 50.182594 | yes | 1.362E-07 | 677.35779 | 9567.0547 |
| EAPPMEKPEVVK | 12 | P62841 | Small ribosomal subunit protein uS19 OS | RPS15 | 17.041 | 2 | 50.182594 | yes | 1.156E-07 | 685.35522 | 9567.0547 |
| EAPPMEKPEVVK | 12 | P62841 | Small ribosomal subunit protein uS19 OS | RPS15 | 17.041 | 2 | 50.182594 | yes | 8.965E-07 | 451.90759 | 9567.0547 |
| EAPPMEKPEVVK | 12 | P62841 | Small ribosomal subunit protein uS19 OS | RPS15 | 17.041 | 2 | 50.182594 | yes | 7.078E-07 | 457.23926 | 9567.0547 |
| EAPPMEKPEVVK | 12 | P62841 | Small ribosomal subunit protein uS19 OS | RPS15 | 17.041 | 3 | 50.182594 | yes | 1.362E-07 | 677.35779 | 9567.0547 |
| EAPPMEKPEVVK | 12 | P62841 | Small ribosomal subunit protein uS19 OS | RPS15 | 17.041 | 3 | 50.182594 | yes | 1.156E-07 | 685.35522 | 9567.0547 |
| EAPPMEKPEVVK | 12 | P62841 | Small ribosomal subunit protein uS19 OS | RPS15 | 17.041 | 3 | 50.182594 | yes | 8.965E-07 | 451.90759 | 9567.0547 |
| EAPPMEKPEVVK | 12 | P62841 | Small ribosomal subunit protein uS19 OS | RPS15 | 17.041 | 3 | 50.182594 | yes | 7.078E-07 | 457.23926 | 9567.0547 |
| KEAPPMEKPEVVK | 13 | P62841 | Small ribosomal subunit protein uS19 OS | RPS15 | 17.041 | 3 | 50.182594 | yes | 1.115E-07 | 494.60593 | 853.33197 |
| KEAPPMEKPEVVK | 13 | P62841 | Small ribosomal subunit protein uS19 OS | RPS15 | 17.041 | 3 | 50.182594 | yes | 1.678E-07 | 741.40527 | 853.33197 |
| KEAPPMEKPEVVK | 13 | P62841 | Small ribosomal subunit protein uS19 OS | RPS15 | 17.041 | 3 | 50.182594 | yes | 9.861E-08 | 499.93756 | 853.33197 |
| KEAPPMEKPEVVK | 13 | P62841 | Small ribosomal subunit protein uS19 OS | RPS15 | 17.041 | 3 | 50.182594 | yes | 1.849E-07 | 749.40271 | 853.33197 |
| KEAPPMEKPEVVK | 13 | P62841 | Small ribosomal subunit protein uS19 OS | RPS15 | 17.041 | 2 | 50.182594 | yes | 1.115E-07 | 494.60593 | 853.33197 |
| KEAPPMEKPEVVK | 13 | P62841 | Small ribosomal subunit protein uS19 OS | RPS15 | 17.041 | 2 | 50.182594 | yes | 1.678E-07 | 741.40527 | 853.33197 |
| KEAPPMEKPEVVK | 13 | P62841 | Small ribosomal subunit protein uS19 OS | RPS15 | 17.041 | 2 | 50.182594 | yes | 9.861E-08 | 499.93756 | 853.33197 |
| KEAPPMEKPEVVK | 13 | P62841 | Small ribosomal subunit protein uS19 OS | RPS15 | 17.041 | 2 | 50.182594 | yes | 1.849E-07 | 749.40271 | 853.33197 |
| TFNQVEIKPEMIGI | 27 | P62841 | Small ribosomal subunit protein uS19 OS | RPS15 | 17.041 | 4 | 50.182594 | yes | 1.027E-07 | 792.91644 | 853.2146 |
| TFNQVEIKPEMIGI | 27 | P62841 | Small ribosomal subunit protein uS19 OS | RPS15 | 17.041 | 4 | 50.182594 | yes | 1.011E-07 | 634.53461 | 853.2146 |
| TFNQVEIKPEMIGI | 27 | P62841 | Small ribosomal subunit protein uS19 OS | RPS15 | 17.041 | 4 | 50.182594 | yes | 1.26E-06 | 796.91516 | 853.2146 |
| TFNQVEIKPEMIGI | 27 | P62841 | Small ribosomal subunit protein uS19 OS | RPS15 | 17.041 | 5 | 50.182594 | yes | 1.027E-07 | 792.91644 | 853.2146 |
| TFNQVEIKPEMIGI | 27 | P62841 | Small ribosomal subunit protein uS19 OS | RPS15 | 17.041 | 5 | 50.182594 | yes | 1.011E-07 | 634.53461 | 853.2146 |
| TFNQVEIKPEMIGI | 27 | P62841 | Small ribosomal subunit protein uS19 OS | RPS15 | 17.041 | 5 | 50.182594 | yes | 1.26E-06 | 796.91516 | 853.2146 |
| PGIGATHSSR | 10 | P62841 | Small ribosomal subunit protein uS19 OS | RPS15 | 17.041 | 2 | 50.182594 | yes | 7.189E-07 | 491.75687 | 192.82469 |
| KQHSLLK | 7 | P62841 | Small ribosomal subunit protein uS19 OS | RPS15 | 17.041 | 2 | 50.182594 | yes | 2.523E-07 | 427.26636 | 119.71042 |
| TFNQVEIKPEMIGI | 24 | P62841 | Small ribosomal subunit protein uS19 OS | RPS15 | 17.041 | 4 | 50.182594 | yes | 1.248E-07 | 711.86237 | 58.796204 |
| PEMIGHYLGEFSIT | 19 | P62841 | Small ribosomal subunit protein uS19 OS | RPS15 | 17.041 | 3 | 50.182594 | yes | 1.345E-07 | 737.05023 | 40.011486 |
| QMVIDVLHPGK | 11 | P62847 | Small ribosomal subunit protein eS24 OS | RPS24 | 15.424 | 2 | 49.72103 | yes | 1.084E-07 | 618.8421 | 12060.904 |
| QMVIDVLHPGK | 11 | P62847 | Small ribosomal subunit protein eS24 OS | RPS24 | 15.424 | 2 | 49.72103 | yes | 6.377E-07 | 626.83954 | 12060.904 |
| KQMVIDVLHPGK | 12 | P62847 | Small ribosomal subunit protein eS24 OS | RPS24 | 15.424 | 2 | 49.72103 | yes | 1.007E-07 | 682.88959 | 620.04602 |
| KQMVIDVLHPGK | 12 | P62847 | Small ribosomal subunit protein eS24 OS | RPS24 | 15.424 | 2 | 49.72103 | yes | 1.941E-07 | 690.88702 | 620.04602 |
| MNDTVTIR | 8 | P62847 | Small ribosomal subunit protein eS24 OS | RPS24 | 15.424 | 2 | 49.72103 | yes | 2.817E-07 | 475.24222 | 342.36005 |
| MNDTVTIR | 8 | P62847 | Small ribosomal subunit protein eS24 OS | RPS24 | 15.424 | 2 | 49.72103 | yes | 1.087E-07 | 496.2475 | 342.36005 |
| TTGFGMIYDSLDY | 15 | P62847 | Small ribosomal subunit protein eS24 OS | RPS24 | 15.424 | 2 | 49.72103 | yes | 6.097E-07 | 841.39252 | 141.59464 |
| LITPAVVSER | 10 | P62851 | Small ribosomal subunit protein eS25 OS | RPS25 | 13.743 | 2 | 46.683308 | yes | 5.36E-07 | 542.82166 | 56784.691 |
| AALQELLSK | 9 | P62851 | Small ribosomal subunit protein eS25 OS | RPS25 | 13.743 | 2 | 46.683308 | yes | 1.905E-07 | 486.78986 | 42734.852 |

| | | | | | | | | | | | |
|---|---|---|---|---|---|---|---|---|---|---|---|
| LNNLVLFDK | 9 | P62851 | Small ribosomal subunit protein eS25 OS | RPS25 | 13.743 | 2 | 46.683308 | yes | 1.561E-07 | 538.31097 | 32119.449 |
| DKLNNLVLFDK | 11 | P62851 | Small ribosomal subunit protein eS25 OS | RPS25 | 13.743 | 2 | 46.683308 | yes | 3.914E-07 | 659.87189 | 10632.265 |
| AQVIYTR | 7 | P62851 | Small ribosomal subunit protein eS25 OS | RPS25 | 13.743 | 2 | 46.683308 | yes | 2.085E-07 | 425.74271 | 5204.3413 |
| ATYDKLCK | 8 | P62851 | Small ribosomal subunit protein eS25 OS | RPS25 | 13.743 | 2 | 46.683308 | yes | 1.497E-07 | 499.75241 | 598.00305 |
| NIVEAAAVR | 9 | P62854 | Small ribosomal subunit protein eS26 OS | RPS26 | 13.016 | 2 | 52.035244 | no | 1.809E-07 | 471.772 | 7238.0601 |
| GHVQPIR | 7 | P62854 | Small ribosomal subunit protein eS26 OS | RPS26 | 13.016 | 2 | 52.035244 | no | 9.823E-08 | 403.7352 | 1869.771 |
| DISEASVFDAYVLI | 15 | P62854 | Small ribosomal subunit protein eS26 OS | RPS26 | 13.016 | 2 | 52.035244 | yes | 1.519E-07 | 827.42218 | 394.12195 |
| FRPAGAAPRPPPK | 13 | P62854 | Small ribosomal subunit protein eS26 OS | RPS26 | 13.016 | 3 | 52.035244 | no | 3.173E-06 | 454.59821 | 238.17671 |
| PAGAAPRPPPKPM | 13 | P62854 | Small ribosomal subunit protein eS26 OS | RPS26 | 13.016 | 2 | 52.035244 | no | 7.335E-07 | 643.85553 | 126.23824 |
| LHYCVSCAIHSK | 12 | P62854 | Small ribosomal subunit protein eS26 OS | RPS26 | 13.016 | 2 | 52.035244 | no | 9.722E-08 | 737.8501 | 120.55975 |
| LYVKLHYCVSCAI | 16 | P62854 | Small ribosomal subunit protein eS26 OS | RPS26 | 13.016 | 3 | 52.035244 | no | 3.191E-07 | 660.0061 | 14.547429 |
| VEFMDDTSR | 9 | P62857 | Small ribosomal subunit protein eS28 OS | RPS28 | 7.842 | 2 | 49.799377 | yes | 7.627E-07 | 550.23987 | 7670.8721 |
| VEFMDDTSR | 9 | P62857 | Small ribosomal subunit protein eS28 OS | RPS28 | 7.842 | 2 | 49.799377 | yes | 1.003E-07 | 558.2373 | 7670.8721 |
| EGDVLTLLESER | 12 | P62857 | Small ribosomal subunit protein eS28 OS | RPS28 | 7.842 | 2 | 49.799377 | yes | 1.573E-07 | 680.85138 | 10458.787 |
| TGSQGQCTQVR | 11 | P62857 | Small ribosomal subunit protein eS28 OS | RPS28 | 7.842 | 2 | 49.799377 | yes | 4.317E-07 | 611.28564 | 44.693577 |
| FVNVVPTFGK | 10 | P62861 | Ubiquitin-like FUBI-ribosomal protein eS | FAU | 14.39 | 2 | 46.8129 | yes | 1.465E-07 | 554.31348 | 66007.922 |
| RFVNVVPTFGK | 11 | P62861 | Ubiquitin-like FUBI-ribosomal protein eS | FAU | 14.39 | 2 | 46.8129 | yes | 1.566E-07 | 632.36407 | 2614.7317 |
| LFVSGACDASAK | 12 | P62873 | Guanine nucleotide-binding protein G(I)/( | GNB1 | 37.377 | 2 | 37.738743 | yes | 5.74E-07 | 613.29773 | 47.157677 |
| MLLAHVDLIEK | 11 | P62875 | DNA-directed RNA polymerases I, II, anc | POLR2L | 7.646 | 2 | 50.932533 | yes | 9.766E-08 | 641.36542 | 321.03638 |
| LLNYAPLEK | 9 | P62875 | DNA-directed RNA polymerases I, II, anc | POLR2L | 7.646 | 2 | 50.932533 | yes | 9.722E-08 | 530.80548 | 251.45401 |
| AAAMDVDTPSGTI | 18 | P62877 | E3 ubiquitin-protein ligase RBX1 OS=Hc | RBX1 | 12.274 | 2 | 50.575386 | yes | 1.318E-07 | 846.38068 | 635.69061 |
| QVCPLDNR | 8 | P62877 | E3 ubiquitin-protein ligase RBX1 OS=Hc | RBX1 | 12.274 | 2 | 50.575386 | yes | 9.759E-08 | 501.24527 | 205.70294 |
| LLVSASQDGK | 10 | P62879 | Guanine nucleotide-binding protein G(I)/( | GNB2 | 37.332 | 2 | 49.30438 | no | 1.047E-07 | 509.28238 | 358.27472 |
| LIIWDSYTTNK | 11 | P62879 | Guanine nucleotide-binding protein G(I)/( | GNB2 | 37.332 | 2 | 49.30438 | no | 1.376E-07 | 677.35608 | 357.83774 |
| VHAIPLR | 7 | P62879 | Guanine nucleotide-binding protein G(I)/( | GNB2 | 37.332 | 2 | 49.30438 | no | 1.058E-07 | 403.25577 | 157.36366 |
| ELPGHTGYLSCCR | 13 | P62879 | Guanine nucleotide-binding protein G(I)/( | GNB2 | 37.332 | 2 | 49.30438 | no | 2.787E-06 | 775.34796 | 68.539352 |
| LVILANNCPALR | 12 | P62888 | Large ribosomal subunit protein eL30 OS | RPL30 | 12.785 | 2 | 49.9376 | yes | 1.607E-07 | 677.38721 | 32599.49 |
| SEIEYYAMLAK | 11 | P62888 | Large ribosomal subunit protein eL30 OS | RPL30 | 12.785 | 2 | 49.9376 | yes | 1.305E-07 | 659.32343 | 1814.2015 |
| SEIEYYAMLAK | 11 | P62888 | Large ribosomal subunit protein eL30 OS | RPL30 | 12.785 | 2 | 49.9376 | yes | 9.975E-08 | 667.32086 | 1814.2015 |
| SLESINSR | 8 | P62888 | Large ribosomal subunit protein eL30 OS | RPL30 | 12.785 | 2 | 49.9376 | yes | 1.292E-07 | 453.23798 | 2421.9253 |
| KSLESINSR | 9 | P62888 | Large ribosomal subunit protein eL30 OS | RPL30 | 12.785 | 2 | 49.9376 | yes | 2.229E-07 | 517.28546 | 1332.7275 |
| SMPEQTGEK | 9 | P62888 | Large ribosomal subunit protein eL30 OS | RPL30 | 12.785 | 2 | 49.9376 | yes | 3.078E-06 | 503.72913 | 514.03662 |
| KSEIEYYAMLAK | 12 | P62888 | Large ribosomal subunit protein eL30 OS | RPL30 | 12.785 | 2 | 49.9376 | yes | 3.277E-07 | 731.36835 | 123.23035 |
| TGVHHYSGNNIEL | 19 | P62888 | Large ribosomal subunit protein eL30 OS | RPL30 | 12.785 | 4 | 49.9376 | yes | 2.709E-06 | 504.49045 | 24.01413 |
| TGVHHYSGNNIEL | 19 | P62888 | Large ribosomal subunit protein eL30 OS | RPL30 | 12.785 | 4 | 49.9376 | yes | 2.982E-05 | 672.31818 | 24.01413 |
| TGVHHYSGNNIEL | 19 | P62888 | Large ribosomal subunit protein eL30 OS | RPL30 | 12.785 | 3 | 49.9376 | yes | 2.709E-06 | 504.49045 | 24.01413 |
| TGVHHYSGNNIEL | 19 | P62888 | Large ribosomal subunit protein eL30 OS | RPL30 | 12.785 | 3 | 49.9376 | yes | 2.982E-05 | 672.31818 | 24.01413 |
| PIPQWIR | 7 | P62891 | Large ribosomal subunit protein eL39 OS | RPL39 | 6.407 | 2 | 49.950794 | yes | 1.108E-07 | 455.26889 | 1862.5962 |
| QNRPIPQWIR | 10 | P62891 | Large ribosomal subunit protein eL39 OS | RPL39 | 6.407 | 2 | 49.950794 | yes | 1.061E-07 | 654.37018 | 756.59143 |

| | | | | | | | | | | |
|---|---|---|---|---|---|---|---|---|---|---|
| QNRPIPQWIR | 10 | P62891 | Large ribosomal subunit protein eL39 OS RPL39 | 6.407 | 2 | 49.950794 | yes | 9.968E-08 | 436.58255 | 756.59143 |
| QNRPIPQWIR | 10 | P62891 | Large ribosomal subunit protein eL39 OS RPL39 | 6.407 | 3 | 49.950794 | yes | 1.061E-07 | 654.37018 | 756.59143 |
| QNRPIPQWIR | 10 | P62891 | Large ribosomal subunit protein eL39 OS RPL39 | 6.407 | 3 | 49.950794 | yes | 9.968E-08 | 436.58255 | 756.59143 |
| EYTINIHK | 8 | P62899 | Large ribosomal subunit protein eL31 OS RPL31 | 14.463 | 2 | 49.913662 | yes | 1.552E-07 | 509.27182 | 29591.631 |
| SAINEVVTR | 9 | P62899 | Large ribosomal subunit protein eL31 OS RPL31 | 14.463 | 2 | 49.913662 | yes | 1.242E-07 | 494.77472 | 27878.637 |
| LYTLVTYVPVTTF | 14 | P62899 | Large ribosomal subunit protein eL31 OS RPL31 | 14.463 | 2 | 49.913662 | yes | 9.984E-08 | 822.966 | 26482.385 |
| EMGTPDVR | 8 | P62899 | Large ribosomal subunit protein eL31 OS RPL31 | 14.463 | 2 | 49.913662 | yes | 1.339E-07 | 452.71329 | 5166.4512 |
| RIHGVGFK | 8 | P62899 | Large ribosomal subunit protein eL31 OS RPL31 | 14.463 | 2 | 49.913662 | yes | 2.611E-07 | 457.27197 | 303.7338 |
| NLQTVNVDEN | 10 | P62899 | Large ribosomal subunit protein eL31 OS RPL31 | 14.463 | 2 | 49.913662 | yes | 1.535E-07 | 573.27527 | 54.120529 |
| AVDIPHMDIEALK | 13 | P62906 | Large ribosomal subunit protein uL1 OS= RPL10A | 24.832 | 2 | 52.50577 | yes | 5.248E-07 | 726.38177 | 7627.1431 |
| AVDIPHMDIEALK | 13 | P62906 | Large ribosomal subunit protein uL1 OS= RPL10A | 24.832 | 2 | 52.50577 | yes | 2.329E-07 | 484.59027 | 7627.1431 |
| AVDIPHMDIEALK | 13 | P62906 | Large ribosomal subunit protein uL1 OS= RPL10A | 24.832 | 2 | 52.50577 | yes | 1.201E-06 | 734.37921 | 7627.1431 |
| AVDIPHMDIEALK | 13 | P62906 | Large ribosomal subunit protein uL1 OS= RPL10A | 24.832 | 3 | 52.50577 | yes | 5.248E-07 | 726.38177 | 7627.1431 |
| AVDIPHMDIEALK | 13 | P62906 | Large ribosomal subunit protein uL1 OS= RPL10A | 24.832 | 3 | 52.50577 | yes | 2.329E-07 | 484.59027 | 7627.1431 |
| AVDIPHMDIEALK | 13 | P62906 | Large ribosomal subunit protein uL1 OS= RPL10A | 24.832 | 3 | 52.50577 | yes | 1.201E-06 | 734.37921 | 7627.1431 |
| KNWQNVR | 7 | P62906 | Large ribosomal subunit protein uL1 OS= RPL10A | 24.832 | 2 | 52.50577 | yes | 9.913E-08 | 472.75668 | 9222.0967 |
| KYDAFLASESLIK | 13 | P62906 | Large ribosomal subunit protein uL1 OS= RPL10A | 24.832 | 2 | 52.50577 | yes | 2.6E-07 | 742.90338 | 3237.1646 |
| KYDAFLASESLIK | 13 | P62906 | Large ribosomal subunit protein uL1 OS= RPL10A | 24.832 | 2 | 52.50577 | yes | 1.225E-07 | 495.60471 | 3237.1646 |
| KYDAFLASESLIK | 13 | P62906 | Large ribosomal subunit protein uL1 OS= RPL10A | 24.832 | 3 | 52.50577 | yes | 2.6E-07 | 742.90338 | 3237.1646 |
| KYDAFLASESLIK | 13 | P62906 | Large ribosomal subunit protein uL1 OS= RPL10A | 24.832 | 3 | 52.50577 | yes | 1.225E-07 | 495.60471 | 3237.1646 |
| FPSLLTHNENMVA | 14 | P62906 | Large ribosomal subunit protein uL1 OS= RPL10A | 24.832 | 2 | 52.50577 | yes | 2.829E-07 | 800.91125 | 2798.8662 |
| FPSLLTHNENMVA | 14 | P62906 | Large ribosomal subunit protein uL1 OS= RPL10A | 24.832 | 2 | 52.50577 | yes | 2.763E-07 | 808.90869 | 2798.8662 |
| DTLYEAVR | 8 | P62906 | Large ribosomal subunit protein uL1 OS= RPL10A | 24.832 | 2 | 52.50577 | yes | 8.894E-07 | 483.74817 | 4822.1689 |
| FLETVELQISLK | 12 | P62906 | Large ribosomal subunit protein uL1 OS= RPL10A | 24.832 | 2 | 52.50577 | yes | 9.722E-08 | 710.40833 | 4199.7847 |
| YDAFLASESLIK | 12 | P62906 | Large ribosomal subunit protein uL1 OS= RPL10A | 24.832 | 2 | 52.50577 | yes | 1.994E-07 | 678.8559 | 4193.7583 |
| KFLETVELQISLK | 13 | P62906 | Large ribosomal subunit protein uL1 OS= RPL10A | 24.832 | 2 | 52.50577 | yes | 1.193E-07 | 774.45581 | 808.83917 |
| KFLETVELQISLK | 13 | P62906 | Large ribosomal subunit protein uL1 OS= RPL10A | 24.832 | 2 | 52.50577 | yes | 2.888E-07 | 516.63965 | 808.83917 |
| KFLETVELQISLK | 13 | P62906 | Large ribosomal subunit protein uL1 OS= RPL10A | 24.832 | 3 | 52.50577 | yes | 1.193E-07 | 774.45581 | 808.83917 |
| KFLETVELQISLK | 13 | P62906 | Large ribosomal subunit protein uL1 OS= RPL10A | 24.832 | 3 | 52.50577 | yes | 2.888E-07 | 516.63965 | 808.83917 |
| AVDIPHMDIEALK | 14 | P62906 | Large ribosomal subunit protein uL1 OS= RPL10A | 24.832 | 2 | 52.50577 | yes | 2.693E-07 | 790.42926 | 876.47791 |
| RFSGTVR | 7 | P62906 | Large ribosomal subunit protein uL1 OS= RPL10A | 24.832 | 2 | 52.50577 | yes | 3.209E-07 | 411.73267 | 683.5011 |
| KVLCLAVAVGHV | 13 | P62906 | Large ribosomal subunit protein uL1 OS= RPL10A | 24.832 | 3 | 52.50577 | yes | 9.934E-08 | 465.28311 | 423.11551 |
| AGKFPSLLTHNEN | 17 | P62906 | Large ribosomal subunit protein uL1 OS= RPL10A | 24.832 | 3 | 52.50577 | yes | 1.679E-07 | 619.66107 | 192.94582 |
| VLCLAVAVGHVK | 12 | P62906 | Large ribosomal subunit protein uL1 OS= RPL10A | 24.832 | 3 | 52.50577 | yes | 3.21E-07 | 422.58481 | 111.37511 |
| FSVCVLGDQQHCI | 16 | P62906 | Large ribosomal subunit protein uL1 OS= RPL10A | 24.832 | 3 | 52.50577 | yes | 9.992E-07 | 631.61371 | 18.905237 |
| HMLPSGFR | 8 | P62910 | Large ribosomal subunit protein eL32 OS RPL32 | 15.86 | 2 | 55.90927 | yes | 1.467E-07 | 472.74219 | 15019.463 |
| HMLPSGFR | 8 | P62910 | Large ribosomal subunit protein eL32 OS RPL32 | 15.86 | 2 | 55.90927 | yes | 3.123E-07 | 480.73962 | 15019.463 |
| ELEVLLMCNK | 10 | P62910 | Large ribosomal subunit protein eL32 OS RPL32 | 15.86 | 2 | 55.90927 | yes | 1.144E-07 | 624.81995 | 13326.044 |
| ELEVLLMCNK | 10 | P62910 | Large ribosomal subunit protein eL32 OS RPL32 | 15.86 | 2 | 55.90927 | yes | 9.722E-08 | 632.81744 | 13326.044 |

| | | | | | | | | | | |
|---|---|---|---|---|---|---|---|---|---|---|
| FLVHNVK | 7 | P62910 | Large ribosomal subunit protein eL32 OS RPL32 | 15.86 | 2 | 55.90927 | yes | 1.692E-07 | 428.75562 | 15669.063 |
| GQILMPNIGYGSNI | 14 | P62910 | Large ribosomal subunit protein eL32 OS RPL32 | 15.86 | 2 | 55.90927 | yes | 2.358E-07 | 746.38483 | 7454.8315 |
| GQILMPNIGYGSNI | 14 | P62910 | Large ribosomal subunit protein eL32 OS RPL32 | 15.86 | 2 | 55.90927 | yes | 1.733E-07 | 754.38232 | 7454.8315 |
| SYCAEIAHNVSSK | 13 | P62910 | Large ribosomal subunit protein eL32 OS RPL32 | 15.86 | 2 | 55.90927 | yes | 9.722E-08 | 733.34064 | 9962.7471 |
| AALRPLVK | 8 | P62910 | Large ribosomal subunit protein eL32 OS RPL32 | 15.86 | 2 | 55.90927 | yes | 1.435E-07 | 434.29236 | 2316.7751 |
| AALRPLVK | 8 | P62910 | Large ribosomal subunit protein eL32 OS RPL32 | 15.86 | 2 | 55.90927 | yes | 1.625E-07 | 455.29764 | 2316.7751 |
| KFLVHNVK | 8 | P62910 | Large ribosomal subunit protein eL32 OS RPL32 | 15.86 | 2 | 55.90927 | yes | 1.776E-07 | 492.8031 | 1913.3341 |
| AALRPLVKPK | 10 | P62910 | Large ribosomal subunit protein eL32 OS RPL32 | 15.86 | 2 | 55.90927 | yes | 1.661E-07 | 546.86621 | 60.276974 |
| YDGIILPGK | 9 | P62913 | Large ribosomal subunit protein uL5 OS= RPL11 | 20.253 | 2 | 48.658527 | yes | 2.536E-07 | 488.27911 | 33437.305 |
| VLEQLTGQTPVFSI | 14 | P62913 | Large ribosomal subunit protein uL5 OS= RPL11 | 20.253 | 2 | 48.658527 | yes | 1.726E-07 | 773.92743 | 30946.443 |
| IAVHCTVR | 8 | P62913 | Large ribosomal subunit protein uL5 OS= RPL11 | 20.253 | 2 | 48.658527 | yes | 1.192E-07 | 478.26074 | 12071.584 |
| LCLNICVGESGDR | 13 | P62913 | Large ribosomal subunit protein uL5 OS= RPL11 | 20.253 | 2 | 48.658527 | yes | 1.118E-07 | 746.84778 | 11776.653 |
| AEEILEK | 7 | P62913 | Large ribosomal subunit protein uL5 OS= RPL11 | 20.253 | 2 | 48.658527 | yes | 1.348E-07 | 416.22656 | 1624.3583 |
| KNNFSDTGNFGFG | 23 | P62913 | Large ribosomal subunit protein uL5 OS= RPL11 | 20.253 | 4 | 48.658527 | yes | 1.429E-07 | 638.56982 | 661.03326 |
| PGFSIADKK | 9 | P62913 | Large ribosomal subunit protein uL5 OS= RPL11 | 20.253 | 2 | 48.658527 | yes | 4.183E-07 | 481.76892 | 449.49591 |
| KLCLNICVGESGDI | 14 | P62913 | Large ribosomal subunit protein uL5 OS= RPL11 | 20.253 | 2 | 48.658527 | yes | 1.166E-07 | 810.89526 | 123.92678 |
| KLCLNICVGESGDI | 14 | P62913 | Large ribosomal subunit protein uL5 OS= RPL11 | 20.253 | 2 | 48.658527 | yes | 5.419E-06 | 540.93262 | 123.92678 |
| KLCLNICVGESGDI | 14 | P62913 | Large ribosomal subunit protein uL5 OS= RPL11 | 20.253 | 3 | 48.658527 | yes | 1.166E-07 | 810.89526 | 123.92678 |
| KLCLNICVGESGDI | 14 | P62913 | Large ribosomal subunit protein uL5 OS= RPL11 | 20.253 | 3 | 48.658527 | yes | 5.419E-06 | 540.93262 | 123.92678 |
| AVVGVVAGGGR | 11 | P62917 | Large ribosomal subunit protein uL2 OS= RPL8 | 28.025 | 2 | 50.09155 | yes | 1.164E-07 | 471.27997 | 28452.527 |
| DIIHDPGR | 8 | P62917 | Large ribosomal subunit protein uL2 OS= RPL8 | 28.025 | 2 | 50.09155 | yes | 1.545E-07 | 461.74069 | 26306.254 |
| IDKPILK | 7 | P62917 | Large ribosomal subunit protein uL2 OS= RPL8 | 28.025 | 2 | 50.09155 | yes | 1.184E-07 | 413.77347 | 14126.894 |
| KVGLIAAR | 8 | P62917 | Large ribosomal subunit protein uL2 OS= RPL8 | 28.025 | 2 | 50.09155 | yes | 1.589E-07 | 414.2767 | 10619.9 |
| ASGNYATVISHNPI | 16 | P62917 | Large ribosomal subunit protein uL2 OS= RPL8 | 28.025 | 2 | 50.09155 | yes | 1.248E-07 | 844.91559 | 2266.9866 |
| ASGNYATVISHNPI | 16 | P62917 | Large ribosomal subunit protein uL2 OS= RPL8 | 28.025 | 2 | 50.09155 | yes | 1.028E-07 | 563.61279 | 2266.9866 |
| ASGNYATVISHNPI | 16 | P62917 | Large ribosomal subunit protein uL2 OS= RPL8 | 28.025 | 3 | 50.09155 | yes | 1.248E-07 | 844.91559 | 2266.9866 |
| ASGNYATVISHNPI | 16 | P62917 | Large ribosomal subunit protein uL2 OS= RPL8 | 28.025 | 3 | 50.09155 | yes | 1.028E-07 | 563.61279 | 2266.9866 |
| TELFIAAEGIHTGQ | 20 | P62917 | Large ribosomal subunit protein uL2 OS= RPL8 | 28.025 | 3 | 50.09155 | yes | 1.424E-07 | 747.70514 | 4402.7935 |
| GVAMNPVEHPFG | 26 | P62917 | Large ribosomal subunit protein uL2 OS= RPL8 | 28.025 | 5 | 50.09155 | yes | 1.104E-07 | 551.47961 | 1104.9458 |
| GVAMNPVEHPFG | 26 | P62917 | Large ribosomal subunit protein uL2 OS= RPL8 | 28.025 | 5 | 50.09155 | yes | 1.738E-07 | 689.09766 | 1104.9458 |
| GVAMNPVEHPFG | 26 | P62917 | Large ribosomal subunit protein uL2 OS= RPL8 | 28.025 | 5 | 50.09155 | yes | 9.904E-08 | 685.09894 | 1104.9458 |
| GVAMNPVEHPFG | 26 | P62917 | Large ribosomal subunit protein uL2 OS= RPL8 | 28.025 | 4 | 50.09155 | yes | 1.104E-07 | 551.47961 | 1104.9458 |
| GVAMNPVEHPFG | 26 | P62917 | Large ribosomal subunit protein uL2 OS= RPL8 | 28.025 | 4 | 50.09155 | yes | 1.738E-07 | 689.09766 | 1104.9458 |
| GVAMNPVEHPFG | 26 | P62917 | Large ribosomal subunit protein uL2 OS= RPL8 | 28.025 | 4 | 50.09155 | yes | 9.904E-08 | 685.09894 | 1104.9458 |
| RTELFIAAEGIHTG | 21 | P62917 | Large ribosomal subunit protein uL2 OS= RPL8 | 28.025 | 3 | 50.09155 | yes | 1.257E-07 | 799.73883 | 568.53406 |
| AQLNIGNVLPVGT | 30 | P62917 | Large ribosomal subunit protein uL2 OS= RPL8 | 28.025 | 4 | 50.09155 | yes | 1.549E-07 | 817.65973 | 541.19934 |
| GVAMNPVEHPFG | 21 | P62917 | Large ribosomal subunit protein uL2 OS= RPL8 | 28.025 | 4 | 50.09155 | yes | 1.996E-07 | 546.51953 | 234.6953 |
| GVAMNPVEHPFG | 21 | P62917 | Large ribosomal subunit protein uL2 OS= RPL8 | 28.025 | 4 | 50.09155 | yes | 1.833E-07 | 728.35693 | 234.6953 |
| GVAMNPVEHPFG | 21 | P62917 | Large ribosomal subunit protein uL2 OS= RPL8 | 28.025 | 3 | 50.09155 | yes | 1.996E-07 | 546.51953 | 234.6953 |

| | | | | | | | | | | | |
|---|---|---|---|---|---|---|---|---|---|---|---|
| GVAMNPVEHPFG( | 21 | P62917 | Large ribosomal subunit protein uL2 OS= | RPL8 | 28.025 | 3 | 50.09155 | yes | 1.833E-07 | 728.35693 | 234.6953 |
| KVISSANR | 8 | P62917 | Large ribosomal subunit protein uL2 OS= | RPL8 | 28.025 | 2 | 50.09155 | yes | 6.801E-07 | 437.75888 | 314.45865 |
| VKLPSGSK | 8 | P62917 | Large ribosomal subunit protein uL2 OS= | RPL8 | 28.025 | 2 | 50.09155 | yes | 5.372E-07 | 408.2529 | 30.638107 |
| FEDENFILK | 9 | P62937 | Peptidyl-prolyl cis-trans isomerase A OS= | PPIA | 18.013 | 2 | 50.64698 | yes | 3.174E-06 | 577.79004 | 1596.5547 |
| IIPGFMCQGGDFTI | 14 | P62937 | Peptidyl-prolyl cis-trans isomerase A OS= | PPIA | 18.013 | 2 | 50.64698 | no | 2.045E-07 | 799.87634 | 1399.2239 |
| VSFELFADK | 9 | P62937 | Peptidyl-prolyl cis-trans isomerase A OS= | PPIA | 18.013 | 2 | 50.64698 | yes | 1.318E-07 | 528.27405 | 896.82813 |
| EGMNIVEAMER | 11 | P62937 | Peptidyl-prolyl cis-trans isomerase A OS= | PPIA | 18.013 | 2 | 50.64698 | yes | 9.732E-08 | 639.79449 | 400.30643 |
| KITIADCGQLE | 11 | P62937 | Peptidyl-prolyl cis-trans isomerase A OS= | PPIA | 18.013 | 2 | 50.64698 | yes | 3.851E-07 | 624.31866 | 100.33738 |
| ECPSDECGAGVFM | 19 | P62979 | Ubiquitin-ribosomal protein eS31 fusion | RPS27A | 17.965 | 3 | 50.79333 | yes | 1.615E-07 | 724.62415 | 8243.376 |
| ECPSDECGAGVFM | 19 | P62979 | Ubiquitin-ribosomal protein eS31 fusion | RPS27A | 17.965 | 3 | 50.79333 | yes | 1.536E-07 | 729.95581 | 8243.376 |
| ESTLHLVLR | 9 | P62979 | Ubiquitin-ribosomal protein eS31 fusion | RPS27A | 17.965 | 2 | 50.79333 | no | 1.113E-07 | 534.31403 | 6466.9941 |
| RECPSDECGAGVF | 20 | P62979 | Ubiquitin-ribosomal protein eS31 fusion | RPS27A | 17.965 | 3 | 50.79333 | yes | 9.722E-08 | 776.65784 | 2343.6191 |
| RECPSDECGAGVF | 20 | P62979 | Ubiquitin-ribosomal protein eS31 fusion | RPS27A | 17.965 | 3 | 50.79333 | yes | 1.084E-07 | 781.9895 | 2343.6191 |
| TLSDYNIQK | 9 | P62979 | Ubiquitin-ribosomal protein eS31 fusion | RPS27A | 17.965 | 2 | 50.79333 | no | 2.685E-06 | 541.27985 | 3174.915 |
| EGIPPDQQR | 9 | P62979 | Ubiquitin-ribosomal protein eS31 fusion | RPS27A | 17.965 | 2 | 50.79333 | no | 3.201E-06 | 520.26196 | 89.328735 |
| HYCGKCCLTYCFN | 18 | P62979 | Ubiquitin-ribosomal protein eS31 fusion | RPS27A | 17.965 | 4 | 50.79333 | yes | 1.868E-06 | 595.75488 | 43.997601 |
| YNCDKMICR | 9 | P62987 | Ubiquitin-ribosomal protein eL40 fusion | UBA52 | 14.729 | 2 | 51.834557 | yes | 9.722E-08 | 630.27014 | 142.18269 |
| ESESAPGDFSLSVI | 14 | P62993 | Growth factor receptor-bound protein 2 O | GRB2 | 25.207 | 2 | 51.882133 | yes | 1.972E-07 | 726.84625 | 752.52728 |
| HDGAFLIR | 8 | P62993 | Growth factor receptor-bound protein 2 O | GRB2 | 25.207 | 2 | 51.882133 | yes | 2.776E-07 | 464.7536 | 519.02875 |
| NQQIFLR | 7 | P62993 | Growth factor receptor-bound protein 2 O | GRB2 | 25.207 | 2 | 51.882133 | yes | 2.983E-07 | 459.76141 | 444.39865 |
| VLNEECDQNWYK | 12 | P62993 | Growth factor receptor-bound protein 2 O | GRB2 | 25.207 | 2 | 51.882133 | yes | 9.85E-08 | 799.3512 | 296.59882 |
| FGNDVQHFK | 9 | P62993 | Growth factor receptor-bound protein 2 O | GRB2 | 25.207 | 2 | 51.882133 | yes | 9.722E-08 | 546.26709 | 266.09058 |
| NYVTPVNR | 8 | P62993 | Growth factor receptor-bound protein 2 O | GRB2 | 25.207 | 2 | 51.882133 | yes | 1.129E-07 | 481.75635 | 249.85739 |
| PHPWFFGK | 8 | P62993 | Growth factor receptor-bound protein 2 O | GRB2 | 25.207 | 2 | 51.882133 | yes | 1.037E-07 | 508.26105 | 201.01755 |
| ATADDELSFK | 10 | P62993 | Growth factor receptor-bound protein 2 O | GRB2 | 25.207 | 2 | 51.882133 | yes | 2.478E-06 | 548.76147 | 172.69582 |
| GACHGQTGMFPR | 12 | P62993 | Growth factor receptor-bound protein 2 O | GRB2 | 25.207 | 2 | 51.882133 | yes | 2.042E-07 | 659.7926 | 89.049637 |
| GFAFVYFENVDD/ | 14 | P62995 | Transformer-2 protein homolog beta OS= | TRA2B | 33.666 | 2 | 51.784477 | yes | 1.02E-07 | 811.38031 | 7541.9341 |
| RPHTPTPGIYMGR | 13 | P62995 | Transformer-2 protein homolog beta OS= | TRA2B | 33.666 | 3 | 51.784477 | yes | 1.174E-07 | 494.92599 | 1656.0602 |
| ANPDPNCCLGVFC | 21 | P62995 | Transformer-2 protein homolog beta OS= | TRA2B | 33.666 | 3 | 51.784477 | yes | 9.722E-08 | 795.37183 | 246.15784 |
| GYDDRDYYSR | 10 | P62995 | Transformer-2 protein homolog beta OS= | TRA2B | 33.666 | 2 | 51.784477 | yes | 9.722E-08 | 655.27582 | 174.26561 |
| ANGMELDGR | 9 | P62995 | Transformer-2 protein homolog beta OS= | TRA2B | 33.666 | 2 | 51.784477 | no | 5.211E-07 | 481.72165 | 115.38696 |
| SPSPYYSR | 8 | P62995 | Transformer-2 protein homolog beta OS= | TRA2B | 33.666 | 2 | 51.784477 | no | 9.892E-08 | 478.72726 | 103.96046 |
| SDSGEQNYGER | 11 | P62995 | Transformer-2 protein homolog beta OS= | TRA2B | 33.666 | 2 | 51.784477 | yes | 9.728E-08 | 642.26038 | 93.052322 |
| AAQDRDQIYR | 10 | P62995 | Transformer-2 protein homolog beta OS= | TRA2B | 33.666 | 2 | 51.784477 | yes | 1.004E-07 | 618.31 | 34.692848 |
| CVVVGDGAVGK | 11 | P63000 | Ras-related C3 botulinum toxin substrate | RAC1 | 21.451 | 2 | 51.667805 | no | 1.597E-07 | 530.77643 | 693.79749 |
| TVFDEAIR | 8 | P63000 | Ras-related C3 botulinum toxin substrate | RAC1 | 21.451 | 2 | 51.667805 | no | 1.707E-07 | 475.75073 | 197.22398 |
| LTPITYPQGLAMA | 14 | P63000 | Ras-related C3 botulinum toxin substrate | RAC1 | 21.451 | 2 | 51.667805 | yes | 9.722E-08 | 752.41565 | 173.81259 |
| AVLCPPPVK | 9 | P63000 | Ras-related C3 botulinum toxin substrate | RAC1 | 21.451 | 2 | 51.667805 | no | 7.549E-07 | 490.78351 | 132.14923 |
| YLECSALTQR | 10 | P63000 | Ras-related C3 botulinum toxin substrate | RAC1 | 21.451 | 2 | 51.667805 | no | 9.722E-08 | 620.80316 | 94.887131 |

| | | | | | | | | | | | |
|---|---|---|---|---|---|---|---|---|---|---|---|
| ITEYLCEPLR | 10 | P63010 | AP-2 complex subunit beta OS=Homo saj | AP2B1 | 104.553 | 2 | 54.31121 | no | 1.041E-07 | 647.32904 | 1399.2607 |
| GLEISGTFTHR | 11 | P63010 | AP-2 complex subunit beta OS=Homo saj | AP2B1 | 104.553 | 2 | 54.31121 | yes | 9.974E-08 | 609.31732 | 1356.2969 |
| DIPNENELQFQIK | 13 | P63010 | AP-2 complex subunit beta OS=Homo saj | AP2B1 | 104.553 | 2 | 54.31121 | yes | 1.601E-07 | 794.4043 | 1070.2242 |
| AAMIWIVGEYAEF | 13 | P63010 | AP-2 complex subunit beta OS=Homo saj | AP2B1 | 104.553 | 2 | 54.31121 | no | 4.815E-07 | 754.88196 | 658.51758 |
| LSHANSAVVLSAV | 14 | P63010 | AP-2 complex subunit beta OS=Homo saj | AP2B1 | 104.553 | 2 | 54.31121 | no | 9.722E-08 | 698.40137 | 328.68259 |
| LSHANSAVVLSAV | 14 | P63010 | AP-2 complex subunit beta OS=Homo saj | AP2B1 | 104.553 | 2 | 54.31121 | no | 1.037E-07 | 465.93668 | 328.68259 |
| LSHANSAVVLSAV | 14 | P63010 | AP-2 complex subunit beta OS=Homo saj | AP2B1 | 104.553 | 3 | 54.31121 | no | 9.722E-08 | 698.40137 | 328.68259 |
| LSHANSAVVLSAV | 14 | P63010 | AP-2 complex subunit beta OS=Homo saj | AP2B1 | 104.553 | 3 | 54.31121 | no | 1.037E-07 | 465.93668 | 328.68259 |
| KGEIFELK | 8 | P63010 | AP-2 complex subunit beta OS=Homo saj | AP2B1 | 104.553 | 2 | 54.31121 | no | 9.722E-08 | 482.27911 | 653.90826 |
| EYATEVDVDFVR | 12 | P63010 | AP-2 complex subunit beta OS=Homo saj | AP2B1 | 104.553 | 2 | 54.31121 | no | 1.37E-07 | 721.84357 | 609.32953 |
| NVEGQDMLYQSLl | 13 | P63010 | AP-2 complex subunit beta OS=Homo saj | AP2B1 | 104.553 | 2 | 54.31121 | no | 1.134E-07 | 762.87177 | 524.00983 |
| DCEDPNPLIR | 10 | P63010 | AP-2 complex subunit beta OS=Homo saj | AP2B1 | 104.553 | 2 | 54.31121 | no | 9.722E-08 | 614.78497 | 396.26675 |
| MEPLNNLQVAVK | 12 | P63010 | AP-2 complex subunit beta OS=Homo saj | AP2B1 | 104.553 | 2 | 54.31121 | no | 1.597E-07 | 678.37122 | 396.11911 |
| SQPDMAIMAVNSI | 15 | P63010 | AP-2 complex subunit beta OS=Homo saj | AP2B1 | 104.553 | 2 | 54.31121 | yes | 9.722E-08 | 819.40491 | 374.05011 |
| KVIAAMTVGK | 10 | P63010 | AP-2 complex subunit beta OS=Homo saj | AP2B1 | 104.553 | 2 | 54.31121 | yes | 8.593E-07 | 509.30991 | 351.76138 |
| VNYVVQEAIVVIR | 13 | P63010 | AP-2 complex subunit beta OS=Homo saj | AP2B1 | 104.553 | 2 | 54.31121 | yes | 2.504E-07 | 751.44049 | 319.89694 |
| IQPGNPNYTLSLK | 13 | P63010 | AP-2 complex subunit beta OS=Homo saj | AP2B1 | 104.553 | 2 | 54.31121 | yes | 1.321E-07 | 722.89337 | 311.64185 |
| CLKDEDPYVR | 10 | P63010 | AP-2 complex subunit beta OS=Homo saj | AP2B1 | 104.553 | 2 | 54.31121 | no | 1.236E-07 | 647.80847 | 266.31818 |
| CVSTLLDLIQTK | 12 | P63010 | AP-2 complex subunit beta OS=Homo saj | AP2B1 | 104.553 | 2 | 54.31121 | no | 9.722E-08 | 695.88416 | 216.51427 |
| YNDPIYVK | 8 | P63010 | AP-2 complex subunit beta OS=Homo saj | AP2B1 | 104.553 | 2 | 54.31121 | no | 1.872E-07 | 506.26093 | 216.40558 |
| DSDYYNMLLK | 10 | P63010 | AP-2 complex subunit beta OS=Homo saj | AP2B1 | 104.553 | 2 | 54.31121 | yes | 1.039E-07 | 631.29211 | 180.86604 |
| ECHLNADTVSSK | 12 | P63010 | AP-2 complex subunit beta OS=Homo saj | AP2B1 | 104.553 | 2 | 54.31121 | yes | 1.14E-07 | 680.81171 | 176.96832 |
| ALQHMTDFAIQFN | 14 | P63010 | AP-2 complex subunit beta OS=Homo saj | AP2B1 | 104.553 | 2 | 54.31121 | yes | 2.193E-07 | 832.41669 | 94.518593 |
| LQNNNVYTIAK | 11 | P63010 | AP-2 complex subunit beta OS=Homo saj | AP2B1 | 104.553 | 2 | 54.31121 | yes | 1.124E-07 | 639.34607 | 93.171791 |
| LASQANIAQVLAE | 15 | P63010 | AP-2 complex subunit beta OS=Homo saj | AP2B1 | 104.553 | 2 | 54.31121 | no | 1.967E-07 | 784.95398 | 85.67543 |
| FLELLPK | 7 | P63010 | AP-2 complex subunit beta OS=Homo saj | AP2B1 | 104.553 | 2 | 54.31121 | yes | 9.225E-07 | 430.26804 | 67.311615 |
| QGHIYMEMNFTNI | 13 | P63010 | AP-2 complex subunit beta OS=Homo saj | AP2B1 | 104.553 | 2 | 54.31121 | yes | 2.166E-07 | 806.86597 | 63.750969 |
| VIAAMTVGK | 9 | P63010 | AP-2 complex subunit beta OS=Homo saj | AP2B1 | 104.553 | 2 | 54.31121 | yes | 2.462E-06 | 445.26242 | 52.826511 |
| EAQSICER | 8 | P63010 | AP-2 complex subunit beta OS=Homo saj | AP2B1 | 104.553 | 2 | 54.31121 | no | 1.029E-06 | 496.72693 | 49.17226 |
| AVWLPAVK | 8 | P63010 | AP-2 complex subunit beta OS=Homo saj | AP2B1 | 104.553 | 2 | 54.31121 | yes | 1.08E-06 | 442.27365 | 16.40947 |
| LVYLYLMNYAK | 11 | P63010 | AP-2 complex subunit beta OS=Homo saj | AP2B1 | 104.553 | 2 | 54.31121 | no | 9.722E-08 | 695.87561 | 14.009979 |
| QVFLATWK | 8 | P63010 | AP-2 complex subunit beta OS=Homo saj | AP2B1 | 104.553 | 2 | 54.31121 | yes | 8.365E-07 | 496.78183 | 10.259858 |
| QADYVPSDQDLLI | 13 | P63092 | Guanine nucleotide-binding protein G(s) : | GNAS | 45.665 | 2 | 50.015285 | yes | 1.826E-07 | 760.37299 | 316.7984 |
| ALWEDEGVR | 9 | P63092 | Guanine nucleotide-binding protein G(s) : | GNAS | 45.665 | 2 | 50.015285 | yes | 9.934E-08 | 537.76434 | 294.59177 |
| IEDYFPEFAR | 10 | P63092 | Guanine nucleotide-binding protein G(s) : | GNAS | 45.665 | 2 | 50.015285 | yes | 1.888E-07 | 643.80621 | 216.88438 |
| LQEALNLFK | 9 | P63092 | Guanine nucleotide-binding protein G(s) : | GNAS | 45.665 | 2 | 50.015285 | yes | 2.918E-07 | 538.31097 | 90.570587 |
| VLTSGIFETK | 10 | P63092 | Guanine nucleotide-binding protein G(s) : | GNAS | 45.665 | 2 | 50.015285 | yes | 8.031E-07 | 547.80823 | 38.788029 |
| DTQLQQIVDK | 10 | P63098 | Calcineurin subunit B type 1 OS=Homo s | PPP3R1 | 19.3 | 2 | 41.806152 | yes | 3.332E-07 | 594.31696 | 68.276688 |
| YLAEVAAGDDKK | 12 | P63104 | 14-3-3 protein zeta/delta OS=Homo sapie | YWHAZ | 27.746 | 2 | 53.899403 | yes | 1.297E-07 | 640.33008 | 909.48126 |

| | | | | | | | | | | | |
|---|---|---|---|---|---|---|---|---|---|---|---|
| SVTEQGAELSNEE | 14 | P63104 | 14-3-3 protein zeta/delta OS=Homo sapie | YWHAZ | 27.746 | 2 | 53.899403 | yes | 9.722E-08 | 774.86047 | 323.87256 |
| YDDMAACMK | 9 | P63104 | 14-3-3 protein zeta/delta OS=Homo sapie | YWHAZ | 27.746 | 2 | 53.899403 | yes | 9.722E-08 | 552.71155 | 259.8703 |
| VVSSIEQK | 8 | P63104 | 14-3-3 protein zeta/delta OS=Homo sapie | YWHAZ | 27.746 | 2 | 53.899403 | yes | 3.191E-07 | 445.25311 | 80.903214 |
| MDKNELVQK | 9 | P63104 | 14-3-3 protein zeta/delta OS=Homo sapie | YWHAZ | 27.746 | 2 | 53.899403 | yes | 4.487E-07 | 573.7948 | 54.206429 |
| VVIFQQEQENK | 11 | P63151 | Serine/threonine-protein phosphatase 2A | PPP2R2A | 51.693 | 2 | 53.793385 | yes | 1.004E-07 | 681.35663 | 409.02707 |
| INLWHLEITDR | 11 | P63151 | Serine/threonine-protein phosphatase 2A | PPP2R2A | 51.693 | 2 | 53.793385 | no | 9.722E-08 | 705.38043 | 198.70837 |
| ILHTAWHPK | 9 | P63151 | Serine/threonine-protein phosphatase 2A | PPP2R2A | 51.693 | 2 | 53.793385 | yes | 1.082E-07 | 551.81146 | 192.78201 |
| NAAQFLLSTNDK | 12 | P63151 | Serine/threonine-protein phosphatase 2A | PPP2R2A | 51.693 | 2 | 53.793385 | yes | 4.704E-07 | 661.34094 | 165.14667 |
| LFEEPEDPSNR | 11 | P63151 | Serine/threonine-protein phosphatase 2A | PPP2R2A | 51.693 | 2 | 53.793385 | no | 7.994E-07 | 666.80695 | 132.88138 |
| PMDLMVEASPR | 11 | P63151 | Serine/threonine-protein phosphatase 2A | PPP2R2A | 51.693 | 2 | 53.793385 | no | 1.068E-07 | 623.30194 | 99.889618 |
| YRDPTTVTTLR | 11 | P63151 | Serine/threonine-protein phosphatase 2A | PPP2R2A | 51.693 | 2 | 53.793385 | yes | 2.926E-07 | 661.85681 | 58.427872 |
| LCSLYENDCIFDK | 13 | P63151 | Serine/threonine-protein phosphatase 2A | PPP2R2A | 51.693 | 2 | 53.793385 | no | 1.557E-07 | 838.86835 | 46.34906 |
| NFGSYVTHETK | 11 | P63167 | Dynein light chain 1, cytoplasmic OS=Ho | DYNLL1 | 10.366 | 2 | 53.582645 | no | 9.809E-08 | 641.80676 | 8932.7539 |
| YNPTWHCIVGR | 11 | P63167 | Dynein light chain 1, cytoplasmic OS=Ho | DYNLL1 | 10.366 | 2 | 53.582645 | no | 1.762E-07 | 701.83789 | 5538.397 |
| NADMSEEMQQDS | 22 | P63167 | Dynein light chain 1, cytoplasmic OS=Ho | DYNLL1 | 10.366 | 3 | 53.582645 | yes | 9.722E-08 | 838.68579 | 137.2663 |
| KYNPTWHCIVGR | 12 | P63167 | Dynein light chain 1, cytoplasmic OS=Ho | DYNLL1 | 10.366 | 2 | 53.582645 | no | 9.722E-08 | 765.88538 | 64.286819 |
| YLYTLVITDK | 10 | P63173 | Large ribosomal subunit protein eL38 OS | RPL38 | 8.218 | 2 | 55.24216 | yes | 1.041E-07 | 614.84485 | 10841.478 |
| QSLPPGLAVK | 10 | P63173 | Large ribosomal subunit protein eL38 OS | RPL38 | 8.218 | 2 | 55.24216 | yes | 3.387E-06 | 505.30566 | 1990.8568 |
| KIEEIKDFLLTAR | 13 | P63173 | Large ribosomal subunit protein eL38 OS | RPL38 | 8.218 | 2 | 55.24216 | yes | 3.781E-07 | 788.45886 | 1904.8857 |
| IEEIKDFLLTAR | 12 | P63173 | Large ribosomal subunit protein eL38 OS | RPL38 | 8.218 | 2 | 55.24216 | yes | 9.722E-08 | 724.41138 | 1583.6208 |
| DFLLTAR | 7 | P63173 | Large ribosomal subunit protein eL38 OS | RPL38 | 8.218 | 2 | 55.24216 | yes | 1.773E-07 | 418.23724 | 148.36813 |
| CSRYLYTLVITDKl | 15 | P63173 | Large ribosomal subunit protein eL38 OS | RPL38 | 8.218 | 3 | 55.24216 | yes | 9.722E-08 | 630.33276 | 64.774452 |
| TDDIPVWDQEFLK | 13 | P63208 | S-phase kinase-associated protein 1 OS=I | SKP1 | 18.658 | 2 | 55.775665 | yes | 2.543E-07 | 803.39337 | 1398.5685 |
| GLLDVTCK | 8 | P63208 | S-phase kinase-associated protein 1 OS=I | SKP1 | 18.658 | 2 | 55.775665 | yes | 1.274E-07 | 453.24167 | 471.90698 |
| VIQWCTHHK | 9 | P63208 | S-phase kinase-associated protein 1 OS=I | SKP1 | 18.658 | 2 | 55.775665 | yes | 1.002E-07 | 604.80328 | 445.51117 |
| NDFTEEEEAQVR | 12 | P63208 | S-phase kinase-associated protein 1 OS=I | SKP1 | 18.658 | 2 | 55.775665 | yes | 9.722E-08 | 733.82336 | 251.36526 |
| KENQWCEEK | 9 | P63208 | S-phase kinase-associated protein 1 OS=I | SKP1 | 18.658 | 2 | 55.775665 | yes | 1.031E-06 | 625.77716 | 222.53149 |
| DDPPPPEDDENKE | 14 | P63208 | S-phase kinase-associated protein 1 OS=I | SKP1 | 18.658 | 2 | 55.775665 | yes | 7.206E-07 | 812.85229 | 136.8916 |
| ENQWCEEK | 8 | P63208 | S-phase kinase-associated protein 1 OS=I | SKP1 | 18.658 | 2 | 55.775665 | yes | 9.722E-08 | 561.72968 | 74.249847 |
| TPEEIRK | 7 | P63208 | S-phase kinase-associated protein 1 OS=I | SKP1 | 18.658 | 2 | 55.775665 | yes | 6.689E-07 | 436.74545 | 39.113689 |
| TYAICGAIR | 9 | P63220 | Small ribosomal subunit protein eS21 OS | RPS21 | 9.112 | 2 | 49.454975 | yes | 1.176E-07 | 512.76587 | 8008.7139 |
| MGESDDSILR | 10 | P63220 | Small ribosomal subunit protein eS21 OS | RPS21 | 9.112 | 2 | 49.454975 | yes | 1.326E-07 | 561.75842 | 3137.6133 |
| MGESDDSILR | 10 | P63220 | Small ribosomal subunit protein eS21 OS | RPS21 | 9.112 | 2 | 49.454975 | yes | 1.029E-07 | 569.75586 | 3137.6133 |
| DHASIQMNVAEVI | 14 | P63220 | Small ribosomal subunit protein eS21 OS | RPS21 | 9.112 | 2 | 49.454975 | yes | 1.649E-06 | 778.87231 | 579.9115 |
| VHLVGIDIFTGK | 12 | P63241 | Eukaryotic translation initiation factor 5A | EIF5A | 16.833 | 2 | 47.716763 | no | 2.453E-07 | 649.87701 | 2602.6333 |
| LPEGDLGK | 8 | P63241 | Eukaryotic translation initiation factor 5A | EIF5A | 16.833 | 2 | 47.716763 | no | 1.526E-07 | 414.72672 | 608.6236 |
| EDLRLPEGDLGK | 12 | P63241 | Eukaryotic translation initiation factor 5A | EIF5A | 16.833 | 2 | 47.716763 | yes | 1.273E-07 | 671.35406 | 290.89124 |
| KNGFVVLK | 8 | P63241 | Eukaryotic translation initiation factor 5A | EIF5A | 16.833 | 2 | 47.716763 | no | 4.547E-07 | 452.78436 | 43.594566 |
| LWDLTTGTTTR | 11 | P63244 | Small ribosomal subunit protein RACK1 | RACK1 | 35.077 | 2 | 51.851986 | yes | 1.106E-07 | 632.83026 | 10224.432 |

| | | | | | | | | | | | |
|---|---|---|---|---|---|---|---|---|---|---|---|
| DGQAMLWDLNEC | 13 | P63244 | Small ribosomal subunit protein RACK1 | RACK1 | 35.077 | 2 | 51.851986 | yes | 9.722E-08 | 738.84302 | 3503.2932 |
| DGQAMLWDLNEC | 13 | P63244 | Small ribosomal subunit protein RACK1 | RACK1 | 35.077 | 2 | 51.851986 | yes | 1.136E-06 | 746.84045 | 3503.2932 |
| VWQVTIGTR | 9 | P63244 | Small ribosomal subunit protein RACK1 | RACK1 | 35.077 | 2 | 51.851986 | yes | 2.095E-07 | 530.3009 | 5769.0127 |
| DVLSVAFSSDNR | 12 | P63244 | Small ribosomal subunit protein RACK1 | RACK1 | 35.077 | 2 | 51.851986 | yes | 1.007E-07 | 655.32275 | 4494.7275 |
| LWNTLGVCK | 9 | P63244 | Small ribosomal subunit protein RACK1 | RACK1 | 35.077 | 2 | 51.851986 | yes | 1.791E-07 | 545.78931 | 4156.9595 |
| YWLCAATGPSIK | 12 | P63244 | Small ribosomal subunit protein RACK1 | RACK1 | 35.077 | 2 | 51.851986 | yes | 9.789E-08 | 683.84485 | 3924.5916 |
| DETNYGIPQR | 10 | P63244 | Small ribosomal subunit protein RACK1 | RACK1 | 35.077 | 2 | 51.851986 | yes | 3.966E-07 | 596.78326 | 3099.6091 |
| IWDLEGK | 7 | P63244 | Small ribosomal subunit protein RACK1 | RACK1 | 35.077 | 2 | 51.851986 | yes | 1.063E-07 | 430.72925 | 1122.0756 |
| QEVISTSSK | 9 | P63244 | Small ribosomal subunit protein RACK1 | RACK1 | 35.077 | 2 | 51.851986 | yes | 1.281E-07 | 489.75876 | 999.05975 |
| IIVDELK | 7 | P63244 | Small ribosomal subunit protein RACK1 | RACK1 | 35.077 | 2 | 51.851986 | yes | 1.863E-07 | 415.2551 | 909.03473 |
| TNHIGHTGYLNTV | 27 | P63244 | Small ribosomal subunit protein RACK1 | RACK1 | 35.077 | 4 | 51.851986 | yes | 9.727E-08 | 686.58307 | 524.16229 |
| VWNLANCK | 8 | P63244 | Small ribosomal subunit protein RACK1 | RACK1 | 35.077 | 2 | 51.851986 | yes | 2.365E-07 | 502.75275 | 425.94296 |
| DKTIIMWK | 8 | P63244 | Small ribosomal subunit protein RACK1 | RACK1 | 35.077 | 2 | 51.851986 | yes | 2.582E-07 | 517.78882 | 219.539 |
| HLYTLDGGDIINAI | 20 | P63244 | Small ribosomal subunit protein RACK1 | RACK1 | 35.077 | 3 | 51.851986 | yes | 9.722E-08 | 759.37579 | 149.2189 |
| ACLLCSLVK | 9 | P63272 | Transcription elongation factor SPT4 OS= | SUPT4H1 | 13.194 | 2 | 46.94243 | yes | 1.436E-07 | 532.28558 | 520.67548 |
| VSNFKPGVYAVSV | 16 | P63272 | Transcription elongation factor SPT4 OS= | SUPT4H1 | 13.194 | 2 | 46.94243 | yes | 1.668E-06 | 840.95703 | 26.082054 |
| DHPFGFVAVPTK | 12 | P63279 | SUMO-conjugating enzyme UBC9 OS=H | UBE2I | 18.007 | 2 | 45.175262 | yes | 1.936E-07 | 657.8457 | 56.6329 |
| QLSESQVK | 8 | P67775 | Serine/threonine-protein phosphatase 2A | PPP2CA | 35.595 | 2 | 45.373405 | yes | 1.87E-07 | 459.74817 | 130.44347 |
| GAEAANVTGPGGV | 19 | P67809 | Y-box-binding protein 1 OS=Homo sapie | YBX1 | 35.925 | 2 | 50.39698 | yes | 9.809E-08 | 848.43665 | 42197.547 |
| FPPYYMR | 7 | P67809 | Y-box-binding protein 1 OS=Homo sapie | YBX1 | 35.925 | 2 | 50.39698 | yes | 2.365E-07 | 487.23364 | 3215.6738 |
| FPPYYMR | 7 | P67809 | Y-box-binding protein 1 OS=Homo sapie | YBX1 | 35.925 | 2 | 50.39698 | yes | 4.528E-07 | 495.23111 | 3215.6738 |
| NEGSESAPEGQAQ | 15 | P67809 | Y-box-binding protein 1 OS=Homo sapie | YBX1 | 35.925 | 2 | 50.39698 | yes | 1.011E-07 | 794.35333 | 2389.168 |
| RPQYSNPPVQGEV | 30 | P67809 | Y-box-binding protein 1 OS=Homo sapie | YBX1 | 35.925 | 4 | 50.39698 | yes | 2.082E-07 | 806.63788 | 1307.7218 |
| RFPPYYMR | 8 | P67809 | Y-box-binding protein 1 OS=Homo sapie | YBX1 | 35.925 | 2 | 50.39698 | yes | 2.639E-06 | 565.28424 | 997.61835 |
| PGTTGSGAGSGGP | 26 | P67809 | Y-box-binding protein 1 OS=Homo sapie | YBX1 | 35.925 | 3 | 50.39698 | yes | 1.542E-07 | 695.66626 | 159.24428 |
| RPQYSNPPVQGEV | 27 | P67809 | Y-box-binding protein 1 OS=Homo sapie | YBX1 | 35.925 | 4 | 50.39698 | yes | 5.917E-07 | 718.58234 | 27.480373 |
| PGTTGSGAGSGGP | 27 | P67809 | Y-box-binding protein 1 OS=Homo sapie | YBX1 | 35.925 | 3 | 50.39698 | yes | 5.907E-07 | 738.36462 | 12.002019 |
| EIPIVHR | 7 | P67812 | Signal peptidase complex catalytic subuni | SEC11A | 20.626 | 2 | 39.945988 | yes | 4.294E-07 | 432.25851 | 70.412903 |
| FNLTGLNEQVPHY | 14 | P67870 | Casein kinase II subunit beta OS=Homo s | CSNK2B | 24.943 | 2 | 56.17845 | yes | 9.722E-08 | 844.43115 | 221.31941 |
| YQQGDFGYCPR | 11 | P67870 | Casein kinase II subunit beta OS=Homo s | CSNK2B | 24.943 | 2 | 56.17845 | yes | 9.722E-08 | 695.7959 | 207.06088 |
| AEGDVAALNR | 10 | P67936 | Tropomyosin alpha-4 chain OS=Homo sa | TPM4 | 28.522 | 2 | 51.616154 | yes | 5.963E-07 | 508.26199 | 390.17041 |
| IQALQQQADEAED | 14 | P67936 | Tropomyosin alpha-4 chain OS=Homo sa | TPM4 | 28.522 | 2 | 51.616154 | yes | 9.722E-08 | 807.88953 | 348.06653 |
| AGLNSLEAVK | 10 | P67936 | Tropomyosin alpha-4 chain OS=Homo sa | TPM4 | 28.522 | 2 | 51.616154 | yes | 1.062E-07 | 522.29022 | 153.76308 |
| EKAEGDVAALNR | 12 | P67936 | Tropomyosin alpha-4 chain OS=Homo sa | TPM4 | 28.522 | 2 | 51.616154 | yes | 1.367E-07 | 636.83075 | 104.70506 |
| IYHPNIDEK | 9 | P68036 | Ubiquitin-conjugating enzyme E2 L3 OS= | UBE2L3 | 17.862 | 2 | 54.948616 | no | 9.722E-08 | 564.78784 | 131.35188 |
| ADLAEEYSK | 9 | P68036 | Ubiquitin-conjugating enzyme E2 L3 OS= | UBE2L3 | 17.862 | 2 | 54.948616 | yes | 1.001E-07 | 513.24292 | 77.893692 |
| GQVCLPVISAENW | 14 | P68036 | Ubiquitin-conjugating enzyme E2 L3 OS= | UBE2L3 | 17.862 | 2 | 54.948616 | no | 5.345E-07 | 800.91125 | 56.92696 |
| LPLQDVYK | 8 | P68104 | Elongation factor 1-alpha 1 OS=Homo sa | EEF1A1 | 50.141 | 2 | 50.932682 | no | 2.293E-07 | 488.27911 | 25895.877 |
| IGGIGTVPVGR | 11 | P68104 | Elongation factor 1-alpha 1 OS=Homo sa | EEF1A1 | 50.141 | 2 | 50.932682 | no | 1.397E-07 | 513.30872 | 18470.473 |

| | | | | | | | | | | | |
|---|---|---|---|---|---|---|---|---|---|---|---|
| STTTGHLIYK | 10 | P68104 | Elongation factor 1-alpha 1 OS=Homo sa | EEF1A1 | 50.141 | 2 | 50.932682 | no | 3.999E-07 | 560.80347 | 12461.793 |
| QTVAVGVIK | 9 | P68104 | Elongation factor 1-alpha 1 OS=Homo sa | EEF1A1 | 50.141 | 2 | 50.932682 | no | 7.823E-07 | 457.78711 | 9176.5566 |
| VETGVLKPGMVV | 24 | P68104 | Elongation factor 1-alpha 1 OS=Homo sa | EEF1A1 | 50.141 | 3 | 50.932682 | yes | 1.159E-07 | 839.13287 | 3791.708 |
| VETGVLKPGMVV | 24 | P68104 | Elongation factor 1-alpha 1 OS=Homo sa | EEF1A1 | 50.141 | 3 | 50.932682 | yes | 9.723E-08 | 844.46448 | 3791.708 |
| THINIVVIGHVDSC | 15 | P68104 | Elongation factor 1-alpha 1 OS=Homo sa | EEF1A1 | 50.141 | 2 | 50.932682 | no | 1.808E-07 | 794.94391 | 2701.1401 |
| THINIVVIGHVDSC | 15 | P68104 | Elongation factor 1-alpha 1 OS=Homo sa | EEF1A1 | 50.141 | 2 | 50.932682 | no | 3.429E-06 | 530.2984 | 2701.1401 |
| THINIVVIGHVDSC | 15 | P68104 | Elongation factor 1-alpha 1 OS=Homo sa | EEF1A1 | 50.141 | 3 | 50.932682 | no | 1.808E-07 | 794.94391 | 2701.1401 |
| THINIVVIGHVDSC | 15 | P68104 | Elongation factor 1-alpha 1 OS=Homo sa | EEF1A1 | 50.141 | 3 | 50.932682 | no | 3.429E-06 | 530.2984 | 2701.1401 |
| RYEEIVK | 7 | P68104 | Elongation factor 1-alpha 1 OS=Homo sa | EEF1A1 | 50.141 | 2 | 50.932682 | yes | 3.266E-07 | 468.76108 | 4277.1074 |
| QLIVGVNK | 8 | P68104 | Elongation factor 1-alpha 1 OS=Homo sa | EEF1A1 | 50.141 | 2 | 50.932682 | no | 2.718E-07 | 435.77402 | 3859.3113 |
| SGDAAIVDMVPGK | 13 | P68104 | Elongation factor 1-alpha 1 OS=Homo sa | EEF1A1 | 50.141 | 2 | 50.932682 | yes | 1.101E-07 | 630.31866 | 1660.9528 |
| SGDAAIVDMVPGK | 13 | P68104 | Elongation factor 1-alpha 1 OS=Homo sa | EEF1A1 | 50.141 | 2 | 50.932682 | yes | 6.647E-06 | 638.3161 | 1660.9528 |
| EHALLAYTLGVK | 12 | P68104 | Elongation factor 1-alpha 1 OS=Homo sa | EEF1A1 | 50.141 | 2 | 50.932682 | no | 9.722E-08 | 657.87445 | 2121.1348 |
| YYVTIIDAPGHR | 12 | P68104 | Elongation factor 1-alpha 1 OS=Homo sa | EEF1A1 | 50.141 | 2 | 50.932682 | yes | 1.001E-07 | 702.86713 | 671.58643 |
| YYVTIIDAPGHR | 12 | P68104 | Elongation factor 1-alpha 1 OS=Homo sa | EEF1A1 | 50.141 | 2 | 50.932682 | yes | 1.301E-07 | 468.91385 | 671.58643 |
| YYVTIIDAPGHR | 12 | P68104 | Elongation factor 1-alpha 1 OS=Homo sa | EEF1A1 | 50.141 | 3 | 50.932682 | yes | 1.001E-07 | 702.86713 | 671.58643 |
| YYVTIIDAPGHR | 12 | P68104 | Elongation factor 1-alpha 1 OS=Homo sa | EEF1A1 | 50.141 | 3 | 50.932682 | yes | 1.301E-07 | 468.91385 | 671.58643 |
| MDSTEPPYSQK | 11 | P68104 | Elongation factor 1-alpha 1 OS=Homo sa | EEF1A1 | 50.141 | 2 | 50.932682 | yes | 1.362E-07 | 641.78461 | 624.77399 |
| EVSTYIK | 7 | P68104 | Elongation factor 1-alpha 1 OS=Homo sa | EEF1A1 | 50.141 | 2 | 50.932682 | yes | 1.091E-07 | 420.2291 | 54.818272 |
| PMCVESFSDYPPL | 15 | P68104 | Elongation factor 1-alpha 1 OS=Homo sa | EEF1A1 | 50.141 | 3 | 50.932682 | yes | 0.00013 | 585.60071 | 19.635147 |
| EIIDLVLDR | 9 | P68363 | Tubulin alpha-1B chain OS=Homo sapier | TUBA1B | 50.152 | 2 | 51.410347 | no | 1.613E-07 | 543.31366 | 12921.14 |
| DVNAAIATIK | 10 | P68363 | Tubulin alpha-1B chain OS=Homo sapier | TUBA1B | 50.152 | 2 | 51.410347 | no | 1.364E-07 | 508.29276 | 6729.2021 |
| YMACCLLYR | 9 | P68363 | Tubulin alpha-1B chain OS=Homo sapier | TUBA1B | 50.152 | 2 | 51.410347 | no | 1.263E-07 | 625.27997 | 3270.6899 |
| YMACCLLYR | 9 | P68363 | Tubulin alpha-1B chain OS=Homo sapier | TUBA1B | 50.152 | 2 | 51.410347 | no | 1.022E-07 | 633.2774 | 3270.6899 |
| QLFHPEQLITGK | 12 | P68363 | Tubulin alpha-1B chain OS=Homo sapier | TUBA1B | 50.152 | 2 | 51.410347 | no | 1.051E-07 | 705.89063 | 2002.4095 |
| QLFHPEQLITGK | 12 | P68363 | Tubulin alpha-1B chain OS=Homo sapier | TUBA1B | 50.152 | 2 | 51.410347 | no | 4.249E-07 | 470.9295 | 2002.4095 |
| QLFHPEQLITGK | 12 | P68363 | Tubulin alpha-1B chain OS=Homo sapier | TUBA1B | 50.152 | 3 | 51.410347 | no | 1.051E-07 | 705.89063 | 2002.4095 |
| QLFHPEQLITGK | 12 | P68363 | Tubulin alpha-1B chain OS=Homo sapier | TUBA1B | 50.152 | 3 | 51.410347 | no | 4.249E-07 | 470.9295 | 2002.4095 |
| SIQFVDWCPTGFK | 13 | P68363 | Tubulin alpha-1B chain OS=Homo sapier | TUBA1B | 50.152 | 2 | 51.410347 | no | 1.527E-07 | 792.87939 | 3950.4504 |
| IHFPLATYAPVISA | 16 | P68363 | Tubulin alpha-1B chain OS=Homo sapier | TUBA1B | 50.152 | 3 | 51.410347 | no | 8.142E-06 | 586.32593 | 3392.4617 |
| EDAANNYAR | 9 | P68363 | Tubulin alpha-1B chain OS=Homo sapier | TUBA1B | 50.152 | 2 | 51.410347 | no | 3.343E-07 | 512.22815 | 2900.1929 |
| AFVHWYVGEGME | 20 | P68363 | Tubulin alpha-1B chain OS=Homo sapier | TUBA1B | 50.152 | 3 | 51.410347 | no | 1.018E-07 | 777.3443 | 1082.4548 |
| PTYTNLNR | 8 | P68363 | Tubulin alpha-1B chain OS=Homo sapier | TUBA1B | 50.152 | 2 | 51.410347 | no | 2.081E-07 | 489.75378 | 1044.0526 |
| FDLMYAK | 7 | P68363 | Tubulin alpha-1B chain OS=Homo sapier | TUBA1B | 50.152 | 2 | 51.410347 | no | 2.407E-07 | 444.22021 | 263.41541 |
| FDLMYAK | 7 | P68363 | Tubulin alpha-1B chain OS=Homo sapier | TUBA1B | 50.152 | 2 | 51.410347 | no | 9.722E-08 | 452.21768 | 263.41541 |
| EDMAALEK | 8 | P68363 | Tubulin alpha-1B chain OS=Homo sapier | TUBA1B | 50.152 | 2 | 51.410347 | no | 1.437E-07 | 453.71548 | 256.31812 |
| AVCMLSNTTAIAE | 17 | P68363 | Tubulin alpha-1B chain OS=Homo sapier | TUBA1B | 50.152 | 3 | 51.410347 | no | 9.722E-08 | 622.30634 | 107.00553 |
| YMACCLLYRGDV | 15 | P68363 | Tubulin alpha-1B chain OS=Homo sapier | TUBA1B | 50.152 | 3 | 51.410347 | no | 1.138E-06 | 615.63336 | 44.038677 |
| HGKYMACCLLYR | 12 | P68363 | Tubulin alpha-1B chain OS=Homo sapier | TUBA1B | 50.152 | 3 | 51.410347 | no | 6.184E-07 | 524.58087 | 34.186989 |

| | | | | | | | | | | | |
|---|---|---|---|---|---|---|---|---|---|---|---|
| EIIDPVLDR | 9 | P68366 | Tubulin alpha-4A chain OS=Homo sapier | TUBA4A | 49.925 | 2 | 51.342426 | yes | 1.07E-07 | 535.29803 | 1425.5645 |
| DVNAAIAAIK | 10 | P68366 | Tubulin alpha-4A chain OS=Homo sapier | TUBA4A | 49.925 | 2 | 51.342426 | yes | 9.722E-08 | 493.28748 | 1135.6995 |
| LAVNMVPFPR | 10 | P68371 | Tubulin beta-4B chain OS=Homo sapiens | TUBB4B | 49.831 | 2 | 53.748657 | no | 1.253E-07 | 572.3208 | 17437.943 |
| LAVNMVPFPR | 10 | P68371 | Tubulin beta-4B chain OS=Homo sapiens | TUBB4B | 49.831 | 2 | 53.748657 | no | 9.722E-08 | 580.31824 | 17437.943 |
| AVLVDLEPGTMDS | 15 | P68371 | Tubulin beta-4B chain OS=Homo sapiens | TUBB4B | 49.831 | 2 | 53.748657 | no | 2.627E-07 | 801.41382 | 16642.361 |
| AVLVDLEPGTMDS | 15 | P68371 | Tubulin beta-4B chain OS=Homo sapiens | TUBB4B | 49.831 | 2 | 53.748657 | no | 1.433E-07 | 809.41125 | 16642.361 |
| EVDEQMLNVQNK | 12 | P68371 | Tubulin beta-4B chain OS=Homo sapiens | TUBB4B | 49.831 | 2 | 53.748657 | no | 1.138E-07 | 723.84833 | 12863.833 |
| EVDEQMLNVQNK | 12 | P68371 | Tubulin beta-4B chain OS=Homo sapiens | TUBB4B | 49.831 | 2 | 53.748657 | no | 9.722E-08 | 731.84576 | 12863.833 |
| NSSYFVEWIPNNV | 14 | P68371 | Tubulin beta-4B chain OS=Homo sapiens | TUBB4B | 49.831 | 2 | 53.748657 | no | 1.346E-07 | 848.9201 | 17520.912 |
| NMMAACDPR | 9 | P68371 | Tubulin beta-4B chain OS=Homo sapiens | TUBB4B | 49.831 | 2 | 53.748657 | no | 3.539E-07 | 533.21735 | 5838.7007 |
| NMMAACDPR | 9 | P68371 | Tubulin beta-4B chain OS=Homo sapiens | TUBB4B | 49.831 | 2 | 53.748657 | no | 2.105E-07 | 541.21478 | 5838.7007 |
| NMMAACDPR | 9 | P68371 | Tubulin beta-4B chain OS=Homo sapiens | TUBB4B | 49.831 | 2 | 53.748657 | no | 2.031E-07 | 541.21478 | 5838.7007 |
| FPGQLNADLR | 10 | P68371 | Tubulin beta-4B chain OS=Homo sapiens | TUBB4B | 49.831 | 2 | 53.748657 | no | 1.849E-06 | 565.80127 | 16021.587 |
| LHFFMPGFAPLTSI | 14 | P68371 | Tubulin beta-4B chain OS=Homo sapiens | TUBB4B | 49.831 | 2 | 53.748657 | no | 1.931E-07 | 810.92139 | 4589.749 |
| LHFFMPGFAPLTSI | 14 | P68371 | Tubulin beta-4B chain OS=Homo sapiens | TUBB4B | 49.831 | 2 | 53.748657 | no | 1.084E-07 | 818.91888 | 4589.749 |
| LHFFMPGFAPLTSI | 14 | P68371 | Tubulin beta-4B chain OS=Homo sapiens | TUBB4B | 49.831 | 2 | 53.748657 | no | 2.363E-07 | 546.28168 | 4589.749 |
| LHFFMPGFAPLTSI | 14 | P68371 | Tubulin beta-4B chain OS=Homo sapiens | TUBB4B | 49.831 | 2 | 53.748657 | no | 1.076E-06 | 540.95001 | 4589.749 |
| LHFFMPGFAPLTSI | 14 | P68371 | Tubulin beta-4B chain OS=Homo sapiens | TUBB4B | 49.831 | 3 | 53.748657 | no | 1.931E-07 | 810.92139 | 4589.749 |
| LHFFMPGFAPLTSI | 14 | P68371 | Tubulin beta-4B chain OS=Homo sapiens | TUBB4B | 49.831 | 3 | 53.748657 | no | 1.084E-07 | 818.91888 | 4589.749 |
| LHFFMPGFAPLTSI | 14 | P68371 | Tubulin beta-4B chain OS=Homo sapiens | TUBB4B | 49.831 | 3 | 53.748657 | no | 2.363E-07 | 546.28168 | 4589.749 |
| LHFFMPGFAPLTSI | 14 | P68371 | Tubulin beta-4B chain OS=Homo sapiens | TUBB4B | 49.831 | 3 | 53.748657 | no | 1.076E-06 | 540.95001 | 4589.749 |
| YLTVAAVFR | 9 | P68371 | Tubulin beta-4B chain OS=Homo sapiens | TUBB4B | 49.831 | 2 | 53.748657 | no | 1.897E-07 | 520.30035 | 13588.66 |
| INVYYNEATGGK | 12 | P68371 | Tubulin beta-4B chain OS=Homo sapiens | TUBB4B | 49.831 | 2 | 53.748657 | yes | 1.331E-07 | 664.8277 | 9995.8398 |
| IMNTFSVVPSPK | 12 | P68371 | Tubulin beta-4B chain OS=Homo sapiens | TUBB4B | 49.831 | 2 | 53.748657 | no | 9.886E-08 | 660.35504 | 4013.5059 |
| IMNTFSVVPSPK | 12 | P68371 | Tubulin beta-4B chain OS=Homo sapiens | TUBB4B | 49.831 | 2 | 53.748657 | no | 3.155E-07 | 668.35248 | 4013.5059 |
| ALTVPELTQQMFD | 15 | P68371 | Tubulin beta-4B chain OS=Homo sapiens | TUBB4B | 49.831 | 2 | 53.748657 | no | 4.062E-07 | 846.43726 | 7853.3833 |
| IREEYPDR | 8 | P68371 | Tubulin beta-4B chain OS=Homo sapiens | TUBB4B | 49.831 | 2 | 53.748657 | no | 1.554E-07 | 539.26984 | 6703.7573 |
| GHYTEGAELVDSV | 18 | P68371 | Tubulin beta-4B chain OS=Homo sapiens | TUBB4B | 49.831 | 3 | 53.748657 | no | 1.018E-07 | 653.66547 | 4296.833 |
| ISEQFTAMFR | 10 | P68371 | Tubulin beta-4B chain OS=Homo sapiens | TUBB4B | 49.831 | 2 | 53.748657 | no | 1.021E-07 | 623.30023 | 916.3233 |
| ISEQFTAMFR | 10 | P68371 | Tubulin beta-4B chain OS=Homo sapiens | TUBB4B | 49.831 | 2 | 53.748657 | no | 1.411E-06 | 615.3028 | 916.3233 |
| KLAVNMVPFPR | 11 | P68371 | Tubulin beta-4B chain OS=Homo sapiens | TUBB4B | 49.831 | 2 | 53.748657 | no | 1.214E-07 | 636.36829 | 1168.6129 |
| FWEVISDEHGIDPT | 27 | P68371 | Tubulin beta-4B chain OS=Homo sapiens | TUBB4B | 49.831 | 4 | 53.748657 | no | 1.973E-07 | 779.86127 | 1130.1747 |
| TAVCDIPPR | 9 | P68371 | Tubulin beta-4B chain OS=Homo sapiens | TUBB4B | 49.831 | 2 | 53.748657 | no | 3.74E-06 | 514.76331 | 776.14539 |
| SGPFGQIFR | 9 | P68371 | Tubulin beta-4B chain OS=Homo sapiens | TUBB4B | 49.831 | 2 | 53.748657 | no | 2.713E-06 | 504.76672 | 137.52295 |
| GGPNIITLADIVK | 13 | P68400 | Casein kinase II subunit alpha OS=Homo | CSNK2A1 | 45.144 | 2 | 54.815975 | no | 9.722E-08 | 655.88757 | 108.25644 |
| VLGTEDLYDYIDK | 13 | P68400 | Casein kinase II subunit alpha OS=Homo | CSNK2A1 | 45.144 | 2 | 54.815975 | yes | 5.839E-07 | 772.37994 | 59.100128 |
| FNDILGR | 7 | P68400 | Casein kinase II subunit alpha OS=Homo | CSNK2A1 | 45.144 | 2 | 54.815975 | no | 4.221E-07 | 417.72705 | 58.919338 |
| FYMYEILK | 8 | P68400 | Casein kinase II subunit alpha OS=Homo | CSNK2A1 | 45.144 | 2 | 54.815975 | no | 2.957E-06 | 553.7832 | 16.583065 |
| EIAQDFK | 7 | P68431 | Histone H3.1 OS=Homo sapiens OX=96( | H3C1 | 15.405 | 2 | 44.468357 | yes | 2.182E-07 | 425.7189 | 256.10181 |

| | | | | | | | | | | |
|---|---|---|---|---|---|---|---|---|---|---|
| MFLSFPTTK | 9 | P69905 | Hemoglobin subunit alpha OS=Homo sap | HBA1 | 15.258 | 2 | 52.647182 | yes | 1.281E-07 | 536.28082 | 1180.8718 |
| MFLSFPTTK | 9 | P69905 | Hemoglobin subunit alpha OS=Homo sap | HBA1 | 15.258 | 2 | 52.647182 | yes | 1.467E-07 | 544.27826 | 1180.8718 |
| FLASVSTVLTSK | 12 | P69905 | Hemoglobin subunit alpha OS=Homo sap | HBA1 | 15.258 | 2 | 52.647182 | yes | 9.722E-08 | 626.86102 | 744.62109 |
| TYFPHFDLSHGSA( | 16 | P69905 | Hemoglobin subunit alpha OS=Homo sap | HBA1 | 15.258 | 3 | 52.647182 | yes | 1.882E-07 | 611.96881 | 348.30682 |
| VGAHAGEYGAEA | 15 | P69905 | Hemoglobin subunit alpha OS=Homo sap | HBA1 | 15.258 | 2 | 52.647182 | yes | 3.308E-06 | 765.37079 | 32.943214 |
| EIQTLLSHK | 9 | P78316 | Nucleolar protein 14 OS=Homo sapiens ( | NOP14 | 97.669 | 2 | 54.487827 | yes | 1.043E-07 | 534.80603 | 1164.9922 |
| NSELLVVSAR | 10 | P78316 | Nucleolar protein 14 OS=Homo sapiens ( | NOP14 | 97.669 | 2 | 54.487827 | yes | 9.927E-08 | 544.30896 | 1071.0306 |
| EDVATWQQSSLSL | 14 | P78316 | Nucleolar protein 14 OS=Homo sapiens ( | NOP14 | 97.669 | 2 | 54.487827 | yes | 1.073E-07 | 810.40485 | 938.02203 |
| HDVGLPGVSR | 10 | P78316 | Nucleolar protein 14 OS=Homo sapiens ( | NOP14 | 97.669 | 2 | 54.487827 | yes | 1.179E-07 | 518.78033 | 846.22125 |
| APTSTEANHIR | 11 | P78316 | Nucleolar protein 14 OS=Homo sapiens ( | NOP14 | 97.669 | 2 | 54.487827 | yes | 9.722E-08 | 598.80457 | 812.06299 |
| GTLSAELTAAHFG | 21 | P78316 | Nucleolar protein 14 OS=Homo sapiens ( | NOP14 | 97.669 | 3 | 54.487827 | yes | 2.638E-07 | 665.35333 | 665.79248 |
| FALEQQR | 7 | P78316 | Nucleolar protein 14 OS=Homo sapiens ( | NOP14 | 97.669 | 2 | 54.487827 | yes | 3.783E-07 | 446.23779 | 649.84467 |
| ANSNPFEVK | 9 | P78316 | Nucleolar protein 14 OS=Homo sapiens ( | NOP14 | 97.669 | 2 | 54.487827 | yes | 9.723E-08 | 503.25363 | 536.54785 |
| MQLSEIMER | 9 | P78316 | Nucleolar protein 14 OS=Homo sapiens ( | NOP14 | 97.669 | 2 | 54.487827 | yes | 1.899E-07 | 568.77557 | 456.60062 |
| EDALELTEK | 9 | P78316 | Nucleolar protein 14 OS=Homo sapiens ( | NOP14 | 97.669 | 2 | 54.487827 | yes | 1.671E-07 | 524.26385 | 413.43893 |
| KDNQFLAR | 8 | P78316 | Nucleolar protein 14 OS=Homo sapiens ( | NOP14 | 97.669 | 2 | 54.487827 | yes | 9.727E-08 | 496.26962 | 360.91077 |
| QLFNSLATQEGEW | 14 | P78316 | Nucleolar protein 14 OS=Homo sapiens ( | NOP14 | 97.669 | 2 | 54.487827 | yes | 1.026E-07 | 825.90973 | 347.6073 |
| HMSADDLNDGFV | 18 | P78316 | Nucleolar protein 14 OS=Homo sapiens ( | NOP14 | 97.669 | 3 | 54.487827 | yes | 2.945E-07 | 688.30933 | 346.21051 |
| ELIEELIAK | 9 | P78316 | Nucleolar protein 14 OS=Homo sapiens ( | NOP14 | 97.669 | 2 | 54.487827 | yes | 2.068E-06 | 529.31061 | 321.26752 |
| KELIEELIAK | 10 | P78316 | Nucleolar protein 14 OS=Homo sapiens ( | NOP14 | 97.669 | 2 | 54.487827 | yes | 4.609E-07 | 593.35809 | 299.09515 |
| SMEEQLLVVER | 11 | P78316 | Nucleolar protein 14 OS=Homo sapiens ( | NOP14 | 97.669 | 2 | 54.487827 | yes | 9.79E-08 | 666.84503 | 258.08652 |
| FGEYNSNMSPEEK | 13 | P78316 | Nucleolar protein 14 OS=Homo sapiens ( | NOP14 | 97.669 | 2 | 54.487827 | yes | 2.103E-07 | 766.32214 | 227.24835 |
| DAMHEMEEMIETI | 13 | P78316 | Nucleolar protein 14 OS=Homo sapiens ( | NOP14 | 97.669 | 2 | 54.487827 | yes | 1.258E-07 | 797.33331 | 206.43938 |
| CPILSLQDVVK | 11 | P78316 | Nucleolar protein 14 OS=Homo sapiens ( | NOP14 | 97.669 | 2 | 54.487827 | yes | 9.722E-08 | 636.35504 | 155.66006 |
| MLGKDEDENVK | 11 | P78316 | Nucleolar protein 14 OS=Homo sapiens ( | NOP14 | 97.669 | 2 | 54.487827 | yes | 9.722E-08 | 639.30573 | 111.26519 |
| ASQGSTLVHPFR | 12 | P78316 | Nucleolar protein 14 OS=Homo sapiens ( | NOP14 | 97.669 | 2 | 54.487827 | yes | 9.722E-08 | 650.34387 | 108.88036 |
| MKTEAELAK | 9 | P78316 | Nucleolar protein 14 OS=Homo sapiens ( | NOP14 | 97.669 | 2 | 54.487827 | yes | 3.171E-07 | 510.77335 | 84.268791 |
| CNHPSLAEGNK | 11 | P78316 | Nucleolar protein 14 OS=Homo sapiens ( | NOP14 | 97.669 | 2 | 54.487827 | yes | 4.097E-07 | 613.78278 | 81.968658 |
| ELGFEMK | 7 | P78316 | Nucleolar protein 14 OS=Homo sapiens ( | NOP14 | 97.669 | 2 | 54.487827 | yes | 1.061E-07 | 427.20984 | 63.427181 |
| QLCRPLTCEK | 10 | P78316 | Nucleolar protein 14 OS=Homo sapiens ( | NOP14 | 97.669 | 2 | 54.487827 | yes | 9.722E-08 | 652.82611 | 57.300079 |
| MNVEEDVQEEQSI | 13 | P78316 | Nucleolar protein 14 OS=Homo sapiens ( | NOP14 | 97.669 | 2 | 54.487827 | yes | 2.639E-07 | 782.84338 | 31.201534 |
| HMSADDLNDGFV | 15 | P78316 | Nucleolar protein 14 OS=Homo sapiens ( | NOP14 | 97.669 | 2 | 54.487827 | yes | 2.035E-07 | 838.88287 | 27.812094 |
| DVYVTTHTPR | 10 | P78317 | E3 ubiquitin-protein ligase RNF4 OS=Ho | RNF4 | 21.319 | 2 | 47.10804 | yes | 1.399E-07 | 594.80402 | 376.58795 |
| NANTCPTCR | 9 | P78317 | E3 ubiquitin-protein ligase RNF4 OS=Ho | RNF4 | 21.319 | 2 | 47.10804 | yes | 5.842E-07 | 547.22931 | 66.222794 |
| YGALPDQGIAK | 11 | P78318 | Immunoglobulin-binding protein 1 OS=H | IGBP1 | 39.222 | 2 | 50.058556 | yes | 9.919E-08 | 566.80347 | 140.44913 |
| NEDLEEIASTDLK | 13 | P78318 | Immunoglobulin-binding protein 1 OS=H | IGBP1 | 39.222 | 2 | 50.058556 | yes | 4.668E-07 | 738.85687 | 123.67789 |
| LPELFETGR | 9 | P78318 | Immunoglobulin-binding protein 1 OS=H | IGBP1 | 39.222 | 2 | 50.058556 | yes | 1.491E-06 | 531.28491 | 113.14381 |
| GEFEHSETR | 9 | P78332 | RNA-binding protein 6 OS=Homo sapien | RBM6 | 128.644 | 2 | 53.906406 | yes | 1.006E-07 | 546.24127 | 8353.6953 |
| GSGTTDLDFR | 10 | P78332 | RNA-binding protein 6 OS=Homo sapien | RBM6 | 128.644 | 2 | 53.906406 | yes | 1.547E-07 | 534.75146 | 7734.1118 |

| | | | | | | | | | | | |
|---|---|---|---|---|---|---|---|---|---|---|---|
| EEILNAFR | 8 | P78332 | RNA-binding protein 6 OS=Homo sapien | RBM6 | 128.644 | 2 | 53.906406 | yes | 6.005E-07 | 496.26401 | 6412.9321 |
| LSGVPEDATK | 10 | P78332 | RNA-binding protein 6 OS=Homo sapien | RBM6 | 128.644 | 2 | 53.906406 | yes | 2.388E-07 | 508.76657 | 6210.3892 |
| QELITYPQPQK | 11 | P78332 | RNA-binding protein 6 OS=Homo sapien | RBM6 | 128.644 | 2 | 53.906406 | yes | 2.54E-07 | 672.86151 | 6207.6665 |
| TSIPAPLEK | 9 | P78332 | RNA-binding protein 6 OS=Homo sapien | RBM6 | 128.644 | 2 | 53.906406 | yes | 5.177E-07 | 478.27658 | 5805.2192 |
| EEQSSDAGLFK | 11 | P78332 | RNA-binding protein 6 OS=Homo sapien | RBM6 | 128.644 | 2 | 53.906406 | yes | 9.722E-08 | 605.78296 | 5245.4092 |
| LPGSQMFGYGQSK | 13 | P78332 | RNA-binding protein 6 OS=Homo sapien | RBM6 | 128.644 | 2 | 53.906406 | yes | 1.375E-07 | 700.33734 | 2581.2439 |
| LPGSQMFGYGQSK | 13 | P78332 | RNA-binding protein 6 OS=Homo sapien | RBM6 | 128.644 | 2 | 53.906406 | yes | 1.003E-07 | 708.33484 | 2581.2439 |
| MVAVNLATGK | 10 | P78332 | RNA-binding protein 6 OS=Homo sapien | RBM6 | 128.644 | 2 | 53.906406 | yes | 1.834E-07 | 502.28387 | 4612.2017 |
| GGEGPGHDFR | 10 | P78332 | RNA-binding protein 6 OS=Homo sapien | RBM6 | 128.644 | 2 | 53.906406 | yes | 1.612E-07 | 514.73083 | 4268.0449 |
| EAPHMNYR | 8 | P78332 | RNA-binding protein 6 OS=Homo sapien | RBM6 | 128.644 | 2 | 53.906406 | yes | 9.919E-08 | 509.23218 | 4197.9297 |
| QSEQELAYLER | 11 | P78332 | RNA-binding protein 6 OS=Homo sapien | RBM6 | 128.644 | 2 | 53.906406 | yes | 1.062E-07 | 683.33588 | 3581.4976 |
| APAEDVFK | 8 | P78332 | RNA-binding protein 6 OS=Homo sapien | RBM6 | 128.644 | 2 | 53.906406 | yes | 1.233E-07 | 438.72672 | 3466.4695 |
| DKDGTQVDFR | 10 | P78332 | RNA-binding protein 6 OS=Homo sapien | RBM6 | 128.644 | 2 | 53.906406 | yes | 4.648E-07 | 590.78326 | 3130.3235 |
| ILQNLDPPFSIDGK | 14 | P78332 | RNA-binding protein 6 OS=Homo sapien | RBM6 | 128.644 | 2 | 53.906406 | yes | 5.357E-07 | 778.91956 | 3007.9238 |
| DAHAVDFR | 8 | P78332 | RNA-binding protein 6 OS=Homo sapien | RBM6 | 128.644 | 2 | 53.906406 | yes | 1.202E-07 | 465.72504 | 2958.1814 |
| HQQLSDLHK | 9 | P78332 | RNA-binding protein 6 OS=Homo sapien | RBM6 | 128.644 | 2 | 53.906406 | no | 1.847E-07 | 553.29108 | 2677.2285 |
| GTYDLDFR | 8 | P78332 | RNA-binding protein 6 OS=Homo sapien | RBM6 | 128.644 | 2 | 53.906406 | yes | 1.02E-07 | 493.73254 | 2635.8748 |
| EMGSCMEFK | 9 | P78332 | RNA-binding protein 6 OS=Homo sapien | RBM6 | 128.644 | 2 | 53.906406 | yes | 9.722E-08 | 559.71942 | 1311.4489 |
| EMGSCMEFK | 9 | P78332 | RNA-binding protein 6 OS=Homo sapien | RBM6 | 128.644 | 2 | 53.906406 | yes | 9.722E-08 | 567.71686 | 1311.4489 |
| GGCVQQATGWR | 11 | P78332 | RNA-binding protein 6 OS=Homo sapien | RBM6 | 128.644 | 2 | 53.906406 | yes | 9.933E-08 | 610.28546 | 2569.2212 |
| TGPMGHTYGFIDL | 21 | P78332 | RNA-binding protein 6 OS=Homo sapien | RBM6 | 128.644 | 3 | 53.906406 | yes | 9.935E-08 | 763.36365 | 1278.9081 |
| TGPMGHTYGFIDL | 21 | P78332 | RNA-binding protein 6 OS=Homo sapien | RBM6 | 128.644 | 3 | 53.906406 | yes | 1.274E-07 | 572.7746 | 1278.9081 |
| TGPMGHTYGFIDL | 21 | P78332 | RNA-binding protein 6 OS=Homo sapien | RBM6 | 128.644 | 4 | 53.906406 | yes | 9.935E-08 | 763.36365 | 1278.9081 |
| TGPMGHTYGFIDL | 21 | P78332 | RNA-binding protein 6 OS=Homo sapien | RBM6 | 128.644 | 4 | 53.906406 | yes | 1.274E-07 | 572.7746 | 1278.9081 |
| EEGGLDFLGR | 10 | P78332 | RNA-binding protein 6 OS=Homo sapien | RBM6 | 128.644 | 2 | 53.906406 | yes | 1.767E-07 | 546.76965 | 2325.8875 |
| DTPHSDFR | 8 | P78332 | RNA-binding protein 6 OS=Homo sapien | RBM6 | 128.644 | 2 | 53.906406 | yes | 1.066E-06 | 487.71997 | 2292.3193 |
| GGDFSSSDFQSR | 12 | P78332 | RNA-binding protein 6 OS=Homo sapien | RBM6 | 128.644 | 2 | 53.906406 | yes | 1.112E-07 | 645.27325 | 1624.6007 |
| DGSHADFR | 8 | P78332 | RNA-binding protein 6 OS=Homo sapien | RBM6 | 128.644 | 2 | 53.906406 | yes | 1.893E-07 | 452.69904 | 1484.2611 |
| DRDTPHSDFR | 10 | P78332 | RNA-binding protein 6 OS=Homo sapien | RBM6 | 128.644 | 3 | 53.906406 | yes | 9.722E-08 | 415.85843 | 677.5672 |
| DRDTPHSDFR | 10 | P78332 | RNA-binding protein 6 OS=Homo sapien | RBM6 | 128.644 | 3 | 53.906406 | yes | 6.084E-07 | 623.284 | 677.5672 |
| DRDTPHSDFR | 10 | P78332 | RNA-binding protein 6 OS=Homo sapien | RBM6 | 128.644 | 2 | 53.906406 | yes | 9.722E-08 | 415.85843 | 677.5672 |
| DRDTPHSDFR | 10 | P78332 | RNA-binding protein 6 OS=Homo sapien | RBM6 | 128.644 | 2 | 53.906406 | yes | 6.084E-07 | 623.284 | 677.5672 |
| GGDGTSMDYR | 10 | P78332 | RNA-binding protein 6 OS=Homo sapien | RBM6 | 128.644 | 2 | 53.906406 | yes | 1.682E-07 | 529.71399 | 622.35461 |
| GGDGTSMDYR | 10 | P78332 | RNA-binding protein 6 OS=Homo sapien | RBM6 | 128.644 | 2 | 53.906406 | yes | 9.722E-08 | 537.71149 | 622.35461 |
| HSGNFPGR | 8 | P78332 | RNA-binding protein 6 OS=Homo sapien | RBM6 | 128.644 | 2 | 53.906406 | yes | 3.108E-07 | 436.2121 | 1061.1752 |
| DIHSGDFR | 8 | P78332 | RNA-binding protein 6 OS=Homo sapien | RBM6 | 128.644 | 2 | 53.906406 | yes | 2.377E-07 | 473.7225 | 1057.0524 |
| LVDKEDIDTSSK | 12 | P78332 | RNA-binding protein 6 OS=Homo sapien | RBM6 | 128.644 | 2 | 53.906406 | yes | 9.722E-08 | 675.34338 | 502.89301 |
| LVDKEDIDTSSK | 12 | P78332 | RNA-binding protein 6 OS=Homo sapien | RBM6 | 128.644 | 2 | 53.906406 | yes | 9.722E-08 | 450.56467 | 502.89301 |
| LVDKEDIDTSSK | 12 | P78332 | RNA-binding protein 6 OS=Homo sapien | RBM6 | 128.644 | 3 | 53.906406 | yes | 9.722E-08 | 675.34338 | 502.89301 |

| | | | | | | | | | | | |
|---|---|---|---|---|---|---|---|---|---|---|---|
| LVDKEDIDTSSK | 12 | P78332 | RNA-binding protein 6 OS=Homo sapien | RBM6 | 128.644 | 3 | 53.906406 | yes | 9.722E-08 | 450.56467 | 502.89301 |
| DRDAHAVDFR | 10 | P78332 | RNA-binding protein 6 OS=Homo sapien | RBM6 | 128.644 | 3 | 53.906406 | yes | 9.722E-08 | 401.19513 | 342.62558 |
| DRDAHAVDFR | 10 | P78332 | RNA-binding protein 6 OS=Homo sapien | RBM6 | 128.644 | 3 | 53.906406 | yes | 8.767E-06 | 601.28906 | 342.62558 |
| DRDAHAVDFR | 10 | P78332 | RNA-binding protein 6 OS=Homo sapien | RBM6 | 128.644 | 2 | 53.906406 | yes | 9.722E-08 | 401.19513 | 342.62558 |
| DRDAHAVDFR | 10 | P78332 | RNA-binding protein 6 OS=Homo sapien | RBM6 | 128.644 | 2 | 53.906406 | yes | 8.767E-06 | 601.28906 | 342.62558 |
| FQENASEGK | 9 | P78332 | RNA-binding protein 6 OS=Homo sapien | RBM6 | 128.644 | 2 | 53.906406 | yes | 1.461E-07 | 505.23291 | 659.8526 |
| KPLPPTVK | 8 | P78332 | RNA-binding protein 6 OS=Homo sapien | RBM6 | 128.644 | 2 | 53.906406 | yes | 5.414E-07 | 440.28674 | 659.32568 |
| DLQDQDYR | 8 | P78332 | RNA-binding protein 6 OS=Homo sapien | RBM6 | 128.644 | 2 | 53.906406 | yes | 2.573E-07 | 526.73578 | 643.18213 |
| TPDGMPVK | 8 | P78332 | RNA-binding protein 6 OS=Homo sapien | RBM6 | 128.644 | 2 | 53.906406 | yes | 9.809E-08 | 422.7153 | 607.41949 |
| DYPPPPLK | 8 | P78332 | RNA-binding protein 6 OS=Homo sapien | RBM6 | 128.644 | 2 | 53.906406 | yes | 3.855E-07 | 463.75275 | 595.90857 |
| DLSDLDFR | 8 | P78332 | RNA-binding protein 6 OS=Homo sapien | RBM6 | 128.644 | 2 | 53.906406 | yes | 3.034E-07 | 490.73782 | 583.49237 |
| KGTGLGYGHPGL/ | 21 | P78332 | RNA-binding protein 6 OS=Homo sapien | RBM6 | 128.644 | 4 | 53.906406 | yes | 1.783E-07 | 519.00537 | 453.60693 |
| QPNQPLRPADKEP | 16 | P78332 | RNA-binding protein 6 OS=Homo sapien | RBM6 | 128.644 | 4 | 53.906406 | yes | 4.02E-07 | 468.7485 | 432.78442 |
| EESTHDHTIERPAF | 18 | P78332 | RNA-binding protein 6 OS=Homo sapien | RBM6 | 128.644 | 4 | 53.906406 | yes | 1.495E-07 | 524.51056 | 306.90097 |
| EGPPMDYR | 8 | P78332 | RNA-binding protein 6 OS=Homo sapien | RBM6 | 128.644 | 2 | 53.906406 | yes | 1.737E-07 | 482.71329 | 214.25826 |
| FAPGWNR | 7 | P78332 | RNA-binding protein 6 OS=Homo sapien | RBM6 | 128.644 | 2 | 53.906406 | yes | 3.594E-07 | 424.21411 | 159.28336 |
| NSDWSSDTNR | 10 | P78332 | RNA-binding protein 6 OS=Homo sapien | RBM6 | 128.644 | 2 | 53.906406 | yes | 3.582E-07 | 591.24451 | 118.01316 |
| LQSFDSPER | 9 | P78332 | RNA-binding protein 6 OS=Homo sapien | RBM6 | 128.644 | 2 | 53.906406 | yes | 1.259E-07 | 539.76184 | 98.851784 |
| KLVDKEDIDTSSK | 13 | P78332 | RNA-binding protein 6 OS=Homo sapien | RBM6 | 128.644 | 2 | 53.906406 | yes | 9.722E-08 | 739.39087 | 97.563217 |
| QPNQPLRPADK | 11 | P78332 | RNA-binding protein 6 OS=Homo sapien | RBM6 | 128.644 | 2 | 53.906406 | yes | 1.061E-07 | 632.34387 | 70.841232 |
| EKLQSFDSPER | 11 | P78332 | RNA-binding protein 6 OS=Homo sapien | RBM6 | 128.644 | 2 | 53.906406 | yes | 9.722E-08 | 668.33063 | 69.4198 |
| KENEEDKLTDWNI | 13 | P78332 | RNA-binding protein 6 OS=Homo sapien | RBM6 | 128.644 | 2 | 53.906406 | yes | 2.182E-07 | 824.89429 | 34.187481 |
| DSLPFDFQGHSGPI | 28 | P78332 | RNA-binding protein 6 OS=Homo sapien | RBM6 | 128.644 | 4 | 53.906406 | yes | 1.365E-06 | 774.60339 | 12.321661 |
| EEIAVWTNLTEPR | 13 | P78337 | Pituitary homeobox 1 OS=Homo sapiens | PITX1 | 34.129 | 2 | 49.723812 | yes | 1.341E-07 | 779.39899 | 1636.6748 |
| SLAPAPLSTK | 10 | P78337 | Pituitary homeobox 1 OS=Homo sapiens | PITX1 | 34.129 | 2 | 49.723812 | yes | 4.865E-07 | 492.78986 | 1610.9888 |
| NQQLDLCK | 8 | P78337 | Pituitary homeobox 1 OS=Homo sapiens | PITX1 | 34.129 | 2 | 49.723812 | yes | 3.296E-07 | 509.75293 | 531.00403 |
| DTCNSSLASLR | 11 | P78337 | Pituitary homeobox 1 OS=Homo sapiens | PITX1 | 34.129 | 2 | 49.723812 | no | 1.053E-07 | 612.2879 | 444.28452 |
| YPDMSMR | 7 | P78337 | Pituitary homeobox 1 OS=Homo sapiens | PITX1 | 34.129 | 2 | 49.723812 | yes | 1.007E-07 | 450.19113 | 182.22328 |
| LPEGLRPPPPPPHD | 23 | P78337 | Pituitary homeobox 1 OS=Homo sapiens | PITX1 | 34.129 | 4 | 49.723812 | yes | 2.102E-07 | 625.5824 | 88.586212 |
| LPEGLRPPPPPPHD | 23 | P78337 | Pituitary homeobox 1 OS=Homo sapiens | PITX1 | 34.129 | 4 | 49.723812 | yes | 5.246E-07 | 500.66739 | 88.586212 |
| LPEGLRPPPPPPHD | 23 | P78337 | Pituitary homeobox 1 OS=Homo sapiens | PITX1 | 34.129 | 5 | 49.723812 | yes | 2.102E-07 | 625.5824 | 88.586212 |
| LPEGLRPPPPPPHD | 23 | P78337 | Pituitary homeobox 1 OS=Homo sapiens | PITX1 | 34.129 | 5 | 49.723812 | yes | 5.246E-07 | 500.66739 | 88.586212 |
| NRYPDMSMR | 9 | P78337 | Pituitary homeobox 1 OS=Homo sapiens | PITX1 | 34.129 | 2 | 49.723812 | yes | 1.432E-06 | 585.26312 | 133.02209 |
| TQTPPLGQTPQLGI | 15 | P78344 | Eukaryotic translation initiation factor 4 g | EIF4G2 | 102.362 | 2 | 53.91595 | yes | 2.166E-07 | 789.94611 | 884.06793 |
| VIILSLDR | 8 | P78344 | Eukaryotic translation initiation factor 4 g | EIF4G2 | 102.362 | 2 | 53.91595 | yes | 1.313E-07 | 464.79492 | 740.13318 |
| FLLQDTVELR | 10 | P78344 | Eukaryotic translation initiation factor 4 g | EIF4G2 | 102.362 | 2 | 53.91595 | yes | 9.81E-08 | 617.34552 | 740.02338 |
| LQDEFENR | 8 | P78344 | Eukaryotic translation initiation factor 4 g | EIF4G2 | 102.362 | 2 | 53.91595 | yes | 5.191E-07 | 525.74615 | 563.24872 |
| ENPLLPEEEEQR | 12 | P78344 | Eukaryotic translation initiation factor 4 g | EIF4G2 | 102.362 | 2 | 53.91595 | yes | 2.035E-07 | 741.85718 | 508.10297 |
| YSSLYAQLCLR | 11 | P78344 | Eukaryotic translation initiation factor 4 g | EIF4G2 | 102.362 | 2 | 53.91595 | yes | 9.885E-08 | 687.34772 | 491.89328 |

| | | | | | | | | | | | |
|---|---|---|---|---|---|---|---|---|---|---|---|
| LDPSPQTIYK | 10 | P78344 | Eukaryotic translation initiation factor 4 g | EIF4G2 | 102.362 | 2 | 53.91595 | yes | 1.01E-07 | 581.31116 | 488.94299 |
| EWLTELFQQSK | 11 | P78344 | Eukaryotic translation initiation factor 4 g | EIF4G2 | 102.362 | 2 | 53.91595 | yes | 9.722E-08 | 704.85901 | 440.79669 |
| SYLAQFAAR | 9 | P78344 | Eukaryotic translation initiation factor 4 g | EIF4G2 | 102.362 | 2 | 53.91595 | yes | 9.722E-08 | 513.77197 | 429.27353 |
| EDITQEFPGK | 10 | P78344 | Eukaryotic translation initiation factor 4 g | EIF4G2 | 102.362 | 2 | 53.91595 | yes | 1.795E-07 | 582.28259 | 406.54919 |
| SLMDQYFAR | 9 | P78344 | Eukaryotic translation initiation factor 4 g | EIF4G2 | 102.362 | 2 | 53.91595 | yes | 9.809E-08 | 565.76862 | 406.34851 |
| SDFFLEGPFMPPR | 13 | P78344 | Eukaryotic translation initiation factor 4 g | EIF4G2 | 102.362 | 2 | 53.91595 | yes | 2.149E-07 | 770.36865 | 332.18372 |
| DLGVFIPAPMAQG | 15 | P78344 | Eukaryotic translation initiation factor 4 g | EIF4G2 | 102.362 | 2 | 53.91595 | yes | 1.318E-07 | 801.91016 | 272.91971 |
| LEVDIPLVK | 9 | P78344 | Eukaryotic translation initiation factor 4 g | EIF4G2 | 102.362 | 2 | 53.91595 | yes | 2.492E-07 | 513.31567 | 267.00537 |
| MLPEIDQNK | 9 | P78344 | Eukaryotic translation initiation factor 4 g | EIF4G2 | 102.362 | 2 | 53.91595 | yes | 1.165E-07 | 544.27625 | 261.2702 |
| TNPPLIQEKPAK | 12 | P78344 | Eukaryotic translation initiation factor 4 g | EIF4G2 | 102.362 | 2 | 53.91595 | yes | 6.039E-07 | 668.38519 | 249.99719 |
| NVDVYDKR | 8 | P78344 | Eukaryotic translation initiation factor 4 g | EIF4G2 | 102.362 | 2 | 53.91595 | yes | 1.809E-07 | 504.75909 | 186.42352 |
| MCSLMLSK | 8 | P78344 | Eukaryotic translation initiation factor 4 g | EIF4G2 | 102.362 | 2 | 53.91595 | yes | 2.087E-07 | 485.23193 | 148.62396 |
| GQLNADEISLR | 11 | P78344 | Eukaryotic translation initiation factor 4 g | EIF4G2 | 102.362 | 2 | 53.91595 | yes | 1.09E-07 | 608.32001 | 118.44421 |
| HFLPEMLSK | 9 | P78344 | Eukaryotic translation initiation factor 4 g | EIF4G2 | 102.362 | 2 | 53.91595 | yes | 3.222E-06 | 551.29169 | 117.76444 |
| LDLIHESILHK | 11 | P78344 | Eukaryotic translation initiation factor 4 g | EIF4G2 | 102.362 | 2 | 53.91595 | yes | 1.343E-07 | 659.37988 | 104.0113 |
| TNPPLIQEK | 9 | P78344 | Eukaryotic translation initiation factor 4 g | EIF4G2 | 102.362 | 2 | 53.91595 | yes | 3.889E-07 | 520.29279 | 74.100792 |
| LCLELLNVGVESK | 13 | P78344 | Eukaryotic translation initiation factor 4 g | EIF4G2 | 102.362 | 2 | 53.91595 | yes | 1.481E-07 | 737.40271 | 73.793091 |
| GVILLIVDK | 9 | P78344 | Eukaryotic translation initiation factor 4 g | EIF4G2 | 102.362 | 2 | 53.91595 | yes | 1.503E-06 | 485.3208 | 56.301788 |
| AFLDNGPK | 8 | P78344 | Eukaryotic translation initiation factor 4 g | EIF4G2 | 102.362 | 2 | 53.91595 | yes | 3.327E-07 | 431.2269 | 29.851677 |
| ASSLISLLK | 9 | P78344 | Eukaryotic translation initiation factor 4 g | EIF4G2 | 102.362 | 2 | 53.91595 | yes | 2.618E-06 | 466.29477 | 20.227345 |
| KGQLNADEISLR | 12 | P78344 | Eukaryotic translation initiation factor 4 g | EIF4G2 | 102.362 | 2 | 53.91595 | yes | 1.672E-06 | 672.36749 | 19.904833 |
| QLAIGVNEVTR | 11 | P78345 | Ribonuclease P protein subunit p38 OS=I | RPP38 | 31.835 | 2 | 55.53771 | yes | 1.059E-07 | 600.34076 | 728.35791 |
| NTTDFVDEVR | 10 | P78345 | Ribonuclease P protein subunit p38 OS=I | RPP38 | 31.835 | 2 | 55.53771 | yes | 9.722E-08 | 598.28314 | 535.78125 |
| TSLNNPYIIR | 10 | P78345 | Ribonuclease P protein subunit p38 OS=I | RPP38 | 31.835 | 2 | 55.53771 | yes | 1.128E-07 | 595.83002 | 473.46945 |
| VPSLSVPWLQDR | 12 | P78345 | Ribonuclease P protein subunit p38 OS=I | RPP38 | 31.835 | 2 | 55.53771 | yes | 9.722E-08 | 698.88281 | 466.53607 |
| QQVSGWTPAHVR | 12 | P78345 | Ribonuclease P protein subunit p38 OS=I | RPP38 | 31.835 | 2 | 55.53771 | yes | 9.723E-08 | 683.35474 | 228.22876 |
| QQVSGWTPAHVR | 12 | P78345 | Ribonuclease P protein subunit p38 OS=I | RPP38 | 31.835 | 2 | 55.53771 | yes | 9.722E-08 | 455.90558 | 228.22876 |
| QQVSGWTPAHVR | 12 | P78345 | Ribonuclease P protein subunit p38 OS=I | RPP38 | 31.835 | 3 | 55.53771 | yes | 9.723E-08 | 683.35474 | 228.22876 |
| QQVSGWTPAHVR | 12 | P78345 | Ribonuclease P protein subunit p38 OS=I | RPP38 | 31.835 | 3 | 55.53771 | yes | 9.722E-08 | 455.90558 | 228.22876 |
| SVPACQVPR | 9 | P78345 | Ribonuclease P protein subunit p38 OS=I | RPP38 | 31.835 | 2 | 55.53771 | yes | 9.728E-08 | 507.26349 | 298.97958 |
| QASVTLQPLK | 10 | P78345 | Ribonuclease P protein subunit p38 OS=I | RPP38 | 31.835 | 2 | 55.53771 | yes | 9.722E-08 | 542.82166 | 227.84488 |
| IAPVIGLK | 8 | P78345 | Ribonuclease P protein subunit p38 OS=I | RPP38 | 31.835 | 2 | 55.53771 | yes | 1.627E-07 | 405.776 | 226.97276 |
| CSIAVDISENLK | 12 | P78345 | Ribonuclease P protein subunit p38 OS=I | RPP38 | 31.835 | 2 | 55.53771 | yes | 9.722E-08 | 674.84247 | 166.81186 |
| SVKPAMITSHLIQI | 18 | P78345 | Ribonuclease P protein subunit p38 OS=I | RPP38 | 31.835 | 3 | 55.53771 | yes | 1.453E-07 | 661.04694 | 111.87996 |
| ELLLVLVCK | 9 | P78345 | Ribonuclease P protein subunit p38 OS=I | RPP38 | 31.835 | 2 | 55.53771 | yes | 3.63E-07 | 543.83319 | 77.237259 |
| KNTTDFVDEVR | 11 | P78345 | Ribonuclease P protein subunit p38 OS=I | RPP38 | 31.835 | 2 | 55.53771 | yes | 2.283E-07 | 662.33063 | 70.775146 |
| KLIPNPNK | 8 | P78345 | Ribonuclease P protein subunit p38 OS=I | RPP38 | 31.835 | 2 | 55.53771 | yes | 2.654E-07 | 462.28726 | 65.930855 |
| ELLDTSFEDLSK | 12 | P78345 | Ribonuclease P protein subunit p38 OS=I | RPP38 | 31.835 | 2 | 55.53771 | yes | 2.002E-06 | 698.84576 | 49.654343 |
| EKCSIAVDISENLK | 14 | P78345 | Ribonuclease P protein subunit p38 OS=I | RPP38 | 31.835 | 2 | 55.53771 | yes | 2.385E-07 | 803.41125 | 32.881519 |

| | | | | | | | | | | | |
|---|---|---|---|---|---|---|---|---|---|---|---|
| AALLHGETR | 9 | P78346 | Ribonuclease P protein subunit p30 OS=I | RPP30 | 29.322 | 2 | 53.59739 | yes | 1.189E-06 | 484.26962 | 501.38928 |
| TAFGIISTVK | 10 | P78346 | Ribonuclease P protein subunit p30 OS=I | RPP30 | 29.322 | 2 | 53.59739 | yes | 1.102E-07 | 518.80548 | 426.1113 |
| RPPINVAIDR | 10 | P78346 | Ribonuclease P protein subunit p30 OS=I | RPP30 | 29.322 | 2 | 53.59739 | yes | 1.96E-07 | 575.8382 | 234.61977 |
| GLAFELVYSPAIK | 13 | P78346 | Ribonuclease P protein subunit p30 OS=I | RPP30 | 29.322 | 2 | 53.59739 | yes | 9.722E-08 | 704.39777 | 164.93892 |
| NVIISSAAER | 10 | P78346 | Ribonuclease P protein subunit p30 OS=I | RPP30 | 29.322 | 2 | 53.59739 | yes | 3.09E-07 | 530.29327 | 121.10494 |
| NVIISSAAERPLEIF | 15 | P78346 | Ribonuclease P protein subunit p30 OS=I | RPP30 | 29.322 | 3 | 53.59739 | yes | 1.838E-07 | 556.65283 | 104.15884 |
| LFHIACTHLDVDL | 21 | P78346 | Ribonuclease P protein subunit p30 OS=I | RPP30 | 29.322 | 3 | 53.59739 | yes | 9.722E-08 | 828.75903 | 78.99482 |
| YTISSALNLMQICK | 14 | P78346 | Ribonuclease P protein subunit p30 OS=I | RPP30 | 29.322 | 2 | 53.59739 | yes | 9.722E-08 | 821.42059 | 76.607788 |
| KQEIEKPVAVSELI | 23 | P78346 | Ribonuclease P protein subunit p30 OS=I | RPP30 | 29.322 | 4 | 53.59739 | yes | 1.071E-07 | 639.36774 | 54.526974 |
| GLVETAAHLGYSV | 23 | P78346 | Ribonuclease P protein subunit p30 OS=I | RPP30 | 29.322 | 3 | 53.59739 | yes | 2.498E-07 | 818.4444 | 32.166542 |
| PSEGDEDCLPASK | 13 | P78346 | Ribonuclease P protein subunit p30 OS=I | RPP30 | 29.322 | 2 | 53.59739 | yes | 1.507E-07 | 702.80103 | 31.448078 |
| KTAFGIISTVK | 11 | P78346 | Ribonuclease P protein subunit p30 OS=I | RPP30 | 29.322 | 2 | 53.59739 | yes | 2.663E-07 | 582.85297 | 20.851095 |
| SPSWYGIPR | 9 | P78347 | General transcription factor II-I OS=Hom | GTF2I | 112.417 | 2 | 52.396595 | yes | 4.922E-06 | 531.77197 | 516.54425 |
| SILSPGGSCGPIK | 13 | P78347 | General transcription factor II-I OS=Hom | GTF2I | 112.417 | 2 | 52.396595 | yes | 3.265E-07 | 636.83447 | 171.72327 |
| FAQALGLTEAVK | 12 | P78347 | General transcription factor II-I OS=Hom | GTF2I | 112.417 | 2 | 52.396595 | yes | 9.722E-08 | 624.35333 | 164.97633 |
| MSVDAVEIETLR | 12 | P78347 | General transcription factor II-I OS=Hom | GTF2I | 112.417 | 2 | 52.396595 | yes | 1.105E-07 | 681.85028 | 132.74942 |
| APSYLEISSMR | 11 | P78347 | General transcription factor II-I OS=Hom | GTF2I | 112.417 | 2 | 52.396595 | yes | 9.722E-08 | 627.31335 | 115.07864 |
| ILDSAEFIK | 9 | P78347 | General transcription factor II-I OS=Hom | GTF2I | 112.417 | 2 | 52.396595 | yes | 1.881E-06 | 518.28967 | 101.31017 |
| HPENYDLATLK | 11 | P78347 | General transcription factor II-I OS=Hom | GTF2I | 112.417 | 2 | 52.396595 | yes | 2.705E-07 | 650.83026 | 86.034187 |
| KPEMFETAIK | 10 | P78347 | General transcription factor II-I OS=Hom | GTF2I | 112.417 | 2 | 52.396595 | yes | 1.596E-07 | 597.31537 | 84.179596 |
| QVEELFER | 8 | P78347 | General transcription factor II-I OS=Hom | GTF2I | 112.417 | 2 | 52.396595 | yes | 5.335E-07 | 525.26672 | 82.639389 |
| VMVTDADR | 8 | P78347 | General transcription factor II-I OS=Hom | GTF2I | 112.417 | 2 | 52.396595 | yes | 1.864E-06 | 453.7211 | 78.378685 |
| EFSFEAWNAK | 10 | P78347 | General transcription factor II-I OS=Hom | GTF2I | 112.417 | 2 | 52.396595 | yes | 9.758E-08 | 614.78528 | 77.57634 |
| MVDQLFCK | 8 | P78347 | General transcription factor II-I OS=Hom | GTF2I | 112.417 | 2 | 52.396595 | yes | 1.552E-07 | 520.74884 | 69.429955 |
| FGEAIGMGFPVK | 12 | P78347 | General transcription factor II-I OS=Hom | GTF2I | 112.417 | 2 | 52.396595 | yes | 8.759E-07 | 626.82336 | 63.765099 |
| FAEALGSTEAK | 11 | P78347 | General transcription factor II-I OS=Hom | GTF2I | 112.417 | 2 | 52.396595 | yes | 1.061E-06 | 562.28516 | 54.046844 |
| EDLQLDKPASGVK | 13 | P78347 | General transcription factor II-I OS=Hom | GTF2I | 112.417 | 2 | 52.396595 | yes | 1.157E-07 | 700.375 | 51.951176 |
| TVEDYFCFCYGK | 12 | P78347 | General transcription factor II-I OS=Hom | GTF2I | 112.417 | 2 | 52.396595 | yes | 5.386E-07 | 794.82599 | 6.0486045 |
| HFTEDIQTR | 9 | P78362 | SRSF protein kinase 2 OS=Homo sapiens | SRPK2 | 77.527 | 2 | 52.487835 | no | 9.992E-08 | 573.78052 | 4450.0137 |
| IADLGNACWVHK | 12 | P78362 | SRSF protein kinase 2 OS=Homo sapiens | SRPK2 | 77.527 | 2 | 52.487835 | no | 1.32E-07 | 692.34552 | 1966.7439 |
| LQEIEELER | 9 | P78362 | SRSF protein kinase 2 OS=Homo sapiens | SRPK2 | 77.527 | 2 | 52.487835 | yes | 9.819E-08 | 579.80371 | 1740.1461 |
| AADLLVNPLDPR | 12 | P78362 | SRSF protein kinase 2 OS=Homo sapiens | SRPK2 | 77.527 | 2 | 52.487835 | yes | 2.672E-07 | 647.36169 | 1397.597 |
| TTGLEEAAEAETA | 14 | P78362 | SRSF protein kinase 2 OS=Homo sapiens | SRPK2 | 77.527 | 2 | 52.487835 | yes | 9.722E-08 | 710.84375 | 1052.7446 |
| SNYQGLPVR | 9 | P78362 | SRSF protein kinase 2 OS=Homo sapiens | SRPK2 | 77.527 | 2 | 52.487835 | yes | 1.109E-07 | 517.2749 | 898.27118 |
| TVSASSTGDLPK | 12 | P78362 | SRSF protein kinase 2 OS=Homo sapiens | SRPK2 | 77.527 | 2 | 52.487835 | yes | 1.43E-07 | 581.80115 | 887.03809 |
| MAAEATEWQK | 10 | P78362 | SRSF protein kinase 2 OS=Homo sapiens | SRPK2 | 77.527 | 2 | 52.487835 | yes | 2.563E-07 | 582.7713 | 536.88904 |
| AGAPPPSGSAVST | 18 | P78362 | SRSF protein kinase 2 OS=Homo sapiens | SRPK2 | 77.527 | 2 | 52.487835 | yes | 3.814E-07 | 825.9259 | 336.47205 |
| IIHTDIKPENILMC | 21 | P78362 | SRSF protein kinase 2 OS=Homo sapiens | SRPK2 | 77.527 | 4 | 52.487835 | yes | 2.127E-07 | 629.57257 | 233.39294 |
| SAQHYTETALDEII | 14 | P78362 | SRSF protein kinase 2 OS=Homo sapiens | SRPK2 | 77.527 | 2 | 52.487835 | yes | 1.194E-07 | 803.39136 | 115.02144 |

| | | | | | | | | | | | |
|---|---|---|---|---|---|---|---|---|---|---|---|
| ASAGECLR | 8 | P78362 | SRSF protein kinase 2 OS=Homo sapiens | SRPK2 | 77.527 | 2 | 52.487835 | yes | 4.396E-06 | 432.20563 | 75.268318 |
| AGAPPPSGSAVST. | 22 | P78362 | SRSF protein kinase 2 OS=Homo sapiens | SRPK2 | 77.527 | 3 | 52.487835 | yes | 2.313E-06 | 682.7041 | 23.025114 |
| DVKPENFLVGR | 11 | P78368 | Casein kinase I isoform gamma-2 OS=Ho | CSNK1G2 | 47.458 | 2 | 46.077602 | yes | 2.454E-07 | 637.34857 | 152.40317 |
| SSGTSSGVLMVGP | 16 | P78368 | Casein kinase I isoform gamma-2 OS=Ho | CSNK1G2 | 47.458 | 2 | 46.077602 | yes | 1.66E-07 | 798.39594 | 62.796116 |
| QHAIHIIDFGLAK | 13 | P78368 | Casein kinase I isoform gamma-2 OS=Ho | CSNK1G2 | 47.458 | 3 | 46.077602 | yes | 4.111E-07 | 488.27701 | 25.423571 |
| DASLMVTNDGATI | 15 | P78371 | T-complex protein 1 subunit beta OS=Ho | CCT2 | 57.489 | 2 | 52.230804 | yes | 9.722E-08 | 774.90051 | 310.68039 |
| ILIANTGMDTDK | 12 | P78371 | T-complex protein 1 subunit beta OS=Ho | CCT2 | 57.489 | 2 | 52.230804 | yes | 1.063E-07 | 646.33173 | 256.90408 |
| QDLMNIAGTTLSS | 14 | P78371 | T-complex protein 1 subunit beta OS=Ho | CCT2 | 57.489 | 2 | 52.230804 | yes | 9.722E-08 | 739.87958 | 224.35095 |
| IHPQTIIAGWR | 11 | P78371 | T-complex protein 1 subunit beta OS=Ho | CCT2 | 57.489 | 2 | 52.230804 | yes | 7.163E-07 | 646.36713 | 208.25182 |
| NIGVDNPAAK | 10 | P78371 | T-complex protein 1 subunit beta OS=Ho | CCT2 | 57.489 | 2 | 52.230804 | yes | 4.075E-07 | 499.76691 | 176.30157 |
| HGINCFINR | 9 | P78371 | T-complex protein 1 subunit beta OS=Ho | CCT2 | 57.489 | 2 | 52.230804 | yes | 1.122E-07 | 565.77985 | 143.31458 |
| EAVAMESYAK | 10 | P78371 | T-complex protein 1 subunit beta OS=Ho | CCT2 | 57.489 | 2 | 52.230804 | yes | 1.301E-07 | 549.76044 | 125.33773 |
| VAEIEHAEK | 9 | P78371 | T-complex protein 1 subunit beta OS=Ho | CCT2 | 57.489 | 2 | 52.230804 | yes | 1.625E-07 | 513.26672 | 110.68792 |
| LALVTGGEIASTFI | 20 | P78371 | T-complex protein 1 subunit beta OS=Ho | CCT2 | 57.489 | 3 | 52.230804 | yes | 1.639E-07 | 699.7124 | 80.57431 |
| GATQQILDEAER | 12 | P78371 | T-complex protein 1 subunit beta OS=Ho | CCT2 | 57.489 | 2 | 52.230804 | yes | 6.816E-07 | 665.8335 | 67.535034 |
| AAHSEGNTTAGLI | 15 | P78371 | T-complex protein 1 subunit beta OS=Ho | CCT2 | 57.489 | 2 | 52.230804 | yes | 2.579E-06 | 765.85193 | 46.641937 |
| EALLSSAVDHGSD | 16 | P78371 | T-complex protein 1 subunit beta OS=Ho | CCT2 | 57.489 | 2 | 52.230804 | yes | 1.422E-07 | 828.90741 | 18.578569 |
| SLHDALCVLAQTV | 14 | P78371 | T-complex protein 1 subunit beta OS=Ho | CCT2 | 57.489 | 2 | 52.230804 | yes | 4.643E-05 | 777.91907 | 0.9289573 |
| SSNPMMVLQLPEF | 13 | P78406 | mRNA export factor RAE1 OS=Homo sa | RAE1 | 40.969 | 2 | 53.75566 | yes | 1.039E-07 | 751.37872 | 596.18048 |
| NAAEELKPR | 9 | P78406 | mRNA export factor RAE1 OS=Homo sa | RAE1 | 40.969 | 2 | 53.75566 | yes | 1.763E-07 | 514.28015 | 404.56961 |
| CWEVQDSGQTIPK | 13 | P78406 | mRNA export factor RAE1 OS=Homo sa | RAE1 | 40.969 | 2 | 53.75566 | yes | 9.722E-08 | 774.36157 | 377.63684 |
| VAIHYINPPNPAK | 13 | P78406 | mRNA export factor RAE1 OS=Homo sa | RAE1 | 40.969 | 2 | 53.75566 | yes | 9.825E-08 | 717.39862 | 173.6138 |
| VAIHYINPPNPAK | 13 | P78406 | mRNA export factor RAE1 OS=Homo sa | RAE1 | 40.969 | 2 | 53.75566 | yes | 3.354E-07 | 478.6015 | 173.6138 |
| VAIHYINPPNPAK | 13 | P78406 | mRNA export factor RAE1 OS=Homo sa | RAE1 | 40.969 | 3 | 53.75566 | yes | 9.825E-08 | 717.39862 | 173.6138 |
| VAIHYINPPNPAK | 13 | P78406 | mRNA export factor RAE1 OS=Homo sa | RAE1 | 40.969 | 3 | 53.75566 | yes | 3.354E-07 | 478.6015 | 173.6138 |
| APNYSCVMTGSW. | 14 | P78406 | mRNA export factor RAE1 OS=Homo sa | RAE1 | 40.969 | 2 | 53.75566 | yes | 1.022E-07 | 808.3476 | 211.78432 |
| MWDLSSNQAIQIA | 20 | P78406 | mRNA export factor RAE1 OS=Homo sa | RAE1 | 40.969 | 3 | 53.75566 | yes | 1.877E-07 | 751.37579 | 82.803223 |
| LWSLAEIATSSDK | 13 | P78411 | Iroquois-class homeodomain protein IRX | IRX5 | 50.361 | 2 | 50.61768 | yes | 1.156E-07 | 710.86957 | 227.93216 |
| LDALQGPPR | 9 | P78411 | Iroquois-class homeodomain protein IRX | IRX5 | 50.361 | 2 | 50.61768 | yes | 9.734E-08 | 483.772 | 67.101456 |
| ETEGSLSDSDFK | 12 | P78411 | Iroquois-class homeodomain protein IRX | IRX5 | 50.361 | 2 | 50.61768 | yes | 9.755E-08 | 657.78845 | 49.698498 |
| TFLPGAGNEVLEL | 14 | P78524 | DENN domain-containing protein 2B OS | DENND2B | 126.485 | 2 | 43.61498 | yes | 7.351E-07 | 758.41193 | 286.32089 |
| GLPQLPSSCYSVDI | 14 | P78524 | DENN domain-containing protein 2B OS | DENND2B | 126.485 | 2 | 43.61498 | yes | 2.517E-07 | 789.88269 | 106.01814 |
| TASFGYLDR | 9 | P78524 | DENN domain-containing protein 2B OS | DENND2B | 126.485 | 2 | 43.61498 | yes | 1.463E-06 | 515.2536 | 42.489101 |
| VTELALTASDR | 11 | P78527 | DNA-dependent protein kinase catalytic s | PRKDC | 469.089 | 2 | 53.533348 | yes | 9.722E-08 | 588.31696 | 394.0929 |
| LQETLSAADR | 10 | P78527 | DNA-dependent protein kinase catalytic s | PRKDC | 469.089 | 2 | 53.533348 | yes | 1.301E-07 | 552.28821 | 382.74591 |
| LGLPGDEVDNK | 11 | P78527 | DNA-dependent protein kinase catalytic s | PRKDC | 469.089 | 2 | 53.533348 | yes | 3.886E-07 | 578.79584 | 353.39545 |
| SLGPPQGEEDSVPI | 14 | P78527 | DNA-dependent protein kinase catalytic s | PRKDC | 469.089 | 2 | 53.533348 | yes | 2.907E-07 | 734.35736 | 352.84418 |
| LGASLAFNNIYR | 12 | P78527 | DNA-dependent protein kinase catalytic s | PRKDC | 469.089 | 2 | 53.533348 | yes | 2.263E-07 | 669.86188 | 318.22455 |
| LNESTFDTQITK | 12 | P78527 | DNA-dependent protein kinase catalytic s | PRKDC | 469.089 | 2 | 53.533348 | yes | 9.722E-08 | 698.85138 | 313.3714 |

| | | | | | | | | | | | |
|---|---|---|---|---|---|---|---|---|---|---|---|
| DILETHLR | 8 | P78527 | DNA-dependent protein kinase catalytic s | PRKDC | 469.089 | 2 | 53.533348 | yes | 1.527E-07 | 498.77728 | 279.33127 |
| GLSSLLCNFTK | 11 | P78527 | DNA-dependent protein kinase catalytic s | PRKDC | 469.089 | 2 | 53.533348 | yes | 3.938E-07 | 620.32373 | 276.91922 |
| VIAGLYQR | 8 | P78527 | DNA-dependent protein kinase catalytic s | PRKDC | 469.089 | 2 | 53.533348 | yes | 1.346E-07 | 460.27164 | 268.9473 |
| NELEIPGQYDGR | 12 | P78527 | DNA-dependent protein kinase catalytic s | PRKDC | 469.089 | 2 | 53.533348 | yes | 3.796E-07 | 695.8335 | 266.33099 |
| LLNTWTNR | 8 | P78527 | DNA-dependent protein kinase catalytic s | PRKDC | 469.089 | 2 | 53.533348 | yes | 2.841E-07 | 509.27744 | 264.33496 |
| FVPLLPGNR | 9 | P78527 | DNA-dependent protein kinase catalytic s | PRKDC | 469.089 | 2 | 53.533348 | yes | 1.656E-07 | 506.80054 | 256.09009 |
| ILELSGSSSEDSEK | 14 | P78527 | DNA-dependent protein kinase catalytic s | PRKDC | 469.089 | 2 | 53.533348 | yes | 9.766E-08 | 740.85431 | 255.90924 |
| YNFPVEVEVPMEI | 13 | P78527 | DNA-dependent protein kinase catalytic s | PRKDC | 469.089 | 2 | 53.533348 | yes | 1.155E-07 | 804.88995 | 250.36566 |
| NLSSNEAISLEEIR | 14 | P78527 | DNA-dependent protein kinase catalytic s | PRKDC | 469.089 | 2 | 53.533348 | yes | 9.722E-08 | 787.90466 | 249.8519 |
| SDPGLLTNTMDVI | 15 | P78527 | DNA-dependent protein kinase catalytic s | PRKDC | 469.089 | 2 | 53.533348 | yes | 1.392E-07 | 818.9162 | 227.02814 |
| LLPAELPAK | 9 | P78527 | DNA-dependent protein kinase catalytic s | PRKDC | 469.089 | 2 | 53.533348 | yes | 1.551E-07 | 476.2973 | 214.09184 |
| SIGEYDVLR | 9 | P78527 | DNA-dependent protein kinase catalytic s | PRKDC | 469.089 | 2 | 53.533348 | yes | 6.91E-07 | 526.27454 | 202.8324 |
| YPEETLSLMTK | 11 | P78527 | DNA-dependent protein kinase catalytic s | PRKDC | 469.089 | 2 | 53.533348 | yes | 1.121E-07 | 656.32867 | 202.7942 |
| NNWEVSALSR | 10 | P78527 | DNA-dependent protein kinase catalytic s | PRKDC | 469.089 | 2 | 53.533348 | yes | 9.722E-08 | 588.29382 | 184.97133 |
| QITQSALLAEAR | 12 | P78527 | DNA-dependent protein kinase catalytic s | PRKDC | 469.089 | 2 | 53.533348 | yes | 9.722E-08 | 650.86462 | 182.66966 |
| HSSLITPLQAVAQI | 14 | P78527 | DNA-dependent protein kinase catalytic s | PRKDC | 469.089 | 2 | 53.533348 | yes | 9.722E-08 | 760.93079 | 85.852074 |
| HSSLITPLQAVAQI | 14 | P78527 | DNA-dependent protein kinase catalytic s | PRKDC | 469.089 | 2 | 53.533348 | yes | 1.211E-07 | 507.62296 | 85.852074 |
| HSSLITPLQAVAQI | 14 | P78527 | DNA-dependent protein kinase catalytic s | PRKDC | 469.089 | 3 | 53.533348 | yes | 9.722E-08 | 760.93079 | 85.852074 |
| HSSLITPLQAVAQI | 14 | P78527 | DNA-dependent protein kinase catalytic s | PRKDC | 469.089 | 3 | 53.533348 | yes | 1.211E-07 | 507.62296 | 85.852074 |
| LATTILQHWK | 10 | P78527 | DNA-dependent protein kinase catalytic s | PRKDC | 469.089 | 2 | 53.533348 | yes | 9.722E-08 | 605.85077 | 170.70955 |
| NFWSHETR | 8 | P78527 | DNA-dependent protein kinase catalytic s | PRKDC | 469.089 | 2 | 53.533348 | yes | 9.722E-08 | 538.74902 | 157.9325 |
| LAAVVSACK | 9 | P78527 | DNA-dependent protein kinase catalytic s | PRKDC | 469.089 | 2 | 53.533348 | yes | 1.12E-07 | 459.75751 | 153.82324 |
| TYSVVPMTSR | 10 | P78527 | DNA-dependent protein kinase catalytic s | PRKDC | 469.089 | 2 | 53.533348 | yes | 4.047E-07 | 570.78955 | 149.12595 |
| LACDVDQVTR | 10 | P78527 | DNA-dependent protein kinase catalytic s | PRKDC | 469.089 | 2 | 53.533348 | yes | 1.526E-07 | 588.78754 | 146.91382 |
| AALSALESFLK | 11 | P78527 | DNA-dependent protein kinase catalytic s | PRKDC | 469.089 | 2 | 53.533348 | yes | 1.002E-07 | 575.32935 | 141.60568 |
| LPLISGFYK | 9 | P78527 | DNA-dependent protein kinase catalytic s | PRKDC | 469.089 | 2 | 53.533348 | yes | 1.074E-07 | 519.30511 | 129.8916 |
| LLEEALLR | 8 | P78527 | DNA-dependent protein kinase catalytic s | PRKDC | 469.089 | 2 | 53.533348 | yes | 4.187E-07 | 478.79239 | 126.29176 |
| MQLVCCR | 7 | P78527 | DNA-dependent protein kinase catalytic s | PRKDC | 469.089 | 2 | 53.533348 | yes | 9.722E-08 | 483.71954 | 124.8122 |
| VCLDIIYK | 8 | P78527 | DNA-dependent protein kinase catalytic s | PRKDC | 469.089 | 2 | 53.533348 | yes | 1.343E-07 | 512.28082 | 120.69009 |
| GGSWIQEINVAEK | 13 | P78527 | DNA-dependent protein kinase catalytic s | PRKDC | 469.089 | 2 | 53.533348 | yes | 1.066E-07 | 715.86737 | 113.315 |
| NCISTVVHQGLIR | 13 | P78527 | DNA-dependent protein kinase catalytic s | PRKDC | 469.089 | 2 | 53.533348 | yes | 9.85E-08 | 748.90375 | 112.30036 |
| FNNYVDCMK | 9 | P78527 | DNA-dependent protein kinase catalytic s | PRKDC | 469.089 | 2 | 53.533348 | yes | 9.722E-08 | 595.75208 | 109.6273 |
| TLCLEVVLCR | 10 | P78527 | DNA-dependent protein kinase catalytic s | PRKDC | 469.089 | 2 | 53.533348 | yes | 9.791E-08 | 631.83344 | 105.87457 |
| SLNSIEFR | 8 | P78527 | DNA-dependent protein kinase catalytic s | PRKDC | 469.089 | 2 | 53.533348 | yes | 2.671E-07 | 483.25616 | 101.89478 |
| SQGCSEQVLTVLK | 13 | P78527 | DNA-dependent protein kinase catalytic s | PRKDC | 469.089 | 2 | 53.533348 | yes | 4.314E-07 | 724.87433 | 99.566994 |
| LPPDVLR | 7 | P78527 | DNA-dependent protein kinase catalytic s | PRKDC | 469.089 | 2 | 53.533348 | yes | 5.625E-07 | 405.24762 | 94.866631 |
| GIFTSEIGTK | 10 | P78527 | DNA-dependent protein kinase catalytic s | PRKDC | 469.089 | 2 | 53.533348 | yes | 2.784E-06 | 526.78479 | 89.074532 |
| LLQDFNR | 7 | P78527 | DNA-dependent protein kinase catalytic s | PRKDC | 469.089 | 2 | 53.533348 | yes | 1.684E-07 | 453.24561 | 86.201973 |
| HECMAPLTALVK | 12 | P78527 | DNA-dependent protein kinase catalytic s | PRKDC | 469.089 | 2 | 53.533348 | yes | 2.167E-07 | 685.35199 | 85.154846 |

| | | | | | | | | | | | |
|---|---|---|---|---|---|---|---|---|---|---|---|
| DQNILLGTTYR | 11 | P78527 | DNA-dependent protein kinase catalytic s | PRKDC | 469.089 | 2 | 53.533348 | yes | 5.566E-07 | 647.34351 | 84.027832 |
| MSTSPEAFLALR | 12 | P78527 | DNA-dependent protein kinase catalytic s | PRKDC | 469.089 | 2 | 53.533348 | yes | 3.433E-07 | 661.84229 | 81.161331 |
| DLCNTHLMR | 9 | P78527 | DNA-dependent protein kinase catalytic s | PRKDC | 469.089 | 2 | 53.533348 | yes | 2.8E-07 | 580.271 | 79.68074 |
| QCLPSLDLSCK | 11 | P78527 | DNA-dependent protein kinase catalytic s | PRKDC | 469.089 | 2 | 53.533348 | yes | 1.252E-06 | 660.81793 | 76.566216 |
| VEGMTELYFQLK | 12 | P78527 | DNA-dependent protein kinase catalytic s | PRKDC | 469.089 | 2 | 53.533348 | yes | 9.722E-08 | 729.37085 | 76.154701 |
| CFGTGAAGNR | 10 | P78527 | DNA-dependent protein kinase catalytic s | PRKDC | 469.089 | 2 | 53.533348 | yes | 1.211E-06 | 505.72726 | 73.776558 |
| ILDVMYSR | 8 | P78527 | DNA-dependent protein kinase catalytic s | PRKDC | 469.089 | 2 | 53.533348 | yes | 1.367E-07 | 498.76279 | 73.109032 |
| NLLTVTSSDEMMK | 13 | P78527 | DNA-dependent protein kinase catalytic s | PRKDC | 469.089 | 2 | 53.533348 | yes | 2.353E-07 | 734.85474 | 65.600403 |
| ECSPWMSDFK | 10 | P78527 | DNA-dependent protein kinase catalytic s | PRKDC | 469.089 | 2 | 53.533348 | yes | 4.166E-07 | 643.76263 | 63.725449 |
| NTCTSVYTK | 9 | P78527 | DNA-dependent protein kinase catalytic s | PRKDC | 469.089 | 2 | 53.533348 | yes | 5.807E-07 | 537.25024 | 61.28886 |
| LSFAVPFR | 8 | P78527 | DNA-dependent protein kinase catalytic s | PRKDC | 469.089 | 2 | 53.533348 | yes | 2.021E-07 | 468.76871 | 60.401859 |
| TETVTSFR | 8 | P78527 | DNA-dependent protein kinase catalytic s | PRKDC | 469.089 | 2 | 53.533348 | yes | 3.604E-07 | 470.74036 | 60.392303 |
| LSLMYAR | 7 | P78527 | DNA-dependent protein kinase catalytic s | PRKDC | 469.089 | 2 | 53.533348 | yes | 6.01E-07 | 427.23364 | 57.776539 |
| SMEEDPQTSR | 10 | P78527 | DNA-dependent protein kinase catalytic s | PRKDC | 469.089 | 2 | 53.533348 | yes | 1.94E-06 | 590.25098 | 57.393978 |
| FLCIFLEK | 8 | P78527 | DNA-dependent protein kinase catalytic s | PRKDC | 469.089 | 2 | 53.533348 | yes | 6.499E-06 | 535.2912 | 53.725376 |
| DPTVHDDVLELEM | 18 | P78527 | DNA-dependent protein kinase catalytic s | PRKDC | 469.089 | 3 | 53.533348 | yes | 1.526E-07 | 714.00031 | 51.903336 |
| AYVPALQMAFK | 11 | P78527 | DNA-dependent protein kinase catalytic s | PRKDC | 469.089 | 2 | 53.533348 | yes | 1.281E-06 | 619.83374 | 49.510872 |
| LPVLAGCLK | 9 | P78527 | DNA-dependent protein kinase catalytic s | PRKDC | 469.089 | 2 | 53.533348 | yes | 3.846E-07 | 485.79135 | 46.556194 |
| NILEESLCELVAK | 13 | P78527 | DNA-dependent protein kinase catalytic s | PRKDC | 469.089 | 2 | 53.533348 | yes | 9.809E-08 | 759.39764 | 46.293148 |
| IPALDLLIK | 9 | P78527 | DNA-dependent protein kinase catalytic s | PRKDC | 469.089 | 2 | 53.533348 | yes | 1.445E-06 | 498.32861 | 43.276188 |
| YAVPSAGLR | 9 | P78527 | DNA-dependent protein kinase catalytic s | PRKDC | 469.089 | 2 | 53.533348 | yes | 5.467E-06 | 467.26126 | 43.001278 |
| AIRPQIDLK | 9 | P78527 | DNA-dependent protein kinase catalytic s | PRKDC | 469.089 | 2 | 53.533348 | yes | 3.218E-07 | 527.3244 | 38.517269 |
| DVDFMYVELIQR | 12 | P78527 | DNA-dependent protein kinase catalytic s | PRKDC | 469.089 | 2 | 53.533348 | yes | 9.722E-08 | 764.37921 | 36.492588 |
| QFINLMLPMK | 10 | P78527 | DNA-dependent protein kinase catalytic s | PRKDC | 469.089 | 2 | 53.533348 | yes | 3.925E-07 | 617.83795 | 34.110615 |
| QDWVDGEPTEAEI | 13 | P78527 | DNA-dependent protein kinase catalytic s | PRKDC | 469.089 | 2 | 53.533348 | yes | 8.616E-07 | 752.33356 | 33.228161 |
| LAGANPAVITCDE | 20 | P78527 | DNA-dependent protein kinase catalytic s | PRKDC | 469.089 | 3 | 53.533348 | yes | 1.835E-06 | 707.70514 | 33.128521 |
| SSFDWLTGSSTDPI | 30 | P78527 | DNA-dependent protein kinase catalytic s | PRKDC | 469.089 | 4 | 53.533348 | yes | 4.897E-07 | 809.38611 | 25.454517 |
| YSCFALFVK | 9 | P78527 | DNA-dependent protein kinase catalytic s | PRKDC | 469.089 | 2 | 53.533348 | yes | 2.905E-06 | 567.78625 | 18.761097 |
| YFEGVSPK | 8 | P78527 | DNA-dependent protein kinase catalytic s | PRKDC | 469.089 | 2 | 53.533348 | yes | 2.495E-06 | 463.73456 | 17.682665 |
| AQEPESGLSEETQV | 15 | P78527 | DNA-dependent protein kinase catalytic s | PRKDC | 469.089 | 2 | 53.533348 | yes | 3.71E-07 | 816.3916 | 16.303398 |
| GKPLPEYHVR | 10 | P78527 | DNA-dependent protein kinase catalytic s | PRKDC | 469.089 | 2 | 53.533348 | yes | 2.536E-07 | 598.33276 | 12.612848 |
| AALEEWLPR | 9 | P78549 | Endonuclease III-like protein 1 OS=Hom | NTHL1 | 34.39 | 2 | 51.003357 | yes | 2.296E-07 | 542.79291 | 668.64227 |
| VAYEGSDSEK | 10 | P78549 | Endonuclease III-like protein 1 OS=Hom | NTHL1 | 34.39 | 2 | 51.003357 | yes | 1.639E-07 | 542.74329 | 276.07529 |
| DAPVDHLGTEHCY | 20 | P78549 | Endonuclease III-like protein 1 OS=Hom | NTHL1 | 34.39 | 3 | 51.003357 | yes | 1.063E-07 | 732.66028 | 136.07927 |
| LIYPVGFWR | 9 | P78549 | Endonuclease III-like protein 1 OS=Hom | NTHL1 | 34.39 | 2 | 51.003357 | yes | 9.722E-08 | 575.82404 | 82.217567 |
| DQVTAGAMQR | 10 | P78549 | Endonuclease III-like protein 1 OS=Hom | NTHL1 | 34.39 | 2 | 51.003357 | yes | 1.423E-06 | 538.76129 | 28.472507 |
| VLAGVVMTTGTD | 14 | P78563 | Double-stranded RNA-specific editase 1 ( | ADARB1 | 80.764 | 2 | 54.532314 | yes | 9.849E-08 | 695.88416 | 528.28546 |
| YDFLSESGESHAK | 13 | P78563 | Double-stranded RNA-specific editase 1 ( | ADARB1 | 80.764 | 2 | 54.532314 | yes | 2.32E-07 | 735.33081 | 421.18372 |
| GLALNDCHAEIISR | 14 | P78563 | Double-stranded RNA-specific editase 1 ( | ADARB1 | 80.764 | 2 | 54.532314 | yes | 9.722E-08 | 784.89612 | 137.49371 |

| | | | | | | | | | | | |
|---|---|---|---|---|---|---|---|---|---|---|---|
| GLALNDCHAEIISR | 14 | P78563 | Double-stranded RNA-specific editase 1 ( | ADARB1 | 80.764 | 2 | 54.532314 | yes | 2.017E-07 | 523.59985 | 137.49371 |
| GLALNDCHAEIISR | 14 | P78563 | Double-stranded RNA-specific editase 1 ( | ADARB1 | 80.764 | 3 | 54.532314 | yes | 9.722E-08 | 784.89612 | 137.49371 |
| GLALNDCHAEIISR | 14 | P78563 | Double-stranded RNA-specific editase 1 ( | ADARB1 | 80.764 | 3 | 54.532314 | yes | 2.017E-07 | 523.59985 | 137.49371 |
| LLTMSCSDK | 9 | P78563 | Double-stranded RNA-specific editase 1 ( | ADARB1 | 80.764 | 2 | 54.532314 | yes | 2.725E-06 | 527.74902 | 240.57378 |
| ITKPNVYHESK | 11 | P78563 | Double-stranded RNA-specific editase 1 ( | ADARB1 | 80.764 | 2 | 54.532314 | yes | 1.473E-07 | 658.35388 | 208.18008 |
| IFSPHEPILEGSR | 13 | P78563 | Double-stranded RNA-specific editase 1 ( | ADARB1 | 80.764 | 2 | 54.532314 | yes | 9.722E-08 | 741.39099 | 188.47562 |
| IESGEGTIPVR | 11 | P78563 | Double-stranded RNA-specific editase 1 ( | ADARB1 | 80.764 | 2 | 54.532314 | yes | 3.545E-07 | 579.31171 | 187.79636 |
| VPSHLLR | 7 | P78563 | Double-stranded RNA-specific editase 1 ( | ADARB1 | 80.764 | 2 | 54.532314 | yes | 2.396E-07 | 411.25323 | 159.88556 |
| PLLSGISNAEAR | 12 | P78563 | Double-stranded RNA-specific editase 1 ( | ADARB1 | 80.764 | 2 | 54.532314 | yes | 1.429E-07 | 614.33826 | 147.07367 |
| KTPGPVLPK | 9 | P78563 | Double-stranded RNA-specific editase 1 ( | ADARB1 | 80.764 | 2 | 54.532314 | yes | 1.173E-07 | 468.79749 | 145.30609 |
| FLYTQLELYLNNK | 13 | P78563 | Double-stranded RNA-specific editase 1 ( | ADARB1 | 80.764 | 2 | 54.532314 | yes | 9.722E-08 | 829.94305 | 130.39093 |
| NPVMILNELRPGL | 14 | P78563 | Double-stranded RNA-specific editase 1 ( | ADARB1 | 80.764 | 2 | 54.532314 | yes | 1.567E-07 | 797.46088 | 124.10106 |
| CINGEYMSDR | 10 | P78563 | Double-stranded RNA-specific editase 1 ( | ADARB1 | 80.764 | 2 | 54.532314 | yes | 2.458E-07 | 622.75537 | 52.385036 |
| NPVMILNELR | 10 | P78563 | Double-stranded RNA-specific editase 1 ( | ADARB1 | 80.764 | 2 | 54.532314 | yes | 9.722E-08 | 599.83429 | 39.46468 |
| ENAGEDPGLAR | 11 | P81605 | Dermcidin OS=Homo sapiens OX=9606 ( | DCD | 11.284 | 2 | 29.135355 | yes | 2.574E-06 | 564.76764 | 37.533024 |
| VPINDVLAEDK | 11 | P82650 | Small ribosomal subunit protein mS22 OS | MRPS22 | 41.281 | 2 | 50.543976 | yes | 1.13E-07 | 606.82715 | 2472.5168 |
| MPPVLEER | 8 | P82650 | Small ribosomal subunit protein mS22 OS | MRPS22 | 41.281 | 2 | 50.543976 | yes | 1.062E-07 | 485.75494 | 1954.0088 |
| LMTQAQLEEATR | 12 | P82650 | Small ribosomal subunit protein mS22 OS | MRPS22 | 41.281 | 2 | 50.543976 | yes | 2.346E-07 | 695.85339 | 1860.0059 |
| IDGLLIDQIQR | 11 | P82650 | Small ribosomal subunit protein mS22 OS | MRPS22 | 41.281 | 2 | 50.543976 | yes | 1.008E-07 | 642.36951 | 1828.4611 |
| MTGLNLQK | 8 | P82650 | Small ribosomal subunit protein mS22 OS | MRPS22 | 41.281 | 2 | 50.543976 | yes | 9.927E-08 | 452.74966 | 807.5918 |
| TYEDIDKR | 8 | P82650 | Small ribosomal subunit protein mS22 OS | MRPS22 | 41.281 | 2 | 50.543976 | yes | 3.07E-07 | 520.25635 | 705.38904 |
| YVFTDISYSIPHR | 13 | P82650 | Small ribosomal subunit protein mS22 OS | MRPS22 | 41.281 | 2 | 50.543976 | yes | 1.994E-07 | 799.40411 | 474.84326 |
| MIQVYFPK | 8 | P82650 | Small ribosomal subunit protein mS22 OS | MRPS22 | 41.281 | 2 | 50.543976 | yes | 9.893E-08 | 513.27808 | 432.93771 |
| GKYDLLR | 7 | P82650 | Small ribosomal subunit protein mS22 OS | MRPS22 | 41.281 | 2 | 50.543976 | yes | 1.962E-07 | 432.75052 | 428.93579 |
| TFKPAIQELKPPTY | 15 | P82650 | Small ribosomal subunit protein mS22 OS | MRPS22 | 41.281 | 3 | 50.543976 | yes | 2.245E-07 | 587.66968 | 184.51424 |
| TFKPAIQELKPPTY | 15 | P82650 | Small ribosomal subunit protein mS22 OS | MRPS22 | 41.281 | 3 | 50.543976 | yes | 4.119E-07 | 441.00409 | 184.51424 |
| TFKPAIQELKPPTY | 15 | P82650 | Small ribosomal subunit protein mS22 OS | MRPS22 | 41.281 | 4 | 50.543976 | yes | 2.245E-07 | 587.66968 | 184.51424 |
| TFKPAIQELKPPTY | 15 | P82650 | Small ribosomal subunit protein mS22 OS | MRPS22 | 41.281 | 4 | 50.543976 | yes | 4.119E-07 | 441.00409 | 184.51424 |
| KIDGLLIDQIQR | 12 | P82650 | Small ribosomal subunit protein mS22 OS | MRPS22 | 41.281 | 2 | 50.543976 | yes | 1.009E-07 | 706.41699 | 337.80533 |
| ILTPIIFK | 8 | P82650 | Small ribosomal subunit protein mS22 OS | MRPS22 | 41.281 | 2 | 50.543976 | yes | 1.381E-07 | 472.81259 | 314.03348 |
| ILEGTETTK | 9 | P82650 | Small ribosomal subunit protein mS22 OS | MRPS22 | 41.281 | 2 | 50.543976 | yes | 1.937E-07 | 496.26895 | 249.66791 |
| DQAAEGINLIK | 11 | P82650 | Small ribosomal subunit protein mS22 OS | MRPS22 | 41.281 | 2 | 50.543976 | yes | 1.36E-07 | 586.31952 | 163.30464 |
| KILTPIIFK | 9 | P82650 | Small ribosomal subunit protein mS22 OS | MRPS22 | 41.281 | 2 | 50.543976 | yes | 9.779E-08 | 536.86011 | 138.23058 |
| YFGGMVWYFVNN | 13 | P82650 | Small ribosomal subunit protein mS22 OS | MRPS22 | 41.281 | 2 | 50.543976 | yes | 3.49E-07 | 812.88446 | 30.882334 |
| LDDDPPPSTLLK | 12 | P82914 | Small ribosomal subunit protein uS15m C | MRPS15 | 29.843 | 2 | 49.430878 | yes | 2.172E-07 | 655.84552 | 743.56885 |
| IVANPEDTR | 9 | P82914 | Small ribosomal subunit protein uS15m C | MRPS15 | 29.843 | 2 | 49.430878 | yes | 1.877E-07 | 507.76437 | 394.30661 |
| SYEEHLEK | 8 | P82914 | Small ribosomal subunit protein uS15m C | MRPS15 | 29.843 | 2 | 49.430878 | yes | 3.353E-07 | 517.7431 | 379.61169 |
| NTNYDVFEK | 9 | P82914 | Small ribosomal subunit protein uS15m C | MRPS15 | 29.843 | 2 | 49.430878 | yes | 1.963E-07 | 565.26166 | 313.68646 |
| LLSLEMANK | 9 | P82914 | Small ribosomal subunit protein uS15m C | MRPS15 | 29.843 | 2 | 49.430878 | yes | 1.416E-06 | 509.78372 | 249.84949 |

| | | | | | | | | | | | |
|---|---|---|---|---|---|---|---|---|---|---|---|
| DYQNVPGIEK | 10 | P82914 | Small ribosomal subunit protein uS15m C | MRPS15 | 29.843 | 2 | 49.430878 | yes | 2.003E-07 | 581.79059 | 249.33403 |
| YLLMSIDQR | 9 | P82914 | Small ribosomal subunit protein uS15m C | MRPS15 | 29.843 | 2 | 49.430878 | yes | 4.826E-07 | 569.79987 | 204.517 |
| VFQETQK | 7 | P82914 | Small ribosomal subunit protein uS15m C | MRPS15 | 29.843 | 2 | 49.430878 | yes | 1.365E-05 | 440.23218 | 95.200012 |
| KIVANPEDTR | 10 | P82914 | Small ribosomal subunit protein uS15m C | MRPS15 | 29.843 | 2 | 49.430878 | yes | 1.032E-07 | 571.81183 | 80.206627 |
| IKQEQFMK | 8 | P82914 | Small ribosomal subunit protein uS15m C | MRPS15 | 29.843 | 2 | 49.430878 | yes | 9.299E-07 | 526.28387 | 72.807404 |
| DSQLYAVDYETLT | 14 | P82930 | Small ribosomal subunit protein mS34 OS | MRPS34 | 25.651 | 2 | 53.575073 | yes | 1.02E-07 | 837.40448 | 1802.6271 |
| LPLFGLGR | 8 | P82930 | Small ribosomal subunit protein mS34 OS | MRPS34 | 25.651 | 2 | 53.575073 | yes | 1.549E-07 | 436.77127 | 1266.1274 |
| MEPWDYPAK | 9 | P82930 | Small ribosomal subunit protein mS34 OS | MRPS34 | 25.651 | 2 | 53.575073 | yes | 2.283E-06 | 568.75769 | 475.43903 |
| VRPDYTAQNLDH( | 14 | P82930 | Small ribosomal subunit protein mS34 OS | MRPS34 | 25.651 | 3 | 53.575073 | yes | 9.722E-08 | 538.6059 | 130.8013 |
| VRPDYTAQNLDH( | 14 | P82930 | Small ribosomal subunit protein mS34 OS | MRPS34 | 25.651 | 3 | 53.575073 | yes | 9.722E-08 | 807.40515 | 130.8013 |
| VRPDYTAQNLDH( | 14 | P82930 | Small ribosomal subunit protein mS34 OS | MRPS34 | 25.651 | 2 | 53.575073 | yes | 9.722E-08 | 538.6059 | 130.8013 |
| VRPDYTAQNLDH( | 14 | P82930 | Small ribosomal subunit protein mS34 OS | MRPS34 | 25.651 | 2 | 53.575073 | yes | 9.722E-08 | 807.40515 | 130.8013 |
| LLQLLGR | 7 | P82930 | Small ribosomal subunit protein mS34 OS | MRPS34 | 25.651 | 2 | 53.575073 | yes | 9.849E-08 | 406.77127 | 171.76952 |
| AWGILTFK | 8 | P82930 | Small ribosomal subunit protein mS34 OS | MRPS34 | 25.651 | 2 | 53.575073 | yes | 8.215E-07 | 468.27109 | 128.13455 |
| EIEHVMYHDWR | 11 | P82930 | Small ribosomal subunit protein mS34 OS | MRPS34 | 25.651 | 2 | 53.575073 | yes | 1.27E-07 | 757.84589 | 45.378681 |
| NGDTSTEEPMLNV | 15 | P82930 | Small ribosomal subunit protein mS34 OS | MRPS34 | 25.651 | 2 | 53.575073 | yes | 9.722E-08 | 845.88867 | 39.277443 |
| ECEGIVPVPLAEK | 13 | P82932 | Small ribosomal subunit protein bS6m OS | MRPS6 | 14.227 | 2 | 48.841312 | yes | 1.211E-07 | 720.87378 | 2514.7651 |
| HPLTQELK | 8 | P82932 | Small ribosomal subunit protein bS6m OS | MRPS6 | 14.227 | 2 | 48.841312 | yes | 1.097E-07 | 483.27435 | 619.30963 |
| YELALILK | 8 | P82932 | Small ribosomal subunit protein bS6m OS | MRPS6 | 14.227 | 2 | 48.841312 | yes | 1.728E-06 | 481.79968 | 283.08908 |
| AMQRPETAATLK | 12 | P82932 | Small ribosomal subunit protein bS6m OS | MRPS6 | 14.227 | 2 | 48.841312 | yes | 1.594E-07 | 658.85321 | 238.96942 |
| IALEFDKDR | 9 | P83111 | Serine beta-lactamase-like protein LACTI | LACTB | 60.694 | 2 | 53.643288 | yes | 3.022E-07 | 553.79565 | 2654.4312 |
| VPCKPETVMR | 10 | P83111 | Serine beta-lactamase-like protein LACTI | LACTB | 60.694 | 2 | 53.643288 | yes | 9.968E-08 | 608.8125 | 990.06348 |
| VPCKPETVMR | 10 | P83111 | Serine beta-lactamase-like protein LACTI | LACTB | 60.694 | 2 | 53.643288 | yes | 1.165E-07 | 406.21075 | 990.06348 |
| VPCKPETVMR | 10 | P83111 | Serine beta-lactamase-like protein LACTI | LACTB | 60.694 | 3 | 53.643288 | yes | 9.968E-08 | 608.8125 | 990.06348 |
| VPCKPETVMR | 10 | P83111 | Serine beta-lactamase-like protein LACTI | LACTB | 60.694 | 3 | 53.643288 | yes | 1.165E-07 | 406.21075 | 990.06348 |
| LVNTPYVDNSYK | 12 | P83111 | Serine beta-lactamase-like protein LACTI | LACTB | 60.694 | 2 | 53.643288 | yes | 1.38E-07 | 706.85645 | 1615.8566 |
| NDFEQGELYLR | 11 | P83111 | Serine beta-lactamase-like protein LACTI | LACTB | 60.694 | 2 | 53.643288 | yes | 9.722E-08 | 692.33063 | 892.54944 |
| EKFENSIESLR | 11 | P83111 | Serine beta-lactamase-like protein LACTI | LACTB | 60.694 | 2 | 53.643288 | yes | 9.728E-08 | 676.34625 | 885.41626 |
| YLDYMQK | 7 | P83111 | Serine beta-lactamase-like protein LACTI | LACTB | 60.694 | 2 | 53.643288 | yes | 9.892E-08 | 480.72839 | 501.91248 |
| KNDFEQGELYLR | 12 | P83111 | Serine beta-lactamase-like protein LACTI | LACTB | 60.694 | 2 | 53.643288 | yes | 3.61E-07 | 756.37805 | 387.3439 |
| HYASHTGGAVGA: | 31 | P83111 | Serine beta-lactamase-like protein LACTI | LACTB | 60.694 | 4 | 53.643288 | yes | 2.152E-07 | 806.41241 | 358.70056 |
| FENSIESLR | 9 | P83111 | Serine beta-lactamase-like protein LACTI | LACTB | 60.694 | 2 | 53.643288 | yes | 3.836E-07 | 547.77747 | 313.21954 |
| IKDEVGAPGIVVG' | 19 | P83111 | Serine beta-lactamase-like protein LACTI | LACTB | 60.694 | 3 | 53.643288 | yes | 9.728E-08 | 613.67889 | 128.91464 |
| ENVAFEQEK | 9 | P83111 | Serine beta-lactamase-like protein LACTI | LACTB | 60.694 | 2 | 53.643288 | yes | 1.23E-07 | 547.26166 | 115.21432 |
| LLISHLSGIR | 10 | P83111 | Serine beta-lactamase-like protein LACTI | LACTB | 60.694 | 2 | 53.643288 | yes | 9.722E-08 | 554.84552 | 89.626152 |
| YAMAWGVVER | 10 | P83111 | Serine beta-lactamase-like protein LACTI | LACTB | 60.694 | 2 | 53.643288 | yes | 4.119E-07 | 599.28967 | 66.199966 |
| HINQIFIR | 8 | P83369 | U7 snRNA-associated Sm-like protein LS | LSM11 | 39.5 | 2 | 50.94598 | yes | 1.014E-07 | 520.80365 | 276.77811 |
| SVPSSLQASAR | 11 | P83369 | U7 snRNA-associated Sm-like protein LS | LSM11 | 39.5 | 2 | 50.94598 | yes | 2.896E-06 | 551.7962 | 123.69089 |
| GVCTGFLVAFDK | 12 | P83369 | U7 snRNA-associated Sm-like protein LS | LSM11 | 39.5 | 2 | 50.94598 | yes | 9.722E-08 | 657.33154 | 72.316757 |

| | | | | | | | | | | | |
|---|---|---|---|---|---|---|---|---|---|---|---|
| SAVEDSTLSR | 10 | P83369 | U7 snRNA-associated Sm-like protein LS | LSM11 | 39.5 | 2 | 50.94598 | yes | 3.669E-05 | 532.76459 | 21.743349 |
| VELCSFSGYK | 10 | P83731 | Large ribosomal subunit protein eL24 OS | RPL24 | 17.779 | 2 | 52.158176 | yes | 9.722E-08 | 595.28156 | 25270.764 |
| AITGASLADIMAK | 13 | P83731 | Large ribosomal subunit protein eL24 OS | RPL24 | 17.779 | 2 | 52.158176 | yes | 2.874E-06 | 631.34467 | 12140.963 |
| AITGASLADIMAK | 13 | P83731 | Large ribosomal subunit protein eL24 OS | RPL24 | 17.779 | 2 | 52.158176 | yes | 9.722E-08 | 639.3421 | 12140.963 |
| QINWTVLYR | 9 | P83731 | Large ribosomal subunit protein eL24 OS | RPL24 | 17.779 | 2 | 52.158176 | yes | 1.092E-07 | 596.82727 | 13699.241 |
| VFQFLNAK | 8 | P83731 | Large ribosomal subunit protein eL24 OS | RPL24 | 17.779 | 2 | 52.158176 | yes | 2.533E-07 | 483.77402 | 13121.037 |
| CESAFLSK | 8 | P83731 | Large ribosomal subunit protein eL24 OS | RPL24 | 17.779 | 2 | 52.158176 | yes | 2.06E-07 | 471.22348 | 6050.6626 |
| GQSEEIQK | 8 | P83731 | Large ribosomal subunit protein eL24 OS | RPL24 | 17.779 | 2 | 52.158176 | yes | 1.719E-06 | 459.72998 | 1008.6769 |
| MKVELCSFSGYK | 12 | P83731 | Large ribosomal subunit protein eL24 OS | RPL24 | 17.779 | 2 | 52.158176 | yes | 1.197E-07 | 724.84924 | 361.91177 |
| MKVELCSFSGYK | 12 | P83731 | Large ribosomal subunit protein eL24 OS | RPL24 | 17.779 | 2 | 52.158176 | yes | 3.827E-07 | 732.84674 | 361.91177 |
| TDGKVFQFLNAK | 12 | P83731 | Large ribosomal subunit protein eL24 OS | RPL24 | 17.779 | 2 | 52.158176 | yes | 5.144E-07 | 684.36951 | 33.650806 |
| QEMVDIIETVYR | 12 | P83876 | Thioredoxin-like protein 4A OS=Homo s | TXNL4A | 16.787 | 2 | 55.46245 | yes | 3.137E-07 | 748.37671 | 119.47569 |
| MDEVLYSIAEK | 11 | P83876 | Thioredoxin-like protein 4A OS=Homo s | TXNL4A | 16.787 | 2 | 55.46245 | yes | 9.722E-08 | 649.32086 | 76.026855 |
| LECVEPNCR | 9 | P83881 | Large ribosomal subunit protein eL42 OS | RPL36A | 12.441 | 2 | 53.547813 | no | 9.722E-08 | 588.76044 | 19286.873 |
| HFELGGDK | 8 | P83881 | Large ribosomal subunit protein eL42 OS | RPL36A | 12.441 | 2 | 53.547813 | no | 1.113E-07 | 451.72195 | 7464.7456 |
| HFELGGDKK | 9 | P83881 | Large ribosomal subunit protein eL42 OS | RPL36A | 12.441 | 2 | 53.547813 | no | 1.864E-07 | 515.76947 | 4602.8462 |
| GKDSLYAQGK | 10 | P83881 | Large ribosomal subunit protein eL42 OS | RPL36A | 12.441 | 2 | 53.547813 | yes | 1.703E-07 | 533.78003 | 3706.3052 |
| QSGYGGQTKPIFR | 13 | P83881 | Large ribosomal subunit protein eL42 OS | RPL36A | 12.441 | 3 | 53.547813 | no | 1.435E-07 | 480.25278 | 672.82617 |
| QSGYGGQTKPIFR | 13 | P83881 | Large ribosomal subunit protein eL42 OS | RPL36A | 12.441 | 3 | 53.547813 | no | 9.819E-08 | 719.87549 | 672.82617 |
| QSGYGGQTKPIFR | 13 | P83881 | Large ribosomal subunit protein eL42 OS | RPL36A | 12.441 | 2 | 53.547813 | no | 1.435E-07 | 480.25278 | 672.82617 |
| QSGYGGQTKPIFR | 13 | P83881 | Large ribosomal subunit protein eL42 OS | RPL36A | 12.441 | 2 | 53.547813 | no | 9.819E-08 | 719.87549 | 672.82617 |
| DSLYAQGK | 8 | P83881 | Large ribosomal subunit protein eL42 OS | RPL36A | 12.441 | 2 | 53.547813 | yes | 1.097E-07 | 441.2218 | 233.21515 |
| KQSGYGGQTKPIF | 14 | P83881 | Large ribosomal subunit protein eL42 OS | RPL36A | 12.441 | 3 | 53.547813 | no | 3.314E-07 | 522.95111 | 106.55657 |
| CKHFELGGDK | 10 | P83881 | Large ribosomal subunit protein eL42 OS | RPL36A | 12.441 | 2 | 53.547813 | no | 9.722E-08 | 595.78479 | 91.742867 |
| NSDEADLVPAK | 11 | P83916 | Chromobox protein homolog 1 OS=Hom | CBX1 | 21.418 | 2 | 45.742462 | yes | 1.823E-07 | 579.78552 | 62.019245 |
| CPQVVISFYEER | 12 | P83916 | Chromobox protein homolog 1 OS=Hom | CBX1 | 21.418 | 2 | 45.742462 | yes | 1.755E-07 | 763.86902 | 61.038021 |
| VLTQMGSPSIR | 11 | P84022 | Mothers against decapentaplegic homolog | SMAD3 | 48.081 | 2 | 54.873993 | yes | 9.722E-08 | 594.82391 | 448.99628 |
| TGQLDELEK | 9 | P84022 | Mothers against decapentaplegic homolog | SMAD3 | 48.081 | 2 | 54.873993 | yes | 1.335E-07 | 516.76404 | 160.46587 |
| AITTQNVNTK | 10 | P84022 | Mothers against decapentaplegic homolog | SMAD3 | 48.081 | 2 | 54.873993 | yes | 3.147E-07 | 545.29858 | 157.84285 |
| AMELCEFAFNMK | 12 | P84022 | Mothers against decapentaplegic homolog | SMAD3 | 48.081 | 2 | 54.873993 | yes | 9.722E-08 | 745.82745 | 131.7226 |
| LGLQHLR | 7 | P84085 | ADP-ribosylation factor 5 OS=Homo sap | ARF5 | 20.53 | 2 | 48.14596 | yes | 1.385E-07 | 418.7587 | 159.40767 |
| MLQEDELR | 8 | P84085 | ADP-ribosylation factor 5 OS=Homo sap | ARF5 | 20.53 | 2 | 48.14596 | yes | 3.071E-07 | 517.25275 | 99.435387 |
| NICFTVWDVGGQI | 14 | P84085 | ADP-ribosylation factor 5 OS=Homo sap | ARF5 | 20.53 | 2 | 48.14596 | yes | 1.192E-07 | 819.88269 | 87.866913 |
| QDMPNAMPVSEL | 15 | P84085 | ADP-ribosylation factor 5 OS=Homo sap | ARF5 | 20.53 | 2 | 48.14596 | yes | 7.622E-07 | 838.3869 | 54.096268 |
| VQESADELQK | 10 | P84085 | ADP-ribosylation factor 5 OS=Homo sap | ARF5 | 20.53 | 2 | 48.14596 | yes | 2.568E-05 | 573.78552 | 21.707403 |
| SHTILLVQPTK | 11 | P84090 | Enhancer of rudimentary homolog OS=H | ERH | 12.259 | 2 | 53.72231 | yes | 1.088E-07 | 639.87445 | 5966.8081 |
| MYEEHLK | 7 | P84090 | Enhancer of rudimentary homolog OS=H | ERH | 12.259 | 2 | 53.72231 | yes | 2.564E-07 | 475.22601 | 4243.1025 |
| ADTQTYQPYNK | 11 | P84090 | Enhancer of rudimentary homolog OS=H | ERH | 12.259 | 2 | 53.72231 | yes | 9.722E-08 | 664.80951 | 242.93633 |
| YLECSALQQDGVI | 13 | P84095 | Rho-related GTP-binding protein RhoG C | RHOG | 21.309 | 2 | 51.200737 | yes | 9.722E-08 | 755.86395 | 284.30255 |

| | | | | | | | | | | | |
|---|---|---|---|---|---|---|---|---|---|---|---|
| AVLNPTPIK | 9 | P84095 | Rho-related GTP-binding protein RhoG ( | RHOG | 21.309 | 2 | 51.200737 | yes | 1.736E-07 | 476.79492 | 213.44537 |
| LLADQAEAR | 9 | P84098 | Large ribosomal subunit protein eL19 OS | RPL19 | 23.466 | 2 | 52.766087 | yes | 1.382E-07 | 493.76691 | 38675.105 |
| ILMEHIHK | 8 | P84098 | Large ribosomal subunit protein eL19 OS | RPL19 | 23.466 | 2 | 52.766087 | yes | 1.466E-07 | 510.78659 | 13687.561 |
| ILMEHIHK | 8 | P84098 | Large ribosomal subunit protein eL19 OS | RPL19 | 23.466 | 2 | 52.766087 | yes | 1.1E-07 | 518.78406 | 13687.561 |
| KLLADQAEAR | 10 | P84098 | Large ribosomal subunit protein eL19 OS | RPL19 | 23.466 | 2 | 52.766087 | yes | 9.722E-08 | 557.81439 | 2303.8948 |
| HMYHSLYLK | 9 | P84098 | Large ribosomal subunit protein eL19 OS | RPL19 | 23.466 | 2 | 52.766087 | yes | 2.987E-07 | 596.30261 | 1069.8246 |
| HMYHSLYLK | 9 | P84098 | Large ribosomal subunit protein eL19 OS | RPL19 | 23.466 | 2 | 52.766087 | yes | 9.722E-08 | 604.30005 | 1069.8246 |
| KVWLDPNETNEIA | 18 | P84098 | Large ribosomal subunit protein eL19 OS | RPL19 | 23.466 | 3 | 52.766087 | yes | 9.926E-08 | 691.0116 | 158.06268 |
| LIKDGLIIR | 9 | P84098 | Large ribosomal subunit protein eL19 OS | RPL19 | 23.466 | 2 | 52.766087 | yes | 5.913E-06 | 520.84497 | 105.83762 |
| VWLDPNETNEIAN | 17 | P84098 | Large ribosomal subunit protein eL19 OS | RPL19 | 23.466 | 3 | 52.766087 | yes | 1.297E-07 | 648.31329 | 38.296898 |
| AFGYYGPLR | 9 | P84103 | Serine/arginine-rich splicing factor 3 OS= | SRSF3 | 19.33 | 2 | 50.30808 | yes | 1.295E-07 | 522.2691 | 34331.477 |
| NPPGFAFVEFEDPI | 14 | P84103 | Serine/arginine-rich splicing factor 3 OS= | SRSF3 | 19.33 | 2 | 50.30808 | no | 1.157E-06 | 811.38593 | 14420.65 |
| VYVGNLGNNGNK | 12 | P84103 | Serine/arginine-rich splicing factor 3 OS= | SRSF3 | 19.33 | 2 | 50.30808 | yes | 9.819E-08 | 624.82019 | 1391.2632 |
| GPPPSWGR | 8 | P84103 | Serine/arginine-rich splicing factor 3 OS= | SRSF3 | 19.33 | 2 | 50.30808 | yes | 3.268E-06 | 427.21939 | 656.76086 |
| VELSNGEK | 8 | P84103 | Serine/arginine-rich splicing factor 3 OS= | SRSF3 | 19.33 | 2 | 50.30808 | yes | 1.449E-07 | 438.22708 | 242.27638 |
| VRVELSNGEK | 10 | P84103 | Serine/arginine-rich splicing factor 3 OS= | SRSF3 | 19.33 | 2 | 50.30808 | yes | 1.473E-06 | 565.81183 | 14.794253 |
| LASVPEYR | 8 | P85037 | Forkhead box protein K1 OS=Homo sapi | FOXK1 | 75.458 | 2 | 51.698948 | yes | 9.722E-08 | 467.75327 | 434.64496 |
| TPFGPLSSR | 9 | P85037 | Forkhead box protein K1 OS=Homo sapi | FOXK1 | 75.458 | 2 | 51.698948 | yes | 1.022E-07 | 481.25873 | 226.1685 |
| IQFTSLYHK | 9 | P85037 | Forkhead box protein K1 OS=Homo sapi | FOXK1 | 75.458 | 2 | 51.698948 | yes | 9.722E-08 | 568.80859 | 176.29799 |
| HLQLSFQEPHFYLI | 14 | P85037 | Forkhead box protein K1 OS=Homo sapi | FOXK1 | 75.458 | 3 | 51.698948 | yes | 1.508E-06 | 605.64941 | 173.63922 |
| SAPASPTHPGLMS | 15 | P85037 | Forkhead box protein K1 OS=Homo sapi | FOXK1 | 75.458 | 2 | 51.698948 | yes | 9.722E-08 | 753.38013 | 169.86684 |
| SGGLQTPECLSR | 12 | P85037 | Forkhead box protein K1 OS=Homo sapi | FOXK1 | 75.458 | 2 | 51.698948 | yes | 1.729E-07 | 652.81677 | 152.29675 |
| NGVFVDGAFQR | 11 | P85037 | Forkhead box protein K1 OS=Homo sapi | FOXK1 | 75.458 | 2 | 51.698948 | yes | 1.078E-06 | 605.3042 | 128.84648 |
| QPSVTIGR | 8 | P85037 | Forkhead box protein K1 OS=Homo sapi | FOXK1 | 75.458 | 2 | 51.698948 | yes | 3.44E-07 | 429.24561 | 57.163551 |
| AASEQQADTSGGI | 16 | P85037 | Forkhead box protein K1 OS=Homo sapi | FOXK1 | 75.458 | 2 | 51.698948 | yes | 9.727E-08 | 774.84229 | 53.20845 |
| GAPALQLPK | 9 | P85037 | Forkhead box protein K1 OS=Homo sapi | FOXK1 | 75.458 | 2 | 51.698948 | yes | 2.879E-06 | 447.77402 | 21.354511 |
| GLPDQELFSLNEG | 15 | P85299 | Proline-rich protein 5 OS=Homo sapiens | PRR5 | 42.754 | 2 | 26.926037 | yes | 3.043E-06 | 837.42828 | 21.787712 |
| TGQQAEPLVVDLI | 13 | P98082 | Disabled homolog 2 OS=Homo sapiens C | DAB2 | 82.448 | 2 | 40.602264 | yes | 3.939E-07 | 699.38538 | 62.632092 |
| QCCDCCGLGLR | 11 | P98095 | Fibulin-2 OS=Homo sapiens OX=9606 G | FBLN2 | 126.573 | 2 | 53.072304 | yes | 9.722E-08 | 699.77271 | 72.044113 |
| QGSVTTFLAK | 10 | P98095 | Fibulin-2 OS=Homo sapiens OX=9606 G | FBLN2 | 126.573 | 2 | 53.072304 | yes | 8.953E-07 | 526.29279 | 40.760193 |
| HQTHGSLLR | 9 | P98160 | Basement membrane-specific heparan sul | HSPG2 | 468.83 | 2 | 47.83542 | yes | 2.05E-07 | 524.78595 | 139.80074 |
| YELGSGLAVLR | 11 | P98160 | Basement membrane-specific heparan sul | HSPG2 | 468.83 | 2 | 47.83542 | yes | 1.248E-07 | 589.3324 | 129.15057 |
| SPAYTLVWTR | 10 | P98160 | Basement membrane-specific heparan sul | HSPG2 | 468.83 | 2 | 47.83542 | yes | 5.917E-07 | 597.31934 | 84.350983 |
| LPQVSPADSGEYV | 15 | P98160 | Basement membrane-specific heparan sul | HSPG2 | 468.83 | 2 | 47.83542 | yes | 3.608E-06 | 839.39868 | 14.577021 |
| TLQLSGSTYCMPM | 26 | P98164 | Low-density lipoprotein receptor-related | LRP2 | 521.958 | 4 | 41.137886 | yes | 3.657E-07 | 757.8299 | 8.7202435 |
| VFPPLTSLPGPVYI | 15 | P98171 | Rho GTPase-activating protein 4 OS=Ho | ARHGAP4 | 105.026 | 2 | 46.5237 | yes | 1.219E-06 | 822.45581 | 156.74202 |
| WQLSEQLR | 8 | P98171 | Rho GTPase-activating protein 4 OS=Ho | ARHGAP4 | 105.026 | 2 | 46.5237 | yes | 2.778E-06 | 530.28271 | 89.494545 |
| LSHIAEDVGR | 10 | P98171 | Rho GTPase-activating protein 4 OS=Ho | ARHGAP4 | 105.026 | 2 | 46.5237 | yes | 1.747E-07 | 548.79089 | 63.991001 |
| VNQLVQTLIVQPD | 14 | P98171 | Rho GTPase-activating protein 4 OS=Ho | ARHGAP4 | 105.026 | 2 | 46.5237 | yes | 2.647E-07 | 811.96484 | 53.571503 |

| | | | | | | | | | | | |
|---|---|---|---|---|---|---|---|---|---|---|---|
| ESAALSEVLAGPL | 16 | P98171 | Rho GTPase-activating protein 4 OS=Hoı | ARHGAP4 | 105.026 | 2 | 46.5237 | yes | 1.537E-07 | 806.43866 | 49.796368 |
| QGLGPASTTSPSPC | 16 | P98171 | Rho GTPase-activating protein 4 OS=Hoı | ARHGAP4 | 105.026 | 2 | 46.5237 | yes | 3.635E-06 | 755.38641 | 13.393791 |
| LDQQTLPLGGR | 11 | P98175 | RNA-binding protein 10 OS=Homo sapie | RBM10 | 103.533 | 2 | 51.478638 | yes | 1.146E-07 | 599.33295 | 3852.7507 |
| DGLGSDNIGSR | 11 | P98175 | RNA-binding protein 10 OS=Homo sapie | RBM10 | 103.533 | 2 | 51.478638 | yes | 2.961E-07 | 545.75983 | 2932.2986 |
| NSFQPISSLR | 10 | P98175 | RNA-binding protein 10 OS=Homo sapie | RBM10 | 103.533 | 2 | 51.478638 | yes | 1.465E-07 | 574.80658 | 2842.6902 |
| LACLLCR | 7 | P98175 | RNA-binding protein 10 OS=Homo sapie | RBM10 | 103.533 | 2 | 51.478638 | no | 1.065E-07 | 453.2384 | 2799.3892 |
| MLPQAATEDDIR | 12 | P98175 | RNA-binding protein 10 OS=Homo sapie | RBM10 | 103.533 | 2 | 51.478638 | yes | 5.253E-07 | 680.33228 | 1129.1918 |
| MLPQAATEDDIR | 12 | P98175 | RNA-binding protein 10 OS=Homo sapie | RBM10 | 103.533 | 2 | 51.478638 | yes | 3.555E-06 | 688.32971 | 1129.1918 |
| TYVPALEQSADGH | 14 | P98175 | RNA-binding protein 10 OS=Homo sapie | RBM10 | 103.533 | 2 | 51.478638 | yes | 1.752E-07 | 758.37555 | 1121.6971 |
| TYVPALEQSADGH | 14 | P98175 | RNA-binding protein 10 OS=Homo sapie | RBM10 | 103.533 | 2 | 51.478638 | yes | 3.079E-06 | 505.91946 | 1121.6971 |
| TYVPALEQSADGH | 14 | P98175 | RNA-binding protein 10 OS=Homo sapie | RBM10 | 103.533 | 3 | 51.478638 | yes | 1.752E-07 | 758.37555 | 1121.6971 |
| TYVPALEQSADGH | 14 | P98175 | RNA-binding protein 10 OS=Homo sapie | RBM10 | 103.533 | 3 | 51.478638 | yes | 3.079E-06 | 505.91946 | 1121.6971 |
| YGIPEPPEPK | 10 | P98175 | RNA-binding protein 10 OS=Homo sapie | RBM10 | 103.533 | 2 | 51.478638 | no | 2.37E-07 | 563.7926 | 1856.921 |
| ESATADAGYAILE | 14 | P98175 | RNA-binding protein 10 OS=Homo sapie | RBM10 | 103.533 | 2 | 51.478638 | yes | 9.722E-08 | 719.85663 | 1759.4408 |
| GSSYGVTSTESYK | 13 | P98175 | RNA-binding protein 10 OS=Homo sapie | RBM10 | 103.533 | 2 | 51.478638 | yes | 9.819E-08 | 683.31207 | 1486.3893 |
| GQLQSHGVQAR | 11 | P98175 | RNA-binding protein 10 OS=Homo sapie | RBM10 | 103.533 | 2 | 51.478638 | yes | 1.125E-07 | 590.81268 | 1311.4838 |
| INEDWLCNK | 9 | P98175 | RNA-binding protein 10 OS=Homo sapie | RBM10 | 103.533 | 2 | 51.478638 | yes | 2.189E-07 | 596.27679 | 1003.4448 |
| VSMHYSDPKPK | 11 | P98175 | RNA-binding protein 10 OS=Homo sapie | RBM10 | 103.533 | 2 | 51.478638 | yes | 1.145E-07 | 644.82135 | 767.62762 |
| QGIVTPIEAQTR | 12 | P98175 | RNA-binding protein 10 OS=Homo sapie | RBM10 | 103.533 | 2 | 51.478638 | yes | 1.011E-07 | 656.86462 | 688.07269 |
| AHLSENELEALEK | 13 | P98175 | RNA-binding protein 10 OS=Homo sapie | RBM10 | 103.533 | 2 | 51.478638 | yes | 1.231E-07 | 741.87537 | 598.75861 |
| ASNIVMLR | 8 | P98175 | RNA-binding protein 10 OS=Homo sapie | RBM10 | 103.533 | 2 | 51.478638 | yes | 1.532E-07 | 452.25766 | 562.39264 |
| CGVQNFK | 7 | P98175 | RNA-binding protein 10 OS=Homo sapie | RBM10 | 103.533 | 2 | 51.478638 | yes | 2.226E-07 | 426.70526 | 558.33875 |
| QHTSMDLPK | 9 | P98175 | RNA-binding protein 10 OS=Homo sapie | RBM10 | 103.533 | 2 | 51.478638 | yes | 6.452E-07 | 528.76074 | 519.63855 |
| TINVEFAK | 8 | P98175 | RNA-binding protein 10 OS=Homo sapie | RBM10 | 103.533 | 2 | 51.478638 | yes | 1.153E-07 | 461.25565 | 443.7413 |
| HSPTGPPGFPR | 11 | P98175 | RNA-binding protein 10 OS=Homo sapie | RBM10 | 103.533 | 2 | 51.478638 | yes | 2.348E-07 | 575.29364 | 207.7403 |
| RESATADAGYAIL | 15 | P98175 | RNA-binding protein 10 OS=Homo sapie | RBM10 | 103.533 | 3 | 51.478638 | yes | 1.003E-07 | 532.27386 | 64.017151 |
| RESATADAGYAIL | 15 | P98175 | RNA-binding protein 10 OS=Homo sapie | RBM10 | 103.533 | 3 | 51.478638 | yes | 4.543E-07 | 797.90723 | 64.017151 |
| RESATADAGYAIL | 15 | P98175 | RNA-binding protein 10 OS=Homo sapie | RBM10 | 103.533 | 2 | 51.478638 | yes | 1.003E-07 | 532.27386 | 64.017151 |
| RESATADAGYAIL | 15 | P98175 | RNA-binding protein 10 OS=Homo sapie | RBM10 | 103.533 | 2 | 51.478638 | yes | 4.543E-07 | 797.90723 | 64.017151 |
| EKYGIPEPPEPK | 12 | P98175 | RNA-binding protein 10 OS=Homo sapie | RBM10 | 103.533 | 2 | 51.478638 | no | 2.335E-06 | 692.36139 | 72.199272 |
| KQGIVTPIEAQTR | 13 | P98175 | RNA-binding protein 10 OS=Homo sapie | RBM10 | 103.533 | 2 | 51.478638 | yes | 9.728E-08 | 720.91211 | 49.006779 |
| DKQTQLNR | 8 | P98175 | RNA-binding protein 10 OS=Homo sapie | RBM10 | 103.533 | 2 | 51.478638 | yes | 1.107E-06 | 501.76999 | 20.411539 |
| PGGYGYGYGR | 10 | P98179 | RNA-binding protein 3 OS=Homo sapien | RBM3 | 17.171 | 2 | 48.904213 | yes | 1.088E-07 | 523.73816 | 2034.391 |
| AMNGESLDGR | 10 | P98179 | RNA-binding protein 3 OS=Homo sapien | RBM3 | 17.171 | 2 | 48.904213 | yes | 1.233E-07 | 525.23767 | 247.15627 |
| GFGFITFTNPEHAS | 18 | P98179 | RNA-binding protein 3 OS=Homo sapien | RBM3 | 17.171 | 3 | 48.904213 | yes | 1.896E-07 | 661.32446 | 79.134979 |
| TGPNLHGLFGR | 11 | P99999 | Cytochrome c OS=Homo sapiens OX=96 | CYCS | 11.749 | 2 | 48.309624 | yes | 1.165E-07 | 584.8147 | 160.53041 |
| EQLTPSQIMSLEK | 13 | Q00059 | Transcription factor A, mitochondrial OS | TFAM | 29.097 | 2 | 52.27366 | yes | 1.645E-07 | 752.38977 | 2998.9688 |
| EQLTPSQIMSLEK | 13 | Q00059 | Transcription factor A, mitochondrial OS | TFAM | 29.097 | 2 | 52.27366 | yes | 9.722E-08 | 760.38727 | 2998.9688 |
| KPVSSYLR | 8 | Q00059 | Transcription factor A, mitochondrial OS | TFAM | 29.097 | 2 | 52.27366 | yes | 9.997E-08 | 475.27692 | 5829.0542 |

| | | | | | | | | | | | |
|---|---|---|---|---|---|---|---|---|---|---|---|
| ELYIQHAK | 8 | Q00059 | Transcription factor A, mitochondrial OS | TFAM | 29.097 | 2 | 52.27366 | yes | 2.537E-07 | 501.27435 | 3152.3279 |
| ELTLLGKPK | 9 | Q00059 | Transcription factor A, mitochondrial OS | TFAM | 29.097 | 2 | 52.27366 | yes | 2.303E-07 | 499.81586 | 2065.6423 |
| SAYNVYVAER | 10 | Q00059 | Transcription factor A, mitochondrial OS | TFAM | 29.097 | 2 | 52.27366 | yes | 9.722E-08 | 586.29077 | 1588.8162 |
| EQLPIFK | 7 | Q00059 | Transcription factor A, mitochondrial OS | TFAM | 29.097 | 2 | 52.27366 | yes | 2.314E-06 | 437.75528 | 685.08356 |
| AEWQVYK | 7 | Q00059 | Transcription factor A, mitochondrial OS | TFAM | 29.097 | 2 | 52.27366 | yes | 2.125E-07 | 462.23471 | 452.33212 |
| KIYQDAYR | 8 | Q00059 | Transcription factor A, mitochondrial OS | TFAM | 29.097 | 2 | 52.27366 | yes | 9.809E-08 | 528.77728 | 155.67462 |
| PVSSYLR | 7 | Q00059 | Transcription factor A, mitochondrial OS | TFAM | 29.097 | 2 | 52.27366 | yes | 5.025E-07 | 411.22943 | 119.23187 |
| KELTLLGK | 8 | Q00059 | Transcription factor A, mitochondrial OS | TFAM | 29.097 | 2 | 52.27366 | yes | 1.007E-07 | 451.28949 | 61.604401 |
| IQTQPGYANTLR | 12 | Q00325 | Solute carrier family 25 member 3 OS=H | SLC25A3 | 40.095 | 2 | 49.408104 | yes | 1.992E-07 | 681.36224 | 6916.7139 |
| FGFYEVFK | 8 | Q00325 | Solute carrier family 25 member 3 OS=H | SLC25A3 | 40.095 | 2 | 49.408104 | yes | 3.51E-07 | 518.76056 | 1947.0743 |
| LPRPPPPEMPESLK | 14 | Q00325 | Solute carrier family 25 member 3 OS=H | SLC25A3 | 40.095 | 3 | 49.408104 | yes | 1.078E-06 | 529.95697 | 520.71576 |
| LPRPPPPEMPESLK | 14 | Q00325 | Solute carrier family 25 member 3 OS=H | SLC25A3 | 40.095 | 3 | 49.408104 | yes | 1.138E-07 | 794.43182 | 520.71576 |
| LPRPPPPEMPESLK | 14 | Q00325 | Solute carrier family 25 member 3 OS=H | SLC25A3 | 40.095 | 2 | 49.408104 | yes | 1.078E-06 | 529.95697 | 520.71576 |
| LPRPPPPEMPESLK | 14 | Q00325 | Solute carrier family 25 member 3 OS=H | SLC25A3 | 40.095 | 2 | 49.408104 | yes | 1.138E-07 | 794.43182 | 520.71576 |
| LPRPPPPEMPESLK | 15 | Q00325 | Solute carrier family 25 member 3 OS=H | SLC25A3 | 40.095 | 3 | 49.408104 | yes | 4.89E-07 | 572.65527 | 584.30658 |
| GVAPLWMR | 8 | Q00325 | Solute carrier family 25 member 3 OS=H | SLC25A3 | 40.095 | 2 | 49.408104 | yes | 1.437E-07 | 465.25491 | 243.90965 |
| GVAPLWMR | 8 | Q00325 | Solute carrier family 25 member 3 OS=H | SLC25A3 | 40.095 | 2 | 49.408104 | yes | 3.279E-07 | 473.25238 | 243.90965 |
| MYKEEGLK | 8 | Q00325 | Solute carrier family 25 member 3 OS=H | SLC25A3 | 40.095 | 2 | 49.408104 | yes | 1.107E-07 | 499.25479 | 466.12811 |
| GIFNGFSVTLK | 11 | Q00325 | Solute carrier family 25 member 3 OS=H | SLC25A3 | 40.095 | 2 | 49.408104 | yes | 1.035E-07 | 591.82953 | 354.55469 |
| FVVPKPR | 7 | Q00325 | Solute carrier family 25 member 3 OS=H | SLC25A3 | 40.095 | 2 | 49.408104 | yes | 1.071E-07 | 421.76599 | 239.10977 |
| ICLEIMQR | 8 | Q00341 | Vigilin OS=Homo sapiens OX=9606 GN | HDLBP | 141.456 | 2 | 55.498333 | yes | 9.722E-08 | 531.77539 | 807.76501 |
| ELQAEQEDR | 9 | Q00341 | Vigilin OS=Homo sapiens OX=9606 GN | HDLBP | 141.456 | 2 | 55.498333 | yes | 9.722E-08 | 559.25964 | 728.54327 |
| MDYVEINIDHK | 11 | Q00341 | Vigilin OS=Homo sapiens OX=9606 GN | HDLBP | 141.456 | 2 | 55.498333 | yes | 1.047E-07 | 688.82941 | 698.24139 |
| HEVLLISAEQDKR | 13 | Q00341 | Vigilin OS=Homo sapiens OX=9606 GN | HDLBP | 141.456 | 3 | 55.498333 | yes | 9.722E-08 | 513.28259 | 324.15076 |
| HEVLLISAEQDKR | 13 | Q00341 | Vigilin OS=Homo sapiens OX=9606 GN | HDLBP | 141.456 | 3 | 55.498333 | yes | 1.099E-07 | 769.42029 | 324.15076 |
| HEVLLISAEQDKR | 13 | Q00341 | Vigilin OS=Homo sapiens OX=9606 GN | HDLBP | 141.456 | 2 | 55.498333 | yes | 9.722E-08 | 513.28259 | 324.15076 |
| HEVLLISAEQDKR | 13 | Q00341 | Vigilin OS=Homo sapiens OX=9606 GN | HDLBP | 141.456 | 2 | 55.498333 | yes | 1.099E-07 | 769.42029 | 324.15076 |
| LQTQASATVAIPK | 13 | Q00341 | Vigilin OS=Homo sapiens OX=9606 GN | HDLBP | 141.456 | 2 | 55.498333 | yes | 9.887E-08 | 664.38263 | 503.30652 |
| IEGDPQGVQQAK | 12 | Q00341 | Vigilin OS=Homo sapiens OX=9606 GN | HDLBP | 141.456 | 2 | 55.498333 | yes | 5.083E-07 | 635.32532 | 467.02722 |
| LGQALTEVYAK | 11 | Q00341 | Vigilin OS=Homo sapiens OX=9606 GN | HDLBP | 141.456 | 2 | 55.498333 | yes | 1.865E-07 | 596.83228 | 458.85934 |
| EQLAQAVAR | 9 | Q00341 | Vigilin OS=Homo sapiens OX=9606 GN | HDLBP | 141.456 | 2 | 55.498333 | yes | 1.071E-06 | 493.2749 | 410.95441 |
| INIPPPSVNR | 10 | Q00341 | Vigilin OS=Homo sapiens OX=9606 GN | HDLBP | 141.456 | 2 | 55.498333 | yes | 1.06E-07 | 553.81946 | 407.07889 |
| DMNQFGEGEQAK | 12 | Q00341 | Vigilin OS=Homo sapiens OX=9606 GN | HDLBP | 141.456 | 2 | 55.498333 | yes | 9.722E-08 | 677.29059 | 326.57275 |
| LHNSLIGTK | 9 | Q00341 | Vigilin OS=Homo sapiens OX=9606 GN | HDLBP | 141.456 | 2 | 55.498333 | yes | 1.885E-07 | 491.78763 | 275.90833 |
| ELLELASR | 8 | Q00341 | Vigilin OS=Homo sapiens OX=9606 GN | HDLBP | 141.456 | 2 | 55.498333 | yes | 9.734E-08 | 465.76639 | 257.30472 |
| DQGLSIMVSGK | 11 | Q00341 | Vigilin OS=Homo sapiens OX=9606 GN | HDLBP | 141.456 | 2 | 55.498333 | yes | 9.722E-08 | 567.7948 | 255.90485 |
| TGAHLELSLAK | 11 | Q00341 | Vigilin OS=Homo sapiens OX=9606 GN | HDLBP | 141.456 | 2 | 55.498333 | yes | 9.722E-08 | 570.32458 | 209.40739 |
| VHIEFTEGEDK | 11 | Q00341 | Vigilin OS=Homo sapiens OX=9606 GN | HDLBP | 141.456 | 2 | 55.498333 | yes | 1.779E-07 | 652.31189 | 206.91417 |
| DLANIAEVEVSIPA | 15 | Q00341 | Vigilin OS=Homo sapiens OX=9606 GN | HDLBP | 141.456 | 2 | 55.498333 | yes | 1.67E-07 | 784.93018 | 205.3353 |

| | | | | | | | | | | | |
|---|---|---|---|---|---|---|---|---|---|---|---|
| LVGEIMQETGTR | 12 | Q00341 | Vigilin OS=Homo sapiens OX=9606 GN | HDLBP | 141.456 | 2 | 55.498333 | yes | 1.082E-07 | 667.34265 | 193.05995 |
| KGAVITQIR | 9 | Q00341 | Vigilin OS=Homo sapiens OX=9606 GN | HDLBP | 141.456 | 2 | 55.498333 | yes | 2.3E-07 | 493.31128 | 190.52721 |
| EIAEEYGGVMVSF | 15 | Q00341 | Vigilin OS=Homo sapiens OX=9606 GN | HDLBP | 141.456 | 2 | 55.498333 | yes | 9.722E-08 | 842.40601 | 167.23438 |
| VIFPAAEDKDQDL | 19 | Q00341 | Vigilin OS=Homo sapiens OX=9606 GN | HDLBP | 141.456 | 3 | 55.498333 | yes | 1.117E-07 | 696.05255 | 138.58388 |
| IQIPRPDDPSNQIK | 14 | Q00341 | Vigilin OS=Homo sapiens OX=9606 GN | HDLBP | 141.456 | 2 | 55.498333 | yes | 3.845E-07 | 810.93884 | 67.553848 |
| IQIPRPDDPSNQIK | 14 | Q00341 | Vigilin OS=Homo sapiens OX=9606 GN | HDLBP | 141.456 | 2 | 55.498333 | yes | 1.049E-06 | 540.96167 | 67.553848 |
| IQIPRPDDPSNQIK | 14 | Q00341 | Vigilin OS=Homo sapiens OX=9606 GN | HDLBP | 141.456 | 3 | 55.498333 | yes | 3.845E-07 | 810.93884 | 67.553848 |
| IQIPRPDDPSNQIK | 14 | Q00341 | Vigilin OS=Homo sapiens OX=9606 GN | HDLBP | 141.456 | 3 | 55.498333 | yes | 1.049E-06 | 540.96167 | 67.553848 |
| QLLHLAEEK | 9 | Q00341 | Vigilin OS=Homo sapiens OX=9606 GN | HDLBP | 141.456 | 2 | 55.498333 | yes | 1.49E-07 | 540.80603 | 120.11218 |
| SDIVQLR | 7 | Q00341 | Vigilin OS=Homo sapiens OX=9606 GN | HDLBP | 141.456 | 2 | 55.498333 | yes | 9.621E-07 | 415.74017 | 106.5006 |
| DAPWTASSSEK | 11 | Q00341 | Vigilin OS=Homo sapiens OX=9606 GN | HDLBP | 141.456 | 2 | 55.498333 | yes | 1.828E-07 | 589.76984 | 93.679497 |
| CDIIIISGR | 9 | Q00341 | Vigilin OS=Homo sapiens OX=9606 GN | HDLBP | 141.456 | 2 | 55.498333 | yes | 6.639E-07 | 523.7868 | 91.484665 |
| SIMEECGGVHIHFI | 26 | Q00341 | Vigilin OS=Homo sapiens OX=9606 GN | HDLBP | 141.456 | 4 | 55.498333 | yes | 1.222E-07 | 703.84027 | 76.122253 |
| RCDIIIISGR | 10 | Q00341 | Vigilin OS=Homo sapiens OX=9606 GN | HDLBP | 141.456 | 2 | 55.498333 | yes | 1.658E-07 | 601.83734 | 64.703934 |
| TTTIAVEVK | 9 | Q00341 | Vigilin OS=Homo sapiens OX=9606 GN | HDLBP | 141.456 | 2 | 55.498333 | yes | 4.942E-07 | 481.28186 | 53.403168 |
| SFTVDIR | 7 | Q00341 | Vigilin OS=Homo sapiens OX=9606 GN | HDLBP | 141.456 | 2 | 55.498333 | yes | 2.858E-06 | 419.2269 | 53.256783 |
| HEVLLISAEQDK | 12 | Q00341 | Vigilin OS=Homo sapiens OX=9606 GN | HDLBP | 141.456 | 2 | 55.498333 | yes | 4.664E-07 | 691.36975 | 52.961979 |
| KTTTIAVEVK | 10 | Q00341 | Vigilin OS=Homo sapiens OX=9606 GN | HDLBP | 141.456 | 2 | 55.498333 | yes | 3.599E-07 | 545.32935 | 52.763901 |
| IQEIIEDLEAQVTLI | 21 | Q00341 | Vigilin OS=Homo sapiens OX=9606 GN | HDLBP | 141.456 | 3 | 55.498333 | yes | 9.9E-08 | 814.09283 | 41.615505 |
| FPDREENAVHSTE | 27 | Q00341 | Vigilin OS=Homo sapiens OX=9606 GN | HDLBP | 141.456 | 4 | 55.498333 | yes | 1.555E-06 | 742.83807 | 39.220009 |
| DKFPEVIINFPDPA | 16 | Q00341 | Vigilin OS=Homo sapiens OX=9606 GN | HDLBP | 141.456 | 3 | 55.498333 | yes | 3.231E-07 | 619.99634 | 20.156782 |
| VIDVGSEWR | 9 | Q00403 | Transcription initiation factor IIB OS=Ho | GTF2B | 34.833 | 2 | 56.20659 | yes | 3.413E-07 | 530.77472 | 723.3988 |
| EIGDIAGVADVTIR | 14 | Q00403 | Transcription initiation factor IIB OS=Ho | GTF2B | 34.833 | 2 | 56.20659 | yes | 9.722E-08 | 714.88831 | 624.11127 |
| APDLFPTDFK | 10 | Q00403 | Transcription initiation factor IIB OS=Ho | GTF2B | 34.833 | 2 | 56.20659 | yes | 4.158E-06 | 575.7926 | 498.69431 |
| FCSNLCLPK | 9 | Q00403 | Transcription initiation factor IIB OS=Ho | GTF2B | 34.833 | 2 | 56.20659 | yes | 9.722E-08 | 569.77283 | 306.94562 |
| AVELDLVPGR | 10 | Q00403 | Transcription initiation factor IIB OS=Ho | GTF2B | 34.833 | 2 | 56.20659 | yes | 9.722E-08 | 534.80603 | 267.9313 |
| QVQMAATHIAR | 11 | Q00403 | Transcription initiation factor IIB OS=Ho | GTF2B | 34.833 | 2 | 56.20659 | yes | 1.027E-06 | 613.32715 | 249.92938 |
| GTGAASFDEFGNS | 14 | Q00403 | Transcription initiation factor IIB OS=Ho | GTF2B | 34.833 | 2 | 56.20659 | yes | 1.091E-07 | 694.30988 | 246.7724 |
| KAVELDLVPGR | 11 | Q00403 | Transcription initiation factor IIB OS=Ho | GTF2B | 34.833 | 2 | 56.20659 | yes | 9.722E-08 | 598.85352 | 162.90665 |
| FDTPVDKLPQL | 11 | Q00403 | Transcription initiation factor IIB OS=Ho | GTF2B | 34.833 | 2 | 56.20659 | yes | 9.722E-08 | 636.84534 | 37.860882 |
| ANDAIASACLYIA( | 15 | Q00403 | Transcription initiation factor IIB OS=Ho | GTF2B | 34.833 | 2 | 56.20659 | yes | 4.491E-07 | 834.89526 | 21.166265 |
| DEAAPLNPGMYS( | 14 | Q00444 | Homeobox protein Hox-C5 OS=Homo sa | HOXC5 | 24.977 | 2 | 51.743427 | yes | 9.722E-08 | 760.85614 | 309.14334 |
| AHPDRPACSAAAA | 20 | Q00444 | Homeobox protein Hox-C5 OS=Homo sa | HOXC5 | 24.977 | 3 | 51.743427 | yes | 1.377E-07 | 656.3186 | 188.80379 |
| LFDVCTVSR | 9 | Q00534 | Cyclin-dependent kinase 6 OS=Homo sap | CDK6 | 36.939 | 2 | 41.095554 | yes | 3.679E-07 | 548.77643 | 59.461342 |
| DLKPQNLLINR | 11 | Q00535 | Cyclin-dependent kinase 5 OS=Homo sap | CDK5 | 33.305 | 2 | 49.339256 | yes | 1.043E-07 | 662.39081 | 204.34698 |
| IGEGTYGTVFK | 11 | Q00535 | Cyclin-dependent kinase 5 OS=Homo sap | CDK5 | 33.305 | 2 | 49.339256 | yes | 1.166E-07 | 586.30334 | 109.80833 |
| LDDDDEGVPSSAL | 14 | Q00535 | Cyclin-dependent kinase 5 OS=Homo sap | CDK5 | 33.305 | 2 | 49.339256 | yes | 1.788E-07 | 744.84424 | 100.16003 |
| PLFPGNDVDDQLK | 13 | Q00535 | Cyclin-dependent kinase 5 OS=Homo sap | CDK5 | 33.305 | 2 | 49.339256 | yes | 2.501E-05 | 729.36719 | 88.990578 |
| GLGFCHSR | 8 | Q00535 | Cyclin-dependent kinase 5 OS=Homo sap | CDK5 | 33.305 | 2 | 49.339256 | yes | 9.134E-07 | 467.22162 | 38.909836 |

| | | | | | | | | | | | |
|---|---|---|---|---|---|---|---|---|---|---|---|
| LADFGLAR | 8 | Q00537 | Cyclin-dependent kinase 17 OS=Homo sa | CDK17 | 59.583 | 2 | 52.93346 | no | 1.076E-07 | 431.74271 | 1203.7883 |
| DLKPQNLLINEK | 12 | Q00537 | Cyclin-dependent kinase 17 OS=Homo sa | CDK17 | 59.583 | 2 | 52.93346 | yes | 9.733E-08 | 712.909 | 132.56671 |
| LQINSPPFDQPMSI | 14 | Q00537 | Cyclin-dependent kinase 17 OS=Homo sa | CDK17 | 59.583 | 2 | 52.93346 | yes | 2.149E-06 | 815.40631 | 88.056099 |
| ISMEDLNK | 8 | Q00537 | Cyclin-dependent kinase 17 OS=Homo sa | CDK17 | 59.583 | 2 | 52.93346 | yes | 5.505E-07 | 475.2366 | 80.671524 |
| LEHEEGAPCTAIR | 13 | Q00537 | Cyclin-dependent kinase 17 OS=Homo sa | CDK17 | 59.583 | 2 | 52.93346 | no | 9.722E-08 | 741.85394 | 78.289886 |
| LTENLVALK | 9 | Q00537 | Cyclin-dependent kinase 17 OS=Homo sa | CDK17 | 59.583 | 2 | 52.93346 | no | 1.362E-07 | 500.80551 | 74.305962 |
| LGEGTYATVYK | 11 | Q00537 | Cyclin-dependent kinase 17 OS=Homo sa | CDK17 | 59.583 | 2 | 52.93346 | no | 1.362E-07 | 601.30859 | 58.025188 |
| ASLSEIGFGK | 10 | Q00537 | Cyclin-dependent kinase 17 OS=Homo sa | CDK17 | 59.583 | 2 | 52.93346 | yes | 8.649E-07 | 504.77167 | 47.216461 |
| LDIVHENLK | 9 | Q00537 | Cyclin-dependent kinase 17 OS=Homo sa | CDK17 | 59.583 | 2 | 52.93346 | yes | 9.722E-08 | 540.80603 | 43.333103 |
| FFFDVGSNK | 9 | Q00577 | Transcriptional activator protein Pur-alph | PURA | 34.911 | 2 | 51.6328 | yes | 2.519E-07 | 530.75854 | 10423.026 |
| KYYMDLK | 7 | Q00577 | Transcriptional activator protein Pur-alph | PURA | 34.911 | 2 | 51.6328 | yes | 1.341E-07 | 480.74658 | 2656.7168 |
| GPGLGSTQGQTIAI | 23 | Q00577 | Transcriptional activator protein Pur-alph | PURA | 34.911 | 3 | 51.6328 | yes | 9.722E-08 | 771.08502 | 1351.9614 |
| VSEVKPTYR | 9 | Q00577 | Transcriptional activator protein Pur-alph | PURA | 34.911 | 2 | 51.6328 | yes | 2.93E-07 | 539.79822 | 1140.4122 |
| LTLSMSVAVEFR | 12 | Q00577 | Transcriptional activator protein Pur-alph | PURA | 34.911 | 2 | 51.6328 | yes | 9.722E-08 | 676.86578 | 305.43607 |
| NSITVPYKVWAK | 12 | Q00577 | Transcriptional activator protein Pur-alph | PURA | 34.911 | 2 | 51.6328 | yes | 9.722E-08 | 703.39557 | 279.46356 |
| NSITVPYK | 8 | Q00577 | Transcriptional activator protein Pur-alph | PURA | 34.911 | 2 | 51.6328 | yes | 2.809E-07 | 461.25565 | 226.81252 |
| FGHTFCK | 7 | Q00577 | Transcriptional activator protein Pur-alph | PURA | 34.911 | 2 | 51.6328 | yes | 1.069E-07 | 448.70779 | 117.6412 |
| RFFFDVGSNK | 10 | Q00577 | Transcriptional activator protein Pur-alph | PURA | 34.911 | 2 | 51.6328 | yes | 1.988E-06 | 608.80908 | 88.982437 |
| LSPVGWVSSSQGK | 13 | Q00587 | Cdc42 effector protein 1 OS=Homo sapie | CDC42EP1 | 40.295 | 2 | 52.988342 | yes | 1.421E-07 | 666.35132 | 366.51028 |
| LTADMISHPLGDFI | 14 | Q00587 | Cdc42 effector protein 1 OS=Homo sapie | CDC42EP1 | 40.295 | 2 | 52.988342 | yes | 9.722E-08 | 786.89557 | 168.96332 |
| LTADMISHPLGDFI | 14 | Q00587 | Cdc42 effector protein 1 OS=Homo sapie | CDC42EP1 | 40.295 | 2 | 52.988342 | yes | 1.093E-07 | 524.9328 | 168.96332 |
| LTADMISHPLGDFI | 14 | Q00587 | Cdc42 effector protein 1 OS=Homo sapie | CDC42EP1 | 40.295 | 3 | 52.988342 | yes | 9.722E-08 | 786.89557 | 168.96332 |
| LTADMISHPLGDFI | 14 | Q00587 | Cdc42 effector protein 1 OS=Homo sapie | CDC42EP1 | 40.295 | 3 | 52.988342 | yes | 1.093E-07 | 524.9328 | 168.96332 |
| ASWESLDEEWR | 11 | Q00587 | Cdc42 effector protein 1 OS=Homo sapie | CDC42EP1 | 40.295 | 2 | 52.988342 | yes | 9.851E-08 | 704.31238 | 177.61743 |
| HELIEFR | 7 | Q00610 | Clathrin heavy chain 1 OS=Homo sapiens | CLTC | 191.615 | 2 | 55.411663 | yes | 1.644E-07 | 472.25345 | 1049.9677 |
| LLYNNVSNFGR | 11 | Q00610 | Clathrin heavy chain 1 OS=Homo sapiens | CLTC | 191.615 | 2 | 55.411663 | yes | 9.892E-08 | 648.83838 | 1008.9628 |
| VIQCFAETGQVQK | 13 | Q00610 | Clathrin heavy chain 1 OS=Homo sapiens | CLTC | 191.615 | 2 | 55.411663 | yes | 9.733E-08 | 754.38232 | 965.06653 |
| HDVVFLITK | 9 | Q00610 | Clathrin heavy chain 1 OS=Homo sapiens | CLTC | 191.615 | 2 | 55.411663 | yes | 2.976E-07 | 536.31348 | 911.13745 |
| IYIDSNNNPER | 11 | Q00610 | Clathrin heavy chain 1 OS=Homo sapiens | CLTC | 191.615 | 2 | 55.411663 | yes | 9.722E-08 | 667.82037 | 897.31653 |
| NNLAGAEELFAR | 12 | Q00610 | Clathrin heavy chain 1 OS=Homo sapiens | CLTC | 191.615 | 2 | 55.411663 | yes | 1.075E-07 | 652.83331 | 897.21692 |
| EHLELFWSR | 9 | Q00610 | Clathrin heavy chain 1 OS=Homo sapiens | CLTC | 191.615 | 2 | 55.411663 | yes | 1.141E-07 | 608.80908 | 847.48395 |
| VANVELYYR | 9 | Q00610 | Clathrin heavy chain 1 OS=Homo sapiens | CLTC | 191.615 | 2 | 55.411663 | yes | 9.722E-08 | 563.79822 | 531.1004 |
| EVCFACVDGK | 10 | Q00610 | Clathrin heavy chain 1 OS=Homo sapiens | CLTC | 191.615 | 2 | 55.411663 | yes | 9.934E-08 | 592.75739 | 527.28326 |
| ALEHFTDLYDIK | 12 | Q00610 | Clathrin heavy chain 1 OS=Homo sapiens | CLTC | 191.615 | 2 | 55.411663 | yes | 1.033E-07 | 732.87207 | 526.73022 |
| WLLLTGISAQQNR | 13 | Q00610 | Clathrin heavy chain 1 OS=Homo sapiens | CLTC | 191.615 | 2 | 55.411663 | yes | 3.256E-07 | 750.4201 | 468.97134 |
| YESLELCR | 8 | Q00610 | Clathrin heavy chain 1 OS=Homo sapiens | CLTC | 191.615 | 2 | 55.411663 | yes | 1.172E-07 | 535.25275 | 419.40991 |
| LASTLVHLGEYQA | 19 | Q00610 | Clathrin heavy chain 1 OS=Homo sapiens | CLTC | 191.615 | 3 | 55.411663 | yes | 1.086E-07 | 657.68134 | 417.36075 |
| VDKLDASESLR | 11 | Q00610 | Clathrin heavy chain 1 OS=Homo sapiens | CLTC | 191.615 | 2 | 55.411663 | yes | 9.722E-08 | 616.8277 | 413.6637 |
| ISGETIFVTAPHEA | 23 | Q00610 | Clathrin heavy chain 1 OS=Homo sapiens | CLTC | 191.615 | 3 | 55.411663 | yes | 1.376E-07 | 785.0885 | 383.19028 |

| | | | | | | | | | | | |
|---|---|---|---|---|---|---|---|---|---|---|---|
| DAMQYASESK | 10 | Q00610 | Clathrin heavy chain 1 OS=Homo sapiens | CLTC | 191.615 | 2 | 55.411663 | yes | 9.722E-08 | 565.24512 | 347.31644 |
| HSSLAGCQIINYR | 13 | Q00610 | Clathrin heavy chain 1 OS=Homo sapiens | CLTC | 191.615 | 3 | 55.411663 | yes | 9.722E-08 | 506.9209 | 158.1424 |
| HSSLAGCQIINYR | 13 | Q00610 | Clathrin heavy chain 1 OS=Homo sapiens | CLTC | 191.615 | 3 | 55.411663 | yes | 9.722E-08 | 759.87775 | 158.1424 |
| HSSLAGCQIINYR | 13 | Q00610 | Clathrin heavy chain 1 OS=Homo sapiens | CLTC | 191.615 | 2 | 55.411663 | yes | 9.722E-08 | 506.9209 | 158.1424 |
| HSSLAGCQIINYR | 13 | Q00610 | Clathrin heavy chain 1 OS=Homo sapiens | CLTC | 191.615 | 2 | 55.411663 | yes | 9.722E-08 | 759.87775 | 158.1424 |
| CNEPAVWSQLAK | 12 | Q00610 | Clathrin heavy chain 1 OS=Homo sapiens | CLTC | 191.615 | 2 | 55.411663 | yes | 1.93E-07 | 701.84283 | 309.2915 |
| SVDPTLALSVYLR | 13 | Q00610 | Clathrin heavy chain 1 OS=Homo sapiens | CLTC | 191.615 | 2 | 55.411663 | yes | 1.296E-07 | 717.40356 | 204.47243 |
| ENPYYDSR | 8 | Q00610 | Clathrin heavy chain 1 OS=Homo sapiens | CLTC | 191.615 | 2 | 55.411663 | yes | 1.999E-07 | 522.2251 | 202.43347 |
| FNALFAQGNYSEA | 15 | Q00610 | Clathrin heavy chain 1 OS=Homo sapiens | CLTC | 191.615 | 2 | 55.411663 | yes | 1.614E-07 | 815.89661 | 178.10976 |
| LDASESLR | 8 | Q00610 | Clathrin heavy chain 1 OS=Homo sapiens | CLTC | 191.615 | 2 | 55.411663 | yes | 2.573E-07 | 445.73254 | 164.67094 |
| VMEYINR | 7 | Q00610 | Clathrin heavy chain 1 OS=Homo sapiens | CLTC | 191.615 | 2 | 55.411663 | yes | 9.722E-08 | 462.73401 | 153.83047 |
| AMLSANIR | 8 | Q00610 | Clathrin heavy chain 1 OS=Homo sapiens | CLTC | 191.615 | 2 | 55.411663 | yes | 3.371E-07 | 438.242 | 128.16216 |
| KDPELWGSVLLES | 17 | Q00610 | Clathrin heavy chain 1 OS=Homo sapiens | CLTC | 191.615 | 3 | 55.411663 | yes | 1.553E-07 | 668.34595 | 102.55042 |
| NLILVVR | 7 | Q00610 | Clathrin heavy chain 1 OS=Homo sapiens | CLTC | 191.615 | 2 | 55.411663 | yes | 1.096E-07 | 413.77908 | 98.781685 |
| KFDVNTSAVQVLI | 21 | Q00610 | Clathrin heavy chain 1 OS=Homo sapiens | CLTC | 191.615 | 3 | 55.411663 | yes | 9.722E-08 | 790.09216 | 90.397507 |
| KFNALFAQGNYSE | 16 | Q00610 | Clathrin heavy chain 1 OS=Homo sapiens | CLTC | 191.615 | 3 | 55.411663 | yes | 9.722E-08 | 586.96515 | 90.17366 |
| RPISADSAIMNPAS | 15 | Q00610 | Clathrin heavy chain 1 OS=Homo sapiens | CLTC | 191.615 | 2 | 55.411663 | yes | 2.851E-07 | 779.40631 | 88.872612 |
| YGYIHLYDLETGT | 19 | Q00610 | Clathrin heavy chain 1 OS=Homo sapiens | CLTC | 191.615 | 3 | 55.411663 | yes | 1.857E-07 | 794.70129 | 80.349762 |
| AHTMTDDVTFWK | 12 | Q00610 | Clathrin heavy chain 1 OS=Homo sapiens | CLTC | 191.615 | 2 | 55.411663 | yes | 9.722E-08 | 726.33484 | 75.685097 |
| VSQPIEGHAASFA | 16 | Q00610 | Clathrin heavy chain 1 OS=Homo sapiens | CLTC | 191.615 | 3 | 55.411663 | yes | 8.339E-06 | 572.96167 | 51.323048 |
| QSVELCK | 7 | Q00610 | Clathrin heavy chain 1 OS=Homo sapiens | CLTC | 191.615 | 2 | 55.411663 | yes | 4.471E-07 | 432.2182 | 40.631554 |
| IHEGCEEPATHNA | 16 | Q00610 | Clathrin heavy chain 1 OS=Homo sapiens | CLTC | 191.615 | 3 | 55.411663 | yes | 8.235E-07 | 592.94928 | 38.303032 |
| QLPLVKPYLR | 10 | Q00610 | Clathrin heavy chain 1 OS=Homo sapiens | CLTC | 191.615 | 3 | 55.411663 | yes | 5.634E-07 | 409.59216 | 34.980358 |
| GQCDLELINVCNE | 18 | Q00610 | Clathrin heavy chain 1 OS=Homo sapiens | CLTC | 191.615 | 3 | 55.411663 | yes | 1.456E-07 | 718.33826 | 28.731371 |
| ASDGSFSLPLKPVI | 22 | Q00653 | Nuclear factor NF-kappa-B p100 subunit | NFKB2 | 96.75 | 3 | 52.883774 | yes | 1.522E-07 | 775.08124 | 706.29376 |
| IEVDLVTHSDPPR | 13 | Q00653 | Nuclear factor NF-kappa-B p100 subunit | NFKB2 | 96.75 | 2 | 52.883774 | yes | 9.81E-08 | 739.38593 | 321.68188 |
| IEVDLVTHSDPPR | 13 | Q00653 | Nuclear factor NF-kappa-B p100 subunit | NFKB2 | 96.75 | 2 | 52.883774 | yes | 1.12E-07 | 493.2597 | 321.68188 |
| IEVDLVTHSDPPR | 13 | Q00653 | Nuclear factor NF-kappa-B p100 subunit | NFKB2 | 96.75 | 3 | 52.883774 | yes | 9.81E-08 | 739.38593 | 321.68188 |
| IEVDLVTHSDPPR | 13 | Q00653 | Nuclear factor NF-kappa-B p100 subunit | NFKB2 | 96.75 | 3 | 52.883774 | yes | 1.12E-07 | 493.2597 | 321.68188 |
| QCSELGICAVSVG | 15 | Q00653 | Nuclear factor NF-kappa-B p100 subunit | NFKB2 | 96.75 | 2 | 52.883774 | yes | 9.722E-08 | 802.89221 | 458.24667 |
| GGDEVYLLCDK | 11 | Q00653 | Nuclear factor NF-kappa-B p100 subunit | NFKB2 | 96.75 | 2 | 52.883774 | yes | 9.809E-08 | 634.79498 | 350.2019 |
| VMDLSIVR | 8 | Q00653 | Nuclear factor NF-kappa-B p100 subunit | NFKB2 | 96.75 | 2 | 52.883774 | yes | 1.422E-07 | 466.76532 | 267.64215 |
| NMMGTMIQK | 9 | Q00653 | Nuclear factor NF-kappa-B p100 subunit | NFKB2 | 96.75 | 2 | 52.883774 | yes | 2.75E-07 | 527.24811 | 245.37018 |
| QYAIVFR | 7 | Q00653 | Nuclear factor NF-kappa-B p100 subunit | NFKB2 | 96.75 | 2 | 52.883774 | yes | 1.475E-07 | 448.75308 | 206.21962 |
| LNSSIVEPK | 9 | Q00653 | Nuclear factor NF-kappa-B p100 subunit | NFKB2 | 96.75 | 2 | 52.883774 | yes | 4.776E-07 | 493.77948 | 191.33937 |
| ICNYEGPAK | 9 | Q00653 | Nuclear factor NF-kappa-B p100 subunit | NFKB2 | 96.75 | 2 | 52.883774 | yes | 1.965E-07 | 526.2475 | 137.78865 |
| ALLDYGVTADAR | 12 | Q00653 | Nuclear factor NF-kappa-B p100 subunit | NFKB2 | 96.75 | 2 | 52.883774 | yes | 4.74E-07 | 632.83026 | 112.71448 |
| QFTYYPLVEDK | 11 | Q00653 | Nuclear factor NF-kappa-B p100 subunit | NFKB2 | 96.75 | 2 | 52.883774 | yes | 9.722E-08 | 701.84808 | 70.333832 |
| IERPVTVFLQLK | 12 | Q00653 | Nuclear factor NF-kappa-B p100 subunit | NFKB2 | 96.75 | 2 | 52.883774 | yes | 2.718E-07 | 721.94012 | 52.346153 |

| | | | | | | | | | | | |
|---|---|---|---|---|---|---|---|---|---|---|---|
| EPAPETADGPYLV | 19 | Q00653 | Nuclear factor NF-kappa-B p100 subunit | NFKB2 | 96.75 | 3 | 52.883774 | yes | 1.595E-06 | 685.02112 | 50.01675 |
| PVTVFLQLK | 9 | Q00653 | Nuclear factor NF-kappa-B p100 subunit | NFKB2 | 96.75 | 2 | 52.883774 | yes | 1.075E-07 | 522.82623 | 45.919128 |
| SRPQGLTEAEQR | 12 | Q00653 | Nuclear factor NF-kappa-B p100 subunit | NFKB2 | 96.75 | 2 | 52.883774 | yes | 4.41E-07 | 686.35242 | 25.128799 |
| YNILGTNTIMDK | 12 | Q00839 | Heterogeneous nuclear ribonucleoprotein | HNRNPU | 90.585 | 2 | 53.21819 | yes | 1.92E-07 | 691.85284 | 32199.197 |
| DLPEHAVLK | 9 | Q00839 | Heterogeneous nuclear ribonucleoprotein | HNRNPU | 90.585 | 2 | 53.21819 | yes | 5.161E-07 | 511.28748 | 26246.371 |
| MCLFAGFQR | 9 | Q00839 | Heterogeneous nuclear ribonucleoprotein | HNRNPU | 90.585 | 2 | 53.21819 | yes | 1.349E-07 | 565.2677 | 10686.013 |
| MCLFAGFQR | 9 | Q00839 | Heterogeneous nuclear ribonucleoprotein | HNRNPU | 90.585 | 2 | 53.21819 | yes | 1.153E-07 | 573.26514 | 10686.013 |
| NFILDQTNVSAAA | 15 | Q00839 | Heterogeneous nuclear ribonucleoprotein | HNRNPU | 90.585 | 2 | 53.21819 | yes | 1.151E-07 | 824.42609 | 20380.064 |
| DIDIHEVR | 8 | Q00839 | Heterogeneous nuclear ribonucleoprotein | HNRNPU | 90.585 | 2 | 53.21819 | yes | 2.695E-07 | 498.75909 | 18111.545 |
| GYFEYIEENK | 10 | Q00839 | Heterogeneous nuclear ribonucleoprotein | HNRNPU | 90.585 | 2 | 53.21819 | yes | 2.455E-07 | 646.29572 | 11181.976 |
| LNTLLQR | 7 | Q00839 | Heterogeneous nuclear ribonucleoprotein | HNRNPU | 90.585 | 2 | 53.21819 | yes | 1.26E-07 | 429.26379 | 6551.335 |
| VSELKEELK | 9 | Q00839 | Heterogeneous nuclear ribonucleoprotein | HNRNPU | 90.585 | 2 | 53.21819 | yes | 9.722E-08 | 537.80573 | 4977.6011 |
| NGQDLGVAFK | 10 | Q00839 | Heterogeneous nuclear ribonucleoprotein | HNRNPU | 90.585 | 2 | 53.21819 | yes | 1.421E-06 | 524.77472 | 3855.8972 |
| LQAALDDEEAGGI | 13 | Q00839 | Heterogeneous nuclear ribonucleoprotein | HNRNPU | 90.585 | 2 | 53.21819 | yes | 1.001E-07 | 672.82312 | 3276.6479 |
| LLEQYKEESK | 10 | Q00839 | Heterogeneous nuclear ribonucleoprotein | HNRNPU | 90.585 | 2 | 53.21819 | yes | 3.02E-07 | 633.83246 | 2515.4182 |
| KAVVVCPK | 8 | Q00839 | Heterogeneous nuclear ribonucleoprotein | HNRNPU | 90.585 | 2 | 53.21819 | yes | 1.465E-07 | 450.77042 | 2230.4338 |
| GKVCFEMK | 8 | Q00839 | Heterogeneous nuclear ribonucleoprotein | HNRNPU | 90.585 | 2 | 53.21819 | yes | 2.294E-07 | 499.74353 | 1695.7892 |
| KMCLFAGFQR | 10 | Q00839 | Heterogeneous nuclear ribonucleoprotein | HNRNPU | 90.585 | 2 | 53.21819 | yes | 1.009E-07 | 629.31519 | 1198.2126 |
| DCEVVMMIGLPG | 15 | Q00839 | Heterogeneous nuclear ribonucleoprotein | HNRNPU | 90.585 | 2 | 53.21819 | yes | 1.102E-07 | 788.88025 | 1022.2296 |
| TCNCETEDYGEK | 12 | Q00839 | Heterogeneous nuclear ribonucleoprotein | HNRNPU | 90.585 | 2 | 53.21819 | yes | 3.979E-07 | 753.2796 | 938.02454 |
| VSELKEELKK | 10 | Q00839 | Heterogeneous nuclear ribonucleoprotein | HNRNPU | 90.585 | 2 | 53.21819 | yes | 9.722E-08 | 601.85315 | 503.13971 |
| FIEIAAR | 7 | Q00839 | Heterogeneous nuclear ribonucleoprotein | HNRNPU | 90.585 | 2 | 53.21819 | yes | 1.976E-07 | 410.23981 | 259.20392 |
| KDCEVVMMIGLP | 16 | Q00839 | Heterogeneous nuclear ribonucleoprotein | HNRNPU | 90.585 | 3 | 53.21819 | yes | 1.744E-07 | 568.95422 | 222.50005 |
| AVVVCPKDEDYK | 12 | Q00839 | Heterogeneous nuclear ribonucleoprotein | HNRNPU | 90.585 | 2 | 53.21819 | yes | 1.075E-07 | 711.85028 | 205.08998 |
| SSGPTSLFAVTVAI | 18 | Q00839 | Heterogeneous nuclear ribonucleoprotein | HNRNPU | 90.585 | 3 | 53.21819 | yes | 1.108E-07 | 572.30896 | 171.2406 |
| PWSQHYHQGYY | 11 | Q00839 | Heterogeneous nuclear ribonucleoprotein | HNRNPU | 90.585 | 2 | 53.21819 | yes | 9.809E-08 | 733.31781 | 114.27835 |
| LQAALDDEEAGGI | 32 | Q00839 | Heterogeneous nuclear ribonucleoprotein | HNRNPU | 90.585 | 4 | 53.21819 | yes | 1.71E-07 | 782.3653 | 76.478539 |
| AVIDLNNR | 8 | Q01081 | Splicing factor U2AF 35 kDa subunit OS | U2AF1 | 27.873 | 2 | 54.021385 | yes | 1.899E-07 | 457.75635 | 6341.1528 |
| DKVNCSFYFK | 10 | Q01081 | Splicing factor U2AF 35 kDa subunit OS | U2AF1 | 27.873 | 2 | 54.021385 | no | 1.616E-07 | 654.30811 | 3054.136 |
| GGFCNFMHLK | 10 | Q01081 | Splicing factor U2AF 35 kDa subunit OS | U2AF1 | 27.873 | 2 | 54.021385 | yes | 1.491E-07 | 605.77844 | 1295.0533 |
| VNCSFYFK | 8 | Q01081 | Splicing factor U2AF 35 kDa subunit OS | U2AF1 | 27.873 | 2 | 54.021385 | no | 2.207E-07 | 532.74713 | 1085.0006 |
| GGFCNFMHLKPIS | 14 | Q01081 | Splicing factor U2AF 35 kDa subunit OS | U2AF1 | 27.873 | 2 | 54.021385 | yes | 1.108E-07 | 832.41339 | 396.40308 |
| NPQNSSQSADGLR | 13 | Q01081 | Splicing factor U2AF 35 kDa subunit OS | U2AF1 | 27.873 | 2 | 54.021385 | yes | 1.143E-07 | 687.32385 | 72.39817 |
| QYEMGECTR | 9 | Q01081 | Splicing factor U2AF 35 kDa subunit OS | U2AF1 | 27.873 | 2 | 54.021385 | no | 9.722E-08 | 587.23682 | 35.055286 |
| EQWANLEQLSAIR | 13 | Q01082 | Spectrin beta chain, non-erythrocytic 1 O | SPTBN1 | 274.61 | 2 | 54.35775 | yes | 2.252E-07 | 779.4046 | 231.89975 |
| TALPAQSAATLPA | 14 | Q01082 | Spectrin beta chain, non-erythrocytic 1 O | SPTBN1 | 274.61 | 2 | 54.35775 | yes | 9.722E-08 | 684.38568 | 158.41121 |
| HLLGVEDLLQK | 11 | Q01082 | Spectrin beta chain, non-erythrocytic 1 O | SPTBN1 | 274.61 | 2 | 54.35775 | yes | 9.722E-08 | 632.86664 | 134.88371 |
| FMELLEPLNER | 11 | Q01082 | Spectrin beta chain, non-erythrocytic 1 O | SPTBN1 | 274.61 | 2 | 54.35775 | yes | 9.722E-08 | 695.85541 | 131.96863 |
| FESLEPEMNNQAS | 14 | Q01082 | Spectrin beta chain, non-erythrocytic 1 O | SPTBN1 | 274.61 | 2 | 54.35775 | yes | 1.692E-07 | 826.37268 | 116.32243 |

| | | | | | | | | | | | |
|---|---|---|---|---|---|---|---|---|---|---|---|
| VLDNAIETEK | 10 | Q01082 | Spectrin beta chain, non-erythrocytic 1 O: | SPTBN1 | 274.61 | 2 | 54.35775 | yes | 1.424E-07 | 566.29822 | 112.88767 |
| DGMAFNALIHK | 11 | Q01082 | Spectrin beta chain, non-erythrocytic 1 O: | SPTBN1 | 274.61 | 2 | 54.35775 | yes | 1.214E-07 | 608.81079 | 111.20228 |
| LAEISDVWEEMK | 12 | Q01082 | Spectrin beta chain, non-erythrocytic 1 O: | SPTBN1 | 274.61 | 2 | 54.35775 | yes | 9.722E-08 | 725.35016 | 107.57101 |
| LEMNLGLQK | 9 | Q01082 | Spectrin beta chain, non-erythrocytic 1 O: | SPTBN1 | 274.61 | 2 | 54.35775 | yes | 2.314E-07 | 523.28918 | 106.78236 |
| ETWLSENQR | 9 | Q01082 | Spectrin beta chain, non-erythrocytic 1 O: | SPTBN1 | 274.61 | 2 | 54.35775 | no | 2.896E-06 | 581.77802 | 91.633827 |
| ALVADSHPESER | 12 | Q01082 | Spectrin beta chain, non-erythrocytic 1 O: | SPTBN1 | 274.61 | 2 | 54.35775 | yes | 1.071E-07 | 655.82037 | 89.478256 |
| MWEVLESTTQTK | 12 | Q01082 | Spectrin beta chain, non-erythrocytic 1 O: | SPTBN1 | 274.61 | 2 | 54.35775 | yes | 1.47E-07 | 726.85559 | 78.909805 |
| LTTLELLEVR | 10 | Q01082 | Spectrin beta chain, non-erythrocytic 1 O: | SPTBN1 | 274.61 | 2 | 54.35775 | yes | 1.32E-07 | 593.85571 | 68.694138 |
| AELFTQSCADLDK | 13 | Q01082 | Spectrin beta chain, non-erythrocytic 1 O: | SPTBN1 | 274.61 | 2 | 54.35775 | yes | 1.846E-07 | 749.34814 | 68.282494 |
| MLTAQDMSYDEA | 13 | Q01082 | Spectrin beta chain, non-erythrocytic 1 O: | SPTBN1 | 274.61 | 2 | 54.35775 | yes | 1.342E-07 | 765.83179 | 65.577698 |
| TQILAASYELHK | 12 | Q01082 | Spectrin beta chain, non-erythrocytic 1 O: | SPTBN1 | 274.61 | 2 | 54.35775 | yes | 9.722E-08 | 687.37482 | 58.979156 |
| TAASGIPYHSEVP\ | 17 | Q01082 | Spectrin beta chain, non-erythrocytic 1 O: | SPTBN1 | 274.61 | 3 | 54.35775 | yes | 3.609E-06 | 585.98071 | 58.875198 |
| LVSDGNINSDR | 11 | Q01082 | Spectrin beta chain, non-erythrocytic 1 O: | SPTBN1 | 274.61 | 2 | 54.35775 | yes | 1.251E-06 | 595.29401 | 52.569828 |
| NDSFTTCIELGK | 12 | Q01082 | Spectrin beta chain, non-erythrocytic 1 O: | SPTBN1 | 274.61 | 2 | 54.35775 | yes | 9.291E-07 | 692.82428 | 51.965912 |
| LLEVLSGER | 9 | Q01082 | Spectrin beta chain, non-erythrocytic 1 O: | SPTBN1 | 274.61 | 2 | 54.35775 | yes | 2.724E-06 | 508.29276 | 43.496422 |
| VIESTQDLGNDLA( | 20 | Q01082 | Spectrin beta chain, non-erythrocytic 1 O: | SPTBN1 | 274.61 | 3 | 54.35775 | yes | 2.071E-06 | 710.7002 | 39.406731 |
| DVEDEILWVGER | 12 | Q01082 | Spectrin beta chain, non-erythrocytic 1 O: | SPTBN1 | 274.61 | 2 | 54.35775 | yes | 2.502E-07 | 730.35681 | 38.781162 |
| EVDDLEQWIAER | 12 | Q01082 | Spectrin beta chain, non-erythrocytic 1 O: | SPTBN1 | 274.61 | 2 | 54.35775 | yes | 1.115E-06 | 751.85974 | 37.536575 |
| LLTQHENIK | 9 | Q01082 | Spectrin beta chain, non-erythrocytic 1 O: | SPTBN1 | 274.61 | 2 | 54.35775 | yes | 1.283E-07 | 548.31146 | 33.817612 |
| EAVCEVALDYK | 11 | Q01082 | Spectrin beta chain, non-erythrocytic 1 O: | SPTBN1 | 274.61 | 2 | 54.35775 | yes | 5.523E-07 | 648.81067 | 31.24379 |
| HQILEQAVEDYAE | 20 | Q01082 | Spectrin beta chain, non-erythrocytic 1 O: | SPTBN1 | 274.61 | 3 | 54.35775 | yes | 1.083E-07 | 780.06067 | 30.765356 |
| SLLDACESR | 9 | Q01082 | Spectrin beta chain, non-erythrocytic 1 O: | SPTBN1 | 274.61 | 2 | 54.35775 | yes | 3.912E-06 | 525.74786 | 30.243469 |
| DALLLWCQMK | 10 | Q01082 | Spectrin beta chain, non-erythrocytic 1 O: | SPTBN1 | 274.61 | 2 | 54.35775 | no | 3.232E-06 | 639.32269 | 29.565094 |
| DLDDFQSWLSR | 11 | Q01082 | Spectrin beta chain, non-erythrocytic 1 O: | SPTBN1 | 274.61 | 2 | 54.35775 | yes | 7.663E-07 | 691.32275 | 24.799564 |
| ELEAENYHDIK | 11 | Q01082 | Spectrin beta chain, non-erythrocytic 1 O: | SPTBN1 | 274.61 | 2 | 54.35775 | yes | 5.631E-07 | 680.82263 | 13.260387 |
| AKDEQSAVSMLK | 12 | Q01082 | Spectrin beta chain, non-erythrocytic 1 O: | SPTBN1 | 274.61 | 2 | 54.35775 | yes | 7.216E-05 | 653.83722 | 11.198324 |
| FEDVVNQSSPK | 11 | Q01085 | Nucleolysin TIAR OS=Homo sapiens OX | TIAL1 | 41.591 | 2 | 50.55414 | yes | 9.771E-08 | 625.30658 | 1334.9263 |
| GYGFVSFYNK | 10 | Q01085 | Nucleolysin TIAR OS=Homo sapiens OX | TIAL1 | 41.591 | 2 | 50.55414 | yes | 4.492E-07 | 591.28491 | 441.97855 |
| FSTHESAAHAIVS\ | 25 | Q01085 | Nucleolysin TIAR OS=Homo sapiens OX | TIAL1 | 41.591 | 4 | 50.55414 | yes | 1.357E-07 | 648.58575 | 84.074371 |
| VEVTEFEDIK | 10 | Q01105 | Protein SET OS=Homo sapiens OX=960( | SET | 33.489 | 2 | 40.776287 | no | 3.847E-07 | 604.80591 | 101.37017 |
| LNEQASEEILK | 11 | Q01105 | Protein SET OS=Homo sapiens OX=960( | SET | 33.489 | 2 | 40.776287 | yes | 2.525E-06 | 637.33533 | 60.66132 |
| VGDVYIPR | 8 | Q01130 | Serine/arginine-rich splicing factor 2 OS= | SRSF2 | 25.477 | 2 | 31.848753 | yes | 1.705E-06 | 459.7558 | 1333.4639 |
| EGSPAPLEPEPGA/ | 17 | Q01167 | Forkhead box protein K2 OS=Homo sapi( | FOXK2 | 69.062 | 2 | 47.86446 | yes | 1.243E-07 | 837.92029 | 51.230686 |
| GAPPLQLPR | 9 | Q01167 | Forkhead box protein K2 OS=Homo sapi( | FOXK2 | 69.062 | 2 | 47.86446 | yes | 7.944E-06 | 474.78491 | 24.663868 |
| ASGMTTLSAELSSI | 14 | Q01196 | Runt-related transcription factor 1 OS=H( | RUNX1 | 48.737 | 2 | 48.258152 | yes | 1.172E-07 | 705.84833 | 90.404648 |
| LSELEQLR | 8 | Q01196 | Runt-related transcription factor 1 OS=H( | RUNX1 | 48.737 | 2 | 48.258152 | yes | 3.256E-07 | 494.2771 | 64.797401 |
| EMNDAAMFYTNR | 12 | Q01518 | Adenylyl cyclase-associated protein 1 OS | CAP1 | 51.902 | 2 | 41.15904 | yes | 5.197E-06 | 731.80811 | 65.27758 |
| INSITVDNCK | 10 | Q01518 | Adenylyl cyclase-associated protein 1 OS | CAP1 | 51.902 | 2 | 41.15904 | yes | 3.648E-07 | 582.28992 | 61.917614 |
| NSLDCEIVSAK | 11 | Q01518 | Adenylyl cyclase-associated protein 1 OS | CAP1 | 51.902 | 2 | 41.15904 | yes | 3.378E-06 | 618.30048 | 46.011002 |

| | | | | | | | | | | | |
|---|---|---|---|---|---|---|---|---|---|---|---|
| LSDLLAPISEQIK | 13 | Q01518 | Adenylyl cyclase-associated protein 1 OS | CAP1 | 51.902 | 2 | 41.15904 | yes | 4.885E-07 | 713.91119 | 34.715569 |
| EEHEVAVLGAPHN | 25 | Q01628 | Interferon-induced transmembrane protein | IFITM3 | 14.633 | 4 | 37.888992 | yes | 6.533E-07 | 668.60425 | 67.69355 |
| MVGDVTGAQAYA | 16 | Q01628 | Interferon-induced transmembrane protein | IFITM3 | 14.633 | 2 | 37.888992 | no | 5.618E-07 | 785.38251 | 21.399755 |
| ALAAPAAEEKEEA | 14 | Q01650 | Large neutral amino acids transporter sma | SLC7A5 | 55.011 | 2 | 45.143974 | yes | 1.947E-07 | 728.37555 | 89.29937 |
| KPELERPIK | 9 | Q01650 | Large neutral amino acids transporter sma | SLC7A5 | 55.011 | 2 | 45.143974 | yes | 3.159E-06 | 555.33752 | 26.902948 |
| LENLGIPEEELLR | 13 | Q01658 | Protein Dr1 OS=Homo sapiens OX=9606 | DR1 | 19.444 | 2 | 46.17973 | yes | 1.629E-07 | 762.91705 | 189.32526 |
| MMLEEQVR | 8 | Q01664 | Transcription factor AP-4 OS=Homo sapi | TFAP4 | 38.726 | 2 | 47.81475 | yes | 5.521E-07 | 518.25171 | 202.36884 |
| MQSINAGFQSLK | 12 | Q01664 | Transcription factor AP-4 OS=Homo sapi | TFAP4 | 38.726 | 2 | 47.81475 | yes | 1.253E-07 | 662.3399 | 178.60452 |
| TLIPHTDGEK | 10 | Q01664 | Transcription factor AP-4 OS=Homo sapi | TFAP4 | 38.726 | 2 | 47.81475 | yes | 3.561E-07 | 555.79309 | 170.85081 |
| AVIVKPVR | 8 | Q01664 | Transcription factor AP-4 OS=Homo sapi | TFAP4 | 38.726 | 2 | 47.81475 | yes | 2.562E-07 | 441.3002 | 168.79201 |
| SLEAHMYPEK | 10 | Q01664 | Transcription factor AP-4 OS=Homo sapi | TFAP4 | 38.726 | 2 | 47.81475 | yes | 3.518E-05 | 602.78699 | 44.958908 |
| MFLPSLGHR | 9 | Q01780 | Exosome complex component 10 OS=Ho | EXOSC10 | 100.832 | 2 | 54.95397 | yes | 3.463E-07 | 529.28424 | 423.84879 |
| VGILLDEASGVNK | 13 | Q01780 | Exosome complex component 10 OS=Ho | EXOSC10 | 100.832 | 2 | 54.95397 | yes | 9.722E-08 | 657.86682 | 410.62146 |
| NQQPVLPAGLQVI | 14 | Q01780 | Exosome complex component 10 OS=Ho | EXOSC10 | 100.832 | 2 | 54.95397 | yes | 1.523E-07 | 744.9303 | 371.69449 |
| AAAEQAISVR | 10 | Q01780 | Exosome complex component 10 OS=Ho | EXOSC10 | 100.832 | 2 | 54.95397 | yes | 4.849E-07 | 508.28018 | 313.25958 |
| TEDFIIDTLELR | 12 | Q01780 | Exosome complex component 10 OS=Ho | EXOSC10 | 100.832 | 2 | 54.95397 | yes | 3.077E-07 | 732.88269 | 187.69168 |
| FALGSVVAVTK | 11 | Q01780 | Exosome complex component 10 OS=Ho | EXOSC10 | 100.832 | 2 | 54.95397 | yes | 4.285E-07 | 546.3266 | 154.46945 |
| NIIRPQLK | 8 | Q01780 | Exosome complex component 10 OS=Ho | EXOSC10 | 100.832 | 2 | 54.95397 | yes | 3.741E-07 | 491.31381 | 153.81438 |
| LYCNVDSNK | 9 | Q01780 | Exosome complex component 10 OS=Ho | EXOSC10 | 100.832 | 2 | 54.95397 | yes | 9.894E-08 | 556.75568 | 131.98068 |
| EFTPYDYSQSDFK | 13 | Q01780 | Exosome complex component 10 OS=Ho | EXOSC10 | 100.832 | 2 | 54.95397 | yes | 4.177E-07 | 813.85156 | 114.49339 |
| VMQYHGCR | 8 | Q01780 | Exosome complex component 10 OS=Ho | EXOSC10 | 100.832 | 2 | 54.95397 | yes | 9.722E-08 | 525.73401 | 110.7114 |
| AQNIMESFENPFR | 13 | Q01780 | Exosome complex component 10 OS=Ho | EXOSC10 | 100.832 | 2 | 54.95397 | yes | 1.163E-07 | 791.86957 | 93.159622 |
| HSLDHLLK | 8 | Q01780 | Exosome complex component 10 OS=Ho | EXOSC10 | 100.832 | 2 | 54.95397 | yes | 5.735E-07 | 481.77454 | 80.510834 |
| IDNSNTPFLPK | 11 | Q01780 | Exosome complex component 10 OS=Ho | EXOSC10 | 100.832 | 2 | 54.95397 | yes | 6.191E-07 | 623.32733 | 72.359825 |
| LLNCQEFAVDLEH | 17 | Q01780 | Exosome complex component 10 OS=Ho | EXOSC10 | 100.832 | 3 | 54.95397 | yes | 1.156E-07 | 711.00568 | 65.766327 |
| VFHGADSDIEWLQ | 14 | Q01780 | Exosome complex component 10 OS=Ho | EXOSC10 | 100.832 | 2 | 54.95397 | yes | 1.862E-07 | 822.90448 | 48.388908 |
| FIKPIFTDESYLELY | 16 | Q01780 | Exosome complex component 10 OS=Ho | EXOSC10 | 100.832 | 3 | 54.95397 | yes | 9.722E-08 | 678.69092 | 45.469505 |
| GNGQPVQLQVVW | 14 | Q01780 | Exosome complex component 10 OS=Ho | EXOSC10 | 100.832 | 2 | 54.95397 | yes | 4.353E-07 | 804.9339 | 44.263435 |
| LLQCMSR | 7 | Q01780 | Exosome complex component 10 OS=Ho | EXOSC10 | 100.832 | 2 | 54.95397 | yes | 1.439E-07 | 454.22806 | 43.510094 |
| QYQLADWR | 8 | Q01780 | Exosome complex component 10 OS=Ho | EXOSC10 | 100.832 | 2 | 54.95397 | yes | 3.066E-06 | 540.26709 | 42.883312 |
| LAQVQVQK | 8 | Q01780 | Exosome complex component 10 OS=Ho | EXOSC10 | 100.832 | 2 | 54.95397 | yes | 9.958E-08 | 457.27692 | 38.162254 |
| REDESYGYVLPNH | 17 | Q01780 | Exosome complex component 10 OS=Ho | EXOSC10 | 100.832 | 3 | 54.95397 | yes | 5.823E-07 | 694.66431 | 37.702103 |
| PIFTDESYLELYR | 13 | Q01780 | Exosome complex component 10 OS=Ho | EXOSC10 | 100.832 | 2 | 54.95397 | yes | 2.001E-07 | 823.40906 | 27.674107 |
| LLYEIQNR | 8 | Q01804 | OTU domain-containing protein 4 OS=Ho | OTUD4 | 124.045 | 2 | 52.572807 | yes | 9.722E-08 | 524.79291 | 956.44391 |
| AHPPTQILNR | 10 | Q01804 | OTU domain-containing protein 4 OS=Ho | OTUD4 | 124.045 | 2 | 52.572807 | yes | 1.702E-06 | 573.82257 | 549.72083 |
| LQHPSGVR | 8 | Q01804 | OTU domain-containing protein 4 OS=Ho | OTUD4 | 124.045 | 2 | 52.572807 | yes | 3.469E-07 | 447.25122 | 425.93964 |
| MACIHYLR | 8 | Q01804 | OTU domain-containing protein 4 OS=Ho | OTUD4 | 124.045 | 2 | 52.572807 | yes | 1.712E-07 | 532.26245 | 413.77148 |
| ESNYFGLSPEER | 12 | Q01804 | OTU domain-containing protein 4 OS=Ho | OTUD4 | 124.045 | 2 | 52.572807 | yes | 9.722E-08 | 714.3255 | 394.74994 |
| APPPESWNTVSGK | 13 | Q01804 | OTU domain-containing protein 4 OS=Ho | OTUD4 | 124.045 | 2 | 52.572807 | yes | 2.394E-07 | 685.34094 | 369.13626 |

| | | | | | | | | | | | |
|---|---|---|---|---|---|---|---|---|---|---|---|
| EDATPMDAYLR | 11 | Q01804 | OTU domain-containing protein 4 OS=H( | OTUD4 | 124.045 | 2 | 52.572807 | yes | 1.064E-07 | 641.2926 | 363.40155 |
| TAADVVSPGANS\ | 16 | Q01804 | OTU domain-containing protein 4 OS=H( | OTUD4 | 124.045 | 2 | 52.572807 | yes | 9.81E-08 | 773.37878 | 326.90845 |
| KDFIIYR | 7 | Q01804 | OTU domain-containing protein 4 OS=H( | OTUD4 | 124.045 | 2 | 52.572807 | yes | 3.241E-07 | 477.77402 | 187.43404 |
| ETVPVELEPK | 10 | Q01804 | OTU domain-containing protein 4 OS=H( | OTUD4 | 124.045 | 2 | 52.572807 | yes | 3.231E-07 | 570.81097 | 183.97153 |
| DCGSVSTVDEFPE. | 15 | Q01804 | OTU domain-containing protein 4 OS=H( | OTUD4 | 124.045 | 2 | 52.572807 | yes | 1.272E-07 | 834.86212 | 158.97549 |
| GELDLSLENLDLSI | 14 | Q01804 | OTU domain-containing protein 4 OS=H( | OTUD4 | 124.045 | 2 | 52.572807 | yes | 9.722E-08 | 773.40393 | 139.87465 |
| SLNPAVYR | 8 | Q01804 | OTU domain-containing protein 4 OS=H( | OTUD4 | 124.045 | 2 | 52.572807 | yes | 1.098E-06 | 460.25345 | 110.96512 |
| QFYNQTYGSR | 10 | Q01804 | OTU domain-containing protein 4 OS=H( | OTUD4 | 124.045 | 2 | 52.572807 | yes | 1.197E-07 | 632.29126 | 79.823685 |
| VEDFDHTSR | 9 | Q01804 | OTU domain-containing protein 4 OS=H( | OTUD4 | 124.045 | 2 | 52.572807 | yes | 3.296E-07 | 553.24908 | 75.503815 |
| FFFNLGVK | 8 | Q01804 | OTU domain-containing protein 4 OS=H( | OTUD4 | 124.045 | 2 | 52.572807 | yes | 1.54E-06 | 486.27109 | 60.435364 |
| RPEPSTLENITDDK | 14 | Q01804 | OTU domain-containing protein 4 OS=H( | OTUD4 | 124.045 | 2 | 52.572807 | yes | 2.171E-07 | 807.9021 | 29.041176 |
| KADTALASIPPVAI | 16 | Q01804 | OTU domain-containing protein 4 OS=H( | OTUD4 | 124.045 | 2 | 52.572807 | yes | 4.131E-05 | 784.43811 | 16.982285 |
| ELVVTQLGYDTR | 12 | Q01813 | ATP-dependent 6-phosphofructokinase, p | PFKP | 85.597 | 2 | 53.54435 | yes | 1.025E-07 | 697.36975 | 1837.8883 |
| NVIFQPVAELK | 11 | Q01813 | ATP-dependent 6-phosphofructokinase, p | PFKP | 85.597 | 2 | 53.54435 | yes | 1.002E-07 | 629.36371 | 1311.2891 |
| YLEEIATQMR | 10 | Q01813 | ATP-dependent 6-phosphofructokinase, p | PFKP | 85.597 | 2 | 53.54435 | yes | 9.722E-08 | 627.31335 | 1066.1763 |
| VTILGHVQR | 9 | Q01813 | ATP-dependent 6-phosphofructokinase, p | PFKP | 85.597 | 2 | 53.54435 | yes | 2.877E-07 | 511.8089 | 897.40619 |
| AACNLLQR | 8 | Q01813 | ATP-dependent 6-phosphofructokinase, p | PFKP | 85.597 | 2 | 53.54435 | yes | 2.746E-07 | 473.25037 | 758.73401 |
| MLAIYDGFDGFAK | 13 | Q01813 | ATP-dependent 6-phosphofructokinase, p | PFKP | 85.597 | 2 | 53.54435 | yes | 9.722E-08 | 724.34998 | 745.91803 |
| SFAGNLNTYK | 10 | Q01813 | ATP-dependent 6-phosphofructokinase, p | PFKP | 85.597 | 2 | 53.54435 | yes | 1.125E-07 | 557.78003 | 625.54279 |
| DLQSNVEHLTEK | 12 | Q01813 | ATP-dependent 6-phosphofructokinase, p | PFKP | 85.597 | 2 | 53.54435 | yes | 1.003E-07 | 706.85443 | 540.62817 |
| FTTDDSICVLGISK | 14 | Q01813 | ATP-dependent 6-phosphofructokinase, p | PFKP | 85.597 | 2 | 53.54435 | yes | 9.722E-08 | 778.38727 | 518.5108 |
| FLEHLSGAGK | 10 | Q01813 | ATP-dependent 6-phosphofructokinase, p | PFKP | 85.597 | 2 | 53.54435 | yes | 1.836E-07 | 529.7851 | 455.90164 |
| VGIADGHR | 8 | Q01813 | ATP-dependent 6-phosphofructokinase, p | PFKP | 85.597 | 2 | 53.54435 | yes | 2.145E-07 | 412.72229 | 434.88385 |
| LPDDQIPK | 8 | Q01813 | ATP-dependent 6-phosphofructokinase, p | PFKP | 85.597 | 2 | 53.54435 | yes | 2.729E-07 | 463.25311 | 343.15155 |
| LNIIIVAEGAIDTQI | 22 | Q01813 | ATP-dependent 6-phosphofructokinase, p | PFKP | 85.597 | 3 | 53.54435 | yes | 1.001E-07 | 789.77582 | 180.68268 |
| RFQDAVR | 7 | Q01813 | ATP-dependent 6-phosphofructokinase, p | PFKP | 85.597 | 2 | 53.54435 | yes | 9.722E-08 | 446.24341 | 167.89369 |
| KEWSGLLEELAR | 12 | Q01813 | ATP-dependent 6-phosphofructokinase, p | PFKP | 85.597 | 2 | 53.54435 | yes | 1.003E-07 | 715.88556 | 103.20368 |
| IIEVVDAIMTTAQS | 17 | Q01813 | ATP-dependent 6-phosphofructokinase, p | PFKP | 85.597 | 3 | 53.54435 | yes | 5.804E-07 | 638.00342 | 92.324684 |
| TNCNVAVINVGAI | 22 | Q01813 | ATP-dependent 6-phosphofructokinase, p | PFKP | 85.597 | 3 | 53.54435 | yes | 1.123E-07 | 724.03339 | 84.374687 |
| EWSGLLEELAR | 11 | Q01813 | ATP-dependent 6-phosphofructokinase, p | PFKP | 85.597 | 2 | 53.54435 | yes | 1.495E-07 | 651.83807 | 80.726212 |
| KFLEHLSGAGK | 11 | Q01813 | ATP-dependent 6-phosphofructokinase, p | PFKP | 85.597 | 2 | 53.54435 | yes | 9.722E-08 | 593.83258 | 69.760658 |
| RLNIIIVAEGAIDT( | 23 | Q01813 | ATP-dependent 6-phosphofructokinase, p | PFKP | 85.597 | 3 | 53.54435 | yes | 1.79E-06 | 841.80957 | 3.2069356 |
| ILAASWPQNR | 10 | Q01831 | DNA repair protein complementing XP-C | XPC | 105.953 | 2 | 52.66501 | yes | 9.722E-08 | 578.31708 | 148.3718 |
| DVDTYYLSNLVK | 12 | Q01831 | DNA repair protein complementing XP-C | XPC | 105.953 | 2 | 52.66501 | yes | 2.548E-07 | 715.36414 | 32.657162 |
| LEFETYLR | 8 | Q01831 | DNA repair protein complementing XP-C | XPC | 105.953 | 2 | 52.66501 | yes | 4.096E-06 | 535.77948 | 31.608536 |
| HMDQPLPTAIGLY | 14 | Q01831 | DNA repair protein complementing XP-C | XPC | 105.953 | 2 | 52.66501 | yes | 4.217E-07 | 792.41614 | 28.219208 |
| GDATVSYEDPPTA | 14 | Q01844 | RNA-binding protein EWS OS=Homo sa | EWSR1 | 68.479 | 2 | 51.129234 | yes | 9.734E-08 | 725.83844 | 30485.691 |
| TGQPMIHIYLDK | 12 | Q01844 | RNA-binding protein EWS OS=Homo sa | EWSR1 | 68.479 | 2 | 51.129234 | yes | 8.563E-07 | 708.37122 | 3631.7632 |
| TGQPMIHIYLDK | 12 | Q01844 | RNA-binding protein EWS OS=Homo sa | EWSR1 | 68.479 | 2 | 51.129234 | yes | 8.785E-05 | 716.36865 | 3631.7632 |

| | | | | | | | | | | | |
|---|---|---|---|---|---|---|---|---|---|---|---|
| GGPGGPGGPGGPM | 15 | Q01844 | RNA-binding protein EWS OS=Homo sa | EWSR1 | 68.479 | 2 | 51.129234 | yes | 1.041E-07 | 604.28546 | 994.35474 |
| RTGQPMIHIYLDK | 13 | Q01844 | RNA-binding protein EWS OS=Homo sa | EWSR1 | 68.479 | 3 | 51.129234 | yes | 9.722E-08 | 524.61694 | 353.63815 |
| RTGQPMIHIYLDK | 13 | Q01844 | RNA-binding protein EWS OS=Homo sa | EWSR1 | 68.479 | 3 | 51.129234 | yes | 2.105E-07 | 786.42175 | 353.63815 |
| RTGQPMIHIYLDK | 13 | Q01844 | RNA-binding protein EWS OS=Homo sa | EWSR1 | 68.479 | 2 | 51.129234 | yes | 9.722E-08 | 524.61694 | 353.63815 |
| RTGQPMIHIYLDK | 13 | Q01844 | RNA-binding protein EWS OS=Homo sa | EWSR1 | 68.479 | 2 | 51.129234 | yes | 2.105E-07 | 786.42175 | 353.63815 |
| GGPGGPPGPLMEQ | 17 | Q01844 | RNA-binding protein EWS OS=Homo sa | EWSR1 | 68.479 | 2 | 51.129234 | yes | 1.301E-07 | 797.87689 | 216.15302 |
| ASPFLECHGR | 10 | Q01955 | Collagen alpha-3(IV) chain OS=Homo sa | COL4A3 | 161.813 | 2 | 46.093525 | yes | 1.655E-07 | 587.2771 | 847.05524 |
| CSVGLMSSVVAPA | 22 | Q01959 | Sodium-dependent dopamine transporter | SLC6A3 | 68.495 | 3 | 36.69171 | yes | 6.713E-07 | 738.71344 | 81.885597 |
| EPCALTLAQR | 10 | Q01968 | Inositol polyphosphate 5-phosphatase OC | OCRL | 104.205 | 2 | 46.799454 | yes | 1.85E-07 | 579.80042 | 124.94695 |
| LCMPDANEVK | 10 | Q01968 | Inositol polyphosphate 5-phosphatase OC | OCRL | 104.205 | 2 | 46.799454 | yes | 4.038E-07 | 588.77301 | 112.15883 |
| AQSQLLVPEQK | 11 | Q01968 | Inositol polyphosphate 5-phosphatase OC | OCRL | 104.205 | 2 | 46.799454 | yes | 1.467E-07 | 620.84845 | 81.522049 |
| AFVDFNEGEIK | 11 | Q01968 | Inositol polyphosphate 5-phosphatase OC | OCRL | 104.205 | 2 | 46.799454 | yes | 1.584E-07 | 634.81152 | 74.718903 |
| VQGDWIR | 7 | Q01968 | Inositol polyphosphate 5-phosphatase OC | OCRL | 104.205 | 2 | 46.799454 | yes | 7.801E-07 | 437.23251 | 35.722107 |
| LIDLEEDSFLEK | 12 | Q01968 | Inositol polyphosphate 5-phosphatase OC | OCRL | 104.205 | 2 | 46.799454 | yes | 5.433E-07 | 725.8692 | 24.586653 |
| SLLQMVPLDEGAS | 21 | Q01968 | Inositol polyphosphate 5-phosphatase OC | OCRL | 104.205 | 3 | 46.799454 | yes | 2.982E-06 | 769.75073 | 17.977434 |
| LLAQLAQECQEQR | 13 | Q01970 | 1-phosphatidylinositol 4,5-bisphosphate p | PLCB3 | 138.799 | 2 | 43.78651 | yes | 2.448E-07 | 793.90137 | 31.929979 |
| EVVLDANTTQFK | 12 | Q01970 | 1-phosphatidylinositol 4,5-bisphosphate p | PLCB3 | 138.799 | 2 | 43.78651 | yes | 4.913E-07 | 682.85645 | 20.929836 |
| QKELEELER | 9 | Q02040 | A-kinase anchor protein 17A OS=Homo s | AKAP17A | 80.736 | 2 | 47.174633 | yes | 1.163E-06 | 587.30914 | 237.51144 |
| SISNWEVMER | 10 | Q02040 | A-kinase anchor protein 17A OS=Homo s | AKAP17A | 80.736 | 2 | 47.174633 | yes | 1.384E-07 | 625.79535 | 112.31116 |
| LQELEQQR | 8 | Q02040 | A-kinase anchor protein 17A OS=Homo s | AKAP17A | 80.736 | 2 | 47.174633 | yes | 6.177E-07 | 522.27765 | 95.335999 |
| LSGFSDILK | 9 | Q02040 | A-kinase anchor protein 17A OS=Homo s | AKAP17A | 80.736 | 2 | 47.174633 | yes | 4.953E-07 | 490.27658 | 65.087769 |
| NVDIPMLDPYR | 11 | Q02040 | A-kinase anchor protein 17A OS=Homo s | AKAP17A | 80.736 | 2 | 47.174633 | yes | 5.522E-07 | 666.83447 | 39.82597 |
| ESGSEKPSEDVLVI | 14 | Q02040 | A-kinase anchor protein 17A OS=Homo s | AKAP17A | 80.736 | 2 | 47.174633 | yes | 8.202E-07 | 752.38049 | 29.870764 |
| SSTTTTPVQSGANY | 16 | Q02086 | Transcription factor Sp2 OS=Homo sapie | SP2 | 64.901 | 2 | 48.51859 | yes | 1.136E-07 | 788.91248 | 43.70882 |
| LNVLANVIR | 9 | Q02218 | 2-oxoglutarate dehydrogenase complex co | OGDH | 115.936 | 2 | 52.211266 | no | 1.34E-07 | 506.31912 | 90.029335 |
| THLTELQR | 8 | Q02218 | 2-oxoglutarate dehydrogenase complex co | OGDH | 115.936 | 2 | 52.211266 | yes | 4.921E-07 | 499.2749 | 82.141922 |
| SSPYPTDVAR | 10 | Q02218 | 2-oxoglutarate dehydrogenase complex co | OGDH | 115.936 | 2 | 52.211266 | no | 3.02E-06 | 546.76965 | 66.613884 |
| DVVVDLVCYR | 10 | Q02218 | 2-oxoglutarate dehydrogenase complex co | OGDH | 115.936 | 2 | 52.211266 | no | 3.665E-07 | 619.31592 | 57.124393 |
| ELEQIFCQFDSK | 12 | Q02218 | 2-oxoglutarate dehydrogenase complex co | OGDH | 115.936 | 2 | 52.211266 | yes | 9.722E-08 | 772.35852 | 53.722885 |
| AEQFYCGDTEGK | 12 | Q02218 | 2-oxoglutarate dehydrogenase complex co | OGDH | 115.936 | 2 | 52.211266 | yes | 6.687E-07 | 702.79047 | 45.722816 |
| KPLIIFTPK | 9 | Q02218 | 2-oxoglutarate dehydrogenase complex co | OGDH | 115.936 | 2 | 52.211266 | no | 5.07E-07 | 528.84442 | 32.382668 |
| LEAADEGSGDVK | 12 | Q02218 | 2-oxoglutarate dehydrogenase complex co | OGDH | 115.936 | 2 | 52.211266 | no | 6.469E-06 | 595.7804 | 7.5554733 |
| ELQSAFNEITK | 11 | Q02224 | Centromere-associated protein E OS=Hor | CENPE | 316.416 | 2 | 53.610855 | yes | 1.176E-05 | 640.33008 | 137.4312 |
| TKDDLATTQSNYK | 13 | Q02224 | Centromere-associated protein E OS=Hor | CENPE | 316.416 | 2 | 53.610855 | yes | 4.185E-06 | 742.86499 | 126.3964 |
| DQLAQLLEEK | 10 | Q02224 | Centromere-associated protein E OS=Hor | CENPE | 316.416 | 2 | 53.610855 | yes | 9.722E-08 | 593.81934 | 106.7537 |
| LQEVLQSESDQLK | 13 | Q02224 | Centromere-associated protein E OS=Hor | CENPE | 316.416 | 2 | 53.610855 | yes | 3.207E-06 | 758.8963 | 67.113701 |
| MNQEIVNLSK | 10 | Q02224 | Centromere-associated protein E OS=Hor | CENPE | 316.416 | 2 | 53.610855 | yes | 1.982E-07 | 588.30811 | 54.524994 |
| IEIEMLGLSK | 10 | Q02224 | Centromere-associated protein E OS=Hor | CENPE | 316.416 | 2 | 53.610855 | yes | 2.748E-06 | 566.81775 | 53.704216 |
| LNLENIETENIR | 12 | Q02224 | Centromere-associated protein E OS=Hor | CENPE | 316.416 | 2 | 53.610855 | yes | 3.74E-07 | 729.38336 | 48.373726 |

| | | | | | | | | | | | |
|---|---|---|---|---|---|---|---|---|---|---|---|
| MTAVNETQEK | 10 | Q02224 | Centromere-associated protein E OS=Hor | CENPE | 316.416 | 2 | 53.610855 | yes | 4.207E-07 | 575.77405 | 44.049885 |
| LSDGQVGGFINYR | 13 | Q02224 | Centromere-associated protein E OS=Hor | CENPE | 316.416 | 2 | 53.610855 | yes | 6.103E-07 | 713.35968 | 38.65379 |
| DLLCNLELEGK | 11 | Q02224 | Centromere-associated protein E OS=Hor | CENPE | 316.416 | 2 | 53.610855 | yes | 5.202E-07 | 652.33173 | 33.35334 |
| QTLFDAETVALDA | 14 | Q02224 | Centromere-associated protein E OS=Hor | CENPE | 316.416 | 2 | 53.610855 | yes | 5.168E-07 | 761.39337 | 30.599039 |
| ALLQEFDNAVLSK | 13 | Q02241 | Kinesin-like protein KIF23 OS=Homo sa | KIF23 | 110.059 | 2 | 54.3365 | yes | 6.491E-07 | 724.39319 | 4066.9216 |
| LQGMVTETTMK | 11 | Q02241 | Kinesin-like protein KIF23 OS=Homo sa | KIF23 | 110.059 | 2 | 54.3365 | yes | 1.355E-07 | 619.80957 | 1874.7626 |
| LQGMVTETTMK | 11 | Q02241 | Kinesin-like protein KIF23 OS=Homo sa | KIF23 | 110.059 | 2 | 54.3365 | yes | 4.436E-07 | 627.80707 | 1874.7626 |
| EAGNINQSLMTLR | 13 | Q02241 | Kinesin-like protein KIF23 OS=Homo sa | KIF23 | 110.059 | 2 | 54.3365 | yes | 1.048E-07 | 723.87213 | 1830.1082 |
| EAGNINQSLMTLR | 13 | Q02241 | Kinesin-like protein KIF23 OS=Homo sa | KIF23 | 110.059 | 2 | 54.3365 | yes | 9.722E-08 | 731.86957 | 1830.1082 |
| AEDYEENLQVMR | 12 | Q02241 | Kinesin-like protein KIF23 OS=Homo sa | KIF23 | 110.059 | 2 | 54.3365 | yes | 1.128E-07 | 748.83795 | 3501.6853 |
| AICGLTPGR | 9 | Q02241 | Kinesin-like protein KIF23 OS=Homo sa | KIF23 | 110.059 | 2 | 54.3365 | yes | 1.731E-07 | 472.75275 | 2712.1204 |
| GGGQSVQFTDIETI | 15 | Q02241 | Kinesin-like protein KIF23 OS=Homo sa | KIF23 | 110.059 | 2 | 54.3365 | yes | 1.002E-07 | 790.40173 | 2570.6882 |
| CSVAVEMR | 8 | Q02241 | Kinesin-like protein KIF23 OS=Homo sa | KIF23 | 110.059 | 2 | 54.3365 | yes | 2.697E-07 | 476.22296 | 2484.1292 |
| DPVGVYCR | 8 | Q02241 | Kinesin-like protein KIF23 OS=Homo sa | KIF23 | 110.059 | 2 | 54.3365 | yes | 1.33E-07 | 483.2291 | 2397.9817 |
| MIVCVNPK | 8 | Q02241 | Kinesin-like protein KIF23 OS=Homo sa | KIF23 | 110.059 | 2 | 54.3365 | yes | 1.131E-07 | 480.75391 | 1041.1006 |
| MIVCVNPK | 8 | Q02241 | Kinesin-like protein KIF23 OS=Homo sa | KIF23 | 110.059 | 2 | 54.3365 | yes | 4.902E-07 | 488.75134 | 1041.1006 |
| TCMDVLR | 7 | Q02241 | Kinesin-like protein KIF23 OS=Homo sa | KIF23 | 110.059 | 2 | 54.3365 | yes | 9.935E-08 | 447.71222 | 1999.8268 |
| TTTIYEEDKR | 10 | Q02241 | Kinesin-like protein KIF23 OS=Homo sa | KIF23 | 110.059 | 2 | 54.3365 | yes | 9.758E-08 | 628.31189 | 1980.6047 |
| QMMIDEFNK | 9 | Q02241 | Kinesin-like protein KIF23 OS=Homo sa | KIF23 | 110.059 | 2 | 54.3365 | yes | 1.234E-07 | 578.26227 | 1962.7303 |
| FAEVTQEVEVAR | 12 | Q02241 | Kinesin-like protein KIF23 OS=Homo sa | KIF23 | 110.059 | 2 | 54.3365 | yes | 9.722E-08 | 689.35406 | 1890.5034 |
| SHSVFNIK | 8 | Q02241 | Kinesin-like protein KIF23 OS=Homo sa | KIF23 | 110.059 | 2 | 54.3365 | yes | 1.374E-07 | 466.25345 | 1740.2869 |
| IANTHLNR | 8 | Q02241 | Kinesin-like protein KIF23 OS=Homo sa | KIF23 | 110.059 | 2 | 54.3365 | yes | 1.173E-07 | 469.76196 | 1715.0442 |
| NGLLFTYGVTGSG | 14 | Q02241 | Kinesin-like protein KIF23 OS=Homo sa | KIF23 | 110.059 | 2 | 54.3365 | yes | 9.919E-08 | 707.37225 | 1703.1798 |
| NLQQELETQNQK | 12 | Q02241 | Kinesin-like protein KIF23 OS=Homo sa | KIF23 | 110.059 | 2 | 54.3365 | yes | 4.827E-07 | 736.87061 | 1407.6237 |
| THTMTGSPGEGGL | 16 | Q02241 | Kinesin-like protein KIF23 OS=Homo sa | KIF23 | 110.059 | 3 | 54.3365 | yes | 1.762E-07 | 537.60333 | 677.53101 |
| THTMTGSPGEGGL | 16 | Q02241 | Kinesin-like protein KIF23 OS=Homo sa | KIF23 | 110.059 | 3 | 54.3365 | yes | 1.286E-07 | 805.90137 | 677.53101 |
| THTMTGSPGEGGL | 16 | Q02241 | Kinesin-like protein KIF23 OS=Homo sa | KIF23 | 110.059 | 2 | 54.3365 | yes | 1.762E-07 | 537.60333 | 677.53101 |
| THTMTGSPGEGGL | 16 | Q02241 | Kinesin-like protein KIF23 OS=Homo sa | KIF23 | 110.059 | 2 | 54.3365 | yes | 1.286E-07 | 805.90137 | 677.53101 |
| QVFGTHTTQK | 10 | Q02241 | Kinesin-like protein KIF23 OS=Homo sa | KIF23 | 110.059 | 2 | 54.3365 | yes | 9.722E-08 | 573.79877 | 1305.1266 |
| LLFQPDQNAPPIR | 13 | Q02241 | Kinesin-like protein KIF23 OS=Homo sa | KIF23 | 110.059 | 2 | 54.3365 | yes | 1.815E-07 | 754.91461 | 1252.6244 |
| IPTYNTPLK | 9 | Q02241 | Kinesin-like protein KIF23 OS=Homo sa | KIF23 | 110.059 | 2 | 54.3365 | yes | 3.584E-07 | 523.79767 | 1252.0958 |
| ENQMYGTNK | 9 | Q02241 | Kinesin-like protein KIF23 OS=Homo sa | KIF23 | 110.059 | 2 | 54.3365 | yes | 1.984E-07 | 542.74005 | 1075.5881 |
| STEEAFEVFWR | 11 | Q02241 | Kinesin-like protein KIF23 OS=Homo sa | KIF23 | 110.059 | 2 | 54.3365 | yes | 1.562E-07 | 700.8277 | 792.95581 |
| WVDHKPASNMQT | 32 | Q02241 | Kinesin-like protein KIF23 OS=Homo sa | KIF23 | 110.059 | 5 | 54.3365 | yes | 9.722E-08 | 717.15839 | 737.17078 |
| GMYWTEGR | 8 | Q02241 | Kinesin-like protein KIF23 OS=Homo sa | KIF23 | 110.059 | 2 | 54.3365 | yes | 3.759E-07 | 500.22128 | 645.81799 |
| QLEMQNK | 7 | Q02241 | Kinesin-like protein KIF23 OS=Homo sa | KIF23 | 110.059 | 2 | 54.3365 | yes | 1.438E-07 | 445.72366 | 362.51077 |
| QVDPEFADMITVQ | 17 | Q02241 | Kinesin-like protein KIF23 OS=Homo sa | KIF23 | 110.059 | 3 | 54.3365 | yes | 1.675E-07 | 686.31665 | 271.81253 |
| LIEALEKR | 8 | Q02241 | Kinesin-like protein KIF23 OS=Homo sa | KIF23 | 110.059 | 2 | 54.3365 | yes | 1.628E-07 | 486.29785 | 269.49854 |
| EVVPTFR | 7 | Q02241 | Kinesin-like protein KIF23 OS=Homo sa | KIF23 | 110.059 | 2 | 54.3365 | yes | 1.357E-06 | 424.23724 | 222.31383 |

| | | | | | | | | | | | |
|---|---|---|---|---|---|---|---|---|---|---|---|
| NEIEIEEDHCGR | 12 | Q02241 | Kinesin-like protein KIF23 OS=Homo sa| | KIF23 | 110.059 | 2 | 54.3365 | yes | 1.446E-07 | 750.82281 | 206.19589 |
| AGSQLGPGYQHH/ | 16 | Q02241 | Kinesin-like protein KIF23 OS=Homo sa| | KIF23 | 110.059 | 2 | 54.3365 | yes | 9.733E-08 | 838.41858 | 197.63229 |
| WEKECER | 7 | Q02241 | Kinesin-like protein KIF23 OS=Homo sa| | KIF23 | 110.059 | 2 | 54.3365 | yes | 1.528E-07 | 518.72949 | 192.69608 |
| NHNMYVAGCTEV | 15 | Q02241 | Kinesin-like protein KIF23 OS=Homo sa| | KIF23 | 110.059 | 3 | 54.3365 | yes | 9.722E-08 | 584.26776 | 158.15047 |
| GSVAGSVGCCSER | 13 | Q02487 | Desmocollin-2 OS=Homo sapiens OX=9( | DSC2 | 99.962 | 2 | 51.4882 | yes | 9.722E-08 | 663.28229 | 73.66407 |
| GTQLSQVEILQR | 12 | Q02535 | DNA-binding protein inhibitor ID-3 OS=' | ID3 | 13 | 2 | 51.224564 | yes | 9.722E-08 | 686.38318 | 60.797768 |
| GCYEAVCCLSER | 12 | Q02535 | DNA-binding protein inhibitor ID-3 OS=' | ID3 | 13 | 2 | 51.224564 | yes | 1.294E-07 | 752.30487 | 29.545219 |
| DLTTAGAVTQCYI | 13 | Q02543 | Large ribosomal subunit protein eL20 OS | RPL18A | 20.763 | 2 | 52.442978 | yes | 1.516E-07 | 728.34845 | 29185.32 |
| CHTPPLYR | 8 | Q02543 | Large ribosomal subunit protein eL20 OS | RPL18A | 20.763 | 2 | 52.442978 | yes | 1.229E-07 | 522.25818 | 22248.229 |
| SSGEIVYCGQVFEI | 14 | Q02543 | Large ribosomal subunit protein eL20 OS | RPL18A | 20.763 | 2 | 52.442978 | yes | 1.555E-07 | 801.87708 | 19476.936 |
| IFAPNHVVAK | 10 | Q02543 | Large ribosomal subunit protein eL20 OS | RPL18A | 20.763 | 2 | 52.442978 | yes | 9.722E-08 | 548.31909 | 15850.342 |
| AHSIQIMK | 8 | Q02543 | Large ribosomal subunit protein eL20 OS | RPL18A | 20.763 | 2 | 52.442978 | yes | 3.029E-07 | 464.25766 | 6028.354 |
| AHSIQIMK | 8 | Q02543 | Large ribosomal subunit protein eL20 OS | RPL18A | 20.763 | 2 | 52.442978 | yes | 3.412E-06 | 472.25513 | 6028.354 |
| IKFPLPHR | 8 | Q02543 | Large ribosomal subunit protein eL20 OS | RPL18A | 20.763 | 2 | 52.442978 | yes | 2.072E-06 | 504.3111 | 10211.448 |
| FWYFVSQLK | 9 | Q02543 | Large ribosomal subunit protein eL20 OS | RPL18A | 20.763 | 2 | 52.442978 | yes | 9.722E-08 | 609.32129 | 6718.0205 |
| NFGIWLR | 7 | Q02543 | Large ribosomal subunit protein eL20 OS | RPL18A | 20.763 | 2 | 52.442978 | yes | 1.193E-05 | 453.25323 | 4036.844 |
| SGTHNMYR | 8 | Q02543 | Large ribosomal subunit protein eL20 OS | RPL18A | 20.763 | 2 | 52.442978 | yes | 2.179E-07 | 483.21652 | 857.32336 |
| VEEIAASK | 8 | Q02543 | Large ribosomal subunit protein eL20 OS | RPL18A | 20.763 | 2 | 52.442978 | yes | 9.096E-07 | 423.73199 | 159.25946 |
| VGELKDDDFEK | 11 | Q02750 | Dual specificity mitogen-activated proteir | MAP2K1 | 43.44 | 2 | 42.664963 | yes | 2.927E-07 | 647.81171 | 42.064445 |
| GEDLTEEEDGGIIR | 14 | Q02790 | Peptidyl-prolyl cis-trans isomerase FKBP | FKBP4 | 51.805 | 2 | 40.31782 | yes | 4.091E-07 | 766.85736 | 53.986847 |
| LDPDMAFCANIR | 12 | Q02809 | Procollagen-lysine,2-oxoglutarate 5-dioxy | PLOD1 | 83.551 | 2 | 54.651325 | yes | 9.722E-08 | 711.82886 | 258.84952 |
| IFQNLDGALDEVV | 15 | Q02809 | Procollagen-lysine,2-oxoglutarate 5-dioxy | PLOD1 | 83.551 | 2 | 54.651325 | yes | 9.722E-08 | 837.45905 | 128.20464 |
| SEDYVDIVQGR | 11 | Q02809 | Procollagen-lysine,2-oxoglutarate 5-dioxy | PLOD1 | 83.551 | 2 | 54.651325 | yes | 1.842E-07 | 640.80951 | 126.33326 |
| LQLNYLGNYIPR | 12 | Q02809 | Procollagen-lysine,2-oxoglutarate 5-dioxy | PLOD1 | 83.551 | 2 | 54.651325 | yes | 6.725E-07 | 732.40393 | 121.00636 |
| IFLDPEKR | 8 | Q02809 | Procollagen-lysine,2-oxoglutarate 5-dioxy | PLOD1 | 83.551 | 2 | 54.651325 | yes | 1.486E-07 | 509.29001 | 107.82211 |
| IQALGLGEDWNVI | 14 | Q02809 | Procollagen-lysine,2-oxoglutarate 5-dioxy | PLOD1 | 83.551 | 2 | 54.651325 | yes | 1.197E-07 | 786.40686 | 102.82874 |
| FLGSGGFIGYAPNI | 16 | Q02809 | Procollagen-lysine,2-oxoglutarate 5-dioxy | PLOD1 | 83.551 | 2 | 54.651325 | yes | 9.722E-08 | 814.42755 | 100.70021 |
| HTLGHLLSLDSYR | 13 | Q02809 | Procollagen-lysine,2-oxoglutarate 5-dioxy | PLOD1 | 83.551 | 3 | 54.651325 | yes | 9.935E-08 | 504.6037 | 78.01326 |
| QQDVFMFLTNR | 11 | Q02809 | Procollagen-lysine,2-oxoglutarate 5-dioxy | PLOD1 | 83.551 | 2 | 54.651325 | yes | 1.001E-07 | 699.84534 | 76.874733 |
| FLLEYIAPMTEK | 12 | Q02809 | Procollagen-lysine,2-oxoglutarate 5-dioxy | PLOD1 | 83.551 | 2 | 54.651325 | yes | 3.016E-07 | 727.88361 | 75.446899 |
| AQFDLAFVVR | 10 | Q02809 | Procollagen-lysine,2-oxoglutarate 5-dioxy | PLOD1 | 83.551 | 2 | 54.651325 | yes | 9.722E-08 | 583.32184 | 43.952076 |
| NVIAPLMTR | 9 | Q02809 | Procollagen-lysine,2-oxoglutarate 5-dioxy | PLOD1 | 83.551 | 2 | 54.651325 | yes | 2.506E-05 | 507.79187 | 39.914749 |
| AIPQLQGYLR | 10 | Q02878 | Large ribosomal subunit protein eL6 OS= | RPL6 | 32.728 | 2 | 53.110615 | yes | 1.795E-07 | 579.83514 | 44192.555 |
| YYPTEDVPR | 9 | Q02878 | Large ribosomal subunit protein eL6 OS= | RPL6 | 32.728 | 2 | 53.110615 | yes | 1.02E-07 | 570.27203 | 24117.975 |
| KPFSQHVR | 8 | Q02878 | Large ribosomal subunit protein eL6 OS= | RPL6 | 32.728 | 2 | 53.110615 | yes | 1.164E-07 | 499.78015 | 18044.311 |
| EKYEITEQR | 9 | Q02878 | Large ribosomal subunit protein eL6 OS= | RPL6 | 32.728 | 2 | 53.110615 | yes | 9.722E-08 | 598.30133 | 15888.983 |
| YEITEQR | 7 | Q02878 | Large ribosomal subunit protein eL6 OS= | RPL6 | 32.728 | 2 | 53.110615 | yes | 2.299E-07 | 469.73254 | 6751.0381 |
| HQEGEIFDTEK | 11 | Q02878 | Large ribosomal subunit protein eL6 OS= | RPL6 | 32.728 | 2 | 53.110615 | yes | 1.165E-07 | 666.80695 | 6306.7397 |
| AVDSQILPK | 9 | Q02878 | Large ribosomal subunit protein eL6 OS= | RPL6 | 32.728 | 2 | 53.110615 | yes | 1.127E-06 | 485.78201 | 4877.5234 |

| | | | | | | | | | | | |
|---|---|---|---|---|---|---|---|---|---|---|---|
| FVIATSTK | 8 | Q02878 | Large ribosomal subunit protein eL6 OS= | RPL6 | 32.728 | 2 | 53.110615 | yes | 1.208E-07 | 433.75275 | 2019.6217 |
| VLATVTKPVGGDI | 13 | Q02878 | Large ribosomal subunit protein eL6 OS= | RPL6 | 32.728 | 2 | 53.110615 | yes | 9.082E-07 | 642.8797 | 501.24323 |
| VLATVTKPVGGDI | 13 | Q02878 | Large ribosomal subunit protein eL6 OS= | RPL6 | 32.728 | 2 | 53.110615 | yes | 2.48E-07 | 428.92224 | 501.24323 |
| VLATVTKPVGGDI | 13 | Q02878 | Large ribosomal subunit protein eL6 OS= | RPL6 | 32.728 | 3 | 53.110615 | yes | 9.082E-07 | 642.8797 | 501.24323 |
| VLATVTKPVGGDI | 13 | Q02878 | Large ribosomal subunit protein eL6 OS= | RPL6 | 32.728 | 3 | 53.110615 | yes | 2.48E-07 | 428.92224 | 501.24323 |
| HLTDAYFK | 8 | Q02878 | Large ribosomal subunit protein eL6 OS= | RPL6 | 32.728 | 2 | 53.110615 | yes | 5.632E-07 | 497.75327 | 656.71838 |
| PFSQHVR | 7 | Q02878 | Large ribosomal subunit protein eL6 OS= | RPL6 | 32.728 | 2 | 53.110615 | yes | 1.98E-07 | 435.73267 | 395.98056 |
| SVFALTNGIYPHK | 13 | Q02878 | Large ribosomal subunit protein eL6 OS= | RPL6 | 32.728 | 2 | 53.110615 | yes | 0.0001831 | 723.89063 | 394.37564 |
| EVTFVPGLYK | 10 | Q02880 | DNA topoisomerase 2-beta OS=Homo sa | TOP2B | 183.268 | 2 | 53.624928 | no | 9.722E-08 | 576.8186 | 3243.095 |
| ELILFSNSDNER | 12 | Q02880 | DNA topoisomerase 2-beta OS=Homo sa | TOP2B | 183.268 | 2 | 53.624928 | no | 9.722E-08 | 718.85443 | 2910.3718 |
| FLYDDNQR | 8 | Q02880 | DNA topoisomerase 2-beta OS=Homo sa | TOP2B | 183.268 | 2 | 53.624928 | no | 9.722E-08 | 535.74872 | 1461.9342 |
| IFDEILVNAADNK | 13 | Q02880 | DNA topoisomerase 2-beta OS=Homo sa | TOP2B | 183.268 | 2 | 53.624928 | no | 1.186E-07 | 731.38281 | 1411.4567 |
| EWLTNFMEDR | 10 | Q02880 | DNA topoisomerase 2-beta OS=Homo sa | TOP2B | 183.268 | 2 | 53.624928 | no | 1.145E-07 | 670.8006 | 1190.7043 |
| SLAVSGLGVIGR | 12 | Q02880 | DNA topoisomerase 2-beta OS=Homo sa | TOP2B | 183.268 | 2 | 53.624928 | yes | 1.441E-07 | 564.84039 | 898.56091 |
| QIMENAEINNIIK | 13 | Q02880 | DNA topoisomerase 2-beta OS=Homo sa | TOP2B | 183.268 | 2 | 53.624928 | no | 1.123E-07 | 765.40326 | 868.25238 |
| EYHTDTTVK | 9 | Q02880 | DNA topoisomerase 2-beta OS=Homo sa | TOP2B | 183.268 | 2 | 53.624928 | no | 1.843E-07 | 547.26166 | 771.93359 |
| ENMTLQPK | 8 | Q02880 | DNA topoisomerase 2-beta OS=Homo sa | TOP2B | 183.268 | 2 | 53.624928 | no | 9.212E-07 | 480.7446 | 680.79321 |
| GIPVVEHK | 8 | Q02880 | DNA topoisomerase 2-beta OS=Homo sa | TOP2B | 183.268 | 2 | 53.624928 | no | 2.049E-07 | 439.75836 | 581.7981 |
| AYDLAGSCR | 9 | Q02880 | DNA topoisomerase 2-beta OS=Homo sa | TOP2B | 183.268 | 2 | 53.624928 | yes | 9.819E-08 | 506.72946 | 523.16827 |
| KVLFTCFK | 8 | Q02880 | DNA topoisomerase 2-beta OS=Homo sa | TOP2B | 183.268 | 2 | 53.624928 | no | 1.206E-07 | 521.79132 | 402.44049 |
| DKLDETGVALK | 11 | Q02880 | DNA topoisomerase 2-beta OS=Homo sa | TOP2B | 183.268 | 2 | 53.624928 | yes | 9.722E-08 | 594.82715 | 389.34271 |
| LQLEETMPSPYGR | 13 | Q02880 | DNA topoisomerase 2-beta OS=Homo sa | TOP2B | 183.268 | 2 | 53.624928 | yes | 1.138E-07 | 760.87433 | 368.48877 |
| LCNIFSTK | 8 | Q02880 | DNA topoisomerase 2-beta OS=Homo sa | TOP2B | 183.268 | 2 | 53.624928 | no | 2.85E-07 | 491.75494 | 365.35352 |
| SIPSLVDGFKPGQI | 14 | Q02880 | DNA topoisomerase 2-beta OS=Homo sa | TOP2B | 183.268 | 2 | 53.624928 | yes | 1.377E-07 | 750.91211 | 347.91104 |
| EYFADMER | 8 | Q02880 | DNA topoisomerase 2-beta OS=Homo sa | TOP2B | 183.268 | 2 | 53.624928 | yes | 1.8E-07 | 530.72388 | 292.83426 |
| LPNYDAR | 7 | Q02880 | DNA topoisomerase 2-beta OS=Homo sa | TOP2B | 183.268 | 2 | 53.624928 | yes | 2.115E-07 | 424.71667 | 279.08963 |
| LDKDIVALMTR | 11 | Q02880 | DNA topoisomerase 2-beta OS=Homo sa | TOP2B | 183.268 | 2 | 53.624928 | yes | 1.113E-07 | 637.86047 | 263.9422 |
| IIPEITAMK | 9 | Q02880 | DNA topoisomerase 2-beta OS=Homo sa | TOP2B | 183.268 | 2 | 53.624928 | yes | 1.933E-07 | 508.29645 | 242.81755 |
| KEWLTNFMEDR | 11 | Q02880 | DNA topoisomerase 2-beta OS=Homo sa | TOP2B | 183.268 | 2 | 53.624928 | no | 9.722E-08 | 734.84808 | 234.82999 |
| VIHELANER | 9 | Q02880 | DNA topoisomerase 2-beta OS=Homo sa | TOP2B | 183.268 | 2 | 53.624928 | yes | 3.301E-07 | 540.79346 | 229.85274 |
| LHGLPEQFLYGTA | 15 | Q02880 | DNA topoisomerase 2-beta OS=Homo sa | TOP2B | 183.268 | 2 | 53.624928 | yes | 1.027E-07 | 837.94611 | 215.94199 |
| EWLVGMLGAESTI | 13 | Q02880 | DNA topoisomerase 2-beta OS=Homo sa | TOP2B | 183.268 | 2 | 53.624928 | yes | 2.024E-07 | 710.86066 | 211.42805 |
| DLIQMLVQR | 9 | Q02880 | DNA topoisomerase 2-beta OS=Homo sa | TOP2B | 183.268 | 2 | 53.624928 | yes | 1.197E-07 | 558.31573 | 202.64915 |
| LLFPAVDDNLLK | 12 | Q02880 | DNA topoisomerase 2-beta OS=Homo sa | TOP2B | 183.268 | 2 | 53.624928 | yes | 1.197E-07 | 679.38995 | 188.83659 |
| EDVLAGMSGK | 10 | Q02880 | DNA topoisomerase 2-beta OS=Homo sa | TOP2B | 183.268 | 2 | 53.624928 | yes | 1.117E-07 | 503.74731 | 179.62009 |
| EKVEELIK | 8 | Q02880 | DNA topoisomerase 2-beta OS=Homo sa | TOP2B | 183.268 | 2 | 53.624928 | yes | 1.259E-07 | 494.28967 | 175.04153 |
| HLTYNDFINK | 10 | Q02880 | DNA topoisomerase 2-beta OS=Homo sa | TOP2B | 183.268 | 2 | 53.624928 | yes | 1.084E-07 | 632.81964 | 154.46777 |
| HVDYVVDQVVGK | 12 | Q02880 | DNA topoisomerase 2-beta OS=Homo sa | TOP2B | 183.268 | 2 | 53.624928 | yes | 9.722E-08 | 679.35913 | 142.11655 |
| NTVEITELPVR | 11 | Q02880 | DNA topoisomerase 2-beta OS=Homo sa | TOP2B | 183.268 | 2 | 53.624928 | yes | 4.013E-07 | 635.8537 | 137.6965 |

| | | | | | | | | | | | | |
|---|---|---|---|---|---|---|---|---|---|---|---|---|
| KYETVQDILK | 10 | Q02880 | DNA topoisomerase 2-beta OS=Homo sa| | TOP2B | 183.268 | 2 | 53.624928 | yes | 1.439E-07 | 618.84534 | 133.8244 |
| KEWLVGMLGAES | 14 | Q02880 | DNA topoisomerase 2-beta OS=Homo sa| | TOP2B | 183.268 | 2 | 53.624928 | yes | 9.722E-08 | 774.90814 | 132.7684 |
| TPALISDYK | 9 | Q02880 | DNA topoisomerase 2-beta OS=Homo sa| | TOP2B | 183.268 | 2 | 53.624928 | yes | 1.855E-07 | 504.27402 | 129.60626 |
| SYDDAESLK | 9 | Q02880 | DNA topoisomerase 2-beta OS=Homo sa| | TOP2B | 183.268 | 2 | 53.624928 | yes | 4.547E-07 | 514.23254 | 125.58586 |
| GFQQISFVNSIATT | 15 | Q02880 | DNA topoisomerase 2-beta OS=Homo sa| | TOP2B | 183.268 | 2 | 53.624928 | yes | 1.349E-07 | 820.93579 | 125.30581 |
| LSYYGLR | 7 | Q02880 | DNA topoisomerase 2-beta OS=Homo sa| | TOP2B | 183.268 | 2 | 53.624928 | yes | 9.722E-08 | 436.23724 | 122.60541 |
| WDVCLTLSEK | 10 | Q02880 | DNA topoisomerase 2-beta OS=Homo sa| | TOP2B | 183.268 | 2 | 53.624928 | yes | 4.348E-07 | 625.80792 | 115.69489 |
| KGDLDTAAVK | 10 | Q02880 | DNA topoisomerase 2-beta OS=Homo sa| | TOP2B | 183.268 | 2 | 53.624928 | yes | 2.492E-07 | 509.28238 | 112.61282 |
| AGVSVKPFQVK | 11 | Q02880 | DNA topoisomerase 2-beta OS=Homo sa| | TOP2B | 183.268 | 2 | 53.624928 | yes | 1.303E-07 | 580.34534 | 109.87638 |
| KSPSDLWK | 8 | Q02880 | DNA topoisomerase 2-beta OS=Homo sa| | TOP2B | 183.268 | 2 | 53.624928 | no | 1.603E-07 | 480.76108 | 100.98638 |
| LAQAEAAGLHK | 11 | Q02880 | DNA topoisomerase 2-beta OS=Homo sa| | TOP2B | 183.268 | 2 | 53.624928 | yes | 4.712E-06 | 554.80908 | 85.712456 |
| HGFLEEFITPIVK | 13 | Q02880 | DNA topoisomerase 2-beta OS=Homo sa| | TOP2B | 183.268 | 2 | 53.624928 | yes | 1.277E-07 | 765.42175 | 71.862419 |
| EIVNNVR | 7 | Q02880 | DNA topoisomerase 2-beta OS=Homo sa| | TOP2B | 183.268 | 2 | 53.624928 | yes | 5.487E-07 | 422.23779 | 68.051041 |
| YGVFPLR | 7 | Q02880 | DNA topoisomerase 2-beta OS=Homo sa| | TOP2B | 183.268 | 2 | 53.624928 | no | 9.174E-07 | 426.24234 | 65.970222 |
| GTIQELGQNQYAV | 22 | Q02880 | DNA topoisomerase 2-beta OS=Homo sa| | TOP2B | 183.268 | 3 | 53.624928 | yes | 1.011E-07 | 808.41156 | 62.453243 |
| LQTTLTCNSMVLF | 20 | Q02880 | DNA topoisomerase 2-beta OS=Homo sa| | TOP2B | 183.268 | 3 | 53.624928 | yes | 2.175E-07 | 790.37555 | 56.967869 |
| HFDGEDYTCITFQI | 18 | Q02880 | DNA topoisomerase 2-beta OS=Homo sa| | TOP2B | 183.268 | 3 | 53.624928 | yes | 4.334E-07 | 724.98962 | 27.782806 |
| EDLAAFVEELDK | 12 | Q02880 | DNA topoisomerase 2-beta OS=Homo sa| | TOP2B | 183.268 | 2 | 53.624928 | yes | 3.385E-06 | 689.84045 | 13.346924 |
| VVDEDPDER | 9 | Q02930 | Cyclic AMP-responsive element-binding | CREB5 | 56.919 | 2 | 45.45191 | yes | 1.846E-07 | 537.24091 | 38.853016 |
| EEGVLTLWR | 9 | Q02978 | Mitochondrial 2-oxoglutarate/malate carri | SLC25A11 | 34.062 | 2 | 50.561993 | yes | 1.049E-07 | 551.79822 | 437.60306 |
| LTGADGTPPGFLLI | 14 | Q02978 | Mitochondrial 2-oxoglutarate/malate carri | SLC25A11 | 34.062 | 2 | 50.561993 | yes | 1.084E-07 | 693.88501 | 307.66705 |
| LGIYTVLFER | 10 | Q02978 | Mitochondrial 2-oxoglutarate/malate carri | SLC25A11 | 34.062 | 2 | 50.561993 | yes | 9.766E-08 | 605.84515 | 243.35818 |
| GCIPTMAR | 8 | Q02978 | Mitochondrial 2-oxoglutarate/malate carri | SLC25A11 | 34.062 | 2 | 50.561993 | yes | 1.606E-07 | 453.22021 | 207.25087 |
| NVFNALIR | 8 | Q02978 | Mitochondrial 2-oxoglutarate/malate carri | SLC25A11 | 34.062 | 2 | 50.561993 | yes | 4.998E-07 | 473.77707 | 182.40707 |
| MIDGKPEYK | 9 | Q02978 | Mitochondrial 2-oxoglutarate/malate carri | SLC25A11 | 34.062 | 2 | 50.561993 | yes | 1.376E-07 | 540.77332 | 119.37602 |
| AVVVNAAQLASY: | 16 | Q02978 | Mitochondrial 2-oxoglutarate/malate carri | SLC25A11 | 34.062 | 2 | 50.561993 | yes | 1.414E-07 | 818.43866 | 109.99137 |
| TSFHALTSILK | 11 | Q02978 | Mitochondrial 2-oxoglutarate/malate carri | SLC25A11 | 34.062 | 2 | 50.561993 | yes | 1.686E-07 | 609.34808 | 79.391853 |
| YEGFFSLWK | 9 | Q02978 | Mitochondrial 2-oxoglutarate/malate carri | SLC25A11 | 34.062 | 2 | 50.561993 | yes | 9.964E-08 | 588.78986 | 51.921474 |
| LGPHTVLTFIFLEQ | 17 | Q02978 | Mitochondrial 2-oxoglutarate/malate carri | SLC25A11 | 34.062 | 3 | 50.561993 | yes | 6.589E-07 | 663.36066 | 13.848723 |
| IDQILESLER | 10 | Q03001 | Dystonin OS=Homo sapiens OX=9606 G | DST | 860.662 | 2 | 55.676228 | yes | 9.81E-08 | 608.33258 | 1091.8939 |
| DLEDPGIDPSVVK | 13 | Q03001 | Dystonin OS=Homo sapiens OX=9606 G | DST | 860.662 | 2 | 55.676228 | yes | 9.722E-08 | 692.35376 | 849.13129 |
| NLLISVQSR | 9 | Q03001 | Dystonin OS=Homo sapiens OX=9606 G | DST | 860.662 | 2 | 55.676228 | yes | 2.608E-07 | 515.30621 | 791.87439 |
| LASMSPIGTDLET\ | 15 | Q03001 | Dystonin OS=Homo sapiens OX=9606 G | DST | 860.662 | 2 | 55.676228 | yes | 4.973E-07 | 781.41071 | 782.73584 |
| LMEWLEESEK | 10 | Q03001 | Dystonin OS=Homo sapiens OX=9606 G | DST | 860.662 | 2 | 55.676228 | yes | 1.008E-07 | 647.30518 | 698.17389 |
| LEMLQQIANR | 10 | Q03001 | Dystonin OS=Homo sapiens OX=9606 G | DST | 860.662 | 2 | 55.676228 | yes | 9.722E-08 | 608.32935 | 680.66638 |
| VEEIDAAILR | 10 | Q03001 | Dystonin OS=Homo sapiens OX=9606 G | DST | 860.662 | 2 | 55.676228 | yes | 1.131E-07 | 564.81659 | 643.07129 |
| LVSDTITQK | 9 | Q03001 | Dystonin OS=Homo sapiens OX=9606 G | DST | 860.662 | 2 | 55.676228 | yes | 2.674E-07 | 502.78476 | 627.77789 |
| LASALAGLIAK | 11 | Q03001 | Dystonin OS=Homo sapiens OX=9606 G | DST | 860.662 | 2 | 55.676228 | yes | 1.905E-07 | 514.32916 | 613.16156 |
| TQALTEVDVPGK | 12 | Q03001 | Dystonin OS=Homo sapiens OX=9606 G | DST | 860.662 | 2 | 55.676228 | yes | 9.848E-08 | 629.33789 | 610.8205 |

| | | | | | | | | | | |
|---|---|---|---|---|---|---|---|---|---|---|
| PLGGLPETAK | 10 | Q03001 | Dystonin OS=Homo sapiens OX=9606 G DST | 860.662 | 2 | 55.676228 | yes | 4.149E-07 | 491.78201 | 574.72052 |
| YVAADTLYSQIK | 12 | Q03001 | Dystonin OS=Homo sapiens OX=9606 G DST | 860.662 | 2 | 55.676228 | yes | 1.147E-07 | 686.36139 | 572.13513 |
| TLEQALQLAR | 10 | Q03001 | Dystonin OS=Homo sapiens OX=9606 G DST | 860.662 | 2 | 55.676228 | yes | 1.375E-07 | 571.83002 | 556.61432 |
| GFHSGEDSGLITTA | 17 | Q03001 | Dystonin OS=Homo sapiens OX=9606 G DST | 860.662 | 3 | 55.676228 | yes | 4.622E-07 | 563.94452 | 276.15784 |
| GFHSGEDSGLITTA | 17 | Q03001 | Dystonin OS=Homo sapiens OX=9606 G DST | 860.662 | 3 | 55.676228 | yes | 2.762E-07 | 845.41321 | 276.15784 |
| GFHSGEDSGLITTA | 17 | Q03001 | Dystonin OS=Homo sapiens OX=9606 G DST | 860.662 | 2 | 55.676228 | yes | 4.622E-07 | 563.94452 | 276.15784 |
| GFHSGEDSGLITTA | 17 | Q03001 | Dystonin OS=Homo sapiens OX=9606 G DST | 860.662 | 2 | 55.676228 | yes | 2.762E-07 | 845.41321 | 276.15784 |
| VAQALCEDLSALV | 14 | Q03001 | Dystonin OS=Homo sapiens OX=9606 G DST | 860.662 | 2 | 55.676228 | yes | 9.833E-08 | 758.90564 | 543.4657 |
| VQMQELSTR | 9 | Q03001 | Dystonin OS=Homo sapiens OX=9606 G DST | 860.662 | 2 | 55.676228 | yes | 9.82E-08 | 546.2793 | 539.3949 |
| VLQEDILLR | 9 | Q03001 | Dystonin OS=Homo sapiens OX=9606 G DST | 860.662 | 2 | 55.676228 | yes | 1.002E-07 | 549.82953 | 525.84277 |
| SIDELNSAWDSLN | 14 | Q03001 | Dystonin OS=Homo sapiens OX=9606 G DST | 860.662 | 2 | 55.676228 | yes | 9.722E-08 | 796.38354 | 519.04675 |
| TLATQLNQQK | 10 | Q03001 | Dystonin OS=Homo sapiens OX=9606 G DST | 860.662 | 2 | 55.676228 | yes | 4.537E-07 | 572.81964 | 511.63644 |
| FFLGNQFGDSQQL | 14 | Q03001 | Dystonin OS=Homo sapiens OX=9606 G DST | 860.662 | 2 | 55.676228 | yes | 2.402E-07 | 828.9101 | 504.3334 |
| SLDSELEIANDPDK | 14 | Q03001 | Dystonin OS=Homo sapiens OX=9606 G DST | 860.662 | 2 | 55.676228 | yes | 4.301E-07 | 773.36755 | 485.75073 |
| LQNVQIALDYLK | 12 | Q03001 | Dystonin OS=Homo sapiens OX=9606 G DST | 860.662 | 2 | 55.676228 | yes | 1.163E-07 | 709.40613 | 482.64429 |
| ISEIQMTAPLK | 11 | Q03001 | Dystonin OS=Homo sapiens OX=9606 G DST | 860.662 | 2 | 55.676228 | yes | 9.722E-08 | 615.84174 | 474.63715 |
| LDDMLSELR | 9 | Q03001 | Dystonin OS=Homo sapiens OX=9606 G DST | 860.662 | 2 | 55.676228 | yes | 9.819E-08 | 546.27368 | 462.90134 |
| DYHAELMR | 8 | Q03001 | Dystonin OS=Homo sapiens OX=9606 G DST | 860.662 | 2 | 55.676228 | yes | 1.478E-07 | 517.73981 | 458.70544 |
| YYQADQLVQR | 10 | Q03001 | Dystonin OS=Homo sapiens OX=9606 G DST | 860.662 | 2 | 55.676228 | yes | 9.722E-08 | 642.32257 | 456.56161 |
| ALEDDIINHNK | 11 | Q03001 | Dystonin OS=Homo sapiens OX=9606 G DST | 860.662 | 2 | 55.676228 | yes | 9.722E-08 | 641.32532 | 441.43906 |
| LLDVMELAEK | 10 | Q03001 | Dystonin OS=Homo sapiens OX=9606 G DST | 860.662 | 2 | 55.676228 | yes | 9.819E-08 | 580.81519 | 436.11765 |
| VDMVTEQLHSK | 11 | Q03001 | Dystonin OS=Homo sapiens OX=9606 G DST | 860.662 | 2 | 55.676228 | yes | 9.722E-08 | 643.8241 | 434.83746 |
| LAQDLVVEASDSK | 13 | Q03001 | Dystonin OS=Homo sapiens OX=9606 G DST | 860.662 | 2 | 55.676228 | yes | 9.722E-08 | 687.85919 | 430.19351 |
| IISQHQAVIIATQSA | 21 | Q03001 | Dystonin OS=Homo sapiens OX=9606 G DST | 860.662 | 3 | 55.676228 | yes | 2.636E-07 | 764.10913 | 419.41821 |
| DDLHESVFR | 9 | Q03001 | Dystonin OS=Homo sapiens OX=9606 G DST | 860.662 | 2 | 55.676228 | yes | 9.728E-08 | 559.26727 | 413.77753 |
| LQITLTR | 7 | Q03001 | Dystonin OS=Homo sapiens OX=9606 G DST | 860.662 | 2 | 55.676228 | yes | 9.978E-08 | 422.76617 | 407.03531 |
| NIMLEQDIAGR | 11 | Q03001 | Dystonin OS=Homo sapiens OX=9606 G DST | 860.662 | 2 | 55.676228 | yes | 9.722E-08 | 630.32428 | 390.37595 |
| LEMSAVADIFDR | 12 | Q03001 | Dystonin OS=Homo sapiens OX=9606 G DST | 860.662 | 2 | 55.676228 | yes | 9.722E-08 | 683.83722 | 377.44473 |
| HSVYDTTNR | 9 | Q03001 | Dystonin OS=Homo sapiens OX=9606 G DST | 860.662 | 2 | 55.676228 | yes | 9.849E-08 | 546.75708 | 374.34308 |
| LSVQDYSTEGLWK | 13 | Q03001 | Dystonin OS=Homo sapiens OX=9606 G DST | 860.662 | 2 | 55.676228 | yes | 1.027E-07 | 763.38031 | 367.97363 |
| VFANEVNSHR | 10 | Q03001 | Dystonin OS=Homo sapiens OX=9606 G DST | 860.662 | 2 | 55.676228 | yes | 9.722E-08 | 586.79401 | 360.36285 |
| CDWIDQAIVK | 10 | Q03001 | Dystonin OS=Homo sapiens OX=9606 G DST | 860.662 | 2 | 55.676228 | yes | 2.024E-07 | 624.30811 | 354.36096 |
| TFPNNPVELK | 10 | Q03001 | Dystonin OS=Homo sapiens OX=9606 G DST | 860.662 | 2 | 55.676228 | yes | 3.109E-07 | 579.81134 | 353.54913 |
| STQYQSLLR | 9 | Q03001 | Dystonin OS=Homo sapiens OX=9606 G DST | 860.662 | 2 | 55.676228 | yes | 9.722E-08 | 548.29327 | 350.74133 |
| LIWDSLEER | 9 | Q03001 | Dystonin OS=Homo sapiens OX=9606 G DST | 860.662 | 2 | 55.676228 | yes | 1.073E-07 | 580.80096 | 341.67078 |
| DLDFDWHK | 8 | Q03001 | Dystonin OS=Homo sapiens OX=9606 G DST | 860.662 | 2 | 55.676228 | yes | 1.169E-07 | 538.24579 | 336.42404 |
| LQGICDLLTQTENI | 14 | Q03001 | Dystonin OS=Homo sapiens OX=9606 G DST | 860.662 | 2 | 55.676228 | yes | 1.142E-07 | 830.9198 | 305.57144 |
| FQVEQIGDNK | 10 | Q03001 | Dystonin OS=Homo sapiens OX=9606 G DST | 860.662 | 2 | 55.676228 | yes | 9.869E-08 | 589.29602 | 301.28583 |
| VHYETALAESEK | 12 | Q03001 | Dystonin OS=Homo sapiens OX=9606 G DST | 860.662 | 2 | 55.676228 | yes | 9.722E-08 | 688.83826 | 297.19962 |

| | | | | | | | | | | |
|---|---|---|---|---|---|---|---|---|---|---|
| LKDDLGTITNK | 11 | Q03001 | Dystonin OS=Homo sapiens OX=9606 G DST | 860.662 | 2 | 55.676228 | yes | 2.512E-07 | 609.34045 | 294.76544 |
| YLTMLQTQQK | 10 | Q03001 | Dystonin OS=Homo sapiens OX=9606 G DST | 860.662 | 2 | 55.676228 | yes | 2.258E-07 | 627.33154 | 294.69238 |
| LCEEEAVIADK | 11 | Q03001 | Dystonin OS=Homo sapiens OX=9606 G DST | 860.662 | 2 | 55.676228 | yes | 9.722E-08 | 638.80811 | 292.35309 |
| SEAYQQQIEMER | 12 | Q03001 | Dystonin OS=Homo sapiens OX=9606 G DST | 860.662 | 2 | 55.676228 | yes | 9.722E-08 | 756.34338 | 290.28912 |
| ENTAYFEFFNDAK | 13 | Q03001 | Dystonin OS=Homo sapiens OX=9606 G DST | 860.662 | 2 | 55.676228 | yes | 9.81E-08 | 798.35425 | 287.8804 |
| QEFIDGILSSK | 11 | Q03001 | Dystonin OS=Homo sapiens OX=9606 G DST | 860.662 | 2 | 55.676228 | yes | 1.101E-07 | 618.82715 | 285.49976 |
| RLEEFYSK | 8 | Q03001 | Dystonin OS=Homo sapiens OX=9606 G DST | 860.662 | 2 | 55.676228 | yes | 9.722E-08 | 536.2771 | 284.57111 |
| AALQLQLNTIK | 11 | Q03001 | Dystonin OS=Homo sapiens OX=9606 G DST | 860.662 | 2 | 55.676228 | yes | 1.015E-07 | 606.86914 | 279.36026 |
| QLSLLDSR | 8 | Q03001 | Dystonin OS=Homo sapiens OX=9606 G DST | 860.662 | 2 | 55.676228 | yes | 9.722E-08 | 466.26401 | 272.05127 |
| LVNCEPIGTQASK | 13 | Q03001 | Dystonin OS=Homo sapiens OX=9606 G DST | 860.662 | 2 | 55.676228 | yes | 1.121E-07 | 708.86121 | 269.06082 |
| TIAEGENLLLK | 11 | Q03001 | Dystonin OS=Homo sapiens OX=9606 G DST | 860.662 | 2 | 55.676228 | yes | 4.643E-07 | 600.84534 | 268.37863 |
| YAEQYSATVK | 10 | Q03001 | Dystonin OS=Homo sapiens OX=9606 G DST | 860.662 | 2 | 55.676228 | yes | 1.049E-07 | 580.28516 | 261.91772 |
| ALYNQYLQFK | 10 | Q03001 | Dystonin OS=Homo sapiens OX=9606 G DST | 860.662 | 2 | 55.676228 | yes | 9.722E-08 | 644.34021 | 261.84576 |
| AIEEFESSLK | 10 | Q03001 | Dystonin OS=Homo sapiens OX=9606 G DST | 860.662 | 2 | 55.676228 | yes | 1.501E-07 | 576.79279 | 261.49448 |
| YITISGNR | 8 | Q03001 | Dystonin OS=Homo sapiens OX=9606 G DST | 860.662 | 2 | 55.676228 | yes | 9.722E-08 | 462.25089 | 260.20889 |
| ALQGQISSQQVAV | 15 | Q03001 | Dystonin OS=Homo sapiens OX=9606 G DST | 860.662 | 2 | 55.676228 | yes | 9.722E-08 | 793.43085 | 260.05923 |
| WDSLTGQLSDR | 11 | Q03001 | Dystonin OS=Homo sapiens OX=9606 G DST | 860.662 | 2 | 55.676228 | yes | 9.722E-08 | 639.30969 | 257.73737 |
| LQTFLETK | 8 | Q03001 | Dystonin OS=Homo sapiens OX=9606 G DST | 860.662 | 2 | 55.676228 | yes | 1.12E-07 | 490.27658 | 257.50797 |
| FHGVLPDDEDALR | 13 | Q03001 | Dystonin OS=Homo sapiens OX=9606 G DST | 860.662 | 2 | 55.676228 | yes | 1.062E-07 | 742.36243 | 251.0127 |
| QTTGDEVLIIQDK | 13 | Q03001 | Dystonin OS=Homo sapiens OX=9606 G DST | 860.662 | 2 | 55.676228 | yes | 1.002E-07 | 730.38556 | 250.6212 |
| LEQDQTSAQLQVQ | 14 | Q03001 | Dystonin OS=Homo sapiens OX=9606 G DST | 860.662 | 2 | 55.676228 | yes | 9.722E-08 | 808.41791 | 247.3233 |
| SFSEDVISHK | 10 | Q03001 | Dystonin OS=Homo sapiens OX=9606 G DST | 860.662 | 2 | 55.676228 | yes | 2.414E-07 | 574.78278 | 244.60657 |
| AGKPPFSK | 8 | Q03001 | Dystonin OS=Homo sapiens OX=9606 G DST | 860.662 | 2 | 55.676228 | yes | 4.042E-07 | 416.23981 | 238.74754 |
| HKDIIDDLVK | 10 | Q03001 | Dystonin OS=Homo sapiens OX=9606 G DST | 860.662 | 2 | 55.676228 | yes | 1.143E-07 | 598.33771 | 237.84015 |
| LIIAMLER | 8 | Q03001 | Dystonin OS=Homo sapiens OX=9606 G DST | 860.662 | 2 | 55.676228 | yes | 9.722E-08 | 479.79135 | 233.41119 |
| DKWDTICGK | 9 | Q03001 | Dystonin OS=Homo sapiens OX=9606 G DST | 860.662 | 2 | 55.676228 | yes | 1.549E-07 | 561.76605 | 232.88181 |
| EQVETLWPWIDK | 12 | Q03001 | Dystonin OS=Homo sapiens OX=9606 G DST | 860.662 | 2 | 55.676228 | yes | 9.849E-08 | 772.39319 | 228.02367 |
| LEDLVQESMEEK | 12 | Q03001 | Dystonin OS=Homo sapiens OX=9606 G DST | 860.662 | 2 | 55.676228 | yes | 9.722E-08 | 725.34253 | 225.68146 |
| SELLQQALCNAK | 12 | Q03001 | Dystonin OS=Homo sapiens OX=9606 G DST | 860.662 | 2 | 55.676228 | yes | 9.722E-08 | 687.8559 | 222.02118 |
| LMSLGDIR | 8 | Q03001 | Dystonin OS=Homo sapiens OX=9606 G DST | 860.662 | 2 | 55.676228 | yes | 5.228E-07 | 452.74966 | 221.36032 |
| EISSHGLPSDK | 11 | Q03001 | Dystonin OS=Homo sapiens OX=9606 G DST | 860.662 | 2 | 55.676228 | yes | 9.722E-08 | 585.29346 | 209.5661 |
| LTHSLQEELEK | 11 | Q03001 | Dystonin OS=Homo sapiens OX=9606 G DST | 860.662 | 2 | 55.676228 | yes | 1.654E-07 | 663.84863 | 104.42435 |
| LTHSLQEELEK | 11 | Q03001 | Dystonin OS=Homo sapiens OX=9606 G DST | 860.662 | 2 | 55.676228 | yes | 2.589E-06 | 442.90152 | 104.42435 |
| LTHSLQEELEK | 11 | Q03001 | Dystonin OS=Homo sapiens OX=9606 G DST | 860.662 | 3 | 55.676228 | yes | 1.654E-07 | 663.84863 | 104.42435 |
| LTHSLQEELEK | 11 | Q03001 | Dystonin OS=Homo sapiens OX=9606 G DST | 860.662 | 3 | 55.676228 | yes | 2.589E-06 | 442.90152 | 104.42435 |
| LLDPEDVDVSSPD | 15 | Q03001 | Dystonin OS=Homo sapiens OX=9606 G DST | 860.662 | 2 | 55.676228 | yes | 9.722E-08 | 829.3938 | 205.11958 |
| IMTACSEEEK | 10 | Q03001 | Dystonin OS=Homo sapiens OX=9606 G DST | 860.662 | 2 | 55.676228 | yes | 1.084E-07 | 599.25995 | 202.83092 |
| SAETNIDQDINNLK | 14 | Q03001 | Dystonin OS=Homo sapiens OX=9606 G DST | 860.662 | 2 | 55.676228 | yes | 1.461E-07 | 787.88647 | 202.47263 |
| TLTAQSHMYEK | 11 | Q03001 | Dystonin OS=Homo sapiens OX=9606 G DST | 860.662 | 2 | 55.676228 | yes | 9.722E-08 | 654.81628 | 199.61977 |

| | | | | | | | | | | |
|---|---|---|---|---|---|---|---|---|---|---|
| DMQGSAQALAEV | 14 | Q03001 | Dystonin OS=Homo sapiens OX=9606 G DST | 860.662 | 2 | 55.676228 | yes | 2.563E-07 | 723.86652 | 199.12135 |
| DAYKPITDADK | 11 | Q03001 | Dystonin OS=Homo sapiens OX=9606 G DST | 860.662 | 2 | 55.676228 | yes | 3.798E-07 | 618.80896 | 193.67747 |
| AAQLQEALLHCGI | 13 | Q03001 | Dystonin OS=Homo sapiens OX=9606 G DST | 860.662 | 2 | 55.676228 | yes | 2.073E-07 | 733.88025 | 186.75323 |
| VELFENLSEK | 10 | Q03001 | Dystonin OS=Homo sapiens OX=9606 G DST | 860.662 | 2 | 55.676228 | yes | 9.722E-08 | 604.3139 | 182.58246 |
| QQVQILLQEFATR | 13 | Q03001 | Dystonin OS=Homo sapiens OX=9606 G DST | 860.662 | 2 | 55.676228 | yes | 9.722E-08 | 787.43848 | 180.91583 |
| WINQHLMK | 8 | Q03001 | Dystonin OS=Homo sapiens OX=9606 G DST | 860.662 | 2 | 55.676228 | yes | 9.722E-08 | 535.28424 | 175.1622 |
| LILAGNALQSDSK | 13 | Q03001 | Dystonin OS=Homo sapiens OX=9606 G DST | 860.662 | 2 | 55.676228 | yes | 9.722E-08 | 665.37225 | 172.508 |
| FILADGASQGMAA | 15 | Q03001 | Dystonin OS=Homo sapiens OX=9606 G DST | 860.662 | 2 | 55.676228 | yes | 9.722E-08 | 777.89032 | 170.24895 |
| DFQAWLDTK | 9 | Q03001 | Dystonin OS=Homo sapiens OX=9606 G DST | 860.662 | 2 | 55.676228 | yes | 1.108E-07 | 562.27454 | 166.06342 |
| HVNDLYEDLR | 10 | Q03001 | Dystonin OS=Homo sapiens OX=9606 G DST | 860.662 | 2 | 55.676228 | yes | 1.998E-07 | 637.31219 | 165.82857 |
| LQGIGHFQNTIR | 12 | Q03001 | Dystonin OS=Homo sapiens OX=9606 G DST | 860.662 | 2 | 55.676228 | yes | 4.265E-07 | 692.37823 | 162.83105 |
| MVAEDNER | 8 | Q03001 | Dystonin OS=Homo sapiens OX=9606 G DST | 860.662 | 2 | 55.676228 | yes | 6.909E-07 | 482.21365 | 158.38301 |
| NYDTICQINSER | 12 | Q03001 | Dystonin OS=Homo sapiens OX=9606 G DST | 860.662 | 2 | 55.676228 | yes | 1.437E-07 | 756.841 | 156.90939 |
| EFSILLQK | 8 | Q03001 | Dystonin OS=Homo sapiens OX=9606 G DST | 860.662 | 2 | 55.676228 | yes | 3.664E-07 | 489.28696 | 156.39462 |
| LRDEIMALR | 9 | Q03001 | Dystonin OS=Homo sapiens OX=9606 G DST | 860.662 | 2 | 55.676228 | yes | 1.182E-07 | 558.81335 | 144.50372 |
| LLQGYHPNDIEK | 12 | Q03001 | Dystonin OS=Homo sapiens OX=9606 G DST | 860.662 | 2 | 55.676228 | yes | 6.71E-07 | 713.86987 | 143.35242 |
| LDVLESLIK | 9 | Q03001 | Dystonin OS=Homo sapiens OX=9606 G DST | 860.662 | 2 | 55.676228 | yes | 2.254E-07 | 515.31317 | 142.32607 |
| ATNELIWLNEK | 11 | Q03001 | Dystonin OS=Homo sapiens OX=9606 G DST | 860.662 | 2 | 55.676228 | yes | 9.722E-08 | 665.8537 | 138.74202 |
| ILTTEQTK | 8 | Q03001 | Dystonin OS=Homo sapiens OX=9606 G DST | 860.662 | 2 | 55.676228 | yes | 2.401E-07 | 467.2662 | 134.05817 |
| CENFTTCWR | 9 | Q03001 | Dystonin OS=Homo sapiens OX=9606 G DST | 860.662 | 2 | 55.676228 | yes | 3.301E-07 | 637.25806 | 133.57741 |
| AADPSSLQSHIPVI | 16 | Q03001 | Dystonin OS=Homo sapiens OX=9606 G DST | 860.662 | 2 | 55.676228 | yes | 9.722E-08 | 839.44397 | 131.2939 |
| EVIPQEIEEVK | 11 | Q03001 | Dystonin OS=Homo sapiens OX=9606 G DST | 860.662 | 2 | 55.676228 | yes | 2.281E-07 | 656.85339 | 130.09598 |
| LEEQIAQHK | 9 | Q03001 | Dystonin OS=Homo sapiens OX=9606 G DST | 860.662 | 2 | 55.676228 | yes | 2.923E-07 | 548.29327 | 128.40384 |
| WQQTPAPTDTEA\ | 14 | Q03001 | Dystonin OS=Homo sapiens OX=9606 G DST | 860.662 | 2 | 55.676228 | yes | 4.661E-07 | 786.38867 | 127.59101 |
| WETVCALSISK | 11 | Q03001 | Dystonin OS=Homo sapiens OX=9606 G DST | 860.662 | 2 | 55.676228 | yes | 1.417E-07 | 647.32904 | 126.4157 |
| TAEVLLDAR | 9 | Q03001 | Dystonin OS=Homo sapiens OX=9606 G DST | 860.662 | 2 | 55.676228 | yes | 5.954E-07 | 494.2771 | 123.20276 |
| SAEREEQEESVYN | 20 | Q03001 | Dystonin OS=Homo sapiens OX=9606 G DST | 860.662 | 3 | 55.676228 | yes | 1.166E-07 | 810.71881 | 120.50925 |
| ALRPEVER | 8 | Q03001 | Dystonin OS=Homo sapiens OX=9606 G DST | 860.662 | 2 | 55.676228 | yes | 1.251E-07 | 485.27744 | 114.19479 |
| WQELNQLTIDR | 11 | Q03001 | Dystonin OS=Homo sapiens OX=9606 G DST | 860.662 | 2 | 55.676228 | yes | 2.841E-07 | 708.36749 | 108.34465 |
| DYELQTMTYR | 10 | Q03001 | Dystonin OS=Homo sapiens OX=9606 G DST | 860.662 | 2 | 55.676228 | yes | 1.611E-07 | 660.30048 | 107.02833 |
| QQQEAAETIR | 10 | Q03001 | Dystonin OS=Homo sapiens OX=9606 G DST | 860.662 | 2 | 55.676228 | yes | 2.688E-07 | 587.29657 | 106.23535 |
| HLHQAVSIGQSLK | 13 | Q03001 | Dystonin OS=Homo sapiens OX=9606 G DST | 860.662 | 2 | 55.676228 | yes | 1.67E-07 | 709.39917 | 103.89934 |
| AQEESSAMMQWL | 14 | Q03001 | Dystonin OS=Homo sapiens OX=9606 G DST | 860.662 | 2 | 55.676228 | yes | 1.477E-07 | 833.88184 | 100.76733 |
| HLEVLHSLLK | 10 | Q03001 | Dystonin OS=Homo sapiens OX=9606 G DST | 860.662 | 2 | 55.676228 | yes | 2.496E-07 | 594.85858 | 97.0634 |
| KSTVEVIK | 8 | Q03001 | Dystonin OS=Homo sapiens OX=9606 G DST | 860.662 | 2 | 55.676228 | yes | 1.42E-07 | 452.27911 | 96.218063 |
| QQLDGALR | 8 | Q03001 | Dystonin OS=Homo sapiens OX=9606 G DST | 860.662 | 2 | 55.676228 | yes | 1.136E-07 | 450.7485 | 95.36338 |
| QFHEAWSK | 8 | Q03001 | Dystonin OS=Homo sapiens OX=9606 G DST | 860.662 | 2 | 55.676228 | yes | 9.81E-08 | 516.74854 | 93.045952 |
| EEEEVAYDWSER | 12 | Q03001 | Dystonin OS=Homo sapiens OX=9606 G DST | 860.662 | 2 | 55.676228 | yes | 1.865E-07 | 771.32318 | 92.549919 |
| HTVQDPLMELK | 11 | Q03001 | Dystonin OS=Homo sapiens OX=9606 G DST | 860.662 | 2 | 55.676228 | yes | 9.722E-08 | 655.84229 | 91.081894 |

| | | | | | | | | | | |
|---|---|---|---|---|---|---|---|---|---|---|
| WQNVHVQIDNR | 11 | Q03001 | Dystonin OS=Homo sapiens OX=9606 G DST | 860.662 | 2 | 55.676228 | yes | 1.658E-07 | 704.85767 | 86.927246 |
| TKVELFENLSEK | 12 | Q03001 | Dystonin OS=Homo sapiens OX=9606 G DST | 860.662 | 2 | 55.676228 | yes | 1.57E-07 | 718.88519 | 80.960625 |
| NECSSVYSK | 9 | Q03001 | Dystonin OS=Homo sapiens OX=9606 G DST | 860.662 | 2 | 55.676228 | yes | 1.05E-07 | 537.23206 | 74.850121 |
| LFNAIIHK | 8 | Q03001 | Dystonin OS=Homo sapiens OX=9606 G DST | 860.662 | 2 | 55.676228 | yes | 1.482E-06 | 478.28983 | 70.911034 |
| DKEVIPQEIEEVK | 13 | Q03001 | Dystonin OS=Homo sapiens OX=9606 G DST | 860.662 | 2 | 55.676228 | yes | 9.81E-08 | 778.41431 | 63.636665 |
| KVPIVQPSFGAEDI | 16 | Q03001 | Dystonin OS=Homo sapiens OX=9606 G DST | 860.662 | 2 | 55.676228 | yes | 9.802E-08 | 842.96704 | 63.142624 |
| QVFHALEDELQK | 12 | Q03001 | Dystonin OS=Homo sapiens OX=9606 G DST | 860.662 | 2 | 55.676228 | yes | 2.854E-07 | 728.87518 | 62.688427 |
| HQVDLAK | 7 | Q03001 | Dystonin OS=Homo sapiens OX=9606 G DST | 860.662 | 2 | 55.676228 | yes | 2.639E-07 | 405.72705 | 62.192776 |
| EELLQYK | 7 | Q03001 | Dystonin OS=Homo sapiens OX=9606 G DST | 860.662 | 2 | 55.676228 | yes | 6.276E-07 | 461.74765 | 61.315815 |
| IFGEDEVELMNWL | 19 | Q03001 | Dystonin OS=Homo sapiens OX=9606 G DST | 860.662 | 3 | 55.676228 | yes | 3.286E-07 | 773.03168 | 60.287491 |
| KHVNDLYEDLR | 11 | Q03001 | Dystonin OS=Homo sapiens OX=9606 G DST | 860.662 | 2 | 55.676228 | yes | 3.043E-06 | 701.35968 | 57.42857 |
| QSSINAMNEK | 10 | Q03001 | Dystonin OS=Homo sapiens OX=9606 G DST | 860.662 | 2 | 55.676228 | yes | 9.722E-08 | 561.26642 | 56.98233 |
| EATDYLR | 7 | Q03001 | Dystonin OS=Homo sapiens OX=9606 G DST | 860.662 | 2 | 55.676228 | yes | 9.722E-08 | 434.21399 | 52.813118 |
| HLDTLHNFVSR | 11 | Q03001 | Dystonin OS=Homo sapiens OX=9606 G DST | 860.662 | 2 | 55.676228 | yes | 1.183E-07 | 669.8493 | 52.640701 |
| KTAEVLLDAR | 10 | Q03001 | Dystonin OS=Homo sapiens OX=9606 G DST | 860.662 | 2 | 55.676228 | yes | 9.722E-08 | 558.32458 | 49.34473 |
| CEQLNLK | 7 | Q03001 | Dystonin OS=Homo sapiens OX=9606 G DST | 860.662 | 2 | 55.676228 | yes | 1.247E-07 | 452.73148 | 47.51202 |
| NNIENLISTLK | 11 | Q03001 | Dystonin OS=Homo sapiens OX=9606 G DST | 860.662 | 2 | 55.676228 | yes | 9.722E-08 | 629.8537 | 45.605042 |
| RFQVEQIGDNK | 11 | Q03001 | Dystonin OS=Homo sapiens OX=9606 G DST | 860.662 | 2 | 55.676228 | yes | 1.074E-07 | 667.34656 | 43.904755 |
| DTQDFIR | 7 | Q03001 | Dystonin OS=Homo sapiens OX=9606 G DST | 860.662 | 2 | 55.676228 | yes | 1.179E-06 | 447.71942 | 43.451775 |
| EKADQLVER | 9 | Q03001 | Dystonin OS=Homo sapiens OX=9606 G DST | 860.662 | 2 | 55.676228 | yes | 1.823E-06 | 544.29077 | 42.617489 |
| VPEGGEGIGANDV | 16 | Q03001 | Dystonin OS=Homo sapiens OX=9606 G DST | 860.662 | 2 | 55.676228 | yes | 9.491E-07 | 785.39136 | 39.645939 |
| FWCDHMSLIVTIK | 13 | Q03001 | Dystonin OS=Homo sapiens OX=9606 G DST | 860.662 | 2 | 55.676228 | yes | 4.144E-07 | 825.41254 | 39.616707 |
| NTQAAEALVK | 10 | Q03001 | Dystonin OS=Homo sapiens OX=9606 G DST | 860.662 | 2 | 55.676228 | yes | 9.587E-07 | 522.78784 | 39.585789 |
| VSLIDMVQR | 9 | Q03001 | Dystonin OS=Homo sapiens OX=9606 G DST | 860.662 | 2 | 55.676228 | yes | 2.482E-07 | 530.79462 | 36.138844 |
| TFTMEILR | 8 | Q03001 | Dystonin OS=Homo sapiens OX=9606 G DST | 860.662 | 2 | 55.676228 | yes | 1.823E-07 | 505.7706 | 35.399574 |
| IATTAEPADK | 10 | Q03001 | Dystonin OS=Homo sapiens OX=9606 G DST | 860.662 | 2 | 55.676228 | yes | 3.877E-06 | 508.76657 | 34.219452 |
| STSVSSQAAQAAS | 23 | Q03001 | Dystonin OS=Homo sapiens OX=9606 G DST | 860.662 | 3 | 55.676228 | yes | 2.559E-07 | 739.04486 | 29.763783 |
| ELDQKEENIK | 10 | Q03001 | Dystonin OS=Homo sapiens OX=9606 G DST | 860.662 | 2 | 55.676228 | yes | 9.722E-08 | 623.3197 | 25.791321 |
| FCLENMTQK | 9 | Q03001 | Dystonin OS=Homo sapiens OX=9606 G DST | 860.662 | 2 | 55.676228 | yes | 5.91E-07 | 585.76776 | 21.576719 |
| ELIAEHKPHIDK | 12 | Q03001 | Dystonin OS=Homo sapiens OX=9606 G DST | 860.662 | 2 | 55.676228 | yes | 9.92E-07 | 715.39355 | 18.038477 |
| LKEFSILLQK | 10 | Q03001 | Dystonin OS=Homo sapiens OX=9606 G DST | 860.662 | 2 | 55.676228 | yes | 2.895E-06 | 609.87646 | 16.858374 |
| DVTELSQYMQEST | 20 | Q03001 | Dystonin OS=Homo sapiens OX=9606 G DST | 860.662 | 3 | 55.676228 | yes | 3.812E-06 | 801.03363 | 12.016084 |
| LTFNNPTTEFR | 11 | Q03111 | Protein ENL OS=Homo sapiens OX=960 MLLT1 | 62.057 | 2 | 53.396645 | yes | 9.722E-08 | 670.33569 | 250.72641 |
| AYTDELVELHR | 11 | Q03111 | Protein ENL OS=Homo sapiens OX=960 MLLT1 | 62.057 | 2 | 53.396645 | yes | 9.722E-08 | 673.34094 | 235.49103 |
| AGGVMVMPEGAD | 16 | Q03111 | Protein ENL OS=Homo sapiens OX=960 MLLT1 | 62.057 | 2 | 53.396645 | yes | 1.11E-07 | 788.87653 | 175.94223 |
| KLQSCLEAVAT | 11 | Q03111 | Protein ENL OS=Homo sapiens OX=960 MLLT1 | 62.057 | 2 | 53.396645 | yes | 1.388E-07 | 610.32117 | 120.6319 |
| VVFWLHDSFPKPR | 13 | Q03111 | Protein ENL OS=Homo sapiens OX=960 MLLT1 | 62.057 | 3 | 53.396645 | yes | 9.722E-08 | 543.29633 | 80.48307 |
| MDNQCTVQVR | 10 | Q03111 | Protein ENL OS=Homo sapiens OX=960 MLLT1 | 62.057 | 2 | 53.396645 | yes | 1.805E-07 | 646.78973 | 59.377407 |
| GPEQCDIQHFVEK | 13 | Q03111 | Protein ENL OS=Homo sapiens OX=960 MLLT1 | 62.057 | 2 | 53.396645 | yes | 1.451E-07 | 793.867 | 36.094261 |

| | | | | | | | | | | | |
|---|---|---|---|---|---|---|---|---|---|---|---|
| APPNALPENLLR | 12 | Q03112 | Histone-lysine N-methyltransferase MEC | MECOM | 138.136 | 2 | 36.8558 | yes | 6.542E-07 | 652.86969 | 85.191292 |
| PPVTPATSQDQPLI | 20 | Q03112 | Histone-lysine N-methyltransferase MEC | MECOM | 138.136 | 3 | 36.8558 | yes | 0.0001138 | 705.01251 | 0.40448 |
| SSPEGPDPPVLTEV | 16 | Q03164 | Histone-lysine N-methyltransferase 2A O | KMT2A | 431.765 | 2 | 55.914978 | yes | 2.494E-06 | 819.91467 | 83.849068 |
| TDLSTTVATPSSGI | 15 | Q03164 | Histone-lysine N-methyltransferase 2A O | KMT2A | 431.765 | 2 | 55.914978 | yes | 9.722E-08 | 739.39087 | 72.157967 |
| VFVDFEGISLR | 11 | Q03164 | Histone-lysine N-methyltransferase 2A O | KMT2A | 431.765 | 2 | 55.914978 | yes | 2.132E-07 | 641.34552 | 51.044338 |
| SSIAGSEDAEPLAP | 24 | Q03164 | Histone-lysine N-methyltransferase 2A O | KMT2A | 431.765 | 4 | 55.914978 | yes | 2.142E-06 | 619.09698 | 34.355106 |
| NSYHPECLGPNYP | 18 | Q03164 | Histone-lysine N-methyltransferase 2A O | KMT2A | 431.765 | 3 | 55.914978 | yes | 2.316E-06 | 701.67029 | 8.6583738 |
| ALYESELADAR | 11 | Q03252 | Lamin-B2 OS=Homo sapiens OX=9606 ( | LMNB2 | 69.949 | 2 | 50.499607 | yes | 9.794E-08 | 619.30658 | 109.77334 |
| GLESDVAELR | 10 | Q03252 | Lamin-B2 OS=Homo sapiens OX=9606 ( | LMNB2 | 69.949 | 2 | 50.499607 | yes | 3.017E-07 | 544.78278 | 89.321899 |
| DLESLFHR | 8 | Q03252 | Lamin-B2 OS=Homo sapiens OX=9606 ( | LMNB2 | 69.949 | 2 | 50.499607 | yes | 5.534E-07 | 508.76163 | 84.537437 |
| MAQALEELR | 9 | Q03252 | Lamin-B2 OS=Homo sapiens OX=9606 ( | LMNB2 | 69.949 | 2 | 50.499607 | yes | 1.776E-07 | 530.77643 | 72.402122 |
| GQSSWGTGESFR | 12 | Q03252 | Lamin-B2 OS=Homo sapiens OX=9606 ( | LMNB2 | 69.949 | 2 | 50.499607 | yes | 4.527E-07 | 649.79163 | 45.324963 |
| LALDMEINAYR | 11 | Q03252 | Lamin-B2 OS=Homo sapiens OX=9606 ( | LMNB2 | 69.949 | 2 | 50.499607 | yes | 2.329E-07 | 654.83447 | 43.744259 |
| LESLSYQLSGLQK | 13 | Q03252 | Lamin-B2 OS=Homo sapiens OX=9606 ( | LMNB2 | 69.949 | 2 | 50.499607 | yes | 6.077E-07 | 733.3985 | 27.254932 |
| ATSSSSGSLSATGR | 14 | Q03252 | Lamin-B2 OS=Homo sapiens OX=9606 ( | LMNB2 | 69.949 | 2 | 50.499607 | yes | 5.572E-07 | 634.8075 | 19.574265 |
| VGGLGVNLTGANI | 13 | Q03468 | DNA excision repair protein ERCC-6 OS | ERCC6 | 168.417 | 2 | 54.764767 | yes | 9.722E-08 | 614.34387 | 464.27148 |
| GAEVNAVTSNR | 11 | Q03468 | DNA excision repair protein ERCC-6 OS | ERCC6 | 168.417 | 2 | 54.764767 | yes | 2.196E-07 | 559.28345 | 358.23007 |
| LLTAGTIEEK | 10 | Q03468 | DNA excision repair protein ERCC-6 OS | ERCC6 | 168.417 | 2 | 54.764767 | yes | 1.388E-07 | 537.80573 | 338.47031 |
| HDAIMDGASPDYV | 21 | Q03468 | DNA excision repair protein ERCC-6 OS | ERCC6 | 168.417 | 3 | 54.764767 | yes | 9.722E-08 | 758.35492 | 312.04138 |
| IILSGSPMQNNLR | 13 | Q03468 | DNA excision repair protein ERCC-6 OS | ERCC6 | 168.417 | 2 | 54.764767 | yes | 9.722E-08 | 721.89282 | 310.58124 |
| LMQDDISR | 8 | Q03468 | DNA excision repair protein ERCC-6 OS | ERCC6 | 168.417 | 2 | 54.764767 | yes | 1.343E-07 | 489.23965 | 302.4801 |
| MIVVESLLK | 9 | Q03468 | DNA excision repair protein ERCC-6 OS | ERCC6 | 168.417 | 2 | 54.764767 | yes | 9.722E-08 | 516.31213 | 222.38113 |
| NPNAAVTLACK | 11 | Q03468 | DNA excision repair protein ERCC-6 OS | ERCC6 | 168.417 | 2 | 54.764767 | yes | 1.845E-07 | 579.80042 | 216.25246 |
| NHLILPER | 8 | Q03468 | DNA excision repair protein ERCC-6 OS | ERCC6 | 168.417 | 2 | 54.764767 | yes | 2.913E-07 | 496.28781 | 213.2645 |
| CLGAVSGVPTWTC | 15 | Q03468 | DNA excision repair protein ERCC-6 OS | ERCC6 | 168.417 | 2 | 54.764767 | yes | 1.004E-07 | 799.3988 | 208.28 |
| NLCTFHR | 7 | Q03468 | DNA excision repair protein ERCC-6 OS | ERCC6 | 168.417 | 2 | 54.764767 | yes | 1.166E-07 | 474.22943 | 129.41223 |
| EQSNDDYVLEK | 11 | Q03468 | DNA excision repair protein ERCC-6 OS | ERCC6 | 168.417 | 2 | 54.764767 | yes | 9.722E-08 | 670.30426 | 125.28088 |
| IQPAFGADHDVPK | 13 | Q03468 | DNA excision repair protein ERCC-6 OS | ERCC6 | 168.417 | 2 | 54.764767 | yes | 1.424E-07 | 697.85681 | 58.487827 |
| IQPAFGADHDVPK | 13 | Q03468 | DNA excision repair protein ERCC-6 OS | ERCC6 | 168.417 | 2 | 54.764767 | yes | 9.11E-06 | 465.57361 | 58.487827 |
| IQPAFGADHDVPK | 13 | Q03468 | DNA excision repair protein ERCC-6 OS | ERCC6 | 168.417 | 3 | 54.764767 | yes | 1.424E-07 | 697.85681 | 58.487827 |
| IQPAFGADHDVPK | 13 | Q03468 | DNA excision repair protein ERCC-6 OS | ERCC6 | 168.417 | 3 | 54.764767 | yes | 9.11E-06 | 465.57361 | 58.487827 |
| KQVTVYR | 7 | Q03468 | DNA excision repair protein ERCC-6 OS | ERCC6 | 168.417 | 2 | 54.764767 | yes | 3.958E-07 | 447.26379 | 114.0637 |
| VYQNFVDSK | 9 | Q03468 | DNA excision repair protein ERCC-6 OS | ERCC6 | 168.417 | 2 | 54.764767 | yes | 1.006E-07 | 550.27454 | 108.72785 |
| EILQEFESK | 9 | Q03468 | DNA excision repair protein ERCC-6 OS | ERCC6 | 168.417 | 2 | 54.764767 | yes | 1.268E-07 | 561.78748 | 94.660324 |
| MDGTTTIASR | 10 | Q03468 | DNA excision repair protein ERCC-6 OS | ERCC6 | 168.417 | 2 | 54.764767 | yes | 8.6E-07 | 526.75568 | 92.857224 |
| ICNHPDLFSGGPK | 13 | Q03468 | DNA excision repair protein ERCC-6 OS | ERCC6 | 168.417 | 3 | 54.764767 | yes | 4.493E-07 | 481.23462 | 38.090572 |
| TSMPSGDESIDEK | 13 | Q03468 | DNA excision repair protein ERCC-6 OS | ERCC6 | 168.417 | 2 | 54.764767 | yes | 3.171E-06 | 698.30084 | 14.536297 |
| TVLVIAHR | 8 | Q03519 | Antigen peptide transporter 2 OS=Homo | TAP2 | 75.664 | 2 | 46.64303 | yes | 1.506E-07 | 454.78745 | 92.752022 |
| DLEQAISQR | 9 | Q04206 | Transcription factor p65 OS=Homo sapie | RELA | 60.22 | 2 | 48.60866 | yes | 1.499E-07 | 530.27509 | 118.909 |

| | | | | | | | | | | | |
|---|---|---|---|---|---|---|---|---|---|---|---|
| LCFQVTVR | 8 | Q04206 | Transcription factor p65 OS=Homo sapie | RELA | 60.22 | 2 | 48.60866 | yes | 2.482E-07 | 511.77621 | 80.51284 |
| TPPYADPSLQAPV | 14 | Q04206 | Transcription factor p65 OS=Homo sapie | RELA | 60.22 | 2 | 48.60866 | yes | 1.123E-07 | 756.3963 | 72.349312 |
| GSFSQADVHR | 10 | Q04206 | Transcription factor p65 OS=Homo sapie | RELA | 60.22 | 2 | 48.60866 | yes | 2.968E-06 | 552.26508 | 57.354809 |
| VPTTEKPTVTVNF | 14 | Q04637 | Eukaryotic translation initiation factor 4 g | EIF4G1 | 175.492 | 3 | 54.1342 | yes | 2.73E-07 | 530.29462 | 8372.585 |
| VPTTEKPTVTVNF | 14 | Q04637 | Eukaryotic translation initiation factor 4 g | EIF4G1 | 175.492 | 3 | 54.1342 | yes | 3.365E-07 | 794.93829 | 8372.585 |
| VPTTEKPTVTVNF | 14 | Q04637 | Eukaryotic translation initiation factor 4 g | EIF4G1 | 175.492 | 2 | 54.1342 | yes | 2.73E-07 | 530.29462 | 8372.585 |
| VPTTEKPTVTVNF | 14 | Q04637 | Eukaryotic translation initiation factor 4 g | EIF4G1 | 175.492 | 2 | 54.1342 | yes | 3.365E-07 | 794.93829 | 8372.585 |
| LTPQMFQQLMK | 11 | Q04637 | Eukaryotic translation initiation factor 4 g | EIF4G1 | 175.492 | 2 | 54.1342 | yes | 1.898E-07 | 682.85687 | 6516.6304 |
| LTPQMFQQLMK | 11 | Q04637 | Eukaryotic translation initiation factor 4 g | EIF4G1 | 175.492 | 2 | 54.1342 | yes | 9.722E-08 | 690.85437 | 6516.6304 |
| LKEELEEAR | 9 | Q04637 | Eukaryotic translation initiation factor 4 g | EIF4G1 | 175.492 | 2 | 54.1342 | yes | 9.722E-08 | 558.7984 | 9071.793 |
| MFFDALYDEDVVI | 13 | Q04637 | Eukaryotic translation initiation factor 4 g | EIF4G1 | 175.492 | 2 | 54.1342 | yes | 1.063E-07 | 796.37109 | 4278.792 |
| MFFDALYDEDVVI | 13 | Q04637 | Eukaryotic translation initiation factor 4 g | EIF4G1 | 175.492 | 2 | 54.1342 | yes | 2.87E-06 | 804.36853 | 4278.792 |
| AALSEEELEK | 10 | Q04637 | Eukaryotic translation initiation factor 4 g | EIF4G1 | 175.492 | 2 | 54.1342 | yes | 1.039E-07 | 559.78241 | 8342.6006 |
| TPLRPLDPTR | 10 | Q04637 | Eukaryotic translation initiation factor 4 g | EIF4G1 | 175.492 | 2 | 54.1342 | yes | 2.307E-07 | 583.33801 | 8085.3149 |
| EITKPLRPLGK | 11 | Q04637 | Eukaryotic translation initiation factor 4 g | EIF4G1 | 175.492 | 3 | 54.1342 | yes | 3.729E-07 | 417.93097 | 3918.0432 |
| EITKPLRPLGK | 11 | Q04637 | Eukaryotic translation initiation factor 4 g | EIF4G1 | 175.492 | 3 | 54.1342 | yes | 2.896E-07 | 626.39282 | 3918.0432 |
| EITKPLRPLGK | 11 | Q04637 | Eukaryotic translation initiation factor 4 g | EIF4G1 | 175.492 | 2 | 54.1342 | yes | 3.729E-07 | 417.93097 | 3918.0432 |
| EITKPLRPLGK | 11 | Q04637 | Eukaryotic translation initiation factor 4 g | EIF4G1 | 175.492 | 2 | 54.1342 | yes | 2.896E-07 | 626.39282 | 3918.0432 |
| AIIEEYLHLNDMK | 13 | Q04637 | Eukaryotic translation initiation factor 4 g | EIF4G1 | 175.492 | 2 | 54.1342 | yes | 1.399E-07 | 794.90564 | 2593.7886 |
| AIIEEYLHLNDMK | 13 | Q04637 | Eukaryotic translation initiation factor 4 g | EIF4G1 | 175.492 | 2 | 54.1342 | yes | 9.924E-08 | 802.90308 | 2593.7886 |
| AIIEEYLHLNDMK | 13 | Q04637 | Eukaryotic translation initiation factor 4 g | EIF4G1 | 175.492 | 2 | 54.1342 | yes | 2.685E-06 | 530.27283 | 2593.7886 |
| AIIEEYLHLNDMK | 13 | Q04637 | Eukaryotic translation initiation factor 4 g | EIF4G1 | 175.492 | 3 | 54.1342 | yes | 1.399E-07 | 794.90564 | 2593.7886 |
| AIIEEYLHLNDMK | 13 | Q04637 | Eukaryotic translation initiation factor 4 g | EIF4G1 | 175.492 | 3 | 54.1342 | yes | 9.924E-08 | 802.90308 | 2593.7886 |
| AIIEEYLHLNDMK | 13 | Q04637 | Eukaryotic translation initiation factor 4 g | EIF4G1 | 175.492 | 3 | 54.1342 | yes | 2.685E-06 | 530.27283 | 2593.7886 |
| QVTQLAIDTEER | 12 | Q04637 | Eukaryotic translation initiation factor 4 g | EIF4G1 | 175.492 | 2 | 54.1342 | yes | 1.532E-07 | 701.86224 | 7468.1675 |
| VEYTLGEESEAPG | 15 | Q04637 | Eukaryotic translation initiation factor 4 g | EIF4G1 | 175.492 | 2 | 54.1342 | yes | 9.722E-08 | 832.89172 | 3525.6289 |
| VEYTLGEESEAPG | 15 | Q04637 | Eukaryotic translation initiation factor 4 g | EIF4G1 | 175.492 | 2 | 54.1342 | yes | 1.062E-07 | 555.59692 | 3525.6289 |
| VEYTLGEESEAPG | 15 | Q04637 | Eukaryotic translation initiation factor 4 g | EIF4G1 | 175.492 | 3 | 54.1342 | yes | 9.722E-08 | 832.89172 | 3525.6289 |
| VEYTLGEESEAPG | 15 | Q04637 | Eukaryotic translation initiation factor 4 g | EIF4G1 | 175.492 | 3 | 54.1342 | yes | 1.062E-07 | 555.59692 | 3525.6289 |
| HGVESTLER | 9 | Q04637 | Eukaryotic translation initiation factor 4 g | EIF4G1 | 175.492 | 2 | 54.1342 | yes | 1.489E-07 | 514.26196 | 6428.9272 |
| DITEEIMSGAR | 11 | Q04637 | Eukaryotic translation initiation factor 4 g | EIF4G1 | 175.492 | 2 | 54.1342 | yes | 9.722E-08 | 611.2926 | 5207.1187 |
| IHNAENIQPGEQK | 13 | Q04637 | Eukaryotic translation initiation factor 4 g | EIF4G1 | 175.492 | 2 | 54.1342 | yes | 1.062E-07 | 739.37335 | 4663.6836 |
| DKDDDEVFEK | 10 | Q04637 | Eukaryotic translation initiation factor 4 g | EIF4G1 | 175.492 | 2 | 54.1342 | yes | 9.722E-08 | 620.2724 | 4460.5854 |
| EDAFYSWESSK | 11 | Q04637 | Eukaryotic translation initiation factor 4 g | EIF4G1 | 175.492 | 2 | 54.1342 | yes | 9.722E-08 | 674.78821 | 4320.9209 |
| AQPPSSAASR | 10 | Q04637 | Eukaryotic translation initiation factor 4 g | EIF4G1 | 175.492 | 2 | 54.1342 | yes | 4.464E-07 | 486.24887 | 4136.6279 |
| EAVGDLLDAFK | 11 | Q04637 | Eukaryotic translation initiation factor 4 g | EIF4G1 | 175.492 | 2 | 54.1342 | yes | 1.352E-07 | 589.30859 | 3771.5127 |
| SDQWKPLNLEEK | 12 | Q04637 | Eukaryotic translation initiation factor 4 g | EIF4G1 | 175.492 | 2 | 54.1342 | yes | 4.925E-07 | 743.88043 | 3736.0413 |
| YLCDEQK | 7 | Q04637 | Eukaryotic translation initiation factor 4 g | EIF4G1 | 175.492 | 2 | 54.1342 | yes | 5.776E-07 | 478.2131 | 3514.0269 |
| IIATVLMTEDIK | 12 | Q04637 | Eukaryotic translation initiation factor 4 g | EIF4G1 | 175.492 | 2 | 54.1342 | yes | 9.962E-08 | 673.88361 | 1754.9745 |

| | | | | | | | | | | | |
|---|---|---|---|---|---|---|---|---|---|---|---|
| IIATVLMTEDIK | 12 | Q04637 | Eukaryotic translation initiation factor 4 g | EIF4G1 | 175.492 | 2 | 54.1342 | yes | 1.117E-07 | 681.8811 | 1754.9745 |
| EAALPPVSPLK | 11 | Q04637 | Eukaryotic translation initiation factor 4 g | EIF4G1 | 175.492 | 2 | 54.1342 | yes | 2.85E-06 | 561.33191 | 3504.917 |
| EMDEAATAEER | 11 | Q04637 | Eukaryotic translation initiation factor 4 g | EIF4G1 | 175.492 | 2 | 54.1342 | yes | 9.919E-08 | 626.26154 | 1522.6578 |
| EMDEAATAEER | 11 | Q04637 | Eukaryotic translation initiation factor 4 g | EIF4G1 | 175.492 | 2 | 54.1342 | yes | 9.722E-08 | 634.25897 | 1522.6578 |
| AALSEEELEKK | 11 | Q04637 | Eukaryotic translation initiation factor 4 g | EIF4G1 | 175.492 | 2 | 54.1342 | yes | 5.523E-07 | 623.8299 | 2837.1526 |
| ALPSEELNR | 9 | Q04637 | Eukaryotic translation initiation factor 4 g | EIF4G1 | 175.492 | 2 | 54.1342 | yes | 2.193E-07 | 514.77222 | 2079.7087 |
| QHFYPSR | 7 | Q04637 | Eukaryotic translation initiation factor 4 g | EIF4G1 | 175.492 | 2 | 54.1342 | yes | 2.647E-07 | 467.73013 | 2011.038 |
| KVGTLWR | 7 | Q04637 | Eukaryotic translation initiation factor 4 g | EIF4G1 | 175.492 | 2 | 54.1342 | yes | 1.358E-07 | 430.26105 | 1958.9038 |
| KIIATVLMTEDIK | 13 | Q04637 | Eukaryotic translation initiation factor 4 g | EIF4G1 | 175.492 | 2 | 54.1342 | yes | 3.126E-07 | 737.93109 | 799.41357 |
| KIIATVLMTEDIK | 13 | Q04637 | Eukaryotic translation initiation factor 4 g | EIF4G1 | 175.492 | 2 | 54.1342 | yes | 9.722E-08 | 745.92859 | 799.41357 |
| PSDAASEAAR | 10 | Q04637 | Eukaryotic translation initiation factor 4 g | EIF4G1 | 175.492 | 2 | 54.1342 | yes | 4.163E-07 | 487.73053 | 1433.1658 |
| GVIDLIFEK | 9 | Q04637 | Eukaryotic translation initiation factor 4 g | EIF4G1 | 175.492 | 2 | 54.1342 | yes | 2.574E-07 | 517.30005 | 1345.9684 |
| GSRPIDTSR | 9 | Q04637 | Eukaryotic translation initiation factor 4 g | EIF4G1 | 175.492 | 2 | 54.1342 | yes | 2.493E-07 | 494.76215 | 948.97394 |
| SQGAIIADR | 9 | Q04637 | Eukaryotic translation initiation factor 4 g | EIF4G1 | 175.492 | 2 | 54.1342 | yes | 2.54E-07 | 465.75378 | 726.4707 |
| KAASLTEDR | 9 | Q04637 | Eukaryotic translation initiation factor 4 g | EIF4G1 | 175.492 | 2 | 54.1342 | yes | 2.41E-07 | 495.76437 | 656.8219 |
| EAEMEEHR | 8 | Q04637 | Eukaryotic translation initiation factor 4 g | EIF4G1 | 175.492 | 2 | 54.1342 | yes | 1.514E-07 | 515.71655 | 427.78372 |
| ERPSQPEGLR | 10 | Q04637 | Eukaryotic translation initiation factor 4 g | EIF4G1 | 175.492 | 2 | 54.1342 | yes | 1.967E-07 | 584.80707 | 409.98779 |
| PGSIDSNNQLFAPC | 16 | Q04637 | Eukaryotic translation initiation factor 4 g | EIF4G1 | 175.492 | 2 | 54.1342 | yes | 1.199E-07 | 815.40259 | 397.22607 |
| ETGEPYR | 7 | Q04637 | Eukaryotic translation initiation factor 4 g | EIF4G1 | 175.492 | 2 | 54.1342 | yes | 1.943E-07 | 426.19833 | 370.49173 |
| GGPPGPPISR | 10 | Q04637 | Eukaryotic translation initiation factor 4 g | EIF4G1 | 175.492 | 2 | 54.1342 | yes | 3.801E-07 | 467.75888 | 368.93402 |
| GSSGGSGAKPSDA | 19 | Q04637 | Eukaryotic translation initiation factor 4 g | EIF4G1 | 175.492 | 2 | 54.1342 | yes | 1.148E-07 | 831.88751 | 179.61566 |
| GSSGGSGAKPSDA | 19 | Q04637 | Eukaryotic translation initiation factor 4 g | EIF4G1 | 175.492 | 2 | 54.1342 | yes | 1.485E-06 | 554.92743 | 179.61566 |
| GSSGGSGAKPSDA | 19 | Q04637 | Eukaryotic translation initiation factor 4 g | EIF4G1 | 175.492 | 3 | 54.1342 | yes | 1.148E-07 | 831.88751 | 179.61566 |
| GSSGGSGAKPSDA | 19 | Q04637 | Eukaryotic translation initiation factor 4 g | EIF4G1 | 175.492 | 3 | 54.1342 | yes | 1.485E-06 | 554.92743 | 179.61566 |
| PATSTLNR | 8 | Q04637 | Eukaryotic translation initiation factor 4 g | EIF4G1 | 175.492 | 2 | 54.1342 | yes | 1.175E-07 | 430.23523 | 290.39978 |
| EFLPEGQDIGAFV/ | 17 | Q04637 | Eukaryotic translation initiation factor 4 g | EIF4G1 | 175.492 | 3 | 54.1342 | yes | 3.648E-07 | 626.64752 | 129.26866 |
| LQGINCGPDFTPSF | 19 | Q04637 | Eukaryotic translation initiation factor 4 g | EIF4G1 | 175.492 | 3 | 54.1342 | yes | 9.754E-08 | 688.67041 | 123.35532 |
| GGPGGELPR | 9 | Q04637 | Eukaryotic translation initiation factor 4 g | EIF4G1 | 175.492 | 2 | 54.1342 | yes | 1.167E-07 | 420.22214 | 107.80488 |
| VQQLMAK | 7 | Q04637 | Eukaryotic translation initiation factor 4 g | EIF4G1 | 175.492 | 2 | 54.1342 | yes | 1.334E-07 | 409.23364 | 100.36609 |
| AASLTEDRDR | 10 | Q04637 | Eukaryotic translation initiation factor 4 g | EIF4G1 | 175.492 | 2 | 54.1342 | yes | 1.161E-06 | 567.28088 | 74.814255 |
| PTVTVNFR | 8 | Q04637 | Eukaryotic translation initiation factor 4 g | EIF4G1 | 175.492 | 2 | 54.1342 | yes | 3.476E-07 | 467.26126 | 70.550522 |
| QKEMDEAATAEEI | 13 | Q04637 | Eukaryotic translation initiation factor 4 g | EIF4G1 | 175.492 | 2 | 54.1342 | yes | 1.481E-07 | 754.33832 | 59.523064 |
| KEAVGDLLDAFK | 12 | Q04637 | Eukaryotic translation initiation factor 4 g | EIF4G1 | 175.492 | 2 | 54.1342 | yes | 9.722E-08 | 653.35608 | 46.253983 |
| RGGPPGPPISR | 11 | Q04637 | Eukaryotic translation initiation factor 4 g | EIF4G1 | 175.492 | 2 | 54.1342 | yes | 8.316E-07 | 545.80945 | 34.789886 |
| CLMALKVPTTEKF | 20 | Q04637 | Eukaryotic translation initiation factor 4 g | EIF4G1 | 175.492 | 3 | 54.1342 | yes | 2.52E-07 | 769.08508 | 34.027931 |
| NCQTLVNLCSR | 11 | Q04721 | Neurogenic locus notch homolog protein | NOTCH2 | 265.405 | 2 | 52.429783 | yes | 9.722E-08 | 682.8241 | 113.77213 |
| MNDGTTPLILAAR | 13 | Q04721 | Neurogenic locus notch homolog protein | NOTCH2 | 265.405 | 2 | 52.429783 | yes | 2.921E-07 | 686.86627 | 89.676468 |
| IGGFTCLCMPGFK | 13 | Q04721 | Neurogenic locus notch homolog protein | NOTCH2 | 265.405 | 2 | 52.429783 | yes | 1.987E-07 | 744.34583 | 60.726994 |
| ILLDHFANR | 9 | Q04721 | Neurogenic locus notch homolog protein | NOTCH2 | 265.405 | 2 | 52.429783 | yes | 1.402E-07 | 549.80634 | 57.583851 |

| | | | | | | | | | | | |
|---|---|---|---|---|---|---|---|---|---|---|---|
| TGEMALHLAAR | 11 | Q04721 | Neurogenic locus notch homolog protein | NOTCH2 | 265.405 | 2 | 52.429783 | yes | 1.095E-07 | 585.30841 | 57.335781 |
| YSCVCSPGFTGQR | 13 | Q04721 | Neurogenic locus notch homolog protein | NOTCH2 | 265.405 | 2 | 52.429783 | yes | 9.722E-08 | 759.82684 | 37.916985 |
| GLLCEENIDDCAR | 13 | Q04721 | Neurogenic locus notch homolog protein | NOTCH2 | 265.405 | 2 | 52.429783 | yes | 3.352E-07 | 782.84015 | 36.29258 |
| VASFSCMCPEGK | 12 | Q04721 | Neurogenic locus notch homolog protein | NOTCH2 | 265.405 | 2 | 52.429783 | yes | 9.819E-08 | 686.78833 | 30.415407 |
| NDAPTPGTSTTPGI | 15 | Q04726 | Transducin-like enhancer protein 3 OS=H | TLE3 | 83.417 | 2 | 30.151556 | yes | 2.27E-06 | 742.87061 | 40.29557 |
| SGDSEVYQLGDVS | 15 | Q04837 | Single-stranded DNA-binding protein, mi | SSBP1 | 17.26 | 2 | 50.934483 | yes | 1.028E-07 | 806.37848 | 375.57565 |
| IDYGEYMDK | 9 | Q04837 | Single-stranded DNA-binding protein, mi | SSBP1 | 17.26 | 2 | 50.934483 | yes | 1.108E-07 | 567.24463 | 168.79887 |
| VGQDPVLR | 8 | Q04837 | Single-stranded DNA-binding protein, mi | SSBP1 | 17.26 | 2 | 50.934483 | yes | 9.722E-08 | 442.25345 | 150.81548 |
| ISVFRPGLR | 9 | Q04837 | Single-stranded DNA-binding protein, mi | SSBP1 | 17.26 | 2 | 50.934483 | yes | 1.249E-07 | 522.81927 | 31.63213 |
| RQATTIIADNIIFLS | 19 | Q04837 | Single-stranded DNA-binding protein, mi | SSBP1 | 17.26 | 3 | 50.934483 | yes | 2.041E-07 | 716.72681 | 22.937683 |
| GGDEIFLLCDK | 11 | Q04864 | Proto-oncogene c-Rel OS=Homo sapiens | REL | 68.52 | 2 | 52.299908 | yes | 9.722E-08 | 633.80536 | 657.15033 |
| GIFSQADVHR | 10 | Q04864 | Proto-oncogene c-Rel OS=Homo sapiens | REL | 68.52 | 2 | 52.299908 | yes | 1.089E-07 | 565.29108 | 418.49088 |
| TYPSIQIMNYYGK | 13 | Q04864 | Proto-oncogene c-Rel OS=Homo sapiens | REL | 68.52 | 2 | 52.299908 | yes | 2.59E-07 | 789.38708 | 267.59683 |
| PGLLGSIGEGR | 11 | Q04864 | Proto-oncogene c-Rel OS=Homo sapiens | REL | 68.52 | 2 | 52.299908 | yes | 2.485E-07 | 528.29584 | 256.73993 |
| SGNTNPLSSFSTR | 13 | Q04864 | Proto-oncogene c-Rel OS=Homo sapiens | REL | 68.52 | 2 | 52.299908 | yes | 1.152E-07 | 684.33112 | 221.20103 |
| FVLNDWEAK | 9 | Q04864 | Proto-oncogene c-Rel OS=Homo sapiens | REL | 68.52 | 2 | 52.299908 | yes | 1.696E-07 | 561.28491 | 199.9837 |
| TLPSNSQGIPPFLR | 14 | Q04864 | Proto-oncogene c-Rel OS=Homo sapiens | REL | 68.52 | 2 | 52.299908 | yes | 5.837E-07 | 763.91992 | 143.37169 |
| DGYYEAEFGQER | 12 | Q04864 | Proto-oncogene c-Rel OS=Homo sapiens | REL | 68.52 | 2 | 52.299908 | yes | 1.221E-07 | 732.30731 | 129.33098 |
| KEPNLFSHDAVVR | 13 | Q04864 | Proto-oncogene c-Rel OS=Homo sapiens | REL | 68.52 | 3 | 52.299908 | yes | 9.722E-08 | 504.6037 | 59.660038 |
| SAGSIPGEHSTDNN | 15 | Q04864 | Proto-oncogene c-Rel OS=Homo sapiens | REL | 68.52 | 2 | 52.299908 | yes | 5.066E-07 | 771.35059 | 44.751736 |
| AITEPVTVK | 9 | Q04864 | Proto-oncogene c-Rel OS=Homo sapiens | REL | 68.52 | 2 | 52.299908 | yes | 1.737E-06 | 479.28439 | 43.15527 |
| AVTELNEPLSNED | 14 | Q04917 | 14-3-3 protein eta OS=Homo sapiens OX | YWHAH | 28.219 | 2 | 46.553154 | yes | 1.53E-07 | 793.88647 | 265.60602 |
| NSVVEASEAAYK | 12 | Q04917 | 14-3-3 protein eta OS=Homo sapiens OX | YWHAH | 28.219 | 2 | 46.553154 | yes | 7.811E-07 | 634.31189 | 182.97169 |
| LLNSDLGELINK | 12 | Q05397 | Focal adhesion kinase 1 OS=Homo sapier | PTK2 | 119.234 | 2 | 51.24259 | yes | 9.722E-08 | 664.87463 | 140.79317 |
| LGDFGLSR | 8 | Q05397 | Focal adhesion kinase 1 OS=Homo sapier | PTK2 | 119.234 | 2 | 51.24259 | no | 2.626E-06 | 432.73233 | 129.43141 |
| AQLSTILEEEK | 11 | Q05397 | Focal adhesion kinase 1 OS=Homo sapier | PTK2 | 119.234 | 2 | 51.24259 | yes | 3.44E-07 | 630.83771 | 128.37807 |
| THLGTGMER | 9 | Q05397 | Focal adhesion kinase 1 OS=Homo sapier | PTK2 | 119.234 | 2 | 51.24259 | yes | 7.989E-06 | 501.24527 | 112.99593 |
| WMAPESINFR | 10 | Q05397 | Focal adhesion kinase 1 OS=Homo sapier | PTK2 | 119.234 | 2 | 51.24259 | yes | 1.741E-07 | 625.80298 | 86.295082 |
| FLQEALTMR | 9 | Q05397 | Focal adhesion kinase 1 OS=Homo sapier | PTK2 | 119.234 | 2 | 51.24259 | yes | 1.083E-07 | 554.79462 | 85.011513 |
| NLLDVIDQAR | 10 | Q05397 | Focal adhesion kinase 1 OS=Homo sapier | PTK2 | 119.234 | 2 | 51.24259 | yes | 2.405E-07 | 578.81964 | 83.336411 |
| IQPAPPEEYVPMV | 14 | Q05397 | Focal adhesion kinase 1 OS=Homo sapier | PTK2 | 119.234 | 2 | 51.24259 | yes | 4.515E-07 | 799.41833 | 47.389149 |
| FFEILSPVYR | 10 | Q05397 | Focal adhesion kinase 1 OS=Homo sapier | PTK2 | 119.234 | 2 | 51.24259 | yes | 6.717E-06 | 635.84515 | 46.938782 |
| TLLATVDETIPLLP | 19 | Q05397 | Focal adhesion kinase 1 OS=Homo sapier | PTK2 | 119.234 | 3 | 51.24259 | yes | 2.554E-07 | 683.3844 | 40.677326 |
| LFPPDDSPLPVSSR | 14 | Q05519 | Serine/arginine-rich splicing factor 11 OS | SRSF11 | 53.543 | 2 | 49.078228 | yes | 1.176E-07 | 763.89612 | 2289.2495 |
| LNHVAAGLVSPSL | 14 | Q05519 | Serine/arginine-rich splicing factor 11 OS | SRSF11 | 53.543 | 2 | 49.078228 | yes | 1.102E-07 | 703.41174 | 941.96265 |
| LMSTVDPK | 8 | Q05519 | Serine/arginine-rich splicing factor 11 OS | SRSF11 | 53.543 | 2 | 49.078228 | yes | 6.707E-07 | 445.73624 | 340.23499 |
| FHDPDSAVVAQHI | 21 | Q05519 | Serine/arginine-rich splicing factor 11 OS | SRSF11 | 53.543 | 3 | 49.078228 | yes | 1.856E-07 | 790.06085 | 66.589661 |
| FHDPDSAVVAQHI | 21 | Q05519 | Serine/arginine-rich splicing factor 11 OS | SRSF11 | 53.543 | 3 | 49.078228 | yes | 3.957E-06 | 592.79749 | 66.589661 |
| FHDPDSAVVAQHI | 21 | Q05519 | Serine/arginine-rich splicing factor 11 OS | SRSF11 | 53.543 | 4 | 49.078228 | yes | 1.856E-07 | 790.06085 | 66.589661 |

| | | | | | | | | | | | |
|---|---|---|---|---|---|---|---|---|---|---|---|
| FHDPDSAVVAQHI | 21 | Q05519 | Serine/arginine-rich splicing factor 11 OS | SRSF11 | 53.543 | 4 | 49.078228 | yes | 3.957E-06 | 592.79749 | 66.589661 |
| EAQSLISAAIEPDK | 14 | Q05519 | Serine/arginine-rich splicing factor 11 OS | SRSF11 | 53.543 | 2 | 49.078228 | yes | 1.071E-07 | 736.38556 | 78.66581 |
| EAQSLISAAIEPDK | 15 | Q05519 | Serine/arginine-rich splicing factor 11 OS | SRSF11 | 53.543 | 2 | 49.078228 | yes | 1.926E-07 | 800.43304 | 76.86879 |
| TLFGFLGK | 8 | Q05519 | Serine/arginine-rich splicing factor 11 OS | SRSF11 | 53.543 | 2 | 49.078228 | yes | 1.823E-07 | 441.75781 | 27.532787 |
| AAVEDNHLLIK | 11 | Q05823 | 2-5A-dependent ribonuclease OS=Homo | RNASEL | 83.533 | 2 | 54.304344 | yes | 9.722E-08 | 611.84314 | 43.709747 |
| SGLQVEDLDR | 10 | Q05932 | Folylpolyglutamate synthase, mitochondr | FPGS | 64.61 | 2 | 47.631187 | yes | 1.288E-07 | 566.28564 | 61.3232 |
| LNIIHVTGTK | 10 | Q05932 | Folylpolyglutamate synthase, mitochondr | FPGS | 64.61 | 2 | 47.631187 | yes | 6.079E-07 | 548.32965 | 49.655994 |
| EVSFDVELPK | 10 | Q06033 | Inter-alpha-trypsin inhibitor heavy chain I | ITIH3 | 99.85 | 2 | 40.580685 | yes | 3.948E-07 | 581.80316 | 388.92722 |
| VTFELTYEELLK | 12 | Q06033 | Inter-alpha-trypsin inhibitor heavy chain I | ITIH3 | 99.85 | 2 | 40.580685 | yes | 7.841E-07 | 742.89777 | 48.695057 |
| TVWQYHFR | 8 | Q06124 | Tyrosine-protein phosphatase non-recepto | PTPN11 | 68.011 | 2 | 47.147026 | yes | 1.39E-07 | 568.78546 | 45.843166 |
| HGSFLVR | 7 | Q06124 | Tyrosine-protein phosphatase non-recepto | PTPN11 | 68.011 | 2 | 47.147026 | yes | 3.55E-06 | 408.22977 | 26.758091 |
| EVLSMDDEIQK | 11 | Q06210 | Glutamine--fructose-6-phosphate aminotr | GFPT1 | 78.807 | 2 | 48.503124 | yes | 2.459E-07 | 653.81342 | 163.79092 |
| EIFEQPESVVNTMI | 14 | Q06210 | Glutamine--fructose-6-phosphate aminotr | GFPT1 | 78.807 | 2 | 48.503124 | yes | 3.119E-07 | 839.90887 | 105.37399 |
| VNFDDYTVNLGGI | 14 | Q06210 | Glutamine--fructose-6-phosphate aminotr | GFPT1 | 78.807 | 2 | 48.503124 | yes | 1.138E-07 | 777.89355 | 93.937469 |
| CQNALQQVVAR | 11 | Q06210 | Glutamine--fructose-6-phosphate aminotr | GFPT1 | 78.807 | 2 | 48.503124 | yes | 7.324E-07 | 643.83533 | 88.851006 |
| SVHFPGQAVGTR | 12 | Q06210 | Glutamine--fructose-6-phosphate aminotr | GFPT1 | 78.807 | 3 | 48.503124 | yes | 2.929E-07 | 419.22293 | 38.693012 |
| SVHFPGQAVGTR | 12 | Q06210 | Glutamine--fructose-6-phosphate aminotr | GFPT1 | 78.807 | 3 | 48.503124 | yes | 0.0001561 | 628.33075 | 38.693012 |
| SVHFPGQAVGTR | 12 | Q06210 | Glutamine--fructose-6-phosphate aminotr | GFPT1 | 78.807 | 2 | 48.503124 | yes | 2.929E-07 | 419.22293 | 38.693012 |
| SVHFPGQAVGTR | 12 | Q06210 | Glutamine--fructose-6-phosphate aminotr | GFPT1 | 78.807 | 2 | 48.503124 | yes | 0.0001561 | 628.33075 | 38.693012 |
| LSTDHIPILYR | 11 | Q06210 | Glutamine--fructose-6-phosphate aminotr | GFPT1 | 78.807 | 2 | 48.503124 | yes | 9.499E-07 | 664.37207 | 48.751083 |
| VDSTTCLFPVEEK | 13 | Q06210 | Glutamine--fructose-6-phosphate aminotr | GFPT1 | 78.807 | 2 | 48.503124 | yes | 1.966E-07 | 762.86615 | 44.495125 |
| GALTVGITNTVGS | 17 | Q06210 | Glutamine--fructose-6-phosphate aminotr | GFPT1 | 78.807 | 2 | 48.503124 | yes | 1.287E-07 | 816.9494 | 31.875393 |
| FGFAESIANQR | 11 | Q06265 | Exosome complex component RRP45 OS | EXOSC9 | 48.95 | 2 | 51.76255 | yes | 2.012E-07 | 620.30945 | 1248.6376 |
| VAEITELILK | 10 | Q06265 | Exosome complex component RRP45 OS | EXOSC9 | 48.95 | 2 | 51.76255 | yes | 1.032E-07 | 564.84735 | 490.32672 |
| APIDTSDVEEK | 11 | Q06265 | Exosome complex component RRP45 OS | EXOSC9 | 48.95 | 2 | 51.76255 | yes | 9.722E-08 | 602.29059 | 452.12878 |
| EICTIQSSGGIMLLI | 15 | Q06265 | Exosome complex component RRP45 OS | EXOSC9 | 48.95 | 2 | 51.76255 | yes | 9.757E-08 | 825.43365 | 440.71872 |
| ETPLSNCER | 9 | Q06265 | Exosome complex component RRP45 OS | EXOSC9 | 48.95 | 2 | 51.76255 | yes | 1.146E-07 | 553.25073 | 233.11136 |
| CIDTESLCVVAGEI | 14 | Q06265 | Exosome complex component RRP45 OS | EXOSC9 | 48.95 | 2 | 51.76255 | yes | 2.107E-07 | 790.86835 | 105.52032 |
| QSDLLVK | 7 | Q06265 | Exosome complex component RRP45 OS | EXOSC9 | 48.95 | 2 | 51.76255 | yes | 6.55E-07 | 401.73709 | 34.434204 |
| MYLCLSQER | 9 | Q06330 | Recombining binding protein suppressor | RBPJ | 55.638 | 2 | 55.070972 | yes | 1.048E-07 | 600.28101 | 552.586 |
| QPVQVPVTLVR | 11 | Q06330 | Recombining binding protein suppressor | RBPJ | 55.638 | 2 | 55.070972 | yes | 9.722E-08 | 618.37714 | 435.18027 |
| IIQFQATPCPK | 11 | Q06330 | Recombining binding protein suppressor | RBPJ | 55.638 | 2 | 55.070972 | yes | 1.124E-07 | 651.84735 | 271.20447 |
| TLYISDSDKR | 10 | Q06330 | Recombining binding protein suppressor | RBPJ | 55.638 | 2 | 55.070972 | yes | 2.398E-07 | 599.30914 | 267.47763 |
| NADLCIASGTK | 11 | Q06330 | Recombining binding protein suppressor | RBPJ | 55.638 | 2 | 55.070972 | yes | 1.179E-07 | 575.28204 | 251.44485 |
| MFYGNSDDIGVFL | 15 | Q06330 | Recombining binding protein suppressor | RBPJ | 55.638 | 2 | 55.070972 | yes | 9.722E-08 | 846.90051 | 247.83342 |
| VWFGDVEAETMY | 13 | Q06330 | Recombining binding protein suppressor | RBPJ | 55.638 | 2 | 55.070972 | no | 9.722E-08 | 801.86652 | 198.21516 |
| LVCSVTGMALPR | 12 | Q06330 | Recombining binding protein suppressor | RBPJ | 55.638 | 2 | 55.070972 | yes | 3.037E-07 | 652.34668 | 119.16454 |
| PHCSAAGAILR | 11 | Q06330 | Recombining binding protein suppressor | RBPJ | 55.638 | 2 | 55.070972 | yes | 2.758E-06 | 576.80078 | 116.97335 |
| GDQTVLILHAK | 11 | Q06330 | Recombining binding protein suppressor | RBPJ | 55.638 | 2 | 55.070972 | yes | 1.154E-07 | 597.8457 | 99.781898 |

| | | | | | | | | | | | |
|---|---|---|---|---|---|---|---|---|---|---|---|
| DGYIHYGQTVK | 11 | Q06330 | Recombining binding protein suppressor ( | RBPJ | 55.638 | 2 | 55.070972 | yes | 4.95E-07 | 640.81714 | 99.646416 |
| HFMLSVK | 7 | Q06330 | Recombining binding protein suppressor ( | RBPJ | 55.638 | 2 | 55.070972 | yes | 9.722E-08 | 431.23621 | 54.355282 |
| KHFMLSVK | 8 | Q06330 | Recombining binding protein suppressor ( | RBPJ | 55.638 | 2 | 55.070972 | yes | 9.85E-08 | 495.28369 | 54.010365 |
| QTALLDADDPVSQ | 16 | Q06330 | Recombining binding protein suppressor ( | RBPJ | 55.638 | 3 | 55.070972 | yes | 1.79E-07 | 584.30383 | 46.33968 |
| VSIDNWCR | 8 | Q06481 | Amyloid beta precursor like protein 2 OS= | APLP2 | 86.956 | 2 | 47.375862 | yes | 1.343E-07 | 525.2453 | 55.088627 |
| VDENMVIDETLDV | 14 | Q06481 | Amyloid beta precursor like protein 2 OS= | APLP2 | 86.956 | 2 | 47.375862 | yes | 1.846E-06 | 810.39526 | 41.447529 |
| FCSDCIVTALR | 11 | Q06587 | E3 ubiquitin-protein ligase RING1 OS=H | RING1 | 42.43 | 2 | 51.061176 | yes | 3.341E-07 | 671.31812 | 310.8678 |
| TWELSLYELHR | 11 | Q06587 | E3 ubiquitin-protein ligase RING1 OS=H | RING1 | 42.43 | 2 | 51.061176 | yes | 4.184E-07 | 723.87244 | 126.1506 |
| TTGNATVDHLSK | 12 | Q06587 | E3 ubiquitin-protein ligase RING1 OS=H | RING1 | 42.43 | 2 | 51.061176 | yes | 9.722E-08 | 622.3175 | 42.262215 |
| FHEQFIVR | 8 | Q06787 | Fragile X messenger ribonucleoprotein 1 | FMR1 | 71.175 | 2 | 53.588226 | yes | 5.553E-07 | 538.28778 | 17082.572 |
| LDVPEDLR | 8 | Q06787 | Fragile X messenger ribonucleoprotein 1 | FMR1 | 71.175 | 2 | 53.588226 | yes | 1.481E-07 | 478.75601 | 9575.5352 |
| AHMLIDMHFR | 10 | Q06787 | Fragile X messenger ribonucleoprotein 1 | FMR1 | 71.175 | 2 | 53.588226 | yes | 9.722E-08 | 635.81281 | 3060.252 |
| AHMLIDMHFR | 10 | Q06787 | Fragile X messenger ribonucleoprotein 1 | FMR1 | 71.175 | 2 | 53.588226 | yes | 1.172E-06 | 643.81024 | 3060.252 |
| QIPFHDVR | 8 | Q06787 | Fragile X messenger ribonucleoprotein 1 | FMR1 | 71.175 | 2 | 53.588226 | yes | 1.876E-07 | 506.27216 | 5819.3384 |
| TTDGSLQIR | 9 | Q06787 | Fragile X messenger ribonucleoprotein 1 | FMR1 | 71.175 | 2 | 53.588226 | yes | 3.538E-07 | 495.76437 | 5591.8545 |
| IKLDVPEDLR | 10 | Q06787 | Fragile X messenger ribonucleoprotein 1 | FMR1 | 71.175 | 2 | 53.588226 | yes | 9.722E-08 | 599.34552 | 5009.4438 |
| EPCCWWLAK | 9 | Q06787 | Fragile X messenger ribonucleoprotein 1 | FMR1 | 71.175 | 2 | 53.588226 | yes | 1.169E-07 | 625.27826 | 3431.2151 |
| KVPGVTAIDLDED | 28 | Q06787 | Fragile X messenger ribonucleoprotein 1 | FMR1 | 71.175 | 4 | 53.588226 | yes | 1.611E-07 | 784.87494 | 3199.3257 |
| SYVTDDGQGMGR | 12 | Q06787 | Fragile X messenger ribonucleoprotein 1 | FMR1 | 71.175 | 2 | 53.588226 | yes | 9.722E-08 | 643.27753 | 1761.7379 |
| TLQNTSSEGSR | 11 | Q06787 | Fragile X messenger ribonucleoprotein 1 | FMR1 | 71.175 | 2 | 53.588226 | yes | 1.095E-07 | 590.28363 | 1213.6587 |
| QIGASSRPPPNR | 12 | Q06787 | Fragile X messenger ribonucleoprotein 1 | FMR1 | 71.175 | 3 | 53.588226 | yes | 9.722E-08 | 427.2337 | 417.76416 |
| QIGASSRPPPNR | 12 | Q06787 | Fragile X messenger ribonucleoprotein 1 | FMR1 | 71.175 | 3 | 53.588226 | yes | 1.574E-07 | 640.34692 | 417.76416 |
| QIGASSRPPPNR | 12 | Q06787 | Fragile X messenger ribonucleoprotein 1 | FMR1 | 71.175 | 2 | 53.588226 | yes | 9.722E-08 | 427.2337 | 417.76416 |
| QIGASSRPPPNR | 12 | Q06787 | Fragile X messenger ribonucleoprotein 1 | FMR1 | 71.175 | 2 | 53.588226 | yes | 1.574E-07 | 640.34692 | 417.76416 |
| VLVASSVVAGESQ | 18 | Q06787 | Fragile X messenger ribonucleoprotein 1 | FMR1 | 71.175 | 3 | 53.588226 | yes | 1.538E-07 | 614.35083 | 657.16931 |
| VGPNAPEEK | 9 | Q06787 | Fragile X messenger ribonucleoprotein 1 | FMR1 | 71.175 | 2 | 53.588226 | yes | 1.629E-07 | 470.74036 | 656.2594 |
| MEELVVEVR | 9 | Q06787 | Fragile X messenger ribonucleoprotein 1 | FMR1 | 71.175 | 2 | 53.588226 | yes | 9.809E-08 | 573.29718 | 445.81079 |
| ANEKEPCCWWLA | 13 | Q06787 | Fragile X messenger ribonucleoprotein 1 | FMR1 | 71.175 | 2 | 53.588226 | yes | 2.425E-07 | 846.38708 | 397.62949 |
| ENSTHFSQPNSTK | 13 | Q06787 | Fragile X messenger ribonucleoprotein 1 | FMR1 | 71.175 | 2 | 53.588226 | yes | 1.112E-07 | 738.83929 | 368.70282 |
| DSIANATVLLDYH | 18 | Q06787 | Fragile X messenger ribonucleoprotein 1 | FMR1 | 71.175 | 3 | 53.588226 | yes | 9.722E-08 | 688.36511 | 147.76372 |
| DINESDEVEVYSR | 13 | Q06787 | Fragile X messenger ribonucleoprotein 1 | FMR1 | 71.175 | 2 | 53.588226 | yes | 1.223E-07 | 777.84955 | 108.05511 |
| VRIEAENEK | 9 | Q06787 | Fragile X messenger ribonucleoprotein 1 | FMR1 | 71.175 | 2 | 53.588226 | yes | 1.547E-07 | 544.29077 | 65.517921 |
| LVQAFQFTDK | 10 | Q06830 | Peroxiredoxin-1 OS=Homo sapiens OX=9 | PRDX1 | 22.111 | 2 | 50.364468 | yes | 4.581E-07 | 598.81915 | 940.40784 |
| ATAVMPDGQFK | 11 | Q06830 | Peroxiredoxin-1 OS=Homo sapiens OX=9 | PRDX1 | 22.111 | 2 | 50.364468 | yes | 1.108E-07 | 582.78955 | 926.2981 |
| QITVNDLPVGR | 11 | Q06830 | Peroxiredoxin-1 OS=Homo sapiens OX=9 | PRDX1 | 22.111 | 2 | 50.364468 | no | 2.89E-07 | 606.34076 | 821.51477 |
| TIAQDYGVLK | 10 | Q06830 | Peroxiredoxin-1 OS=Homo sapiens OX=9 | PRDX1 | 22.111 | 2 | 50.364468 | yes | 9.81E-08 | 554.30585 | 529.21802 |
| HGEVCPAGWKPG | 22 | Q06830 | Peroxiredoxin-1 OS=Homo sapiens OX=9 | PRDX1 | 22.111 | 4 | 50.364468 | yes | 9.963E-08 | 602.30225 | 361.13983 |
| QGGLGPMNIPLVS | 16 | Q06830 | Peroxiredoxin-1 OS=Homo sapiens OX=9 | PRDX1 | 22.111 | 2 | 50.364468 | yes | 9.974E-08 | 811.93219 | 260.89383 |
| IGHPAPNFK | 9 | Q06830 | Peroxiredoxin-1 OS=Homo sapiens OX=9 | PRDX1 | 22.111 | 2 | 50.364468 | yes | 1.354E-07 | 490.76926 | 255.34937 |

| | | | | | | | | | | | |
|---|---|---|---|---|---|---|---|---|---|---|---|
| GLFIIDDK | 8 | Q06830 | Peroxiredoxin-1 OS=Homo sapiens OX=9 | PRDX1 | 22.111 | 2 | 50.364468 | no | 2.864E-07 | 460.75803 | 239.02808 |
| LNCQVIGASVDSH | 24 | Q06830 | Peroxiredoxin-1 OS=Homo sapiens OX=9 | PRDX1 | 22.111 | 4 | 50.364468 | yes | 1.417E-07 | 689.08765 | 67.97628 |
| DISLSDYK | 8 | Q06830 | Peroxiredoxin-1 OS=Homo sapiens OX=9 | PRDX1 | 22.111 | 2 | 50.364468 | yes | 5.026E-07 | 470.73474 | 32.987881 |
| KLNCQVIGASVDS | 25 | Q06830 | Peroxiredoxin-1 OS=Homo sapiens OX=9 | PRDX1 | 22.111 | 4 | 50.364468 | yes | 7.309E-07 | 721.11139 | 22.237206 |
| SKEYFSK | 7 | Q06830 | Peroxiredoxin-1 OS=Homo sapiens OX=9 | PRDX1 | 22.111 | 2 | 50.364468 | yes | 3.497E-05 | 444.72672 | 10.278951 |
| GCGTVLLSGPR | 11 | Q07020 | Large ribosomal subunit protein eL18 OS | RPL18 | 21.635 | 2 | 49.821934 | yes | 4.039E-07 | 558.79517 | 42360.539 |
| TAVVVGTITDDVR | 13 | Q07020 | Large ribosomal subunit protein eL18 OS | RPL18 | 21.635 | 2 | 49.821934 | yes | 1.491E-07 | 673.36975 | 32001.504 |
| TNSTFNQVVLK | 11 | Q07020 | Large ribosomal subunit protein eL18 OS | RPL18 | 21.635 | 2 | 49.821934 | yes | 6.104E-07 | 625.84058 | 17102.984 |
| SQDIYLR | 7 | Q07020 | Large ribosomal subunit protein eL18 OS | RPL18 | 21.635 | 2 | 49.821934 | yes | 1.844E-07 | 447.73761 | 16462.314 |
| ILTFDQLALDSPK | 13 | Q07020 | Large ribosomal subunit protein eL18 OS | RPL18 | 21.635 | 2 | 49.821934 | yes | 1.002E-07 | 730.90338 | 6133.7441 |
| TNRPPLSLSR | 10 | Q07020 | Large ribosomal subunit protein eL18 OS | RPL18 | 21.635 | 2 | 49.821934 | yes | 1.152E-07 | 570.82782 | 4791.3481 |
| RTNSTFNQVVLK | 12 | Q07020 | Large ribosomal subunit protein eL18 OS | RPL18 | 21.635 | 2 | 49.821934 | yes | 1.405E-07 | 703.89117 | 1123.6709 |
| APGTPHSHTKPYV | 14 | Q07020 | Large ribosomal subunit protein eL18 OS | RPL18 | 21.635 | 3 | 49.821934 | yes | 1.374E-07 | 516.60742 | 346.1676 |
| APGTPHSHTKPYV | 14 | Q07020 | Large ribosomal subunit protein eL18 OS | RPL18 | 21.635 | 3 | 49.821934 | yes | 1.224E-06 | 774.40753 | 346.1676 |
| APGTPHSHTKPYV | 14 | Q07020 | Large ribosomal subunit protein eL18 OS | RPL18 | 21.635 | 2 | 49.821934 | yes | 1.374E-07 | 516.60742 | 346.1676 |
| APGTPHSHTKPYV | 14 | Q07020 | Large ribosomal subunit protein eL18 OS | RPL18 | 21.635 | 2 | 49.821934 | yes | 1.224E-06 | 774.40753 | 346.1676 |
| LKVCALR | 7 | Q07020 | Large ribosomal subunit protein eL18 OS | RPL18 | 21.635 | 2 | 49.821934 | yes | 2.367E-07 | 430.26276 | 432.11325 |
| EVSFQSTGESEWK | 13 | Q07021 | Complement component 1 Q subcompone | C1QBP | 31.363 | 2 | 49.69021 | yes | 1.01E-07 | 757.34393 | 1234.8916 |
| MSGGWELELNGTI | 15 | Q07021 | Complement component 1 Q subcompone | C1QBP | 31.363 | 2 | 49.69021 | yes | 1.292E-07 | 811.37994 | 747.34589 |
| AFVDFLSDEIKEEF | 14 | Q07021 | Complement component 1 Q subcompone | C1QBP | 31.363 | 3 | 49.69021 | yes | 1.318E-07 | 566.61755 | 192.72041 |
| VEEQEPELTSTPNI | 20 | Q07021 | Complement component 1 Q subcompone | C1QBP | 31.363 | 3 | 49.69021 | yes | 5.183E-07 | 763.06165 | 34.658009 |
| LALQALTEK | 9 | Q07065 | Cytoskeleton-associated protein 4 OS=Hc | CKAP4 | 66.023 | 2 | 51.7158 | yes | 1.02E-07 | 493.79767 | 1904.6521 |
| SVGELPSTVESLQI | 14 | Q07065 | Cytoskeleton-associated protein 4 OS=Hc | CKAP4 | 66.023 | 2 | 51.7158 | yes | 1.122E-07 | 737.39337 | 1709.7994 |
| ASVSQVEADLK | 11 | Q07065 | Cytoskeleton-associated protein 4 OS=Hc | CKAP4 | 66.023 | 2 | 51.7158 | yes | 1.002E-07 | 573.80371 | 1352.1575 |
| DLSDGIHVVK | 10 | Q07065 | Cytoskeleton-associated protein 4 OS=Hc | CKAP4 | 66.023 | 2 | 51.7158 | yes | 9.729E-08 | 541.79565 | 1314.3755 |
| SINDNIAIFTEVQK | 14 | Q07065 | Cytoskeleton-associated protein 4 OS=Hc | CKAP4 | 66.023 | 2 | 51.7158 | yes | 1.456E-07 | 796.41992 | 1215.6632 |
| LQHVEDGVLSMQ | 17 | Q07065 | Cytoskeleton-associated protein 4 OS=Hc | CKAP4 | 66.023 | 3 | 51.7158 | yes | 9.722E-08 | 613.98425 | 898.69 |
| DFTSLENTVEER | 12 | Q07065 | Cytoskeleton-associated protein 4 OS=Hc | CKAP4 | 66.023 | 2 | 51.7158 | yes | 2.111E-07 | 720.33606 | 716.32355 |
| EWDMEALR | 8 | Q07065 | Cytoskeleton-associated protein 4 OS=Hc | CKAP4 | 66.023 | 2 | 51.7158 | yes | 2.127E-07 | 525.23969 | 609.31287 |
| SLGETQLVLYGDV | 18 | Q07065 | Cytoskeleton-associated protein 4 OS=Hc | CKAP4 | 66.023 | 3 | 51.7158 | yes | 1.245E-07 | 683.70026 | 536.0918 |
| LQNEILK | 7 | Q07065 | Cytoskeleton-associated protein 4 OS=Hc | CKAP4 | 66.023 | 2 | 51.7158 | yes | 2.608E-07 | 429.25818 | 475.7247 |
| EELGQGLQGVEQK | 13 | Q07065 | Cytoskeleton-associated protein 4 OS=Hc | CKAP4 | 66.023 | 2 | 51.7158 | yes | 3.154E-07 | 707.86224 | 418.81131 |
| LSSLDNLK | 8 | Q07065 | Cytoskeleton-associated protein 4 OS=Hc | CKAP4 | 66.023 | 2 | 51.7158 | yes | 9.722E-08 | 445.25311 | 360.47531 |
| QTESLESLLSK | 11 | Q07065 | Cytoskeleton-associated protein 4 OS=Hc | CKAP4 | 66.023 | 2 | 51.7158 | yes | 1.328E-07 | 617.8299 | 344.62469 |
| HSEAFEALQQK | 11 | Q07065 | Cytoskeleton-associated protein 4 OS=Hc | CKAP4 | 66.023 | 2 | 51.7158 | yes | 1.062E-07 | 644.32001 | 327.05386 |
| GAHPSGGADDVAI | 13 | Q07065 | Cytoskeleton-associated protein 4 OS=Hc | CKAP4 | 66.023 | 2 | 51.7158 | yes | 5.647E-07 | 591.28088 | 293.95059 |
| LPPQDFLDR | 9 | Q07065 | Cytoskeleton-associated protein 4 OS=Hc | CKAP4 | 66.023 | 2 | 51.7158 | yes | 3.346E-06 | 550.79041 | 293.26816 |
| VASLEESEGNK | 11 | Q07065 | Cytoskeleton-associated protein 4 OS=Hc | CKAP4 | 66.023 | 2 | 51.7158 | yes | 2.158E-07 | 581.78296 | 229.63644 |
| IETNENNLESAK | 12 | Q07065 | Cytoskeleton-associated protein 4 OS=Hc | CKAP4 | 66.023 | 2 | 51.7158 | yes | 3.568E-07 | 681.33081 | 229.05154 |

| | | | | | | | | | | | |
|---|---|---|---|---|---|---|---|---|---|---|---|
| TAVDSLVAYSVK | 12 | Q07065 | Cytoskeleton-associated protein 4 OS=Hc | CKAP4 | 66.023 | 2 | 51.7158 | yes | 1.398E-07 | 626.84283 | 156.74834 |
| GLLDDLR | 7 | Q07065 | Cytoskeleton-associated protein 4 OS=Hc | CKAP4 | 66.023 | 2 | 51.7158 | yes | 1.785E-07 | 401.2269 | 111.43217 |
| VASLEESEGNKQD | 15 | Q07065 | Cytoskeleton-associated protein 4 OS=Hc | CKAP4 | 66.023 | 2 | 51.7158 | yes | 1.118E-07 | 823.91522 | 80.264725 |
| HEEQPAPGYDTHC | 14 | Q07157 | Tight junction protein ZO-1 OS=Homo sa | TJP1 | 195.46 | 3 | 54.241085 | yes | 3.46E-07 | 531.90619 | 1463.0544 |
| PSGAPIIGPK | 10 | Q07157 | Tight junction protein ZO-1 OS=Homo sa | TJP1 | 195.46 | 2 | 54.241085 | yes | 2.049E-07 | 468.7793 | 1170.1626 |
| DGNIQEGDVVLK | 12 | Q07157 | Tight junction protein ZO-1 OS=Homo sa | TJP1 | 195.46 | 2 | 54.241085 | yes | 9.722E-08 | 643.83295 | 891.80792 |
| AEQLASVQYTLPK | 13 | Q07157 | Tight junction protein ZO-1 OS=Homo sa | TJP1 | 195.46 | 2 | 54.241085 | yes | 9.722E-08 | 724.39319 | 831.72528 |
| EISQDSLAAR | 10 | Q07157 | Tight junction protein ZO-1 OS=Homo sa | TJP1 | 195.46 | 2 | 54.241085 | yes | 9.722E-08 | 545.2804 | 762.9729 |
| VNNVDFTNIIR | 11 | Q07157 | Tight junction protein ZO-1 OS=Homo sa | TJP1 | 195.46 | 2 | 54.241085 | yes | 1.142E-07 | 652.8515 | 748.38171 |
| GKPPEADGVDR | 11 | Q07157 | Tight junction protein ZO-1 OS=Homo sa | TJP1 | 195.46 | 2 | 54.241085 | yes | 1.426E-07 | 570.78583 | 712.46521 |
| FEEPAPLSYDSR | 12 | Q07157 | Tight junction protein ZO-1 OS=Homo sa | TJP1 | 195.46 | 2 | 54.241085 | yes | 9.789E-08 | 705.83044 | 642.18237 |
| ISKPGAVSTPVK | 12 | Q07157 | Tight junction protein ZO-1 OS=Homo sa | TJP1 | 195.46 | 2 | 54.241085 | yes | 2.897E-07 | 592.3559 | 609.52783 |
| EEPDIYQIAK | 10 | Q07157 | Tight junction protein ZO-1 OS=Homo sa | TJP1 | 195.46 | 2 | 54.241085 | yes | 1.765E-07 | 603.30603 | 583.50537 |
| LRPEAQPHPSAGPI | 14 | Q07157 | Tight junction protein ZO-1 OS=Homo sa | TJP1 | 195.46 | 3 | 54.241085 | yes | 9.722E-08 | 495.60379 | 275.81024 |
| LRPEAQPHPSAGPI | 14 | Q07157 | Tight junction protein ZO-1 OS=Homo sa | TJP1 | 195.46 | 3 | 54.241085 | yes | 3.62E-07 | 742.90204 | 275.81024 |
| LRPEAQPHPSAGPI | 14 | Q07157 | Tight junction protein ZO-1 OS=Homo sa | TJP1 | 195.46 | 2 | 54.241085 | yes | 9.722E-08 | 495.60379 | 275.81024 |
| LRPEAQPHPSAGPI | 14 | Q07157 | Tight junction protein ZO-1 OS=Homo sa | TJP1 | 195.46 | 2 | 54.241085 | yes | 3.62E-07 | 742.90204 | 275.81024 |
| APGFGFGIAISGGR | 14 | Q07157 | Tight junction protein ZO-1 OS=Homo sa | TJP1 | 195.46 | 2 | 54.241085 | yes | 1.348E-07 | 653.84875 | 535.65063 |
| GGPAEGQLQENDI | 13 | Q07157 | Tight junction protein ZO-1 OS=Homo sa | TJP1 | 195.46 | 2 | 54.241085 | yes | 9.965E-08 | 685.81836 | 532.58759 |
| ESPYGLSFNK | 10 | Q07157 | Tight junction protein ZO-1 OS=Homo sa | TJP1 | 195.46 | 2 | 54.241085 | yes | 1.457E-07 | 571.27985 | 529.39105 |
| EDLSAQPVQTK | 11 | Q07157 | Tight junction protein ZO-1 OS=Homo sa | TJP1 | 195.46 | 2 | 54.241085 | yes | 5.867E-07 | 608.31439 | 514.70502 |
| GEEVTILAQK | 10 | Q07157 | Tight junction protein ZO-1 OS=Homo sa | TJP1 | 195.46 | 2 | 54.241085 | yes | 1.973E-07 | 544.30334 | 474.16748 |
| KDPYPEEMMR | 10 | Q07157 | Tight junction protein ZO-1 OS=Homo sa | TJP1 | 195.46 | 2 | 54.241085 | yes | 9.722E-08 | 648.29156 | 428.54431 |
| DLEQPTYR | 8 | Q07157 | Tight junction protein ZO-1 OS=Homo sa | TJP1 | 195.46 | 2 | 54.241085 | yes | 1.57E-07 | 511.2511 | 420.67233 |
| IVESDVGDSFYIR | 13 | Q07157 | Tight junction protein ZO-1 OS=Homo sa | TJP1 | 195.46 | 2 | 54.241085 | yes | 9.722E-08 | 750.37244 | 370.38208 |
| DNSILPPLDK | 10 | Q07157 | Tight junction protein ZO-1 OS=Homo sa | TJP1 | 195.46 | 2 | 54.241085 | yes | 9.722E-08 | 556.30334 | 357.41321 |
| EGLEEGDQILR | 11 | Q07157 | Tight junction protein ZO-1 OS=Homo sa | TJP1 | 195.46 | 2 | 54.241085 | yes | 1.246E-07 | 629.81732 | 355.81116 |
| KNEEYGLR | 8 | Q07157 | Tight junction protein ZO-1 OS=Homo sa | TJP1 | 195.46 | 2 | 54.241085 | yes | 1.584E-07 | 504.75909 | 350.4306 |
| LASHIFVK | 8 | Q07157 | Tight junction protein ZO-1 OS=Homo sa | TJP1 | 195.46 | 2 | 54.241085 | yes | 2.667E-07 | 457.77655 | 288.71219 |
| ADGATSDDLDLHI | 15 | Q07157 | Tight junction protein ZO-1 OS=Homo sa | TJP1 | 195.46 | 2 | 54.241085 | yes | 1.42E-07 | 808.34515 | 268.41782 |
| SVASSQPAKPTK | 12 | Q07157 | Tight junction protein ZO-1 OS=Homo sa | TJP1 | 195.46 | 2 | 54.241085 | yes | 3.323E-07 | 600.83276 | 263.20087 |
| YESSSYTDQFSR | 12 | Q07157 | Tight junction protein ZO-1 OS=Homo sa | TJP1 | 195.46 | 2 | 54.241085 | yes | 9.722E-08 | 735.31262 | 244.0204 |
| SHSLAQPPEFDSG\ | 25 | Q07157 | Tight junction protein ZO-1 OS=Homo sa | TJP1 | 195.46 | 4 | 54.241085 | yes | 1.166E-07 | 685.34094 | 226.183 |
| YQINNISTVPK | 11 | Q07157 | Tight junction protein ZO-1 OS=Homo sa | TJP1 | 195.46 | 2 | 54.241085 | yes | 1.222E-07 | 638.84845 | 145.28105 |
| FEEPAPLSYDSRPI | 14 | Q07157 | Tight junction protein ZO-1 OS=Homo sa | TJP1 | 195.46 | 3 | 54.241085 | yes | 9.722E-08 | 555.27399 | 144.72102 |
| LRPEAQPHPSAGPI | 19 | Q07157 | Tight junction protein ZO-1 OS=Homo sa | TJP1 | 195.46 | 4 | 54.241085 | yes | 1.421E-07 | 500.01953 | 137.51263 |
| THFEYEK | 7 | Q07157 | Tight junction protein ZO-1 OS=Homo sa | TJP1 | 195.46 | 2 | 54.241085 | yes | 1.353E-07 | 477.2218 | 133.70496 |
| DNPHFQSGETSIVI | 19 | Q07157 | Tight junction protein ZO-1 OS=Homo sa | TJP1 | 195.46 | 3 | 54.241085 | yes | 1.041E-07 | 696.0199 | 126.42644 |
| TVTPAYNR | 8 | Q07157 | Tight junction protein ZO-1 OS=Homo sa | TJP1 | 195.46 | 2 | 54.241085 | yes | 9.722E-08 | 461.24307 | 98.73233 |

| | | | | | | | | | | | |
|---|---|---|---|---|---|---|---|---|---|---|---|
| HALLDVTPNAVDI | 13 | Q07157 | Tight junction protein ZO-1 OS=Homo sa | TJP1 | 195.46 | 3 | 54.241085 | yes | 9.722E-08 | 474.25629 | 49.1763 |
| HALLDVTPNAVDI | 13 | Q07157 | Tight junction protein ZO-1 OS=Homo sa | TJP1 | 195.46 | 3 | 54.241085 | yes | 3.284E-07 | 710.8808 | 49.1763 |
| HALLDVTPNAVDI | 13 | Q07157 | Tight junction protein ZO-1 OS=Homo sa | TJP1 | 195.46 | 2 | 54.241085 | yes | 9.722E-08 | 474.25629 | 49.1763 |
| HALLDVTPNAVDI | 13 | Q07157 | Tight junction protein ZO-1 OS=Homo sa | TJP1 | 195.46 | 2 | 54.241085 | yes | 3.284E-07 | 710.8808 | 49.1763 |
| PVTIFGPIADVAR | 13 | Q07157 | Tight junction protein ZO-1 OS=Homo sa | TJP1 | 195.46 | 2 | 54.241085 | yes | 1.152E-07 | 678.3877 | 95.512283 |
| EAGFLRPVTIFGPI | 19 | Q07157 | Tight junction protein ZO-1 OS=Homo sa | TJP1 | 195.46 | 3 | 54.241085 | yes | 1.276E-07 | 677.04584 | 95.242737 |
| VVDTLYNGK | 9 | Q07157 | Tight junction protein ZO-1 OS=Homo sa | TJP1 | 195.46 | 2 | 54.241085 | yes | 1.06E-07 | 504.77167 | 89.909889 |
| QYFEQYSR | 8 | Q07157 | Tight junction protein ZO-1 OS=Homo sa | TJP1 | 195.46 | 2 | 54.241085 | yes | 1.272E-07 | 560.75653 | 83.559677 |
| TVEEVTVER | 9 | Q07157 | Tight junction protein ZO-1 OS=Homo sa | TJP1 | 195.46 | 2 | 54.241085 | yes | 3.868E-07 | 531.27728 | 74.096054 |
| EKGETLLSPLVMC | 19 | Q07157 | Tight junction protein ZO-1 OS=Homo sa | TJP1 | 195.46 | 3 | 54.241085 | yes | 1.612E-07 | 689.36395 | 72.392929 |
| DQEPSLSSHVDPTI | 14 | Q07157 | Tight junction protein ZO-1 OS=Homo sa | TJP1 | 195.46 | 2 | 54.241085 | yes | 4.368E-07 | 770.36792 | 70.681076 |
| QLSYFDR | 7 | Q07157 | Tight junction protein ZO-1 OS=Homo sa | TJP1 | 195.46 | 2 | 54.241085 | yes | 7.824E-07 | 464.7298 | 68.150719 |
| LGSWLAIR | 8 | Q07157 | Tight junction protein ZO-1 OS=Homo sa | TJP1 | 195.46 | 2 | 54.241085 | yes | 9.722E-08 | 458.27417 | 64.347855 |
| QTPSLPEPK | 9 | Q07157 | Tight junction protein ZO-1 OS=Homo sa | TJP1 | 195.46 | 2 | 54.241085 | yes | 2.587E-06 | 498.77167 | 55.943909 |
| INGTVTENMSLTD | 15 | Q07157 | Tight junction protein ZO-1 OS=Homo sa | TJP1 | 195.46 | 2 | 54.241085 | yes | 1.859E-07 | 797.39307 | 47.384071 |
| KGDSVGLR | 8 | Q07157 | Tight junction protein ZO-1 OS=Homo sa | TJP1 | 195.46 | 2 | 54.241085 | no | 9.722E-08 | 416.23779 | 45.925781 |
| VAMVNGVSMDNV | 22 | Q07157 | Tight junction protein ZO-1 OS=Homo sa | TJP1 | 195.46 | 3 | 54.241085 | yes | 1.539E-07 | 805.73511 | 43.352085 |
| QPAVSHPGHRPDK | 28 | Q07157 | Tight junction protein ZO-1 OS=Homo sa | TJP1 | 195.46 | 4 | 54.241085 | yes | 7.981E-07 | 807.41248 | 34.626789 |
| PTSQNQFSEHDK | 12 | Q07157 | Tight junction protein ZO-1 OS=Homo sa | TJP1 | 195.46 | 2 | 54.241085 | yes | 2.348E-07 | 709.32074 | 25.613249 |
| TFHTIGFCPYGPR | 13 | Q07352 | mRNA decay activator protein ZFP36L1 | ZFP36L1 | 36.314 | 2 | 50.733616 | no | 9.727E-08 | 776.87189 | 469.30295 |
| CHFIHNAEER | 10 | Q07352 | mRNA decay activator protein ZFP36L1 | ZFP36L1 | 36.314 | 2 | 50.733616 | yes | 1.066E-07 | 656.7962 | 214.8886 |
| DLSADRPR | 8 | Q07352 | mRNA decay activator protein ZFP36L1 | ZFP36L1 | 36.314 | 2 | 50.733616 | yes | 1.967E-07 | 465.24359 | 186.48094 |
| HSVTLPSSK | 9 | Q07352 | mRNA decay activator protein ZFP36L1 | ZFP36L1 | 36.314 | 2 | 50.733616 | yes | 1.993E-07 | 478.26401 | 115.15399 |
| AVGTPAGGGFPR | 12 | Q07352 | mRNA decay activator protein ZFP36L1 | ZFP36L1 | 36.314 | 2 | 50.733616 | yes | 3.886E-06 | 543.78815 | 110.31109 |
| FHQNQLLSSLK | 11 | Q07352 | mRNA decay activator protein ZFP36L1 | ZFP36L1 | 36.314 | 2 | 50.733616 | yes | 2.69E-06 | 657.86188 | 14.8328 |
| SGSMDPSGAHPSV | 14 | Q07666 | KH domain-containing, RNA-binding, sig | KHDRBS1 | 48.228 | 2 | 49.904346 | yes | 2.011E-07 | 692.81732 | 21592.338 |
| SGSMDPSGAHPSV | 14 | Q07666 | KH domain-containing, RNA-binding, sig | KHDRBS1 | 48.228 | 2 | 49.904346 | yes | 1.215E-07 | 462.21399 | 21592.338 |
| SGSMDPSGAHPSV | 14 | Q07666 | KH domain-containing, RNA-binding, sig | KHDRBS1 | 48.228 | 2 | 49.904346 | yes | 2.466E-07 | 467.54562 | 21592.338 |
| SGSMDPSGAHPSV | 14 | Q07666 | KH domain-containing, RNA-binding, sig | KHDRBS1 | 48.228 | 2 | 49.904346 | yes | 9.991E-08 | 700.81482 | 21592.338 |
| SGSMDPSGAHPSV | 14 | Q07666 | KH domain-containing, RNA-binding, sig | KHDRBS1 | 48.228 | 3 | 49.904346 | yes | 2.011E-07 | 692.81732 | 21592.338 |
| SGSMDPSGAHPSV | 14 | Q07666 | KH domain-containing, RNA-binding, sig | KHDRBS1 | 48.228 | 3 | 49.904346 | yes | 1.215E-07 | 462.21399 | 21592.338 |
| SGSMDPSGAHPSV | 14 | Q07666 | KH domain-containing, RNA-binding, sig | KHDRBS1 | 48.228 | 3 | 49.904346 | yes | 2.466E-07 | 467.54562 | 21592.338 |
| SGSMDPSGAHPSV | 14 | Q07666 | KH domain-containing, RNA-binding, sig | KHDRBS1 | 48.228 | 3 | 49.904346 | yes | 9.991E-08 | 700.81482 | 21592.338 |
| ILGPQGNTIK | 10 | Q07666 | KH domain-containing, RNA-binding, sig | KHDRBS1 | 48.228 | 2 | 49.904346 | yes | 2.111E-07 | 520.80859 | 67100.156 |
| YLPELMAEK | 9 | Q07666 | KH domain-containing, RNA-binding, sig | KHDRBS1 | 48.228 | 2 | 49.904346 | no | 4.905E-07 | 547.28357 | 33539.555 |
| YLPELMAEK | 9 | Q07666 | KH domain-containing, RNA-binding, sig | KHDRBS1 | 48.228 | 2 | 49.904346 | no | 1.059E-06 | 555.28101 | 33539.555 |
| QPPLPHR | 7 | Q07666 | KH domain-containing, RNA-binding, sig | KHDRBS1 | 48.228 | 2 | 49.904346 | yes | 3.077E-07 | 422.74304 | 14329.706 |
| LQEETGAK | 8 | Q07666 | KH domain-containing, RNA-binding, sig | KHDRBS1 | 48.228 | 2 | 49.904346 | no | 1.555E-07 | 438.22708 | 11100.022 |
| AKEEELR | 7 | Q07666 | KH domain-containing, RNA-binding, sig | KHDRBS1 | 48.228 | 2 | 49.904346 | no | 1.265E-07 | 437.73508 | 10331.652 |

| | | | | | | | | | | | |
|---|---|---|---|---|---|---|---|---|---|---|---|
| DDEENYLDLFSHK | 13 | Q07666 | KH domain-containing, RNA-binding, sig | KHDRBS1 | 48.228 | 2 | 49.904346 | yes | 1.87E-07 | 812.85992 | 8705.4443 |
| GVPPPPTVR | 9 | Q07666 | KH domain-containing, RNA-binding, sig | KHDRBS1 | 48.228 | 2 | 49.904346 | yes | 3.752E-07 | 460.27164 | 5888.1201 |
| GAAPPPPPVPR | 11 | Q07666 | KH domain-containing, RNA-binding, sig | KHDRBS1 | 48.228 | 2 | 49.904346 | yes | 1.797E-07 | 528.30347 | 5224.6191 |
| RLQEETGAK | 9 | Q07666 | KH domain-containing, RNA-binding, sig | KHDRBS1 | 48.228 | 2 | 49.904346 | no | 4.355E-07 | 516.27765 | 1958.7806 |
| SSGRSGSMDPSGA | 18 | Q07666 | KH domain-containing, RNA-binding, sig | KHDRBS1 | 48.228 | 4 | 49.904346 | yes | 4.265E-07 | 447.7077 | 763.47443 |
| APPARPVK | 8 | Q07666 | KH domain-containing, RNA-binding, sig | KHDRBS1 | 48.228 | 2 | 49.904346 | yes | 1.777E-06 | 418.26105 | 440.3476 |
| LCQENQWLR | 9 | Q07866 | Kinesin light chain 1 OS=Homo sapiens ( | KLC1 | 65.31 | 2 | 46.362556 | no | 4.659E-06 | 623.80347 | 46.426064 |
| VAEVLNDPENMEI | 13 | Q07866 | Kinesin light chain 1 OS=Homo sapiens ( | KLC1 | 65.31 | 2 | 46.362556 | yes | 1.58E-07 | 744.35596 | 35.479584 |
| DGTGVVEFVR | 10 | Q07955 | Serine/arginine-rich splicing factor 1 OS= | SRSF1 | 27.745 | 2 | 53.158012 | yes | 1.525E-07 | 539.78003 | 15874.48 |
| SHEGETAYIR | 10 | Q07955 | Serine/arginine-rich splicing factor 1 OS= | SRSF1 | 27.745 | 2 | 53.158012 | yes | 5.561E-07 | 581.77802 | 7274.4683 |
| IYVGNLPPDIR | 11 | Q07955 | Serine/arginine-rich splicing factor 1 OS= | SRSF1 | 27.745 | 2 | 53.158012 | yes | 9.722E-08 | 628.85352 | 6712.355 |
| EAGDVCYADVYR | 12 | Q07955 | Serine/arginine-rich splicing factor 1 OS= | SRSF1 | 27.745 | 2 | 53.158012 | yes | 9.722E-08 | 709.30627 | 6364.4067 |
| KEDMTYAVR | 9 | Q07955 | Serine/arginine-rich splicing factor 1 OS= | SRSF1 | 27.745 | 2 | 53.158012 | yes | 1.012E-07 | 556.77386 | 2487.0803 |
| KEDMTYAVR | 9 | Q07955 | Serine/arginine-rich splicing factor 1 OS= | SRSF1 | 27.745 | 2 | 53.158012 | yes | 4.573E-07 | 564.7713 | 2487.0803 |
| VVVSGLPPSGSWQ | 16 | Q07955 | Serine/arginine-rich splicing factor 1 OS= | SRSF1 | 27.745 | 2 | 53.158012 | yes | 8.948E-07 | 834.95142 | 4765.582 |
| GGPPFAFVEFEDPI | 14 | Q07955 | Serine/arginine-rich splicing factor 1 OS= | SRSF1 | 27.745 | 2 | 53.158012 | yes | 2.234E-06 | 782.87518 | 4217.519 |
| TKDIEDVFYK | 10 | Q07955 | Serine/arginine-rich splicing factor 1 OS= | SRSF1 | 27.745 | 2 | 53.158012 | yes | 1.013E-07 | 629.32172 | 1837.1621 |
| DGYDYDGYR | 9 | Q07955 | Serine/arginine-rich splicing factor 1 OS= | SRSF1 | 27.745 | 2 | 53.158012 | yes | 1.213E-07 | 562.21997 | 1518.722 |
| LRVEFPR | 7 | Q07955 | Serine/arginine-rich splicing factor 1 OS= | SRSF1 | 27.745 | 2 | 53.158012 | no | 3.153E-06 | 458.77179 | 1409.6343 |
| EDMTYAVR | 8 | Q07955 | Serine/arginine-rich splicing factor 1 OS= | SRSF1 | 27.745 | 2 | 53.158012 | yes | 1.461E-07 | 492.72638 | 914.22076 |
| DIEDVFYK | 8 | Q07955 | Serine/arginine-rich splicing factor 1 OS= | SRSF1 | 27.745 | 2 | 53.158012 | yes | 1.211E-07 | 514.75037 | 528.54303 |
| RGGPPFAFVEFEDI | 15 | Q07955 | Serine/arginine-rich splicing factor 1 OS= | SRSF1 | 27.745 | 3 | 53.158012 | yes | 9.722E-08 | 574.28625 | 219.91367 |
| TVPWLENR | 8 | Q08043 | Alpha-actinin-3 OS=Homo sapiens OX=9 | ACTN3 | 103.242 | 2 | 27.540052 | yes | 2.934E-06 | 507.772 | 1808.8347 |
| VSFWTIHPSANR | 12 | Q08050 | Forkhead box protein M1 OS=Homo sapi | FOXM1 | 84.283 | 2 | 52.04427 | yes | 1.014E-07 | 707.86493 | 153.58221 |
| CVSEMLVIQHR | 11 | Q08050 | Forkhead box protein M1 OS=Homo sapi | FOXM1 | 84.283 | 2 | 52.04427 | yes | 9.722E-08 | 686.34723 | 128.16002 |
| ENCHLEQR | 8 | Q08050 | Forkhead box protein M1 OS=Homo sapi | FOXM1 | 84.283 | 2 | 52.04427 | yes | 3.119E-07 | 543.24329 | 92.532204 |
| RPNPELR | 7 | Q08050 | Forkhead box protein M1 OS=Homo sapi | FOXM1 | 84.283 | 2 | 52.04427 | yes | 3.238E-06 | 441.25122 | 77.669823 |
| HNLSLHDMFVR | 11 | Q08050 | Forkhead box protein M1 OS=Homo sapi | FOXM1 | 84.283 | 3 | 52.04427 | yes | 9.759E-08 | 456.89957 | 46.405716 |
| QEMEEKENCHLEC | 14 | Q08050 | Forkhead box protein M1 OS=Homo sapi | FOXM1 | 84.283 | 4 | 52.04427 | yes | 4.478E-06 | 469.70447 | 42.551235 |
| RTEVTLETLGPK | 12 | Q08050 | Forkhead box protein M1 OS=Homo sapi | FOXM1 | 84.283 | 2 | 52.04427 | yes | 5.133E-07 | 672.38007 | 33.840668 |
| LVEDKPGSR | 9 | Q08170 | Serine/arginine-rich splicing factor 4 OS= | SRSF4 | 56.679 | 2 | 52.843777 | yes | 9.793E-08 | 500.77472 | 397.0275 |
| NEGVIEFVSYSDM | 14 | Q08170 | Serine/arginine-rich splicing factor 4 OS= | SRSF4 | 56.679 | 2 | 52.843777 | yes | 1.295E-07 | 809.37689 | 293.0062 |
| SKPNLPSESR | 10 | Q08170 | Serine/arginine-rich splicing factor 4 OS= | SRSF4 | 56.679 | 2 | 52.843777 | yes | 1.263E-07 | 557.7962 | 149.33774 |
| NGYGFVEFDDLR | 12 | Q08170 | Serine/arginine-rich splicing factor 4 OS= | SRSF4 | 56.679 | 2 | 52.843777 | yes | 9.722E-08 | 716.33063 | 147.17494 |
| YLFLGDYVDR | 10 | Q08209 | Protein phosphatase 3 catalytic subunit al | PPP3CA | 58.688 | 2 | 49.715565 | no | 1.094E-07 | 630.81659 | 1303.5897 |
| LFEVGGSPANTR | 12 | Q08209 | Protein phosphatase 3 catalytic subunit al | PPP3CA | 58.688 | 2 | 49.715565 | no | 2.893E-07 | 624.32257 | 132.11934 |
| IITEGASILR | 10 | Q08209 | Protein phosphatase 3 catalytic subunit al | PPP3CA | 58.688 | 2 | 49.715565 | yes | 1.008E-07 | 536.82166 | 97.860489 |
| DAMPSDANLNSIN | 14 | Q08209 | Protein phosphatase 3 catalytic subunit al | PPP3CA | 58.688 | 2 | 49.715565 | yes | 3.287E-06 | 745.3512 | 85.281052 |
| GLTPTGMLPSGVL | 17 | Q08209 | Protein phosphatase 3 catalytic subunit al | PPP3CA | 58.688 | 2 | 49.715565 | yes | 1.909E-06 | 786.4267 | 49.001884 |

| | | | | | | | | | | | |
|---|---|---|---|---|---|---|---|---|---|---|---|
| EVFDNDGKPR | 10 | Q08209 | Protein phosphatase 3 catalytic subunit al | PPP3CA | 58.688 | 2 | 49.715565 | yes | 2.276E-07 | 588.78583 | 31.857025 |
| LAQFEPSQR | 9 | Q08211 | ATP-dependent RNA helicase A OS=Hor | DHX9 | 140.959 | 2 | 51.98221 | yes | 9.722E-08 | 538.28015 | 10567.463 |
| DVVQAYPEVR | 10 | Q08211 | ATP-dependent RNA helicase A OS=Hor | DHX9 | 140.959 | 2 | 51.98221 | yes | 1.885E-07 | 588.3064 | 9389.6475 |
| AAECNIVVTQPR | 12 | Q08211 | ATP-dependent RNA helicase A OS=Hor | DHX9 | 140.959 | 2 | 51.98221 | yes | 9.722E-08 | 679.34827 | 8066.9399 |
| VFDPVPVGVTK | 11 | Q08211 | ATP-dependent RNA helicase A OS=Hor | DHX9 | 140.959 | 2 | 51.98221 | yes | 9.81E-08 | 579.33191 | 6714.9106 |
| LGGIGQFLAK | 10 | Q08211 | ATP-dependent RNA helicase A OS=Hor | DHX9 | 140.959 | 2 | 51.98221 | yes | 2.66E-07 | 502.30038 | 6591.6978 |
| LAAQSCALSLVR | 12 | Q08211 | ATP-dependent RNA helicase A OS=Hor | DHX9 | 140.959 | 2 | 51.98221 | yes | 9.898E-08 | 644.85571 | 6411.0947 |
| AAMEALVVEVTK | 12 | Q08211 | ATP-dependent RNA helicase A OS=Hor | DHX9 | 140.959 | 2 | 51.98221 | yes | 9.722E-08 | 630.84705 | 3154.2341 |
| AAMEALVVEVTK | 12 | Q08211 | ATP-dependent RNA helicase A OS=Hor | DHX9 | 140.959 | 2 | 51.98221 | yes | 2.065E-07 | 638.84448 | 3154.2341 |
| KVFDPVPVGVTK | 12 | Q08211 | ATP-dependent RNA helicase A OS=Hor | DHX9 | 140.959 | 2 | 51.98221 | yes | 1.122E-07 | 643.37933 | 5675.6636 |
| QPAIISQLDPVNER | 14 | Q08211 | ATP-dependent RNA helicase A OS=Hor | DHX9 | 140.959 | 2 | 51.98221 | yes | 1.146E-07 | 790.42554 | 5515.8682 |
| YQILPLHSQIPR | 12 | Q08211 | ATP-dependent RNA helicase A OS=Hor | DHX9 | 140.959 | 3 | 51.98221 | yes | 2.95E-07 | 488.94891 | 2757.4512 |
| YQILPLHSQIPR | 12 | Q08211 | ATP-dependent RNA helicase A OS=Hor | DHX9 | 140.959 | 3 | 51.98221 | yes | 1.218E-07 | 732.91974 | 2757.4512 |
| YQILPLHSQIPR | 12 | Q08211 | ATP-dependent RNA helicase A OS=Hor | DHX9 | 140.959 | 2 | 51.98221 | yes | 2.95E-07 | 488.94891 | 2757.4512 |
| YQILPLHSQIPR | 12 | Q08211 | ATP-dependent RNA helicase A OS=Hor | DHX9 | 140.959 | 2 | 51.98221 | yes | 1.218E-07 | 732.91974 | 2757.4512 |
| AIEPPPLDAVIEAE | 18 | Q08211 | ATP-dependent RNA helicase A OS=Hor | DHX9 | 140.959 | 3 | 51.98221 | yes | 2.066E-07 | 657.6897 | 5076.105 |
| ISAVSVAER | 9 | Q08211 | ATP-dependent RNA helicase A OS=Hor | DHX9 | 140.959 | 2 | 51.98221 | yes | 1.331E-07 | 466.26401 | 4996.6523 |
| GISHVIVDEIHER | 13 | Q08211 | ATP-dependent RNA helicase A OS=Hor | DHX9 | 140.959 | 2 | 51.98221 | yes | 3.588E-07 | 752.39935 | 4962.7134 |
| MGGEEAEIR | 9 | Q08211 | ATP-dependent RNA helicase A OS=Hor | DHX9 | 140.959 | 2 | 51.98221 | yes | 1.883E-07 | 496.22931 | 4582.9404 |
| YTQVGPDHNR | 10 | Q08211 | ATP-dependent RNA helicase A OS=Hor | DHX9 | 140.959 | 2 | 51.98221 | yes | 2.291E-07 | 593.78363 | 4554.8413 |
| FESILPR | 7 | Q08211 | ATP-dependent RNA helicase A OS=Hor | DHX9 | 140.959 | 2 | 51.98221 | yes | 7.239E-07 | 431.24509 | 3918.9551 |
| LSMSQLNEK | 9 | Q08211 | ATP-dependent RNA helicase A OS=Hor | DHX9 | 140.959 | 2 | 51.98221 | yes | 1.05E-07 | 525.26843 | 3914.47 |
| LETHMTPEMFR | 11 | Q08211 | ATP-dependent RNA helicase A OS=Hor | DHX9 | 140.959 | 2 | 51.98221 | yes | 3.496E-07 | 696.32593 | 3834.5742 |
| NFLYAWCGK | 9 | Q08211 | ATP-dependent RNA helicase A OS=Hor | DHX9 | 140.959 | 2 | 51.98221 | yes | 1.784E-07 | 579.77368 | 3788.5352 |
| YGDGPRPPK | 9 | Q08211 | ATP-dependent RNA helicase A OS=Hor | DHX9 | 140.959 | 2 | 51.98221 | yes | 1.134E-07 | 493.75635 | 3120.9133 |
| SSVNCPFSSQDMK | 13 | Q08211 | ATP-dependent RNA helicase A OS=Hor | DHX9 | 140.959 | 2 | 51.98221 | yes | 1.251E-07 | 743.81866 | 1350.9451 |
| SSVNCPFSSQDMK | 13 | Q08211 | ATP-dependent RNA helicase A OS=Hor | DHX9 | 140.959 | 2 | 51.98221 | yes | 4.372E-07 | 751.81616 | 1350.9451 |
| DFVNYLVR | 8 | Q08211 | ATP-dependent RNA helicase A OS=Hor | DHX9 | 140.959 | 2 | 51.98221 | yes | 1.456E-07 | 513.27435 | 2186.5581 |
| TPLHEIALSIK | 11 | Q08211 | ATP-dependent RNA helicase A OS=Hor | DHX9 | 140.959 | 2 | 51.98221 | yes | 1.236E-07 | 611.36371 | 881.41351 |
| TPLHEIALSIK | 11 | Q08211 | ATP-dependent RNA helicase A OS=Hor | DHX9 | 140.959 | 2 | 51.98221 | yes | 2.064E-07 | 407.91156 | 881.41351 |
| TPLHEIALSIK | 11 | Q08211 | ATP-dependent RNA helicase A OS=Hor | DHX9 | 140.959 | 3 | 51.98221 | yes | 1.236E-07 | 611.36371 | 881.41351 |
| TPLHEIALSIK | 11 | Q08211 | ATP-dependent RNA helicase A OS=Hor | DHX9 | 140.959 | 3 | 51.98221 | yes | 2.064E-07 | 407.91156 | 881.41351 |
| PSAAGINLMIGSTF | 14 | Q08211 | ATP-dependent RNA helicase A OS=Hor | DHX9 | 140.959 | 2 | 51.98221 | yes | 1.311E-07 | 694.37177 | 828.22156 |
| PSAAGINLMIGSTF | 14 | Q08211 | ATP-dependent RNA helicase A OS=Hor | DHX9 | 140.959 | 2 | 51.98221 | yes | 3.432E-06 | 702.3692 | 828.22156 |
| LNQYFQK | 7 | Q08211 | ATP-dependent RNA helicase A OS=Hor | DHX9 | 140.959 | 2 | 51.98221 | yes | 2.215E-07 | 470.74799 | 1253.4769 |
| QLYHLGVVEAYS( | 16 | Q08211 | ATP-dependent RNA helicase A OS=Hor | DHX9 | 140.959 | 3 | 51.98221 | yes | 3.654E-07 | 593.32098 | 1087.8679 |
| LQISHEAAACITGL | 15 | Q08211 | ATP-dependent RNA helicase A OS=Hor | DHX9 | 140.959 | 2 | 51.98221 | yes | 9.809E-08 | 820.43286 | 535.93903 |
| LQISHEAAACITGL | 15 | Q08211 | ATP-dependent RNA helicase A OS=Hor | DHX9 | 140.959 | 2 | 51.98221 | yes | 6.513E-07 | 547.29102 | 535.93903 |
| LQISHEAAACITGL | 15 | Q08211 | ATP-dependent RNA helicase A OS=Hor | DHX9 | 140.959 | 3 | 51.98221 | yes | 9.809E-08 | 820.43286 | 535.93903 |

| | | | | | | | | | | | |
|---|---|---|---|---|---|---|---|---|---|---|---|
| LQISHEAAACITGL | 15 | Q08211 | ATP-dependent RNA helicase A OS=Hor | DHX9 | 140.959 | 3 | 51.98221 | yes | 6.513E-07 | 547.29102 | 535.93903 |
| EGETVEPYK | 9 | Q08211 | ATP-dependent RNA helicase A OS=Hor | DHX9 | 140.959 | 2 | 51.98221 | yes | 1.112E-07 | 526.25073 | 1001.5797 |
| VRPGFCFHLCSR | 12 | Q08211 | ATP-dependent RNA helicase A OS=Hor | DHX9 | 140.959 | 3 | 51.98221 | yes | 1.112E-07 | 512.58344 | 670.24585 |
| YPSPFFVFGEK | 11 | Q08211 | ATP-dependent RNA helicase A OS=Hor | DHX9 | 140.959 | 2 | 51.98221 | yes | 1.078E-07 | 659.32935 | 657.89917 |
| MTPSYEIR | 8 | Q08211 | ATP-dependent RNA helicase A OS=Hor | DHX9 | 140.959 | 2 | 51.98221 | yes | 9.722E-08 | 498.7446 | 555.91449 |
| LNMATLR | 7 | Q08211 | ATP-dependent RNA helicase A OS=Hor | DHX9 | 140.959 | 2 | 51.98221 | yes | 1.519E-07 | 409.73129 | 447.75537 |
| KILTTEGR | 8 | Q08211 | ATP-dependent RNA helicase A OS=Hor | DHX9 | 140.959 | 2 | 51.98221 | yes | 1.503E-07 | 459.27435 | 302.02603 |
| KMTPSYEIR | 9 | Q08211 | ATP-dependent RNA helicase A OS=Hor | DHX9 | 140.959 | 2 | 51.98221 | yes | 5.675E-07 | 562.79205 | 267.02457 |
| KEGETVEPYK | 10 | Q08211 | ATP-dependent RNA helicase A OS=Hor | DHX9 | 140.959 | 2 | 51.98221 | yes | 9.885E-08 | 590.29822 | 195.02408 |
| AGYGAGVGGGYR | 12 | Q08211 | ATP-dependent RNA helicase A OS=Hor | DHX9 | 140.959 | 2 | 51.98221 | yes | 1.041E-07 | 542.76215 | 170.24721 |
| SCGYSVR | 7 | Q08211 | ATP-dependent RNA helicase A OS=Hor | DHX9 | 140.959 | 2 | 51.98221 | yes | 1.029E-07 | 414.68707 | 145.70628 |
| SFIAEMTIYIK | 11 | Q08211 | ATP-dependent RNA helicase A OS=Hor | DHX9 | 140.959 | 2 | 51.98221 | yes | 6.81E-07 | 658.35193 | 97.389526 |
| RISAVSVAER | 10 | Q08211 | ATP-dependent RNA helicase A OS=Hor | DHX9 | 140.959 | 2 | 51.98221 | yes | 9.722E-08 | 544.31458 | 50.676929 |
| KFESEILEAISQNS' | 19 | Q08211 | ATP-dependent RNA helicase A OS=Hor | DHX9 | 140.959 | 3 | 51.98221 | yes | 1.996E-06 | 725.73883 | 33.353401 |
| PHASIMFCTVGVL | 15 | Q08211 | ATP-dependent RNA helicase A OS=Hor | DHX9 | 140.959 | 3 | 51.98221 | yes | 3.491E-07 | 567.63733 | 15.898046 |
| EQFLQQK | 7 | Q08378 | Golgin subfamily A member 3 OS=Homo | GOLGA3 | 167.355 | 2 | 20.493107 | yes | 4.446E-06 | 460.74545 | 147.90251 |
| AAELWGEQAEAR | 12 | Q08379 | Golgin subfamily A member 2 OS=Homo | GOLGA2 | 113.087 | 2 | 53.87067 | yes | 1.354E-07 | 665.82294 | 483.8645 |
| AGMQLNLEELQK | 12 | Q08379 | Golgin subfamily A member 2 OS=Homo | GOLGA2 | 113.087 | 2 | 53.87067 | yes | 9.722E-08 | 687.35828 | 253.18472 |
| LLELQELVLR | 10 | Q08379 | Golgin subfamily A member 2 OS=Homo | GOLGA2 | 113.087 | 2 | 53.87067 | yes | 1.217E-07 | 613.37933 | 244.48048 |
| VEELEHR | 7 | Q08379 | Golgin subfamily A member 2 OS=Homo | GOLGA2 | 113.087 | 2 | 53.87067 | yes | 3.915E-07 | 456.2327 | 230.6257 |
| EQLAELQSGFVK | 12 | Q08379 | Golgin subfamily A member 2 OS=Homo | GOLGA2 | 113.087 | 2 | 53.87067 | yes | 9.733E-08 | 674.85901 | 229.48785 |
| TFSSTESLR | 9 | Q08379 | Golgin subfamily A member 2 OS=Homo | GOLGA2 | 113.087 | 2 | 53.87067 | yes | 1.234E-06 | 514.25635 | 199.14546 |
| AELQTALAHTQHA | 15 | Q08379 | Golgin subfamily A member 2 OS=Homo | GOLGA2 | 113.087 | 3 | 53.87067 | yes | 1.392E-07 | 539.95337 | 97.732758 |
| AELQTALAHTQHA | 15 | Q08379 | Golgin subfamily A member 2 OS=Homo | GOLGA2 | 113.087 | 3 | 53.87067 | yes | 1.024E-06 | 809.42645 | 97.732758 |
| AELQTALAHTQHA | 15 | Q08379 | Golgin subfamily A member 2 OS=Homo | GOLGA2 | 113.087 | 2 | 53.87067 | yes | 1.392E-07 | 539.95337 | 97.732758 |
| AELQTALAHTQHA | 15 | Q08379 | Golgin subfamily A member 2 OS=Homo | GOLGA2 | 113.087 | 2 | 53.87067 | yes | 1.024E-06 | 809.42645 | 97.732758 |
| ALSAVSTQQK | 10 | Q08379 | Golgin subfamily A member 2 OS=Homo | GOLGA2 | 113.087 | 2 | 53.87067 | yes | 2.024E-07 | 516.78784 | 192.12166 |
| SQEAQSLQQQR | 11 | Q08379 | Golgin subfamily A member 2 OS=Homo | GOLGA2 | 113.087 | 2 | 53.87067 | yes | 9.722E-08 | 651.82349 | 184.02104 |
| LAHLLASAQK | 10 | Q08379 | Golgin subfamily A member 2 OS=Homo | GOLGA2 | 113.087 | 2 | 53.87067 | yes | 1.21E-07 | 526.31659 | 178.16736 |
| QELQETQER | 9 | Q08379 | Golgin subfamily A member 2 OS=Homo | GOLGA2 | 113.087 | 2 | 53.87067 | yes | 9.722E-08 | 580.78076 | 177.20786 |
| LTNENMEITSALQ! | 19 | Q08379 | Golgin subfamily A member 2 OS=Homo | GOLGA2 | 113.087 | 3 | 53.87067 | yes | 1.909E-07 | 724.69153 | 160.53308 |
| FMELMQEK | 8 | Q08379 | Golgin subfamily A member 2 OS=Homo | GOLGA2 | 113.087 | 2 | 53.87067 | yes | 1.004E-07 | 528.24866 | 158.95435 |
| VQELETSLAELR | 12 | Q08379 | Golgin subfamily A member 2 OS=Homo | GOLGA2 | 113.087 | 2 | 53.87067 | yes | 9.722E-08 | 694.375 | 153.09828 |
| DQYLGHLQQYVA | 21 | Q08379 | Golgin subfamily A member 2 OS=Homo | GOLGA2 | 113.087 | 3 | 53.87067 | yes | 1.073E-07 | 828.41156 | 64.748322 |
| EKEEYISR | 8 | Q08379 | Golgin subfamily A member 2 OS=Homo | GOLGA2 | 113.087 | 2 | 53.87067 | yes | 6.447E-07 | 527.26422 | 49.683796 |
| EPEAAAPAPGTGG | 22 | Q08379 | Golgin subfamily A member 2 OS=Homo | GOLGA2 | 113.087 | 3 | 53.87067 | yes | 1.61E-07 | 722.6554 | 48.680794 |
| AQLEAHLGQVME! | 15 | Q08379 | Golgin subfamily A member 2 OS=Homo | GOLGA2 | 113.087 | 2 | 53.87067 | yes | 1.727E-07 | 834.4303 | 44.843964 |
| LEAATQQNQQLR | 12 | Q08379 | Golgin subfamily A member 2 OS=Homo | GOLGA2 | 113.087 | 2 | 53.87067 | yes | 1.376E-07 | 700.36804 | 31.900013 |
| DNPTAQQIMQLLR | 13 | Q08379 | Golgin subfamily A member 2 OS=Homo | GOLGA2 | 113.087 | 2 | 53.87067 | yes | 3.303E-07 | 764.40106 | 22.651676 |

| | | | | | | | | | | | |
|---|---|---|---|---|---|---|---|---|---|---|---|
| YSSDYFQAPSDYR | 13 | Q08380 | Galectin-3-binding protein OS=Homo sap | LGALS3BI | 65.331 | 2 | 50.84147 | yes | 9.722E-08 | 799.84155 | 178.09877 |
| STHTLDLSR | 9 | Q08380 | Galectin-3-binding protein OS=Homo sap | LGALS3BI | 65.331 | 2 | 50.84147 | yes | 2.205E-07 | 515.26984 | 83.00856 |
| SDLAVPSELALLK | 13 | Q08380 | Galectin-3-binding protein OS=Homo sap | LGALS3BI | 65.331 | 2 | 50.84147 | yes | 1.591E-06 | 678.39264 | 67.084633 |
| FDEISFVNFAR | 11 | Q08945 | FACT complex subunit SSRP1 OS=Hom | SSRP1 | 81.075 | 2 | 55.424683 | yes | 1.363E-07 | 672.83276 | 263.79944 |
| IPYTTVLR | 8 | Q08945 | FACT complex subunit SSRP1 OS=Hom | SSRP1 | 81.075 | 2 | 55.424683 | yes | 1.811E-07 | 481.78711 | 231.10245 |
| ADVIQATGDAICIF | 15 | Q08945 | FACT complex subunit SSRP1 OS=Hom | SSRP1 | 81.075 | 2 | 55.424683 | yes | 1.01E-07 | 825.41943 | 226.38542 |
| IYPTFLHLHGK | 11 | Q08945 | FACT complex subunit SSRP1 OS=Hom | SSRP1 | 81.075 | 2 | 55.424683 | yes | 3.046E-07 | 663.37189 | 207.29649 |
| ASSGLLYPLER | 11 | Q08945 | FACT complex subunit SSRP1 OS=Hom | SSRP1 | 81.075 | 2 | 55.424683 | yes | 2.37E-07 | 603.3299 | 198.64407 |
| GFIYVHKPPVHIR | 13 | Q08945 | FACT complex subunit SSRP1 OS=Hom | SSRP1 | 81.075 | 3 | 55.424683 | yes | 9.722E-08 | 521.63666 | 96.225433 |
| GFIYVHKPPVHIR | 13 | Q08945 | FACT complex subunit SSRP1 OS=Hom | SSRP1 | 81.075 | 3 | 55.424683 | yes | 9.938E-08 | 781.95135 | 96.225433 |
| GFIYVHKPPVHIR | 13 | Q08945 | FACT complex subunit SSRP1 OS=Hom | SSRP1 | 81.075 | 2 | 55.424683 | yes | 9.722E-08 | 521.63666 | 96.225433 |
| GFIYVHKPPVHIR | 13 | Q08945 | FACT complex subunit SSRP1 OS=Hom | SSRP1 | 81.075 | 2 | 55.424683 | yes | 9.938E-08 | 781.95135 | 96.225433 |
| SDHPGISITDLSK | 13 | Q08945 | FACT complex subunit SSRP1 OS=Hom | SSRP1 | 81.075 | 2 | 55.424683 | yes | 9.722E-08 | 685.3515 | 190.88147 |
| ELQCLTPR | 8 | Q08945 | FACT complex subunit SSRP1 OS=Hom | SSRP1 | 81.075 | 2 | 55.424683 | yes | 3.334E-06 | 508.76331 | 132.23065 |
| LFDFVNAK | 8 | Q08945 | FACT complex subunit SSRP1 OS=Hom | SSRP1 | 81.075 | 2 | 55.424683 | yes | 5.655E-07 | 477.25818 | 119.05316 |
| QGTQYTFSSIER | 12 | Q08945 | FACT complex subunit SSRP1 OS=Hom | SSRP1 | 81.075 | 2 | 55.424683 | yes | 4.833E-07 | 708.84131 | 111.62019 |
| LFLLPHK | 7 | Q08945 | FACT complex subunit SSRP1 OS=Hom | SSRP1 | 81.075 | 2 | 55.424683 | yes | 1.878E-06 | 434.27618 | 84.103683 |
| NMSGSLYEMVSR | 12 | Q08945 | FACT complex subunit SSRP1 OS=Hom | SSRP1 | 81.075 | 2 | 55.424683 | yes | 9.722E-08 | 687.31305 | 61.259899 |
| NEVTLEFHQNDDA | 21 | Q08945 | FACT complex subunit SSRP1 OS=Hom | SSRP1 | 81.075 | 3 | 55.424683 | yes | 1.346E-07 | 825.72009 | 38.313965 |
| MIEGLYK | 7 | Q08AD1 | Calmodulin-regulated spectrin-associated | CAMSAP2 | 168.089 | 2 | 53.89426 | yes | 1.032E-07 | 427.22803 | 572.70831 |
| FLQDYDIR | 8 | Q08AD1 | Calmodulin-regulated spectrin-associated | CAMSAP2 | 168.089 | 2 | 53.89426 | yes | 1.121E-07 | 535.26929 | 378.01364 |
| TLSASVDAITIHSH | 19 | Q08AD1 | Calmodulin-regulated spectrin-associated | CAMSAP2 | 168.089 | 3 | 53.89426 | yes | 5.888E-07 | 703.37604 | 366.86731 |
| DYTVSLDSDMDD | 15 | Q08AD1 | Calmodulin-regulated spectrin-associated | CAMSAP2 | 168.089 | 2 | 53.89426 | yes | 2.653E-07 | 831.34601 | 337.30017 |
| EALSPCPSTVSTK | 13 | Q08AD1 | Calmodulin-regulated spectrin-associated | CAMSAP2 | 168.089 | 2 | 53.89426 | yes | 1.234E-07 | 688.83997 | 336.81857 |
| SFVCFGDDGEPQL | 14 | Q08AD1 | Calmodulin-regulated spectrin-associated | CAMSAP2 | 168.089 | 2 | 53.89426 | yes | 9.722E-08 | 799.86139 | 293.2041 |
| SESVEGFLSPSR | 12 | Q08AD1 | Calmodulin-regulated spectrin-associated | CAMSAP2 | 168.089 | 2 | 53.89426 | yes | 3.338E-07 | 647.81732 | 287.69803 |
| FSPSQVPIQTR | 11 | Q08AD1 | Calmodulin-regulated spectrin-associated | CAMSAP2 | 168.089 | 2 | 53.89426 | yes | 9.734E-08 | 630.34076 | 228.59618 |
| IACNLAWLVAK | 11 | Q08AD1 | Calmodulin-regulated spectrin-associated | CAMSAP2 | 168.089 | 2 | 53.89426 | yes | 1.659E-07 | 629.85248 | 219.40248 |
| VCCGFFFK | 8 | Q08AD1 | Calmodulin-regulated spectrin-associated | CAMSAP2 | 168.089 | 2 | 53.89426 | yes | 1.596E-07 | 532.73822 | 209.03879 |
| LMEDMDTVIKPR | 12 | Q08AD1 | Calmodulin-regulated spectrin-associated | CAMSAP2 | 168.089 | 2 | 53.89426 | yes | 9.722E-08 | 724.3678 | 100.32085 |
| LMEDMDTVIKPR | 12 | Q08AD1 | Calmodulin-regulated spectrin-associated | CAMSAP2 | 168.089 | 2 | 53.89426 | yes | 4.578E-07 | 483.24762 | 100.32085 |
| LMEDMDTVIKPR | 12 | Q08AD1 | Calmodulin-regulated spectrin-associated | CAMSAP2 | 168.089 | 3 | 53.89426 | yes | 9.722E-08 | 724.3678 | 100.32085 |
| LMEDMDTVIKPR | 12 | Q08AD1 | Calmodulin-regulated spectrin-associated | CAMSAP2 | 168.089 | 3 | 53.89426 | yes | 4.578E-07 | 483.24762 | 100.32085 |
| ESMENPQAK | 9 | Q08AD1 | Calmodulin-regulated spectrin-associated | CAMSAP2 | 168.089 | 2 | 53.89426 | yes | 3.602E-07 | 517.23456 | 199.50813 |
| KGDGISPLR | 9 | Q08AD1 | Calmodulin-regulated spectrin-associated | CAMSAP2 | 168.089 | 2 | 53.89426 | yes | 1.989E-07 | 471.772 | 165.74677 |
| DMPSIPVLNAAK | 12 | Q08AD1 | Calmodulin-regulated spectrin-associated | CAMSAP2 | 168.089 | 2 | 53.89426 | yes | 9.722E-08 | 628.33936 | 164.83862 |
| LSLQQEMLMQMR | 12 | Q08AD1 | Calmodulin-regulated spectrin-associated | CAMSAP2 | 168.089 | 2 | 53.89426 | yes | 9.722E-08 | 754.37512 | 162.63687 |
| GPPVSSLSLASLNT | 24 | Q08AD1 | Calmodulin-regulated spectrin-associated | CAMSAP2 | 168.089 | 3 | 53.89426 | yes | 2.64E-07 | 785.0592 | 140.93065 |
| LKEHHTVEAPGGC | 14 | Q08AD1 | Calmodulin-regulated spectrin-associated | CAMSAP2 | 168.089 | 3 | 53.89426 | yes | 1.025E-07 | 510.93896 | 67.205894 |

| | | | | | | | | | | | |
|---|---|---|---|---|---|---|---|---|---|---|---|
| LKEHHTVEAPGGQ | 14 | Q08AD1 | Calmodulin-regulated spectrin-associated | CAMSAP2 | 168.089 | 3 | 53.89426 | yes | 3.18E-07 | 765.90479 | 67.205894 |
| LKEHHTVEAPGGQ | 14 | Q08AD1 | Calmodulin-regulated spectrin-associated | CAMSAP2 | 168.089 | 2 | 53.89426 | yes | 1.025E-07 | 510.93896 | 67.205894 |
| LKEHHTVEAPGGQ | 14 | Q08AD1 | Calmodulin-regulated spectrin-associated | CAMSAP2 | 168.089 | 2 | 53.89426 | yes | 3.18E-07 | 765.90479 | 67.205894 |
| VVRPQGAEPVK | 11 | Q08AD1 | Calmodulin-regulated spectrin-associated | CAMSAP2 | 168.089 | 2 | 53.89426 | yes | 3.747E-07 | 590.34583 | 118.19431 |
| LNQSSPDNVTDTK | 13 | Q08AD1 | Calmodulin-regulated spectrin-associated | CAMSAP2 | 168.089 | 2 | 53.89426 | yes | 9.722E-08 | 709.84155 | 98.650505 |
| EHHTVEAPGGQK | 12 | Q08AD1 | Calmodulin-regulated spectrin-associated | CAMSAP2 | 168.089 | 3 | 53.89426 | yes | 4.939E-07 | 430.54596 | 75.692963 |
| HIIQNALAHCCLAC | 15 | Q08AD1 | Calmodulin-regulated spectrin-associated | CAMSAP2 | 168.089 | 3 | 53.89426 | yes | 9.722E-08 | 569.29236 | 68.927681 |
| QQLEAEMEHK | 10 | Q08AD1 | Calmodulin-regulated spectrin-associated | CAMSAP2 | 168.089 | 2 | 53.89426 | yes | 9.722E-08 | 621.79279 | 57.722168 |
| NVLDSSSDFPSSGI | 28 | Q08AD1 | Calmodulin-regulated spectrin-associated | CAMSAP2 | 168.089 | 4 | 53.89426 | yes | 2.261E-06 | 750.09851 | 51.94899 |
| PLLGHDAVIQALA | 15 | Q08AD1 | Calmodulin-regulated spectrin-associated | CAMSAP2 | 168.089 | 3 | 53.89426 | yes | 2.509E-07 | 525.30688 | 51.661018 |
| QFSHIPAK | 8 | Q08AD1 | Calmodulin-regulated spectrin-associated | CAMSAP2 | 168.089 | 2 | 53.89426 | yes | 2.302E-06 | 464.25598 | 46.455284 |
| ILEEMEK | 7 | Q08AD1 | Calmodulin-regulated spectrin-associated | CAMSAP2 | 168.089 | 2 | 53.89426 | yes | 5.551E-07 | 446.22824 | 43.256107 |
| SAHEIILDFIR | 11 | Q08AE8 | Protein spire homolog 1 OS=Homo sapiei | SPIRE1 | 85.544 | 2 | 29.654474 | yes | 2.424E-06 | 657.36426 | 22.719164 |
| ETMDVLLEISR | 11 | Q08AG7 | Mitotic-spindle organizing protein 1 OS= | MZT1 | 8.479 | 2 | 51.04523 | yes | 9.722E-08 | 653.3396 | 856.58191 |
| ETMDVLLEISR | 11 | Q08AG7 | Mitotic-spindle organizing protein 1 OS= | MZT1 | 8.479 | 2 | 51.04523 | yes | 1.108E-05 | 661.33704 | 856.58191 |
| FYALDPSFPR | 10 | Q08J23 | RNA cytosine C(5)-methyltransferase NS | NSUN2 | 86.471 | 2 | 55.23669 | yes | 1.687E-06 | 606.80603 | 5299.8506 |
| ILCDVPCSGDGTM | 14 | Q08J23 | RNA cytosine C(5)-methyltransferase NS | NSUN2 | 86.471 | 2 | 55.23669 | yes | 2.368E-07 | 790.84692 | 5057.9722 |
| ILLTQENPFFR | 11 | Q08J23 | RNA cytosine C(5)-methyltransferase NS | NSUN2 | 86.471 | 2 | 55.23669 | yes | 1.195E-07 | 689.37988 | 4321.8862 |
| WMPGITQWK | 9 | Q08J23 | RNA cytosine C(5)-methyltransferase NS | NSUN2 | 86.471 | 2 | 55.23669 | yes | 1.994E-07 | 573.79187 | 2739.6104 |
| ILDMCAAPGSK | 11 | Q08J23 | RNA cytosine C(5)-methyltransferase NS | NSUN2 | 86.471 | 2 | 55.23669 | yes | 9.722E-08 | 581.78339 | 2585.1538 |
| ELEDLEVDGQK | 11 | Q08J23 | RNA cytosine C(5)-methyltransferase NS | NSUN2 | 86.471 | 2 | 55.23669 | yes | 1.808E-07 | 637.80914 | 2558.8638 |
| LFEHYYQELK | 10 | Q08J23 | RNA cytosine C(5)-methyltransferase NS | NSUN2 | 86.471 | 2 | 55.23669 | yes | 1.345E-07 | 685.34296 | 1070.0979 |
| LFEHYYQELK | 10 | Q08J23 | RNA cytosine C(5)-methyltransferase NS | NSUN2 | 86.471 | 2 | 55.23669 | yes | 9.722E-08 | 457.23108 | 1070.0979 |
| LFEHYYQELK | 10 | Q08J23 | RNA cytosine C(5)-methyltransferase NS | NSUN2 | 86.471 | 3 | 55.23669 | yes | 1.345E-07 | 685.34296 | 1070.0979 |
| LFEHYYQELK | 10 | Q08J23 | RNA cytosine C(5)-methyltransferase NS | NSUN2 | 86.471 | 3 | 55.23669 | yes | 9.722E-08 | 457.23108 | 1070.0979 |
| CYLLVHQAK | 9 | Q08J23 | RNA cytosine C(5)-methyltransferase NS | NSUN2 | 86.471 | 2 | 55.23669 | yes | 9.881E-08 | 566.30261 | 1711.4156 |
| LQIDVDGR | 8 | Q08J23 | RNA cytosine C(5)-methyltransferase NS | NSUN2 | 86.471 | 2 | 55.23669 | yes | 1.422E-07 | 458.24835 | 1566.1521 |
| RGEAGWEGGYPEI | 15 | Q08J23 | RNA cytosine C(5)-methyltransferase NS | NSUN2 | 86.471 | 3 | 55.23669 | yes | 1.077E-07 | 549.94238 | 685.8316 |
| RGEAGWEGGYPEI | 15 | Q08J23 | RNA cytosine C(5)-methyltransferase NS | NSUN2 | 86.471 | 3 | 55.23669 | yes | 2.387E-07 | 824.40991 | 685.8316 |
| RGEAGWEGGYPEI | 15 | Q08J23 | RNA cytosine C(5)-methyltransferase NS | NSUN2 | 86.471 | 2 | 55.23669 | yes | 1.077E-07 | 549.94238 | 685.8316 |
| RGEAGWEGGYPEI | 15 | Q08J23 | RNA cytosine C(5)-methyltransferase NS | NSUN2 | 86.471 | 2 | 55.23669 | yes | 2.387E-07 | 824.40991 | 685.8316 |
| IITVSMEDVK | 10 | Q08J23 | RNA cytosine C(5)-methyltransferase NS | NSUN2 | 86.471 | 2 | 55.23669 | yes | 2.226E-07 | 567.80737 | 1158.5642 |
| KEILFYDR | 8 | Q08J23 | RNA cytosine C(5)-methyltransferase NS | NSUN2 | 86.471 | 2 | 55.23669 | yes | 2.117E-07 | 542.29529 | 1031.9312 |
| LQAMHLER | 8 | Q08J23 | RNA cytosine C(5)-methyltransferase NS | NSUN2 | 86.471 | 2 | 55.23669 | yes | 1.612E-07 | 499.26602 | 1025.7314 |
| EILHCLK | 7 | Q08J23 | RNA cytosine C(5)-methyltransferase NS | NSUN2 | 86.471 | 2 | 55.23669 | yes | 9.809E-08 | 456.75223 | 963.80585 |
| EILFYDR | 7 | Q08J23 | RNA cytosine C(5)-methyltransferase NS | NSUN2 | 86.471 | 2 | 55.23669 | yes | 4.715E-07 | 478.2478 | 874.40948 |
| GAEQLAEGGR | 10 | Q08J23 | RNA cytosine C(5)-methyltransferase NS | NSUN2 | 86.471 | 2 | 55.23669 | yes | 1.096E-07 | 494.24634 | 798.117 |
| LSSETYSQAK | 10 | Q08J23 | RNA cytosine C(5)-methyltransferase NS | NSUN2 | 86.471 | 2 | 55.23669 | yes | 9.722E-08 | 557.27478 | 728.46259 |
| NNSGEEFDCAFR | 12 | Q08J23 | RNA cytosine C(5)-methyltransferase NS | NSUN2 | 86.471 | 2 | 55.23669 | yes | 1.022E-07 | 723.29114 | 689.87292 |

| | | | | | | | | | | | |
|---|---|---|---|---|---|---|---|---|---|---|---|
| MMGLEVLGEK | 10 | Q08J23 | RNA cytosine C(5)-methyltransferase NS | NSUN2 | 86.471 | 2 | 55.23669 | yes | 1.118E-07 | 553.78284 | 613.7594 |
| NVLLNNSEK | 9 | Q08J23 | RNA cytosine C(5)-methyltransferase NS | NSUN2 | 86.471 | 2 | 55.23669 | yes | 2.177E-07 | 515.78003 | 588.92603 |
| GEAGWEGGYPEIV | 14 | Q08J23 | RNA cytosine C(5)-methyltransferase NS | NSUN2 | 86.471 | 2 | 55.23669 | yes | 9.722E-08 | 746.35938 | 477.45389 |
| KLSSETYSQAK | 11 | Q08J23 | RNA cytosine C(5)-methyltransferase NS | NSUN2 | 86.471 | 2 | 55.23669 | yes | 9.938E-08 | 621.32227 | 289.24902 |
| EPLPATLR | 8 | Q08J23 | RNA cytosine C(5)-methyltransferase NS | NSUN2 | 86.471 | 2 | 55.23669 | yes | 3.24E-06 | 448.76364 | 283.98889 |
| WTTLNSLQLHGLC | 15 | Q08J23 | RNA cytosine C(5)-methyltransferase NS | NSUN2 | 86.471 | 3 | 55.23669 | yes | 9.722E-08 | 594.00031 | 184.72528 |
| ESTQLSPADLTEGI | 14 | Q08J23 | RNA cytosine C(5)-methyltransferase NS | NSUN2 | 86.471 | 2 | 55.23669 | yes | 1.579E-07 | 738.36481 | 179.49612 |
| HTQIRPTMFPPKDI | 16 | Q08J23 | RNA cytosine C(5)-methyltransferase NS | NSUN2 | 86.471 | 4 | 55.23669 | yes | 3.165E-06 | 481.25427 | 80.04332 |
| HTQIRPTMFPPKDI | 16 | Q08J23 | RNA cytosine C(5)-methyltransferase NS | NSUN2 | 86.471 | 4 | 55.23669 | yes | 1.588E-05 | 641.33661 | 80.04332 |
| HTQIRPTMFPPKDI | 16 | Q08J23 | RNA cytosine C(5)-methyltransferase NS | NSUN2 | 86.471 | 3 | 55.23669 | yes | 3.165E-06 | 481.25427 | 80.04332 |
| HTQIRPTMFPPKDI | 16 | Q08J23 | RNA cytosine C(5)-methyltransferase NS | NSUN2 | 86.471 | 3 | 55.23669 | yes | 1.588E-05 | 641.33661 | 80.04332 |
| KDGVCGPPPSK | 11 | Q08J23 | RNA cytosine C(5)-methyltransferase NS | NSUN2 | 86.471 | 2 | 55.23669 | yes | 1.924E-07 | 571.28717 | 157.99249 |
| HTQIRPTMFPPK | 12 | Q08J23 | RNA cytosine C(5)-methyltransferase NS | NSUN2 | 86.471 | 3 | 55.23669 | yes | 3.806E-06 | 484.93085 | 99.340446 |
| FHQFLVSETESGNI | 16 | Q08J23 | RNA cytosine C(5)-methyltransferase NS | NSUN2 | 86.471 | 3 | 55.23669 | yes | 1.169E-07 | 617.63922 | 69.260933 |
| QLYMVSK | 7 | Q08J23 | RNA cytosine C(5)-methyltransferase NS | NSUN2 | 86.471 | 2 | 55.23669 | yes | 2.147E-07 | 434.73349 | 45.633106 |
| ILPHHQNTGGFFV/ | 18 | Q08J23 | RNA cytosine C(5)-methyltransferase NS | NSUN2 | 86.471 | 3 | 55.23669 | yes | 1.611E-06 | 659.70715 | 39.068703 |
| KWTTLNSLQLHGI | 16 | Q08J23 | RNA cytosine C(5)-methyltransferase NS | NSUN2 | 86.471 | 3 | 55.23669 | yes | 5.476E-07 | 636.69867 | 18.542803 |
| LMIWDTR | 7 | Q09028 | Histone-binding protein RBBP4 OS=Hon | RBBP4 | 47.656 | 2 | 51.32987 | no | 1.483E-07 | 467.74438 | 1140.2603 |
| TPSSDVLVFDYTK | 13 | Q09028 | Histone-binding protein RBBP4 OS=Hon | RBBP4 | 47.656 | 2 | 51.32987 | no | 9.722E-08 | 736.36938 | 1004.723 |
| YMPQNPCIIATK | 12 | Q09028 | Histone-binding protein RBBP4 OS=Hon | RBBP4 | 47.656 | 2 | 51.32987 | yes | 9.722E-08 | 718.35724 | 790.75641 |
| RLNVWDLSK | 9 | Q09028 | Histone-binding protein RBBP4 OS=Hon | RBBP4 | 47.656 | 2 | 51.32987 | no | 1.427E-07 | 565.81946 | 627.49487 |
| EAAFDDAVEER | 11 | Q09028 | Histone-binding protein RBBP4 OS=Hon | RBBP4 | 47.656 | 2 | 51.32987 | yes | 1.767E-07 | 626.27802 | 421.90753 |
| INHEGEVNR | 9 | Q09028 | Histone-binding protein RBBP4 OS=Hon | RBBP4 | 47.656 | 2 | 51.32987 | no | 3.873E-07 | 534.26508 | 338.18256 |
| ADKEAAFDDAVEI | 14 | Q09028 | Histone-binding protein RBBP4 OS=Hon | RBBP4 | 47.656 | 2 | 51.32987 | yes | 9.975E-08 | 804.36285 | 58.087955 |
| VINEEYK | 7 | Q09028 | Histone-binding protein RBBP4 OS=Hon | RBBP4 | 47.656 | 2 | 51.32987 | no | 9.722E-08 | 447.73199 | 45.842491 |
| IFANTESYLK | 10 | Q09161 | Nuclear cap-binding protein subunit 1 OS | NCBP1 | 91.84 | 2 | 53.222572 | yes | 1.38E-07 | 593.31116 | 14219.01 |
| ANNYNEAVYLVR | 12 | Q09161 | Nuclear cap-binding protein subunit 1 OS | NCBP1 | 91.84 | 2 | 53.222572 | yes | 3.714E-07 | 713.35968 | 10595.089 |
| ATNDEIFSILK | 11 | Q09161 | Nuclear cap-binding protein subunit 1 OS | NCBP1 | 91.84 | 2 | 53.222572 | yes | 1.113E-07 | 625.83496 | 9009.9619 |
| LDTMNTTCVDR | 11 | Q09161 | Nuclear cap-binding protein subunit 1 OS | NCBP1 | 91.84 | 2 | 53.222572 | yes | 9.722E-08 | 663.29486 | 3583.6677 |
| LDTMNTTCVDR | 11 | Q09161 | Nuclear cap-binding protein subunit 1 OS | NCBP1 | 91.84 | 2 | 53.222572 | yes | 1.041E-07 | 671.2923 | 3583.6677 |
| NHPQMIAVLVDK | 12 | Q09161 | Nuclear cap-binding protein subunit 1 OS | NCBP1 | 91.84 | 2 | 53.222572 | yes | 3.756E-07 | 682.8714 | 3566.7715 |
| NHPQMIAVLVDK | 12 | Q09161 | Nuclear cap-binding protein subunit 1 OS | NCBP1 | 91.84 | 2 | 53.222572 | yes | 2.815E-07 | 690.86884 | 3566.7715 |
| TCAAQLVSYPGK | 12 | Q09161 | Nuclear cap-binding protein subunit 1 OS | NCBP1 | 91.84 | 2 | 53.222572 | yes | 6.406E-07 | 647.82666 | 4492.4214 |
| SFSHSFSALAK | 11 | Q09161 | Nuclear cap-binding protein subunit 1 OS | NCBP1 | 91.84 | 2 | 53.222572 | yes | 1.232E-07 | 591.30109 | 3406.1074 |
| KTCAAQLVSYPGI | 13 | Q09161 | Nuclear cap-binding protein subunit 1 OS | NCBP1 | 91.84 | 2 | 53.222572 | yes | 1.069E-07 | 711.87415 | 817.18854 |
| KTCAAQLVSYPGI | 13 | Q09161 | Nuclear cap-binding protein subunit 1 OS | NCBP1 | 91.84 | 2 | 53.222572 | yes | 1.456E-07 | 474.91852 | 817.18854 |
| KTCAAQLVSYPGI | 13 | Q09161 | Nuclear cap-binding protein subunit 1 OS | NCBP1 | 91.84 | 3 | 53.222572 | yes | 1.069E-07 | 711.87415 | 817.18854 |
| KTCAAQLVSYPGI | 13 | Q09161 | Nuclear cap-binding protein subunit 1 OS | NCBP1 | 91.84 | 3 | 53.222572 | yes | 1.456E-07 | 474.91852 | 817.18854 |
| FVIEENLHCIIK | 12 | Q09161 | Nuclear cap-binding protein subunit 1 OS | NCBP1 | 91.84 | 2 | 53.222572 | yes | 2.369E-07 | 757.9054 | 1308.7473 |

| | | | | | | | | | | | |
|---|---|---|---|---|---|---|---|---|---|---|---|
| CETDGTSVLTPWY | 14 | Q09161 | Nuclear cap-binding protein subunit 1 OS | NCBP1 | 91.84 | 2 | 53.222572 | yes | 1.107E-07 | 828.88232 | 1153.7672 |
| KDGVLEEQIER | 11 | Q09161 | Nuclear cap-binding protein subunit 1 OS | NCBP1 | 91.84 | 2 | 53.222572 | yes | 2.678E-07 | 658.34625 | 795.13177 |
| FIMILTEHLVR | 11 | Q09161 | Nuclear cap-binding protein subunit 1 OS | NCBP1 | 91.84 | 2 | 53.222572 | yes | 9.722E-08 | 694.39197 | 142.00496 |
| FIMILTEHLVR | 11 | Q09161 | Nuclear cap-binding protein subunit 1 OS | NCBP1 | 91.84 | 2 | 53.222572 | yes | 9.722E-08 | 686.39447 | 142.00496 |
| FIMILTEHLVR | 11 | Q09161 | Nuclear cap-binding protein subunit 1 OS | NCBP1 | 91.84 | 2 | 53.222572 | yes | 2.988E-07 | 457.9321 | 142.00496 |
| FIMILTEHLVR | 11 | Q09161 | Nuclear cap-binding protein subunit 1 OS | NCBP1 | 91.84 | 2 | 53.222572 | yes | 4.419E-07 | 463.26373 | 142.00496 |
| FIMILTEHLVR | 11 | Q09161 | Nuclear cap-binding protein subunit 1 OS | NCBP1 | 91.84 | 3 | 53.222572 | yes | 9.722E-08 | 694.39197 | 142.00496 |
| FIMILTEHLVR | 11 | Q09161 | Nuclear cap-binding protein subunit 1 OS | NCBP1 | 91.84 | 3 | 53.222572 | yes | 9.722E-08 | 686.39447 | 142.00496 |
| FIMILTEHLVR | 11 | Q09161 | Nuclear cap-binding protein subunit 1 OS | NCBP1 | 91.84 | 3 | 53.222572 | yes | 2.988E-07 | 457.9321 | 142.00496 |
| FIMILTEHLVR | 11 | Q09161 | Nuclear cap-binding protein subunit 1 OS | NCBP1 | 91.84 | 3 | 53.222572 | yes | 4.419E-07 | 463.26373 | 142.00496 |
| VMFEVWR | 7 | Q09161 | Nuclear cap-binding protein subunit 1 OS | NCBP1 | 91.84 | 2 | 53.222572 | yes | 1.202E-07 | 483.74692 | 182.72807 |
| VMFEVWR | 7 | Q09161 | Nuclear cap-binding protein subunit 1 OS | NCBP1 | 91.84 | 2 | 53.222572 | yes | 1.794E-06 | 491.74438 | 182.72807 |
| NYNFGGEFVEAMI | 14 | Q09161 | Nuclear cap-binding protein subunit 1 OS | NCBP1 | 91.84 | 2 | 53.222572 | yes | 1.055E-07 | 823.88519 | 145.0135 |
| NYNFGGEFVEAMI | 14 | Q09161 | Nuclear cap-binding protein subunit 1 OS | NCBP1 | 91.84 | 2 | 53.222572 | yes | 3.692E-06 | 831.88269 | 145.0135 |
| LLCTVAR | 7 | Q09161 | Nuclear cap-binding protein subunit 1 OS | NCBP1 | 91.84 | 2 | 53.222572 | yes | 4.574E-07 | 416.73911 | 175.78833 |
| LFVWEILHSTIR | 12 | Q09161 | Nuclear cap-binding protein subunit 1 OS | NCBP1 | 91.84 | 2 | 53.222572 | yes | 5.906E-07 | 757.42993 | 130.9767 |
| YGDESSNSLPGHS | 23 | Q09161 | Nuclear cap-binding protein subunit 1 OS | NCBP1 | 91.84 | 3 | 53.222572 | yes | 5.438E-07 | 808.06177 | 114.07964 |
| LVQAIFPTPDPAAI | 15 | Q09472 | Histone acetyltransferase p300 OS=Homo | EP300 | 264.161 | 2 | 40.408566 | yes | 4.04E-07 | 790.95599 | 82.759354 |
| QALMPTLEALYR | 12 | Q09472 | Histone acetyltransferase p300 OS=Homo | EP300 | 264.161 | 2 | 40.408566 | yes | 4.6E-07 | 703.37903 | 29.918716 |
| AEGPEVDVNLPK | 12 | Q09666 | Neuroblast differentiation-associated prot | AHNAK | 629.102 | 2 | 54.324913 | yes | 1.002E-07 | 634.33008 | 873.11621 |
| ISMPDLDLNLK | 11 | Q09666 | Neuroblast differentiation-associated prot | AHNAK | 629.102 | 2 | 54.324913 | yes | 1.069E-07 | 629.83923 | 534.75201 |
| ISMPDFDLHLK | 11 | Q09666 | Neuroblast differentiation-associated prot | AHNAK | 629.102 | 2 | 54.324913 | yes | 2.466E-07 | 658.33936 | 411.5014 |
| GEGPDVDVNLPK | 12 | Q09666 | Neuroblast differentiation-associated prot | AHNAK | 629.102 | 2 | 54.324913 | yes | 9.809E-08 | 620.31439 | 363.40524 |
| MPEMHFK | 7 | Q09666 | Neuroblast differentiation-associated prot | AHNAK | 629.102 | 2 | 54.324913 | yes | 1.139E-07 | 460.21185 | 309.21591 |
| VKGDVDVSLPK | 11 | Q09666 | Neuroblast differentiation-associated prot | AHNAK | 629.102 | 2 | 54.324913 | yes | 2.407E-07 | 578.83228 | 280.62848 |
| VDVDVPDVNIEGP | 16 | Q09666 | Neuroblast differentiation-associated prot | AHNAK | 629.102 | 2 | 54.324913 | yes | 1.89E-07 | 841.4176 | 273.57437 |
| ISMPEVDLNLK | 11 | Q09666 | Neuroblast differentiation-associated prot | AHNAK | 629.102 | 2 | 54.324913 | yes | 2.14E-07 | 629.83923 | 225.34872 |
| VDIDVPDVNIEGPI | 16 | Q09666 | Neuroblast differentiation-associated prot | AHNAK | 629.102 | 2 | 54.324913 | yes | 1.36E-07 | 848.42542 | 209.60849 |
| ISMPDVDLHMK | 11 | Q09666 | Neuroblast differentiation-associated prot | AHNAK | 629.102 | 2 | 54.324913 | yes | 3.631E-06 | 643.31757 | 168.48151 |
| VGVEVPDVNIEGP | 16 | Q09666 | Neuroblast differentiation-associated prot | AHNAK | 629.102 | 2 | 54.324913 | yes | 1.006E-07 | 819.42267 | 161.19733 |
| ISMPDVGLNLK | 11 | Q09666 | Neuroblast differentiation-associated prot | AHNAK | 629.102 | 2 | 54.324913 | yes | 1E-07 | 593.82867 | 158.69653 |
| VEGDMQVPDLDI | 13 | Q09666 | Neuroblast differentiation-associated prot | AHNAK | 629.102 | 2 | 54.324913 | yes | 1.468E-07 | 729.8609 | 152.7256 |
| MPEMNIK | 7 | Q09666 | Neuroblast differentiation-associated prot | AHNAK | 629.102 | 2 | 54.324913 | yes | 1.467E-07 | 431.7117 | 152.00197 |
| GGVQVPAVDISSS | 17 | Q09666 | Neuroblast differentiation-associated prot | AHNAK | 629.102 | 2 | 54.324913 | yes | 3.238E-07 | 799.92847 | 145.26221 |
| VDINAPDVDVR | 11 | Q09666 | Neuroblast differentiation-associated prot | AHNAK | 629.102 | 2 | 54.324913 | yes | 2.374E-06 | 606.81458 | 137.79703 |
| VKGDMDVSVPK | 11 | Q09666 | Neuroblast differentiation-associated prot | AHNAK | 629.102 | 2 | 54.324913 | yes | 3.677E-07 | 587.81049 | 136.98062 |
| LNVGAPDVTLR | 11 | Q09666 | Neuroblast differentiation-associated prot | AHNAK | 629.102 | 2 | 54.324913 | yes | 1.009E-07 | 577.83002 | 136.28612 |
| VSMPDVELNLK | 11 | Q09666 | Neuroblast differentiation-associated prot | AHNAK | 629.102 | 2 | 54.324913 | yes | 1.045E-07 | 622.83136 | 135.7943 |
| ISMPDVDFNLK | 11 | Q09666 | Neuroblast differentiation-associated prot | AHNAK | 629.102 | 2 | 54.324913 | yes | 2.704E-07 | 639.82355 | 132.67448 |

| | | | | | | | | | | |
|---|---|---|---|---|---|---|---|---|---|---|
| ISMPDLDLHLK | 11 | Q09666 | Neuroblast differentiation-associated prot AHNAK | 629.102 | 2 | 54.324913 | yes | 1.211E-07 | 641.34723 | 119.55106 |
| ISMPDVDLNLK | 11 | Q09666 | Neuroblast differentiation-associated prot AHNAK | 629.102 | 2 | 54.324913 | yes | 3.59E-07 | 622.83136 | 118.529 |
| VNVEAPDVNLEGI | 16 | Q09666 | Neuroblast differentiation-associated prot AHNAK | 629.102 | 2 | 54.324913 | yes | 1.338E-07 | 805.92285 | 116.20471 |
| SSGCDVNLPGVNV | 14 | Q09666 | Neuroblast differentiation-associated prot AHNAK | 629.102 | 2 | 54.324913 | yes | 1.354E-07 | 723.35626 | 114.49003 |
| VDIEGPDVNIEGPI | 16 | Q09666 | Neuroblast differentiation-associated prot AHNAK | 629.102 | 2 | 54.324913 | yes | 2.307E-07 | 834.40979 | 108.39632 |
| LEGPDVSLK | 9 | Q09666 | Neuroblast differentiation-associated prot AHNAK | 629.102 | 2 | 54.324913 | yes | 1.775E-07 | 479.2662 | 107.84433 |
| LQVTMPGIK | 9 | Q09666 | Neuroblast differentiation-associated prot AHNAK | 629.102 | 2 | 54.324913 | yes | 1.557E-07 | 493.78879 | 107.12895 |
| ISMPDVNLNLK | 11 | Q09666 | Neuroblast differentiation-associated prot AHNAK | 629.102 | 2 | 54.324913 | yes | 4.031E-07 | 622.33936 | 105.04452 |
| ISAPNVDFNLEGPI | 14 | Q09666 | Neuroblast differentiation-associated prot AHNAK | 629.102 | 2 | 54.324913 | yes | 5.918E-07 | 750.88831 | 101.24516 |
| GDLDIAGPNLEGD | 15 | Q09666 | Neuroblast differentiation-associated prot AHNAK | 629.102 | 2 | 54.324913 | yes | 2.394E-07 | 780.88068 | 100.18005 |
| APDVDVNIAGPDA | 16 | Q09666 | Neuroblast differentiation-associated prot AHNAK | 629.102 | 2 | 54.324913 | yes | 2.058E-07 | 783.41211 | 99.737556 |
| GDLDASVPSMK | 11 | Q09666 | Neuroblast differentiation-associated prot AHNAK | 629.102 | 2 | 54.324913 | yes | 2.804E-07 | 560.27118 | 98.626152 |
| ISMQDVDLSLGSPI | 14 | Q09666 | Neuroblast differentiation-associated prot AHNAK | 629.102 | 2 | 54.324913 | yes | 9.722E-08 | 745.38196 | 97.519661 |
| VDINAPDVEVQGK | 13 | Q09666 | Neuroblast differentiation-associated prot AHNAK | 629.102 | 2 | 54.324913 | yes | 1.107E-07 | 692.35938 | 94.455185 |
| MSLPDVDLDLK | 11 | Q09666 | Neuroblast differentiation-associated prot AHNAK | 629.102 | 2 | 54.324913 | yes | 1.144E-07 | 623.32343 | 93.780029 |
| VDIDAPDVSIEGPI | 16 | Q09666 | Neuroblast differentiation-associated prot AHNAK | 629.102 | 2 | 54.324913 | yes | 3.012E-06 | 820.9043 | 92.959442 |
| ISMPDIDLNLTGPK | 14 | Q09666 | Neuroblast differentiation-associated prot AHNAK | 629.102 | 2 | 54.324913 | yes | 2.104E-07 | 757.40015 | 92.653427 |
| GPGVDLPSVNLSM | 15 | Q09666 | Neuroblast differentiation-associated prot AHNAK | 629.102 | 2 | 54.324913 | yes | 1.123E-06 | 755.90033 | 88.784721 |
| MPEMHFR | 7 | Q09666 | Neuroblast differentiation-associated prot AHNAK | 629.102 | 2 | 54.324913 | yes | 1.808E-07 | 474.21494 | 85.7724 |
| VKGEYDMTVPK | 11 | Q09666 | Neuroblast differentiation-associated prot AHNAK | 629.102 | 2 | 54.324913 | yes | 5.504E-07 | 633.82355 | 84.247444 |
| GEGPEVDMNLPK | 12 | Q09666 | Neuroblast differentiation-associated prot AHNAK | 629.102 | 2 | 54.324913 | yes | 3.252E-07 | 643.30829 | 83.764343 |
| GEGPDVDVSLPK | 12 | Q09666 | Neuroblast differentiation-associated prot AHNAK | 629.102 | 2 | 54.324913 | yes | 4.031E-07 | 606.80896 | 81.041183 |
| GEGPDVDVTLPK | 12 | Q09666 | Neuroblast differentiation-associated prot AHNAK | 629.102 | 2 | 54.324913 | yes | 2.03E-07 | 613.81677 | 76.707703 |
| GPQITGPSLEGDLC | 16 | Q09666 | Neuroblast differentiation-associated prot AHNAK | 629.102 | 2 | 54.324913 | yes | 3.571E-07 | 791.4278 | 76.036911 |
| SPEPGQTWTR | 10 | Q09666 | Neuroblast differentiation-associated prot AHNAK | 629.102 | 2 | 54.324913 | yes | 1.856E-06 | 579.78052 | 75.871315 |
| AEGPDVAVDLPK | 12 | Q09666 | Neuroblast differentiation-associated prot AHNAK | 629.102 | 2 | 54.324913 | yes | 1.02E-07 | 605.81934 | 75.797195 |
| GEVDVSLANVEGI | 15 | Q09666 | Neuroblast differentiation-associated prot AHNAK | 629.102 | 2 | 54.324913 | yes | 3.974E-07 | 772.89374 | 74.545242 |
| VDIEAPDVSLEGPI | 16 | Q09666 | Neuroblast differentiation-associated prot AHNAK | 629.102 | 2 | 54.324913 | yes | 1.617E-07 | 827.91217 | 72.743057 |
| TPEMIIQKPK | 10 | Q09666 | Neuroblast differentiation-associated prot AHNAK | 629.102 | 2 | 54.324913 | yes | 9.819E-08 | 592.83899 | 71.370544 |
| VKGDVDVSVPK | 11 | Q09666 | Neuroblast differentiation-associated prot AHNAK | 629.102 | 2 | 54.324913 | yes | 2.042E-07 | 571.8244 | 67.634453 |
| GEGPEFDVNLSK | 12 | Q09666 | Neuroblast differentiation-associated prot AHNAK | 629.102 | 2 | 54.324913 | yes | 9.722E-08 | 646.31189 | 62.531654 |
| IKGDVDVTGPK | 11 | Q09666 | Neuroblast differentiation-associated prot AHNAK | 629.102 | 2 | 54.324913 | yes | 3.929E-07 | 564.81659 | 62.420094 |
| MDVNVGDIDIEGP | 16 | Q09666 | Neuroblast differentiation-associated prot AHNAK | 629.102 | 2 | 54.324913 | yes | 9.902E-08 | 844.39581 | 61.644726 |
| GEGPEVDVTLPK | 12 | Q09666 | Neuroblast differentiation-associated prot AHNAK | 629.102 | 2 | 54.324913 | yes | 2.321E-07 | 620.82465 | 59.81559 |
| LPTGQISGPEIK | 12 | Q09666 | Neuroblast differentiation-associated prot AHNAK | 629.102 | 2 | 54.324913 | yes | 9.722E-08 | 620.35083 | 58.465458 |
| FGTFGGLGSK | 10 | Q09666 | Neuroblast differentiation-associated prot AHNAK | 629.102 | 2 | 54.324913 | yes | 5.925E-06 | 485.75327 | 58.305084 |
| VDTNAPDLSLEGP | 16 | Q09666 | Neuroblast differentiation-associated prot AHNAK | 629.102 | 2 | 54.324913 | yes | 9.815E-08 | 821.40198 | 58.078754 |
| APNISMPDVDLNL | 14 | Q09666 | Neuroblast differentiation-associated prot AHNAK | 629.102 | 2 | 54.324913 | yes | 7.92E-07 | 763.89777 | 57.736298 |
| GEGPDVHMTLPK | 12 | Q09666 | Neuroblast differentiation-associated prot AHNAK | 629.102 | 2 | 54.324913 | yes | 4.841E-07 | 640.81879 | 56.826313 |

| | | | | | | | | | | | |
|---|---|---|---|---|---|---|---|---|---|---|---|
| ISMSEVDLNVAAP | 14 | Q09666 | Neuroblast differentiation-associated prot | AHNAK | 629.102 | 2 | 54.324913 | yes | 1.987E-07 | 737.38452 | 55.629265 |
| ISMPDFDLNLK | 11 | Q09666 | Neuroblast differentiation-associated prot | AHNAK | 629.102 | 2 | 54.324913 | yes | 2.866E-06 | 646.83136 | 53.400326 |
| VKGDVDISLPK | 11 | Q09666 | Neuroblast differentiation-associated prot | AHNAK | 629.102 | 2 | 54.324913 | yes | 4.264E-07 | 585.84009 | 51.842022 |
| GDVDVSLPEVEGE | 15 | Q09666 | Neuroblast differentiation-associated prot | AHNAK | 629.102 | 2 | 54.324913 | yes | 2.067E-06 | 802.37964 | 50.823009 |
| SPQISMSDIDLNLK | 14 | Q09666 | Neuroblast differentiation-associated prot | AHNAK | 629.102 | 2 | 54.324913 | yes | 9.722E-08 | 780.90051 | 50.252483 |
| VSAPGVQGDVK | 11 | Q09666 | Neuroblast differentiation-associated prot | AHNAK | 629.102 | 2 | 54.324913 | yes | 2.756E-05 | 528.78784 | 50.120083 |
| EMGVDVHFPK | 10 | Q09666 | Neuroblast differentiation-associated prot | AHNAK | 629.102 | 2 | 54.324913 | yes | 7.224E-07 | 579.78424 | 48.125336 |
| MPFLSISSPK | 10 | Q09666 | Neuroblast differentiation-associated prot | AHNAK | 629.102 | 2 | 54.324913 | yes | 2.581E-07 | 553.79938 | 46.414848 |
| MPDLHLK | 7 | Q09666 | Neuroblast differentiation-associated prot | AHNAK | 629.102 | 2 | 54.324913 | yes | 2.961E-07 | 427.23364 | 45.237221 |
| VTAPDVDLHLK | 11 | Q09666 | Neuroblast differentiation-associated prot | AHNAK | 629.102 | 2 | 54.324913 | yes | 6.165E-07 | 604.33771 | 40.366356 |
| VDISAPDVDVHGP | 18 | Q09666 | Neuroblast differentiation-associated prot | AHNAK | 629.102 | 3 | 54.324913 | yes | 9.967E-08 | 667.33392 | 39.367512 |
| MPDMHVNMPK | 10 | Q09666 | Neuroblast differentiation-associated prot | AHNAK | 629.102 | 2 | 54.324913 | yes | 9.181E-07 | 600.27216 | 38.577148 |
| MPSMNIQTHK | 10 | Q09666 | Neuroblast differentiation-associated prot | AHNAK | 629.102 | 2 | 54.324913 | yes | 2.568E-05 | 593.789 | 36.829662 |
| AVEVQGPSLESGD | 16 | Q09666 | Neuroblast differentiation-associated prot | AHNAK | 629.102 | 2 | 54.324913 | yes | 4.684E-07 | 805.39447 | 36.293465 |
| VDVEGPDVNIEGP | 16 | Q09666 | Neuroblast differentiation-associated prot | AHNAK | 629.102 | 2 | 54.324913 | yes | 2.522E-07 | 827.40198 | 33.076031 |
| KPDIDITGPK | 10 | Q09666 | Neuroblast differentiation-associated prot | AHNAK | 629.102 | 2 | 54.324913 | yes | 3.238E-06 | 542.30585 | 31.024885 |
| ISIPDVDLDLK | 11 | Q09666 | Neuroblast differentiation-associated prot | AHNAK | 629.102 | 2 | 54.324913 | yes | 2.551E-06 | 614.34521 | 29.790234 |
| VGIQLPEVELSVST | 15 | Q09666 | Neuroblast differentiation-associated prot | AHNAK | 629.102 | 2 | 54.324913 | yes | 5.503E-07 | 799.95361 | 25.071394 |
| LPSGSGAASPTGSA | 18 | Q09666 | Neuroblast differentiation-associated prot | AHNAK | 629.102 | 2 | 54.324913 | yes | 8.974E-07 | 821.9234 | 19.279289 |
| ELLLPNWQGSGSH | 20 | Q09666 | Neuroblast differentiation-associated prot | AHNAK | 629.102 | 3 | 54.324913 | yes | 1.867E-06 | 726.3869 | 18.954651 |
| SAVDVVFQAQK | 11 | Q0PNE2 | Elongator complex protein 6 OS=Homo s | ELP6 | 29.794 | 2 | 37.381336 | yes | 3.286E-06 | 596.32202 | 52.818405 |
| GQLVFLEGLK | 10 | Q0PNE2 | Elongator complex protein 6 OS=Homo s | ELP6 | 29.794 | 2 | 37.381336 | yes | 6.043E-07 | 552.3266 | 51.592545 |
| LTLLCDAK | 8 | Q0PNE2 | Elongator complex protein 6 OS=Homo s | ELP6 | 29.794 | 2 | 37.381336 | yes | 4.486E-06 | 467.25732 | 25.621887 |
| DPVATFLETCR | 11 | Q0VG06 | Fanconi anemia core complex-associated | FAAP100 | 93.434 | 2 | 42.90573 | yes | 2.819E-07 | 654.81628 | 43.661808 |
| WYLENVYPELR | 11 | Q10471 | Polypeptide N-acetylgalactosaminyltransf | GALNT2 | 64.733 | 2 | 54.283756 | yes | 1.804E-07 | 741.37482 | 463.67175 |
| NFYYAAVPSAR | 11 | Q10471 | Polypeptide N-acetylgalactosaminyltransf | GALNT2 | 64.733 | 2 | 54.283756 | yes | 1.283E-07 | 629.81439 | 342.01071 |
| WDYMTPEQR | 9 | Q10471 | Polypeptide N-acetylgalactosaminyltransf | GALNT2 | 64.733 | 2 | 54.283756 | yes | 9.722E-08 | 613.26898 | 246.27908 |
| HVGSNLCLDSR | 11 | Q10471 | Polypeptide N-acetylgalactosaminyltransf | GALNT2 | 64.733 | 2 | 54.283756 | yes | 9.722E-08 | 629.30383 | 208.81932 |
| NVPYGNIQSR | 10 | Q10471 | Polypeptide N-acetylgalactosaminyltransf | GALNT2 | 64.733 | 2 | 54.283756 | yes | 2.908E-07 | 574.29639 | 186.51022 |
| GGFDWNLVFK | 10 | Q10471 | Polypeptide N-acetylgalactosaminyltransf | GALNT2 | 64.733 | 2 | 54.283756 | yes | 9.9E-08 | 591.80072 | 157.69394 |
| HMDLCLTVVDR | 11 | Q10471 | Polypeptide N-acetylgalactosaminyltransf | GALNT2 | 64.733 | 2 | 54.283756 | yes | 2.926E-07 | 679.83142 | 130.41989 |
| FYFEELGK | 8 | Q10471 | Polypeptide N-acetylgalactosaminyltransf | GALNT2 | 64.733 | 2 | 54.283756 | yes | 9.819E-08 | 516.75549 | 127.94404 |
| TPMIAGGLFVMDK | 13 | Q10471 | Polypeptide N-acetylgalactosaminyltransf | GALNT2 | 64.733 | 2 | 54.283756 | yes | 1.097E-07 | 690.35669 | 107.04419 |
| LSCKPFK | 7 | Q10471 | Polypeptide N-acetylgalactosaminyltransf | GALNT2 | 64.733 | 2 | 54.283756 | yes | 1.466E-07 | 440.24149 | 71.867249 |
| AQSMETLPPGK | 11 | Q10471 | Polypeptide N-acetylgalactosaminyltransf | GALNT2 | 64.733 | 2 | 54.283756 | yes | 3.353E-05 | 579.7948 | 60.095081 |
| KEDWNEIDPIK | 11 | Q10471 | Polypeptide N-acetylgalactosaminyltransf | GALNT2 | 64.733 | 2 | 54.283756 | yes | 3.014E-06 | 693.84863 | 54.007755 |
| QKWEQIEGNSK | 11 | Q10471 | Polypeptide N-acetylgalactosaminyltransf | GALNT2 | 64.733 | 2 | 54.283756 | yes | 1.007E-06 | 673.83856 | 30.858179 |
| VDLPATSVVITFHN | 17 | Q10471 | Polypeptide N-acetylgalactosaminyltransf | GALNT2 | 64.733 | 3 | 54.283756 | yes | 4.507E-07 | 623.66699 | 23.143051 |
| QGNPVAPIK | 9 | Q10471 | Polypeptide N-acetylgalactosaminyltransf | GALNT2 | 64.733 | 2 | 54.283756 | yes | 5.195E-07 | 462.26907 | 20.107056 |

| | | | | | | | | | | | |
|---|---|---|---|---|---|---|---|---|---|---|---|
| TPTMAGGLFSIDR | 13 | Q10472 | Polypeptide N-acetylgalactosaminyltransf | GALNT1 | 64.219 | 2 | 50.630363 | no | 9.733E-08 | 683.34521 | 68.031105 |
| HYFSLGEIR | 9 | Q10472 | Polypeptide N-acetylgalactosaminyltransf | GALNT1 | 64.219 | 2 | 50.630363 | yes | 2.286E-06 | 561.29053 | 35.545815 |
| SQQWLLR | 7 | Q10472 | Polypeptide N-acetylgalactosaminyltransf | GALNT1 | 64.219 | 2 | 50.630363 | no | 5.163E-07 | 465.76141 | 31.188457 |
| VNYVVQEAIVVIK | 13 | Q10567 | AP-1 complex subunit beta-1 OS=Homo : | AP1B1 | 104.607 | 2 | 56.434254 | yes | 9.722E-08 | 737.43744 | 50.986397 |
| ALQVMTDFAIQFN | 14 | Q10567 | AP-1 complex subunit beta-1 OS=Homo : | AP1B1 | 104.607 | 2 | 56.434254 | yes | 0.0003002 | 827.4245 | 27.503069 |
| EVLLVALGSR | 10 | Q10570 | Cleavage and polyadenylation specificity | CPSF1 | 160.884 | 2 | 55.789536 | yes | 9.934E-08 | 528.82422 | 389.31415 |
| YIVQVSPLGIR | 11 | Q10570 | Cleavage and polyadenylation specificity | CPSF1 | 160.884 | 2 | 55.789536 | yes | 9.722E-08 | 622.8717 | 344.14218 |
| LSVVEYDPGTHDL | 14 | Q10570 | Cleavage and polyadenylation specificity | CPSF1 | 160.884 | 2 | 55.789536 | yes | 9.722E-08 | 786.89886 | 266.43091 |
| VYAVATSTNTPCA | 14 | Q10570 | Cleavage and polyadenylation specificity | CPSF1 | 160.884 | 2 | 55.789536 | yes | 9.722E-08 | 755.86957 | 218.1989 |
| NVLDGELLNR | 10 | Q10570 | Cleavage and polyadenylation specificity | CPSF1 | 160.884 | 2 | 55.789536 | yes | 1.937E-07 | 571.81183 | 200.8073 |
| DLSGMFTTESR | 11 | Q10570 | Cleavage and polyadenylation specificity | CPSF1 | 160.884 | 2 | 55.789536 | yes | 9.722E-08 | 622.28479 | 153.47186 |
| VITLCLYR | 8 | Q10570 | Cleavage and polyadenylation specificity | CPSF1 | 160.884 | 2 | 55.789536 | yes | 1.079E-07 | 519.29425 | 142.01578 |
| HGFLILSR | 8 | Q10570 | Cleavage and polyadenylation specificity | CPSF1 | 160.884 | 2 | 55.789536 | yes | 2.119E-07 | 471.77963 | 134.31207 |
| DGFVQNVHTPR | 11 | Q10570 | Cleavage and polyadenylation specificity | CPSF1 | 160.884 | 2 | 55.789536 | yes | 9.722E-08 | 635.32037 | 91.915367 |
| TLSLHYFEEPELR | 13 | Q10570 | Cleavage and polyadenylation specificity | CPSF1 | 160.884 | 2 | 55.789536 | yes | 1.308E-07 | 817.41467 | 63.241394 |
| NLVVAGTSQLYVY | 14 | Q10570 | Cleavage and polyadenylation specificity | CPSF1 | 160.884 | 2 | 55.789536 | yes | 3.722E-06 | 791.93304 | 54.599316 |
| NFILAADVMK | 10 | Q10570 | Cleavage and polyadenylation specificity | CPSF1 | 160.884 | 2 | 55.789536 | yes | 1.074E-07 | 561.30481 | 52.250553 |
| IFLWSLR | 7 | Q10570 | Cleavage and polyadenylation specificity | CPSF1 | 160.884 | 2 | 55.789536 | yes | 4.647E-07 | 467.77908 | 19.686686 |
| LLLPHPSR | 8 | Q12769 | Nuclear pore complex protein Nup160 OS | NUP160 | 162.122 | 2 | 56.140762 | yes | 2.37E-07 | 466.78745 | 238.85162 |
| GFNPAQPLNIR | 11 | Q12769 | Nuclear pore complex protein Nup160 OS | NUP160 | 162.122 | 2 | 56.140762 | yes | 9.722E-08 | 613.83563 | 169.00975 |
| LLTGWMPTAIR | 11 | Q12769 | Nuclear pore complex protein Nup160 OS | NUP160 | 162.122 | 2 | 56.140762 | yes | 9.722E-08 | 629.85248 | 163.44591 |
| NPIHAIGLLIR | 11 | Q12769 | Nuclear pore complex protein Nup160 OS | NUP160 | 162.122 | 2 | 56.140762 | yes | 9.722E-08 | 608.87988 | 75.359413 |
| NPIHAIGLLIR | 11 | Q12769 | Nuclear pore complex protein Nup160 OS | NUP160 | 162.122 | 2 | 56.140762 | yes | 0.0001563 | 406.25568 | 75.359413 |
| NPIHAIGLLIR | 11 | Q12769 | Nuclear pore complex protein Nup160 OS | NUP160 | 162.122 | 3 | 56.140762 | yes | 9.722E-08 | 608.87988 | 75.359413 |
| NPIHAIGLLIR | 11 | Q12769 | Nuclear pore complex protein Nup160 OS | NUP160 | 162.122 | 3 | 56.140762 | yes | 0.0001563 | 406.25568 | 75.359413 |
| KVDAAELLR | 9 | Q12769 | Nuclear pore complex protein Nup160 OS | NUP160 | 162.122 | 2 | 56.140762 | yes | 3.165E-07 | 507.80075 | 148.8011 |
| LLSTYLER | 8 | Q12769 | Nuclear pore complex protein Nup160 OS | NUP160 | 162.122 | 2 | 56.140762 | yes | 2.636E-07 | 497.78201 | 145.00616 |
| LGDAVIWGTGQLF | 22 | Q12769 | Nuclear pore complex protein Nup160 OS | NUP160 | 162.122 | 3 | 56.140762 | yes | 1.895E-07 | 822.76776 | 144.95816 |
| QLVVVLCER | 9 | Q12769 | Nuclear pore complex protein Nup160 OS | NUP160 | 162.122 | 2 | 56.140762 | yes | 9.722E-08 | 558.31573 | 128.10648 |
| NLDLSWSELK | 10 | Q12769 | Nuclear pore complex protein Nup160 OS | NUP160 | 162.122 | 2 | 56.140762 | yes | 9.722E-08 | 602.81403 | 93.191154 |
| NLQQEFWCK | 9 | Q12769 | Nuclear pore complex protein Nup160 OS | NUP160 | 162.122 | 2 | 56.140762 | yes | 9.194E-07 | 626.7926 | 87.070488 |
| EMDYETEVEMEK | 12 | Q12769 | Nuclear pore complex protein Nup160 OS | NUP160 | 162.122 | 2 | 56.140762 | yes | 1.031E-07 | 766.81018 | 85.290268 |
| AGTVMFEYGMR | 11 | Q12769 | Nuclear pore complex protein Nup160 OS | NUP160 | 162.122 | 2 | 56.140762 | yes | 9.903E-08 | 631.28882 | 80.955414 |
| CYLVTGEGQK | 10 | Q12769 | Nuclear pore complex protein Nup160 OS | NUP160 | 162.122 | 2 | 56.140762 | yes | 4.238E-07 | 577.77917 | 70.220177 |
| SFVELSGAER | 10 | Q12769 | Nuclear pore complex protein Nup160 OS | NUP160 | 162.122 | 2 | 56.140762 | yes | 1.265E-07 | 547.77747 | 65.412613 |
| YSESAGGFYYVES | 15 | Q12769 | Nuclear pore complex protein Nup160 OS | NUP160 | 162.122 | 2 | 56.140762 | yes | 2.244E-06 | 822.36481 | 57.583771 |
| VIILMLTNQTVHR | 13 | Q12769 | Nuclear pore complex protein Nup160 OS | NUP160 | 162.122 | 2 | 56.140762 | yes | 9.729E-08 | 769.44781 | 34.294685 |
| VSPTSVPTTPR | 11 | Q12772 | Sterol regulatory element-binding protein | SREBF2 | 123.688 | 2 | 52.79069 | yes | 2.943E-07 | 571.31421 | 278.1283 |
| ATAILLACR | 9 | Q12772 | Sterol regulatory element-binding protein | SREBF2 | 123.688 | 2 | 52.79069 | yes | 6.87E-07 | 494.78406 | 108.83971 |

| | | | | | | | | | | | |
|---|---|---|---|---|---|---|---|---|---|---|---|
| ATPATEAGFEDEA | 14 | Q12772 | Sterol regulatory element-binding protein | SREBF2 | 123.688 | 2 | 52.79069 | yes | 1.384E-07 | 718.83063 | 71.008705 |
| ALEVTESPLVK | 11 | Q12772 | Sterol regulatory element-binding protein | SREBF2 | 123.688 | 2 | 52.79069 | yes | 1.395E-07 | 593.3399 | 50.689392 |
| AVLLAEAAR | 9 | Q12772 | Sterol regulatory element-binding protein | SREBF2 | 123.688 | 2 | 52.79069 | yes | 9.382E-07 | 457.27692 | 38.119995 |
| TDSLVLTTLK | 10 | Q12772 | Sterol regulatory element-binding protein | SREBF2 | 123.688 | 2 | 52.79069 | yes | 9.722E-08 | 545.82135 | 34.905685 |
| SCNDCQQMIVK | 11 | Q12772 | Sterol regulatory element-binding protein | SREBF2 | 123.688 | 2 | 52.79069 | yes | 1.101E-07 | 691.79669 | 24.063175 |
| ELQVPEENR | 9 | Q12774 | Rho guanine nucleotide exchange factor 5 | ARHGEF5 | 176.8 | 2 | 55.559402 | no | 2.471E-07 | 557.2804 | 261.29608 |
| ALVSSESYLQR | 11 | Q12774 | Rho guanine nucleotide exchange factor 5 | ARHGEF5 | 176.8 | 2 | 55.559402 | yes | 1.163E-07 | 626.83026 | 208.08954 |
| HPGPSDTVVFR | 11 | Q12774 | Rho guanine nucleotide exchange factor 5 | ARHGEF5 | 176.8 | 2 | 55.559402 | yes | 3.498E-06 | 606.31201 | 174.5191 |
| IFPLISQSR | 9 | Q12774 | Rho guanine nucleotide exchange factor 5 | ARHGEF5 | 176.8 | 2 | 55.559402 | yes | 4.677E-07 | 530.8111 | 168.09662 |
| GPLPQASDPAVAR | 13 | Q12774 | Rho guanine nucleotide exchange factor 5 | ARHGEF5 | 176.8 | 2 | 55.559402 | yes | 1.321E-06 | 639.84369 | 151.72162 |
| LSATQQSELR | 10 | Q12774 | Rho guanine nucleotide exchange factor 5 | ARHGEF5 | 176.8 | 2 | 55.559402 | no | 4.329E-07 | 566.80145 | 137.79323 |
| LESLSETPGPSSPR | 14 | Q12774 | Rho guanine nucleotide exchange factor 5 | ARHGEF5 | 176.8 | 2 | 55.559402 | yes | 8.476E-07 | 728.86755 | 100.48198 |
| AHHALEQLIR | 10 | Q12774 | Rho guanine nucleotide exchange factor 5 | ARHGEF5 | 176.8 | 2 | 55.559402 | yes | 3.076E-07 | 594.33582 | 75.444504 |
| QNTQGAQAEFLFR | 13 | Q12774 | Rho guanine nucleotide exchange factor 5 | ARHGEF5 | 176.8 | 2 | 55.559402 | yes | 1.923E-07 | 755.37585 | 72.691711 |
| SGELTALEFSASPC | 16 | Q12774 | Rho guanine nucleotide exchange factor 5 | ARHGEF5 | 176.8 | 2 | 55.559402 | yes | 9.977E-08 | 817.92285 | 64.386574 |
| STESFTSTSR | 10 | Q12774 | Rho guanine nucleotide exchange factor 5 | ARHGEF5 | 176.8 | 2 | 55.559402 | yes | 1.05E-07 | 551.75421 | 56.275547 |
| LLLQNILK | 8 | Q12774 | Rho guanine nucleotide exchange factor 5 | ARHGEF5 | 176.8 | 2 | 55.559402 | yes | 1.823E-07 | 477.82095 | 42.122963 |
| NSTVLLSMTHEDQ | 14 | Q12774 | Rho guanine nucleotide exchange factor 5 | ARHGEF5 | 176.8 | 2 | 55.559402 | yes | 9.722E-08 | 801.89325 | 39.170837 |
| RTEELIYLSQK | 11 | Q12774 | Rho guanine nucleotide exchange factor 5 | ARHGEF5 | 176.8 | 2 | 55.559402 | yes | 4.954E-07 | 690.38007 | 32.65366 |
| DYSTVSASPTALST | 16 | Q12774 | Rho guanine nucleotide exchange factor 5 | ARHGEF5 | 176.8 | 2 | 55.559402 | yes | 7.043E-07 | 820.92255 | 24.986155 |
| DEAGVSEHPEAPA | 14 | Q12774 | Rho guanine nucleotide exchange factor 5 | ARHGEF5 | 176.8 | 2 | 55.559402 | yes | 1.003E-07 | 732.83929 | 23.213615 |
| LPSDLDGMFIER | 12 | Q12778 | Forkhead box protein O1 OS=Homo sapi | FOXO1 | 69.662 | 2 | 53.8806 | yes | 9.722E-08 | 696.84503 | 450.61661 |
| LDCDMESIIR | 10 | Q12778 | Forkhead box protein O1 OS=Homo sapi | FOXO1 | 69.662 | 2 | 53.8806 | yes | 9.975E-08 | 626.289 | 237.84456 |
| NAWGNLSYADLIT | 14 | Q12778 | Forkhead box protein O1 OS=Homo sapi | FOXO1 | 69.662 | 2 | 53.8806 | no | 1.461E-07 | 783.40155 | 224.13048 |
| MGLLHQEK | 8 | Q12778 | Forkhead box protein O1 OS=Homo sapi | FOXO1 | 69.662 | 2 | 53.8806 | yes | 2.747E-06 | 478.25513 | 159.7849 |
| SSWWMLNPEGGK | 12 | Q12778 | Forkhead box protein O1 OS=Homo sapi | FOXO1 | 69.662 | 2 | 53.8806 | no | 9.722E-08 | 696.32428 | 114.93811 |
| TSSNASTISGR | 11 | Q12778 | Forkhead box protein O1 OS=Homo sapi | FOXO1 | 69.662 | 2 | 53.8806 | yes | 2.174E-07 | 540.76764 | 85.550362 |
| DKGDSNSSAGWK | 12 | Q12778 | Forkhead box protein O1 OS=Homo sapi | FOXO1 | 69.662 | 2 | 53.8806 | no | 1.34E-07 | 626.28363 | 57.886475 |
| GTQLLSSGSDGLV | 14 | Q12788 | Transducin beta-like protein 3 OS=Homo | TBL3 | 89.035 | 2 | 53.800903 | yes | 9.722E-08 | 681.36719 | 420.76703 |
| LLLFSSATDAAIR | 13 | Q12788 | Transducin beta-like protein 3 OS=Homo | TBL3 | 89.035 | 2 | 53.800903 | yes | 9.722E-08 | 689.39044 | 404.50912 |
| EAPEELLAYEGVR | 13 | Q12788 | Transducin beta-like protein 3 OS=Homo | TBL3 | 89.035 | 2 | 53.800903 | yes | 2.114E-07 | 738.37244 | 244.38011 |
| LDDHALTGASDSR | 13 | Q12788 | Transducin beta-like protein 3 OS=Homo | TBL3 | 89.035 | 2 | 53.800903 | yes | 9.81E-08 | 679.32074 | 190.48874 |
| ESFLVTGSQDCTV | 14 | Q12788 | Transducin beta-like protein 3 OS=Homo | TBL3 | 89.035 | 2 | 53.800903 | yes | 3.751E-07 | 785.87451 | 177.06712 |
| LPVPAAAPTPWET | 15 | Q12788 | Transducin beta-like protein 3 OS=Homo | TBL3 | 89.035 | 2 | 53.800903 | yes | 9.722E-08 | 807.93555 | 82.428619 |
| LPVPAAAPTPWET | 15 | Q12788 | Transducin beta-like protein 3 OS=Homo | TBL3 | 89.035 | 2 | 53.800903 | yes | 3.869E-07 | 538.95947 | 82.428619 |
| LPVPAAAPTPWET | 15 | Q12788 | Transducin beta-like protein 3 OS=Homo | TBL3 | 89.035 | 3 | 53.800903 | yes | 9.722E-08 | 807.93555 | 82.428619 |
| LPVPAAAPTPWET | 15 | Q12788 | Transducin beta-like protein 3 OS=Homo | TBL3 | 89.035 | 3 | 53.800903 | yes | 3.869E-07 | 538.95947 | 82.428619 |
| LWALQDFSCLK | 11 | Q12788 | Transducin beta-like protein 3 OS=Homo | TBL3 | 89.035 | 2 | 53.800903 | yes | 1.564E-07 | 690.85266 | 153.64722 |
| VNILEVASGAVLR | 13 | Q12788 | Transducin beta-like protein 3 OS=Homo | TBL3 | 89.035 | 2 | 53.800903 | yes | 2.409E-07 | 670.89844 | 131.75905 |

| | | | | | | | | | | | |
|---|---|---|---|---|---|---|---|---|---|---|---|
| VWSLQDR | 7 | Q12788 | Transducin beta-like protein 3 OS=Homo | TBL3 | 89.035 | 2 | 53.800903 | yes | 3.564E-07 | 452.23779 | 105.18948 |
| AALEALLPYTER | 12 | Q12788 | Transducin beta-like protein 3 OS=Homo | TBL3 | 89.035 | 2 | 53.800903 | yes | 2.889E-07 | 673.86938 | 95.659157 |
| FCVTWNTNSR | 10 | Q12788 | Transducin beta-like protein 3 OS=Homo | TBL3 | 89.035 | 2 | 53.800903 | yes | 3.46E-07 | 642.79315 | 84.292419 |
| QFAGYSEEVLDVR | 13 | Q12788 | Transducin beta-like protein 3 OS=Homo | TBL3 | 89.035 | 2 | 53.800903 | yes | 1.841E-07 | 756.87012 | 67.200363 |
| LWALPQCQLLGVI | 17 | Q12788 | Transducin beta-like protein 3 OS=Homo | TBL3 | 89.035 | 3 | 53.800903 | yes | 2.2E-07 | 661.35248 | 14.194944 |
| ALGLAISLDRPHTV | 22 | Q12788 | Transducin beta-like protein 3 OS=Homo | TBL3 | 89.035 | 3 | 53.800903 | yes | 9.191E-07 | 786.47241 | 8.5822086 |
| NYHPIVVPGLGR | 12 | Q12789 | General transcription factor 3C polypeptic | GTF3C1 | 238.875 | 2 | 56.326996 | yes | 9.722E-08 | 661.37238 | 394.30148 |
| PGIPESSLLR | 10 | Q12789 | General transcription factor 3C polypeptic | GTF3C1 | 238.875 | 2 | 56.326996 | yes | 1.001E-07 | 534.80603 | 315.24222 |
| ALIGQEGDPDLK | 12 | Q12789 | General transcription factor 3C polypeptic | GTF3C1 | 238.875 | 2 | 56.326996 | yes | 9.722E-08 | 628.33008 | 314.96362 |
| SKYDILMEK | 9 | Q12789 | General transcription factor 3C polypeptic | GTF3C1 | 238.875 | 2 | 56.326996 | yes | 1.588E-07 | 563.79425 | 304.10724 |
| LIIVASQAMR | 10 | Q12789 | General transcription factor 3C polypeptic | GTF3C1 | 238.875 | 2 | 56.326996 | yes | 9.722E-08 | 551.32605 | 299.28119 |
| GYYSPGIVSTR | 11 | Q12789 | General transcription factor 3C polypeptic | GTF3C1 | 238.875 | 2 | 56.326996 | yes | 9.722E-08 | 600.3064 | 298.27423 |
| TVPPVDIVFER | 11 | Q12789 | General transcription factor 3C polypeptic | GTF3C1 | 238.875 | 2 | 56.326996 | yes | 3.384E-07 | 636.35333 | 289.41129 |
| IASNVLNTK | 9 | Q12789 | General transcription factor 3C polypeptic | GTF3C1 | 238.875 | 2 | 56.326996 | yes | 3.503E-07 | 480.27966 | 227.21561 |
| CTMVEAFDR | 9 | Q12789 | General transcription factor 3C polypeptic | GTF3C1 | 238.875 | 2 | 56.326996 | yes | 9.722E-08 | 564.74426 | 201.05411 |
| WQGELQR | 7 | Q12789 | General transcription factor 3C polypeptic | GTF3C1 | 238.875 | 2 | 56.326996 | yes | 3.824E-07 | 458.73541 | 195.35695 |
| ALVGDFMNR | 9 | Q12789 | General transcription factor 3C polypeptic | GTF3C1 | 238.875 | 2 | 56.326996 | yes | 6.662E-07 | 511.75803 | 190.56735 |
| YIFSVVENLQR | 11 | Q12789 | General transcription factor 3C polypeptic | GTF3C1 | 238.875 | 2 | 56.326996 | yes | 1.949E-07 | 684.36951 | 189.65987 |
| LIESLFTIQK | 10 | Q12789 | General transcription factor 3C polypeptic | GTF3C1 | 238.875 | 2 | 56.326996 | yes | 1.063E-07 | 596.35284 | 181.65594 |
| LPDFSYCILER | 11 | Q12789 | General transcription factor 3C polypeptic | GTF3C1 | 238.875 | 2 | 56.326996 | yes | 1.094E-07 | 706.8476 | 179.21585 |
| YISCVFAEESDLSR | 14 | Q12789 | General transcription factor 3C polypeptic | GTF3C1 | 238.875 | 2 | 56.326996 | yes | 3.126E-07 | 838.38525 | 163.6929 |
| YEEIDLETGILESR | 14 | Q12789 | General transcription factor 3C polypeptic | GTF3C1 | 238.875 | 2 | 56.326996 | yes | 9.722E-08 | 833.91217 | 162.90196 |
| DMLTQTYDLIER | 12 | Q12789 | General transcription factor 3C polypeptic | GTF3C1 | 238.875 | 2 | 56.326996 | yes | 2.615E-07 | 749.36633 | 149.85399 |
| NLIIEAVTNLR | 11 | Q12789 | General transcription factor 3C polypeptic | GTF3C1 | 238.875 | 2 | 56.326996 | yes | 3.196E-07 | 628.37207 | 145.95642 |
| RDPVALEDVYPIH | 19 | Q12789 | General transcription factor 3C polypeptic | GTF3C1 | 238.875 | 3 | 56.326996 | yes | 1.344E-07 | 751.3963 | 121.06666 |
| TNHIETLGK | 9 | Q12789 | General transcription factor 3C polypeptic | GTF3C1 | 238.875 | 2 | 56.326996 | yes | 2.433E-07 | 506.77472 | 88.666382 |
| LYQYMLNAGLAK | 12 | Q12789 | General transcription factor 3C polypeptic | GTF3C1 | 238.875 | 2 | 56.326996 | yes | 1.265E-06 | 692.86829 | 86.049217 |
| VVDGHLNLPVCK | 12 | Q12789 | General transcription factor 3C polypeptic | GTF3C1 | 238.875 | 2 | 56.326996 | yes | 1.158E-07 | 675.86353 | 84.221916 |
| VVNLHPLK | 8 | Q12789 | General transcription factor 3C polypeptic | GTF3C1 | 238.875 | 2 | 56.326996 | yes | 3.321E-07 | 460.28983 | 83.432121 |
| SSRPFER | 7 | Q12789 | General transcription factor 3C polypeptic | GTF3C1 | 238.875 | 2 | 56.326996 | yes | 3.7E-07 | 439.72757 | 82.370796 |
| GPFVTWQVVR | 10 | Q12789 | General transcription factor 3C polypeptic | GTF3C1 | 238.875 | 2 | 56.326996 | yes | 1.275E-07 | 594.82983 | 71.006134 |
| LQEIHPECGPCK | 12 | Q12789 | General transcription factor 3C polypeptic | GTF3C1 | 238.875 | 2 | 56.326996 | yes | 9.722E-08 | 734.3396 | 68.712692 |
| LSVMLSTR | 8 | Q12789 | General transcription factor 3C polypeptic | GTF3C1 | 238.875 | 2 | 56.326996 | yes | 2.692E-07 | 453.75751 | 64.601433 |
| VCLAEVYQDK | 10 | Q12789 | General transcription factor 3C polypeptic | GTF3C1 | 238.875 | 2 | 56.326996 | yes | 9.722E-08 | 612.80011 | 56.944641 |
| KLIIVASQAMR | 11 | Q12789 | General transcription factor 3C polypeptic | GTF3C1 | 238.875 | 2 | 56.326996 | yes | 1.876E-07 | 615.37354 | 52.505653 |
| SMEVKPAQASHTN | 18 | Q12789 | General transcription factor 3C polypeptic | GTF3C1 | 238.875 | 3 | 56.326996 | yes | 4.111E-07 | 692.68365 | 41.240906 |
| YIPPIPVHR | 9 | Q12789 | General transcription factor 3C polypeptic | GTF3C1 | 238.875 | 2 | 56.326996 | yes | 5.089E-07 | 546.32166 | 39.936832 |
| ERPDLQLQDR | 10 | Q12789 | General transcription factor 3C polypeptic | GTF3C1 | 238.875 | 2 | 56.326996 | yes | 9.722E-08 | 635.33093 | 39.882824 |
| FQDKDQVFIFLK | 12 | Q12789 | General transcription factor 3C polypeptic | GTF3C1 | 238.875 | 2 | 56.326996 | yes | 5.062E-07 | 764.41394 | 8.8192177 |

| | | | | | | | | | | |
|---|---|---|---|---|---|---|---|---|---|---|
| EDVSLHGYK | 9 | Q12792 | Twinfilin-1 OS=Homo sapiens OX=9606 TWF1 | 40.283 | 2 | 52.899204 | yes | 1.004E-07 | 524.25891 | 121.50897 |
| INEVQTDVGVDTK | 13 | Q12792 | Twinfilin-1 OS=Homo sapiens OX=9606 TWF1 | 40.283 | 2 | 52.899204 | yes | 9.722E-08 | 709.36212 | 107.14711 |
| QLQMDVIR | 8 | Q12792 | Twinfilin-1 OS=Homo sapiens OX=9606 TWF1 | 40.283 | 2 | 52.899204 | yes | 9.722E-08 | 501.77368 | 101.81477 |
| HQTLQGVAFPISR | 13 | Q12792 | Twinfilin-1 OS=Homo sapiens OX=9606 TWF1 | 40.283 | 2 | 52.899204 | yes | 1.002E-07 | 727.39917 | 84.944267 |
| QPCYILFR | 8 | Q12792 | Twinfilin-1 OS=Homo sapiens OX=9606 TWF1 | 40.283 | 2 | 52.899204 | yes | 2.968E-07 | 548.78406 | 84.730957 |
| NEIIILANTTNTELK | 15 | Q12792 | Twinfilin-1 OS=Homo sapiens OX=9606 TWF1 | 40.283 | 2 | 52.899204 | yes | 9.722E-08 | 843.96729 | 70.624886 |
| QLNYVQLEIDIK | 12 | Q12792 | Twinfilin-1 OS=Homo sapiens OX=9606 TWF1 | 40.283 | 2 | 52.899204 | yes | 4.682E-06 | 738.40887 | 28.329945 |
| GLFLPEDENLR | 11 | Q12797 | Aspartyl/asparaginyl beta-hydroxylase OS ASPH | 85.863 | 2 | 54.01147 | yes | 1.392E-07 | 651.83807 | 7288.2168 |
| GSLLTLQR | 8 | Q12797 | Aspartyl/asparaginyl beta-hydroxylase OS ASPH | 85.863 | 2 | 54.01147 | yes | 9.728E-08 | 444.26907 | 3591.9304 |
| MHLGLVIPK | 9 | Q12797 | Aspartyl/asparaginyl beta-hydroxylase OS ASPH | 85.863 | 2 | 54.01147 | yes | 5.796E-07 | 504.30716 | 3432.7656 |
| FYFHLGDAMQR | 11 | Q12797 | Aspartyl/asparaginyl beta-hydroxylase OS ASPH | 85.863 | 2 | 54.01147 | yes | 1.088E-07 | 692.82697 | 1699.4824 |
| FYFHLGDAMQR | 11 | Q12797 | Aspartyl/asparaginyl beta-hydroxylase OS ASPH | 85.863 | 2 | 54.01147 | yes | 9.722E-08 | 700.8244 | 1699.4824 |
| IEEAVNAFK | 9 | Q12797 | Aspartyl/asparaginyl beta-hydroxylase OS ASPH | 85.863 | 2 | 54.01147 | yes | 1.605E-07 | 510.77167 | 1662.3103 |
| VHYGFILK | 8 | Q12797 | Aspartyl/asparaginyl beta-hydroxylase OS ASPH | 85.863 | 2 | 54.01147 | yes | 2.664E-07 | 488.78436 | 1658.9128 |
| AQPWWTPK | 8 | Q12797 | Aspartyl/asparaginyl beta-hydroxylase OS ASPH | 85.863 | 2 | 54.01147 | yes | 4.355E-07 | 507.26379 | 1623.0332 |
| KYPQSPR | 7 | Q12797 | Aspartyl/asparaginyl beta-hydroxylase OS ASPH | 85.863 | 2 | 54.01147 | yes | 1.602E-06 | 438.24033 | 1596.8978 |
| GDWSQFTLWQQG | 13 | Q12797 | Aspartyl/asparaginyl beta-hydroxylase OS ASPH | 85.863 | 2 | 54.01147 | yes | 9.722E-08 | 804.88135 | 1463.5963 |
| IAESIPYLK | 9 | Q12797 | Aspartyl/asparaginyl beta-hydroxylase OS ASPH | 85.863 | 2 | 54.01147 | yes | 4.82E-07 | 517.30005 | 1434.2688 |
| GHFASVWQR | 9 | Q12797 | Aspartyl/asparaginyl beta-hydroxylase OS ASPH | 85.863 | 2 | 54.01147 | yes | 1.726E-07 | 544.27521 | 1421.3978 |
| AQCEDDLAEK | 10 | Q12797 | Aspartyl/asparaginyl beta-hydroxylase OS ASPH | 85.863 | 2 | 54.01147 | yes | 9.722E-08 | 589.75336 | 1033.881 |
| TCTLLEK | 7 | Q12797 | Aspartyl/asparaginyl beta-hydroxylase OS ASPH | 85.863 | 2 | 54.01147 | yes | 9.722E-08 | 432.72839 | 766.25854 |
| GKIEEAVNAFK | 11 | Q12797 | Aspartyl/asparaginyl beta-hydroxylase OS ASPH | 85.863 | 2 | 54.01147 | yes | 1.002E-07 | 603.3299 | 727.13403 |
| QQFLGHMR | 8 | Q12797 | Aspartyl/asparaginyl beta-hydroxylase OS ASPH | 85.863 | 2 | 54.01147 | yes | 9.722E-08 | 508.75836 | 479.20551 |
| ETGYTELVK | 9 | Q12797 | Aspartyl/asparaginyl beta-hydroxylase OS ASPH | 85.863 | 2 | 54.01147 | yes | 6.259E-07 | 520.26892 | 433.31314 |
| FPETTGCR | 8 | Q12797 | Aspartyl/asparaginyl beta-hydroxylase OS ASPH | 85.863 | 2 | 54.01147 | yes | 2.652E-07 | 484.21872 | 288.55798 |
| SLYNVNGLK | 9 | Q12797 | Aspartyl/asparaginyl beta-hydroxylase OS ASPH | 85.863 | 2 | 54.01147 | yes | 1.612E-07 | 504.27966 | 275.00177 |
| YSIMHPGTHVWPH | 20 | Q12797 | Aspartyl/asparaginyl beta-hydroxylase OS ASPH | 85.863 | 4 | 54.01147 | yes | 9.722E-08 | 587.77576 | 144.11021 |
| WYELGHK | 7 | Q12797 | Aspartyl/asparaginyl beta-hydroxylase OS ASPH | 85.863 | 2 | 54.01147 | yes | 4.038E-07 | 466.73486 | 135.1037 |
| KVYEEVLSVTPND | 17 | Q12797 | Aspartyl/asparaginyl beta-hydroxylase OS ASPH | 85.863 | 3 | 54.01147 | yes | 9.722E-08 | 632.66315 | 121.63779 |
| LLNKFDK | 7 | Q12797 | Aspartyl/asparaginyl beta-hydroxylase OS ASPH | 85.863 | 2 | 54.01147 | yes | 9.722E-08 | 439.26074 | 58.4333 |
| DEGLAVMDK | 9 | Q12797 | Aspartyl/asparaginyl beta-hydroxylase OS ASPH | 85.863 | 2 | 54.01147 | yes | 1.823E-07 | 489.23404 | 51.993687 |
| AELDAAEK | 8 | Q12797 | Aspartyl/asparaginyl beta-hydroxylase OS ASPH | 85.863 | 2 | 54.01147 | yes | 3.181E-06 | 423.71381 | 43.125404 |
| RGHFASVWQR | 10 | Q12797 | Aspartyl/asparaginyl beta-hydroxylase OS ASPH | 85.863 | 3 | 54.01147 | yes | 3.204E-07 | 415.2196 | 25.149174 |
| LFTNFSGADLLK | 12 | Q12800 | Alpha-globin transcription factor CP2 OS TFCP2 | 57.256 | 2 | 52.918804 | yes | 1.187E-07 | 663.35864 | 755.15027 |
| LTIYVCQESLQLR | 13 | Q12800 | Alpha-globin transcription factor CP2 OS TFCP2 | 57.256 | 2 | 52.918804 | yes | 9.722E-08 | 811.93219 | 476.95642 |
| DDVIQICGPADGIR | 14 | Q12800 | Alpha-globin transcription factor CP2 OS TFCP2 | 57.256 | 2 | 52.918804 | yes | 1.499E-07 | 764.87482 | 385.21088 |
| VQIDTFK | 7 | Q12800 | Alpha-globin transcription factor CP2 OS TFCP2 | 57.256 | 2 | 52.918804 | no | 2.222E-07 | 425.73709 | 173.65292 |
| AETNDSYHIILK | 12 | Q12800 | Alpha-globin transcription factor CP2 OS TFCP2 | 57.256 | 2 | 52.918804 | yes | 4.435E-07 | 702.36188 | 147.21596 |
| LQYTEHQQLEGWI | 13 | Q12800 | Alpha-globin transcription factor CP2 OS TFCP2 | 57.256 | 2 | 52.918804 | no | 2.164E-07 | 844.41296 | 19.780369 |

| | | | | | | | | | | | |
|---|---|---|---|---|---|---|---|---|---|---|---|
| LQYTEHQQLEGWI | 13 | Q12800 | Alpha-globin transcription factor CP2 OS | TFCP2 | 57.256 | 2 | 52.918804 | no | 4.562E-06 | 563.27777 | 19.780369 |
| LQYTEHQQLEGWI | 13 | Q12800 | Alpha-globin transcription factor CP2 OS | TFCP2 | 57.256 | 3 | 52.918804 | no | 2.164E-07 | 844.41296 | 19.780369 |
| LQYTEHQQLEGWI | 13 | Q12800 | Alpha-globin transcription factor CP2 OS | TFCP2 | 57.256 | 3 | 52.918804 | no | 4.562E-06 | 563.27777 | 19.780369 |
| NAVLGLPVALQDI | 13 | Q12802 | A-kinase anchor protein 13 OS=Homo saj | AKAP13 | 307.551 | 2 | 54.60787 | yes | 1.003E-07 | 669.39301 | 337.4509 |
| ATLALDSVLTEEG | 14 | Q12802 | A-kinase anchor protein 13 OS=Homo saj | AKAP13 | 307.551 | 2 | 54.60787 | yes | 1.202E-07 | 723.88794 | 276.74716 |
| PSSLIEQEK | 9 | Q12802 | A-kinase anchor protein 13 OS=Homo saj | AKAP13 | 307.551 | 2 | 54.60787 | yes | 2.403E-07 | 515.77441 | 240.93199 |
| LCDNIVSESESTTA | 15 | Q12802 | A-kinase anchor protein 13 OS=Homo saj | AKAP13 | 307.551 | 2 | 54.60787 | yes | 1.073E-07 | 841.38849 | 232.5654 |
| ETLMHFAVR | 9 | Q12802 | A-kinase anchor protein 13 OS=Homo saj | AKAP13 | 307.551 | 2 | 54.60787 | yes | 9.722E-08 | 552.28693 | 205.72519 |
| IVDAVIEQVK | 10 | Q12802 | A-kinase anchor protein 13 OS=Homo saj | AKAP13 | 307.551 | 2 | 54.60787 | yes | 1.209E-07 | 557.32935 | 185.37309 |
| ALQLSNSPGASSAI | 16 | Q12802 | A-kinase anchor protein 13 OS=Homo saj | AKAP13 | 307.551 | 2 | 54.60787 | yes | 1.072E-07 | 795.92792 | 179.67656 |
| NAASDAEMNHR | 11 | Q12802 | A-kinase anchor protein 13 OS=Homo saj | AKAP13 | 307.551 | 2 | 54.60787 | yes | 1.297E-07 | 608.26221 | 164.80519 |
| YIFASLDQK | 9 | Q12802 | A-kinase anchor protein 13 OS=Homo saj | AKAP13 | 307.551 | 2 | 54.60787 | yes | 6.34E-07 | 542.78729 | 154.95918 |
| YPVLFQR | 7 | Q12802 | A-kinase anchor protein 13 OS=Homo saj | AKAP13 | 307.551 | 2 | 54.60787 | yes | 1.974E-07 | 461.76089 | 154.63512 |
| LGIPECILLVTQR | 13 | Q12802 | A-kinase anchor protein 13 OS=Homo saj | AKAP13 | 307.551 | 2 | 54.60787 | yes | 1.192E-07 | 756.43433 | 154.61862 |
| ILLLLEEK | 8 | Q12802 | A-kinase anchor protein 13 OS=Homo saj | AKAP13 | 307.551 | 2 | 54.60787 | yes | 1.441E-07 | 485.81281 | 148.52403 |
| GSLQAHDTSSLPT | 17 | Q12802 | A-kinase anchor protein 13 OS=Homo saj | AKAP13 | 307.551 | 3 | 54.60787 | yes | 4.446E-07 | 604.98059 | 133.32037 |
| LFGLTKPK | 8 | Q12802 | A-kinase anchor protein 13 OS=Homo saj | AKAP13 | 307.551 | 2 | 54.60787 | yes | 9.722E-08 | 452.28674 | 113.85209 |
| DLCQVSHPHTK | 11 | Q12802 | A-kinase anchor protein 13 OS=Homo saj | AKAP13 | 307.551 | 2 | 54.60787 | yes | 1.021E-07 | 661.31952 | 109.37438 |
| SGSLDSELSVSPK | 13 | Q12802 | A-kinase anchor protein 13 OS=Homo saj | AKAP13 | 307.551 | 2 | 54.60787 | yes | 4.166E-07 | 653.33026 | 105.31431 |
| DDALSFVPSQK | 11 | Q12802 | A-kinase anchor protein 13 OS=Homo saj | AKAP13 | 307.551 | 2 | 54.60787 | yes | 2.32E-07 | 603.80371 | 101.38009 |
| DAYTCANCSAFVI | 14 | Q12802 | A-kinase anchor protein 13 OS=Homo saj | AKAP13 | 307.551 | 2 | 54.60787 | yes | 3.587E-07 | 822.35071 | 100.47618 |
| GTYQYDLER | 9 | Q12802 | A-kinase anchor protein 13 OS=Homo saj | AKAP13 | 307.551 | 2 | 54.60787 | yes | 8.284E-07 | 572.76709 | 96.788208 |
| HSSHGSDVSLSQII | 18 | Q12802 | A-kinase anchor protein 13 OS=Homo saj | AKAP13 | 307.551 | 4 | 54.60787 | yes | 1.267E-07 | 491.25925 | 45.152786 |
| HSSHGSDVSLSQII | 18 | Q12802 | A-kinase anchor protein 13 OS=Homo saj | AKAP13 | 307.551 | 4 | 54.60787 | yes | 4.209E-07 | 654.67657 | 45.152786 |
| HSSHGSDVSLSQII | 18 | Q12802 | A-kinase anchor protein 13 OS=Homo saj | AKAP13 | 307.551 | 3 | 54.60787 | yes | 1.267E-07 | 491.25925 | 45.152786 |
| HSSHGSDVSLSQII | 18 | Q12802 | A-kinase anchor protein 13 OS=Homo saj | AKAP13 | 307.551 | 3 | 54.60787 | yes | 4.209E-07 | 654.67657 | 45.152786 |
| GLESAFTEK | 9 | Q12802 | A-kinase anchor protein 13 OS=Homo saj | AKAP13 | 307.551 | 2 | 54.60787 | yes | 2.078E-07 | 491.24802 | 79.148552 |
| FLSHSTDSLNK | 11 | Q12802 | A-kinase anchor protein 13 OS=Homo saj | AKAP13 | 307.551 | 2 | 54.60787 | yes | 1.149E-07 | 624.81458 | 76.790642 |
| GTEGLSSCGNR | 11 | Q12802 | A-kinase anchor protein 13 OS=Homo saj | AKAP13 | 307.551 | 2 | 54.60787 | yes | 3.408E-07 | 569.25128 | 76.278969 |
| LPTEWNVLGTDQS | 20 | Q12802 | A-kinase anchor protein 13 OS=Homo saj | AKAP13 | 307.551 | 3 | 54.60787 | yes | 2E-06 | 736.03442 | 70.563019 |
| GTATPELHTATDY | 14 | Q12802 | A-kinase anchor protein 13 OS=Homo saj | AKAP13 | 307.551 | 2 | 54.60787 | yes | 9.722E-08 | 766.87061 | 57.99939 |
| LMNIQQQLMK | 10 | Q12802 | A-kinase anchor protein 13 OS=Homo saj | AKAP13 | 307.551 | 2 | 54.60787 | yes | 2.319E-06 | 623.83594 | 49.723358 |
| EEEVQQGQQDLEI | 13 | Q12802 | A-kinase anchor protein 13 OS=Homo saj | AKAP13 | 307.551 | 2 | 54.60787 | yes | 1.14E-06 | 780.36285 | 37.477959 |
| NVLQGGESTK | 10 | Q12802 | A-kinase anchor protein 13 OS=Homo saj | AKAP13 | 307.551 | 2 | 54.60787 | yes | 3.617E-07 | 516.76965 | 35.726158 |
| SSGMPTDQESLSS( | 20 | Q12802 | A-kinase anchor protein 13 OS=Homo saj | AKAP13 | 307.551 | 3 | 54.60787 | yes | 4.388E-07 | 688.98828 | 32.201427 |
| GGEKEEGDDGQDI | 14 | Q12802 | A-kinase anchor protein 13 OS=Homo saj | AKAP13 | 307.551 | 2 | 54.60787 | yes | 3.389E-07 | 752.82916 | 16.807741 |
| ADQVCINLR | 9 | Q12805 | EGF-containing fibulin-like extracellular | EFEMP1 | 54.641 | 2 | 54.666378 | yes | 1.249E-07 | 544.77948 | 316.78616 |
| DIDECDIVPDACK | 13 | Q12805 | EGF-containing fibulin-like extracellular | EFEMP1 | 54.641 | 2 | 54.666378 | yes | 9.809E-08 | 775.3291 | 276.4241 |
| CVCPVSNAMCR | 11 | Q12805 | EGF-containing fibulin-like extracellular | EFEMP1 | 54.641 | 2 | 54.666378 | yes | 9.722E-08 | 677.28015 | 248.43488 |

| | | | | | | | | | | | |
|---|---|---|---|---|---|---|---|---|---|---|---|
| LNCEDIDECR | 10 | Q12805 | EGF-containing fibulin-like extracellular | EFEMP1 | 54.641 | 2 | 54.666378 | yes | 9.722E-08 | 662.2688 | 128.3795 |
| IPSNPSHR | 8 | Q12805 | EGF-containing fibulin-like extracellular | EFEMP1 | 54.641 | 2 | 54.666378 | yes | 6.426E-07 | 454.24084 | 116.91299 |
| GSFACQCPPGYQK | 13 | Q12805 | EGF-containing fibulin-like extracellular | EFEMP1 | 54.641 | 2 | 54.666378 | yes | 1.67E-07 | 750.32391 | 65.276382 |
| SGNENGEFYLR | 11 | Q12805 | EGF-containing fibulin-like extracellular | EFEMP1 | 54.641 | 2 | 54.666378 | yes | 2.526E-06 | 643.29401 | 57.646782 |
| FSCMCPQGYQVVI | 13 | Q12805 | EGF-containing fibulin-like extracellular | EFEMP1 | 54.641 | 2 | 54.666378 | yes | 9.728E-08 | 816.35999 | 51.37233 |
| QTSPVSAMLVLVK | 13 | Q12805 | EGF-containing fibulin-like extracellular | EFEMP1 | 54.641 | 2 | 54.666378 | yes | 5.379E-05 | 686.89703 | 14.877378 |
| FTPMPSTPR | 9 | Q12815 | Tastin OS=Homo sapiens OX=9606 GN= | TROAP | 83.858 | 2 | 52.573586 | yes | 5.755E-07 | 517.26038 | 1280.696 |
| EVVTHSDEGGVAS | 19 | Q12815 | Tastin OS=Homo sapiens OX=9606 GN= | TROAP | 83.858 | 3 | 52.573586 | yes | 9.727E-08 | 642.32898 | 341.78125 |
| PSFQELR | 7 | Q12815 | Tastin OS=Homo sapiens OX=9606 GN= | TROAP | 83.858 | 2 | 52.573586 | yes | 2.021E-07 | 438.73233 | 318.30295 |
| WVQKPPLNIQR | 11 | Q12815 | Tastin OS=Homo sapiens OX=9606 GN= | TROAP | 83.858 | 2 | 52.573586 | yes | 1.257E-07 | 689.90131 | 142.34364 |
| WVQKPPLNIQR | 11 | Q12815 | Tastin OS=Homo sapiens OX=9606 GN= | TROAP | 83.858 | 2 | 52.573586 | yes | 1.194E-07 | 460.26999 | 142.34364 |
| WVQKPPLNIQR | 11 | Q12815 | Tastin OS=Homo sapiens OX=9606 GN= | TROAP | 83.858 | 3 | 52.573586 | yes | 1.257E-07 | 689.90131 | 142.34364 |
| WVQKPPLNIQR | 11 | Q12815 | Tastin OS=Homo sapiens OX=9606 GN= | TROAP | 83.858 | 3 | 52.573586 | yes | 1.194E-07 | 460.26999 | 142.34364 |
| LFDQESCIR | 9 | Q12815 | Tastin OS=Homo sapiens OX=9606 GN= | TROAP | 83.858 | 2 | 52.573586 | yes | 9.722E-08 | 584.27679 | 268.63022 |
| VQQAQWLR | 8 | Q12815 | Tastin OS=Homo sapiens OX=9606 GN= | TROAP | 83.858 | 2 | 52.573586 | yes | 2.099E-07 | 514.78546 | 263.57684 |
| LDDECAFYTSR | 11 | Q12815 | Tastin OS=Homo sapiens OX=9606 GN= | TROAP | 83.858 | 2 | 52.573586 | yes | 9.722E-08 | 688.79297 | 139.95642 |
| IGILQQLLR | 9 | Q12815 | Tastin OS=Homo sapiens OX=9606 GN= | TROAP | 83.858 | 2 | 52.573586 | yes | 1.02E-07 | 527.34259 | 134.94714 |
| IEPEIPESSR | 10 | Q12815 | Tastin OS=Homo sapiens OX=9606 GN= | TROAP | 83.858 | 2 | 52.573586 | yes | 4.215E-07 | 578.79584 | 134.71985 |
| LVGISQPR | 8 | Q12815 | Tastin OS=Homo sapiens OX=9606 GN= | TROAP | 83.858 | 2 | 52.573586 | yes | 3.607E-07 | 435.26379 | 129.02206 |
| SLEGSGKPPVATPS | 20 | Q12815 | Tastin OS=Homo sapiens OX=9606 GN= | TROAP | 83.858 | 3 | 52.573586 | yes | 2.495E-07 | 659.00458 | 88.892784 |
| PLESYCR | 7 | Q12815 | Tastin OS=Homo sapiens OX=9606 GN= | TROAP | 83.858 | 2 | 52.573586 | yes | 9.819E-08 | 462.71582 | 88.234741 |
| VPSPGPPTLTSYSV | 16 | Q12815 | Tastin OS=Homo sapiens OX=9606 GN= | TROAP | 83.858 | 2 | 52.573586 | yes | 3.255E-06 | 835.95923 | 65.272133 |
| VELGASEPCTLEHI | 14 | Q12815 | Tastin OS=Homo sapiens OX=9606 GN= | TROAP | 83.858 | 2 | 52.573586 | yes | 2.928E-07 | 799.38556 | 51.30014 |
| RLTVQPK | 7 | Q12815 | Tastin OS=Homo sapiens OX=9606 GN= | TROAP | 83.858 | 2 | 52.573586 | yes | 7.77E-07 | 421.26636 | 43.13028 |
| SCLTAIHCFHEAR | 13 | Q12815 | Tastin OS=Homo sapiens OX=9606 GN= | TROAP | 83.858 | 2 | 52.573586 | yes | 9.722E-08 | 801.3692 | 42.656483 |
| NPLEELRPSPR | 11 | Q12815 | Tastin OS=Homo sapiens OX=9606 GN= | TROAP | 83.858 | 3 | 52.573586 | yes | 2.614E-06 | 436.57373 | 27.640821 |
| AVSISTEPPTYLR | 13 | Q12824 | SWI/SNF-related matrix-associated actin- | SMARCB1 | 44.142 | 2 | 50.38329 | yes | 9.81E-08 | 717.38538 | 228.14905 |
| LDMEIDGQK | 9 | Q12824 | SWI/SNF-related matrix-associated actin- | SMARCB1 | 44.142 | 2 | 50.38329 | yes | 4.621E-07 | 524.75262 | 110.45512 |
| DAFTWNMNEK | 10 | Q12824 | SWI/SNF-related matrix-associated actin- | SMARCB1 | 44.142 | 2 | 50.38329 | yes | 5.502E-07 | 628.27423 | 34.098537 |
| EQLRAEILK | 9 | Q12830 | Nucleosome-remodeling factor subunit B | BPTF | 338.263 | 2 | 52.830555 | yes | 8.054E-07 | 550.32715 | 2136.5867 |
| VMVAPISGSVTTG | 15 | Q12830 | Nucleosome-remodeling factor subunit B | BPTF | 338.263 | 2 | 52.830555 | yes | 2.408E-07 | 724.3949 | 70.674828 |
| VGSPATVTFQQNK | 13 | Q12830 | Nucleosome-remodeling factor subunit B | BPTF | 338.263 | 2 | 52.830555 | yes | 4.067E-07 | 688.86206 | 69.198013 |
| WAAAQAEVAPK | 11 | Q12830 | Nucleosome-remodeling factor subunit B | BPTF | 338.263 | 2 | 52.830555 | yes | 6.412E-06 | 571.30365 | 64.02153 |
| PALDIWPYPSPR | 12 | Q12830 | Nucleosome-remodeling factor subunit B | BPTF | 338.263 | 2 | 52.830555 | yes | 3.897E-07 | 706.37207 | 59.757835 |
| SQQVAAAAHEAN | 13 | Q12830 | Nucleosome-remodeling factor subunit B | BPTF | 338.263 | 2 | 52.830555 | yes | 9.722E-08 | 662.83386 | 57.128292 |
| IICPIGVPETPK | 12 | Q12830 | Nucleosome-remodeling factor subunit B | BPTF | 338.263 | 2 | 52.830555 | yes | 2.551E-06 | 662.37067 | 47.698406 |
| EPMDLATMEER | 11 | Q12830 | Nucleosome-remodeling factor subunit B | BPTF | 338.263 | 2 | 52.830555 | yes | 3.022E-06 | 661.29175 | 45.076084 |
| NLSESPVITK | 10 | Q12830 | Nucleosome-remodeling factor subunit B | BPTF | 338.263 | 2 | 52.830555 | yes | 5.663E-07 | 544.30334 | 37.775238 |
| VLTISTLR | 8 | Q12830 | Nucleosome-remodeling factor subunit B | BPTF | 338.263 | 2 | 52.830555 | yes | 3.042E-06 | 451.78711 | 33.067657 |

| | | | | | | | | | | | |
|---|---|---|---|---|---|---|---|---|---|---|---|
| LIIEEDTENENEK | 13 | Q12830 | Nucleosome-remodeling factor subunit B | BPTF | 338.263 | 2 | 52.830555 | yes | 3.16E-06 | 788.37286 | 25.70994 |
| ILDAPEIR | 8 | Q12834 | Cell division cycle protein 20 homolog O | CDC20 | 54.723 | 2 | 53.969025 | yes | 1.875E-07 | 463.76892 | 843.40167 |
| SSLIHQGIR | 9 | Q12834 | Cell division cycle protein 20 homolog O | CDC20 | 54.723 | 2 | 53.969025 | yes | 5.567E-07 | 505.79071 | 687.72998 |
| SAAQMEVASFLLS | 14 | Q12834 | Cell division cycle protein 20 homolog O | CDC20 | 54.723 | 2 | 53.969025 | yes | 1.261E-07 | 741.38708 | 618.21863 |
| CFELDPAR | 8 | Q12834 | Cell division cycle protein 20 homolog O | CDC20 | 54.723 | 2 | 53.969025 | yes | 1.407E-07 | 504.23438 | 484.22299 |
| VQTTPSKPGGDR | 12 | Q12834 | Cell division cycle protein 20 homolog O | CDC20 | 54.723 | 2 | 53.969025 | yes | 9.789E-08 | 621.8255 | 228.267 |
| VQTTPSKPGGDR | 12 | Q12834 | Cell division cycle protein 20 homolog O | CDC20 | 54.723 | 2 | 53.969025 | yes | 4.402E-07 | 414.88608 | 228.267 |
| VQTTPSKPGGDR | 12 | Q12834 | Cell division cycle protein 20 homolog O | CDC20 | 54.723 | 3 | 53.969025 | yes | 9.789E-08 | 621.8255 | 228.267 |
| VQTTPSKPGGDR | 12 | Q12834 | Cell division cycle protein 20 homolog O | CDC20 | 54.723 | 3 | 53.969025 | yes | 4.402E-07 | 414.88608 | 228.267 |
| EAAGPAPSPMR | 11 | Q12834 | Cell division cycle protein 20 homolog O | CDC20 | 54.723 | 2 | 53.969025 | yes | 1.998E-07 | 542.26624 | 255.17062 |
| YIPSLPDR | 8 | Q12834 | Cell division cycle protein 20 homolog O | CDC20 | 54.723 | 2 | 53.969025 | yes | 1.345E-07 | 480.76108 | 217.75919 |
| AWALNLNGFDVE | 15 | Q12834 | Cell division cycle protein 20 homolog O | CDC20 | 54.723 | 2 | 53.969025 | yes | 1.071E-07 | 838.91754 | 81.580772 |
| VAEHHVATLSGHS | 20 | Q12834 | Cell division cycle protein 20 homolog O | CDC20 | 54.723 | 4 | 53.969025 | yes | 1.561E-07 | 547.52356 | 69.517464 |
| ENQPENSQTPTK | 12 | Q12834 | Cell division cycle protein 20 homolog O | CDC20 | 54.723 | 2 | 53.969025 | yes | 9.722E-08 | 686.82062 | 45.001648 |
| VLSLTMSPDGATV | 23 | Q12834 | Cell division cycle protein 20 homolog O | CDC20 | 54.723 | 3 | 53.969025 | yes | 3.598E-07 | 759.38611 | 43.361736 |
| SSPVVNDGVVR | 11 | Q12849 | G-rich sequence factor 1 OS=Homo sapie | GRSF1 | 53.127 | 2 | 51.04695 | yes | 1.361E-07 | 564.80402 | 2626.418 |
| YIELFLNSCPK | 11 | Q12849 | G-rich sequence factor 1 OS=Homo sapie | GRSF1 | 53.127 | 2 | 51.04695 | yes | 1.083E-07 | 692.35248 | 1505.3391 |
| ITMEYSSSGK | 10 | Q12849 | G-rich sequence factor 1 OS=Homo sapie | GRSF1 | 53.127 | 2 | 51.04695 | yes | 1.001E-07 | 551.75787 | 1056.7372 |
| LEEEVDDVFLIR | 12 | Q12849 | G-rich sequence factor 1 OS=Homo sapie | GRSF1 | 53.127 | 2 | 51.04695 | yes | 1.248E-07 | 738.88269 | 741.36389 |
| RGDALIEMESEQD | 16 | Q12849 | G-rich sequence factor 1 OS=Homo sapie | GRSF1 | 53.127 | 3 | 51.04695 | yes | 1.147E-07 | 616.63165 | 705.82678 |
| ASLLPQSLAAAAA | 17 | Q12849 | G-rich sequence factor 1 OS=Homo sapie | GRSF1 | 53.127 | 2 | 51.04695 | yes | 9.902E-08 | 818.97266 | 651.37726 |
| GLPYSCNEK | 9 | Q12849 | G-rich sequence factor 1 OS=Homo sapie | GRSF1 | 53.127 | 2 | 51.04695 | yes | 2.355E-07 | 534.24493 | 525.74438 |
| GDALIEMESEQDV | 15 | Q12849 | G-rich sequence factor 1 OS=Homo sapie | GRSF1 | 53.127 | 2 | 51.04695 | yes | 9.722E-08 | 846.39325 | 348.7522 |
| YIEIFPSR | 8 | Q12849 | G-rich sequence factor 1 OS=Homo sapie | GRSF1 | 53.127 | 2 | 51.04695 | yes | 2.131E-07 | 512.77673 | 268.63107 |
| LPEAADFGTTSSLI | 19 | Q12849 | G-rich sequence factor 1 OS=Homo sapie | GRSF1 | 53.127 | 3 | 51.04695 | yes | 2.119E-07 | 706.01422 | 248.82512 |
| NGENGIHFLLNR | 12 | Q12849 | G-rich sequence factor 1 OS=Homo sapie | GRSF1 | 53.127 | 2 | 51.04695 | yes | 2.082E-07 | 692.36005 | 104.21935 |
| KIASFPTAK | 9 | Q12849 | G-rich sequence factor 1 OS=Homo sapie | GRSF1 | 53.127 | 2 | 51.04695 | yes | 7.206E-07 | 481.78711 | 102.7315 |
| IASFPTAK | 8 | Q12849 | G-rich sequence factor 1 OS=Homo sapie | GRSF1 | 53.127 | 2 | 51.04695 | yes | 9.722E-08 | 417.73962 | 43.325302 |
| WVLGALLR | 8 | Q12849 | G-rich sequence factor 1 OS=Homo sapie | GRSF1 | 53.127 | 2 | 51.04695 | yes | 1.75E-07 | 464.29236 | 41.520874 |
| GIPLESTDGER | 11 | Q12857 | Nuclear factor 1 A-type OS=Homo sapier | NFIA | 55.945 | 2 | 50.97993 | no | 1.763E-07 | 587.29095 | 4256.1753 |
| EDFVLTVTGK | 10 | Q12857 | Nuclear factor 1 A-type OS=Homo sapier | NFIA | 55.945 | 2 | 50.97993 | no | 1.108E-07 | 554.79785 | 872.81738 |
| YHPQETLK | 8 | Q12857 | Nuclear factor 1 A-type OS=Homo sapier | NFIA | 55.945 | 2 | 50.97993 | yes | 1.787E-07 | 508.26401 | 578.38336 |
| SLPSTSSTSSTK | 12 | Q12857 | Nuclear factor 1 A-type OS=Homo sapier | NFIA | 55.945 | 2 | 50.97993 | yes | 1.352E-07 | 591.79608 | 499.04263 |
| VHNPFLPTPMLPPI | 21 | Q12857 | Nuclear factor 1 A-type OS=Homo sapier | NFIA | 55.945 | 3 | 50.97993 | yes | 1.377E-07 | 768.41125 | 456.37225 |
| DELLSEKPEVK | 11 | Q12857 | Nuclear factor 1 A-type OS=Homo sapier | NFIA | 55.945 | 2 | 50.97993 | yes | 9.81E-08 | 643.84552 | 376.76913 |
| AVKDELLSEKPEV | 14 | Q12857 | Nuclear factor 1 A-type OS=Homo sapier | NFIA | 55.945 | 3 | 50.97993 | yes | 9.722E-08 | 528.96631 | 187.06398 |
| AVKDELLSEKPEV | 14 | Q12857 | Nuclear factor 1 A-type OS=Homo sapier | NFIA | 55.945 | 3 | 50.97993 | yes | 5.586E-07 | 792.9458 | 187.06398 |
| AVKDELLSEKPEV | 14 | Q12857 | Nuclear factor 1 A-type OS=Homo sapier | NFIA | 55.945 | 2 | 50.97993 | yes | 9.722E-08 | 528.96631 | 187.06398 |
| AVKDELLSEKPEV | 14 | Q12857 | Nuclear factor 1 A-type OS=Homo sapier | NFIA | 55.945 | 2 | 50.97993 | yes | 5.586E-07 | 792.9458 | 187.06398 |

| | | | | | | | | | | | |
|---|---|---|---|---|---|---|---|---|---|---|---|
| DIRPEYR | 7 | Q12857 | Nuclear factor 1 A-type OS=Homo sapier | NFIA | 55.945 | 2 | 50.97993 | yes | 1.562E-07 | 474.7485 | 312.03705 |
| KPPCCVLSNPDQK | 13 | Q12857 | Nuclear factor 1 A-type OS=Homo sapier | NFIA | 55.945 | 2 | 50.97993 | no | 1.116E-07 | 771.87378 | 216.21799 |
| SGFSSPSPSQTSSL( | 23 | Q12857 | Nuclear factor 1 A-type OS=Homo sapier | NFIA | 55.945 | 3 | 50.97993 | yes | 1.073E-07 | 802.04736 | 169.24498 |
| AFAYTWFNLQAR | 12 | Q12857 | Nuclear factor 1 A-type OS=Homo sapier | NFIA | 55.945 | 2 | 50.97993 | no | 1.832E-07 | 744.37512 | 160.54277 |
| ASPHATPSTLHFPT | 31 | Q12857 | Nuclear factor 1 A-type OS=Homo sapier | NFIA | 55.945 | 4 | 50.97993 | yes | 3.343E-07 | 838.9361 | 138.37459 |
| DKPLPPLLAR | 10 | Q12873 | Chromodomain-helicase-DNA-binding pr | CHD3 | 226.592 | 2 | 55.778 | no | 9.722E-08 | 560.34784 | 523.63531 |
| MMLTHLVVR | 9 | Q12873 | Chromodomain-helicase-DNA-binding pr | CHD3 | 226.592 | 2 | 55.778 | no | 9.819E-08 | 550.30939 | 434.09274 |
| LLEQALVIEEQLR | 13 | Q12873 | Chromodomain-helicase-DNA-binding pr | CHD3 | 226.592 | 2 | 55.778 | no | 1.061E-07 | 777.44849 | 247.54832 |
| VGGNIEVLGFNTR | 13 | Q12873 | Chromodomain-helicase-DNA-binding pr | CHD3 | 226.592 | 2 | 55.778 | yes | 9.919E-08 | 688.37006 | 245.13011 |
| VLIFSQMTK | 9 | Q12873 | Chromodomain-helicase-DNA-binding pr | CHD3 | 226.592 | 2 | 55.778 | no | 3.075E-07 | 533.80188 | 176.84445 |
| WSMPELMPDPSAI | 15 | Q12873 | Chromodomain-helicase-DNA-binding pr | CHD3 | 226.592 | 2 | 55.778 | yes | 1.355E-07 | 845.87622 | 130.84454 |
| IINHSVDK | 8 | Q12873 | Chromodomain-helicase-DNA-binding pr | CHD3 | 226.592 | 2 | 55.778 | yes | 1.241E-07 | 463.25873 | 116.61709 |
| WSCPHCEK | 8 | Q12873 | Chromodomain-helicase-DNA-binding pr | CHD3 | 226.592 | 2 | 55.778 | no | 1.679E-07 | 552.22351 | 94.607697 |
| AYVSLFMR | 8 | Q12873 | Chromodomain-helicase-DNA-binding pr | CHD3 | 226.592 | 2 | 55.778 | no | 2.346E-07 | 493.76004 | 82.397499 |
| ENEFSFEDNAIK | 12 | Q12873 | Chromodomain-helicase-DNA-binding pr | CHD3 | 226.592 | 2 | 55.778 | yes | 9.722E-08 | 721.82532 | 80.371017 |
| LPSGAYEGGALIK | 13 | Q12873 | Chromodomain-helicase-DNA-binding pr | CHD3 | 226.592 | 2 | 55.778 | yes | 5.253E-05 | 638.35083 | 77.775414 |
| ELIMGEDPAQPR | 12 | Q12873 | Chromodomain-helicase-DNA-binding pr | CHD3 | 226.592 | 2 | 55.778 | yes | 1.041E-07 | 678.33484 | 75.8582 |
| AFLNAVMR | 8 | Q12873 | Chromodomain-helicase-DNA-binding pr | CHD3 | 226.592 | 2 | 55.778 | yes | 9.819E-08 | 461.25238 | 72.896843 |
| AAYLNLSQEPAHP | 20 | Q12873 | Chromodomain-helicase-DNA-binding pr | CHD3 | 226.592 | 3 | 55.778 | yes | 2.636E-07 | 721.03723 | 65.730171 |
| LLPSALGVK | 9 | Q12873 | Chromodomain-helicase-DNA-binding pr | CHD3 | 226.592 | 2 | 55.778 | yes | 3.158E-06 | 449.29202 | 64.147194 |
| LNEIWHR | 7 | Q12873 | Chromodomain-helicase-DNA-binding pr | CHD3 | 226.592 | 2 | 55.778 | yes | 4.251E-07 | 484.25906 | 59.935738 |
| FGTEELFK | 8 | Q12873 | Chromodomain-helicase-DNA-binding pr | CHD3 | 226.592 | 2 | 55.778 | no | 1.419E-06 | 485.74765 | 39.758778 |
| HLCEPGADGSETF. | 19 | Q12873 | Chromodomain-helicase-DNA-binding pr | CHD3 | 226.592 | 3 | 55.778 | yes | 2.451E-06 | 672.30231 | 39.325996 |
| GNYHYLVK | 8 | Q12873 | Chromodomain-helicase-DNA-binding pr | CHD3 | 226.592 | 2 | 55.778 | yes | 7.347E-07 | 497.26126 | 38.086552 |
| HHYEQQQEDLAR | 12 | Q12873 | Chromodomain-helicase-DNA-binding pr | CHD3 | 226.592 | 2 | 55.778 | no | 4.379E-07 | 777.35822 | 33.706448 |
| EEDKIEEIER | 10 | Q12873 | Chromodomain-helicase-DNA-binding pr | CHD3 | 226.592 | 2 | 55.778 | yes | 3.566E-07 | 645.31464 | 33.126205 |
| EFSANNPFK | 9 | Q12873 | Chromodomain-helicase-DNA-binding pr | CHD3 | 226.592 | 2 | 55.778 | no | 1.398E-05 | 527.2536 | 13.786035 |
| WACLVVDEAHR | 11 | Q12873 | Chromodomain-helicase-DNA-binding pr | CHD3 | 226.592 | 3 | 55.778 | yes | 0.0001257 | 452.55566 | 3.6859384 |
| VKDPHYAEMEEK | 12 | Q12873 | Chromodomain-helicase-DNA-binding pr | CHD3 | 226.592 | 3 | 55.778 | yes | 3.687E-06 | 492.56601 | 3.6728861 |
| METILEQQR | 9 | Q12874 | Splicing factor 3A subunit 3 OS=Homo sa | SF3A3 | 58.849 | 2 | 49.747543 | yes | 2.921E-07 | 595.29773 | 711.93732 |
| YMEVSGNLR | 9 | Q12874 | Splicing factor 3A subunit 3 OS=Homo sa | SF3A3 | 58.849 | 2 | 49.747543 | yes | 1.153E-07 | 534.76074 | 426.3349 |
| VKPLQDQNELFGK | 13 | Q12874 | Splicing factor 3A subunit 3 OS=Homo sa | SF3A3 | 58.849 | 2 | 49.747543 | yes | 4.045E-07 | 758.41193 | 406.86005 |
| DLYDDKDGLR | 10 | Q12874 | Splicing factor 3A subunit 3 OS=Homo sa | SF3A3 | 58.849 | 2 | 49.747543 | yes | 1.078E-07 | 605.29095 | 301.71664 |
| YLDLHDCYLK | 10 | Q12874 | Splicing factor 3A subunit 3 OS=Homo sa | SF3A3 | 58.849 | 2 | 49.747543 | yes | 1.006E-07 | 670.32117 | 215.77014 |
| IQAEFEK | 7 | Q12874 | Splicing factor 3A subunit 3 OS=Homo sa | SF3A3 | 58.849 | 2 | 49.747543 | yes | 2.601E-07 | 432.72672 | 199.6573 |
| TYEDLKR | 7 | Q12874 | Splicing factor 3A subunit 3 OS=Homo sa | SF3A3 | 58.849 | 2 | 49.747543 | yes | 2.195E-07 | 462.74289 | 168.76988 |
| SLESLDTSLFAK | 12 | Q12874 | Splicing factor 3A subunit 3 OS=Homo sa | SF3A3 | 58.849 | 2 | 49.747543 | yes | 1.66E-07 | 655.84552 | 150.73582 |
| SALLALGLK | 9 | Q12874 | Splicing factor 3A subunit 3 OS=Homo sa | SF3A3 | 58.849 | 2 | 49.747543 | yes | 5.241E-07 | 443.29202 | 72.638397 |
| LALVISELEGK | 11 | Q12899 | Tripartite motif-containing protein 26 OS | TRIM26 | 62.166 | 2 | 53.77441 | yes | 1.727E-07 | 586.35028 | 1368.6675 |

| | | | | | | | | | | | |
|---|---|---|---|---|---|---|---|---|---|---|---|
| GEADILAALK | 10 | Q12899 | Tripartite motif-containing protein 26 OS | TRIM26 | 62.166 | 2 | 53.77441 | yes | 1.102E-07 | 500.78729 | 990.44666 |
| LLCVMCR | 7 | Q12899 | Tripartite motif-containing protein 26 OS | TRIM26 | 62.166 | 2 | 53.77441 | yes | 1.475E-07 | 476.23227 | 801.474 |
| LEQELTEGR | 9 | Q12899 | Tripartite motif-containing protein 26 OS | TRIM26 | 62.166 | 2 | 53.77441 | yes | 1.962E-07 | 537.7749 | 800.15094 |
| FWVGKPIAR | 9 | Q12899 | Tripartite motif-containing protein 26 OS | TRIM26 | 62.166 | 2 | 53.77441 | yes | 9.81E-08 | 537.31635 | 705.50031 |
| AQQPAAELMQDTI | 13 | Q12899 | Tripartite motif-containing protein 26 OS | TRIM26 | 62.166 | 2 | 53.77441 | yes | 1.394E-07 | 729.85394 | 626.44421 |
| VYWEVEVER | 9 | Q12899 | Tripartite motif-containing protein 26 OS | TRIM26 | 62.166 | 2 | 53.77441 | yes | 9.809E-08 | 604.80096 | 476.88782 |
| DPVTIDCGHVFCR | 13 | Q12899 | Tripartite motif-containing protein 26 OS | TRIM26 | 62.166 | 2 | 53.77441 | yes | 9.722E-08 | 788.35577 | 237.25175 |
| DPVTIDCGHVFCR | 13 | Q12899 | Tripartite motif-containing protein 26 OS | TRIM26 | 62.166 | 2 | 53.77441 | yes | 2.037E-06 | 525.90625 | 237.25175 |
| DPVTIDCGHVFCR | 13 | Q12899 | Tripartite motif-containing protein 26 OS | TRIM26 | 62.166 | 3 | 53.77441 | yes | 9.722E-08 | 788.35577 | 237.25175 |
| DPVTIDCGHVFCR | 13 | Q12899 | Tripartite motif-containing protein 26 OS | TRIM26 | 62.166 | 3 | 53.77441 | yes | 2.037E-06 | 525.90625 | 237.25175 |
| ILNHLSTLR | 9 | Q12899 | Tripartite motif-containing protein 26 OS | TRIM26 | 62.166 | 2 | 53.77441 | yes | 1.113E-07 | 533.82202 | 360.21149 |
| LHYYCEDDGK | 10 | Q12899 | Tripartite motif-containing protein 26 OS | TRIM26 | 62.166 | 2 | 53.77441 | yes | 5.791E-07 | 650.26917 | 349.36642 |
| QYIVAEFEQGHQF | 15 | Q12899 | Tripartite motif-containing protein 26 OS | TRIM26 | 62.166 | 3 | 53.77441 | yes | 1.463E-07 | 622.31622 | 258.28656 |
| CVTYTSLYK | 9 | Q12899 | Tripartite motif-containing protein 26 OS | TRIM26 | 62.166 | 2 | 53.77441 | yes | 1.122E-07 | 567.77863 | 197.50745 |
| EEHLLEQLAK | 10 | Q12899 | Tripartite motif-containing protein 26 OS | TRIM26 | 62.166 | 2 | 53.77441 | yes | 9.722E-08 | 605.32733 | 158.92593 |
| EREEHLLEQLAK | 12 | Q12899 | Tripartite motif-containing protein 26 OS | TRIM26 | 62.166 | 2 | 53.77441 | yes | 6.952E-07 | 747.89917 | 64.837845 |
| LSSSGIWANTSPEA | 23 | Q12899 | Tripartite motif-containing protein 26 OS | TRIM26 | 62.166 | 3 | 53.77441 | yes | 3.625E-07 | 833.77118 | 21.002598 |
| IGCIITAR | 8 | Q12904 | Aminoacyl tRNA synthase complex-intera | AIMP1 | 34.353 | 2 | 51.594357 | yes | 1.06E-07 | 452.25766 | 469.33218 |
| KHPDADSLYVEEV | 21 | Q12904 | Aminoacyl tRNA synthase complex-intera | AIMP1 | 34.353 | 3 | 51.594357 | yes | 9.728E-08 | 780.38867 | 197.41512 |
| QQSIAGSADSKPID | 17 | Q12904 | Aminoacyl tRNA synthase complex-intera | AIMP1 | 34.353 | 3 | 51.594357 | yes | 1.012E-07 | 586.97089 | 137.68121 |
| AILQATLR | 8 | Q12904 | Aminoacyl tRNA synthase complex-intera | AIMP1 | 34.353 | 2 | 51.594357 | yes | 9.842E-08 | 443.27945 | 92.848083 |
| MVILLCNLKPAK | 12 | Q12904 | Aminoacyl tRNA synthase complex-intera | AIMP1 | 34.353 | 2 | 51.594357 | yes | 9.722E-08 | 700.4118 | 71.267548 |
| GVLSQAMVMCAS | 16 | Q12904 | Aminoacyl tRNA synthase complex-intera | AIMP1 | 34.353 | 2 | 51.594357 | yes | 3.817E-07 | 847.89917 | 27.903133 |
| ILITTVPPNLR | 11 | Q12905 | Interleukin enhancer-binding factor 2 OS | ILF2 | 43.063 | 2 | 53.122826 | yes | 1.91E-06 | 618.88733 | 13078.82 |
| WFEENASQSTVK | 12 | Q12905 | Interleukin enhancer-binding factor 2 OS | ILF2 | 43.063 | 2 | 53.122826 | yes | 4.866E-07 | 713.33588 | 13039.563 |
| QPLALNVAYR | 10 | Q12905 | Interleukin enhancer-binding factor 2 OS | ILF2 | 43.063 | 2 | 53.122826 | yes | 1.071E-07 | 572.82727 | 11944.861 |
| ILSHGGFR | 8 | Q12905 | Interleukin enhancer-binding factor 2 OS | ILF2 | 43.063 | 2 | 53.122826 | yes | 1.071E-07 | 443.74832 | 10514.006 |
| VLQSALAAIR | 10 | Q12905 | Interleukin enhancer-binding factor 2 OS | ILF2 | 43.063 | 2 | 53.122826 | yes | 1.318E-07 | 521.3244 | 3749.0449 |
| KLDPELHLDIK | 11 | Q12905 | Interleukin enhancer-binding factor 2 OS | ILF2 | 43.063 | 2 | 53.122826 | yes | 1.001E-07 | 660.8797 | 3503.0022 |
| LDPELHLDIK | 10 | Q12905 | Interleukin enhancer-binding factor 2 OS | ILF2 | 43.063 | 2 | 53.122826 | yes | 1.102E-07 | 596.83228 | 1112.3812 |
| NQDLAPNSAEQAS | 20 | Q12905 | Interleukin enhancer-binding factor 2 OS | ILF2 | 43.063 | 3 | 53.122826 | yes | 2.153E-07 | 700.37079 | 335.48859 |
| AYEKPPEK | 8 | Q12905 | Interleukin enhancer-binding factor 2 OS | ILF2 | 43.063 | 2 | 53.122826 | yes | 1.021E-07 | 481.25311 | 269.2709 |
| ILPTLEAVAALGNI | 14 | Q12905 | Interleukin enhancer-binding factor 2 OS | ILF2 | 43.063 | 2 | 53.122826 | yes | 8.245E-07 | 705.42175 | 184.30621 |
| PAPDETSFSEALLK | 14 | Q12905 | Interleukin enhancer-binding factor 2 OS | ILF2 | 43.063 | 2 | 53.122826 | yes | 2.779E-07 | 752.88013 | 157.74838 |
| KGTMTTGHNVAD | 18 | Q12905 | Interleukin enhancer-binding factor 2 OS | ILF2 | 43.063 | 3 | 53.122826 | yes | 9.722E-08 | 633.02405 | 126.38478 |
| RNQDLAPNSAEQA | 21 | Q12905 | Interleukin enhancer-binding factor 2 OS | ILF2 | 43.063 | 3 | 53.122826 | yes | 2.757E-06 | 752.40454 | 7.3261166 |
| LAAFGQLHK | 9 | Q12906 | Interleukin enhancer-binding factor 3 OS | ILF3 | 95.339 | 2 | 52.167088 | yes | 1.467E-07 | 492.78491 | 17948.83 |
| FVMEVEVDGQK | 11 | Q12906 | Interleukin enhancer-binding factor 3 OS | ILF3 | 95.339 | 2 | 52.167088 | no | 1.379E-07 | 640.81317 | 8252.627 |
| FVMEVEVDGQK | 11 | Q12906 | Interleukin enhancer-binding factor 3 OS | ILF3 | 95.339 | 2 | 52.167088 | no | 2.199E-06 | 648.81067 | 8252.627 |

| | | | | | | | | | | | |
|---|---|---|---|---|---|---|---|---|---|---|---|
| LFPDTPLALDANK | 13 | Q12906 | Interleukin enhancer-binding factor 3 OS= | ILF3 | 95.339 | 2 | 52.167088 | yes | 1.317E-07 | 707.88245 | 14928.371 |
| VLGMDPLPSK | 10 | Q12906 | Interleukin enhancer-binding factor 3 OS= | ILF3 | 95.339 | 2 | 52.167088 | yes | 3.656E-07 | 528.79156 | 6959.1865 |
| VLGMDPLPSK | 10 | Q12906 | Interleukin enhancer-binding factor 3 OS= | ILF3 | 95.339 | 2 | 52.167088 | yes | 1.062E-07 | 536.789 | 6959.1865 |
| EKPTTALLDK | 10 | Q12906 | Interleukin enhancer-binding factor 3 OS= | ILF3 | 95.339 | 2 | 52.167088 | yes | 1.713E-07 | 558.31897 | 13617.06 |
| VLQDMGLPTGAE( | 14 | Q12906 | Interleukin enhancer-binding factor 3 OS= | ILF3 | 95.339 | 2 | 52.167088 | yes | 1.269E-07 | 722.36664 | 6240.1597 |
| VLQDMGLPTGAE( | 14 | Q12906 | Interleukin enhancer-binding factor 3 OS= | ILF3 | 95.339 | 2 | 52.167088 | yes | 2.299E-07 | 730.36414 | 6240.1597 |
| EATDAIGHLDR | 11 | Q12906 | Interleukin enhancer-binding factor 3 OS= | ILF3 | 95.339 | 2 | 52.167088 | yes | 3.432E-07 | 599.29657 | 12186.701 |
| YELISETGGSHDK | 13 | Q12906 | Interleukin enhancer-binding factor 3 OS= | ILF3 | 95.339 | 2 | 52.167088 | no | 1.021E-07 | 718.33862 | 5762.7329 |
| YELISETGGSHDK | 13 | Q12906 | Interleukin enhancer-binding factor 3 OS= | ILF3 | 95.339 | 2 | 52.167088 | no | 1.849E-07 | 479.22818 | 5762.7329 |
| YELISETGGSHDK | 13 | Q12906 | Interleukin enhancer-binding factor 3 OS= | ILF3 | 95.339 | 3 | 52.167088 | no | 1.021E-07 | 718.33862 | 5762.7329 |
| YELISETGGSHDK | 13 | Q12906 | Interleukin enhancer-binding factor 3 OS= | ILF3 | 95.339 | 3 | 52.167088 | no | 1.849E-07 | 479.22818 | 5762.7329 |
| AVSDWIDEQEK | 11 | Q12906 | Interleukin enhancer-binding factor 3 OS= | ILF3 | 95.339 | 2 | 52.167088 | yes | 1.451E-07 | 660.30933 | 11315.424 |
| EDITQSAQHALR | 12 | Q12906 | Interleukin enhancer-binding factor 3 OS= | ILF3 | 95.339 | 2 | 52.167088 | yes | 9.723E-08 | 684.84692 | 8439.3984 |
| LNQLKPGLQYK | 11 | Q12906 | Interleukin enhancer-binding factor 3 OS= | ILF3 | 95.339 | 2 | 52.167088 | yes | 1.857E-07 | 651.38245 | 3611.8 |
| LNQLKPGLQYK | 11 | Q12906 | Interleukin enhancer-binding factor 3 OS= | ILF3 | 95.339 | 2 | 52.167088 | yes | 9.976E-08 | 434.59073 | 3611.8 |
| LNQLKPGLQYK | 11 | Q12906 | Interleukin enhancer-binding factor 3 OS= | ILF3 | 95.339 | 3 | 52.167088 | yes | 1.857E-07 | 651.38245 | 3611.8 |
| LNQLKPGLQYK | 11 | Q12906 | Interleukin enhancer-binding factor 3 OS= | ILF3 | 95.339 | 3 | 52.167088 | yes | 9.976E-08 | 434.59073 | 3611.8 |
| SCVIVIR | 7 | Q12906 | Interleukin enhancer-binding factor 3 OS= | ILF3 | 95.339 | 2 | 52.167088 | yes | 2.124E-07 | 423.74692 | 6849.6533 |
| GWPLELLCEK | 10 | Q12906 | Interleukin enhancer-binding factor 3 OS= | ILF3 | 95.339 | 2 | 52.167088 | yes | 1.003E-07 | 622.8208 | 6596.52 |
| CLAALASLR | 9 | Q12906 | Interleukin enhancer-binding factor 3 OS= | ILF3 | 95.339 | 2 | 52.167088 | yes | 3.12E-07 | 487.77621 | 5408.7656 |
| NPVMELNEK | 9 | Q12906 | Interleukin enhancer-binding factor 3 OS= | ILF3 | 95.339 | 2 | 52.167088 | no | 1.026E-07 | 537.26843 | 4672.7227 |
| EPPLSLTIHLTSPV' | 16 | Q12906 | Interleukin enhancer-binding factor 3 OS= | ILF3 | 95.339 | 3 | 52.167088 | yes | 1.182E-07 | 587.00861 | 3555.5254 |
| PMGAGEALR | 9 | Q12906 | Interleukin enhancer-binding factor 3 OS= | ILF3 | 95.339 | 2 | 52.167088 | yes | 9.818E-08 | 451.23166 | 2482.1819 |
| AYAALAALEK | 10 | Q12906 | Interleukin enhancer-binding factor 3 OS= | ILF3 | 95.339 | 2 | 52.167088 | yes | 1.995E-07 | 510.78986 | 2244.696 |
| AEPPQAMNALMR | 12 | Q12906 | Interleukin enhancer-binding factor 3 OS= | ILF3 | 95.339 | 2 | 52.167088 | yes | 1.115E-07 | 664.82611 | 1483.6049 |
| VADNLAIQLAAVT | 16 | Q12906 | Interleukin enhancer-binding factor 3 OS= | ILF3 | 95.339 | 2 | 52.167088 | yes | 9.722E-08 | 835.9516 | 1450.9005 |
| VPTWGPLR | 8 | Q12906 | Interleukin enhancer-binding factor 3 OS= | ILF3 | 95.339 | 2 | 52.167088 | yes | 1.638E-07 | 463.26636 | 1102.8499 |
| RPMEEDGEEK | 10 | Q12906 | Interleukin enhancer-binding factor 3 OS= | ILF3 | 95.339 | 2 | 52.167088 | yes | 4.24E-07 | 610.2666 | 1052.8959 |
| IFVNDDR | 7 | Q12906 | Interleukin enhancer-binding factor 3 OS= | ILF3 | 95.339 | 2 | 52.167088 | yes | 4.087E-07 | 439.72195 | 873.97333 |
| LFPDTPLALDANK | 14 | Q12906 | Interleukin enhancer-binding factor 3 OS= | ILF3 | 95.339 | 2 | 52.167088 | yes | 9.322E-07 | 771.92993 | 576.24744 |
| YELISETGGSHDKI | 14 | Q12906 | Interleukin enhancer-binding factor 3 OS= | ILF3 | 95.339 | 2 | 52.167088 | no | 1.211E-07 | 796.38916 | 571.87866 |
| SIGTANRPMGAGE | 16 | Q12906 | Interleukin enhancer-binding factor 3 OS= | ILF3 | 95.339 | 2 | 52.167088 | yes | 1.874E-07 | 800.91486 | 312.47382 |
| GDLDLELVLLCK | 12 | Q12906 | Interleukin enhancer-binding factor 3 OS= | ILF3 | 95.339 | 2 | 52.167088 | yes | 1.004E-07 | 694.37872 | 310.36987 |
| VLECLASGIVMPD | 23 | Q12906 | Interleukin enhancer-binding factor 3 OS= | ILF3 | 95.339 | 3 | 52.167088 | yes | 9.722E-08 | 837.39191 | 248.62317 |
| YEILQSVDDAAIVI | 15 | Q12906 | Interleukin enhancer-binding factor 3 OS= | ILF3 | 95.339 | 2 | 52.167088 | yes | 9.809E-08 | 838.95892 | 116.55814 |
| PAVVAPAPVVEA' | 34 | Q12906 | Interleukin enhancer-binding factor 3 OS= | ILF3 | 95.339 | 4 | 52.167088 | yes | 9.722E-08 | 822.93951 | 90.507477 |
| NADHSMNYQYR | 11 | Q12906 | Interleukin enhancer-binding factor 3 OS= | ILF3 | 95.339 | 2 | 52.167088 | yes | 1.423E-07 | 699.79639 | 71.933929 |
| GVVDSEDIPLNLSI | 14 | Q12931 | Heat shock protein 75 kDa, mitochondrial | TRAP1 | 80.11 | 2 | 51.60084 | yes | 9.722E-08 | 757.39648 | 3058.5854 |
| AQLLQPTLEINPR | 13 | Q12931 | Heat shock protein 75 kDa, mitochondrial | TRAP1 | 80.11 | 2 | 51.60084 | yes | 1.192E-07 | 746.92773 | 116.46415 |

| | | | | | | | | | | | |
|---|---|---|---|---|---|---|---|---|---|---|---|
| ETEELMAWMR | 10 | Q12931 | Heat shock protein 75 kDa, mitochondrial | TRAP1 | 80.11 | 2 | 51.60084 | yes | 1.377E-07 | 648.29156 | 45.224743 |
| AFLDALQNQAEAS | 15 | Q12931 | Heat shock protein 75 kDa, mitochondrial | TRAP1 | 80.11 | 2 | 51.60084 | yes | 2.75E-07 | 796.89935 | 26.341711 |
| DLAMADLEQK | 10 | Q12933 | TNF receptor-associated factor 2 OS=Hoi | TRAF2 | 55.86 | 2 | 52.827908 | yes | 1.236E-07 | 567.27899 | 166.61836 |
| TATFENIVCVLNR | 13 | Q12933 | TNF receptor-associated factor 2 OS=Hoi | TRAF2 | 55.86 | 2 | 52.827908 | yes | 9.722E-08 | 768.89557 | 130.93237 |
| IPAIFSPAFYTSR | 13 | Q12933 | TNF receptor-associated factor 2 OS=Hoi | TRAF2 | 55.86 | 2 | 52.827908 | yes | 2.18E-07 | 735.39301 | 52.650021 |
| EDLSVGLPR | 9 | Q12947 | Forkhead box protein F2 OS=Homo sapie | FOXF2 | 45.994 | 2 | 50.30997 | yes | 2.32E-07 | 493.26929 | 1664.6671 |
| HNLSLNECFIK | 11 | Q12947 | Forkhead box protein F2 OS=Homo sapie | FOXF2 | 45.994 | 2 | 50.30997 | no | 9.819E-08 | 687.84534 | 353.47281 |
| TTEGGPPPAPLR | 12 | Q12947 | Forkhead box protein F2 OS=Homo sapie | FOXF2 | 45.994 | 2 | 50.30997 | yes | 1.121E-06 | 617.82495 | 46.485821 |
| YQHHSTPVCDR | 11 | Q12947 | Forkhead box protein F2 OS=Homo sapie | FOXF2 | 45.994 | 2 | 50.30997 | yes | 1.846E-07 | 700.31219 | 40.972191 |
| AYGPYTPQPQPK | 12 | Q12948 | Forkhead box protein C1 OS=Homo sapie | FOXC1 | 56.789 | 2 | 52.439457 | yes | 1.439E-07 | 673.84058 | 1562.0034 |
| GSPQSAAAELSSGI | 23 | Q12948 | Forkhead box protein C1 OS=Homo sapie | FOXC1 | 56.789 | 3 | 52.439457 | yes | 1.164E-07 | 697.02216 | 644.01691 |
| TSGAFVYDCSK | 11 | Q12948 | Forkhead box protein C1 OS=Homo sapie | FOXC1 | 56.789 | 2 | 52.439457 | yes | 1.146E-07 | 617.77405 | 581.30334 |
| EMFESQR | 7 | Q12948 | Forkhead box protein C1 OS=Homo sapie | FOXC1 | 56.789 | 2 | 52.439457 | yes | 3.662E-07 | 463.70547 | 272.70923 |
| TSGAFVYDCSKF | 12 | Q12948 | Forkhead box protein C1 OS=Homo sapie | FOXC1 | 56.789 | 2 | 52.439457 | yes | 9.722E-08 | 691.30829 | 121.12382 |
| QPPPAPPEQADGN | 23 | Q12948 | Forkhead box protein C1 OS=Homo sapie | FOXC1 | 56.789 | 3 | 52.439457 | yes | 1.44E-07 | 772.05731 | 54.593761 |
| ALHLLEEYR | 9 | Q12959 | Disks large homolog 1 OS=Homo sapiens | DLG1 | 100.456 | 2 | 51.58941 | yes | 9.729E-08 | 572.31146 | 198.01511 |
| HCILDVSGNAIK | 12 | Q12959 | Disks large homolog 1 OS=Homo sapiens | DLG1 | 100.456 | 2 | 51.58941 | no | 1.473E-07 | 663.84534 | 194.2157 |
| INDDLISEFPDK | 12 | Q12959 | Disks large homolog 1 OS=Homo sapiens | DLG1 | 100.456 | 2 | 51.58941 | no | 1.954E-07 | 703.34589 | 157.039 |
| LLAVNNVCLEEVT | 22 | Q12959 | Disks large homolog 1 OS=Homo sapiens | DLG1 | 100.456 | 3 | 51.58941 | yes | 9.849E-08 | 818.09656 | 150.60194 |
| EQMMNSSISSGSG | 16 | Q12959 | Disks large homolog 1 OS=Homo sapiens | DLG1 | 100.456 | 2 | 51.58941 | yes | 9.722E-08 | 835.87726 | 117.24072 |
| DYHFVTSR | 8 | Q12959 | Disks large homolog 1 OS=Homo sapiens | DLG1 | 100.456 | 2 | 51.58941 | yes | 1.443E-07 | 512.74597 | 107.12377 |
| AVLGDDEITR | 10 | Q12959 | Disks large homolog 1 OS=Homo sapiens | DLG1 | 100.456 | 2 | 51.58941 | yes | 1.627E-05 | 544.78278 | 65.669258 |
| SMENIMEMNK | 10 | Q12959 | Disks large homolog 1 OS=Homo sapiens | DLG1 | 100.456 | 2 | 51.58941 | yes | 4.573E-07 | 613.76434 | 65.194878 |
| AASHEQAAAALK | 12 | Q12959 | Disks large homolog 1 OS=Homo sapiens | DLG1 | 100.456 | 2 | 51.58941 | yes | 1.937E-07 | 584.30945 | 56.355171 |
| VSQTPESLDFLK | 12 | Q12965 | Unconventional myosin-Ie OS=Homo sap | MYO1E | 127.062 | 2 | 52.925304 | yes | 1.305E-07 | 682.35883 | 1607.5435 |
| VLQVSIGPGLPK | 12 | Q12965 | Unconventional myosin-Ie OS=Homo sap | MYO1E | 127.062 | 2 | 52.925304 | yes | 1.219E-07 | 604.37408 | 1391.5884 |
| HQVEYLGLK | 9 | Q12965 | Unconventional myosin-Ie OS=Homo sap | MYO1E | 127.062 | 2 | 52.925304 | no | 6.271E-07 | 543.80072 | 1095.1652 |
| AEQEEYVQEGIR | 12 | Q12965 | Unconventional myosin-Ie OS=Homo sap | MYO1E | 127.062 | 2 | 52.925304 | no | 9.722E-08 | 725.84406 | 942.54083 |
| VFDFLVDSINK | 11 | Q12965 | Unconventional myosin-Ie OS=Homo sap | MYO1E | 127.062 | 2 | 52.925304 | yes | 1.004E-07 | 648.84534 | 890.64038 |
| IVCDLIENK | 9 | Q12965 | Unconventional myosin-Ie OS=Homo sap | MYO1E | 127.062 | 2 | 52.925304 | yes | 1.104E-07 | 552.29187 | 803.42444 |
| ITENSIVENLK | 11 | Q12965 | Unconventional myosin-Ie OS=Homo sap | MYO1E | 127.062 | 2 | 52.925304 | yes | 9.81E-08 | 630.3457 | 641.02789 |
| NQYVPYPHAPGSC | 14 | Q12965 | Unconventional myosin-Ie OS=Homo sap | MYO1E | 127.062 | 3 | 52.925304 | yes | 9.722E-08 | 538.59882 | 320.05637 |
| NQYVPYPHAPGSC | 14 | Q12965 | Unconventional myosin-Ie OS=Homo sap | MYO1E | 127.062 | 3 | 52.925304 | yes | 9.722E-08 | 807.39459 | 320.05637 |
| NQYVPYPHAPGSC | 14 | Q12965 | Unconventional myosin-Ie OS=Homo sap | MYO1E | 127.062 | 2 | 52.925304 | yes | 9.722E-08 | 538.59882 | 320.05637 |
| NQYVPYPHAPGSC | 14 | Q12965 | Unconventional myosin-Ie OS=Homo sap | MYO1E | 127.062 | 2 | 52.925304 | yes | 9.722E-08 | 807.39459 | 320.05637 |
| VSYDMDGFCER | 11 | Q12965 | Unconventional myosin-Ie OS=Homo sap | MYO1E | 127.062 | 2 | 52.925304 | yes | 9.722E-08 | 689.77374 | 606.24023 |
| SLFPENLQADK | 11 | Q12965 | Unconventional myosin-Ie OS=Homo sap | MYO1E | 127.062 | 2 | 52.925304 | yes | 3.184E-07 | 631.32477 | 550.20123 |
| FSNTLELK | 8 | Q12965 | Unconventional myosin-Ie OS=Homo sap | MYO1E | 127.062 | 2 | 52.925304 | yes | 1.576E-07 | 476.26093 | 547.56549 |
| QANDLVSTLMK | 11 | Q12965 | Unconventional myosin-Ie OS=Homo sap | MYO1E | 127.062 | 2 | 52.925304 | yes | 1.063E-07 | 610.32117 | 531.69409 |

| | | | | | | | | | | | |
|---|---|---|---|---|---|---|---|---|---|---|---|
| WTPIEYFNNK | 10 | Q12965 | Unconventional myosin-Ie OS=Homo sap | MYO1E | 127.062 | 2 | 52.925304 | yes | 2.298E-07 | 656.32202 | 480.83563 |
| EKIDFADTVTK | 11 | Q12965 | Unconventional myosin-Ie OS=Homo sap | MYO1E | 127.062 | 2 | 52.925304 | yes | 9.819E-08 | 633.83246 | 459.56781 |
| APESLFLLEEMR | 12 | Q12965 | Unconventional myosin-Ie OS=Homo sap | MYO1E | 127.062 | 2 | 52.925304 | yes | 3.684E-07 | 717.86853 | 411.1481 |
| CTPHYIR | 7 | Q12965 | Unconventional myosin-Ie OS=Homo sap | MYO1E | 127.062 | 2 | 52.925304 | no | 1.688E-07 | 473.73181 | 407.8252 |
| ATWPSWQGEEK | 11 | Q12965 | Unconventional myosin-Ie OS=Homo sap | MYO1E | 127.062 | 2 | 52.925304 | yes | 1.861E-07 | 659.80676 | 326.23468 |
| EEASDLLLNK | 10 | Q12965 | Unconventional myosin-Ie OS=Homo sap | MYO1E | 127.062 | 2 | 52.925304 | yes | 3.159E-07 | 566.29822 | 291.56595 |
| TEFLSLLAK | 9 | Q12965 | Unconventional myosin-Ie OS=Homo sap | MYO1E | 127.062 | 2 | 52.925304 | yes | 4.078E-07 | 511.30005 | 289.52469 |
| QVQFHQGFGDLA | 19 | Q12965 | Unconventional myosin-Ie OS=Homo sap | MYO1E | 127.062 | 3 | 52.925304 | yes | 9.81E-08 | 705.04449 | 281.41751 |
| ENQCVIISGESGA | 15 | Q12965 | Unconventional myosin-Ie OS=Homo sap | MYO1E | 127.062 | 2 | 52.925304 | yes | 9.809E-08 | 774.86975 | 281.02539 |
| HSGVDDMVLLSK | 12 | Q12965 | Unconventional myosin-Ie OS=Homo sap | MYO1E | 127.062 | 2 | 52.925304 | yes | 9.892E-08 | 650.83191 | 275.32559 |
| PKPQVPQCK | 9 | Q12965 | Unconventional myosin-Ie OS=Homo sap | MYO1E | 127.062 | 2 | 52.925304 | yes | 3.526E-07 | 541.2948 | 216.69803 |
| CLYLIGR | 7 | Q12965 | Unconventional myosin-Ie OS=Homo sap | MYO1E | 127.062 | 2 | 52.925304 | yes | 9.819E-08 | 447.74692 | 188.8129 |
| SFHIFYQLIEGASA | 17 | Q12965 | Unconventional myosin-Ie OS=Homo sap | MYO1E | 127.062 | 3 | 52.925304 | yes | 1.091E-07 | 656.66693 | 159.10905 |
| ISNFLLEK | 8 | Q12965 | Unconventional myosin-Ie OS=Homo sap | MYO1E | 127.062 | 2 | 52.925304 | no | 5.17E-07 | 482.27911 | 155.55678 |
| YIMSYISR | 8 | Q12965 | Unconventional myosin-Ie OS=Homo sap | MYO1E | 127.062 | 2 | 52.925304 | yes | 1.01E-07 | 516.76276 | 141.74403 |
| GVYQYHWQSHN | 13 | Q12965 | Unconventional myosin-Ie OS=Homo sap | MYO1E | 127.062 | 2 | 52.925304 | yes | 1.51E-07 | 823.39716 | 79.446213 |
| RDLLLTPK | 8 | Q12965 | Unconventional myosin-Ie OS=Homo sap | MYO1E | 127.062 | 2 | 52.925304 | yes | 1.006E-06 | 478.30038 | 63.09906 |
| EDPSGWWTGR | 10 | Q12965 | Unconventional myosin-Ie OS=Homo sap | MYO1E | 127.062 | 2 | 52.925304 | yes | 1.83E-07 | 595.76489 | 39.340351 |
| SLYTSMARPPLPR | 13 | Q12965 | Unconventional myosin-Ie OS=Homo sap | MYO1E | 127.062 | 3 | 52.925304 | yes | 4.771E-06 | 496.9379 | 34.033699 |
| QGVLHLLQSVNM | 21 | Q12965 | Unconventional myosin-Ie OS=Homo sap | MYO1E | 127.062 | 3 | 52.925304 | yes | 3.226E-07 | 795.73749 | 29.75275 |
| VPDQGAAGVR | 10 | Q12965 | Unconventional myosin-Ie OS=Homo sap | MYO1E | 127.062 | 2 | 52.925304 | yes | 2.797E-05 | 485.25925 | 25.893623 |
| DMSQISVSQGAG | 15 | Q12968 | Nuclear factor of activated T-cells, cytopl | NFATC3 | 115.594 | 2 | 53.160786 | yes | 2.697E-07 | 761.36993 | 1350.798 |
| DHLYLPLEPSYR | 12 | Q12968 | Nuclear factor of activated T-cells, cytopl | NFATC3 | 115.594 | 2 | 53.160786 | yes | 1.741E-07 | 751.88556 | 1212.8451 |
| ESSLSPSPASSISSR | 15 | Q12968 | Nuclear factor of activated T-cells, cytopl | NFATC3 | 115.594 | 2 | 53.160786 | yes | 1.09E-07 | 746.36792 | 1074.7411 |
| LVFGEDGAPAPPP | 17 | Q12968 | Nuclear factor of activated T-cells, cytopl | NFATC3 | 115.594 | 2 | 53.160786 | yes | 9.722E-08 | 832.42554 | 865.50629 |
| SSVTDENWLSPR | 12 | Q12968 | Nuclear factor of activated T-cells, cytopl | NFATC3 | 115.594 | 2 | 53.160786 | yes | 1.579E-07 | 695.8335 | 712.16547 |
| FTYTPVLMK | 9 | Q12968 | Nuclear factor of activated T-cells, cytopl | NFATC3 | 115.594 | 2 | 53.160786 | yes | 1.104E-07 | 550.29645 | 579.12793 |
| ETSIDDGLGSQYPI | 15 | Q12968 | Nuclear factor of activated T-cells, cytopl | NFATC3 | 115.594 | 2 | 53.160786 | yes | 2.132E-07 | 811.89905 | 533.0434 |
| SAQELPHIEK | 10 | Q12968 | Nuclear factor of activated T-cells, cytopl | NFATC3 | 115.594 | 2 | 53.160786 | yes | 1.248E-07 | 576.3064 | 418.88297 |
| TVATASQEIIAST | 15 | Q12968 | Nuclear factor of activated T-cells, cytopl | NFATC3 | 115.594 | 2 | 53.160786 | yes | 9.722E-08 | 766.93018 | 303.05679 |
| LELCSDDQGSLSP | 15 | Q12968 | Nuclear factor of activated T-cells, cytopl | NFATC3 | 115.594 | 2 | 53.160786 | yes | 9.722E-08 | 824.3858 | 246.8065 |
| VHIPQPSGK | 9 | Q12968 | Nuclear factor of activated T-cells, cytopl | NFATC3 | 115.594 | 2 | 53.160786 | yes | 4.3E-07 | 481.77454 | 210.76419 |
| NYEGTCEIPESK | 12 | Q12968 | Nuclear factor of activated T-cells, cytopl | NFATC3 | 115.594 | 2 | 53.160786 | yes | 9.722E-08 | 713.8114 | 185.37543 |
| TTANCGAHDELD | 14 | Q12968 | Nuclear factor of activated T-cells, cytopl | NFATC3 | 115.594 | 2 | 53.160786 | yes | 1.122E-07 | 810.85156 | 155.72351 |
| PKPGHTPIFR | 10 | Q12968 | Nuclear factor of activated T-cells, cytopl | NFATC3 | 115.594 | 2 | 53.160786 | yes | 1.69E-07 | 575.33002 | 143.01302 |
| TSEDQAAILPGK | 12 | Q12968 | Nuclear factor of activated T-cells, cytopl | NFATC3 | 115.594 | 2 | 53.160786 | yes | 1.104E-07 | 615.32227 | 114.9268 |
| KTSEDQAAILPGK | 13 | Q12968 | Nuclear factor of activated T-cells, cytopl | NFATC3 | 115.594 | 2 | 53.160786 | yes | 1.117E-06 | 679.36975 | 78.330429 |
| QAPLPSPESLDLG | 14 | Q12968 | Nuclear factor of activated T-cells, cytopl | NFATC3 | 115.594 | 2 | 53.160786 | yes | 1.701E-07 | 740.39374 | 76.403366 |
| HSSAEVCYAGSLS | 27 | Q12968 | Nuclear factor of activated T-cells, cytopl | NFATC3 | 115.594 | 4 | 53.160786 | yes | 2.954E-06 | 710.08478 | 48.40794 |

| | | | | | | | | | | | |
|---|---|---|---|---|---|---|---|---|---|---|---|
| GQIQLDPQTVETK | 13 | Q12979 | Active breakpoint cluster region-related p | ABR | 97.598 | 2 | 27.830484 | yes | 2.878E-06 | 728.88574 | 23.302753 |
| EAYDSLCTSGVVR | 13 | Q12980 | GATOR1 complex protein NPRL3 OS=H | NPRL3 | 63.605 | 2 | 41.298576 | yes | 3.579E-07 | 728.84045 | 32.662529 |
| SVCPPTTLTGVLEI | 14 | Q12986 | Transcriptional repressor NF-X1 OS=Hor | NFX1 | 124.395 | 2 | 53.986103 | yes | 2.563E-07 | 765.40326 | 1707.1866 |
| FNTDAAEFIPQEK | 13 | Q12986 | Transcriptional repressor NF-X1 OS=Hor | NFX1 | 124.395 | 2 | 53.986103 | yes | 7.363E-07 | 755.36462 | 1669.2545 |
| YECMVCCELVR | 11 | Q12986 | Transcriptional repressor NF-X1 OS=Hor | NFX1 | 124.395 | 2 | 53.986103 | yes | 1.312E-07 | 759.81403 | 1507.7974 |
| SDGFGDFSCLK | 11 | Q12986 | Transcriptional repressor NF-X1 OS=Hor | NFX1 | 124.395 | 2 | 53.986103 | yes | 1.022E-07 | 616.76624 | 1223.5177 |
| ITDMQLGGSVEISI | 14 | Q12986 | Transcriptional repressor NF-X1 OS=Hor | NFX1 | 124.395 | 2 | 53.986103 | yes | 1.004E-07 | 739.38196 | 1022.1446 |
| DVLCGTDVGK | 10 | Q12986 | Transcriptional repressor NF-X1 OS=Hor | NFX1 | 124.395 | 2 | 53.986103 | yes | 1.84E-07 | 532.25806 | 889.78351 |
| KGVLDGYGAR | 10 | Q12986 | Transcriptional repressor NF-X1 OS=Hor | NFX1 | 124.395 | 2 | 53.986103 | yes | 2.253E-07 | 518.28271 | 857.50665 |
| LECDEECSALER | 12 | Q12986 | Transcriptional repressor NF-X1 OS=Hor | NFX1 | 124.395 | 2 | 53.986103 | yes | 1.096E-07 | 755.81104 | 848.14124 |
| IIHDLAQVYGLES\ | 21 | Q12986 | Transcriptional repressor NF-X1 OS=Hor | NFX1 | 124.395 | 3 | 53.986103 | yes | 1.003E-07 | 788.39703 | 790.88373 |
| EMVICSEASSTYQI | 14 | Q12986 | Transcriptional repressor NF-X1 OS=Hor | NFX1 | 124.395 | 2 | 53.986103 | yes | 9.722E-08 | 830.8689 | 640.07794 |
| ATQFVYSYGR | 10 | Q12986 | Transcriptional repressor NF-X1 OS=Hor | NFX1 | 124.395 | 2 | 53.986103 | yes | 1.362E-07 | 596.29327 | 587.90833 |
| CPLICGR | 7 | Q12986 | Transcriptional repressor NF-X1 OS=Hor | NFX1 | 124.395 | 2 | 53.986103 | yes | 1.684E-07 | 438.21494 | 553.35144 |
| SHGLQNQPWQK | 11 | Q12986 | Transcriptional repressor NF-X1 OS=Hor | NFX1 | 124.395 | 2 | 53.986103 | yes | 1.537E-07 | 661.83362 | 529.4856 |
| ELPCTSLK | 8 | Q12986 | Transcriptional repressor NF-X1 OS=Hor | NFX1 | 124.395 | 2 | 53.986103 | yes | 2.779E-07 | 474.24695 | 494.87167 |
| TSVISCRCSFR | 11 | Q12986 | Transcriptional repressor NF-X1 OS=Hor | NFX1 | 124.395 | 2 | 53.986103 | yes | 1.262E-05 | 686.82666 | 443.20428 |
| STCDSENLAVINK | 13 | Q12986 | Transcriptional repressor NF-X1 OS=Hor | NFX1 | 124.395 | 2 | 53.986103 | yes | 9.722E-08 | 725.84576 | 442.10327 |
| VCGKPLPCGSLDF | 19 | Q12986 | Transcriptional repressor NF-X1 OS=Hor | NFX1 | 124.395 | 3 | 53.986103 | yes | 2.226E-07 | 740.01996 | 408.42679 |
| SEDATFMCDK | 10 | Q12986 | Transcriptional repressor NF-X1 OS=Hor | NFX1 | 124.395 | 2 | 53.986103 | yes | 1.004E-07 | 602.23645 | 370.44489 |
| TCMDPVPSCGK | 11 | Q12986 | Transcriptional repressor NF-X1 OS=Hor | NFX1 | 124.395 | 2 | 53.986103 | yes | 9.722E-08 | 626.26196 | 355.40182 |
| CNEICCVDK | 9 | Q12986 | Transcriptional repressor NF-X1 OS=Hor | NFX1 | 124.395 | 2 | 53.986103 | yes | 9.722E-08 | 599.23846 | 345.33173 |
| DDLNERPAK | 9 | Q12986 | Transcriptional repressor NF-X1 OS=Hor | NFX1 | 124.395 | 2 | 53.986103 | yes | 2.093E-07 | 529.26727 | 316.82724 |
| AEAPPVSGTFK | 11 | Q12986 | Transcriptional repressor NF-X1 OS=Hor | NFX1 | 124.395 | 2 | 53.986103 | yes | 1.192E-07 | 573.29547 | 243.71802 |
| TTPKPEDAGPESTI | 29 | Q12986 | Transcriptional repressor NF-X1 OS=Hor | NFX1 | 124.395 | 4 | 53.986103 | yes | 8.62E-07 | 753.36517 | 243.2345 |
| CPPCTFLTQK | 10 | Q12986 | Transcriptional repressor NF-X1 OS=Hor | NFX1 | 124.395 | 2 | 53.986103 | yes | 9.809E-08 | 626.29663 | 241.82591 |
| RPPWEVEGAR | 10 | Q12986 | Transcriptional repressor NF-X1 OS=Hor | NFX1 | 124.395 | 2 | 53.986103 | yes | 1.143E-07 | 598.81219 | 205.23303 |
| EMETLVEAVNK | 11 | Q12986 | Transcriptional repressor NF-X1 OS=Hor | NFX1 | 124.395 | 2 | 53.986103 | yes | 9.722E-08 | 631.81848 | 191.21695 |
| GVLDGYGAR | 9 | Q12986 | Transcriptional repressor NF-X1 OS=Hor | NFX1 | 124.395 | 2 | 53.986103 | yes | 2.693E-07 | 454.23523 | 189.25482 |
| GECLVDEPCK | 10 | Q12986 | Transcriptional repressor NF-X1 OS=Hor | NFX1 | 124.395 | 2 | 53.986103 | yes | 2.76E-07 | 603.76013 | 184.55223 |
| LCHEGDCGPCSR | 12 | Q12986 | Transcriptional repressor NF-X1 OS=Hor | NFX1 | 124.395 | 2 | 53.986103 | yes | 1.03E-07 | 724.27917 | 183.92293 |
| KATQFVYSYGR | 11 | Q12986 | Transcriptional repressor NF-X1 OS=Hor | NFX1 | 124.395 | 2 | 53.986103 | yes | 9.722E-08 | 660.34076 | 179.10292 |
| SEDATFMCDKR | 11 | Q12986 | Transcriptional repressor NF-X1 OS=Hor | NFX1 | 124.395 | 2 | 53.986103 | yes | 9.722E-08 | 680.28705 | 157.25929 |
| NPGSSNLQK | 9 | Q12986 | Transcriptional repressor NF-X1 OS=Hor | NFX1 | 124.395 | 2 | 53.986103 | yes | 3.304E-07 | 472.74344 | 146.02791 |
| NSGLNCGTQR | 10 | Q12986 | Transcriptional repressor NF-X1 OS=Hor | NFX1 | 124.395 | 2 | 53.986103 | yes | 1.992E-07 | 553.75403 | 143.33014 |
| SPASQADGQSGWI | 13 | Q12986 | Transcriptional repressor NF-X1 OS=Hor | NFX1 | 124.395 | 2 | 53.986103 | yes | 9.722E-08 | 673.8078 | 135.97234 |
| CGNHTCSQVCHP( | 20 | Q12986 | Transcriptional repressor NF-X1 OS=Hor | NFX1 | 124.395 | 4 | 53.986103 | yes | 1.011E-07 | 628.50421 | 125.31121 |
| VELQCECGR | 9 | Q12986 | Transcriptional repressor NF-X1 OS=Hor | NFX1 | 124.395 | 2 | 53.986103 | yes | 9.722E-08 | 575.75262 | 120.58219 |
| KTCMDPVPSCGK | 12 | Q12986 | Transcriptional repressor NF-X1 OS=Hor | NFX1 | 124.395 | 2 | 53.986103 | yes | 5.326E-07 | 690.30945 | 110.77755 |

| | | | | | | | | | | | |
|---|---|---|---|---|---|---|---|---|---|---|---|
| LKCEWSNR | 8 | Q12986 | Transcriptional repressor NF-X1 OS=Hor | NFX1 | 124.395 | 2 | 53.986103 | yes | 9.722E-08 | 546.76636 | 108.61436 |
| KAQSLAEQTSDTA | 20 | Q12986 | Transcriptional repressor NF-X1 OS=Hor | NFX1 | 124.395 | 3 | 53.986103 | yes | 1.269E-07 | 694.00995 | 95.089493 |
| NYSSPPPCHLSR | 12 | Q12986 | Transcriptional repressor NF-X1 OS=Hor | NFX1 | 124.395 | 2 | 53.986103 | yes | 3.081E-07 | 707.83026 | 85.409821 |
| ADCGHPCMAPCH | 23 | Q12986 | Transcriptional repressor NF-X1 OS=Hor | NFX1 | 124.395 | 4 | 53.986103 | yes | 3.323E-07 | 651.01123 | 79.854912 |
| SQQTSFQSSPCNK | 13 | Q12986 | Transcriptional repressor NF-X1 OS=Hor | NFX1 | 124.395 | 2 | 53.986103 | yes | 1.183E-07 | 749.83319 | 66.022598 |
| NNMGPIPK | 8 | Q12986 | Transcriptional repressor NF-X1 OS=Hor | NFX1 | 124.395 | 2 | 53.986103 | yes | 1.905E-07 | 435.72873 | 50.297169 |
| SHSFPPMNR | 9 | Q12986 | Transcriptional repressor NF-X1 OS=Hor | NFX1 | 124.395 | 2 | 53.986103 | yes | 2.417E-05 | 536.7533 | 50.030792 |
| SNIPCHLVDISCGL | 26 | Q12986 | Transcriptional repressor NF-X1 OS=Hor | NFX1 | 124.395 | 4 | 53.986103 | yes | 3.398E-07 | 731.83905 | 39.992096 |
| VTAPVWSCQSCYI | 21 | Q12986 | Transcriptional repressor NF-X1 OS=Hor | NFX1 | 124.395 | 4 | 53.986103 | yes | 3.154E-06 | 652.30798 | 5.2511382 |
| NEIPHSCGEVCR | 12 | Q12986 | Transcriptional repressor NF-X1 OS=Hor | NFX1 | 124.395 | 2 | 53.986103 | yes | 6.113E-06 | 729.31665 | 0.30336 |
| FMDLYPCSASELK | 13 | Q12996 | Cleavage stimulation factor subunit 3 OS= | CSTF3 | 82.922 | 2 | 53.89546 | yes | 1.396E-07 | 780.8573 | 181.25497 |
| MAQAYDFALDK | 11 | Q12996 | Cleavage stimulation factor subunit 3 OS= | CSTF3 | 82.922 | 2 | 53.89546 | yes | 2.552E-07 | 636.80011 | 130.52696 |
| GVEAVGSYAENQI | 13 | Q12996 | Cleavage stimulation factor subunit 3 OS= | CSTF3 | 82.922 | 2 | 53.89546 | yes | 9.722E-08 | 690.33112 | 103.17329 |
| VLHIDLWK | 8 | Q12996 | Cleavage stimulation factor subunit 3 OS= | CSTF3 | 82.922 | 2 | 53.89546 | yes | 1.248E-07 | 512.30292 | 72.382599 |
| IPNTVEEAVR | 10 | Q12996 | Cleavage stimulation factor subunit 3 OS= | CSTF3 | 82.922 | 2 | 53.89546 | yes | 4.566E-06 | 564.3064 | 45.873203 |
| MMADEALGSGLV | 14 | Q13011 | Delta(3,5)-Delta(2,4)-dienoyl-CoA isome | ECH1 | 35.817 | 2 | 34.695194 | yes | 9.583E-07 | 718.84723 | 64.23114 |
| ELLESLPLSK | 10 | Q13042 | Cell division cycle protein 16 homolog O | CDC16 | 71.656 | 2 | 52.308395 | yes | 1.022E-07 | 564.82916 | 284.05923 |
| ANELFYLSHK | 10 | Q13042 | Cell division cycle protein 16 homolog O | CDC16 | 71.656 | 2 | 52.308395 | yes | 9.722E-08 | 611.31677 | 145.38998 |
| IYDALDNR | 8 | Q13042 | Cell division cycle protein 16 homolog O | CDC16 | 71.656 | 2 | 52.308395 | yes | 1.778E-07 | 490.24582 | 132.26711 |
| QTAEETGLTPLETS | 15 | Q13042 | Cell division cycle protein 16 homolog O | CDC16 | 71.656 | 2 | 52.308395 | yes | 9.722E-08 | 816.90741 | 102.19762 |
| NIISPPWDFR | 10 | Q13042 | Cell division cycle protein 16 homolog O | CDC16 | 71.656 | 2 | 52.308395 | yes | 5.47E-07 | 622.82477 | 56.816586 |
| DPSSDWEMSQSSI | 14 | Q13042 | Cell division cycle protein 16 homolog O | CDC16 | 71.656 | 2 | 52.308395 | yes | 2.679E-07 | 798.84595 | 42.325325 |
| WFLDALEK | 8 | Q13042 | Cell division cycle protein 16 homolog O | CDC16 | 71.656 | 2 | 52.308395 | yes | 7.277E-07 | 511.2713 | 27.621908 |
| GVDLSGNDFK | 10 | Q13045 | Protein flightless-1 homolog OS=Homo s | FLII | 144.752 | 2 | 55.136467 | yes | 3.713E-07 | 526.25635 | 733.47107 |
| LLQSLLDTR | 9 | Q13045 | Protein flightless-1 homolog OS=Homo s | FLII | 144.752 | 2 | 55.136467 | yes | 1.948E-07 | 529.81384 | 555.55615 |
| VPECLYTLPSLR | 12 | Q13045 | Protein flightless-1 homolog OS=Homo s | FLII | 144.752 | 2 | 55.136467 | yes | 9.722E-08 | 724.38434 | 377.24527 |
| NQLTSLPSAICK | 12 | Q13045 | Protein flightless-1 homolog OS=Homo s | FLII | 144.752 | 2 | 55.136467 | yes | 9.722E-08 | 666.35303 | 371.34866 |
| AVQGAQQPSLYQI | 14 | Q13045 | Protein flightless-1 homolog OS=Homo s | FLII | 144.752 | 2 | 55.136467 | yes | 2.634E-07 | 779.92047 | 295.61224 |
| LDFDGLPSGIGK | 12 | Q13045 | Protein flightless-1 homolog OS=Homo s | FLII | 144.752 | 2 | 55.136467 | yes | 8.427E-07 | 609.8219 | 239.75061 |
| SLEGTEAQVFK | 11 | Q13045 | Protein flightless-1 homolog OS=Homo s | FLII | 144.752 | 2 | 55.136467 | yes | 1.801E-07 | 604.81152 | 196.66014 |
| EGEEATAEAEEK | 12 | Q13045 | Protein flightless-1 homolog OS=Homo s | FLII | 144.752 | 2 | 55.136467 | yes | 9.722E-08 | 646.77808 | 185.53703 |
| FVFLLDR | 7 | Q13045 | Protein flightless-1 homolog OS=Homo s | FLII | 144.752 | 2 | 55.136467 | yes | 3.807E-07 | 455.26328 | 122.19708 |
| YWVPVEYEEEEK | 12 | Q13045 | Protein flightless-1 homolog OS=Homo s | FLII | 144.752 | 2 | 55.136467 | yes | 9.722E-08 | 800.36432 | 94.682434 |
| ADLTALFLPR | 10 | Q13045 | Protein flightless-1 homolog OS=Homo s | FLII | 144.752 | 2 | 55.136467 | yes | 1.653E-07 | 558.82422 | 88.531044 |
| ACQVYIQHMR | 10 | Q13045 | Protein flightless-1 homolog OS=Homo s | FLII | 144.752 | 2 | 55.136467 | yes | 1.363E-07 | 653.31317 | 75.114174 |
| NAEAVLQSPGLSG | 14 | Q13045 | Protein flightless-1 homolog OS=Homo s | FLII | 144.752 | 2 | 55.136467 | yes | 1.762E-07 | 685.86737 | 56.739616 |
| LAGASPATVAAAA | 20 | Q13045 | Protein flightless-1 homolog OS=Homo s | FLII | 144.752 | 2 | 55.136467 | yes | 2.4E-07 | 819.94409 | 50.223392 |
| CFHAWSAFCK | 10 | Q13045 | Protein flightless-1 homolog OS=Homo s | FLII | 144.752 | 2 | 55.136467 | yes | 1.152E-07 | 657.28137 | 34.782623 |
| KFESLFPGK | 9 | Q13045 | Protein flightless-1 homolog OS=Homo s | FLII | 144.752 | 2 | 55.136467 | yes | 2.523E-07 | 526.79242 | 27.313766 |

| | | | | | | | | | | |
|---|---|---|---|---|---|---|---|---|---|---|
| LEHLSVSHNNLTT | 25 | Q13045 | Protein flightless-1 homolog OS=Homo s | FLII | 144.752 | 4 | 55.136467 | yes | 3.979E-07 | 689.36877 | 23.7306 |
| ELTLQDVELLK | 11 | Q13049 | E3 ubiquitin-protein ligase TRIM32 OS= | TRIM32 | 71.989 | 2 | 54.333374 | yes | 9.722E-08 | 650.87158 | 67.131516 |
| EMDMSPEEVVASI | 14 | Q13049 | E3 ubiquitin-protein ligase TRIM32 OS= | TRIM32 | 71.989 | 2 | 54.333374 | yes | 4.394E-07 | 788.85272 | 52.479019 |
| GPEAASNIQQCLFI | 15 | Q13049 | E3 ubiquitin-protein ligase TRIM32 OS= | TRIM32 | 71.989 | 2 | 54.333374 | yes | 4.881E-06 | 838.42725 | 51.352715 |
| GWAPFPVAPVPAF | 15 | Q13064 | Probable E3 ubiquitin-protein ligase mak | MKRN3 | 55.645 | 2 | 38.78602 | yes | 6.06E-07 | 807.94879 | 56.15057 |
| FCYYASR | 7 | Q13064 | Probable E3 ubiquitin-protein ligase mak | MKRN3 | 55.645 | 2 | 38.78602 | yes | 4.987E-07 | 483.71054 | 33.879169 |
| VWKPQLFER | 9 | Q13084 | Large ribosomal subunit protein bL28m C | MRPL28 | 30.157 | 2 | 47.408047 | yes | 5.393E-07 | 601.83765 | 333.56628 |
| VWKPQLFER | 9 | Q13084 | Large ribosomal subunit protein bL28m C | MRPL28 | 30.157 | 2 | 47.408047 | yes | 1.612E-07 | 401.56088 | 333.56628 |
| VWKPQLFER | 9 | Q13084 | Large ribosomal subunit protein bL28m C | MRPL28 | 30.157 | 3 | 47.408047 | yes | 5.393E-07 | 601.83765 | 333.56628 |
| VWKPQLFER | 9 | Q13084 | Large ribosomal subunit protein bL28m C | MRPL28 | 30.157 | 3 | 47.408047 | yes | 1.612E-07 | 401.56088 | 333.56628 |
| TPTPVHYR | 8 | Q13084 | Large ribosomal subunit protein bL28m C | MRPL28 | 30.157 | 2 | 47.408047 | yes | 3.503E-07 | 485.75888 | 197.73824 |
| LPGHYLR | 7 | Q13084 | Large ribosomal subunit protein bL28m C | MRPL28 | 30.157 | 2 | 47.408047 | yes | 1.337E-07 | 428.24542 | 173.17807 |
| EFYSEILDK | 9 | Q13084 | Large ribosomal subunit protein bL28m C | MRPL28 | 30.157 | 2 | 47.408047 | yes | 1.982E-07 | 572.28204 | 149.50218 |
| KVWKPQLFER | 10 | Q13084 | Large ribosomal subunit protein bL28m C | MRPL28 | 30.157 | 3 | 47.408047 | yes | 2.222E-07 | 444.25919 | 86.169785 |
| LLEEKDPVPLFK | 12 | Q13084 | Large ribosomal subunit protein bL28m C | MRPL28 | 30.157 | 2 | 47.408047 | yes | 5.319E-07 | 714.41089 | 76.037537 |
| QDPQLHPEDPER | 12 | Q13084 | Large ribosomal subunit protein bL28m C | MRPL28 | 30.157 | 3 | 47.408047 | yes | 3.967E-07 | 487.56366 | 32.905533 |
| QDPQLHPEDPER | 12 | Q13084 | Large ribosomal subunit protein bL28m C | MRPL28 | 30.157 | 3 | 47.408047 | yes | 4.79E-07 | 730.84186 | 32.905533 |
| QDPQLHPEDPER | 12 | Q13084 | Large ribosomal subunit protein bL28m C | MRPL28 | 30.157 | 2 | 47.408047 | yes | 3.967E-07 | 487.56366 | 32.905533 |
| QDPQLHPEDPER | 12 | Q13084 | Large ribosomal subunit protein bL28m C | MRPL28 | 30.157 | 2 | 47.408047 | yes | 4.79E-07 | 730.84186 | 32.905533 |
| FTVTVTMR | 8 | Q13084 | Large ribosomal subunit protein bL28m C | MRPL28 | 30.157 | 2 | 47.408047 | yes | 2.718E-07 | 477.75751 | 35.659882 |
| LGGIPVGVVAVET | 14 | Q13085 | Acetyl-CoA carboxylase 1 OS=Homo sap | ACACA | 265.554 | 2 | 47.490887 | yes | 1.318E-07 | 683.90625 | 106.83433 |
| FVVMVTPEDLK | 11 | Q13085 | Acetyl-CoA carboxylase 1 OS=Homo sap | ACACA | 265.554 | 2 | 47.490887 | no | 1.559E-07 | 639.34412 | 70.181641 |
| GSVLEPEGTVEIK | 13 | Q13085 | Acetyl-CoA carboxylase 1 OS=Homo sap | ACACA | 265.554 | 2 | 47.490887 | yes | 3.081E-07 | 679.36414 | 59.53587 |
| DFTVASPAEFVTR | 13 | Q13085 | Acetyl-CoA carboxylase 1 OS=Homo sap | ACACA | 265.554 | 2 | 47.490887 | no | 5.235E-07 | 720.36188 | 57.957897 |
| MAALEVYVR | 9 | Q13085 | Acetyl-CoA carboxylase 1 OS=Homo sap | ACACA | 265.554 | 2 | 47.490887 | yes | 2.154E-06 | 526.28387 | 33.776733 |
| IGLAEEIR | 8 | Q13085 | Acetyl-CoA carboxylase 1 OS=Homo sap | ACACA | 265.554 | 2 | 47.490887 | yes | 2.812E-06 | 450.76108 | 26.198212 |
| DIIVIGNDITYR | 12 | Q13085 | Acetyl-CoA carboxylase 1 OS=Homo sap | ACACA | 265.554 | 2 | 47.490887 | yes | 6.015E-06 | 696.38007 | 12.642496 |
| LFLELEPQVR | 10 | Q13098 | COP9 signalosome complex subunit 1 OS | GPS1 | 55.537 | 2 | 39.984978 | yes | 4.271E-07 | 622.3559 | 79.640274 |
| CAAGLAELAAR | 11 | Q13098 | COP9 signalosome complex subunit 1 OS | GPS1 | 55.537 | 2 | 39.984978 | yes | 6.79E-07 | 551.78729 | 50.48838 |
| LAFPGETLSDIPCK | 14 | Q13111 | Chromatin assembly factor 1 subunit A O | CHAF1A | 106.91 | 2 | 50.432503 | yes | 2.748E-07 | 774.39233 | 421.34964 |
| NADIFNSDVVIVEI | 14 | Q13111 | Chromatin assembly factor 1 subunit A O | CHAF1A | 106.91 | 2 | 50.432503 | yes | 9.809E-08 | 795.90973 | 190.45784 |
| EHMVLAPR | 8 | Q13111 | Chromatin assembly factor 1 subunit A O | CHAF1A | 106.91 | 2 | 50.432503 | yes | 1.614E-05 | 476.75528 | 158.93288 |
| LISENSVYEK | 10 | Q13111 | Chromatin assembly factor 1 subunit A O | CHAF1A | 106.91 | 2 | 50.432503 | yes | 1.816E-07 | 591.30603 | 148.06531 |
| TEEEGVGCGGAGF | 13 | Q13111 | Chromatin assembly factor 1 subunit A O | CHAF1A | 106.91 | 2 | 50.432503 | yes | 1.741E-07 | 639.77258 | 61.416721 |
| DRPAFPVK | 8 | Q13111 | Chromatin assembly factor 1 subunit A O | CHAF1A | 106.91 | 2 | 50.432503 | yes | 3.733E-07 | 465.26379 | 40.639149 |
| EQDSWSEAGGILF | 14 | Q13111 | Chromatin assembly factor 1 subunit A O | CHAF1A | 106.91 | 2 | 50.432503 | yes | 2.156E-07 | 783.87537 | 33.062977 |
| DELGIPLK | 8 | Q13112 | Chromatin assembly factor 1 subunit B O | CHAF1B | 61.493 | 2 | 51.40039 | yes | 1.43E-06 | 442.75803 | 1091.2668 |
| LASAGVDTNVR | 11 | Q13112 | Chromatin assembly factor 1 subunit B O | CHAF1B | 61.493 | 2 | 51.40039 | yes | 2.426E-07 | 551.7962 | 185.21045 |
| TLQPSSQNTK | 10 | Q13112 | Chromatin assembly factor 1 subunit B O | CHAF1B | 61.493 | 2 | 51.40039 | yes | 2.018E-06 | 552.28821 | 113.75372 |

| | | | | | | | | | | | |
|---|---|---|---|---|---|---|---|---|---|---|---|
| EPVYSLDFQHGTA | 15 | Q13112 | Chromatin assembly factor 1 subunit B O | CHAF1B | 61.493 | 2 | 51.40039 | yes | 2.396E-07 | 838.90497 | 109.26862 |
| VITCEIAWHNK | 11 | Q13112 | Chromatin assembly factor 1 subunit B O | CHAF1B | 61.493 | 2 | 51.40039 | yes | 9.722E-08 | 685.8479 | 107.37187 |
| MLSGIGAEGEAR | 12 | Q13112 | Chromatin assembly factor 1 subunit B O | CHAF1B | 61.493 | 2 | 51.40039 | yes | 9.862E-07 | 595.79535 | 90.984856 |
| RPIAHLPCPGK | 11 | Q13112 | Chromatin assembly factor 1 subunit B O | CHAF1B | 61.493 | 3 | 51.40039 | yes | 1.212E-06 | 415.901 | 25.888477 |
| RPIAHLPCPGK | 11 | Q13112 | Chromatin assembly factor 1 subunit B O | CHAF1B | 61.493 | 3 | 51.40039 | yes | 4.536E-06 | 623.3479 | 25.888477 |
| RPIAHLPCPGK | 11 | Q13112 | Chromatin assembly factor 1 subunit B O | CHAF1B | 61.493 | 2 | 51.40039 | yes | 1.212E-06 | 415.901 | 25.888477 |
| RPIAHLPCPGK | 11 | Q13112 | Chromatin assembly factor 1 subunit B O | CHAF1B | 61.493 | 2 | 51.40039 | yes | 4.536E-06 | 623.3479 | 25.888477 |
| DPPSITPAVK | 10 | Q13112 | Chromatin assembly factor 1 subunit B O | CHAF1B | 61.493 | 2 | 51.40039 | yes | 3.838E-07 | 512.78729 | 51.286064 |
| LSPIAPAPGFSPSA | 16 | Q13118 | Krueppel-like factor 10 OS=Homo sapien | KLF10 | 52.556 | 2 | 49.755863 | yes | 5.687E-07 | 755.91687 | 429.98834 |
| THLNVEAAR | 9 | Q13118 | Krueppel-like factor 10 OS=Homo sapien | KLF10 | 52.556 | 2 | 49.755863 | yes | 5.381E-07 | 505.77252 | 216.99792 |
| MEMISERPK | 9 | Q13118 | Krueppel-like factor 10 OS=Homo sapien | KLF10 | 52.556 | 2 | 49.755863 | yes | 3.011E-06 | 560.77808 | 154.09061 |
| VTPQIDSSR | 9 | Q13118 | Krueppel-like factor 10 OS=Homo sapien | KLF10 | 52.556 | 2 | 49.755863 | yes | 2.45E-07 | 501.76437 | 118.63293 |
| KLPNWQMEVSK | 11 | Q13118 | Krueppel-like factor 10 OS=Homo sapien | KLF10 | 52.556 | 2 | 49.755863 | yes | 4.665E-07 | 680.35809 | 116.06493 |
| AASILNYQNNSFR | 13 | Q13118 | Krueppel-like factor 10 OS=Homo sapien | KLF10 | 52.556 | 2 | 49.755863 | yes | 3.449E-07 | 749.37585 | 112.07354 |
| KFACPMCDR | 9 | Q13118 | Krueppel-like factor 10 OS=Homo sapien | KLF10 | 52.556 | 2 | 49.755863 | yes | 1.005E-07 | 592.75409 | 90.1381 |
| SQPAPVSPQQK | 11 | Q13118 | Krueppel-like factor 10 OS=Homo sapien | KLF10 | 52.556 | 2 | 49.755863 | yes | 1.229E-06 | 583.81183 | 76.268463 |
| NTVADVDEK | 9 | Q13118 | Krueppel-like factor 10 OS=Homo sapien | KLF10 | 52.556 | 2 | 49.755863 | yes | 4.377E-07 | 495.74054 | 50.012028 |
| FACPMCDR | 8 | Q13118 | Krueppel-like factor 10 OS=Homo sapien | KLF10 | 52.556 | 2 | 49.755863 | yes | 1.675E-07 | 528.7066 | 13.531666 |
| LTQILSYLR | 9 | Q13123 | Protein Red OS=Homo sapiens OX=9606 | IK | 65.603 | 2 | 52.315388 | yes | 1.376E-07 | 553.83203 | 1145.3585 |
| AVGPTAEADK | 10 | Q13123 | Protein Red OS=Homo sapiens OX=9606 | IK | 65.603 | 2 | 52.315388 | yes | 1.768E-07 | 479.74564 | 895.02576 |
| FAGSAGWEGTESL | 14 | Q13123 | Protein Red OS=Homo sapiens OX=9606 | IK | 65.603 | 2 | 52.315388 | yes | 1.022E-07 | 720.34369 | 690.05957 |
| NELFLPGR | 8 | Q13123 | Protein Red OS=Homo sapiens OX=9606 | IK | 65.603 | 2 | 52.315388 | yes | 3.401E-07 | 473.26126 | 615.09253 |
| GLDFALLQK | 9 | Q13123 | Protein Red OS=Homo sapiens OX=9606 | IK | 65.603 | 2 | 52.315388 | yes | 9.819E-08 | 502.79239 | 510.02548 |
| AAFQYGIK | 8 | Q13123 | Protein Red OS=Homo sapiens OX=9606 | IK | 65.603 | 2 | 52.315388 | yes | 1.856E-07 | 449.24509 | 266.38937 |
| VDDEPMDVDK | 10 | Q13123 | Protein Red OS=Homo sapiens OX=9606 | IK | 65.603 | 2 | 52.315388 | yes | 1.127E-07 | 581.75024 | 217.237 |
| HSYFEKPK | 8 | Q13123 | Protein Red OS=Homo sapiens OX=9606 | IK | 65.603 | 2 | 52.315388 | yes | 9.722E-08 | 518.26654 | 177.83485 |
| KMEADGVEVK | 10 | Q13123 | Protein Red OS=Homo sapiens OX=9606 | IK | 65.603 | 2 | 52.315388 | yes | 1.34E-07 | 553.28156 | 168.73048 |
| QLIQESK | 7 | Q13123 | Protein Red OS=Homo sapiens OX=9606 | IK | 65.603 | 2 | 52.315388 | yes | 2.912E-07 | 423.23999 | 154.42747 |
| KLLMTPR | 7 | Q13123 | Protein Red OS=Homo sapiens OX=9606 | IK | 65.603 | 2 | 52.315388 | yes | 4.63E-07 | 429.76514 | 107.16637 |
| DGVNKDYEETELI | 20 | Q13123 | Protein Red OS=Homo sapiens OX=9606 | IK | 65.603 | 3 | 52.315388 | yes | 1.127E-07 | 773.36426 | 100.52721 |
| LTNEDFR | 7 | Q13123 | Protein Red OS=Homo sapiens OX=9606 | IK | 65.603 | 2 | 52.315388 | yes | 3.001E-07 | 447.71942 | 87.135742 |
| FLGGDMEHTHLVI | 13 | Q13123 | Protein Red OS=Homo sapiens OX=9606 | IK | 65.603 | 2 | 52.315388 | yes | 1.031E-07 | 742.37177 | 78.949158 |
| EKEEEELMEKPQK | 13 | Q13123 | Protein Red OS=Homo sapiens OX=9606 | IK | 65.603 | 2 | 52.315388 | yes | 1.609E-07 | 823.90076 | 77.869347 |
| MEADGVEVK | 9 | Q13123 | Protein Red OS=Homo sapiens OX=9606 | IK | 65.603 | 2 | 52.315388 | yes | 6.063E-05 | 489.23404 | 44.392132 |
| VDDEPMDVDKGP | 16 | Q13123 | Protein Red OS=Homo sapiens OX=9606 | IK | 65.603 | 2 | 52.315388 | yes | 9.722E-08 | 845.38544 | 36.496513 |
| VPFLVAETPR | 10 | Q13131 | 5'-AMP-activated protein kinase catalytic | PRKAA1 | 64.01 | 2 | 54.7212 | yes | 9.722E-08 | 564.82422 | 281.93213 |
| HMLQVDPMK | 9 | Q13131 | 5'-AMP-activated protein kinase catalytic | PRKAA1 | 64.01 | 2 | 54.7212 | yes | 2.868E-06 | 549.77539 | 119.19482 |
| SIDDEITEAK | 10 | Q13131 | 5'-AMP-activated protein kinase catalytic | PRKAA1 | 64.01 | 2 | 54.7212 | yes | 1.385E-07 | 560.77203 | 94.60041 |
| KNPVTSTYSK | 10 | Q13131 | 5'-AMP-activated protein kinase catalytic | PRKAA1 | 64.01 | 2 | 54.7212 | yes | 2.337E-07 | 562.80096 | 53.29351 |

| | | | | | | | | | | | |
|---|---|---|---|---|---|---|---|---|---|---|---|
| PGSHTIEFFEMCAN | 17 | Q13131 | 5'-AMP-activated protein kinase catalytic | PRKAA1 | 64.01 | 3 | 54.7212 | yes | 1.36E-06 | 665.32184 | 34.886353 |
| HTLDELNPQK | 10 | Q13131 | 5'-AMP-activated protein kinase catalytic | PRKAA1 | 64.01 | 2 | 54.7212 | yes | 1.002E-07 | 597.80933 | 28.726025 |
| GLAAGSAETLPAN | 15 | Q13136 | Liprin-alpha-1 OS=Homo sapiens OX=96 | PPFIA1 | 135.779 | 2 | 57.615223 | yes | 9.722E-08 | 737.88605 | 382.10754 |
| VIELQEIISK | 10 | Q13136 | Liprin-alpha-1 OS=Homo sapiens OX=96 | PPFIA1 | 135.779 | 2 | 57.615223 | yes | 9.722E-08 | 586.35028 | 273.38602 |
| QLNTALPQEFAAL | 15 | Q13136 | Liprin-alpha-1 OS=Homo sapiens OX=96 | PPFIA1 | 135.779 | 2 | 57.615223 | yes | 1.777E-07 | 822.95142 | 155.0817 |
| SYFMECLVDAR | 11 | Q13136 | Liprin-alpha-1 OS=Homo sapiens OX=96 | PPFIA1 | 135.779 | 2 | 57.615223 | no | 1.003E-07 | 695.81012 | 147.33414 |
| VQTLNEQDWER | 11 | Q13136 | Liprin-alpha-1 OS=Homo sapiens OX=96 | PPFIA1 | 135.779 | 2 | 57.615223 | yes | 1.389E-07 | 709.33893 | 121.90079 |
| SGAIMSALSDTEIQ | 15 | Q13136 | Liprin-alpha-1 OS=Homo sapiens OX=96 | PPFIA1 | 135.779 | 2 | 57.615223 | no | 2.138E-07 | 789.89325 | 113.39423 |
| DVLVWSNDR | 9 | Q13136 | Liprin-alpha-1 OS=Homo sapiens OX=96 | PPFIA1 | 135.779 | 2 | 57.615223 | no | 1.692E-07 | 552.27765 | 107.2394 |
| EIGISNPLHR | 10 | Q13136 | Liprin-alpha-1 OS=Homo sapiens OX=96 | PPFIA1 | 135.779 | 2 | 57.615223 | no | 9.838E-08 | 568.31458 | 104.52737 |
| LGVMTLLPPSR | 11 | Q13136 | Liprin-alpha-1 OS=Homo sapiens OX=96 | PPFIA1 | 135.779 | 2 | 57.615223 | yes | 2.441E-06 | 592.34698 | 104.34021 |
| EFNNLLVMGTDR | 12 | Q13136 | Liprin-alpha-1 OS=Homo sapiens OX=96 | PPFIA1 | 135.779 | 2 | 57.615223 | yes | 4.098E-07 | 704.84808 | 92.806114 |
| GALHTVSHEDIR | 12 | Q13136 | Liprin-alpha-1 OS=Homo sapiens OX=96 | PPFIA1 | 135.779 | 2 | 57.615223 | yes | 1.093E-07 | 667.84418 | 90.128342 |
| VTSSMSSPSMQPK | 13 | Q13136 | Liprin-alpha-1 OS=Homo sapiens OX=96 | PPFIA1 | 135.779 | 2 | 57.615223 | yes | 1.506E-07 | 683.82068 | 82.156937 |
| KHELLEEAR | 9 | Q13136 | Liprin-alpha-1 OS=Homo sapiens OX=96 | PPFIA1 | 135.779 | 2 | 57.615223 | no | 1.261E-07 | 562.80658 | 74.915787 |
| CSLLEEELGATHK | 13 | Q13136 | Liprin-alpha-1 OS=Homo sapiens OX=96 | PPFIA1 | 135.779 | 2 | 57.615223 | yes | 9.722E-08 | 743.86395 | 73.26461 |
| MQMDGNVSGTQR | 12 | Q13136 | Liprin-alpha-1 OS=Homo sapiens OX=96 | PPFIA1 | 135.779 | 2 | 57.615223 | yes | 9.722E-08 | 662.29266 | 61.257725 |
| VGSGSLDNLGR | 11 | Q13136 | Liprin-alpha-1 OS=Homo sapiens OX=96 | PPFIA1 | 135.779 | 2 | 57.615223 | yes | 9.389E-07 | 537.78052 | 57.649323 |
| AELDHMR | 7 | Q13136 | Liprin-alpha-1 OS=Homo sapiens OX=96 | PPFIA1 | 135.779 | 2 | 57.615223 | yes | 1.577E-07 | 436.20816 | 57.151997 |
| QAQSPAGVSSEVE | 16 | Q13136 | Liprin-alpha-1 OS=Homo sapiens OX=96 | PPFIA1 | 135.779 | 2 | 57.615223 | yes | 7.125E-07 | 814.92816 | 16.01297 |
| LSENEIICNALQR | 13 | Q13137 | Calcium-binding and coiled-coil domain- | CALCOCO | 52.254 | 2 | 54.127773 | yes | 1E-07 | 780.39594 | 261.78857 |
| VDQLQAQLSTQEK | 13 | Q13137 | Calcium-binding and coiled-coil domain- | CALCOCO | 52.254 | 2 | 54.127773 | yes | 9.722E-08 | 744.38867 | 152.65219 |
| ENDHLFLSLTEQR | 13 | Q13137 | Calcium-binding and coiled-coil domain- | CALCOCO | 52.254 | 2 | 54.127773 | yes | 1.022E-07 | 801.39954 | 72.31929 |
| QQELMDENFDLSK | 13 | Q13137 | Calcium-binding and coiled-coil domain- | CALCOCO | 52.254 | 2 | 54.127773 | yes | 9.722E-08 | 798.86414 | 46.724854 |
| KQEELETLQSINK | 13 | Q13137 | Calcium-binding and coiled-coil domain- | CALCOCO | 52.254 | 2 | 54.127773 | yes | 2.761E-06 | 780.41742 | 23.458725 |
| VFQNEVLGTLQR | 12 | Q13144 | Translation initiation factor eIF-2B subun | EIF2B5 | 80.38 | 2 | 48.110863 | yes | 2.169E-07 | 702.38568 | 113.28191 |
| GYNPAEVGAAGK | 12 | Q13144 | Translation initiation factor eIF-2B subun | EIF2B5 | 80.38 | 2 | 48.110863 | yes | 2.253E-06 | 567.2829 | 96.804733 |
| AAPVVAPPGVVVS | 14 | Q13144 | Translation initiation factor eIF-2B subun | EIF2B5 | 80.38 | 2 | 48.110863 | yes | 1.197E-07 | 680.901 | 64.549995 |
| PTSLNVVR | 8 | Q13144 | Translation initiation factor eIF-2B subun | EIF2B5 | 80.38 | 2 | 48.110863 | yes | 2.024E-07 | 443.26126 | 57.582371 |
| YAYNISLK | 8 | Q13144 | Translation initiation factor eIF-2B subun | EIF2B5 | 80.38 | 2 | 48.110863 | yes | 2.947E-07 | 486.26346 | 41.605827 |
| FTEYETQVK | 9 | Q13148 | TAR DNA-binding protein 43 OS=Homo | TARDBP | 44.74 | 2 | 52.327873 | yes | 1.459E-07 | 572.77966 | 2029.8872 |
| GISVHISNAEPK | 12 | Q13148 | TAR DNA-binding protein 43 OS=Homo | TARDBP | 44.74 | 2 | 52.327873 | yes | 9.722E-08 | 626.33826 | 692.67871 |
| GISVHISNAEPK | 12 | Q13148 | TAR DNA-binding protein 43 OS=Homo | TARDBP | 44.74 | 2 | 52.327873 | yes | 9.778E-08 | 417.89456 | 692.67871 |
| GISVHISNAEPK | 12 | Q13148 | TAR DNA-binding protein 43 OS=Homo | TARDBP | 44.74 | 3 | 52.327873 | yes | 9.722E-08 | 626.33826 | 692.67871 |
| GISVHISNAEPK | 12 | Q13148 | TAR DNA-binding protein 43 OS=Homo | TARDBP | 44.74 | 3 | 52.327873 | yes | 9.778E-08 | 417.89456 | 692.67871 |
| CTEDMTEDELR | 11 | Q13148 | TAR DNA-binding protein 43 OS=Homo | TARDBP | 44.74 | 2 | 52.327873 | yes | 9.722E-08 | 699.77924 | 1056.9091 |
| QSQDEPLR | 8 | Q13148 | TAR DNA-binding protein 43 OS=Homo | TARDBP | 44.74 | 2 | 52.327873 | yes | 1.188E-07 | 486.74088 | 491.14117 |
| NPVSQCMR | 8 | Q13148 | TAR DNA-binding protein 43 OS=Homo | TARDBP | 44.74 | 2 | 52.327873 | yes | 1.004E-07 | 496.22604 | 418.33966 |
| KMDETDASSAVK | 12 | Q13148 | TAR DNA-binding protein 43 OS=Homo | TARDBP | 44.74 | 2 | 52.327873 | yes | 9.722E-08 | 641.30322 | 376.71341 |

| | | | | | | | | | | | |
|---|---|---|---|---|---|---|---|---|---|---|---|
| MDETDASSAVK | 11 | Q13148 | TAR DNA-binding protein 43 OS=Homo | TARDBP | 44.74 | 2 | 52.327873 | yes | 9.722E-08 | 577.25574 | 147.41386 |
| LFIGGLNVQTSESC | 16 | Q13151 | Heterogeneous nuclear ribonucleoprotein | HNRNPA0 | 30.841 | 2 | 46.857925 | yes | 1.456E-07 | 845.95978 | 15756.626 |
| MENSQLCK | 8 | Q13151 | Heterogeneous nuclear ribonucleoprotein | HNRNPA0 | 30.841 | 2 | 46.857925 | yes | 5.148E-07 | 526.23096 | 664.72937 |
| RGFGFVYFQNHDA | 16 | Q13151 | Heterogeneous nuclear ribonucleoprotein | HNRNPA0 | 30.841 | 3 | 46.857925 | yes | 1.873E-07 | 624.63232 | 474.1785 |
| GHFEAFGTLTDCV | 21 | Q13151 | Heterogeneous nuclear ribonucleoprotein | HNRNPA0 | 30.841 | 3 | 46.857925 | yes | 2.972E-07 | 773.71942 | 131.22142 |
| GDVAEGDLIEHFS | 20 | Q13151 | Heterogeneous nuclear ribonucleoprotein | HNRNPA0 | 30.841 | 3 | 46.857925 | yes | 6.847E-07 | 726.68317 | 102.57954 |
| SCENLAPFNTALK | 13 | Q13155 | Aminoacyl tRNA synthase complex-intera | AIMP2 | 35.349 | 2 | 50.333664 | yes | 1.283E-07 | 732.86121 | 549.84387 |
| LLCEHFR | 7 | Q13155 | Aminoacyl tRNA synthase complex-intera | AIMP2 | 35.349 | 2 | 50.333664 | yes | 1.147E-07 | 487.74747 | 542.88501 |
| VLSTVHTHSSVK | 12 | Q13155 | Aminoacyl tRNA synthase complex-intera | AIMP2 | 35.349 | 2 | 50.333664 | yes | 9.819E-08 | 647.85931 | 149.99699 |
| CFGEQNK | 7 | Q13155 | Aminoacyl tRNA synthase complex-intera | AIMP2 | 35.349 | 2 | 50.333664 | yes | 1.003E-07 | 441.69235 | 49.907173 |
| QITLNDLPVGR | 11 | Q13162 | Peroxiredoxin-4 OS=Homo sapiens OX=9 | PRDX4 | 30.54 | 2 | 54.09539 | yes | 9.722E-08 | 613.34857 | 172.46486 |
| IPLLSDLTHQISK | 13 | Q13162 | Peroxiredoxin-4 OS=Homo sapiens OX=9 | PRDX4 | 30.54 | 2 | 54.09539 | yes | 1.32E-07 | 732.92468 | 113.14713 |
| DYGVYLEDSGHTI | 14 | Q13162 | Peroxiredoxin-4 OS=Homo sapiens OX=9 | PRDX4 | 30.54 | 2 | 54.09539 | yes | 1.034E-07 | 812.88373 | 101.77184 |
| LVQAFQYTDK | 10 | Q13162 | Peroxiredoxin-4 OS=Homo sapiens OX=9 | PRDX4 | 30.54 | 2 | 54.09539 | yes | 4.367E-06 | 606.81659 | 32.540871 |
| SLSDSESDDSK | 11 | Q13185 | Chromobox protein homolog 3 OS=Homo | CBX3 | 20.812 | 2 | 53.200565 | yes | 9.722E-08 | 585.24384 | 414.36194 |
| KVEEAEPEEFVVE | 14 | Q13185 | Chromobox protein homolog 3 OS=Homo | CBX3 | 20.812 | 2 | 53.200565 | yes | 1.045E-07 | 831.41705 | 327.21777 |
| VEEAEPEEFVVEK | 13 | Q13185 | Chromobox protein homolog 3 OS=Homo | CBX3 | 20.812 | 2 | 53.200565 | yes | 1.672E-07 | 767.36957 | 251.48804 |
| WKDSDEADLVLA | 13 | Q13185 | Chromobox protein homolog 3 OS=Homo | CBX3 | 20.812 | 2 | 53.200565 | yes | 9.722E-08 | 745.38031 | 243.745 |
| LTWHSCPEDEAQ | 12 | Q13185 | Chromobox protein homolog 3 OS=Homo | CBX3 | 20.812 | 2 | 53.200565 | yes | 1.018E-07 | 736.80914 | 178.53261 |
| DSDEADLVLAK | 11 | Q13185 | Chromobox protein homolog 3 OS=Homo | CBX3 | 20.812 | 2 | 53.200565 | yes | 9.722E-08 | 588.29315 | 126.7423 |
| LADFGVAGQLTDT | 16 | Q13188 | Serine/threonine-protein kinase 3 OS=Ho | STK3 | 56.302 | 2 | 52.144325 | no | 9.742E-08 | 819.41382 | 219.5694 |
| AIFMIPTNPPPTFR | 14 | Q13188 | Serine/threonine-protein kinase 3 OS=Ho | STK3 | 56.302 | 2 | 52.144325 | no | 9.728E-08 | 801.42908 | 199.56363 |
| AGNILLNTEGHAK | 13 | Q13188 | Serine/threonine-protein kinase 3 OS=Ho | STK3 | 56.302 | 2 | 52.144325 | no | 2.081E-07 | 669.36224 | 167.91853 |
| ALDPMMER | 8 | Q13188 | Serine/threonine-protein kinase 3 OS=Ho | STK3 | 56.302 | 2 | 52.144325 | yes | 3.209E-07 | 481.72534 | 93.902191 |
| LGEGSYGSVFK | 11 | Q13188 | Serine/threonine-protein kinase 3 OS=Ho | STK3 | 56.302 | 2 | 52.144325 | yes | 9.722E-08 | 572.28766 | 65.705742 |
| GLEYLHFMR | 9 | Q13188 | Serine/threonine-protein kinase 3 OS=Ho | STK3 | 56.302 | 2 | 52.144325 | no | 4.978E-07 | 583.2948 | 63.125668 |
| AVEIEELTYIIK | 12 | Q13190 | Syntaxin-5 OS=Homo sapiens OX=9606 | STX5 | 39.673 | 2 | 45.266438 | yes | 4.913E-07 | 710.90033 | 52.987076 |
| LASMSNDFK | 9 | Q13190 | Syntaxin-5 OS=Homo sapiens OX=9606 | STX5 | 39.673 | 2 | 45.266438 | yes | 1.903E-07 | 506.74203 | 27.519739 |
| DPPMPLEAWCPR | 12 | Q13191 | E3 ubiquitin-protein ligase CBL-B OS=H | CBLB | 109.45 | 2 | 36.995804 | yes | 6.402E-07 | 734.83923 | 42.717102 |
| FGGSGSQVDSAR | 12 | Q13200 | 26S proteasome non-ATPase regulatory s | PSMD2 | 100.2 | 2 | 45.042225 | yes | 1.983E-07 | 584.27307 | 68.272003 |
| EWQELDDAEK | 10 | Q13200 | 26S proteasome non-ATPase regulatory s | PSMD2 | 100.2 | 2 | 45.042225 | yes | 4.019E-07 | 631.7804 | 48.107635 |
| TITGFQTHTTPVLL | 19 | Q13200 | 26S proteasome non-ATPase regulatory s | PSMD2 | 100.2 | 3 | 45.042225 | yes | 7.791E-07 | 693.70422 | 39.52853 |
| MLVTFDEELRPLP | 17 | Q13200 | 26S proteasome non-ATPase regulatory s | PSMD2 | 100.2 | 3 | 45.042225 | yes | 7.991E-07 | 667.69946 | 31.588167 |
| AAVLFATDIAAR | 12 | Q13206 | Probable ATP-dependent RNA helicase D | DDX10 | 100.888 | 2 | 50.368668 | yes | 5.901E-07 | 609.8457 | 103.01666 |
| NPEYVWVHEK | 10 | Q13206 | Probable ATP-dependent RNA helicase D | DDX10 | 100.888 | 2 | 50.368668 | yes | 1.398E-07 | 650.81964 | 98.103264 |
| MEVYNEFVR | 9 | Q13206 | Probable ATP-dependent RNA helicase D | DDX10 | 100.888 | 2 | 50.368668 | yes | 9.81E-08 | 593.78168 | 93.310471 |
| QTLLFSATQTK | 11 | Q13206 | Probable ATP-dependent RNA helicase D | DDX10 | 100.888 | 2 | 50.368668 | yes | 1.728E-07 | 619.34296 | 87.511353 |
| ELAYQTFEVLR | 11 | Q13206 | Probable ATP-dependent RNA helicase D | DDX10 | 100.888 | 2 | 50.368668 | yes | 9.819E-08 | 684.86151 | 87.014709 |
| AMVQQLLQK | 9 | Q13206 | Probable ATP-dependent RNA helicase D | DDX10 | 100.888 | 2 | 50.368668 | yes | 5.103E-07 | 529.80499 | 81.2658 |

| | | | | | | | | | | | |
|---|---|---|---|---|---|---|---|---|---|---|---|
| INNINILVCTPGR | 13 | Q13206 | Probable ATP-dependent RNA helicase D | DDX10 | 100.888 | 2 | 50.368668 | yes | 3.944E-07 | 742.40613 | 56.508816 |
| APSLTNDEVEEFR | 13 | Q13206 | Probable ATP-dependent RNA helicase D | DDX10 | 100.888 | 2 | 50.368668 | yes | 1.929E-07 | 753.85718 | 53.676941 |
| GSQAPSLPNTSEA( | 15 | Q13206 | Probable ATP-dependent RNA helicase D | DDX10 | 100.888 | 2 | 50.368668 | yes | 2.072E-06 | 757.87592 | 52.951233 |
| LRPGVSILALHGR | 13 | Q13206 | Probable ATP-dependent RNA helicase D | DDX10 | 100.888 | 3 | 50.368668 | yes | 5.483E-07 | 463.621 | 47.308037 |
| HNVFGLDLK | 9 | Q13206 | Probable ATP-dependent RNA helicase D | DDX10 | 100.888 | 2 | 50.368668 | yes | 4.499E-06 | 521.78766 | 46.984283 |
| SIVFFSSCK | 9 | Q13206 | Probable ATP-dependent RNA helicase D | DDX10 | 100.888 | 2 | 50.368668 | yes | 3.294E-06 | 537.76807 | 42.866371 |
| QTIGLALQGK | 10 | Q13206 | Probable ATP-dependent RNA helicase D | DDX10 | 100.888 | 2 | 50.368668 | yes | 2.698E-07 | 514.80859 | 34.143089 |
| ISVLYSFLR | 9 | Q13206 | Probable ATP-dependent RNA helicase D | DDX10 | 100.888 | 2 | 50.368668 | yes | 4.325E-07 | 549.32129 | 29.714157 |
| KLESILAQDQDLK | 13 | Q13206 | Probable ATP-dependent RNA helicase D | DDX10 | 100.888 | 2 | 50.368668 | yes | 9.839E-08 | 750.91705 | 17.853668 |
| DFPDDVIQFAR | 11 | Q13214 | Semaphorin-3B OS=Homo sapiens OX=9 | SEMA3B | 83.122 | 2 | 48.061604 | yes | 9.194E-06 | 661.82239 | 336.18024 |
| THLLACGTGAFHP | 23 | Q13214 | Semaphorin-3B OS=Homo sapiens OX=9 | SEMA3B | 83.122 | 4 | 48.061604 | yes | 3.103E-07 | 635.30865 | 128.18027 |
| SAVAQIALHR | 10 | Q13214 | Semaphorin-3B OS=Homo sapiens OX=9 | SEMA3B | 83.122 | 2 | 48.061604 | yes | 3.309E-07 | 533.31183 | 123.98534 |
| TCCYQALLVDEER | 13 | Q13214 | Semaphorin-3B OS=Homo sapiens OX=9 | SEMA3B | 83.122 | 2 | 48.061604 | yes | 1.393E-07 | 828.87146 | 122.04281 |
| ETAVEAAPALGR | 12 | Q13214 | Semaphorin-3B OS=Homo sapiens OX=9 | SEMA3B | 83.122 | 2 | 48.061604 | yes | 1.97E-07 | 592.81714 | 105.6619 |
| VCTECCLAR | 9 | Q13214 | Semaphorin-3B OS=Homo sapiens OX=9 | SEMA3B | 83.122 | 2 | 48.061604 | yes | 1.819E-07 | 584.74902 | 94.305428 |
| LAWPAPVEWR | 10 | Q13214 | Semaphorin-3B OS=Homo sapiens OX=9 | SEMA3B | 83.122 | 2 | 48.061604 | yes | 2.534E-07 | 612.82983 | 92.403603 |
| DIGTECMNFVK | 11 | Q13214 | Semaphorin-3B OS=Homo sapiens OX=9 | SEMA3B | 83.122 | 2 | 48.061604 | yes | 1.488E-06 | 657.29688 | 59.672897 |
| DPYCAWDGVACT | 13 | Q13214 | Semaphorin-3B OS=Homo sapiens OX=9 | SEMA3B | 83.122 | 2 | 48.061604 | yes | 1.206E-07 | 785.82428 | 58.576588 |
| AEEAAPAAPPGPK | 13 | Q13214 | Semaphorin-3B OS=Homo sapiens OX=9 | SEMA3B | 83.122 | 2 | 48.061604 | yes | 2.47E-07 | 603.31171 | 58.360294 |
| VAAADGHYDVLF | 22 | Q13214 | Semaphorin-3B OS=Homo sapiens OX=9 | SEMA3B | 83.122 | 3 | 48.061604 | yes | 3.604E-07 | 754.39862 | 28.708969 |
| IYVGNLPTDVR | 11 | Q13242 | Serine/arginine-rich splicing factor 9 OS= | SRSF9 | 25.543 | 2 | 50.778442 | yes | 1.219E-07 | 623.84314 | 5846.8569 |
| EAGDVCYADVQK | 12 | Q13242 | Serine/arginine-rich splicing factor 9 OS= | SRSF9 | 25.543 | 2 | 50.778442 | yes | 1.021E-07 | 677.80084 | 3384.3787 |
| DGVGMVEYLR | 10 | Q13242 | Serine/arginine-rich splicing factor 9 OS= | SRSF9 | 25.543 | 2 | 50.778442 | yes | 1.855E-07 | 569.78168 | 2902.7256 |
| KEDMEYALR | 9 | Q13242 | Serine/arginine-rich splicing factor 9 OS= | SRSF9 | 25.543 | 2 | 50.778442 | yes | 1.088E-07 | 577.77917 | 1538.429 |
| SHEGETSYIR | 10 | Q13242 | Serine/arginine-rich splicing factor 9 OS= | SRSF9 | 25.543 | 2 | 50.778442 | yes | 9.722E-08 | 589.77545 | 1055.7926 |
| VLVSGLPPSGSWQ | 16 | Q13242 | Serine/arginine-rich splicing factor 9 OS= | SRSF9 | 25.543 | 2 | 50.778442 | yes | 2.823E-06 | 841.95923 | 903.59674 |
| EKDLEDLFYK | 10 | Q13242 | Serine/arginine-rich splicing factor 9 OS= | SRSF9 | 25.543 | 2 | 50.778442 | yes | 1.162E-07 | 650.32697 | 757.8988 |
| GSPHYFSPFRPY | 12 | Q13242 | Serine/arginine-rich splicing factor 9 OS= | SRSF9 | 25.543 | 2 | 50.778442 | yes | 3.089E-07 | 727.84619 | 512.59412 |
| DLEDLFYK | 8 | Q13242 | Serine/arginine-rich splicing factor 9 OS= | SRSF9 | 25.543 | 2 | 50.778442 | yes | 5.642E-07 | 521.75824 | 83.350029 |
| STSYGYSR | 8 | Q13242 | Serine/arginine-rich splicing factor 9 OS= | SRSF9 | 25.543 | 2 | 50.778442 | yes | 4.751E-07 | 460.70905 | 45.717323 |
| VTIEHAR | 7 | Q13243 | Serine/arginine-rich splicing factor 5 OS= | SRSF5 | 31.264 | 2 | 51.983536 | yes | 9.927E-08 | 413.23251 | 3192.2783 |
| DADDAVYELDGK | 12 | Q13243 | Serine/arginine-rich splicing factor 5 OS= | SRSF5 | 31.264 | 2 | 51.983536 | yes | 9.722E-08 | 655.79095 | 2496.7332 |
| LNEGVVEFASYGI | 15 | Q13243 | Serine/arginine-rich splicing factor 5 OS= | SRSF5 | 31.264 | 2 | 51.983536 | yes | 1.102E-07 | 820.91193 | 1353.7172 |
| QAGEVTFADAHRI | 14 | Q13243 | Serine/arginine-rich splicing factor 5 OS= | SRSF5 | 31.264 | 2 | 51.983536 | yes | 9.722E-08 | 763.88916 | 214.82887 |
| QAGEVTFADAHRI | 14 | Q13243 | Serine/arginine-rich splicing factor 5 OS= | SRSF5 | 31.264 | 2 | 51.983536 | yes | 1.39E-07 | 509.59518 | 214.82887 |
| QAGEVTFADAHRI | 14 | Q13243 | Serine/arginine-rich splicing factor 5 OS= | SRSF5 | 31.264 | 3 | 51.983536 | yes | 9.722E-08 | 763.88916 | 214.82887 |
| QAGEVTFADAHRI | 14 | Q13243 | Serine/arginine-rich splicing factor 5 OS= | SRSF5 | 31.264 | 3 | 51.983536 | yes | 1.39E-07 | 509.59518 | 214.82887 |
| GFGFVEFEDPR | 11 | Q13243 | Serine/arginine-rich splicing factor 5 OS= | SRSF5 | 31.264 | 2 | 51.983536 | yes | 2.063E-07 | 650.30383 | 363.04651 |
| QAGEVTFADAHR | 12 | Q13243 | Serine/arginine-rich splicing factor 5 OS= | SRSF5 | 31.264 | 2 | 51.983536 | yes | 1.164E-07 | 651.31531 | 170.381 |

| | | | | | | | | | | | |
|---|---|---|---|---|---|---|---|---|---|---|---|
| VSWQDLK | 7 | Q13243 | Serine/arginine-rich splicing factor 5 OS= | SRSF5 | 31.264 | 2 | 51.983536 | yes | 1.313E-07 | 438.23471 | 64.097061 |
| LIVENLSSR | 9 | Q13247 | Serine/arginine-rich splicing factor 6 OS= | SRSF6 | 39.587 | 2 | 55.46661 | no | 1.584E-07 | 515.79822 | 15135.058 |
| TNEGVIEFR | 9 | Q13247 | Serine/arginine-rich splicing factor 6 OS= | SRSF6 | 39.587 | 2 | 55.46661 | yes | 2.033E-07 | 532.77222 | 2401.1699 |
| VIVEHAR | 7 | Q13247 | Serine/arginine-rich splicing factor 6 OS= | SRSF6 | 39.587 | 2 | 55.46661 | no | 6.127E-07 | 412.24286 | 2351.96 |
| LIEDKPR | 7 | Q13247 | Serine/arginine-rich splicing factor 6 OS= | SRSF6 | 39.587 | 2 | 55.46661 | yes | 1.407E-07 | 435.7558 | 2184.4861 |
| NGYGFVEFEDSR | 12 | Q13247 | Serine/arginine-rich splicing factor 6 OS= | SRSF6 | 39.587 | 2 | 55.46661 | yes | 1.26E-07 | 710.31238 | 1100.7462 |
| DKYGPPVR | 8 | Q13247 | Serine/arginine-rich splicing factor 6 OS= | SRSF6 | 39.587 | 2 | 55.46661 | yes | 9.722E-08 | 466.25345 | 605.23529 |
| DADDAVYELNGK | 12 | Q13247 | Serine/arginine-rich splicing factor 6 OS= | SRSF6 | 39.587 | 2 | 55.46661 | no | 9.809E-08 | 655.29895 | 494.77109 |
| QAGEVTYADAHK | 12 | Q13247 | Serine/arginine-rich splicing factor 6 OS= | SRSF6 | 39.587 | 2 | 55.46661 | no | 3.609E-07 | 645.30969 | 241.36516 |
| QAGEVTYADAHK | 12 | Q13247 | Serine/arginine-rich splicing factor 6 OS= | SRSF6 | 39.587 | 2 | 55.46661 | no | 4.18E-07 | 430.54221 | 241.36516 |
| QAGEVTYADAHK | 12 | Q13247 | Serine/arginine-rich splicing factor 6 OS= | SRSF6 | 39.587 | 3 | 55.46661 | no | 3.609E-07 | 645.30969 | 241.36516 |
| QAGEVTYADAHK | 12 | Q13247 | Serine/arginine-rich splicing factor 6 OS= | SRSF6 | 39.587 | 3 | 55.46661 | no | 4.18E-07 | 430.54221 | 241.36516 |
| CSWQDLK | 7 | Q13247 | Serine/arginine-rich splicing factor 6 OS= | SRSF6 | 39.587 | 2 | 55.46661 | no | 1.017E-07 | 468.71582 | 429.54974 |
| SNSPLPVPPSK | 11 | Q13247 | Serine/arginine-rich splicing factor 6 OS= | SRSF6 | 39.587 | 2 | 55.46661 | yes | 2.883E-07 | 561.81134 | 108.1767 |
| DGYSYGSR | 8 | Q13247 | Serine/arginine-rich splicing factor 6 OS= | SRSF6 | 39.587 | 2 | 55.46661 | yes | 1.002E-06 | 452.69342 | 80.133286 |
| ALDKLDGTEINGR | 13 | Q13247 | Serine/arginine-rich splicing factor 6 OS= | SRSF6 | 39.587 | 2 | 55.46661 | yes | 9.722E-08 | 701.37024 | 77.065834 |
| LDLDLTADSQPPV | 15 | Q13263 | Transcription intermediary factor 1-beta ( | TRIM28 | 88.55 | 2 | 53.022953 | yes | 9.908E-08 | 829.93542 | 1183.6815 |
| MIVDPVEPHGEMI | 13 | Q13263 | Transcription intermediary factor 1-beta ( | TRIM28 | 88.55 | 2 | 53.022953 | yes | 9.722E-08 | 741.35999 | 379.89008 |
| MIVDPVEPHGEMI | 13 | Q13263 | Transcription intermediary factor 1-beta ( | TRIM28 | 88.55 | 2 | 53.022953 | yes | 1.3E-07 | 494.57574 | 379.89008 |
| MIVDPVEPHGEMI | 13 | Q13263 | Transcription intermediary factor 1-beta ( | TRIM28 | 88.55 | 3 | 53.022953 | yes | 9.722E-08 | 741.35999 | 379.89008 |
| MIVDPVEPHGEMI | 13 | Q13263 | Transcription intermediary factor 1-beta ( | TRIM28 | 88.55 | 3 | 53.022953 | yes | 1.3E-07 | 494.57574 | 379.89008 |
| ADVQSIIGLQR | 11 | Q13263 | Transcription intermediary factor 1-beta ( | TRIM28 | 88.55 | 2 | 53.022953 | yes | 9.862E-08 | 600.34076 | 697.24854 |
| AASAAAASAAAAS | 30 | Q13263 | Transcription intermediary factor 1-beta ( | TRIM28 | 88.55 | 3 | 53.022953 | yes | 1.432E-07 | 810.37738 | 665.48444 |
| DIVENYFMR | 9 | Q13263 | Transcription intermediary factor 1-beta ( | TRIM28 | 88.55 | 2 | 53.022953 | yes | 5.058E-07 | 593.78168 | 523.42114 |
| TVYCNVHK | 8 | Q13263 | Transcription intermediary factor 1-beta ( | TRIM28 | 88.55 | 2 | 53.022953 | yes | 2.069E-07 | 510.75021 | 508.16379 |
| SGEGEVSGLMR | 11 | Q13263 | Transcription intermediary factor 1-beta ( | TRIM28 | 88.55 | 2 | 53.022953 | yes | 9.722E-08 | 561.26642 | 433.81708 |
| VLVNDAQK | 8 | Q13263 | Transcription intermediary factor 1-beta ( | TRIM28 | 88.55 | 2 | 53.022953 | yes | 4.089E-07 | 443.75327 | 314.21307 |
| MNEAFGDTK | 9 | Q13263 | Transcription intermediary factor 1-beta ( | TRIM28 | 88.55 | 2 | 53.022953 | yes | 9.722E-08 | 506.72385 | 241.12091 |
| FQWDLNAWTK | 10 | Q13263 | Transcription intermediary factor 1-beta ( | TRIM28 | 88.55 | 2 | 53.022953 | yes | 9.722E-08 | 654.8222 | 237.35739 |
| MAILQIMK | 8 | Q13263 | Transcription intermediary factor 1-beta ( | TRIM28 | 88.55 | 2 | 53.022953 | yes | 1.859E-07 | 474.27448 | 201.3042 |
| LIYFQLHR | 8 | Q13263 | Transcription intermediary factor 1-beta ( | TRIM28 | 88.55 | 2 | 53.022953 | yes | 1.065E-07 | 545.31384 | 196.47409 |
| DHQYQFLEDAVR | 12 | Q13263 | Transcription intermediary factor 1-beta ( | TRIM28 | 88.55 | 2 | 53.022953 | yes | 9.722E-08 | 760.86005 | 193.97499 |
| PGTNSTGPAPMAP | 15 | Q13263 | Transcription intermediary factor 1-beta ( | TRIM28 | 88.55 | 2 | 53.022953 | yes | 2.407E-07 | 725.85901 | 29.834639 |
| KLLASLVK | 8 | Q13263 | Transcription intermediary factor 1-beta ( | TRIM28 | 88.55 | 2 | 53.022953 | yes | 2.158E-06 | 436.3024 | 16.86112 |
| DYPDEVINFMR | 11 | Q13275 | Semaphorin-3F OS=Homo sapiens OX=9 | SEMA3F | 88.382 | 2 | 56.093685 | yes | 1.11E-07 | 699.82153 | 142.30313 |
| VQLHVLGR | 8 | Q13275 | Semaphorin-3F OS=Homo sapiens OX=9 | SEMA3F | 88.382 | 2 | 56.093685 | yes | 1.368E-07 | 461.28506 | 84.993179 |
| LTTIAVDQVDAAD | 15 | Q13275 | Semaphorin-3F OS=Homo sapiens OX=9 | SEMA3F | 88.382 | 2 | 56.093685 | yes | 9.722E-08 | 772.89935 | 74.64267 |
| TEQGLLLR | 8 | Q13275 | Semaphorin-3F OS=Homo sapiens OX=9 | SEMA3F | 88.382 | 2 | 56.093685 | yes | 1.857E-07 | 465.27435 | 70.295044 |
| FYVHNDIFR | 9 | Q13283 | Ras GTPase-activating protein-binding pr | G3BP1 | 52.165 | 2 | 51.26468 | yes | 1.225E-07 | 605.80383 | 43052.684 |

| | | | | | | | | | | | |
|---|---|---|---|---|---|---|---|---|---|---|---|
| FYVHNDIFR | 9 | Q13283 | Ras GTPase-activating protein-binding pr | G3BP1 | 52.165 | 2 | 51.26468 | yes | 1.981E-07 | 404.20499 | 43052.684 |
| FYVHNDIFR | 9 | Q13283 | Ras GTPase-activating protein-binding pr | G3BP1 | 52.165 | 3 | 51.26468 | yes | 1.225E-07 | 605.80383 | 43052.684 |
| FYVHNDIFR | 9 | Q13283 | Ras GTPase-activating protein-binding pr | G3BP1 | 52.165 | 3 | 51.26468 | yes | 1.981E-07 | 404.20499 | 43052.684 |
| HPDSHQLFIGNLPI | 18 | Q13283 | Ras GTPase-activating protein-binding pr | G3BP1 | 52.165 | 3 | 51.26468 | yes | 9.722E-08 | 695.01672 | 21552.676 |
| HPDSHQLFIGNLPI | 18 | Q13283 | Ras GTPase-activating protein-binding pr | G3BP1 | 52.165 | 3 | 51.26468 | yes | 2.693E-07 | 521.51434 | 21552.676 |
| HPDSHQLFIGNLPI | 18 | Q13283 | Ras GTPase-activating protein-binding pr | G3BP1 | 52.165 | 4 | 51.26468 | yes | 9.722E-08 | 695.01672 | 21552.676 |
| HPDSHQLFIGNLPI | 18 | Q13283 | Ras GTPase-activating protein-binding pr | G3BP1 | 52.165 | 4 | 51.26468 | yes | 2.693E-07 | 521.51434 | 21552.676 |
| EAGEQGDIEPR | 11 | Q13283 | Ras GTPase-activating protein-binding pr | G3BP1 | 52.165 | 2 | 51.26468 | yes | 1.369E-07 | 600.7782 | 38776.211 |
| TFSWASVTSK | 10 | Q13283 | Ras GTPase-activating protein-binding pr | G3BP1 | 52.165 | 2 | 51.26468 | yes | 1.456E-07 | 557.28241 | 35919.813 |
| QYYTLLNQAPDMI | 15 | Q13283 | Ras GTPase-activating protein-binding pr | G3BP1 | 52.165 | 3 | 51.26468 | yes | 1.209E-07 | 621.64545 | 17501.391 |
| QYYTLLNQAPDMI | 15 | Q13283 | Ras GTPase-activating protein-binding pr | G3BP1 | 52.165 | 3 | 51.26468 | yes | 1.393E-07 | 626.97711 | 17501.391 |
| VMSQNFTNCHTK | 12 | Q13283 | Ras GTPase-activating protein-binding pr | G3BP1 | 52.165 | 2 | 51.26468 | yes | 5.79E-07 | 733.82941 | 8044.7036 |
| VMSQNFTNCHTK | 12 | Q13283 | Ras GTPase-activating protein-binding pr | G3BP1 | 52.165 | 2 | 51.26468 | yes | 1.03E-07 | 741.82684 | 8044.7036 |
| VLSNRPIMFR | 10 | Q13283 | Ras GTPase-activating protein-binding pr | G3BP1 | 52.165 | 2 | 51.26468 | yes | 1.332E-07 | 616.85022 | 6053.8906 |
| VLSNRPIMFR | 10 | Q13283 | Ras GTPase-activating protein-binding pr | G3BP1 | 52.165 | 2 | 51.26468 | yes | 1.893E-07 | 411.56924 | 6053.8906 |
| VLSNRPIMFR | 10 | Q13283 | Ras GTPase-activating protein-binding pr | G3BP1 | 52.165 | 3 | 51.26468 | yes | 1.332E-07 | 616.85022 | 6053.8906 |
| VLSNRPIMFR | 10 | Q13283 | Ras GTPase-activating protein-binding pr | G3BP1 | 52.165 | 3 | 51.26468 | yes | 1.893E-07 | 411.56924 | 6053.8906 |
| VPASQPRPESK | 11 | Q13283 | Ras GTPase-activating protein-binding pr | G3BP1 | 52.165 | 2 | 51.26468 | yes | 1.499E-07 | 598.32513 | 11006.474 |
| NLPPSGAVPVTGII | 19 | Q13283 | Ras GTPase-activating protein-binding pr | G3BP1 | 52.165 | 3 | 51.26468 | yes | 2.624E-07 | 627.03149 | 10548.027 |
| PADAVYGQK | 9 | Q13283 | Ras GTPase-activating protein-binding pr | G3BP1 | 52.165 | 2 | 51.26468 | yes | 7.21E-07 | 474.74289 | 7309.1685 |
| HVDAHATLNDGV | 28 | Q13283 | Ras GTPase-activating protein-binding pr | G3BP1 | 52.165 | 4 | 51.26468 | yes | 1.99E-07 | 747.13855 | 3632.0791 |
| HVDAHATLNDGV | 28 | Q13283 | Ras GTPase-activating protein-binding pr | G3BP1 | 52.165 | 4 | 51.26468 | yes | 9.727E-08 | 751.13727 | 3632.0791 |
| PESKPESQIPPQRP | 16 | Q13283 | Ras GTPase-activating protein-binding pr | G3BP1 | 52.165 | 4 | 51.26468 | yes | 1.301E-06 | 469.25244 | 1478.7894 |
| RFMQTFVLAPEGS | 17 | Q13283 | Ras GTPase-activating protein-binding pr | G3BP1 | 52.165 | 3 | 51.26468 | yes | 9.722E-08 | 632.33301 | 658.10889 |
| RFMQTFVLAPEGS | 17 | Q13283 | Ras GTPase-activating protein-binding pr | G3BP1 | 52.165 | 3 | 51.26468 | yes | 1.995E-06 | 637.66461 | 658.10889 |
| NSSYVHGGLDSNC | 14 | Q13283 | Ras GTPase-activating protein-binding pr | G3BP1 | 52.165 | 2 | 51.26468 | yes | 1.114E-07 | 717.83405 | 604.52295 |
| HPDSHQLFIGNLPI | 22 | Q13283 | Ras GTPase-activating protein-binding pr | G3BP1 | 52.165 | 4 | 51.26468 | yes | 5.505E-07 | 635.82776 | 198.57274 |
| HPDSHQLFIGNLPI | 22 | Q13283 | Ras GTPase-activating protein-binding pr | G3BP1 | 52.165 | 4 | 51.26468 | yes | 2.323E-07 | 847.43457 | 198.57274 |
| HPDSHQLFIGNLPI | 22 | Q13283 | Ras GTPase-activating protein-binding pr | G3BP1 | 52.165 | 4 | 51.26468 | yes | 1.877E-06 | 508.86368 | 198.57274 |
| HPDSHQLFIGNLPI | 22 | Q13283 | Ras GTPase-activating protein-binding pr | G3BP1 | 52.165 | 3 | 51.26468 | yes | 5.505E-07 | 635.82776 | 198.57274 |
| HPDSHQLFIGNLPI | 22 | Q13283 | Ras GTPase-activating protein-binding pr | G3BP1 | 52.165 | 3 | 51.26468 | yes | 2.323E-07 | 847.43457 | 198.57274 |
| HPDSHQLFIGNLPI | 22 | Q13283 | Ras GTPase-activating protein-binding pr | G3BP1 | 52.165 | 3 | 51.26468 | yes | 1.877E-06 | 508.86368 | 198.57274 |
| HPDSHQLFIGNLPI | 22 | Q13283 | Ras GTPase-activating protein-binding pr | G3BP1 | 52.165 | 5 | 51.26468 | yes | 5.505E-07 | 635.82776 | 198.57274 |
| HPDSHQLFIGNLPI | 22 | Q13283 | Ras GTPase-activating protein-binding pr | G3BP1 | 52.165 | 5 | 51.26468 | yes | 2.323E-07 | 847.43457 | 198.57274 |
| HPDSHQLFIGNLPI | 22 | Q13283 | Ras GTPase-activating protein-binding pr | G3BP1 | 52.165 | 5 | 51.26468 | yes | 1.877E-06 | 508.86368 | 198.57274 |
| INIPPQR | 7 | Q13283 | Ras GTPase-activating protein-binding pr | G3BP1 | 52.165 | 2 | 51.26468 | yes | 9.795E-08 | 419.2507 | 446.20117 |
| KVMSQNFTNCHTI | 13 | Q13283 | Ras GTPase-activating protein-binding pr | G3BP1 | 52.165 | 2 | 51.26468 | yes | 1.163E-07 | 797.87689 | 206.22937 |
| KVMSQNFTNCHTI | 13 | Q13283 | Ras GTPase-activating protein-binding pr | G3BP1 | 52.165 | 2 | 51.26468 | yes | 2.985E-07 | 805.87433 | 206.22937 |
| FMQTFVLAPEGSV | 16 | Q13283 | Ras GTPase-activating protein-binding pr | G3BP1 | 52.165 | 3 | 51.26468 | yes | 1.381E-07 | 580.29926 | 245.51874 |

| | | | | | | | | | | | |
|---|---|---|---|---|---|---|---|---|---|---|---|
| VPASQPRPESKPES | 20 | Q13283 | Ras GTPase-activating protein-binding pr | G3BP1 | 52.165 | 4 | 51.26468 | yes | 1.521E-07 | 557.79999 | 241.31154 |
| DFFQSYGNVVELR | 13 | Q13283 | Ras GTPase-activating protein-binding pr | G3BP1 | 52.165 | 2 | 51.26468 | yes | 1.227E-06 | 787.38593 | 119.40784 |
| PESQIPPQRPQR | 12 | Q13283 | Ras GTPase-activating protein-binding pr | G3BP1 | 52.165 | 2 | 51.26468 | yes | 4.513E-06 | 716.88641 | 109.10628 |
| GGMVQKPGFGVG | 13 | Q13283 | Ras GTPase-activating protein-binding pr | G3BP1 | 52.165 | 2 | 51.26468 | yes | 9.722E-08 | 645.34277 | 72.367859 |
| LPNFGFVVFDDSE | 17 | Q13283 | Ras GTPase-activating protein-binding pr | G3BP1 | 52.165 | 3 | 51.26468 | yes | 4.145E-07 | 646.65961 | 49.408192 |
| PESKPESQIPPQR | 13 | Q13283 | Ras GTPase-activating protein-binding pr | G3BP1 | 52.165 | 2 | 51.26468 | yes | 8.23E-06 | 746.89136 | 39.059505 |
| SELKDFFQSYGNV | 17 | Q13283 | Ras GTPase-activating protein-binding pr | G3BP1 | 52.165 | 3 | 51.26468 | yes | 1.014E-07 | 677.67755 | 37.881813 |
| LQNLSLEGLR | 10 | Q13309 | S-phase kinase-associated protein 2 OS=I | SKP2 | 47.761 | 2 | 44.85049 | yes | 2.046E-07 | 571.83002 | 70.377892 |
| FSPAGPVLSIR | 11 | Q13310 | Polyadenylate-binding protein 4 OS=Hon | PABPC4 | 70.783 | 2 | 53.658638 | yes | 1.063E-07 | 572.32965 | 93284.781 |
| EFSPFGSITSAK | 12 | Q13310 | Polyadenylate-binding protein 4 OS=Hon | PABPC4 | 70.783 | 2 | 53.658638 | yes | 1.54E-07 | 635.81934 | 65108.574 |
| NFGEEVDDESLK | 12 | Q13310 | Polyadenylate-binding protein 4 OS=Hon | PABPC4 | 70.783 | 2 | 53.658638 | yes | 9.722E-08 | 691.30951 | 27012.629 |
| GFGFVSYEK | 9 | Q13310 | Polyadenylate-binding protein 4 OS=Hon | PABPC4 | 70.783 | 2 | 53.658638 | yes | 2.589E-07 | 517.25311 | 23436.467 |
| SGVGNVFIK | 9 | Q13310 | Polyadenylate-binding protein 4 OS=Hon | PABPC4 | 70.783 | 2 | 53.658638 | no | 1.602E-07 | 460.76364 | 20728.521 |
| NLDDTIDDEK | 10 | Q13310 | Polyadenylate-binding protein 4 OS=Hon | PABPC4 | 70.783 | 2 | 53.658638 | no | 1.699E-07 | 589.26459 | 20063.139 |
| AHLTNQYMQR | 10 | Q13310 | Polyadenylate-binding protein 4 OS=Hon | PABPC4 | 70.783 | 2 | 53.658638 | yes | 1.149E-07 | 631.30896 | 7143.5176 |
| AHLTNQYMQR | 10 | Q13310 | Polyadenylate-binding protein 4 OS=Hon | PABPC4 | 70.783 | 2 | 53.658638 | yes | 9.796E-08 | 639.3064 | 7143.5176 |
| YQGVNLYIK | 9 | Q13310 | Polyadenylate-binding protein 4 OS=Hon | PABPC4 | 70.783 | 2 | 53.658638 | yes | 1.561E-07 | 549.3031 | 10963.883 |
| IVGSKPLYVALAQ | 14 | Q13310 | Polyadenylate-binding protein 4 OS=Hon | PABPC4 | 70.783 | 3 | 53.658638 | yes | 1.26E-07 | 505.63995 | 5174.6895 |
| IVGSKPLYVALAQ | 14 | Q13310 | Polyadenylate-binding protein 4 OS=Hon | PABPC4 | 70.783 | 3 | 53.658638 | yes | 3.95E-07 | 757.9563 | 5174.6895 |
| IVGSKPLYVALAQ | 14 | Q13310 | Polyadenylate-binding protein 4 OS=Hon | PABPC4 | 70.783 | 2 | 53.658638 | yes | 1.26E-07 | 505.63995 | 5174.6895 |
| IVGSKPLYVALAQ | 14 | Q13310 | Polyadenylate-binding protein 4 OS=Hon | PABPC4 | 70.783 | 2 | 53.658638 | yes | 3.95E-07 | 757.9563 | 5174.6895 |
| SKVDEAVAVLQAI | 16 | Q13310 | Polyadenylate-binding protein 4 OS=Hon | PABPC4 | 70.783 | 3 | 53.658638 | yes | 1.059E-07 | 568.31268 | 9605.2568 |
| KSGVGNVFIK | 10 | Q13310 | Polyadenylate-binding protein 4 OS=Hon | PABPC4 | 70.783 | 2 | 53.658638 | no | 1.592E-07 | 524.8111 | 7549.4995 |
| KEFSPFGSITSAK | 13 | Q13310 | Polyadenylate-binding protein 4 OS=Hon | PABPC4 | 70.783 | 2 | 53.658638 | yes | 1.022E-07 | 699.86682 | 3449.7695 |
| KEFSPFGSITSAK | 13 | Q13310 | Polyadenylate-binding protein 4 OS=Hon | PABPC4 | 70.783 | 2 | 53.658638 | yes | 8.739E-06 | 466.91364 | 3449.7695 |
| KEFSPFGSITSAK | 13 | Q13310 | Polyadenylate-binding protein 4 OS=Hon | PABPC4 | 70.783 | 3 | 53.658638 | yes | 1.022E-07 | 699.86682 | 3449.7695 |
| KEFSPFGSITSAK | 13 | Q13310 | Polyadenylate-binding protein 4 OS=Hon | PABPC4 | 70.783 | 3 | 53.658638 | yes | 8.739E-06 | 466.91364 | 3449.7695 |
| KAHLTNQYMQR | 11 | Q13310 | Polyadenylate-binding protein 4 OS=Hon | PABPC4 | 70.783 | 2 | 53.658638 | yes | 2.213E-07 | 695.35645 | 2276.9702 |
| KAHLTNQYMQR | 11 | Q13310 | Polyadenylate-binding protein 4 OS=Hon | PABPC4 | 70.783 | 2 | 53.658638 | yes | 9.722E-08 | 703.35388 | 2276.9702 |
| LFPLIQTMHSNLA( | 15 | Q13310 | Polyadenylate-binding protein 4 OS=Hon | PABPC4 | 70.783 | 2 | 53.658638 | yes | 2.176E-07 | 835.45837 | 2205.8765 |
| LFPLIQTMHSNLA( | 15 | Q13310 | Polyadenylate-binding protein 4 OS=Hon | PABPC4 | 70.783 | 2 | 53.658638 | yes | 1.673E-07 | 843.45581 | 2205.8765 |
| SPHPAIQPLQAPQI | 38 | Q13310 | Polyadenylate-binding protein 4 OS=Hon | PABPC4 | 70.783 | 5 | 53.658638 | yes | 9.722E-08 | 793.41675 | 1448.0042 |
| NLDDTIDDEKLR | 12 | Q13310 | Polyadenylate-binding protein 4 OS=Hon | PABPC4 | 70.783 | 2 | 53.658638 | no | 5.522E-07 | 723.85718 | 604.77783 |
| NLDDTIDDEKLR | 12 | Q13310 | Polyadenylate-binding protein 4 OS=Hon | PABPC4 | 70.783 | 2 | 53.658638 | no | 4.491E-07 | 482.9072 | 604.77783 |
| NLDDTIDDEKLR | 12 | Q13310 | Polyadenylate-binding protein 4 OS=Hon | PABPC4 | 70.783 | 3 | 53.658638 | no | 5.522E-07 | 723.85718 | 604.77783 |
| NLDDTIDDEKLR | 12 | Q13310 | Polyadenylate-binding protein 4 OS=Hon | PABPC4 | 70.783 | 3 | 53.658638 | no | 4.491E-07 | 482.9072 | 604.77783 |
| VMLEDGR | 7 | Q13310 | Polyadenylate-binding protein 4 OS=Hon | PABPC4 | 70.783 | 2 | 53.658638 | yes | 1.613E-06 | 410.20511 | 998.91705 |
| LAMDFGGAGAAQ | 34 | Q13310 | Polyadenylate-binding protein 4 OS=Hon | PABPC4 | 70.783 | 4 | 53.658638 | yes | 1.341E-07 | 837.15424 | 546.02649 |
| VDEAVAVLQAHH | 14 | Q13310 | Polyadenylate-binding protein 4 OS=Hon | PABPC4 | 70.783 | 2 | 53.658638 | yes | 1.005E-07 | 744.40192 | 240.08049 |

| | | | | | | | | | | | |
|---|---|---|---|---|---|---|---|---|---|---|---|
| VDEAVAVLQAHH | 14 | Q13310 | Polyadenylate-binding protein 4 OS=Hon | PABPC4 | 70.783 | 2 | 53.658638 | yes | 8.96E-07 | 496.6037 | 240.08049 |
| VDEAVAVLQAHH | 14 | Q13310 | Polyadenylate-binding protein 4 OS=Hon | PABPC4 | 70.783 | 3 | 53.658638 | yes | 1.005E-07 | 744.40192 | 240.08049 |
| VDEAVAVLQAHH | 14 | Q13310 | Polyadenylate-binding protein 4 OS=Hon | PABPC4 | 70.783 | 3 | 53.658638 | yes | 8.96E-07 | 496.6037 | 240.08049 |
| HLAPTGSECPDR | 12 | Q13310 | Polyadenylate-binding protein 4 OS=Hon | PABPC4 | 70.783 | 2 | 53.658638 | yes | 1.157E-07 | 670.30658 | 446.12561 |
| PQGFQGMPSAIR | 12 | Q13310 | Polyadenylate-binding protein 4 OS=Hon | PABPC4 | 70.783 | 2 | 53.658638 | yes | 1.133E-07 | 644.82697 | 430.8381 |
| EREAELGAK | 9 | Q13310 | Polyadenylate-binding protein 4 OS=Hon | PABPC4 | 70.783 | 2 | 53.658638 | yes | 2.808E-07 | 501.76437 | 143.58449 |
| NFGEEVDDESLKE | 20 | Q13310 | Polyadenylate-binding protein 4 OS=Hon | PABPC4 | 70.783 | 3 | 53.658638 | yes | 2.526E-07 | 773.3656 | 49.158897 |
| SQPVHILAVR | 10 | Q13322 | Growth factor receptor-bound protein 10 | GRB10 | 67.232 | 2 | 37.713707 | yes | 5.761E-07 | 560.33527 | 54.99472 |
| LECPETDCEK | 10 | Q13325 | Interferon-induced protein with tetratrico | IFIT5 | 55.847 | 2 | 45.998974 | yes | 1.682E-07 | 640.76031 | 71.38649 |
| GHFGPINSVAFHPI | 16 | Q13347 | Eukaryotic translation initiation factor 3 s | EIF3I | 36.502 | 3 | 53.446426 | yes | 1.768E-07 | 560.6145 | 917.84875 |
| GHFGPINSVAFHPI | 16 | Q13347 | Eukaryotic translation initiation factor 3 s | EIF3I | 36.502 | 3 | 53.446426 | yes | 3.84E-07 | 840.41809 | 917.84875 |
| GHFGPINSVAFHPI | 16 | Q13347 | Eukaryotic translation initiation factor 3 s | EIF3I | 36.502 | 2 | 53.446426 | yes | 1.768E-07 | 560.6145 | 917.84875 |
| GHFGPINSVAFHPI | 16 | Q13347 | Eukaryotic translation initiation factor 3 s | EIF3I | 36.502 | 2 | 53.446426 | yes | 3.84E-07 | 840.41809 | 917.84875 |
| MKPILLQGHER | 11 | Q13347 | Eukaryotic translation initiation factor 3 s | EIF3I | 36.502 | 3 | 53.446426 | yes | 1.973E-07 | 441.25183 | 892.18115 |
| MKPILLQGHER | 11 | Q13347 | Eukaryotic translation initiation factor 3 s | EIF3I | 36.502 | 3 | 53.446426 | yes | 1.025E-07 | 661.37408 | 892.18115 |
| MKPILLQGHER | 11 | Q13347 | Eukaryotic translation initiation factor 3 s | EIF3I | 36.502 | 2 | 53.446426 | yes | 1.973E-07 | 441.25183 | 892.18115 |
| MKPILLQGHER | 11 | Q13347 | Eukaryotic translation initiation factor 3 s | EIF3I | 36.502 | 2 | 53.446426 | yes | 1.025E-07 | 661.37408 | 892.18115 |
| QINDIQLSR | 9 | Q13347 | Eukaryotic translation initiation factor 3 s | EIF3I | 36.502 | 2 | 53.446426 | yes | 2.859E-07 | 543.79877 | 820.83508 |
| DPSQIDNNEPYMK | 13 | Q13347 | Eukaryotic translation initiation factor 3 s | EIF3I | 36.502 | 2 | 53.446426 | yes | 2.16E-07 | 775.8432 | 664.36938 |
| SGEVLVNVK | 9 | Q13347 | Eukaryotic translation initiation factor 3 s | EIF3I | 36.502 | 2 | 53.446426 | yes | 2.975E-06 | 472.7742 | 649.47858 |
| LWDCETGK | 8 | Q13347 | Eukaryotic translation initiation factor 3 s | EIF3I | 36.502 | 2 | 53.446426 | yes | 2.518E-07 | 504.72638 | 466.40903 |
| FFHLAFEEEFGR | 12 | Q13347 | Eukaryotic translation initiation factor 3 s | EIF3I | 36.502 | 2 | 53.446426 | yes | 9.904E-08 | 764.86462 | 446.27588 |
| HVLTGSADNSCR | 12 | Q13347 | Eukaryotic translation initiation factor 3 s | EIF3I | 36.502 | 2 | 53.446426 | yes | 1.029E-07 | 658.8042 | 393.34781 |
| SYSSGGEDGYVR | 12 | Q13347 | Eukaryotic translation initiation factor 3 s | EIF3I | 36.502 | 2 | 53.446426 | yes | 9.795E-08 | 638.77563 | 384.98074 |
| DMTMFVTASK | 10 | Q13347 | Eukaryotic translation initiation factor 3 s | EIF3I | 36.502 | 2 | 53.446426 | yes | 9.722E-08 | 565.76465 | 332.69827 |
| DPIVNVWYSVNGI | 14 | Q13347 | Eukaryotic translation initiation factor 3 s | EIF3I | 36.502 | 2 | 53.446426 | yes | 9.722E-08 | 824.40991 | 276.43106 |
| EGDLLFTVAK | 10 | Q13347 | Eukaryotic translation initiation factor 3 s | EIF3I | 36.502 | 2 | 53.446426 | yes | 1.603E-06 | 546.80042 | 142.16373 |
| DLLTDEPFSR | 10 | Q13356 | RING-type E3 ubiquitin-protein ligase PF | PPIL2 | 58.824 | 2 | 52.9886 | yes | 1.022E-07 | 596.79584 | 612.82318 |
| QDIITLQDPTNLDK | 14 | Q13356 | RING-type E3 ubiquitin-protein ligase PF | PPIL2 | 58.824 | 2 | 52.9886 | yes | 9.722E-08 | 807.42267 | 548.27777 |
| IIDPDEEK | 8 | Q13356 | RING-type E3 ubiquitin-protein ligase PF | PPIL2 | 58.824 | 2 | 52.9886 | yes | 1.944E-07 | 479.74002 | 467.59125 |
| FNVSNFYHVK | 10 | Q13356 | RING-type E3 ubiquitin-protein ligase PF | PPIL2 | 58.824 | 2 | 52.9886 | yes | 1.118E-07 | 627.81696 | 456.06891 |
| AAEEEPSTSATVPI | 16 | Q13356 | RING-type E3 ubiquitin-protein ligase PF | PPIL2 | 58.824 | 2 | 52.9886 | yes | 9.722E-08 | 817.88251 | 453.67853 |
| SQFFITFR | 8 | Q13356 | RING-type E3 ubiquitin-protein ligase PF | PPIL2 | 58.824 | 2 | 52.9886 | yes | 3.999E-07 | 523.27692 | 317.83978 |
| ETLQELYK | 8 | Q13356 | RING-type E3 ubiquitin-protein ligase PF | PPIL2 | 58.824 | 2 | 52.9886 | yes | 1.524E-07 | 512.27148 | 264.59277 |
| TCENFIR | 7 | Q13356 | RING-type E3 ubiquitin-protein ligase PF | PPIL2 | 58.824 | 2 | 52.9886 | yes | 1.67E-07 | 470.22128 | 186.4623 |
| LNAAHYSTGK | 10 | Q13356 | RING-type E3 ubiquitin-protein ligase PF | PPIL2 | 58.824 | 2 | 52.9886 | yes | 2.065E-07 | 531.27234 | 183.5321 |
| QDPSYYLK | 8 | Q13356 | RING-type E3 ubiquitin-protein ligase PF | PPIL2 | 58.824 | 2 | 52.9886 | yes | 3.812E-05 | 507.25055 | 180.75645 |
| GILSMANSGPNSN | 14 | Q13356 | RING-type E3 ubiquitin-protein ligase PF | PPIL2 | 58.824 | 2 | 52.9886 | yes | 1.061E-07 | 709.34625 | 159.35117 |
| SSQPQAGSQGPQT | 15 | Q13356 | RING-type E3 ubiquitin-protein ligase PF | PPIL2 | 58.824 | 2 | 52.9886 | yes | 1.022E-07 | 788.37915 | 144.62637 |

| | | | | | | | | | | | |
|---|---|---|---|---|---|---|---|---|---|---|---|
| KPDLPQTNFR | 10 | Q13356 | RING-type E3 ubiquitin-protein ligase PF | PPIL2 | 58.824 | 2 | 52.9886 | yes | 1.295E-07 | 608.32764 | 112.10645 |
| HYYDGTIFHR | 10 | Q13356 | RING-type E3 ubiquitin-protein ligase PF | PPIL2 | 58.824 | 2 | 52.9886 | yes | 1.414E-07 | 654.80963 | 61.142426 |
| GDEILAATMK | 10 | Q13356 | RING-type E3 ubiquitin-protein ligase PF | PPIL2 | 58.824 | 2 | 52.9886 | yes | 1.494E-06 | 524.77081 | 52.141693 |
| IDATTVFVDPYEEA | 22 | Q13356 | RING-type E3 ubiquitin-protein ligase PF | PPIL2 | 58.824 | 3 | 52.9886 | yes | 3.691E-06 | 827.73083 | 44.95755 |
| EFKGDEILAATMK | 13 | Q13356 | RING-type E3 ubiquitin-protein ligase PF | PPIL2 | 58.824 | 2 | 52.9886 | yes | 1.47E-07 | 726.87378 | 42.673813 |
| LFMEMNQK | 8 | Q13362 | Serine/threonine-protein phosphatase 2A | PPP2R5C | 61.061 | 2 | 41.74472 | no | 3.36E-07 | 520.74884 | 48.022758 |
| VLLPLHK | 7 | Q13362 | Serine/threonine-protein phosphatase 2A | PPP2R5C | 61.061 | 2 | 41.74472 | no | 1.559E-06 | 410.27618 | 40.7766 |
| AALSEMVEYITHN | 14 | Q13362 | Serine/threonine-protein phosphatase 2A | PPP2R5C | 61.061 | 2 | 41.74472 | yes | 4.248E-07 | 817.40375 | 25.019394 |
| DIPYHNR | 7 | Q13370 | cGMP-inhibited 3',5'-cyclic phosphodiest | PDE3B | 124.334 | 2 | 21.298742 | yes | 4.182E-06 | 457.72757 | 45.275581 |
| EVLVLTSVR | 9 | Q13371 | Phosducin-like protein OS=Homo sapiens | PDCL | 34.282 | 2 | 42.29697 | yes | 3.099E-07 | 508.31094 | 46.303093 |
| FGSYAAIQENALA | 14 | Q13393 | Phospholipase D1 OS=Homo sapiens OX | PLD1 | 124.185 | 2 | 51.869682 | yes | 2.805E-07 | 741.883 | 87.997429 |
| GENSILGQLK | 10 | Q13393 | Phospholipase D1 OS=Homo sapiens OX | PLD1 | 124.185 | 2 | 51.869682 | yes | 8.713E-07 | 529.79565 | 42.22578 |
| NEPVQNLPIQK | 11 | Q13393 | Phospholipase D1 OS=Homo sapiens OX | PLD1 | 124.185 | 2 | 51.869682 | yes | 3.638E-06 | 640.35388 | 40.762539 |
| DFINKPVLAK | 10 | Q13393 | Phospholipase D1 OS=Homo sapiens OX | PLD1 | 124.185 | 2 | 51.869682 | yes | 9.722E-08 | 572.83984 | 30.68877 |
| SLLLLGNER | 9 | Q13395 | Probable methyltransferase TARBP1 OS= | TARBP1 | 181.675 | 2 | 50.106205 | yes | 3.602E-07 | 507.80075 | 142.80641 |
| AVWGGLAAPGAS | 17 | Q13395 | Probable methyltransferase TARBP1 OS= | TARBP1 | 181.675 | 2 | 50.106205 | yes | 1.345E-07 | 763.90735 | 108.78128 |
| SPGQPIGSCSPLGL | 15 | Q13395 | Probable methyltransferase TARBP1 OS= | TARBP1 | 181.675 | 2 | 50.106205 | yes | 3.846E-07 | 749.39014 | 105.52851 |
| TVQAGLGQADAL | 14 | Q13395 | Probable methyltransferase TARBP1 OS= | TARBP1 | 181.675 | 2 | 50.106205 | yes | 1.215E-07 | 700.87823 | 83.796532 |
| LLPVLVQCGGAAL | 14 | Q13395 | Probable methyltransferase TARBP1 OS= | TARBP1 | 181.675 | 2 | 50.106205 | yes | 8.18E-07 | 733.9292 | 81.819725 |
| ALLGALCQGEASA | 15 | Q13395 | Probable methyltransferase TARBP1 OS= | TARBP1 | 181.675 | 2 | 50.106205 | yes | 9.902E-08 | 773.38812 | 76.896561 |
| LLVLSALAEK | 10 | Q13395 | Probable methyltransferase TARBP1 OS= | TARBP1 | 181.675 | 2 | 50.106205 | yes | 2.672E-07 | 528.83679 | 56.690475 |
| LAGRPQLAAALAE | 17 | Q13395 | Probable methyltransferase TARBP1 OS= | TARBP1 | 181.675 | 3 | 50.106205 | yes | 3.413E-07 | 584.00385 | 55.771606 |
| LSGLIEDEWITIDK | 14 | Q13395 | Probable methyltransferase TARBP1 OS= | TARBP1 | 181.675 | 2 | 50.106205 | yes | 4.627E-07 | 816.42999 | 28.084232 |
| WTGMIIGPPR | 10 | Q13404 | Ubiquitin-conjugating enzyme E2 variant | UBE2V1 | 16.495 | 2 | 42.71778 | yes | 2.903E-07 | 564.30511 | 27.144182 |
| FVESVDEYQFVER | 13 | Q13405 | Large ribosomal subunit protein mL49 OS | MRPL49 | 19.199 | 2 | 51.11326 | yes | 1.05E-07 | 823.88849 | 1131.9559 |
| MHNIPVYK | 8 | Q13405 | Large ribosomal subunit protein mL49 OS | MRPL49 | 19.199 | 2 | 51.11326 | yes | 5.101E-06 | 501.26547 | 1081.6331 |
| DPPPNLPYFVR | 11 | Q13405 | Large ribosomal subunit protein mL49 OS | MRPL49 | 19.199 | 2 | 51.11326 | yes | 9.722E-08 | 657.8457 | 843.53174 |
| TPVTQVNEVTGTL | 14 | Q13405 | Large ribosomal subunit protein mL49 OS | MRPL49 | 19.199 | 2 | 51.11326 | yes | 1.15E-07 | 757.91229 | 713.50208 |
| KVEGDIWALQK | 11 | Q13405 | Large ribosomal subunit protein mL49 OS | MRPL49 | 19.199 | 2 | 51.11326 | yes | 9.722E-08 | 643.85876 | 570.81458 |
| GYFDQELK | 8 | Q13405 | Large ribosomal subunit protein mL49 OS | MRPL49 | 19.199 | 2 | 51.11326 | yes | 3.982E-07 | 500.24274 | 363.21466 |
| VEGDIWALQK | 10 | Q13405 | Large ribosomal subunit protein mL49 OS | MRPL49 | 19.199 | 2 | 51.11326 | yes | 1.365E-07 | 579.81134 | 125.97234 |
| DITHGNR | 7 | Q13405 | Large ribosomal subunit protein mL49 OS | MRPL49 | 19.199 | 2 | 51.11326 | yes | 1.152E-07 | 406.7041 | 35.272537 |
| EAVAPVQEESDLE | 14 | Q13409 | Cytoplasmic dynein 1 intermediate chain | DYNC1I2 | 71.457 | 2 | 47.723072 | yes | 1.272E-07 | 772.37793 | 37.901634 |
| AFEDDAIQLVAR | 12 | Q13415 | Origin recognition complex subunit 1 OS | ORC1 | 97.35 | 2 | 51.9224 | yes | 2.966E-07 | 674.34882 | 111.89996 |
| LLLVEPSR | 8 | Q13415 | Origin recognition complex subunit 1 OS | ORC1 | 97.35 | 2 | 51.9224 | yes | 2.707E-07 | 463.78711 | 91.666756 |
| SAESPSWTPAEHV | 15 | Q13415 | Origin recognition complex subunit 1 OS | ORC1 | 97.35 | 2 | 51.9224 | yes | 1.878E-07 | 798.88629 | 86.590027 |
| LQTLSPALK | 9 | Q13415 | Origin recognition complex subunit 1 OS | ORC1 | 97.35 | 2 | 51.9224 | yes | 3.037E-07 | 485.80023 | 81.963898 |
| TATVHEVIR | 9 | Q13415 | Origin recognition complex subunit 1 OS | ORC1 | 97.35 | 2 | 51.9224 | yes | 9.723E-08 | 513.29053 | 74.00251 |
| FCEVPACK | 8 | Q13415 | Origin recognition complex subunit 1 OS | ORC1 | 97.35 | 2 | 51.9224 | yes | 9.722E-08 | 505.72534 | 63.339546 |

| | | | | | | | | | | | |
|---|---|---|---|---|---|---|---|---|---|---|---|
| LLELFEDDSDPPPK | 14 | Q13415 | Origin recognition complex subunit 1 OS= | ORC1 | 97.35 | 2 | 51.9224 | yes | 4.567E-07 | 807.8985 | 57.644028 |
| CAAPQIR | 7 | Q13415 | Origin recognition complex subunit 1 OS= | ORC1 | 97.35 | 2 | 51.9224 | yes | 4.574E-07 | 408.21326 | 32.401909 |
| QTPTHPLTPR | 10 | Q13415 | Origin recognition complex subunit 1 OS= | ORC1 | 97.35 | 2 | 51.9224 | yes | 2.578E-06 | 574.31458 | 22.827507 |
| ETGLSYTEDDKK | 12 | Q13415 | Origin recognition complex subunit 1 OS= | ORC1 | 97.35 | 2 | 51.9224 | yes | 2.985E-06 | 693.3252 | 11.037281 |
| EAFLVNSDLTLR | 12 | Q13416 | Origin recognition complex subunit 2 OS= | ORC2 | 65.972 | 2 | 48.260185 | yes | 1.172E-07 | 689.37225 | 197.02919 |
| VSPSFSAELK | 10 | Q13416 | Origin recognition complex subunit 2 OS= | ORC2 | 65.972 | 2 | 48.260185 | yes | 1.366E-06 | 532.78479 | 64.052574 |
| AQLLVNPK | 8 | Q13416 | Origin recognition complex subunit 2 OS= | ORC2 | 65.972 | 2 | 48.260185 | yes | 4.613E-07 | 441.77402 | 50.69738 |
| DQNYVEIMGR | 10 | Q13416 | Origin recognition complex subunit 2 OS= | ORC2 | 65.972 | 2 | 48.260185 | yes | 3.018E-06 | 612.78754 | 24.187536 |
| SVMIDEDMTAR | 11 | Q13418 | Integrin-linked protein kinase OS=Homo | ILK | 51.42 | 2 | 47.210014 | yes | 2.249E-07 | 634.2865 | 142.23042 |
| MGQNLNR | 7 | Q13418 | Integrin-linked protein kinase OS=Homo | ILK | 51.42 | 2 | 47.210014 | yes | 2.834E-07 | 416.70834 | 99.191269 |
| EVPFADLSNMEIG | 15 | Q13418 | Integrin-linked protein kinase OS=Homo | ILK | 51.42 | 2 | 47.210014 | yes | 5.469E-07 | 840.90222 | 86.383423 |
| FDMIVPILEK | 10 | Q13418 | Integrin-linked protein kinase OS=Homo | ILK | 51.42 | 2 | 47.210014 | yes | 1.376E-07 | 602.83594 | 66.142754 |
| MYAPAWVAPEAL | 14 | Q13418 | Integrin-linked protein kinase OS=Homo | ILK | 51.42 | 2 | 47.210014 | yes | 1.813E-07 | 787.9054 | 60.726749 |
| PVLMAVTEK | 9 | Q13425 | Beta-2-syntrophin OS=Homo sapiens OX | SNTB2 | 57.95 | 2 | 47.317028 | yes | 2.773E-07 | 494.28079 | 106.99575 |
| QEAGGLGISIK | 11 | Q13425 | Beta-2-syntrophin OS=Homo sapiens OX | SNTB2 | 57.95 | 2 | 47.317028 | yes | 3.126E-07 | 536.80347 | 67.239067 |
| DLLLYDCMPWTR | 12 | Q13425 | Beta-2-syntrophin OS=Homo sapiens OX | SNTB2 | 57.95 | 2 | 47.317028 | yes | 1.355E-07 | 791.87329 | 55.063866 |
| QGIEMHLFR | 9 | Q13425 | Beta-2-syntrophin OS=Homo sapiens OX | SNTB2 | 57.95 | 2 | 47.317028 | yes | 2.21E-06 | 565.79242 | 47.630653 |
| AGLVELLLR | 9 | Q13425 | Beta-2-syntrophin OS=Homo sapiens OX | SNTB2 | 57.95 | 2 | 47.317028 | yes | 2.758E-06 | 492.31604 | 19.944391 |
| ILSCGELIPK | 10 | Q13427 | Peptidyl-prolyl cis-trans isomerase G OS= | PPIG | 88.618 | 2 | 51.92363 | yes | 7.303E-07 | 565.31793 | 203.84964 |
| VVFELFSDVCPK | 12 | Q13427 | Peptidyl-prolyl cis-trans isomerase G OS= | PPIG | 88.618 | 2 | 51.92363 | yes | 1.456E-07 | 720.3656 | 145.89597 |
| EFLLSMANR | 9 | Q13427 | Peptidyl-prolyl cis-trans isomerase G OS= | PPIG | 88.618 | 2 | 51.92363 | yes | 9.722E-08 | 540.77893 | 137.54062 |
| CFFDIAINNQPAGR | 14 | Q13427 | Peptidyl-prolyl cis-trans isomerase G OS= | PPIG | 88.618 | 2 | 51.92363 | yes | 1.761E-07 | 811.89081 | 119.55175 |
| TDAASKPFAEVR | 12 | Q13427 | Peptidyl-prolyl cis-trans isomerase G OS= | PPIG | 88.618 | 2 | 51.92363 | yes | 6.239E-07 | 646.33569 | 115.67506 |
| VLTELLEQER | 10 | Q13428 | Treacle protein OS=Homo sapiens OX=9 | TCOF1 | 152.107 | 2 | 53.56694 | yes | 9.85E-08 | 615.34045 | 925.12659 |
| TNVVTMPTAHPR | 12 | Q13428 | Treacle protein OS=Homo sapiens OX=9 | TCOF1 | 152.107 | 2 | 53.56694 | yes | 9.722E-08 | 662.34552 | 633.36005 |
| GSLGQGTAPVLPG | 14 | Q13428 | Treacle protein OS=Homo sapiens OX=9 | TCOF1 | 152.107 | 2 | 53.56694 | yes | 9.722E-08 | 641.36169 | 513.41077 |
| LPEVQQATK | 9 | Q13428 | Treacle protein OS=Homo sapiens OX=9 | TCOF1 | 152.107 | 2 | 53.56694 | yes | 1.048E-07 | 507.28494 | 450.4339 |
| VGPATPSAQVGK | 12 | Q13428 | Treacle protein OS=Homo sapiens OX=9 | TCOF1 | 152.107 | 2 | 53.56694 | yes | 2.412E-07 | 556.30896 | 209.43802 |
| ASEAQPPVAR | 10 | Q13428 | Treacle protein OS=Homo sapiens OX=9 | TCOF1 | 152.107 | 2 | 53.56694 | yes | 1.694E-07 | 513.27234 | 200.2782 |
| LDSSPSVSSTLAAK | 14 | Q13428 | Treacle protein OS=Homo sapiens OX=9 | TCOF1 | 152.107 | 2 | 53.56694 | yes | 1.585E-07 | 681.85919 | 198.15569 |
| NPQNSTVLAR | 10 | Q13428 | Treacle protein OS=Homo sapiens OX=9 | TCOF1 | 152.107 | 2 | 53.56694 | yes | 1.176E-07 | 550.29639 | 186.69453 |
| TSQVGAASAPAK | 12 | Q13428 | Treacle protein OS=Homo sapiens OX=9 | TCOF1 | 152.107 | 2 | 53.56694 | yes | 1.83E-07 | 544.29077 | 159.14908 |
| TGPAAAQVQVGK | 12 | Q13428 | Treacle protein OS=Homo sapiens OX=9 | TCOF1 | 152.107 | 2 | 53.56694 | yes | 1.137E-07 | 563.81439 | 111.77573 |
| DEPEEELQK | 9 | Q13428 | Treacle protein OS=Homo sapiens OX=9 | TCOF1 | 152.107 | 2 | 53.56694 | yes | 1.457E-06 | 558.75641 | 103.73248 |
| PASTMGMGPLGK | 12 | Q13428 | Treacle protein OS=Homo sapiens OX=9 | TCOF1 | 152.107 | 2 | 53.56694 | yes | 1.296E-07 | 573.78595 | 75.663132 |
| SLGNILQAK | 9 | Q13428 | Treacle protein OS=Homo sapiens OX=9 | TCOF1 | 152.107 | 2 | 53.56694 | yes | 1.634E-07 | 472.2822 | 64.230675 |
| GMGTVEGGDQSN | 14 | Q13428 | Treacle protein OS=Homo sapiens OX=9 | TCOF1 | 152.107 | 2 | 53.56694 | yes | 6.211E-07 | 688.80914 | 37.769684 |
| KNPASLPLTQAAL | 14 | Q13428 | Treacle protein OS=Homo sapiens OX=9 | TCOF1 | 152.107 | 2 | 53.56694 | yes | 1.367E-06 | 726.43268 | 36.139832 |
| KTGPAAAQVQVG | 13 | Q13428 | Treacle protein OS=Homo sapiens OX=9 | TCOF1 | 152.107 | 2 | 53.56694 | yes | 2.394E-07 | 627.86188 | 11.126393 |

| | | | | | | | | | | | |
|---|---|---|---|---|---|---|---|---|---|---|---|
| LAEIGAPIQGNR | 12 | Q13435 | Splicing factor 3B subunit 2 OS=Homo sa | SF3B2 | 100.228 | 2 | 54.704933 | yes | 9.862E-08 | 619.84619 | 3418.1978 |
| AAVLLEQER | 9 | Q13435 | Splicing factor 3B subunit 2 OS=Homo sa | SF3B2 | 100.228 | 2 | 54.704933 | yes | 9.722E-08 | 514.79041 | 1953.2813 |
| VGEPVALSEEER | 12 | Q13435 | Splicing factor 3B subunit 2 OS=Homo sa | SF3B2 | 100.228 | 2 | 54.704933 | yes | 9.758E-08 | 657.83044 | 1547.8684 |
| ISLGMPVGPNAHK | 13 | Q13435 | Splicing factor 3B subunit 2 OS=Homo sa | SF3B2 | 100.228 | 2 | 54.704933 | yes | 1.697E-07 | 660.85828 | 1449.848 |
| EDFSDMVAEHAAI | 13 | Q13435 | Splicing factor 3B subunit 2 OS=Homo sa | SF3B2 | 100.228 | 2 | 54.704933 | yes | 1.596E-07 | 725.3194 | 1417.5236 |
| EMDDPSVGPK | 10 | Q13435 | Splicing factor 3B subunit 2 OS=Homo sa | SF3B2 | 100.228 | 2 | 54.704933 | yes | 1.093E-07 | 537.74225 | 1217.1058 |
| KPGDLSDELR | 10 | Q13435 | Splicing factor 3B subunit 2 OS=Homo sa | SF3B2 | 100.228 | 2 | 54.704933 | yes | 1.913E-07 | 565.29602 | 1177.8245 |
| YGPPPSYPNLK | 11 | Q13435 | Splicing factor 3B subunit 2 OS=Homo sa | SF3B2 | 100.228 | 2 | 54.704933 | yes | 5.838E-07 | 616.81915 | 1129.7662 |
| LHDAFFK | 7 | Q13435 | Splicing factor 3B subunit 2 OS=Homo sa | SF3B2 | 100.228 | 2 | 54.704933 | yes | 6.04E-07 | 439.23196 | 677.04181 |
| TATVGGAMMGST | 23 | Q13435 | Splicing factor 3B subunit 2 OS=Homo sa | SF3B2 | 100.228 | 3 | 54.704933 | yes | 1.37E-07 | 802.03033 | 617.58771 |
| QTGIVLNR | 8 | Q13435 | Splicing factor 3B subunit 2 OS=Homo sa | SF3B2 | 100.228 | 2 | 54.704933 | yes | 3.66E-07 | 450.76672 | 511.70392 |
| LTIHGDLYYEGK | 12 | Q13435 | Splicing factor 3B subunit 2 OS=Homo sa | SF3B2 | 100.228 | 2 | 54.704933 | yes | 9.81E-08 | 704.85901 | 447.84177 |
| QLVARPDVVEMH | 20 | Q13435 | Splicing factor 3B subunit 2 OS=Homo sa | SF3B2 | 100.228 | 3 | 54.704933 | yes | 1.714E-07 | 750.05121 | 389.37463 |
| VPPPWLIAMQR | 11 | Q13435 | Splicing factor 3B subunit 2 OS=Homo sa | SF3B2 | 100.228 | 2 | 54.704933 | yes | 9.722E-08 | 654.36829 | 379.74738 |
| EHELLEQQK | 9 | Q13435 | Splicing factor 3B subunit 2 OS=Homo sa | SF3B2 | 100.228 | 2 | 54.704933 | yes | 6.327E-07 | 577.29602 | 346.41705 |
| GIEKPPFELPDFIK | 14 | Q13435 | Splicing factor 3B subunit 2 OS=Homo sa | SF3B2 | 100.228 | 2 | 54.704933 | yes | 1.386E-07 | 815.448 | 171.90849 |
| GIEKPPFELPDFIK | 14 | Q13435 | Splicing factor 3B subunit 2 OS=Homo sa | SF3B2 | 100.228 | 2 | 54.704933 | yes | 4.014E-06 | 543.96771 | 171.90849 |
| GIEKPPFELPDFIK | 14 | Q13435 | Splicing factor 3B subunit 2 OS=Homo sa | SF3B2 | 100.228 | 3 | 54.704933 | yes | 1.386E-07 | 815.448 | 171.90849 |
| GIEKPPFELPDFIK | 14 | Q13435 | Splicing factor 3B subunit 2 OS=Homo sa | SF3B2 | 100.228 | 3 | 54.704933 | yes | 4.014E-06 | 543.96771 | 171.90849 |
| EQQAQVEKEDFSI | 21 | Q13435 | Splicing factor 3B subunit 2 OS=Homo sa | SF3B2 | 100.228 | 3 | 54.704933 | yes | 9.722E-08 | 797.36914 | 282.54001 |
| IDIDYQK | 7 | Q13435 | Splicing factor 3B subunit 2 OS=Homo sa | SF3B2 | 100.228 | 2 | 54.704933 | yes | 9.722E-08 | 447.73199 | 225.5744 |
| ATEHPEPPK | 9 | Q13435 | Splicing factor 3B subunit 2 OS=Homo sa | SF3B2 | 100.228 | 2 | 54.704933 | yes | 2.151E-07 | 524.25891 | 176.12207 |
| TEEEEIDR | 8 | Q13435 | Splicing factor 3B subunit 2 OS=Homo sa | SF3B2 | 100.228 | 2 | 54.704933 | yes | 3.252E-07 | 510.72763 | 124.50736 |
| LAQQQAALLMQQ | 15 | Q13435 | Splicing factor 3B subunit 2 OS=Homo sa | SF3B2 | 100.228 | 3 | 54.704933 | yes | 2.978E-07 | 586.30518 | 103.5742 |
| GPPPPPGDENR | 11 | Q13435 | Splicing factor 3B subunit 2 OS=Homo sa | SF3B2 | 100.228 | 2 | 54.704933 | yes | 3.16E-07 | 566.77271 | 99.498299 |
| KIEEAMDGSETPQ | 21 | Q13435 | Splicing factor 3B subunit 2 OS=Homo sa | SF3B2 | 100.228 | 3 | 54.704933 | yes | 9.722E-08 | 788.06641 | 92.567856 |
| PDVVEMHDVTAQ | 15 | Q13435 | Splicing factor 3B subunit 2 OS=Homo sa | SF3B2 | 100.228 | 2 | 54.704933 | yes | 1.385E-07 | 840.8985 | 33.647682 |
| RGIEKPPFELPDFII | 15 | Q13435 | Splicing factor 3B subunit 2 OS=Homo sa | SF3B2 | 100.228 | 3 | 54.704933 | yes | 6.211E-07 | 596.00146 | 33.245255 |
| EEQCILYLGPR | 11 | Q13451 | Peptidyl-prolyl cis-trans isomerase FKBP | FKBP5 | 51.213 | 2 | 42.292908 | yes | 3.101E-07 | 689.34521 | 108.52377 |
| LQISMCQK | 8 | Q13451 | Peptidyl-prolyl cis-trans isomerase FKBP | FKBP5 | 51.213 | 2 | 42.292908 | yes | 1.425E-06 | 504.25427 | 45.891411 |
| YTGMLETVR | 9 | Q13459 | Unconventional myosin-IXb OS=Homo s | MYO9B | 243.401 | 2 | 53.797485 | no | 9.81E-08 | 535.271 | 391.38809 |
| VALLEDVNR | 9 | Q13459 | Unconventional myosin-IXb OS=Homo s | MYO9B | 243.401 | 2 | 53.797485 | yes | 2.338E-07 | 514.79041 | 359.11777 |
| VSPPAPGSAPETPE | 16 | Q13459 | Unconventional myosin-IXb OS=Homo s | MYO9B | 243.401 | 2 | 53.797485 | yes | 3.891E-07 | 789.88593 | 311.00824 |
| ATGAALTPTEER | 12 | Q13459 | Unconventional myosin-IXb OS=Homo s | MYO9B | 243.401 | 2 | 53.797485 | yes | 1.439E-07 | 608.81201 | 272.99557 |
| GLEAPSGQQHR | 11 | Q13459 | Unconventional myosin-IXb OS=Homo s | MYO9B | 243.401 | 2 | 53.797485 | yes | 9.722E-08 | 590.29688 | 199.00142 |
| AQPAAETTDGER | 12 | Q13459 | Unconventional myosin-IXb OS=Homo s | MYO9B | 243.401 | 2 | 53.797485 | yes | 1.87E-06 | 623.28894 | 191.88103 |
| LGFSSPYEGVLNK | 13 | Q13459 | Unconventional myosin-IXb OS=Homo s | MYO9B | 243.401 | 2 | 53.797485 | yes | 9.722E-08 | 705.86682 | 167.20444 |
| FIQVSYLESGIVR | 13 | Q13459 | Unconventional myosin-IXb OS=Homo s | MYO9B | 243.401 | 2 | 53.797485 | yes | 2.759E-07 | 755.91687 | 159.89951 |
| AAGMSSPGAQSHI | 18 | Q13459 | Unconventional myosin-IXb OS=Homo s | MYO9B | 243.401 | 3 | 53.797485 | yes | 1.464E-07 | 607.95648 | 154.11584 |

| | | | | | | | | | | | |
|---|---|---|---|---|---|---|---|---|---|---|---|
| PDIVALLR | 8 | Q13459 | Unconventional myosin-IXb OS=Homo s | MYO9B | 243.401 | 2 | 53.797485 | no | 1.055E-07 | 448.78183 | 142.04564 |
| FLPIYNPK | 8 | Q13459 | Unconventional myosin-IXb OS=Homo s | MYO9B | 243.401 | 2 | 53.797485 | no | 2.837E-07 | 496.28421 | 140.90855 |
| QALQETLHR | 9 | Q13459 | Unconventional myosin-IXb OS=Homo s | MYO9B | 243.401 | 2 | 53.797485 | yes | 9.722E-08 | 548.29889 | 122.80458 |
| QAMEMVGFLPATI | 13 | Q13459 | Unconventional myosin-IXb OS=Homo s | MYO9B | 243.401 | 2 | 53.797485 | yes | 2.455E-07 | 711.85962 | 119.32809 |
| EILVEVLTK | 9 | Q13459 | Unconventional myosin-IXb OS=Homo s | MYO9B | 243.401 | 2 | 53.797485 | yes | 1.078E-07 | 522.32098 | 107.4112 |
| LENFPIHAITGVLK | 14 | Q13459 | Unconventional myosin-IXb OS=Homo s | MYO9B | 243.401 | 2 | 53.797485 | yes | 1.35E-07 | 776.4483 | 97.673347 |
| TILGAGPVLEAFGI | 16 | Q13459 | Unconventional myosin-IXb OS=Homo s | MYO9B | 243.401 | 2 | 53.797485 | yes | 1.019E-07 | 779.43542 | 95.355553 |
| ITTCVEMLIK | 10 | Q13459 | Unconventional myosin-IXb OS=Homo s | MYO9B | 243.401 | 2 | 53.797485 | yes | 1.053E-07 | 604.32489 | 94.074837 |
| DLHNQMIK | 8 | Q13459 | Unconventional myosin-IXb OS=Homo s | MYO9B | 243.401 | 2 | 53.797485 | yes | 3.568E-07 | 499.75803 | 88.246994 |
| SLLQSLSR | 8 | Q13459 | Unconventional myosin-IXb OS=Homo s | MYO9B | 243.401 | 2 | 53.797485 | yes | 3.94E-07 | 452.26654 | 86.216347 |
| VNQCIVISGESGSC | 15 | Q13459 | Unconventional myosin-IXb OS=Homo s | MYO9B | 243.401 | 2 | 53.797485 | yes | 2.456E-07 | 767.88013 | 77.045486 |
| MYENQQLGK | 9 | Q13459 | Unconventional myosin-IXb OS=Homo s | MYO9B | 243.401 | 2 | 53.797485 | yes | 2.628E-07 | 555.76605 | 74.900627 |
| ALLCSVCK | 8 | Q13459 | Unconventional myosin-IXb OS=Homo s | MYO9B | 243.401 | 2 | 53.797485 | yes | 6.077E-07 | 475.74353 | 70.262123 |
| EVISTLLEK | 9 | Q13459 | Unconventional myosin-IXb OS=Homo s | MYO9B | 243.401 | 2 | 53.797485 | yes | 2.863E-07 | 516.3028 | 57.173416 |
| CYVLVEVK | 8 | Q13459 | Unconventional myosin-IXb OS=Homo s | MYO9B | 243.401 | 2 | 53.797485 | yes | 3.936E-06 | 505.27298 | 55.802139 |
| LASAVELWR | 9 | Q13459 | Unconventional myosin-IXb OS=Homo s | MYO9B | 243.401 | 2 | 53.797485 | yes | 1.664E-07 | 522.79547 | 53.702778 |
| QPTEQPQAMAVGI | 13 | Q13459 | Unconventional myosin-IXb OS=Homo s | MYO9B | 243.401 | 2 | 53.797485 | yes | 3.917E-07 | 692.84808 | 50.497681 |
| TSFSTSDVSK | 10 | Q13459 | Unconventional myosin-IXb OS=Homo s | MYO9B | 243.401 | 2 | 53.797485 | yes | 5.787E-07 | 529.75366 | 46.375046 |
| AQDEHPQEDGYYI | 18 | Q13459 | Unconventional myosin-IXb OS=Homo s | MYO9B | 243.401 | 3 | 53.797485 | yes | 1.522E-07 | 746.6748 | 41.821987 |
| LIFHLVK | 7 | Q13459 | Unconventional myosin-IXb OS=Homo s | MYO9B | 243.401 | 2 | 53.797485 | yes | 1.594E-07 | 435.284 | 41.085968 |
| YVHMQLVAQATA | 14 | Q13459 | Unconventional myosin-IXb OS=Homo s | MYO9B | 243.401 | 2 | 53.797485 | yes | 1.984E-07 | 794.91687 | 40.439682 |
| TPIESLFIEATEK | 13 | Q13459 | Unconventional myosin-IXb OS=Homo s | MYO9B | 243.401 | 2 | 53.797485 | yes | 9.722E-08 | 739.39288 | 40.078373 |
| AAVTIQACWR | 10 | Q13459 | Unconventional myosin-IXb OS=Homo s | MYO9B | 243.401 | 2 | 53.797485 | yes | 4.486E-06 | 588.30316 | 32.933228 |
| LIISMTLHDR | 10 | Q13459 | Unconventional myosin-IXb OS=Homo s | MYO9B | 243.401 | 2 | 53.797485 | yes | 8.231E-07 | 599.83429 | 26.881157 |
| QALQTDPAAVK | 11 | Q13459 | Unconventional myosin-IXb OS=Homo s | MYO9B | 243.401 | 2 | 53.797485 | yes | 0.0002419 | 571.31421 | 24.94931 |
| HNLTLNDCFVK | 11 | Q13461 | Forkhead box protein E3 OS=Homo sapi | FOXE3 | 33.235 | 2 | 25.993137 | yes | 3.194E-06 | 680.83752 | 483.2319 |
| ATIDCAGILK | 10 | Q13469 | Nuclear factor of activated T-cells, cytopl | NFATC2 | 100.146 | 2 | 46.076874 | yes | 1.66E-07 | 531.28662 | 94.95546 |
| VLEIPLEPK | 9 | Q13469 | Nuclear factor of activated T-cells, cytopl | NFATC2 | 100.146 | 2 | 46.076874 | yes | 4.744E-07 | 519.31567 | 37.825119 |
| FTLPVPGFEGYR | 12 | Q13469 | Nuclear factor of activated T-cells, cytopl | NFATC2 | 100.146 | 2 | 46.076874 | yes | 2.746E-07 | 691.85876 | 21.831215 |
| ILGGFAPEHK | 10 | Q13470 | Non-receptor tyrosine-protein kinase TNI | TNK1 | 72.468 | 2 | 53.688488 | yes | 5.527E-07 | 534.79547 | 216.4823 |
| AAALSGGLLSDPE | 16 | Q13470 | Non-receptor tyrosine-protein kinase TNI | TNK1 | 72.468 | 2 | 53.688488 | yes | 9.899E-08 | 799.43085 | 154.58032 |
| VDQLFHLSSR | 10 | Q13470 | Non-receptor tyrosine-protein kinase TNI | TNK1 | 72.468 | 2 | 53.688488 | yes | 9.722E-08 | 601.31982 | 148.40387 |
| VADFGLVR | 8 | Q13470 | Non-receptor tyrosine-protein kinase TNI | TNK1 | 72.468 | 2 | 53.688488 | yes | 1.171E-07 | 438.75052 | 146.813 |
| PIPYAWCAPESLR | 13 | Q13470 | Non-receptor tyrosine-protein kinase TNI | TNK1 | 72.468 | 2 | 53.688488 | yes | 3.535E-07 | 780.38739 | 143.13631 |
| GELLGSGCFGVVH | 14 | Q13470 | Non-receptor tyrosine-protein kinase TNI | TNK1 | 72.468 | 2 | 53.688488 | yes | 3.712E-07 | 744.37482 | 116.76199 |
| VGPEGPMGTELGI | 16 | Q13470 | Non-receptor tyrosine-protein kinase TNI | TNK1 | 72.468 | 2 | 53.688488 | yes | 2.173E-07 | 837.91144 | 59.235947 |
| MLPEAGSLWLLK | 12 | Q13470 | Non-receptor tyrosine-protein kinase TNI | TNK1 | 72.468 | 2 | 53.688488 | yes | 1.856E-06 | 679.38104 | 48.63575 |
| ALYSLALR | 8 | Q13470 | Non-receptor tyrosine-protein kinase TNI | TNK1 | 72.468 | 2 | 53.688488 | yes | 4.954E-07 | 453.77402 | 43.255291 |
| SLESVLSLGPR | 11 | Q13470 | Non-receptor tyrosine-protein kinase TNI | TNK1 | 72.468 | 2 | 53.688488 | yes | 3.733E-07 | 579.3299 | 31.447882 |

| | | | | | | | | | | | |
|---|---|---|---|---|---|---|---|---|---|---|---|
| LYTQGYISYPR | 11 | Q13472 | DNA topoisomerase 3-alpha OS=Homo s | TOP3A | 112.373 | 2 | 53.93602 | no | 2.642E-07 | 680.84845 | 1739.5872 |
| SAVWLPDSVLEAS | 14 | Q13472 | DNA topoisomerase 3-alpha OS=Homo s | TOP3A | 112.373 | 2 | 53.93602 | yes | 1.229E-07 | 765.40155 | 470.21036 |
| AIQAFVPEIFHR | 12 | Q13472 | DNA topoisomerase 3-alpha OS=Homo s | TOP3A | 112.373 | 2 | 53.93602 | yes | 9.901E-08 | 714.39331 | 311.69272 |
| QCQALVIWTDCDI | 13 | Q13472 | DNA topoisomerase 3-alpha OS=Homo s | TOP3A | 112.373 | 2 | 53.93602 | yes | 9.723E-08 | 832.87958 | 195.70494 |
| SDQAHPPIHPTK | 12 | Q13472 | DNA topoisomerase 3-alpha OS=Homo s | TOP3A | 112.373 | 2 | 53.93602 | yes | 9.722E-08 | 664.34131 | 167.44615 |
| YCPENFVDIK | 10 | Q13472 | DNA topoisomerase 3-alpha OS=Homo s | TOP3A | 112.373 | 2 | 53.93602 | yes | 1.171E-07 | 642.80011 | 167.12158 |
| WGAFAQSILER | 11 | Q13472 | DNA topoisomerase 3-alpha OS=Homo s | TOP3A | 112.373 | 2 | 53.93602 | yes | 9.722E-08 | 639.33551 | 146.69937 |
| TETNIFPR | 8 | Q13472 | DNA topoisomerase 3-alpha OS=Homo s | TOP3A | 112.373 | 2 | 53.93602 | yes | 3.834E-07 | 489.25616 | 119.50688 |
| EGENIGFEIIHVCK | 14 | Q13472 | DNA topoisomerase 3-alpha OS=Homo s | TOP3A | 112.373 | 2 | 53.93602 | yes | 1.071E-07 | 822.90613 | 112.82732 |
| HGIGTDATHAEHII | 17 | Q13472 | DNA topoisomerase 3-alpha OS=Homo s | TOP3A | 112.373 | 3 | 53.93602 | yes | 9.935E-08 | 610.64288 | 69.910667 |
| TACENLTEPDQR | 12 | Q13472 | DNA topoisomerase 3-alpha OS=Homo s | TOP3A | 112.373 | 2 | 53.93602 | yes | 9.722E-08 | 717.31989 | 63.463272 |
| QVFIEAVAK | 9 | Q13472 | DNA topoisomerase 3-alpha OS=Homo s | TOP3A | 112.373 | 2 | 53.93602 | yes | 2.489E-06 | 502.79239 | 57.338444 |
| WRPQALDTVELEI | 13 | Q13472 | DNA topoisomerase 3-alpha OS=Homo s | TOP3A | 112.373 | 2 | 53.93602 | yes | 1.062E-07 | 792.92267 | 29.610472 |
| DSSVCPVCQPHPV | 15 | Q13472 | DNA topoisomerase 3-alpha OS=Homo s | TOP3A | 112.373 | 3 | 53.93602 | yes | 7.956E-07 | 600.94336 | 10.730743 |
| LTPVSPESSSTEEK | 14 | Q13501 | Sequestosome-1 OS=Homo sapiens OX=! | SQSTM1 | 47.688 | 2 | 50.797516 | yes | 4.464E-07 | 745.86469 | 1415.0013 |
| NYDIGAALDTIQY! | 15 | Q13501 | Sequestosome-1 OS=Homo sapiens OX=! | SQSTM1 | 47.688 | 2 | 50.797516 | yes | 1.691E-07 | 836.41486 | 1284.376 |
| LAFPSPFGHLSEGI | 18 | Q13501 | Sequestosome-1 OS=Homo sapiens OX=! | SQSTM1 | 47.688 | 3 | 50.797516 | yes | 2.937E-07 | 658.32697 | 164.61177 |
| DHRPPCAQEAPR | 12 | Q13501 | Sequestosome-1 OS=Homo sapiens OX=! | SQSTM1 | 47.688 | 2 | 50.797516 | yes | 1.361E-07 | 717.33875 | 137.20897 |
| VAALFPALRPGGF | 18 | Q13501 | Sequestosome-1 OS=Homo sapiens OX=! | SQSTM1 | 47.688 | 3 | 50.797516 | yes | 9.722E-08 | 657.69525 | 116.5304 |
| EAALYPHLPPEAD | 15 | Q13501 | Sequestosome-1 OS=Homo sapiens OX=! | SQSTM1 | 47.688 | 2 | 50.797516 | yes | 1.073E-06 | 838.42554 | 115.72459 |
| VAALFPALR | 9 | Q13501 | Sequestosome-1 OS=Homo sapiens OX=! | SQSTM1 | 47.688 | 2 | 50.797516 | yes | 5.029E-07 | 479.29764 | 91.898018 |
| PGGFQAHYR | 9 | Q13501 | Sequestosome-1 OS=Homo sapiens OX=! | SQSTM1 | 47.688 | 2 | 50.797516 | yes | 9.903E-07 | 516.75415 | 51.491837 |
| IQSALADIAQSQLI | 14 | Q13503 | Mediator of RNA polymerase II transcrip | MED21 | 15.565 | 2 | 53.15923 | yes | 9.722E-08 | 743.41718 | 121.62696 |
| ECLTLLSQR | 9 | Q13505 | Metaxin-1 OS=Homo sapiens OX=9606 ( | MTX1 | 51.463 | 2 | 40.778316 | yes | 3.846E-07 | 560.29498 | 72.963394 |
| ISVYYNEASSHK | 12 | Q13509 | Tubulin beta-3 chain OS=Homo sapiens ( | TUBB3 | 50.433 | 2 | 51.695362 | yes | 9.722E-08 | 699.33844 | 399.54791 |
| SGAFGHLFRPDNF | 26 | Q13509 | Tubulin beta-3 chain OS=Homo sapiens ( | TUBB3 | 50.433 | 4 | 51.695362 | yes | 9.037E-07 | 699.59137 | 127.05933 |
| SGAFGHLFR | 9 | Q13509 | Tubulin beta-3 chain OS=Homo sapiens ( | TUBB3 | 50.433 | 2 | 51.695362 | yes | 1.708E-07 | 496.25906 | 114.69651 |
| LHFFMPGFAPLTA | 14 | Q13509 | Tubulin beta-3 chain OS=Homo sapiens ( | TUBB3 | 50.433 | 2 | 51.695362 | yes | 1.405E-07 | 802.92395 | 26.141182 |
| GKLELVDNK | 9 | Q13523 | Serine/threonine-protein kinase PRP4 hor | PRPF4B | 116.987 | 2 | 48.583736 | yes | 1.35E-07 | 508.29276 | 178.17163 |
| ENVDTFEASVK | 11 | Q13523 | Serine/threonine-protein kinase PRP4 hor | PRPF4B | 116.987 | 2 | 48.583736 | yes | 1.127E-07 | 619.79858 | 111.39839 |
| LDDLALLEDLEK | 12 | Q13523 | Serine/threonine-protein kinase PRP4 hor | PRPF4B | 116.987 | 2 | 48.583736 | yes | 2.557E-07 | 693.87177 | 99.6035 |
| LNDADPDDKFHCI | 14 | Q13523 | Serine/threonine-protein kinase PRP4 hor | PRPF4B | 116.987 | 3 | 48.583736 | yes | 1.198E-07 | 572.59839 | 96.711227 |
| DLLADLIGCQR | 11 | Q13523 | Serine/threonine-protein kinase PRP4 hor | PRPF4B | 116.987 | 2 | 48.583736 | yes | 1.202E-07 | 637.33209 | 94.201599 |
| AELDNELMEGK | 11 | Q13523 | Serine/threonine-protein kinase PRP4 hor | PRPF4B | 116.987 | 2 | 48.583736 | yes | 3.885E-07 | 624.79248 | 89.679749 |
| SYSQQLFLALK | 11 | Q13523 | Serine/threonine-protein kinase PRP4 hor | PRPF4B | 116.987 | 2 | 48.583736 | yes | 2.385E-07 | 649.36115 | 56.618679 |
| LAQALHQCSR | 10 | Q13546 | Receptor-interacting serine/threonine-pro | RIPK1 | 75.931 | 2 | 52.964695 | yes | 5.308E-07 | 592.30365 | 141.88988 |
| MQSLQLDCVAVP! | 15 | Q13546 | Receptor-interacting serine/threonine-pro | RIPK1 | 75.931 | 2 | 52.964695 | yes | 9.722E-08 | 845.9162 | 120.39107 |
| IADLGLASFK | 10 | Q13546 | Receptor-interacting serine/threonine-pro | RIPK1 | 75.931 | 2 | 52.964695 | yes | 4.255E-07 | 517.79767 | 117.45184 |
| VSLCFHR | 7 | Q13546 | Receptor-interacting serine/threonine-pro | RIPK1 | 75.931 | 2 | 52.964695 | yes | 1.804E-07 | 459.73434 | 104.03516 |

| | | | | | | | | | | | |
|---|---|---|---|---|---|---|---|---|---|---|---|
| YSLVMEYMEK | 10 | Q13546 | Receptor-interacting serine/threonine-pro | RIPK1 | 75.931 | 2 | 52.964695 | yes | 1.346E-07 | 646.79871 | 101.07269 |
| PTFPGIEEK | 9 | Q13546 | Receptor-interacting serine/threonine-pro | RIPK1 | 75.931 | 2 | 52.964695 | yes | 5.618E-06 | 509.2662 | 87.553749 |
| AEMSTPLSVK | 10 | Q13546 | Receptor-interacting serine/threonine-pro | RIPK1 | 75.931 | 2 | 52.964695 | yes | 3.892E-06 | 531.77863 | 80.608841 |
| LLGVIIEEGK | 10 | Q13546 | Receptor-interacting serine/threonine-pro | RIPK1 | 75.931 | 2 | 52.964695 | yes | 3.705E-07 | 535.82642 | 55.491661 |
| VMTVSFHK | 8 | Q13547 | Histone deacetylase 1 OS=Homo sapiens | HDAC1 | 55.103 | 2 | 54.76191 | no | 8.588E-07 | 474.75223 | 620.82422 |
| YGEYFPGTGDLR | 12 | Q13547 | Histone deacetylase 1 OS=Homo sapiens | HDAC1 | 55.103 | 2 | 54.76191 | no | 1.847E-07 | 687.81989 | 422.47864 |
| YYAVNYPLR | 9 | Q13547 | Histone deacetylase 1 OS=Homo sapiens | HDAC1 | 55.103 | 2 | 54.76191 | yes | 9.722E-08 | 579.80072 | 321.99048 |
| LGCFNLTIK | 9 | Q13547 | Histone deacetylase 1 OS=Homo sapiens | HDAC1 | 55.103 | 2 | 54.76191 | yes | 1.456E-07 | 533.29169 | 234.30644 |
| MTHNLLLNYGLYI | 13 | Q13547 | Histone deacetylase 1 OS=Homo sapiens | HDAC1 | 55.103 | 2 | 54.76191 | no | 9.722E-08 | 804.42175 | 108.47415 |
| MTHNLLLNYGLYI | 13 | Q13547 | Histone deacetylase 1 OS=Homo sapiens | HDAC1 | 55.103 | 2 | 54.76191 | no | 3.144E-06 | 536.61694 | 108.47415 |
| MTHNLLLNYGLYI | 13 | Q13547 | Histone deacetylase 1 OS=Homo sapiens | HDAC1 | 55.103 | 3 | 54.76191 | no | 9.722E-08 | 804.42175 | 108.47415 |
| MTHNLLLNYGLYI | 13 | Q13547 | Histone deacetylase 1 OS=Homo sapiens | HDAC1 | 55.103 | 3 | 54.76191 | no | 3.144E-06 | 536.61694 | 108.47415 |
| YHSDDYIK | 8 | Q13547 | Histone deacetylase 1 OS=Homo sapiens | HDAC1 | 55.103 | 2 | 54.76191 | yes | 1.334E-07 | 520.73779 | 165.95602 |
| MEIYRPHK | 8 | Q13547 | Histone deacetylase 1 OS=Homo sapiens | HDAC1 | 55.103 | 2 | 54.76191 | no | 1.462E-07 | 537.28168 | 112.78809 |
| QQTDIAVNWAGGI | 17 | Q13547 | Histone deacetylase 1 OS=Homo sapiens | HDAC1 | 55.103 | 3 | 54.76191 | yes | 6.145E-07 | 615.98334 | 100.40048 |
| SIRPDNMSEYSK | 12 | Q13547 | Histone deacetylase 1 OS=Homo sapiens | HDAC1 | 55.103 | 2 | 54.76191 | no | 9.722E-08 | 713.83521 | 97.867821 |
| VLYIDIDIHHGDGV | 23 | Q13547 | Histone deacetylase 1 OS=Homo sapiens | HDAC1 | 55.103 | 4 | 54.76191 | no | 1.967E-07 | 670.32379 | 66.327293 |
| KMEIYRPHK | 9 | Q13547 | Histone deacetylase 1 OS=Homo sapiens | HDAC1 | 55.103 | 2 | 54.76191 | no | 2.435E-07 | 601.32916 | 59.655117 |
| DGIDDESYEAIFKF | 18 | Q13547 | Histone deacetylase 1 OS=Homo sapiens | HDAC1 | 55.103 | 3 | 54.76191 | yes | 1.374E-06 | 681.99084 | 38.609066 |
| DLKPENLLLASK | 12 | Q13554 | Calcium/calmodulin-dependent protein ki | CAMK2B | 72.678 | 2 | 53.201942 | no | 2.739E-07 | 670.89282 | 403.25281 |
| STVASMMHR | 9 | Q13554 | Calcium/calmodulin-dependent protein ki | CAMK2B | 72.678 | 2 | 53.201942 | no | 1.138E-07 | 510.2417 | 306.03482 |
| LTQYIDGQGR | 10 | Q13554 | Calcium/calmodulin-dependent protein ki | CAMK2B | 72.678 | 2 | 53.201942 | no | 9.722E-08 | 575.7962 | 248.88251 |
| GAILTTMLATR | 11 | Q13554 | Calcium/calmodulin-dependent protein ki | CAMK2B | 72.678 | 2 | 53.201942 | no | 1.337E-07 | 574.3288 | 182.38971 |
| NLINQMLTINPAK | 13 | Q13554 | Calcium/calmodulin-dependent protein ki | CAMK2B | 72.678 | 2 | 53.201942 | no | 9.722E-08 | 735.41089 | 130.00864 |
| HPWVCQR | 7 | Q13554 | Calcium/calmodulin-dependent protein ki | CAMK2B | 72.678 | 2 | 53.201942 | no | 4.381E-07 | 491.73743 | 39.094322 |
| GAILTTMLVSR | 11 | Q13555 | Calcium/calmodulin-dependent protein ki | CAMK2G | 62.608 | 2 | 50.704536 | yes | 1.117E-07 | 581.33667 | 318.27057 |
| QETVECLR | 8 | Q13555 | Calcium/calmodulin-dependent protein ki | CAMK2G | 62.608 | 2 | 50.704536 | yes | 3.143E-07 | 517.75043 | 171.02902 |
| FTDDYQLFEELGK | 13 | Q13555 | Calcium/calmodulin-dependent protein ki | CAMK2G | 62.608 | 2 | 50.704536 | yes | 9.728E-08 | 802.87756 | 147.58824 |
| FYFENLLSK | 9 | Q13555 | Calcium/calmodulin-dependent protein ki | CAMK2G | 62.608 | 2 | 50.704536 | yes | 4.074E-07 | 580.80298 | 118.61928 |
| FTDEYQLFEELGK | 13 | Q13557 | Calcium/calmodulin-dependent protein ki | CAMK2D | 56.37 | 2 | 51.42871 | yes | 1.603E-07 | 809.88538 | 85.808556 |
| LTQYMDGSGMPK | 12 | Q13557 | Calcium/calmodulin-dependent protein ki | CAMK2D | 56.37 | 2 | 51.42871 | yes | 9.722E-08 | 664.30469 | 60.42757 |
| MLTINPAK | 8 | Q13557 | Calcium/calmodulin-dependent protein ki | CAMK2D | 56.37 | 2 | 51.42871 | yes | 1.885E-07 | 444.25458 | 31.205851 |
| FYFENALSK | 9 | Q13557 | Calcium/calmodulin-dependent protein ki | CAMK2D | 56.37 | 2 | 51.42871 | yes | 5.298E-05 | 559.77948 | 9.4874897 |
| NLEFGLQPHK | 10 | Q13569 | G/T mismatch-specific thymine DNA gly | TDG | 46.054 | 2 | 50.37085 | yes | 1.157E-07 | 591.81696 | 1056.8755 |
| YGIGFTNMVER | 11 | Q13569 | G/T mismatch-specific thymine DNA gly | TDG | 46.054 | 2 | 50.37085 | yes | 9.81E-08 | 643.81354 | 507.08264 |
| FNGVSEAELLTK | 12 | Q13569 | G/T mismatch-specific thymine DNA gly | TDG | 46.054 | 2 | 50.37085 | yes | 2.711E-07 | 654.3457 | 152.38354 |
| LAEALYIADR | 10 | Q13573 | SNW domain-containing protein 1 OS=H | SNW1 | 61.495 | 2 | 55.204365 | yes | 1.022E-07 | 567.81134 | 1959.5707 |
| YTPSQQGVAFNSG | 15 | Q13573 | SNW domain-containing protein 1 OS=H | SNW1 | 61.495 | 2 | 55.204365 | yes | 1.528E-07 | 777.88098 | 1458.8907 |
| MSNALAIQVDSEG | 14 | Q13573 | SNW domain-containing protein 1 OS=H | SNW1 | 61.495 | 2 | 55.204365 | yes | 9.81E-08 | 731.86395 | 1182.3007 |

| | | | | | | | | | | | |
|---|---|---|---|---|---|---|---|---|---|---|---|
| DISEVIALGVPNPR | 14 | Q13573 | SNW domain-containing protein 1 OS=H | SNW1 | 61.495 | 2 | 55.204365 | yes | 9.722E-08 | 740.41193 | 855.56122 |
| DMAQSIYRPSK | 11 | Q13573 | SNW domain-containing protein 1 OS=H | SNW1 | 61.495 | 2 | 55.204365 | yes | 9.722E-08 | 648.32428 | 412.4534 |
| DMAQSIYRPSK | 11 | Q13573 | SNW domain-containing protein 1 OS=H | SNW1 | 61.495 | 2 | 55.204365 | yes | 1.13E-06 | 656.32172 | 412.4534 |
| IPPCISNWK | 9 | Q13573 | SNW domain-containing protein 1 OS=H | SNW1 | 61.495 | 2 | 55.204365 | yes | 3.196E-06 | 557.78931 | 758.12042 |
| LAPAQYIR | 8 | Q13573 | SNW domain-containing protein 1 OS=H | SNW1 | 61.495 | 2 | 55.204365 | yes | 1.747E-07 | 466.27164 | 630.59882 |
| IKYDAIAR | 8 | Q13573 | SNW domain-containing protein 1 OS=H | SNW1 | 61.495 | 2 | 55.204365 | yes | 9.722E-08 | 475.27692 | 559.73956 |
| GLQTVHINENFAK | 13 | Q13573 | SNW domain-containing protein 1 OS=H | SNW1 | 61.495 | 2 | 55.204365 | yes | 2.845E-07 | 735.88861 | 248.2868 |
| GLQTVHINENFAK | 13 | Q13573 | SNW domain-containing protein 1 OS=H | SNW1 | 61.495 | 2 | 55.204365 | yes | 3.397E-07 | 490.92816 | 248.2868 |
| GLQTVHINENFAK | 13 | Q13573 | SNW domain-containing protein 1 OS=H | SNW1 | 61.495 | 3 | 55.204365 | yes | 2.845E-07 | 735.88861 | 248.2868 |
| GLQTVHINENFAK | 13 | Q13573 | SNW domain-containing protein 1 OS=H | SNW1 | 61.495 | 3 | 55.204365 | yes | 3.397E-07 | 490.92816 | 248.2868 |
| TSNEVQYDQR | 10 | Q13573 | SNW domain-containing protein 1 OS=H | SNW1 | 61.495 | 2 | 55.204365 | yes | 4.618E-07 | 620.28363 | 323.54657 |
| FVPDKEFSGSDR | 12 | Q13573 | SNW domain-containing protein 1 OS=H | SNW1 | 61.495 | 2 | 55.204365 | yes | 1.136E-07 | 692.33063 | 323.38763 |
| REPPPYGYR | 9 | Q13573 | SNW domain-containing protein 1 OS=H | SNW1 | 61.495 | 2 | 55.204365 | yes | 9.64E-07 | 567.78815 | 151.92 |
| QTSLVSSR | 8 | Q13573 | SNW domain-containing protein 1 OS=H | SNW1 | 61.495 | 2 | 55.204365 | yes | 9.722E-08 | 439.24051 | 93.766685 |
| DPMEPPR | 7 | Q13573 | SNW domain-containing protein 1 OS=H | SNW1 | 61.495 | 2 | 55.204365 | yes | 1.384E-07 | 421.19727 | 75.841568 |
| NLDKDMYGDDLE | 14 | Q13573 | SNW domain-containing protein 1 OS=H | SNW1 | 61.495 | 2 | 55.204365 | yes | 9.722E-08 | 827.8725 | 51.669323 |
| DMAQSIYR | 8 | Q13573 | SNW domain-containing protein 1 OS=H | SNW1 | 61.495 | 2 | 55.204365 | yes | 5.546E-06 | 492.23438 | 32.569519 |
| TLNWGGGYTDEP | 15 | Q13574 | Diacylglycerol kinase zeta OS=Homo sap | DGKZ | 103.981 | 2 | 52.508305 | no | 9.722E-08 | 812.38611 | 85.3358 |
| AQDTELAAYLENI | 13 | Q13574 | Diacylglycerol kinase zeta OS=Homo sap | DGKZ | 103.981 | 2 | 52.508305 | yes | 9.975E-08 | 747.36517 | 64.79734 |
| PLLVFVNPK | 9 | Q13574 | Diacylglycerol kinase zeta OS=Homo sap | DGKZ | 103.981 | 2 | 52.508305 | no | 3.201E-07 | 513.82092 | 64.189156 |
| TLLHHAVSTGSK | 12 | Q13574 | Diacylglycerol kinase zeta OS=Homo sap | DGKZ | 103.981 | 2 | 52.508305 | yes | 3.305E-07 | 625.84619 | 18.838623 |
| YGPLSGVNVVYD | 14 | Q13595 | Transformer-2 protein homolog alpha OS | TRA2A | 32.689 | 2 | 52.786396 | yes | 1.418E-07 | 783.89917 | 2786.0239 |
| AHTPTPGIYMGR | 12 | Q13595 | Transformer-2 protein homolog alpha OS | TRA2A | 32.689 | 2 | 52.786396 | yes | 1.624E-07 | 650.82697 | 1163.5192 |
| GFAFVYFER | 9 | Q13595 | Transformer-2 protein homolog alpha OS | TRA2A | 32.689 | 2 | 52.786396 | yes | 1.677E-07 | 568.28217 | 1004.6196 |
| SDVEENNFEGR | 11 | Q13595 | Transformer-2 protein homolog alpha OS | TRA2A | 32.689 | 2 | 52.786396 | yes | 9.722E-08 | 669.28387 | 158.59634 |
| YEDYDYR | 7 | Q13595 | Transformer-2 protein homolog alpha OS | TRA2A | 32.689 | 2 | 52.786396 | yes | 2.297E-05 | 512.20636 | 72.499512 |
| ANPDPNTCLGVFG | 21 | Q13595 | Transformer-2 protein homolog alpha OS | TRA2A | 32.689 | 3 | 52.786396 | yes | 9.722E-08 | 775.71082 | 48.055344 |
| LGALTAEEIALK | 12 | Q13601 | KRR1 small subunit processome compon | KRR1 | 43.665 | 2 | 51.743843 | yes | 1.048E-07 | 614.86102 | 709.8576 |
| NIHPIYNIK | 9 | Q13601 | KRR1 small subunit processome compon | KRR1 | 43.665 | 2 | 51.743843 | yes | 9.935E-08 | 556.31659 | 484.59262 |
| ECWPLVQK | 8 | Q13601 | KRR1 small subunit processome compon | KRR1 | 43.665 | 2 | 51.743843 | yes | 1.582E-07 | 530.26825 | 412.76291 |
| KVVLDTMK | 8 | Q13601 | KRR1 small subunit processome compon | KRR1 | 43.665 | 2 | 51.743843 | yes | 1.032E-07 | 467.27551 | 285.61829 |
| TFDPYIIIR | 9 | Q13601 | KRR1 small subunit processome compon | KRR1 | 43.665 | 2 | 51.743843 | yes | 9.722E-08 | 569.31879 | 223.97374 |
| ELASGEYFLK | 10 | Q13601 | KRR1 small subunit processome compon | KRR1 | 43.665 | 2 | 51.743843 | yes | 4.149E-07 | 578.79785 | 169.6586 |
| ALNEHHVNATLDI | 24 | Q13601 | KRR1 small subunit processome compon | KRR1 | 43.665 | 4 | 51.743843 | yes | 4.486E-07 | 664.32825 | 68.080444 |
| LQPSVSEQELLR | 12 | Q13608 | Peroxisomal ATPase PEX6 OS=Homo sa | PEX6 | 104.062 | 2 | 51.82304 | yes | 2.134E-07 | 699.883 | 577.36194 |
| GSCSLLPGPPFAR | 13 | Q13608 | Peroxisomal ATPase PEX6 OS=Homo sa | PEX6 | 104.062 | 2 | 51.82304 | yes | 1.536E-07 | 679.8479 | 546.46515 |
| LLALGSGAWVR | 11 | Q13608 | Peroxisomal ATPase PEX6 OS=Homo sa | PEX6 | 104.062 | 2 | 51.82304 | yes | 1.346E-07 | 571.83765 | 499.00406 |
| LQGVLGGTGDSLC | 16 | Q13608 | Peroxisomal ATPase PEX6 OS=Homo sa | PEX6 | 104.062 | 2 | 51.82304 | yes | 9.722E-08 | 758.40991 | 450.44348 |
| LVFVGANEDR | 10 | Q13608 | Peroxisomal ATPase PEX6 OS=Homo sa | PEX6 | 104.062 | 2 | 51.82304 | yes | 9.722E-08 | 560.29327 | 420.78497 |

| | | | | | | | | | | | |
|---|---|---|---|---|---|---|---|---|---|---|---|
| LQPGGALLTGTSS' | 17 | Q13608 | Peroxisomal ATPase PEX6 OS=Homo sa | PEX6 | 104.062 | 2 | 51.82304 | yes | 1.667E-07 | 841.99359 | 295.27377 |
| VQVLEPR | 7 | Q13608 | Peroxisomal ATPase PEX6 OS=Homo sa | PEX6 | 104.062 | 2 | 51.82304 | yes | 1.059E-07 | 420.75052 | 231.25824 |
| SGLLLHGPPGTGK | 13 | Q13608 | Peroxisomal ATPase PEX6 OS=Homo sa | PEX6 | 104.062 | 2 | 51.82304 | yes | 1.007E-07 | 617.35114 | 207.94011 |
| LQAIFSR | 7 | Q13608 | Peroxisomal ATPase PEX6 OS=Homo sa | PEX6 | 104.062 | 2 | 51.82304 | yes | 4.68E-07 | 417.74524 | 144.01793 |
| YLEGSIAPEDK | 11 | Q13608 | Peroxisomal ATPase PEX6 OS=Homo sa | PEX6 | 104.062 | 2 | 51.82304 | yes | 9.722E-08 | 611.30353 | 139.58743 |
| ALTAHLPLGQEVN | 19 | Q13608 | Peroxisomal ATPase PEX6 OS=Homo sa | PEX6 | 104.062 | 3 | 51.82304 | yes | 9.809E-08 | 672.3847 | 124.54022 |
| DRDGLGEDAR | 10 | Q13608 | Peroxisomal ATPase PEX6 OS=Homo sa | PEX6 | 104.062 | 2 | 51.82304 | yes | 3.185E-06 | 552.25745 | 115.12105 |
| AVATECSLTFLSVI | 14 | Q13608 | Peroxisomal ATPase PEX6 OS=Homo sa | PEX6 | 104.062 | 2 | 51.82304 | yes | 1.172E-07 | 763.40015 | 79.54232 |
| PALQGLLGPGTR | 12 | Q13608 | Peroxisomal ATPase PEX6 OS=Homo sa | PEX6 | 104.062 | 2 | 51.82304 | yes | 7.037E-05 | 590.34583 | 57.492779 |
| GCLVTASADK | 10 | Q13610 | Periodic tryptophan protein 1 homolog O | PWP1 | 55.828 | 2 | 52.946167 | yes | 9.75E-08 | 511.25278 | 1462.4825 |
| QVTCVAWVR | 9 | Q13610 | Periodic tryptophan protein 1 homolog O | PWP1 | 55.828 | 2 | 52.946167 | yes | 1.375E-07 | 559.79242 | 1022.7755 |
| NSSISGPFGSR | 11 | Q13610 | Periodic tryptophan protein 1 homolog O | PWP1 | 55.828 | 2 | 52.946167 | yes | 1.197E-07 | 554.77271 | 759.2627 |
| SVALYDCR | 8 | Q13610 | Periodic tryptophan protein 1 homolog O | PWP1 | 55.828 | 2 | 52.946167 | yes | 2.123E-07 | 492.23438 | 622.85107 |
| PAASLAVHTDK | 11 | Q13610 | Periodic tryptophan protein 1 homolog O | PWP1 | 55.828 | 2 | 52.946167 | yes | 1.46E-07 | 555.30109 | 422.64691 |
| ETPDKVELSK | 10 | Q13610 | Periodic tryptophan protein 1 homolog O | PWP1 | 55.828 | 2 | 52.946167 | yes | 9.789E-08 | 573.30603 | 362.16736 |
| IWDILGDRPSLVHS | 15 | Q13610 | Periodic tryptophan protein 1 homolog O | PWP1 | 55.828 | 3 | 52.946167 | yes | 2.025E-07 | 588.65656 | 260.4838 |
| PSDNLIVCGR | 10 | Q13610 | Periodic tryptophan protein 1 homolog O | PWP1 | 55.828 | 2 | 52.946167 | yes | 1.06E-07 | 565.78479 | 201.6761 |
| SDKPIFTLNAHNDI | 25 | Q13610 | Periodic tryptophan protein 1 homolog O | PWP1 | 55.828 | 4 | 52.946167 | yes | 1.886E-07 | 686.35406 | 188.45798 |
| EPLEDGDPEDDR | 12 | Q13610 | Periodic tryptophan protein 1 homolog O | PWP1 | 55.828 | 2 | 52.946167 | yes | 1.374E-07 | 693.78644 | 146.13087 |
| NVLASASADNTVI | 32 | Q13610 | Periodic tryptophan protein 1 homolog O | PWP1 | 55.828 | 4 | 52.946167 | yes | 9.722E-08 | 824.68024 | 91.556602 |
| IWDILGDR | 8 | Q13610 | Periodic tryptophan protein 1 homolog O | PWP1 | 55.828 | 2 | 52.946167 | yes | 3.422E-07 | 494.26654 | 65.68222 |
| VQTLQFHPFEAQT | 21 | Q13610 | Periodic tryptophan protein 1 homolog O | PWP1 | 55.828 | 3 | 52.946167 | yes | 3.923E-07 | 803.74103 | 55.212337 |
| MLLAGAVR | 8 | Q13613 | Myotubularin-related protein 1 OS=Hom | MTMR1 | 74.678 | 2 | 49.132084 | yes | 7.573E-07 | 415.74948 | 162.35318 |
| LGIFENLNK | 9 | Q13613 | Myotubularin-related protein 1 OS=Hom | MTMR1 | 74.678 | 2 | 49.132084 | yes | 1.065E-07 | 524.29529 | 122.94913 |
| EIVYPSIDEAR | 11 | Q13613 | Myotubularin-related protein 1 OS=Hom | MTMR1 | 74.678 | 2 | 49.132084 | yes | 1.349E-06 | 646.33008 | 105.9587 |
| EWISFGHR | 8 | Q13613 | Myotubularin-related protein 1 OS=Hom | MTMR1 | 74.678 | 2 | 49.132084 | yes | 2.221E-06 | 516.25653 | 81.710709 |
| CSQPLVGPNDK | 11 | Q13613 | Myotubularin-related protein 1 OS=Hom | MTMR1 | 74.678 | 2 | 49.132084 | yes | 5.159E-06 | 607.79535 | 58.785301 |
| TSVVVHCSDGWD | 13 | Q13613 | Myotubularin-related protein 1 OS=Hom | MTMR1 | 74.678 | 2 | 49.132084 | no | 1.908E-05 | 759.34369 | 28.420959 |
| LYDPLLEYR | 9 | Q13614 | Myotubularin-related protein 2 OS=Hom | MTMR2 | 73.382 | 2 | 51.473804 | yes | 1.097E-07 | 591.31366 | 166.93942 |
| YLQAIMDSNAQSF | 14 | Q13614 | Myotubularin-related protein 2 OS=Hom | MTMR2 | 73.382 | 2 | 51.473804 | yes | 9.722E-08 | 803.38812 | 155.1058 |
| GENSYGLETVCK | 12 | Q13614 | Myotubularin-related protein 2 OS=Hom | MTMR2 | 73.382 | 2 | 51.473804 | yes | 9.722E-08 | 678.80865 | 88.168961 |
| IFIFDAR | 7 | Q13614 | Myotubularin-related protein 2 OS=Hom | MTMR2 | 73.382 | 2 | 51.473804 | yes | 7.639E-07 | 441.24762 | 58.7388 |
| MKPQEPIHNR | 10 | Q13614 | Myotubularin-related protein 2 OS=Hom | MTMR2 | 73.382 | 3 | 51.473804 | yes | 1.373E-07 | 417.22052 | 34.426556 |
| GEDSLEVPVEQFR | 13 | Q13615 | Myotubularin-related protein 3 OS=Hom | MTMR3 | 133.619 | 2 | 48.820812 | yes | 1.101E-07 | 752.86755 | 79.190094 |
| GQTPGGAQFVGLE | 16 | Q13616 | Cullin-1 OS=Homo sapiens OX=9606 G | CUL1 | 89.679 | 2 | 46.519753 | yes | 3.988E-07 | 832.93579 | 72.686829 |
| ESFESQFLADTER | 13 | Q13616 | Cullin-1 OS=Homo sapiens OX=9606 G | CUL1 | 89.679 | 2 | 46.519753 | yes | 1.537E-07 | 779.85461 | 61.395458 |
| LLIQAAIVR | 9 | Q13616 | Cullin-1 OS=Homo sapiens OX=9606 G | CUL1 | 89.679 | 2 | 46.519753 | yes | 3.963E-07 | 498.83185 | 47.241463 |
| VLNGICAYLNR | 11 | Q13616 | Cullin-1 OS=Homo sapiens OX=9606 G | CUL1 | 89.679 | 2 | 46.519753 | yes | 5.936E-07 | 646.84265 | 34.690113 |
| IQDGLGELK | 9 | Q13616 | Cullin-1 OS=Homo sapiens OX=9606 G | CUL1 | 89.679 | 2 | 46.519753 | yes | 2.784E-06 | 486.77167 | 34.636524 |

| | | | | | | | | | | | |
|---|---|---|---|---|---|---|---|---|---|---|---|
| MFQDIGVSK | 9 | Q13616 | Cullin-1 OS=Homo sapiens OX=9606 GN | CUL1 | 89.679 | 2 | 46.519753 | yes | 1.223E-05 | 512.76025 | 28.94376 |
| ALTSVVNYR | 9 | Q13617 | Cullin-2 OS=Homo sapiens OX=9606 GN | CUL2 | 86.984 | 2 | 45.804173 | yes | 1.736E-07 | 511.7851 | 84.011551 |
| AVVMLEYVER | 10 | Q13617 | Cullin-2 OS=Homo sapiens OX=9606 GN | CUL2 | 86.984 | 2 | 45.804173 | yes | 8.09E-07 | 604.8208 | 62.339859 |
| VLTTGYWPTQSAT | 15 | Q13618 | Cullin-3 OS=Homo sapiens OX=9606 GN | CUL3 | 88.931 | 2 | 54.12789 | yes | 2.45E-07 | 825.4303 | 128.82681 |
| GLTEQEVETILDK | 13 | Q13618 | Cullin-3 OS=Homo sapiens OX=9606 GN | CUL3 | 88.931 | 2 | 54.12789 | yes | 9.722E-08 | 737.88538 | 103.68624 |
| FLLESFNNDR | 10 | Q13618 | Cullin-3 OS=Homo sapiens OX=9606 GN | CUL3 | 88.931 | 2 | 54.12789 | yes | 1.112E-06 | 627.80933 | 57.321152 |
| EVVTEHLINK | 10 | Q13618 | Cullin-3 OS=Homo sapiens OX=9606 GN | CUL3 | 88.931 | 2 | 54.12789 | yes | 3.432E-07 | 591.3299 | 49.670883 |
| MQHNVLVAEVTQ | 15 | Q13618 | Cullin-3 OS=Homo sapiens OX=9606 GN | CUL3 | 88.931 | 3 | 54.12789 | yes | 3.121E-07 | 579.98206 | 42.94949 |
| DILMYMDR | 8 | Q13618 | Cullin-3 OS=Homo sapiens OX=9606 GN | CUL3 | 88.931 | 2 | 54.12789 | yes | 3.037E-06 | 528.74628 | 26.701994 |
| TMCECMSSYLR | 11 | Q13618 | Cullin-3 OS=Homo sapiens OX=9606 GN | CUL3 | 88.931 | 2 | 54.12789 | yes | 2.961E-07 | 719.2851 | 17.362564 |
| DIMVHFK | 7 | Q13619 | Cullin-4A OS=Homo sapiens OX=9606 ( | CUL4A | 87.681 | 2 | 47.567657 | yes | 2.949E-07 | 445.23364 | 133.65598 |
| MATGIEDSELR | 11 | Q13619 | Cullin-4A OS=Homo sapiens OX=9606 ( | CUL4A | 87.681 | 2 | 47.567657 | yes | 3.245E-07 | 611.2926 | 90.243126 |
| QYQIDAAIVR | 10 | Q13619 | Cullin-4A OS=Homo sapiens OX=9606 ( | CUL4A | 87.681 | 2 | 47.567657 | no | 2.537E-07 | 588.8222 | 79.102287 |
| TLQSLACGK | 9 | Q13619 | Cullin-4A OS=Homo sapiens OX=9606 ( | CUL4A | 87.681 | 2 | 47.567657 | no | 4.983E-07 | 489.25787 | 62.097462 |
| VITYLDHSTQKPLI | 19 | Q13619 | Cullin-4A OS=Homo sapiens OX=9606 ( | CUL4A | 87.681 | 3 | 47.567657 | yes | 2.642E-07 | 739.06458 | 59.848591 |
| KGSFSALVGR | 10 | Q13619 | Cullin-4A OS=Homo sapiens OX=9606 ( | CUL4A | 87.681 | 2 | 47.567657 | yes | 2.387E-06 | 511.29309 | 56.62455 |
| LPDNYTQDTWR | 11 | Q13619 | Cullin-4A OS=Homo sapiens OX=9606 ( | CUL4A | 87.681 | 2 | 47.567657 | yes | 1.176E-06 | 704.82825 | 51.613388 |
| ESFETFINK | 9 | Q13619 | Cullin-4A OS=Homo sapiens OX=9606 ( | CUL4A | 87.681 | 2 | 47.567657 | yes | 7.443E-07 | 557.77441 | 45.566498 |
| EDSLDSVLFLK | 11 | Q13619 | Cullin-4A OS=Homo sapiens OX=9606 ( | CUL4A | 87.681 | 2 | 47.567657 | no | 1.302E-07 | 633.33484 | 43.792412 |
| GLDHLLDENR | 10 | Q13619 | Cullin-4A OS=Homo sapiens OX=9606 ( | CUL4A | 87.681 | 2 | 47.567657 | yes | 5.597E-07 | 591.29907 | 42.203484 |
| DKDMVQDLLDFK | 12 | Q13619 | Cullin-4A OS=Homo sapiens OX=9606 ( | CUL4A | 87.681 | 2 | 47.567657 | yes | 3.085E-06 | 733.8634 | 28.627253 |
| QLLGEHLTAILQK | 13 | Q13619 | Cullin-4A OS=Homo sapiens OX=9606 ( | CUL4A | 87.681 | 2 | 47.567657 | no | 1.441E-07 | 732.43268 | 17.499897 |
| NQSDADLEALR | 11 | Q13625 | Apoptosis-stimulating of p53 protein 2 OS | TP53BP2 | 125.617 | 2 | 47.775932 | yes | 1.26E-07 | 616.29932 | 47.183384 |
| MDLTLAELQEMAS | 14 | Q13625 | Apoptosis-stimulating of p53 protein 2 OS | TP53BP2 | 125.617 | 2 | 47.775932 | yes | 3.079E-07 | 804.39203 | 22.215185 |
| IVDFGSSCQLGQR | 13 | Q13627 | Dual specificity tyrosine-phosphorylation | DYRK1A | 85.584 | 2 | 53.53628 | no | 9.722E-08 | 733.85645 | 472.58719 |
| YYIVHLK | 7 | Q13627 | Dual specificity tyrosine-phosphorylation | DYRK1A | 85.584 | 2 | 53.53628 | no | 9.81E-08 | 468.27109 | 430.2868 |
| IYQYIQSR | 8 | Q13627 | Dual specificity tyrosine-phosphorylation | DYRK1A | 85.584 | 2 | 53.53628 | no | 9.722E-08 | 535.7851 | 352.44968 |
| HINEVYYAK | 9 | Q13627 | Dual specificity tyrosine-phosphorylation | DYRK1A | 85.584 | 2 | 53.53628 | no | 9.722E-08 | 568.79041 | 210.30415 |
| GVSLNLTR | 8 | Q13627 | Dual specificity tyrosine-phosphorylation | DYRK1A | 85.584 | 2 | 53.53628 | no | 1.105E-07 | 430.25345 | 207.91618 |
| VEQEWVAIK | 9 | Q13627 | Dual specificity tyrosine-phosphorylation | DYRK1A | 85.584 | 2 | 53.53628 | yes | 2.024E-07 | 551.3006 | 201.80025 |
| MLDYDPK | 7 | Q13627 | Dual specificity tyrosine-phosphorylation | DYRK1A | 85.584 | 2 | 53.53628 | yes | 9.722E-08 | 441.20731 | 147.83325 |
| LPDGTWNLK | 9 | Q13627 | Dual specificity tyrosine-phosphorylation | DYRK1A | 85.584 | 2 | 53.53628 | yes | 2.879E-06 | 522.27966 | 141.2785 |
| KLSVDLIK | 8 | Q13627 | Dual specificity tyrosine-phosphorylation | DYRK1A | 85.584 | 2 | 53.53628 | no | 6.786E-07 | 458.2973 | 132.68854 |
| LHNILGVETGGPGG | 15 | Q13627 | Dual specificity tyrosine-phosphorylation | DYRK1A | 85.584 | 2 | 53.53628 | yes | 7.188E-07 | 738.89954 | 116.4323 |
| HSGGHFTAAVQAN | 23 | Q13627 | Dual specificity tyrosine-phosphorylation | DYRK1A | 85.584 | 4 | 53.53628 | yes | 2.035E-07 | 631.53674 | 44.798882 |
| HSGGHFTAAVQAN | 23 | Q13627 | Dual specificity tyrosine-phosphorylation | DYRK1A | 85.584 | 4 | 53.53628 | yes | 4.626E-06 | 841.71326 | 44.798882 |
| HSGGHFTAAVQAN | 23 | Q13627 | Dual specificity tyrosine-phosphorylation | DYRK1A | 85.584 | 3 | 53.53628 | yes | 2.035E-07 | 631.53674 | 44.798882 |
| HSGGHFTAAVQAN | 23 | Q13627 | Dual specificity tyrosine-phosphorylation | DYRK1A | 85.584 | 3 | 53.53628 | yes | 4.626E-06 | 841.71326 | 44.798882 |
| LLELMNK | 7 | Q13627 | Dual specificity tyrosine-phosphorylation | DYRK1A | 85.584 | 2 | 53.53628 | yes | 2.119E-07 | 430.74915 | 86.077263 |

| | | | | | | | | | | | |
|---|---|---|---|---|---|---|---|---|---|---|---|
| DPATAPLR | 8 | Q13627 | Dual specificity tyrosine-phosphorylation | DYRK1A | 85.584 | 2 | 53.53628 | yes | 3.34E-07 | 420.73233 | 75.238678 |
| TLTQGGVTYR | 10 | Q13643 | Four and a half LIM domains protein 3 O | FHL3 | 31.192 | 2 | 46.374073 | yes | 4.004E-07 | 548.29327 | 358.93256 |
| ELFYEDR | 7 | Q13643 | Four and a half LIM domains protein 3 O | FHL3 | 31.192 | 2 | 46.374073 | yes | 2.635E-07 | 486.22708 | 328.78131 |
| YVSFEDR | 7 | Q13643 | Four and a half LIM domains protein 3 O | FHL3 | 31.192 | 2 | 46.374073 | yes | 1.577E-07 | 458.21399 | 327.53052 |
| GAHYCVPCYENK | 12 | Q13643 | Four and a half LIM domains protein 3 O | FHL3 | 31.192 | 2 | 46.374073 | yes | 5.927E-07 | 749.3161 | 196.82065 |
| HFHEGCFR | 8 | Q13643 | Four and a half LIM domains protein 3 O | FHL3 | 31.192 | 2 | 46.374073 | yes | 3.423E-07 | 545.23779 | 113.01636 |
| RPIVGLGGGK | 10 | Q13643 | Four and a half LIM domains protein 3 O | FHL3 | 31.192 | 2 | 46.374073 | yes | 5.797E-07 | 477.29819 | 84.357285 |
| CNESLYGR | 8 | Q13643 | Four and a half LIM domains protein 3 O | FHL3 | 31.192 | 2 | 46.374073 | yes | 5.658E-07 | 499.72165 | 36.065044 |
| AGPEPQELQGIR | 12 | Q13671 | Ras and Rab interactor 1 OS=Homo sapie | RIN1 | 84.099 | 2 | 51.33079 | yes | 1.561E-07 | 647.84113 | 88.748039 |
| TEPPGTFLVR | 10 | Q13671 | Ras and Rab interactor 1 OS=Homo sapie | RIN1 | 84.099 | 2 | 51.33079 | yes | 5.43E-07 | 558.80603 | 60.011536 |
| GPAGGPVLPQLK | 12 | Q13671 | Ras and Rab interactor 1 OS=Homo sapie | RIN1 | 84.099 | 2 | 51.33079 | yes | 4.328E-06 | 567.33752 | 50.028667 |
| LPTTGYLVYR | 10 | Q13671 | Ras and Rab interactor 1 OS=Homo sapie | RIN1 | 84.099 | 2 | 51.33079 | yes | 9.722E-08 | 591.82953 | 48.527069 |
| VAYQDPSSGCTSK | 13 | Q13671 | Ras and Rab interactor 1 OS=Homo sapie | RIN1 | 84.099 | 2 | 51.33079 | yes | 1.584E-07 | 700.31152 | 31.925009 |
| TNTLAVTGGEDDK | 13 | Q13685 | Angio-associated migratory cell protein C | AAMP | 46.751 | 2 | 45.175056 | yes | 2.053E-07 | 660.8175 | 75.753151 |
| LSDGELLFECAGH | 14 | Q13685 | Angio-associated migratory cell protein C | AAMP | 46.751 | 2 | 45.175056 | yes | 2.645E-07 | 788.3772 | 61.819416 |
| DASLVVTTSGDHK | 13 | Q13685 | Angio-associated migratory cell protein C | AAMP | 46.751 | 2 | 45.175056 | yes | 1.936E-07 | 665.33588 | 53.915474 |
| TFQGPNCPATCGR | 13 | Q13685 | Angio-associated migratory cell protein C | AAMP | 46.751 | 2 | 45.175056 | yes | 3.339E-07 | 733.31921 | 33.064053 |
| GHTAEILDFALSK | 13 | Q13685 | Angio-associated migratory cell protein C | AAMP | 46.751 | 2 | 45.175056 | yes | 3.075E-07 | 701.37225 | 29.601238 |
| VPPEFLVQR | 9 | Q13724 | Mannosyl-oligosaccharide glucosidase OS | MOGS | 91.918 | 2 | 51.977863 | yes | 2.052E-07 | 542.8111 | 926.30432 |
| FTLLPPTSPGDTAP | 15 | Q13724 | Mannosyl-oligosaccharide glucosidase OS | MOGS | 91.918 | 2 | 51.977863 | yes | 3.427E-07 | 771.41412 | 498.50574 |
| LLDPTSSR | 8 | Q13724 | Mannosyl-oligosaccharide glucosidase OS | MOGS | 91.918 | 2 | 51.977863 | yes | 9.759E-08 | 444.74289 | 321.22446 |
| EQILGDEAR | 9 | Q13724 | Mannosyl-oligosaccharide glucosidase OS | MOGS | 91.918 | 2 | 51.977863 | yes | 2.388E-06 | 515.76184 | 307.20447 |
| SLAASSSFYGQR | 12 | Q13724 | Mannosyl-oligosaccharide glucosidase OS | MOGS | 91.918 | 2 | 51.977863 | yes | 9.722E-08 | 637.31219 | 259.51862 |
| PHVYFGMK | 8 | Q13724 | Mannosyl-oligosaccharide glucosidase OS | MOGS | 91.918 | 2 | 51.977863 | yes | 2.896E-07 | 489.74692 | 221.04764 |
| TLPSGLDDYPR | 11 | Q13724 | Mannosyl-oligosaccharide glucosidase OS | MOGS | 91.918 | 2 | 51.977863 | yes | 9.54E-07 | 617.30914 | 213.25926 |
| GRDPALPTLLNPK | 13 | Q13724 | Mannosyl-oligosaccharide glucosidase OS | MOGS | 91.918 | 2 | 51.977863 | yes | 1.3E-07 | 696.40393 | 121.01156 |
| NSEHDPPYWR | 10 | Q13724 | Mannosyl-oligosaccharide glucosidase OS | MOGS | 91.918 | 2 | 51.977863 | yes | 1.029E-06 | 650.78888 | 120.62695 |
| CWVALGAR | 8 | Q13724 | Mannosyl-oligosaccharide glucosidase OS | MOGS | 91.918 | 2 | 51.977863 | yes | 1.899E-07 | 466.74219 | 95.213951 |
| SPKPLLTGLMWAC | 22 | Q13724 | Mannosyl-oligosaccharide glucosidase OS | MOGS | 91.918 | 3 | 51.977863 | yes | 2.638E-07 | 770.41559 | 75.976624 |
| WVLAWYR | 7 | Q13724 | Mannosyl-oligosaccharide glucosidase OS | MOGS | 91.918 | 2 | 51.977863 | yes | 1.531E-07 | 497.26889 | 69.882629 |
| AVQLKPRPPQGLV | 14 | Q13724 | Mannosyl-oligosaccharide glucosidase OS | MOGS | 91.918 | 3 | 51.977863 | yes | 2.945E-07 | 520.32288 | 59.838547 |
| HLWSPFGLR | 9 | Q13724 | Mannosyl-oligosaccharide glucosidase OS | MOGS | 91.918 | 2 | 51.977863 | yes | 5.67E-07 | 556.80365 | 52.457569 |
| EVLLPEVGAK | 10 | Q13724 | Mannosyl-oligosaccharide glucosidase OS | MOGS | 91.918 | 2 | 51.977863 | yes | 4.818E-07 | 527.81079 | 48.61301 |
| IQEVGEITNLR | 11 | Q13751 | Laminin subunit beta-3 OS=Homo sapien | LAMB3 | 129.573 | 2 | 53.673073 | yes | 9.722E-08 | 636.35132 | 333.14114 |
| MNEIQAIAAR | 10 | Q13751 | Laminin subunit beta-3 OS=Homo sapien | LAMB3 | 129.573 | 2 | 53.673073 | yes | 1.988E-07 | 558.79517 | 149.99806 |
| AGGAFLMAGQVA | 16 | Q13751 | Laminin subunit beta-3 OS=Homo sapien | LAMB3 | 129.573 | 2 | 53.673073 | yes | 9.722E-08 | 809.92212 | 137.1221 |
| LEMSSLPDLTPTFN | 15 | Q13751 | Laminin subunit beta-3 OS=Homo sapien | LAMB3 | 129.573 | 2 | 53.673073 | yes | 3.548E-06 | 846.92926 | 108.19653 |
| GACYPPVGDLLVC | 14 | Q13751 | Laminin subunit beta-3 OS=Homo sapien | LAMB3 | 129.573 | 2 | 53.673073 | yes | 1.41E-06 | 737.37958 | 99.030548 |
| LPNVDLVLSQTK | 12 | Q13751 | Laminin subunit beta-3 OS=Homo sapien | LAMB3 | 129.573 | 2 | 53.673073 | yes | 9.722E-08 | 663.88501 | 96.932297 |

| | | | | | | | | | | | |
|---|---|---|---|---|---|---|---|---|---|---|---|
| LLIQQVR | 7 | Q13751 | Laminin subunit beta-3 OS=Homo sapien | LAMB3 | 129.573 | 2 | 53.673073 | yes | 1.12E-07 | 435.28198 | 92.723488 |
| CLCLPNVVGPK | 11 | Q13751 | Laminin subunit beta-3 OS=Homo sapien | LAMB3 | 129.573 | 2 | 53.673073 | yes | 1.076E-07 | 628.82813 | 83.948875 |
| EGFGGLMCSAAAI | 14 | Q13751 | Laminin subunit beta-3 OS=Homo sapien | LAMB3 | 129.573 | 2 | 53.673073 | yes | 1.29E-07 | 720.3421 | 71.777672 |
| LGQSSMLGEQGAI | 13 | Q13751 | Laminin subunit beta-3 OS=Homo sapien | LAMB3 | 129.573 | 2 | 53.673073 | yes | 1.124E-07 | 667.33008 | 59.347191 |
| VAEVQQVLRPAEI | 13 | Q13751 | Laminin subunit beta-3 OS=Homo sapien | LAMB3 | 129.573 | 2 | 53.673073 | yes | 9.722E-08 | 733.91992 | 42.730114 |
| AAEESASQIQSSA( | 15 | Q13751 | Laminin subunit beta-3 OS=Homo sapien | LAMB3 | 129.573 | 2 | 53.673073 | yes | 1.797E-07 | 781.8739 | 39.042259 |
| VENVASSSGPMR | 12 | Q13751 | Laminin subunit beta-3 OS=Homo sapien | LAMB3 | 129.573 | 2 | 53.673073 | yes | 1.682E-07 | 617.29822 | 24.509167 |
| LMAEIQDLK | 9 | Q13769 | THO complex subunit 5 homolog OS=Ho | THOC5 | 78.508 | 2 | 57.3749 | yes | 9.722E-08 | 530.789 | 203.48935 |
| YTCQELQR | 8 | Q13769 | THO complex subunit 5 homolog OS=Ho | THOC5 | 78.508 | 2 | 57.3749 | yes | 1.147E-07 | 549.25586 | 189.79227 |
| AMEGEVNVCYK | 11 | Q13769 | THO complex subunit 5 homolog OS=Ho | THOC5 | 78.508 | 2 | 57.3749 | yes | 9.722E-08 | 650.289 | 129.16989 |
| LGGLHFPK | 8 | Q13769 | THO complex subunit 5 homolog OS=Ho | THOC5 | 78.508 | 2 | 57.3749 | yes | 5.323E-07 | 434.75562 | 105.9277 |
| YYSEEAEVDLR | 11 | Q13769 | THO complex subunit 5 homolog OS=Ho | THOC5 | 78.508 | 2 | 57.3749 | yes | 1.509E-07 | 687.31464 | 82.408195 |
| IQSCVHFMTLK | 11 | Q13769 | THO complex subunit 5 homolog OS=Ho | THOC5 | 78.508 | 2 | 57.3749 | yes | 9.722E-08 | 682.34668 | 66.269562 |
| KTPNPANQYQFDI | 13 | Q13769 | THO complex subunit 5 homolog OS=Ho | THOC5 | 78.508 | 2 | 57.3749 | yes | 2.669E-06 | 775.88354 | 33.087254 |
| AQIHSCISR | 9 | Q13772 | Nuclear receptor coactivator 4 OS=Homo | NCOA4 | 69.727 | 2 | 44.973057 | yes | 4.294E-07 | 536.27185 | 119.03656 |
| GVEIENLGNLK | 11 | Q13772 | Nuclear receptor coactivator 4 OS=Homo | NCOA4 | 69.727 | 2 | 44.973057 | yes | 2.007E-07 | 593.32733 | 82.000587 |
| CLNDHLEAK | 9 | Q13772 | Nuclear receptor coactivator 4 OS=Homo | NCOA4 | 69.727 | 2 | 44.973057 | yes | 8.858E-07 | 550.26367 | 30.910883 |
| ITTTVGLNIGTVD\ | 16 | Q13795 | ADP-ribosylation factor-related protein 1 | ARFRP1 | 22.615 | 2 | 34.226242 | yes | 1.052E-06 | 794.45123 | 18.217407 |
| DLTGVQNLR | 9 | Q13813 | Spectrin alpha chain, non-erythrocytic 1 ( | SPTAN1 | 284.54 | 2 | 54.9534 | yes | 4.362E-07 | 508.28018 | 149.57498 |
| LLEAQSHFR | 9 | Q13813 | Spectrin alpha chain, non-erythrocytic 1 ( | SPTAN1 | 284.54 | 2 | 54.9534 | yes | 9.722E-08 | 550.79602 | 144.44212 |
| HALLEADVAAHQI | 14 | Q13813 | Spectrin alpha chain, non-erythrocytic 1 ( | SPTAN1 | 284.54 | 3 | 54.9534 | yes | 4.74E-07 | 515.93048 | 143.90543 |
| GVIDMGNSLIER | 12 | Q13813 | Spectrin alpha chain, non-erythrocytic 1 ( | SPTAN1 | 284.54 | 2 | 54.9534 | yes | 9.722E-08 | 652.33734 | 137.91327 |
| KLDPAQSASR | 10 | Q13813 | Spectrin alpha chain, non-erythrocytic 1 ( | SPTAN1 | 284.54 | 2 | 54.9534 | yes | 1.589E-06 | 536.79089 | 136.56979 |
| VLETAEDIQER | 11 | Q13813 | Spectrin alpha chain, non-erythrocytic 1 ( | SPTAN1 | 284.54 | 2 | 54.9534 | yes | 1.208E-07 | 651.83044 | 124.44379 |
| ELPTAFDYVEFTR | 13 | Q13813 | Spectrin alpha chain, non-erythrocytic 1 ( | SPTAN1 | 284.54 | 2 | 54.9534 | yes | 9.81E-08 | 794.38812 | 118.96677 |
| EELYQNLTR | 9 | Q13813 | Spectrin alpha chain, non-erythrocytic 1 ( | SPTAN1 | 284.54 | 2 | 54.9534 | yes | 3.431E-07 | 583.29602 | 116.17912 |
| LDILDQER | 8 | Q13813 | Spectrin alpha chain, non-erythrocytic 1 ( | SPTAN1 | 284.54 | 2 | 54.9534 | yes | 4.176E-07 | 501.26672 | 106.78214 |
| GACAGSEDAVK | 11 | Q13813 | Spectrin alpha chain, non-erythrocytic 1 ( | SPTAN1 | 284.54 | 2 | 54.9534 | yes | 3.705E-07 | 532.73749 | 106.51344 |
| LAQFVEHWK | 9 | Q13813 | Spectrin alpha chain, non-erythrocytic 1 ( | SPTAN1 | 284.54 | 2 | 54.9534 | yes | 9.722E-08 | 579.30872 | 98.325218 |
| GEIDAHEDSFK | 11 | Q13813 | Spectrin alpha chain, non-erythrocytic 1 ( | SPTAN1 | 284.54 | 2 | 54.9534 | yes | 7.557E-07 | 624.28058 | 95.507561 |
| VNSLGETAER | 10 | Q13813 | Spectrin alpha chain, non-erythrocytic 1 ( | SPTAN1 | 284.54 | 2 | 54.9534 | yes | 6.521E-07 | 538.27252 | 85.742455 |
| LIDVNHYAK | 9 | Q13813 | Spectrin alpha chain, non-erythrocytic 1 ( | SPTAN1 | 284.54 | 2 | 54.9534 | yes | 2.036E-07 | 536.79291 | 85.390938 |
| MNEVISLWK | 9 | Q13813 | Spectrin alpha chain, non-erythrocytic 1 ( | SPTAN1 | 284.54 | 2 | 54.9534 | yes | 9.722E-08 | 560.297 | 84.059196 |
| DLASVNNLLK | 10 | Q13813 | Spectrin alpha chain, non-erythrocytic 1 ( | SPTAN1 | 284.54 | 2 | 54.9534 | yes | 9.004E-07 | 543.81134 | 83.352539 |
| QQVAPTDDETGK | 12 | Q13813 | Spectrin alpha chain, non-erythrocytic 1 ( | SPTAN1 | 284.54 | 2 | 54.9534 | yes | 1.97E-07 | 644.80444 | 80.75647 |
| KQEALVAR | 8 | Q13813 | Spectrin alpha chain, non-erythrocytic 1 ( | SPTAN1 | 284.54 | 2 | 54.9534 | yes | 3.478E-07 | 457.77454 | 78.342484 |
| LGDSHDLQR | 9 | Q13813 | Spectrin alpha chain, non-erythrocytic 1 ( | SPTAN1 | 284.54 | 2 | 54.9534 | yes | 3.506E-07 | 520.75958 | 75.09684 |
| EKEPIAASTNR | 11 | Q13813 | Spectrin alpha chain, non-erythrocytic 1 ( | SPTAN1 | 284.54 | 2 | 54.9534 | yes | 9.722E-08 | 608.32001 | 72.540771 |
| ECEDVMDWINDK | 12 | Q13813 | Spectrin alpha chain, non-erythrocytic 1 ( | SPTAN1 | 284.54 | 2 | 54.9534 | yes | 1.873E-07 | 777.31598 | 68.568588 |

| | | | | | | | | | | | |
|---|---|---|---|---|---|---|---|---|---|---|---|
| LLVGSEDYGR | 10 | Q13813 | Spectrin alpha chain, non-erythrocytic 1 ( | SPTAN1 | 284.54 | 2 | 54.9534 | yes | 5.876E-07 | 554.78528 | 67.337074 |
| DVDEIEAWISEK | 12 | Q13813 | Spectrin alpha chain, non-erythrocytic 1 ( | SPTAN1 | 284.54 | 2 | 54.9534 | yes | 1.705E-07 | 717.34338 | 66.848083 |
| DVTGAEALLER | 11 | Q13813 | Spectrin alpha chain, non-erythrocytic 1 ( | SPTAN1 | 284.54 | 2 | 54.9534 | yes | 9.722E-08 | 587.30914 | 62.433296 |
| ENLLEEQGSIALR | 13 | Q13813 | Spectrin alpha chain, non-erythrocytic 1 ( | SPTAN1 | 284.54 | 2 | 54.9534 | yes | 1.685E-07 | 736.39117 | 59.432884 |
| LIQEQHPEEELIK | 13 | Q13813 | Spectrin alpha chain, non-erythrocytic 1 ( | SPTAN1 | 284.54 | 2 | 54.9534 | yes | 9.722E-08 | 803.4278 | 51.229561 |
| HQLLEADISAHED | 14 | Q13813 | Spectrin alpha chain, non-erythrocytic 1 ( | SPTAN1 | 284.54 | 3 | 54.9534 | yes | 9.819E-08 | 545.26917 | 47.895145 |
| QKLDILDQER | 10 | Q13813 | Spectrin alpha chain, non-erythrocytic 1 ( | SPTAN1 | 284.54 | 2 | 54.9534 | yes | 6.388E-07 | 629.34351 | 46.131252 |
| DLTSWVTEMK | 10 | Q13813 | Spectrin alpha chain, non-erythrocytic 1 ( | SPTAN1 | 284.54 | 2 | 54.9534 | yes | 1.764E-06 | 605.29462 | 41.458923 |
| DMDDEESWIK | 10 | Q13813 | Spectrin alpha chain, non-erythrocytic 1 ( | SPTAN1 | 284.54 | 2 | 54.9534 | yes | 3.654E-07 | 634.26099 | 38.286366 |
| LQQSHPLSATQIQ | 15 | Q13813 | Spectrin alpha chain, non-erythrocytic 1 ( | SPTAN1 | 284.54 | 2 | 54.9534 | yes | 5.397E-07 | 839.46777 | 36.983868 |
| AQLADSFHLQQFF | 14 | Q13813 | Spectrin alpha chain, non-erythrocytic 1 ( | SPTAN1 | 284.54 | 3 | 54.9534 | yes | 2.427E-06 | 569.95825 | 35.59383 |
| LDENSAFLQFNWK | 13 | Q13813 | Spectrin alpha chain, non-erythrocytic 1 ( | SPTAN1 | 284.54 | 2 | 54.9534 | yes | 1.406E-05 | 806.39374 | 35.200642 |
| ALSSEGKPYVTK | 12 | Q13813 | Spectrin alpha chain, non-erythrocytic 1 ( | SPTAN1 | 284.54 | 2 | 54.9534 | yes | 3.046E-07 | 640.34827 | 32.744377 |
| HQAFEAELHANAI | 14 | Q13813 | Spectrin alpha chain, non-erythrocytic 1 ( | SPTAN1 | 284.54 | 3 | 54.9534 | yes | 2.117E-07 | 536.92194 | 31.126421 |
| EAIVTSEELGQDLI | 21 | Q13813 | Spectrin alpha chain, non-erythrocytic 1 ( | SPTAN1 | 284.54 | 3 | 54.9534 | yes | 4.361E-07 | 789.40771 | 28.408657 |
| MQHNLEQQIQAR | 12 | Q13813 | Spectrin alpha chain, non-erythrocytic 1 ( | SPTAN1 | 284.54 | 2 | 54.9534 | yes | 2.742E-07 | 748.37537 | 26.861855 |
| LAALADQWQFLV | 14 | Q13813 | Spectrin alpha chain, non-erythrocytic 1 ( | SPTAN1 | 284.54 | 2 | 54.9534 | yes | 1.393E-05 | 815.95123 | 24.722992 |
| KVEDLFLTFAK | 11 | Q13813 | Spectrin alpha chain, non-erythrocytic 1 ( | SPTAN1 | 284.54 | 2 | 54.9534 | yes | 4.58E-07 | 655.8714 | 24.147577 |
| ELVLALYDYQEK | 12 | Q13813 | Spectrin alpha chain, non-erythrocytic 1 ( | SPTAN1 | 284.54 | 2 | 54.9534 | yes | 0.0001298 | 742.38757 | 18.760612 |
| DLSSVQTLLTK | 11 | Q13813 | Spectrin alpha chain, non-erythrocytic 1 ( | SPTAN1 | 284.54 | 2 | 54.9534 | yes | 6.846E-07 | 602.84283 | 18.262312 |
| VIDSSDVVVQVLD | 15 | Q13823 | Nucleolar GTP-binding protein 2 OS=Ho | GNL2 | 83.655 | 2 | 53.099552 | yes | 1.464E-07 | 807.93848 | 1331.9655 |
| MVLNDWQR | 8 | Q13823 | Nucleolar GTP-binding protein 2 OS=Ho | GNL2 | 83.655 | 2 | 53.099552 | yes | 1.984E-07 | 531.26349 | 1176.5209 |
| LPMSLLHDR | 9 | Q13823 | Nucleolar GTP-binding protein 2 OS=Ho | GNL2 | 83.655 | 2 | 53.099552 | yes | 1.659E-07 | 541.2948 | 1075.2671 |
| IFAKPEEQR | 9 | Q13823 | Nucleolar GTP-binding protein 2 OS=Ho | GNL2 | 83.655 | 2 | 53.099552 | yes | 9.722E-08 | 559.30365 | 835.11322 |
| AKPEYISK | 8 | Q13823 | Nucleolar GTP-binding protein 2 OS=Ho | GNL2 | 83.655 | 2 | 53.099552 | yes | 2.45E-07 | 468.26346 | 645.89288 |
| SPHIETYLK | 9 | Q13823 | Nucleolar GTP-binding protein 2 OS=Ho | GNL2 | 83.655 | 2 | 53.099552 | yes | 9.722E-08 | 544.29279 | 629.27014 |
| SPEDHIGAVLER | 12 | Q13823 | Nucleolar GTP-binding protein 2 OS=Ho | GNL2 | 83.655 | 2 | 53.099552 | yes | 1.521E-07 | 661.83856 | 611.74677 |
| TLEEDVDDR | 9 | Q13823 | Nucleolar GTP-binding protein 2 OS=Ho | GNL2 | 83.655 | 2 | 53.099552 | yes | 9.81E-08 | 546.24622 | 590.86328 |
| VWQYITLMR | 9 | Q13823 | Nucleolar GTP-binding protein 2 OS=Ho | GNL2 | 83.655 | 2 | 53.099552 | yes | 1.004E-07 | 605.32605 | 538.4425 |
| IDSWENAEDFLEK | 13 | Q13823 | Nucleolar GTP-binding protein 2 OS=Ho | GNL2 | 83.655 | 2 | 53.099552 | yes | 1.14E-07 | 798.36481 | 377.26907 |
| CDLVPTWATK | 10 | Q13823 | Nucleolar GTP-binding protein 2 OS=Ho | GNL2 | 83.655 | 2 | 53.099552 | yes | 1.107E-07 | 595.79736 | 366.30295 |
| NEAQEEIYK | 9 | Q13823 | Nucleolar GTP-binding protein 2 OS=Ho | GNL2 | 83.655 | 2 | 53.099552 | yes | 1.811E-07 | 562.26691 | 352.50586 |
| GGEPDLQTVGK | 11 | Q13823 | Nucleolar GTP-binding protein 2 OS=Ho | GNL2 | 83.655 | 2 | 53.099552 | yes | 4.886E-07 | 550.78278 | 322.87378 |
| HLIFVLNK | 8 | Q13823 | Nucleolar GTP-binding protein 2 OS=Ho | GNL2 | 83.655 | 2 | 53.099552 | yes | 5.48E-07 | 492.30548 | 285.47467 |
| QISVGFIGYPNVGK | 14 | Q13823 | Nucleolar GTP-binding protein 2 OS=Ho | GNL2 | 83.655 | 2 | 53.099552 | yes | 2.177E-07 | 739.90375 | 249.19196 |
| IWGELYK | 7 | Q13823 | Nucleolar GTP-binding protein 2 OS=Ho | GNL2 | 83.655 | 2 | 53.099552 | yes | 2.013E-07 | 454.74747 | 187.75749 |
| DLVTEDTGVR | 10 | Q13823 | Nucleolar GTP-binding protein 2 OS=Ho | GNL2 | 83.655 | 2 | 53.099552 | yes | 5.358E-07 | 552.78021 | 140.64391 |
| KTLEEDVDDR | 10 | Q13823 | Nucleolar GTP-binding protein 2 OS=Ho | GNL2 | 83.655 | 2 | 53.099552 | yes | 9.722E-08 | 610.2937 | 130.41269 |
| FQEEMDTVMK | 10 | Q13823 | Nucleolar GTP-binding protein 2 OS=Ho | GNL2 | 83.655 | 2 | 53.099552 | yes | 1.664E-07 | 629.27814 | 126.64424 |

| | | | | | | | | | | | |
|---|---|---|---|---|---|---|---|---|---|---|---|
| YYETHNVK | 8 | Q13823 | Nucleolar GTP-binding protein 2 OS=Ho | GNL2 | 83.655 | 2 | 53.099552 | yes | 1.535E-07 | 527.2536 | 110.1112 |
| RIWGELYK | 8 | Q13823 | Nucleolar GTP-binding protein 2 OS=Ho | GNL2 | 83.655 | 2 | 53.099552 | yes | 1.155E-06 | 532.79803 | 98.59314 |
| KVCNVAPIAGETK | 13 | Q13823 | Nucleolar GTP-binding protein 2 OS=Ho | GNL2 | 83.655 | 2 | 53.099552 | yes | 1.105E-07 | 693.87415 | 70.924034 |
| GAFIQLLR | 8 | Q13823 | Nucleolar GTP-binding protein 2 OS=Ho | GNL2 | 83.655 | 2 | 53.099552 | yes | 4.235E-06 | 459.28198 | 67.364334 |
| DRDLVTEDTGVR | 12 | Q13823 | Nucleolar GTP-binding protein 2 OS=Ho | GNL2 | 83.655 | 2 | 53.099552 | yes | 1.073E-07 | 688.34424 | 51.472309 |
| VCNVAPIAGETK | 12 | Q13823 | Nucleolar GTP-binding protein 2 OS=Ho | GNL2 | 83.655 | 2 | 53.099552 | yes | 9.727E-08 | 629.82666 | 39.000462 |
| PLQYQSTVASGTV | 15 | Q13823 | Nucleolar GTP-binding protein 2 OS=Ho | GNL2 | 83.655 | 2 | 53.099552 | yes | 9.894E-08 | 789.41772 | 35.681976 |
| NEAQEEIYKK | 10 | Q13823 | Nucleolar GTP-binding protein 2 OS=Ho | GNL2 | 83.655 | 2 | 53.099552 | yes | 9.722E-08 | 626.31439 | 32.553535 |
| IIKPLQYQSTVASG | 18 | Q13823 | Nucleolar GTP-binding protein 2 OS=Ho | GNL2 | 83.655 | 3 | 53.099552 | yes | 1.002E-07 | 644.70197 | 29.155859 |
| HFILDECDK | 9 | Q13838 | Spliceosome RNA helicase DDX39B OS= | DDX39B | 48.992 | 2 | 45.48601 | yes | 1.836E-07 | 588.7713 | 140.19614 |
| KNCPHIVVGTPGR | 13 | Q13838 | Spliceosome RNA helicase DDX39B OS= | DDX39B | 48.992 | 3 | 45.48601 | yes | 9.417E-05 | 478.92599 | 20.110443 |
| VAVFFGGLSIK | 11 | Q13838 | Spliceosome RNA helicase DDX39B OS= | DDX39B | 48.992 | 2 | 45.48601 | yes | 2.362E-07 | 569.33698 | 14.854727 |
| LGQGVLVQVSPSL | 15 | Q13868 | Exosome complex component RRP4 OS= | EXOSC2 | 32.789 | 2 | 50.45422 | yes | 1.783E-07 | 762.46143 | 1046.6968 |
| YIGEVGDIVVGR | 12 | Q13868 | Exosome complex component RRP4 OS= | EXOSC2 | 32.789 | 2 | 50.45422 | yes | 9.809E-08 | 638.84845 | 420.2034 |
| NCIISLVTQR | 10 | Q13868 | Exosome complex component RRP4 OS= | EXOSC2 | 32.789 | 2 | 50.45422 | yes | 9.809E-08 | 602.32935 | 366.81622 |
| HLVVPGDTITTDT( | 17 | Q13868 | Exosome complex component RRP4 OS= | EXOSC2 | 32.789 | 3 | 50.45422 | yes | 1.157E-07 | 620.64886 | 62.685589 |
| DILKPEIMEEIVME | 16 | Q13868 | Exosome complex component RRP4 OS= | EXOSC2 | 32.789 | 3 | 50.45422 | yes | 6.555E-07 | 649.33728 | 18.635729 |
| AILVDLEPGTMDS' | 15 | Q13885 | Tubulin beta-2A chain OS=Homo sapiens | TUBB2A | 49.907 | 2 | 49.915455 | no | 4.184E-07 | 808.42163 | 32142.586 |
| AILVDLEPGTMDS' | 15 | Q13885 | Tubulin beta-2A chain OS=Homo sapiens | TUBB2A | 49.907 | 2 | 49.915455 | no | 1.361E-07 | 816.41907 | 32142.586 |
| FWEVISDEHGIDPT | 27 | Q13885 | Tubulin beta-2A chain OS=Homo sapiens | TUBB2A | 49.907 | 4 | 49.915455 | no | 3.499E-06 | 776.35736 | 3399.1489 |
| INVYYNEAAGNK | 12 | Q13885 | Tubulin beta-2A chain OS=Homo sapiens | TUBB2A | 49.907 | 2 | 49.915455 | yes | 9.982E-08 | 678.33313 | 571.63605 |
| YLTVAAIFR | 9 | Q13885 | Tubulin beta-2A chain OS=Homo sapiens | TUBB2A | 49.907 | 2 | 49.915455 | no | 1.008E-07 | 527.30823 | 221.01607 |
| SDHLALHMK | 9 | Q13887 | Krueppel-like factor 5 OS=Homo sapiens | KLF5 | 50.792 | 2 | 50.745907 | no | 1.22E-07 | 526.2713 | 389.26022 |
| PFQCGVCNR | 9 | Q13887 | Krueppel-like factor 5 OS=Homo sapiens | KLF5 | 50.792 | 2 | 50.745907 | yes | 9.723E-08 | 569.25006 | 116.45845 |
| HLYVVVDGSR | 10 | Q13888 | General transcription factor IIH subunit 2 | GTF2H2 | 44.419 | 2 | 53.342255 | no | 9.722E-08 | 572.80908 | 333.2645 |
| VSVIGLSAEVR | 11 | Q13888 | General transcription factor IIH subunit 2 | GTF2H2 | 44.419 | 2 | 53.342255 | no | 2.034E-07 | 565.3324 | 312.88257 |
| YCELPVECK | 9 | Q13888 | General transcription factor IIH subunit 2 | GTF2H2 | 44.419 | 2 | 53.342255 | no | 2.447E-07 | 599.26758 | 144.65337 |
| VCTVLAR | 7 | Q13888 | General transcription factor IIH subunit 2 | GTF2H2 | 44.419 | 2 | 53.342255 | no | 2.827E-07 | 409.73129 | 144.42 |
| FCYGCQGELK | 10 | Q13888 | General transcription factor IIH subunit 2 | GTF2H2 | 44.419 | 2 | 53.342255 | no | 9.9E-08 | 631.27063 | 112.04401 |
| ICGLTLVSAPHLAI | 14 | Q13888 | General transcription factor IIH subunit 2 | GTF2H2 | 44.419 | 3 | 53.342255 | no | 9.722E-08 | 503.28531 | 87.915314 |
| ATIEDILFK | 9 | Q13888 | General transcription factor IIH subunit 2 | GTF2H2 | 44.419 | 2 | 53.342255 | no | 5.045E-07 | 525.29749 | 76.658264 |
| TMEDQDLKPNR | 11 | Q13888 | General transcription factor IIH subunit 2 | GTF2H2 | 44.419 | 2 | 53.342255 | no | 2.206E-07 | 673.82208 | 76.028984 |
| ELLTHHVSPPPASS | 22 | Q13888 | General transcription factor IIH subunit 2 | GTF2H2 | 44.419 | 4 | 53.342255 | yes | 1.246E-05 | 601.80359 | 37.503227 |
| ALCYIHR | 7 | Q13889 | General transcription factor IIH subunit 3 | GTF2H3 | 34.379 | 2 | 40.03395 | yes | 7.229E-07 | 466.74219 | 241.32201 |
| GQHTETLLAGSLA | 14 | Q13889 | General transcription factor IIH subunit 3 | GTF2H3 | 34.379 | 2 | 40.03395 | yes | 4.245E-07 | 713.38843 | 35.820202 |
| ISLPPVLK | 8 | Q13889 | General transcription factor IIH subunit 3 | GTF2H3 | 34.379 | 2 | 40.03395 | yes | 4.143E-06 | 433.78912 | 25.452885 |
| EAIIVGSIITK | 11 | Q13895 | Bystin OS=Homo sapiens OX=9606 GN= | BYSL | 49.602 | 2 | 51.771873 | yes | 9.722E-08 | 572.35284 | 5331.4565 |
| GILIPLCESGTCTLI | 15 | Q13895 | Bystin OS=Homo sapiens OX=9606 GN= | BYSL | 49.602 | 2 | 51.771873 | yes | 1.806E-07 | 845.43677 | 4636.7319 |
| LQPHPQLSPEIR | 12 | Q13895 | Bystin OS=Homo sapiens OX=9606 GN= | BYSL | 49.602 | 2 | 51.771873 | yes | 3.62E-07 | 707.89368 | 1632.2301 |

| | | | | | | | | | | | |
|---|---|---|---|---|---|---|---|---|---|---|---|
| LQPHPQLSPEIR | 12 | Q13895 | Bystin OS=Homo sapiens OX=9606 GN= | BYSL | 49.602 | 2 | 51.771873 | yes | 1.188E-07 | 472.26489 | 1632.2301 |
| LQPHPQLSPEIR | 12 | Q13895 | Bystin OS=Homo sapiens OX=9606 GN= | BYSL | 49.602 | 3 | 51.771873 | yes | 3.62E-07 | 707.89368 | 1632.2301 |
| LQPHPQLSPEIR | 12 | Q13895 | Bystin OS=Homo sapiens OX=9606 GN= | BYSL | 49.602 | 3 | 51.771873 | yes | 1.188E-07 | 472.26489 | 1632.2301 |
| GTGEAEEEYVGPR | 13 | Q13895 | Bystin OS=Homo sapiens OX=9606 GN= | BYSL | 49.602 | 2 | 51.771873 | yes | 9.722E-08 | 697.31512 | 3222.8472 |
| TLADIIMEK | 9 | Q13895 | Bystin OS=Homo sapiens OX=9606 GN= | BYSL | 49.602 | 2 | 51.771873 | yes | 1.102E-07 | 517.28357 | 2632.3237 |
| HAPLADQILAGNA | 15 | Q13895 | Bystin OS=Homo sapiens OX=9606 GN= | BYSL | 49.602 | 2 | 51.771873 | yes | 1.512E-07 | 773.42847 | 1265.2053 |
| HAPLADQILAGNA | 15 | Q13895 | Bystin OS=Homo sapiens OX=9606 GN= | BYSL | 49.602 | 2 | 51.771873 | yes | 1.284E-07 | 515.95471 | 1265.2053 |
| HAPLADQILAGNA | 15 | Q13895 | Bystin OS=Homo sapiens OX=9606 GN= | BYSL | 49.602 | 3 | 51.771873 | yes | 1.512E-07 | 773.42847 | 1265.2053 |
| HAPLADQILAGNA | 15 | Q13895 | Bystin OS=Homo sapiens OX=9606 GN= | BYSL | 49.602 | 3 | 51.771873 | yes | 1.284E-07 | 515.95471 | 1265.2053 |
| FYNLVLLPR | 9 | Q13895 | Bystin OS=Homo sapiens OX=9606 GN= | BYSL | 49.602 | 2 | 51.771873 | yes | 3.097E-07 | 567.83716 | 1767.8474 |
| VRDDVAEYK | 9 | Q13895 | Bystin OS=Homo sapiens OX=9606 GN= | BYSL | 49.602 | 2 | 51.771873 | yes | 3.825E-07 | 547.77747 | 878.7334 |
| AATMTAAGHHAE | 21 | Q13895 | Bystin OS=Homo sapiens OX=9606 GN= | BYSL | 49.602 | 4 | 51.771873 | yes | 9.722E-08 | 552.26031 | 575.9386 |
| CSIPVLHSSAAMLI | 14 | Q13895 | Bystin OS=Homo sapiens OX=9606 GN= | BYSL | 49.602 | 2 | 51.771873 | yes | 1.554E-07 | 757.39691 | 249.56352 |
| CSIPVLHSSAAMLI | 14 | Q13895 | Bystin OS=Homo sapiens OX=9606 GN= | BYSL | 49.602 | 2 | 51.771873 | yes | 1.398E-07 | 505.26703 | 249.56352 |
| CSIPVLHSSAAMLI | 14 | Q13895 | Bystin OS=Homo sapiens OX=9606 GN= | BYSL | 49.602 | 3 | 51.771873 | yes | 1.554E-07 | 757.39691 | 249.56352 |
| CSIPVLHSSAAMLI | 14 | Q13895 | Bystin OS=Homo sapiens OX=9606 GN= | BYSL | 49.602 | 3 | 51.771873 | yes | 1.398E-07 | 505.26703 | 249.56352 |
| QQQEELEAEHGTC | 20 | Q13895 | Bystin OS=Homo sapiens OX=9606 GN= | BYSL | 49.602 | 4 | 51.771873 | yes | 9.722E-08 | 548.51483 | 493.23099 |
| ALFKPGAWFK | 10 | Q13895 | Bystin OS=Homo sapiens OX=9606 GN= | BYSL | 49.602 | 2 | 51.771873 | yes | 2.396E-07 | 582.83185 | 366.3555 |
| YKADLATDQK | 10 | Q13895 | Bystin OS=Homo sapiens OX=9606 GN= | BYSL | 49.602 | 2 | 51.771873 | yes | 1.266E-06 | 576.7984 | 238.30479 |
| KYALPYR | 7 | Q13895 | Bystin OS=Homo sapiens OX=9606 GN= | BYSL | 49.602 | 2 | 51.771873 | yes | 1.913E-07 | 455.76089 | 218.45016 |
| RTLADIIMEK | 10 | Q13895 | Bystin OS=Homo sapiens OX=9606 GN= | BYSL | 49.602 | 2 | 51.771873 | yes | 5.483E-07 | 595.33411 | 72.359985 |
| AIEMFMNK | 8 | Q13895 | Bystin OS=Homo sapiens OX=9606 GN= | BYSL | 49.602 | 2 | 51.771873 | yes | 2.413E-06 | 500.23553 | 33.877079 |
| NVTLNPDPNEIK | 12 | Q13907 | Isopentenyl-diphosphate Delta-isomerase | IDI1 | 26.32 | 2 | 53.1882 | yes | 9.722E-08 | 677.35406 | 147.61166 |
| AFSVFLFNTENK | 12 | Q13907 | Isopentenyl-diphosphate Delta-isomerase | IDI1 | 26.32 | 2 | 53.1882 | yes | 1.391E-07 | 708.86151 | 93.096382 |
| KNVTLNPDPNEIK | 13 | Q13907 | Isopentenyl-diphosphate Delta-isomerase | IDI1 | 26.32 | 2 | 53.1882 | yes | 9.722E-08 | 741.40155 | 86.585739 |
| SKFENEEFFR | 10 | Q13951 | Core-binding factor subunit beta OS=Hor | CBFB | 21.509 | 2 | 41.21404 | yes | 3.62E-07 | 666.81458 | 47.097179 |
| VSQLLAVTGK | 10 | Q13951 | Core-binding factor subunit beta OS=Hor | CBFB | 21.509 | 2 | 41.21404 | yes | 5.13E-06 | 508.31094 | 39.24976 |
| LDGMGCLEFDEEF | 13 | Q13951 | Core-binding factor subunit beta OS=Hor | CBFB | 21.509 | 2 | 41.21404 | yes | 1.197E-06 | 785.82922 | 15.395439 |
| TNTPQGVLPSSQLI | 14 | Q14004 | Cyclin-dependent kinase 13 OS=Homo sa | CDK13 | 164.924 | 2 | 51.16935 | yes | 9.722E-08 | 735.40155 | 162.18634 |
| TKPPLQVTK | 9 | Q14004 | Cyclin-dependent kinase 13 OS=Homo sa | CDK13 | 164.924 | 2 | 51.16935 | yes | 8.103E-07 | 506.31351 | 113.26158 |
| TKPLTPSIGAK | 11 | Q14004 | Cyclin-dependent kinase 13 OS=Homo sa | CDK13 | 164.924 | 2 | 51.16935 | yes | 4.484E-07 | 556.83734 | 92.541756 |
| CTAEQALQCEFLR | 13 | Q14004 | Cyclin-dependent kinase 13 OS=Homo sa | CDK13 | 164.924 | 2 | 51.16935 | yes | 1.22E-07 | 813.37415 | 67.261208 |
| GGDVSPSPYSSSSV | 15 | Q14004 | Cyclin-dependent kinase 13 OS=Homo sa | CDK13 | 164.924 | 2 | 51.16935 | yes | 5.113E-07 | 784.85242 | 34.704407 |
| GNTETSASASQTN | 16 | Q14004 | Cyclin-dependent kinase 13 OS=Homo sa | CDK13 | 164.924 | 2 | 51.16935 | yes | 2.736E-06 | 816.38464 | 12.074403 |
| LLPDTINSINLDR | 13 | Q14008 | Cytoskeleton-associated protein 5 OS=Hc | CKAP5 | 225.495 | 2 | 53.69977 | yes | 1.365E-07 | 742.40936 | 1233.7615 |
| TSAQVVLDGLVDI | 13 | Q14008 | Cytoskeleton-associated protein 5 OS=Hc | CKAP5 | 225.495 | 2 | 53.69977 | yes | 1.165E-07 | 672.87207 | 931.47607 |
| GEAVQEELLK | 10 | Q14008 | Cytoskeleton-associated protein 5 OS=Hc | CKAP5 | 225.495 | 2 | 53.69977 | yes | 3.554E-07 | 558.30078 | 631.11163 |
| TTGEVVSGVVSK | 12 | Q14008 | Cytoskeleton-associated protein 5 OS=Hc | CKAP5 | 225.495 | 2 | 53.69977 | yes | 9.722E-08 | 581.81934 | 604.28491 |
| YSDADIEPFLK | 11 | Q14008 | Cytoskeleton-associated protein 5 OS=Hc | CKAP5 | 225.495 | 2 | 53.69977 | yes | 9.722E-08 | 649.31915 | 603.57251 |

| | | | | | | | | | | | |
|---|---|---|---|---|---|---|---|---|---|---|---|
| EGLDEVAGIINDAI | 14 | Q14008 | Cytoskeleton-associated protein 5 OS=Hc | CKAP5 | 225.495 | 2 | 53.69977 | yes | 9.722E-08 | 722.36993 | 590.32312 |
| IEDLEEGQQVIR | 12 | Q14008 | Cytoskeleton-associated protein 5 OS=Hc | CKAP5 | 225.495 | 2 | 53.69977 | yes | 1.025E-07 | 714.87012 | 588.81177 |
| EFGFSGLNVK | 10 | Q14008 | Cytoskeleton-associated protein 5 OS=Hc | CKAP5 | 225.495 | 2 | 53.69977 | yes | 9.722E-08 | 549.28491 | 579.34082 |
| MCLVYPASK | 9 | Q14008 | Cytoskeleton-associated protein 5 OS=Hc | CKAP5 | 225.495 | 2 | 53.69977 | yes | 9.729E-08 | 534.76447 | 550.50824 |
| TEMPCQALVR | 10 | Q14008 | Cytoskeleton-associated protein 5 OS=Hc | CKAP5 | 225.495 | 2 | 53.69977 | yes | 1.874E-07 | 602.79431 | 481.56522 |
| LDDIFEPVLIPEPK | 14 | Q14008 | Cytoskeleton-associated protein 5 OS=Hc | CKAP5 | 225.495 | 2 | 53.69977 | yes | 2.049E-07 | 812.94525 | 443.70361 |
| HCTASTLPK | 9 | Q14008 | Cytoskeleton-associated protein 5 OS=Hc | CKAP5 | 225.495 | 2 | 53.69977 | yes | 3.709E-07 | 507.75549 | 431.26184 |
| LACMEEFQK | 9 | Q14008 | Cytoskeleton-associated protein 5 OS=Hc | CKAP5 | 225.495 | 2 | 53.69977 | yes | 1.253E-07 | 578.26227 | 430.53552 |
| EGLAELYEYK | 10 | Q14008 | Cytoskeleton-associated protein 5 OS=Hc | CKAP5 | 225.495 | 2 | 53.69977 | yes | 9.722E-08 | 607.8006 | 412.81409 |
| DQVLAMLEK | 9 | Q14008 | Cytoskeleton-associated protein 5 OS=Hc | CKAP5 | 225.495 | 2 | 53.69977 | yes | 1.001E-07 | 523.78119 | 378.39935 |
| QQTSLFIAR | 9 | Q14008 | Cytoskeleton-associated protein 5 OS=Hc | CKAP5 | 225.495 | 2 | 53.69977 | yes | 1.047E-07 | 532.2984 | 374.38065 |
| CEMPELVQHK | 10 | Q14008 | Cytoskeleton-associated protein 5 OS=Hc | CKAP5 | 225.495 | 2 | 53.69977 | yes | 1.092E-07 | 635.79956 | 345.89212 |
| NESELEAHLCR | 11 | Q14008 | Cytoskeleton-associated protein 5 OS=Hc | CKAP5 | 225.495 | 2 | 53.69977 | yes | 9.722E-08 | 679.31189 | 317.74863 |
| MQGQSPPAPTR | 11 | Q14008 | Cytoskeleton-associated protein 5 OS=Hc | CKAP5 | 225.495 | 2 | 53.69977 | yes | 1.161E-07 | 585.29022 | 311.22549 |
| TLLHTLCK | 8 | Q14008 | Cytoskeleton-associated protein 5 OS=Hc | CKAP5 | 225.495 | 2 | 53.69977 | yes | 9.732E-08 | 493.27859 | 310.58975 |
| TALAATNPAVR | 11 | Q14008 | Cytoskeleton-associated protein 5 OS=Hc | CKAP5 | 225.495 | 2 | 53.69977 | yes | 1.053E-07 | 542.80908 | 292.94522 |
| NSSQFFQSYVER | 12 | Q14008 | Cytoskeleton-associated protein 5 OS=Hc | CKAP5 | 225.495 | 2 | 53.69977 | yes | 1.082E-07 | 746.3468 | 287.15433 |
| EIAVHIGDR | 9 | Q14008 | Cytoskeleton-associated protein 5 OS=Hc | CKAP5 | 225.495 | 2 | 53.69977 | yes | 2.048E-07 | 505.2749 | 286.70102 |
| ALAVMVDHLESEI | 13 | Q14008 | Cytoskeleton-associated protein 5 OS=Hc | CKAP5 | 225.495 | 2 | 53.69977 | yes | 1.119E-07 | 721.3714 | 283.40512 |
| HINDSAPEVR | 10 | Q14008 | Cytoskeleton-associated protein 5 OS=Hc | CKAP5 | 225.495 | 2 | 53.69977 | yes | 1.993E-07 | 569.28601 | 274.76913 |
| QELLGWLAEK | 10 | Q14008 | Cytoskeleton-associated protein 5 OS=Hc | CKAP5 | 225.495 | 2 | 53.69977 | yes | 1.597E-07 | 593.82697 | 273.83395 |
| WNFTTPR | 7 | Q14008 | Cytoskeleton-associated protein 5 OS=Hc | CKAP5 | 225.495 | 2 | 53.69977 | yes | 4.638E-07 | 461.23251 | 247.43272 |
| FVTDSNAVVQLK | 12 | Q14008 | Cytoskeleton-associated protein 5 OS=Hc | CKAP5 | 225.495 | 2 | 53.69977 | yes | 1.06E-07 | 660.86151 | 246.8924 |
| LIAVEIYR | 8 | Q14008 | Cytoskeleton-associated protein 5 OS=Hc | CKAP5 | 225.495 | 2 | 53.69977 | yes | 1.245E-07 | 488.79492 | 245.23952 |
| ILDHLTMIDNK | 11 | Q14008 | Cytoskeleton-associated protein 5 OS=Hc | CKAP5 | 225.495 | 2 | 53.69977 | yes | 1.043E-07 | 656.8501 | 236.16478 |
| LEAGDYADLVK | 11 | Q14008 | Cytoskeleton-associated protein 5 OS=Hc | CKAP5 | 225.495 | 2 | 53.69977 | yes | 9.902E-08 | 597.30603 | 229.58363 |
| ETNFQVMQMK | 10 | Q14008 | Cytoskeleton-associated protein 5 OS=Hc | CKAP5 | 225.495 | 2 | 53.69977 | yes | 9.722E-08 | 628.29413 | 218.71323 |
| LSGYEEALK | 9 | Q14008 | Cytoskeleton-associated protein 5 OS=Hc | CKAP5 | 225.495 | 2 | 53.69977 | yes | 7.156E-07 | 505.26367 | 206.11497 |
| NQSETLNWLSNAI | 14 | Q14008 | Cytoskeleton-associated protein 5 OS=Hc | CKAP5 | 225.495 | 2 | 53.69977 | yes | 9.722E-08 | 809.41522 | 196.50481 |
| VNMPAKPAPPTK | 12 | Q14008 | Cytoskeleton-associated protein 5 OS=Hc | CKAP5 | 225.495 | 2 | 53.69977 | yes | 1.001E-07 | 625.84991 | 193.92967 |
| LIGNLSEK | 8 | Q14008 | Cytoskeleton-associated protein 5 OS=Hc | CKAP5 | 225.495 | 2 | 53.69977 | yes | 3.123E-07 | 437.25565 | 170.57043 |
| LHIVALIAQK | 10 | Q14008 | Cytoskeleton-associated protein 5 OS=Hc | CKAP5 | 225.495 | 2 | 53.69977 | yes | 9.819E-08 | 553.35822 | 157.74255 |
| SMSGHPEAAQMV | 13 | Q14008 | Cytoskeleton-associated protein 5 OS=Hc | CKAP5 | 225.495 | 2 | 53.69977 | yes | 1.46E-07 | 700.8241 | 157.68095 |
| ELEEEWVK | 8 | Q14008 | Cytoskeleton-associated protein 5 OS=Hc | CKAP5 | 225.495 | 2 | 53.69977 | yes | 1.527E-07 | 531.26111 | 154.97134 |
| AVNPFLADVDK | 11 | Q14008 | Cytoskeleton-associated protein 5 OS=Hc | CKAP5 | 225.495 | 2 | 53.69977 | yes | 2.625E-07 | 594.81659 | 154.89897 |
| DMSMLEER | 8 | Q14008 | Cytoskeleton-associated protein 5 OS=Hc | CKAP5 | 225.495 | 2 | 53.69977 | yes | 1.175E-07 | 505.71771 | 137.08821 |
| LEKDEIIK | 8 | Q14008 | Cytoskeleton-associated protein 5 OS=Hc | CKAP5 | 225.495 | 2 | 53.69977 | yes | 1.11E-07 | 494.28967 | 134.40872 |
| FFDTNTSVLMK | 11 | Q14008 | Cytoskeleton-associated protein 5 OS=Hc | CKAP5 | 225.495 | 2 | 53.69977 | yes | 1.012E-07 | 651.82355 | 130.22867 |
| EALESVEVLIK | 11 | Q14008 | Cytoskeleton-associated protein 5 OS=Hc | CKAP5 | 225.495 | 2 | 53.69977 | yes | 9.722E-08 | 615.35303 | 114.25072 |

| | | | | | | | | | | | |
|---|---|---|---|---|---|---|---|---|---|---|---|
| PAAPGGAGNTGTK | 13 | Q14008 | Cytoskeleton-associated protein 5 OS=Hc | CKAP5 | 225.495 | 2 | 53.69977 | yes | 5.413E-07 | 549.78052 | 107.37443 |
| PAVPTVASSTDML | 16 | Q14008 | Cytoskeleton-associated protein 5 OS=Hc | CKAP5 | 225.495 | 3 | 53.69977 | yes | 6.998E-07 | 547.61523 | 105.89466 |
| CLTGLAVGLR | 10 | Q14008 | Cytoskeleton-associated protein 5 OS=Hc | CKAP5 | 225.495 | 2 | 53.69977 | yes | 9.722E-08 | 530.30261 | 105.66978 |
| LIYNTHMADEK | 11 | Q14008 | Cytoskeleton-associated protein 5 OS=Hc | CKAP5 | 225.495 | 2 | 53.69977 | yes | 1.05E-07 | 667.8241 | 65.303444 |
| EGVIGCLDLILK | 12 | Q14008 | Cytoskeleton-associated protein 5 OS=Hc | CKAP5 | 225.495 | 2 | 53.69977 | yes | 6.26E-07 | 665.37598 | 49.706108 |
| KFVTDSNAVVQLK | 13 | Q14008 | Cytoskeleton-associated protein 5 OS=Hc | CKAP5 | 225.495 | 2 | 53.69977 | yes | 4.357E-07 | 724.909 | 44.872684 |
| DAAFEALGTALK | 12 | Q14008 | Cytoskeleton-associated protein 5 OS=Hc | CKAP5 | 225.495 | 2 | 53.69977 | yes | 6.486E-07 | 603.8219 | 44.081955 |
| TSLKEDEDK | 9 | Q14008 | Cytoskeleton-associated protein 5 OS=Hc | CKAP5 | 225.495 | 2 | 53.69977 | yes | 3.897E-06 | 532.75897 | 41.34549 |
| KEALESVEVLIK | 12 | Q14008 | Cytoskeleton-associated protein 5 OS=Hc | CKAP5 | 225.495 | 2 | 53.69977 | yes | 9.722E-08 | 679.40051 | 33.633965 |
| EQHQHSDLDSNQT | 33 | Q14008 | Cytoskeleton-associated protein 5 OS=Hc | CKAP5 | 225.495 | 5 | 53.69977 | yes | 1.388E-07 | 711.73035 | 19.708029 |
| DTNVMLVALAAK | 12 | Q14008 | Cytoskeleton-associated protein 5 OS=Hc | CKAP5 | 225.495 | 2 | 53.69977 | yes | 2.765E-06 | 623.34723 | 18.689692 |
| KFGQYAGHVVPTI | 16 | Q14008 | Cytoskeleton-associated protein 5 OS=Hc | CKAP5 | 225.495 | 3 | 53.69977 | yes | 4.951E-07 | 596.33319 | 14.803185 |
| GFGFVTFENIDDAI | 14 | Q14011 | Cold-inducible RNA-binding protein OS= | CIRBP | 18.648 | 2 | 54.870667 | yes | 9.722E-08 | 780.37244 | 4223.3955 |
| YGQISEVVVVK | 11 | Q14011 | Cold-inducible RNA-binding protein OS= | CIRBP | 18.648 | 2 | 54.870667 | yes | 9.722E-08 | 610.8479 | 2740.2844 |
| DSYDSYATHNE | 11 | Q14011 | Cold-inducible RNA-binding protein OS= | CIRBP | 18.648 | 2 | 54.870667 | yes | 9.722E-08 | 651.24945 | 146.9911 |
| FGEVVDCTLK | 10 | Q14103 | Heterogeneous nuclear ribonucleoprotein | HNRNPD | 38.435 | 2 | 52.232037 | yes | 1.542E-07 | 584.28937 | 19433.527 |
| MFIGGLSWDTTK | 12 | Q14103 | Heterogeneous nuclear ribonucleoprotein | HNRNPD | 38.435 | 2 | 52.232037 | yes | 1.152E-07 | 678.33685 | 8920.7656 |
| MFIGGLSWDTTK | 12 | Q14103 | Heterogeneous nuclear ribonucleoprotein | HNRNPD | 38.435 | 2 | 52.232037 | yes | 9.722E-08 | 686.33429 | 8920.7656 |
| IFVGGLSPDTPEEK | 14 | Q14103 | Heterogeneous nuclear ribonucleoprotein | HNRNPD | 38.435 | 2 | 52.232037 | yes | 1.15E-07 | 744.88269 | 15231.458 |
| KYHNVGLSK | 9 | Q14103 | Heterogeneous nuclear ribonucleoprotein | HNRNPD | 38.435 | 2 | 52.232037 | yes | 1.389E-07 | 523.29309 | 2624.5251 |
| RGFCFITFK | 9 | Q14103 | Heterogeneous nuclear ribonucleoprotein | HNRNPD | 38.435 | 2 | 52.232037 | yes | 1.637E-07 | 588.30511 | 496.94891 |
| GFCFITFK | 8 | Q14103 | Heterogeneous nuclear ribonucleoprotein | HNRNPD | 38.435 | 2 | 52.232037 | yes | 3.253E-07 | 510.25458 | 121.82283 |
| YHNVGLSK | 8 | Q14103 | Heterogeneous nuclear ribonucleoprotein | HNRNPD | 38.435 | 2 | 52.232037 | yes | 2.844E-07 | 459.24561 | 68.84877 |
| EYFGGFGEVESIEL | 19 | Q14103 | Heterogeneous nuclear ribonucleoprotein | HNRNPD | 38.435 | 3 | 52.232037 | yes | 1.455E-07 | 720.99799 | 49.318981 |
| EQYQQQQQWGSR | 12 | Q14103 | Heterogeneous nuclear ribonucleoprotein | HNRNPD | 38.435 | 2 | 52.232037 | yes | 9.722E-08 | 783.35822 | 46.774818 |
| KIFVGGLSPDTPEE | 15 | Q14103 | Heterogeneous nuclear ribonucleoprotein | HNRNPD | 38.435 | 2 | 52.232037 | yes | 2.755E-06 | 808.93018 | 23.000837 |
| ADLFGEELER | 10 | Q14106 | Protein Tob2 OS=Homo sapiens OX=960 | TOB2 | 36.633 | 2 | 43.78197 | yes | 2.676E-07 | 589.78802 | 92.517021 |
| CVHIGEMVDPVVI | 17 | Q14106 | Protein Tob2 OS=Homo sapiens OX=960 | TOB2 | 36.633 | 3 | 43.78197 | yes | 2.45E-07 | 622.98657 | 44.593323 |
| SAQPITFTTASFAA | 16 | Q14106 | Protein Tob2 OS=Homo sapiens OX=960 | TOB2 | 36.633 | 2 | 43.78197 | yes | 2.897E-07 | 821.4278 | 20.897146 |
| MVYWTDVAGR | 10 | Q14112 | Nidogen-2 OS=Homo sapiens OX=9606 | NID2 | 151.254 | 2 | 21.384537 | yes | 4.156E-06 | 599.28967 | 49.29488 |
| SPSLIFTNR | 9 | Q14114 | Low-density lipoprotein receptor-related | LRP8 | 105.634 | 2 | 56.430897 | yes | 1.551E-07 | 517.7851 | 191.77719 |
| SMNFDNPVYR | 10 | Q14114 | Low-density lipoprotein receptor-related | LRP8 | 105.634 | 2 | 56.430897 | yes | 9.722E-08 | 621.78223 | 79.237862 |
| GFMYWSDWGDQ | 13 | Q14114 | Low-density lipoprotein receptor-related | LRP8 | 105.634 | 2 | 56.430897 | yes | 9.759E-08 | 795.83771 | 55.13147 |
| ELFVLPGEPR | 10 | Q14114 | Low-density lipoprotein receptor-related | LRP8 | 105.634 | 2 | 56.430897 | yes | 7.794E-05 | 578.82166 | 51.634953 |
| IYWCDLSYR | 9 | Q14114 | Low-density lipoprotein receptor-related | LRP8 | 105.634 | 2 | 56.430897 | yes | 4.641E-07 | 638.29498 | 44.717964 |
| TSFVCTYCSK | 10 | Q14119 | Vascular endothelial zinc finger 1 OS=Hc | VEZF1 | 56.932 | 2 | 52.381443 | yes | 9.722E-08 | 626.77045 | 151.0009 |
| CQTCTAAFATK | 11 | Q14119 | Vascular endothelial zinc finger 1 OS=Hc | VEZF1 | 56.932 | 2 | 52.381443 | yes | 9.722E-08 | 629.78137 | 131.26956 |
| FLDDLGLK | 8 | Q14126 | Desmoglein-2 OS=Homo sapiens OX=96 | DSG2 | 122.294 | 2 | 37.053814 | yes | 6.574E-07 | 460.75803 | 47.07679 |
| AASYTEEDENHTA | 14 | Q14126 | Desmoglein-2 OS=Homo sapiens OX=96 | DSG2 | 122.294 | 2 | 37.053814 | yes | 6.346E-07 | 783.33936 | 21.690105 |

| | | | | | | | | | | | |
|---|---|---|---|---|---|---|---|---|---|---|---|
| ILDVNDNIPVVENI | 14 | Q14126 | Desmoglein-2 OS=Homo sapiens OX=96 | DSG2 | 122.294 | 2 | 37.053814 | yes | 1.156E-05 | 791.4278 | 20.647436 |
| KPPVTFAEK | 9 | Q14134 | Tripartite motif-containing protein 29 OS | TRIM29 | 65.835 | 2 | 52.785328 | yes | 4.001E-07 | 508.79239 | 752.7901 |
| IIEIEDEAEK | 10 | Q14134 | Tripartite motif-containing protein 29 OS | TRIM29 | 65.835 | 2 | 52.785328 | yes | 9.722E-08 | 594.80334 | 457.65329 |
| SLGSALKPGEGR | 12 | Q14134 | Tripartite motif-containing protein 29 OS | TRIM29 | 65.835 | 2 | 52.785328 | yes | 9.722E-08 | 586.32513 | 349.45889 |
| AILEQNFR | 8 | Q14134 | Tripartite motif-containing protein 29 OS | TRIM29 | 65.835 | 2 | 52.785328 | yes | 1.027E-07 | 495.772 | 346.52463 |
| SPYAGLQLGAAK | 12 | Q14134 | Tripartite motif-containing protein 29 OS | TRIM29 | 65.835 | 2 | 52.785328 | yes | 9.722E-08 | 588.32458 | 263.66876 |
| DDLLNVCMR | 9 | Q14134 | Tripartite motif-containing protein 29 OS | TRIM29 | 65.835 | 2 | 52.785328 | yes | 1.136E-07 | 568.26538 | 254.53426 |
| NHSTVTVEEAK | 11 | Q14134 | Tripartite motif-containing protein 29 OS | TRIM29 | 65.835 | 2 | 52.785328 | yes | 9.722E-08 | 607.8042 | 252.87737 |
| SALFAGNEWR | 10 | Q14134 | Tripartite motif-containing protein 29 OS | TRIM29 | 65.835 | 2 | 52.785328 | yes | 2.4E-07 | 575.78564 | 248.54785 |
| EQDAVDQVK | 9 | Q14134 | Tripartite motif-containing protein 29 OS | TRIM29 | 65.835 | 2 | 52.785328 | yes | 1.897E-07 | 516.25385 | 196.62944 |
| YSMYLTPK | 8 | Q14134 | Tripartite motif-containing protein 29 OS | TRIM29 | 65.835 | 2 | 52.785328 | yes | 3.881E-07 | 501.75189 | 131.83861 |
| NFNNLYGTK | 9 | Q14134 | Tripartite motif-containing protein 29 OS | TRIM29 | 65.835 | 2 | 52.785328 | yes | 1.138E-07 | 535.76691 | 129.46571 |
| SGSEEVLCDSCIGN | 15 | Q14134 | Tripartite motif-containing protein 29 OS | TRIM29 | 65.835 | 2 | 52.785328 | yes | 9.728E-08 | 827.85602 | 89.845726 |
| RPIIQFVESGDDK | 13 | Q14134 | Tripartite motif-containing protein 29 OS | TRIM29 | 65.835 | 2 | 52.785328 | yes | 9.722E-08 | 752.39374 | 56.687469 |
| KPTVSIMEPGETR | 13 | Q14134 | Tripartite motif-containing protein 29 OS | TRIM29 | 65.835 | 2 | 52.785328 | yes | 1.136E-07 | 722.87683 | 49.88726 |
| ETELSLQK | 8 | Q14134 | Tripartite motif-containing protein 29 OS | TRIM29 | 65.835 | 2 | 52.785328 | yes | 5.15E-06 | 474.25583 | 46.87294 |
| SCLVCQASFCELHI | 24 | Q14134 | Tripartite motif-containing protein 29 OS | TRIM29 | 65.835 | 4 | 52.785328 | yes | 4.361E-07 | 708.34509 | 36.398952 |
| LTDEQVALVR | 10 | Q14137 | Ribosome biogenesis protein BOP1 OS=I | BOP1 | 83.63 | 2 | 51.23584 | yes | 3.307E-07 | 572.32202 | 407.21945 |
| PFLLVASQR | 9 | Q14137 | Ribosome biogenesis protein BOP1 OS=I | BOP1 | 83.63 | 2 | 51.23584 | yes | 1.318E-07 | 515.80585 | 357.05008 |
| TAAPSVRPEK | 10 | Q14137 | Ribosome biogenesis protein BOP1 OS=I | BOP1 | 83.63 | 2 | 51.23584 | yes | 3.582E-07 | 528.29584 | 308.50961 |
| SFIPSLVEK | 9 | Q14137 | Ribosome biogenesis protein BOP1 OS=I | BOP1 | 83.63 | 2 | 51.23584 | yes | 5.404E-07 | 510.29224 | 230.24049 |
| MGWIQPR | 7 | Q14137 | Ribosome biogenesis protein BOP1 OS=I | BOP1 | 83.63 | 2 | 51.23584 | yes | 4.202E-07 | 444.23145 | 194.12386 |
| CLDLYLCPR | 9 | Q14137 | Ribosome biogenesis protein BOP1 OS=I | BOP1 | 83.63 | 2 | 51.23584 | yes | 1.012E-07 | 605.29138 | 173.12119 |
| VNVDPEDLIPK | 11 | Q14137 | Ribosome biogenesis protein BOP1 OS=I | BOP1 | 83.63 | 2 | 51.23584 | yes | 1.764E-07 | 619.83496 | 165.7336 |
| TTEEQVQASTPCPI | 14 | Q14137 | Ribosome biogenesis protein BOP1 OS=I | BOP1 | 83.63 | 2 | 51.23584 | yes | 9.722E-08 | 802.37268 | 80.582558 |
| LWEVATAR | 8 | Q14137 | Ribosome biogenesis protein BOP1 OS=I | BOP1 | 83.63 | 2 | 51.23584 | yes | 3.225E-06 | 473.26126 | 71.119644 |
| MDDPDYWR | 8 | Q14137 | Ribosome biogenesis protein BOP1 OS=I | BOP1 | 83.63 | 2 | 51.23584 | yes | 9.722E-08 | 549.2215 | 62.617603 |
| TRDELDQFLDK | 11 | Q14137 | Ribosome biogenesis protein BOP1 OS=I | BOP1 | 83.63 | 2 | 51.23584 | yes | 2.059E-07 | 690.34369 | 61.311779 |
| LAWEQQEPGER | 11 | Q14137 | Ribosome biogenesis protein BOP1 OS=I | BOP1 | 83.63 | 2 | 51.23584 | yes | 3.708E-07 | 671.82294 | 51.800095 |
| WLEASEEER | 9 | Q14137 | Ribosome biogenesis protein BOP1 OS=I | BOP1 | 83.63 | 2 | 51.23584 | yes | 2.474E-07 | 574.76459 | 48.934074 |
| STLMDTLFNTK | 11 | Q14141 | Septin-6 OS=Homo sapiens OX=9606 GI | SEPTIN6 | 49.717 | 2 | 48.51518 | no | 1.137E-07 | 635.82104 | 70.742004 |
| LTIVSTVGFGDQIN | 15 | Q14141 | Septin-6 OS=Homo sapiens OX=9606 GI | SEPTIN6 | 49.717 | 2 | 48.51518 | yes | 3.403E-07 | 796.43811 | 61.959595 |
| LWEGAISIPR | 10 | Q14142 | Tripartite motif-containing protein 14 OS | TRIM14 | 49.773 | 2 | 51.526024 | yes | 1.824E-07 | 571.32184 | 632.96558 |
| EQLDIMNDLSNR | 12 | Q14142 | Tripartite motif-containing protein 14 OS | TRIM14 | 49.773 | 2 | 51.526024 | yes | 1.539E-07 | 724.34595 | 520.02966 |
| ATFQEPLYPALR | 12 | Q14142 | Tripartite motif-containing protein 14 OS | TRIM14 | 49.773 | 2 | 51.526024 | yes | 2.691E-07 | 703.37738 | 491.84079 |
| CGLLGSLGPVPVL | 14 | Q14142 | Tripartite motif-containing protein 14 OS | TRIM14 | 49.773 | 2 | 51.526024 | yes | 1.031E-07 | 719.41595 | 301.23578 |
| TPTLDPDTMHAR | 12 | Q14142 | Tripartite motif-containing protein 14 OS | TRIM14 | 49.773 | 2 | 51.526024 | yes | 9.722E-08 | 677.82465 | 298.96777 |
| LLLDEEEALAK | 11 | Q14142 | Tripartite motif-containing protein 14 OS | TRIM14 | 49.773 | 2 | 51.526024 | yes | 9.722E-08 | 622.34265 | 181.21013 |
| VWSISQEPDPVQR | 13 | Q14142 | Tripartite motif-containing protein 14 OS | TRIM14 | 49.773 | 2 | 51.526024 | yes | 1.004E-07 | 770.89136 | 168.75443 |

| | | | | | | | | | | | |
|---|---|---|---|---|---|---|---|---|---|---|---|
| SELVEGCGWR | 10 | Q14142 | Tripartite motif-containing protein 14 OS | TRIM14 | 49.773 | 2 | 51.526024 | yes | 1.004E-07 | 596.77441 | 125.85864 |
| CVCALCPVLGAHI | 13 | Q14142 | Tripartite motif-containing protein 14 OS | TRIM14 | 49.773 | 2 | 51.526024 | yes | 7.525E-07 | 756.86523 | 121.22707 |
| GHPVGLALEAAVI | 16 | Q14142 | Tripartite motif-containing protein 14 OS | TRIM14 | 49.773 | 3 | 51.526024 | yes | 5.541E-07 | 542.64227 | 73.869179 |
| LSQECLK | 7 | Q14142 | Tripartite motif-containing protein 14 OS | TRIM14 | 49.773 | 2 | 51.526024 | yes | 1.862E-07 | 439.22601 | 61.51815 |
| VAELFCR | 7 | Q14142 | Tripartite motif-containing protein 14 OS | TRIM14 | 49.773 | 2 | 51.526024 | yes | 3.53E-07 | 447.72873 | 54.475246 |
| CEILQSDSR | 9 | Q14145 | Kelch-like ECH-associated protein 1 OS= | KEAP1 | 69.667 | 2 | 56.54318 | yes | 6.715E-07 | 554.25861 | 327.19006 |
| VVLASSSPVFK | 11 | Q14145 | Kelch-like ECH-associated protein 1 OS= | KEAP1 | 69.667 | 2 | 56.54318 | yes | 9.722E-08 | 567.33191 | 308.39551 |
| SALGITVHQGR | 11 | Q14145 | Kelch-like ECH-associated protein 1 OS= | KEAP1 | 69.667 | 2 | 56.54318 | yes | 9.722E-08 | 569.82001 | 202.2822 |
| FYVQALLR | 8 | Q14145 | Kelch-like ECH-associated protein 1 OS= | KEAP1 | 69.667 | 2 | 56.54318 | yes | 1.234E-07 | 505.2951 | 188.40005 |
| LIYTAGGYFR | 10 | Q14145 | Kelch-like ECH-associated protein 1 OS= | KEAP1 | 69.667 | 2 | 56.54318 | yes | 1.153E-07 | 580.80859 | 183.72986 |
| TFSYTLEDHTK | 11 | Q14145 | Kelch-like ECH-associated protein 1 OS= | KEAP1 | 69.667 | 2 | 56.54318 | yes | 9.722E-08 | 671.3197 | 182.40178 |
| LLYAVGGFDGTNI | 13 | Q14145 | Kelch-like ECH-associated protein 1 OS= | KEAP1 | 69.667 | 2 | 56.54318 | yes | 1.234E-07 | 691.85681 | 171.22067 |
| MITAMNTIR | 9 | Q14145 | Kelch-like ECH-associated protein 1 OS= | KEAP1 | 69.667 | 2 | 56.54318 | yes | 1.868E-06 | 525.77539 | 161.78978 |
| SGVGVAVTMEPCI | 13 | Q14145 | Kelch-like ECH-associated protein 1 OS= | KEAP1 | 69.667 | 2 | 56.54318 | yes | 1.003E-07 | 681.82886 | 143.89293 |
| EQGMEVVSIEGIHI | 15 | Q14145 | Kelch-like ECH-associated protein 1 OS= | KEAP1 | 69.667 | 3 | 56.54318 | yes | 2.161E-07 | 551.61523 | 72.440582 |
| LADLQVPR | 8 | Q14145 | Kelch-like ECH-associated protein 1 OS= | KEAP1 | 69.667 | 2 | 56.54318 | yes | 3.582E-07 | 456.26907 | 135.87892 |
| QAFGIMNELR | 10 | Q14145 | Kelch-like ECH-associated protein 1 OS= | KEAP1 | 69.667 | 2 | 56.54318 | yes | 1.073E-07 | 589.80298 | 132.52029 |
| IGVGVAVLNR | 10 | Q14145 | Kelch-like ECH-associated protein 1 OS= | KEAP1 | 69.667 | 2 | 56.54318 | yes | 1.397E-07 | 499.31128 | 127.66567 |
| LNSAECYYPER | 11 | Q14145 | Kelch-like ECH-associated protein 1 OS= | KEAP1 | 69.667 | 2 | 56.54318 | yes | 9.722E-08 | 701.30884 | 118.569 |
| LSQQLCDVTLQVK | 13 | Q14145 | Kelch-like ECH-associated protein 1 OS= | KEAP1 | 69.667 | 2 | 56.54318 | yes | 9.722E-08 | 766.41107 | 115.41024 |
| EYIYMHFGEVAK | 12 | Q14145 | Kelch-like ECH-associated protein 1 OS= | KEAP1 | 69.667 | 2 | 56.54318 | yes | 9.722E-08 | 743.85541 | 83.600937 |
| SGVGVAVTMEPCI | 14 | Q14145 | Kelch-like ECH-associated protein 1 OS= | KEAP1 | 69.667 | 2 | 56.54318 | yes | 9.722E-08 | 745.87634 | 75.212463 |
| YQDAPAAQFMAH | 13 | Q14145 | Kelch-like ECH-associated protein 1 OS= | KEAP1 | 69.667 | 2 | 56.54318 | yes | 0.0001095 | 739.34827 | 20.805819 |
| AALPIAQYGNR | 11 | Q14147 | Probable ATP-dependent RNA helicase D | DHX34 | 128.12 | 2 | 52.713608 | yes | 4.24E-07 | 587.32239 | 181.3149 |
| INLSVLGPATR | 11 | Q14147 | Probable ATP-dependent RNA helicase D | DHX34 | 128.12 | 2 | 52.713608 | yes | 9.722E-08 | 570.84039 | 158.65079 |
| AGPAPPGASDGVD | 18 | Q14147 | Probable ATP-dependent RNA helicase D | DHX34 | 128.12 | 2 | 52.713608 | yes | 7.058E-07 | 847.42322 | 128.94644 |
| LQEFWISQASAEQ | 14 | Q14147 | Probable ATP-dependent RNA helicase D | DHX34 | 128.12 | 2 | 52.713608 | yes | 1.063E-07 | 846.92065 | 122.73553 |
| VALDSLVLQMK | 11 | Q14147 | Probable ATP-dependent RNA helicase D | DHX34 | 128.12 | 2 | 52.713608 | yes | 9.722E-08 | 608.85211 | 101.28761 |
| HSIPALADLPR | 11 | Q14147 | Probable ATP-dependent RNA helicase D | DHX34 | 128.12 | 2 | 52.713608 | yes | 6.822E-07 | 595.33801 | 100.65136 |
| HDLAQLQAAASSA | 18 | Q14147 | Probable ATP-dependent RNA helicase D | DHX34 | 128.12 | 3 | 52.713608 | yes | 9.722E-08 | 627.98529 | 91.402115 |
| EHQVVVVAGDTG | 15 | Q14147 | Probable ATP-dependent RNA helicase D | DHX34 | 128.12 | 3 | 52.713608 | yes | 5.419E-07 | 519.25598 | 42.000267 |
| EHQVVVVAGDTG | 15 | Q14147 | Probable ATP-dependent RNA helicase D | DHX34 | 128.12 | 3 | 52.713608 | yes | 9.809E-08 | 778.38031 | 42.000267 |
| EHQVVVVAGDTG | 15 | Q14147 | Probable ATP-dependent RNA helicase D | DHX34 | 128.12 | 2 | 52.713608 | yes | 5.419E-07 | 519.25598 | 42.000267 |
| EHQVVVVAGDTG | 15 | Q14147 | Probable ATP-dependent RNA helicase D | DHX34 | 128.12 | 2 | 52.713608 | yes | 9.809E-08 | 778.38031 | 42.000267 |
| SAQSSPECAAAR | 12 | Q14147 | Probable ATP-dependent RNA helicase D | DHX34 | 128.12 | 2 | 52.713608 | yes | 3.502E-07 | 617.77765 | 68.868866 |
| EPSLPQYEVLIVDI | 18 | Q14147 | Probable ATP-dependent RNA helicase D | DHX34 | 128.12 | 3 | 52.713608 | yes | 3.272E-07 | 718.03552 | 68.14785 |
| LYEMANLR | 8 | Q14147 | Probable ATP-dependent RNA helicase D | DHX34 | 128.12 | 2 | 52.713608 | yes | 3.824E-07 | 505.26041 | 66.300629 |
| VLESIDHK | 8 | Q14147 | Probable ATP-dependent RNA helicase D | DHX34 | 128.12 | 2 | 52.713608 | yes | 3.404E-06 | 470.75854 | 44.923962 |
| WDWNCPETR | 9 | Q14147 | Probable ATP-dependent RNA helicase D | DHX34 | 128.12 | 2 | 52.713608 | yes | 3.516E-07 | 632.26422 | 40.244423 |

| | | | | | | | | | | | |
|---|---|---|---|---|---|---|---|---|---|---|---|
| ALLHYLDFGQK | 11 | Q14147 | Probable ATP-dependent RNA helicase D | DHX34 | 128.12 | 2 | 52.713608 | yes | 2.068E-06 | 652.85352 | 27.208698 |
| HLHNDFLLGVLQR | 13 | Q14147 | Probable ATP-dependent RNA helicase D | DHX34 | 128.12 | 3 | 52.713608 | yes | 3.694E-06 | 521.29144 | 22.042007 |
| FWTFFER | 7 | Q14147 | Probable ATP-dependent RNA helicase D | DHX34 | 128.12 | 2 | 52.713608 | yes | 4.602E-07 | 516.75055 | 21.489676 |
| ILDILGETCK | 10 | Q14151 | Scaffold attachment factor B2 OS=Homo | SAFB2 | 107.474 | 2 | 53.02662 | no | 1.202E-07 | 581.31281 | 1973.9969 |
| LEAFHER | 7 | Q14151 | Scaffold attachment factor B2 OS=Homo | SAFB2 | 107.474 | 2 | 53.02662 | yes | 9.722E-08 | 451.22995 | 1229.2719 |
| NTLETSSLNFK | 11 | Q14151 | Scaffold attachment factor B2 OS=Homo | SAFB2 | 107.474 | 2 | 53.02662 | yes | 1.421E-06 | 627.32227 | 902.54974 |
| LQLECQR | 7 | Q14151 | Scaffold attachment factor B2 OS=Homo | SAFB2 | 107.474 | 2 | 53.02662 | yes | 9.722E-08 | 473.74237 | 843.94843 |
| SKGEPVISVK | 10 | Q14151 | Scaffold attachment factor B2 OS=Homo | SAFB2 | 107.474 | 2 | 53.02662 | yes | 1.707E-07 | 522.30841 | 736.93616 |
| FHDFDHR | 7 | Q14151 | Scaffold attachment factor B2 OS=Homo | SAFB2 | 107.474 | 2 | 53.02662 | no | 7.498E-07 | 487.21738 | 647.04205 |
| DWGEHNQR | 8 | Q14151 | Scaffold attachment factor B2 OS=Homo | SAFB2 | 107.474 | 2 | 53.02662 | yes | 1.202E-07 | 521.22845 | 516.36993 |
| NLWVSGLSSTTR | 12 | Q14151 | Scaffold attachment factor B2 OS=Homo | SAFB2 | 107.474 | 2 | 53.02662 | yes | 1.198E-07 | 660.84894 | 470.81769 |
| GQYQDHAIDR | 10 | Q14151 | Scaffold attachment factor B2 OS=Homo | SAFB2 | 107.474 | 2 | 53.02662 | yes | 1.263E-07 | 601.78107 | 454.78088 |
| DGWGGYGSDKR | 11 | Q14151 | Scaffold attachment factor B2 OS=Homo | SAFB2 | 107.474 | 2 | 53.02662 | no | 1.043E-07 | 599.26782 | 438.54263 |
| RPYDLDR | 7 | Q14151 | Scaffold attachment factor B2 OS=Homo | SAFB2 | 107.474 | 2 | 53.02662 | no | 3.786E-07 | 467.74069 | 408.85181 |
| DQDELKPGPTNR | 12 | Q14151 | Scaffold attachment factor B2 OS=Homo | SAFB2 | 107.474 | 2 | 53.02662 | yes | 1.147E-07 | 685.33893 | 399.59784 |
| QLPAQPPEHAVDC | 17 | Q14151 | Scaffold attachment factor B2 OS=Homo | SAFB2 | 107.474 | 3 | 53.02662 | yes | 2.911E-06 | 607.30396 | 383.15634 |
| DIKPIIK | 7 | Q14151 | Scaffold attachment factor B2 OS=Homo | SAFB2 | 107.474 | 2 | 53.02662 | yes | 2.2E-07 | 413.77347 | 312.15115 |
| ADFPRPDHR | 9 | Q14151 | Scaffold attachment factor B2 OS=Homo | SAFB2 | 107.474 | 2 | 53.02662 | yes | 4.24E-07 | 555.77557 | 308.56927 |
| EYVAAQLR | 8 | Q14151 | Scaffold attachment factor B2 OS=Homo | SAFB2 | 107.474 | 2 | 53.02662 | yes | 3.481E-07 | 475.25873 | 247.96048 |
| FDFDACNEVPPAP | 14 | Q14151 | Scaffold attachment factor B2 OS=Homo | SAFB2 | 107.474 | 2 | 53.02662 | no | 1.673E-07 | 803.86395 | 176.00854 |
| GLSGPSGPGHMAS | 14 | Q14151 | Scaffold attachment factor B2 OS=Homo | SAFB2 | 107.474 | 2 | 53.02662 | yes | 9.729E-08 | 655.81714 | 165.12521 |
| AWQGAMDAGAAS | 13 | Q14151 | Scaffold attachment factor B2 OS=Homo | SAFB2 | 107.474 | 2 | 53.02662 | yes | 9.722E-08 | 646.29602 | 150.22057 |
| RDDAYWPEGK | 10 | Q14151 | Scaffold attachment factor B2 OS=Homo | SAFB2 | 107.474 | 2 | 53.02662 | yes | 3.747E-07 | 618.78583 | 148.01208 |
| DIKPIIKDEK | 10 | Q14151 | Scaffold attachment factor B2 OS=Homo | SAFB2 | 107.474 | 2 | 53.02662 | yes | 9.898E-08 | 599.85571 | 137.32973 |
| CYGFVTMSTSDEA | 15 | Q14151 | Scaffold attachment factor B2 OS=Homo | SAFB2 | 107.474 | 2 | 53.02662 | yes | 2.177E-07 | 848.86328 | 123.01243 |
| KFDFDACNEVPPA | 15 | Q14151 | Scaffold attachment factor B2 OS=Homo | SAFB2 | 107.474 | 3 | 53.02662 | no | 1.37E-07 | 578.94336 | 117.95313 |
| ADSLLAVVK | 9 | Q14151 | Scaffold attachment factor B2 OS=Homo | SAFB2 | 107.474 | 2 | 53.02662 | no | 1.8E-07 | 458.27911 | 104.93459 |
| VPHPHPHPPPYPH | 16 | Q14151 | Scaffold attachment factor B2 OS=Homo | SAFB2 | 107.474 | 4 | 53.02662 | yes | 9.863E-07 | 478.99841 | 84.284462 |
| DDAYWPEGK | 9 | Q14151 | Scaffold attachment factor B2 OS=Homo | SAFB2 | 107.474 | 2 | 53.02662 | yes | 1.745E-07 | 540.73529 | 66.474747 |
| REPAEQPGDGER | 12 | Q14151 | Scaffold attachment factor B2 OS=Homo | SAFB2 | 107.474 | 2 | 53.02662 | no | 5.099E-07 | 670.81311 | 51.514839 |
| IGLINDMVR | 9 | Q14152 | Eukaryotic translation initiation factor 3 s | EIF3A | 166.57 | 2 | 53.891575 | yes | 1.063E-07 | 515.78931 | 3162.6384 |
| MHLSQIQR | 8 | Q14152 | Eukaryotic translation initiation factor 3 s | EIF3A | 166.57 | 2 | 53.891575 | yes | 1.375E-07 | 506.77145 | 2682.8389 |
| LESLNIQR | 8 | Q14152 | Eukaryotic translation initiation factor 3 s | EIF3A | 166.57 | 2 | 53.891575 | yes | 1.589E-07 | 486.77728 | 2276.4565 |
| RLEEIPLIK | 9 | Q14152 | Eukaryotic translation initiation factor 3 s | EIF3A | 166.57 | 2 | 53.891575 | yes | 1.855E-07 | 555.84772 | 2170.9976 |
| FCLQYTR | 7 | Q14152 | Eukaryotic translation initiation factor 3 s | EIF3A | 166.57 | 2 | 53.891575 | yes | 1.794E-07 | 494.23947 | 2090.7495 |
| KIDYFER | 7 | Q14152 | Eukaryotic translation initiation factor 3 s | EIF3A | 166.57 | 2 | 53.891575 | yes | 1.025E-07 | 485.75327 | 1895.0304 |
| DMDLWEQQEEER | 12 | Q14152 | Eukaryotic translation initiation factor 3 s | EIF3A | 166.57 | 2 | 53.891575 | yes | 9.722E-08 | 804.33575 | 1737.3844 |
| LLDMDGIIVEK | 11 | Q14152 | Eukaryotic translation initiation factor 3 s | EIF3A | 166.57 | 2 | 53.891575 | yes | 9.722E-08 | 623.34161 | 1476.3387 |
| ITTMQLER | 8 | Q14152 | Eukaryotic translation initiation factor 3 s | EIF3A | 166.57 | 2 | 53.891575 | yes | 1.726E-07 | 496.26569 | 1430.1077 |

| | | | | | | | | | | | |
|---|---|---|---|---|---|---|---|---|---|---|---|
| KLEEVER | 7 | Q14152 | Eukaryotic translation initiation factor 3 s | EIF3A | 166.57 | 2 | 53.891575 | yes | 9.722E-08 | 451.75073 | 1428.4808 |
| QCLDLLR | 7 | Q14152 | Eukaryotic translation initiation factor 3 s | EIF3A | 166.57 | 2 | 53.891575 | yes | 1.798E-07 | 459.24728 | 1379.9105 |
| LATLLGLQAPPTR | 13 | Q14152 | Eukaryotic translation initiation factor 3 s | EIF3A | 166.57 | 2 | 53.891575 | yes | 9.722E-08 | 675.90881 | 1281.3988 |
| YLELCVDLR | 9 | Q14152 | Eukaryotic translation initiation factor 3 s | EIF3A | 166.57 | 2 | 53.891575 | yes | 5.406E-07 | 590.80518 | 1160.1281 |
| HCDLQVR | 7 | Q14152 | Eukaryotic translation initiation factor 3 s | EIF3A | 166.57 | 2 | 53.891575 | yes | 1.001E-07 | 464.2269 | 1116.2207 |
| NICQQVNIK | 9 | Q14152 | Eukaryotic translation initiation factor 3 s | EIF3A | 166.57 | 2 | 53.891575 | yes | 9.722E-08 | 558.79517 | 1057.1504 |
| VLLATLSIPITPER | 14 | Q14152 | Eukaryotic translation initiation factor 3 s | EIF3A | 166.57 | 2 | 53.891575 | yes | 5.579E-07 | 761.96381 | 1046.7653 |
| IHEPIMLK | 8 | Q14152 | Eukaryotic translation initiation factor 3 s | EIF3A | 166.57 | 2 | 53.891575 | yes | 1.638E-07 | 490.78351 | 1023.3425 |
| KLCDNLR | 7 | Q14152 | Eukaryotic translation initiation factor 3 s | EIF3A | 166.57 | 2 | 53.891575 | yes | 9.722E-08 | 459.74493 | 902.99109 |
| AVEDIHGLFSLSK | 13 | Q14152 | Eukaryotic translation initiation factor 3 s | EIF3A | 166.57 | 2 | 53.891575 | yes | 1.545E-07 | 708.38007 | 902.612 |
| RAEEQMLK | 8 | Q14152 | Eukaryotic translation initiation factor 3 s | EIF3A | 166.57 | 2 | 53.891575 | yes | 2.289E-06 | 502.76331 | 813.34985 |
| EKEELEQR | 8 | Q14152 | Eukaryotic translation initiation factor 3 s | EIF3A | 166.57 | 2 | 53.891575 | yes | 1.355E-07 | 530.76709 | 790.2323 |
| PAYFQRPENALK | 12 | Q14152 | Eukaryotic translation initiation factor 3 s | EIF3A | 166.57 | 3 | 53.891575 | yes | 3.034E-07 | 478.58939 | 375.00464 |
| PAYFQRPENALK | 12 | Q14152 | Eukaryotic translation initiation factor 3 s | EIF3A | 166.57 | 3 | 53.891575 | yes | 9.91E-08 | 717.38043 | 375.00464 |
| PAYFQRPENALK | 12 | Q14152 | Eukaryotic translation initiation factor 3 s | EIF3A | 166.57 | 2 | 53.891575 | yes | 3.034E-07 | 478.58939 | 375.00464 |
| PAYFQRPENALK | 12 | Q14152 | Eukaryotic translation initiation factor 3 s | EIF3A | 166.57 | 2 | 53.891575 | yes | 9.91E-08 | 717.38043 | 375.00464 |
| ILQEHEQIK | 9 | Q14152 | Eukaryotic translation initiation factor 3 s | EIF3A | 166.57 | 2 | 53.891575 | yes | 1.233E-07 | 569.31677 | 747.52429 |
| TLSFGSDLNYATR | 13 | Q14152 | Eukaryotic translation initiation factor 3 s | EIF3A | 166.57 | 2 | 53.891575 | yes | 1.022E-07 | 722.85699 | 740.27832 |
| ELEIEER | 7 | Q14152 | Eukaryotic translation initiation factor 3 s | EIF3A | 166.57 | 2 | 53.891575 | yes | 2.314E-07 | 459.23236 | 706.59827 |
| ANEFLEVGK | 9 | Q14152 | Eukaryotic translation initiation factor 3 s | EIF3A | 166.57 | 2 | 53.891575 | yes | 2.482E-07 | 503.76382 | 593.58234 |
| FLWESYR | 7 | Q14152 | Eukaryotic translation initiation factor 3 s | EIF3A | 166.57 | 2 | 53.891575 | yes | 2.236E-07 | 500.74799 | 586.67432 |
| LYHDIAQQAFK | 11 | Q14152 | Eukaryotic translation initiation factor 3 s | EIF3A | 166.57 | 2 | 53.891575 | yes | 1.114E-07 | 667.34857 | 483.45261 |
| KGPEADSEWR | 10 | Q14152 | Eukaryotic translation initiation factor 3 s | EIF3A | 166.57 | 2 | 53.891575 | yes | 1.04E-07 | 587.77802 | 458.39172 |
| ALEVIKPAHILQEK | 14 | Q14152 | Eukaryotic translation initiation factor 3 s | EIF3A | 166.57 | 2 | 53.891575 | yes | 9.722E-08 | 794.97467 | 440.7265 |
| KQPALDVLYDVM | 13 | Q14152 | Eukaryotic translation initiation factor 3 s | EIF3A | 166.57 | 2 | 53.891575 | yes | 9.722E-08 | 760.41309 | 386.03729 |
| EGLYQYK | 7 | Q14152 | Eukaryotic translation initiation factor 3 s | EIF3A | 166.57 | 2 | 53.891575 | yes | 9.819E-08 | 450.72672 | 378.98499 |
| HHNQSTAINLNNP | 23 | Q14152 | Eukaryotic translation initiation factor 3 s | EIF3A | 166.57 | 4 | 53.891575 | yes | 1.22E-07 | 665.31641 | 338.72772 |
| NETDEDGWTTVR | 12 | Q14152 | Eukaryotic translation initiation factor 3 s | EIF3A | 166.57 | 2 | 53.891575 | yes | 9.722E-08 | 711.81024 | 326.73141 |
| FNVLQYVVPEVK | 12 | Q14152 | Eukaryotic translation initiation factor 3 s | EIF3A | 166.57 | 2 | 53.891575 | yes | 6.684E-07 | 717.9032 | 309.31207 |
| LEEIPLIK | 8 | Q14152 | Eukaryotic translation initiation factor 3 s | EIF3A | 166.57 | 2 | 53.891575 | yes | 2.471E-06 | 477.79715 | 278.37692 |
| NQLTAMSSVLAK | 12 | Q14152 | Eukaryotic translation initiation factor 3 s | EIF3A | 166.57 | 2 | 53.891575 | yes | 1.027E-07 | 631.84229 | 244.46234 |
| RANEFLEVGK | 10 | Q14152 | Eukaryotic translation initiation factor 3 s | EIF3A | 166.57 | 2 | 53.891575 | yes | 1.583E-06 | 581.81439 | 237.51297 |
| EEQHQLAVTAYLK | 13 | Q14152 | Eukaryotic translation initiation factor 3 s | EIF3A | 166.57 | 2 | 53.891575 | yes | 9.722E-08 | 765.40155 | 222.45801 |
| QSVYEEK | 7 | Q14152 | Eukaryotic translation initiation factor 3 s | EIF3A | 166.57 | 2 | 53.891575 | yes | 9.722E-08 | 441.71381 | 216.42476 |
| SGNALFHASTLHR | 13 | Q14152 | Eukaryotic translation initiation factor 3 s | EIF3A | 166.57 | 2 | 53.891575 | yes | 1.177E-07 | 705.86548 | 194.83823 |
| PLVKPGGWR | 9 | Q14152 | Eukaryotic translation initiation factor 3 s | EIF3A | 166.57 | 2 | 53.891575 | yes | 1.751E-07 | 505.30072 | 188.8295 |
| GMDDDRGPR | 9 | Q14152 | Eukaryotic translation initiation factor 3 s | EIF3A | 166.57 | 2 | 53.891575 | yes | 1.139E-07 | 509.72217 | 177.70279 |
| RVPPPALSR | 9 | Q14152 | Eukaryotic translation initiation factor 3 s | EIF3A | 166.57 | 2 | 53.891575 | yes | 7.059E-07 | 496.80362 | 169.46297 |
| KNLTQDEMQR | 10 | Q14152 | Eukaryotic translation initiation factor 3 s | EIF3A | 166.57 | 2 | 53.891575 | yes | 9.722E-08 | 631.81152 | 148.68695 |

| | | | | | | | | | | |
|---|---|---|---|---|---|---|---|---|---|---|
| PGPWRPLVK | 9 | Q14152 | Eukaryotic translation initiation factor 3 s | EIF3A | 166.57 | 2 | 53.891575 | yes | 1.828E-07 | 525.31635 | 138.08331 |
| GADDDRPSWR | 10 | Q14152 | Eukaryotic translation initiation factor 3 s | EIF3A | 166.57 | 2 | 53.891575 | yes | 1.021E-07 | 587.76544 | 121.82175 |
| QVEQLEK | 7 | Q14152 | Eukaryotic translation initiation factor 3 s | EIF3A | 166.57 | 2 | 53.891575 | yes | 6.446E-07 | 437.23746 | 106.39382 |
| TKNETDEDGWTT\ | 14 | Q14152 | Eukaryotic translation initiation factor 3 s | EIF3A | 166.57 | 2 | 53.891575 | yes | 9.722E-08 | 826.38153 | 105.12308 |
| QPALDVLYDVMK | 12 | Q14152 | Eukaryotic translation initiation factor 3 s | EIF3A | 166.57 | 2 | 53.891575 | yes | 1.315E-07 | 696.3656 | 101.33674 |
| GLDDDRGPWR | 10 | Q14152 | Eukaryotic translation initiation factor 3 s | EIF3A | 166.57 | 2 | 53.891575 | yes | 1.02E-07 | 593.78363 | 94.592041 |
| SAYEEQR | 7 | Q14152 | Eukaryotic translation initiation factor 3 s | EIF3A | 166.57 | 2 | 53.891575 | yes | 2.503E-06 | 441.70123 | 89.064316 |
| RLGDSSLSR | 9 | Q14152 | Eukaryotic translation initiation factor 3 s | EIF3A | 166.57 | 2 | 53.891575 | yes | 4.205E-07 | 495.76999 | 51.879295 |
| ILQEHEQIKK | 10 | Q14152 | Eukaryotic translation initiation factor 3 s | EIF3A | 166.57 | 2 | 53.891575 | yes | 2.814E-07 | 633.36426 | 42.031685 |
| LLLTPWVK | 8 | Q14152 | Eukaryotic translation initiation factor 3 s | EIF3A | 166.57 | 2 | 53.891575 | yes | 4.858E-07 | 485.31021 | 30.621052 |
| KPPKPQLMANYYI | 14 | Q14152 | Eukaryotic translation initiation factor 3 s | EIF3A | 166.57 | 2 | 53.891575 | yes | 1.567E-07 | 846.4505 | 28.180984 |
| RLATLLGLQAPPTI | 14 | Q14152 | Eukaryotic translation initiation factor 3 s | EIF3A | 166.57 | 3 | 53.891575 | yes | 6.74E-07 | 502.97534 | 26.735722 |
| NLTQDEMQR | 9 | Q14152 | Eukaryotic translation initiation factor 3 s | EIF3A | 166.57 | 2 | 53.891575 | yes | 2.01E-05 | 567.76404 | 25.581059 |
| EAVTACAVTSLIK | 13 | Q14153 | Protein FAM53B OS=Homo sapiens OX= | FAM53B | 45.768 | 2 | 48.831554 | yes | 3.869E-07 | 681.86853 | 212.74969 |
| AWTALLSASGPGC | 14 | Q14153 | Protein FAM53B OS=Homo sapiens OX= | FAM53B | 45.768 | 2 | 48.831554 | yes | 1.1E-07 | 672.35693 | 172.99573 |
| SLSFSDEMSSCR | 12 | Q14153 | Protein FAM53B OS=Homo sapiens OX= | FAM53B | 45.768 | 2 | 48.831554 | yes | 1.099E-07 | 703.28973 | 151.82152 |
| DGSLASNPYSGDL | 15 | Q14157 | Ubiquitin-associated protein 2-like OS=H | UBAP2L | 114.535 | 2 | 51.811913 | yes | 1.076E-07 | 762.86249 | 7970.9351 |
| GFGDVGEAK | 9 | Q14157 | Ubiquitin-associated protein 2-like OS=H | UBAP2L | 114.535 | 2 | 51.811913 | yes | 1.145E-07 | 440.21399 | 2923.8704 |
| IFTASNVSSVPLPA | 24 | Q14157 | Ubiquitin-associated protein 2-like OS=H | UBAP2L | 114.535 | 3 | 51.811913 | yes | 1.35E-07 | 824.7746 | 2366.7524 |
| NPSDSAVHSPFTK | 13 | Q14157 | Ubiquitin-associated protein 2-like OS=H | UBAP2L | 114.535 | 2 | 51.811913 | yes | 1.009E-07 | 693.83606 | 2163.3718 |
| QLIDITGK | 8 | Q14157 | Ubiquitin-associated protein 2-like OS=H | UBAP2L | 114.535 | 2 | 51.811913 | yes | 1.256E-07 | 444.26346 | 1896.2355 |
| GGSTTGSQFLEQFI | 14 | Q14157 | Ubiquitin-associated protein 2-like OS=H | UBAP2L | 114.535 | 2 | 51.811913 | yes | 9.9E-08 | 743.86224 | 1249.3093 |
| GQENGLDGTK | 10 | Q14157 | Ubiquitin-associated protein 2-like OS=H | UBAP2L | 114.535 | 2 | 51.811913 | yes | 1.104E-07 | 509.74362 | 992.42841 |
| QRPQATAEQIR | 11 | Q14157 | Ubiquitin-associated protein 2-like OS=H | UBAP2L | 114.535 | 3 | 51.811913 | yes | 2.23E-07 | 433.23721 | 495.61765 |
| QRPQATAEQIR | 11 | Q14157 | Ubiquitin-associated protein 2-like OS=H | UBAP2L | 114.535 | 3 | 51.811913 | yes | 1.098E-07 | 649.35217 | 495.61765 |
| QRPQATAEQIR | 11 | Q14157 | Ubiquitin-associated protein 2-like OS=H | UBAP2L | 114.535 | 2 | 51.811913 | yes | 2.23E-07 | 433.23721 | 495.61765 |
| QRPQATAEQIR | 11 | Q14157 | Ubiquitin-associated protein 2-like OS=H | UBAP2L | 114.535 | 2 | 51.811913 | yes | 1.098E-07 | 649.35217 | 495.61765 |
| IDLAVLLGK | 9 | Q14157 | Ubiquitin-associated protein 2-like OS=H | UBAP2L | 114.535 | 2 | 51.811913 | yes | 1.225E-07 | 471.30515 | 672.08557 |
| SQTSSIPQKPQTNK | 14 | Q14157 | Ubiquitin-associated protein 2-like OS=H | UBAP2L | 114.535 | 2 | 51.811913 | yes | 9.951E-08 | 772.40735 | 586.552 |
| RYPSSISSSPQK | 12 | Q14157 | Ubiquitin-associated protein 2-like OS=H | UBAP2L | 114.535 | 2 | 51.811913 | yes | 1.259E-07 | 668.84644 | 413.17932 |
| YPSSISSSPQK | 11 | Q14157 | Ubiquitin-associated protein 2-like OS=H | UBAP2L | 114.535 | 2 | 51.811913 | yes | 9.722E-08 | 590.79584 | 226.69736 |
| MMTSVGTNR | 9 | Q14157 | Ubiquitin-associated protein 2-like OS=H | UBAP2L | 114.535 | 2 | 51.811913 | yes | 2.466E-07 | 519.73895 | 147.74101 |
| QAFTPSSTMMEVF | 17 | Q14157 | Ubiquitin-associated protein 2-like OS=H | UBAP2L | 114.535 | 3 | 51.811913 | yes | 1.124E-07 | 658.64978 | 124.7298 |
| QTAISQPASGNTFS | 24 | Q14157 | Ubiquitin-associated protein 2-like OS=H | UBAP2L | 114.535 | 4 | 51.811913 | yes | 1.104E-07 | 629.80597 | 111.82923 |
| AINVLLEGNPDTH: | 20 | Q14157 | Ubiquitin-associated protein 2-like OS=H | UBAP2L | 114.535 | 3 | 51.811913 | yes | 2.331E-07 | 737.36853 | 64.290146 |
| LGLSDNEIQR | 10 | Q14160 | Protein scribble homolog OS=Homo sapi | SCRIB | 174.915 | 2 | 51.62047 | no | 9.722E-08 | 572.80145 | 54.944424 |
| LPLLPPESPGPLR | 13 | Q14160 | Protein scribble homolog OS=Homo sapi | SCRIB | 174.915 | 2 | 51.62047 | yes | 1.189E-06 | 693.41119 | 37.968128 |
| AISTPETPLTK | 11 | Q14181 | DNA polymerase alpha subunit B OS=Hc | POLA2 | 65.948 | 2 | 49.376667 | yes | 1.039E-07 | 579.32428 | 119.31653 |
| VLGCPEALTGSYK | 13 | Q14181 | DNA polymerase alpha subunit B OS=Hc | POLA2 | 65.948 | 2 | 49.376667 | yes | 1.036E-06 | 697.85284 | 109.05138 |

| | | | | | | | | | | | |
|---|---|---|---|---|---|---|---|---|---|---|---|
| QSPCIAVQVVR | 11 | Q14181 | DNA polymerase alpha subunit B OS=Hc | POLA2 | 65.948 | 2 | 49.376667 | yes | 3.96E-07 | 628.84265 | 30.721252 |
| DVLGCVCVNPGR | 12 | Q14181 | DNA polymerase alpha subunit B OS=Hc | POLA2 | 65.948 | 2 | 49.376667 | yes | 8.409E-07 | 673.32117 | 28.244732 |
| ALEPYVTEMAQG | 30 | Q14186 | Transcription factor Dp-1 OS=Homo sapi | TFDP1 | 45.071 | 4 | 46.610287 | yes | 2.62E-06 | 758.62238 | 156.05017 |
| VFIDQNLSPGK | 11 | Q14186 | Transcription factor Dp-1 OS=Homo sapi | TFDP1 | 45.071 | 2 | 46.610287 | yes | 1.515E-07 | 609.3299 | 112.86025 |
| AGVGIEGDQWK | 11 | Q14191 | Bifunctional 3'-5' exonuclease/ATP-deper | WRN | 162.461 | 2 | 54.754375 | yes | 1.287E-07 | 580.29077 | 169.18388 |
| MTQQVTGELR | 10 | Q14191 | Bifunctional 3'-5' exonuclease/ATP-deper | WRN | 162.461 | 2 | 54.754375 | yes | 1.339E-07 | 581.79785 | 162.20724 |
| NLEILDDTVQR | 11 | Q14191 | Bifunctional 3'-5' exonuclease/ATP-deper | WRN | 162.461 | 2 | 54.754375 | yes | 9.722E-08 | 658.34625 | 134.56674 |
| LLSAVDILGEK | 11 | Q14191 | Bifunctional 3'-5' exonuclease/ATP-deper | WRN | 162.461 | 2 | 54.754375 | yes | 2.574E-07 | 579.34247 | 117.28773 |
| VIHSVLEER | 9 | Q14191 | Bifunctional 3'-5' exonuclease/ATP-deper | WRN | 162.461 | 2 | 54.754375 | yes | 2.638E-07 | 541.30365 | 110.06898 |
| DISENLYSLR | 10 | Q14191 | Bifunctional 3'-5' exonuclease/ATP-deper | WRN | 162.461 | 2 | 54.754375 | yes | 9.809E-08 | 605.30914 | 107.25234 |
| EEEILLSDMNK | 11 | Q14191 | Bifunctional 3'-5' exonuclease/ATP-deper | WRN | 162.461 | 2 | 54.754375 | yes | 9.722E-08 | 660.82123 | 106.75789 |
| VALIQLCVSESK | 12 | Q14191 | Bifunctional 3'-5' exonuclease/ATP-deper | WRN | 162.461 | 2 | 54.754375 | yes | 1.233E-07 | 673.87103 | 102.66409 |
| QLTSISEEVMDLAI | 14 | Q14191 | Bifunctional 3'-5' exonuclease/ATP-deper | WRN | 162.461 | 2 | 54.754375 | yes | 1.008E-06 | 782.40039 | 94.75795 |
| QIILSHFEDK | 10 | Q14191 | Bifunctional 3'-5' exonuclease/ATP-deper | WRN | 162.461 | 2 | 54.754375 | yes | 2.885E-07 | 615.3299 | 80.784233 |
| QLITEGFLVEVSR | 13 | Q14191 | Bifunctional 3'-5' exonuclease/ATP-deper | WRN | 162.461 | 2 | 54.754375 | yes | 9.722E-08 | 745.91431 | 80.518349 |
| SLCFQYPPVYVGK | 13 | Q14191 | Bifunctional 3'-5' exonuclease/ATP-deper | WRN | 162.461 | 2 | 54.754375 | yes | 1.341E-07 | 779.39215 | 63.384922 |
| CTETWSLNSLVK | 12 | Q14191 | Bifunctional 3'-5' exonuclease/ATP-deper | WRN | 162.461 | 2 | 54.754375 | yes | 9.722E-08 | 719.35577 | 59.670029 |
| ASLGIMGTEK | 10 | Q14191 | Bifunctional 3'-5' exonuclease/ATP-deper | WRN | 162.461 | 2 | 54.754375 | yes | 4.151E-07 | 503.7655 | 55.326958 |
| LYSYKPCDK | 9 | Q14191 | Bifunctional 3'-5' exonuclease/ATP-deper | WRN | 162.461 | 2 | 54.754375 | yes | 2.385E-07 | 587.28406 | 53.707737 |
| MDVPPAILATNK | 12 | Q14191 | Bifunctional 3'-5' exonuclease/ATP-deper | WRN | 162.461 | 2 | 54.754375 | yes | 2.868E-06 | 635.34723 | 48.700394 |
| QVIHYGAPK | 9 | Q14191 | Bifunctional 3'-5' exonuclease/ATP-deper | WRN | 162.461 | 2 | 54.754375 | yes | 2.905E-06 | 506.78235 | 35.388298 |
| RLPVWFAK | 8 | Q14191 | Bifunctional 3'-5' exonuclease/ATP-deper | WRN | 162.461 | 2 | 54.754375 | yes | 1.086E-06 | 508.80563 | 33.914658 |
| EHEVLIHVEDETW | 22 | Q14191 | Bifunctional 3'-5' exonuclease/ATP-deper | WRN | 162.461 | 4 | 54.754375 | yes | 1.003E-06 | 657.32507 | 19.263708 |
| RDNVAVMATGYC | 13 | Q14191 | Bifunctional 3'-5' exonuclease/ATP-deper | WRN | 162.461 | 2 | 54.754375 | yes | 4.607E-06 | 691.34827 | 13.268061 |
| DNQNFCVPCYEK | 12 | Q14192 | Four and a half LIM domains protein 2 O | FHL2 | 32.193 | 2 | 52.432514 | yes | 9.893E-08 | 787.32416 | 1346.8329 |
| ECFVCTACR | 9 | Q14192 | Four and a half LIM domains protein 2 O | FHL2 | 32.193 | 2 | 52.432514 | yes | 9.722E-08 | 601.74121 | 936.71332 |
| CQQPIGTK | 8 | Q14192 | Four and a half LIM domains protein 2 O | FHL2 | 32.193 | 2 | 52.432514 | yes | 4.368E-07 | 466.23694 | 570.72186 |
| CAGCTNPISGLGG | 15 | Q14192 | Four and a half LIM domains protein 2 O | FHL2 | 32.193 | 2 | 52.432514 | yes | 3.387E-07 | 746.84778 | 555.87152 |
| CSLSLVGR | 8 | Q14192 | Four and a half LIM domains protein 2 O | FHL2 | 32.193 | 2 | 52.432514 | yes | 1.342E-07 | 446.23947 | 514.46155 |
| KPITTGGVTYR | 11 | Q14192 | Four and a half LIM domains protein 2 O | FHL2 | 32.193 | 2 | 52.432514 | yes | 3.142E-07 | 596.83789 | 374.59192 |
| DDILCPDCGK | 10 | Q14192 | Four and a half LIM domains protein 2 O | FHL2 | 32.193 | 2 | 52.432514 | yes | 1.113E-07 | 596.75226 | 323.52838 |
| QHAMQCVQCK | 10 | Q14192 | Four and a half LIM domains protein 2 O | FHL2 | 32.193 | 2 | 52.432514 | yes | 9.85E-08 | 645.28101 | 294.53314 |
| GSSWHETCFICHR | 13 | Q14192 | Four and a half LIM domains protein 2 O | FHL2 | 32.193 | 2 | 52.432514 | yes | 3.309E-07 | 838.85645 | 74.157059 |
| FDCHHCNESLFGK | 13 | Q14192 | Four and a half LIM domains protein 2 O | FHL2 | 32.193 | 2 | 52.432514 | yes | 2.592E-07 | 825.84302 | 59.344028 |
| QIGENLIVPGGVK | 13 | Q14194 | Dihydropyrimidinase-related protein 1 OS | CRMP1 | 62.184 | 2 | 46.886864 | yes | 1.448E-07 | 662.38519 | 35.005371 |
| LGELILTSESSR | 12 | Q14197 | Large ribosomal subunit protein mL62 OS | MRPL58 | 23.631 | 2 | 51.623577 | yes | 1.27E-07 | 652.85645 | 143.97504 |
| DMITEASQTPK | 11 | Q14197 | Large ribosomal subunit protein mL62 OS | MRPL58 | 23.631 | 2 | 51.623577 | yes | 7.065E-07 | 610.79498 | 108.07385 |
| LYPESQGSDTAWF | 13 | Q14197 | Large ribosomal subunit protein mL62 OS | MRPL58 | 23.631 | 2 | 51.623577 | yes | 5.092E-05 | 755.35205 | 85.983459 |
| NLADCLQK | 8 | Q14197 | Large ribosomal subunit protein mL62 OS | MRPL58 | 23.631 | 2 | 51.623577 | yes | 3.211E-05 | 481.24222 | 84.301903 |

| | | | | | | | | | | | |
|---|---|---|---|---|---|---|---|---|---|---|---|
| LTISYCR | 7 | Q14197 | Large ribosomal subunit protein mL62 OS | MRPL58 | 23.631 | 2 | 51.623577 | yes | 1.356E-07 | 456.73401 | 52.168762 |
| FHLATAEWIAEPV | 14 | Q14197 | Large ribosomal subunit protein mL62 OS | MRPL58 | 23.631 | 3 | 51.623577 | yes | 9.722E-08 | 547.2912 | 22.662251 |
| LEETQALLR | 9 | Q14203 | Dynactin subunit 1 OS=Homo sapiens OX | DCTN1 | 141.695 | 2 | 48.86733 | yes | 2.337E-06 | 536.80347 | 48.251213 |
| WVGVILDEAK | 10 | Q14203 | Dynactin subunit 1 OS=Homo sapiens OX | DCTN1 | 141.695 | 2 | 48.86733 | yes | 1.093E-07 | 565.31622 | 43.195778 |
| LAETVFNFQEK | 11 | Q14204 | Cytoplasmic dynein 1 heavy chain 1 OS= | DYNC1H1 | 532.408 | 2 | 49.269394 | yes | 1.648E-06 | 663.34045 | 236.77681 |
| LQGEFQLR | 8 | Q14204 | Cytoplasmic dynein 1 heavy chain 1 OS= | DYNC1H1 | 532.408 | 2 | 49.269394 | yes | 1.186E-05 | 495.772 | 78.827423 |
| VLLTTQGVDMISK | 13 | Q14204 | Cytoplasmic dynein 1 heavy chain 1 OS= | DYNC1H1 | 532.408 | 2 | 49.269394 | yes | 1.05E-07 | 702.89197 | 48.030319 |
| VFEEDALSWEDK | 12 | Q14204 | Cytoplasmic dynein 1 heavy chain 1 OS= | DYNC1H1 | 532.408 | 2 | 49.269394 | yes | 1.344E-07 | 734.33557 | 41.861866 |
| SLLQALNEVK | 10 | Q14204 | Cytoplasmic dynein 1 heavy chain 1 OS= | DYNC1H1 | 532.408 | 2 | 49.269394 | yes | 2.286E-06 | 557.82697 | 40.918556 |
| VQVALEELQDLK | 12 | Q14204 | Cytoplasmic dynein 1 heavy chain 1 OS= | DYNC1H1 | 532.408 | 2 | 49.269394 | yes | 1.682E-06 | 692.88776 | 33.894184 |
| YATLATVSR | 9 | Q14204 | Cytoplasmic dynein 1 heavy chain 1 OS= | DYNC1H1 | 532.408 | 2 | 49.269394 | yes | 3.081E-07 | 491.27182 | 30.509516 |
| MVVLSLPR | 8 | Q14204 | Cytoplasmic dynein 1 heavy chain 1 OS= | DYNC1H1 | 532.408 | 2 | 49.269394 | yes | 1.313E-06 | 457.77823 | 26.223389 |
| INEWLTLVEK | 10 | Q14204 | Cytoplasmic dynein 1 heavy chain 1 OS= | DYNC1H1 | 532.408 | 2 | 49.269394 | yes | 2.208E-07 | 622.8479 | 24.500593 |
| TLINELVK | 8 | Q14204 | Cytoplasmic dynein 1 heavy chain 1 OS= | DYNC1H1 | 532.408 | 2 | 49.269394 | yes | 3.662E-07 | 465.28696 | 24.222614 |
| LNTQEIFDDWAR | 12 | Q14204 | Cytoplasmic dynein 1 heavy chain 1 OS= | DYNC1H1 | 532.408 | 2 | 49.269394 | yes | 5.689E-05 | 754.36243 | 19.485119 |
| VAQTGQDLK | 9 | Q14232 | Translation initiation factor eIF-2B subun | EIF2B1 | 33.713 | 2 | 52.50658 | yes | 3.916E-07 | 480.26144 | 244.67299 |
| FISLASLEYSDYSK | 14 | Q14232 | Translation initiation factor eIF-2B subun | EIF2B1 | 33.713 | 2 | 52.50658 | yes | 1.434E-07 | 811.90106 | 237.75714 |
| DKGETIQGLR | 10 | Q14232 | Translation initiation factor eIF-2B subun | EIF2B1 | 33.713 | 2 | 52.50658 | yes | 9.722E-08 | 558.80402 | 185.51463 |
| IGTNQMAVCAK | 11 | Q14232 | Translation initiation factor eIF-2B subun | EIF2B1 | 33.713 | 2 | 52.50658 | yes | 1.013E-07 | 596.79431 | 145.06415 |
| IADLCHTFIK | 10 | Q14232 | Translation initiation factor eIF-2B subun | EIF2B1 | 33.713 | 2 | 52.50658 | yes | 2.172E-07 | 609.32098 | 136.2285 |
| FSVYVTESQPDLS( | 15 | Q14232 | Translation initiation factor eIF-2B subun | EIF2B1 | 33.713 | 2 | 52.50658 | yes | 2.512E-07 | 828.90942 | 101.64102 |
| LFPLNQQDVPDK | 12 | Q14232 | Translation initiation factor eIF-2B subun | EIF2B1 | 33.713 | 2 | 52.50658 | yes | 2.353E-07 | 707.37225 | 38.722008 |
| TLLEFLK | 7 | Q14232 | Translation initiation factor eIF-2B subun | EIF2B1 | 33.713 | 2 | 52.50658 | yes | 8.135E-07 | 432.26547 | 31.603941 |
| EDPDMASAVAAIR | 13 | Q14232 | Translation initiation factor eIF-2B subun | EIF2B1 | 33.713 | 2 | 52.50658 | yes | 2.21E-06 | 673.32446 | 17.233273 |
| VLITTDLLAR | 10 | Q14240 | Eukaryotic initiation factor 4A-II OS=Ho | EIF4A2 | 46.403 | 2 | 52.39321 | no | 1.222E-07 | 557.84515 | 7910.0459 |
| KVDWLTEK | 8 | Q14240 | Eukaryotic initiation factor 4A-II OS=Ho | EIF4A2 | 46.403 | 2 | 52.39321 | no | 1.939E-07 | 509.78201 | 6993.3403 |
| GYDVIAQAQSGTC | 14 | Q14240 | Eukaryotic initiation factor 4A-II OS=Ho | EIF4A2 | 46.403 | 2 | 52.39321 | no | 9.795E-08 | 697.84918 | 3681.4785 |
| GIYAYGFEKPSAIQ | 16 | Q14240 | Eukaryotic initiation factor 4A-II OS=Ho | EIF4A2 | 46.403 | 3 | 52.39321 | no | 1.08E-07 | 609.98444 | 2287.1296 |
| QFYINVER | 8 | Q14240 | Eukaryotic initiation factor 4A-II OS=Ho | EIF4A2 | 46.403 | 2 | 52.39321 | no | 1.403E-07 | 534.77728 | 2221.072 |
| GIYAYGFEK | 9 | Q14240 | Eukaryotic initiation factor 4A-II OS=Ho | EIF4A2 | 46.403 | 2 | 52.39321 | no | 2.239E-07 | 524.26093 | 1943.9846 |
| MFVLDEADEMLSI | 13 | Q14240 | Eukaryotic initiation factor 4A-II OS=Ho | EIF4A2 | 46.403 | 2 | 52.39321 | no | 1.283E-07 | 778.36017 | 591.29431 |
| MFVLDEADEMLSI | 13 | Q14240 | Eukaryotic initiation factor 4A-II OS=Ho | EIF4A2 | 46.403 | 2 | 52.39321 | no | 9.722E-08 | 786.35767 | 591.29431 |
| LQAEAPHIVVGTP( | 15 | Q14240 | Eukaryotic initiation factor 4A-II OS=Ho | EIF4A2 | 46.403 | 3 | 52.39321 | yes | 1.394E-07 | 515.62299 | 426.94702 |
| LQAEAPHIVVGTP( | 15 | Q14240 | Eukaryotic initiation factor 4A-II OS=Ho | EIF4A2 | 46.403 | 3 | 52.39321 | yes | 1.007E-07 | 772.93079 | 426.94702 |
| LQAEAPHIVVGTP( | 15 | Q14240 | Eukaryotic initiation factor 4A-II OS=Ho | EIF4A2 | 46.403 | 2 | 52.39321 | yes | 1.394E-07 | 515.62299 | 426.94702 |
| LQAEAPHIVVGTP( | 15 | Q14240 | Eukaryotic initiation factor 4A-II OS=Ho | EIF4A2 | 46.403 | 2 | 52.39321 | yes | 1.007E-07 | 772.93079 | 426.94702 |
| DQIYEIFQK | 9 | Q14240 | Eukaryotic initiation factor 4A-II OS=Ho | EIF4A2 | 46.403 | 2 | 52.39321 | yes | 9.893E-08 | 592.30334 | 743.95935 |
| ELAQQIQK | 8 | Q14240 | Eukaryotic initiation factor 4A-II OS=Ho | EIF4A2 | 46.403 | 2 | 52.39321 | no | 2.331E-06 | 479.27182 | 659.51935 |
| VFDMLNR | 7 | Q14240 | Eukaryotic initiation factor 4A-II OS=Ho | EIF4A2 | 46.403 | 2 | 52.39321 | no | 1.063E-07 | 447.72873 | 506.36099 |

| | | | | | | | | | | | |
|---|---|---|---|---|---|---|---|---|---|---|---|
| GVAINFVTEEDKR | 13 | Q14240 | Eukaryotic initiation factor 4A-II OS=Ho | EIF4A2 | 46.403 | 2 | 52.39321 | yes | 9.722E-08 | 739.38593 | 322.08826 |
| KEELTLEGIK | 10 | Q14240 | Eukaryotic initiation factor 4A-II OS=Ho | EIF4A2 | 46.403 | 2 | 52.39321 | yes | 9.722E-08 | 580.33209 | 277.54645 |
| DFTVSALHGDMD( | 14 | Q14240 | Eukaryotic initiation factor 4A-II OS=Ho | EIF4A2 | 46.403 | 2 | 52.39321 | yes | 9.722E-08 | 782.35901 | 218.47824 |
| ETQALVLAPTR | 11 | Q14240 | Eukaryotic initiation factor 4A-II OS=Ho | EIF4A2 | 46.403 | 2 | 52.39321 | yes | 2.112E-07 | 599.84314 | 176.95174 |
| VILALGDYMGATC | 23 | Q14240 | Eukaryotic initiation factor 4A-II OS=Ho | EIF4A2 | 46.403 | 3 | 52.39321 | yes | 1.115E-07 | 817.06433 | 144.46545 |
| GIDVQQVSLVINY | 19 | Q14240 | Eukaryotic initiation factor 4A-II OS=Ho | EIF4A2 | 46.403 | 3 | 52.39321 | no | 9.927E-08 | 715.38306 | 101.1827 |
| AIIPCIK | 7 | Q14240 | Eukaryotic initiation factor 4A-II OS=Ho | EIF4A2 | 46.403 | 2 | 52.39321 | yes | 1.081E-06 | 407.7464 | 39.485439 |
| KGVAINFVTEEDK | 14 | Q14240 | Eukaryotic initiation factor 4A-II OS=Ho | EIF4A2 | 46.403 | 2 | 52.39321 | yes | 9.914E-08 | 803.43341 | 39.242374 |
| ERDVIMR | 7 | Q14240 | Eukaryotic initiation factor 4A-II OS=Ho | EIF4A2 | 46.403 | 2 | 52.39321 | no | 3.362E-07 | 459.74493 | 24.609549 |
| FGTGGAAVPEK | 11 | Q14241 | Elongin-A OS=Homo sapiens OX=9606 ( | ELOA | 89.909 | 2 | 53.630894 | yes | 1.539E-07 | 517.26929 | 881.68982 |
| SEKPAGADLAK | 11 | Q14241 | Elongin-A OS=Homo sapiens OX=9606 ( | ELOA | 89.909 | 2 | 53.630894 | yes | 1.027E-07 | 543.79309 | 637.38373 |
| GHSNAFQDR | 9 | Q14241 | Elongin-A OS=Homo sapiens OX=9606 ( | ELOA | 89.909 | 2 | 53.630894 | yes | 1.988E-07 | 516.23633 | 418.49957 |
| NAEPDEQDFEK | 11 | Q14241 | Elongin-A OS=Homo sapiens OX=9606 ( | ELOA | 89.909 | 2 | 53.630894 | yes | 1.07E-07 | 661.28076 | 410.6524 |
| CTPDQLYR | 8 | Q14241 | Elongin-A OS=Homo sapiens OX=9606 ( | ELOA | 89.909 | 2 | 53.630894 | yes | 1.972E-07 | 526.74512 | 398.56586 |
| MMTLHQQCIR | 10 | Q14241 | Elongin-A OS=Homo sapiens OX=9606 ( | ELOA | 89.909 | 2 | 53.630894 | yes | 1.149E-07 | 659.31488 | 166.34171 |
| MMTLHQQCIR | 10 | Q14241 | Elongin-A OS=Homo sapiens OX=9606 ( | ELOA | 89.909 | 2 | 53.630894 | yes | 1.483E-07 | 439.879 | 166.34171 |
| MMTLHQQCIR | 10 | Q14241 | Elongin-A OS=Homo sapiens OX=9606 ( | ELOA | 89.909 | 3 | 53.630894 | yes | 1.149E-07 | 659.31488 | 166.34171 |
| MMTLHQQCIR | 10 | Q14241 | Elongin-A OS=Homo sapiens OX=9606 ( | ELOA | 89.909 | 3 | 53.630894 | yes | 1.483E-07 | 439.879 | 166.34171 |
| QGLDSFDTGK | 10 | Q14241 | Elongin-A OS=Homo sapiens OX=9606 ( | ELOA | 89.909 | 2 | 53.630894 | yes | 1.006E-07 | 534.25385 | 263.92892 |
| KLVPVER | 7 | Q14241 | Elongin-A OS=Homo sapiens OX=9606 ( | ELOA | 89.909 | 2 | 53.630894 | yes | 1.123E-07 | 420.76871 | 220.86682 |
| MAFVNSVAKPPR | 12 | Q14241 | Elongin-A OS=Homo sapiens OX=9606 ( | ELOA | 89.909 | 2 | 53.630894 | yes | 9.722E-08 | 658.86084 | 100.6777 |
| MAFVNSVAKPPR | 12 | Q14241 | Elongin-A OS=Homo sapiens OX=9606 ( | ELOA | 89.909 | 2 | 53.630894 | yes | 1.164E-07 | 439.57629 | 100.6777 |
| MAFVNSVAKPPR | 12 | Q14241 | Elongin-A OS=Homo sapiens OX=9606 ( | ELOA | 89.909 | 3 | 53.630894 | yes | 9.722E-08 | 658.86084 | 100.6777 |
| MAFVNSVAKPPR | 12 | Q14241 | Elongin-A OS=Homo sapiens OX=9606 ( | ELOA | 89.909 | 3 | 53.630894 | yes | 1.164E-07 | 439.57629 | 100.6777 |
| NIQFAHANKPK | 11 | Q14241 | Elongin-A OS=Homo sapiens OX=9606 ( | ELOA | 89.909 | 2 | 53.630894 | yes | 1.288E-07 | 634.34894 | 166.40826 |
| KHEHVGSFAR | 10 | Q14241 | Elongin-A OS=Homo sapiens OX=9606 ( | ELOA | 89.909 | 2 | 53.630894 | yes | 1.059E-07 | 584.3045 | 123.9001 |
| LAANPDPK | 8 | Q14241 | Elongin-A OS=Homo sapiens OX=9606 ( | ELOA | 89.909 | 2 | 53.630894 | yes | 1.467E-07 | 413.2269 | 106.46914 |
| LSELERPHK | 9 | Q14241 | Elongin-A OS=Homo sapiens OX=9606 ( | ELOA | 89.909 | 2 | 53.630894 | yes | 9.722E-08 | 554.80908 | 97.355675 |
| EEEMEGDYQETW | 13 | Q14241 | Elongin-A OS=Homo sapiens OX=9606 ( | ELOA | 89.909 | 2 | 53.630894 | yes | 9.79E-08 | 837.33539 | 89.861023 |
| EKPPSSGVK | 9 | Q14241 | Elongin-A OS=Homo sapiens OX=9606 ( | ELOA | 89.909 | 2 | 53.630894 | yes | 4.357E-07 | 464.75854 | 77.083054 |
| KLSELERPHK | 10 | Q14241 | Elongin-A OS=Homo sapiens OX=9606 ( | ELOA | 89.909 | 2 | 53.630894 | yes | 4.38E-07 | 618.85657 | 46.671047 |
| AELGAGGDGHR | 11 | Q14244 | Ensconsin OS=Homo sapiens OX=9606 ( | MAP7 | 84.052 | 2 | 53.13809 | yes | 2.011E-07 | 541.2547 | 1347.0425 |
| LFVTPPEGSSR | 11 | Q14244 | Ensconsin OS=Homo sapiens OX=9606 ( | MAP7 | 84.052 | 2 | 53.13809 | yes | 1.028E-07 | 595.31421 | 923.76532 |
| LSSSSATLLNSPDR | 14 | Q14244 | Ensconsin OS=Homo sapiens OX=9606 ( | MAP7 | 84.052 | 2 | 53.13809 | yes | 9.722E-08 | 724.37299 | 853.99982 |
| VTVESTPDLEK | 11 | Q14244 | Ensconsin OS=Homo sapiens OX=9606 ( | MAP7 | 84.052 | 2 | 53.13809 | yes | 1.405E-07 | 609.31665 | 576.17645 |
| VEEATVEER | 9 | Q14244 | Ensconsin OS=Homo sapiens OX=9606 ( | MAP7 | 84.052 | 2 | 53.13809 | yes | 9.722E-08 | 531.25909 | 573.27417 |
| ENVLFLTSGTR | 11 | Q14244 | Ensconsin OS=Homo sapiens OX=9606 ( | MAP7 | 84.052 | 2 | 53.13809 | yes | 9.722E-08 | 618.83276 | 567.03326 |
| SLPHLPGTPR | 10 | Q14244 | Ensconsin OS=Homo sapiens OX=9606 ( | MAP7 | 84.052 | 2 | 53.13809 | yes | 1.063E-07 | 537.80634 | 523.31946 |
| LLTPTHSFLAR | 11 | Q14244 | Ensconsin OS=Homo sapiens OX=9606 ( | MAP7 | 84.052 | 2 | 53.13809 | yes | 9.722E-08 | 628.36151 | 519.85413 |

| | | | | | | | | | | |
|---|---|---|---|---|---|---|---|---|---|---|
| IIHGTASYK | 9 | Q14244 | Ensconsin OS=Homo sapiens OX=9606 ( | MAP7 | 84.052 | 2 | 53.13809 | yes | 9.99E-08 | 495.27435 | 481.16089 |
| LQLSPWESSVVNR | 13 | Q14244 | Ensconsin OS=Homo sapiens OX=9606 ( | MAP7 | 84.052 | 2 | 53.13809 | yes | 1.292E-07 | 757.90173 | 477.6756 |
| TSAGTTDPEEATR | 13 | Q14244 | Ensconsin OS=Homo sapiens OX=9606 ( | MAP7 | 84.052 | 2 | 53.13809 | yes | 9.862E-08 | 668.30475 | 415.81674 |
| RLEAEQAR | 8 | Q14244 | Ensconsin OS=Homo sapiens OX=9606 ( | MAP7 | 84.052 | 2 | 53.13809 | yes | 2.173E-07 | 486.76471 | 358.77536 |
| YVDPVISK | 8 | Q14244 | Ensconsin OS=Homo sapiens OX=9606 ( | MAP7 | 84.052 | 2 | 53.13809 | yes | 3.296E-07 | 460.75803 | 299.74878 |
| PASAISGQNNNHS( | 24 | Q14244 | Ensconsin OS=Homo sapiens OX=9606 ( | MAP7 | 84.052 | 4 | 53.13809 | yes | 1.563E-07 | 617.56635 | 223.25861 |
| EQEELER | 7 | Q14244 | Ensconsin OS=Homo sapiens OX=9606 ( | MAP7 | 84.052 | 2 | 53.13809 | yes | 1.7E-07 | 466.71964 | 222.9823 |
| REQEELER | 8 | Q14244 | Ensconsin OS=Homo sapiens OX=9606 ( | MAP7 | 84.052 | 2 | 53.13809 | yes | 2.299E-05 | 544.7702 | 67.54451 |
| VANEPSLK | 8 | Q14244 | Ensconsin OS=Homo sapiens OX=9606 ( | MAP7 | 84.052 | 2 | 53.13809 | yes | 1.291E-06 | 429.23999 | 35.683968 |
| YGLFPANYVELR | 12 | Q14247 | Src substrate cortactin OS=Homo sapiens | CTTN | 61.587 | 2 | 53.79213 | yes | 4.708E-07 | 721.37738 | 7581.3994 |
| VDQSAVGFEYQGI | 13 | Q14247 | Src substrate cortactin OS=Homo sapiens | CTTN | 61.587 | 2 | 53.79213 | yes | 9.722E-08 | 714.34369 | 5949.8857 |
| LPSSPVYEDAASFI | 14 | Q14247 | Src substrate cortactin OS=Homo sapiens | CTTN | 61.587 | 2 | 53.79213 | yes | 7.02E-07 | 755.87482 | 5111.2578 |
| QDSAAVGFDYK | 11 | Q14247 | Src substrate cortactin OS=Homo sapiens | CTTN | 61.587 | 2 | 53.79213 | yes | 1.128E-07 | 600.78021 | 2941.5935 |
| LQLHESQK | 8 | Q14247 | Src substrate cortactin OS=Homo sapiens | CTTN | 61.587 | 2 | 53.79213 | yes | 1.145E-07 | 491.76944 | 2470.1584 |
| ENVFQEHQTLK | 11 | Q14247 | Src substrate cortactin OS=Homo sapiens | CTTN | 61.587 | 2 | 53.79213 | yes | 9.722E-08 | 686.84644 | 2381.3599 |
| SAVGFDYQGK | 10 | Q14247 | Src substrate cortactin OS=Homo sapiens | CTTN | 61.587 | 2 | 53.79213 | yes | 1.846E-07 | 536.25891 | 2302.8364 |
| SAVGHEYQSK | 10 | Q14247 | Src substrate cortactin OS=Homo sapiens | CTTN | 61.587 | 2 | 53.79213 | yes | 2.441E-07 | 553.26727 | 2183.6433 |
| CALGWDHQEK | 10 | Q14247 | Src substrate cortactin OS=Homo sapiens | CTTN | 61.587 | 2 | 53.79213 | yes | 9.849E-08 | 622.27985 | 1933.6382 |
| TVPVEAVTSK | 10 | Q14247 | Src substrate cortactin OS=Homo sapiens | CTTN | 61.587 | 2 | 53.79213 | yes | 4.163E-07 | 515.7926 | 1708.8729 |
| EKELETGPK | 9 | Q14247 | Src substrate cortactin OS=Homo sapiens | CTTN | 61.587 | 2 | 53.79213 | yes | 9.722E-08 | 515.77441 | 1020.0092 |
| SAVGFEYQGK | 10 | Q14247 | Src substrate cortactin OS=Homo sapiens | CTTN | 61.587 | 2 | 53.79213 | yes | 9.722E-08 | 543.26672 | 916.72083 |
| KLEEQAR | 7 | Q14247 | Src substrate cortactin OS=Homo sapiens | CTTN | 61.587 | 2 | 53.79213 | yes | 1.525E-07 | 437.24307 | 560.05072 |
| FGVQSER | 7 | Q14247 | Src substrate cortactin OS=Homo sapiens | CTTN | 61.587 | 2 | 53.79213 | yes | 9.722E-08 | 411.70886 | 460.72418 |
| ANFENLAK | 8 | Q14247 | Src substrate cortactin OS=Homo sapiens | CTTN | 61.587 | 2 | 53.79213 | yes | 2.162E-07 | 453.73761 | 330.64465 |
| HCSQVDSVR | 9 | Q14247 | Src substrate cortactin OS=Homo sapiens | CTTN | 61.587 | 2 | 53.79213 | yes | 7.061E-07 | 544.2511 | 281.66837 |
| FGVQMDR | 7 | Q14247 | Src substrate cortactin OS=Homo sapiens | CTTN | 61.587 | 2 | 53.79213 | yes | 3.292E-07 | 434.70273 | 29.27087 |
| FLVPIESIER | 10 | Q14249 | Endonuclease G, mitochondrial OS=Hom | ENDOG | 32.621 | 2 | 48.78191 | yes | 1.557E-07 | 601.84259 | 269.32837 |
| EDDSVHAYHR | 10 | Q14249 | Endonuclease G, mitochondrial OS=Hom | ENDOG | 32.621 | 2 | 48.78191 | yes | 1.105E-07 | 614.77069 | 164.39299 |
| YGLPGLAQLK | 10 | Q14249 | Endonuclease G, mitochondrial OS=Hom | ENDOG | 32.621 | 2 | 48.78191 | yes | 4.449E-07 | 530.31348 | 125.05276 |
| VLILEAAGGQIELR | 14 | Q14249 | Endonuclease G, mitochondrial OS=Hom | ENDOG | 32.621 | 2 | 48.78191 | yes | 6.706E-07 | 741.43793 | 46.54171 |
| NHVAVPTHFFK | 11 | Q14249 | Endonuclease G, mitochondrial OS=Hom | ENDOG | 32.621 | 2 | 48.78191 | yes | 1.173E-07 | 648.84601 | 39.507561 |
| DADIGVAEAER | 11 | Q14254 | Flotillin-2 OS=Homo sapiens OX=9606 ( | FLOT2 | 47.065 | 2 | 53.966488 | yes | 9.722E-08 | 573.27527 | 180.54068 |
| NVVLQTLEGHLR | 12 | Q14254 | Flotillin-2 OS=Homo sapiens OX=9606 ( | FLOT2 | 47.065 | 2 | 53.966488 | yes | 9.722E-08 | 689.89368 | 110.21114 |
| ISLEIMTLQPR | 11 | Q14254 | Flotillin-2 OS=Homo sapiens OX=9606 ( | FLOT2 | 47.065 | 2 | 53.966488 | yes | 1.957E-07 | 650.86829 | 76.587395 |
| SAFSEEVNIK | 10 | Q14254 | Flotillin-2 OS=Homo sapiens OX=9606 ( | FLOT2 | 47.065 | 2 | 53.966488 | yes | 3.969E-07 | 562.28516 | 54.945503 |
| TAEAQLAYELQGA | 14 | Q14254 | Flotillin-2 OS=Homo sapiens OX=9606 ( | FLOT2 | 47.065 | 2 | 53.966488 | yes | 4.168E-07 | 760.88879 | 43.204014 |
| VDYLSSLGK | 9 | Q14254 | Flotillin-2 OS=Homo sapiens OX=9606 ( | FLOT2 | 47.065 | 2 | 53.966488 | yes | 8.092E-07 | 491.2662 | 17.31205 |
| AEELHYPLGER | 11 | Q14257 | Reticulocalbin-2 OS=Homo sapiens OX= | RCN2 | 36.877 | 2 | 54.649014 | yes | 1.003E-07 | 657.32788 | 1124.4446 |
| QQFVEYDK | 8 | Q14257 | Reticulocalbin-2 OS=Homo sapiens OX= | RCN2 | 36.877 | 2 | 54.649014 | yes | 2.6E-07 | 528.75348 | 1091.5294 |

| | | | | | | | | | | | |
|---|---|---|---|---|---|---|---|---|---|---|---|
| QLHDDYFYHDEL | 12 | Q14257 | Reticulocalbin-2 OS=Homo sapiens OX= | RCN2 | 36.877 | 2 | 54.649014 | yes | 9.722E-08 | 797.84406 | 546.20044 |
| RLQAIIK | 7 | Q14257 | Reticulocalbin-2 OS=Homo sapiens OX= | RCN2 | 36.877 | 2 | 54.649014 | yes | 1.41E-07 | 421.28455 | 244.67474 |
| FVNDYDKDNDGR | 12 | Q14257 | Reticulocalbin-2 OS=Homo sapiens OX= | RCN2 | 36.877 | 2 | 54.649014 | yes | 9.722E-08 | 729.31824 | 177.16788 |
| KPPPVPALPSK | 11 | Q14258 | E3 ubiquitin/ISG15 ligase TRIM25 OS=I | TRIM25 | 70.974 | 2 | 54.724762 | yes | 1.164E-07 | 565.85022 | 5535.3081 |
| ALLDASETTSTR | 12 | Q14258 | E3 ubiquitin/ISG15 ligase TRIM25 OS=I | TRIM25 | 70.974 | 2 | 54.724762 | yes | 9.722E-08 | 632.82263 | 5180.7358 |
| GISTKPVYIPEVEL | 17 | Q14258 | E3 ubiquitin/ISG15 ligase TRIM25 OS=I | TRIM25 | 70.974 | 3 | 54.724762 | yes | 3.445E-07 | 642.02283 | 2371.571 |
| GISTKPVYIPEVEL | 17 | Q14258 | E3 ubiquitin/ISG15 ligase TRIM25 OS=I | TRIM25 | 70.974 | 3 | 54.724762 | yes | 3.069E-06 | 481.76892 | 2371.571 |
| GISTKPVYIPEVEL | 17 | Q14258 | E3 ubiquitin/ISG15 ligase TRIM25 OS=I | TRIM25 | 70.974 | 4 | 54.724762 | yes | 3.445E-07 | 642.02283 | 2371.571 |
| GISTKPVYIPEVEL | 17 | Q14258 | E3 ubiquitin/ISG15 ligase TRIM25 OS=I | TRIM25 | 70.974 | 4 | 54.724762 | yes | 3.069E-06 | 481.76892 | 2371.571 |
| LPTFGAPEQLVDLI | 14 | Q14258 | E3 ubiquitin/ISG15 ligase TRIM25 OS=I | TRIM25 | 70.974 | 2 | 54.724762 | yes | 5.413E-07 | 764.4245 | 3286.1108 |
| RDEFEFLEK | 9 | Q14258 | E3 ubiquitin/ISG15 ligase TRIM25 OS=I | TRIM25 | 70.974 | 2 | 54.724762 | yes | 2.282E-07 | 606.7984 | 3077.3533 |
| NSASWCVEWFNTI | 13 | Q14258 | E3 ubiquitin/ISG15 ligase TRIM25 OS=I | TRIM25 | 70.974 | 2 | 54.724762 | yes | 9.722E-08 | 814.86176 | 2270.4141 |
| GIHYWEVELQK | 11 | Q14258 | E3 ubiquitin/ISG15 ligase TRIM25 OS=I | TRIM25 | 70.974 | 2 | 54.724762 | yes | 1.512E-07 | 701.36169 | 965.24084 |
| GIHYWEVELQK | 11 | Q14258 | E3 ubiquitin/ISG15 ligase TRIM25 OS=I | TRIM25 | 70.974 | 2 | 54.724762 | yes | 1.949E-07 | 467.91022 | 965.24084 |
| GIHYWEVELQK | 11 | Q14258 | E3 ubiquitin/ISG15 ligase TRIM25 OS=I | TRIM25 | 70.974 | 3 | 54.724762 | yes | 1.512E-07 | 701.36169 | 965.24084 |
| GIHYWEVELQK | 11 | Q14258 | E3 ubiquitin/ISG15 ligase TRIM25 OS=I | TRIM25 | 70.974 | 3 | 54.724762 | yes | 1.949E-07 | 467.91022 | 965.24084 |
| ISAWHNNVEK | 10 | Q14258 | E3 ubiquitin/ISG15 ligase TRIM25 OS=I | TRIM25 | 70.974 | 2 | 54.724762 | yes | 9.722E-08 | 599.3042 | 1851.8821 |
| EEIEQSLTK | 9 | Q14258 | E3 ubiquitin/ISG15 ligase TRIM25 OS=I | TRIM25 | 70.974 | 2 | 54.724762 | yes | 3.477E-07 | 538.77716 | 1586.9028 |
| VLETFLAK | 8 | Q14258 | E3 ubiquitin/ISG15 ligase TRIM25 OS=I | TRIM25 | 70.974 | 2 | 54.724762 | yes | 1.248E-07 | 460.77621 | 1538.0991 |
| GIHQSTIDLK | 10 | Q14258 | E3 ubiquitin/ISG15 ligase TRIM25 OS=I | TRIM25 | 70.974 | 2 | 54.724762 | yes | 1.022E-07 | 556.30896 | 1336.1125 |
| EPPADVWTPPAR | 12 | Q14258 | E3 ubiquitin/ISG15 ligase TRIM25 OS=I | TRIM25 | 70.974 | 2 | 54.724762 | yes | 5.626E-07 | 668.33826 | 1237.9333 |
| PVYIPEVELNHK | 12 | Q14258 | E3 ubiquitin/ISG15 ligase TRIM25 OS=I | TRIM25 | 70.974 | 2 | 54.724762 | yes | 9.795E-08 | 719.39044 | 830.04718 |
| ATSSHPNSTSLK | 12 | Q14258 | E3 ubiquitin/ISG15 ligase TRIM25 OS=I | TRIM25 | 70.974 | 2 | 54.724762 | yes | 1.344E-07 | 615.30969 | 707.59424 |
| QAGLEAAAK | 9 | Q14258 | E3 ubiquitin/ISG15 ligase TRIM25 OS=I | TRIM25 | 70.974 | 2 | 54.724762 | yes | 1.354E-07 | 429.73761 | 570.90216 |
| SRPELLEYYIK | 11 | Q14258 | E3 ubiquitin/ISG15 ligase TRIM25 OS=I | TRIM25 | 70.974 | 2 | 54.724762 | yes | 1.163E-07 | 705.88501 | 441.38666 |
| KSEIQTLK | 8 | Q14258 | E3 ubiquitin/ISG15 ligase TRIM25 OS=I | TRIM25 | 70.974 | 2 | 54.724762 | yes | 1.611E-07 | 473.78201 | 410.24365 |
| VALSECYTVASVA | 20 | Q14258 | E3 ubiquitin/ISG15 ligase TRIM25 OS=I | TRIM25 | 70.974 | 3 | 54.724762 | yes | 1.214E-07 | 763.36102 | 402.27109 |
| LTVMYSQINGASR | 13 | Q14258 | E3 ubiquitin/ISG15 ligase TRIM25 OS=I | TRIM25 | 70.974 | 2 | 54.724762 | yes | 9.722E-08 | 720.3692 | 280.33484 |
| FDTIYQILLK | 10 | Q14258 | E3 ubiquitin/ISG15 ligase TRIM25 OS=I | TRIM25 | 70.974 | 2 | 54.724762 | yes | 9.664E-07 | 627.36066 | 275.44138 |
| KVEQLQQEYTEMI | 13 | Q14258 | E3 ubiquitin/ISG15 ligase TRIM25 OS=I | TRIM25 | 70.974 | 2 | 54.724762 | yes | 1.436E-07 | 827.41125 | 245.04555 |
| VEQLQQEYTEMK | 12 | Q14258 | E3 ubiquitin/ISG15 ligase TRIM25 OS=I | TRIM25 | 70.974 | 2 | 54.724762 | yes | 9.722E-08 | 763.36377 | 160.57571 |
| VILDYNTAHNK | 11 | Q14258 | E3 ubiquitin/ISG15 ligase TRIM25 OS=I | TRIM25 | 70.974 | 2 | 54.724762 | yes | 9.722E-08 | 644.33826 | 123.17249 |
| KGIHYWEVELQK | 12 | Q14258 | E3 ubiquitin/ISG15 ligase TRIM25 OS=I | TRIM25 | 70.974 | 2 | 54.724762 | yes | 3.431E-07 | 765.40918 | 22.189373 |
| SCPQGQAASSATT | 14 | Q14296 | Fas-activated serine/threonine kinase OS= | FASTK | 61.105 | 2 | 47.00477 | yes | 3.152E-07 | 711.3255 | 340.44083 |
| GGQGLEAALSCPR | 13 | Q14296 | Fas-activated serine/threonine kinase OS= | FASTK | 61.105 | 2 | 47.00477 | yes | 1.422E-07 | 658.32477 | 283.67285 |
| VLLSAQTSPAR | 11 | Q14296 | Fas-activated serine/threonine kinase OS= | FASTK | 61.105 | 2 | 47.00477 | yes | 2.894E-07 | 571.83002 | 184.68959 |
| DIVAEGLR | 8 | Q14296 | Fas-activated serine/threonine kinase OS= | FASTK | 61.105 | 2 | 47.00477 | yes | 2.314E-07 | 436.74545 | 125.16656 |
| AHHYSVALR | 9 | Q14296 | Fas-activated serine/threonine kinase OS= | FASTK | 61.105 | 2 | 47.00477 | yes | 2.844E-07 | 527.28308 | 71.646278 |
| CLNNLAASQLK | 11 | Q14318 | Peptidyl-prolyl cis-trans isomerase FKBP | FKBP8 | 44.562 | 2 | 56.427032 | yes | 9.722E-08 | 616.32678 | 122.45136 |

| | | | | | | | | | | | |
|---|---|---|---|---|---|---|---|---|---|---|---|
| ADFVLAANSYDL/ | 15 | Q14318 | Peptidyl-prolyl cis-trans isomerase FKBP | FKBP8 | 44.562 | 2 | 56.427032 | yes | 1.251E-07 | 805.92487 | 99.187294 |
| SCSLVLEHQPDNII | 14 | Q14318 | Peptidyl-prolyl cis-trans isomerase FKBP | FKBP8 | 44.562 | 2 | 56.427032 | yes | 1.66E-07 | 820.40906 | 59.437424 |
| YLGINSDGLVVGR | 13 | Q14331 | Protein FRG1 OS=Homo sapiens OX=96 | FRG1 | 29.173 | 2 | 51.57267 | yes | 9.927E-08 | 681.87244 | 2089.6501 |
| GTYIHALDNGLFTI | 19 | Q14331 | Protein FRG1 OS=Homo sapiens OX=96 | FRG1 | 29.173 | 3 | 51.57267 | yes | 9.919E-08 | 675.68994 | 1453.566 |
| MALLASNSCFIR | 12 | Q14331 | Protein FRG1 OS=Homo sapiens OX=96 | FRG1 | 29.173 | 2 | 51.57267 | yes | 9.849E-08 | 691.84961 | 385.46558 |
| CNEAGDIEAK | 10 | Q14331 | Protein FRG1 OS=Homo sapiens OX=96 | FRG1 | 29.173 | 2 | 51.57267 | yes | 1.668E-07 | 553.7428 | 361.33591 |
| DGFLHETLLDR | 11 | Q14331 | Protein FRG1 OS=Homo sapiens OX=96 | FRG1 | 29.173 | 2 | 51.57267 | yes | 9.976E-08 | 658.33569 | 262.40997 |
| KFQSFQDHK | 9 | Q14331 | Protein FRG1 OS=Homo sapiens OX=96 | FRG1 | 29.173 | 2 | 51.57267 | yes | 3.343E-07 | 582.79346 | 156.86722 |
| KDGFLHETLLDR | 12 | Q14331 | Protein FRG1 OS=Homo sapiens OX=96 | FRG1 | 29.173 | 2 | 51.57267 | yes | 9.722E-08 | 722.38318 | 122.22293 |
| QCEINYVK | 8 | Q14331 | Protein FRG1 OS=Homo sapiens OX=96 | FRG1 | 29.173 | 2 | 51.57267 | yes | 8.867E-06 | 527.25531 | 65.167458 |
| TVLELQYVLDK | 11 | Q14444 | Caprin-1 OS=Homo sapiens OX=9606 G | CAPRIN1 | 78.367 | 2 | 51.453556 | yes | 9.79E-08 | 660.87408 | 76060.43 |
| YQEVTNNLEFAK | 12 | Q14444 | Caprin-1 OS=Homo sapiens OX=9606 G | CAPRIN1 | 78.367 | 2 | 51.453556 | yes | 1.75E-07 | 728.35938 | 53510.633 |
| SFMALSQDIQK | 11 | Q14444 | Caprin-1 OS=Homo sapiens OX=9606 G | CAPRIN1 | 78.367 | 2 | 51.453556 | yes | 1.413E-07 | 634.32117 | 23163.285 |
| SFMALSQDIQK | 11 | Q14444 | Caprin-1 OS=Homo sapiens OX=9606 G | CAPRIN1 | 78.367 | 2 | 51.453556 | yes | 1.537E-07 | 642.31866 | 23163.285 |
| QFMAETQFTSGEK | 13 | Q14444 | Caprin-1 OS=Homo sapiens OX=9606 G | CAPRIN1 | 78.367 | 2 | 51.453556 | yes | 9.722E-08 | 752.34283 | 19097.061 |
| QFMAETQFTSGEK | 13 | Q14444 | Caprin-1 OS=Homo sapiens OX=9606 G | CAPRIN1 | 78.367 | 2 | 51.453556 | yes | 9.809E-08 | 760.34033 | 19097.061 |
| LNQDQLDAVSK | 11 | Q14444 | Caprin-1 OS=Homo sapiens OX=9606 G | CAPRIN1 | 78.367 | 2 | 51.453556 | yes | 1.088E-07 | 615.81989 | 35735.031 |
| LGDDEVR | 7 | Q14444 | Caprin-1 OS=Homo sapiens OX=9606 G | CAPRIN1 | 78.367 | 2 | 51.453556 | yes | 1.481E-07 | 402.19833 | 34792.832 |
| LDDYQER | 7 | Q14444 | Caprin-1 OS=Homo sapiens OX=9606 G | CAPRIN1 | 78.367 | 2 | 51.453556 | yes | 1.106E-07 | 469.71432 | 31866.898 |
| EKPVCGTTYK | 10 | Q14444 | Caprin-1 OS=Homo sapiens OX=9606 G | CAPRIN1 | 78.367 | 2 | 51.453556 | yes | 1.202E-07 | 591.7948 | 25746.518 |
| QILGVIDK | 8 | Q14444 | Caprin-1 OS=Homo sapiens OX=9606 G | CAPRIN1 | 78.367 | 2 | 51.453556 | yes | 1.305E-07 | 443.27383 | 23306.258 |
| GKLDDYQER | 9 | Q14444 | Caprin-1 OS=Homo sapiens OX=9606 G | CAPRIN1 | 78.367 | 2 | 51.453556 | yes | 9.722E-08 | 562.27252 | 13242.137 |
| DGYQQNFK | 8 | Q14444 | Caprin-1 OS=Homo sapiens OX=9606 G | CAPRIN1 | 78.367 | 2 | 51.453556 | yes | 2.065E-07 | 500.23016 | 10482.419 |
| QILGVIDKK | 9 | Q14444 | Caprin-1 OS=Homo sapiens OX=9606 G | CAPRIN1 | 78.367 | 2 | 51.453556 | yes | 1.024E-07 | 507.32132 | 3827.9167 |
| LNEQYEHASIHLW | 19 | Q14444 | Caprin-1 OS=Homo sapiens OX=9606 G | CAPRIN1 | 78.367 | 4 | 51.453556 | yes | 1.562E-07 | 574.54059 | 319.02975 |
| LNEQYEHASIHLW | 19 | Q14444 | Caprin-1 OS=Homo sapiens OX=9606 G | CAPRIN1 | 78.367 | 4 | 51.453556 | yes | 4.23E-07 | 765.71832 | 319.02975 |
| LNEQYEHASIHLW | 19 | Q14444 | Caprin-1 OS=Homo sapiens OX=9606 G | CAPRIN1 | 78.367 | 3 | 51.453556 | yes | 1.562E-07 | 574.54059 | 319.02975 |
| LNEQYEHASIHLW | 19 | Q14444 | Caprin-1 OS=Homo sapiens OX=9606 G | CAPRIN1 | 78.367 | 3 | 51.453556 | yes | 4.23E-07 | 765.71832 | 319.02975 |
| DYSGYQR | 7 | Q14444 | Caprin-1 OS=Homo sapiens OX=9606 G | CAPRIN1 | 78.367 | 2 | 51.453556 | yes | 1.12E-07 | 444.69595 | 372.02588 |
| SNQPYYNSR | 9 | Q14444 | Caprin-1 OS=Homo sapiens OX=9606 G | CAPRIN1 | 78.367 | 2 | 51.453556 | yes | 2.196E-07 | 564.75708 | 136.20079 |
| PVCGTTYK | 8 | Q14444 | Caprin-1 OS=Homo sapiens OX=9606 G | CAPRIN1 | 78.367 | 2 | 51.453556 | yes | 9.927E-08 | 463.22601 | 84.836029 |
| IFCSEDEQSR | 10 | Q14451 | Growth factor receptor-bound protein 7 O | GRB7 | 59.682 | 2 | 49.453197 | yes | 1.029E-07 | 635.77203 | 307.99173 |
| SQPLLIPTTGR | 11 | Q14451 | Growth factor receptor-bound protein 7 O | GRB7 | 59.682 | 2 | 49.453197 | yes | 5.011E-07 | 591.8457 | 269.85446 |
| LLINDFGR | 8 | Q14493 | Histone RNA hairpin-binding protein OS | SLBP | 31.286 | 2 | 51.06094 | yes | 1.967E-07 | 474.26907 | 1088.4257 |
| CSDWASAVEEDEI | 14 | Q14493 | Histone RNA hairpin-binding protein OS | SLBP | 31.286 | 2 | 51.06094 | yes | 1.354E-07 | 842.83252 | 759.35248 |
| KLLINDFGR | 9 | Q14493 | Histone RNA hairpin-binding protein OS | SLBP | 31.286 | 2 | 51.06094 | yes | 1.152E-07 | 538.31659 | 400.81708 |
| RSWDQQIK | 8 | Q14493 | Histone RNA hairpin-binding protein OS | SLBP | 31.286 | 2 | 51.06094 | yes | 1.006E-07 | 530.78033 | 176.51874 |
| HMDSQVEDEFDLI | 21 | Q14493 | Histone RNA hairpin-binding protein OS | SLBP | 31.286 | 3 | 51.06094 | yes | 9.722E-08 | 845.37732 | 135.56451 |
| VLGVPIIVQASQAI | 15 | Q14498 | RNA-binding protein 39 OS=Homo sapie | RBM39 | 59.38 | 2 | 51.97482 | yes | 5.459E-07 | 776.45886 | 3957.3406 |

| | | | | | | | | | | | |
|---|---|---|---|---|---|---|---|---|---|---|---|
| GYGFITFSDSECAK | 14 | Q14498 | RNA-binding protein 39 OS=Homo sapie | RBM39 | 59.38 | 2 | 51.97482 | yes | 1.417E-07 | 791.34814 | 3445.7554 |
| IESIQLMMDSETGI | 14 | Q14498 | RNA-binding protein 39 OS=Homo sapie | RBM39 | 59.38 | 2 | 51.97482 | yes | 2.046E-07 | 805.38165 | 886.03046 |
| IESIQLMMDSETGI | 14 | Q14498 | RNA-binding protein 39 OS=Homo sapie | RBM39 | 59.38 | 2 | 51.97482 | yes | 3.944E-07 | 813.37909 | 886.03046 |
| IESIQLMMDSETGI | 14 | Q14498 | RNA-binding protein 39 OS=Homo sapie | RBM39 | 59.38 | 2 | 51.97482 | yes | 3.401E-07 | 813.37909 | 886.03046 |
| IGLPHSIK | 8 | Q14498 | RNA-binding protein 39 OS=Homo sapie | RBM39 | 59.38 | 2 | 51.97482 | yes | 2.369E-07 | 432.76871 | 1047.7161 |
| NSAQGNVYVK | 10 | Q14498 | RNA-binding protein 39 OS=Homo sapie | RBM39 | 59.38 | 2 | 51.97482 | yes | 2.616E-07 | 540.27765 | 959.59375 |
| TVFCMQLAAR | 10 | Q14498 | RNA-binding protein 39 OS=Homo sapie | RBM39 | 59.38 | 2 | 51.97482 | no | 9.722E-08 | 598.79938 | 909.90118 |
| DLEEFFSTVGK | 11 | Q14498 | RNA-binding protein 39 OS=Homo sapie | RBM39 | 59.38 | 2 | 51.97482 | yes | 9.728E-08 | 636.31134 | 783.24451 |
| TGIDLGTTGR | 10 | Q14498 | RNA-binding protein 39 OS=Homo sapie | RBM39 | 59.38 | 2 | 51.97482 | yes | 1.093E-07 | 495.76437 | 680.30981 |
| HGGVIHIYVDK | 11 | Q14498 | RNA-binding protein 39 OS=Homo sapie | RBM39 | 59.38 | 2 | 51.97482 | yes | 1.155E-07 | 619.33801 | 326.24896 |
| HGGVIHIYVDK | 11 | Q14498 | RNA-binding protein 39 OS=Homo sapie | RBM39 | 59.38 | 2 | 51.97482 | yes | 2.04E-07 | 413.22778 | 326.24896 |
| HGGVIHIYVDK | 11 | Q14498 | RNA-binding protein 39 OS=Homo sapie | RBM39 | 59.38 | 3 | 51.97482 | yes | 1.155E-07 | 619.33801 | 326.24896 |
| HGGVIHIYVDK | 11 | Q14498 | RNA-binding protein 39 OS=Homo sapie | RBM39 | 59.38 | 3 | 51.97482 | yes | 2.04E-07 | 413.22778 | 326.24896 |
| AAAMANNLQK | 10 | Q14498 | RNA-binding protein 39 OS=Homo sapie | RBM39 | 59.38 | 2 | 51.97482 | yes | 1.607E-07 | 516.26874 | 606.96594 |
| CPSIAAAIAAVNAI | 17 | Q14498 | RNA-binding protein 39 OS=Homo sapie | RBM39 | 59.38 | 3 | 51.97482 | yes | 1.144E-07 | 564.63849 | 365.26312 |
| EPIDNLTPEER | 11 | Q14498 | RNA-binding protein 39 OS=Homo sapie | RBM39 | 59.38 | 2 | 51.97482 | yes | 1.406E-07 | 656.82263 | 241.39964 |
| ALEQLNGFELAGR | 13 | Q14498 | RNA-binding protein 39 OS=Homo sapie | RBM39 | 59.38 | 2 | 51.97482 | no | 2.033E-07 | 709.37537 | 234.88445 |
| VVDLDPCLSK | 10 | Q14517 | Protocadherin Fat 1 OS=Homo sapiens O | FAT1 | 506.273 | 2 | 55.668854 | yes | 1.136E-07 | 573.29718 | 459.03555 |
| LIMQISATDADIR | 13 | Q14517 | Protocadherin Fat 1 OS=Homo sapiens O | FAT1 | 506.273 | 2 | 55.668854 | yes | 9.723E-08 | 723.8847 | 383.65787 |
| TSGVLSTTGTPFDI | 14 | Q14517 | Protocadherin Fat 1 OS=Homo sapiens O | FAT1 | 506.273 | 2 | 55.668854 | yes | 9.722E-08 | 719.86224 | 327.15732 |
| EEPQLFSTVVVK | 12 | Q14517 | Protocadherin Fat 1 OS=Homo sapiens O | FAT1 | 506.273 | 2 | 55.668854 | yes | 1.26E-07 | 688.37701 | 326.9585 |
| STLPTGIQVK | 10 | Q14517 | Protocadherin Fat 1 OS=Homo sapiens O | FAT1 | 506.273 | 2 | 55.668854 | yes | 3.274E-07 | 522.30841 | 322.91672 |
| VDTEVGHVIR | 10 | Q14517 | Protocadherin Fat 1 OS=Homo sapiens O | FAT1 | 506.273 | 2 | 55.668854 | yes | 9.722E-08 | 562.80658 | 288.9902 |
| EEQAVYHLLVR | 11 | Q14517 | Protocadherin Fat 1 OS=Homo sapiens O | FAT1 | 506.273 | 2 | 55.668854 | yes | 9.722E-08 | 678.86713 | 272.34689 |
| LSFVTPR | 7 | Q14517 | Protocadherin Fat 1 OS=Homo sapiens O | FAT1 | 506.273 | 2 | 55.668854 | yes | 3.548E-07 | 410.23981 | 268.79538 |
| LDPATGSLYTSEK | 13 | Q14517 | Protocadherin Fat 1 OS=Homo sapiens O | FAT1 | 506.273 | 2 | 55.668854 | yes | 6.308E-07 | 691.34589 | 264.56119 |
| GSPPMSEITSVR | 12 | Q14517 | Protocadherin Fat 1 OS=Homo sapiens O | FAT1 | 506.273 | 2 | 55.668854 | yes | 1.832E-07 | 630.81628 | 258.01007 |
| FTQDVYSAVVK | 11 | Q14517 | Protocadherin Fat 1 OS=Homo sapiens O | FAT1 | 506.273 | 2 | 55.668854 | yes | 9.722E-08 | 628.82971 | 253.46294 |
| VSFHSLR | 7 | Q14517 | Protocadherin Fat 1 OS=Homo sapiens O | FAT1 | 506.273 | 2 | 55.668854 | yes | 2.656E-07 | 423.23505 | 245.35513 |
| MGVYITHPAWEVI | 13 | Q14517 | Protocadherin Fat 1 OS=Homo sapiens O | FAT1 | 506.273 | 2 | 55.668854 | yes | 9.722E-08 | 779.89539 | 118.31189 |
| MGVYITHPAWEVI | 13 | Q14517 | Protocadherin Fat 1 OS=Homo sapiens O | FAT1 | 506.273 | 2 | 55.668854 | yes | 1.998E-07 | 520.26599 | 118.31189 |
| MGVYITHPAWEVI | 13 | Q14517 | Protocadherin Fat 1 OS=Homo sapiens O | FAT1 | 506.273 | 3 | 55.668854 | yes | 9.722E-08 | 779.89539 | 118.31189 |
| MGVYITHPAWEVI | 13 | Q14517 | Protocadherin Fat 1 OS=Homo sapiens O | FAT1 | 506.273 | 3 | 55.668854 | yes | 1.998E-07 | 520.26599 | 118.31189 |
| LVNLQCEETGVAK | 13 | Q14517 | Protocadherin Fat 1 OS=Homo sapiens O | FAT1 | 506.273 | 2 | 55.668854 | yes | 1.368E-07 | 730.87433 | 226.21478 |
| HFVIDSATGIITLSN | 18 | Q14517 | Protocadherin Fat 1 OS=Homo sapiens O | FAT1 | 506.273 | 3 | 55.668854 | yes | 1.013E-07 | 665.36542 | 225.1525 |
| TLYSDTIPEDVLPC | 15 | Q14517 | Protocadherin Fat 1 OS=Homo sapiens O | FAT1 | 506.273 | 2 | 55.668854 | yes | 9.722E-08 | 824.42743 | 225.11259 |
| ELQAVYTLSLK | 11 | Q14517 | Protocadherin Fat 1 OS=Homo sapiens O | FAT1 | 506.273 | 2 | 55.668854 | yes | 1.489E-07 | 632.86102 | 223.75272 |
| IGEETGVIETSDR | 13 | Q14517 | Protocadherin Fat 1 OS=Homo sapiens O | FAT1 | 506.273 | 2 | 55.668854 | yes | 9.722E-08 | 703.34393 | 221.77649 |
| IFVTIADNASPK | 12 | Q14517 | Protocadherin Fat 1 OS=Homo sapiens O | FAT1 | 506.273 | 2 | 55.668854 | yes | 1.446E-07 | 638.35083 | 205.15752 |

| | | | | | | | | | | | |
|---|---|---|---|---|---|---|---|---|---|---|---|
| ITSGNPQGFFSIHPI | 15 | Q14517 | Protocadherin Fat 1 OS=Homo sapiens O | FAT1 | 506.273 | 2 | 55.668854 | yes | 9.722E-08 | 815.42285 | 97.496742 |
| ITSGNPQGFFSIHPI | 15 | Q14517 | Protocadherin Fat 1 OS=Homo sapiens O | FAT1 | 506.273 | 2 | 55.668854 | yes | 3.025E-07 | 543.95099 | 97.496742 |
| ITSGNPQGFFSIHPI | 15 | Q14517 | Protocadherin Fat 1 OS=Homo sapiens O | FAT1 | 506.273 | 3 | 55.668854 | yes | 9.722E-08 | 815.42285 | 97.496742 |
| ITSGNPQGFFSIHPI | 15 | Q14517 | Protocadherin Fat 1 OS=Homo sapiens O | FAT1 | 506.273 | 3 | 55.668854 | yes | 3.025E-07 | 543.95099 | 97.496742 |
| LNPDTGELK | 9 | Q14517 | Protocadherin Fat 1 OS=Homo sapiens O | FAT1 | 506.273 | 2 | 55.668854 | yes | 1.001E-06 | 493.76129 | 191.90446 |
| YEVNIGSSAAK | 11 | Q14517 | Protocadherin Fat 1 OS=Homo sapiens O | FAT1 | 506.273 | 2 | 55.668854 | yes | 1.57E-06 | 569.79059 | 191.72922 |
| LYGSSGISSMAK | 12 | Q14517 | Protocadherin Fat 1 OS=Homo sapiens O | FAT1 | 506.273 | 2 | 55.668854 | yes | 1.32E-07 | 600.80011 | 179.45755 |
| VIHVTSPQFK | 10 | Q14517 | Protocadherin Fat 1 OS=Homo sapiens O | FAT1 | 506.273 | 2 | 55.668854 | yes | 1.017E-07 | 578.32965 | 165.82349 |
| IVSGDSENLFK | 11 | Q14517 | Protocadherin Fat 1 OS=Homo sapiens O | FAT1 | 506.273 | 2 | 55.668854 | yes | 7.996E-07 | 604.81152 | 158.47626 |
| SNQAEYDLMVK | 11 | Q14517 | Protocadherin Fat 1 OS=Homo sapiens O | FAT1 | 506.273 | 2 | 55.668854 | yes | 9.722E-08 | 649.30829 | 152.56825 |
| VLDANDNSPVCEI | 13 | Q14517 | Protocadherin Fat 1 OS=Homo sapiens O | FAT1 | 506.273 | 2 | 55.668854 | yes | 9.809E-08 | 730.83795 | 134.01381 |
| LHIEWISK | 8 | Q14517 | Protocadherin Fat 1 OS=Homo sapiens O | FAT1 | 506.273 | 2 | 55.668854 | yes | 8.026E-07 | 513.29254 | 124.24384 |
| INCEGTIPR | 9 | Q14517 | Protocadherin Fat 1 OS=Homo sapiens O | FAT1 | 506.273 | 2 | 55.668854 | yes | 2.683E-07 | 530.26624 | 114.84797 |
| VQATDADAGLNR | 12 | Q14517 | Protocadherin Fat 1 OS=Homo sapiens O | FAT1 | 506.273 | 2 | 55.668854 | yes | 9.722E-08 | 615.80731 | 110.11593 |
| PLYEASLLLPTYK | 13 | Q14517 | Protocadherin Fat 1 OS=Homo sapiens O | FAT1 | 506.273 | 2 | 55.668854 | yes | 9.722E-08 | 754.42395 | 102.12984 |
| GGNTAILNR | 9 | Q14517 | Protocadherin Fat 1 OS=Homo sapiens O | FAT1 | 506.273 | 2 | 55.668854 | yes | 4.348E-07 | 458.25397 | 92.129822 |
| WYQFSILAR | 9 | Q14517 | Protocadherin Fat 1 OS=Homo sapiens O | FAT1 | 506.273 | 2 | 55.668854 | yes | 1.474E-07 | 592.31659 | 91.4291 |
| VSVDESVMSTHST | 15 | Q14517 | Protocadherin Fat 1 OS=Homo sapiens O | FAT1 | 506.273 | 2 | 55.668854 | yes | 9.809E-08 | 803.38049 | 90.877495 |
| TYVGHPVK | 8 | Q14517 | Protocadherin Fat 1 OS=Homo sapiens O | FAT1 | 506.273 | 2 | 55.668854 | yes | 7.905E-07 | 450.75052 | 89.308891 |
| LCAGLDCPWK | 10 | Q14517 | Protocadherin Fat 1 OS=Homo sapiens O | FAT1 | 506.273 | 2 | 55.668854 | yes | 1.455E-07 | 610.28357 | 79.945099 |
| DHYLLQVK | 8 | Q14517 | Protocadherin Fat 1 OS=Homo sapiens O | FAT1 | 506.273 | 2 | 55.668854 | yes | 6.184E-07 | 508.2822 | 78.40094 |
| TLDYEQSR | 8 | Q14517 | Protocadherin Fat 1 OS=Homo sapiens O | FAT1 | 506.273 | 2 | 55.668854 | yes | 9.722E-08 | 506.24072 | 74.379898 |
| SNAEITYTLLGSGA | 16 | Q14517 | Protocadherin Fat 1 OS=Homo sapiens O | FAT1 | 506.273 | 2 | 55.668854 | yes | 7.707E-07 | 827.42017 | 72.07814 |
| ILSPLDR | 7 | Q14517 | Protocadherin Fat 1 OS=Homo sapiens O | FAT1 | 506.273 | 2 | 55.668854 | yes | 1.512E-07 | 407.24509 | 67.786118 |
| GQIFTLEK | 8 | Q14517 | Protocadherin Fat 1 OS=Homo sapiens O | FAT1 | 506.273 | 2 | 55.668854 | yes | 2.343E-06 | 468.26346 | 65.391495 |
| DMPAAGSLGSSSR | 13 | Q14517 | Protocadherin Fat 1 OS=Homo sapiens O | FAT1 | 506.273 | 2 | 55.668854 | yes | 1.771E-07 | 618.2879 | 65.391472 |
| AIPAYSHLR | 9 | Q14517 | Protocadherin Fat 1 OS=Homo sapiens O | FAT1 | 506.273 | 2 | 55.668854 | yes | 1.675E-07 | 514.28778 | 60.479939 |
| LVLDQVHTASGTA | 18 | Q14517 | Protocadherin Fat 1 OS=Homo sapiens O | FAT1 | 506.273 | 3 | 55.668854 | yes | 3.016E-07 | 603.33527 | 60.34343 |
| NSFEGSAIPEHPEF | 26 | Q14517 | Protocadherin Fat 1 OS=Homo sapiens O | FAT1 | 506.273 | 4 | 55.668854 | yes | 8.221E-07 | 728.08643 | 58.616547 |
| GTDYSILEIHHGR | 13 | Q14517 | Protocadherin Fat 1 OS=Homo sapiens O | FAT1 | 506.273 | 2 | 55.668854 | yes | 9.862E-08 | 749.37585 | 52.12368 |
| FAGLTSVK | 8 | Q14517 | Protocadherin Fat 1 OS=Homo sapiens O | FAT1 | 506.273 | 2 | 55.668854 | yes | 1.121E-07 | 411.73962 | 43.913914 |
| AEHSGTVLYSLVK | 13 | Q14517 | Protocadherin Fat 1 OS=Homo sapiens O | FAT1 | 506.273 | 2 | 55.668854 | yes | 9.722E-08 | 702.38007 | 30.118851 |
| DKGTPPQFSSVK | 12 | Q14517 | Protocadherin Fat 1 OS=Homo sapiens O | FAT1 | 506.273 | 2 | 55.668854 | yes | 3.033E-05 | 645.83807 | 26.822908 |
| EKDHYLLQVK | 10 | Q14517 | Protocadherin Fat 1 OS=Homo sapiens O | FAT1 | 506.273 | 2 | 55.668854 | yes | 5.426E-07 | 636.85095 | 25.451136 |
| TLNLDNYVFFGGI | 15 | Q14517 | Protocadherin Fat 1 OS=Homo sapiens O | FAT1 | 506.273 | 3 | 55.668854 | yes | 8.907E-07 | 589.30554 | 19.840727 |
| GKPVLELPER | 10 | Q14527 | Helicase-like transcription factor OS=Hoı | HLTF | 113.929 | 2 | 53.076084 | yes | 1.431E-07 | 569.33496 | 1078.3499 |
| AVLDLESER | 9 | Q14527 | Helicase-like transcription factor OS=Hoı | HLTF | 113.929 | 2 | 53.076084 | yes | 9.788E-08 | 516.27203 | 1039.1918 |
| LSYPTFFPR | 9 | Q14527 | Helicase-like transcription factor OS=Hoı | HLTF | 113.929 | 2 | 53.076084 | yes | 9.722E-08 | 564.29785 | 769.16156 |
| AGGVGLNLSAASI | 13 | Q14527 | Helicase-like transcription factor OS=Hoı | HLTF | 113.929 | 2 | 53.076084 | yes | 2.111E-07 | 586.82275 | 703.82849 |

| | | | | | | | | | | | |
|---|---|---|---|---|---|---|---|---|---|---|---|
| TLGFNLESGWGSG | 14 | Q14527 | Helicase-like transcription factor OS=Hor | HLTF | 113.929 | 2 | 53.076084 | yes | 1.163E-07 | 740.86261 | 629.59021 |
| ELAAGAFGTK | 10 | Q14527 | Helicase-like transcription factor OS=Hor | HLTF | 113.929 | 2 | 53.076084 | yes | 3.344E-07 | 482.75854 | 617.06097 |
| LKPFIDR | 7 | Q14527 | Helicase-like transcription factor OS=Hor | HLTF | 113.929 | 2 | 53.076084 | yes | 2.687E-07 | 444.76871 | 587.15759 |
| VILDEGHAIR | 10 | Q14527 | Helicase-like transcription factor OS=Hor | HLTF | 113.929 | 2 | 53.076084 | yes | 1.26E-07 | 561.81696 | 451.1597 |
| QALAWMVSR | 9 | Q14527 | Helicase-like transcription factor OS=Hor | HLTF | 113.929 | 2 | 53.076084 | yes | 9.722E-08 | 531.28168 | 426.37134 |
| IREPALLSK | 9 | Q14527 | Helicase-like transcription factor OS=Hor | HLTF | 113.929 | 2 | 53.076084 | yes | 1.121E-07 | 513.81897 | 411.82935 |
| GDSPLHSIR | 9 | Q14527 | Helicase-like transcription factor OS=Hor | HLTF | 113.929 | 2 | 53.076084 | yes | 3.462E-07 | 491.25925 | 358.94086 |
| WVLTGTPIQNSLK | 13 | Q14527 | Helicase-like transcription factor OS=Hor | HLTF | 113.929 | 2 | 53.076084 | yes | 4.749E-07 | 728.91156 | 344.77951 |
| MSELSSSRPK | 10 | Q14527 | Helicase-like transcription factor OS=Hor | HLTF | 113.929 | 2 | 53.076084 | yes | 1.353E-07 | 561.28461 | 336.09509 |
| CSEQPSISDIK | 11 | Q14527 | Helicase-like transcription factor OS=Hor | HLTF | 113.929 | 2 | 53.076084 | yes | 1.272E-07 | 632.29791 | 312.85828 |
| INALMHALTDLR | 12 | Q14527 | Helicase-like transcription factor OS=Hor | HLTF | 113.929 | 2 | 53.076084 | yes | 1.806E-07 | 684.37683 | 262.21304 |
| EYNVNDDSMK | 10 | Q14527 | Helicase-like transcription factor OS=Hor | HLTF | 113.929 | 2 | 53.076084 | yes | 9.927E-08 | 607.75336 | 240.75775 |
| THEMEPAEAIETPI | 19 | Q14527 | Helicase-like transcription factor OS=Hor | HLTF | 113.929 | 3 | 53.076084 | yes | 6.689E-07 | 724.36487 | 169.35748 |
| VFIQHITLSDEER | 13 | Q14527 | Helicase-like transcription factor OS=Hor | HLTF | 113.929 | 2 | 53.076084 | yes | 9.722E-08 | 793.91229 | 104.4209 |
| PVTMGDEGGLR | 11 | Q14527 | Helicase-like transcription factor OS=Hor | HLTF | 113.929 | 2 | 53.076084 | yes | 1.168E-07 | 566.27679 | 100.2202 |
| KSDMEWTSSSK | 11 | Q14527 | Helicase-like transcription factor OS=Hor | HLTF | 113.929 | 2 | 53.076084 | yes | 1.105E-07 | 643.2901 | 82.218475 |
| RLQSLIK | 7 | Q14527 | Helicase-like transcription factor OS=Hor | HLTF | 113.929 | 2 | 53.076084 | yes | 4.254E-07 | 429.28198 | 70.168655 |
| TIQRPVTMGDEGG | 15 | Q14527 | Helicase-like transcription factor OS=Hor | HLTF | 113.929 | 2 | 53.076084 | yes | 9.722E-08 | 815.42249 | 51.753723 |
| LFEDLKEDDKTHE | 29 | Q14527 | Helicase-like transcription factor OS=Hor | HLTF | 113.929 | 5 | 53.076084 | yes | 9.911E-08 | 681.54034 | 43.776947 |
| QDIVLTTYNILTHI | 18 | Q14527 | Helicase-like transcription factor OS=Hor | HLTF | 113.929 | 3 | 53.076084 | yes | 1.078E-07 | 699.02838 | 39.679668 |
| KVFIQHITLSDEER | 14 | Q14527 | Helicase-like transcription factor OS=Hor | HLTF | 113.929 | 3 | 53.076084 | yes | 5.59E-06 | 572.30896 | 21.734203 |
| GQDIFIIQTIPR | 12 | Q14558 | Phosphoribosyl pyrophosphate synthase-a | PRPSAP1 | 39.394 | 2 | 55.239334 | yes | 9.722E-08 | 700.89844 | 87.591675 |
| LIEESSVDEVVVTI | 22 | Q14558 | Phosphoribosyl pyrophosphate synthase-a | PRPSAP1 | 39.394 | 3 | 55.239334 | yes | 3.458E-07 | 817.75861 | 78.114105 |
| VFSANSTAACTEL | 15 | Q14558 | Phosphoribosyl pyrophosphate synthase-a | PRPSAP1 | 39.394 | 2 | 55.239334 | yes | 5.564E-07 | 785.38251 | 32.0942 |
| NIIGVIPYFPYSK | 13 | Q14558 | Phosphoribosyl pyrophosphate synthase-a | PRPSAP1 | 39.394 | 2 | 55.239334 | yes | 6.182E-07 | 755.91882 | 30.656857 |
| LIVTSATLDAVK | 12 | Q14562 | ATP-dependent RNA helicase DHX8 OS | DHX8 | 139.315 | 2 | 53.76238 | yes | 5.195E-07 | 615.86884 | 328.67151 |
| ELEALMPSAAGQE | 14 | Q14562 | ATP-dependent RNA helicase DHX8 OS | DHX8 | 139.315 | 2 | 53.76238 | yes | 1.14E-07 | 737.36633 | 243.48184 |
| DEMLTTNVPEIQR | 13 | Q14562 | ATP-dependent RNA helicase DHX8 OS | DHX8 | 139.315 | 2 | 53.76238 | yes | 9.722E-08 | 773.38251 | 199.97192 |
| TNLASTVLSLK | 11 | Q14562 | ATP-dependent RNA helicase DHX8 OS | DHX8 | 139.315 | 2 | 53.76238 | yes | 3.989E-07 | 573.84009 | 172.80473 |
| TYPVEILYTK | 10 | Q14562 | ATP-dependent RNA helicase DHX8 OS | DHX8 | 139.315 | 2 | 53.76238 | yes | 9.722E-08 | 613.83698 | 167.26917 |
| GIGMMPNDIPEWK | 13 | Q14562 | ATP-dependent RNA helicase DHX8 OS | DHX8 | 139.315 | 2 | 53.76238 | yes | 2.523E-07 | 744.35468 | 144.78473 |
| NLVGETNEETSMR | 13 | Q14562 | ATP-dependent RNA helicase DHX8 OS | DHX8 | 139.315 | 2 | 53.76238 | yes | 9.722E-08 | 740.34082 | 144.55325 |
| FEDCTSPETVIK | 12 | Q14562 | ATP-dependent RNA helicase DHX8 OS | DHX8 | 139.315 | 2 | 53.76238 | yes | 1.344E-06 | 713.33197 | 134.21509 |
| EAEMDSIPMGLNK | 13 | Q14562 | ATP-dependent RNA helicase DHX8 OS | DHX8 | 139.315 | 2 | 53.76238 | yes | 1.543E-07 | 717.8338 | 130.69115 |
| LEYLSLVSK | 9 | Q14562 | ATP-dependent RNA helicase DHX8 OS | DHX8 | 139.315 | 2 | 53.76238 | yes | 1.37E-07 | 526.30536 | 123.37526 |
| MAEFPLEPMLCK | 12 | Q14562 | ATP-dependent RNA helicase DHX8 OS | DHX8 | 139.315 | 2 | 53.76238 | yes | 1.047E-07 | 733.34802 | 117.29695 |
| IFDPAPPGSR | 10 | Q14562 | ATP-dependent RNA helicase DHX8 OS | DHX8 | 139.315 | 2 | 53.76238 | yes | 6.371E-07 | 528.77728 | 116.26266 |
| TMLDEDDVK | 9 | Q14562 | ATP-dependent RNA helicase DHX8 OS | DHX8 | 139.315 | 2 | 53.76238 | yes | 4.428E-07 | 533.24207 | 109.60692 |
| HWVDPLPDAEGR | 12 | Q14562 | ATP-dependent RNA helicase DHX8 OS | DHX8 | 139.315 | 2 | 53.76238 | yes | 1.04E-07 | 696.33875 | 105.65083 |

| | | | | | | | | | | | |
|---|---|---|---|---|---|---|---|---|---|---|---|
| LIQTMRPPAKPSTS | 15 | Q14562 | ATP-dependent RNA helicase DHX8 OS | DHX8 | 139.315 | 3 | 53.76238 | yes | 3.325E-06 | 552.31512 | 80.587776 |
| TIHTDVLFGLLK | 12 | Q14562 | ATP-dependent RNA helicase DHX8 OS | DHX8 | 139.315 | 2 | 53.76238 | yes | 9.722E-08 | 678.89795 | 67.202965 |
| WEGLVHISELR | 11 | Q14562 | ATP-dependent RNA helicase DHX8 OS | DHX8 | 139.315 | 2 | 53.76238 | yes | 4.443E-06 | 669.86188 | 39.960567 |
| EAFAIVSVPVSEIR | 14 | Q14573 | Inositol 1,4,5-trisphosphate receptor type | ITPR3 | 304.107 | 2 | 52.256 | yes | 2.598E-07 | 758.92212 | 76.899353 |
| YSLGPSLR | 8 | Q14573 | Inositol 1,4,5-trisphosphate receptor type | ITPR3 | 304.107 | 2 | 52.256 | yes | 3.422E-07 | 446.74799 | 63.945744 |
| LQDIITALEER | 11 | Q14573 | Inositol 1,4,5-trisphosphate receptor type | ITPR3 | 304.107 | 2 | 52.256 | yes | 9.722E-08 | 650.85901 | 62.30677 |
| LFTTTEQDEQGSK | 13 | Q14573 | Inositol 1,4,5-trisphosphate receptor type | ITPR3 | 304.107 | 2 | 52.256 | yes | 1.028E-07 | 742.34918 | 32.883457 |
| ISYLLSVFK | 9 | Q14573 | Inositol 1,4,5-trisphosphate receptor type | ITPR3 | 304.107 | 2 | 52.256 | yes | 3.679E-06 | 535.31824 | 27.78495 |
| LLLALMESR | 9 | Q14573 | Inositol 1,4,5-trisphosphate receptor type | ITPR3 | 304.107 | 2 | 52.256 | yes | 4.267E-05 | 523.30737 | 17.881737 |
| SQLIIHQR | 8 | Q14587 | Zinc finger protein 268 OS=Homo sapien | ZNF268 | 108.374 | 2 | 48.473316 | no | 4.243E-07 | 497.79324 | 706.15845 |
| SNLIVHQR | 8 | Q14587 | Zinc finger protein 268 OS=Homo sapien | ZNF268 | 108.374 | 2 | 48.473316 | no | 1.143E-07 | 483.77762 | 88.095146 |
| TASIWVPPLQER | 12 | Q14587 | Zinc finger protein 268 OS=Homo sapien | ZNF268 | 108.374 | 2 | 48.473316 | yes | 1.241E-06 | 698.88281 | 27.46143 |
| VKPYDCPECGK | 11 | Q14592 | Zinc finger protein 460 OS=Homo sapien | ZNF460 | 63.665 | 2 | 45.778595 | yes | 1.743E-07 | 676.80231 | 135.98288 |
| SNFVLHNK | 8 | Q14592 | Zinc finger protein 460 OS=Homo sapien | ZNF460 | 63.665 | 2 | 45.778595 | yes | 3.037E-06 | 479.75888 | 126.33195 |
| KPFACSECGK | 10 | Q14592 | Zinc finger protein 460 OS=Homo sapien | ZNF460 | 63.665 | 2 | 45.778595 | yes | 2.661E-07 | 592.26538 | 76.385841 |
| HFSIHTGEKPYEC\ | 18 | Q14592 | Zinc finger protein 460 OS=Homo sapien | ZNF460 | 63.665 | 4 | 45.778595 | no | 6.618E-07 | 545.24902 | 55.364758 |
| PFVCNECGK | 9 | Q14592 | Zinc finger protein 460 OS=Homo sapien | ZNF460 | 63.665 | 2 | 45.778595 | no | 3.241E-07 | 555.73895 | 42.96946 |
| FAHTVVTSR | 9 | Q14624 | Inter-alpha-trypsin inhibitor heavy chain 1 | ITIH4 | 103.358 | 2 | 51.562862 | yes | 1.268E-07 | 509.27744 | 543.97015 |
| FKPTLSQQQK | 10 | Q14624 | Inter-alpha-trypsin inhibitor heavy chain 1 | ITIH4 | 103.358 | 2 | 51.562862 | yes | 1.688E-07 | 602.83789 | 409.24768 |
| YIFHNFMER | 9 | Q14624 | Inter-alpha-trypsin inhibitor heavy chain 1 | ITIH4 | 103.358 | 2 | 51.562862 | yes | 9.722E-08 | 628.79767 | 363.28607 |
| VRPQQLVK | 8 | Q14624 | Inter-alpha-trypsin inhibitor heavy chain 1 | ITIH4 | 103.358 | 2 | 51.562862 | yes | 3.451E-07 | 484.306 | 310.49628 |
| LQDIVSALEDR | 11 | Q14643 | Inositol 1,4,5-trisphosphate receptor type | ITPR1 | 313.93 | 2 | 45.53162 | yes | 2.535E-07 | 629.83551 | 69.76033 |
| DLQNFLR | 7 | Q14643 | Inositol 1,4,5-trisphosphate receptor type | ITPR1 | 313.93 | 2 | 45.53162 | no | 3.893E-07 | 453.24561 | 49.631744 |
| LLLAIMESR | 9 | Q14643 | Inositol 1,4,5-trisphosphate receptor type | ITPR1 | 313.93 | 2 | 45.53162 | no | 1.029E-06 | 523.30737 | 44.337841 |
| LVTNLSGQLSELK | 13 | Q14643 | Inositol 1,4,5-trisphosphate receptor type | ITPR1 | 313.93 | 2 | 45.53162 | yes | 3.76E-07 | 701.401 | 31.957682 |
| SEDLFNEMNWQK | 12 | Q14643 | Inositol 1,4,5-trisphosphate receptor type | ITPR1 | 313.93 | 2 | 45.53162 | yes | 1.822E-07 | 770.84045 | 24.192799 |
| EIYTSNHMWK | 10 | Q14643 | Inositol 1,4,5-trisphosphate receptor type | ITPR1 | 313.93 | 2 | 45.53162 | yes | 2.906E-06 | 654.80573 | 22.519119 |
| SLKPDDLGSLR | 11 | Q14644 | Ras GTPase-activating protein 3 OS=Hor | RASA3 | 95.699 | 2 | 51.237827 | yes | 1.308E-06 | 600.83276 | 140.77591 |
| VDLWNASNLK | 10 | Q14644 | Ras GTPase-activating protein 3 OS=Hor | RASA3 | 95.699 | 2 | 51.237827 | yes | 3.249E-07 | 580.30896 | 89.01918 |
| NFLDLISSSGR | 11 | Q14644 | Ras GTPase-activating protein 3 OS=Hor | RASA3 | 95.699 | 2 | 51.237827 | yes | 1.173E-07 | 604.81714 | 80.167984 |
| ALYIQANNCVEAK | 13 | Q14644 | Ras GTPase-activating protein 3 OS=Hor | RASA3 | 95.699 | 2 | 51.237827 | yes | 9.722E-08 | 747.37451 | 42.747131 |
| EKQEAAVPLVR | 11 | Q14644 | Ras GTPase-activating protein 3 OS=Hor | RASA3 | 95.699 | 2 | 51.237827 | yes | 3.527E-06 | 620.35645 | 22.630508 |
| QEAAVPLVR | 9 | Q14644 | Ras GTPase-activating protein 3 OS=Hor | RASA3 | 95.699 | 2 | 51.237827 | yes | 3.599E-06 | 491.78763 | 20.813961 |
| SLNPPLTIVR | 10 | Q14669 | E3 ubiquitin-protein ligase TRIP12 OS=I | TRIP12 | 220.434 | 2 | 53.507042 | yes | 9.722E-08 | 555.33752 | 232.43011 |
| SSAVVVDAIPVFLI | 15 | Q14669 | E3 ubiquitin-protein ligase TRIP12 OS=I | TRIP12 | 220.434 | 2 | 53.507042 | yes | 1.009E-07 | 787.44543 | 214.42842 |
| QAESVMQDLGSSI | 13 | Q14669 | E3 ubiquitin-protein ligase TRIP12 OS=I | TRIP12 | 220.434 | 2 | 53.507042 | yes | 1.572E-07 | 704.33026 | 171.23793 |
| SVYHLEDIVR | 10 | Q14669 | E3 ubiquitin-protein ligase TRIP12 OS=I | TRIP12 | 220.434 | 2 | 53.507042 | yes | 9.722E-08 | 615.82751 | 170.64365 |
| VEPVGNAPLLALV | 15 | Q14669 | E3 ubiquitin-protein ligase TRIP12 OS=I | TRIP12 | 220.434 | 2 | 53.507042 | yes | 1.183E-07 | 778.96161 | 72.196594 |
| VEPVGNAPLLALV | 15 | Q14669 | E3 ubiquitin-protein ligase TRIP12 OS=I | TRIP12 | 220.434 | 2 | 53.507042 | yes | 2.607E-07 | 519.64349 | 72.196594 |

| | | | | | | | | | | | |
|---|---|---|---|---|---|---|---|---|---|---|---|
| VEPVGNAPLLALV | 15 | Q14669 | E3 ubiquitin-protein ligase TRIP12 OS=I | TRIP12 | 220.434 | 3 | 53.507042 | yes | 1.183E-07 | 778.96161 | 72.196594 |
| VEPVGNAPLLALV | 15 | Q14669 | E3 ubiquitin-protein ligase TRIP12 OS=I | TRIP12 | 220.434 | 3 | 53.507042 | yes | 2.607E-07 | 519.64349 | 72.196594 |
| MSQLFHR | 7 | Q14669 | E3 ubiquitin-protein ligase TRIP12 OS=I | TRIP12 | 220.434 | 2 | 53.507042 | yes | 1.337E-07 | 459.73434 | 141.76585 |
| LPDYSSIEIMR | 11 | Q14669 | E3 ubiquitin-protein ligase TRIP12 OS=I | TRIP12 | 220.434 | 2 | 53.507042 | yes | 1.197E-07 | 662.33429 | 139.4724 |
| DDSLDLSPQGR | 11 | Q14669 | E3 ubiquitin-protein ligase TRIP12 OS=I | TRIP12 | 220.434 | 2 | 53.507042 | yes | 3.533E-07 | 601.78601 | 129.64207 |
| EGIFAVDTMLK | 11 | Q14669 | E3 ubiquitin-protein ligase TRIP12 OS=I | TRIP12 | 220.434 | 2 | 53.507042 | yes | 9.722E-08 | 612.32068 | 124.52802 |
| HLDQEQQLK | 9 | Q14669 | E3 ubiquitin-protein ligase TRIP12 OS=I | TRIP12 | 220.434 | 2 | 53.507042 | yes | 3.618E-06 | 569.7962 | 119.7733 |
| IIYFADAELLK | 11 | Q14669 | E3 ubiquitin-protein ligase TRIP12 OS=I | TRIP12 | 220.434 | 2 | 53.507042 | yes | 9.722E-08 | 648.36591 | 96.830345 |
| EIIPTSEFINSK | 12 | Q14669 | E3 ubiquitin-protein ligase TRIP12 OS=I | TRIP12 | 220.434 | 2 | 53.507042 | yes | 1.1E-06 | 689.36664 | 76.096039 |
| QLLLYLTSK | 9 | Q14669 | E3 ubiquitin-protein ligase TRIP12 OS=I | TRIP12 | 220.434 | 2 | 53.507042 | yes | 4.442E-07 | 539.82898 | 72.287048 |
| FFNTHQLK | 8 | Q14669 | E3 ubiquitin-protein ligase TRIP12 OS=I | TRIP12 | 220.434 | 2 | 53.507042 | yes | 1.456E-07 | 517.77454 | 70.909119 |
| THTIWYKPVR | 10 | Q14669 | E3 ubiquitin-protein ligase TRIP12 OS=I | TRIP12 | 220.434 | 2 | 53.507042 | yes | 5.353E-07 | 650.86169 | 35.275642 |
| THTIWYKPVR | 10 | Q14669 | E3 ubiquitin-protein ligase TRIP12 OS=I | TRIP12 | 220.434 | 2 | 53.507042 | yes | 6.762E-07 | 434.24353 | 35.275642 |
| THTIWYKPVR | 10 | Q14669 | E3 ubiquitin-protein ligase TRIP12 OS=I | TRIP12 | 220.434 | 3 | 53.507042 | yes | 5.353E-07 | 650.86169 | 35.275642 |
| THTIWYKPVR | 10 | Q14669 | E3 ubiquitin-protein ligase TRIP12 OS=I | TRIP12 | 220.434 | 3 | 53.507042 | yes | 6.762E-07 | 434.24353 | 35.275642 |
| SETSKPGPSGLQAI | 14 | Q14669 | E3 ubiquitin-protein ligase TRIP12 OS=I | TRIP12 | 220.434 | 2 | 53.507042 | yes | 1.108E-07 | 693.86481 | 65.659744 |
| NHAVSSHIASMLS | 18 | Q14669 | E3 ubiquitin-protein ligase TRIP12 OS=I | TRIP12 | 220.434 | 3 | 53.507042 | yes | 3.182E-05 | 642.32306 | 52.425034 |
| SESPPAELPSLR | 12 | Q14669 | E3 ubiquitin-protein ligase TRIP12 OS=I | TRIP12 | 220.434 | 2 | 53.507042 | yes | 4.053E-07 | 641.83551 | 47.911331 |
| SSFLASLNPK | 10 | Q14669 | E3 ubiquitin-protein ligase TRIP12 OS=I | TRIP12 | 220.434 | 2 | 53.507042 | yes | 9.325E-07 | 532.29279 | 42.246822 |
| ALTYMMEALPR | 11 | Q14669 | E3 ubiquitin-protein ligase TRIP12 OS=I | TRIP12 | 220.434 | 2 | 53.507042 | yes | 1.768E-07 | 648.32794 | 32.41312 |
| DQYANYVVQK | 10 | Q14671 | Pumilio homolog 1 OS=Homo sapiens O | PUM1 | 126.474 | 2 | 54.666813 | no | 9.722E-08 | 614.30383 | 3128.9761 |
| YISAAPGAEAK | 11 | Q14671 | Pumilio homolog 1 OS=Homo sapiens O | PUM1 | 126.474 | 2 | 54.666813 | no | 1.165E-07 | 539.28241 | 2914.5776 |
| GNVLVLSQHK | 10 | Q14671 | Pumilio homolog 1 OS=Homo sapiens O | PUM1 | 126.474 | 2 | 54.666813 | yes | 6.376E-07 | 547.81946 | 2456.6553 |
| SQDDAMVDYFFQ | 13 | Q14671 | Pumilio homolog 1 OS=Homo sapiens O | PUM1 | 126.474 | 2 | 54.666813 | yes | 9.916E-08 | 811.3512 | 1201.8174 |
| SQDDAMVDYFFQ | 13 | Q14671 | Pumilio homolog 1 OS=Homo sapiens O | PUM1 | 126.474 | 2 | 54.666813 | yes | 3.471E-07 | 819.34863 | 1201.8174 |
| GIFLGDQWR | 9 | Q14671 | Pumilio homolog 1 OS=Homo sapiens O | PUM1 | 126.474 | 2 | 54.666813 | yes | 9.722E-08 | 546.28528 | 2167.4236 |
| FASNVVEK | 8 | Q14671 | Pumilio homolog 1 OS=Homo sapiens O | PUM1 | 126.474 | 2 | 54.666813 | no | 9.849E-08 | 447.23999 | 2078.6016 |
| FFEFGSLEQK | 10 | Q14671 | Pumilio homolog 1 OS=Homo sapiens O | PUM1 | 126.474 | 2 | 54.666813 | yes | 6.429E-07 | 616.30334 | 1677.563 |
| DSLTGSSDLYK | 11 | Q14671 | Pumilio homolog 1 OS=Homo sapiens O | PUM1 | 126.474 | 2 | 54.666813 | yes | 1.799E-07 | 593.28534 | 1476.6934 |
| GQVFALSTHPYGC | 14 | Q14671 | Pumilio homolog 1 OS=Homo sapiens O | PUM1 | 126.474 | 2 | 54.666813 | yes | 9.81E-08 | 796.88556 | 721.11951 |
| GQVFALSTHPYGC | 14 | Q14671 | Pumilio homolog 1 OS=Homo sapiens O | PUM1 | 126.474 | 2 | 54.666813 | yes | 1.897E-07 | 531.59277 | 721.11951 |
| GQVFALSTHPYGC | 14 | Q14671 | Pumilio homolog 1 OS=Homo sapiens O | PUM1 | 126.474 | 3 | 54.666813 | yes | 9.81E-08 | 796.88556 | 721.11951 |
| GQVFALSTHPYGC | 14 | Q14671 | Pumilio homolog 1 OS=Homo sapiens O | PUM1 | 126.474 | 3 | 54.666813 | yes | 1.897E-07 | 531.59277 | 721.11951 |
| YGMSDVMPSGR | 11 | Q14671 | Pumilio homolog 1 OS=Homo sapiens O | PUM1 | 126.474 | 2 | 54.666813 | yes | 1.083E-07 | 600.26282 | 1007.8868 |
| YPNLQLR | 7 | Q14671 | Pumilio homolog 1 OS=Homo sapiens O | PUM1 | 126.474 | 2 | 54.666813 | yes | 1.853E-07 | 452.25598 | 963.3302 |
| ELDGHVLK | 8 | Q14671 | Pumilio homolog 1 OS=Homo sapiens O | PUM1 | 126.474 | 2 | 54.666813 | no | 1.499E-07 | 455.75327 | 769.94403 |
| GHVLSLALQMYG | 14 | Q14671 | Pumilio homolog 1 OS=Homo sapiens O | PUM1 | 126.474 | 2 | 54.666813 | yes | 1.003E-07 | 802.90546 | 371.77518 |
| GHVLSLALQMYG | 14 | Q14671 | Pumilio homolog 1 OS=Homo sapiens O | PUM1 | 126.474 | 2 | 54.666813 | yes | 1.828E-07 | 535.60602 | 371.77518 |
| GHVLSLALQMYG | 14 | Q14671 | Pumilio homolog 1 OS=Homo sapiens O | PUM1 | 126.474 | 3 | 54.666813 | yes | 1.003E-07 | 802.90546 | 371.77518 |

| | | | | | | | | | | | |
|---|---|---|---|---|---|---|---|---|---|---|---|
| GHVLSLALQMYG( | 14 | Q14671 | Pumilio homolog 1 OS=Homo sapiens O: | PUM1 | 126.474 | 3 | 54.666813 | yes | 1.828E-07 | 535.60602 | 371.77518 |
| AMGEQLLPGK | 10 | Q14671 | Pumilio homolog 1 OS=Homo sapiens O: | PUM1 | 126.474 | 2 | 54.666813 | yes | 9.892E-08 | 522.28131 | 697.44336 |
| AVLWQDSFSPHLK | 13 | Q14671 | Pumilio homolog 1 OS=Homo sapiens O: | PUM1 | 126.474 | 2 | 54.666813 | yes | 1.004E-07 | 764.40137 | 628.88232 |
| MIDVAEPGQR | 10 | Q14671 | Pumilio homolog 1 OS=Homo sapiens O: | PUM1 | 126.474 | 2 | 54.666813 | yes | 9.79E-08 | 558.2793 | 585.93781 |
| WPTGDNIHAEHQ\ | 14 | Q14671 | Pumilio homolog 1 OS=Homo sapiens O: | PUM1 | 126.474 | 3 | 54.666813 | yes | 1.733E-07 | 553.93774 | 195.08264 |
| WPTGDNIHAEHQ\ | 14 | Q14671 | Pumilio homolog 1 OS=Homo sapiens O: | PUM1 | 126.474 | 3 | 54.666813 | yes | 2.908E-07 | 830.40295 | 195.08264 |
| WPTGDNIHAEHQ\ | 14 | Q14671 | Pumilio homolog 1 OS=Homo sapiens O: | PUM1 | 126.474 | 2 | 54.666813 | yes | 1.733E-07 | 553.93774 | 195.08264 |
| WPTGDNIHAEHQ\ | 14 | Q14671 | Pumilio homolog 1 OS=Homo sapiens O: | PUM1 | 126.474 | 2 | 54.666813 | yes | 2.908E-07 | 830.40295 | 195.08264 |
| LGDLKEEGDVMD] | 13 | Q14671 | Pumilio homolog 1 OS=Homo sapiens O: | PUM1 | 126.474 | 2 | 54.666813 | yes | 1.934E-07 | 724.85052 | 268.91589 |
| RPGQSFHVNSEVN | 19 | Q14671 | Pumilio homolog 1 OS=Homo sapiens O: | PUM1 | 126.474 | 4 | 54.666813 | yes | 9.722E-08 | 528.27515 | 268.00314 |
| RDSLTGSSDLYK | 12 | Q14671 | Pumilio homolog 1 OS=Homo sapiens O: | PUM1 | 126.474 | 2 | 54.666813 | yes | 5.157E-07 | 671.33588 | 139.27657 |
| QHGEQLGGGGSG( | 20 | Q14671 | Pumilio homolog 1 OS=Homo sapiens O: | PUM1 | 126.474 | 3 | 54.666813 | yes | 9.722E-08 | 620.9455 | 70.870476 |
| AMGEQLLPGKK | 11 | Q14671 | Pumilio homolog 1 OS=Homo sapiens O: | PUM1 | 126.474 | 2 | 54.666813 | yes | 9.722E-08 | 586.3288 | 66.236664 |
| KFWETDESSK | 10 | Q14671 | Pumilio homolog 1 OS=Homo sapiens O: | PUM1 | 126.474 | 2 | 54.666813 | yes | 1.578E-07 | 628.79333 | 48.726475 |
| HRWPTGDNIHAEI | 16 | Q14671 | Pumilio homolog 1 OS=Homo sapiens O: | PUM1 | 126.474 | 4 | 54.666813 | yes | 1.387E-06 | 488.99512 | 45.351223 |
| LLQVGEEGPQAV/ | 14 | Q14674 | Separin OS=Homo sapiens OX=9606 GN | ESPL1 | 233.176 | 2 | 52.151882 | yes | 9.722E-08 | 719.89862 | 317.96002 |
| GSSSGLLSPQR | 11 | Q14674 | Separin OS=Homo sapiens OX=9606 GN | ESPL1 | 233.176 | 2 | 52.151882 | yes | 3.828E-07 | 544.78839 | 255.35435 |
| FLDGQAVLR | 9 | Q14674 | Separin OS=Homo sapiens OX=9606 GN | ESPL1 | 233.176 | 2 | 52.151882 | yes | 2.077E-06 | 509.78763 | 217.38333 |
| DSPPVSVQIPTGQ! | 15 | Q14674 | Separin OS=Homo sapiens OX=9606 GN | ESPL1 | 233.176 | 2 | 52.151882 | yes | 1.122E-07 | 783.90973 | 212.55396 |
| NAAAQGSPEATLR | 13 | Q14674 | Separin OS=Homo sapiens OX=9606 GN | ESPL1 | 233.176 | 2 | 52.151882 | yes | 9.989E-08 | 643.32837 | 210.3998 |
| WQVLSEVLSCSEK | 13 | Q14674 | Separin OS=Homo sapiens OX=9606 GN | ESPL1 | 233.176 | 2 | 52.151882 | yes | 2.336E-07 | 782.88745 | 196.87611 |
| EFLSNPPAGFPSSR | 14 | Q14674 | Separin OS=Homo sapiens OX=9606 GN | ESPL1 | 233.176 | 2 | 52.151882 | yes | 1.557E-07 | 753.3728 | 196.80385 |
| GPALELVATVAK | 12 | Q14674 | Separin OS=Homo sapiens OX=9606 GN | ESPL1 | 233.176 | 2 | 52.151882 | yes | 9.819E-08 | 584.85046 | 182.2684 |
| LQELLQDCGWK | 11 | Q14674 | Separin OS=Homo sapiens OX=9606 GN | ESPL1 | 233.176 | 2 | 52.151882 | yes | 1.175E-07 | 695.34521 | 157.92094 |
| EYGASPVLSQGVD | 15 | Q14674 | Separin OS=Homo sapiens OX=9606 GN | ESPL1 | 233.176 | 2 | 52.151882 | yes | 4.26E-07 | 787.8941 | 144.02641 |
| SMEAPSPPLR | 10 | Q14674 | Separin OS=Homo sapiens OX=9606 GN | ESPL1 | 233.176 | 2 | 52.151882 | yes | 3.295E-06 | 542.77643 | 139.7511 |
| AQELLNEAVGR | 11 | Q14674 | Separin OS=Homo sapiens OX=9606 GN | ESPL1 | 233.176 | 2 | 52.151882 | yes | 2.508E-07 | 600.32257 | 135.54359 |
| FLALCLGHR | 9 | Q14674 | Separin OS=Homo sapiens OX=9606 GN | ESPL1 | 233.176 | 2 | 52.151882 | yes | 2.804E-07 | 543.79749 | 120.34684 |
| EPGPIAPSTNSSPV. | 16 | Q14674 | Separin OS=Homo sapiens OX=9606 GN | ESPL1 | 233.176 | 2 | 52.151882 | yes | 6.847E-07 | 797.4278 | 90.409691 |
| SVLNEFDAIQK | 11 | Q14674 | Separin OS=Homo sapiens OX=9606 GN | ESPL1 | 233.176 | 2 | 52.151882 | yes | 1.736E-06 | 632.33258 | 74.288261 |
| AVAAHQLFDASGI | 29 | Q14674 | Separin OS=Homo sapiens OX=9606 GN | ESPL1 | 233.176 | 4 | 52.151882 | yes | 1.467E-07 | 767.87317 | 60.327293 |
| EAAPQDYEAVAR | 12 | Q14674 | Separin OS=Homo sapiens OX=9606 GN | ESPL1 | 233.176 | 2 | 52.151882 | yes | 7.301E-06 | 660.31494 | 54.071602 |
| ALCLLELTLEHCR | 13 | Q14674 | Separin OS=Homo sapiens OX=9606 GN | ESPL1 | 233.176 | 2 | 52.151882 | yes | 1.488E-07 | 814.4184 | 48.916664 |
| QACDAILR | 8 | Q14674 | Separin OS=Homo sapiens OX=9606 GN | ESPL1 | 233.176 | 2 | 52.151882 | yes | 4.038E-07 | 473.74237 | 47.836147 |
| VNFGTLLSSQK | 11 | Q14674 | Separin OS=Homo sapiens OX=9606 GN | ESPL1 | 233.176 | 2 | 52.151882 | yes | 1.402E-07 | 597.3299 | 43.787827 |
| CLDQALALWK | 10 | Q14674 | Separin OS=Homo sapiens OX=9606 GN | ESPL1 | 233.176 | 2 | 52.151882 | yes | 5.686E-07 | 609.32098 | 42.586861 |
| WVLVTAGVR | 9 | Q14674 | Separin OS=Homo sapiens OX=9606 GN | ESPL1 | 233.176 | 2 | 52.151882 | yes | 1.79E-07 | 500.80054 | 35.258339 |
| HQLLTHLHR | 9 | Q14674 | Separin OS=Homo sapiens OX=9606 GN | ESPL1 | 233.176 | 2 | 52.151882 | yes | 9.819E-08 | 577.83069 | 33.875198 |
| LQGLTVPSNSHLV | 18 | Q14674 | Separin OS=Homo sapiens OX=9606 GN | ESPL1 | 233.176 | 3 | 52.151882 | yes | 3.847E-07 | 645.04205 | 6.0539188 |

| | | | | | | | | | | | |
|---|---|---|---|---|---|---|---|---|---|---|---|
| GAGNGVVPAGVIL | 15 | Q14676 | Mediator of DNA damage checkpoint pro | MDC1 | 226.667 | 2 | 49.891594 | yes | 1.989E-07 | 704.89899 | 106.92005 |
| GPLTVEETPR | 10 | Q14676 | Mediator of DNA damage checkpoint pro | MDC1 | 226.667 | 2 | 49.891594 | yes | 2.219E-07 | 549.79309 | 87.177536 |
| TPETLVPTAPK | 11 | Q14676 | Mediator of DNA damage checkpoint pro | MDC1 | 226.667 | 2 | 49.891594 | yes | 1E-07 | 577.32678 | 84.836151 |
| VLFTGVVDAR | 10 | Q14676 | Mediator of DNA damage checkpoint pro | MDC1 | 226.667 | 2 | 49.891594 | yes | 4.442E-07 | 538.80859 | 78.642654 |
| NFGFSLQDALSR | 12 | Q14676 | Mediator of DNA damage checkpoint pro | MDC1 | 226.667 | 2 | 49.891594 | yes | 2.654E-06 | 677.84113 | 75.092789 |
| MPDCSVALPFPSIS | 15 | Q14676 | Mediator of DNA damage checkpoint pro | MDC1 | 226.667 | 2 | 49.891594 | yes | 1.089E-07 | 824.90729 | 57.743011 |
| LHIFSGAHGPEK | 12 | Q14676 | Mediator of DNA damage checkpoint pro | MDC1 | 226.667 | 2 | 49.891594 | yes | 1.236E-07 | 646.84094 | 56.245525 |
| FTPELQPK | 8 | Q14676 | Mediator of DNA damage checkpoint pro | MDC1 | 226.667 | 2 | 49.891594 | yes | 1.179E-06 | 480.26346 | 49.201878 |
| FLCALGR | 7 | Q14676 | Mediator of DNA damage checkpoint pro | MDC1 | 226.667 | 2 | 49.891594 | yes | 2.053E-06 | 418.72601 | 35.261269 |
| NQLVTPEPTSR | 11 | Q14676 | Mediator of DNA damage checkpoint pro | MDC1 | 226.667 | 2 | 49.891594 | yes | 3.437E-07 | 621.32788 | 34.040817 |
| SQASTTVDINTQVI | 15 | Q14676 | Mediator of DNA damage checkpoint pro | MDC1 | 226.667 | 2 | 49.891594 | yes | 1.741E-07 | 810.90741 | 26.497667 |
| IGSTIDDTISK | 11 | Q14677 | Clathrin interactor 1 OS=Homo sapiens C | CLINT1 | 68.26 | 2 | 53.12677 | yes | 1.715E-07 | 575.30353 | 82.691925 |
| ELVEFAQDDDR | 11 | Q14677 | Clathrin interactor 1 OS=Homo sapiens C | CLINT1 | 68.26 | 2 | 53.12677 | yes | 3.347E-07 | 668.80444 | 82.647133 |
| ATNVVMNYSEIES | 14 | Q14677 | Clathrin interactor 1 OS=Homo sapiens C | CLINT1 | 68.26 | 2 | 53.12677 | yes | 4.892E-07 | 792.88232 | 80.81797 |
| TIDLGAAAHYTGD | 14 | Q14677 | Clathrin interactor 1 OS=Homo sapiens C | CLINT1 | 68.26 | 2 | 53.12677 | yes | 9.722E-08 | 716.85699 | 49.003651 |
| GEFKDEEETVTTK | 13 | Q14677 | Clathrin interactor 1 OS=Homo sapiens C | CLINT1 | 68.26 | 2 | 53.12677 | yes | 9.722E-08 | 756.85687 | 43.956684 |
| DKYVGVSSDSVGC | 15 | Q14677 | Clathrin interactor 1 OS=Homo sapiens C | CLINT1 | 68.26 | 2 | 53.12677 | yes | 1.425E-07 | 786.88629 | 35.84782 |
| HIHITQATETTTTR | 14 | Q14677 | Clathrin interactor 1 OS=Homo sapiens C | CLINT1 | 68.26 | 2 | 53.12677 | yes | 3.341E-06 | 805.41827 | 11.521435 |
| GMAEGHHAVNIEC | 15 | Q14678 | KN motif and ankyrin repeat domain-cont | KANK1 | 147.29 | 3 | 44.131413 | yes | 2.823E-06 | 521.59631 | 72.726357 |
| AASQINVCGVR | 11 | Q14678 | KN motif and ankyrin repeat domain-cont | KANK1 | 147.29 | 2 | 44.131413 | yes | 3.888E-07 | 587.80347 | 72.07798 |
| FCLNTLQHEWFR | 12 | Q14678 | KN motif and ankyrin repeat domain-cont | KANK1 | 147.29 | 3 | 44.131413 | yes | 2.31E-07 | 550.93304 | 50.03933 |
| ITGNYLGYTCK | 11 | Q14678 | KN motif and ankyrin repeat domain-cont | KANK1 | 147.29 | 2 | 44.131413 | yes | 6.991E-07 | 645.31335 | 21.037361 |
| MLSACNLLK | 9 | Q14678 | KN motif and ankyrin repeat domain-cont | KANK1 | 147.29 | 2 | 44.131413 | yes | 8.225E-06 | 525.27771 | 19.865629 |
| LSSIQLGQSEK | 11 | Q14679 | Tubulin monoglutamylase TTLL4 OS=Hc | TTLL4 | 133.379 | 2 | 53.05184 | yes | 4.267E-07 | 595.32477 | 181.02948 |
| IYLFSDGLVR | 10 | Q14679 | Tubulin monoglutamylase TTLL4 OS=Hc | TTLL4 | 133.379 | 2 | 53.05184 | yes | 9.722E-08 | 591.82953 | 149.54932 |
| TIISSEPYVTSLLK | 14 | Q14679 | Tubulin monoglutamylase TTLL4 OS=Hc | TTLL4 | 133.379 | 2 | 53.05184 | yes | 2.389E-07 | 775.93744 | 136.31685 |
| ILVEMEDEFSR | 11 | Q14679 | Tubulin monoglutamylase TTLL4 OS=Hc | TTLL4 | 133.379 | 2 | 53.05184 | yes | 9.722E-08 | 684.32922 | 130.06982 |
| ALWNYLSQK | 9 | Q14679 | Tubulin monoglutamylase TTLL4 OS=Hc | TTLL4 | 133.379 | 2 | 53.05184 | yes | 3.352E-07 | 561.80072 | 117.99295 |
| GIGIQVIHK | 9 | Q14679 | Tubulin monoglutamylase TTLL4 OS=Hc | TTLL4 | 133.379 | 2 | 53.05184 | yes | 1.359E-07 | 482.80054 | 91.592705 |
| EPSLSTQTLPVIK | 13 | Q14679 | Tubulin monoglutamylase TTLL4 OS=Hc | TTLL4 | 133.379 | 2 | 53.05184 | yes | 6.409E-07 | 706.90338 | 75.186646 |
| YFNILTTQWEQK | 12 | Q14679 | Tubulin monoglutamylase TTLL4 OS=Hc | TTLL4 | 133.379 | 2 | 53.05184 | yes | 9.722E-08 | 785.89862 | 49.824783 |
| FMHLTNYSVNK | 11 | Q14679 | Tubulin monoglutamylase TTLL4 OS=Hc | TTLL4 | 133.379 | 2 | 53.05184 | yes | 6.505E-07 | 677.33466 | 43.035019 |
| SCMEILTK | 8 | Q14679 | Tubulin monoglutamylase TTLL4 OS=Hc | TTLL4 | 133.379 | 2 | 53.05184 | yes | 8.452E-07 | 491.24081 | 6.53617 |
| SNNWSLEDVTASI | 14 | Q14680 | Maternal embryonic leucine zipper kinase | MELK | 74.643 | 2 | 51.925964 | yes | 1.57E-07 | 783.35754 | 373.68558 |
| LHYNVTTTR | 9 | Q14680 | Maternal embryonic leucine zipper kinase | MELK | 74.643 | 2 | 51.925964 | yes | 2.982E-07 | 552.79346 | 368.89551 |
| LVEDILSSCK | 10 | Q14680 | Maternal embryonic leucine zipper kinase | MELK | 74.643 | 2 | 51.925964 | yes | 9.722E-08 | 582.30249 | 323.34894 |
| LMTGVISPER | 10 | Q14680 | Maternal embryonic leucine zipper kinase | MELK | 74.643 | 2 | 51.925964 | yes | 2.368E-07 | 551.79987 | 237.88319 |
| SLTPALCR | 8 | Q14680 | Maternal embryonic leucine zipper kinase | MELK | 74.643 | 2 | 51.925964 | yes | 1.691E-07 | 459.24728 | 235.7933 |
| MKDYDELLK | 9 | Q14680 | Maternal embryonic leucine zipper kinase | MELK | 74.643 | 2 | 51.925964 | yes | 2.466E-07 | 577.79175 | 234.06047 |

| | | | | | | | | | | | |
|---|---|---|---|---|---|---|---|---|---|---|---|
| LIDFGLCAK | 9 | Q14680 | Maternal embryonic leucine zipper kinase | MELK | 74.643 | 2 | 51.925964 | yes | 9.897E-08 | 518.77844 | 152.03598 |
| EILTTPNR | 8 | Q14680 | Maternal embryonic leucine zipper kinase | MELK | 74.643 | 2 | 51.925964 | yes | 2.964E-07 | 472.26401 | 135.01154 |
| LKGDAWVYK | 9 | Q14680 | Maternal embryonic leucine zipper kinase | MELK | 74.643 | 2 | 51.925964 | yes | 2.648E-07 | 540.29785 | 116.1653 |
| NEEYFMFPEPK | 11 | Q14680 | Maternal embryonic leucine zipper kinase | MELK | 74.643 | 2 | 51.925964 | yes | 2.952E-07 | 715.81848 | 111.35294 |
| IKTEIEALK | 9 | Q14680 | Maternal embryonic leucine zipper kinase | MELK | 74.643 | 2 | 51.925964 | yes | 2.189E-07 | 522.8186 | 106.22974 |
| VITVLTR | 7 | Q14680 | Maternal embryonic leucine zipper kinase | MELK | 74.643 | 2 | 51.925964 | yes | 8.387E-07 | 401.26328 | 72.582916 |
| YWTESNGVESK | 11 | Q14680 | Maternal embryonic leucine zipper kinase | MELK | 74.643 | 2 | 51.925964 | yes | 3.076E-06 | 650.2962 | 70.513664 |
| HVDFVQK | 7 | Q14680 | Maternal embryonic leucine zipper kinase | MELK | 74.643 | 2 | 51.925964 | yes | 2.242E-07 | 436.73486 | 57.595539 |
| GKYDVPK | 7 | Q14680 | Maternal embryonic leucine zipper kinase | MELK | 74.643 | 2 | 51.925964 | yes | 4.858E-06 | 403.72397 | 48.653748 |
| YYELHETIGTGGF | 15 | Q14680 | Maternal embryonic leucine zipper kinase | MELK | 74.643 | 2 | 51.925964 | yes | 9.722E-08 | 843.41211 | 46.496803 |
| SGVISGGASDLK | 12 | Q14683 | Structural maintenance of chromosomes p | SMC1A | 143.233 | 2 | 53.793556 | yes | 1.308E-07 | 545.79059 | 286.35751 |
| ELNQVMEQLGDA | 13 | Q14683 | Structural maintenance of chromosomes p | SMC1A | 143.233 | 2 | 53.793556 | yes | 9.722E-08 | 751.867 | 272.24539 |
| TALFEEISR | 9 | Q14683 | Structural maintenance of chromosomes p | SMC1A | 143.233 | 2 | 53.793556 | yes | 1.756E-07 | 533.28241 | 270.79507 |
| LKDEVVR | 7 | Q14683 | Structural maintenance of chromosomes p | SMC1A | 143.233 | 2 | 53.793556 | yes | 1.467E-07 | 429.7558 | 229.0927 |
| AQVQLQLFK | 9 | Q14683 | Structural maintenance of chromosomes p | SMC1A | 143.233 | 2 | 53.793556 | yes | 5.333E-07 | 537.81897 | 219.24904 |
| TVALDGTLFQK | 11 | Q14683 | Structural maintenance of chromosomes p | SMC1A | 143.233 | 2 | 53.793556 | yes | 4.498E-07 | 596.83228 | 204.52948 |
| LIDLCQPTQK | 10 | Q14683 | Structural maintenance of chromosomes p | SMC1A | 143.233 | 2 | 53.793556 | yes | 3.174E-07 | 608.32373 | 203.77145 |
| AFVSMVYSEEGAI | 15 | Q14683 | Structural maintenance of chromosomes p | SMC1A | 143.233 | 2 | 53.793556 | yes | 2.224E-07 | 845.37488 | 191.59006 |
| LGIQLDFEK | 9 | Q14683 | Structural maintenance of chromosomes p | SMC1A | 143.233 | 2 | 53.793556 | yes | 9.722E-08 | 531.7951 | 188.6489 |
| LNEQQSVLQR | 10 | Q14683 | Structural maintenance of chromosomes p | SMC1A | 143.233 | 2 | 53.793556 | yes | 9.722E-08 | 607.828 | 164.35545 |
| RIDEINK | 7 | Q14683 | Structural maintenance of chromosomes p | SMC1A | 143.233 | 2 | 53.793556 | yes | 1.381E-06 | 444.25089 | 149.25902 |
| VIVGGSSEYK | 10 | Q14683 | Structural maintenance of chromosomes p | SMC1A | 143.233 | 2 | 53.793556 | yes | 1.081E-07 | 519.77692 | 137.44118 |
| DLTLEENQVK | 10 | Q14683 | Structural maintenance of chromosomes p | SMC1A | 143.233 | 2 | 53.793556 | yes | 9.722E-08 | 594.80896 | 127.00136 |
| VLTFDLTK | 8 | Q14683 | Structural maintenance of chromosomes p | SMC1A | 143.233 | 2 | 53.793556 | yes | 1.283E-07 | 468.77368 | 123.45644 |
| LYHNEVEIEK | 10 | Q14683 | Structural maintenance of chromosomes p | SMC1A | 143.233 | 2 | 53.793556 | yes | 2.041E-07 | 637.32477 | 108.59232 |
| LESELANFGPR | 11 | Q14683 | Structural maintenance of chromosomes p | SMC1A | 143.233 | 2 | 53.793556 | yes | 2.952E-07 | 616.81714 | 106.86285 |
| LIEIENFK | 8 | Q14683 | Structural maintenance of chromosomes p | SMC1A | 143.233 | 2 | 53.793556 | yes | 3.846E-07 | 503.28439 | 104.8737 |
| VHMWEQTVK | 9 | Q14683 | Structural maintenance of chromosomes p | SMC1A | 143.233 | 2 | 53.793556 | yes | 3.054E-07 | 579.29224 | 98.92202 |
| DLIHGAPVGK | 10 | Q14683 | Structural maintenance of chromosomes p | SMC1A | 143.233 | 2 | 53.793556 | yes | 4.935E-07 | 503.78763 | 85.800797 |
| KGDMDELEK | 9 | Q14683 | Structural maintenance of chromosomes p | SMC1A | 143.233 | 2 | 53.793556 | yes | 1.596E-07 | 532.75006 | 44.114632 |
| GEPETFLPLDYLE | 20 | Q14683 | Structural maintenance of chromosomes p | SMC1A | 143.233 | 3 | 53.793556 | yes | 3.839E-06 | 774.05798 | 30.58021 |
| VVQLHEYSEELEK | 13 | Q14683 | Structural maintenance of chromosomes p | SMC1A | 143.233 | 2 | 53.793556 | yes | 4.46E-07 | 801.90411 | 28.287529 |
| AEEDTQFNYHR | 11 | Q14683 | Structural maintenance of chromosomes p | SMC1A | 143.233 | 2 | 53.793556 | yes | 9.722E-08 | 705.30768 | 15.437717 |
| SILVSPTGPSR | 11 | Q14684 | Ribosomal RNA processing protein 1 hon | RRP1B | 84.428 | 2 | 51.89273 | yes | 9.722E-08 | 557.31677 | 803.49341 |
| DHTLVQTIAR | 10 | Q14684 | Ribosomal RNA processing protein 1 hon | RRP1B | 84.428 | 2 | 51.89273 | yes | 8.113E-07 | 577.31982 | 453.86993 |
| AGPGSLELCGLPS | 15 | Q14684 | Ribosomal RNA processing protein 1 hon | RRP1B | 84.428 | 2 | 51.89273 | yes | 1.234E-07 | 757.38757 | 364.54041 |
| APAMQPAEIQFAQ | 14 | Q14684 | Ribosomal RNA processing protein 1 hon | RRP1B | 84.428 | 2 | 51.89273 | yes | 2.042E-07 | 779.39575 | 291.36118 |
| KVTFGLNR | 8 | Q14684 | Ribosomal RNA processing protein 1 hon | RRP1B | 84.428 | 2 | 51.89273 | yes | 9.722E-08 | 467.77707 | 240.64101 |
| ELLADQNLK | 9 | Q14684 | Ribosomal RNA processing protein 1 hon | RRP1B | 84.428 | 2 | 51.89273 | yes | 9.722E-08 | 522.29022 | 238.591 |

| | | | | | | | | | | | |
|---|---|---|---|---|---|---|---|---|---|---|---|
| ETGGFSQEELLK | 12 | Q14684 | Ribosomal RNA processing protein 1 hon | RRP1B | 84.428 | 2 | 51.89273 | yes | 1.57E-07 | 669.33282 | 235.53183 |
| KPLTTTPR | 8 | Q14684 | Ribosomal RNA processing protein 1 hon | RRP1B | 84.428 | 2 | 51.89273 | yes | 1.301E-07 | 457.27692 | 224.80351 |
| TPTSSPASSPLVAK | 14 | Q14684 | Ribosomal RNA processing protein 1 hon | RRP1B | 84.428 | 2 | 51.89273 | yes | 1.831E-07 | 671.86426 | 148.79303 |
| FDTPFLPK | 8 | Q14684 | Ribosomal RNA processing protein 1 hon | RRP1B | 84.428 | 2 | 51.89273 | yes | 2.937E-07 | 482.76056 | 146.93788 |
| SSTATHPPGPAVQ | 16 | Q14684 | Ribosomal RNA processing protein 1 hon | RRP1B | 84.428 | 2 | 51.89273 | yes | 1.048E-07 | 802.92316 | 140.8905 |
| NTPHFNR | 7 | Q14684 | Ribosomal RNA processing protein 1 hon | RRP1B | 84.428 | 2 | 51.89273 | yes | 3.427E-07 | 443.21994 | 135.77405 |
| GSPTGGAQLLK | 11 | Q14684 | Ribosomal RNA processing protein 1 hon | RRP1B | 84.428 | 2 | 51.89273 | yes | 5.276E-07 | 514.79041 | 105.55757 |
| VTFGLNR | 7 | Q14684 | Ribosomal RNA processing protein 1 hon | RRP1B | 84.428 | 2 | 51.89273 | yes | 3.102E-07 | 403.72958 | 89.123306 |
| VAFDPEQKPLHGV | 15 | Q14684 | Ribosomal RNA processing protein 1 hon | RRP1B | 84.428 | 2 | 51.89273 | yes | 1.984E-07 | 839.46979 | 26.368813 |
| VAFDPEQKPLHGV | 15 | Q14684 | Ribosomal RNA processing protein 1 hon | RRP1B | 84.428 | 2 | 51.89273 | yes | 1.186E-05 | 559.9823 | 26.368813 |
| VAFDPEQKPLHGV | 15 | Q14684 | Ribosomal RNA processing protein 1 hon | RRP1B | 84.428 | 3 | 51.89273 | yes | 1.984E-07 | 839.46979 | 26.368813 |
| VAFDPEQKPLHGV | 15 | Q14684 | Ribosomal RNA processing protein 1 hon | RRP1B | 84.428 | 3 | 51.89273 | yes | 1.186E-05 | 559.9823 | 26.368813 |
| VFLDVLMK | 8 | Q14684 | Ribosomal RNA processing protein 1 hon | RRP1B | 84.428 | 2 | 51.89273 | yes | 4.306E-07 | 482.78043 | 52.215908 |
| VVVLNCEPSK | 10 | Q14690 | Protein RRP5 homolog OS=Homo sapien | PDCD11 | 208.701 | 2 | 55.520275 | yes | 9.722E-08 | 572.80518 | 308.37811 |
| VFYTGQVVK | 9 | Q14690 | Protein RRP5 homolog OS=Homo sapien | PDCD11 | 208.701 | 2 | 55.520275 | yes | 3.494E-07 | 520.79242 | 285.38419 |
| ACILCVHPR | 9 | Q14690 | Protein RRP5 homolog OS=Homo sapien | PDCD11 | 208.701 | 2 | 55.520275 | yes | 1.511E-07 | 563.28644 | 281.49167 |
| AINIGQLVDVK | 11 | Q14690 | Protein RRP5 homolog OS=Homo sapien | PDCD11 | 208.701 | 2 | 55.520275 | yes | 9.722E-08 | 585.34808 | 275.91287 |
| WLEVEIAPDIR | 11 | Q14690 | Protein RRP5 homolog OS=Homo sapien | PDCD11 | 208.701 | 2 | 55.520275 | yes | 9.722E-08 | 670.86407 | 253.34279 |
| TSIIEAQYLR | 10 | Q14690 | Protein RRP5 homolog OS=Homo sapien | PDCD11 | 208.701 | 2 | 55.520275 | yes | 2.095E-07 | 597.3299 | 217.95358 |
| LQVGQGVSLTLK | 12 | Q14690 | Protein RRP5 homolog OS=Homo sapien | PDCD11 | 208.701 | 2 | 55.520275 | yes | 9.722E-08 | 621.87445 | 185.00365 |
| FAQLEFQLGDAER | 13 | Q14690 | Protein RRP5 homolog OS=Homo sapien | PDCD11 | 208.701 | 2 | 55.520275 | yes | 1.411E-07 | 762.37805 | 181.03166 |
| VLLCDPEAK | 9 | Q14690 | Protein RRP5 homolog OS=Homo sapien | PDCD11 | 208.701 | 2 | 55.520275 | yes | 1.418E-07 | 522.77332 | 169.96182 |
| IEEALMDPGR | 10 | Q14690 | Protein RRP5 homolog OS=Homo sapien | PDCD11 | 208.701 | 2 | 55.520275 | yes | 2.733E-06 | 565.77917 | 164.71442 |
| VLCLSQSEGR | 10 | Q14690 | Protein RRP5 homolog OS=Homo sapien | PDCD11 | 208.701 | 2 | 55.520275 | yes | 2.007E-07 | 574.79004 | 158.6461 |
| VTPNEGLTVSFPF( | 15 | Q14690 | Protein RRP5 homolog OS=Homo sapien | PDCD11 | 208.701 | 2 | 55.520275 | yes | 8.499E-07 | 796.91956 | 152.36118 |
| YHDIEPGAVVK | 11 | Q14690 | Protein RRP5 homolog OS=Homo sapien | PDCD11 | 208.701 | 2 | 55.520275 | yes | 1.267E-07 | 614.32202 | 149.10942 |
| FQEAGELYNR | 10 | Q14690 | Protein RRP5 homolog OS=Homo sapien | PDCD11 | 208.701 | 2 | 55.520275 | yes | 2.081E-07 | 613.79364 | 147.1219 |
| LEEGEVAMGR | 10 | Q14690 | Protein RRP5 homolog OS=Homo sapien | PDCD11 | 208.701 | 2 | 55.520275 | yes | 2.638E-07 | 545.76349 | 141.37009 |
| VGQYLNCIVEK | 11 | Q14690 | Protein RRP5 homolog OS=Homo sapien | PDCD11 | 208.701 | 2 | 55.520275 | yes | 9.722E-08 | 661.84229 | 140.91232 |
| LGPSVVGLAR | 10 | Q14690 | Protein RRP5 homolog OS=Homo sapien | PDCD11 | 208.701 | 2 | 55.520275 | yes | 1.997E-07 | 484.798 | 124.01382 |
| DGLEVAVLPHNIR | 13 | Q14690 | Protein RRP5 homolog OS=Homo sapien | PDCD11 | 208.701 | 2 | 55.520275 | yes | 1.083E-07 | 716.89899 | 113.6678 |
| PIFLQPGR | 8 | Q14690 | Protein RRP5 homolog OS=Homo sapien | PDCD11 | 208.701 | 2 | 55.520275 | yes | 4.051E-07 | 464.27417 | 96.1996 |
| VVHLSLRPIFLQPC | 15 | Q14690 | Protein RRP5 homolog OS=Homo sapien | PDCD11 | 208.701 | 3 | 55.520275 | yes | 1.848E-07 | 578.01752 | 87.976219 |
| YHIGDEVK | 8 | Q14690 | Protein RRP5 homolog OS=Homo sapien | PDCD11 | 208.701 | 2 | 55.520275 | yes | 1.263E-07 | 480.74289 | 86.362076 |
| LPVITCYADAK | 11 | Q14690 | Protein RRP5 homolog OS=Homo sapien | PDCD11 | 208.701 | 2 | 55.520275 | yes | 2.117E-07 | 625.82611 | 72.633476 |
| QYQAGQTVTCFLK | 13 | Q14690 | Protein RRP5 homolog OS=Homo sapien | PDCD11 | 208.701 | 2 | 55.520275 | yes | 9.758E-08 | 772.38232 | 67.839546 |
| KVVILNVDLLK | 11 | Q14690 | Protein RRP5 homolog OS=Homo sapien | PDCD11 | 208.701 | 2 | 55.520275 | yes | 1.21E-07 | 627.41321 | 62.929619 |
| AIFENTLSTYPK | 12 | Q14690 | Protein RRP5 homolog OS=Homo sapien | PDCD11 | 208.701 | 2 | 55.520275 | yes | 2.067E-07 | 692.36139 | 62.375664 |
| LNEQVTQEQPLK | 12 | Q14690 | Protein RRP5 homolog OS=Homo sapien | PDCD11 | 208.701 | 2 | 55.520275 | yes | 1.108E-07 | 713.88043 | 60.016903 |

| | | | | | | | | | | | |
|---|---|---|---|---|---|---|---|---|---|---|---|
| ALECLPSK | 8 | Q14690 | Protein RRP5 homolog OS=Homo sapien | PDCD11 | 208.701 | 2 | 55.520275 | yes | 3.428E-07 | 459.24167 | 55.805298 |
| FVTSTSDHFVEGQ | 18 | Q14690 | Protein RRP5 homolog OS=Homo sapien | PDCD11 | 208.701 | 3 | 55.520275 | yes | 3.332E-06 | 641.98639 | 37.263103 |
| KPALVSTVEGGQD | 15 | Q14690 | Protein RRP5 homolog OS=Homo sapien | PDCD11 | 208.701 | 2 | 55.520275 | yes | 1.052E-07 | 763.41467 | 34.539165 |
| QPESADDFDR | 10 | Q14690 | Protein RRP5 homolog OS=Homo sapien | PDCD11 | 208.701 | 2 | 55.520275 | yes | 3.74E-06 | 590.24927 | 33.606579 |
| AFQQSVEQDNLFD | 22 | Q14690 | Protein RRP5 homolog OS=Homo sapien | PDCD11 | 208.701 | 3 | 55.520275 | yes | 3.383E-06 | 825.05035 | 30.219755 |
| AGTYFSNQAVR | 11 | Q14690 | Protein RRP5 homolog OS=Homo sapien | PDCD11 | 208.701 | 2 | 55.520275 | yes | 5.857E-05 | 607.30164 | 14.706586 |
| GSEHQAIVQHLEK | 13 | Q14690 | Protein RRP5 homolog OS=Homo sapien | PDCD11 | 208.701 | 2 | 55.520275 | yes | 3.714E-07 | 738.38367 | 14.381871 |
| LLQDLQLGDEEDA | 14 | Q14692 | Ribosome biogenesis protein BMS1 hom | BMS1 | 145.808 | 2 | 54.833363 | yes | 1.421E-07 | 807.9021 | 1747.783 |
| LVYAPLSGVGGVI | 16 | Q14692 | Ribosome biogenesis protein BMS1 hom | BMS1 | 145.808 | 2 | 54.833363 | yes | 1.84E-07 | 825.95868 | 1292.3376 |
| STLIQCLIR | 9 | Q14692 | Ribosome biogenesis protein BMS1 hom | BMS1 | 145.808 | 2 | 54.833363 | yes | 2.658E-07 | 552.31573 | 1151.2137 |
| MEDLTNPEDIR | 11 | Q14692 | Ribosome biogenesis protein BMS1 hom | BMS1 | 145.808 | 2 | 54.833363 | yes | 1.74E-07 | 666.80865 | 824.94543 |
| IMGVLTHLDSFK | 12 | Q14692 | Ribosome biogenesis protein BMS1 hom | BMS1 | 145.808 | 2 | 54.833363 | yes | 1.345E-07 | 680.86829 | 319.76102 |
| IMGVLTHLDSFK | 12 | Q14692 | Ribosome biogenesis protein BMS1 hom | BMS1 | 145.808 | 2 | 54.833363 | yes | 1.6E-07 | 454.24796 | 319.76102 |
| IMGVLTHLDSFK | 12 | Q14692 | Ribosome biogenesis protein BMS1 hom | BMS1 | 145.808 | 3 | 54.833363 | yes | 1.345E-07 | 680.86829 | 319.76102 |
| IMGVLTHLDSFK | 12 | Q14692 | Ribosome biogenesis protein BMS1 hom | BMS1 | 145.808 | 3 | 54.833363 | yes | 1.6E-07 | 454.24796 | 319.76102 |
| VQYEGFRPGMYV | 13 | Q14692 | Ribosome biogenesis protein BMS1 hom | BMS1 | 145.808 | 3 | 54.833363 | yes | 1.556E-07 | 534.60126 | 303.21835 |
| VQYEGFRPGMYV | 13 | Q14692 | Ribosome biogenesis protein BMS1 hom | BMS1 | 145.808 | 3 | 54.833363 | yes | 1.91E-07 | 801.39832 | 303.21835 |
| VQYEGFRPGMYV | 13 | Q14692 | Ribosome biogenesis protein BMS1 hom | BMS1 | 145.808 | 2 | 54.833363 | yes | 1.556E-07 | 534.60126 | 303.21835 |
| VQYEGFRPGMYV | 13 | Q14692 | Ribosome biogenesis protein BMS1 hom | BMS1 | 145.808 | 2 | 54.833363 | yes | 1.91E-07 | 801.39832 | 303.21835 |
| GMFNSALEVAK | 11 | Q14692 | Ribosome biogenesis protein BMS1 hom | BMS1 | 145.808 | 2 | 54.833363 | yes | 1.211E-07 | 583.79736 | 598.25128 |
| AFAVQSAVR | 9 | Q14692 | Ribosome biogenesis protein BMS1 hom | BMS1 | 145.808 | 2 | 54.833363 | yes | 1.324E-07 | 474.76672 | 568.06836 |
| VSLYGYLR | 8 | Q14692 | Ribosome biogenesis protein BMS1 hom | BMS1 | 145.808 | 2 | 54.833363 | yes | 9.722E-08 | 485.77145 | 537.98297 |
| AEFEDQDDEAR | 11 | Q14692 | Ribosome biogenesis protein BMS1 hom | BMS1 | 145.808 | 2 | 54.833363 | yes | 9.722E-08 | 662.76801 | 497.55264 |
| IAATGVVLDLDK | 12 | Q14692 | Ribosome biogenesis protein BMS1 hom | BMS1 | 145.808 | 2 | 54.833363 | yes | 1.006E-07 | 607.85315 | 468.80988 |
| SGPNTQNEDIEK | 12 | Q14692 | Ribosome biogenesis protein BMS1 hom | BMS1 | 145.808 | 2 | 54.833363 | yes | 9.819E-08 | 666.30731 | 447.75394 |
| FWTEVYPGAK | 10 | Q14692 | Ribosome biogenesis protein BMS1 hom | BMS1 | 145.808 | 2 | 54.833363 | yes | 9.722E-08 | 599.3006 | 380.99643 |
| AGLSPANCQSDR | 12 | Q14692 | Ribosome biogenesis protein BMS1 hom | BMS1 | 145.808 | 2 | 54.833363 | yes | 9.903E-08 | 638.29095 | 341.24982 |
| LSKPSQVSSGQK | 12 | Q14692 | Ribosome biogenesis protein BMS1 hom | BMS1 | 145.808 | 2 | 54.833363 | yes | 1.823E-07 | 623.34351 | 309.10724 |
| DSLYKPILR | 9 | Q14692 | Ribosome biogenesis protein BMS1 hom | BMS1 | 145.808 | 2 | 54.833363 | yes | 9.766E-08 | 552.82422 | 296.02637 |
| WKEDLSR | 7 | Q14692 | Ribosome biogenesis protein BMS1 hom | BMS1 | 145.808 | 2 | 54.833363 | yes | 1.276E-07 | 467.24307 | 265.22787 |
| KVSLYGYLR | 9 | Q14692 | Ribosome biogenesis protein BMS1 hom | BMS1 | 145.808 | 2 | 54.833363 | yes | 1.521E-07 | 549.81897 | 252.69034 |
| LLMSDIVFMR | 10 | Q14692 | Ribosome biogenesis protein BMS1 hom | BMS1 | 145.808 | 2 | 54.833363 | yes | 1.079E-07 | 612.82758 | 244.76567 |
| VLAEDEELYGDFE | 22 | Q14692 | Ribosome biogenesis protein BMS1 hom | BMS1 | 145.808 | 3 | 54.833363 | yes | 1.728E-07 | 841.71851 | 209.96828 |
| QQQAAPNLR | 9 | Q14692 | Ribosome biogenesis protein BMS1 hom | BMS1 | 145.808 | 2 | 54.833363 | yes | 9.722E-08 | 513.27795 | 149.57111 |
| KAAEAFLR | 8 | Q14692 | Ribosome biogenesis protein BMS1 hom | BMS1 | 145.808 | 2 | 54.833363 | yes | 1.783E-07 | 453.26379 | 140.98039 |
| QMDLNTGR | 8 | Q14692 | Ribosome biogenesis protein BMS1 hom | BMS1 | 145.808 | 2 | 54.833363 | yes | 1.004E-07 | 467.72418 | 132.69476 |
| HFNSLHIPK | 9 | Q14692 | Ribosome biogenesis protein BMS1 hom | BMS1 | 145.808 | 2 | 54.833363 | yes | 1.674E-06 | 546.80109 | 102.75439 |
| FQTIPLYYIEDHNC | 15 | Q14692 | Ribosome biogenesis protein BMS1 hom | BMS1 | 145.808 | 3 | 54.833363 | yes | 4.632E-07 | 622.64423 | 97.639038 |
| FLVEAPHDWDLEI | 19 | Q14692 | Ribosome biogenesis protein BMS1 hom | BMS1 | 145.808 | 3 | 54.833363 | yes | 2.976E-07 | 767.37207 | 68.634026 |

| | | | | | | | | | | | |
|---|---|---|---|---|---|---|---|---|---|---|---|
| FRPLTWQTSHPYII | 17 | Q14692 | Ribosome biogenesis protein BMS1 hom | BMS1 | 145.808 | 3 | 54.833363 | yes | 5.86E-07 | 701.03741 | 60.602501 |
| KHHIPVVDR | 9 | Q14692 | Ribosome biogenesis protein BMS1 hom | BMS1 | 145.808 | 2 | 54.833363 | yes | 5.147E-07 | 550.81982 | 50.292637 |
| VQYEGFR | 7 | Q14692 | Ribosome biogenesis protein BMS1 hom | BMS1 | 145.808 | 2 | 54.833363 | yes | 3.689E-07 | 449.72452 | 41.060551 |
| DPIIFSVGWR | 10 | Q14692 | Ribosome biogenesis protein BMS1 hom | BMS1 | 145.808 | 2 | 54.833363 | yes | 4.549E-06 | 595.32184 | 33.166702 |
| KILALLDALSTVHS | 16 | Q14692 | Ribosome biogenesis protein BMS1 hom | BMS1 | 145.808 | 3 | 54.833363 | yes | 2.633E-07 | 579.68048 | 32.981155 |
| VIISDIDGTITR | 12 | Q14693 | Phosphatidate phosphatase LPIN1 OS=H | LPIN1 | 98.665 | 2 | 45.395977 | yes | 1.009E-06 | 651.86682 | 98.587257 |
| GYLHWVNER | 9 | Q14693 | Phosphatidate phosphatase LPIN1 OS=H | LPIN1 | 98.665 | 2 | 45.395977 | yes | 1.862E-07 | 587.29364 | 67.594391 |
| SDTLGHILPTLGK | 13 | Q14693 | Phosphatidate phosphatase LPIN1 OS=H | LPIN1 | 98.665 | 2 | 45.395977 | yes | 3.62E-07 | 676.38263 | 53.449673 |
| EGLVPVSEDPVAII | 14 | Q14694 | Ubiquitin carboxyl-terminal hydrolase 10 | USP10 | 87.134 | 2 | 52.436764 | yes | 2.154E-07 | 726.90088 | 40474.305 |
| NIEYPVDLEISK | 12 | Q14694 | Ubiquitin carboxyl-terminal hydrolase 10 | USP10 | 87.134 | 2 | 52.436764 | yes | 1.259E-07 | 710.37195 | 27798.717 |
| TVQDALESLVAR | 12 | Q14694 | Ubiquitin carboxyl-terminal hydrolase 10 | USP10 | 87.134 | 2 | 52.436764 | yes | 1.113E-07 | 651.35663 | 22994.262 |
| YSPPAISPLVSEK | 13 | Q14694 | Ubiquitin carboxyl-terminal hydrolase 10 | USP10 | 87.134 | 2 | 52.436764 | yes | 6.017E-07 | 694.37701 | 21316.51 |
| PGAAFEPTYIYR | 12 | Q14694 | Ubiquitin carboxyl-terminal hydrolase 10 | USP10 | 87.134 | 2 | 52.436764 | yes | 1.477E-07 | 692.84845 | 10906.356 |
| VINQYQVVKPTAE | 14 | Q14694 | Ubiquitin carboxyl-terminal hydrolase 10 | USP10 | 87.134 | 2 | 52.436764 | yes | 1.224E-07 | 822.95703 | 4943.0713 |
| VINQYQVVKPTAE | 14 | Q14694 | Ubiquitin carboxyl-terminal hydrolase 10 | USP10 | 87.134 | 2 | 52.436764 | yes | 1.889E-07 | 548.97375 | 4943.0713 |
| VINQYQVVKPTAE | 14 | Q14694 | Ubiquitin carboxyl-terminal hydrolase 10 | USP10 | 87.134 | 3 | 52.436764 | yes | 1.224E-07 | 822.95703 | 4943.0713 |
| VINQYQVVKPTAE | 14 | Q14694 | Ubiquitin carboxyl-terminal hydrolase 10 | USP10 | 87.134 | 3 | 52.436764 | yes | 1.889E-07 | 548.97375 | 4943.0713 |
| ESVQGYTTK | 9 | Q14694 | Ubiquitin carboxyl-terminal hydrolase 10 | USP10 | 87.134 | 2 | 52.436764 | yes | 3.464E-07 | 506.75092 | 8705.6943 |
| LPDGQEYQR | 9 | Q14694 | Ubiquitin carboxyl-terminal hydrolase 10 | USP10 | 87.134 | 2 | 52.436764 | yes | 1.46E-07 | 553.26727 | 6620.2578 |
| TAYLLYYR | 8 | Q14694 | Ubiquitin carboxyl-terminal hydrolase 10 | USP10 | 87.134 | 2 | 52.436764 | yes | 1.943E-07 | 531.78455 | 5400.2305 |
| LLSPSNEK | 8 | Q14694 | Ubiquitin carboxyl-terminal hydrolase 10 | USP10 | 87.134 | 2 | 52.436764 | yes | 2.064E-07 | 444.24527 | 4535.0249 |
| TKQEVEISR | 9 | Q14694 | Ubiquitin carboxyl-terminal hydrolase 10 | USP10 | 87.134 | 2 | 52.436764 | yes | 1.29E-07 | 545.29858 | 3999.9736 |
| FIPLYSK | 7 | Q14694 | Ubiquitin carboxyl-terminal hydrolase 10 | USP10 | 87.134 | 2 | 52.436764 | yes | 1.207E-07 | 434.25238 | 3849.8801 |
| SVVYQQSSK | 9 | Q14694 | Ubiquitin carboxyl-terminal hydrolase 10 | USP10 | 87.134 | 2 | 52.436764 | yes | 1.636E-07 | 513.26672 | 3839.7839 |
| QADFVQTPITGIFG | 18 | Q14694 | Ubiquitin carboxyl-terminal hydrolase 10 | USP10 | 87.134 | 3 | 52.436764 | yes | 2.382E-06 | 653.01453 | 3257.2542 |
| SWASLFHDSK | 10 | Q14694 | Ubiquitin carboxyl-terminal hydrolase 10 | USP10 | 87.134 | 2 | 52.436764 | yes | 1.149E-07 | 589.28546 | 3141.6523 |
| PSSSSPVAYVETK | 13 | Q14694 | Ubiquitin carboxyl-terminal hydrolase 10 | USP10 | 87.134 | 2 | 52.436764 | yes | 9.809E-08 | 676.34064 | 2492.5508 |
| RPPGYYSYLK | 10 | Q14694 | Ubiquitin carboxyl-terminal hydrolase 10 | USP10 | 87.134 | 2 | 52.436764 | yes | 2.442E-07 | 622.32715 | 2280.9077 |
| SWASLFHDSKPSS | 23 | Q14694 | Ubiquitin carboxyl-terminal hydrolase 10 | USP10 | 87.134 | 4 | 52.436764 | yes | 5.044E-07 | 628.31042 | 2178.9912 |
| IAELLENVTLIHKP | 20 | Q14694 | Ubiquitin carboxyl-terminal hydrolase 10 | USP10 | 87.134 | 4 | 52.436764 | yes | 1.653E-07 | 568.33624 | 2043.5044 |
| IAELLENVTLIHK | 13 | Q14694 | Ubiquitin carboxyl-terminal hydrolase 10 | USP10 | 87.134 | 2 | 52.436764 | yes | 9.734E-08 | 746.94031 | 989.44159 |
| IAELLENVTLIHK | 13 | Q14694 | Ubiquitin carboxyl-terminal hydrolase 10 | USP10 | 87.134 | 2 | 52.436764 | yes | 1.85E-06 | 498.29596 | 989.44159 |
| IAELLENVTLIHK | 13 | Q14694 | Ubiquitin carboxyl-terminal hydrolase 10 | USP10 | 87.134 | 3 | 52.436764 | yes | 9.734E-08 | 746.94031 | 989.44159 |
| IAELLENVTLIHK | 13 | Q14694 | Ubiquitin carboxyl-terminal hydrolase 10 | USP10 | 87.134 | 3 | 52.436764 | yes | 1.85E-06 | 498.29596 | 989.44159 |
| QEVEISR | 7 | Q14694 | Ubiquitin carboxyl-terminal hydrolase 10 | USP10 | 87.134 | 2 | 52.436764 | yes | 2.244E-07 | 430.72726 | 1968.606 |
| LPPVLVLHLK | 10 | Q14694 | Ubiquitin carboxyl-terminal hydrolase 10 | USP10 | 87.134 | 2 | 52.436764 | yes | 1.008E-07 | 564.87878 | 1720.5016 |
| KLLSPSNEK | 9 | Q14694 | Ubiquitin carboxyl-terminal hydrolase 10 | USP10 | 87.134 | 2 | 52.436764 | yes | 1.066E-07 | 508.29276 | 1520.5736 |
| VINQYQVVK | 9 | Q14694 | Ubiquitin carboxyl-terminal hydrolase 10 | USP10 | 87.134 | 2 | 52.436764 | yes | 9.728E-08 | 545.81641 | 1478.3866 |
| PCTSTPMIDSFVR | 13 | Q14694 | Ubiquitin carboxyl-terminal hydrolase 10 | USP10 | 87.134 | 2 | 52.436764 | yes | 9.722E-08 | 755.85504 | 1294.7478 |

| | | | | | | | | | | | |
|---|---|---|---|---|---|---|---|---|---|---|---|
| LMNEFTNMPVPPK | 15 | Q14694 | Ubiquitin carboxyl-terminal hydrolase 10 | USP10 | 87.134 | 3 | 52.436764 | yes | 2.906E-06 | 590.97253 | 900.64331 |
| KRPPGYYSYLK | 11 | Q14694 | Ubiquitin carboxyl-terminal hydrolase 10 | USP10 | 87.134 | 3 | 52.436764 | yes | 1.285E-06 | 457.91882 | 394.18628 |
| KRPPGYYSYLK | 11 | Q14694 | Ubiquitin carboxyl-terminal hydrolase 10 | USP10 | 87.134 | 3 | 52.436764 | yes | 4.163E-07 | 686.37463 | 394.18628 |
| KRPPGYYSYLK | 11 | Q14694 | Ubiquitin carboxyl-terminal hydrolase 10 | USP10 | 87.134 | 2 | 52.436764 | yes | 1.285E-06 | 457.91882 | 394.18628 |
| KRPPGYYSYLK | 11 | Q14694 | Ubiquitin carboxyl-terminal hydrolase 10 | USP10 | 87.134 | 2 | 52.436764 | yes | 4.163E-07 | 686.37463 | 394.18628 |
| SSVELPPYSGTVLC | 21 | Q14694 | Ubiquitin carboxyl-terminal hydrolase 10 | USP10 | 87.134 | 3 | 52.436764 | yes | 1.197E-07 | 736.69995 | 176.1922 |
| ELLSPGVK | 8 | Q14694 | Ubiquitin carboxyl-terminal hydrolase 10 | USP10 | 87.134 | 2 | 52.436764 | yes | 1.434E-07 | 421.75275 | 171.43587 |
| LMNEFTNMPVPPK | 13 | Q14694 | Ubiquitin carboxyl-terminal hydrolase 10 | USP10 | 87.134 | 2 | 52.436764 | yes | 7.039E-07 | 759.37817 | 150.33139 |
| DIRPGAAFEPTYIY | 15 | Q14694 | Ubiquitin carboxyl-terminal hydrolase 10 | USP10 | 87.134 | 3 | 52.436764 | yes | 4.33E-07 | 590.30542 | 63.653721 |
| PPGYYSYLK | 9 | Q14694 | Ubiquitin carboxyl-terminal hydrolase 10 | USP10 | 87.134 | 2 | 52.436764 | yes | 9.722E-08 | 544.27655 | 54.167076 |
| ITPDGITK | 8 | Q14694 | Ubiquitin carboxyl-terminal hydrolase 10 | USP10 | 87.134 | 2 | 52.436764 | yes | 4.951E-07 | 422.74237 | 47.459637 |
| LDLLEDR | 7 | Q14697 | Neutral alpha-glucosidase AB OS=Homo | GANAB | 106.874 | 2 | 36.63444 | yes | 6.775E-07 | 437.23746 | 16.670938 |
| VAVFDTGLSEK | 11 | Q14703 | Membrane-bound transcription factor site | MBTPS1 | 117.749 | 2 | 39.69869 | yes | 4.431E-07 | 583.30859 | 49.341988 |
| VVEGTPLIDGR | 11 | Q14739 | Delta(14)-sterol reductase LBR OS=Hom | LBR | 70.704 | 2 | 52.48181 | yes | 9.722E-08 | 578.32202 | 1488.0762 |
| FADGEVVR | 8 | Q14739 | Delta(14)-sterol reductase LBR OS=Hom | LBR | 70.704 | 2 | 52.48181 | yes | 1.503E-07 | 446.7298 | 699.70709 |
| ENDIKPLTSFR | 11 | Q14739 | Delta(14)-sterol reductase LBR OS=Hom | LBR | 70.704 | 2 | 52.48181 | yes | 9.722E-08 | 660.35132 | 604.22217 |
| YKDGTELELK | 10 | Q14739 | Delta(14)-sterol reductase LBR OS=Hom | LBR | 70.704 | 2 | 52.48181 | yes | 6.189E-07 | 598.3139 | 513.91101 |
| TFEVTPIR | 8 | Q14739 | Delta(14)-sterol reductase LBR OS=Hom | LBR | 70.704 | 2 | 52.48181 | yes | 1.331E-06 | 481.76892 | 237.99968 |
| KFADGEVVR | 9 | Q14739 | Delta(14)-sterol reductase LBR OS=Hom | LBR | 70.704 | 2 | 52.48181 | yes | 5.983E-07 | 510.77728 | 153.03961 |
| YNGEPEHIER | 10 | Q14739 | Delta(14)-sterol reductase LBR OS=Hom | LBR | 70.704 | 2 | 52.48181 | yes | 9.974E-08 | 622.28876 | 152.57977 |
| ELPPGVEELLNK | 12 | Q14764 | Major vault protein OS=Homo sapiens O. | MVP | 99.327 | 2 | 53.965622 | yes | 4.972E-07 | 669.3692 | 364.28259 |
| LFSVPDFVGDACK | 13 | Q14764 | Major vault protein OS=Homo sapiens O. | MVP | 99.327 | 2 | 53.965622 | yes | 9.722E-08 | 727.85284 | 266.11661 |
| ILDQSEAEK | 9 | Q14764 | Major vault protein OS=Homo sapiens O. | MVP | 99.327 | 2 | 53.965622 | yes | 2.205E-07 | 516.76404 | 236.20573 |
| SLQPLAPR | 8 | Q14764 | Major vault protein OS=Homo sapiens O. | MVP | 99.327 | 2 | 53.965622 | yes | 1.902E-07 | 441.26379 | 214.79402 |
| IEGEGSVLQAK | 11 | Q14764 | Major vault protein OS=Homo sapiens O. | MVP | 99.327 | 2 | 53.965622 | yes | 5.877E-07 | 565.80621 | 210.80089 |
| DLAVAGPEMQVK | 12 | Q14764 | Major vault protein OS=Homo sapiens O. | MVP | 99.327 | 2 | 53.965622 | yes | 9.722E-08 | 629.32904 | 177.62274 |
| HADLEIR | 7 | Q14764 | Major vault protein OS=Homo sapiens O. | MVP | 99.327 | 2 | 53.965622 | yes | 5.057E-07 | 427.22995 | 159.377 |
| ELELVYAR | 8 | Q14764 | Major vault protein OS=Homo sapiens O. | MVP | 99.327 | 2 | 53.965622 | yes | 1.437E-07 | 496.7742 | 132.05241 |
| AQALAIETEAELQI | 14 | Q14764 | Major vault protein OS=Homo sapiens O. | MVP | 99.327 | 2 | 53.965622 | yes | 5.796E-07 | 771.90973 | 125.24625 |
| ALQPLEEGEDEEK | 13 | Q14764 | Major vault protein OS=Homo sapiens O. | MVP | 99.327 | 2 | 53.965622 | yes | 6.807E-07 | 743.849 | 124.29684 |
| QMTEAIGPSTIR | 12 | Q14764 | Major vault protein OS=Homo sapiens O. | MVP | 99.327 | 2 | 53.965622 | yes | 9.722E-08 | 652.33734 | 115.81601 |
| HYCTVANPVSR | 11 | Q14764 | Major vault protein OS=Homo sapiens O. | MVP | 99.327 | 2 | 53.965622 | yes | 1.155E-06 | 652.31421 | 73.261581 |
| GPLEYVPSAK | 10 | Q14764 | Major vault protein OS=Homo sapiens O. | MVP | 99.327 | 2 | 53.965622 | yes | 6.397E-07 | 530.78729 | 64.63546 |
| VSHQAGDHWLIR | 12 | Q14764 | Major vault protein OS=Homo sapiens O. | MVP | 99.327 | 2 | 53.965622 | yes | 4.771E-07 | 709.86804 | 56.449524 |
| LLQSLGLK | 8 | Q14764 | Major vault protein OS=Homo sapiens O. | MVP | 99.327 | 2 | 53.965622 | yes | 3.157E-07 | 436.28421 | 46.476585 |
| VPHNAAVQVYDY | 13 | Q14764 | Major vault protein OS=Homo sapiens O. | MVP | 99.327 | 2 | 53.965622 | yes | 8.248E-07 | 766.38623 | 37.522663 |
| ALLDFEDK | 8 | Q14764 | Major vault protein OS=Homo sapiens O. | MVP | 99.327 | 2 | 53.965622 | yes | 1.925E-06 | 475.74512 | 33.91227 |
| LKEPDAPSK | 9 | Q14781 | Chromobox protein homolog 2 OS=Hom | CBX2 | 56.081 | 2 | 47.64179 | yes | 2.42E-07 | 492.77167 | 173.59875 |
| ESPTSVGFFNLR | 12 | Q14781 | Chromobox protein homolog 2 OS=Hom | CBX2 | 56.081 | 2 | 47.64179 | yes | 1.286E-07 | 677.34351 | 87.829117 |

| | | | | | | | | | | | |
|---|---|---|---|---|---|---|---|---|---|---|---|
| NGMPGVGLLAR | 11 | Q14781 | Chromobox protein homolog 2 OS=Homo | CBX2 | 56.081 | 2 | 47.64179 | yes | 4.307E-07 | 542.80023 | 81.908112 |
| LLLAFQK | 7 | Q14781 | Chromobox protein homolog 2 OS=Homo | CBX2 | 56.081 | 2 | 47.64179 | yes | 2.117E-07 | 416.76819 | 44.307961 |
| LPPPLSAPVAGLA | 16 | Q14781 | Chromobox protein homolog 2 OS=Homo | CBX2 | 56.081 | 2 | 47.64179 | yes | 6.844E-07 | 757.96887 | 24.154245 |
| LTAMSSCSR | 9 | Q14781 | Chromobox protein homolog 2 OS=Homo | CBX2 | 56.081 | 2 | 47.64179 | yes | 2.924E-06 | 506.73114 | 16.193657 |
| QLQIQVR | 7 | Q14789 | Golgin subfamily B member 1 OS=Homo | GOLGB1 | 376.02 | 2 | 31.62268 | yes | 1.78E-06 | 442.76926 | 49.825802 |
| VAAENALSVAEEQ | 15 | Q14789 | Golgin subfamily B member 1 OS=Homo | GOLGB1 | 376.02 | 2 | 31.62268 | yes | 2.245E-06 | 800.42047 | 31.472153 |
| GMDSCSLEIANWR | 13 | Q14807 | Kinesin-like protein KIF22 OS=Homo sa | KIF22 | 73.263 | 2 | 54.50266 | yes | 9.722E-08 | 769.83997 | 128.87556 |
| ESGAINTSLFVLGK | 14 | Q14807 | Kinesin-like protein KIF22 OS=Homo sa | KIF22 | 73.263 | 2 | 54.50266 | yes | 9.722E-08 | 718.39319 | 123.45253 |
| FYLDTVSALNFAA | 14 | Q14807 | Kinesin-like protein KIF22 OS=Homo sa | KIF22 | 73.263 | 2 | 54.50266 | yes | 1.339E-07 | 794.41193 | 115.54774 |
| THTMLGSPEQPGV | 16 | Q14807 | Kinesin-like protein KIF22 OS=Homo sa | KIF22 | 73.263 | 3 | 54.50266 | yes | 2.793E-07 | 573.96643 | 104.03036 |
| YQFDAFYGER | 10 | Q14807 | Kinesin-like protein KIF22 OS=Homo sa | KIF22 | 73.263 | 2 | 54.50266 | yes | 7.975E-07 | 648.28821 | 97.386444 |
| LLASQGSQGAPLL | 17 | Q14807 | Kinesin-like protein KIF22 OS=Homo sa | KIF22 | 73.263 | 2 | 54.50266 | yes | 1E-07 | 834.46997 | 88.379333 |
| HFLPASR | 7 | Q14807 | Kinesin-like protein KIF22 OS=Homo sa | KIF22 | 73.263 | 2 | 54.50266 | yes | 1.52E-07 | 414.22977 | 87.999046 |
| ALMDLLQLTR | 10 | Q14807 | Kinesin-like protein KIF22 OS=Homo sa | KIF22 | 73.263 | 2 | 54.50266 | yes | 2.052E-07 | 587.33667 | 76.537903 |
| ILDLLNEGSAR | 11 | Q14807 | Kinesin-like protein KIF22 OS=Homo sa | KIF22 | 73.263 | 2 | 54.50266 | yes | 5.924E-07 | 600.83276 | 43.233078 |
| ELLGPPEAK | 9 | Q14807 | Kinesin-like protein KIF22 OS=Homo sa | KIF22 | 73.263 | 2 | 54.50266 | yes | 2.448E-06 | 477.26874 | 37.355312 |
| VGGNIEVLGFNAR | 13 | Q14839 | Chromodomain-helicase-DNA-binding pr | CHD4 | 218.005 | 2 | 51.318935 | no | 9.722E-08 | 673.36475 | 348.04373 |
| ENEFSFEDNAIR | 12 | Q14839 | Chromodomain-helicase-DNA-binding pr | CHD4 | 218.005 | 2 | 51.318935 | no | 9.918E-08 | 735.82843 | 313.98026 |
| IGVMSLIR | 8 | Q14839 | Chromodomain-helicase-DNA-binding pr | CHD4 | 218.005 | 2 | 51.318935 | yes | 3.028E-06 | 444.77042 | 283.55295 |
| SAIDLTPIVVEDK | 13 | Q14839 | Chromodomain-helicase-DNA-binding pr | CHD4 | 218.005 | 2 | 51.318935 | yes | 9.722E-08 | 700.38757 | 221.81764 |
| VVVEPPEGEEK | 11 | Q14839 | Chromodomain-helicase-DNA-binding pr | CHD4 | 218.005 | 2 | 51.318935 | yes | 1.114E-06 | 606.31134 | 217.86308 |
| WQDIQNDPR | 9 | Q14839 | Chromodomain-helicase-DNA-binding pr | CHD4 | 218.005 | 2 | 51.318935 | no | 1.461E-07 | 586.2782 | 167.56206 |
| WSMPELAEVEENI | 14 | Q14839 | Chromodomain-helicase-DNA-binding pr | CHD4 | 218.005 | 2 | 51.318935 | yes | 9.722E-08 | 845.41125 | 166.40889 |
| GHVHYLIK | 8 | Q14839 | Chromodomain-helicase-DNA-binding pr | CHD4 | 218.005 | 2 | 51.318935 | yes | 2.947E-06 | 483.77963 | 152.60031 |
| AYHMVCLDPDME | 13 | Q14839 | Chromodomain-helicase-DNA-binding pr | CHD4 | 218.005 | 2 | 51.318935 | yes | 1.457E-07 | 804.84637 | 150.32933 |
| IDGGITGNMR | 10 | Q14839 | Chromodomain-helicase-DNA-binding pr | CHD4 | 218.005 | 2 | 51.318935 | yes | 3.175E-07 | 517.25836 | 137.43518 |
| TYEIWHR | 7 | Q14839 | Chromodomain-helicase-DNA-binding pr | CHD4 | 218.005 | 2 | 51.318935 | yes | 3.907E-07 | 502.75107 | 128.29105 |
| LHDMLGPHMLR | 11 | Q14839 | Chromodomain-helicase-DNA-binding pr | CHD4 | 218.005 | 3 | 51.318935 | yes | 3.367E-07 | 440.56189 | 55.959301 |
| LHDMLGPHMLR | 11 | Q14839 | Chromodomain-helicase-DNA-binding pr | CHD4 | 218.005 | 3 | 51.318935 | yes | 1.99E-06 | 660.33917 | 55.959301 |
| LHDMLGPHMLR | 11 | Q14839 | Chromodomain-helicase-DNA-binding pr | CHD4 | 218.005 | 2 | 51.318935 | yes | 3.367E-07 | 440.56189 | 55.959301 |
| LHDMLGPHMLR | 11 | Q14839 | Chromodomain-helicase-DNA-binding pr | CHD4 | 218.005 | 2 | 51.318935 | yes | 1.99E-06 | 660.33917 | 55.959301 |
| AFLNAIMR | 8 | Q14839 | Chromodomain-helicase-DNA-binding pr | CHD4 | 218.005 | 2 | 51.318935 | no | 4.713E-07 | 468.26019 | 105.02509 |
| APEPTPQQVAQQQ | 13 | Q14839 | Chromodomain-helicase-DNA-binding pr | CHD4 | 218.005 | 2 | 51.318935 | yes | 6.396E-07 | 711.35461 | 96.750381 |
| QLEELLSDMK | 10 | Q14839 | Chromodomain-helicase-DNA-binding pr | CHD4 | 218.005 | 2 | 51.318935 | yes | 1.144E-07 | 603.30774 | 92.097748 |
| KLERPPETPTVDPT | 16 | Q14839 | Chromodomain-helicase-DNA-binding pr | CHD4 | 218.005 | 3 | 51.318935 | yes | 4.967E-07 | 603.00354 | 84.233109 |
| EGIQWEAK | 8 | Q14839 | Chromodomain-helicase-DNA-binding pr | CHD4 | 218.005 | 2 | 51.318935 | yes | 3.371E-07 | 480.74289 | 58.742321 |
| CCNHPYLFPVAAM | 17 | Q14839 | Chromodomain-helicase-DNA-binding pr | CHD4 | 218.005 | 3 | 51.318935 | yes | 2.363E-07 | 668.97583 | 52.627853 |
| MMMVLGAK | 8 | Q14839 | Chromodomain-helicase-DNA-binding pr | CHD4 | 218.005 | 2 | 51.318935 | yes | 1.647E-06 | 440.72629 | 46.105732 |
| EGEDSSVIHYDDK | 13 | Q14839 | Chromodomain-helicase-DNA-binding pr | CHD4 | 218.005 | 2 | 51.318935 | yes | 5.503E-07 | 747.32318 | 41.854885 |

| | | | | | | | | | | | |
|---|---|---|---|---|---|---|---|---|---|---|---|
| MPNGMYDGSALII | 13 | Q14839 | Chromodomain-helicase-DNA-binding pr | CHD4 | 218.005 | 2 | 51.318935 | yes | 7.999E-07 | 712.83667 | 34.157204 |
| GAADVEKVEEK | 11 | Q14839 | Chromodomain-helicase-DNA-binding pr | CHD4 | 218.005 | 2 | 51.318935 | yes | 2.485E-06 | 587.80115 | 30.811264 |
| EPAAPVSIQR | 10 | Q14847 | LIM and SH3 domain protein 1 OS=Hom | LASP1 | 29.718 | 2 | 53.356102 | yes | 2.385E-07 | 534.29584 | 532.74023 |
| GFSVVADTPELQR | 13 | Q14847 | LIM and SH3 domain protein 1 OS=Hom | LASP1 | 29.718 | 2 | 53.356102 | yes | 9.722E-08 | 709.86737 | 472.78125 |
| QSFTMVADTPENL | 14 | Q14847 | LIM and SH3 domain protein 1 OS=Hom | LASP1 | 29.718 | 2 | 53.356102 | yes | 1.675E-07 | 804.88794 | 435.07635 |
| YHEEFEK | 7 | Q14847 | LIM and SH3 domain protein 1 OS=Hom | LASP1 | 29.718 | 2 | 53.356102 | yes | 2.945E-07 | 491.21927 | 304.87222 |
| YKEEFEK | 7 | Q14847 | LIM and SH3 domain protein 1 OS=Hom | LASP1 | 29.718 | 2 | 53.356102 | yes | 2.296E-07 | 486.73727 | 301.71973 |
| TQDQISNIK | 9 | Q14847 | LIM and SH3 domain protein 1 OS=Hom | LASP1 | 29.718 | 2 | 53.356102 | yes | 1.527E-07 | 523.77747 | 242.01288 |
| IVYPTEK | 7 | Q14847 | LIM and SH3 domain protein 1 OS=Hom | LASP1 | 29.718 | 2 | 53.356102 | yes | 1.3E-07 | 425.23947 | 226.14482 |
| QQSELQSQVR | 10 | Q14847 | LIM and SH3 domain protein 1 OS=Hom | LASP1 | 29.718 | 2 | 53.356102 | yes | 9.722E-08 | 601.80981 | 214.52188 |
| TGDTGMLPANYVI | 15 | Q14847 | LIM and SH3 domain protein 1 OS=Hom | LASP1 | 29.718 | 2 | 53.356102 | yes | 9.722E-08 | 776.37158 | 47.385609 |
| DIRPEFR | 7 | Q14938 | Nuclear factor 1 X-type OS=Homo sapier | NFIX | 55.099 | 2 | 50.84622 | yes | 2.502E-07 | 466.75107 | 530.28137 |
| AVKDELLGEKPEII | 14 | Q14938 | Nuclear factor 1 X-type OS=Homo sapier | NFIX | 55.099 | 3 | 50.84622 | yes | 8.669E-07 | 523.63464 | 259.41467 |
| AVKDELLGEKPEII | 14 | Q14938 | Nuclear factor 1 X-type OS=Homo sapier | NFIX | 55.099 | 3 | 50.84622 | yes | 4.925E-07 | 784.94836 | 259.41467 |
| AVKDELLGEKPEII | 14 | Q14938 | Nuclear factor 1 X-type OS=Homo sapier | NFIX | 55.099 | 2 | 50.84622 | yes | 8.669E-07 | 523.63464 | 259.41467 |
| AVKDELLGEKPEII | 14 | Q14938 | Nuclear factor 1 X-type OS=Homo sapier | NFIX | 55.099 | 2 | 50.84622 | yes | 4.925E-07 | 784.94836 | 259.41467 |
| EDFVLTITGK | 10 | Q14938 | Nuclear factor 1 X-type OS=Homo sapier | NFIX | 55.099 | 2 | 50.84622 | yes | 9.766E-08 | 561.80573 | 454.45871 |
| KDIRPEFR | 8 | Q14938 | Nuclear factor 1 X-type OS=Homo sapier | NFIX | 55.099 | 2 | 50.84622 | yes | 1.066E-07 | 530.79852 | 116.25983 |
| LDFCSALSSQGSSI | 15 | Q14938 | Nuclear factor 1 X-type OS=Homo sapier | NFIX | 55.099 | 2 | 50.84622 | yes | 9.722E-08 | 806.37518 | 113.84687 |
| AFSYTWFNLQAR | 12 | Q14938 | Nuclear factor 1 X-type OS=Homo sapier | NFIX | 55.099 | 2 | 50.84622 | yes | 1.012E-07 | 752.37262 | 112.78938 |
| VPGSFLLPPPPPVA | 15 | Q14938 | Nuclear factor 1 X-type OS=Homo sapier | NFIX | 55.099 | 2 | 50.84622 | yes | 2.031E-07 | 772.45337 | 79.382538 |
| IVFLLVGHK | 9 | Q14964 | Ras-related protein Rab-39A OS=Homo s | RAB39A | 25.007 | 2 | 35.907623 | yes | 7.656E-07 | 513.32892 | 36.933662 |
| TALLPSDSVFAEEI | 14 | Q14966 | Zinc finger protein 638 OS=Homo sapien | ZNF638 | 220.625 | 2 | 58.713104 | yes | 1.627E-07 | 767.89099 | 445.2735 |
| ALEDVVQR | 8 | Q14966 | Zinc finger protein 638 OS=Homo sapien | ZNF638 | 220.625 | 2 | 58.713104 | yes | 1.373E-07 | 465.25616 | 345.12888 |
| YGYTEDPLEVR | 11 | Q14966 | Zinc finger protein 638 OS=Homo sapien | ZNF638 | 220.625 | 2 | 58.713104 | yes | 3.011E-07 | 671.3197 | 314.87772 |
| AVEIVTSTSAAK | 12 | Q14966 | Zinc finger protein 638 OS=Homo sapien | ZNF638 | 220.625 | 2 | 58.713104 | yes | 9.722E-08 | 588.82715 | 286.1105 |
| DILILSSHK | 9 | Q14966 | Zinc finger protein 638 OS=Homo sapien | ZNF638 | 220.625 | 2 | 58.713104 | yes | 1.054E-07 | 513.3031 | 272.47784 |
| MNVQVTQHR | 9 | Q14966 | Zinc finger protein 638 OS=Homo sapien | ZNF638 | 220.625 | 2 | 58.713104 | yes | 9.722E-08 | 556.7851 | 211.80049 |
| FKPTQSSLTR | 10 | Q14966 | Zinc finger protein 638 OS=Homo sapien | ZNF638 | 220.625 | 2 | 58.713104 | yes | 1.567E-07 | 582.8222 | 204.00714 |
| NVPFSELNLK | 10 | Q14966 | Zinc finger protein 638 OS=Homo sapien | ZNF638 | 220.625 | 2 | 58.713104 | yes | 5.183E-07 | 580.81915 | 172.45056 |
| YIETTPLTIK | 10 | Q14966 | Zinc finger protein 638 OS=Homo sapien | ZNF638 | 220.625 | 2 | 58.713104 | yes | 2.252E-07 | 589.83698 | 166.31277 |
| ENDPEANIDTIYDI | 14 | Q14966 | Zinc finger protein 638 OS=Homo sapien | ZNF638 | 220.625 | 2 | 58.713104 | yes | 9.722E-08 | 832.87354 | 165.14218 |
| DVTVLSVAEEQDL | 15 | Q14966 | Zinc finger protein 638 OS=Homo sapien | ZNF638 | 220.625 | 2 | 58.713104 | yes | 1.396E-07 | 829.94598 | 163.76047 |
| GYSVEEVYDLAK | 12 | Q14966 | Zinc finger protein 638 OS=Homo sapien | ZNF638 | 220.625 | 2 | 58.713104 | yes | 4.177E-07 | 686.83521 | 162.03087 |
| VNDVLIVPYR | 10 | Q14966 | Zinc finger protein 638 OS=Homo sapien | ZNF638 | 220.625 | 2 | 58.713104 | yes | 2.971E-07 | 594.34277 | 156.24709 |
| EAYLEMEFK | 9 | Q14966 | Zinc finger protein 638 OS=Homo sapien | ZNF638 | 220.625 | 2 | 58.713104 | yes | 9.722E-08 | 580.27063 | 145.97089 |
| NYQSQADIPIR | 11 | Q14966 | Zinc finger protein 638 OS=Homo sapien | ZNF638 | 220.625 | 2 | 58.713104 | yes | 9.722E-08 | 652.83331 | 139.52951 |
| QSSVTQVTEQSPK | 13 | Q14966 | Zinc finger protein 638 OS=Homo sapien | ZNF638 | 220.625 | 2 | 58.713104 | yes | 9.722E-08 | 709.85974 | 135.45134 |
| NLEDDTLSECK | 11 | Q14966 | Zinc finger protein 638 OS=Homo sapien | ZNF638 | 220.625 | 2 | 58.713104 | yes | 1.006E-07 | 662.29028 | 125.18522 |

| | | | | | | | | | | | |
|---|---|---|---|---|---|---|---|---|---|---|---|
| MDLQIGTEK | 9 | Q14966 | Zinc finger protein 638 OS=Homo sapien | ZNF638 | 220.625 | 2 | 58.713104 | yes | 1.191E-07 | 517.763 | 116.88577 |
| SDSNLGGHSIR | 11 | Q14966 | Zinc finger protein 638 OS=Homo sapien | ZNF638 | 220.625 | 2 | 58.713104 | yes | 9.722E-08 | 571.78107 | 98.852478 |
| WDDEPHISASVAV | 14 | Q14966 | Zinc finger protein 638 OS=Homo sapien | ZNF638 | 220.625 | 2 | 58.713104 | yes | 3.012E-07 | 777.38336 | 98.361076 |
| MSGLHISGGQSVL | 17 | Q14966 | Zinc finger protein 638 OS=Homo sapien | ZNF638 | 220.625 | 3 | 58.713104 | yes | 1.962E-07 | 584.98193 | 98.211899 |
| IPHEPVINSSNVHV | 17 | Q14966 | Zinc finger protein 638 OS=Homo sapien | ZNF638 | 220.625 | 3 | 58.713104 | yes | 1.44E-07 | 614.65875 | 88.528854 |
| VDHHVFISNR | 10 | Q14966 | Zinc finger protein 638 OS=Homo sapien | ZNF638 | 220.625 | 2 | 58.713104 | yes | 9.722E-08 | 612.31763 | 68.281738 |
| EKLDFHEAQQK | 11 | Q14966 | Zinc finger protein 638 OS=Homo sapien | ZNF638 | 220.625 | 2 | 58.713104 | yes | 9.722E-08 | 686.84644 | 45.01408 |
| DSNKPVTIPENSEI | 15 | Q14966 | Zinc finger protein 638 OS=Homo sapien | ZNF638 | 220.625 | 2 | 58.713104 | yes | 2.277E-07 | 835.93341 | 39.419331 |
| SFFSVESGTK | 10 | Q14966 | Zinc finger protein 638 OS=Homo sapien | ZNF638 | 220.625 | 2 | 58.713104 | yes | 3.636E-07 | 544.7666 | 36.177273 |
| RLPNLPSQSR | 10 | Q14966 | Zinc finger protein 638 OS=Homo sapien | ZNF638 | 220.625 | 2 | 58.713104 | yes | 2.108E-07 | 584.33325 | 32.549614 |
| DEEAIFITLVK | 11 | Q14966 | Zinc finger protein 638 OS=Homo sapien | ZNF638 | 220.625 | 2 | 58.713104 | yes | 1.027E-07 | 639.35303 | 16.65193 |
| KALEDVVQR | 9 | Q14966 | Zinc finger protein 638 OS=Homo sapien | ZNF638 | 220.625 | 2 | 58.713104 | yes | 4.664E-06 | 529.30365 | 15.496145 |
| WLAIDANAR | 9 | Q14974 | Importin subunit beta-1 OS=Homo sapier | KPNB1 | 97.171 | 2 | 53.34343 | yes | 2.068E-07 | 515.27747 | 375.65674 |
| VLANPGNSQVAR | 12 | Q14974 | Importin subunit beta-1 OS=Homo sapier | KPNB1 | 97.171 | 2 | 53.34343 | yes | 9.722E-08 | 613.336 | 360.13339 |
| SDYDMVDYLNELI | 13 | Q14974 | Importin subunit beta-1 OS=Homo sapier | KPNB1 | 97.171 | 2 | 53.34343 | yes | 1.584E-07 | 816.86414 | 342.93307 |
| TTLVIMER | 8 | Q14974 | Importin subunit beta-1 OS=Homo sapier | KPNB1 | 97.171 | 2 | 53.34343 | yes | 2.908E-07 | 481.7706 | 316.30643 |
| SSAYESLMEIVK | 12 | Q14974 | Importin subunit beta-1 OS=Homo sapier | KPNB1 | 97.171 | 2 | 53.34343 | yes | 9.722E-08 | 678.83942 | 311.56671 |
| ESCLEAYTGIVQGI | 15 | Q14974 | Importin subunit beta-1 OS=Homo sapier | KPNB1 | 97.171 | 2 | 53.34343 | yes | 1.098E-07 | 834.41907 | 218.56938 |
| PMIHELLTEGR | 11 | Q14974 | Importin subunit beta-1 OS=Homo sapier | KPNB1 | 97.171 | 2 | 53.34343 | yes | 1.158E-07 | 648.34247 | 129.63702 |
| LAATNALLNSLEF | 15 | Q14974 | Importin subunit beta-1 OS=Homo sapier | KPNB1 | 97.171 | 2 | 53.34343 | yes | 9.722E-08 | 803.44598 | 122.40543 |
| VAALQNLVK | 9 | Q14974 | Importin subunit beta-1 OS=Homo sapier | KPNB1 | 97.171 | 2 | 53.34343 | yes | 5.955E-07 | 478.30038 | 90.396576 |
| LLETTDRPDGHQN | 16 | Q14974 | Importin subunit beta-1 OS=Homo sapier | KPNB1 | 97.171 | 3 | 53.34343 | yes | 1.161E-07 | 626.98541 | 62.88924 |
| TLATWATK | 8 | Q14974 | Importin subunit beta-1 OS=Homo sapier | KPNB1 | 97.171 | 2 | 53.34343 | yes | 2.778E-06 | 446.25037 | 50.020851 |
| NYVLQTLGTETYR | 13 | Q14974 | Importin subunit beta-1 OS=Homo sapier | KPNB1 | 97.171 | 2 | 53.34343 | yes | 2.475E-07 | 779.39899 | 48.889748 |
| SPAVKPAAAPK | 11 | Q14978 | Nucleolar and coiled-body phosphoprotei | NOLC1 | 73.604 | 2 | 50.4609 | yes | 1.106E-07 | 518.8111 | 142.46835 |
| GGSISVQVNSIK | 12 | Q14978 | Nucleolar and coiled-body phosphoprotei | NOLC1 | 73.604 | 2 | 50.4609 | yes | 8.627E-07 | 594.83276 | 142.01843 |
| DNQLSEVANK | 10 | Q14978 | Nucleolar and coiled-body phosphoprotei | NOLC1 | 73.604 | 2 | 50.4609 | yes | 3.196E-06 | 559.27783 | 102.69616 |
| VAGGAAPSKPASA | 14 | Q14978 | Nucleolar and coiled-body phosphoprotei | NOLC1 | 73.604 | 2 | 50.4609 | yes | 9.809E-08 | 606.34076 | 68.36451 |
| NKPGPYSSVPPPSA | 18 | Q14978 | Nucleolar and coiled-body phosphoprotei | NOLC1 | 73.604 | 3 | 50.4609 | yes | 1.298E-06 | 606.32458 | 60.295685 |
| KLQANGPVAK | 10 | Q14978 | Nucleolar and coiled-body phosphoprotei | NOLC1 | 73.604 | 2 | 50.4609 | yes | 1.086E-07 | 513.30872 | 32.241085 |
| VREEEIEVDSR | 11 | Q14978 | Nucleolar and coiled-body phosphoprotei | NOLC1 | 73.604 | 2 | 50.4609 | yes | 3.343E-06 | 680.83881 | 30.806023 |
| PSLSLGTITDEEMK | 14 | Q14980 | Nuclear mitotic apparatus protein 1 OS=H | NUMA1 | 238.26 | 2 | 50.317505 | yes | 1.331E-07 | 760.87927 | 110.46691 |
| LLQAETASNSAR | 12 | Q14980 | Nuclear mitotic apparatus protein 1 OS=H | NUMA1 | 238.26 | 2 | 50.317505 | yes | 2.461E-07 | 630.83075 | 70.271698 |
| LDFVCSFLQK | 10 | Q14980 | Nuclear mitotic apparatus protein 1 OS=H | NUMA1 | 238.26 | 2 | 50.317505 | yes | 1.833E-07 | 628.8208 | 64.354141 |
| DSAQTSVTQAQR | 12 | Q14980 | Nuclear mitotic apparatus protein 1 OS=H | NUMA1 | 238.26 | 2 | 50.317505 | yes | 1.007E-07 | 646.31549 | 60.455986 |
| LTAQVEQLEVFQR | 13 | Q14980 | Nuclear mitotic apparatus protein 1 OS=H | NUMA1 | 238.26 | 2 | 50.317505 | yes | 1.163E-07 | 780.92267 | 57.531639 |
| DQLQEQLQALK | 11 | Q14980 | Nuclear mitotic apparatus protein 1 OS=H | NUMA1 | 238.26 | 2 | 50.317505 | yes | 2.204E-07 | 657.35663 | 55.426922 |
| QFLEVELDQAR | 11 | Q14980 | Nuclear mitotic apparatus protein 1 OS=H | NUMA1 | 238.26 | 2 | 50.317505 | yes | 2.902E-07 | 674.34882 | 54.540859 |
| EAEQMGNELER | 11 | Q14980 | Nuclear mitotic apparatus protein 1 OS=H | NUMA1 | 238.26 | 2 | 50.317505 | yes | 2.074E-06 | 653.29059 | 53.538513 |

| | | | | | | | | | | | |
|---|---|---|---|---|---|---|---|---|---|---|---|
| ELSAALQDK | 9 | Q14980 | Nuclear mitotic apparatus protein 1 OS=H | NUMA1 | 238.26 | 2 | 50.317505 | yes | 1.548E-06 | 487.76129 | 49.405369 |
| EELEQASQAHGAR | 13 | Q14980 | Nuclear mitotic apparatus protein 1 OS=H | NUMA1 | 238.26 | 2 | 50.317505 | yes | 9.819E-08 | 713.33948 | 47.814011 |
| ELGELIPLR | 9 | Q14980 | Nuclear mitotic apparatus protein 1 OS=H | NUMA1 | 238.26 | 2 | 50.317505 | yes | 2.613E-07 | 520.31097 | 41.377087 |
| SLEAQVAHADQQI | 14 | Q14980 | Nuclear mitotic apparatus protein 1 OS=H | NUMA1 | 238.26 | 2 | 50.317505 | yes | 2.414E-07 | 783.40515 | 30.613104 |
| INQLSEENGDLSFK | 14 | Q14980 | Nuclear mitotic apparatus protein 1 OS=H | NUMA1 | 238.26 | 2 | 50.317505 | yes | 6.239E-07 | 797.39136 | 25.286093 |
| ATSSTQSLAR | 10 | Q14980 | Nuclear mitotic apparatus protein 1 OS=H | NUMA1 | 238.26 | 2 | 50.317505 | yes | 2.784E-06 | 511.26727 | 24.501884 |
| QQNQELQEQLR | 11 | Q14980 | Nuclear mitotic apparatus protein 1 OS=H | NUMA1 | 238.26 | 2 | 50.317505 | yes | 2.311E-07 | 707.35767 | 18.156166 |
| HLCQQLQAEQAA | 15 | Q14980 | Nuclear mitotic apparatus protein 1 OS=H | NUMA1 | 238.26 | 3 | 50.317505 | yes | 0.0001935 | 575.61768 | 15.037603 |
| FSGMVLLCK | 9 | Q14995 | Nuclear receptor subfamily 1 group D me | NR1D2 | 64.644 | 2 | 47.927887 | yes | 1.231E-07 | 527.77484 | 78.705215 |
| MLIEMQSAMK | 10 | Q14995 | Nuclear receptor subfamily 1 group D me | NR1D2 | 64.644 | 2 | 47.927887 | yes | 1.353E-07 | 591.28998 | 78.062325 |
| SSSPPSSDFAK | 11 | Q14995 | Nuclear receptor subfamily 1 group D me | NR1D2 | 64.644 | 2 | 47.927887 | yes | 2.755E-05 | 555.25909 | 31.956802 |
| SLNLPVGSSQVLV | 15 | Q14997 | Proteasome activator complex subunit 4 C | PSME4 | 211.334 | 2 | 44.966877 | yes | 2.009E-07 | 783.4541 | 58.44529 |
| VPLGPGLHAYPDE | 16 | Q14999 | Cullin-7 OS=Homo sapiens OX=9606 GN | CUL7 | 191.161 | 3 | 38.309483 | yes | 2.935E-06 | 582.98944 | 42.168991 |
| VKPLLLQLQR | 10 | Q14999 | Cullin-7 OS=Homo sapiens OX=9606 GN | CUL7 | 191.161 | 2 | 38.309483 | yes | 5.313E-07 | 604.39789 | 32.612442 |
| VSAVDELAMATEI | 13 | Q149N8 | E3 ubiquitin-protein ligase SHPRH OS=H | SHPRH | 193.08 | 2 | 46.555763 | yes | 8.956E-07 | 696.3454 | 77.475006 |
| EETVGIFSPLSVK | 13 | Q149N8 | E3 ubiquitin-protein ligase SHPRH OS=H | SHPRH | 193.08 | 2 | 46.555763 | yes | 1.53E-07 | 703.38232 | 70.891205 |
| WCISGTPVQR | 10 | Q149N8 | E3 ubiquitin-protein ligase SHPRH OS=H | SHPRH | 193.08 | 2 | 46.555763 | yes | 2.443E-07 | 602.3006 | 49.957104 |
| VGIYLLEAGLAK | 12 | Q149N8 | E3 ubiquitin-protein ligase SHPRH OS=H | SHPRH | 193.08 | 2 | 46.555763 | yes | 6.243E-07 | 623.8739 | 43.498348 |
| ADELFTEYESK | 11 | Q149N8 | E3 ubiquitin-protein ligase SHPRH OS=H | SHPRH | 193.08 | 2 | 46.555763 | yes | 6.872E-07 | 666.30371 | 29.806038 |
| QDALTLPEGK | 10 | Q149N8 | E3 ubiquitin-protein ligase SHPRH OS=H | SHPRH | 193.08 | 2 | 46.555763 | yes | 3.211E-06 | 536.28766 | 26.071671 |
| ILEDTQFVDGYK | 12 | Q14C86 | GTPase-activating protein and VPS9 dom | GAPVD1 | 164.98 | 2 | 37.491978 | yes | 5.944E-07 | 714.35626 | 47.152725 |
| NLLFSGSPAVTMT | 16 | Q14CB8 | Rho GTPase-activating protein 19 OS=Ho | ARHGAP1 | 55.757 | 2 | 52.291348 | yes | 9.722E-08 | 846.44287 | 96.301277 |
| TNIPDKDR | 8 | Q14CB8 | Rho GTPase-activating protein 19 OS=Ho | ARHGAP1 | 55.757 | 2 | 52.291348 | yes | 8.629E-07 | 479.75125 | 86.50679 |
| GQPIIFNPDFFVEK | 14 | Q14CB8 | Rho GTPase-activating protein 19 OS=Ho | ARHGAP1 | 55.757 | 2 | 52.291348 | yes | 3.129E-07 | 825.92993 | 62.45723 |
| IADLMQFDDK | 10 | Q14CB8 | Rho GTPase-activating protein 19 OS=Ho | ARHGAP1 | 55.757 | 2 | 52.291348 | yes | 1.402E-06 | 598.2868 | 40.542645 |
| NVTANDLQENITK | 13 | Q14CB8 | Rho GTPase-activating protein 19 OS=Ho | ARHGAP1 | 55.757 | 2 | 52.291348 | yes | 2.932E-06 | 730.37299 | 26.354738 |
| SHLSLVGTASGLG | 16 | Q14CW9 | Ataxin-7-like protein 3 OS=Homo sapiens | ATXN7L3 | 38.651 | 2 | 41.492462 | yes | 3.482E-07 | 764.40991 | 25.136385 |
| IGLTNLLGAR | 10 | Q14CZ7 | FAST kinase domain-containing protein 3 | FASTKD3 | 75.69 | 2 | 51.855892 | yes | 9.722E-08 | 514.31659 | 191.63278 |
| ELVEYLQR | 8 | Q14CZ7 | FAST kinase domain-containing protein 3 | FASTKD3 | 75.69 | 2 | 51.855892 | yes | 9.722E-08 | 525.28491 | 126.4155 |
| ESHLDTLSR | 9 | Q14CZ7 | FAST kinase domain-containing protein 3 | FASTKD3 | 75.69 | 2 | 51.855892 | yes | 5.195E-07 | 529.26727 | 117.72747 |
| LNYLPPNASALFR | 13 | Q14CZ7 | FAST kinase domain-containing protein 3 | FASTKD3 | 75.69 | 2 | 51.855892 | yes | 5.442E-06 | 738.40393 | 85.623993 |
| IFKPLFLQR | 9 | Q14CZ7 | FAST kinase domain-containing protein 3 | FASTKD3 | 75.69 | 2 | 51.855892 | yes | 1.451E-06 | 581.36078 | 58.992035 |
| ILQACTEK | 8 | Q14CZ7 | FAST kinase domain-containing protein 3 | FASTKD3 | 75.69 | 2 | 51.855892 | yes | 3.319E-07 | 481.75241 | 33.075851 |
| NEESQMFYR | 9 | Q14CZ7 | FAST kinase domain-containing protein 3 | FASTKD3 | 75.69 | 2 | 51.855892 | yes | 1.328E-07 | 602.25861 | 27.176504 |
| VDEHQTFLNK | 10 | Q14CZ7 | FAST kinase domain-containing protein 3 | FASTKD3 | 75.69 | 2 | 51.855892 | yes | 3.322E-06 | 615.80933 | 26.692848 |
| VFDLQFSTDSPR | 12 | Q15003 | Condensin complex subunit 2 OS=Homo | NCAPH | 82.564 | 2 | 41.624653 | yes | 3.418E-07 | 706.34625 | 83.481232 |
| YAGGNPVCVRPTI | 14 | Q15004 | PCNA-associated factor OS=Homo sapier | PCLAF | 11.987 | 2 | 48.303265 | yes | 1.301E-07 | 758.39044 | 43.629303 |
| YAGGNPVCVRPTI | 14 | Q15004 | PCNA-associated factor OS=Homo sapier | PCLAF | 11.987 | 2 | 48.303265 | yes | 1.053E-06 | 505.92941 | 43.629303 |
| YAGGNPVCVRPTI | 14 | Q15004 | PCNA-associated factor OS=Homo sapier | PCLAF | 11.987 | 3 | 48.303265 | yes | 1.301E-07 | 758.39044 | 43.629303 |

| | | | | | | | | | | |
|---|---|---|---|---|---|---|---|---|---|---|
| YAGGNPVCVRPTI | 14 | Q15004 | PCNA-associated factor OS=Homo sapie | PCLAF | 11.987 | 3 | 48.303265 | yes | 1.053E-06 | 505.92941 | 43.629303 |
| DSEKENQIPEEAG | 19 | Q15004 | PCNA-associated factor OS=Homo sapie | PCLAF | 11.987 | 3 | 48.303265 | yes | 7.059E-07 | 658.97986 | 31.012083 |
| ENQIPEEAGSSGL | 15 | Q15004 | PCNA-associated factor OS=Homo sapie | PCLAF | 11.987 | 2 | 48.303265 | yes | 1.166E-07 | 758.36792 | 27.104641 |
| VLGSSTSATNSTS | 17 | Q15004 | PCNA-associated factor OS=Homo sapie | PCLAF | 11.987 | 2 | 48.303265 | yes | 1.108E-06 | 820.90796 | 15.672243 |
| SIFLVAHR | 8 | Q15005 | Signal peptidase complex subunit 2 OS=I | SPCS2 | 25.003 | 2 | 38.500874 | yes | 5.177E-07 | 471.77963 | 86.00515 |
| QVQQPSVAQLR | 11 | Q15007 | Pre-mRNA-splicing regulator WTAP OS= | WTAP | 44.244 | 2 | 51.459232 | yes | 1.995E-07 | 627.35168 | 185.88782 |
| YTDLNSNDVTGLI | 13 | Q15007 | Pre-mRNA-splicing regulator WTAP OS= | WTAP | 44.244 | 2 | 51.459232 | yes | 9.722E-08 | 734.35736 | 174.31204 |
| MLIQENQELGR | 11 | Q15007 | Pre-mRNA-splicing regulator WTAP OS= | WTAP | 44.244 | 2 | 51.459232 | yes | 9.967E-08 | 665.84283 | 135.80461 |
| IAQLEAELALQK | 12 | Q15007 | Pre-mRNA-splicing regulator WTAP OS= | WTAP | 44.244 | 2 | 51.459232 | yes | 9.722E-08 | 663.88501 | 60.976582 |
| GAEILEVLHSLPA | 15 | Q15008 | 26S proteasome non-ATPase regulatory s | PSMD6 | 45.532 | 2 | 52.07096 | yes | 9.722E-08 | 802.46198 | 20.73295 |
| NVDAILEEYANCK | 13 | Q15014 | Mortality factor 4-like protein 2 OS=Hon | MORF4L2 | 32.308 | 2 | 54.998566 | yes | 9.776E-08 | 769.86139 | 1846.4379 |
| FERPQYAEILLAHI | 29 | Q15014 | Mortality factor 4-like protein 2 OS=Hon | MORF4L2 | 32.308 | 4 | 54.998566 | yes | 1.169E-07 | 830.68158 | 585.22772 |
| FERPQYAEILLAHI | 29 | Q15014 | Mortality factor 4-like protein 2 OS=Hon | MORF4L2 | 32.308 | 4 | 54.998566 | yes | 9.722E-08 | 664.7467 | 585.22772 |
| FERPQYAEILLAHI | 29 | Q15014 | Mortality factor 4-like protein 2 OS=Hon | MORF4L2 | 32.308 | 5 | 54.998566 | yes | 1.169E-07 | 830.68158 | 585.22772 |
| FERPQYAEILLAHI | 29 | Q15014 | Mortality factor 4-like protein 2 OS=Hon | MORF4L2 | 32.308 | 5 | 54.998566 | yes | 9.722E-08 | 664.7467 | 585.22772 |
| NSASLFTASDYK | 12 | Q15014 | Mortality factor 4-like protein 2 OS=Hon | MORF4L2 | 32.308 | 2 | 54.998566 | yes | 1.009E-07 | 652.31189 | 858.39246 |
| GQQSAEEENFK | 11 | Q15014 | Mortality factor 4-like protein 2 OS=Hon | MORF4L2 | 32.308 | 2 | 54.998566 | yes | 1.037E-07 | 633.78351 | 778.00903 |
| EYAVNEVVAGIK | 12 | Q15014 | Mortality factor 4-like protein 2 OS=Hon | MORF4L2 | 32.308 | 2 | 54.998566 | no | 1.347E-07 | 646.34827 | 774.87573 |
| ADPTVESEEAFK | 12 | Q15014 | Mortality factor 4-like protein 2 OS=Hon | MORF4L2 | 32.308 | 2 | 54.998566 | yes | 1.185E-07 | 661.80914 | 745.67096 |
| SAENPPSGSVR | 11 | Q15014 | Mortality factor 4-like protein 2 OS=Hon | MORF4L2 | 32.308 | 2 | 54.998566 | yes | 1.122E-07 | 550.7702 | 454.52344 |
| NLEPALPGR | 9 | Q15014 | Mortality factor 4-like protein 2 OS=Hon | MORF4L2 | 32.308 | 2 | 54.998566 | yes | 1.457E-07 | 483.772 | 393.82138 |
| IGAMLAYTPLDEK | 13 | Q15014 | Mortality factor 4-like protein 2 OS=Hon | MORF4L2 | 32.308 | 2 | 54.998566 | no | 9.722E-08 | 719.36835 | 151.65056 |
| IGAMLAYTPLDEK | 13 | Q15014 | Mortality factor 4-like protein 2 OS=Hon | MORF4L2 | 32.308 | 2 | 54.998566 | no | 9.722E-08 | 711.37085 | 151.65056 |
| VASAEYHR | 8 | Q15014 | Mortality factor 4-like protein 2 OS=Hon | MORF4L2 | 32.308 | 2 | 54.998566 | yes | 1.187E-07 | 466.73288 | 156.44455 |
| QLFQLPAK | 8 | Q15014 | Mortality factor 4-like protein 2 OS=Hon | MORF4L2 | 32.308 | 2 | 54.998566 | yes | 1.215E-07 | 472.78183 | 65.129837 |
| KNVDAILEEYANC | 14 | Q15014 | Mortality factor 4-like protein 2 OS=Hon | MORF4L2 | 32.308 | 2 | 54.998566 | yes | 3.921E-07 | 833.90887 | 30.600405 |
| LYPWGVVEVENPI | 19 | Q15019 | Septin-2 OS=Homo sapiens OX=9606 GI | SEPTIN2 | 41.488 | 3 | 51.579678 | yes | 1.612E-07 | 762.37939 | 78.331581 |
| ILDEIEEHNIK | 11 | Q15019 | Septin-2 OS=Homo sapiens OX=9606 GI | SEPTIN2 | 41.488 | 2 | 51.579678 | yes | 9.722E-08 | 676.85645 | 75.249786 |
| TIISYIDEQFER | 12 | Q15019 | Septin-2 OS=Homo sapiens OX=9606 GI | SEPTIN2 | 41.488 | 2 | 51.579678 | yes | 2.56E-07 | 757.38031 | 71.512535 |
| TVQIEASTVEIEER | 14 | Q15019 | Septin-2 OS=Homo sapiens OX=9606 GI | SEPTIN2 | 41.488 | 2 | 51.579678 | yes | 2.588E-06 | 802.41229 | 34.221008 |
| IQLIFER | 7 | Q15020 | Squamous cell carcinoma antigen recogni | SART3 | 109.935 | 2 | 55.599365 | yes | 2.147E-07 | 459.77402 | 553.9234 |
| DLVLSVHNR | 9 | Q15020 | Squamous cell carcinoma antigen recogni | SART3 | 109.935 | 2 | 55.599365 | yes | 2.088E-07 | 526.79602 | 292.99582 |
| PMFVSPCVDK | 10 | Q15020 | Squamous cell carcinoma antigen recogni | SART3 | 109.935 | 2 | 55.599365 | yes | 4.837E-07 | 590.28046 | 254.06204 |
| SALQALEMDR | 10 | Q15020 | Squamous cell carcinoma antigen recogni | SART3 | 109.935 | 2 | 55.599365 | yes | 1.514E-07 | 567.28461 | 244.88556 |
| EAALVQQEEEK | 11 | Q15020 | Squamous cell carcinoma antigen recogni | SART3 | 109.935 | 2 | 55.599365 | yes | 1.015E-07 | 637.31714 | 242.72751 |
| TEGSLEDWDIAVQ | 14 | Q15020 | Squamous cell carcinoma antigen recogni | SART3 | 109.935 | 2 | 55.599365 | yes | 9.722E-08 | 795.88593 | 202.21159 |
| GLALWEAYR | 9 | Q15020 | Squamous cell carcinoma antigen recogni | SART3 | 109.935 | 2 | 55.599365 | yes | 2.748E-07 | 539.78766 | 188.13914 |
| YSQYLDR | 7 | Q15020 | Squamous cell carcinoma antigen recogni | SART3 | 109.935 | 2 | 55.599365 | yes | 9.722E-08 | 472.72726 | 186.62279 |
| NCPWTVALWSR | 11 | Q15020 | Squamous cell carcinoma antigen recogni | SART3 | 109.935 | 2 | 55.599365 | yes | 9.81E-08 | 695.34027 | 175.36435 |

| | | | | | | | | | | | |
|---|---|---|---|---|---|---|---|---|---|---|---|
| YLLAMER | 7 | Q15020 | Squamous cell carcinoma antigen recogni | SART3 | 109.935 | 2 | 55.599365 | yes | 2.891E-07 | 448.23895 | 163.72816 |
| HGVDHQVISVTFE | 14 | Q15020 | Squamous cell carcinoma antigen recogni | SART3 | 109.935 | 3 | 55.599365 | yes | 1.823E-07 | 532.61072 | 81.103355 |
| HGVDHQVISVTFE | 14 | Q15020 | Squamous cell carcinoma antigen recogni | SART3 | 109.935 | 3 | 55.599365 | yes | 2.886E-06 | 798.41248 | 81.103355 |
| HGVDHQVISVTFE | 14 | Q15020 | Squamous cell carcinoma antigen recogni | SART3 | 109.935 | 2 | 55.599365 | yes | 1.823E-07 | 532.61072 | 81.103355 |
| HGVDHQVISVTFE | 14 | Q15020 | Squamous cell carcinoma antigen recogni | SART3 | 109.935 | 2 | 55.599365 | yes | 2.886E-06 | 798.41248 | 81.103355 |
| EHVYDLFEK | 9 | Q15020 | Squamous cell carcinoma antigen recogni | SART3 | 109.935 | 2 | 55.599365 | yes | 4.194E-07 | 590.28766 | 138.20523 |
| TQLSLLPR | 8 | Q15020 | Squamous cell carcinoma antigen recogni | SART3 | 109.935 | 2 | 55.599365 | yes | 2.678E-07 | 464.28473 | 138.05223 |
| ATAAETSASEPEAI | 16 | Q15020 | Squamous cell carcinoma antigen recogni | SART3 | 109.935 | 2 | 55.599365 | yes | 1.075E-07 | 810.86542 | 127.87118 |
| LAEYQAYIDFEMK | 13 | Q15020 | Squamous cell carcinoma antigen recogni | SART3 | 109.935 | 2 | 55.599365 | yes | 9.722E-08 | 810.88434 | 123.26345 |
| ELWDSIMTR | 9 | Q15020 | Squamous cell carcinoma antigen recogni | SART3 | 109.935 | 2 | 55.599365 | yes | 9.727E-08 | 575.78168 | 119.74844 |
| YKPYEEALLQAEA | 15 | Q15020 | Squamous cell carcinoma antigen recogni | SART3 | 109.935 | 3 | 55.599365 | yes | 1.39E-07 | 593.30884 | 86.78904 |
| CAAVDVEPPSK | 11 | Q15020 | Squamous cell carcinoma antigen recogni | SART3 | 109.935 | 2 | 55.599365 | yes | 1.079E-06 | 586.78442 | 84.29361 |
| VAISNPPQR | 9 | Q15020 | Squamous cell carcinoma antigen recogni | SART3 | 109.935 | 2 | 55.599365 | yes | 1.025E-07 | 491.27744 | 71.727936 |
| KAPGGPMLLPQTY | 16 | Q15020 | Squamous cell carcinoma antigen recogni | SART3 | 109.935 | 3 | 55.599365 | yes | 1.123E-07 | 552.96783 | 68.524071 |
| EFESAIVEAAR | 11 | Q15020 | Squamous cell carcinoma antigen recogni | SART3 | 109.935 | 2 | 55.599365 | yes | 2.627E-07 | 611.30914 | 51.950127 |
| MSNADFAK | 8 | Q15020 | Squamous cell carcinoma antigen recogni | SART3 | 109.935 | 2 | 55.599365 | yes | 2.49E-07 | 442.20255 | 51.614449 |
| DLAYCVSQLPLTE | 14 | Q15021 | Condensin complex subunit 1 OS=Homo | NCAPD2 | 157.183 | 2 | 51.279743 | yes | 1.022E-07 | 832.91925 | 113.17638 |
| VACCPLER | 8 | Q15021 | Condensin complex subunit 1 OS=Homo | NCAPD2 | 157.183 | 2 | 51.279743 | yes | 1.768E-07 | 502.73624 | 83.606766 |
| SNLMVATGDLAIR | 13 | Q15021 | Condensin complex subunit 1 OS=Homo | NCAPD2 | 157.183 | 2 | 51.279743 | yes | 1.157E-07 | 680.86627 | 79.368874 |
| AIIDEFEQK | 9 | Q15021 | Condensin complex subunit 1 OS=Homo | NCAPD2 | 157.183 | 2 | 51.279743 | yes | 4.772E-07 | 546.78223 | 79.279526 |
| EAITHLLGVALTR | 13 | Q15021 | Condensin complex subunit 1 OS=Homo | NCAPD2 | 157.183 | 2 | 51.279743 | yes | 9.722E-08 | 697.41174 | 62.529263 |
| ELEIGQAGSQR | 11 | Q15021 | Condensin complex subunit 1 OS=Homo | NCAPD2 | 157.183 | 2 | 51.279743 | yes | 2.581E-07 | 594.30438 | 51.085754 |
| KITEAIGIISK | 11 | Q15021 | Condensin complex subunit 1 OS=Homo | NCAPD2 | 157.183 | 2 | 51.279743 | yes | 5.576E-07 | 586.86609 | 39.603062 |
| MLDNFDCFGDK | 11 | Q15021 | Condensin complex subunit 1 OS=Homo | NCAPD2 | 157.183 | 2 | 51.279743 | yes | 9.85E-08 | 681.27869 | 24.529882 |
| LLFTMLEK | 8 | Q15021 | Condensin complex subunit 1 OS=Homo | NCAPD2 | 157.183 | 2 | 51.279743 | yes | 3.209E-06 | 497.78574 | 14.517859 |
| NLIAPIFLHR | 10 | Q15022 | Polycomb protein SUZ12 OS=Homo sapi | SUZ12 | 83.055 | 2 | 52.657356 | yes | 9.722E-08 | 597.36133 | 406.69586 |
| TFVAQMTVFDK | 11 | Q15022 | Polycomb protein SUZ12 OS=Homo sapi | SUZ12 | 83.055 | 2 | 52.657356 | yes | 1.03E-07 | 643.82611 | 206.05417 |
| FIFNYVYHPK | 10 | Q15022 | Polycomb protein SUZ12 OS=Homo sapi | SUZ12 | 83.055 | 2 | 52.657356 | yes | 9.722E-08 | 664.34534 | 177.6778 |
| LWNLHVMK | 8 | Q15022 | Polycomb protein SUZ12 OS=Homo sapi | SUZ12 | 83.055 | 2 | 52.657356 | yes | 4.955E-07 | 520.78912 | 175.2883 |
| ALETDSVSGVSK | 12 | Q15022 | Polycomb protein SUZ12 OS=Homo sapi | SUZ12 | 83.055 | 2 | 52.657356 | yes | 2.818E-07 | 596.8064 | 146.8707 |
| IFYQFLYNNNTR | 12 | Q15022 | Polycomb protein SUZ12 OS=Homo sapi | SUZ12 | 83.055 | 2 | 52.657356 | yes | 9.722E-08 | 796.89642 | 133.24438 |
| ESLTTDLQTR | 10 | Q15022 | Polycomb protein SUZ12 OS=Homo sapi | SUZ12 | 83.055 | 2 | 52.657356 | yes | 1.096E-06 | 582.29877 | 85.51561 |
| KLYSLLK | 7 | Q15022 | Polycomb protein SUZ12 OS=Homo sapi | SUZ12 | 83.055 | 2 | 52.657356 | yes | 1.292E-07 | 432.78131 | 84.651207 |
| SYSLLFR | 7 | Q15022 | Polycomb protein SUZ12 OS=Homo sapi | SUZ12 | 83.055 | 2 | 52.657356 | yes | 4.155E-07 | 443.24509 | 32.591225 |
| LSVENVPCIVTLCI | 14 | Q15024 | Exosome complex component RRP42 OS | EXOSC7 | 31.822 | 2 | 51.225506 | yes | 1.109E-07 | 816.42841 | 1260.2773 |
| ASVTLSEAEK | 10 | Q15024 | Exosome complex component RRP42 OS | EXOSC7 | 31.822 | 2 | 51.225506 | yes | 2.451E-07 | 538.77716 | 520.26593 |
| GGDDLGTEIANTL | 15 | Q15024 | Exosome complex component RRP42 OS | EXOSC7 | 31.822 | 2 | 51.225506 | yes | 1.142E-07 | 797.88904 | 402.85193 |
| VYIVHGVQEDLR | 12 | Q15024 | Exosome complex component RRP42 OS | EXOSC7 | 31.822 | 2 | 51.225506 | yes | 2.471E-07 | 714.38568 | 370.76065 |
| LGHTDILVGVK | 11 | Q15024 | Exosome complex component RRP42 OS | EXOSC7 | 31.822 | 2 | 51.225506 | yes | 1.002E-07 | 576.34277 | 230.99486 |

| | | | | | | | | | | | |
|---|---|---|---|---|---|---|---|---|---|---|---|
| DIELSDDPYDCIR | 13 | Q15024 | Exosome complex component RRP42 OS | EXOSC7 | 31.822 | 2 | 51.225506 | yes | 9.722E-08 | 805.85376 | 97.84462 |
| VPEVVALGAAEK | 12 | Q15025 | TNFAIP3-interacting protein 1 OS=Hom | TNIP1 | 71.865 | 2 | 56.08142 | yes | 1.179E-07 | 591.84009 | 291.46271 |
| IFEEDFQR | 8 | Q15025 | TNFAIP3-interacting protein 1 OS=Hom | TNIP1 | 71.865 | 2 | 56.08142 | yes | 1.112E-07 | 542.25891 | 191.33582 |
| EDGNLMLHLQR | 11 | Q15025 | TNFAIP3-interacting protein 1 OS=Hom | TNIP1 | 71.865 | 2 | 56.08142 | yes | 9.722E-08 | 663.33514 | 141.31244 |
| LQAQVTLSNAQLK | 13 | Q15025 | TNFAIP3-interacting protein 1 OS=Hom | TNIP1 | 71.865 | 2 | 56.08142 | yes | 9.722E-08 | 707.40662 | 138.55989 |
| YLQDQLSPLTR | 11 | Q15025 | TNFAIP3-interacting protein 1 OS=Hom | TNIP1 | 71.865 | 2 | 56.08142 | yes | 2.438E-07 | 667.35913 | 126.7617 |
| TSILQTLCEQLR | 12 | Q15025 | TNFAIP3-interacting protein 1 OS=Hom | TNIP1 | 71.865 | 2 | 56.08142 | yes | 4.793E-07 | 731.39014 | 68.06147 |
| AVAGQQQASVTA | 14 | Q15025 | TNFAIP3-interacting protein 1 OS=Hom | TNIP1 | 71.865 | 2 | 56.08142 | yes | 9.722E-08 | 658.35187 | 52.067429 |
| KQELVTQNELLK | 12 | Q15025 | TNFAIP3-interacting protein 1 OS=Hom | TNIP1 | 71.865 | 2 | 56.08142 | yes | 1.118E-07 | 721.91431 | 34.180611 |
| IAVEPVNPSELPK | 13 | Q15029 | 116 kDa U5 small nuclear ribonucleoprot | EFTUD2 | 109.436 | 2 | 55.093933 | yes | 1.59E-07 | 696.89032 | 767.10791 |
| ILAQVVGDVDTSL | 15 | Q15029 | 116 kDa U5 small nuclear ribonucleoprot | EFTUD2 | 109.436 | 2 | 55.093933 | yes | 1.013E-07 | 791.9436 | 623.66028 |
| ILDAVVAQEPLHR | 13 | Q15029 | 116 kDa U5 small nuclear ribonucleoprot | EFTUD2 | 109.436 | 2 | 55.093933 | yes | 1.032E-07 | 730.91461 | 309.55057 |
| ILDAVVAQEPLHR | 13 | Q15029 | 116 kDa U5 small nuclear ribonucleoprot | EFTUD2 | 109.436 | 2 | 55.093933 | yes | 1.528E-06 | 487.61218 | 309.55057 |
| ILDAVVAQEPLHR | 13 | Q15029 | 116 kDa U5 small nuclear ribonucleoprot | EFTUD2 | 109.436 | 3 | 55.093933 | yes | 1.032E-07 | 730.91461 | 309.55057 |
| ILDAVVAQEPLHR | 13 | Q15029 | 116 kDa U5 small nuclear ribonucleoprot | EFTUD2 | 109.436 | 3 | 55.093933 | yes | 1.528E-06 | 487.61218 | 309.55057 |
| DSIVQGFQWGTR | 12 | Q15029 | 116 kDa U5 small nuclear ribonucleoprot | EFTUD2 | 109.436 | 2 | 55.093933 | yes | 1.634E-07 | 697.34656 | 578.57574 |
| LPPTDAYYK | 9 | Q15029 | 116 kDa U5 small nuclear ribonucleoprot | EFTUD2 | 109.436 | 2 | 55.093933 | yes | 1.217E-06 | 534.27405 | 547.3045 |
| EGPLCDELIR | 10 | Q15029 | 116 kDa U5 small nuclear ribonucleoprot | EFTUD2 | 109.436 | 2 | 55.093933 | yes | 1.032E-07 | 601.29773 | 539.74713 |
| ITMIAEPLEK | 10 | Q15029 | 116 kDa U5 small nuclear ribonucleoprot | EFTUD2 | 109.436 | 2 | 55.093933 | yes | 1.028E-07 | 572.81775 | 377.47012 |
| VLSGTIHAGQPVK | 13 | Q15029 | 116 kDa U5 small nuclear ribonucleoprot | EFTUD2 | 109.436 | 2 | 55.093933 | yes | 9.79E-08 | 653.8775 | 335.40616 |
| LWGDIYFNPK | 10 | Q15029 | 116 kDa U5 small nuclear ribonucleoprot | EFTUD2 | 109.436 | 2 | 55.093933 | yes | 9.722E-08 | 626.82166 | 182.84163 |
| LAVTVCINK | 9 | Q15029 | 116 kDa U5 small nuclear ribonucleoprot | EFTUD2 | 109.436 | 2 | 55.093933 | yes | 1.993E-07 | 509.29172 | 155.85713 |
| YDWDLLAAR | 9 | Q15029 | 116 kDa U5 small nuclear ribonucleoprot | EFTUD2 | 109.436 | 2 | 55.093933 | yes | 1.037E-07 | 561.78253 | 144.24808 |
| STPVTVVLPDTK | 12 | Q15029 | 116 kDa U5 small nuclear ribonucleoprot | EFTUD2 | 109.436 | 2 | 55.093933 | yes | 1.49E-07 | 628.85846 | 100.82927 |
| GNEEAQIFRPLK | 12 | Q15029 | 116 kDa U5 small nuclear ribonucleoprot | EFTUD2 | 109.436 | 2 | 55.093933 | yes | 9.722E-08 | 701.37787 | 88.632202 |
| CFAETPNKK | 9 | Q15029 | 116 kDa U5 small nuclear ribonucleoprot | EFTUD2 | 109.436 | 2 | 55.093933 | yes | 1.871E-07 | 547.76862 | 87.470276 |
| GLSEDVSISK | 10 | Q15029 | 116 kDa U5 small nuclear ribonucleoprot | EFTUD2 | 109.436 | 2 | 55.093933 | yes | 4.047E-07 | 517.77185 | 80.037994 |
| LGEFFQTK | 8 | Q15029 | 116 kDa U5 small nuclear ribonucleoprot | EFTUD2 | 109.436 | 2 | 55.093933 | yes | 2.203E-07 | 485.25565 | 75.241333 |
| KGLSEDVSISK | 11 | Q15029 | 116 kDa U5 small nuclear ribonucleoprot | EFTUD2 | 109.436 | 2 | 55.093933 | yes | 3.323E-07 | 581.81934 | 71.368065 |
| FFGEFTGFVDMCV | 20 | Q15029 | 116 kDa U5 small nuclear ribonucleoprot | EFTUD2 | 109.436 | 3 | 55.093933 | yes | 1.991E-06 | 781.70276 | 56.723843 |
| GHVTQDAPIPGSPI | 18 | Q15029 | 116 kDa U5 small nuclear ribonucleoprot | EFTUD2 | 109.436 | 3 | 55.093933 | yes | 1.154E-06 | 632.00714 | 50.724617 |
| PLEPQPAPHLAR | 12 | Q15029 | 116 kDa U5 small nuclear ribonucleoprot | EFTUD2 | 109.436 | 3 | 55.093933 | yes | 5.047E-07 | 442.58234 | 47.680882 |
| NVTLCGHLHHGK | 12 | Q15029 | 116 kDa U5 small nuclear ribonucleoprot | EFTUD2 | 109.436 | 2 | 55.093933 | yes | 4.156E-07 | 686.84875 | 38.239857 |
| KFFGEFTGFVDMC | 21 | Q15029 | 116 kDa U5 small nuclear ribonucleoprot | EFTUD2 | 109.436 | 3 | 55.093933 | yes | 2.457E-06 | 824.40106 | 17.274145 |
| MLVVGGIDR | 9 | Q15046 | Lysine--tRNA ligase OS=Homo sapiens ( | KARS1 | 68.049 | 2 | 51.09865 | yes | 2.657E-07 | 480.27078 | 805.46771 |
| IAPELYHK | 8 | Q15046 | Lysine--tRNA ligase OS=Homo sapiens ( | KARS1 | 68.049 | 2 | 51.09865 | yes | 1.296E-07 | 485.77145 | 423.68243 |
| ILDDICVAK | 9 | Q15046 | Lysine--tRNA ligase OS=Homo sapiens ( | KARS1 | 68.049 | 2 | 51.09865 | yes | 2.801E-07 | 523.78119 | 337.73224 |
| VAMFLTDSNNIK | 12 | Q15046 | Lysine--tRNA ligase OS=Homo sapiens ( | KARS1 | 68.049 | 2 | 51.09865 | yes | 1.05E-07 | 676.8476 | 242.29568 |
| SQAIHQLK | 8 | Q15046 | Lysine--tRNA ligase OS=Homo sapiens ( | KARS1 | 68.049 | 2 | 51.09865 | yes | 7.426E-07 | 462.76672 | 153.48732 |

| | | | | | | | | | | |
|---|---|---|---|---|---|---|---|---|---|---|
| INMVEELEK | 9 | Q15046 | Lysine--tRNA ligase OS=Homo sapiens ( | KARS1 | 68.049 | 2 | 51.09865 | yes | 1.161E-07 | 552.78394 | 131.11119 |
| LQVMANSR | 8 | Q15046 | Lysine--tRNA ligase OS=Homo sapiens ( | KARS1 | 68.049 | 2 | 51.09865 | yes | 3.584E-07 | 459.74493 | 107.23012 |
| SEEEFIHINNK | 11 | Q15046 | Lysine--tRNA ligase OS=Homo sapiens ( | KARS1 | 68.049 | 2 | 51.09865 | yes | 9.722E-08 | 680.33063 | 97.895973 |
| AVECPPPR | 8 | Q15046 | Lysine--tRNA ligase OS=Homo sapiens ( | KARS1 | 68.049 | 2 | 51.09865 | yes | 3.173E-07 | 463.23166 | 78.783981 |
| LPETNLFETEETR | 13 | Q15046 | Lysine--tRNA ligase OS=Homo sapiens ( | KARS1 | 68.049 | 2 | 51.09865 | yes | 1.741E-07 | 789.88593 | 32.282513 |
| RGDIIGVQGNPGK | 13 | Q15046 | Lysine--tRNA ligase OS=Homo sapiens ( | KARS1 | 68.049 | 2 | 51.09865 | yes | 1.964E-06 | 655.86243 | 28.267572 |
| KPQLDVTR | 8 | Q15050 | Ribosome biogenesis regulatory protein h | RRS1 | 41.194 | 2 | 52.700047 | yes | 1.758E-07 | 478.77982 | 331.32108 |
| FQPLFGDFAAEK | 12 | Q15050 | Ribosome biogenesis regulatory protein h | RRS1 | 41.194 | 2 | 52.700047 | yes | 1.157E-07 | 685.34296 | 280.37939 |
| MQLPSAAGLHPTC | 17 | Q15050 | Ribosome biogenesis regulatory protein h | RRS1 | 41.194 | 3 | 52.700047 | yes | 9.723E-08 | 587.30182 | 247.11763 |
| MNSGPPGLGGK | 11 | Q15050 | Ribosome biogenesis regulatory protein h | RRS1 | 41.194 | 2 | 52.700047 | yes | 2.439E-07 | 507.75549 | 234.916 |
| TNLVWDEVSGQW | 13 | Q15050 | Ribosome biogenesis regulatory protein h | RRS1 | 41.194 | 2 | 52.700047 | yes | 9.722E-08 | 795.38898 | 181.01965 |
| KFQPLFGDFAAEK | 13 | Q15050 | Ribosome biogenesis regulatory protein h | RRS1 | 41.194 | 2 | 52.700047 | yes | 2.494E-06 | 749.39044 | 127.94485 |
| CAGPTPEAELQAL | 15 | Q15050 | Ribosome biogenesis regulatory protein h | RRS1 | 41.194 | 2 | 52.700047 | yes | 1.112E-07 | 792.39594 | 120.01131 |
| VEEAIVAR | 8 | Q15050 | Ribosome biogenesis regulatory protein h | RRS1 | 41.194 | 2 | 52.700047 | yes | 5.366E-07 | 443.75327 | 107.72538 |
| NQLELLR | 7 | Q15050 | Ribosome biogenesis regulatory protein h | RRS1 | 41.194 | 2 | 52.700047 | yes | 3.065E-07 | 443.26126 | 105.53142 |
| KNQLELLR | 8 | Q15050 | Ribosome biogenesis regulatory protein h | RRS1 | 41.194 | 2 | 52.700047 | yes | 1.457E-07 | 507.30875 | 76.15448 |
| IVEQPTVSVTEPK | 13 | Q15054 | DNA polymerase delta subunit 3 OS=Hor | POLD3 | 51.401 | 2 | 50.88794 | yes | 9.722E-08 | 713.89301 | 167.54314 |
| DSGPLFNTDYDILI | 14 | Q15054 | DNA polymerase delta subunit 3 OS=Hor | POLD3 | 51.401 | 2 | 50.88794 | yes | 1.2E-07 | 799.39087 | 105.86668 |
| QVSITGFFQR | 10 | Q15054 | DNA polymerase delta subunit 3 OS=Hor | POLD3 | 51.401 | 2 | 50.88794 | yes | 7.442E-07 | 591.81696 | 93.098251 |
| FSAIQCAAAVPR | 12 | Q15054 | DNA polymerase delta subunit 3 OS=Hor | POLD3 | 51.401 | 2 | 50.88794 | yes | 1.433E-06 | 645.83478 | 44.581909 |
| EALTYDGALLGDF | 13 | Q15056 | Eukaryotic translation initiation factor 4H | EIF4H | 27.386 | 2 | 46.100346 | yes | 1.653E-07 | 697.3515 | 162.17163 |
| NFGGTVSIIPVGEA | 15 | Q15058 | Kinesin-like protein KIF14 OS=Homo sa | KIF14 | 186.492 | 2 | 55.509743 | yes | 1.253E-07 | 744.90649 | 814.15582 |
| FCEDLFSQVAR | 11 | Q15058 | Kinesin-like protein KIF14 OS=Homo sa | KIF14 | 186.492 | 2 | 55.509743 | yes | 9.722E-08 | 686.32172 | 801.49774 |
| LAAPLLER | 8 | Q15058 | Kinesin-like protein KIF14 OS=Homo sa | KIF14 | 186.492 | 2 | 55.509743 | yes | 1.037E-07 | 441.77402 | 800.19977 |
| TADMPLTPNPVGF | 13 | Q15058 | Kinesin-like protein KIF14 OS=Homo sa | KIF14 | 186.492 | 2 | 55.509743 | yes | 9.722E-08 | 684.85065 | 662.61102 |
| ESSLLVSELEDTTE | 15 | Q15058 | Kinesin-like protein KIF14 OS=Homo sa | KIF14 | 186.492 | 2 | 55.509743 | yes | 1.09E-07 | 840.41473 | 636.79968 |
| ILLDSGLEK | 9 | Q15058 | Kinesin-like protein KIF14 OS=Homo sa | KIF14 | 186.492 | 2 | 55.509743 | yes | 1.47E-07 | 494.28967 | 625.19983 |
| HILEITVLR | 9 | Q15058 | Kinesin-like protein KIF14 OS=Homo sa | KIF14 | 186.492 | 2 | 55.509743 | yes | 2.726E-07 | 547.34003 | 520.15338 |
| VISALSEQANQR | 12 | Q15058 | Kinesin-like protein KIF14 OS=Homo sa | KIF14 | 186.492 | 2 | 55.509743 | yes | 1.994E-07 | 658.35187 | 442.46701 |
| ELQHELFR | 8 | Q15058 | Kinesin-like protein KIF14 OS=Homo sa | KIF14 | 186.492 | 2 | 55.509743 | yes | 2.049E-07 | 536.28271 | 442.45551 |
| ILEALETEK | 9 | Q15058 | Kinesin-like protein KIF14 OS=Homo sa | KIF14 | 186.492 | 2 | 55.509743 | yes | 1.073E-07 | 523.29242 | 440.83249 |
| LGISTFWSLEK | 11 | Q15058 | Kinesin-like protein KIF14 OS=Homo sa | KIF14 | 186.492 | 2 | 55.509743 | yes | 1.063E-07 | 640.8479 | 413.31265 |
| NAIQIVQQAVK | 11 | Q15058 | Kinesin-like protein KIF14 OS=Homo sa | KIF14 | 186.492 | 2 | 55.509743 | yes | 9.722E-08 | 606.35895 | 386.37845 |
| EMAQQELSSQK | 11 | Q15058 | Kinesin-like protein KIF14 OS=Homo sa | KIF14 | 186.492 | 2 | 55.509743 | yes | 9.902E-08 | 639.80334 | 314.81845 |
| EVQILQQNR | 9 | Q15058 | Kinesin-like protein KIF14 OS=Homo sa | KIF14 | 186.492 | 2 | 55.509743 | yes | 9.722E-08 | 564.31201 | 310.43173 |
| QEITSLR | 7 | Q15058 | Kinesin-like protein KIF14 OS=Homo sa | KIF14 | 186.492 | 2 | 55.509743 | yes | 6.259E-07 | 423.73761 | 302.95859 |
| EITVEHPDTK | 10 | Q15058 | Kinesin-like protein KIF14 OS=Homo sa | KIF14 | 186.492 | 2 | 55.509743 | yes | 1.012E-07 | 584.79584 | 284.64014 |
| YVGQLAVLK | 9 | Q15058 | Kinesin-like protein KIF14 OS=Homo sa | KIF14 | 186.492 | 2 | 55.509743 | yes | 2.153E-07 | 495.80276 | 270.58945 |
| VENSQVTVAVR | 11 | Q15058 | Kinesin-like protein KIF14 OS=Homo sa | KIF14 | 186.492 | 2 | 55.509743 | yes | 9.722E-08 | 601.33038 | 252.79559 |

| | | | | | | | | | | | |
|---|---|---|---|---|---|---|---|---|---|---|---|
| FNHPVEVQK | 9 | Q15058 | Kinesin-like protein KIF14 OS=Homo saj | KIF14 | 186.492 | 2 | 55.509743 | yes | 1.019E-07 | 549.29053 | 245.77513 |
| SHSVFTLVMTQTK | 13 | Q15058 | Kinesin-like protein KIF14 OS=Homo saj | KIF14 | 186.492 | 2 | 55.509743 | yes | 9.722E-08 | 739.88721 | 218.77393 |
| NELLMAQR | 8 | Q15058 | Kinesin-like protein KIF14 OS=Homo saj | KIF14 | 186.492 | 2 | 55.509743 | yes | 1.299E-07 | 487.75803 | 209.6908 |
| VTHIVHQELESLAI | 14 | Q15058 | Kinesin-like protein KIF14 OS=Homo saj | KIF14 | 186.492 | 2 | 55.509743 | yes | 9.809E-08 | 802.44373 | 91.984711 |
| VTHIVHQELESLAI | 14 | Q15058 | Kinesin-like protein KIF14 OS=Homo saj | KIF14 | 186.492 | 2 | 55.509743 | yes | 1.084E-07 | 535.29828 | 91.984711 |
| VTHIVHQELESLAI | 14 | Q15058 | Kinesin-like protein KIF14 OS=Homo saj | KIF14 | 186.492 | 3 | 55.509743 | yes | 9.809E-08 | 802.44373 | 91.984711 |
| VTHIVHQELESLAI | 14 | Q15058 | Kinesin-like protein KIF14 OS=Homo saj | KIF14 | 186.492 | 3 | 55.509743 | yes | 1.084E-07 | 535.29828 | 91.984711 |
| SLENIFAESK | 10 | Q15058 | Kinesin-like protein KIF14 OS=Homo saj | KIF14 | 186.492 | 2 | 55.509743 | yes | 2.134E-07 | 569.29297 | 177.2742 |
| SDMSECENDDPLL | 14 | Q15058 | Kinesin-like protein KIF14 OS=Homo saj | KIF14 | 186.492 | 2 | 55.509743 | yes | 1.483E-07 | 840.84564 | 146.10701 |
| INLIDLAGSER | 11 | Q15058 | Kinesin-like protein KIF14 OS=Homo saj | KIF14 | 186.492 | 2 | 55.509743 | yes | 9.722E-08 | 600.83276 | 134.20505 |
| CSLPQLK | 7 | Q15058 | Kinesin-like protein KIF14 OS=Homo saj | KIF14 | 186.492 | 2 | 55.509743 | yes | 1.117E-07 | 423.23111 | 132.09181 |
| LSMMIQEANAISSI | 14 | Q15058 | Kinesin-like protein KIF14 OS=Homo saj | KIF14 | 186.492 | 2 | 55.509743 | yes | 1.187E-07 | 761.88385 | 124.92831 |
| IHDLLVCK | 8 | Q15058 | Kinesin-like protein KIF14 OS=Homo saj | KIF14 | 186.492 | 2 | 55.509743 | yes | 4.293E-07 | 499.27859 | 103.16392 |
| AKEEMMQGIQIAK | 13 | Q15058 | Kinesin-like protein KIF14 OS=Homo saj | KIF14 | 186.492 | 2 | 55.509743 | yes | 9.722E-08 | 738.8811 | 96.882156 |
| TEFVEGEEHDHR | 12 | Q15058 | Kinesin-like protein KIF14 OS=Homo saj | KIF14 | 186.492 | 2 | 55.509743 | yes | 1.282E-07 | 742.82367 | 86.13987 |
| NLHSSHSSGLMDK | 13 | Q15058 | Kinesin-like protein KIF14 OS=Homo saj | KIF14 | 186.492 | 2 | 55.509743 | yes | 1.416E-07 | 706.83301 | 76.560692 |
| SVFIPYR | 7 | Q15058 | Kinesin-like protein KIF14 OS=Homo saj | KIF14 | 186.492 | 2 | 55.509743 | yes | 7.567E-07 | 441.24762 | 74.627579 |
| ISGCLHDIQVHPIK | 14 | Q15058 | Kinesin-like protein KIF14 OS=Homo saj | KIF14 | 186.492 | 3 | 55.509743 | yes | 2.593E-06 | 539.62408 | 34.704319 |
| ISGCLHDIQVHPIK | 14 | Q15058 | Kinesin-like protein KIF14 OS=Homo saj | KIF14 | 186.492 | 3 | 55.509743 | yes | 1.365E-07 | 808.9325 | 34.704319 |
| ISGCLHDIQVHPIK | 14 | Q15058 | Kinesin-like protein KIF14 OS=Homo saj | KIF14 | 186.492 | 2 | 55.509743 | yes | 2.593E-06 | 539.62408 | 34.704319 |
| ISGCLHDIQVHPIK | 14 | Q15058 | Kinesin-like protein KIF14 OS=Homo saj | KIF14 | 186.492 | 2 | 55.509743 | yes | 1.365E-07 | 808.9325 | 34.704319 |
| QALEDHSIR | 9 | Q15058 | Kinesin-like protein KIF14 OS=Homo saj | KIF14 | 186.492 | 2 | 55.509743 | yes | 1.344E-07 | 534.77527 | 65.824692 |
| QHLEQEIYVNK | 11 | Q15058 | Kinesin-like protein KIF14 OS=Homo saj | KIF14 | 186.492 | 2 | 55.509743 | yes | 1.22E-07 | 700.86206 | 56.818359 |
| YKPNSSHDIQLSG' | 24 | Q15058 | Kinesin-like protein KIF14 OS=Homo saj | KIF14 | 186.492 | 4 | 55.509743 | yes | 9.722E-08 | 678.59192 | 54.089661 |
| AIEIIISALK | 10 | Q15058 | Kinesin-like protein KIF14 OS=Homo saj | KIF14 | 186.492 | 2 | 55.509743 | yes | 1.492E-07 | 535.8446 | 40.196198 |
| TNAMGLIR | 8 | Q15058 | Kinesin-like protein KIF14 OS=Homo saj | KIF14 | 186.492 | 2 | 55.509743 | yes | 4.388E-06 | 438.242 | 32.351776 |
| IYNGLFAISK | 10 | Q15058 | Kinesin-like protein KIF14 OS=Homo saj | KIF14 | 186.492 | 2 | 55.509743 | yes | 1.962E-07 | 563.31879 | 26.440599 |
| TFTVQTTWSSMK | 12 | Q15058 | Kinesin-like protein KIF14 OS=Homo saj | KIF14 | 186.492 | 2 | 55.509743 | yes | 3.514E-06 | 708.84503 | 22.259533 |
| RLEMETLATK | 10 | Q15058 | Kinesin-like protein KIF14 OS=Homo saj | KIF14 | 186.492 | 2 | 55.509743 | yes | 5.118E-07 | 596.32373 | 20.493429 |
| KTADMPLTPNPVC | 14 | Q15058 | Kinesin-like protein KIF14 OS=Homo saj | KIF14 | 186.492 | 2 | 55.509743 | yes | 1.139E-06 | 748.89813 | 8.0815887 |
| LILLITQVTASEK | 13 | Q15061 | WD repeat-containing protein 43 OS=Ho | WDR43 | 74.891 | 2 | 52.127686 | yes | 9.722E-08 | 714.93726 | 379.48685 |
| EPHMCLVR | 8 | Q15061 | WD repeat-containing protein 43 OS=Ho | WDR43 | 74.891 | 2 | 52.127686 | yes | 1.852E-07 | 521.25208 | 350.8114 |
| LLNVWQVR | 8 | Q15061 | WD repeat-containing protein 43 OS=Ho | WDR43 | 74.891 | 2 | 52.127686 | yes | 1.013E-07 | 514.30603 | 350.37161 |
| LGAMDIDTHK | 10 | Q15061 | WD repeat-containing protein 43 OS=Ho | WDR43 | 74.891 | 2 | 52.127686 | yes | 1.133E-07 | 550.77386 | 319.01013 |
| LWVLETK | 7 | Q15061 | WD repeat-containing protein 43 OS=Ho | WDR43 | 74.891 | 2 | 52.127686 | yes | 2.122E-07 | 444.76309 | 252.68495 |
| HIVEWNVQTCK | 11 | Q15061 | WD repeat-containing protein 43 OS=Ho | WDR43 | 74.891 | 2 | 52.127686 | yes | 1.092E-07 | 707.35083 | 218.77399 |
| TPVMNSEAK | 9 | Q15061 | WD repeat-containing protein 43 OS=Ho | WDR43 | 74.891 | 2 | 52.127686 | yes | 1.022E-07 | 488.74203 | 203.55971 |
| GDNSSVSSLCISPD | 16 | Q15061 | WD repeat-containing protein 43 OS=Ho | WDR43 | 74.891 | 2 | 52.127686 | yes | 3.349E-07 | 811.86993 | 162.73752 |
| LHQEYVPSAHLSG | 24 | Q15061 | WD repeat-containing protein 43 OS=Ho | WDR43 | 74.891 | 4 | 52.127686 | yes | 1.015E-07 | 682.07977 | 128.2189 |

| | | | | | | | | | | | |
|---|---|---|---|---|---|---|---|---|---|---|---|
| SGGNEVSIEER | 11 | Q15061 | WD repeat-containing protein 43 OS=Ho | WDR43 | 74.891 | 2 | 52.127686 | yes | 9.722E-08 | 588.7782 | 122.21399 |
| PIPILAAGFCSDK | 13 | Q15061 | WD repeat-containing protein 43 OS=Ho | WDR43 | 74.891 | 2 | 52.127686 | yes | 2.766E-07 | 694.86578 | 57.056862 |
| STPKPIPILAAGFC | 17 | Q15061 | WD repeat-containing protein 43 OS=Ho | WDR43 | 74.891 | 3 | 52.127686 | yes | 2.075E-06 | 601.32208 | 22.866873 |
| TTVLYECCPGYMI | 13 | Q15063 | Periostin OS=Homo sapiens OX=9606 G | POSTN | 93.314 | 2 | 46.787025 | yes | 1.47E-07 | 825.35968 | 66.859962 |
| ASSEMALYQK | 10 | Q15070 | Mitochondrial inner membrane protein O | OXA1L | 48.548 | 2 | 50.167263 | yes | 9.873E-08 | 564.27368 | 179.93719 |
| LAGDHIEYYK | 10 | Q15070 | Mitochondrial inner membrane protein O | OXA1L | 48.548 | 2 | 50.167263 | yes | 3.189E-07 | 604.80096 | 107.04997 |
| EGFLESFK | 8 | Q15070 | Mitochondrial inner membrane protein O | OXA1L | 48.548 | 2 | 50.167263 | yes | 1.617E-07 | 478.73984 | 80.835922 |
| NQLELAAR | 8 | Q15070 | Mitochondrial inner membrane protein O | OXA1L | 48.548 | 2 | 50.167263 | yes | 1.107E-07 | 457.75635 | 58.693291 |
| IHNHLPEIQK | 10 | Q15070 | Mitochondrial inner membrane protein O | OXA1L | 48.548 | 2 | 50.167263 | yes | 3.099E-06 | 614.84351 | 24.128614 |
| DNPPNIPSSSSKPK | 14 | Q15070 | Mitochondrial inner membrane protein O | OXA1L | 48.548 | 2 | 50.167263 | yes | 4.055E-07 | 734.37555 | 23.566275 |
| TSLIVHVR | 8 | Q15072 | Zinc finger protein OZF OS=Homo sapie | ZNF146 | 33.308 | 2 | 48.574028 | yes | 3.615E-07 | 462.78491 | 217.91844 |
| LFECNECGK | 9 | Q15072 | Zinc finger protein OZF OS=Homo sapie | ZNF146 | 33.308 | 2 | 48.574028 | yes | 1.128E-07 | 578.7417 | 177.20679 |
| SNLTEHEHFHTR | 12 | Q15072 | Zinc finger protein OZF OS=Homo sapie | ZNF146 | 33.308 | 2 | 48.574028 | yes | 2.525E-07 | 754.35547 | 19.850437 |
| GSTAPVGGGAFPT | 17 | Q15084 | Protein disulfide-isomerase A6 OS=Hom | PDIA6 | 48.122 | 2 | 53.125927 | yes | 1.554E-07 | 808.42554 | 228.66983 |
| GSFSEQGINEFLR | 13 | Q15084 | Protein disulfide-isomerase A6 OS=Hom | PDIA6 | 48.122 | 2 | 53.125927 | yes | 9.722E-08 | 742.36243 | 177.05441 |
| LAAVDATVNQVL | 15 | Q15084 | Protein disulfide-isomerase A6 OS=Hom | PDIA6 | 48.122 | 2 | 53.125927 | yes | 9.722E-08 | 764.4281 | 110.8474 |
| DVIELTDDSFDK | 12 | Q15084 | Protein disulfide-isomerase A6 OS=Hom | PDIA6 | 48.122 | 2 | 53.125927 | yes | 9.722E-08 | 698.82758 | 98.710571 |
| GESPVDYDGGR | 11 | Q15084 | Protein disulfide-isomerase A6 OS=Hom | PDIA6 | 48.122 | 2 | 53.125927 | yes | 1.396E-07 | 576.25183 | 78.815979 |
| AIYDFTDTVIR | 11 | Q15118 | [Pyruvate dehydrogenase (acetyl-transfer | PDK1 | 49.245 | 2 | 33.496086 | yes | 1.221E-06 | 657.34045 | 26.056808 |
| VMVALGEEDLSIK | 13 | Q15119 | [Pyruvate dehydrogenase (acetyl-transfer | PDK2 | 46.154 | 2 | 49.40109 | yes | 1.036E-07 | 702.37616 | 41.829853 |
| ISDLGGGVPLR | 11 | Q15120 | [Pyruvate dehydrogenase (acetyl-transfer | PDK3 | 46.939 | 2 | 54.33882 | yes | 2.539E-07 | 542.31146 | 289.41531 |
| EVNLLPDNLLNR | 12 | Q15120 | [Pyruvate dehydrogenase (acetyl-transfer | PDK3 | 46.939 | 2 | 54.33882 | yes | 9.722E-08 | 705.39099 | 178.64638 |
| AAPLAGFGYGLPI | 15 | Q15120 | [Pyruvate dehydrogenase (acetyl-transfer | PDK3 | 46.939 | 2 | 54.33882 | yes | 2.332E-07 | 745.41174 | 131.55779 |
| QFLDFGR | 7 | Q15120 | [Pyruvate dehydrogenase (acetyl-transfer | PDK3 | 46.939 | 2 | 54.33882 | yes | 9.722E-08 | 441.72705 | 114.36597 |
| LFNYMYSTAPR | 11 | Q15120 | [Pyruvate dehydrogenase (acetyl-transfer | PDK3 | 46.939 | 2 | 54.33882 | no | 1.362E-07 | 681.82916 | 75.626991 |
| HIGSIDPTCNVAD | 16 | Q15120 | [Pyruvate dehydrogenase (acetyl-transfer | PDK3 | 46.939 | 3 | 54.33882 | yes | 3.253E-06 | 575.62604 | 33.072765 |
| TSYMFLR | 7 | Q15120 | [Pyruvate dehydrogenase (acetyl-transfer | PDK3 | 46.939 | 2 | 54.33882 | yes | 1.215E-06 | 459.23111 | 31.511332 |
| HNDVVPTMAQGV | 16 | Q15120 | [Pyruvate dehydrogenase (acetyl-transfer | PDK3 | 46.939 | 3 | 54.33882 | yes | 1.16E-06 | 600.96979 | 28.594833 |
| AAVVPLGTWR | 10 | Q15125 | 3-beta-hydroxysteroid-Delta(8),Delta(7)-i | EBP | 26.353 | 2 | 54.104485 | yes | 1.124E-06 | 535.31128 | 298.82755 |
| HLTHAQSTLDAK | 12 | Q15125 | 3-beta-hydroxysteroid-Delta(8),Delta(7)-i | EBP | 26.353 | 2 | 54.104485 | yes | 9.722E-08 | 661.34656 | 70.2593 |
| VPLDEALQR | 9 | Q15149 | Plectin OS=Homo sapiens OX=9606 GN | PLEC | 531.791 | 2 | 57.015724 | yes | 2.318E-07 | 520.79041 | 2272.0361 |
| VPLDVACAR | 9 | Q15149 | Plectin OS=Homo sapiens OX=9606 GN | PLEC | 531.791 | 2 | 57.015724 | yes | 7.212E-07 | 500.76584 | 2036.2064 |
| LLLWSQR | 7 | Q15149 | Plectin OS=Homo sapiens OX=9606 GN | PLEC | 531.791 | 2 | 57.015724 | yes | 1.742E-07 | 458.27417 | 1779.3923 |
| WQAVLAQTDVR | 11 | Q15149 | Plectin OS=Homo sapiens OX=9606 GN | PLEC | 531.791 | 2 | 57.015724 | yes | 1.022E-07 | 643.84619 | 1730.7605 |
| VTLVQTLEIQR | 11 | Q15149 | Plectin OS=Homo sapiens OX=9606 GN | PLEC | 531.791 | 2 | 57.015724 | yes | 4.301E-07 | 650.38519 | 1629.0048 |
| VPVDVAYR | 8 | Q15149 | Plectin OS=Homo sapiens OX=9606 GN | PLEC | 531.791 | 2 | 57.015724 | no | 1.117E-07 | 459.7558 | 1597.5481 |
| EAQAVPATLPELE | 16 | Q15149 | Plectin OS=Homo sapiens OX=9606 GN | PLEC | 531.791 | 2 | 57.015724 | yes | 2.866E-07 | 834.44617 | 1583.7389 |
| GGELVYTDSEAR | 12 | Q15149 | Plectin OS=Homo sapiens OX=9606 GN | PLEC | 531.791 | 2 | 57.015724 | yes | 9.722E-08 | 648.80695 | 1500.5314 |
| VVIVDPETGK | 10 | Q15149 | Plectin OS=Homo sapiens OX=9606 GN | PLEC | 531.791 | 2 | 57.015724 | yes | 1.974E-07 | 528.80042 | 1426.7175 |

| | | | | | | | | | | |
|---|---|---|---|---|---|---|---|---|---|---|
| LLFNDVQTLK | 10 | Q15149 | Plectin OS=Homo sapiens OX=9606 GN= PLEC | 531.791 | 2 | 57.015724 | yes | 1.241E-07 | 595.84259 | 1411.7322 |
| VLADPSDDTK | 10 | Q15149 | Plectin OS=Homo sapiens OX=9606 GN= PLEC | 531.791 | 2 | 57.015724 | no | 2.321E-07 | 530.76147 | 1390.9146 |
| VLALPEPSPAAPTI | 15 | Q15149 | Plectin OS=Homo sapiens OX=9606 GN= PLEC | 531.791 | 2 | 57.015724 | yes | 1.102E-07 | 766.44574 | 1356.4946 |
| LQLEATER | 8 | Q15149 | Plectin OS=Homo sapiens OX=9606 GN= PLEC | 531.791 | 2 | 57.015724 | yes | 2.405E-07 | 480.26144 | 1327.9572 |
| LQNVQIALDYLR | 12 | Q15149 | Plectin OS=Homo sapiens OX=9606 GN= PLEC | 531.791 | 2 | 57.015724 | yes | 1.017E-07 | 723.40918 | 1295.4338 |
| AGLVGPEFHEK | 11 | Q15149 | Plectin OS=Homo sapiens OX=9606 GN= PLEC | 531.791 | 2 | 57.015724 | yes | 9.722E-08 | 592.30896 | 1206.8608 |
| NLLDEELQR | 9 | Q15149 | Plectin OS=Homo sapiens OX=9606 GN= PLEC | 531.791 | 2 | 57.015724 | yes | 1.441E-07 | 565.29602 | 1204.6306 |
| AQAEAQQPTFDAI | 14 | Q15149 | Plectin OS=Homo sapiens OX=9606 GN= PLEC | 531.791 | 2 | 57.015724 | yes | 1.321E-07 | 773.38641 | 1147.5146 |
| LQLEACETR | 9 | Q15149 | Plectin OS=Homo sapiens OX=9606 GN= PLEC | 531.791 | 2 | 57.015724 | yes | 1.223E-07 | 560.27679 | 1109.2906 |
| LFNAIIHR | 8 | Q15149 | Plectin OS=Homo sapiens OX=9606 GN= PLEC | 531.791 | 2 | 57.015724 | yes | 2.16E-07 | 492.29291 | 1096.8695 |
| GGAEGELQALR | 11 | Q15149 | Plectin OS=Homo sapiens OX=9606 GN= PLEC | 531.791 | 2 | 57.015724 | yes | 1.13E-07 | 550.78839 | 1091.8148 |
| HISDLYEDLR | 10 | Q15149 | Plectin OS=Homo sapiens OX=9606 GN= PLEC | 531.791 | 2 | 57.015724 | yes | 2.664E-07 | 630.81458 | 1062.3651 |
| EQLQLLEEQHR | 11 | Q15149 | Plectin OS=Homo sapiens OX=9606 GN= PLEC | 531.791 | 2 | 57.015724 | yes | 1.145E-07 | 711.87042 | 1036.8026 |
| GLVEDTLR | 8 | Q15149 | Plectin OS=Homo sapiens OX=9606 GN= PLEC | 531.791 | 2 | 57.015724 | yes | 2.342E-07 | 451.75073 | 1029.0494 |
| ELIPTEEALR | 10 | Q15149 | Plectin OS=Homo sapiens OX=9606 GN= PLEC | 531.791 | 2 | 57.015724 | yes | 2.47E-07 | 585.8219 | 1017.4734 |
| GIYQSLEGAVQAG | 16 | Q15149 | Plectin OS=Homo sapiens OX=9606 GN= PLEC | 531.791 | 2 | 57.015724 | yes | 1.065E-07 | 831.44647 | 1015.9321 |
| SLLAWQSLR | 9 | Q15149 | Plectin OS=Homo sapiens OX=9606 GN= PLEC | 531.791 | 2 | 57.015724 | yes | 1.716E-07 | 537.30872 | 1011.8304 |
| LLDAQLATGGIVD | 15 | Q15149 | Plectin OS=Homo sapiens OX=9606 GN= PLEC | 531.791 | 2 | 57.015724 | yes | 9.722E-08 | 769.93048 | 990.94354 |
| VSITEAMHR | 9 | Q15149 | Plectin OS=Homo sapiens OX=9606 GN= PLEC | 531.791 | 2 | 57.015724 | yes | 1.528E-07 | 522.26874 | 964.00769 |
| SQVMDEATALQLI | 13 | Q15149 | Plectin OS=Homo sapiens OX=9606 GN= PLEC | 531.791 | 2 | 57.015724 | yes | 9.85E-08 | 731.37195 | 963.25677 |
| LEQLFQDEVAK | 11 | Q15149 | Plectin OS=Homo sapiens OX=9606 GN= PLEC | 531.791 | 2 | 57.015724 | yes | 1.549E-07 | 660.3457 | 962.60315 |
| LVASMEEAR | 9 | Q15149 | Plectin OS=Homo sapiens OX=9606 GN= PLEC | 531.791 | 2 | 57.015724 | yes | 2.836E-07 | 503.25531 | 960.95209 |
| GYFSEEMNR | 9 | Q15149 | Plectin OS=Homo sapiens OX=9606 GN= PLEC | 531.791 | 2 | 57.015724 | yes | 9.722E-08 | 566.74005 | 957.41187 |
| LAEVEAALEK | 10 | Q15149 | Plectin OS=Homo sapiens OX=9606 GN= PLEC | 531.791 | 2 | 57.015724 | yes | 3.684E-07 | 536.79785 | 955.69446 |
| LAEDEAFQR | 9 | Q15149 | Plectin OS=Homo sapiens OX=9606 GN= PLEC | 531.791 | 2 | 57.015724 | yes | 2.39E-07 | 539.76184 | 937.95355 |
| LLDAQLSTGGIVDI | 16 | Q15149 | Plectin OS=Homo sapiens OX=9606 GN= PLEC | 531.791 | 2 | 57.015724 | yes | 1.084E-07 | 807.44086 | 931.89575 |
| APVPASELLASGV | 16 | Q15149 | Plectin OS=Homo sapiens OX=9606 GN= PLEC | 531.791 | 2 | 57.015724 | yes | 2.81E-07 | 783.94611 | 930.37994 |
| GYFDEEMNR | 9 | Q15149 | Plectin OS=Homo sapiens OX=9606 GN= PLEC | 531.791 | 2 | 57.015724 | no | 9.722E-08 | 580.73749 | 920.36682 |
| GTQGAEEVLR | 10 | Q15149 | Plectin OS=Homo sapiens OX=9606 GN= PLEC | 531.791 | 2 | 57.015724 | yes | 1.043E-07 | 530.27509 | 902.59106 |
| GLVGPELHDR | 10 | Q15149 | Plectin OS=Homo sapiens OX=9606 GN= PLEC | 531.791 | 2 | 57.015724 | yes | 1.953E-07 | 546.79346 | 887.64905 |
| LSVAAQEAAR | 10 | Q15149 | Plectin OS=Homo sapiens OX=9606 GN= PLEC | 531.791 | 2 | 57.015724 | yes | 1.362E-07 | 508.28018 | 882.84955 |
| AQLEPVASPAK | 11 | Q15149 | Plectin OS=Homo sapiens OX=9606 GN= PLEC | 531.791 | 2 | 57.015724 | yes | 9.722E-08 | 555.81134 | 877.14478 |
| LPLLAVCDYK | 10 | Q15149 | Plectin OS=Homo sapiens OX=9606 GN= PLEC | 531.791 | 2 | 57.015724 | yes | 1.183E-07 | 596.32574 | 875.00348 |
| LSVYAALQR | 9 | Q15149 | Plectin OS=Homo sapiens OX=9606 GN= PLEC | 531.791 | 2 | 57.015724 | yes | 5.259E-07 | 510.79547 | 872.04736 |
| AELELELGR | 9 | Q15149 | Plectin OS=Homo sapiens OX=9606 GN= PLEC | 531.791 | 2 | 57.015724 | yes | 1.205E-07 | 515.28241 | 870.23218 |
| AQVEQELTTLR | 11 | Q15149 | Plectin OS=Homo sapiens OX=9606 GN= PLEC | 531.791 | 2 | 57.015724 | yes | 1.073E-07 | 644.34882 | 858.8667 |
| SLQEEHVAVAQLF | 13 | Q15149 | Plectin OS=Homo sapiens OX=9606 GN= PLEC | 531.791 | 2 | 57.015724 | yes | 9.79E-08 | 740.39935 | 428.16782 |
| SLQEEHVAVAQLF | 13 | Q15149 | Plectin OS=Homo sapiens OX=9606 GN= PLEC | 531.791 | 2 | 57.015724 | yes | 1.709E-07 | 493.93533 | 428.16782 |
| SLQEEHVAVAQLF | 13 | Q15149 | Plectin OS=Homo sapiens OX=9606 GN= PLEC | 531.791 | 3 | 57.015724 | yes | 9.79E-08 | 740.39935 | 428.16782 |

| | | | | | | | | | | |
|---|---|---|---|---|---|---|---|---|---|---|
| SLQEEHVAVAQLF | 13 | Q15149 | Plectin OS=Homo sapiens OX=9606 GN= PLEC | 531.791 | 3 | 57.015724 | yes | 1.709E-07 | 493.93533 | 428.16782 |
| SQVEEELFSVR | 11 | Q15149 | Plectin OS=Homo sapiens OX=9606 GN= PLEC | 531.791 | 2 | 57.015724 | yes | 9.809E-08 | 661.83295 | 853.86621 |
| GLLSAEVAR | 9 | Q15149 | Plectin OS=Homo sapiens OX=9606 GN= PLEC | 531.791 | 2 | 57.015724 | yes | 1.562E-07 | 458.26654 | 840.75317 |
| EGLTSIEEVTK | 11 | Q15149 | Plectin OS=Homo sapiens OX=9606 GN= PLEC | 531.791 | 2 | 57.015724 | yes | 1.002E-07 | 603.31665 | 834.69122 |
| AFCGFEDPR | 9 | Q15149 | Plectin OS=Homo sapiens OX=9606 GN= PLEC | 531.791 | 2 | 57.015724 | yes | 1.542E-07 | 549.7373 | 834.24481 |
| AALAHSEEVTASQ | 18 | Q15149 | Plectin OS=Homo sapiens OX=9606 GN= PLEC | 531.791 | 3 | 57.015724 | yes | 9.722E-08 | 595.31104 | 829.6283 |
| QQGLASYDYVR | 11 | Q15149 | Plectin OS=Homo sapiens OX=9606 GN= PLEC | 531.791 | 2 | 57.015724 | yes | 1.265E-07 | 650.32001 | 802.34808 |
| LTAEDLFEAR | 10 | Q15149 | Plectin OS=Homo sapiens OX=9606 GN= PLEC | 531.791 | 2 | 57.015724 | yes | 1.202E-07 | 582.7984 | 796.14783 |
| LHVAILER | 8 | Q15149 | Plectin OS=Homo sapiens OX=9606 GN= PLEC | 531.791 | 2 | 57.015724 | yes | 1.1E-07 | 475.79272 | 795.55811 |
| SIQEELQQLR | 10 | Q15149 | Plectin OS=Homo sapiens OX=9606 GN= PLEC | 531.791 | 2 | 57.015724 | yes | 9.722E-08 | 622.33569 | 782.92139 |
| ELYQQLQR | 8 | Q15149 | Plectin OS=Homo sapiens OX=9606 GN= PLEC | 531.791 | 2 | 57.015724 | yes | 3.637E-07 | 539.28802 | 755.95917 |
| FAEQTLR | 7 | Q15149 | Plectin OS=Homo sapiens OX=9606 GN= PLEC | 531.791 | 2 | 57.015724 | yes | 2.421E-07 | 432.73233 | 727.54913 |
| HPQGEQMYR | 9 | Q15149 | Plectin OS=Homo sapiens OX=9606 GN= PLEC | 531.791 | 2 | 57.015724 | yes | 1.192E-07 | 573.26147 | 724.64691 |
| MQAVQEATR | 9 | Q15149 | Plectin OS=Homo sapiens OX=9606 GN= PLEC | 531.791 | 2 | 57.015724 | yes | 1.047E-07 | 517.25836 | 708.80695 |
| QLAEEDLAQQR | 11 | Q15149 | Plectin OS=Homo sapiens OX=9606 GN= PLEC | 531.791 | 2 | 57.015724 | yes | 9.903E-08 | 650.82825 | 705.7226 |
| QLLEEELAR | 9 | Q15149 | Plectin OS=Homo sapiens OX=9606 GN= PLEC | 531.791 | 2 | 57.015724 | yes | 9.722E-08 | 550.80096 | 705.68787 |
| LQEAGILSAEELQF | 14 | Q15149 | Plectin OS=Homo sapiens OX=9606 GN= PLEC | 531.791 | 2 | 57.015724 | yes | 2.193E-07 | 778.91754 | 701.00244 |
| GLHQSIEEFR | 10 | Q15149 | Plectin OS=Homo sapiens OX=9606 GN= PLEC | 531.791 | 2 | 57.015724 | yes | 1.934E-07 | 608.30945 | 699.77142 |
| AQLMADFQAGR | 11 | Q15149 | Plectin OS=Homo sapiens OX=9606 GN= PLEC | 531.791 | 2 | 57.015724 | yes | 9.909E-08 | 604.29803 | 696.55396 |
| EMSVYEAYR | 9 | Q15149 | Plectin OS=Homo sapiens OX=9606 GN= PLEC | 531.791 | 2 | 57.015724 | yes | 1.483E-07 | 574.25806 | 676.48602 |
| LEDLLQDAQDEK | 12 | Q15149 | Plectin OS=Homo sapiens OX=9606 GN= PLEC | 531.791 | 2 | 57.015724 | yes | 9.722E-08 | 708.84625 | 674.69952 |
| VSIYEAMR | 8 | Q15149 | Plectin OS=Homo sapiens OX=9606 GN= PLEC | 531.791 | 2 | 57.015724 | yes | 1.655E-07 | 484.74713 | 651.94012 |
| LLEAAAQSTK | 10 | Q15149 | Plectin OS=Homo sapiens OX=9606 GN= PLEC | 531.791 | 2 | 57.015724 | yes | 9.722E-08 | 516.29022 | 651.19666 |
| TLKPEEQR | 8 | Q15149 | Plectin OS=Homo sapiens OX=9606 GN= PLEC | 531.791 | 2 | 57.015724 | yes | 2.386E-07 | 500.77472 | 646.39453 |
| LYVHEAVK | 8 | Q15149 | Plectin OS=Homo sapiens OX=9606 GN= PLEC | 531.791 | 2 | 57.015724 | yes | 1.459E-07 | 479.77145 | 639.61707 |
| EGVVGPELHHK | 11 | Q15149 | Plectin OS=Homo sapiens OX=9606 GN= PLEC | 531.791 | 2 | 57.015724 | yes | 1.113E-07 | 601.31982 | 599.00671 |
| VPDVQDGVR | 9 | Q15149 | Plectin OS=Homo sapiens OX=9606 GN= PLEC | 531.791 | 2 | 57.015724 | yes | 1.836E-07 | 492.75909 | 578.55115 |
| LQAEEAER | 8 | Q15149 | Plectin OS=Homo sapiens OX=9606 GN= PLEC | 531.791 | 2 | 57.015724 | yes | 1.296E-07 | 473.23544 | 574.95282 |
| SLVPAAELLESR | 12 | Q15149 | Plectin OS=Homo sapiens OX=9606 GN= PLEC | 531.791 | 2 | 57.015724 | yes | 1.796E-07 | 642.86151 | 570.36316 |
| SYVDPSTDER | 10 | Q15149 | Plectin OS=Homo sapiens OX=9606 GN= PLEC | 531.791 | 2 | 57.015724 | yes | 1.033E-07 | 584.75946 | 568.96765 |
| AGVVGPELHEQLL | 17 | Q15149 | Plectin OS=Homo sapiens OX=9606 GN= PLEC | 531.791 | 3 | 57.015724 | yes | 1.066E-07 | 592.98785 | 559.35077 |
| LQLEETDHQK | 10 | Q15149 | Plectin OS=Homo sapiens OX=9606 GN= PLEC | 531.791 | 2 | 57.015724 | yes | 9.722E-08 | 620.81201 | 558.34796 |
| IISLETYNLLR | 11 | Q15149 | Plectin OS=Homo sapiens OX=9606 GN= PLEC | 531.791 | 2 | 57.015724 | yes | 1.046E-07 | 667.88757 | 547.9173 |
| TPVEVPVGGFK | 11 | Q15149 | Plectin OS=Homo sapiens OX=9606 GN= PLEC | 531.791 | 2 | 57.015724 | yes | 9.722E-08 | 565.31622 | 545.68213 |
| AEMEVLLASK | 10 | Q15149 | Plectin OS=Homo sapiens OX=9606 GN= PLEC | 531.791 | 2 | 57.015724 | yes | 1.707E-07 | 545.79425 | 543.867 |
| LPVEVAYK | 8 | Q15149 | Plectin OS=Homo sapiens OX=9606 GN= PLEC | 531.791 | 2 | 57.015724 | yes | 3.988E-07 | 459.76837 | 537.16748 |
| LLEAQIATGGIIDPI | 19 | Q15149 | Plectin OS=Homo sapiens OX=9606 GN= PLEC | 531.791 | 3 | 57.015724 | yes | 1.418E-07 | 683.69189 | 533.32135 |
| QAEVELASR | 9 | Q15149 | Plectin OS=Homo sapiens OX=9606 GN= PLEC | 531.791 | 2 | 57.015724 | yes | 1.499E-07 | 501.76437 | 525.72162 |
| MGIVGPEFK | 9 | Q15149 | Plectin OS=Homo sapiens OX=9606 GN= PLEC | 531.791 | 2 | 57.015724 | yes | 2.97E-07 | 489.25986 | 524.94104 |

| | | | | | | | | | | |
|---|---|---|---|---|---|---|---|---|---|---|
| QITMEELVR | 9 | Q15149 | Plectin OS=Homo sapiens OX=9606 GN= PLEC | 531.791 | 2 | 57.015724 | yes | 9.722E-08 | 559.79736 | 518.46826 |
| VPPGYHPLDVEK | 12 | Q15149 | Plectin OS=Homo sapiens OX=9606 GN= PLEC | 531.791 | 2 | 57.015724 | yes | 9.844E-07 | 675.85626 | 257.77328 |
| VPPGYHPLDVEK | 12 | Q15149 | Plectin OS=Homo sapiens OX=9606 GN= PLEC | 531.791 | 2 | 57.015724 | yes | 1.543E-06 | 450.90659 | 257.77328 |
| VPPGYHPLDVEK | 12 | Q15149 | Plectin OS=Homo sapiens OX=9606 GN= PLEC | 531.791 | 3 | 57.015724 | yes | 9.844E-07 | 675.85626 | 257.77328 |
| VPPGYHPLDVEK | 12 | Q15149 | Plectin OS=Homo sapiens OX=9606 GN= PLEC | 531.791 | 3 | 57.015724 | yes | 1.543E-06 | 450.90659 | 257.77328 |
| QLAEGTAQQR | 10 | Q15149 | Plectin OS=Homo sapiens OX=9606 GN= PLEC | 531.791 | 2 | 57.015724 | yes | 1.082E-07 | 551.28601 | 491.1611 |
| LAAIGEATR | 9 | Q15149 | Plectin OS=Homo sapiens OX=9606 GN= PLEC | 531.791 | 2 | 57.015724 | yes | 9.934E-08 | 451.25873 | 487.72787 |
| QLQLAQEAAQK | 11 | Q15149 | Plectin OS=Homo sapiens OX=9606 GN= PLEC | 531.791 | 2 | 57.015724 | yes | 1.72E-07 | 614.33826 | 473.82013 |
| MQEEVVR | 7 | Q15149 | Plectin OS=Homo sapiens OX=9606 GN= PLEC | 531.791 | 2 | 57.015724 | yes | 4.873E-07 | 445.72366 | 452.05566 |
| LQAEEVAQQK | 10 | Q15149 | Plectin OS=Homo sapiens OX=9606 GN= PLEC | 531.791 | 2 | 57.015724 | yes | 1.241E-07 | 572.30383 | 443.87668 |
| SELELTLGK | 9 | Q15149 | Plectin OS=Homo sapiens OX=9606 GN= PLEC | 531.791 | 2 | 57.015724 | yes | 1.493E-07 | 495.27933 | 434.08588 |
| ELAEEAAR | 8 | Q15149 | Plectin OS=Homo sapiens OX=9606 GN= PLEC | 531.791 | 2 | 57.015724 | yes | 9.819E-08 | 444.7247 | 428.03983 |
| LKTEAEIALK | 10 | Q15149 | Plectin OS=Homo sapiens OX=9606 GN= PLEC | 531.791 | 2 | 57.015724 | yes | 2.539E-07 | 558.33716 | 423.83728 |
| ESADPLGAWLQDA | 14 | Q15149 | Plectin OS=Homo sapiens OX=9606 GN= PLEC | 531.791 | 2 | 57.015724 | yes | 1.622E-07 | 764.87317 | 413.67804 |
| ALQALEELR | 9 | Q15149 | Plectin OS=Homo sapiens OX=9606 GN= PLEC | 531.791 | 2 | 57.015724 | yes | 1.467E-07 | 521.79822 | 404.42639 |
| ENAAYFQFFSDVR | 13 | Q15149 | Plectin OS=Homo sapiens OX=9606 GN= PLEC | 531.791 | 2 | 57.015724 | yes | 2.041E-07 | 797.37024 | 384.66904 |
| FLQEEAEK | 8 | Q15149 | Plectin OS=Homo sapiens OX=9606 GN= PLEC | 531.791 | 2 | 57.015724 | yes | 1.356E-07 | 497.24802 | 380.02905 |
| GCLDEETSR | 9 | Q15149 | Plectin OS=Homo sapiens OX=9606 GN= PLEC | 531.791 | 2 | 57.015724 | yes | 6.302E-07 | 533.72711 | 373.73233 |
| LSYTQLLR | 8 | Q15149 | Plectin OS=Homo sapiens OX=9606 GN= PLEC | 531.791 | 2 | 57.015724 | yes | 2.295E-07 | 497.29001 | 371.79361 |
| HKPLLIDMNK | 10 | Q15149 | Plectin OS=Homo sapiens OX=9606 GN= PLEC | 531.791 | 2 | 57.015724 | yes | 5.319E-07 | 604.8446 | 370.63895 |
| QLAAEEER | 8 | Q15149 | Plectin OS=Homo sapiens OX=9606 GN= PLEC | 531.791 | 2 | 57.015724 | yes | 4.271E-07 | 473.23544 | 370.1813 |
| LTVDEAVR | 8 | Q15149 | Plectin OS=Homo sapiens OX=9606 GN= PLEC | 531.791 | 2 | 57.015724 | yes | 2.05E-07 | 451.75073 | 365.62857 |
| AEAELLQQQK | 10 | Q15149 | Plectin OS=Homo sapiens OX=9606 GN= PLEC | 531.791 | 2 | 57.015724 | yes | 2.085E-07 | 579.31171 | 360.70316 |
| LTVNEAVK | 8 | Q15149 | Plectin OS=Homo sapiens OX=9606 GN= PLEC | 531.791 | 2 | 57.015724 | yes | 1E-07 | 437.25565 | 360.16815 |
| QTLSIYQALK | 10 | Q15149 | Plectin OS=Homo sapiens OX=9606 GN= PLEC | 531.791 | 2 | 57.015724 | yes | 9.722E-08 | 582.83478 | 357.75046 |
| KESYSALMR | 9 | Q15149 | Plectin OS=Homo sapiens OX=9606 GN= PLEC | 531.791 | 2 | 57.015724 | yes | 1.58E-07 | 542.77643 | 356.32727 |
| IIITVVEEQEQK | 12 | Q15149 | Plectin OS=Homo sapiens OX=9606 GN= PLEC | 531.791 | 2 | 57.015724 | yes | 9.968E-08 | 714.90088 | 354.79352 |
| AVVQLKPR | 8 | Q15149 | Plectin OS=Homo sapiens OX=9606 GN= PLEC | 531.791 | 2 | 57.015724 | yes | 2.881E-07 | 455.79526 | 352.02402 |
| LSVYQAMK | 8 | Q15149 | Plectin OS=Homo sapiens OX=9606 GN= PLEC | 531.791 | 2 | 57.015724 | yes | 2.115E-07 | 470.25204 | 349.69794 |
| QYDIDDAIAK | 10 | Q15149 | Plectin OS=Homo sapiens OX=9606 GN= PLEC | 531.791 | 2 | 57.015724 | yes | 4.891E-07 | 576.28259 | 348.45218 |
| SLAQAEAEK | 9 | Q15149 | Plectin OS=Homo sapiens OX=9606 GN= PLEC | 531.791 | 2 | 57.015724 | yes | 2.325E-06 | 473.74564 | 346.0094 |
| LGFHLPLEVAYQR | 13 | Q15149 | Plectin OS=Homo sapiens OX=9606 GN= PLEC | 531.791 | 2 | 57.015724 | yes | 2.482E-07 | 771.92499 | 345.71423 |
| GWLYYEAGQR | 10 | Q15149 | Plectin OS=Homo sapiens OX=9606 GN= PLEC | 531.791 | 2 | 57.015724 | yes | 3.055E-07 | 621.79877 | 339.8403 |
| QLAEEDAAR | 9 | Q15149 | Plectin OS=Homo sapiens OX=9606 GN= PLEC | 531.791 | 2 | 57.015724 | yes | 1.558E-07 | 501.74619 | 328.01166 |
| FPVTDAVNK | 9 | Q15149 | Plectin OS=Homo sapiens OX=9606 GN= PLEC | 531.791 | 2 | 57.015724 | yes | 1.845E-07 | 495.76639 | 318.0831 |
| QEQALLEEIER | 11 | Q15149 | Plectin OS=Homo sapiens OX=9606 GN= PLEC | 531.791 | 2 | 57.015724 | yes | 9.722E-08 | 679.3515 | 315.87656 |
| DPYTGQSVSLFQA | 15 | Q15149 | Plectin OS=Homo sapiens OX=9606 GN= PLEC | 531.791 | 2 | 57.015724 | yes | 1.022E-07 | 827.4278 | 310.55847 |
| NDDIADGNPK | 10 | Q15149 | Plectin OS=Homo sapiens OX=9606 GN= PLEC | 531.791 | 2 | 57.015724 | yes | 3.355E-07 | 529.74109 | 305.32584 |
| DSQDAGGFGPEDR | 13 | Q15149 | Plectin OS=Homo sapiens OX=9606 GN= PLEC | 531.791 | 2 | 57.015724 | yes | 9.722E-08 | 675.78149 | 304.15457 |

| | | | | | | | | | | |
|---|---|---|---|---|---|---|---|---|---|---|
| QEELYSELQAR | 11 | Q15149 | Plectin OS=Homo sapiens OX=9606 GN= PLEC | 531.791 | 2 | 57.015724 | yes | 1.92E-07 | 683.33588 | 302.66226 |
| LSIYNALK | 8 | Q15149 | Plectin OS=Homo sapiens OX=9606 GN= PLEC | 531.791 | 2 | 57.015724 | yes | 1.048E-07 | 461.27383 | 299.02237 |
| EAIAELER | 8 | Q15149 | Plectin OS=Homo sapiens OX=9606 GN= PLEC | 531.791 | 2 | 57.015724 | yes | 2.886E-07 | 465.74817 | 288.40506 |
| SMVEEGTGLR | 10 | Q15149 | Plectin OS=Homo sapiens OX=9606 GN= PLEC | 531.791 | 2 | 57.015724 | yes | 1.076E-07 | 539.76349 | 286.91504 |
| LKAEAELLQQQK | 12 | Q15149 | Plectin OS=Homo sapiens OX=9606 GN= PLEC | 531.791 | 2 | 57.015724 | yes | 9.823E-08 | 699.90118 | 286.7616 |
| QLEMSAEAER | 10 | Q15149 | Plectin OS=Homo sapiens OX=9606 GN= PLEC | 531.791 | 2 | 57.015724 | yes | 1.386E-07 | 582.27167 | 275.55093 |
| RQELEAELAK | 10 | Q15149 | Plectin OS=Homo sapiens OX=9606 GN= PLEC | 531.791 | 2 | 57.015724 | yes | 1.701E-07 | 593.82495 | 262.86786 |
| SDEGQLSPATR | 11 | Q15149 | Plectin OS=Homo sapiens OX=9606 GN= PLEC | 531.791 | 2 | 57.015724 | yes | 9.722E-08 | 580.78076 | 262.1062 |
| GPLPTEEQR | 9 | Q15149 | Plectin OS=Homo sapiens OX=9606 GN= PLEC | 531.791 | 2 | 57.015724 | yes | 2.503E-06 | 513.76434 | 261.64505 |
| ELMWLNEK | 8 | Q15149 | Plectin OS=Homo sapiens OX=9606 GN= PLEC | 531.791 | 2 | 57.015724 | yes | 9.722E-08 | 531.76807 | 256.53162 |
| ELAEQELEK | 9 | Q15149 | Plectin OS=Homo sapiens OX=9606 GN= PLEC | 531.791 | 2 | 57.015724 | yes | 1.211E-07 | 544.77716 | 256.30014 |
| QQEELLAEENQR | 12 | Q15149 | Plectin OS=Homo sapiens OX=9606 GN= PLEC | 531.791 | 2 | 57.015724 | yes | 1.432E-07 | 743.86023 | 255.84274 |
| AALEEVER | 8 | Q15149 | Plectin OS=Homo sapiens OX=9606 GN= PLEC | 531.791 | 2 | 57.015724 | yes | 1.532E-07 | 458.74036 | 252.91005 |
| MVEGYQGLR | 9 | Q15149 | Plectin OS=Homo sapiens OX=9606 GN= PLEC | 531.791 | 2 | 57.015724 | yes | 3.131E-07 | 526.76331 | 242.6263 |
| VPVDVAYQR | 9 | Q15149 | Plectin OS=Homo sapiens OX=9606 GN= PLEC | 531.791 | 2 | 57.015724 | no | 2.646E-07 | 523.7851 | 235.55266 |
| DYELQLVTYK | 10 | Q15149 | Plectin OS=Homo sapiens OX=9606 GN= PLEC | 531.791 | 2 | 57.015724 | yes | 9.722E-08 | 636.32953 | 230.49953 |
| LAAEQELIR | 9 | Q15149 | Plectin OS=Homo sapiens OX=9606 GN= PLEC | 531.791 | 2 | 57.015724 | yes | 2.044E-07 | 521.79822 | 215.41669 |
| SLSAIYLEK | 9 | Q15149 | Plectin OS=Homo sapiens OX=9606 GN= PLEC | 531.791 | 2 | 57.015724 | yes | 1.866E-07 | 512.28967 | 214.80814 |
| SKEQAELEAAR | 11 | Q15149 | Plectin OS=Homo sapiens OX=9606 GN= PLEC | 531.791 | 2 | 57.015724 | yes | 1.049E-07 | 616.3175 | 207.87799 |
| MSAAQALK | 8 | Q15149 | Plectin OS=Homo sapiens OX=9606 GN= PLEC | 531.791 | 2 | 57.015724 | yes | 3.108E-07 | 410.2233 | 207.57175 |
| VDGAEWGVDLPS | 21 | Q15149 | Plectin OS=Homo sapiens OX=9606 GN= PLEC | 531.791 | 3 | 57.015724 | yes | 2.886E-07 | 741.36609 | 205.14728 |
| DVAEVDTVR | 9 | Q15149 | Plectin OS=Homo sapiens OX=9606 GN= PLEC | 531.791 | 2 | 57.015724 | yes | 2.035E-07 | 502.25638 | 203.18169 |
| NLVDNITGQR | 10 | Q15149 | Plectin OS=Homo sapiens OX=9606 GN= PLEC | 531.791 | 2 | 57.015724 | yes | 1.107E-07 | 565.30164 | 199.43066 |
| SLESLHSFVAAT | 14 | Q15149 | Plectin OS=Homo sapiens OX=9606 GN= PLEC | 531.791 | 2 | 57.015724 | yes | 1.213E-07 | 730.89081 | 198.04636 |
| CVEDPETGLR | 10 | Q15149 | Plectin OS=Homo sapiens OX=9606 GN= PLEC | 531.791 | 2 | 57.015724 | yes | 1.828E-07 | 588.27167 | 196.7215 |
| AGVAAPATQVAQ | 20 | Q15149 | Plectin OS=Homo sapiens OX=9606 GN= PLEC | 531.791 | 3 | 57.015724 | yes | 9.722E-08 | 665.70422 | 195.62834 |
| RPELEDSTLR | 10 | Q15149 | Plectin OS=Homo sapiens OX=9606 GN= PLEC | 531.791 | 2 | 57.015724 | yes | 2.383E-07 | 608.32001 | 193.54956 |
| GANVIAGVWLEE | 16 | Q15149 | Plectin OS=Homo sapiens OX=9606 GN= PLEC | 531.791 | 2 | 57.015724 | yes | 9.722E-08 | 821.43341 | 176.06213 |
| VAGMLMPR | 8 | Q15149 | Plectin OS=Homo sapiens OX=9606 GN= PLEC | 531.791 | 2 | 57.015724 | yes | 3.052E-07 | 437.7355 | 174.66426 |
| LPLDKEPAR | 9 | Q15149 | Plectin OS=Homo sapiens OX=9606 GN= PLEC | 531.791 | 2 | 57.015724 | yes | 1.104E-07 | 519.80072 | 166.54657 |
| RVVIVDPETGK | 11 | Q15149 | Plectin OS=Homo sapiens OX=9606 GN= PLEC | 531.791 | 2 | 57.015724 | yes | 1.139E-07 | 606.85095 | 163.10396 |
| RQEQIQAMPLADS | 17 | Q15149 | Plectin OS=Homo sapiens OX=9606 GN= PLEC | 531.791 | 3 | 57.015724 | yes | 2.204E-07 | 647.67053 | 151.18842 |
| LAQGHTTVDELA | 13 | Q15149 | Plectin OS=Homo sapiens OX=9606 GN= PLEC | 531.791 | 2 | 57.015724 | yes | 9.722E-08 | 705.87042 | 150.70084 |
| LISLFQAMK | 9 | Q15149 | Plectin OS=Homo sapiens OX=9606 GN= PLEC | 531.791 | 2 | 57.015724 | yes | 3.824E-07 | 525.80444 | 147.7196 |
| QQQQMEQER | 9 | Q15149 | Plectin OS=Homo sapiens OX=9606 GN= PLEC | 531.791 | 2 | 57.015724 | yes | 2.93E-07 | 602.7724 | 141.98351 |
| QSSEAEIQAK | 10 | Q15149 | Plectin OS=Homo sapiens OX=9606 GN= PLEC | 531.791 | 2 | 57.015724 | yes | 9.854E-08 | 545.7724 | 138.87552 |
| AKLEQLFQDEVA | 13 | Q15149 | Plectin OS=Homo sapiens OX=9606 GN= PLEC | 531.791 | 2 | 57.015724 | yes | 9.722E-08 | 759.91174 | 69.053726 |
| AKLEQLFQDEVA | 13 | Q15149 | Plectin OS=Homo sapiens OX=9606 GN= PLEC | 531.791 | 2 | 57.015724 | yes | 4.914E-06 | 506.9436 | 69.053726 |
| AKLEQLFQDEVA | 13 | Q15149 | Plectin OS=Homo sapiens OX=9606 GN= PLEC | 531.791 | 3 | 57.015724 | yes | 9.722E-08 | 759.91174 | 69.053726 |

| | | | | | | | | | | |
|---|---|---|---|---|---|---|---|---|---|---|
| AKLEQLFQDEVAK | 13 | Q15149 | Plectin OS=Homo sapiens OX=9606 GN= PLEC | 531.791 | 3 | 57.015724 | yes | 4.914E-06 | 506.9436 | 69.053726 |
| LLEAQIATGGVIDI | 19 | Q15149 | Plectin OS=Homo sapiens OX=9606 GN= PLEC | 531.791 | 3 | 57.015724 | no | 1.062E-07 | 671.70074 | 136.196 |
| EEAAVDAQQQK | 11 | Q15149 | Plectin OS=Homo sapiens OX=9606 GN= PLEC | 531.791 | 2 | 57.015724 | yes | 1.049E-06 | 608.79382 | 125.93382 |
| SAEAELQSK | 9 | Q15149 | Plectin OS=Homo sapiens OX=9606 GN= PLEC | 531.791 | 2 | 57.015724 | yes | 1.496E-07 | 481.7431 | 117.66267 |
| RQVEEEILALK | 11 | Q15149 | Plectin OS=Homo sapiens OX=9606 GN= PLEC | 531.791 | 2 | 57.015724 | yes | 1.908E-07 | 664.38263 | 116.86418 |
| RYASGSSASLGGP | 18 | Q15149 | Plectin OS=Homo sapiens OX=9606 GN= PLEC | 531.791 | 2 | 57.015724 | yes | 1.326E-07 | 833.90521 | 108.47706 |
| KGLLSAEVAR | 10 | Q15149 | Plectin OS=Homo sapiens OX=9606 GN= PLEC | 531.791 | 2 | 57.015724 | yes | 9.722E-08 | 522.31403 | 108.23389 |
| EEEEVGFDWSDR | 12 | Q15149 | Plectin OS=Homo sapiens OX=9606 GN= PLEC | 531.791 | 2 | 57.015724 | yes | 1.313E-07 | 749.31006 | 104.91476 |
| KGWLYYEAGQR | 11 | Q15149 | Plectin OS=Homo sapiens OX=9606 GN= PLEC | 531.791 | 2 | 57.015724 | yes | 9.722E-08 | 685.84619 | 103.25593 |
| QYLHLPPEIVPASL | 16 | Q15149 | Plectin OS=Homo sapiens OX=9606 GN= PLEC | 531.791 | 3 | 57.015724 | yes | 2.739E-06 | 621.01587 | 87.62133 |
| QVEEEILALK | 10 | Q15149 | Plectin OS=Homo sapiens OX=9606 GN= PLEC | 531.791 | 2 | 57.015724 | yes | 1.161E-06 | 586.33209 | 87.560402 |
| KELIPTEEALR | 11 | Q15149 | Plectin OS=Homo sapiens OX=9606 GN= PLEC | 531.791 | 2 | 57.015724 | yes | 3.146E-07 | 649.86938 | 87.326355 |
| CDNFTSSWR | 9 | Q15149 | Plectin OS=Homo sapiens OX=9606 GN= PLEC | 531.791 | 2 | 57.015724 | yes | 1.717E-07 | 586.7431 | 82.300781 |
| TELATQEK | 8 | Q15149 | Plectin OS=Homo sapiens OX=9606 GN= PLEC | 531.791 | 2 | 57.015724 | yes | 3.446E-07 | 460.2402 | 78.764465 |
| GRLPLLAVCDYK | 12 | Q15149 | Plectin OS=Homo sapiens OX=9606 GN= PLEC | 531.791 | 2 | 57.015724 | yes | 1.065E-07 | 702.88702 | 77.058289 |
| AQAEVEGLGK | 10 | Q15149 | Plectin OS=Homo sapiens OX=9606 GN= PLEC | 531.791 | 2 | 57.015724 | yes | 2.73E-06 | 501.26672 | 75.266693 |
| LKAEATEAAR | 10 | Q15149 | Plectin OS=Homo sapiens OX=9606 GN= PLEC | 531.791 | 2 | 57.015724 | yes | 2.159E-06 | 530.29327 | 75.244797 |
| LEAQHQALVTLWI | 20 | Q15149 | Plectin OS=Homo sapiens OX=9606 GN= PLEC | 531.791 | 3 | 57.015724 | yes | 1.073E-07 | 799.7547 | 73.77169 |
| EQAELEAAR | 9 | Q15149 | Plectin OS=Homo sapiens OX=9606 GN= PLEC | 531.791 | 2 | 57.015724 | yes | 7.323E-07 | 508.754 | 69.169395 |
| QEELQQLEQQR | 11 | Q15149 | Plectin OS=Homo sapiens OX=9606 GN= PLEC | 531.791 | 2 | 57.015724 | yes | 9.722E-08 | 714.85748 | 68.005714 |
| SLAAEEEAAR | 10 | Q15149 | Plectin OS=Homo sapiens OX=9606 GN= PLEC | 531.791 | 2 | 57.015724 | yes | 2.782E-07 | 523.75928 | 64.646553 |
| DDGTGQLLLPLSD | 15 | Q15149 | Plectin OS=Homo sapiens OX=9606 GN= PLEC | 531.791 | 2 | 57.015724 | yes | 2.058E-07 | 785.90723 | 64.038162 |
| QAADAEMEK | 9 | Q15149 | Plectin OS=Homo sapiens OX=9606 GN= PLEC | 531.791 | 2 | 57.015724 | yes | 1.703E-06 | 496.72131 | 58.514954 |
| YASGSSASLGGPE | 17 | Q15149 | Plectin OS=Homo sapiens OX=9606 GN= PLEC | 531.791 | 2 | 57.015724 | yes | 1.961E-07 | 755.85461 | 51.791782 |
| LDLQYAK | 7 | Q15149 | Plectin OS=Homo sapiens OX=9606 GN= PLEC | 531.791 | 2 | 57.015724 | yes | 2.992E-07 | 425.73709 | 49.813057 |
| QVQVALETAQR | 11 | Q15149 | Plectin OS=Homo sapiens OX=9606 GN= PLEC | 531.791 | 2 | 57.015724 | yes | 9.269E-07 | 621.84369 | 48.181587 |
| RDVQLIR | 7 | Q15149 | Plectin OS=Homo sapiens OX=9606 GN= PLEC | 531.791 | 2 | 57.015724 | yes | 1.009E-06 | 450.27469 | 46.679459 |
| KASDSELER | 9 | Q15149 | Plectin OS=Homo sapiens OX=9606 GN= PLEC | 531.791 | 2 | 57.015724 | yes | 2.611E-06 | 517.75928 | 45.096451 |
| QALHSLELHYQAF | 15 | Q15149 | Plectin OS=Homo sapiens OX=9606 GN= PLEC | 531.791 | 3 | 57.015724 | yes | 9.728E-08 | 609.32843 | 44.160694 |
| ESYSALMR | 8 | Q15149 | Plectin OS=Homo sapiens OX=9606 GN= PLEC | 531.791 | 2 | 57.015724 | yes | 1.65E-07 | 478.72894 | 40.731384 |
| SSIAGLLLK | 9 | Q15149 | Plectin OS=Homo sapiens OX=9606 GN= PLEC | 531.791 | 2 | 57.015724 | yes | 1.297E-07 | 451.28949 | 35.44342 |
| RLTVNEAVK | 9 | Q15149 | Plectin OS=Homo sapiens OX=9606 GN= PLEC | 531.791 | 2 | 57.015724 | yes | 2.108E-05 | 515.30621 | 35.313942 |
| AIYEVLFR | 8 | Q15149 | Plectin OS=Homo sapiens OX=9606 GN= PLEC | 531.791 | 2 | 57.015724 | yes | 3.554E-06 | 505.78711 | 35.125328 |
| CRPDQLTGLSLLPI | 17 | Q15149 | Plectin OS=Homo sapiens OX=9606 GN= PLEC | 531.791 | 3 | 57.015724 | yes | 5.634E-06 | 643.01544 | 31.137014 |
| ATVSAPFGK | 9 | Q15149 | Plectin OS=Homo sapiens OX=9606 GN= PLEC | 531.791 | 2 | 57.015724 | yes | 5.416E-07 | 439.24252 | 27.972084 |
| KQEELQQLEQQR | 12 | Q15149 | Plectin OS=Homo sapiens OX=9606 GN= PLEC | 531.791 | 2 | 57.015724 | yes | 3.537E-07 | 778.90497 | 25.276186 |
| QSAEEQAQAR | 10 | Q15149 | Plectin OS=Homo sapiens OX=9606 GN= PLEC | 531.791 | 2 | 57.015724 | yes | 2.983E-06 | 559.26526 | 22.346853 |
| DTHDQLSEPSEVR | 13 | Q15149 | Plectin OS=Homo sapiens OX=9606 GN= PLEC | 531.791 | 2 | 57.015724 | yes | 1.064E-07 | 756.84985 | 15.120356 |
| GDECQLVGPAQPS | 16 | Q15149 | Plectin OS=Homo sapiens OX=9606 GN= PLEC | 531.791 | 3 | 57.015724 | yes | 0.0001099 | 603.61768 | 1.1901586 |

| | | | | | | | | | | | |
|---|---|---|---|---|---|---|---|---|---|---|---|
| LLGELHTLR | 9 | Q15154 | Pericentriolar material 1 protein OS=Hon | PCM1 | 228.545 | 2 | 54.20475 | yes | 2.207E-07 | 526.31659 | 193.43875 |
| INFSDLDQR | 9 | Q15154 | Pericentriolar material 1 protein OS=Hon | PCM1 | 228.545 | 2 | 54.20475 | yes | 5.825E-07 | 554.27509 | 135.46323 |
| MLQQQEQLR | 9 | Q15154 | Pericentriolar material 1 protein OS=Hon | PCM1 | 228.545 | 2 | 54.20475 | yes | 2.459E-07 | 587.30585 | 129.86751 |
| ALYALQDIVSR | 11 | Q15154 | Pericentriolar material 1 protein OS=Hon | PCM1 | 228.545 | 2 | 54.20475 | yes | 2.307E-07 | 624.85095 | 121.58841 |
| LTHLIDHLK | 9 | Q15154 | Pericentriolar material 1 protein OS=Hon | PCM1 | 228.545 | 2 | 54.20475 | yes | 1.082E-07 | 545.3244 | 102.57476 |
| LMQDLDNNSITVK | 13 | Q15154 | Pericentriolar material 1 protein OS=Hon | PCM1 | 228.545 | 2 | 54.20475 | yes | 9.722E-08 | 745.87958 | 94.158104 |
| WVSELSYVEEK | 11 | Q15154 | Pericentriolar material 1 protein OS=Hon | PCM1 | 228.545 | 2 | 54.20475 | yes | 2.128E-07 | 684.83771 | 90.55471 |
| QLGSILQDSLAK | 12 | Q15154 | Pericentriolar material 1 protein OS=Hon | PCM1 | 228.545 | 2 | 54.20475 | yes | 1.011E-07 | 636.86151 | 89.16967 |
| GEPAMESSQIVSR | 13 | Q15154 | Pericentriolar material 1 protein OS=Hon | PCM1 | 228.545 | 2 | 54.20475 | yes | 3.731E-07 | 695.83521 | 81.891251 |
| MVLTLTQQNDESK | 13 | Q15154 | Pericentriolar material 1 protein OS=Hon | PCM1 | 228.545 | 2 | 54.20475 | yes | 1.666E-07 | 753.87708 | 70.054764 |
| QMLNELMR | 8 | Q15154 | Pericentriolar material 1 protein OS=Hon | PCM1 | 228.545 | 2 | 54.20475 | yes | 9.722E-08 | 517.7597 | 68.632881 |
| KLPEEEVESSR | 11 | Q15154 | Pericentriolar material 1 protein OS=Hon | PCM1 | 228.545 | 2 | 54.20475 | yes | 5.942E-07 | 651.83044 | 66.66378 |
| IQALQTACPDLQLS | 24 | Q15154 | Pericentriolar material 1 protein OS=Hon | PCM1 | 228.545 | 3 | 54.20475 | yes | 2.14E-06 | 848.42462 | 44.03894 |
| EQWQEQINQLK | 11 | Q15154 | Pericentriolar material 1 protein OS=Hon | PCM1 | 228.545 | 2 | 54.20475 | yes | 9.722E-08 | 722.36499 | 43.789589 |
| LPEMEPLVPR | 10 | Q15154 | Pericentriolar material 1 protein OS=Hon | PCM1 | 228.545 | 2 | 54.20475 | yes | 3.853E-06 | 590.82336 | 43.140404 |
| QAALLALQHK | 10 | Q15154 | Pericentriolar material 1 protein OS=Hon | PCM1 | 228.545 | 2 | 54.20475 | yes | 9.311E-07 | 546.82983 | 34.06263 |
| FLESPDFQPSIAK | 13 | Q15172 | Serine/threonine-protein phosphatase 2A | PPP2R5A | 56.194 | 2 | 50.55735 | yes | 8.778E-07 | 739.87994 | 126.78888 |
| ATLNELVEYVSTN | 14 | Q15172 | Serine/threonine-protein phosphatase 2A | PPP2R5A | 56.194 | 2 | 50.55735 | yes | 5.567E-07 | 804.91504 | 93.694206 |
| KIEEPLFK | 8 | Q15172 | Serine/threonine-protein phosphatase 2A | PPP2R5A | 56.194 | 2 | 50.55735 | yes | 1.731E-06 | 502.29477 | 65.116959 |
| SQGSQAELHPLPQ | 15 | Q15172 | Serine/threonine-protein phosphatase 2A | PPP2R5A | 56.194 | 2 | 50.55735 | yes | 9.766E-08 | 816.93884 | 55.942951 |
| LFDDLTSSYK | 10 | Q15172 | Serine/threonine-protein phosphatase 2A | PPP2R5A | 56.194 | 2 | 50.55735 | yes | 1.261E-07 | 594.79279 | 52.737259 |
| CVSSSHFQVAER | 12 | Q15172 | Serine/threonine-protein phosphatase 2A | PPP2R5A | 56.194 | 2 | 50.55735 | yes | 1.189E-07 | 703.8277 | 25.183018 |
| SILCCLR | 7 | Q15185 | Prostaglandin E synthase 3 OS=Homo saj | PTGES3 | 18.698 | 2 | 53.653645 | yes | 1.037E-07 | 461.23587 | 467.15579 |
| LTFSCLGGSDNFK | 13 | Q15185 | Prostaglandin E synthase 3 OS=Homo saj | PTGES3 | 18.698 | 2 | 53.653645 | yes | 9.722E-08 | 723.34009 | 241.6481 |
| KGESGQSWPR | 10 | Q15185 | Prostaglandin E synthase 3 OS=Homo saj | PTGES3 | 18.698 | 2 | 53.653645 | yes | 1.623E-07 | 566.2807 | 56.966579 |
| LNWLSVDFNNWK | 12 | Q15185 | Prostaglandin E synthase 3 OS=Homo saj | PTGES3 | 18.698 | 2 | 53.653645 | yes | 9.722E-08 | 768.38568 | 52.125778 |
| LSDFGLCTGLK | 11 | Q15208 | Serine/threonine-protein kinase 38 OS=H | STK38 | 54.191 | 2 | 43.7623 | yes | 2.456E-07 | 605.81049 | 92.261909 |
| VELDNMPLR | 9 | Q15233 | Non-POU domain-containing octamer-bir | NONO | 54.232 | 2 | 51.91537 | yes | 1.385E-07 | 543.78424 | 32362.859 |
| VELDNMPLR | 9 | Q15233 | Non-POU domain-containing octamer-bir | NONO | 54.232 | 2 | 51.91537 | yes | 1.02E-07 | 551.78168 | 32362.859 |
| FAQPGSFEYEYAM | 14 | Q15233 | Non-POU domain-containing octamer-bir | NONO | 54.232 | 2 | 51.91537 | yes | 2.653E-07 | 848.3772 | 51386.551 |
| AGEVFIHK | 8 | Q15233 | Non-POU domain-containing octamer-bir | NONO | 54.232 | 2 | 51.91537 | yes | 3.211E-06 | 450.75052 | 44562.531 |
| MGQMAMGGAMG | 15 | Q15233 | Non-POU domain-containing octamer-bir | NONO | 54.232 | 2 | 51.91537 | yes | 1.142E-07 | 769.83838 | 11207.936 |
| MGQMAMGGAMG | 15 | Q15233 | Non-POU domain-containing octamer-bir | NONO | 54.232 | 2 | 51.91537 | yes | 9.722E-08 | 777.83582 | 11207.936 |
| MGQMAMGGAMG | 15 | Q15233 | Non-POU domain-containing octamer-bir | NONO | 54.232 | 2 | 51.91537 | yes | 1.109E-07 | 777.83582 | 11207.936 |
| MGQMAMGGAMG | 15 | Q15233 | Non-POU domain-containing octamer-bir | NONO | 54.232 | 2 | 51.91537 | yes | 1.455E-07 | 777.83582 | 11207.936 |
| MGQMAMGGAMG | 15 | Q15233 | Non-POU domain-containing octamer-bir | NONO | 54.232 | 2 | 51.91537 | yes | 1.12E-07 | 777.83582 | 11207.936 |
| MGQMAMGGAMG | 15 | Q15233 | Non-POU domain-containing octamer-bir | NONO | 54.232 | 2 | 51.91537 | yes | 2.923E-07 | 513.56134 | 11207.936 |
| MGQMAMGGAMG | 15 | Q15233 | Non-POU domain-containing octamer-bir | NONO | 54.232 | 2 | 51.91537 | yes | 9.722E-08 | 785.83331 | 11207.936 |
| MGQMAMGGAMG | 15 | Q15233 | Non-POU domain-containing octamer-bir | NONO | 54.232 | 3 | 51.91537 | yes | 1.142E-07 | 769.83838 | 11207.936 |

| | | | | | | | | | | | |
|---|---|---|---|---|---|---|---|---|---|---|---|
| MGQMAMGGAMG | 15 | Q15233 | Non-POU domain-containing octamer-bir | NONO | 54.232 | 3 | 51.91537 | yes | 9.722E-08 | 777.83582 | 11207.936 |
| MGQMAMGGAMG | 15 | Q15233 | Non-POU domain-containing octamer-bir | NONO | 54.232 | 3 | 51.91537 | yes | 1.109E-07 | 777.83582 | 11207.936 |
| MGQMAMGGAMG | 15 | Q15233 | Non-POU domain-containing octamer-bir | NONO | 54.232 | 3 | 51.91537 | yes | 1.455E-07 | 777.83582 | 11207.936 |
| MGQMAMGGAMG | 15 | Q15233 | Non-POU domain-containing octamer-bir | NONO | 54.232 | 3 | 51.91537 | yes | 1.12E-07 | 777.83582 | 11207.936 |
| MGQMAMGGAMG | 15 | Q15233 | Non-POU domain-containing octamer-bir | NONO | 54.232 | 3 | 51.91537 | yes | 2.923E-07 | 513.56134 | 11207.936 |
| MGQMAMGGAMG | 15 | Q15233 | Non-POU domain-containing octamer-bir | NONO | 54.232 | 3 | 51.91537 | yes | 9.722E-08 | 785.83331 | 11207.936 |
| EKLEMEMEAAR | 11 | Q15233 | Non-POU domain-containing octamer-bir | NONO | 54.232 | 2 | 51.91537 | yes | 1.084E-07 | 668.81543 | 11478.931 |
| EKLEMEMEAAR | 11 | Q15233 | Non-POU domain-containing octamer-bir | NONO | 54.232 | 2 | 51.91537 | yes | 9.722E-08 | 676.81287 | 11478.931 |
| EKLEMEMEAAR | 11 | Q15233 | Non-POU domain-containing octamer-bir | NONO | 54.232 | 2 | 51.91537 | yes | 1.095E-07 | 676.81287 | 11478.931 |
| EKLEMEMEAAR | 11 | Q15233 | Non-POU domain-containing octamer-bir | NONO | 54.232 | 2 | 51.91537 | yes | 2.258E-07 | 684.8103 | 11478.931 |
| AAPGAEFAPNK | 11 | Q15233 | Non-POU domain-containing octamer-bir | NONO | 54.232 | 2 | 51.91537 | yes | 2.29E-07 | 536.77472 | 23334.092 |
| AAPGAEFAPNKR | 12 | Q15233 | Non-POU domain-containing octamer-bir | NONO | 54.232 | 2 | 51.91537 | yes | 1.272E-07 | 614.82526 | 22648.443 |
| FACHSASLTVR | 11 | Q15233 | Non-POU domain-containing octamer-bir | NONO | 54.232 | 3 | 51.91537 | yes | 2.337E-07 | 416.87665 | 10745.348 |
| FACHSASLTVR | 11 | Q15233 | Non-POU domain-containing octamer-bir | NONO | 54.232 | 3 | 51.91537 | yes | 1.691E-07 | 624.81134 | 10745.348 |
| FACHSASLTVR | 11 | Q15233 | Non-POU domain-containing octamer-bir | NONO | 54.232 | 2 | 51.91537 | yes | 2.337E-07 | 416.87665 | 10745.348 |
| FACHSASLTVR | 11 | Q15233 | Non-POU domain-containing octamer-bir | NONO | 54.232 | 2 | 51.91537 | yes | 1.691E-07 | 624.81134 | 10745.348 |
| HEHQVMLMR | 9 | Q15233 | Non-POU domain-containing octamer-bir | NONO | 54.232 | 2 | 51.91537 | yes | 9.92E-08 | 590.78931 | 5152.981 |
| HEHQVMLMR | 9 | Q15233 | Non-POU domain-containing octamer-bir | NONO | 54.232 | 2 | 51.91537 | yes | 1.632E-07 | 598.7868 | 5152.981 |
| HEHQVMLMR | 9 | Q15233 | Non-POU domain-containing octamer-bir | NONO | 54.232 | 2 | 51.91537 | yes | 8.161E-07 | 598.7868 | 5152.981 |
| AVVIVDDR | 8 | Q15233 | Non-POU domain-containing octamer-bir | NONO | 54.232 | 2 | 51.91537 | no | 1.701E-06 | 443.75327 | 14884.907 |
| RMEELHNQEVQK | 12 | Q15233 | Non-POU domain-containing octamer-bir | NONO | 54.232 | 3 | 51.91537 | yes | 1.305E-07 | 514.25604 | 4355.6484 |
| RMEELHNQEVQK | 12 | Q15233 | Non-POU domain-containing octamer-bir | NONO | 54.232 | 3 | 51.91537 | yes | 1.113E-07 | 770.88049 | 4355.6484 |
| RMEELHNQEVQK | 12 | Q15233 | Non-POU domain-containing octamer-bir | NONO | 54.232 | 3 | 51.91537 | yes | 1.001E-07 | 519.58771 | 4355.6484 |
| RMEELHNQEVQK | 12 | Q15233 | Non-POU domain-containing octamer-bir | NONO | 54.232 | 3 | 51.91537 | yes | 1.066E-07 | 778.87793 | 4355.6484 |
| RMEELHNQEVQK | 12 | Q15233 | Non-POU domain-containing octamer-bir | NONO | 54.232 | 2 | 51.91537 | yes | 1.305E-07 | 514.25604 | 4355.6484 |
| RMEELHNQEVQK | 12 | Q15233 | Non-POU domain-containing octamer-bir | NONO | 54.232 | 2 | 51.91537 | yes | 1.113E-07 | 770.88049 | 4355.6484 |
| RMEELHNQEVQK | 12 | Q15233 | Non-POU domain-containing octamer-bir | NONO | 54.232 | 2 | 51.91537 | yes | 1.001E-07 | 519.58771 | 4355.6484 |
| RMEELHNQEVQK | 12 | Q15233 | Non-POU domain-containing octamer-bir | NONO | 54.232 | 2 | 51.91537 | yes | 1.066E-07 | 778.87793 | 4355.6484 |
| FGQAATMEGIGAI( | 22 | Q15233 | Non-POU domain-containing octamer-bir | NONO | 54.232 | 3 | 51.91537 | yes | 1.084E-07 | 721.69324 | 6428.7832 |
| FGQAATMEGIGAI( | 22 | Q15233 | Non-POU domain-containing octamer-bir | NONO | 54.232 | 3 | 51.91537 | yes | 1.235E-07 | 727.0249 | 6428.7832 |
| DKGFGFIR | 8 | Q15233 | Non-POU domain-containing octamer-bir | NONO | 54.232 | 2 | 51.91537 | yes | 4.156E-07 | 470.25598 | 12083.071 |
| QQEEMMR | 7 | Q15233 | Non-POU domain-containing octamer-bir | NONO | 54.232 | 2 | 51.91537 | yes | 9.934E-08 | 476.20477 | 3908.7666 |
| QQEEMMR | 7 | Q15233 | Non-POU domain-containing octamer-bir | NONO | 54.232 | 2 | 51.91537 | yes | 1.412E-07 | 484.20224 | 3908.7666 |
| QQEEMMR | 7 | Q15233 | Non-POU domain-containing octamer-bir | NONO | 54.232 | 2 | 51.91537 | yes | 9.968E-08 | 484.20224 | 3908.7666 |
| MEELHNQEVQK | 11 | Q15233 | Non-POU domain-containing octamer-bir | NONO | 54.232 | 2 | 51.91537 | yes | 9.722E-08 | 692.8299 | 3264.0916 |
| MEELHNQEVQK | 11 | Q15233 | Non-POU domain-containing octamer-bir | NONO | 54.232 | 2 | 51.91537 | yes | 4.714E-07 | 462.22235 | 3264.0916 |
| MEELHNQEVQK | 11 | Q15233 | Non-POU domain-containing octamer-bir | NONO | 54.232 | 2 | 51.91537 | yes | 9.722E-08 | 700.82739 | 3264.0916 |
| MEELHNQEVQK | 11 | Q15233 | Non-POU domain-containing octamer-bir | NONO | 54.232 | 3 | 51.91537 | yes | 9.722E-08 | 692.8299 | 3264.0916 |
| MEELHNQEVQK | 11 | Q15233 | Non-POU domain-containing octamer-bir | NONO | 54.232 | 3 | 51.91537 | yes | 4.714E-07 | 462.22235 | 3264.0916 |

| | | | | | | | | | | | |
|---|---|---|---|---|---|---|---|---|---|---|---|
| MEELHNQEVQK | 11 | Q15233 | Non-POU domain-containing octamer-bi | NONO | 54.232 | 3 | 51.91537 | yes | 9.722E-08 | 700.82739 | 3264.0916 |
| LEMEMEAAR | 9 | Q15233 | Non-POU domain-containing octamer-bi | NONO | 54.232 | 2 | 51.91537 | yes | 2.59E-07 | 540.24664 | 3694.491 |
| LEMEMEAAR | 9 | Q15233 | Non-POU domain-containing octamer-bi | NONO | 54.232 | 2 | 51.91537 | yes | 4.705E-06 | 548.24408 | 3694.491 |
| QQQDQVDR | 8 | Q15233 | Non-POU domain-containing octamer-bi | NONO | 54.232 | 2 | 51.91537 | yes | 6.495E-07 | 508.74142 | 5865.4951 |
| GIVEFSGKPAAR | 12 | Q15233 | Non-POU domain-containing octamer-bi | NONO | 54.232 | 2 | 51.91537 | yes | 9.722E-08 | 616.34332 | 2049.6243 |
| GIVEFSGKPAAR | 12 | Q15233 | Non-POU domain-containing octamer-bi | NONO | 54.232 | 2 | 51.91537 | yes | 1.895E-06 | 411.23129 | 2049.6243 |
| GIVEFSGKPAAR | 12 | Q15233 | Non-POU domain-containing octamer-bi | NONO | 54.232 | 3 | 51.91537 | yes | 9.722E-08 | 616.34332 | 2049.6243 |
| GIVEFSGKPAAR | 12 | Q15233 | Non-POU domain-containing octamer-bi | NONO | 54.232 | 3 | 51.91537 | yes | 1.895E-06 | 411.23129 | 2049.6243 |
| CSEGSFLLTTFPR | 13 | Q15233 | Non-POU domain-containing octamer-bi | NONO | 54.232 | 2 | 51.91537 | yes | 3.297E-06 | 757.86902 | 3080.6907 |
| ALIEMEK | 7 | Q15233 | Non-POU domain-containing octamer-bi | NONO | 54.232 | 2 | 51.91537 | yes | 2.164E-07 | 417.22549 | 557.20227 |
| ALIEMEK | 7 | Q15233 | Non-POU domain-containing octamer-bi | NONO | 54.232 | 2 | 51.91537 | yes | 1.022E-07 | 425.22296 | 557.20227 |
| RQQEEMMR | 8 | Q15233 | Non-POU domain-containing octamer-bi | NONO | 54.232 | 2 | 51.91537 | yes | 1.082E-07 | 554.25531 | 802.84424 |
| GIVEFSGK | 8 | Q15233 | Non-POU domain-containing octamer-bi | NONO | 54.232 | 2 | 51.91537 | yes | 1.805E-07 | 418.72925 | 53.641769 |
| EGVVDIYNCVK | 11 | Q15262 | Receptor-type tyrosine-protein phosphata | PTPRK | 162.103 | 2 | 49.678 | yes | 1.011E-07 | 648.31866 | 93.793282 |
| ESEETIIQTDEDVP | 20 | Q15262 | Receptor-type tyrosine-protein phosphata | PTPRK | 162.103 | 3 | 49.678 | yes | 1.909E-06 | 728.03021 | 35.452507 |
| LQAEDCSIACLPR | 13 | Q15262 | Receptor-type tyrosine-protein phosphata | PTPRK | 162.103 | 2 | 49.678 | yes | 1.475E-07 | 766.8634 | 23.072224 |
| FSNLLGTVYR | 10 | Q15269 | Periodic tryptophan protein 2 homolog O | PWP2 | 102.453 | 2 | 54.251377 | yes | 9.842E-08 | 585.31934 | 326.54794 |
| FVCIYHVR | 8 | Q15269 | Periodic tryptophan protein 2 homolog O | PWP2 | 102.453 | 2 | 54.251377 | yes | 1.446E-07 | 547.28424 | 290.63046 |
| LLMLHGQK | 8 | Q15269 | Periodic tryptophan protein 2 homolog O | PWP2 | 102.453 | 2 | 54.251377 | yes | 4.14E-07 | 470.27585 | 204.88191 |
| CVGLSPDGR | 9 | Q15269 | Periodic tryptophan protein 2 homolog O | PWP2 | 102.453 | 2 | 54.251377 | yes | 2.115E-06 | 480.73199 | 172.81668 |
| EGDFNNLTAAAFI | 14 | Q15269 | Periodic tryptophan protein 2 homolog O | PWP2 | 102.453 | 2 | 54.251377 | yes | 9.722E-08 | 767.86786 | 100.45606 |
| VLEFLASSFEVSR | 13 | Q15269 | Periodic tryptophan protein 2 homolog O | PWP2 | 102.453 | 2 | 54.251377 | yes | 4.743E-07 | 742.39319 | 69.185905 |
| LVQEALEAVPR | 11 | Q15269 | Periodic tryptophan protein 2 homolog O | PWP2 | 102.453 | 2 | 54.251377 | yes | 1.612E-07 | 612.85095 | 60.411877 |
| SVLASASWDK | 10 | Q15269 | Periodic tryptophan protein 2 homolog O | PWP2 | 102.453 | 2 | 54.251377 | yes | 1.21E-06 | 532.27454 | 56.0975 |
| EFNAFVLDK | 9 | Q15269 | Periodic tryptophan protein 2 homolog O | PWP2 | 102.453 | 2 | 54.251377 | yes | 4.761E-07 | 541.77948 | 44.626301 |
| GSVHSVSFSPDGR | 13 | Q15269 | Periodic tryptophan protein 2 homolog O | PWP2 | 102.453 | 2 | 54.251377 | yes | 5.226E-07 | 666.32056 | 30.492897 |
| LQIWDTAGQER | 11 | Q15286 | Ras-related protein Rab-35 OS=Homo sa | RAB35 | 23.026 | 2 | 48.82093 | no | 1.101E-07 | 658.83331 | 985.34625 |
| LLIIGDSGVGK | 11 | Q15286 | Ras-related protein Rab-35 OS=Homo sa | RAB35 | 23.026 | 2 | 48.82093 | yes | 1.257E-07 | 536.32404 | 185.3934 |
| FAGQMGIQLFETS | 15 | Q15286 | Ras-related protein Rab-35 OS=Homo sa | RAB35 | 23.026 | 2 | 48.82093 | yes | 4.247E-07 | 814.41107 | 63.050877 |
| DYDHLFK | 7 | Q15286 | Ras-related protein Rab-35 OS=Homo sa | RAB35 | 23.026 | 2 | 48.82093 | yes | 6.363E-07 | 469.22433 | 59.335693 |
| GTHGVIVVYDVTS | 21 | Q15286 | Ras-related protein Rab-35 OS=Homo sa | RAB35 | 23.026 | 3 | 48.82093 | yes | 2.187E-06 | 741.05481 | 27.032162 |
| MIDMPVER | 8 | Q15287 | RNA-binding protein with serine-rich do | RNPS1 | 34.209 | 2 | 42.04331 | yes | 3.535E-07 | 495.741 | 1339.8016 |
| DHIMEIFSTYGK | 12 | Q15287 | RNA-binding protein with serine-rich do | RNPS1 | 34.209 | 2 | 42.04331 | yes | 3.22E-07 | 720.84503 | 667.48145 |
| MLPPPPMWR | 9 | Q15287 | RNA-binding protein with serine-rich do | RNPS1 | 34.209 | 2 | 42.04331 | yes | 2.846E-06 | 562.79083 | 351.44846 |
| HMDGGQIDGQEIT | 27 | Q15287 | RNA-binding protein with serine-rich do | RNPS1 | 34.209 | 4 | 42.04331 | yes | 2.208E-06 | 728.37244 | 280.28485 |
| TDSQDLVASFR | 11 | Q15291 | Retinoblastoma-binding protein 5 OS=H | RBBP5 | 59.153 | 2 | 43.806686 | yes | 2.439E-07 | 619.8042 | 54.892895 |
| TFDQLTPDESK | 11 | Q15293 | Reticulocalbin-1 OS=Homo sapiens OX= | RCN1 | 38.89 | 2 | 51.587276 | yes | 1.717E-07 | 640.80389 | 15645.879 |
| LDKDEIR | 7 | Q15293 | Reticulocalbin-1 OS=Homo sapiens OX= | RCN1 | 38.89 | 2 | 51.587276 | yes | 1.697E-07 | 444.74289 | 8865.7803 |
| HWILPQDYDHAQ | 16 | Q15293 | Reticulocalbin-1 OS=Homo sapiens OX= | RCN1 | 38.89 | 3 | 51.587276 | yes | 1.856E-07 | 650.64661 | 7661.4155 |

| | | | | | | | | | | |
|---|---|---|---|---|---|---|---|---|---|---|
| HLVYESDK | 8 | Q15293 | Reticulocalbin-1 OS=Homo sapiens OX= | RCN1 | 38.89 | 2 | 51.587276 | yes | 1.094E-07 | 495.74817 | 6462.0313 |
| IDNDGDGFVTTEE | 15 | Q15293 | Reticulocalbin-1 OS=Homo sapiens OX= | RCN1 | 38.89 | 2 | 51.587276 | yes | 1.26E-07 | 826.88611 | 5272.4121 |
| YIFDNVAK | 8 | Q15293 | Reticulocalbin-1 OS=Homo sapiens OX= | RCN1 | 38.89 | 2 | 51.587276 | yes | 3.55E-07 | 485.25565 | 4066.677 |
| VVRPDSELGERPP | 29 | Q15293 | Reticulocalbin-1 OS=Homo sapiens OX= | RCN1 | 38.89 | 5 | 51.587276 | yes | 4.228E-07 | 672.72437 | 1869.4875 |
| VVRPDSELGERPP | 29 | Q15293 | Reticulocalbin-1 OS=Homo sapiens OX= | RCN1 | 38.89 | 5 | 51.587276 | yes | 3.976E-07 | 840.65363 | 1869.4875 |
| VVRPDSELGERPP | 29 | Q15293 | Reticulocalbin-1 OS=Homo sapiens OX= | RCN1 | 38.89 | 4 | 51.587276 | yes | 4.228E-07 | 672.72437 | 1869.4875 |
| VVRPDSELGERPP | 29 | Q15293 | Reticulocalbin-1 OS=Homo sapiens OX= | RCN1 | 38.89 | 4 | 51.587276 | yes | 3.976E-07 | 840.65363 | 1869.4875 |
| AADLNGDLTATR | 12 | Q15293 | Reticulocalbin-1 OS=Homo sapiens OX= | RCN1 | 38.89 | 2 | 51.587276 | yes | 9.903E-08 | 609.30969 | 3687.4272 |
| EIVVLETLEDIDK | 13 | Q15293 | Reticulocalbin-1 OS=Homo sapiens OX= | RCN1 | 38.89 | 2 | 51.587276 | yes | 9.722E-08 | 758.41125 | 3468.7717 |
| EQFNEFR | 7 | Q15293 | Reticulocalbin-1 OS=Homo sapiens OX= | RCN1 | 38.89 | 2 | 51.587276 | yes | 1.357E-07 | 485.22488 | 2287.8262 |
| ISWEEYK | 7 | Q15293 | Reticulocalbin-1 OS=Homo sapiens OX= | RCN1 | 38.89 | 2 | 51.587276 | yes | 1.382E-07 | 477.73199 | 1782.6533 |
| RYIFDNVAK | 9 | Q15293 | Reticulocalbin-1 OS=Homo sapiens OX= | RCN1 | 38.89 | 2 | 51.587276 | yes | 1.883E-07 | 563.30621 | 723.64087 |
| DGKLDKDEIR | 10 | Q15293 | Reticulocalbin-1 OS=Homo sapiens OX= | RCN1 | 38.89 | 2 | 51.587276 | yes | 1.219E-07 | 594.81458 | 256.58725 |
| DDKISWEEYK | 10 | Q15293 | Reticulocalbin-1 OS=Homo sapiens OX= | RCN1 | 38.89 | 2 | 51.587276 | yes | 9.779E-08 | 656.8064 | 25.963558 |
| TGPLPWPAGFQPR | 13 | Q15334 | Lethal(2) giant larvae protein homolog 1 ( | LLGL1 | 115.419 | 2 | 53.066475 | yes | 3.072E-07 | 712.37769 | 264.52698 |
| APVVAIAVLDGR | 12 | Q15334 | Lethal(2) giant larvae protein homolog 1 ( | LLGL1 | 115.419 | 2 | 53.066475 | yes | 9.722E-08 | 590.85608 | 202.28085 |
| DLAQAPDMQGGH | 23 | Q15334 | Lethal(2) giant larvae protein homolog 1 ( | LLGL1 | 115.419 | 3 | 53.066475 | yes | 2.753E-07 | 819.06891 | 112.69084 |
| AETLVTLDFTSR | 12 | Q15334 | Lethal(2) giant larvae protein homolog 1 ( | LLGL1 | 115.419 | 2 | 53.066475 | yes | 4.226E-07 | 676.85645 | 99.86068 |
| SADDSLSGVVR | 11 | Q15334 | Lethal(2) giant larvae protein homolog 1 ( | LLGL1 | 115.419 | 2 | 53.066475 | yes | 1.284E-07 | 553.27783 | 93.273132 |
| GLLLTGHEDGTVR | 13 | Q15334 | Lethal(2) giant larvae protein homolog 1 ( | LLGL1 | 115.419 | 2 | 53.066475 | yes | 9.722E-08 | 684.36749 | 46.380318 |
| EDISGIASCVFTR | 13 | Q15334 | Lethal(2) giant larvae protein homolog 1 ( | LLGL1 | 115.419 | 2 | 53.066475 | yes | 9.722E-08 | 727.85083 | 42.193359 |
| RPEQAVEAVLGK | 12 | Q15334 | Lethal(2) giant larvae protein homolog 1 ( | LLGL1 | 115.419 | 2 | 53.066475 | yes | 4.162E-07 | 648.86713 | 32.024364 |
| HGQGFYLISPSEFE | 15 | Q15334 | Lethal(2) giant larvae protein homolog 1 ( | LLGL1 | 115.419 | 3 | 53.066475 | yes | 4.374E-05 | 589.6214 | 12.1852 |
| STQESLTAGGTDL | 14 | Q15345 | Leucine-rich repeat-containing protein 41 | LRRC41 | 88.65 | 2 | 53.784164 | yes | 9.722E-08 | 704.35175 | 157.24355 |
| DEGSLLLGSR | 10 | Q15345 | Leucine-rich repeat-containing protein 41 | LRRC41 | 88.65 | 2 | 53.784164 | yes | 2.876E-07 | 523.77747 | 142.66841 |
| FSPLLHSSR | 9 | Q15345 | Leucine-rich repeat-containing protein 41 | LRRC41 | 88.65 | 2 | 53.784164 | yes | 9.722E-08 | 522.28528 | 103.58039 |
| GAQPAPLLCSVLK | 13 | Q15345 | Leucine-rich repeat-containing protein 41 | LRRC41 | 88.65 | 2 | 53.784164 | yes | 1.161E-07 | 677.38159 | 90.420799 |
| HLLFSDVAAQQSL | 14 | Q15345 | Leucine-rich repeat-containing protein 41 | LRRC41 | 88.65 | 3 | 53.784164 | yes | 3.901E-07 | 528.95459 | 42.771339 |
| HLLFSDVAAQQSL | 14 | Q15345 | Leucine-rich repeat-containing protein 41 | LRRC41 | 88.65 | 3 | 53.784164 | yes | 4.559E-07 | 792.92828 | 42.771339 |
| HLLFSDVAAQQSL | 14 | Q15345 | Leucine-rich repeat-containing protein 41 | LRRC41 | 88.65 | 2 | 53.784164 | yes | 3.901E-07 | 528.95459 | 42.771339 |
| HLLFSDVAAQQSL | 14 | Q15345 | Leucine-rich repeat-containing protein 41 | LRRC41 | 88.65 | 2 | 53.784164 | yes | 4.559E-07 | 792.92828 | 42.771339 |
| ICDETANATSTLK | 13 | Q15361 | Transcription termination factor 1 OS=Hc | TTF1 | 103.052 | 2 | 39.77104 | yes | 4.391E-07 | 712.34033 | 38.54916 |
| AITIAGVPQSVTEC | 16 | Q15365 | Poly(rC)-binding protein 1 OS=Homo sap | PCBP1 | 37.498 | 2 | 54.531372 | yes | 3.268E-07 | 836.95056 | 7858.6045 |
| LVVPATQCGSLIGI | 14 | Q15365 | Poly(rC)-binding protein 1 OS=Homo sap | PCBP1 | 37.498 | 2 | 54.531372 | yes | 3.376E-07 | 721.9054 | 7258.1123 |
| QGANINEIR | 9 | Q15365 | Poly(rC)-binding protein 1 OS=Homo sap | PCBP1 | 37.498 | 2 | 54.531372 | yes | 1.715E-07 | 507.76999 | 3064.9934 |
| IANPVEGSSGR | 11 | Q15365 | Poly(rC)-binding protein 1 OS=Homo sap | PCBP1 | 37.498 | 2 | 54.531372 | yes | 9.722E-08 | 543.78052 | 2710.1313 |
| IITLTGPTNAIFK | 13 | Q15365 | Poly(rC)-binding protein 1 OS=Homo sap | PCBP1 | 37.498 | 2 | 54.531372 | yes | 1.33E-07 | 694.91101 | 2534.3352 |
| QQSHFAMMHGGT | 25 | Q15365 | Poly(rC)-binding protein 1 OS=Homo sap | PCBP1 | 37.498 | 4 | 54.531372 | yes | 9.766E-08 | 652.2995 | 112.26071 |
| QQSHFAMMHGGT | 25 | Q15365 | Poly(rC)-binding protein 1 OS=Homo sap | PCBP1 | 37.498 | 4 | 54.531372 | yes | 3.365E-07 | 656.29822 | 112.26071 |

| | | | | | | | | | | | |
|---|---|---|---|---|---|---|---|---|---|---|---|
| QQSHFAMMHGGT | 25 | Q15365 | Poly(rC)-binding protein 1 OS=Homo sa[ | PCBP1 | 37.498 | 4 | 54.531372 | yes | 3.944E-07 | 656.29822 | 112.26071 |
| QICLVMLETLSQSI | 17 | Q15365 | Poly(rC)-binding protein 1 OS=Homo sa[ | PCBP1 | 37.498 | 3 | 54.531372 | yes | 1.43E-07 | 654.00452 | 150.27135 |
| LEEDINSSMTNST/ | 23 | Q15365 | Poly(rC)-binding protein 1 OS=Homo sa[ | PCBP1 | 37.498 | 3 | 54.531372 | yes | 8.585E-07 | 830.41486 | 102.3662 |
| VMTIPYQPMPASS | 23 | Q15365 | Poly(rC)-binding protein 1 OS=Homo sa[ | PCBP1 | 37.498 | 3 | 54.531372 | yes | 9.722E-08 | 825.73248 | 81.921204 |
| AFAMIIDKLEEDIN | 31 | Q15365 | Poly(rC)-binding protein 1 OS=Homo sa[ | PCBP1 | 37.498 | 4 | 54.531372 | yes | 3.67E-05 | 845.43127 | 19.089087 |
| IITLAGPTNAIFK | 13 | Q15366 | Poly(rC)-binding protein 2 OS=Homo sa[ | PCBP2 | 38.581 | 2 | 52.2427 | yes | 5.024E-07 | 679.90576 | 14763.858 |
| LVVPASQCGSLIGI | 14 | Q15366 | Poly(rC)-binding protein 2 OS=Homo sa[ | PCBP2 | 38.581 | 2 | 52.2427 | no | 1.587E-07 | 714.89758 | 14595.388 |
| GVTIPYRPK | 9 | Q15366 | Poly(rC)-binding protein 2 OS=Homo sa[ | PCBP2 | 38.581 | 2 | 52.2427 | yes | 2.314E-07 | 515.80585 | 11421.484 |
| IANPVEGSTDR | 11 | Q15366 | Poly(rC)-binding protein 2 OS=Homo sa[ | PCBP2 | 38.581 | 2 | 52.2427 | yes | 9.766E-08 | 579.79114 | 7239.7388 |
| INISEGNCPER | 11 | Q15366 | Poly(rC)-binding protein 2 OS=Homo sa[ | PCBP2 | 38.581 | 2 | 52.2427 | no | 1.039E-07 | 644.80115 | 4474.1733 |
| QMSGAQIK | 8 | Q15366 | Poly(rC)-binding protein 2 OS=Homo sa[ | PCBP2 | 38.581 | 2 | 52.2427 | no | 1.339E-07 | 431.7262 | 1527.2291 |
| LHQLAMQQSHFPN | 31 | Q15366 | Poly(rC)-binding protein 2 OS=Homo sa[ | PCBP2 | 38.581 | 4 | 52.2427 | yes | 1.16E-07 | 846.40405 | 432.76956 |
| LHQLAMQQSHFPN | 31 | Q15366 | Poly(rC)-binding protein 2 OS=Homo sa[ | PCBP2 | 38.581 | 4 | 52.2427 | yes | 9.722E-08 | 677.32471 | 432.76956 |
| LHQLAMQQSHFPN | 31 | Q15366 | Poly(rC)-binding protein 2 OS=Homo sa[ | PCBP2 | 38.581 | 5 | 52.2427 | yes | 1.16E-07 | 846.40405 | 432.76956 |
| LHQLAMQQSHFPN | 31 | Q15366 | Poly(rC)-binding protein 2 OS=Homo sa[ | PCBP2 | 38.581 | 5 | 52.2427 | yes | 9.722E-08 | 677.32471 | 432.76956 |
| PSSSPVIFAGGQDF | 14 | Q15366 | Poly(rC)-binding protein 2 OS=Homo sa[ | PCBP2 | 38.581 | 2 | 52.2427 | yes | 2.475E-07 | 709.35718 | 744.64331 |
| PKPSSSPVIFAGGQ | 16 | Q15366 | Poly(rC)-binding protein 2 OS=Homo sa[ | PCBP2 | 38.581 | 2 | 52.2427 | yes | 9.81E-08 | 821.93103 | 108.41911 |
| QICVVMLETLSQSI | 16 | Q15366 | Poly(rC)-binding protein 2 OS=Homo sa[ | PCBP2 | 38.581 | 3 | 52.2427 | yes | 3.291E-07 | 610.65485 | 89.273293 |
| LEEDISSSMTNSTA | 23 | Q15366 | Poly(rC)-binding protein 2 OS=Homo sa[ | PCBP2 | 38.581 | 3 | 52.2427 | yes | 1.974E-07 | 821.41125 | 28.349163 |
| INEIRQMSGAQIK | 13 | Q15366 | Poly(rC)-binding protein 2 OS=Homo sa[ | PCBP2 | 38.581 | 2 | 52.2427 | no | 8.965E-05 | 752.40106 | 0.4044838 |
| EIPSHVLSK | 9 | Q15369 | Elongin-C OS=Homo sapiens OX=9606 ( | ELOC | 12.474 | 2 | 51.81894 | yes | 2.163E-07 | 505.28748 | 1370.4669 |
| EHALTSGTIK | 10 | Q15369 | Elongin-C OS=Homo sapiens OX=9606 ( | ELOC | 12.474 | 2 | 51.81894 | yes | 9.722E-08 | 528.78784 | 848.29407 |
| VCMYFTYK | 8 | Q15369 | Elongin-C OS=Homo sapiens OX=9606 ( | ELOC | 12.474 | 2 | 51.81894 | yes | 9.873E-08 | 556.25116 | 406.87665 |
| TYGGCEGPDAMY | 14 | Q15369 | Elongin-C OS=Homo sapiens OX=9606 ( | ELOC | 12.474 | 2 | 51.81894 | yes | 9.722E-08 | 774.32886 | 405.62357 |
| LISSDGHEFIVK | 12 | Q15369 | Elongin-C OS=Homo sapiens OX=9606 ( | ELOC | 12.474 | 3 | 51.81894 | yes | 3.207E-07 | 448.91013 | 67.262154 |
| PQAPATVGLAFR | 12 | Q15370 | Elongin-B OS=Homo sapiens OX=9606 ( | ELOB | 13.133 | 2 | 53.06479 | yes | 5.925E-07 | 614.34583 | 1607.1548 |
| TLGECGFTSQTAR | 13 | Q15370 | Elongin-B OS=Homo sapiens OX=9606 ( | ELOB | 13.133 | 2 | 53.06479 | yes | 9.722E-08 | 714.33282 | 1097.3619 |
| LYKDDQLLDDGK | 12 | Q15370 | Elongin-B OS=Homo sapiens OX=9606 ( | ELOB | 13.133 | 2 | 53.06479 | yes | 9.819E-08 | 711.85919 | 363.80563 |
| ESSTVFELK | 9 | Q15370 | Elongin-B OS=Homo sapiens OX=9606 ( | ELOB | 13.133 | 2 | 53.06479 | yes | 9.722E-08 | 520.26892 | 333.78055 |
| RIVEGILK | 8 | Q15370 | Elongin-B OS=Homo sapiens OX=9606 ( | ELOB | 13.133 | 2 | 53.06479 | yes | 4.508E-07 | 464.30292 | 125.02469 |
| DDQLLDDGK | 9 | Q15370 | Elongin-B OS=Homo sapiens OX=9606 ( | ELOB | 13.133 | 2 | 53.06479 | yes | 1.579E-07 | 509.73801 | 87.78302 |
| TTIFTDAK | 8 | Q15370 | Elongin-B OS=Homo sapiens OX=9606 ( | ELOB | 13.133 | 2 | 53.06479 | yes | 3.657E-06 | 448.73984 | 27.689346 |
| VQIPIMLVGNK | 11 | Q15382 | GTP-binding protein Rheb OS=Homo sa[ | RHEB | 20.498 | 2 | 44.41067 | yes | 2.84E-07 | 606.36267 | 235.94537 |
| ALAESWNAAFLES | 16 | Q15382 | GTP-binding protein Rheb OS=Homo sa[ | RHEB | 20.498 | 2 | 44.41067 | yes | 2.204E-07 | 847.92285 | 72.300766 |
| ENQTAVDVFR | 10 | Q15382 | GTP-binding protein Rheb OS=Homo sa[ | RHEB | 20.498 | 2 | 44.41067 | yes | 3.145E-06 | 589.79364 | 61.016823 |
| LPSSIEYSDLSR | 12 | Q15386 | Ubiquitin-protein ligase E3C OS=Homo s | UBE3C | 123.923 | 2 | 47.488445 | yes | 1.078E-05 | 683.84607 | 116.26872 |
| LPEFYDETLLR | 11 | Q15386 | Ubiquitin-protein ligase E3C OS=Homo s | UBE3C | 123.923 | 2 | 47.488445 | yes | 1.318E-07 | 698.36139 | 37.598713 |
| LPTISQR | 7 | Q15388 | Mitochondrial import receptor subunit TC | TOMM20 | 16.298 | 2 | 47.22336 | yes | 3.484E-07 | 407.74271 | 2883.4429 |
| IVSAQSLAEDDVE | 13 | Q15388 | Mitochondrial import receptor subunit TC | TOMM20 | 16.298 | 2 | 47.22336 | yes | 1.441E-07 | 688.33301 | 235.45497 |

| | | | | | | | | | | | |
|---|---|---|---|---|---|---|---|---|---|---|---|
| RSDPNFK | 7 | Q15388 | Mitochondrial import receptor subunit TC | TOMM20 | 16.298 | 2 | 47.22336 | yes | 1.375E-07 | 432.22214 | 108.30417 |
| EGLEYIPLR | 9 | Q15392 | Delta(24)-sterol reductase OS=Homo sapi | DHCR24 | 60.102 | 2 | 51.811905 | yes | 9.819E-08 | 545.3006 | 727.43823 |
| PGWLTVSLR | 9 | Q15392 | Delta(24)-sterol reductase OS=Homo sapi | DHCR24 | 60.102 | 2 | 51.811905 | yes | 9.722E-08 | 514.79803 | 308.35324 |
| LGCQDAFPEVYDK | 13 | Q15392 | Delta(24)-sterol reductase OS=Homo sapi | DHCR24 | 60.102 | 2 | 51.811905 | yes | 1.339E-07 | 771.35065 | 262.3251 |
| HVENYLK | 7 | Q15392 | Delta(24)-sterol reductase OS=Homo sapi | DHCR24 | 60.102 | 2 | 51.811905 | yes | 5.681E-07 | 451.74017 | 116.21707 |
| LNSIGNYYK | 9 | Q15392 | Delta(24)-sterol reductase OS=Homo sapi | DHCR24 | 60.102 | 2 | 51.811905 | yes | 3.141E-05 | 536.2771 | 77.857811 |
| GLEAICAK | 8 | Q15392 | Delta(24)-sterol reductase OS=Homo sapi | DHCR24 | 60.102 | 2 | 51.811905 | yes | 2.292E-07 | 431.22858 | 52.550453 |
| YLFGWMVPPK | 10 | Q15392 | Delta(24)-sterol reductase OS=Homo sapi | DHCR24 | 60.102 | 2 | 51.811905 | yes | 1.641E-07 | 619.32556 | 50.609573 |
| TVLDPVTGDLSDT | 14 | Q15393 | Splicing factor 3B subunit 3 OS=Homo sa | SF3B3 | 135.578 | 2 | 52.569668 | yes | 1.607E-07 | 744.88068 | 1817.3752 |
| DYIVVGSDSGR | 11 | Q15393 | Splicing factor 3B subunit 3 OS=Homo sa | SF3B3 | 135.578 | 2 | 52.569668 | yes | 1.558E-07 | 584.28564 | 1601.6437 |
| GMIFVCSATHK | 11 | Q15393 | Splicing factor 3B subunit 3 OS=Homo sa | SF3B3 | 135.578 | 2 | 52.569668 | yes | 9.722E-08 | 625.80463 | 1481.2645 |
| MFLYNLTLQR | 10 | Q15393 | Splicing factor 3B subunit 3 OS=Homo sa | SF3B3 | 135.578 | 2 | 52.569668 | yes | 1.084E-07 | 649.84991 | 739.37823 |
| MFLYNLTLQR | 10 | Q15393 | Splicing factor 3B subunit 3 OS=Homo sa | SF3B3 | 135.578 | 2 | 52.569668 | yes | 4.134E-07 | 657.84735 | 739.37823 |
| MQGQEAVLAMSS | 13 | Q15393 | Splicing factor 3B subunit 3 OS=Homo sa | SF3B3 | 135.578 | 2 | 52.569668 | yes | 1.123E-07 | 704.3396 | 1325.3436 |
| HIANYISGIQTIGHI | 15 | Q15393 | Splicing factor 3B subunit 3 OS=Homo sa | SF3B3 | 135.578 | 3 | 52.569668 | yes | 1.188E-07 | 560.63739 | 590.06366 |
| HIANYISGIQTIGHI | 15 | Q15393 | Splicing factor 3B subunit 3 OS=Homo sa | SF3B3 | 135.578 | 3 | 52.569668 | yes | 2.044E-07 | 840.45245 | 590.06366 |
| HIANYISGIQTIGHI | 15 | Q15393 | Splicing factor 3B subunit 3 OS=Homo sa | SF3B3 | 135.578 | 2 | 52.569668 | yes | 1.188E-07 | 560.63739 | 590.06366 |
| HIANYISGIQTIGHI | 15 | Q15393 | Splicing factor 3B subunit 3 OS=Homo sa | SF3B3 | 135.578 | 2 | 52.569668 | yes | 2.044E-07 | 840.45245 | 590.06366 |
| IVILEYQPSK | 10 | Q15393 | Splicing factor 3B subunit 3 OS=Homo sa | SF3B3 | 135.578 | 2 | 52.569668 | yes | 2.962E-07 | 595.34497 | 899.60022 |
| IVPGQFLAVDPK | 12 | Q15393 | Splicing factor 3B subunit 3 OS=Homo sa | SF3B3 | 135.578 | 2 | 52.569668 | yes | 9.934E-08 | 642.37152 | 886.58984 |
| FLAVGLVDNTVR | 12 | Q15393 | Splicing factor 3B subunit 3 OS=Homo sa | SF3B3 | 135.578 | 2 | 52.569668 | yes | 2.34E-07 | 652.37207 | 874.74811 |
| IHQETFGK | 8 | Q15393 | Splicing factor 3B subunit 3 OS=Homo sa | SF3B3 | 135.578 | 2 | 52.569668 | yes | 2.247E-07 | 480.25089 | 784.31036 |
| SVAGGFVYTYK | 11 | Q15393 | Splicing factor 3B subunit 3 OS=Homo sa | SF3B3 | 135.578 | 2 | 52.569668 | yes | 1.285E-07 | 596.30585 | 756.58752 |
| TPVEEVPAAIAPFC | 16 | Q15393 | Splicing factor 3B subunit 3 OS=Homo sa | SF3B3 | 135.578 | 2 | 52.569668 | yes | 1.171E-07 | 841.44904 | 721.92151 |
| ILELLRPDPNTGK | 13 | Q15393 | Splicing factor 3B subunit 3 OS=Homo sa | SF3B3 | 135.578 | 2 | 52.569668 | yes | 1.788E-07 | 733.4223 | 688.25238 |
| LVYILNR | 7 | Q15393 | Splicing factor 3B subunit 3 OS=Homo sa | SF3B3 | 135.578 | 2 | 52.569668 | yes | 9.722E-08 | 445.77655 | 578.31287 |
| SWLSYSYQSR | 10 | Q15393 | Splicing factor 3B subunit 3 OS=Homo sa | SF3B3 | 135.578 | 2 | 52.569668 | yes | 1.025E-07 | 638.80145 | 536.57605 |
| NFGDQPDIR | 9 | Q15393 | Splicing factor 3B subunit 3 OS=Homo sa | SF3B3 | 135.578 | 2 | 52.569668 | yes | 3.069E-06 | 531.25415 | 407.76285 |
| SEHPPLCGR | 9 | Q15393 | Splicing factor 3B subunit 3 OS=Homo sa | SF3B3 | 135.578 | 2 | 52.569668 | yes | 8.083E-07 | 526.75073 | 401.25812 |
| ATGISFAIHGNFSG | 16 | Q15393 | Splicing factor 3B subunit 3 OS=Homo sa | SF3B3 | 135.578 | 2 | 52.569668 | yes | 9.722E-08 | 804.41248 | 181.52975 |
| ATGISFAIHGNFSG | 16 | Q15393 | Splicing factor 3B subunit 3 OS=Homo sa | SF3B3 | 135.578 | 2 | 52.569668 | yes | 1.36E-07 | 536.61072 | 181.52975 |
| ATGISFAIHGNFSG | 16 | Q15393 | Splicing factor 3B subunit 3 OS=Homo sa | SF3B3 | 135.578 | 3 | 52.569668 | yes | 9.722E-08 | 804.41248 | 181.52975 |
| ATGISFAIHGNFSG | 16 | Q15393 | Splicing factor 3B subunit 3 OS=Homo sa | SF3B3 | 135.578 | 3 | 52.569668 | yes | 1.36E-07 | 536.61072 | 181.52975 |
| QQEIVVSR | 8 | Q15393 | Splicing factor 3B subunit 3 OS=Homo sa | SF3B3 | 135.578 | 2 | 52.569668 | yes | 2.685E-07 | 479.76944 | 314.82629 |
| YFDTVPVAAAMC | 15 | Q15393 | Splicing factor 3B subunit 3 OS=Homo sa | SF3B3 | 135.578 | 2 | 52.569668 | yes | 9.722E-08 | 842.92548 | 303.14798 |
| VIVSDVQESFIWVI | 14 | Q15393 | Splicing factor 3B subunit 3 OS=Homo sa | SF3B3 | 135.578 | 2 | 52.569668 | yes | 1.05E-07 | 838.95398 | 278.73389 |
| KQQMAEEMVEAA | 17 | Q15393 | Splicing factor 3B subunit 3 OS=Homo sa | SF3B3 | 135.578 | 3 | 52.569668 | yes | 1.001E-07 | 650.95593 | 173.05228 |
| ITLETDEDMVTEIR | 14 | Q15393 | Splicing factor 3B subunit 3 OS=Homo sa | SF3B3 | 135.578 | 2 | 52.569668 | yes | 9.81E-08 | 832.90601 | 167.81512 |
| RNDLDDPER | 9 | Q15393 | Splicing factor 3B subunit 3 OS=Homo sa | SF3B3 | 135.578 | 2 | 52.569668 | yes | 2.715E-07 | 565.26526 | 164.81223 |

| | | | | | | | | | | | |
|---|---|---|---|---|---|---|---|---|---|---|---|
| AGNGQWASVIR | 11 | Q15393 | Splicing factor 3B subunit 3 OS=Homo sa | SF3B3 | 135.578 | 2 | 52.569668 | yes | 3.094E-07 | 579.80438 | 164.4153 |
| KFVIHPESNNLIIIE | 26 | Q15393 | Splicing factor 3B subunit 3 OS=Homo sa | SF3B3 | 135.578 | 4 | 52.569668 | yes | 2.128E-07 | 750.13837 | 124.19605 |
| SYYFPVK | 7 | Q15393 | Splicing factor 3B subunit 3 OS=Homo sa | SF3B3 | 135.578 | 2 | 52.569668 | yes | 1.197E-07 | 452.23419 | 121.64005 |
| DLILNPR | 7 | Q15393 | Splicing factor 3B subunit 3 OS=Homo sa | SF3B3 | 135.578 | 2 | 52.569668 | yes | 2.654E-07 | 420.75052 | 98.641617 |
| AEVIMNYHVGETV | 18 | Q15393 | Splicing factor 3B subunit 3 OS=Homo sa | SF3B3 | 135.578 | 3 | 52.569668 | yes | 1.949E-07 | 677.69086 | 96.438171 |
| NDLDDPER | 8 | Q15393 | Splicing factor 3B subunit 3 OS=Homo sa | SF3B3 | 135.578 | 2 | 52.569668 | yes | 1.267E-06 | 487.21469 | 78.012154 |
| VHTLLTVEVFGVII | 14 | Q15393 | Splicing factor 3B subunit 3 OS=Homo sa | SF3B3 | 135.578 | 2 | 52.569668 | yes | 1.205E-07 | 791.96942 | 33.341545 |
| TIAFAHDSTR | 10 | Q15397 | Pumilio homolog 3 OS=Homo sapiens OX | PUM3 | 73.585 | 2 | 50.287144 | yes | 1.36E-07 | 559.78308 | 199.97122 |
| PQIAEIIR | 8 | Q15397 | Pumilio homolog 3 OS=Homo sapiens OX | PUM3 | 73.585 | 2 | 50.287144 | yes | 1.429E-07 | 470.28473 | 158.20422 |
| LELIMDEMK | 9 | Q15397 | Pumilio homolog 3 OS=Homo sapiens OX | PUM3 | 73.585 | 2 | 50.287144 | yes | 9.819E-08 | 561.28271 | 150.34795 |
| TNYDIVVR | 8 | Q15397 | Pumilio homolog 3 OS=Homo sapiens OX | PUM3 | 73.585 | 2 | 50.287144 | yes | 6.064E-07 | 490.26401 | 146.58177 |
| SLIPTLEK | 8 | Q15397 | Pumilio homolog 3 OS=Homo sapiens OX | PUM3 | 73.585 | 2 | 50.287144 | yes | 3.886E-07 | 450.77368 | 135.83206 |
| SEMIEAIR | 8 | Q15397 | Pumilio homolog 3 OS=Homo sapiens OX | PUM3 | 73.585 | 2 | 50.287144 | yes | 1.785E-07 | 474.7446 | 108.92997 |
| QMWEILR | 7 | Q15397 | Pumilio homolog 3 OS=Homo sapiens OX | PUM3 | 73.585 | 2 | 50.287144 | yes | 1.266E-07 | 488.25766 | 100.17834 |
| GIEILLEK | 8 | Q15397 | Pumilio homolog 3 OS=Homo sapiens OX | PUM3 | 73.585 | 2 | 50.287144 | yes | 2.319E-07 | 457.78149 | 64.441338 |
| VLEVQPEK | 8 | Q15397 | Pumilio homolog 3 OS=Homo sapiens OX | PUM3 | 73.585 | 2 | 50.287144 | yes | 3.151E-06 | 471.26874 | 57.23576 |
| VLLYLLSPR | 9 | Q15397 | Pumilio homolog 3 OS=Homo sapiens OX | PUM3 | 73.585 | 2 | 50.287144 | yes | 2.938E-06 | 537.33954 | 42.071819 |
| EAVVYLAHTHDGA | 14 | Q15397 | Pumilio homolog 3 OS=Homo sapiens OX | PUM3 | 73.585 | 3 | 50.287144 | yes | 8.924E-07 | 513.59521 | 19.832907 |
| EAVVYLAHTHDGA | 14 | Q15397 | Pumilio homolog 3 OS=Homo sapiens OX | PUM3 | 73.585 | 3 | 50.287144 | yes | 3.406E-07 | 769.88916 | 19.832907 |
| EAVVYLAHTHDGA | 14 | Q15397 | Pumilio homolog 3 OS=Homo sapiens OX | PUM3 | 73.585 | 2 | 50.287144 | yes | 8.924E-07 | 513.59521 | 19.832907 |
| EAVVYLAHTHDGA | 14 | Q15397 | Pumilio homolog 3 OS=Homo sapiens OX | PUM3 | 73.585 | 2 | 50.287144 | yes | 3.406E-07 | 769.88916 | 19.832907 |
| EASFVLHTIGK | 11 | Q15418 | Ribosomal protein S6 kinase alpha-1 OS= | RPS6KA1 | 82.723 | 2 | 37.185375 | yes | 6.219E-07 | 601.3324 | 49.312988 |
| NFWVSGLSSTTR | 12 | Q15424 | Scaffold attachment factor B1 OS=Homo | SAFB | 102.642 | 2 | 52.76869 | yes | 9.81E-08 | 677.84113 | 2470.1448 |
| YHSDFNR | 7 | Q15424 | Scaffold attachment factor B1 OS=Homo | SAFB | 102.642 | 2 | 52.76869 | yes | 1.59E-07 | 469.70938 | 944.90955 |
| AETLSGLGDSGAA | 28 | Q15424 | Scaffold attachment factor B1 OS=Homo | SAFB | 102.642 | 3 | 52.76869 | yes | 1.622E-07 | 846.40356 | 852.16699 |
| MQAQWER | 7 | Q15424 | Scaffold attachment factor B1 OS=Homo | SAFB | 102.642 | 2 | 52.76869 | yes | 3.553E-07 | 474.72144 | 823.05865 |
| SWQGTADGGMMI | 13 | Q15424 | Scaffold attachment factor B1 OS=Homo | SAFB | 102.642 | 2 | 52.76869 | yes | 1.02E-07 | 706.2901 | 769.46619 |
| ELPEQLQEHAIEDR | 14 | Q15424 | Scaffold attachment factor B1 OS=Homo | SAFB | 102.642 | 2 | 52.76869 | yes | 3.819E-07 | 839.91779 | 584.58472 |
| YPDHSVDR | 8 | Q15424 | Scaffold attachment factor B1 OS=Homo | SAFB | 102.642 | 2 | 52.76869 | yes | 3.902E-07 | 494.72778 | 442.5137 |
| CYGFVTMSTAEEA | 15 | Q15424 | Scaffold attachment factor B1 OS=Homo | SAFB | 102.642 | 2 | 52.76869 | yes | 9.722E-08 | 847.87366 | 391.99802 |
| ELVEGEMK | 8 | Q15424 | Scaffold attachment factor B1 OS=Homo | SAFB | 102.642 | 2 | 52.76869 | yes | 1.256E-07 | 467.73114 | 274.85889 |
| MSSPEDDSDTKR | 12 | Q15424 | Scaffold attachment factor B1 OS=Homo | SAFB | 102.642 | 2 | 52.76869 | yes | 1.041E-07 | 684.29083 | 272.52167 |
| DDAYWPEAK | 9 | Q15424 | Scaffold attachment factor B1 OS=Homo | SAFB | 102.642 | 2 | 52.76869 | yes | 1.122E-07 | 547.7431 | 251.38696 |
| AIEDEGGNPDEIEI | 21 | Q15424 | Scaffold attachment factor B1 OS=Homo | SAFB | 102.642 | 3 | 52.76869 | yes | 1.776E-07 | 749.02039 | 244.08243 |
| RDDAYWPEAK | 10 | Q15424 | Scaffold attachment factor B1 OS=Homo | SAFB | 102.642 | 2 | 52.76869 | yes | 9.722E-08 | 625.79364 | 217.82515 |
| ERLEIAR | 7 | Q15424 | Scaffold attachment factor B1 OS=Homo | SAFB | 102.642 | 2 | 52.76869 | yes | 1.305E-07 | 443.75888 | 78.391541 |
| NQDATVYVGGLD | 14 | Q15427 | Splicing factor 3B subunit 4 OS=Homo sa | SF3B4 | 44.386 | 2 | 52.120697 | yes | 9.722E-08 | 754.86499 | 513.58698 |
| LLAAQNPLSQADR | 13 | Q15427 | Splicing factor 3B subunit 4 OS=Homo sa | SF3B4 | 44.386 | 2 | 52.120697 | yes | 9.722E-08 | 698.8808 | 116.66661 |
| FMSAYEQR | 8 | Q15428 | Splicing factor 3A subunit 2 OS=Homo sa | SF3A2 | 49.256 | 2 | 52.841854 | yes | 1.662E-07 | 516.23438 | 324.4942 |

| | | | | | | | | | | | |
|---|---|---|---|---|---|---|---|---|---|---|---|
| QFFLQFHFK | 9 | Q15428 | Splicing factor 3A subunit 2 OS=Homo sa | SF3A2 | 49.256 | 2 | 52.841854 | yes | 1.348E-07 | 621.3269 | 239.093 |
| IEPPDRR | 7 | Q15428 | Splicing factor 3A subunit 2 OS=Homo sa | SF3A2 | 49.256 | 2 | 52.841854 | yes | 2.813E-06 | 441.74323 | 236.17915 |
| NHLGSYECK | 9 | Q15428 | Splicing factor 3A subunit 2 OS=Homo sa | SF3A2 | 49.256 | 2 | 52.841854 | yes | 2.543E-06 | 554.24805 | 206.23384 |
| FWTHWNR | 7 | Q15428 | Splicing factor 3A subunit 2 OS=Homo sa | SF3A2 | 49.256 | 2 | 52.841854 | yes | 1.657E-07 | 523.7514 | 186.22198 |
| LCLTLHNNEGSYL | 19 | Q15428 | Splicing factor 3A subunit 2 OS=Homo sa | SF3A2 | 49.256 | 3 | 52.841854 | yes | 9.722E-08 | 719.35663 | 165.01453 |
| TGSGGVASSSESNI | 14 | Q15428 | Splicing factor 3A subunit 2 OS=Homo sa | SF3A2 | 49.256 | 2 | 52.841854 | yes | 5.061E-07 | 648.29474 | 131.8791 |
| MEKPPAPPSLPAG | 18 | Q15428 | Splicing factor 3A subunit 2 OS=Homo sa | SF3A2 | 49.256 | 3 | 52.841854 | yes | 1.034E-07 | 590.65881 | 83.937416 |
| QLALETIDINKDPY | 17 | Q15428 | Splicing factor 3A subunit 2 OS=Homo sa | SF3A2 | 49.256 | 3 | 52.841854 | yes | 7.341E-07 | 680.35547 | 82.151726 |
| EAPAQPAPEK | 10 | Q15428 | Splicing factor 3A subunit 2 OS=Homo sa | SF3A2 | 49.256 | 2 | 52.841854 | yes | 1.211E-06 | 519.26672 | 81.561127 |
| AVLNPLCQVDYR | 12 | Q15436 | Protein transport protein Sec23A OS=Hoi | SEC23A | 86.161 | 2 | 55.882965 | no | 1.78E-07 | 724.37177 | 325.97012 |
| SGYQDMPEYENFI | 13 | Q15436 | Protein transport protein Sec23A OS=Hoi | SEC23A | 86.161 | 2 | 55.882965 | yes | 2.082E-07 | 818.34082 | 137.12657 |
| AETEEGPDVLR | 11 | Q15436 | Protein transport protein Sec23A OS=Hoi | SEC23A | 86.161 | 2 | 55.882965 | yes | 9.903E-08 | 608.2962 | 85.10257 |
| YIDTEHGGSQAR | 12 | Q15436 | Protein transport protein Sec23A OS=Hoi | SEC23A | 86.161 | 2 | 55.882965 | yes | 4.835E-07 | 667.31018 | 79.417915 |
| HLLQAPVDDAQEI | 17 | Q15436 | Protein transport protein Sec23A OS=Hoi | SEC23A | 86.161 | 3 | 55.882965 | yes | 3.699E-07 | 648.00952 | 76.275436 |
| MVQVHELGCEGIS | 14 | Q15436 | Protein transport protein Sec23A OS=Hoi | SEC23A | 86.161 | 2 | 55.882965 | yes | 9.722E-08 | 793.8869 | 64.496536 |
| FSWNVWPSSR | 10 | Q15436 | Protein transport protein Sec23A OS=Hoi | SEC23A | 86.161 | 2 | 55.882965 | no | 1.589E-07 | 633.3067 | 41.328281 |
| LWACNFCYQR | 10 | Q15436 | Protein transport protein Sec23A OS=Hoi | SEC23A | 86.161 | 2 | 55.882965 | yes | 3.65E-07 | 709.31061 | 28.476028 |
| DPWPVPQGK | 9 | Q15436 | Protein transport protein Sec23A OS=Hoi | SEC23A | 86.161 | 2 | 55.882965 | yes | 5.86E-07 | 512.26654 | 18.820576 |
| PAMPMQQAR | 9 | Q15437 | Protein transport protein Sec23B OS=Hoi | SEC23B | 86.48 | 2 | 54.86003 | yes | 8.442E-07 | 515.25208 | 295.86423 |
| AGYQDMPEYENFI | 13 | Q15437 | Protein transport protein Sec23B OS=Hoi | SEC23B | 86.48 | 2 | 54.86003 | yes | 1.066E-07 | 796.34033 | 182.89888 |
| YINTEHGGSQAR | 12 | Q15437 | Protein transport protein Sec23B OS=Hoi | SEC23B | 86.48 | 2 | 54.86003 | yes | 9.722E-08 | 666.81818 | 134.65437 |
| HYEMLANR | 8 | Q15437 | Protein transport protein Sec23B OS=Hoi | SEC23B | 86.48 | 2 | 54.86003 | yes | 1.779E-07 | 517.2478 | 134.07323 |
| HIEAAFDQEAAAV | 17 | Q15437 | Protein transport protein Sec23B OS=Hoi | SEC23B | 86.48 | 3 | 54.86003 | yes | 4.026E-07 | 614.97711 | 123.14107 |
| IAGAIGPCVSLNVK | 14 | Q15437 | Protein transport protein Sec23B OS=Hoi | SEC23B | 86.48 | 2 | 54.86003 | yes | 9.722E-08 | 699.89233 | 112.25784 |
| MVQVHELSCEGIS | 14 | Q15437 | Protein transport protein Sec23B OS=Hoi | SEC23B | 86.48 | 2 | 54.86003 | yes | 1.046E-07 | 808.89221 | 107.6169 |
| PAQPQEHPFASSR | 13 | Q15437 | Protein transport protein Sec23B OS=Hoi | SEC23B | 86.48 | 3 | 54.86003 | yes | 4.105E-07 | 484.57239 | 47.061531 |
| PAQPQEHPFASSR | 13 | Q15437 | Protein transport protein Sec23B OS=Hoi | SEC23B | 86.48 | 3 | 54.86003 | yes | 3.363E-07 | 726.35492 | 47.061531 |
| PAQPQEHPFASSR | 13 | Q15437 | Protein transport protein Sec23B OS=Hoi | SEC23B | 86.48 | 2 | 54.86003 | yes | 4.105E-07 | 484.57239 | 47.061531 |
| PAQPQEHPFASSR | 13 | Q15437 | Protein transport protein Sec23B OS=Hoi | SEC23B | 86.48 | 2 | 54.86003 | yes | 3.363E-07 | 726.35492 | 47.061531 |
| HLLQAPLDDAQEI | 17 | Q15437 | Protein transport protein Sec23B OS=Hoi | SEC23B | 86.48 | 3 | 54.86003 | yes | 9.722E-08 | 644.34967 | 93.059364 |
| QIQDMLGLTK | 10 | Q15437 | Protein transport protein Sec23B OS=Hoi | SEC23B | 86.48 | 2 | 54.86003 | yes | 1.152E-07 | 573.81299 | 90.171768 |
| FLQPVHK | 7 | Q15437 | Protein transport protein Sec23B OS=Hoi | SEC23B | 86.48 | 2 | 54.86003 | yes | 2.619E-07 | 434.75562 | 74.16922 |
| MAFGATLDVK | 10 | Q15437 | Protein transport protein Sec23B OS=Hoi | SEC23B | 86.48 | 2 | 54.86003 | yes | 1.534E-07 | 526.77588 | 57.063915 |
| NWADVQSQLR | 10 | Q15437 | Protein transport protein Sec23B OS=Hoi | SEC23B | 86.48 | 2 | 54.86003 | yes | 1.712E-06 | 608.80707 | 35.122952 |
| LWACNFCFQR | 10 | Q15437 | Protein transport protein Sec23B OS=Hoi | SEC23B | 86.48 | 2 | 54.86003 | yes | 1.57E-06 | 701.31317 | 26.554033 |
| VQAQVIQETIVPK | 13 | Q15459 | Splicing factor 3A subunit 1 OS=Homo sa | SF3A1 | 88.887 | 2 | 53.499405 | yes | 9.722E-08 | 726.92468 | 984.79523 |
| EVLDQVCYR | 9 | Q15459 | Splicing factor 3A subunit 1 OS=Homo sa | SF3A1 | 88.887 | 2 | 53.499405 | yes | 1.341E-07 | 591.28461 | 477.26035 |
| LQYEGIFIK | 9 | Q15459 | Splicing factor 3A subunit 1 OS=Homo sa | SF3A1 | 88.887 | 2 | 53.499405 | yes | 1.835E-07 | 555.81335 | 378.03079 |
| VQVPNMQDK | 9 | Q15459 | Splicing factor 3A subunit 1 OS=Homo sa | SF3A1 | 88.887 | 2 | 53.499405 | yes | 1.078E-07 | 529.76862 | 376.04102 |

| | | | | | | | | | | | |
|---|---|---|---|---|---|---|---|---|---|---|---|
| TEDSLMPEEEFLR | 13 | Q15459 | Splicing factor 3A subunit 1 OS=Homo sa | SF3A1 | 88.887 | 2 | 53.499405 | yes | 1.456E-07 | 798.36652 | 314.51846 |
| TDIFGVEETAIGK | 13 | Q15459 | Splicing factor 3A subunit 1 OS=Homo sa | SF3A1 | 88.887 | 2 | 53.499405 | yes | 9.722E-08 | 690.35626 | 272.70392 |
| AQEPSAAIPK | 10 | Q15459 | Splicing factor 3A subunit 1 OS=Homo sa | SF3A1 | 88.887 | 2 | 53.499405 | yes | 2.452E-07 | 506.2771 | 252.89886 |
| QFLTQLMQK | 9 | Q15459 | Splicing factor 3A subunit 1 OS=Homo sa | SF3A1 | 88.887 | 2 | 53.499405 | yes | 1.444E-07 | 568.81024 | 235.35846 |
| VTWDGHSGSMAR | 12 | Q15459 | Splicing factor 3A subunit 1 OS=Homo sa | SF3A1 | 88.887 | 2 | 53.499405 | yes | 4.085E-07 | 652.29602 | 180.13976 |
| EDSAPSKPVVGIIY | 20 | Q15459 | Splicing factor 3A subunit 1 OS=Homo sa | SF3A1 | 88.887 | 3 | 53.499405 | yes | 2.315E-07 | 717.38794 | 168.94344 |
| RTDIFGVEETAIGK | 14 | Q15459 | Splicing factor 3A subunit 1 OS=Homo sa | SF3A1 | 88.887 | 2 | 53.499405 | yes | 7.907E-07 | 768.40686 | 55.940533 |
| NKGPVSIK | 8 | Q15459 | Splicing factor 3A subunit 1 OS=Homo sa | SF3A1 | 88.887 | 2 | 53.499405 | yes | 9.722E-08 | 421.75836 | 35.34137 |
| DSNSLAYYNMAN | 21 | Q15459 | Splicing factor 3A subunit 1 OS=Homo sa | SF3A1 | 88.887 | 3 | 53.499405 | yes | 1.086E-06 | 755.71594 | 4.939096 |
| HDYSLSLR | 8 | Q15464 | SH2 domain-containing adapter protein B | SHB | 55.043 | 2 | 49.970444 | yes | 2.869E-07 | 495.75378 | 207.49529 |
| IMTEFQR | 7 | Q15464 | SH2 domain-containing adapter protein B | SHB | 55.043 | 2 | 49.970444 | yes | 5.532E-07 | 462.73401 | 113.26546 |
| SNQGFMHMK | 9 | Q15464 | SH2 domain-containing adapter protein B | SHB | 55.043 | 2 | 49.970444 | yes | 9.962E-08 | 540.2417 | 104.34262 |
| GAEHLSLLYPVAV | 14 | Q15464 | SH2 domain-containing adapter protein B | SHB | 55.043 | 2 | 49.970444 | yes | 1.459E-07 | 762.9303 | 60.653572 |
| GESAGYMEPYEA | 14 | Q15464 | SH2 domain-containing adapter protein B | SHB | 55.043 | 2 | 49.970444 | yes | 1.104E-07 | 794.34082 | 62.143082 |
| VDPAVPLEK | 9 | Q15464 | SH2 domain-containing adapter protein B | SHB | 55.043 | 2 | 49.970444 | yes | 6.913E-07 | 484.27658 | 60.21513 |
| EPEFYECFPCDGK | 13 | Q15468 | SCL-interrupting locus protein OS=Homo | STIL | 142.955 | 2 | 23.178812 | yes | 3.7E-06 | 839.3316 | 16.12764 |
| TIWDGEETSYCFK | 13 | Q15475 | Homeobox protein SIX1 OS=Homo sapie | SIX1 | 32.211 | 2 | 34.841866 | yes | 9.313E-07 | 818.35339 | 31.485125 |
| DLLPGEYVQMAG | 13 | Q15477 | Superkiller complex protein 2 OS=Homo | SKIC2 | 137.756 | 2 | 54.632507 | yes | 1.084E-07 | 724.86377 | 365.15903 |
| SMLYSGSDVIR | 11 | Q15477 | Superkiller complex protein 2 OS=Homo | SKIC2 | 137.756 | 2 | 54.632507 | yes | 9.722E-08 | 614.30554 | 352.98093 |
| METAATLLR | 9 | Q15477 | Superkiller complex protein 2 OS=Homo | SKIC2 | 137.756 | 2 | 54.632507 | yes | 1.141E-07 | 503.2735 | 309.88354 |
| PSQLQSQFR | 9 | Q15477 | Superkiller complex protein 2 OS=Homo | SKIC2 | 137.756 | 2 | 54.632507 | yes | 3.695E-07 | 545.78564 | 256.34186 |
| GLDPTGTVILLCK | 13 | Q15477 | Superkiller complex protein 2 OS=Homo | SKIC2 | 137.756 | 2 | 54.632507 | yes | 9.722E-08 | 693.88672 | 190.70824 |
| QIYVISTVTR | 10 | Q15477 | Superkiller complex protein 2 OS=Homo | SKIC2 | 137.756 | 2 | 54.632507 | yes | 4.369E-07 | 590.34021 | 189.78745 |
| LQPGDMAAITTK | 12 | Q15477 | Superkiller complex protein 2 OS=Homo | SKIC2 | 137.756 | 2 | 54.632507 | yes | 1.587E-06 | 623.32904 | 185.29617 |
| TQGELFLLLDSR | 12 | Q15477 | Superkiller complex protein 2 OS=Homo | SKIC2 | 137.756 | 2 | 54.632507 | yes | 1.179E-07 | 696.38007 | 129.30327 |
| EEAEEEIDFEK | 11 | Q15477 | Superkiller complex protein 2 OS=Homo | SKIC2 | 137.756 | 2 | 54.632507 | yes | 9.722E-08 | 684.29608 | 125.92003 |
| LVGEPVLGAK | 10 | Q15477 | Superkiller complex protein 2 OS=Homo | SKIC2 | 137.756 | 2 | 54.632507 | yes | 2.453E-07 | 491.80023 | 122.23138 |
| QAILHLER | 8 | Q15477 | Superkiller complex protein 2 OS=Homo | SKIC2 | 137.756 | 2 | 54.632507 | yes | 1.689E-07 | 490.28781 | 108.33601 |
| SEIHLFLQR | 9 | Q15477 | Superkiller complex protein 2 OS=Homo | SKIC2 | 137.756 | 2 | 54.632507 | yes | 5.029E-07 | 571.81946 | 95.17524 |
| AHEQALAELTK | 11 | Q15477 | Superkiller complex protein 2 OS=Homo | SKIC2 | 137.756 | 2 | 54.632507 | yes | 9.877E-08 | 605.82495 | 91.413315 |
| VPEMADLHR | 9 | Q15477 | Superkiller complex protein 2 OS=Homo | SKIC2 | 137.756 | 2 | 54.632507 | yes | 4.927E-07 | 534.26874 | 84.727112 |
| EIVEMLFSR | 9 | Q15477 | Superkiller complex protein 2 OS=Homo | SKIC2 | 137.756 | 2 | 54.632507 | yes | 2.296E-06 | 562.29443 | 77.267181 |
| GDTVSASPCSAPL | 15 | Q15477 | Superkiller complex protein 2 OS=Homo | SKIC2 | 137.756 | 2 | 54.632507 | yes | 1.042E-07 | 744.85919 | 71.980133 |
| HPTTGQILGYK | 11 | Q15477 | Superkiller complex protein 2 OS=Homo | SKIC2 | 137.756 | 2 | 54.632507 | yes | 6.617E-07 | 607.83002 | 71.364792 |
| LFLPEGPCDHTVV | 14 | Q15477 | Superkiller complex protein 2 OS=Homo | SKIC2 | 137.756 | 2 | 54.632507 | yes | 1.084E-07 | 806.41364 | 45.589836 |
| GLGVHHSGILPILK | 14 | Q15477 | Superkiller complex protein 2 OS=Homo | SKIC2 | 137.756 | 3 | 54.632507 | yes | 2.755E-07 | 480.96103 | 20.644083 |
| GLGVHHSGILPILK | 14 | Q15477 | Superkiller complex protein 2 OS=Homo | SKIC2 | 137.756 | 3 | 54.632507 | yes | 1.135E-06 | 720.93793 | 20.644083 |
| GLGVHHSGILPILK | 14 | Q15477 | Superkiller complex protein 2 OS=Homo | SKIC2 | 137.756 | 2 | 54.632507 | yes | 2.755E-07 | 480.96103 | 20.644083 |
| GLGVHHSGILPILK | 14 | Q15477 | Superkiller complex protein 2 OS=Homo | SKIC2 | 137.756 | 2 | 54.632507 | yes | 1.135E-06 | 720.93793 | 20.644083 |

| | | | | | | | | | | | |
|---|---|---|---|---|---|---|---|---|---|---|---|
| ASSLEDLVLK | 10 | Q15477 | Superkiller complex protein 2 OS=Homo | SKIC2 | 137.756 | 2 | 54.632507 | yes | 3.388E-06 | 537.80573 | 37.708084 |
| SFSEFPSR | 8 | Q15477 | Superkiller complex protein 2 OS=Homo | SKIC2 | 137.756 | 2 | 54.632507 | yes | 9.46E-07 | 478.72726 | 35.661266 |
| HDSVFVAAHTSAC | 14 | Q15477 | Superkiller complex protein 2 OS=Homo | SKIC2 | 137.756 | 2 | 54.632507 | yes | 9.997E-08 | 713.8573 | 29.294348 |
| KLEELIQGAQCVH | 16 | Q15477 | Superkiller complex protein 2 OS=Homo | SKIC2 | 137.756 | 3 | 54.632507 | yes | 9.061E-07 | 622.32813 | 19.705229 |
| GKGEITPAAIQK | 12 | Q15532 | Protein SSXT OS=Homo sapiens OX=96 | SS18 | 45.93 | 2 | 48.875156 | yes | 1.092E-07 | 606.85095 | 411.54422 |
| GEITPAAIQK | 10 | Q15532 | Protein SSXT OS=Homo sapiens OX=96 | SS18 | 45.93 | 2 | 48.875156 | yes | 2.855E-07 | 514.29279 | 69.979637 |
| PYGYDQGQYGNY | 14 | Q15532 | Protein SSXT OS=Homo sapiens OX=96 | SS18 | 45.93 | 2 | 48.875156 | yes | 4.636E-06 | 840.84985 | 48.093513 |
| LLSTDAEAVSTR | 12 | Q15545 | Transcription initiation factor TFIID subu | TAF7 | 40.26 | 2 | 45.46261 | yes | 1.842E-07 | 631.83295 | 70.271637 |
| LVDLPCVMESLK | 12 | Q15545 | Transcription initiation factor TFIID subu | TAF7 | 40.26 | 2 | 45.46261 | yes | 7.543E-06 | 702.36731 | 64.610039 |
| WEIIAEDETK | 10 | Q15545 | Transcription initiation factor TFIID subu | TAF7 | 40.26 | 2 | 45.46261 | yes | 3.352E-06 | 617.30353 | 36.676502 |
| DLVLPTQALPASP/ | 16 | Q15554 | Telomeric repeat-binding factor 2 OS=Hc | TERF2 | 59.594 | 2 | 53.994244 | yes | 3.464E-07 | 817.4798 | 255.42761 |
| WTVEESEWVK | 10 | Q15554 | Telomeric repeat-binding factor 2 OS=Hc | TERF2 | 59.594 | 2 | 53.994244 | yes | 2.231E-07 | 646.81152 | 202.97498 |
| FYFHEALR | 8 | Q15554 | Telomeric repeat-binding factor 2 OS=Hc | TERF2 | 59.594 | 2 | 53.994244 | yes | 1.007E-07 | 541.77454 | 200.09444 |
| HEPGLGGPAER | 11 | Q15554 | Telomeric repeat-binding factor 2 OS=Hc | TERF2 | 59.594 | 2 | 53.994244 | yes | 7.047E-06 | 560.2807 | 146.10501 |
| TEFTLTEAVVESSI | 14 | Q15554 | Telomeric repeat-binding factor 2 OS=Hc | TERF2 | 59.594 | 2 | 53.994244 | yes | 6.298E-07 | 784.89374 | 130.52315 |
| NPPTTIGMMTLK | 12 | Q15554 | Telomeric repeat-binding factor 2 OS=Hc | TERF2 | 59.594 | 2 | 53.994244 | yes | 9.722E-08 | 652.34106 | 129.72406 |
| FLESHLDDAEPYLI | 18 | Q15554 | Telomeric repeat-binding factor 2 OS=Hc | TERF2 | 59.594 | 3 | 53.994244 | yes | 3.54E-06 | 698.34686 | 98.539116 |
| EAAVIICIK | 9 | Q15554 | Telomeric repeat-binding factor 2 OS=Hc | TERF2 | 59.594 | 2 | 53.994244 | yes | 2.526E-06 | 508.79407 | 90.361031 |
| VMQCLSR | 7 | Q15554 | Telomeric repeat-binding factor 2 OS=Hc | TERF2 | 59.594 | 2 | 53.994244 | yes | 1.129E-06 | 447.22021 | 67.587532 |
| TLSGAQDSEAAFA | 14 | Q15554 | Telomeric repeat-binding factor 2 OS=Hc | TERF2 | 59.594 | 2 | 53.994244 | yes | 9.722E-08 | 698.34119 | 54.105881 |
| AAGAGTAGPASGI | 17 | Q15554 | Telomeric repeat-binding factor 2 OS=Hc | TERF2 | 59.594 | 2 | 53.994244 | yes | 2.123E-07 | 719.37585 | 39.683998 |
| NDLLNIIR | 8 | Q15554 | Telomeric repeat-binding factor 2 OS=Hc | TERF2 | 59.594 | 2 | 53.994244 | yes | 2.462E-06 | 485.78763 | 31.841776 |
| DIMQALLVR | 9 | Q15554 | Telomeric repeat-binding factor 2 OS=Hc | TERF2 | 59.594 | 2 | 53.994244 | yes | 3.701E-07 | 529.80499 | 20.461418 |
| LSAQIEECIFR | 11 | Q15560 | Transcription elongation factor A protein | TCEA2 | 33.601 | 2 | 32.35384 | yes | 1.543E-06 | 683.34521 | 20.322367 |
| GPSNAFFLVK | 10 | Q15561 | Transcriptional enhancer factor TEF-3 OS | TEAD4 | 48.329 | 2 | 43.36937 | yes | 2.62E-07 | 540.29785 | 33.092537 |
| DWLYEHR | 7 | Q15583 | Homeobox protein TGIF1 OS=Homo sap | TGIF1 | 43.013 | 2 | 33.44105 | yes | 3.297E-06 | 509.74069 | 34.53204 |
| PSSPGSVLAR | 10 | Q15583 | Homeobox protein TGIF1 OS=Homo sap | TGIF1 | 43.013 | 2 | 33.44105 | yes | 1.235E-06 | 485.76944 | 34.175396 |
| QLTCTEEMAQR | 11 | Q15599 | Na(+)/H(+) exchange regulatory cofactor | NHERF2 | 37.414 | 2 | 48.096672 | yes | 1.2E-07 | 683.80811 | 33.755703 |
| FINFQLR | 7 | Q15629 | Translocating chain-associated membrane | TRAM1 | 43.072 | 2 | 55.428112 | yes | 2.494E-07 | 469.26636 | 442.9968 |
| EHSAFQAPAVK | 11 | Q15629 | Translocating chain-associated membrane | TRAM1 | 43.072 | 2 | 55.428112 | yes | 3.576E-07 | 592.80658 | 414.22549 |
| LDFSTGNFNVLAV | 14 | Q15629 | Translocating chain-associated membrane | TRAM1 | 43.072 | 2 | 55.428112 | yes | 9.722E-08 | 776.90955 | 167.55783 |
| LSVNSVTAGDYSR | 13 | Q15631 | Translin OS=Homo sapiens OX=9606 GI | TSN | 26.183 | 2 | 56.51124 | yes | 1.813E-07 | 684.84131 | 240.40607 |
| EAVTEILGIEPDR | 13 | Q15631 | Translin OS=Homo sapiens OX=9606 GI | TSN | 26.183 | 2 | 56.51124 | yes | 1.292E-07 | 721.38031 | 231.1769 |
| KVEEVVYDLSIR | 12 | Q15631 | Translin OS=Homo sapiens OX=9606 GI | TSN | 26.183 | 2 | 56.51124 | yes | 9.722E-08 | 725.401 | 144.04497 |
| VVQSLEQTAR | 10 | Q15631 | Translin OS=Homo sapiens OX=9606 GI | TSN | 26.183 | 2 | 56.51124 | yes | 1.427E-07 | 565.81183 | 137.9588 |
| KVVQSLEQTAR | 11 | Q15631 | Translin OS=Homo sapiens OX=9606 GI | TSN | 26.183 | 2 | 56.51124 | yes | 1.301E-07 | 629.85931 | 56.591724 |
| EILTLLQGVHQGA | 20 | Q15631 | Translin OS=Homo sapiens OX=9606 GI | TSN | 26.183 | 3 | 56.51124 | yes | 6.852E-07 | 722.06354 | 35.710564 |
| TPISLLQEYGTR | 12 | Q15633 | RISC-loading complex subunit TARBP2 | TARBP2 | 39.04 | 2 | 54.35935 | yes | 1.02E-07 | 689.37225 | 1788.6174 |
| SCSLGSLGALGPA | 16 | Q15633 | RISC-loading complex subunit TARBP2 | TARBP2 | 39.04 | 2 | 54.35935 | yes | 1.188E-07 | 833.37891 | 1368.0111 |

| | | | | | | | | | | | |
|---|---|---|---|---|---|---|---|---|---|---|---|
| VHTVPLDAR | 9 | Q15633 | RISC-loading complex subunit TARBP2 | TARBP2 | 39.04 | 2 | 54.35935 | yes | 2.572E-07 | 504.28528 | 1206.337 |
| GPGCTWDSLR | 10 | Q15633 | RISC-loading complex subunit TARBP2 | TARBP2 | 39.04 | 2 | 54.35935 | yes | 1.11E-07 | 574.76129 | 800.35376 |
| FIEIGSGTSK | 10 | Q15633 | RISC-loading complex subunit TARBP2 | TARBP2 | 39.04 | 2 | 54.35935 | yes | 2.41E-07 | 519.77692 | 797.9043 |
| VTVGDTSCTGQGF | 15 | Q15633 | RISC-loading complex subunit TARBP2 | TARBP2 | 39.04 | 2 | 54.35935 | yes | 1.576E-07 | 747.34863 | 287.02124 |
| TPVYDLLK | 8 | Q15633 | RISC-loading complex subunit TARBP2 | TARBP2 | 39.04 | 2 | 54.35935 | yes | 1.086E-07 | 474.77368 | 276.18265 |
| EFTMTCR | 7 | Q15633 | RISC-loading complex subunit TARBP2 | TARBP2 | 39.04 | 2 | 54.35935 | yes | 9.722E-08 | 472.70187 | 270.1185 |
| AEGQAHQPNFTFR | 13 | Q15633 | RISC-loading complex subunit TARBP2 | TARBP2 | 39.04 | 2 | 54.35935 | yes | 9.722E-08 | 751.86041 | 123.32047 |
| AEGQAHQPNFTFR | 13 | Q15633 | RISC-loading complex subunit TARBP2 | TARBP2 | 39.04 | 2 | 54.35935 | yes | 1.243E-07 | 501.57602 | 123.32047 |
| AEGQAHQPNFTFR | 13 | Q15633 | RISC-loading complex subunit TARBP2 | TARBP2 | 39.04 | 3 | 54.35935 | yes | 9.722E-08 | 751.86041 | 123.32047 |
| AEGQAHQPNFTFR | 13 | Q15633 | RISC-loading complex subunit TARBP2 | TARBP2 | 39.04 | 3 | 54.35935 | yes | 1.243E-07 | 501.57602 | 123.32047 |
| LPEYTVTQESGPA | 15 | Q15633 | RISC-loading complex subunit TARBP2 | TARBP2 | 39.04 | 2 | 54.35935 | yes | 1.851E-07 | 842.91809 | 175.97862 |
| KEFTMTCR | 8 | Q15633 | RISC-loading complex subunit TARBP2 | TARBP2 | 39.04 | 2 | 54.35935 | yes | 2.456E-05 | 536.74933 | 22.765877 |
| TVIPGMPTVIPPGL | 16 | Q15637 | Splicing factor 1 OS=Homo sapiens OX= | SF1 | 68.33 | 2 | 50.626144 | yes | 1.441E-07 | 824.97638 | 8639.04 |
| HNLITEMVALNPD | 26 | Q15637 | Splicing factor 1 OS=Homo sapiens OX= | SF1 | 68.33 | 4 | 50.626144 | yes | 1.159E-07 | 734.63385 | 4277.8745 |
| HNLITEMVALNPD | 26 | Q15637 | Splicing factor 1 OS=Homo sapiens OX= | SF1 | 68.33 | 4 | 50.626144 | yes | 1.35E-07 | 587.90851 | 4277.8745 |
| HNLITEMVALNPD | 26 | Q15637 | Splicing factor 1 OS=Homo sapiens OX= | SF1 | 68.33 | 5 | 50.626144 | yes | 1.159E-07 | 734.63385 | 4277.8745 |
| HNLITEMVALNPD | 26 | Q15637 | Splicing factor 1 OS=Homo sapiens OX= | SF1 | 68.33 | 5 | 50.626144 | yes | 1.35E-07 | 587.90851 | 4277.8745 |
| ATGANATPLDFPS | 14 | Q15637 | Splicing factor 1 OS=Homo sapiens OX= | SF1 | 68.33 | 2 | 50.626144 | yes | 9.894E-08 | 716.35938 | 5904.7573 |
| AYIVQLQIEDLTR | 13 | Q15637 | Splicing factor 1 OS=Homo sapiens OX= | SF1 | 68.33 | 2 | 50.626144 | yes | 9.733E-08 | 781.43286 | 3733.8252 |
| SITNTTVCTK | 10 | Q15637 | Splicing factor 1 OS=Homo sapiens OX= | SF1 | 68.33 | 2 | 50.626144 | yes | 1.168E-07 | 562.78442 | 2584.8613 |
| CGGAGHIASDCK | 12 | Q15637 | Splicing factor 1 OS=Homo sapiens OX= | SF1 | 68.33 | 2 | 50.626144 | yes | 1.214E-07 | 616.76099 | 819.04779 |
| CGGAGHIASDCK | 12 | Q15637 | Splicing factor 1 OS=Homo sapiens OX= | SF1 | 68.33 | 2 | 50.626144 | yes | 2.165E-07 | 411.50974 | 819.04779 |
| CGGAGHIASDCK | 12 | Q15637 | Splicing factor 1 OS=Homo sapiens OX= | SF1 | 68.33 | 3 | 50.626144 | yes | 1.214E-07 | 616.76099 | 819.04779 |
| CGGAGHIASDCK | 12 | Q15637 | Splicing factor 1 OS=Homo sapiens OX= | SF1 | 68.33 | 3 | 50.626144 | yes | 2.165E-07 | 411.50974 | 819.04779 |
| QGIETPEDQNDLR | 13 | Q15637 | Splicing factor 1 OS=Homo sapiens OX= | SF1 | 68.33 | 2 | 50.626144 | yes | 1.974E-07 | 757.85773 | 1634.7861 |
| WNQDTMEQK | 9 | Q15637 | Splicing factor 1 OS=Homo sapiens OX= | SF1 | 68.33 | 2 | 50.626144 | yes | 3.843E-06 | 590.25861 | 682.62683 |
| WNQDTMEQK | 9 | Q15637 | Splicing factor 1 OS=Homo sapiens OX= | SF1 | 68.33 | 2 | 50.626144 | yes | 2.223E-05 | 598.25604 | 682.62683 |
| KAVEQIR | 7 | Q15637 | Splicing factor 1 OS=Homo sapiens OX= | SF1 | 68.33 | 2 | 50.626144 | yes | 2.642E-07 | 422.25598 | 867.1709 |
| ILRPWQSSETR | 11 | Q15637 | Splicing factor 1 OS=Homo sapiens OX= | SF1 | 68.33 | 2 | 50.626144 | yes | 1.538E-07 | 686.87024 | 590.07892 |
| FQRPGDPQSAQDK | 13 | Q15637 | Splicing factor 1 OS=Homo sapiens OX= | SF1 | 68.33 | 2 | 50.626144 | yes | 3.25E-07 | 737.35767 | 193.60568 |
| TGDLGIPPNPEDR | 13 | Q15637 | Splicing factor 1 OS=Homo sapiens OX= | SF1 | 68.33 | 2 | 50.626144 | yes | 5.709E-05 | 690.84131 | 172.40483 |
| QGIETPEDQNDLR | 14 | Q15637 | Splicing factor 1 OS=Homo sapiens OX= | SF1 | 68.33 | 2 | 50.626144 | yes | 5.064E-07 | 821.90521 | 37.29126 |
| EGGEGYVPTSYLR | 13 | Q15642 | Cdc42-interacting protein 4 OS=Homo sa | TRIP10 | 68.353 | 2 | 48.334423 | yes | 1.163E-07 | 714.34369 | 82.311722 |
| LSLLLNDISR | 10 | Q15645 | Pachytene checkpoint protein 2 homolog | TRIP13 | 48.551 | 2 | 55.865757 | yes | 2.439E-07 | 572.34021 | 273.68021 |
| NVNSNLITWNR | 11 | Q15645 | Pachytene checkpoint protein 2 homolog | TRIP13 | 48.551 | 2 | 55.865757 | yes | 9.722E-08 | 665.84674 | 164.96059 |
| VVNAVLTQIDQIK | 13 | Q15645 | Pachytene checkpoint protein 2 homolog | TRIP13 | 48.551 | 2 | 55.865757 | yes | 1.719E-07 | 720.92468 | 150.2708 |
| KEDINLSVR | 9 | Q15645 | Pachytene checkpoint protein 2 homolog | TRIP13 | 48.551 | 2 | 55.865757 | yes | 1.114E-07 | 537.30109 | 140.14043 |
| QYIGPPSAAAIFK | 13 | Q15645 | Pachytene checkpoint protein 2 homolog | TRIP13 | 48.551 | 2 | 55.865757 | yes | 9.722E-08 | 681.87445 | 138.61987 |
| QALPCVAESPTVH | 19 | Q15645 | Pachytene checkpoint protein 2 homolog | TRIP13 | 48.551 | 3 | 55.865757 | yes | 3.237E-07 | 719.7005 | 65.352638 |

| | | | | | | | | | | | |
|---|---|---|---|---|---|---|---|---|---|---|---|
| IDVAFVDR | 8 | Q15645 | Pachytene checkpoint protein 2 homolog | TRIP13 | 48.551 | 2 | 55.865757 | yes | 1.131E-07 | 467.75327 | 57.732548 |
| VVLLHGPPGTGK | 12 | Q15645 | Pachytene checkpoint protein 2 homolog | TRIP13 | 48.551 | 2 | 55.865757 | yes | 1.666E-07 | 587.85077 | 48.131287 |
| NVQSVSIIDTELK | 13 | Q15645 | Pachytene checkpoint protein 2 homolog | TRIP13 | 48.551 | 2 | 55.865757 | yes | 3.789E-07 | 723.39594 | 42.500454 |
| HSNVVILTTSNITE | 15 | Q15645 | Pachytene checkpoint protein 2 homolog | TRIP13 | 48.551 | 2 | 55.865757 | yes | 1.041E-07 | 828.45178 | 35.331532 |
| VPDALVFTIQTR | 12 | Q15646 | 2'-5'-oligoadenylate synthase-like protein | OASL | 59.227 | 2 | 52.407322 | yes | 9.722E-08 | 680.38519 | 1369.0518 |
| LSFQVPGSER | 10 | Q15646 | 2'-5'-oligoadenylate synthase-like protein | OASL | 59.227 | 2 | 52.407322 | yes | 9.722E-08 | 560.29327 | 1091.8127 |
| GTAEPITVTIVPAY | 15 | Q15646 | 2'-5'-oligoadenylate synthase-like protein | OASL | 59.227 | 2 | 52.407322 | yes | 1.194E-07 | 794.44067 | 1069.2615 |
| DIHLTVEQR | 9 | Q15646 | 2'-5'-oligoadenylate synthase-like protein | OASL | 59.227 | 2 | 52.407322 | yes | 1.043E-07 | 555.79877 | 773.83716 |
| QQIEDQQGLPK | 11 | Q15646 | 2'-5'-oligoadenylate synthase-like protein | OASL | 59.227 | 2 | 52.407322 | yes | 9.722E-08 | 642.33313 | 606.16156 |
| EEVLDAVR | 8 | Q15646 | 2'-5'-oligoadenylate synthase-like protein | OASL | 59.227 | 2 | 52.407322 | yes | 1.602E-07 | 465.74817 | 272.16425 |
| HWYQQYVK | 8 | Q15646 | 2'-5'-oligoadenylate synthase-like protein | OASL | 59.227 | 2 | 52.407322 | yes | 1.155E-07 | 576.28528 | 265.14133 |
| KKGEALFPAS | 10 | Q15646 | 2'-5'-oligoadenylate synthase-like protein | OASL | 59.227 | 2 | 52.407322 | yes | 4.883E-07 | 524.29529 | 204.58073 |
| TVEEFLR | 7 | Q15646 | 2'-5'-oligoadenylate synthase-like protein | OASL | 59.227 | 2 | 52.407322 | yes | 1.606E-06 | 447.23999 | 161.28091 |
| EWKEEVLDAVR | 11 | Q15646 | 2'-5'-oligoadenylate synthase-like protein | OASL | 59.227 | 3 | 52.407322 | yes | 9.722E-08 | 458.57352 | 58.230492 |
| EWKEEVLDAVR | 11 | Q15646 | 2'-5'-oligoadenylate synthase-like protein | OASL | 59.227 | 3 | 52.407322 | yes | 2.516E-07 | 687.35663 | 58.230492 |
| EWKEEVLDAVR | 11 | Q15646 | 2'-5'-oligoadenylate synthase-like protein | OASL | 59.227 | 2 | 52.407322 | yes | 9.722E-08 | 458.57352 | 58.230492 |
| EWKEEVLDAVR | 11 | Q15646 | 2'-5'-oligoadenylate synthase-like protein | OASL | 59.227 | 2 | 52.407322 | yes | 2.516E-07 | 687.35663 | 58.230492 |
| VGSFGNGTVLR | 11 | Q15646 | 2'-5'-oligoadenylate synthase-like protein | OASL | 59.227 | 2 | 52.407322 | yes | 1.879E-07 | 553.80127 | 104.02827 |
| ALMQELYSTPASR | 13 | Q15646 | 2'-5'-oligoadenylate synthase-like protein | OASL | 59.227 | 2 | 52.407322 | yes | 3.595E-06 | 754.87433 | 78.847923 |
| ALGPSLPNSQPPPE | 21 | Q15646 | 2'-5'-oligoadenylate synthase-like protein | OASL | 59.227 | 3 | 52.407322 | yes | 3.986E-07 | 736.07544 | 24.489595 |
| YYVSPSDLLDDK | 12 | Q15648 | Mediator of RNA polymerase II transcrip | MED1 | 168.479 | 2 | 52.033566 | yes | 1.1E-07 | 707.84045 | 226.28656 |
| GLSDALICTDDFIA | 15 | Q15648 | Mediator of RNA polymerase II transcrip | MED1 | 168.479 | 2 | 52.033566 | yes | 2.082E-07 | 819.90582 | 156.49934 |
| TASPIILHENNVSR | 14 | Q15648 | Mediator of RNA polymerase II transcrip | MED1 | 168.479 | 2 | 52.033566 | yes | 9.722E-08 | 775.91791 | 149.39619 |
| SSSGLGSSGSLSQK | 14 | Q15648 | Mediator of RNA polymerase II transcrip | MED1 | 168.479 | 2 | 52.033566 | yes | 1.04E-07 | 641.31769 | 112.81435 |
| SLQGTLVSK | 9 | Q15648 | Mediator of RNA polymerase II transcrip | MED1 | 168.479 | 2 | 52.033566 | yes | 1.675E-07 | 466.7742 | 98.780861 |
| GLVNLYNLPGDNK | 13 | Q15648 | Mediator of RNA polymerase II transcrip | MED1 | 168.479 | 2 | 52.033566 | yes | 1.001E-07 | 708.87769 | 79.216621 |
| MYLALQSLEQDLS | 14 | Q15648 | Mediator of RNA polymerase II transcrip | MED1 | 168.479 | 2 | 52.033566 | yes | 9.722E-08 | 819.92401 | 61.621372 |
| KPSLTAVIDK | 10 | Q15648 | Mediator of RNA polymerase II transcrip | MED1 | 168.479 | 2 | 52.033566 | yes | 2.012E-07 | 536.32404 | 58.520069 |
| SQTPPGVATPPIPK | 14 | Q15648 | Mediator of RNA polymerase II transcrip | MED1 | 168.479 | 2 | 52.033566 | yes | 2.021E-07 | 695.39044 | 42.834698 |
| VSTSGSSVDSSK | 12 | Q15648 | Mediator of RNA polymerase II transcrip | MED1 | 168.479 | 2 | 52.033566 | yes | 1.012E-06 | 570.77258 | 35.053524 |
| LPIAPLIMGSHPVD | 16 | Q15648 | Mediator of RNA polymerase II transcrip | MED1 | 168.479 | 3 | 52.033566 | yes | 4.45E-06 | 567.9834 | 32.215927 |
| SLLLNPHLR | 9 | Q15649 | Zinc finger HIT domain-containing protei | ZNHIT3 | 17.608 | 2 | 53.228043 | yes | 1.736E-07 | 531.82458 | 341.28528 |
| VPYCSVVCFR | 10 | Q15649 | Zinc finger HIT domain-containing protei | ZNHIT3 | 17.608 | 2 | 53.228043 | yes | 1.108E-07 | 643.80463 | 234.0119 |
| CSTVVCVICLEKPI | 14 | Q15649 | Zinc finger HIT domain-containing protei | ZNHIT3 | 17.608 | 2 | 53.228043 | yes | 9.722E-08 | 846.92773 | 159.90009 |
| QLMVNLDQGEDK | 12 | Q15649 | Zinc finger HIT domain-containing protei | ZNHIT3 | 17.608 | 2 | 53.228043 | yes | 1.166E-07 | 695.33759 | 83.607841 |
| NLGESATLR | 9 | Q15649 | Zinc finger HIT domain-containing protei | ZNHIT3 | 17.608 | 2 | 53.228043 | yes | 3.435E-06 | 480.75909 | 71.854004 |
| DLLPHQELR | 9 | Q15650 | Activating signal cointegrator 1 OS=Hom | TRIP4 | 66.147 | 2 | 52.162853 | yes | 8.594E-07 | 560.80908 | 6942.2808 |
| KVTIDFAGR | 9 | Q15650 | Activating signal cointegrator 1 OS=Hom | TRIP4 | 66.147 | 2 | 52.162853 | yes | 1.112E-07 | 503.78763 | 4790.3398 |
| LINNCLICGR | 10 | Q15650 | Activating signal cointegrator 1 OS=Hom | TRIP4 | 66.147 | 2 | 52.162853 | yes | 1.324E-07 | 616.81555 | 4672.126 |

| | | | | | | | | | | | |
|---|---|---|---|---|---|---|---|---|---|---|---|
| EYVTDLLQGNEGK | 13 | Q15650 | Activating signal cointegrator 1 OS=Hom | TRIP4 | 66.147 | 2 | 52.162853 | yes | 1.006E-07 | 733.36212 | 4390.1099 |
| GQFIEELITK | 10 | Q15650 | Activating signal cointegrator 1 OS=Hom | TRIP4 | 66.147 | 2 | 52.162853 | yes | 1.391E-07 | 589.32678 | 1851.3048 |
| HPCDCLGQK | 9 | Q15650 | Activating signal cointegrator 1 OS=Hom | TRIP4 | 66.147 | 2 | 52.162853 | yes | 1.104E-07 | 557.74207 | 1752.3414 |
| AVAGAVSGEPLVH | 20 | Q15650 | Activating signal cointegrator 1 OS=Hom | TRIP4 | 66.147 | 3 | 52.162853 | yes | 1.283E-07 | 741.04425 | 1109.6654 |
| SWYTPHR | 7 | Q15650 | Activating signal cointegrator 1 OS=Hom | TRIP4 | 66.147 | 2 | 52.162853 | yes | 2.866E-06 | 473.73013 | 1105.106 |
| FVNLYTR | 7 | Q15650 | Activating signal cointegrator 1 OS=Hom | TRIP4 | 66.147 | 2 | 52.162853 | yes | 1.52E-07 | 456.75052 | 991.52917 |
| KGQFIEELITK | 11 | Q15650 | Activating signal cointegrator 1 OS=Hom | TRIP4 | 66.147 | 2 | 52.162853 | yes | 9.81E-08 | 653.37427 | 955.33691 |
| KPSPQEVSELQAT | 15 | Q15650 | Activating signal cointegrator 1 OS=Hom | TRIP4 | 66.147 | 3 | 52.162853 | yes | 9.722E-08 | 578.30035 | 867.74829 |
| NPQEMVVK | 8 | Q15650 | Activating signal cointegrator 1 OS=Hom | TRIP4 | 66.147 | 2 | 52.162853 | yes | 2.013E-07 | 472.74713 | 524.60254 |
| ILEEENSLAEYHSR | 14 | Q15650 | Activating signal cointegrator 1 OS=Hom | TRIP4 | 66.147 | 2 | 52.162853 | yes | 1.011E-07 | 845.40759 | 512.31378 |
| KDEILDGQK | 9 | Q15650 | Activating signal cointegrator 1 OS=Hom | TRIP4 | 66.147 | 2 | 52.162853 | yes | 1.55E-07 | 523.27985 | 408.72513 |
| KLMSGVENSGK | 11 | Q15650 | Activating signal cointegrator 1 OS=Hom | TRIP4 | 66.147 | 2 | 52.162853 | yes | 9.723E-08 | 575.30023 | 347.64978 |
| LAVLLPGR | 8 | Q15650 | Activating signal cointegrator 1 OS=Hom | TRIP4 | 66.147 | 2 | 52.162853 | yes | 2.093E-07 | 419.77908 | 209.47485 |
| VTIDFAGR | 8 | Q15650 | Activating signal cointegrator 1 OS=Hom | TRIP4 | 66.147 | 2 | 52.162853 | yes | 2.253E-07 | 439.74017 | 116.57788 |
| DEILDGQK | 8 | Q15650 | Activating signal cointegrator 1 OS=Hom | TRIP4 | 66.147 | 2 | 52.162853 | yes | 1.038E-06 | 459.23236 | 101.06192 |
| TQVIDDESDYFASI | 21 | Q15650 | Activating signal cointegrator 1 OS=Hom | TRIP4 | 66.147 | 3 | 52.162853 | yes | 9.818E-08 | 816.69946 | 58.949169 |
| LWIAATAK | 8 | Q15650 | Activating signal cointegrator 1 OS=Hom | TRIP4 | 66.147 | 2 | 52.162853 | yes | 3.411E-05 | 437.26328 | 33.11681 |
| EQFPDISQESDSPF | 19 | Q15650 | Activating signal cointegrator 1 OS=Hom | TRIP4 | 66.147 | 3 | 52.162853 | yes | 8.019E-07 | 758.35254 | 24.900799 |
| ILQESIDVAPFTTK | 14 | Q15652 | Probable JmjC domain-containing histone | JMJD1C | 284.525 | 2 | 44.812443 | yes | 2.175E-07 | 781.42725 | 51.85704 |
| CGFVVCLDCYK | 11 | Q15652 | Probable JmjC domain-containing histone | JMJD1C | 284.525 | 2 | 44.812443 | yes | 6.84E-07 | 710.80652 | 44.735378 |
| EFWDGFEEVSK | 11 | Q15652 | Probable JmjC domain-containing histone | JMJD1C | 284.525 | 2 | 44.812443 | yes | 4.99E-07 | 686.8064 | 41.220043 |
| LLEEYGVR | 8 | Q15652 | Probable JmjC domain-containing histone | JMJD1C | 284.525 | 2 | 44.812443 | yes | 1.097E-06 | 489.76639 | 37.425323 |
| ATYVNSQATGER | 12 | Q15652 | Probable JmjC domain-containing histone | JMJD1C | 284.525 | 2 | 44.812443 | yes | 2.06E-07 | 648.81256 | 25.732401 |
| LKDWPSGEDFK | 11 | Q15652 | Probable JmjC domain-containing histone | JMJD1C | 284.525 | 2 | 44.812443 | yes | 3.492E-07 | 661.32477 | 24.308489 |
| CATCSQPILDR | 11 | Q15654 | Thyroid receptor-interacting protein 6 OS | TRIP6 | 50.288 | 2 | 53.949047 | yes | 9.722E-08 | 660.80536 | 865.72644 |
| GQHFYAVER | 9 | Q15654 | Thyroid receptor-interacting protein 6 OS | TRIP6 | 50.288 | 2 | 53.949047 | yes | 1.231E-07 | 553.77252 | 762.16431 |
| AYCEGCYVATLEK | 13 | Q15654 | Thyroid receptor-interacting protein 6 OS | TRIP6 | 50.288 | 2 | 53.949047 | yes | 9.768E-08 | 782.34454 | 696.47119 |
| QAYEPPPPPAYR | 12 | Q15654 | Thyroid receptor-interacting protein 6 OS | TRIP6 | 50.288 | 2 | 53.949047 | yes | 1.002E-07 | 693.34607 | 573.33417 |
| VFHVGCFVCSTCR | 13 | Q15654 | Thyroid receptor-interacting protein 6 OS | TRIP6 | 50.288 | 2 | 53.949047 | yes | 4.568E-07 | 814.86017 | 243.98399 |
| GASQASGPLPGPH | 19 | Q15654 | Thyroid receptor-interacting protein 6 OS | TRIP6 | 50.288 | 3 | 53.949047 | yes | 1.588E-07 | 614.99182 | 33.766529 |
| GLDGIPFTVDATSC | 23 | Q15654 | Thyroid receptor-interacting protein 6 OS | TRIP6 | 50.288 | 4 | 53.949047 | yes | 6.6E-06 | 657.81824 | 14.011365 |
| WVELEKPLK | 9 | Q15678 | Tyrosine-protein phosphatase non-recepto | PTPN14 | 135.262 | 2 | 40.5381 | yes | 3.972E-07 | 571.33441 | 39.543522 |
| LTVEDLEK | 8 | Q15691 | Microtubule-associated protein RP/EB fa | MAPRE1 | 30 | 2 | 49.494358 | yes | 3.507E-07 | 473.75821 | 37.331795 |
| QGQETAVAPSLVA | 20 | Q15691 | Microtubule-associated protein RP/EB fa | MAPRE1 | 30 | 3 | 49.494358 | yes | 1.025E-07 | 673.71265 | 24.920618 |
| YGQIQQAK | 8 | Q15699 | ALX homeobox protein 1 OS=Homo sapi | ALX1 | 36.962 | 2 | 49.04871 | yes | 4.335E-07 | 468.25089 | 215.64981 |
| TTFTSLQLEELEK | 13 | Q15699 | ALX homeobox protein 1 OS=Homo sapi | ALX1 | 36.962 | 2 | 49.04871 | yes | 1.074E-07 | 769.90106 | 140.08394 |
| VEGQPLHTELNR | 12 | Q15699 | ALX homeobox protein 1 OS=Homo sapi | ALX1 | 36.962 | 2 | 49.04871 | yes | 1.215E-07 | 696.86511 | 106.31898 |
| SLFSSIGEVESAK | 13 | Q15717 | ELAV-like protein 1 OS=Homo sapiens ( | ELAVL1 | 36.092 | 2 | 49.153114 | yes | 1.53E-07 | 677.34845 | 43919.938 |
| VLVDQTTGLSR | 11 | Q15717 | ELAV-like protein 1 OS=Homo sapiens ( | ELAVL1 | 36.092 | 2 | 49.153114 | yes | 1.727E-07 | 594.83276 | 24869.355 |

| | | | | | | | | | | | |
|---|---|---|---|---|---|---|---|---|---|---|---|
| DANLYISGLPR | 11 | Q15717 | ELAV-like protein 1 OS=Homo sapiens ( | ELAVL1 | 36.092 | 2 | 49.153114 | yes | 1.149E-07 | 609.82751 | 20992.834 |
| FGGPVHHQAQR | 11 | Q15717 | ELAV-like protein 1 OS=Homo sapiens ( | ELAVL1 | 36.092 | 3 | 49.153114 | yes | 1.101E-07 | 411.87936 | 6912.3564 |
| FGGPVHHQAQR | 11 | Q15717 | ELAV-like protein 1 OS=Homo sapiens ( | ELAVL1 | 36.092 | 3 | 49.153114 | yes | 1.274E-07 | 617.31543 | 6912.3564 |
| FGGPVHHQAQR | 11 | Q15717 | ELAV-like protein 1 OS=Homo sapiens ( | ELAVL1 | 36.092 | 2 | 49.153114 | yes | 1.101E-07 | 411.87936 | 6912.3564 |
| FGGPVHHQAQR | 11 | Q15717 | ELAV-like protein 1 OS=Homo sapiens ( | ELAVL1 | 36.092 | 2 | 49.153114 | yes | 1.274E-07 | 617.31543 | 6912.3564 |
| FAANPNQNK | 9 | Q15717 | ELAV-like protein 1 OS=Homo sapiens ( | ELAVL1 | 36.092 | 2 | 49.153114 | yes | 5.689E-07 | 502.25143 | 12310.389 |
| NVALLSQLYHSPA | 14 | Q15717 | ELAV-like protein 1 OS=Homo sapiens ( | ELAVL1 | 36.092 | 2 | 49.153114 | yes | 2.635E-07 | 784.93079 | 3641.8071 |
| DVEDMFSR | 8 | Q15717 | ELAV-like protein 1 OS=Homo sapiens ( | ELAVL1 | 36.092 | 2 | 49.153114 | yes | 1.063E-07 | 499.71603 | 1826.0702 |
| VAGHSLGYGFVN | 17 | Q15717 | ELAV-like protein 1 OS=Homo sapiens ( | ELAVL1 | 36.092 | 3 | 49.153114 | yes | 3.16E-07 | 594.97729 | 705.51343 |
| SNGYEDHMAEDC | 13 | Q15717 | ELAV-like protein 1 OS=Homo sapiens ( | ELAVL1 | 36.092 | 2 | 49.153114 | yes | 5.488E-07 | 813.30139 | 169.05499 |
| RFGGPVHHQAQR | 12 | Q15717 | ELAV-like protein 1 OS=Homo sapiens ( | ELAVL1 | 36.092 | 3 | 49.153114 | yes | 3.568E-06 | 463.91309 | 7.6266146 |
| LIDFGLAR | 8 | Q15746 | Myosin light chain kinase, smooth muscle | MYLK | 210.716 | 2 | 42.29987 | yes | 3.097E-07 | 452.76617 | 74.155617 |
| IEDMAVSLIR | 10 | Q15776 | Zinc finger protein with KRAB and SCA | ZKSCAN8 | 65.816 | 2 | 47.75468 | yes | 3.272E-07 | 573.81299 | 123.82643 |
| AFSQNTGLILHQR | 13 | Q15776 | Zinc finger protein with KRAB and SCA | ZKSCAN8 | 65.816 | 2 | 47.75468 | yes | 1.896E-07 | 742.90204 | 56.884567 |
| SALLSHQDIHNK | 12 | Q15776 | Zinc finger protein with KRAB and SCA | ZKSCAN8 | 65.816 | 2 | 47.75468 | yes | 1.265E-07 | 681.85986 | 27.768919 |
| FCLGLLSNVNR | 11 | Q15796 | Mothers against decapentaplegic homolog | SMAD2 | 52.307 | 2 | 53.14539 | no | 1.051E-07 | 646.84265 | 855.0976 |
| VETPVLPPVLVPR | 13 | Q15796 | Mothers against decapentaplegic homolog | SMAD2 | 52.307 | 2 | 53.14539 | no | 1.648E-07 | 708.43469 | 376.83359 |
| GLPHVIYCR | 9 | Q15796 | Mothers against decapentaplegic homolog | SMAD2 | 52.307 | 2 | 53.14539 | no | 1.071E-07 | 557.79492 | 305.4472 |
| VLTQMGSPSVR | 11 | Q15796 | Mothers against decapentaplegic homolog | SMAD2 | 52.307 | 2 | 53.14539 | yes | 9.722E-08 | 587.8161 | 279.85367 |
| AITTQNCNTK | 10 | Q15796 | Mothers against decapentaplegic homolog | SMAD2 | 52.307 | 2 | 53.14539 | yes | 2.086E-07 | 575.77966 | 201.25662 |
| AIENCEYAFNLK | 12 | Q15796 | Mothers against decapentaplegic homolog | SMAD2 | 52.307 | 2 | 53.14539 | yes | 9.722E-08 | 736.34796 | 123.21124 |
| DEVCVNPYHYQR | 12 | Q15796 | Mothers against decapentaplegic homolog | SMAD2 | 52.307 | 2 | 53.14539 | no | 1.597E-07 | 790.35156 | 50.009876 |
| VESPVLPPVLVPR | 13 | Q15797 | Mothers against decapentaplegic homolog | SMAD1 | 52.261 | 2 | 24.579813 | yes | 3.428E-06 | 701.42682 | 28.284817 |
| DLCCFLDSER | 10 | Q15834 | Coiled-coil domain-containing protein 85 | CCDC85B | 22.091 | 2 | 45.47847 | yes | 8.608E-07 | 657.77631 | 67.415024 |
| QWQLFGTQASR | 11 | Q15834 | Coiled-coil domain-containing protein 85 | CCDC85B | 22.091 | 2 | 45.47847 | yes | 1.838E-07 | 661.336 | 33.897655 |
| ALLPLELQDDGSD | 15 | Q15906 | Vacuolar protein sorting-associated protei | VPS72 | 40.595 | 2 | 51.639248 | yes | 1.706E-07 | 814.90991 | 269.44803 |
| DPVTDIPYATAR | 12 | Q15906 | Vacuolar protein sorting-associated protei | VPS72 | 40.595 | 2 | 51.639248 | yes | 9.722E-08 | 659.83551 | 178.01941 |
| EVCPVTHR | 8 | Q15906 | Vacuolar protein sorting-associated protei | VPS72 | 40.595 | 2 | 51.639248 | yes | 1.483E-07 | 499.24783 | 147.72679 |
| PLTQEELLR | 9 | Q15906 | Vacuolar protein sorting-associated protei | VPS72 | 40.595 | 2 | 51.639248 | yes | 1.61E-07 | 549.81134 | 100.29855 |
| AYKEPLK | 7 | Q15906 | Vacuolar protein sorting-associated protei | VPS72 | 40.595 | 2 | 51.639248 | yes | 9.722E-07 | 424.74747 | 41.969273 |
| EVCPVTHRPALYR | 13 | Q15906 | Vacuolar protein sorting-associated protei | VPS72 | 40.595 | 3 | 51.639248 | yes | 8.43E-05 | 533.28046 | 17.499138 |
| EFAAALTAER | 10 | Q15910 | Histone-lysine N-methyltransferase EZH2 | EZH2 | 85.364 | 2 | 54.041103 | yes | 1.732E-07 | 539.78003 | 321.08884 |
| TEILNQEWK | 9 | Q15910 | Histone-lysine N-methyltransferase EZH2 | EZH2 | 85.364 | 2 | 54.041103 | yes | 6.875E-07 | 580.80096 | 221.78352 |
| AIQTGEELFFDYR | 13 | Q15910 | Histone-lysine N-methyltransferase EZH2 | EZH2 | 85.364 | 2 | 54.041103 | yes | 2.292E-07 | 794.88574 | 155.97791 |
| IQPVHILTSVSSLR | 14 | Q15910 | Histone-lysine N-methyltransferase EZH2 | EZH2 | 85.364 | 2 | 54.041103 | yes | 9.977E-07 | 775.45667 | 126.56842 |
| IFEAISSMFPDK | 12 | Q15910 | Histone-lysine N-methyltransferase EZH2 | EZH2 | 85.364 | 2 | 54.041103 | yes | 1.3E-07 | 692.84448 | 99.489395 |
| LWAAHCR | 7 | Q15910 | Histone-lysine N-methyltransferase EZH2 | EZH2 | 85.364 | 2 | 54.041103 | no | 4.097E-07 | 457.22668 | 50.686146 |
| FANHSVNPNCYAI | 13 | Q15910 | Histone-lysine N-methyltransferase EZH2 | EZH2 | 85.364 | 2 | 54.041103 | no | 9.722E-08 | 761.34882 | 31.162491 |
| ECDPDLCLTCGAA | 19 | Q15910 | Histone-lysine N-methyltransferase EZH2 | EZH2 | 85.364 | 3 | 54.041103 | yes | 3.355E-06 | 750.6347 | 27.660816 |

| | | | | | | | | | | | |
|---|---|---|---|---|---|---|---|---|---|---|---|
| NEFISEYCGEIISQI | 20 | Q15910 | Histone-lysine N-methyltransferase EZH2 | EZH2 | 85.364 | 3 | 54.041103 | yes | 5.689E-06 | 811.03253 | 25.197918 |
| FEDYLNAESR | 10 | Q16181 | Septin-7 OS=Homo sapiens OX=9606 GI | SEPTIN7 | 50.68 | 2 | 50.240395 | yes | 9.839E-08 | 622.28314 | 117.5696 |
| DVTNNVHYENYR | 12 | Q16181 | Septin-7 OS=Homo sapiens OX=9606 GI | SEPTIN7 | 50.68 | 2 | 50.240395 | yes | 3.354E-07 | 762.34729 | 18.955313 |
| SQSAAVTPSSTTSS | 16 | Q16186 | Proteasomal ubiquitin receptor ADRM1 ( | ADRM1 | 42.154 | 2 | 46.09743 | yes | 1.654E-07 | 784.38153 | 32.41935 |
| DISMEIDSPENMM | 14 | Q16204 | Coiled-coil domain-containing protein 6 ( | CCDC6 | 53.291 | 2 | 43.55287 | yes | 2.542E-07 | 834.3573 | 27.893858 |
| NIHNDELMGIR | 11 | Q16206 | Ecto-NOX disulfide-thiol exchanger 2 OS | ENOX2 | 70.083 | 2 | 54.410915 | yes | 1.222E-07 | 656.32733 | 352.13821 |
| FCGFEGLK | 8 | Q16206 | Ecto-NOX disulfide-thiol exchanger 2 OS | ENOX2 | 70.083 | 2 | 54.410915 | yes | 9.722E-08 | 479.22858 | 259.53647 |
| ALYLSGYR | 8 | Q16206 | Ecto-NOX disulfide-thiol exchanger 2 OS | ENOX2 | 70.083 | 2 | 54.410915 | yes | 1.679E-07 | 471.7558 | 237.55115 |
| ICTSDVECLMGR | 12 | Q16206 | Ecto-NOX disulfide-thiol exchanger 2 OS | ENOX2 | 70.083 | 2 | 54.410915 | yes | 9.722E-08 | 720.81763 | 234.42244 |
| EDDPNKEQQLK | 11 | Q16206 | Ecto-NOX disulfide-thiol exchanger 2 OS | ENOX2 | 70.083 | 2 | 54.410915 | yes | 1.295E-07 | 672.3255 | 219.97673 |
| FAEEYMVDK | 9 | Q16206 | Ecto-NOX disulfide-thiol exchanger 2 OS | ENOX2 | 70.083 | 2 | 54.410915 | yes | 9.723E-08 | 566.255 | 192.60716 |
| LCASNQDSEYPLE | 14 | Q16206 | Ecto-NOX disulfide-thiol exchanger 2 OS | ENOX2 | 70.083 | 2 | 54.410915 | yes | 9.818E-08 | 827.37488 | 143.57227 |
| ETEESALVSQAEA | 15 | Q16206 | Ecto-NOX disulfide-thiol exchanger 2 OS | ENOX2 | 70.083 | 2 | 54.410915 | yes | 3.541E-07 | 802.9043 | 143.24564 |
| ETEESALVSQAEA | 22 | Q16206 | Ecto-NOX disulfide-thiol exchanger 2 OS | ENOX2 | 70.083 | 3 | 54.410915 | yes | 1.497E-07 | 816.72937 | 140.0797 |
| DDLYEWECK | 9 | Q16206 | Ecto-NOX disulfide-thiol exchanger 2 OS | ENOX2 | 70.083 | 2 | 54.410915 | yes | 2.89E-07 | 629.25824 | 137.87038 |
| LLQQALQGMQQH | 15 | Q16206 | Ecto-NOX disulfide-thiol exchanger 2 OS | ENOX2 | 70.083 | 3 | 54.410915 | yes | 9.819E-08 | 583.6662 | 122.03735 |
| AWDHFTK | 7 | Q16206 | Ecto-NOX disulfide-thiol exchanger 2 OS | ENOX2 | 70.083 | 2 | 54.410915 | yes | 1.043E-07 | 452.71924 | 101.77489 |
| QEMTGVGASLEK | 12 | Q16206 | Ecto-NOX disulfide-thiol exchanger 2 OS | ENOX2 | 70.083 | 2 | 54.410915 | yes | 9.722E-08 | 625.30829 | 90.304565 |
| TQIALEPEQILR | 12 | Q16342 | Programmed cell death protein 2 OS=Hor | PDCD2 | 38.593 | 2 | 49.319923 | yes | 1.045E-07 | 705.90118 | 309.76825 |
| EQPCCAGLR | 9 | Q16342 | Programmed cell death protein 2 OS=Hor | PDCD2 | 38.593 | 2 | 49.319923 | yes | 1.716E-07 | 545.74207 | 138.01868 |
| EHQTLDWR | 8 | Q16342 | Programmed cell death protein 2 OS=Hor | PDCD2 | 38.593 | 2 | 49.319923 | yes | 1.073E-07 | 542.76215 | 117.81633 |
| PVELGFAESAPAW | 14 | Q16342 | Programmed cell death protein 2 OS=Hor | PDCD2 | 38.593 | 2 | 49.319923 | yes | 1.076E-07 | 765.39099 | 96.514885 |
| DIPDCPCGAK | 10 | Q16342 | Programmed cell death protein 2 OS=Hor | PDCD2 | 38.593 | 2 | 49.319923 | yes | 1.049E-07 | 566.7417 | 32.449047 |
| VFLTVPSLSSTAEE | 15 | Q16363 | Laminin subunit alpha-4 OS=Homo sapie | LAMA4 | 202.525 | 2 | 50.701576 | yes | 9.728E-08 | 804.42999 | 76.59967 |
| LLLQIPSTIR | 10 | Q16394 | Exostosin-1 OS=Homo sapiens OX=9606 | EXT1 | 86.255 | 2 | 56.405254 | yes | 1.006E-07 | 577.36877 | 266.14267 |
| IVLTTLEIIQDR | 12 | Q16394 | Exostosin-1 OS=Homo sapiens OX=9606 | EXT1 | 86.255 | 2 | 56.405254 | yes | 9.722E-08 | 707.41919 | 205.08165 |
| FTAVIHAVTPLVS( | 20 | Q16394 | Exostosin-1 OS=Homo sapiens OX=9606 | EXT1 | 86.255 | 3 | 56.405254 | yes | 9.809E-08 | 712.41229 | 145.53384 |
| MESCFDFTLCK | 11 | Q16394 | Exostosin-1 OS=Homo sapiens OX=9606 | EXT1 | 86.255 | 2 | 56.405254 | yes | 9.722E-08 | 719.29602 | 62.187244 |
| WPATAVPVVVIEC | 16 | Q16394 | Exostosin-1 OS=Homo sapiens OX=9606 | EXT1 | 86.255 | 2 | 56.405254 | yes | 1.923E-07 | 841.46161 | 51.217056 |
| ILTLSQIGR | 9 | Q16401 | 26S proteasome non-ATPase regulatory s | PSMD5 | 56.196 | 2 | 44.886303 | yes | 2.035E-07 | 500.81113 | 47.140244 |
| SLPFSENVSAVQK | 13 | Q16513 | Serine/threonine-protein kinase N2 OS=H | PKN2 | 112.035 | 2 | 53.742626 | yes | 1.629E-07 | 703.36975 | 270.13565 |
| IPQLAPPASDSTVT | 15 | Q16513 | Serine/threonine-protein kinase N2 OS=H | PKN2 | 112.035 | 2 | 53.742626 | yes | 2.713E-07 | 762.91705 | 246.56209 |
| VKPPFIPTIR | 10 | Q16513 | Serine/threonine-protein kinase N2 OS=H | PKN2 | 112.035 | 2 | 53.742626 | yes | 9.556E-07 | 584.36609 | 185.90895 |
| PAALTGTLEVR | 11 | Q16513 | Serine/threonine-protein kinase N2 OS=H | PKN2 | 112.035 | 2 | 53.742626 | yes | 2.492E-07 | 564.32458 | 172.81804 |
| LDFDLEPEPPPAPP | 15 | Q16513 | Serine/threonine-protein kinase N2 OS=H | PKN2 | 112.035 | 2 | 53.742626 | yes | 1.05E-07 | 845.4278 | 141.12393 |
| PVISPLELR | 9 | Q16513 | Serine/threonine-protein kinase N2 OS=H | PKN2 | 112.035 | 2 | 53.742626 | yes | 1.836E-07 | 512.31348 | 132.28918 |
| FLSTEAISIMR | 11 | Q16513 | Serine/threonine-protein kinase N2 OS=H | PKN2 | 112.035 | 2 | 53.742626 | yes | 1.703E-07 | 634.33936 | 118.58148 |
| ATSVALPGWSPSE | 15 | Q16513 | Serine/threonine-protein kinase N2 OS=H | PKN2 | 112.035 | 2 | 53.742626 | yes | 1.568E-07 | 779.89661 | 110.49586 |
| ASSLGEIDESSELR | 14 | Q16513 | Serine/threonine-protein kinase N2 OS=H | PKN2 | 112.035 | 2 | 53.742626 | yes | 1.055E-07 | 746.85992 | 107.82471 |

| | | | | | | | | | | | |
|---|---|---|---|---|---|---|---|---|---|---|---|
| LMGCQDILENVPG | 14 | Q16513 | Serine/threonine-protein kinase N2 OS=H | PKN2 | 112.035 | 2 | 53.742626 | yes | 2.577E-07 | 801.39233 | 99.263542 |
| LDFSDTMVQQK | 11 | Q16513 | Serine/threonine-protein kinase N2 OS=H | PKN2 | 112.035 | 2 | 53.742626 | yes | 2.42E-07 | 656.3161 | 91.240135 |
| LDNLLLDTEGFVK | 13 | Q16513 | Serine/threonine-protein kinase N2 OS=H | PKN2 | 112.035 | 2 | 53.742626 | yes | 9.742E-08 | 738.90088 | 78.283676 |
| IEFAVAEGAK | 10 | Q16513 | Serine/threonine-protein kinase N2 OS=H | PKN2 | 112.035 | 2 | 53.742626 | yes | 2.124E-07 | 517.77948 | 74.422501 |
| GEILLTELQGDSR | 13 | Q16513 | Serine/threonine-protein kinase N2 OS=H | PKN2 | 112.035 | 2 | 53.742626 | yes | 4.098E-06 | 715.87793 | 71.169754 |
| LEDFLDNQR | 9 | Q16513 | Serine/threonine-protein kinase N2 OS=H | PKN2 | 112.035 | 2 | 53.742626 | yes | 9.722E-08 | 575.2804 | 65.541092 |
| DEVDSLMCEK | 10 | Q16513 | Serine/threonine-protein kinase N2 OS=H | PKN2 | 112.035 | 2 | 53.742626 | yes | 9.722E-08 | 613.25739 | 57.516884 |
| SLAYVDNILK | 10 | Q16513 | Serine/threonine-protein kinase N2 OS=H | PKN2 | 112.035 | 2 | 53.742626 | yes | 1.667E-07 | 568.32153 | 56.687561 |
| ILSEEEQEMFR | 11 | Q16513 | Serine/threonine-protein kinase N2 OS=H | PKN2 | 112.035 | 2 | 53.742626 | yes | 9.722E-08 | 705.83209 | 41.938595 |
| LIDWSALMDK | 10 | Q16513 | Serine/threonine-protein kinase N2 OS=H | PKN2 | 112.035 | 2 | 53.742626 | yes | 6.113E-07 | 596.30756 | 37.417019 |
| GFGYGQGAGALVI | 15 | Q16527 | Cysteine and glycine-rich protein 2 OS=H | CSRP2 | 20.954 | 2 | 53.313225 | yes | 1.539E-07 | 716.85205 | 337.11768 |
| GYGYGQGAGTLN | 15 | Q16527 | Cysteine and glycine-rich protein 2 OS=H | CSRP2 | 20.954 | 2 | 53.313225 | yes | 9.722E-08 | 780.349 | 180.65787 |
| IIGAGKPWHK | 10 | Q16527 | Cysteine and glycine-rich protein 2 OS=H | CSRP2 | 20.954 | 2 | 53.313225 | yes | 1.674E-07 | 553.82709 | 136.96561 |
| CCFLCMVCR | 9 | Q16527 | Cysteine and glycine-rich protein 2 OS=H | CSRP2 | 20.954 | 2 | 53.313225 | yes | 1.965E-07 | 653.25513 | 135.59918 |
| CGDSVYAAEK | 10 | Q16527 | Cysteine and glycine-rich protein 2 OS=H | CSRP2 | 20.954 | 2 | 53.313225 | yes | 9.722E-08 | 550.23987 | 41.305862 |
| TVYHAEEVQCDGI | 13 | Q16527 | Cysteine and glycine-rich protein 2 OS=H | CSRP2 | 20.954 | 2 | 53.313225 | yes | 9.733E-08 | 782.34644 | 19.551315 |
| KNLDSTTVAIHDE | 18 | Q16527 | Cysteine and glycine-rich protein 2 OS=H | CSRP2 | 20.954 | 3 | 53.313225 | yes | 1.354E-06 | 712.68079 | 15.607388 |
| LVFSNVNLK | 9 | Q16531 | DNA damage-binding protein 1 OS=Hom | DDB1 | 126.968 | 2 | 52.36534 | yes | 1.708E-06 | 517.30566 | 372.53171 |
| LLASINSTVR | 10 | Q16531 | DNA damage-binding protein 1 OS=Hom | DDB1 | 126.968 | 2 | 52.36534 | yes | 1.233E-07 | 537.31934 | 315.07092 |
| VTLGTQPTVLR | 11 | Q16531 | DNA damage-binding protein 1 OS=Hom | DDB1 | 126.968 | 2 | 52.36534 | yes | 1.006E-07 | 592.85352 | 294.59888 |
| LPSFELLHK | 9 | Q16531 | DNA damage-binding protein 1 OS=Hom | DDB1 | 126.968 | 2 | 52.36534 | yes | 9.976E-08 | 542.31348 | 268.82538 |
| QGQGQLVTCSGAI | 14 | Q16531 | DNA damage-binding protein 1 OS=Hom | DDB1 | 126.968 | 2 | 52.36534 | yes | 9.722E-08 | 740.86426 | 198.57916 |
| YLLGDMEGR | 9 | Q16531 | DNA damage-binding protein 1 OS=Hom | DDB1 | 126.968 | 2 | 52.36534 | yes | 5.061E-07 | 527.25531 | 192.381 |
| LEELHVIDVK | 10 | Q16531 | DNA damage-binding protein 1 OS=Hom | DDB1 | 126.968 | 2 | 52.36534 | yes | 9.722E-08 | 597.84009 | 172.40573 |
| ETDDTLVLSFVGQ | 15 | Q16531 | DNA damage-binding protein 1 OS=Hom | DDB1 | 126.968 | 2 | 52.36534 | yes | 2.294E-06 | 840.9256 | 159.05153 |
| YLAIAPPIIK | 10 | Q16531 | DNA damage-binding protein 1 OS=Hom | DDB1 | 126.968 | 2 | 52.36534 | yes | 1.301E-07 | 549.84973 | 144.83482 |
| PSETGIIGIIDPECR | 15 | Q16531 | DNA damage-binding protein 1 OS=Hom | DDB1 | 126.968 | 2 | 52.36534 | yes | 9.722E-08 | 828.91669 | 142.85538 |
| YNACILEYK | 9 | Q16531 | DNA damage-binding protein 1 OS=Hom | DDB1 | 126.968 | 2 | 52.36534 | yes | 1.009E-07 | 587.28406 | 124.30693 |
| TVPLYESPR | 9 | Q16531 | DNA damage-binding protein 1 OS=Hom | DDB1 | 126.968 | 2 | 52.36534 | yes | 1.348E-06 | 531.28491 | 111.55038 |
| LYEWTTEK | 8 | Q16531 | DNA damage-binding protein 1 OS=Hom | DDB1 | 126.968 | 2 | 52.36534 | yes | 1.612E-07 | 535.26367 | 81.630898 |
| EMLGGEIIPR | 10 | Q16531 | DNA damage-binding protein 1 OS=Hom | DDB1 | 126.968 | 2 | 52.36534 | yes | 1.004E-07 | 557.79987 | 75.212288 |
| QSGESIDIITR | 11 | Q16531 | DNA damage-binding protein 1 OS=Hom | DDB1 | 126.968 | 2 | 52.36534 | yes | 5.018E-07 | 609.81989 | 72.080391 |
| IAVMELFR | 8 | Q16531 | DNA damage-binding protein 1 OS=Hom | DDB1 | 126.968 | 2 | 52.36534 | yes | 4.217E-07 | 489.7757 | 71.201164 |
| ALYYLQIHPQELR | 13 | Q16531 | DNA damage-binding protein 1 OS=Hom | DDB1 | 126.968 | 3 | 52.36534 | yes | 1.836E-07 | 548.63501 | 70.822899 |
| PMEGNFEEIAR | 11 | Q16531 | DNA damage-binding protein 1 OS=Hom | DDB1 | 126.968 | 2 | 52.36534 | yes | 2.818E-07 | 646.8006 | 67.400398 |
| SVLLLAYK | 8 | Q16531 | DNA damage-binding protein 1 OS=Hom | DDB1 | 126.968 | 2 | 52.36534 | yes | 1.121E-07 | 453.78659 | 56.399151 |
| SVLLLAYKPMEGN | 19 | Q16531 | DNA damage-binding protein 1 OS=Hom | DDB1 | 126.968 | 3 | 52.36534 | yes | 9.81E-08 | 727.3855 | 53.050014 |
| TKGDFILVGDLMR | 13 | Q16531 | DNA damage-binding protein 1 OS=Hom | DDB1 | 126.968 | 2 | 52.36534 | yes | 9.722E-08 | 732.89758 | 48.338207 |
| IEVQDTSGGTTALI | 14 | Q16531 | DNA damage-binding protein 1 OS=Hom | DDB1 | 126.968 | 2 | 52.36534 | yes | 5.317E-07 | 724.37299 | 46.90583 |

| | | | | | | | | | | | |
|---|---|---|---|---|---|---|---|---|---|---|---|
| EATADDLIK | 9 | Q16531 | DNA damage-binding protein 1 OS=Hom | DDB1 | 126.968 | 2 | 52.36534 | yes | 3.573E-06 | 488.2533 | 40.841259 |
| LYDGLFK | 7 | Q16531 | DNA damage-binding protein 1 OS=Hom | DDB1 | 126.968 | 2 | 52.36534 | yes | 1.463E-07 | 428.23419 | 27.758659 |
| KVTLGTQPTVLR | 12 | Q16531 | DNA damage-binding protein 1 OS=Hom | DDB1 | 126.968 | 2 | 52.36534 | yes | 3.168E-06 | 656.901 | 22.424095 |
| DVPSSEQPELFLK | 13 | Q16537 | Serine/threonine-protein phosphatase 2A | PPP2R5E | 54.7 | 2 | 51.1793 | yes | 1.025E-07 | 744.88269 | 592.91113 |
| GCLTEQTYPEVVR | 13 | Q16537 | Serine/threonine-protein phosphatase 2A | PPP2R5E | 54.7 | 2 | 51.1793 | yes | 1.022E-07 | 776.3772 | 579.58582 |
| FLESQEFQPSIAK | 13 | Q16537 | Serine/threonine-protein phosphatase 2A | PPP2R5E | 54.7 | 2 | 51.1793 | yes | 9.722E-08 | 762.39069 | 386.19431 |
| KLEDLELK | 8 | Q16537 | Serine/threonine-protein phosphatase 2A | PPP2R5E | 54.7 | 2 | 51.1793 | yes | 9.722E-08 | 494.28967 | 329.79593 |
| SQGKPIELTPLPLL | 15 | Q16537 | Serine/threonine-protein phosphatase 2A | PPP2R5E | 54.7 | 2 | 51.1793 | yes | 9.722E-08 | 817.49799 | 95.165054 |
| SQGKPIELTPLPLL | 15 | Q16537 | Serine/threonine-protein phosphatase 2A | PPP2R5E | 54.7 | 2 | 51.1793 | yes | 9.722E-08 | 545.33441 | 95.165054 |
| SQGKPIELTPLPLL | 15 | Q16537 | Serine/threonine-protein phosphatase 2A | PPP2R5E | 54.7 | 3 | 51.1793 | yes | 9.722E-08 | 817.49799 | 95.165054 |
| SQGKPIELTPLPLL | 15 | Q16537 | Serine/threonine-protein phosphatase 2A | PPP2R5E | 54.7 | 3 | 51.1793 | yes | 9.722E-08 | 545.33441 | 95.165054 |
| MVSCNIFR | 8 | Q16537 | Serine/threonine-protein phosphatase 2A | PPP2R5E | 54.7 | 2 | 51.1793 | yes | 2.916E-07 | 513.74658 | 189.76598 |
| STLNELVDYITISR | 14 | Q16537 | Serine/threonine-protein phosphatase 2A | PPP2R5E | 54.7 | 2 | 51.1793 | yes | 1.082E-07 | 812.43304 | 188.91316 |
| QINNIFLR | 8 | Q16537 | Serine/threonine-protein phosphatase 2A | PPP2R5E | 54.7 | 2 | 51.1793 | no | 2.524E-07 | 509.29562 | 155.01727 |
| CVSSPHFQVAER | 12 | Q16537 | Serine/threonine-protein phosphatase 2A | PPP2R5E | 54.7 | 3 | 51.1793 | no | 1.819E-07 | 472.89447 | 70.719543 |
| CVSSPHFQVAER | 12 | Q16537 | Serine/threonine-protein phosphatase 2A | PPP2R5E | 54.7 | 3 | 51.1793 | no | 1.802E-07 | 708.83807 | 70.719543 |
| CVSSPHFQVAER | 12 | Q16537 | Serine/threonine-protein phosphatase 2A | PPP2R5E | 54.7 | 2 | 51.1793 | no | 1.819E-07 | 472.89447 | 70.719543 |
| CVSSPHFQVAER | 12 | Q16537 | Serine/threonine-protein phosphatase 2A | PPP2R5E | 54.7 | 2 | 51.1793 | no | 1.802E-07 | 708.83807 | 70.719543 |
| IQEPLFK | 7 | Q16537 | Serine/threonine-protein phosphatase 2A | PPP2R5E | 54.7 | 2 | 51.1793 | no | 1.085E-06 | 437.75528 | 114.30957 |
| DPSLTEPVIR | 10 | Q16537 | Serine/threonine-protein phosphatase 2A | PPP2R5E | 54.7 | 2 | 51.1793 | yes | 1.684E-07 | 563.80878 | 92.027496 |
| PGVAQPEDTVQFR | 13 | Q16540 | Large ribosomal subunit protein uL23m C | MRPL23 | 17.782 | 2 | 51.022717 | yes | 1.879E-07 | 722.36499 | 956.6355 |
| NYLEGIYNVPVAA | 15 | Q16540 | Large ribosomal subunit protein uL23m C | MRPL23 | 17.782 | 2 | 51.022717 | yes | 9.722E-08 | 839.4516 | 937.62109 |
| IPMEMTR | 7 | Q16540 | Large ribosomal subunit protein uL23m C | MRPL23 | 17.782 | 2 | 51.022717 | yes | 1.625E-07 | 439.21716 | 879.77026 |
| NVVYPLYR | 8 | Q16540 | Large ribosomal subunit protein uL23m C | MRPL23 | 17.782 | 2 | 51.022717 | yes | 1.555E-07 | 512.28473 | 541.40442 |
| TNFFIQLVR | 9 | Q16540 | Large ribosomal subunit protein uL23m C | MRPL23 | 17.782 | 2 | 51.022717 | yes | 2.15E-07 | 569.3244 | 241.68207 |
| LQAEAQQLR | 9 | Q16543 | Hsp90 co-chaperone Cdc37 OS=Homo sa | CDC37 | 44.469 | 2 | 53.5954 | yes | 9.722E-08 | 528.79346 | 1395.535 |
| SMPWNVDTLSK | 11 | Q16543 | Hsp90 co-chaperone Cdc37 OS=Homo sa | CDC37 | 44.469 | 2 | 53.5954 | yes | 1.003E-07 | 639.31335 | 901.93616 |
| TEEDSEEVR | 9 | Q16543 | Hsp90 co-chaperone Cdc37 OS=Homo sa | CDC37 | 44.469 | 2 | 53.5954 | yes | 2.535E-07 | 547.23584 | 641.08746 |
| DVQMLQDAISK | 11 | Q16543 | Hsp90 co-chaperone Cdc37 OS=Homo sa | CDC37 | 44.469 | 2 | 53.5954 | yes | 1.248E-07 | 624.31866 | 529.24316 |
| SMVNTKPEK | 9 | Q16543 | Hsp90 co-chaperone Cdc37 OS=Homo sa | CDC37 | 44.469 | 2 | 53.5954 | yes | 3.056E-07 | 517.271 | 262.78653 |
| CIDSGLWVPNSK | 12 | Q16543 | Hsp90 co-chaperone Cdc37 OS=Homo sa | CDC37 | 44.469 | 2 | 53.5954 | yes | 4.197E-07 | 688.33734 | 211.62277 |
| LGPGGLDPVEVYE | 21 | Q16543 | Hsp90 co-chaperone Cdc37 OS=Homo sa | CDC37 | 44.469 | 3 | 53.5954 | yes | 3.346E-07 | 757.05811 | 14.031735 |
| EMFEDTVEER | 10 | Q16576 | Histone-binding protein RBBP7 OS=Hon | RBBP7 | 47.821 | 2 | 54.90253 | yes | 9.722E-08 | 642.77429 | 936.67169 |
| GEFGGFGSVTGK | 12 | Q16576 | Histone-binding protein RBBP7 OS=Hon | RBBP7 | 47.821 | 2 | 54.90253 | yes | 3.457E-07 | 571.77747 | 147.88803 |
| LHTFESHKDEIFQV | 31 | Q16576 | Histone-binding protein RBBP7 OS=Hon | RBBP7 | 47.821 | 5 | 54.90253 | yes | 9.722E-08 | 724.55304 | 121.7802 |
| FTVQMPTSQSPAV | 14 | Q16594 | Transcription initiation factor TFIID subu | TAF9 | 28.974 | 2 | 51.75503 | yes | 9.722E-08 | 760.89252 | 103.14095 |
| VINQMLEFAFR | 11 | Q16594 | Transcription initiation factor TFIID subu | TAF9 | 28.974 | 2 | 51.75503 | no | 2.261E-06 | 684.36066 | 31.83786 |
| LFVMEESTER | 10 | Q16600 | Zinc finger protein 239 OS=Homo sapien | ZNF239 | 51.591 | 2 | 54.54765 | yes | 1.067E-07 | 620.79755 | 469.10294 |
| NFSQSSELLLHQR | 13 | Q16600 | Zinc finger protein 239 OS=Homo sapien | ZNF239 | 51.591 | 2 | 54.54765 | yes | 9.722E-08 | 779.90228 | 360.21674 |

| | | | | | | | | | | | |
|---|---|---|---|---|---|---|---|---|---|---|---|
| ESLDPIDCNCK | 11 | Q16600 | Zinc finger protein 239 OS=Homo sapien | ZNF239 | 51.591 | 2 | 54.54765 | yes | 9.722E-08 | 675.78687 | 291.00684 |
| GESCSENLQVK | 11 | Q16600 | Zinc finger protein 239 OS=Homo sapien | ZNF239 | 51.591 | 2 | 54.54765 | yes | 1.019E-07 | 625.78772 | 249.00531 |
| ILNTSPDGHPYEK | 13 | Q16600 | Zinc finger protein 239 OS=Homo sapien | ZNF239 | 51.591 | 2 | 54.54765 | yes | 1.246E-07 | 735.86481 | 155.64076 |
| GFSQSSNLHIHR | 12 | Q16600 | Zinc finger protein 239 OS=Homo sapien | ZNF239 | 51.591 | 2 | 54.54765 | yes | 9.722E-08 | 691.84985 | 100.51827 |
| GFSQSSNLHIHQR | 13 | Q16600 | Zinc finger protein 239 OS=Homo sapien | ZNF239 | 51.591 | 2 | 54.54765 | yes | 9.722E-08 | 755.87909 | 39.184273 |
| APYPNYDR | 8 | Q16610 | Extracellular matrix protein 1 OS=Homo | ECM1 | 60.675 | 2 | 50.830147 | yes | 1.035E-06 | 498.2327 | 4745.1787 |
| FCEAEFSVK | 9 | Q16610 | Extracellular matrix protein 1 OS=Homo | ECM1 | 60.675 | 2 | 50.830147 | yes | 1.108E-07 | 558.75513 | 4711.7754 |
| RAPYPNYDR | 9 | Q16610 | Extracellular matrix protein 1 OS=Homo | ECM1 | 60.675 | 2 | 50.830147 | yes | 9.722E-08 | 576.28326 | 212.91241 |
| CCDLPFPEQACCA | 17 | Q16610 | Extracellular matrix protein 1 OS=Homo | ECM1 | 60.675 | 3 | 50.830147 | yes | 1.743E-07 | 714.94604 | 112.23273 |
| VYVGNLGTGAGK | 12 | Q16629 | Serine/arginine-rich splicing factor 7 OS= | SRSF7 | 27.367 | 2 | 47.513596 | yes | 1.442E-07 | 568.30896 | 6490.6426 |
| VELSTGMPR | 9 | Q16629 | Serine/arginine-rich splicing factor 7 OS= | SRSF7 | 27.367 | 2 | 47.513596 | yes | 2.256E-06 | 495.25787 | 2686.7632 |
| VELSTGMPR | 9 | Q16629 | Serine/arginine-rich splicing factor 7 OS= | SRSF7 | 27.367 | 2 | 47.513596 | yes | 2.894E-07 | 503.25531 | 2686.7632 |
| RPFDPNDR | 8 | Q16629 | Serine/arginine-rich splicing factor 7 OS= | SRSF7 | 27.367 | 2 | 47.513596 | yes | 1.705E-07 | 508.74905 | 3504.3308 |
| AFSYYGPLR | 9 | Q16629 | Serine/arginine-rich splicing factor 7 OS= | SRSF7 | 27.367 | 2 | 47.513596 | yes | 1.637E-07 | 537.27435 | 3281.0662 |
| CYECGEK | 7 | Q16629 | Serine/arginine-rich splicing factor 7 OS= | SRSF7 | 27.367 | 2 | 47.513596 | yes | 1.313E-07 | 473.17569 | 37.649002 |
| GAAPNVVYTYTGI | 13 | Q16630 | Cleavage and polyadenylation specificity | CPSF6 | 59.21 | 2 | 51.55124 | yes | 9.849E-08 | 670.84589 | 3609.855 |
| QFLSQFEMQSR | 11 | Q16630 | Cleavage and polyadenylation specificity | CPSF6 | 59.21 | 2 | 51.55124 | yes | 9.839E-08 | 700.83502 | 1244.1011 |
| GFALVGVGSEASS | 14 | Q16630 | Cleavage and polyadenylation specificity | CPSF6 | 59.21 | 2 | 51.55124 | yes | 2.702E-07 | 654.84332 | 1236.6809 |
| TPLSEAEFEEIMNF | 14 | Q16630 | Cleavage and polyadenylation specificity | CPSF6 | 59.21 | 2 | 51.55124 | yes | 1.345E-07 | 833.39307 | 1184.4478 |
| AISSSAISR | 9 | Q16630 | Cleavage and polyadenylation specificity | CPSF6 | 59.21 | 2 | 51.55124 | yes | 4.065E-07 | 446.24835 | 477.32001 |
| KLMDLLPK | 8 | Q16630 | Cleavage and polyadenylation specificity | CPSF6 | 59.21 | 2 | 51.55124 | yes | 2.154E-07 | 479.2937 | 367.61877 |
| ELHGQNPVVTPCN | 14 | Q16630 | Cleavage and polyadenylation specificity | CPSF6 | 59.21 | 2 | 51.55124 | yes | 9.722E-08 | 796.89612 | 276.76895 |
| LMDLLPK | 7 | Q16630 | Cleavage and polyadenylation specificity | CPSF6 | 59.21 | 2 | 51.55124 | yes | 3.131E-07 | 415.24625 | 144.42857 |
| ETCVVVYTGYGNI | 13 | Q16637 | Survival motor neuron protein OS=Homo | SMN1 | 31.849 | 2 | 54.161377 | yes | 1.017E-07 | 759.35626 | 3143.8347 |
| LGPGKPGLK | 9 | Q16637 | Survival motor neuron protein OS=Homo | SMN1 | 31.849 | 2 | 54.161377 | yes | 1.577E-07 | 433.77655 | 2658.0547 |
| NTAASLQQWK | 10 | Q16637 | Survival motor neuron protein OS=Homo | SMN1 | 31.849 | 2 | 54.161377 | yes | 1.067E-07 | 573.79877 | 1984.6412 |
| KNTAASLQQWK | 11 | Q16637 | Survival motor neuron protein OS=Homo | SMN1 | 31.849 | 2 | 54.161377 | yes | 2.113E-07 | 637.84619 | 518.13251 |
| NGDICETSGKPK | 12 | Q16637 | Survival motor neuron protein OS=Homo | SMN1 | 31.849 | 2 | 54.161377 | yes | 1.006E-07 | 653.30884 | 415.0509 |
| CSAIWSEDGCIYPA | 22 | Q16637 | Survival motor neuron protein OS=Homo | SMN1 | 31.849 | 3 | 54.161377 | yes | 9.722E-08 | 835.38727 | 92.795258 |
| AGVSFSGHR | 9 | Q16643 | Drebrin OS=Homo sapiens OX=9606 GN | DBN1 | 71.43 | 2 | 53.574276 | yes | 2.009E-07 | 480.23831 | 351.7616 |
| EQFWEQAK | 8 | Q16643 | Drebrin OS=Homo sapiens OX=9606 GN | DBN1 | 71.43 | 2 | 53.574276 | yes | 4.488E-07 | 533.2536 | 170.34172 |
| VMYGFCSVK | 9 | Q16643 | Drebrin OS=Homo sapiens OX=9606 GN | DBN1 | 71.43 | 2 | 53.574276 | yes | 9.722E-08 | 545.75665 | 96.646545 |
| MAPTPIPTR | 9 | Q16643 | Drebrin OS=Homo sapiens OX=9606 GN | DBN1 | 71.43 | 2 | 53.574276 | yes | 3.263E-07 | 492.27078 | 46.832455 |
| QQTLEAEEAK | 10 | Q16643 | Drebrin OS=Homo sapiens OX=9606 GN | DBN1 | 71.43 | 2 | 53.574276 | yes | 5.791E-07 | 573.78552 | 36.691288 |
| KSESEVEEAAAIIA | 21 | Q16643 | Drebrin OS=Homo sapiens OX=9606 GN | DBN1 | 71.43 | 3 | 53.574276 | yes | 1.505E-07 | 770.72766 | 31.670773 |
| LELLAAYEEVIR | 12 | Q16643 | Drebrin OS=Homo sapiens OX=9606 GN | DBN1 | 71.43 | 2 | 53.574276 | yes | 4.533E-05 | 709.89813 | 22.200424 |
| KQQTLEAEEAK | 11 | Q16643 | Drebrin OS=Homo sapiens OX=9606 GN | DBN1 | 71.43 | 2 | 53.574276 | yes | 3.177E-05 | 637.83295 | 20.527012 |
| VFGAAPLENVVR | 12 | Q16656 | Nuclear respiratory factor 1 OS=Homo sa | NRF1 | 53.542 | 2 | 50.20482 | yes | 9.85E-08 | 636.35895 | 226.67139 |
| FLIVAHDDGR | 10 | Q16658 | Fascin OS=Homo sapiens OX=9606 GN= | FSCN1 | 54.53 | 2 | 51.101276 | yes | 9.722E-08 | 571.80127 | 135.8309 |

| | | | | | | | | | | | |
|---|---|---|---|---|---|---|---|---|---|---|---|
| PADEIAVDR | 9 | Q16658 | Fascin OS=Homo sapiens OX=9606 GN= | FSCN1 | 54.53 | 2 | 51.101276 | yes | 9.904E-08 | 493.2511 | 111.83834 |
| YSVQTADHR | 9 | Q16658 | Fascin OS=Homo sapiens OX=9606 GN= | FSCN1 | 54.53 | 2 | 51.101276 | yes | 4.886E-07 | 538.75958 | 90.543053 |
| WSLQSEAHR | 9 | Q16658 | Fascin OS=Homo sapiens OX=9606 GN= | FSCN1 | 54.53 | 2 | 51.101276 | yes | 3.904E-07 | 557.27545 | 87.317307 |
| LSCFAQTVSPAEK | 13 | Q16658 | Fascin OS=Homo sapiens OX=9606 GN= | FSCN1 | 54.53 | 2 | 51.101276 | yes | 1.103E-06 | 719.35577 | 44.290855 |
| IGDFGLAR | 8 | Q16659 | Mitogen-activated protein kinase 6 OS=H | MAPK6 | 82.681 | 2 | 49.95333 | no | 9.968E-08 | 424.73486 | 258.96841 |
| LNDLNSSVSQLELI | 14 | Q16659 | Mitogen-activated protein kinase 6 OS=H | MAPK6 | 82.681 | 2 | 49.95333 | yes | 1.051E-07 | 780.41742 | 18.151558 |
| TTIYEIQDDR | 10 | Q16666 | Gamma-interferon-inducible protein 16 O | IFI16 | 88.256 | 2 | 53.27042 | yes | 1.025E-07 | 627.30408 | 3293.2561 |
| GEFTYYEIQDNTGI | 14 | Q16666 | Gamma-interferon-inducible protein 16 O | IFI16 | 88.256 | 2 | 53.27042 | yes | 2.837E-07 | 832.87555 | 2752.2979 |
| IQIADLMEEK | 10 | Q16666 | Gamma-interferon-inducible protein 16 O | IFI16 | 88.256 | 2 | 53.27042 | no | 1.101E-07 | 595.3103 | 2430.147 |
| IIAIANYVCR | 10 | Q16666 | Gamma-interferon-inducible protein 16 O | IFI16 | 88.256 | 2 | 53.27042 | yes | 1.062E-07 | 596.82898 | 1817.0659 |
| LTCFELAPK | 9 | Q16666 | Gamma-interferon-inducible protein 16 O | IFI16 | 88.256 | 2 | 53.27042 | yes | 2.829E-07 | 539.78369 | 1641.6368 |
| MEVVVHGR | 8 | Q16666 | Gamma-interferon-inducible protein 16 O | IFI16 | 88.256 | 2 | 53.27042 | yes | 1.4E-07 | 463.74747 | 1498.5934 |
| INQLCSQTK | 9 | Q16666 | Gamma-interferon-inducible protein 16 O | IFI16 | 88.256 | 2 | 53.27042 | yes | 1.901E-07 | 546.2793 | 1467.5924 |
| GLEVINDYHFR | 11 | Q16666 | Gamma-interferon-inducible protein 16 O | IFI16 | 88.256 | 2 | 53.27042 | no | 9.758E-08 | 681.84369 | 1415.4235 |
| SVIHSHIK | 8 | Q16666 | Gamma-interferon-inducible protein 16 O | IFI16 | 88.256 | 2 | 53.27042 | yes | 9.722E-08 | 460.76926 | 1386.0808 |
| EVDATSPAPSTSST | 16 | Q16666 | Gamma-interferon-inducible protein 16 O | IFI16 | 88.256 | 2 | 53.27042 | yes | 1.362E-07 | 788.88867 | 1201.6964 |
| IMFHATVATQTQF | 17 | Q16666 | Gamma-interferon-inducible protein 16 O | IFI16 | 88.256 | 3 | 53.27042 | yes | 1.247E-07 | 669.34875 | 544.46155 |
| IMFHATVATQTQF | 17 | Q16666 | Gamma-interferon-inducible protein 16 O | IFI16 | 88.256 | 3 | 53.27042 | yes | 8.695E-07 | 502.26337 | 544.46155 |
| IMFHATVATQTQF | 17 | Q16666 | Gamma-interferon-inducible protein 16 O | IFI16 | 88.256 | 4 | 53.27042 | yes | 1.247E-07 | 669.34875 | 544.46155 |
| IMFHATVATQTQF | 17 | Q16666 | Gamma-interferon-inducible protein 16 O | IFI16 | 88.256 | 4 | 53.27042 | yes | 8.695E-07 | 502.26337 | 544.46155 |
| MFHATVATENEVI | 14 | Q16666 | Gamma-interferon-inducible protein 16 O | IFI16 | 88.256 | 2 | 53.27042 | yes | 9.935E-08 | 826.3985 | 900.54401 |
| SLLSNDLK | 8 | Q16666 | Gamma-interferon-inducible protein 16 O | IFI16 | 88.256 | 2 | 53.27042 | no | 2.496E-07 | 445.25311 | 736.49323 |
| QLPYPSEASTTFPE | 18 | Q16666 | Gamma-interferon-inducible protein 16 O | IFI16 | 88.256 | 3 | 53.27042 | yes | 1.504E-07 | 687.34094 | 589.66309 |
| LISEMHSFIQIK | 12 | Q16666 | Gamma-interferon-inducible protein 16 O | IFI16 | 88.256 | 2 | 53.27042 | yes | 1.218E-07 | 723.39471 | 553.35864 |
| LTTINCEEGDKLK | 13 | Q16666 | Gamma-interferon-inducible protein 16 O | IFI16 | 88.256 | 2 | 53.27042 | yes | 9.722E-08 | 760.88489 | 458.65213 |
| FRGDAGLGK | 9 | Q16666 | Gamma-interferon-inducible protein 16 O | IFI16 | 88.256 | 2 | 53.27042 | yes | 2.951E-07 | 460.75107 | 242.05856 |
| KIIAIANYVCR | 11 | Q16666 | Gamma-interferon-inducible protein 16 O | IFI16 | 88.256 | 2 | 53.27042 | yes | 6.397E-07 | 660.87646 | 233.87691 |
| LTTINCEEGDK | 11 | Q16666 | Gamma-interferon-inducible protein 16 O | IFI16 | 88.256 | 2 | 53.27042 | yes | 1.781E-07 | 640.29535 | 210.50566 |
| QASGNIVYGVFMI | 15 | Q16666 | Gamma-interferon-inducible protein 16 O | IFI16 | 88.256 | 3 | 53.27042 | yes | 9.722E-08 | 555.29236 | 85.165756 |
| QASGNIVYGVFMI | 15 | Q16666 | Gamma-interferon-inducible protein 16 O | IFI16 | 88.256 | 3 | 53.27042 | yes | 3.261E-07 | 832.43488 | 85.165756 |
| QASGNIVYGVFMI | 15 | Q16666 | Gamma-interferon-inducible protein 16 O | IFI16 | 88.256 | 2 | 53.27042 | yes | 9.722E-08 | 555.29236 | 85.165756 |
| QASGNIVYGVFMI | 15 | Q16666 | Gamma-interferon-inducible protein 16 O | IFI16 | 88.256 | 2 | 53.27042 | yes | 3.261E-07 | 832.43488 | 85.165756 |
| LQLFCFR | 7 | Q16666 | Gamma-interferon-inducible protein 16 O | IFI16 | 88.256 | 2 | 53.27042 | yes | 1.025E-07 | 492.26019 | 134.61186 |
| PFEYETPEMEK | 11 | Q16666 | Gamma-interferon-inducible protein 16 O | IFI16 | 88.256 | 2 | 53.27042 | yes | 1.879E-07 | 700.30792 | 67.898239 |
| VLSTTKPFEYETPE | 17 | Q16666 | Gamma-interferon-inducible protein 16 O | IFI16 | 88.256 | 3 | 53.27042 | yes | 4.562E-06 | 676.99933 | 30.792393 |
| VFNIDLK | 7 | Q16666 | Gamma-interferon-inducible protein 16 O | IFI16 | 88.256 | 2 | 53.27042 | yes | 4.671E-07 | 424.74747 | 23.053236 |
| LIVVTDSCGR | 10 | Q16670 | Zinc finger and SCAN domain-containing | ZSCAN26 | 55.254 | 2 | 49.612385 | yes | 4.203E-07 | 560.29498 | 211.23283 |
| ILVEEMAPLK | 10 | Q16670 | Zinc finger and SCAN domain-containing | ZSCAN26 | 55.254 | 2 | 49.612385 | yes | 1.11E-07 | 571.82813 | 91.281029 |
| AFSLTSDLIR | 10 | Q16670 | Zinc finger and SCAN domain-containing | ZSCAN26 | 55.254 | 2 | 49.612385 | yes | 1.018E-07 | 561.81134 | 56.567307 |

| | | | | | | | | | | | |
|---|---|---|---|---|---|---|---|---|---|---|---|
| VQAELDQVVGR | 11 | Q16678 | Cytochrome P450 1B1 OS=Homo sapien: | CYP1B1 | 60.846 | 2 | 52.602875 | yes | 9.722E-08 | 607.33038 | 4552.3096 |
| GSADGAFLDPR | 11 | Q16678 | Cytochrome P450 1B1 OS=Homo sapien: | CYP1B1 | 60.846 | 2 | 52.602875 | yes | 5.905E-07 | 553.26727 | 3083.3013 |
| NFSNFILDK | 9 | Q16678 | Cytochrome P450 1B1 OS=Homo sapien: | CYP1B1 | 60.846 | 2 | 52.602875 | yes | 4.976E-06 | 549.28491 | 2386.7817 |
| HCESLRPGAAPR | 12 | Q16678 | Cytochrome P450 1B1 OS=Homo sapien: | CYP1B1 | 60.846 | 3 | 52.602875 | yes | 9.81E-08 | 450.89468 | 976.71924 |
| HCESLRPGAAPR | 12 | Q16678 | Cytochrome P450 1B1 OS=Homo sapien: | CYP1B1 | 60.846 | 3 | 52.602875 | yes | 2.323E-07 | 675.83838 | 976.71924 |
| HCESLRPGAAPR | 12 | Q16678 | Cytochrome P450 1B1 OS=Homo sapien: | CYP1B1 | 60.846 | 2 | 52.602875 | yes | 9.81E-08 | 450.89468 | 976.71924 |
| HCESLRPGAAPR | 12 | Q16678 | Cytochrome P450 1B1 OS=Homo sapien: | CYP1B1 | 60.846 | 2 | 52.602875 | yes | 2.323E-07 | 675.83838 | 976.71924 |
| LGSCPIVVLNGER | 13 | Q16678 | Cytochrome P450 1B1 OS=Homo sapien: | CYP1B1 | 60.846 | 2 | 52.602875 | yes | 4.118E-07 | 707.37958 | 919.05292 |
| CIGEELSK | 8 | Q16678 | Cytochrome P450 1B1 OS=Homo sapien: | CYP1B1 | 60.846 | 2 | 52.602875 | yes | 1.259E-06 | 468.22876 | 861.89813 |
| YSHDDPEFR | 9 | Q16678 | Cytochrome P450 1B1 OS=Homo sapien: | CYP1B1 | 60.846 | 2 | 52.602875 | yes | 9.722E-08 | 583.24908 | 822.83398 |
| YGDVFQIR | 8 | Q16678 | Cytochrome P450 1B1 OS=Homo sapien: | CYP1B1 | 60.846 | 2 | 52.602875 | yes | 1.401E-07 | 499.25873 | 696.51508 |
| ELLSHNEEFGR | 11 | Q16678 | Cytochrome P450 1B1 OS=Homo sapien: | CYP1B1 | 60.846 | 2 | 52.602875 | yes | 9.823E-08 | 665.82294 | 619.81964 |
| FLDKDGLINK | 10 | Q16678 | Cytochrome P450 1B1 OS=Homo sapien: | CYP1B1 | 60.846 | 2 | 52.602875 | yes | 3.139E-07 | 581.82697 | 597.78546 |
| MNFSYGLTIK | 10 | Q16678 | Cytochrome P450 1B1 OS=Homo sapien: | CYP1B1 | 60.846 | 2 | 52.602875 | yes | 9.722E-08 | 587.30225 | 437.55115 |
| QVLEGHVLSEAR | 12 | Q16678 | Cytochrome P450 1B1 OS=Homo sapien: | CYP1B1 | 60.846 | 2 | 52.602875 | yes | 1.05E-07 | 669.36224 | 210.19743 |
| QVLEGHVLSEAR | 12 | Q16678 | Cytochrome P450 1B1 OS=Homo sapien: | CYP1B1 | 60.846 | 2 | 52.602875 | yes | 1.834E-07 | 446.57724 | 210.19743 |
| QVLEGHVLSEAR | 12 | Q16678 | Cytochrome P450 1B1 OS=Homo sapien: | CYP1B1 | 60.846 | 3 | 52.602875 | yes | 1.05E-07 | 669.36224 | 210.19743 |
| QVLEGHVLSEAR | 12 | Q16678 | Cytochrome P450 1B1 OS=Homo sapien: | CYP1B1 | 60.846 | 3 | 52.602875 | yes | 1.834E-07 | 446.57724 | 210.19743 |
| EFEQLNR | 7 | Q16678 | Cytochrome P450 1B1 OS=Homo sapien: | CYP1B1 | 60.846 | 2 | 52.602875 | yes | 1.345E-07 | 468.2327 | 412.69159 |
| DMMDAFILSAEK | 12 | Q16678 | Cytochrome P450 1B1 OS=Homo sapien: | CYP1B1 | 60.846 | 2 | 52.602875 | yes | 9.818E-08 | 685.82019 | 235.78601 |
| SMAFGHYSEHWK | 12 | Q16678 | Cytochrome P450 1B1 OS=Homo sapien: | CYP1B1 | 60.846 | 2 | 52.602875 | yes | 1.286E-07 | 740.32733 | 210.25334 |
| FSSFVPVTIPHATT | 26 | Q16678 | Cytochrome P450 1B1 OS=Homo sapien: | CYP1B1 | 60.846 | 4 | 52.602875 | yes | 1.087E-07 | 696.87341 | 156.71181 |
| RYGDVFQIR | 9 | Q16678 | Cytochrome P450 1B1 OS=Homo sapien: | CYP1B1 | 60.846 | 2 | 52.602875 | yes | 2.397E-07 | 577.30927 | 119.38652 |
| WPNPENFDPAR | 11 | Q16678 | Cytochrome P450 1B1 OS=Homo sapien: | CYP1B1 | 60.846 | 2 | 52.602875 | yes | 7.421E-07 | 671.81238 | 74.723984 |
| MNFSYGLTIKPK | 12 | Q16678 | Cytochrome P450 1B1 OS=Homo sapien: | CYP1B1 | 60.846 | 2 | 52.602875 | yes | 2.636E-07 | 699.87616 | 26.482508 |
| VAFITGGGTGLGK | 13 | Q16698 | 2,4-dienoyl-CoA reductase [(3E)-enoyl-C | DECR1 | 36.068 | 2 | 33.6753 | yes | 1.177E-06 | 589.3324 | 68.771439 |
| LLAENNEIISNIR | 13 | Q16706 | Alpha-mannosidase 2 OS=Homo sapiens | MAN2A1 | 131.141 | 2 | 51.716915 | yes | 1.083E-07 | 749.91486 | 179.63152 |
| YLVVYNPLEQDR | 12 | Q16706 | Alpha-mannosidase 2 OS=Homo sapiens | MAN2A1 | 131.141 | 2 | 51.716915 | yes | 3.474E-07 | 754.89081 | 172.00648 |
| MLLDQYR | 7 | Q16706 | Alpha-mannosidase 2 OS=Homo sapiens | MAN2A1 | 131.141 | 2 | 51.716915 | yes | 1.37E-06 | 469.74185 | 112.64899 |
| GTGLFCSTTQGK | 12 | Q16706 | Alpha-mannosidase 2 OS=Homo sapiens | MAN2A1 | 131.141 | 2 | 51.716915 | yes | 1.809E-07 | 628.8006 | 96.042694 |
| FLSSSLYTALTEAR | 14 | Q16706 | Alpha-mannosidase 2 OS=Homo sapiens | MAN2A1 | 131.141 | 2 | 51.716915 | yes | 1.2E-07 | 779.90924 | 73.964813 |
| DSVINLSESVEDGI | 15 | Q16706 | Alpha-mannosidase 2 OS=Homo sapiens | MAN2A1 | 131.141 | 2 | 51.716915 | yes | 9.81E-08 | 794.88867 | 62.431324 |
| VLLAPLGDDFR | 11 | Q16706 | Alpha-mannosidase 2 OS=Homo sapiens | MAN2A1 | 131.141 | 2 | 51.716915 | yes | 9.722E-08 | 608.34021 | 59.640785 |
| NKVEDSGIFTIK | 12 | Q16706 | Alpha-mannosidase 2 OS=Homo sapiens | MAN2A1 | 131.141 | 2 | 51.716915 | yes | 1.397E-07 | 675.86682 | 42.543098 |
| WWDIIDIQK | 9 | Q16706 | Alpha-mannosidase 2 OS=Homo sapiens | MAN2A1 | 131.141 | 2 | 51.716915 | yes | 1.234E-07 | 608.82166 | 41.057827 |
| LFHSLMVLEK | 10 | Q16706 | Alpha-mannosidase 2 OS=Homo sapiens | MAN2A1 | 131.141 | 2 | 51.716915 | yes | 3.051E-06 | 608.84155 | 36.882874 |
| TFNDYFR | 7 | Q16706 | Alpha-mannosidase 2 OS=Homo sapiens | MAN2A1 | 131.141 | 2 | 51.716915 | yes | 9.895E-07 | 481.72195 | 35.19994 |
| AHIPPLGLK | 9 | Q16706 | Alpha-mannosidase 2 OS=Homo sapiens | MAN2A1 | 131.141 | 2 | 51.716915 | yes | 9.571E-07 | 473.29764 | 29.839373 |
| VLDASWYSPGTR | 12 | Q16762 | Thiosulfate sulfurtransferase OS=Homo s | TST | 33.429 | 2 | 49.984764 | yes | 9.938E-08 | 676.33569 | 145.21107 |

| | | | | | | | | | | | |
|---|---|---|---|---|---|---|---|---|---|---|---|
| VHQVLYR | 7 | Q16762 | Thiosulfate sulfurtransferase OS=Homo s | TST | 33.429 | 2 | 49.984764 | yes | 1.142E-07 | 457.76398 | 108.67397 |
| TYEQVLENLESK | 12 | Q16762 | Thiosulfate sulfurtransferase OS=Homo s | TST | 33.429 | 2 | 49.984764 | yes | 1.87E-07 | 726.86444 | 101.58833 |
| FLGTEPEPDAVGL | 19 | Q16762 | Thiosulfate sulfurtransferase OS=Homo s | TST | 33.429 | 3 | 49.984764 | yes | 2.69E-07 | 670.66907 | 45.300228 |
| HVPGASFFDIEECF | 14 | Q16762 | Thiosulfate sulfurtransferase OS=Homo s | TST | 33.429 | 3 | 49.984764 | yes | 2.078E-06 | 555.25598 | 28.032415 |
| LLLENYEEYAAR | 12 | Q16763 | Ubiquitin-conjugating enzyme E2 S OS=l | UBE2S | 23.846 | 2 | 46.11005 | yes | 1.325E-06 | 742.375 | 239.2511 |
| EVTTLTADPPDGIK | 14 | Q16763 | Ubiquitin-conjugating enzyme E2 S OS=l | UBE2S | 23.846 | 2 | 46.11005 | yes | 1.65E-07 | 728.88013 | 78.757095 |
| DWTAELGIR | 9 | Q16763 | Ubiquitin-conjugating enzyme E2 S OS=l | UBE2S | 23.846 | 2 | 46.11005 | yes | 3.495E-07 | 530.77472 | 28.139471 |
| IPQAIAQLSK | 10 | Q16795 | NADH dehydrogenase [ubiquinone] 1 alp | NDUFA9 | 42.51 | 2 | 54.86535 | yes | 1.884E-07 | 534.82422 | 1000.6248 |
| MGSQVIIPYR | 10 | Q16795 | NADH dehydrogenase [ubiquinone] 1 alp | NDUFA9 | 42.51 | 2 | 54.86535 | yes | 1.936E-07 | 582.31573 | 803.46332 |
| FGPIPLGSLGWK | 12 | Q16795 | NADH dehydrogenase [ubiquinone] 1 alp | NDUFA9 | 42.51 | 2 | 54.86535 | yes | 1.737E-07 | 636.36096 | 518.02118 |
| YVVNHLGR | 8 | Q16795 | NADH dehydrogenase [ubiquinone] 1 alp | NDUFA9 | 42.51 | 2 | 54.86535 | yes | 6.846E-07 | 479.26688 | 492.0694 |
| YIFAVAHR | 8 | Q16795 | NADH dehydrogenase [ubiquinone] 1 alp | NDUFA9 | 42.51 | 2 | 54.86535 | yes | 1.128E-07 | 488.77179 | 315.6225 |
| NFDFEDVFVK | 10 | Q16795 | NADH dehydrogenase [ubiquinone] 1 alp | NDUFA9 | 42.51 | 2 | 54.86535 | yes | 1.004E-07 | 630.30078 | 296.07217 |
| WLSAEIEDVKPAK | 13 | Q16795 | NADH dehydrogenase [ubiquinone] 1 alp | NDUFA9 | 42.51 | 2 | 54.86535 | yes | 1.141E-07 | 743.401 | 262.17877 |
| QPVYVVDVSK | 10 | Q16795 | NADH dehydrogenase [ubiquinone] 1 alp | NDUFA9 | 42.51 | 2 | 54.86535 | yes | 1.006E-07 | 567.31366 | 230.73843 |
| FLNSFASMHR | 10 | Q16795 | NADH dehydrogenase [ubiquinone] 1 alp | NDUFA9 | 42.51 | 2 | 54.86535 | yes | 9.722E-08 | 605.29529 | 194.65398 |
| VVQHSNVVINLIG | 14 | Q16795 | NADH dehydrogenase [ubiquinone] 1 alp | NDUFA9 | 42.51 | 2 | 54.86535 | yes | 9.722E-08 | 774.45447 | 192.86057 |
| YDIMHLR | 7 | Q16795 | NADH dehydrogenase [ubiquinone] 1 alp | NDUFA9 | 42.51 | 2 | 54.86535 | yes | 1.664E-07 | 474.242 | 151.10605 |
| LPHLPGLEDLGIQA | 20 | Q16795 | NADH dehydrogenase [ubiquinone] 1 alp | NDUFA9 | 42.51 | 3 | 54.86535 | yes | 4.195E-07 | 718.74384 | 148.1409 |
| FIHVSHLNANIK | 12 | Q16795 | NADH dehydrogenase [ubiquinone] 1 alp | NDUFA9 | 42.51 | 2 | 54.86535 | yes | 9.722E-08 | 696.89093 | 124.33312 |
| SFAFVGPSR | 9 | Q16795 | NADH dehydrogenase [ubiquinone] 1 alp | NDUFA9 | 42.51 | 2 | 54.86535 | yes | 3.525E-07 | 484.25345 | 78.129456 |
| YLLFHLVK | 8 | Q16795 | NADH dehydrogenase [ubiquinone] 1 alp | NDUFA9 | 42.51 | 2 | 54.86535 | yes | 1.283E-07 | 516.81567 | 52.134026 |
| MHITDMK | 7 | Q16795 | NADH dehydrogenase [ubiquinone] 1 alp | NDUFA9 | 42.51 | 2 | 54.86535 | yes | 8.038E-06 | 438.20932 | 50.594307 |
| CVGAELGLDCPGF | 13 | Q16831 | Uridine phosphorylase 1 OS=Homo sapie | UPP1 | 33.935 | 2 | 49.389233 | yes | 1.037E-07 | 702.32391 | 40.017647 |
| DYPNICAGTDR | 11 | Q16831 | Uridine phosphorylase 1 OS=Homo sapie | UPP1 | 33.935 | 2 | 49.389233 | yes | 1.472E-06 | 641.28003 | 28.150169 |
| AEFEQIVLGK | 10 | Q16831 | Uridine phosphorylase 1 OS=Homo sapie | UPP1 | 33.935 | 2 | 49.389233 | yes | 3.774E-06 | 567.31366 | 24.135164 |
| GVAYDVPNPVFLE | 15 | Q16850 | Lanosterol 14-alpha demethylase OS=Ho | CYP51A1 | 57.279 | 2 | 47.008587 | yes | 1.422E-07 | 838.43811 | 57.804737 |
| STIQTAEALR | 10 | Q16875 | 6-phosphofructo-2-kinase/fructose-2,6-bi | PFKFB3 | 59.61 | 2 | 49.932102 | yes | 2.189E-07 | 545.29858 | 158.94023 |
| EGGQIAVFDATNT | 15 | Q16875 | 6-phosphofructo-2-kinase/fructose-2,6-bi | PFKFB3 | 59.61 | 2 | 49.932102 | yes | 9.975E-08 | 790.38916 | 126.4469 |
| LTNSPTVIVMVGL | 16 | Q16875 | 6-phosphofructo-2-kinase/fructose-2,6-bi | PFKFB3 | 59.61 | 2 | 49.932102 | yes | 2.601E-07 | 834.47931 | 42.905678 |
| YPTGESYQDLVQR | 13 | Q16875 | 6-phosphofructo-2-kinase/fructose-2,6-bi | PFKFB3 | 59.61 | 2 | 49.932102 | yes | 1.613E-07 | 778.37299 | 26.511444 |
| LSEQELQFR | 9 | Q16891 | MICOS complex subunit MIC60 OS=Ho | IMMT | 83.678 | 2 | 50.35089 | yes | 9.817E-08 | 575.29858 | 139.3107 |
| ELDSITPEVLPGWI | 14 | Q16891 | MICOS complex subunit MIC60 OS=Ho | IMMT | 83.678 | 2 | 50.35089 | yes | 1.742E-07 | 792.41943 | 131.12769 |
| AVDEAADALLK | 11 | Q16891 | MICOS complex subunit MIC60 OS=Ho | IMMT | 83.678 | 2 | 50.35089 | yes | 1.014E-06 | 558.30078 | 123.71997 |
| VAMIDETR | 8 | Q16891 | MICOS complex subunit MIC60 OS=Ho | IMMT | 83.678 | 2 | 50.35089 | yes | 1.001E-07 | 467.73676 | 111.1967 |
| SEIQAEQDR | 9 | Q16891 | MICOS complex subunit MIC60 OS=Ho | IMMT | 83.678 | 2 | 50.35089 | yes | 2.338E-07 | 538.25433 | 88.169662 |
| VVSQYHELVVQAI | 13 | Q16891 | MICOS complex subunit MIC60 OS=Ho | IMMT | 83.678 | 2 | 50.35089 | yes | 1.263E-07 | 764.41754 | 68.611816 |
| AAMDNSEIAGEK | 12 | Q16891 | MICOS complex subunit MIC60 OS=Ho | IMMT | 83.678 | 2 | 50.35089 | yes | 1.34E-07 | 618.28229 | 44.130077 |
| AHQLWLSVEALK | 12 | Q16891 | MICOS complex subunit MIC60 OS=Ho | IMMT | 83.678 | 2 | 50.35089 | yes | 5.252E-07 | 697.89319 | 34.287052 |

| | | | | | | | | | | | |
|---|---|---|---|---|---|---|---|---|---|---|---|
| YSTSGSSGLTTGK | 13 | Q16891 | MICOS complex subunit MIC60 OS=Hoı | IMMT | 83.678 | 2 | 50.35089 | yes | 4.863E-06 | 623.30151 | 30.362492 |
| QHITLALEK | 9 | Q16891 | MICOS complex subunit MIC60 OS=Hoı | IMMT | 83.678 | 2 | 50.35089 | yes | 5.351E-07 | 526.80859 | 29.59005 |
| VNTSVPYSVSNTV | 15 | Q17RS7 | Flap endonuclease GEN homolog 1 OS=I | GEN1 | 102.885 | 2 | 47.799866 | yes | 1.256E-07 | 797.4278 | 148.18298 |
| DLPGIPLQNESR | 12 | Q17RS7 | Flap endonuclease GEN homolog 1 OS=I | GEN1 | 102.885 | 2 | 47.799866 | yes | 5.527E-07 | 669.85425 | 90.513977 |
| GDQLLQEDYK | 10 | Q17RS7 | Flap endonuclease GEN homolog 1 OS=I | GEN1 | 102.885 | 2 | 47.799866 | yes | 2.314E-07 | 604.79333 | 50.77879 |
| ENNSGTCLDSPLPI | 15 | Q17RS7 | Flap endonuclease GEN homolog 1 OS=I | GEN1 | 102.885 | 2 | 47.799866 | yes | 1.294E-07 | 836.90161 | 39.462318 |
| GGPGRTGLAGAPC | 18 | Q17RW2 | Collagen alpha-1(XXIV) chain OS=Hom | COL24A1 | 175.497 | 3 | 20.088293 | yes | 4.592E-06 | 515.95471 | 100.8071 |
| FGPLIVDWPHK | 11 | Q17RY0 | Cytoplasmic polyadenylation element-bin | CPEB4 | 80.152 | 2 | 54.99952 | yes | 9.722E-08 | 654.85858 | 2242.3875 |
| SWMEDSLNR | 9 | Q17RY0 | Cytoplasmic polyadenylation element-bin | CPEB4 | 80.152 | 2 | 54.99952 | yes | 2.161E-07 | 569.2533 | 979.35699 |
| GLNGGITPLNSISPI | 16 | Q17RY0 | Cytoplasmic polyadenylation element-bin | CPEB4 | 80.152 | 2 | 54.99952 | yes | 4.844E-07 | 790.95398 | 918.43164 |
| NFASNHIQLQK | 11 | Q17RY0 | Cytoplasmic polyadenylation element-bin | CPEB4 | 80.152 | 2 | 54.99952 | yes | 9.722E-08 | 650.34387 | 562.09924 |
| FVQLQHGEIDK | 11 | Q17RY0 | Cytoplasmic polyadenylation element-bin | CPEB4 | 80.152 | 2 | 54.99952 | yes | 1.008E-07 | 657.34607 | 256.34671 |
| FVQLQHGEIDK | 11 | Q17RY0 | Cytoplasmic polyadenylation element-bin | CPEB4 | 80.152 | 2 | 54.99952 | yes | 5.388E-07 | 438.56647 | 256.34671 |
| FVQLQHGEIDK | 11 | Q17RY0 | Cytoplasmic polyadenylation element-bin | CPEB4 | 80.152 | 3 | 54.99952 | yes | 1.008E-07 | 657.34607 | 256.34671 |
| FVQLQHGEIDK | 11 | Q17RY0 | Cytoplasmic polyadenylation element-bin | CPEB4 | 80.152 | 3 | 54.99952 | yes | 5.388E-07 | 438.56647 | 256.34671 |
| ADNIFPFPDRPR | 12 | Q17RY0 | Cytoplasmic polyadenylation element-bin | CPEB4 | 80.152 | 2 | 54.99952 | yes | 9.722E-08 | 722.87024 | 275.60541 |
| FVQLQHGEIDKR | 12 | Q17RY0 | Cytoplasmic polyadenylation element-bin | CPEB4 | 80.152 | 2 | 54.99952 | yes | 9.862E-08 | 735.39661 | 254.39919 |
| SQQQEQQDPLEK | 12 | Q17RY0 | Cytoplasmic polyadenylation element-bin | CPEB4 | 80.152 | 2 | 54.99952 | yes | 9.722E-08 | 729.34698 | 222.28415 |
| SPASPHPPPFTHR | 13 | Q17RY0 | Cytoplasmic polyadenylation element-bin | CPEB4 | 80.152 | 2 | 54.99952 | yes | 9.723E-08 | 714.36255 | 153.85484 |
| RFGPLIVDWPHK | 12 | Q17RY0 | Cytoplasmic polyadenylation element-bin | CPEB4 | 80.152 | 2 | 54.99952 | yes | 1.116E-07 | 732.90918 | 29.899952 |
| RFGPLIVDWPHK | 12 | Q17RY0 | Cytoplasmic polyadenylation element-bin | CPEB4 | 80.152 | 2 | 54.99952 | yes | 8.578E-07 | 488.94186 | 29.899952 |
| RFGPLIVDWPHK | 12 | Q17RY0 | Cytoplasmic polyadenylation element-bin | CPEB4 | 80.152 | 3 | 54.99952 | yes | 1.116E-07 | 732.90918 | 29.899952 |
| RFGPLIVDWPHK | 12 | Q17RY0 | Cytoplasmic polyadenylation element-bin | CPEB4 | 80.152 | 3 | 54.99952 | yes | 8.578E-07 | 488.94186 | 29.899952 |
| LLQMPSVVNYSGI | 14 | Q27J81 | Inverted formin-2 OS=Homo sapiens OX | INF2 | 135.625 | 2 | 55.794502 | yes | 1.558E-07 | 788.92944 | 93.355721 |
| AQLVLAACESLLT | 15 | Q27J81 | Inverted formin-2 OS=Homo sapiens OX | INF2 | 135.625 | 2 | 55.794502 | yes | 9.722E-08 | 816.44287 | 69.201813 |
| LQASISAFR | 9 | Q27J81 | Inverted formin-2 OS=Homo sapiens OX | INF2 | 135.625 | 2 | 55.794502 | yes | 5.826E-07 | 496.77982 | 67.730812 |
| SHDPEEAIR | 9 | Q29RF7 | Sister chromatid cohesion protein PDS5 I | PDS5A | 150.831 | 2 | 53.02922 | no | 1.966E-06 | 527.25165 | 120.22239 |
| TVTAAGAENIQQK | 13 | Q29RF7 | Sister chromatid cohesion protein PDS5 I | PDS5A | 150.831 | 2 | 53.02922 | yes | 6.122E-07 | 665.85168 | 77.796936 |
| IETDLPQIR | 9 | Q29RF7 | Sister chromatid cohesion protein PDS5 I | PDS5A | 150.831 | 2 | 53.02922 | yes | 2.903E-07 | 542.80347 | 76.743584 |
| HDVIVTIITAAK | 12 | Q29RF7 | Sister chromatid cohesion protein PDS5 I | PDS5A | 150.831 | 2 | 53.02922 | yes | 9.722E-08 | 640.88226 | 74.023148 |
| EVQLAQIFEPLSR | 13 | Q29RF7 | Sister chromatid cohesion protein PDS5 I | PDS5A | 150.831 | 2 | 53.02922 | yes | 3.878E-07 | 765.41974 | 43.242096 |
| PATALCGVVSADC | 14 | Q29RF7 | Sister chromatid cohesion protein PDS5 I | PDS5A | 150.831 | 2 | 53.02922 | yes | 1.487E-07 | 673.34265 | 42.381222 |
| AAVGQESPGGLEA | 17 | Q29RF7 | Sister chromatid cohesion protein PDS5 I | PDS5A | 150.831 | 2 | 53.02922 | yes | 2.258E-07 | 778.38916 | 39.765896 |
| VAEAAIQIFR | 10 | Q29RF7 | Sister chromatid cohesion protein PDS5 I | PDS5A | 150.831 | 2 | 53.02922 | yes | 3.583E-06 | 559.32184 | 25.238972 |
| SPTTVLFTSSPIK | 13 | Q2KHR2 | DNA-binding protein RFX7 OS=Homo s | RFX7 | 146.896 | 2 | 48.650066 | yes | 3.247E-07 | 689.38483 | 164.07108 |
| GLGMNNLPSNYTA | 14 | Q2KHR2 | DNA-binding protein RFX7 OS=Homo s | RFX7 | 146.896 | 2 | 48.650066 | yes | 2.482E-07 | 754.36975 | 126.22077 |
| NLSGSTLYPVSNIP | 15 | Q2KHR2 | DNA-binding protein RFX7 OS=Homo s | RFX7 | 146.896 | 2 | 48.650066 | yes | 2.411E-07 | 809.43341 | 123.96442 |
| LEGSVFLLDSDSK | 13 | Q2KHR2 | DNA-binding protein RFX7 OS=Homo s | RFX7 | 146.896 | 2 | 48.650066 | yes | 1.527E-07 | 705.36157 | 115.31467 |
| LAQTTPVDSALGS | 15 | Q2KHR2 | DNA-binding protein RFX7 OS=Homo s | RFX7 | 146.896 | 2 | 48.650066 | yes | 4.333E-07 | 751.8941 | 115.21695 |

| | | | | | | | | | | | |
|---|---|---|---|---|---|---|---|---|---|---|---|
| YCYSGLR | 7 | Q2KHR2 | DNA-binding protein RFX7 OS=Homo sa | RFX7 | 146.896 | 2 | 48.650066 | yes | 2.864E-06 | 459.71054 | 91.894081 |
| TEGSTAGAQIPSK | 13 | Q2KHR2 | DNA-binding protein RFX7 OS=Homo sa | RFX7 | 146.896 | 2 | 48.650066 | yes | 3.23E-06 | 623.81732 | 62.837826 |
| QEGFAVPAPLDNK | 13 | Q2KHR2 | DNA-binding protein RFX7 OS=Homo sa | RFX7 | 146.896 | 2 | 48.650066 | yes | 1.423E-06 | 693.35663 | 61.769245 |
| NTLEEHPETSLPK | 13 | Q2KHR2 | DNA-binding protein RFX7 OS=Homo sa | RFX7 | 146.896 | 2 | 48.650066 | yes | 1.12E-07 | 747.87537 | 58.887135 |
| KITNVLLSK | 9 | Q2KHR2 | DNA-binding protein RFX7 OS=Homo sa | RFX7 | 146.896 | 2 | 48.650066 | yes | 4.873E-07 | 508.32913 | 45.239201 |
| SSSAGGTSAVEVK | 13 | Q2KHR2 | DNA-binding protein RFX7 OS=Homo sa | RFX7 | 146.896 | 2 | 48.650066 | yes | 1.139E-07 | 590.2962 | 35.301548 |
| VLSQPFDTVLELAI | 14 | Q2KHR2 | DNA-binding protein RFX7 OS=Homo sa | RFX7 | 146.896 | 2 | 48.650066 | yes | 1.143E-05 | 794.44067 | 28.789715 |
| ELQAEPGLPGAAA | 14 | Q2M1P5 | Kinesin-like protein KIF7 OS=Homo sapi | KIF7 | 150.588 | 2 | 53.293213 | yes | 1.592E-07 | 690.36749 | 162.58464 |
| EAGAELLTEVNR | 12 | Q2M1P5 | Kinesin-like protein KIF7 OS=Homo sapi | KIF7 | 150.588 | 2 | 53.293213 | yes | 9.722E-08 | 651.33844 | 114.17781 |
| LSYLSSSETR | 10 | Q2M1P5 | Kinesin-like protein KIF7 OS=Homo sapi | KIF7 | 150.588 | 2 | 53.293213 | yes | 1.605E-07 | 571.78802 | 81.217209 |
| ACTDAAYSLLR | 11 | Q2M1P5 | Kinesin-like protein KIF7 OS=Homo sapi | KIF7 | 150.588 | 2 | 53.293213 | yes | 5.276E-07 | 620.80316 | 76.783821 |
| RLEAEMSK | 8 | Q2M1P5 | Kinesin-like protein KIF7 OS=Homo sapi | KIF7 | 150.588 | 2 | 53.293213 | yes | 6.403E-06 | 482.25003 | 66.192627 |
| QGSAQSQQQIR | 11 | Q2M1P5 | Kinesin-like protein KIF7 OS=Homo sapi | KIF7 | 150.588 | 2 | 53.293213 | yes | 3.604E-06 | 615.81293 | 45.883572 |
| VAAAQSQVQVLK | 12 | Q2M1P5 | Kinesin-like protein KIF7 OS=Homo sapi | KIF7 | 150.588 | 2 | 53.293213 | yes | 9.722E-08 | 621.36426 | 42.350086 |
| VRPLLPK | 7 | Q2M1P5 | Kinesin-like protein KIF7 OS=Homo sapi | KIF7 | 150.588 | 2 | 53.293213 | yes | 4.319E-07 | 411.78162 | 33.922947 |
| LTDPIPTTETSIAPF | 15 | Q2M2I8 | AP2-associated protein kinase 1 OS=Hon | AAK1 | 103.885 | 2 | 51.63785 | yes | 9.722E-08 | 806.43304 | 129.88725 |
| LTAEELLNK | 9 | Q2M2I8 | AP2-associated protein kinase 1 OS=Hon | AAK1 | 103.885 | 2 | 51.63785 | yes | 1.658E-06 | 515.7926 | 70.058487 |
| VQTTPPPAVQGQK | 13 | Q2M2I8 | AP2-associated protein kinase 1 OS=Hon | AAK1 | 103.885 | 2 | 51.63785 | yes | 5.648E-07 | 675.87244 | 44.227978 |
| GHYVLCDFGSATN | 14 | Q2M2I8 | AP2-associated protein kinase 1 OS=Hon | AAK1 | 103.885 | 2 | 51.63785 | yes | 1.15E-07 | 784.86176 | 43.995628 |
| STQLLQAAAAEAS | 16 | Q2M2I8 | AP2-associated protein kinase 1 OS=Hon | AAK1 | 103.885 | 2 | 51.63785 | yes | 1.561E-06 | 808.4361 | 40.848499 |
| LGEPSEIDQR | 10 | Q2M389 | WASH complex subunit 4 OS=Homo sap | WASHC4 | 136.404 | 2 | 50.854916 | yes | 2.493E-07 | 572.28564 | 100.06091 |
| KLDLISELR | 9 | Q2M389 | WASH complex subunit 4 OS=Homo sap | WASHC4 | 136.404 | 2 | 50.854916 | yes | 9.722E-08 | 543.82953 | 41.783916 |
| MLVDVFAPEFR | 11 | Q2M389 | WASH complex subunit 4 OS=Homo sap | WASHC4 | 136.404 | 2 | 50.854916 | yes | 1.131E-06 | 662.34192 | 27.040621 |
| ELSMPGFIQK | 10 | Q2NKJ3 | CST complex subunit CTC1 OS=Homo sa | CTC1 | 134.609 | 2 | 45.875767 | yes | 2.116E-06 | 575.30225 | 98.721565 |
| LQTPSSFPTLATLK | 14 | Q2NKJ3 | CST complex subunit CTC1 OS=Homo sa | CTC1 | 134.609 | 2 | 45.875767 | yes | 1.717E-07 | 752.4245 | 38.314915 |
| DDEVSEDEEADQL | 14 | Q2T9J0 | Peroxisomal leader peptide-processing pr | TYSND1 | 59.309 | 2 | 47.46486 | yes | 1.324E-07 | 825.34229 | 31.318047 |
| GGSWAAPSLPSGV | 14 | Q2TAK8 | PWWP domain-containing DNA repair fa | PWWP3A | 78.637 | 2 | 45.74996 | yes | 2.11E-06 | 671.34912 | 100.98562 |
| YLQGVYQEVGAK | 12 | Q2TAK8 | PWWP domain-containing DNA repair fa | PWWP3A | 78.637 | 2 | 45.74996 | yes | 2.74E-07 | 677.8537 | 93.296791 |
| GSPEEPVVGCPLG | 15 | Q2TAK8 | PWWP domain-containing DNA repair fa | PWWP3A | 78.637 | 2 | 45.74996 | yes | 1.752E-07 | 791.38812 | 30.652775 |
| YIHLENLLAR | 10 | Q2TAY7 | WD40 repeat-containing protein SMU1 C | SMU1 | 57.544 | 2 | 52.431225 | yes | 1.174E-07 | 621.3537 | 1245.678 |
| YIHLENLLAR | 10 | Q2TAY7 | WD40 repeat-containing protein SMU1 C | SMU1 | 57.544 | 2 | 52.431225 | yes | 3.61E-07 | 414.57153 | 1245.678 |
| YIHLENLLAR | 10 | Q2TAY7 | WD40 repeat-containing protein SMU1 C | SMU1 | 57.544 | 3 | 52.431225 | yes | 1.174E-07 | 621.3537 | 1245.678 |
| YIHLENLLAR | 10 | Q2TAY7 | WD40 repeat-containing protein SMU1 C | SMU1 | 57.544 | 3 | 52.431225 | yes | 3.61E-07 | 414.57153 | 1245.678 |
| EGGDFVCCALSPR | 13 | Q2TAY7 | WD40 repeat-containing protein SMU1 C | SMU1 | 57.544 | 2 | 52.431225 | yes | 1.02E-07 | 734.32141 | 2400.2476 |
| DTEMLATGAQDGI | 13 | Q2TAY7 | WD40 repeat-containing protein SMU1 C | SMU1 | 57.544 | 2 | 52.431225 | yes | 9.742E-08 | 668.80609 | 2124.4365 |
| LMALLGQALK | 10 | Q2TAY7 | WD40 repeat-containing protein SMU1 C | SMU1 | 57.544 | 2 | 52.431225 | yes | 1.371E-07 | 529.32556 | 2085.7476 |
| DSSQILSASFDQTII | 15 | Q2TAY7 | WD40 repeat-containing protein SMU1 C | SMU1 | 57.544 | 2 | 52.431225 | yes | 1.05E-07 | 834.41541 | 2035.5205 |
| SIEIESSDVIR | 11 | Q2TAY7 | WD40 repeat-containing protein SMU1 C | SMU1 | 57.544 | 2 | 52.431225 | yes | 9.934E-08 | 645.33282 | 886.53381 |
| DVEEEKFPTQLSR | 13 | Q2TAY7 | WD40 repeat-containing protein SMU1 C | SMU1 | 57.544 | 2 | 52.431225 | yes | 9.722E-08 | 789.39392 | 881.67719 |

| | | | | | | | | | | |
|---|---|---|---|---|---|---|---|---|---|---|
| NPEHFVVCNR | 10 | Q2TAY7 | WD40 repeat-containing protein SMU1 C | SMU1 | 57.544 | 2 | 52.431225 | yes | 9.927E-08 | 636.30115 | 410.19043 |
| NPEHFVVCNR | 10 | Q2TAY7 | WD40 repeat-containing protein SMU1 C | SMU1 | 57.544 | 2 | 52.431225 | yes | 1.232E-07 | 424.5365 | 410.19043 |
| NPEHFVVCNR | 10 | Q2TAY7 | WD40 repeat-containing protein SMU1 C | SMU1 | 57.544 | 3 | 52.431225 | yes | 9.927E-08 | 636.30115 | 410.19043 |
| NPEHFVVCNR | 10 | Q2TAY7 | WD40 repeat-containing protein SMU1 C | SMU1 | 57.544 | 3 | 52.431225 | yes | 1.232E-07 | 424.5365 | 410.19043 |
| SNTVVIMNMQGQ | 15 | Q2TAY7 | WD40 repeat-containing protein SMU1 C | SMU1 | 57.544 | 2 | 52.431225 | yes | 1.301E-07 | 845.44238 | 803.02856 |
| TTECSNTFK | 9 | Q2TAY7 | WD40 repeat-containing protein SMU1 C | SMU1 | 57.544 | 2 | 52.431225 | yes | 1.12E-07 | 544.23987 | 669.80157 |
| QTDPMIMLK | 9 | Q2TAY7 | WD40 repeat-containing protein SMU1 C | SMU1 | 57.544 | 2 | 52.431225 | yes | 9.935E-08 | 538.77759 | 475.12683 |
| GVTCLSFSK | 9 | Q2TAY7 | WD40 repeat-containing protein SMU1 C | SMU1 | 57.544 | 2 | 52.431225 | yes | 9.722E-08 | 499.75241 | 390.79898 |
| GHSSFVNEATFTQ | 28 | Q2TAY7 | WD40 repeat-containing protein SMU1 C | SMU1 | 57.544 | 4 | 52.431225 | yes | 1.22E-07 | 739.59937 | 354.80273 |
| LIMQYLK | 7 | Q2TAY7 | WD40 repeat-containing protein SMU1 C | SMU1 | 57.544 | 2 | 52.431225 | yes | 3.824E-07 | 454.76733 | 295.82837 |
| DVIGIAHHPHQNL | 24 | Q2TAY7 | WD40 repeat-containing protein SMU1 C | SMU1 | 57.544 | 4 | 52.431225 | yes | 1.602E-07 | 661.09875 | 187.34415 |
| AAIAQALAGEVSV | 18 | Q2TAY7 | WD40 repeat-containing protein SMU1 C | SMU1 | 57.544 | 3 | 52.431225 | yes | 1.06E-06 | 579.32825 | 59.82674 |
| FPTQLSR | 7 | Q2TAY7 | WD40 repeat-containing protein SMU1 C | SMU1 | 57.544 | 2 | 52.431225 | yes | 4.679E-06 | 424.73486 | 16.319508 |
| VYLCLPNVR | 9 | Q2TBE0 | CWF19-like protein 2 OS=Homo sapiens | CWF19L2 | 103.787 | 2 | 51.876762 | yes | 5.633E-07 | 567.31042 | 202.01672 |
| EIIGGMLDIEPR | 12 | Q2TBE0 | CWF19-like protein 2 OS=Homo sapiens | CWF19L2 | 103.787 | 2 | 51.876762 | yes | 5.219E-06 | 671.85541 | 134.5618 |
| HLIVAIGVK | 9 | Q2TBE0 | CWF19-like protein 2 OS=Homo sapiens | CWF19L2 | 103.787 | 2 | 51.876762 | yes | 3.354E-07 | 475.31329 | 91.539543 |
| EVGDMAPIYFK | 11 | Q2TBE0 | CWF19-like protein 2 OS=Homo sapiens | CWF19L2 | 103.787 | 2 | 51.876762 | yes | 2.44E-06 | 635.31281 | 87.907669 |
| NFEDIVAER | 9 | Q2TBE0 | CWF19-like protein 2 OS=Homo sapiens | CWF19L2 | 103.787 | 2 | 51.876762 | yes | 1.112E-07 | 546.76965 | 82.276772 |
| LGEEEENQR | 9 | Q2TBE0 | CWF19-like protein 2 OS=Homo sapiens | CWF19L2 | 103.787 | 2 | 51.876762 | yes | 3.903E-07 | 552.25183 | 77.997635 |
| AEMMGNMELAEQ | 14 | Q2TBE0 | CWF19-like protein 2 OS=Homo sapiens | CWF19L2 | 103.787 | 2 | 51.876762 | yes | 3.533E-07 | 797.86731 | 74.682175 |
| CLYCFDSSQFPK | 12 | Q2TBE0 | CWF19-like protein 2 OS=Homo sapiens | CWF19L2 | 103.787 | 2 | 51.876762 | yes | 9.722E-08 | 776.33398 | 67.608612 |
| AIMESDEEWSMNI | 13 | Q2TBE0 | CWF19-like protein 2 OS=Homo sapiens | CWF19L2 | 103.787 | 2 | 51.876762 | yes | 9.722E-08 | 785.3316 | 50.63575 |
| ETITQIPK | 8 | Q2TBE0 | CWF19-like protein 2 OS=Homo sapiens | CWF19L2 | 103.787 | 2 | 51.876762 | yes | 4.083E-07 | 465.26874 | 36.969444 |
| KPTYSDK | 7 | Q2TBE0 | CWF19-like protein 2 OS=Homo sapiens | CWF19L2 | 103.787 | 2 | 51.876762 | yes | 2.811E-06 | 419.7189 | 27.078604 |
| LMTNLALGR | 9 | Q2VPK5 | Cytoplasmic tRNA 2-thiolation protein 2 | CTU2 | 56.108 | 2 | 51.094017 | yes | 1.45E-07 | 494.78406 | 238.08798 |
| HGDVVVVR | 8 | Q2VPK5 | Cytoplasmic tRNA 2-thiolation protein 2 | CTU2 | 56.108 | 2 | 51.094017 | yes | 3.334E-07 | 440.7536 | 95.095016 |
| LFSVPSVFTPAVD | 15 | Q2VPK5 | Cytoplasmic tRNA 2-thiolation protein 2 | CTU2 | 56.108 | 2 | 51.094017 | yes | 4.174E-07 | 804.43762 | 90.300514 |
| EAQPVVVIR | 9 | Q2VPK5 | Cytoplasmic tRNA 2-thiolation protein 2 | CTU2 | 56.108 | 2 | 51.094017 | yes | 9.722E-08 | 505.80328 | 71.050789 |
| GAFLAWDTGFSDI | 14 | Q2VPK5 | Cytoplasmic tRNA 2-thiolation protein 2 | CTU2 | 56.108 | 2 | 51.094017 | yes | 3.942E-07 | 786.35986 | 40.933437 |
| EVPAHSTEVGR | 11 | Q32MZ4 | Leucine-rich repeat flightless-interacting | LRRFIP1 | 89.254 | 2 | 57.619232 | yes | 1.132E-07 | 591.29907 | 227.14713 |
| EIDCLSPEAQK | 11 | Q32MZ4 | Leucine-rich repeat flightless-interacting | LRRFIP1 | 89.254 | 2 | 57.619232 | yes | 9.862E-08 | 645.30573 | 166.23979 |
| TNFMYQVDTLK | 11 | Q32MZ4 | Leucine-rich repeat flightless-interacting | LRRFIP1 | 89.254 | 2 | 57.619232 | yes | 1.052E-07 | 680.33429 | 151.19376 |
| KSPVPVETLK | 10 | Q32MZ4 | Leucine-rich repeat flightless-interacting | LRRFIP1 | 89.254 | 2 | 57.619232 | yes | 9.722E-08 | 549.33191 | 145.03603 |
| ELTYQNTDLSEIKI | 19 | Q32MZ4 | Leucine-rich repeat flightless-interacting | LRRFIP1 | 89.254 | 3 | 57.619232 | yes | 1.209E-07 | 766.04456 | 141.311 |
| IDGATQSSPAEPK | 13 | Q32MZ4 | Leucine-rich repeat flightless-interacting | LRRFIP1 | 89.254 | 2 | 57.619232 | yes | 7.155E-07 | 650.82263 | 82.579933 |
| QREEMLEK | 8 | Q32MZ4 | Leucine-rich repeat flightless-interacting | LRRFIP1 | 89.254 | 2 | 57.619232 | yes | 9.722E-08 | 531.76605 | 57.723621 |
| AGGEELDEGVAK | 12 | Q32MZ4 | Leucine-rich repeat flightless-interacting | LRRFIP1 | 89.254 | 2 | 57.619232 | yes | 1.216E-07 | 587.78296 | 55.160561 |
| SAVEAQNEVTENI | 14 | Q32MZ4 | Leucine-rich repeat flightless-interacting | LRRFIP1 | 89.254 | 2 | 57.619232 | yes | 9.722E-08 | 758.36792 | 45.690525 |
| EFTNQEAAEPK | 11 | Q32MZ4 | Leucine-rich repeat flightless-interacting | LRRFIP1 | 89.254 | 2 | 57.619232 | yes | 2.818E-06 | 632.2962 | 17.36422 |

| | | | | | | | | | | | |
|---|---|---|---|---|---|---|---|---|---|---|---|
| INNIDNMYR | 9 | Q32P41 | tRNA (guanine(37)-N1)-methyltransferas | TRMT5 | 58.257 | 2 | 33.595093 | yes | 1.197E-06 | 584.77441 | 260.30194 |
| LQEIGLGAFER | 11 | Q32P44 | Echinoderm microtubule-associated prote | EML3 | 95.197 | 2 | 52.423393 | yes | 1.063E-07 | 616.83533 | 325.24854 |
| VLGAGGAGPAPAT | 16 | Q32P44 | Echinoderm microtubule-associated prote | EML3 | 95.197 | 2 | 52.423393 | yes | 1.294E-07 | 689.87549 | 244.20671 |
| STNDSVLAVGFNP | 14 | Q32P44 | Echinoderm microtubule-associated prote | EML3 | 95.197 | 2 | 52.423393 | yes | 1.133E-07 | 738.87573 | 215.06522 |
| CLAVHPDGVR | 10 | Q32P44 | Echinoderm microtubule-associated prote | EML3 | 95.197 | 2 | 52.423393 | yes | 1.121E-07 | 562.28748 | 145.40901 |
| SNYNLEGISVK | 11 | Q32P44 | Echinoderm microtubule-associated prote | EML3 | 95.197 | 2 | 52.423393 | yes | 9.722E-08 | 612.31696 | 135.96112 |
| MDGAAGPGDGPA | 13 | Q32P44 | Echinoderm microtubule-associated prote | EML3 | 95.197 | 2 | 52.423393 | yes | 1.328E-07 | 607.26697 | 124.44542 |
| WLVLDTETR | 9 | Q32P44 | Echinoderm microtubule-associated prote | EML3 | 95.197 | 2 | 52.423393 | yes | 9.722E-08 | 566.80347 | 122.8877 |
| VVAVADDFCK | 10 | Q32P44 | Echinoderm microtubule-associated prote | EML3 | 95.197 | 2 | 52.423393 | yes | 1.43E-07 | 562.27625 | 92.367325 |
| GTQTETEVELK | 11 | Q32P44 | Echinoderm microtubule-associated prote | EML3 | 95.197 | 2 | 52.423393 | yes | 1.142E-06 | 617.81171 | 57.823242 |
| VQEQEMELVK | 10 | Q32P44 | Echinoderm microtubule-associated prote | EML3 | 95.197 | 2 | 52.423393 | yes | 4.963E-07 | 616.81317 | 42.205524 |
| GGKDPLSSPGGPG | 15 | Q32P44 | Echinoderm microtubule-associated prote | EML3 | 95.197 | 2 | 52.423393 | yes | 4.736E-07 | 684.84692 | 28.354557 |
| SLSELYLGNNPGLI | 14 | Q38SD2 | Leucine-rich repeat serine/threonine-prot | LRRK1 | 225.393 | 2 | 50.129128 | yes | 4.664E-07 | 766.90698 | 75.513489 |
| YFCGVPSPLR | 10 | Q38SD2 | Leucine-rich repeat serine/threonine-prot | LRRK1 | 225.393 | 2 | 50.129128 | yes | 2.752E-07 | 598.30005 | 71.429108 |
| VSCQLQVQR | 9 | Q38SD2 | Leucine-rich repeat serine/threonine-prot | LRRK1 | 225.393 | 2 | 50.129128 | yes | 3.385E-07 | 559.29279 | 48.433323 |
| APQVVHGEATIR | 12 | Q38SD2 | Leucine-rich repeat serine/threonine-prot | LRRK1 | 225.393 | 2 | 50.129128 | yes | 9.893E-08 | 639.35168 | 42.322346 |
| LPLYAAIK | 8 | Q38SD2 | Leucine-rich repeat serine/threonine-prot | LRRK1 | 225.393 | 2 | 50.129128 | yes | 9.66E-07 | 444.78131 | 42.122551 |
| LSDYGISR | 8 | Q38SD2 | Leucine-rich repeat serine/threonine-prot | LRRK1 | 225.393 | 2 | 50.129128 | yes | 6.907E-07 | 455.73508 | 40.452946 |
| ELTPHGVLVDAAV | 16 | Q38SD2 | Leucine-rich repeat serine/threonine-prot | LRRK1 | 225.393 | 3 | 50.129128 | yes | 9.966E-08 | 540.3103 | 35.378529 |
| PVLGQPEEVQFR | 12 | Q38SD2 | Leucine-rich repeat serine/threonine-prot | LRRK1 | 225.393 | 2 | 50.129128 | yes | 2.733E-07 | 699.87244 | 32.831406 |
| SGNEDIAIFLLR | 12 | Q38SD2 | Leucine-rich repeat serine/threonine-prot | LRRK1 | 225.393 | 2 | 50.129128 | yes | 4.317E-07 | 674.367 | 24.042696 |
| TSNNIEDDTILK | 12 | Q3B7T1 | Erythroid differentiation-related factor 1 ( | EDRF1 | 138.529 | 2 | 36.16026 | yes | 9.793E-07 | 681.841 | 53.656532 |
| TLSGALDIMVR | 11 | Q3B7T1 | Erythroid differentiation-related factor 1 ( | EDRF1 | 138.529 | 2 | 36.16026 | yes | 7.325E-07 | 588.32629 | 49.840294 |
| LLSIFESR | 8 | Q3B7T1 | Erythroid differentiation-related factor 1 ( | EDRF1 | 138.529 | 2 | 36.16026 | yes | 2.012E-06 | 482.77673 | 43.454269 |
| WETLLTVR | 8 | Q3KR37 | Protein Aster-B OS=Homo sapiens OX=9 | GRAMD1E | 85.401 | 2 | 30.249723 | yes | 2.239E-06 | 509.29001 | 98.724335 |
| HPQLFER | 7 | Q3L8U1 | Chromodomain-helicase-DNA-binding pr | CHD9 | 326.022 | 2 | 52.132442 | yes | 2.4E-07 | 463.74579 | 500.39191 |
| NLLVYGWGR | 9 | Q3L8U1 | Chromodomain-helicase-DNA-binding pr | CHD9 | 326.022 | 2 | 52.132442 | no | 3.815E-07 | 539.29565 | 489.67157 |
| TDISLDDPNFWQK | 13 | Q3L8U1 | Chromodomain-helicase-DNA-binding pr | CHD9 | 326.022 | 2 | 52.132442 | no | 1.138E-07 | 789.87537 | 290.79825 |
| VLIFSQMVR | 9 | Q3L8U1 | Chromodomain-helicase-DNA-binding pr | CHD9 | 326.022 | 2 | 52.132442 | no | 1.375E-07 | 546.81537 | 278.03275 |
| FSKPDSDR | 8 | Q3L8U1 | Chromodomain-helicase-DNA-binding pr | CHD9 | 326.022 | 2 | 52.132442 | no | 8.332E-07 | 476.23016 | 263.53888 |
| EETIIEVELTNIQK | 14 | Q3L8U1 | Chromodomain-helicase-DNA-binding pr | CHD9 | 326.022 | 2 | 52.132442 | no | 1.061E-07 | 829.94598 | 237.70966 |
| CVIIDEAHR | 9 | Q3L8U1 | Chromodomain-helicase-DNA-binding pr | CHD9 | 326.022 | 2 | 52.132442 | no | 1.621E-07 | 556.77948 | 194.47249 |
| VVSTFGVVFDPDR | 13 | Q3L8U1 | Chromodomain-helicase-DNA-binding pr | CHD9 | 326.022 | 2 | 52.132442 | yes | 9.723E-08 | 719.37225 | 194.14088 |
| ADPALCFLER | 10 | Q3L8U1 | Chromodomain-helicase-DNA-binding pr | CHD9 | 326.022 | 2 | 52.132442 | no | 1.327E-07 | 596.29498 | 158.9449 |
| LQNTQVR | 7 | Q3L8U1 | Chromodomain-helicase-DNA-binding pr | CHD9 | 326.022 | 2 | 52.132442 | yes | 1.751E-07 | 429.74323 | 135.12698 |
| VMSPENFPTASVE | 15 | Q3L8U1 | Chromodomain-helicase-DNA-binding pr | CHD9 | 326.022 | 2 | 52.132442 | yes | 1.418E-07 | 796.88489 | 130.30133 |
| NCILADEMGLGK | 12 | Q3L8U1 | Chromodomain-helicase-DNA-binding pr | CHD9 | 326.022 | 2 | 52.132442 | no | 2.31E-07 | 660.81793 | 105.37444 |
| NFMNVSGPHR | 10 | Q3L8U1 | Chromodomain-helicase-DNA-binding pr | CHD9 | 326.022 | 2 | 52.132442 | yes | 9.722E-08 | 579.77728 | 101.60852 |
| KEIEDLLR | 8 | Q3L8U1 | Chromodomain-helicase-DNA-binding pr | CHD9 | 326.022 | 2 | 52.132442 | no | 2.311E-07 | 508.29276 | 97.152435 |

| | | | | | | | | | | | |
|---|---|---|---|---|---|---|---|---|---|---|---|
| AVLQSMSGR | 9 | Q3L8U1 | Chromodomain-helicase-DNA-binding pı | CHD9 | 326.022 | 2 | 52.132442 | no | 1.612E-07 | 474.75021 | 85.610023 |
| FVFLLCTR | 8 | Q3L8U1 | Chromodomain-helicase-DNA-binding pı | CHD9 | 326.022 | 2 | 52.132442 | no | 9.722E-08 | 528.28894 | 82.694496 |
| SYDANTVASFYTT | 14 | Q3L8U1 | Chromodomain-helicase-DNA-binding pı | CHD9 | 326.022 | 2 | 52.132442 | yes | 1.997E-07 | 784.36737 | 79.829918 |
| TEPLTPNPASK | 11 | Q3L8U1 | Chromodomain-helicase-DNA-binding pı | CHD9 | 326.022 | 2 | 52.132442 | yes | 9.562E-07 | 577.80621 | 77.050766 |
| FCEEDIDQILLR | 12 | Q3L8U1 | Chromodomain-helicase-DNA-binding pı | CHD9 | 326.022 | 2 | 52.132442 | no | 1.188E-07 | 775.87958 | 72.106171 |
| ILTGPVVR | 8 | Q3L8U1 | Chromodomain-helicase-DNA-binding pı | CHD9 | 326.022 | 2 | 52.132442 | yes | 1.366E-06 | 427.77655 | 70.101982 |
| LDVNSLTGEER | 11 | Q3L8U1 | Chromodomain-helicase-DNA-binding pı | CHD9 | 326.022 | 2 | 52.132442 | yes | 3.271E-07 | 616.80951 | 68.090363 |
| TDYHILR | 7 | Q3L8U1 | Chromodomain-helicase-DNA-binding pı | CHD9 | 326.022 | 2 | 52.132442 | yes | 1.679E-05 | 459.24561 | 67.104935 |
| KAEIDIEAISGR | 12 | Q3L8U1 | Chromodomain-helicase-DNA-binding pı | CHD9 | 326.022 | 2 | 52.132442 | yes | 3.184E-07 | 651.35663 | 65.011826 |
| ELQNHLGLSAPVP | 14 | Q3L8U1 | Chromodomain-helicase-DNA-binding pı | CHD9 | 326.022 | 3 | 52.132442 | yes | 1.414E-07 | 510.95108 | 60.374779 |
| TQTSSFDIQK | 10 | Q3L8U1 | Chromodomain-helicase-DNA-binding pı | CHD9 | 326.022 | 2 | 52.132442 | yes | 5.512E-07 | 577.78802 | 44.855347 |
| NNLNPGQNSLSQS | 14 | Q3L8U1 | Chromodomain-helicase-DNA-binding pı | CHD9 | 326.022 | 2 | 52.132442 | yes | 9.722E-08 | 750.87372 | 41.1866 |
| NVEGVDIFFFNR | 12 | Q3L8U1 | Chromodomain-helicase-DNA-binding pı | CHD9 | 326.022 | 2 | 52.132442 | yes | 7.211E-06 | 728.86462 | 31.236414 |
| PSDMAQTQLQSQ | 14 | Q3L8U1 | Chromodomain-helicase-DNA-binding pı | CHD9 | 326.022 | 2 | 52.132442 | yes | 5.623E-07 | 780.87537 | 16.500805 |
| TETPPPLASLNVSK | 14 | Q3MHD2 | Protein LSM12 OS=Homo sapiens OX=9 | LSM12 | 21.701 | 2 | 52.933693 | yes | 1.354E-06 | 727.3985 | 46453.793 |
| LQGEVVAFDYQSK | 13 | Q3MHD2 | Protein LSM12 OS=Homo sapiens OX=9 | LSM12 | 21.701 | 2 | 52.933693 | yes | 1.001E-07 | 742.375 | 33760.445 |
| EGSALSHVR | 9 | Q3MHD2 | Protein LSM12 OS=Homo sapiens OX=9 | LSM12 | 21.701 | 2 | 52.933693 | yes | 3.406E-07 | 478.25143 | 1953.6147 |
| CPSSSGKPNHADII | 30 | Q3MHD2 | Protein LSM12 OS=Homo sapiens OX=9 | LSM12 | 21.701 | 4 | 52.933693 | yes | 5.381E-07 | 846.18317 | 402.36841 |
| NIVVMEEVVITPPY | 20 | Q3MHD2 | Protein LSM12 OS=Homo sapiens OX=9 | LSM12 | 21.701 | 3 | 52.933693 | yes | 1.313E-07 | 787.73248 | 82.567932 |
| NIVVMEEVVITPPY | 20 | Q3MHD2 | Protein LSM12 OS=Homo sapiens OX=9 | LSM12 | 21.701 | 3 | 52.933693 | yes | 4.532E-07 | 793.06409 | 82.567932 |
| GKEGSALSHVR | 11 | Q3MHD2 | Protein LSM12 OS=Homo sapiens OX=9 | LSM12 | 21.701 | 2 | 52.933693 | yes | 9.722E-08 | 570.80963 | 52.335106 |
| TSELSEAPQK | 10 | Q3V6T2 | Girdin OS=Homo sapiens OX=9606 GN= | CCDC88A | 216.042 | 2 | 31.330397 | yes | 1.877E-06 | 545.27478 | 32.115135 |
| IEALEQENSELER | 13 | Q3V6T2 | Girdin OS=Homo sapiens OX=9606 GN= | CCDC88A | 216.042 | 2 | 31.330397 | yes | 7.416E-06 | 780.38104 | 31.842796 |
| EAESCDCLQGFQL | 32 | Q3ZCM7 | Tubulin beta-8 chain OS=Homo sapiens ( | TUBB8 | 49.776 | 4 | 47.92939 | yes | 1.23E-07 | 828.63898 | 208.63918 |
| LEQEEQQR | 8 | Q3ZCQ8 | Mitochondrial import inner membrane tra | TIMM50 | 39.647 | 2 | 51.5412 | yes | 9.722E-08 | 530.2569 | 363.64542 |
| QNLFLGSLTSR | 11 | Q3ZCQ8 | Mitochondrial import inner membrane tra | TIMM50 | 39.647 | 2 | 51.5412 | yes | 1.484E-06 | 618.34076 | 360.68533 |
| TIALNGVEDVR | 11 | Q3ZCQ8 | Mitochondrial import inner membrane tra | TIMM50 | 39.647 | 2 | 51.5412 | yes | 4.067E-07 | 593.82495 | 155.21654 |
| RAPDQAAEIGSR | 12 | Q3ZCQ8 | Mitochondrial import inner membrane tra | TIMM50 | 39.647 | 2 | 51.5412 | yes | 3.42E-07 | 635.82855 | 31.542015 |
| TSFFLGEVGNAAK | 13 | Q49A26 | Cytokine-like nuclear factor N-PAC OS= | GLYR1 | 60.548 | 2 | 56.187763 | yes | 1.52E-07 | 670.84589 | 427.41235 |
| DLTIPESSTVK | 11 | Q49A26 | Cytokine-like nuclear factor N-PAC OS= | GLYR1 | 60.548 | 2 | 56.187763 | yes | 2.224E-07 | 595.31915 | 382.88254 |
| WQPTASEPVK | 10 | Q49A26 | Cytokine-like nuclear factor N-PAC OS= | GLYR1 | 60.548 | 2 | 56.187763 | yes | 2.964E-07 | 571.79565 | 325.78342 |
| FQQAVDAVEEFLR | 13 | Q49A26 | Cytokine-like nuclear factor N-PAC OS= | GLYR1 | 60.548 | 2 | 56.187763 | yes | 1.061E-07 | 776.39374 | 291.21429 |
| FFGTEDHAWIK | 11 | Q49A26 | Cytokine-like nuclear factor N-PAC OS= | GLYR1 | 60.548 | 2 | 56.187763 | yes | 1.811E-07 | 675.82751 | 162.51053 |
| GMMAGPMAAFK | 11 | Q49A26 | Cytokine-like nuclear factor N-PAC OS= | GLYR1 | 60.548 | 2 | 56.187763 | yes | 9.722E-08 | 556.25848 | 111.01855 |
| DLVLGPSGVLQGI | 14 | Q49A26 | Cytokine-like nuclear factor N-PAC OS= | GLYR1 | 60.548 | 2 | 56.187763 | yes | 9.722E-08 | 712.41699 | 110.18336 |
| KLSLSEGK | 8 | Q49A26 | Cytokine-like nuclear factor N-PAC OS= | GLYR1 | 60.548 | 2 | 56.187763 | yes | 3.057E-06 | 431.25565 | 107.49975 |
| CDLFIQEGAR | 10 | Q49A26 | Cytokine-like nuclear factor N-PAC OS= | GLYR1 | 60.548 | 2 | 56.187763 | yes | 6.959E-07 | 604.79004 | 99.679192 |
| MGHTVTVWNR | 10 | Q49A26 | Cytokine-like nuclear factor N-PAC OS= | GLYR1 | 60.548 | 2 | 56.187763 | yes | 9.722E-08 | 600.80078 | 90.406487 |
| ALDQSDNDMSAV | 14 | Q49A26 | Cytokine-like nuclear factor N-PAC OS= | GLYR1 | 60.548 | 2 | 56.187763 | yes | 9.722E-08 | 792.85156 | 51.277565 |

| | | | | | | | | | | | |
|---|---|---|---|---|---|---|---|---|---|---|---|
| AATTQLVSNSGLA | 17 | Q49A88 | Coiled-coil domain-containing protein 14 | CCDC14 | 106.302 | 2 | 46.92068 | yes | 3.04E-06 | 802.4361 | 69.769592 |
| ALVAEQEDSEIQR | 13 | Q49A88 | Coiled-coil domain-containing protein 14 | CCDC14 | 106.302 | 2 | 46.92068 | yes | 5.052E-07 | 744.37042 | 54.83757 |
| LLSDLSVDSAR | 11 | Q49A88 | Coiled-coil domain-containing protein 14 | CCDC14 | 106.302 | 2 | 46.92068 | yes | 1.441E-07 | 588.31696 | 53.686321 |
| GMEEASAPGIISAL | 16 | Q49A88 | Coiled-coil domain-containing protein 14 | CCDC14 | 106.302 | 2 | 46.92068 | yes | 5.615E-07 | 780.90051 | 49.127228 |
| EGISIAQIR | 9 | Q49AM1 | Transcription termination factor 2, mitoch | MTERF2 | 44.414 | 2 | 43.318607 | yes | 2.64E-07 | 493.7851 | 38.977825 |
| CTMPDEPLVR | 10 | Q4AC94 | C2 domain-containing protein 3 OS=Hom | C2CD3 | 260.389 | 2 | 49.452248 | yes | 1.029E-07 | 609.28632 | 87.428108 |
| ALPELNQDMTCTK | 13 | Q4AC94 | C2 domain-containing protein 3 OS=Hom | C2CD3 | 260.389 | 2 | 49.452248 | yes | 3.624E-07 | 760.85779 | 81.594368 |
| TALITEVVR | 9 | Q4AC94 | C2 domain-containing protein 3 OS=Hom | C2CD3 | 260.389 | 2 | 49.452248 | yes | 2.355E-07 | 501.30313 | 77.289444 |
| ELLLDPQQTLVFK | 13 | Q4AC94 | C2 domain-containing protein 3 OS=Hom | C2CD3 | 260.389 | 2 | 49.452248 | yes | 1.897E-07 | 772.44012 | 56.466164 |
| VTMELITDNK | 10 | Q4AC94 | C2 domain-containing protein 3 OS=Hom | C2CD3 | 260.389 | 2 | 49.452248 | yes | 4.574E-05 | 582.30249 | 43.408508 |
| TLTVSGVYPLFGR | 13 | Q4AC94 | C2 domain-containing protein 3 OS=Hom | C2CD3 | 260.389 | 2 | 49.452248 | yes | 1.915E-07 | 705.39301 | 33.115368 |
| MDEGEPATVTISTI | 15 | Q4AC94 | C2 domain-containing protein 3 OS=Hom | C2CD3 | 260.389 | 2 | 49.452248 | yes | 3.343E-06 | 802.38525 | 19.490423 |
| EGPSCPSPGPFCR | 13 | Q4AC94 | C2 domain-containing protein 3 OS=Hom | C2CD3 | 260.389 | 2 | 49.452248 | yes | 1.682E-06 | 724.30829 | 10.233788 |
| IISTEPLPGR | 10 | Q4G0J3 | La-related protein 7 OS=Homo sapiens O | LARP7 | 66.899 | 2 | 44.485783 | yes | 2.519E-07 | 541.81384 | 224.49449 |
| VNATGPQFVSGVI | 15 | Q4G0J3 | La-related protein 7 OS=Homo sapiens O | LARP7 | 66.899 | 2 | 44.485783 | yes | 4.873E-07 | 758.43011 | 127.12154 |
| GFAFVEFETK | 10 | Q4G0J3 | La-related protein 7 OS=Homo sapiens O | LARP7 | 66.899 | 2 | 44.485783 | yes | 2.685E-07 | 587.7926 | 104.45493 |
| AIEFLNNPPEEAPR | 14 | Q4G0J3 | La-related protein 7 OS=Homo sapiens O | LARP7 | 66.899 | 2 | 44.485783 | yes | 2.176E-07 | 798.90448 | 99.223228 |
| TVYYVELLPK | 9 | Q4G0J3 | La-related protein 7 OS=Homo sapiens O | LARP7 | 66.899 | 2 | 44.485783 | yes | 5.062E-06 | 531.31567 | 31.662529 |
| PGTTVASGVTAR | 12 | Q4KMP7 | TBC1 domain family member 10B OS=H | TBC1D10E | 87.199 | 2 | 54.74044 | yes | 9.722E-08 | 558.80402 | 108.70546 |
| SCQGMYETMEQLI | 13 | Q4KMP7 | TBC1 domain family member 10B OS=H | TBC1D10E | 87.199 | 2 | 54.74044 | yes | 9.809E-08 | 816.84436 | 49.764133 |
| WLDMFSNWDK | 10 | Q4KMP7 | TBC1 domain family member 10B OS=H | TBC1D10E | 87.199 | 2 | 54.74044 | yes | 9.722E-08 | 671.30023 | 29.33774 |
| SILGLGQAR | 9 | Q4VC44 | FLYWCH-type zinc finger-containing pro | FLYWCH1 | 80.109 | 2 | 48.669086 | yes | 1.117E-07 | 457.77454 | 760.24231 |
| HLPDILVINCEVNS | 16 | Q504Q3 | PAN2-PAN3 deadenylation complex cata | PAN2 | 135.368 | 3 | 55.946075 | yes | 1.478E-07 | 613.32074 | 200.03075 |
| LTGLACDR | 8 | Q504Q3 | PAN2-PAN3 deadenylation complex cata | PAN2 | 135.368 | 2 | 55.946075 | yes | 4.807E-07 | 453.2291 | 192.64207 |
| VGFIGYAPNPR | 11 | Q504Q3 | PAN2-PAN3 deadenylation complex cata | PAN2 | 135.368 | 2 | 55.946075 | yes | 1.019E-07 | 595.81946 | 192.51526 |
| FFFCGHTSGK | 10 | Q504Q3 | PAN2-PAN3 deadenylation complex cata | PAN2 | 135.368 | 2 | 55.946075 | yes | 1.548E-07 | 594.26874 | 189.48956 |
| YAVETPGVTIMR | 12 | Q504Q3 | PAN2-PAN3 deadenylation complex cata | PAN2 | 135.368 | 2 | 55.946075 | yes | 2.257E-07 | 668.8501 | 146.84978 |
| NYDFAQVLK | 9 | Q504Q3 | PAN2-PAN3 deadenylation complex cata | PAN2 | 135.368 | 2 | 55.946075 | yes | 1.299E-07 | 549.28491 | 130.74661 |
| VPAILYYVK | 9 | Q504Q3 | PAN2-PAN3 deadenylation complex cata | PAN2 | 135.368 | 2 | 55.946075 | yes | 4.613E-07 | 533.3208 | 117.9445 |
| GDPCQGNNFLR | 11 | Q504Q3 | PAN2-PAN3 deadenylation complex cata | PAN2 | 135.368 | 2 | 55.946075 | yes | 1.148E-07 | 639.28821 | 112.42524 |
| AITPLQVHVDPAFI | 15 | Q504Q3 | PAN2-PAN3 deadenylation complex cata | PAN2 | 135.368 | 2 | 55.946075 | yes | 1.063E-07 | 838.97778 | 107.34663 |
| MQAEVAFK | 8 | Q504Q3 | PAN2-PAN3 deadenylation complex cata | PAN2 | 135.368 | 2 | 55.946075 | yes | 1.824E-06 | 462.23639 | 95.821823 |
| KYLELSK | 7 | Q504Q3 | PAN2-PAN3 deadenylation complex cata | PAN2 | 135.368 | 2 | 55.946075 | yes | 2.175E-06 | 440.76056 | 72.461639 |
| LGLEDFDFK | 9 | Q504Q3 | PAN2-PAN3 deadenylation complex cata | PAN2 | 135.368 | 2 | 55.946075 | yes | 1.678E-07 | 542.27148 | 65.856781 |
| HEAVQFDMNWK | 11 | Q504Q3 | PAN2-PAN3 deadenylation complex cata | PAN2 | 135.368 | 2 | 55.946075 | yes | 8.412E-07 | 702.8219 | 61.182072 |
| RAPPVDAEILR | 11 | Q504Q3 | PAN2-PAN3 deadenylation complex cata | PAN2 | 135.368 | 2 | 55.946075 | yes | 1.003E-07 | 618.85657 | 53.105881 |
| YSSFQVNGSDDIR | 13 | Q504Q3 | PAN2-PAN3 deadenylation complex cata | PAN2 | 135.368 | 2 | 55.946075 | yes | 3.901E-07 | 744.34167 | 52.743732 |
| CLIQNHLCQK | 10 | Q504Q3 | PAN2-PAN3 deadenylation complex cata | PAN2 | 135.368 | 2 | 55.946075 | yes | 2.942E-07 | 657.32629 | 36.249783 |
| EEEPHLHMVSK | 11 | Q504Q3 | PAN2-PAN3 deadenylation complex cata | PAN2 | 135.368 | 2 | 55.946075 | yes | 9.722E-08 | 668.32172 | 19.83292 |

| | | | | | | | | | | | |
|---|---|---|---|---|---|---|---|---|---|---|---|
| GPLLEEQALTK | 11 | Q52LJ0 | Protein FAM98B OS=Homo sapiens OX= | FAM98B | 45.548 | 2 | 53.941376 | yes | 9.722E-08 | 599.83752 | 416.97614 |
| SLCNLEESITSAGR | 14 | Q52LJ0 | Protein FAM98B OS=Homo sapiens OX= | FAM98B | 45.548 | 2 | 53.941376 | yes | 9.796E-08 | 768.86975 | 382.67462 |
| TTITMAHLLAAR | 12 | Q52LJ0 | Protein FAM98B OS=Homo sapiens OX= | FAM98B | 45.548 | 2 | 53.941376 | yes | 9.935E-08 | 649.86609 | 231.32918 |
| INDALSCEYECR | 12 | Q52LJ0 | Protein FAM98B OS=Homo sapiens OX= | FAM98B | 45.548 | 2 | 53.941376 | yes | 9.722E-08 | 765.32159 | 109.82997 |
| EMACPYSVLISGD | 15 | Q52LJ0 | Protein FAM98B OS=Homo sapiens OX= | FAM98B | 45.548 | 2 | 53.941376 | yes | 3.891E-07 | 841.91003 | 106.92273 |
| MDLNSEQAEQLEI | 13 | Q52LJ0 | Protein FAM98B OS=Homo sapiens OX= | FAM98B | 45.548 | 2 | 53.941376 | yes | 9.722E-08 | 781.85938 | 59.240559 |
| APQLVGQFIAR | 11 | Q53EL6 | Programmed cell death protein 4 OS=Hor | PDCD4 | 51.736 | 2 | 55.512383 | yes | 3.8E-07 | 600.34839 | 526.65198 |
| DLPELALDTPR | 11 | Q53EL6 | Programmed cell death protein 4 OS=Hor | PDCD4 | 51.736 | 2 | 55.512383 | yes | 9.722E-08 | 620.33258 | 319.68552 |
| SSTITVDQMK | 10 | Q53EL6 | Programmed cell death protein 4 OS=Hor | PDCD4 | 51.736 | 2 | 55.512383 | yes | 1.994E-07 | 555.27899 | 153.04901 |
| GDSVSDSGSDALR | 13 | Q53EL6 | Programmed cell death protein 4 OS=Hor | PDCD4 | 51.736 | 2 | 55.512383 | yes | 9.722E-08 | 633.28387 | 95.582169 |
| ATVLLSMSK | 9 | Q53EL6 | Programmed cell death protein 4 OS=Hor | PDCD4 | 51.736 | 2 | 55.512383 | yes | 1.078E-06 | 475.27298 | 50.939392 |
| DLLPAAQYCCR | 11 | Q53EP0 | Fibronectin type III domain-containing pr | FNDC3B | 132.889 | 2 | 52.517292 | yes | 9.722E-08 | 683.81573 | 842.2713 |
| LECAAAGPQSLK | 12 | Q53EP0 | Fibronectin type III domain-containing pr | FNDC3B | 132.889 | 2 | 52.517292 | yes | 9.722E-08 | 622.81879 | 637.66394 |
| LCPAMGYTFR | 10 | Q53EP0 | Fibronectin type III domain-containing pr | FNDC3B | 132.889 | 2 | 52.517292 | yes | 1.755E-07 | 608.28607 | 524.10162 |
| LNSPPSSIYK | 10 | Q53EP0 | Fibronectin type III domain-containing pr | FNDC3B | 132.889 | 2 | 52.517292 | yes | 1.084E-07 | 553.29803 | 450.41022 |
| LTEFTCYSFR | 10 | Q53EP0 | Fibronectin type III domain-containing pr | FNDC3B | 132.889 | 2 | 52.517292 | yes | 9.722E-08 | 662.30554 | 423.9649 |
| IIYSGEELECNLK | 13 | Q53EP0 | Fibronectin type III domain-containing pr | FNDC3B | 132.889 | 2 | 52.517292 | yes | 1.152E-07 | 784.38727 | 422.54868 |
| DLLPETTYR | 9 | Q53EP0 | Fibronectin type III domain-containing pr | FNDC3B | 132.889 | 2 | 52.517292 | yes | 1.12E-07 | 554.28766 | 418.95126 |
| GPVTSHGFSVK | 11 | Q53EP0 | Fibronectin type III domain-containing pr | FNDC3B | 132.889 | 2 | 52.517292 | yes | 9.722E-08 | 558.29584 | 351.35162 |
| YTGEDLTCTVK | 11 | Q53EP0 | Fibronectin type III domain-containing pr | FNDC3B | 132.889 | 2 | 52.517292 | yes | 1.556E-07 | 643.80029 | 342.2485 |
| LEWGEDEESLELI | 20 | Q53EP0 | Fibronectin type III domain-containing pr | FNDC3B | 132.889 | 3 | 52.517292 | yes | 2.627E-07 | 798.03613 | 282.55612 |
| QCFFGSQK | 8 | Q53EP0 | Fibronectin type III domain-containing pr | FNDC3B | 132.889 | 2 | 52.517292 | yes | 2.74E-07 | 501.2291 | 160.98346 |
| ITNYLLEWDEGKR | 13 | Q53EP0 | Fibronectin type III domain-containing pr | FNDC3B | 132.889 | 2 | 52.517292 | yes | 1.138E-07 | 818.9201 | 147.11217 |
| VYAMYNSVK | 9 | Q53EP0 | Fibronectin type III domain-containing pr | FNDC3B | 132.889 | 2 | 52.517292 | yes | 4.982E-07 | 537.76807 | 124.49284 |
| SNDSDLQEYELEV | 14 | Q53EP0 | Fibronectin type III domain-containing pr | FNDC3B | 132.889 | 2 | 52.517292 | yes | 1.898E-07 | 834.88361 | 117.94249 |
| ITNYLLEWDEGK | 12 | Q53EP0 | Fibronectin type III domain-containing pr | FNDC3B | 132.889 | 2 | 52.517292 | yes | 1.118E-07 | 740.86957 | 111.83923 |
| SEVMLTGDMGSLI | 16 | Q53EP0 | Fibronectin type III domain-containing pr | FNDC3B | 132.889 | 2 | 52.517292 | yes | 2.311E-07 | 847.88422 | 77.904175 |
| CLDTSQELSGAFSI | 22 | Q53EP0 | Fibronectin type III domain-containing pr | FNDC3B | 132.889 | 3 | 52.517292 | yes | 4.921E-07 | 795.38989 | 31.109991 |
| VPAIEENLLDDK | 12 | Q53ET0 | CREB-regulated transcription coactivator | CRTC2 | 73.302 | 2 | 44.38749 | yes | 2.212E-07 | 678.35626 | 174.79845 |
| TMAWGNFPAEK | 11 | Q53ET0 | CREB-regulated transcription coactivator | CRTC2 | 73.302 | 2 | 44.38749 | yes | 7.881E-07 | 626.29498 | 124.46306 |
| GGILDGEMDPK | 11 | Q53ET0 | CREB-regulated transcription coactivator | CRTC2 | 73.302 | 2 | 44.38749 | yes | 3.199E-06 | 566.27118 | 45.69664 |
| NQITQLESLK | 10 | Q53EZ4 | Centrosomal protein of 55 kDa OS=Hom | CEP55 | 54.179 | 2 | 51.87812 | yes | 1.161E-07 | 587.32733 | 1271.1127 |
| NQQWLVYDQQR | 11 | Q53EZ4 | Centrosomal protein of 55 kDa OS=Hom | CEP55 | 54.179 | 2 | 51.87812 | yes | 2.667E-07 | 739.36279 | 1140.2262 |
| TSVDEITSGK | 10 | Q53EZ4 | Centrosomal protein of 55 kDa OS=Hom | CEP55 | 54.179 | 2 | 51.87812 | yes | 1.938E-07 | 518.76147 | 559.34473 |
| QQLSAATSR | 9 | Q53EZ4 | Centrosomal protein of 55 kDa OS=Hom | CEP55 | 54.179 | 2 | 51.87812 | yes | 2.759E-07 | 481.25671 | 475.2868 |
| LKGEIAHLK | 9 | Q53EZ4 | Centrosomal protein of 55 kDa OS=Hom | CEP55 | 54.179 | 2 | 51.87812 | yes | 1.496E-07 | 504.81366 | 427.2753 |
| SPTAALNESLVECI | 15 | Q53EZ4 | Centrosomal protein of 55 kDa OS=Hom | CEP55 | 54.179 | 2 | 51.87812 | yes | 1.829E-07 | 808.40344 | 344.96707 |
| RADVQHLEDDR | 11 | Q53EZ4 | Centrosomal protein of 55 kDa OS=Hom | CEP55 | 54.179 | 3 | 51.87812 | yes | 1.002E-07 | 451.88837 | 150.90005 |
| RADVQHLEDDR | 11 | Q53EZ4 | Centrosomal protein of 55 kDa OS=Hom | CEP55 | 54.179 | 3 | 51.87812 | yes | 6.811E-07 | 677.32892 | 150.90005 |

| | | | | | | | | | | | |
|---|---|---|---|---|---|---|---|---|---|---|---|
| RADVQHLEDDR | 11 | Q53EZ4 | Centrosomal protein of 55 kDa OS=Hom | CEP55 | 54.179 | 2 | 51.87812 | yes | 1.002E-07 | 451.88837 | 150.90005 |
| RADVQHLEDDR | 11 | Q53EZ4 | Centrosomal protein of 55 kDa OS=Hom | CEP55 | 54.179 | 2 | 51.87812 | yes | 6.811E-07 | 677.32892 | 150.90005 |
| KTETAAHSLPQQT | 14 | Q53EZ4 | Centrosomal protein of 55 kDa OS=Hom | CEP55 | 54.179 | 3 | 51.87812 | yes | 9.722E-08 | 513.94238 | 150.45357 |
| KTETAAHSLPQQT | 14 | Q53EZ4 | Centrosomal protein of 55 kDa OS=Hom | CEP55 | 54.179 | 3 | 51.87812 | yes | 8.531E-07 | 770.40991 | 150.45357 |
| KTETAAHSLPQQT | 14 | Q53EZ4 | Centrosomal protein of 55 kDa OS=Hom | CEP55 | 54.179 | 2 | 51.87812 | yes | 9.722E-08 | 513.94238 | 150.45357 |
| KTETAAHSLPQQT | 14 | Q53EZ4 | Centrosomal protein of 55 kDa OS=Hom | CEP55 | 54.179 | 2 | 51.87812 | yes | 8.531E-07 | 770.40991 | 150.45357 |
| DLLVHVEYCSK | 11 | Q53EZ4 | Centrosomal protein of 55 kDa OS=Hom | CEP55 | 54.179 | 2 | 51.87812 | yes | 9.722E-08 | 681.83972 | 222.65054 |
| ALSEEKDVLK | 10 | Q53EZ4 | Centrosomal protein of 55 kDa OS=Hom | CEP55 | 54.179 | 2 | 51.87812 | yes | 9.819E-08 | 566.31641 | 212.24326 |
| LREENDIAR | 9 | Q53EZ4 | Centrosomal protein of 55 kDa OS=Hom | CEP55 | 54.179 | 2 | 51.87812 | yes | 1.048E-07 | 558.29382 | 197.17941 |
| TETAAHSLPQQTK | 13 | Q53EZ4 | Centrosomal protein of 55 kDa OS=Hom | CEP55 | 54.179 | 2 | 51.87812 | yes | 1.021E-07 | 706.36243 | 147.95866 |
| EVHNLNQLLYSQF | 13 | Q53EZ4 | Centrosomal protein of 55 kDa OS=Hom | CEP55 | 54.179 | 2 | 51.87812 | yes | 9.81E-08 | 807.42334 | 136.59183 |
| ADVQHLEDDR | 10 | Q53EZ4 | Centrosomal protein of 55 kDa OS=Hom | CEP55 | 54.179 | 2 | 51.87812 | yes | 2.944E-07 | 599.27838 | 136.23734 |
| KPESEGYLQEEK | 12 | Q53EZ4 | Centrosomal protein of 55 kDa OS=Hom | CEP55 | 54.179 | 2 | 51.87812 | yes | 2.563E-07 | 718.84882 | 114.15769 |
| CYNDLLASAK | 10 | Q53EZ4 | Centrosomal protein of 55 kDa OS=Hom | CEP55 | 54.179 | 2 | 51.87812 | yes | 1.746E-07 | 577.77917 | 73.034775 |
| NAYQLTEK | 8 | Q53EZ4 | Centrosomal protein of 55 kDa OS=Hom | CEP55 | 54.179 | 2 | 51.87812 | yes | 1.397E-07 | 483.74817 | 67.390694 |
| QLHEFAITEPLVTF | 21 | Q53EZ4 | Centrosomal protein of 55 kDa OS=Hom | CEP55 | 54.179 | 3 | 51.87812 | yes | 4.51E-07 | 820.41156 | 66.234589 |
| AGSFITGIDVTSK | 13 | Q53F19 | Nuclear cap-binding protein subunit 3 OS | NCBP3 | 70.593 | 2 | 51.70569 | yes | 1.292E-07 | 648.3457 | 4579.894 |
| ALIGDDVGLTSYK | 13 | Q53F19 | Nuclear cap-binding protein subunit 3 OS | NCBP3 | 70.593 | 2 | 51.70569 | yes | 9.728E-08 | 676.35883 | 3495.4275 |
| AFSSNPVVR | 9 | Q53F19 | Nuclear cap-binding protein subunit 3 OS | NCBP3 | 70.593 | 2 | 51.70569 | yes | 1.001E-07 | 488.76416 | 2373.917 |
| MTMYADEVESQL | 13 | Q53F19 | Nuclear cap-binding protein subunit 3 OS | NCBP3 | 70.593 | 2 | 51.70569 | yes | 1.937E-07 | 772.85217 | 1142.8005 |
| MTMYADEVESQL | 13 | Q53F19 | Nuclear cap-binding protein subunit 3 OS | NCBP3 | 70.593 | 2 | 51.70569 | yes | 4.1E-07 | 780.84967 | 1142.8005 |
| ALINMSSLPAQDK | 13 | Q53F19 | Nuclear cap-binding protein subunit 3 OS | NCBP3 | 70.593 | 2 | 51.70569 | yes | 9.722E-08 | 694.36615 | 2023.4324 |
| SEVNLAQR | 8 | Q53F19 | Nuclear cap-binding protein subunit 3 OS | NCBP3 | 70.593 | 2 | 51.70569 | yes | 1.837E-07 | 458.74597 | 1879.6537 |
| LDNLPSLQIEVSR | 13 | Q53F19 | Nuclear cap-binding protein subunit 3 OS | NCBP3 | 70.593 | 2 | 51.70569 | yes | 1.382E-07 | 742.40936 | 1732.87 |
| ADSVSSSNIK | 10 | Q53F19 | Nuclear cap-binding protein subunit 3 OS | NCBP3 | 70.593 | 2 | 51.70569 | yes | 5.406E-07 | 504.25381 | 1649.0135 |
| YGNPNYGGMK | 10 | Q53F19 | Nuclear cap-binding protein subunit 3 OS | NCBP3 | 70.593 | 2 | 51.70569 | yes | 1.219E-07 | 550.74512 | 1020.7418 |
| VVVEYHEELPALK | 13 | Q53F19 | Nuclear cap-binding protein subunit 3 OS | NCBP3 | 70.593 | 2 | 51.70569 | yes | 9.09E-07 | 763.41669 | 413.00583 |
| VVVEYHEELPALK | 13 | Q53F19 | Nuclear cap-binding protein subunit 3 OS | NCBP3 | 70.593 | 2 | 51.70569 | yes | 8.595E-07 | 509.28021 | 413.00583 |
| VVVEYHEELPALK | 13 | Q53F19 | Nuclear cap-binding protein subunit 3 OS | NCBP3 | 70.593 | 3 | 51.70569 | yes | 9.09E-07 | 763.41669 | 413.00583 |
| VVVEYHEELPALK | 13 | Q53F19 | Nuclear cap-binding protein subunit 3 OS | NCBP3 | 70.593 | 3 | 51.70569 | yes | 8.595E-07 | 509.28021 | 413.00583 |
| APGAEEDDSELQR | 13 | Q53F19 | Nuclear cap-binding protein subunit 3 OS | NCBP3 | 70.593 | 2 | 51.70569 | yes | 9.787E-08 | 708.81549 | 729.87323 |
| ELIPDTSR | 8 | Q53F19 | Nuclear cap-binding protein subunit 3 OS | NCBP3 | 70.593 | 2 | 51.70569 | yes | 3.525E-07 | 465.74817 | 714.13745 |
| GILSNSWK | 8 | Q53F19 | Nuclear cap-binding protein subunit 3 OS | NCBP3 | 70.593 | 2 | 51.70569 | yes | 2.354E-07 | 452.74799 | 250.03493 |
| RSQYYMK | 7 | Q53F19 | Nuclear cap-binding protein subunit 3 OS | NCBP3 | 70.593 | 2 | 51.70569 | yes | 5.855E-07 | 488.23947 | 139.46147 |
| MISTPSPK | 8 | Q53F19 | Nuclear cap-binding protein subunit 3 OS | NCBP3 | 70.593 | 2 | 51.70569 | yes | 2.13E-06 | 430.73096 | 105.53455 |
| YVASVSLHDR | 10 | Q53G59 | Kelch-like protein 12 OS=Homo sapiens | KLHL12 | 63.278 | 2 | 54.229286 | yes | 9.722E-08 | 573.79877 | 383.95471 |
| IYVIGGYDGR | 10 | Q53G59 | Kelch-like protein 12 OS=Homo sapiens | KLHL12 | 63.278 | 2 | 54.229286 | yes | 1.379E-07 | 556.79041 | 371.52353 |
| YITDVIDAEPFIR | 13 | Q53G59 | Kelch-like protein 12 OS=Homo sapiens | KLHL12 | 63.278 | 2 | 54.229286 | yes | 9.893E-08 | 776.40631 | 296.29865 |
| CDAGVCVLR | 9 | Q53G59 | Kelch-like protein 12 OS=Homo sapiens | KLHL12 | 63.278 | 2 | 54.229286 | yes | 8.025E-07 | 525.24695 | 271.8335 |

| | | | | | | | | | | | |
|---|---|---|---|---|---|---|---|---|---|---|---|
| CYVGATVLR | 9 | Q53G59 | Kelch-like protein 12 OS=Homo sapiens | KLHL12 | 63.278 | 2 | 54.229286 | yes | 4.477E-07 | 519.77368 | 220.94304 |
| DIMTNTHAK | 9 | Q53G59 | Kelch-like protein 12 OS=Homo sapiens | KLHL12 | 63.278 | 2 | 54.229286 | yes | 1.674E-07 | 515.75293 | 218.70335 |
| TQEWSFLPSITR | 12 | Q53G59 | Kelch-like protein 12 OS=Homo sapiens | KLHL12 | 63.278 | 2 | 54.229286 | yes | 1.073E-07 | 732.87769 | 196.30141 |
| EESLPNLLQYVR | 12 | Q53G59 | Kelch-like protein 12 OS=Homo sapiens | KLHL12 | 63.278 | 2 | 54.229286 | yes | 9.723E-08 | 730.89081 | 180.57845 |
| MPLLTPR | 7 | Q53G59 | Kelch-like protein 12 OS=Homo sapiens | KLHL12 | 63.278 | 2 | 54.229286 | yes | 1.027E-07 | 414.24402 | 129.28572 |
| TDSWTTVTSMTTP | 14 | Q53G59 | Kelch-like protein 12 OS=Homo sapiens | KLHL12 | 63.278 | 2 | 54.229286 | yes | 9.722E-08 | 792.37213 | 101.23681 |
| HFPEVVQHEEFILI | 21 | Q53G59 | Kelch-like protein 12 OS=Homo sapiens | KLHL12 | 63.278 | 3 | 54.229286 | yes | 1.043E-07 | 832.09192 | 46.09145 |
| DFAETHNCVDLM( | 21 | Q53G59 | Kelch-like protein 12 OS=Homo sapiens | KLHL12 | 63.278 | 3 | 54.229286 | yes | 9.975E-08 | 814.0351 | 35.818371 |
| LSLFPASPR | 9 | Q53GA4 | Pleckstrin homology-like domain family A | PHLDA2 | 17.093 | 2 | 33.521664 | yes | 1.215E-06 | 494.28473 | 568.03857 |
| TIAVDFASEDIYDK | 14 | Q53GQ0 | Very-long-chain 3-oxoacyl-CoA reductase | HSD17B12 | 34.325 | 2 | 55.187622 | yes | 9.722E-08 | 793.88287 | 131.79695 |
| MTQLVLPGMVER | 12 | Q53GQ0 | Very-long-chain 3-oxoacyl-CoA reductase | HSD17B12 | 34.325 | 2 | 55.187622 | yes | 1.723E-07 | 687.36761 | 125.23355 |
| MININILSVCK | 11 | Q53GQ0 | Very-long-chain 3-oxoacyl-CoA reductase | HSD17B12 | 34.325 | 2 | 55.187622 | yes | 3.048E-06 | 652.85687 | 56.479614 |
| ISYSLFTALR | 10 | Q53GQ0 | Very-long-chain 3-oxoacyl-CoA reductase | HSD17B12 | 34.325 | 2 | 55.187622 | yes | 8.935E-07 | 585.82953 | 33.212429 |
| KPTLDKPSPETFVI | 14 | Q53GQ0 | Very-long-chain 3-oxoacyl-CoA reductase | HSD17B12 | 34.325 | 3 | 55.187622 | yes | 9.284E-07 | 529.6311 | 31.251825 |
| AATIPVSQISTIAGS | 16 | Q53GS7 | mRNA export factor GLE1 OS=Homo sap | GLE1 | 79.837 | 2 | 52.680534 | yes | 2.23E-06 | 772.43811 | 539.59656 |
| VDLAAFQTR | 9 | Q53GS7 | mRNA export factor GLE1 OS=Homo sap | GLE1 | 79.837 | 2 | 52.680534 | yes | 1.581E-07 | 510.77728 | 509.62335 |
| LWQEEQER | 8 | Q53GS7 | mRNA export factor GLE1 OS=Homo sap | GLE1 | 79.837 | 2 | 52.680534 | yes | 4.49E-07 | 559.26727 | 345.30743 |
| EGMALEDYQR | 10 | Q53GS7 | mRNA export factor GLE1 OS=Homo sap | GLE1 | 79.837 | 2 | 52.680534 | yes | 4.156E-07 | 606.27167 | 340.95474 |
| LYAAIIQLR | 9 | Q53GS7 | mRNA export factor GLE1 OS=Homo sap | GLE1 | 79.837 | 2 | 52.680534 | yes | 1.796E-07 | 530.82928 | 309.05597 |
| MYELVHR | 7 | Q53GS7 | mRNA export factor GLE1 OS=Homo sap | GLE1 | 79.837 | 2 | 52.680534 | yes | 1.88E-07 | 474.242 | 254.31392 |
| IEAITSSGQMGSFII | 15 | Q53GS7 | mRNA export factor GLE1 OS=Homo sap | GLE1 | 79.837 | 2 | 52.680534 | yes | 6.541E-07 | 798.90613 | 224.61671 |
| QYQVQFWK | 8 | Q53GS7 | mRNA export factor GLE1 OS=Homo sap | GLE1 | 79.837 | 2 | 52.680534 | yes | 1.076E-07 | 563.78766 | 170.75192 |
| IHSLLSGKPVQSG( | 15 | Q53GS7 | mRNA export factor GLE1 OS=Homo sap | GLE1 | 79.837 | 3 | 52.680534 | yes | 4.79E-07 | 512.62659 | 68.563644 |
| IHSLLSGKPVQSG( | 15 | Q53GS7 | mRNA export factor GLE1 OS=Homo sap | GLE1 | 79.837 | 3 | 52.680534 | yes | 2.305E-06 | 768.43628 | 68.563644 |
| IHSLLSGKPVQSG( | 15 | Q53GS7 | mRNA export factor GLE1 OS=Homo sap | GLE1 | 79.837 | 2 | 52.680534 | yes | 4.79E-07 | 512.62659 | 68.563644 |
| IHSLLSGKPVQSG( | 15 | Q53GS7 | mRNA export factor GLE1 OS=Homo sap | GLE1 | 79.837 | 2 | 52.680534 | yes | 2.305E-06 | 768.43628 | 68.563644 |
| VQALSEMASEQLK | 13 | Q53GS7 | mRNA export factor GLE1 OS=Homo sap | GLE1 | 79.837 | 2 | 52.680534 | yes | 9.722E-08 | 717.3689 | 134.38704 |
| GFLTSSFWR | 9 | Q53GS7 | mRNA export factor GLE1 OS=Homo sap | GLE1 | 79.837 | 2 | 52.680534 | yes | 9.722E-08 | 550.77985 | 122.65694 |
| GNQLCSLISGIIR | 13 | Q53GS7 | mRNA export factor GLE1 OS=Homo sap | GLE1 | 79.837 | 2 | 52.680534 | yes | 2.83E-07 | 715.89282 | 44.734268 |
| LKEIFDK | 7 | Q53GS7 | mRNA export factor GLE1 OS=Homo sap | GLE1 | 79.837 | 2 | 52.680534 | yes | 4.344E-06 | 446.76056 | 43.898674 |
| RQDEEEAQVK | 10 | Q53GS7 | mRNA export factor GLE1 OS=Homo sap | GLE1 | 79.837 | 2 | 52.680534 | yes | 1.301E-07 | 616.29932 | 29.725563 |
| KVQALSEMASEQI | 14 | Q53GS7 | mRNA export factor GLE1 OS=Homo sap | GLE1 | 79.837 | 2 | 52.680534 | yes | 4.203E-07 | 781.41632 | 22.622066 |
| LQEAQMQQGPEAI | 14 | Q53GS7 | mRNA export factor GLE1 OS=Homo sap | GLE1 | 79.837 | 2 | 52.680534 | yes | 1.156E-07 | 797.88574 | 20.064423 |
| TIVTDVFQGSMR | 12 | Q53GS9 | Ubiquitin carboxyl-terminal hydrolase 39 | USP39 | 65.381 | 2 | 46.77647 | yes | 3.074E-07 | 677.34521 | 466.79834 |
| EYLSEEVQAVHK | 12 | Q53GS9 | Ubiquitin carboxyl-terminal hydrolase 39 | USP39 | 65.381 | 2 | 46.77647 | yes | 5.671E-07 | 716.35938 | 225.53056 |
| SVLDFDFEK | 9 | Q53GS9 | Ubiquitin carboxyl-terminal hydrolase 39 | USP39 | 65.381 | 2 | 46.77647 | yes | 1.634E-07 | 550.26892 | 198.78905 |
| NYFLEEDNYK | 10 | Q53GS9 | Ubiquitin carboxyl-terminal hydrolase 39 | USP39 | 65.381 | 2 | 46.77647 | yes | 1.473E-07 | 667.79858 | 193.54103 |
| IHVLHHGTGK | 10 | Q53GS9 | Ubiquitin carboxyl-terminal hydrolase 39 | USP39 | 65.381 | 2 | 46.77647 | yes | 1.625E-07 | 549.81201 | 92.044868 |
| KLPHPDLPAEEK | 12 | Q53GS9 | Ubiquitin carboxyl-terminal hydrolase 39 | USP39 | 65.381 | 3 | 46.77647 | yes | 4.566E-06 | 458.58563 | 44.027397 |

| | | | | | | | | | | | |
|---|---|---|---|---|---|---|---|---|---|---|---|
| KLPHPDLPAEEK | 12 | Q53GS9 | Ubiquitin carboxyl-terminal hydrolase 39 | USP39 | 65.381 | 3 | 46.77647 | yes | 2.754E-07 | 687.37482 | 44.027397 |
| KLPHPDLPAEEK | 12 | Q53GS9 | Ubiquitin carboxyl-terminal hydrolase 39 | USP39 | 65.381 | 2 | 46.77647 | yes | 4.566E-06 | 458.58563 | 44.027397 |
| KLPHPDLPAEEK | 12 | Q53GS9 | Ubiquitin carboxyl-terminal hydrolase 39 | USP39 | 65.381 | 2 | 46.77647 | yes | 2.754E-07 | 687.37482 | 44.027397 |
| EVDEDSEPER | 10 | Q53GS9 | Ubiquitin carboxyl-terminal hydrolase 39 | USP39 | 65.381 | 2 | 46.77647 | yes | 5.112E-06 | 602.75183 | 14.534603 |
| MGMPSSLAHR | 10 | Q53H96 | Pyrroline-5-carboxylate reductase 3 OS=I | PYCR3 | 28.664 | 2 | 44.3703 | yes | 2.219E-07 | 543.76276 | 145.48396 |
| EIFLTVPVGGGESI | 15 | Q53HL2 | Borealin OS=Homo sapiens OX=9606 GI | CDCA8 | 31.323 | 2 | 50.36234 | yes | 3.524E-07 | 787.43286 | 442.57034 |
| EVDNLYNIEILR | 12 | Q53HL2 | Borealin OS=Homo sapiens OX=9606 GI | CDCA8 | 31.323 | 2 | 50.36234 | yes | 1.214E-07 | 745.89612 | 264.45633 |
| LAQICSSIR | 9 | Q53HL2 | Borealin OS=Homo sapiens OX=9606 GI | CDCA8 | 31.323 | 2 | 50.36234 | yes | 9.81E-08 | 524.28442 | 220.85521 |
| LLASDLQR | 8 | Q53HL2 | Borealin OS=Homo sapiens OX=9606 GI | CDCA8 | 31.323 | 2 | 50.36234 | yes | 2.294E-07 | 458.26654 | 196.26129 |
| PTPGLTPR | 8 | Q53HL2 | Borealin OS=Homo sapiens OX=9606 GI | CDCA8 | 31.323 | 2 | 50.36234 | yes | 4.868E-06 | 419.74271 | 165.14856 |
| LTAEAIQTPLK | 11 | Q53HL2 | Borealin OS=Homo sapiens OX=9606 GI | CDCA8 | 31.323 | 2 | 50.36234 | yes | 2.147E-07 | 592.8479 | 162.14964 |
| GIGGMDGTQQQIF | 14 | Q53R41 | FAST kinase domain-containing protein 1 | FASTKD1 | 97.411 | 2 | 50.64144 | yes | 3.66E-07 | 740.36664 | 48.834244 |
| VDFECILDK | 9 | Q53R41 | FAST kinase domain-containing protein 1 | FASTKD1 | 97.411 | 2 | 50.64144 | yes | 9.733E-08 | 569.77606 | 30.074711 |
| DKPWFDFR | 8 | Q53T94 | TATA box-binding protein-associated fac | TAF1B | 68.832 | 2 | 40.290504 | yes | 7.818E-07 | 555.77197 | 51.890709 |
| NSLYWLSTQK | 10 | Q53T94 | TATA box-binding protein-associated fac | TAF1B | 68.832 | 2 | 40.290504 | yes | 4.105E-07 | 620.32202 | 24.758892 |
| GLGGLLEVPEQPR | 13 | Q562E7 | WD repeat-containing protein 81 OS=Ho | WDR81 | 211.697 | 2 | 54.763298 | yes | 1.084E-07 | 682.88025 | 119.84372 |
| GSPACPSLLR | 10 | Q562E7 | WD repeat-containing protein 81 OS=Ho | WDR81 | 211.697 | 2 | 54.763298 | yes | 3.319E-06 | 529.27661 | 64.93763 |
| LLVQTIQETTGR | 12 | Q562E7 | WD repeat-containing protein 81 OS=Ho | WDR81 | 211.697 | 2 | 54.763298 | yes | 3.035E-07 | 679.88556 | 55.576054 |
| SLVLDWVHGR | 10 | Q562E7 | WD repeat-containing protein 81 OS=Ho | WDR81 | 211.697 | 2 | 54.763298 | yes | 9.722E-08 | 591.32495 | 40.344353 |
| ELQIPALSTR | 10 | Q562F6 | Shugoshin 2 OS=Homo sapiens OX=960 | SGO2 | 144.74 | 2 | 52.488472 | yes | 9.174E-07 | 564.32458 | 143.84152 |
| MTLQTGFEQGDR | 12 | Q562F6 | Shugoshin 2 OS=Homo sapiens OX=960 | SGO2 | 144.74 | 2 | 52.488472 | yes | 9.722E-08 | 691.82208 | 141.95935 |
| LNFENTFLR | 9 | Q562F6 | Shugoshin 2 OS=Homo sapiens OX=960 | SGO2 | 144.74 | 2 | 52.488472 | yes | 4.695E-07 | 577.30365 | 122.32107 |
| ILNNSSIFK | 9 | Q562F6 | Shugoshin 2 OS=Homo sapiens OX=960 | SGO2 | 144.74 | 2 | 52.488472 | yes | 1.83E-07 | 518.29529 | 100.64026 |
| SHSLFLTQK | 9 | Q562F6 | Shugoshin 2 OS=Homo sapiens OX=960 | SGO2 | 144.74 | 2 | 52.488472 | yes | 6.87E-07 | 530.79291 | 98.060951 |
| TQNVLGLQK | 9 | Q562F6 | Shugoshin 2 OS=Homo sapiens OX=960 | SGO2 | 144.74 | 2 | 52.488472 | yes | 2.761E-07 | 500.79294 | 95.585846 |
| SEDLSSER | 8 | Q562F6 | Shugoshin 2 OS=Homo sapiens OX=960 | SGO2 | 144.74 | 2 | 52.488472 | yes | 1.647E-06 | 461.70926 | 85.048477 |
| QDLPGLSSESAR | 12 | Q562F6 | Shugoshin 2 OS=Homo sapiens OX=960 | SGO2 | 144.74 | 2 | 52.488472 | yes | 1.877E-07 | 630.31494 | 84.668884 |
| ESCDQILDSYK | 11 | Q562F6 | Shugoshin 2 OS=Homo sapiens OX=960 | SGO2 | 144.74 | 2 | 52.488472 | yes | 1.807E-07 | 679.30066 | 60.532711 |
| DNENQCDYR | 9 | Q562F6 | Shugoshin 2 OS=Homo sapiens OX=960 | SGO2 | 144.74 | 2 | 52.488472 | yes | 9.722E-08 | 607.23053 | 38.787498 |
| DNFFSLTPK | 9 | Q562F6 | Shugoshin 2 OS=Homo sapiens OX=960 | SGO2 | 144.74 | 2 | 52.488472 | yes | 5.063E-07 | 534.77167 | 34.299141 |
| DQLIIDAGQK | 10 | Q56NI9 | N-acetyltransferase ESCO2 OS=Homo sa | ESCO2 | 68.307 | 2 | 54.969795 | yes | 1.63E-07 | 550.80096 | 257.31183 |
| VVGCLIAEPIK | 11 | Q56NI9 | N-acetyltransferase ESCO2 OS=Homo sa | ESCO2 | 68.307 | 2 | 54.969795 | yes | 9.745E-08 | 599.84686 | 211.6214 |
| LPSANQGSPFK | 11 | Q56NI9 | N-acetyltransferase ESCO2 OS=Homo sa | ESCO2 | 68.307 | 2 | 54.969795 | yes | 1.26E-07 | 573.30109 | 163.49847 |
| WYLNPLER | 8 | Q56NI9 | N-acetyltransferase ESCO2 OS=Homo sa | ESCO2 | 68.307 | 2 | 54.969795 | yes | 2.909E-07 | 545.78766 | 156.71989 |
| TFLFISDEK | 9 | Q56NI9 | N-acetyltransferase ESCO2 OS=Homo sa | ESCO2 | 68.307 | 2 | 54.969795 | yes | 1.504E-07 | 550.28711 | 112.27164 |
| SSLENEPSLGR | 11 | Q56NI9 | N-acetyltransferase ESCO2 OS=Homo sa | ESCO2 | 68.307 | 2 | 54.969795 | yes | 1.074E-07 | 594.79639 | 93.586952 |
| SALSTVSFYNQNK | 13 | Q56NI9 | N-acetyltransferase ESCO2 OS=Homo sa | ESCO2 | 68.307 | 2 | 54.969795 | yes | 2.122E-06 | 729.86481 | 92.468895 |
| SEVIEDSDVETVSI | 15 | Q56NI9 | N-acetyltransferase ESCO2 OS=Homo sa | ESCO2 | 68.307 | 2 | 54.969795 | yes | 2.961E-07 | 833.38873 | 55.119877 |
| IVLVLPHDPSFAIK | 14 | Q56NI9 | N-acetyltransferase ESCO2 OS=Homo sa | ESCO2 | 68.307 | 2 | 54.969795 | yes | 9.722E-08 | 774.96106 | 50.598736 |

| | | | | | | | | | | | |
|---|---|---|---|---|---|---|---|---|---|---|---|
| NEAFSSEDSLGENI | 14 | Q56NI9 | N-acetyltransferase ESCO2 OS=Homo sa | ESCO2 | 68.307 | 2 | 54.969795 | yes | 2.024E-07 | 763.83392 | 32.864357 |
| FYTQLDAAQMR | 11 | Q58A45 | PAN2-PAN3 deadenylation complex subı | PAN3 | 95.613 | 2 | 53.03331 | yes | 9.81E-08 | 672.32428 | 798.49597 |
| SSNFGYITSCYK | 12 | Q58A45 | PAN2-PAN3 deadenylation complex subı | PAN3 | 95.613 | 2 | 53.03331 | yes | 9.722E-08 | 713.81903 | 512.66663 |
| AMELVTINYSSDL | 14 | Q58A45 | PAN2-PAN3 deadenylation complex subı | PAN3 | 95.613 | 2 | 53.03331 | yes | 1.418E-07 | 792.40289 | 410.24615 |
| NDVIEEDLAK | 10 | Q58A45 | PAN2-PAN3 deadenylation complex subı | PAN3 | 95.613 | 2 | 53.03331 | yes | 9.809E-08 | 573.28784 | 406.79126 |
| SVLVVTYSDLK | 11 | Q58A45 | PAN2-PAN3 deadenylation complex subı | PAN3 | 95.613 | 2 | 53.03331 | yes | 1.931E-07 | 612.34772 | 345.9462 |
| ANAPSFFMADELR | 13 | Q58A45 | PAN2-PAN3 deadenylation complex subı | PAN3 | 95.613 | 2 | 53.03331 | yes | 9.722E-08 | 734.84808 | 266.9588 |
| TIHTAGLACR | 10 | Q58A45 | PAN2-PAN3 deadenylation complex subı | PAN3 | 95.613 | 2 | 53.03331 | yes | 9.722E-08 | 550.28748 | 218.38887 |
| HFNDPNADAYFTK | 13 | Q58A45 | PAN2-PAN3 deadenylation complex subı | PAN3 | 95.613 | 2 | 53.03331 | yes | 1.663E-07 | 770.3468 | 201.52779 |
| VVLALACNSLAGI | 15 | Q58A45 | PAN2-PAN3 deadenylation complex subı | PAN3 | 95.613 | 2 | 53.03331 | yes | 9.974E-08 | 792.94812 | 194.26001 |
| WGQHEGPLPR | 10 | Q58A45 | PAN2-PAN3 deadenylation complex subı | PAN3 | 95.613 | 2 | 53.03331 | yes | 1.23E-07 | 588.79907 | 181.97908 |
| ITPHTSPAPR | 10 | Q58A45 | PAN2-PAN3 deadenylation complex subı | PAN3 | 95.613 | 2 | 53.03331 | yes | 9.927E-07 | 538.79602 | 173.62549 |
| CMVLVDMWK | 9 | Q58A45 | PAN2-PAN3 deadenylation complex subı | PAN3 | 95.613 | 2 | 53.03331 | yes | 9.722E-08 | 591.27942 | 168.66495 |
| DPTWSETGDR | 10 | Q58A45 | PAN2-PAN3 deadenylation complex subı | PAN3 | 95.613 | 2 | 53.03331 | yes | 9.733E-08 | 582.25183 | 161.78064 |
| IQHSNIVTLR | 10 | Q58A45 | PAN2-PAN3 deadenylation complex subı | PAN3 | 95.613 | 2 | 53.03331 | yes | 2.958E-07 | 590.84351 | 132.75383 |
| NPIGCLADR | 9 | Q58A45 | PAN2-PAN3 deadenylation complex subı | PAN3 | 95.613 | 2 | 53.03331 | yes | 4.618E-07 | 508.25311 | 126.70283 |
| DDLPYCLR | 8 | Q58A45 | PAN2-PAN3 deadenylation complex subı | PAN3 | 95.613 | 2 | 53.03331 | yes | 3.023E-07 | 526.2475 | 114.56069 |
| LGMLEER | 7 | Q58A45 | PAN2-PAN3 deadenylation complex subı | PAN3 | 95.613 | 2 | 53.03331 | yes | 1.885E-07 | 424.22073 | 86.16301 |
| SGVPINMVWWNR | 12 | Q58A45 | PAN2-PAN3 deadenylation complex subı | PAN3 | 95.613 | 2 | 53.03331 | yes | 2.552E-06 | 729.86914 | 62.036652 |
| LGTINERPEFQK | 12 | Q58A45 | PAN2-PAN3 deadenylation complex subı | PAN3 | 95.613 | 2 | 53.03331 | yes | 6.133E-07 | 716.38318 | 59.733467 |
| MTNSSSSPSLLNDS | 16 | Q58A45 | PAN2-PAN3 deadenylation complex subı | PAN3 | 95.613 | 2 | 53.03331 | yes | 1.012E-07 | 819.88562 | 55.528824 |
| KPDLGDPGTGAAA | 26 | Q58A45 | PAN2-PAN3 deadenylation complex subı | PAN3 | 95.613 | 3 | 53.03331 | yes | 4.057E-07 | 742.02448 | 52.30265 |
| YALLLQELAR | 10 | Q58EX7 | Puratrophin-1 OS=Homo sapiens OX=96 | PLEKHG4 | 130.803 | 2 | 45.65657 | yes | 3.311E-06 | 595.35059 | 45.335186 |
| DTFVLQASSLAIK | 13 | Q58EX7 | Puratrophin-1 OS=Homo sapiens OX=96 | PLEKHG4 | 130.803 | 2 | 45.65657 | yes | 2.054E-07 | 696.89032 | 37.658138 |
| EALTWAEEGQR | 11 | Q58EX7 | Puratrophin-1 OS=Homo sapiens OX=96 | PLEKHG4 | 130.803 | 2 | 45.65657 | yes | 1.783E-07 | 645.30969 | 28.228397 |
| ELYPVEFR | 8 | Q5BJF2 | Sigma intracellular receptor 2 OS=Homo | TMEM97 | 20.848 | 2 | 45.087307 | yes | 1.967E-07 | 526.77417 | 133.45387 |
| VTDLVNQQQTLEI | 14 | Q5BJF6 | Outer dense fiber protein 2 OS=Homo saı | ODF2 | 95.402 | 2 | 58.523544 | yes | 9.722E-08 | 822.9256 | 152.77245 |
| LMEQQGALLK | 10 | Q5BJF6 | Outer dense fiber protein 2 OS=Homo saı | ODF2 | 95.402 | 2 | 58.523544 | yes | 9.013E-07 | 565.81555 | 116.64217 |
| ILDLETQLSR | 10 | Q5BJF6 | Outer dense fiber protein 2 OS=Homo saı | ODF2 | 95.402 | 2 | 58.523544 | yes | 2.566E-07 | 594.33514 | 93.288902 |
| LEADEVAAQLER | 12 | Q5BJF6 | Outer dense fiber protein 2 OS=Homo saı | ODF2 | 95.402 | 2 | 58.523544 | yes | 9.722E-08 | 672.34369 | 91.188866 |
| NYEGMIDNYK | 10 | Q5BJF6 | Outer dense fiber protein 2 OS=Homo saı | ODF2 | 95.402 | 2 | 58.523544 | yes | 2.035E-07 | 623.77405 | 78.231239 |
| QMTCTDINTLTR | 12 | Q5BJF6 | Outer dense fiber protein 2 OS=Homo saı | ODF2 | 95.402 | 2 | 58.523544 | yes | 6.418E-07 | 727.34235 | 68.661583 |
| SMQNYVQFLK | 10 | Q5BJF6 | Outer dense fiber protein 2 OS=Homo saı | ODF2 | 95.402 | 2 | 58.523544 | yes | 1.028E-07 | 629.31842 | 64.020676 |
| ASFAPMEDK | 9 | Q5BJF6 | Outer dense fiber protein 2 OS=Homo saı | ODF2 | 95.402 | 2 | 58.523544 | yes | 1.276E-06 | 498.22876 | 64.009178 |
| GHLQAQLR | 8 | Q5BJF6 | Outer dense fiber protein 2 OS=Homo saı | ODF2 | 95.402 | 2 | 58.523544 | yes | 1.108E-07 | 461.7645 | 54.727936 |
| AEVEAIMEQLK | 11 | Q5BJF6 | Outer dense fiber protein 2 OS=Homo saı | ODF2 | 95.402 | 2 | 58.523544 | yes | 1.179E-07 | 630.82886 | 50.709625 |
| LAECQDQLQGYEI | 13 | Q5BJF6 | Outer dense fiber protein 2 OS=Homo saı | ODF2 | 95.402 | 2 | 58.523544 | yes | 7.04E-07 | 805.36737 | 43.836239 |
| LVEAEMDGAAAA | 13 | Q5BJF6 | Outer dense fiber protein 2 OS=Homo saı | ODF2 | 95.402 | 2 | 58.523544 | yes | 9.769E-07 | 638.3161 | 43.069347 |
| LEQSESTNR | 9 | Q5BJF6 | Outer dense fiber protein 2 OS=Homo saı | ODF2 | 95.402 | 2 | 58.523544 | yes | 2.256E-07 | 532.25433 | 40.998894 |

| | | | | | | | | | | |
|---|---|---|---|---|---|---|---|---|---|---|
| KLEATSAQNIEFL( | 18 | Q5BJF6 | Outer dense fiber protein 2 OS=Homo sa| | ODF2 | 95.402 | 3 | 58.523544 | yes | 4.109E-07 | 668.37726 | 18.880121 |
| ILADAAAEGVPVR | 13 | Q5BJH7 | Protein YIF1B OS=Homo sapiens OX=9( | YIF1B | 34.436 | 2 | 46.083546 | yes | 1.658E-07 | 641.36169 | 45.424168 |
| SGYGFNEPEQSR | 12 | Q5BKZ1 | DBIRD complex subunit ZNF326 OS=H( | ZNF326 | 65.654 | 2 | 51.838253 | yes | 9.722E-08 | 685.80219 | 1109.3271 |
| MDFEDDYTHSACI | 13 | Q5BKZ1 | DBIRD complex subunit ZNF326 OS=H( | ZNF326 | 65.654 | 2 | 51.838253 | yes | 1.079E-07 | 844.8194 | 462.00497 |
| FGPYESYDSR | 10 | Q5BKZ1 | DBIRD complex subunit ZNF326 OS=H( | ZNF326 | 65.654 | 2 | 51.838253 | yes | 1.81E-07 | 610.76459 | 456.5957 |
| VVMEFLHECMVN | 13 | Q5BKZ1 | DBIRD complex subunit ZNF326 OS=H( | ZNF326 | 65.654 | 2 | 51.838253 | yes | 9.722E-08 | 818.38824 | 223.29707 |
| VVMEFLHECMVN | 13 | Q5BKZ1 | DBIRD complex subunit ZNF326 OS=H( | ZNF326 | 65.654 | 2 | 51.838253 | yes | 3.972E-07 | 545.92792 | 223.29707 |
| VVMEFLHECMVN | 13 | Q5BKZ1 | DBIRD complex subunit ZNF326 OS=H( | ZNF326 | 65.654 | 3 | 51.838253 | yes | 9.722E-08 | 818.38824 | 223.29707 |
| VVMEFLHECMVN | 13 | Q5BKZ1 | DBIRD complex subunit ZNF326 OS=H( | ZNF326 | 65.654 | 3 | 51.838253 | yes | 3.972E-07 | 545.92792 | 223.29707 |
| NSLDSFGGR | 9 | Q5BKZ1 | DBIRD complex subunit ZNF326 OS=H( | ZNF326 | 65.654 | 2 | 51.838253 | yes | 1.938E-07 | 476.72778 | 381.93848 |
| LRPGFMEDR | 9 | Q5BKZ1 | DBIRD complex subunit ZNF326 OS=H( | ZNF326 | 65.654 | 2 | 51.838253 | yes | 3.152E-06 | 560.78204 | 304.70596 |
| PGIVVDYQNK | 10 | Q5BKZ1 | DBIRD complex subunit ZNF326 OS=H( | ZNF326 | 65.654 | 2 | 51.838253 | yes | 1.366E-07 | 566.80347 | 262.12665 |
| VETVHCSACSVYI | 25 | Q5BKZ1 | DBIRD complex subunit ZNF326 OS=H( | ZNF326 | 65.654 | 4 | 51.838253 | yes | 5.261E-07 | 713.35608 | 155.08583 |
| MAFTCSFCK | 9 | Q5BKZ1 | DBIRD complex subunit ZNF326 OS=H( | ZNF326 | 65.654 | 2 | 51.838253 | yes | 9.722E-08 | 576.23773 | 154.47635 |
| QQTNNQTEVVK | 11 | Q5BKZ1 | DBIRD complex subunit ZNF326 OS=H( | ZNF326 | 65.654 | 2 | 51.838253 | yes | 9.81E-08 | 644.82825 | 117.8148 |
| NTYQGFNGMDR | 11 | Q5BKZ1 | DBIRD complex subunit ZNF326 OS=H( | ZNF326 | 65.654 | 2 | 51.838253 | yes | 1.876E-07 | 651.78021 | 113.19694 |
| DIELHLESSSHQET | 20 | Q5BKZ1 | DBIRD complex subunit ZNF326 OS=H( | ZNF326 | 65.654 | 3 | 51.838253 | yes | 1.986E-07 | 787.05566 | 78.510201 |
| DYGPGSYGGMDR | 12 | Q5BKZ1 | DBIRD complex subunit ZNF326 OS=H( | ZNF326 | 65.654 | 2 | 51.838253 | yes | 8.645E-07 | 637.75897 | 70.952942 |
| STNVTVAAAR | 10 | Q5BKZ1 | DBIRD complex subunit ZNF326 OS=H( | ZNF326 | 65.654 | 2 | 51.838253 | yes | 4.914E-07 | 495.27237 | 61.879089 |
| DYGHGSYGGQR | 11 | Q5BKZ1 | DBIRD complex subunit ZNF326 OS=H( | ZNF326 | 65.654 | 2 | 51.838253 | yes | 4.243E-06 | 598.75763 | 60.036366 |
| ALLAANEEEDR | 11 | Q5C9Z4 | Nucleolar MIF4G domain-containing pro | NOM1 | 96.258 | 2 | 51.01603 | yes | 1.198E-07 | 615.8017 | 437.72131 |
| NAGSGSETQLR | 11 | Q5C9Z4 | Nucleolar MIF4G domain-containing pro | NOM1 | 96.258 | 2 | 51.01603 | yes | 1.061E-07 | 560.27307 | 325.73422 |
| VSWDSVLSAEQT( | 14 | Q5C9Z4 | Nucleolar MIF4G domain-containing pro | NOM1 | 96.258 | 2 | 51.01603 | yes | 4.317E-07 | 767.87842 | 161.43204 |
| ELITEAQTK | 9 | Q5C9Z4 | Nucleolar MIF4G domain-containing pro | NOM1 | 96.258 | 2 | 51.01603 | yes | 7.68E-07 | 516.78223 | 124.31925 |
| KIPGYDPEPVEK | 12 | Q5C9Z4 | Nucleolar MIF4G domain-containing pro | NOM1 | 96.258 | 2 | 51.01603 | yes | 9.722E-08 | 686.36139 | 69.55407 |
| KFDAIYK | 7 | Q5C9Z4 | Nucleolar MIF4G domain-containing pro | NOM1 | 96.258 | 2 | 51.01603 | yes | 4.362E-07 | 442.74747 | 56.556675 |
| QLVGTVSSK | 9 | Q5C9Z4 | Nucleolar MIF4G domain-containing pro | NOM1 | 96.258 | 2 | 51.01603 | yes | 3.418E-06 | 459.76639 | 55.961281 |
| TRPSAAATAAAR | 12 | Q5C9Z4 | Nucleolar MIF4G domain-containing pro | NOM1 | 96.258 | 2 | 51.01603 | yes | 2.295E-07 | 572.31506 | 50.750217 |
| LFISHFLLK | 9 | Q5C9Z4 | Nucleolar MIF4G domain-containing pro | NOM1 | 96.258 | 2 | 51.01603 | yes | 2.518E-07 | 559.34204 | 40.636131 |
| VVEFSELDKPR | 11 | Q5C9Z4 | Nucleolar MIF4G domain-containing pro | NOM1 | 96.258 | 3 | 51.01603 | yes | 4.63E-07 | 440.23822 | 38.975513 |
| FMLETMLALK | 10 | Q5C9Z4 | Nucleolar MIF4G domain-containing pro | NOM1 | 96.258 | 2 | 51.01603 | yes | 1.338E-06 | 598.82452 | 31.390362 |
| TAGPEQGPGLGGR | 13 | Q5C9Z4 | Nucleolar MIF4G domain-containing pro | NOM1 | 96.258 | 2 | 51.01603 | yes | 4.018E-05 | 598.80457 | 19.532269 |
| LATLFESYK | 9 | Q5D0E6 | DALR anticodon-binding domain-contair | DALRD3 | 59.363 | 2 | 51.511868 | yes | 9.722E-08 | 536.28967 | 89.271416 |
| VLLHCPALR | 9 | Q5D0E6 | DALR anticodon-binding domain-contair | DALRD3 | 59.363 | 2 | 51.511868 | yes | 4.2E-07 | 539.81311 | 69.577377 |
| TLSPGILGR | 9 | Q5D0E6 | DALR anticodon-binding domain-contair | DALRD3 | 59.363 | 2 | 51.511868 | yes | 4.12E-07 | 457.27692 | 39.93679 |
| LGVQADTNTVLGI | 16 | Q5D1E8 | Endoribonuclease ZC3H12A OS=Homo s | ZC3H12A | 65.7 | 2 | 49.856102 | yes | 1.001E-07 | 828.96198 | 82.617622 |
| APNLEPPLPEEEK | 13 | Q5D1E8 | Endoribonuclease ZC3H12A OS=Homo s | ZC3H12A | 65.7 | 2 | 49.856102 | yes | 1.809E-07 | 731.87482 | 58.110252 |
| QEGLTQTYAPSGR | 13 | Q5D1E8 | Endoribonuclease ZC3H12A OS=Homo s | ZC3H12A | 65.7 | 2 | 49.856102 | yes | 1.316E-07 | 704.3468 | 51.559837 |
| ASQLGIFISK | 10 | Q5EBL8 | PDZ domain-containing protein 11 OS=H | PDZD11 | 16.132 | 2 | 43.819614 | yes | 2.434E-07 | 532.31097 | 166.87186 |

| | | | | | | | | | | | |
|---|---|---|---|---|---|---|---|---|---|---|---|
| KPPGAQLGFNIR | 12 | Q5EBL8 | PDZ domain-containing protein 11 OS=H | PDZD11 | 16.132 | 2 | 43.819614 | yes | 3.297E-06 | 649.37238 | 98.718918 |
| EQSWMTMANLEK | 12 | Q5F1R6 | DnaJ homolog subfamily C member 21 O | DNAJC21 | 62.028 | 2 | 54.093006 | yes | 1.047E-07 | 734.33398 | 820.56677 |
| DASEEELKK | 9 | Q5F1R6 | DnaJ homolog subfamily C member 21 O | DNAJC21 | 62.028 | 2 | 54.093006 | yes | 2.11E-07 | 524.76147 | 678.63 |
| VDPEDTNLNQDSA | 14 | Q5F1R6 | DnaJ homolog subfamily C member 21 O | DNAJC21 | 62.028 | 2 | 54.093006 | yes | 1.613E-07 | 773.35498 | 674.59302 |
| CHYEALGVR | 9 | Q5F1R6 | DnaJ homolog subfamily C member 21 O | DNAJC21 | 62.028 | 2 | 54.093006 | yes | 9.892E-08 | 552.76636 | 627.39069 |
| LVEEQNAEK | 9 | Q5F1R6 | DnaJ homolog subfamily C member 21 O | DNAJC21 | 62.028 | 2 | 54.093006 | yes | 1.423E-07 | 530.26947 | 497.05185 |
| EKNELVR | 7 | Q5F1R6 | DnaJ homolog subfamily C member 21 O | DNAJC21 | 62.028 | 2 | 54.093006 | yes | 4.29E-07 | 444.25089 | 173.32451 |
| GFYTVYR | 7 | Q5F1R6 | DnaJ homolog subfamily C member 21 O | DNAJC21 | 62.028 | 2 | 54.093006 | yes | 1.915E-06 | 453.22943 | 132.05429 |
| ELQEMEAR | 8 | Q5F1R6 | DnaJ homolog subfamily C member 21 O | DNAJC21 | 62.028 | 2 | 54.093006 | yes | 1.602E-07 | 503.23712 | 127.39308 |
| APSSSSLNSATSSQ | 16 | Q5F1R6 | DnaJ homolog subfamily C member 21 O | DNAJC21 | 62.028 | 2 | 54.093006 | yes | 9.722E-08 | 769.86829 | 86.440887 |
| KLVEEQNAEK | 10 | Q5F1R6 | DnaJ homolog subfamily C member 21 O | DNAJC21 | 62.028 | 2 | 54.093006 | yes | 9.722E-08 | 594.31696 | 69.003258 |
| EAQDIILQLR | 10 | Q5FBB7 | Shugoshin 1 OS=Homo sapiens OX=960 | SGO1 | 64.19 | 2 | 30.242344 | yes | 2.242E-06 | 599.84314 | 29.928768 |
| ILEAWEMNEK | 10 | Q5H9R7 | Serine/threonine-protein phosphatase 6 re | PPP6R3 | 97.67 | 2 | 50.848446 | yes | 3.418E-06 | 631.80792 | 267.80426 |
| LVEIVHPSQEEDR | 13 | Q5H9R7 | Serine/threonine-protein phosphatase 6 re | PPP6R3 | 97.67 | 2 | 50.848446 | yes | 9.722E-08 | 775.8941 | 90.982529 |
| LVEIVHPSQEEDR | 13 | Q5H9R7 | Serine/threonine-protein phosphatase 6 re | PPP6R3 | 97.67 | 2 | 50.848446 | yes | 3.763E-07 | 517.59851 | 90.982529 |
| LVEIVHPSQEEDR | 13 | Q5H9R7 | Serine/threonine-protein phosphatase 6 re | PPP6R3 | 97.67 | 3 | 50.848446 | yes | 9.722E-08 | 775.8941 | 90.982529 |
| LVEIVHPSQEEDR | 13 | Q5H9R7 | Serine/threonine-protein phosphatase 6 re | PPP6R3 | 97.67 | 3 | 50.848446 | yes | 3.763E-07 | 517.59851 | 90.982529 |
| HGYMGHLTR | 9 | Q5H9R7 | Serine/threonine-protein phosphatase 6 re | PPP6R3 | 97.67 | 2 | 50.848446 | yes | 5.132E-07 | 536.26129 | 172.89885 |
| LGEDESLLMK | 10 | Q5H9R7 | Serine/threonine-protein phosphatase 6 re | PPP6R3 | 97.67 | 2 | 50.848446 | yes | 5.596E-07 | 567.78918 | 159.26172 |
| LLTCIEPPQPR | 11 | Q5H9R7 | Serine/threonine-protein phosphatase 6 re | PPP6R3 | 97.67 | 2 | 50.848446 | yes | 2.664E-06 | 662.35809 | 158.54004 |
| VLSILISR | 8 | Q5H9R7 | Serine/threonine-protein phosphatase 6 re | PPP6R3 | 97.67 | 2 | 50.848446 | yes | 1.018E-07 | 450.79749 | 117.40585 |
| LSTSQDAACK | 10 | Q5H9R7 | Serine/threonine-protein phosphatase 6 re | PPP6R3 | 97.67 | 2 | 50.848446 | yes | 1.258E-07 | 540.75311 | 113.77326 |
| ELMDEEDVLQECI | 13 | Q5H9R7 | Serine/threonine-protein phosphatase 6 re | PPP6R3 | 97.67 | 2 | 50.848446 | yes | 1.058E-07 | 819.35529 | 112.60625 |
| TTWGVLDPPVGN | 14 | Q5H9R7 | Serine/threonine-protein phosphatase 6 re | PPP6R3 | 97.67 | 2 | 50.848446 | yes | 4.723E-07 | 756.89392 | 109.97404 |
| QDLFEPSSANTEDI | 14 | Q5H9R7 | Serine/threonine-protein phosphatase 6 re | PPP6R3 | 97.67 | 2 | 50.848446 | yes | 5.136E-07 | 790.85736 | 91.887512 |
| IANCIVHSTDK | 11 | Q5H9R7 | Serine/threonine-protein phosphatase 6 re | PPP6R3 | 97.67 | 2 | 50.848446 | yes | 1.597E-07 | 629.31647 | 85.420822 |
| GPNSALVQQLIK | 12 | Q5H9R7 | Serine/threonine-protein phosphatase 6 re | PPP6R3 | 97.67 | 2 | 50.848446 | yes | 9.79E-08 | 634.37207 | 65.563141 |
| QDVLNWLNEEK | 11 | Q5H9R7 | Serine/threonine-protein phosphatase 6 re | PPP6R3 | 97.67 | 2 | 50.848446 | yes | 1.338E-07 | 694.34625 | 61.790405 |
| RPTFEGHIEICPPGI | 24 | Q5H9R7 | Serine/threonine-protein phosphatase 6 re | PPP6R3 | 97.67 | 4 | 50.848446 | yes | 2.778E-06 | 678.56763 | 46.943943 |
| WETFCTSSLGETN | 14 | Q5H9R7 | Serine/threonine-protein phosphatase 6 re | PPP6R3 | 97.67 | 2 | 50.848446 | yes | 1.046E-07 | 830.36957 | 46.738495 |
| KLIEFLLK | 8 | Q5H9R7 | Serine/threonine-protein phosphatase 6 re | PPP6R3 | 97.67 | 2 | 50.848446 | yes | 3.31E-06 | 502.33115 | 31.91223 |
| FADQDDIGNVSFD | 14 | Q5H9R7 | Serine/threonine-protein phosphatase 6 re | PPP6R3 | 97.67 | 2 | 50.848446 | yes | 3.269E-06 | 799.85773 | 26.386833 |
| VLVLGDSGVGK | 11 | Q5HYI8 | Rab-like protein 3 OS=Homo sapiens OX | RABL3 | 26.423 | 2 | 48.788246 | yes | 1.104E-07 | 522.30841 | 170.46077 |
| AASALYACTK | 10 | Q5HYJ3 | Protein FAM76B OS=Homo sapiens OX= | FAM76B | 38.708 | 2 | 52.272717 | yes | 9.722E-08 | 549.26843 | 395.53445 |
| ISNLSPEEEQGLWI | 14 | Q5HYJ3 | Protein FAM76B OS=Homo sapiens OX= | FAM76B | 38.708 | 2 | 52.272717 | yes | 1.448E-07 | 815.40961 | 223.77315 |
| ETVEQLQAK | 9 | Q5HYJ3 | Protein FAM76B OS=Homo sapiens OX= | FAM76B | 38.708 | 2 | 52.272717 | yes | 2.378E-06 | 523.27985 | 186.7749 |
| SSATIQNETPK | 11 | Q5HYJ3 | Protein FAM76B OS=Homo sapiens OX= | FAM76B | 38.708 | 2 | 52.272717 | yes | 9.722E-08 | 588.29877 | 124.93519 |
| ADFQYQESNLR | 11 | Q5HYJ3 | Protein FAM76B OS=Homo sapiens OX= | FAM76B | 38.708 | 2 | 52.272717 | yes | 1.122E-07 | 685.82037 | 115.2194 |
| LLCWLCTLSYK | 11 | Q5HYJ3 | Protein FAM76B OS=Homo sapiens OX= | FAM76B | 38.708 | 2 | 52.272717 | no | 1.783E-07 | 728.87 | 47.354996 |

| | | | | | | | | | | | |
|---|---|---|---|---|---|---|---|---|---|---|---|
| SLGSSHSNSSSSSL' | 17 | Q5HYJ3 | Protein FAM76B OS=Homo sapiens OX= | FAM76B | 38.708 | 2 | 52.272717 | yes | 1.1E-07 | 847.89502 | 19.866112 |
| LELYAQATEALLK | 13 | Q5JPH6 | Probable glutamate--tRNA ligase, mitoch | EARS2 | 58.689 | 2 | 45.252605 | yes | 1.908E-07 | 731.91119 | 96.559044 |
| DVLNPVYVER | 10 | Q5JPH6 | Probable glutamate--tRNA ligase, mitoch | EARS2 | 58.689 | 2 | 45.252605 | yes | 7.452E-07 | 602.32202 | 78.484795 |
| GGPAGPYQQSQR | 12 | Q5JPH6 | Probable glutamate--tRNA ligase, mitoch | EARS2 | 58.689 | 2 | 45.252605 | yes | 1.472E-06 | 623.30219 | 27.339539 |
| HEVASVEGDPVIM | 14 | Q5JPH6 | Probable glutamate--tRNA ligase, mitoch | EARS2 | 58.689 | 2 | 45.252605 | yes | 3.151E-06 | 755.88214 | 12.01816 |
| VYQVTEQQISEK | 12 | Q5JRA6 | Transport and Golgi organization protein | MIA3 | 213.703 | 2 | 50.3329 | yes | 9.819E-08 | 726.37244 | 59.476746 |
| ELLPHSSGR | 9 | Q5JST6 | EF-hand domain-containing family memb | EFHC2 | 87.397 | 2 | 35.02357 | yes | 8.995E-07 | 498.26706 | 86.918556 |
| APGESGPQLAR | 11 | Q5JTC6 | APC membrane recruitment protein 1 OS | AMER1 | 124.03 | 2 | 38.72254 | yes | 5.029E-07 | 541.78308 | 49.251427 |
| LLPEEYHR | 8 | Q5JTH9 | RRP12-like protein OS=Homo sapiens O | RRP12 | 143.703 | 2 | 54.652313 | yes | 2.417E-07 | 528.77728 | 1648.2466 |
| AMDLAQAGSTVES | 14 | Q5JTH9 | RRP12-like protein OS=Homo sapiens O | RRP12 | 143.703 | 2 | 54.652313 | yes | 1.123E-07 | 704.34283 | 817.20129 |
| AMDLAQAGSTVES | 14 | Q5JTH9 | RRP12-like protein OS=Homo sapiens O | RRP12 | 143.703 | 2 | 54.652313 | yes | 9.722E-08 | 712.34033 | 817.20129 |
| VLDPASSDFTR | 11 | Q5JTH9 | RRP12-like protein OS=Homo sapiens O | RRP12 | 143.703 | 2 | 54.652313 | yes | 1.063E-07 | 604.30133 | 1609.0676 |
| AHINLVR | 7 | Q5JTH9 | RRP12-like protein OS=Homo sapiens O | RRP12 | 143.703 | 2 | 54.652313 | yes | 9.722E-08 | 411.75085 | 1119.8561 |
| LHNELQSGSLR | 11 | Q5JTH9 | RRP12-like protein OS=Homo sapiens O | RRP12 | 143.703 | 2 | 54.652313 | yes | 1.152E-07 | 627.3335 | 997.04059 |
| RVPSPVLIK | 9 | Q5JTH9 | RRP12-like protein OS=Homo sapiens O | RRP12 | 143.703 | 2 | 54.652313 | yes | 4.987E-07 | 504.83185 | 873.69952 |
| FWESNSAAHK | 10 | Q5JTH9 | RRP12-like protein OS=Homo sapiens O | RRP12 | 143.703 | 2 | 54.652313 | yes | 1.315E-07 | 588.77527 | 620.85675 |
| FCIQEIEK | 8 | Q5JTH9 | RRP12-like protein OS=Homo sapiens O | RRP12 | 143.703 | 2 | 54.652313 | yes | 1.248E-07 | 533.7655 | 513.96613 |
| SCSETLLR | 8 | Q5JTH9 | RRP12-like protein OS=Homo sapiens O | RRP12 | 143.703 | 2 | 54.652313 | yes | 2.293E-07 | 483.23965 | 476.81863 |
| AAQHGVCSVLK | 11 | Q5JTH9 | RRP12-like protein OS=Homo sapiens O | RRP12 | 143.703 | 2 | 54.652313 | yes | 1.112E-07 | 585.30841 | 476.79977 |
| EGGGDEPLNFLDP | 14 | Q5JTH9 | RRP12-like protein OS=Homo sapiens O | RRP12 | 143.703 | 2 | 54.652313 | yes | 9.722E-08 | 744.35425 | 438.61566 |
| CLQSLCDLR | 9 | Q5JTH9 | RRP12-like protein OS=Homo sapiens O | RRP12 | 143.703 | 2 | 54.652313 | yes | 2.375E-07 | 582.77863 | 409.63477 |
| HVQLVMEAIGK | 11 | Q5JTH9 | RRP12-like protein OS=Homo sapiens O | RRP12 | 143.703 | 2 | 54.652313 | yes | 9.722E-08 | 612.8421 | 367.18756 |
| CLLHIVR | 7 | Q5JTH9 | RRP12-like protein OS=Homo sapiens O | RRP12 | 143.703 | 2 | 54.652313 | yes | 4.388E-07 | 455.76819 | 340.56122 |
| KFGFELVK | 8 | Q5JTH9 | RRP12-like protein OS=Homo sapiens O | RRP12 | 143.703 | 2 | 54.652313 | yes | 2.295E-07 | 484.28421 | 310.12418 |
| EATTTLHMLTLLK | 13 | Q5JTH9 | RRP12-like protein OS=Homo sapiens O | RRP12 | 143.703 | 2 | 54.652313 | yes | 1.269E-07 | 736.41309 | 277.94064 |
| APAHHPAAISTAK | 13 | Q5JTH9 | RRP12-like protein OS=Homo sapiens O | RRP12 | 143.703 | 3 | 54.652313 | yes | 1.624E-07 | 424.56668 | 135.42892 |
| APAHHPAAISTAK | 13 | Q5JTH9 | RRP12-like protein OS=Homo sapiens O | RRP12 | 143.703 | 3 | 54.652313 | yes | 9.81E-08 | 636.34637 | 135.42892 |
| APAHHPAAISTAK | 13 | Q5JTH9 | RRP12-like protein OS=Homo sapiens O | RRP12 | 143.703 | 2 | 54.652313 | yes | 1.624E-07 | 424.56668 | 135.42892 |
| APAHHPAAISTAK | 13 | Q5JTH9 | RRP12-like protein OS=Homo sapiens O | RRP12 | 143.703 | 2 | 54.652313 | yes | 9.81E-08 | 636.34637 | 135.42892 |
| GHSSDSNPAICR | 12 | Q5JTH9 | RRP12-like protein OS=Homo sapiens O | RRP12 | 143.703 | 2 | 54.652313 | yes | 9.722E-08 | 650.78857 | 227.51863 |
| VLEEVCASPQGPG | 26 | Q5JTH9 | RRP12-like protein OS=Homo sapiens O | RRP12 | 143.703 | 4 | 54.652313 | yes | 2.664E-07 | 713.62195 | 175.4549 |
| AVEEGLTYK | 9 | Q5JTH9 | RRP12-like protein OS=Homo sapiens O | RRP12 | 143.703 | 2 | 54.652313 | yes | 4.735E-07 | 505.26367 | 173.35764 |
| GRPDPYAYIPLNR | 13 | Q5JTH9 | RRP12-like protein OS=Homo sapiens O | RRP12 | 143.703 | 2 | 54.652313 | yes | 1.274E-07 | 766.40442 | 164.97209 |
| PTDVAISFK | 9 | Q5JTH9 | RRP12-like protein OS=Homo sapiens O | RRP12 | 143.703 | 2 | 54.652313 | yes | 1.083E-07 | 489.26874 | 132.90257 |
| LQWDLGLGHLPR | 12 | Q5JTH9 | RRP12-like protein OS=Homo sapiens O | RRP12 | 143.703 | 2 | 54.652313 | yes | 1.122E-07 | 702.89093 | 115.91614 |
| TLGMAISERPDLR | 13 | Q5JTH9 | RRP12-like protein OS=Homo sapiens O | RRP12 | 143.703 | 2 | 54.652313 | yes | 2.453E-05 | 729.89032 | 100.56356 |
| TLGMAISER | 9 | Q5JTH9 | RRP12-like protein OS=Homo sapiens O | RRP12 | 143.703 | 2 | 54.652313 | yes | 4.12E-06 | 489.25787 | 97.35952 |
| SWLLPVIR | 8 | Q5JTH9 | RRP12-like protein OS=Homo sapiens O | RRP12 | 143.703 | 2 | 54.652313 | yes | 2.312E-06 | 492.30548 | 89.299889 |
| LYSTIRPYLESK | 12 | Q5JTH9 | RRP12-like protein OS=Homo sapiens O | RRP12 | 143.703 | 2 | 54.652313 | yes | 0.0001282 | 735.40356 | 87.655403 |

| | | | | | | | | | | | |
|---|---|---|---|---|---|---|---|---|---|---|---|
| TLLDSLR | 7 | Q5JTH9 | RRP12-like protein OS=Homo sapiens O: | RRP12 | 143.703 | 2 | 54.652313 | yes | 1.685E-06 | 409.24252 | 58.728405 |
| VAVTVMDVAHLA | 13 | Q5JTH9 | RRP12-like protein OS=Homo sapiens O: | RRP12 | 143.703 | 3 | 54.652313 | yes | 1.584E-07 | 451.92349 | 53.211639 |
| CLSEQIADAYSSFF | 14 | Q5JTV8 | Torsin-1A-interacting protein 1 OS=Hom | TOR1AIP1 | 66.249 | 2 | 39.114418 | yes | 4.781E-07 | 823.87756 | 21.262922 |
| SQPAILLLTAAR | 12 | Q5JTV8 | Torsin-1A-interacting protein 1 OS=Hom | TOR1AIP1 | 66.249 | 2 | 39.114418 | yes | 3.72E-06 | 627.38245 | 17.500031 |
| SEYQWITSPSVK | 12 | Q5JTW2 | Centrosomal protein of 78 kDa OS=Homo | CEP78 | 76.397 | 2 | 53.6817 | yes | 9.722E-08 | 712.85645 | 250.88875 |
| EGVLDFNADR | 10 | Q5JTW2 | Centrosomal protein of 78 kDa OS=Homo | CEP78 | 76.397 | 2 | 53.6817 | yes | 9.722E-08 | 568.27252 | 218.88622 |
| TIILGSGHK | 9 | Q5JTW2 | Centrosomal protein of 78 kDa OS=Homo | CEP78 | 76.397 | 2 | 53.6817 | yes | 4.691E-07 | 463.27692 | 208.13705 |
| FHAFLDLLK | 9 | Q5JTW2 | Centrosomal protein of 78 kDa OS=Homo | CEP78 | 76.397 | 2 | 53.6817 | yes | 1.122E-07 | 552.31604 | 152.56462 |
| ALDLQQCGLTNEC | 15 | Q5JTW2 | Centrosomal protein of 78 kDa OS=Homo | CEP78 | 76.397 | 2 | 53.6817 | yes | 2.578E-07 | 809.39868 | 109.73133 |
| QESFEGFIAR | 10 | Q5JTW2 | Centrosomal protein of 78 kDa OS=Homo | CEP78 | 76.397 | 2 | 53.6817 | yes | 9.722E-08 | 592.29077 | 78.173317 |
| ALEDEKPEPK | 10 | Q5JTW2 | Centrosomal protein of 78 kDa OS=Homo | CEP78 | 76.397 | 2 | 53.6817 | yes | 2.581E-07 | 578.29822 | 60.95702 |
| KALEDEKPEPK | 11 | Q5JTW2 | Centrosomal protein of 78 kDa OS=Homo | CEP78 | 76.397 | 2 | 53.6817 | yes | 3.804E-07 | 642.3457 | 27.628616 |
| GCLSISSVLK | 10 | Q5JTW2 | Centrosomal protein of 78 kDa OS=Homo | CEP78 | 76.397 | 2 | 53.6817 | yes | 2.522E-06 | 532.29443 | 26.056137 |
| TGAVGDLVIIGDR | 13 | Q5JTZ9 | Alanine--tRNA ligase, mitochondrial OS= | AARS2 | 107.341 | 2 | 48.631416 | yes | 1.121E-07 | 643.35913 | 97.983002 |
| ELGQSLAQEVK | 11 | Q5JTZ9 | Alanine--tRNA ligase, mitochondrial OS= | AARS2 | 107.341 | 2 | 48.631416 | yes | 4.099E-06 | 601.32477 | 62.088463 |
| PIFLGTVDPR | 10 | Q5JTZ9 | Alanine--tRNA ligase, mitochondrial OS= | AARS2 | 107.341 | 2 | 48.631416 | yes | 5.71E-07 | 557.81641 | 58.054966 |
| AGLDPDLETR | 10 | Q5JTZ9 | Alanine--tRNA ligase, mitochondrial OS= | AARS2 | 107.341 | 2 | 48.631416 | yes | 5.117E-05 | 543.7749 | 55.917175 |
| AGQEDVLFPVAR | 12 | Q5JTZ9 | Alanine--tRNA ligase, mitochondrial OS= | AARS2 | 107.341 | 2 | 48.631416 | yes | 2.323E-07 | 651.34607 | 45.094635 |
| HVDTGMGLER | 10 | Q5JTZ9 | Alanine--tRNA ligase, mitochondrial OS= | AARS2 | 107.341 | 2 | 48.631416 | yes | 5.191E-06 | 557.7691 | 37.894806 |
| TNFYAEQGGQASI | 14 | Q5JTZ9 | Alanine--tRNA ligase, mitochondrial OS= | AARS2 | 107.341 | 2 | 48.631416 | yes | 5.923E-07 | 772.34222 | 24.043743 |
| LWISYFDGDPK | 11 | Q5JTZ9 | Alanine--tRNA ligase, mitochondrial OS= | AARS2 | 107.341 | 2 | 48.631416 | yes | 4.162E-07 | 670.82971 | 12.851949 |
| CQTVIVQSFLR | 11 | Q5JVF3 | PCI domain-containing protein 2 OS=Ho | PCID2 | 46.03 | 2 | 52.22744 | yes | 1.058E-07 | 675.86353 | 414.08286 |
| HEAFFIR | 7 | Q5JVF3 | PCI domain-containing protein 2 OS=Ho | PCID2 | 46.03 | 2 | 52.22744 | yes | 1.577E-07 | 460.24286 | 378.19397 |
| LQMASPEEK | 9 | Q5JVF3 | PCI domain-containing protein 2 OS=Ho | PCID2 | 46.03 | 2 | 52.22744 | yes | 1.2E-07 | 516.75513 | 326.27676 |
| MLLGHMPTVELLI | 13 | Q5JVF3 | PCI domain-containing protein 2 OS=Ho | PCID2 | 46.03 | 2 | 52.22744 | yes | 9.722E-08 | 741.41455 | 140.83368 |
| MLLGHMPTVELLI | 13 | Q5JVF3 | PCI domain-containing protein 2 OS=Ho | PCID2 | 46.03 | 2 | 52.22744 | yes | 6.039E-07 | 494.61212 | 140.83368 |
| MLLGHMPTVELLI | 13 | Q5JVF3 | PCI domain-containing protein 2 OS=Ho | PCID2 | 46.03 | 3 | 52.22744 | yes | 9.722E-08 | 741.41455 | 140.83368 |
| MLLGHMPTVELLI | 13 | Q5JVF3 | PCI domain-containing protein 2 OS=Ho | PCID2 | 46.03 | 3 | 52.22744 | yes | 6.039E-07 | 494.61212 | 140.83368 |
| DGASCAELVSFK | 12 | Q5JVF3 | PCI domain-containing protein 2 OS=Ho | PCID2 | 46.03 | 2 | 52.22744 | yes | 9.722E-08 | 642.30048 | 208.14384 |
| YHLMQFAEVTR | 11 | Q5JVF3 | PCI domain-containing protein 2 OS=Ho | PCID2 | 46.03 | 2 | 52.22744 | yes | 9.722E-08 | 697.8479 | 187.07037 |
| CQQVLEPPYDEMI | 18 | Q5JVF3 | PCI domain-containing protein 2 OS=Ho | PCID2 | 46.03 | 3 | 52.22744 | yes | 6.673E-07 | 735.34698 | 186.9379 |
| KVYLLLK | 7 | Q5JVF3 | PCI domain-containing protein 2 OS=Ho | PCID2 | 46.03 | 2 | 52.22744 | yes | 9.722E-08 | 438.7995 | 108.48261 |
| CGIFLILEK | 9 | Q5JVF3 | PCI domain-containing protein 2 OS=Ho | PCID2 | 46.03 | 2 | 52.22744 | yes | 9.403E-07 | 546.80975 | 86.324768 |
| AVSEGNLLLLHEA | 16 | Q5JVF3 | PCI domain-containing protein 2 OS=Ho | PCID2 | 46.03 | 3 | 52.22744 | yes | 9.722E-08 | 559.98932 | 82.612587 |
| VFANNADQQLVK | 12 | Q5JVF3 | PCI domain-containing protein 2 OS=Ho | PCID2 | 46.03 | 2 | 52.22744 | yes | 1.467E-07 | 673.85681 | 40.01955 |
| CTYAVGNHDFIEA | 15 | Q5JVF3 | PCI domain-containing protein 2 OS=Ho | PCID2 | 46.03 | 3 | 52.22744 | yes | 0.0001426 | 596.60675 | 32.172005 |
| THQLSLDAFLVAL | 14 | Q5JVF3 | PCI domain-containing protein 2 OS=Ho | PCID2 | 46.03 | 2 | 52.22744 | yes | 7.775E-07 | 778.44574 | 21.377321 |
| VFDAFDQR | 8 | Q5JVS0 | Intracellular hyaluronan-binding protein 4 | HABP4 | 45.786 | 2 | 48.609814 | yes | 1.123E-07 | 499.24051 | 308.89783 |
| LFPVPVDPGTPCR | 13 | Q5JXC2 | Migration and invasion-inhibitory protein | MIIP | 42.825 | 2 | 39.98867 | yes | 4.269E-07 | 727.87665 | 51.719551 |

| | | | | | | | | | | | |
|---|---|---|---|---|---|---|---|---|---|---|---|
| NSFKGQYK | 8 | Q5K651 | Sterile alpha motif domain-containing prc | SAMD9 | 184.282 | 2 | 19.66184 | yes | 4.759E-06 | 486.25089 | 135.03137 |
| LMNLLQER | 8 | Q5M775 | Cytospin-B OS=Homo sapiens OX=9606 | SPECC1 | 118.585 | 2 | 45.638004 | yes | 1.789E-07 | 508.78149 | 62.731895 |
| TATAGAISELTESR | 14 | Q5M775 | Cytospin-B OS=Homo sapiens OX=9606 | SPECC1 | 118.585 | 2 | 45.638004 | yes | 6.742E-05 | 703.85974 | 34.756477 |
| LALALENEGYIK | 12 | Q5MIZ7 | Serine/threonine-protein phosphatase 4 re | PPP4R3B | 97.459 | 2 | 32.730648 | yes | 1.43E-06 | 667.37177 | 54.092457 |
| AESDGSLLLESK | 12 | Q5MIZ7 | Serine/threonine-protein phosphatase 4 re | PPP4R3B | 97.459 | 2 | 32.730648 | yes | 4.011E-06 | 624.81952 | 17.212076 |
| MGLLSPSGIGGVSI | 15 | Q5PRF9 | Protein Smaug homolog 2 OS=Homo sap | SAMD4B | 75.484 | 2 | 54.958347 | yes | 4.565E-07 | 714.38739 | 467.31921 |
| NTFQEDGSGMK | 11 | Q5PRF9 | Protein Smaug homolog 2 OS=Homo sap | SAMD4B | 75.484 | 2 | 54.958347 | yes | 9.722E-08 | 607.26135 | 351.54187 |
| NALALWLSHLEEF | 13 | Q5PRF9 | Protein Smaug homolog 2 OS=Homo sap | SAMD4B | 75.484 | 2 | 54.958347 | yes | 9.722E-08 | 776.41754 | 214.05585 |
| HALTSPSLGGQGR | 13 | Q5PRF9 | Protein Smaug homolog 2 OS=Homo sap | SAMD4B | 75.484 | 2 | 54.958347 | yes | 9.728E-08 | 640.83893 | 173.81763 |
| VVSLLLSHLPLLQI | 20 | Q5PRF9 | Protein Smaug homolog 2 OS=Homo sap | SAMD4B | 75.484 | 3 | 54.958347 | yes | 1.14E-07 | 710.41583 | 120.29839 |
| AALEMQNYR | 9 | Q5PRF9 | Protein Smaug homolog 2 OS=Homo sap | SAMD4B | 75.484 | 2 | 54.958347 | yes | 8.374E-07 | 548.26624 | 76.021713 |
| CLTHEAFTETQK | 12 | Q5PRF9 | Protein Smaug homolog 2 OS=Homo sap | SAMD4B | 75.484 | 2 | 54.958347 | yes | 9.722E-08 | 732.84302 | 69.988571 |
| LTSSSIDPGLSIK | 13 | Q5QJE6 | Deoxynucleotidyltransferase terminal-inte | DNTTIP2 | 84.47 | 2 | 52.39512 | yes | 1.058E-07 | 659.36664 | 1714.8651 |
| LYTSAPNTSQGK | 12 | Q5QJE6 | Deoxynucleotidyltransferase terminal-inte | DNTTIP2 | 84.47 | 2 | 52.39512 | yes | 1.076E-07 | 633.81989 | 1215.9915 |
| APEMTNELK | 9 | Q5QJE6 | Deoxynucleotidyltransferase terminal-inte | DNTTIP2 | 84.47 | 2 | 52.39512 | yes | 2.757E-07 | 516.75513 | 1183.2551 |
| SLTQTFATVEVGR | 13 | Q5QJE6 | Deoxynucleotidyltransferase terminal-inte | DNTTIP2 | 84.47 | 2 | 52.39512 | yes | 9.722E-08 | 704.87518 | 1161.3893 |
| QILIACSPVSSVR | 13 | Q5QJE6 | Deoxynucleotidyltransferase terminal-inte | DNTTIP2 | 84.47 | 2 | 52.39512 | yes | 1.017E-07 | 715.3952 | 1098.1362 |
| QLGGLYINFNADK | 13 | Q5QJE6 | Deoxynucleotidyltransferase terminal-inte | DNTTIP2 | 84.47 | 2 | 52.39512 | yes | 9.823E-08 | 726.87769 | 926.85968 |
| ASIQAASAESSGQI | 14 | Q5QJE6 | Deoxynucleotidyltransferase terminal-inte | DNTTIP2 | 84.47 | 2 | 52.39512 | yes | 9.722E-08 | 667.83093 | 711.34247 |
| TIVEELLADSEFR | 13 | Q5QJE6 | Deoxynucleotidyltransferase terminal-inte | DNTTIP2 | 84.47 | 2 | 52.39512 | yes | 3.366E-07 | 761.39337 | 651.04932 |
| QIVGTPVNSEDSD | 15 | Q5QJE6 | Deoxynucleotidyltransferase terminal-inte | DNTTIP2 | 84.47 | 2 | 52.39512 | yes | 1.283E-07 | 809.38934 | 548.51447 |
| AVITPDFEK | 9 | Q5QJE6 | Deoxynucleotidyltransferase terminal-inte | DNTTIP2 | 84.47 | 2 | 52.39512 | yes | 1.865E-07 | 510.27402 | 490.76563 |
| NFYLEEEDK | 9 | Q5QJE6 | Deoxynucleotidyltransferase terminal-inte | DNTTIP2 | 84.47 | 2 | 52.39512 | yes | 1.751E-07 | 593.76678 | 340.83896 |
| YSEIMAEK | 8 | Q5QJE6 | Deoxynucleotidyltransferase terminal-inte | DNTTIP2 | 84.47 | 2 | 52.39512 | yes | 1.19E-07 | 485.73114 | 248.95439 |
| KYSEIMAEK | 9 | Q5QJE6 | Deoxynucleotidyltransferase terminal-inte | DNTTIP2 | 84.47 | 2 | 52.39512 | yes | 1.387E-07 | 549.77863 | 195.60834 |
| DLDEDANGITDEG | 14 | Q5QJE6 | Deoxynucleotidyltransferase terminal-inte | DNTTIP2 | 84.47 | 2 | 52.39512 | yes | 9.722E-08 | 746.32593 | 133.83046 |
| TAGDGWFGMK | 10 | Q5QJE6 | Deoxynucleotidyltransferase terminal-inte | DNTTIP2 | 84.47 | 2 | 52.39512 | yes | 3.797E-07 | 535.24219 | 68.869583 |
| ESYTEEIVSEAESH | 20 | Q5QJE6 | Deoxynucleotidyltransferase terminal-inte | DNTTIP2 | 84.47 | 3 | 52.39512 | yes | 9.722E-08 | 737.01337 | 64.306358 |
| NHCVPPYSESK | 11 | Q5QJE6 | Deoxynucleotidyltransferase terminal-inte | DNTTIP2 | 84.47 | 2 | 52.39512 | yes | 1.165E-06 | 659.29822 | 23.803455 |
| QCSELVDCLPK | 11 | Q5QP82 | DDB1- and CUL4-associated factor 10 O | DCAF10 | 60.583 | 2 | 54.044662 | yes | 3.228E-06 | 674.81543 | 64.927635 |
| SLEVGSYPILR | 11 | Q5QP82 | DDB1- and CUL4-associated factor 10 O | DCAF10 | 60.583 | 2 | 54.044662 | yes | 9.722E-08 | 617.34552 | 59.163872 |
| NIEYDTNTR | 9 | Q5QP82 | DDB1- and CUL4-associated factor 10 O | DCAF10 | 60.583 | 2 | 54.044662 | yes | 2.247E-07 | 563.26221 | 20.065454 |
| TTSSSDLTTSSSSS( | 17 | Q5QP82 | DDB1- and CUL4-associated factor 10 O | DCAF10 | 60.583 | 2 | 54.044662 | yes | 8.575E-07 | 829.37921 | 13.688244 |
| GGGPAGAGGEAP/ | 16 | Q5RKV6 | Exosome complex component MTR3 OS= | EXOSC6 | 28.236 | 2 | 49.666256 | yes | 1.76E-07 | 654.83636 | 733.51453 |
| LYPVLQQSLVR | 11 | Q5RKV6 | Exosome complex component MTR3 OS= | EXOSC6 | 28.236 | 2 | 49.666256 | yes | 2.781E-07 | 658.39026 | 649.00421 |
| GSAYLEAGGTK | 11 | Q5RKV6 | Exosome complex component MTR3 OS= | EXOSC6 | 28.236 | 2 | 49.666256 | yes | 3.136E-07 | 527.26422 | 188.96877 |
| VLCAVSGPR | 9 | Q5RKV6 | Exosome complex component MTR3 OS= | EXOSC6 | 28.236 | 2 | 49.666256 | yes | 5.539E-07 | 479.76056 | 174.69136 |
| LGLEGCQR | 8 | Q5RKV6 | Exosome complex component MTR3 OS= | EXOSC6 | 28.236 | 2 | 49.666256 | yes | 5.861E-07 | 466.73456 | 149.6651 |
| LRPVYAR | 7 | Q5RKV6 | Exosome complex component MTR3 OS= | EXOSC6 | 28.236 | 2 | 49.666256 | yes | 1.108E-07 | 437.76651 | 124.64086 |

| | | | | | | | | | | | |
|---|---|---|---|---|---|---|---|---|---|---|---|
| ELALALQEALEPA | 15 | Q5RKV6 | Exosome complex component MTR3 OS= | EXOSC6 | 28.236 | 2 | 49.666256 | yes | 1.012E-07 | 811.95923 | 58.349697 |
| ALSLGTIPSLTR | 12 | Q5SNT2 | Transmembrane protein 201 OS=Homo sa | TMEM201 | 72.236 | 2 | 49.215405 | yes | 2.676E-07 | 614.86664 | 110.95119 |
| ALLLSHQFK | 9 | Q5SNT2 | Transmembrane protein 201 OS=Homo sa | TMEM201 | 72.236 | 2 | 49.215405 | yes | 2.868E-07 | 528.81366 | 63.395535 |
| QLAAFAPR | 8 | Q5SNT2 | Transmembrane protein 201 OS=Homo sa | TMEM201 | 72.236 | 2 | 49.215405 | yes | 3.634E-05 | 437.2507 | 19.117519 |
| EADQTHAQNFSSA | 15 | Q5SNT2 | Transmembrane protein 201 OS=Homo sa | TMEM201 | 72.236 | 2 | 49.215405 | yes | 1.058E-07 | 816.88428 | 10.609901 |
| ALLLVPASVNCLR | 13 | Q5SRE5 | Nucleoporin NUP188 OS=Homo sapiens | NUP188 | 196.043 | 2 | 45.88148 | yes | 3.455E-07 | 713.41595 | 106.6385 |
| SLDAPSWPGVYR | 12 | Q5SRE5 | Nucleoporin NUP188 OS=Homo sapiens | NUP188 | 196.043 | 2 | 45.88148 | yes | 2.996E-07 | 674.33826 | 71.447372 |
| HTLNPEETSSVVR | 13 | Q5SRE5 | Nucleoporin NUP188 OS=Homo sapiens | NUP188 | 196.043 | 2 | 45.88148 | yes | 2.234E-07 | 734.87317 | 59.571262 |
| LLYPLGGQTNLR | 12 | Q5SRE5 | Nucleoporin NUP188 OS=Homo sapiens | NUP188 | 196.043 | 2 | 45.88148 | yes | 9.756E-06 | 672.88531 | 45.60408 |
| IADYYYEER | 9 | Q5SRE5 | Nucleoporin NUP188 OS=Homo sapiens | NUP188 | 196.043 | 2 | 45.88148 | yes | 1.715E-07 | 611.27478 | 43.564137 |
| HLVDETMDPFVDI | 13 | Q5SRE5 | Nucleoporin NUP188 OS=Homo sapiens | NUP188 | 196.043 | 2 | 45.88148 | yes | 4.934E-07 | 787.36938 | 26.118681 |
| TRPSSTVIK | 9 | Q5SSJ5 | Heterochromatin protein 1-binding protei | HP1BP3 | 61.207 | 2 | 49.659332 | yes | 1.623E-07 | 494.79294 | 153.40276 |
| YPSLELER | 8 | Q5SSJ5 | Heterochromatin protein 1-binding protei | HP1BP3 | 61.207 | 2 | 49.659332 | yes | 1.725E-07 | 503.76382 | 144.1924 |
| ATDTSQGELVHPK | 13 | Q5SSJ5 | Heterochromatin protein 1-binding protei | HP1BP3 | 61.207 | 2 | 49.659332 | yes | 1.013E-07 | 712.85443 | 142.53889 |
| SSAVDPEPQVK | 11 | Q5SSJ5 | Heterochromatin protein 1-binding protei | HP1BP3 | 61.207 | 2 | 49.659332 | yes | 3.582E-07 | 578.79584 | 126.70634 |
| KYVSQYYPK | 9 | Q5SSJ5 | Heterochromatin protein 1-binding protei | HP1BP3 | 61.207 | 2 | 49.659332 | yes | 6.42E-06 | 588.30841 | 94.993553 |
| LEDVLPLAFTR | 11 | Q5SSJ5 | Heterochromatin protein 1-binding protei | HP1BP3 | 61.207 | 2 | 49.659332 | yes | 2.007E-07 | 637.36115 | 63.062202 |
| GASGSFVVVQK | 11 | Q5SSJ5 | Heterochromatin protein 1-binding protei | HP1BP3 | 61.207 | 2 | 49.659332 | yes | 5.272E-07 | 539.79822 | 29.098148 |
| LSSLQLELSK | 10 | Q5ST30 | Valine--tRNA ligase, mitochondrial OS=I | VARS2 | 118.49 | 2 | 44.695614 | yes | 7.554E-07 | 559.32678 | 62.844772 |
| VQEVVQVLR | 9 | Q5ST30 | Valine--tRNA ligase, mitochondrial OS=I | VARS2 | 118.49 | 2 | 44.695614 | yes | 2.101E-07 | 535.32184 | 57.471638 |
| ATLEEHLR | 8 | Q5SW79 | Centrosomal protein of 170 kDa OS=Hon | CEP170 | 175.293 | 2 | 53.246346 | yes | 5.397E-07 | 484.76163 | 921.91632 |
| VADAATEVQHK | 11 | Q5SW79 | Centrosomal protein of 170 kDa OS=Hon | CEP170 | 175.293 | 2 | 53.246346 | yes | 1.183E-06 | 584.80145 | 547.25433 |
| TLPQLPNEEK | 10 | Q5SW79 | Centrosomal protein of 170 kDa OS=Hon | CEP170 | 175.293 | 2 | 53.246346 | yes | 2.459E-07 | 584.81403 | 467.48123 |
| AMDISAMPR | 9 | Q5SW79 | Centrosomal protein of 170 kDa OS=Hon | CEP170 | 175.293 | 2 | 53.246346 | yes | 5.146E-07 | 496.23862 | 330.57056 |
| DDCELMLQSR | 10 | Q5SW79 | Centrosomal protein of 170 kDa OS=Hon | CEP170 | 175.293 | 2 | 53.246346 | yes | 1.485E-07 | 633.77631 | 330.34894 |
| EPSYFEIPTK | 10 | Q5SW79 | Centrosomal protein of 170 kDa OS=Hon | CEP170 | 175.293 | 2 | 53.246346 | no | 3.456E-07 | 605.80316 | 310.74121 |
| IPEQTYITLK | 10 | Q5SW79 | Centrosomal protein of 170 kDa OS=Hon | CEP170 | 175.293 | 2 | 53.246346 | yes | 5.075E-07 | 603.34247 | 286.7402 |
| LQSAGSAMPTSSSI | 15 | Q5SW79 | Centrosomal protein of 170 kDa OS=Hon | CEP170 | 175.293 | 2 | 53.246346 | yes | 5.199E-07 | 749.86395 | 262.02881 |
| TLLHLGSSAPGK | 12 | Q5SW79 | Centrosomal protein of 170 kDa OS=Hon | CEP170 | 175.293 | 2 | 53.246346 | yes | 9.722E-08 | 590.83789 | 249.13255 |
| DWALSSAAAVME | 14 | Q5SW79 | Centrosomal protein of 170 kDa OS=Hon | CEP170 | 175.293 | 2 | 53.246346 | yes | 6.871E-07 | 768.36157 | 189.08893 |
| EFQQPSQITESTIHI | 19 | Q5SW79 | Centrosomal protein of 170 kDa OS=Hon | CEP170 | 175.293 | 3 | 53.246346 | yes | 3.929E-07 | 738.37433 | 180.1059 |
| QDTELQEK | 8 | Q5SW79 | Centrosomal protein of 170 kDa OS=Hon | CEP170 | 175.293 | 2 | 53.246346 | yes | 2.538E-06 | 495.74054 | 178.37218 |
| LGEASDSELADAD | 14 | Q5SW79 | Centrosomal protein of 170 kDa OS=Hon | CEP170 | 175.293 | 2 | 53.246346 | yes | 9.722E-08 | 710.82556 | 168.24103 |
| VFDESLNFR | 9 | Q5SW79 | Centrosomal protein of 170 kDa OS=Hon | CEP170 | 175.293 | 2 | 53.246346 | yes | 2.242E-07 | 563.78003 | 146.47057 |
| IKEQEDYIR | 9 | Q5SW79 | Centrosomal protein of 170 kDa OS=Hon | CEP170 | 175.293 | 2 | 53.246346 | no | 1.884E-07 | 597.31171 | 145.87938 |
| TPLTSADEHVHSK | 13 | Q5SW79 | Centrosomal protein of 170 kDa OS=Hon | CEP170 | 175.293 | 2 | 53.246346 | yes | 9.722E-08 | 711.35461 | 139.57065 |
| TSSMEISSILQELK | 14 | Q5SW79 | Centrosomal protein of 170 kDa OS=Hon | CEP170 | 175.293 | 2 | 53.246346 | no | 1E-07 | 783.4082 | 138.71321 |
| ETPTQVYQK | 9 | Q5SW79 | Centrosomal protein of 170 kDa OS=Hon | CEP170 | 175.293 | 2 | 53.246346 | yes | 1.348E-07 | 547.27985 | 137.70938 |
| FTIQLQLSQK | 10 | Q5SW79 | Centrosomal protein of 170 kDa OS=Hon | CEP170 | 175.293 | 2 | 53.246346 | yes | 1.361E-07 | 603.34808 | 134.76158 |

| | | | | | | | | | | | |
|---|---|---|---|---|---|---|---|---|---|---|---|
| IDLLAQPR | 8 | Q5SW79 | Centrosomal protein of 170 kDa OS=Hon | CEP170 | 175.293 | 2 | 53.246346 | yes | 2.964E-07 | 463.27692 | 127.3035 |
| QHAVINYDASTDE | 17 | Q5SW79 | Centrosomal protein of 170 kDa OS=Hon | CEP170 | 175.293 | 3 | 53.246346 | yes | 9.722E-08 | 647.32178 | 83.271332 |
| ALHPAAVSAAAEF | 28 | Q5SW79 | Centrosomal protein of 170 kDa OS=Hon | CEP170 | 175.293 | 4 | 53.246346 | no | 1.404E-07 | 751.1073 | 83.028687 |
| QTSSTPSSLALTSA | 16 | Q5SW79 | Centrosomal protein of 170 kDa OS=Hon | CEP170 | 175.293 | 2 | 53.246346 | yes | 9.722E-08 | 797.40759 | 80.12059 |
| TSPVVSGSSSK | 11 | Q5SW79 | Centrosomal protein of 170 kDa OS=Hon | CEP170 | 175.293 | 2 | 53.246346 | yes | 1.053E-07 | 518.26947 | 73.400368 |
| ISQDLALIAR | 10 | Q5SW79 | Centrosomal protein of 170 kDa OS=Hon | CEP170 | 175.293 | 2 | 53.246346 | no | 1.4E-06 | 550.32715 | 52.55616 |
| TDEGPDTPSYNR | 12 | Q5SW79 | Centrosomal protein of 170 kDa OS=Hon | CEP170 | 175.293 | 2 | 53.246346 | yes | 1.879E-07 | 676.29169 | 44.11964 |
| SETDKETSLVK | 11 | Q5SW79 | Centrosomal protein of 170 kDa OS=Hon | CEP170 | 175.293 | 2 | 53.246346 | yes | 1.526E-07 | 618.81952 | 44.079556 |
| LRAESEVPIVK | 11 | Q5SW79 | Centrosomal protein of 170 kDa OS=Hon | CEP170 | 175.293 | 2 | 53.246346 | no | 5.42E-07 | 620.86664 | 43.181541 |
| ARPQLILSR | 9 | Q5SY16 | Polynucleotide 5'-hydroxyl-kinase NOL9 | NOL9 | 79.323 | 2 | 50.636837 | yes | 6.518E-07 | 527.33002 | 60.198074 |
| NNYENYIDIVK | 11 | Q5SY16 | Polynucleotide 5'-hydroxyl-kinase NOL9 | NOL9 | 79.323 | 2 | 50.636837 | yes | 9.733E-08 | 692.84082 | 38.958752 |
| LAAFADALEFADE | 15 | Q5SY16 | Polynucleotide 5'-hydroxyl-kinase NOL9 | NOL9 | 79.323 | 2 | 50.636837 | yes | 1.018E-06 | 820.39612 | 18.440596 |
| MILEIQSMQGK | 11 | Q5SZL2 | Centrosomal protein of 85 kDa-like OS=I | CEP85L | 91.809 | 2 | 51.320007 | yes | 1.609E-07 | 639.33325 | 97.04187 |
| ALCSILNQR | 9 | Q5SZL2 | Centrosomal protein of 85 kDa-like OS=I | CEP85L | 91.809 | 2 | 51.320007 | yes | 2.201E-06 | 537.78986 | 77.371956 |
| IIDSQQDEIDR | 11 | Q5SZL2 | Centrosomal protein of 85 kDa-like OS=I | CEP85L | 91.809 | 2 | 51.320007 | yes | 2.326E-07 | 666.3255 | 64.515312 |
| NLQEALIDTEK | 11 | Q5SZL2 | Centrosomal protein of 85 kDa-like OS=I | CEP85L | 91.809 | 2 | 51.320007 | yes | 8.46E-07 | 637.33533 | 63.817734 |
| LSDVCQLR | 8 | Q5SZL2 | Centrosomal protein of 85 kDa-like OS=I | CEP85L | 91.809 | 2 | 51.320007 | yes | 6.271E-07 | 495.75549 | 61.917503 |
| EVLQLNEFLK | 10 | Q5SZL2 | Centrosomal protein of 85 kDa-like OS=I | CEP85L | 91.809 | 2 | 51.320007 | yes | 1.163E-07 | 616.8479 | 35.062469 |
| EPNLSLLLGIR | 11 | Q5SZL2 | Centrosomal protein of 85 kDa-like OS=I | CEP85L | 91.809 | 2 | 51.320007 | yes | 9.722E-08 | 612.86914 | 18.28632 |
| HTQWGIDFLER | 11 | Q5T0N5 | Formin-binding protein 1-like OS=Homo | FNBP1L | 70.066 | 2 | 54.89607 | yes | 9.934E-08 | 701.34912 | 197.33176 |
| EVVAEEMAHR | 10 | Q5T0N5 | Formin-binding protein 1-like OS=Homo | FNBP1L | 70.066 | 2 | 54.89607 | yes | 1.806E-07 | 585.78223 | 172.61113 |
| LAETMNNIDR | 10 | Q5T0N5 | Formin-binding protein 1-like OS=Homo | FNBP1L | 70.066 | 2 | 54.89607 | yes | 1.108E-07 | 588.78754 | 154.91663 |
| NPQMGDPGSLQPK | 13 | Q5T0N5 | Formin-binding protein 1-like OS=Homo | FNBP1L | 70.066 | 2 | 54.89607 | yes | 9.722E-08 | 684.83246 | 120.30643 |
| HFYVVIPQIYK | 11 | Q5T0N5 | Formin-binding protein 1-like OS=Homo | FNBP1L | 70.066 | 2 | 54.89607 | yes | 1.045E-07 | 703.8952 | 63.338871 |
| AQQYLDMCWK | 10 | Q5T0N5 | Formin-binding protein 1-like OS=Homo | FNBP1L | 70.066 | 2 | 54.89607 | yes | 5.812E-07 | 671.79956 | 40.816444 |
| TSAVSSPLLDQQR | 13 | Q5T200 | Zinc finger CCCH domain-containing prc | ZC3H13 | 196.636 | 2 | 52.643486 | yes | 1.027E-07 | 701.37024 | 181.8824 |
| TGNCLYGNTCR | 11 | Q5T200 | Zinc finger CCCH domain-containing prc | ZC3H13 | 196.636 | 2 | 52.643486 | yes | 1.265E-07 | 658.27954 | 156.43077 |
| FVHGPSPR | 8 | Q5T200 | Zinc finger CCCH domain-containing prc | ZC3H13 | 196.636 | 2 | 52.643486 | yes | 1.319E-06 | 448.74051 | 117.08312 |
| GRVPELPEK | 9 | Q5T200 | Zinc finger CCCH domain-containing prc | ZC3H13 | 196.636 | 2 | 52.643486 | yes | 9.722E-08 | 512.79291 | 100.27391 |
| AQDIIGHHQSEDR | 13 | Q5T200 | Zinc finger CCCH domain-containing prc | ZC3H13 | 196.636 | 2 | 52.643486 | yes | 9.722E-08 | 753.35822 | 27.653923 |
| EYEQDQSSSR | 10 | Q5T200 | Zinc finger CCCH domain-containing prc | ZC3H13 | 196.636 | 2 | 52.643486 | yes | 1.073E-07 | 614.75745 | 26.064095 |
| ILQYLECPQYLR | 12 | Q5T280 | Putative methyltransferase C9orf114 OS= | SPOUT1 | 42.009 | 2 | 51.77564 | yes | 1.028E-07 | 798.41614 | 1416.4225 |
| TYLAGQIAR | 9 | Q5T280 | Putative methyltransferase C9orf114 OS= | SPOUT1 | 42.009 | 2 | 51.77564 | yes | 3.65E-07 | 496.77982 | 561.495 |
| AGLYWGYTVR | 10 | Q5T280 | Putative methyltransferase C9orf114 OS= | SPOUT1 | 42.009 | 2 | 51.77564 | yes | 9.796E-08 | 593.30621 | 503.1322 |
| GSDVASAQLPNFR | 13 | Q5T280 | Putative methyltransferase C9orf114 OS= | SPOUT1 | 42.009 | 2 | 51.77564 | yes | 9.782E-08 | 681.34406 | 332.47159 |
| KGQACVQLAR | 10 | Q5T280 | Putative methyltransferase C9orf114 OS= | SPOUT1 | 42.009 | 2 | 51.77564 | yes | 1.207E-07 | 565.80859 | 136.49969 |
| TVEGEFTGVGK | 11 | Q5T280 | Putative methyltransferase C9orf114 OS= | SPOUT1 | 42.009 | 2 | 51.77564 | yes | 1.576E-07 | 562.28516 | 117.47671 |
| RLEEEEAAAEK | 11 | Q5T280 | Putative methyltransferase C9orf114 OS= | SPOUT1 | 42.009 | 2 | 51.77564 | yes | 3.654E-07 | 637.81476 | 92.421379 |
| HQDLQFAGLLNPL | 20 | Q5T280 | Putative methyltransferase C9orf114 OS= | SPOUT1 | 42.009 | 3 | 51.77564 | yes | 1.135E-07 | 776.05518 | 85.146942 |

| | | | | | | | | | | | |
|---|---|---|---|---|---|---|---|---|---|---|---|
| GQACVQLAR | 9 | Q5T280 | Putative methyltransferase C9orf114 OS= | SPOUT1 | 42.009 | 2 | 51.77564 | yes | 9.722E-08 | 501.76111 | 84.034561 |
| LEEEEAAAEKEDR | 13 | Q5T280 | Putative methyltransferase C9orf114 OS= | SPOUT1 | 42.009 | 2 | 51.77564 | yes | 3.257E-07 | 759.84955 | 12.794674 |
| LLISDTQHYGVTV | 18 | Q5T2T1 | MAGUK p55 subfamily member 7 OS=H | MPP7 | 65.524 | 3 | 55.113613 | yes | 1.499E-07 | 680.02875 | 434.90103 |
| SGLIHVGDELR | 11 | Q5T2T1 | MAGUK p55 subfamily member 7 OS=H | MPP7 | 65.524 | 2 | 55.113613 | yes | 9.722E-08 | 598.32513 | 384.93274 |
| ALLSVHDTVAQK | 12 | Q5T2T1 | MAGUK p55 subfamily member 7 OS=H | MPP7 | 65.524 | 2 | 55.113613 | yes | 1.003E-07 | 641.36169 | 329.89398 |
| NNYYGTSIDSVR | 12 | Q5T2T1 | MAGUK p55 subfamily member 7 OS=H | MPP7 | 65.524 | 2 | 55.113613 | yes | 2.161E-07 | 694.82568 | 181.53552 |
| RPEILVQPLK | 10 | Q5T2T1 | MAGUK p55 subfamily member 7 OS=H | MPP7 | 65.524 | 2 | 55.113613 | yes | 1.155E-07 | 596.87427 | 154.88666 |
| ALFDYNPNEDK | 11 | Q5T2T1 | MAGUK p55 subfamily member 7 OS=H | MPP7 | 65.524 | 2 | 55.113613 | yes | 2.124E-07 | 663.30408 | 114.53552 |
| NREPLGATIK | 10 | Q5T2T1 | MAGUK p55 subfamily member 7 OS=H | MPP7 | 65.524 | 2 | 55.113613 | yes | 1.506E-07 | 549.81696 | 95.620064 |
| VCLLDVQPHTVK | 12 | Q5T2T1 | MAGUK p55 subfamily member 7 OS=H | MPP7 | 65.524 | 2 | 55.113613 | yes | 9.722E-08 | 704.88446 | 90.667953 |
| KDEQTGAIIVAR | 12 | Q5T2T1 | MAGUK p55 subfamily member 7 OS=H | MPP7 | 65.524 | 2 | 55.113613 | yes | 9.722E-08 | 650.86462 | 88.041428 |
| SQESDGVEYIFISK | 14 | Q5T2T1 | MAGUK p55 subfamily member 7 OS=H | MPP7 | 65.524 | 2 | 55.113613 | yes | 9.722E-08 | 801.38831 | 87.584 |
| LVVLVGPVGVGLI | 16 | Q5T2T1 | MAGUK p55 subfamily member 7 OS=H | MPP7 | 65.524 | 2 | 55.113613 | yes | 1.169E-07 | 803.50055 | 58.653748 |
| EVNGIPVEDK | 10 | Q5T2T1 | MAGUK p55 subfamily member 7 OS=H | MPP7 | 65.524 | 2 | 55.113613 | yes | 6.516E-05 | 550.28516 | 30.52491 |
| RPEEIIQILAQSQG | 19 | Q5T2T1 | MAGUK p55 subfamily member 7 OS=H | MPP7 | 65.524 | 3 | 55.113613 | yes | 2.082E-07 | 714.73541 | 14.616962 |
| GCYIGQELTAR | 11 | Q5T440 | Putative transferase CAF17, mitochondria | IBA57 | 38.155 | 2 | 50.248478 | yes | 9.838E-08 | 634.30859 | 207.89926 |
| LLTQDEGPALVPG | 15 | Q5T440 | Putative transferase CAF17, mitochondria | IBA57 | 38.155 | 2 | 50.248478 | yes | 1.038E-07 | 761.91486 | 128.43713 |
| VTVEPHPELR | 10 | Q5T440 | Putative transferase CAF17, mitochondria | IBA57 | 38.155 | 2 | 50.248478 | yes | 5.437E-07 | 588.8222 | 51.282169 |
| QLQLQNQELLR | 11 | Q5T5P2 | Sickle tail protein homolog OS=Homo sa | KIAA1217 | 214.116 | 2 | 55.885704 | yes | 1.248E-07 | 691.89117 | 60.354328 |
| DVEDGAFLLR | 10 | Q5T5P2 | Sickle tail protein homolog OS=Homo sa | KIAA1217 | 214.116 | 2 | 55.885704 | yes | 5.693E-07 | 567.79309 | 53.972923 |
| QLAALTQAIR | 10 | Q5T5P2 | Sickle tail protein homolog OS=Homo sa | KIAA1217 | 214.116 | 2 | 55.885704 | yes | 5.422E-07 | 542.82727 | 51.818462 |
| GEFPTLQNK | 9 | Q5T5P2 | Sickle tail protein homolog OS=Homo sa | KIAA1217 | 214.116 | 2 | 55.885704 | yes | 1.582E-07 | 517.26929 | 46.587753 |
| GTDAAQAAQYMA | 15 | Q5T5P2 | Sickle tail protein homolog OS=Homo sa | KIAA1217 | 214.116 | 2 | 55.885704 | yes | 9.722E-08 | 793.35291 | 41.210316 |
| SMTDVLTMLR | 10 | Q5T5P2 | Sickle tail protein homolog OS=Homo sa | KIAA1217 | 214.116 | 2 | 55.885704 | yes | 3.809E-07 | 583.79901 | 31.695101 |
| SLEPSGILLK | 10 | Q5T5U3 | Rho GTPase-activating protein 21 OS=H | ARHGAP2 | 217.463 | 2 | 46.24468 | yes | 1.805E-07 | 528.8186 | 113.74471 |
| LSPEVQSVAESK | 12 | Q5T5U3 | Rho GTPase-activating protein 21 OS=H | ARHGAP2 | 217.463 | 2 | 46.24468 | yes | 1.698E-06 | 637.33533 | 102.31638 |
| NFQTTCGMSLPR | 12 | Q5T5U3 | Rho GTPase-activating protein 21 OS=H | ARHGAP2 | 217.463 | 2 | 46.24468 | yes | 2.205E-06 | 706.32648 | 89.979256 |
| VVDLLSNR | 8 | Q5T5U3 | Rho GTPase-activating protein 21 OS=H | ARHGAP2 | 217.463 | 2 | 46.24468 | yes | 2.548E-07 | 458.26654 | 87.017967 |
| YADFIEANR | 9 | Q5T5U3 | Rho GTPase-activating protein 21 OS=H | ARHGAP2 | 217.463 | 2 | 46.24468 | yes | 2.151E-07 | 549.76434 | 85.576263 |
| STGSLLTPTR | 10 | Q5T5U3 | Rho GTPase-activating protein 21 OS=H | ARHGAP2 | 217.463 | 2 | 46.24468 | yes | 8.759E-07 | 516.78784 | 78.156525 |
| GLEYTGIYR | 9 | Q5T5U3 | Rho GTPase-activating protein 21 OS=H | ARHGAP2 | 217.463 | 2 | 46.24468 | yes | 1.726E-07 | 536.2771 | 77.171738 |
| LIECDTLSR | 9 | Q5T5U3 | Rho GTPase-activating protein 21 OS=H | ARHGAP2 | 217.463 | 2 | 46.24468 | yes | 3.401E-07 | 553.77917 | 65.216591 |
| GSNFTVAPSVVNS | 16 | Q5T5U3 | Rho GTPase-activating protein 21 OS=H | ARHGAP2 | 217.463 | 2 | 46.24468 | yes | 2.522E-07 | 832.40533 | 62.626762 |
| EGWLHFR | 7 | Q5T5U3 | Rho GTPase-activating protein 21 OS=H | ARHGAP2 | 217.463 | 2 | 46.24468 | yes | 2.909E-07 | 472.74051 | 58.297131 |
| TSQGFGFTLR | 10 | Q5T5U3 | Rho GTPase-activating protein 21 OS=H | ARHGAP2 | 217.463 | 2 | 46.24468 | yes | 7.06E-07 | 557.28802 | 56.155037 |
| SSHLEETGSDSGTI | 25 | Q5T5U3 | Rho GTPase-activating protein 21 OS=H | ARHGAP2 | 217.463 | 3 | 46.24468 | yes | 1.611E-07 | 841.07196 | 43.855614 |
| QQTSTPVLTQPGR | 13 | Q5T5U3 | Rho GTPase-activating protein 21 OS=H | ARHGAP2 | 217.463 | 2 | 46.24468 | yes | 2.825E-07 | 706.87823 | 39.762577 |
| LSETPGSK | 8 | Q5T5U3 | Rho GTPase-activating protein 21 OS=H | ARHGAP2 | 217.463 | 2 | 46.24468 | yes | 1.676E-05 | 409.71634 | 36.097363 |
| KLPEPLFTNDK | 11 | Q5T5U3 | Rho GTPase-activating protein 21 OS=H | ARHGAP2 | 217.463 | 2 | 46.24468 | yes | 1.64E-06 | 651.35864 | 31.117054 |

| | | | | | | | | | | | |
|---|---|---|---|---|---|---|---|---|---|---|---|
| WSGFTEQDDR | 10 | Q5T5U3 | Rho GTPase-activating protein 21 OS=H | ARHGAP2 | 217.463 | 2 | 46.24468 | yes | 8.842E-07 | 620.76508 | 29.481596 |
| DLNVISSLLK | 10 | Q5T5U3 | Rho GTPase-activating protein 21 OS=H | ARHGAP2 | 217.463 | 2 | 46.24468 | yes | 4.792E-07 | 551.32935 | 22.997408 |
| SNTAVCVCNESVR | 13 | Q5T5U3 | Rho GTPase-activating protein 21 OS=H | ARHGAP2 | 217.463 | 2 | 46.24468 | yes | 2.379E-07 | 748.33502 | 16.067842 |
| EIVQQSLSVGNFA | 15 | Q5T5X7 | BEN domain-containing protein 3 OS=H | BEND3 | 94.475 | 2 | 50.50177 | yes | 9.793E-08 | 809.93103 | 48.834473 |
| LQEDMKVK | 8 | Q5T5Y3 | Calmodulin-regulated spectrin-associated | CAMSAP1 | 177.972 | 2 | 52.365776 | yes | 1.965E-06 | 503.7655 | 371.04565 |
| LLLSSELYCR | 10 | Q5T5Y3 | Calmodulin-regulated spectrin-associated | CAMSAP1 | 177.972 | 2 | 52.365776 | yes | 4.041E-07 | 627.33154 | 354.80844 |
| APVHFVEPLSPTG | 18 | Q5T5Y3 | Calmodulin-regulated spectrin-associated | CAMSAP1 | 177.972 | 3 | 52.365776 | yes | 1.068E-07 | 624.33557 | 225.7106 |
| ASLIEVDLSDLK | 12 | Q5T5Y3 | Calmodulin-regulated spectrin-associated | CAMSAP1 | 177.972 | 2 | 52.365776 | yes | 9.81E-08 | 651.86121 | 218.17183 |
| DNIPEDLRDPFYV | 27 | Q5T5Y3 | Calmodulin-regulated spectrin-associated | CAMSAP1 | 177.972 | 4 | 52.365776 | yes | 2.349E-07 | 825.16614 | 178.00244 |
| GTAAFSPSHPLLPL | 15 | Q5T5Y3 | Calmodulin-regulated spectrin-associated | CAMSAP1 | 177.972 | 2 | 52.365776 | yes | 9.722E-08 | 782.43573 | 177.80101 |
| LNETISTLQQAILK | 14 | Q5T5Y3 | Calmodulin-regulated spectrin-associated | CAMSAP1 | 177.972 | 2 | 52.365776 | yes | 9.722E-08 | 786.4538 | 163.77893 |
| ESVPVEEPLR | 10 | Q5T5Y3 | Calmodulin-regulated spectrin-associated | CAMSAP1 | 177.972 | 2 | 52.365776 | yes | 1.487E-07 | 577.80621 | 154.54257 |
| IAANLQWICAK | 11 | Q5T5Y3 | Calmodulin-regulated spectrin-associated | CAMSAP1 | 177.972 | 2 | 52.365776 | yes | 9.722E-08 | 644.34753 | 110.12613 |
| VESMEALPILSR | 12 | Q5T5Y3 | Calmodulin-regulated spectrin-associated | CAMSAP1 | 177.972 | 2 | 52.365776 | yes | 5.099E-07 | 672.86322 | 107.86085 |
| TMSVSVDALTIHN | 19 | Q5T5Y3 | Calmodulin-regulated spectrin-associated | CAMSAP1 | 177.972 | 3 | 52.365776 | yes | 3.098E-07 | 726.37726 | 101.84035 |
| AAFLHVVK | 8 | Q5T5Y3 | Calmodulin-regulated spectrin-associated | CAMSAP1 | 177.972 | 2 | 52.365776 | yes | 2.966E-06 | 442.77127 | 73.680008 |
| ISQQQEQLLMK | 11 | Q5T5Y3 | Calmodulin-regulated spectrin-associated | CAMSAP1 | 177.972 | 2 | 52.365776 | yes | 9.722E-08 | 673.36084 | 71.770538 |
| QFNLIPAK | 8 | Q5T5Y3 | Calmodulin-regulated spectrin-associated | CAMSAP1 | 177.972 | 2 | 52.365776 | yes | 8.333E-07 | 465.77402 | 68.052765 |
| NSILEELEK | 9 | Q5T5Y3 | Calmodulin-regulated spectrin-associated | CAMSAP1 | 177.972 | 2 | 52.365776 | yes | 1.975E-07 | 537.78748 | 65.95137 |
| QQLLESPAHQK | 11 | Q5T5Y3 | Calmodulin-regulated spectrin-associated | CAMSAP1 | 177.972 | 2 | 52.365776 | yes | 2.008E-07 | 639.84369 | 65.777336 |
| ESLAFAQQHK | 10 | Q5T5Y3 | Calmodulin-regulated spectrin-associated | CAMSAP1 | 177.972 | 2 | 52.365776 | yes | 1.312E-05 | 579.79877 | 65.52784 |
| SIQGEDIPDQR | 11 | Q5T5Y3 | Calmodulin-regulated spectrin-associated | CAMSAP1 | 177.972 | 2 | 52.365776 | yes | 9.82E-08 | 629.30713 | 65.157761 |
| HYLHPEEPEYLGK | 13 | Q5T5Y3 | Calmodulin-regulated spectrin-associated | CAMSAP1 | 177.972 | 3 | 52.365776 | yes | 6.103E-07 | 537.93158 | 51.284412 |
| SSRPPVPISNATK | 13 | Q5T5Y3 | Calmodulin-regulated spectrin-associated | CAMSAP1 | 177.972 | 2 | 52.365776 | yes | 3.858E-07 | 677.37787 | 34.458542 |
| LDDICLK | 7 | Q5T5Y3 | Calmodulin-regulated spectrin-associated | CAMSAP1 | 177.972 | 2 | 52.365776 | yes | 7.797E-07 | 438.72839 | 31.317575 |
| EFHLGTLNK | 9 | Q5T6F0 | DDB1- and CUL4-associated factor 12 O | DCAF12 | 50.518 | 2 | 50.376232 | yes | 2.281E-07 | 529.7851 | 141.88246 |
| VFASQWLNHR | 10 | Q5T6F0 | DDB1- and CUL4-associated factor 12 O | DCAF12 | 50.518 | 2 | 50.376232 | yes | 9.81E-08 | 629.328 | 76.054016 |
| VPVYAHITHK | 10 | Q5T6F0 | DDB1- and CUL4-associated factor 12 O | DCAF12 | 50.518 | 2 | 50.376232 | yes | 2.349E-07 | 582.82983 | 48.948715 |
| SQPEPSPVLSQLSQ | 15 | Q5T6F2 | Ubiquitin-associated protein 2 OS=Homo | UBAP2 | 117.116 | 2 | 50.95546 | yes | 5.96E-07 | 826.93378 | 4335.0117 |
| LLQLPSTTIENISVS | 22 | Q5T6F2 | Ubiquitin-associated protein 2 OS=Homo | UBAP2 | 117.116 | 3 | 50.95546 | yes | 1.035E-06 | 810.45148 | 4295.9912 |
| DGSLANNPYPGDV | 15 | Q5T6F2 | Ubiquitin-associated protein 2 OS=Homo | UBAP2 | 117.116 | 2 | 50.95546 | yes | 1.342E-07 | 774.37042 | 2869.3684 |
| MANITSSQILDQLK | 14 | Q5T6F2 | Ubiquitin-associated protein 2 OS=Homo | UBAP2 | 117.116 | 2 | 50.95546 | yes | 1.149E-07 | 781.41632 | 980.99786 |
| MANITSSQILDQLK | 14 | Q5T6F2 | Ubiquitin-associated protein 2 OS=Homo | UBAP2 | 117.116 | 2 | 50.95546 | yes | 1.722E-07 | 789.41382 | 980.99786 |
| ESTPGDSPSTVNK | 13 | Q5T6F2 | Ubiquitin-associated protein 2 OS=Homo | UBAP2 | 117.116 | 2 | 50.95546 | yes | 1.659E-07 | 659.80969 | 1389.0647 |
| SQQTLDTPK | 9 | Q5T6F2 | Ubiquitin-associated protein 2 OS=Homo | UBAP2 | 117.116 | 2 | 50.95546 | yes | 2.368E-07 | 509.26419 | 1028.1086 |
| QVVQATAEQMR | 11 | Q5T6F2 | Ubiquitin-associated protein 2 OS=Homo | UBAP2 | 117.116 | 2 | 50.95546 | yes | 1.192E-07 | 630.8219 | 433.65405 |
| QVVQATAEQMR | 11 | Q5T6F2 | Ubiquitin-associated protein 2 OS=Homo | UBAP2 | 117.116 | 2 | 50.95546 | yes | 7.305E-07 | 638.81934 | 433.65405 |
| EKPQISAAQSTQPQ | 15 | Q5T6F2 | Ubiquitin-associated protein 2 OS=Homo | UBAP2 | 117.116 | 2 | 50.95546 | yes | 1.003E-07 | 820.93378 | 188.72937 |
| EKPQISAAQSTQPQ | 15 | Q5T6F2 | Ubiquitin-associated protein 2 OS=Homo | UBAP2 | 117.116 | 2 | 50.95546 | yes | 1.722E-07 | 547.62494 | 188.72937 |

| | | | | | | | | | | | |
|---|---|---|---|---|---|---|---|---|---|---|---|
| EKPQISAAQSTQP( | 15 | Q5T6F2 | Ubiquitin-associated protein 2 OS=Homo | UBAP2 | 117.116 | 3 | 50.95546 | yes | 1.003E-07 | 820.93378 | 188.72937 |
| EKPQISAAQSTQP( | 15 | Q5T6F2 | Ubiquitin-associated protein 2 OS=Homo | UBAP2 | 117.116 | 3 | 50.95546 | yes | 1.722E-07 | 547.62494 | 188.72937 |
| MMTSVSSDHCR | 11 | Q5T6F2 | Ubiquitin-associated protein 2 OS=Homo | UBAP2 | 117.116 | 2 | 50.95546 | yes | 1.023E-07 | 676.77319 | 267.8143 |
| PAYGNSPYWTN | 11 | Q5T6F2 | Ubiquitin-associated protein 2 OS=Homo | UBAP2 | 117.116 | 2 | 50.95546 | yes | 2.291E-07 | 635.2804 | 154.87701 |
| LAQVIFDK | 8 | Q5T6F2 | Ubiquitin-associated protein 2 OS=Homo | UBAP2 | 117.116 | 2 | 50.95546 | yes | 9.722E-08 | 467.27383 | 99.862198 |
| NDSDFEAK | 8 | Q5T6F2 | Ubiquitin-associated protein 2 OS=Homo | UBAP2 | 117.116 | 2 | 50.95546 | yes | 2.977E-05 | 463.19852 | 37.990978 |
| LTEGQYVLCR | 10 | Q5T6S3 | PHD finger protein 19 OS=Homo sapiens | PHF19 | 65.591 | 2 | 50.359745 | yes | 9.81E-08 | 619.81354 | 285.83835 |
| GLLPNENSASSELI | 14 | Q5T6S3 | PHD finger protein 19 OS=Homo sapiens | PHF19 | 65.591 | 2 | 50.359745 | yes | 3.777E-07 | 743.87842 | 118.08671 |
| LTSTPVTDR | 9 | Q5T6S3 | PHD finger protein 19 OS=Homo sapiens | PHF19 | 65.591 | 2 | 50.359745 | yes | 5.45E-07 | 495.26672 | 105.52253 |
| SKPGLLPHEFQQQ | 14 | Q5T6S3 | PHD finger protein 19 OS=Homo sapiens | PHF19 | 65.591 | 3 | 50.359745 | yes | 1.452E-07 | 546.2984 | 100.11299 |
| QSCLVTFEDNSK | 12 | Q5T6S3 | PHD finger protein 19 OS=Homo sapiens | PHF19 | 65.591 | 2 | 50.359745 | yes | 3.208E-07 | 714.32721 | 83.457275 |
| WTDGLYYLGK | 10 | Q5T6S3 | PHD finger protein 19 OS=Homo sapiens | PHF19 | 65.591 | 2 | 50.359745 | yes | 2.426E-07 | 608.30585 | 82.418526 |
| TSGPLNEILICGK | 13 | Q5T6S3 | PHD finger protein 19 OS=Homo sapiens | PHF19 | 65.591 | 2 | 50.359745 | yes | 1.438E-07 | 701.37396 | 70.128006 |
| CIFALAVR | 8 | Q5T6S3 | PHD finger protein 19 OS=Homo sapiens | PHF19 | 65.591 | 2 | 50.359745 | yes | 4.314E-07 | 475.26804 | 70.02533 |
| SSITNYFGAAGR | 12 | Q5T6S3 | PHD finger protein 19 OS=Homo sapiens | PHF19 | 65.591 | 2 | 50.359745 | yes | 2.958E-07 | 622.30695 | 54.272873 |
| DIQHAGVPGEEPK | 13 | Q5T6S3 | PHD finger protein 19 OS=Homo sapiens | PHF19 | 65.591 | 2 | 50.359745 | yes | 3.154E-06 | 688.84387 | 46.805916 |
| GPHLLNALNSYK | 12 | Q5T6S3 | PHD finger protein 19 OS=Homo sapiens | PHF19 | 65.591 | 2 | 50.359745 | yes | 3.704E-07 | 663.86188 | 44.703384 |
| DSYGATSHLPNK | 12 | Q5T6S3 | PHD finger protein 19 OS=Homo sapiens | PHF19 | 65.591 | 2 | 50.359745 | yes | 1.896E-06 | 645.30969 | 24.093445 |
| PLPSFCPPR | 9 | Q5T749 | Keratinocyte proline-rich protein OS=Ho | KPRP | 64.136 | 2 | 32.435627 | yes | 1.518E-06 | 535.77625 | 124.45901 |
| LGFQQSLTVAAPG | 16 | Q5T7B8 | Kinesin-like protein KIF24 OS=Homo sa | KIF24 | 151.904 | 2 | 49.735344 | yes | 1.006E-07 | 816.93884 | 55.531357 |
| WEVLQQEQR | 9 | Q5T7V8 | RAB6-interacting golgin OS=Homo sapie | GORAB | 42.266 | 2 | 49.786743 | yes | 1.003E-07 | 608.30945 | 90.530594 |
| FSYFPSYVDK | 10 | Q5T7W7 | Thiosulfate sulfurtransferase/rhodanese-li | TSTD2 | 58.263 | 2 | 37.335125 | yes | 6.084E-07 | 626.79785 | 51.721996 |
| LFVFDER | 7 | Q5T7W7 | Thiosulfate sulfurtransferase/rhodanese-li | TSTD2 | 58.263 | 2 | 37.335125 | yes | 3.703E-06 | 463.24252 | 30.739855 |
| LKEYEAAVEQLK | 12 | Q5T9A4 | ATPase family AAA domain-containing p | ATAD3B | 72.573 | 2 | 53.3161 | yes | 1.084E-07 | 710.88776 | 867.49628 |
| LLSRPQDVLEGVV | 21 | Q5T9A4 | ATPase family AAA domain-containing p | ATAD3B | 72.573 | 3 | 53.3161 | no | 3.359E-07 | 760.09705 | 720.0293 |
| HILLYGPPGTGK | 12 | Q5T9A4 | ATPase family AAA domain-containing p | ATAD3B | 72.573 | 2 | 53.3161 | no | 9.722E-08 | 626.85608 | 531.77332 |
| EYEAAVEQLK | 10 | Q5T9A4 | ATPase family AAA domain-containing p | ATAD3B | 72.573 | 2 | 53.3161 | yes | 3.839E-07 | 590.29822 | 522.76221 |
| LHFDNCVLKPATE | 15 | Q5T9A4 | ATPase family AAA domain-containing p | ATAD3B | 72.573 | 3 | 53.3161 | yes | 9.722E-08 | 576.96277 | 227.92163 |
| PATEGKR | 7 | Q5T9A4 | ATPase family AAA domain-containing p | ATAD3B | 72.573 | 1 | 53.3161 | no | 2.663E-07 | 758.41553 | 106.19146 |
| EALNLAQMQEQTI | 20 | Q5T9A4 | ATPase family AAA domain-containing p | ATAD3B | 72.573 | 3 | 53.3161 | yes | 1.843E-07 | 777.39337 | 74.411774 |
| WQALPALR | 8 | Q5TAQ9 | DDB1- and CUL4-associated factor 8 OS | DCAF8 | 66.853 | 2 | 40.61242 | yes | 1.503E-06 | 477.77963 | 40.368584 |
| IWAPTAEASTELT( | 16 | Q5TAQ9 | DDB1- and CUL4-associated factor 8 OS | DCAF8 | 66.853 | 2 | 40.61242 | yes | 9.751E-06 | 844.44873 | 27.187706 |
| GTWLASGSDDLK | 12 | Q5TAQ9 | DDB1- and CUL4-associated factor 8 OS | DCAF8 | 66.853 | 2 | 40.61242 | yes | 3.933E-07 | 625.30658 | 16.028036 |
| ELSQCNCIDLSK | 12 | Q5TAX3 | Terminal uridylyltransferase 4 OS=Homo | TUT4 | 185.166 | 2 | 53.81501 | yes | 9.722E-08 | 733.83435 | 212.28635 |
| IAIEDPFSVK | 10 | Q5TAX3 | Terminal uridylyltransferase 4 OS=Homo | TUT4 | 185.166 | 2 | 53.81501 | yes | 3.628E-07 | 559.80823 | 186.93634 |
| LYGSSLTR | 8 | Q5TAX3 | Terminal uridylyltransferase 4 OS=Homo | TUT4 | 185.166 | 2 | 53.81501 | yes | 2.05E-06 | 448.74545 | 175.48788 |
| SPLALETPNR | 10 | Q5TAX3 | Terminal uridylyltransferase 4 OS=Homo | TUT4 | 185.166 | 2 | 53.81501 | yes | 9.903E-08 | 549.30109 | 162.53058 |
| YFACPQTK | 8 | Q5TAX3 | Terminal uridylyltransferase 4 OS=Homo | TUT4 | 185.166 | 2 | 53.81501 | yes | 2.954E-07 | 507.73929 | 148.53844 |
| VITTFLPECSLR | 12 | Q5TAX3 | Terminal uridylyltransferase 4 OS=Homo | TUT4 | 185.166 | 2 | 53.81501 | yes | 1.231E-07 | 718.38434 | 146.10535 |

| | | | | | | | | | | | |
|---|---|---|---|---|---|---|---|---|---|---|---|
| IQDILTR | 7 | Q5TAX3 | Terminal uridylyltransferase 4 OS=Homo | TUT4 | 185.166 | 2 | 53.81501 | yes | 4.174E-06 | 429.7558 | 135.8118 |
| SPEYTNCR | 8 | Q5TAX3 | Terminal uridylyltransferase 4 OS=Homo | TUT4 | 185.166 | 2 | 53.81501 | yes | 1.658E-07 | 513.71912 | 104.19611 |
| NVLYVDVESDFH/ | 14 | Q5TAX3 | Terminal uridylyltransferase 4 OS=Homo | TUT4 | 185.166 | 2 | 53.81501 | yes | 9.722E-08 | 818.4043 | 95.758797 |
| MNHPDLLIK | 9 | Q5TAX3 | Terminal uridylyltransferase 4 OS=Homo | TUT4 | 185.166 | 2 | 53.81501 | yes | 3.229E-07 | 540.79718 | 84.828445 |
| MLATYAAIDPR | 11 | Q5TAX3 | Terminal uridylyltransferase 4 OS=Homo | TUT4 | 185.166 | 2 | 53.81501 | yes | 1.734E-07 | 611.31842 | 82.78157 |
| EIIENLAK | 8 | Q5TAX3 | Terminal uridylyltransferase 4 OS=Homo | TUT4 | 185.166 | 2 | 53.81501 | yes | 4.641E-07 | 465.26874 | 65.392143 |
| AVQVIGNQTLK | 11 | Q5TAX3 | Terminal uridylyltransferase 4 OS=Homo | TUT4 | 185.166 | 2 | 53.81501 | yes | 1.664E-07 | 585.8457 | 61.67905 |
| EAEYFFDSR | 9 | Q5TAX3 | Terminal uridylyltransferase 4 OS=Homo | TUT4 | 185.166 | 2 | 53.81501 | yes | 2.192E-05 | 582.25385 | 56.154892 |
| VQYLGYTMK | 9 | Q5TAX3 | Terminal uridylyltransferase 4 OS=Homo | TUT4 | 185.166 | 2 | 53.81501 | yes | 4.054E-07 | 551.78369 | 47.521629 |
| EQILIGLEK | 9 | Q5TAX3 | Terminal uridylyltransferase 4 OS=Homo | TUT4 | 185.166 | 2 | 53.81501 | yes | 3.656E-07 | 521.81079 | 38.868595 |
| QEIVEEMSK | 9 | Q5TAX3 | Terminal uridylyltransferase 4 OS=Homo | TUT4 | 185.166 | 2 | 53.81501 | yes | 5.257E-07 | 546.76569 | 35.784424 |
| ELSQEDVEEVWR | 12 | Q5TB30 | DEP domain-containing protein 1A OS=I | DEPDC1 | 92.96 | 2 | 54.470818 | yes | 9.722E-08 | 759.85718 | 152.65498 |
| QQTIQLLR | 8 | Q5TB30 | DEP domain-containing protein 1A OS=I | DEPDC1 | 92.96 | 2 | 54.470818 | yes | 1.022E-07 | 500.3009 | 120.97651 |
| LQISNPGFQER | 11 | Q5TB30 | DEP domain-containing protein 1A OS=I | DEPDC1 | 92.96 | 2 | 54.470818 | yes | 1.735E-07 | 644.83588 | 72.444992 |
| SNDMNNPTYVGFI | 14 | Q5TB30 | DEP domain-containing protein 1A OS=I | DEPDC1 | 92.96 | 2 | 54.470818 | yes | 9.722E-08 | 822.35956 | 62.435253 |
| SQSLLLR | 7 | Q5TB30 | DEP domain-containing protein 1A OS=I | DEPDC1 | 92.96 | 2 | 54.470818 | yes | 3.031E-06 | 408.75052 | 26.364746 |
| QPMLILR | 7 | Q5TB30 | DEP domain-containing protein 1A OS=I | DEPDC1 | 92.96 | 2 | 54.470818 | yes | 4.987E-07 | 435.76514 | 20.998327 |
| STECLLLSLLHR | 12 | Q5TB30 | DEP domain-containing protein 1A OS=I | DEPDC1 | 92.96 | 2 | 54.470818 | yes | 1.004E-06 | 721.3952 | 19.82135 |
| WGSENVDDNNQL | 14 | Q5TB30 | DEP domain-containing protein 1A OS=I | DEPDC1 | 92.96 | 2 | 54.470818 | yes | 1.813E-07 | 847.3819 | 16.892656 |
| EIMAQLEER | 9 | Q5TC82 | Roquin-1 OS=Homo sapiens OX=9606 G | RC3H1 | 125.737 | 2 | 52.2435 | yes | 1.15E-07 | 559.77917 | 1214.7275 |
| GVGLNSTTQSVLS | 14 | Q5TC82 | Roquin-1 OS=Homo sapiens OX=9606 G | RC3H1 | 125.737 | 2 | 52.2435 | yes | 9.722E-08 | 709.88354 | 1020.7163 |
| VYPAPSYTR | 9 | Q5TC82 | Roquin-1 OS=Homo sapiens OX=9606 G | RC3H1 | 125.737 | 2 | 52.2435 | yes | 1.238E-07 | 527.27185 | 734.28735 |
| GTDPSYDSSLKPGI | 14 | Q5TC82 | Roquin-1 OS=Homo sapiens OX=9606 G | RC3H1 | 125.737 | 2 | 52.2435 | yes | 1.084E-07 | 726.35425 | 640.52484 |
| ISMSEVASHGK | 11 | Q5TC82 | Roquin-1 OS=Homo sapiens OX=9606 G | RC3H1 | 125.737 | 2 | 52.2435 | yes | 9.722E-08 | 573.28461 | 374.11276 |
| WPGMISSEQLSLEI | 19 | Q5TC82 | Roquin-1 OS=Homo sapiens OX=9606 G | RC3H1 | 125.737 | 3 | 52.2435 | yes | 1.501E-07 | 747.04407 | 336.00504 |
| ELSMENQCSLDMI | 13 | Q5TC82 | Roquin-1 OS=Homo sapiens OX=9606 G | RC3H1 | 125.737 | 2 | 52.2435 | yes | 9.722E-08 | 792.83875 | 132.22772 |
| GASCTFAHSQEEL | 15 | Q5TC82 | Roquin-1 OS=Homo sapiens OX=9606 G | RC3H1 | 125.737 | 2 | 52.2435 | yes | 9.722E-08 | 847.37793 | 117.20197 |
| LPPPPQPHPSLDEL | 16 | Q5TC82 | Roquin-1 OS=Homo sapiens OX=9606 G | RC3H1 | 125.737 | 4 | 52.2435 | yes | 4.719E-07 | 458.24689 | 115.68195 |
| PLQMVPR | 7 | Q5TC82 | Roquin-1 OS=Homo sapiens OX=9606 G | RC3H1 | 125.737 | 2 | 52.2435 | yes | 1.28E-06 | 420.74164 | 94.226097 |
| DVVAAGSVEMMN | 16 | Q5TC82 | Roquin-1 OS=Homo sapiens OX=9606 G | RC3H1 | 125.737 | 2 | 52.2435 | yes | 1.94E-07 | 833.39478 | 79.918816 |
| KPISLGCGHTVCK | 13 | Q5TC82 | Roquin-1 OS=Homo sapiens OX=9606 G | RC3H1 | 125.737 | 2 | 52.2435 | yes | 1.136E-07 | 728.8736 | 50.575996 |
| KCVEELALYLKPL | 17 | Q5TC82 | Roquin-1 OS=Homo sapiens OX=9606 G | RC3H1 | 125.737 | 3 | 52.2435 | yes | 2.145E-07 | 659.69946 | 31.480579 |
| WTLGGMVNR | 9 | Q5TCZ1 | SH3 and PX domain-containing protein 2 | SH3PXD2/ | 125.29 | 2 | 52.293217 | yes | 1.084E-07 | 517.26599 | 249.25142 |
| AASQGSDSPLLPA( | 15 | Q5TCZ1 | SH3 and PX domain-containing protein 2 | SH3PXD2/ | 125.29 | 2 | 52.293217 | yes | 2.554E-07 | 749.38641 | 169.70653 |
| NSIPVSPVRPK | 11 | Q5TCZ1 | SH3 and PX domain-containing protein 2 | SH3PXD2/ | 125.29 | 2 | 52.293217 | yes | 1.596E-07 | 597.3537 | 159.14618 |
| AQISSPNLR | 9 | Q5TCZ1 | SH3 and PX domain-containing protein 2 | SH3PXD2/ | 125.29 | 2 | 52.293217 | yes | 4.742E-07 | 493.2749 | 159.01259 |
| VQALNTVNQSK | 11 | Q5TCZ1 | SH3 and PX domain-containing protein 2 | SH3PXD2/ | 125.29 | 2 | 52.293217 | yes | 9.927E-08 | 601.33038 | 152.47681 |
| EGWAPASYIDK | 11 | Q5TCZ1 | SH3 and PX domain-containing protein 2 | SH3PXD2/ | 125.29 | 2 | 52.293217 | yes | 9.722E-08 | 618.7984 | 107.70502 |
| AKDDLPTR | 8 | Q5TCZ1 | SH3 and PX domain-containing protein 2 | SH3PXD2/ | 125.29 | 2 | 52.293217 | yes | 1.239E-06 | 458.24835 | 83.819443 |

| | | | | | | | | | | | |
|---|---|---|---|---|---|---|---|---|---|---|---|
| QESGWWYVR | 9 | Q5TCZ1 | SH3 and PX domain-containing protein 2 | SH3PXD2/ | 125.29 | 2 | 52.293217 | yes | 1.948E-07 | 605.78564 | 70.942596 |
| SQFIHNNLK | 9 | Q5TCZ1 | SH3 and PX domain-containing protein 2 | SH3PXD2/ | 125.29 | 2 | 52.293217 | yes | 1.671E-07 | 550.79602 | 63.169609 |
| NSGGWWYVQIGE | 13 | Q5TCZ1 | SH3 and PX domain-containing protein 2 | SH3PXD2/ | 125.29 | 2 | 52.293217 | yes | 4.657E-07 | 762.36749 | 38.972088 |
| DADANAGLTSCPR | 13 | Q5TCZ1 | SH3 and PX domain-containing protein 2 | SH3PXD2/ | 125.29 | 2 | 52.293217 | yes | 1.001E-07 | 674.30151 | 24.624342 |
| DSLLPLQDTAYR | 12 | Q5TGY3 | Transcription factor Gibbin OS=Homo sa | AHDC1 | 168.35 | 2 | 49.98555 | yes | 3.924E-07 | 696.36188 | 227.10287 |
| VSAEPTPLLK | 10 | Q5TGY3 | Transcription factor Gibbin OS=Homo sa | AHDC1 | 168.35 | 2 | 49.98555 | yes | 1.073E-07 | 527.81079 | 123.94727 |
| VSSLSSSLSVEGK | 13 | Q5TGY3 | Transcription factor Gibbin OS=Homo sa | AHDC1 | 168.35 | 2 | 49.98555 | yes | 1.104E-07 | 640.34064 | 116.88996 |
| NLLLGPGKPK | 10 | Q5TGY3 | Transcription factor Gibbin OS=Homo sa | AHDC1 | 168.35 | 2 | 49.98555 | yes | 9.938E-08 | 518.82928 | 116.58456 |
| AFASTGLESGASGI | 14 | Q5TGY3 | Transcription factor Gibbin OS=Homo sa | AHDC1 | 168.35 | 2 | 49.98555 | yes | 1.037E-07 | 655.82037 | 105.2621 |
| QFAGASQWPFR | 11 | Q5TGY3 | Transcription factor Gibbin OS=Homo sa | AHDC1 | 168.35 | 2 | 49.98555 | yes | 3.179E-07 | 647.82001 | 91.297806 |
| AYSPTCSPTLGFK | 13 | Q5TGY3 | Transcription factor Gibbin OS=Homo sa | AHDC1 | 168.35 | 2 | 49.98555 | yes | 1.326E-07 | 714.84503 | 83.629417 |
| QNLFTGYFR | 9 | Q5TGY3 | Transcription factor Gibbin OS=Homo sa | AHDC1 | 168.35 | 2 | 49.98555 | yes | 8.453E-07 | 573.29053 | 59.679474 |
| ASGTYAGPPTSAL | 17 | Q5TGY3 | Transcription factor Gibbin OS=Homo sa | AHDC1 | 168.35 | 2 | 49.98555 | yes | 2.898E-07 | 822.92065 | 52.51688 |
| LASPQPSYAADAN | 16 | Q5TGY3 | Transcription factor Gibbin OS=Homo sa | AHDC1 | 168.35 | 2 | 49.98555 | yes | 1.066E-07 | 817.88647 | 38.5443 |
| ASTVSPGGYMVPK | 13 | Q5TGY3 | Transcription factor Gibbin OS=Homo sa | AHDC1 | 168.35 | 2 | 49.98555 | yes | 2.835E-06 | 647.32904 | 31.808506 |
| LAACEPLK | 8 | Q5TGY3 | Transcription factor Gibbin OS=Homo sa | AHDC1 | 168.35 | 2 | 49.98555 | yes | 0.0002107 | 451.24423 | 10.826408 |
| LTAILLQIK | 9 | Q5TKA1 | Protein lin-9 homolog OS=Homo sapiens | LIN9 | 61.946 | 2 | 40.42041 | yes | 6.402E-07 | 506.84189 | 33.691486 |
| CSSAFFEEER | 10 | Q5TKA1 | Protein lin-9 homolog OS=Homo sapiens | LIN9 | 61.946 | 2 | 40.42041 | yes | 4.035E-07 | 631.26135 | 23.93643 |
| GEEPIFVVTGR | 11 | Q5U5Q3 | RNA-binding E3 ubiquitin-protein ligase | MEX3C | 69.367 | 2 | 39.180893 | yes | 4.74E-07 | 602.32202 | 87.997757 |
| TIGDLSTLTASEIK | 14 | Q5UIP0 | Telomere-associated protein RIF1 OS=H | RIF1 | 274.466 | 2 | 51.328777 | yes | 1.109E-07 | 724.89575 | 190.79945 |
| LIEQAPIQMGEEA | 15 | Q5UIP0 | Telomere-associated protein RIF1 OS=H | RIF1 | 274.466 | 2 | 51.328777 | yes | 2.36E-07 | 842.44037 | 163.18707 |
| KIFATLTR | 8 | Q5UIP0 | Telomere-associated protein RIF1 OS=H | RIF1 | 274.466 | 2 | 51.328777 | yes | 1.642E-07 | 475.2951 | 137.14104 |
| LIAEQTLQENLIEK | 14 | Q5UIP0 | Telomere-associated protein RIF1 OS=H | RIF1 | 274.466 | 2 | 51.328777 | yes | 9.722E-08 | 821.45654 | 121.24888 |
| CVWSPLASPSTSIL | 15 | Q5UIP0 | Telomere-associated protein RIF1 OS=H | RIF1 | 274.466 | 2 | 51.328777 | yes | 9.809E-08 | 823.43457 | 115.04523 |
| PLALFYENSK | 10 | Q5UIP0 | Telomere-associated protein RIF1 OS=H | RIF1 | 274.466 | 2 | 51.328777 | yes | 2.067E-07 | 591.31366 | 108.29387 |
| EKPGSEVLTLLLK | 13 | Q5UIP0 | Telomere-associated protein RIF1 OS=H | RIF1 | 274.466 | 2 | 51.328777 | yes | 2.424E-07 | 713.92944 | 107.58084 |
| VMMLVYPEELKP | 18 | Q5UIP0 | Telomere-associated protein RIF1 OS=H | RIF1 | 274.466 | 3 | 51.328777 | yes | 1.133E-07 | 697.05273 | 73.524635 |
| IGISDISSLSEK | 12 | Q5UIP0 | Telomere-associated protein RIF1 OS=H | RIF1 | 274.466 | 2 | 51.328777 | yes | 1.387E-07 | 624.83771 | 68.114265 |
| GATALEMGMPLLI | 15 | Q5UIP0 | Telomere-associated protein RIF1 OS=H | RIF1 | 274.466 | 2 | 51.328777 | yes | 1.002E-07 | 786.92804 | 42.507252 |
| DSFDNCSLGESSK | 13 | Q5UIP0 | Telomere-associated protein RIF1 OS=H | RIF1 | 274.466 | 2 | 51.328777 | yes | 3.98E-07 | 723.29608 | 36.114212 |
| HANTLITAVHDSF | 18 | Q5UIP0 | Telomere-associated protein RIF1 OS=H | RIF1 | 274.466 | 3 | 51.328777 | yes | 1.804E-05 | 627.33899 | 35.434795 |
| VSFADPIYQAGLA | 19 | Q5UIP0 | Telomere-associated protein RIF1 OS=H | RIF1 | 274.466 | 3 | 51.328777 | yes | 2.352E-07 | 708.02362 | 34.009228 |
| NTENNDVEISETK | 13 | Q5UIP0 | Telomere-associated protein RIF1 OS=H | RIF1 | 274.466 | 2 | 51.328777 | yes | 1.585E-06 | 746.84174 | 21.698275 |
| GASSPYGAPGTPR | 13 | Q5UIP0 | Telomere-associated protein RIF1 OS=H | RIF1 | 274.466 | 2 | 51.328777 | yes | 1.329E-06 | 609.29907 | 20.80077 |
| LSCMANSVIK | 10 | Q5UIP0 | Telomere-associated protein RIF1 OS=H | RIF1 | 274.466 | 2 | 51.328777 | yes | 3.139E-06 | 561.78595 | 20.140993 |
| NVITQWEAK | 9 | Q5VSY0 | G kinase-anchoring protein 1 OS=Homo s | GKAP1 | 42.079 | 2 | 58.47071 | yes | 3.891E-07 | 544.79041 | 166.00784 |
| DRPLTVSLK | 9 | Q5VSY0 | G kinase-anchoring protein 1 OS=Homo s | GKAP1 | 42.079 | 2 | 58.47071 | yes | 1.425E-06 | 514.80859 | 133.73653 |
| AEILLQVDESQSIK | 14 | Q5VSY0 | G kinase-anchoring protein 1 OS=Homo s | GKAP1 | 42.079 | 2 | 58.47071 | yes | 3.961E-07 | 786.92761 | 107.17288 |
| ASAVLSSVPTTASI | 14 | Q5VSY0 | G kinase-anchoring protein 1 OS=Homo s | GKAP1 | 42.079 | 2 | 58.47071 | yes | 2.587E-06 | 694.87262 | 89.963631 |

| | | | | | | | | | | | |
|---|---|---|---|---|---|---|---|---|---|---|---|
| EQQQSEANELR | 11 | Q5VSY0 | G kinase-anchoring protein 1 OS=Homo s | GKAP1 | 42.079 | 2 | 58.47071 | yes | 9.722E-08 | 666.31293 | 71.503464 |
| MLQEGEMK | 8 | Q5VSY0 | G kinase-anchoring protein 1 OS=Homo s | GKAP1 | 42.079 | 2 | 58.47071 | yes | 1.326E-07 | 483.22516 | 61.003635 |
| HYDTDEVR | 8 | Q5VT06 | Centrosome-associated protein 350 OS=H | CEP350 | 350.931 | 2 | 54.24917 | yes | 3.176E-07 | 517.73053 | 125.49136 |
| ELNMPFSGGQDSF | 15 | Q5VT06 | Centrosome-associated protein 350 OS=H | CEP350 | 350.931 | 2 | 54.24917 | yes | 4.55E-07 | 822.37213 | 102.91784 |
| ATSPGPTDLETR | 12 | Q5VT06 | Centrosome-associated protein 350 OS=H | CEP350 | 350.931 | 2 | 54.24917 | yes | 3.606E-07 | 622.80951 | 78.965965 |
| YLDDSWVNAPISK | 13 | Q5VT06 | Centrosome-associated protein 350 OS=H | CEP350 | 350.931 | 2 | 54.24917 | yes | 9.722E-08 | 754.375 | 78.494957 |
| VLIGNVQPGILR | 12 | Q5VT06 | Centrosome-associated protein 350 OS=H | CEP350 | 350.931 | 2 | 54.24917 | yes | 1.073E-07 | 639.89825 | 77.815041 |
| ITANEDPPVISK | 12 | Q5VT06 | Centrosome-associated protein 350 OS=H | CEP350 | 350.931 | 2 | 54.24917 | yes | 9.722E-08 | 642.3457 | 57.101055 |
| DTIDVLNQISEK | 12 | Q5VT06 | Centrosome-associated protein 350 OS=H | CEP350 | 350.931 | 2 | 54.24917 | yes | 1.153E-07 | 687.85919 | 32.299305 |
| VWQEAEFQNMSR | 12 | Q5VT06 | Centrosome-associated protein 350 OS=H | CEP350 | 350.931 | 2 | 54.24917 | yes | 6.474E-07 | 762.84863 | 31.44153 |
| TGSSHLISTSSWR | 13 | Q5VT06 | Centrosome-associated protein 350 OS=H | CEP350 | 350.931 | 2 | 54.24917 | yes | 4.043E-06 | 709.8548 | 31.049179 |
| LSVNNMAHDTDPI | 13 | Q5VT06 | Centrosome-associated protein 350 OS=H | CEP350 | 350.931 | 2 | 54.24917 | yes | 1.464E-07 | 721.34064 | 13.900656 |
| WVGVLSELHK | 10 | Q5VT25 | Serine/threonine-protein kinase MRCK al | CDC42BPA | 197.307 | 2 | 55.638523 | yes | 1.34E-07 | 584.32965 | 140.16074 |
| CSILMLADTENEK | 13 | Q5VT25 | Serine/threonine-protein kinase MRCK al | CDC42BPA | 197.307 | 2 | 55.638523 | yes | 9.722E-08 | 762.35767 | 128.29044 |
| ASQPSVVISQVIDM | 15 | Q5VT25 | Serine/threonine-protein kinase MRCK al | CDC42BPA | 197.307 | 2 | 55.638523 | yes | 2.284E-07 | 815.43506 | 124.35649 |
| LFLYDIAEGK | 10 | Q5VT25 | Serine/threonine-protein kinase MRCK al | CDC42BPA | 197.307 | 2 | 55.638523 | yes | 3.659E-06 | 584.81604 | 117.70438 |
| FQFPAQVTDVSEN | 15 | Q5VT25 | Serine/threonine-protein kinase MRCK al | CDC42BPA | 197.307 | 2 | 55.638523 | yes | 6.233E-07 | 840.91504 | 116.81143 |
| TTQAAAIIDHER | 12 | Q5VT25 | Serine/threonine-protein kinase MRCK al | CDC42BPA | 197.307 | 2 | 55.638523 | yes | 9.722E-08 | 663.34406 | 105.50075 |
| SIFYEEELSK | 10 | Q5VT25 | Serine/threonine-protein kinase MRCK al | CDC42BPA | 197.307 | 2 | 55.638523 | yes | 9.722E-08 | 622.80591 | 100.90552 |
| TLDNNLATEAYER | 13 | Q5VT25 | Serine/threonine-protein kinase MRCK al | CDC42BPA | 197.307 | 2 | 55.638523 | yes | 7.294E-07 | 755.36261 | 80.408005 |
| SMSASSGLSAR | 11 | Q5VT25 | Serine/threonine-protein kinase MRCK al | CDC42BPA | 197.307 | 2 | 55.638523 | yes | 1.475E-06 | 527.2533 | 79.284477 |
| LELQSALDAEIR | 12 | Q5VT25 | Serine/threonine-protein kinase MRCK al | CDC42BPA | 197.307 | 2 | 55.638523 | yes | 9.974E-08 | 679.36975 | 78.905556 |
| ALAIVCDFK | 9 | Q5VT25 | Serine/threonine-protein kinase MRCK al | CDC42BPA | 197.307 | 2 | 55.638523 | yes | 1.139E-06 | 518.77844 | 71.201271 |
| GIGTAYEGHVR | 11 | Q5VT25 | Serine/threonine-protein kinase MRCK al | CDC42BPA | 197.307 | 2 | 55.638523 | yes | 1.242E-07 | 580.29639 | 66.016907 |
| IALGNEEGLFVVHT | 16 | Q5VT25 | Serine/threonine-protein kinase MRCK al | CDC42BPA | 197.307 | 3 | 55.638523 | yes | 9.722E-08 | 575.98932 | 39.122322 |
| NVYPEEMIVALR | 12 | Q5VT52 | Regulation of nuclear pre-mRNA domain | RPRD2 | 156.02 | 2 | 50.84136 | yes | 6.011E-06 | 717.37646 | 39.726299 |
| VSSSCLDLPDSTEE | 15 | Q5VT52 | Regulation of nuclear pre-mRNA domain | RPRD2 | 156.02 | 2 | 50.84136 | yes | 9.722E-08 | 833.87744 | 35.796928 |
| ITEDSAVTTFEALK | 14 | Q5VTB9 | E3 ubiquitin-protein ligase RNF220 OS=I | RNF220 | 62.766 | 2 | 48.37122 | yes | 1.157E-07 | 762.89325 | 1099.9987 |
| ATTLLEGGFR | 10 | Q5VTB9 | E3 ubiquitin-protein ligase RNF220 OS=I | RNF220 | 62.766 | 2 | 48.37122 | yes | 1.693E-07 | 532.79041 | 421.37805 |
| GSGFIMCSGK | 10 | Q5VTB9 | E3 ubiquitin-protein ligase RNF220 OS=I | RNF220 | 62.766 | 2 | 48.37122 | yes | 1.557E-07 | 522.23608 | 283.15875 |
| GAVLNGGPPSTR | 12 | Q5VTB9 | E3 ubiquitin-protein ligase RNF220 OS=I | RNF220 | 62.766 | 2 | 48.37122 | yes | 5.58E-07 | 563.3042 | 88.446098 |
| MLQDLLSDR | 9 | Q5VU43 | Myomegalin OS=Homo sapiens OX=960 | PDE4DIP | 265.103 | 2 | 47.457684 | yes | 1.415E-07 | 545.78168 | 118.46249 |
| LNEALQAER | 9 | Q5VU43 | Myomegalin OS=Homo sapiens OX=960 | PDE4DIP | 265.103 | 2 | 47.457684 | yes | 1.386E-07 | 522.27765 | 102.3077 |
| LTQEVLLLR | 9 | Q5VU43 | Myomegalin OS=Homo sapiens OX=960 | PDE4DIP | 265.103 | 2 | 47.457684 | yes | 1.326E-07 | 542.83984 | 76.27002 |
| EVEDLSATLLCK | 12 | Q5VU43 | Myomegalin OS=Homo sapiens OX=960 | PDE4DIP | 265.103 | 2 | 47.457684 | yes | 5.239E-07 | 689.35016 | 54.507507 |
| LNEMSYELK | 9 | Q5VU43 | Myomegalin OS=Homo sapiens OX=960 | PDE4DIP | 265.103 | 2 | 47.457684 | yes | 4.081E-07 | 563.77606 | 44.743046 |
| TLQVELEGAQVLR | 13 | Q5VU43 | Myomegalin OS=Homo sapiens OX=960 | PDE4DIP | 265.103 | 2 | 47.457684 | yes | 2.346E-07 | 728.41193 | 40.396114 |
| IQQTEATNK | 9 | Q5VU43 | Myomegalin OS=Homo sapiens OX=960 | PDE4DIP | 265.103 | 2 | 47.457684 | yes | 9.459E-05 | 516.76965 | 23.474869 |
| FLDPEFR | 7 | Q5VUA4 | Zinc finger protein 318 OS=Homo sapien | ZNF318 | 251.113 | 2 | 54.551445 | yes | 1.992E-07 | 462.23471 | 274.6181 |

| | | | | | | | | | | |
|---|---|---|---|---|---|---|---|---|---|---|
| ILSMLADSTSTQEI | 14 | Q5VUA4 | Zinc finger protein 318 OS=Homo sapien | ZNF318 | 251.113 | 2 | 54.551445 | yes | 9.722E-08 | 762.3847 | 230.0788 |
| PAPLNTFLSIK | 11 | Q5VUA4 | Zinc finger protein 318 OS=Homo sapien | ZNF318 | 251.113 | 2 | 54.551445 | yes | 2.229E-07 | 600.85297 | 223.0912 |
| AFGGEEVILK | 10 | Q5VUA4 | Zinc finger protein 318 OS=Homo sapien | ZNF318 | 251.113 | 2 | 54.551445 | yes | 2.895E-07 | 531.7951 | 173.56276 |
| TLVLDFETEGER | 12 | Q5VUA4 | Zinc finger protein 318 OS=Homo sapien | ZNF318 | 251.113 | 2 | 54.551445 | yes | 2.467E-07 | 704.85138 | 173.01427 |
| LVEIHLR | 7 | Q5VUA4 | Zinc finger protein 318 OS=Homo sapien | ZNF318 | 251.113 | 2 | 54.551445 | yes | 1.167E-07 | 440.27417 | 166.52902 |
| ASSLPSSAPAVK | 12 | Q5VUA4 | Zinc finger protein 318 OS=Homo sapien | ZNF318 | 251.113 | 2 | 54.551445 | yes | 2.982E-07 | 557.80878 | 139.37981 |
| QAGLAVVLETER | 12 | Q5VUA4 | Zinc finger protein 318 OS=Homo sapien | ZNF318 | 251.113 | 2 | 54.551445 | yes | 1.66E-07 | 643.35913 | 124.3745 |
| ESSADLLLPPDIISI | 15 | Q5VUA4 | Zinc finger protein 318 OS=Homo sapien | ZNF318 | 251.113 | 2 | 54.551445 | yes | 6.451E-07 | 799.43781 | 121.09691 |
| LDSFLSEAR | 9 | Q5VUA4 | Zinc finger protein 318 OS=Homo sapien | ZNF318 | 251.113 | 2 | 54.551445 | yes | 5.843E-07 | 519.26672 | 120.20592 |
| TGLTENVDR | 9 | Q5VUA4 | Zinc finger protein 318 OS=Homo sapien | ZNF318 | 251.113 | 2 | 54.551445 | yes | 2.303E-07 | 502.754 | 119.57761 |
| YVDENPLYEER | 11 | Q5VUA4 | Zinc finger protein 318 OS=Homo sapien | ZNF318 | 251.113 | 2 | 54.551445 | yes | 1.054E-07 | 713.82788 | 118.1814 |
| LESLEETNPEYAK | 13 | Q5VUA4 | Zinc finger protein 318 OS=Homo sapien | ZNF318 | 251.113 | 2 | 54.551445 | yes | 9.893E-08 | 761.86719 | 108.04243 |
| ESEIETNTELK | 11 | Q5VUA4 | Zinc finger protein 318 OS=Homo sapien | ZNF318 | 251.113 | 2 | 54.551445 | yes | 2.413E-06 | 646.81445 | 98.386589 |
| SFPDIEDEEK | 10 | Q5VUA4 | Zinc finger protein 318 OS=Homo sapien | ZNF318 | 251.113 | 2 | 54.551445 | yes | 7.08E-07 | 604.76953 | 90.095444 |
| ALEDFEATDLK | 11 | Q5VUA4 | Zinc finger protein 318 OS=Homo sapien | ZNF318 | 251.113 | 2 | 54.551445 | yes | 9.088E-06 | 626.30878 | 90.000069 |
| ASQDGSGFSR | 10 | Q5VUA4 | Zinc finger protein 318 OS=Homo sapien | ZNF318 | 251.113 | 2 | 54.551445 | yes | 3.742E-07 | 506.22815 | 80.189415 |
| LSPQACSFTK | 10 | Q5VUA4 | Zinc finger protein 318 OS=Homo sapien | ZNF318 | 251.113 | 2 | 54.551445 | yes | 7.804E-07 | 569.78168 | 71.744156 |
| QESLGLELK | 9 | Q5VUA4 | Zinc finger protein 318 OS=Homo sapien | ZNF318 | 251.113 | 2 | 54.551445 | yes | 9.131E-07 | 508.78476 | 70.179947 |
| DLYDIFYSSGGK | 12 | Q5VUA4 | Zinc finger protein 318 OS=Homo sapien | ZNF318 | 251.113 | 2 | 54.551445 | yes | 9.722E-08 | 682.82208 | 70.061043 |
| HTDFLLPHER | 10 | Q5VUA4 | Zinc finger protein 318 OS=Homo sapien | ZNF318 | 251.113 | 2 | 54.551445 | yes | 1.02E-07 | 632.82526 | 63.055248 |
| WSVVEHVGPK | 10 | Q5VUA4 | Zinc finger protein 318 OS=Homo sapien | ZNF318 | 251.113 | 2 | 54.551445 | yes | 2.774E-06 | 569.30621 | 54.143013 |
| STGSTYGFLQPLTI | 14 | Q5VUA4 | Zinc finger protein 318 OS=Homo sapien | ZNF318 | 251.113 | 2 | 54.551445 | yes | 1.73E-06 | 764.39374 | 49.103676 |
| LEDQLSEGR | 9 | Q5VUA4 | Zinc finger protein 318 OS=Homo sapien | ZNF318 | 251.113 | 2 | 54.551445 | yes | 2.359E-06 | 523.75928 | 44.853054 |
| SSGTTAKPLPVVK | 13 | Q5VUA4 | Zinc finger protein 318 OS=Homo sapien | ZNF318 | 251.113 | 2 | 54.551445 | yes | 1.034E-07 | 642.8797 | 43.898064 |
| IVLAGCVPQAQPR | 13 | Q5VV42 | Threonylcarbamoyladenosine tRNA meth | CDKAL1 | 65.112 | 2 | 50.60784 | yes | 9.742E-08 | 704.89008 | 216.46896 |
| GNLASYPIDELVDI | 14 | Q5VV42 | Threonylcarbamoyladenosine tRNA meth | CDKAL1 | 65.112 | 2 | 50.60784 | yes | 4.656E-07 | 781.39648 | 182.20934 |
| LVEVIPEGAMLR | 12 | Q5VV42 | Threonylcarbamoyladenosine tRNA meth | CDKAL1 | 65.112 | 2 | 50.60784 | yes | 9.927E-08 | 663.87616 | 177.08275 |
| NPAEDHFR | 8 | Q5VV42 | Threonylcarbamoyladenosine tRNA meth | CDKAL1 | 65.112 | 2 | 50.60784 | yes | 6.685E-07 | 493.22794 | 79.10482 |
| VVEVVEETIK | 10 | Q5VV42 | Threonylcarbamoyladenosine tRNA meth | CDKAL1 | 65.112 | 2 | 50.60784 | yes | 3.697E-06 | 572.8266 | 59.249367 |
| GLSIIGVQQIDR | 12 | Q5VV42 | Threonylcarbamoyladenosine tRNA meth | CDKAL1 | 65.112 | 2 | 50.60784 | yes | 1.639E-06 | 649.875 | 53.387066 |
| MVEVDIYESGK | 11 | Q5VV42 | Threonylcarbamoyladenosine tRNA meth | CDKAL1 | 65.112 | 2 | 50.60784 | yes | 5.457E-06 | 635.30518 | 49.875793 |
| MEQVPAQVK | 9 | Q5VV42 | Threonylcarbamoyladenosine tRNA meth | CDKAL1 | 65.112 | 2 | 50.60784 | yes | 3.153E-06 | 515.2735 | 46.175575 |
| IIQLLGTTPR | 10 | Q5VW36 | Focadhesin OS=Homo sapiens OX=9606 | FOCAD | 200.059 | 2 | 54.173508 | yes | 2.285E-07 | 556.34534 | 296.55539 |
| LYLVSQGR | 8 | Q5VW36 | Focadhesin OS=Homo sapiens OX=9606 | FOCAD | 200.059 | 2 | 54.173508 | yes | 1.016E-06 | 468.26907 | 157.62183 |
| VHVAALLR | 8 | Q5VW36 | Focadhesin OS=Homo sapiens OX=9606 | FOCAD | 200.059 | 2 | 54.173508 | yes | 9.722E-08 | 439.78217 | 145.31206 |
| AYHAILQGR | 9 | Q5VW36 | Focadhesin OS=Homo sapiens OX=9606 | FOCAD | 200.059 | 2 | 54.173508 | yes | 9.809E-08 | 514.78546 | 133.743 |
| GQNLHQILK | 9 | Q5VW36 | Focadhesin OS=Homo sapiens OX=9606 | FOCAD | 200.059 | 2 | 54.173508 | yes | 1.204E-07 | 525.80634 | 131.91574 |
| CLLEMTDDDANR | 12 | Q5VW36 | Focadhesin OS=Homo sapiens OX=9606 | FOCAD | 200.059 | 2 | 54.173508 | yes | 1.385E-07 | 726.80829 | 127.20345 |
| TVAGIPNFILK | 11 | Q5VW36 | Focadhesin OS=Homo sapiens OX=9606 | FOCAD | 200.059 | 2 | 54.173508 | yes | 2.266E-07 | 586.85553 | 120.32588 |

| | | | | | | | | | | | |
|---|---|---|---|---|---|---|---|---|---|---|---|
| DMLTDEITK | 9 | Q5VW36 | Focadhesin OS=Homo sapiens OX=9606 | FOCAD | 200.059 | 2 | 54.173508 | yes | 6.865E-05 | 533.26025 | 104.95728 |
| LGELELQLK | 9 | Q5VW36 | Focadhesin OS=Homo sapiens OX=9606 | FOCAD | 200.059 | 2 | 54.173508 | yes | 8.398E-07 | 521.81079 | 104.33358 |
| VYPELQR | 7 | Q5VW36 | Focadhesin OS=Homo sapiens OX=9606 | FOCAD | 200.059 | 2 | 54.173508 | yes | 4.095E-07 | 452.74799 | 102.33432 |
| LLTSLWEK | 8 | Q5VW36 | Focadhesin OS=Homo sapiens OX=9606 | FOCAD | 200.059 | 2 | 54.173508 | yes | 2.067E-07 | 495.28696 | 102.31855 |
| SYSGENTASAIAR | 13 | Q5VW36 | Focadhesin OS=Homo sapiens OX=9606 | FOCAD | 200.059 | 2 | 54.173508 | yes | 2.704E-07 | 663.81787 | 91.949448 |
| FMAVSDVPSLSVC | 14 | Q5VW36 | Focadhesin OS=Homo sapiens OX=9606 | FOCAD | 200.059 | 2 | 54.173508 | yes | 4.241E-07 | 718.87634 | 90.434258 |
| LLSVTEDQK | 9 | Q5VW36 | Focadhesin OS=Homo sapiens OX=9606 | FOCAD | 200.059 | 2 | 54.173508 | yes | 9.057E-07 | 516.78223 | 77.913544 |
| IVSHANTGVLK | 11 | Q5VW36 | Focadhesin OS=Homo sapiens OX=9606 | FOCAD | 200.059 | 2 | 54.173508 | yes | 6.219E-07 | 569.83258 | 64.160652 |
| SEAIQTSHFQGR | 12 | Q5VW36 | Focadhesin OS=Homo sapiens OX=9606 | FOCAD | 200.059 | 2 | 54.173508 | yes | 9.722E-08 | 680.83386 | 62.739586 |
| STWNALSPK | 9 | Q5VW36 | Focadhesin OS=Homo sapiens OX=9606 | FOCAD | 200.059 | 2 | 54.173508 | yes | 3.588E-07 | 502.26401 | 49.997238 |
| LSDISGQEMNLLLI | 15 | Q5VW36 | Focadhesin OS=Homo sapiens OX=9606 | FOCAD | 200.059 | 2 | 54.173508 | yes | 6.621E-07 | 846.43896 | 29.110966 |
| VDILDAAVQTK | 11 | Q5VWN6 | Protein TASOR 2 OS=Homo sapiens OX | TASOR2 | 268.843 | 2 | 39.6165 | yes | 4.48E-07 | 586.82971 | 74.712654 |
| LDLEDSYTLR | 10 | Q5VWN6 | Protein TASOR 2 OS=Homo sapiens OX | TASOR2 | 268.843 | 2 | 39.6165 | yes | 4.797E-07 | 612.80896 | 44.04549 |
| LPSGFDLIPPAEK | 13 | Q5VWN6 | Protein TASOR 2 OS=Homo sapiens OX | TASOR2 | 268.843 | 2 | 39.6165 | yes | 4.465E-06 | 692.37958 | 35.321026 |
| FAVLLTDKPTIPR | 13 | Q5VWN6 | Protein TASOR 2 OS=Homo sapiens OX | TASOR2 | 268.843 | 3 | 39.6165 | yes | 8.14E-07 | 490.96082 | 11.525296 |
| YLQDALGEFIK | 11 | Q5VWQ8 | Disabled homolog 2-interacting protein O | DAB2IP | 131.626 | 2 | 49.299637 | yes | 1.537E-07 | 648.84534 | 128.42181 |
| ATPVNLAGLATVF | 13 | Q5VWQ8 | Disabled homolog 2-interacting protein O | DAB2IP | 131.626 | 2 | 49.299637 | yes | 1.521E-07 | 641.8775 | 123.37296 |
| NSYLGLVSLPAAS | 17 | Q5VWQ8 | Disabled homolog 2-interacting protein O | DAB2IP | 131.626 | 2 | 49.299637 | yes | 1.047E-07 | 837.96234 | 97.352623 |
| LGSFSTAAEELAR | 13 | Q5VWQ8 | Disabled homolog 2-interacting protein O | DAB2IP | 131.626 | 2 | 49.299637 | yes | 1.08E-06 | 676.34625 | 91.743896 |
| KLEEYETLFK | 10 | Q5VWQ8 | Disabled homolog 2-interacting protein O | DAB2IP | 131.626 | 2 | 49.299637 | yes | 2.602E-07 | 650.34521 | 80.098053 |
| WYPVVTPNPK | 10 | Q5VWQ8 | Disabled homolog 2-interacting protein O | DAB2IP | 131.626 | 2 | 49.299637 | yes | 2.126E-07 | 600.82422 | 43.047924 |
| GGISNVWFDR | 10 | Q5VWZ2 | Lysophospholipase-like protein 1 OS=Ho | LYPLAL1 | 26.317 | 2 | 45.886063 | yes | 1.714E-07 | 575.78564 | 69.911751 |
| VMELLVHLNK | 10 | Q5VYK3 | Proteasome adapter and scaffold protein I | ECPAS | 204.291 | 2 | 43.236034 | yes | 2.673E-07 | 598.34698 | 24.069988 |
| FYTEEFDFK | 9 | Q5VYS8 | Terminal uridylyltransferase 7 OS=Homo | TUT7 | 171.23 | 2 | 53.762115 | no | 1.291E-07 | 613.27423 | 224.61736 |
| LPDCSLR | 7 | Q5VYS8 | Terminal uridylyltransferase 7 OS=Homo | TUT7 | 171.23 | 2 | 53.762115 | yes | 4.598E-07 | 430.71838 | 195.86148 |
| IAIEDPYSVK | 10 | Q5VYS8 | Terminal uridylyltransferase 7 OS=Homo | TUT7 | 171.23 | 2 | 53.762115 | yes | 3.463E-07 | 567.80573 | 182.10492 |
| ELLGLQQAEER | 11 | Q5VYS8 | Terminal uridylyltransferase 7 OS=Homo | TUT7 | 171.23 | 2 | 53.762115 | yes | 1.026E-07 | 643.34094 | 180.01495 |
| DFSPTIIEDQAR | 12 | Q5VYS8 | Terminal uridylyltransferase 7 OS=Homo | TUT7 | 171.23 | 2 | 53.762115 | yes | 2.703E-07 | 696.34369 | 170.25819 |
| IQLEPLPPLTPK | 12 | Q5VYS8 | Terminal uridylyltransferase 7 OS=Homo | TUT7 | 171.23 | 2 | 53.762115 | yes | 2.894E-07 | 673.40814 | 169.48828 |
| SSLLKPLNAITCISI | 18 | Q5VYS8 | Terminal uridylyltransferase 7 OS=Homo | TUT7 | 171.23 | 3 | 53.762115 | yes | 9.81E-08 | 666.69446 | 129.70663 |
| LCDVLIESIAFAHK | 14 | Q5VYS8 | Terminal uridylyltransferase 7 OS=Homo | TUT7 | 171.23 | 3 | 53.762115 | yes | 1.618E-07 | 539.28857 | 63.583038 |
| LCDVLIESIAFAHK | 14 | Q5VYS8 | Terminal uridylyltransferase 7 OS=Homo | TUT7 | 171.23 | 3 | 53.762115 | yes | 1.063E-07 | 808.42926 | 63.583038 |
| LCDVLIESIAFAHK | 14 | Q5VYS8 | Terminal uridylyltransferase 7 OS=Homo | TUT7 | 171.23 | 2 | 53.762115 | yes | 1.618E-07 | 539.28857 | 63.583038 |
| LCDVLIESIAFAHK | 14 | Q5VYS8 | Terminal uridylyltransferase 7 OS=Homo | TUT7 | 171.23 | 2 | 53.762115 | yes | 1.063E-07 | 808.42926 | 63.583038 |
| VPVVVCR | 7 | Q5VYS8 | Terminal uridylyltransferase 7 OS=Homo | TUT7 | 171.23 | 2 | 53.762115 | no | 9.722E-08 | 414.74164 | 126.84878 |
| IMENVFQHK | 9 | Q5VYS8 | Terminal uridylyltransferase 7 OS=Homo | TUT7 | 171.23 | 2 | 53.762115 | yes | 9.723E-08 | 573.29224 | 118.25246 |
| NPPVIPVLQEIYK | 13 | Q5VYS8 | Terminal uridylyltransferase 7 OS=Homo | TUT7 | 171.23 | 2 | 53.762115 | yes | 1.127E-07 | 755.43744 | 108.58081 |
| EHVISIR | 7 | Q5VYS8 | Terminal uridylyltransferase 7 OS=Homo | TUT7 | 171.23 | 2 | 53.762115 | yes | 9.722E-08 | 427.24814 | 98.579659 |
| SPTVVCSLCK | 10 | Q5VYS8 | Terminal uridylyltransferase 7 OS=Homo | TUT7 | 171.23 | 2 | 53.762115 | yes | 8.487E-07 | 575.78339 | 98.026825 |

| | | | | | | | | | | | |
|---|---|---|---|---|---|---|---|---|---|---|---|
| DQEDALNQR | 9 | Q5VYS8 | Terminal uridylyltransferase 7 OS=Homo | TUT7 | 171.23 | 2 | 53.762115 | yes | 7.862E-07 | 544.75201 | 96.605774 |
| GQVSLILDVK | 10 | Q5VYS8 | Terminal uridylyltransferase 7 OS=Homo | TUT7 | 171.23 | 2 | 53.762115 | yes | 9.722E-08 | 536.32404 | 89.105232 |
| RVFGIPVK | 8 | Q5VYS8 | Terminal uridylyltransferase 7 OS=Homo | TUT7 | 171.23 | 2 | 53.762115 | yes | 7.946E-07 | 458.29236 | 81.58461 |
| EVINHHPDVQTK | 12 | Q5VYS8 | Terminal uridylyltransferase 7 OS=Homo | TUT7 | 171.23 | 3 | 53.762115 | yes | 2.322E-07 | 472.91251 | 77.975449 |
| LGNFNLQDIEK | 11 | Q5VYS8 | Terminal uridylyltransferase 7 OS=Homo | TUT7 | 171.23 | 2 | 53.762115 | yes | 9.903E-08 | 645.83807 | 67.058113 |
| LYGSSCSR | 8 | Q5VYS8 | Terminal uridylyltransferase 7 OS=Homo | TUT7 | 171.23 | 2 | 53.762115 | yes | 2.261E-06 | 465.21091 | 66.339897 |
| EDKPIQCTPQK | 11 | Q5VYS8 | Terminal uridylyltransferase 7 OS=Homo | TUT7 | 171.23 | 2 | 53.762115 | yes | 3.456E-07 | 672.33484 | 52.74007 |
| VSAGNENACLTTK | 13 | Q5VYS8 | Terminal uridylyltransferase 7 OS=Homo | TUT7 | 171.23 | 2 | 53.762115 | yes | 1.328E-07 | 682.82739 | 44.572811 |
| MCDIGDASR | 9 | Q5VYS8 | Terminal uridylyltransferase 7 OS=Homo | TUT7 | 171.23 | 2 | 53.762115 | yes | 4.565E-07 | 512.71295 | 32.61557 |
| DVVIWEHTDSAAC | 19 | Q5VYS8 | Terminal uridylyltransferase 7 OS=Homo | TUT7 | 171.23 | 3 | 53.762115 | yes | 3.956E-07 | 672.32874 | 28.921013 |
| GPCAVSSNPYAFK | 13 | Q5VYS8 | Terminal uridylyltransferase 7 OS=Homo | TUT7 | 171.23 | 2 | 53.762115 | yes | 5.871E-07 | 699.32953 | 24.005079 |
| ALLVELNK | 8 | Q5VYS8 | Terminal uridylyltransferase 7 OS=Homo | TUT7 | 171.23 | 2 | 53.762115 | yes | 2.92E-06 | 450.28165 | 16.184336 |
| LIPGCEVILATPYG | 15 | Q5VZ89 | DENN domain-containing protein 4C OS | DENND4C | 212.711 | 2 | 44.445847 | yes | 2.191E-07 | 829.95038 | 66.132774 |
| DTGLAFFDDCIEK | 13 | Q5VZ89 | DENN domain-containing protein 4C OS | DENND4C | 212.711 | 2 | 44.445847 | yes | 4.194E-07 | 765.84265 | 26.333611 |
| LGELLHDK | 8 | Q5VZE5 | N-alpha-acetyltransferase 35, NatC auxili | NAA35 | 83.639 | 2 | 46.726078 | yes | 2.301E-07 | 462.76108 | 99.512932 |
| TMVAFDMDGK | 10 | Q5VZE5 | N-alpha-acetyltransferase 35, NatC auxili | NAA35 | 83.639 | 2 | 46.726078 | yes | 1.604E-07 | 557.74902 | 39.407269 |
| SLLQTTFLVDNK | 12 | Q5VZE5 | N-alpha-acetyltransferase 35, NatC auxili | NAA35 | 83.639 | 2 | 46.726078 | yes | 1.486E-07 | 689.88245 | 36.160957 |
| YLHPPTHLK | 9 | Q5VZF2 | Muscleblind-like protein 2 OS=Homo sap | MBNL2 | 40.518 | 2 | 51.36519 | yes | 1.164E-07 | 553.31128 | 388.2713 |
| TDKLEVCR | 8 | Q5VZF2 | Muscleblind-like protein 2 OS=Homo sap | MBNL2 | 40.518 | 2 | 51.36519 | yes | 1.042E-07 | 510.76077 | 348.22714 |
| AAQHQANQAAVA | 35 | Q5VZF2 | Muscleblind-like protein 2 OS=Homo sap | MBNL2 | 40.518 | 4 | 51.36519 | yes | 1.156E-07 | 841.19513 | 98.392227 |
| NGMECQESALR | 11 | Q5VZF2 | Muscleblind-like protein 2 OS=Homo sap | MBNL2 | 40.518 | 2 | 51.36519 | yes | 9.722E-08 | 647.77936 | 43.670052 |
| SSGFISELPSEEGK | 14 | Q5VZK9 | F-actin-uncapping protein LRRC16A OS | CARMIL1 | 151.557 | 2 | 46.828274 | yes | 3.184E-07 | 733.85413 | 100.79114 |
| VLVLTSCR | 8 | Q5VZK9 | F-actin-uncapping protein LRRC16A OS | CARMIL1 | 151.557 | 2 | 46.828274 | yes | 5.985E-07 | 474.27078 | 71.106094 |
| SIITVEELTEIER | 13 | Q5VZK9 | F-actin-uncapping protein LRRC16A OS | CARMIL1 | 151.557 | 2 | 46.828274 | yes | 1.462E-07 | 766.41431 | 67.813919 |
| LEELVLENAGLR | 12 | Q5VZK9 | F-actin-uncapping protein LRRC16A OS | CARMIL1 | 151.557 | 2 | 46.828274 | yes | 6.222E-07 | 678.38007 | 66.651665 |
| TLPQQESLEIELAE | 19 | Q5VZK9 | F-actin-uncapping protein LRRC16A OS | CARMIL1 | 151.557 | 3 | 46.828274 | yes | 5.492E-07 | 727.7265 | 47.963165 |
| GVSLDLSNCELR | 12 | Q5VZK9 | F-actin-uncapping protein LRRC16A OS | CARMIL1 | 151.557 | 2 | 46.828274 | yes | 8.914E-06 | 681.83771 | 46.576473 |
| MDLVQEDQR | 9 | Q5VZM2 | Ras-related GTP-binding protein B OS=H | RRAGB | 43.251 | 2 | 53.67933 | yes | 3.328E-06 | 567.26642 | 55.083664 |
| LAASFQSMEVR | 11 | Q5VZM2 | Ras-related GTP-binding protein B OS=H | RRAGB | 43.251 | 2 | 53.67933 | yes | 9.722E-08 | 619.81354 | 29.113171 |
| NLSLESQIK | 9 | Q5W0B1 | ORC ubiquitin ligase 1 OS=Homo sapien | OBI1 | 81.117 | 2 | 54.320618 | yes | 4.796E-06 | 516.29022 | 224.70149 |
| VPITPENPCK | 10 | Q5W0B1 | ORC ubiquitin ligase 1 OS=Homo sapien | OBI1 | 81.117 | 2 | 54.320618 | yes | 2.427E-07 | 577.79736 | 153.80077 |
| EAMNSICQTALSA | 16 | Q5W0B1 | ORC ubiquitin ligase 1 OS=Homo sapien | OBI1 | 81.117 | 2 | 54.320618 | yes | 1.204E-07 | 848.38745 | 141.00029 |
| FAVAALQSK | 9 | Q5W0B1 | ORC ubiquitin ligase 1 OS=Homo sapien | OBI1 | 81.117 | 2 | 54.320618 | yes | 1.984E-07 | 467.77145 | 119.32097 |
| ISELDSMMSESDN | 15 | Q5W0B1 | ORC ubiquitin ligase 1 OS=Homo sapien | OBI1 | 81.117 | 2 | 54.320618 | yes | 9.722E-08 | 836.85565 | 108.38594 |
| EYEDEIDCLQK | 11 | Q5W0B1 | ORC ubiquitin ligase 1 OS=Homo sapien | OBI1 | 81.117 | 2 | 54.320618 | yes | 9.722E-08 | 721.31122 | 108.36859 |
| TSSEASMDAAYLD | 14 | Q5W0B1 | ORC ubiquitin ligase 1 OS=Homo sapien | OBI1 | 81.117 | 2 | 54.320618 | yes | 9.722E-08 | 744.82977 | 97.892303 |
| HLVTDNPSK | 9 | Q5W0B1 | ORC ubiquitin ligase 1 OS=Homo sapien | OBI1 | 81.117 | 2 | 54.320618 | yes | 3.4E-06 | 505.76691 | 96.063629 |
| YIEELESQVAQLK | 13 | Q5W0B1 | ORC ubiquitin ligase 1 OS=Homo sapien | OBI1 | 81.117 | 2 | 54.320618 | yes | 1.332E-07 | 775.40906 | 59.280445 |
| IQSSLSSASPSK | 12 | Q5W0B1 | ORC ubiquitin ligase 1 OS=Homo sapien | OBI1 | 81.117 | 2 | 54.320618 | yes | 1.415E-07 | 596.31439 | 57.681728 |

| | | | | | | | | | | | |
|---|---|---|---|---|---|---|---|---|---|---|---|
| KSECFSSPK | 9 | Q5W0B1 | ORC ubiquitin ligase 1 OS=Homo sapien | OBI1 | 81.117 | 2 | 54.320618 | yes | 5.86E-07 | 535.25275 | 51.321346 |
| AAALSGGGGPGA( | 16 | Q5XG87 | Terminal nucleotidyltransferase 4A OS=I | TENT4A | 84.651 | 2 | 51.437405 | yes | 9.722E-08 | 669.34967 | 265.15646 |
| ASTYGLNYLLSGS | 14 | Q5XG87 | Terminal nucleotidyltransferase 4A OS=I | TENT4A | 84.651 | 2 | 51.437405 | yes | 5.003E-07 | 751.38593 | 129.78438 |
| LTDQETEVK | 9 | Q5XG87 | Terminal nucleotidyltransferase 4A OS=I | TENT4A | 84.651 | 2 | 51.437405 | yes | 1.685E-07 | 531.76929 | 90.145813 |
| VTQEVIDYR | 9 | Q5XG87 | Terminal nucleotidyltransferase 4A OS=I | TENT4A | 84.651 | 2 | 51.437405 | yes | 2.582E-07 | 561.79309 | 82.309044 |
| EPESPYGQR | 9 | Q5XG87 | Terminal nucleotidyltransferase 4A OS=I | TENT4A | 84.651 | 2 | 51.437405 | yes | 8.293E-07 | 531.74615 | 71.69368 |
| VDISFNMETGVR | 12 | Q5XG87 | Terminal nucleotidyltransferase 4A OS=I | TENT4A | 84.651 | 2 | 51.437405 | yes | 1.11E-07 | 684.33484 | 51.846767 |
| LLQDISEAR | 9 | Q63HN8 | E3 ubiquitin-protein ligase RNF213 OS= | RNF213 | 591.408 | 2 | 52.815144 | yes | 3.286E-07 | 522.78784 | 147.34727 |
| LIAVADSVLTK | 11 | Q63HN8 | E3 ubiquitin-protein ligase RNF213 OS= | RNF213 | 591.408 | 2 | 52.815144 | yes | 1.219E-06 | 565.34497 | 146.17203 |
| LLSLVDSAGQR | 11 | Q63HN8 | E3 ubiquitin-protein ligase RNF213 OS= | RNF213 | 591.408 | 2 | 52.815144 | yes | 9.722E-08 | 579.82751 | 133.75536 |
| LLNFDTELSTK | 11 | Q63HN8 | E3 ubiquitin-protein ligase RNF213 OS= | RNF213 | 591.408 | 2 | 52.815144 | yes | 9.722E-08 | 640.84027 | 125.33045 |
| FATSLVDNSVPLL | 14 | Q63HN8 | E3 ubiquitin-protein ligase RNF213 OS= | RNF213 | 591.408 | 2 | 52.815144 | yes | 9.81E-08 | 766.42755 | 121.09775 |
| YVDLGLGTHR | 10 | Q63HN8 | E3 ubiquitin-protein ligase RNF213 OS= | RNF213 | 591.408 | 2 | 52.815144 | yes | 9.722E-08 | 565.80127 | 109.64844 |
| ILANVTEDAK | 10 | Q63HN8 | E3 ubiquitin-protein ligase RNF213 OS= | RNF213 | 591.408 | 2 | 52.815144 | yes | 2.935E-07 | 537.29547 | 89.328613 |
| AADFLSEPEGGPE | 16 | Q63HN8 | E3 ubiquitin-protein ligase RNF213 OS= | RNF213 | 591.408 | 2 | 52.815144 | yes | 1.226E-07 | 824.87982 | 88.800163 |
| VLVEGIVCISK | 11 | Q63HN8 | E3 ubiquitin-protein ligase RNF213 OS= | RNF213 | 591.408 | 2 | 52.815144 | yes | 1.183E-07 | 608.85211 | 84.938057 |
| VITEVLCASQGFM | 14 | Q63HN8 | E3 ubiquitin-protein ligase RNF213 OS= | RNF213 | 591.408 | 2 | 52.815144 | yes | 1.209E-07 | 805.90509 | 80.217026 |
| LLLDEITR | 8 | Q63HN8 | E3 ubiquitin-protein ligase RNF213 OS= | RNF213 | 591.408 | 2 | 52.815144 | yes | 2.054E-06 | 486.78986 | 80.083275 |
| TEDAAQELLLPES | 14 | Q63HN8 | E3 ubiquitin-protein ligase RNF213 OS= | RNF213 | 591.408 | 2 | 52.815144 | yes | 5.718E-07 | 772.39612 | 76.202072 |
| LCVGIVASER | 10 | Q63HN8 | E3 ubiquitin-protein ligase RNF213 OS= | RNF213 | 591.408 | 2 | 52.815144 | yes | 1.283E-07 | 552.29755 | 71.978424 |
| STDFLPVDCPVR | 12 | Q63HN8 | E3 ubiquitin-protein ligase RNF213 OS= | RNF213 | 591.408 | 2 | 52.815144 | yes | 9.722E-08 | 703.34265 | 66.923866 |
| ETVQEFDLEK | 10 | Q63HN8 | E3 ubiquitin-protein ligase RNF213 OS= | RNF213 | 591.408 | 2 | 52.815144 | yes | 3.87E-07 | 619.30096 | 65.422691 |
| ESLLGDMEWR | 10 | Q63HN8 | E3 ubiquitin-protein ligase RNF213 OS= | RNF213 | 591.408 | 2 | 52.815144 | yes | 1.582E-07 | 618.28992 | 64.970848 |
| FVDIFQQTPLGR | 12 | Q63HN8 | E3 ubiquitin-protein ligase RNF213 OS= | RNF213 | 591.408 | 2 | 52.815144 | yes | 3.947E-06 | 710.88281 | 59.291824 |
| VGFVGISNWALDF | 15 | Q63HN8 | E3 ubiquitin-protein ligase RNF213 OS= | RNF213 | 591.408 | 2 | 52.815144 | yes | 2.383E-07 | 787.4223 | 56.523438 |
| ILIFQTDFEDGIR | 13 | Q63HN8 | E3 ubiquitin-protein ligase RNF213 OS= | RNF213 | 591.408 | 2 | 52.815144 | yes | 9.759E-08 | 783.91174 | 53.137138 |
| CGIPVIIMGETGCG | 15 | Q63HN8 | E3 ubiquitin-protein ligase RNF213 OS= | RNF213 | 591.408 | 2 | 52.815144 | yes | 3.778E-07 | 796.38568 | 52.764946 |
| DQEYTQLCR | 9 | Q63HN8 | E3 ubiquitin-protein ligase RNF213 OS= | RNF213 | 591.408 | 2 | 52.815144 | yes | 1.968E-07 | 606.76935 | 49.47776 |
| HVVPLPDGK | 9 | Q63HN8 | E3 ubiquitin-protein ligase RNF213 OS= | RNF213 | 591.408 | 2 | 52.815144 | yes | 1.139E-06 | 481.27692 | 48.794788 |
| ETLDQINQELIQAK | 14 | Q63HN8 | E3 ubiquitin-protein ligase RNF213 OS= | RNF213 | 591.408 | 2 | 52.815144 | yes | 9.722E-08 | 821.93597 | 45.172363 |
| EVIDMELSALR | 11 | Q63HN8 | E3 ubiquitin-protein ligase RNF213 OS= | RNF213 | 591.408 | 2 | 52.815144 | yes | 9.919E-08 | 638.33429 | 43.6003 |
| VMNHITDGPR | 10 | Q63HN8 | E3 ubiquitin-protein ligase RNF213 OS= | RNF213 | 591.408 | 2 | 52.815144 | yes | 1.318E-07 | 570.28491 | 32.378433 |
| FGIVLSAVITK | 11 | Q63HN8 | E3 ubiquitin-protein ligase RNF213 OS= | RNF213 | 591.408 | 2 | 52.815144 | yes | 5.036E-07 | 574.35791 | 25.393709 |
| ITVEYLQHIVEQK | 13 | Q63HN8 | E3 ubiquitin-protein ligase RNF213 OS= | RNF213 | 591.408 | 2 | 52.815144 | yes | 1.653E-07 | 800.44067 | 17.010223 |
| ALTEELHQK | 9 | Q63HN8 | E3 ubiquitin-protein ligase RNF213 OS= | RNF213 | 591.408 | 2 | 52.815144 | yes | 4.948E-07 | 534.78784 | 16.119881 |
| YLLVLTK | 7 | Q63HN8 | E3 ubiquitin-protein ligase RNF213 OS= | RNF213 | 591.408 | 2 | 52.815144 | yes | 3.889E-06 | 425.27585 | 8.5822315 |
| SPTSAPTQR | 9 | Q63HR2 | Tensin-2 OS=Homo sapiens OX=9606 G | TNS2 | 152.581 | 2 | 52.37645 | yes | 5.253E-07 | 472.74344 | 129.28639 |
| LSPGEALPPVSQA( | 17 | Q63HR2 | Tensin-2 OS=Homo sapiens OX=9606 G | TNS2 | 152.581 | 2 | 52.37645 | yes | 6.017E-07 | 804.93323 | 123.09966 |
| ISAGADQALATLT | 15 | Q63HR2 | Tensin-2 OS=Homo sapiens OX=9606 G | TNS2 | 152.581 | 2 | 52.37645 | yes | 9.722E-08 | 759.90088 | 103.69669 |

| | | | | | | | | | | | |
|---|---|---|---|---|---|---|---|---|---|---|---|
| LALPTAALYGLR | 12 | Q63HR2 | Tensin-2 OS=Homo sapiens OX=9606 G | TNS2 | 152.581 | 2 | 52.37645 | yes | 1.066E-07 | 629.87952 | 99.232475 |
| VTSACQALPPVEL | 14 | Q63HR2 | Tensin-2 OS=Homo sapiens OX=9606 G | TNS2 | 152.581 | 2 | 52.37645 | yes | 4.888E-07 | 770.91125 | 98.330521 |
| VTIDGTGVSCR | 11 | Q63HR2 | Tensin-2 OS=Homo sapiens OX=9606 G | TNS2 | 152.581 | 2 | 52.37645 | yes | 6.347E-07 | 582.78754 | 97.362129 |
| SFSLDPLMER | 10 | Q63HR2 | Tensin-2 OS=Homo sapiens OX=9606 G | TNS2 | 152.581 | 2 | 52.37645 | yes | 9.319E-07 | 597.7948 | 78.36702 |
| VATPPPSAQPWK | 12 | Q63HR2 | Tensin-2 OS=Homo sapiens OX=9606 G | TNS2 | 152.581 | 2 | 52.37645 | yes | 9.722E-08 | 639.8457 | 68.727058 |
| ILAAAFPARPDEQI | 14 | Q63HR2 | Tensin-2 OS=Homo sapiens OX=9606 G | TNS2 | 152.581 | 3 | 52.37645 | yes | 5.028E-07 | 518.95111 | 67.4963 |
| DQLDEAWTDER | 11 | Q63HR2 | Tensin-2 OS=Homo sapiens OX=9606 G | TNS2 | 152.581 | 2 | 52.37645 | yes | 9.904E-08 | 689.2995 | 63.75679 |
| PATGDFGYR | 9 | Q63HR2 | Tensin-2 OS=Homo sapiens OX=9606 G | TNS2 | 152.581 | 2 | 52.37645 | yes | 1.946E-07 | 492.2327 | 59.843861 |
| PTPAVVHFK | 9 | Q63HR2 | Tensin-2 OS=Homo sapiens OX=9606 G | TNS2 | 152.581 | 2 | 52.37645 | yes | 9.345E-07 | 498.28726 | 39.771549 |
| ASSAALSCSPR | 11 | Q63HR2 | Tensin-2 OS=Homo sapiens OX=9606 G | TNS2 | 152.581 | 2 | 52.37645 | yes | 3.635E-07 | 553.7666 | 39.54089 |
| DSHSFQGAYGLAL | 14 | Q63HR2 | Tensin-2 OS=Homo sapiens OX=9606 G | TNS2 | 152.581 | 2 | 52.37645 | yes | 3.579E-06 | 747.3728 | 19.095289 |
| AQSLEPYGTGLR | 12 | Q63ZY3 | KN motif and ankyrin repeat domain-cont | KANK2 | 91.175 | 2 | 51.85112 | yes | 3.329E-07 | 646.33569 | 169.85724 |
| VPTQEPTHR | 9 | Q63ZY3 | KN motif and ankyrin repeat domain-cont | KANK2 | 91.175 | 2 | 51.85112 | yes | 2.112E-07 | 532.77783 | 174.24701 |
| YLDNPNALTER | 11 | Q63ZY3 | KN motif and ankyrin repeat domain-cont | KANK2 | 91.175 | 2 | 51.85112 | yes | 4.48E-07 | 653.32532 | 112.95802 |
| ALQELQAAQAR | 11 | Q63ZY3 | KN motif and ankyrin repeat domain-cont | KANK2 | 91.175 | 2 | 51.85112 | yes | 9.722E-08 | 599.83057 | 101.35517 |
| VPSVAEAPQLR | 11 | Q63ZY3 | KN motif and ankyrin repeat domain-cont | KANK2 | 91.175 | 2 | 51.85112 | yes | 1.82E-07 | 583.83002 | 82.558762 |
| VAVLETQLK | 9 | Q63ZY3 | KN motif and ankyrin repeat domain-cont | KANK2 | 91.175 | 2 | 51.85112 | yes | 3.733E-07 | 500.80551 | 61.689838 |
| LPLAQLGR | 8 | Q64LD2 | WD repeat-containing protein 25 OS=Ho | WDR25 | 60.162 | 2 | 53.79872 | yes | 9.722E-08 | 434.27417 | 129.42982 |
| CEDCVVPYTPR | 11 | Q64LD2 | WD repeat-containing protein 25 OS=Ho | WDR25 | 60.162 | 2 | 53.79872 | yes | 4.054E-07 | 698.30524 | 117.83393 |
| EGSEFLSSTDASTR | 14 | Q64LD2 | WD repeat-containing protein 25 OS=Ho | WDR25 | 60.162 | 2 | 53.79872 | yes | 9.722E-08 | 743.83643 | 96.833992 |
| ISNQIFHER | 9 | Q64LD2 | WD repeat-containing protein 25 OS=Ho | WDR25 | 60.162 | 2 | 53.79872 | yes | 2.247E-07 | 572.29889 | 67.610153 |
| TIIAWDFR | 8 | Q64LD2 | WD repeat-containing protein 25 OS=Ho | WDR25 | 60.162 | 2 | 53.79872 | yes | 1.153E-06 | 511.27692 | 62.173141 |
| SHMLLSTSMDK | 11 | Q64LD2 | WD repeat-containing protein 25 OS=Ho | WDR25 | 60.162 | 2 | 53.79872 | yes | 0.0001121 | 625.29938 | 39.459129 |
| NDYLQYLDMK | 10 | Q659A1 | Little elongation complex subunit 2 OS=I | ICE2 | 110.012 | 2 | 50.28625 | yes | 9.819E-08 | 651.80536 | 53.453201 |
| KFDPVGEILK | 10 | Q659A1 | Little elongation complex subunit 2 OS=I | ICE2 | 110.012 | 2 | 50.28625 | yes | 1.376E-07 | 573.33191 | 52.475124 |
| WVPLHLDVVR | 10 | Q659C4 | La-related protein 1B OS=Homo sapiens | LARP1B | 105.323 | 2 | 50.02499 | yes | 1.215E-07 | 617.35876 | 246.3334 |
| WVPLHLDVVR | 10 | Q659C4 | La-related protein 1B OS=Homo sapiens | LARP1B | 105.323 | 2 | 50.02499 | yes | 4.481E-07 | 411.90826 | 246.3334 |
| WVPLHLDVVR | 10 | Q659C4 | La-related protein 1B OS=Homo sapiens | LARP1B | 105.323 | 3 | 50.02499 | yes | 1.215E-07 | 617.35876 | 246.3334 |
| WVPLHLDVVR | 10 | Q659C4 | La-related protein 1B OS=Homo sapiens | LARP1B | 105.323 | 3 | 50.02499 | yes | 4.481E-07 | 411.90826 | 246.3334 |
| VDPPISDEFGR | 11 | Q659C4 | La-related protein 1B OS=Homo sapiens | LARP1B | 105.323 | 2 | 50.02499 | yes | 1.301E-07 | 616.30133 | 217.41539 |
| DSTEVEIVDEK | 11 | Q659C4 | La-related protein 1B OS=Homo sapiens | LARP1B | 105.323 | 2 | 50.02499 | yes | 1.003E-07 | 632.30115 | 209.08012 |
| QIEYYFSVENLER | 13 | Q659C4 | La-related protein 1B OS=Homo sapiens | LARP1B | 105.323 | 2 | 50.02499 | yes | 1.735E-07 | 845.40961 | 158.88089 |
| ILIVTQTPPYVK | 12 | Q659C4 | La-related protein 1B OS=Homo sapiens | LARP1B | 105.323 | 2 | 50.02499 | yes | 1.154E-07 | 686.41595 | 92.763779 |
| LQEYLCSFK | 9 | Q659C4 | La-related protein 1B OS=Homo sapiens | LARP1B | 105.323 | 2 | 50.02499 | yes | 9.928E-08 | 594.29187 | 86.491531 |
| RDDQDDVSSVR | 11 | Q659C4 | La-related protein 1B OS=Homo sapiens | LARP1B | 105.323 | 2 | 50.02499 | yes | 4.424E-07 | 646.2973 | 55.80875 |
| NLLLDPASHK | 10 | Q66K74 | Microtubule-associated protein 1S OS=H | MAP1S | 112.212 | 2 | 56.65682 | yes | 2.605E-07 | 554.31146 | 141.16 |
| ALCYVISGQDQR | 12 | Q66K74 | Microtubule-associated protein 1S OS=H | MAP1S | 112.212 | 2 | 56.65682 | yes | 9.988E-08 | 705.3457 | 129.54417 |
| EPVVTPQDLEGPG | 14 | Q66K74 | Microtubule-associated protein 1S OS=H | MAP1S | 112.212 | 2 | 56.65682 | yes | 9.722E-08 | 747.38336 | 96.675064 |
| GDNLETLVLLNPS | 15 | Q66K74 | Microtubule-associated protein 1S OS=H | MAP1S | 112.212 | 2 | 56.65682 | yes | 9.722E-08 | 814.43048 | 73.561333 |

| | | | | | | | | | | | |
|---|---|---|---|---|---|---|---|---|---|---|---|
| HSATFSSIVK | 10 | Q66K74 | Microtubule-associated protein 1S OS=H | MAP1S | 112.212 | 2 | 56.65682 | yes | 2.446E-07 | 538.79041 | 44.166069 |
| SEVAAGGGSWDD | 13 | Q66K74 | Microtubule-associated protein 1S OS=H | MAP1S | 112.212 | 2 | 56.65682 | yes | 4.41E-07 | 653.78656 | 38.999329 |
| GDGEVLEEIVTK | 12 | Q66PJ3 | ADP-ribosylation factor-like protein 6-int | ARL6IP4 | 26.375 | 2 | 50.56461 | yes | 3.015E-07 | 644.83521 | 1208.7561 |
| KVVDPETGR | 9 | Q66PJ3 | ADP-ribosylation factor-like protein 6-int | ARL6IP4 | 26.375 | 2 | 50.56461 | yes | 1.037E-07 | 500.77472 | 355.79913 |
| EKAEAQQVEALPC | 21 | Q66PJ3 | ADP-ribosylation factor-like protein 6-int | ARL6IP4 | 26.375 | 4 | 50.56461 | yes | 3.975E-07 | 598.05286 | 142.50615 |
| GDCLAFQMR | 9 | Q66PJ3 | ADP-ribosylation factor-like protein 6-int | ARL6IP4 | 26.375 | 2 | 50.56461 | yes | 9.766E-08 | 549.24695 | 81.405785 |
| GGILTSTGFYK | 11 | Q68CP9 | AT-rich interactive domain-containing pr | ARID2 | 197.391 | 2 | 36.939857 | yes | 2.97E-06 | 572.30585 | 60.519672 |
| GMEILGNLCK | 10 | Q68CP9 | AT-rich interactive domain-containing pr | ARID2 | 197.391 | 2 | 36.939857 | yes | 6.457E-07 | 567.78595 | 42.170704 |
| MEAENLASVIDAR | 13 | Q68CQ4 | U3 small nucleolar RNA-associated prote | UTP25 | 87.056 | 2 | 52.398777 | yes | 1.534E-07 | 709.85083 | 269.49298 |
| IGVAILQR | 8 | Q68CQ4 | U3 small nucleolar RNA-associated prote | UTP25 | 87.056 | 2 | 52.398777 | yes | 2.945E-07 | 435.28198 | 186.5694 |
| FGVGDDDDFR | 10 | Q68CQ4 | U3 small nucleolar RNA-associated prote | UTP25 | 87.056 | 2 | 52.398777 | yes | 1.084E-07 | 571.74109 | 146.5412 |
| SSSPFTPLQK | 10 | Q68CQ4 | U3 small nucleolar RNA-associated prote | UTP25 | 87.056 | 2 | 52.398777 | yes | 2.818E-07 | 546.29022 | 96.110725 |
| NVHLFITGEK | 10 | Q68CQ4 | U3 small nucleolar RNA-associated prote | UTP25 | 87.056 | 2 | 52.398777 | yes | 9.722E-08 | 579.31934 | 95.715927 |
| DFGEEHPFYDR | 11 | Q68CQ4 | U3 small nucleolar RNA-associated prote | UTP25 | 87.056 | 2 | 52.398777 | yes | 9.722E-08 | 706.29932 | 94.462135 |
| GEEATWTCTVLYS | 14 | Q68CQ4 | U3 small nucleolar RNA-associated prote | UTP25 | 87.056 | 2 | 52.398777 | yes | 3.449E-07 | 822.88232 | 92.737923 |
| RPEDYEAVFVGNI | 18 | Q68CQ4 | U3 small nucleolar RNA-associated prote | UTP25 | 87.056 | 3 | 52.398777 | yes | 3.373E-07 | 727.0116 | 82.16024 |
| NVPMTGSISHVLV | 21 | Q68CQ4 | U3 small nucleolar RNA-associated prote | UTP25 | 87.056 | 3 | 52.398777 | yes | 1.49E-07 | 787.09497 | 77.765724 |
| YCVNMQGQVAVF | 12 | Q68CQ4 | U3 small nucleolar RNA-associated prote | UTP25 | 87.056 | 2 | 52.398777 | yes | 7.645E-07 | 712.84229 | 75.605858 |
| SQSQLLNTLTK | 11 | Q68CQ4 | U3 small nucleolar RNA-associated prote | UTP25 | 87.056 | 2 | 52.398777 | yes | 4.742E-07 | 616.84589 | 67.993523 |
| TNSQFLSGPQK | 11 | Q68CQ4 | U3 small nucleolar RNA-associated prote | UTP25 | 87.056 | 2 | 52.398777 | yes | 1.409E-07 | 603.80933 | 66.606674 |
| MWSLNNWSK | 9 | Q68CQ4 | U3 small nucleolar RNA-associated prote | UTP25 | 87.056 | 2 | 52.398777 | yes | 2.412E-07 | 583.27661 | 60.043945 |
| AAQMLQSNK | 9 | Q68CQ4 | U3 small nucleolar RNA-associated prote | UTP25 | 87.056 | 2 | 52.398777 | yes | 7.575E-06 | 495.75549 | 55.589233 |
| SLHLQKPLESTWT | 14 | Q68CQ4 | U3 small nucleolar RNA-associated prote | UTP25 | 87.056 | 3 | 52.398777 | yes | 3.246E-06 | 556.64203 | 41.201416 |
| DLFYPER | 7 | Q68CQ4 | U3 small nucleolar RNA-associated prote | UTP25 | 87.056 | 2 | 52.398777 | yes | 8.193E-06 | 470.23218 | 37.565811 |
| ELFLIMNSYR | 10 | Q68CQ4 | U3 small nucleolar RNA-associated prote | UTP25 | 87.056 | 2 | 52.398777 | yes | 6.861E-07 | 643.33411 | 28.254049 |
| VLIVVPFR | 8 | Q68CQ4 | U3 small nucleolar RNA-associated prote | UTP25 | 87.056 | 2 | 52.398777 | yes | 3.858E-06 | 471.81039 | 27.458227 |
| NLENVIQSQR | 10 | Q68CZ1 | Protein fantom OS=Homo sapiens OX=9( | RPGRIP1L | 151.202 | 2 | 43.546604 | yes | 3.708E-06 | 600.82019 | 94.680336 |
| LPPASSVSTLVLAI | 15 | Q68CZ1 | Protein fantom OS=Homo sapiens OX=9( | RPGRIP1L | 151.202 | 2 | 43.546604 | yes | 2.544E-07 | 754.44574 | 64.894409 |
| VNVPLISLAHDR | 12 | Q68CZ1 | Protein fantom OS=Homo sapiens OX=9( | RPGRIP1L | 151.202 | 2 | 43.546604 | yes | 2.358E-05 | 667.383 | 39.910431 |
| ENEIELSLLQLR | 12 | Q68CZ1 | Protein fantom OS=Homo sapiens OX=9( | RPGRIP1L | 151.202 | 2 | 43.546604 | yes | 2.114E-06 | 728.90393 | 33.588177 |
| IFCTASLIGTK | 11 | Q68CZ1 | Protein fantom OS=Homo sapiens OX=9( | RPGRIP1L | 151.202 | 2 | 43.546604 | yes | 1.854E-06 | 605.82867 | 25.907194 |
| ANENAGLQECPR | 12 | Q68CZ1 | Protein fantom OS=Homo sapiens OX=9( | RPGRIP1L | 151.202 | 2 | 43.546604 | yes | 6.443E-07 | 679.80951 | 15.057085 |
| ASEAASPLPDSPGI | 15 | Q68CZ2 | Tensin-3 OS=Homo sapiens OX=9606 G | TNS3 | 155.266 | 2 | 41.738083 | yes | 1.331E-06 | 721.34393 | 112.61644 |
| PMVSLLGSGR | 10 | Q68CZ2 | Tensin-3 OS=Homo sapiens OX=9606 G | TNS3 | 155.266 | 2 | 41.738083 | yes | 7.771E-07 | 508.78149 | 107.70544 |
| VATPPPSVLQLNK | 13 | Q68CZ2 | Tensin-3 OS=Homo sapiens OX=9606 G | TNS3 | 155.266 | 2 | 41.738083 | yes | 3.363E-07 | 682.40082 | 81.78627 |
| VELVFSATPEK | 11 | Q68CZ2 | Tensin-3 OS=Homo sapiens OX=9606 G | TNS3 | 155.266 | 2 | 41.738083 | yes | 5.09E-07 | 610.33209 | 62.382351 |
| PTGSPLSAEFSGTR | 14 | Q68CZ2 | Tensin-3 OS=Homo sapiens OX=9606 G | TNS3 | 155.266 | 2 | 41.738083 | yes | 3.569E-07 | 703.84918 | 43.793755 |
| CPADNPGLVQAQI | 14 | Q68CZ2 | Tensin-3 OS=Homo sapiens OX=9606 G | TNS3 | 155.266 | 2 | 41.738083 | yes | 3.5E-07 | 761.87518 | 37.83984 |
| ICIVIEPAQLLK | 12 | Q68CZ2 | Tensin-3 OS=Homo sapiens OX=9606 G | TNS3 | 155.266 | 2 | 41.738083 | yes | 0.0001158 | 698.91528 | 11.624566 |

| | | | | | | | | | | | |
|---|---|---|---|---|---|---|---|---|---|---|---|
| DIVENLETQSK | 11 | Q68CZ6 | HAUS augmin-like complex subunit 3 OS | HAUS3 | 69.651 | 2 | 45.032852 | yes | 1.243E-06 | 638.32501 | 45.079765 |
| ENAQLLNMPVVK | 12 | Q68CZ6 | HAUS augmin-like complex subunit 3 OS | HAUS3 | 69.651 | 2 | 45.032852 | yes | 1.986E-07 | 678.37122 | 41.350563 |
| WAEESLHSLTSK | 12 | Q68CZ6 | HAUS augmin-like complex subunit 3 OS | HAUS3 | 69.651 | 2 | 45.032852 | yes | 1.065E-06 | 694.34625 | 35.879986 |
| GDFDLQIAK | 9 | Q68CZ6 | HAUS augmin-like complex subunit 3 OS | HAUS3 | 69.651 | 2 | 45.032852 | yes | 3.187E-07 | 503.76382 | 33.496464 |
| CQLMASVTSHK | 11 | Q68CZ6 | HAUS augmin-like complex subunit 3 OS | HAUS3 | 69.651 | 2 | 45.032852 | yes | 3.217E-06 | 631.30499 | 31.946676 |
| SGKPILEGAALDEA | 16 | Q68CZ6 | HAUS augmin-like complex subunit 3 OS | HAUS3 | 69.651 | 2 | 45.032852 | yes | 1.033E-06 | 806.45123 | 14.556362 |
| GMPLPHAEK | 9 | Q68D10 | Protein SPT2 homolog OS=Homo sapiens | SPTY2D1 | 75.599 | 2 | 52.48447 | yes | 2.329E-07 | 490.25513 | 749.86871 |
| LGMQEDLEEMR | 11 | Q68D10 | Protein SPT2 homolog OS=Homo sapiens | SPTY2D1 | 75.599 | 2 | 52.48447 | yes | 1.149E-07 | 675.80505 | 504.56512 |
| YKDESDYALR | 10 | Q68D10 | Protein SPT2 homolog OS=Homo sapiens | SPTY2D1 | 75.599 | 2 | 52.48447 | yes | 2.411E-07 | 630.29877 | 401.164 |
| EIFGYDR | 7 | Q68D10 | Protein SPT2 homolog OS=Homo sapiens | SPTY2D1 | 75.599 | 2 | 52.48447 | yes | 4.089E-07 | 450.21652 | 400.29315 |
| CTVVSETISSK | 11 | Q68D10 | Protein SPT2 homolog OS=Homo sapiens | SPTY2D1 | 75.599 | 2 | 52.48447 | yes | 9.722E-08 | 605.80286 | 369.77902 |
| PISGSVSSAR | 10 | Q68D10 | Protein SPT2 homolog OS=Homo sapiens | SPTY2D1 | 75.599 | 2 | 52.48447 | yes | 1.286E-07 | 480.75909 | 361.33334 |
| SAPPPMNFTDLLR | 13 | Q68D10 | Protein SPT2 homolog OS=Homo sapiens | SPTY2D1 | 75.599 | 2 | 52.48447 | yes | 6.366E-07 | 729.87415 | 346.11777 |
| YSLAVGPPK | 9 | Q68D10 | Protein SPT2 homolog OS=Homo sapiens | SPTY2D1 | 75.599 | 2 | 52.48447 | yes | 9.728E-08 | 466.26602 | 331.48584 |
| LPFPTGYK | 8 | Q68D10 | Protein SPT2 homolog OS=Homo sapiens | SPTY2D1 | 75.599 | 2 | 52.48447 | yes | 2.438E-07 | 461.75528 | 320.26291 |
| GPGRPVSSPHELR | 13 | Q68D10 | Protein SPT2 homolog OS=Homo sapiens | SPTY2D1 | 75.599 | 3 | 52.48447 | yes | 1.358E-07 | 463.58463 | 119.3504 |
| GPGRPVSSPHELR | 13 | Q68D10 | Protein SPT2 homolog OS=Homo sapiens | SPTY2D1 | 75.599 | 3 | 52.48447 | yes | 9.218E-07 | 694.87329 | 119.3504 |
| GPGRPVSSPHELR | 13 | Q68D10 | Protein SPT2 homolog OS=Homo sapiens | SPTY2D1 | 75.599 | 2 | 52.48447 | yes | 1.358E-07 | 463.58463 | 119.3504 |
| GPGRPVSSPHELR | 13 | Q68D10 | Protein SPT2 homolog OS=Homo sapiens | SPTY2D1 | 75.599 | 2 | 52.48447 | yes | 9.218E-07 | 694.87329 | 119.3504 |
| PVSSPHELR | 9 | Q68D10 | Protein SPT2 homolog OS=Homo sapiens | SPTY2D1 | 75.599 | 2 | 52.48447 | yes | 7.038E-07 | 511.2749 | 192.41376 |
| TVSNSVPGRPVSSI | 30 | Q68D10 | Protein SPT2 homolog OS=Homo sapiens | SPTY2D1 | 75.599 | 4 | 52.48447 | yes | 3.642E-07 | 731.14771 | 186.70552 |
| PVSSLGPGQTVSSS | 21 | Q68D10 | Protein SPT2 homolog OS=Homo sapiens | SPTY2D1 | 75.599 | 3 | 52.48447 | yes | 1.004E-07 | 675.37225 | 147.51341 |
| GQGVNNVPK | 9 | Q68D10 | Protein SPT2 homolog OS=Homo sapiens | SPTY2D1 | 75.599 | 2 | 52.48447 | yes | 1.532E-07 | 456.7485 | 118.87079 |
| HLALSSSK | 8 | Q68D10 | Protein SPT2 homolog OS=Homo sapiens | SPTY2D1 | 75.599 | 2 | 52.48447 | yes | 8.462E-07 | 421.74017 | 118.86195 |
| PHSSTSSPSVPK | 12 | Q68D10 | Protein SPT2 homolog OS=Homo sapiens | SPTY2D1 | 75.599 | 2 | 52.48447 | yes | 5.25E-07 | 605.80676 | 114.27402 |
| EILMIASK | 8 | Q68D10 | Protein SPT2 homolog OS=Homo sapiens | SPTY2D1 | 75.599 | 2 | 52.48447 | yes | 3.535E-07 | 452.76224 | 92.727066 |
| PTVSSGPVPR | 10 | Q68D10 | Protein SPT2 homolog OS=Homo sapiens | SPTY2D1 | 75.599 | 2 | 52.48447 | yes | 2.017E-07 | 498.77728 | 80.357582 |
| SEERPMTAEELR | 12 | Q68D10 | Protein SPT2 homolog OS=Homo sapiens | SPTY2D1 | 75.599 | 2 | 52.48447 | yes | 3.045E-07 | 724.34595 | 55.448635 |
| GVQSAAVQAFLK | 12 | Q68D10 | Protein SPT2 homolog OS=Homo sapiens | SPTY2D1 | 75.599 | 2 | 52.48447 | yes | 1.16E-06 | 609.8457 | 53.715771 |
| RPVSGLGPPGR | 11 | Q68D10 | Protein SPT2 homolog OS=Homo sapiens | SPTY2D1 | 75.599 | 2 | 52.48447 | yes | 1.06E-07 | 546.81726 | 45.451946 |
| AGSGNSSQPSLR | 12 | Q68D10 | Protein SPT2 homolog OS=Homo sapiens | SPTY2D1 | 75.599 | 2 | 52.48447 | yes | 1.128E-07 | 580.78638 | 40.431957 |
| KQFEPVEIK | 9 | Q68D10 | Protein SPT2 homolog OS=Homo sapiens | SPTY2D1 | 75.599 | 2 | 52.48447 | yes | 1.026E-06 | 559.31622 | 38.799541 |
| RYSLAVGPPK | 10 | Q68D10 | Protein SPT2 homolog OS=Homo sapiens | SPTY2D1 | 75.599 | 2 | 52.48447 | yes | 9.009E-05 | 544.31659 | 17.535786 |
| QYLSTPDSQSLR | 12 | Q68E01 | Integrator complex subunit 3 OS=Homo s | INTS3 | 118.07 | 2 | 49.56043 | yes | 3.387E-05 | 697.84918 | 84.455063 |
| FMECLMIGR | 9 | Q68E01 | Integrator complex subunit 3 OS=Homo s | INTS3 | 118.07 | 2 | 49.56043 | yes | 1.021E-07 | 578.76923 | 53.655827 |
| LLLSTSLDAK | 10 | Q68E01 | Integrator complex subunit 3 OS=Homo s | INTS3 | 118.07 | 2 | 49.56043 | yes | 1.163E-07 | 530.81604 | 52.541389 |
| EVDFCISLLR | 10 | Q68E01 | Integrator complex subunit 3 OS=Homo s | INTS3 | 118.07 | 2 | 49.56043 | yes | 5.971E-07 | 626.32373 | 43.680138 |
| EANDALNAYVCK | 12 | Q68E01 | Integrator complex subunit 3 OS=Homo s | INTS3 | 118.07 | 2 | 49.56043 | yes | 2.318E-07 | 684.31665 | 41.983551 |
| SGVLGADGVCMT | 15 | Q68E01 | Integrator complex subunit 3 OS=Homo s | INTS3 | 118.07 | 2 | 49.56043 | yes | 2.618E-07 | 786.86456 | 39.460003 |

| | | | | | | | | | | | |
|---|---|---|---|---|---|---|---|---|---|---|---|
| YQDWFQR | 7 | Q68E01 | Integrator complex subunit 3 OS=Homo s | INTS3 | 118.07 | 2 | 49.56043 | yes | 1.713E-06 | 521.74066 | 33.13868 |
| LNSILTQPWR | 10 | Q69YN4 | Protein virilizer homolog OS=Homo sapi | VIRMA | 202.025 | 2 | 53.58344 | yes | 1.541E-07 | 614.34583 | 157.9469 |
| FPCVVYINEVR | 11 | Q69YN4 | Protein virilizer homolog OS=Homo sapi | VIRMA | 202.025 | 2 | 53.58344 | yes | 9.722E-08 | 698.35809 | 152.91751 |
| VVTAGSAILQK | 11 | Q69YN4 | Protein virilizer homolog OS=Homo sapi | VIRMA | 202.025 | 2 | 53.58344 | yes | 3.159E-07 | 543.82953 | 95.694611 |
| VDLIELSEK | 9 | Q69YN4 | Protein virilizer homolog OS=Homo sapi | VIRMA | 202.025 | 2 | 53.58344 | yes | 2.185E-07 | 523.29242 | 84.798027 |
| QYTSQTTR | 8 | Q69YN4 | Protein virilizer homolog OS=Homo sapi | VIRMA | 202.025 | 2 | 53.58344 | yes | 1.901E-07 | 492.74088 | 66.677399 |
| VNTDGLVLR | 9 | Q69YN4 | Protein virilizer homolog OS=Homo sapi | VIRMA | 202.025 | 2 | 53.58344 | yes | 4.542E-07 | 493.7851 | 58.389511 |
| LVLEHSK | 7 | Q69YN4 | Protein virilizer homolog OS=Homo sapi | VIRMA | 202.025 | 2 | 53.58344 | yes | 3.556E-07 | 413.24509 | 43.88102 |
| HPSAEQSSHIDVVI | 14 | Q69YN4 | Protein virilizer homolog OS=Homo sapi | VIRMA | 202.025 | 3 | 53.58344 | yes | 9.623E-07 | 521.26208 | 21.638187 |
| HPSAEQSSHIDVVI | 14 | Q69YN4 | Protein virilizer homolog OS=Homo sapi | VIRMA | 202.025 | 3 | 53.58344 | yes | 3.105E-06 | 781.38953 | 21.638187 |
| HPSAEQSSHIDVVI | 14 | Q69YN4 | Protein virilizer homolog OS=Homo sapi | VIRMA | 202.025 | 2 | 53.58344 | yes | 9.623E-07 | 521.26208 | 21.638187 |
| HPSAEQSSHIDVVI | 14 | Q69YN4 | Protein virilizer homolog OS=Homo sapi | VIRMA | 202.025 | 2 | 53.58344 | yes | 3.105E-06 | 781.38953 | 21.638187 |
| LTELLDLYR | 9 | Q69YN4 | Protein virilizer homolog OS=Homo sapi | VIRMA | 202.025 | 2 | 53.58344 | yes | 4.224E-07 | 568.32153 | 42.984253 |
| EVVPQDGIPPPK | 12 | Q69YN4 | Protein virilizer homolog OS=Homo sapi | VIRMA | 202.025 | 2 | 53.58344 | yes | 1.122E-07 | 638.35083 | 39.890163 |
| NSSALHSLLK | 10 | Q69YN4 | Protein virilizer homolog OS=Homo sapi | VIRMA | 202.025 | 2 | 53.58344 | yes | 3.754E-06 | 535.30365 | 29.046635 |
| QPIALNVR | 8 | Q69YN4 | Protein virilizer homolog OS=Homo sapi | VIRMA | 202.025 | 2 | 53.58344 | yes | 1.085E-06 | 455.77707 | 25.706257 |
| LVKPGTAASLSK | 12 | Q69YQ0 | Cytospin-A OS=Homo sapiens OX=9606 | SPECC1L | 124.544 | 2 | 40.826332 | yes | 3.82E-07 | 586.3559 | 32.73024 |
| RPNYGEIPVQEHLI | 15 | Q69YQ0 | Cytospin-A OS=Homo sapiens OX=9606 | SPECC1L | 124.544 | 3 | 40.826332 | yes | 5.112E-06 | 607.6637 | 18.451965 |
| TSLGGDAPEEPEK | 13 | Q6DN12 | Multiple C2 and transmembrane domain-c | MCTP2 | 99.596 | 2 | 50.95695 | yes | 2.918E-07 | 665.31207 | 35.332893 |
| VDISALPLK | 9 | Q6DN12 | Multiple C2 and transmembrane domain-c | MCTP2 | 99.596 | 2 | 50.95695 | yes | 1.269E-05 | 478.29477 | 19.758879 |
| NVSGGSMTEMFV( | 15 | Q6DN12 | Multiple C2 and transmembrane domain-c | MCTP2 | 99.596 | 2 | 50.95695 | yes | 8.406E-07 | 814.39453 | 16.736292 |
| TRPLLINLSK | 10 | Q6DN12 | Multiple C2 and transmembrane domain-c | MCTP2 | 99.596 | 2 | 50.95695 | yes | 9.722E-08 | 577.86639 | 16.728037 |
| VLINFNAPNPQDR | 13 | Q6DN90 | IQ motif and SEC7 domain-containing pr | IQSEC1 | 108.315 | 2 | 53.32779 | yes | 1.448E-07 | 749.39404 | 512.29993 |
| SLAESIDDALNCR | 13 | Q6DN90 | IQ motif and SEC7 domain-containing pr | IQSEC1 | 108.315 | 2 | 53.32779 | yes | 9.722E-08 | 732.34338 | 371.92087 |
| YCICNPGVVR | 10 | Q6DN90 | IQ motif and SEC7 domain-containing pr | IQSEC1 | 108.315 | 2 | 53.32779 | yes | 9.722E-08 | 619.29443 | 348.52396 |
| EMLMGIYER | 9 | Q6DN90 | IQ motif and SEC7 domain-containing pr | IQSEC1 | 108.315 | 2 | 53.32779 | yes | 1.022E-07 | 571.27264 | 267.80933 |
| LIEAFSQR | 8 | Q6DN90 | IQ motif and SEC7 domain-containing pr | IQSEC1 | 108.315 | 2 | 53.32779 | no | 3.705E-07 | 482.26654 | 261.96869 |
| KPIGSLHPGLGCVI | 19 | Q6DN90 | IQ motif and SEC7 domain-containing pr | IQSEC1 | 108.315 | 4 | 53.32779 | yes | 1.017E-07 | 510.28992 | 111.18719 |
| KPIGSLHPGLGCVI | 19 | Q6DN90 | IQ motif and SEC7 domain-containing pr | IQSEC1 | 108.315 | 4 | 53.32779 | yes | 1.189E-07 | 680.05078 | 111.18719 |
| KPIGSLHPGLGCVI | 19 | Q6DN90 | IQ motif and SEC7 domain-containing pr | IQSEC1 | 108.315 | 3 | 53.32779 | yes | 1.017E-07 | 510.28992 | 111.18719 |
| KPIGSLHPGLGCVI | 19 | Q6DN90 | IQ motif and SEC7 domain-containing pr | IQSEC1 | 108.315 | 3 | 53.32779 | yes | 1.189E-07 | 680.05078 | 111.18719 |
| ESIAEVQEMEK | 11 | Q6DN90 | IQ motif and SEC7 domain-containing pr | IQSEC1 | 108.315 | 2 | 53.32779 | no | 1.861E-07 | 646.80554 | 208.67973 |
| LQHSTSILR | 9 | Q6DN90 | IQ motif and SEC7 domain-containing pr | IQSEC1 | 108.315 | 2 | 53.32779 | yes | 1.851E-07 | 527.80383 | 190.65112 |
| GVDDGEDIPR | 10 | Q6DN90 | IQ motif and SEC7 domain-containing pr | IQSEC1 | 108.315 | 2 | 53.32779 | yes | 2.147E-07 | 536.7489 | 185.38708 |
| AGGAAPDYWALA | 14 | Q6DN90 | IQ motif and SEC7 domain-containing pr | IQSEC1 | 108.315 | 2 | 53.32779 | yes | 2.287E-07 | 714.35693 | 146.26501 |
| LTSSVPGADIK | 11 | Q6DN90 | IQ motif and SEC7 domain-containing pr | IQSEC1 | 108.315 | 2 | 53.32779 | yes | 1.967E-07 | 544.30334 | 135.90674 |
| ACLDDSYASGEGL | 14 | Q6DN90 | IQ motif and SEC7 domain-containing pr | IQSEC1 | 108.315 | 2 | 53.32779 | yes | 9.722E-08 | 743.32996 | 117.6548 |
| LFEVPDPNKPQK | 12 | Q6DN90 | IQ motif and SEC7 domain-containing pr | IQSEC1 | 108.315 | 2 | 53.32779 | yes | 9.722E-08 | 706.38263 | 111.48689 |
| GVQYLIER | 8 | Q6DN90 | IQ motif and SEC7 domain-containing pr | IQSEC1 | 108.315 | 2 | 53.32779 | yes | 2.657E-07 | 489.27435 | 111.03044 |

| | | | | | | | | | | | |
|---|---|---|---|---|---|---|---|---|---|---|---|
| NSWDSPAFSNDVI | 14 | Q6DN90 | IQ motif and SEC7 domain-containing pr | IQSEC1 | 108.315 | 2 | 53.32779 | yes | 9.722E-08 | 804.3761 | 101.51263 |
| QVEMLER | 7 | Q6DN90 | IQ motif and SEC7 domain-containing pr | IQSEC1 | 108.315 | 2 | 53.32779 | yes | 2.511E-07 | 452.73148 | 97.543152 |
| IGLNLFNK | 8 | Q6DN90 | IQ motif and SEC7 domain-containing pr | IQSEC1 | 108.315 | 2 | 53.32779 | no | 8.043E-07 | 459.77402 | 93.226341 |
| QMIGEFLGNR | 10 | Q6DN90 | IQ motif and SEC7 domain-containing pr | IQSEC1 | 108.315 | 2 | 53.32779 | no | 1.043E-07 | 582.79517 | 92.840996 |
| MQFSFEGPEK | 10 | Q6DN90 | IQ motif and SEC7 domain-containing pr | IQSEC1 | 108.315 | 2 | 53.32779 | yes | 9.722E-08 | 600.27368 | 76.964294 |
| VLLSDSNLHDA | 11 | Q6FI81 | Anamorsin OS=Homo sapiens OX=9606 | CIAPIN1 | 33.583 | 2 | 55.008034 | yes | 1.207E-07 | 592.30133 | 551.60187 |
| EPLTPEEVQSVR | 12 | Q6FI81 | Anamorsin OS=Homo sapiens OX=9606 | CIAPIN1 | 33.583 | 2 | 55.008034 | yes | 1.026E-07 | 692.35938 | 473.39706 |
| KPNFEVGSSR | 10 | Q6FI81 | Anamorsin OS=Homo sapiens OX=9606 | CIAPIN1 | 33.583 | 2 | 55.008034 | yes | 3.612E-07 | 560.79089 | 318.05081 |
| LQALTGNEGR | 10 | Q6FI81 | Anamorsin OS=Homo sapiens OX=9606 | CIAPIN1 | 33.583 | 2 | 55.008034 | yes | 1.351E-07 | 529.78308 | 268.36682 |
| EPVETAVDNNSK | 12 | Q6FI81 | Anamorsin OS=Homo sapiens OX=9606 | CIAPIN1 | 33.583 | 2 | 55.008034 | yes | 9.722E-08 | 651.81226 | 253.4951 |
| SACGNCYLGDAFI | 13 | Q6FI81 | Anamorsin OS=Homo sapiens OX=9606 | CIAPIN1 | 33.583 | 2 | 55.008034 | yes | 1.001E-07 | 745.81122 | 196.43346 |
| NCTCGLAEELEK | 12 | Q6FI81 | Anamorsin OS=Homo sapiens OX=9606 | CIAPIN1 | 33.583 | 2 | 55.008034 | yes | 1.002E-07 | 712.31323 | 153.82193 |
| LCSALTLSGLVEVI | 14 | Q6FI81 | Anamorsin OS=Homo sapiens OX=9606 | CIAPIN1 | 33.583 | 2 | 55.008034 | yes | 9.722E-08 | 745.41833 | 152.79471 |
| KPDPASLR | 8 | Q6FI81 | Anamorsin OS=Homo sapiens OX=9606 | CIAPIN1 | 33.583 | 2 | 55.008034 | yes | 1.377E-07 | 442.25345 | 108.64216 |
| SSPSVKPAVDPAA | 15 | Q6FI81 | Anamorsin OS=Homo sapiens OX=9606 | CIAPIN1 | 33.583 | 2 | 55.008034 | yes | 9.722E-08 | 712.89081 | 94.342125 |
| EHLGHESDNLLFV | 18 | Q6FI81 | Anamorsin OS=Homo sapiens OX=9606 | CIAPIN1 | 33.583 | 3 | 55.008034 | yes | 9.722E-08 | 679.68481 | 50.491478 |
| ILRPGGCLFLK | 11 | Q6FI81 | Anamorsin OS=Homo sapiens OX=9606 | CIAPIN1 | 33.583 | 2 | 55.008034 | yes | 1.424E-07 | 637.3761 | 49.165722 |
| CASCPYLGMPAFK | 13 | Q6FI81 | Anamorsin OS=Homo sapiens OX=9606 | CIAPIN1 | 33.583 | 2 | 55.008034 | yes | 5.491E-07 | 751.33545 | 34.486359 |
| VGLTGFECR | 9 | Q6FIF0 | AN1-type zinc finger protein 6 OS=Homo | ZFAND6 | 22.556 | 2 | 43.02191 | yes | 2.765E-07 | 519.75549 | 73.667229 |
| EQAVIPLTDSEYK | 13 | Q6GQQ9 | OTU domain-containing protein 7B OS=I | OTUD7B | 92.526 | 2 | 53.97894 | yes | 1.823E-07 | 746.88013 | 252.29832 |
| YSQEVMQSLSILR | 13 | Q6GQQ9 | OTU domain-containing protein 7B OS=I | OTUD7B | 92.526 | 2 | 53.97894 | yes | 1.225E-06 | 777.40326 | 60.456387 |
| LLPLHFAVDPGK | 12 | Q6GQQ9 | OTU domain-containing protein 7B OS=I | OTUD7B | 92.526 | 2 | 53.97894 | no | 4.934E-07 | 653.87952 | 59.604961 |
| ESGLVYTEDEWQI | 13 | Q6GQQ9 | OTU domain-containing protein 7B OS=I | OTUD7B | 92.526 | 2 | 53.97894 | yes | 9.722E-08 | 792.36481 | 58.003086 |
| LNWWVSVDPTSQ | 13 | Q6GQQ9 | OTU domain-containing protein 7B OS=I | OTUD7B | 92.526 | 2 | 53.97894 | yes | 4.319E-07 | 794.39935 | 34.452415 |
| PSGGGVHCQEPR | 12 | Q6GQQ9 | OTU domain-containing protein 7B OS=I | OTUD7B | 92.526 | 3 | 53.97894 | yes | 1.783E-07 | 427.53152 | 33.489685 |
| HQYQEEMIQR | 10 | Q6GQQ9 | OTU domain-containing protein 7B OS=I | OTUD7B | 92.526 | 2 | 53.97894 | yes | 2.657E-07 | 681.31696 | 31.353893 |
| LASVILSLEVK | 11 | Q6GQQ9 | OTU domain-containing protein 7B OS=I | OTUD7B | 92.526 | 2 | 53.97894 | yes | 0.0001121 | 586.36847 | 6.8136911 |
| TLLVYDMNLR | 10 | Q6I9Y2 | THO complex subunit 7 homolog OS=Ho | THOC7 | 23.743 | 2 | 53.763756 | yes | 1.286E-07 | 619.33411 | 207.87799 |
| MLSTLSQCEFSMG | 14 | Q6I9Y2 | THO complex subunit 7 homolog OS=Ho | THOC7 | 23.743 | 2 | 53.763756 | yes | 1.146E-07 | 809.86731 | 103.48857 |
| EIECSIAGAHEK | 12 | Q6I9Y2 | THO complex subunit 7 homolog OS=Ho | THOC7 | 23.743 | 2 | 53.763756 | yes | 4.424E-07 | 672.31665 | 95.821312 |
| ELEHLSHIK | 9 | Q6I9Y2 | THO complex subunit 7 homolog OS=Ho | THOC7 | 23.743 | 2 | 53.763756 | yes | 1.02E-07 | 553.30365 | 90.55175 |
| LLIDGDGAGDDR | 12 | Q6I9Y2 | THO complex subunit 7 homolog OS=Ho | THOC7 | 23.743 | 2 | 53.763756 | yes | 9.722E-08 | 608.79382 | 71.004501 |
| STSLETQDDDNIR | 13 | Q6IA86 | Elongator complex protein 2 OS=Homo s | ELP2 | 92.501 | 2 | 30.255466 | yes | 2.238E-06 | 747.33936 | 63.285774 |
| LLLDPSSPPTK | 11 | Q6IAA8 | Ragulator complex protein LAMTOR1 O | LAMTOR1 | 17.745 | 2 | 47.064083 | yes | 1.41E-07 | 584.33459 | 105.11781 |
| YGVMDTTTAQGR | 12 | Q6IAN0 | Dehydrogenase/reductase SDR family me | DHRS7B | 35.12 | 2 | 57.41321 | yes | 2.109E-07 | 650.30353 | 179.6277 |
| VMETNYFGPVALT | 14 | Q6IAN0 | Dehydrogenase/reductase SDR family me | DHRS7B | 35.12 | 2 | 57.41321 | yes | 1.446E-07 | 785.40271 | 138.71916 |
| HATQAFFDCLR | 11 | Q6IAN0 | Dehydrogenase/reductase SDR family me | DHRS7B | 35.12 | 2 | 57.41321 | yes | 9.722E-08 | 683.32208 | 78.462036 |
| QGHIVAISSIQGK | 13 | Q6IAN0 | Dehydrogenase/reductase SDR family me | DHRS7B | 35.12 | 2 | 57.41321 | yes | 9.722E-08 | 669.38043 | 49.515446 |
| SPVEVAQDVLAAV | 15 | Q6IAN0 | Dehydrogenase/reductase SDR family me | DHRS7B | 35.12 | 2 | 57.41321 | yes | 5.994E-06 | 741.91174 | 36.185471 |

| | | | | | | | | | | | |
|---|---|---|---|---|---|---|---|---|---|---|---|
| NGGALEELIR | 10 | Q6IAN0 | Dehydrogenase/reductase SDR family me | DHRS7B | 35.12 | 2 | 57.41321 | yes | 1.432E-06 | 536.29327 | 28.815868 |
| ALLPSMIK | 8 | Q6IAN0 | Dehydrogenase/reductase SDR family me | DHRS7B | 35.12 | 2 | 57.41321 | yes | 2.652E-06 | 436.76733 | 6.1844115 |
| QLSSVQEDR | 9 | Q6IBW4 | Condensin-2 complex subunit H2 OS=Hc | NCAPH2 | 68.227 | 2 | 50.76261 | yes | 1.091E-07 | 531.26471 | 124.15668 |
| TEEQPMEVSVCR | 12 | Q6IBW4 | Condensin-2 complex subunit H2 OS=Hc | NCAPH2 | 68.227 | 2 | 50.76261 | yes | 9.723E-08 | 732.82654 | 112.99097 |
| SPQQSAALPR | 10 | Q6IBW4 | Condensin-2 complex subunit H2 OS=Hc | NCAPH2 | 68.227 | 2 | 50.76261 | yes | 2.21E-07 | 527.78564 | 102.62698 |
| MNTCVPHPR | 9 | Q6IBW4 | Condensin-2 complex subunit H2 OS=Hc | NCAPH2 | 68.227 | 2 | 50.76261 | yes | 2.314E-07 | 556.26044 | 85.906143 |
| FVQETELSQR | 10 | Q6IBW4 | Condensin-2 complex subunit H2 OS=Hc | NCAPH2 | 68.227 | 2 | 50.76261 | yes | 3.878E-07 | 618.81458 | 72.081764 |
| NNNPLYSR | 8 | Q6IBW4 | Condensin-2 complex subunit H2 OS=Hc | NCAPH2 | 68.227 | 2 | 50.76261 | yes | 3.572E-06 | 489.24359 | 70.940598 |
| EGAPEPASCVK | 11 | Q6IBW4 | Condensin-2 complex subunit H2 OS=Hc | NCAPH2 | 68.227 | 2 | 50.76261 | yes | 2.255E-06 | 572.7688 | 64.329338 |
| TCALGVQPK | 9 | Q6IE81 | Protein Jade-1 OS=Homo sapiens OX=96 | JADE1 | 95.534 | 2 | 48.650677 | no | 1.12E-07 | 487.26041 | 53.082371 |
| FVCDLTQADR | 10 | Q6IEG0 | U11/U12 small nuclear ribonucleoprotein | SNRNP48 | 39.966 | 2 | 49.57527 | yes | 1.02E-07 | 612.78754 | 92.559357 |
| QSGGSYLDAECSR | 13 | Q6IEG0 | U11/U12 small nuclear ribonucleoprotein | SNRNP48 | 39.966 | 2 | 49.57527 | yes | 1.125E-06 | 715.30426 | 15.580627 |
| LLNSGYFESIPVPK | 14 | Q6IMN6 | Caprin-2 OS=Homo sapiens OX=9606 G | CAPRIN2 | 125.925 | 2 | 48.078438 | yes | 1.203E-07 | 782.4245 | 63.556408 |
| EYSGAPYSQR | 10 | Q6IMN6 | Caprin-2 OS=Homo sapiens OX=9606 G | CAPRIN2 | 125.925 | 2 | 48.078438 | yes | 4.238E-07 | 579.26471 | 57.519703 |
| DVPKPVHQPVGSS | 18 | Q6IMN6 | Caprin-2 OS=Homo sapiens OX=9606 G | CAPRIN2 | 125.925 | 3 | 48.078438 | yes | 4.036E-07 | 625.0108 | 51.686428 |
| QGEEQPWEADYA | 13 | Q6IMN6 | Caprin-2 OS=Homo sapiens OX=9606 G | CAPRIN2 | 125.925 | 2 | 48.078438 | yes | 3.36E-06 | 789.8446 | 25.973736 |
| HSLTSQSQISPK | 12 | Q6IMN6 | Caprin-2 OS=Homo sapiens OX=9606 G | CAPRIN2 | 125.925 | 2 | 48.078438 | yes | 1.908E-07 | 656.84644 | 23.321171 |
| LLLDDLVPTSR | 11 | Q6IN84 | rRNA methyltransferase 1, mitochondrial | MRM1 | 38.639 | 2 | 51.413364 | yes | 9.722E-08 | 621.35864 | 1124.5811 |
| QLLQDPQEPSAR | 12 | Q6IN84 | rRNA methyltransferase 1, mitochondrial | MRM1 | 38.639 | 2 | 51.413364 | yes | 1.612E-07 | 691.35718 | 512.58887 |
| DIPVLRPR | 8 | Q6IN84 | rRNA methyltransferase 1, mitochondrial | MRM1 | 38.639 | 2 | 51.413364 | yes | 4.241E-07 | 483.29819 | 390.15729 |
| NSCPLTPVVSK | 11 | Q6IN84 | rRNA methyltransferase 1, mitochondrial | MRM1 | 38.639 | 2 | 51.413364 | yes | 3.072E-06 | 601.31592 | 370.9053 |
| KGFPTEGER | 9 | Q6IN84 | rRNA methyltransferase 1, mitochondrial | MRM1 | 38.639 | 2 | 51.413364 | yes | 3.013E-06 | 510.75909 | 144.12917 |
| SAHFLGVDK | 9 | Q6IN84 | rRNA methyltransferase 1, mitochondrial | MRM1 | 38.639 | 2 | 51.413364 | yes | 2.271E-07 | 487.25873 | 124.11961 |
| RNSCPLTPVVSK | 12 | Q6IN84 | rRNA methyltransferase 1, mitochondrial | MRM1 | 38.639 | 2 | 51.413364 | yes | 6.046E-07 | 679.36646 | 26.698854 |
| YQVHQGVCMEVS | 17 | Q6IN84 | rRNA methyltransferase 1, mitochondrial | MRM1 | 38.639 | 4 | 51.413364 | yes | 1.673E-06 | 514.76086 | 21.321617 |
| HGERPGGEELSR | 12 | Q6IN84 | rRNA methyltransferase 1, mitochondrial | MRM1 | 38.639 | 2 | 51.413364 | yes | 2.938E-06 | 662.32367 | 18.370312 |
| TALQEQLLQR | 10 | Q6IPM2 | IQ domain-containing protein E OS=Hon | IQCE | 77.298 | 2 | 46.293518 | yes | 1.599E-07 | 600.34076 | 40.585186 |
| MADNLLAAADK | 11 | Q6IQ16 | Speckle-type POZ protein-like OS=Homc | SPOPL | 44.648 | 2 | 47.83982 | yes | 3.729E-07 | 566.78699 | 60.436859 |
| LAEDLGNLWENTI | 13 | Q6IQ16 | Speckle-type POZ protein-like OS=Homc | SPOPL | 44.648 | 2 | 47.83982 | yes | 2.646E-07 | 765.88098 | 45.232285 |
| FTDCSFFVR | 9 | Q6IQ16 | Speckle-type POZ protein-like OS=Homc | SPOPL | 44.648 | 2 | 47.83982 | yes | 1.248E-07 | 589.76862 | 45.091774 |
| QQPSALFAR | 9 | Q6IQ19 | Centriole, cilia and spindle-associated prc | CCSAP | 30.217 | 2 | 52.405735 | yes | 3.908E-07 | 509.27744 | 347.96667 |
| ASSSENPWMTEYN | 14 | Q6IQ19 | Centriole, cilia and spindle-associated prc | CCSAP | 30.217 | 2 | 52.405735 | yes | 9.722E-08 | 844.85577 | 94.085594 |
| WEEYGPCYR | 9 | Q6IQ19 | Centriole, cilia and spindle-associated prc | CCSAP | 30.217 | 2 | 52.405735 | yes | 2.792E-07 | 630.26111 | 62.145824 |
| HPFALYGWGEK | 11 | Q6IQ19 | Centriole, cilia and spindle-associated prc | CCSAP | 30.217 | 2 | 52.405735 | yes | 3.29E-06 | 652.82477 | 46.323498 |
| SVDISLGDSPR | 11 | Q6IQ23 | Pleckstrin homology domain-containing f | PLEKHA7 | 127.135 | 2 | 45.841373 | yes | 9.821E-07 | 573.29346 | 95.065277 |
| SLPLDQTLPR | 10 | Q6IQ23 | Pleckstrin homology domain-containing f | PLEKHA7 | 127.135 | 2 | 45.841373 | yes | 1.727E-07 | 570.32458 | 81.934608 |
| LQESQPGNAVIDK | 13 | Q6IQ49 | Splicing regulator SDE2 OS=Homo sapie | SDE2 | 49.742 | 2 | 48.667137 | yes | 1.117E-07 | 699.86481 | 306.0827 |
| EPIEEEPTGAGLNK | 14 | Q6IQ49 | Splicing regulator SDE2 OS=Homo sapie | SDE2 | 49.742 | 2 | 48.667137 | yes | 1.939E-07 | 742.36737 | 267.12369 |
| EQIDPALFAK | 10 | Q6IQ49 | Splicing regulator SDE2 OS=Homo sapie | SDE2 | 49.742 | 2 | 48.667137 | yes | 3.039E-07 | 566.30585 | 249.01245 |

| | | | | | | | | | | | |
|---|---|---|---|---|---|---|---|---|---|---|---|
| MVSAEISENR | 10 | Q6IQ49 | Splicing regulator SDE2 OS=Homo sapie | SDE2 | 49.742 | 2 | 48.667137 | yes | 1.743E-07 | 568.27423 | 243.60283 |
| VAEVAPEER | 9 | Q6IQ49 | Splicing regulator SDE2 OS=Homo sapie | SDE2 | 49.742 | 2 | 48.667137 | yes | 7.163E-07 | 500.25891 | 82.725555 |
| CELMALGLK | 9 | Q6IQ49 | Splicing regulator SDE2 OS=Homo sapie | SDE2 | 49.742 | 2 | 48.667137 | yes | 1.759E-07 | 517.77228 | 54.749344 |
| HILEDSCAELGESK | 14 | Q6IQ49 | Splicing regulator SDE2 OS=Homo sapie | SDE2 | 49.742 | 2 | 48.667137 | yes | 4.923E-07 | 794.36957 | 42.342705 |
| VASLDYLGTVAAF | 13 | Q6KC79 | Nipped-B-like protein OS=Homo sapiens | NIPBL | 316.052 | 2 | 50.80475 | yes | 9.722E-08 | 668.367 | 115.37901 |
| EIETTGQIMHR | 11 | Q6KC79 | Nipped-B-like protein OS=Homo sapiens | NIPBL | 316.052 | 2 | 50.80475 | yes | 9.85E-08 | 657.82715 | 91.476257 |
| IAEEVNCLLACR | 12 | Q6KC79 | Nipped-B-like protein OS=Homo sapiens | NIPBL | 316.052 | 2 | 50.80475 | yes | 4.213E-07 | 724.35522 | 71.365715 |
| LTPEEMMDSSTFK | 13 | Q6KC79 | Nipped-B-like protein OS=Homo sapiens | NIPBL | 316.052 | 2 | 50.80475 | yes | 6.15E-07 | 758.33893 | 69.924545 |
| YAGFIHMK | 8 | Q6KC79 | Nipped-B-like protein OS=Homo sapiens | NIPBL | 316.052 | 2 | 50.80475 | yes | 1.101E-06 | 483.74692 | 49.991737 |
| LSLDDVQK | 8 | Q6KC79 | Nipped-B-like protein OS=Homo sapiens | NIPBL | 316.052 | 2 | 50.80475 | yes | 3.351E-06 | 459.25055 | 49.948788 |
| HFDFDLEDFK | 10 | Q6KC79 | Nipped-B-like protein OS=Homo sapiens | NIPBL | 316.052 | 2 | 50.80475 | yes | 1.937E-07 | 656.79584 | 45.721355 |
| CLSEVVAVDPSILA | 15 | Q6KC79 | Nipped-B-like protein OS=Homo sapiens | NIPBL | 316.052 | 2 | 50.80475 | yes | 2.881E-07 | 814.93744 | 42.915195 |
| IGLVEDLNK | 9 | Q6KC79 | Nipped-B-like protein OS=Homo sapiens | NIPBL | 316.052 | 2 | 50.80475 | yes | 1.17E-05 | 500.78729 | 38.494442 |
| VDSQASITQDSDSI | 15 | Q6KC79 | Nipped-B-like protein OS=Homo sapiens | NIPBL | 316.052 | 2 | 50.80475 | yes | 1.495E-07 | 797.38373 | 37.551414 |
| IRPQLMVK | 8 | Q6KC79 | Nipped-B-like protein OS=Homo sapiens | NIPBL | 316.052 | 2 | 50.80475 | yes | 7.378E-07 | 492.80478 | 30.268448 |
| GAKPVVVLQK | 10 | Q6KC79 | Nipped-B-like protein OS=Homo sapiens | NIPBL | 316.052 | 2 | 50.80475 | yes | 1.507E-06 | 519.83716 | 21.94062 |
| LDPHGGGLLSSK | 12 | Q6KC79 | Nipped-B-like protein OS=Homo sapiens | NIPBL | 316.052 | 2 | 50.80475 | yes | 2.311E-07 | 590.81964 | 15.101349 |
| NPPAFTELQLPR | 12 | Q6L8Q7 | 2',5'-phosphodiesterase 12 OS=Homo sap | PDE12 | 67.352 | 2 | 35.872513 | yes | 7.702E-07 | 691.875 | 50.362011 |
| VTVLNYASPITAV: | 15 | Q6MZQ0 | Proline-rich protein 5-like OS=Homo sap | PRR5L | 40.836 | 2 | 25.87967 | yes | 3.212E-06 | 795.94611 | 24.010571 |
| SFSGPTYTLAR | 11 | Q6MZQ0 | Proline-rich protein 5-like OS=Homo sap | PRR5L | 40.836 | 2 | 25.87967 | yes | 5.156E-06 | 600.3064 | 11.137654 |
| SMVLLLHSQR | 10 | Q6N022 | Teneurin-4 OS=Homo sapiens OX=9606 | TENM4 | 307.957 | 2 | 24.467054 | yes | 3.449E-06 | 592.33441 | 22.745352 |
| AEADEIFQELEK | 12 | Q6NS38 | DNA oxidative demethylase ALKBH2 O: | ALKBH2 | 29.323 | 2 | 54.856255 | yes | 9.722E-08 | 711.34338 | 293.9747 |
| AEGLDCSYTVLFG | 14 | Q6NS38 | DNA oxidative demethylase ALKBH2 O: | ALKBH2 | 29.323 | 2 | 54.856255 | yes | 9.722E-08 | 780.37415 | 251.30762 |
| PWIPVLER | 8 | Q6NS38 | DNA oxidative demethylase ALKBH2 O: | ALKBH2 | 29.323 | 2 | 54.856255 | yes | 2.736E-07 | 505.2951 | 158.99731 |
| EVEYFTGALAR | 11 | Q6NS38 | DNA oxidative demethylase ALKBH2 O: | ALKBH2 | 29.323 | 2 | 54.856255 | yes | 2.215E-07 | 628.31952 | 155.34148 |
| ELELVACGLER | 11 | Q6NSJ5 | Volume-regulated anion channel subunit | LRRC8E | 90.248 | 2 | 45.27262 | yes | 1.599E-06 | 644.83191 | 87.578659 |
| LETLPSQLGLCSGI | 15 | Q6NSJ5 | Volume-regulated anion channel subunit | LRRC8E | 90.248 | 2 | 45.27262 | yes | 9.268E-07 | 822.44287 | 78.987518 |
| LEALPEELGNCGG | 15 | Q6NSJ5 | Volume-regulated anion channel subunit | LRRC8E | 90.248 | 2 | 45.27262 | yes | 1.901E-07 | 800.40601 | 66.014145 |
| FAVFLSEVSESR | 12 | Q6NSJ5 | Volume-regulated anion channel subunit | LRRC8E | 90.248 | 2 | 45.27262 | yes | 1.006E-06 | 685.8512 | 28.515331 |
| PTAAVQLPR | 9 | Q6NTE8 | MRN complex-interacting protein OS=H | MRNIP | 37.744 | 2 | 54.377037 | yes | 2.66E-07 | 476.78235 | 242.81198 |
| ELWSPIQQVTATS | 15 | Q6NTE8 | MRN complex-interacting protein OS=H | MRNIP | 37.744 | 2 | 54.377037 | yes | 1.541E-07 | 837.93848 | 154.91687 |
| LNLLQGQVSELPL | 14 | Q6NTE8 | MRN complex-interacting protein OS=H | MRNIP | 37.744 | 2 | 54.377037 | yes | 9.722E-08 | 790.46198 | 98.422562 |
| FSQDLPR | 7 | Q6NTE8 | MRN complex-interacting protein OS=H | MRNIP | 37.744 | 2 | 54.377037 | yes | 1.415E-07 | 431.72452 | 66.346619 |
| GVQDSGGSEVAW | 16 | Q6NTE8 | MRN complex-interacting protein OS=H | MRNIP | 37.744 | 2 | 54.377037 | yes | 1.752E-07 | 801.38135 | 37.05101 |
| YETLFQALDR | 10 | Q6NUK1 | Mitochondrial adenyl nucleotide antiport | SLC25A24 | 53.355 | 2 | 53.55179 | yes | 9.722E-08 | 628.31952 | 116.69197 |
| QLLAGGIAGAVSR | 13 | Q6NUK1 | Mitochondrial adenyl nucleotide antiport | SLC25A24 | 53.355 | 2 | 53.55179 | yes | 9.722E-08 | 606.85657 | 113.86339 |
| NLGIPLGQDAEEK | 13 | Q6NUK1 | Mitochondrial adenyl nucleotide antiport | SLC25A24 | 53.355 | 2 | 53.55179 | yes | 2.434E-06 | 692.35938 | 78.614914 |
| TGQYSGIYDCAK | 12 | Q6NUK1 | Mitochondrial adenyl nucleotide antiport | SLC25A24 | 53.355 | 2 | 53.55179 | yes | 6.986E-07 | 681.80334 | 50.693768 |
| NGDGVVDIGELQE | 16 | Q6NUK1 | Mitochondrial adenyl nucleotide antiport | SLC25A24 | 53.355 | 2 | 53.55179 | yes | 3.946E-07 | 835.92084 | 41.78299 |

| | | | | | | | | | | | |
|---|---|---|---|---|---|---|---|---|---|---|---|
| LDFEEFMK | 8 | Q6NUK1 | Mitochondrial adenyl nucleotide antiport | SLC25A24 | 53.355 | 2 | 53.55179 | yes | 4.737E-07 | 529.74677 | 26.851242 |
| LTNQELLR | 8 | Q6NUQ4 | Transmembrane protein 214 OS=Homo sa | TMEM214 | 77.151 | 2 | 50.590096 | yes | 2.97E-06 | 493.7851 | 574.6665 |
| GLLQQVQGPR | 10 | Q6NUQ4 | Transmembrane protein 214 OS=Homo sa | TMEM214 | 77.151 | 2 | 50.590096 | yes | 1.527E-07 | 548.31708 | 131.84923 |
| SSGFLPASQQACA | 14 | Q6NUQ4 | Transmembrane protein 214 OS=Homo sa | TMEM214 | 77.151 | 2 | 50.590096 | yes | 9.757E-08 | 726.35101 | 106.46052 |
| SQSVFSGNPSIWLI | 14 | Q6NUQ4 | Transmembrane protein 214 OS=Homo sa | TMEM214 | 77.151 | 2 | 50.590096 | yes | 2.2E-07 | 775.40411 | 91.006767 |
| SLQETIQSLK | 10 | Q6NUQ4 | Transmembrane protein 214 OS=Homo sa | TMEM214 | 77.151 | 2 | 50.590096 | yes | 9.625E-07 | 573.8219 | 74.689316 |
| SHSSFQASLTGR | 12 | Q6NUQ4 | Transmembrane protein 214 OS=Homo sa | TMEM214 | 77.151 | 2 | 50.590096 | yes | 4.122E-07 | 639.31531 | 41.999252 |
| LLLMHPNLTK | 10 | Q6NUQ4 | Transmembrane protein 214 OS=Homo sa | TMEM214 | 77.151 | 2 | 50.590096 | yes | 1.685E-07 | 590.34955 | 37.455673 |
| YYIEYMESTR | 10 | Q6NYC1 | Bifunctional arginine demethylase and lys | JMJD6 | 46.462 | 2 | 50.258537 | yes | 9.825E-08 | 677.80286 | 84.693192 |
| TCSMVGNGDTTS( | 19 | Q6NYC1 | Bifunctional arginine demethylase and lys | JMJD6 | 46.462 | 3 | 50.258537 | yes | 5.349E-05 | 693.27509 | 0.5056047 |
| ETAELSETLTR | 11 | Q6NYC8 | Phostensin OS=Homo sapiens OX=9606 | PPP1R18 | 67.943 | 2 | 42.75781 | yes | 2.888E-07 | 625.31714 | 64.088058 |
| GMKPTEGWK | 9 | Q6NYC8 | Phostensin OS=Homo sapiens OX=9606 | PPP1R18 | 67.943 | 2 | 42.75781 | yes | 7.856E-07 | 517.26038 | 58.893169 |
| LGLTEAHK | 8 | Q6NYC8 | Phostensin OS=Homo sapiens OX=9606 | PPP1R18 | 67.943 | 2 | 42.75781 | yes | 3.013E-06 | 434.74799 | 54.077278 |
| VPPFTFHVK | 9 | Q6NZI2 | Caveolae-associated protein 1 OS=Homo | CAVIN1 | 43.477 | 2 | 46.883747 | yes | 3.483E-06 | 536.30292 | 230.9429 |
| VMIYQDEVK | 9 | Q6NZI2 | Caveolae-associated protein 1 OS=Homo | CAVIN1 | 43.477 | 2 | 46.883747 | yes | 1.464E-07 | 562.78644 | 224.2735 |
| KLEVNEAELLR | 11 | Q6NZI2 | Caveolae-associated protein 1 OS=Homo | CAVIN1 | 43.477 | 2 | 46.883747 | yes | 1.449E-07 | 657.37482 | 131.16756 |
| EGQVEVLK | 8 | Q6NZI2 | Caveolae-associated protein 1 OS=Homo | CAVIN1 | 43.477 | 2 | 46.883747 | yes | 5.891E-07 | 451.25311 | 71.725357 |
| GTPPLTPSDSPQTR | 14 | Q6NZY4 | Zinc finger CCHC domain-containing prc | ZCCHC8 | 78.578 | 2 | 49.95927 | yes | 1.476E-07 | 727.36774 | 373.35046 |
| GTPPPVFTPPLPK | 13 | Q6NZY4 | Zinc finger CCHC domain-containing prc | ZCCHC8 | 78.578 | 2 | 49.95927 | yes | 1.112E-07 | 674.38721 | 290.08252 |
| IFGSIPMQACQQK | 13 | Q6NZY4 | Zinc finger CCHC domain-containing prc | ZCCHC8 | 78.578 | 2 | 49.95927 | yes | 9.968E-08 | 754.37341 | 284.50232 |
| LVNYPGFNISTPR | 13 | Q6NZY4 | Zinc finger CCHC domain-containing prc | ZCCHC8 | 78.578 | 2 | 49.95927 | yes | 3.418E-07 | 739.39355 | 255.6564 |
| QTLDEPEVPEIFTK | 14 | Q6NZY4 | Zinc finger CCHC domain-containing prc | ZCCHC8 | 78.578 | 2 | 49.95927 | yes | 1.122E-07 | 823.41962 | 123.15883 |
| PSGILVNDTK | 10 | Q6NZY4 | Zinc finger CCHC domain-containing prc | ZCCHC8 | 78.578 | 2 | 49.95927 | yes | 1.76E-07 | 522.29022 | 94.898201 |
| SLPPFIYR | 8 | Q6NZY4 | Zinc finger CCHC domain-containing prc | ZCCHC8 | 78.578 | 2 | 49.95927 | yes | 1.072E-07 | 496.78183 | 72.365578 |
| EAELENSGLALYD | 15 | Q6NZY4 | Zinc finger CCHC domain-containing prc | ZCCHC8 | 78.578 | 2 | 49.95927 | yes | 1.496E-07 | 804.89124 | 65.336395 |
| YHQVFSHIVSLEG( | 19 | Q6NZY4 | Zinc finger CCHC domain-containing prc | ZCCHC8 | 78.578 | 3 | 49.95927 | yes | 1.465E-07 | 747.72693 | 44.103737 |
| LEHIDGVNLTVPV | 14 | Q6P087 | Mitochondrial mRNA pseudouridine synt | RPUSD3 | 38.461 | 3 | 49.547745 | yes | 1.357E-07 | 511.95935 | 755.74738 |
| LEHIDGVNLTVPV | 14 | Q6P087 | Mitochondrial mRNA pseudouridine synt | RPUSD3 | 38.461 | 3 | 49.547745 | yes | 1.022E-07 | 767.43542 | 755.74738 |
| LEHIDGVNLTVPV | 14 | Q6P087 | Mitochondrial mRNA pseudouridine synt | RPUSD3 | 38.461 | 2 | 49.547745 | yes | 1.357E-07 | 511.95935 | 755.74738 |
| LEHIDGVNLTVPV | 14 | Q6P087 | Mitochondrial mRNA pseudouridine synt | RPUSD3 | 38.461 | 2 | 49.547745 | yes | 1.022E-07 | 767.43542 | 755.74738 |
| TLQCLGLR | 8 | Q6P087 | Mitochondrial mRNA pseudouridine synt | RPUSD3 | 38.461 | 2 | 49.547745 | yes | 1.043E-07 | 480.7684 | 1172.3953 |
| FLLPAENNKPQR | 12 | Q6P087 | Mitochondrial mRNA pseudouridine synt | RPUSD3 | 38.461 | 2 | 49.547745 | yes | 2.082E-07 | 713.89368 | 1034.429 |
| QVLDEALLR | 9 | Q6P087 | Mitochondrial mRNA pseudouridine synt | RPUSD3 | 38.461 | 2 | 49.547745 | yes | 1.941E-07 | 528.80603 | 869.7301 |
| EQELQVVR | 8 | Q6P087 | Mitochondrial mRNA pseudouridine synt | RPUSD3 | 38.461 | 2 | 49.547745 | yes | 1.284E-07 | 500.77472 | 863.90082 |
| SGPLGDQPFAGLL | 15 | Q6P087 | Mitochondrial mRNA pseudouridine synt | RPUSD3 | 38.461 | 2 | 49.547745 | yes | 5.35E-07 | 748.909 | 539.16168 |
| EELVDALR | 8 | Q6P087 | Mitochondrial mRNA pseudouridine synt | RPUSD3 | 38.461 | 2 | 49.547745 | yes | 1.986E-07 | 472.75601 | 497.33575 |
| LHLTPSQAAQLPLI | 19 | Q6P087 | Mitochondrial mRNA pseudouridine synt | RPUSD3 | 38.461 | 3 | 49.547745 | yes | 2.193E-07 | 728.08337 | 424.91821 |
| KGPLVTLNKPQGL | 18 | Q6P087 | Mitochondrial mRNA pseudouridine synt | RPUSD3 | 38.461 | 3 | 49.547745 | yes | 2.169E-07 | 616.37531 | 112.51326 |
| GAGESGEEDPFPSI | 14 | Q6P0Q8 | Microtubule-associated serine/threonine-p | MAST2 | 196.436 | 2 | 53.53771 | yes | 2.731E-07 | 717.81024 | 257.79965 |

| | | | | | | | | | | | |
|---|---|---|---|---|---|---|---|---|---|---|---|
| NIGALPVDMVR | 11 | Q6P0Q8 | Microtubule-associated serine/threonine-p | MAST2 | 196.436 | 2 | 53.53771 | yes | 9.722E-08 | 592.82642 | 160.1983 |
| DLKPDNLLITSMG | 16 | Q6P0Q8 | Microtubule-associated serine/threonine-p | MAST2 | 196.436 | 3 | 53.53771 | no | 1.522E-07 | 598.99762 | 157.01389 |
| NQIQQAFVER | 10 | Q6P0Q8 | Microtubule-associated serine/threonine-p | MAST2 | 196.436 | 2 | 53.53771 | no | 1.084E-07 | 616.82275 | 154.6528 |
| LLQDAHER | 8 | Q6P0Q8 | Microtubule-associated serine/threonine-p | MAST2 | 196.436 | 2 | 53.53771 | no | 2.974E-07 | 491.25925 | 145.55505 |
| EQDVVTGVSPLLF | 14 | Q6P0Q8 | Microtubule-associated serine/threonine-p | MAST2 | 196.436 | 2 | 53.53771 | yes | 9.722E-08 | 780.42505 | 131.84177 |
| LLHQNPLER | 9 | Q6P0Q8 | Microtubule-associated serine/threonine-p | MAST2 | 196.436 | 2 | 53.53771 | yes | 1.274E-07 | 560.31708 | 125.45704 |
| LAAALAASEK | 10 | Q6P0Q8 | Microtubule-associated serine/threonine-p | MAST2 | 196.436 | 2 | 53.53771 | yes | 9.823E-08 | 472.7742 | 113.94818 |
| NQSLGQSAPSLTA( | 16 | Q6P0Q8 | Microtubule-associated serine/threonine-p | MAST2 | 196.436 | 2 | 53.53771 | yes | 9.968E-08 | 786.42303 | 107.91005 |
| GTSQLAEGATAK | 12 | Q6P0Q8 | Microtubule-associated serine/threonine-p | MAST2 | 196.436 | 2 | 53.53771 | yes | 4.422E-07 | 567.29346 | 106.2915 |
| YIVSQLGLTR | 10 | Q6P0Q8 | Microtubule-associated serine/threonine-p | MAST2 | 196.436 | 2 | 53.53771 | yes | 2.466E-07 | 575.33496 | 102.73457 |
| VAISTTPLENTSIK | 14 | Q6P0Q8 | Microtubule-associated serine/threonine-p | MAST2 | 196.436 | 2 | 53.53771 | yes | 1.044E-07 | 737.41156 | 98.708801 |
| EVSPLEVVGAR | 11 | Q6P0Q8 | Microtubule-associated serine/threonine-p | MAST2 | 196.436 | 2 | 53.53771 | yes | 2.372E-07 | 578.32202 | 87.885513 |
| FPEGPEEASSTLR | 13 | Q6P0Q8 | Microtubule-associated serine/threonine-p | MAST2 | 196.436 | 2 | 53.53771 | yes | 7.743E-07 | 710.34119 | 32.145409 |
| DRPQPPPPDR | 10 | Q6P0Q8 | Microtubule-associated serine/threonine-p | MAST2 | 196.436 | 2 | 53.53771 | yes | 3.797E-06 | 587.80182 | 28.442385 |
| TPSEEDFETIK | 11 | Q6P0Q8 | Microtubule-associated serine/threonine-p | MAST2 | 196.436 | 2 | 53.53771 | yes | 2.093E-07 | 648.30371 | 20.932924 |
| LMIIIAR | 7 | Q6P0Q8 | Microtubule-associated serine/threonine-p | MAST2 | 196.436 | 2 | 53.53771 | yes | 1.619E-06 | 415.27005 | 20.002306 |
| KLMIIIAR | 8 | Q6P0Q8 | Microtubule-associated serine/threonine-p | MAST2 | 196.436 | 2 | 53.53771 | yes | 2.42E-07 | 479.31754 | 19.007454 |
| QLSQDDCK | 8 | Q6P0Q8 | Microtubule-associated serine/threonine-p | MAST2 | 196.436 | 2 | 53.53771 | yes | 2.082E-07 | 497.21893 | 17.102852 |
| CSGLLDAPR | 9 | Q6P0Q8 | Microtubule-associated serine/threonine-p | MAST2 | 196.436 | 2 | 53.53771 | yes | 3.607E-06 | 494.74765 | 16.26272 |
| ISAISVAER | 9 | Q6P158 | Putative ATP-dependent RNA helicase D | DHX57 | 155.605 | 2 | 53.984962 | no | 1.122E-07 | 473.27182 | 7937.5903 |
| VPLEQLCLR | 9 | Q6P158 | Putative ATP-dependent RNA helicase D | DHX57 | 155.605 | 2 | 53.984962 | yes | 1.02E-07 | 564.31573 | 2944.9299 |
| LLYCTTGVLLR | 11 | Q6P158 | Putative ATP-dependent RNA helicase D | DHX57 | 155.605 | 2 | 53.984962 | no | 1.087E-07 | 654.87085 | 1725.7625 |
| VGLTVGYQIR | 10 | Q6P158 | Putative ATP-dependent RNA helicase D | DHX57 | 155.605 | 2 | 53.984962 | yes | 1.448E-07 | 553.32184 | 1452.5685 |
| DLGALTPDER | 10 | Q6P158 | Putative ATP-dependent RNA helicase D | DHX57 | 155.605 | 2 | 53.984962 | yes | 1.351E-07 | 543.7749 | 1428.8188 |
| YSDPPVNFLPVPSI | 14 | Q6P158 | Putative ATP-dependent RNA helicase D | DHX57 | 155.605 | 2 | 53.984962 | yes | 1.024E-07 | 794.41193 | 1391.0846 |
| QFQSILQER | 9 | Q6P158 | Putative ATP-dependent RNA helicase D | DHX57 | 155.605 | 2 | 53.984962 | yes | 1.067E-07 | 574.80658 | 1044.9119 |
| YVLQDGSPYMR | 11 | Q6P158 | Putative ATP-dependent RNA helicase D | DHX57 | 155.605 | 2 | 53.984962 | yes | 9.781E-08 | 664.81879 | 1011.9102 |
| TAFEEVEEDLR | 11 | Q6P158 | Putative ATP-dependent RNA helicase D | DHX57 | 155.605 | 2 | 53.984962 | yes | 9.722E-08 | 669.31464 | 843.43036 |
| VANIICTQPR | 10 | Q6P158 | Putative ATP-dependent RNA helicase D | DHX57 | 155.605 | 2 | 53.984962 | yes | 1.108E-07 | 586.31622 | 841.19702 |
| DAVPDQQLDFK | 11 | Q6P158 | Putative ATP-dependent RNA helicase D | DHX57 | 155.605 | 2 | 53.984962 | yes | 1.401E-07 | 638.31439 | 782.42139 |
| VLQEMASLK | 9 | Q6P158 | Putative ATP-dependent RNA helicase D | DHX57 | 155.605 | 2 | 53.984962 | yes | 9.722E-08 | 509.78372 | 565.99805 |
| NPSIDLCTCPR | 11 | Q6P158 | Putative ATP-dependent RNA helicase D | DHX57 | 155.605 | 2 | 53.984962 | yes | 1.043E-07 | 666.80536 | 545.55579 |
| QQLPEIQR | 8 | Q6P158 | Putative ATP-dependent RNA helicase D | DHX57 | 155.605 | 2 | 53.984962 | yes | 1.231E-07 | 506.28271 | 539.46283 |
| GEFVVSLDDGWIR | 13 | Q6P158 | Putative ATP-dependent RNA helicase D | DHX57 | 155.605 | 2 | 53.984962 | yes | 1.908E-07 | 746.87518 | 484.87738 |
| QSLPAWEER | 9 | Q6P158 | Putative ATP-dependent RNA helicase D | DHX57 | 155.605 | 2 | 53.984962 | yes | 1.527E-07 | 558.27765 | 453.88077 |
| LSLHLQDQDSVK | 12 | Q6P158 | Putative ATP-dependent RNA helicase D | DHX57 | 155.605 | 2 | 53.984962 | yes | 9.722E-08 | 691.86737 | 442.95703 |
| NVQENSLEICK | 11 | Q6P158 | Putative ATP-dependent RNA helicase D | DHX57 | 155.605 | 2 | 53.984962 | yes | 9.722E-08 | 667.32446 | 381.43402 |
| YGFNTER | 7 | Q6P158 | Putative ATP-dependent RNA helicase D | DHX57 | 155.605 | 2 | 53.984962 | yes | 2.454E-07 | 443.70633 | 333.94611 |
| QFTELLSDIGFAR | 13 | Q6P158 | Putative ATP-dependent RNA helicase D | DHX57 | 155.605 | 2 | 53.984962 | yes | 9.722E-08 | 748.89081 | 323.65808 |

| | | | | | | | | | | | |
|---|---|---|---|---|---|---|---|---|---|---|---|
| LIEPPHTDSLR | 11 | Q6P158 | Putative ATP-dependent RNA helicase D | DHX57 | 155.605 | 2 | 53.984962 | yes | 4.831E-07 | 639.34607 | 158.29965 |
| LIEPPHTDSLR | 11 | Q6P158 | Putative ATP-dependent RNA helicase D | DHX57 | 155.605 | 2 | 53.984962 | yes | 4.398E-07 | 426.56647 | 158.29965 |
| LIEPPHTDSLR | 11 | Q6P158 | Putative ATP-dependent RNA helicase D | DHX57 | 155.605 | 3 | 53.984962 | yes | 4.831E-07 | 639.34607 | 158.29965 |
| LIEPPHTDSLR | 11 | Q6P158 | Putative ATP-dependent RNA helicase D | DHX57 | 155.605 | 3 | 53.984962 | yes | 4.398E-07 | 426.56647 | 158.29965 |
| ETILNLLR | 8 | Q6P158 | Putative ATP-dependent RNA helicase D | DHX57 | 155.605 | 2 | 53.984962 | yes | 1.815E-07 | 486.29785 | 191.05879 |
| HQVVVISGMTGC( | 14 | Q6P158 | Putative ATP-dependent RNA helicase D | DHX57 | 155.605 | 2 | 53.984962 | yes | 9.722E-08 | 736.87103 | 89.55864 |
| HQVVVISGMTGC( | 14 | Q6P158 | Putative ATP-dependent RNA helicase D | DHX57 | 155.605 | 2 | 53.984962 | yes | 1.067E-07 | 491.58313 | 89.55864 |
| HQVVVISGMTGC( | 14 | Q6P158 | Putative ATP-dependent RNA helicase D | DHX57 | 155.605 | 3 | 53.984962 | yes | 9.722E-08 | 736.87103 | 89.55864 |
| HQVVVISGMTGC( | 14 | Q6P158 | Putative ATP-dependent RNA helicase D | DHX57 | 155.605 | 3 | 53.984962 | yes | 1.067E-07 | 491.58313 | 89.55864 |
| LTPLGYHLASLPVI | 16 | Q6P158 | Putative ATP-dependent RNA helicase D | DHX57 | 155.605 | 3 | 53.984962 | yes | 9.722E-08 | 584.33319 | 171.03479 |
| CELDQLLQDK | 10 | Q6P158 | Putative ATP-dependent RNA helicase D | DHX57 | 155.605 | 2 | 53.984962 | yes | 5.168E-07 | 631.30829 | 167.98126 |
| RYDWQAK | 7 | Q6P158 | Putative ATP-dependent RNA helicase D | DHX57 | 155.605 | 2 | 53.984962 | yes | 1.036E-07 | 483.74323 | 164.32219 |
| IKNPSIDLCTCPR | 13 | Q6P158 | Putative ATP-dependent RNA helicase D | DHX57 | 155.605 | 2 | 53.984962 | yes | 3.274E-07 | 787.3949 | 161.71233 |
| FVAASHQVAELVI | 13 | Q6P158 | Putative ATP-dependent RNA helicase D | DHX57 | 155.605 | 3 | 53.984962 | yes | 1.267E-07 | 466.9295 | 146.59697 |
| TMSIMDFEK | 9 | Q6P158 | Putative ATP-dependent RNA helicase D | DHX57 | 155.605 | 2 | 53.984962 | yes | 9.997E-08 | 551.2514 | 114.25274 |
| RLEGDTALQGVSI | 22 | Q6P158 | Putative ATP-dependent RNA helicase D | DHX57 | 155.605 | 3 | 53.984962 | yes | 1.699E-06 | 825.09796 | 113.46768 |
| CVIHPLHSSLSSEE | 28 | Q6P158 | Putative ATP-dependent RNA helicase D | DHX57 | 155.605 | 4 | 53.984962 | yes | 1.31E-07 | 762.15112 | 90.272545 |
| LEGDTALQGVSHI | 21 | Q6P158 | Putative ATP-dependent RNA helicase D | DHX57 | 155.605 | 3 | 53.984962 | yes | 2.236E-07 | 773.06427 | 84.859314 |
| HFDSPYLLYHEK | 12 | Q6P158 | Putative ATP-dependent RNA helicase D | DHX57 | 155.605 | 2 | 53.984962 | yes | 1.075E-07 | 774.87769 | 71.832336 |
| KHQVVVISGMTG( | 15 | Q6P158 | Putative ATP-dependent RNA helicase D | DHX57 | 155.605 | 2 | 53.984962 | yes | 1.007E-07 | 800.91852 | 35.598595 |
| KHQVVVISGMTG( | 15 | Q6P158 | Putative ATP-dependent RNA helicase D | DHX57 | 155.605 | 2 | 53.984962 | yes | 0.0001773 | 534.28143 | 35.598595 |
| KHQVVVISGMTG( | 15 | Q6P158 | Putative ATP-dependent RNA helicase D | DHX57 | 155.605 | 3 | 53.984962 | yes | 1.007E-07 | 800.91852 | 35.598595 |
| KHQVVVISGMTG( | 15 | Q6P158 | Putative ATP-dependent RNA helicase D | DHX57 | 155.605 | 3 | 53.984962 | yes | 0.0001773 | 534.28143 | 35.598595 |
| QEEAFALK | 8 | Q6P158 | Putative ATP-dependent RNA helicase D | DHX57 | 155.605 | 2 | 53.984962 | yes | 1.535E-06 | 468.24527 | 63.52874 |
| RISAISVAER | 10 | Q6P158 | Putative ATP-dependent RNA helicase D | DHX57 | 155.605 | 2 | 53.984962 | no | 4.151E-07 | 551.32239 | 43.731762 |
| KLEFAFANSDYLA | 19 | Q6P158 | Putative ATP-dependent RNA helicase D | DHX57 | 155.605 | 3 | 53.984962 | yes | 4.284E-06 | 735.72455 | 19.218441 |
| DLQEQDADAGSEI | 13 | Q6P158 | Putative ATP-dependent RNA helicase D | DHX57 | 155.605 | 2 | 53.984962 | yes | 1.464E-07 | 717.31061 | 13.519943 |
| SAPLEIGLQR | 10 | Q6P1J9 | Parafibromin OS=Homo sapiens OX=960 | CDC73 | 60.577 | 2 | 53.98572 | yes | 1.536E-07 | 542.31146 | 913.39923 |
| EGIVQTEQIR | 10 | Q6P1J9 | Parafibromin OS=Homo sapiens OX=960 | CDC73 | 60.577 | 2 | 53.98572 | yes | 2.748E-07 | 586.81714 | 427.63593 |
| VVAVFVQGPAWQ | 14 | Q6P1J9 | Parafibromin OS=Homo sapiens OX=960 | CDC73 | 60.577 | 2 | 53.98572 | yes | 1.512E-07 | 788.43774 | 360.14362 |
| SFVDAEVDVTR | 11 | Q6P1J9 | Parafibromin OS=Homo sapiens OX=960 | CDC73 | 60.577 | 2 | 53.98572 | yes | 1.041E-07 | 619.30658 | 309.50323 |
| PAPNAAPVDPTLR | 13 | Q6P1J9 | Parafibromin OS=Homo sapiens OX=960 | CDC73 | 60.577 | 2 | 53.98572 | yes | 9.722E-08 | 659.85931 | 298.47214 |
| IEDEECVR | 8 | Q6P1J9 | Parafibromin OS=Homo sapiens OX=960 | CDC73 | 60.577 | 2 | 53.98572 | yes | 1.236E-07 | 525.23206 | 279.62607 |
| AADEVLAEAK | 10 | Q6P1J9 | Parafibromin OS=Homo sapiens OX=960 | CDC73 | 60.577 | 2 | 53.98572 | yes | 1.146E-07 | 508.76657 | 261.96997 |
| AATENIPVVR | 10 | Q6P1J9 | Parafibromin OS=Homo sapiens OX=960 | CDC73 | 60.577 | 2 | 53.98572 | yes | 9.809E-08 | 535.30365 | 213.69615 |
| TDLDDDITALK | 11 | Q6P1J9 | Parafibromin OS=Homo sapiens OX=960 | CDC73 | 60.577 | 2 | 53.98572 | yes | 9.722E-08 | 610.30627 | 210.92529 |
| GDEVIFGEFSWPK | 13 | Q6P1J9 | Parafibromin OS=Homo sapiens OX=960 | CDC73 | 60.577 | 2 | 53.98572 | yes | 9.809E-08 | 755.86426 | 181.67355 |
| LMPQDWDR | 8 | Q6P1J9 | Parafibromin OS=Homo sapiens OX=960 | CDC73 | 60.577 | 2 | 53.98572 | yes | 9.722E-08 | 530.74768 | 180.4178 |
| IDTMGTYHGMTLI | 13 | Q6P1J9 | Parafibromin OS=Homo sapiens OX=960 | CDC73 | 60.577 | 2 | 53.98572 | yes | 9.722E-08 | 734.35217 | 140.44356 |

| | | | | | | | | | | | |
|---|---|---|---|---|---|---|---|---|---|---|---|
| TNYVVWGTGK | 10 | Q6P1J9 | Parafibromin OS=Homo sapiens OX=960 | CDC73 | 60.577 | 2 | 53.98572 | yes | 2.45E-07 | 562.79041 | 121.19014 |
| WDVTVLELSYHK | 12 | Q6P1J9 | Parafibromin OS=Homo sapiens OX=960 | CDC73 | 60.577 | 2 | 53.98572 | yes | 1.318E-07 | 745.38794 | 116.46644 |
| FWETLDR | 7 | Q6P1J9 | Parafibromin OS=Homo sapiens OX=960 | CDC73 | 60.577 | 2 | 53.98572 | yes | 8.878E-07 | 483.73761 | 112.37337 |
| TKQPIPAAYNR | 11 | Q6P1J9 | Parafibromin OS=Homo sapiens OX=960 | CDC73 | 60.577 | 2 | 53.98572 | yes | 1.045E-07 | 629.84875 | 107.43283 |
| SLSEAMSVEK | 10 | Q6P1J9 | Parafibromin OS=Homo sapiens OX=960 | CDC73 | 60.577 | 2 | 53.98572 | yes | 4.633E-07 | 540.76569 | 87.528084 |
| NIFAILQSVK | 10 | Q6P1J9 | Parafibromin OS=Homo sapiens OX=960 | CDC73 | 60.577 | 2 | 53.98572 | yes | 7.176E-07 | 566.83984 | 68.655342 |
| KTQTPAAQPVPR | 12 | Q6P1J9 | Parafibromin OS=Homo sapiens OX=960 | CDC73 | 60.577 | 2 | 53.98572 | yes | 1.478E-07 | 647.36731 | 67.892487 |
| KDQMQPGGTAISV | 18 | Q6P1J9 | Parafibromin OS=Homo sapiens OX=960 | CDC73 | 60.577 | 3 | 53.98572 | yes | 6.58E-07 | 650.00342 | 45.966499 |
| TQTPAAQPVPR | 11 | Q6P1J9 | Parafibromin OS=Homo sapiens OX=960 | CDC73 | 60.577 | 2 | 53.98572 | yes | 1.18E-07 | 583.31982 | 44.931187 |
| PVSQARPPPNQK | 12 | Q6P1J9 | Parafibromin OS=Homo sapiens OX=960 | CDC73 | 60.577 | 3 | 53.98572 | yes | 1.466E-07 | 440.24573 | 34.541504 |
| LLDTMVDTFLQK | 12 | Q6P1K2 | Polyamine-modulated factor 1 OS=Homo | PMF1 | 23.34 | 2 | 46.546272 | yes | 1.532E-07 | 712.37872 | 96.050461 |
| FIAQLQTSIR | 10 | Q6P1K2 | Polyamine-modulated factor 1 OS=Homo | PMF1 | 23.34 | 2 | 46.546272 | yes | 4.3E-07 | 588.84039 | 38.259899 |
| TSPTVATQTGASA | 17 | Q6P1L5 | Protein FAM117B OS=Homo sapiens OX | FAM117B | 61.968 | 2 | 51.347294 | yes | 9.722E-08 | 818.91046 | 23.308994 |
| APPPLVQR | 8 | Q6P1L5 | Protein FAM117B OS=Homo sapiens OX | FAM117B | 61.968 | 2 | 51.347294 | yes | 1.222E-05 | 439.26636 | 9.2907267 |
| PLLPTPDLTLK | 11 | Q6P1L5 | Protein FAM117B OS=Homo sapiens OX | FAM117B | 61.968 | 2 | 51.347294 | yes | 4.549E-07 | 604.36847 | 9.1867809 |
| LGVEAALINTNLR | 13 | Q6P1M0 | Long-chain fatty acid transport protein 4 ( | SLC27A4 | 72.065 | 2 | 52.464603 | yes | 9.722E-08 | 692.40137 | 231.81076 |
| DPLFYLDAQK | 10 | Q6P1M0 | Long-chain fatty acid transport protein 4 ( | SLC27A4 | 72.065 | 2 | 52.464603 | yes | 1.486E-07 | 605.31116 | 138.14954 |
| DALLHCLTTSR | 11 | Q6P1M0 | Long-chain fatty acid transport protein 4 ( | SLC27A4 | 72.065 | 2 | 52.464603 | yes | 1.077E-06 | 643.82971 | 90.862961 |
| GENVSTTEVEGTL | 15 | Q6P1M0 | Long-chain fatty acid transport protein 4 ( | SLC27A4 | 72.065 | 2 | 52.464603 | yes | 2.396E-07 | 789.88391 | 74.271347 |
| GLASGDVAAIFME | 15 | Q6P1M0 | Long-chain fatty acid transport protein 4 ( | SLC27A4 | 72.065 | 2 | 52.464603 | yes | 1.489E-07 | 775.88519 | 58.100914 |
| QSIWTNFSSR | 10 | Q6P1M0 | Long-chain fatty acid transport protein 4 ( | SLC27A4 | 72.065 | 2 | 52.464603 | yes | 2.439E-07 | 613.30164 | 56.63409 |
| ILSFVYPIR | 9 | Q6P1M0 | Long-chain fatty acid transport protein 4 ( | SLC27A4 | 72.065 | 2 | 52.464603 | yes | 2.706E-07 | 554.33167 | 43.905575 |
| TVPILFASTVR | 11 | Q6P1M0 | Long-chain fatty acid transport protein 4 ( | SLC27A4 | 72.065 | 2 | 52.464603 | yes | 7.021E-07 | 602.35846 | 43.450497 |
| SLVPYGPFPCK | 11 | Q6P1M3 | LLGL scribble cell polarity complex com | LLGL2 | 113.448 | 2 | 54.39029 | yes | 1.519E-07 | 632.82336 | 337.65906 |
| FWDASGVCLR | 10 | Q6P1M3 | LLGL scribble cell polarity complex com | LLGL2 | 113.448 | 2 | 54.39029 | yes | 1.164E-07 | 605.78729 | 276.43906 |
| TISSDAVLQR | 10 | Q6P1M3 | LLGL scribble cell polarity complex com | LLGL2 | 113.448 | 2 | 54.39029 | yes | 1.863E-07 | 545.29858 | 276.03867 |
| VGSFDPYSDDPR | 12 | Q6P1M3 | LLGL scribble cell polarity complex com | LLGL2 | 113.448 | 2 | 54.39029 | yes | 9.926E-08 | 677.79913 | 273.22165 |
| EDVSGIASCVFTK | 13 | Q6P1M3 | LLGL scribble cell polarity complex com | LLGL2 | 113.448 | 2 | 54.39029 | yes | 9.722E-08 | 706.83997 | 265.4704 |
| VSVAHFGSR | 9 | Q6P1M3 | LLGL scribble cell polarity complex com | LLGL2 | 113.448 | 2 | 54.39029 | yes | 2.041E-07 | 480.2565 | 187.88589 |
| LTALEGSR | 8 | Q6P1M3 | LLGL scribble cell polarity complex com | LLGL2 | 113.448 | 2 | 54.39029 | yes | 3.172E-07 | 423.73761 | 135.98531 |
| SAEDSFTGFVR | 11 | Q6P1M3 | LLGL scribble cell polarity complex com | LLGL2 | 113.448 | 2 | 54.39029 | yes | 1.489E-07 | 608.28564 | 122.25588 |
| GGASELQEDESFT | 15 | Q6P1M3 | LLGL scribble cell polarity complex com | LLGL2 | 113.448 | 2 | 54.39029 | yes | 2.886E-07 | 819.88391 | 116.6467 |
| ILWLTTR | 7 | Q6P1M3 | LLGL scribble cell polarity complex com | LLGL2 | 113.448 | 2 | 54.39029 | yes | 1.486E-06 | 451.77655 | 111.96688 |
| QPGLVMER | 8 | Q6P1M3 | LLGL scribble cell polarity complex com | LLGL2 | 113.448 | 2 | 54.39029 | yes | 1.189E-07 | 465.24728 | 97.681396 |
| SPDMQGSHQLLVV | 19 | Q6P1M3 | LLGL scribble cell polarity complex com | LLGL2 | 113.448 | 3 | 54.39029 | yes | 1.034E-07 | 720.35278 | 89.837349 |
| VFEMVEALQEHPF | 13 | Q6P1M3 | LLGL scribble cell polarity complex com | LLGL2 | 113.448 | 2 | 54.39029 | yes | 1.192E-07 | 792.89557 | 37.635658 |
| VFEMVEALQEHPF | 13 | Q6P1M3 | LLGL scribble cell polarity complex com | LLGL2 | 113.448 | 2 | 54.39029 | yes | 2.655E-06 | 528.9328 | 37.635658 |
| VFEMVEALQEHPF | 13 | Q6P1M3 | LLGL scribble cell polarity complex com | LLGL2 | 113.448 | 3 | 54.39029 | yes | 1.192E-07 | 792.89557 | 37.635658 |
| VFEMVEALQEHPF | 13 | Q6P1M3 | LLGL scribble cell polarity complex com | LLGL2 | 113.448 | 3 | 54.39029 | yes | 2.655E-06 | 528.9328 | 37.635658 |

| | | | | | | | | | | |
|---|---|---|---|---|---|---|---|---|---|---|
| TLYFADTYLK | 10 | Q6P1M3 | LLGL scribble cell polarity complex com | LLGL2 | 113.448 | 2 | 54.39029 | yes | 1.174E-07 | 617.82135 | 59.361946 |
| FLRPGHDPVR | 10 | Q6P1M3 | LLGL scribble cell polarity complex com | LLGL2 | 113.448 | 2 | 54.39029 | yes | 9.722E-08 | 597.33057 | 59.206047 |
| GLVVIWDLQGSR | 12 | Q6P1M3 | LLGL scribble cell polarity complex com | LLGL2 | 113.448 | 2 | 54.39029 | yes | 9.722E-08 | 671.8775 | 46.308437 |
| EIQLMHR | 7 | Q6P1M3 | LLGL scribble cell polarity complex com | LLGL2 | 113.448 | 2 | 54.39029 | yes | 4.972E-07 | 463.74747 | 28.883993 |
| LALYQTAIESAR | 12 | Q6P1N0 | Coiled-coil and C2 domain-containing pr | CC2D1A | 104.063 | 2 | 51.775887 | yes | 3.503E-06 | 668.367 | 1215.0946 |
| AQLEQGGVGIR | 11 | Q6P1N0 | Coiled-coil and C2 domain-containing pr | CC2D1A | 104.063 | 2 | 51.775887 | yes | 9.885E-08 | 564.31201 | 295.4621 |
| NLVESELQR | 9 | Q6P1N0 | Coiled-coil and C2 domain-containing pr | CC2D1A | 104.063 | 2 | 51.775887 | yes | 3.649E-07 | 544.29077 | 256.8306 |
| LDALEIACEVR | 11 | Q6P1N0 | Coiled-coil and C2 domain-containing pr | CC2D1A | 104.063 | 2 | 51.775887 | yes | 9.722E-08 | 644.83191 | 230.61087 |
| TLLEALEQR | 9 | Q6P1N0 | Coiled-coil and C2 domain-containing pr | CC2D1A | 104.063 | 2 | 51.775887 | yes | 1.004E-07 | 536.80347 | 199.47556 |
| GLEPMLEASR | 10 | Q6P1N0 | Coiled-coil and C2 domain-containing pr | CC2D1A | 104.063 | 2 | 51.775887 | yes | 5.181E-07 | 551.78168 | 130.06931 |
| AQQQLAFLEGR | 11 | Q6P1N0 | Coiled-coil and C2 domain-containing pr | CC2D1A | 104.063 | 2 | 51.775887 | yes | 1.166E-07 | 630.83838 | 119.77368 |
| QLQFYTEAAR | 10 | Q6P1N0 | Coiled-coil and C2 domain-containing pr | CC2D1A | 104.063 | 2 | 51.775887 | yes | 1.002E-07 | 613.81183 | 117.26394 |
| TLENLLASIR | 10 | Q6P1N0 | Coiled-coil and C2 domain-containing pr | CC2D1A | 104.063 | 2 | 51.775887 | yes | 9.722E-08 | 565.3324 | 111.99041 |
| APPVPAPAR | 9 | Q6P1N0 | Coiled-coil and C2 domain-containing pr | CC2D1A | 104.063 | 2 | 51.775887 | yes | 5.278E-07 | 438.25851 | 70.268188 |
| SFDAVLEALSR | 11 | Q6P1N0 | Coiled-coil and C2 domain-containing pr | CC2D1A | 104.063 | 2 | 51.775887 | yes | 4.388E-07 | 604.31952 | 70.147156 |
| QLLQAALR | 8 | Q6P1N0 | Coiled-coil and C2 domain-containing pr | CC2D1A | 104.063 | 2 | 51.775887 | yes | 1.899E-07 | 456.78491 | 50.491272 |
| MASLCMR | 7 | Q6P1N0 | Coiled-coil and C2 domain-containing pr | CC2D1A | 104.063 | 2 | 51.775887 | yes | 6.305E-07 | 434.69553 | 35.962784 |
| VIPLISDAGLR | 11 | Q6P1Q0 | LETM1 domain-containing protein 1 OS= | LETMD1 | 41.791 | 2 | 50.507298 | yes | 1.055E-07 | 577.35059 | 347.33029 |
| FHQLPYR | 7 | Q6P1Q0 | LETM1 domain-containing protein 1 OS= | LETMD1 | 41.791 | 2 | 50.507298 | yes | 6.103E-06 | 480.75613 | 317.20856 |
| LGIGQLTAQEVK | 12 | Q6P1Q0 | LETM1 domain-containing protein 1 OS= | LETMD1 | 41.791 | 2 | 50.507298 | yes | 9.79E-08 | 628.86407 | 304.55685 |
| GLNSTHIGEDR | 11 | Q6P1Q0 | LETM1 domain-containing protein 1 OS= | LETMD1 | 41.791 | 2 | 50.507298 | yes | 1.616E-07 | 599.79419 | 118.68696 |
| THTTVIHQLDK | 11 | Q6P1Q0 | LETM1 domain-containing protein 1 OS= | LETMD1 | 41.791 | 2 | 50.507298 | yes | 1.478E-07 | 646.8515 | 90.617065 |
| QSHPEIISYLEK | 12 | Q6P1Q0 | LETM1 domain-containing protein 1 OS= | LETMD1 | 41.791 | 2 | 50.507298 | yes | 1.022E-07 | 722.37756 | 53.951859 |
| GTHPAIHDILALR | 13 | Q6P1Q0 | LETM1 domain-containing protein 1 OS= | LETMD1 | 41.791 | 2 | 50.507298 | yes | 5.23E-06 | 707.40167 | 17.339153 |
| TLDNLNPDVR | 10 | Q6P1X5 | Transcription initiation factor TFIID subu | TAF2 | 136.972 | 2 | 51.71588 | yes | 9.789E-08 | 578.80145 | 338.07834 |
| IAALEAVVDYTK | 12 | Q6P1X5 | Transcription initiation factor TFIID subu | TAF2 | 136.972 | 2 | 51.71588 | yes | 1.874E-07 | 646.85846 | 329.91156 |
| LLSLASTASSQK | 12 | Q6P1X5 | Transcription initiation factor TFIID subu | TAF2 | 136.972 | 2 | 51.71588 | yes | 9.722E-08 | 603.34045 | 304.17914 |
| IDPDMSVLR | 9 | Q6P1X5 | Transcription initiation factor TFIID subu | TAF2 | 136.972 | 2 | 51.71588 | yes | 1.219E-07 | 523.271 | 280.24371 |
| IANSMVSTWTGPP | 16 | Q6P1X5 | Transcription initiation factor TFIID subu | TAF2 | 136.972 | 2 | 51.71588 | yes | 9.903E-08 | 845.91821 | 248.1629 |
| HTITVSCLR | 9 | Q6P1X5 | Transcription initiation factor TFIID subu | TAF2 | 136.972 | 2 | 51.71588 | yes | 1.199E-07 | 543.78986 | 233.9256 |
| IHINFSLDQPK | 11 | Q6P1X5 | Transcription initiation factor TFIID subu | TAF2 | 136.972 | 2 | 51.71588 | yes | 2.606E-07 | 656.35645 | 222.67717 |
| MSWSDEWVLK | 10 | Q6P1X5 | Transcription initiation factor TFIID subu | TAF2 | 136.972 | 2 | 51.71588 | yes | 9.722E-08 | 640.80261 | 183.94415 |
| TFGVNEYR | 8 | Q6P1X5 | Transcription initiation factor TFIID subu | TAF2 | 136.972 | 2 | 51.71588 | yes | 3.363E-07 | 493.24051 | 182.4942 |
| LILEEITR | 8 | Q6P1X5 | Transcription initiation factor TFIID subu | TAF2 | 136.972 | 2 | 51.71588 | yes | 1.189E-07 | 493.79767 | 161.19489 |
| MSACFCLAK | 9 | Q6P1X5 | Transcription initiation factor TFIID subu | TAF2 | 136.972 | 2 | 51.71588 | yes | 3.445E-07 | 544.2403 | 110.19034 |
| DVVAQQESILALE | 14 | Q6P1X5 | Transcription initiation factor TFIID subu | TAF2 | 136.972 | 2 | 51.71588 | yes | 9.722E-08 | 771.9223 | 105.59575 |
| GGLHFVVPSVEGS | 17 | Q6P1X5 | Transcription initiation factor TFIID subu | TAF2 | 136.972 | 3 | 51.71588 | yes | 3.203E-07 | 591.29803 | 99.529831 |
| NGHVPSDPALFK | 12 | Q6P1X5 | Transcription initiation factor TFIID subu | TAF2 | 136.972 | 2 | 51.71588 | yes | 5.73E-07 | 641.33295 | 89.28421 |
| NVLELEIK | 8 | Q6P1X5 | Transcription initiation factor TFIID subu | TAF2 | 136.972 | 2 | 51.71588 | yes | 2.635E-06 | 479.28439 | 71.901962 |

| | | | | | | | | | | | |
|---|---|---|---|---|---|---|---|---|---|---|---|
| FSDNYYR | 7 | Q6P1X5 | Transcription initiation factor TFIID subu | TAF2 | 136.972 | 2 | 51.71588 | yes | 5.226E-07 | 482.71161 | 62.453392 |
| FQSHMWSQMLVS | 18 | Q6P1X5 | Transcription initiation factor TFIID subu | TAF2 | 136.972 | 3 | 51.71588 | yes | 2.782E-07 | 705.34479 | 26.256157 |
| LMNSGTSHDWR | 11 | Q6P1X5 | Transcription initiation factor TFIID subu | TAF2 | 136.972 | 2 | 51.71588 | yes | 8.931E-05 | 652.29602 | 22.239109 |
| LFQAPCQR | 8 | Q6P2C8 | Mediator of RNA polymerase II transcript | MED27 | 35.432 | 2 | 48.966034 | yes | 6.583E-07 | 510.25818 | 121.16502 |
| TPLYSQLLQAYK | 12 | Q6P2C8 | Mediator of RNA polymerase II transcript | MED27 | 35.432 | 2 | 48.966034 | yes | 1.083E-07 | 712.89282 | 59.809036 |
| VTDHATTALLHY( | 23 | Q6P2C8 | Mediator of RNA polymerase II transcript | MED27 | 35.432 | 4 | 48.966034 | yes | 8.546E-07 | 651.84552 | 46.080738 |
| FLQDGLPPTWR | 11 | Q6P2C8 | Mediator of RNA polymerase II transcript | MED27 | 35.432 | 2 | 48.966034 | yes | 4.73E-06 | 665.35114 | 37.021492 |
| SANQMGVSAK | 10 | Q6P2C8 | Mediator of RNA polymerase II transcript | MED27 | 35.432 | 2 | 48.966034 | yes | 6.927E-06 | 496.74512 | 36.607059 |
| FLITGADQNR | 10 | Q6P2E9 | Enhancer of mRNA-decapping protein 4 ( | EDC4 | 151.662 | 2 | 53.970535 | yes | 9.722E-08 | 567.79877 | 4355.0034 |
| EAFQSVVLPAFEK | 13 | Q6P2E9 | Enhancer of mRNA-decapping protein 4 ( | EDC4 | 151.662 | 2 | 53.970535 | yes | 1.06E-07 | 732.89032 | 3512.2585 |
| AAADTLQGPMQA | 15 | Q6P2E9 | Enhancer of mRNA-decapping protein 4 ( | EDC4 | 151.662 | 2 | 53.970535 | yes | 6.286E-07 | 782.38281 | 2746.3013 |
| EPVLAQLR | 8 | Q6P2E9 | Enhancer of mRNA-decapping protein 4 ( | EDC4 | 151.662 | 2 | 53.970535 | yes | 1.742E-07 | 463.27692 | 2585.281 |
| LCTQLEGLQSTVT( | 18 | Q6P2E9 | Enhancer of mRNA-decapping protein 4 ( | EDC4 | 151.662 | 3 | 53.970535 | yes | 1.884E-07 | 676.67749 | 2425.7463 |
| FWQIYIEGQDEPR | 13 | Q6P2E9 | Enhancer of mRNA-decapping protein 4 ( | EDC4 | 151.662 | 2 | 53.970535 | yes | 1.505E-07 | 840.90448 | 2059.8689 |
| EQQAAVTSSIMQA | 15 | Q6P2E9 | Enhancer of mRNA-decapping protein 4 ( | EDC4 | 151.662 | 2 | 53.970535 | yes | 3.051E-07 | 825.90051 | 2046.4911 |
| HTEVLPAEEENDS | 34 | Q6P2E9 | Enhancer of mRNA-decapping protein 4 ( | EDC4 | 151.662 | 4 | 53.970535 | yes | 1.857E-07 | 844.90448 | 1982.3905 |
| GLLPGLLPAPADK | 13 | Q6P2E9 | Enhancer of mRNA-decapping protein 4 ( | EDC4 | 151.662 | 2 | 53.970535 | yes | 3.023E-07 | 631.37933 | 1948.4935 |
| ETCSTLAESPR | 11 | Q6P2E9 | Enhancer of mRNA-decapping protein 4 ( | EDC4 | 151.662 | 2 | 53.970535 | yes | 2.675E-07 | 625.78772 | 1780.5171 |
| AEVWDLDMLR | 10 | Q6P2E9 | Enhancer of mRNA-decapping protein 4 ( | EDC4 | 151.662 | 2 | 53.970535 | yes | 1.102E-07 | 624.30811 | 766.55579 |
| AEVWDLDMLR | 10 | Q6P2E9 | Enhancer of mRNA-decapping protein 4 ( | EDC4 | 151.662 | 2 | 53.970535 | yes | 8.619E-07 | 632.30554 | 766.55579 |
| LSYLEEAVMHLDI | 19 | Q6P2E9 | Enhancer of mRNA-decapping protein 4 ( | EDC4 | 151.662 | 3 | 53.970535 | yes | 1.271E-07 | 742.7002 | 1502.127 |
| IVELPAPADFLSLS | 20 | Q6P2E9 | Enhancer of mRNA-decapping protein 4 ( | EDC4 | 151.662 | 3 | 53.970535 | yes | 3.168E-07 | 714.72797 | 1389.5754 |
| LFCVHTK | 7 | Q6P2E9 | Enhancer of mRNA-decapping protein 4 ( | EDC4 | 151.662 | 2 | 53.970535 | yes | 1.987E-07 | 452.73911 | 1273.635 |
| HSQEELLQR | 9 | Q6P2E9 | Enhancer of mRNA-decapping protein 4 ( | EDC4 | 151.662 | 2 | 53.970535 | yes | 1.116E-07 | 570.29382 | 1219.8496 |
| TMPPINLQEK | 10 | Q6P2E9 | Enhancer of mRNA-decapping protein 4 ( | EDC4 | 151.662 | 2 | 53.970535 | yes | 2.553E-07 | 585.81299 | 1063.7998 |
| EVEIVASSDSSISSI | 15 | Q6P2E9 | Enhancer of mRNA-decapping protein 4 ( | EDC4 | 151.662 | 2 | 53.970535 | yes | 1.003E-07 | 769.38324 | 973.06708 |
| LFQFLQAEPHNSL( | 15 | Q6P2E9 | Enhancer of mRNA-decapping protein 4 ( | EDC4 | 151.662 | 3 | 53.970535 | yes | 9.722E-08 | 576.97375 | 889.77332 |
| SSHSTWPVDVSQI | 14 | Q6P2E9 | Enhancer of mRNA-decapping protein 4 ( | EDC4 | 151.662 | 2 | 53.970535 | yes | 1.244E-07 | 785.89661 | 863.56555 |
| ALQDVQIR | 8 | Q6P2E9 | Enhancer of mRNA-decapping protein 4 ( | EDC4 | 151.662 | 2 | 53.970535 | yes | 1.074E-07 | 471.772 | 816.6991 |
| KQDPDVPFWR | 10 | Q6P2E9 | Enhancer of mRNA-decapping protein 4 ( | EDC4 | 151.662 | 2 | 53.970535 | yes | 1.728E-07 | 644.32764 | 749.79041 |
| AEVQHQLHVAVG | 24 | Q6P2E9 | Enhancer of mRNA-decapping protein 4 ( | EDC4 | 151.662 | 4 | 53.970535 | yes | 2.378E-07 | 660.86084 | 542.28656 |
| QPEGTPLNHFR | 11 | Q6P2E9 | Enhancer of mRNA-decapping protein 4 ( | EDC4 | 151.662 | 2 | 53.970535 | yes | 1.683E-07 | 648.32819 | 325.45901 |
| RIVELPAPADFLSL | 21 | Q6P2E9 | Enhancer of mRNA-decapping protein 4 ( | EDC4 | 151.662 | 3 | 53.970535 | yes | 1.911E-07 | 766.76166 | 183.20593 |
| VISVSTSER | 9 | Q6P2E9 | Enhancer of mRNA-decapping protein 4 ( | EDC4 | 151.662 | 2 | 53.970535 | yes | 2.216E-06 | 489.26672 | 165.92302 |
| IQEEILVHIR | 10 | Q6P2E9 | Enhancer of mRNA-decapping protein 4 ( | EDC4 | 151.662 | 2 | 53.970535 | yes | 9.722E-08 | 625.36676 | 151.815 |
| GHSTCLSEGALSPI | 28 | Q6P2E9 | Enhancer of mRNA-decapping protein 4 ( | EDC4 | 151.662 | 4 | 53.970535 | yes | 9.722E-08 | 708.33826 | 132.9902 |
| PLSCLLFCDNHK | 12 | Q6P2E9 | Enhancer of mRNA-decapping protein 4 ( | EDC4 | 151.662 | 2 | 53.970535 | yes | 2.214E-07 | 752.35779 | 111.99293 |
| LDRPAGGPSAESPI | 14 | Q6P2E9 | Enhancer of mRNA-decapping protein 4 ( | EDC4 | 151.662 | 2 | 53.970535 | yes | 2.311E-07 | 705.36023 | 101.76242 |
| SLEPMAGQLSNSV | 16 | Q6P2E9 | Enhancer of mRNA-decapping protein 4 ( | EDC4 | 151.662 | 2 | 53.970535 | yes | 2.058E-07 | 816.91669 | 100.59607 |

| | | | | | | | | | | | |
|---|---|---|---|---|---|---|---|---|---|---|---|
| RLSLMLHGLVTPS | 15 | Q6P2E9 | Enhancer of mRNA-decapping protein 4 ( | EDC4 | 151.662 | 2 | 53.970535 | yes | 9.157E-07 | 817.47656 | 47.537201 |
| DHMGSVMAQVR | 11 | Q6P2E9 | Enhancer of mRNA-decapping protein 4 ( | EDC4 | 151.662 | 2 | 53.970535 | yes | 1.244E-06 | 615.78955 | 46.601219 |
| LQLDGSLTMSSSG | 20 | Q6P2E9 | Enhancer of mRNA-decapping protein 4 ( | EDC4 | 151.662 | 3 | 53.970535 | yes | 1.297E-06 | 679.00897 | 35.341103 |
| CLHEWKPHDGR | 11 | Q6P2E9 | Enhancer of mRNA-decapping protein 4 ( | EDC4 | 151.662 | 2 | 53.970535 | yes | 3.035E-06 | 717.83838 | 30.297539 |
| PAPFGDVCLLR | 11 | Q6P2H3 | Centrosomal protein of 85 kDa OS=Homo | CEP85 | 85.64 | 2 | 54.944122 | yes | 1.025E-07 | 622.82642 | 991.74805 |
| LSASEVEVQLIR | 12 | Q6P2H3 | Centrosomal protein of 85 kDa OS=Homo | CEP85 | 85.64 | 2 | 54.944122 | yes | 2.953E-07 | 672.38007 | 406.68649 |
| NLTLQEHLR | 9 | Q6P2H3 | Centrosomal protein of 85 kDa OS=Homo | CEP85 | 85.64 | 2 | 54.944122 | yes | 9.722E-08 | 562.31458 | 380.69656 |
| ELSVQNQDLIEK | 12 | Q6P2H3 | Centrosomal protein of 85 kDa OS=Homo | CEP85 | 85.64 | 2 | 54.944122 | yes | 9.968E-08 | 708.37244 | 373.42697 |
| KLEEVQQLR | 9 | Q6P2H3 | Centrosomal protein of 85 kDa OS=Homo | CEP85 | 85.64 | 2 | 54.944122 | yes | 1.355E-07 | 571.83002 | 212.65186 |
| YLADLPTLEDHQK | 13 | Q6P2H3 | Centrosomal protein of 85 kDa OS=Homo | CEP85 | 85.64 | 2 | 54.944122 | yes | 1.974E-07 | 771.89355 | 175.54362 |
| EIQLESLR | 8 | Q6P2H3 | Centrosomal protein of 85 kDa OS=Homo | CEP85 | 85.64 | 2 | 54.944122 | yes | 7.856E-07 | 494.2771 | 169.13646 |
| EAEFSSAGHSLQD | 14 | Q6P2H3 | Centrosomal protein of 85 kDa OS=Homo | CEP85 | 85.64 | 2 | 54.944122 | yes | 9.722E-08 | 753.34698 | 146.75266 |
| DSALQQLR | 8 | Q6P2H3 | Centrosomal protein of 85 kDa OS=Homo | CEP85 | 85.64 | 2 | 54.944122 | yes | 2.443E-07 | 465.75378 | 138.8593 |
| QQLELIR | 7 | Q6P2H3 | Centrosomal protein of 85 kDa OS=Homo | CEP85 | 85.64 | 2 | 54.944122 | yes | 1.064E-07 | 450.26907 | 104.69445 |
| LPLSGLAESVGMT | 14 | Q6P2H3 | Centrosomal protein of 85 kDa OS=Homo | CEP85 | 85.64 | 2 | 54.944122 | yes | 9.722E-08 | 715.88721 | 100.52325 |
| SQVQSLEQEVAQE | 21 | Q6P2H3 | Centrosomal protein of 85 kDa OS=Homo | CEP85 | 85.64 | 3 | 54.944122 | yes | 9.722E-08 | 773.04706 | 32.658527 |
| KHISQLEQK | 9 | Q6P2H3 | Centrosomal protein of 85 kDa OS=Homo | CEP85 | 85.64 | 2 | 54.944122 | yes | 8.234E-07 | 555.81696 | 27.821501 |
| EQSCELSTCR | 10 | Q6P2H3 | Centrosomal protein of 85 kDa OS=Homo | CEP85 | 85.64 | 2 | 54.944122 | yes | 1.738E-07 | 635.26355 | 15.47488 |
| IDLTLLNR | 8 | Q6P2Q9 | Pre-mRNA-processing-splicing factor 8 C | PRPF8 | 273.601 | 2 | 54.528656 | yes | 1.28E-07 | 479.29001 | 681.91882 |
| GYLPSHYER | 9 | Q6P2Q9 | Pre-mRNA-processing-splicing factor 8 C | PRPF8 | 273.601 | 2 | 54.528656 | yes | 1.473E-07 | 561.27234 | 506.52078 |
| VQMLLSDR | 8 | Q6P2Q9 | Pre-mRNA-processing-splicing factor 8 C | PRPF8 | 273.601 | 2 | 54.528656 | yes | 1.177E-07 | 481.26041 | 450.56091 |
| LTPSGYEWGR | 10 | Q6P2Q9 | Pre-mRNA-processing-splicing factor 8 C | PRPF8 | 273.601 | 2 | 54.528656 | yes | 2.353E-07 | 583.28546 | 434.38449 |
| SLPVEEQPK | 9 | Q6P2Q9 | Pre-mRNA-processing-splicing factor 8 C | PRPF8 | 273.601 | 2 | 54.528656 | yes | 2.533E-07 | 513.77692 | 428.67487 |
| WLGNLLAR | 8 | Q6P2Q9 | Pre-mRNA-processing-splicing factor 8 C | PRPF8 | 273.601 | 2 | 54.528656 | yes | 2.39E-07 | 471.77963 | 428.65286 |
| TNHIYVSSDDIK | 12 | Q6P2Q9 | Pre-mRNA-processing-splicing factor 8 C | PRPF8 | 273.601 | 2 | 54.528656 | yes | 9.722E-08 | 696.34369 | 399.38669 |
| LDWVEVGLQVCR | 12 | Q6P2Q9 | Pre-mRNA-processing-splicing factor 8 C | PRPF8 | 273.601 | 2 | 54.528656 | yes | 9.722E-08 | 737.37958 | 398.1564 |
| AISAANLHLR | 10 | Q6P2Q9 | Pre-mRNA-processing-splicing factor 8 C | PRPF8 | 273.601 | 2 | 54.528656 | yes | 1.779E-07 | 533.31183 | 377.23236 |
| DIILGMEISAPSQQ | 15 | Q6P2Q9 | Pre-mRNA-processing-splicing factor 8 C | PRPF8 | 273.601 | 2 | 54.528656 | yes | 2.225E-07 | 829.43256 | 376.84766 |
| IMADNPSWDGEK | 12 | Q6P2Q9 | Pre-mRNA-processing-splicing factor 8 C | PRPF8 | 273.601 | 2 | 54.528656 | yes | 1.026E-07 | 681.80334 | 354.38162 |
| LVVDSHVQYR | 10 | Q6P2Q9 | Pre-mRNA-processing-splicing factor 8 C | PRPF8 | 273.601 | 2 | 54.528656 | yes | 9.722E-08 | 608.32764 | 347.90628 |
| LLENMPMPWEQIF | 13 | Q6P2Q9 | Pre-mRNA-processing-splicing factor 8 C | PRPF8 | 273.601 | 2 | 54.528656 | yes | 9.819E-08 | 828.91547 | 322.66144 |
| FTLWWSPTINR | 11 | Q6P2Q9 | Pre-mRNA-processing-splicing factor 8 C | PRPF8 | 273.601 | 2 | 54.528656 | yes | 7.847E-07 | 710.87225 | 313.121 |
| TILQHLSEAWR | 11 | Q6P2Q9 | Pre-mRNA-processing-splicing factor 8 C | PRPF8 | 273.601 | 2 | 54.528656 | yes | 9.722E-08 | 677.36731 | 280.18417 |
| EQSQLTATQTR | 11 | Q6P2Q9 | Pre-mRNA-processing-splicing factor 8 C | PRPF8 | 273.601 | 2 | 54.528656 | yes | 1.219E-07 | 631.82037 | 277.51837 |
| YWIDIQLR | 8 | Q6P2Q9 | Pre-mRNA-processing-splicing factor 8 C | PRPF8 | 273.601 | 2 | 54.528656 | yes | 1.018E-07 | 553.80328 | 253.79602 |
| YELQLANPK | 9 | Q6P2Q9 | Pre-mRNA-processing-splicing factor 8 C | PRPF8 | 273.601 | 2 | 54.528656 | yes | 1.057E-06 | 538.29279 | 252.01096 |
| FGFVDAQK | 8 | Q6P2Q9 | Pre-mRNA-processing-splicing factor 8 C | PRPF8 | 273.601 | 2 | 54.528656 | yes | 1.02E-07 | 456.23471 | 221.67863 |
| NNVNVASLTQSEI | 14 | Q6P2Q9 | Pre-mRNA-processing-splicing factor 8 C | PRPF8 | 273.601 | 2 | 54.528656 | yes | 1.183E-07 | 772.90497 | 217.58469 |
| IIHTSVWAGQK | 11 | Q6P2Q9 | Pre-mRNA-processing-splicing factor 8 C | PRPF8 | 273.601 | 2 | 54.528656 | yes | 9.722E-08 | 620.34583 | 213.1965 |

| | | | | | | | | | | | |
|---|---|---|---|---|---|---|---|---|---|---|---|
| ETGYTYILPK | 10 | Q6P2Q9 | Pre-mRNA-processing-splicing factor 8 C | PRPF8 | 273.601 | 2 | 54.528656 | yes | 9.722E-08 | 592.81354 | 206.27319 |
| YMPHAVLK | 8 | Q6P2Q9 | Pre-mRNA-processing-splicing factor 8 C | PRPF8 | 273.601 | 2 | 54.528656 | yes | 8.149E-07 | 479.76257 | 203.66721 |
| QTDVGITHFR | 10 | Q6P2Q9 | Pre-mRNA-processing-splicing factor 8 C | PRPF8 | 273.601 | 2 | 54.528656 | yes | 9.722E-08 | 587.3042 | 198.30827 |
| GSELQLPFQACLK | 13 | Q6P2Q9 | Pre-mRNA-processing-splicing factor 8 C | PRPF8 | 273.601 | 2 | 54.528656 | yes | 2.938E-06 | 745.88721 | 193.10782 |
| FSPIPFPPLSYK | 12 | Q6P2Q9 | Pre-mRNA-processing-splicing factor 8 C | PRPF8 | 273.601 | 2 | 54.528656 | yes | 3.817E-07 | 696.88171 | 186.30544 |
| ANPALYVLR | 9 | Q6P2Q9 | Pre-mRNA-processing-splicing factor 8 C | PRPF8 | 273.601 | 2 | 54.528656 | yes | 3.224E-07 | 508.798 | 176.90385 |
| DGVWNLQNEVTK | 12 | Q6P2Q9 | Pre-mRNA-processing-splicing factor 8 C | PRPF8 | 273.601 | 2 | 54.528656 | yes | 9.722E-08 | 701.85168 | 161.54756 |
| TSYEEFTHK | 9 | Q6P2Q9 | Pre-mRNA-processing-splicing factor 8 C | PRPF8 | 273.601 | 2 | 54.528656 | yes | 1.562E-06 | 571.26166 | 157.31003 |
| LIVDHNIADYMTA | 14 | Q6P2Q9 | Pre-mRNA-processing-splicing factor 8 C | PRPF8 | 273.601 | 2 | 54.528656 | yes | 9.722E-08 | 802.41107 | 156.71593 |
| QILMASGSTTFTK | 13 | Q6P2Q9 | Pre-mRNA-processing-splicing factor 8 C | PRPF8 | 273.601 | 2 | 54.528656 | yes | 1.074E-07 | 692.86066 | 156.5451 |
| VILKPDK | 7 | Q6P2Q9 | Pre-mRNA-processing-splicing factor 8 C | PRPF8 | 273.601 | 2 | 54.528656 | yes | 7.408E-07 | 406.76566 | 153.14136 |
| NNVVINYK | 8 | Q6P2Q9 | Pre-mRNA-processing-splicing factor 8 C | PRPF8 | 273.601 | 2 | 54.528656 | yes | 1.61E-06 | 482.26654 | 146.21176 |
| ISLIQIFR | 8 | Q6P2Q9 | Pre-mRNA-processing-splicing factor 8 C | PRPF8 | 273.601 | 2 | 54.528656 | yes | 9.722E-08 | 495.31076 | 143.55627 |
| KWQQLQAK | 8 | Q6P2Q9 | Pre-mRNA-processing-splicing factor 8 C | PRPF8 | 273.601 | 2 | 54.528656 | yes | 9.722E-08 | 515.29565 | 134.88785 |
| PADTEPPPLLVYK | 13 | Q6P2Q9 | Pre-mRNA-processing-splicing factor 8 C | PRPF8 | 273.601 | 2 | 54.528656 | yes | 2.941E-07 | 720.39264 | 124.63657 |
| EMEPLGWIHTQPN | 28 | Q6P2Q9 | Pre-mRNA-processing-splicing factor 8 C | PRPF8 | 273.601 | 4 | 54.528656 | yes | 2.641E-07 | 793.38574 | 116.32654 |
| QNPFWWTHQR | 10 | Q6P2Q9 | Pre-mRNA-processing-splicing factor 8 C | PRPF8 | 273.601 | 2 | 54.528656 | yes | 9.722E-08 | 700.33636 | 112.66246 |
| YYVLNALK | 8 | Q6P2Q9 | Pre-mRNA-processing-splicing factor 8 C | PRPF8 | 273.601 | 2 | 54.528656 | yes | 9.722E-08 | 492.28165 | 105.82526 |
| WGDYDSHDIER | 11 | Q6P2Q9 | Pre-mRNA-processing-splicing factor 8 C | PRPF8 | 273.601 | 2 | 54.528656 | yes | 9.722E-08 | 696.79437 | 105.43188 |
| TAEEVAALIR | 10 | Q6P2Q9 | Pre-mRNA-processing-splicing factor 8 C | PRPF8 | 273.601 | 2 | 54.528656 | yes | 2.255E-06 | 536.80347 | 102.2981 |
| GITPLLER | 8 | Q6P2Q9 | Pre-mRNA-processing-splicing factor 8 C | PRPF8 | 273.601 | 2 | 54.528656 | yes | 8.847E-07 | 449.77145 | 101.30173 |
| VWAEYALK | 8 | Q6P2Q9 | Pre-mRNA-processing-splicing factor 8 C | PRPF8 | 273.601 | 2 | 54.528656 | yes | 1.592E-06 | 490.26602 | 98.6269 |
| VPGLPTPIENMILR | 14 | Q6P2Q9 | Pre-mRNA-processing-splicing factor 8 C | PRPF8 | 273.601 | 2 | 54.528656 | yes | 3.768E-07 | 775.44214 | 96.937057 |
| ALNMAIPGGPK | 11 | Q6P2Q9 | Pre-mRNA-processing-splicing factor 8 C | PRPF8 | 273.601 | 2 | 54.528656 | yes | 2.036E-07 | 534.79718 | 96.770706 |
| DLILADYGK | 9 | Q6P2Q9 | Pre-mRNA-processing-splicing factor 8 C | PRPF8 | 273.601 | 2 | 54.528656 | yes | 1.754E-07 | 504.27402 | 89.844818 |
| EDMPPEHVR | 9 | Q6P2Q9 | Pre-mRNA-processing-splicing factor 8 C | PRPF8 | 273.601 | 2 | 54.528656 | yes | 1.001E-06 | 555.25586 | 77.732742 |
| FGNAFHLCR | 9 | Q6P2Q9 | Pre-mRNA-processing-splicing factor 8 C | PRPF8 | 273.601 | 2 | 54.528656 | yes | 3.541E-07 | 561.2691 | 76.102974 |
| LTLEDLEDSWDR | 12 | Q6P2Q9 | Pre-mRNA-processing-splicing factor 8 C | PRPF8 | 273.601 | 2 | 54.528656 | yes | 9.722E-08 | 746.35175 | 72.247238 |
| PLIQQAMAK | 9 | Q6P2Q9 | Pre-mRNA-processing-splicing factor 8 C | PRPF8 | 273.601 | 2 | 54.528656 | yes | 3.277E-07 | 500.28641 | 69.499832 |
| EFYHEVHR | 8 | Q6P2Q9 | Pre-mRNA-processing-splicing factor 8 C | PRPF8 | 273.601 | 2 | 54.528656 | yes | 2.905E-07 | 558.76471 | 55.874794 |
| FPPVVFYTPK | 10 | Q6P2Q9 | Pre-mRNA-processing-splicing factor 8 C | PRPF8 | 273.601 | 2 | 54.528656 | yes | 6.576E-07 | 597.83154 | 49.54055 |
| EELGLIEQAYDNPI | 19 | Q6P2Q9 | Pre-mRNA-processing-splicing factor 8 C | PRPF8 | 273.601 | 3 | 54.528656 | yes | 5.548E-07 | 728.69043 | 42.980522 |
| ALDIPLVK | 8 | Q6P2Q9 | Pre-mRNA-processing-splicing factor 8 C | PRPF8 | 273.601 | 2 | 54.528656 | yes | 1.709E-07 | 434.77875 | 26.424488 |
| YGLTGPWIK | 9 | Q6P4A7 | Sideroflexin-4 OS=Homo sapiens OX=96 | SFXN4 | 37.999 | 2 | 49.64179 | yes | 3.103E-07 | 517.78711 | 207.00136 |
| SLATVHPDSSNLIP | 15 | Q6P4A7 | Sideroflexin-4 OS=Homo sapiens OX=96 | SFXN4 | 37.999 | 2 | 49.64179 | yes | 4.46E-07 | 789.92792 | 141.96111 |
| SYTCKPLER | 9 | Q6P4A7 | Sideroflexin-4 OS=Homo sapiens OX=96 | SFXN4 | 37.999 | 2 | 49.64179 | yes | 1.641E-07 | 577.28717 | 113.10223 |
| DAVPAFIEPNVR | 12 | Q6P4A7 | Sideroflexin-4 OS=Homo sapiens OX=96 | SFXN4 | 37.999 | 2 | 49.64179 | yes | 1.014E-07 | 664.35388 | 112.04459 |
| KNPGSLWILK | 10 | Q6P4A7 | Sideroflexin-4 OS=Homo sapiens OX=96 | SFXN4 | 37.999 | 2 | 49.64179 | yes | 5.553E-07 | 578.34784 | 58.677212 |
| SLEQEEETQPGR | 12 | Q6P4A7 | Sideroflexin-4 OS=Homo sapiens OX=96 | SFXN4 | 37.999 | 2 | 49.64179 | yes | 4.021E-07 | 722.83118 | 32.238033 |

| | | | | | | | | | | | |
|---|---|---|---|---|---|---|---|---|---|---|---|
| LYLPQEPTSEETPR | 14 | Q6P5Z2 | Serine/threonine-protein kinase N3 OS=H | PKN3 | 99.421 | 2 | 53.944496 | yes | 1.041E-07 | 830.41486 | 596.99567 |
| TTNWQALLAR | 10 | Q6P5Z2 | Serine/threonine-protein kinase N3 OS=H | PKN3 | 99.421 | 2 | 53.944496 | yes | 1.11E-07 | 587.32239 | 328.3515 |
| LHVEAAVAEGAK | 12 | Q6P5Z2 | Serine/threonine-protein kinase N3 OS=H | PKN3 | 99.421 | 2 | 53.944496 | yes | 1.41E-07 | 597.82751 | 303.74033 |
| SPAAALASSPSEGV | 16 | Q6P5Z2 | Serine/threonine-protein kinase N3 OS=H | PKN3 | 99.421 | 2 | 53.944496 | yes | 1.017E-07 | 800.40991 | 284.14627 |
| ELEIGVHWR | 9 | Q6P5Z2 | Serine/threonine-protein kinase N3 OS=H | PKN3 | 99.421 | 2 | 53.944496 | yes | 1.055E-07 | 569.80383 | 258.19952 |
| LLGCEQLLTAVPG | 14 | Q6P5Z2 | Serine/threonine-protein kinase N3 OS=H | PKN3 | 99.421 | 2 | 53.944496 | yes | 9.722E-08 | 763.92163 | 181.76772 |
| LGAGEQDAEEIK | 12 | Q6P5Z2 | Serine/threonine-protein kinase N3 OS=H | PKN3 | 99.421 | 2 | 53.944496 | yes | 9.722E-08 | 630.30933 | 176.20703 |
| ALAEAQAQLQESS | 15 | Q6P5Z2 | Serine/threonine-protein kinase N3 OS=H | PKN3 | 99.421 | 2 | 53.944496 | yes | 1.423E-07 | 801.4101 | 163.1077 |
| QLCGVAFLR | 9 | Q6P5Z2 | Serine/threonine-protein kinase N3 OS=H | PKN3 | 99.421 | 2 | 53.944496 | yes | 9.722E-08 | 532.28949 | 135.60526 |
| LLAAAQQMLR | 10 | Q6P5Z2 | Serine/threonine-protein kinase N3 OS=H | PKN3 | 99.421 | 2 | 53.944496 | yes | 9.722E-08 | 557.82373 | 131.04874 |
| DEIESLYCEK | 10 | Q6P5Z2 | Serine/threonine-protein kinase N3 OS=H | PKN3 | 99.421 | 2 | 53.944496 | yes | 1.41E-07 | 643.28448 | 111.30889 |
| RLEQLHGELR | 10 | Q6P5Z2 | Serine/threonine-protein kinase N3 OS=H | PKN3 | 99.421 | 3 | 53.944496 | yes | 9.722E-08 | 417.57031 | 101.5918 |
| QLHVELK | 7 | Q6P5Z2 | Serine/threonine-protein kinase N3 OS=H | PKN3 | 99.421 | 2 | 53.944496 | yes | 8.353E-07 | 433.75836 | 85.240135 |
| VSEPVLSLPVR | 11 | Q6P6C2 | RNA demethylase ALKBH5 OS=Homo s | ALKBH5 | 44.256 | 2 | 52.105564 | yes | 9.722E-08 | 598.3559 | 1143.9744 |
| SVLLPTHR | 8 | Q6P6C2 | RNA demethylase ALKBH5 OS=Homo s | ALKBH5 | 44.256 | 2 | 52.105564 | yes | 9.722E-08 | 461.77707 | 858.24323 |
| LFSQDECAK | 9 | Q6P6C2 | RNA demethylase ALKBH5 OS=Homo s | ALKBH5 | 44.256 | 2 | 52.105564 | yes | 9.066E-07 | 549.25024 | 488.60632 |
| SLSSSVLPPSYASD | 15 | Q6P6C2 | RNA demethylase ALKBH5 OS=Homo s | ALKBH5 | 44.256 | 2 | 52.105564 | yes | 2.801E-07 | 783.39392 | 463.74881 |
| GLYNEHTVDR | 10 | Q6P6C2 | RNA demethylase ALKBH5 OS=Homo s | ALKBH5 | 44.256 | 2 | 52.105564 | yes | 4.507E-07 | 602.29126 | 221.52509 |
| GSFSSENYWR | 10 | Q6P6C2 | RNA demethylase ALKBH5 OS=Homo s | ALKBH5 | 44.256 | 2 | 52.105564 | yes | 1.004E-07 | 616.7702 | 144.02699 |
| SDYEEQQLQK | 10 | Q6P6C2 | RNA demethylase ALKBH5 OS=Homo s | ALKBH5 | 44.256 | 2 | 52.105564 | yes | 9.722E-08 | 634.2937 | 121.42975 |
| AAASGYTDLR | 10 | Q6P6C2 | RNA demethylase ALKBH5 OS=Homo s | ALKBH5 | 44.256 | 2 | 52.105564 | yes | 2.039E-06 | 533.76184 | 80.542946 |
| ILEMDKEENR | 10 | Q6P6C2 | RNA demethylase ALKBH5 OS=Homo s | ALKBH5 | 44.256 | 2 | 52.105564 | yes | 1.102E-06 | 638.81372 | 63.396954 |
| LYPPGDVDEIPEW | 21 | Q6P6C2 | RNA demethylase ALKBH5 OS=Homo s | ALKBH5 | 44.256 | 3 | 52.105564 | yes | 0.0001297 | 825.7688 | 32.185646 |
| MLEHALR | 7 | Q6P9F0 | Coiled-coil domain-containing protein 62 | CCDC62 | 77.749 | 2 | 23.137278 | yes | 3.709E-06 | 435.23672 | 27.877399 |
| FDLALAATEAR | 11 | Q6PD62 | RNA polymerase-associated protein CTR | CTR9 | 133.503 | 2 | 51.401604 | yes | 9.722E-08 | 589.31421 | 153.11313 |
| ILGSLYAASEDQEI | 14 | Q6PD62 | RNA polymerase-associated protein CTR | CTR9 | 133.503 | 2 | 51.401604 | yes | 3.934E-07 | 762.38306 | 59.967323 |
| NILMFTGETEATK | 13 | Q6PD62 | RNA polymerase-associated protein CTR | CTR9 | 133.503 | 2 | 51.401604 | yes | 1.173E-06 | 727.8634 | 58.370651 |
| KYFLASLDR | 9 | Q6PD62 | RNA polymerase-associated protein CTR | CTR9 | 133.503 | 2 | 51.401604 | yes | 9.601E-07 | 556.80859 | 58.054489 |
| TNPGCPAEVR | 10 | Q6PD62 | RNA polymerase-associated protein CTR | CTR9 | 133.503 | 2 | 51.401604 | yes | 1.755E-06 | 550.76129 | 53.228901 |
| WLVQNGEK | 8 | Q6PF05 | Tetratricopeptide repeat protein 23-like O | TTC23L | 40.837 | 2 | 45.655148 | yes | 1.783E-07 | 487.25873 | 181.12907 |
| SSSQLPIPSAIPR | 13 | Q6PGN9 | Proline/serine-rich coiled-coil protein 1 C | PSRC1 | 38.797 | 2 | 51.382626 | yes | 1.013E-07 | 676.88025 | 592.36609 |
| SDPNAVAPAPQGV | 14 | Q6PGN9 | Proline/serine-rich coiled-coil protein 1 C | PSRC1 | 38.797 | 2 | 51.382626 | yes | 1.344E-07 | 689.8573 | 535.55896 |
| LAAQLEQCALQDF | 13 | Q6PGN9 | Proline/serine-rich coiled-coil protein 1 C | PSRC1 | 38.797 | 2 | 51.382626 | yes | 9.722E-08 | 758.38281 | 352.89642 |
| LEEILDEANR | 10 | Q6PGN9 | Proline/serine-rich coiled-coil protein 1 C | PSRC1 | 38.797 | 2 | 51.382626 | yes | 1.462E-06 | 601.30658 | 311.60535 |
| STPSPSSLTPR | 11 | Q6PGN9 | Proline/serine-rich coiled-coil protein 1 C | PSRC1 | 38.797 | 2 | 51.382626 | yes | 1.252E-06 | 565.29602 | 255.46797 |
| SVLAPQPSTSNSQI | 14 | Q6PGN9 | Proline/serine-rich coiled-coil protein 1 C | PSRC1 | 38.797 | 2 | 51.382626 | yes | 1.269E-07 | 736.3786 | 245.34264 |
| LSLGPLSPEK | 10 | Q6PGN9 | Proline/serine-rich coiled-coil protein 1 C | PSRC1 | 38.797 | 2 | 51.382626 | yes | 3.218E-07 | 520.80298 | 168.05107 |
| SPASSPLTR | 9 | Q6PGN9 | Proline/serine-rich coiled-coil protein 1 C | PSRC1 | 38.797 | 2 | 51.382626 | yes | 9.989E-08 | 458.24835 | 153.8499 |
| ALSIVESEQDK | 11 | Q6PGP7 | Superkiller complex protein 3 OS=Homo | SKIC3 | 175.487 | 2 | 56.58588 | yes | 3.87E-07 | 609.81427 | 243.31955 |

| | | | | | | | | | | | |
|---|---|---|---|---|---|---|---|---|---|---|---|
| LSDYDSSEEAIR | 12 | Q6PGP7 | Superkiller complex protein 3 OS=Homo | SKIC3 | 175.487 | 2 | 56.58588 | yes | 9.722E-08 | 692.815 | 226.1142 |
| IVDNLGASGNSLY | 15 | Q6PGP7 | Superkiller complex protein 3 OS=Homo | SKIC3 | 175.487 | 2 | 56.58588 | yes | 9.809E-08 | 803.91284 | 193.80612 |
| FFENYNQSLEK | 11 | Q6PGP7 | Superkiller complex protein 3 OS=Homo | SKIC3 | 175.487 | 2 | 56.58588 | yes | 2.348E-07 | 709.83295 | 191.20879 |
| VVYQPGYPK | 9 | Q6PGP7 | Superkiller complex protein 3 OS=Homo | SKIC3 | 175.487 | 2 | 56.58588 | yes | 1.435E-06 | 525.78455 | 186.99319 |
| VAAIQQILGK | 10 | Q6PGP7 | Superkiller complex protein 3 OS=Homo | SKIC3 | 175.487 | 2 | 56.58588 | yes | 1.305E-07 | 520.82678 | 154.77563 |
| AALVDYLDGK | 10 | Q6PGP7 | Superkiller complex protein 3 OS=Homo | SKIC3 | 175.487 | 2 | 56.58588 | yes | 2.06E-06 | 532.78479 | 147.42204 |
| LLDLYESVDK | 10 | Q6PGP7 | Superkiller complex protein 3 OS=Homo | SKIC3 | 175.487 | 2 | 56.58588 | yes | 1.229E-07 | 597.81628 | 140.91446 |
| SGPGLIGLGIK | 11 | Q6PGP7 | Superkiller complex protein 3 OS=Homo | SKIC3 | 175.487 | 2 | 56.58588 | yes | 1.17E-07 | 506.31351 | 128.41772 |
| VVAQYAQR | 8 | Q6PGP7 | Superkiller complex protein 3 OS=Homo | SKIC3 | 175.487 | 2 | 56.58588 | yes | 3.07E-07 | 467.75888 | 96.784401 |
| SNPDQPAVILLLR | 13 | Q6PGP7 | Superkiller complex protein 3 OS=Homo | SKIC3 | 175.487 | 2 | 56.58588 | yes | 9.722E-08 | 718.41699 | 88.985703 |
| NELLHLGGR | 9 | Q6PGP7 | Superkiller complex protein 3 OS=Homo | SKIC3 | 175.487 | 2 | 56.58588 | yes | 3.398E-07 | 504.7829 | 88.205696 |
| ISEAETLCTK | 10 | Q6PGP7 | Superkiller complex protein 3 OS=Homo | SKIC3 | 175.487 | 2 | 56.58588 | yes | 3.437E-07 | 576.28424 | 78.960999 |
| AHILTALAITEYK | 13 | Q6PGP7 | Superkiller complex protein 3 OS=Homo | SKIC3 | 175.487 | 2 | 56.58588 | yes | 9.722E-08 | 722.41394 | 36.128044 |
| AHILTALAITEYK | 13 | Q6PGP7 | Superkiller complex protein 3 OS=Homo | SKIC3 | 175.487 | 2 | 56.58588 | yes | 3.336E-07 | 481.94504 | 36.128044 |
| AHILTALAITEYK | 13 | Q6PGP7 | Superkiller complex protein 3 OS=Homo | SKIC3 | 175.487 | 3 | 56.58588 | yes | 9.722E-08 | 722.41394 | 36.128044 |
| AHILTALAITEYK | 13 | Q6PGP7 | Superkiller complex protein 3 OS=Homo | SKIC3 | 175.487 | 3 | 56.58588 | yes | 3.336E-07 | 481.94504 | 36.128044 |
| ASELNPESIYSVFK | 14 | Q6PGP7 | Superkiller complex protein 3 OS=Homo | SKIC3 | 175.487 | 2 | 56.58588 | yes | 1.876E-07 | 792.40125 | 66.729118 |
| KNELLHLGGR | 10 | Q6PGP7 | Superkiller complex protein 3 OS=Homo | SKIC3 | 175.487 | 2 | 56.58588 | yes | 3.72E-07 | 568.83038 | 62.092911 |
| AGQHSQAVADLQ. | 16 | Q6PGP7 | Superkiller complex protein 3 OS=Homo | SKIC3 | 175.487 | 3 | 56.58588 | yes | 9.722E-08 | 545.95691 | 52.073101 |
| VNVHVLGVLLGQI | 13 | Q6PGP7 | Superkiller complex protein 3 OS=Homo | SKIC3 | 175.487 | 2 | 56.58588 | yes | 1.903E-07 | 688.42462 | 48.781677 |
| ALLYTAVNQLAM( | 21 | Q6PGP7 | Superkiller complex protein 3 OS=Homo | SKIC3 | 175.487 | 3 | 56.58588 | yes | 3.656E-07 | 733.35968 | 36.92992 |
| EAVAQYQMIIK | 11 | Q6PGP7 | Superkiller complex protein 3 OS=Homo | SKIC3 | 175.487 | 2 | 56.58588 | yes | 4.853E-07 | 647.34723 | 34.924393 |
| AALLSPGDPAIWA | 24 | Q6PGP7 | Superkiller complex protein 3 OS=Homo | SKIC3 | 175.487 | 3 | 56.58588 | yes | 2.617E-07 | 817.73138 | 26.124313 |
| DDYELIDVLVNNA | 14 | Q6PGP7 | Superkiller complex protein 3 OS=Homo | SKIC3 | 175.487 | 2 | 56.58588 | yes | 3.163E-05 | 810.90942 | 25.722677 |
| IPSEDHQVLYR | 11 | Q6PGP7 | Superkiller complex protein 3 OS=Homo | SKIC3 | 175.487 | 2 | 56.58588 | yes | 2.414E-06 | 678.84894 | 17.146748 |
| LICLVTGSPSIR | 12 | Q6PI48 | Aspartate--tRNA ligase, mitochondrial O! | DARS2 | 73.563 | 2 | 41.240612 | yes | 3.607E-07 | 658.37378 | 66.018799 |
| WALLSCR | 7 | Q6PID8 | Kelch domain-containing protein 10 OS= | KLHDC10 | 49.098 | 2 | 39.86967 | yes | 4.336E-07 | 453.23672 | 46.111687 |
| LNLQTFQWVK | 10 | Q6PID8 | Kelch domain-containing protein 10 OS= | KLHDC10 | 49.098 | 2 | 39.86967 | yes | 7.791E-07 | 638.85608 | 32.11414 |
| IVQELPQLLDAR | 12 | Q6PIU2 | Neutral cholesterol ester hydrolase 1 OS= | NCEH1 | 45.808 | 2 | 39.074417 | yes | 4.804E-07 | 697.90375 | 153.29082 |
| VTDTDFDGVEVR | 12 | Q6PIU2 | Neutral cholesterol ester hydrolase 1 OS= | NCEH1 | 45.808 | 2 | 39.074417 | yes | 7.493E-05 | 676.82007 | 48.816837 |
| EIQLLFPQSR | 10 | Q6PIY7 | Poly(A) RNA polymerase GLD2 OS=Hoi | TENT2 | 56.028 | 2 | 48.175655 | yes | 1.187E-07 | 615.8457 | 213.53606 |
| LSGYIERPQLIR | 12 | Q6PIY7 | Poly(A) RNA polymerase GLD2 OS=Hoi | TENT2 | 56.028 | 3 | 48.175655 | yes | 2.801E-07 | 482.28055 | 113.54562 |
| YSMPPLFHTHYVP | 17 | Q6PIY7 | Poly(A) RNA polymerase GLD2 OS=Hoi | TENT2 | 56.028 | 3 | 48.175655 | yes | 1.388E-07 | 691.35223 | 105.30296 |
| VRPLVLVIK | 9 | Q6PIY7 | Poly(A) RNA polymerase GLD2 OS=Hoi | TENT2 | 56.028 | 2 | 48.175655 | yes | 2.013E-07 | 518.86572 | 77.924545 |
| EITLPEAK | 8 | Q6PIY7 | Poly(A) RNA polymerase GLD2 OS=Hoi | TENT2 | 56.028 | 2 | 48.175655 | yes | 3.3E-07 | 450.75546 | 20.492064 |
| QASLSQSIR | 9 | Q6PJF5 | Inactive rhomboid protein 2 OS=Homo sa | RHBDF2 | 96.687 | 2 | 46.85072 | yes | 1.456E-07 | 495.27237 | 92.15818 |
| SVLDATGQR | 9 | Q6PJF5 | Inactive rhomboid protein 2 OS=Homo sa | RHBDF2 | 96.687 | 2 | 46.85072 | yes | 6.852E-06 | 473.75125 | 41.207504 |
| GSPNTASAEATLPI | 14 | Q6PJG6 | BRCA1-associated ATM activator 1 OS= | BRAT1 | 88.12 | 2 | 43.242535 | yes | 2.671E-07 | 686.3468 | 55.682152 |
| VTQALNVLTTTFG | 14 | Q6PJG6 | BRCA1-associated ATM activator 1 OS= | BRAT1 | 88.12 | 2 | 43.242535 | yes | 3.603E-07 | 760.92517 | 49.225571 |

| | | | | | | | | | | | |
|---|---|---|---|---|---|---|---|---|---|---|---|
| STLAESSDHVEK | 12 | Q6PJG6 | BRCA1-associated ATM activator 1 OS= | BRAT1 | 88.12 | 2 | 43.242535 | yes | 8.581E-06 | 651.81226 | 21.324419 |
| KPSTAVEYLAAHL | 15 | Q6PJI9 | GATOR2 complex protein WDR59 OS=I | WDR59 | 109.794 | 3 | 45.957947 | yes | 1.692E-07 | 538.96649 | 290.9704 |
| WSSENVVVEFR | 11 | Q6PJI9 | GATOR2 complex protein WDR59 OS=I | WDR59 | 109.794 | 2 | 45.957947 | yes | 2.541E-07 | 676.33569 | 91.089027 |
| YLNILPCQVPVWK | 13 | Q6PJI9 | GATOR2 complex protein WDR59 OS=I | WDR59 | 109.794 | 2 | 45.957947 | yes | 2.716E-07 | 815.4447 | 55.880886 |
| KPTVALSAVAGAS | 16 | Q6PJI9 | GATOR2 complex protein WDR59 OS=I | WDR59 | 109.794 | 3 | 45.957947 | yes | 6.048E-06 | 509.63489 | 38.408676 |
| LLDPANTQQFDDF | 14 | Q6PJI9 | GATOR2 complex protein WDR59 OS=I | WDR59 | 109.794 | 2 | 45.957947 | yes | 2.266E-06 | 826.40173 | 31.573652 |
| NAASALLVGR | 10 | Q6PJI9 | GATOR2 complex protein WDR59 OS=I | WDR59 | 109.794 | 2 | 45.957947 | yes | 9.592E-07 | 486.28528 | 26.601913 |
| NANCLATSHDGD\ | 14 | Q6PJI9 | GATOR2 complex protein WDR59 OS=I | WDR59 | 109.794 | 2 | 45.957947 | yes | 2.964E-07 | 765.34167 | 26.18926 |
| IIPTVNVGTWK | 11 | Q6PJP8 | DNA cross-link repair 1A protein OS=Ho | DCLRE1A | 116.4 | 2 | 46.941612 | yes | 1.192E-06 | 614.35846 | 54.940815 |
| SGVLCSLEER | 10 | Q6PJP8 | DNA cross-link repair 1A protein OS=Ho | DCLRE1A | 116.4 | 2 | 46.941612 | yes | 1.861E-06 | 575.28204 | 53.887772 |
| FVQWLKPQK | 9 | Q6PJP8 | DNA cross-link repair 1A protein OS=Ho | DCLRE1A | 116.4 | 2 | 46.941612 | yes | 1.436E-07 | 587.34259 | 44.822388 |
| RIPVLSPK | 8 | Q6PJT7 | Zinc finger CCCH domain-containing prc | ZC3H14 | 82.876 | 2 | 51.400917 | yes | 3.209E-07 | 455.29764 | 110.46824 |
| KLPVVSSVVK | 10 | Q6PJT7 | Zinc finger CCCH domain-containing prc | ZC3H14 | 82.876 | 2 | 51.400917 | yes | 3.963E-07 | 528.34479 | 106.98938 |
| PAVAPPAPPSSSQI | 16 | Q6PJT7 | Zinc finger CCCH domain-containing prc | ZC3H14 | 82.876 | 2 | 51.400917 | yes | 1.374E-07 | 817.91962 | 73.298477 |
| LMSTVKPLR | 9 | Q6PJT7 | Zinc finger CCCH domain-containing prc | ZC3H14 | 82.876 | 2 | 51.400917 | yes | 3.063E-07 | 522.81537 | 68.883202 |
| MSMEDENFR | 9 | Q6PJT7 | Zinc finger CCCH domain-containing prc | ZC3H14 | 82.876 | 2 | 51.400917 | yes | 9.722E-08 | 579.73132 | 55.438976 |
| QTLPVAPR | 8 | Q6PJT7 | Zinc finger CCCH domain-containing prc | ZC3H14 | 82.876 | 2 | 51.400917 | yes | 1.174E-06 | 441.26379 | 54.895969 |
| TTNYSTVPQK | 10 | Q6PJT7 | Zinc finger CCCH domain-containing prc | ZC3H14 | 82.876 | 2 | 51.400917 | yes | 1.737E-07 | 569.79059 | 35.099754 |
| TGSISSSVSVPAKP | 16 | Q6PJT7 | Zinc finger CCCH domain-containing prc | ZC3H14 | 82.876 | 2 | 51.400917 | yes | 1.062E-07 | 801.42828 | 25.677961 |
| LQLAQTEACQDV/ | 14 | Q6PJW8 | Consortin OS=Homo sapiens OX=9606 ( | CNST | 79.597 | 2 | 51.798344 | yes | 9.722E-08 | 801.89886 | 1223.1584 |
| TEPLISPGCDR | 11 | Q6PJW8 | Consortin OS=Homo sapiens OX=9606 ( | CNST | 79.597 | 2 | 51.798344 | yes | 1.326E-07 | 622.8006 | 474.96884 |
| TCLGTESSK | 9 | Q6PJW8 | Consortin OS=Homo sapiens OX=9606 ( | CNST | 79.597 | 2 | 51.798344 | yes | 1.127E-07 | 491.72913 | 245.86343 |
| CSTQLLVSEDPK | 12 | Q6PJW8 | Consortin OS=Homo sapiens OX=9606 ( | CNST | 79.597 | 2 | 51.798344 | yes | 9.722E-08 | 688.83997 | 213.19525 |
| ICATHQDPLLSK | 12 | Q6PJW8 | Consortin OS=Homo sapiens OX=9606 ( | CNST | 79.597 | 2 | 51.798344 | yes | 1E-07 | 691.85846 | 194.34276 |
| GFNGEDFER | 9 | Q6PJW8 | Consortin OS=Homo sapiens OX=9606 ( | CNST | 79.597 | 2 | 51.798344 | yes | 1.732E-07 | 535.73053 | 117.479 |
| RNDTVGDHPAQM | 18 | Q6PJW8 | Consortin OS=Homo sapiens OX=9606 ( | CNST | 79.597 | 4 | 51.798344 | yes | 1.041E-07 | 512.25378 | 96.095497 |
| KCSTQLLVSEDPK | 13 | Q6PJW8 | Consortin OS=Homo sapiens OX=9606 ( | CNST | 79.597 | 2 | 51.798344 | yes | 9.722E-08 | 752.88745 | 59.11377 |
| SKTEPLISPGCDR | 13 | Q6PJW8 | Consortin OS=Homo sapiens OX=9606 ( | CNST | 79.597 | 2 | 51.798344 | yes | 9.914E-08 | 730.36414 | 33.135006 |
| LFGAPEPSTIAR | 12 | Q6PKG0 | La-related protein 1 OS=Homo sapiens O | LARP1 | 123.511 | 2 | 51.095512 | yes | 1.665E-07 | 629.84314 | 33663.762 |
| FQQVPTDALANK | 12 | Q6PKG0 | La-related protein 1 OS=Homo sapiens O | LARP1 | 123.511 | 2 | 51.095512 | yes | 2.025E-07 | 666.35132 | 31271.799 |
| SLPTTVPESPNYR | 13 | Q6PKG0 | La-related protein 1 OS=Homo sapiens O | LARP1 | 123.511 | 2 | 51.095512 | yes | 1.887E-07 | 730.87262 | 23915.334 |
| FEPEYSQIK | 9 | Q6PKG0 | La-related protein 1 OS=Homo sapiens O | LARP1 | 123.511 | 2 | 51.095512 | yes | 1.995E-07 | 570.78223 | 20230.162 |
| WVPLQIDMKPEVI | 14 | Q6PKG0 | La-related protein 1 OS=Homo sapiens O | LARP1 | 123.511 | 3 | 51.095512 | yes | 1.202E-07 | 569.97986 | 9334.5254 |
| WVPLQIDMKPEVI | 14 | Q6PKG0 | La-related protein 1 OS=Homo sapiens O | LARP1 | 123.511 | 3 | 51.095512 | yes | 2.145E-07 | 575.31152 | 9334.5254 |
| LGIGQSQEMNTLF | 14 | Q6PKG0 | La-related protein 1 OS=Homo sapiens O | LARP1 | 123.511 | 2 | 51.095512 | no | 1.532E-07 | 797.40631 | 7284.1895 |
| LGIGQSQEMNTLF | 14 | Q6PKG0 | La-related protein 1 OS=Homo sapiens O | LARP1 | 123.511 | 2 | 51.095512 | no | 3.218E-07 | 805.40375 | 7284.1895 |
| YGLECLFR | 8 | Q6PKG0 | La-related protein 1 OS=Homo sapiens O | LARP1 | 123.511 | 2 | 51.095512 | no | 1.339E-07 | 529.26038 | 12606.517 |
| DYEAGQLYGLEK | 12 | Q6PKG0 | La-related protein 1 OS=Homo sapiens O | LARP1 | 123.511 | 2 | 51.095512 | yes | 1.083E-07 | 693.33282 | 12521.731 |
| QIEYYFSVDNLER | 13 | Q6PKG0 | La-related protein 1 OS=Homo sapiens O | LARP1 | 123.511 | 2 | 51.095512 | yes | 1.429E-07 | 838.40173 | 10865.257 |

| | | | | | | | | | | |
|---|---|---|---|---|---|---|---|---|---|---|
| KFDGVEGPR | 9 | Q6PKG0 | La-related protein 1 OS=Homo sapiens O | LARP1 | 123.511 | 2 | 51.095512 | yes | 3.101E-06 | 502.76163 | 10502.866 |
| HSSNPPLESHVGW | 18 | Q6PKG0 | La-related protein 1 OS=Homo sapiens O | LARP1 | 123.511 | 3 | 51.095512 | yes | 1.26E-07 | 678.98651 | 9807.8857 |
| FQHPSHELLK | 10 | Q6PKG0 | La-related protein 1 OS=Homo sapiens O | LARP1 | 123.511 | 2 | 51.095512 | no | 6.475E-07 | 618.3302 | 4732.418 |
| FQHPSHELLK | 10 | Q6PKG0 | La-related protein 1 OS=Homo sapiens O | LARP1 | 123.511 | 2 | 51.095512 | no | 1.531E-07 | 412.55591 | 4732.418 |
| FQHPSHELLK | 10 | Q6PKG0 | La-related protein 1 OS=Homo sapiens O | LARP1 | 123.511 | 3 | 51.095512 | no | 6.475E-07 | 618.3302 | 4732.418 |
| FQHPSHELLK | 10 | Q6PKG0 | La-related protein 1 OS=Homo sapiens O | LARP1 | 123.511 | 3 | 51.095512 | no | 1.531E-07 | 412.55591 | 4732.418 |
| VVEIVDEK | 8 | Q6PKG0 | La-related protein 1 OS=Homo sapiens O | LARP1 | 123.511 | 2 | 51.095512 | yes | 1.296E-07 | 465.76077 | 9013.4014 |
| VDPPMGEEGNHK | 12 | Q6PKG0 | La-related protein 1 OS=Homo sapiens O | LARP1 | 123.511 | 2 | 51.095512 | yes | 4.626E-07 | 655.29572 | 2902.1624 |
| VDPPMGEEGNHK | 12 | Q6PKG0 | La-related protein 1 OS=Homo sapiens O | LARP1 | 123.511 | 2 | 51.095512 | yes | 2.189E-07 | 663.29315 | 2902.1624 |
| QLALEDAK | 8 | Q6PKG0 | La-related protein 1 OS=Homo sapiens O | LARP1 | 123.511 | 2 | 51.095512 | yes | 1.767E-07 | 444.24527 | 5545.2705 |
| LQEYLGK | 7 | Q6PKG0 | La-related protein 1 OS=Homo sapiens O | LARP1 | 123.511 | 2 | 51.095512 | yes | 1.045E-07 | 425.73709 | 5228.6133 |
| KVNMISR | 7 | Q6PKG0 | La-related protein 1 OS=Homo sapiens O | LARP1 | 123.511 | 2 | 51.095512 | yes | 9.819E-08 | 424.24457 | 2860.9106 |
| TEEVSNLK | 8 | Q6PKG0 | La-related protein 1 OS=Homo sapiens O | LARP1 | 123.511 | 2 | 51.095512 | yes | 1.232E-07 | 460.2402 | 2760.3105 |
| FDGVEGPR | 8 | Q6PKG0 | La-related protein 1 OS=Homo sapiens O | LARP1 | 123.511 | 2 | 51.095512 | yes | 6.816E-07 | 438.71414 | 2756.5144 |
| QEVENFK | 7 | Q6PKG0 | La-related protein 1 OS=Homo sapiens O | LARP1 | 123.511 | 2 | 51.095512 | no | 1.839E-07 | 447.2218 | 2587.5544 |
| YYSYGLEK | 8 | Q6PKG0 | La-related protein 1 OS=Homo sapiens O | LARP1 | 123.511 | 2 | 51.095512 | yes | 1.867E-07 | 511.74512 | 2533.1497 |
| LASRPTRPPEPR | 12 | Q6PKG0 | La-related protein 1 OS=Homo sapiens O | LARP1 | 123.511 | 3 | 51.095512 | yes | 2.232E-07 | 459.59674 | 1187.3982 |
| LASRPTRPPEPR | 12 | Q6PKG0 | La-related protein 1 OS=Homo sapiens O | LARP1 | 123.511 | 3 | 51.095512 | yes | 1.328E-07 | 688.89148 | 1187.3982 |
| LASRPTRPPEPR | 12 | Q6PKG0 | La-related protein 1 OS=Homo sapiens O | LARP1 | 123.511 | 2 | 51.095512 | yes | 2.232E-07 | 459.59674 | 1187.3982 |
| LASRPTRPPEPR | 12 | Q6PKG0 | La-related protein 1 OS=Homo sapiens O | LARP1 | 123.511 | 2 | 51.095512 | yes | 1.328E-07 | 688.89148 | 1187.3982 |
| DFVEAPPPK | 9 | Q6PKG0 | La-related protein 1 OS=Homo sapiens O | LARP1 | 123.511 | 2 | 51.095512 | yes | 2.682E-07 | 500.26093 | 1896.7491 |
| DFQEETVK | 8 | Q6PKG0 | La-related protein 1 OS=Homo sapiens O | LARP1 | 123.511 | 2 | 51.095512 | yes | 1.57E-07 | 498.23764 | 1770.2277 |
| PTRPPEPR | 8 | Q6PKG0 | La-related protein 1 OS=Homo sapiens O | LARP1 | 123.511 | 2 | 51.095512 | yes | 1.028E-05 | 475.26434 | 1766.5027 |
| WTSQHSNTQTLGK | 13 | Q6PKG0 | La-related protein 1 OS=Homo sapiens O | LARP1 | 123.511 | 2 | 51.095512 | yes | 1.006E-07 | 744.36548 | 1491.1539 |
| VDPPMGEEGNHKI | 13 | Q6PKG0 | La-related protein 1 OS=Homo sapiens O | LARP1 | 123.511 | 2 | 51.095512 | yes | 1.517E-07 | 733.34625 | 1255.1863 |
| ETESAPGSPR | 10 | Q6PKG0 | La-related protein 1 OS=Homo sapiens O | LARP1 | 123.511 | 2 | 51.095512 | yes | 2.373E-07 | 515.74365 | 1103.6271 |
| RDFVEAPPPK | 10 | Q6PKG0 | La-related protein 1 OS=Homo sapiens O | LARP1 | 123.511 | 2 | 51.095512 | yes | 3.92E-07 | 578.31146 | 917.68109 |
| FRLDIFK | 7 | Q6PKG0 | La-related protein 1 OS=Homo sapiens O | LARP1 | 123.511 | 2 | 51.095512 | yes | 4.989E-07 | 469.77655 | 821.31647 |
| KMYEEFK | 7 | Q6PKG0 | La-related protein 1 OS=Homo sapiens O | LARP1 | 123.511 | 2 | 51.095512 | yes | 3.08E-06 | 487.73624 | 500.05161 |
| GEPGPNDVR | 9 | Q6PKG0 | La-related protein 1 OS=Homo sapiens O | LARP1 | 123.511 | 2 | 51.095512 | yes | 4.274E-07 | 470.72778 | 490.56799 |
| NLDIDPK | 7 | Q6PKG0 | La-related protein 1 OS=Homo sapiens O | LARP1 | 123.511 | 2 | 51.095512 | yes | 1.846E-07 | 407.7189 | 342.35086 |
| KMDADGFLPITLIA | 18 | Q6PKG0 | La-related protein 1 OS=Homo sapiens O | LARP1 | 123.511 | 3 | 51.095512 | yes | 1.913E-07 | 678.02765 | 152.2142 |
| KMDADGFLPITLIA | 18 | Q6PKG0 | La-related protein 1 OS=Homo sapiens O | LARP1 | 123.511 | 3 | 51.095512 | yes | 9.722E-08 | 683.35931 | 152.2142 |
| ENGFTQHVYHK | 11 | Q6PKG0 | La-related protein 1 OS=Homo sapiens O | LARP1 | 123.511 | 2 | 51.095512 | yes | 9.722E-08 | 680.32562 | 288.24991 |
| ILIVTQTPHYMR | 12 | Q6PKG0 | La-related protein 1 OS=Homo sapiens O | LARP1 | 123.511 | 2 | 51.095512 | yes | 1.181E-07 | 744.40558 | 55.687492 |
| ILIVTQTPHYMR | 12 | Q6PKG0 | La-related protein 1 OS=Homo sapiens O | LARP1 | 123.511 | 2 | 51.095512 | yes | 2.124E-07 | 496.60614 | 55.687492 |
| ILIVTQTPHYMR | 12 | Q6PKG0 | La-related protein 1 OS=Homo sapiens O | LARP1 | 123.511 | 2 | 51.095512 | yes | 1.18E-05 | 736.40814 | 55.687492 |
| ILIVTQTPHYMR | 12 | Q6PKG0 | La-related protein 1 OS=Homo sapiens O | LARP1 | 123.511 | 3 | 51.095512 | yes | 1.181E-07 | 744.40558 | 55.687492 |
| ILIVTQTPHYMR | 12 | Q6PKG0 | La-related protein 1 OS=Homo sapiens O | LARP1 | 123.511 | 3 | 51.095512 | yes | 2.124E-07 | 496.60614 | 55.687492 |

| | | | | | | | | | | | |
|---|---|---|---|---|---|---|---|---|---|---|---|
| ILIVTQTPHYMR | 12 | Q6PKG0 | La-related protein 1 OS=Homo sapiens O | LARP1 | 123.511 | 3 | 51.095512 | yes | 1.18E-05 | 736.40814 | 55.687492 |
| SVQPQSHKPQPTR | 13 | Q6PKG0 | La-related protein 1 OS=Homo sapiens O | LARP1 | 123.511 | 3 | 51.095512 | yes | 1.508E-07 | 497.26718 | 140.69405 |
| QEVENFKK | 8 | Q6PKG0 | La-related protein 1 OS=Homo sapiens O | LARP1 | 123.511 | 2 | 51.095512 | no | 9.729E-08 | 511.26929 | 136.42952 |
| DSKVVEIVDEK | 11 | Q6PKG0 | La-related protein 1 OS=Homo sapiens O | LARP1 | 123.511 | 2 | 51.095512 | yes | 9.759E-08 | 630.83771 | 87.859337 |
| PAAMISQPPTPPTC | 18 | Q6PKG0 | La-related protein 1 OS=Homo sapiens O | LARP1 | 123.511 | 3 | 51.095512 | yes | 3.814E-07 | 615.66107 | 39.628014 |
| DFQEETVKDYEAC | 20 | Q6PKG0 | La-related protein 1 OS=Homo sapiens O | LARP1 | 123.511 | 3 | 51.095512 | yes | 1.15E-06 | 788.04102 | 12.248859 |
| LITNTAEAVLQK | 12 | Q6PL18 | ATPase family AAA domain-containing p | ATAD2 | 158.555 | 2 | 40.374107 | yes | 4.059E-07 | 650.8772 | 58.200916 |
| GCLFYGPPGTGK | 12 | Q6PL18 | ATPase family AAA domain-containing p | ATAD2 | 158.555 | 2 | 40.374107 | no | 8.806E-07 | 627.3028 | 41.002205 |
| LSEDLMEFR | 9 | Q6Q0C0 | E3 ubiquitin-protein ligase TRAF7 OS=H | TRAF7 | 74.61 | 2 | 52.255074 | yes | 9.919E-08 | 570.27368 | 329.224 |
| VWDIVGTELK | 10 | Q6Q0C0 | E3 ubiquitin-protein ligase TRAF7 OS=H | TRAF7 | 74.61 | 2 | 52.255074 | yes | 9.838E-08 | 580.32153 | 319.61487 |
| DPVITTCGHTFCR | 13 | Q6Q0C0 | E3 ubiquitin-protein ligase TRAF7 OS=H | TRAF7 | 74.61 | 2 | 52.255074 | yes | 9.722E-08 | 782.35577 | 195.03972 |
| FDVLDENQSK | 10 | Q6Q0C0 | E3 ubiquitin-protein ligase TRAF7 OS=H | TRAF7 | 74.61 | 2 | 52.255074 | yes | 2.35E-07 | 597.78552 | 193.43279 |
| CPNNPSCPPLLR | 12 | Q6Q0C0 | E3 ubiquitin-protein ligase TRAF7 OS=H | TRAF7 | 74.61 | 2 | 52.255074 | yes | 9.85E-08 | 712.84229 | 173.47728 |
| EFLQQTDDR | 9 | Q6Q0C0 | E3 ubiquitin-protein ligase TRAF7 OS=H | TRAF7 | 74.61 | 2 | 52.255074 | yes | 2.898E-07 | 576.27002 | 158.22841 |
| SLHSESSMSLR | 11 | Q6Q0C0 | E3 ubiquitin-protein ligase TRAF7 OS=H | TRAF7 | 74.61 | 2 | 52.255074 | yes | 1.792E-07 | 617.29822 | 150.02972 |
| ELTGLNHWVR | 10 | Q6Q0C0 | E3 ubiquitin-protein ligase TRAF7 OS=H | TRAF7 | 74.61 | 2 | 52.255074 | yes | 4.119E-07 | 612.82782 | 137.29616 |
| FHEMHVALAQK | 11 | Q6Q0C0 | E3 ubiquitin-protein ligase TRAF7 OS=H | TRAF7 | 74.61 | 2 | 52.255074 | yes | 2.98E-06 | 655.83734 | 135.37105 |
| VWDTCTTYK | 9 | Q6Q0C0 | E3 ubiquitin-protein ligase TRAF7 OS=H | TRAF7 | 74.61 | 2 | 52.255074 | yes | 2.748E-07 | 587.26587 | 106.57202 |
| LCCQLCCSVFK | 11 | Q6Q0C0 | E3 ubiquitin-protein ligase TRAF7 OS=H | TRAF7 | 74.61 | 2 | 52.255074 | yes | 5.675E-07 | 737.81909 | 95.326706 |
| DEEDSMPPISTPR | 13 | Q6Q0C0 | E3 ubiquitin-protein ligase TRAF7 OS=H | TRAF7 | 74.61 | 2 | 52.255074 | yes | 8.296E-06 | 737.32996 | 94.069412 |
| TLTGHVGTVYALA | 22 | Q6Q0C0 | E3 ubiquitin-protein ligase TRAF7 OS=H | TRAF7 | 74.61 | 3 | 52.255074 | yes | 4.533E-07 | 758.07764 | 89.779129 |
| TLEGHDGIVLALC | 18 | Q6Q0C0 | E3 ubiquitin-protein ligase TRAF7 OS=H | TRAF7 | 74.61 | 3 | 52.255074 | yes | 9.809E-08 | 662.00452 | 78.142609 |
| MNLEAHLK | 8 | Q6Q0C0 | E3 ubiquitin-protein ligase TRAF7 OS=H | TRAF7 | 74.61 | 2 | 52.255074 | yes | 3.132E-07 | 478.25513 | 67.984566 |
| DASMLNDELSHIN | 15 | Q6Q0C0 | E3 ubiquitin-protein ligase TRAF7 OS=H | TRAF7 | 74.61 | 2 | 52.255074 | yes | 1.083E-07 | 843.39923 | 66.084709 |
| DQEIAFLR | 8 | Q6Q0C0 | E3 ubiquitin-protein ligase TRAF7 OS=H | TRAF7 | 74.61 | 2 | 52.255074 | yes | 2.115E-07 | 496.26401 | 65.135323 |
| TPSSSSTLAYSPR | 13 | Q6Q0C0 | E3 ubiquitin-protein ligase TRAF7 OS=H | TRAF7 | 74.61 | 2 | 52.255074 | yes | 4.97E-07 | 677.33588 | 58.260471 |
| ILAPYTDFHYR | 11 | Q6R327 | Rapamycin-insensitive companion of mT( | RICTOR | 192.218 | 2 | 54.754807 | yes | 9.722E-08 | 698.35645 | 323.60126 |
| ILTAATDACR | 10 | Q6R327 | Rapamycin-insensitive companion of mT( | RICTOR | 192.218 | 2 | 54.754807 | yes | 1.28E-07 | 546.2793 | 282.76144 |
| YLIQDSSILQK | 11 | Q6R327 | Rapamycin-insensitive companion of mT( | RICTOR | 192.218 | 2 | 54.754807 | yes | 9.722E-08 | 654.36389 | 266.19702 |
| LYANLDLDFAK | 11 | Q6R327 | Rapamycin-insensitive companion of mT( | RICTOR | 192.218 | 2 | 54.754807 | yes | 1.082E-07 | 641.83752 | 260.24084 |
| LSDGFVAAEAK | 11 | Q6R327 | Rapamycin-insensitive companion of mT( | RICTOR | 192.218 | 2 | 54.754807 | yes | 8.389E-07 | 554.28766 | 213.67886 |
| LTVSSLDYSR | 10 | Q6R327 | Rapamycin-insensitive companion of mT( | RICTOR | 192.218 | 2 | 54.754807 | yes | 1.037E-07 | 570.7984 | 210.12988 |
| NDSGEENVPLDLT | 14 | Q6R327 | Rapamycin-insensitive companion of mT( | RICTOR | 192.218 | 2 | 54.754807 | yes | 2.221E-07 | 779.87079 | 194.43936 |
| GFSYLNER | 8 | Q6R327 | Rapamycin-insensitive companion of mT( | RICTOR | 192.218 | 2 | 54.754807 | yes | 1.453E-07 | 493.24051 | 185.25262 |
| NVIDCQLSR | 9 | Q6R327 | Rapamycin-insensitive companion of mT( | RICTOR | 192.218 | 2 | 54.754807 | yes | 1.102E-07 | 552.77698 | 155.23183 |
| TLQLETSFMGNK | 12 | Q6R327 | Rapamycin-insensitive companion of mT( | RICTOR | 192.218 | 2 | 54.754807 | yes | 3.566E-07 | 684.84503 | 153.43402 |
| HSPDTAEGQLK | 11 | Q6R327 | Rapamycin-insensitive companion of mT( | RICTOR | 192.218 | 2 | 54.754807 | yes | 3.617E-07 | 591.79114 | 151.69032 |
| DVLFTDTITMK | 11 | Q6R327 | Rapamycin-insensitive companion of mT( | RICTOR | 192.218 | 2 | 54.754807 | yes | 9.722E-08 | 642.33124 | 142.90329 |
| QCEVLSIR | 8 | Q6R327 | Rapamycin-insensitive companion of mT( | RICTOR | 192.218 | 2 | 54.754807 | yes | 2.156E-07 | 502.76331 | 130.99561 |

| | | | | | | | | | | | |
|---|---|---|---|---|---|---|---|---|---|---|---|
| LKVDYLIAR | 9 | Q6R327 | Rapamycin-insensitive companion of mT( | RICTOR | 192.218 | 2 | 54.754807 | yes | 9.722E-08 | 545.83459 | 130.79041 |
| DTEEALLINLR | 11 | Q6R327 | Rapamycin-insensitive companion of mT( | RICTOR | 192.218 | 2 | 54.754807 | yes | 9.722E-08 | 643.8512 | 120.24651 |
| CIDIQQSNEVER | 12 | Q6R327 | Rapamycin-insensitive companion of mT( | RICTOR | 192.218 | 2 | 54.754807 | yes | 2.08E-07 | 745.84882 | 114.16816 |
| PALSHLGDK | 9 | Q6R327 | Rapamycin-insensitive companion of mT( | RICTOR | 192.218 | 2 | 54.754807 | yes | 2.804E-06 | 469.25873 | 107.83546 |
| SLADCNFSYTSSR | 13 | Q6R327 | Rapamycin-insensitive companion of mT( | RICTOR | 192.218 | 2 | 54.754807 | yes | 9.722E-08 | 754.32776 | 99.479607 |
| LVINLSSSVSTK | 12 | Q6R327 | Rapamycin-insensitive companion of mT( | RICTOR | 192.218 | 2 | 54.754807 | yes | 9.722E-08 | 624.36389 | 90.710365 |
| AFAHDAGGLPSGT | 18 | Q6R327 | Rapamycin-insensitive companion of mT( | RICTOR | 192.218 | 3 | 54.754807 | yes | 2.743E-07 | 552.28979 | 87.417625 |
| LALLNEAK | 8 | Q6R327 | Rapamycin-insensitive companion of mT( | RICTOR | 192.218 | 2 | 54.754807 | yes | 2.226E-07 | 436.26602 | 76.060013 |
| KPVDGDNYVR | 10 | Q6R327 | Rapamycin-insensitive companion of mT( | RICTOR | 192.218 | 2 | 54.754807 | yes | 1.704E-07 | 581.7962 | 69.91198 |
| DIVQWLNASSGMI | 13 | Q6R327 | Rapamycin-insensitive companion of mT( | RICTOR | 192.218 | 2 | 54.754807 | yes | 2.532E-07 | 724.86377 | 58.789127 |
| MGIIATFR | 8 | Q6R327 | Rapamycin-insensitive companion of mT( | RICTOR | 192.218 | 2 | 54.754807 | yes | 4.038E-07 | 454.75476 | 58.654301 |
| NSFHLLR | 7 | Q6R327 | Rapamycin-insensitive companion of mT( | RICTOR | 192.218 | 2 | 54.754807 | yes | 2.295E-06 | 443.74832 | 50.631207 |
| STELLLGVK | 9 | Q6R327 | Rapamycin-insensitive companion of mT( | RICTOR | 192.218 | 2 | 54.754807 | yes | 1.986E-07 | 480.29224 | 49.346329 |
| ILNSLTLPNK | 10 | Q6R327 | Rapamycin-insensitive companion of mT( | RICTOR | 192.218 | 2 | 54.754807 | yes | 9.722E-08 | 556.83734 | 47.060444 |
| ANLHALIQMK | 10 | Q6R327 | Rapamycin-insensitive companion of mT( | RICTOR | 192.218 | 2 | 54.754807 | yes | 9.722E-08 | 569.82373 | 41.357155 |
| NQDHLLK | 7 | Q6R327 | Rapamycin-insensitive companion of mT( | RICTOR | 192.218 | 2 | 54.754807 | yes | 1.442E-07 | 434.23779 | 40.52079 |
| CHETGLLTIK | 10 | Q6R327 | Rapamycin-insensitive companion of mT( | RICTOR | 192.218 | 2 | 54.754807 | yes | 4.969E-06 | 586.31061 | 35.156277 |
| ASAALNCLK | 9 | Q6R327 | Rapamycin-insensitive companion of mT( | RICTOR | 192.218 | 2 | 54.754807 | yes | 9.56E-07 | 474.25256 | 34.61367 |
| TIPDDTPMCR | 10 | Q6R327 | Rapamycin-insensitive companion of mT( | RICTOR | 192.218 | 2 | 54.754807 | yes | 2.49E-07 | 603.26813 | 25.340036 |
| IWDLQGSEEPVFR | 13 | Q6RFH5 | WD repeat-containing protein 74 OS=Ho | WDR74 | 42.442 | 2 | 53.35835 | yes | 1.466E-07 | 788.39374 | 594.69531 |
| LSGLEQPQGALQT | 14 | Q6RFH5 | WD repeat-containing protein 74 OS=Ho | WDR74 | 42.442 | 2 | 53.35835 | yes | 1.063E-07 | 749.4046 | 516.55164 |
| SQLNCLLLSGR | 11 | Q6RFH5 | WD repeat-containing protein 74 OS=Ho | WDR74 | 42.442 | 2 | 53.35835 | yes | 1.145E-07 | 630.84009 | 437.23837 |
| NDWLDLR | 7 | Q6RFH5 | WD repeat-containing protein 74 OS=Ho | WDR74 | 42.442 | 2 | 53.35835 | yes | 1.329E-07 | 466.23523 | 156.84642 |
| PLLASCGLDR | 10 | Q6RFH5 | WD repeat-containing protein 74 OS=Ho | WDR74 | 42.442 | 2 | 53.35835 | yes | 1.566E-07 | 551.28967 | 156.57013 |
| DTSSDPLLELR | 11 | Q6RFH5 | WD repeat-containing protein 74 OS=Ho | WDR74 | 42.442 | 2 | 53.35835 | yes | 1.338E-07 | 623.3197 | 155.75731 |
| LVTCTGYHQVR | 11 | Q6RFH5 | WD repeat-containing protein 74 OS=Ho | WDR74 | 42.442 | 2 | 53.35835 | yes | 1.948E-07 | 667.33771 | 129.49246 |
| QAANFTAGGQPR | 12 | Q6RFH5 | WD repeat-containing protein 74 OS=Ho | WDR74 | 42.442 | 2 | 53.35835 | yes | 1.934E-07 | 609.30475 | 121.99384 |
| WNHVWVGTETGI | 14 | Q6RFH5 | WD repeat-containing protein 74 OS=Ho | WDR74 | 42.442 | 2 | 53.35835 | yes | 9.722E-08 | 820.43317 | 71.74836 |
| DKDTSSDPLLELR | 13 | Q6RFH5 | WD repeat-containing protein 74 OS=Ho | WDR74 | 42.442 | 2 | 53.35835 | yes | 4.27E-07 | 744.88068 | 68.099998 |
| QDPAHPHVVATG( | 14 | Q6RFH5 | WD repeat-containing protein 74 OS=Ho | WDR74 | 42.442 | 2 | 53.35835 | yes | 3.384E-06 | 707.3653 | 56.532272 |
| VYDPASPQR | 9 | Q6RFH5 | WD repeat-containing protein 74 OS=Ho | WDR74 | 42.442 | 2 | 53.35835 | yes | 3.121E-07 | 516.75909 | 47.260452 |
| HCPGGEGMFR | 10 | Q6RFH5 | WD repeat-containing protein 74 OS=Ho | WDR74 | 42.442 | 2 | 53.35835 | yes | 4.58E-07 | 574.24219 | 45.496567 |
| HFSTEDGIFQGQR | 13 | Q6RFH5 | WD repeat-containing protein 74 OS=Ho | WDR74 | 42.442 | 2 | 53.35835 | yes | 2.749E-07 | 761.35767 | 41.544613 |
| SFSAMEDDQR | 10 | Q6SJ93 | Serine protease FAM111B OS=Homo sap | FAM111B | 84.675 | 2 | 53.038067 | yes | 9.722E-08 | 593.24567 | 390.13773 |
| VSNENLDYAILK | 12 | Q6SJ93 | Serine protease FAM111B OS=Homo sap | FAM111B | 84.675 | 2 | 53.038067 | yes | 4.083E-07 | 689.86426 | 322.85672 |
| IDGCTVIPLNER | 12 | Q6SJ93 | Serine protease FAM111B OS=Homo sap | FAM111B | 84.675 | 2 | 53.038067 | yes | 1.123E-07 | 693.8559 | 243.51483 |
| YAINLDVQK | 9 | Q6SJ93 | Serine protease FAM111B OS=Homo sap | FAM111B | 84.675 | 2 | 53.038067 | yes | 2.838E-06 | 532.29279 | 225.79825 |
| GETIEGALCK | 10 | Q6SJ93 | Serine protease FAM111B OS=Homo sap | FAM111B | 84.675 | 2 | 53.038067 | yes | 1.385E-07 | 539.26587 | 223.85881 |
| TIDGHINLGMPLK | 13 | Q6SJ93 | Serine protease FAM111B OS=Homo sap | FAM111B | 84.675 | 2 | 53.038067 | yes | 9.722E-08 | 704.88446 | 161.93134 |

| | | | | | | | | | | | |
|---|---|---|---|---|---|---|---|---|---|---|---|
| INELHEK | 7 | Q6SJ93 | Serine protease FAM111B OS=Homo sap | FAM111B | 84.675 | 2 | 53.038067 | yes | 3.436E-07 | 441.73761 | 129.60826 |
| SDIGEFEWK | 9 | Q6SJ93 | Serine protease FAM111B OS=Homo sap | FAM111B | 84.675 | 2 | 53.038067 | yes | 3.376E-07 | 555.75873 | 127.07848 |
| NYYFCSLPR | 9 | Q6SJ93 | Serine protease FAM111B OS=Homo sap | FAM111B | 84.675 | 2 | 53.038067 | yes | 2.26E-07 | 610.28186 | 95.624573 |
| LCIYALK | 7 | Q6SJ93 | Serine protease FAM111B OS=Homo sap | FAM111B | 84.675 | 2 | 53.038067 | yes | 2.167E-07 | 440.75168 | 86.500206 |
| QSMVDEVSGK | 10 | Q6SJ93 | Serine protease FAM111B OS=Homo sap | FAM111B | 84.675 | 2 | 53.038067 | yes | 1.191E-05 | 540.25549 | 73.600708 |
| QISPQPSTGLIYLIC | 22 | Q6SJ93 | Serine protease FAM111B OS=Homo sap | FAM111B | 84.675 | 3 | 53.038067 | yes | 1.87E-07 | 792.76892 | 66.550858 |
| LETYDEEK | 8 | Q6SJ93 | Serine protease FAM111B OS=Homo sap | FAM111B | 84.675 | 2 | 53.038067 | yes | 1.392E-07 | 513.73492 | 56.326248 |
| VLEMDISK | 8 | Q6SJ93 | Serine protease FAM111B OS=Homo sap | FAM111B | 84.675 | 2 | 53.038067 | yes | 1.972E-07 | 467.74933 | 52.926838 |
| QCENPNMECILFH | 19 | Q6SJ93 | Serine protease FAM111B OS=Homo sap | FAM111B | 84.675 | 3 | 53.038067 | yes | 7.368E-07 | 763.03088 | 36.18103 |
| VQGLLEEEAAAR | 12 | Q6SPF0 | Sterile alpha motif domain-containing prc | SAMD1 | 56.052 | 2 | 53.39961 | yes | 1.004E-07 | 643.34094 | 200.64331 |
| TDVLTGLSIR | 10 | Q6SPF0 | Sterile alpha motif domain-containing prc | SAMD1 | 56.052 | 2 | 53.39961 | yes | 9.722E-08 | 537.81134 | 74.848129 |
| LGALALPR | 8 | Q6SPF0 | Sterile alpha motif domain-containing prc | SAMD1 | 56.052 | 2 | 53.39961 | yes | 5.754E-07 | 405.76343 | 5.6998191 |
| TTEDLFAVIHR | 11 | Q6T4R5 | Actin remodeling regulator NHS OS=Hor | NHS | 179.136 | 2 | 56.507065 | yes | 1.304E-07 | 651.34607 | 300.47354 |
| LPLDFANTPSR | 11 | Q6T4R5 | Actin remodeling regulator NHS OS=Hor | NHS | 179.136 | 2 | 56.507065 | yes | 1.907E-07 | 615.82751 | 286.50473 |
| NCAFPTEGFQR | 11 | Q6T4R5 | Actin remodeling regulator NHS OS=Hor | NHS | 179.136 | 2 | 56.507065 | yes | 2.766E-07 | 663.7984 | 278.29474 |
| ATTPSLPSVDNEFI | 14 | Q6T4R5 | Actin remodeling regulator NHS OS=Hor | NHS | 179.136 | 2 | 56.507065 | yes | 2.227E-07 | 753.37775 | 265.1069 |
| SSLQQPSLK | 9 | Q6T4R5 | Actin remodeling regulator NHS OS=Hor | NHS | 179.136 | 2 | 56.507065 | yes | 9.938E-08 | 494.2771 | 159.0491 |
| SNPPPSLAITPTILK | 15 | Q6T4R5 | Actin remodeling regulator NHS OS=Hor | NHS | 179.136 | 2 | 56.507065 | yes | 9.531E-07 | 774.95343 | 151.4705 |
| AEETQGNVDEASL | 14 | Q6T4R5 | Actin remodeling regulator NHS OS=Hor | NHS | 179.136 | 2 | 56.507065 | yes | 9.722E-08 | 745.85211 | 138.45573 |
| NPIHNIPSTLDK | 12 | Q6T4R5 | Actin remodeling regulator NHS OS=Hor | NHS | 179.136 | 2 | 56.507065 | yes | 1.642E-07 | 674.86462 | 108.20613 |
| MENANLPTK | 9 | Q6T4R5 | Actin remodeling regulator NHS OS=Hor | NHS | 179.136 | 2 | 56.507065 | yes | 2.122E-07 | 509.25531 | 106.27637 |
| VQQEIDSDESPVAI | 14 | Q6T4R5 | Actin remodeling regulator NHS OS=Hor | NHS | 179.136 | 2 | 56.507065 | yes | 2.512E-07 | 786.87866 | 97.309837 |
| QEPSWINQSEQGII | 14 | Q6T4R5 | Actin remodeling regulator NHS OS=Hor | NHS | 179.136 | 2 | 56.507065 | yes | 9.722E-08 | 822.40485 | 88.823181 |
| DLDEVEAPGPEEP. | 15 | Q6T4R5 | Actin remodeling regulator NHS OS=Hor | NHS | 179.136 | 2 | 56.507065 | yes | 9.722E-08 | 812.37848 | 86.407539 |
| TETEPIPENTPTK | 13 | Q6T4R5 | Actin remodeling regulator NHS OS=Hor | NHS | 179.136 | 2 | 56.507065 | yes | 1.482E-07 | 728.86194 | 52.56472 |
| ALPLPTPEEK | 10 | Q6T4R5 | Actin remodeling regulator NHS OS=Hor | NHS | 179.136 | 2 | 56.507065 | yes | 2.205E-06 | 547.80823 | 50.405724 |
| HACSLFQELESDIC | 18 | Q6T4R5 | Actin remodeling regulator NHS OS=Hor | NHS | 179.136 | 3 | 56.507065 | yes | 3.451E-07 | 728.68823 | 49.341675 |
| QEAVPVSNLDIESI | 14 | Q6T4R5 | Actin remodeling regulator NHS OS=Hor | NHS | 179.136 | 2 | 56.507065 | yes | 4.074E-07 | 764.8963 | 46.109058 |
| VNAEGFSSK | 9 | Q6T4R5 | Actin remodeling regulator NHS OS=Hor | NHS | 179.136 | 2 | 56.507065 | yes | 5.765E-05 | 469.73254 | 35.555882 |
| VGAKPSAYEEGES | 20 | Q6T4R5 | Actin remodeling regulator NHS OS=Hor | NHS | 179.136 | 3 | 56.507065 | yes | 2.183E-06 | 722.00293 | 14.945372 |
| DIELSPEAQAK | 11 | Q6UB35 | Monofunctional C1-tetrahydrofolate syntl | MTHFD1L | 105.79 | 2 | 55.914925 | yes | 1.345E-07 | 600.80896 | 323.90588 |
| LQPLSPVPSDIEISI | 15 | Q6UB35 | Monofunctional C1-tetrahydrofolate syntl | MTHFD1L | 105.79 | 2 | 55.914925 | yes | 1.423E-07 | 825.95667 | 311.69635 |
| DVDGVTDINLGK | 12 | Q6UB35 | Monofunctional C1-tetrahydrofolate syntl | MTHFD1L | 105.79 | 2 | 55.914925 | yes | 2.313E-07 | 623.3197 | 303.96445 |
| EVLSLLQEK | 9 | Q6UB35 | Monofunctional C1-tetrahydrofolate syntl | MTHFD1L | 105.79 | 2 | 55.914925 | yes | 1.145E-07 | 529.80823 | 222.56943 |
| YVLVAGITPTPLGI | 16 | Q6UB35 | Monofunctional C1-tetrahydrofolate syntl | MTHFD1L | 105.79 | 2 | 55.914925 | yes | 2.98E-07 | 807.95868 | 221.82266 |
| TDPSTLTEEEVSK | 13 | Q6UB35 | Monofunctional C1-tetrahydrofolate syntl | MTHFD1L | 105.79 | 2 | 55.914925 | yes | 2.097E-07 | 718.34357 | 182.07693 |
| LDIDPSTITWQR | 12 | Q6UB35 | Monofunctional C1-tetrahydrofolate syntl | MTHFD1L | 105.79 | 2 | 55.914925 | yes | 9.722E-08 | 722.87518 | 144.80167 |
| IQNMVSSGR | 9 | Q6UB35 | Monofunctional C1-tetrahydrofolate syntl | MTHFD1L | 105.79 | 2 | 55.914925 | yes | 1.557E-07 | 496.25311 | 120.22061 |
| EFSEIQLAR | 9 | Q6UB35 | Monofunctional C1-tetrahydrofolate syntl | MTHFD1L | 105.79 | 2 | 55.914925 | yes | 1.531E-07 | 546.78784 | 99.479698 |

| | | | | | | | | | | |
|---|---|---|---|---|---|---|---|---|---|---|
| EIGLLADEIEIYGK | 14 | Q6UB35 | Monofunctional C1-tetrahydrofolate syntl | MTHFD1L | 105.79 | 2 | 55.914925 | yes | 9.722E-08 | 781.91925 | 89.271973 |
| MHGGGPSVTAGV] | 15 | Q6UB35 | Monofunctional C1-tetrahydrofolate syntl | MTHFD1L | 105.79 | 2 | 55.914925 | yes | 9.722E-08 | 704.37427 | 81.921951 |
| VLNALKPEK | 9 | Q6UB35 | Monofunctional C1-tetrahydrofolate syntl | MTHFD1L | 105.79 | 2 | 55.914925 | yes | 1.509E-07 | 506.31351 | 72.809006 |
| KITIGQGNTEK | 11 | Q6UB35 | Monofunctional C1-tetrahydrofolate syntl | MTHFD1L | 105.79 | 2 | 55.914925 | yes | 1.657E-07 | 594.83276 | 60.003143 |
| ILHENTQTDK | 10 | Q6UB35 | Monofunctional C1-tetrahydrofolate syntl | MTHFD1L | 105.79 | 2 | 55.914925 | yes | 5.059E-07 | 599.80676 | 34.182964 |
| VQELDPAGHK | 10 | Q6UB99 | Ankyrin repeat domain-containing proteir | ANKRD11 | 297.914 | 2 | 49.558628 | yes | 2.392E-06 | 547.28546 | 59.792305 |
| APLPPVPAEK | 10 | Q6UB99 | Ankyrin repeat domain-containing proteir | ANKRD11 | 297.914 | 2 | 49.558628 | yes | 2.932E-06 | 509.80023 | 52.787155 |
| MGEASDLPR | 9 | Q6UB99 | Ankyrin repeat domain-containing proteir | ANKRD11 | 297.914 | 2 | 49.558628 | yes | 1.961E-06 | 488.23184 | 45.065586 |
| EDSAFDEYCNK | 11 | Q6UB99 | Ankyrin repeat domain-containing proteir | ANKRD11 | 297.914 | 2 | 49.558628 | yes | 6.704E-07 | 689.26678 | 32.607323 |
| ANPYFEYLQIR | 11 | Q6UB99 | Ankyrin repeat domain-containing proteir | ANKRD11 | 297.914 | 2 | 49.558628 | yes | 1.499E-06 | 707.36169 | 29.611658 |
| DFAGWTALHEACI | 14 | Q6UB99 | Ankyrin repeat domain-containing proteir | ANKRD11 | 297.914 | 2 | 49.558628 | yes | 1.021E-07 | 824.37024 | 29.194515 |
| DWLAGPHMK | 9 | Q6UB99 | Ankyrin repeat domain-containing proteir | ANKRD11 | 297.914 | 2 | 49.558628 | yes | 2.906E-06 | 527.76056 | 28.95232 |
| GQSDVLLSTTSK | 12 | Q6ULP2 | Aftiphilin OS=Homo sapiens OX=9606 C | AFTPH | 102.113 | 2 | 36.299595 | yes | 7.154E-07 | 618.32751 | 30.687847 |
| QTSDNLSEECQLA | 14 | Q6ULP2 | Aftiphilin OS=Homo sapiens OX=9606 C | AFTPH | 102.113 | 2 | 36.299595 | yes | 1.779E-05 | 825.87305 | 9.381258 |
| MGLEVIPVTSTTNI | 14 | Q6UN15 | Pre-mRNA 3'-end-processing factor FIP1 | FIP1L1 | 66.527 | 2 | 53.53003 | yes | 1.197E-07 | 745.40015 | 1982.3409 |
| AEFTSPPSLFK | 11 | Q6UN15 | Pre-mRNA 3'-end-processing factor FIP1 | FIP1L1 | 66.527 | 2 | 53.53003 | yes | 9.722E-08 | 612.31897 | 399.84268 |
| ANENSNIQVLSER | 13 | Q6UN15 | Pre-mRNA 3'-end-processing factor FIP1 | FIP1L1 | 66.527 | 2 | 53.53003 | yes | 9.796E-08 | 737.36823 | 370.78717 |
| QWDYYAR | 7 | Q6UN15 | Pre-mRNA 3'-end-processing factor FIP1 | FIP1L1 | 66.527 | 2 | 53.53003 | yes | 3.158E-07 | 501.22742 | 153.12613 |
| RLPGAIDVIGQTIT | 17 | Q6UN15 | Pre-mRNA 3'-end-processing factor FIP1 | FIP1L1 | 66.527 | 3 | 53.53003 | yes | 3.316E-07 | 604.02301 | 140.879 |
| LPGAIDVIGQTITIS | 16 | Q6UN15 | Pre-mRNA 3'-end-processing factor FIP1 | FIP1L1 | 66.527 | 2 | 53.53003 | yes | 1.05E-07 | 827.48035 | 79.191223 |
| SNPSIQATLNK | 11 | Q6UUV7 | CREB-regulated transcription coactivator | CRTC3 | 66.96 | 2 | 36.07994 | yes | 7.426E-07 | 586.81714 | 81.004471 |
| VIITGDAFVPGER | 13 | Q6UWP7 | Lysocardiolipin acyltransferase 1 OS=Ho | LCLAT1 | 48.921 | 2 | 51.149334 | yes | 9.722E-08 | 687.37482 | 313.03723 |
| YEYVLHPR | 8 | Q6UWP7 | Lysocardiolipin acyltransferase 1 OS=Ho | LCLAT1 | 48.921 | 2 | 51.149334 | yes | 1.396E-07 | 538.77985 | 295.40915 |
| TTGFTFVVDR | 10 | Q6UWP7 | Lysocardiolipin acyltransferase 1 OS=Ho | LCLAT1 | 48.921 | 2 | 51.149334 | yes | 1.257E-07 | 571.79565 | 228.30969 |
| YPIDTLPTSK | 10 | Q6UWP7 | Lysocardiolipin acyltransferase 1 OS=Ho | LCLAT1 | 48.921 | 2 | 51.149334 | yes | 1.197E-07 | 567.80573 | 224.0925 |
| EDLQLWCHK | 9 | Q6UWP7 | Lysocardiolipin acyltransferase 1 OS=Ho | LCLAT1 | 48.921 | 2 | 51.149334 | yes | 1.058E-07 | 614.7926 | 76.42865 |
| EIHFHVHR | 8 | Q6UWP7 | Lysocardiolipin acyltransferase 1 OS=Ho | LCLAT1 | 48.921 | 2 | 51.149334 | yes | 1.684E-07 | 537.7832 | 49.270119 |
| DPQENIFLCQALR | 13 | Q6UWZ7 | BRCA1-A complex subunit Abraxas 1 OS | ABRAXAS | 46.663 | 2 | 50.981728 | yes | 1.077E-07 | 802.3985 | 538.50671 |
| INEMYASLQEELK | 13 | Q6UWZ7 | BRCA1-A complex subunit Abraxas 1 OS | ABRAXAS | 46.663 | 2 | 50.981728 | yes | 1.113E-07 | 784.38727 | 528.16162 |
| HSDQIMTFR | 9 | Q6UWZ7 | BRCA1-A complex subunit Abraxas 1 OS | ABRAXAS | 46.663 | 2 | 50.981728 | yes | 1.026E-07 | 567.77167 | 368.25052 |
| FFEEDGSLK | 9 | Q6UWZ7 | BRCA1-A complex subunit Abraxas 1 OS | ABRAXAS | 46.663 | 2 | 50.981728 | yes | 9.823E-08 | 536.2533 | 304.97614 |
| LEHSLYKPQK | 10 | Q6UWZ7 | BRCA1-A complex subunit Abraxas 1 OS | ABRAXAS | 46.663 | 2 | 50.981728 | yes | 1.166E-07 | 621.8457 | 241.15729 |
| LLDTQDKR | 8 | Q6UWZ7 | BRCA1-A complex subunit Abraxas 1 OS | ABRAXAS | 46.663 | 2 | 50.981728 | yes | 1.035E-07 | 494.77472 | 233.42523 |
| TVSGSCMSTGFSR | 13 | Q6UWZ7 | BRCA1-A complex subunit Abraxas 1 OS | ABRAXAS | 46.663 | 2 | 50.981728 | yes | 9.722E-08 | 688.80029 | 192.04594 |
| MSSPETDEEIEK | 12 | Q6UWZ7 | BRCA1-A complex subunit Abraxas 1 OS | ABRAXAS | 46.663 | 2 | 50.981728 | yes | 1.101E-07 | 697.80322 | 164.32085 |
| KVEDSEQAVDK | 11 | Q6UWZ7 | BRCA1-A complex subunit Abraxas 1 OS | ABRAXAS | 46.663 | 2 | 50.981728 | yes | 9.722E-08 | 624.30933 | 107.89033 |
| NSITDSQMDDVEV | 21 | Q6UWZ7 | BRCA1-A complex subunit Abraxas 1 OS | ABRAXAS | 46.663 | 3 | 50.981728 | yes | 4.922E-06 | 810.38446 | 9.0306435 |
| VTGDTVYNMLR | 11 | Q6UX04 | Spliceosome-associated protein CWC27 h | CWC27 | 53.848 | 2 | 34.27532 | yes | 1.042E-06 | 634.81879 | 34.575386 |
| KYGVDLIR | 8 | Q6UXN9 | WD repeat-containing protein 82 OS=Ho | WDR82 | 35.08 | 2 | 51.87907 | yes | 9.819E-08 | 482.28473 | 2037.7863 |

| | | | | | | | | | | | |
|---|---|---|---|---|---|---|---|---|---|---|---|
| YLSLHDNK | 8 | Q6UXN9 | WD repeat-containing protein 82 OS=Ho | WDR82 | 35.08 | 2 | 51.87907 | yes | 2.542E-07 | 495.25616 | 1505.5487 |
| IHVWNGESGIK | 11 | Q6UXN9 | WD repeat-containing protein 82 OS=Ho | WDR82 | 35.08 | 2 | 51.87907 | yes | 9.722E-08 | 620.32764 | 943.2334 |
| LILISTNGSFIR | 12 | Q6UXN9 | WD repeat-containing protein 82 OS=Ho | WDR82 | 35.08 | 2 | 51.87907 | yes | 1.039E-07 | 667.39557 | 734.53564 |
| HTGPITCLQFNPK | 13 | Q6UXN9 | WD repeat-containing protein 82 OS=Ho | WDR82 | 35.08 | 2 | 51.87907 | yes | 1.066E-07 | 756.88501 | 363.026 |
| HTGPITCLQFNPK | 13 | Q6UXN9 | WD repeat-containing protein 82 OS=Ho | WDR82 | 35.08 | 2 | 51.87907 | yes | 2.062E-07 | 504.92578 | 363.026 |
| HTGPITCLQFNPK | 13 | Q6UXN9 | WD repeat-containing protein 82 OS=Ho | WDR82 | 35.08 | 3 | 51.87907 | yes | 1.066E-07 | 756.88501 | 363.026 |
| HTGPITCLQFNPK | 13 | Q6UXN9 | WD repeat-containing protein 82 OS=Ho | WDR82 | 35.08 | 3 | 51.87907 | yes | 2.062E-07 | 504.92578 | 363.026 |
| GVVMHTFGGYAN | 14 | Q6UXN9 | WD repeat-containing protein 82 OS=Ho | WDR82 | 35.08 | 2 | 51.87907 | yes | 2.813E-07 | 734.35608 | 303.81119 |
| GVVMHTFGGYAN | 14 | Q6UXN9 | WD repeat-containing protein 82 OS=Ho | WDR82 | 35.08 | 2 | 51.87907 | yes | 1.166E-07 | 489.90649 | 303.81119 |
| GVVMHTFGGYAN | 14 | Q6UXN9 | WD repeat-containing protein 82 OS=Ho | WDR82 | 35.08 | 3 | 51.87907 | yes | 2.813E-07 | 734.35608 | 303.81119 |
| GVVMHTFGGYAN | 14 | Q6UXN9 | WD repeat-containing protein 82 OS=Ho | WDR82 | 35.08 | 3 | 51.87907 | yes | 1.166E-07 | 489.90649 | 303.81119 |
| TCEWTGLK | 8 | Q6UXN9 | WD repeat-containing protein 82 OS=Ho | WDR82 | 35.08 | 2 | 51.87907 | yes | 1.367E-07 | 497.73676 | 583.57361 |
| YTHAANTVVYSSN | 14 | Q6UXN9 | WD repeat-containing protein 82 OS=Ho | WDR82 | 35.08 | 2 | 51.87907 | yes | 9.722E-08 | 777.88098 | 518.64911 |
| SPNCQGLMHLQGI | 13 | Q6UXN9 | WD repeat-containing protein 82 OS=Ho | WDR82 | 35.08 | 2 | 51.87907 | yes | 3.665E-07 | 735.35303 | 308.17606 |
| YFPGHSK | 7 | Q6UXN9 | WD repeat-containing protein 82 OS=Ho | WDR82 | 35.08 | 2 | 51.87907 | yes | 2.127E-07 | 418.2085 | 299.686 |
| SFDKGPFATFK | 11 | Q6UXN9 | WD repeat-containing protein 82 OS=Ho | WDR82 | 35.08 | 2 | 51.87907 | yes | 2.212E-07 | 622.81915 | 269.27625 |
| YGVDLIR | 7 | Q6UXN9 | WD repeat-containing protein 82 OS=Ho | WDR82 | 35.08 | 2 | 51.87907 | yes | 1.543E-07 | 418.23724 | 242.52193 |
| LTDSVLR | 7 | Q6UXN9 | WD repeat-containing protein 82 OS=Ho | WDR82 | 35.08 | 2 | 51.87907 | yes | 9.85E-08 | 402.23471 | 185.51379 |
| AVTLEASFTPDSQI | 23 | Q6UXN9 | WD repeat-containing protein 82 OS=Ho | WDR82 | 35.08 | 3 | 51.87907 | yes | 2.306E-05 | 815.0614 | 37.415424 |
| VAAVGGQSYLSSI | 15 | Q6VY07 | Phosphofurin acidic cluster sorting protei | PACS1 | 104.899 | 2 | 43.815308 | yes | 2.436E-07 | 760.92517 | 50.018646 |
| HLIPQESR | 8 | Q6W2J9 | BCL-6 corepressor OS=Homo sapiens OX | BCOR | 192.189 | 2 | 54.92092 | yes | 2.076E-07 | 490.26962 | 260.56284 |
| NAGETLLQR | 9 | Q6W2J9 | BCL-6 corepressor OS=Homo sapiens OX | BCOR | 192.189 | 2 | 54.92092 | no | 5.777E-07 | 501.27237 | 205.08856 |
| KEVTQATQPEAIP( | 25 | Q6W2J9 | BCL-6 corepressor OS=Homo sapiens OX | BCOR | 192.189 | 4 | 54.92092 | yes | 1.878E-07 | 681.35559 | 141.20346 |
| FLTDYLNDLQGR | 12 | Q6W2J9 | BCL-6 corepressor OS=Homo sapiens OX | BCOR | 192.189 | 2 | 54.92092 | yes | 1.794E-07 | 727.86737 | 128.82927 |
| AGSGLVLSGSEIPK | 14 | Q6W2J9 | BCL-6 corepressor OS=Homo sapiens OX | BCOR | 192.189 | 2 | 54.92092 | yes | 1.399E-07 | 657.86682 | 67.158829 |
| DNAGYCALHEAC | 14 | Q6W2J9 | BCL-6 corepressor OS=Homo sapiens OX | BCOR | 192.189 | 2 | 54.92092 | yes | 9.722E-08 | 804.33813 | 40.235901 |
| QVSASLLFSCSK | 12 | Q6W2J9 | BCL-6 corepressor OS=Homo sapiens OX | BCOR | 192.189 | 2 | 54.92092 | yes | 2.043E-06 | 663.83972 | 39.111439 |
| VESGYFSLEK | 10 | Q6WCQ1 | Myosin phosphatase Rho-interacting prot | MPRIP | 116.534 | 2 | 47.884483 | yes | 1.238E-07 | 579.78748 | 61.2813 |
| STEPSVTPDLLNFK | 14 | Q6WCQ1 | Myosin phosphatase Rho-interacting prot | MPRIP | 116.534 | 2 | 47.884483 | yes | 7.8E-07 | 774.40125 | 24.957773 |
| ELEVLSEQYSQK | 12 | Q6WCQ1 | Myosin phosphatase Rho-interacting prot | MPRIP | 116.534 | 2 | 47.884483 | yes | 0.0002858 | 726.86444 | 8.1779652 |
| VMEETPNILR | 10 | Q6WKZ4 | Rab11 family-interacting protein 1 OS=H | RAB11FIP | 137.168 | 2 | 48.21992 | yes | 1.714E-07 | 601.31592 | 124.48775 |
| ELEDYIDNLLVR | 12 | Q6WKZ4 | Rab11 family-interacting protein 1 OS=H | RAB11FIP | 137.168 | 2 | 48.21992 | no | 1.21E-07 | 746.38812 | 116.31643 |
| NNMTASMFDLSM | 13 | Q6WKZ4 | Rab11 family-interacting protein 1 OS=H | RAB11FIP | 137.168 | 2 | 48.21992 | no | 1.179E-07 | 745.32782 | 69.71962 |
| TPPGIPAPSEPLLCI | 15 | Q6XE24 | RNA-binding motif, single-stranded-inter | RBMS3 | 47.84 | 2 | 46.28416 | yes | 1.602E-07 | 788.92383 | 89.121193 |
| VEETMALPQEGSL | 15 | Q6XZF7 | Dynamin-binding protein OS=Homo sapi | DNMBP | 177.348 | 2 | 48.530544 | yes | 1.135E-07 | 815.90887 | 121.74659 |
| DLSLYLQHIR | 10 | Q6XZF7 | Dynamin-binding protein OS=Homo sapi | DNMBP | 177.348 | 2 | 48.530544 | yes | 2.307E-06 | 629.35114 | 58.525051 |
| HRPTCETLEKEGP( | 18 | Q6XZF7 | Dynamin-binding protein OS=Homo sapi | DNMBP | 177.348 | 4 | 48.530544 | yes | 8.862E-05 | 527.74945 | 9.9290915 |
| SLLEIQQEEAR | 11 | Q6Y7W6 | GRB10-interacting GYF protein 2 OS=H | GIGYF2 | 150.07 | 2 | 52.15949 | yes | 9.959E-08 | 658.34625 | 2709.2778 |
| ISDQNIIPSVTR | 12 | Q6Y7W6 | GRB10-interacting GYF protein 2 OS=H | GIGYF2 | 150.07 | 2 | 52.15949 | yes | 8.326E-07 | 671.86987 | 2410.2563 |

| | | | | | | | | | | | |
|---|---|---|---|---|---|---|---|---|---|---|---|
| ADPSLLGFSVNAS: | 16 | Q6Y7W6 | GRB10-interacting GYF protein 2 OS=H( | GIGYF2 | 150.07 | 2 | 52.15949 | yes | 1.002E-07 | 825.4101 | 1658.9523 |
| ALQQQQQQQQK | 12 | Q6Y7W6 | GRB10-interacting GYF protein 2 OS=H( | GIGYF2 | 150.07 | 2 | 52.15949 | yes | 2.801E-07 | 742.38422 | 1640.7545 |
| RFEFDFR | 7 | Q6Y7W6 | GRB10-interacting GYF protein 2 OS=H( | GIGYF2 | 150.07 | 2 | 52.15949 | yes | 1.223E-07 | 508.75107 | 1038.7144 |
| GVSIPLMHEAMQK | 13 | Q6Y7W6 | GRB10-interacting GYF protein 2 OS=H( | GIGYF2 | 150.07 | 2 | 52.15949 | yes | 1.535E-07 | 720.87054 | 1028.1896 |
| ACDESFQPLGDIM | 14 | Q6Y7W6 | GRB10-interacting GYF protein 2 OS=H( | GIGYF2 | 150.07 | 2 | 52.15949 | yes | 1.163E-07 | 805.8631 | 976.1626 |
| EVESPYEVHDYIR | 13 | Q6Y7W6 | GRB10-interacting GYF protein 2 OS=H( | GIGYF2 | 150.07 | 2 | 52.15949 | yes | 1.004E-07 | 818.38611 | 453.71667 |
| EVESPYEVHDYIR | 13 | Q6Y7W6 | GRB10-interacting GYF protein 2 OS=H( | GIGYF2 | 150.07 | 2 | 52.15949 | yes | 1.393E-07 | 545.92651 | 453.71667 |
| EVESPYEVHDYIR | 13 | Q6Y7W6 | GRB10-interacting GYF protein 2 OS=H( | GIGYF2 | 150.07 | 3 | 52.15949 | yes | 1.004E-07 | 818.38611 | 453.71667 |
| EVESPYEVHDYIR | 13 | Q6Y7W6 | GRB10-interacting GYF protein 2 OS=H( | GIGYF2 | 150.07 | 3 | 52.15949 | yes | 1.393E-07 | 545.92651 | 453.71667 |
| AYLGDTSEAK | 10 | Q6Y7W6 | GRB10-interacting GYF protein 2 OS=H( | GIGYF2 | 150.07 | 2 | 52.15949 | yes | 1.532E-07 | 527.75623 | 792.07922 |
| QQQQEALR | 8 | Q6Y7W6 | GRB10-interacting GYF protein 2 OS=H( | GIGYF2 | 150.07 | 2 | 52.15949 | yes | 1.81E-07 | 500.76215 | 737.36285 |
| QQQEEALR | 8 | Q6Y7W6 | GRB10-interacting GYF protein 2 OS=H( | GIGYF2 | 150.07 | 2 | 52.15949 | yes | 1.531E-07 | 501.25415 | 694.73413 |
| MVAYLQDSALDD | 14 | Q6Y7W6 | GRB10-interacting GYF protein 2 OS=H( | GIGYF2 | 150.07 | 2 | 52.15949 | yes | 1.229E-07 | 813.37744 | 676.33441 |
| NSNMGFWDDAVK | 12 | Q6Y7W6 | GRB10-interacting GYF protein 2 OS=H( | GIGYF2 | 150.07 | 2 | 52.15949 | yes | 1.017E-07 | 692.30353 | 662.68628 |
| WRPHSPDGPR | 10 | Q6Y7W6 | GRB10-interacting GYF protein 2 OS=H( | GIGYF2 | 150.07 | 3 | 52.15949 | yes | 9.722E-08 | 402.20386 | 129.63907 |
| WRPHSPDGPR | 10 | Q6Y7W6 | GRB10-interacting GYF protein 2 OS=H( | GIGYF2 | 150.07 | 3 | 52.15949 | yes | 1.192E-07 | 602.80212 | 129.63907 |
| WRPHSPDGPR | 10 | Q6Y7W6 | GRB10-interacting GYF protein 2 OS=H( | GIGYF2 | 150.07 | 2 | 52.15949 | yes | 9.722E-08 | 402.20386 | 129.63907 |
| WRPHSPDGPR | 10 | Q6Y7W6 | GRB10-interacting GYF protein 2 OS=H( | GIGYF2 | 150.07 | 2 | 52.15949 | yes | 1.192E-07 | 602.80212 | 129.63907 |
| LSGWGNVSK | 9 | Q6Y7W6 | GRB10-interacting GYF protein 2 OS=H( | GIGYF2 | 150.07 | 2 | 52.15949 | yes | 9.722E-08 | 474.25089 | 220.79509 |
| MENSLPAK | 8 | Q6Y7W6 | GRB10-interacting GYF protein 2 OS=H( | GIGYF2 | 150.07 | 2 | 52.15949 | yes | 3.929E-07 | 445.22601 | 219.08124 |
| KQEELLR | 7 | Q6Y7W6 | GRB10-interacting GYF protein 2 OS=H( | GIGYF2 | 150.07 | 2 | 52.15949 | yes | 2.061E-07 | 458.26654 | 216.18086 |
| NFSMSVNSAAVLF | 13 | Q6Y7W6 | GRB10-interacting GYF protein 2 OS=H( | GIGYF2 | 150.07 | 2 | 52.15949 | yes | 1.485E-07 | 698.35608 | 197.55914 |
| EEMLALFLK | 9 | Q6Y7W6 | GRB10-interacting GYF protein 2 OS=H( | GIGYF2 | 150.07 | 2 | 52.15949 | yes | 1.85E-06 | 547.30176 | 180.80888 |
| SGSGSIDDDR | 10 | Q6Y7W6 | GRB10-interacting GYF protein 2 OS=H( | GIGYF2 | 150.07 | 2 | 52.15949 | yes | 5.876E-07 | 504.71506 | 60.208958 |
| RQQEEILR | 8 | Q6Y7W6 | GRB10-interacting GYF protein 2 OS=H( | GIGYF2 | 150.07 | 2 | 52.15949 | yes | 1.367E-07 | 536.29889 | 56.933907 |
| LAGCTVFITGASR | 13 | Q6YN16 | Hydroxysteroid dehydrogenase-like prote | HSDL2 | 45.395 | 2 | 46.65746 | yes | 1.503E-07 | 676.85321 | 163.68646 |
| SGAVEETFR | 9 | Q6YN16 | Hydroxysteroid dehydrogenase-like prote | HSDL2 | 45.395 | 2 | 46.65746 | yes | 3.313E-07 | 498.24326 | 69.486557 |
| SFTGNFVIDENILK | 14 | Q6YN16 | Hydroxysteroid dehydrogenase-like prote | HSDL2 | 45.395 | 2 | 46.65746 | yes | 2.683E-07 | 798.91705 | 60.710861 |
| KVDIIADAAYSIFQ | 17 | Q6YN16 | Hydroxysteroid dehydrogenase-like prote | HSDL2 | 45.395 | 3 | 46.65746 | yes | 5.051E-07 | 636.35944 | 38.552792 |
| HQRIHTGERPYQC | 18 | Q6ZMS7 | Zinc finger protein 783 OS=Homo sapien | ZNF783 | 61.015 | 4 | 21.872648 | yes | 4.017E-06 | 570.771 | 6.1375723 |
| GEEPVFVVTGR | 11 | Q6ZN04 | RNA-binding protein MEX3B OS=Homo | MEX3B | 58.833 | 2 | 50.012283 | yes | 1.007E-07 | 595.31421 | 233.7576 |
| PSSLFADLER | 10 | Q6ZN04 | RNA-binding protein MEX3B OS=Homo | MEX3B | 58.833 | 2 | 50.012283 | yes | 1.145E-07 | 567.79309 | 153.80359 |
| IQQQTHTYIVTPSR | 14 | Q6ZN04 | RNA-binding protein MEX3B OS=Homo | MEX3B | 58.833 | 3 | 50.012283 | no | 3.875E-07 | 557.96527 | 111.65482 |
| PTPSITPTPGR | 11 | Q6ZN04 | RNA-binding protein MEX3B OS=Homo | MEX3B | 58.833 | 2 | 50.012283 | yes | 1.04E-07 | 562.30896 | 82.012779 |
| EEIEAHIALR | 10 | Q6ZN04 | RNA-binding protein MEX3B OS=Homo | MEX3B | 58.833 | 2 | 50.012283 | yes | 9.935E-08 | 590.81964 | 36.038311 |
| VVGLVVGPK | 9 | Q6ZN04 | RNA-binding protein MEX3B OS=Homo | MEX3B | 58.833 | 2 | 50.012283 | no | 1.348E-06 | 434.28674 | 19.012104 |
| DTALLLDPNIYR | 12 | Q6ZN18 | Zinc finger protein AEBP2 OS=Homo sa | AEBP2 | 54.467 | 2 | 38.730633 | yes | 1.265E-06 | 702.38007 | 147.12808 |
| VYNTPSTSQSWLQ | 14 | Q6ZN18 | Zinc finger protein AEBP2 OS=Homo sa | AEBP2 | 54.467 | 2 | 38.730633 | yes | 5.025E-07 | 833.91284 | 119.06317 |
| HVPTHFSQQNSSK | 13 | Q6ZN18 | Zinc finger protein AEBP2 OS=Homo sa | AEBP2 | 54.467 | 3 | 38.730633 | yes | 2.992E-06 | 499.57953 | 34.373039 |

| | | | | | | | | | | | |
|---|---|---|---|---|---|---|---|---|---|---|---|
| MMAPQEAVPAEP! | 15 | Q6ZN55 | Zinc finger protein 574 OS=Homo sapien | ZNF574 | 98.901 | 2 | 53.4161 | yes | 1.735E-07 | 791.88385 | 688.22675 |
| MMAPQEAVPAEP! | 15 | Q6ZN55 | Zinc finger protein 574 OS=Homo sapien | ZNF574 | 98.901 | 2 | 53.4161 | yes | 3.154E-06 | 799.88129 | 688.22675 |
| MMAPQEAVPAEP! | 15 | Q6ZN55 | Zinc finger protein 574 OS=Homo sapien | ZNF574 | 98.901 | 2 | 53.4161 | yes | 7.967E-07 | 799.88129 | 688.22675 |
| VAVEHSYR | 8 | Q6ZN55 | Zinc finger protein 574 OS=Homo sapien | ZNF574 | 98.901 | 2 | 53.4161 | yes | 2.026E-07 | 480.7485 | 1145.1133 |
| LFSTETSLQVHR | 12 | Q6ZN55 | Zinc finger protein 574 OS=Homo sapien | ZNF574 | 98.901 | 2 | 53.4161 | yes | 9.722E-08 | 709.37537 | 956.3653 |
| PFNSPANLAR | 10 | Q6ZN55 | Zinc finger protein 574 OS=Homo sapien | ZNF574 | 98.901 | 2 | 53.4161 | yes | 1.734E-07 | 543.78815 | 912.51233 |
| APAPVVLGSPVVL | 21 | Q6ZN55 | Zinc finger protein 574 OS=Homo sapien | ZNF574 | 98.901 | 3 | 53.4161 | yes | 1.182E-07 | 661.05798 | 861.55664 |
| CAAAAAQAPR | 10 | Q6ZN55 | Zinc finger protein 574 OS=Homo sapien | ZNF574 | 98.901 | 2 | 53.4161 | yes | 2.958E-07 | 493.74545 | 810.85217 |
| LVHAQHFPYR | 10 | Q6ZN55 | Zinc finger protein 574 OS=Homo sapien | ZNF574 | 98.901 | 2 | 53.4161 | yes | 1.696E-07 | 634.33838 | 373.20383 |
| LVHAQHFPYR | 10 | Q6ZN55 | Zinc finger protein 574 OS=Homo sapien | ZNF574 | 98.901 | 2 | 53.4161 | yes | 3.556E-07 | 423.228 | 373.20383 |
| LVHAQHFPYR | 10 | Q6ZN55 | Zinc finger protein 574 OS=Homo sapien | ZNF574 | 98.901 | 3 | 53.4161 | yes | 1.696E-07 | 634.33838 | 373.20383 |
| LVHAQHFPYR | 10 | Q6ZN55 | Zinc finger protein 574 OS=Homo sapien | ZNF574 | 98.901 | 3 | 53.4161 | yes | 3.556E-07 | 423.228 | 373.20383 |
| LQAHEAAHAAAG | 19 | Q6ZN55 | Zinc finger protein 574 OS=Homo sapien | ZNF574 | 98.901 | 3 | 53.4161 | yes | 9.819E-08 | 614.32697 | 676.45599 |
| CPSCGAAFPSSLR | 13 | Q6ZN55 | Zinc finger protein 574 OS=Homo sapien | ZNF574 | 98.901 | 2 | 53.4161 | yes | 1.429E-07 | 705.31866 | 619.77197 |
| AFTQSSTLR | 9 | Q6ZN55 | Zinc finger protein 574 OS=Homo sapien | ZNF574 | 98.901 | 2 | 53.4161 | yes | 2.223E-07 | 505.76691 | 618.05951 |
| PYSCPDCGK | 9 | Q6ZN55 | Zinc finger protein 574 OS=Homo sapien | ZNF574 | 98.901 | 2 | 53.4161 | yes | 2.018E-07 | 542.21533 | 331.90723 |
| LLEVHQLVVHAGI | 13 | Q6ZN55 | Zinc finger protein 574 OS=Homo sapien | ZNF574 | 98.901 | 3 | 53.4161 | yes | 1.076E-07 | 490.95618 | 159.1581 |
| LLEVHQLVVHAGI | 13 | Q6ZN55 | Zinc finger protein 574 OS=Homo sapien | ZNF574 | 98.901 | 3 | 53.4161 | yes | 1.103E-07 | 735.9306 | 159.1581 |
| LLEVHQLVVHAGI | 13 | Q6ZN55 | Zinc finger protein 574 OS=Homo sapien | ZNF574 | 98.901 | 2 | 53.4161 | yes | 1.076E-07 | 490.95618 | 159.1581 |
| LLEVHQLVVHAGI | 13 | Q6ZN55 | Zinc finger protein 574 OS=Homo sapien | ZNF574 | 98.901 | 2 | 53.4161 | yes | 1.103E-07 | 735.9306 | 159.1581 |
| ALFASQELWLNHF | 13 | Q6ZN55 | Zinc finger protein 574 OS=Homo sapien | ZNF574 | 98.901 | 2 | 53.4161 | yes | 9.722E-08 | 792.91772 | 283.72998 |
| AHTPNPLHSCPCG | 14 | Q6ZN55 | Zinc finger protein 574 OS=Homo sapien | ZNF574 | 98.901 | 3 | 53.4161 | yes | 1.295E-07 | 525.91003 | 113.77634 |
| AHTPNPLHSCPCG | 14 | Q6ZN55 | Zinc finger protein 574 OS=Homo sapien | ZNF574 | 98.901 | 3 | 53.4161 | yes | 1.645E-07 | 788.36139 | 113.77634 |
| AHTPNPLHSCPCG | 14 | Q6ZN55 | Zinc finger protein 574 OS=Homo sapien | ZNF574 | 98.901 | 2 | 53.4161 | yes | 1.295E-07 | 525.91003 | 113.77634 |
| AHTPNPLHSCPCG | 14 | Q6ZN55 | Zinc finger protein 574 OS=Homo sapien | ZNF574 | 98.901 | 2 | 53.4161 | yes | 1.645E-07 | 788.36139 | 113.77634 |
| YHHTGEYPYK | 10 | Q6ZN55 | Zinc finger protein 574 OS=Homo sapien | ZNF574 | 98.901 | 2 | 53.4161 | yes | 1.172E-07 | 647.7962 | 174.92976 |
| PFACEVCGK | 9 | Q6ZN55 | Zinc finger protein 574 OS=Homo sapien | ZNF574 | 98.901 | 2 | 53.4161 | yes | 1.617E-07 | 534.23608 | 171.97537 |
| RFECGTCGK | 9 | Q6ZN55 | Zinc finger protein 574 OS=Homo sapien | ZNF574 | 98.901 | 2 | 53.4161 | yes | 1.429E-07 | 557.74207 | 169.50531 |
| LTHTGERPYR | 10 | Q6ZN55 | Zinc finger protein 574 OS=Homo sapien | ZNF574 | 98.901 | 3 | 53.4161 | yes | 4.336E-07 | 410.55103 | 137.03674 |
| KLFSTETSLQVHR | 13 | Q6ZN55 | Zinc finger protein 574 OS=Homo sapien | ZNF574 | 98.901 | 2 | 53.4161 | yes | 2.236E-07 | 773.42285 | 25.0243 |
| TTSSAVTETGPPAI | 16 | Q6ZNA4 | E3 ubiquitin-protein ligase Arkadia OS=I | RNF111 | 108.862 | 2 | 53.667255 | yes | 9.722E-08 | 801.89325 | 107.85033 |
| ISTVIQPLR | 9 | Q6ZNA4 | E3 ubiquitin-protein ligase Arkadia OS=I | RNF111 | 108.862 | 2 | 53.667255 | yes | 1.076E-07 | 513.81897 | 82.281822 |
| HSQTILNAK | 9 | Q6ZNA4 | E3 ubiquitin-protein ligase Arkadia OS=I | RNF111 | 108.862 | 2 | 53.667255 | yes | 4.973E-07 | 506.28271 | 26.335934 |
| GFHPDPEALK | 10 | Q6ZQX7 | Protein LIAT1 OS=Homo sapiens OX=9( | LIAT1 | 49.656 | 2 | 36.089848 | yes | 7.413E-07 | 555.78253 | 76.807053 |
| IPYNAVLDLVNNK | 13 | Q6ZRP7 | Sulfhydryl oxidase 2 OS=Homo sapiens ( | QSOX2 | 77.529 | 2 | 44.768734 | yes | 3.357E-06 | 736.909 | 61.007759 |
| AGEDAVWVLDSG | 15 | Q6ZRP7 | Sulfhydryl oxidase 2 OS=Homo sapiens ( | QSOX2 | 77.529 | 2 | 44.768734 | yes | 2.076E-07 | 780.88629 | 43.682735 |
| ECGEHFEEMAK | 11 | Q6ZRP7 | Sulfhydryl oxidase 2 OS=Homo sapiens ( | QSOX2 | 77.529 | 2 | 44.768734 | yes | 2.988E-07 | 683.77374 | 16.955177 |
| LAVDALAEGGSEA | 16 | Q6ZRV2 | Protein FAM83H OS=Homo sapiens OX= | FAM83H | 127.123 | 2 | 51.853878 | yes | 2.128E-07 | 804.89685 | 201.1225 |
| APAGDLLPSAFR | 12 | Q6ZRV2 | Protein FAM83H OS=Homo sapiens OX= | FAM83H | 127.123 | 2 | 51.853878 | yes | 1.463E-07 | 607.83002 | 151.29698 |

| | | | | | | | | | | | |
|---|---|---|---|---|---|---|---|---|---|---|---|
| AGFADPDDFTLGA | 16 | Q6ZRV2 | Protein FAM83H OS=Homo sapiens OX= | FAM83H | 127.123 | 2 | 51.853878 | yes | 3.189E-07 | 803.87842 | 93.423134 |
| HSFATEGAGAVEN | 18 | Q6ZRV2 | Protein FAM83H OS=Homo sapiens OX= | FAM83H | 127.123 | 3 | 51.853878 | yes | 9.722E-08 | 602.62372 | 78.859917 |
| EEGLGFPSFLDPDI | 14 | Q6ZRV2 | Protein FAM83H OS=Homo sapiens OX= | FAM83H | 127.123 | 2 | 51.853878 | yes | 5.725E-07 | 789.87537 | 68.836189 |
| HGSDPAFAPGPR | 12 | Q6ZRV2 | Protein FAM83H OS=Homo sapiens OX= | FAM83H | 127.123 | 2 | 51.853878 | yes | 4.632E-07 | 604.79401 | 63.771954 |
| EGPEEPGLAK | 10 | Q6ZRV2 | Protein FAM83H OS=Homo sapiens OX= | FAM83H | 127.123 | 2 | 51.853878 | yes | 2.52E-07 | 513.75873 | 59.046791 |
| LSSATANALYSSN | 15 | Q6ZRV2 | Protein FAM83H OS=Homo sapiens OX= | FAM83H | 127.123 | 2 | 51.853878 | yes | 1.638E-07 | 784.40735 | 57.682819 |
| HLRPPQYVTR | 10 | Q6ZRV2 | Protein FAM83H OS=Homo sapiens OX= | FAM83H | 127.123 | 3 | 51.853878 | yes | 7.85E-07 | 422.90704 | 57.380981 |
| TVAGPTYYCR | 10 | Q6ZRV2 | Protein FAM83H OS=Homo sapiens OX= | FAM83H | 127.123 | 2 | 51.853878 | yes | 1.361E-06 | 594.2793 | 32.383698 |
| LSLGQGDSTEAAT | 16 | Q6ZRV2 | Protein FAM83H OS=Homo sapiens OX= | FAM83H | 127.123 | 2 | 51.853878 | yes | 1.127E-07 | 832.39209 | 32.237083 |
| FDPENPQTLR | 10 | Q6ZRY4 | RNA-binding protein with multiple splici | RBPMS2 | 22.497 | 2 | 50.89207 | yes | 9.722E-08 | 608.80145 | 813.36243 |
| WYPSSDTTQQGW | 13 | Q6ZRY4 | RNA-binding protein with multiple splici | RBPMS2 | 22.497 | 2 | 50.89207 | yes | 3.134E-07 | 792.35986 | 181.24162 |
| TLFVSGLPVDIKPF | 14 | Q6ZRY4 | RNA-binding protein with multiple splici | RBPMS2 | 22.497 | 3 | 50.89207 | yes | 2.203E-07 | 514.63983 | 42.055759 |
| TLFVSGLPVDIKPF | 14 | Q6ZRY4 | RNA-binding protein with multiple splici | RBPMS2 | 22.497 | 3 | 50.89207 | yes | 5.083E-07 | 771.45612 | 42.055759 |
| TLFVSGLPVDIKPF | 14 | Q6ZRY4 | RNA-binding protein with multiple splici | RBPMS2 | 22.497 | 2 | 50.89207 | yes | 2.203E-07 | 514.63983 | 42.055759 |
| TLFVSGLPVDIKPF | 14 | Q6ZRY4 | RNA-binding protein with multiple splici | RBPMS2 | 22.497 | 2 | 50.89207 | yes | 5.083E-07 | 771.45612 | 42.055759 |
| TKNMDDNTFAMA | 31 | Q6ZSY5 | Protein phosphatase 1 regulatory subunit | PPP1R3F | 82.798 | 4 | 26.592459 | yes | 3.098E-06 | 845.91168 | 49.937653 |
| LLLQQNK | 7 | Q6ZSZ5 | Rho guanine nucleotide exchange factor 1 | ARHGEF1 | 151.643 | 2 | 37.710545 | yes | 1.108E-06 | 428.76617 | 84.722878 |
| DQLIAQSLLEK | 11 | Q6ZSZ5 | Rho guanine nucleotide exchange factor 1 | ARHGEF1 | 151.643 | 2 | 37.710545 | yes | 5.764E-07 | 629.35608 | 79.718781 |
| FYDYEANMPDR | 11 | Q6ZUT1 | Uncharacterized protein NKAPD1 OS=H | NKAPD1 | 34.11 | 2 | 51.80664 | yes | 9.722E-08 | 710.79553 | 89.572945 |
| DLCCLLQVDGEAF | 13 | Q6ZW31 | Rho GTPase-activating protein SYDE1 O | SYDE1 | 79.793 | 2 | 53.725395 | yes | 9.722E-08 | 774.8609 | 969.04718 |
| ALVLAWDPGVR | 11 | Q6ZW31 | Rho GTPase-activating protein SYDE1 O | SYDE1 | 79.793 | 2 | 53.725395 | yes | 1.076E-07 | 598.84296 | 882.90674 |
| GLLSCLPDVER | 11 | Q6ZW31 | Rho GTPase-activating protein SYDE1 O | SYDE1 | 79.793 | 2 | 53.725395 | yes | 9.819E-08 | 629.82666 | 808.90118 |
| ELPTPLITQPLYK | 13 | Q6ZW31 | Rho GTPase-activating protein SYDE1 O | SYDE1 | 79.793 | 2 | 53.725395 | yes | 1.186E-07 | 756.93726 | 742.3913 |
| SSGPGLASAVDFK | 13 | Q6ZW31 | Rho GTPase-activating protein SYDE1 O | SYDE1 | 79.793 | 2 | 53.725395 | yes | 1.143E-07 | 618.31696 | 685.10248 |
| GCQAQQLAVR | 10 | Q6ZW31 | Rho GTPase-activating protein SYDE1 O | SYDE1 | 79.793 | 2 | 53.725395 | yes | 1E-07 | 565.79041 | 579.64368 |
| ATLTLLLDHLR | 11 | Q6ZW31 | Rho GTPase-activating protein SYDE1 O | SYDE1 | 79.793 | 2 | 53.725395 | yes | 1.001E-07 | 633.38245 | 429.1604 |
| APPAALWGR | 9 | Q6ZW31 | Rho GTPase-activating protein SYDE1 O | SYDE1 | 79.793 | 2 | 53.725395 | yes | 4.603E-07 | 469.76398 | 379.39545 |
| RQPPLHLPLADPE | 19 | Q6ZW31 | Rho GTPase-activating protein SYDE1 O | SYDE1 | 79.793 | 4 | 53.725395 | yes | 1.673E-07 | 548.56689 | 350.38785 |
| LEPQGLLYAK | 10 | Q6ZW31 | Rho GTPase-activating protein SYDE1 O | SYDE1 | 79.793 | 2 | 53.725395 | yes | 1.478E-07 | 566.32404 | 330.6188 |
| LDHTFHLELEAAR | 13 | Q6ZW31 | Rho GTPase-activating protein SYDE1 O | SYDE1 | 79.793 | 3 | 53.725395 | yes | 2.005E-07 | 517.9353 | 155.37701 |
| LDHTFHLELEAAR | 13 | Q6ZW31 | Rho GTPase-activating protein SYDE1 O | SYDE1 | 79.793 | 3 | 53.725395 | yes | 2.69E-07 | 776.39935 | 155.37701 |
| LDHTFHLELEAAR | 13 | Q6ZW31 | Rho GTPase-activating protein SYDE1 O | SYDE1 | 79.793 | 2 | 53.725395 | yes | 2.005E-07 | 517.9353 | 155.37701 |
| LDHTFHLELEAAR | 13 | Q6ZW31 | Rho GTPase-activating protein SYDE1 O | SYDE1 | 79.793 | 2 | 53.725395 | yes | 2.69E-07 | 776.39935 | 155.37701 |
| LSLHLYGLGGLRP | 20 | Q6ZW31 | Rho GTPase-activating protein SYDE1 O | SYDE1 | 79.793 | 3 | 53.725395 | yes | 1.021E-07 | 682.7251 | 202.88025 |
| VVLEAMAR | 8 | Q6ZW31 | Rho GTPase-activating protein SYDE1 O | SYDE1 | 79.793 | 2 | 53.725395 | yes | 4.735E-07 | 444.75223 | 169.60571 |
| LSLHLYGLGGLR | 12 | Q6ZW31 | Rho GTPase-activating protein SYDE1 O | SYDE1 | 79.793 | 2 | 53.725395 | yes | 9.722E-08 | 649.88263 | 153.10637 |
| ERPPGQVPLIIQK | 13 | Q6ZW31 | Rho GTPase-activating protein SYDE1 O | SYDE1 | 79.793 | 2 | 53.725395 | yes | 2.166E-07 | 737.94067 | 130.21074 |
| LVSSFHAYNR | 10 | Q6ZW31 | Rho GTPase-activating protein SYDE1 O | SYDE1 | 79.793 | 2 | 53.725395 | yes | 2.08E-07 | 597.3067 | 109.00247 |
| CVGQIER | 7 | Q6ZW31 | Rho GTPase-activating protein SYDE1 O | SYDE1 | 79.793 | 2 | 53.725395 | yes | 1.123E-07 | 431.216 | 75.792274 |

| | | | | | | | | | | | |
|---|---|---|---|---|---|---|---|---|---|---|---|
| QSPDVAPYLRPK | 12 | Q6ZW31 | Rho GTPase-activating protein SYDE1 O | SYDE1 | 79.793 | 2 | 53.725395 | yes | 9.722E-08 | 685.875 | 63.016945 |
| GTVSFGDFVQVAF | 13 | Q6ZWJ1 | Syntaxin-binding protein 4 OS=Homo sap | STXBP4 | 61.662 | 2 | 37.93605 | yes | 5.58E-07 | 691.85681 | 40.383606 |
| ILEMDPSNVQGK | 12 | Q6ZXV5 | Protein O-mannosyl-transferase TMTC3 ( | TMTC3 | 104.009 | 2 | 49.618195 | yes | 4.018E-07 | 665.83722 | 221.35956 |
| DLHPSTPLK | 9 | Q6ZXV5 | Protein O-mannosyl-transferase TMTC3 ( | TMTC3 | 104.009 | 2 | 49.618195 | yes | 1.017E-07 | 504.27966 | 115.7467 |
| ISSSSFIEPIFPTSK | 15 | Q6ZXV5 | Protein O-mannosyl-transferase TMTC3 ( | TMTC3 | 104.009 | 2 | 49.618195 | yes | 2.846E-06 | 820.4325 | 58.51807 |
| GVLIECDPAMK | 11 | Q6ZYL4 | General transcription factor IIH subunit 5 | GTF2H5 | 8.054 | 2 | 48.415592 | yes | 1.545E-07 | 616.80432 | 321.0834 |
| VGELMDQNAFSLT | 15 | Q6ZYL4 | General transcription factor IIH subunit 5 | GTF2H5 | 8.054 | 2 | 48.415592 | yes | 1.782E-07 | 840.91669 | 66.383415 |
| QFLLYLDESNALG | 14 | Q6ZYL4 | General transcription factor IIH subunit 5 | GTF2H5 | 8.054 | 2 | 48.415592 | yes | 1.151E-07 | 805.92487 | 54.932072 |
| VPATNLILGEGR | 12 | Q709F0 | Acyl-CoA dehydrogenase family member | ACAD11 | 87.265 | 2 | 52.515465 | yes | 3.824E-07 | 620.35645 | 300.418 |
| MAGIAQGVYSR | 11 | Q709F0 | Acyl-CoA dehydrogenase family member | ACAD11 | 87.265 | 2 | 52.515465 | yes | 9.722E-08 | 576.79517 | 174.75462 |
| LDNIVFHPK | 9 | Q709F0 | Acyl-CoA dehydrogenase family member | ACAD11 | 87.265 | 2 | 52.515465 | yes | 1.437E-07 | 541.80328 | 133.23341 |
| ALQIMCER | 8 | Q709F0 | Acyl-CoA dehydrogenase family member | ACAD11 | 87.265 | 2 | 52.515465 | yes | 2.625E-07 | 510.75189 | 104.43875 |
| ALFSIGFPVPK | 11 | Q709F0 | Acyl-CoA dehydrogenase family member | ACAD11 | 87.265 | 2 | 52.515465 | yes | 2.292E-07 | 588.34479 | 52.080753 |
| AAHSMDTLGSAGA | 14 | Q709F0 | Acyl-CoA dehydrogenase family member | ACAD11 | 87.265 | 2 | 52.515465 | yes | 8.161E-07 | 658.81677 | 46.469868 |
| GFQTYVLR | 8 | Q709F0 | Acyl-CoA dehydrogenase family member | ACAD11 | 87.265 | 2 | 52.515465 | yes | 2.995E-06 | 492.26907 | 38.095795 |
| WGKPLVIDK | 9 | Q709F0 | Acyl-CoA dehydrogenase family member | ACAD11 | 87.265 | 2 | 52.515465 | yes | 1.005E-06 | 528.31604 | 24.589142 |
| IGVAVPLSGTVAR | 13 | Q70CQ4 | Ubiquitin carboxyl-terminal hydrolase 31 | USP31 | 146.652 | 2 | 48.55891 | yes | 3.458E-07 | 620.37463 | 108.86005 |
| IVLLVCNR | 8 | Q70CQ4 | Ubiquitin carboxyl-terminal hydrolase 31 | USP31 | 146.652 | 2 | 48.55891 | yes | 1.13E-07 | 493.7944 | 70.945755 |
| PASVTSAASSR | 11 | Q70CQ4 | Ubiquitin carboxyl-terminal hydrolase 31 | USP31 | 146.652 | 2 | 48.55891 | yes | 2.184E-06 | 517.26727 | 66.519073 |
| APLAVMEGVFK | 11 | Q70CQ4 | Ubiquitin carboxyl-terminal hydrolase 31 | USP31 | 146.652 | 2 | 48.55891 | yes | 3.239E-06 | 581.32043 | 52.334633 |
| ALWTLEYTPQHSR | 13 | Q70CQ4 | Ubiquitin carboxyl-terminal hydrolase 31 | USP31 | 146.652 | 2 | 48.55891 | yes | 2.253E-07 | 801.40717 | 36.259911 |
| MFGAWLGELDK | 11 | Q70E73 | Ras-associated and pleckstrin homology c | RAPH1 | 135.256 | 2 | 51.10168 | yes | 9.722E-08 | 633.81299 | 63.264698 |
| VAVVNPQPQQWS | 13 | Q70E73 | Ras-associated and pleckstrin homology c | RAPH1 | 135.256 | 2 | 51.10168 | yes | 1.73E-07 | 740.89899 | 61.150017 |
| YISTTALCNEVER | 13 | Q70EK8 | Inactive ubiquitin carboxyl-terminal hydr | USP53 | 120.807 | 2 | 45.66878 | yes | 1.779E-07 | 778.37469 | 47.150032 |
| WVFFDDANVK | 10 | Q70EK8 | Inactive ubiquitin carboxyl-terminal hydr | USP53 | 120.807 | 2 | 45.66878 | yes | 1.451E-06 | 620.80347 | 44.130714 |
| FSEWLNIENSER | 12 | Q70EK8 | Inactive ubiquitin carboxyl-terminal hydr | USP53 | 120.807 | 2 | 45.66878 | yes | 3.322E-06 | 762.35986 | 38.324532 |
| VYQPGSMLSLAPT | 14 | Q70EK8 | Inactive ubiquitin carboxyl-terminal hydr | USP53 | 120.807 | 2 | 45.66878 | yes | 4.112E-05 | 746.3974 | 36.453037 |
| LLTSVEVDNIEPSA | 17 | Q70EL1 | Inactive ubiquitin carboxyl-terminal hydr | USP54 | 187.389 | 3 | 48.176563 | yes | 3.249E-07 | 643.0022 | 92.592621 |
| FMDLDELQNQGR | 12 | Q70EL1 | Inactive ubiquitin carboxyl-terminal hydr | USP54 | 187.389 | 2 | 48.176563 | yes | 1.187E-07 | 733.34064 | 57.38953 |
| ESLNIDSIFSK | 11 | Q70EL1 | Inactive ubiquitin carboxyl-terminal hydr | USP54 | 187.389 | 2 | 48.176563 | yes | 3.804E-07 | 626.82465 | 34.730434 |
| SVQQFLAMCDR | 11 | Q70EL1 | Inactive ubiquitin carboxyl-terminal hydr | USP54 | 187.389 | 2 | 48.176563 | yes | 5.424E-07 | 677.81573 | 31.138987 |
| DWEIESTSSESK | 12 | Q70EL1 | Inactive ubiquitin carboxyl-terminal hydr | USP54 | 187.389 | 2 | 48.176563 | yes | 1.136E-06 | 699.30701 | 17.426699 |
| SNVALPANSEDGC | 14 | Q70EL4 | Ubiquitin carboxyl-terminal hydrolase 43 | USP43 | 122.81 | 2 | 52.77018 | yes | 3.103E-06 | 693.83405 | 77.857254 |
| EYTPQLSAEFK | 11 | Q70EL4 | Ubiquitin carboxyl-terminal hydrolase 43 | USP43 | 122.81 | 2 | 52.77018 | yes | 9.722E-08 | 656.82465 | 75.867699 |
| FSLSLHSESK | 10 | Q70EL4 | Ubiquitin carboxyl-terminal hydrolase 43 | USP43 | 122.81 | 2 | 52.77018 | yes | 1.188E-07 | 567.79309 | 58.12817 |
| STSPEAGLGPWPSY | 15 | Q70EL4 | Ubiquitin carboxyl-terminal hydrolase 43 | USP43 | 122.81 | 2 | 52.77018 | yes | 8.754E-07 | 800.39374 | 54.002914 |
| SEAPVQNLGSLFSI | 15 | Q70EL4 | Ubiquitin carboxyl-terminal hydrolase 43 | USP43 | 122.81 | 2 | 52.77018 | yes | 2.502E-07 | 809.43341 | 52.172226 |
| AVSWAQLQQSILS | 14 | Q70EL4 | Ubiquitin carboxyl-terminal hydrolase 43 | USP43 | 122.81 | 2 | 52.77018 | yes | 2.752E-07 | 779.93304 | 42.433723 |
| GTSELDRPLQGTLT | 17 | Q70EL4 | Ubiquitin carboxyl-terminal hydrolase 43 | USP43 | 122.81 | 3 | 52.77018 | yes | 8.001E-07 | 624.01794 | 26.051256 |

| | | | | | | | | | | | |
|---|---|---|---|---|---|---|---|---|---|---|---|
| TLLQQLQK | 8 | Q70SY1 | Cyclic AMP-responsive element-binding | CREB3L2 | 57.415 | 2 | 49.270134 | no | 1.05E-07 | 486.29785 | 54.341961 |
| WYLSTDFPSTSIK | 13 | Q70SY1 | Cyclic AMP-responsive element-binding | CREB3L2 | 57.415 | 2 | 49.270134 | yes | 4.868E-07 | 772.88519 | 49.645241 |
| LETNEFQQLQSK | 12 | Q70UQ0 | Inhibitor of nuclear factor kappa-B kinase | IKBIP | 39.309 | 2 | 49.156494 | yes | 1.063E-07 | 732.87012 | 92.529594 |
| VENQYQLLK | 9 | Q70UQ0 | Inhibitor of nuclear factor kappa-B kinase | IKBIP | 39.309 | 2 | 49.156494 | yes | 3.203E-07 | 567.81134 | 65.709702 |
| VPTTLAEYCIK | 11 | Q712K3 | Ubiquitin-conjugating enzyme E2 R2 OS= | UBE2R2 | 27.166 | 2 | 39.454887 | yes | 9.442E-07 | 647.83923 | 164.92661 |
| SLQEEPVEGFR | 11 | Q712K3 | Ubiquitin-conjugating enzyme E2 R2 OS= | UBE2R2 | 27.166 | 2 | 39.454887 | yes | 4.575E-07 | 645.81989 | 116.96008 |
| ENLAIESQSK | 10 | Q71F23 | Centromere protein U OS=Homo sapiens | CENPU | 47.522 | 2 | 48.787315 | yes | 1.237E-06 | 559.78802 | 100.11559 |
| SSDTSGNEASEIES | 16 | Q71F23 | Centromere protein U OS=Homo sapiens | CENPU | 47.522 | 2 | 48.787315 | yes | 1.043E-06 | 820.36829 | 82.2743 |
| ELNIVLPEFEK | 11 | Q71F23 | Centromere protein U OS=Homo sapiens | CENPU | 47.522 | 2 | 48.787315 | yes | 7.368E-07 | 665.86627 | 76.305725 |
| TGPLSAQPSVEK | 12 | Q71F23 | Centromere protein U OS=Homo sapiens | CENPU | 47.522 | 2 | 48.787315 | yes | 1.104E-07 | 607.32477 | 66.75621 |
| EPSSVLVQPLR | 11 | Q71RC2 | La-related protein 4 OS=Homo sapiens O | LARP4 | 80.596 | 2 | 53.689156 | yes | 9.722E-08 | 612.85095 | 20880.285 |
| MPGELVLENR | 10 | Q71RC2 | La-related protein 4 OS=Homo sapiens O | LARP4 | 80.596 | 2 | 53.689156 | yes | 9.722E-08 | 579.3028 | 8617.5723 |
| MPGELVLENR | 10 | Q71RC2 | La-related protein 4 OS=Homo sapiens O | LARP4 | 80.596 | 2 | 53.689156 | yes | 1.003E-06 | 587.30023 | 8617.5723 |
| EIPETTPIEEVK | 12 | Q71RC2 | La-related protein 4 OS=Homo sapiens O | LARP4 | 80.596 | 2 | 53.689156 | yes | 1.292E-06 | 692.86395 | 14605.556 |
| SSPMVQVDEK | 10 | Q71RC2 | La-related protein 4 OS=Homo sapiens O | LARP4 | 80.596 | 2 | 53.689156 | yes | 1.21E-07 | 560.27118 | 6865.4707 |
| SSGGSEHSTEGSVS | 22 | Q71RC2 | La-related protein 4 OS=Homo sapiens O | LARP4 | 80.596 | 3 | 53.689156 | yes | 3.94E-07 | 720.99658 | 6428.937 |
| MLLFVEQVASK | 11 | Q71RC2 | La-related protein 4 OS=Homo sapiens O | LARP4 | 80.596 | 2 | 53.689156 | yes | 9.722E-08 | 632.85211 | 2507.5166 |
| MLLFVEQVASK | 11 | Q71RC2 | La-related protein 4 OS=Homo sapiens O | LARP4 | 80.596 | 2 | 53.689156 | yes | 9.722E-08 | 640.84961 | 2507.5166 |
| QLEFCFSR | 8 | Q71RC2 | La-related protein 4 OS=Homo sapiens O | LARP4 | 80.596 | 2 | 53.689156 | yes | 1.061E-07 | 543.75549 | 3174.2285 |
| LSYAEVCQKPPK | 12 | Q71RC2 | La-related protein 4 OS=Homo sapiens O | LARP4 | 80.596 | 2 | 53.689156 | yes | 1.452E-06 | 710.36865 | 2532.3955 |
| TNAAAMNMGR | 10 | Q71RC2 | La-related protein 4 OS=Homo sapiens O | LARP4 | 80.596 | 2 | 53.689156 | yes | 1.524E-07 | 518.73676 | 726.97144 |
| TNAAAMNMGR | 10 | Q71RC2 | La-related protein 4 OS=Homo sapiens O | LARP4 | 80.596 | 2 | 53.689156 | yes | 6.261E-06 | 526.73425 | 726.97144 |
| TNAAAMNMGR | 10 | Q71RC2 | La-related protein 4 OS=Homo sapiens O | LARP4 | 80.596 | 2 | 53.689156 | yes | 1.061E-06 | 526.73425 | 726.97144 |
| HNPTVTGHQEQTY | 16 | Q71RC2 | La-related protein 4 OS=Homo sapiens O | LARP4 | 80.596 | 3 | 53.689156 | yes | 6.297E-07 | 627.64648 | 2141.3625 |
| ASTASPCNNNINA | 23 | Q71RC2 | La-related protein 4 OS=Homo sapiens O | LARP4 | 80.596 | 3 | 53.689156 | yes | 1.622E-07 | 790.7204 | 1733.4137 |
| TFQGKPIMAR | 10 | Q71RC2 | La-related protein 4 OS=Homo sapiens O | LARP4 | 80.596 | 2 | 53.689156 | yes | 5.115E-07 | 574.81586 | 1493.6306 |
| NEDNGAPENSVEK | 21 | Q71RC2 | La-related protein 4 OS=Homo sapiens O | LARP4 | 80.596 | 4 | 53.689156 | yes | 1.652E-07 | 587.5282 | 1307.3396 |
| GTGLNPNAK | 9 | Q71RC2 | La-related protein 4 OS=Homo sapiens O | LARP4 | 80.596 | 2 | 53.689156 | yes | 1.161E-07 | 436.23523 | 852.72931 |
| KLSYAEVCQKPPK | 13 | Q71RC2 | La-related protein 4 OS=Homo sapiens O | LARP4 | 80.596 | 2 | 53.689156 | yes | 5.281E-07 | 774.41614 | 299.17392 |
| KLSYAEVCQKPPK | 13 | Q71RC2 | La-related protein 4 OS=Homo sapiens O | LARP4 | 80.596 | 2 | 53.689156 | yes | 8.248E-07 | 516.61322 | 299.17392 |
| KLSYAEVCQKPPK | 13 | Q71RC2 | La-related protein 4 OS=Homo sapiens O | LARP4 | 80.596 | 3 | 53.689156 | yes | 5.281E-07 | 774.41614 | 299.17392 |
| KLSYAEVCQKPPK | 13 | Q71RC2 | La-related protein 4 OS=Homo sapiens O | LARP4 | 80.596 | 3 | 53.689156 | yes | 8.248E-07 | 516.61322 | 299.17392 |
| DLIEDSSVQK | 10 | Q71RC2 | La-related protein 4 OS=Homo sapiens O | LARP4 | 80.596 | 2 | 53.689156 | yes | 2.761E-07 | 567.28784 | 470.71231 |
| ISRPHPSTAESK | 12 | Q71RC2 | La-related protein 4 OS=Homo sapiens O | LARP4 | 80.596 | 3 | 53.689156 | yes | 1.13E-07 | 437.23349 | 162.79816 |
| ISRPHPSTAESK | 12 | Q71RC2 | La-related protein 4 OS=Homo sapiens O | LARP4 | 80.596 | 3 | 53.689156 | yes | 1.163E-07 | 655.34656 | 162.79816 |
| ISRPHPSTAESK | 12 | Q71RC2 | La-related protein 4 OS=Homo sapiens O | LARP4 | 80.596 | 2 | 53.689156 | yes | 1.13E-07 | 437.23349 | 162.79816 |
| ISRPHPSTAESK | 12 | Q71RC2 | La-related protein 4 OS=Homo sapiens O | LARP4 | 80.596 | 2 | 53.689156 | yes | 1.163E-07 | 655.34656 | 162.79816 |
| EQYVPPR | 7 | Q71RC2 | La-related protein 4 OS=Homo sapiens O | LARP4 | 80.596 | 2 | 53.689156 | yes | 1.991E-07 | 444.73233 | 270.35687 |
| KLTTDPDLILEVLF | 14 | Q71RC2 | La-related protein 4 OS=Homo sapiens O | LARP4 | 80.596 | 2 | 53.689156 | yes | 1.753E-07 | 813.47723 | 177.8503 |

| | | | | | | | | | | | |
|---|---|---|---|---|---|---|---|---|---|---|---|
| LSYAEVCQKPPKE | 23 | Q71RC2 | La-related protein 4 OS=Homo sapiens O | LARP4 | 80.596 | 4 | 53.689156 | yes | 9.722E-08 | 657.10718 | 110.37576 |
| AINTFFAK | 8 | Q71RC2 | La-related protein 4 OS=Homo sapiens O | LARP4 | 80.596 | 2 | 53.689156 | yes | 1.411E-07 | 456.2529 | 105.49936 |
| RCIVILR | 7 | Q71RC2 | La-related protein 4 OS=Homo sapiens O | LARP4 | 80.596 | 2 | 53.689156 | yes | 2.124E-07 | 465.28931 | 39.661446 |
| ETSTLQVEQNGDY | 15 | Q71RC2 | La-related protein 4 OS=Homo sapiens O | LARP4 | 80.596 | 2 | 53.689156 | yes | 1.318E-07 | 848.89227 | 31.475189 |
| LTEGCSFR | 8 | Q71UM5 | Ribosomal protein eS27-like OS=Homo s | RPS27L | 9.478 | 2 | 50.72201 | no | 1.003E-07 | 485.22656 | 17430.59 |
| LVQSPNSYFMDVK | 13 | Q71UM5 | Ribosomal protein eS27-like OS=Homo s | RPS27L | 9.478 | 2 | 50.72201 | no | 2.021E-07 | 764.37921 | 4713.106 |
| LVQSPNSYFMDVK | 13 | Q71UM5 | Ribosomal protein eS27-like OS=Homo s | RPS27L | 9.478 | 2 | 50.72201 | no | 1.069E-07 | 772.37671 | 4713.106 |
| DLLHPSLEEEK | 11 | Q71UM5 | Ribosomal protein eS27-like OS=Homo s | RPS27L | 9.478 | 2 | 50.72201 | yes | 1.742E-07 | 655.33533 | 3321.9094 |
| DLLHPSLEEEK | 11 | Q71UM5 | Ribosomal protein eS27-like OS=Homo s | RPS27L | 9.478 | 2 | 50.72201 | yes | 1.064E-07 | 437.22598 | 3321.9094 |
| DLLHPSLEEEK | 11 | Q71UM5 | Ribosomal protein eS27-like OS=Homo s | RPS27L | 9.478 | 3 | 50.72201 | yes | 1.742E-07 | 655.33533 | 3321.9094 |
| DLLHPSLEEEK | 11 | Q71UM5 | Ribosomal protein eS27-like OS=Homo s | RPS27L | 9.478 | 3 | 50.72201 | yes | 1.064E-07 | 437.22598 | 3321.9094 |
| DLLHPSLEEEKK | 12 | Q71UM5 | Ribosomal protein eS27-like OS=Homo s | RPS27L | 9.478 | 2 | 50.72201 | yes | 9.727E-08 | 719.38281 | 317.84137 |
| SSMCSAPSSSSSAK | 14 | Q765P7 | Protein MTSS 2 OS=Homo sapiens OX=9 | MTSS2 | 79.93 | 2 | 39.218914 | yes | 1.313E-05 | 687.28723 | 70.144745 |
| LPPASEQVIK | 10 | Q765P7 | Protein MTSS 2 OS=Homo sapiens OX=9 | MTSS2 | 79.93 | 2 | 39.218914 | yes | 2.7E-05 | 541.31622 | 51.828129 |
| TPTVPDSPGYMGP | 15 | Q765P7 | Protein MTSS 2 OS=Homo sapiens OX=9 | MTSS2 | 79.93 | 2 | 39.218914 | yes | 4.717E-07 | 788.3772 | 44.965107 |
| NSNIAQNYR | 9 | Q765P7 | Protein MTSS 2 OS=Homo sapiens OX=9 | MTSS2 | 79.93 | 2 | 39.218914 | yes | 1.981E-06 | 540.26508 | 44.41753 |
| SSYPIWEDFNSK | 12 | Q765P7 | Protein MTSS 2 OS=Homo sapiens OX=9 | MTSS2 | 79.93 | 2 | 39.218914 | yes | 1.045E-06 | 736.83826 | 21.163202 |
| VGSHEQPSGATLQ | 14 | Q765P7 | Protein MTSS 2 OS=Homo sapiens OX=9 | MTSS2 | 79.93 | 2 | 39.218914 | yes | 3.359E-06 | 733.87097 | 18.275011 |
| FGEVSDVEIITR | 12 | Q76FK4 | Nucleolar protein 8 OS=Homo sapiens O. | NOL8 | 131.617 | 2 | 45.58002 | yes | 1.806E-07 | 682.85645 | 71.788223 |
| VLPVLHLK | 8 | Q76FK4 | Nucleolar protein 8 OS=Homo sapiens O. | NOL8 | 131.617 | 2 | 45.58002 | yes | 1.27E-06 | 459.81039 | 60.889465 |
| LQGSDLFWR | 9 | Q76FK4 | Nucleolar protein 8 OS=Homo sapiens O. | NOL8 | 131.617 | 2 | 45.58002 | yes | 8.701E-07 | 561.29053 | 36.800148 |
| TGGVDFHMK | 9 | Q76FK4 | Nucleolar protein 8 OS=Homo sapiens O. | NOL8 | 131.617 | 2 | 45.58002 | yes | 2.703E-07 | 496.23694 | 31.403545 |
| EMYYNIAMDLK | 11 | Q76FK4 | Nucleolar protein 8 OS=Homo sapiens O. | NOL8 | 131.617 | 2 | 45.58002 | yes | 4.496E-06 | 695.82269 | 21.11455 |
| NSTEFSQMEK | 10 | Q76FK4 | Nucleolar protein 8 OS=Homo sapiens O. | NOL8 | 131.617 | 2 | 45.58002 | yes | 5.028E-07 | 600.76373 | 16.526234 |
| KVPLVSPDSSR | 11 | Q76M96 | Coiled-coil domain-containing protein 80 | CCDC80 | 108.174 | 2 | 51.538506 | yes | 1.536E-07 | 592.83533 | 209.96686 |
| STLEEPNLQPLQR | 13 | Q76M96 | Coiled-coil domain-containing protein 80 | CCDC80 | 108.174 | 2 | 51.538506 | yes | 4.759E-07 | 762.90448 | 149.44601 |
| LEAMYEVIDQGPII | 14 | Q76M96 | Coiled-coil domain-containing protein 80 | CCDC80 | 108.174 | 2 | 51.538506 | yes | 9.968E-08 | 817.41632 | 104.08041 |
| DDVYCELAER | 10 | Q76M96 | Coiled-coil domain-containing protein 80 | CCDC80 | 108.174 | 2 | 51.538506 | yes | 1.81E-07 | 635.27441 | 99.332138 |
| QSLENFLSR | 9 | Q76M96 | Coiled-coil domain-containing protein 80 | CCDC80 | 108.174 | 2 | 51.538506 | yes | 2.816E-06 | 547.28546 | 91.129715 |
| VVDDEDLVDQR | 11 | Q76M96 | Coiled-coil domain-containing protein 80 | CCDC80 | 108.174 | 2 | 51.538506 | yes | 9.728E-08 | 651.81226 | 90.831444 |
| SVADLLGSFEGK | 12 | Q76M96 | Coiled-coil domain-containing protein 80 | CCDC80 | 108.174 | 2 | 51.538506 | yes | 9.722E-08 | 611.81934 | 83.843666 |
| DEYLESFCK | 9 | Q76M96 | Coiled-coil domain-containing protein 80 | CCDC80 | 108.174 | 2 | 51.538506 | yes | 1.12E-07 | 595.75537 | 69.77906 |
| ILSNEYEEK | 9 | Q76M96 | Coiled-coil domain-containing protein 80 | CCDC80 | 108.174 | 2 | 51.538506 | yes | 8.33E-07 | 562.77716 | 63.158695 |
| LLLITAPK | 8 | Q76M96 | Coiled-coil domain-containing protein 80 | CCDC80 | 108.174 | 2 | 51.538506 | yes | 8.334E-07 | 434.79694 | 39.447163 |
| QYYEVPITMK | 10 | Q76M96 | Coiled-coil domain-containing protein 80 | CCDC80 | 108.174 | 2 | 51.538506 | yes | 8.239E-06 | 636.32068 | 37.467865 |
| HLDMGSPQLR | 10 | Q76N32 | Centrosomal protein of 68 kDa OS=Homo | CEP68 | 81.103 | 2 | 38.178646 | yes | 5.404E-07 | 577.29279 | 62.726273 |
| VFYVDHVNR | 9 | Q76N89 | E3 ubiquitin-protein ligase HECW1 OS=H | HECW1 | 179.554 | 2 | 47.414074 | yes | 3.39E-06 | 574.79602 | 123.19033 |
| EEPLEEEATTQSR | 13 | Q76N89 | E3 ubiquitin-protein ligase HECW1 OS=H | HECW1 | 179.554 | 2 | 47.414074 | yes | 2.408E-06 | 759.84955 | 66.334534 |
| GMFFNPDPYLK | 11 | Q76N89 | E3 ubiquitin-protein ligase HECW1 OS=H | HECW1 | 179.554 | 2 | 47.414074 | no | 1.443E-07 | 664.8208 | 45.419842 |

| | | | | | | | | | | | |
|---|---|---|---|---|---|---|---|---|---|---|---|
| EGSLSPVNSQK | 11 | Q76N89 | E3 ubiquitin-protein ligase HECW1 OS=] | HECW1 | 179.554 | 2 | 47.414074 | yes | 7.042E-07 | 573.29346 | 29.812456 |
| ITLLLQSPAVK | 11 | Q76N89 | E3 ubiquitin-protein ligase HECW1 OS=] | HECW1 | 179.554 | 2 | 47.414074 | yes | 1.335E-07 | 591.87646 | 28.497379 |
| LIICNVTGIAHR | 12 | Q7KZ85 | Transcription elongation factor SPT6 OS= | SUPT6H | 199.073 | 2 | 36.38546 | yes | 7.051E-07 | 683.88483 | 38.241577 |
| LGTLSPAFSTR | 11 | Q7KZF4 | Staphylococcal nuclease domain-containi | SND1 | 101.997 | 2 | 53.978302 | yes | 6.564E-07 | 575.31677 | 586.80249 |
| VITEYLNAQESAK | 13 | Q7KZF4 | Staphylococcal nuclease domain-containi | SND1 | 101.997 | 2 | 53.978302 | yes | 9.996E-08 | 733.38031 | 503.07178 |
| MVLSGCAIIVR | 11 | Q7KZF4 | Staphylococcal nuclease domain-containi | SND1 | 101.997 | 2 | 53.978302 | yes | 1.084E-07 | 609.8385 | 437.30219 |
| IHVFYIDYGNR | 11 | Q7KZF4 | Staphylococcal nuclease domain-containi | SND1 | 101.997 | 2 | 53.978302 | yes | 9.722E-08 | 698.85406 | 402.62033 |
| FVDGEWYR | 8 | Q7KZF4 | Staphylococcal nuclease domain-containi | SND1 | 101.997 | 2 | 53.978302 | yes | 2.673E-07 | 536.24835 | 396.5448 |
| YTIENPR | 7 | Q7KZF4 | Staphylococcal nuclease domain-containi | SND1 | 101.997 | 2 | 53.978302 | yes | 3.097E-07 | 446.7298 | 356.70975 |
| EGLVMVEVR | 9 | Q7KZF4 | Staphylococcal nuclease domain-containi | SND1 | 101.997 | 2 | 53.978302 | yes | 9.722E-08 | 516.28131 | 353.89719 |
| QINLSNIR | 8 | Q7KZF4 | Staphylococcal nuclease domain-containi | SND1 | 101.997 | 2 | 53.978302 | yes | 1.355E-07 | 479.27744 | 246.05472 |
| SEAVVEYVFSGSR | 13 | Q7KZF4 | Staphylococcal nuclease domain-containi | SND1 | 101.997 | 2 | 53.978302 | yes | 5.362E-07 | 715.3515 | 231.6992 |
| VMQVLNADAIVV | 13 | Q7KZF4 | Staphylococcal nuclease domain-containi | SND1 | 101.997 | 2 | 53.978302 | yes | 9.722E-08 | 700.40253 | 222.7724 |
| EVEVEVESMDK | 11 | Q7KZF4 | Staphylococcal nuclease domain-containi | SND1 | 101.997 | 2 | 53.978302 | yes | 1.05E-07 | 647.29755 | 202.68738 |
| QFLPFLQR | 8 | Q7KZF4 | Staphylococcal nuclease domain-containi | SND1 | 101.997 | 2 | 53.978302 | yes | 5.977E-07 | 524.80054 | 164.11952 |
| CVDWSIAVYTR | 11 | Q7KZF4 | Staphylococcal nuclease domain-containi | SND1 | 101.997 | 2 | 53.978302 | yes | 2.437E-07 | 685.33209 | 157.25876 |
| TDAVDSVVR | 9 | Q7KZF4 | Staphylococcal nuclease domain-containi | SND1 | 101.997 | 2 | 53.978302 | yes | 2.236E-07 | 481.2511 | 155.54575 |
| EYGMIYLGK | 9 | Q7KZF4 | Staphylococcal nuclease domain-containi | SND1 | 101.997 | 2 | 53.978302 | yes | 2.295E-07 | 537.27045 | 150.24117 |
| EVCFTIENK | 9 | Q7KZF4 | Staphylococcal nuclease domain-containi | SND1 | 101.997 | 2 | 53.978302 | yes | 2.362E-07 | 570.27368 | 146.60927 |
| SSHYDELLAAEAR | 13 | Q7KZF4 | Staphylococcal nuclease domain-containi | SND1 | 101.997 | 2 | 53.978302 | yes | 9.729E-08 | 731.35205 | 129.57188 |
| PASPATETVPAFSE | 15 | Q7KZF4 | Staphylococcal nuclease domain-containi | SND1 | 101.997 | 2 | 53.978302 | yes | 9.722E-08 | 780.38867 | 114.64366 |
| ADDADEFGYSR | 11 | Q7KZF4 | Staphylococcal nuclease domain-containi | SND1 | 101.997 | 2 | 53.978302 | yes | 9.722E-08 | 623.25458 | 111.06696 |
| SLLSAEEAAK | 10 | Q7KZF4 | Staphylococcal nuclease domain-containi | SND1 | 101.997 | 2 | 53.978302 | yes | 5.399E-07 | 509.77438 | 111.00038 |
| VHFTAER | 7 | Q7KZF4 | Staphylococcal nuclease domain-containi | SND1 | 101.997 | 2 | 53.978302 | yes | 2.237E-06 | 430.22467 | 94.204643 |
| TIHLSSIRPPR | 11 | Q7KZF4 | Staphylococcal nuclease domain-containi | SND1 | 101.997 | 3 | 53.978302 | yes | 9.052E-07 | 426.25433 | 91.25795 |
| EADGSETPEPFAAI | 16 | Q7KZF4 | Staphylococcal nuclease domain-containi | SND1 | 101.997 | 2 | 53.978302 | yes | 1.335E-07 | 824.87048 | 90.087769 |
| PVNAIIEHVR | 10 | Q7KZF4 | Staphylococcal nuclease domain-containi | SND1 | 101.997 | 2 | 53.978302 | yes | 1.664E-07 | 574.33276 | 83.375046 |
| HFVDSHHQKPVN/ | 19 | Q7KZF4 | Staphylococcal nuclease domain-containi | SND1 | 101.997 | 4 | 53.978302 | yes | 2.145E-07 | 566.55151 | 71.745461 |
| DTPDEPWAFPAR | 12 | Q7KZF4 | Staphylococcal nuclease domain-containi | SND1 | 101.997 | 2 | 53.978302 | yes | 6.905E-07 | 701.32532 | 38.119705 |
| VPASPLPGLER | 11 | Q7KZI7 | Serine/threonine-protein kinase MARK2 ( | MARK2 | 87.911 | 2 | 52.93043 | yes | 1.021E-07 | 568.32715 | 1739.7098 |
| TPLPTLNER | 9 | Q7KZI7 | Serine/threonine-protein kinase MARK2 ( | MARK2 | 87.911 | 2 | 52.93043 | yes | 2.363E-07 | 520.79041 | 1557.8245 |
| EEIQDSLVGQR | 11 | Q7KZI7 | Serine/threonine-protein kinase MARK2 ( | MARK2 | 87.911 | 2 | 52.93043 | yes | 9.722E-08 | 637.32275 | 1115.1099 |
| STFHAGQLR | 9 | Q7KZI7 | Serine/threonine-protein kinase MARK2 ( | MARK2 | 87.911 | 2 | 52.93043 | yes | 1.172E-07 | 508.76724 | 716.23871 |
| SSELEGDTITLKPR | 14 | Q7KZI7 | Serine/threonine-protein kinase MARK2 ( | MARK2 | 87.911 | 2 | 52.93043 | yes | 9.722E-08 | 773.40961 | 690.89374 |
| TTSSMEPNEMMR | 12 | Q7KZI7 | Serine/threonine-protein kinase MARK2 ( | MARK2 | 87.911 | 2 | 52.93043 | yes | 9.722E-08 | 707.29401 | 518.45166 |
| VLNHPNIVK | 9 | Q7KZI7 | Serine/threonine-protein kinase MARK2 ( | MARK2 | 87.911 | 2 | 52.93043 | yes | 2.514E-07 | 517.31128 | 427.72031 |
| IADFGFSNEFTFGN | 15 | Q7KZI7 | Serine/threonine-protein kinase MARK2 ( | MARK2 | 87.911 | 2 | 52.93043 | yes | 9.918E-08 | 847.39648 | 369.95099 |
| TQLNSSSLQK | 10 | Q7KZI7 | Serine/threonine-protein kinase MARK2 ( | MARK2 | 87.911 | 2 | 52.93043 | yes | 9.722E-08 | 553.29602 | 349.83334 |
| DTEQPTLGHLDSK | 13 | Q7KZI7 | Serine/threonine-protein kinase MARK2 ( | MARK2 | 87.911 | 2 | 52.93043 | yes | 9.975E-08 | 720.85187 | 308.14523 |

| | | | | | | | | | | | |
|---|---|---|---|---|---|---|---|---|---|---|---|
| SMSASVHPNK | 10 | Q7KZI7 | Serine/threonine-protein kinase MARK2 ( | MARK2 | 87.911 | 2 | 52.93043 | yes | 3.346E-07 | 529.25836 | 297.45227 |
| VLDANSCQSELHE | 14 | Q7KZI7 | Serine/threonine-protein kinase MARK2 ( | MARK2 | 87.911 | 2 | 52.93043 | yes | 9.722E-08 | 815.38049 | 221.18964 |
| RGASGSIFSK | 10 | Q7KZI7 | Serine/threonine-protein kinase MARK2 ( | MARK2 | 87.911 | 2 | 52.93043 | yes | 1.475E-07 | 505.2749 | 163.94888 |
| TELMVSMGYTR | 11 | Q7KZI7 | Serine/threonine-protein kinase MARK2 ( | MARK2 | 87.911 | 2 | 52.93043 | yes | 9.722E-08 | 644.30725 | 153.32253 |
| KFLILNPSK | 9 | Q7KZI7 | Serine/threonine-protein kinase MARK2 ( | MARK2 | 87.911 | 2 | 52.93043 | yes | 1.022E-07 | 530.33167 | 118.16864 |
| PSADLTNSSAPSPS | 16 | Q7KZI7 | Serine/threonine-protein kinase MARK2 ( | MARK2 | 87.911 | 2 | 52.93043 | yes | 9.722E-08 | 798.38666 | 117.75952 |
| WMNVGHEDDELK | 25 | Q7KZI7 | Serine/threonine-protein kinase MARK2 ( | MARK2 | 87.911 | 4 | 52.93043 | yes | 2.753E-07 | 761.36304 | 88.77475 |
| ISGTSMAFK | 9 | Q7KZI7 | Serine/threonine-protein kinase MARK2 ( | MARK2 | 87.911 | 2 | 52.93043 | yes | 1.006E-07 | 471.24167 | 65.133636 |
| FLILNPSK | 8 | Q7KZI7 | Serine/threonine-protein kinase MARK2 ( | MARK2 | 87.911 | 2 | 52.93043 | yes | 4.512E-06 | 466.28421 | 56.316315 |
| VETLRPHVVGSGG | 16 | Q7KZI7 | Serine/threonine-protein kinase MARK2 ( | MARK2 | 87.911 | 2 | 52.93043 | yes | 1.327E-06 | 832.93939 | 4.1586843 |
| YQQFPEFK | 8 | Q7KZN9 | Cytochrome c oxidase assembly protein C | COX15 | 46.031 | 2 | 47.899265 | yes | 1.692E-07 | 543.76636 | 98.939651 |
| ILNHDMTLTEFK | 12 | Q7KZN9 | Cytochrome c oxidase assembly protein C | COX15 | 46.031 | 2 | 47.899265 | yes | 1.236E-07 | 731.37396 | 63.835281 |
| LLVATSVAAR | 10 | Q7L014 | Probable ATP-dependent RNA helicase D | DDX46 | 117.363 | 2 | 53.564266 | yes | 9.722E-08 | 500.81113 | 456.68747 |
| FDETEQALANER | 12 | Q7L014 | Probable ATP-dependent RNA helicase D | DDX46 | 117.363 | 2 | 53.564266 | yes | 9.722E-08 | 711.82843 | 286.27252 |
| GAEIIVCTPGR | 11 | Q7L014 | Probable ATP-dependent RNA helicase D | DDX46 | 117.363 | 2 | 53.564266 | yes | 1.088E-07 | 586.80823 | 249.80942 |
| GGTILAPTVSAK | 12 | Q7L014 | Probable ATP-dependent RNA helicase D | DDX46 | 117.363 | 2 | 53.564266 | yes | 9.722E-08 | 557.82697 | 227.16252 |
| NFYVEVPELAK | 11 | Q7L014 | Probable ATP-dependent RNA helicase D | DDX46 | 117.363 | 2 | 53.564266 | yes | 1.032E-07 | 654.84534 | 225.13797 |
| SWVQCGISMK | 10 | Q7L014 | Probable ATP-dependent RNA helicase D | DDX46 | 117.363 | 2 | 53.564266 | yes | 9.872E-08 | 598.28357 | 146.8038 |
| ILSKPIEVQVGGR | 13 | Q7L014 | Probable ATP-dependent RNA helicase D | DDX46 | 117.363 | 3 | 53.564266 | yes | 1.156E-07 | 465.94879 | 69.115479 |
| ILSKPIEVQVGGR | 13 | Q7L014 | Probable ATP-dependent RNA helicase D | DDX46 | 117.363 | 3 | 53.564266 | yes | 2.198E-07 | 698.41956 | 69.115479 |
| ILSKPIEVQVGGR | 13 | Q7L014 | Probable ATP-dependent RNA helicase D | DDX46 | 117.363 | 2 | 53.564266 | yes | 1.156E-07 | 465.94879 | 69.115479 |
| ILSKPIEVQVGGR | 13 | Q7L014 | Probable ATP-dependent RNA helicase D | DDX46 | 117.363 | 2 | 53.564266 | yes | 2.198E-07 | 698.41956 | 69.115479 |
| LLEPVDHGK | 9 | Q7L014 | Probable ATP-dependent RNA helicase D | DDX46 | 117.363 | 2 | 53.564266 | yes | 4.398E-07 | 504.27966 | 130.70372 |
| ISEYSEAAITIR | 12 | Q7L014 | Probable ATP-dependent RNA helicase D | DDX46 | 117.363 | 2 | 53.564266 | yes | 2.488E-07 | 676.85645 | 129.47794 |
| LNYVPLEK | 8 | Q7L014 | Probable ATP-dependent RNA helicase D | DDX46 | 117.363 | 2 | 53.564266 | yes | 3.436E-07 | 488.27911 | 90.017044 |
| MIDMLAANSGR | 11 | Q7L014 | Probable ATP-dependent RNA helicase D | DDX46 | 117.363 | 2 | 53.564266 | yes | 4.241E-07 | 589.78644 | 89.864845 |
| MSQEEVNVFR | 10 | Q7L014 | Probable ATP-dependent RNA helicase D | DDX46 | 117.363 | 2 | 53.564266 | yes | 1.557E-07 | 619.79535 | 89.164177 |
| MFDMGFEPQVMR | 12 | Q7L014 | Probable ATP-dependent RNA helicase D | DDX46 | 117.363 | 2 | 53.564266 | yes | 9.722E-08 | 744.32764 | 81.155785 |
| IYLAIESANELAVQ | 15 | Q7L014 | Probable ATP-dependent RNA helicase D | DDX46 | 117.363 | 2 | 53.564266 | yes | 9.722E-08 | 831.45905 | 79.2118 |
| LEMEGITVK | 9 | Q7L014 | Probable ATP-dependent RNA helicase D | DDX46 | 117.363 | 2 | 53.564266 | yes | 6.406E-07 | 510.27573 | 71.988472 |
| AMENIGELK | 9 | Q7L014 | Probable ATP-dependent RNA helicase D | DDX46 | 117.363 | 2 | 53.564266 | yes | 6.321E-06 | 502.75769 | 53.795616 |
| IVDNVRPDR | 9 | Q7L014 | Probable ATP-dependent RNA helicase D | DDX46 | 117.363 | 2 | 53.564266 | yes | 1.897E-07 | 542.29889 | 52.179546 |
| SVVCSDVEQQVIV | 19 | Q7L014 | Probable ATP-dependent RNA helicase D | DDX46 | 117.363 | 3 | 53.564266 | yes | 3.383E-07 | 740.04724 | 49.59412 |
| AMENIGELKK | 10 | Q7L014 | Probable ATP-dependent RNA helicase D | DDX46 | 117.363 | 2 | 53.564266 | yes | 2.877E-07 | 566.80518 | 39.65654 |
| KLLEPVDHGK | 10 | Q7L014 | Probable ATP-dependent RNA helicase D | DDX46 | 117.363 | 2 | 53.564266 | yes | 3.3E-06 | 568.32715 | 36.606682 |
| IEYEPFR | 7 | Q7L014 | Probable ATP-dependent RNA helicase D | DDX46 | 117.363 | 2 | 53.564266 | yes | 9.918E-05 | 477.23999 | 30.922028 |
| IPAVTYPK | 8 | Q7L0Y3 | tRNA methyltransferase 10 homolog C O | TRMT10C | 47.347 | 2 | 54.322487 | yes | 1.851E-07 | 444.76309 | 942.75684 |
| VYVIGSFVDK | 10 | Q7L0Y3 | tRNA methyltransferase 10 homolog C O | TRMT10C | 47.347 | 2 | 54.322487 | yes | 3.54E-07 | 563.81079 | 788.84192 |
| SMQPGTSLAK | 10 | Q7L0Y3 | tRNA methyltransferase 10 homolog C O | TRMT10C | 47.347 | 2 | 54.322487 | yes | 1.377E-07 | 510.26315 | 602.16278 |

| | | | | | | | | | | | |
|---|---|---|---|---|---|---|---|---|---|---|---|
| LNLATECLPLDK | 12 | Q7L0Y3 | tRNA methyltransferase 10 homolog C O | TRMT10C | 47.347 | 2 | 54.322487 | yes | 4.934E-07 | 693.86853 | 489.22186 |
| SSVQEECVSTISSS | 15 | Q7L0Y3 | tRNA methyltransferase 10 homolog C O | TRMT10C | 47.347 | 2 | 54.322487 | yes | 9.722E-08 | 814.37762 | 426.14398 |
| NLTLDQMIR | 9 | Q7L0Y3 | tRNA methyltransferase 10 homolog C O | TRMT10C | 47.347 | 2 | 54.322487 | yes | 9.722E-08 | 552.29755 | 361.54953 |
| LLLTSTEK | 8 | Q7L0Y3 | tRNA methyltransferase 10 homolog C O | TRMT10C | 47.347 | 2 | 54.322487 | yes | 9.722E-08 | 452.77112 | 346.80194 |
| NESTPPSEELELDK | 14 | Q7L0Y3 | tRNA methyltransferase 10 homolog C O | TRMT10C | 47.347 | 2 | 54.322487 | yes | 9.722E-08 | 794.37286 | 330.60736 |
| EFIEMWR | 7 | Q7L0Y3 | tRNA methyltransferase 10 homolog C O | TRMT10C | 47.347 | 2 | 54.322487 | yes | 1.631E-07 | 505.74185 | 328.36047 |
| TLMECVSNTAK | 11 | Q7L0Y3 | tRNA methyltransferase 10 homolog C O | TRMT10C | 47.347 | 2 | 54.322487 | yes | 9.722E-08 | 627.29688 | 317.00977 |
| LLETTEEDK | 9 | Q7L0Y3 | tRNA methyltransferase 10 homolog C O | TRMT10C | 47.347 | 2 | 54.322487 | yes | 2.017E-07 | 539.26917 | 291.28711 |
| YLQWEIGNK | 9 | Q7L0Y3 | tRNA methyltransferase 10 homolog C O | TRMT10C | 47.347 | 2 | 54.322487 | yes | 1.301E-06 | 575.79822 | 291.21671 |
| SHVDLFPK | 8 | Q7L0Y3 | tRNA methyltransferase 10 homolog C O | TRMT10C | 47.347 | 2 | 54.322487 | yes | 6.563E-07 | 471.7558 | 249.63065 |
| LLETTEEDKQK | 11 | Q7L0Y3 | tRNA methyltransferase 10 homolog C O | TRMT10C | 47.347 | 2 | 54.322487 | yes | 9.722E-08 | 667.34589 | 177.90225 |
| NVDPFHIYFCNLK | 13 | Q7L0Y3 | tRNA methyltransferase 10 homolog C O | TRMT10C | 47.347 | 2 | 54.322487 | yes | 1.154E-07 | 833.90594 | 79.317039 |
| NVDPFHIYFCNLK | 13 | Q7L0Y3 | tRNA methyltransferase 10 homolog C O | TRMT10C | 47.347 | 2 | 54.322487 | yes | 1.047E-07 | 556.27307 | 79.317039 |
| NVDPFHIYFCNLK | 13 | Q7L0Y3 | tRNA methyltransferase 10 homolog C O | TRMT10C | 47.347 | 3 | 54.322487 | yes | 1.154E-07 | 833.90594 | 79.317039 |
| NVDPFHIYFCNLK | 13 | Q7L0Y3 | tRNA methyltransferase 10 homolog C O | TRMT10C | 47.347 | 3 | 54.322487 | yes | 1.047E-07 | 556.27307 | 79.317039 |
| DEDPLAATR | 9 | Q7L0Y3 | tRNA methyltransferase 10 homolog C O | TRMT10C | 47.347 | 2 | 54.322487 | yes | 6.113E-07 | 494.24072 | 133.36754 |
| NNGNWQEALQFV | 14 | Q7L0Y3 | tRNA methyltransferase 10 homolog C O | TRMT10C | 47.347 | 2 | 54.322487 | yes | 9.722E-08 | 822.9101 | 63.968746 |
| AEVLSEEPILK | 11 | Q7L1Q6 | eIF5-mimic protein 2 OS=Homo sapiens ( | BZW1 | 48.044 | 2 | 50.516415 | yes | 9.79E-08 | 614.34521 | 155.06883 |
| DINAVAASLR | 10 | Q7L1Q6 | eIF5-mimic protein 2 OS=Homo sapiens ( | BZW1 | 48.044 | 2 | 50.516415 | yes | 5.676E-07 | 515.28802 | 123.33503 |
| EGVSAAFAVK | 10 | Q7L1Q6 | eIF5-mimic protein 2 OS=Homo sapiens ( | BZW1 | 48.044 | 2 | 50.516415 | yes | 2.429E-07 | 489.76639 | 83.486023 |
| LMELFPANK | 9 | Q7L1Q6 | eIF5-mimic protein 2 OS=Homo sapiens ( | BZW1 | 48.044 | 2 | 50.516415 | yes | 1.184E-07 | 531.78625 | 78.996948 |
| FGVFLSEVSENK | 12 | Q7L1W4 | Volume-regulated anion channel subunit | LRRC8D | 98.202 | 2 | 50.34878 | yes | 1.373E-07 | 678.3457 | 97.209625 |
| VGQLSQLTQLELK | 13 | Q7L1W4 | Volume-regulated anion channel subunit | LRRC8D | 98.202 | 2 | 50.34878 | yes | 9.818E-08 | 728.92212 | 81.588341 |
| ELYLIGNLNSENNI | 14 | Q7L1W4 | Volume-regulated anion channel subunit | LRRC8D | 98.202 | 2 | 50.34878 | yes | 2.253E-07 | 810.91504 | 66.003471 |
| MIGLESLR | 8 | Q7L1W4 | Volume-regulated anion channel subunit | LRRC8D | 98.202 | 2 | 50.34878 | yes | 3.976E-07 | 459.75751 | 65.013893 |
| TIEEIISFQHLK | 12 | Q7L1W4 | Volume-regulated anion channel subunit | LRRC8D | 98.202 | 2 | 50.34878 | yes | 1.776E-07 | 729.40356 | 55.953995 |
| LESLPVAVFSLQK | 13 | Q7L1W4 | Volume-regulated anion channel subunit | LRRC8D | 98.202 | 2 | 50.34878 | yes | 5.781E-07 | 715.91632 | 34.087967 |
| ALSETACEDSEENI | 14 | Q7L1W4 | Volume-regulated anion channel subunit | LRRC8D | 98.202 | 2 | 50.34878 | yes | 1.078E-07 | 791.83051 | 11.451086 |
| LQSDDILPLGK | 11 | Q7L2E3 | ATP-dependent RNA helicase DHX30 OS | DHX30 | 133.939 | 2 | 54.992085 | yes | 3.855E-07 | 599.83752 | 13391.272 |
| AIFQQPPVGVR | 11 | Q7L2E3 | ATP-dependent RNA helicase DHX30 OS | DHX30 | 133.939 | 2 | 54.992085 | yes | 9.728E-08 | 606.34839 | 12815.488 |
| IPQLLLER | 8 | Q7L2E3 | ATP-dependent RNA helicase DHX30 OS | DHX30 | 133.939 | 2 | 54.992085 | yes | 2.729E-07 | 491.3082 | 11084.514 |
| ATISLSDSDLLR | 12 | Q7L2E3 | ATP-dependent RNA helicase DHX30 OS | DHX30 | 133.939 | 2 | 54.992085 | yes | 4.544E-07 | 645.84863 | 9873.6631 |
| GSSFEMTDDDSAI | 14 | Q7L2E3 | ATP-dependent RNA helicase DHX30 OS | DHX30 | 133.939 | 2 | 54.992085 | yes | 2.417E-07 | 765.82245 | 9661.0537 |
| EYLTTLGQR | 9 | Q7L2E3 | ATP-dependent RNA helicase DHX30 OS | DHX30 | 133.939 | 2 | 54.992085 | yes | 1.427E-07 | 540.78784 | 9329.4375 |
| LVLMSATGDNER | 12 | Q7L2E3 | ATP-dependent RNA helicase DHX30 OS | DHX30 | 133.939 | 2 | 54.992085 | yes | 1.532E-07 | 653.32703 | 4462.8706 |
| LVLMSATGDNER | 12 | Q7L2E3 | ATP-dependent RNA helicase DHX30 OS | DHX30 | 133.939 | 2 | 54.992085 | yes | 1.701E-06 | 661.32446 | 4462.8706 |
| LQEALGMHESK | 11 | Q7L2E3 | ATP-dependent RNA helicase DHX30 OS | DHX30 | 133.939 | 2 | 54.992085 | yes | 1.165E-07 | 621.81097 | 3987.1226 |
| LQEALGMHESK | 11 | Q7L2E3 | ATP-dependent RNA helicase DHX30 OS | DHX30 | 133.939 | 2 | 54.992085 | yes | 1.104E-07 | 629.80847 | 3987.1226 |
| YFGGCPVIK | 9 | Q7L2E3 | ATP-dependent RNA helicase DHX30 OS | DHX30 | 133.939 | 2 | 54.992085 | yes | 1.251E-07 | 520.76532 | 7133.9551 |

| | | | | | | | | | | | |
|---|---|---|---|---|---|---|---|---|---|---|---|
| CNVIITQPR | 9 | Q7L2E3 | ATP-dependent RNA helicase DHX30 O: | DHX30 | 133.939 | 2 | 54.992085 | yes | 1.002E-07 | 550.79767 | 6980.6123 |
| LSQSLLELWR | 10 | Q7L2E3 | ATP-dependent RNA helicase DHX30 O: | DHX30 | 133.939 | 2 | 54.992085 | yes | 9.722E-08 | 622.85352 | 6907.2305 |
| TPLENLVLQAK | 11 | Q7L2E3 | ATP-dependent RNA helicase DHX30 O: | DHX30 | 133.939 | 2 | 54.992085 | yes | 2.624E-06 | 613.36115 | 6005.3291 |
| GPCGSFDVR | 9 | Q7L2E3 | ATP-dependent RNA helicase DHX30 O: | DHX30 | 133.939 | 2 | 54.992085 | yes | 1.31E-07 | 497.72418 | 5707.1465 |
| VIQIATSSSTAK | 12 | Q7L2E3 | ATP-dependent RNA helicase DHX30 O: | DHX30 | 133.939 | 2 | 54.992085 | yes | 9.917E-08 | 603.34045 | 5183.8857 |
| EHYLEDILAK | 10 | Q7L2E3 | ATP-dependent RNA helicase DHX30 O: | DHX30 | 133.939 | 2 | 54.992085 | yes | 9.722E-08 | 615.8219 | 4802.7056 |
| VSHELGPSLR | 10 | Q7L2E3 | ATP-dependent RNA helicase DHX30 O: | DHX30 | 133.939 | 2 | 54.992085 | yes | 1.222E-07 | 547.80127 | 4760.5244 |
| DPFSSSLQNR | 10 | Q7L2E3 | ATP-dependent RNA helicase DHX30 O: | DHX30 | 133.939 | 2 | 54.992085 | yes | 1.531E-07 | 575.77802 | 4581.0156 |
| VSCLETVWVSR | 11 | Q7L2E3 | ATP-dependent RNA helicase DHX30 O: | DHX30 | 133.939 | 2 | 54.992085 | yes | 9.722E-08 | 668.3399 | 4516.9707 |
| ALLSHDSGSDHLA | 16 | Q7L2E3 | ATP-dependent RNA helicase DHX30 O: | DHX30 | 133.939 | 3 | 54.992085 | yes | 9.722E-08 | 575.62866 | 2246.845 |
| ALLSHDSGSDHLA | 16 | Q7L2E3 | ATP-dependent RNA helicase DHX30 O: | DHX30 | 133.939 | 3 | 54.992085 | yes | 1.156E-07 | 431.97333 | 2246.845 |
| ALLSHDSGSDHLA | 16 | Q7L2E3 | ATP-dependent RNA helicase DHX30 O: | DHX30 | 133.939 | 4 | 54.992085 | yes | 9.722E-08 | 575.62866 | 2246.845 |
| ALLSHDSGSDHLA | 16 | Q7L2E3 | ATP-dependent RNA helicase DHX30 O: | DHX30 | 133.939 | 4 | 54.992085 | yes | 1.156E-07 | 431.97333 | 2246.845 |
| NLLNSVIGR | 9 | Q7L2E3 | ATP-dependent RNA helicase DHX30 O: | DHX30 | 133.939 | 2 | 54.992085 | yes | 1.822E-07 | 493.29309 | 4146.5454 |
| ISAVSVAQR | 9 | Q7L2E3 | ATP-dependent RNA helicase DHX30 O: | DHX30 | 133.939 | 2 | 54.992085 | yes | 1.873E-07 | 465.772 | 4121.4927 |
| CQSGFAYHLFPR | 12 | Q7L2E3 | ATP-dependent RNA helicase DHX30 O: | DHX30 | 133.939 | 2 | 54.992085 | yes | 1.064E-07 | 741.85095 | 2043.4971 |
| CQSGFAYHLFPR | 12 | Q7L2E3 | ATP-dependent RNA helicase DHX30 O: | DHX30 | 133.939 | 2 | 54.992085 | yes | 2.913E-07 | 494.90308 | 2043.4971 |
| CQSGFAYHLFPR | 12 | Q7L2E3 | ATP-dependent RNA helicase DHX30 O: | DHX30 | 133.939 | 3 | 54.992085 | yes | 1.064E-07 | 741.85095 | 2043.4971 |
| CQSGFAYHLFPR | 12 | Q7L2E3 | ATP-dependent RNA helicase DHX30 O: | DHX30 | 133.939 | 3 | 54.992085 | yes | 2.913E-07 | 494.90308 | 2043.4971 |
| NLMQFHTVGTK | 11 | Q7L2E3 | ATP-dependent RNA helicase DHX30 O: | DHX30 | 133.939 | 2 | 54.992085 | yes | 1.702E-07 | 638.32935 | 1746.1912 |
| NLMQFHTVGTK | 11 | Q7L2E3 | ATP-dependent RNA helicase DHX30 O: | DHX30 | 133.939 | 2 | 54.992085 | yes | 2.15E-07 | 646.32678 | 1746.1912 |
| DSGPLSDPITGK | 12 | Q7L2E3 | ATP-dependent RNA helicase DHX30 O: | DHX30 | 133.939 | 2 | 54.992085 | yes | 1.112E-07 | 593.80115 | 3453.9766 |
| AVAGWEEVLR | 10 | Q7L2E3 | ATP-dependent RNA helicase DHX30 O: | DHX30 | 133.939 | 2 | 54.992085 | yes | 1.01E-07 | 565.30365 | 3225.4167 |
| QAAAAACQLFK | 11 | Q7L2E3 | ATP-dependent RNA helicase DHX30 O: | DHX30 | 133.939 | 2 | 54.992085 | yes | 9.819E-08 | 589.80298 | 3068.2712 |
| LQSNPSLEGVSHV | 21 | Q7L2E3 | ATP-dependent RNA helicase DHX30 O: | DHX30 | 133.939 | 3 | 54.992085 | yes | 1.029E-06 | 782.06793 | 3034.6599 |
| DSSQVHPLAVLLL | 21 | Q7L2E3 | ATP-dependent RNA helicase DHX30 O: | DHX30 | 133.939 | 3 | 54.992085 | yes | 9.722E-08 | 762.41315 | 1493.377 |
| DSSQVHPLAVLLL | 21 | Q7L2E3 | ATP-dependent RNA helicase DHX30 O: | DHX30 | 133.939 | 3 | 54.992085 | yes | 1.576E-07 | 572.06165 | 1493.377 |
| DSSQVHPLAVLLL | 21 | Q7L2E3 | ATP-dependent RNA helicase DHX30 O: | DHX30 | 133.939 | 4 | 54.992085 | yes | 9.722E-08 | 762.41315 | 1493.377 |
| DSSQVHPLAVLLL | 21 | Q7L2E3 | ATP-dependent RNA helicase DHX30 O: | DHX30 | 133.939 | 4 | 54.992085 | yes | 1.576E-07 | 572.06165 | 1493.377 |
| SGNILLHK | 8 | Q7L2E3 | ATP-dependent RNA helicase DHX30 O: | DHX30 | 133.939 | 2 | 54.992085 | yes | 4.587E-07 | 441.26379 | 2924.8301 |
| ALTQFPLPK | 9 | Q7L2E3 | ATP-dependent RNA helicase DHX30 O: | DHX30 | 133.939 | 2 | 54.992085 | yes | 1.034E-06 | 507.80276 | 2856.5037 |
| LAHISTDPR | 9 | Q7L2E3 | ATP-dependent RNA helicase DHX30 O: | DHX30 | 133.939 | 2 | 54.992085 | yes | 2.771E-06 | 505.2749 | 2853.3511 |
| RPCTIQVPEPILR | 13 | Q7L2E3 | ATP-dependent RNA helicase DHX30 O: | DHX30 | 133.939 | 3 | 54.992085 | yes | 1.361E-07 | 526.96436 | 1312.5507 |
| RPCTIQVPEPILR | 13 | Q7L2E3 | ATP-dependent RNA helicase DHX30 O: | DHX30 | 133.939 | 3 | 54.992085 | yes | 1.11E-07 | 789.94287 | 1312.5507 |
| RPCTIQVPEPILR | 13 | Q7L2E3 | ATP-dependent RNA helicase DHX30 O: | DHX30 | 133.939 | 2 | 54.992085 | yes | 1.361E-07 | 526.96436 | 1312.5507 |
| RPCTIQVPEPILR | 13 | Q7L2E3 | ATP-dependent RNA helicase DHX30 O: | DHX30 | 133.939 | 2 | 54.992085 | yes | 1.11E-07 | 789.94287 | 1312.5507 |
| VPGFMYPVK | 9 | Q7L2E3 | ATP-dependent RNA helicase DHX30 O: | DHX30 | 133.939 | 2 | 54.992085 | yes | 3.265E-07 | 519.27808 | 2527.0393 |
| SVEVEGYGSK | 10 | Q7L2E3 | ATP-dependent RNA helicase DHX30 O: | DHX30 | 133.939 | 2 | 54.992085 | yes | 2.02E-07 | 527.75623 | 2373.0981 |
| FKPNSVTYR | 9 | Q7L2E3 | ATP-dependent RNA helicase DHX30 O: | DHX30 | 133.939 | 2 | 54.992085 | yes | 1.696E-07 | 556.2984 | 2166.092 |

| | | | | | | | | | | | |
|---|---|---|---|---|---|---|---|---|---|---|---|
| FIHGLIK | 7 | Q7L2E3 | ATP-dependent RNA helicase DHX30 O | DHX30 | 133.939 | 2 | 54.992085 | yes | 1.8E-07 | 414.26053 | 1642.2971 |
| GVLMAGLYPNLIQ | 15 | Q7L2E3 | ATP-dependent RNA helicase DHX30 O | DHX30 | 133.939 | 2 | 54.992085 | yes | 9.722E-08 | 822.46869 | 1564.4326 |
| LVYVHTNGPK | 10 | Q7L2E3 | ATP-dependent RNA helicase DHX30 O | DHX30 | 133.939 | 2 | 54.992085 | yes | 9.722E-08 | 564.31403 | 1430.2192 |
| LESKPPSR | 8 | Q7L2E3 | ATP-dependent RNA helicase DHX30 O | DHX30 | 133.939 | 2 | 54.992085 | yes | 9.722E-08 | 457.25873 | 1376.652 |
| KIETFLNHYPVESS | 21 | Q7L2E3 | ATP-dependent RNA helicase DHX30 O | DHX30 | 133.939 | 4 | 54.992085 | yes | 1.014E-07 | 633.08386 | 1300.1797 |
| QLNPESIR | 8 | Q7L2E3 | ATP-dependent RNA helicase DHX30 O | DHX30 | 133.939 | 2 | 54.992085 | yes | 2.297E-07 | 478.76163 | 963.5307 |
| AIVLAAIFR | 9 | Q7L2E3 | ATP-dependent RNA helicase DHX30 O | DHX30 | 133.939 | 2 | 54.992085 | yes | 9.722E-08 | 487.31329 | 960.79974 |
| QFSENIYEAFLVGI | 14 | Q7L2E3 | ATP-dependent RNA helicase DHX30 O | DHX30 | 133.939 | 2 | 54.992085 | yes | 5.894E-07 | 822.91705 | 764.9024 |
| NELFDAAK | 8 | Q7L2E3 | ATP-dependent RNA helicase DHX30 O | DHX30 | 133.939 | 2 | 54.992085 | yes | 1.836E-07 | 454.22961 | 749.57275 |
| PYVPLLEAEEVR | 12 | Q7L2E3 | ATP-dependent RNA helicase DHX30 O | DHX30 | 133.939 | 2 | 54.992085 | yes | 1.198E-06 | 707.88245 | 707.21759 |
| DKLVYVHTNGPK | 12 | Q7L2E3 | ATP-dependent RNA helicase DHX30 O | DHX30 | 133.939 | 2 | 54.992085 | yes | 9.722E-08 | 685.875 | 184.28217 |
| DKLVYVHTNGPK | 12 | Q7L2E3 | ATP-dependent RNA helicase DHX30 O | DHX30 | 133.939 | 2 | 54.992085 | yes | 0.0002964 | 457.58575 | 184.28217 |
| DKLVYVHTNGPK | 12 | Q7L2E3 | ATP-dependent RNA helicase DHX30 O | DHX30 | 133.939 | 3 | 54.992085 | yes | 9.722E-08 | 685.875 | 184.28217 |
| DKLVYVHTNGPK | 12 | Q7L2E3 | ATP-dependent RNA helicase DHX30 O | DHX30 | 133.939 | 3 | 54.992085 | yes | 0.0002964 | 457.58575 | 184.28217 |
| DTILNAIEQHPVVV | 23 | Q7L2E3 | ATP-dependent RNA helicase DHX30 O | DHX30 | 133.939 | 3 | 54.992085 | yes | 1.483E-07 | 808.41266 | 357.74661 |
| CLHPLLVVVSCLT | 14 | Q7L2E3 | ATP-dependent RNA helicase DHX30 O | DHX30 | 133.939 | 2 | 54.992085 | yes | 1.829E-07 | 833.96021 | 169.41652 |
| CLHPLLVVVSCLT | 14 | Q7L2E3 | ATP-dependent RNA helicase DHX30 O | DHX30 | 133.939 | 2 | 54.992085 | yes | 1.38E-07 | 556.3092 | 169.41652 |
| CLHPLLVVVSCLT | 14 | Q7L2E3 | ATP-dependent RNA helicase DHX30 O | DHX30 | 133.939 | 3 | 54.992085 | yes | 1.829E-07 | 833.96021 | 169.41652 |
| CLHPLLVVVSCLT | 14 | Q7L2E3 | ATP-dependent RNA helicase DHX30 O | DHX30 | 133.939 | 3 | 54.992085 | yes | 1.38E-07 | 556.3092 | 169.41652 |
| KVTLHIK | 7 | Q7L2E3 | ATP-dependent RNA helicase DHX30 O | DHX30 | 133.939 | 2 | 54.992085 | yes | 2.177E-07 | 419.77908 | 319.1994 |
| KLQSNPSLEGVSH | 22 | Q7L2E3 | ATP-dependent RNA helicase DHX30 O | DHX30 | 133.939 | 3 | 54.992085 | yes | 1.373E-07 | 824.76624 | 317.45578 |
| LPPMCVNPTPGGT | 16 | Q7L2E3 | ATP-dependent RNA helicase DHX30 O | DHX30 | 133.939 | 2 | 54.992085 | yes | 6.745E-07 | 848.92908 | 296.78973 |
| IETFLNHYPVESSV | 20 | Q7L2E3 | ATP-dependent RNA helicase DHX30 O | DHX30 | 133.939 | 3 | 54.992085 | yes | 9.722E-08 | 801.07776 | 276.75586 |
| ELGETQR | 7 | Q7L2E3 | ATP-dependent RNA helicase DHX30 O | DHX30 | 133.939 | 2 | 54.992085 | yes | 9.722E-08 | 416.71161 | 186.09523 |
| SNGSVFVR | 8 | Q7L2E3 | ATP-dependent RNA helicase DHX30 O | DHX30 | 133.939 | 2 | 54.992085 | yes | 1.122E-07 | 433.22995 | 110.10333 |
| RISAVSVAQR | 10 | Q7L2E3 | ATP-dependent RNA helicase DHX30 O | DHX30 | 133.939 | 2 | 54.992085 | yes | 3.334E-07 | 543.82257 | 82.53598 |
| RNVGFQVR | 8 | Q7L2E3 | ATP-dependent RNA helicase DHX30 O | DHX30 | 133.939 | 2 | 54.992085 | yes | 4.989E-07 | 488.27777 | 78.830124 |
| AVDSPNIK | 8 | Q7L2E3 | ATP-dependent RNA helicase DHX30 O | DHX30 | 133.939 | 2 | 54.992085 | yes | 1.772E-07 | 422.23218 | 53.884159 |
| MVPFQVPEILR | 11 | Q7L2E3 | ATP-dependent RNA helicase DHX30 O | DHX30 | 133.939 | 2 | 54.992085 | yes | 2.195E-07 | 664.87341 | 52.047981 |
| WLTYFMAVK | 9 | Q7L2E3 | ATP-dependent RNA helicase DHX30 O | DHX30 | 133.939 | 2 | 54.992085 | yes | 7.336E-07 | 587.80188 | 15.507224 |
| WISDWNLTTEK | 11 | Q7L2H7 | Eukaryotic translation initiation factor 3 s | EIF3M | 42.503 | 2 | 52.038918 | yes | 1.565E-07 | 696.84332 | 2609.1758 |
| LLYEALVDCK | 10 | Q7L2H7 | Eukaryotic translation initiation factor 3 s | EIF3M | 42.503 | 2 | 52.038918 | yes | 9.722E-08 | 612.32068 | 2561.0032 |
| LLTFMGMAVENK | 12 | Q7L2H7 | Eukaryotic translation initiation factor 3 s | EIF3M | 42.503 | 2 | 52.038918 | yes | 1.019E-07 | 677.34888 | 1181.5408 |
| YTVYCSLIK | 9 | Q7L2H7 | Eukaryotic translation initiation factor 3 s | EIF3M | 42.503 | 2 | 52.038918 | yes | 1.665E-07 | 573.79681 | 1093.6892 |
| EGERPSLR | 8 | Q7L2H7 | Eukaryotic translation initiation factor 3 s | EIF3M | 42.503 | 2 | 52.038918 | yes | 1.516E-07 | 472.25143 | 400.85849 |
| QEALIESLCEK | 11 | Q7L2H7 | Eukaryotic translation initiation factor 3 s | EIF3M | 42.503 | 2 | 52.038918 | yes | 4.322E-07 | 660.32922 | 194.17751 |
| ALKDPNAFLFDHL | 20 | Q7L2H7 | Eukaryotic translation initiation factor 3 s | EIF3M | 42.503 | 3 | 52.038918 | yes | 1.692E-07 | 760.77533 | 126.60651 |
| VAASCGAIQYIPTE | 19 | Q7L2H7 | Eukaryotic translation initiation factor 3 s | EIF3M | 42.503 | 3 | 52.038918 | yes | 9.722E-08 | 697.68951 | 103.27708 |
| KWISDWNLTTEK | 12 | Q7L2H7 | Eukaryotic translation initiation factor 3 s | EIF3M | 42.503 | 2 | 52.038918 | yes | 2.159E-07 | 760.89081 | 72.218651 |

| | | | | | | | | | | | |
|---|---|---|---|---|---|---|---|---|---|---|---|
| DFIDSLGLLHEQNI | 16 | Q7L2H7 | Eukaryotic translation initiation factor 3 s | EIF3M | 42.503 | 3 | 52.038918 | yes | 4.362E-07 | 610.97333 | 22.645987 |
| MVGLDIDSR | 9 | Q7L2J0 | 7SK snRNA methylphosphate capping en | MEPCE | 74.356 | 2 | 55.153065 | yes | 9.722E-08 | 503.25531 | 565.68677 |
| VLKPEWFR | 8 | Q7L2J0 | 7SK snRNA methylphosphate capping en | MEPCE | 74.356 | 2 | 55.153065 | yes | 3.68E-07 | 537.80835 | 536.81244 |
| AAPPDVGEER | 10 | Q7L2J0 | 7SK snRNA methylphosphate capping en | MEPCE | 74.356 | 2 | 55.153065 | yes | 3.026E-07 | 520.75397 | 406.18024 |
| HYLSEELR | 8 | Q7L2J0 | 7SK snRNA methylphosphate capping en | MEPCE | 74.356 | 2 | 55.153065 | yes | 1.893E-07 | 523.76691 | 348.20151 |
| GRDPVEILIPK | 11 | Q7L2J0 | 7SK snRNA methylphosphate capping en | MEPCE | 74.356 | 2 | 55.153065 | yes | 1.003E-07 | 618.86914 | 126.3423 |
| EPFLVPAPPPPLK | 13 | Q7L2J0 | 7SK snRNA methylphosphate capping en | MEPCE | 74.356 | 2 | 55.153065 | yes | 3.394E-06 | 701.41064 | 104.22978 |
| SCFPASLTASR | 11 | Q7L2J0 | 7SK snRNA methylphosphate capping en | MEPCE | 74.356 | 2 | 55.153065 | yes | 4.009E-06 | 598.79004 | 93.291725 |
| CAPSAGSPAAAVC | 14 | Q7L2J0 | 7SK snRNA methylphosphate capping en | MEPCE | 74.356 | 2 | 55.153065 | yes | 9.722E-08 | 636.31171 | 63.821548 |
| SMSTFIEEAYK | 11 | Q7L2Z9 | Centromere protein Q OS=Homo sapiens | CENPQ | 30.596 | 2 | 34.094627 | yes | 3.285E-06 | 653.30518 | 42.415775 |
| IQILASEVEEEEER | 14 | Q7L2Z9 | Centromere protein Q OS=Homo sapiens | CENPQ | 30.596 | 2 | 34.094627 | yes | 1.08E-06 | 837.41504 | 33.310841 |
| GMGLGFTSSMR | 11 | Q7L4I2 | Arginine/serine-rich coiled-coil protein 2 | RSRC2 | 50.56 | 2 | 51.34421 | yes | 9.722E-08 | 572.26788 | 473.21069 |
| NTAMDAQEALAR | 12 | Q7L4I2 | Arginine/serine-rich coiled-coil protein 2 | RSRC2 | 50.56 | 2 | 51.34421 | yes | 9.722E-08 | 645.80896 | 400.30093 |
| QQEEVFR | 7 | Q7L4I2 | Arginine/serine-rich coiled-coil protein 2 | RSRC2 | 50.56 | 2 | 51.34421 | yes | 8.929E-07 | 468.2327 | 231.85265 |
| NLDAQYEMAR | 10 | Q7L4I2 | Arginine/serine-rich coiled-coil protein 2 | RSRC2 | 50.56 | 2 | 51.34421 | yes | 1.914E-07 | 605.77966 | 125.46198 |
| SQSAEIWEK | 9 | Q7L4I2 | Arginine/serine-rich coiled-coil protein 2 | RSRC2 | 50.56 | 2 | 51.34421 | yes | 2.488E-06 | 539.26422 | 106.12646 |
| EQSEVSVSPR | 10 | Q7L4I2 | Arginine/serine-rich coiled-coil protein 2 | RSRC2 | 50.56 | 2 | 51.34421 | yes | 1.783E-07 | 559.27783 | 27.54072 |
| LGTPQQIAIAR | 11 | Q7L576 | Cytoplasmic FMR1-interacting protein 1 | CYFIP1 | 145.183 | 2 | 53.656467 | no | 1.104E-07 | 584.34583 | 574.77362 |
| TVEVLEPEVTK | 11 | Q7L576 | Cytoplasmic FMR1-interacting protein 1 | CYFIP1 | 145.183 | 2 | 53.656467 | no | 2.542E-07 | 622.34265 | 369.91418 |
| HVQLLGR | 7 | Q7L576 | Cytoplasmic FMR1-interacting protein 1 | CYFIP1 | 145.183 | 2 | 53.656467 | no | 1.003E-07 | 411.75085 | 316.37711 |
| AVGPSSTQLYMVI | 13 | Q7L576 | Cytoplasmic FMR1-interacting protein 1 | CYFIP1 | 145.183 | 2 | 53.656467 | yes | 1.31E-07 | 704.86627 | 308.97897 |
| FISELAR | 7 | Q7L576 | Cytoplasmic FMR1-interacting protein 1 | CYFIP1 | 145.183 | 2 | 53.656467 | no | 2.651E-07 | 418.23724 | 268.05707 |
| TVCFQNLR | 8 | Q7L576 | Cytoplasmic FMR1-interacting protein 1 | CYFIP1 | 145.183 | 2 | 53.656467 | yes | 3.938E-07 | 519.26349 | 258.05042 |
| TVLPFSQEFQR | 11 | Q7L576 | Cytoplasmic FMR1-interacting protein 1 | CYFIP1 | 145.183 | 2 | 53.656467 | yes | 1.035E-07 | 676.35388 | 247.0141 |
| SGDGEGTPVEHVR | 13 | Q7L576 | Cytoplasmic FMR1-interacting protein 1 | CYFIP1 | 145.183 | 2 | 53.656467 | yes | 9.722E-08 | 670.31549 | 207.58098 |
| SFLDDPIWR | 9 | Q7L576 | Cytoplasmic FMR1-interacting protein 1 | CYFIP1 | 145.183 | 2 | 53.656467 | yes | 1.094E-06 | 574.79041 | 203.83253 |
| ALNLAYSSIYGSYI | 14 | Q7L576 | Cytoplasmic FMR1-interacting protein 1 | CYFIP1 | 145.183 | 2 | 53.656467 | yes | 9.975E-08 | 789.40155 | 202.7146 |
| LADQIFAYYK | 10 | Q7L576 | Cytoplasmic FMR1-interacting protein 1 | CYFIP1 | 145.183 | 2 | 53.656467 | no | 1.154E-07 | 616.32153 | 200.05304 |
| SSLEGPTILDIEK | 13 | Q7L576 | Cytoplasmic FMR1-interacting protein 1 | CYFIP1 | 145.183 | 2 | 53.656467 | yes | 1.825E-07 | 701.3772 | 183.27431 |
| VMAGSLLLDK | 10 | Q7L576 | Cytoplasmic FMR1-interacting protein 1 | CYFIP1 | 145.183 | 2 | 53.656467 | yes | 2.828E-07 | 523.79938 | 174.69443 |
| LMNFMYFQR | 9 | Q7L576 | Cytoplasmic FMR1-interacting protein 1 | CYFIP1 | 145.183 | 2 | 53.656467 | yes | 2.031E-07 | 625.29645 | 163.48184 |
| MESVFNHAIR | 10 | Q7L576 | Cytoplasmic FMR1-interacting protein 1 | CYFIP1 | 145.183 | 2 | 53.656467 | yes | 9.722E-08 | 602.3006 | 140.32904 |
| YAPLHLVPLIER | 12 | Q7L576 | Cytoplasmic FMR1-interacting protein 1 | CYFIP1 | 145.183 | 2 | 53.656467 | no | 1.26E-07 | 710.91919 | 116.59542 |
| CFQPPIHQSLASS | 13 | Q7L576 | Cytoplasmic FMR1-interacting protein 1 | CYFIP1 | 145.183 | 2 | 53.656467 | yes | 4.792E-07 | 736.35358 | 82.834618 |
| YSNSEVVTGSGR | 12 | Q7L576 | Cytoplasmic FMR1-interacting protein 1 | CYFIP1 | 145.183 | 2 | 53.656467 | yes | 5.19E-07 | 628.29932 | 69.701271 |
| HTVYAALQDFSQV | 16 | Q7L576 | Cytoplasmic FMR1-interacting protein 1 | CYFIP1 | 145.183 | 3 | 53.656467 | yes | 9.809E-08 | 616.99164 | 61.352627 |
| MYLTPSEK | 8 | Q7L576 | Cytoplasmic FMR1-interacting protein 1 | CYFIP1 | 145.183 | 2 | 53.656467 | no | 9.104E-07 | 484.74152 | 24.939939 |
| GTNLIIQETR | 10 | Q7L590 | Protein MCM10 homolog OS=Homo sap | MCM10 | 98.184 | 2 | 54.585815 | yes | 9.809E-08 | 572.81964 | 499.4989 |
| MTSAPSQPLQTISR | 14 | Q7L590 | Protein MCM10 homolog OS=Homo sap | MCM10 | 98.184 | 2 | 54.585815 | yes | 2.01E-07 | 758.89307 | 281.20941 |

| | | | | | | | | | | | |
|---|---|---|---|---|---|---|---|---|---|---|---|
| LEGAPATMTPK | 11 | Q7L590 | Protein MCM10 homolog OS=Homo sap | MCM10 | 98.184 | 2 | 54.585815 | yes | 1.334E-06 | 558.29187 | 210.24905 |
| VLIMGEALDLGTC | 14 | Q7L590 | Protein MCM10 homolog OS=Homo sap | MCM10 | 98.184 | 2 | 54.585815 | yes | 9.722E-08 | 760.39655 | 186.10719 |
| IQTTLSNLVVK | 11 | Q7L590 | Protein MCM10 homolog OS=Homo sap | MCM10 | 98.184 | 2 | 54.585815 | yes | 1.509E-07 | 608.36902 | 171.23195 |
| ELMDLPTCGAR | 11 | Q7L590 | Protein MCM10 homolog OS=Homo sap | MCM10 | 98.184 | 2 | 54.585815 | yes | 2.913E-06 | 631.797 | 129.08772 |
| ATASGIMGSPK | 11 | Q7L590 | Protein MCM10 homolog OS=Homo sap | MCM10 | 98.184 | 2 | 54.585815 | yes | 1.43E-07 | 510.26315 | 116.62836 |
| TNEELQEELR | 10 | Q7L590 | Protein MCM10 homolog OS=Homo sap | MCM10 | 98.184 | 2 | 54.585815 | yes | 1.42E-07 | 630.80695 | 97.335464 |
| TCAYTHFK | 8 | Q7L590 | Protein MCM10 homolog OS=Homo sap | MCM10 | 98.184 | 2 | 54.585815 | yes | 8.201E-07 | 514.23694 | 84.384705 |
| ADLQSTFSGGR | 11 | Q7L590 | Protein MCM10 homolog OS=Homo sap | MCM10 | 98.184 | 2 | 54.585815 | yes | 3E-07 | 569.77802 | 64.544739 |
| EQLAYLESEEFQK | 13 | Q7L590 | Protein MCM10 homolog OS=Homo sap | MCM10 | 98.184 | 2 | 54.585815 | yes | 9.722E-08 | 807.38831 | 53.697075 |
| EAEAEMQER | 9 | Q7L590 | Protein MCM10 homolog OS=Homo sap | MCM10 | 98.184 | 2 | 54.585815 | yes | 6.562E-07 | 546.73492 | 51.586918 |
| YFEPLVK | 7 | Q7L590 | Protein MCM10 homolog OS=Homo sap | MCM10 | 98.184 | 2 | 54.585815 | yes | 3.302E-07 | 448.24982 | 50.239212 |
| IGGETLLPR | 9 | Q7L590 | Protein MCM10 homolog OS=Homo sap | MCM10 | 98.184 | 2 | 54.585815 | yes | 1.277E-07 | 478.2822 | 23.389645 |
| NGEPCTQTVNLR | 12 | Q7L590 | Protein MCM10 homolog OS=Homo sap | MCM10 | 98.184 | 2 | 54.585815 | yes | 8.065E-06 | 694.83301 | 22.522291 |
| LSEIQALTLTK | 11 | Q7L592 | Protein arginine methyltransferase NDUF | NDUFAF7 | 49.238 | 2 | 36.36617 | yes | 7.073E-07 | 608.86102 | 38.651127 |
| MLIQQFR | 7 | Q7L5Y9 | E3 ubiquitin-protein transferase MAEA C | MAEA | 45.288 | 2 | 44.400642 | yes | 3.348E-06 | 468.26019 | 43.605656 |
| IQEFIELIR | 9 | Q7L5Y9 | E3 ubiquitin-protein transferase MAEA C | MAEA | 45.288 | 2 | 44.400642 | yes | 2.207E-07 | 580.83734 | 34.101978 |
| INTATTVADLK | 11 | Q7L622 | G2/M phase-specific E3 ubiquitin-protein | G2E3 | 80.505 | 2 | 49.649506 | yes | 1.823E-07 | 573.8219 | 98.514122 |
| DILGYHVIQR | 10 | Q7L622 | G2/M phase-specific E3 ubiquitin-protein | G2E3 | 80.505 | 2 | 49.649506 | yes | 1.014E-07 | 607.33801 | 80.278656 |
| GIIYNSGEFQK | 11 | Q7L622 | G2/M phase-specific E3 ubiquitin-protein | G2E3 | 80.505 | 2 | 49.649506 | yes | 7.76E-07 | 628.31952 | 45.230751 |
| AGNILLTEPGQVK | 13 | Q7L7X3 | Serine/threonine-protein kinase TAO1 OS | TAOK1 | 116.071 | 2 | 51.10542 | no | 1.736E-07 | 670.38263 | 172.22328 |
| NFVDSCLQK | 9 | Q7L7X3 | Serine/threonine-protein kinase TAO1 OS | TAOK1 | 116.071 | 2 | 51.10542 | no | 2.46E-07 | 555.76605 | 126.84174 |
| EIEAFDSESMR | 11 | Q7L7X3 | Serine/threonine-protein kinase TAO1 OS | TAOK1 | 116.071 | 2 | 51.10542 | yes | 2.085E-07 | 657.28754 | 93.27314 |
| DLEHAMLLR | 9 | Q7L7X3 | Serine/threonine-protein kinase TAO1 OS | TAOK1 | 116.071 | 2 | 51.10542 | yes | 1.161E-07 | 549.29224 | 92.819504 |
| LDEAQEAECQVLK | 13 | Q7L7X3 | Serine/threonine-protein kinase TAO1 OS | TAOK1 | 116.071 | 2 | 51.10542 | yes | 9.722E-08 | 766.8667 | 80.206726 |
| EHFATIR | 7 | Q7L7X3 | Serine/threonine-protein kinase TAO1 OS | TAOK1 | 116.071 | 2 | 51.10542 | yes | 1.716E-06 | 437.23251 | 71.289223 |
| ELNSFLESQK | 10 | Q7L7X3 | Serine/threonine-protein kinase TAO1 OS | TAOK1 | 116.071 | 2 | 51.10542 | yes | 4.084E-06 | 597.80371 | 64.056717 |
| IEEEMLALQNER | 12 | Q7L7X3 | Serine/threonine-protein kinase TAO1 OS | TAOK1 | 116.071 | 2 | 51.10542 | yes | 2.521E-07 | 737.86395 | 61.437508 |
| QYLELECR | 8 | Q7L7X3 | Serine/threonine-protein kinase TAO1 OS | TAOK1 | 116.071 | 2 | 51.10542 | yes | 1.101E-06 | 555.76605 | 58.652462 |
| VMEETPSILR | 10 | Q7L804 | Rab11 family-interacting protein 2 OS=H | RAB11FIP2 | 58.28 | 2 | 53.93772 | yes | 9.722E-08 | 587.81049 | 118.93327 |
| KPFLLGPQR | 9 | Q7L804 | Rab11 family-interacting protein 2 OS=H | RAB11FIP2 | 58.28 | 2 | 53.93772 | yes | 2.514E-07 | 528.32166 | 100.92371 |
| SGTNDTYTIIQLGK | 14 | Q7L804 | Rab11 family-interacting protein 2 OS=H | RAB11FIP2 | 58.28 | 2 | 53.93772 | yes | 1.147E-07 | 755.89099 | 46.475601 |
| QNDPFTNVTASLP | 15 | Q7L804 | Rab11 family-interacting protein 2 OS=H | RAB11FIP2 | 58.28 | 2 | 53.93772 | yes | 9.722E-08 | 830.42047 | 42.50547 |
| AAVPLGGFLCNVA | 15 | Q7L8L6 | FAST kinase domain-containing protein 5 | FASTKD5 | 86.575 | 2 | 49.09007 | yes | 1.069E-07 | 766.40051 | 128.44302 |
| LAVQFTNR | 8 | Q7L8L6 | FAST kinase domain-containing protein 5 | FASTKD5 | 86.575 | 2 | 49.09007 | yes | 1.539E-07 | 474.76672 | 118.09789 |
| SSTNLSEFVMR | 11 | Q7L8L6 | FAST kinase domain-containing protein 5 | FASTKD5 | 86.575 | 2 | 49.09007 | yes | 2.337E-07 | 635.80847 | 38.552917 |
| LAFLHEK | 7 | Q7L8L6 | FAST kinase domain-containing protein 5 | FASTKD5 | 86.575 | 2 | 49.09007 | yes | 5.575E-07 | 429.24762 | 32.987518 |
| LEHVGVSLTDDLM | 15 | Q7L8L6 | FAST kinase domain-containing protein 5 | FASTKD5 | 86.575 | 2 | 49.09007 | yes | 2.788E-06 | 835.92456 | 28.107031 |
| VFLQLRPEYR | 10 | Q7L8L6 | FAST kinase domain-containing protein 5 | FASTKD5 | 86.575 | 2 | 49.09007 | yes | 3.548E-07 | 660.87476 | 18.391747 |
| NQAVNFLR | 8 | Q7LBR1 | Charged multivesicular body protein 1b C | CHMP1B | 22.11 | 2 | 47.021904 | yes | 1.418E-07 | 481.26434 | 288.18726 |

| | | | | | | | | | | | |
|---|---|---|---|---|---|---|---|---|---|---|---|
| NNPVMSLQDQVR | 12 | Q7LGA3 | Heparan sulfate 2-O-sulfotransferase 1 OS | HS2ST1 | 41.882 | 2 | 52.197487 | yes | 9.79E-08 | 700.8512 | 568.80109 |
| LQQSDIWK | 8 | Q7LGA3 | Heparan sulfate 2-O-sulfotransferase 1 OS | HS2ST1 | 41.882 | 2 | 52.197487 | yes | 9.819E-08 | 509.27182 | 541.65051 |
| LVSYYYFLR | 9 | Q7LGA3 | Heparan sulfate 2-O-sulfotransferase 1 OS | HS2ST1 | 41.882 | 2 | 52.197487 | yes | 1.028E-07 | 612.3266 | 488.26096 |
| KKPIYINVIR | 10 | Q7LGA3 | Heparan sulfate 2-O-sulfotransferase 1 OS | HS2ST1 | 41.882 | 2 | 52.197487 | yes | 9.722E-08 | 622.39789 | 91.062218 |
| KKPIYINVIR | 10 | Q7LGA3 | Heparan sulfate 2-O-sulfotransferase 1 OS | HS2ST1 | 41.882 | 2 | 52.197487 | yes | 5.057E-07 | 415.2677 | 91.062218 |
| KKPIYINVIR | 10 | Q7LGA3 | Heparan sulfate 2-O-sulfotransferase 1 OS | HS2ST1 | 41.882 | 3 | 52.197487 | yes | 9.722E-08 | 622.39789 | 91.062218 |
| KKPIYINVIR | 10 | Q7LGA3 | Heparan sulfate 2-O-sulfotransferase 1 OS | HS2ST1 | 41.882 | 3 | 52.197487 | yes | 5.057E-07 | 415.2677 | 91.062218 |
| FGDDYRPGLR | 10 | Q7LGA3 | Heparan sulfate 2-O-sulfotransferase 1 OS | HS2ST1 | 41.882 | 2 | 52.197487 | yes | 9.722E-08 | 598.29639 | 143.76389 |
| KPIYINVIR | 9 | Q7LGA3 | Heparan sulfate 2-O-sulfotransferase 1 OS | HS2ST1 | 41.882 | 2 | 52.197487 | yes | 1.354E-07 | 558.3504 | 128.39003 |
| EMKPGFYHGHVS | 18 | Q7LGA3 | Heparan sulfate 2-O-sulfotransferase 1 OS | HS2ST1 | 41.882 | 3 | 52.197487 | yes | 4.198E-07 | 709.34363 | 57.569977 |
| SLCEDPVCTPFVK | 13 | Q7LGC8 | Carbohydrate sulfotransferase 3 OS=Hom | CHST3 | 54.707 | 2 | 47.59003 | yes | 1.298E-07 | 776.36273 | 70.815163 |
| LAQVVQAACGPA | 14 | Q7LGC8 | Carbohydrate sulfotransferase 3 OS=Hom | CHST3 | 54.707 | 2 | 47.59003 | yes | 4.365E-06 | 736.37903 | 65.252693 |
| MAVEDHLK | 8 | Q7RTP6 | [F-actin]-monooxygenase MICAL3 OS=H | MICAL3 | 224.295 | 2 | 42.751427 | yes | 3.97E-07 | 471.73929 | 71.32058 |
| LPALEGTLQPQK | 12 | Q7RTP6 | [F-actin]-monooxygenase MICAL3 OS=H | MICAL3 | 224.295 | 2 | 42.751427 | yes | 2.89E-07 | 647.87189 | 61.990788 |
| ELGISPIMTGK | 11 | Q7RTP6 | [F-actin]-monooxygenase MICAL3 OS=H | MICAL3 | 224.295 | 2 | 42.751427 | yes | 2.413E-06 | 573.31537 | 59.252655 |
| HHPDLIFCR | 9 | Q7RTV0 | PHD finger-like domain-containing prote | PHF5A | 12.406 | 2 | 49.786953 | yes | 1.128E-07 | 597.79547 | 976.25781 |
| TDLFYER | 7 | Q7RTV0 | PHD finger-like domain-containing prote | PHF5A | 12.406 | 2 | 49.786953 | yes | 1.003E-07 | 472.22961 | 183.85786 |
| KQAGVAIGR | 9 | Q7RTV0 | PHD finger-like domain-containing prote | PHF5A | 12.406 | 2 | 49.786953 | yes | 1.913E-06 | 450.27469 | 38.193333 |
| VIVDFAGSSK | 10 | Q7Z2E3 | Aprataxin OS=Homo sapiens OX=9606 C | APTX | 40.741 | 2 | 52.359985 | yes | 2.064E-07 | 511.77948 | 457.76157 |
| NPGLETHR | 8 | Q7Z2E3 | Aprataxin OS=Homo sapiens OX=9606 C | APTX | 40.741 | 2 | 52.359985 | yes | 2.565E-07 | 462.23831 | 398.57062 |
| EHLELLK | 7 | Q7Z2E3 | Aprataxin OS=Homo sapiens OX=9606 C | APTX | 40.741 | 2 | 52.359985 | yes | 2.123E-07 | 441.25818 | 356.81848 |
| LPHLEAVVIGR | 11 | Q7Z2E3 | Aprataxin OS=Homo sapiens OX=9606 C | APTX | 40.741 | 2 | 52.359985 | yes | 9.722E-08 | 602.36407 | 337.28577 |
| ESLGHWSQGLK | 11 | Q7Z2E3 | Aprataxin OS=Homo sapiens OX=9606 C | APTX | 40.741 | 2 | 52.359985 | yes | 1.34E-07 | 621.31732 | 320.30353 |
| QVGVNPTSIDSVV | 16 | Q7Z2E3 | Aprataxin OS=Homo sapiens OX=9606 C | APTX | 40.741 | 2 | 52.359985 | yes | 1.014E-07 | 806.94885 | 201.7524 |
| DEQVVVIK | 8 | Q7Z2E3 | Aprataxin OS=Homo sapiens OX=9606 C | APTX | 40.741 | 2 | 52.359985 | yes | 5.307E-06 | 465.26874 | 16.895935 |
| CHECQQLLPSIPQI | 15 | Q7Z2E3 | Aprataxin OS=Homo sapiens OX=9606 C | APTX | 40.741 | 3 | 52.359985 | yes | 2.938E-06 | 617.64655 | 16.593699 |
| FVVLETGGEAGITI | 14 | Q7Z2W4 | Zinc finger CCCH-type antiviral protein 1 | ZC3HAV1 | 101.432 | 2 | 51.536686 | yes | 1.338E-07 | 724.89081 | 10959.836 |
| ASLEDAPVDDLTR | 13 | Q7Z2W4 | Zinc finger CCCH-type antiviral protein 1 | ZC3HAV1 | 101.432 | 2 | 51.536686 | yes | 1.023E-07 | 701.34644 | 10168.819 |
| TVFSPTLPAAR | 11 | Q7Z2W4 | Zinc finger CCCH-type antiviral protein 1 | ZC3HAV1 | 101.432 | 2 | 51.536686 | yes | 1.136E-07 | 580.32715 | 7679.7026 |
| SSLGSLQTPEAVTT | 15 | Q7Z2W4 | Zinc finger CCCH-type antiviral protein 1 | ZC3HAV1 | 101.432 | 2 | 51.536686 | yes | 2.941E-07 | 773.90723 | 7059.3765 |
| VMSCDSFPIR | 10 | Q7Z2W4 | Zinc finger CCCH-type antiviral protein 1 | ZC3HAV1 | 101.432 | 2 | 51.536686 | yes | 1.002E-07 | 606.28101 | 2949.6323 |
| VMSCDSFPIR | 10 | Q7Z2W4 | Zinc finger CCCH-type antiviral protein 1 | ZC3HAV1 | 101.432 | 2 | 51.536686 | yes | 2.438E-07 | 614.27844 | 2949.6323 |
| ILCAHGGR | 8 | Q7Z2W4 | Zinc finger CCCH-type antiviral protein 1 | ZC3HAV1 | 101.432 | 2 | 51.536686 | yes | 1.398E-07 | 442.23196 | 4316.6851 |
| LHICDHFTR | 9 | Q7Z2W4 | Zinc finger CCCH-type antiviral protein 1 | ZC3HAV1 | 101.432 | 2 | 51.536686 | yes | 1.066E-07 | 599.79291 | 3532.1965 |
| KFTYLGSQDR | 10 | Q7Z2W4 | Zinc finger CCCH-type antiviral protein 1 | ZC3HAV1 | 101.432 | 2 | 51.536686 | yes | 2.651E-07 | 607.81183 | 3435.5747 |
| VHFHLPYR | 8 | Q7Z2W4 | Zinc finger CCCH-type antiviral protein 1 | ZC3HAV1 | 101.432 | 2 | 51.536686 | no | 1.974E-07 | 534.79053 | 3316.7595 |
| YSHEVLSEENFK | 12 | Q7Z2W4 | Zinc finger CCCH-type antiviral protein 1 | ZC3HAV1 | 101.432 | 2 | 51.536686 | yes | 9.901E-08 | 741.349 | 1655.0032 |
| YSHEVLSEENFK | 12 | Q7Z2W4 | Zinc finger CCCH-type antiviral protein 1 | ZC3HAV1 | 101.432 | 2 | 51.536686 | yes | 1.815E-07 | 494.56842 | 1655.0032 |
| YSHEVLSEENFK | 12 | Q7Z2W4 | Zinc finger CCCH-type antiviral protein 1 | ZC3HAV1 | 101.432 | 3 | 51.536686 | yes | 9.901E-08 | 741.349 | 1655.0032 |

| | | | | | | | | | | |
|---|---|---|---|---|---|---|---|---|---|---|
| YSHEVLSEENFK | 12 | Q7Z2W4 | Zinc finger CCCH-type antiviral protein 1ZC3HAV1 | 101.432 | 3 | 51.536686 | yes | 1.815E-07 | 494.56842 | 1655.0032 |
| GTGLLSSDYR | 10 | Q7Z2W4 | Zinc finger CCCH-type antiviral protein 1ZC3HAV1 | 101.432 | 2 | 51.536686 | yes | 9.722E-08 | 534.76965 | 2973.9722 |
| SNPSVFVIFQK | 11 | Q7Z2W4 | Zinc finger CCCH-type antiviral protein 1ZC3HAV1 | 101.432 | 2 | 51.536686 | yes | 9.893E-08 | 633.34808 | 2842.7339 |
| SLTSWTNDQGAR | 12 | Q7Z2W4 | Zinc finger CCCH-type antiviral protein 1ZC3HAV1 | 101.432 | 2 | 51.536686 | yes | 1.06E-07 | 668.31799 | 2766.4646 |
| ADPEVCCFITK | 11 | Q7Z2W4 | Zinc finger CCCH-type antiviral protein 1ZC3HAV1 | 101.432 | 2 | 51.536686 | yes | 1.043E-07 | 691.30994 | 2752.251 |
| SCTPSPDQISHR | 12 | Q7Z2W4 | Zinc finger CCCH-type antiviral protein 1ZC3HAV1 | 101.432 | 2 | 51.536686 | yes | 1.427E-07 | 692.81732 | 2750.0239 |
| TWTDFEHMETIEK | 13 | Q7Z2W4 | Zinc finger CCCH-type antiviral protein 1ZC3HAV1 | 101.432 | 2 | 51.536686 | yes | 3.885E-07 | 833.87451 | 1225.4983 |
| TWTDFEHMETIEK | 13 | Q7Z2W4 | Zinc finger CCCH-type antiviral protein 1ZC3HAV1 | 101.432 | 2 | 51.536686 | yes | 1.998E-07 | 556.25208 | 1225.4983 |
| TWTDFEHMETIEK | 13 | Q7Z2W4 | Zinc finger CCCH-type antiviral protein 1ZC3HAV1 | 101.432 | 3 | 51.536686 | yes | 3.885E-07 | 833.87451 | 1225.4983 |
| TWTDFEHMETIEK | 13 | Q7Z2W4 | Zinc finger CCCH-type antiviral protein 1ZC3HAV1 | 101.432 | 3 | 51.536686 | yes | 1.998E-07 | 556.25208 | 1225.4983 |
| LSEIHHLHPEYVR | 13 | Q7Z2W4 | Zinc finger CCCH-type antiviral protein 1ZC3HAV1 | 101.432 | 3 | 51.536686 | yes | 1.202E-07 | 543.95471 | 1126.4697 |
| LSEIHHLHPEYVR | 13 | Q7Z2W4 | Zinc finger CCCH-type antiviral protein 1ZC3HAV1 | 101.432 | 3 | 51.536686 | yes | 1.084E-07 | 815.42847 | 1126.4697 |
| LSEIHHLHPEYVR | 13 | Q7Z2W4 | Zinc finger CCCH-type antiviral protein 1ZC3HAV1 | 101.432 | 2 | 51.536686 | yes | 1.202E-07 | 543.95471 | 1126.4697 |
| LSEIHHLHPEYVR | 13 | Q7Z2W4 | Zinc finger CCCH-type antiviral protein 1ZC3HAV1 | 101.432 | 2 | 51.536686 | yes | 1.084E-07 | 815.42847 | 1126.4697 |
| KGTGLLSSDYR | 11 | Q7Z2W4 | Zinc finger CCCH-type antiviral protein 1ZC3HAV1 | 101.432 | 2 | 51.536686 | yes | 9.894E-08 | 598.81714 | 1914.6993 |
| NESGTWIQYGEEK | 13 | Q7Z2W4 | Zinc finger CCCH-type antiviral protein 1ZC3HAV1 | 101.432 | 2 | 51.536686 | yes | 8.294E-07 | 770.84937 | 1678.7218 |
| VDDHDSEEICLDH | 16 | Q7Z2W4 | Zinc finger CCCH-type antiviral protein 1ZC3HAV1 | 101.432 | 3 | 51.536686 | yes | 9.722E-08 | 662.284 | 1366.8682 |
| FTYLGSQDR | 9 | Q7Z2W4 | Zinc finger CCCH-type antiviral protein 1ZC3HAV1 | 101.432 | 2 | 51.536686 | yes | 9.722E-08 | 543.76434 | 1325.7261 |
| KVLAIMR | 7 | Q7Z2W4 | Zinc finger CCCH-type antiviral protein 1ZC3HAV1 | 101.432 | 2 | 51.536686 | yes | 1.292E-07 | 415.76767 | 1237.0037 |
| IENSELLDK | 9 | Q7Z2W4 | Zinc finger CCCH-type antiviral protein 1ZC3HAV1 | 101.432 | 2 | 51.536686 | yes | 2.286E-07 | 530.77966 | 1108.0012 |
| LLFYATSR | 8 | Q7Z2W4 | Zinc finger CCCH-type antiviral protein 1ZC3HAV1 | 101.432 | 2 | 51.536686 | yes | 3.255E-07 | 485.77145 | 1045.161 |
| TSSVSLTATFR | 11 | Q7Z2W4 | Zinc finger CCCH-type antiviral protein 1ZC3HAV1 | 101.432 | 2 | 51.536686 | yes | 1.311E-07 | 585.31171 | 955.922 |
| IADDADPR | 8 | Q7Z2W4 | Zinc finger CCCH-type antiviral protein 1ZC3HAV1 | 101.432 | 2 | 51.536686 | yes | 4.394E-07 | 436.70905 | 931.17535 |
| ATDLGGTSQAGTS | 15 | Q7Z2W4 | Zinc finger CCCH-type antiviral protein 1ZC3HAV1 | 101.432 | 2 | 51.536686 | yes | 4.824E-07 | 725.35004 | 738.69592 |
| KIENSELLDK | 10 | Q7Z2W4 | Zinc finger CCCH-type antiviral protein 1ZC3HAV1 | 101.432 | 2 | 51.536686 | yes | 9.81E-08 | 594.82715 | 732.841 |
| IQDAGPASR | 9 | Q7Z2W4 | Zinc finger CCCH-type antiviral protein 1ZC3HAV1 | 101.432 | 2 | 51.536686 | yes | 4.989E-07 | 457.73816 | 567.28931 |
| NCPYDAK | 7 | Q7Z2W4 | Zinc finger CCCH-type antiviral protein 1ZC3HAV1 | 101.432 | 2 | 51.536686 | yes | 2.134E-07 | 434.18689 | 559.836 |
| NHELSGLNK | 9 | Q7Z2W4 | Zinc finger CCCH-type antiviral protein 1ZC3HAV1 | 101.432 | 2 | 51.536686 | yes | 1.397E-05 | 506.26453 | 319.31631 |
| PQEDFCFLSSK | 11 | Q7Z2W4 | Zinc finger CCCH-type antiviral protein 1ZC3HAV1 | 101.432 | 2 | 51.536686 | yes | 1.18E-07 | 679.30829 | 298.07513 |
| WQMLIGK | 7 | Q7Z2W4 | Zinc finger CCCH-type antiviral protein 1ZC3HAV1 | 101.432 | 2 | 51.536686 | yes | 1.563E-07 | 438.24402 | 277.60602 |
| PANSVFTTK | 9 | Q7Z2W4 | Zinc finger CCCH-type antiviral protein 1ZC3HAV1 | 101.432 | 2 | 51.536686 | yes | 1.005E-07 | 482.75854 | 274.23932 |
| QQICNQQPPCSR | 12 | Q7Z2W4 | Zinc finger CCCH-type antiviral protein 1ZC3HAV1 | 101.432 | 2 | 51.536686 | yes | 1E-07 | 758.3432 | 198.18359 |
| KTVFSPTLPAAR | 12 | Q7Z2W4 | Zinc finger CCCH-type antiviral protein 1ZC3HAV1 | 101.432 | 2 | 51.536686 | yes | 1.827E-07 | 644.37463 | 183.51425 |
| SHNLMDR | 7 | Q7Z2W4 | Zinc finger CCCH-type antiviral protein 1ZC3HAV1 | 101.432 | 2 | 51.536686 | yes | 3.375E-07 | 436.70578 | 134.84059 |
| PCDNLHLCK | 9 | Q7Z2W4 | Zinc finger CCCH-type antiviral protein 1ZC3HAV1 | 101.432 | 2 | 51.536686 | yes | 8.446E-07 | 578.7655 | 131.88844 |
| RLSTPSSVTKPANS | 19 | Q7Z2W4 | Zinc finger CCCH-type antiviral protein 1ZC3HAV1 | 101.432 | 3 | 51.536686 | yes | 1.499E-07 | 674.37238 | 118.1718 |
| GYCNPGIHLCSVG | 20 | Q7Z2W4 | Zinc finger CCCH-type antiviral protein 1ZC3HAV1 | 101.432 | 3 | 51.536686 | yes | 1.035E-06 | 772.36139 | 114.30462 |
| PTFVPQWYVQQM | 13 | Q7Z2W4 | Zinc finger CCCH-type antiviral protein 1ZC3HAV1 | 101.432 | 2 | 51.536686 | yes | 9.729E-08 | 826.4187 | 104.63805 |
| RPTFVPQWYVQQI | 14 | Q7Z2W4 | Zinc finger CCCH-type antiviral protein 1ZC3HAV1 | 101.432 | 3 | 51.536686 | yes | 5.753E-07 | 603.31525 | 67.461983 |

| | | | | | | | | | | |
|---|---|---|---|---|---|---|---|---|---|---|
| GVVPFQAGSR | 10 | Q7Z2W4 | Zinc finger CCCH-type antiviral protein 1 | ZC3HAV1 | 101.432 | 2 | 51.536686 | yes | 2.897E-06 | 509.27744 | 55.907684 |
| CNYSQSER | 8 | Q7Z2W4 | Zinc finger CCCH-type antiviral protein 1 | ZC3HAV1 | 101.432 | 2 | 51.536686 | yes | 2.281E-06 | 522.21417 | 43.910885 |
| RLSTPSSVTK | 10 | Q7Z2W4 | Zinc finger CCCH-type antiviral protein 1 | ZC3HAV1 | 101.432 | 2 | 51.536686 | yes | 4.156E-07 | 538.30896 | 42.916374 |
| LSTPSSVTK | 9 | Q7Z2W4 | Zinc finger CCCH-type antiviral protein 1 | ZC3HAV1 | 101.432 | 2 | 51.536686 | yes | 1.559E-06 | 460.25839 | 33.899178 |
| IENSELLDKFTWK | 13 | Q7Z2W4 | Zinc finger CCCH-type antiviral protein 1 | ZC3HAV1 | 101.432 | 2 | 51.536686 | yes | 3.306E-07 | 811.92487 | 18.723272 |
| VMLLLDTAGGAAI | 13 | Q7Z2Z1 | Treslin OS=Homo sapiens OX=9606 GN= | TICRR | 210.857 | 2 | 53.039272 | yes | 9.903E-08 | 644.35809 | 406.91748 |
| LLTYLSCR | 8 | Q7Z2Z1 | Treslin OS=Homo sapiens OX=9606 GN= | TICRR | 210.857 | 2 | 53.039272 | yes | 3.472E-06 | 513.27606 | 159.59818 |
| NTLDSEVPAAYQT | 15 | Q7Z2Z1 | Treslin OS=Homo sapiens OX=9606 GN= | TICRR | 210.857 | 2 | 53.039272 | yes | 9.728E-08 | 817.40704 | 120.96243 |
| SNVLSVEEGEGLR | 13 | Q7Z2Z1 | Treslin OS=Homo sapiens OX=9606 GN= | TICRR | 210.857 | 2 | 53.039272 | yes | 3.114E-06 | 694.85443 | 116.75803 |
| LFTSPLCDVSK | 11 | Q7Z2Z1 | Treslin OS=Homo sapiens OX=9606 GN= | TICRR | 210.857 | 2 | 53.039272 | yes | 3.321E-07 | 633.82355 | 105.87097 |
| SWEDFEEELEAR | 12 | Q7Z2Z1 | Treslin OS=Homo sapiens OX=9606 GN= | TICRR | 210.857 | 2 | 53.039272 | yes | 5.543E-07 | 770.33356 | 94.014771 |
| MEGMLFLPVEAGI | 13 | Q7Z2Z1 | Treslin OS=Homo sapiens OX=9606 GN= | TICRR | 210.857 | 2 | 53.039272 | yes | 2.629E-06 | 711.362 | 65.073051 |
| NLFNQELLSPSK | 12 | Q7Z2Z1 | Treslin OS=Homo sapiens OX=9606 GN= | TICRR | 210.857 | 2 | 53.039272 | yes | 9.722E-08 | 695.37225 | 62.850029 |
| SHSVSAVDGLEDK | 13 | Q7Z2Z1 | Treslin OS=Homo sapiens OX=9606 GN= | TICRR | 210.857 | 2 | 53.039272 | yes | 9.819E-08 | 672.3255 | 62.338627 |
| AEEASSWGQFGLS | 15 | Q7Z2Z1 | Treslin OS=Homo sapiens OX=9606 GN= | TICRR | 210.857 | 2 | 53.039272 | yes | 1.417E-07 | 806.37354 | 54.076672 |
| ELEVNCLNQVK | 11 | Q7Z2Z1 | Treslin OS=Homo sapiens OX=9606 GN= | TICRR | 210.857 | 2 | 53.039272 | yes | 9.808E-08 | 673.34265 | 45.233334 |
| EESTIAHQEDSK | 12 | Q7Z2Z1 | Treslin OS=Homo sapiens OX=9606 GN= | TICRR | 210.857 | 2 | 53.039272 | yes | 1.423E-07 | 687.31262 | 36.967136 |
| HTQAGEGTSLETK | 13 | Q7Z2Z1 | Treslin OS=Homo sapiens OX=9606 GN= | TICRR | 210.857 | 2 | 53.039272 | yes | 3.635E-06 | 679.83093 | 33.804432 |
| THSASFYSVSQPK | 13 | Q7Z2Z1 | Treslin OS=Homo sapiens OX=9606 GN= | TICRR | 210.857 | 2 | 53.039272 | yes | 7.522E-06 | 719.85168 | 21.615566 |
| NIGIITHYK | 9 | Q7Z333 | Probable helicase senataxin OS=Homo sa | SETX | 302.88 | 2 | 55.56374 | yes | 4.456E-07 | 529.80328 | 382.92374 |
| ELIPFTEMTNASEK | 14 | Q7Z333 | Probable helicase senataxin OS=Homo sa | SETX | 302.88 | 2 | 55.56374 | yes | 2.02E-07 | 805.39252 | 189.20477 |
| ENFYQLQVR | 9 | Q7Z333 | Probable helicase senataxin OS=Homo sa | SETX | 302.88 | 2 | 55.56374 | yes | 1.227E-07 | 598.80658 | 187.10959 |
| VNELCVEALCR | 11 | Q7Z333 | Probable helicase senataxin OS=Homo sa | SETX | 302.88 | 2 | 55.56374 | yes | 1.699E-07 | 681.82886 | 178.71864 |
| LIELIHGK | 8 | Q7Z333 | Probable helicase senataxin OS=Homo sa | SETX | 302.88 | 2 | 55.56374 | yes | 9.722E-08 | 461.78964 | 151.73181 |
| LLTDSSTDALEK | 12 | Q7Z333 | Probable helicase senataxin OS=Homo sa | SETX | 302.88 | 2 | 55.56374 | yes | 1.452E-07 | 646.83264 | 151.39876 |
| TIVGLLYR | 8 | Q7Z333 | Probable helicase senataxin OS=Homo sa | SETX | 302.88 | 2 | 55.56374 | yes | 6.719E-07 | 467.78964 | 136.31114 |
| LISPQNLSVR | 10 | Q7Z333 | Probable helicase senataxin OS=Homo sa | SETX | 302.88 | 2 | 55.56374 | yes | 3.426E-05 | 563.83258 | 127.76614 |
| LCTCTEPIR | 9 | Q7Z333 | Probable helicase senataxin OS=Homo sa | SETX | 302.88 | 2 | 55.56374 | yes | 1.102E-07 | 575.27319 | 122.80996 |
| DLLTTTSER | 9 | Q7Z333 | Probable helicase senataxin OS=Homo sa | SETX | 302.88 | 2 | 55.56374 | yes | 1.035E-06 | 518.26947 | 98.981743 |
| DNDSYINVQEIK | 12 | Q7Z333 | Probable helicase senataxin OS=Homo sa | SETX | 302.88 | 2 | 55.56374 | yes | 1.06E-07 | 719.34644 | 91.251099 |
| FSLDSQVNHR | 10 | Q7Z333 | Probable helicase senataxin OS=Homo sa | SETX | 302.88 | 2 | 55.56374 | yes | 1.727E-07 | 601.79926 | 86.119064 |
| SLDYVAQLR | 9 | Q7Z333 | Probable helicase senataxin OS=Homo sa | SETX | 302.88 | 2 | 55.56374 | yes | 1.001E-07 | 532.79041 | 82.485115 |
| LLTSQELQMQR | 11 | Q7Z333 | Probable helicase senataxin OS=Homo sa | SETX | 302.88 | 2 | 55.56374 | yes | 2.625E-07 | 673.85846 | 73.561516 |
| AADEDVFR | 8 | Q7Z333 | Probable helicase senataxin OS=Homo sa | SETX | 302.88 | 2 | 55.56374 | yes | 4.898E-07 | 461.71689 | 66.872253 |
| DLDKEFDR | 8 | Q7Z333 | Probable helicase senataxin OS=Homo sa | SETX | 302.88 | 2 | 55.56374 | yes | 8.082E-07 | 519.24854 | 62.994347 |
| LSDCESTDVK | 10 | Q7Z333 | Probable helicase senataxin OS=Homo sa | SETX | 302.88 | 2 | 55.56374 | yes | 9.722E-08 | 577.25574 | 57.013512 |
| EGYQLGQQSLCK | 12 | Q7Z333 | Probable helicase senataxin OS=Homo sa | SETX | 302.88 | 2 | 55.56374 | yes | 3.226E-06 | 705.83771 | 56.170795 |
| EFLDYQLDELSR | 12 | Q7Z333 | Probable helicase senataxin OS=Homo sa | SETX | 302.88 | 2 | 55.56374 | yes | 1.557E-07 | 764.36993 | 54.461311 |
| AVLNPNPMDFCTK | 13 | Q7Z333 | Probable helicase senataxin OS=Homo sa | SETX | 302.88 | 2 | 55.56374 | yes | 1.667E-07 | 753.8576 | 53.974457 |

| | | | | | | | | | | | |
|---|---|---|---|---|---|---|---|---|---|---|---|
| SSNYLSCR | 8 | Q7Z333 | Probable helicase senataxin OS=Homo sa | SETX | 302.88 | 2 | 55.56374 | yes | 4.003E-07 | 493.72165 | 52.391827 |
| YPYLLLHER | 9 | Q7Z333 | Probable helicase senataxin OS=Homo sa | SETX | 302.88 | 2 | 55.56374 | yes | 1.03E-07 | 602.32965 | 51.564419 |
| SLETSSALSPSLK | 13 | Q7Z333 | Probable helicase senataxin OS=Homo sa | SETX | 302.88 | 2 | 55.56374 | yes | 2.201E-06 | 660.35626 | 43.097469 |
| AQEYGYDQSMMA | 13 | Q7Z333 | Probable helicase senataxin OS=Homo sa | SETX | 302.88 | 2 | 55.56374 | yes | 9.722E-08 | 775.32416 | 39.961227 |
| MEQANCSFQVFDI | 13 | Q7Z333 | Probable helicase senataxin OS=Homo sa | SETX | 302.88 | 2 | 55.56374 | yes | 2.689E-07 | 802.3476 | 38.249672 |
| VPLLEILK | 8 | Q7Z333 | Probable helicase senataxin OS=Homo sa | SETX | 302.88 | 2 | 55.56374 | yes | 2.283E-07 | 462.81006 | 34.252663 |
| VSTSNEDFSLK | 11 | Q7Z333 | Probable helicase senataxin OS=Homo sa | SETX | 302.88 | 2 | 55.56374 | yes | 3.872E-07 | 613.79858 | 23.25808 |
| LNELMSQIR | 9 | Q7Z3B4 | Nucleoporin p54 OS=Homo sapiens OX= | NUP54 | 55.436 | 2 | 45.48889 | yes | 2.938E-06 | 552.29755 | 63.880577 |
| SGYAIQADEEQLR | 13 | Q7Z3B4 | Nucleoporin p54 OS=Homo sapiens OX= | NUP54 | 55.436 | 2 | 45.48889 | yes | 1.835E-07 | 740.35736 | 41.866718 |
| SAYLSQHR | 8 | Q7Z3I7 | Zinc finger protein 572 OS=Homo sapien | ZNF572 | 61.239 | 2 | 44.593853 | yes | 8.892E-07 | 481.24615 | 69.04686 |
| SFGLSSHLIR | 10 | Q7Z3I7 | Zinc finger protein 572 OS=Homo sapien | ZNF572 | 61.239 | 2 | 44.593853 | yes | 2.703E-07 | 558.81165 | 66.083824 |
| LLVSDSNSFMER | 12 | Q7Z3I7 | Zinc finger protein 572 OS=Homo sapien | ZNF572 | 61.239 | 2 | 44.593853 | yes | 2.138E-07 | 699.34009 | 33.880341 |
| VTSSIPVFDLQDG( | 15 | Q7Z3K3 | Pogo transposable element with ZNF don | POGZ | 155.344 | 2 | 44.62504 | yes | 3.462E-07 | 795.90973 | 181.88202 |
| ISYEWAVR | 8 | Q7Z3K3 | Pogo transposable element with ZNF don | POGZ | 155.344 | 2 | 44.62504 | yes | 5.608E-07 | 512.26654 | 74.288673 |
| LAEWVLTQR | 9 | Q7Z3K3 | Pogo transposable element with ZNF don | POGZ | 155.344 | 2 | 44.62504 | yes | 2.126E-07 | 558.31403 | 45.899097 |
| NMYPPPSFPTNK | 12 | Q7Z3K3 | Pogo transposable element with ZNF don | POGZ | 155.344 | 2 | 44.62504 | yes | 4.102E-06 | 696.83447 | 41.947308 |
| LIMLVDDFYYGR | 12 | Q7Z3K3 | Pogo transposable element with ZNF don | POGZ | 155.344 | 2 | 44.62504 | yes | 4.931E-06 | 752.87885 | 28.504343 |
| VPEPNENVGDAV( | 15 | Q7Z3K3 | Pogo transposable element with ZNF don | POGZ | 155.344 | 2 | 44.62504 | yes | 1.186E-06 | 798.89685 | 24.682568 |
| VVLFALCCNTEQA | 18 | Q7Z3K3 | Pogo transposable element with ZNF don | POGZ | 155.344 | 3 | 44.62504 | yes | 0.0001069 | 722.34705 | 18.062115 |
| TLEEEEMMDEGK | 12 | Q7Z3K6 | Mesoderm induction early response prote | MIER3 | 61.437 | 2 | 43.382828 | yes | 2.615E-07 | 720.79706 | 45.046185 |
| DFHLIQK | 7 | Q7Z3K6 | Mesoderm induction early response prote | MIER3 | 61.437 | 2 | 43.382828 | yes | 4.092E-07 | 450.75052 | 19.3913 |
| EQQNDISSELQNR | 13 | Q7Z3T8 | Zinc finger FYVE domain-containing pro | ZFYVE16 | 168.904 | 2 | 43.369694 | yes | 3.762E-07 | 780.86609 | 47.865211 |
| STIPVQQGLPTSK | 13 | Q7Z3T8 | Zinc finger FYVE domain-containing pro | ZFYVE16 | 168.904 | 2 | 43.369694 | yes | 2.62E-07 | 678.38007 | 45.209705 |
| LPDFSFQEDK | 10 | Q7Z3T8 | Zinc finger FYVE domain-containing pro | ZFYVE16 | 168.904 | 2 | 43.369694 | yes | 6.204E-07 | 613.29041 | 36.405468 |
| LLLGLTSADGK | 11 | Q7Z401 | C-myc promoter-binding protein OS=Hor | DENND4A | 209.245 | 2 | 50.691086 | yes | 4.233E-07 | 544.32153 | 104.88644 |
| NVVLGVTQFK | 10 | Q7Z401 | C-myc promoter-binding protein OS=Hor | DENND4A | 209.245 | 2 | 50.691086 | yes | 1.677E-06 | 552.82422 | 89.435585 |
| EPITDVSVIIK | 11 | Q7Z401 | C-myc promoter-binding protein OS=Hor | DENND4A | 209.245 | 2 | 50.691086 | yes | 4.342E-07 | 607.35553 | 80.893349 |
| STRPNTLDIGKPPL | 15 | Q7Z401 | C-myc promoter-binding protein OS=Hor | DENND4A | 209.245 | 3 | 50.691086 | yes | 9.728E-08 | 555.65295 | 54.21032 |
| RLDLPSNLPGLILS | 20 | Q7Z401 | C-myc promoter-binding protein OS=Hor | DENND4A | 209.245 | 3 | 50.691086 | yes | 5.804E-07 | 755.06696 | 41.175697 |
| INQEATPGDIVEK | 13 | Q7Z401 | C-myc promoter-binding protein OS=Hor | DENND4A | 209.245 | 2 | 50.691086 | yes | 8.701E-07 | 707.36462 | 27.702312 |
| QSLLVKELQSLAV | 16 | Q7Z403 | Transmembrane channel-like protein 6 O | TMC6 | 90.046 | 3 | 34.97258 | yes | 9.084E-07 | 595.01941 | 212.14302 |
| LVLDQLR | 7 | Q7Z406 | Myosin-14 OS=Homo sapiens OX=9606 | MYH14 | 227.871 | 2 | 41.424465 | yes | 3.516E-07 | 428.76617 | 382.8136 |
| GKPAVGDMLR | 10 | Q7Z417 | FMR1-interacting protein NUFIP2 OS=H | NUFIP2 | 76.122 | 2 | 52.361546 | yes | 1.631E-07 | 522.28693 | 27080.551 |
| GKPAVGDMLR | 10 | Q7Z417 | FMR1-interacting protein NUFIP2 OS=H | NUFIP2 | 76.122 | 2 | 52.361546 | yes | 9.722E-08 | 530.28442 | 27080.551 |
| ADTSSQGALVFLS | 14 | Q7Z417 | FMR1-interacting protein NUFIP2 OS=H | NUFIP2 | 76.122 | 2 | 52.361546 | yes | 1.334E-07 | 712.375 | 36284.523 |
| TSPQVLGSILK | 11 | Q7Z417 | FMR1-interacting protein NUFIP2 OS=H | NUFIP2 | 76.122 | 2 | 52.361546 | yes | 1.459E-07 | 571.84259 | 34500.129 |
| EMESIWNLQK | 10 | Q7Z417 | FMR1-interacting protein NUFIP2 OS=H | NUFIP2 | 76.122 | 2 | 52.361546 | yes | 9.728E-08 | 639.31335 | 15580.319 |
| EMESIWNLQK | 10 | Q7Z417 | FMR1-interacting protein NUFIP2 OS=H | NUFIP2 | 76.122 | 2 | 52.361546 | yes | 9.733E-08 | 647.31079 | 15580.319 |
| AAIVYHTK | 8 | Q7Z417 | FMR1-interacting protein NUFIP2 OS=H | NUFIP2 | 76.122 | 2 | 52.361546 | yes | 1.33E-07 | 451.75836 | 22833.672 |

| | | | | | | | | | | |
|---|---|---|---|---|---|---|---|---|---|---|
| IMQQETSVPTLK | 12 | Q7Z417 | FMR1-interacting protein NUFIP2 OS=H | NUFIP2 | 76.122 | 2 | 52.361546 | yes | 1.865E-07 | 687.86853 | 10615.798 |
| IMQQETSVPTLK | 12 | Q7Z417 | FMR1-interacting protein NUFIP2 OS=H | NUFIP2 | 76.122 | 2 | 52.361546 | yes | 9.722E-08 | 695.86597 | 10615.798 |
| LSQVPMSALK | 10 | Q7Z417 | FMR1-interacting protein NUFIP2 OS=H | NUFIP2 | 76.122 | 2 | 52.361546 | yes | 1.552E-07 | 537.30481 | 10602.536 |
| LSQVPMSALK | 10 | Q7Z417 | FMR1-interacting protein NUFIP2 OS=H | NUFIP2 | 76.122 | 2 | 52.361546 | yes | 2.616E-07 | 545.30225 | 10602.536 |
| NDSWGSFDLR | 10 | Q7Z417 | FMR1-interacting protein NUFIP2 OS=H | NUFIP2 | 76.122 | 2 | 52.361546 | yes | 4.377E-07 | 598.7702 | 17993.604 |
| VDGSKPIWK | 9 | Q7Z417 | FMR1-interacting protein NUFIP2 OS=H | NUFIP2 | 76.122 | 2 | 52.361546 | yes | 1.279E-07 | 515.29004 | 15596.936 |
| IITYNEAMDSPDQ | 13 | Q7Z417 | FMR1-interacting protein NUFIP2 OS=H | NUFIP2 | 76.122 | 2 | 52.361546 | yes | 9.722E-08 | 748.83234 | 13030.881 |
| QGLETFKPDYSEQ | 14 | Q7Z417 | FMR1-interacting protein NUFIP2 OS=H | NUFIP2 | 76.122 | 3 | 52.361546 | yes | 1.017E-07 | 557.2738 | 4525.3203 |
| QGLETFKPDYSEQ | 14 | Q7Z417 | FMR1-interacting protein NUFIP2 OS=H | NUFIP2 | 76.122 | 3 | 52.361546 | yes | 9.572E-07 | 835.40704 | 4525.3203 |
| QGLETFKPDYSEQ | 14 | Q7Z417 | FMR1-interacting protein NUFIP2 OS=H | NUFIP2 | 76.122 | 2 | 52.361546 | yes | 1.017E-07 | 557.2738 | 4525.3203 |
| QGLETFKPDYSEQ | 14 | Q7Z417 | FMR1-interacting protein NUFIP2 OS=H | NUFIP2 | 76.122 | 2 | 52.361546 | yes | 9.572E-07 | 835.40704 | 4525.3203 |
| GCENLNIVQDK | 11 | Q7Z417 | FMR1-interacting protein NUFIP2 OS=H | NUFIP2 | 76.122 | 2 | 52.361546 | yes | 1.047E-07 | 645.31134 | 8758.7568 |
| YETGPGGTSR | 10 | Q7Z417 | FMR1-interacting protein NUFIP2 OS=H | NUFIP2 | 76.122 | 2 | 52.361546 | yes | 1.506E-07 | 512.73834 | 4155.1016 |
| TGYGELNGNAGEI | 13 | Q7Z417 | FMR1-interacting protein NUFIP2 OS=H | NUFIP2 | 76.122 | 2 | 52.361546 | yes | 9.722E-08 | 669.30768 | 3069.4814 |
| RIITYNEAMDSPD( | 14 | Q7Z417 | FMR1-interacting protein NUFIP2 OS=H | NUFIP2 | 76.122 | 2 | 52.361546 | yes | 1.035E-07 | 826.88287 | 2309.9915 |
| HTLQQHQETPK | 11 | Q7Z417 | FMR1-interacting protein NUFIP2 OS=H | NUFIP2 | 76.122 | 2 | 52.361546 | yes | 3.206E-07 | 673.84418 | 1981.2794 |
| VLNGNQQVVDTSI | 14 | Q7Z417 | FMR1-interacting protein NUFIP2 OS=H | NUFIP2 | 76.122 | 2 | 52.361546 | yes | 1.7E-07 | 757.91229 | 1339.3171 |
| PPPVFPVDNSSAK | 13 | Q7Z417 | FMR1-interacting protein NUFIP2 OS=H | NUFIP2 | 76.122 | 2 | 52.361546 | yes | 1.254E-07 | 677.8537 | 883.6192 |
| PAVGDMLR | 8 | Q7Z417 | FMR1-interacting protein NUFIP2 OS=H | NUFIP2 | 76.122 | 2 | 52.361546 | yes | 2.782E-07 | 429.72873 | 826.39685 |
| NLSSDEATNPISR | 13 | Q7Z417 | FMR1-interacting protein NUFIP2 OS=H | NUFIP2 | 76.122 | 2 | 52.361546 | yes | 1.835E-06 | 702.34167 | 790.04492 |
| KFDDRPK | 7 | Q7Z417 | FMR1-interacting protein NUFIP2 OS=H | NUFIP2 | 76.122 | 2 | 52.361546 | yes | 1.841E-07 | 453.24561 | 778.5885 |
| GADNDGSGSESGY | 17 | Q7Z417 | FMR1-interacting protein NUFIP2 OS=H | NUFIP2 | 76.122 | 2 | 52.361546 | yes | 1.055E-07 | 821.84497 | 415.59323 |
| RTSPQVLGSILK | 12 | Q7Z417 | FMR1-interacting protein NUFIP2 OS=H | NUFIP2 | 76.122 | 2 | 52.361546 | yes | 1.623E-07 | 649.89319 | 322.03711 |
| SGTTSESGALSLEP | 22 | Q7Z417 | FMR1-interacting protein NUFIP2 OS=H | NUFIP2 | 76.122 | 3 | 52.361546 | yes | 1.086E-07 | 738.70099 | 271.89377 |
| EDSWTLFK | 8 | Q7Z417 | FMR1-interacting protein NUFIP2 OS=H | NUFIP2 | 76.122 | 2 | 52.361546 | yes | 1.014E-07 | 513.25055 | 197.37663 |
| MGADLLGSVQAK | 12 | Q7Z460 | CLIP-associating protein 1 OS=Homo saı | CLASP1 | 169.451 | 2 | 56.232708 | no | 1.066E-07 | 595.31592 | 1498.5222 |
| GIHDADSEAR | 10 | Q7Z460 | CLIP-associating protein 1 OS=Homo saı | CLASP1 | 169.451 | 2 | 56.232708 | yes | 1.603E-07 | 535.7467 | 1427.5374 |
| IITWTTEPK | 9 | Q7Z460 | CLIP-associating protein 1 OS=Homo saı | CLASP1 | 169.451 | 2 | 56.232708 | yes | 1.423E-07 | 544.80298 | 1246.6134 |
| EGAGAVDEEDFIK | 13 | Q7Z460 | CLIP-associating protein 1 OS=Homo saı | CLASP1 | 169.451 | 2 | 56.232708 | yes | 2.324E-07 | 690.31989 | 1137.2825 |
| IMATSGVVAVR | 11 | Q7Z460 | CLIP-associating protein 1 OS=Homo saı | CLASP1 | 169.451 | 2 | 56.232708 | yes | 9.727E-08 | 552.31573 | 1133.5597 |
| DAAINSLVEIYR | 12 | Q7Z460 | CLIP-associating protein 1 OS=Homo saı | CLASP1 | 169.451 | 2 | 56.232708 | yes | 1.575E-07 | 682.36444 | 1132.6628 |
| AFDDVPVVQIYSS | 14 | Q7Z460 | CLIP-associating protein 1 OS=Homo saı | CLASP1 | 169.451 | 2 | 56.232708 | yes | 1.774E-07 | 798.40686 | 1054.688 |
| AQIGTVLPSLIDR | 13 | Q7Z460 | CLIP-associating protein 1 OS=Homo saı | CLASP1 | 169.451 | 2 | 56.232708 | yes | 9.81E-08 | 691.90375 | 1048.7021 |
| EAVFDDDMEQLR | 12 | Q7Z460 | CLIP-associating protein 1 OS=Homo saı | CLASP1 | 169.451 | 2 | 56.232708 | yes | 9.722E-08 | 734.32465 | 508.96262 |
| EAVFDDDMEQLR | 12 | Q7Z460 | CLIP-associating protein 1 OS=Homo saı | CLASP1 | 169.451 | 2 | 56.232708 | yes | 3.169E-07 | 742.32214 | 508.96262 |
| LLGSGYGGLTGGS | 15 | Q7Z460 | CLIP-associating protein 1 OS=Homo saı | CLASP1 | 169.451 | 2 | 56.232708 | yes | 1.196E-07 | 691.35718 | 964.86761 |
| SVSTTGSLQR | 10 | Q7Z460 | CLIP-associating protein 1 OS=Homo saı | CLASP1 | 169.451 | 2 | 56.232708 | yes | 9.872E-08 | 518.27509 | 889.65875 |
| HISVLAETIK | 10 | Q7Z460 | CLIP-associating protein 1 OS=Homo saı | CLASP1 | 169.451 | 2 | 56.232708 | yes | 1.858E-07 | 555.82953 | 858.74835 |
| MESCLAQVLQK | 11 | Q7Z460 | CLIP-associating protein 1 OS=Homo saı | CLASP1 | 169.451 | 2 | 56.232708 | yes | 6.285E-07 | 653.82831 | 811.60278 |

| | | | | | | | | | | | |
|---|---|---|---|---|---|---|---|---|---|---|---|
| LCEIFTR | 7 | Q7Z460 | CLIP-associating protein 1 OS=Homo sa| CLASP1 | 169.451 | 2 | 56.232708 | no | 1.355E-07 | 469.74185 | 744.33765 |
| TSLLNTQPPR | 10 | Q7Z460 | CLIP-associating protein 1 OS=Homo sa| CLASP1 | 169.451 | 2 | 56.232708 | yes | 9.722E-08 | 563.81439 | 693.75476 |
| EDSLGVWEEHFK | 12 | Q7Z460 | CLIP-associating protein 1 OS=Homo sa| CLASP1 | 169.451 | 2 | 56.232708 | yes | 9.722E-08 | 738.34369 | 685.81934 |
| SDIDVNAAASAK | 12 | Q7Z460 | CLIP-associating protein 1 OS=Homo sa| CLASP1 | 169.451 | 2 | 56.232708 | yes | 1.636E-07 | 581.29095 | 683.97302 |
| KGALLELLK | 9 | Q7Z460 | CLIP-associating protein 1 OS=Homo sa| CLASP1 | 169.451 | 2 | 56.232708 | yes | 1.566E-07 | 492.82623 | 614.59167 |
| FIVDQTQTPNLK | 12 | Q7Z460 | CLIP-associating protein 1 OS=Homo sa| CLASP1 | 169.451 | 2 | 56.232708 | yes | 9.722E-08 | 702.38007 | 503.81763 |
| NSDSIVSLPQSDR | 13 | Q7Z460 | CLIP-associating protein 1 OS=Homo sa| CLASP1 | 169.451 | 2 | 56.232708 | yes | 9.722E-08 | 709.34955 | 500.17044 |
| FGLGQPGR | 8 | Q7Z460 | CLIP-associating protein 1 OS=Homo sa| CLASP1 | 169.451 | 2 | 56.232708 | yes | 1.082E-07 | 416.22723 | 474.8028 |
| DVPIDHSDLVADL | 15 | Q7Z460 | CLIP-associating protein 1 OS=Homo sa| CLASP1 | 169.451 | 2 | 56.232708 | yes | 1.105E-07 | 825.44086 | 396.41293 |
| EGLLGLQNLLK | 11 | Q7Z460 | CLIP-associating protein 1 OS=Homo sa| CLASP1 | 169.451 | 2 | 56.232708 | no | 1.493E-07 | 599.36371 | 367.1315 |
| DGGAASPATEGR | 12 | Q7Z460 | CLIP-associating protein 1 OS=Homo sa| CLASP1 | 169.451 | 2 | 56.232708 | yes | 2.146E-07 | 544.75201 | 354.47812 |
| QSSGSATNVASTP | 16 | Q7Z460 | CLIP-associating protein 1 OS=Homo sa| CLASP1 | 169.451 | 2 | 56.232708 | yes | 1.802E-07 | 796.36896 | 350.56915 |
| LIPVITSNCTSK | 12 | Q7Z460 | CLIP-associating protein 1 OS=Homo sa| CLASP1 | 169.451 | 2 | 56.232708 | yes | 1.12E-07 | 666.86322 | 319.07602 |
| LLNLYIK | 7 | Q7Z460 | CLIP-associating protein 1 OS=Homo sa| CLASP1 | 169.451 | 2 | 56.232708 | no | 2.468E-07 | 438.78131 | 313.42545 |
| SQEDLNEPIKR | 11 | Q7Z460 | CLIP-associating protein 1 OS=Homo sa| CLASP1 | 169.451 | 2 | 56.232708 | yes | 9.722E-08 | 664.84387 | 309.54544 |
| YIESLAR | 7 | Q7Z460 | CLIP-associating protein 1 OS=Homo sa| CLASP1 | 169.451 | 2 | 56.232708 | yes | 2.033E-07 | 426.23471 | 304.26422 |
| EQDQTLLLK | 9 | Q7Z460 | CLIP-associating protein 1 OS=Homo sa| CLASP1 | 169.451 | 2 | 56.232708 | yes | 1.165E-07 | 544.30334 | 285.05682 |
| SADLEHDQTMLD | 13 | Q7Z460 | CLIP-associating protein 1 OS=Homo sa| CLASP1 | 169.451 | 2 | 56.232708 | yes | 1.53E-07 | 751.8432 | 281.80185 |
| NGGIPHYLR | 9 | Q7Z460 | CLIP-associating protein 1 OS=Homo sa| CLASP1 | 169.451 | 2 | 56.232708 | yes | 1.417E-07 | 513.77759 | 257.54373 |
| NSSNTSVGSPSNT | 16 | Q7Z460 | CLIP-associating protein 1 OS=Homo sa| CLASP1 | 169.451 | 2 | 56.232708 | yes | 9.722E-08 | 789.37933 | 253.30759 |
| MFADPHSK | 8 | Q7Z460 | CLIP-associating protein 1 OS=Homo sa| CLASP1 | 169.451 | 2 | 56.232708 | yes | 2.933E-07 | 466.71838 | 245.76038 |
| KEGLLGLQNLLK | 12 | Q7Z460 | CLIP-associating protein 1 OS=Homo sa| CLASP1 | 169.451 | 2 | 56.232708 | no | 9.722E-08 | 663.41119 | 245.75696 |
| ECDIVSR | 7 | Q7Z460 | CLIP-associating protein 1 OS=Homo sa| CLASP1 | 169.451 | 2 | 56.232708 | yes | 7.363E-07 | 439.70547 | 176.71866 |
| SQEDLNEPIK | 10 | Q7Z460 | CLIP-associating protein 1 OS=Homo sa| CLASP1 | 169.451 | 2 | 56.232708 | yes | 1.022E-07 | 586.79333 | 154.39259 |
| KECDIVSR | 8 | Q7Z460 | CLIP-associating protein 1 OS=Homo sa| CLASP1 | 169.451 | 2 | 56.232708 | yes | 1.423E-06 | 503.75293 | 98.897392 |
| IPGSVNAMR | 9 | Q7Z460 | CLIP-associating protein 1 OS=Homo sa| CLASP1 | 169.451 | 2 | 56.232708 | yes | 2.062E-06 | 472.75275 | 90.827766 |
| LQVGQELIDYFSD | 14 | Q7Z460 | CLIP-associating protein 1 OS=Homo sa| CLASP1 | 169.451 | 2 | 56.232708 | yes | 9.722E-08 | 827.9198 | 89.164307 |
| LGSSTLGSK | 9 | Q7Z460 | CLIP-associating protein 1 OS=Homo sa| CLASP1 | 169.451 | 2 | 56.232708 | yes | 2.646E-07 | 425.23746 | 80.684082 |
| RLGSSTLGSK | 10 | Q7Z460 | CLIP-associating protein 1 OS=Homo sa| CLASP1 | 169.451 | 2 | 56.232708 | yes | 8.084E-07 | 503.28799 | 72.26915 |
| QMDPTDFVNSSET | 14 | Q7Z460 | CLIP-associating protein 1 OS=Homo sa| CLASP1 | 169.451 | 2 | 56.232708 | yes | 9.722E-08 | 813.85687 | 65.501717 |
| NYAELTIMK | 9 | Q7Z460 | CLIP-associating protein 1 OS=Homo sa| CLASP1 | 169.451 | 2 | 56.232708 | yes | 6.955E-07 | 541.78119 | 64.029236 |
| AAEEAASTLASSI | 20 | Q7Z460 | CLIP-associating protein 1 OS=Homo sa| CLASP1 | 169.451 | 3 | 56.232708 | yes | 1.461E-07 | 705.01251 | 63.570568 |
| SSSSSQESLNR | 11 | Q7Z460 | CLIP-associating protein 1 OS=Homo sa| CLASP1 | 169.451 | 2 | 56.232708 | no | 2.069E-07 | 591.27325 | 38.354012 |
| EILSDDKHDWEQ | 13 | Q7Z460 | CLIP-associating protein 1 OS=Homo sa| CLASP1 | 169.451 | 2 | 56.232708 | yes | 4.245E-07 | 835.89209 | 23.96769 |
| SLLLAGAAEYDN | 18 | Q7Z460 | CLIP-associating protein 1 OS=Homo sa| CLASP1 | 169.451 | 3 | 56.232708 | yes | 4.179E-07 | 689.02155 | 13.930777 |
| SSSSSQESLNRPLS | 16 | Q7Z460 | CLIP-associating protein 1 OS=Homo sa| CLASP1 | 169.451 | 2 | 56.232708 | yes | 7.973E-07 | 839.42377 | 10.675946 |
| PATAAAAAAGSR | 12 | Q7Z478 | ATP-dependent RNA helicase DHX29 O | DHX29 | 155.236 | 2 | 52.186733 | yes | 9.722E-08 | 507.76999 | 1079.5269 |
| VLIDSVLR | 8 | Q7Z478 | ATP-dependent RNA helicase DHX29 O | DHX29 | 155.236 | 2 | 52.186733 | yes | 9.819E-08 | 457.78711 | 648.34235 |
| VVVVAGETGSGK | 12 | Q7Z478 | ATP-dependent RNA helicase DHX29 O | DHX29 | 155.236 | 2 | 52.186733 | yes | 9.722E-08 | 551.80878 | 643.39587 |

| | | | | | | | | | | | |
|---|---|---|---|---|---|---|---|---|---|---|---|
| IIGVINEHK | 9 | Q7Z478 | ATP-dependent RNA helicase DHX29 O! | DHX29 | 155.236 | 2 | 52.186733 | yes | 9.978E-08 | 511.80328 | 498.98679 |
| LACIVETAQGK | 11 | Q7Z478 | ATP-dependent RNA helicase DHX29 O! | DHX29 | 155.236 | 2 | 52.186733 | yes | 1.28E-07 | 595.31592 | 472.21954 |
| LTPLGQHLAALPV | 16 | Q7Z478 | ATP-dependent RNA helicase DHX29 O! | DHX29 | 155.236 | 3 | 52.186733 | yes | 9.85E-08 | 557.66992 | 368.37973 |
| CNLGSPEDFLSK | 12 | Q7Z478 | ATP-dependent RNA helicase DHX29 O! | DHX29 | 155.236 | 2 | 52.186733 | yes | 1.536E-07 | 683.81903 | 329.62201 |
| VPLEELCLHIMK | 12 | Q7Z478 | ATP-dependent RNA helicase DHX29 O! | DHX29 | 155.236 | 2 | 52.186733 | yes | 1.004E-07 | 741.39636 | 291.14926 |
| AVLVAGLYDNVG | 13 | Q7Z478 | ATP-dependent RNA helicase DHX29 O! | DHX29 | 155.236 | 2 | 52.186733 | yes | 1.74E-06 | 659.87189 | 289.12198 |
| NSLCGYQIR | 9 | Q7Z478 | ATP-dependent RNA helicase DHX29 O! | DHX29 | 155.236 | 2 | 52.186733 | yes | 5.365E-07 | 555.77167 | 262.55591 |
| ASQTLSFQEIALLK | 14 | Q7Z478 | ATP-dependent RNA helicase DHX29 O! | DHX29 | 155.236 | 2 | 52.186733 | yes | 1.532E-07 | 774.93524 | 233.47733 |
| AQVHPSSVNR | 10 | Q7Z478 | ATP-dependent RNA helicase DHX29 O! | DHX29 | 155.236 | 2 | 52.186733 | yes | 1.408E-07 | 547.7887 | 233.10512 |
| TSLLTLEDVK | 10 | Q7Z478 | ATP-dependent RNA helicase DHX29 O! | DHX29 | 155.236 | 2 | 52.186733 | yes | 2.376E-07 | 559.81879 | 175.23576 |
| SPFTTPIGR | 9 | Q7Z478 | ATP-dependent RNA helicase DHX29 O! | DHX29 | 155.236 | 2 | 52.186733 | yes | 2.684E-07 | 488.26654 | 167.5316 |
| IGACELNEPK | 10 | Q7Z478 | ATP-dependent RNA helicase DHX29 O! | DHX29 | 155.236 | 2 | 52.186733 | yes | 2.675E-07 | 565.77917 | 157.01596 |
| YLHLAAK | 7 | Q7Z478 | ATP-dependent RNA helicase DHX29 O! | DHX29 | 155.236 | 2 | 52.186733 | yes | 7.726E-07 | 408.24234 | 137.75523 |
| SLEEEEKFDPNER | 13 | Q7Z478 | ATP-dependent RNA helicase DHX29 O! | DHX29 | 155.236 | 2 | 52.186733 | yes | 1.437E-07 | 811.37067 | 132.99777 |
| LQEDGLLSNVSHV | 21 | Q7Z478 | ATP-dependent RNA helicase DHX29 O! | DHX29 | 155.236 | 3 | 52.186733 | yes | 1.021E-07 | 796.74329 | 128.21831 |
| FSTYFTHCPILR | 12 | Q7Z478 | ATP-dependent RNA helicase DHX29 O! | DHX29 | 155.236 | 2 | 52.186733 | yes | 9.722E-08 | 771.38214 | 126.73928 |
| DVWLEWSDAEK | 11 | Q7Z478 | ATP-dependent RNA helicase DHX29 O! | DHX29 | 155.236 | 2 | 52.186733 | yes | 2.515E-07 | 689.3197 | 125.06967 |
| SAEAGIAGEAQSK | 13 | Q7Z478 | ATP-dependent RNA helicase DHX29 O! | DHX29 | 155.236 | 2 | 52.186733 | yes | 3.046E-07 | 609.8017 | 119.87708 |
| TQHAILYMNPHK | 12 | Q7Z478 | ATP-dependent RNA helicase DHX29 O! | DHX29 | 155.236 | 2 | 52.186733 | yes | 1.49E-07 | 726.87445 | 97.464081 |
| SALAMADSDHLTI | 21 | Q7Z478 | ATP-dependent RNA helicase DHX29 O! | DHX29 | 155.236 | 3 | 52.186733 | yes | 1.012E-07 | 780.71582 | 96.315033 |
| DLQTHGWLLYQEI | 13 | Q7Z478 | ATP-dependent RNA helicase DHX29 O! | DHX29 | 155.236 | 2 | 52.186733 | yes | 2.158E-07 | 815.91486 | 85.437317 |
| QFLIDWVR | 8 | Q7Z478 | ATP-dependent RNA helicase DHX29 O! | DHX29 | 155.236 | 2 | 52.186733 | yes | 1.967E-07 | 538.79803 | 84.124252 |
| EMETLEDHPVFNP | 16 | Q7Z478 | ATP-dependent RNA helicase DHX29 O! | DHX29 | 155.236 | 3 | 52.186733 | yes | 9.911E-08 | 629.95868 | 59.667332 |
| KIGACELNEPK | 11 | Q7Z478 | ATP-dependent RNA helicase DHX29 O! | DHX29 | 155.236 | 2 | 52.186733 | yes | 5.253E-07 | 629.82666 | 42.392513 |
| KLQEDGLLSNVSH | 22 | Q7Z478 | ATP-dependent RNA helicase DHX29 O! | DHX29 | 155.236 | 3 | 52.186733 | yes | 1.029E-06 | 839.44165 | 24.71055 |
| YAEQQNEEEK | 10 | Q7Z478 | ATP-dependent RNA helicase DHX29 O! | DHX29 | 155.236 | 2 | 52.186733 | yes | 2.974E-06 | 634.27551 | 17.322466 |
| IKDDLTTIR | 9 | Q7Z4H7 | HAUS augmin-like complex subunit 6 OS | HAUS6 | 108.622 | 2 | 56.61076 | yes | 1.194E-07 | 537.81134 | 310.85107 |
| TPENLITEIR | 10 | Q7Z4H7 | HAUS augmin-like complex subunit 6 OS | HAUS6 | 108.622 | 2 | 56.61076 | yes | 9.722E-08 | 593.32733 | 309.99921 |
| SECIGLENQIK | 11 | Q7Z4H7 | HAUS augmin-like complex subunit 6 OS | HAUS6 | 108.622 | 2 | 56.61076 | yes | 1.224E-07 | 645.82153 | 152.08279 |
| TTIACSLGELPNLK | 14 | Q7Z4H7 | HAUS augmin-like complex subunit 6 OS | HAUS6 | 108.622 | 2 | 56.61076 | yes | 2.783E-07 | 758.90564 | 150.67943 |
| METFSPAVGNR | 11 | Q7Z4H7 | HAUS augmin-like complex subunit 6 OS | HAUS6 | 108.622 | 2 | 56.61076 | yes | 9.722E-08 | 604.79004 | 120.07598 |
| VISDCECVPQK | 11 | Q7Z4H7 | HAUS augmin-like complex subunit 6 OS | HAUS6 | 108.622 | 2 | 56.61076 | yes | 9.722E-08 | 667.80762 | 117.94929 |
| QTTPESDFNLQALI | 14 | Q7Z4H7 | HAUS augmin-like complex subunit 6 OS | HAUS6 | 108.622 | 2 | 56.61076 | yes | 6.606E-07 | 810.40485 | 93.675743 |
| FCWPPFDQK | 9 | Q7Z4H7 | HAUS augmin-like complex subunit 6 OS | HAUS6 | 108.622 | 2 | 56.61076 | yes | 1.019E-06 | 612.77893 | 60.433033 |
| IDVMGGSEEEFMK | 13 | Q7Z4H7 | HAUS augmin-like complex subunit 6 OS | HAUS6 | 108.622 | 2 | 56.61076 | yes | 9.935E-08 | 736.32581 | 57.967079 |
| QDLECLAFTK | 10 | Q7Z4H7 | HAUS augmin-like complex subunit 6 OS | HAUS6 | 108.622 | 2 | 56.61076 | yes | 2.124E-07 | 612.80011 | 50.266262 |
| HCCEWIK | 7 | Q7Z4H7 | HAUS augmin-like complex subunit 6 OS | HAUS6 | 108.622 | 2 | 56.61076 | yes | 6.076E-06 | 516.72314 | 24.035896 |
| AMQETVDALR | 10 | Q7Z4S6 | Kinesin-like protein KIF21A OS=Homo s | KIF21A | 187.18 | 2 | 48.644203 | yes | 1.12E-07 | 567.28461 | 86.308289 |
| LQMELMEYK | 9 | Q7Z4S6 | Kinesin-like protein KIF21A OS=Homo s | KIF21A | 187.18 | 2 | 48.644203 | no | 1.826E-06 | 592.78815 | 53.199089 |

| | | | | | | | | | | | |
|---|---|---|---|---|---|---|---|---|---|---|---|
| ITQLVSDQANHVL | 15 | Q7Z4S6 | Kinesin-like protein KIF21A OS=Homo s | KIF21A | 187.18 | 2 | 48.644203 | yes | 2.035E-07 | 832.95758 | 29.857418 |
| MTISNMEADMNR | 12 | Q7Z4S6 | Kinesin-like protein KIF21A OS=Homo s | KIF21A | 187.18 | 2 | 48.644203 | yes | 9.927E-05 | 706.802 | 8.9665756 |
| INHLPAETR | 9 | Q7Z569 | BRCA1-associated protein OS=Homo sap | BRAP | 67.305 | 2 | 50.120605 | yes | 2.237E-07 | 525.78815 | 152.52502 |
| ANQVLLQNK | 9 | Q7Z569 | BRCA1-associated protein OS=Homo sap | BRAP | 67.305 | 2 | 50.120605 | yes | 3.778E-07 | 514.2984 | 134.57965 |
| FVAPFNEVIEQMK | 13 | Q7Z569 | BRCA1-associated protein OS=Homo sap | BRAP | 67.305 | 2 | 50.120605 | yes | 1.355E-07 | 776.3974 | 131.21642 |
| DSTPNQYMVLIK | 12 | Q7Z569 | BRCA1-associated protein OS=Homo sap | BRAP | 67.305 | 2 | 50.120605 | yes | 1.219E-07 | 704.86066 | 125.67432 |
| DLQITEIQEQLR | 12 | Q7Z569 | BRCA1-associated protein OS=Homo sap | BRAP | 67.305 | 2 | 50.120605 | yes | 1.182E-07 | 743.39899 | 123.08212 |
| TTLASAVACLEGK | 13 | Q7Z569 | BRCA1-associated protein OS=Homo sap | BRAP | 67.305 | 2 | 50.120605 | yes | 1.512E-07 | 660.84503 | 69.62468 |
| CTVCLER | 7 | Q7Z569 | BRCA1-associated protein OS=Homo sap | BRAP | 67.305 | 2 | 50.120605 | yes | 4.682E-07 | 469.21515 | 65.967545 |
| VWDYAGDNYVHI | 12 | Q7Z569 | BRCA1-associated protein OS=Homo sap | BRAP | 67.305 | 2 | 50.120605 | yes | 9.897E-08 | 747.84167 | 63.27475 |
| DVMFYLETQQK | 11 | Q7Z569 | BRCA1-associated protein OS=Homo sap | BRAP | 67.305 | 2 | 50.120605 | yes | 2.698E-06 | 701.3396 | 33.59901 |
| WDDTTCPVCR | 10 | Q7Z569 | BRCA1-associated protein OS=Homo sap | BRAP | 67.305 | 2 | 50.120605 | yes | 6.661E-07 | 655.26862 | 20.649645 |
| ALPTQQGLQGR | 11 | Q7Z5J4 | Retinoic acid-induced protein 1 OS=Hom | RAI1 | 203.353 | 2 | 43.68368 | yes | 3.617E-07 | 584.82526 | 198.39203 |
| QYAEQGAQVPFR | 12 | Q7Z5J4 | Retinoic acid-induced protein 1 OS=Hom | RAI1 | 203.353 | 2 | 43.68368 | yes | 2.489E-07 | 697.34656 | 87.372993 |
| QPSQAGLSCDR | 11 | Q7Z5J4 | Retinoic acid-induced protein 1 OS=Hom | RAI1 | 203.353 | 2 | 43.68368 | yes | 5.066E-07 | 609.78021 | 23.838772 |
| LMDGTSQALAK | 11 | Q7Z5K2 | Wings apart-like protein homolog OS=Hc | WAPL | 132.946 | 2 | 51.14725 | yes | 4.809E-07 | 567.7948 | 115.5709 |
| IVEDDASISSCNK | 13 | Q7Z5K2 | Wings apart-like protein homolog OS=Hc | WAPL | 132.946 | 2 | 51.14725 | yes | 1.715E-07 | 719.32996 | 79.250198 |
| CLSVISLATK | 10 | Q7Z5K2 | Wings apart-like protein homolog OS=Hc | WAPL | 132.946 | 2 | 51.14725 | yes | 3.448E-07 | 546.31012 | 71.71595 |
| LLGGLDHIVDK | 11 | Q7Z5K2 | Wings apart-like protein homolog OS=Hc | WAPL | 132.946 | 2 | 51.14725 | yes | 9.722E-08 | 590.34021 | 64.47789 |
| ECVDHLSR | 8 | Q7Z5K2 | Wings apart-like protein homolog OS=Hc | WAPL | 132.946 | 2 | 51.14725 | yes | 3.148E-07 | 508.23492 | 63.515144 |
| ISHVVVEDTVVSD | 14 | Q7Z5K2 | Wings apart-like protein homolog OS=Hc | WAPL | 132.946 | 2 | 51.14725 | yes | 1.136E-07 | 763.90668 | 56.63068 |
| ASLDLMIR | 8 | Q7Z5K2 | Wings apart-like protein homolog OS=Hc | WAPL | 132.946 | 2 | 51.14725 | yes | 6.378E-07 | 459.75751 | 39.456078 |
| AHGMVAMVFK | 10 | Q7Z5K2 | Wings apart-like protein homolog OS=Hc | WAPL | 132.946 | 2 | 51.14725 | yes | 9.381E-06 | 545.78046 | 34.23365 |
| TIFVGGVPR | 9 | Q7Z5Q1 | Cytoplasmic polyadenylation element-bin | CPEB2 | 64.945 | 2 | 50.45121 | no | 1.209E-07 | 473.27945 | 1873.1101 |
| LYLCVSSPTIK | 11 | Q7Z5Q1 | Cytoplasmic polyadenylation element-bin | CPEB2 | 64.945 | 2 | 50.45121 | no | 6.97E-07 | 640.84961 | 1068.2661 |
| AVELAMIMDR | 10 | Q7Z5Q1 | Cytoplasmic polyadenylation element-bin | CPEB2 | 64.945 | 2 | 50.45121 | no | 1.027E-07 | 574.79376 | 463.60425 |
| AVELAMIMDR | 10 | Q7Z5Q1 | Cytoplasmic polyadenylation element-bin | CPEB2 | 64.945 | 2 | 50.45121 | no | 3.005E-07 | 582.7912 | 463.60425 |
| FGPLVVDWPHK | 11 | Q7Z5Q1 | Cytoplasmic polyadenylation element-bin | CPEB2 | 64.945 | 2 | 50.45121 | no | 1.43E-07 | 647.85077 | 653.97845 |
| EFHKPLVK | 8 | Q7Z5Q1 | Cytoplasmic polyadenylation element-bin | CPEB2 | 64.945 | 2 | 50.45121 | no | 1.383E-07 | 499.2951 | 599.95978 |
| KTIFVGGVPR | 10 | Q7Z5Q1 | Cytoplasmic polyadenylation element-bin | CPEB2 | 64.945 | 2 | 50.45121 | no | 1.507E-07 | 537.3269 | 186.95311 |
| FVQLQHGDIDK | 11 | Q7Z5Q1 | Cytoplasmic polyadenylation element-bin | CPEB2 | 64.945 | 2 | 50.45121 | yes | 1.347E-07 | 650.33826 | 103.02214 |
| KPFSGNVIAPPK | 12 | Q7Z5Q1 | Cytoplasmic polyadenylation element-bin | CPEB2 | 64.945 | 2 | 50.45121 | yes | 9.809E-08 | 627.86389 | 65.267654 |
| SPVSPQLQQQHQA | 22 | Q7Z5Q1 | Cytoplasmic polyadenylation element-bin | CPEB2 | 64.945 | 3 | 50.45121 | yes | 4.415E-07 | 802.08752 | 60.861916 |
| NSYNHHQPLLK | 11 | Q7Z5Q1 | Cytoplasmic polyadenylation element-bin | CPEB2 | 64.945 | 2 | 50.45121 | yes | 1.342E-06 | 675.8493 | 29.982862 |
| LLCCDIDPVLVK | 12 | Q7Z5W3 | RNA 5'-monophosphate methyltransferas | BCDIN3D | 33.201 | 2 | 41.46892 | yes | 3.494E-07 | 722.88055 | 40.213207 |
| VQDFLQR | 7 | Q7Z628 | Neuroepithelial cell-transforming gene 1 | NET1 | 67.741 | 2 | 54.919556 | no | 1.945E-07 | 453.24561 | 223.76918 |
| VLLCHGELR | 9 | Q7Z628 | Neuroepithelial cell-transforming gene 1 | NET1 | 67.741 | 2 | 54.919556 | yes | 2.441E-07 | 548.80023 | 129.23576 |
| EKDDDVVSLSSLD | 15 | Q7Z628 | Neuroepithelial cell-transforming gene 1 | NET1 | 67.741 | 2 | 54.919556 | yes | 9.722E-08 | 831.92523 | 41.357941 |
| CGSGMQMAEDSK | 12 | Q7Z628 | Neuroepithelial cell-transforming gene 1 | NET1 | 67.741 | 2 | 54.919556 | yes | 2.563E-06 | 650.75195 | 18.539137 |

| | | | | | | | | | | | |
|---|---|---|---|---|---|---|---|---|---|---|---|
| EYLEGSNLITK | 11 | Q7Z6B7 | SLIT-ROBO Rho GTPase-activating prot | SRGAP1 | 124.265 | 2 | 53.093147 | yes | 2.028E-07 | 633.83246 | 486.40848 |
| VLNELYTVMK | 10 | Q7Z6B7 | SLIT-ROBO Rho GTPase-activating prot | SRGAP1 | 124.265 | 2 | 53.093147 | yes | 9.809E-08 | 605.33105 | 452.54083 |
| SGDPVFHIR | 9 | Q7Z6B7 | SLIT-ROBO Rho GTPase-activating prot | SRGAP1 | 124.265 | 2 | 53.093147 | yes | 6.135E-07 | 514.26959 | 425.06595 |
| FMQISEDSTR | 10 | Q7Z6B7 | SLIT-ROBO Rho GTPase-activating prot | SRGAP1 | 124.265 | 2 | 53.093147 | yes | 5.641E-07 | 607.27954 | 327.03162 |
| HAPDVVLDTLEQV | 14 | Q7Z6B7 | SLIT-ROBO Rho GTPase-activating prot | SRGAP1 | 124.265 | 2 | 53.093147 | yes | 1.796E-07 | 782.42249 | 130.24937 |
| HAPDVVLDTLEQV | 14 | Q7Z6B7 | SLIT-ROBO Rho GTPase-activating prot | SRGAP1 | 124.265 | 2 | 53.093147 | yes | 1.315E-07 | 521.95074 | 130.24937 |
| HAPDVVLDTLEQV | 14 | Q7Z6B7 | SLIT-ROBO Rho GTPase-activating prot | SRGAP1 | 124.265 | 3 | 53.093147 | yes | 1.796E-07 | 782.42249 | 130.24937 |
| HAPDVVLDTLEQV | 14 | Q7Z6B7 | SLIT-ROBO Rho GTPase-activating prot | SRGAP1 | 124.265 | 3 | 53.093147 | yes | 1.315E-07 | 521.95074 | 130.24937 |
| DSGQVIPLIVESCII | 15 | Q7Z6B7 | SLIT-ROBO Rho GTPase-activating prot | SRGAP1 | 124.265 | 2 | 53.093147 | yes | 1.452E-07 | 843.44818 | 251.47412 |
| KIDSPPIR | 8 | Q7Z6B7 | SLIT-ROBO Rho GTPase-activating prot | SRGAP1 | 124.265 | 2 | 53.093147 | yes | 9.722E-08 | 463.27692 | 244.13184 |
| EIAFQLHEDLMK | 12 | Q7Z6B7 | SLIT-ROBO Rho GTPase-activating prot | SRGAP1 | 124.265 | 2 | 53.093147 | yes | 6.623E-07 | 737.37396 | 220.50726 |
| ELDGPVYEK | 9 | Q7Z6B7 | SLIT-ROBO Rho GTPase-activating prot | SRGAP1 | 124.265 | 2 | 53.093147 | yes | 2.166E-07 | 525.26111 | 204.3381 |
| GASLLLYHR | 9 | Q7Z6B7 | SLIT-ROBO Rho GTPase-activating prot | SRGAP1 | 124.265 | 2 | 53.093147 | no | 1.992E-07 | 515.29565 | 195.65056 |
| TYLSAEYNLETSR | 13 | Q7Z6B7 | SLIT-ROBO Rho GTPase-activating prot | SRGAP1 | 124.265 | 2 | 53.093147 | no | 9.722E-08 | 773.8728 | 169.32892 |
| TNPTIGPAPPPQGP | 17 | Q7Z6B7 | SLIT-ROBO Rho GTPase-activating prot | SRGAP1 | 124.265 | 2 | 53.093147 | yes | 9.722E-08 | 844.4361 | 157.06685 |
| TIIIHHETIFPDAK | 14 | Q7Z6B7 | SLIT-ROBO Rho GTPase-activating prot | SRGAP1 | 124.265 | 3 | 53.093147 | yes | 1.26E-07 | 545.63489 | 72.114342 |
| TIIIHHETIFPDAK | 14 | Q7Z6B7 | SLIT-ROBO Rho GTPase-activating prot | SRGAP1 | 124.265 | 3 | 53.093147 | yes | 1.503E-07 | 817.94867 | 72.114342 |
| TIIIHHETIFPDAK | 14 | Q7Z6B7 | SLIT-ROBO Rho GTPase-activating prot | SRGAP1 | 124.265 | 2 | 53.093147 | yes | 1.26E-07 | 545.63489 | 72.114342 |
| TIIIHHETIFPDAK | 14 | Q7Z6B7 | SLIT-ROBO Rho GTPase-activating prot | SRGAP1 | 124.265 | 2 | 53.093147 | yes | 1.503E-07 | 817.94867 | 72.114342 |
| HPDGYLAR | 8 | Q7Z6B7 | SLIT-ROBO Rho GTPase-activating prot | SRGAP1 | 124.265 | 2 | 53.093147 | yes | 3.933E-07 | 464.73541 | 142.617 |
| FNDLISCIR | 9 | Q7Z6B7 | SLIT-ROBO Rho GTPase-activating prot | SRGAP1 | 124.265 | 2 | 53.093147 | yes | 1.529E-07 | 569.28967 | 132.80771 |
| STVSETYLSKPSIA | 15 | Q7Z6B7 | SLIT-ROBO Rho GTPase-activating prot | SRGAP1 | 124.265 | 2 | 53.093147 | yes | 6.811E-07 | 805.93542 | 115.5978 |
| AQLVEQQK | 8 | Q7Z6B7 | SLIT-ROBO Rho GTPase-activating prot | SRGAP1 | 124.265 | 2 | 53.093147 | yes | 3.393E-07 | 472.26401 | 111.42368 |
| ANQQETEQFYFMI | 13 | Q7Z6B7 | SLIT-ROBO Rho GTPase-activating prot | SRGAP1 | 124.265 | 2 | 53.093147 | yes | 1.234E-07 | 832.37469 | 95.276703 |
| ADSEASSGPVTED | 14 | Q7Z6B7 | SLIT-ROBO Rho GTPase-activating prot | SRGAP1 | 124.265 | 2 | 53.093147 | yes | 9.722E-08 | 696.80988 | 82.044174 |
| IDNLYER | 7 | Q7Z6B7 | SLIT-ROBO Rho GTPase-activating prot | SRGAP1 | 124.265 | 2 | 53.093147 | yes | 3.834E-07 | 461.73508 | 64.985008 |
| TYHMYHAESISAE | 15 | Q7Z6B7 | SLIT-ROBO Rho GTPase-activating prot | SRGAP1 | 124.265 | 3 | 53.093147 | no | 1.452E-07 | 585.26654 | 47.356186 |
| KAEIETEYSR | 10 | Q7Z6B7 | SLIT-ROBO Rho GTPase-activating prot | SRGAP1 | 124.265 | 2 | 53.093147 | yes | 1.06E-07 | 613.30658 | 41.005196 |
| KLLLTLPR | 8 | Q7Z6B7 | SLIT-ROBO Rho GTPase-activating prot | SRGAP1 | 124.265 | 2 | 53.093147 | yes | 5.641E-07 | 477.32895 | 27.483284 |
| STSSSSDTMSTFK | 13 | Q7Z6B7 | SLIT-ROBO Rho GTPase-activating prot | SRGAP1 | 124.265 | 2 | 53.093147 | yes | 1.125E-07 | 683.29559 | 26.678793 |
| FINLYGLQHQGIFI | 14 | Q7Z6B7 | SLIT-ROBO Rho GTPase-activating prot | SRGAP1 | 124.265 | 3 | 53.093147 | yes | 0.0002135 | 569.31061 | 26.622286 |
| DHATLSDIYLNNV | 16 | Q7Z6B7 | SLIT-ROBO Rho GTPase-activating prot | SRGAP1 | 124.265 | 3 | 53.093147 | yes | 3.511E-06 | 625.65247 | 17.409264 |
| AEYMTTRPPNVPP | 17 | Q7Z6B7 | SLIT-ROBO Rho GTPase-activating prot | SRGAP1 | 124.265 | 3 | 53.093147 | yes | 3.698E-06 | 652.01202 | 12.926477 |
| YAIPTIDAEAYAIG | 15 | Q7Z6E9 | E3 ubiquitin-protein ligase RBBP6 OS=F | RBBP6 | 201.565 | 2 | 51.703552 | yes | 1.405E-07 | 798.41943 | 291.11118 |
| LGYLVSPPQQIR | 12 | Q7Z6E9 | E3 ubiquitin-protein ligase RBBP6 OS=F | RBBP6 | 201.565 | 2 | 51.703552 | yes | 9.722E-08 | 685.89319 | 258.09 |
| FLPLNIR | 7 | Q7Z6E9 | E3 ubiquitin-protein ligase RBBP6 OS=F | RBBP6 | 201.565 | 2 | 51.703552 | yes | 1.766E-06 | 436.77127 | 253.13582 |
| AIDDSSASISLAQL | 16 | Q7Z6E9 | E3 ubiquitin-protein ligase RBBP6 OS=F | RBBP6 | 201.565 | 2 | 51.703552 | yes | 1.084E-07 | 810.42798 | 252.26926 |
| AADCDLQITNAQT | 14 | Q7Z6E9 | E3 ubiquitin-protein ligase RBBP6 OS=F | RBBP6 | 201.565 | 2 | 51.703552 | yes | 2.605E-07 | 774.86975 | 232.873 |
| RIPIGGVK | 8 | Q7Z6E9 | E3 ubiquitin-protein ligase RBBP6 OS=F | RBBP6 | 201.565 | 2 | 51.703552 | yes | 1.257E-06 | 420.2767 | 224.63106 |

| | | | | | | | | | | | |
|---|---|---|---|---|---|---|---|---|---|---|---|
| LLYILNPPETQVEK | 14 | Q7Z6E9 | E3 ubiquitin-protein ligase RBBP6 OS=H | RBBP6 | 201.565 | 2 | 51.703552 | yes | 5.569E-07 | 828.96399 | 195.23752 |
| EDYVGGQSHR | 10 | Q7Z6E9 | E3 ubiquitin-protein ligase RBBP6 OS=H | RBBP6 | 201.565 | 2 | 51.703552 | yes | 2.038E-07 | 574.25995 | 151.25931 |
| TDSLFVLPSR | 10 | Q7Z6E9 | E3 ubiquitin-protein ligase RBBP6 OS=H | RBBP6 | 201.565 | 2 | 51.703552 | yes | 1.202E-07 | 567.81134 | 119.9737 |
| STQPISSVGK | 10 | Q7Z6E9 | E3 ubiquitin-protein ligase RBBP6 OS=H | RBBP6 | 201.565 | 2 | 51.703552 | yes | 1.049E-07 | 502.27457 | 104.84513 |
| GTSSIAITALMEEK | 14 | Q7Z6E9 | E3 ubiquitin-protein ligase RBBP6 OS=H | RBBP6 | 201.565 | 2 | 51.703552 | yes | 9.729E-08 | 725.87653 | 84.080032 |
| IGSTENISNTK | 11 | Q7Z6E9 | E3 ubiquitin-protein ligase RBBP6 OS=H | RBBP6 | 201.565 | 2 | 51.703552 | yes | 1.084E-07 | 582.29877 | 82.731064 |
| DEPMDAESITFK | 12 | Q7Z6E9 | E3 ubiquitin-protein ligase RBBP6 OS=H | RBBP6 | 201.565 | 2 | 51.703552 | yes | 2.933E-07 | 691.81085 | 68.055145 |
| GVDKDFESSSMK | 12 | Q7Z6E9 | E3 ubiquitin-protein ligase RBBP6 OS=H | RBBP6 | 201.565 | 2 | 51.703552 | yes | 3.255E-07 | 665.30322 | 60.240105 |
| DVSHEIIQHEVK | 12 | Q7Z6E9 | E3 ubiquitin-protein ligase RBBP6 OS=H | RBBP6 | 201.565 | 2 | 51.703552 | yes | 1.596E-07 | 717.3728 | 45.431469 |
| LDEFTNDFAK | 10 | Q7Z6E9 | E3 ubiquitin-protein ligase RBBP6 OS=H | RBBP6 | 201.565 | 2 | 51.703552 | yes | 1.128E-07 | 600.28259 | 42.691875 |
| NVSTKPSNIVK | 11 | Q7Z6E9 | E3 ubiquitin-protein ligase RBBP6 OS=H | RBBP6 | 201.565 | 2 | 51.703552 | yes | 1.02E-07 | 593.84314 | 28.820091 |
| ENIVKPAK | 8 | Q7Z6E9 | E3 ubiquitin-protein ligase RBBP6 OS=H | RBBP6 | 201.565 | 2 | 51.703552 | yes | 3.561E-07 | 449.77145 | 28.775608 |
| ILPAAALASEHSK | 13 | Q7Z6E9 | E3 ubiquitin-protein ligase RBBP6 OS=H | RBBP6 | 201.565 | 3 | 51.703552 | yes | 9.722E-08 | 436.58212 | 22.389709 |
| TALLESDEHTCPTC | 28 | Q7Z6E9 | E3 ubiquitin-protein ligase RBBP6 OS=H | RBBP6 | 201.565 | 4 | 51.703552 | yes | 5.216E-06 | 784.86322 | 8.3638554 |
| LPIMQMR | 7 | Q7Z6M4 | Transcription termination factor 4, mitoch | MTERF4 | 43.958 | 2 | 56.314377 | yes | 1.402E-07 | 444.74335 | 104.76057 |
| ILHSCPSVLR | 10 | Q7Z6M4 | Transcription termination factor 4, mitoch | MTERF4 | 43.958 | 2 | 56.314377 | yes | 9.722E-08 | 591.3266 | 104.32214 |
| VLYCCPEIFTMR | 12 | Q7Z6M4 | Transcription termination factor 4, mitoch | MTERF4 | 43.958 | 2 | 56.314377 | yes | 9.722E-08 | 794.86969 | 92.377922 |
| QGTPVVQGSLELE | 14 | Q7Z6M4 | Transcription termination factor 4, mitoch | MTERF4 | 43.958 | 2 | 56.314377 | yes | 1.167E-07 | 756.90448 | 73.549835 |
| GQTQIPNPLLK | 11 | Q7Z6M4 | Transcription termination factor 4, mitoch | MTERF4 | 43.958 | 2 | 56.314377 | yes | 3.763E-07 | 604.85352 | 66.780029 |
| VSEAEFLAR | 9 | Q7Z6M4 | Transcription termination factor 4, mitoch | MTERF4 | 43.958 | 2 | 56.314377 | yes | 6.859E-07 | 511.26929 | 61.775589 |
| QQDINDTVR | 9 | Q7Z6M4 | Transcription termination factor 4, mitoch | MTERF4 | 43.958 | 2 | 56.314377 | yes | 4.314E-07 | 544.7702 | 26.808153 |
| LTTASNGGVIEELS | 17 | Q7Z6M4 | Transcription termination factor 4, mitoch | MTERF4 | 43.958 | 3 | 56.314377 | yes | 2.707E-06 | 602.64026 | 8.8896351 |
| FLQFVTGTSK | 10 | Q7Z6Z7 | E3 ubiquitin-protein ligase HUWE1 OS= | HUWE1 | 481.891 | 2 | 36.33549 | yes | 6.846E-06 | 564.30841 | 30.416269 |
| LGSSGLGSASSIQA | 17 | Q7Z6Z7 | E3 ubiquitin-protein ligase HUWE1 OS= | HUWE1 | 481.891 | 2 | 36.33549 | yes | 3.033E-05 | 780.92065 | 29.482735 |
| VLQVIIQLR | 9 | Q7Z6Z7 | E3 ubiquitin-protein ligase HUWE1 OS= | HUWE1 | 481.891 | 2 | 36.33549 | yes | 2.683E-06 | 541.35822 | 26.308268 |
| ACVSMLGVPVDPI | 20 | Q7Z6Z7 | E3 ubiquitin-protein ligase HUWE1 OS= | HUWE1 | 481.891 | 3 | 36.33549 | yes | 4.288E-06 | 718.03448 | 20.332363 |
| GNDTPLALESTNT | 15 | Q7Z6Z7 | E3 ubiquitin-protein ligase HUWE1 OS= | HUWE1 | 481.891 | 2 | 36.33549 | yes | 7.111E-07 | 795.38629 | 12.968059 |
| SYSEDDIHR | 9 | Q7Z739 | YTH domain-containing family protein 3 | YTHDF3 | 63.862 | 2 | 50.704792 | no | 9.728E-08 | 561.24652 | 7513.0703 |
| GNVGIGGSAVPPP | 16 | Q7Z739 | YTH domain-containing family protein 3 | YTHDF3 | 63.862 | 2 | 50.704792 | yes | 8.737E-07 | 730.41699 | 3900.7905 |
| AITDGQAGFGNDT | 16 | Q7Z739 | YTH domain-containing family protein 3 | YTHDF3 | 63.862 | 2 | 50.704792 | yes | 1.376E-07 | 797.88904 | 3565.5598 |
| PTSWAAIAR | 9 | Q7Z739 | YTH domain-containing family protein 3 | YTHDF3 | 63.862 | 2 | 50.704792 | yes | 1.034E-07 | 486.76672 | 3286.8596 |
| VPGISSIEQGMTGL | 15 | Q7Z739 | YTH domain-containing family protein 3 | YTHDF3 | 63.862 | 2 | 50.704792 | yes | 5.471E-07 | 758.90564 | 1639.0337 |
| VPGISSIEQGMTGL | 15 | Q7Z739 | YTH domain-containing family protein 3 | YTHDF3 | 63.862 | 2 | 50.704792 | yes | 1.612E-07 | 766.90308 | 1639.0337 |
| YSIWCSTEHGNK | 12 | Q7Z739 | YTH domain-containing family protein 3 | YTHDF3 | 63.862 | 2 | 50.704792 | no | 1.002E-07 | 741.32751 | 3193.4712 |
| DTQEVPLEK | 9 | Q7Z739 | YTH domain-containing family protein 3 | YTHDF3 | 63.862 | 2 | 50.704792 | no | 2.134E-07 | 529.77185 | 2930.8416 |
| IGGDLTAAVTK | 11 | Q7Z739 | YTH domain-containing family protein 3 | YTHDF3 | 63.862 | 2 | 50.704792 | yes | 2.772E-07 | 523.29803 | 2468.376 |
| RLDAAYR | 7 | Q7Z739 | YTH domain-containing family protein 3 | YTHDF3 | 63.862 | 2 | 50.704792 | no | 1.927E-07 | 432.73795 | 1971.7218 |
| SATSVDQRPK | 10 | Q7Z739 | YTH domain-containing family protein 3 | YTHDF3 | 63.862 | 2 | 50.704792 | yes | 1.066E-07 | 565.79364 | 1823.6575 |
| HNMNIGTWDEK | 11 | Q7Z739 | YTH domain-containing family protein 3 | YTHDF3 | 63.862 | 2 | 50.704792 | yes | 2.95E-07 | 672.80371 | 676.95227 |

| | | | | | | | | | | | |
|---|---|---|---|---|---|---|---|---|---|---|---|
| HNMNIGTWDEK | 11 | Q7Z739 | YTH domain-containing family protein 3 | YTHDF3 | 63.862 | 2 | 50.704792 | yes | 3.448E-06 | 680.80115 | 676.95227 |
| AINNYNPK | 8 | Q7Z739 | YTH domain-containing family protein 3 | YTHDF3 | 63.862 | 2 | 50.704792 | yes | 2.529E-07 | 467.24307 | 743.11261 |
| HTTSIFDDFAHYEI | 14 | Q7Z739 | YTH domain-containing family protein 3 | YTHDF3 | 63.862 | 3 | 50.704792 | no | 9.935E-08 | 570.93011 | 648.29767 |
| RQEEEEAMR | 9 | Q7Z739 | YTH domain-containing family protein 3 | YTHDF3 | 63.862 | 2 | 50.704792 | yes | 1.236E-07 | 589.26697 | 355.46249 |
| QEEEEAMR | 8 | Q7Z739 | YTH domain-containing family protein 3 | YTHDF3 | 63.862 | 2 | 50.704792 | yes | 1.3E-07 | 511.2164 | 277.18338 |
| DVPNNQLR | 8 | Q7Z739 | YTH domain-containing family protein 3 | YTHDF3 | 63.862 | 2 | 50.704792 | no | 5.849E-06 | 478.25143 | 196.6209 |
| VSVQNGSIHQK | 11 | Q7Z739 | YTH domain-containing family protein 3 | YTHDF3 | 63.862 | 2 | 50.704792 | yes | 1.001E-07 | 598.82275 | 121.4741 |
| YSIWCSTEHGNKR | 13 | Q7Z739 | YTH domain-containing family protein 3 | YTHDF3 | 63.862 | 2 | 50.704792 | no | 5.641E-07 | 819.37811 | 37.018333 |
| DFDWNLK | 7 | Q7Z739 | YTH domain-containing family protein 3 | YTHDF3 | 63.862 | 2 | 50.704792 | no | 4.928E-07 | 469.22433 | 10.33147 |
| MVITAPPVTNQPV | 16 | Q7Z7C8 | Transcription initiation factor TFIID subu | TAF8 | 34.263 | 2 | 42.586742 | yes | 2.961E-07 | 846.97131 | 176.39291 |
| EIAAVFQDNR | 10 | Q7Z7H8 | Large ribosomal subunit protein uL10m C | MRPL10 | 29.283 | 2 | 49.180084 | yes | 1.71E-07 | 581.7962 | 533.2359 |
| MIAVCQNVALSAI | 15 | Q7Z7H8 | Large ribosomal subunit protein uL10m C | MRPL10 | 29.283 | 2 | 49.180084 | yes | 1.061E-07 | 824.90527 | 163.85025 |
| VFPNQVLKPFLED | 15 | Q7Z7H8 | Large ribosomal subunit protein uL10m C | MRPL10 | 29.283 | 3 | 49.180084 | yes | 1.388E-07 | 587.65759 | 136.13243 |
| QGFINYSK | 8 | Q7Z7H8 | Large ribosomal subunit protein uL10m C | MRPL10 | 29.283 | 2 | 49.180084 | yes | 1.118E-07 | 478.74545 | 47.229298 |
| CGVQSFYTPR | 10 | Q7Z7K6 | Centromere protein V OS=Homo sapiens | CENPV | 29.946 | 2 | 50.777348 | yes | 9.809E-08 | 607.78479 | 519.87836 |
| LLLDTFEYQGLVK | 13 | Q7Z7K6 | Centromere protein V OS=Homo sapiens | CENPV | 29.946 | 2 | 50.777348 | yes | 1.017E-07 | 769.92688 | 441.88809 |
| HFIVPASR | 8 | Q7Z7K6 | Centromere protein V OS=Homo sapiens | CENPV | 29.946 | 2 | 50.777348 | yes | 9.734E-08 | 463.76398 | 363.92905 |
| SQAVEKPPSEKPR | 13 | Q7Z7K6 | Centromere protein V OS=Homo sapiens | CENPV | 29.946 | 2 | 50.777348 | yes | 1.677E-06 | 726.89392 | 118.37781 |
| GAEHITTYTFNTHI | 14 | Q7Z7K6 | Centromere protein V OS=Homo sapiens | CENPV | 29.946 | 2 | 50.777348 | yes | 9.722E-08 | 810.39429 | 118.18649 |
| SMVTEEFNGSDWI | 14 | Q7Z7K6 | Centromere protein V OS=Homo sapiens | CENPV | 29.946 | 2 | 50.777348 | yes | 2.638E-07 | 829.85376 | 12.352293 |
| LLFVPCSR | 8 | Q86SF2 | N-acetylgalactosaminyltransferase 7 OS=I | GALNT7 | 75.39 | 2 | 46.188023 | yes | 1.626E-07 | 496.27332 | 100.9649 |
| GFETAYCIDSMGK | 13 | Q86SF2 | N-acetylgalactosaminyltransferase 7 OS=I | GALNT7 | 75.39 | 2 | 46.188023 | yes | 2.089E-07 | 739.81818 | 41.616482 |
| AKPLVLGPEFK | 11 | Q86SF2 | N-acetylgalactosaminyltransferase 7 OS=I | GALNT7 | 75.39 | 2 | 46.188023 | yes | 3.777E-07 | 599.86334 | 33.360249 |
| IYYMVAPSPEAMF | 13 | Q86SQ0 | Pleckstrin homology-like domain family I | PHLDB2 | 142.159 | 2 | 54.41084 | yes | 2.688E-07 | 764.37036 | 174.52844 |
| EGLYLSDTLPR | 11 | Q86SQ0 | Pleckstrin homology-like domain family I | PHLDB2 | 142.159 | 2 | 54.41084 | yes | 9.722E-08 | 632.33258 | 166.94006 |
| ELDMECALLDGEC | 14 | Q86SQ0 | Pleckstrin homology-like domain family I | PHLDB2 | 142.159 | 2 | 54.41084 | yes | 9.968E-08 | 825.87109 | 159.44041 |
| HFEDLEFQQLEHE | 15 | Q86SQ0 | Pleckstrin homology-like domain family I | PHLDB2 | 142.159 | 3 | 54.41084 | yes | 9.722E-08 | 648.63428 | 154.54869 |
| SQELAAMEETR | 11 | Q86SQ0 | Pleckstrin homology-like domain family I | PHLDB2 | 142.159 | 2 | 54.41084 | yes | 4.395E-07 | 632.79553 | 140.04999 |
| ENLTQQLLR | 9 | Q86SQ0 | Pleckstrin homology-like domain family I | PHLDB2 | 142.159 | 2 | 54.41084 | yes | 2.605E-07 | 557.81439 | 121.62349 |
| ELASENINLR | 10 | Q86SQ0 | Pleckstrin homology-like domain family I | PHLDB2 | 142.159 | 2 | 54.41084 | yes | 1.163E-07 | 579.80933 | 114.36441 |
| GFPVNPNTLK | 10 | Q86SQ0 | Pleckstrin homology-like domain family I | PHLDB2 | 142.159 | 2 | 54.41084 | yes | 3.062E-07 | 543.80072 | 104.10257 |
| SPSPLGTSVR | 10 | Q86SQ0 | Pleckstrin homology-like domain family I | PHLDB2 | 142.159 | 2 | 54.41084 | yes | 1.188E-06 | 500.77472 | 101.63212 |
| ADFDHYTGR | 9 | Q86SQ0 | Pleckstrin homology-like domain family I | PHLDB2 | 142.159 | 2 | 54.41084 | yes | 1.766E-06 | 541.23853 | 101.40247 |
| DDLMDYHR | 8 | Q86SQ0 | Pleckstrin homology-like domain family I | PHLDB2 | 142.159 | 2 | 54.41084 | yes | 3.133E-07 | 532.72693 | 100.00266 |
| LSTGTTVEDVQK | 12 | Q86SQ0 | Pleckstrin homology-like domain family I | PHLDB2 | 142.159 | 2 | 54.41084 | yes | 5.835E-07 | 639.33282 | 91.641083 |
| SGAASMPSSPK | 11 | Q86SQ0 | Pleckstrin homology-like domain family I | PHLDB2 | 142.159 | 2 | 54.41084 | yes | 1.684E-07 | 510.24493 | 82.975418 |
| TPPPPSSTFPK | 11 | Q86SQ0 | Pleckstrin homology-like domain family I | PHLDB2 | 142.159 | 2 | 54.41084 | yes | 1.276E-07 | 578.30585 | 82.429916 |
| TQLDNCPESMR | 11 | Q86SQ0 | Pleckstrin homology-like domain family I | PHLDB2 | 142.159 | 2 | 54.41084 | yes | 1.165E-07 | 675.79248 | 57.628769 |
| MSIQDSLALQPK | 12 | Q86SQ0 | Pleckstrin homology-like domain family I | PHLDB2 | 142.159 | 2 | 54.41084 | yes | 9.722E-08 | 665.85541 | 53.563431 |

| | | | | | | | | | | | |
|---|---|---|---|---|---|---|---|---|---|---|---|
| SHDNVYSLGGLEC | 14 | Q86SQ0 | Pleckstrin homology-like domain family I | PHLDB2 | 142.159 | 2 | 54.41084 | yes | 2.249E-07 | 752.36298 | 43.30146 |
| EKENLCNLEK | 10 | Q86SQ0 | Pleckstrin homology-like domain family I | PHLDB2 | 142.159 | 2 | 54.41084 | yes | 7.143E-06 | 638.81372 | 31.182905 |
| SPNPLLTFSVK | 11 | Q86SQ0 | Pleckstrin homology-like domain family I | PHLDB2 | 142.159 | 2 | 54.41084 | yes | 9.722E-08 | 601.84259 | 30.462818 |
| VCQSLINVEGK | 11 | Q86T03 | Type 1 phosphatidylinositol 4,5-bisphosp | PIP4P1 | 29.47 | 2 | 33.713676 | yes | 1.168E-06 | 623.82666 | 35.577995 |
| FIAELGVPLSQVK | 13 | Q86T24 | Transcriptional regulator Kaiso OS=Hom | ZBTB33 | 74.485 | 2 | 31.367937 | yes | 1.865E-06 | 700.91101 | 44.84301 |
| NVVLRPSGAK | 10 | Q86T82 | Ubiquitin carboxyl-terminal hydrolase 37 | USP37 | 110.17 | 2 | 38.395695 | yes | 3.311E-06 | 520.81421 | 10.62308 |
| VLILHLK | 7 | Q86T82 | Ubiquitin carboxyl-terminal hydrolase 37 | USP37 | 110.17 | 2 | 38.395695 | yes | 5.251E-07 | 418.29184 | 7.1222034 |
| MVIENELEDPAIM | 14 | Q86TB9 | Protein PAT1 homolog 1 OS=Homo sapi | PATL1 | 86.851 | 2 | 54.00473 | yes | 9.722E-08 | 830.40765 | 690.9574 |
| MVIENELEDPAIM | 14 | Q86TB9 | Protein PAT1 homolog 1 OS=Homo sapi | PATL1 | 86.851 | 2 | 54.00473 | yes | 1.089E-06 | 838.40515 | 690.9574 |
| MVIENELEDPAIM | 14 | Q86TB9 | Protein PAT1 homolog 1 OS=Homo sapi | PATL1 | 86.851 | 2 | 54.00473 | yes | 1.89E-07 | 838.40515 | 690.9574 |
| ICSMYDNLR | 9 | Q86TB9 | Protein PAT1 homolog 1 OS=Homo sapi | PATL1 | 86.851 | 2 | 54.00473 | yes | 9.722E-08 | 586.26538 | 1103.5013 |
| LSAAEEIQGDGPK | 13 | Q86TB9 | Protein PAT1 homolog 1 OS=Homo sapi | PATL1 | 86.851 | 2 | 54.00473 | yes | 9.722E-08 | 657.83044 | 1069.653 |
| MIDAVVTSR | 9 | Q86TB9 | Protein PAT1 homolog 1 OS=Homo sapi | PATL1 | 86.851 | 2 | 54.00473 | yes | 1.01E-07 | 496.26569 | 935.42291 |
| STSPIIGSPPVR | 12 | Q86TB9 | Protein PAT1 homolog 1 OS=Homo sapi | PATL1 | 86.851 | 2 | 54.00473 | yes | 3.707E-07 | 605.84314 | 832.5022 |
| KDPYANLMLQR | 11 | Q86TB9 | Protein PAT1 homolog 1 OS=Homo sapi | PATL1 | 86.851 | 2 | 54.00473 | yes | 9.722E-08 | 674.85571 | 706.12952 |
| LTVSSVNNPR | 10 | Q86TB9 | Protein PAT1 homolog 1 OS=Homo sapi | PATL1 | 86.851 | 2 | 54.00473 | yes | 4.353E-07 | 543.79877 | 702.94165 |
| TYSLLLDVEDYER | 13 | Q86TB9 | Protein PAT1 homolog 1 OS=Homo sapi | PATL1 | 86.851 | 2 | 54.00473 | yes | 9.849E-08 | 808.39612 | 680.99341 |
| LQLVQGIR | 8 | Q86TB9 | Protein PAT1 homolog 1 OS=Homo sapi | PATL1 | 86.851 | 2 | 54.00473 | yes | 2.592E-07 | 463.79272 | 544.5885 |
| LEHAYKPVQFEGS | 16 | Q86TB9 | Protein PAT1 homolog 1 OS=Homo sapi | PATL1 | 86.851 | 3 | 54.00473 | yes | 3.053E-07 | 601.65271 | 320.52432 |
| LAELEEK | 7 | Q86TB9 | Protein PAT1 homolog 1 OS=Homo sapi | PATL1 | 86.851 | 2 | 54.00473 | yes | 3.189E-07 | 416.22656 | 203.56581 |
| AQLLGGAQLQPGF | 13 | Q86TB9 | Protein PAT1 homolog 1 OS=Homo sapi | PATL1 | 86.851 | 2 | 54.00473 | yes | 1.317E-07 | 654.87274 | 115.91089 |
| LNLLETK | 7 | Q86TB9 | Protein PAT1 homolog 1 OS=Homo sapi | PATL1 | 86.851 | 2 | 54.00473 | yes | 9.722E-08 | 415.75275 | 94.633965 |
| LPVAVNEQTGNGI | 14 | Q86TB9 | Protein PAT1 homolog 1 OS=Homo sapi | PATL1 | 86.851 | 2 | 54.00473 | yes | 2.994E-05 | 742.3786 | 34.747437 |
| PSDDHFVQIMCIR | 13 | Q86TB9 | Protein PAT1 homolog 1 OS=Homo sapi | PATL1 | 86.851 | 2 | 54.00473 | yes | 1.463E-06 | 809.37927 | 28.801224 |
| KTLVIIEK | 8 | Q86TB9 | Protein PAT1 homolog 1 OS=Homo sapi | PATL1 | 86.851 | 2 | 54.00473 | yes | 3.155E-06 | 472.31296 | 27.761612 |
| GKLPGQERPSDDH | 21 | Q86TB9 | Protein PAT1 homolog 1 OS=Homo sapi | PATL1 | 86.851 | 4 | 54.00473 | yes | 3.41E-06 | 621.5625 | 17.414587 |
| TLVIIEK | 7 | Q86TB9 | Protein PAT1 homolog 1 OS=Homo sapi | PATL1 | 86.851 | 2 | 54.00473 | yes | 4E-07 | 408.26547 | 16.258224 |
| EGTLIEDSPDFR | 12 | Q86TC9 | Myopalladin OS=Homo sapiens OX=960 | MYPN | 145.258 | 2 | 53.077774 | yes | 3.266E-07 | 689.82788 | 134.9467 |
| LMNEIEFR | 8 | Q86TC9 | Myopalladin OS=Homo sapiens OX=960 | MYPN | 145.258 | 2 | 53.077774 | yes | 1.541E-07 | 526.26569 | 115.09112 |
| VTEGSPVTFTCK | 12 | Q86TC9 | Myopalladin OS=Homo sapiens OX=960 | MYPN | 145.258 | 2 | 53.077774 | yes | 9.722E-08 | 663.32391 | 111.14027 |
| SAPSMPSQGLAK | 12 | Q86TC9 | Myopalladin OS=Homo sapiens OX=960 | MYPN | 145.258 | 2 | 53.077774 | yes | 5.501E-07 | 587.30023 | 73.935905 |
| KAPVILEK | 8 | Q86TC9 | Myopalladin OS=Homo sapiens OX=960 | MYPN | 145.258 | 2 | 53.077774 | yes | 1.722E-07 | 449.29202 | 65.201874 |
| CIAPIFDK | 8 | Q86TC9 | Myopalladin OS=Homo sapiens OX=960 | MYPN | 145.258 | 2 | 53.077774 | yes | 1.14E-07 | 482.25204 | 49.514599 |
| DKEPLQER | 8 | Q86TC9 | Myopalladin OS=Homo sapiens OX=960 | MYPN | 145.258 | 2 | 53.077774 | yes | 3.493E-07 | 507.76437 | 38.757381 |
| TGQNSFSLELSVV | 15 | Q86TC9 | Myopalladin OS=Homo sapiens OX=960 | MYPN | 145.258 | 2 | 53.077774 | yes | 4.051E-07 | 790.41992 | 32.104893 |
| SIVFDSEYCR | 10 | Q86TG7 | Retrotransposon-derived protein PEG10 ( | PEG10 | 80.173 | 2 | 54.648796 | yes | 7.523E-07 | 638.28735 | 3976.2808 |
| LTEENTTLR | 9 | Q86TG7 | Retrotransposon-derived protein PEG10 ( | PEG10 | 80.173 | 2 | 54.648796 | yes | 1.66E-07 | 538.78278 | 3515.4158 |
| HVFEDPQR | 8 | Q86TG7 | Retrotransposon-derived protein PEG10 ( | PEG10 | 80.173 | 2 | 54.648796 | yes | 1.696E-07 | 514.2514 | 1786.4016 |
| APNNFTIQNQYPR | 13 | Q86TG7 | Retrotransposon-derived protein PEG10 ( | PEG10 | 80.173 | 2 | 54.648796 | yes | 1.11E-07 | 781.88916 | 1412.1344 |

| | | | | | | | | | | | |
|---|---|---|---|---|---|---|---|---|---|---|---|
| LQVSYDCR | 8 | Q86TG7 | Retrotransposon-derived protein PEG10 ( | PEG10 | 80.173 | 2 | 54.648796 | yes | 2.474E-07 | 520.74512 | 950.96313 |
| VCFVTSMMTGR | 11 | Q86TG7 | Retrotransposon-derived protein PEG10 ( | PEG10 | 80.173 | 2 | 54.648796 | yes | 1.276E-07 | 644.79596 | 833.26837 |
| DELSEEINNLR | 11 | Q86TG7 | Retrotransposon-derived protein PEG10 ( | PEG10 | 80.173 | 2 | 54.648796 | yes | 9.742E-08 | 666.3255 | 692.06189 |
| PIASGPVVHETHDI | 22 | Q86TG7 | Retrotransposon-derived protein PEG10 ( | PEG10 | 80.173 | 4 | 54.648796 | yes | 1.281E-06 | 595.06195 | 133.39532 |
| PIASGPVVHETHDI | 22 | Q86TG7 | Retrotransposon-derived protein PEG10 ( | PEG10 | 80.173 | 4 | 54.648796 | yes | 9.722E-08 | 793.08014 | 133.39532 |
| PIASGPVVHETHDI | 22 | Q86TG7 | Retrotransposon-derived protein PEG10 ( | PEG10 | 80.173 | 3 | 54.648796 | yes | 1.281E-06 | 595.06195 | 133.39532 |
| PIASGPVVHETHDI | 22 | Q86TG7 | Retrotransposon-derived protein PEG10 ( | PEG10 | 80.173 | 3 | 54.648796 | yes | 9.722E-08 | 793.08014 | 133.39532 |
| SPQDDASSPHLQV | 22 | Q86TG7 | Retrotransposon-derived protein PEG10 ( | PEG10 | 80.173 | 4 | 54.648796 | yes | 1.719E-07 | 607.31158 | 130.8925 |
| SPQDDASSPHLQV | 22 | Q86TG7 | Retrotransposon-derived protein PEG10 ( | PEG10 | 80.173 | 4 | 54.648796 | yes | 1.224E-07 | 809.41302 | 130.8925 |
| SPQDDASSPHLQV | 22 | Q86TG7 | Retrotransposon-derived protein PEG10 ( | PEG10 | 80.173 | 3 | 54.648796 | yes | 1.719E-07 | 607.31158 | 130.8925 |
| SPQDDASSPHLQV | 22 | Q86TG7 | Retrotransposon-derived protein PEG10 ( | PEG10 | 80.173 | 3 | 54.648796 | yes | 1.224E-07 | 809.41302 | 130.8925 |
| QSEENNNLQSQVC | 14 | Q86TG7 | Retrotransposon-derived protein PEG10 ( | PEG10 | 80.173 | 2 | 54.648796 | yes | 9.722E-08 | 823.39246 | 235.79962 |
| KLNLCLYCGTGGH | 21 | Q86TG7 | Retrotransposon-derived protein PEG10 ( | PEG10 | 80.173 | 3 | 54.648796 | yes | 1.414E-07 | 804.70129 | 191.96208 |
| ALVLPHIASHHQV | 23 | Q86TG7 | Retrotransposon-derived protein PEG10 ( | PEG10 | 80.173 | 4 | 54.648796 | yes | 3.113E-06 | 601.07385 | 178.30309 |
| IKDWPILVEAIDGF | 14 | Q86TG7 | Retrotransposon-derived protein PEG10 ( | PEG10 | 80.173 | 2 | 54.648796 | yes | 9.729E-08 | 812.95648 | 172.09387 |
| RDELSEEINNLR | 12 | Q86TG7 | Retrotransposon-derived protein PEG10 ( | PEG10 | 80.173 | 2 | 54.648796 | yes | 9.722E-08 | 744.3761 | 82.977356 |
| IKDWPILVEAIDGF | 36 | Q86TG7 | Retrotransposon-derived protein PEG10 ( | PEG10 | 80.173 | 5 | 54.648796 | yes | 1.006E-07 | 797.42859 | 77.941986 |
| LVDPHIEMIPGAHS | 32 | Q86TG7 | Retrotransposon-derived protein PEG10 ( | PEG10 | 80.173 | 5 | 54.648796 | yes | 2.627E-07 | 691.5517 | 40.573669 |
| LNLCLYCGTGGHY | 20 | Q86TG7 | Retrotransposon-derived protein PEG10 ( | PEG10 | 80.173 | 3 | 54.648796 | yes | 5.999E-07 | 762.00299 | 9.2670975 |
| AHVDMYVR | 8 | Q86TJ2 | Transcriptional adapter 2-beta OS=Homo | TADA2B | 48.471 | 2 | 50.671425 | yes | 3.189E-06 | 495.74493 | 104.13079 |
| EFDDLFENMHK | 11 | Q86TJ2 | Transcriptional adapter 2-beta OS=Homo | TADA2B | 48.471 | 2 | 50.671425 | yes | 9.728E-08 | 712.81116 | 74.032516 |
| VLCSSLNLSPAR | 12 | Q86TJ2 | Transcriptional adapter 2-beta OS=Homo | TADA2B | 48.471 | 2 | 50.671425 | yes | 1.028E-06 | 658.85321 | 64.360817 |
| ILNFLTESGWISR | 13 | Q86TJ2 | Transcriptional adapter 2-beta OS=Homo | TADA2B | 48.471 | 2 | 50.671425 | yes | 3.201E-06 | 768.41449 | 29.139391 |
| GYGFITFSDSECAF | 14 | Q86U06 | Probable RNA-binding protein 23 OS=H | RBM23 | 48.731 | 2 | 49.372196 | yes | 2.856E-07 | 805.3512 | 148.15604 |
| SPPLATGYR | 9 | Q86U06 | Probable RNA-binding protein 23 OS=H | RBM23 | 48.731 | 2 | 49.372196 | yes | 1.862E-07 | 481.25873 | 141.84169 |
| LAAMANNLQK | 10 | Q86U06 | Probable RNA-binding protein 23 OS=H | RBM23 | 48.731 | 2 | 49.372196 | yes | 3.566E-07 | 537.29224 | 131.79172 |
| LLGVPIIVQASQAE | 15 | Q86U06 | Probable RNA-binding protein 23 OS=H | RBM23 | 48.731 | 2 | 49.372196 | yes | 3.735E-07 | 783.46667 | 82.014084 |
| DLEDFFSAVGK | 11 | Q86U06 | Probable RNA-binding protein 23 OS=H | RBM23 | 48.731 | 2 | 49.372196 | yes | 1.04E-07 | 614.29822 | 57.096531 |
| IDNIVLMK | 8 | Q86U06 | Probable RNA-binding protein 23 OS=H | RBM23 | 48.731 | 2 | 49.372196 | yes | 0.0002638 | 473.27551 | 27.659369 |
| TSLALDESLFR | 11 | Q86U42 | Polyadenylate-binding protein 2 OS=Hon | PABPN1 | 32.75 | 2 | 49.000744 | yes | 1.078E-07 | 626.33258 | 2212.1106 |
| GFAYIEFSDK | 10 | Q86U42 | Polyadenylate-binding protein 2 OS=Hon | PABPN1 | 32.75 | 2 | 49.000744 | yes | 1.517E-07 | 588.78223 | 608.51276 |
| ELQNEVEK | 8 | Q86U42 | Polyadenylate-binding protein 2 OS=Hon | PABPN1 | 32.75 | 2 | 49.000744 | yes | 3.268E-07 | 494.75092 | 486.3959 |
| MVAPPAEPTR | 10 | Q86U70 | LIM domain-binding protein 1 OS=Homo | LDB1 | 46.533 | 2 | 52.28215 | no | 3.884E-07 | 534.77893 | 350.10031 |
| PMFTQVCVEGR | 11 | Q86U70 | LIM domain-binding protein 1 OS=Homo | LDB1 | 46.533 | 2 | 52.28215 | yes | 9.722E-08 | 662.31287 | 90.804901 |
| CGLSNSTLNYLR | 12 | Q86U70 | LIM domain-binding protein 1 OS=Homo | LDB1 | 46.533 | 2 | 52.28215 | yes | 1.897E-07 | 699.3457 | 59.980103 |
| SVGCACPGCSSK | 12 | Q86U70 | LIM domain-binding protein 1 OS=Homo | LDB1 | 46.533 | 2 | 52.28215 | yes | 1.759E-07 | 656.25995 | 39.512821 |
| LLCELFIR | 8 | Q86U86 | Protein polybromo-1 OS=Homo sapiens ( | PBRM1 | 192.948 | 2 | 40.986515 | yes | 3.736E-07 | 532.30206 | 99.802574 |
| SELPDYYLTIK | 11 | Q86U86 | Protein polybromo-1 OS=Homo sapiens ( | PBRM1 | 192.948 | 2 | 40.986515 | yes | 1.533E-06 | 671.35046 | 51.767971 |
| MPISLQQIR | 9 | Q86U86 | Protein polybromo-1 OS=Homo sapiens ( | PBRM1 | 192.948 | 2 | 40.986515 | yes | 5.178E-07 | 543.31042 | 36.275845 |

| | | | | | | | | | | | |
|---|---|---|---|---|---|---|---|---|---|---|---|
| LWDLYLR | 7 | Q86U86 | Protein polybromo-1 OS=Homo sapiens ( | PBRM1 | 192.948 | 2 | 40.986515 | yes | 1.228E-06 | 489.77402 | 25.765995 |
| LGSTVVDLSVPGK | 13 | Q86U90 | Threonylcarbamoyl-AMP synthase OS=H | YRDC | 29.328 | 2 | 33.38609 | yes | 1.249E-06 | 636.36389 | 65.137238 |
| PLAVCLGR | 8 | Q86U90 | Threonylcarbamoyl-AMP synthase OS=H | YRDC | 29.328 | 2 | 33.38609 | yes | 2.861E-06 | 443.25238 | 62.101578 |
| LPGSGAVQAASPE | 14 | Q86U90 | Threonylcarbamoyl-AMP synthase OS=H | YRDC | 29.328 | 2 | 33.38609 | yes | 3.543E-06 | 670.35187 | 50.305264 |
| TVEVAEGEAVR | 11 | Q86UE4 | Protein LYRIC OS=Homo sapiens OX=9 | MTDH | 63.837 | 2 | 54.67671 | yes | 1.146E-07 | 580.30133 | 5224.9189 |
| WNSVSPASAGK | 11 | Q86UE4 | Protein LYRIC OS=Homo sapiens OX=9 | MTDH | 63.837 | 2 | 54.67671 | yes | 2.093E-07 | 552.27765 | 2326.5854 |
| EDLPVNTSK | 9 | Q86UE4 | Protein LYRIC OS=Homo sapiens OX=9 | MTDH | 63.837 | 2 | 54.67671 | yes | 1.897E-07 | 501.75876 | 1974.2355 |
| TISTSDPAEVLVK | 13 | Q86UE4 | Protein LYRIC OS=Homo sapiens OX=9 | MTDH | 63.837 | 2 | 54.67671 | yes | 1.657E-07 | 680.37195 | 1847.8931 |
| TELGLDLGLEPK | 12 | Q86UE4 | Protein LYRIC OS=Homo sapiens OX=9 | MTDH | 63.837 | 2 | 54.67671 | yes | 3.11E-07 | 642.8559 | 1386.5486 |
| SSSQVPPILQETDK | 14 | Q86UE4 | Protein LYRIC OS=Homo sapiens OX=9 | MTDH | 63.837 | 2 | 54.67671 | yes | 2.847E-06 | 764.8963 | 1151.8361 |
| LSSQISAGEEK | 11 | Q86UE4 | Protein LYRIC OS=Homo sapiens OX=9 | MTDH | 63.837 | 2 | 54.67671 | yes | 9.722E-08 | 574.79333 | 1113.7936 |
| EMLSVGLGFLR | 11 | Q86UE4 | Protein LYRIC OS=Homo sapiens OX=9 | MTDH | 63.837 | 2 | 54.67671 | yes | 4.893E-07 | 611.33667 | 388.62146 |
| EMLSVGLGFLR | 11 | Q86UE4 | Protein LYRIC OS=Homo sapiens OX=9 | MTDH | 63.837 | 2 | 54.67671 | yes | 9.722E-08 | 619.33411 | 388.62146 |
| GDSHLNVQVSNFK | 13 | Q86UE4 | Protein LYRIC OS=Homo sapiens OX=9 | MTDH | 63.837 | 3 | 54.67671 | yes | 9.722E-08 | 482.24417 | 329.56592 |
| GDSHLNVQVSNFK | 13 | Q86UE4 | Protein LYRIC OS=Homo sapiens OX=9 | MTDH | 63.837 | 3 | 54.67671 | yes | 1.238E-07 | 722.86261 | 329.56592 |
| GDSHLNVQVSNFK | 13 | Q86UE4 | Protein LYRIC OS=Homo sapiens OX=9 | MTDH | 63.837 | 2 | 54.67671 | yes | 9.722E-08 | 482.24417 | 329.56592 |
| GDSHLNVQVSNFK | 13 | Q86UE4 | Protein LYRIC OS=Homo sapiens OX=9 | MTDH | 63.837 | 2 | 54.67671 | yes | 1.238E-07 | 722.86261 | 329.56592 |
| SQEPIPDDQK | 10 | Q86UE4 | Protein LYRIC OS=Homo sapiens OX=9 | MTDH | 63.837 | 2 | 54.67671 | yes | 9.722E-08 | 578.77765 | 151.83835 |
| SETSWESPK | 9 | Q86UE4 | Protein LYRIC OS=Homo sapiens OX=9 | MTDH | 63.837 | 2 | 54.67671 | yes | 1.09E-07 | 525.74054 | 147.89302 |
| QNSVPPSQTK | 10 | Q86UE4 | Protein LYRIC OS=Homo sapiens OX=9 | MTDH | 63.837 | 2 | 54.67671 | yes | 3.296E-06 | 543.2829 | 97.49778 |
| DDDFPSLQAIAR | 12 | Q86UK7 | E3 ubiquitin-protein ligase ZNF598 OS=I | ZNF598 | 98.638 | 2 | 53.84738 | yes | 1.41E-07 | 674.33063 | 1008.0638 |
| DLLGENFQK | 9 | Q86UK7 | E3 ubiquitin-protein ligase ZNF598 OS=I | ZNF598 | 98.638 | 2 | 53.84738 | yes | 1.952E-07 | 532.27454 | 952.16888 |
| NLQLIQSIR | 9 | Q86UK7 | E3 ubiquitin-protein ligase ZNF598 OS=I | ZNF598 | 98.638 | 2 | 53.84738 | yes | 1.902E-07 | 542.82727 | 812.75415 |
| SAWQATTQQAGL | 15 | Q86UK7 | E3 ubiquitin-protein ligase ZNF598 OS=I | ZNF598 | 98.638 | 2 | 53.84738 | yes | 2.379E-07 | 846.89154 | 683.35187 |
| MPPPPGFSAVVLL | 14 | Q86UK7 | E3 ubiquitin-protein ligase ZNF598 OS=I | ZNF598 | 98.638 | 2 | 53.84738 | yes | 1.456E-07 | 726.91779 | 668.71918 |
| QGLISAAQYYK | 11 | Q86UK7 | E3 ubiquitin-protein ligase ZNF598 OS=I | ZNF598 | 98.638 | 2 | 53.84738 | yes | 2.16E-07 | 621.3299 | 525.42755 |
| KYDIYFADGK | 10 | Q86UK7 | E3 ubiquitin-protein ligase ZNF598 OS=I | ZNF598 | 98.638 | 2 | 53.84738 | yes | 2.647E-07 | 610.30334 | 468.38751 |
| HFLCEEGR | 8 | Q86UK7 | E3 ubiquitin-protein ligase ZNF598 OS=I | ZNF598 | 98.638 | 2 | 53.84738 | yes | 1.121E-07 | 524.23749 | 447.92599 |
| CSTEQFTHAFR | 11 | Q86UK7 | E3 ubiquitin-protein ligase ZNF598 OS=I | ZNF598 | 98.638 | 2 | 53.84738 | yes | 9.742E-08 | 692.30914 | 334.48227 |
| HIDLQFSYAPR | 11 | Q86UK7 | E3 ubiquitin-protein ligase ZNF598 OS=I | ZNF598 | 98.638 | 2 | 53.84738 | yes | 9.722E-08 | 673.84619 | 334.258 |
| QLLQHECPR | 9 | Q86UK7 | E3 ubiquitin-protein ligase ZNF598 OS=I | ZNF598 | 98.638 | 2 | 53.84738 | yes | 9.916E-08 | 590.79822 | 296.56079 |
| QHELFCCR | 8 | Q86UK7 | E3 ubiquitin-protein ligase ZNF598 OS=I | ZNF598 | 98.638 | 2 | 53.84738 | yes | 1.014E-07 | 575.25006 | 295.05804 |
| CDHPVCYR | 8 | Q86UK7 | E3 ubiquitin-protein ligase ZNF598 OS=I | ZNF598 | 98.638 | 2 | 53.84738 | yes | 9.722E-08 | 553.72894 | 266.45615 |
| DFLQSDEAR | 9 | Q86UK7 | E3 ubiquitin-protein ligase ZNF598 OS=I | ZNF598 | 98.638 | 2 | 53.84738 | yes | 2.247E-07 | 540.75146 | 228.14481 |
| ASVAAQQQEEAR | 12 | Q86UK7 | E3 ubiquitin-protein ligase ZNF598 OS=I | ZNF598 | 98.638 | 2 | 53.84738 | yes | 1.087E-07 | 644.31799 | 164.63971 |
| KLPAFATIPIHQLQ | 17 | Q86UK7 | E3 ubiquitin-protein ligase ZNF598 OS=I | ZNF598 | 98.638 | 3 | 53.84738 | yes | 9.757E-08 | 657.71069 | 158.0258 |
| NEGVVGGEDYEE | 15 | Q86UK7 | E3 ubiquitin-protein ligase ZNF598 OS=I | ZNF598 | 98.638 | 2 | 53.84738 | yes | 9.722E-08 | 833.86316 | 136.12408 |
| VGSEKPGTTLPQP | 32 | Q86UK7 | E3 ubiquitin-protein ligase ZNF598 OS=I | ZNF598 | 98.638 | 4 | 53.84738 | yes | 1.087E-06 | 795.6593 | 113.51236 |
| EKHFLCEEGR | 10 | Q86UK7 | E3 ubiquitin-protein ligase ZNF598 OS=I | ZNF598 | 98.638 | 2 | 53.84738 | yes | 1.088E-07 | 652.80621 | 42.722233 |

| | | | | | | | | | | | |
|---|---|---|---|---|---|---|---|---|---|---|---|
| QQELLSAHTDFCN | 14 | Q86UK7 | E3 ubiquitin-protein ligase ZNF598 OS=l | ZNF598 | 98.638 | 3 | 53.84738 | yes | 1.593E-07 | 573.60199 | 31.810263 |
| VCPTCQQVLAHGI | 24 | Q86UK7 | E3 ubiquitin-protein ligase ZNF598 OS=l | ZNF598 | 98.638 | 4 | 53.84738 | yes | 7.651E-07 | 654.31451 | 28.015326 |
| KVAQPPLSAQATC | 19 | Q86UK7 | E3 ubiquitin-protein ligase ZNF598 OS=l | ZNF598 | 98.638 | 3 | 53.84738 | yes | 9.56E-07 | 632.00952 | 15.602623 |
| EADEDAVQFANR | 12 | Q86UL3 | Glycerol-3-phosphate acyltransferase 4 O | GPAT4 | 52.072 | 2 | 52.41138 | yes | 9.722E-08 | 682.8075 | 78.331993 |
| EQMEAEIAHLK | 11 | Q86UP2 | Kinectin OS=Homo sapiens OX=9606 Gl | KTN1 | 156.276 | 2 | 54.640892 | yes | 9.722E-08 | 649.8241 | 2275.3044 |
| VQELQNLLK | 9 | Q86UP2 | Kinectin OS=Homo sapiens OX=9606 Gl | KTN1 | 156.276 | 2 | 54.640892 | yes | 1.454E-07 | 542.82166 | 2033.2303 |
| ENEVQSLHSK | 10 | Q86UP2 | Kinectin OS=Homo sapiens OX=9606 Gl | KTN1 | 156.276 | 2 | 54.640892 | yes | 9.722E-08 | 585.79114 | 1873.2781 |
| GELTTLIHQLQEK | 13 | Q86UP2 | Kinectin OS=Homo sapiens OX=9606 Gl | KTN1 | 156.276 | 2 | 54.640892 | yes | 1.154E-07 | 755.41718 | 1854.0737 |
| LQTLVSEQPNK | 11 | Q86UP2 | Kinectin OS=Homo sapiens OX=9606 Gl | KTN1 | 156.276 | 2 | 54.640892 | yes | 9.722E-08 | 628.84589 | 1803.0166 |
| QQQVEAVELEAK | 12 | Q86UP2 | Kinectin OS=Homo sapiens OX=9606 Gl | KTN1 | 156.276 | 2 | 54.640892 | yes | 9.722E-08 | 686.35938 | 1726.5773 |
| AQQSLELIQSK | 11 | Q86UP2 | Kinectin OS=Homo sapiens OX=9606 Gl | KTN1 | 156.276 | 2 | 54.640892 | yes | 1.022E-07 | 622.84589 | 1703.8634 |
| LMQLMESEQK | 10 | Q86UP2 | Kinectin OS=Homo sapiens OX=9606 Gl | KTN1 | 156.276 | 2 | 54.640892 | yes | 2.086E-07 | 618.80176 | 1634.5587 |
| WLQDLQEENESLk | 13 | Q86UP2 | Kinectin OS=Homo sapiens OX=9606 Gl | KTN1 | 156.276 | 2 | 54.640892 | yes | 9.926E-08 | 816.39923 | 1548.5198 |
| VQLQEAER | 8 | Q86UP2 | Kinectin OS=Homo sapiens OX=9606 Gl | KTN1 | 156.276 | 2 | 54.640892 | yes | 1.041E-07 | 486.75909 | 1545.5714 |
| LQALANEQAAAAl | 17 | Q86UP2 | Kinectin OS=Homo sapiens OX=9606 Gl | KTN1 | 156.276 | 3 | 54.640892 | yes | 1.073E-07 | 602.98303 | 1412.9371 |
| EEIGNVQLEK | 10 | Q86UP2 | Kinectin OS=Homo sapiens OX=9606 Gl | KTN1 | 156.276 | 2 | 54.640892 | yes | 4.054E-07 | 579.80371 | 1286.954 |
| TQLLQDVQDENK | 12 | Q86UP2 | Kinectin OS=Homo sapiens OX=9606 Gl | KTN1 | 156.276 | 2 | 54.640892 | yes | 9.919E-08 | 715.85974 | 1131.1661 |
| WEEVQSYIR | 9 | Q86UP2 | Kinectin OS=Homo sapiens OX=9606 Gl | KTN1 | 156.276 | 2 | 54.640892 | yes | 1.16E-07 | 605.29858 | 1055.54 |
| ECMAGTSGSEEVk | 13 | Q86UP2 | Kinectin OS=Homo sapiens OX=9606 Gl | KTN1 | 156.276 | 2 | 54.640892 | yes | 9.722E-08 | 692.78961 | 852.94501 |
| DAVSNTTNQLESK | 13 | Q86UP2 | Kinectin OS=Homo sapiens OX=9606 Gl | KTN1 | 156.276 | 2 | 54.640892 | yes | 1.34E-07 | 703.84155 | 798.87598 |
| LLEEQLQHEISNK | 13 | Q86UP2 | Kinectin OS=Homo sapiens OX=9606 Gl | KTN1 | 156.276 | 2 | 54.640892 | yes | 1.004E-07 | 790.91754 | 795.11444 |
| LVNELTEK | 8 | Q86UP2 | Kinectin OS=Homo sapiens OX=9606 Gl | KTN1 | 156.276 | 2 | 54.640892 | yes | 2.212E-07 | 473.2662 | 778.10828 |
| QTEDSLASER | 10 | Q86UP2 | Kinectin OS=Homo sapiens OX=9606 Gl | KTN1 | 156.276 | 2 | 54.640892 | yes | 9.722E-08 | 568.26495 | 709.34314 |
| DIQNMNFLLK | 10 | Q86UP2 | Kinectin OS=Homo sapiens OX=9606 Gl | KTN1 | 156.276 | 2 | 54.640892 | yes | 9.722E-08 | 618.32629 | 670.2453 |
| EISGLWNELDSLK | 13 | Q86UP2 | Kinectin OS=Homo sapiens OX=9606 Gl | KTN1 | 156.276 | 2 | 54.640892 | yes | 1.003E-07 | 752.38812 | 644.76654 |
| FKDFLLSLK | 9 | Q86UP2 | Kinectin OS=Homo sapiens OX=9606 Gl | KTN1 | 156.276 | 2 | 54.640892 | yes | 1.328E-07 | 555.83154 | 626.70929 |
| LQQEEVQK | 8 | Q86UP2 | Kinectin OS=Homo sapiens OX=9606 Gl | KTN1 | 156.276 | 2 | 54.640892 | yes | 1.212E-06 | 501.26672 | 587.44757 |
| EKEHYQVLE | 9 | Q86UP2 | Kinectin OS=Homo sapiens OX=9606 Gl | KTN1 | 156.276 | 2 | 54.640892 | yes | 1.198E-07 | 587.79059 | 584.51691 |
| RLEAMLK | 7 | Q86UP2 | Kinectin OS=Homo sapiens OX=9606 Gl | KTN1 | 156.276 | 2 | 54.640892 | yes | 1.482E-07 | 430.75476 | 565.58063 |
| EREHLEMELEK | 11 | Q86UP2 | Kinectin OS=Homo sapiens OX=9606 Gl | KTN1 | 156.276 | 3 | 54.640892 | yes | 9.722E-08 | 481.56964 | 278.82083 |
| EREHLEMELEK | 11 | Q86UP2 | Kinectin OS=Homo sapiens OX=9606 Gl | KTN1 | 156.276 | 3 | 54.640892 | yes | 2.176E-07 | 721.85083 | 278.82083 |
| EREHLEMELEK | 11 | Q86UP2 | Kinectin OS=Homo sapiens OX=9606 Gl | KTN1 | 156.276 | 2 | 54.640892 | yes | 9.722E-08 | 481.56964 | 278.82083 |
| EREHLEMELEK | 11 | Q86UP2 | Kinectin OS=Homo sapiens OX=9606 Gl | KTN1 | 156.276 | 2 | 54.640892 | yes | 2.176E-07 | 721.85083 | 278.82083 |
| VETLMVPSK | 9 | Q86UP2 | Kinectin OS=Homo sapiens OX=9606 Gl | KTN1 | 156.276 | 2 | 54.640892 | yes | 1.29E-07 | 502.27826 | 553.87012 |
| GKEEQMNTMK | 10 | Q86UP2 | Kinectin OS=Homo sapiens OX=9606 Gl | KTN1 | 156.276 | 2 | 54.640892 | yes | 3.313E-07 | 598.27594 | 501.07166 |
| QLTQEMMTEK | 10 | Q86UP2 | Kinectin OS=Homo sapiens OX=9606 Gl | KTN1 | 156.276 | 2 | 54.640892 | yes | 1.018E-07 | 619.79138 | 498.81845 |
| QEALPLHQETK | 11 | Q86UP2 | Kinectin OS=Homo sapiens OX=9606 Gl | KTN1 | 156.276 | 2 | 54.640892 | yes | 9.722E-08 | 647.34351 | 247.54044 |
| QEALPLHQETK | 11 | Q86UP2 | Kinectin OS=Homo sapiens OX=9606 Gl | KTN1 | 156.276 | 2 | 54.640892 | yes | 2.167E-07 | 431.8981 | 247.54044 |
| QEALPLHQETK | 11 | Q86UP2 | Kinectin OS=Homo sapiens OX=9606 Gl | KTN1 | 156.276 | 3 | 54.640892 | yes | 9.722E-08 | 647.34351 | 247.54044 |

| | | | | | | | | | | | |
|---|---|---|---|---|---|---|---|---|---|---|---|
| QEALPLHQETK | 11 | Q86UP2 | Kinectin OS=Homo sapiens OX=9606 Gl | KTN1 | 156.276 | 3 | 54.640892 | yes | 2.167E-07 | 431.8981 | 247.54044 |
| EHNVFQNK | 8 | Q86UP2 | Kinectin OS=Homo sapiens OX=9606 Gl | KTN1 | 156.276 | 2 | 54.640892 | yes | 2.532E-07 | 508.25143 | 477.82767 |
| DLLTELQK | 8 | Q86UP2 | Kinectin OS=Homo sapiens OX=9606 Gl | KTN1 | 156.276 | 2 | 54.640892 | yes | 2.97E-07 | 480.27402 | 467.35147 |
| ALKEEIGNVQLEK | 13 | Q86UP2 | Kinectin OS=Homo sapiens OX=9606 Gl | KTN1 | 156.276 | 2 | 54.640892 | yes | 9.926E-08 | 735.91174 | 461.19528 |
| MQQSVYVK | 8 | Q86UP2 | Kinectin OS=Homo sapiens OX=9606 Gl | KTN1 | 156.276 | 2 | 54.640892 | yes | 1.546E-07 | 491.75494 | 431.49844 |
| AQQLSITSK | 9 | Q86UP2 | Kinectin OS=Homo sapiens OX=9606 Gl | KTN1 | 156.276 | 2 | 54.640892 | yes | 2.828E-07 | 488.2771 | 419.75464 |
| KVEPVPVTK | 9 | Q86UP2 | Kinectin OS=Homo sapiens OX=9606 Gl | KTN1 | 156.276 | 2 | 54.640892 | yes | 1.106E-07 | 498.80804 | 386.12372 |
| KVETLMVPSK | 10 | Q86UP2 | Kinectin OS=Homo sapiens OX=9606 Gl | KTN1 | 156.276 | 2 | 54.640892 | yes | 9.722E-08 | 566.32574 | 331.55212 |
| AQLNETLTK | 9 | Q86UP2 | Kinectin OS=Homo sapiens OX=9606 Gl | KTN1 | 156.276 | 2 | 54.640892 | yes | 1.248E-07 | 509.28238 | 329.85849 |
| EEELNAIR | 8 | Q86UP2 | Kinectin OS=Homo sapiens OX=9606 Gl | KTN1 | 156.276 | 2 | 54.640892 | yes | 2.867E-07 | 487.2511 | 323.59464 |
| SVLAETEGILQK | 12 | Q86UP2 | Kinectin OS=Homo sapiens OX=9606 Gl | KTN1 | 156.276 | 2 | 54.640892 | yes | 9.722E-08 | 644.36139 | 320.64401 |
| LDDSYSEAVR | 10 | Q86UP2 | Kinectin OS=Homo sapiens OX=9606 Gl | KTN1 | 156.276 | 2 | 54.640892 | yes | 9.902E-08 | 577.76984 | 314.63556 |
| TAEHEAAQQDLQS | 14 | Q86UP2 | Kinectin OS=Homo sapiens OX=9606 Gl | KTN1 | 156.276 | 2 | 54.640892 | yes | 1.001E-07 | 778.37097 | 299.72894 |
| QNDQVSFASLVEE | 16 | Q86UP2 | Kinectin OS=Homo sapiens OX=9606 Gl | KTN1 | 156.276 | 3 | 54.640892 | yes | 1.137E-07 | 612.323 | 294.56128 |
| QPTPPSEAAASK | 12 | Q86UP2 | Kinectin OS=Homo sapiens OX=9606 Gl | KTN1 | 156.276 | 2 | 54.640892 | yes | 3.033E-07 | 592.30133 | 283.49759 |
| EHLEMELEK | 9 | Q86UP2 | Kinectin OS=Homo sapiens OX=9606 Gl | KTN1 | 156.276 | 2 | 54.640892 | yes | 2.082E-07 | 579.27899 | 268.38123 |
| QKPVLEEQVIK | 11 | Q86UP2 | Kinectin OS=Homo sapiens OX=9606 Gl | KTN1 | 156.276 | 2 | 54.640892 | yes | 1.224E-07 | 655.88757 | 242.13193 |
| QNEELNLLK | 9 | Q86UP2 | Kinectin OS=Homo sapiens OX=9606 Gl | KTN1 | 156.276 | 2 | 54.640892 | yes | 1.794E-07 | 550.80096 | 223.99265 |
| NWEAMEALASTEI | 13 | Q86UP2 | Kinectin OS=Homo sapiens OX=9606 Gl | KTN1 | 156.276 | 2 | 54.640892 | yes | 9.722E-08 | 740.34283 | 205.8838 |
| KLDDSYSEAVR | 11 | Q86UP2 | Kinectin OS=Homo sapiens OX=9606 Gl | KTN1 | 156.276 | 2 | 54.640892 | yes | 2.149E-07 | 641.81732 | 194.53896 |
| AHVQEVAQHNLK | 12 | Q86UP2 | Kinectin OS=Homo sapiens OX=9606 Gl | KTN1 | 156.276 | 2 | 54.640892 | yes | 2.255E-07 | 687.36786 | 181.97723 |
| DVVEQMEK | 8 | Q86UP2 | Kinectin OS=Homo sapiens OX=9606 Gl | KTN1 | 156.276 | 2 | 54.640892 | yes | 3.937E-07 | 489.23404 | 128.7984 |
| SVEELLEAELLK | 12 | Q86UP2 | Kinectin OS=Homo sapiens OX=9606 Gl | KTN1 | 156.276 | 2 | 54.640892 | yes | 9.722E-08 | 686.88214 | 118.57755 |
| AQIQQFHSQIAAQ | 25 | Q86UP2 | Kinectin OS=Homo sapiens OX=9606 Gl | KTN1 | 156.276 | 4 | 54.640892 | yes | 9.722E-08 | 684.60822 | 91.664383 |
| SENKDIENLR | 10 | Q86UP2 | Kinectin OS=Homo sapiens OX=9606 Gl | KTN1 | 156.276 | 2 | 54.640892 | yes | 2.192E-07 | 609.30969 | 79.87677 |
| RQEALPLHQETK | 12 | Q86UP2 | Kinectin OS=Homo sapiens OX=9606 Gl | KTN1 | 156.276 | 2 | 54.640892 | yes | 1.027E-05 | 725.39404 | 40.080196 |
| KNAEQAATQLK | 11 | Q86UP2 | Kinectin OS=Homo sapiens OX=9606 Gl | KTN1 | 156.276 | 2 | 54.640892 | yes | 1.712E-06 | 601.33038 | 32.105072 |
| NAEQAATQLK | 10 | Q86UP2 | Kinectin OS=Homo sapiens OX=9606 Gl | KTN1 | 156.276 | 2 | 54.640892 | yes | 3.289E-06 | 537.2829 | 32.017822 |
| DKLLAAVK | 8 | Q86UP2 | Kinectin OS=Homo sapiens OX=9606 Gl | KTN1 | 156.276 | 2 | 54.640892 | yes | 3.443E-07 | 429.27637 | 28.48321 |
| AKECMAGTSGSEE | 15 | Q86UP2 | Kinectin OS=Homo sapiens OX=9606 Gl | KTN1 | 156.276 | 2 | 54.640892 | yes | 9.722E-08 | 792.35565 | 14.734079 |
| SFMCTVCGR | 9 | Q86UQ0 | Zinc finger protein 589 OS=Homo sapien | ZNF589 | 41.19 | 2 | 47.939518 | yes | 1.229E-07 | 559.23303 | 154.4908 |
| EVMLENLR | 8 | Q86UQ0 | Zinc finger protein 589 OS=Homo sapien | ZNF589 | 41.19 | 2 | 47.939518 | yes | 2.588E-07 | 502.26569 | 71.577766 |
| IGMETFPAVAEK | 12 | Q86US8 | Telomerase-binding protein EST1A OS=l | SMG6 | 160.462 | 2 | 54.72668 | yes | 1.165E-07 | 646.83136 | 1282.5931 |
| YVSVAPVPDTMGl | 13 | Q86US8 | Telomerase-binding protein EST1A OS=l | SMG6 | 160.462 | 2 | 54.72668 | yes | 9.926E-08 | 682.34998 | 1204.4285 |
| SLAASNPILTAK | 12 | Q86US8 | Telomerase-binding protein EST1A OS=l | SMG6 | 160.462 | 2 | 54.72668 | yes | 4.117E-06 | 593.34552 | 1017.0804 |
| EVVLLTDDR | 9 | Q86US8 | Telomerase-binding protein EST1A OS=l | SMG6 | 160.462 | 2 | 54.72668 | yes | 1.261E-07 | 530.28766 | 934.90186 |
| DPNVENPEQIR | 11 | Q86US8 | Telomerase-binding protein EST1A OS=l | SMG6 | 160.462 | 2 | 54.72668 | yes | 1.559E-07 | 655.82037 | 775.2583 |
| RPDLEIYKPGLSR | 13 | Q86US8 | Telomerase-binding protein EST1A OS=l | SMG6 | 160.462 | 3 | 54.72668 | yes | 3.55E-07 | 515.2912 | 732.17194 |
| YALISAQR | 8 | Q86US8 | Telomerase-binding protein EST1A OS=l | SMG6 | 160.462 | 2 | 54.72668 | yes | 1.077E-07 | 461.26126 | 690.78955 |

| | | | | | | | | | | | |
|---|---|---|---|---|---|---|---|---|---|---|---|
| VTFDAEAMNK | 10 | Q86US8 | Telomerase-binding protein EST1A OS=I | SMG6 | 160.462 | 2 | 54.72668 | yes | 9.722E-08 | 563.26587 | 631.36182 |
| CMICQGDIAR | 10 | Q86US8 | Telomerase-binding protein EST1A OS=I | SMG6 | 160.462 | 2 | 54.72668 | yes | 1.058E-07 | 612.27014 | 616.7533 |
| AELLQLYER | 9 | Q86US8 | Telomerase-binding protein EST1A OS=I | SMG6 | 160.462 | 2 | 54.72668 | yes | 9.722E-08 | 567.81134 | 601.58398 |
| LEDYMDGLAIR | 11 | Q86US8 | Telomerase-binding protein EST1A OS=I | SMG6 | 160.462 | 2 | 54.72668 | yes | 1.145E-07 | 648.31866 | 598.1322 |
| GNELESIAFR | 10 | Q86US8 | Telomerase-binding protein EST1A OS=I | SMG6 | 160.462 | 2 | 54.72668 | yes | 9.913E-08 | 568.29077 | 451.83252 |
| GILATLAPQAGSR | 13 | Q86US8 | Telomerase-binding protein EST1A OS=I | SMG6 | 160.462 | 2 | 54.72668 | yes | 2.15E-07 | 627.86188 | 353.1008 |
| LEIWIHPSHPR | 11 | Q86US8 | Telomerase-binding protein EST1A OS=I | SMG6 | 160.462 | 2 | 54.72668 | yes | 9.722E-08 | 692.87787 | 340.52164 |
| IQAVLEDHSQMR | 12 | Q86US8 | Telomerase-binding protein EST1A OS=I | SMG6 | 160.462 | 2 | 54.72668 | yes | 9.722E-08 | 713.85901 | 321.10315 |
| LWDPNNPDQKPAI | 14 | Q86US8 | Telomerase-binding protein EST1A OS=I | SMG6 | 160.462 | 2 | 54.72668 | yes | 1.451E-07 | 818.4281 | 159.76031 |
| LWDPNNPDQKPAI | 14 | Q86US8 | Telomerase-binding protein EST1A OS=I | SMG6 | 160.462 | 2 | 54.72668 | yes | 9.722E-08 | 545.95447 | 159.76031 |
| LWDPNNPDQKPAI | 14 | Q86US8 | Telomerase-binding protein EST1A OS=I | SMG6 | 160.462 | 3 | 54.72668 | yes | 1.451E-07 | 818.4281 | 159.76031 |
| LWDPNNPDQKPAI | 14 | Q86US8 | Telomerase-binding protein EST1A OS=I | SMG6 | 160.462 | 3 | 54.72668 | yes | 9.722E-08 | 545.95447 | 159.76031 |
| QMELEIR | 7 | Q86US8 | Telomerase-binding protein EST1A OS=I | SMG6 | 160.462 | 2 | 54.72668 | yes | 4.923E-07 | 459.73929 | 318.40363 |
| ESLMSLFEETK | 11 | Q86US8 | Telomerase-binding protein EST1A OS=I | SMG6 | 160.462 | 2 | 54.72668 | yes | 9.722E-08 | 657.3183 | 314.29141 |
| SIEFLEQR | 8 | Q86US8 | Telomerase-binding protein EST1A OS=I | SMG6 | 160.462 | 2 | 54.72668 | yes | 3.125E-07 | 511.26929 | 308.25153 |
| NLQQQELHR | 9 | Q86US8 | Telomerase-binding protein EST1A OS=I | SMG6 | 160.462 | 2 | 54.72668 | yes | 1.257E-07 | 583.30725 | 281.45203 |
| VSSFVPDLK | 9 | Q86US8 | Telomerase-binding protein EST1A OS=I | SMG6 | 160.462 | 2 | 54.72668 | yes | 1.279E-07 | 496.27658 | 204.43651 |
| KLDAVYYYMR | 10 | Q86US8 | Telomerase-binding protein EST1A OS=I | SMG6 | 160.462 | 2 | 54.72668 | yes | 9.722E-08 | 661.33411 | 186.47632 |
| ESLMSLFEETKR | 12 | Q86US8 | Telomerase-binding protein EST1A OS=I | SMG6 | 160.462 | 2 | 54.72668 | yes | 9.722E-08 | 735.3689 | 141.92844 |
| FILSFLHAHGK | 11 | Q86US8 | Telomerase-binding protein EST1A OS=I | SMG6 | 160.462 | 2 | 54.72668 | yes | 3.376E-07 | 635.35876 | 135.23856 |
| NAFYQVIEK | 9 | Q86US8 | Telomerase-binding protein EST1A OS=I | SMG6 | 160.462 | 2 | 54.72668 | yes | 1.646E-07 | 556.29279 | 110.06546 |
| QAQASYYK | 8 | Q86US8 | Telomerase-binding protein EST1A OS=I | SMG6 | 160.462 | 2 | 54.72668 | yes | 7.386E-05 | 479.73508 | 85.220123 |
| QHEEFDLSPDQWI | 13 | Q86US8 | Telomerase-binding protein EST1A OS=I | SMG6 | 160.462 | 2 | 54.72668 | yes | 1.573E-07 | 843.87897 | 59.734737 |
| FKLEDYMDGLAIR | 13 | Q86US8 | Telomerase-binding protein EST1A OS=I | SMG6 | 160.462 | 2 | 54.72668 | yes | 1.895E-07 | 785.90033 | 54.14064 |
| KPDLQIYQPGR | 11 | Q86US8 | Telomerase-binding protein EST1A OS=I | SMG6 | 160.462 | 2 | 54.72668 | yes | 1.689E-07 | 657.86188 | 52.307598 |
| PYNQLALLAVYTI | 13 | Q86US8 | Telomerase-binding protein EST1A OS=I | SMG6 | 160.462 | 2 | 54.72668 | yes | 9.862E-08 | 761.4248 | 48.175049 |
| FQNSDNPYYYPR | 12 | Q86US8 | Telomerase-binding protein EST1A OS=I | SMG6 | 160.462 | 2 | 54.72668 | yes | 9.722E-08 | 782.3468 | 42.899246 |
| IDEPDGLGPR | 10 | Q86US8 | Telomerase-binding protein EST1A OS=I | SMG6 | 160.462 | 2 | 54.72668 | yes | 1.821E-07 | 534.76965 | 41.99263 |
| LLPQVEAAR | 9 | Q86UT8 | Centrosomal AT-AC splicing factor OS=I | CENATAC | 37.975 | 2 | 50.514538 | yes | 9.79E-08 | 498.79547 | 149.19513 |
| VGANFDHSSR | 10 | Q86UT8 | Centrosomal AT-AC splicing factor OS=I | CENATAC | 37.975 | 2 | 50.514538 | yes | 3.902E-07 | 545.25726 | 80.977791 |
| YVPEHER | 7 | Q86UT8 | Centrosomal AT-AC splicing factor OS=I | CENATAC | 37.975 | 2 | 50.514538 | yes | 2.71E-06 | 465.22742 | 62.120094 |
| LQGEVLALEEER | 12 | Q86UU1 | Pleckstrin homology-like domain family I | PHLDB1 | 151.162 | 2 | 52.707302 | yes | 9.722E-08 | 693.36719 | 100.61057 |
| LFEDLEFQQLER | 12 | Q86UU1 | Pleckstrin homology-like domain family I | PHLDB1 | 151.162 | 2 | 52.707302 | yes | 5.591E-07 | 783.89355 | 50.337109 |
| IGTLQDRPPSPFR | 13 | Q86UU1 | Pleckstrin homology-like domain family I | PHLDB1 | 151.162 | 3 | 52.707302 | yes | 1.316E-06 | 495.27203 | 42.013626 |
| EEPTTTLPSMGETI | 14 | Q86UW6 | NEDD4-binding protein 2 OS=Homo sap | N4BP2 | 198.802 | 2 | 54.325413 | yes | 3.044E-06 | 760.86108 | 135.79385 |
| TGLSEPNLEIGTNI | 15 | Q86UW6 | NEDD4-binding protein 2 OS=Homo sap | N4BP2 | 198.802 | 2 | 54.325413 | yes | 2.119E-07 | 794.39667 | 79.857094 |
| EIMSEEIALQEK | 12 | Q86UW6 | NEDD4-binding protein 2 OS=Homo sap | N4BP2 | 198.802 | 2 | 54.325413 | yes | 9.722E-08 | 710.35541 | 64.76207 |
| SSPGEIVEER | 10 | Q86UW6 | NEDD4-binding protein 2 OS=Homo sap | N4BP2 | 198.802 | 2 | 54.325413 | yes | 8.122E-07 | 551.7724 | 64.160934 |
| SDVQEAIPYR | 10 | Q86UW6 | NEDD4-binding protein 2 OS=Homo sap | N4BP2 | 198.802 | 2 | 54.325413 | yes | 9.722E-08 | 589.29602 | 53.545784 |

| | | | | | | | | | | | |
|---|---|---|---|---|---|---|---|---|---|---|---|
| TVVAQEFVHQNEI | 21 | Q86UW6 | NEDD4-binding protein 2 OS=Homo sap | N4BP2 | 198.802 | 3 | 54.325413 | yes | 4.99E-07 | 785.05591 | 52.199745 |
| ISISDSIK | 8 | Q86UW6 | NEDD4-binding protein 2 OS=Homo sap | N4BP2 | 198.802 | 2 | 54.325413 | yes | 5.699E-07 | 431.74765 | 49.188301 |
| LCEDTEFENFQK | 12 | Q86UW6 | NEDD4-binding protein 2 OS=Homo sap | N4BP2 | 198.802 | 2 | 54.325413 | yes | 9.722E-08 | 780.33777 | 42.056904 |
| SQCDDASEPLNSY | 14 | Q86UW6 | NEDD4-binding protein 2 OS=Homo sap | N4BP2 | 198.802 | 2 | 54.325413 | yes | 9.722E-08 | 807.34106 | 41.629265 |
| IELCAYSCEDR | 11 | Q86UW6 | NEDD4-binding protein 2 OS=Homo sap | N4BP2 | 198.802 | 2 | 54.325413 | yes | 1.301E-07 | 708.30011 | 41.390232 |
| MLEHYQR | 7 | Q86UW6 | NEDD4-binding protein 2 OS=Homo sap | N4BP2 | 198.802 | 2 | 54.325413 | yes | 2.021E-07 | 488.73709 | 37.884922 |
| HLEFTEEK | 8 | Q86UW6 | NEDD4-binding protein 2 OS=Homo sap | N4BP2 | 198.802 | 2 | 54.325413 | yes | 3.558E-05 | 516.75348 | 31.066305 |
| QCYLPDNAQGR | 11 | Q86UW9 | Probable E3 ubiquitin-protein ligase DTX | DTX2 | 67.247 | 2 | 47.00511 | yes | 2.139E-07 | 661.30133 | 60.57209 |
| KVLELLK | 7 | Q86UW9 | Probable E3 ubiquitin-protein ligase DTX | DTX2 | 67.247 | 2 | 47.00511 | yes | 1.422E-07 | 421.78912 | 24.491013 |
| IEDGISSTLPSK | 12 | Q86V48 | Leucine zipper protein 1 OS=Homo sapie | LUZP1 | 120.275 | 2 | 57.385548 | yes | 1.703E-07 | 623.8299 | 223.24284 |
| SLTLSFVSER | 10 | Q86V48 | Leucine zipper protein 1 OS=Homo sapie | LUZP1 | 120.275 | 2 | 57.385548 | yes | 3.045E-07 | 569.80878 | 222.70825 |
| EGVNQPAAVVME | 21 | Q86V48 | Leucine zipper protein 1 OS=Homo sapie | LUZP1 | 120.275 | 3 | 57.385548 | yes | 9.722E-08 | 746.02692 | 216.29323 |
| QVSSPSFTNR | 10 | Q86V48 | Leucine zipper protein 1 OS=Homo sapie | LUZP1 | 120.275 | 2 | 57.385548 | yes | 8.487E-07 | 561.78052 | 200.20573 |
| LLASQLEEIK | 10 | Q86V48 | Leucine zipper protein 1 OS=Homo sapie | LUZP1 | 120.275 | 2 | 57.385548 | yes | 9.722E-08 | 572.33459 | 189.73625 |
| IMGGSGTETTLEK | 13 | Q86V48 | Leucine zipper protein 1 OS=Homo sapie | LUZP1 | 120.275 | 2 | 57.385548 | yes | 9.722E-08 | 662.32666 | 186.13695 |
| LVNTTITPEPEPK | 13 | Q86V48 | Leucine zipper protein 1 OS=Homo sapie | LUZP1 | 120.275 | 2 | 57.385548 | yes | 9.722E-08 | 719.89301 | 183.99593 |
| SSIIIKPSDPVER | 13 | Q86V48 | Leucine zipper protein 1 OS=Homo sapie | LUZP1 | 120.275 | 2 | 57.385548 | yes | 1.181E-07 | 720.90649 | 183.10367 |
| TQSSLTVSEVLTR | 13 | Q86V48 | Leucine zipper protein 1 OS=Homo sapie | LUZP1 | 120.275 | 2 | 57.385548 | yes | 9.722E-08 | 710.88574 | 182.32257 |
| SCLFEQGTR | 9 | Q86V48 | Leucine zipper protein 1 OS=Homo sapie | LUZP1 | 120.275 | 2 | 57.385548 | yes | 1.146E-07 | 549.25586 | 180.34071 |
| SSTDFSELEQPR | 12 | Q86V48 | Leucine zipper protein 1 OS=Homo sapie | LUZP1 | 120.275 | 2 | 57.385548 | yes | 9.855E-08 | 698.323 | 160.81294 |
| AEDELLDLQDK | 11 | Q86V48 | Leucine zipper protein 1 OS=Homo sapie | LUZP1 | 120.275 | 2 | 57.385548 | yes | 1.113E-07 | 644.81702 | 127.91467 |
| TEQSLASELEK | 11 | Q86V48 | Leucine zipper protein 1 OS=Homo sapie | LUZP1 | 120.275 | 2 | 57.385548 | yes | 3.138E-07 | 617.81171 | 122.5141 |
| ISSELEMLR | 9 | Q86V48 | Leucine zipper protein 1 OS=Homo sapie | LUZP1 | 120.275 | 2 | 57.385548 | yes | 9.839E-08 | 539.28406 | 121.11794 |
| EFALNNENYSLSN | 14 | Q86V48 | Leucine zipper protein 1 OS=Homo sapie | LUZP1 | 120.275 | 2 | 57.385548 | yes | 9.722E-08 | 835.89209 | 118.15062 |
| NSHAPPAETIR | 11 | Q86V48 | Leucine zipper protein 1 OS=Homo sapie | LUZP1 | 120.275 | 2 | 57.385548 | yes | 1.242E-07 | 596.80707 | 105.28684 |
| VTSSITIYPSDSSSP | 16 | Q86V48 | Leucine zipper protein 1 OS=Homo sapie | LUZP1 | 120.275 | 2 | 57.385548 | yes | 6.926E-07 | 848.92303 | 91.047958 |
| HDITLQLAEAER | 12 | Q86V48 | Leucine zipper protein 1 OS=Homo sapie | LUZP1 | 120.275 | 2 | 57.385548 | yes | 2.965E-07 | 698.36499 | 83.542175 |
| YPPAAQEHSK | 10 | Q86V48 | Leucine zipper protein 1 OS=Homo sapie | LUZP1 | 120.275 | 2 | 57.385548 | yes | 4.02E-07 | 564.27765 | 79.742767 |
| SHSAPSEVGFSDAI | 14 | Q86V48 | Leucine zipper protein 1 OS=Homo sapie | LUZP1 | 120.275 | 2 | 57.385548 | yes | 1.346E-07 | 723.83405 | 74.681404 |
| NNDLQDNYLSEQI | 14 | Q86V48 | Leucine zipper protein 1 OS=Homo sapie | LUZP1 | 120.275 | 2 | 57.385548 | yes | 1.288E-07 | 847.88446 | 65.40741 |
| RLDELEEATK | 10 | Q86V48 | Leucine zipper protein 1 OS=Homo sapie | LUZP1 | 120.275 | 2 | 57.385548 | yes | 2.902E-06 | 602.31439 | 60.513649 |
| EKLEEEENLTR | 11 | Q86V48 | Leucine zipper protein 1 OS=Homo sapie | LUZP1 | 120.275 | 2 | 57.385548 | yes | 1.675E-07 | 695.34644 | 52.826252 |
| MDMSLDDIIK | 10 | Q86V81 | THO complex subunit 4 OS=Homo sapie | ALYREF | 26.888 | 2 | 53.585743 | yes | 9.789E-08 | 590.78302 | 6187.4946 |
| MDMSLDDIIK | 10 | Q86V81 | THO complex subunit 4 OS=Homo sapie | ALYREF | 26.888 | 2 | 53.585743 | yes | 9.722E-08 | 598.78052 | 6187.4946 |
| MDMSLDDIIK | 10 | Q86V81 | THO complex subunit 4 OS=Homo sapie | ALYREF | 26.888 | 2 | 53.585743 | yes | 4.815E-07 | 598.78052 | 6187.4946 |
| SLGTADVHFER | 11 | Q86V81 | THO complex subunit 4 OS=Homo sapie | ALYREF | 26.888 | 2 | 53.585743 | yes | 9.722E-08 | 616.30695 | 7460.9219 |
| QYNGVPLDGR | 10 | Q86V81 | THO complex subunit 4 OS=Homo sapie | ALYREF | 26.888 | 2 | 53.585743 | yes | 9.903E-08 | 559.78308 | 3887.1584 |
| AAVHYDR | 7 | Q86V81 | THO complex subunit 4 OS=Homo sapie | ALYREF | 26.888 | 2 | 53.585743 | yes | 3.359E-07 | 416.20901 | 3349.4487 |
| WQHDLFDSGFGG | 21 | Q86V81 | THO complex subunit 4 OS=Homo sapie | ALYREF | 26.888 | 3 | 53.585743 | yes | 1.109E-07 | 707.99231 | 2993.7842 |

| | | | | | | | | | | | |
|---|---|---|---|---|---|---|---|---|---|---|---|
| QLPDKWQHDLFD: | 26 | Q86V81 | THO complex subunit 4 OS=Homo sapie | ALYREF | 26.888 | 4 | 53.585743 | yes | 9.789E-08 | 676.57538 | 1884.7848 |
| QQLSAEELDAQLD | 18 | Q86V81 | THO complex subunit 4 OS=Homo sapie | ALYREF | 26.888 | 3 | 53.585743 | yes | 1.096E-07 | 678.99573 | 689.1167 |
| PMNIQLVTSQIDA( | 15 | Q86V81 | THO complex subunit 4 OS=Homo sapie | ALYREF | 26.888 | 3 | 53.585743 | yes | 2.907E-07 | 577.30157 | 105.78966 |
| PMNIQLVTSQIDA( | 15 | Q86V81 | THO complex subunit 4 OS=Homo sapie | ALYREF | 26.888 | 3 | 53.585743 | yes | 1.276E-06 | 571.96991 | 105.78966 |
| ADKMDMSLDDIIK | 13 | Q86V81 | THO complex subunit 4 OS=Homo sapie | ALYREF | 26.888 | 2 | 53.585743 | yes | 1.57E-07 | 768.86786 | 85.103294 |
| MEKAANARNHDD | 13 | Q86VI3 | Ras GTPase-activating-like protein IQGA | IQGAP3 | 184.699 | 2 | 56.85035 | yes | 2.005E-05 | 772.35516 | 822.987 |
| YHLLLVAAK | 9 | Q86VI3 | Ras GTPase-activating-like protein IQGA | IQGAP3 | 184.699 | 2 | 56.85035 | yes | 6.802E-07 | 514.3186 | 46.604801 |
| NWINQTEAQTGQF | 13 | Q86VI3 | Ras GTPase-activating-like protein IQGA | IQGAP3 | 184.699 | 2 | 56.85035 | yes | 6.158E-06 | 773.37384 | 35.865932 |
| FEGLEADADDSNT | 14 | Q86VI3 | Ras GTPase-activating-like protein IQGA | IQGAP3 | 184.699 | 2 | 56.85035 | yes | 1.819E-06 | 770.33154 | 33.001133 |
| GDQEQAMLHAVQ | 13 | Q86VI3 | Ras GTPase-activating-like protein IQGA | IQGAP3 | 184.699 | 2 | 56.85035 | yes | 9.722E-08 | 741.85956 | 29.480501 |
| EEVQAGVAAANTI | 13 | Q86VI3 | Ras GTPase-activating-like protein IQGA | IQGAP3 | 184.699 | 2 | 56.85035 | yes | 5.044E-06 | 644.33063 | 18.055904 |
| AQELGLVELLEK | 12 | Q86VI3 | Ras GTPase-activating-like protein IQGA | IQGAP3 | 184.699 | 2 | 56.85035 | yes | 1.503E-07 | 671.38483 | 9.9373932 |
| QACTAQTPEPLR | 12 | Q86VI3 | Ras GTPase-activating-like protein IQGA | IQGAP3 | 184.699 | 2 | 56.85035 | yes | 1.796E-05 | 686.33789 | 5.0248365 |
| FTVTIGEDER | 10 | Q86VM9 | Zinc finger CCCH domain-containing prc | ZC3H18 | 106.379 | 2 | 53.990685 | yes | 1.003E-07 | 583.78802 | 461.41733 |
| GNCTWGMNCR | 10 | Q86VM9 | Zinc finger CCCH domain-containing prc | ZC3H18 | 106.379 | 2 | 53.990685 | yes | 1.021E-07 | 628.24188 | 291.08292 |
| SQDQDSEVNELSR | 13 | Q86VM9 | Zinc finger CCCH domain-containing prc | ZC3H18 | 106.379 | 2 | 53.990685 | yes | 9.722E-08 | 753.83698 | 201.75681 |
| TTAPVPEPTKPGDI | 15 | Q86VM9 | Zinc finger CCCH domain-containing prc | ZC3H18 | 106.379 | 2 | 53.990685 | yes | 1.222E-07 | 781.91229 | 183.76747 |
| EAATTGPQVK | 10 | Q86VM9 | Zinc finger CCCH domain-containing prc | ZC3H18 | 106.379 | 2 | 53.990685 | yes | 1.076E-07 | 501.26672 | 167.62564 |
| AGAEDDEEKGEGT | 15 | Q86VM9 | Zinc finger CCCH domain-containing prc | ZC3H18 | 106.379 | 3 | 53.990685 | yes | 9.935E-08 | 520.89728 | 69.810425 |
| AGAEDDEEKGEGT | 15 | Q86VM9 | Zinc finger CCCH domain-containing prc | ZC3H18 | 106.379 | 3 | 53.990685 | yes | 2.096E-06 | 780.84229 | 69.810425 |
| AGAEDDEEKGEGT | 15 | Q86VM9 | Zinc finger CCCH domain-containing prc | ZC3H18 | 106.379 | 2 | 53.990685 | yes | 9.935E-08 | 520.89728 | 69.810425 |
| AGAEDDEEKGEGT | 15 | Q86VM9 | Zinc finger CCCH domain-containing prc | ZC3H18 | 106.379 | 2 | 53.990685 | yes | 2.096E-06 | 780.84229 | 69.810425 |
| KPAPPPAPPQATK | 13 | Q86VM9 | Zinc finger CCCH domain-containing prc | ZC3H18 | 106.379 | 2 | 53.990685 | yes | 1.547E-07 | 650.37463 | 136.28647 |
| EFDKENEVFR | 10 | Q86VM9 | Zinc finger CCCH domain-containing prc | ZC3H18 | 106.379 | 2 | 53.990685 | yes | 9.722E-08 | 656.81201 | 104.41154 |
| TKGEPAPPPGK | 11 | Q86VM9 | Zinc finger CCCH domain-containing prc | ZC3H18 | 106.379 | 2 | 53.990685 | yes | 3.054E-07 | 539.79822 | 98.17894 |
| GGQYENFR | 8 | Q86VM9 | Zinc finger CCCH domain-containing prc | ZC3H18 | 106.379 | 2 | 53.990685 | yes | 1.541E-06 | 485.7225 | 93.196266 |
| ELSDAVNVR | 9 | Q86VP1 | Tax1-binding protein 1 OS=Homo sapien | TAX1BP1 | 90.878 | 2 | 53.018074 | yes | 2.741E-07 | 501.76437 | 112.18241 |
| LQLCLAEK | 8 | Q86VP1 | Tax1-binding protein 1 OS=Homo sapien | TAX1BP1 | 90.878 | 2 | 53.018074 | yes | 1.58E-07 | 487.7706 | 94.683533 |
| LELAEVQDNYK | 11 | Q86VP1 | Tax1-binding protein 1 OS=Homo sapien | TAX1BP1 | 90.878 | 2 | 53.018074 | yes | 9.722E-08 | 661.33533 | 92.586624 |
| LMSEVQTLK | 9 | Q86VP1 | Tax1-binding protein 1 OS=Homo sapien | TAX1BP1 | 90.878 | 2 | 53.018074 | yes | 4.302E-07 | 524.789 | 89.777351 |
| QLADAVAELK | 10 | Q86VP1 | Tax1-binding protein 1 OS=Homo sapien | TAX1BP1 | 90.878 | 2 | 53.018074 | yes | 5.223E-07 | 529.29803 | 89.106689 |
| LQMAADHYK | 9 | Q86VP1 | Tax1-binding protein 1 OS=Homo sapien | TAX1BP1 | 90.878 | 2 | 53.018074 | yes | 9.935E-08 | 538.76331 | 88.052567 |
| GLTEVTQSLK | 10 | Q86VP1 | Tax1-binding protein 1 OS=Homo sapien | TAX1BP1 | 90.878 | 2 | 53.018074 | yes | 5.1E-07 | 538.30334 | 69.853516 |
| LSDQSANNNNVFT | 14 | Q86VP1 | Tax1-binding protein 1 OS=Homo sapien | TAX1BP1 | 90.878 | 2 | 53.018074 | yes | 9.722E-08 | 776.37354 | 50.42688 |
| CDQLQAEQK | 9 | Q86VP1 | Tax1-binding protein 1 OS=Homo sapien | TAX1BP1 | 90.878 | 2 | 53.018074 | yes | 3.089E-06 | 560.25861 | 43.051834 |
| ALTLIAGSPLK | 11 | Q86VP6 | Cullin-associated NEDD8-dissociated prc | CAND1 | 136.376 | 2 | 38.54582 | yes | 5.146E-07 | 542.34222 | 61.964386 |
| VQAALVWER | 9 | Q86VR2 | Reticulophagy regulator 3 OS=Homo sap | RETREG3 | 51.397 | 2 | 42.51663 | yes | 2.991E-07 | 536.3009 | 103.55911 |
| ALVEVLGPYEPLL | 15 | Q86VR2 | Reticulophagy regulator 3 OS=Homo sap | RETREG3 | 51.397 | 2 | 42.51663 | yes | 4.85E-07 | 828.47198 | 63.157478 |
| LASTGSGQSFLAR | 13 | Q86VS8 | Protein Hook homolog 3 OS=Homo sapie | HOOK3 | 83.126 | 2 | 46.91828 | yes | 2.274E-07 | 647.84113 | 62.023388 |

| | | | | | | | | | | | |
|---|---|---|---|---|---|---|---|---|---|---|---|
| IEELQEALR | 9 | Q86VS8 | Protein Hook homolog 3 OS=Homo sapie | HOOK3 | 83.126 | 2 | 46.91828 | yes | 3.151E-06 | 550.80096 | 60.247429 |
| QNQGAAPEIQALK | 13 | Q86VS8 | Protein Hook homolog 3 OS=Homo sapie | HOOK3 | 83.126 | 2 | 46.91828 | yes | 1.441E-07 | 684.36749 | 26.61112 |
| SSTPTTIFPIR | 11 | Q86W42 | THO complex subunit 6 homolog OS=Ho | THOC6 | 37.535 | 2 | 50.914536 | yes | 9.722E-08 | 610.33771 | 246.75175 |
| VLTAAGNSCR | 10 | Q86W42 | THO complex subunit 6 homolog OS=Ho | THOC6 | 37.535 | 2 | 50.914536 | yes | 1.46E-07 | 524.76385 | 174.79382 |
| HLLSAGDGEVK | 11 | Q86W42 | THO complex subunit 6 homolog OS=Ho | THOC6 | 37.535 | 2 | 50.914536 | yes | 9.789E-08 | 563.29858 | 131.4875 |
| SPEVLSGGEDGAV | 14 | Q86W42 | THO complex subunit 6 homolog OS=Ho | THOC6 | 37.535 | 2 | 50.914536 | yes | 1.338E-07 | 686.83881 | 92.244568 |
| VDVFTNLGYR | 10 | Q86W42 | THO complex subunit 6 homolog OS=Ho | THOC6 | 37.535 | 2 | 50.914536 | yes | 1.023E-07 | 592.30896 | 69.408676 |
| GHTDYIHCLALR | 12 | Q86W42 | THO complex subunit 6 homolog OS=Ho | THOC6 | 37.535 | 2 | 50.914536 | yes | 6.291E-07 | 728.36169 | 44.502911 |
| CVNQWQLSGELK | 12 | Q86W42 | THO complex subunit 6 homolog OS=Ho | THOC6 | 37.535 | 2 | 50.914536 | yes | 1.585E-07 | 731.36139 | 42.686039 |
| AVPLAVPLGQTEV | 19 | Q86W42 | THO complex subunit 6 homolog OS=Ho | THOC6 | 37.535 | 3 | 50.914536 | yes | 1.12E-07 | 679.72125 | 35.889244 |
| APEDVIQALEEK | 12 | Q86W50 | RNA N6-adenosine-methyltransferase MI | METTL16 | 63.622 | 2 | 49.348583 | yes | 1.042E-07 | 671.34845 | 155.13371 |
| LPGVAGQYLFK | 11 | Q86W50 | RNA N6-adenosine-methyltransferase MI | METTL16 | 63.622 | 2 | 49.348583 | yes | 1.982E-07 | 596.83984 | 147.09245 |
| DKPPDFAYLASK | 12 | Q86W50 | RNA N6-adenosine-methyltransferase MI | METTL16 | 63.622 | 2 | 49.348583 | yes | 2.941E-07 | 676.34827 | 146.25337 |
| ALTCTLLR | 8 | Q86W50 | RNA N6-adenosine-methyltransferase MI | METTL16 | 63.622 | 2 | 49.348583 | yes | 1.42E-07 | 474.27078 | 106.3964 |
| NVEQNNLSDLIK | 12 | Q86W50 | RNA N6-adenosine-methyltransferase MI | METTL16 | 63.622 | 2 | 49.348583 | yes | 1.325E-06 | 693.86481 | 81.163589 |
| TLLMDALK | 8 | Q86W50 | RNA N6-adenosine-methyltransferase MI | METTL16 | 63.622 | 2 | 49.348583 | yes | 3.076E-07 | 452.76224 | 54.82003 |
| VTYTEFCQGR | 10 | Q86W50 | RNA N6-adenosine-methyltransferase MI | METTL16 | 63.622 | 2 | 49.348583 | yes | 5.342E-07 | 630.78754 | 28.244797 |
| DLMNQLCTYIR | 11 | Q86W50 | RNA N6-adenosine-methyltransferase MI | METTL16 | 63.622 | 2 | 49.348583 | yes | 4.11E-07 | 713.84448 | 24.87122 |
| WLGTPIEEMR | 10 | Q86W56 | Poly(ADP-ribose) glycohydrolase OS=Hc | PARG | 111.111 | 2 | 53.182323 | yes | 2.46E-07 | 616.31061 | 198.33061 |
| LENVSQLSLDK | 11 | Q86W56 | Poly(ADP-ribose) glycohydrolase OS=Hc | PARG | 111.111 | 2 | 53.182323 | yes | 1.192E-07 | 623.33789 | 170.17572 |
| LPGNISSLNVECR | 13 | Q86W56 | Poly(ADP-ribose) glycohydrolase OS=Hc | PARG | 111.111 | 2 | 53.182323 | yes | 1.208E-07 | 729.87213 | 104.3471 |
| KLTVGDVYK | 9 | Q86W56 | Poly(ADP-ribose) glycohydrolase OS=Hc | PARG | 111.111 | 2 | 53.182323 | yes | 3.203E-07 | 511.79767 | 96.272614 |
| SEYSSYPDINFNR | 13 | Q86W56 | Poly(ADP-ribose) glycohydrolase OS=Hc | PARG | 111.111 | 2 | 53.182323 | yes | 9.722E-08 | 796.3548 | 95.287895 |
| NCSTPGPDIK | 10 | Q86W56 | Poly(ADP-ribose) glycohydrolase OS=Hc | PARG | 111.111 | 2 | 53.182323 | yes | 1.827E-07 | 544.75568 | 92.762726 |
| ITDHFMR | 7 | Q86W56 | Poly(ADP-ribose) glycohydrolase OS=Hc | PARG | 111.111 | 2 | 53.182323 | yes | 1.806E-07 | 460.22635 | 78.915161 |
| DAHVQFR | 7 | Q86W56 | Poly(ADP-ribose) glycohydrolase OS=Hc | PARG | 111.111 | 2 | 53.182323 | yes | 4.786E-07 | 436.72229 | 74.666122 |
| QSLEDFPEWER | 11 | Q86W56 | Poly(ADP-ribose) glycohydrolase OS=Hc | PARG | 111.111 | 2 | 53.182323 | yes | 9.722E-08 | 718.32806 | 41.324905 |
| WELIQTALLNK | 11 | Q86W56 | Poly(ADP-ribose) glycohydrolase OS=Hc | PARG | 111.111 | 2 | 53.182323 | yes | 9.722E-08 | 664.88226 | 39.415302 |
| DVVYFTFGDSELM | 14 | Q86W56 | Poly(ADP-ribose) glycohydrolase OS=Hc | PARG | 111.111 | 2 | 53.182323 | yes | 6.556E-07 | 839.8927 | 35.085194 |
| PSANHTVTIR | 10 | Q86W56 | Poly(ADP-ribose) glycohydrolase OS=Hc | PARG | 111.111 | 2 | 53.182323 | yes | 9.722E-08 | 548.29889 | 27.736046 |
| AGEVPKPFPTHYK | 13 | Q86W56 | Poly(ADP-ribose) glycohydrolase OS=Hc | PARG | 111.111 | 2 | 53.182323 | yes | 2.916E-07 | 735.89063 | 20.758831 |
| CTEIVAIDALHFR | 13 | Q86W56 | Poly(ADP-ribose) glycohydrolase OS=Hc | PARG | 111.111 | 2 | 53.182323 | yes | 5.185E-07 | 772.89813 | 13.844667 |
| WGAATTSPAASDA | 14 | Q86W56 | Poly(ADP-ribose) glycohydrolase OS=Hc | PARG | 111.111 | 2 | 53.182323 | yes | 6.188E-06 | 681.32587 | 13.34695 |
| GLLEMMETDEK | 11 | Q86W92 | Liprin-beta-1 OS=Homo sapiens OX=960 | PPFIBP1 | 114.025 | 2 | 54.790653 | yes | 1.001E-07 | 648.29651 | 1445.6343 |
| SVDLAEYAPNLR | 12 | Q86W92 | Liprin-beta-1 OS=Homo sapiens OX=960 | PPFIBP1 | 114.025 | 2 | 54.790653 | no | 3.759E-07 | 674.34882 | 1250.5315 |
| DTEGLIQEINDLR | 13 | Q86W92 | Liprin-beta-1 OS=Homo sapiens OX=960 | PPFIBP1 | 114.025 | 2 | 54.790653 | yes | 9.839E-08 | 758.38611 | 1245.7043 |
| ALHLVEDLR | 9 | Q86W92 | Liprin-beta-1 OS=Homo sapiens OX=960 | PPFIBP1 | 114.025 | 2 | 54.790653 | yes | 3.796E-07 | 533.30621 | 1067.6078 |
| WLDDIGLPQYK | 11 | Q86W92 | Liprin-beta-1 OS=Homo sapiens OX=960 | PPFIBP1 | 114.025 | 2 | 54.790653 | no | 1.006E-07 | 674.35083 | 978.23309 |
| DLGQSNSDLDMPF | 15 | Q86W92 | Liprin-beta-1 OS=Homo sapiens OX=960 | PPFIBP1 | 114.025 | 2 | 54.790653 | yes | 1.348E-07 | 819.37744 | 958.23877 |

| | | | | | | | | | | | |
|---|---|---|---|---|---|---|---|---|---|---|---|
| LAFSNFGNLR | 10 | Q86W92 | Liprin-beta-1 OS=Homo sapiens OX=96( | PPFIBP1 | 114.025 | 2 | 54.790653 | yes | 1.131E-07 | 569.80383 | 919.15936 |
| DLEFCLEEHR | 10 | Q86W92 | Liprin-beta-1 OS=Homo sapiens OX=96( | PPFIBP1 | 114.025 | 2 | 54.790653 | yes | 9.722E-08 | 674.30353 | 809.45929 |
| LDFNWVTR | 8 | Q86W92 | Liprin-beta-1 OS=Homo sapiens OX=96( | PPFIBP1 | 114.025 | 2 | 54.790653 | yes | 3.021E-06 | 525.77197 | 796.02362 |
| HLATHFNLLIGAE/ | 18 | Q86W92 | Liprin-beta-1 OS=Homo sapiens OX=96( | PPFIBP1 | 114.025 | 3 | 54.790653 | yes | 9.986E-08 | 676.69733 | 337.95154 |
| HLATHFNLLIGAE/ | 18 | Q86W92 | Liprin-beta-1 OS=Homo sapiens OX=96( | PPFIBP1 | 114.025 | 3 | 54.790653 | yes | 1.934E-07 | 507.77481 | 337.95154 |
| HLATHFNLLIGAE/ | 18 | Q86W92 | Liprin-beta-1 OS=Homo sapiens OX=96( | PPFIBP1 | 114.025 | 4 | 54.790653 | yes | 9.986E-08 | 676.69733 | 337.95154 |
| HLATHFNLLIGAE/ | 18 | Q86W92 | Liprin-beta-1 OS=Homo sapiens OX=96( | PPFIBP1 | 114.025 | 4 | 54.790653 | yes | 1.934E-07 | 507.77481 | 337.95154 |
| VSEMDSER | 8 | Q86W92 | Liprin-beta-1 OS=Homo sapiens OX=96( | PPFIBP1 | 114.025 | 2 | 54.790653 | yes | 1.341E-07 | 476.70566 | 552.16272 |
| GFKPGLDMR | 9 | Q86W92 | Liprin-beta-1 OS=Homo sapiens OX=96( | PPFIBP1 | 114.025 | 2 | 54.790653 | yes | 4.731E-07 | 510.7684 | 478.04138 |
| VVSVLHHLSIK | 11 | Q86W92 | Liprin-beta-1 OS=Homo sapiens OX=96( | PPFIBP1 | 114.025 | 2 | 54.790653 | yes | 9.823E-08 | 616.3797 | 425.51123 |
| LDLMAEISNLK | 11 | Q86W92 | Liprin-beta-1 OS=Homo sapiens OX=96( | PPFIBP1 | 114.025 | 2 | 54.790653 | yes | 1.393E-07 | 623.83923 | 414.91577 |
| DRLDYEDK | 8 | Q86W92 | Liprin-beta-1 OS=Homo sapiens OX=96( | PPFIBP1 | 114.025 | 2 | 54.790653 | yes | 1.974E-07 | 527.24603 | 259.54578 |
| GSGVHGGLMVLEI | 14 | Q86W92 | Liprin-beta-1 OS=Homo sapiens OX=96( | PPFIBP1 | 114.025 | 2 | 54.790653 | yes | 1.346E-07 | 704.87189 | 233.88699 |
| MKGEGVEIVDR | 11 | Q86W92 | Liprin-beta-1 OS=Homo sapiens OX=96( | PPFIBP1 | 114.025 | 2 | 54.790653 | yes | 1.152E-07 | 616.81879 | 216.47292 |
| ETAEHLDLAGASS | 14 | Q86W92 | Liprin-beta-1 OS=Homo sapiens OX=96( | PPFIBP1 | 114.025 | 2 | 54.790653 | yes | 1.437E-07 | 728.85498 | 184.72968 |
| LLTSSLQK | 8 | Q86W92 | Liprin-beta-1 OS=Homo sapiens OX=96( | PPFIBP1 | 114.025 | 2 | 54.790653 | yes | 1.401E-07 | 445.2713 | 180.22916 |
| INNFEPNCLR | 10 | Q86W92 | Liprin-beta-1 OS=Homo sapiens OX=96( | PPFIBP1 | 114.025 | 2 | 54.790653 | yes | 9.927E-08 | 638.80878 | 178.98618 |
| HWIASGQTLLQAS | 19 | Q86W92 | Liprin-beta-1 OS=Homo sapiens OX=96( | PPFIBP1 | 114.025 | 3 | 54.790653 | yes | 9.722E-08 | 718.37103 | 176.90631 |
| MQDTVVLAQGK | 11 | Q86W92 | Liprin-beta-1 OS=Homo sapiens OX=96( | PPFIBP1 | 114.025 | 2 | 54.790653 | yes | 4.038E-07 | 595.31592 | 166.36024 |
| KLQLALQALGSEE | 19 | Q86W92 | Liprin-beta-1 OS=Homo sapiens OX=96( | PPFIBP1 | 114.025 | 3 | 54.790653 | yes | 4.712E-07 | 689.36743 | 152.56133 |
| AVESLMAANEEK | 12 | Q86W92 | Liprin-beta-1 OS=Homo sapiens OX=96( | PPFIBP1 | 114.025 | 2 | 54.790653 | yes | 9.909E-08 | 646.31354 | 131.0638 |
| LEQMEDSEGTVR | 12 | Q86W92 | Liprin-beta-1 OS=Homo sapiens OX=96( | PPFIBP1 | 114.025 | 2 | 54.790653 | yes | 9.722E-08 | 697.31683 | 124.28603 |
| LYEEDDLDR | 9 | Q86W92 | Liprin-beta-1 OS=Homo sapiens OX=96( | PPFIBP1 | 114.025 | 2 | 54.790653 | yes | 2.818E-07 | 584.26184 | 122.85764 |
| KGFKPGLDMR | 10 | Q86W92 | Liprin-beta-1 OS=Homo sapiens OX=96( | PPFIBP1 | 114.025 | 2 | 54.790653 | yes | 3.471E-07 | 574.81586 | 107.23923 |
| NSPFQIPPPSPDSK | 14 | Q86W92 | Liprin-beta-1 OS=Homo sapiens OX=96( | PPFIBP1 | 114.025 | 2 | 54.790653 | yes | 3.29E-06 | 755.88043 | 92.430237 |
| TASAPNLAETEK | 12 | Q86W92 | Liprin-beta-1 OS=Homo sapiens OX=96( | PPFIBP1 | 114.025 | 2 | 54.790653 | yes | 2.906E-06 | 616.31189 | 84.963791 |
| AVESLMAANEEKI | 14 | Q86W92 | Liprin-beta-1 OS=Homo sapiens OX=96( | PPFIBP1 | 114.025 | 2 | 54.790653 | yes | 9.723E-08 | 781.87756 | 83.571625 |
| MLHYMTVDDLLS | 14 | Q86W92 | Liprin-beta-1 OS=Homo sapiens OX=96( | PPFIBP1 | 114.025 | 2 | 54.790653 | yes | 9.722E-08 | 839.93079 | 65.734444 |
| ETSDGEKETIQK | 12 | Q86W92 | Liprin-beta-1 OS=Homo sapiens OX=96( | PPFIBP1 | 114.025 | 2 | 54.790653 | yes | 9.722E-08 | 682.83063 | 35.884583 |
| KMQDTVVLAQGK | 12 | Q86W92 | Liprin-beta-1 OS=Homo sapiens OX=96( | PPFIBP1 | 114.025 | 2 | 54.790653 | yes | 2.912E-07 | 659.3634 | 32.823986 |
| RPSDENTIAPSEVC | 15 | Q86W92 | Liprin-beta-1 OS=Homo sapiens OX=96( | PPFIBP1 | 114.025 | 2 | 54.790653 | yes | 1.093E-07 | 835.92084 | 29.23999 |
| VAEGQHKEAKLDI | 13 | Q86WA8 | Lon protease homolog 2, peroxisomal OS | LONP2 | 94.617 | 2 | 59.25273 | yes | 2.856E-06 | 740.89697 | 3952.8337 |
| MEIIQVPGYTQEEI | 14 | Q86WA8 | Lon protease homolog 2, peroxisomal OS | LONP2 | 94.617 | 2 | 59.25273 | yes | 3.022E-07 | 832.91364 | 531.32123 |
| GPILCFVGPPGVGI | 14 | Q86WA8 | Lon protease homolog 2, peroxisomal OS | LONP2 | 94.617 | 2 | 59.25273 | yes | 9.722E-08 | 699.38409 | 317.44788 |
| LQILDAVSLEER | 12 | Q86WA8 | Lon protease homolog 2, peroxisomal OS | LONP2 | 94.617 | 2 | 59.25273 | yes | 1.294E-07 | 693.38538 | 281.35403 |
| SDVAMTGEITLR | 12 | Q86WA8 | Lon protease homolog 2, peroxisomal OS | LONP2 | 94.617 | 2 | 59.25273 | yes | 1.961E-07 | 646.82941 | 268.56885 |
| VLEYLAVR | 8 | Q86WA8 | Lon protease homolog 2, peroxisomal OS | LONP2 | 94.617 | 2 | 59.25273 | yes | 4.166E-07 | 481.78711 | 235.37984 |
| IALGGVCDQSDIR | 13 | Q86WA8 | Lon protease homolog 2, peroxisomal OS | LONP2 | 94.617 | 2 | 59.25273 | yes | 9.722E-08 | 702.35101 | 208.92238 |
| MPQSMPEYALTR | 12 | Q86WA8 | Lon protease homolog 2, peroxisomal OS | LONP2 | 94.617 | 2 | 59.25273 | yes | 1.099E-07 | 712.33905 | 192.27405 |

| | | | | | | | | | | | |
|---|---|---|---|---|---|---|---|---|---|---|---|
| ESAHLAISWLR | 11 | Q86WA8 | Lon protease homolog 2, peroxisomal OS | LONP2 | 94.617 | 2 | 59.25273 | yes | 9.722E-08 | 641.84875 | 186.1597 |
| FQIVQVLK | 8 | Q86WA8 | Lon protease homolog 2, peroxisomal OS | LONP2 | 94.617 | 2 | 59.25273 | yes | 4.359E-06 | 487.8053 | 184.48309 |
| GLVLPVGGIK | 10 | Q86WA8 | Lon protease homolog 2, peroxisomal OS | LONP2 | 94.617 | 2 | 59.25273 | yes | 3.413E-07 | 476.81314 | 178.00128 |
| MTIPLLVR | 8 | Q86WA8 | Lon protease homolog 2, peroxisomal OS | LONP2 | 94.617 | 2 | 59.25273 | yes | 2.311E-07 | 471.79388 | 165.22079 |
| ILLDNDHYAMEK | 12 | Q86WA8 | Lon protease homolog 2, peroxisomal OS | LONP2 | 94.617 | 2 | 59.25273 | yes | 9.722E-08 | 731.35577 | 103.73075 |
| LEEFPNTCK | 9 | Q86WA8 | Lon protease homolog 2, peroxisomal OS | LONP2 | 94.617 | 2 | 59.25273 | yes | 2.644E-06 | 569.26587 | 39.630669 |
| VIAIRPIR | 8 | Q86WA8 | Lon protease homolog 2, peroxisomal OS | LONP2 | 94.617 | 2 | 59.25273 | yes | 2.138E-07 | 469.31891 | 37.28056 |
| EKPYPIAEVEQLDI | 14 | Q86WA8 | Lon protease homolog 2, peroxisomal OS | LONP2 | 94.617 | 2 | 59.25273 | yes | 2.789E-06 | 843.93848 | 16.802891 |
| AVLTILLAHK | 10 | Q86WB0 | Zinc finger C3HC-type protein 1 OS=Ho | ZC3HC1 | 55.262 | 2 | 40.130863 | yes | 4.194E-07 | 539.85278 | 8.6109056 |
| EFQVISEQLQK | 11 | Q86WP2 | Vasculin OS=Homo sapiens OX=9606 G | GPBP1 | 53.34 | 2 | 55.68248 | yes | 2.273E-07 | 674.85901 | 771.90674 |
| PAAPPTKPTQWK | 12 | Q86WP2 | Vasculin OS=Homo sapiens OX=9606 G | GPBP1 | 53.34 | 3 | 55.68248 | yes | 1.242E-07 | 441.24695 | 65.645752 |
| PAAPPTKPTQWK | 12 | Q86WP2 | Vasculin OS=Homo sapiens OX=9606 G | GPBP1 | 53.34 | 3 | 55.68248 | yes | 3.14E-06 | 661.36676 | 65.645752 |
| PAAPPTKPTQWK | 12 | Q86WP2 | Vasculin OS=Homo sapiens OX=9606 G | GPBP1 | 53.34 | 2 | 55.68248 | yes | 1.242E-07 | 441.24695 | 65.645752 |
| PAAPPTKPTQWK | 12 | Q86WP2 | Vasculin OS=Homo sapiens OX=9606 G | GPBP1 | 53.34 | 2 | 55.68248 | yes | 3.14E-06 | 661.36676 | 65.645752 |
| HSENFAWTENR | 11 | Q86WP2 | Vasculin OS=Homo sapiens OX=9606 G | GPBP1 | 53.34 | 2 | 55.68248 | yes | 9.722E-08 | 695.81036 | 78.262367 |
| STVAESVSQQILR | 13 | Q86WX3 | Active regulator of SIRT1 OS=Homo sap | RPS19BP1 | 15.434 | 2 | 46.961323 | yes | 1.431E-07 | 709.38593 | 166.34123 |
| LQTLVGALR | 9 | Q86X02 | Cerebellar degeneration-related protein 2- | CDR2L | 53.011 | 2 | 47.93377 | yes | 7.492E-07 | 485.80585 | 182.41695 |
| DLELTNHR | 8 | Q86X02 | Cerebellar degeneration-related protein 2- | CDR2L | 53.011 | 2 | 47.93377 | yes | 2.107E-07 | 499.25671 | 112.77466 |
| VYEQLDLTAR | 10 | Q86X02 | Cerebellar degeneration-related protein 2- | CDR2L | 53.011 | 2 | 47.93377 | yes | 4.296E-07 | 604.31952 | 87.899284 |
| IHGLTETIER | 10 | Q86X02 | Cerebellar degeneration-related protein 2- | CDR2L | 53.011 | 2 | 47.93377 | yes | 1.23E-07 | 584.81964 | 58.376392 |
| EVDEQYHALLEK | 12 | Q86X02 | Cerebellar degeneration-related protein 2- | CDR2L | 53.011 | 2 | 47.93377 | yes | 1.606E-07 | 737.36462 | 42.60014 |
| QLCEMEACR | 9 | Q86X02 | Cerebellar degeneration-related protein 2- | CDR2L | 53.011 | 2 | 47.93377 | yes | 3.517E-07 | 598.74646 | 37.323906 |
| TCLIPNQNVK | 10 | Q86X52 | Chondroitin sulfate synthase 1 OS=Homo | CHSY1 | 91.785 | 2 | 51.845997 | yes | 2.023E-07 | 593.8161 | 839.17908 |
| ADMQILPVSGEFSI | 14 | Q86X52 | Chondroitin sulfate synthase 1 OS=Homo | CHSY1 | 91.785 | 2 | 51.845997 | yes | 1.42E-07 | 775.38757 | 552.55798 |
| YLYSAVDGQPPR | 12 | Q86X52 | Chondroitin sulfate synthase 1 OS=Homo | CHSY1 | 91.785 | 2 | 51.845997 | yes | 9.722E-08 | 683.34351 | 450.81479 |
| LVVLLFNSDSNPD | 14 | Q86X52 | Chondroitin sulfate synthase 1 OS=Homo | CHSY1 | 91.785 | 2 | 51.845997 | yes | 9.894E-08 | 780.91705 | 412.41568 |
| NYGFGITCIYK | 11 | Q86X52 | Chondroitin sulfate synthase 1 OS=Homo | CHSY1 | 91.785 | 2 | 51.845997 | yes | 1.663E-07 | 668.32373 | 376.86856 |
| IHQAITLHPNK | 11 | Q86X52 | Chondroitin sulfate synthase 1 OS=Homo | CHSY1 | 91.785 | 2 | 51.845997 | yes | 1.128E-07 | 636.36456 | 294.69858 |
| LHSYMLSR | 8 | Q86X52 | Chondroitin sulfate synthase 1 OS=Homo | CHSY1 | 91.785 | 2 | 51.845997 | yes | 9.823E-08 | 503.76056 | 228.29849 |
| INILIPLSGR | 10 | Q86X52 | Chondroitin sulfate synthase 1 OS=Homo | CHSY1 | 91.785 | 2 | 51.845997 | yes | 2.283E-07 | 548.34784 | 180.33377 |
| EDLQLGIPPSFMR | 13 | Q86X52 | Chondroitin sulfate synthase 1 OS=Homo | CHSY1 | 91.785 | 2 | 51.845997 | yes | 1.052E-06 | 751.88721 | 161.76469 |
| ADDDVYIK | 8 | Q86X52 | Chondroitin sulfate synthase 1 OS=Homo | CHSY1 | 91.785 | 2 | 51.845997 | no | 1.902E-07 | 469.72693 | 131.27533 |
| VPSDNHFAFTQK | 12 | Q86X52 | Chondroitin sulfate synthase 1 OS=Homo | CHSY1 | 91.785 | 2 | 51.845997 | yes | 4.347E-06 | 695.84113 | 114.0304 |
| LVPFQLPGSK | 10 | Q86X52 | Chondroitin sulfate synthase 1 OS=Homo | CHSY1 | 91.785 | 2 | 51.845997 | yes | 9.722E-08 | 543.32129 | 111.73972 |
| EMYTTHEDVEVGI | 13 | Q86X52 | Chondroitin sulfate synthase 1 OS=Homo | CHSY1 | 91.785 | 2 | 51.845997 | no | 1.553E-07 | 783.34863 | 79.198708 |
| KINILIPLSGR | 11 | Q86X52 | Chondroitin sulfate synthase 1 OS=Homo | CHSY1 | 91.785 | 2 | 51.845997 | yes | 9.722E-08 | 612.39532 | 51.365395 |
| FMGNFEK | 7 | Q86X52 | Chondroitin sulfate synthase 1 OS=Homo | CHSY1 | 91.785 | 2 | 51.845997 | yes | 2.896E-07 | 436.70218 | 25.490128 |
| NFLFVGVMTAQK | 12 | Q86X52 | Chondroitin sulfate synthase 1 OS=Homo | CHSY1 | 91.785 | 2 | 51.845997 | yes | 6.635E-07 | 677.86304 | 24.399321 |
| YMHDHYLDK | 9 | Q86X52 | Chondroitin sulfate synthase 1 OS=Homo | CHSY1 | 91.785 | 2 | 51.845997 | no | 2.277E-06 | 611.27148 | 20.529099 |

| | | | | | | | | | | | |
|---|---|---|---|---|---|---|---|---|---|---|---|
| GAAVDEYFR | 9 | Q86X55 | Histone-arginine methyltransferase CARM | CARM1 | 65.854 | 2 | 48.521564 | yes | 1.136E-07 | 514.24579 | 103.83267 |
| CLFQSPLFAK | 10 | Q86X55 | Histone-arginine methyltransferase CARM | CARM1 | 65.854 | 2 | 48.521564 | yes | 3.651E-06 | 605.81805 | 51.980495 |
| AGDTLSGTCLLIAN | 15 | Q86X55 | Histone-arginine methyltransferase CARM | CARM1 | 65.854 | 2 | 48.521564 | yes | 1.252E-07 | 767.4007 | 51.104313 |
| MLESYLHAK | 9 | Q86X55 | Histone-arginine methyltransferase CARM | CARM1 | 65.854 | 2 | 48.521564 | yes | 1.812E-07 | 546.28131 | 42.624886 |
| LLTIGDANGEIQR | 13 | Q86X55 | Histone-arginine methyltransferase CARM | CARM1 | 65.854 | 2 | 48.521564 | yes | 3.837E-06 | 700.38062 | 30.60951 |
| HSLLELER | 8 | Q86X67 | NAD(P)H pyrophosphatase NUDT13, mi | NUDT13 | 39.689 | 2 | 32.345387 | yes | 1.546E-06 | 498.77728 | 87.925529 |
| SHGTDLYR | 8 | Q86X95 | Corepressor interacting with RBPJ 1 OS= | CIR1 | 52.314 | 2 | 50.505962 | yes | 1.232E-07 | 474.73032 | 349.00934 |
| LLMGDER | 7 | Q86X95 | Corepressor interacting with RBPJ 1 OS= | CIR1 | 52.314 | 2 | 50.505962 | yes | 2.645E-07 | 417.21292 | 143.27798 |
| LLMGDER | 7 | Q86X95 | Corepressor interacting with RBPJ 1 OS= | CIR1 | 52.314 | 2 | 50.505962 | yes | 5.788E-06 | 425.21039 | 143.27798 |
| DQPFGIQVR | 9 | Q86X95 | Corepressor interacting with RBPJ 1 OS= | CIR1 | 52.314 | 2 | 50.505962 | yes | 7.982E-07 | 530.28271 | 199.46054 |
| NGLNFMYEAPPGA | 14 | Q86X95 | Corepressor interacting with RBPJ 1 OS= | CIR1 | 52.314 | 2 | 50.505962 | yes | 5.275E-07 | 754.86377 | 108.78988 |
| QEELMQQYLK | 10 | Q86X95 | Corepressor interacting with RBPJ 1 OS= | CIR1 | 52.314 | 2 | 50.505962 | yes | 7.531E-07 | 655.32648 | 65.553604 |
| KQEELMQQYLK | 11 | Q86X95 | Corepressor interacting with RBPJ 1 OS= | CIR1 | 52.314 | 2 | 50.505962 | yes | 9.79E-08 | 719.37396 | 47.4981 |
| EITGSSLIQK | 10 | Q86XI2 | Condensin-2 complex subunit G2 OS=Ho | NCAPG2 | 130.961 | 2 | 52.356674 | yes | 3.768E-07 | 538.30334 | 260.60873 |
| STGILGVCK | 9 | Q86XI2 | Condensin-2 complex subunit G2 OS=Ho | NCAPG2 | 130.961 | 2 | 52.356674 | yes | 2.083E-07 | 467.75494 | 129.24757 |
| CLPMILDNK | 9 | Q86XI2 | Condensin-2 complex subunit G2 OS=Ho | NCAPG2 | 130.961 | 2 | 52.356674 | yes | 1.77E-06 | 552.28302 | 124.35616 |
| FASVLPEYLK | 10 | Q86XI2 | Condensin-2 complex subunit G2 OS=Ho | NCAPG2 | 130.961 | 2 | 52.356674 | yes | 1.459E-07 | 583.82642 | 96.67659 |
| VTGELAFDTSSAD | 15 | Q86XI2 | Condensin-2 complex subunit G2 OS=Ho | NCAPG2 | 130.961 | 2 | 52.356674 | yes | 3.525E-07 | 784.38354 | 91.556686 |
| ADLESLLQTPGGK | 13 | Q86XI2 | Condensin-2 complex subunit G2 OS=Ho | NCAPG2 | 130.961 | 2 | 52.356674 | yes | 1.064E-07 | 664.85645 | 90.177719 |
| NQLQGLQK | 8 | Q86XI2 | Condensin-2 complex subunit G2 OS=Ho | NCAPG2 | 130.961 | 2 | 52.356674 | yes | 1.908E-07 | 464.76416 | 64.067764 |
| FLYESSSR | 8 | Q86XI2 | Condensin-2 complex subunit G2 OS=Ho | NCAPG2 | 130.961 | 2 | 52.356674 | yes | 2.972E-07 | 494.74036 | 60.208858 |
| ADLESLLQTPGGK | 15 | Q86XI2 | Condensin-2 complex subunit G2 OS=Ho | NCAPG2 | 130.961 | 2 | 52.356674 | yes | 9.722E-08 | 791.43341 | 47.69104 |
| VAFVDMLLK | 9 | Q86XI2 | Condensin-2 complex subunit G2 OS=Ho | NCAPG2 | 130.961 | 2 | 52.356674 | yes | 9.809E-08 | 518.29901 | 40.734486 |
| HCLNACIQR | 9 | Q86XI2 | Condensin-2 complex subunit G2 OS=Ho | NCAPG2 | 130.961 | 2 | 52.356674 | yes | 3.249E-07 | 586.27661 | 27.307354 |
| HIAEDPPCSCSHR | 13 | Q86XJ1 | GAS2-like protein 3 OS=Homo sapiens C | GAS2L3 | 75.215 | 3 | 49.12685 | yes | 1.893E-07 | 522.55762 | 160.73616 |
| FSIEYLSEGR | 10 | Q86XJ1 | GAS2-like protein 3 OS=Homo sapiens C | GAS2L3 | 75.215 | 2 | 49.12685 | yes | 3.93E-07 | 600.7984 | 152.38904 |
| TPLSIVSLPQSSTK | 14 | Q86XJ1 | GAS2-like protein 3 OS=Homo sapiens C | GAS2L3 | 75.215 | 2 | 49.12685 | yes | 1.065E-07 | 729.41412 | 144.32564 |
| ANIPVRPK | 8 | Q86XJ1 | GAS2-like protein 3 OS=Homo sapiens C | GAS2L3 | 75.215 | 2 | 49.12685 | yes | 4.777E-07 | 447.77963 | 94.306084 |
| YGVEPPVLVK | 10 | Q86XJ1 | GAS2-like protein 3 OS=Homo sapiens C | GAS2L3 | 75.215 | 2 | 49.12685 | yes | 1.178E-06 | 550.82117 | 61.917538 |
| QVYLCLLEIGR | 11 | Q86XJ1 | GAS2-like protein 3 OS=Homo sapiens C | GAS2L3 | 75.215 | 2 | 49.12685 | yes | 6.41E-07 | 682.37378 | 40.694168 |
| IFSYLLEQDLCR | 12 | Q86XK2 | F-box only protein 11 OS=Homo sapiens | FBXO11 | 103.585 | 2 | 51.59584 | yes | 3.074E-07 | 778.89252 | 180.40398 |
| FSELANDPILWK | 12 | Q86XK2 | F-box only protein 11 OS=Homo sapiens | FBXO11 | 103.585 | 2 | 51.59584 | yes | 9.722E-08 | 716.8772 | 99.68457 |
| IMNNQDAIEK | 10 | Q86XK2 | F-box only protein 11 OS=Homo sapiens | FBXO11 | 103.585 | 2 | 51.59584 | yes | 2.081E-07 | 588.28992 | 81.207016 |
| ESFQQLYK | 8 | Q86XK2 | F-box only protein 11 OS=Homo sapiens | FBXO11 | 103.585 | 2 | 51.59584 | yes | 3.241E-07 | 521.76385 | 66.890144 |
| GLLEENDIFR | 10 | Q86XK2 | F-box only protein 11 OS=Homo sapiens | FBXO11 | 103.585 | 2 | 51.59584 | yes | 2.585E-07 | 603.31171 | 60.374409 |
| FGGLFLASGVNVT | 15 | Q86XK2 | F-box only protein 11 OS=Homo sapiens | FBXO11 | 103.585 | 2 | 51.59584 | yes | 1.66E-07 | 770.91327 | 53.59491 |
| YSPVHGSQLR | 10 | Q86XL3 | Ankyrin repeat and LEM domain-contain | ANKLE2 | 104.115 | 2 | 52.044384 | yes | 9.722E-08 | 572.29889 | 62.153851 |
| LAQALLEQGGR | 11 | Q86XL3 | Ankyrin repeat and LEM domain-contain | ANKLE2 | 104.115 | 2 | 52.044384 | yes | 1.54E-07 | 578.32764 | 58.311253 |
| LEEYLTQQEIGK | 12 | Q86XL3 | Ankyrin repeat and LEM domain-contain | ANKLE2 | 104.115 | 2 | 52.044384 | yes | 1.402E-07 | 725.87482 | 41.66156 |

| | | | | | | | | | | | |
|---|---|---|---|---|---|---|---|---|---|---|---|
| YVVDLYLNTPDK | 12 | Q86XL3 | Ankyrin repeat and LEM domain-contain | ANKLE2 | 104.115 | 2 | 52.044384 | yes | 2.108E-07 | 720.37445 | 39.170506 |
| RPPTATAGLR | 10 | Q86XN8 | RNA-binding protein MEX3D OS=Homo | MEX3D | 64.883 | 2 | 51.208244 | yes | 4.332E-06 | 520.30402 | 335.95297 |
| GEEPVFIVTGR | 11 | Q86XN8 | RNA-binding protein MEX3D OS=Homo | MEX3D | 64.883 | 2 | 51.208244 | yes | 1.042E-07 | 602.32202 | 233.39635 |
| THTYIVTPGR | 10 | Q86XN8 | RNA-binding protein MEX3D OS=Homo | MEX3D | 64.883 | 2 | 51.208244 | yes | 9.722E-08 | 572.80908 | 224.01787 |
| HSPTLPEPGGLR | 12 | Q86XN8 | RNA-binding protein MEX3D OS=Homo | MEX3D | 64.883 | 2 | 51.208244 | yes | 5.322E-07 | 630.83838 | 120.69775 |
| LELPLSR | 7 | Q86XN8 | RNA-binding protein MEX3D OS=Homo | MEX3D | 64.883 | 2 | 51.208244 | yes | 4.253E-07 | 414.2529 | 30.743259 |
| SEPECPACR | 9 | Q86XN8 | RNA-binding protein MEX3D OS=Homo | MEX3D | 64.883 | 2 | 51.208244 | yes | 8.672E-07 | 553.22369 | 21.347294 |
| VSYLVFDEADR | 11 | Q86XP3 | ATP-dependent RNA helicase DDX42 OS | DDX42 | 102.976 | 2 | 47.89675 | yes | 1.461E-07 | 657.32227 | 211.05191 |
| LNIGGGGLGYR | 11 | Q86XP3 | ATP-dependent RNA helicase DDX42 OS | DDX42 | 102.976 | 2 | 47.89675 | yes | 1.741E-07 | 538.79602 | 173.86485 |
| SAPPQLPSFYK | 11 | Q86XP3 | ATP-dependent RNA helicase DDX42 OS | DDX42 | 102.976 | 2 | 47.89675 | yes | 1.931E-07 | 617.82697 | 113.42495 |
| DIPVLVATDVAAR | 13 | Q86XP3 | ATP-dependent RNA helicase DDX42 OS | DDX42 | 102.976 | 2 | 47.89675 | yes | 1.236E-07 | 670.38263 | 91.758804 |
| DSNFAGDLVR | 10 | Q86XP3 | ATP-dependent RNA helicase DDX42 OS | DDX42 | 102.976 | 2 | 47.89675 | yes | 4.406E-07 | 547.26727 | 80.235321 |
| MFDMGFEYQVR | 11 | Q86XP3 | ATP-dependent RNA helicase DDX42 OS | DDX42 | 102.976 | 2 | 47.89675 | yes | 2.656E-07 | 711.81268 | 66.329811 |
| HGDGYRHPESSSR | 13 | Q86XP3 | ATP-dependent RNA helicase DDX42 OS | DDX42 | 102.976 | 3 | 47.89675 | yes | 7.973E-07 | 495.55902 | 51.094521 |
| GVAYTLLTPK | 10 | Q86XP3 | ATP-dependent RNA helicase DDX42 OS | DDX42 | 102.976 | 2 | 47.89675 | yes | 6.388E-07 | 531.81335 | 50.966 |
| SVAVYGGGSMWE | 15 | Q86XP3 | ATP-dependent RNA helicase DDX42 OS | DDX42 | 102.976 | 2 | 47.89675 | yes | 3.137E-06 | 785.37195 | 47.977585 |
| QTLLFSATFR | 10 | Q86XP3 | ATP-dependent RNA helicase DDX42 OS | DDX42 | 102.976 | 2 | 47.89675 | yes | 1.919E-06 | 592.32715 | 43.776943 |
| SHFVAASLSNQK | 12 | Q86XP3 | ATP-dependent RNA helicase DDX42 OS | DDX42 | 102.976 | 2 | 47.89675 | yes | 0.0001403 | 644.83588 | 17.78327 |
| LIQESPTVELK | 11 | Q86XR8 | Centrosomal protein of 57 kDa OS=Homo | CEP57 | 57.089 | 2 | 46.093666 | yes | 3.107E-06 | 628.85846 | 93.007843 |
| LDLLEQEYNK | 10 | Q86XR8 | Centrosomal protein of 57 kDa OS=Homo | CEP57 | 57.089 | 2 | 46.093666 | yes | 3.097E-07 | 632.82465 | 65.997971 |
| VINSIPLAK | 9 | Q86XR8 | Centrosomal protein of 57 kDa OS=Homo | CEP57 | 57.089 | 2 | 46.093666 | yes | 1.655E-07 | 477.80276 | 49.801895 |
| LCLGDMPFVAGK | 12 | Q86XR8 | Centrosomal protein of 57 kDa OS=Homo | CEP57 | 57.089 | 2 | 46.093666 | yes | 5.108E-05 | 654.32794 | 37.208733 |
| YKLPIPFPEGK | 11 | Q86Y07 | Serine/threonine-protein kinase VRK2 OS | VRK2 | 58.141 | 2 | 51.80869 | yes | 1.162E-07 | 644.86865 | 672.95044 |
| LPWEQNLK | 8 | Q86Y07 | Serine/threonine-protein kinase VRK2 OS | VRK2 | 58.141 | 2 | 51.80869 | yes | 9.722E-08 | 514.28217 | 672.22296 |
| SDVEILGYCMLR | 12 | Q86Y07 | Serine/threonine-protein kinase VRK2 OS | VRK2 | 58.141 | 2 | 51.80869 | yes | 1.306E-07 | 728.35217 | 598.39288 |
| TNLLDELPQSVLK | 13 | Q86Y07 | Serine/threonine-protein kinase VRK2 OS | VRK2 | 58.141 | 2 | 51.80869 | yes | 1.006E-07 | 735.41412 | 589.46851 |
| VLDDMEGNQWVI | 14 | Q86Y07 | Serine/threonine-protein kinase VRK2 OS | VRK2 | 58.141 | 2 | 51.80869 | yes | 1.638E-07 | 802.39288 | 537.41492 |
| LIGLMNNEAAQES | 15 | Q86Y07 | Serine/threonine-protein kinase VRK2 OS | VRK2 | 58.141 | 2 | 51.80869 | yes | 2.366E-07 | 823.91199 | 382.77118 |
| VEYQENGPLFSEL | 14 | Q86Y07 | Serine/threonine-protein kinase VRK2 OS | VRK2 | 58.141 | 2 | 51.80869 | yes | 1.022E-07 | 826.91193 | 170.13226 |
| QYQENPR | 7 | Q86Y07 | Serine/threonine-protein kinase VRK2 OS | VRK2 | 58.141 | 2 | 51.80869 | yes | 4.502E-07 | 467.7225 | 139.01418 |
| GQSINVHTPNSQK | 13 | Q86Y07 | Serine/threonine-protein kinase VRK2 OS | VRK2 | 58.141 | 2 | 51.80869 | yes | 2.894E-07 | 705.36023 | 110.84038 |
| LPIPFPEGK | 9 | Q86Y07 | Serine/threonine-protein kinase VRK2 OS | VRK2 | 58.141 | 2 | 51.80869 | yes | 1.17E-07 | 499.28949 | 46.423649 |
| KIGSGGFGLIYLAF | 21 | Q86Y07 | Serine/threonine-protein kinase VRK2 OS | VRK2 | 58.141 | 3 | 51.80869 | yes | 3.456E-06 | 746.41583 | 18.396469 |
| KPPSRQK | 7 | Q86Y38 | Xylosyltransferase 1 OS=Homo sapiens C | XYLT1 | 107.57 | 2 | 40.922104 | yes | 3.769E-07 | 420.75613 | 73.924866 |
| VTLNPPGTFLEGV | 15 | Q86Y39 | NADH dehydrogenase [ubiquinone] 1 alp | NDUFA11 | 14.853 | 2 | 41.842915 | yes | 9.043E-07 | 771.92993 | 90.914391 |
| LEGWEVFAK | 9 | Q86Y39 | NADH dehydrogenase [ubiquinone] 1 alp | NDUFA11 | 14.853 | 2 | 41.842915 | yes | 3.314E-07 | 539.78204 | 81.477272 |
| VPGLTQAVQVAK | 12 | Q86Y91 | Kinesin-like protein KIF18B OS=Homo s | KIF18B | 93.011 | 2 | 50.559097 | yes | 9.766E-08 | 605.86133 | 194.38069 |
| QSFLPCLR | 8 | Q86Y91 | Kinesin-like protein KIF18B OS=Homo s | KIF18B | 93.011 | 2 | 50.559097 | yes | 1.028E-07 | 510.7684 | 80.19606 |
| ALQEESLGMEAQV | 15 | Q86Y91 | Kinesin-like protein KIF18B OS=Homo s | KIF18B | 93.011 | 2 | 50.559097 | yes | 2.951E-07 | 845.40924 | 75.410637 |

| | | | | | | | | | | | |
|---|---|---|---|---|---|---|---|---|---|---|---|
| WGGTHDGPK | 9 | Q86Y91 | Kinesin-like protein KIF18B OS=Homo s | KIF18B | 93.011 | 2 | 50.559097 | yes | 3.637E-06 | 477.72504 | 57.53772 |
| GSLPDTQPSQGPST | 16 | Q86Y91 | Kinesin-like protein KIF18B OS=Homo s | KIF18B | 93.011 | 2 | 50.559097 | yes | 6.604E-07 | 798.89685 | 32.661407 |
| AVEDSTLQVVVR | 12 | Q86Y91 | Kinesin-like protein KIF18B OS=Homo s | KIF18B | 93.011 | 2 | 50.559097 | yes | 1E-07 | 658.36444 | 26.858402 |
| RPVVQVVDER | 10 | Q86Y91 | Kinesin-like protein KIF18B OS=Homo s | KIF18B | 93.011 | 2 | 50.559097 | yes | 1.576E-07 | 598.84094 | 11.456659 |
| VIANQQQVLQR | 11 | Q86YD1 | Prostate tumor-overexpressed gene 1 prot | PTOV1 | 46.869 | 2 | 51.73035 | yes | 9.722E-08 | 648.87274 | 80.794426 |
| VLMLLYSSK | 9 | Q86YD1 | Prostate tumor-overexpressed gene 1 prot | PTOV1 | 46.869 | 2 | 51.73035 | no | 4.448E-07 | 527.30426 | 45.652954 |
| VTPEPGAGPTQGL | 15 | Q86YP4 | Transcriptional repressor p66-alpha OS=I | GATAD2A | 68.063 | 2 | 53.659588 | yes | 2.021E-07 | 746.90955 | 283.90427 |
| GEGLVGDGPVDM | 13 | Q86YP4 | Transcriptional repressor p66-alpha OS=I | GATAD2A | 68.063 | 2 | 53.659588 | yes | 6.174E-07 | 651.31134 | 165.96605 |
| LLQQGTAPAQAK | 12 | Q86YP4 | Transcriptional repressor p66-alpha OS=I | GATAD2A | 68.063 | 2 | 53.659588 | yes | 1.403E-07 | 613.34857 | 141.92595 |
| GLLASDLNTDGDM | 14 | Q86YP4 | Transcriptional repressor p66-alpha OS=I | GATAD2A | 68.063 | 2 | 53.659588 | yes | 2.277E-06 | 739.3512 | 122.37366 |
| LGPQASSQVVMPI | 16 | Q86YP4 | Transcriptional repressor p66-alpha OS=I | GATAD2A | 68.063 | 2 | 53.659588 | yes | 1.371E-07 | 839.96906 | 103.31013 |
| ALQQEQEIEQR | 11 | Q86YP4 | Transcriptional repressor p66-alpha OS=I | GATAD2A | 68.063 | 2 | 53.659588 | no | 5.157E-07 | 686.3468 | 101.16071 |
| ASVPSVQIQGQR | 12 | Q86YP4 | Transcriptional repressor p66-alpha OS=I | GATAD2A | 68.063 | 2 | 53.659588 | yes | 9.722E-08 | 635.34912 | 74.248238 |
| LQNSASATALVSR | 13 | Q86YP4 | Transcriptional repressor p66-alpha OS=I | GATAD2A | 68.063 | 2 | 53.659588 | yes | 1.376E-07 | 659.35968 | 65.773193 |
| PSLQTSSAR | 9 | Q86YP4 | Transcriptional repressor p66-alpha OS=I | GATAD2A | 68.063 | 2 | 53.659588 | yes | 1.02E-07 | 473.75125 | 59.653748 |
| MSAATVLSR | 9 | Q86YP4 | Transcriptional repressor p66-alpha OS=I | GATAD2A | 68.063 | 2 | 53.659588 | yes | 3.03E-06 | 468.25256 | 49.000237 |
| IIQQGLIR | 8 | Q86YP4 | Transcriptional repressor p66-alpha OS=I | GATAD2A | 68.063 | 2 | 53.659588 | yes | 4.867E-07 | 470.80054 | 37.727226 |
| EATAQKPTGSVGS | 23 | Q86YP4 | Transcriptional repressor p66-alpha OS=I | GATAD2A | 68.063 | 3 | 53.659588 | yes | 2.562E-07 | 765.41656 | 29.587294 |
| EQTMCPVCLDR | 11 | Q86YT6 | E3 ubiquitin-protein ligase MIB1 OS=Ho | MIB1 | 110.136 | 2 | 52.826057 | yes | 9.722E-08 | 704.8045 | 194.31335 |
| VGFEGMSDLK | 10 | Q86YT6 | E3 ubiquitin-protein ligase MIB1 OS=Ho | MIB1 | 110.136 | 2 | 52.826057 | yes | 3.674E-07 | 541.763 | 156.35371 |
| VQQIYSDSDLK | 11 | Q86YT6 | E3 ubiquitin-protein ligase MIB1 OS=Ho | MIB1 | 110.136 | 2 | 52.826057 | yes | 1.303E-07 | 648.32751 | 78.470169 |
| ILDSAPTGIK | 10 | Q86YT6 | E3 ubiquitin-protein ligase MIB1 OS=Ho | MIB1 | 110.136 | 2 | 52.826057 | yes | 6.172E-07 | 507.79514 | 73.236916 |
| AVHHAAFGDEGA | 19 | Q86YT6 | E3 ubiquitin-protein ligase MIB1 OS=Ho | MIB1 | 110.136 | 3 | 52.826057 | yes | 9.722E-08 | 676.68518 | 71.112442 |
| NTLIMGLGTQGAE | 14 | Q86YT6 | E3 ubiquitin-protein ligase MIB1 OS=Ho | MIB1 | 110.136 | 2 | 52.826057 | yes | 2.421E-06 | 716.87683 | 69.585045 |
| WTFNPAVLTK | 10 | Q86YT6 | E3 ubiquitin-protein ligase MIB1 OS=Ho | MIB1 | 110.136 | 2 | 52.826057 | yes | 3.176E-06 | 588.82422 | 69.001251 |
| MSECPICR | 8 | Q86YT6 | E3 ubiquitin-protein ligase MIB1 OS=Ho | MIB1 | 110.136 | 2 | 52.826057 | no | 4.348E-07 | 526.71973 | 66.603767 |
| LQQQLQDIK | 9 | Q86YT6 | E3 ubiquitin-protein ligase MIB1 OS=Ho | MIB1 | 110.136 | 2 | 52.826057 | yes | 9.722E-08 | 557.31677 | 47.175625 |
| VASAGSAISNASGI | 15 | Q86YT6 | E3 ubiquitin-protein ligase MIB1 OS=Ho | MIB1 | 110.136 | 2 | 52.826057 | yes | 1.339E-07 | 688.84186 | 46.467701 |
| QDGGEGHVGTVR | 12 | Q86YT6 | E3 ubiquitin-protein ligase MIB1 OS=Ho | MIB1 | 110.136 | 2 | 52.826057 | yes | 3.777E-07 | 606.29181 | 42.51432 |
| VLQCLLWGPR | 10 | Q86YV5 | Inactive tyrosine-protein kinase PRAG1 ( | PRAG1 | 149.624 | 2 | 39.48397 | yes | 4.558E-07 | 621.34479 | 45.820568 |
| ALDAGLALGPSSPI | 16 | Q86YV9 | BLOC-2 complex member HPS6 OS=Ho | HPS6 | 82.975 | 2 | 49.81115 | yes | 1.002E-07 | 775.94867 | 94.243385 |
| ELVAGDSAVR | 10 | Q86YV9 | BLOC-2 complex member HPS6 OS=Ho | HPS6 | 82.975 | 2 | 49.81115 | yes | 4.278E-07 | 508.77219 | 93.216385 |
| AVLQAVGQLVQK | 12 | Q86YV9 | BLOC-2 complex member HPS6 OS=Ho | HPS6 | 82.975 | 2 | 49.81115 | yes | 1.066E-07 | 627.38245 | 62.320709 |
| LLLAEFAQHR | 10 | Q86YV9 | BLOC-2 complex member HPS6 OS=Ho | HPS6 | 82.975 | 2 | 49.81115 | yes | 4.629E-07 | 599.34058 | 51.845554 |
| TLEPSGEASTSLGR | 14 | Q86YV9 | BLOC-2 complex member HPS6 OS=Ho | HPS6 | 82.975 | 2 | 49.81115 | yes | 3.804E-06 | 702.85187 | 46.366524 |
| RIPLFSR | 7 | Q86YW0 | 1-phosphatidylinositol 4,5-bisphosphate p | PLCZ1 | 70.412 | 2 | 46.84867 | yes | 1.457E-07 | 444.77432 | 410.77817 |
| GPYESGSGHSSGL( | 16 | Q86YZ3 | Hornerin OS=Homo sapiens OX=9606 G | HRNR | 282.39 | 3 | 39.159214 | yes | 4.753E-07 | 528.90985 | 99.295937 |
| FILHIPSEER | 10 | Q8IU60 | m7GpppN-mRNA hydrolase OS=Homo s | DCP2 | 48.424 | 2 | 51.94708 | yes | 1.375E-07 | 620.83789 | 714.91229 |
| LYIIPGIPK | 9 | Q8IU60 | m7GpppN-mRNA hydrolase OS=Homo s | DCP2 | 48.424 | 2 | 51.94708 | yes | 2.089E-07 | 507.32333 | 312.29938 |

| | | | | | | | | | | | |
|---|---|---|---|---|---|---|---|---|---|---|---|
| EVFEETGFDIK | 11 | Q8IU60 | m7GpppN-mRNA hydrolase OS=Homo s | DCP2 | 48.424 | 2 | 51.94708 | yes | 9.722E-08 | 657.31665 | 229.65973 |
| FDHNAIMK | 8 | Q8IU60 | m7GpppN-mRNA hydrolase OS=Homo s | DCP2 | 48.424 | 2 | 51.94708 | yes | 4.897E-07 | 488.23947 | 191.75996 |
| QYQDSPNQK | 9 | Q8IU60 | m7GpppN-mRNA hydrolase OS=Homo s | DCP2 | 48.424 | 2 | 51.94708 | yes | 9.809E-08 | 554.2569 | 82.09668 |
| NIEWFSIEK | 9 | Q8IU60 | m7GpppN-mRNA hydrolase OS=Homo s | DCP2 | 48.424 | 2 | 51.94708 | yes | 4.134E-06 | 583.29803 | 16.228924 |
| LTVTDFFQK | 9 | Q8IUC4 | Rhophilin-2 OS=Homo sapiens OX=960( | RHPN2 | 76.994 | 2 | 55.2612 | yes | 9.722E-08 | 549.7951 | 79.326599 |
| TDALLPAAPQPLEI | 14 | Q8IUC4 | Rhophilin-2 OS=Homo sapiens OX=960( | RHPN2 | 76.994 | 2 | 55.2612 | yes | 1.659E-07 | 753.41412 | 39.123337 |
| AAILQTEVDALR | 12 | Q8IUD2 | ELKS/Rab6-interacting/CAST family mei | ERC1 | 128.087 | 2 | 52.379414 | no | 1.331E-07 | 650.367 | 469.47037 |
| DNTIMDLQTQLK | 12 | Q8IUD2 | ELKS/Rab6-interacting/CAST family mei | ERC1 | 128.087 | 2 | 52.379414 | yes | 9.753E-08 | 710.36102 | 315.1741 |
| LETLTNQFSDSK | 12 | Q8IUD2 | ELKS/Rab6-interacting/CAST family mei | ERC1 | 128.087 | 2 | 52.379414 | yes | 9.722E-08 | 691.84357 | 238.77298 |
| SAQMLEEAR | 9 | Q8IUD2 | ELKS/Rab6-interacting/CAST family mei | ERC1 | 128.087 | 2 | 52.379414 | yes | 1.003E-07 | 517.75043 | 191.24352 |
| EMVLAQEESAR | 11 | Q8IUD2 | ELKS/Rab6-interacting/CAST family mei | ERC1 | 128.087 | 2 | 52.379414 | yes | 1.262E-07 | 631.80591 | 180.38298 |
| DLNQLFQQDSSSR | 13 | Q8IUD2 | ELKS/Rab6-interacting/CAST family mei | ERC1 | 128.087 | 2 | 52.379414 | yes | 9.727E-08 | 769.36572 | 177.82991 |
| EHASSLASSGLK | 12 | Q8IUD2 | ELKS/Rab6-interacting/CAST family mei | ERC1 | 128.087 | 2 | 52.379414 | yes | 1.043E-07 | 593.80676 | 162.82622 |
| AHEAALEAR | 9 | Q8IUD2 | ELKS/Rab6-interacting/CAST family mei | ERC1 | 128.087 | 2 | 52.379414 | yes | 1.212E-07 | 484.25143 | 159.10701 |
| ETHLTNLR | 8 | Q8IUD2 | ELKS/Rab6-interacting/CAST family mei | ERC1 | 128.087 | 2 | 52.379414 | yes | 1.912E-07 | 492.26706 | 144.63136 |
| VSLLQGDLSEK | 11 | Q8IUD2 | ELKS/Rab6-interacting/CAST family mei | ERC1 | 128.087 | 2 | 52.379414 | yes | 4.048E-07 | 594.82715 | 139.00044 |
| KIENLQEQLR | 10 | Q8IUD2 | ELKS/Rab6-interacting/CAST family mei | ERC1 | 128.087 | 2 | 52.379414 | no | 9.722E-08 | 635.85931 | 132.87527 |
| AAGLQAEIGQVK | 12 | Q8IUD2 | ELKS/Rab6-interacting/CAST family mei | ERC1 | 128.087 | 2 | 52.379414 | yes | 8.241E-07 | 592.83533 | 131.97484 |
| QVEELLMAMEK | 11 | Q8IUD2 | ELKS/Rab6-interacting/CAST family mei | ERC1 | 128.087 | 2 | 52.379414 | yes | 1.084E-07 | 660.83051 | 121.68351 |
| GQLQDELEK | 9 | Q8IUD2 | ELKS/Rab6-interacting/CAST family mei | ERC1 | 128.087 | 2 | 52.379414 | yes | 3.161E-07 | 530.26947 | 118.04539 |
| QEALLAAISEK | 11 | Q8IUD2 | ELKS/Rab6-interacting/CAST family mei | ERC1 | 128.087 | 2 | 52.379414 | no | 4.658E-07 | 586.82971 | 109.89156 |
| IEELEEALR | 9 | Q8IUD2 | ELKS/Rab6-interacting/CAST family mei | ERC1 | 128.087 | 2 | 52.379414 | yes | 3.265E-07 | 551.29297 | 100.12456 |
| LLEMLQSK | 8 | Q8IUD2 | ELKS/Rab6-interacting/CAST family mei | ERC1 | 128.087 | 2 | 52.379414 | no | 8.763E-07 | 481.27298 | 75.968285 |
| KDTELLALQTK | 11 | Q8IUD2 | ELKS/Rab6-interacting/CAST family mei | ERC1 | 128.087 | 2 | 52.379414 | yes | 1.171E-07 | 630.36389 | 69.123688 |
| LSSTQQSLAEK | 11 | Q8IUD2 | ELKS/Rab6-interacting/CAST family mei | ERC1 | 128.087 | 2 | 52.379414 | yes | 4.245E-07 | 596.31439 | 67.824394 |
| VKQELESMK | 9 | Q8IUD2 | ELKS/Rab6-interacting/CAST family mei | ERC1 | 128.087 | 2 | 52.379414 | yes | 9.722E-08 | 546.29187 | 66.515305 |
| TLEIALEQK | 9 | Q8IUD2 | ELKS/Rab6-interacting/CAST family mei | ERC1 | 128.087 | 2 | 52.379414 | yes | 1.79E-06 | 522.80042 | 62.10083 |
| IENLQEQLR | 9 | Q8IUD2 | ELKS/Rab6-interacting/CAST family mei | ERC1 | 128.087 | 2 | 52.379414 | no | 6.678E-07 | 571.81183 | 61.536221 |
| TFWSPELK | 8 | Q8IUD2 | ELKS/Rab6-interacting/CAST family mei | ERC1 | 128.087 | 2 | 52.379414 | no | 1.988E-07 | 504.26346 | 60.696194 |
| LMADNYEDDHFK | 12 | Q8IUD2 | ELKS/Rab6-interacting/CAST family mei | ERC1 | 128.087 | 2 | 52.379414 | yes | 1.335E-07 | 749.3194 | 48.586792 |
| QIQDMAEEK | 9 | Q8IUD2 | ELKS/Rab6-interacting/CAST family mei | ERC1 | 128.087 | 2 | 52.379414 | yes | 9.796E-08 | 546.25549 | 40.684353 |
| ALQTVIEMK | 9 | Q8IUD2 | ELKS/Rab6-interacting/CAST family mei | ERC1 | 128.087 | 2 | 52.379414 | no | 2.24E-05 | 516.79156 | 39.855556 |
| EKQEEIDNYK | 10 | Q8IUD2 | ELKS/Rab6-interacting/CAST family mei | ERC1 | 128.087 | 2 | 52.379414 | yes | 4.616E-07 | 648.30933 | 19.542793 |
| HLQLAVR | 7 | Q8IUE6 | Histone H2A type 2-B OS=Homo sapiens | H2AC21 | 13.996 | 2 | 45.45327 | yes | 1.845E-07 | 418.7587 | 851.16815 |
| LYHPTVPLAR | 10 | Q8IUF8 | Ribosomal oxygenase 2 OS=Homo sapier | RIOX2 | 52.801 | 2 | 52.39705 | yes | 1.425E-07 | 583.83765 | 727.69598 |
| FKDELWR | 7 | Q8IUF8 | Ribosomal oxygenase 2 OS=Homo sapier | RIOX2 | 52.801 | 2 | 52.39705 | yes | 4.213E-07 | 497.26126 | 502.45105 |
| ATIQFHQPQR | 10 | Q8IUF8 | Ribosomal oxygenase 2 OS=Homo sapier | RIOX2 | 52.801 | 2 | 52.39705 | yes | 9.722E-08 | 613.32544 | 354.00754 |
| AHFLQLR | 7 | Q8IUF8 | Ribosomal oxygenase 2 OS=Homo sapier | RIOX2 | 52.801 | 2 | 52.39705 | yes | 1.584E-07 | 442.7587 | 291.24936 |
| FPLSHLDALK | 10 | Q8IUF8 | Ribosomal oxygenase 2 OS=Homo sapier | RIOX2 | 52.801 | 2 | 52.39705 | yes | 1.417E-07 | 570.82422 | 289.23132 |

| | | | | | | | | | | | |
|---|---|---|---|---|---|---|---|---|---|---|---|
| QIWNSPAISVK | 11 | Q8IUF8 | Ribosomal oxygenase 2 OS=Homo sapier | RIOX2 | 52.801 | 2 | 52.39705 | yes | 9.722E-08 | 621.8457 | 253.98216 |
| DDPALATYYGSLF | 14 | Q8IUF8 | Ribosomal oxygenase 2 OS=Homo sapier | RIOX2 | 52.801 | 2 | 52.39705 | yes | 9.728E-08 | 780.88269 | 220.4565 |
| QLLLQVESTTVAT | 14 | Q8IUF8 | Ribosomal oxygenase 2 OS=Homo sapier | RIOX2 | 52.801 | 2 | 52.39705 | yes | 2.969E-07 | 779.9436 | 202.64911 |
| EFWEQKPLLIQR | 12 | Q8IUF8 | Ribosomal oxygenase 2 OS=Homo sapier | RIOX2 | 52.801 | 2 | 52.39705 | yes | 9.722E-08 | 793.93811 | 200.86852 |
| VMLADSPR | 8 | Q8IUH3 | RNA-binding protein 45 OS=Homo sapie | RBM45 | 53.503 | 2 | 43.54849 | yes | 2.543E-07 | 444.73401 | 327.01501 |
| YTPESVLR | 8 | Q8IUH3 | RNA-binding protein 45 OS=Homo sapie | RBM45 | 53.503 | 2 | 43.54849 | yes | 5.285E-07 | 482.75854 | 174.13011 |
| TLQAEVPLSDCVQ | 14 | Q8IUR6 | CREB3 regulatory factor OS=Homo sapie | CREBRF | 72.149 | 2 | 49.808723 | yes | 1.003E-07 | 794.40601 | 91.863899 |
| DMGIVDILHK | 10 | Q8IUR7 | Armadillo repeat-containing protein 8 OS | ARMC8 | 75.51 | 2 | 45.902073 | yes | 1.709E-07 | 570.80774 | 88.943771 |
| IIETENMMDR | 10 | Q8IUR7 | Armadillo repeat-containing protein 8 OS | ARMC8 | 75.51 | 2 | 45.902073 | yes | 3.63E-07 | 626.289 | 27.127453 |
| LLSDSDLNVLMK | 12 | Q8IUR7 | Armadillo repeat-containing protein 8 OS | ARMC8 | 75.51 | 2 | 45.902073 | yes | 5.706E-06 | 674.36304 | 23.219688 |
| HSNVNLTIFTAR | 12 | Q8IUX4 | DNA dC->dU-editing enzyme APOBEC- | APOBEC3I | 45.021 | 2 | 52.87171 | no | 1.005E-07 | 686.87024 | 996.21741 |
| FDDNYAFLHR | 10 | Q8IUX4 | DNA dC->dU-editing enzyme APOBEC- | APOBEC3I | 45.021 | 2 | 52.87171 | yes | 9.722E-08 | 649.30164 | 415.15463 |
| SLSQEGASVEIMG | 15 | Q8IUX4 | DNA dC->dU-editing enzyme APOBEC- | APOBEC3I | 45.021 | 2 | 52.87171 | yes | 9.722E-08 | 799.89014 | 183.14415 |
| YNFLFLDSK | 9 | Q8IUX4 | DNA dC->dU-editing enzyme APOBEC- | APOBEC3I | 45.021 | 2 | 52.87171 | yes | 1.002E-07 | 573.7951 | 114.32005 |
| RNTVWLCYEVK | 11 | Q8IUX4 | DNA dC->dU-editing enzyme APOBEC- | APOBEC3I | 45.021 | 2 | 52.87171 | no | 2.149E-07 | 734.37427 | 57.386108 |
| NESWLCFTMEVVI | 13 | Q8IUX4 | DNA dC->dU-editing enzyme APOBEC- | APOBEC3I | 45.021 | 2 | 52.87171 | yes | 9.964E-05 | 821.88385 | 16.185776 |
| MLQDGCELR | 9 | Q8IV48 | 3'-5' exoribonuclease 1 OS=Homo sapien | ERI1 | 40.064 | 2 | 52.777676 | yes | 1.033E-07 | 561.25751 | 629.25372 |
| ADTFPQVLK | 9 | Q8IV48 | 3'-5' exoribonuclease 1 OS=Homo sapien | ERI1 | 40.064 | 2 | 52.777676 | yes | 5.929E-07 | 509.78201 | 615.77295 |
| YSLLTDGSWDMSI | 13 | Q8IV48 | 3'-5' exoribonuclease 1 OS=Homo sapien | ERI1 | 40.064 | 2 | 52.777676 | yes | 9.722E-08 | 751.84521 | 511.12021 |
| FITSSASDFSDPVY | 15 | Q8IV48 | 3'-5' exoribonuclease 1 OS=Homo sapien | ERI1 | 40.064 | 2 | 52.777676 | yes | 1.017E-07 | 832.39612 | 431.09891 |
| FLNIQCQLSR | 10 | Q8IV48 | 3'-5' exoribonuclease 1 OS=Homo sapien | ERI1 | 40.064 | 2 | 52.777676 | yes | 9.722E-08 | 639.83478 | 410.01727 |
| EIAITNGCINR | 11 | Q8IV48 | 3'-5' exoribonuclease 1 OS=Homo sapien | ERI1 | 40.064 | 2 | 52.777676 | yes | 1.002E-07 | 630.8219 | 173.76913 |
| MHAGQLMSVSSSI | 27 | Q8IV48 | 3'-5' exoribonuclease 1 OS=Homo sapien | ERI1 | 40.064 | 4 | 52.777676 | yes | 6.162E-07 | 733.86212 | 70.506142 |
| GLESVDSEALAR | 12 | Q8IVF7 | Formin-like protein 3 OS=Homo sapiens | FMNL3 | 117.213 | 2 | 42.70404 | yes | 3.792E-07 | 623.81732 | 38.249359 |
| MPMPEPCELEER | 12 | Q8IVF7 | Formin-like protein 3 OS=Homo sapiens | FMNL3 | 117.213 | 2 | 42.70404 | yes | 7.241E-07 | 759.32532 | 36.543232 |
| LLDLENENMMR | 11 | Q8IVF7 | Formin-like protein 3 OS=Homo sapiens | FMNL3 | 117.213 | 2 | 42.70404 | yes | 2.91E-07 | 689.32867 | 29.24202 |
| LGLEEFLQK | 9 | Q8IVF7 | Formin-like protein 3 OS=Homo sapiens | FMNL3 | 117.213 | 2 | 42.70404 | yes | 1.484E-06 | 538.80298 | 26.840275 |
| LPVAIPAHK | 9 | Q8IVH8 | Mitogen-activated protein kinase kinase k | MAP4K3 | 101.316 | 2 | 53.71192 | yes | 9.722E-08 | 473.29764 | 393.03497 |
| SNEVTQEISDSTR | 13 | Q8IVH8 | Mitogen-activated protein kinase kinase k | MAP4K3 | 101.316 | 2 | 53.71192 | yes | 1.004E-07 | 733.34192 | 189.75084 |
| SVEEELHQR | 9 | Q8IVH8 | Mitogen-activated protein kinase kinase k | MAP4K3 | 101.316 | 2 | 53.71192 | yes | 1.05E-07 | 563.77802 | 149.86966 |
| LLQHPFVTQHLTR | 13 | Q8IVH8 | Mitogen-activated protein kinase kinase k | MAP4K3 | 101.316 | 2 | 53.71192 | yes | 9.846E-08 | 795.44916 | 57.441299 |
| LLQHPFVTQHLTR | 13 | Q8IVH8 | Mitogen-activated protein kinase kinase k | MAP4K3 | 101.316 | 2 | 53.71192 | yes | 9.722E-08 | 530.63519 | 57.441299 |
| LLQHPFVTQHLTR | 13 | Q8IVH8 | Mitogen-activated protein kinase kinase k | MAP4K3 | 101.316 | 3 | 53.71192 | yes | 9.846E-08 | 795.44916 | 57.441299 |
| LLQHPFVTQHLTR | 13 | Q8IVH8 | Mitogen-activated protein kinase kinase k | MAP4K3 | 101.316 | 3 | 53.71192 | yes | 9.722E-08 | 530.63519 | 57.441299 |
| DTILVCLDCCIK | 12 | Q8IVH8 | Mitogen-activated protein kinase kinase k | MAP4K3 | 101.316 | 2 | 53.71192 | yes | 3.768E-07 | 755.35913 | 109.35183 |
| SEITFGQVK | 9 | Q8IVH8 | Mitogen-activated protein kinase kinase k | MAP4K3 | 101.316 | 2 | 53.71192 | yes | 2.038E-07 | 504.77167 | 79.019394 |
| IGSGTYGDVYK | 11 | Q8IVH8 | Mitogen-activated protein kinase kinase k | MAP4K3 | 101.316 | 2 | 53.71192 | yes | 1.899E-06 | 580.28516 | 66.307732 |
| ETLQGLYYLHSK | 12 | Q8IVH8 | Mitogen-activated protein kinase kinase k | MAP4K3 | 101.316 | 2 | 53.71192 | yes | 2.591E-07 | 726.38007 | 27.073477 |
| LADFGVSAQITATI | 16 | Q8IVH8 | Mitogen-activated protein kinase kinase k | MAP4K3 | 101.316 | 2 | 53.71192 | yes | 6.375E-07 | 803.44598 | 18.064081 |

| | | | | | | | | | | | |
|---|---|---|---|---|---|---|---|---|---|---|---|
| TTYPSQIPLK | 10 | Q8IVL1 | Neuron navigator 2 OS=Homo sapiens O | NAV2 | 268.168 | 2 | 48.756878 | yes | 2.519E-06 | 574.32153 | 89.400139 |
| LFLSCPIDVDGSR | 13 | Q8IVL1 | Neuron navigator 2 OS=Homo sapiens O | NAV2 | 268.168 | 2 | 48.756878 | yes | 1.108E-07 | 739.86902 | 78.633583 |
| TALGSSLPGLVNQ | 16 | Q8IVL1 | Neuron navigator 2 OS=Homo sapiens O | NAV2 | 268.168 | 2 | 48.756878 | yes | 5.136E-07 | 800.93066 | 61.184681 |
| IVVSFQEEMK | 10 | Q8IVL1 | Neuron navigator 2 OS=Homo sapiens O | NAV2 | 268.168 | 2 | 48.756878 | yes | 1.456E-07 | 605.31281 | 56.742077 |
| APSQVSISASPR | 12 | Q8IVL1 | Neuron navigator 2 OS=Homo sapiens O | NAV2 | 268.168 | 2 | 48.756878 | yes | 2.112E-07 | 600.32257 | 50.588894 |
| SSTSSIDSNISSK | 13 | Q8IVL1 | Neuron navigator 2 OS=Homo sapiens O | NAV2 | 268.168 | 2 | 48.756878 | yes | 2.294E-07 | 656.815 | 44.19532 |
| LSEYIVLR | 8 | Q8IVL1 | Neuron navigator 2 OS=Homo sapiens O | NAV2 | 268.168 | 2 | 48.756878 | yes | 4.122E-07 | 496.79239 | 38.235485 |
| TTLSESPLSSPAAS | 16 | Q8IVL1 | Neuron navigator 2 OS=Homo sapiens O | NAV2 | 268.168 | 2 | 48.756878 | yes | 8.199E-06 | 786.90942 | 23.04154 |
| LEALSSAHQLDQL | 14 | Q8IVL1 | Neuron navigator 2 OS=Homo sapiens O | NAV2 | 268.168 | 2 | 48.756878 | yes | 1.122E-07 | 790.92316 | 16.800684 |
| DQEWYDAEIAR | 11 | Q8IVM0 | Coiled-coil domain-containing protein 50 | CCDC50 | 35.822 | 2 | 40.590508 | yes | 3.944E-07 | 698.31238 | 68.840179 |
| LLAQQLVSPVK | 11 | Q8IVS2 | Malonyl-CoA-acyl carrier protein transac | MCAT | 42.962 | 2 | 52.35375 | yes | 9.85E-08 | 598.37408 | 1747.4131 |
| LMEPAVEPLTQAL | 14 | Q8IVS2 | Malonyl-CoA-acyl carrier protein transac | MCAT | 42.962 | 2 | 52.35375 | yes | 3.503E-07 | 770.42621 | 598.95502 |
| LMEPAVEPLTQAL | 14 | Q8IVS2 | Malonyl-CoA-acyl carrier protein transac | MCAT | 42.962 | 2 | 52.35375 | yes | 8.143E-06 | 778.42365 | 598.95502 |
| FNFACLEAR | 9 | Q8IVS2 | Malonyl-CoA-acyl carrier protein transac | MCAT | 42.962 | 2 | 52.35375 | yes | 1.59E-07 | 564.26874 | 1007.4698 |
| KPLVSVYSNVHAI | 14 | Q8IVS2 | Malonyl-CoA-acyl carrier protein transac | MCAT | 42.962 | 3 | 52.35375 | yes | 2.606E-07 | 536.29858 | 76.76371 |
| KPLVSVYSNVHAI | 14 | Q8IVS2 | Malonyl-CoA-acyl carrier protein transac | MCAT | 42.962 | 3 | 52.35375 | yes | 9.722E-08 | 803.94427 | 76.76371 |
| KPLVSVYSNVHAI | 14 | Q8IVS2 | Malonyl-CoA-acyl carrier protein transac | MCAT | 42.962 | 2 | 52.35375 | yes | 2.606E-07 | 536.29858 | 76.76371 |
| KPLVSVYSNVHAI | 14 | Q8IVS2 | Malonyl-CoA-acyl carrier protein transac | MCAT | 42.962 | 2 | 52.35375 | yes | 9.722E-08 | 803.94427 | 76.76371 |
| GFPQTFEVGPGR | 12 | Q8IVS2 | Malonyl-CoA-acyl carrier protein transac | MCAT | 42.962 | 2 | 52.35375 | yes | 3.301E-07 | 646.32513 | 147.31587 |
| GLLNYPR | 7 | Q8IVS2 | Malonyl-CoA-acyl carrier protein transac | MCAT | 42.962 | 2 | 52.35375 | yes | 1.915E-07 | 416.73743 | 104.92267 |
| TVHCQPAIFVASL | 18 | Q8IVS2 | Malonyl-CoA-acyl carrier protein transac | MCAT | 42.962 | 3 | 52.35375 | yes | 9.722E-08 | 647.68024 | 102.11696 |
| MPGQCSVLLFPGQ | 22 | Q8IVS2 | Malonyl-CoA-acyl carrier protein transac | MCAT | 42.962 | 3 | 52.35375 | yes | 1.374E-07 | 769.04651 | 75.298477 |
| VISGHQEALR | 10 | Q8IVS2 | Malonyl-CoA-acyl carrier protein transac | MCAT | 42.962 | 2 | 52.35375 | yes | 3.125E-07 | 555.3067 | 45.143288 |
| WEQTMHAIYER | 11 | Q8IVS2 | Malonyl-CoA-acyl carrier protein transac | MCAT | 42.962 | 2 | 52.35375 | yes | 1.683E-07 | 732.34045 | 21.924929 |
| YPILGIPQAHR | 11 | Q8IVT2 | Mitotic interactor and substrate of PLK1 ( | MISP | 75.357 | 2 | 52.031467 | yes | 9.722E-08 | 632.86169 | 265.80725 |
| ALSSDSILSPAPDA | 15 | Q8IVT2 | Mitotic interactor and substrate of PLK1 ( | MISP | 75.357 | 2 | 52.031467 | yes | 9.722E-08 | 750.38867 | 195.37549 |
| STPLEENVVDR | 11 | Q8IVT2 | Mitotic interactor and substrate of PLK1 ( | MISP | 75.357 | 2 | 52.031467 | yes | 9.722E-08 | 629.81732 | 179.51567 |
| EQIDFLAAR | 9 | Q8IVT2 | Mitotic interactor and substrate of PLK1 ( | MISP | 75.357 | 2 | 52.031467 | yes | 3.98E-07 | 531.78253 | 158.75128 |
| SQSSDLLER | 9 | Q8IVT2 | Mitotic interactor and substrate of PLK1 ( | MISP | 75.357 | 2 | 52.031467 | yes | 6.188E-07 | 517.75928 | 154.05936 |
| QQFLSLEQANK | 11 | Q8IVT2 | Mitotic interactor and substrate of PLK1 ( | MISP | 75.357 | 2 | 52.031467 | yes | 3.155E-07 | 653.34351 | 103.63043 |
| WAVIQGQAVR | 10 | Q8IVT2 | Mitotic interactor and substrate of PLK1 ( | MISP | 75.357 | 2 | 52.031467 | yes | 3.748E-06 | 564.31964 | 36.763138 |
| PSSLSTAEAK | 10 | Q8IVT2 | Mitotic interactor and substrate of PLK1 ( | MISP | 75.357 | 2 | 52.031467 | yes | 5.269E-06 | 495.75876 | 35.538784 |
| GTPAGTTPGASQA | 15 | Q8IVT2 | Mitotic interactor and substrate of PLK1 ( | MISP | 75.357 | 2 | 52.031467 | yes | 3.698E-07 | 670.84387 | 19.030941 |
| EDSSWSSLDAR | 11 | Q8IVT5 | Kinase suppressor of Ras 1 OS=Homo sa | KSR1 | 102.16 | 2 | 47.358624 | yes | 1.542E-07 | 626.77563 | 132.49268 |
| LIDISIGSLR | 10 | Q8IVT5 | Kinase suppressor of Ras 1 OS=Homo sa | KSR1 | 102.16 | 2 | 47.358624 | yes | 2.305E-07 | 543.82953 | 125.04012 |
| FGLGVLESSNPK | 12 | Q8IVT5 | Kinase suppressor of Ras 1 OS=Homo sa | KSR1 | 102.16 | 2 | 47.358624 | yes | 4.228E-07 | 624.33514 | 114.96632 |
| LQYALTCLR | 9 | Q8IVT5 | Kinase suppressor of Ras 1 OS=Homo sa | KSR1 | 102.16 | 2 | 47.358624 | yes | 1.487E-07 | 569.30786 | 107.03788 |
| AAEPHFGTLPK | 11 | Q8IVT5 | Kinase suppressor of Ras 1 OS=Homo sa | KSR1 | 102.16 | 2 | 47.358624 | yes | 4.443E-07 | 584.31146 | 87.082581 |
| TPELNSYPR | 9 | Q8IVT5 | Kinase suppressor of Ras 1 OS=Homo sa | KSR1 | 102.16 | 2 | 47.358624 | yes | 3.548E-06 | 538.77222 | 72.653938 |

| | | | | | | | | | | |
|---|---|---|---|---|---|---|---|---|---|---|
| ISFLPLTR | 8 | Q8IVT5 | Kinase suppressor of Ras 1 OS=Homo sa| KSR1 | 102.16 | 2 | 47.358624 | yes | 7.707E-07 | 473.78964 | 55.376652 |
| ALQQCGQLQK | 10 | Q8IVT5 | Kinase suppressor of Ras 1 OS=Homo sa| KSR1 | 102.16 | 2 | 47.358624 | yes | 1.346E-07 | 587.30585 | 51.882671 |
| LLEMDGHNQDHL | 13 | Q8IVT5 | Kinase suppressor of Ras 1 OS=Homo sa| KSR1 | 102.16 | 2 | 47.358624 | yes | 4.21E-05 | 775.375 | 7.598825 |
| VQWGVFVPR | 9 | Q8IWA0 | WD repeat-containing protein 75 OS=Ho | WDR75 | 94.499 | 2 | 57.184666 | yes | 2.97E-07 | 544.30603 | 410.59723 |
| VWILTDDSDIYK | 12 | Q8IWA0 | WD repeat-containing protein 75 OS=Ho | WDR75 | 94.499 | 2 | 57.184666 | yes | 1.019E-07 | 734.37195 | 271.62799 |
| ELSFVLDYINQSPK | 14 | Q8IWA0 | WD repeat-containing protein 75 OS=Ho | WDR75 | 94.499 | 2 | 57.184666 | yes | 1.36E-07 | 826.93018 | 196.61073 |
| SEQPTLVTASK | 11 | Q8IWA0 | WD repeat-containing protein 75 OS=Ho | WDR75 | 94.499 | 2 | 57.184666 | yes | 1.128E-07 | 580.81152 | 107.65837 |
| YIFCVSGDFVK | 11 | Q8IWA0 | WD repeat-containing protein 75 OS=Ho | WDR75 | 94.499 | 2 | 57.184666 | yes | 9.722E-08 | 667.82611 | 105.31223 |
| TQGFILNTK | 9 | Q8IWA0 | WD repeat-containing protein 75 OS=Ho | WDR75 | 94.499 | 2 | 57.184666 | yes | 1.525E-06 | 511.28748 | 69.996765 |
| CGGSELNFR | 9 | Q8IWA0 | WD repeat-containing protein 75 OS=Ho | WDR75 | 94.499 | 2 | 57.184666 | yes | 8.739E-07 | 520.23492 | 63.503231 |
| EKPDIFQLVSVK | 12 | Q8IWA0 | WD repeat-containing protein 75 OS=Ho | WDR75 | 94.499 | 2 | 57.184666 | yes | 2.616E-07 | 701.90063 | 47.578468 |
| VQDTSNTGLGEDI | 19 | Q8IWA0 | WD repeat-containing protein 75 OS=Ho | WDR75 | 94.499 | 3 | 57.184666 | yes | 2.068E-07 | 685.68329 | 32.693085 |
| CIAFGNEGVYVAA | 15 | Q8IWA0 | WD repeat-containing protein 75 OS=Ho | WDR75 | 94.499 | 2 | 57.184666 | yes | 4.508E-07 | 813.40887 | 31.462811 |
| KIDYSWIAAL | 10 | Q8IWA0 | WD repeat-containing protein 75 OS=Ho | WDR75 | 94.499 | 2 | 57.184666 | yes | 4.278E-07 | 590.32404 | 12.752155 |
| LVFLEDGTSQVR | 12 | Q8IWC1 | MAP7 domain-containing protein 3 OS=I | MAP7D3 | 98.43 | 2 | 54.52364 | yes | 1.265E-07 | 682.36444 | 1302.274 |
| LEQGTSALIR | 10 | Q8IWC1 | MAP7 domain-containing protein 3 OS=I | MAP7D3 | 98.43 | 2 | 54.52364 | yes | 9.728E-08 | 544.30896 | 759.14984 |
| IMLQNLQER | 9 | Q8IWC1 | MAP7 domain-containing protein 3 OS=I | MAP7D3 | 98.43 | 2 | 54.52364 | yes | 1.014E-07 | 572.81079 | 746.91858 |
| GWLDQEDQEAPL( | 14 | Q8IWC1 | MAP7 domain-containing protein 3 OS=I | MAP7D3 | 98.43 | 2 | 54.52364 | yes | 9.722E-08 | 828.89685 | 517.74921 |
| NCPPSPLPLISK | 12 | Q8IWC1 | MAP7 domain-containing protein 3 OS=I | MAP7D3 | 98.43 | 2 | 54.52364 | yes | 3.74E-07 | 661.86047 | 409.38361 |
| MVAAANEIAK | 10 | Q8IWC1 | MAP7 domain-containing protein 3 OS=I | MAP7D3 | 98.43 | 2 | 54.52364 | yes | 2.97E-07 | 509.2735 | 376.53296 |
| STAGIMNAEAATK | 13 | Q8IWC1 | MAP7 domain-containing protein 3 OS=I | MAP7D3 | 98.43 | 2 | 54.52364 | yes | 2.311E-07 | 632.81372 | 349.78677 |
| CEALSQR | 7 | Q8IWC1 | MAP7 domain-containing protein 3 OS=I | MAP7D3 | 98.43 | 2 | 54.52364 | yes | 4.329E-07 | 432.20563 | 332.02957 |
| ESMEASPEAMVK | 12 | Q8IWC1 | MAP7 domain-containing protein 3 OS=I | MAP7D3 | 98.43 | 2 | 54.52364 | yes | 9.722E-08 | 654.79413 | 329.85611 |
| STFKPVIDGSMLK | 13 | Q8IWC1 | MAP7 domain-containing protein 3 OS=I | MAP7D3 | 98.43 | 2 | 54.52364 | yes | 1.689E-07 | 711.88672 | 309.25253 |
| VTGVTNYVMQYV | 18 | Q8IWC1 | MAP7 domain-containing protein 3 OS=I | MAP7D3 | 98.43 | 3 | 54.52364 | yes | 1.171E-07 | 695.37848 | 306.58505 |
| TSLEASMEASPK | 12 | Q8IWC1 | MAP7 domain-containing protein 3 OS=I | MAP7D3 | 98.43 | 2 | 54.52364 | yes | 9.722E-08 | 625.80029 | 298.43875 |
| SEGFHDILPK | 10 | Q8IWC1 | MAP7 domain-containing protein 3 OS=I | MAP7D3 | 98.43 | 2 | 54.52364 | yes | 3.003E-07 | 571.79565 | 296.3302 |
| LADDYQQK | 8 | Q8IWC1 | MAP7 domain-containing protein 3 OS=I | MAP7D3 | 98.43 | 2 | 54.52364 | yes | 1.499E-07 | 490.73782 | 228.74992 |
| QMPLSSAGLQNSV | 15 | Q8IWC1 | MAP7 domain-containing protein 3 OS=I | MAP7D3 | 98.43 | 2 | 54.52364 | yes | 9.722E-08 | 765.90088 | 210.24399 |
| QSPQTSFPYK | 10 | Q8IWC1 | MAP7 domain-containing protein 3 OS=I | MAP7D3 | 98.43 | 2 | 54.52364 | yes | 9.722E-08 | 591.79309 | 186.53465 |
| WSWGGSAMANSE | 14 | Q8IWC1 | MAP7 domain-containing protein 3 OS=I | MAP7D3 | 98.43 | 2 | 54.52364 | yes | 1.495E-07 | 749.32501 | 180.82013 |
| LSSYTECYK | 9 | Q8IWC1 | MAP7 domain-containing protein 3 OS=I | MAP7D3 | 98.43 | 2 | 54.52364 | yes | 1.658E-07 | 575.75787 | 145.01109 |
| PVIDGSMLK | 9 | Q8IWC1 | MAP7 domain-containing protein 3 OS=I | MAP7D3 | 98.43 | 2 | 54.52364 | yes | 3.661E-07 | 480.26517 | 143.21381 |
| RIEEIMK | 7 | Q8IWC1 | MAP7 domain-containing protein 3 OS=I | MAP7D3 | 98.43 | 2 | 54.52364 | yes | 2.154E-07 | 459.75751 | 142.24173 |
| IMPIQHTLSVQSAS | 18 | Q8IWC1 | MAP7 domain-containing protein 3 OS=I | MAP7D3 | 98.43 | 3 | 54.52364 | yes | 1.519E-07 | 643.01544 | 102.78847 |
| HMIYEESGNK | 10 | Q8IWC1 | MAP7 domain-containing protein 3 OS=I | MAP7D3 | 98.43 | 2 | 54.52364 | yes | 1.166E-07 | 604.27423 | 74.55854 |
| MPFNNAK | 7 | Q8IWC1 | MAP7 domain-containing protein 3 OS=I | MAP7D3 | 98.43 | 2 | 54.52364 | yes | 3.144E-07 | 411.20236 | 73.394684 |
| AVMFPMSTMK | 10 | Q8IWC1 | MAP7 domain-containing protein 3 OS=I | MAP7D3 | 98.43 | 2 | 54.52364 | yes | 4.339E-07 | 571.77399 | 42.587353 |
| DTSIQEVVSR | 10 | Q8IWC1 | MAP7 domain-containing protein 3 OS=I | MAP7D3 | 98.43 | 2 | 54.52364 | yes | 7.847E-07 | 567.29346 | 41.196381 |

| | | | | | | | | | | | |
|---|---|---|---|---|---|---|---|---|---|---|---|
| SILPYPVSPK | 10 | Q8IWI9 | MAX gene-associated protein OS=Homo | MGA | 336.16 | 2 | 30.77474 | yes | 4.334E-06 | 550.82117 | 851.99652 |
| LDEYLENEGK | 10 | Q8IWI9 | MAX gene-associated protein OS=Homo | MGA | 336.16 | 2 | 30.77474 | yes | 4.648E-05 | 605.28534 | 52.383858 |
| YWITGLDSNLK | 11 | Q8IWI9 | MAX gene-associated protein OS=Homo | MGA | 336.16 | 2 | 30.77474 | yes | 3.303E-06 | 655.34296 | 34.099895 |
| TKPIHTIIR | 9 | Q8IWI9 | MAX gene-associated protein OS=Homo | MGA | 336.16 | 2 | 30.77474 | yes | 2.065E-06 | 539.84021 | 20.995245 |
| LQLIAFR | 7 | Q8IWK6 | Adhesion G protein-coupled receptor A3 | ADGRA3 | 146.151 | 2 | 43.12752 | yes | 2.72E-07 | 430.77127 | 172.93562 |
| AEFGDFDIK | 9 | Q8IWS0 | PHD finger protein 6 OS=Homo sapiens ( | PHF6 | 41.291 | 2 | 51.473843 | yes | 1.411E-07 | 521.24799 | 3391.9116 |
| YIENMSR | 7 | Q8IWS0 | PHD finger protein 6 OS=Homo sapiens ( | PHF6 | 41.291 | 2 | 51.473843 | yes | 1.614E-07 | 456.71582 | 1581.574 |
| LHIFNAK | 7 | Q8IWS0 | PHD finger protein 6 OS=Homo sapiens ( | PHF6 | 41.291 | 2 | 51.473843 | yes | 1.493E-07 | 421.7478 | 1411.76 |
| DKECGQLLISENQI | 14 | Q8IWS0 | PHD finger protein 6 OS=Homo sapiens ( | PHF6 | 41.291 | 2 | 51.473843 | yes | 1.031E-07 | 831.4118 | 922.8714 |
| TVLQEIK | 7 | Q8IWS0 | PHD finger protein 6 OS=Homo sapiens ( | PHF6 | 41.291 | 2 | 51.473843 | yes | 1.248E-07 | 415.75275 | 690.37122 |
| ECGQLLISENQK | 12 | Q8IWS0 | PHD finger protein 6 OS=Homo sapiens ( | PHF6 | 41.291 | 2 | 51.473843 | yes | 9.888E-08 | 709.85083 | 683.49158 |
| TYHYHCALHDK | 11 | Q8IWS0 | PHD finger protein 6 OS=Homo sapiens ( | PHF6 | 41.291 | 2 | 51.473843 | yes | 1.083E-07 | 722.82495 | 479.23785 |
| TYHYHCGVQDK | 11 | Q8IWS0 | PHD finger protein 6 OS=Homo sapiens ( | PHF6 | 41.291 | 2 | 51.473843 | yes | 9.722E-08 | 704.30914 | 473.58551 |
| CGFCHVGEEENEA | 14 | Q8IWS0 | PHD finger protein 6 OS=Homo sapiens ( | PHF6 | 41.291 | 2 | 51.473843 | yes | 1.894E-07 | 847.33832 | 24.042561 |
| CGFCHVGEEENEA | 14 | Q8IWS0 | PHD finger protein 6 OS=Homo sapiens ( | PHF6 | 41.291 | 2 | 51.473843 | yes | 3.526E-07 | 565.22797 | 24.042561 |
| CGFCHVGEEENEA | 14 | Q8IWS0 | PHD finger protein 6 OS=Homo sapiens ( | PHF6 | 41.291 | 3 | 51.473843 | yes | 1.894E-07 | 847.33832 | 24.042561 |
| CGFCHVGEEENEA | 14 | Q8IWS0 | PHD finger protein 6 OS=Homo sapiens ( | PHF6 | 41.291 | 3 | 51.473843 | yes | 3.526E-07 | 565.22797 | 24.042561 |
| TAHNSEADLEESFI | 24 | Q8IWS0 | PHD finger protein 6 OS=Homo sapiens ( | PHF6 | 41.291 | 4 | 51.473843 | yes | 1.112E-07 | 675.05322 | 43.417461 |
| FAVFLSEVSENK | 12 | Q8IWT6 | Volume-regulated anion channel subunit | LRRC8A | 94.199 | 2 | 52.750107 | no | 1.114E-07 | 685.35352 | 155.36644 |
| VGELTNLTQIELR | 13 | Q8IWT6 | Volume-regulated anion channel subunit | LRRC8A | 94.199 | 2 | 52.750107 | yes | 9.911E-07 | 743.41718 | 123.29055 |
| MANLTELELIR | 11 | Q8IWT6 | Volume-regulated anion channel subunit | LRRC8A | 94.199 | 2 | 52.750107 | yes | 9.722E-08 | 651.85791 | 99.389458 |
| CFDSPWTTR | 9 | Q8IWT6 | Volume-regulated anion channel subunit | LRRC8A | 94.199 | 2 | 52.750107 | no | 9.918E-08 | 585.25586 | 90.165253 |
| IETLPPELFQCR | 12 | Q8IWT6 | Volume-regulated anion channel subunit | LRRC8A | 94.199 | 2 | 52.750107 | yes | 3.404E-07 | 751.88721 | 72.098076 |
| LEHFVSILLK | 10 | Q8IWT6 | Volume-regulated anion channel subunit | LRRC8A | 94.199 | 2 | 52.750107 | yes | 8.788E-05 | 599.86334 | 56.917717 |
| YFADTQPAYR | 10 | Q8IWT6 | Volume-regulated anion channel subunit | LRRC8A | 94.199 | 2 | 52.750107 | yes | 2.584E-05 | 616.29077 | 46.8256 |
| EESSYSDIPDVK | 12 | Q8IWT6 | Volume-regulated anion channel subunit | LRRC8A | 94.199 | 2 | 52.750107 | yes | 2.945E-07 | 684.81189 | 42.132645 |
| LIVLNSLK | 8 | Q8IWT6 | Volume-regulated anion channel subunit | LRRC8A | 94.199 | 2 | 52.750107 | yes | 3.07E-06 | 450.29987 | 16.581558 |
| EMLTTFGTATYK | 12 | Q8IWV8 | E3 ubiquitin-protein ligase UBR2 OS=Ho | UBR2 | 200.539 | 2 | 46.668102 | yes | 1.5E-07 | 681.83411 | 72.962883 |
| PSSSVQGTGLIR | 12 | Q8IWW6 | Rho GTPase-activating protein 12 OS=Ho | ARHGAP1 | 96.255 | 2 | 50.983757 | yes | 9.818E-08 | 601.33038 | 89.623131 |
| DQVFGSNLANLCQ | 14 | Q8IWW6 | Rho GTPase-activating protein 12 OS=Ho | ARHGAP1 | 96.255 | 2 | 50.983757 | yes | 1.415E-07 | 811.39117 | 69.7733 |
| YGLLNVTK | 8 | Q8IWW6 | Rho GTPase-activating protein 12 OS=Ho | ARHGAP1 | 96.255 | 2 | 50.983757 | yes | 9.722E-08 | 454.26602 | 64.774506 |
| VSSIDSSEQK | 10 | Q8IWW6 | Rho GTPase-activating protein 12 OS=Ho | ARHGAP1 | 96.255 | 2 | 50.983757 | yes | 1.859E-05 | 540.2644 | 48.995678 |
| VSGNLAVIQK | 10 | Q8IWW6 | Rho GTPase-activating protein 12 OS=Ho | ARHGAP1 | 96.255 | 2 | 50.983757 | yes | 2.536E-07 | 514.80859 | 43.307613 |
| IMQSLHLQR | 9 | Q8IWW6 | Rho GTPase-activating protein 12 OS=Ho | ARHGAP1 | 96.255 | 2 | 50.983757 | yes | 1.541E-06 | 563.31348 | 33.272427 |
| LPPPMPPSER | 10 | Q8IWX8 | Calcium homeostasis endoplasmic reticul | CHERP | 103.703 | 2 | 52.211014 | yes | 5.31E-07 | 560.79462 | 3618.1455 |
| GVGVALDDPYENY | 14 | Q8IWX8 | Calcium homeostasis endoplasmic reticul | CHERP | 103.703 | 2 | 52.211014 | yes | 1.163E-07 | 784.37299 | 3462.0105 |
| LALEQQQLICK | 11 | Q8IWX8 | Calcium homeostasis endoplasmic reticul | CHERP | 103.703 | 2 | 52.211014 | yes | 2.97E-07 | 672.37122 | 3095.084 |
| MGWSGSGGLGAK | 12 | Q8IWX8 | Calcium homeostasis endoplasmic reticul | CHERP | 103.703 | 2 | 52.211014 | yes | 2.759E-07 | 554.26624 | 1533.0908 |
| MGWSGSGGLGAK | 12 | Q8IWX8 | Calcium homeostasis endoplasmic reticul | CHERP | 103.703 | 2 | 52.211014 | yes | 2.577E-06 | 562.26367 | 1533.0908 |

| | | | | | | | | | | | |
|---|---|---|---|---|---|---|---|---|---|---|---|
| ITADGAHFELR | 11 | Q8IWX8 | Calcium homeostasis endoplasmic reticul | CHERP | 103.703 | 2 | 52.211014 | yes | 1.823E-07 | 615.31732 | 1088.7181 |
| ITADGAHFELR | 11 | Q8IWX8 | Calcium homeostasis endoplasmic reticul | CHERP | 103.703 | 2 | 52.211014 | yes | 9.722E-08 | 410.5473 | 1088.7181 |
| ITADGAHFELR | 11 | Q8IWX8 | Calcium homeostasis endoplasmic reticul | CHERP | 103.703 | 3 | 52.211014 | yes | 1.823E-07 | 615.31732 | 1088.7181 |
| ITADGAHFELR | 11 | Q8IWX8 | Calcium homeostasis endoplasmic reticul | CHERP | 103.703 | 3 | 52.211014 | yes | 9.722E-08 | 410.5473 | 1088.7181 |
| EQGIQDPIK | 9 | Q8IWX8 | Calcium homeostasis endoplasmic reticul | CHERP | 103.703 | 2 | 52.211014 | yes | 1.197E-07 | 514.27454 | 2092.6541 |
| LEDHEYKPLDPK | 12 | Q8IWX8 | Calcium homeostasis endoplasmic reticul | CHERP | 103.703 | 2 | 52.211014 | yes | 1.564E-07 | 742.375 | 372.40872 |
| LEDHEYKPLDPK | 12 | Q8IWX8 | Calcium homeostasis endoplasmic reticul | CHERP | 103.703 | 2 | 52.211014 | yes | 9.722E-08 | 495.25244 | 372.40872 |
| LEDHEYKPLDPK | 12 | Q8IWX8 | Calcium homeostasis endoplasmic reticul | CHERP | 103.703 | 3 | 52.211014 | yes | 1.564E-07 | 742.375 | 372.40872 |
| LEDHEYKPLDPK | 12 | Q8IWX8 | Calcium homeostasis endoplasmic reticul | CHERP | 103.703 | 3 | 52.211014 | yes | 9.722E-08 | 495.25244 | 372.40872 |
| LLQLWEK | 7 | Q8IWX8 | Calcium homeostasis endoplasmic reticul | CHERP | 103.703 | 2 | 52.211014 | yes | 1.003E-07 | 465.27637 | 723.9682 |
| GHQMLVK | 7 | Q8IWX8 | Calcium homeostasis endoplasmic reticul | CHERP | 103.703 | 2 | 52.211014 | yes | 3.203E-07 | 406.72601 | 653.37994 |
| DKWDQYK | 7 | Q8IWX8 | Calcium homeostasis endoplasmic reticul | CHERP | 103.703 | 2 | 52.211014 | yes | 1.128E-07 | 491.73508 | 496.4953 |
| SPPHCELMAGHLR | 13 | Q8IWX8 | Calcium homeostasis endoplasmic reticul | CHERP | 103.703 | 2 | 52.211014 | yes | 9.722E-08 | 752.86102 | 349.13589 |
| SYSFIAR | 7 | Q8IWX8 | Calcium homeostasis endoplasmic reticul | CHERP | 103.703 | 2 | 52.211014 | yes | 2.035E-07 | 422.22159 | 316.50577 |
| NWMFSNAK | 8 | Q8IWX8 | Calcium homeostasis endoplasmic reticul | CHERP | 103.703 | 2 | 52.211014 | yes | 1.362E-07 | 499.23166 | 292.57901 |
| MEMPLPPDDQELI | 13 | Q8IWX8 | Calcium homeostasis endoplasmic reticul | CHERP | 103.703 | 2 | 52.211014 | yes | 1.369E-07 | 806.87091 | 167.29068 |
| ELLAALQK | 8 | Q8IWX8 | Calcium homeostasis endoplasmic reticul | CHERP | 103.703 | 2 | 52.211014 | yes | 2.292E-07 | 443.27383 | 163.41313 |
| VVVPIYCTSFLAVI | 20 | Q8IWX8 | Calcium homeostasis endoplasmic reticul | CHERP | 103.703 | 3 | 52.211014 | yes | 4.746E-07 | 785.07513 | 142.6543 |
| LHLIYLINDVLHH( | 16 | Q8IWX8 | Calcium homeostasis endoplasmic reticul | CHERP | 103.703 | 3 | 52.211014 | yes | 9.722E-08 | 682.03516 | 120.35213 |
| VVVPIYCTSFLAVI | 17 | Q8IWX8 | Calcium homeostasis endoplasmic reticul | CHERP | 103.703 | 3 | 52.211014 | yes | 4.759E-07 | 657.00439 | 26.897806 |
| AFADPEVLR | 9 | Q8IWZ3 | Ankyrin repeat and KH domain-containin | ANKHD1 | 269.458 | 2 | 55.120026 | no | 1.772E-07 | 509.27182 | 1795.4987 |
| ILLNAGAEINSR | 12 | Q8IWZ3 | Ankyrin repeat and KH domain-containin | ANKHD1 | 269.458 | 2 | 55.120026 | no | 4.507E-07 | 635.85931 | 1163.3749 |
| TVDPETQAR | 9 | Q8IWZ3 | Ankyrin repeat and KH domain-containin | ANKHD1 | 269.458 | 2 | 55.120026 | no | 4.318E-07 | 508.754 | 1124.3691 |
| GADVNAPPVPSSR | 13 | Q8IWZ3 | Ankyrin repeat and KH domain-containin | ANKHD1 | 269.458 | 2 | 55.120026 | no | 9.722E-08 | 633.8255 | 1058.5747 |
| FCELLIHR | 8 | Q8IWZ3 | Ankyrin repeat and KH domain-containin | ANKHD1 | 269.458 | 2 | 55.120026 | yes | 1.219E-07 | 544.28949 | 1053.8141 |
| VFLQGPAPVGTPS | 16 | Q8IWZ3 | Ankyrin repeat and KH domain-containin | ANKHD1 | 269.458 | 2 | 55.120026 | yes | 1.321E-07 | 843.95172 | 1002.7322 |
| SLELIQGEPLNK | 12 | Q8IWZ3 | Ankyrin repeat and KH domain-containin | ANKHD1 | 269.458 | 2 | 55.120026 | yes | 9.722E-08 | 670.87463 | 906.19763 |
| LSVPASVVSR | 10 | Q8IWZ3 | Ankyrin repeat and KH domain-containin | ANKHD1 | 269.458 | 2 | 55.120026 | yes | 7.717E-07 | 507.80075 | 836.4057 |
| NTALTLACFQGR | 12 | Q8IWZ3 | Ankyrin repeat and KH domain-containin | ANKHD1 | 269.458 | 2 | 55.120026 | no | 1.404E-07 | 676.34296 | 781.18475 |
| ESALTLACYK | 10 | Q8IWZ3 | Ankyrin repeat and KH domain-containin | ANKHD1 | 269.458 | 2 | 55.120026 | no | 1.535E-07 | 578.28937 | 753.77826 |
| DTPLSLACSGGR | 12 | Q8IWZ3 | Ankyrin repeat and KH domain-containin | ANKHD1 | 269.458 | 2 | 55.120026 | no | 9.722E-08 | 617.29822 | 684.54083 |
| AEVVSLLLDR | 10 | Q8IWZ3 | Ankyrin repeat and KH domain-containin | ANKHD1 | 269.458 | 2 | 55.120026 | yes | 9.809E-08 | 557.82697 | 655.93616 |
| TSQENSPALLGVQ | 14 | Q8IWZ3 | Ankyrin repeat and KH domain-containin | ANKHD1 | 269.458 | 2 | 55.120026 | yes | 1.066E-07 | 736.39117 | 603.82672 |
| YIATITDK | 8 | Q8IWZ3 | Ankyrin repeat and KH domain-containin | ANKHD1 | 269.458 | 2 | 55.120026 | no | 2.081E-07 | 462.75546 | 531.89764 |
| LEALLEAAGIGK | 12 | Q8IWZ3 | Ankyrin repeat and KH domain-containin | ANKHD1 | 269.458 | 2 | 55.120026 | no | 1.065E-07 | 592.8479 | 509.91608 |
| SLAEACSDGDVNA | 15 | Q8IWZ3 | Ankyrin repeat and KH domain-containin | ANKHD1 | 269.458 | 2 | 55.120026 | yes | 9.809E-08 | 782.35699 | 495.62469 |
| QEVVDLLLAR | 10 | Q8IWZ3 | Ankyrin repeat and KH domain-containin | ANKHD1 | 269.458 | 2 | 55.120026 | yes | 9.722E-08 | 578.34021 | 443.5488 |
| SQELNFVMDVNSS | 14 | Q8IWZ3 | Ankyrin repeat and KH domain-containin | ANKHD1 | 269.458 | 2 | 55.120026 | yes | 9.722E-08 | 799.37994 | 440.36627 |
| DGSTMLIEAAK | 11 | Q8IWZ3 | Ankyrin repeat and KH domain-containin | ANKHD1 | 269.458 | 2 | 55.120026 | no | 1.166E-07 | 568.2868 | 432.60049 |

| | | | | | | | | | | |
|---|---|---|---|---|---|---|---|---|---|---|
| KLLDEGR | 7 | Q8IWZ3 | Ankyrin repeat and KH domain-containin | ANKHD1 | 269.458 | 2 | 55.120026 | yes | 1.602E-07 | 415.74017 | 367.54349 |
| GGDIEAQSER | 10 | Q8IWZ3 | Ankyrin repeat and KH domain-containin | ANKHD1 | 269.458 | 2 | 55.120026 | yes | 3.423E-07 | 531.24652 | 351.24359 |
| EVNQFPSDIECMR | 13 | Q8IWZ3 | Ankyrin repeat and KH domain-containin | ANKHD1 | 269.458 | 2 | 55.120026 | yes | 1.715E-07 | 812.85834 | 314.20706 |
| GHLDMVR | 7 | Q8IWZ3 | Ankyrin repeat and KH domain-containin | ANKHD1 | 269.458 | 2 | 55.120026 | yes | 9.722E-08 | 414.21326 | 271.85968 |
| TVSLPLSSPNIK | 12 | Q8IWZ3 | Ankyrin repeat and KH domain-containin | ANKHD1 | 269.458 | 2 | 55.120026 | yes | 4.603E-06 | 628.36646 | 257.86835 |
| GGCNITAIQDVTG/ | 20 | Q8IWZ3 | Ankyrin repeat and KH domain-containin | ANKHD1 | 269.458 | 3 | 55.120026 | yes | 2.197E-07 | 695.34076 | 225.0173 |
| ITPLMSAFR | 9 | Q8IWZ3 | Ankyrin repeat and KH domain-containin | ANKHD1 | 269.458 | 2 | 55.120026 | yes | 1.166E-07 | 518.28644 | 208.18866 |
| CHQCVETIVK | 10 | Q8IWZ3 | Ankyrin repeat and KH domain-containin | ANKHD1 | 269.458 | 2 | 55.120026 | yes | 5.595E-06 | 637.30499 | 199.98753 |
| AGHLCTVQFLISK | 13 | Q8IWZ3 | Ankyrin repeat and KH domain-containin | ANKHD1 | 269.458 | 2 | 55.120026 | yes | 9.722E-08 | 737.39777 | 196.39613 |
| KLSVPASVVSR | 11 | Q8IWZ3 | Ankyrin repeat and KH domain-containin | ANKHD1 | 269.458 | 2 | 55.120026 | yes | 2.877E-07 | 571.84821 | 190.13814 |
| IPPPIGTER | 9 | Q8IWZ3 | Ankyrin repeat and KH domain-containin | ANKHD1 | 269.458 | 2 | 55.120026 | yes | 1.531E-06 | 490.2822 | 166.35626 |
| MESFSAVPPTK | 11 | Q8IWZ3 | Ankyrin repeat and KH domain-containin | ANKHD1 | 269.458 | 2 | 55.120026 | yes | 4.337E-07 | 597.29718 | 130.89857 |
| VLLDHGAGINTHS | 17 | Q8IWZ3 | Ankyrin repeat and KH domain-containin | ANKHD1 | 269.458 | 3 | 55.120026 | yes | 9.722E-08 | 617.98309 | 103.14684 |
| LKDGSTMLIEAAK | 13 | Q8IWZ3 | Ankyrin repeat and KH domain-containin | ANKHD1 | 269.458 | 2 | 55.120026 | no | 1.347E-07 | 688.87634 | 101.00851 |
| YAVQLINALIQDP/ | 15 | Q8IWZ3 | Ankyrin repeat and KH domain-containin | ANKHD1 | 269.458 | 2 | 55.120026 | yes | 9.722E-08 | 828.9696 | 62.672279 |
| LEGEVTPNSLSTSY | 15 | Q8IWZ3 | Ankyrin repeat and KH domain-containin | ANKHD1 | 269.458 | 2 | 55.120026 | yes | 1.522E-07 | 812.90686 | 56.690422 |
| ANVVTTPSTNR | 11 | Q8IWZ3 | Ankyrin repeat and KH domain-containin | ANKHD1 | 269.458 | 2 | 55.120026 | yes | 4.295E-07 | 580.30695 | 49.101856 |
| VVQYLVK | 7 | Q8IWZ3 | Ankyrin repeat and KH domain-containin | ANKHD1 | 269.458 | 2 | 55.120026 | yes | 1.87E-07 | 424.76566 | 46.867569 |
| DTVSLHQQCSHR | 12 | Q8IWZ3 | Ankyrin repeat and KH domain-containin | ANKHD1 | 269.458 | 2 | 55.120026 | yes | 1.701E-07 | 734.34149 | 40.88438 |
| ATANNDHTVVSL/ | 35 | Q8IWZ3 | Ankyrin repeat and KH domain-containin | ANKHD1 | 269.458 | 5 | 55.120026 | yes | 1.073E-07 | 718.36853 | 40.361858 |
| QLFACVPK | 8 | Q8IWZ3 | Ankyrin repeat and KH domain-containin | ANKHD1 | 269.458 | 2 | 55.120026 | yes | 1.392E-06 | 481.76004 | 38.996143 |
| TDEMHTALMEAC | 21 | Q8IWZ3 | Ankyrin repeat and KH domain-containin | ANKHD1 | 269.458 | 3 | 55.120026 | no | 1.554E-06 | 801.67804 | 38.680851 |
| ELEDLIPK | 8 | Q8IWZ3 | Ankyrin repeat and KH domain-containin | ANKHD1 | 269.458 | 2 | 55.120026 | yes | 4.748E-07 | 478.76859 | 37.824467 |
| TGLGLASLPGPVK | 13 | Q8IWZ8 | SURP and G-patch domain-containing pr | SUGP1 | 72.471 | 2 | 54.414005 | yes | 9.722E-08 | 605.36371 | 75.696747 |
| AMQDMQLLWEK | 11 | Q8IWZ8 | SURP and G-patch domain-containing pr | SUGP1 | 72.471 | 2 | 54.414005 | yes | 1.768E-07 | 696.83612 | 63.604816 |
| DNPAFAFLHDK | 11 | Q8IWZ8 | SURP and G-patch domain-containing pr | SUGP1 | 72.471 | 2 | 54.414005 | yes | 5.483E-07 | 637.81183 | 60.388336 |
| ASSTGSFTAPDPGI | 15 | Q8IWZ8 | SURP and G-patch domain-containing pr | SUGP1 | 72.471 | 2 | 54.414005 | yes | 2.694E-07 | 718.35681 | 57.307571 |
| DVDASPSPLSVQD | 15 | Q8IWZ8 | SURP and G-patch domain-containing pr | SUGP1 | 72.471 | 2 | 54.414005 | yes | 2.041E-07 | 785.90161 | 25.885714 |
| EPVSVGTPSEGEGI | 30 | Q8IX01 | SURP and G-patch domain-containing pr | SUGP2 | 120.208 | 4 | 54.533707 | yes | 3.898E-05 | 798.12061 | 805.66357 |
| APSSLSDAVPQR | 12 | Q8IX01 | SURP and G-patch domain-containing pr | SUGP2 | 120.208 | 2 | 54.533707 | yes | 9.722E-08 | 614.32001 | 394.11194 |
| ALNIVDQEGSLLGI | 14 | Q8IX01 | SURP and G-patch domain-containing pr | SUGP2 | 120.208 | 2 | 54.533707 | yes | 1.165E-07 | 728.90393 | 371.50195 |
| VVGTIDQLVK | 10 | Q8IX01 | SURP and G-patch domain-containing pr | SUGP2 | 120.208 | 2 | 54.533707 | yes | 5.881E-07 | 536.32404 | 336.69244 |
| ATTGTQTLLSSGTI | 14 | Q8IX01 | SURP and G-patch domain-containing pr | SUGP2 | 120.208 | 2 | 54.533707 | yes | 1.079E-07 | 697.36774 | 323.78989 |
| DLEFSHSDSR | 10 | Q8IX01 | SURP and G-patch domain-containing pr | SUGP2 | 120.208 | 2 | 54.533707 | yes | 9.722E-08 | 596.76508 | 197.09531 |
| SSNPSISDDSYFR | 13 | Q8IX01 | SURP and G-patch domain-containing pr | SUGP2 | 120.208 | 2 | 54.533707 | yes | 9.722E-08 | 737.82587 | 190.6993 |
| KIPTVNR | 7 | Q8IX01 | SURP and G-patch domain-containing pr | SUGP2 | 120.208 | 2 | 54.533707 | yes | 1.211E-06 | 414.25851 | 162.63657 |
| WAGFHTIK | 8 | Q8IX01 | SURP and G-patch domain-containing pr | SUGP2 | 120.208 | 2 | 54.533707 | yes | 9.722E-08 | 480.25851 | 130.74532 |
| VIEGSLSPK | 9 | Q8IX01 | SURP and G-patch domain-containing pr | SUGP2 | 120.208 | 2 | 54.533707 | yes | 5.589E-07 | 465.26874 | 120.74086 |
| NCFFEIIKPFDK | 12 | Q8IX01 | SURP and G-patch domain-containing pr | SUGP2 | 120.208 | 2 | 54.533707 | yes | 1.155E-07 | 779.39215 | 117.08907 |

| | | | | | | | | | | | |
|---|---|---|---|---|---|---|---|---|---|---|---|
| AWLVSSGCPLQVI | 13 | Q8IX01 | SURP and G-patch domain-containing pr | SUGP2 | 120.208 | 2 | 54.533707 | yes | 1.128E-07 | 722.88446 | 111.68829 |
| DYDVDHPGEADS' | 15 | Q8IX01 | SURP and G-patch domain-containing pr | SUGP2 | 120.208 | 2 | 54.533707 | yes | 2.184E-07 | 844.38153 | 106.8457 |
| YSLSGSVAHSR | 11 | Q8IX01 | SURP and G-patch domain-containing pr | SUGP2 | 120.208 | 2 | 54.533707 | yes | 1.151E-07 | 582.29382 | 86.887543 |
| EPVSVGTPSEGEGI | 22 | Q8IX01 | SURP and G-patch domain-containing pr | SUGP2 | 120.208 | 3 | 54.533707 | yes | 1.26E-06 | 727.34155 | 78.54641 |
| ITQETFDAVLQEK | 13 | Q8IX01 | SURP and G-patch domain-containing pr | SUGP2 | 120.208 | 2 | 54.533707 | yes | 2.254E-07 | 761.39337 | 73.648468 |
| DLDFAQQK | 8 | Q8IX01 | SURP and G-patch domain-containing pr | SUGP2 | 120.208 | 2 | 54.533707 | yes | 5.205E-06 | 482.74036 | 68.428825 |
| GETQGLLTAK | 10 | Q8IX01 | SURP and G-patch domain-containing pr | SUGP2 | 120.208 | 2 | 54.533707 | yes | 6.876E-07 | 509.28238 | 63.943024 |
| NLGFQMLQK | 9 | Q8IX01 | SURP and G-patch domain-containing pr | SUGP2 | 120.208 | 2 | 54.533707 | yes | 5.448E-07 | 539.78931 | 59.411354 |
| EDTFDVFR | 8 | Q8IX01 | SURP and G-patch domain-containing pr | SUGP2 | 120.208 | 2 | 54.533707 | yes | 2.161E-07 | 514.73779 | 56.193497 |
| QAPGLSQAK | 9 | Q8IX01 | SURP and G-patch domain-containing pr | SUGP2 | 120.208 | 2 | 54.533707 | yes | 1.413E-07 | 450.25089 | 55.22636 |
| IPLGLDLK | 8 | Q8IX01 | SURP and G-patch domain-containing pr | SUGP2 | 120.208 | 2 | 54.533707 | yes | 3.514E-06 | 434.77875 | 34.505856 |
| RVIEGSLSPK | 10 | Q8IX01 | SURP and G-patch domain-containing pr | SUGP2 | 120.208 | 2 | 54.533707 | yes | 1.431E-05 | 543.31934 | 22.375877 |
| DFCFFTIK | 8 | Q8IX01 | SURP and G-patch domain-containing pr | SUGP2 | 120.208 | 2 | 54.533707 | yes | 9.836E-07 | 539.25732 | 19.048576 |
| TQMITINR | 8 | Q8IX12 | Cell division cycle and apoptosis regulato | CCAR1 | 132.822 | 2 | 53.96509 | yes | 4.237E-07 | 488.76584 | 666.54346 |
| SCALAEDPQELR | 12 | Q8IX12 | Cell division cycle and apoptosis regulato | CCAR1 | 132.822 | 2 | 53.96509 | yes | 2.485E-07 | 694.82739 | 646.43176 |
| RLDPPSR | 7 | Q8IX12 | Cell division cycle and apoptosis regulato | CCAR1 | 132.822 | 2 | 53.96509 | yes | 3.108E-07 | 420.73795 | 619.73865 |
| EISTPTHWSK | 10 | Q8IX12 | Cell division cycle and apoptosis regulato | CCAR1 | 132.822 | 2 | 53.96509 | yes | 3.348E-07 | 593.29858 | 511.11035 |
| TILNLENSNK | 10 | Q8IX12 | Cell division cycle and apoptosis regulato | CCAR1 | 132.822 | 2 | 53.96509 | yes | 1.942E-07 | 573.31171 | 476.38486 |
| VLVEATYNPNMPI | 14 | Q8IX12 | Cell division cycle and apoptosis regulato | CCAR1 | 132.822 | 2 | 53.96509 | yes | 9.722E-08 | 811.91595 | 411.93091 |
| AGLLQPPVR | 9 | Q8IX12 | Cell division cycle and apoptosis regulato | CCAR1 | 132.822 | 2 | 53.96509 | yes | 1.077E-07 | 475.79272 | 401.47583 |
| NLSTVMDEIHTVL | 14 | Q8IX12 | Cell division cycle and apoptosis regulato | CCAR1 | 132.822 | 2 | 53.96509 | yes | 9.722E-08 | 800.42419 | 321.24545 |
| DLSQLQENLK | 10 | Q8IX12 | Cell division cycle and apoptosis regulato | CCAR1 | 132.822 | 2 | 53.96509 | yes | 9.722E-08 | 594.31696 | 298.79898 |
| LQLLEEK | 7 | Q8IX12 | Cell division cycle and apoptosis regulato | CCAR1 | 132.822 | 2 | 53.96509 | yes | 1.096E-07 | 436.75803 | 196.25995 |
| SEWETLSR | 8 | Q8IX12 | Cell division cycle and apoptosis regulato | CCAR1 | 132.822 | 2 | 53.96509 | yes | 2.005E-07 | 504.24326 | 183.27965 |
| YILPDEPAIIVHPN' | 18 | Q8IX12 | Cell division cycle and apoptosis regulato | CCAR1 | 132.822 | 3 | 53.96509 | yes | 1.022E-07 | 683.03857 | 153.6922 |
| KDLSQLQENLK | 11 | Q8IX12 | Cell division cycle and apoptosis regulato | CCAR1 | 132.822 | 2 | 53.96509 | yes | 9.722E-08 | 658.36444 | 43.009373 |
| SLASTLDCETAR | 12 | Q8IX90 | Spindle and kinetochore-associated protei | SKA3 | 46.36 | 2 | 54.758198 | yes | 9.722E-08 | 662.31409 | 103.248 |
| MDDFECVTPK | 10 | Q8IX90 | Spindle and kinetochore-associated protei | SKA3 | 46.36 | 2 | 54.758198 | yes | 2.343E-07 | 621.26245 | 45.481873 |
| YNSNLATPIAIK | 12 | Q8IX90 | Spindle and kinetochore-associated protei | SKA3 | 46.36 | 2 | 54.758198 | yes | 4.488E-06 | 652.86407 | 36.955143 |
| NSILFLNSLDAK | 12 | Q8IXB1 | DnaJ homolog subfamily C member 10 O | DNAJC10 | 91.08 | 2 | 53.061893 | yes | 9.722E-08 | 667.86938 | 106.2973 |
| ESLVSFAMQHVR | 12 | Q8IXB1 | DnaJ homolog subfamily C member 10 O | DNAJC10 | 91.08 | 2 | 53.061893 | yes | 9.722E-08 | 702.35864 | 79.726952 |
| AYEVLKDEDLR | 11 | Q8IXB1 | DnaJ homolog subfamily C member 10 O | DNAJC10 | 91.08 | 2 | 53.061893 | yes | 1.977E-07 | 675.84863 | 73.332283 |
| GVNSYPSLFIFR | 12 | Q8IXB1 | DnaJ homolog subfamily C member 10 O | DNAJC10 | 91.08 | 2 | 53.061893 | yes | 2.114E-07 | 700.37207 | 68.506104 |
| IGAVNCGDDR | 10 | Q8IXB1 | DnaJ homolog subfamily C member 10 O | DNAJC10 | 91.08 | 2 | 53.061893 | yes | 3.563E-07 | 538.7431 | 64.110123 |
| NFQEEQINTR | 10 | Q8IXB1 | DnaJ homolog subfamily C member 10 O | DNAJC10 | 91.08 | 2 | 53.061893 | yes | 1.536E-06 | 639.80731 | 56.836212 |
| ASNLLYGQLK | 10 | Q8IXB1 | DnaJ homolog subfamily C member 10 O | DNAJC10 | 91.08 | 2 | 53.061893 | yes | 5.19E-07 | 553.81384 | 52.014755 |
| ILYDILAFAK | 10 | Q8IXB1 | DnaJ homolog subfamily C member 10 O | DNAJC10 | 91.08 | 2 | 53.061893 | yes | 3.059E-06 | 583.8446 | 19.660898 |
| MFTSMDPPPVELII | 14 | Q8IXH7 | Negative elongation factor C/D OS=Hom | NELFCD | 66.247 | 2 | 58.119667 | yes | 9.722E-08 | 816.90985 | 206.67532 |
| TSLATILDGGEENI | 16 | Q8IXH7 | Negative elongation factor C/D OS=Hom | NELFCD | 66.247 | 2 | 58.119667 | yes | 3.812E-07 | 845.43073 | 178.26089 |

| | | | | | | | | | | | |
|---|---|---|---|---|---|---|---|---|---|---|---|
| WVDWTVSEPR | 10 | Q8IXH7 | Negative elongation factor C/D OS=Hom | NELFCD | 66.247 | 2 | 58.119667 | yes | 1.312E-07 | 637.81183 | 133.34685 |
| FPVVAMGVLK | 10 | Q8IXH7 | Negative elongation factor C/D OS=Hom | NELFCD | 66.247 | 2 | 58.119667 | yes | 4.25E-07 | 530.81482 | 110.91994 |
| LDTDISLIR | 9 | Q8IXH7 | Negative elongation factor C/D OS=Hom | NELFCD | 66.247 | 2 | 58.119667 | yes | 2.58E-07 | 523.29803 | 83.580879 |
| GHDASQITLALGT. | 19 | Q8IXH7 | Negative elongation factor C/D OS=Hom | NELFCD | 66.247 | 3 | 58.119667 | yes | 9.722E-08 | 643.66565 | 59.749336 |
| GALNPADITVLFK | 13 | Q8IXH7 | Negative elongation factor C/D OS=Hom | NELFCD | 66.247 | 2 | 58.119667 | yes | 4.314E-07 | 679.88757 | 30.994987 |
| AVETVHNLCCNEI | 14 | Q8IXH7 | Negative elongation factor C/D OS=Hom | NELFCD | 66.247 | 2 | 58.119667 | yes | 3.732E-07 | 844.37994 | 16.045256 |
| DCALSPDELK | 10 | Q8IXI2 | Mitochondrial Rho GTPase 1 OS=Homo | RHOT1 | 70.785 | 2 | 53.154827 | yes | 9.722E-08 | 574.26862 | 115.01282 |
| QEYSISPTDFCR | 12 | Q8IXI2 | Mitochondrial Rho GTPase 1 OS=Homo | RHOT1 | 70.785 | 2 | 53.154827 | yes | 9.819E-08 | 751.83264 | 102.045 |
| HISDGVADSGLTLl | 14 | Q8IXI2 | Mitochondrial Rho GTPase 1 OS=Homo | RHOT1 | 70.785 | 2 | 53.154827 | yes | 5.838E-07 | 706.87262 | 82.458275 |
| WIPLINER | 8 | Q8IXI2 | Mitochondrial Rho GTPase 1 OS=Homo | RHOT1 | 70.785 | 2 | 53.154827 | yes | 1.244E-07 | 520.79803 | 68.913902 |
| AEEITIPADVTPER | 14 | Q8IXI2 | Mitochondrial Rho GTPase 1 OS=Homo | RHOT1 | 70.785 | 2 | 53.154827 | yes | 2.516E-07 | 770.8963 | 59.078716 |
| DLVGMGHHFLPSI | 16 | Q8IXK0 | Polyhomeotic-like protein 2 OS=Homo sa | PHC2 | 90.714 | 3 | 56.43404 | yes | 1.018E-07 | 588.96277 | 163.16833 |
| AQEIDGQALLLLK | 13 | Q8IXK0 | Polyhomeotic-like protein 2 OS=Homo sa | PHC2 | 90.714 | 2 | 56.43404 | no | 9.722E-08 | 706.41138 | 146.05606 |
| VGLFHSDR | 8 | Q8IXK0 | Polyhomeotic-like protein 2 OS=Homo sa | PHC2 | 90.714 | 2 | 56.43404 | yes | 1.68E-07 | 465.74323 | 63.115063 |
| FCSMACAK | 8 | Q8IXK0 | Polyhomeotic-like protein 2 OS=Homo sa | PHC2 | 90.714 | 2 | 56.43404 | yes | 1.415E-06 | 487.69827 | 56.274361 |
| DLELPDMHMR | 10 | Q8IXK0 | Polyhomeotic-like protein 2 OS=Homo sa | PHC2 | 90.714 | 2 | 56.43404 | yes | 9.723E-08 | 628.79175 | 51.679066 |
| DGTFDPDNLEK | 11 | Q8IXM3 | Large ribosomal subunit protein mL41 OS | MRPL41 | 15.383 | 2 | 51.842762 | yes | 1.54E-07 | 625.7804 | 1172.7587 |
| LFQLYPR | 7 | Q8IXM3 | Large ribosomal subunit protein mL41 OS | MRPL41 | 15.383 | 2 | 51.842762 | yes | 3.072E-07 | 468.76871 | 565.36078 |
| YGFEPTQEGK | 10 | Q8IXM3 | Large ribosomal subunit protein mL41 OS | MRPL41 | 15.383 | 2 | 51.842762 | yes | 9.722E-08 | 578.26947 | 542.5611 |
| GIGFLTSGWR | 10 | Q8IXM3 | Large ribosomal subunit protein mL41 OS | MRPL41 | 15.383 | 2 | 51.842762 | yes | 2.259E-07 | 547.29309 | 486.64105 |
| TELEIEGAR | 9 | Q8IXQ6 | Protein mono-ADP-ribosyltransferase PA | PARP9 | 96.344 | 2 | 53.32466 | yes | 1.811E-07 | 509.26419 | 211.52686 |
| VSAGEIAVTGAGR | 13 | Q8IXQ6 | Protein mono-ADP-ribosyltransferase PA | PARP9 | 96.344 | 2 | 53.32466 | yes | 1.253E-07 | 594.32257 | 203.80359 |
| IDNEVLMAAFQR | 12 | Q8IXQ6 | Protein mono-ADP-ribosyltransferase PA | PARP9 | 96.344 | 2 | 53.32466 | yes | 1.064E-07 | 703.85846 | 143.46899 |
| CPVPPTQELLDQK | 13 | Q8IXQ6 | Protein mono-ADP-ribosyltransferase PA | PARP9 | 96.344 | 2 | 53.32466 | yes | 1.548E-06 | 762.88995 | 133.39009 |
| SLGQWTIQQQK | 11 | Q8IXQ6 | Protein mono-ADP-ribosyltransferase PA | PARP9 | 96.344 | 2 | 53.32466 | yes | 9.513E-07 | 658.8515 | 131.39299 |
| YGAGIYFTK | 9 | Q8IXQ6 | Protein mono-ADP-ribosyltransferase PA | PARP9 | 96.344 | 2 | 53.32466 | yes | 2.874E-07 | 510.26346 | 117.87808 |
| SILQQAGVEMK | 11 | Q8IXQ6 | Protein mono-ADP-ribosyltransferase PA | PARP9 | 96.344 | 2 | 53.32466 | yes | 9.722E-08 | 602.32373 | 97.237381 |
| QIIHAVGPR | 9 | Q8IXQ6 | Protein mono-ADP-ribosyltransferase PA | PARP9 | 96.344 | 2 | 53.32466 | yes | 2.092E-07 | 495.79581 | 90.598137 |
| VQEEMAR | 7 | Q8IXQ6 | Protein mono-ADP-ribosyltransferase PA | PARP9 | 96.344 | 2 | 53.32466 | yes | 4.244E-07 | 431.70801 | 79.719795 |
| MYSTPCDPK | 9 | Q8IXQ6 | Protein mono-ADP-ribosyltransferase PA | PARP9 | 96.344 | 2 | 53.32466 | yes | 6.375E-07 | 549.73334 | 68.567345 |
| MLSLNNYSVPQST | 14 | Q8IXQ6 | Protein mono-ADP-ribosyltransferase PA | PARP9 | 96.344 | 2 | 53.32466 | yes | 1.606E-07 | 805.40375 | 65.290855 |
| AGGFEIQEESK | 11 | Q8IXQ6 | Protein mono-ADP-ribosyltransferase PA | PARP9 | 96.344 | 2 | 53.32466 | yes | 4.442E-07 | 597.78552 | 37.350666 |
| TSSVSITEIISPGR | 14 | Q8IXQ6 | Protein mono-ADP-ribosyltransferase PA | PARP9 | 96.344 | 2 | 53.32466 | yes | 3.851E-06 | 723.89355 | 32.231972 |
| CGLQVLK | 7 | Q8IXQ6 | Protein mono-ADP-ribosyltransferase PA | PARP9 | 96.344 | 2 | 53.32466 | yes | 1.824E-06 | 409.23364 | 16.257755 |
| LCGYPLCQK | 9 | Q8IXW5 | Putative RNA polymerase II subunit B1 C | RPAP2 | 69.509 | 2 | 51.070232 | yes | 1.308E-07 | 569.77283 | 116.37199 |
| FLYGQNYASVCLK | 20 | Q8IXW5 | Putative RNA polymerase II subunit B1 C | RPAP2 | 69.509 | 3 | 51.070232 | yes | 3.631E-07 | 763.06458 | 64.44281 |
| SVSSSVQVCPEVG | 14 | Q8IXW5 | Putative RNA polymerase II subunit B1 C | RPAP2 | 69.509 | 2 | 51.070232 | yes | 9.722E-08 | 731.86395 | 56.922173 |
| TSDIDNPSHFEK | 12 | Q8IXW5 | Putative RNA polymerase II subunit B1 C | RPAP2 | 69.509 | 2 | 51.070232 | yes | 2.187E-06 | 695.31769 | 43.18602 |
| VYDITER | 7 | Q8IXW5 | Putative RNA polymerase II subunit B1 C | RPAP2 | 69.509 | 2 | 51.070232 | yes | 1.648E-07 | 448.22961 | 41.84341 |

| | | | | | | | | | | | |
|---|---|---|---|---|---|---|---|---|---|---|---|
| ESQNSLDESLPFR | 13 | Q8IXW5 | Putative RNA polymerase II subunit B1 C | RPAP2 | 69.509 | 2 | 51.070232 | yes | 1.032E-07 | 761.36261 | 36.836571 |
| DATCELPLQK | 10 | Q8IXW5 | Putative RNA polymerase II subunit B1 C | RPAP2 | 69.509 | 2 | 51.070232 | yes | 8.5E-07 | 587.79224 | 36.369286 |
| FITPAHYSDVVDEI | 14 | Q8IXW5 | Putative RNA polymerase II subunit B1 C | RPAP2 | 69.509 | 2 | 51.070232 | yes | 3.801E-07 | 824.90192 | 19.548952 |
| GSGTAIKPLPSYEN | 16 | Q8IXW5 | Putative RNA polymerase II subunit B1 C | RPAP2 | 69.509 | 2 | 51.070232 | yes | 6.109E-07 | 837.95667 | 18.401194 |
| GTCSVEDPLLVCQ | 14 | Q8IXZ2 | Zinc finger CCCH domain-containing prc | ZC3H3 | 101.942 | 2 | 41.25173 | yes | 4.41E-07 | 803.38422 | 63.171764 |
| LLQGLIDDYK | 10 | Q8IXZ2 | Zinc finger CCCH domain-containing prc | ZC3H3 | 101.942 | 2 | 41.25173 | yes | 3.602E-07 | 589.32678 | 54.30743 |
| LFPSPGLPTR | 10 | Q8IY17 | Patatin-like phospholipase domain-contai | PNPLA6 | 150.954 | 2 | 52.589718 | yes | 1.186E-07 | 542.8111 | 384.32059 |
| VPDMAEIQSR | 10 | Q8IY17 | Patatin-like phospholipase domain-contai | PNPLA6 | 150.954 | 2 | 52.589718 | yes | 1.552E-07 | 573.28461 | 371.84509 |
| LGASALDSIQEFR | 13 | Q8IY17 | Patatin-like phospholipase domain-contai | PNPLA6 | 150.954 | 2 | 52.589718 | yes | 9.722E-08 | 703.86737 | 341.25641 |
| IEDPSLLNSR | 10 | Q8IY17 | Patatin-like phospholipase domain-contai | PNPLA6 | 150.954 | 2 | 52.589718 | yes | 1.254E-07 | 572.30383 | 303.23941 |
| FDQIYDVGYQYGK | 13 | Q8IY17 | Patatin-like phospholipase domain-contai | PNPLA6 | 150.954 | 2 | 52.589718 | yes | 9.722E-08 | 798.37244 | 255.74831 |
| CVSMPGDISGLQG | 16 | Q8IY17 | Patatin-like phospholipase domain-contai | PNPLA6 | 150.954 | 2 | 52.589718 | yes | 9.938E-08 | 815.88745 | 202.00891 |
| LSGWLAQQEDAHI | 13 | Q8IY17 | Patatin-like phospholipase domain-contai | PNPLA6 | 150.954 | 2 | 52.589718 | yes | 9.722E-08 | 755.87347 | 183.76401 |
| LPEGTLGHIK | 10 | Q8IY17 | Patatin-like phospholipase domain-contai | PNPLA6 | 150.954 | 2 | 52.589718 | yes | 1.526E-07 | 532.80859 | 179.25433 |
| SDFYEIMR | 8 | Q8IY17 | Patatin-like phospholipase domain-contai | PNPLA6 | 150.954 | 2 | 52.589718 | yes | 2.733E-07 | 530.74207 | 174.32684 |
| SDFDMAYER | 9 | Q8IY17 | Patatin-like phospholipase domain-contai | PNPLA6 | 150.954 | 2 | 52.589718 | yes | 1.702E-07 | 567.23206 | 169.12395 |
| DGFQDVLAPGEGS | 16 | Q8IY17 | Patatin-like phospholipase domain-contai | PNPLA6 | 150.954 | 2 | 52.589718 | yes | 2.555E-07 | 788.37354 | 147.29411 |
| GCSHIGVLK | 9 | Q8IY17 | Patatin-like phospholipase domain-contai | PNPLA6 | 150.954 | 2 | 52.589718 | yes | 2.328E-06 | 485.76056 | 130.82341 |
| QLVLLHR | 7 | Q8IY17 | Patatin-like phospholipase domain-contai | PNPLA6 | 150.954 | 2 | 52.589718 | yes | 9.722E-08 | 439.78217 | 122.92681 |
| MVSTSATDEPR | 11 | Q8IY17 | Patatin-like phospholipase domain-contai | PNPLA6 | 150.954 | 2 | 52.589718 | yes | 1.027E-07 | 597.27698 | 119.89077 |
| QTSSIFEAAK | 10 | Q8IY17 | Patatin-like phospholipase domain-contai | PNPLA6 | 150.954 | 2 | 52.589718 | yes | 2.666E-06 | 541.27985 | 114.51543 |
| LIHLLSQK | 8 | Q8IY17 | Patatin-like phospholipase domain-contai | PNPLA6 | 150.954 | 2 | 52.589718 | yes | 1.452E-07 | 476.30292 | 95.937378 |
| VLTGNTIALVLGG | 17 | Q8IY17 | Patatin-like phospholipase domain-contai | PNPLA6 | 150.954 | 2 | 52.589718 | yes | 1.811E-07 | 784.95959 | 95.5168 |
| VVQIIMVR | 8 | Q8IY17 | Patatin-like phospholipase domain-contai | PNPLA6 | 150.954 | 2 | 52.589718 | no | 1.134E-07 | 479.29932 | 91.829338 |
| LGQGDYVFR | 9 | Q8IY17 | Patatin-like phospholipase domain-contai | PNPLA6 | 150.954 | 2 | 52.589718 | yes | 2.782E-07 | 527.76947 | 80.31588 |
| AVFGGWSR | 8 | Q8IY17 | Patatin-like phospholipase domain-contai | PNPLA6 | 150.954 | 2 | 52.589718 | yes | 4.818E-07 | 440.22723 | 72.209457 |
| LPVEAFSAVFTK | 12 | Q8IY17 | Patatin-like phospholipase domain-contai | PNPLA6 | 150.954 | 2 | 52.589718 | yes | 1.218E-07 | 654.86353 | 68.944778 |
| VLGHFEKPLFLELC | 15 | Q8IY17 | Patatin-like phospholipase domain-contai | PNPLA6 | 150.954 | 3 | 52.589718 | yes | 1.153E-07 | 620.00629 | 61.659439 |
| TVEWLNMR | 8 | Q8IY17 | Patatin-like phospholipase domain-contai | PNPLA6 | 150.954 | 2 | 52.589718 | no | 1.591E-07 | 524.76587 | 61.299862 |
| PLFLELCR | 8 | Q8IY17 | Patatin-like phospholipase domain-contai | PNPLA6 | 150.954 | 2 | 52.589718 | yes | 7.997E-07 | 524.28644 | 27.013847 |
| VSQSTSSLVDTSVS | 20 | Q8IY17 | Patatin-like phospholipase domain-contai | PNPLA6 | 150.954 | 3 | 52.589718 | yes | 2.842E-06 | 689.02216 | 5.2292409 |
| LQAVDSGLIALR | 12 | Q8IY18 | Structural maintenance of chromosomes p | SMC5 | 128.807 | 2 | 52.9948 | yes | 9.893E-08 | 628.37207 | 342.3811 |
| VFEMVVNTACK | 11 | Q8IY18 | Structural maintenance of chromosomes p | SMC5 | 128.807 | 2 | 52.9948 | yes | 1.656E-07 | 649.31757 | 277.74329 |
| LLQNLPYSEK | 10 | Q8IY18 | Structural maintenance of chromosomes p | SMC5 | 128.807 | 2 | 52.9948 | yes | 1E-07 | 602.83228 | 260.45544 |
| ENTSQYFFITPK | 12 | Q8IY18 | Structural maintenance of chromosomes p | SMC5 | 128.807 | 2 | 52.9948 | yes | 1.58E-07 | 737.86426 | 259.21713 |
| LVTELTNLIK | 10 | Q8IY18 | Structural maintenance of chromosomes p | SMC5 | 128.807 | 2 | 52.9948 | yes | 9.722E-08 | 572.35284 | 257.20255 |
| ALCEGEIIDK | 10 | Q8IY18 | Structural maintenance of chromosomes p | SMC5 | 128.807 | 2 | 52.9948 | yes | 9.888E-08 | 574.2868 | 256.44766 |
| MIEDLQNELK | 10 | Q8IY18 | Structural maintenance of chromosomes p | SMC5 | 128.807 | 2 | 52.9948 | yes | 9.722E-08 | 616.81317 | 205.39207 |
| SIGPPEMHK | 9 | Q8IY18 | Structural maintenance of chromosomes p | SMC5 | 128.807 | 2 | 52.9948 | yes | 1.317E-06 | 498.25256 | 185.80119 |

| | | | | | | | | | | | |
|---|---|---|---|---|---|---|---|---|---|---|---|
| KVELDQYR | 8 | Q8IY18 | Structural maintenance of chromosomes p | SMC5 | 128.807 | 2 | 52.9948 | yes | 6.695E-07 | 525.78253 | 182.89656 |
| IELLEATEK | 9 | Q8IY18 | Structural maintenance of chromosomes p | SMC5 | 128.807 | 2 | 52.9948 | yes | 1.416E-07 | 523.29242 | 160.12236 |
| VDLILQNTTVISEK | 14 | Q8IY18 | Structural maintenance of chromosomes p | SMC5 | 128.807 | 2 | 52.9948 | yes | 9.722E-08 | 786.9458 | 140.08749 |
| ICTSLHIQK | 9 | Q8IY18 | Structural maintenance of chromosomes p | SMC5 | 128.807 | 2 | 52.9948 | yes | 1.073E-07 | 550.30005 | 126.06621 |
| NQSFWFINK | 9 | Q8IY18 | Structural maintenance of chromosomes p | SMC5 | 128.807 | 2 | 52.9948 | yes | 3.876E-06 | 592.2984 | 120.18819 |
| QYGFFSYLR | 9 | Q8IY18 | Structural maintenance of chromosomes p | SMC5 | 128.807 | 2 | 52.9948 | yes | 1.272E-07 | 590.79291 | 95.204689 |
| LESDYMAASSQLR | 13 | Q8IY18 | Structural maintenance of chromosomes p | SMC5 | 128.807 | 2 | 52.9948 | yes | 9.722E-08 | 735.84833 | 84.019272 |
| GMVEIELFR | 9 | Q8IY18 | Structural maintenance of chromosomes p | SMC5 | 128.807 | 2 | 52.9948 | yes | 1.632E-07 | 547.28918 | 82.562363 |
| SSIVCAICLGLAGK | 14 | Q8IY18 | Structural maintenance of chromosomes p | SMC5 | 128.807 | 2 | 52.9948 | yes | 8.361E-07 | 724.88361 | 80.650658 |
| EEEIEQLTEELK | 12 | Q8IY18 | Structural maintenance of chromosomes p | SMC5 | 128.807 | 2 | 52.9948 | yes | 1.027E-06 | 745.36707 | 73.805367 |
| TSTPSPQPSK | 10 | Q8IY18 | Structural maintenance of chromosomes p | SMC5 | 128.807 | 2 | 52.9948 | yes | 2.669E-06 | 515.26422 | 66.382133 |
| HIEELQQALIVK | 12 | Q8IY18 | Structural maintenance of chromosomes p | SMC5 | 128.807 | 2 | 52.9948 | yes | 3.161E-07 | 710.91156 | 59.849464 |
| HLDLIEMLEAK | 11 | Q8IY18 | Structural maintenance of chromosomes p | SMC5 | 128.807 | 2 | 52.9948 | yes | 2.361E-05 | 656.35248 | 44.893337 |
| DTYDAVLWLR | 10 | Q8IY18 | Structural maintenance of chromosomes p | SMC5 | 128.807 | 2 | 52.9948 | yes | 9.926E-08 | 626.32202 | 39.853195 |
| LTEQHFIELDENR | 13 | Q8IY18 | Structural maintenance of chromosomes p | SMC5 | 128.807 | 2 | 52.9948 | yes | 3.849E-07 | 822.40485 | 15.965342 |
| VAQFLTVTVDLEQ | 14 | Q8IY18 | Structural maintenance of chromosomes p | SMC5 | 128.807 | 2 | 52.9948 | yes | 5.217E-06 | 809.9436 | 9.5736504 |
| ALVNQVAATVQN | 13 | Q8IY21 | Probable ATP-dependent RNA helicase D | DDX60 | 197.854 | 2 | 52.961742 | yes | 1.062E-07 | 692.38879 | 230.66237 |
| VQDFIPDTWQR | 11 | Q8IY21 | Probable ATP-dependent RNA helicase D | DDX60 | 197.854 | 2 | 52.961742 | yes | 9.722E-08 | 702.84894 | 167.69249 |
| LGAVENAAESVST | 16 | Q8IY21 | Probable ATP-dependent RNA helicase D | DDX60 | 197.854 | 2 | 52.961742 | yes | 9.722E-08 | 818.43304 | 167.47348 |
| NIAFYYENENVK | 12 | Q8IY21 | Probable ATP-dependent RNA helicase D | DDX60 | 197.854 | 2 | 52.961742 | yes | 1.4E-07 | 752.35938 | 161.00032 |
| TALILHLQK | 9 | Q8IY21 | Probable ATP-dependent RNA helicase D | DDX60 | 197.854 | 2 | 52.961742 | yes | 1.297E-07 | 518.82928 | 155.5616 |
| VFCVTSCSLSLR | 12 | Q8IY21 | Probable ATP-dependent RNA helicase D | DDX60 | 197.854 | 2 | 52.961742 | yes | 2.459E-07 | 714.85236 | 150.64728 |
| YSELLQEDDR | 10 | Q8IY21 | Probable ATP-dependent RNA helicase D | DDX60 | 197.854 | 2 | 52.961742 | yes | 6.862E-07 | 634.2937 | 145.21626 |
| NLEIPQDCTYADQ | 14 | Q8IY21 | Probable ATP-dependent RNA helicase D | DDX60 | 197.854 | 2 | 52.961742 | yes | 3.094E-07 | 847.88812 | 144.3756 |
| LSALAPLFGSLK | 12 | Q8IY21 | Probable ATP-dependent RNA helicase D | DDX60 | 197.854 | 2 | 52.961742 | yes | 2.754E-07 | 608.86865 | 140.87567 |
| IMEDFTTFLR | 10 | Q8IY21 | Probable ATP-dependent RNA helicase D | DDX60 | 197.854 | 2 | 52.961742 | yes | 9.722E-08 | 636.8183 | 127.66716 |
| WNNITEEAHK | 10 | Q8IY21 | Probable ATP-dependent RNA helicase D | DDX60 | 197.854 | 2 | 52.961742 | yes | 3.473E-07 | 621.29907 | 127.21818 |
| VITSHWAEDMKPI | 17 | Q8IY21 | Probable ATP-dependent RNA helicase D | DDX60 | 197.854 | 3 | 52.961742 | yes | 3.835E-07 | 676.35327 | 126.13075 |
| IIENNTASK | 9 | Q8IY21 | Probable ATP-dependent RNA helicase D | DDX60 | 197.854 | 2 | 52.961742 | yes | 2.259E-07 | 495.26672 | 124.47688 |
| SDDPIVTSLVK | 11 | Q8IY21 | Probable ATP-dependent RNA helicase D | DDX60 | 197.854 | 2 | 52.961742 | yes | 9.752E-08 | 587.32172 | 121.33261 |
| ESDDGVVVYVAPT | 14 | Q8IY21 | Probable ATP-dependent RNA helicase D | DDX60 | 197.854 | 2 | 52.961742 | yes | 1.303E-07 | 739.87231 | 120.45745 |
| NLPSGEVLCGVFT | 14 | Q8IY21 | Probable ATP-dependent RNA helicase D | DDX60 | 197.854 | 2 | 52.961742 | yes | 3.405E-07 | 774.89557 | 111.06 |
| DAYTTLLNQLER | 12 | Q8IY21 | Probable ATP-dependent RNA helicase D | DDX60 | 197.854 | 2 | 52.961742 | yes | 3.298E-07 | 718.87262 | 105.55723 |
| HGSLIGLVQDNR | 12 | Q8IY21 | Probable ATP-dependent RNA helicase D | DDX60 | 197.854 | 2 | 52.961742 | yes | 3.106E-06 | 654.85455 | 72.257851 |
| YLVDLISK | 8 | Q8IY21 | Probable ATP-dependent RNA helicase D | DDX60 | 197.854 | 2 | 52.961742 | yes | 3.274E-07 | 475.78149 | 65.451988 |
| HFSQDVMEK | 9 | Q8IY21 | Probable ATP-dependent RNA helicase D | DDX60 | 197.854 | 2 | 52.961742 | yes | 3.194E-06 | 560.75824 | 52.475216 |
| LQVEMVGLTACLI | 13 | Q8IY21 | Probable ATP-dependent RNA helicase D | DDX60 | 197.854 | 2 | 52.961742 | yes | 2.612E-07 | 731.39386 | 49.158264 |
| GLHLNLGDTIMK | 12 | Q8IY21 | Probable ATP-dependent RNA helicase D | DDX60 | 197.854 | 2 | 52.961742 | yes | 9.819E-08 | 656.35809 | 41.269604 |
| DFEVGQPFPLR | 11 | Q8IY21 | Probable ATP-dependent RNA helicase D | DDX60 | 197.854 | 2 | 52.961742 | yes | 5.689E-06 | 652.83533 | 36.585312 |

| | | | | | | | | | | | |
|---|---|---|---|---|---|---|---|---|---|---|---|
| GHFPLSITLVLR | 12 | Q8IY21 | Probable ATP-dependent RNA helicase D | DDX60 | 197.854 | 2 | 52.961742 | yes | 1.79E-06 | 676.90607 | 32.043602 |
| MSLNAYALDFYK | 12 | Q8IY21 | Probable ATP-dependent RNA helicase D | DDX60 | 197.854 | 2 | 52.961742 | yes | 3.01E-07 | 718.34998 | 30.135798 |
| FQDAHFEFYQSK | 12 | Q8IY21 | Probable ATP-dependent RNA helicase D | DDX60 | 197.854 | 2 | 52.961742 | yes | 1.144E-07 | 773.85168 | 28.813858 |
| IIVTQTIK | 8 | Q8IY21 | Probable ATP-dependent RNA helicase D | DDX60 | 197.854 | 2 | 52.961742 | yes | 1.577E-07 | 458.2973 | 26.017397 |
| WCEYFILR | 8 | Q8IY21 | Probable ATP-dependent RNA helicase D | DDX60 | 197.854 | 2 | 52.961742 | yes | 4.159E-07 | 593.78931 | 25.805206 |
| QLVEILFR | 8 | Q8IY21 | Probable ATP-dependent RNA helicase D | DDX60 | 197.854 | 2 | 52.961742 | yes | 6.099E-06 | 509.3082 | 19.512363 |
| VLDPEVTQQTIELI | 14 | Q8IY31 | Intraflagellar transport protein 20 homolo | IFT20 | 15.281 | 2 | 50.798428 | yes | 9.722E-08 | 806.94324 | 60.830311 |
| LSCPITALGR | 10 | Q8IY37 | Probable ATP-dependent RNA helicase D | DHX37 | 129.546 | 2 | 56.685753 | yes | 4.974E-07 | 544.30005 | 1195.3445 |
| SCLLSSPGTMLK | 12 | Q8IY37 | Probable ATP-dependent RNA helicase D | DHX37 | 129.546 | 2 | 56.685753 | yes | 9.722E-08 | 647.33069 | 1145.8596 |
| VVIIDEAHER | 10 | Q8IY37 | Probable ATP-dependent RNA helicase D | DHX37 | 129.546 | 2 | 56.685753 | yes | 9.722E-08 | 590.81964 | 1102.1727 |
| MQPPTESQVTYLR | 13 | Q8IY37 | Probable ATP-dependent RNA helicase D | DHX37 | 129.546 | 2 | 56.685753 | yes | 1.08E-07 | 775.38757 | 1101.9833 |
| PAVFIPVNR | 9 | Q8IY37 | Probable ATP-dependent RNA helicase D | DHX37 | 129.546 | 2 | 56.685753 | yes | 1.084E-07 | 506.80054 | 1026.3566 |
| TMATFPVAPR | 10 | Q8IY37 | Probable ATP-dependent RNA helicase D | DHX37 | 129.546 | 2 | 56.685753 | yes | 3.25E-07 | 545.78931 | 858.75586 |
| VEDFTQNPR | 9 | Q8IY37 | Probable ATP-dependent RNA helicase D | DHX37 | 129.546 | 2 | 56.685753 | yes | 9.722E-08 | 553.26727 | 851.02466 |
| FLLEGQVFR | 9 | Q8IY37 | Probable ATP-dependent RNA helicase D | DHX37 | 129.546 | 2 | 56.685753 | yes | 1.284E-07 | 554.8111 | 842.04193 |
| LSEVQASEAEMR | 12 | Q8IY37 | Probable ATP-dependent RNA helicase D | DHX37 | 129.546 | 2 | 56.685753 | yes | 9.722E-08 | 675.3219 | 400.86395 |
| LSEVQASEAEMR | 12 | Q8IY37 | Probable ATP-dependent RNA helicase D | DHX37 | 129.546 | 2 | 56.685753 | yes | 3.519E-06 | 683.3194 | 400.86395 |
| TWAGQGASLK | 10 | Q8IY37 | Probable ATP-dependent RNA helicase D | DHX37 | 129.546 | 2 | 56.685753 | yes | 3.591E-07 | 509.76944 | 524.50226 |
| QFPVTVHFNK | 10 | Q8IY37 | Probable ATP-dependent RNA helicase D | DHX37 | 129.546 | 2 | 56.685753 | yes | 1.032E-07 | 608.82727 | 517.35504 |
| ISSLSGAHR | 9 | Q8IY37 | Probable ATP-dependent RNA helicase D | DHX37 | 129.546 | 2 | 56.685753 | yes | 1.444E-07 | 464.25397 | 512.82355 |
| RPVEDLILQMK | 11 | Q8IY37 | Probable ATP-dependent RNA helicase D | DHX37 | 129.546 | 2 | 56.685753 | yes | 1.786E-07 | 671.38159 | 497.19168 |
| GVDASNALVLPGK | 13 | Q8IY37 | Probable ATP-dependent RNA helicase D | DHX37 | 129.546 | 2 | 56.685753 | yes | 9.819E-08 | 620.84845 | 451.82727 |
| VTGVSSFR | 8 | Q8IY37 | Probable ATP-dependent RNA helicase D | DHX37 | 129.546 | 2 | 56.685753 | yes | 2.193E-07 | 426.73233 | 416.65411 |
| YEGNVTEETR | 10 | Q8IY37 | Probable ATP-dependent RNA helicase D | DHX37 | 129.546 | 2 | 56.685753 | yes | 1.006E-07 | 599.27277 | 371.4277 |
| TPLEDYSGECFR | 12 | Q8IY37 | Probable ATP-dependent RNA helicase D | DHX37 | 129.546 | 2 | 56.685753 | yes | 1.136E-07 | 737.3194 | 347.98517 |
| LLIMSATLR | 9 | Q8IY37 | Probable ATP-dependent RNA helicase D | DHX37 | 129.546 | 2 | 56.685753 | yes | 1.087E-07 | 509.30991 | 336.26895 |
| VAAVAMSQR | 9 | Q8IY37 | Probable ATP-dependent RNA helicase D | DHX37 | 129.546 | 2 | 56.685753 | yes | 1.363E-07 | 466.75275 | 306.8157 |
| VVSYQIR | 7 | Q8IY37 | Probable ATP-dependent RNA helicase D | DHX37 | 129.546 | 2 | 56.685753 | yes | 8.386E-07 | 432.75052 | 300.70071 |
| VTWVSQASADQR | 12 | Q8IY37 | Probable ATP-dependent RNA helicase D | DHX37 | 129.546 | 2 | 56.685753 | yes | 9.938E-08 | 674.33624 | 250.85666 |
| EKADEVVAPGQEI | 13 | Q8IY37 | Probable ATP-dependent RNA helicase D | DHX37 | 129.546 | 2 | 56.685753 | yes | 9.722E-08 | 700.35681 | 199.76564 |
| SPEMQEER | 8 | Q8IY37 | Probable ATP-dependent RNA helicase D | DHX37 | 129.546 | 2 | 56.685753 | yes | 1.214E-06 | 503.21893 | 193.27699 |
| FMTDGVLLK | 9 | Q8IY37 | Probable ATP-dependent RNA helicase D | DHX37 | 129.546 | 2 | 56.685753 | yes | 4.044E-07 | 512.28082 | 171.89085 |
| QAQVFKPPPEGTR | 13 | Q8IY37 | Probable ATP-dependent RNA helicase D | DHX37 | 129.546 | 2 | 56.685753 | yes | 1.47E-06 | 727.89117 | 146.7861 |
| ADCHEALLAAWK | 12 | Q8IY37 | Probable ATP-dependent RNA helicase D | DHX37 | 129.546 | 2 | 56.685753 | yes | 9.758E-08 | 692.83752 | 144.98933 |
| VQSEEMLEDK | 10 | Q8IY37 | Probable ATP-dependent RNA helicase D | DHX37 | 129.546 | 2 | 56.685753 | yes | 2.15E-07 | 604.27917 | 130.16782 |
| TPLLDDPVFIHPSS | 18 | Q8IY37 | Probable ATP-dependent RNA helicase D | DHX37 | 129.546 | 3 | 56.685753 | yes | 9.722E-08 | 675.70673 | 107.86673 |
| GPPEPPPVQLELEI | 15 | Q8IY37 | Probable ATP-dependent RNA helicase D | DHX37 | 129.546 | 2 | 56.685753 | yes | 8.395E-07 | 822.92761 | 94.630142 |
| QIVTAGLGDHLAR | 13 | Q8IY37 | Probable ATP-dependent RNA helicase D | DHX37 | 129.546 | 3 | 56.685753 | yes | 1.357E-07 | 450.92111 | 83.51339 |
| DDDQKDSVEEMR | 12 | Q8IY37 | Probable ATP-dependent RNA helicase D | DHX37 | 129.546 | 2 | 56.685753 | yes | 3.626E-07 | 733.80682 | 74.717239 |

| | | | | | | | | | | | |
|---|---|---|---|---|---|---|---|---|---|---|---|
| RVQSEEMLEDK | 11 | Q8IY37 | Probable ATP-dependent RNA helicase D | DHX37 | 129.546 | 2 | 56.685753 | yes | 7.978E-07 | 682.32971 | 60.914803 |
| GPPEPPPVQLELEI | 19 | Q8IY37 | Probable ATP-dependent RNA helicase D | DHX37 | 129.546 | 3 | 56.685753 | yes | 3.362E-06 | 701.37207 | 52.184715 |
| VQSEEMLEDKWR | 12 | Q8IY37 | Probable ATP-dependent RNA helicase D | DHX37 | 129.546 | 2 | 56.685753 | yes | 3.84E-06 | 775.36938 | 48.008686 |
| ADEVVAPGQEK | 11 | Q8IY37 | Probable ATP-dependent RNA helicase D | DHX37 | 129.546 | 2 | 56.685753 | yes | 4.245E-07 | 571.78802 | 47.527611 |
| YLLAEYCEWLPQA | 20 | Q8IY37 | Probable ATP-dependent RNA helicase D | DHX37 | 129.546 | 3 | 56.685753 | yes | 1.953E-07 | 836.06421 | 35.023567 |
| QQLPQTPPSCLK | 12 | Q8IY81 | pre-rRNA 2'-O-ribose RNA methyltransfe | FTSJ3 | 96.559 | 2 | 51.508873 | yes | 5.157E-07 | 698.86627 | 487.22717 |
| TEIMSPLYQDEAPI | 14 | Q8IY81 | pre-rRNA 2'-O-ribose RNA methyltransfe | FTSJ3 | 96.559 | 2 | 51.508873 | yes | 1.317E-07 | 811.39252 | 343.69815 |
| AVLQEEQANLWFS | 14 | Q8IY81 | pre-rRNA 2'-O-ribose RNA methyltransfe | FTSJ3 | 96.559 | 2 | 51.508873 | yes | 1.238E-07 | 831.92792 | 317.50989 |
| ILDPEGLALGAVIA | 17 | Q8IY81 | pre-rRNA 2'-O-ribose RNA methyltransfe | FTSJ3 | 96.559 | 2 | 51.508873 | yes | 9.722E-08 | 827.47473 | 232.68117 |
| AEGYAEGDLTLYH | 14 | Q8IY81 | pre-rRNA 2'-O-ribose RNA methyltransfe | FTSJ3 | 96.559 | 2 | 51.508873 | yes | 9.823E-08 | 797.87842 | 227.92381 |
| DLIDNSFNR | 9 | Q8IY81 | pre-rRNA 2'-O-ribose RNA methyltransfe | FTSJ3 | 96.559 | 2 | 51.508873 | yes | 9.893E-08 | 547.26727 | 219.23921 |
| LACDFLAR | 8 | Q8IY81 | pre-rRNA 2'-O-ribose RNA methyltransfe | FTSJ3 | 96.559 | 2 | 51.508873 | yes | 9.85E-08 | 483.24728 | 177.63336 |
| RFQFLQK | 7 | Q8IY81 | pre-rRNA 2'-O-ribose RNA methyltransfe | FTSJ3 | 96.559 | 2 | 51.508873 | yes | 1.946E-07 | 483.77963 | 160.07242 |
| TSVTDFLR | 8 | Q8IY81 | pre-rRNA 2'-O-ribose RNA methyltransfe | FTSJ3 | 96.559 | 2 | 51.508873 | yes | 2.998E-06 | 469.75073 | 135.23289 |
| AEAVVNTVDISER | 13 | Q8IY81 | pre-rRNA 2'-O-ribose RNA methyltransfe | FTSJ3 | 96.559 | 2 | 51.508873 | yes | 2.142E-06 | 701.86224 | 121.60444 |
| DKFYHLAK | 8 | Q8IY81 | pre-rRNA 2'-O-ribose RNA methyltransfe | FTSJ3 | 96.559 | 2 | 51.508873 | yes | 9.722E-08 | 511.27692 | 117.88072 |
| HESAEIFVVCQGFI | 18 | Q8IY81 | pre-rRNA 2'-O-ribose RNA methyltransfe | FTSJ3 | 96.559 | 3 | 51.508873 | yes | 9.819E-08 | 683.0033 | 112.04078 |
| AQEVAELK | 8 | Q8IY81 | pre-rRNA 2'-O-ribose RNA methyltransfe | FTSJ3 | 96.559 | 2 | 51.508873 | yes | 3.715E-07 | 444.24527 | 111.64683 |
| HESAEIFVVCQGFI | 22 | Q8IY81 | pre-rRNA 2'-O-ribose RNA methyltransfe | FTSJ3 | 96.559 | 3 | 51.508873 | yes | 3.941E-07 | 826.07739 | 58.878128 |
| KAEAVVNTVDISE | 14 | Q8IY81 | pre-rRNA 2'-O-ribose RNA methyltransfe | FTSJ3 | 96.559 | 2 | 51.508873 | yes | 2.153E-07 | 765.90973 | 54.493671 |
| VQATKPQASR | 10 | Q8IY81 | pre-rRNA 2'-O-ribose RNA methyltransfe | FTSJ3 | 96.559 | 2 | 51.508873 | yes | 7.241E-07 | 543.3067 | 40.610085 |
| LREEEER | 7 | Q8IY82 | Dynein regulatory complex subunit 7 OS= | DRC7 | 103.498 | 2 | 45.217422 | yes | 1.921E-07 | 480.74088 | 88.495987 |
| VLLGDVSTEAAR | 12 | Q8IY92 | Structure-specific endonuclease subunit S | SLX4 | 200.012 | 2 | 45.39417 | yes | 3.974E-07 | 615.83807 | 139.43672 |
| GAPWLFCDR | 9 | Q8IY92 | Structure-specific endonuclease subunit S | SLX4 | 200.012 | 2 | 45.39417 | yes | 1.863E-07 | 561.26349 | 81.208992 |
| AENFQELLR | 9 | Q8IY92 | Structure-specific endonuclease subunit S | SLX4 | 200.012 | 2 | 45.39417 | yes | 2.808E-07 | 560.29327 | 64.95192 |
| GLFFCQICQK | 10 | Q8IY92 | Structure-specific endonuclease subunit S | SLX4 | 200.012 | 2 | 45.39417 | yes | 1.981E-07 | 650.8125 | 44.522354 |
| SPGLLDTTPIR | 11 | Q8IY92 | Structure-specific endonuclease subunit S | SLX4 | 200.012 | 2 | 45.39417 | yes | 4.753E-06 | 585.3299 | 33.58593 |
| LESAFSER | 8 | Q8IY92 | Structure-specific endonuclease subunit S | SLX4 | 200.012 | 2 | 45.39417 | yes | 2.188E-07 | 469.73254 | 27.447582 |
| QFVGETQSR | 9 | Q8IYA6 | Cytoskeleton-associated protein 2-like OS | CKAP2L | 83.587 | 2 | 53.412487 | yes | 1.806E-07 | 526.26196 | 236.00432 |
| VGPGPTAAAAVEE | 14 | Q8IYA6 | Cytoskeleton-associated protein 2-like OS | CKAP2L | 83.587 | 2 | 53.412487 | yes | 1.281E-07 | 662.84644 | 179.92615 |
| VVLNILQDSNR | 11 | Q8IYA6 | Cytoskeleton-associated protein 2-like OS | CKAP2L | 83.587 | 2 | 53.412487 | yes | 2.284E-07 | 635.85931 | 164.8613 |
| TYPSLLQGEYNNR | 13 | Q8IYA6 | Cytoskeleton-associated protein 2-like OS | CKAP2L | 83.587 | 2 | 53.412487 | yes | 1.796E-07 | 777.88098 | 92.045761 |
| GTFDVIGLYEEAIK | 14 | Q8IYA6 | Cytoskeleton-associated protein 2-like OS | CKAP2L | 83.587 | 2 | 53.412487 | yes | 1.209E-07 | 777.90613 | 76.27861 |
| LQPRPPNTAGSQK | 15 | Q8IYA6 | Cytoskeleton-associated protein 2-like OS | CKAP2L | 83.587 | 3 | 53.412487 | yes | 4.829E-07 | 540.30566 | 73.739662 |
| MVGPGPTAAAAV | 15 | Q8IYA6 | Cytoskeleton-associated protein 2-like OS | CKAP2L | 83.587 | 3 | 53.412487 | yes | 1.553E-06 | 485.91354 | 60.838417 |
| AEQHNYPGIK | 10 | Q8IYA6 | Cytoskeleton-associated protein 2-like OS | CKAP2L | 83.587 | 2 | 53.412487 | yes | 3.146E-06 | 578.79089 | 41.225872 |
| KVVLNILQDSNR | 12 | Q8IYA6 | Cytoskeleton-associated protein 2-like OS | CKAP2L | 83.587 | 2 | 53.412487 | yes | 3.089E-05 | 699.9068 | 39.770435 |
| SIEKEEEEK | 9 | Q8IYA6 | Cytoskeleton-associated protein 2-like OS | CKAP2L | 83.587 | 2 | 53.412487 | yes | 9.722E-08 | 560.77203 | 39.323051 |
| LTSECVSSNPYSK | 13 | Q8IYA6 | Cytoskeleton-associated protein 2-like OS | CKAP2L | 83.587 | 2 | 53.412487 | yes | 5.516E-07 | 736.34033 | 29.729321 |

| | | | | | | | | | | | |
|---|---|---|---|---|---|---|---|---|---|---|---|
| KLQEYLAAK | 9 | Q8IYA6 | Cytoskeleton-associated protein 2-like OS | CKAP2L | 83.587 | 2 | 53.412487 | yes | 1.285E-07 | 532.31097 | 24.695265 |
| KPVGSLNIEQLK | 12 | Q8IYA6 | Cytoskeleton-associated protein 2-like OS | CKAP2L | 83.587 | 2 | 53.412487 | yes | 7.654E-07 | 663.39301 | 23.308571 |
| VNLEVIKPWITK | 12 | Q8IYB3 | Serine/arginine repetitive matrix protein 1 | SRRM1 | 102.336 | 2 | 51.12637 | yes | 1.106E-07 | 720.43469 | 409.67004 |
| VNLEVIKPWITK | 12 | Q8IYB3 | Serine/arginine repetitive matrix protein 1 | SRRM1 | 102.336 | 2 | 51.12637 | yes | 1.038E-07 | 480.62555 | 409.67004 |
| VNLEVIKPWITK | 12 | Q8IYB3 | Serine/arginine repetitive matrix protein 1 | SRRM1 | 102.336 | 3 | 51.12637 | yes | 1.106E-07 | 720.43469 | 409.67004 |
| VNLEVIKPWITK | 12 | Q8IYB3 | Serine/arginine repetitive matrix protein 1 | SRRM1 | 102.336 | 3 | 51.12637 | yes | 1.038E-07 | 480.62555 | 409.67004 |
| VPKPEPIPEPK | 11 | Q8IYB3 | Serine/arginine repetitive matrix protein 1 | SRRM1 | 102.336 | 2 | 51.12637 | yes | 5.75E-07 | 615.85828 | 378.86938 |
| VPKPEPIPEPK | 11 | Q8IYB3 | Serine/arginine repetitive matrix protein 1 | SRRM1 | 102.336 | 2 | 51.12637 | yes | 6.287E-07 | 410.90793 | 378.86938 |
| VPKPEPIPEPK | 11 | Q8IYB3 | Serine/arginine repetitive matrix protein 1 | SRRM1 | 102.336 | 3 | 51.12637 | yes | 5.75E-07 | 615.85828 | 378.86938 |
| VPKPEPIPEPK | 11 | Q8IYB3 | Serine/arginine repetitive matrix protein 1 | SRRM1 | 102.336 | 3 | 51.12637 | yes | 6.287E-07 | 410.90793 | 378.86938 |
| MAAADSVQQR | 10 | Q8IYB3 | Serine/arginine repetitive matrix protein 1 | SRRM1 | 102.336 | 2 | 51.12637 | yes | 2.144E-07 | 538.76129 | 357.58185 |
| EPSVQEATSTSDIL | 15 | Q8IYB3 | Serine/arginine repetitive matrix protein 1 | SRRM1 | 102.336 | 2 | 51.12637 | yes | 9.722E-08 | 802.9043 | 305.931 |
| SVSGSPEPAAK | 11 | Q8IYB3 | Serine/arginine repetitive matrix protein 1 | SRRM1 | 102.336 | 2 | 51.12637 | yes | 1.375E-07 | 515.26422 | 254.17612 |
| RMPPPPR | 7 | Q8IYB3 | Serine/arginine repetitive matrix protein 1 | SRRM1 | 102.336 | 2 | 51.12637 | yes | 1.07E-06 | 425.73944 | 134.19975 |
| AASPSPQSVR | 10 | Q8IYB3 | Serine/arginine repetitive matrix protein 1 | SRRM1 | 102.336 | 2 | 51.12637 | yes | 1.177E-07 | 500.26453 | 66.515564 |
| KVELSESEEDK | 11 | Q8IYB3 | Serine/arginine repetitive matrix protein 1 | SRRM1 | 102.336 | 2 | 51.12637 | yes | 4.594E-07 | 646.81445 | 42.078785 |
| EKTPELPEPSVK | 12 | Q8IYB3 | Serine/arginine repetitive matrix protein 1 | SRRM1 | 102.336 | 2 | 51.12637 | yes | 3.264E-06 | 677.36664 | 30.538355 |
| APQTSSSPPPVR | 12 | Q8IYB3 | Serine/arginine repetitive matrix protein 1 | SRRM1 | 102.336 | 2 | 51.12637 | yes | 1.004E-07 | 612.32257 | 27.98402 |
| FADVLVHR | 8 | Q8IYB7 | DIS3-like exonuclease 2 OS=Homo sapie | DIS3L2 | 99.28 | 2 | 53.922115 | yes | 3.535E-07 | 478.76926 | 284.44272 |
| DCIFTIDPSTAR | 12 | Q8IYB7 | DIS3-like exonuclease 2 OS=Homo sapie | DIS3L2 | 99.28 | 2 | 53.922115 | yes | 1.354E-07 | 698.33228 | 277.48749 |
| DYANTLFICR | 10 | Q8IYB7 | DIS3-like exonuclease 2 OS=Homo sapie | DIS3L2 | 99.28 | 2 | 53.922115 | yes | 9.722E-08 | 636.80573 | 178.42084 |
| AFPEQALLR | 9 | Q8IYB7 | DIS3-like exonuclease 2 OS=Homo sapie | DIS3L2 | 99.28 | 2 | 53.922115 | yes | 1.96E-07 | 522.79547 | 163.95938 |
| DIFIDGVVAR | 10 | Q8IYB7 | DIS3-like exonuclease 2 OS=Homo sapie | DIS3L2 | 99.28 | 2 | 53.922115 | yes | 3.877E-07 | 552.80603 | 162.48244 |
| EDCNFALGQLAK | 12 | Q8IYB7 | DIS3-like exonuclease 2 OS=Homo sapie | DIS3L2 | 99.28 | 2 | 53.922115 | yes | 1.648E-07 | 683.32703 | 158.50481 |
| LLPEEHWK | 8 | Q8IYB7 | DIS3-like exonuclease 2 OS=Homo sapie | DIS3L2 | 99.28 | 2 | 53.922115 | yes | 7.87E-07 | 526.28217 | 152.98434 |
| ILDEWFGR | 8 | Q8IYB7 | DIS3-like exonuclease 2 OS=Homo sapie | DIS3L2 | 99.28 | 2 | 53.922115 | yes | 4.506E-07 | 518.26654 | 132.18382 |
| LLAAALGYR | 9 | Q8IYB7 | DIS3-like exonuclease 2 OS=Homo sapie | DIS3L2 | 99.28 | 2 | 53.922115 | yes | 3.698E-07 | 474.28726 | 120.4174 |
| LSVCVSEK | 8 | Q8IYB7 | DIS3-like exonuclease 2 OS=Homo sapie | DIS3L2 | 99.28 | 2 | 53.922115 | yes | 2.765E-07 | 461.23914 | 113.8476 |
| QAFDVLVLR | 9 | Q8IYB7 | DIS3-like exonuclease 2 OS=Homo sapie | DIS3L2 | 99.28 | 2 | 53.922115 | yes | 2.165E-07 | 530.8111 | 106.07428 |
| GTLIQGVLR | 9 | Q8IYB7 | DIS3-like exonuclease 2 OS=Homo sapie | DIS3L2 | 99.28 | 2 | 53.922115 | yes | 9.733E-08 | 478.798 | 94.095062 |
| ESGPLESEAMVMC | 16 | Q8IYB7 | DIS3-like exonuclease 2 OS=Homo sapie | DIS3L2 | 99.28 | 2 | 53.922115 | yes | 2.585E-07 | 845.92316 | 69.438286 |
| HYALNVPLYTHFT | 17 | Q8IYB7 | DIS3-like exonuclease 2 OS=Homo sapie | DIS3L2 | 99.28 | 3 | 53.922115 | yes | 9.722E-08 | 677.02667 | 68.156143 |
| GVSAVAGPHDIGA | 18 | Q8IYB7 | DIS3-like exonuclease 2 OS=Homo sapie | DIS3L2 | 99.28 | 2 | 53.922115 | yes | 9.722E-08 | 817.91028 | 61.489105 |
| YALFSPSDHR | 10 | Q8IYB7 | DIS3-like exonuclease 2 OS=Homo sapie | DIS3L2 | 99.28 | 2 | 53.922115 | yes | 9.828E-07 | 596.79089 | 59.736595 |
| IYCNALALR | 9 | Q8IYB7 | DIS3-like exonuclease 2 OS=Homo sapie | DIS3L2 | 99.28 | 2 | 53.922115 | yes | 1.979E-07 | 547.2948 | 52.925991 |
| ATSVYLVQK | 9 | Q8IYB7 | DIS3-like exonuclease 2 OS=Homo sapie | DIS3L2 | 99.28 | 2 | 53.922115 | yes | 1.637E-06 | 504.78986 | 45.64621 |
| LDMAPDTLQK | 10 | Q8IYB7 | DIS3-like exonuclease 2 OS=Homo sapie | DIS3L2 | 99.28 | 2 | 53.922115 | yes | 0.0002 | 566.28937 | 38.095924 |
| SLTQTFGDDK | 10 | Q8IYB7 | DIS3-like exonuclease 2 OS=Homo sapie | DIS3L2 | 99.28 | 2 | 53.922115 | yes | 4.106E-06 | 556.26691 | 35.783348 |
| EDVSEGLKR | 9 | Q8IYB7 | DIS3-like exonuclease 2 OS=Homo sapie | DIS3L2 | 99.28 | 2 | 53.922115 | yes | 6.801E-07 | 516.76965 | 34.962799 |

| | | | | | | | | | | | |
|---|---|---|---|---|---|---|---|---|---|---|---|
| LVQQGLLTPDMLI | 13 | Q8IYB8 | ATP-dependent RNA helicase SUPV3L1. | SUPV3L1 | 87.992 | 2 | 55.734184 | yes | 1.325E-07 | 728.41565 | 708.33557 |
| LGADYGLDAR | 10 | Q8IYB8 | ATP-dependent RNA helicase SUPV3L1. | SUPV3L1 | 87.992 | 2 | 55.734184 | yes | 2.2E-07 | 525.76434 | 610.69836 |
| YVFCTAPINK | 10 | Q8IYB8 | ATP-dependent RNA helicase SUPV3L1. | SUPV3L1 | 87.992 | 2 | 55.734184 | yes | 1.197E-07 | 606.80774 | 535.815 |
| LLNLEGFPSGSQSI | 14 | Q8IYB8 | ATP-dependent RNA helicase SUPV3L1. | SUPV3L1 | 87.992 | 2 | 55.734184 | yes | 2.379E-07 | 752.89136 | 493.72217 |
| LLAHEIFEK | 9 | Q8IYB8 | ATP-dependent RNA helicase SUPV3L1. | SUPV3L1 | 87.992 | 2 | 55.734184 | yes | 9.722E-08 | 550.31097 | 466.19821 |
| QIFPVLDCK | 9 | Q8IYB8 | ATP-dependent RNA helicase SUPV3L1. | SUPV3L1 | 87.992 | 2 | 55.734184 | yes | 1.022E-07 | 560.297 | 377.3483 |
| IPPNWYPDAR | 10 | Q8IYB8 | ATP-dependent RNA helicase SUPV3L1. | SUPV3L1 | 87.992 | 2 | 55.734184 | yes | 4.139E-07 | 614.80908 | 341.03122 |
| NEPLTFAWLR | 10 | Q8IYB8 | ATP-dependent RNA helicase SUPV3L1. | SUPV3L1 | 87.992 | 2 | 55.734184 | yes | 1.499E-07 | 623.83258 | 316.67163 |
| FMDMFPDASLIR | 12 | Q8IYB8 | ATP-dependent RNA helicase SUPV3L1. | SUPV3L1 | 87.992 | 2 | 55.734184 | yes | 1.612E-07 | 721.84399 | 250.03819 |
| KFNDPNDPCK | 10 | Q8IYB8 | ATP-dependent RNA helicase SUPV3L1. | SUPV3L1 | 87.992 | 2 | 55.734184 | yes | 9.893E-08 | 617.77966 | 160.43704 |
| RPVDPIR | 7 | Q8IYB8 | ATP-dependent RNA helicase SUPV3L1. | SUPV3L1 | 87.992 | 2 | 55.734184 | yes | 1.171E-06 | 426.75613 | 97.863747 |
| SNAAGVPCDLVTC | 16 | Q8IYB8 | ATP-dependent RNA helicase SUPV3L1. | SUPV3L1 | 87.992 | 2 | 55.734184 | yes | 2.347E-06 | 837.89124 | 89.433426 |
| EWMTQQTEHNK | 11 | Q8IYB8 | ATP-dependent RNA helicase SUPV3L1. | SUPV3L1 | 87.992 | 2 | 55.734184 | yes | 9.722E-08 | 716.3197 | 75.333702 |
| KQPFVCSSLLQFAI | 14 | Q8IYB8 | ATP-dependent RNA helicase SUPV3L1. | SUPV3L1 | 87.992 | 3 | 55.734184 | yes | 9.819E-08 | 560.96783 | 65.809326 |
| GLESAVIYGSLPPC | 16 | Q8IYB8 | ATP-dependent RNA helicase SUPV3L1. | SUPV3L1 | 87.992 | 2 | 55.734184 | yes | 1.545E-07 | 794.93268 | 55.427597 |
| QPFVCSSLLQFAR | 13 | Q8IYB8 | ATP-dependent RNA helicase SUPV3L1. | SUPV3L1 | 87.992 | 2 | 55.734184 | yes | 9.79E-08 | 776.9007 | 35.48941 |
| FSAELIQHIPLSLR | 14 | Q8IYB8 | ATP-dependent RNA helicase SUPV3L1. | SUPV3L1 | 87.992 | 3 | 55.734184 | yes | 4.107E-06 | 541.97876 | 4.1702914 |
| PFVPNVHAAEFVP | 17 | Q8IYD1 | Eukaryotic peptide chain release factor G | GSPT2 | 68.884 | 3 | 35.882477 | yes | 7.69E-07 | 643.01233 | 22.109385 |
| TLFQTWGSSISR | 12 | Q8IYD8 | Fanconi anemia group M protein OS=Ho | FANCM | 232.192 | 2 | 54.68813 | yes | 9.722E-08 | 691.85681 | 145.79408 |
| CLVIDEAHK | 9 | Q8IYD8 | Fanconi anemia group M protein OS=Ho | FANCM | 232.192 | 2 | 54.68813 | yes | 5.782E-07 | 542.77643 | 128.74124 |
| ILALSATPGSDIK | 13 | Q8IYD8 | Fanconi anemia group M protein OS=Ho | FANCM | 232.192 | 2 | 54.68813 | yes | 4.561E-07 | 643.37177 | 116.19633 |
| ALGNYAYCQVVR | 12 | Q8IYD8 | Fanconi anemia group M protein OS=Ho | FANCM | 232.192 | 2 | 54.68813 | yes | 9.722E-08 | 707.35083 | 112.78313 |
| ESNIAVNPSTVK | 12 | Q8IYD8 | Fanconi anemia group M protein OS=Ho | FANCM | 232.192 | 2 | 54.68813 | yes | 2.874E-07 | 629.83551 | 83.870003 |
| SEDSPDILTYSHER | 14 | Q8IYD8 | Fanconi anemia group M protein OS=Ho | FANCM | 232.192 | 2 | 54.68813 | yes | 1.227E-07 | 824.8761 | 74.323853 |
| STSANGISAIQQGD | 15 | Q8IYD8 | Fanconi anemia group M protein OS=Ho | FANCM | 232.192 | 2 | 54.68813 | yes | 1.418E-06 | 738.8681 | 71.945313 |
| IVIILSEGR | 9 | Q8IYD8 | Fanconi anemia group M protein OS=Ho | FANCM | 232.192 | 2 | 54.68813 | yes | 9.722E-08 | 500.31351 | 66.310905 |
| DWFLSEEEFK | 10 | Q8IYD8 | Fanconi anemia group M protein OS=Ho | FANCM | 232.192 | 2 | 54.68813 | yes | 1.819E-07 | 665.30353 | 64.675163 |
| LIVPLGEELAAIQK | 14 | Q8IYD8 | Fanconi anemia group M protein OS=Ho | FANCM | 232.192 | 2 | 54.68813 | yes | 2.371E-07 | 747.4505 | 56.269958 |
| TYIQILESFAR | 11 | Q8IYD8 | Fanconi anemia group M protein OS=Ho | FANCM | 232.192 | 2 | 54.68813 | yes | 2.001E-07 | 670.86407 | 53.448486 |
| SQSEMLNSVNK | 11 | Q8IYD8 | Fanconi anemia group M protein OS=Ho | FANCM | 232.192 | 2 | 54.68813 | yes | 6.121E-07 | 618.7981 | 45.643734 |
| NESNNLASDTFITI | 15 | Q8IYD8 | Fanconi anemia group M protein OS=Ho | FANCM | 232.192 | 2 | 54.68813 | yes | 3.532E-07 | 845.90521 | 45.264004 |
| VMTFVGHASGK | 11 | Q8IYD8 | Fanconi anemia group M protein OS=Ho | FANCM | 232.192 | 2 | 54.68813 | yes | 1.381E-05 | 567.29224 | 21.887068 |
| FCCAVFAK | 8 | Q8IYM2 | Ribonuclease SLFN12 OS=Homo sapiens | SLFN12 | 66.973 | 2 | 44.012215 | no | 3.158E-07 | 501.73044 | 76.48761 |
| ALAGVFFDR | 9 | Q8IYM2 | Ribonuclease SLFN12 OS=Homo sapiens | SLFN12 | 66.973 | 2 | 44.012215 | yes | 2.356E-07 | 498.26907 | 65.454475 |
| AEMSDLDDLER | 11 | Q8IYM2 | Ribonuclease SLFN12 OS=Homo sapiens | SLFN12 | 66.973 | 2 | 44.012215 | yes | 6.118E-07 | 647.28497 | 52.779804 |
| GYSTQTALTLK | 11 | Q8IYM2 | Ribonuclease SLFN12 OS=Homo sapiens | SLFN12 | 66.973 | 2 | 44.012215 | yes | 2.639E-06 | 591.8219 | 12.804629 |
| YQLETISENIR | 11 | Q8IYT2 | Cap-specific mRNA (nucleoside-2'-O-)-m | CMTR2 | 88.12 | 2 | 54.080894 | yes | 9.722E-08 | 683.35406 | 548.83704 |
| MHIPLEVR | 8 | Q8IYT2 | Cap-specific mRNA (nucleoside-2'-O-)-m | CMTR2 | 88.12 | 2 | 54.080894 | yes | 1.847E-07 | 497.77875 | 270.52042 |
| ALFPHHVIPDSFLK | 14 | Q8IYT2 | Cap-specific mRNA (nucleoside-2'-O-)-m | CMTR2 | 88.12 | 2 | 54.080894 | yes | 2.772E-07 | 810.94849 | 82.951874 |

| | | | | | | | | | | | |
|---|---|---|---|---|---|---|---|---|---|---|---|
| ALFPHHVIPDSFLK | 14 | Q8IYT2 | Cap-specific mRNA (nucleoside-2'-O-)-m | CMTR2 | 88.12 | 2 | 54.080894 | yes | 2.258E-07 | 540.96808 | 82.951874 |
| ALFPHHVIPDSFLK | 14 | Q8IYT2 | Cap-specific mRNA (nucleoside-2'-O-)-m | CMTR2 | 88.12 | 3 | 54.080894 | yes | 2.772E-07 | 810.94849 | 82.951874 |
| ALFPHHVIPDSFLK | 14 | Q8IYT2 | Cap-specific mRNA (nucleoside-2'-O-)-m | CMTR2 | 88.12 | 3 | 54.080894 | yes | 2.258E-07 | 540.96808 | 82.951874 |
| SLGGAFNLDSK | 11 | Q8IYT2 | Cap-specific mRNA (nucleoside-2'-O-)-m | CMTR2 | 88.12 | 2 | 54.080894 | yes | 2.03E-06 | 554.78528 | 154.61888 |
| EAIHPLLSK | 9 | Q8IYT2 | Cap-specific mRNA (nucleoside-2'-O-)-m | CMTR2 | 88.12 | 2 | 54.080894 | yes | 9.722E-08 | 504.29785 | 129.11568 |
| MTLNFGTEMK | 10 | Q8IYT2 | Cap-specific mRNA (nucleoside-2'-O-)-m | CMTR2 | 88.12 | 2 | 54.080894 | yes | 2.084E-07 | 586.27795 | 106.74768 |
| CLLVGFSTLR | 10 | Q8IYT2 | Cap-specific mRNA (nucleoside-2'-O-)-m | CMTR2 | 88.12 | 2 | 54.080894 | yes | 9.722E-08 | 583.32355 | 98.049774 |
| EREEIINSLQLQN | 13 | Q8IYT2 | Cap-specific mRNA (nucleoside-2'-O-)-m | CMTR2 | 88.12 | 2 | 54.080894 | yes | 1.238E-07 | 793.41266 | 82.229248 |
| DCAIQYFMQK | 10 | Q8IYT2 | Cap-specific mRNA (nucleoside-2'-O-)-m | CMTR2 | 88.12 | 2 | 54.080894 | yes | 1.018E-07 | 652.29413 | 79.733665 |
| FITFVCPTSSDPLR | 14 | Q8IYT2 | Cap-specific mRNA (nucleoside-2'-O-)-m | CMTR2 | 88.12 | 2 | 54.080894 | yes | 9.742E-08 | 820.41107 | 46.77166 |
| SVNAELCTQAWCI | 13 | Q8IYT2 | Cap-specific mRNA (nucleoside-2'-O-)-m | CMTR2 | 88.12 | 2 | 54.080894 | yes | 2.526E-07 | 783.85559 | 45.205002 |
| MLEALSWK | 8 | Q8IYT2 | Cap-specific mRNA (nucleoside-2'-O-)-m | CMTR2 | 88.12 | 2 | 54.080894 | yes | 6.011E-07 | 489.25986 | 41.043652 |
| GLLLYGPPGTGK | 12 | Q8IYT4 | Katanin p60 ATPase-containing subunit A | KATNAL2 | 61.253 | 2 | 36.783504 | yes | 6.618E-07 | 586.83734 | 76.207802 |
| TNETGYQEAIVK | 12 | Q8IYU8 | Calcium uptake protein 2, mitochondrial ( | MICU2 | 49.667 | 2 | 52.928207 | yes | 9.722E-08 | 676.83826 | 311.51019 |
| LSAGESISLDEFK | 13 | Q8IYU8 | Calcium uptake protein 2, mitochondrial ( | MICU2 | 49.667 | 2 | 52.928207 | yes | 1.697E-07 | 698.35376 | 297.41904 |
| MLDTDGNEMIEK | 12 | Q8IYU8 | Calcium uptake protein 2, mitochondrial ( | MICU2 | 49.667 | 2 | 52.928207 | yes | 9.722E-08 | 698.31018 | 276.40747 |
| EPEINTTLQMR | 11 | Q8IYU8 | Calcium uptake protein 2, mitochondrial ( | MICU2 | 49.667 | 2 | 52.928207 | yes | 9.722E-08 | 666.33484 | 187.79027 |
| NVEHGIIYIGK | 11 | Q8IYU8 | Calcium uptake protein 2, mitochondrial ( | MICU2 | 49.667 | 2 | 52.928207 | yes | 1.682E-06 | 621.8457 | 155.59959 |
| FMQFSSLEHEGEY | 18 | Q8IYU8 | Calcium uptake protein 2, mitochondrial ( | MICU2 | 49.667 | 3 | 52.928207 | yes | 1.065E-07 | 751.33105 | 54.822906 |
| ELCVLSHER | 9 | Q8IYW5 | E3 ubiquitin-protein ligase RNF168 OS=] | RNF168 | 65.02 | 2 | 54.264076 | yes | 4.976E-07 | 571.78479 | 2042.3673 |
| LLSKPGELR | 9 | Q8IYW5 | E3 ubiquitin-protein ligase RNF168 OS=] | RNF168 | 65.02 | 2 | 54.264076 | yes | 1.783E-07 | 506.81113 | 937.89471 |
| GSPDEYHLR | 9 | Q8IYW5 | E3 ubiquitin-protein ligase RNF168 OS=] | RNF168 | 65.02 | 2 | 54.264076 | yes | 9.728E-08 | 537.25415 | 784.62598 |
| LLALQLQK | 8 | Q8IYW5 | E3 ubiquitin-protein ligase RNF168 OS=] | RNF168 | 65.02 | 2 | 54.264076 | yes | 2.725E-07 | 463.8053 | 676.78583 |
| ETAVMPCGR | 9 | Q8IYW5 | E3 ubiquitin-protein ligase RNF168 OS=] | RNF168 | 65.02 | 2 | 54.264076 | yes | 2.776E-07 | 510.7337 | 592.46948 |
| SAHSLQPSISQK | 12 | Q8IYW5 | E3 ubiquitin-protein ligase RNF168 OS=] | RNF168 | 65.02 | 2 | 54.264076 | yes | 9.722E-08 | 641.84113 | 258.47412 |
| SAHSLQPSISQK | 12 | Q8IYW5 | E3 ubiquitin-protein ligase RNF168 OS=] | RNF168 | 65.02 | 2 | 54.264076 | yes | 9.722E-08 | 428.22986 | 258.47412 |
| SAHSLQPSISQK | 12 | Q8IYW5 | E3 ubiquitin-protein ligase RNF168 OS=] | RNF168 | 65.02 | 3 | 54.264076 | yes | 9.722E-08 | 641.84113 | 258.47412 |
| SAHSLQPSISQK | 12 | Q8IYW5 | E3 ubiquitin-protein ligase RNF168 OS=] | RNF168 | 65.02 | 3 | 54.264076 | yes | 9.722E-08 | 428.22986 | 258.47412 |
| LLAEEEEEEKR | 11 | Q8IYW5 | E3 ubiquitin-protein ligase RNF168 OS=] | RNF168 | 65.02 | 2 | 54.264076 | yes | 9.938E-08 | 687.841 | 291.64856 |
| SVFQMFQR | 8 | Q8IYW5 | E3 ubiquitin-protein ligase RNF168 OS=] | RNF168 | 65.02 | 2 | 54.264076 | yes | 1.352E-07 | 521.76056 | 228.80312 |
| NQESSFEAVK | 10 | Q8IYW5 | E3 ubiquitin-protein ligase RNF168 OS=] | RNF168 | 65.02 | 2 | 54.264076 | yes | 4.91E-07 | 569.7724 | 196.26917 |
| ASLCCPFCR | 9 | Q8IYW5 | E3 ubiquitin-protein ligase RNF168 OS=] | RNF168 | 65.02 | 2 | 54.264076 | yes | 9.722E-08 | 585.74628 | 196.17776 |
| EYEEEISK | 8 | Q8IYW5 | E3 ubiquitin-protein ligase RNF168 OS=] | RNF168 | 65.02 | 2 | 54.264076 | yes | 2.193E-07 | 513.73492 | 172.56374 |
| ELCVLSHERPK | 11 | Q8IYW5 | E3 ubiquitin-protein ligase RNF168 OS=] | RNF168 | 65.02 | 2 | 54.264076 | yes | 9.823E-08 | 684.35864 | 157.04504 |
| EVDKEQMVPNR | 11 | Q8IYW5 | E3 ubiquitin-protein ligase RNF168 OS=] | RNF168 | 65.02 | 2 | 54.264076 | yes | 9.722E-08 | 672.83246 | 152.3736 |
| SQFGSASHSEAVQ | 16 | Q8IYW5 | E3 ubiquitin-protein ligase RNF168 OS=] | RNF168 | 65.02 | 3 | 54.264076 | yes | 2.978E-07 | 573.60791 | 75.42691 |
| EQMVPNR | 7 | Q8IYW5 | E3 ubiquitin-protein ligase RNF168 OS=] | RNF168 | 65.02 | 2 | 54.264076 | yes | 2.475E-07 | 437.216 | 60.301418 |
| PCFQSTVEK | 9 | Q8IYW5 | E3 ubiquitin-protein ligase RNF168 OS=] | RNF168 | 65.02 | 2 | 54.264076 | yes | 1.312E-07 | 548.26062 | 56.01823 |
| KNQESSFEAVK | 11 | Q8IYW5 | E3 ubiquitin-protein ligase RNF168 OS=] | RNF168 | 65.02 | 2 | 54.264076 | yes | 4.684E-07 | 633.81989 | 28.913771 |

| | | | | | | | | | | | |
|---|---|---|---|---|---|---|---|---|---|---|---|
| AHVAELER | 8 | Q8IZ52 | Chondroitin sulfate synthase 2 OS=Homo | CHPF | 85.468 | 2 | 55.311867 | yes | 2.13E-06 | 462.7485 | 441.81189 |
| VPWLSVQTAAPSP | 15 | Q8IZ52 | Chondroitin sulfate synthase 2 OS=Homo | CHPF | 85.468 | 2 | 55.311867 | yes | 1.022E-07 | 811.45667 | 172.78883 |
| LSEDLYHR | 8 | Q8IZ52 | Chondroitin sulfate synthase 2 OS=Homo | CHPF | 85.468 | 2 | 55.311867 | yes | 1.214E-07 | 516.75909 | 153.74873 |
| VLPYHPAQPGQAA | 14 | Q8IZ52 | Chondroitin sulfate synthase 2 OS=Homo | CHPF | 85.468 | 2 | 55.311867 | yes | 4.064E-07 | 738.90155 | 112.59871 |
| VEILPVPYVTEASP | 14 | Q8IZ52 | Chondroitin sulfate synthase 2 OS=Homo | CHPF | 85.468 | 2 | 55.311867 | yes | 5.345E-07 | 786.93524 | 94.083847 |
| VAHADVFAPVK | 11 | Q8IZ52 | Chondroitin sulfate synthase 2 OS=Homo | CHPF | 85.468 | 2 | 55.311867 | yes | 1.048E-07 | 577.32184 | 91.499184 |
| PQDFIGGEPTPGR | 13 | Q8IZ52 | Chondroitin sulfate synthase 2 OS=Homo | CHPF | 85.468 | 2 | 55.311867 | yes | 1.823E-07 | 685.83856 | 75.905945 |
| SALTAHPVR | 9 | Q8IZ52 | Chondroitin sulfate synthase 2 OS=Homo | CHPF | 85.468 | 2 | 55.311867 | yes | 1.742E-06 | 476.27216 | 67.249672 |
| MLLQQLR | 7 | Q8IZ52 | Chondroitin sulfate synthase 2 OS=Homo | CHPF | 85.468 | 2 | 55.311867 | yes | 3.142E-07 | 451.26804 | 63.083038 |
| CLQSVLEGLGSR | 12 | Q8IZ52 | Chondroitin sulfate synthase 2 OS=Homo | CHPF | 85.468 | 2 | 55.311867 | yes | 9.722E-08 | 659.84283 | 44.925201 |
| AVEPALLQR | 9 | Q8IZ52 | Chondroitin sulfate synthase 2 OS=Homo | CHPF | 85.468 | 2 | 55.311867 | yes | 1.798E-07 | 498.79547 | 23.341072 |
| LMDLLSK | 7 | Q8IZ52 | Chondroitin sulfate synthase 2 OS=Homo | CHPF | 85.468 | 2 | 55.311867 | yes | 1.844E-06 | 410.23587 | 8.4378824 |
| AEDLVPTLVSR | 11 | Q8IZ69 | tRNA (uracil-5-)-methyltransferase homo | TRMT2A | 68.726 | 2 | 50.01245 | yes | 1.96E-07 | 600.33514 | 1713.8024 |
| VIGVELCPEAVED. | 15 | Q8IZ69 | tRNA (uracil-5-)-methyltransferase homo | TRMT2A | 68.726 | 2 | 50.01245 | yes | 1.031E-07 | 828.91669 | 911.05237 |
| LLYVSCNPR | 9 | Q8IZ69 | tRNA (uracil-5-)-methyltransferase homo | TRMT2A | 68.726 | 2 | 50.01245 | yes | 1.377E-07 | 561.29224 | 720.53357 |
| LSPEELAELK | 10 | Q8IZ69 | tRNA (uracil-5-)-methyltransferase homo | TRMT2A | 68.726 | 2 | 50.01245 | yes | 1.376E-07 | 564.81097 | 684.15796 |
| AAMGNFVDLCR | 11 | Q8IZ69 | tRNA (uracil-5-)-methyltransferase homo | TRMT2A | 68.726 | 2 | 50.01245 | yes | 1.354E-07 | 627.29193 | 598.85059 |
| GIPFRPVK | 8 | Q8IZ69 | tRNA (uracil-5-)-methyltransferase homo | TRMT2A | 68.726 | 2 | 50.01245 | yes | 9.935E-08 | 457.28455 | 593.84955 |
| LFGQPPCAFVTFR | 13 | Q8IZ69 | tRNA (uracil-5-)-methyltransferase homo | TRMT2A | 68.726 | 2 | 50.01245 | yes | 1.164E-07 | 770.39246 | 475.26074 |
| TSLAQHFTAGPGR | 13 | Q8IZ69 | tRNA (uracil-5-)-methyltransferase homo | TRMT2A | 68.726 | 3 | 50.01245 | yes | 1.832E-07 | 448.23358 | 174.19865 |
| TSLAQHFTAGPGR | 13 | Q8IZ69 | tRNA (uracil-5-)-methyltransferase homo | TRMT2A | 68.726 | 3 | 50.01245 | yes | 1.055E-07 | 671.84674 | 174.19865 |
| TSLAQHFTAGPGR | 13 | Q8IZ69 | tRNA (uracil-5-)-methyltransferase homo | TRMT2A | 68.726 | 2 | 50.01245 | yes | 1.832E-07 | 448.23358 | 174.19865 |
| TSLAQHFTAGPGR | 13 | Q8IZ69 | tRNA (uracil-5-)-methyltransferase homo | TRMT2A | 68.726 | 2 | 50.01245 | yes | 1.055E-07 | 671.84674 | 174.19865 |
| ACCPLEGVR | 9 | Q8IZ69 | tRNA (uracil-5-)-methyltransferase homo | TRMT2A | 68.726 | 2 | 50.01245 | yes | 3.925E-07 | 531.24695 | 245.18903 |
| FGLQPHK | 7 | Q8IZ69 | tRNA (uracil-5-)-methyltransferase homo | TRMT2A | 68.726 | 2 | 50.01245 | yes | 1.36E-07 | 413.73215 | 164.74167 |
| CIHEDLLGLTFR | 12 | Q8IZ69 | tRNA (uracil-5-)-methyltransferase homo | TRMT2A | 68.726 | 2 | 50.01245 | yes | 9.975E-08 | 737.37958 | 145.52141 |
| DDLFTSEIFK | 10 | Q8IZ69 | tRNA (uracil-5-)-methyltransferase homo | TRMT2A | 68.726 | 2 | 50.01245 | yes | 1.051E-07 | 607.8006 | 144.40828 |
| VLHGALWK | 8 | Q8IZ69 | tRNA (uracil-5-)-methyltransferase homo | TRMT2A | 68.726 | 2 | 50.01245 | yes | 9.126E-07 | 462.2767 | 112.55596 |
| QLECEQVLQK | 10 | Q8IZ69 | tRNA (uracil-5-)-methyltransferase homo | TRMT2A | 68.726 | 2 | 50.01245 | yes | 1.036E-07 | 637.8241 | 111.1639 |
| AFQEFIR | 7 | Q8IZ69 | tRNA (uracil-5-)-methyltransferase homo | TRMT2A | 68.726 | 2 | 50.01245 | yes | 2.739E-07 | 455.74271 | 83.903664 |
| LELQNVPR | 8 | Q8IZ69 | tRNA (uracil-5-)-methyltransferase homo | TRMT2A | 68.726 | 2 | 50.01245 | yes | 4.332E-07 | 484.77982 | 77.214401 |
| GGTCAVAAPFDTV | 20 | Q8IZ69 | tRNA (uracil-5-)-methyltransferase homo | TRMT2A | 68.726 | 3 | 50.01245 | yes | 7.321E-07 | 681.33856 | 59.17292 |
| HQAMAIAYFHPQK | 13 | Q8IZ69 | tRNA (uracil-5-)-methyltransferase homo | TRMT2A | 68.726 | 3 | 50.01245 | yes | 1.063E-07 | 514.59424 | 56.030659 |
| KQLECEQVLQK | 11 | Q8IZ69 | tRNA (uracil-5-)-methyltransferase homo | TRMT2A | 68.726 | 2 | 50.01245 | yes | 1.686E-07 | 701.87158 | 42.45723 |
| QEGESEPPVTR | 11 | Q8IZ69 | tRNA (uracil-5-)-methyltransferase homo | TRMT2A | 68.726 | 2 | 50.01245 | yes | 4.609E-07 | 614.79382 | 21.575766 |
| VSCLSEELIASR | 12 | Q8IZE3 | Protein-associating with the carboxyl-tern | SCYL3 | 82.858 | 2 | 55.631287 | yes | 9.722E-08 | 682.34796 | 28.900433 |
| LGVSGYLVSR | 10 | Q8IZH2 | 5'-3' exoribonuclease 1 OS=Homo sapiens | XRN1 | 194.108 | 2 | 53.128323 | yes | 2.662E-07 | 525.80072 | 1199.9841 |
| MQYFQPTIQEK | 11 | Q8IZH2 | 5'-3' exoribonuclease 1 OS=Homo sapiens | XRN1 | 194.108 | 2 | 53.128323 | yes | 9.729E-08 | 706.8476 | 976.48596 |
| FYLEEPPGTQK | 11 | Q8IZH2 | 5'-3' exoribonuclease 1 OS=Homo sapiens | XRN1 | 194.108 | 2 | 53.128323 | yes | 9.81E-08 | 654.82715 | 918.12146 |

| | | | | | | | | | | | |
|---|---|---|---|---|---|---|---|---|---|---|---|
| GHPVSTLSR | 9 | Q8IZH2 | 5'-3' exoribonuclease 1 OS=Homo sapien: | XRN1 | 194.108 | 2 | 53.128323 | yes | 5.555E-07 | 477.26178 | 802.0177 |
| IDGSNTVDHK | 10 | Q8IZH2 | 5'-3' exoribonuclease 1 OS=Homo sapien: | XRN1 | 194.108 | 2 | 53.128323 | yes | 1.001E-07 | 543.26471 | 783.10211 |
| SSCDLQILDAAIVE | 15 | Q8IZH2 | 5'-3' exoribonuclease 1 OS=Homo sapien: | XRN1 | 194.108 | 2 | 53.128323 | yes | 9.722E-08 | 831.42438 | 775.6283 |
| YLNEAAGVAAEE/ | 14 | Q8IZH2 | 5'-3' exoribonuclease 1 OS=Homo sapien: | XRN1 | 194.108 | 2 | 53.128323 | yes | 9.722E-08 | 732.35986 | 735.98169 |
| SVFVNWPHLEEAF | 13 | Q8IZH2 | 5'-3' exoribonuclease 1 OS=Homo sapien: | XRN1 | 194.108 | 2 | 53.128323 | yes | 9.809E-08 | 792.40192 | 287.64185 |
| SVFVNWPHLEEAF | 13 | Q8IZH2 | 5'-3' exoribonuclease 1 OS=Homo sapien: | XRN1 | 194.108 | 2 | 53.128323 | yes | 9.722E-08 | 528.6037 | 287.64185 |
| SVFVNWPHLEEAF | 13 | Q8IZH2 | 5'-3' exoribonuclease 1 OS=Homo sapien: | XRN1 | 194.108 | 3 | 53.128323 | yes | 9.809E-08 | 792.40192 | 287.64185 |
| SVFVNWPHLEEAF | 13 | Q8IZH2 | 5'-3' exoribonuclease 1 OS=Homo sapien: | XRN1 | 194.108 | 3 | 53.128323 | yes | 9.722E-08 | 528.6037 | 287.64185 |
| EYIDYEFSVLK | 11 | Q8IZH2 | 5'-3' exoribonuclease 1 OS=Homo sapien: | XRN1 | 194.108 | 2 | 53.128323 | yes | 1.075E-07 | 703.3479 | 558.3269 |
| YNPGYVLASR | 10 | Q8IZH2 | 5'-3' exoribonuclease 1 OS=Homo sapien: | XRN1 | 194.108 | 2 | 53.128323 | yes | 9.722E-08 | 570.29584 | 515.36542 |
| SGVQVFQQSSR | 11 | Q8IZH2 | 5'-3' exoribonuclease 1 OS=Homo sapien: | XRN1 | 194.108 | 2 | 53.128323 | yes | 3.458E-07 | 611.81238 | 401.74246 |
| VQEIITWLK | 9 | Q8IZH2 | 5'-3' exoribonuclease 1 OS=Homo sapien: | XRN1 | 194.108 | 2 | 53.128323 | yes | 1.229E-07 | 565.33441 | 384.43835 |
| EHFSEVFVDLK | 11 | Q8IZH2 | 5'-3' exoribonuclease 1 OS=Homo sapien: | XRN1 | 194.108 | 2 | 53.128323 | yes | 9.722E-08 | 675.34045 | 361.19913 |
| ALYFCGFPTLK | 11 | Q8IZH2 | 5'-3' exoribonuclease 1 OS=Homo sapien: | XRN1 | 194.108 | 2 | 53.128323 | yes | 9.722E-08 | 658.83899 | 358.61563 |
| VVAVSDGETK | 10 | Q8IZH2 | 5'-3' exoribonuclease 1 OS=Homo sapien: | XRN1 | 194.108 | 2 | 53.128323 | yes | 9.917E-08 | 502.76657 | 357.74765 |
| LPTSALVNLSHGSI | 14 | Q8IZH2 | 5'-3' exoribonuclease 1 OS=Homo sapien: | XRN1 | 194.108 | 2 | 53.128323 | yes | 1.328E-07 | 726.40192 | 328.80688 |
| DTEFIYPSPWPEK | 13 | Q8IZH2 | 5'-3' exoribonuclease 1 OS=Homo sapien: | XRN1 | 194.108 | 2 | 53.128323 | yes | 7.338E-07 | 804.88269 | 322.0809 |
| MPQTMTVCQVK | 11 | Q8IZH2 | 5'-3' exoribonuclease 1 OS=Homo sapien: | XRN1 | 194.108 | 2 | 53.128323 | yes | 9.722E-08 | 661.81689 | 311.24002 |
| IHFELGK | 7 | Q8IZH2 | 5'-3' exoribonuclease 1 OS=Homo sapien: | XRN1 | 194.108 | 2 | 53.128323 | yes | 2.807E-07 | 422.23981 | 309.8284 |
| APELFSYIAK | 10 | Q8IZH2 | 5'-3' exoribonuclease 1 OS=Homo sapien: | XRN1 | 194.108 | 2 | 53.128323 | yes | 1.186E-07 | 569.81079 | 299.34949 |
| QIANEIPVSSNR | 12 | Q8IZH2 | 5'-3' exoribonuclease 1 OS=Homo sapien: | XRN1 | 194.108 | 2 | 53.128323 | yes | 1.085E-07 | 664.35187 | 282.9783 |
| VIFSIPCEPNLDAL | 20 | Q8IZH2 | 5'-3' exoribonuclease 1 OS=Homo sapien: | XRN1 | 194.108 | 3 | 53.128323 | yes | 1.022E-07 | 779.4071 | 260.12564 |
| IISLDAWR | 8 | Q8IZH2 | 5'-3' exoribonuclease 1 OS=Homo sapien: | XRN1 | 194.108 | 2 | 53.128323 | yes | 2.205E-07 | 487.27692 | 258.07056 |
| LAVNFGVSKPSE | 12 | Q8IZH2 | 5'-3' exoribonuclease 1 OS=Homo sapien: | XRN1 | 194.108 | 2 | 53.128323 | yes | 1.151E-07 | 624.33514 | 234.00804 |
| RDEYGLPSQPK | 11 | Q8IZH2 | 5'-3' exoribonuclease 1 OS=Homo sapien: | XRN1 | 194.108 | 2 | 53.128323 | yes | 2.997E-07 | 645.32788 | 222.63652 |
| PLEQQHGVIPDR | 12 | Q8IZH2 | 5'-3' exoribonuclease 1 OS=Homo sapien: | XRN1 | 194.108 | 3 | 53.128323 | yes | 1.399E-07 | 463.58087 | 109.0743 |
| PLEQQHGVIPDR | 12 | Q8IZH2 | 5'-3' exoribonuclease 1 OS=Homo sapien: | XRN1 | 194.108 | 3 | 53.128323 | yes | 9.893E-08 | 694.86768 | 109.0743 |
| PLEQQHGVIPDR | 12 | Q8IZH2 | 5'-3' exoribonuclease 1 OS=Homo sapien: | XRN1 | 194.108 | 2 | 53.128323 | yes | 1.399E-07 | 463.58087 | 109.0743 |
| PLEQQHGVIPDR | 12 | Q8IZH2 | 5'-3' exoribonuclease 1 OS=Homo sapien: | XRN1 | 194.108 | 2 | 53.128323 | yes | 9.893E-08 | 694.86768 | 109.0743 |
| VTVKPHLLYR | 10 | Q8IZH2 | 5'-3' exoribonuclease 1 OS=Homo sapien: | XRN1 | 194.108 | 3 | 53.128323 | yes | 5.436E-07 | 409.25204 | 197.95258 |
| QVVPFVYQTIVK | 12 | Q8IZH2 | 5'-3' exoribonuclease 1 OS=Homo sapien: | XRN1 | 194.108 | 2 | 53.128323 | yes | 1.189E-07 | 710.91357 | 191.58833 |
| GQENSLCWTALDI | 13 | Q8IZH2 | 5'-3' exoribonuclease 1 OS=Homo sapien: | XRN1 | 194.108 | 2 | 53.128323 | yes | 1.434E-07 | 761.35376 | 169.69637 |
| YPCLSEVVK | 9 | Q8IZH2 | 5'-3' exoribonuclease 1 OS=Homo sapien: | XRN1 | 194.108 | 2 | 53.128323 | yes | 1.893E-07 | 547.78119 | 130.92514 |
| DAEFCLFDR | 9 | Q8IZH2 | 5'-3' exoribonuclease 1 OS=Homo sapien: | XRN1 | 194.108 | 2 | 53.128323 | yes | 1.29E-07 | 586.75568 | 125.0667 |
| NEEVPGYTK | 9 | Q8IZH2 | 5'-3' exoribonuclease 1 OS=Homo sapien: | XRN1 | 194.108 | 2 | 53.128323 | yes | 2.165E-07 | 518.75092 | 120.82921 |
| TLDDLFPLR | 9 | Q8IZH2 | 5'-3' exoribonuclease 1 OS=Homo sapien: | XRN1 | 194.108 | 2 | 53.128323 | yes | 9.722E-08 | 545.3006 | 105.82365 |
| FTGSIFIGR | 9 | Q8IZH2 | 5'-3' exoribonuclease 1 OS=Homo sapien: | XRN1 | 194.108 | 2 | 53.128323 | yes | 6.078E-07 | 499.27692 | 93.12236 |
| LLEAMETCNHSLK | 13 | Q8IZH2 | 5'-3' exoribonuclease 1 OS=Homo sapien: | XRN1 | 194.108 | 2 | 53.128323 | yes | 1.165E-07 | 773.3736 | 35.302002 |
| LLEAMETCNHSLK | 13 | Q8IZH2 | 5'-3' exoribonuclease 1 OS=Homo sapien: | XRN1 | 194.108 | 2 | 53.128323 | yes | 5.489E-07 | 515.91815 | 35.302002 |

| | | | | | | | | | | | |
|---|---|---|---|---|---|---|---|---|---|---|---|
| LLEAMETCNHSLK | 13 | Q8IZH2 | 5'-3' exoribonuclease 1 OS=Homo sapien: | XRN1 | 194.108 | 3 | 53.128323 | yes | 1.165E-07 | 773.3736 | 35.302002 |
| LLEAMETCNHSLK | 13 | Q8IZH2 | 5'-3' exoribonuclease 1 OS=Homo sapien: | XRN1 | 194.108 | 3 | 53.128323 | yes | 5.489E-07 | 515.91815 | 35.302002 |
| KVFFMAVDGVAP | 13 | Q8IZH2 | 5'-3' exoribonuclease 1 OS=Homo sapien: | XRN1 | 194.108 | 2 | 53.128323 | yes | 1.769E-07 | 718.88959 | 63.765312 |
| KNEEVPGYTK | 10 | Q8IZH2 | 5'-3' exoribonuclease 1 OS=Homo sapien: | XRN1 | 194.108 | 2 | 53.128323 | yes | 1.463E-06 | 582.7984 | 59.234856 |
| ENFSVPVGLR | 10 | Q8IZH2 | 5'-3' exoribonuclease 1 OS=Homo sapien: | XRN1 | 194.108 | 2 | 53.128323 | yes | 2.348E-07 | 559.30365 | 57.710678 |
| KLAVNFGVSKPSE | 13 | Q8IZH2 | 5'-3' exoribonuclease 1 OS=Homo sapien: | XRN1 | 194.108 | 2 | 53.128323 | yes | 2.016E-07 | 688.38263 | 45.889286 |
| TILMCGPLIK | 10 | Q8IZL8 | Proline-, glutamic acid- and leucine-rich p | PELP1 | 119.7 | 2 | 49.050343 | yes | 1.819E-07 | 573.32471 | 168.14609 |
| LPSLGAGFSQGLK | 13 | Q8IZL8 | Proline-, glutamic acid- and leucine-rich p | PELP1 | 119.7 | 2 | 49.050343 | yes | 1.073E-07 | 637.85876 | 155.33383 |
| YAAQLPALFR | 10 | Q8IZL8 | Proline-, glutamic acid- and leucine-rich p | PELP1 | 119.7 | 2 | 49.050343 | yes | 1.143E-07 | 575.3244 | 125.88802 |
| LLPQVLNSWSIGR | 13 | Q8IZL8 | Proline-, glutamic acid- and leucine-rich p | PELP1 | 119.7 | 2 | 49.050343 | yes | 3.857E-07 | 741.92499 | 89.712227 |
| LLPGPENHR | 9 | Q8IZL8 | Proline-, glutamic acid- and leucine-rich p | PELP1 | 119.7 | 2 | 49.050343 | yes | 6.891E-07 | 516.7829 | 59.010185 |
| KPIDYTVLDDVGH | 16 | Q8IZP0 | Abl interactor 1 OS=Homo sapiens OX=9 | ABI1 | 55.081 | 3 | 50.86511 | yes | 9.722E-08 | 585.98071 | 386.55652 |
| ALIESYQNLTR | 11 | Q8IZP0 | Abl interactor 1 OS=Homo sapiens OX=9 | ABI1 | 55.081 | 2 | 50.86511 | yes | 5.475E-07 | 654.35132 | 53.429062 |
| LLWNPDIAAEYR | 12 | Q8IZT6 | Abnormal spindle-like microcephaly-asso | ASPM | 409.801 | 2 | 53.709 | yes | 1.202E-07 | 730.88025 | 195.70367 |
| VAITEHTEVR | 10 | Q8IZT6 | Abnormal spindle-like microcephaly-asso | ASPM | 409.801 | 2 | 53.709 | yes | 1.184E-06 | 577.81183 | 116.46317 |
| STVILQALVR | 10 | Q8IZT6 | Abnormal spindle-like microcephaly-asso | ASPM | 409.801 | 2 | 53.709 | yes | 3.926E-07 | 550.34534 | 101.92582 |
| AATLIQSFYR | 10 | Q8IZT6 | Abnormal spindle-like microcephaly-asso | ASPM | 409.801 | 2 | 53.709 | yes | 1.488E-07 | 585.31934 | 96.611916 |
| AVISLQSYFR | 10 | Q8IZT6 | Abnormal spindle-like microcephaly-asso | ASPM | 409.801 | 2 | 53.709 | yes | 2.896E-07 | 592.32715 | 83.829926 |
| AAVVLQNHYR | 10 | Q8IZT6 | Abnormal spindle-like microcephaly-asso | ASPM | 409.801 | 2 | 53.709 | yes | 2.408E-07 | 585.82257 | 83.648682 |
| CLNSAVGEHEK | 11 | Q8IZT6 | Abnormal spindle-like microcephaly-asso | ASPM | 409.801 | 2 | 53.709 | yes | 9.849E-08 | 622.29041 | 81.329918 |
| DSGSFEQYSENIK | 13 | Q8IZT6 | Abnormal spindle-like microcephaly-asso | ASPM | 409.801 | 2 | 53.709 | yes | 4.235E-07 | 752.33356 | 80.671707 |
| SSTLIIQSMFR | 11 | Q8IZT6 | Abnormal spindle-like microcephaly-asso | ASPM | 409.801 | 2 | 53.709 | yes | 1.758E-07 | 641.84485 | 79.833984 |
| NSSISIPFIPETPVR | 15 | Q8IZT6 | Abnormal spindle-like microcephaly-asso | ASPM | 409.801 | 2 | 53.709 | yes | 6.742E-07 | 828.95142 | 78.418129 |
| KLEIEIEAR | 9 | Q8IZT6 | Abnormal spindle-like microcephaly-asso | ASPM | 409.801 | 2 | 53.709 | yes | 1.628E-07 | 550.81915 | 77.599686 |
| YYSIILQSR | 9 | Q8IZT6 | Abnormal spindle-like microcephaly-asso | ASPM | 409.801 | 2 | 53.709 | yes | 1.008E-07 | 571.81384 | 75.826958 |
| AAVIIQTHFR | 10 | Q8IZT6 | Abnormal spindle-like microcephaly-asso | ASPM | 409.801 | 2 | 53.709 | yes | 1.388E-07 | 578.33527 | 74.975044 |
| TMELLTQNWDLSI | 13 | Q8IZT6 | Abnormal spindle-like microcephaly-asso | ASPM | 409.801 | 2 | 53.709 | yes | 1.13E-07 | 789.89526 | 74.674942 |
| AAALFIQQCYR | 11 | Q8IZT6 | Abnormal spindle-like microcephaly-asso | ASPM | 409.801 | 2 | 53.709 | yes | 1.054E-07 | 670.84265 | 74.549171 |
| TQAVICIQSYYR | 12 | Q8IZT6 | Abnormal spindle-like microcephaly-asso | ASPM | 409.801 | 2 | 53.709 | yes | 2.537E-06 | 751.37701 | 72.104233 |
| AAVISLQSAYR | 11 | Q8IZT6 | Abnormal spindle-like microcephaly-asso | ASPM | 409.801 | 2 | 53.709 | yes | 9.722E-08 | 589.83002 | 70.31839 |
| ASAIIIQR | 8 | Q8IZT6 | Abnormal spindle-like microcephaly-asso | ASPM | 409.801 | 2 | 53.709 | yes | 2.514E-07 | 436.27164 | 67.181862 |
| AAVLIQATFR | 10 | Q8IZT6 | Abnormal spindle-like microcephaly-asso | ASPM | 409.801 | 2 | 53.709 | yes | 1.272E-06 | 545.3244 | 59.386616 |
| MHNVDIVLQVLK | 12 | Q8IZT6 | Abnormal spindle-like microcephaly-asso | ASPM | 409.801 | 2 | 53.709 | yes | 1.413E-07 | 704.90271 | 57.212631 |
| LYQFLHLR | 8 | Q8IZT6 | Abnormal spindle-like microcephaly-asso | ASPM | 409.801 | 2 | 53.709 | yes | 1.682E-07 | 545.31384 | 48.779255 |
| ELLNVHSANVSK | 12 | Q8IZT6 | Abnormal spindle-like microcephaly-asso | ASPM | 409.801 | 2 | 53.709 | yes | 9.722E-08 | 655.85681 | 47.88147 |
| IVISVNWTPLK | 11 | Q8IZT6 | Abnormal spindle-like microcephaly-asso | ASPM | 409.801 | 2 | 53.709 | yes | 9.992E-08 | 635.3819 | 46.636436 |
| HQAILLGNAEEQK | 13 | Q8IZT6 | Abnormal spindle-like microcephaly-asso | ASPM | 409.801 | 2 | 53.709 | yes | 1.142E-07 | 725.88605 | 43.36882 |
| DNSQPVHLESTIAI | 19 | Q8IZT6 | Abnormal spindle-like microcephaly-asso | ASPM | 409.801 | 3 | 53.709 | yes | 1.751E-07 | 737.0343 | 42.860695 |
| TALALHYLLTYK | 12 | Q8IZT6 | Abnormal spindle-like microcephaly-asso | ASPM | 409.801 | 2 | 53.709 | yes | 5.414E-07 | 703.90576 | 40.306732 |

| | | | | | | | | | | | |
|---|---|---|---|---|---|---|---|---|---|---|---|
| MAAIVNQSALCCY | 14 | Q8IZT6 | Abnormal spindle-like microcephaly-asso | ASPM | 409.801 | 2 | 53.709 | yes | 2.897E-06 | 828.88635 | 34.124359 |
| ILSPDSFIK | 9 | Q8IZT6 | Abnormal spindle-like microcephaly-asso | ASPM | 409.801 | 2 | 53.709 | yes | 3.637E-07 | 510.29224 | 26.829685 |
| VNAALVIQK | 9 | Q8IZT6 | Abnormal spindle-like microcephaly-asso | ASPM | 409.801 | 2 | 53.709 | yes | 2.595E-05 | 478.30038 | 20.540442 |
| AFDHENTSELYK | 12 | Q8IZT6 | Abnormal spindle-like microcephaly-asso | ASPM | 409.801 | 2 | 53.709 | yes | 1.279E-07 | 727.33331 | 19.432665 |
| EEYMQMR | 7 | Q8IZT6 | Abnormal spindle-like microcephaly-asso | ASPM | 409.801 | 2 | 53.709 | yes | 5.479E-05 | 493.70715 | 15.682796 |
| ALCDLGPWVK | 10 | Q8N0U8 | Vitamin K epoxide reductase complex sul | VKORC1L | 19.836 | 2 | 42.3769 | yes | 3.06E-07 | 579.80243 | 56.782089 |
| HLLMSQQTLR | 10 | Q8N0V3 | Putative ribosome-binding factor A, mito | RBFA | 38.36 | 2 | 50.99893 | yes | 9.722E-08 | 613.83734 | 197.51306 |
| VSLTPDFSACR | 11 | Q8N0V3 | Putative ribosome-binding factor A, mito | RBFA | 38.36 | 2 | 50.99893 | yes | 6.228E-07 | 626.80316 | 180.75351 |
| NVPPIVFVQDK | 11 | Q8N0V3 | Putative ribosome-binding factor A, mito | RBFA | 38.36 | 2 | 50.99893 | yes | 8.531E-07 | 628.3559 | 70.280983 |
| ENQEGAANVNVR | 12 | Q8N0X7 | Spartin OS=Homo sapiens OX=9606 GN | SPART | 72.834 | 2 | 46.912373 | yes | 1.443E-07 | 650.81567 | 76.743622 |
| GLATSLQNDLQEV | 15 | Q8N0X7 | Spartin OS=Homo sapiens OX=9606 GN | SPART | 72.834 | 2 | 46.912373 | yes | 3.8E-07 | 806.93066 | 34.374504 |
| CIVNNVSAETVQT | 15 | Q8N0X7 | Spartin OS=Homo sapiens OX=9606 GN | SPART | 72.834 | 2 | 46.912373 | yes | 2.041E-07 | 845.43304 | 32.783939 |
| TSLKPAHLLR | 10 | Q8N0Y2 | Zinc finger protein 444 OS=Homo sapien | ZNF444 | 35.204 | 2 | 43.270298 | yes | 2.945E-07 | 568.35095 | 31.398413 |
| PFACWECGK | 9 | Q8N0Y2 | Zinc finger protein 444 OS=Homo sapien | ZNF444 | 35.204 | 2 | 43.270298 | yes | 2.66E-07 | 577.74152 | 29.609058 |
| LSQVPDPPDNMNI | 15 | Q8N0Z3 | Spindle and centriole-associated protein 1 | SPICE1 | 96.265 | 2 | 39.55058 | yes | 3.774E-06 | 819.91144 | 59.806568 |
| FSPLQDVLR | 9 | Q8N0Z3 | Spindle and centriole-associated protein 1 | SPICE1 | 96.265 | 2 | 39.55058 | yes | 4.517E-07 | 537.80072 | 56.072124 |
| EEVAEFPQEELPVI | 14 | Q8N0Z3 | Spindle and centriole-associated protein 1 | SPICE1 | 96.265 | 2 | 39.55058 | yes | 5.78E-07 | 822.4118 | 44.320107 |
| TNESLIQR | 8 | Q8N0Z3 | Spindle and centriole-associated protein 1 | SPICE1 | 96.265 | 2 | 39.55058 | yes | 5.251E-06 | 480.75909 | 32.893867 |
| GVEAIPEGSHIK | 12 | Q8N108 | Mesoderm induction early response prote | MIER1 | 57.984 | 2 | 49.279602 | yes | 1.068E-07 | 618.83276 | 103.86 |
| LLDESESAASSR | 12 | Q8N108 | Mesoderm induction early response prote | MIER1 | 57.984 | 2 | 49.279602 | yes | 2.703E-07 | 632.80444 | 49.78355 |
| EELSVWTEEECR | 12 | Q8N108 | Mesoderm induction early response prote | MIER1 | 57.984 | 2 | 49.279602 | yes | 1.049E-07 | 783.8407 | 47.040169 |
| ALETIGANLQK | 11 | Q8N122 | Regulatory-associated protein of mTOR ( | RPTOR | 149.038 | 2 | 52.48334 | yes | 4.138E-07 | 579.3299 | 193.47749 |
| LDDQIFLNR | 9 | Q8N122 | Regulatory-associated protein of mTOR ( | RPTOR | 149.038 | 2 | 52.48334 | yes | 1.427E-07 | 567.30109 | 132.19164 |
| LYSLLSDPIPEVR | 13 | Q8N122 | Regulatory-associated protein of mTOR ( | RPTOR | 149.038 | 2 | 52.48334 | yes | 9.728E-08 | 751.41669 | 103.96136 |
| LECWIDPLSMGPQ | 14 | Q8N122 | Regulatory-associated protein of mTOR ( | RPTOR | 149.038 | 2 | 52.48334 | yes | 1.874E-07 | 837.40491 | 89.972878 |
| ILAVDSSCQADLV | 14 | Q8N122 | Regulatory-associated protein of mTOR ( | RPTOR | 149.038 | 2 | 52.48334 | yes | 9.722E-08 | 759.89526 | 75.902206 |
| SLIVAGLGDGSIR | 13 | Q8N122 | Regulatory-associated protein of mTOR ( | RPTOR | 149.038 | 2 | 52.48334 | yes | 1.122E-07 | 629.36169 | 58.494705 |
| MPESVNVLQIVK | 12 | Q8N122 | Regulatory-associated protein of mTOR ( | RPTOR | 149.038 | 2 | 52.48334 | yes | 2.18E-07 | 678.88141 | 40.096756 |
| YYDGFMGQR | 9 | Q8N122 | Regulatory-associated protein of mTOR ( | RPTOR | 149.038 | 2 | 52.48334 | yes | 4.483E-06 | 568.74512 | 34.406406 |
| LETVTEVIR | 9 | Q8N157 | Jouberin OS=Homo sapiens OX=9606 GI | AHI1 | 137.116 | 2 | 49.812588 | yes | 1.318E-06 | 530.30585 | 113.43314 |
| SDFMISHPMVK | 11 | Q8N157 | Jouberin OS=Homo sapiens OX=9606 GI | AHI1 | 137.116 | 2 | 49.812588 | yes | 1.002E-07 | 646.31232 | 34.993252 |
| IHVVDEHTGQYVK | 13 | Q8N157 | Jouberin OS=Homo sapiens OX=9606 GI | AHI1 | 137.116 | 2 | 49.812588 | yes | 1.993E-07 | 762.89392 | 18.336018 |
| ETTSDDVSAANTN | 16 | Q8N157 | Jouberin OS=Homo sapiens OX=9606 GI | AHI1 | 137.116 | 2 | 49.812588 | yes | 3.258E-06 | 840.38959 | 13.481524 |
| FAEFQYLQPGPPR | 13 | Q8N163 | Cell cycle and apoptosis regulator protein | CCAR2 | 102.902 | 2 | 52.991516 | yes | 2.67E-07 | 775.39355 | 432.02579 |
| VLLLSSPGLEELYI | 14 | Q8N163 | Cell cycle and apoptosis regulator protein | CCAR2 | 102.902 | 2 | 52.991516 | yes | 9.722E-08 | 794.95087 | 230.50848 |
| VVTQNICQYR | 10 | Q8N163 | Cell cycle and apoptosis regulator protein | CCAR2 | 102.902 | 2 | 52.991516 | yes | 1.148E-07 | 640.8244 | 220.66945 |
| SPAPPLLHVAALG | 15 | Q8N163 | Cell cycle and apoptosis regulator protein | CCAR2 | 102.902 | 2 | 52.991516 | yes | 1.581E-07 | 749.94067 | 104.89792 |
| SPAPPLLHVAALG | 15 | Q8N163 | Cell cycle and apoptosis regulator protein | CCAR2 | 102.902 | 2 | 52.991516 | yes | 2.047E-05 | 500.2962 | 104.89792 |
| SPAPPLLHVAALG | 15 | Q8N163 | Cell cycle and apoptosis regulator protein | CCAR2 | 102.902 | 3 | 52.991516 | yes | 1.581E-07 | 749.94067 | 104.89792 |

| | | | | | | | | | | | |
|---|---|---|---|---|---|---|---|---|---|---|---|
| SPAPPLLHVAALG | 15 | Q8N163 | Cell cycle and apoptosis regulator protein | CCAR2 | 102.902 | 3 | 52.991516 | yes | 2.047E-05 | 500.2962 | 104.89792 |
| AAYNPGQAVPWN | 15 | Q8N163 | Cell cycle and apoptosis regulator protein | CCAR2 | 102.902 | 2 | 52.991516 | yes | 4.762E-07 | 793.40973 | 201.50513 |
| VQTLSNQPLLK | 11 | Q8N163 | Cell cycle and apoptosis regulator protein | CCAR2 | 102.902 | 2 | 52.991516 | yes | 1.735E-07 | 620.86664 | 171.39032 |
| ETPEHPLK | 8 | Q8N163 | Cell cycle and apoptosis regulator protein | CCAR2 | 102.902 | 2 | 52.991516 | yes | 1.061E-06 | 475.75073 | 144.39928 |
| LTPLQLEIQR | 10 | Q8N163 | Cell cycle and apoptosis regulator protein | CCAR2 | 102.902 | 2 | 52.991516 | yes | 1.362E-07 | 605.86133 | 126.76581 |
| TLAAEMQELR | 10 | Q8N163 | Cell cycle and apoptosis regulator protein | CCAR2 | 102.902 | 2 | 52.991516 | yes | 9.722E-08 | 581.30023 | 99.462875 |
| DDGEEEFAGAK | 11 | Q8N163 | Cell cycle and apoptosis regulator protein | CCAR2 | 102.902 | 2 | 52.991516 | yes | 9.722E-08 | 584.24365 | 77.16494 |
| ILLTLGIR | 8 | Q8N163 | Cell cycle and apoptosis regulator protein | CCAR2 | 102.902 | 2 | 52.991516 | yes | 2.725E-07 | 449.80783 | 60.116196 |
| IHTLELK | 7 | Q8N163 | Cell cycle and apoptosis regulator protein | CCAR2 | 102.902 | 2 | 52.991516 | yes | 1.661E-05 | 427.26074 | 54.103832 |
| ADSWVEKEEPAPS | 14 | Q8N163 | Cell cycle and apoptosis regulator protein | CCAR2 | 102.902 | 2 | 52.991516 | yes | 1.493E-06 | 779.85461 | 53.612312 |
| CAQAQTGIDLSGC | 15 | Q8N163 | Cell cycle and apoptosis regulator protein | CCAR2 | 102.902 | 2 | 52.991516 | yes | 9.722E-08 | 805.36908 | 52.685146 |
| FSATEVTNK | 9 | Q8N163 | Cell cycle and apoptosis regulator protein | CCAR2 | 102.902 | 2 | 52.991516 | yes | 4.544E-06 | 498.75348 | 48.55072 |
| EEEAIKEEVVK | 11 | Q8N163 | Cell cycle and apoptosis regulator protein | CCAR2 | 102.902 | 2 | 52.991516 | yes | 1.828E-07 | 651.84302 | 20.763226 |
| VVSPPEPEKEEAA | 14 | Q8N163 | Cell cycle and apoptosis regulator protein | CCAR2 | 102.902 | 2 | 52.991516 | yes | 2.143E-06 | 755.39337 | 20.513601 |
| YFNPSGEPLELDSI | 14 | Q8N1B3 | Cyclin-Q OS=Homo sapiens OX=9606 G | CCNQ | 28.369 | 2 | 46.662872 | yes | 1.059E-06 | 812.38611 | 92.193901 |
| SIPIATACTIYHK | 13 | Q8N1B3 | Cyclin-Q OS=Homo sapiens OX=9606 G | CCNQ | 28.369 | 2 | 46.662872 | yes | 1.501E-07 | 737.88977 | 64.976028 |
| LSGLDVSFCDK | 11 | Q8N1E6 | F-box/LRR-repeat protein 14 OS=Homo : | FBXL14 | 45.887 | 2 | 35.436832 | yes | 8.327E-07 | 620.79755 | 105.04183 |
| LVPLNQESVEER | 12 | Q8N1F7 | Nuclear pore complex protein Nup93 OS= | NUP93 | 93.489 | 2 | 56.76819 | yes | 1.414E-07 | 706.87262 | 370.90469 |
| ESMLVEWEQVK | 11 | Q8N1F7 | Nuclear pore complex protein Nup93 OS= | NUP93 | 93.489 | 2 | 56.76819 | yes | 1.272E-07 | 689.3396 | 276.21289 |
| LLSPVVPQISAPQS | 16 | Q8N1F7 | Nuclear pore complex protein Nup93 OS= | NUP93 | 93.489 | 2 | 56.76819 | yes | 1.272E-07 | 839.48035 | 276.15277 |
| QMTDVLLTPATDA | 15 | Q8N1F7 | Nuclear pore complex protein Nup93 OS= | NUP93 | 93.489 | 2 | 56.76819 | yes | 1.195E-07 | 808.93182 | 243.08737 |
| SSLDNIEMAYAR | 12 | Q8N1F7 | Nuclear pore complex protein Nup93 OS= | NUP93 | 93.489 | 2 | 56.76819 | yes | 9.722E-08 | 685.32446 | 193.63174 |
| NLQEIQQAGER | 11 | Q8N1F7 | Nuclear pore complex protein Nup93 OS= | NUP93 | 93.489 | 2 | 56.76819 | yes | 1.71E-07 | 643.32837 | 176.04204 |
| GLDISHISQR | 10 | Q8N1F7 | Nuclear pore complex protein Nup93 OS= | NUP93 | 93.489 | 2 | 56.76819 | yes | 3.017E-07 | 563.3042 | 161.39819 |
| DNALLSAIEESR | 12 | Q8N1F7 | Nuclear pore complex protein Nup93 OS= | NUP93 | 93.489 | 2 | 56.76819 | yes | 9.722E-08 | 659.33588 | 151.68135 |
| CVSELVIESR | 10 | Q8N1F7 | Nuclear pore complex protein Nup93 OS= | NUP93 | 93.489 | 2 | 56.76819 | yes | 1.284E-07 | 596.30554 | 120.61662 |
| SISDMWTMVK | 10 | Q8N1F7 | Nuclear pore complex protein Nup93 OS= | NUP93 | 93.489 | 2 | 56.76819 | yes | 1.238E-07 | 599.28577 | 95.724319 |
| TFGMAEEYHR | 10 | Q8N1F7 | Nuclear pore complex protein Nup93 OS= | NUP93 | 93.489 | 2 | 56.76819 | yes | 9.722E-08 | 620.77441 | 89.215561 |
| LTLSQFQK | 8 | Q8N1F7 | Nuclear pore complex protein Nup93 OS= | NUP93 | 93.489 | 2 | 56.76819 | yes | 9.722E-08 | 482.77673 | 83.15583 |
| CGDLLAASQVVNI | 13 | Q8N1F7 | Nuclear pore complex protein Nup93 OS= | NUP93 | 93.489 | 2 | 56.76819 | yes | 1.745E-07 | 701.85901 | 81.320168 |
| QALAYLEQSYK | 11 | Q8N1F7 | Nuclear pore complex protein Nup93 OS= | NUP93 | 93.489 | 2 | 56.76819 | yes | 1.065E-07 | 657.34045 | 81.040062 |
| TLITFAGMIPYR | 12 | Q8N1F7 | Nuclear pore complex protein Nup93 OS= | NUP93 | 93.489 | 2 | 56.76819 | yes | 9.722E-08 | 691.87866 | 39.983616 |
| EALQYFYFLR | 10 | Q8N1F7 | Nuclear pore complex protein Nup93 OS= | NUP93 | 93.489 | 2 | 56.76819 | yes | 4.658E-07 | 675.34808 | 32.739471 |
| AVYCIIGR | 8 | Q8N1F7 | Nuclear pore complex protein Nup93 OS= | NUP93 | 93.489 | 2 | 56.76819 | yes | 1.785E-07 | 476.25766 | 31.149151 |
| FTSDTKPIINK | 11 | Q8N1F7 | Nuclear pore complex protein Nup93 OS= | NUP93 | 93.489 | 2 | 56.76819 | yes | 9.722E-08 | 632.35083 | 21.417658 |
| CDVTDNQSEVADI | 13 | Q8N1F7 | Nuclear pore complex protein Nup93 OS= | NUP93 | 93.489 | 2 | 56.76819 | yes | 2.021E-07 | 740.81464 | 18.420813 |
| VVSVQLGDGTR | 11 | Q8N1G0 | Zinc finger protein 687 OS=Homo sapien | ZNF687 | 129.529 | 2 | 44.208004 | yes | 4.414E-07 | 565.81183 | 97.866119 |
| TATGPSTGGGTVIS | 15 | Q8N1G0 | Zinc finger protein 687 OS=Homo sapien | ZNF687 | 129.529 | 2 | 44.208004 | yes | 2.281E-07 | 681.35461 | 45.95607 |
| GLAPSAVPGLSK | 12 | Q8N1G1 | RNA exonuclease 1 homolog OS=Homo : | REXO1 | 131.51 | 2 | 49.32742 | yes | 1.043E-07 | 548.82166 | 229.62036 |

| | | | | | | | | | | | |
|---|---|---|---|---|---|---|---|---|---|---|---|
| GLSGLTTLFPGQK | 13 | Q8N1G1 | RNA exonuclease 1 homolog OS=Homo : | REXO1 | 131.51 | 2 | 49.32742 | yes | 2.139E-07 | 659.87189 | 38.152538 |
| EGLEAEGGGLR | 11 | Q8N1G1 | RNA exonuclease 1 homolog OS=Homo : | REXO1 | 131.51 | 2 | 49.32742 | yes | 4.789E-06 | 544.27252 | 29.759289 |
| SVFDVLDGEEMR | 12 | Q8N1G2 | Cap-specific mRNA (nucleoside-2'-O-)-m | CMTR1 | 95.321 | 2 | 55.048355 | yes | 9.722E-08 | 698.82428 | 718.43909 |
| LWGIPDQAR | 9 | Q8N1G2 | Cap-specific mRNA (nucleoside-2'-O-)-m | CMTR1 | 95.321 | 2 | 55.048355 | yes | 1.675E-07 | 528.28528 | 659.01654 |
| IRPVFDYR | 8 | Q8N1G2 | Cap-specific mRNA (nucleoside-2'-O-)-m | CMTR1 | 95.321 | 2 | 55.048355 | yes | 2.636E-07 | 533.29565 | 351.42099 |
| NFVLDNTDR | 9 | Q8N1G2 | Cap-specific mRNA (nucleoside-2'-O-)-m | CMTR1 | 95.321 | 2 | 55.048355 | yes | 1.47E-07 | 547.26727 | 351.07297 |
| ADSLVEGTSSR | 11 | Q8N1G2 | Cap-specific mRNA (nucleoside-2'-O-)-m | CMTR1 | 95.321 | 2 | 55.048355 | yes | 9.722E-08 | 561.27527 | 294.3306 |
| IHAFVQDTTLSEPF | 14 | Q8N1G2 | Cap-specific mRNA (nucleoside-2'-O-)-m | CMTR1 | 95.321 | 2 | 55.048355 | yes | 4.498E-07 | 807.41772 | 144.31197 |
| IHAFVQDTTLSEPF | 14 | Q8N1G2 | Cap-specific mRNA (nucleoside-2'-O-)-m | CMTR1 | 95.321 | 2 | 55.048355 | yes | 4.819E-07 | 538.61426 | 144.31197 |
| IHAFVQDTTLSEPF | 14 | Q8N1G2 | Cap-specific mRNA (nucleoside-2'-O-)-m | CMTR1 | 95.321 | 3 | 55.048355 | yes | 4.498E-07 | 807.41772 | 144.31197 |
| IHAFVQDTTLSEPF | 14 | Q8N1G2 | Cap-specific mRNA (nucleoside-2'-O-)-m | CMTR1 | 95.321 | 3 | 55.048355 | yes | 4.819E-07 | 538.61426 | 144.31197 |
| GDHEFTDYMIR | 11 | Q8N1G2 | Cap-specific mRNA (nucleoside-2'-O-)-m | CMTR1 | 95.321 | 2 | 55.048355 | yes | 9.722E-08 | 692.30353 | 248.66597 |
| AVSKPSRPDMNPI | 14 | Q8N1G2 | Cap-specific mRNA (nucleoside-2'-O-)-m | CMTR1 | 95.321 | 3 | 55.048355 | yes | 9.722E-08 | 523.28394 | 238.67903 |
| EDVLSFIQMHR | 11 | Q8N1G2 | Cap-specific mRNA (nucleoside-2'-O-)-m | CMTR1 | 95.321 | 2 | 55.048355 | yes | 5.057E-07 | 687.84534 | 195.28485 |
| ANPYEMIR | 8 | Q8N1G2 | Cap-specific mRNA (nucleoside-2'-O-)-m | CMTR1 | 95.321 | 2 | 55.048355 | yes | 1.036E-07 | 497.24475 | 186.1256 |
| HFVPMGLYIVR | 11 | Q8N1G2 | Cap-specific mRNA (nucleoside-2'-O-)-m | CMTR1 | 95.321 | 2 | 55.048355 | yes | 5.224E-07 | 666.36829 | 185.68091 |
| SQIYTWDGR | 9 | Q8N1G2 | Cap-specific mRNA (nucleoside-2'-O-)-m | CMTR1 | 95.321 | 2 | 55.048355 | yes | 1.001E-07 | 563.26984 | 173.0079 |
| YSMYNSVSQK | 10 | Q8N1G2 | Cap-specific mRNA (nucleoside-2'-O-)-m | CMTR1 | 95.321 | 2 | 55.048355 | yes | 9.722E-08 | 603.77661 | 138.72549 |
| GFDQELNVDWR | 11 | Q8N1G2 | Cap-specific mRNA (nucleoside-2'-O-)-m | CMTR1 | 95.321 | 2 | 55.048355 | yes | 9.722E-08 | 689.82294 | 114.37169 |
| MANMDFVFDR | 10 | Q8N1G2 | Cap-specific mRNA (nucleoside-2'-O-)-m | CMTR1 | 95.321 | 2 | 55.048355 | yes | 9.722E-08 | 623.27319 | 108.24335 |
| TVNEPWTMGFSK | 12 | Q8N1G2 | Cap-specific mRNA (nucleoside-2'-O-)-m | CMTR1 | 95.321 | 2 | 55.048355 | yes | 9.722E-08 | 698.83191 | 99.18602 |
| GVFFLNR | 7 | Q8N1G2 | Cap-specific mRNA (nucleoside-2'-O-)-m | CMTR1 | 95.321 | 2 | 55.048355 | yes | 9.722E-08 | 426.73996 | 92.56118 |
| DSYGKPLVK | 9 | Q8N1G2 | Cap-specific mRNA (nucleoside-2'-O-)-m | CMTR1 | 95.321 | 2 | 55.048355 | yes | 1.296E-07 | 503.78201 | 91.689224 |
| DYLFAVNIK | 9 | Q8N1G2 | Cap-specific mRNA (nucleoside-2'-O-)-m | CMTR1 | 95.321 | 2 | 55.048355 | yes | 4.749E-07 | 541.79767 | 84.104805 |
| LFWEWGDGIR | 10 | Q8N1G2 | Cap-specific mRNA (nucleoside-2'-O-)-m | CMTR1 | 95.321 | 2 | 55.048355 | yes | 1.204E-07 | 639.81696 | 77.899124 |
| KDIVEASSQK | 10 | Q8N1G2 | Cap-specific mRNA (nucleoside-2'-O-)-m | CMTR1 | 95.321 | 2 | 55.048355 | yes | 2.908E-07 | 552.7984 | 75.730209 |
| TDPECTAPIK | 10 | Q8N1G2 | Cap-specific mRNA (nucleoside-2'-O-)-m | CMTR1 | 95.321 | 2 | 55.048355 | yes | 2.419E-07 | 566.27118 | 74.741997 |
| VCLFKPITSR | 10 | Q8N1G2 | Cap-specific mRNA (nucleoside-2'-O-)-m | CMTR1 | 95.321 | 2 | 55.048355 | yes | 3.75E-07 | 610.8446 | 71.116356 |
| QHSSDSFDDAFK | 12 | Q8N1G2 | Cap-specific mRNA (nucleoside-2'-O-)-m | CMTR1 | 95.321 | 2 | 55.048355 | yes | 9.722E-08 | 692.29419 | 62.836258 |
| RTDPECTAPIK | 11 | Q8N1G2 | Cap-specific mRNA (nucleoside-2'-O-)-m | CMTR1 | 95.321 | 2 | 55.048355 | yes | 9.893E-08 | 644.32172 | 52.176434 |
| SNESHCSLQIK | 11 | Q8N1G2 | Cap-specific mRNA (nucleoside-2'-O-)-m | CMTR1 | 95.321 | 2 | 55.048355 | yes | 9.722E-08 | 651.80896 | 31.218565 |
| ASTTSLSGSDSETE | 16 | Q8N1G2 | Cap-specific mRNA (nucleoside-2'-O-)-m | CMTR1 | 95.321 | 2 | 55.048355 | yes | 2.609E-07 | 778.84973 | 13.206422 |
| GIPECILLVTQR | 12 | Q8N1W1 | Rho guanine nucleotide exchange factor 2 | ARHGEF28 | 191.891 | 2 | 44.300873 | yes | 8.117E-07 | 699.89233 | 45.44912 |
| ESLLHLAMR | 9 | Q8N1W1 | Rho guanine nucleotide exchange factor 2 | ARHGEF28 | 191.891 | 2 | 44.300873 | yes | 9.986E-07 | 535.2948 | 24.66354 |
| AQQSLLGHWK | 10 | Q8N1W1 | Rho guanine nucleotide exchange factor 2 | ARHGEF28 | 191.891 | 2 | 44.300873 | yes | 5.86E-05 | 584.31708 | 17.568972 |
| LVEDVTNFQGR | 11 | Q8N1W1 | Rho guanine nucleotide exchange factor 2 | ARHGEF28 | 191.891 | 2 | 44.300873 | yes | 2.247E-07 | 639.32788 | 13.135096 |
| AVQLLHQR | 8 | Q8N2A8 | Mitochondrial cardiolipin hydrolase OS=I | PLD6 | 28.273 | 2 | 50.536324 | yes | 9.088E-06 | 482.78796 | 103.12122 |
| LFLEEFER | 8 | Q8N2A8 | Mitochondrial cardiolipin hydrolase OS=I | PLD6 | 28.273 | 2 | 50.536324 | yes | 9.783E-08 | 541.77948 | 85.064148 |
| ENVLITEDDEYVR | 13 | Q8N2A8 | Mitochondrial cardiolipin hydrolase OS=I | PLD6 | 28.273 | 2 | 50.536324 | yes | 4.013E-07 | 797.88336 | 56.928455 |

| | | | | | | | | | | | |
|---|---|---|---|---|---|---|---|---|---|---|---|
| GWGDSVGTPSER | 12 | Q8N2K0 | Lysophosphatidylserine lipase ABHD12 ( | ABHD12 | 45.097 | 2 | 50.574482 | yes | 1.145E-07 | 624.28619 | 119.73746 |
| VPYFIDLK | 8 | Q8N2K0 | Lysophosphatidylserine lipase ABHD12 ( | ABHD12 | 45.097 | 2 | 50.574482 | yes | 3.354E-07 | 497.78403 | 87.196716 |
| TEPVALEHER | 10 | Q8N2K0 | Lysophosphatidylserine lipase ABHD12 ( | ABHD12 | 45.097 | 2 | 50.574482 | yes | 9.759E-08 | 590.80145 | 82.792252 |
| QATTYGMADGDF | 14 | Q8N2M8 | CLK4-associating serine/arginine rich prc | CLASRP | 77.162 | 2 | 38.487076 | yes | 5.187E-07 | 766.34595 | 66.588058 |
| CVCPAGFR | 8 | Q8N2S1 | Latent-transforming growth factor beta-bi | LTBP4 | 173.436 | 2 | 53.2591 | yes | 9.489E-07 | 483.71783 | 68.371246 |
| VPPPCDLGR | 9 | Q8N2S1 | Latent-transforming growth factor beta-bi | LTBP4 | 173.436 | 2 | 53.2591 | yes | 3.841E-06 | 505.75803 | 60.206932 |
| AGPDLASCLDVDE | 15 | Q8N2S1 | Latent-transforming growth factor beta-bi | LTBP4 | 173.436 | 2 | 53.2591 | yes | 9.722E-08 | 839.36401 | 59.581738 |
| SEEIAQINLER | 11 | Q8N2Z9 | Centromere protein S OS=Homo sapiens | CENPS | 15.893 | 2 | 54.201897 | yes | 9.722E-08 | 651.33844 | 358.49582 |
| MEEEAETEEQQR | 12 | Q8N2Z9 | Centromere protein S OS=Homo sapiens | CENPS | 15.893 | 2 | 54.201897 | yes | 9.722E-08 | 775.81738 | 167.77185 |
| QTIAAISELTFR | 12 | Q8N2Z9 | Centromere protein S OS=Homo sapiens | CENPS | 15.893 | 2 | 54.201897 | yes | 9.722E-08 | 675.37482 | 76.76738 |
| SVGDVALSQIVR | 12 | Q8N3C0 | Activating signal cointegrator 1 complex | ASCC3 | 251.461 | 2 | 56.30131 | yes | 1.558E-07 | 622.35388 | 10084.021 |
| GPVLESIVAR | 10 | Q8N3C0 | Activating signal cointegrator 1 complex | ASCC3 | 251.461 | 2 | 56.30131 | yes | 6.248E-07 | 520.80859 | 8905.8799 |
| AVILVHDIK | 9 | Q8N3C0 | Activating signal cointegrator 1 complex | ASCC3 | 251.461 | 2 | 56.30131 | yes | 4.387E-07 | 504.31604 | 8011.3115 |
| ADLIVTTPEK | 10 | Q8N3C0 | Activating signal cointegrator 1 complex | ASCC3 | 251.461 | 2 | 56.30131 | yes | 4.963E-07 | 543.80573 | 7102.2759 |
| LLILDEVHLLHEDI | 14 | Q8N3C0 | Activating signal cointegrator 1 complex | ASCC3 | 251.461 | 3 | 56.30131 | yes | 1.42E-07 | 572.32104 | 3372.9524 |
| LLILDEVHLLHEDI | 14 | Q8N3C0 | Activating signal cointegrator 1 complex | ASCC3 | 251.461 | 3 | 56.30131 | yes | 4.666E-07 | 429.49261 | 3372.9524 |
| LLILDEVHLLHEDI | 14 | Q8N3C0 | Activating signal cointegrator 1 complex | ASCC3 | 251.461 | 4 | 56.30131 | yes | 1.42E-07 | 572.32104 | 3372.9524 |
| LLILDEVHLLHEDI | 14 | Q8N3C0 | Activating signal cointegrator 1 complex | ASCC3 | 251.461 | 4 | 56.30131 | yes | 4.666E-07 | 429.49261 | 3372.9524 |
| IVGLSTALANAR | 12 | Q8N3C0 | Activating signal cointegrator 1 complex | ASCC3 | 251.461 | 2 | 56.30131 | yes | 1.266E-07 | 593.35114 | 6614.3711 |
| EQLVIEVGR | 9 | Q8N3C0 | Activating signal cointegrator 1 complex | ASCC3 | 251.461 | 2 | 56.30131 | yes | 5.722E-07 | 521.79822 | 6431.5586 |
| INILLQTYISR | 11 | Q8N3C0 | Activating signal cointegrator 1 complex | ASCC3 | 251.461 | 2 | 56.30131 | yes | 1.037E-07 | 667.39557 | 4716.0977 |
| SIEPLTYGR | 9 | Q8N3C0 | Activating signal cointegrator 1 complex | ASCC3 | 251.461 | 2 | 56.30131 | yes | 1.916E-07 | 518.2771 | 4713.5664 |
| TVEELFVNCK | 10 | Q8N3C0 | Activating signal cointegrator 1 complex | ASCC3 | 251.461 | 2 | 56.30131 | yes | 3.871E-07 | 619.80792 | 4518.8145 |
| FLSHLIEK | 8 | Q8N3C0 | Activating signal cointegrator 1 complex | ASCC3 | 251.461 | 2 | 56.30131 | yes | 1.279E-07 | 493.78711 | 4484.311 |
| TQMLVTTPEK | 10 | Q8N3C0 | Activating signal cointegrator 1 complex | ASCC3 | 251.461 | 2 | 56.30131 | yes | 9.723E-08 | 574.30499 | 4312.2524 |
| LWGWASPLR | 9 | Q8N3C0 | Activating signal cointegrator 1 complex | ASCC3 | 251.461 | 2 | 56.30131 | yes | 1.821E-07 | 543.29816 | 4217.0166 |
| TGYFSSTDLGR | 11 | Q8N3C0 | Activating signal cointegrator 1 complex | ASCC3 | 251.461 | 2 | 56.30131 | yes | 9.722E-08 | 602.28564 | 3946.6245 |
| LSHYLTLLTQR | 11 | Q8N3C0 | Activating signal cointegrator 1 complex | ASCC3 | 251.461 | 2 | 56.30131 | yes | 5.143E-07 | 672.88531 | 3601.8665 |
| IISHFSQDDLTALV | 19 | Q8N3C0 | Activating signal cointegrator 1 complex | ASCC3 | 251.461 | 3 | 56.30131 | yes | 1.059E-07 | 726.03668 | 3533.4912 |
| TEGDIFAIVSK | 11 | Q8N3C0 | Activating signal cointegrator 1 complex | ASCC3 | 251.461 | 2 | 56.30131 | yes | 9.809E-08 | 590.31641 | 3444.7073 |
| GPVLEVIVSR | 10 | Q8N3C0 | Activating signal cointegrator 1 complex | ASCC3 | 251.461 | 2 | 56.30131 | yes | 2.284E-07 | 534.82422 | 3438.9358 |
| QWLNMDER | 8 | Q8N3C0 | Activating signal cointegrator 1 complex | ASCC3 | 251.461 | 2 | 56.30131 | yes | 1.204E-07 | 546.25055 | 3395.0037 |
| TVAAELAIFR | 10 | Q8N3C0 | Activating signal cointegrator 1 complex | ASCC3 | 251.461 | 2 | 56.30131 | yes | 1.853E-07 | 545.81641 | 3356.4321 |
| IVYVAPMK | 8 | Q8N3C0 | Activating signal cointegrator 1 complex | ASCC3 | 251.461 | 2 | 56.30131 | yes | 1.113E-07 | 460.76733 | 3126.2375 |
| FQALQDNCK | 9 | Q8N3C0 | Activating signal cointegrator 1 complex | ASCC3 | 251.461 | 2 | 56.30131 | yes | 9.823E-08 | 562.26367 | 3118.0132 |
| TNFISSHTEKPVR | 13 | Q8N3C0 | Activating signal cointegrator 1 complex | ASCC3 | 251.461 | 3 | 56.30131 | yes | 8.388E-07 | 505.93533 | 1526.205 |
| TNFISSHTEKPVR | 13 | Q8N3C0 | Activating signal cointegrator 1 complex | ASCC3 | 251.461 | 3 | 56.30131 | yes | 3.906E-07 | 758.39935 | 1526.205 |
| TNFISSHTEKPVR | 13 | Q8N3C0 | Activating signal cointegrator 1 complex | ASCC3 | 251.461 | 2 | 56.30131 | yes | 8.388E-07 | 505.93533 | 1526.205 |
| TNFISSHTEKPVR | 13 | Q8N3C0 | Activating signal cointegrator 1 complex | ASCC3 | 251.461 | 2 | 56.30131 | yes | 3.906E-07 | 758.39935 | 1526.205 |

| | | | | | | | | | | |
|---|---|---|---|---|---|---|---|---|---|---|
| LIMNPSYYNLGDV | 20 | Q8N3C0 | Activating signal cointegrator 1 complex : ASCC3 | 251.461 | 3 | 56.30131 | yes | 4.599E-07 | 756.0318 | 2633.2292 |
| EQALLNAR | 8 | Q8N3C0 | Activating signal cointegrator 1 complex : ASCC3 | 251.461 | 2 | 56.30131 | yes | 1.294E-07 | 457.75635 | 2601.5254 |
| AYQIDPTLR | 9 | Q8N3C0 | Activating signal cointegrator 1 complex : ASCC3 | 251.461 | 2 | 56.30131 | yes | 2.184E-07 | 538.79041 | 2484.8914 |
| FLNSSNDHR | 9 | Q8N3C0 | Activating signal cointegrator 1 complex : ASCC3 | 251.461 | 2 | 56.30131 | yes | 1.625E-07 | 545.25726 | 2432.8599 |
| IASYYYLK | 8 | Q8N3C0 | Activating signal cointegrator 1 complex : ASCC3 | 251.461 | 2 | 56.30131 | yes | 2.796E-07 | 510.77368 | 2432.3645 |
| AMLPCPDYDTDTK | 13 | Q8N3C0 | Activating signal cointegrator 1 complex : ASCC3 | 251.461 | 2 | 56.30131 | yes | 2.358E-07 | 763.82874 | 1199.8018 |
| AMLPCPDYDTDTK | 13 | Q8N3C0 | Activating signal cointegrator 1 complex : ASCC3 | 251.461 | 2 | 56.30131 | yes | 7.366E-07 | 771.82617 | 1199.8018 |
| MQSINEDLK | 9 | Q8N3C0 | Activating signal cointegrator 1 complex : ASCC3 | 251.461 | 2 | 56.30131 | yes | 5.319E-07 | 539.26587 | 2327.812 |
| ANPLAYGISHK | 11 | Q8N3C0 | Activating signal cointegrator 1 complex : ASCC3 | 251.461 | 2 | 56.30131 | yes | 1.704E-07 | 585.81696 | 2312.7195 |
| ALAAEMTDYFSR | 12 | Q8N3C0 | Activating signal cointegrator 1 complex : ASCC3 | 251.461 | 2 | 56.30131 | yes | 9.809E-08 | 687.82153 | 1122.9934 |
| ALAAEMTDYFSR | 12 | Q8N3C0 | Activating signal cointegrator 1 complex : ASCC3 | 251.461 | 2 | 56.30131 | yes | 3.844E-05 | 695.81903 | 1122.9934 |
| PVLIFVSSR | 9 | Q8N3C0 | Activating signal cointegrator 1 complex : ASCC3 | 251.461 | 2 | 56.30131 | yes | 3.533E-07 | 509.3082 | 2243.8794 |
| TVLDQALR | 8 | Q8N3C0 | Activating signal cointegrator 1 complex : ASCC3 | 251.461 | 2 | 56.30131 | yes | 1.73E-07 | 458.26654 | 2180.4106 |
| DHVFSSMVESELH | 16 | Q8N3C0 | Activating signal cointegrator 1 complex : ASCC3 | 251.461 | 3 | 56.30131 | yes | 9.918E-08 | 596.28583 | 2180.0713 |
| WPTMTYR | 7 | Q8N3C0 | Activating signal cointegrator 1 complex : ASCC3 | 251.461 | 2 | 56.30131 | yes | 2.087E-07 | 477.72873 | 1038.549 |
| WPTMTYR | 7 | Q8N3C0 | Activating signal cointegrator 1 complex : ASCC3 | 251.461 | 2 | 56.30131 | yes | 1.013E-07 | 485.7262 | 1038.549 |
| SKDEGWFLILGEV | 16 | Q8N3C0 | Activating signal cointegrator 1 complex : ASCC3 | 251.461 | 3 | 56.30131 | yes | 3.944E-07 | 631.33527 | 2042.7651 |
| VSDSLTDLALK | 11 | Q8N3C0 | Activating signal cointegrator 1 complex : ASCC3 | 251.461 | 2 | 56.30131 | yes | 1.381E-07 | 581.32172 | 1862.1936 |
| EMENIIATVR | 10 | Q8N3C0 | Activating signal cointegrator 1 complex : ASCC3 | 251.461 | 2 | 56.30131 | yes | 1.21E-07 | 588.30811 | 1825.7574 |
| LHEQVLDLGLTWK | 13 | Q8N3C0 | Activating signal cointegrator 1 complex : ASCC3 | 251.461 | 2 | 56.30131 | yes | 1.393E-07 | 776.43011 | 1811.0077 |
| FLNEHLQEACTPE | 19 | Q8N3C0 | Activating signal cointegrator 1 complex : ASCC3 | 251.461 | 3 | 56.30131 | yes | 9.722E-08 | 761.3877 | 905.42328 |
| FLNEHLQEACTPE | 19 | Q8N3C0 | Activating signal cointegrator 1 complex : ASCC3 | 251.461 | 3 | 56.30131 | yes | 9.79E-08 | 571.2926 | 905.42328 |
| FLNEHLQEACTPE | 19 | Q8N3C0 | Activating signal cointegrator 1 complex : ASCC3 | 251.461 | 4 | 56.30131 | yes | 9.722E-08 | 761.3877 | 905.42328 |
| FLNEHLQEACTPE | 19 | Q8N3C0 | Activating signal cointegrator 1 complex : ASCC3 | 251.461 | 4 | 56.30131 | yes | 9.79E-08 | 571.2926 | 905.42328 |
| MASMNKPAFQAIR | 13 | Q8N3C0 | Activating signal cointegrator 1 complex : ASCC3 | 251.461 | 2 | 56.30131 | yes | 2.451E-07 | 732.87616 | 584.40588 |
| MASMNKPAFQAIR | 13 | Q8N3C0 | Activating signal cointegrator 1 complex : ASCC3 | 251.461 | 2 | 56.30131 | yes | 2.14E-07 | 488.91986 | 584.40588 |
| MASMNKPAFQAIR | 13 | Q8N3C0 | Activating signal cointegrator 1 complex : ASCC3 | 251.461 | 2 | 56.30131 | yes | 5.828E-07 | 740.8736 | 584.40588 |
| MASMNKPAFQAIR | 13 | Q8N3C0 | Activating signal cointegrator 1 complex : ASCC3 | 251.461 | 3 | 56.30131 | yes | 2.451E-07 | 732.87616 | 584.40588 |
| MASMNKPAFQAIR | 13 | Q8N3C0 | Activating signal cointegrator 1 complex : ASCC3 | 251.461 | 3 | 56.30131 | yes | 2.14E-07 | 488.91986 | 584.40588 |
| MASMNKPAFQAIR | 13 | Q8N3C0 | Activating signal cointegrator 1 complex : ASCC3 | 251.461 | 3 | 56.30131 | yes | 5.828E-07 | 740.8736 | 584.40588 |
| RLEPLGIIVK | 10 | Q8N3C0 | Activating signal cointegrator 1 complex : ASCC3 | 251.461 | 2 | 56.30131 | yes | 1.197E-07 | 569.37134 | 1747.7122 |
| HNEDHMNSELAK | 12 | Q8N3C0 | Activating signal cointegrator 1 complex : ASCC3 | 251.461 | 3 | 56.30131 | yes | 1.647E-07 | 475.54562 | 819.50586 |
| HNEDHMNSELAK | 12 | Q8N3C0 | Activating signal cointegrator 1 complex : ASCC3 | 251.461 | 3 | 56.30131 | yes | 1.004E-07 | 712.81482 | 819.50586 |
| HNEDHMNSELAK | 12 | Q8N3C0 | Activating signal cointegrator 1 complex : ASCC3 | 251.461 | 2 | 56.30131 | yes | 1.647E-07 | 475.54562 | 819.50586 |
| HNEDHMNSELAK | 12 | Q8N3C0 | Activating signal cointegrator 1 complex : ASCC3 | 251.461 | 2 | 56.30131 | yes | 1.004E-07 | 712.81482 | 819.50586 |
| QFSILPPHILTR | 12 | Q8N3C0 | Activating signal cointegrator 1 complex : ASCC3 | 251.461 | 2 | 56.30131 | yes | 9.809E-08 | 711.41681 | 772.95386 |
| QFSILPPHILTR | 12 | Q8N3C0 | Activating signal cointegrator 1 complex : ASCC3 | 251.461 | 2 | 56.30131 | yes | 3.003E-06 | 474.61365 | 772.95386 |
| QFSILPPHILTR | 12 | Q8N3C0 | Activating signal cointegrator 1 complex : ASCC3 | 251.461 | 3 | 56.30131 | yes | 9.809E-08 | 711.41681 | 772.95386 |
| QFSILPPHILTR | 12 | Q8N3C0 | Activating signal cointegrator 1 complex : ASCC3 | 251.461 | 3 | 56.30131 | yes | 3.003E-06 | 474.61365 | 772.95386 |

| | | | | | | | | | | |
|---|---|---|---|---|---|---|---|---|---|---|
| QDNYNEEVADLK | 12 | Q8N3C0 | Activating signal cointegrator 1 complex : ASCC3 | 251.461 | 2 | 56.30131 | yes | 9.722E-08 | 719.32825 | 1469.9628 |
| QCVHQIPSVMME/ | 20 | Q8N3C0 | Activating signal cointegrator 1 complex : ASCC3 | 251.461 | 3 | 56.30131 | yes | 1.317E-07 | 775.72192 | 1397.9397 |
| TAMSLIER | 8 | Q8N3C0 | Activating signal cointegrator 1 complex : ASCC3 | 251.461 | 2 | 56.30131 | yes | 1.19E-07 | 460.74713 | 1348.5714 |
| GSWDDLVEGHNE | 23 | Q8N3C0 | Activating signal cointegrator 1 complex : ASCC3 | 251.461 | 3 | 56.30131 | yes | 9.938E-08 | 843.74536 | 1316.2222 |
| GKPESCAVTPR | 11 | Q8N3C0 | Activating signal cointegrator 1 complex : ASCC3 | 251.461 | 2 | 56.30131 | yes | 1.448E-07 | 601.30334 | 1311.4354 |
| QMGLFNFR | 8 | Q8N3C0 | Activating signal cointegrator 1 complex : ASCC3 | 251.461 | 2 | 56.30131 | yes | 6.776E-07 | 506.75528 | 1250.8516 |
| AGHQVMVFVHAR | 12 | Q8N3C0 | Activating signal cointegrator 1 complex : ASCC3 | 251.461 | 2 | 56.30131 | yes | 9.927E-08 | 676.3562 | 1022.8771 |
| VIELTGDVTPDMK | 13 | Q8N3C0 | Activating signal cointegrator 1 complex : ASCC3 | 251.461 | 2 | 56.30131 | yes | 9.722E-08 | 709.36578 | 487.4726 |
| VIELTGDVTPDMK | 13 | Q8N3C0 | Activating signal cointegrator 1 complex : ASCC3 | 251.461 | 2 | 56.30131 | yes | 1.534E-06 | 717.36322 | 487.4726 |
| TNIAMLTVLHEIR | 13 | Q8N3C0 | Activating signal cointegrator 1 complex : ASCC3 | 251.461 | 2 | 56.30131 | yes | 9.722E-08 | 755.92413 | 457.33771 |
| TNIAMLTVLHEIR | 13 | Q8N3C0 | Activating signal cointegrator 1 complex : ASCC3 | 251.461 | 2 | 56.30131 | yes | 1.613E-07 | 504.28519 | 457.33771 |
| TNIAMLTVLHEIR | 13 | Q8N3C0 | Activating signal cointegrator 1 complex : ASCC3 | 251.461 | 3 | 56.30131 | yes | 9.722E-08 | 755.92413 | 457.33771 |
| TNIAMLTVLHEIR | 13 | Q8N3C0 | Activating signal cointegrator 1 complex : ASCC3 | 251.461 | 3 | 56.30131 | yes | 1.613E-07 | 504.28519 | 457.33771 |
| TNGSFLWCEVEK | 12 | Q8N3C0 | Activating signal cointegrator 1 complex : ASCC3 | 251.461 | 2 | 56.30131 | yes | 9.758E-08 | 735.34009 | 899.51965 |
| VLVCTATLAWGV | 22 | Q8N3C0 | Activating signal cointegrator 1 complex : ASCC3 | 251.461 | 3 | 56.30131 | yes | 1.116E-07 | 782.78375 | 865.6109 |
| TASHYYIK | 8 | Q8N3C0 | Activating signal cointegrator 1 complex : ASCC3 | 251.461 | 2 | 56.30131 | yes | 6.964E-07 | 491.75327 | 863.46649 |
| WISYTYLYVR | 10 | Q8N3C0 | Activating signal cointegrator 1 complex : ASCC3 | 251.461 | 2 | 56.30131 | yes | 1.39E-07 | 682.3559 | 860.02728 |
| AGRPQFDDQGK | 11 | Q8N3C0 | Activating signal cointegrator 1 complex : ASCC3 | 251.461 | 2 | 56.30131 | yes | 2.506E-07 | 609.79675 | 852.54749 |
| AGRPQFDK | 8 | Q8N3C0 | Activating signal cointegrator 1 complex : ASCC3 | 251.461 | 2 | 56.30131 | yes | 2.566E-07 | 459.74323 | 808.0874 |
| LHADQEYVLQVSI | 15 | Q8N3C0 | Activating signal cointegrator 1 complex : ASCC3 | 251.461 | 3 | 56.30131 | yes | 9.722E-08 | 600.3197 | 785.63928 |
| AEEFDQIK | 8 | Q8N3C0 | Activating signal cointegrator 1 complex : ASCC3 | 251.461 | 2 | 56.30131 | yes | 9.967E-08 | 490.2402 | 708.71411 |
| NCGHIPFFFPTQGH | 21 | Q8N3C0 | Activating signal cointegrator 1 complex : ASCC3 | 251.461 | 4 | 56.30131 | yes | 1.262E-07 | 620.04816 | 351.85938 |
| NCGHIPFFFPTQGH | 21 | Q8N3C0 | Activating signal cointegrator 1 complex : ASCC3 | 251.461 | 4 | 56.30131 | yes | 2.812E-07 | 826.39508 | 351.85938 |
| NCGHIPFFFPTQGH | 21 | Q8N3C0 | Activating signal cointegrator 1 complex : ASCC3 | 251.461 | 3 | 56.30131 | yes | 1.262E-07 | 620.04816 | 351.85938 |
| NCGHIPFFFPTQGH | 21 | Q8N3C0 | Activating signal cointegrator 1 complex : ASCC3 | 251.461 | 3 | 56.30131 | yes | 2.812E-07 | 826.39508 | 351.85938 |
| FRPVPLGQTFLGIK | 14 | Q8N3C0 | Activating signal cointegrator 1 complex : ASCC3 | 251.461 | 2 | 56.30131 | yes | 9.722E-08 | 786.96667 | 315.21701 |
| FRPVPLGQTFLGIK | 14 | Q8N3C0 | Activating signal cointegrator 1 complex : ASCC3 | 251.461 | 2 | 56.30131 | yes | 1.208E-07 | 524.98022 | 315.21701 |
| FRPVPLGQTFLGIK | 14 | Q8N3C0 | Activating signal cointegrator 1 complex : ASCC3 | 251.461 | 3 | 56.30131 | yes | 9.722E-08 | 786.96667 | 315.21701 |
| FRPVPLGQTFLGIK | 14 | Q8N3C0 | Activating signal cointegrator 1 complex : ASCC3 | 251.461 | 3 | 56.30131 | yes | 1.208E-07 | 524.98022 | 315.21701 |
| DLADWLNIK | 9 | Q8N3C0 | Activating signal cointegrator 1 complex : ASCC3 | 251.461 | 2 | 56.30131 | yes | 1E-07 | 544.29279 | 628.15332 |
| QIVGTDNGR | 9 | Q8N3C0 | Activating signal cointegrator 1 complex : ASCC3 | 251.461 | 2 | 56.30131 | yes | 3.49E-07 | 480.24887 | 610.35614 |
| AVYIAPLK | 8 | Q8N3C0 | Activating signal cointegrator 1 complex : ASCC3 | 251.461 | 2 | 56.30131 | yes | 1.311E-07 | 437.77347 | 598.20685 |
| IHYPHVYDSQAEA | 15 | Q8N3C0 | Activating signal cointegrator 1 complex : ASCC3 | 251.461 | 3 | 56.30131 | yes | 9.722E-08 | 596.95062 | 597.93988 |
| KVIELTGDVTPDM | 14 | Q8N3C0 | Activating signal cointegrator 1 complex : ASCC3 | 251.461 | 2 | 56.30131 | yes | 1.165E-07 | 773.41327 | 284.56943 |
| KVIELTGDVTPDM | 14 | Q8N3C0 | Activating signal cointegrator 1 complex : ASCC3 | 251.461 | 2 | 56.30131 | yes | 1.018E-07 | 515.94458 | 284.56943 |
| KVIELTGDVTPDM | 14 | Q8N3C0 | Activating signal cointegrator 1 complex : ASCC3 | 251.461 | 3 | 56.30131 | yes | 1.165E-07 | 773.41327 | 284.56943 |
| KVIELTGDVTPDM | 14 | Q8N3C0 | Activating signal cointegrator 1 complex : ASCC3 | 251.461 | 3 | 56.30131 | yes | 1.018E-07 | 515.94458 | 284.56943 |
| PVPLEVHIQGFPG( | 19 | Q8N3C0 | Activating signal cointegrator 1 complex : ASCC3 | 251.461 | 4 | 56.30131 | yes | 1.141E-07 | 558.53491 | 560.26355 |
| ELTGDMQLSK | 10 | Q8N3C0 | Activating signal cointegrator 1 complex : ASCC3 | 251.461 | 2 | 56.30131 | yes | 1.069E-07 | 561.27899 | 509.68631 |

| | | | | | | | | | | | |
|---|---|---|---|---|---|---|---|---|---|---|---|
| YLNSTLK | 7 | Q8N3C0 | Activating signal cointegrator 1 complex : | ASCC3 | 251.461 | 2 | 56.30131 | yes | 9.777E-08 | 419.73709 | 503.99048 |
| MILPEGIQR | 9 | Q8N3C0 | Activating signal cointegrator 1 complex : | ASCC3 | 251.461 | 2 | 56.30131 | yes | 5.068E-07 | 528.79718 | 503.00262 |
| ALFEIALR | 8 | Q8N3C0 | Activating signal cointegrator 1 complex : | ASCC3 | 251.461 | 2 | 56.30131 | yes | 1.792E-07 | 466.78183 | 431.23471 |
| NLVENLFSNGHIK | 13 | Q8N3C0 | Activating signal cointegrator 1 complex : | ASCC3 | 251.461 | 2 | 56.30131 | yes | 1.007E-07 | 742.89642 | 400.33148 |
| SHSPAKPVLIFVSS | 15 | Q8N3C0 | Activating signal cointegrator 1 complex : | ASCC3 | 251.461 | 2 | 56.30131 | yes | 1.81E-07 | 812.9621 | 368.94193 |
| QVESTQSMIR | 10 | Q8N3C0 | Activating signal cointegrator 1 complex : | ASCC3 | 251.461 | 2 | 56.30131 | yes | 1.013E-07 | 589.79535 | 364.1189 |
| KAGEDLEVSEGLM | 18 | Q8N3C0 | Activating signal cointegrator 1 complex : | ASCC3 | 251.461 | 3 | 56.30131 | yes | 1.277E-07 | 675.64832 | 313.7774 |
| HPPHTELLDLQPLI | 21 | Q8N3C0 | Activating signal cointegrator 1 complex : | ASCC3 | 251.461 | 4 | 56.30131 | yes | 3.69E-07 | 588.32013 | 308.71191 |
| AHLLLQAHLSR | 11 | Q8N3C0 | Activating signal cointegrator 1 complex : | ASCC3 | 251.461 | 2 | 56.30131 | yes | 9.722E-08 | 629.87256 | 136.91684 |
| AHLLLQAHLSR | 11 | Q8N3C0 | Activating signal cointegrator 1 complex : | ASCC3 | 251.461 | 2 | 56.30131 | yes | 3.423E-07 | 420.25079 | 136.91684 |
| AHLLLQAHLSR | 11 | Q8N3C0 | Activating signal cointegrator 1 complex : | ASCC3 | 251.461 | 3 | 56.30131 | yes | 9.722E-08 | 629.87256 | 136.91684 |
| AHLLLQAHLSR | 11 | Q8N3C0 | Activating signal cointegrator 1 complex : | ASCC3 | 251.461 | 3 | 56.30131 | yes | 3.423E-07 | 420.25079 | 136.91684 |
| KTVEELFVNCK | 11 | Q8N3C0 | Activating signal cointegrator 1 complex : | ASCC3 | 251.461 | 2 | 56.30131 | yes | 9.722E-08 | 683.85541 | 244.51224 |
| QHFQQGVIK | 9 | Q8N3C0 | Activating signal cointegrator 1 complex : | ASCC3 | 251.461 | 2 | 56.30131 | yes | 2.565E-07 | 542.79852 | 236.27901 |
| TNFISSHTEK | 10 | Q8N3C0 | Activating signal cointegrator 1 complex : | ASCC3 | 251.461 | 2 | 56.30131 | yes | 9.722E-08 | 582.28821 | 232.95499 |
| KDEIGHILHHVNIC | 16 | Q8N3C0 | Activating signal cointegrator 1 complex : | ASCC3 | 251.461 | 3 | 56.30131 | yes | 9.919E-08 | 608.34772 | 226.67807 |
| LEPLGIIVK | 9 | Q8N3C0 | Activating signal cointegrator 1 complex : | ASCC3 | 251.461 | 2 | 56.30131 | yes | 1.329E-07 | 491.3208 | 179.8143 |
| PVPLGQTFLGIK | 12 | Q8N3C0 | Activating signal cointegrator 1 complex : | ASCC3 | 251.461 | 2 | 56.30131 | yes | 3.492E-07 | 635.3819 | 175.87347 |
| QMFGPFPSSSATA | 20 | Q8N3C0 | Activating signal cointegrator 1 complex : | ASCC3 | 251.461 | 3 | 56.30131 | yes | 9.722E-08 | 705.65094 | 170.66325 |
| DEIGHILHHVNIGL | 15 | Q8N3C0 | Activating signal cointegrator 1 complex : | ASCC3 | 251.461 | 3 | 56.30131 | yes | 1.49E-07 | 565.64941 | 83.282005 |
| DEIGHILHHVNIGL | 15 | Q8N3C0 | Activating signal cointegrator 1 complex : | ASCC3 | 251.461 | 3 | 56.30131 | yes | 1.625E-07 | 847.97046 | 83.282005 |
| DEIGHILHHVNIGL | 15 | Q8N3C0 | Activating signal cointegrator 1 complex : | ASCC3 | 251.461 | 2 | 56.30131 | yes | 1.49E-07 | 565.64941 | 83.282005 |
| DEIGHILHHVNIGL | 15 | Q8N3C0 | Activating signal cointegrator 1 complex : | ASCC3 | 251.461 | 2 | 56.30131 | yes | 1.625E-07 | 847.97046 | 83.282005 |
| PSVRPVPLEVHIQC | 23 | Q8N3C0 | Activating signal cointegrator 1 complex : | ASCC3 | 251.461 | 4 | 56.30131 | yes | 2.877E-06 | 668.34851 | 120.40708 |
| RLIMNPSYYNLGD | 21 | Q8N3C0 | Activating signal cointegrator 1 complex : | ASCC3 | 251.461 | 3 | 56.30131 | yes | 1.534E-07 | 808.06549 | 99.023087 |
| VFNKYPTSK | 9 | Q8N3C0 | Activating signal cointegrator 1 complex : | ASCC3 | 251.461 | 2 | 56.30131 | yes | 9.722E-08 | 542.29529 | 89.839783 |
| NHHVASLSFYTPE | 17 | Q8N3C0 | Activating signal cointegrator 1 complex : | ASCC3 | 251.461 | 4 | 56.30131 | yes | 1.019E-07 | 481.99707 | 44.61068 |
| NHHVASLSFYTPE | 17 | Q8N3C0 | Activating signal cointegrator 1 complex : | ASCC3 | 251.461 | 4 | 56.30131 | yes | 7.341E-07 | 642.32697 | 44.61068 |
| NHHVASLSFYTPE | 17 | Q8N3C0 | Activating signal cointegrator 1 complex : | ASCC3 | 251.461 | 3 | 56.30131 | yes | 1.019E-07 | 481.99707 | 44.61068 |
| NHHVASLSFYTPE | 17 | Q8N3C0 | Activating signal cointegrator 1 complex : | ASCC3 | 251.461 | 3 | 56.30131 | yes | 7.341E-07 | 642.32697 | 44.61068 |
| KWKPIMK | 7 | Q8N3C0 | Activating signal cointegrator 1 complex : | ASCC3 | 251.461 | 2 | 56.30131 | yes | 9.182E-07 | 465.78333 | 77.677475 |
| RWPTMTYR | 8 | Q8N3C0 | Activating signal cointegrator 1 complex : | ASCC3 | 251.461 | 2 | 56.30131 | yes | 1.152E-06 | 555.7793 | 60.715439 |
| PAFQAIR | 7 | Q8N3C0 | Activating signal cointegrator 1 complex : | ASCC3 | 251.461 | 2 | 56.30131 | yes | 9.722E-08 | 401.73215 | 53.210892 |
| FGEGIIITTHDK | 12 | Q8N3C0 | Activating signal cointegrator 1 complex : | ASCC3 | 251.461 | 2 | 56.30131 | yes | 2.465E-07 | 665.8537 | 50.411129 |
| ELFPDGFSIHHAGM | 16 | Q8N3C0 | Activating signal cointegrator 1 complex : | ASCC3 | 251.461 | 4 | 56.30131 | yes | 1.997E-07 | 457.48062 | 44.001888 |
| KSVGDVALSQIVR | 13 | Q8N3C0 | Activating signal cointegrator 1 complex : | ASCC3 | 251.461 | 2 | 56.30131 | yes | 1.96E-07 | 686.40137 | 42.565067 |
| NLDLAAPTAEEAQ | 14 | Q8N3E9 | 1-phosphatidylinositol 4,5-bisphosphate p | PLCD3 | 89.259 | 2 | 51.21271 | yes | 9.722E-08 | 749.87842 | 126.19313 |
| LEGAEIEEFLR | 11 | Q8N3E9 | 1-phosphatidylinositol 4,5-bisphosphate p | PLCD3 | 89.259 | 2 | 51.21271 | yes | 5.014E-07 | 653.33789 | 82.101883 |
| LQEDGLSVWFQR | 12 | Q8N3E9 | 1-phosphatidylinositol 4,5-bisphosphate p | PLCD3 | 89.259 | 2 | 51.21271 | yes | 1.364E-07 | 739.37537 | 70.232162 |

| | | | | | | | | | | | |
|---|---|---|---|---|---|---|---|---|---|---|---|
| FGGAFAPAR | 9 | Q8N3E9 | 1-phosphatidylinositol 4,5-bisphosphate p | PLCD3 | 89.259 | 2 | 51.21271 | yes | 2.91E-06 | 447.23505 | 51.216522 |
| APELALVR | 8 | Q8N3E9 | 1-phosphatidylinositol 4,5-bisphosphate p | PLCD3 | 89.259 | 2 | 51.21271 | yes | 6.177E-07 | 434.76617 | 38.973114 |
| VEQMPQASPGLAI | 14 | Q8N3F8 | MICAL-like protein 1 OS=Homo sapiens | MICALL1 | 93.442 | 2 | 44.92922 | yes | 2.021E-07 | 740.88245 | 21.50309 |
| NLYGTSIDSVR | 11 | Q8N3R9 | Protein PALS1 OS=Homo sapiens OX=9 | PALS1 | 77.294 | 2 | 49.697937 | yes | 1.009E-07 | 612.81458 | 91.537956 |
| ICLLSLR | 7 | Q8N3R9 | Protein PALS1 OS=Homo sapiens OX=9 | PALS1 | 77.294 | 2 | 49.697937 | yes | 1.115E-06 | 437.76257 | 44.770473 |
| SQLIDELADK | 10 | Q8N3U4 | Cohesin subunit SA-2 OS=Homo sapiens | STAG2 | 141.327 | 2 | 48.70127 | yes | 1.113E-07 | 566.29822 | 71.480568 |
| HTSTLAAMK | 9 | Q8N3U4 | Cohesin subunit SA-2 OS=Homo sapiens | STAG2 | 141.327 | 2 | 48.70127 | no | 1.546E-06 | 480.25256 | 56.564842 |
| MYSDAFLNDSYLK | 13 | Q8N3U4 | Cohesin subunit SA-2 OS=Homo sapiens | STAG2 | 141.327 | 2 | 48.70127 | no | 1.216E-07 | 783.8609 | 54.429893 |
| HMQNSEIIR | 9 | Q8N3U4 | Cohesin subunit SA-2 OS=Homo sapiens | STAG2 | 141.327 | 2 | 48.70127 | yes | 1.719E-06 | 564.28491 | 24.467731 |
| LELLLQK | 7 | Q8N3U4 | Cohesin subunit SA-2 OS=Homo sapiens | STAG2 | 141.327 | 2 | 48.70127 | no | 8.13E-05 | 428.77875 | 5.8581591 |
| SDVKPVIQWLK | 11 | Q8N441 | Fibroblast growth factor receptor-like 1 O | FGFRL1 | 54.537 | 2 | 42.733646 | yes | 2.896E-07 | 656.88483 | 181.07922 |
| FVVLPTGDVWSR | 12 | Q8N441 | Fibroblast growth factor receptor-like 1 O | FGFRL1 | 54.537 | 2 | 42.733646 | yes | 2.695E-06 | 688.37207 | 55.796112 |
| EDAGVYVCK | 9 | Q8N441 | Fibroblast growth factor receptor-like 1 O | FGFRL1 | 54.537 | 2 | 42.733646 | yes | 1.84E-06 | 520.7395 | 53.923916 |
| DDQALTRPEAAEP | 14 | Q8N441 | Fibroblast growth factor receptor-like 1 O | FGFRL1 | 54.537 | 2 | 42.733646 | yes | 3.911E-06 | 784.88678 | 14.264911 |
| TSSIATALNTSGAC | 17 | Q8N4C8 | Misshapen-like kinase 1 OS=Homo sapie | MINK1 | 149.822 | 2 | 51.34788 | yes | 9.407E-07 | 775.89209 | 76.246651 |
| TLDEAPRPPK | 10 | Q8N4C8 | Misshapen-like kinase 1 OS=Homo sapie | MINK1 | 149.822 | 2 | 51.34788 | yes | 2.112E-06 | 562.30896 | 34.022713 |
| VMDVTEDEEEEIK | 20 | Q8N4C8 | Misshapen-like kinase 1 OS=Homo sapie | MINK1 | 149.822 | 3 | 51.34788 | yes | 2.837E-07 | 808.05029 | 30.827427 |
| QQQQLQQQQQR | 11 | Q8N4C8 | Misshapen-like kinase 1 OS=Homo sapie | MINK1 | 149.822 | 2 | 51.34788 | yes | 9.722E-08 | 720.86877 | 25.126207 |
| VNILVPGAGLGR | 12 | Q8N4J0 | Carnosine N-methyltransferase OS=Homo | CARNMT1 | 47.186 | 2 | 52.515015 | yes | 9.722E-08 | 583.3562 | 98.062592 |
| DACYQPIIK | 9 | Q8N4J0 | Carnosine N-methyltransferase OS=Homo | CARNMT1 | 47.186 | 2 | 52.515015 | yes | 5.945E-07 | 554.27881 | 45.958687 |
| GSDCVDQFR | 9 | Q8N4Q1 | Mitochondrial intermembrane space impo | CHCHD4 | 15.997 | 2 | 49.005615 | yes | 1.078E-07 | 542.22986 | 61.069 |
| ALGPGGELEPER | 12 | Q8N511 | Transmembrane protein 199 OS=Homo sa | TMEM199 | 23.13 | 2 | 50.036602 | yes | 9.927E-08 | 612.81458 | 59.813862 |
| HGGTLSDLGK | 10 | Q8N511 | Transmembrane protein 199 OS=Homo sa | TMEM199 | 23.13 | 2 | 50.036602 | yes | 1.881E-05 | 492.75909 | 26.477478 |
| NYICDECGQTFK | 12 | Q8N554 | Zinc finger protein 276 OS=Homo sapien | ZNF276 | 67.219 | 2 | 46.678318 | yes | 1.498E-07 | 767.81866 | 25.965481 |
| THIVSIPLR | 9 | Q8N556 | Actin filament-associated protein 1 OS=H | AFAP1 | 80.725 | 2 | 53.88132 | yes | 2.097E-07 | 518.31909 | 238.32054 |
| GCEVIPGLDSK | 11 | Q8N556 | Actin filament-associated protein 1 OS=H | AFAP1 | 80.725 | 2 | 53.88132 | yes | 9.722E-08 | 587.79224 | 118.40652 |
| AVITNILLR | 9 | Q8N556 | Actin filament-associated protein 1 OS=H | AFAP1 | 80.725 | 2 | 53.88132 | yes | 1.462E-07 | 506.82932 | 91.644745 |
| KPQAILEEK | 9 | Q8N556 | Actin filament-associated protein 1 OS=H | AFAP1 | 80.725 | 2 | 53.88132 | yes | 4.801E-06 | 528.30841 | 43.953629 |
| ICAFLLR | 7 | Q8N556 | Actin filament-associated protein 1 OS=H | AFAP1 | 80.725 | 2 | 53.88132 | no | 1.715E-07 | 446.75729 | 38.833717 |
| GLVFAISSDR | 10 | Q8N568 | Serine/threonine-protein kinase DCLK2 ( | DCLK2 | 83.606 | 2 | 40.088814 | yes | 4.217E-07 | 532.79041 | 35.69532 |
| SLSDNVNLPQGVR | 13 | Q8N568 | Serine/threonine-protein kinase DCLK2 ( | DCLK2 | 83.606 | 2 | 40.088814 | yes | 3.241E-06 | 699.87042 | 7.9858828 |
| EAITEAVEAPAAA | 14 | Q8N5A5 | Zinc finger CCCH-type with G patch dom | ZGPAT | 57.359 | 2 | 52.46999 | yes | 9.819E-08 | 699.86481 | 399.25256 |
| VLYLYPTHK | 9 | Q8N5A5 | Zinc finger CCCH-type with G patch dom | ZGPAT | 57.359 | 2 | 52.46999 | yes | 5.141E-07 | 567.32129 | 324.09482 |
| SLDQCVETLQK | 11 | Q8N5A5 | Zinc finger CCCH-type with G patch dom | ZGPAT | 57.359 | 2 | 52.46999 | yes | 9.722E-08 | 660.82684 | 183.11539 |
| LQGQAPGALEAGA | 18 | Q8N5A5 | Zinc finger CCCH-type with G patch dom | ZGPAT | 57.359 | 2 | 52.46999 | yes | 1.253E-07 | 817.93408 | 151.49599 |
| AQEAGLQQEQR | 11 | Q8N5A5 | Zinc finger CCCH-type with G patch dom | ZGPAT | 57.359 | 2 | 52.46999 | yes | 1.171E-07 | 629.31274 | 104.96259 |
| NVFDFLNEK | 9 | Q8N5A5 | Zinc finger CCCH-type with G patch dom | ZGPAT | 57.359 | 2 | 52.46999 | yes | 1.381E-07 | 563.28241 | 94.343483 |
| HSVASAQLQEK | 11 | Q8N5A5 | Zinc finger CCCH-type with G patch dom | ZGPAT | 57.359 | 2 | 52.46999 | yes | 9.722E-08 | 599.31476 | 92.303207 |
| ELIELTEASLVSVR | 14 | Q8N5A5 | Zinc finger CCCH-type with G patch dom | ZGPAT | 57.359 | 2 | 52.46999 | yes | 9.858E-08 | 779.93799 | 82.297798 |

| | | | | | | | | | | | |
|---|---|---|---|---|---|---|---|---|---|---|---|
| EAVVEGDGILPPLI | 14 | Q8N5A5 | Zinc finger CCCH-type with G patch dom | ZGPAT | 57.359 | 2 | 52.46999 | yes | 9.809E-08 | 732.90649 | 80.724213 |
| SIQEALAR | 8 | Q8N5A5 | Zinc finger CCCH-type with G patch dom | ZGPAT | 57.359 | 2 | 52.46999 | yes | 6.46E-07 | 444.25089 | 60.793404 |
| AQLQQVELALGA( | 23 | Q8N5A5 | Zinc finger CCCH-type with G patch dom | ZGPAT | 57.359 | 3 | 52.46999 | yes | 1.091E-07 | 804.75037 | 46.463272 |
| QEDAEYQAFR | 10 | Q8N5A5 | Zinc finger CCCH-type with G patch dom | ZGPAT | 57.359 | 2 | 52.46999 | yes | 1.503E-07 | 628.78076 | 46.03857 |
| ITDVDNGYYTVK | 12 | Q8N5A5 | Zinc finger CCCH-type with G patch dom | ZGPAT | 57.359 | 2 | 52.46999 | yes | 5.819E-07 | 694.34064 | 27.635368 |
| ALASFNQEER | 10 | Q8N5F7 | NF-kappa-B-activating protein OS=Homo | NKAP | 47.139 | 2 | 55.50078 | no | 1.222E-07 | 582.78583 | 271.255 |
| AEEPSDLIGPEAPK | 14 | Q8N5F7 | NF-kappa-B-activating protein OS=Homo | NKAP | 47.139 | 2 | 55.50078 | yes | 9.722E-08 | 726.86444 | 252.9868 |
| ESQEEFLENPWK | 12 | Q8N5F7 | NF-kappa-B-activating protein OS=Homo | NKAP | 47.139 | 2 | 55.50078 | yes | 9.722E-08 | 768.35425 | 112.33416 |
| PYGSDKPWPSLLD | 14 | Q8N5F7 | NF-kappa-B-activating protein OS=Homo | NKAP | 47.139 | 3 | 55.50078 | yes | 9.919E-08 | 534.9436 | 55.044903 |
| PYGSDKPWPSLLD | 14 | Q8N5F7 | NF-kappa-B-activating protein OS=Homo | NKAP | 47.139 | 3 | 55.50078 | yes | 1.478E-07 | 801.91174 | 55.044903 |
| PYGSDKPWPSLLD | 14 | Q8N5F7 | NF-kappa-B-activating protein OS=Homo | NKAP | 47.139 | 2 | 55.50078 | yes | 9.919E-08 | 534.9436 | 55.044903 |
| PYGSDKPWPSLLD | 14 | Q8N5F7 | NF-kappa-B-activating protein OS=Homo | NKAP | 47.139 | 2 | 55.50078 | yes | 1.478E-07 | 801.91174 | 55.044903 |
| IGELGAPEVWGLS | 15 | Q8N5F7 | NF-kappa-B-activating protein OS=Homo | NKAP | 47.139 | 2 | 55.50078 | yes | 1.036E-06 | 776.92212 | 95.418846 |
| AGIVQEDVQPPGL | 14 | Q8N5G0 | Small integral membrane protein 20 OS=l | SMIM20 | 7.703 | 2 | 34.27967 | yes | 1.041E-06 | 725.89862 | 100.07994 |
| LNNDLVGSTENLL | 14 | Q8N5G2 | Macoilin OS=Homo sapiens OX=9606 G | MACO1 | 76.178 | 2 | 48.92316 | yes | 1.725E-07 | 765.41211 | 342.87189 |
| NASGVVNSSPR | 11 | Q8N5G2 | Macoilin OS=Homo sapiens OX=9606 G | MACO1 | 76.178 | 2 | 48.92316 | yes | 4.045E-07 | 544.2782 | 201.77751 |
| SQISSLSSTER | 11 | Q8N5G2 | Macoilin OS=Homo sapiens OX=9606 G | MACO1 | 76.178 | 2 | 48.92316 | yes | 1.94E-07 | 597.8017 | 186.34613 |
| QLEIAQGQILQKD( | 17 | Q8N5G2 | Macoilin OS=Homo sapiens OX=9606 G | MACO1 | 76.178 | 3 | 48.92316 | yes | 3.404E-07 | 661.36877 | 129.64026 |
| LDLFSALGDAK | 11 | Q8N5G2 | Macoilin OS=Homo sapiens OX=9606 G | MACO1 | 76.178 | 2 | 48.92316 | yes | 1.611E-07 | 575.31116 | 116.9844 |
| QENELLQNK | 9 | Q8N5G2 | Macoilin OS=Homo sapiens OX=9606 G | MACO1 | 76.178 | 2 | 48.92316 | yes | 6.931E-07 | 558.28821 | 93.593559 |
| NFHVDLCR | 8 | Q8N5G2 | Macoilin OS=Homo sapiens OX=9606 G | MACO1 | 76.178 | 2 | 48.92316 | yes | 1.086E-07 | 530.7533 | 89.124458 |
| DTEVLMSALSAM( | 15 | Q8N5G2 | Macoilin OS=Homo sapiens OX=9606 G | MACO1 | 76.178 | 2 | 48.92316 | yes | 1.912E-07 | 819.88928 | 84.297516 |
| KLSTTLPEIEYR | 12 | Q8N5G2 | Macoilin OS=Homo sapiens OX=9606 G | MACO1 | 76.178 | 2 | 48.92316 | yes | 3.302E-07 | 725.401 | 33.926678 |
| FVETSPSGLDPNA! | 21 | Q8N5G2 | Macoilin OS=Homo sapiens OX=9606 G | MACO1 | 76.178 | 3 | 48.92316 | yes | 5.843E-07 | 759.73022 | 14.663674 |
| TGSVVLTASTDLR | 13 | Q8N5I2 | Arrestin domain-containing protein 1 OS= | ARRDC1 | 45.982 | 2 | 51.850033 | yes | 1.087E-07 | 660.36188 | 278.96112 |
| VTCIGSCGVSNK | 12 | Q8N5I2 | Arrestin domain-containing protein 1 OS= | ARRDC1 | 45.982 | 2 | 51.850033 | yes | 9.935E-08 | 641.29993 | 225.03827 |
| VQLFEISLSHGR | 12 | Q8N5I2 | Arrestin domain-containing protein 1 OS= | ARRDC1 | 45.982 | 2 | 51.850033 | yes | 9.722E-08 | 693.38043 | 202.66277 |
| DTSPVVASLLQK | 12 | Q8N5I2 | Arrestin domain-containing protein 1 OS= | ARRDC1 | 45.982 | 2 | 51.850033 | yes | 3.364E-07 | 629.35608 | 164.38084 |
| VVYSPGEPLAGTV | 14 | Q8N5I2 | Arrestin domain-containing protein 1 OS= | ARRDC1 | 45.982 | 2 | 51.850033 | yes | 2.895E-07 | 722.89337 | 155.60159 |
| TIAEVEGAGVK | 11 | Q8N5I2 | Arrestin domain-containing protein 1 OS= | ARRDC1 | 45.982 | 2 | 51.850033 | yes | 1.603E-07 | 537.29547 | 87.539169 |
| GYVVGQALQLHA | 32 | Q8N5I2 | Arrestin domain-containing protein 1 OS= | ARRDC1 | 45.982 | 4 | 51.850033 | yes | 5.297E-06 | 838.69403 | 13.20079 |
| VVNEINIEDLCLTK | 14 | Q8N5K1 | CDGSH iron-sulfur domain-containing pı | CISD2 | 15.279 | 2 | 49.261497 | yes | 1.051E-07 | 830.43475 | 148.75725 |
| ELQQYIQAR | 9 | Q8N5P1 | Zinc finger CCCH domain-containing prc | ZC3H8 | 33.576 | 2 | 51.560726 | yes | 1.026E-07 | 574.80658 | 949.93414 |
| FYVQGYCTR | 9 | Q8N5P1 | Zinc finger CCCH domain-containing prc | ZC3H8 | 33.576 | 2 | 51.560726 | yes | 9.872E-08 | 597.27405 | 596.20654 |
| IKLECEQIPK | 10 | Q8N5P1 | Zinc finger CCCH domain-containing prc | ZC3H8 | 33.576 | 2 | 51.560726 | yes | 1.015E-07 | 629.34723 | 579.76501 |
| EMANAAQPEESTK | 13 | Q8N5P1 | Zinc finger CCCH domain-containing prc | ZC3H8 | 33.576 | 2 | 51.560726 | yes | 9.722E-08 | 703.31683 | 238.52211 |
| EMANAAQPEESTK | 13 | Q8N5P1 | Zinc finger CCCH domain-containing prc | ZC3H8 | 33.576 | 2 | 51.560726 | yes | 3.619E-07 | 711.31427 | 238.52211 |
| LECEQIPK | 8 | Q8N5P1 | Zinc finger CCCH domain-containing prc | ZC3H8 | 33.576 | 2 | 51.560726 | yes | 9.722E-08 | 508.75769 | 384.40219 |
| FYHTGTK | 7 | Q8N5P1 | Zinc finger CCCH domain-containing prc | ZC3H8 | 33.576 | 2 | 51.560726 | yes | 1.261E-07 | 427.21378 | 376.69598 |

| | | | | | | | | | | | |
|---|---|---|---|---|---|---|---|---|---|---|---|
| CYQGEYCK | 8 | Q8N5P1 | Zinc finger CCCH domain-containing prc | ZC3H8 | 33.576 | 2 | 51.560726 | yes | 9.766E-08 | 554.21533 | 296.729 |
| QQHLSQAFINQHT | 16 | Q8N5P1 | Zinc finger CCCH domain-containing prc | ZC3H8 | 33.576 | 3 | 51.560726 | yes | 1.055E-07 | 645.99762 | 112.66778 |
| LDTPATSDPLSEEK | 14 | Q8N5U6 | E3 ubiquitin-protein ligase RNF10 OS=H | RNF10 | 89.927 | 2 | 55.302162 | yes | 1.829E-07 | 751.86469 | 450.24265 |
| VALEQQLAEEK | 11 | Q8N5U6 | E3 ubiquitin-protein ligase RNF10 OS=H | RNF10 | 89.927 | 2 | 55.302162 | yes | 9.722E-08 | 629.33789 | 280.30856 |
| EEALSGLAGSR | 11 | Q8N5U6 | E3 ubiquitin-protein ligase RNF10 OS=H | RNF10 | 89.927 | 2 | 55.302162 | yes | 2.646E-07 | 545.2804 | 155.43272 |
| ESVFQPR | 7 | Q8N5U6 | E3 ubiquitin-protein ligase RNF10 OS=H | RNF10 | 89.927 | 2 | 55.302162 | yes | 1.166E-07 | 431.72452 | 125.2776 |
| SVVATESHQYVV( | 22 | Q8N5U6 | E3 ubiquitin-protein ligase RNF10 OS=H | RNF10 | 89.927 | 3 | 55.302162 | yes | 3.113E-07 | 813.07489 | 106.97896 |
| AEFSPAQFSGPK | 12 | Q8N5U6 | E3 ubiquitin-protein ligase RNF10 OS=H | RNF10 | 89.927 | 2 | 55.302162 | yes | 1.421E-06 | 633.31171 | 97.319069 |
| ETLEMFSDDIEK | 12 | Q8N5U6 | E3 ubiquitin-protein ligase RNF10 OS=H | RNF10 | 89.927 | 2 | 55.302162 | yes | 9.722E-08 | 728.82922 | 79.183144 |
| PLSSPNAAATASDI | 16 | Q8N5U6 | E3 ubiquitin-protein ligase RNF10 OS=H | RNF10 | 89.927 | 2 | 55.302162 | yes | 6.79E-07 | 788.36957 | 12.784971 |
| EGSAVFAGFEGR | 12 | Q8N5Y2 | Male-specific lethal 3 homolog OS=Hom | MSL3 | 59.824 | 2 | 51.10841 | yes | 9.722E-08 | 613.79364 | 143.70026 |
| LSESSASPQPK | 11 | Q8N5Y2 | Male-specific lethal 3 homolog OS=Hom | MSL3 | 59.824 | 2 | 51.10841 | yes | 2.606E-07 | 565.78802 | 67.790672 |
| HFDLFLR | 7 | Q8N5Y2 | Male-specific lethal 3 homolog OS=Hom | MSL3 | 59.824 | 2 | 51.10841 | yes | 4.013E-07 | 474.25851 | 49.623463 |
| KAEPEALQSLR | 11 | Q8N5Y2 | Male-specific lethal 3 homolog OS=Hom | MSL3 | 59.824 | 2 | 51.10841 | yes | 5.122E-07 | 621.34607 | 33.29594 |
| YYACHLCGR | 9 | Q8N680 | Zinc finger and BTB domain-containing p | ZBTB2 | 57.337 | 2 | 47.115433 | yes | 1.398E-07 | 600.25787 | 48.491669 |
| ATPSENLVPSSAR | 13 | Q8N684 | Cleavage and polyadenylation specificity | CPSF7 | 52.051 | 2 | 52.491398 | yes | 1.106E-07 | 664.84387 | 4745.0059 |
| TPAILYTYSGLR | 12 | Q8N684 | Cleavage and polyadenylation specificity | CPSF7 | 52.051 | 2 | 52.491398 | yes | 1.493E-07 | 677.87189 | 4466.5972 |
| SYSVGASGSSSR | 12 | Q8N684 | Cleavage and polyadenylation specificity | CPSF7 | 52.051 | 2 | 52.491398 | yes | 9.722E-08 | 572.76508 | 2876.1931 |
| GYAEVVVASENS\ | 15 | Q8N684 | Cleavage and polyadenylation specificity | CPSF7 | 52.051 | 2 | 52.491398 | yes | 9.819E-08 | 794.90192 | 2725.8335 |
| DLLHNEDR | 8 | Q8N684 | Cleavage and polyadenylation specificity | CPSF7 | 52.051 | 2 | 52.491398 | yes | 1.294E-07 | 506.24634 | 2127.9397 |
| SIGVYDVVELK | 11 | Q8N684 | Cleavage and polyadenylation specificity | CPSF7 | 52.051 | 2 | 52.491398 | yes | 9.759E-08 | 611.3399 | 1987.0211 |
| DCLHGIEAK | 9 | Q8N684 | Cleavage and polyadenylation specificity | CPSF7 | 52.051 | 2 | 52.491398 | yes | 1.532E-07 | 521.75293 | 1409.7218 |
| HDDYFQER | 8 | Q8N684 | Cleavage and polyadenylation specificity | CPSF7 | 52.051 | 2 | 52.491398 | yes | 1.148E-07 | 555.23596 | 1064.4473 |
| QNLSQFEAQAR | 11 | Q8N684 | Cleavage and polyadenylation specificity | CPSF7 | 52.051 | 2 | 52.491398 | yes | 1.2E-07 | 646.32312 | 750.93982 |
| LLELLPGK | 8 | Q8N684 | Cleavage and polyadenylation specificity | CPSF7 | 52.051 | 2 | 52.491398 | yes | 1.733E-07 | 441.78659 | 730.22174 |
| VDVRPATR | 8 | Q8N684 | Cleavage and polyadenylation specificity | CPSF7 | 52.051 | 2 | 52.491398 | yes | 4.305E-07 | 457.26434 | 534.75989 |
| SSSTEPPPPVR | 11 | Q8N684 | Cleavage and polyadenylation specificity | CPSF7 | 52.051 | 2 | 52.491398 | yes | 3.354E-06 | 577.29602 | 102.22559 |
| GWGGEDVHLYR | 11 | Q8N6G5 | Chondroitin sulfate N-acetylgalactosamin | CSGALNA | 62.573 | 2 | 42.362743 | yes | 3.067E-07 | 644.80707 | 54.546432 |
| IAEAEIENLTGAR | 13 | Q8N6M0 | Deubiquitinase OTUD6B OS=Homo sapi | OTUD6B | 33.813 | 2 | 42.629906 | yes | 2.943E-07 | 693.86481 | 70.007317 |
| LITVALHR | 8 | Q8N6R0 | eEF1A lysine and N-terminal methyltrans | METTL13 | 78.768 | 2 | 55.15188 | yes | 1.726E-07 | 461.79526 | 495.21222 |
| ETEWLFGMDEGR | 12 | Q8N6R0 | eEF1A lysine and N-terminal methyltrans | METTL13 | 78.768 | 2 | 55.15188 | yes | 9.723E-08 | 735.3219 | 425.51282 |
| GTLDAVLTDEEEK | 13 | Q8N6R0 | eEF1A lysine and N-terminal methyltrans | METTL13 | 78.768 | 2 | 55.15188 | yes | 9.722E-08 | 710.34613 | 423.81198 |
| QQYAWLCSQLR | 11 | Q8N6R0 | eEF1A lysine and N-terminal methyltrans | METTL13 | 78.768 | 2 | 55.15188 | yes | 1.313E-07 | 726.85626 | 307.32349 |
| AMIAGLALLR | 10 | Q8N6R0 | eEF1A lysine and N-terminal methyltrans | METTL13 | 78.768 | 2 | 55.15188 | yes | 1.344E-07 | 514.81787 | 182.0652 |
| LATPELLETAQALI | 15 | Q8N6R0 | eEF1A lysine and N-terminal methyltrans | METTL13 | 78.768 | 2 | 55.15188 | yes | 3.001E-07 | 827.95416 | 146.51559 |
| QLAASAGFR | 9 | Q8N6R0 | eEF1A lysine and N-terminal methyltrans | METTL13 | 78.768 | 2 | 55.15188 | yes | 9.722E-08 | 460.75107 | 117.2058 |
| YTLHVVDSPTVKF | 15 | Q8N6R0 | eEF1A lysine and N-terminal methyltrans | METTL13 | 78.768 | 3 | 55.15188 | yes | 1.173E-07 | 566.97729 | 110.03969 |
| DNHFAIFIIPQGR | 13 | Q8N6R0 | eEF1A lysine and N-terminal methyltrans | METTL13 | 78.768 | 2 | 55.15188 | yes | 9.722E-08 | 764.40698 | 103.43365 |
| VHIADGLDYIASL/ | 20 | Q8N6R0 | eEF1A lysine and N-terminal methyltrans | METTL13 | 78.768 | 3 | 55.15188 | yes | 9.728E-08 | 662.34106 | 92.46347 |

| | | | | | | | | | | | |
|---|---|---|---|---|---|---|---|---|---|---|---|
| DIVNIDISEVVIK | 13 | Q8N6R0 | eEF1A lysine and N-terminal methyltrans | METTL13 | 78.768 | 2 | 55.15188 | yes | 1.022E-07 | 728.9165 | 78.255539 |
| EFGSVDYWEK | 10 | Q8N6R0 | eEF1A lysine and N-terminal methyltrans | METTL13 | 78.768 | 2 | 55.15188 | yes | 2.566E-07 | 630.28259 | 74.464134 |
| AVFPLLYVR | 9 | Q8N6R0 | eEF1A lysine and N-terminal methyltrans | METTL13 | 78.768 | 2 | 55.15188 | yes | 3.035E-07 | 539.32642 | 49.411491 |
| PCYDVIMFDVDSK | 13 | Q8N6R0 | eEF1A lysine and N-terminal methyltrans | METTL13 | 78.768 | 2 | 55.15188 | yes | 9.722E-08 | 794.85474 | 43.899097 |
| GWDDTYVLSDML | 13 | Q8N6R0 | eEF1A lysine and N-terminal methyltrans | METTL13 | 78.768 | 2 | 55.15188 | yes | 3.353E-07 | 771.8609 | 43.408886 |
| FRPVPGSALQIFEL | 20 | Q8N6R0 | eEF1A lysine and N-terminal methyltrans | METTL13 | 78.768 | 3 | 55.15188 | yes | 2.666E-07 | 782.73895 | 35.823578 |
| IFDDVSSGVSQLAS | 15 | Q8N6T3 | ADP-ribosylation factor GTPase-activatir | ARFGAP1 | 44.668 | 2 | 50.067696 | yes | 9.918E-08 | 776.8963 | 491.01266 |
| DKVVALAEGR | 10 | Q8N6T3 | ADP-ribosylation factor GTPase-activatir | ARFGAP1 | 44.668 | 2 | 50.067696 | yes | 1.754E-07 | 529.30365 | 413.02478 |
| GLGVHLSFVR | 10 | Q8N6T3 | ADP-ribosylation factor GTPase-activatir | ARFGAP1 | 44.668 | 2 | 50.067696 | yes | 1.26E-07 | 542.81671 | 352.7005 |
| VSGQPQSVTASSD | 14 | Q8N6T3 | ADP-ribosylation factor GTPase-activatir | ARFGAP1 | 44.668 | 2 | 50.067696 | yes | 2.114E-07 | 695.84406 | 124.12835 |
| YVGFGNTPPPQK | 12 | Q8N6T3 | ADP-ribosylation factor GTPase-activatir | ARFGAP1 | 44.668 | 2 | 50.067696 | yes | 1.271E-07 | 652.83533 | 114.72081 |
| ASELGHSLNENVL | 19 | Q8N6T3 | ADP-ribosylation factor GTPase-activatir | ARFGAP1 | 44.668 | 3 | 50.067696 | yes | 2.164E-07 | 688.6955 | 66.609833 |
| SSDSWEVWGSAST | 15 | Q8N6T3 | ADP-ribosylation factor GTPase-activatir | ARFGAP1 | 44.668 | 2 | 50.067696 | yes | 1.087E-07 | 834.86609 | 53.766884 |
| LVIVNLQPTK | 10 | Q8N6T7 | NAD-dependent protein deacylase sirtuin | SIRT6 | 39.119 | 2 | 54.1344 | yes | 1.001E-07 | 562.85553 | 196.72821 |
| FDTTFESAR | 9 | Q8N6T7 | NAD-dependent protein deacylase sirtuin | SIRT6 | 39.119 | 2 | 54.1344 | yes | 9.849E-08 | 537.24854 | 193.97937 |
| DTILDWEDSLPDR | 13 | Q8N6T7 | NAD-dependent protein deacylase sirtuin | SIRT6 | 39.119 | 2 | 54.1344 | yes | 9.722E-08 | 787.8703 | 160.27171 |
| HLGLEIPAWDGPR | 13 | Q8N6T7 | NAD-dependent protein deacylase sirtuin | SIRT6 | 39.119 | 2 | 54.1344 | yes | 3.04E-07 | 730.88586 | 138.56598 |
| IHGYVDEVMTR | 11 | Q8N6T7 | NAD-dependent protein deacylase sirtuin | SIRT6 | 39.119 | 2 | 54.1344 | yes | 9.722E-08 | 660.32428 | 107.43855 |
| PSGNLPLATK | 10 | Q8N6T7 | NAD-dependent protein deacylase sirtuin | SIRT6 | 39.119 | 2 | 54.1344 | yes | 2.986E-06 | 499.28748 | 94.742218 |
| GPHGVWTMEER | 11 | Q8N6T7 | NAD-dependent protein deacylase sirtuin | SIRT6 | 39.119 | 2 | 54.1344 | yes | 1.073E-07 | 649.80096 | 91.4767 |
| SVNYAAGLSPYAI | 14 | Q8N6T7 | NAD-dependent protein deacylase sirtuin | SIRT6 | 39.119 | 2 | 54.1344 | yes | 1.77E-05 | 749.36462 | 57.209084 |
| ERPTSPAPHRPPK | 13 | Q8N6T7 | NAD-dependent protein deacylase sirtuin | SIRT6 | 39.119 | 3 | 54.1344 | yes | 1.022E-07 | 490.60391 | 45.243763 |
| SFSQSSSLSR | 10 | Q8N720 | Zinc finger protein 655 OS=Homo sapien | ZNF655 | 57.407 | 2 | 54.211536 | yes | 2.723E-07 | 543.26471 | 182.43149 |
| SFSLDSTIDADQR | 13 | Q8N720 | Zinc finger protein 655 OS=Homo sapien | ZNF655 | 57.407 | 2 | 54.211536 | yes | 9.722E-08 | 727.84155 | 137.32362 |
| IFHQSSALTR | 10 | Q8N720 | Zinc finger protein 655 OS=Homo sapien | ZNF655 | 57.407 | 2 | 54.211536 | yes | 9.722E-08 | 580.31458 | 100.40744 |
| ISEEVHSYK | 9 | Q8N720 | Zinc finger protein 655 OS=Homo sapien | ZNF655 | 57.407 | 2 | 54.211536 | yes | 4.595E-07 | 546.27203 | 94.6231 |
| AYEYDEYGLAYIK | 13 | Q8N720 | Zinc finger protein 655 OS=Homo sapien | ZNF655 | 57.407 | 2 | 54.211536 | yes | 3.908E-07 | 799.37469 | 39.399734 |
| LKHDITQVPETR | 12 | Q8N720 | Zinc finger protein 655 OS=Homo sapien | ZNF655 | 57.407 | 2 | 54.211536 | yes | 2.318E-07 | 718.89642 | 16.965618 |
| VFVVHIPR | 8 | Q8N726 | Tumor suppressor ARF OS=Homo sapien | CDKN2A | 13.904 | 2 | 45.333702 | yes | 2.87E-06 | 483.79782 | 385.39337 |
| LGQQPLPR | 8 | Q8N726 | Tumor suppressor ARF OS=Homo sapien | CDKN2A | 13.904 | 2 | 45.333702 | yes | 1.881E-07 | 454.76926 | 383.45285 |
| EGDGVYYNELETF | 13 | Q8N7H5 | RNA polymerase II-associated factor 1 hc | PAF1 | 59.976 | 2 | 49.81617 | yes | 1.127E-07 | 772.8468 | 70.754059 |
| LLEEEIQAPTSSK | 13 | Q8N7H5 | RNA polymerase II-associated factor 1 hc | PAF1 | 59.976 | 2 | 49.81617 | yes | 1.503E-07 | 722.88013 | 64.334274 |
| VTPVEVMPVFPDF | 14 | Q8N7H5 | RNA polymerase II-associated factor 1 hc | PAF1 | 59.976 | 2 | 49.81617 | yes | 4.79E-07 | 802.92328 | 58.230724 |
| QQFTEEEIYK | 10 | Q8N7H5 | RNA polymerase II-associated factor 1 hc | PAF1 | 59.976 | 2 | 49.81617 | yes | 1.002E-07 | 657.81427 | 36.30159 |
| GGSPAVTLLISEK | 13 | Q8N884 | Cyclic GMP-AMP synthase OS=Homo sa | CGAS | 58.815 | 2 | 50.41073 | yes | 1.281E-07 | 636.36389 | 906.49023 |
| GVVDHLLLR | 9 | Q8N884 | Cyclic GMP-AMP synthase OS=Homo sa | CGAS | 58.815 | 2 | 50.41073 | yes | 2.134E-07 | 511.31128 | 788.00964 |
| ISAPNEFDVMFK | 12 | Q8N884 | Cyclic GMP-AMP synthase OS=Homo sa | CGAS | 58.815 | 2 | 50.41073 | yes | 1.451E-07 | 699.3421 | 718.99384 |
| LSFSHIEK | 8 | Q8N884 | Cyclic GMP-AMP synthase OS=Homo sa | CGAS | 58.815 | 2 | 50.41073 | yes | 1.107E-07 | 480.76108 | 531.25891 |
| IQLEEYSNTR | 10 | Q8N884 | Cyclic GMP-AMP synthase OS=Homo sa | CGAS | 58.815 | 2 | 50.41073 | yes | 1.043E-07 | 626.81201 | 489.78433 |

| | | | | | | | | | | | |
|---|---|---|---|---|---|---|---|---|---|---|---|
| DDISTAAGMVK | 11 | Q8N884 | Cyclic GMP-AMP synthase OS=Homo sa | CGAS | 58.815 | 2 | 50.41073 | yes | 1.542E-07 | 554.27118 | 270.42017 |
| EGNGFQEETWR | 11 | Q8N884 | Cyclic GMP-AMP synthase OS=Homo sa | CGAS | 58.815 | 2 | 50.41073 | yes | 9.809E-08 | 676.79694 | 239.83449 |
| IQNWLSAK | 8 | Q8N884 | Cyclic GMP-AMP synthase OS=Homo sa | CGAS | 58.815 | 2 | 50.41073 | yes | 2.985E-07 | 480.26907 | 206.41539 |
| SSWPASTQEGLR | 12 | Q8N884 | Cyclic GMP-AMP synthase OS=Homo sa | CGAS | 58.815 | 2 | 50.41073 | yes | 4.264E-07 | 659.82294 | 184.39359 |
| ISVDITLALESK | 12 | Q8N884 | Cyclic GMP-AMP synthase OS=Homo sa | CGAS | 58.815 | 2 | 50.41073 | yes | 1.052E-07 | 644.87158 | 131.83661 |
| FSSYHVK | 7 | Q8N884 | Cyclic GMP-AMP synthase OS=Homo sa | CGAS | 58.815 | 2 | 50.41073 | yes | 1.533E-07 | 434.22159 | 121.24112 |
| EEINDIKDTDVIMK | 14 | Q8N884 | Cyclic GMP-AMP synthase OS=Homo sa | CGAS | 58.815 | 2 | 50.41073 | yes | 3.583E-07 | 831.91638 | 69.824844 |
| LKPFYLVPK | 9 | Q8N884 | Cyclic GMP-AMP synthase OS=Homo sa | CGAS | 58.815 | 2 | 50.41073 | yes | 1.314E-07 | 552.84442 | 57.037933 |
| GVGLLNTGSYYEF | 15 | Q8N884 | Cyclic GMP-AMP synthase OS=Homo sa | CGAS | 58.815 | 2 | 50.41073 | yes | 1.444E-07 | 818.9201 | 56.727142 |
| SAPDTQERPPVR | 12 | Q8N884 | Cyclic GMP-AMP synthase OS=Homo sa | CGAS | 58.815 | 2 | 50.41073 | yes | 1.257E-07 | 676.84949 | 32.777927 |
| LADQSGLDVIR | 11 | Q8N983 | Large ribosomal subunit protein mL43 OS | MRPL43 | 23.431 | 2 | 52.359516 | yes | 9.722E-08 | 593.82495 | 1002.1136 |
| SVEEISTLVQK | 11 | Q8N983 | Large ribosomal subunit protein mL43 OS | MRPL43 | 23.431 | 2 | 52.359516 | yes | 1.189E-07 | 616.84027 | 839.14526 |
| YVQQLQR | 7 | Q8N983 | Large ribosomal subunit protein mL43 OS | MRPL43 | 23.431 | 2 | 52.359516 | yes | 9.849E-08 | 467.75888 | 169.65308 |
| EVIDFAR | 7 | Q8N983 | Large ribosomal subunit protein mL43 OS | MRPL43 | 23.431 | 2 | 52.359516 | yes | 3.888E-07 | 425.2269 | 131.09764 |
| EVQDPAPAQDTGI | 14 | Q8N983 | Large ribosomal subunit protein mL43 OS | MRPL43 | 23.431 | 2 | 52.359516 | yes | 2.469E-07 | 748.87061 | 84.839989 |
| VLATETVSHK | 10 | Q8N999 | RNA ligase 1 OS=Homo sapiens OX=96 | RLIG1 | 37.491 | 2 | 53.6285 | yes | 9.722E-08 | 542.80347 | 134.2408 |
| IDVLLDTFR | 9 | Q8N9N2 | Activating signal cointegrator 1 complex | ASCC1 | 45.51 | 2 | 50.591988 | yes | 1.563E-07 | 546.30841 | 5933.937 |
| TSISIPKPGQDGEIV | 21 | Q8N9N2 | Activating signal cointegrator 1 complex | ASCC1 | 45.51 | 3 | 50.591988 | yes | 1.062E-06 | 745.06934 | 3779.9465 |
| FQASGLIVK | 9 | Q8N9N2 | Activating signal cointegrator 1 complex | ASCC1 | 45.51 | 2 | 50.591988 | yes | 3.01E-07 | 481.78711 | 2274.3032 |
| LHATVMNTLFR | 11 | Q8N9N2 | Activating signal cointegrator 1 complex | ASCC1 | 45.51 | 2 | 50.591988 | yes | 1.372E-07 | 651.85297 | 2200.8735 |
| YNLYTAEGK | 9 | Q8N9N2 | Activating signal cointegrator 1 complex | ASCC1 | 45.51 | 2 | 50.591988 | yes | 3.338E-07 | 529.76129 | 2166.7637 |
| FQEEVLAK | 8 | Q8N9N2 | Activating signal cointegrator 1 complex | ASCC1 | 45.51 | 2 | 50.591988 | yes | 3.224E-07 | 482.26093 | 1431.7354 |
| LQELVDR | 7 | Q8N9N2 | Activating signal cointegrator 1 complex | ASCC1 | 45.51 | 2 | 50.591988 | yes | 1.133E-07 | 436.74545 | 1321.7137 |
| MEVLRPQLIR | 10 | Q8N9N2 | Activating signal cointegrator 1 complex | ASCC1 | 45.51 | 2 | 50.591988 | yes | 1.296E-07 | 648.87646 | 966.3244 |
| KIEMETK | 7 | Q8N9N2 | Activating signal cointegrator 1 complex | ASCC1 | 45.51 | 2 | 50.591988 | yes | 9.757E-08 | 439.73624 | 865.75903 |
| PGQDGEIVITGQHI | 14 | Q8N9N2 | Activating signal cointegrator 1 complex | ASCC1 | 45.51 | 3 | 50.591988 | yes | 1.03E-07 | 502.92682 | 360.07208 |
| PGQDGEIVITGQHI | 14 | Q8N9N2 | Activating signal cointegrator 1 complex | ASCC1 | 45.51 | 3 | 50.591988 | yes | 1.335E-07 | 753.8866 | 360.07208 |
| PGQDGEIVITGQHI | 14 | Q8N9N2 | Activating signal cointegrator 1 complex | ASCC1 | 45.51 | 2 | 50.591988 | yes | 1.03E-07 | 502.92682 | 360.07208 |
| PGQDGEIVITGQHI | 14 | Q8N9N2 | Activating signal cointegrator 1 complex | ASCC1 | 45.51 | 2 | 50.591988 | yes | 1.335E-07 | 753.8866 | 360.07208 |
| EEFINDISGGK | 11 | Q8N9N2 | Activating signal cointegrator 1 complex | ASCC1 | 45.51 | 2 | 50.591988 | yes | 2.331E-07 | 604.79333 | 100.1875 |
| CVVPQTTVILNND | 14 | Q8N9N5 | Protein BANP OS=Homo sapiens OX=96 | BANP | 56.495 | 2 | 44.599308 | yes | 2.136E-07 | 814.92487 | 75.551323 |
| EVQAVSNLSGQGK | 13 | Q8N9N5 | Protein BANP OS=Homo sapiens OX=96 | BANP | 56.495 | 2 | 44.599308 | yes | 1E-06 | 658.84387 | 22.403782 |
| CGYPGHLTFECR | 12 | Q8N9Q2 | Protein SREK1IP1 OS=Homo sapiens OX | SREK1IP1 | 18.177 | 3 | 50.035217 | yes | 9.927E-08 | 499.55182 | 159.17409 |
| CGYPGHLTFECR | 12 | Q8N9Q2 | Protein SREK1IP1 OS=Homo sapiens OX | SREK1IP1 | 18.177 | 3 | 50.035217 | yes | 1.438E-07 | 748.8241 | 159.17409 |
| CGYPGHLTFECR | 12 | Q8N9Q2 | Protein SREK1IP1 OS=Homo sapiens OX | SREK1IP1 | 18.177 | 2 | 50.035217 | yes | 9.927E-08 | 499.55182 | 159.17409 |
| CGYPGHLTFECR | 12 | Q8N9Q2 | Protein SREK1IP1 OS=Homo sapiens OX | SREK1IP1 | 18.177 | 2 | 50.035217 | yes | 1.438E-07 | 748.8241 | 159.17409 |
| HSSTPNSSEFSR | 12 | Q8N9Q2 | Protein SREK1IP1 OS=Homo sapiens OX | SREK1IP1 | 18.177 | 2 | 50.035217 | yes | 1.443E-07 | 668.2998 | 87.8461 |
| KCGYPGHLTFECR | 13 | Q8N9Q2 | Protein SREK1IP1 OS=Homo sapiens OX | SREK1IP1 | 18.177 | 3 | 50.035217 | yes | 1.127E-07 | 542.25012 | 45.87682 |
| NYPESYPQLPR | 11 | Q8N9R8 | Protein SCAI OS=Homo sapiens OX=96 | SCAI | 70.4 | 2 | 48.875744 | yes | 5.191E-07 | 682.33569 | 140.86578 |

| | | | | | | | | | | | |
|---|---|---|---|---|---|---|---|---|---|---|---|
| FSELTVDMFR | 10 | Q8N9R8 | Protein SCAI OS=Homo sapiens OX=96( | SCAI | 70.4 | 2 | 48.875744 | yes | 1.821E-07 | 622.80261 | 68.983719 |
| EDRPELVVK | 9 | Q8N9R8 | Protein SCAI OS=Homo sapiens OX=96( | SCAI | 70.4 | 2 | 48.875744 | yes | 4.001E-07 | 542.80347 | 44.646667 |
| DIAQLLTHSR | 10 | Q8N9R8 | Protein SCAI OS=Homo sapiens OX=96( | SCAI | 70.4 | 2 | 48.875744 | yes | 2.401E-07 | 577.31982 | 41.952137 |
| DETVENPHLQK | 11 | Q8N9R8 | Protein SCAI OS=Homo sapiens OX=96( | SCAI | 70.4 | 2 | 48.875744 | yes | 1.092E-07 | 655.32275 | 41.172466 |
| PVFVPDVALIANR | 13 | Q8NAT1 | Protein O-linked-mannose beta-1,4-N-ace | POMGNT2 | 66.616 | 2 | 48.675022 | yes | 1.116E-07 | 705.90881 | 87.078476 |
| VDIPSLIQTIR | 11 | Q8NAT1 | Protein O-linked-mannose beta-1,4-N-ace | POMGNT2 | 66.616 | 2 | 48.675022 | yes | 7.337E-07 | 627.87445 | 82.839348 |
| QATEPAPALR | 10 | Q8NAT1 | Protein O-linked-mannose beta-1,4-N-ace | POMGNT2 | 66.616 | 2 | 48.675022 | yes | 1.81E-06 | 527.28802 | 39.198059 |
| NPEWLFR | 7 | Q8NAT1 | Protein O-linked-mannose beta-1,4-N-ace | POMGNT2 | 66.616 | 2 | 48.675022 | yes | 8.422E-07 | 481.24814 | 2.9337754 |
| VCDIILPR | 8 | Q8NAV1 | Pre-mRNA-splicing factor 38A OS=Hom | PRPF38A | 37.477 | 2 | 55.186596 | yes | 5.054E-07 | 493.27859 | 306.6517 |
| MLQIQPEK | 8 | Q8NAV1 | Pre-mRNA-splicing factor 38A OS=Hom | PRPF38A | 37.477 | 2 | 55.186596 | yes | 3.561E-07 | 493.7706 | 121.53558 |
| LTGTAIDCYK | 10 | Q8NAV1 | Pre-mRNA-splicing factor 38A OS=Hom | PRPF38A | 37.477 | 2 | 55.186596 | yes | 1.128E-07 | 571.28156 | 114.09283 |
| EECFGLTAELVVD | 14 | Q8NAV1 | Pre-mRNA-splicing factor 38A OS=Hom | PRPF38A | 37.477 | 2 | 55.186596 | yes | 1.28E-07 | 805.39252 | 110.05984 |
| YLEPLYNDYR | 10 | Q8NAV1 | Pre-mRNA-splicing factor 38A OS=Hom | PRPF38A | 37.477 | 2 | 55.186596 | yes | 2.129E-06 | 673.32477 | 109.75629 |
| MLGALYMR | 8 | Q8NAV1 | Pre-mRNA-splicing factor 38A OS=Hom | PRPF38A | 37.477 | 2 | 55.186596 | yes | 3.251E-07 | 477.74863 | 85.516167 |
| YVLEEAEQLEPR | 12 | Q8NAV1 | Pre-mRNA-splicing factor 38A OS=Hom | PRPF38A | 37.477 | 2 | 55.186596 | yes | 9.722E-08 | 738.37244 | 63.406967 |
| ELSLLLQVEQR | 11 | Q8NB16 | Mixed lineage kinase domain-like protein | MLKL | 54.48 | 2 | 52.442234 | yes | 9.722E-08 | 664.38263 | 95.479996 |
| LQAGSIAIVR | 10 | Q8NB16 | Mixed lineage kinase domain-like protein | MLKL | 54.48 | 2 | 52.442234 | yes | 2.598E-07 | 514.31659 | 90.841377 |
| KFESPNILR | 9 | Q8NB16 | Mixed lineage kinase domain-like protein | MLKL | 54.48 | 2 | 52.442234 | yes | 4.366E-06 | 552.31403 | 36.366863 |
| LAGFELR | 7 | Q8NB16 | Mixed lineage kinase domain-like protein | MLKL | 54.48 | 2 | 52.442234 | yes | 6.392E-07 | 403.23196 | 36.341953 |
| CIPHIIVR | 8 | Q8NB78 | Lysine-specific histone demethylase 2 OS | KDM1B | 92.099 | 2 | 46.31476 | yes | 1.593E-07 | 504.29459 | 65.329414 |
| SVIIIGAGPAGLAA | 16 | Q8NB78 | Lysine-specific histone demethylase 2 OS | KDM1B | 92.099 | 2 | 46.31476 | yes | 2.96E-06 | 718.9328 | 50.22776 |
| IALQFPYR | 8 | Q8NB78 | Lysine-specific histone demethylase 2 OS | KDM1B | 92.099 | 2 | 46.31476 | yes | 4.679E-07 | 504.28726 | 21.577423 |
| ASFDHSPDSLPLR | 13 | Q8NB78 | Lysine-specific histone demethylase 2 OS | KDM1B | 92.099 | 3 | 46.31476 | yes | 4.895E-06 | 481.24051 | 20.444099 |
| IIYVPLPDAATR | 12 | Q8NB90 | ATPase family gene 2 protein homolog A | AFG2A | 97.905 | 2 | 52.30717 | yes | 1.101E-07 | 664.88226 | 955.08105 |
| GVLLYGPPGTGK | 12 | Q8NB90 | ATPase family gene 2 protein homolog A | AFG2A | 97.905 | 2 | 52.30717 | no | 1.002E-07 | 579.82953 | 730.99884 |
| EIEIGVPNAQDR | 12 | Q8NB90 | ATPase family gene 2 protein homolog A | AFG2A | 97.905 | 2 | 52.30717 | yes | 9.722E-08 | 670.84387 | 416.93671 |
| VTYDMIGGLSSQL | 14 | Q8NB90 | ATPase family gene 2 protein homolog A | AFG2A | 97.905 | 2 | 52.30717 | yes | 2.029E-07 | 756.39233 | 358.36737 |
| GVLLYGPPGCSK | 12 | Q8NB90 | ATPase family gene 2 protein homolog A | AFG2A | 97.905 | 2 | 52.30717 | yes | 3.033E-07 | 624.32629 | 297.62708 |
| VLCNEAGLCALR | 12 | Q8NB90 | ATPase family gene 2 protein homolog A | AFG2A | 97.905 | 2 | 52.30717 | yes | 2.035E-07 | 688.34467 | 270.23734 |
| LEQAVEWPLK | 10 | Q8NB90 | ATPase family gene 2 protein homolog A | AFG2A | 97.905 | 2 | 52.30717 | yes | 9.722E-08 | 606.83478 | 171.62534 |
| FYEDYQEK | 8 | Q8NB90 | ATPase family gene 2 protein homolog A | AFG2A | 97.905 | 2 | 52.30717 | yes | 4.416E-07 | 561.24292 | 143.95586 |
| GSSLGAGNVADR | 12 | Q8NB90 | ATPase family gene 2 protein homolog A | AFG2A | 97.905 | 2 | 52.30717 | yes | 1.025E-07 | 552.27563 | 143.11765 |
| HFTQALSTVTPR | 12 | Q8NB90 | ATPase family gene 2 protein homolog A | AFG2A | 97.905 | 2 | 52.30717 | yes | 9.722E-08 | 679.36475 | 113.89027 |
| HPESFIR | 7 | Q8NB90 | ATPase family gene 2 protein homolog A | AFG2A | 97.905 | 2 | 52.30717 | yes | 1.707E-07 | 443.23251 | 111.23061 |
| DFLQAMNDIRPSA | 15 | Q8NB90 | ATPase family gene 2 protein homolog A | AFG2A | 97.905 | 3 | 52.30717 | yes | 1.535E-06 | 588.95551 | 88.162582 |
| STPYKPIDDR | 10 | Q8NB90 | ATPase family gene 2 protein homolog A | AFG2A | 97.905 | 2 | 52.30717 | yes | 4.353E-07 | 596.30383 | 82.301048 |
| VGLSEMAQK | 9 | Q8NB90 | ATPase family gene 2 protein homolog A | AFG2A | 97.905 | 2 | 52.30717 | yes | 2.815E-06 | 481.75241 | 50.784878 |
| QIFAEATLR | 9 | Q8NB90 | ATPase family gene 2 protein homolog A | AFG2A | 97.905 | 2 | 52.30717 | yes | 3.944E-06 | 524.79291 | 46.175022 |
| ALANESGLNFLAIF | 14 | Q8NB90 | ATPase family gene 2 protein homolog A | AFG2A | 97.905 | 2 | 52.30717 | yes | 9.331E-07 | 730.909 | 16.99544 |

| | | | | | | | | | | | |
|---|---|---|---|---|---|---|---|---|---|---|---|
| GPLATGLHPDLDL | 16 | Q8NBF1 | Zinc finger protein GLIS1 OS=Homo sap | GLIS1 | 65.977 | 3 | 48.9166 | yes | 1.087E-07 | 543.63 | 41.981857 |
| LQQDVLQFQK | 10 | Q8NBJ4 | Golgi membrane protein 1 OS=Homo sap | GOLM1 | 45.334 | 2 | 50.924484 | yes | 2.08E-07 | 623.84314 | 1650.214 |
| FSYDLSQCINQMK | 13 | Q8NBJ4 | Golgi membrane protein 1 OS=Homo sap | GOLM1 | 45.334 | 2 | 50.924484 | yes | 9.722E-08 | 817.37109 | 409.44446 |
| WSPESPLQAPR | 11 | Q8NBJ5 | Procollagen galactosyltransferase 1 OS=F | COLGALT | 71.637 | 2 | 56.286686 | yes | 1.01E-07 | 634.32513 | 272.88684 |
| NAAHALPTTLGAL | 15 | Q8NBJ5 | Procollagen galactosyltransferase 1 OS=F | COLGALT | 71.637 | 2 | 56.286686 | yes | 9.722E-08 | 767.92047 | 142.00554 |
| ALQAQEIECR | 10 | Q8NBJ5 | Procollagen galactosyltransferase 1 OS=F | COLGALT | 71.637 | 2 | 56.286686 | yes | 1.417E-07 | 609.30078 | 114.62424 |
| NSDVLQSPLDSAA | 14 | Q8NBJ5 | Procollagen galactosyltransferase 1 OS=F | COLGALT | 71.637 | 2 | 56.286686 | yes | 9.722E-08 | 736.87061 | 100.46599 |
| KLLAAEPLSK | 10 | Q8NBJ5 | Procollagen galactosyltransferase 1 OS=F | COLGALT | 71.637 | 2 | 56.286686 | yes | 1.061E-07 | 535.33441 | 53.391903 |
| EEYGFLPVPLR | 11 | Q8NBJ5 | Procollagen galactosyltransferase 1 OS=F | COLGALT | 71.637 | 2 | 56.286686 | yes | 1.987E-07 | 660.35333 | 42.562447 |
| IGMGATLDIQR | 11 | Q8NBM4 | Ubiquitin-associated domain-containing p | UBAC2 | 38.964 | 2 | 52.55301 | yes | 9.722E-08 | 587.8161 | 218.6395 |
| MFTSTGSSGLYK | 12 | Q8NBM4 | Ubiquitin-associated domain-containing p | UBAC2 | 38.964 | 2 | 52.55301 | yes | 1.877E-07 | 639.80536 | 145.72113 |
| LMEMGFSR | 8 | Q8NBM4 | Ubiquitin-associated domain-containing p | UBAC2 | 38.964 | 2 | 52.55301 | yes | 1.313E-07 | 485.72787 | 125.98485 |
| QLMFSQFAQGR | 11 | Q8NBM4 | Ubiquitin-associated domain-containing p | UBAC2 | 38.964 | 2 | 52.55301 | yes | 9.722E-08 | 656.82697 | 113.39916 |
| TVIVTGANTGIGK | 13 | Q8NBN7 | Retinol dehydrogenase 13 OS=Homo sap | RDH13 | 35.933 | 2 | 28.525663 | yes | 2.728E-06 | 615.85626 | 45.980827 |
| HGLEETAAK | 9 | Q8NBQ5 | Estradiol 17-beta-dehydrogenase 11 OS=l | HSD17B11 | 32.964 | 2 | 51.852077 | yes | 3.036E-06 | 478.24582 | 144.82216 |
| TLTDELAALQITGV | 15 | Q8NBQ5 | Estradiol 17-beta-dehydrogenase 11 OS=l | HSD17B11 | 32.964 | 2 | 51.852077 | yes | 9.722E-08 | 786.9458 | 140.53648 |
| LMHGILTEQK | 10 | Q8NBQ5 | Estradiol 17-beta-dehydrogenase 11 OS=l | HSD17B11 | 32.964 | 2 | 51.852077 | yes | 1.781E-07 | 585.32098 | 125.75346 |
| FAAVGFHK | 8 | Q8NBQ5 | Estradiol 17-beta-dehydrogenase 11 OS=l | HSD17B11 | 32.964 | 2 | 51.852077 | yes | 1.332E-07 | 438.73996 | 112.44617 |
| LTAYEFAK | 8 | Q8NBQ5 | Estradiol 17-beta-dehydrogenase 11 OS=l | HSD17B11 | 32.964 | 2 | 51.852077 | yes | 4.677E-07 | 471.75018 | 40.187653 |
| TLAPTWEELSK | 11 | Q8NBS9 | Thioredoxin domain-containing protein 5 | TXNDC5 | 47.629 | 2 | 42.4601 | yes | 3.019E-07 | 637.83496 | 76.122269 |
| GYPTLLWFR | 9 | Q8NBS9 | Thioredoxin domain-containing protein 5 | TXNDC5 | 47.629 | 2 | 42.4601 | yes | 0.0001267 | 576.81366 | 3.335531 |
| ILDLMEGR | 8 | Q8NBT0 | POC1 centriolar protein homolog A OS=l | POC1A | 45.009 | 2 | 49.596973 | yes | 1.418E-07 | 473.75494 | 146.64494 |
| AAPCAEDPSLER | 12 | Q8NBT0 | POC1 centriolar protein homolog A OS=l | POC1A | 45.009 | 2 | 49.596973 | yes | 2.408E-07 | 679.30627 | 124.16479 |
| SNFDIVDHGEVTK | 13 | Q8NBT0 | POC1 centriolar protein homolog A OS=l | POC1A | 45.009 | 2 | 49.596973 | yes | 5.31E-07 | 730.85443 | 77.775986 |
| IWVPNVK | 7 | Q8NBT0 | POC1 centriolar protein homolog A OS=l | POC1A | 45.009 | 2 | 49.596973 | yes | 1.019E-07 | 428.258 | 75.748444 |
| QCLENQQLIMQR | 12 | Q8NBT0 | POC1 centriolar protein homolog A OS=l | POC1A | 45.009 | 2 | 49.596973 | yes | 3.986E-07 | 780.8847 | 60.319435 |
| DAVTCVDFSINTK | 13 | Q8NBT0 | POC1 centriolar protein homolog A OS=l | POC1A | 45.009 | 2 | 49.596973 | yes | 2.873E-07 | 735.35065 | 35.627789 |
| NEVVGLIFR | 9 | Q8NBU5 | Outer mitochondrial transmembrane helix | ATAD1 | 40.745 | 2 | 45.520485 | yes | 6.261E-07 | 523.80328 | 70.213722 |
| FINLQPSTLTDK | 12 | Q8NBU5 | Outer mitochondrial transmembrane helix | ATAD1 | 40.745 | 2 | 45.520485 | yes | 1.577E-05 | 688.87463 | 65.363708 |
| VHAEAFSR | 8 | Q8NBU5 | Outer mitochondrial transmembrane helix | ATAD1 | 40.745 | 2 | 45.520485 | yes | 5.336E-07 | 458.73541 | 58.410343 |
| WMVDAIDPTR | 10 | Q8NBU5 | Outer mitochondrial transmembrane helix | ATAD1 | 40.745 | 2 | 45.520485 | yes | 6.508E-07 | 602.29498 | 33.434525 |
| DAAFQNVLTHVCI | 14 | Q8NBU5 | Outer mitochondrial transmembrane helix | ATAD1 | 40.745 | 2 | 45.520485 | yes | 1.824E-07 | 801.88269 | 23.41045 |
| MPPPTILSTVPR | 12 | Q8NBZ0 | INO80 complex subunit E OS=Homo sap | INO80E | 26.479 | 2 | 31.772406 | yes | 1.73E-06 | 654.87085 | 25.580057 |
| SAAQAAAQTNSN/ | 16 | Q8NC51 | SERPINE1 mRNA-binding protein 1 OS= | SERBP1 | 44.966 | 2 | 54.09681 | yes | 9.722E-08 | 730.85803 | 11853.494 |
| EMTLDEWK | 8 | Q8NC51 | SERPINE1 mRNA-binding protein 1 OS= | SERBP1 | 44.966 | 2 | 54.09681 | yes | 2.74E-07 | 526.24188 | 8443.3252 |
| EETQPPVALK | 10 | Q8NC51 | SERPINE1 mRNA-binding protein 1 OS= | SERBP1 | 44.966 | 2 | 54.09681 | yes | 1.596E-07 | 556.30334 | 7771.1704 |
| SSFSHYSGLK | 10 | Q8NC51 | SERPINE1 mRNA-binding protein 1 OS= | SERBP1 | 44.966 | 2 | 54.09681 | yes | 1.019E-07 | 556.77222 | 6937.7695 |
| KNPLPPSVGVVDK | 13 | Q8NC51 | SERPINE1 mRNA-binding protein 1 OS= | SERBP1 | 44.966 | 2 | 54.09681 | yes | 2.625E-06 | 675.39301 | 5712.1641 |
| RFEKPLEEK | 9 | Q8NC51 | SERPINE1 mRNA-binding protein 1 OS= | SERBP1 | 44.966 | 2 | 54.09681 | yes | 7.519E-07 | 588.32458 | 3185.5862 |

| | | | | | | | | | | | |
|---|---|---|---|---|---|---|---|---|---|---|---|
| FEKPLEEK | 8 | Q8NC51 | SERPINE1 mRNA-binding protein 1 OS= | SERBP1 | 44.966 | 2 | 54.09681 | yes | 4.415E-07 | 510.27402 | 3075.814 |
| EAGGGGVGGPGAI | 13 | Q8NC51 | SERPINE1 mRNA-binding protein 1 OS= | SERBP1 | 44.966 | 2 | 54.09681 | yes | 1.165E-07 | 507.25415 | 2453.5542 |
| GGSGSHNWGTVK | 12 | Q8NC51 | SERPINE1 mRNA-binding protein 1 OS= | SERBP1 | 44.966 | 2 | 54.09681 | yes | 9.722E-08 | 593.78363 | 2410.3254 |
| AKVEFNIR | 8 | Q8NC51 | SERPINE1 mRNA-binding protein 1 OS= | SERBP1 | 44.966 | 2 | 54.09681 | yes | 2.164E-07 | 488.78235 | 2313.033 |
| KPNEGADGQWK | 11 | Q8NC51 | SERPINE1 mRNA-binding protein 1 OS= | SERBP1 | 44.966 | 2 | 54.09681 | yes | 9.722E-08 | 615.29907 | 2284.0608 |
| PGHLQEGFGCVVT | 15 | Q8NC51 | SERPINE1 mRNA-binding protein 1 OS= | SERBP1 | 44.966 | 2 | 54.09681 | yes | 1.126E-07 | 835.90704 | 908.31934 |
| PGHLQEGFGCVVT | 15 | Q8NC51 | SERPINE1 mRNA-binding protein 1 OS= | SERBP1 | 44.966 | 2 | 54.09681 | yes | 9.938E-08 | 557.60712 | 908.31934 |
| PGHLQEGFGCVVT | 15 | Q8NC51 | SERPINE1 mRNA-binding protein 1 OS= | SERBP1 | 44.966 | 3 | 54.09681 | yes | 1.126E-07 | 835.90704 | 908.31934 |
| PGHLQEGFGCVVT | 15 | Q8NC51 | SERPINE1 mRNA-binding protein 1 OS= | SERBP1 | 44.966 | 3 | 54.09681 | yes | 9.938E-08 | 557.60712 | 908.31934 |
| NPLPPSVGVVDK | 12 | Q8NC51 | SERPINE1 mRNA-binding protein 1 OS= | SERBP1 | 44.966 | 2 | 54.09681 | yes | 2.445E-07 | 611.34552 | 1601.7605 |
| KEETQPPVALK | 11 | Q8NC51 | SERPINE1 mRNA-binding protein 1 OS= | SERBP1 | 44.966 | 2 | 54.09681 | yes | 3.63E-07 | 620.35083 | 1536.5472 |
| GEGGEFSVDR | 10 | Q8NC51 | SERPINE1 mRNA-binding protein 1 OS= | SERBP1 | 44.966 | 2 | 54.09681 | yes | 1.265E-07 | 526.73578 | 1224.1519 |
| ENEVEEVK | 8 | Q8NC51 | SERPINE1 mRNA-binding protein 1 OS= | SERBP1 | 44.966 | 2 | 54.09681 | yes | 9.722E-08 | 488.23511 | 1074.0647 |
| RPDQQLQGEGK | 11 | Q8NC51 | SERPINE1 mRNA-binding protein 1 OS= | SERBP1 | 44.966 | 2 | 54.09681 | yes | 1.001E-07 | 628.32312 | 1071.2491 |
| KGFVLHK | 7 | Q8NC51 | SERPINE1 mRNA-binding protein 1 OS= | SERBP1 | 44.966 | 2 | 54.09681 | yes | 1.503E-07 | 414.75815 | 659.96368 |
| KNPLPPSVGVVDK | 14 | Q8NC51 | SERPINE1 mRNA-binding protein 1 OS= | SERBP1 | 44.966 | 3 | 54.09681 | yes | 1.961E-06 | 493.29608 | 321.98871 |
| KNPLPPSVGVVDK | 14 | Q8NC51 | SERPINE1 mRNA-binding protein 1 OS= | SERBP1 | 44.966 | 3 | 54.09681 | yes | 4.655E-07 | 739.44049 | 321.98871 |
| KNPLPPSVGVVDK | 14 | Q8NC51 | SERPINE1 mRNA-binding protein 1 OS= | SERBP1 | 44.966 | 2 | 54.09681 | yes | 1.961E-06 | 493.29608 | 321.98871 |
| KNPLPPSVGVVDK | 14 | Q8NC51 | SERPINE1 mRNA-binding protein 1 OS= | SERBP1 | 44.966 | 2 | 54.09681 | yes | 4.655E-07 | 739.44049 | 321.98871 |
| DELTESPK | 8 | Q8NC51 | SERPINE1 mRNA-binding protein 1 OS= | SERBP1 | 44.966 | 2 | 54.09681 | yes | 1.154E-07 | 459.72437 | 507.3786 |
| GEGGEFSVDRPIID | 15 | Q8NC51 | SERPINE1 mRNA-binding protein 1 OS= | SERBP1 | 44.966 | 2 | 54.09681 | yes | 1.296E-07 | 823.91028 | 300.59372 |
| EETQPPVALKK | 11 | Q8NC51 | SERPINE1 mRNA-binding protein 1 OS= | SERBP1 | 44.966 | 2 | 54.09681 | yes | 1.425E-07 | 620.35083 | 143.49791 |
| RGGSGSHNWGTVI | 13 | Q8NC51 | SERPINE1 mRNA-binding protein 1 OS= | SERBP1 | 44.966 | 3 | 54.09681 | yes | 2.286E-07 | 448.22522 | 66.816826 |
| RGGSGSHNWGTVI | 13 | Q8NC51 | SERPINE1 mRNA-binding protein 1 OS= | SERBP1 | 44.966 | 3 | 54.09681 | yes | 4.996E-07 | 671.83417 | 66.816826 |
| RGGSGSHNWGTVI | 13 | Q8NC51 | SERPINE1 mRNA-binding protein 1 OS= | SERBP1 | 44.966 | 2 | 54.09681 | yes | 2.286E-07 | 448.22522 | 66.816826 |
| RGGSGSHNWGTVI | 13 | Q8NC51 | SERPINE1 mRNA-binding protein 1 OS= | SERBP1 | 44.966 | 2 | 54.09681 | yes | 4.996E-07 | 671.83417 | 66.816826 |
| ENEVEEVKEEGPK | 13 | Q8NC51 | SERPINE1 mRNA-binding protein 1 OS= | SERBP1 | 44.966 | 3 | 54.09681 | yes | 2.799E-07 | 505.91061 | 48.493259 |
| VVCLESAHPR | 10 | Q8NC56 | LEM domain-containing protein 2 OS=H | LEMD2 | 56.976 | 2 | 48.411465 | yes | 2.637E-07 | 584.3006 | 255.11015 |
| VAGEDMLVWR | 10 | Q8NC56 | LEM domain-containing protein 2 OS=H | LEMD2 | 56.976 | 2 | 48.411465 | yes | 1.152E-07 | 588.29755 | 212.25771 |
| GEDQSELVTTVDK | 13 | Q8NC56 | LEM domain-containing protein 2 OS=H | LEMD2 | 56.976 | 2 | 48.411465 | yes | 1.192E-07 | 710.84375 | 148.98207 |
| LPSSLLGPDPR | 11 | Q8NC56 | LEM domain-containing protein 2 OS=H | LEMD2 | 56.976 | 2 | 48.411465 | yes | 1.456E-07 | 576.32458 | 95.391273 |
| LLPVDCER | 8 | Q8NC56 | LEM domain-containing protein 2 OS=H | LEMD2 | 56.976 | 2 | 48.411465 | yes | 2.16E-07 | 501.25787 | 86.097229 |
| FEAALTWILSSNK | 13 | Q8NC56 | LEM domain-containing protein 2 OS=H | LEMD2 | 56.976 | 2 | 48.411465 | yes | 2.259E-07 | 740.39575 | 39.102409 |
| AVEFLASNESR | 11 | Q8NC56 | LEM domain-containing protein 2 OS=H | LEMD2 | 56.976 | 2 | 48.411465 | yes | 3.528E-06 | 611.80676 | 24.707233 |
| HAGHTLLQIPMGG | 14 | Q8NC60 | Nitric oxide-associated protein 1 OS=Hor | NOA1 | 78.458 | 2 | 54.056362 | yes | 3.435E-07 | 730.39557 | 833.42236 |
| HAGHTLLQIPMGG | 14 | Q8NC60 | Nitric oxide-associated protein 1 OS=Hor | NOA1 | 78.458 | 2 | 54.056362 | yes | 2.135E-05 | 487.26611 | 833.42236 |
| HAGHTLLQIPMGG | 14 | Q8NC60 | Nitric oxide-associated protein 1 OS=Hor | NOA1 | 78.458 | 3 | 54.056362 | yes | 3.435E-07 | 730.39557 | 833.42236 |
| HAGHTLLQIPMGG | 14 | Q8NC60 | Nitric oxide-associated protein 1 OS=Hor | NOA1 | 78.458 | 3 | 54.056362 | yes | 2.135E-05 | 487.26611 | 833.42236 |
| EVNIVLPTQSIVPR | 14 | Q8NC60 | Nitric oxide-associated protein 1 OS=Hor | NOA1 | 78.458 | 2 | 54.056362 | yes | 9.722E-08 | 782.95648 | 935.96826 |

| | | | | | | | | | | | |
|---|---|---|---|---|---|---|---|---|---|---|---|
| VDLLPQDAPGYR | 12 | Q8NC60 | Nitric oxide-associated protein 1 OS=Hor | NOA1 | 78.458 | 2 | 54.056362 | yes | 1.117E-07 | 672.35132 | 401.08783 |
| KPPSLMYNVR | 10 | Q8NC60 | Nitric oxide-associated protein 1 OS=Hor | NOA1 | 78.458 | 2 | 54.056362 | yes | 4.184E-07 | 602.82898 | 371.54974 |
| TFLYSEEQK | 9 | Q8NC60 | Nitric oxide-associated protein 1 OS=Hor | NOA1 | 78.458 | 2 | 54.056362 | yes | 3.107E-07 | 572.77966 | 354.59232 |
| EGLGASEAVADIK | 13 | Q8NC60 | Nitric oxide-associated protein 1 OS=Hor | NOA1 | 78.458 | 2 | 54.056362 | yes | 1.503E-07 | 630.32751 | 284.97345 |
| DAHWFYDTPGITK | 13 | Q8NC60 | Nitric oxide-associated protein 1 OS=Hor | NOA1 | 78.458 | 2 | 54.056362 | yes | 1.151E-07 | 775.86737 | 236.83527 |
| TAEADGGLAR | 10 | Q8NC60 | Nitric oxide-associated protein 1 OS=Hor | NOA1 | 78.458 | 2 | 54.056362 | yes | 9.722E-08 | 480.74088 | 224.16275 |
| GDVYLVGATNAG | 13 | Q8NC60 | Nitric oxide-associated protein 1 OS=Hor | NOA1 | 78.458 | 2 | 54.056362 | yes | 6.321E-07 | 632.83026 | 169.56122 |
| EQYLELVSAALR | 12 | Q8NC60 | Nitric oxide-associated protein 1 OS=Hor | NOA1 | 78.458 | 2 | 54.056362 | yes | 9.722E-08 | 696.38007 | 148.38934 |
| FSSAGWVSVTPNF | 14 | Q8NC60 | Nitric oxide-associated protein 1 OS=Hor | NOA1 | 78.458 | 2 | 54.056362 | yes | 2.874E-07 | 763.88556 | 95.877823 |
| FPICNPTPYR | 10 | Q8NC60 | Nitric oxide-associated protein 1 OS=Hor | NOA1 | 78.458 | 2 | 54.056362 | yes | 9.722E-08 | 632.81079 | 89.485817 |
| ENCILNLLTEK | 11 | Q8NC60 | Nitric oxide-associated protein 1 OS=Hor | NOA1 | 78.458 | 2 | 54.056362 | yes | 1.031E-07 | 673.85284 | 66.329018 |
| ATISPWPGTTLNLI | 15 | Q8NC60 | Nitric oxide-associated protein 1 OS=Hor | NOA1 | 78.458 | 2 | 54.056362 | yes | 1.555E-07 | 806.45886 | 53.238113 |
| TGYGVEELISALQI | 14 | Q8NC60 | Nitric oxide-associated protein 1 OS=Hor | NOA1 | 78.458 | 2 | 54.056362 | yes | 5.222E-06 | 768.40686 | 34.798473 |
| LCAWLVSELR | 10 | Q8NCA5 | Protein FAM98A OS=Homo sapiens OX= | FAM98A | 55.273 | 2 | 53.321453 | yes | 1.435E-07 | 623.83429 | 59422.855 |
| TTISVAHLLAAR | 12 | Q8NCA5 | Protein FAM98A OS=Homo sapiens OX= | FAM98A | 55.273 | 2 | 53.321453 | yes | 1.004E-07 | 626.87225 | 16962.947 |
| TTISVAHLLAAR | 12 | Q8NCA5 | Protein FAM98A OS=Homo sapiens OX= | FAM98A | 55.273 | 2 | 53.321453 | yes | 1.861E-07 | 418.25058 | 16962.947 |
| TTISVAHLLAAR | 12 | Q8NCA5 | Protein FAM98A OS=Homo sapiens OX= | FAM98A | 55.273 | 3 | 53.321453 | yes | 1.004E-07 | 626.87225 | 16962.947 |
| TTISVAHLLAAR | 12 | Q8NCA5 | Protein FAM98A OS=Homo sapiens OX= | FAM98A | 55.273 | 3 | 53.321453 | yes | 1.861E-07 | 418.25058 | 16962.947 |
| AQEGGGSEVFQEL | 14 | Q8NCA5 | Protein FAM98A OS=Homo sapiens OX= | FAM98A | 55.273 | 2 | 53.321453 | yes | 1.258E-06 | 739.85974 | 18171.605 |
| VPPNHVGKPLLK | 12 | Q8NCA5 | Protein FAM98A OS=Homo sapiens OX= | FAM98A | 55.273 | 2 | 53.321453 | yes | 1.244E-07 | 649.90082 | 8803.4854 |
| VPPNHVGKPLLK | 12 | Q8NCA5 | Protein FAM98A OS=Homo sapiens OX= | FAM98A | 55.273 | 2 | 53.321453 | yes | 2.02E-07 | 433.60297 | 8803.4854 |
| VPPNHVGKPLLK | 12 | Q8NCA5 | Protein FAM98A OS=Homo sapiens OX= | FAM98A | 55.273 | 3 | 53.321453 | yes | 1.244E-07 | 649.90082 | 8803.4854 |
| VPPNHVGKPLLK | 12 | Q8NCA5 | Protein FAM98A OS=Homo sapiens OX= | FAM98A | 55.273 | 3 | 53.321453 | yes | 2.02E-07 | 433.60297 | 8803.4854 |
| LDVTVQSFGWSDI | 13 | Q8NCA5 | Protein FAM98A OS=Homo sapiens OX= | FAM98A | 55.273 | 2 | 53.321453 | no | 1.517E-07 | 755.37024 | 17335.613 |
| MLCVNAPPK | 9 | Q8NCA5 | Protein FAM98A OS=Homo sapiens OX= | FAM98A | 55.273 | 2 | 53.321453 | yes | 3.494E-06 | 515.26465 | 5021.4722 |
| MLCVNAPPK | 9 | Q8NCA5 | Protein FAM98A OS=Homo sapiens OX= | FAM98A | 55.273 | 2 | 53.321453 | yes | 1.118E-06 | 523.26208 | 5021.4722 |
| KAQEGGGSEVFQE | 15 | Q8NCA5 | Protein FAM98A OS=Homo sapiens OX= | FAM98A | 55.273 | 3 | 53.321453 | yes | 1.536E-07 | 536.27386 | 4955.5552 |
| KAQEGGGSEVFQE | 15 | Q8NCA5 | Protein FAM98A OS=Homo sapiens OX= | FAM98A | 55.273 | 3 | 53.321453 | yes | 9.728E-08 | 803.90723 | 4955.5552 |
| KAQEGGGSEVFQE | 15 | Q8NCA5 | Protein FAM98A OS=Homo sapiens OX= | FAM98A | 55.273 | 2 | 53.321453 | yes | 1.536E-07 | 536.27386 | 4955.5552 |
| KAQEGGGSEVFQE | 15 | Q8NCA5 | Protein FAM98A OS=Homo sapiens OX= | FAM98A | 55.273 | 2 | 53.321453 | yes | 9.728E-08 | 803.90723 | 4955.5552 |
| GPLLEDGALSQAV | 24 | Q8NCA5 | Protein FAM98A OS=Homo sapiens OX= | FAM98A | 55.273 | 3 | 53.321453 | yes | 1.004E-07 | 778.06042 | 8995.7002 |
| KPMGPAHWEK | 10 | Q8NCA5 | Protein FAM98A OS=Homo sapiens OX= | FAM98A | 55.273 | 2 | 53.321453 | yes | 9.722E-08 | 590.80023 | 4053.283 |
| KPMGPAHWEK | 10 | Q8NCA5 | Protein FAM98A OS=Homo sapiens OX= | FAM98A | 55.273 | 2 | 53.321453 | yes | 9.988E-08 | 598.79767 | 4053.283 |
| LKETLAK | 7 | Q8NCA5 | Protein FAM98A OS=Homo sapiens OX= | FAM98A | 55.273 | 2 | 53.321453 | yes | 3.184E-07 | 401.75528 | 3832.0964 |
| RLDVTVQSFGWSI | 14 | Q8NCA5 | Protein FAM98A OS=Homo sapiens OX= | FAM98A | 55.273 | 3 | 53.321453 | no | 1.102E-07 | 555.94965 | 863.37561 |
| RLDVTVQSFGWSI | 14 | Q8NCA5 | Protein FAM98A OS=Homo sapiens OX= | FAM98A | 55.273 | 3 | 53.321453 | no | 1.722E-07 | 833.42084 | 863.37561 |
| RLDVTVQSFGWSI | 14 | Q8NCA5 | Protein FAM98A OS=Homo sapiens OX= | FAM98A | 55.273 | 2 | 53.321453 | no | 1.102E-07 | 555.94965 | 863.37561 |
| RLDVTVQSFGWSI | 14 | Q8NCA5 | Protein FAM98A OS=Homo sapiens OX= | FAM98A | 55.273 | 2 | 53.321453 | no | 1.722E-07 | 833.42084 | 863.37561 |
| AGQGGGWGGR | 10 | Q8NCA5 | Protein FAM98A OS=Homo sapiens OX= | FAM98A | 55.273 | 2 | 53.321453 | yes | 2.821E-07 | 451.715 | 309.45047 |

| | | | | | | | | | | | |
|---|---|---|---|---|---|---|---|---|---|---|---|
| GICIALGMSK | 10 | Q8NCA5 | Protein FAM98A OS=Homo sapiens OX= | FAM98A | 55.273 | 2 | 53.321453 | yes | 1.711E-07 | 525.27771 | 291.48679 |
| PMGPAHWEK | 9 | Q8NCA5 | Protein FAM98A OS=Homo sapiens OX= | FAM98A | 55.273 | 2 | 53.321453 | yes | 2.111E-07 | 526.75275 | 192.71321 |
| MLCVNAPPKK | 10 | Q8NCA5 | Protein FAM98A OS=Homo sapiens OX= | FAM98A | 55.273 | 2 | 53.321453 | yes | 6.801E-07 | 579.31213 | 57.661335 |
| GGRPNEIEPPPPEM | 19 | Q8NCA5 | Protein FAM98A OS=Homo sapiens OX= | FAM98A | 55.273 | 3 | 53.321453 | no | 3.775E-06 | 719.35797 | 40.857044 |
| LPSSPLGCR | 9 | Q8NCD3 | Holliday junction recognition protein OS= | HJURP | 83.54 | 2 | 42.032635 | yes | 5.309E-07 | 493.75803 | 274.03513 |
| GGPASPGGLQGLE | 15 | Q8NCD3 | Holliday junction recognition protein OS= | HJURP | 83.54 | 2 | 42.032635 | yes | 3.226E-07 | 698.86261 | 225.29298 |
| ENFIPCSEPVK | 11 | Q8NCD3 | Holliday junction recognition protein OS= | HJURP | 83.54 | 2 | 42.032635 | yes | 4.4E-07 | 660.31866 | 130.85867 |
| AEAVFHAPK | 9 | Q8NCE0 | tRNA-splicing endonuclease subunit Sen2 | TSEN2 | 53.247 | 2 | 36.830044 | yes | 6.569E-07 | 506.26654 | 51.781525 |
| YGTDLLLYR | 9 | Q8NCE0 | tRNA-splicing endonuclease subunit Sen2 | TSEN2 | 53.247 | 2 | 36.830044 | yes | 2.399E-06 | 557.3006 | 37.071835 |
| LLRYLSVK | 8 | Q8NCE2 | Myotubularin-related protein 14 OS=Hon | MTMR14 | 72.204 | 2 | 22.864922 | yes | 3.771E-06 | 496.31857 | 764.10382 |
| SPTFAGGLFSISK | 13 | Q8NCL4 | Polypeptide N-acetylgalactosaminyltransf | GALNT6 | 71.159 | 2 | 50.463562 | yes | 9.359E-07 | 656.35083 | 181.51131 |
| DEEWELAQDQLIR | 13 | Q8NCL4 | Polypeptide N-acetylgalactosaminyltransf | GALNT6 | 71.159 | 2 | 50.463562 | yes | 9.809E-08 | 822.89685 | 109.51986 |
| SFGDISER | 8 | Q8NCL4 | Polypeptide N-acetylgalactosaminyltransf | GALNT6 | 71.159 | 2 | 50.463562 | yes | 6.463E-07 | 455.71689 | 74.831917 |
| NSGSGTCLTSQDK | 13 | Q8NCL4 | Polypeptide N-acetylgalactosaminyltransf | GALNT6 | 71.159 | 2 | 50.463562 | yes | 1.419E-07 | 677.79883 | 31.341372 |
| EKPLVAVNTR | 10 | Q8NCN4 | E3 ubiquitin-protein ligase RNF169 OS=I | RNF169 | 77.195 | 2 | 51.87644 | yes | 9.722E-08 | 563.83258 | 205.84218 |
| AAPAEPDFIFR | 11 | Q8NCN4 | E3 ubiquitin-protein ligase RNF169 OS=I | RNF169 | 77.195 | 2 | 51.87644 | yes | 2.353E-06 | 617.31677 | 129.66911 |
| ATLTSLAPEMGEE | 24 | Q8NCN4 | E3 ubiquitin-protein ligase RNF169 OS=I | RNF169 | 77.195 | 3 | 51.87644 | yes | 3.516E-07 | 819.74408 | 68.016403 |
| QLALQLQR | 8 | Q8NCN4 | E3 ubiquitin-protein ligase RNF169 OS=I | RNF169 | 77.195 | 2 | 51.87644 | yes | 1.888E-06 | 485.29562 | 63.623684 |
| EQFEGLGSTPDAK | 13 | Q8NCN4 | E3 ubiquitin-protein ligase RNF169 OS=I | RNF169 | 77.195 | 2 | 51.87644 | yes | 4.904E-07 | 689.82788 | 62.376957 |
| DDGQADSEVLGE( | 15 | Q8NCN4 | E3 ubiquitin-protein ligase RNF169 OS=I | RNF169 | 77.195 | 2 | 51.87644 | yes | 1.596E-07 | 811.34155 | 53.185043 |
| VVLAACSDFFR | 11 | Q8NCP5 | Zinc finger and BTB domain-containing p | ZBTB44 | 63.848 | 2 | 24.643824 | yes | 3.417E-06 | 642.82391 | 22.10239 |
| HQDPGDPGPGIR | 12 | Q8ND04 | Nonsense-mediated mRNA decay factor S | SMG8 | 109.684 | 2 | 48.390343 | yes | 7.663E-07 | 623.30219 | 79.873199 |
| VLEGFLDIDTK | 11 | Q8ND04 | Nonsense-mediated mRNA decay factor S | SMG8 | 109.684 | 2 | 48.390343 | yes | 1.154E-07 | 625.33734 | 65.486092 |
| EDEICVVGIFGK | 12 | Q8ND04 | Nonsense-mediated mRNA decay factor S | SMG8 | 109.684 | 2 | 48.390343 | yes | 2.758E-06 | 683.3396 | 62.678196 |
| AANYDFYQLLEEK | 13 | Q8ND04 | Nonsense-mediated mRNA decay factor S | SMG8 | 109.684 | 2 | 48.390343 | yes | 4.885E-07 | 802.38556 | 59.576958 |
| SFFDNISCDDNR | 12 | Q8ND56 | Protein LSM14 homolog A OS=Homo sa | LSM14A | 50.53 | 2 | 52.11502 | yes | 9.722E-08 | 745.30426 | 3138.5195 |
| DFDFESANAQFNK | 13 | Q8ND56 | Protein LSM14 homolog A OS=Homo sa | LSM14A | 50.53 | 2 | 52.11502 | yes | 1.033E-07 | 766.83624 | 2277.7417 |
| SSPQLDPLR | 9 | Q8ND56 | Protein LSM14 homolog A OS=Homo sa | LSM14A | 50.53 | 2 | 52.11502 | yes | 1.388E-07 | 506.77472 | 1576.351 |
| EFADFEYR | 8 | Q8ND56 | Protein LSM14 homolog A OS=Homo sa | LSM14A | 50.53 | 2 | 52.11502 | yes | 1.6E-07 | 538.73779 | 1456.1282 |
| RPTWAEER | 8 | Q8ND56 | Protein LSM14 homolog A OS=Homo sa | LSM14A | 50.53 | 2 | 52.11502 | yes | 3.592E-07 | 522.76471 | 677.66827 |
| VSRPENEQLR | 10 | Q8ND56 | Protein LSM14 homolog A OS=Homo sa | LSM14A | 50.53 | 2 | 52.11502 | yes | 1.899E-07 | 614.32562 | 458.93546 |
| SSPQLDPLRK | 10 | Q8ND56 | Protein LSM14 homolog A OS=Homo sa | LSM14A | 50.53 | 2 | 52.11502 | yes | 1.222E-07 | 570.8222 | 314.10074 |
| SPVSTRPLPSASQK | 14 | Q8ND56 | Protein LSM14 homolog A OS=Homo sa | LSM14A | 50.53 | 3 | 52.11502 | yes | 2.787E-06 | 485.60358 | 83.436493 |
| SPVSTRPLPSASQK | 14 | Q8ND56 | Protein LSM14 homolog A OS=Homo sa | LSM14A | 50.53 | 3 | 52.11502 | yes | 1.101E-07 | 727.90173 | 83.436493 |
| SPVSTRPLPSASQK | 14 | Q8ND56 | Protein LSM14 homolog A OS=Homo sa | LSM14A | 50.53 | 2 | 52.11502 | yes | 2.787E-06 | 485.60358 | 83.436493 |
| SPVSTRPLPSASQK | 14 | Q8ND56 | Protein LSM14 homolog A OS=Homo sa | LSM14A | 50.53 | 2 | 52.11502 | yes | 1.101E-07 | 727.90173 | 83.436493 |
| SFGTEDRPTDR | 11 | Q8ND56 | Protein LSM14 homolog A OS=Homo sa | LSM14A | 50.53 | 2 | 52.11502 | no | 9.722E-08 | 640.79694 | 135.06136 |
| PLPSASQK | 8 | Q8ND56 | Protein LSM14 homolog A OS=Homo sa | LSM14A | 50.53 | 2 | 52.11502 | yes | 3.214E-07 | 414.23471 | 112.63634 |
| AIMLGAKPPK | 10 | Q8NDD1 | Uncharacterized protein C1orf131 OS=H | C1orf131 | 32.611 | 2 | 50.13145 | yes | 1.02E-07 | 513.31244 | 448.737 |

| | | | | | | | | | | | |
|---|---|---|---|---|---|---|---|---|---|---|---|
| LAQETDIFK | 9 | Q8NDD1 | Uncharacterized protein C1orf131 OS=H | C1orf131 | 32.611 | 2 | 50.13145 | yes | 2.533E-07 | 532.78479 | 327.50601 |
| EQVEVVEFHSNK | 12 | Q8NDD1 | Uncharacterized protein C1orf131 OS=H | C1orf131 | 32.611 | 2 | 50.13145 | yes | 1.163E-07 | 722.85699 | 279.84283 |
| NGTLILSPVDIK | 12 | Q8NDD1 | Uncharacterized protein C1orf131 OS=H | C1orf131 | 32.611 | 2 | 50.13145 | yes | 1.037E-07 | 635.37427 | 183.871 |
| HCAPSGTPTGPEIL | 24 | Q8NDD1 | Uncharacterized protein C1orf131 OS=H | C1orf131 | 32.611 | 3 | 50.13145 | yes | 9.893E-08 | 786.74225 | 180.46277 |
| DVDTQEFNLEK | 11 | Q8NDD1 | Uncharacterized protein C1orf131 OS=H | C1orf131 | 32.611 | 2 | 50.13145 | yes | 3.591E-06 | 669.31464 | 51.027893 |
| ADGGGVVYSGTP | 14 | Q8NDF8 | Terminal nucleotidyltransferase 4B OS=F | TENT4B | 63.267 | 2 | 52.31079 | yes | 2.138E-07 | 697.34094 | 148.03221 |
| VTDEVATYR | 9 | Q8NDF8 | Terminal nucleotidyltransferase 4B OS=F | TENT4B | 63.267 | 2 | 52.31079 | yes | 1.952E-07 | 527.26422 | 127.03088 |
| VGSQDVSLESSQA | 16 | Q8NDF8 | Terminal nucleotidyltransferase 4B OS=F | TENT4B | 63.267 | 2 | 52.31079 | yes | 9.722E-08 | 795.9021 | 125.68785 |
| FQNLDMAHR | 9 | Q8NDT2 | Putative RNA-binding protein 15B OS=H | RBM15B | 97.206 | 2 | 51.17597 | yes | 4.655E-07 | 566.27185 | 159.74133 |
| QAAGVISLPVGGS | 14 | Q8NDT2 | Putative RNA-binding protein 15B OS=H | RBM15B | 97.206 | 2 | 51.17597 | yes | 9.722E-08 | 642.36951 | 82.082031 |
| ALDYYGLYDDR | 11 | Q8NDT2 | Putative RNA-binding protein 15B OS=H | RBM15B | 97.206 | 2 | 51.17597 | yes | 1.091E-07 | 682.31189 | 71.706367 |
| YGIIEEVVIK | 10 | Q8NDT2 | Putative RNA-binding protein 15B OS=H | RBM15B | 97.206 | 2 | 51.17597 | yes | 1.346E-07 | 581.83954 | 70.501923 |
| VAYVNFR | 7 | Q8NDT2 | Putative RNA-binding protein 15B OS=H | RBM15B | 97.206 | 2 | 51.17597 | yes | 9.028E-07 | 434.73743 | 44.529221 |
| TTEAEPKPLEEPK | 13 | Q8NDT2 | Putative RNA-binding protein 15B OS=H | RBM15B | 97.206 | 2 | 51.17597 | yes | 1.221E-07 | 734.88013 | 34.868607 |
| VEPVYLR | 7 | Q8NDT2 | Putative RNA-binding protein 15B OS=H | RBM15B | 97.206 | 2 | 51.17597 | yes | 4.779E-07 | 438.2529 | 34.672493 |
| GQGGAYAFLK | 10 | Q8NDT2 | Putative RNA-binding protein 15B OS=H | RBM15B | 97.206 | 2 | 51.17597 | yes | 1.365E-06 | 506.26654 | 25.882235 |
| LSHTPELGR | 9 | Q8NDT2 | Putative RNA-binding protein 15B OS=H | RBM15B | 97.206 | 2 | 51.17597 | yes | 7.019E-05 | 505.2749 | 16.331308 |
| NLTPQIDGSTLR | 12 | Q8NDV7 | Trinucleotide repeat-containing gene 6A | TNRC6A | 210.298 | 2 | 53.689587 | no | 1.584E-07 | 657.85425 | 1497.8459 |
| MDLSGGMLQDK | 11 | Q8NDV7 | Trinucleotide repeat-containing gene 6A | TNRC6A | 210.298 | 2 | 53.689587 | yes | 1.035E-07 | 597.77832 | 1284.5875 |
| SVSGWNELGK | 10 | Q8NDV7 | Trinucleotide repeat-containing gene 6A | TNRC6A | 210.298 | 2 | 53.689587 | yes | 1.133E-07 | 538.77222 | 727.06439 |
| VLSNSGWGQTPIK | 13 | Q8NDV7 | Trinucleotide repeat-containing gene 6A | TNRC6A | 210.298 | 2 | 53.689587 | yes | 1.921E-07 | 693.87244 | 692.65491 |
| YTPGSSWGESSG | 14 | Q8NDV7 | Trinucleotide repeat-containing gene 6A | TNRC6A | 210.298 | 2 | 53.689587 | yes | 1.192E-07 | 729.31824 | 657.19275 |
| HSLNIGDYNR | 10 | Q8NDV7 | Trinucleotide repeat-containing gene 6A | TNRC6A | 210.298 | 2 | 53.689587 | yes | 1.076E-07 | 594.79144 | 555.39343 |
| IGGGWGNSDAR | 11 | Q8NDV7 | Trinucleotide repeat-containing gene 6A | TNRC6A | 210.298 | 2 | 53.689587 | yes | 4.575E-07 | 545.25726 | 484.37363 |
| DSPEENVQSNK | 11 | Q8NDV7 | Trinucleotide repeat-containing gene 6A | TNRC6A | 210.298 | 2 | 53.689587 | yes | 7.495E-07 | 623.78094 | 440.06241 |
| HGAISSGFR | 9 | Q8NDV7 | Trinucleotide repeat-containing gene 6A | TNRC6A | 210.298 | 2 | 53.689587 | yes | 5.627E-07 | 466.24084 | 295.63846 |
| LTWSPGSVTNTSL | 19 | Q8NDV7 | Trinucleotide repeat-containing gene 6A | TNRC6A | 210.298 | 3 | 53.689587 | yes | 9.722E-08 | 709.7005 | 292.97168 |
| NITIMASGNTGGE | 14 | Q8NDV7 | Trinucleotide repeat-containing gene 6A | TNRC6A | 210.298 | 2 | 53.689587 | yes | 6.222E-07 | 696.84302 | 274.87811 |
| NAVVSDLSEK | 10 | Q8NDV7 | Trinucleotide repeat-containing gene 6A | TNRC6A | 210.298 | 2 | 53.689587 | yes | 2.576E-07 | 531.27728 | 242.63385 |
| QNTAWDTETSPR | 12 | Q8NDV7 | Trinucleotide repeat-containing gene 6A | TNRC6A | 210.298 | 2 | 53.689587 | yes | 9.722E-08 | 703.32074 | 218.7863 |
| NITAPSRPPPGLTG | 28 | Q8NDV7 | Trinucleotide repeat-containing gene 6A | TNRC6A | 210.298 | 4 | 53.689587 | yes | 1.665E-07 | 750.15308 | 210.53908 |
| QFSNISFSR | 9 | Q8NDV7 | Trinucleotide repeat-containing gene 6A | TNRC6A | 210.298 | 2 | 53.689587 | yes | 1.143E-07 | 543.27234 | 197.09248 |
| GPGSRPQISK | 10 | Q8NDV7 | Trinucleotide repeat-containing gene 6A | TNRC6A | 210.298 | 2 | 53.689587 | yes | 9.753E-08 | 513.78815 | 172.07893 |
| SNQSLGWGDSSK | 12 | Q8NDV7 | Trinucleotide repeat-containing gene 6A | TNRC6A | 210.298 | 2 | 53.689587 | yes | 9.722E-08 | 633.29144 | 143.92361 |
| PLSVQQQMMQQS | 13 | Q8NDV7 | Trinucleotide repeat-containing gene 6A | TNRC6A | 210.298 | 2 | 53.689587 | yes | 2.666E-05 | 780.8847 | 133.56978 |
| MEIDDGTSAWGD | 15 | Q8NDV7 | Trinucleotide repeat-containing gene 6A | TNRC6A | 210.298 | 2 | 53.689587 | yes | 1.009E-07 | 804.84595 | 122.44585 |
| IDQHTLLQSIVNR | 13 | Q8NDV7 | Trinucleotide repeat-containing gene 6A | TNRC6A | 210.298 | 2 | 53.689587 | yes | 1.065E-07 | 768.92828 | 118.80135 |
| DLVQEEEQLMEE | 13 | Q8NDV7 | Trinucleotide repeat-containing gene 6A | TNRC6A | 210.298 | 2 | 53.689587 | yes | 5.615E-07 | 810.37708 | 110.40832 |
| AAWNDSQK | 8 | Q8NDV7 | Trinucleotide repeat-containing gene 6A | TNRC6A | 210.298 | 2 | 53.689587 | yes | 1.004E-07 | 460.21704 | 109.89969 |

| | | | | | | | | | | |
|---|---|---|---|---|---|---|---|---|---|---|
| LGSLDCSHSFSSR | 13 | Q8NDV7 | Trinucleotide repeat-containing gene 6A I | TNRC6A | 210.298 | 2 | 53.689587 | yes | 1.627E-07 | 726.83044 | 97.060104 |
| QEEAWINPFVK | 11 | Q8NDV7 | Trinucleotide repeat-containing gene 6A I | TNRC6A | 210.298 | 2 | 53.689587 | yes | 9.722E-08 | 680.84845 | 93.750595 |
| LLAQQQR | 7 | Q8NDV7 | Trinucleotide repeat-containing gene 6A I | TNRC6A | 210.298 | 2 | 53.689587 | yes | 2.635E-07 | 428.7536 | 90.349884 |
| NNHWGEANK | 9 | Q8NDV7 | Trinucleotide repeat-containing gene 6A I | TNRC6A | 210.298 | 2 | 53.689587 | yes | 1.804E-07 | 535.24414 | 85.992271 |
| ITNWLVLK | 8 | Q8NDV7 | Trinucleotide repeat-containing gene 6A I | TNRC6A | 210.298 | 2 | 53.689587 | yes | 1.328E-07 | 493.8053 | 71.002853 |
| QLDPNLLVK | 9 | Q8NDV7 | Trinucleotide repeat-containing gene 6A I | TNRC6A | 210.298 | 2 | 53.689587 | yes | 9.722E-08 | 520.31097 | 45.161373 |
| TLCMQHGPLITFHI | 24 | Q8NDV7 | Trinucleotide repeat-containing gene 6A I | TNRC6A | 210.298 | 4 | 53.689587 | yes | 1.314E-07 | 685.61383 | 24.771816 |
| GPVSSTSDSSTNCI | 14 | Q8NDV7 | Trinucleotide repeat-containing gene 6A I | TNRC6A | 210.298 | 2 | 53.689587 | yes | 1.309E-07 | 713.80939 | 24.476555 |
| TGGTVESDGSTES | 16 | Q8NDV7 | Trinucleotide repeat-containing gene 6A I | TNRC6A | 210.298 | 2 | 53.689587 | yes | 5.305E-07 | 770.83972 | 19.031029 |
| CDLLQAMPR | 9 | Q8NDZ4 | Divergent protein kinase domain 2A OS= | DIPK2A | 49.482 | 2 | 52.613766 | yes | 1.994E-07 | 552.27045 | 229.38629 |
| VIIVDAENVLVAD | 14 | Q8NDZ4 | Divergent protein kinase domain 2A OS= | DIPK2A | 49.482 | 2 | 52.613766 | yes | 9.722E-08 | 749.42981 | 226.75148 |
| LEALLDECANPK | 12 | Q8NDZ4 | Divergent protein kinase domain 2A OS= | DIPK2A | 49.482 | 2 | 52.613766 | yes | 1.49E-07 | 686.84247 | 172.6855 |
| ELAQLDQSICK | 11 | Q8NDZ4 | Divergent protein kinase domain 2A OS= | DIPK2A | 49.482 | 2 | 52.613766 | yes | 1.589E-07 | 652.82941 | 148.47403 |
| GTSGGLLHDPPSEI | 16 | Q8NDZ4 | Divergent protein kinase domain 2A OS= | DIPK2A | 49.482 | 2 | 52.613766 | yes | 4.983E-07 | 789.90973 | 78.071693 |
| NVYFAQYGEPR | 11 | Q8NDZ4 | Divergent protein kinase domain 2A OS= | DIPK2A | 49.482 | 2 | 52.613766 | yes | 2.041E-07 | 672.32257 | 64.602852 |
| VIIVDAENVLVAD | 15 | Q8NDZ4 | Divergent protein kinase domain 2A OS= | DIPK2A | 49.482 | 3 | 52.613766 | yes | 9.722E-08 | 551.98932 | 29.743217 |
| VIIVDAENVLVAD | 15 | Q8NDZ4 | Divergent protein kinase domain 2A OS= | DIPK2A | 49.482 | 3 | 52.613766 | yes | 7.784E-07 | 827.48035 | 29.743217 |
| VIIVDAENVLVAD | 15 | Q8NDZ4 | Divergent protein kinase domain 2A OS= | DIPK2A | 49.482 | 2 | 52.613766 | yes | 9.722E-08 | 551.98932 | 29.743217 |
| VIIVDAENVLVAD | 15 | Q8NDZ4 | Divergent protein kinase domain 2A OS= | DIPK2A | 49.482 | 2 | 52.613766 | yes | 7.784E-07 | 827.48035 | 29.743217 |
| LLDFLNVK | 8 | Q8NDZ4 | Divergent protein kinase domain 2A OS= | DIPK2A | 49.482 | 2 | 52.613766 | yes | 4.619E-07 | 481.28949 | 36.261684 |
| LLTPEAVEGWSDL | 20 | Q8NDZ4 | Divergent protein kinase domain 2A OS= | DIPK2A | 49.482 | 3 | 52.613766 | yes | 1.519E-07 | 765.37933 | 24.30191 |
| TDNGNNLLPFQDR | 13 | Q8NE35 | Cytoplasmic polyadenylation element-bin | CPEB3 | 76.015 | 2 | 45.730064 | yes | 2.566E-07 | 752.36298 | 217.51271 |
| SAAAAYGHQPIMT | 15 | Q8NE35 | Cytoplasmic polyadenylation element-bin | CPEB3 | 76.015 | 2 | 45.730064 | yes | 1.759E-07 | 766.87994 | 37.098083 |
| SPASPSQAPYAQR | 13 | Q8NE35 | Cytoplasmic polyadenylation element-bin | CPEB3 | 76.015 | 2 | 45.730064 | yes | 6.491E-07 | 680.33624 | 26.628641 |
| MEETPTEYLQR | 11 | Q8NE71 | ATP-binding cassette sub-family F memb | ABCF1 | 95.926 | 2 | 53.07616 | yes | 1.006E-07 | 698.82428 | 2420.6467 |
| MEETPTEYLQR | 11 | Q8NE71 | ATP-binding cassette sub-family F memb | ABCF1 | 95.926 | 2 | 53.07616 | yes | 4.646E-07 | 706.82178 | 2420.6467 |
| VVFAELACR | 9 | Q8NE71 | ATP-binding cassette sub-family F memb | ABCF1 | 95.926 | 2 | 53.07616 | yes | 2.147E-07 | 532.78149 | 3410.9207 |
| IGFFNQQYAEQLR | 13 | Q8NE71 | ATP-binding cassette sub-family F memb | ABCF1 | 95.926 | 2 | 53.07616 | yes | 1.418E-07 | 807.40717 | 3343.3938 |
| GFNLPYQDAR | 10 | Q8NE71 | ATP-binding cassette sub-family F memb | ABCF1 | 95.926 | 2 | 53.07616 | yes | 5.551E-07 | 590.79089 | 2893.2097 |
| NLDFGIDMDSR | 11 | Q8NE71 | ATP-binding cassette sub-family F memb | ABCF1 | 95.926 | 2 | 53.07616 | yes | 2.892E-07 | 641.79022 | 2859.4375 |
| GAVIVVSHDAR | 11 | Q8NE71 | ATP-binding cassette sub-family F memb | ABCF1 | 95.926 | 2 | 53.07616 | yes | 1.032E-07 | 562.31458 | 2524.0977 |
| ADDPYAHLSK | 10 | Q8NE71 | ATP-binding cassette sub-family F memb | ABCF1 | 95.926 | 2 | 53.07616 | yes | 1.467E-07 | 558.76965 | 1908.567 |
| EVLEALGEVMVSI | 13 | Q8NE71 | ATP-binding cassette sub-family F memb | ABCF1 | 95.926 | 2 | 53.07616 | yes | 9.854E-08 | 716.37921 | 1885.9105 |
| AVSEEQQPALK | 11 | Q8NE71 | ATP-binding cassette sub-family F memb | ABCF1 | 95.926 | 2 | 53.07616 | yes | 9.91E-08 | 600.31696 | 1862.8025 |
| LTPTHGEMR | 9 | Q8NE71 | ATP-binding cassette sub-family F memb | ABCF1 | 95.926 | 2 | 53.07616 | yes | 2.545E-07 | 521.26093 | 847.13916 |
| LTPTHGEMR | 9 | Q8NE71 | ATP-binding cassette sub-family F memb | ABCF1 | 95.926 | 2 | 53.07616 | yes | 4.744E-07 | 529.25836 | 847.13916 |
| FAALDNEEEDKEE | 17 | Q8NE71 | ATP-binding cassette sub-family F memb | ABCF1 | 95.926 | 3 | 53.07616 | yes | 6.252E-07 | 674.65649 | 1613.0825 |
| NQDEESQEAPELL | 14 | Q8NE71 | ATP-binding cassette sub-family F memb | ABCF1 | 95.926 | 2 | 53.07616 | yes | 1.054E-07 | 815.38373 | 1521.4946 |
| TFFEELAVEDK | 11 | Q8NE71 | ATP-binding cassette sub-family F memb | ABCF1 | 95.926 | 2 | 53.07616 | yes | 1.108E-07 | 664.32446 | 1156.8651 |

| | | | | | | | | | | | |
|---|---|---|---|---|---|---|---|---|---|---|---|
| FSISAHGK | 8 | Q8NE71 | ATP-binding cassette sub-family F memb | ABCF1 | 95.926 | 2 | 53.07616 | yes | 2.428E-07 | 423.72705 | 1069.3578 |
| DVDDDGEEK | 9 | Q8NE71 | ATP-binding cassette sub-family F memb | ABCF1 | 95.926 | 2 | 53.07616 | yes | 5.945E-07 | 511.20145 | 746.78424 |
| ICIVGPNGVGK | 11 | Q8NE71 | ATP-binding cassette sub-family F memb | ABCF1 | 95.926 | 2 | 53.07616 | yes | 9.79E-08 | 557.30786 | 502.48624 |
| ELFVNADLYIVAG | 14 | Q8NE71 | ATP-binding cassette sub-family F memb | ABCF1 | 95.926 | 2 | 53.07616 | yes | 1.055E-07 | 790.42755 | 395.45758 |
| QAMLENASDIK | 11 | Q8NE71 | ATP-binding cassette sub-family F memb | ABCF1 | 95.926 | 2 | 53.07616 | yes | 2.268E-07 | 610.30298 | 165.87274 |
| KNQDEESQEAPEL | 15 | Q8NE71 | ATP-binding cassette sub-family F memb | ABCF1 | 95.926 | 3 | 53.07616 | yes | 1.606E-07 | 586.62323 | 121.21835 |
| KLSVPTSDEEDEV | 19 | Q8NE71 | ATP-binding cassette sub-family F memb | ABCF1 | 95.926 | 3 | 53.07616 | yes | 1.166E-07 | 698.69531 | 115.69086 |
| ILAGLGFDPEMQN | 14 | Q8NE71 | ATP-binding cassette sub-family F memb | ABCF1 | 95.926 | 2 | 53.07616 | yes | 3.134E-07 | 780.89557 | 108.08144 |
| STLLLLLTGK | 10 | Q8NE71 | ATP-binding cassette sub-family F memb | ABCF1 | 95.926 | 2 | 53.07616 | yes | 0.0001533 | 529.8446 | 73.108452 |
| KTFFEELAVEDK | 12 | Q8NE71 | ATP-binding cassette sub-family F memb | ABCF1 | 95.926 | 2 | 53.07616 | yes | 9.722E-08 | 728.37195 | 67.025917 |
| LSVPTSDEEDEVP | 18 | Q8NE71 | ATP-binding cassette sub-family F memb | ABCF1 | 95.926 | 3 | 53.07616 | yes | 3.999E-07 | 655.99695 | 66.799675 |
| TLLIVSHDQGFLDI | 26 | Q8NE71 | ATP-binding cassette sub-family F memb | ABCF1 | 95.926 | 4 | 53.07616 | yes | 1.22E-07 | 749.38019 | 59.224407 |
| DVDDDGEEKELM | 14 | Q8NE71 | ATP-binding cassette sub-family F memb | ABCF1 | 95.926 | 2 | 53.07616 | yes | 3.182E-07 | 840.35687 | 38.654629 |
| ATGAAAAEAK | 10 | Q8NE71 | ATP-binding cassette sub-family F memb | ABCF1 | 95.926 | 2 | 53.07616 | yes | 0.0001313 | 430.72726 | 11.666981 |
| FGLESHAHTIQICK | 14 | Q8NE71 | ATP-binding cassette sub-family F memb | ABCF1 | 95.926 | 2 | 53.07616 | yes | 2.779E-06 | 820.91431 | 4.6802568 |
| LRDPLQVHLPLR | 12 | Q8NE86 | Calcium uniporter protein, mitochondrial | MCU | 39.867 | 3 | 46.214413 | yes | 1.619E-07 | 486.29642 | 153.14287 |
| DAIAQAEMDLK | 11 | Q8NE86 | Calcium uniporter protein, mitochondrial | MCU | 39.867 | 2 | 46.214413 | yes | 4.891E-07 | 602.79755 | 67.202263 |
| NLQEQVMAVTAQ | 14 | Q8NEJ9 | Neuroguidin OS=Homo sapiens OX=960 | NGDN | 35.895 | 2 | 52.546604 | yes | 9.722E-08 | 779.9165 | 180.2915 |
| TAVTGSLSENDPL | 14 | Q8NEJ9 | Neuroguidin OS=Homo sapiens OX=960 | NGDN | 35.895 | 2 | 52.546604 | yes | 3.642E-07 | 730.37299 | 169.37921 |
| INYEESMMVR | 10 | Q8NEJ9 | Neuroguidin OS=Homo sapiens OX=960 | NGDN | 35.895 | 2 | 52.546604 | yes | 9.728E-08 | 636.29156 | 162.78403 |
| EQYSDAPEEIR | 11 | Q8NEJ9 | Neuroguidin OS=Homo sapiens OX=960 | NGDN | 35.895 | 2 | 52.546604 | yes | 1.859E-07 | 668.80444 | 160.23032 |
| LVPVHYDETEAER | 13 | Q8NEJ9 | Neuroguidin OS=Homo sapiens OX=960 | NGDN | 35.895 | 2 | 52.546604 | yes | 1.264E-07 | 779.3808 | 144.88995 |
| ASGGSLQGHDAVI | 14 | Q8NEJ9 | Neuroguidin OS=Homo sapiens OX=960 | NGDN | 35.895 | 2 | 52.546604 | yes | 1.933E-07 | 684.35492 | 48.866535 |
| DNALVELDHR | 10 | Q8NEL9 | Phospholipase DDHD1 OS=Homo sapier | DDHD1 | 100.436 | 2 | 39.983227 | yes | 1.29E-06 | 591.29907 | 85.659073 |
| LYEQLLQK | 8 | Q8NEL9 | Phospholipase DDHD1 OS=Homo sapier | DDHD1 | 100.436 | 2 | 39.983227 | yes | 4.272E-07 | 517.79767 | 44.465069 |
| LYSLFCSR | 8 | Q8NEL9 | Phospholipase DDHD1 OS=Homo sapier | DDHD1 | 100.436 | 2 | 39.983227 | yes | 2.628E-06 | 523.26038 | 37.464138 |
| HYGESITNIGK | 11 | Q8NEL9 | Phospholipase DDHD1 OS=Homo sapier | DDHD1 | 100.436 | 2 | 39.983227 | yes | 3.42E-06 | 609.80933 | 35.114544 |
| QAAGMSQETFEEK | 13 | Q8NEM0 | Microcephalin OS=Homo sapiens OX=96 | MCPH1 | 92.85 | 2 | 53.848885 | yes | 9.722E-08 | 728.32465 | 46.146729 |
| SDVCISSLVLK | 11 | Q8NEM0 | Microcephalin OS=Homo sapiens OX=96 | MCPH1 | 92.85 | 2 | 53.848885 | yes | 5.532E-07 | 610.83136 | 44.684238 |
| ATSSCVTSAPEEAI | 15 | Q8NEM0 | Microcephalin OS=Homo sapiens OX=96 | MCPH1 | 92.85 | 2 | 53.848885 | yes | 2.167E-07 | 789.87506 | 43.230511 |
| TSIFEMSDFSCVGK | 14 | Q8NEM0 | Microcephalin OS=Homo sapiens OX=96 | MCPH1 | 92.85 | 2 | 53.848885 | yes | 9.722E-08 | 804.35767 | 35.529305 |
| TFTTQLVDMGAK | 12 | Q8NEM0 | Microcephalin OS=Homo sapiens OX=96 | MCPH1 | 92.85 | 2 | 53.848885 | yes | 9.722E-08 | 656.33429 | 31.01741 |
| TVDITNFTAK | 10 | Q8NEM0 | Microcephalin OS=Homo sapiens OX=96 | MCPH1 | 92.85 | 2 | 53.848885 | yes | 9.722E-08 | 555.29547 | 21.653694 |
| VVNQYQELVQNE | 14 | Q8NEM7 | Transcription factor SPT20 homolog OS= | SUPT20H | 85.79 | 2 | 42.118168 | yes | 3.184E-07 | 831.42828 | 27.150124 |
| SPCNLAIPSEVDVE | 15 | Q8NEM7 | Transcription factor SPT20 homolog OS= | SUPT20H | 85.79 | 2 | 42.118168 | yes | 5.932E-07 | 829.40869 | 26.044876 |
| LIVAPDADAR | 10 | Q8NEW0 | Zinc transporter 7 OS=Homo sapiens OX | SLC30A7 | 41.626 | 2 | 44.57663 | yes | 2.144E-07 | 520.79041 | 224.95828 |
| DNDAFSYGYVR | 11 | Q8NEW0 | Zinc transporter 7 OS=Homo sapiens OX | SLC30A7 | 41.626 | 2 | 44.57663 | yes | 2.04E-06 | 653.78857 | 26.388626 |
| DTLPWPSAQQDQS | 14 | Q8NEY1 | Neuron navigator 1 OS=Homo sapiens O | NAV1 | 202.472 | 2 | 53.857594 | yes | 1.1E-07 | 800.88373 | 434.09143 |
| AVALDSDNISLK | 12 | Q8NEY1 | Neuron navigator 1 OS=Homo sapiens O | NAV1 | 202.472 | 2 | 53.857594 | yes | 3.316E-07 | 623.33789 | 422.59863 |

| | | | | | | | | | | | |
|---|---|---|---|---|---|---|---|---|---|---|---|
| SLTNLSFLTDSEK | 13 | Q8NEY1 | Neuron navigator 1 OS=Homo sapiens O. | NAV1 | 202.472 | 2 | 53.857594 | yes | 1.017E-07 | 727.87231 | 328.54056 |
| VPGGPPASNLR | 11 | Q8NEY1 | Neuron navigator 1 OS=Homo sapiens O. | NAV1 | 202.472 | 2 | 53.857594 | yes | 9.405E-07 | 532.79602 | 321.47852 |
| TVQNVLDLR | 9 | Q8NEY1 | Neuron navigator 1 OS=Homo sapiens O. | NAV1 | 202.472 | 2 | 53.857594 | yes | 4.158E-07 | 529.30365 | 234.47018 |
| GPYAEVKPLSK | 11 | Q8NEY1 | Neuron navigator 1 OS=Homo sapiens O. | NAV1 | 202.472 | 2 | 53.857594 | yes | 9.722E-08 | 594.83478 | 216.59572 |
| LQAGDAPSVGGSC | 14 | Q8NEY1 | Neuron navigator 1 OS=Homo sapiens O. | NAV1 | 202.472 | 2 | 53.857594 | yes | 9.727E-08 | 687.82513 | 209.84091 |
| YQLQSQEETK | 10 | Q8NEY1 | Neuron navigator 1 OS=Homo sapiens O. | NAV1 | 202.472 | 2 | 53.857594 | yes | 1.418E-07 | 627.30408 | 209.75449 |
| TTPAPVNQTDR | 11 | Q8NEY1 | Neuron navigator 1 OS=Homo sapiens O. | NAV1 | 202.472 | 2 | 53.857594 | yes | 1.128E-07 | 600.30438 | 195.43825 |
| GQLTNIVSPTAATT | 16 | Q8NEY1 | Neuron navigator 1 OS=Homo sapiens O. | NAV1 | 202.472 | 2 | 53.857594 | yes | 1.684E-07 | 813.94409 | 188.50665 |
| HLAETAEEK | 9 | Q8NEY1 | Neuron navigator 1 OS=Homo sapiens O. | NAV1 | 202.472 | 2 | 53.857594 | yes | 2.684E-06 | 514.25635 | 165.57272 |
| LAEYLVER | 8 | Q8NEY1 | Neuron navigator 1 OS=Homo sapiens O. | NAV1 | 202.472 | 2 | 53.857594 | yes | 4.391E-07 | 496.7742 | 163.84465 |
| CPYIIGTTNQPVK | 13 | Q8NEY1 | Neuron navigator 1 OS=Homo sapiens O. | NAV1 | 202.472 | 2 | 53.857594 | yes | 1.11E-07 | 745.88721 | 155.88562 |
| LQEAANYIESPDR | 13 | Q8NEY1 | Neuron navigator 1 OS=Homo sapiens O. | NAV1 | 202.472 | 2 | 53.857594 | yes | 3.921E-07 | 753.36517 | 137.86101 |
| GVNNISVSLK | 10 | Q8NEY1 | Neuron navigator 1 OS=Homo sapiens O. | NAV1 | 202.472 | 2 | 53.857594 | yes | 1.973E-07 | 515.79822 | 127.38228 |
| SGYMSDSDLMGK | 12 | Q8NEY1 | Neuron navigator 1 OS=Homo sapiens O. | NAV1 | 202.472 | 2 | 53.857594 | yes | 1.847E-07 | 645.77069 | 124.93936 |
| MDPASTLGLSTESI | 23 | Q8NEY1 | Neuron navigator 1 OS=Homo sapiens O. | NAV1 | 202.472 | 3 | 53.857594 | yes | 1.088E-07 | 810.73718 | 108.78699 |
| SFVKPPSLANLDK | 13 | Q8NEY1 | Neuron navigator 1 OS=Homo sapiens O. | NAV1 | 202.472 | 2 | 53.857594 | yes | 2.634E-06 | 708.39825 | 102.88782 |
| KLEYDSGSLK | 10 | Q8NEY1 | Neuron navigator 1 OS=Homo sapiens O. | NAV1 | 202.472 | 2 | 53.857594 | yes | 9.722E-08 | 570.30078 | 102.50104 |
| TPLAPLAPNLGK | 12 | Q8NEY1 | Neuron navigator 1 OS=Homo sapiens O. | NAV1 | 202.472 | 2 | 53.857594 | yes | 3.804E-07 | 596.35846 | 100.44555 |
| KPISLGHPGSLK | 12 | Q8NEY1 | Neuron navigator 1 OS=Homo sapiens O. | NAV1 | 202.472 | 2 | 53.857594 | yes | 1.061E-07 | 617.36932 | 88.575768 |
| VLDAEPPEMPPCR | 13 | Q8NEY1 | Neuron navigator 1 OS=Homo sapiens O. | NAV1 | 202.472 | 2 | 53.857594 | yes | 2.244E-06 | 755.85504 | 88.441368 |
| QQEFFLGCSK | 10 | Q8NEY1 | Neuron navigator 1 OS=Homo sapiens O. | NAV1 | 202.472 | 2 | 53.857594 | yes | 9.926E-08 | 622.29242 | 83.500854 |
| LQHGSTETASPSIK | 14 | Q8NEY1 | Neuron navigator 1 OS=Homo sapiens O. | NAV1 | 202.472 | 2 | 53.857594 | yes | 1.662E-07 | 728.37555 | 73.818954 |
| LAELPPTPLR | 10 | Q8NEY1 | Neuron navigator 1 OS=Homo sapiens O. | NAV1 | 202.472 | 2 | 53.857594 | yes | 2.058E-07 | 553.83203 | 73.569756 |
| LEALNSAHQLDQL | 14 | Q8NEY1 | Neuron navigator 1 OS=Homo sapiens O. | NAV1 | 202.472 | 2 | 53.857594 | yes | 1.745E-07 | 804.42865 | 55.090191 |
| AHYSHTMPMR | 10 | Q8NEY1 | Neuron navigator 1 OS=Homo sapiens O. | NAV1 | 202.472 | 2 | 53.857594 | yes | 5.211E-07 | 615.77899 | 49.955067 |
| MLDEAVFQVFK | 11 | Q8NEY1 | Neuron navigator 1 OS=Homo sapiens O. | NAV1 | 202.472 | 2 | 53.857594 | yes | 1.051E-07 | 663.84174 | 49.089394 |
| SNSIPTHEAAFELY | 30 | Q8NEY1 | Neuron navigator 1 OS=Homo sapiens O. | NAV1 | 202.472 | 4 | 53.857594 | yes | 2.644E-07 | 802.89496 | 45.14962 |
| SEGTPAWYMHGEI | 13 | Q8NEY1 | Neuron navigator 1 OS=Homo sapiens O. | NAV1 | 202.472 | 2 | 53.857594 | yes | 1.042E-06 | 760.83301 | 44.006496 |
| PVSSSIDPSLLSTK | 14 | Q8NEY1 | Neuron navigator 1 OS=Homo sapiens O. | NAV1 | 202.472 | 2 | 53.857594 | yes | 4.904E-06 | 715.8905 | 43.84507 |
| TPLAPLAPNLGKPS | 15 | Q8NEY1 | Neuron navigator 1 OS=Homo sapiens O. | NAV1 | 202.472 | 2 | 53.857594 | yes | 2.388E-07 | 766.45135 | 32.375774 |
| LFHANEEEEPEKK | 13 | Q8NEY1 | Neuron navigator 1 OS=Homo sapiens O. | NAV1 | 202.472 | 2 | 53.857594 | yes | 3.796E-07 | 800.38611 | 28.796713 |
| GGPRPVSSSIDPSL | 18 | Q8NEY1 | Neuron navigator 1 OS=Homo sapiens O. | NAV1 | 202.472 | 3 | 53.857594 | yes | 2.116E-06 | 599.995 | 10.892677 |
| VLDWVPK | 7 | Q8NEY1 | Neuron navigator 1 OS=Homo sapiens O. | NAV1 | 202.472 | 2 | 53.857594 | yes | 5.458E-06 | 428.75 | 9.3676319 |
| PMMQHYISLLLK | 12 | Q8NEY1 | Neuron navigator 1 OS=Homo sapiens O. | NAV1 | 202.472 | 2 | 53.857594 | yes | 4.942E-05 | 737.40143 | 1.8152268 |
| SFYSSHYAR | 9 | Q8NEY8 | Periphilin-1 OS=Homo sapiens OX=9606 | PPHLN1 | 52.738 | 2 | 53.319622 | yes | 1.022E-07 | 559.25671 | 846.17767 |
| VLDKPSR | 7 | Q8NEY8 | Periphilin-1 OS=Homo sapiens OX=9606 | PPHLN1 | 52.738 | 2 | 53.319622 | yes | 2.484E-07 | 407.74271 | 682.12213 |
| ERPVQSLK | 8 | Q8NEY8 | Periphilin-1 OS=Homo sapiens OX=9606 | PPHLN1 | 52.738 | 2 | 53.319622 | yes | 1.691E-07 | 478.77982 | 509.13757 |
| YYSHVDYR | 8 | Q8NEY8 | Periphilin-1 OS=Homo sapiens OX=9606 | PPHLN1 | 52.738 | 2 | 53.319622 | yes | 9.722E-08 | 551.75128 | 492.93051 |
| SYSFHQSQHR | 10 | Q8NEY8 | Periphilin-1 OS=Homo sapiens OX=9606 | PPHLN1 | 52.738 | 2 | 53.319622 | yes | 1.043E-07 | 638.79449 | 332.76523 |

| | | | | | | | | | | | |
|---|---|---|---|---|---|---|---|---|---|---|---|
| SVRPGASYK | 9 | Q8NEY8 | Periphilin-1 OS=Homo sapiens OX=960( | PPHLN1 | 52.738 | 2 | 53.319622 | yes | 6.393E-07 | 482.76416 | 223.20981 |
| MLIEKDPSLEK | 11 | Q8NEY8 | Periphilin-1 OS=Homo sapiens OX=960( | PPHLN1 | 52.738 | 2 | 53.319622 | yes | 9.722E-08 | 651.85229 | 212.87547 |
| DTSPSSGSAVSSSK | 14 | Q8NEY8 | Periphilin-1 OS=Homo sapiens OX=960( | PPHLN1 | 52.738 | 2 | 53.319622 | yes | 9.722E-08 | 648.79932 | 169.70251 |
| QNLHEIESAGQTW | 20 | Q8NEY8 | Periphilin-1 OS=Homo sapiens OX=960( | PPHLN1 | 52.738 | 3 | 53.319622 | yes | 5.123E-07 | 773.06097 | 143.34506 |
| EIEQVYR | 7 | Q8NEY8 | Periphilin-1 OS=Homo sapiens OX=960( | PPHLN1 | 52.738 | 2 | 53.319622 | yes | 9.722E-08 | 468.74289 | 141.85413 |
| TKEIEQVYR | 9 | Q8NEY8 | Periphilin-1 OS=Homo sapiens OX=960( | PPHLN1 | 52.738 | 2 | 53.319622 | yes | 2.617E-07 | 583.31421 | 108.24348 |
| TTGAPDITWGMLK | 13 | Q8NEY8 | Periphilin-1 OS=Homo sapiens OX=960( | PPHLN1 | 52.738 | 2 | 53.319622 | yes | 3.256E-07 | 695.85541 | 83.897499 |
| QDCETFGMVVK | 11 | Q8NEY8 | Periphilin-1 OS=Homo sapiens OX=960( | PPHLN1 | 52.738 | 2 | 53.319622 | yes | 3.989E-07 | 657.29688 | 73.535088 |
| KPPLLDR | 7 | Q8NEY8 | Periphilin-1 OS=Homo sapiens OX=960( | PPHLN1 | 52.738 | 2 | 53.319622 | yes | 9.722E-08 | 419.76089 | 61.482166 |
| VVAEELENVR | 10 | Q8NEZ5 | F-box only protein 22 OS=Homo sapiens | FBXO22 | 44.509 | 2 | 48.820656 | yes | 1.101E-07 | 579.31171 | 106.00315 |
| STFVLSNLAEVVE | 14 | Q8NEZ5 | F-box only protein 22 OS=Homo sapiens | FBXO22 | 44.509 | 2 | 48.820656 | yes | 1.608E-07 | 782.42249 | 48.937378 |
| VVLVFGYNCCK | 11 | Q8NEZ5 | F-box only protein 22 OS=Homo sapiens | FBXO22 | 44.509 | 2 | 48.820656 | yes | 2.182E-06 | 679.83344 | 13.457096 |
| ITFADFHR | 8 | Q8NF37 | Lysophosphatidylcholine acyltransferase | LPCAT1 | 59.152 | 2 | 50.683308 | yes | 1.591E-07 | 503.75888 | 132.56787 |
| TLDTIQLAFK | 10 | Q8NF37 | Lysophosphatidylcholine acyltransferase | LPCAT1 | 59.152 | 2 | 50.683308 | yes | 6.003E-07 | 575.32935 | 99.794846 |
| ECVVALSVVCR | 11 | Q8NF37 | Lysophosphatidylcholine acyltransferase | LPCAT1 | 59.152 | 2 | 50.683308 | yes | 3.953E-07 | 646.32849 | 57.777729 |
| LPADTCLLEFAR | 12 | Q8NF37 | Lysophosphatidylcholine acyltransferase | LPCAT1 | 59.152 | 2 | 50.683308 | yes | 9.728E-08 | 703.36084 | 57.068726 |
| KVVDFLLK | 8 | Q8NF37 | Lysophosphatidylcholine acyltransferase | LPCAT1 | 59.152 | 2 | 50.683308 | yes | 2.093E-07 | 481.30768 | 23.603611 |
| AFHEIYR | 7 | Q8NF91 | Nesprin-1 OS=Homo sapiens OX=9606 ( | SYNE1 | 1011.086 | 2 | 39.04974 | yes | 4.82E-07 | 468.24033 | 163.18472 |
| RMQNLNR | 7 | Q8NF91 | Nesprin-1 OS=Homo sapiens OX=9606 ( | SYNE1 | 1011.086 | 2 | 39.04974 | yes | 3.306E-06 | 466.24817 | 84.979065 |
| ALGGEGWEK | 9 | Q8NFF5 | FAD synthase OS=Homo sapiens OX=96 | FLAD1 | 65.266 | 2 | 47.93255 | yes | 2.554E-06 | 473.73508 | 98.996346 |
| FPLVSVR | 7 | Q8NFF5 | FAD synthase OS=Homo sapiens OX=96 | FLAD1 | 65.266 | 2 | 47.93255 | yes | 3.936E-06 | 409.25015 | 47.580513 |
| PAYLLENEEEER | 12 | Q8NFF5 | FAD synthase OS=Homo sapiens OX=96 | FLAD1 | 65.266 | 2 | 47.93255 | yes | 1.23E-07 | 746.35175 | 39.554619 |
| HPQLEAVLMGTR | 12 | Q8NFF5 | FAD synthase OS=Homo sapiens OX=96 | FLAD1 | 65.266 | 2 | 47.93255 | yes | 1.69E-06 | 676.36121 | 26.119814 |
| VFEAEQFGCPQR | 12 | Q8NFG4 | Folliculin OS=Homo sapiens OX=9606 ( | FLCN | 64.474 | 2 | 42.438244 | yes | 3.03E-07 | 734.33789 | 21.469122 |
| IEITSLLPSR | 10 | Q8NFH3 | Nucleoporin Nup43 OS=Homo sapiens O | NUP43 | 42.151 | 2 | 41.518654 | yes | 4.262E-07 | 564.83478 | 111.96663 |
| VPLHCVDR | 8 | Q8NFH3 | Nucleoporin Nup43 OS=Homo sapiens O | NUP43 | 42.151 | 2 | 41.518654 | yes | 3.468E-07 | 498.25818 | 50.121143 |
| TLPLCVIGGDHK | 12 | Q8NFH4 | Nucleoporin Nup37 OS=Homo sapiens O | NUP37 | 36.708 | 2 | 47.352512 | yes | 1.348E-07 | 655.35028 | 181.43198 |
| TLGTPTQPGSTPR | 13 | Q8NFH5 | Nucleoporin NUP35 OS=Homo sapiens ( | NUP35 | 34.774 | 2 | 51.766922 | yes | 9.722E-08 | 656.84644 | 76.298462 |
| CALSSPSLAFTPPII | 15 | Q8NFH5 | Nucleoporin NUP35 OS=Homo sapiens ( | NUP35 | 34.774 | 2 | 51.766922 | yes | 7.2E-07 | 794.92383 | 52.976231 |
| ASTSDYQVISDR | 12 | Q8NFH5 | Nucleoporin NUP35 OS=Homo sapiens ( | NUP35 | 34.774 | 2 | 51.766922 | yes | 5.413E-07 | 671.31769 | 26.777924 |
| TNVNVFSELSAPR | 13 | Q8NFJ5 | Retinoic acid-induced protein 3 OS=Hom | GPRC5A | 40.252 | 2 | 45.72248 | yes | 1.762E-07 | 717.3728 | 334.62354 |
| AHAWPSPYK | 9 | Q8NFJ5 | Retinoic acid-induced protein 3 OS=Hom | GPRC5A | 40.252 | 2 | 45.72248 | yes | 3.425E-06 | 528.76672 | 277.25067 |
| SFYLDLPAGK | 10 | Q8NFT6 | Protein DBF4 homolog B OS=Homo sapi | DBF4B | 67.243 | 2 | 47.16963 | yes | 1.385E-07 | 555.7951 | 51.469135 |
| IIAQLSHSFADIPFQ | 20 | Q8NFT6 | Protein DBF4 homolog B OS=Homo sapi | DBF4B | 67.243 | 3 | 47.16963 | yes | 3.964E-07 | 727.73199 | 49.654503 |
| SFPEISFLGPK | 11 | Q8NFT6 | Protein DBF4 homolog B OS=Homo sapi | DBF4B | 67.243 | 2 | 47.16963 | yes | 3.37E-06 | 611.32935 | 29.759945 |
| EVTEPLNLNPAK | 12 | Q8NFW8 | N-acylneuraminate cytidylyltransferase O | CMAS | 48.38 | 2 | 50.517868 | yes | 1.159E-07 | 662.85901 | 3733.7395 |
| VGLSGAPADACST | 16 | Q8NFW8 | N-acylneuraminate cytidylyltransferase O | CMAS | 48.38 | 2 | 50.517868 | yes | 2.348E-07 | 766.87231 | 2823.3 |
| EEGYDSVFSVVR | 12 | Q8NFW8 | N-acylneuraminate cytidylyltransferase O | CMAS | 48.38 | 2 | 50.517868 | yes | 1.043E-07 | 693.83044 | 2509.0237 |
| MAYYEMR | 7 | Q8NFW8 | N-acylneuraminate cytidylyltransferase O | CMAS | 48.38 | 2 | 50.517868 | yes | 1.125E-07 | 482.20679 | 1270.6155 |

| | | | | | | | | | | | |
|---|---|---|---|---|---|---|---|---|---|---|---|
| LAVVDEWR | 8 | Q8NFW8 | N-acylneuraminate cytidylyltransferase O | CMAS | 48.38 | 2 | 50.517868 | yes | 2.082E-07 | 494.26654 | 1212.3324 |
| GVEKPPHLAALIL/ | 15 | Q8NFW8 | N-acylneuraminate cytidylyltransferase O | CMAS | 48.38 | 3 | 50.517868 | yes | 9.962E-08 | 528.99097 | 550.26202 |
| GVEKPPHLAALIL/ | 15 | Q8NFW8 | N-acylneuraminate cytidylyltransferase O | CMAS | 48.38 | 3 | 50.517868 | yes | 5.089E-07 | 792.98285 | 550.26202 |
| GVEKPPHLAALIL/ | 15 | Q8NFW8 | N-acylneuraminate cytidylyltransferase O | CMAS | 48.38 | 2 | 50.517868 | yes | 9.962E-08 | 528.99097 | 550.26202 |
| GVEKPPHLAALIL/ | 15 | Q8NFW8 | N-acylneuraminate cytidylyltransferase O | CMAS | 48.38 | 2 | 50.517868 | yes | 5.089E-07 | 792.98285 | 550.26202 |
| HLIEMGYLQGGK | 12 | Q8NFW8 | N-acylneuraminate cytidylyltransferase O | CMAS | 48.38 | 2 | 50.517868 | yes | 1.455E-07 | 673.35028 | 1001.2841 |
| EIISYDVK | 8 | Q8NFW8 | N-acylneuraminate cytidylyltransferase O | CMAS | 48.38 | 2 | 50.517868 | yes | 1.503E-07 | 483.76077 | 991.32855 |
| QFGAQVHR | 8 | Q8NFW8 | N-acylneuraminate cytidylyltransferase O | CMAS | 48.38 | 2 | 50.517868 | yes | 2.687E-07 | 471.74884 | 913.27722 |
| EFAEHICLLMEK | 12 | Q8NFW8 | N-acylneuraminate cytidylyltransferase O | CMAS | 48.38 | 2 | 50.517868 | yes | 9.789E-08 | 760.3678 | 295.60733 |
| EFAEHICLLMEK | 12 | Q8NFW8 | N-acylneuraminate cytidylyltransferase O | CMAS | 48.38 | 2 | 50.517868 | yes | 1.541E-07 | 507.24762 | 295.60733 |
| EFAEHICLLMEK | 12 | Q8NFW8 | N-acylneuraminate cytidylyltransferase O | CMAS | 48.38 | 3 | 50.517868 | yes | 9.789E-08 | 760.3678 | 295.60733 |
| EFAEHICLLMEK | 12 | Q8NFW8 | N-acylneuraminate cytidylyltransferase O | CMAS | 48.38 | 3 | 50.517868 | yes | 1.541E-07 | 507.24762 | 295.60733 |
| GAATSVSNPR | 10 | Q8NFW8 | N-acylneuraminate cytidylyltransferase O | CMAS | 48.38 | 2 | 50.517868 | yes | 1.185E-06 | 480.24887 | 427.72861 |
| DAIGISLLK | 9 | Q8NFW8 | N-acylneuraminate cytidylyltransferase O | CMAS | 48.38 | 2 | 50.517868 | yes | 7.009E-07 | 465.28696 | 247.25189 |
| MEVSVSDK | 8 | Q8NFW8 | N-acylneuraminate cytidylyltransferase O | CMAS | 48.38 | 2 | 50.517868 | yes | 3.796E-07 | 447.71548 | 119.19921 |
| VASLLHQVSWR | 11 | Q8NFZ5 | TNFAIP3-interacting protein 2 OS=Homo | TNIP2 | 48.7 | 2 | 49.900066 | yes | 1.236E-07 | 648.36456 | 127.79203 |
| VQMLEQQILAYK | 12 | Q8NFZ5 | TNFAIP3-interacting protein 2 OS=Homo | TNIP2 | 48.7 | 2 | 49.900066 | yes | 2.32E-07 | 732.39996 | 79.008659 |
| EKEVVLLR | 8 | Q8NFZ5 | TNFAIP3-interacting protein 2 OS=Homo | TNIP2 | 48.7 | 2 | 49.900066 | yes | 2.818E-07 | 493.30566 | 23.049295 |
| VTHVEDLNAK | 10 | Q8NFZ5 | TNFAIP3-interacting protein 2 OS=Homo | TNIP2 | 48.7 | 2 | 49.900066 | yes | 9.995E-08 | 563.29858 | 19.686798 |
| EKLQDGR | 7 | Q8NG31 | Kinetochore scaffold 1 OS=Homo sapiens | KNL1 | 265.392 | 2 | 44.887688 | yes | 5.157E-07 | 423.22742 | 36.386089 |
| EQTLAQIDFMQK | 12 | Q8NG31 | Kinetochore scaffold 1 OS=Homo sapiens | KNL1 | 265.392 | 2 | 44.887688 | yes | 4.574E-07 | 726.36359 | 36.014908 |
| AQVESCQLNNR | 11 | Q8NG31 | Kinetochore scaffold 1 OS=Homo sapiens | KNL1 | 265.392 | 2 | 44.887688 | yes | 1.785E-06 | 659.81207 | 22.218084 |
| DNSCVQEIAEK | 11 | Q8NG31 | Kinetochore scaffold 1 OS=Homo sapiens | KNL1 | 265.392 | 2 | 44.887688 | yes | 2.035E-07 | 646.79297 | 20.227118 |
| LAGLDITDASLR | 12 | Q8NHM5 | Lysine-specific demethylase 2B OS=Hom | KDM2B | 152.615 | 2 | 55.562626 | yes | 9.819E-08 | 622.84589 | 133.65814 |
| DSLTEINLSDCNK | 13 | Q8NHM5 | Lysine-specific demethylase 2B OS=Hom | KDM2B | 152.615 | 2 | 55.562626 | yes | 1.292E-07 | 754.84851 | 94.669571 |
| MALANKPLR | 9 | Q8NHM5 | Lysine-specific demethylase 2B OS=Hom | KDM2B | 152.615 | 2 | 55.562626 | yes | 1.431E-07 | 507.29987 | 92.631096 |
| TLDVQWVEGLK | 11 | Q8NHM5 | Lysine-specific demethylase 2B OS=Hom | KDM2B | 152.615 | 2 | 55.562626 | yes | 9.722E-08 | 644.35083 | 79.223198 |
| VTDQCLSFFK | 10 | Q8NHM5 | Lysine-specific demethylase 2B OS=Hom | KDM2B | 152.615 | 2 | 55.562626 | yes | 9.834E-08 | 622.80261 | 55.551537 |
| LQSGILVCTDVMA | 14 | Q8NHQ9 | ATP-dependent RNA helicase DDX55 OS | DDX55 | 68.547 | 2 | 54.808567 | yes | 1.018E-07 | 781.90509 | 941.34167 |
| AEGLDLASCVR | 11 | Q8NHQ9 | ATP-dependent RNA helicase DDX55 OS | DDX55 | 68.547 | 2 | 54.808567 | yes | 1.115E-07 | 595.79535 | 905.04852 |
| HECNLIFR | 8 | Q8NHQ9 | ATP-dependent RNA helicase DDX55 OS | DDX55 | 68.547 | 2 | 54.808567 | yes | 9.722E-08 | 544.76892 | 707.565 |
| LENYYMVCK | 9 | Q8NHQ9 | ATP-dependent RNA helicase DDX55 OS | DDX55 | 68.547 | 2 | 54.808567 | yes | 1.651E-07 | 610.27795 | 621.49628 |
| IMCIHGK | 7 | Q8NHQ9 | ATP-dependent RNA helicase DDX55 OS | DDX55 | 68.547 | 2 | 54.808567 | yes | 1.318E-07 | 429.71985 | 569.19861 |
| SQVGAIIITPTR | 12 | Q8NHQ9 | ATP-dependent RNA helicase DDX55 OS | DDX55 | 68.547 | 2 | 54.808567 | yes | 1.031E-07 | 628.37207 | 525.53558 |
| SLDVLVLDEADR | 12 | Q8NHQ9 | ATP-dependent RNA helicase DDX55 OS | DDX55 | 68.547 | 2 | 54.808567 | yes | 9.722E-08 | 672.85388 | 483.32187 |
| QQGGNIIVATPGR | 13 | Q8NHQ9 | ATP-dependent RNA helicase DDX55 OS | DDX55 | 68.547 | 2 | 54.808567 | yes | 1.26E-07 | 655.86243 | 415.22617 |
| DVAAEAVTGSGK | 12 | Q8NHQ9 | ATP-dependent RNA helicase DDX55 OS | DDX55 | 68.547 | 2 | 54.808567 | yes | 1.012E-07 | 552.78021 | 327.40991 |
| DLDFASLAR | 9 | Q8NHQ9 | ATP-dependent RNA helicase DDX55 OS | DDX55 | 68.547 | 2 | 54.808567 | yes | 1.893E-07 | 504.26144 | 241.66446 |
| NPGEDVER | 8 | Q8NHQ9 | ATP-dependent RNA helicase DDX55 OS | DDX55 | 68.547 | 2 | 54.808567 | yes | 1.603E-07 | 458.21198 | 216.88536 |

| | | | | | | | | | | | |
|---|---|---|---|---|---|---|---|---|---|---|---|
| FNQLVHFLR | 9 | Q8NHQ9 | ATP-dependent RNA helicase DDX55 O: | DDX55 | 68.547 | 2 | 54.808567 | yes | 1.393E-07 | 587.33002 | 206.92438 |
| CPLQEMKPQR | 10 | Q8NHQ9 | ATP-dependent RNA helicase DDX55 O: | DDX55 | 68.547 | 2 | 54.808567 | yes | 9.722E-08 | 643.82086 | 200.11331 |
| ITEEEFEK | 8 | Q8NHQ9 | ATP-dependent RNA helicase DDX55 O: | DDX55 | 68.547 | 2 | 54.808567 | yes | 2.853E-07 | 512.7453 | 196.50496 |
| KAEGLDLASCVR | 12 | Q8NHQ9 | ATP-dependent RNA helicase DDX55 O: | DDX55 | 68.547 | 2 | 54.808567 | yes | 1.784E-07 | 659.84283 | 100.70105 |
| TVDLGISDLEDDC | 13 | Q8NHQ9 | ATP-dependent RNA helicase DDX55 O: | DDX55 | 68.547 | 2 | 54.808567 | yes | 9.722E-08 | 726.31396 | 99.001045 |
| KSQVGAIIITPTR | 13 | Q8NHQ9 | ATP-dependent RNA helicase DDX55 O: | DDX55 | 68.547 | 3 | 54.808567 | yes | 1.021E-07 | 461.94879 | 43.481045 |
| KSQVGAIIITPTR | 13 | Q8NHQ9 | ATP-dependent RNA helicase DDX55 O: | DDX55 | 68.547 | 3 | 54.808567 | yes | 2.224E-07 | 692.41956 | 43.481045 |
| KSQVGAIIITPTR | 13 | Q8NHQ9 | ATP-dependent RNA helicase DDX55 O: | DDX55 | 68.547 | 2 | 54.808567 | yes | 1.021E-07 | 461.94879 | 43.481045 |
| KSQVGAIIITPTR | 13 | Q8NHQ9 | ATP-dependent RNA helicase DDX55 O: | DDX55 | 68.547 | 2 | 54.808567 | yes | 2.224E-07 | 692.41956 | 43.481045 |
| QNAPLTSIQIR | 11 | Q8NHV4 | Protein NEDD1 OS=Homo sapiens OX=9 | NEDD1 | 71.967 | 2 | 52.712994 | yes | 9.722E-08 | 620.85406 | 714.79657 |
| YSVNEGLVAEIER | 13 | Q8NHV4 | Protein NEDD1 OS=Homo sapiens OX=9 | NEDD1 | 71.967 | 2 | 52.712994 | yes | 9.722E-08 | 739.87793 | 640.26501 |
| IILYDTSSK | 9 | Q8NHV4 | Protein NEDD1 OS=Homo sapiens OX=9 | NEDD1 | 71.967 | 2 | 52.712994 | yes | 1.35E-07 | 520.28711 | 391.32642 |
| IWDASSMTLVDK | 12 | Q8NHV4 | Protein NEDD1 OS=Homo sapiens OX=9 | NEDD1 | 71.967 | 2 | 52.712994 | yes | 1.096E-07 | 683.3396 | 329.14771 |
| IASSVTAGVASSLS | 16 | Q8NHV4 | Protein NEDD1 OS=Homo sapiens OX=9 | NEDD1 | 71.967 | 2 | 52.712994 | yes | 9.722E-08 | 753.90411 | 314.39978 |
| SSLGDMFSPIR | 11 | Q8NHV4 | Protein NEDD1 OS=Homo sapiens OX=9 | NEDD1 | 71.967 | 2 | 52.712994 | yes | 9.722E-08 | 605.30023 | 305.76752 |
| CKPVPLLELAEGQ | 14 | Q8NHV4 | Protein NEDD1 OS=Homo sapiens OX=9 | NEDD1 | 71.967 | 2 | 52.712994 | yes | 4.038E-07 | 791.43707 | 235.72128 |
| NQDFSSFDDTGK | 12 | Q8NHV4 | Protein NEDD1 OS=Homo sapiens OX=9 | NEDD1 | 71.967 | 2 | 52.712994 | yes | 9.722E-08 | 680.78619 | 177.48796 |
| DIVNLQVEMIK | 11 | Q8NHV4 | Protein NEDD1 OS=Homo sapiens OX=9 | NEDD1 | 71.967 | 2 | 52.712994 | yes | 9.809E-08 | 651.36029 | 134.97102 |
| FASSGDDIK | 9 | Q8NHV4 | Protein NEDD1 OS=Homo sapiens OX=9 | NEDD1 | 71.967 | 2 | 52.712994 | yes | 9.722E-08 | 470.22455 | 106.53294 |
| QFHMQLNEMHSL | 15 | Q8NHV4 | Protein NEDD1 OS=Homo sapiens OX=9 | NEDD1 | 71.967 | 3 | 52.712994 | yes | 9.848E-08 | 638.31006 | 65.29792 |
| IADSIGNNR | 9 | Q8NHV4 | Protein NEDD1 OS=Homo sapiens OX=9 | NEDD1 | 71.967 | 2 | 52.712994 | yes | 1.304E-07 | 480.24887 | 50.227459 |
| LDDIEEFENIR | 11 | Q8NHZ8 | Anaphase-promoting complex subunit CD | CDC26 | 9.777 | 2 | 47.069134 | yes | 1.408E-07 | 696.83569 | 343.44174 |
| VENTLNIHEDLDK | 13 | Q8NI08 | Nuclear receptor coactivator 7 OS=Homo | NCOA7 | 106.162 | 2 | 31.885077 | yes | 1.281E-05 | 770.38611 | 278.95737 |
| LAYSTLEHGTSLK | 13 | Q8NI08 | Nuclear receptor coactivator 7 OS=Homo | NCOA7 | 106.162 | 2 | 31.885077 | yes | 1.693E-06 | 710.37756 | 50.449253 |
| EGPQVSENLQK | 11 | Q8NI08 | Nuclear receptor coactivator 7 OS=Homo | NCOA7 | 106.162 | 2 | 31.885077 | yes | 7.459E-05 | 614.81201 | 24.167227 |
| SNSCSTFNNDILSK | 14 | Q8NI08 | Nuclear receptor coactivator 7 OS=Homo | NCOA7 | 106.162 | 2 | 31.885077 | yes | 1.96E-06 | 793.85938 | 18.99161 |
| SASLDSPVLLVIK | 13 | Q8NI08 | Nuclear receptor coactivator 7 OS=Homo | NCOA7 | 106.162 | 2 | 31.885077 | yes | 3.507E-06 | 671.40302 | 9.1366558 |
| VLNLGQALER | 10 | Q8NI27 | THO complex subunit 2 OS=Homo sapie | THOC2 | 182.775 | 2 | 53.9944 | yes | 1.128E-07 | 556.82477 | 443.03714 |
| AEGGYFGQIR | 10 | Q8NI27 | THO complex subunit 2 OS=Homo sapie | THOC2 | 182.775 | 2 | 53.9944 | yes | 3.249E-07 | 549.27234 | 439.59174 |
| ADQLDYENFR | 10 | Q8NI27 | THO complex subunit 2 OS=Homo sapie | THOC2 | 182.775 | 2 | 53.9944 | yes | 9.722E-08 | 635.78857 | 421.06195 |
| FVELVHQQK | 9 | Q8NI27 | THO complex subunit 2 OS=Homo sapie | THOC2 | 182.775 | 2 | 53.9944 | yes | 9.733E-08 | 564.31403 | 353.32745 |
| GEFLHLCR | 8 | Q8NI27 | THO complex subunit 2 OS=Homo sapie | THOC2 | 182.775 | 2 | 53.9944 | yes | 1.39E-07 | 516.25818 | 305.58783 |
| GSPVNALQNK | 10 | Q8NI27 | THO complex subunit 2 OS=Homo sapie | THOC2 | 182.775 | 2 | 53.9944 | yes | 2.165E-07 | 514.28015 | 250.77469 |
| LDPETLESLGLIK | 13 | Q8NI27 | THO complex subunit 2 OS=Homo sapie | THOC2 | 182.775 | 2 | 53.9944 | yes | 3.464E-07 | 714.40326 | 231.48422 |
| YDNLITPVVDSLK | 13 | Q8NI27 | THO complex subunit 2 OS=Homo sapie | THOC2 | 182.775 | 2 | 53.9944 | yes | 1.151E-07 | 738.90088 | 170.79948 |
| LIHITIEPLYR | 11 | Q8NI27 | THO complex subunit 2 OS=Homo sapie | THOC2 | 182.775 | 2 | 53.9944 | yes | 9.722E-08 | 684.40588 | 74.2976 |
| LIHITIEPLYR | 11 | Q8NI27 | THO complex subunit 2 OS=Homo sapie | THOC2 | 182.775 | 2 | 53.9944 | yes | 7.854E-07 | 456.60635 | 74.2976 |
| LIHITIEPLYR | 11 | Q8NI27 | THO complex subunit 2 OS=Homo sapie | THOC2 | 182.775 | 3 | 53.9944 | yes | 9.722E-08 | 684.40588 | 74.2976 |
| LIHITIEPLYR | 11 | Q8NI27 | THO complex subunit 2 OS=Homo sapie | THOC2 | 182.775 | 3 | 53.9944 | yes | 7.854E-07 | 456.60635 | 74.2976 |

| | | | | | | | | | | | |
|---|---|---|---|---|---|---|---|---|---|---|---|
| FLCCMLETVTR | 11 | Q8NI27 | THO complex subunit 2 OS=Homo sapie | THOC2 | 182.775 | 2 | 53.9944 | yes | 3.115E-07 | 715.33545 | 139.11943 |
| ECGNYPGFLTILR | 13 | Q8NI27 | THO complex subunit 2 OS=Homo sapie | THOC2 | 182.775 | 2 | 53.9944 | yes | 2.712E-07 | 770.38483 | 133.53481 |
| FLQLCIFPR | 9 | Q8NI27 | THO complex subunit 2 OS=Homo sapie | THOC2 | 182.775 | 2 | 53.9944 | yes | 5.782E-07 | 597.32861 | 121.81663 |
| LTMVVLSSEK | 10 | Q8NI27 | THO complex subunit 2 OS=Homo sapie | THOC2 | 182.775 | 2 | 53.9944 | yes | 2.436E-07 | 553.80994 | 92.814552 |
| RPDLYALAMGYS( | 16 | Q8NI27 | THO complex subunit 2 OS=Homo sapie | THOC2 | 182.775 | 3 | 53.9944 | yes | 1.014E-06 | 594.97839 | 88.178329 |
| CIFSAIDAVYCAR | 13 | Q8NI27 | THO complex subunit 2 OS=Homo sapie | THOC2 | 182.775 | 2 | 53.9944 | yes | 1.835E-06 | 773.36304 | 79.19529 |
| NETYNSHPLLVK | 12 | Q8NI27 | THO complex subunit 2 OS=Homo sapie | THOC2 | 182.775 | 2 | 53.9944 | yes | 9.722E-08 | 707.86987 | 71.364006 |
| TPNFSTLLCYDR | 12 | Q8NI27 | THO complex subunit 2 OS=Homo sapie | THOC2 | 182.775 | 2 | 53.9944 | yes | 9.383E-07 | 743.85339 | 59.646729 |
| ILPWYPK | 7 | Q8NI27 | THO complex subunit 2 OS=Homo sapie | THOC2 | 182.775 | 2 | 53.9944 | yes | 2.346E-06 | 458.76819 | 47.761818 |
| YITSCEMVMAPVH | 23 | Q8NI27 | THO complex subunit 2 OS=Homo sapie | THOC2 | 182.775 | 4 | 53.9944 | yes | 3.751E-07 | 647.57336 | 43.341751 |
| EKTPATTPEAR | 11 | Q8NI27 | THO complex subunit 2 OS=Homo sapie | THOC2 | 182.775 | 2 | 53.9944 | yes | 4.157E-05 | 600.81458 | 33.553165 |
| ALGLFSNDIPHVVI | 14 | Q8NI36 | WD repeat-containing protein 36 OS=Ho | WDR36 | 105.323 | 2 | 56.116543 | yes | 9.722E-08 | 769.42792 | 243.90176 |
| ALGLFSNDIPHVVI | 14 | Q8NI36 | WD repeat-containing protein 36 OS=Ho | WDR36 | 105.323 | 2 | 56.116543 | yes | 9.722E-08 | 513.28772 | 243.90176 |
| ALGLFSNDIPHVVI | 14 | Q8NI36 | WD repeat-containing protein 36 OS=Ho | WDR36 | 105.323 | 3 | 56.116543 | yes | 9.722E-08 | 769.42792 | 243.90176 |
| ALGLFSNDIPHVVI | 14 | Q8NI36 | WD repeat-containing protein 36 OS=Ho | WDR36 | 105.323 | 3 | 56.116543 | yes | 9.722E-08 | 513.28772 | 243.90176 |
| IWIFDGPTGEGR | 12 | Q8NI36 | WD repeat-containing protein 36 OS=Ho | WDR36 | 105.323 | 2 | 56.116543 | yes | 9.722E-08 | 674.33826 | 406.11456 |
| QDWGPITSISFR | 12 | Q8NI36 | WD repeat-containing protein 36 OS=Ho | WDR36 | 105.323 | 2 | 56.116543 | yes | 9.722E-08 | 703.85681 | 333.03802 |
| LLYTFPGWK | 9 | Q8NI36 | WD repeat-containing protein 36 OS=Ho | WDR36 | 105.323 | 2 | 56.116543 | yes | 9.722E-08 | 562.81061 | 253.95938 |
| ESGPSGIETELR | 12 | Q8NI36 | WD repeat-containing protein 36 OS=Ho | WDR36 | 105.323 | 2 | 56.116543 | yes | 1.527E-07 | 637.81476 | 228.72531 |
| NAHSTAIAGLTFLI | 15 | Q8NI36 | WD repeat-containing protein 36 OS=Ho | WDR36 | 105.323 | 3 | 56.116543 | yes | 1.101E-07 | 536.95837 | 217.24667 |
| SLGHGLINK | 9 | Q8NI36 | WD repeat-containing protein 36 OS=Ho | WDR36 | 105.323 | 2 | 56.116543 | yes | 2.9E-07 | 469.77454 | 189.69977 |
| VVNLGVLAQK | 10 | Q8NI36 | WD repeat-containing protein 36 OS=Ho | WDR36 | 105.323 | 2 | 56.116543 | yes | 1.34E-07 | 520.82678 | 189.39946 |
| SFHTYDVQK | 9 | Q8NI36 | WD repeat-containing protein 36 OS=Ho | WDR36 | 105.323 | 2 | 56.116543 | yes | 9.723E-08 | 562.77222 | 175.64015 |
| WLISAAMDCSIR | 12 | Q8NI36 | WD repeat-containing protein 36 OS=Ho | WDR36 | 105.323 | 2 | 56.116543 | yes | 9.819E-08 | 711.84705 | 165.88412 |
| FYVTTCVGK | 9 | Q8NI36 | WD repeat-containing protein 36 OS=Ho | WDR36 | 105.323 | 2 | 56.116543 | yes | 2.343E-07 | 537.76807 | 161.62329 |
| MGHSAPLTNIR | 11 | Q8NI36 | WD repeat-containing protein 36 OS=Ho | WDR36 | 105.323 | 2 | 56.116543 | yes | 5.694E-07 | 598.81384 | 144.12997 |
| EIVHTFK | 7 | Q8NI36 | WD repeat-containing protein 36 OS=Ho | WDR36 | 105.323 | 2 | 56.116543 | yes | 9.722E-08 | 437.24509 | 131.30695 |
| ISAILHPSTYLNK | 13 | Q8NI36 | WD repeat-containing protein 36 OS=Ho | WDR36 | 105.323 | 2 | 56.116543 | yes | 1.548E-06 | 728.91156 | 121.0294 |
| LSCSTWNYQK | 10 | Q8NI36 | WD repeat-containing protein 36 OS=Ho | WDR36 | 105.323 | 2 | 56.116543 | yes | 1.041E-07 | 643.79535 | 92.847099 |
| ILIHSVSLSSSPNIM | 19 | Q8NI36 | WD repeat-containing protein 36 OS=Ho | WDR36 | 105.323 | 3 | 56.116543 | yes | 9.85E-08 | 706.40149 | 41.279053 |
| ILIHSVSLSSSPNIM | 19 | Q8NI36 | WD repeat-containing protein 36 OS=Ho | WDR36 | 105.323 | 3 | 56.116543 | yes | 6.137E-07 | 530.05292 | 41.279053 |
| ILIHSVSLSSSPNIM | 19 | Q8NI36 | WD repeat-containing protein 36 OS=Ho | WDR36 | 105.323 | 4 | 56.116543 | yes | 9.85E-08 | 706.40149 | 41.279053 |
| ILIHSVSLSSSPNIM | 19 | Q8NI36 | WD repeat-containing protein 36 OS=Ho | WDR36 | 105.323 | 4 | 56.116543 | yes | 6.137E-07 | 530.05292 | 41.279053 |
| GLQNTMSVR | 9 | Q8NI36 | WD repeat-containing protein 36 OS=Ho | WDR36 | 105.323 | 2 | 56.116543 | yes | 3.109E-06 | 503.26093 | 62.960648 |
| NLLNLDVIK | 9 | Q8NI36 | WD repeat-containing protein 36 OS=Ho | WDR36 | 105.323 | 2 | 56.116543 | yes | 3.3E-06 | 521.31879 | 62.408455 |
| ESDWDGIIACHQG | 14 | Q8NI36 | WD repeat-containing protein 36 OS=Ho | WDR36 | 105.323 | 2 | 56.116543 | yes | 1.288E-07 | 808.36212 | 51.014469 |
| YDTALNLLK | 9 | Q8NI36 | WD repeat-containing protein 36 OS=Ho | WDR36 | 105.323 | 2 | 56.116543 | yes | 5.478E-07 | 525.7951 | 49.029846 |
| DYGVDPSQFSGTL | 15 | Q8NI37 | Protein phosphatase PTC7 homolog OS=] | PPTC7 | 32.646 | 2 | 44.446724 | yes | 2.191E-07 | 836.88544 | 81.387169 |
| VALLDFGATR | 10 | Q8NI60 | Atypical kinase COQ8A, mitochondrial C | COQ8A | 71.951 | 2 | 49.391735 | yes | 1.441E-07 | 531.80072 | 154.43024 |

| | | | | | | | | | | | |
|---|---|---|---|---|---|---|---|---|---|---|---|
| ELALECDYQR | 10 | Q8NI60 | Atypical kinase COQ8A, mitochondrial C | COQ8A | 71.951 | 2 | 49.391735 | yes | 1.344E-07 | 648.7981 | 104.79799 |
| AVLGSSPFLSEAN/ | 16 | Q8NI60 | Atypical kinase COQ8A, mitochondrial C | COQ8A | 71.951 | 2 | 49.391735 | yes | 1.037E-07 | 824.42047 | 88.393829 |
| TLNNDLGPNWR | 11 | Q8NI60 | Atypical kinase COQ8A, mitochondrial C | COQ8A | 71.951 | 2 | 49.391735 | yes | 3.441E-07 | 650.32562 | 81.665344 |
| LVPPPEETYSLHR | 13 | Q8NI60 | Atypical kinase COQ8A, mitochondrial C | COQ8A | 71.951 | 2 | 49.391735 | yes | 3.758E-07 | 769.40411 | 80.227798 |
| MGGSFLICSK | 10 | Q8NI60 | Atypical kinase COQ8A, mitochondrial C | COQ8A | 71.951 | 2 | 49.391735 | yes | 2.349E-07 | 550.2674 | 54.991478 |
| SFTDLYIQIIR | 11 | Q8NI60 | Atypical kinase COQ8A, mitochondrial C | COQ8A | 71.951 | 2 | 49.391735 | yes | 4.683E-07 | 684.8797 | 53.075665 |
| DKLEYFEERPFAA | 23 | Q8NI60 | Atypical kinase COQ8A, mitochondrial C | COQ8A | 71.951 | 4 | 49.391735 | yes | 3.038E-07 | 662.59613 | 36.395634 |
| DLLVNSGPLAVR | 12 | Q8NI77 | Kinesin-like protein KIF18A OS=Homo s | KIF18A | 102.281 | 2 | 57.086296 | yes | 1.164E-07 | 627.36426 | 445.39035 |
| MSLIDLAGSER | 11 | Q8NI77 | Kinesin-like protein KIF18A OS=Homo s | KIF18A | 102.281 | 2 | 57.086296 | no | 2.072E-07 | 596.30554 | 305.33698 |
| ELQPIVYTPEDCR | 13 | Q8NI77 | Kinesin-like protein KIF18A OS=Homo s | KIF18A | 102.281 | 2 | 57.086296 | yes | 9.849E-08 | 810.39032 | 132.56589 |
| EIEHLVER | 8 | Q8NI77 | Kinesin-like protein KIF18A OS=Homo s | KIF18A | 102.281 | 2 | 57.086296 | yes | 2.513E-07 | 512.77472 | 123.64381 |
| LMISNPQEK | 9 | Q8NI77 | Kinesin-like protein KIF18A OS=Homo s | KIF18A | 102.281 | 2 | 57.086296 | yes | 2.377E-07 | 530.27881 | 123.33887 |
| VVVWADQTAEQP | 13 | Q8NI77 | Kinesin-like protein KIF18A OS=Homo s | KIF18A | 102.281 | 2 | 57.086296 | yes | 9.722E-08 | 735.883 | 105.4939 |
| TASINQNVR | 9 | Q8NI77 | Kinesin-like protein KIF18A OS=Homo s | KIF18A | 102.281 | 2 | 57.086296 | yes | 3.232E-06 | 501.76999 | 84.711479 |
| MLCEVAIPHNR | 11 | Q8NI77 | Kinesin-like protein KIF18A OS=Homo s | KIF18A | 102.281 | 2 | 57.086296 | yes | 9.722E-08 | 670.33411 | 82.686203 |
| AFTNENDQAK | 10 | Q8NI77 | Kinesin-like protein KIF18A OS=Homo s | KIF18A | 102.281 | 2 | 57.086296 | yes | 2.402E-07 | 569.26221 | 81.2509 |
| EAGLSNAAFESDF | 14 | Q8NI77 | Kinesin-like protein KIF18A OS=Homo s | KIF18A | 102.281 | 2 | 57.086296 | yes | 2.314E-07 | 743.34644 | 72.301178 |
| HILVFDPK | 8 | Q8NI77 | Kinesin-like protein KIF18A OS=Homo s | KIF18A | 102.281 | 2 | 57.086296 | yes | 2.779E-05 | 484.78183 | 32.690243 |
| FQEILNCLFQNR | 12 | Q8NI77 | Kinesin-like protein KIF18A OS=Homo s | KIF18A | 102.281 | 2 | 57.086296 | yes | 1.309E-07 | 791.39575 | 28.878727 |
| SPPSQSVQLNDSLS | 15 | Q8NI77 | Kinesin-like protein KIF18A OS=Homo s | KIF18A | 102.281 | 2 | 57.086296 | yes | 4.208E-07 | 793.90466 | 24.402946 |
| QDNSSEK | 7 | Q8NI77 | Kinesin-like protein KIF18A OS=Homo s | KIF18A | 102.281 | 2 | 57.086296 | yes | 4.928E-05 | 404.17758 | 6.4242444 |
| EFLAHAPTK | 9 | Q8TA86 | Retinitis pigmentosa 9 protein OS=Homo | RP9 | 26.107 | 2 | 48.11915 | yes | 1.668E-07 | 507.27435 | 497.13177 |
| EPPEQELQR | 9 | Q8TA86 | Retinitis pigmentosa 9 protein OS=Homo | RP9 | 26.107 | 2 | 48.11915 | yes | 1.716E-07 | 563.2804 | 130.02832 |
| HDAQQLQQLK | 10 | Q8TA86 | Retinitis pigmentosa 9 protein OS=Homo | RP9 | 26.107 | 2 | 48.11915 | yes | 2.204E-06 | 604.82275 | 62.105621 |
| EDETKPEDCIPDVI | 20 | Q8TA86 | Retinitis pigmentosa 9 protein OS=Homo | RP9 | 26.107 | 4 | 48.11915 | yes | 2.497E-07 | 577.75909 | 51.275394 |
| TGDKECPFFIK | 11 | Q8TA86 | Retinitis pigmentosa 9 protein OS=Homo | RP9 | 26.107 | 2 | 48.11915 | yes | 1.197E-07 | 671.32904 | 34.763348 |
| DFTIYNPNLLTASK | 14 | Q8TAA9 | Vang-like protein 1 OS=Homo sapiens O | VANGL1 | 59.975 | 2 | 48.509705 | yes | 2.237E-06 | 798.91705 | 51.829697 |
| AALVVLENYYK | 11 | Q8TAA9 | Vang-like protein 1 OS=Homo sapiens O | VANGL1 | 59.975 | 2 | 48.509705 | yes | 1.138E-07 | 641.85571 | 42.166443 |
| DGDVVLPAGVVV | 13 | Q8TAD8 | Smad nuclear-interacting protein 1 OS=H | SNIP1 | 45.778 | 2 | 53.559803 | yes | 1.137E-07 | 634.36646 | 932.13867 |
| IADIPIDHPSCSK | 13 | Q8TAD8 | Smad nuclear-interacting protein 1 OS=H | SNIP1 | 45.778 | 2 | 53.559803 | yes | 1.675E-07 | 726.86121 | 182.51784 |
| IADIPIDHPSCSK | 13 | Q8TAD8 | Smad nuclear-interacting protein 1 OS=H | SNIP1 | 45.778 | 2 | 53.559803 | yes | 3.285E-07 | 484.90991 | 182.51784 |
| IADIPIDHPSCSK | 13 | Q8TAD8 | Smad nuclear-interacting protein 1 OS=H | SNIP1 | 45.778 | 3 | 53.559803 | yes | 1.675E-07 | 726.86121 | 182.51784 |
| IADIPIDHPSCSK | 13 | Q8TAD8 | Smad nuclear-interacting protein 1 OS=H | SNIP1 | 45.778 | 3 | 53.559803 | yes | 3.285E-07 | 484.90991 | 182.51784 |
| QHAVFQYR | 8 | Q8TAD8 | Smad nuclear-interacting protein 1 OS=H | SNIP1 | 45.778 | 2 | 53.559803 | yes | 4.02E-07 | 524.76978 | 333.2652 |
| EKPSFELSGALLEI | 19 | Q8TAD8 | Smad nuclear-interacting protein 1 OS=H | SNIP1 | 45.778 | 3 | 53.559803 | yes | 1.39E-07 | 718.69531 | 243.39835 |
| LSPEVAPPAHR | 11 | Q8TAD8 | Smad nuclear-interacting protein 1 OS=H | SNIP1 | 45.778 | 2 | 53.559803 | yes | 1.362E-07 | 587.32239 | 124.83456 |
| NDVGGGGSESQEI | 16 | Q8TAD8 | Smad nuclear-interacting protein 1 OS=H | SNIP1 | 45.778 | 2 | 53.559803 | yes | 9.722E-08 | 800.88171 | 88.068298 |
| QSAYLLGR | 8 | Q8TAD8 | Smad nuclear-interacting protein 1 OS=H | SNIP1 | 45.778 | 2 | 53.559803 | yes | 9.722E-08 | 454.25345 | 84.244972 |
| MPQMIVNWQQQC | 13 | Q8TAE8 | Large ribosomal subunit protein mL64 OS | GADD45G | 25.384 | 2 | 52.293446 | yes | 1.497E-07 | 843.91376 | 611.98718 |

| | | | | | | | | | | | |
|---|---|---|---|---|---|---|---|---|---|---|---|
| WPDPEDLLTPR | 11 | Q8TAE8 | Large ribosomal subunit protein mL64 OS | GADD45G | 25.384 | 2 | 52.293446 | yes | 9.722E-08 | 669.83807 | 268.86218 |
| FQELLQDLEK | 10 | Q8TAE8 | Large ribosomal subunit protein mL64 OS | GADD45G | 25.384 | 2 | 52.293446 | yes | 1.634E-07 | 631.83496 | 177.06235 |
| EQHIAECMAK | 10 | Q8TAE8 | Large ribosomal subunit protein mL64 OS | GADD45G | 25.384 | 2 | 52.293446 | yes | 9.722E-08 | 608.77606 | 121.32484 |
| LSASDMLQVR | 10 | Q8TAF3 | WD repeat-containing protein 48 OS=Ho | WDR48 | 76.211 | 2 | 53.94751 | yes | 1.727E-07 | 560.29498 | 193.6044 |
| DKELVASAGLDR | 12 | Q8TAF3 | WD repeat-containing protein 48 OS=Ho | WDR48 | 76.211 | 2 | 53.94751 | yes | 9.722E-08 | 637.34094 | 107.72092 |
| SGGDLTLHYR | 10 | Q8TAF3 | WD repeat-containing protein 48 OS=Ho | WDR48 | 76.211 | 2 | 53.94751 | yes | 3.974E-07 | 559.78308 | 89.984222 |
| DAGFSSPDGSDPK | 13 | Q8TAF3 | WD repeat-containing protein 48 OS=Ho | WDR48 | 76.211 | 2 | 53.94751 | yes | 1.276E-07 | 640.27551 | 58.284073 |
| DGTQCLSGSSDGT | 15 | Q8TAF3 | WD repeat-containing protein 48 OS=Ho | WDR48 | 76.211 | 2 | 53.94751 | yes | 3.843E-07 | 777.34668 | 57.912216 |
| DTNNNVAYWDVL | 13 | Q8TAF3 | WD repeat-containing protein 48 OS=Ho | WDR48 | 76.211 | 2 | 53.94751 | yes | 2.898E-07 | 776.37555 | 42.163555 |
| TLISASSDTTVK | 12 | Q8TAF3 | WD repeat-containing protein 48 OS=Ho | WDR48 | 76.211 | 2 | 53.94751 | yes | 3.173E-06 | 611.8299 | 41.103867 |
| NGVNALQLDPALN | 14 | Q8TAF3 | WD repeat-containing protein 48 OS=Ho | WDR48 | 76.211 | 2 | 53.94751 | yes | 1.788E-06 | 747.90479 | 27.479256 |
| KVQVSYVIR | 9 | Q8TAF3 | WD repeat-containing protein 48 OS=Ho | WDR48 | 76.211 | 2 | 53.94751 | yes | 4.042E-05 | 546.33221 | 18.970261 |
| GGASIIQCHILNDK | 14 | Q8TAF3 | WD repeat-containing protein 48 OS=Ho | WDR48 | 76.211 | 2 | 53.94751 | yes | 2.858E-05 | 763.39319 | 14.60726 |
| SSLALGLELR | 10 | Q8TAP8 | Protein phosphatase 1 regulatory subunit | PPP1R35 | 27.954 | 2 | 49.029007 | yes | 4.924E-07 | 529.81384 | 184.24353 |
| APVPEPGLDLSLSI | 22 | Q8TAP8 | Protein phosphatase 1 regulatory subunit | PPP1R35 | 27.954 | 3 | 49.029007 | yes | 9.813E-07 | 775.74475 | 117.51778 |
| MMMGCGESELK | 11 | Q8TAP8 | Protein phosphatase 1 regulatory subunit | PPP1R35 | 27.954 | 2 | 49.029007 | yes | 1.075E-07 | 657.76349 | 68.159103 |
| APVPEPGLDLSLSI | 15 | Q8TAP8 | Protein phosphatase 1 regulatory subunit | PPP1R35 | 27.954 | 2 | 49.029007 | yes | 1.135E-06 | 774.42505 | 37.984283 |
| MSNELENYFK | 10 | Q8TAP9 | M-phase-specific PLK1-interacting protei | MPLKIP | 19.147 | 2 | 45.32781 | yes | 1.883E-07 | 637.78973 | 51.841389 |
| FGSPSPGGYPGSYS | 15 | Q8TAP9 | M-phase-specific PLK1-interacting protei | MPLKIP | 19.147 | 2 | 45.32781 | yes | 7.092E-06 | 758.3468 | 18.735727 |
| MKEEVPTALVEAI | 15 | Q8TAQ2 | SWI/SNF complex subunit SMARCC2 O | SMARCC2 | 132.88 | 3 | 43.72119 | yes | 5.037E-07 | 570.30652 | 84.304062 |
| PNIFLCPEIEPK | 12 | Q8TAQ2 | SWI/SNF complex subunit SMARCC2 O | SMARCC2 | 132.88 | 2 | 43.72119 | yes | 2.473E-07 | 728.87885 | 71.668823 |
| NLAGDVCAIMR | 11 | Q8TAQ2 | SWI/SNF complex subunit SMARCC2 O | SMARCC2 | 132.88 | 2 | 43.72119 | yes | 7.997E-07 | 610.29974 | 52.997234 |
| EKPTDMQNFGLR | 12 | Q8TAQ2 | SWI/SNF complex subunit SMARCC2 O | SMARCC2 | 132.88 | 2 | 43.72119 | yes | 1.275E-06 | 718.35358 | 30.219631 |
| VDHVMDSVAR | 10 | Q8TAT5 | Endonuclease 8-like 3 OS=Homo sapiens | NEIL3 | 67.77 | 2 | 46.606552 | yes | 1.516E-07 | 564.77698 | 83.548462 |
| ESSSEQYVPDVFY | 14 | Q8TAT6 | Nuclear protein localization protein 4 hon | NPLOC4 | 68.121 | 2 | 49.942623 | yes | 9.974E-08 | 839.38574 | 76.920502 |
| LSPDGHFGSK | 10 | Q8TAT6 | Nuclear protein localization protein 4 hon | NPLOC4 | 68.121 | 2 | 49.942623 | yes | 1.148E-07 | 522.75909 | 39.074978 |
| RSSMSSCGSSGYFS | 30 | Q8TB45 | DEP domain-containing mTOR-interactin | DEPTOR | 46.294 | 4 | 33.96801 | yes | 1.109E-06 | 788.6087 | 238.45337 |
| FVSFPTQVLAK | 11 | Q8TB61 | Adenosine 3'-phospho 5'-phosphosulfate t | SLC35B2 | 47.515 | 2 | 50.059864 | yes | 9.919E-08 | 618.85297 | 252.47885 |
| ASDEVPLAPR | 10 | Q8TB61 | Adenosine 3'-phospho 5'-phosphosulfate t | SLC35B2 | 47.515 | 2 | 50.059864 | yes | 2.785E-07 | 527.78003 | 165.92841 |
| TEAAETTPMWQAI | 14 | Q8TB61 | Adenosine 3'-phospho 5'-phosphosulfate t | SLC35B2 | 47.515 | 2 | 50.059864 | yes | 2.129E-05 | 788.88745 | 55.137104 |
| FFEFGSLDQK | 10 | Q8TB72 | Pumilio homolog 2 OS=Homo sapiens O | PUM2 | 114.216 | 2 | 53.928562 | yes | 3.145E-07 | 609.29547 | 1431.1444 |
| FWEPDDSTK | 9 | Q8TB72 | Pumilio homolog 2 OS=Homo sapiens O | PUM2 | 114.216 | 2 | 53.928562 | yes | 1.509E-07 | 562.74835 | 1043.2189 |
| VLALSQHK | 8 | Q8TB72 | Pumilio homolog 2 OS=Homo sapiens O | PUM2 | 114.216 | 2 | 53.928562 | yes | 9.722E-08 | 448.27164 | 999.61926 |
| FPNLQLR | 7 | Q8TB72 | Pumilio homolog 2 OS=Homo sapiens O | PUM2 | 114.216 | 2 | 53.928562 | yes | 3.733E-07 | 444.25851 | 976.90912 |
| ESLSTSSDLYK | 11 | Q8TB72 | Pumilio homolog 2 OS=Homo sapiens O | PUM2 | 114.216 | 2 | 53.928562 | yes | 9.722E-08 | 615.29846 | 942.29395 |
| MIDMAEPAQR | 10 | Q8TB72 | Pumilio homolog 2 OS=Homo sapiens O | PUM2 | 114.216 | 2 | 53.928562 | yes | 9.722E-08 | 581.27319 | 874.71881 |
| GQVFVLSTHPYGC | 14 | Q8TB72 | Pumilio homolog 2 OS=Homo sapiens O | PUM2 | 114.216 | 2 | 53.928562 | yes | 9.728E-08 | 810.90118 | 253.72954 |
| GQVFVLSTHPYGC | 14 | Q8TB72 | Pumilio homolog 2 OS=Homo sapiens O | PUM2 | 114.216 | 2 | 53.928562 | yes | 4.887E-07 | 540.93658 | 253.72954 |
| GQVFVLSTHPYGC | 14 | Q8TB72 | Pumilio homolog 2 OS=Homo sapiens O | PUM2 | 114.216 | 3 | 53.928562 | yes | 9.728E-08 | 810.90118 | 253.72954 |

| | | | | | | | | | | | |
|---|---|---|---|---|---|---|---|---|---|---|---|
| GQVFVLSTHPYG( | 14 | Q8TB72 | Pumilio homolog 2 OS=Homo sapiens O) | PUM2 | 114.216 | 3 | 53.928562 | yes | 4.887E-07 | 540.93658 | 253.72954 |
| GHVLPLALQMYG( | 14 | Q8TB72 | Pumilio homolog 2 OS=Homo sapiens O) | PUM2 | 114.216 | 2 | 53.928562 | yes | 9.722E-08 | 807.91577 | 209.95721 |
| GHVLPLALQMYG( | 14 | Q8TB72 | Pumilio homolog 2 OS=Homo sapiens O) | PUM2 | 114.216 | 2 | 53.928562 | yes | 1.816E-07 | 538.94629 | 209.95721 |
| GHVLPLALQMYG( | 14 | Q8TB72 | Pumilio homolog 2 OS=Homo sapiens O) | PUM2 | 114.216 | 3 | 53.928562 | yes | 9.722E-08 | 807.91577 | 209.95721 |
| GHVLPLALQMYG( | 14 | Q8TB72 | Pumilio homolog 2 OS=Homo sapiens O) | PUM2 | 114.216 | 3 | 53.928562 | yes | 1.816E-07 | 538.94629 | 209.95721 |
| GMGELLPTK | 9 | Q8TB72 | Pumilio homolog 2 OS=Homo sapiens O) | PUM2 | 114.216 | 2 | 53.928562 | yes | 1.511E-07 | 473.25732 | 381.85422 |
| QASPTEVVER | 10 | Q8TB72 | Pumilio homolog 2 OS=Homo sapiens O) | PUM2 | 114.216 | 2 | 53.928562 | yes | 9.722E-08 | 558.28821 | 312.83209 |
| ETAWGASHHSMS( | 19 | Q8TB72 | Pumilio homolog 2 OS=Homo sapiens O) | PUM2 | 114.216 | 4 | 53.928562 | yes | 9.728E-08 | 539.00586 | 197.46381 |
| MNHDFQALALESI | 13 | Q8TB72 | Pumilio homolog 2 OS=Homo sapiens O) | PUM2 | 114.216 | 2 | 53.928562 | yes | 9.722E-08 | 787.375 | 168.36943 |
| GLPNGMDADCK | 11 | Q8TB72 | Pumilio homolog 2 OS=Homo sapiens O) | PUM2 | 114.216 | 2 | 53.928562 | yes | 1.577E-07 | 589.25244 | 107.59425 |
| SGQGFHGNSEVNA | 18 | Q8TB72 | Pumilio homolog 2 OS=Homo sapiens O) | PUM2 | 114.216 | 3 | 53.928562 | yes | 1.261E-07 | 623.97827 | 75.979172 |
| KFWEPDDSTK | 10 | Q8TB72 | Pumilio homolog 2 OS=Homo sapiens O) | PUM2 | 114.216 | 2 | 53.928562 | yes | 1.714E-07 | 626.79584 | 70.775475 |
| NSPDLGPIGGPPN( | 16 | Q8TB72 | Pumilio homolog 2 OS=Homo sapiens O) | PUM2 | 114.216 | 2 | 53.928562 | yes | 1.511E-07 | 768.37976 | 68.982872 |
| GMISTQPGSTPLAS | 16 | Q8TBC3 | SH3KBP1-binding protein 1 OS=Homo s | SHKBP1 | 76.344 | 2 | 48.4298 | yes | 1.15E-07 | 811.41632 | 31.00292 |
| AVYVLDEPEPAIR | 13 | Q8TBE0 | Bromo adjacent homology domain-contai | BAHD1 | 84.653 | 2 | 47.745808 | yes | 4.644E-07 | 736.39319 | 39.136147 |
| LASLNAAAFLK | 11 | Q8TBE0 | Bromo adjacent homology domain-contai | BAHD1 | 84.653 | 2 | 47.745808 | yes | 1.267E-07 | 559.83203 | 27.69978 |
| SVEEDYVTNIR | 11 | Q8TBM8 | DnaJ homolog subfamily B member 14 O | DNAJB14 | 42.516 | 2 | 52.26407 | yes | 2.461E-07 | 662.82263 | 193.64404 |
| MQTENLGVVYYV | 14 | Q8TBM8 | DnaJ homolog subfamily B member 14 O | DNAJB14 | 42.516 | 2 | 52.26407 | yes | 1.711E-07 | 829.41632 | 91.413574 |
| IGNAYAVLSNPEK | 13 | Q8TBM8 | DnaJ homolog subfamily B member 14 O | DNAJB14 | 42.516 | 2 | 52.26407 | yes | 9.722E-08 | 688.36444 | 89.681679 |
| NYYEVLGVTK | 10 | Q8TBM8 | DnaJ homolog subfamily B member 14 O | DNAJB14 | 42.516 | 2 | 52.26407 | yes | 1.076E-07 | 593.31116 | 82.976501 |
| DQVDGVLSINK | 11 | Q8TBM8 | DnaJ homolog subfamily B member 14 O | DNAJB14 | 42.516 | 2 | 52.26407 | yes | 9.881E-08 | 594.31696 | 58.662338 |
| EYCPQVFR | 8 | Q8TBX8 | Phosphatidylinositol 5-phosphate 4-kinas | PIP4K2C | 47.3 | 2 | 49.92883 | yes | 3.977E-07 | 549.75549 | 127.92178 |
| VSVDNEDSYMLVI | 14 | Q8TBX8 | Phosphatidylinositol 5-phosphate 4-kinas | PIP4K2C | 47.3 | 2 | 49.92883 | yes | 9.975E-08 | 829.38165 | 62.073509 |
| VNNHLFHR | 8 | Q8TBX8 | Phosphatidylinositol 5-phosphate 4-kinas | PIP4K2C | 47.3 | 2 | 49.92883 | yes | 1.108E-07 | 518.77539 | 51.478786 |
| FLISYDR | 7 | Q8TBX8 | Phosphatidylinositol 5-phosphate 4-kinas | PIP4K2C | 47.3 | 2 | 49.92883 | yes | 2.705E-07 | 457.24252 | 37.845543 |
| HGAGAEISTVHPE( | 17 | Q8TBX8 | Phosphatidylinositol 5-phosphate 4-kinas | PIP4K2C | 47.3 | 3 | 49.92883 | yes | 2.815E-07 | 598.96381 | 36.555702 |
| VTNYIFDSLR | 10 | Q8TC07 | TBC1 domain family member 15 OS=Ho | TBC1D15 | 79.491 | 2 | 43.87734 | yes | 2.41E-07 | 614.32202 | 70.636848 |
| INQQEEPGFEVITR | 14 | Q8TC07 | TBC1 domain family member 15 OS=Ho | TBC1D15 | 79.491 | 2 | 43.87734 | yes | 7.47E-07 | 830.42047 | 60.641502 |
| NDSPTQIPVSSDV( | 15 | Q8TC07 | TBC1 domain family member 15 OS=Ho | TBC1D15 | 79.491 | 2 | 43.87734 | yes | 1.327E-06 | 837.89124 | 41.801586 |
| TQLIQLSTLLR | 11 | Q8TC07 | TBC1 domain family member 15 OS=Ho | TBC1D15 | 79.491 | 2 | 43.87734 | yes | 2.546E-07 | 643.39557 | 31.4007 |
| TNDQDGLISGILR | 13 | Q8TC07 | TBC1 domain family member 15 OS=Ho | TBC1D15 | 79.491 | 2 | 43.87734 | yes | 2.484E-07 | 701.37024 | 27.474298 |
| LHVDFAQAR | 9 | Q8TC92 | Ecto-NOX disulfide-thiol exchanger 1 OS | ENOX1 | 73.349 | 2 | 54.826736 | no | 1.039E-07 | 528.7829 | 2204.438 |
| WQLDAYR | 7 | Q8TC92 | Ecto-NOX disulfide-thiol exchanger 1 OS | ENOX1 | 73.349 | 2 | 54.826736 | no | 3.27E-07 | 476.23779 | 1099.6273 |
| NAQSEQLMGIR | 11 | Q8TC92 | Ecto-NOX disulfide-thiol exchanger 1 OS | ENOX1 | 73.349 | 2 | 54.826736 | yes | 9.81E-08 | 623.81409 | 614.48315 |
| QEFTGVGATLEK | 12 | Q8TC92 | Ecto-NOX disulfide-thiol exchanger 1 OS | ENOX1 | 73.349 | 2 | 54.826736 | yes | 1.017E-07 | 640.33008 | 521.63171 |
| AIYLSGYR | 8 | Q8TC92 | Ecto-NOX disulfide-thiol exchanger 1 OS | ENOX1 | 73.349 | 2 | 54.826736 | yes | 9.722E-08 | 471.7558 | 456.13242 |
| LCAFEGIK | 8 | Q8TC92 | Ecto-NOX disulfide-thiol exchanger 1 OS | ENOX1 | 73.349 | 2 | 54.826736 | yes | 1.165E-07 | 469.24423 | 388.59088 |
| FAEEFMVDK | 9 | Q8TC92 | Ecto-NOX disulfide-thiol exchanger 1 OS | ENOX1 | 73.349 | 2 | 54.826736 | yes | 9.722E-08 | 558.25751 | 340.14847 |
| AWDHFSK | 7 | Q8TC92 | Ecto-NOX disulfide-thiol exchanger 1 OS | ENOX1 | 73.349 | 2 | 54.826736 | yes | 9.383E-07 | 445.7114 | 336.36362 |

| | | | | | | | | | | | |
|---|---|---|---|---|---|---|---|---|---|---|---|
| DDFYEWECK | 9 | Q8TC92 | Ecto-NOX disulfide-thiol exchanger 1 OS | ENOX1 | 73.349 | 2 | 54.826736 | yes | 1.031E-07 | 646.25043 | 187.88667 |
| KNIDIWR | 7 | Q8TC92 | Ecto-NOX disulfide-thiol exchanger 1 OS | ENOX1 | 73.349 | 2 | 54.826736 | yes | 4.199E-07 | 472.76926 | 174.23157 |
| EEQSHTQALLK | 11 | Q8TC92 | Ecto-NOX disulfide-thiol exchanger 1 OS | ENOX1 | 73.349 | 2 | 54.826736 | yes | 1.096E-07 | 642.33313 | 139.09546 |
| VDESALAAQAYAI | 14 | Q8TC92 | Ecto-NOX disulfide-thiol exchanger 1 OS | ENOX1 | 73.349 | 2 | 54.826736 | yes | 9.722E-08 | 725.38281 | 120.89808 |
| VLQEQLK | 7 | Q8TC92 | Ecto-NOX disulfide-thiol exchanger 1 OS | ENOX1 | 73.349 | 2 | 54.826736 | no | 4.177E-07 | 429.25818 | 68.250221 |
| ISANEIEMLLMR | 12 | Q8TC92 | Ecto-NOX disulfide-thiol exchanger 1 OS | ENOX1 | 73.349 | 2 | 54.826736 | yes | 8.204E-06 | 710.37036 | 7.5273924 |
| ESDYFTPQGEFR | 12 | Q8TCJ2 | Dolichyl-diphosphooligosaccharide--prot | STT3B | 93.675 | 2 | 53.94544 | yes | 9.722E-08 | 738.3255 | 418.63077 |
| AGSPTLLNCLMYK | 13 | Q8TCJ2 | Dolichyl-diphosphooligosaccharide--prot | STT3B | 93.675 | 2 | 53.94544 | yes | 1.411E-07 | 734.37036 | 350.52322 |
| FYSLWDTGYAK | 11 | Q8TCJ2 | Dolichyl-diphosphooligosaccharide--prot | STT3B | 93.675 | 2 | 53.94544 | yes | 1.152E-07 | 675.8219 | 253.57179 |
| HLEEAFTSEHWLV | 14 | Q8TCJ2 | Dolichyl-diphosphooligosaccharide--prot | STT3B | 93.675 | 3 | 53.94544 | yes | 1.313E-07 | 585.2934 | 178.62206 |
| EAYFWLR | 7 | Q8TCJ2 | Dolichyl-diphosphooligosaccharide--prot | STT3B | 93.675 | 2 | 53.94544 | yes | 1.487E-07 | 492.75052 | 133.82198 |
| FGEMQLDFR | 9 | Q8TCJ2 | Dolichyl-diphosphooligosaccharide--prot | STT3B | 93.675 | 2 | 53.94544 | yes | 9.722E-08 | 571.76862 | 133.68793 |
| TEEGLGPNIK | 10 | Q8TCJ2 | Dolichyl-diphosphooligosaccharide--prot | STT3B | 93.675 | 2 | 53.94544 | yes | 5.362E-06 | 529.27985 | 109.28088 |
| VTNIFPK | 7 | Q8TCJ2 | Dolichyl-diphosphooligosaccharide--prot | STT3B | 93.675 | 2 | 53.94544 | yes | 1.46E-06 | 409.74216 | 34.002449 |
| NILDDFR | 7 | Q8TCJ2 | Dolichyl-diphosphooligosaccharide--prot | STT3B | 93.675 | 2 | 53.94544 | yes | 4.022E-07 | 446.7298 | 31.646336 |
| AWYPLGR | 7 | Q8TCJ2 | Dolichyl-diphosphooligosaccharide--prot | STT3B | 93.675 | 2 | 53.94544 | no | 3.182E-06 | 431.73215 | 30.127161 |
| LFTPSPEIVK | 10 | Q8TCS8 | Polyribonucleotide nucleotidyltransferase | PNPT1 | 85.951 | 2 | 54.83167 | yes | 3.076E-07 | 565.82642 | 754.31616 |
| IGIIDGEYVVNPTR | 14 | Q8TCS8 | Polyribonucleotide nucleotidyltransferase | PNPT1 | 85.951 | 2 | 54.83167 | yes | 9.722E-08 | 773.41718 | 697.04333 |
| VLQSPATTVVR | 11 | Q8TCS8 | Polyribonucleotide nucleotidyltransferase | PNPT1 | 85.951 | 2 | 54.83167 | yes | 1.073E-07 | 585.8457 | 413.49142 |
| IPTNYLR | 7 | Q8TCS8 | Polyribonucleotide nucleotidyltransferase | PNPT1 | 85.951 | 2 | 54.83167 | yes | 1.011E-07 | 438.75052 | 406.88644 |
| YTQQIIQGIQQLVK | 14 | Q8TCS8 | Polyribonucleotide nucleotidyltransferase | PNPT1 | 85.951 | 2 | 54.83167 | yes | 9.723E-08 | 830.47504 | 351.97189 |
| FVGPGGYNLK | 10 | Q8TCS8 | Polyribonucleotide nucleotidyltransferase | PNPT1 | 85.951 | 2 | 54.83167 | yes | 1.118E-06 | 526.28217 | 308.96646 |
| ALYPVIPR | 8 | Q8TCS8 | Polyribonucleotide nucleotidyltransferase | PNPT1 | 85.951 | 2 | 54.83167 | yes | 1.724E-07 | 464.78436 | 303.1662 |
| IVMEAIQQASVAK | 13 | Q8TCS8 | Polyribonucleotide nucleotidyltransferase | PNPT1 | 85.951 | 2 | 54.83167 | yes | 9.722E-08 | 694.38434 | 223.14134 |
| NVSCEVDMFK | 10 | Q8TCS8 | Polyribonucleotide nucleotidyltransferase | PNPT1 | 85.951 | 2 | 54.83167 | yes | 1.381E-07 | 614.77045 | 188.25806 |
| TLHGSALFQR | 10 | Q8TCS8 | Polyribonucleotide nucleotidyltransferase | PNPT1 | 85.951 | 2 | 54.83167 | yes | 6.198E-07 | 565.30927 | 166.47937 |
| LYAVFTDYEHDK | 12 | Q8TCS8 | Polyribonucleotide nucleotidyltransferase | PNPT1 | 85.951 | 2 | 54.83167 | yes | 9.928E-08 | 750.85388 | 149.05768 |
| GITALQADIK | 10 | Q8TCS8 | Polyribonucleotide nucleotidyltransferase | PNPT1 | 85.951 | 2 | 54.83167 | yes | 2.284E-07 | 515.3006 | 129.40944 |
| TDPEKGEIEDYR | 12 | Q8TCS8 | Polyribonucleotide nucleotidyltransferase | PNPT1 | 85.951 | 2 | 54.83167 | yes | 1.51E-07 | 726.33606 | 100.37463 |
| HPTALGLEVGQEI( | 16 | Q8TCS8 | Polyribonucleotide nucleotidyltransferase | PNPT1 | 85.951 | 3 | 54.83167 | yes | 4.059E-07 | 573.65271 | 72.172569 |
| KVLQSPATTVVR | 12 | Q8TCS8 | Polyribonucleotide nucleotidyltransferase | PNPT1 | 85.951 | 2 | 54.83167 | yes | 9.722E-08 | 649.89319 | 60.334976 |
| EMSSSTLNLVVAG | 16 | Q8TCS8 | Polyribonucleotide nucleotidyltransferase | PNPT1 | 85.951 | 2 | 54.83167 | yes | 9.722E-08 | 802.42163 | 56.791603 |
| LDTEEQLK | 8 | Q8TCS8 | Polyribonucleotide nucleotidyltransferase | PNPT1 | 85.951 | 2 | 54.83167 | yes | 2.847E-06 | 488.2533 | 51.860325 |
| ENGPVVETVQVPI | 15 | Q8TCS8 | Polyribonucleotide nucleotidyltransferase | PNPT1 | 85.951 | 2 | 54.83167 | yes | 7.522E-07 | 798.43561 | 25.94672 |
| DISDISFIQSK | 11 | Q8TCU4 | Centrosome-associated protein ALMS1 C | ALMS1 | 461.063 | 2 | 43.778084 | yes | 2.45E-07 | 626.82465 | 45.758629 |
| LYSSITNQQR | 10 | Q8TCU4 | Centrosome-associated protein ALMS1 C | ALMS1 | 461.063 | 2 | 43.778084 | yes | 7.386E-07 | 605.31476 | 42.728714 |
| VSPVSIPAEQK | 11 | Q8TCU4 | Centrosome-associated protein ALMS1 C | ALMS1 | 461.063 | 2 | 43.778084 | yes | 4.337E-05 | 577.8244 | 40.967369 |
| GIENWEFISSTTVR | 14 | Q8TCU4 | Centrosome-associated protein ALMS1 C | ALMS1 | 461.063 | 2 | 43.778084 | yes | 1.356E-06 | 819.90973 | 33.907223 |
| VSMALEETLR | 10 | Q8TCU4 | Centrosome-associated protein ALMS1 C | ALMS1 | 461.063 | 2 | 43.778084 | yes | 9.11E-07 | 574.80261 | 27.394411 |

| | | | | | | | | | | | |
|---|---|---|---|---|---|---|---|---|---|---|---|
| LVVLSTVGVPGTG | 14 | Q8TCY9 | Up-regulator of cell proliferation OS=Hoı | URGCP | 104.988 | 2 | 49.927456 | yes | 9.976E-08 | 663.9032 | 273.37451 |
| CLLENIR | 7 | Q8TCY9 | Up-regulator of cell proliferation OS=Hoı | URGCP | 104.988 | 2 | 49.927456 | yes | 3.316E-07 | 459.24728 | 120.5292 |
| STLLNTMFGLR | 11 | Q8TCY9 | Up-regulator of cell proliferation OS=Hoı | URGCP | 104.988 | 2 | 49.927456 | yes | 2.011E-07 | 626.83954 | 103.10677 |
| GDIGSHWLQFK | 11 | Q8TCY9 | Up-regulator of cell proliferation OS=Hoı | URGCP | 104.988 | 2 | 49.927456 | yes | 3.825E-07 | 644.32764 | 100.57285 |
| EDSVVLSR | 8 | Q8TCY9 | Up-regulator of cell proliferation OS=Hoı | URGCP | 104.988 | 2 | 49.927456 | yes | 3.363E-07 | 452.74036 | 78.497345 |
| ALAGLAFCDPEK | 12 | Q8TCY9 | Up-regulator of cell proliferation OS=Hoı | URGCP | 104.988 | 2 | 49.927456 | yes | 3.532E-06 | 646.32117 | 76.485229 |
| VSVEDMAHAAR | 11 | Q8TCY9 | Up-regulator of cell proliferation OS=Hoı | URGCP | 104.988 | 2 | 49.927456 | yes | 9.267E-07 | 593.28766 | 69.430519 |
| IDHSHVLVK | 9 | Q8TCY9 | Up-regulator of cell proliferation OS=Hoı | URGCP | 104.988 | 2 | 49.927456 | yes | 1.899E-07 | 524.3009 | 61.602539 |
| WMEWGLAR | 8 | Q8TCY9 | Up-regulator of cell proliferation OS=Hoı | URGCP | 104.988 | 2 | 49.927456 | yes | 2.695E-07 | 524.75531 | 56.647751 |
| VSSTDSDSFVK | 11 | Q8TCY9 | Up-regulator of cell proliferation OS=Hoı | URGCP | 104.988 | 2 | 49.927456 | yes | 9.298E-07 | 586.27753 | 27.883089 |
| EMEGDDCEFR | 10 | Q8TCY9 | Up-regulator of cell proliferation OS=Hoı | URGCP | 104.988 | 2 | 49.927456 | yes | 1.608E-07 | 644.23444 | 21.82534 |
| ISLLLHQR | 8 | Q8TD17 | Zinc finger protein 398 OS=Homo sapien | ZNF398 | 71.312 | 2 | 41.099648 | yes | 3.677E-07 | 490.306 | 29.199919 |
| TFDATNPLNLCVK | 13 | Q8TD19 | Serine/threonine-protein kinase Nek9 OS: | NEK9 | 107.169 | 2 | 47.280125 | yes | 1.363E-07 | 746.87683 | 85.276062 |
| VTLLNAPTK | 9 | Q8TD19 | Serine/threonine-protein kinase Nek9 OS: | NEK9 | 107.169 | 2 | 47.280125 | yes | 3.238E-06 | 478.79239 | 59.770302 |
| LGINLLGGPLGGK | 13 | Q8TD19 | Serine/threonine-protein kinase Nek9 OS: | NEK9 | 107.169 | 2 | 47.280125 | yes | 1.69E-07 | 604.8717 | 51.750694 |
| SSTVTEAPIAVVTS | 15 | Q8TD19 | Serine/threonine-protein kinase Nek9 OS: | NEK9 | 107.169 | 2 | 47.280125 | yes | 1.4E-06 | 759.41211 | 41.305664 |
| EVYSWGCGEYGR | 12 | Q8TD19 | Serine/threonine-protein kinase Nek9 OS: | NEK9 | 107.169 | 2 | 47.280125 | yes | 8.39E-07 | 731.80646 | 36.239082 |
| GWHTILIVEK | 10 | Q8TD19 | Serine/threonine-protein kinase Nek9 OS: | NEK9 | 107.169 | 2 | 47.280125 | yes | 6.286E-06 | 598.34534 | 36.102142 |
| SLGKERK | 7 | Q8TDB6 | E3 ubiquitin-protein ligase DTX3L OS=I | DTX3L | 83.555 | 2 | 55.660145 | yes | 6.183E-07 | 409.24814 | 1823.0208 |
| GGELTLLGTQDDIS | 17 | Q8TDB6 | E3 ubiquitin-protein ligase DTX3L OS=I | DTX3L | 83.555 | 2 | 55.660145 | yes | 3.936E-07 | 844.93872 | 561.9646 |
| VCGDFQDIER | 10 | Q8TDB6 | E3 ubiquitin-protein ligase DTX3L OS=I | DTX3L | 83.555 | 2 | 55.660145 | yes | 4.487E-07 | 619.77716 | 411.12625 |
| ESFASEFQK | 9 | Q8TDB6 | E3 ubiquitin-protein ligase DTX3L OS=I | DTX3L | 83.555 | 2 | 55.660145 | yes | 1.993E-07 | 536.75092 | 359.19485 |
| GICVICMDTISNK | 13 | Q8TDB6 | E3 ubiquitin-protein ligase DTX3L OS=I | DTX3L | 83.555 | 2 | 55.660145 | yes | 3.418E-07 | 755.85675 | 345.73648 |
| AYITTLCPSIR | 11 | Q8TDB6 | E3 ubiquitin-protein ligase DTX3L OS=I | DTX3L | 83.555 | 2 | 55.660145 | yes | 6.824E-07 | 647.84485 | 247.92857 |
| HPNVHFVLNQESM | 24 | Q8TDB6 | E3 ubiquitin-protein ligase DTX3L OS=I | DTX3L | 83.555 | 4 | 55.660145 | yes | 2.89E-07 | 675.1037 | 232.60962 |
| TCILFESK | 8 | Q8TDB6 | E3 ubiquitin-protein ligase DTX3L OS=I | DTX3L | 83.555 | 2 | 55.660145 | yes | 2.201E-07 | 499.25479 | 192.48268 |
| LIFTVGYSR | 9 | Q8TDB6 | E3 ubiquitin-protein ligase DTX3L OS=I | DTX3L | 83.555 | 2 | 55.660145 | yes | 2.167E-07 | 528.29785 | 192.22412 |
| GNQPEGSMVFTVS | 14 | Q8TDB6 | E3 ubiquitin-protein ligase DTX3L OS=I | DTX3L | 83.555 | 2 | 55.660145 | yes | 1.066E-07 | 754.86176 | 191.1351 |
| QQFSPSMTER | 10 | Q8TDB6 | E3 ubiquitin-protein ligase DTX3L OS=I | DTX3L | 83.555 | 2 | 55.660145 | yes | 1.34E-07 | 605.77966 | 162.32004 |
| AGIQTEEHPNPGK | 13 | Q8TDB6 | E3 ubiquitin-protein ligase DTX3L OS=I | DTX3L | 83.555 | 2 | 55.660145 | yes | 9.722E-08 | 689.34149 | 158.03848 |
| GSVSSEASELDKK | 13 | Q8TDB6 | E3 ubiquitin-protein ligase DTX3L OS=I | DTX3L | 83.555 | 2 | 55.660145 | yes | 9.722E-08 | 668.83319 | 107.61836 |
| GSVSSEASELDK | 12 | Q8TDB6 | E3 ubiquitin-protein ligase DTX3L OS=I | DTX3L | 83.555 | 2 | 55.660145 | yes | 9.722E-08 | 604.78571 | 79.505829 |
| PICPTCQTSYGIQK | 14 | Q8TDB6 | E3 ubiquitin-protein ligase DTX3L OS=I | DTX3L | 83.555 | 2 | 55.660145 | yes | 9.728E-08 | 826.89221 | 68.096794 |
| QELNHQFTK | 9 | Q8TDB6 | E3 ubiquitin-protein ligase DTX3L OS=I | DTX3L | 83.555 | 2 | 55.660145 | yes | 1.182E-07 | 572.79089 | 42.739849 |
| KLESYFQSSK | 10 | Q8TDB6 | E3 ubiquitin-protein ligase DTX3L OS=I | DTX3L | 83.555 | 2 | 55.660145 | yes | 3.512E-06 | 608.81403 | 26.1684 |
| HEFCAPCINK | 10 | Q8TDB6 | E3 ubiquitin-protein ligase DTX3L OS=I | DTX3L | 83.555 | 2 | 55.660145 | yes | 1.458E-06 | 638.28406 | 23.282187 |
| CSTLIVTDLAAR | 12 | Q8TDD1 | ATP-dependent RNA helicase DDX54 O: | DDX54 | 98.595 | 2 | 53.149876 | yes | 1.091E-07 | 660.35303 | 556.66138 |
| DQEFYIPYRPK | 11 | Q8TDD1 | ATP-dependent RNA helicase DDX54 O: | DDX54 | 98.595 | 2 | 53.149876 | yes | 1.276E-07 | 728.367 | 547.2359 |
| AGLTEPVLIR | 10 | Q8TDD1 | ATP-dependent RNA helicase DDX54 O: | DDX54 | 98.595 | 2 | 53.149876 | yes | 1.025E-07 | 534.82422 | 440.37473 |

| | | | | | | | | | | | |
|---|---|---|---|---|---|---|---|---|---|---|---|
| FEEEELQR | 8 | Q8TDD1 | ATP-dependent RNA helicase DDX54 OS | DDX54 | 98.595 | 2 | 53.149876 | yes | 2.633E-07 | 540.25385 | 375.4425 |
| LQSVEYVVFDEAD | 14 | Q8TDD1 | ATP-dependent RNA helicase DDX54 OS | DDX54 | 98.595 | 2 | 53.149876 | yes | 1.442E-07 | 835.40704 | 275.18326 |
| EPSGVAGVDGML( | 14 | Q8TDD1 | ATP-dependent RNA helicase DDX54 OS | DDX54 | 98.595 | 2 | 53.149876 | yes | 1.879E-07 | 672.83246 | 233.16853 |
| VQELQQGAFGR | 11 | Q8TDD1 | ATP-dependent RNA helicase DDX54 OS | DDX54 | 98.595 | 2 | 53.149876 | yes | 9.722E-08 | 616.82275 | 197.96986 |
| ELALQTLK | 8 | Q8TDD1 | ATP-dependent RNA helicase DDX54 OS | DDX54 | 98.595 | 2 | 53.149876 | yes | 2.226E-07 | 458.27911 | 184.68031 |
| VSCAHIYSALDPTA | 15 | Q8TDD1 | ATP-dependent RNA helicase DDX54 OS | DDX54 | 98.595 | 2 | 53.149876 | yes | 9.722E-08 | 830.90924 | 91.797516 |
| VSCAHIYSALDPTA | 15 | Q8TDD1 | ATP-dependent RNA helicase DDX54 OS | DDX54 | 98.595 | 2 | 53.149876 | yes | 2.48E-07 | 554.27527 | 91.797516 |
| VSCAHIYSALDPTA | 15 | Q8TDD1 | ATP-dependent RNA helicase DDX54 OS | DDX54 | 98.595 | 3 | 53.149876 | yes | 9.722E-08 | 830.90924 | 91.797516 |
| VSCAHIYSALDPTA | 15 | Q8TDD1 | ATP-dependent RNA helicase DDX54 OS | DDX54 | 98.595 | 3 | 53.149876 | yes | 2.48E-07 | 554.27527 | 91.797516 |
| FVGQSGQEDK | 10 | Q8TDD1 | ATP-dependent RNA helicase DDX54 OS | DDX54 | 98.595 | 2 | 53.149876 | yes | 1.022E-07 | 547.75928 | 176.09038 |
| LVHVAVEMSLK | 11 | Q8TDD1 | ATP-dependent RNA helicase DDX54 OS | DDX54 | 98.595 | 2 | 53.149876 | yes | 1.065E-07 | 613.35229 | 152.998 |
| SGGFQSMGLSYPV | 15 | Q8TDD1 | ATP-dependent RNA helicase DDX54 OS | DDX54 | 98.595 | 2 | 53.149876 | yes | 1.02E-07 | 802.89252 | 135.43141 |
| ATIFEINASSR | 11 | Q8TDD1 | ATP-dependent RNA helicase DDX54 OS | DDX54 | 98.595 | 2 | 53.149876 | yes | 1.498E-07 | 604.81714 | 109.42278 |
| VADNAQQQYVR | 11 | Q8TDD1 | ATP-dependent RNA helicase DDX54 OS | DDX54 | 98.595 | 2 | 53.149876 | yes | 9.722E-08 | 646.32312 | 99.309219 |
| LLVEFAR | 7 | Q8TDD1 | ATP-dependent RNA helicase DDX54 OS | DDX54 | 98.595 | 2 | 53.149876 | yes | 2.878E-07 | 424.25543 | 70.474167 |
| SRPAPSPESIK | 11 | Q8TDD1 | ATP-dependent RNA helicase DDX54 OS | DDX54 | 98.595 | 2 | 53.149876 | yes | 1.076E-07 | 584.81964 | 58.165016 |
| IAVNPASLGER | 11 | Q8TDM6 | Disks large homolog 5 OS=Homo sapiens | DLG5 | 213.868 | 2 | 51.578773 | yes | 1.847E-07 | 563.81439 | 174.96767 |
| LQTEVELAESK | 11 | Q8TDM6 | Disks large homolog 5 OS=Homo sapiens | DLG5 | 213.868 | 2 | 51.578773 | yes | 1.829E-06 | 623.8299 | 146.6423 |
| VENLSIQLR | 9 | Q8TDM6 | Disks large homolog 5 OS=Homo sapiens | DLG5 | 213.868 | 2 | 51.578773 | yes | 3.715E-07 | 536.31146 | 142.42976 |
| AVSELAEALR | 10 | Q8TDM6 | Disks large homolog 5 OS=Homo sapiens | DLG5 | 213.868 | 2 | 51.578773 | yes | 3.548E-07 | 529.79565 | 141.37724 |
| LKDDVDMLR | 9 | Q8TDM6 | Disks large homolog 5 OS=Homo sapiens | DLG5 | 213.868 | 2 | 51.578773 | yes | 4.877E-07 | 552.78955 | 127.69421 |
| LSLDLSHR | 8 | Q8TDM6 | Disks large homolog 5 OS=Homo sapiens | DLG5 | 213.868 | 2 | 51.578773 | yes | 1.641E-07 | 470.76416 | 110.76541 |
| SLTPSTTVSSILR | 13 | Q8TDM6 | Disks large homolog 5 OS=Homo sapiens | DLG5 | 213.868 | 2 | 51.578773 | yes | 5.483E-07 | 681.38538 | 103.56271 |
| GLPGDSFYIR | 10 | Q8TDM6 | Disks large homolog 5 OS=Homo sapiens | DLG5 | 213.868 | 2 | 51.578773 | yes | 1.071E-07 | 562.79041 | 98.72773 |
| EGSLAVGDR | 9 | Q8TDM6 | Disks large homolog 5 OS=Homo sapiens | DLG5 | 213.868 | 2 | 51.578773 | yes | 1.108E-07 | 452.23016 | 89.814888 |
| TLCDNLR | 7 | Q8TDM6 | Disks large homolog 5 OS=Homo sapiens | DLG5 | 213.868 | 2 | 51.578773 | yes | 1.831E-07 | 446.22128 | 86.474182 |
| LAFATHGTAFDK | 12 | Q8TDM6 | Disks large homolog 5 OS=Homo sapiens | DLG5 | 213.868 | 2 | 51.578773 | yes | 9.81E-08 | 639.82751 | 84.942741 |
| GSEPLGISIVSGEK | 14 | Q8TDM6 | Disks large homolog 5 OS=Homo sapiens | DLG5 | 213.868 | 2 | 51.578773 | yes | 9.722E-08 | 686.86957 | 79.22744 |
| SCQDSLTLSLLK | 12 | Q8TDM6 | Disks large homolog 5 OS=Homo sapiens | DLG5 | 213.868 | 2 | 51.578773 | yes | 1.119E-07 | 682.85815 | 75.615913 |
| GFGLVDVR | 8 | Q8TDM6 | Disks large homolog 5 OS=Homo sapiens | DLG5 | 213.868 | 2 | 51.578773 | yes | 3.558E-06 | 431.74271 | 70.199059 |
| TVEEVYVEMLKPI | 13 | Q8TDM6 | Disks large homolog 5 OS=Homo sapiens | DLG5 | 213.868 | 2 | 51.578773 | yes | 1.292E-07 | 796.92126 | 49.401642 |
| AANEEMEALR | 10 | Q8TDM6 | Disks large homolog 5 OS=Homo sapiens | DLG5 | 213.868 | 2 | 51.578773 | yes | 1.002E-07 | 567.26642 | 48.267849 |
| VVTPLHINLSGQK | 13 | Q8TDM6 | Disks large homolog 5 OS=Homo sapiens | DLG5 | 213.868 | 2 | 51.578773 | yes | 6.796E-07 | 703.41174 | 44.826687 |
| SCDEDLTSQK | 10 | Q8TDM6 | Disks large homolog 5 OS=Homo sapiens | DLG5 | 213.868 | 2 | 51.578773 | yes | 4.866E-07 | 591.75079 | 42.670319 |
| SLNECESLLR | 10 | Q8TDM6 | Disks large homolog 5 OS=Homo sapiens | DLG5 | 213.868 | 2 | 51.578773 | yes | 2.353E-06 | 610.8006 | 39.91909 |
| HSNPPLYPSRPSV( | 18 | Q8TDM6 | Disks large homolog 5 OS=Homo sapiens | DLG5 | 213.868 | 3 | 51.578773 | yes | 1.23E-06 | 654.34991 | 25.60145 |
| LYDTAMDK | 8 | Q8TDM6 | Disks large homolog 5 OS=Homo sapiens | DLG5 | 213.868 | 2 | 51.578773 | yes | 4.017E-06 | 478.72333 | 23.430815 |
| TCSDYSEMR | 9 | Q8TDM6 | Disks large homolog 5 OS=Homo sapiens | DLG5 | 213.868 | 2 | 51.578773 | yes | 1.478E-07 | 574.72101 | 19.430498 |
| LAPVVIPAQFLEE( | 15 | Q8TDM6 | Disks large homolog 5 OS=Homo sapiens | DLG5 | 213.868 | 2 | 51.578773 | yes | 2.219E-06 | 841.4798 | 15.963203 |

| | | | | | | | | | | | |
|---|---|---|---|---|---|---|---|---|---|---|---|
| TNELLQIIGK | 10 | Q8TDR0 | TRAF3-interacting protein 1 OS=Homo s | TRAF3IP1 | 78.632 | 2 | 42.404316 | yes | 3.046E-07 | 564.83478 | 62.550354 |
| DIKPANVFITATG\ | 16 | Q8TDX7 | Serine/threonine-protein kinase Nek7 OS= | NEK7 | 34.551 | 2 | 51.503315 | no | 9.722E-08 | 836.98523 | 182.99252 |
| AACLLDGVPVALK | 13 | Q8TDX7 | Serine/threonine-protein kinase Nek7 OS= | NEK7 | 34.551 | 2 | 51.503315 | yes | 3.339E-07 | 663.87616 | 126.27788 |
| GQFSEVYR | 8 | Q8TDX7 | Serine/threonine-protein kinase Nek7 OS= | NEK7 | 34.551 | 2 | 51.503315 | yes | 2.292E-07 | 493.24051 | 107.4793 |
| MNLYSLCK | 8 | Q8TDX7 | Serine/threonine-protein kinase Nek7 OS= | NEK7 | 34.551 | 2 | 51.503315 | yes | 1.195E-07 | 514.74884 | 96.086075 |
| VQIFDLMDAK | 10 | Q8TDX7 | Serine/threonine-protein kinase Nek7 OS= | NEK7 | 34.551 | 2 | 51.503315 | yes | 1.462E-07 | 590.30756 | 44.269104 |
| QLVNMCINPDPEK | 13 | Q8TDX7 | Serine/threonine-protein kinase Nek7 OS= | NEK7 | 34.551 | 2 | 51.503315 | yes | 5.153E-07 | 779.3736 | 36.394978 |
| RPDVTYVYDVAK | 12 | Q8TDX7 | Serine/threonine-protein kinase Nek7 OS= | NEK7 | 34.551 | 2 | 51.503315 | yes | 7.449E-07 | 713.37225 | 27.570049 |
| KVQIFDLMDAK | 11 | Q8TDX7 | Serine/threonine-protein kinase Nek7 OS= | NEK7 | 34.551 | 2 | 51.503315 | yes | 2.841E-06 | 654.35504 | 25.579233 |
| IPLLECLTR | 9 | Q8TDY2 | RB1-inducible coiled-coil protein 1 OS=I | RB1CC1 | 183.091 | 2 | 34.35565 | yes | 1.025E-06 | 557.81805 | 28.338139 |
| TEEAFPGGPLGALI | 14 | Q8TE02 | Elongator complex protein 5 OS=Homo s | ELP5 | 34.842 | 2 | 31.360514 | yes | 1.867E-06 | 707.86987 | 72.818398 |
| DDILINRPAK | 10 | Q8TED0 | U3 small nucleolar RNA-associated prote | UTP15 | 58.416 | 2 | 53.53823 | yes | 1.102E-07 | 577.83002 | 393.49817 |
| TPEITVSIIK | 10 | Q8TED0 | U3 small nucleolar RNA-associated prote | UTP15 | 58.416 | 2 | 53.53823 | yes | 2.716E-07 | 550.83173 | 271.27811 |
| ITQDTLYWNNYK | 12 | Q8TED0 | U3 small nucleolar RNA-associated prote | UTP15 | 58.416 | 2 | 53.53823 | yes | 9.722E-08 | 779.88043 | 269.19662 |
| LLSGSLDR | 8 | Q8TED0 | U3 small nucleolar RNA-associated prote | UTP15 | 58.416 | 2 | 53.53823 | yes | 2.13E-07 | 430.74545 | 226.66049 |
| TVTCLCLSSSGQR | 13 | Q8TED0 | U3 small nucleolar RNA-associated prote | UTP15 | 58.416 | 2 | 53.53823 | yes | 1.198E-07 | 734.84778 | 224.56839 |
| VLDPTCTIK | 9 | Q8TED0 | U3 small nucleolar RNA-associated prote | UTP15 | 58.416 | 2 | 53.53823 | yes | 1.068E-07 | 523.78119 | 221.99719 |
| YHVVSGADDYTV | 13 | Q8TED0 | U3 small nucleolar RNA-associated prote | UTP15 | 58.416 | 2 | 53.53823 | yes | 1.466E-07 | 727.3515 | 191.72522 |
| FLLLQGLVEK | 10 | Q8TED0 | U3 small nucleolar RNA-associated prote | UTP15 | 58.416 | 2 | 53.53823 | yes | 1.754E-07 | 580.35791 | 150.60799 |
| AVHTVDFTADK | 11 | Q8TED0 | U3 small nucleolar RNA-associated prote | UTP15 | 58.416 | 2 | 53.53823 | yes | 1.376E-07 | 602.30383 | 121.45032 |
| GVLANALAGR | 10 | Q8TED0 | U3 small nucleolar RNA-associated prote | UTP15 | 58.416 | 2 | 53.53823 | yes | 3.217E-07 | 471.27997 | 100.87437 |
| LWDIPNSK | 8 | Q8TED0 | U3 small nucleolar RNA-associated prote | UTP15 | 58.416 | 2 | 53.53823 | yes | 1.916E-07 | 486.76108 | 57.194191 |
| LNPDLFITGSYDHT | 16 | Q8TED0 | U3 small nucleolar RNA-associated prote | UTP15 | 58.416 | 3 | 53.53823 | yes | 2.732E-07 | 607.31238 | 50.911972 |
| YLVNPEGQVVK | 11 | Q8TED1 | Probable glutathione peroxidase 8 OS=Hc | GPX8 | 23.882 | 2 | 51.595707 | yes | 2.352E-07 | 623.34552 | 902.02356 |
| ILGSEGEPAFR | 11 | Q8TED1 | Probable glutathione peroxidase 8 OS=Hc | GPX8 | 23.882 | 2 | 51.595707 | yes | 1.089E-06 | 588.3064 | 368.267 |
| NYGVTFPIFHK | 11 | Q8TED1 | Probable glutathione peroxidase 8 OS=Hc | GPX8 | 23.882 | 2 | 51.595707 | yes | 1.118E-07 | 661.84821 | 307.01447 |
| INSFYAFEVK | 10 | Q8TED1 | Probable glutathione peroxidase 8 OS=Hc | GPX8 | 23.882 | 2 | 51.595707 | yes | 9.722E-08 | 609.31366 | 296.0296 |
| PDIAALVR | 8 | Q8TED1 | Probable glutathione peroxidase 8 OS=Hc | GPX8 | 23.882 | 2 | 51.595707 | yes | 1.075E-07 | 427.75836 | 78.422203 |
| APVEVAPCADVV7 | 14 | Q8TEP8 | Centrosomal protein of 192 kDa OS=Hon | CEP192 | 279.112 | 2 | 42.754295 | yes | 2.889E-07 | 742.38232 | 48.458515 |
| LQLSFQDDDSISR | 13 | Q8TEP8 | Centrosomal protein of 192 kDa OS=Hon | CEP192 | 279.112 | 2 | 42.754295 | yes | 2.026E-06 | 762.37042 | 39.467323 |
| NEPLSLPELTK | 11 | Q8TEQ6 | Gem-associated protein 5 OS=Homo sapi | GEMIN5 | 168.59 | 2 | 52.181217 | yes | 5.54E-07 | 620.84283 | 3117.8267 |
| VIGELVGHTER | 11 | Q8TEQ6 | Gem-associated protein 5 OS=Homo sapi | GEMIN5 | 168.59 | 2 | 52.181217 | yes | 9.722E-08 | 605.33295 | 1551.0287 |
| VIGELVGHTER | 11 | Q8TEQ6 | Gem-associated protein 5 OS=Homo sapi | GEMIN5 | 168.59 | 2 | 52.181217 | yes | 9.809E-08 | 403.89105 | 1551.0287 |
| VIGELVGHTER | 11 | Q8TEQ6 | Gem-associated protein 5 OS=Homo sapi | GEMIN5 | 168.59 | 3 | 52.181217 | yes | 9.722E-08 | 605.33295 | 1551.0287 |
| VIGELVGHTER | 11 | Q8TEQ6 | Gem-associated protein 5 OS=Homo sapi | GEMIN5 | 168.59 | 3 | 52.181217 | yes | 9.809E-08 | 403.89105 | 1551.0287 |
| FHLGLFTDR | 9 | Q8TEQ6 | Gem-associated protein 5 OS=Homo sapi | GEMIN5 | 168.59 | 2 | 52.181217 | yes | 1.382E-06 | 553.29309 | 2839.2461 |
| VGPGAGESPGTPPI | 15 | Q8TEQ6 | Gem-associated protein 5 OS=Homo sapi | GEMIN5 | 168.59 | 2 | 52.181217 | yes | 9.935E-08 | 713.35968 | 2778.969 |
| GVLQTAAER | 9 | Q8TEQ6 | Gem-associated protein 5 OS=Homo sapi | GEMIN5 | 168.59 | 2 | 52.181217 | yes | 1.506E-07 | 472.76163 | 2079.7244 |
| IVFNLCPLQTEDDI | 14 | Q8TEQ6 | Gem-associated protein 5 OS=Homo sapi | GEMIN5 | 168.59 | 2 | 52.181217 | yes | 1.491E-07 | 846.41907 | 1988.7606 |

| | | | | | | | | | | | |
|---|---|---|---|---|---|---|---|---|---|---|---|
| EEPLCNFR | 8 | Q8TEQ6 | Gem-associated protein 5 OS=Homo sapi | GEMIN5 | 168.59 | 2 | 52.181217 | yes | 1.452E-07 | 532.74512 | 1685.1128 |
| EGCLAFGTDDGK | 12 | Q8TEQ6 | Gem-associated protein 5 OS=Homo sapi | GEMIN5 | 168.59 | 2 | 52.181217 | yes | 9.81E-08 | 635.27441 | 1629.3956 |
| EPELPCGLAPAVSI | 14 | Q8TEQ6 | Gem-associated protein 5 OS=Homo sapi | GEMIN5 | 168.59 | 2 | 52.181217 | yes | 1.665E-07 | 748.38232 | 1622.6637 |
| CYLGATCAYDAAI | 13 | Q8TEQ6 | Gem-associated protein 5 OS=Homo sapi | GEMIN5 | 168.59 | 2 | 52.181217 | yes | 9.819E-08 | 732.3183 | 1527.0568 |
| VTALCWHPTK | 10 | Q8TEQ6 | Gem-associated protein 5 OS=Homo sapi | GEMIN5 | 168.59 | 2 | 52.181217 | yes | 1.569E-07 | 606.81335 | 1229.6536 |
| ELNEDVSADVEER | 13 | Q8TEQ6 | Gem-associated protein 5 OS=Homo sapi | GEMIN5 | 168.59 | 2 | 52.181217 | yes | 9.722E-08 | 752.84174 | 1204.4106 |
| LRPEDPVLK | 9 | Q8TEQ6 | Gem-associated protein 5 OS=Homo sapi | GEMIN5 | 168.59 | 2 | 52.181217 | yes | 1.858E-07 | 533.81641 | 1186.0453 |
| VYEAVELLK | 9 | Q8TEQ6 | Gem-associated protein 5 OS=Homo sapi | GEMIN5 | 168.59 | 2 | 52.181217 | yes | 2.848E-07 | 532.30536 | 1130.1328 |
| QLLLSTSMDR | 10 | Q8TEQ6 | Gem-associated protein 5 OS=Homo sapi | GEMIN5 | 168.59 | 2 | 52.181217 | yes | 1.03E-07 | 582.30811 | 1123.3468 |
| CSDAVPGGLFGFA | 15 | Q8TEQ6 | Gem-associated protein 5 OS=Homo sapi | GEMIN5 | 168.59 | 2 | 52.181217 | yes | 4.888E-07 | 762.86682 | 1033.6345 |
| NDSQHLFIEPR | 11 | Q8TEQ6 | Gem-associated protein 5 OS=Homo sapi | GEMIN5 | 168.59 | 2 | 52.181217 | yes | 1.249E-07 | 678.33875 | 1012.8319 |
| EKPETLIK | 8 | Q8TEQ6 | Gem-associated protein 5 OS=Homo sapi | GEMIN5 | 168.59 | 2 | 52.181217 | yes | 6.937E-07 | 479.28439 | 985.47675 |
| EPVICTPVSSGFEK | 14 | Q8TEQ6 | Gem-associated protein 5 OS=Homo sapi | GEMIN5 | 168.59 | 2 | 52.181217 | yes | 2.213E-07 | 775.38196 | 920.18567 |
| LPVHTEISWK | 10 | Q8TEQ6 | Gem-associated protein 5 OS=Homo sapi | GEMIN5 | 168.59 | 2 | 52.181217 | yes | 9.766E-08 | 605.33496 | 413.92438 |
| LPVHTEISWK | 10 | Q8TEQ6 | Gem-associated protein 5 OS=Homo sapi | GEMIN5 | 168.59 | 2 | 52.181217 | yes | 3.823E-07 | 403.8924 | 413.92438 |
| LPVHTEISWK | 10 | Q8TEQ6 | Gem-associated protein 5 OS=Homo sapi | GEMIN5 | 168.59 | 3 | 52.181217 | yes | 9.766E-08 | 605.33496 | 413.92438 |
| LPVHTEISWK | 10 | Q8TEQ6 | Gem-associated protein 5 OS=Homo sapi | GEMIN5 | 168.59 | 3 | 52.181217 | yes | 3.823E-07 | 403.8924 | 413.92438 |
| YPSATNNTPAK | 11 | Q8TEQ6 | Gem-associated protein 5 OS=Homo sapi | GEMIN5 | 168.59 | 2 | 52.181217 | yes | 3.63E-07 | 582.28821 | 802.8009 |
| DGHYAVAAK | 9 | Q8TEQ6 | Gem-associated protein 5 OS=Homo sapi | GEMIN5 | 168.59 | 2 | 52.181217 | yes | 1.279E-07 | 466.23523 | 751.69684 |
| TVVTEHALHQHTI | 21 | Q8TEQ6 | Gem-associated protein 5 OS=Homo sapi | GEMIN5 | 168.59 | 4 | 52.181217 | yes | 1.83E-07 | 613.32263 | 341.34875 |
| TVVTEHALHQHTI | 21 | Q8TEQ6 | Gem-associated protein 5 OS=Homo sapi | GEMIN5 | 168.59 | 4 | 52.181217 | yes | 1.145E-07 | 817.4278 | 341.34875 |
| TVVTEHALHQHTI | 21 | Q8TEQ6 | Gem-associated protein 5 OS=Homo sapi | GEMIN5 | 168.59 | 3 | 52.181217 | yes | 1.83E-07 | 613.32263 | 341.34875 |
| TVVTEHALHQHTI | 21 | Q8TEQ6 | Gem-associated protein 5 OS=Homo sapi | GEMIN5 | 168.59 | 3 | 52.181217 | yes | 1.145E-07 | 817.4278 | 341.34875 |
| ITSVAWSPHHDGR | 13 | Q8TEQ6 | Gem-associated protein 5 OS=Homo sapi | GEMIN5 | 168.59 | 2 | 52.181217 | yes | 5.269E-07 | 731.86292 | 330.89899 |
| ITSVAWSPHHDGR | 13 | Q8TEQ6 | Gem-associated protein 5 OS=Homo sapi | GEMIN5 | 168.59 | 2 | 52.181217 | yes | 2.629E-07 | 488.24438 | 330.89899 |
| ITSVAWSPHHDGR | 13 | Q8TEQ6 | Gem-associated protein 5 OS=Homo sapi | GEMIN5 | 168.59 | 3 | 52.181217 | yes | 5.269E-07 | 731.86292 | 330.89899 |
| ITSVAWSPHHDGR | 13 | Q8TEQ6 | Gem-associated protein 5 OS=Homo sapi | GEMIN5 | 168.59 | 3 | 52.181217 | yes | 2.629E-07 | 488.24438 | 330.89899 |
| TVAEVQETLAEMI | 14 | Q8TEQ6 | Gem-associated protein 5 OS=Homo sapi | GEMIN5 | 168.59 | 2 | 52.181217 | yes | 3.074E-07 | 795.41382 | 608.40387 |
| QLCFQDQYVK | 10 | Q8TEQ6 | Gem-associated protein 5 OS=Homo sapi | GEMIN5 | 168.59 | 2 | 52.181217 | yes | 1.071E-07 | 664.81879 | 605.87274 |
| AASHLLSIHK | 10 | Q8TEQ6 | Gem-associated protein 5 OS=Homo sapi | GEMIN5 | 168.59 | 2 | 52.181217 | yes | 9.903E-08 | 538.81421 | 595.11182 |
| VWNTLSIK | 8 | Q8TEQ6 | Gem-associated protein 5 OS=Homo sapi | GEMIN5 | 168.59 | 2 | 52.181217 | yes | 1.241E-07 | 480.7793 | 496.72968 |
| LSGEAFDINK | 10 | Q8TEQ6 | Gem-associated protein 5 OS=Homo sapi | GEMIN5 | 168.59 | 2 | 52.181217 | yes | 1.067E-07 | 547.27985 | 485.78137 |
| LTEEGER | 7 | Q8TEQ6 | Gem-associated protein 5 OS=Homo sapi | GEMIN5 | 168.59 | 2 | 52.181217 | yes | 1.103E-07 | 417.20361 | 470.57156 |
| GVVFCYWFNR | 10 | Q8TEQ6 | Gem-associated protein 5 OS=Homo sapi | GEMIN5 | 168.59 | 2 | 52.181217 | yes | 9.722E-08 | 674.31879 | 456.61111 |
| LGDHQSPATPAFK | 13 | Q8TEQ6 | Gem-associated protein 5 OS=Homo sapi | GEMIN5 | 168.59 | 2 | 52.181217 | yes | 1.215E-07 | 684.84894 | 454.66803 |
| DLYLSWGTVLER | 12 | Q8TEQ6 | Gem-associated protein 5 OS=Homo sapi | GEMIN5 | 168.59 | 2 | 52.181217 | yes | 9.722E-08 | 726.38007 | 401.83688 |
| YKLPVHTEISWK | 12 | Q8TEQ6 | Gem-associated protein 5 OS=Homo sapi | GEMIN5 | 168.59 | 2 | 52.181217 | yes | 1.709E-07 | 750.91412 | 155.9012 |
| YKLPVHTEISWK | 12 | Q8TEQ6 | Gem-associated protein 5 OS=Homo sapi | GEMIN5 | 168.59 | 2 | 52.181217 | yes | 1.01E-07 | 500.94516 | 155.9012 |
| YKLPVHTEISWK | 12 | Q8TEQ6 | Gem-associated protein 5 OS=Homo sapi | GEMIN5 | 168.59 | 3 | 52.181217 | yes | 1.709E-07 | 750.91412 | 155.9012 |

| | | | | | | | | | | | |
|---|---|---|---|---|---|---|---|---|---|---|---|
| YKLPVHTEISWK | 12 | Q8TEQ6 | Gem-associated protein 5 OS=Homo sapi | GEMIN5 | 168.59 | 3 | 52.181217 | yes | 1.01E-07 | 500.94516 | 155.9012 |
| LICTIQQHHK | 10 | Q8TEQ6 | Gem-associated protein 5 OS=Homo sapi | GEMIN5 | 168.59 | 2 | 52.181217 | yes | 9.722E-08 | 639.34277 | 298.65546 |
| VKDLIVSGDEK | 11 | Q8TEQ6 | Gem-associated protein 5 OS=Homo sapi | GEMIN5 | 168.59 | 2 | 52.181217 | yes | 1.022E-07 | 601.83496 | 295.56509 |
| SLEAFFLYGR | 10 | Q8TEQ6 | Gem-associated protein 5 OS=Homo sapi | GEMIN5 | 168.59 | 2 | 52.181217 | yes | 1.771E-07 | 601.81384 | 276.84229 |
| VSGFTFSHHPGQY | 26 | Q8TEQ6 | Gem-associated protein 5 OS=Homo sapi | GEMIN5 | 168.59 | 4 | 52.181217 | yes | 1.111E-07 | 703.82123 | 233.0948 |
| GCYLATGSK | 9 | Q8TEQ6 | Gem-associated protein 5 OS=Homo sapi | GEMIN5 | 168.59 | 2 | 52.181217 | yes | 9.834E-08 | 478.72894 | 212.5659 |
| MIDIEGK | 7 | Q8TEQ6 | Gem-associated protein 5 OS=Homo sapi | GEMIN5 | 168.59 | 2 | 52.181217 | yes | 1.56E-07 | 403.20984 | 209.80598 |
| DLIVSGDEK | 9 | Q8TEQ6 | Gem-associated protein 5 OS=Homo sapi | GEMIN5 | 168.59 | 2 | 52.181217 | yes | 1.395E-07 | 488.2533 | 164.61218 |
| YTLFSASSEGQNH | 15 | Q8TEQ6 | Gem-associated protein 5 OS=Homo sapi | GEMIN5 | 168.59 | 2 | 52.181217 | yes | 9.722E-08 | 842.38971 | 136.52989 |
| LESIDGNEEESMK | 13 | Q8TEQ6 | Gem-associated protein 5 OS=Homo sapi | GEMIN5 | 168.59 | 2 | 52.181217 | yes | 3.107E-07 | 740.82721 | 123.85799 |
| RGGGIDPTVK | 10 | Q8TEQ6 | Gem-associated protein 5 OS=Homo sapi | GEMIN5 | 168.59 | 2 | 52.181217 | yes | 6.738E-07 | 500.28271 | 83.758774 |
| DGIVVIIDISK | 11 | Q8TEQ6 | Gem-associated protein 5 OS=Homo sapi | GEMIN5 | 168.59 | 2 | 52.181217 | yes | 5.003E-07 | 586.35028 | 66.479584 |
| EELHQDCLVLATA | 14 | Q8TEQ6 | Gem-associated protein 5 OS=Homo sapi | GEMIN5 | 168.59 | 2 | 52.181217 | yes | 2.668E-07 | 813.91144 | 29.993736 |
| ALALDLGSPAALR | 13 | Q8TER5 | Rho guanine nucleotide exchange factor 4 | ARHGEF4( | 164.658 | 2 | 53.640575 | yes | 9.722E-08 | 634.37207 | 403.00546 |
| LSTCLLSAPSGIQR | 14 | Q8TER5 | Rho guanine nucleotide exchange factor 4 | ARHGEF4( | 164.658 | 2 | 53.640575 | yes | 9.722E-08 | 751.90338 | 309.42026 |
| ALQQPLEQLTR | 11 | Q8TER5 | Rho guanine nucleotide exchange factor 4 | ARHGEF4( | 164.658 | 2 | 53.640575 | yes | 1.038E-07 | 648.86713 | 233.81151 |
| EAYTLQATSPEIK | 13 | Q8TER5 | Rho guanine nucleotide exchange factor 4 | ARHGEF4( | 164.658 | 2 | 53.640575 | yes | 9.826E-08 | 725.87482 | 215.12668 |
| VLDIFEQR | 8 | Q8TER5 | Rho guanine nucleotide exchange factor 4 | ARHGEF4( | 164.658 | 2 | 53.640575 | yes | 1.161E-06 | 510.27966 | 204.58282 |
| AFSAEVQER | 9 | Q8TER5 | Rho guanine nucleotide exchange factor 4 | ARHGEF4( | 164.658 | 2 | 53.640575 | yes | 1.332E-06 | 518.75653 | 165.52058 |
| ALGAAVQLLR | 10 | Q8TER5 | Rho guanine nucleotide exchange factor 4 | ARHGEF4( | 164.658 | 2 | 53.640575 | yes | 9.723E-08 | 506.31912 | 151.03113 |
| GSMEAGPYLPR | 11 | Q8TER5 | Rho guanine nucleotide exchange factor 4 | ARHGEF4( | 164.658 | 2 | 53.640575 | yes | 1.517E-07 | 589.28717 | 147.12634 |
| GLEVTSTEVVDR | 12 | Q8TER5 | Rho guanine nucleotide exchange factor 4 | ARHGEF4( | 164.658 | 2 | 53.640575 | yes | 3.926E-07 | 652.83826 | 126.85727 |
| GTWAAALSAR | 10 | Q8TER5 | Rho guanine nucleotide exchange factor 4 | ARHGEF4( | 164.658 | 2 | 53.640575 | yes | 2.113E-07 | 502.26962 | 123.11462 |
| LEQVESGLHR | 10 | Q8TER5 | Rho guanine nucleotide exchange factor 4 | ARHGEF4( | 164.658 | 2 | 53.640575 | yes | 9.722E-08 | 584.30945 | 120.544 |
| GPDGPWGVGTPR | 12 | Q8TER5 | Rho guanine nucleotide exchange factor 4 | ARHGEF4( | 164.658 | 2 | 53.640575 | yes | 1.182E-07 | 598.29639 | 104.99402 |
| LPWAELICPR | 10 | Q8TER5 | Rho guanine nucleotide exchange factor 4 | ARHGEF4( | 164.658 | 2 | 53.640575 | yes | 5.072E-07 | 627.83679 | 91.994644 |
| GSPTDAEGSPGLSI | 14 | Q8TER5 | Rho guanine nucleotide exchange factor 4 | ARHGEF4( | 164.658 | 2 | 53.640575 | yes | 1.146E-07 | 665.81531 | 91.420235 |
| EAGPELSSECR | 11 | Q8TER5 | Rho guanine nucleotide exchange factor 4 | ARHGEF4( | 164.658 | 2 | 53.640575 | yes | 2.412E-07 | 617.77203 | 77.770439 |
| ELQGCATHPLR | 11 | Q8TER5 | Rho guanine nucleotide exchange factor 4 | ARHGEF4( | 164.658 | 2 | 53.640575 | yes | 2.461E-07 | 641.32208 | 69.618484 |
| YTLDFLVPAK | 10 | Q8TER5 | Rho guanine nucleotide exchange factor 4 | ARHGEF4( | 164.658 | 2 | 53.640575 | yes | 3.903E-07 | 583.82642 | 46.36536 |
| EQGQLLHR | 8 | Q8TER5 | Rho guanine nucleotide exchange factor 4 | ARHGEF4( | 164.658 | 2 | 53.640575 | yes | 1.076E-06 | 490.76724 | 45.528976 |
| DPFTVICGR | 9 | Q8TER5 | Rho guanine nucleotide exchange factor 4 | ARHGEF4( | 164.658 | 2 | 53.640575 | yes | 3.686E-06 | 532.76331 | 26.542889 |
| LSNLHVQQQEQR | 12 | Q8TER5 | Rho guanine nucleotide exchange factor 4 | ARHGEF4( | 164.658 | 2 | 53.640575 | yes | 1.445E-07 | 740.38678 | 17.829872 |
| SWDFLNSYR | 9 | Q8TEU7 | Rap guanine nucleotide exchange factor 6 | RAPGEF6 | 179.423 | 2 | 50.89196 | yes | 3.062E-06 | 594.27765 | 415.21948 |
| GFGIFVEGVEPGSI | 14 | Q8TEU7 | Rap guanine nucleotide exchange factor 6 | RAPGEF6 | 179.423 | 2 | 50.89196 | yes | 1.311E-07 | 711.86682 | 220.30559 |
| VLGTTEEISGK | 11 | Q8TEU7 | Rap guanine nucleotide exchange factor 6 | RAPGEF6 | 179.423 | 2 | 50.89196 | yes | 7.208E-07 | 567.30603 | 152.06325 |
| VLESTPAESSEGLI | 16 | Q8TEU7 | Rap guanine nucleotide exchange factor 6 | RAPGEF6 | 179.423 | 2 | 50.89196 | yes | 1.237E-06 | 829.90961 | 91.618546 |
| VSQVLQVPAVNLI | 16 | Q8TEU7 | Rap guanine nucleotide exchange factor 6 | RAPGEF6 | 179.423 | 3 | 50.89196 | yes | 9.722E-08 | 590.68433 | 70.636398 |
| DMTFLHEGNDSK | 12 | Q8TEU7 | Rap guanine nucleotide exchange factor 6 | RAPGEF6 | 179.423 | 2 | 50.89196 | yes | 2.393E-07 | 697.30627 | 51.258461 |

| | | | | | | | | | | | |
|---|---|---|---|---|---|---|---|---|---|---|---|
| NNTHLALTVK | 10 | Q8TEU7 | Rap guanine nucleotide exchange factor 6 | RAPGEF6 | 179.423 | 2 | 50.89196 | yes | 1.239E-06 | 555.81696 | 32.874561 |
| VFSLIQQAVTR | 11 | Q8TEW0 | Partitioning defective 3 homolog OS=Ho | PARD3 | 151.424 | 2 | 54.04887 | yes | 9.728E-08 | 631.36676 | 735.10828 |
| DPNYWIQVHR | 10 | Q8TEW0 | Partitioning defective 3 homolog OS=Ho | PARD3 | 151.424 | 2 | 54.04887 | yes | 9.722E-08 | 664.33075 | 481.72922 |
| ANMPLHVR | 8 | Q8TEW0 | Partitioning defective 3 homolog OS=Ho | PARD3 | 151.424 | 2 | 54.04887 | yes | 1.637E-07 | 469.25546 | 429.66425 |
| AEDEDIVLTPDGTI | 14 | Q8TEW0 | Partitioning defective 3 homolog OS=Ho | PARD3 | 151.424 | 2 | 54.04887 | yes | 9.927E-08 | 765.86774 | 409.00101 |
| FSPDSQYIDNR | 11 | Q8TEW0 | Partitioning defective 3 homolog OS=Ho | PARD3 | 151.424 | 2 | 54.04887 | yes | 9.722E-08 | 671.30713 | 405.54858 |
| GMIQLIVAR | 9 | Q8TEW0 | Partitioning defective 3 homolog OS=Ho | PARD3 | 151.424 | 2 | 54.04887 | yes | 1.429E-07 | 500.80225 | 334.07629 |
| AEHENLFR | 8 | Q8TEW0 | Partitioning defective 3 homolog OS=Ho | PARD3 | 151.424 | 2 | 54.04887 | yes | 9.849E-08 | 508.25143 | 318.1026 |
| VVVPCGDGHMK | 11 | Q8TEW0 | Partitioning defective 3 homolog OS=Ho | PARD3 | 151.424 | 2 | 54.04887 | yes | 1.727E-06 | 599.789 | 318.04605 |
| TNQDAMETLR | 10 | Q8TEW0 | Partitioning defective 3 homolog OS=Ho | PARD3 | 151.424 | 2 | 54.04887 | yes | 9.722E-08 | 589.77716 | 301.38812 |
| MEGTVSLLVFR | 11 | Q8TEW0 | Partitioning defective 3 homolog OS=Ho | PARD3 | 151.424 | 2 | 54.04887 | yes | 9.722E-08 | 626.34192 | 278.92444 |
| EQYEQLSQSEK | 11 | Q8TEW0 | Partitioning defective 3 homolog OS=Ho | PARD3 | 151.424 | 2 | 54.04887 | yes | 1.385E-07 | 684.8175 | 278.16397 |
| VMLETQELLR | 10 | Q8TEW0 | Partitioning defective 3 homolog OS=Ho | PARD3 | 151.424 | 2 | 54.04887 | yes | 3.718E-07 | 616.33936 | 271.49356 |
| IQESFTSEEER | 11 | Q8TEW0 | Partitioning defective 3 homolog OS=Ho | PARD3 | 151.424 | 2 | 54.04887 | yes | 9.722E-08 | 677.80969 | 254.68913 |
| SSLSASHPMVGK | 12 | Q8TEW0 | Partitioning defective 3 homolog OS=Ho | PARD3 | 151.424 | 2 | 54.04887 | yes | 9.809E-08 | 600.80573 | 211.66216 |
| LIEVNGVDLVGK | 12 | Q8TEW0 | Partitioning defective 3 homolog OS=Ho | PARD3 | 151.424 | 2 | 54.04887 | yes | 2.167E-07 | 628.36646 | 207.33466 |
| SVNSAGLHTVQR | 12 | Q8TEW0 | Partitioning defective 3 homolog OS=Ho | PARD3 | 151.424 | 2 | 54.04887 | yes | 9.722E-08 | 634.83893 | 196.81447 |
| TPIIWFHVVPAANI | 14 | Q8TEW0 | Partitioning defective 3 homolog OS=Ho | PARD3 | 151.424 | 2 | 54.04887 | yes | 9.809E-08 | 796.95099 | 176.39453 |
| TYSFEQPWPNAR | 12 | Q8TEW0 | Partitioning defective 3 homolog OS=Ho | PARD3 | 151.424 | 2 | 54.04887 | yes | 2.152E-07 | 748.35187 | 173.03809 |
| QEDAFHPR | 8 | Q8TEW0 | Partitioning defective 3 homolog OS=Ho | PARD3 | 151.424 | 2 | 54.04887 | yes | 2.522E-07 | 500.23578 | 166.64021 |
| VQDPSYAPPK | 10 | Q8TEW0 | Partitioning defective 3 homolog OS=Ho | PARD3 | 151.424 | 2 | 54.04887 | yes | 9.722E-08 | 551.28241 | 164.39951 |
| QFSDASQLDFVK | 12 | Q8TEW0 | Partitioning defective 3 homolog OS=Ho | PARD3 | 151.424 | 2 | 54.04887 | yes | 1.961E-07 | 692.84082 | 134.85817 |
| SQEEVVSLLR | 10 | Q8TEW0 | Partitioning defective 3 homolog OS=Ho | PARD3 | 151.424 | 2 | 54.04887 | yes | 9.722E-08 | 580.31952 | 129.18109 |
| LNTVDDQK | 8 | Q8TEW0 | Partitioning defective 3 homolog OS=Ho | PARD3 | 151.424 | 2 | 54.04887 | yes | 9.278E-07 | 466.73782 | 125.13693 |
| SMDLGIADETK | 11 | Q8TEW0 | Partitioning defective 3 homolog OS=Ho | PARD3 | 151.424 | 2 | 54.04887 | yes | 9.722E-08 | 590.28174 | 123.3549 |
| DVGPSLGLK | 9 | Q8TEW0 | Partitioning defective 3 homolog OS=Ho | PARD3 | 151.424 | 2 | 54.04887 | yes | 1.137E-06 | 443.25565 | 117.95786 |
| ELNAEPSQMQIPK | 13 | Q8TEW0 | Partitioning defective 3 homolog OS=Ho | PARD3 | 151.424 | 2 | 54.04887 | yes | 3.418E-07 | 742.87433 | 107.41091 |
| EGHMMDALYAQV | 13 | Q8TEW0 | Partitioning defective 3 homolog OS=Ho | PARD3 | 151.424 | 2 | 54.04887 | yes | 1.103E-07 | 746.84979 | 105.69489 |
| HSVSVEVQMQR | 11 | Q8TEW0 | Partitioning defective 3 homolog OS=Ho | PARD3 | 151.424 | 2 | 54.04887 | yes | 1.991E-07 | 650.32733 | 96.554054 |
| VTVCFGR | 7 | Q8TEW0 | Partitioning defective 3 homolog OS=Ho | PARD3 | 151.424 | 2 | 54.04887 | no | 2.107E-07 | 419.71564 | 88.053711 |
| VNDQLIAVNGESL | 16 | Q8TEW0 | Partitioning defective 3 homolog OS=Ho | PARD3 | 151.424 | 2 | 54.04887 | yes | 4.714E-07 | 835.45959 | 80.170486 |
| FEQAQHMFR | 9 | Q8TEW0 | Partitioning defective 3 homolog OS=Ho | PARD3 | 151.424 | 2 | 54.04887 | yes | 1.879E-07 | 597.27966 | 62.04974 |
| ENHADLGIFVK | 11 | Q8TEW0 | Partitioning defective 3 homolog OS=Ho | PARD3 | 151.424 | 2 | 54.04887 | yes | 5.123E-07 | 621.82751 | 57.599873 |
| KGTEGLGFSITSR | 13 | Q8TEW0 | Partitioning defective 3 homolog OS=Ho | PARD3 | 151.424 | 2 | 54.04887 | yes | 1.402E-06 | 676.86206 | 32.754356 |
| LNHPPEQIDSHSR | 13 | Q8TEW0 | Partitioning defective 3 homolog OS=Ho | PARD3 | 151.424 | 2 | 54.04887 | yes | 5.254E-07 | 765.3764 | 23.100704 |
| SLILTALQR | 9 | Q8TEX9 | Importin-4 OS=Homo sapiens OX=9606 | IPO4 | 118.716 | 2 | 41.548317 | yes | 3.592E-07 | 507.81894 | 111.29486 |
| VVPSYMQAVNR | 11 | Q8TEX9 | Importin-4 OS=Homo sapiens OX=9606 | IPO4 | 118.716 | 2 | 41.548317 | yes | 4.408E-06 | 632.32935 | 106.79599 |
| ELLLPDTER | 9 | Q8TEX9 | Importin-4 OS=Homo sapiens OX=9606 | IPO4 | 118.716 | 2 | 41.548317 | yes | 3.454E-07 | 543.29547 | 83.733284 |
| LLMASPTR | 8 | Q8TEX9 | Importin-4 OS=Homo sapiens OX=9606 | IPO4 | 118.716 | 2 | 41.548317 | yes | 1.868E-06 | 444.75223 | 74.920586 |

| | | | | | | | | | | | |
|---|---|---|---|---|---|---|---|---|---|---|---|
| LLECPHLNVR | 10 | Q8TEX9 | Importin-4 OS=Homo sapiens OX=9606 | IPO4 | 118.716 | 2 | 41.548317 | yes | 1.732E-06 | 625.83734 | 53.705887 |
| LAELCGVLK | 9 | Q8TEX9 | Importin-4 OS=Homo sapiens OX=9606 | IPO4 | 118.716 | 2 | 41.548317 | yes | 2.747E-06 | 501.78625 | 19.880175 |
| ATEQLQIVLR | 10 | Q8TEX9 | Importin-4 OS=Homo sapiens OX=9606 | IPO4 | 118.716 | 2 | 41.548317 | yes | 1.136E-06 | 585.8457 | 17.641777 |
| EQNSLSLLEAR | 11 | Q8TF01 | Arginine/serine-rich protein PNISR OS=I | PNISR | 92.577 | 2 | 38.007965 | yes | 5.527E-07 | 630.33313 | 72.60257 |
| EADGDVVNEK | 10 | Q8TF01 | Arginine/serine-rich protein PNISR OS=I | PNISR | 92.577 | 2 | 38.007965 | yes | 3.731E-06 | 538.24872 | 49.545097 |
| IVDALNEFR | 9 | Q8TF40 | Folliculin-interacting protein 1 OS=Homo | FNIP1 | 130.556 | 2 | 38.850853 | yes | 5.255E-07 | 538.79041 | 91.19223 |
| LETVVCTGSVPVD | 14 | Q8TF40 | Folliculin-interacting protein 1 OS=Homo | FNIP1 | 130.556 | 2 | 38.850853 | yes | 4.948E-07 | 752.38977 | 37.199509 |
| FAADFAAEAASK | 12 | Q8TF42 | Ubiquitin-associated and SH3 domain-co | UBASH3B | 72.696 | 2 | 50.900906 | yes | 4.049E-07 | 599.79059 | 126.62689 |
| DAPITVFGCMQAR | 13 | Q8TF42 | Ubiquitin-associated and SH3 domain-co | UBASH3B | 72.696 | 2 | 50.900906 | yes | 9.722E-08 | 733.34998 | 110.93961 |
| VDALGEALQTTVS | 14 | Q8TF42 | Ubiquitin-associated and SH3 domain-co | UBASH3B | 72.696 | 2 | 50.900906 | yes | 2.843E-07 | 730.39117 | 86.615753 |
| LVVSESYDTYISR | 13 | Q8TF42 | Ubiquitin-associated and SH3 domain-co | UBASH3B | 72.696 | 2 | 50.900906 | yes | 2.587E-07 | 766.38556 | 50.579391 |
| LAQNIDVK | 8 | Q8TF42 | Ubiquitin-associated and SH3 domain-co | UBASH3B | 72.696 | 2 | 50.900906 | yes | 1.62E-06 | 450.76108 | 33.127243 |
| YWLSQCFDAK | 10 | Q8TF42 | Ubiquitin-associated and SH3 domain-co | UBASH3B | 72.696 | 2 | 50.900906 | yes | 6.891E-06 | 659.30023 | 25.088135 |
| ILVTPWDYR | 9 | Q8TF46 | DIS3-like exonuclease 1 OS=Homo sapie | DIS3L | 120.788 | 2 | 54.36108 | yes | 2.318E-07 | 581.81641 | 449.56586 |
| ISTQQAETLQDFR | 13 | Q8TF46 | DIS3-like exonuclease 1 OS=Homo sapie | DIS3L | 120.788 | 2 | 54.36108 | yes | 1.624E-07 | 768.88629 | 339.98846 |
| YSDIVVHR | 8 | Q8TF46 | DIS3-like exonuclease 1 OS=Homo sapie | DIS3L | 120.788 | 2 | 54.36108 | yes | 5.239E-07 | 494.76416 | 338.55115 |
| IWESFPHQALLR | 12 | Q8TF46 | DIS3-like exonuclease 1 OS=Homo sapie | DIS3L | 120.788 | 2 | 54.36108 | yes | 9.722E-08 | 748.90405 | 219.41289 |
| YIQGILNVNK | 10 | Q8TF46 | DIS3-like exonuclease 1 OS=Homo sapie | DIS3L | 120.788 | 2 | 54.36108 | yes | 9.722E-08 | 581.33496 | 188.62766 |
| GCEDVDDTLSVR | 12 | Q8TF46 | DIS3-like exonuclease 1 OS=Homo sapie | DIS3L | 120.788 | 2 | 54.36108 | yes | 9.722E-08 | 683.30121 | 158.12248 |
| IPNTELIHQSSPLLI | 15 | Q8TF46 | DIS3-like exonuclease 1 OS=Homo sapie | DIS3L | 120.788 | 3 | 54.36108 | yes | 3.358E-06 | 563.98932 | 66.72789 |
| IPNTELIHQSSPLLI | 15 | Q8TF46 | DIS3-like exonuclease 1 OS=Homo sapie | DIS3L | 120.788 | 3 | 54.36108 | yes | 5.352E-07 | 845.48035 | 66.72789 |
| IPNTELIHQSSPLLI | 15 | Q8TF46 | DIS3-like exonuclease 1 OS=Homo sapie | DIS3L | 120.788 | 2 | 54.36108 | yes | 3.358E-06 | 563.98932 | 66.72789 |
| IPNTELIHQSSPLLI | 15 | Q8TF46 | DIS3-like exonuclease 1 OS=Homo sapie | DIS3L | 120.788 | 2 | 54.36108 | yes | 5.352E-07 | 845.48035 | 66.72789 |
| YTHFTSPIR | 9 | Q8TF46 | DIS3-like exonuclease 1 OS=Homo sapie | DIS3L | 120.788 | 2 | 54.36108 | yes | 2.324E-07 | 561.29053 | 127.92302 |
| YAVSIMWELDK | 11 | Q8TF46 | DIS3-like exonuclease 1 OS=Homo sapie | DIS3L | 120.788 | 2 | 54.36108 | yes | 1.113E-07 | 677.83923 | 127.46381 |
| LEELVWAIGK | 10 | Q8TF46 | DIS3-like exonuclease 1 OS=Homo sapie | DIS3L | 120.788 | 2 | 54.36108 | yes | 3.49E-07 | 579.33191 | 112.33537 |
| CISDGVIYSIR | 11 | Q8TF46 | DIS3-like exonuclease 1 OS=Homo sapie | DIS3L | 120.788 | 2 | 54.36108 | yes | 4.491E-07 | 641.82666 | 102.95544 |
| NIHDLIPK | 8 | Q8TF46 | DIS3-like exonuclease 1 OS=Homo sapie | DIS3L | 120.788 | 2 | 54.36108 | yes | 2.888E-07 | 475.27692 | 100.81821 |
| SHLVFSIDPK | 10 | Q8TF46 | DIS3-like exonuclease 1 OS=Homo sapie | DIS3L | 120.788 | 2 | 54.36108 | yes | 1.467E-07 | 571.81384 | 65.198433 |
| VNIIQEEYQEYR | 12 | Q8TF46 | DIS3-like exonuclease 1 OS=Homo sapie | DIS3L | 120.788 | 2 | 54.36108 | yes | 9.722E-08 | 792.38867 | 63.824165 |
| TVALCENDCDDK | 12 | Q8TF46 | DIS3-like exonuclease 1 OS=Homo sapie | DIS3L | 120.788 | 2 | 54.36108 | yes | 1.491E-07 | 720.29248 | 59.21386 |
| NYLDNFWPDLK | 11 | Q8TF46 | DIS3-like exonuclease 1 OS=Homo sapie | DIS3L | 120.788 | 2 | 54.36108 | yes | 1.802E-07 | 712.84589 | 57.058578 |
| DSDLVSDILIHGMI | 14 | Q8TF46 | DIS3-like exonuclease 1 OS=Homo sapie | DIS3L | 120.788 | 2 | 54.36108 | yes | 9.757E-08 | 771.89526 | 53.076706 |
| QSTELFQCMYFK | 12 | Q8TF46 | DIS3-like exonuclease 1 OS=Homo sapie | DIS3L | 120.788 | 2 | 54.36108 | yes | 2.125E-07 | 791.35742 | 50.427231 |
| AAHELCDSILQSR | 13 | Q8TF46 | DIS3-like exonuclease 1 OS=Homo sapie | DIS3L | 120.788 | 2 | 54.36108 | yes | 1.175E-07 | 750.36719 | 36.640434 |
| SIHGDVVVVELLP | 14 | Q8TF46 | DIS3-like exonuclease 1 OS=Homo sapie | DIS3L | 120.788 | 2 | 54.36108 | yes | 2.436E-07 | 752.94031 | 21.828537 |
| AASLEQHLR | 9 | Q8TF50 | Zinc finger protein 526 OS=Homo sapien | ZNF526 | 73.622 | 2 | 46.281483 | yes | 1.997E-07 | 512.78033 | 90.228973 |
| YLCVDCGR | 8 | Q8TF50 | Zinc finger protein 526 OS=Homo sapien | ZNF526 | 73.622 | 2 | 46.281483 | yes | 1.602E-07 | 521.72589 | 59.938892 |
| TLPAGVLR | 8 | Q8TF63 | Dendritic cell nuclear protein 1 OS=Hom | DCANP1 | 26.704 | 2 | 42.26485 | yes | 3.115E-07 | 413.76089 | 130.68881 |

| | | | | | | | | | | | |
|---|---|---|---|---|---|---|---|---|---|---|---|
| QQSLPSLCSTSDPI | 25 | Q8TF72 | Protein Shroom3 OS=Homo sapiens OX= | SHROOM3 | 216.858 | 4 | 53.571724 | yes | 1.757E-06 | 643.06042 | 356.57999 |
| SAFSSLQNIPESLR | 14 | Q8TF72 | Protein Shroom3 OS=Homo sapiens OX= | SHROOM3 | 216.858 | 2 | 53.571724 | yes | 9.722E-08 | 774.90448 | 135.61479 |
| VLGATSFR | 8 | Q8TF72 | Protein Shroom3 OS=Homo sapiens OX= | SHROOM3 | 216.858 | 2 | 53.571724 | yes | 1.275E-06 | 425.74271 | 106.0632 |
| QFQQSALADYIQR | 13 | Q8TF72 | Protein Shroom3 OS=Homo sapiens OX= | SHROOM3 | 216.858 | 2 | 53.571724 | yes | 1.216E-07 | 784.39679 | 93.853561 |
| QDVLLGQDSGFGI | 15 | Q8TF72 | Protein Shroom3 OS=Homo sapiens OX= | SHROOM3 | 216.858 | 2 | 53.571724 | yes | 1.313E-07 | 788.42249 | 86.418159 |
| HMPGAAHVVGSQ | 17 | Q8TF72 | Protein Shroom3 OS=Homo sapiens OX= | SHROOM3 | 216.858 | 3 | 53.571724 | yes | 1.692E-07 | 573.63 | 86.41407 |
| EGLGQSLSGNFGK | 13 | Q8TF72 | Protein Shroom3 OS=Homo sapiens OX= | SHROOM3 | 216.858 | 2 | 53.571724 | yes | 1.58E-07 | 647.32532 | 80.559677 |
| SMSAEDLLER | 10 | Q8TF72 | Protein Shroom3 OS=Homo sapiens OX= | SHROOM3 | 216.858 | 2 | 53.571724 | yes | 1.432E-07 | 575.77405 | 73.325882 |
| ESGYIAPQGACNK | 13 | Q8TF72 | Protein Shroom3 OS=Homo sapiens OX= | SHROOM3 | 216.858 | 2 | 53.571724 | yes | 2.175E-07 | 697.82208 | 69.734589 |
| KAELIGSLTHK | 11 | Q8TF72 | Protein Shroom3 OS=Homo sapiens OX= | SHROOM3 | 216.858 | 2 | 53.571724 | yes | 1.022E-07 | 598.85352 | 51.836159 |
| SDVLAGPVHVR | 11 | Q8TF72 | Protein Shroom3 OS=Homo sapiens OX= | SHROOM3 | 216.858 | 2 | 53.571724 | yes | 4.238E-07 | 575.32239 | 46.23362 |
| HSVTPAEGDLAR | 12 | Q8TF72 | Protein Shroom3 OS=Homo sapiens OX= | SHROOM3 | 216.858 | 2 | 53.571724 | yes | 6.941E-07 | 626.81769 | 45.897758 |
| SPEASASASPHTPF | 14 | Q8TF72 | Protein Shroom3 OS=Homo sapiens OX= | SHROOM3 | 216.858 | 2 | 53.571724 | yes | 1.7E-07 | 697.83661 | 37.568996 |
| SLADILDPDSR | 11 | Q8TF72 | Protein Shroom3 OS=Homo sapiens OX= | SHROOM3 | 216.858 | 2 | 53.571724 | yes | 4.79E-07 | 601.30658 | 27.648706 |
| TVSTSSTSGNDFEI | 16 | Q8TF72 | Protein Shroom3 OS=Homo sapiens OX= | SHROOM3 | 216.858 | 2 | 53.571724 | yes | 4.4E-07 | 845.3761 | 8.1186304 |
| ELSLLSGEVCNR | 12 | Q8TF76 | Serine/threonine-protein kinase haspin OS | HASPIN | 88.496 | 2 | 24.854322 | yes | 3.38E-06 | 688.84558 | 24.716591 |
| DLALQLLHK | 9 | Q8WTP8 | Apoptosis-enhancing nuclease OS=Homo | AEN | 36.351 | 2 | 52.098015 | yes | 9.722E-08 | 525.81897 | 122.59228 |
| SLWTCPEDR | 9 | Q8WTP8 | Apoptosis-enhancing nuclease OS=Homo | AEN | 36.351 | 2 | 52.098015 | yes | 2.376E-07 | 582.26111 | 56.50631 |
| CVAIDCEMVGTGI | 14 | Q8WTP8 | Apoptosis-enhancing nuclease OS=Homo | AEN | 36.351 | 2 | 52.098015 | yes | 3.924E-07 | 782.84949 | 50.414352 |
| LVEVQWEQQEAR | 12 | Q8WTP8 | Apoptosis-enhancing nuclease OS=Homo | AEN | 36.351 | 2 | 52.098015 | yes | 2.054E-07 | 757.88354 | 48.647141 |
| KPIPLENPK | 9 | Q8WTT2 | Nucleolar complex protein 3 homolog OS | NOC3L | 92.548 | 2 | 55.66121 | yes | 4.623E-07 | 518.31348 | 184.16927 |
| EFEEGLVSQYK | 11 | Q8WTT2 | Nucleolar complex protein 3 homolog OS | NOC3L | 92.548 | 2 | 55.66121 | yes | 1.671E-07 | 664.82208 | 130.48431 |
| FAAHLIAGAPSEGS | 23 | Q8WTT2 | Nucleolar complex protein 3 homolog OS | NOC3L | 92.548 | 3 | 55.66121 | yes | 2.082E-07 | 760.40961 | 109.17001 |
| IIDKYEK | 7 | Q8WTT2 | Nucleolar complex protein 3 homolog OS | NOC3L | 92.548 | 2 | 55.66121 | yes | 9.722E-08 | 454.75803 | 108.93928 |
| LGQASLGVIK | 10 | Q8WTT2 | Nucleolar complex protein 3 homolog OS | NOC3L | 92.548 | 2 | 55.66121 | yes | 3.46E-07 | 493.30566 | 102.45191 |
| LISEMCCEAVK | 11 | Q8WTT2 | Nucleolar complex protein 3 homolog OS | NOC3L | 92.548 | 2 | 55.66121 | yes | 2.283E-07 | 670.3064 | 98.617844 |
| ELIHLLPIK | 9 | Q8WTT2 | Nucleolar complex protein 3 homolog OS | NOC3L | 92.548 | 2 | 55.66121 | yes | 1.061E-07 | 538.34735 | 75.159668 |
| DLSSSEPVHAK | 11 | Q8WTT2 | Nucleolar complex protein 3 homolog OS | NOC3L | 92.548 | 2 | 55.66121 | yes | 9.722E-08 | 585.29346 | 48.418507 |
| ELREAEASESTEK | 13 | Q8WTT2 | Nucleolar complex protein 3 homolog OS | NOC3L | 92.548 | 2 | 55.66121 | yes | 4.44E-05 | 739.85211 | 48.209068 |
| DVDETGITVASLEI | 14 | Q8WU90 | Zinc finger CCCH domain-containing prc | ZC3H15 | 48.603 | 2 | 51.32171 | yes | 1.006E-07 | 752.87811 | 1207.5243 |
| FGQQNPR | 7 | Q8WU90 | Zinc finger CCCH domain-containing prc | ZC3H15 | 48.603 | 2 | 51.32171 | yes | 2.13E-07 | 423.71448 | 806.52899 |
| EDEISLEDLIER | 12 | Q8WU90 | Zinc finger CCCH domain-containing prc | ZC3H15 | 48.603 | 2 | 51.32171 | yes | 1.284E-07 | 730.85938 | 426.71005 |
| SDLEEDNER | 9 | Q8WU90 | Zinc finger CCCH domain-containing prc | ZC3H15 | 48.603 | 2 | 51.32171 | yes | 9.722E-08 | 553.73346 | 414.43597 |
| KLEEVVNK | 8 | Q8WU90 | Zinc finger CCCH domain-containing prc | ZC3H15 | 48.603 | 2 | 51.32171 | yes | 1.968E-07 | 479.78201 | 407.41205 |
| HFLEAIENNK | 10 | Q8WU90 | Zinc finger CCCH domain-containing prc | ZC3H15 | 48.603 | 2 | 51.32171 | yes | 9.722E-08 | 607.81183 | 271.19434 |
| FSTYTSDKDENK | 12 | Q8WU90 | Zinc finger CCCH domain-containing prc | ZC3H15 | 48.603 | 2 | 51.32171 | yes | 9.722E-08 | 717.82281 | 269.90918 |
| ELQELNELFK | 10 | Q8WU90 | Zinc finger CCCH domain-containing prc | ZC3H15 | 48.603 | 2 | 51.32171 | yes | 3.057E-07 | 631.83496 | 238.37311 |
| LEQDMER | 7 | Q8WU90 | Zinc finger CCCH domain-containing prc | ZC3H15 | 48.603 | 2 | 51.32171 | yes | 1.106E-07 | 460.71075 | 118.50998 |
| ELQELNELFKPVV | 17 | Q8WU90 | Zinc finger CCCH domain-containing prc | ZC3H15 | 48.603 | 3 | 51.32171 | yes | 1.24E-07 | 652.69818 | 83.919991 |

| | | | | | | | | | | | |
|---|---|---|---|---|---|---|---|---|---|---|---|
| DTMDNWDEK | 9 | Q8WU90 | Zinc finger CCCH domain-containing prc | ZC3H15 | 48.603 | 2 | 51.32171 | yes | 3.287E-05 | 577.22693 | 26.942497 |
| LEQPDLSSEMSK | 12 | Q8WUA4 | General transcription factor 3C polypeptic | GTF3C2 | 100.68 | 2 | 53.295807 | yes | 9.722E-08 | 682.3241 | 764.80511 |
| EGLPEDGTLYR | 11 | Q8WUA4 | General transcription factor 3C polypeptic | GTF3C2 | 100.68 | 2 | 53.295807 | yes | 9.722E-08 | 625.30658 | 671.80017 |
| GLASPLGHR | 9 | Q8WUA4 | General transcription factor 3C polypeptic | GTF3C2 | 100.68 | 2 | 53.295807 | yes | 1.474E-07 | 454.25906 | 633.74664 |
| FCPSGAWELPGTP | 14 | Q8WUA4 | General transcription factor 3C polypeptic | GTF3C2 | 100.68 | 2 | 53.295807 | yes | 1.167E-07 | 787.87463 | 519.76508 |
| FSSITAHPER | 10 | Q8WUA4 | General transcription factor 3C polypeptic | GTF3C2 | 100.68 | 2 | 53.295807 | yes | 9.834E-08 | 572.79089 | 518.8725 |
| LQLEAIHK | 8 | Q8WUA4 | General transcription factor 3C polypeptic | GTF3C2 | 100.68 | 2 | 53.295807 | yes | 1.859E-07 | 476.28473 | 414.03458 |
| LGLLALACSDGK | 12 | Q8WUA4 | General transcription factor 3C polypeptic | GTF3C2 | 100.68 | 2 | 53.295807 | yes | 9.722E-08 | 609.33154 | 266.55798 |
| AHFNAMFQPSSPT | 14 | Q8WUA4 | General transcription factor 3C polypeptic | GTF3C2 | 100.68 | 3 | 53.295807 | yes | 1.853E-07 | 530.9209 | 110.18175 |
| AHFNAMFQPSSPT | 14 | Q8WUA4 | General transcription factor 3C polypeptic | GTF3C2 | 100.68 | 3 | 53.295807 | yes | 5.497E-07 | 795.87775 | 110.18175 |
| AHFNAMFQPSSPT | 14 | Q8WUA4 | General transcription factor 3C polypeptic | GTF3C2 | 100.68 | 2 | 53.295807 | yes | 1.853E-07 | 530.9209 | 110.18175 |
| AHFNAMFQPSSPT | 14 | Q8WUA4 | General transcription factor 3C polypeptic | GTF3C2 | 100.68 | 2 | 53.295807 | yes | 5.497E-07 | 795.87775 | 110.18175 |
| ANSHFLVSAGSDR | 13 | Q8WUA4 | General transcription factor 3C polypeptic | GTF3C2 | 100.68 | 2 | 53.295807 | yes | 9.722E-08 | 680.83386 | 181.00871 |
| RPYEPINSIK | 10 | Q8WUA4 | General transcription factor 3C polypeptic | GTF3C2 | 100.68 | 2 | 53.295807 | yes | 1.677E-07 | 608.83789 | 99.415482 |
| DLDRPESQSPK | 11 | Q8WUA4 | General transcription factor 3C polypeptic | GTF3C2 | 100.68 | 2 | 53.295807 | yes | 3.073E-07 | 636.31494 | 91.267731 |
| TYTETVNHHYLLF | 26 | Q8WUA4 | General transcription factor 3C polypeptic | GTF3C2 | 100.68 | 4 | 53.295807 | yes | 2.097E-07 | 784.6347 | 70.146759 |
| SKAELLLLK | 9 | Q8WUA4 | General transcription factor 3C polypeptic | GTF3C2 | 100.68 | 2 | 53.295807 | yes | 9.268E-07 | 507.83151 | 64.201561 |
| MQEGEGHSQLCLI | 14 | Q8WUA4 | General transcription factor 3C polypeptic | GTF3C2 | 100.68 | 2 | 53.295807 | yes | 9.722E-08 | 830.36432 | 24.214781 |
| QPAACPGGEEVDC | 16 | Q8WUA4 | General transcription factor 3C polypeptic | GTF3C2 | 100.68 | 2 | 53.295807 | yes | 9.722E-08 | 805.86499 | 24.163702 |
| VIPNAICGICLK | 12 | Q8WUB8 | PHD finger protein 10 OS=Homo sapiens | PHF10 | 56.051 | 2 | 48.624737 | yes | 1.852E-06 | 679.37018 | 148.39482 |
| KYPDLER | 7 | Q8WUB8 | PHD finger protein 10 OS=Homo sapiens | PHF10 | 56.051 | 2 | 48.624737 | yes | 1.875E-06 | 460.74545 | 147.4543 |
| WICDCCQR | 8 | Q8WUB8 | PHD finger protein 10 OS=Homo sapiens | PHF10 | 56.051 | 2 | 48.624737 | yes | 1.122E-07 | 599.23352 | 88.788467 |
| EYSQMQQQNTQK | 12 | Q8WUB8 | PHD finger protein 10 OS=Homo sapiens | PHF10 | 56.051 | 2 | 48.624737 | yes | 4.633E-07 | 756.841 | 49.891914 |
| SDEVIDLMIK | 10 | Q8WUB8 | PHD finger protein 10 OS=Homo sapiens | PHF10 | 56.051 | 2 | 48.624737 | yes | 1.679E-06 | 581.80481 | 44.137558 |
| HAEYSVILQEK | 11 | Q8WUB8 | PHD finger protein 10 OS=Homo sapiens | PHF10 | 56.051 | 2 | 48.624737 | yes | 1.379E-07 | 658.84589 | 44.113667 |
| AMLPGSPLFTR | 11 | Q8WUF5 | RelA-associated inhibitor OS=Homo sapi | PPP1R13L | 89.092 | 2 | 53.528854 | yes | 8.993E-07 | 595.32355 | 64.681793 |
| PASPSLQLLPWR | 12 | Q8WUF5 | RelA-associated inhibitor OS=Homo sapi | PPP1R13L | 89.092 | 2 | 53.528854 | yes | 9.722E-08 | 682.88788 | 29.336527 |
| QLVQDENVR | 9 | Q8WUK0 | Phosphatidylglycerophosphatase and prot | PTPMT1 | 22.844 | 2 | 51.5008 | yes | 1.48E-07 | 550.78839 | 198.03532 |
| WSPEEAVR | 8 | Q8WUK0 | Phosphatidylglycerophosphatase and prot | PTPMT1 | 22.844 | 2 | 51.5008 | yes | 2.507E-07 | 487.24051 | 188.3606 |
| FLCNSSQEWK | 10 | Q8WUK0 | Phosphatidylglycerophosphatase and prot | PTPMT1 | 22.844 | 2 | 51.5008 | yes | 1.272E-07 | 649.79535 | 100.14537 |
| GVITMNEEYETR | 12 | Q8WUK0 | Phosphatidylglycerophosphatase and prot | PTPMT1 | 22.844 | 2 | 51.5008 | yes | 9.722E-08 | 721.33502 | 77.996674 |
| LGVEQLR | 7 | Q8WUK0 | Phosphatidylglycerophosphatase and prot | PTPMT1 | 22.844 | 2 | 51.5008 | yes | 5.847E-07 | 407.74271 | 27.787624 |
| YQSLGQCVYVHCl | 13 | Q8WUK0 | Phosphatidylglycerophosphatase and prot | PTPMT1 | 22.844 | 2 | 51.5008 | yes | 3.859E-07 | 821.37927 | 26.544443 |
| PGQLDVLK | 8 | Q8WUK0 | Phosphatidylglycerophosphatase and prot | PTPMT1 | 22.844 | 2 | 51.5008 | yes | 7.65E-06 | 435.25818 | 16.886831 |
| VAYPQADSNLR | 11 | Q8WUM0 | Nuclear pore complex protein Nup133 OS | NUP133 | 128.98 | 2 | 53.333565 | yes | 9.722E-08 | 617.31476 | 163.72731 |
| DLLQADQLGSLK | 12 | Q8WUM0 | Nuclear pore complex protein Nup133 OS | NUP133 | 128.98 | 2 | 53.333565 | yes | 9.722E-08 | 650.85901 | 146.64537 |
| DGIQLSEYLPEVK | 13 | Q8WUM0 | Nuclear pore complex protein Nup133 OS | NUP133 | 128.98 | 2 | 53.333565 | yes | 9.722E-08 | 745.8905 | 136.42274 |
| HAYSWDINR | 9 | Q8WUM0 | Nuclear pore complex protein Nup133 OS | NUP133 | 128.98 | 2 | 53.333565 | yes | 3.774E-06 | 581.27545 | 104.74033 |
| DMLQAASHYR | 10 | Q8WUM0 | Nuclear pore complex protein Nup133 OS | NUP133 | 128.98 | 2 | 53.333565 | yes | 1.05E-07 | 596.28241 | 93.35965 |

| | | | | | | | | | | | |
|---|---|---|---|---|---|---|---|---|---|---|---|
| NSGLVSITSR | 10 | Q8WUM0 | Nuclear pore complex protein Nup133 OS | NUP133 | 128.98 | 2 | 53.333565 | yes | 3.275E-06 | 517.28546 | 67.828819 |
| SNPYFEFVLK | 10 | Q8WUM0 | Nuclear pore complex protein Nup133 OS | NUP133 | 128.98 | 2 | 53.333565 | yes | 2.738E-07 | 622.32153 | 47.406715 |
| WELDDSSEK | 9 | Q8WUM0 | Nuclear pore complex protein Nup133 OS | NUP133 | 128.98 | 2 | 53.333565 | yes | 3.08E-06 | 554.74329 | 38.248283 |
| LLLCEHAEK | 9 | Q8WUM0 | Nuclear pore complex protein Nup133 OS | NUP133 | 128.98 | 2 | 53.333565 | yes | 2.459E-07 | 556.79205 | 27.719273 |
| LANQAADYFGDAI | 14 | Q8WUM4 | Programmed cell death 6-interacting prote | PDCD6IP | 96.024 | 2 | 54.294525 | yes | 9.722E-08 | 765.86481 | 410.66553 |
| ELPELLQR | 8 | Q8WUM4 | Programmed cell death 6-interacting prote | PDCD6IP | 96.024 | 2 | 54.294525 | yes | 3.478E-07 | 499.28748 | 310.22919 |
| SLLSNLDEVK | 10 | Q8WUM4 | Programmed cell death 6-interacting prote | PDCD6IP | 96.024 | 2 | 54.294525 | yes | 1.513E-07 | 559.30859 | 289.97656 |
| LLDEEEATDNDLR | 13 | Q8WUM4 | Programmed cell death 6-interacting prote | PDCD6IP | 96.024 | 2 | 54.294525 | yes | 9.722E-08 | 766.85736 | 224.14676 |
| KFGEEIAR | 8 | Q8WUM4 | Programmed cell death 6-interacting prote | PDCD6IP | 96.024 | 2 | 54.294525 | yes | 5.702E-07 | 475.25873 | 216.03813 |
| FYNELTEILVR | 11 | Q8WUM4 | Programmed cell death 6-interacting prote | PDCD6IP | 96.024 | 2 | 54.294525 | yes | 1.376E-07 | 698.8772 | 166.03304 |
| NIQVSHQEFSK | 11 | Q8WUM4 | Programmed cell death 6-interacting prote | PDCD6IP | 96.024 | 2 | 54.294525 | yes | 9.722E-08 | 658.83331 | 155.54449 |
| TMQGSEVVNVLK | 12 | Q8WUM4 | Programmed cell death 6-interacting prote | PDCD6IP | 96.024 | 2 | 54.294525 | yes | 1.443E-07 | 652.8476 | 129.16922 |
| EVFPVLAAK | 9 | Q8WUM4 | Programmed cell death 6-interacting prote | PDCD6IP | 96.024 | 2 | 54.294525 | yes | 3.314E-07 | 487.28949 | 125.48476 |
| HYQFASGAFLHIK | 13 | Q8WUM4 | Programmed cell death 6-interacting prote | PDCD6IP | 96.024 | 3 | 54.294525 | yes | 9.722E-08 | 506.93326 | 117.57512 |
| HEGALETLLR | 10 | Q8WUM4 | Programmed cell death 6-interacting prote | PDCD6IP | 96.024 | 2 | 54.294525 | yes | 1.88E-07 | 569.81439 | 116.87946 |
| YYDQICSIEPK | 11 | Q8WUM4 | Programmed cell death 6-interacting prote | PDCD6IP | 96.024 | 2 | 54.294525 | yes | 9.722E-08 | 708.32922 | 115.24043 |
| LALASLGYEK | 10 | Q8WUM4 | Programmed cell death 6-interacting prote | PDCD6IP | 96.024 | 2 | 54.294525 | yes | 1.871E-07 | 532.80298 | 94.616066 |
| TPSNELYKPLR | 11 | Q8WUM4 | Programmed cell death 6-interacting prote | PDCD6IP | 96.024 | 2 | 54.294525 | yes | 9.722E-08 | 659.36169 | 43.034439 |
| KDNDFIYHDR | 10 | Q8WUM4 | Programmed cell death 6-interacting prote | PDCD6IP | 96.024 | 2 | 54.294525 | yes | 9.974E-08 | 661.80981 | 33.373005 |
| STPVNVPISQK | 11 | Q8WUM4 | Programmed cell death 6-interacting prote | PDCD6IP | 96.024 | 2 | 54.294525 | yes | 5.848E-06 | 585.3299 | 24.328287 |
| TSEVDLAKPLVK | 12 | Q8WUM4 | Programmed cell death 6-interacting prote | PDCD6IP | 96.024 | 2 | 54.294525 | yes | 1.641E-06 | 650.37958 | 19.629665 |
| AAAAALSQQQSLC | 15 | Q8WUQ7 | Splicing factor Cactin OS=Homo sapiens | CACTIN | 88.702 | 2 | 52.298824 | yes | 1.12E-07 | 786.41046 | 259.08975 |
| STPEYFLEACADN | 14 | Q8WUQ7 | Splicing factor Cactin OS=Homo sapiens | CACTIN | 88.702 | 2 | 52.298824 | yes | 9.722E-08 | 822.86414 | 222.5502 |
| EQELEMLQR | 9 | Q8WUQ7 | Splicing factor Cactin OS=Homo sapiens | CACTIN | 88.702 | 2 | 52.298824 | yes | 2.321E-06 | 588.28992 | 198.19931 |
| EGVNASVSSDVQS | 16 | Q8WUQ7 | Splicing factor Cactin OS=Homo sapiens | CACTIN | 88.702 | 2 | 52.298824 | yes | 2.144E-07 | 826.90991 | 176.74829 |
| DMTTITEDEISK | 12 | Q8WUQ7 | Splicing factor Cactin OS=Homo sapiens | CACTIN | 88.702 | 2 | 52.298824 | yes | 9.722E-08 | 691.82141 | 155.76305 |
| GISHLEEK | 8 | Q8WUQ7 | Splicing factor Cactin OS=Homo sapiens | CACTIN | 88.702 | 2 | 52.298824 | yes | 1.256E-06 | 456.74289 | 101.22414 |
| VHTGFEWNK | 9 | Q8WUQ7 | Splicing factor Cactin OS=Homo sapiens | CACTIN | 88.702 | 2 | 52.298824 | yes | 8.128E-07 | 559.2749 | 78.830025 |
| SSPFPTVPTRPEK | 13 | Q8WUU5 | GATA zinc finger domain-containing pro | GATAD1 | 28.691 | 2 | 45.40665 | yes | 1.859E-07 | 721.88556 | 71.46418 |
| EYIEIITSSIK | 11 | Q8WUW1 | Protein BRICK1 OS=Homo sapiens OX= | BRK1 | 8.745 | 2 | 52.83515 | yes | 1.001E-07 | 648.35828 | 229.03781 |
| RIEYIEAR | 8 | Q8WUW1 | Protein BRICK1 OS=Homo sapiens OX= | BRK1 | 8.745 | 2 | 52.83515 | yes | 5.6E-07 | 525.29053 | 200.09262 |
| IADFLNSFDMSCR | 13 | Q8WUW1 | Protein BRICK1 OS=Homo sapiens OX= | BRK1 | 8.745 | 2 | 52.83515 | yes | 9.722E-08 | 788.35016 | 191.55313 |
| EIHQDWANR | 9 | Q8WUW1 | Protein BRICK1 OS=Homo sapiens OX= | BRK1 | 8.745 | 2 | 52.83515 | yes | 9.722E-08 | 584.77832 | 155.696 |
| VLLTNVVTGHR | 11 | Q8WV16 | DDB1- and CUL4-associated factor 4 OS | DCAF4 | 55.694 | 2 | 55.805218 | yes | 9.722E-08 | 604.85913 | 119.00247 |
| SMDPSALASDR | 11 | Q8WV16 | DDB1- and CUL4-associated factor 4 OS | DCAF4 | 55.694 | 2 | 55.805218 | yes | 4.226E-06 | 575.26385 | 88.275032 |
| FNLILADTNSDR | 12 | Q8WV16 | DDB1- and CUL4-associated factor 4 OS | DCAF4 | 55.694 | 2 | 55.805218 | yes | 1.114E-07 | 689.85168 | 84.076431 |
| YGIINLQSLK | 10 | Q8WV16 | DDB1- and CUL4-associated factor 4 OS | DCAF4 | 55.694 | 2 | 55.805218 | yes | 1.813E-07 | 574.83734 | 72.521523 |
| AILEMEQYIR | 10 | Q8WV22 | Non-structural maintenance of chromosor | NSMCE1 | 30.856 | 2 | 47.412815 | yes | 2.904E-07 | 633.33154 | 286.55457 |
| MGVMTDVHR | 9 | Q8WV22 | Non-structural maintenance of chromosor | NSMCE1 | 30.856 | 2 | 47.412815 | yes | 6.399E-07 | 523.24951 | 235.56831 |

| | | | | | | | | | | | |
|---|---|---|---|---|---|---|---|---|---|---|---|
| MATDFAENELDLF | 14 | Q8WV22 | Non-structural maintenance of chromosor | NSMCE1 | 30.856 | 2 | 47.412815 | yes | 1.336E-07 | 836.38782 | 109.97691 |
| IAETYSIEMGPR | 12 | Q8WV41 | Sorting nexin-33 OS=Homo sapiens OX= | SNX33 | 65.265 | 2 | 48.52322 | yes | 1.136E-07 | 683.83722 | 241.56152 |
| GETGLFPASYVEI\ | 15 | Q8WV41 | Sorting nexin-33 OS=Homo sapiens OX= | SNX33 | 65.265 | 2 | 48.52322 | yes | 3.364E-07 | 819.4303 | 219.83342 |
| FEEDFIEK | 8 | Q8WV41 | Sorting nexin-33 OS=Homo sapiens OX= | SNX33 | 65.265 | 2 | 48.52322 | yes | 1.194E-06 | 528.74786 | 139.5974 |
| HMMQNYLR | 8 | Q8WV41 | Sorting nexin-33 OS=Homo sapiens OX= | SNX33 | 65.265 | 2 | 48.52322 | yes | 3.984E-07 | 546.75751 | 91.801537 |
| LTPTHAASPVYR | 12 | Q8WV41 | Sorting nexin-33 OS=Homo sapiens OX= | SNX33 | 65.265 | 2 | 48.52322 | yes | 7.014E-07 | 656.85406 | 59.65649 |
| TYEAIGEMFAEQP | 14 | Q8WV41 | Sorting nexin-33 OS=Homo sapiens OX= | SNX33 | 65.265 | 2 | 48.52322 | yes | 4.267E-07 | 807.37946 | 58.943935 |
| LLAEAQER | 8 | Q8WV44 | E3 ubiquitin-protein ligase TRIM41 OS= | TRIM41 | 71.67 | 2 | 52.07658 | yes | 9.722E-08 | 465.25616 | 594.74304 |
| FSADCCVLGAQGI | 14 | Q8WV44 | E3 ubiquitin-protein ligase TRIM41 OS= | TRIM41 | 71.67 | 2 | 52.07658 | yes | 1.12E-07 | 794.35577 | 587.14008 |
| FLAEEQAGLER | 11 | Q8WV44 | E3 ubiquitin-protein ligase TRIM41 OS= | TRIM41 | 71.67 | 2 | 52.07658 | yes | 1.188E-07 | 631.82239 | 454.29462 |
| YQAQSSTEQTLLS | 19 | Q8WV44 | E3 ubiquitin-protein ligase TRIM41 OS= | TRIM41 | 71.67 | 3 | 52.07658 | yes | 3.037E-07 | 717.3623 | 431.66275 |
| LHLPQQPLLQR | 11 | Q8WV44 | E3 ubiquitin-protein ligase TRIM41 OS= | TRIM41 | 71.67 | 2 | 52.07658 | yes | 1.496E-07 | 671.90131 | 273.66748 |
| EMHEAQLGR | 9 | Q8WV44 | E3 ubiquitin-protein ligase TRIM41 OS= | TRIM41 | 71.67 | 2 | 52.07658 | yes | 3.056E-07 | 535.75604 | 268.5466 |
| LQGHVEPLR | 9 | Q8WV44 | E3 ubiquitin-protein ligase TRIM41 OS= | TRIM41 | 71.67 | 2 | 52.07658 | yes | 9.975E-08 | 524.79852 | 222.65063 |
| VTDQGICPK | 9 | Q8WV44 | E3 ubiquitin-protein ligase TRIM41 OS= | TRIM41 | 71.67 | 2 | 52.07658 | yes | 7.848E-07 | 509.25531 | 194.76241 |
| HYWEVEVGGR | 10 | Q8WV44 | E3 ubiquitin-protein ligase TRIM41 OS= | TRIM41 | 71.67 | 2 | 52.07658 | yes | 1.223E-07 | 616.29639 | 194.52161 |
| FGVYLDYEAGR | 11 | Q8WV44 | E3 ubiquitin-protein ligase TRIM41 OS= | TRIM41 | 71.67 | 2 | 52.07658 | yes | 1.047E-06 | 645.31171 | 133.46309 |
| LAEQAAQLSR | 10 | Q8WV44 | E3 ubiquitin-protein ligase TRIM41 OS= | TRIM41 | 71.67 | 2 | 52.07658 | yes | 1.414E-07 | 543.79877 | 113.24599 |
| SELAAVASEFGR | 12 | Q8WV44 | E3 ubiquitin-protein ligase TRIM41 OS= | TRIM41 | 71.67 | 2 | 52.07658 | yes | 4.038E-07 | 618.81458 | 68.3284 |
| RLHLPQQPLLQR | 12 | Q8WV44 | E3 ubiquitin-protein ligase TRIM41 OS= | TRIM41 | 71.67 | 3 | 52.07658 | yes | 3.2E-06 | 500.30368 | 21.589996 |
| GLQEIEESLR | 10 | Q8WV92 | MIT domain-containing protein 1 OS=Hc | MITD1 | 29.315 | 2 | 52.742935 | yes | 9.819E-08 | 587.30914 | 193.67165 |
| QDPQSTAAATVLK | 13 | Q8WV92 | MIT domain-containing protein 1 OS=Hc | MITD1 | 29.315 | 2 | 52.742935 | yes | 9.722E-08 | 665.35406 | 134.48317 |
| SHGVLLEVQYSSS | 17 | Q8WV92 | MIT domain-containing protein 1 OS=Hc | MITD1 | 29.315 | 3 | 52.742935 | yes | 1.036E-07 | 642.99377 | 116.96449 |
| HTHQLYNFLR | 10 | Q8WV92 | MIT domain-containing protein 1 OS=Hc | MITD1 | 29.315 | 3 | 52.742935 | yes | 9.722E-08 | 443.5668 | 93.090912 |
| AVELDSESR | 9 | Q8WV92 | MIT domain-containing protein 1 OS=Hc | MITD1 | 29.315 | 2 | 52.742935 | yes | 4.664E-06 | 503.246 | 72.356346 |
| FSLGYCDFDLRPC | 23 | Q8WV92 | MIT domain-containing protein 1 OS=Hc | MITD1 | 29.315 | 4 | 52.742935 | yes | 5.362E-06 | 715.08228 | 4.1107035 |
| VALLCGPPGLGK | 12 | Q8WVB6 | Chromosome transmission fidelity proteir | CHTF18 | 107.383 | 2 | 52.395245 | yes | 1.281E-07 | 591.33917 | 183.83684 |
| LSDTLHSLR | 9 | Q8WVB6 | Chromosome transmission fidelity proteir | CHTF18 | 107.383 | 2 | 52.395245 | yes | 1.088E-07 | 521.28802 | 107.53395 |
| ITFPSSQQEAQNR | 13 | Q8WVB6 | Chromosome transmission fidelity proteir | CHTF18 | 107.383 | 2 | 52.395245 | yes | 2.426E-07 | 753.37079 | 104.4527 |
| ACINTLQFLYSR | 12 | Q8WVB6 | Chromosome transmission fidelity proteir | CHTF18 | 107.383 | 2 | 52.395245 | yes | 3.488E-06 | 743.37958 | 100.28894 |
| ADPGVLAALCEK | 12 | Q8WVB6 | Chromosome transmission fidelity proteir | CHTF18 | 107.383 | 2 | 52.395245 | yes | 1.446E-07 | 622.32117 | 94.215179 |
| RPPILEDYVHVTS | 18 | Q8WVB6 | Chromosome transmission fidelity proteir | CHTF18 | 107.383 | 3 | 52.395245 | yes | 4.03E-07 | 690.03223 | 72.446701 |
| LWDLVVFGHER | 11 | Q8WVB6 | Chromosome transmission fidelity proteir | CHTF18 | 107.383 | 2 | 52.395245 | yes | 9.722E-08 | 685.86444 | 52.849079 |
| QVDADLQPAGSLI | 17 | Q8WVB6 | Chromosome transmission fidelity proteir | CHTF18 | 107.383 | 3 | 52.395245 | yes | 9.28E-07 | 591.30768 | 50.363659 |
| SHEQVLEEMLEAC | 21 | Q8WVB6 | Chromosome transmission fidelity proteir | CHTF18 | 107.383 | 3 | 52.395245 | yes | 3.577E-07 | 798.39703 | 22.984396 |
| VNTDLGNDLYFVI | 13 | Q8WVC0 | RNA polymerase-associated protein LEO | LEO1 | 75.405 | 2 | 43.93118 | yes | 2.389E-07 | 749.38281 | 58.997692 |
| APLQGDHNHLFIR | 13 | Q8WVC0 | RNA polymerase-associated protein LEO | LEO1 | 75.405 | 3 | 43.93118 | yes | 1.557E-06 | 506.60391 | 58.733051 |
| ALDLENIMR | 9 | Q8WVD3 | E3 ubiquitin-protein ligase RNF138 OS= | RNF138 | 28.193 | 2 | 52.943 | yes | 1.031E-07 | 537.78424 | 364.49597 |
| NFVSHLNQR | 9 | Q8WVD3 | E3 ubiquitin-protein ligase RNF138 OS= | RNF138 | 28.193 | 2 | 52.943 | yes | 9.722E-08 | 557.79126 | 364.24194 |

| | | | | | | | | | | | |
|---|---|---|---|---|---|---|---|---|---|---|---|
| CPLCQESNFTR | 11 | Q8WVD3 | E3 ubiquitin-protein ligase RNF138 OS=] | RNF138 | 28.193 | 2 | 52.943 | yes | 9.722E-08 | 706.30829 | 342.81201 |
| ESGAHCPLCR | 10 | Q8WVD3 | E3 ubiquitin-protein ligase RNF138 OS=] | RNF138 | 28.193 | 2 | 52.943 | yes | 2.04E-07 | 593.75824 | 201.71773 |
| KCFLTAMR | 8 | Q8WVD3 | E3 ubiquitin-protein ligase RNF138 OS=] | RNF138 | 28.193 | 2 | 52.943 | yes | 3.369E-07 | 513.76477 | 124.15504 |
| TTACQHVFCR | 10 | Q8WVD3 | E3 ubiquitin-protein ligase RNF138 OS=] | RNF138 | 28.193 | 2 | 52.943 | yes | 1.248E-07 | 640.28717 | 123.04868 |
| LMGFASFDSTK | 11 | Q8WVK2 | U4/U6.U5 small nuclear ribonucleoprotei | SNRNP27 | 18.861 | 2 | 56.79916 | yes | 9.722E-08 | 602.28937 | 118.38117 |
| VDGSVNAYAINVS | 15 | Q8WVK2 | U4/U6.U5 small nuclear ribonucleoprotei | SNRNP27 | 18.861 | 2 | 56.79916 | yes | 9.722E-08 | 782.90192 | 76.089645 |
| TEEEIEMMK | 9 | Q8WVK2 | U4/U6.U5 small nuclear ribonucleoprotei | SNRNP27 | 18.861 | 2 | 56.79916 | yes | 2.972E-06 | 570.25159 | 69.267387 |
| GGFNRPLDFIA | 11 | Q8WVK2 | U4/U6.U5 small nuclear ribonucleoprotei | SNRNP27 | 18.861 | 2 | 56.79916 | yes | 6.09E-07 | 603.81696 | 36.18259 |
| AESDLDYIQYR | 11 | Q8WVK7 | Spindle and kinetochore-associated protei | SKA2 | 14.189 | 2 | 40.40297 | yes | 4.043E-07 | 686.82263 | 107.32784 |
| MLFPEAQR | 8 | Q8WVM0 | Dimethyladenosine transferase 1, mitochc | TFB1M | 39.543 | 2 | 51.084778 | yes | 4.874E-07 | 496.25513 | 1408.7874 |
| FIPGLQMLSDAAP( | 15 | Q8WVM0 | Dimethyladenosine transferase 1, mitochc | TFB1M | 39.543 | 2 | 51.084778 | yes | 1.199E-07 | 772.91071 | 1004.1657 |
| IVHGDVLTFK | 10 | Q8WVM0 | Dimethyladenosine transferase 1, mitochc | TFB1M | 39.543 | 2 | 51.084778 | yes | 9.722E-08 | 564.82422 | 687.41125 |
| TQMTLTFQK | 9 | Q8WVM0 | Dimethyladenosine transferase 1, mitochc | TFB1M | 39.543 | 2 | 51.084778 | yes | 9.722E-08 | 549.28662 | 618.89716 |
| QLSISHFK | 8 | Q8WVM0 | Dimethyladenosine transferase 1, mitochc | TFB1M | 39.543 | 2 | 51.084778 | yes | 1.055E-07 | 480.26907 | 425.18658 |
| LSVMAQYLCNVR | 12 | Q8WVM0 | Dimethyladenosine transferase 1, mitochc | TFB1M | 39.543 | 2 | 51.084778 | yes | 9.991E-08 | 727.36816 | 414.61902 |
| VVQNVFQFR | 9 | Q8WVM0 | Dimethyladenosine transferase 1, mitochc | TFB1M | 39.543 | 2 | 51.084778 | yes | 1.316E-07 | 568.81421 | 409.01355 |
| WLENISCR | 8 | Q8WVM0 | Dimethyladenosine transferase 1, mitochc | TFB1M | 39.543 | 2 | 51.084778 | yes | 1.165E-07 | 539.26093 | 346.09909 |
| LLELADIDPTLRPR | 14 | Q8WVM0 | Dimethyladenosine transferase 1, mitochc | TFB1M | 39.543 | 3 | 51.084778 | yes | 1.555E-07 | 541.3139 | 271.13062 |
| SLCDVYR | 7 | Q8WVM0 | Dimethyladenosine transferase 1, mitochc | TFB1M | 39.543 | 2 | 51.084778 | yes | 9.722E-08 | 456.71582 | 181.30009 |
| HIFTIPGQAFVPKP | 30 | Q8WVM0 | Dimethyladenosine transferase 1, mitochc | TFB1M | 39.543 | 4 | 51.084778 | yes | 1.555E-07 | 828.46423 | 148.09399 |
| LPPLPTIR | 8 | Q8WVM0 | Dimethyladenosine transferase 1, mitochc | TFB1M | 39.543 | 2 | 51.084778 | yes | 1.777E-07 | 453.79221 | 146.00464 |
| SILNADVAELLVV | 15 | Q8WVM0 | Dimethyladenosine transferase 1, mitochc | TFB1M | 39.543 | 2 | 51.084778 | yes | 4.454E-07 | 806.96143 | 28.305359 |
| HLWAIQSGGSK | 11 | Q8WVS4 | Cytoplasmic dynein 2 intermediate chain | DYNC2I1 | 122.571 | 2 | 42.242138 | yes | 3.126E-07 | 592.31458 | 114.22617 |
| EKLEELPLAQK | 11 | Q8WVS4 | Cytoplasmic dynein 2 intermediate chain | DYNC2I1 | 122.571 | 2 | 42.242138 | yes | 3.177E-06 | 649.37177 | 30.947811 |
| TLSGLLEWECK | 11 | Q8WVV9 | Heterogeneous nuclear ribonucleoprotein | HNRNPLL | 60.084 | 2 | 51.785767 | yes | 9.732E-08 | 668.33429 | 2195.7705 |
| VSVSPVVHVR | 10 | Q8WVV9 | Heterogeneous nuclear ribonucleoprotein | HNRNPLL | 60.084 | 2 | 51.785767 | yes | 1.122E-07 | 539.82202 | 2092.7927 |
| QAILGEHPSSFR | 12 | Q8WVV9 | Heterogeneous nuclear ribonucleoprotein | HNRNPLL | 60.084 | 2 | 51.785767 | yes | 9.722E-08 | 671.34912 | 835.61365 |
| QAILGEHPSSFR | 12 | Q8WVV9 | Heterogeneous nuclear ribonucleoprotein | HNRNPLL | 60.084 | 2 | 51.785767 | yes | 1.422E-07 | 447.90186 | 835.61365 |
| QAILGEHPSSFR | 12 | Q8WVV9 | Heterogeneous nuclear ribonucleoprotein | HNRNPLL | 60.084 | 3 | 51.785767 | yes | 9.722E-08 | 671.34912 | 835.61365 |
| QAILGEHPSSFR | 12 | Q8WVV9 | Heterogeneous nuclear ribonucleoprotein | HNRNPLL | 60.084 | 3 | 51.785767 | yes | 1.422E-07 | 447.90186 | 835.61365 |
| QALVEFENIDSAK | 13 | Q8WVV9 | Heterogeneous nuclear ribonucleoprotein | HNRNPLL | 60.084 | 2 | 51.785767 | yes | 1.025E-07 | 732.37244 | 1251.0427 |
| LCFSTSSHL | 9 | Q8WVV9 | Heterogeneous nuclear ribonucleoprotein | HNRNPLL | 60.084 | 2 | 51.785767 | yes | 2.504E-07 | 526.2475 | 1161.1354 |
| RLNVCVSK | 8 | Q8WVV9 | Heterogeneous nuclear ribonucleoprotein | HNRNPLL | 60.084 | 2 | 51.785767 | yes | 6.017E-07 | 488.27383 | 806.85425 |
| AVTHLNNVK | 9 | Q8WVV9 | Heterogeneous nuclear ribonucleoprotein | HNRNPLL | 60.084 | 2 | 51.785767 | yes | 2.859E-06 | 498.28528 | 453.81381 |
| SFSQPEAGGSHHK | 13 | Q8WVV9 | Heterogeneous nuclear ribonucleoprotein | HNRNPLL | 60.084 | 2 | 51.785767 | yes | 2.202E-07 | 684.81818 | 389.53421 |
| LCNDHEVLTFIK | 12 | Q8WVV9 | Heterogeneous nuclear ribonucleoprotein | HNRNPLL | 60.084 | 3 | 51.785767 | yes | 4.614E-07 | 496.92203 | 92.654602 |
| LCNDHEVLTFIK | 12 | Q8WVV9 | Heterogeneous nuclear ribonucleoprotein | HNRNPLL | 60.084 | 3 | 51.785767 | yes | 1.977E-07 | 744.87939 | 92.654602 |
| LCNDHEVLTFIK | 12 | Q8WVV9 | Heterogeneous nuclear ribonucleoprotein | HNRNPLL | 60.084 | 2 | 51.785767 | yes | 4.614E-07 | 496.92203 | 92.654602 |
| LCNDHEVLTFIK | 12 | Q8WVV9 | Heterogeneous nuclear ribonucleoprotein | HNRNPLL | 60.084 | 2 | 51.785767 | yes | 1.977E-07 | 744.87939 | 92.654602 |

| | | | | | | | | | | | |
|---|---|---|---|---|---|---|---|---|---|---|---|
| VFDAKPSAK | 9 | Q8WVV9 | Heterogeneous nuclear ribonucleoprotein | HNRNPLL | 60.084 | 2 | 51.785767 | yes | 1.625E-07 | 481.76892 | 158.61075 |
| HDGYGSHGPLLPL | 16 | Q8WVV9 | Heterogeneous nuclear ribonucleoprotein | HNRNPLL | 60.084 | 3 | 51.785767 | yes | 3.248E-07 | 568.29352 | 77.568718 |
| HDGYGSHGPLLPL | 16 | Q8WVV9 | Heterogeneous nuclear ribonucleoprotein | HNRNPLL | 60.084 | 3 | 51.785767 | yes | 4.605E-06 | 426.47195 | 77.568718 |
| HDGYGSHGPLLPL | 16 | Q8WVV9 | Heterogeneous nuclear ribonucleoprotein | HNRNPLL | 60.084 | 4 | 51.785767 | yes | 3.248E-07 | 568.29352 | 77.568718 |
| HDGYGSHGPLLPL | 16 | Q8WVV9 | Heterogeneous nuclear ribonucleoprotein | HNRNPLL | 60.084 | 4 | 51.785767 | yes | 4.605E-06 | 426.47195 | 77.568718 |
| NDNDSWDYTKPY | 15 | Q8WVV9 | Heterogeneous nuclear ribonucleoprotein | HNRNPLL | 60.084 | 3 | 51.785767 | yes | 1.323E-07 | 615.27972 | 115.92349 |
| VPNGSNPYTLK | 11 | Q8WVV9 | Heterogeneous nuclear ribonucleoprotein | HNRNPLL | 60.084 | 2 | 51.785767 | yes | 4.583E-07 | 595.31421 | 92.47345 |
| ITRPGNTDDPSGGI | 15 | Q8WVV9 | Heterogeneous nuclear ribonucleoprotein | HNRNPLL | 60.084 | 2 | 51.785767 | yes | 2.206E-07 | 764.87115 | 43.205154 |
| IIAINSELTQPK | 12 | Q8WVX9 | Fatty acyl-CoA reductase 1 OS=Homo sa | FAR1 | 59.357 | 2 | 50.73845 | yes | 2.177E-07 | 663.88501 | 730.96136 |
| ALAEYVVQQEGAI | 13 | Q8WVX9 | Fatty acyl-CoA reductase 1 OS=Homo sa | FAR1 | 59.357 | 2 | 50.73845 | yes | 9.963E-08 | 703.36975 | 407.72055 |
| YVLNEEMSGLPA | 14 | Q8WVX9 | Fatty acyl-CoA reductase 1 OS=Homo sa | FAR1 | 59.357 | 2 | 50.73845 | yes | 1.431E-07 | 775.38757 | 250.04929 |
| DAVQLNVIATR | 11 | Q8WVX9 | Fatty acyl-CoA reductase 1 OS=Homo sa | FAR1 | 59.357 | 2 | 50.73845 | yes | 9.723E-08 | 600.34076 | 187.84903 |
| QLILLAQQMK | 10 | Q8WVX9 | Fatty acyl-CoA reductase 1 OS=Homo sa | FAR1 | 59.357 | 2 | 50.73845 | yes | 1.276E-07 | 593.35486 | 179.33717 |
| VNSVYVLVR | 9 | Q8WVX9 | Fatty acyl-CoA reductase 1 OS=Homo sa | FAR1 | 59.357 | 2 | 50.73845 | yes | 1.083E-07 | 524.8111 | 100.79154 |
| NVLLTGATGFLGK | 13 | Q8WVX9 | Fatty acyl-CoA reductase 1 OS=Homo sa | FAR1 | 59.357 | 2 | 50.73845 | yes | 1.515E-07 | 645.87445 | 70.378403 |
| LRDENPDFR | 9 | Q8WVX9 | Fatty acyl-CoA reductase 1 OS=Homo sa | FAR1 | 59.357 | 2 | 50.73845 | yes | 9.819E-08 | 581.28601 | 67.400833 |
| FGFAIGSQTTK | 11 | Q8WW12 | PEST proteolytic signal-containing nucle | PCNP | 18.925 | 2 | 48.91823 | yes | 1.85E-07 | 578.80347 | 93.557739 |
| ETVPTLAPK | 9 | Q8WW12 | PEST proteolytic signal-containing nucle | PCNP | 18.925 | 2 | 48.91823 | yes | 2.562E-06 | 478.27658 | 54.20562 |
| SAEEEAADLPTKP | 15 | Q8WW12 | PEST proteolytic signal-containing nucle | PCNP | 18.925 | 2 | 48.91823 | yes | 1.087E-07 | 793.89905 | 45.630455 |
| DTPTSAGPNSFNK | 13 | Q8WW12 | PEST proteolytic signal-containing nucle | PCNP | 18.925 | 2 | 48.91823 | yes | 1.722E-07 | 668.31238 | 39.125969 |
| EFSITEFSEGAK | 12 | Q8WWC4 | m-AAA protease-interacting protein 1, mi | MAIP1 | 32.545 | 2 | 49.807377 | yes | 1.065E-07 | 672.81952 | 370.14435 |
| QLLSASYEFQR | 11 | Q8WWC4 | m-AAA protease-interacting protein 1, mi | MAIP1 | 32.545 | 2 | 49.807377 | yes | 1.679E-07 | 671.34351 | 241.08701 |
| VTSLPDNHK | 9 | Q8WWC4 | m-AAA protease-interacting protein 1, mi | MAIP1 | 32.545 | 2 | 49.807377 | yes | 3.81E-07 | 505.76691 | 179.98997 |
| EFTQGVKPDWTIA | 14 | Q8WWC4 | m-AAA protease-interacting protein 1, mi | MAIP1 | 32.545 | 2 | 49.807377 | yes | 1.003E-07 | 824.4281 | 69.727104 |
| EVLHALK | 7 | Q8WWC4 | m-AAA protease-interacting protein 1, mi | MAIP1 | 32.545 | 2 | 49.807377 | yes | 1.317E-07 | 405.24762 | 58.609409 |
| LLAEAGMPSPEWT | 14 | Q8WWH5 | Pseudouridylate synthase TRUB1 OS=Hc | TRUB1 | 37.253 | 2 | 51.59504 | yes | 9.722E-08 | 765.38708 | 122.36972 |
| AAAAVVAAAAR | 11 | Q8WWH5 | Pseudouridylate synthase TRUB1 OS=Hc | TRUB1 | 37.253 | 2 | 51.59504 | yes | 9.722E-08 | 471.27997 | 97.706947 |
| GPTSAELLNR | 10 | Q8WWH5 | Pseudouridylate synthase TRUB1 OS=Hc | TRUB1 | 37.253 | 2 | 51.59504 | yes | 3.368E-07 | 529.28546 | 67.644966 |
| ITQEDIEGILQK | 12 | Q8WWH5 | Pseudouridylate synthase TRUB1 OS=Hc | TRUB1 | 37.253 | 2 | 51.59504 | yes | 1.132E-07 | 693.87738 | 62.51561 |
| ELSSCANVLELTR | 13 | Q8WWH5 | Pseudouridylate synthase TRUB1 OS=Hc | TRUB1 | 37.253 | 2 | 51.59504 | yes | 1.011E-07 | 746.3772 | 48.799389 |
| EPVSLPGIMR | 10 | Q8WWI1 | LIM domain only protein 7 OS=Homo sa | LMO7 | 192.696 | 2 | 51.695183 | yes | 2.361E-07 | 549.80243 | 333.22318 |
| SMSDVSAEDVQNI | 14 | Q8WWI1 | LIM domain only protein 7 OS=Homo sa | LMO7 | 192.696 | 2 | 51.695183 | yes | 3.528E-07 | 775.85938 | 312.82581 |
| MYSFDDVLEEGK | 12 | Q8WWI1 | LIM domain only protein 7 OS=Homo sa | LMO7 | 192.696 | 2 | 51.695183 | yes | 9.722E-08 | 716.81866 | 240.6403 |
| SQDQFSDMR | 9 | Q8WWI1 | LIM domain only protein 7 OS=Homo sa | LMO7 | 192.696 | 2 | 51.695183 | yes | 1.022E-07 | 557.23511 | 201.27419 |
| QQILQEMR | 8 | Q8WWI1 | LIM domain only protein 7 OS=Homo sa | LMO7 | 192.696 | 2 | 51.695183 | yes | 1.412E-06 | 523.27661 | 133.91888 |
| KPQDQLVIER | 10 | Q8WWI1 | LIM domain only protein 7 OS=Homo sa | LMO7 | 192.696 | 2 | 51.695183 | yes | 9.722E-08 | 613.34857 | 120.10564 |
| LTSVVTPR | 8 | Q8WWI1 | LIM domain only protein 7 OS=Homo sa | LMO7 | 192.696 | 2 | 51.695183 | yes | 8.248E-07 | 436.76364 | 106.14858 |
| WVWDQEEER | 9 | Q8WWI1 | LIM domain only protein 7 OS=Homo sa | LMO7 | 192.696 | 2 | 51.695183 | yes | 1.182E-07 | 638.78326 | 75.733238 |
| SLTSCSSDITLR | 12 | Q8WWI1 | LIM domain only protein 7 OS=Homo sa | LMO7 | 192.696 | 2 | 51.695183 | yes | 2.45E-07 | 670.32971 | 73.222534 |

| | | | | | | | | | | | |
|---|---|---|---|---|---|---|---|---|---|---|---|
| WIDATSGIYNSEK | 13 | Q8WWI1 | LIM domain only protein 7 OS=Homo sa | LMO7 | 192.696 | 2 | 51.695183 | yes | 1.248E-07 | 742.35681 | 66.603943 |
| SHSPSASQSGSQLI | 14 | Q8WWI1 | LIM domain only protein 7 OS=Homo sa | LMO7 | 192.696 | 2 | 51.695183 | yes | 1.46E-07 | 714.84491 | 66.466621 |
| WQDDLAK | 7 | Q8WWI1 | LIM domain only protein 7 OS=Homo sa | LMO7 | 192.696 | 2 | 51.695183 | yes | 1.5E-07 | 438.21652 | 63.95583 |
| ICSYCNNILGK | 11 | Q8WWI1 | LIM domain only protein 7 OS=Homo sa | LMO7 | 192.696 | 2 | 51.695183 | yes | 6.827E-07 | 671.31812 | 61.052715 |
| AFENLLGQALTK | 12 | Q8WWI1 | LIM domain only protein 7 OS=Homo sa | LMO7 | 192.696 | 2 | 51.695183 | yes | 1.071E-07 | 652.86407 | 56.869896 |
| RGESLDNLDSPR | 12 | Q8WWI1 | LIM domain only protein 7 OS=Homo sa | LMO7 | 192.696 | 2 | 51.695183 | yes | 2.627E-05 | 679.83661 | 46.814846 |
| EVAATEEDVTR | 11 | Q8WWI1 | LIM domain only protein 7 OS=Homo sa | LMO7 | 192.696 | 2 | 51.695183 | yes | 9.723E-08 | 610.2937 | 46.370838 |
| STTELDDYSTNK | 12 | Q8WWI1 | LIM domain only protein 7 OS=Homo sa | LMO7 | 192.696 | 2 | 51.695183 | yes | 1.906E-07 | 687.30701 | 35.859394 |
| RQPQEEVVHEDQ( | 14 | Q8WWI1 | LIM domain only protein 7 OS=Homo sa | LMO7 | 192.696 | 3 | 51.695183 | yes | 7.121E-07 | 560.27631 | 35.123287 |
| TEEASSGFLPGDR | 13 | Q8WWI1 | LIM domain only protein 7 OS=Homo sa | LMO7 | 192.696 | 2 | 51.695183 | yes | 1.078E-07 | 683.31769 | 34.476372 |
| TSTTGVATTQSPTI | 15 | Q8WWI1 | LIM domain only protein 7 OS=Homo sa | LMO7 | 192.696 | 2 | 51.695183 | yes | 9.758E-08 | 752.88373 | 26.537685 |
| VQFDGTNSAFK | 11 | Q8WWK9 | Cytoskeleton-associated protein 2 OS=Hc | CKAP2 | 76.987 | 2 | 52.173138 | yes | 1.609E-07 | 607.29602 | 542.22656 |
| YNVSTTPYLQSVK | 13 | Q8WWK9 | Cytoskeleton-associated protein 2 OS=Hc | CKAP2 | 76.987 | 2 | 52.173138 | yes | 9.722E-08 | 750.39069 | 460.71692 |
| AILAGAQPIEEMR | 13 | Q8WWK9 | Cytoskeleton-associated protein 2 OS=Hc | CKAP2 | 76.987 | 2 | 52.173138 | yes | 9.766E-08 | 699.87415 | 435.16357 |
| KPVLGSYR | 8 | Q8WWK9 | Cytoskeleton-associated protein 2 OS=Hc | CKAP2 | 76.987 | 2 | 52.173138 | yes | 9.722E-08 | 460.27164 | 321.54431 |
| ANLGENMEK | 9 | Q8WWK9 | Cytoskeleton-associated protein 2 OS=Hc | CKAP2 | 76.987 | 2 | 52.173138 | yes | 1.08E-07 | 503.23712 | 164.64792 |
| ATKPQPVNTSSVT | 15 | Q8WWK9 | Cytoskeleton-associated protein 2 OS=Hc | CKAP2 | 76.987 | 2 | 52.173138 | yes | 2.454E-07 | 778.93579 | 131.3177 |
| LKEHLLR | 7 | Q8WWK9 | Cytoskeleton-associated protein 2 OS=Hc | CKAP2 | 76.987 | 2 | 52.173138 | yes | 1.783E-07 | 454.78745 | 117.80478 |
| HTIVDILTMK | 10 | Q8WWK9 | Cytoskeleton-associated protein 2 OS=Hc | CKAP2 | 76.987 | 2 | 52.173138 | yes | 9.722E-08 | 585.83118 | 83.907593 |
| NLLFQDCEK | 9 | Q8WWK9 | Cytoskeleton-associated protein 2 OS=Hc | CKAP2 | 76.987 | 2 | 52.173138 | yes | 1.458E-06 | 583.77917 | 73.285339 |
| FVSTTSQNTQLVR | 17 | Q8WWK9 | Cytoskeleton-associated protein 2 OS=Hc | CKAP2 | 76.987 | 3 | 52.173138 | yes | 5.673E-07 | 648.69354 | 55.160789 |
| EVSIEDTGVDVDP | 15 | Q8WWK9 | Cytoskeleton-associated protein 2 OS=Hc | CKAP2 | 76.987 | 2 | 52.173138 | yes | 1.018E-07 | 816.38599 | 53.642937 |
| KVQFDGTNSAFK | 12 | Q8WWK9 | Cytoskeleton-associated protein 2 OS=Hc | CKAP2 | 76.987 | 2 | 52.173138 | yes | 2.41E-07 | 671.34351 | 38.726051 |
| VVTSEDQVQEGTK | 13 | Q8WWK9 | Cytoskeleton-associated protein 2 OS=Hc | CKAP2 | 76.987 | 2 | 52.173138 | yes | 9.722E-08 | 710.35175 | 19.434925 |
| TSSSQGIIR | 9 | Q8WWK9 | Cytoskeleton-associated protein 2 OS=Hc | CKAP2 | 76.987 | 2 | 52.173138 | yes | 0.0002513 | 474.75909 | 14.573507 |
| HCIPLKPSNELTNS | 24 | Q8WWK9 | Cytoskeleton-associated protein 2 OS=Hc | CKAP2 | 76.987 | 4 | 52.173138 | yes | 4.22E-06 | 682.8667 | 11.723705 |
| FELAVDAVHR | 10 | Q8WWM7 | Ataxin-2-like protein OS=Homo sapiens ( | ATXN2L | 113.374 | 2 | 53.3485 | yes | 9.85E-08 | 578.80908 | 47917.941 |
| TLEPQELAR | 9 | Q8WWM7 | Ataxin-2-like protein OS=Homo sapiens ( | ATXN2L | 113.374 | 2 | 53.3485 | yes | 1.555E-07 | 528.78784 | 43892.266 |
| ISLAPTDVK | 9 | Q8WWM7 | Ataxin-2-like protein OS=Homo sapiens ( | ATXN2L | 113.374 | 2 | 53.3485 | yes | 2.565E-07 | 472.27658 | 38665.805 |
| FQLEELR | 7 | Q8WWM7 | Ataxin-2-like protein OS=Homo sapiens ( | ATXN2L | 113.374 | 2 | 53.3485 | yes | 1.484E-06 | 467.75327 | 31716.891 |
| AEGLQVGQDAR | 11 | Q8WWM7 | Ataxin-2-like protein OS=Homo sapiens ( | ATXN2L | 113.374 | 2 | 53.3485 | yes | 9.722E-08 | 572.29126 | 25907.986 |
| VPGLQNEQK | 9 | Q8WWM7 | Ataxin-2-like protein OS=Homo sapiens ( | ATXN2L | 113.374 | 2 | 53.3485 | yes | 1.841E-07 | 506.77472 | 24383.469 |
| YIPLPQR | 7 | Q8WWM7 | Ataxin-2-like protein OS=Homo sapiens ( | ATXN2L | 113.374 | 2 | 53.3485 | yes | 1.273E-07 | 443.76089 | 23681.861 |
| EFSLAGGIWHGR | 12 | Q8WWM7 | Ataxin-2-like protein OS=Homo sapiens ( | ATXN2L | 113.374 | 2 | 53.3485 | yes | 1.224E-07 | 665.33856 | 22654.959 |
| EIESSPQYR | 9 | Q8WWM7 | Ataxin-2-like protein OS=Homo sapiens ( | ATXN2L | 113.374 | 2 | 53.3485 | yes | 1.188E-07 | 554.76709 | 18203.943 |
| FNEENYGVK | 9 | Q8WWM7 | Ataxin-2-like protein OS=Homo sapiens ( | ATXN2L | 113.374 | 2 | 53.3485 | yes | 1.065E-07 | 550.25635 | 12370.792 |
| NVDFNYATK | 9 | Q8WWM7 | Ataxin-2-like protein OS=Homo sapiens ( | ATXN2L | 113.374 | 2 | 53.3485 | yes | 2.697E-07 | 536.25891 | 12357.808 |
| FQLEELRK | 8 | Q8WWM7 | Ataxin-2-like protein OS=Homo sapiens ( | ATXN2L | 113.374 | 2 | 53.3485 | yes | 1.804E-07 | 531.80072 | 10356.393 |
| FTDSAIAMNSK | 11 | Q8WWM7 | Ataxin-2-like protein OS=Homo sapiens ( | ATXN2L | 113.374 | 2 | 53.3485 | yes | 9.722E-08 | 592.78442 | 4704.9434 |

| | | | | | | | | | | | |
|---|---|---|---|---|---|---|---|---|---|---|---|
| FTDSAIAMNSK | 11 | Q8WWM7 | Ataxin-2-like protein OS=Homo sapiens ( | ATXN2L | 113.374 | 2 | 53.3485 | yes | 1.957E-07 | 600.78192 | 4704.9434 |
| REDIVDTMVFKPS | 21 | Q8WWM7 | Ataxin-2-like protein OS=Homo sapiens ( | ATXN2L | 113.374 | 4 | 53.3485 | yes | 1.11E-07 | 634.32782 | 2939.7656 |
| REDIVDTMVFKPS | 21 | Q8WWM7 | Ataxin-2-like protein OS=Homo sapiens ( | ATXN2L | 113.374 | 4 | 53.3485 | yes | 1.086E-07 | 638.32654 | 2939.7656 |
| REDIVDTMVFKPS | 21 | Q8WWM7 | Ataxin-2-like protein OS=Homo sapiens ( | ATXN2L | 113.374 | 4 | 53.3485 | yes | 1.421E-07 | 507.6637 | 2939.7656 |
| REDIVDTMVFKPS | 21 | Q8WWM7 | Ataxin-2-like protein OS=Homo sapiens ( | ATXN2L | 113.374 | 4 | 53.3485 | yes | 6.435E-07 | 638.32654 | 2939.7656 |
| REDIVDTMVFKPS | 21 | Q8WWM7 | Ataxin-2-like protein OS=Homo sapiens ( | ATXN2L | 113.374 | 5 | 53.3485 | yes | 1.11E-07 | 634.32782 | 2939.7656 |
| REDIVDTMVFKPS | 21 | Q8WWM7 | Ataxin-2-like protein OS=Homo sapiens ( | ATXN2L | 113.374 | 5 | 53.3485 | yes | 1.086E-07 | 638.32654 | 2939.7656 |
| REDIVDTMVFKPS | 21 | Q8WWM7 | Ataxin-2-like protein OS=Homo sapiens ( | ATXN2L | 113.374 | 5 | 53.3485 | yes | 1.421E-07 | 507.6637 | 2939.7656 |
| REDIVDTMVFKPS | 21 | Q8WWM7 | Ataxin-2-like protein OS=Homo sapiens ( | ATXN2L | 113.374 | 5 | 53.3485 | yes | 6.435E-07 | 638.32654 | 2939.7656 |
| EFNPTKPLLSVNK | 13 | Q8WWM7 | Ataxin-2-like protein OS=Homo sapiens ( | ATXN2L | 113.374 | 2 | 53.3485 | yes | 1.043E-07 | 743.91687 | 3191.5276 |
| EFNPTKPLLSVNK | 13 | Q8WWM7 | Ataxin-2-like protein OS=Homo sapiens ( | ATXN2L | 113.374 | 2 | 53.3485 | yes | 1.947E-07 | 496.28033 | 3191.5276 |
| EFNPTKPLLSVNK | 13 | Q8WWM7 | Ataxin-2-like protein OS=Homo sapiens ( | ATXN2L | 113.374 | 3 | 53.3485 | yes | 1.043E-07 | 743.91687 | 3191.5276 |
| EFNPTKPLLSVNK | 13 | Q8WWM7 | Ataxin-2-like protein OS=Homo sapiens ( | ATXN2L | 113.374 | 3 | 53.3485 | yes | 1.947E-07 | 496.28033 | 3191.5276 |
| MLHFLTAVVGSTC | 16 | Q8WWM7 | Ataxin-2-like protein OS=Homo sapiens ( | ATXN2L | 113.374 | 3 | 53.3485 | yes | 9.722E-08 | 593.30402 | 3002.1182 |
| MLHFLTAVVGSTC | 16 | Q8WWM7 | Ataxin-2-like protein OS=Homo sapiens ( | ATXN2L | 113.374 | 3 | 53.3485 | yes | 1.467E-07 | 598.63568 | 3002.1182 |
| DKFTDSAIAMNSK | 13 | Q8WWM7 | Ataxin-2-like protein OS=Homo sapiens ( | ATXN2L | 113.374 | 2 | 53.3485 | yes | 1.113E-07 | 714.3454 | 1889.7861 |
| DKFTDSAIAMNSK | 13 | Q8WWM7 | Ataxin-2-like protein OS=Homo sapiens ( | ATXN2L | 113.374 | 2 | 53.3485 | yes | 4.415E-07 | 722.34283 | 1889.7861 |
| DKFTDSAIAMNSK | 13 | Q8WWM7 | Ataxin-2-like protein OS=Homo sapiens ( | ATXN2L | 113.374 | 2 | 53.3485 | yes | 9.722E-08 | 476.56601 | 1889.7861 |
| DKFTDSAIAMNSK | 13 | Q8WWM7 | Ataxin-2-like protein OS=Homo sapiens ( | ATXN2L | 113.374 | 3 | 53.3485 | yes | 1.113E-07 | 714.3454 | 1889.7861 |
| DKFTDSAIAMNSK | 13 | Q8WWM7 | Ataxin-2-like protein OS=Homo sapiens ( | ATXN2L | 113.374 | 3 | 53.3485 | yes | 4.415E-07 | 722.34283 | 1889.7861 |
| DKFTDSAIAMNSK | 13 | Q8WWM7 | Ataxin-2-like protein OS=Homo sapiens ( | ATXN2L | 113.374 | 3 | 53.3485 | yes | 9.722E-08 | 476.56601 | 1889.7861 |
| RGAEGILAPQPPPF | 30 | Q8WWM7 | Ataxin-2-like protein OS=Homo sapiens ( | ATXN2L | 113.374 | 5 | 53.3485 | yes | 1.874E-07 | 612.32758 | 2681.1975 |
| RGAEGILAPQPPPF | 30 | Q8WWM7 | Ataxin-2-like protein OS=Homo sapiens ( | ATXN2L | 113.374 | 5 | 53.3485 | yes | 6.376E-07 | 765.15771 | 2681.1975 |
| RGAEGILAPQPPPF | 30 | Q8WWM7 | Ataxin-2-like protein OS=Homo sapiens ( | ATXN2L | 113.374 | 4 | 53.3485 | yes | 1.874E-07 | 612.32758 | 2681.1975 |
| RGAEGILAPQPPPF | 30 | Q8WWM7 | Ataxin-2-like protein OS=Homo sapiens ( | ATXN2L | 113.374 | 4 | 53.3485 | yes | 6.376E-07 | 765.15771 | 2681.1975 |
| PSDVMLVHFR | 10 | Q8WWM7 | Ataxin-2-like protein OS=Homo sapiens ( | ATXN2L | 113.374 | 2 | 53.3485 | yes | 6.602E-07 | 600.81335 | 5182.1973 |
| REDIVDTMVFK | 11 | Q8WWM7 | Ataxin-2-like protein OS=Homo sapiens ( | ATXN2L | 113.374 | 2 | 53.3485 | yes | 2.916E-07 | 676.8476 | 4896.8184 |
| PSGETSVPPPPAVC | 15 | Q8WWM7 | Ataxin-2-like protein OS=Homo sapiens ( | ATXN2L | 113.374 | 2 | 53.3485 | yes | 1.762E-07 | 724.38062 | 3941.3728 |
| RFQLEELR | 8 | Q8WWM7 | Ataxin-2-like protein OS=Homo sapiens ( | ATXN2L | 113.374 | 2 | 53.3485 | yes | 2.219E-07 | 545.80383 | 2704.5876 |
| EDIVDTMVFKPSD | 20 | Q8WWM7 | Ataxin-2-like protein OS=Homo sapiens ( | ATXN2L | 113.374 | 3 | 53.3485 | yes | 9.722E-08 | 793.40094 | 1676.536 |
| RGAEGILAPQPPPF | 20 | Q8WWM7 | Ataxin-2-like protein OS=Homo sapiens ( | ATXN2L | 113.374 | 4 | 53.3485 | yes | 2.02E-07 | 552.29236 | 1279.4783 |
| IAMENDDGRTEEE | 14 | Q8WWM7 | Ataxin-2-like protein OS=Homo sapiens ( | ATXN2L | 113.374 | 2 | 53.3485 | yes | 1.019E-07 | 818.85962 | 622.83319 |
| IAMENDDGRTEEE | 14 | Q8WWM7 | Ataxin-2-like protein OS=Homo sapiens ( | ATXN2L | 113.374 | 2 | 53.3485 | yes | 3.573E-07 | 826.85706 | 622.83319 |
| NGTTYEGIFK | 10 | Q8WWM7 | Ataxin-2-like protein OS=Homo sapiens ( | ATXN2L | 113.374 | 2 | 53.3485 | yes | 9.923E-08 | 565.27985 | 1191.252 |
| GEDKDEGPVAEQV | 14 | Q8WWM7 | Ataxin-2-like protein OS=Homo sapiens ( | ATXN2L | 113.374 | 2 | 53.3485 | yes | 1.083E-07 | 750.86249 | 591.80481 |
| GEDKDEGPVAEQV | 14 | Q8WWM7 | Ataxin-2-like protein OS=Homo sapiens ( | ATXN2L | 113.374 | 2 | 53.3485 | yes | 4.184E-07 | 500.91074 | 591.80481 |
| GEDKDEGPVAEQV | 14 | Q8WWM7 | Ataxin-2-like protein OS=Homo sapiens ( | ATXN2L | 113.374 | 3 | 53.3485 | yes | 1.083E-07 | 750.86249 | 591.80481 |
| GEDKDEGPVAEQV | 14 | Q8WWM7 | Ataxin-2-like protein OS=Homo sapiens ( | ATXN2L | 113.374 | 3 | 53.3485 | yes | 4.184E-07 | 500.91074 | 591.80481 |
| PGLSSLPPR | 9 | Q8WWM7 | Ataxin-2-like protein OS=Homo sapiens ( | ATXN2L | 113.374 | 2 | 53.3485 | yes | 1.859E-07 | 462.26907 | 828.08722 |

| | | | | | | | | | | |
|---|---|---|---|---|---|---|---|---|---|---|
| IAMENDDGR | 9 | Q8WWM7 | Ataxin-2-like protein OS=Homo sapiens ( | ATXN2L | 113.374 | 2 | 53.3485 | yes | 2.067E-07 | 510.7244 | 807.85254 |
| KFGAQFK | 7 | Q8WWM7 | Ataxin-2-like protein OS=Homo sapiens ( | ATXN2L | 113.374 | 2 | 53.3485 | yes | 2.066E-07 | 413.23453 | 572.32654 |
| EDIVDTMVFK | 10 | Q8WWM7 | Ataxin-2-like protein OS=Homo sapiens ( | ATXN2L | 113.374 | 2 | 53.3485 | yes | 9.722E-08 | 598.797 | 517.71057 |
| STSTPTSPGPR | 11 | Q8WWM7 | Ataxin-2-like protein OS=Homo sapiens ( | ATXN2L | 113.374 | 2 | 53.3485 | yes | 2.007E-07 | 544.27252 | 504.68259 |
| GPHHLDNSSPGPG | 17 | Q8WWM7 | Ataxin-2-like protein OS=Homo sapiens ( | ATXN2L | 113.374 | 4 | 53.3485 | yes | 1.122E-07 | 429.45276 | 237.27493 |
| GPHHLDNSSPGPG | 17 | Q8WWM7 | Ataxin-2-like protein OS=Homo sapiens ( | ATXN2L | 113.374 | 4 | 53.3485 | yes | 0.000306 | 572.26788 | 237.27493 |
| GPHHLDNSSPGPG | 17 | Q8WWM7 | Ataxin-2-like protein OS=Homo sapiens ( | ATXN2L | 113.374 | 3 | 53.3485 | yes | 1.122E-07 | 429.45276 | 237.27493 |
| GPHHLDNSSPGPG | 17 | Q8WWM7 | Ataxin-2-like protein OS=Homo sapiens ( | ATXN2L | 113.374 | 3 | 53.3485 | yes | 0.000306 | 572.26788 | 237.27493 |
| ILKEEPK | 7 | Q8WWM7 | Ataxin-2-like protein OS=Homo sapiens ( | ATXN2L | 113.374 | 2 | 53.3485 | yes | 1.318E-07 | 428.76056 | 247.53464 |
| TLSSPSNRPSGETS | 23 | Q8WWM7 | Ataxin-2-like protein OS=Homo sapiens ( | ATXN2L | 113.374 | 3 | 53.3485 | yes | 2.224E-07 | 764.06439 | 180.98685 |
| DNSEEFR | 7 | Q8WWM7 | Ataxin-2-like protein OS=Homo sapiens ( | ATXN2L | 113.374 | 2 | 53.3485 | yes | 9.722E-08 | 448.69086 | 175.81891 |
| GAEGILAPQPPPPQ | 29 | Q8WWM7 | Ataxin-2-like protein OS=Homo sapiens ( | ATXN2L | 113.374 | 4 | 53.3485 | yes | 9.571E-07 | 726.13239 | 27.309914 |
| SFDLLTPHR | 9 | Q8WWN8 | Arf-GAP with Rho-GAP domain, ANK re | ARAP3 | 169.845 | 2 | 41.423294 | yes | 3.517E-07 | 543.29053 | 75.637146 |
| GVIPLTAIEMTR | 12 | Q8WWN8 | Arf-GAP with Rho-GAP domain, ANK re | ARAP3 | 169.845 | 2 | 41.423294 | yes | 2.4E-06 | 650.86829 | 64.671631 |
| SLDQPFLSK | 9 | Q8WWN8 | Arf-GAP with Rho-GAP domain, ANK re | ARAP3 | 169.845 | 2 | 41.423294 | yes | 2.928E-06 | 517.77948 | 28.158869 |
| QPVDLLEYPDYR | 12 | Q8WWQ0 | PH-interacting protein OS=Homo sapiens | PHIP | 206.69 | 2 | 47.0586 | yes | 2.027E-06 | 754.375 | 81.381859 |
| SILLDMATR | 9 | Q8WWQ0 | PH-interacting protein OS=Homo sapiens | PHIP | 206.69 | 2 | 47.0586 | no | 6.356E-07 | 510.28134 | 67.075562 |
| TFNEPGSPIVK | 11 | Q8WWQ0 | PH-interacting protein OS=Homo sapiens | PHIP | 206.69 | 2 | 47.0586 | yes | 1.411E-07 | 594.81659 | 66.728867 |
| DIIDTPMDFATVR | 13 | Q8WWQ0 | PH-interacting protein OS=Homo sapiens | PHIP | 206.69 | 2 | 47.0586 | yes | 8.561E-07 | 747.3689 | 44.141201 |
| LDADLLVPASVK | 12 | Q8WWQ0 | PH-interacting protein OS=Homo sapiens | PHIP | 206.69 | 2 | 47.0586 | yes | 5.291E-07 | 620.86102 | 43.812843 |
| VVVPELSAGVASR | 13 | Q8WWQ0 | PH-interacting protein OS=Homo sapiens | PHIP | 206.69 | 2 | 47.0586 | yes | 1.525E-05 | 642.36951 | 35.504982 |
| VLFSAGHDGNVIV | 18 | Q8WWQ0 | PH-interacting protein OS=Homo sapiens | PHIP | 206.69 | 3 | 47.0586 | yes | 3.57E-07 | 657.01453 | 32.064613 |
| ILKPQLK | 7 | Q8WWQ0 | PH-interacting protein OS=Homo sapiens | PHIP | 206.69 | 2 | 47.0586 | yes | 5.336E-07 | 420.28928 | 9.5632324 |
| ASEVMGPVEAAPI | 15 | Q8WWY3 | U4/U6 small nuclear ribonucleoprotein Pı | PRPF31 | 55.457 | 2 | 53.103413 | yes | 1.223E-07 | 803.38251 | 790.83923 |
| IYEYVESR | 8 | Q8WWY3 | U4/U6 small nuclear ribonucleoprotein Pı | PRPF31 | 55.457 | 2 | 53.103413 | yes | 1.596E-07 | 529.76129 | 612.34753 |
| IMGVAGGLTNLSK | 13 | Q8WWY3 | U4/U6 small nuclear ribonucleoprotein Pı | PRPF31 | 55.457 | 2 | 53.103413 | yes | 2.753E-07 | 630.85266 | 407.96942 |
| LGLTEIR | 7 | Q8WWY3 | U4/U6 small nuclear ribonucleoprotein Pı | PRPF31 | 55.457 | 2 | 53.103413 | yes | 3.103E-07 | 401.24509 | 375.30737 |
| WQEPPPVK | 8 | Q8WWY3 | U4/U6 small nuclear ribonucleoprotein Pı | PRPF31 | 55.457 | 2 | 53.103413 | yes | 2.403E-07 | 490.76364 | 251.79137 |
| QVKPLPAPLDGQR | 13 | Q8WWY3 | U4/U6 small nuclear ribonucleoprotein Pı | PRPF31 | 55.457 | 2 | 53.103413 | yes | 1.024E-07 | 709.90936 | 124.59556 |
| QVKPLPAPLDGQR | 13 | Q8WWY3 | U4/U6 small nuclear ribonucleoprotein Pı | PRPF31 | 55.457 | 2 | 53.103413 | yes | 1.013E-07 | 473.60867 | 124.59556 |
| QVKPLPAPLDGQR | 13 | Q8WWY3 | U4/U6 small nuclear ribonucleoprotein Pı | PRPF31 | 55.457 | 3 | 53.103413 | yes | 1.024E-07 | 709.90936 | 124.59556 |
| QVKPLPAPLDGQR | 13 | Q8WWY3 | U4/U6 small nuclear ribonucleoprotein Pı | PRPF31 | 55.457 | 3 | 53.103413 | yes | 1.013E-07 | 473.60867 | 124.59556 |
| YFSSMAEFLK | 10 | Q8WWY3 | U4/U6 small nuclear ribonucleoprotein Pı | PRPF31 | 55.457 | 2 | 53.103413 | yes | 9.926E-08 | 611.79425 | 230.3672 |
| VDSFHESTEGK | 11 | Q8WWY3 | U4/U6 small nuclear ribonucleoprotein Pı | PRPF31 | 55.457 | 2 | 53.103413 | yes | 9.804E-08 | 618.28058 | 197.3349 |
| LEEACDMALELNA | 15 | Q8WWY3 | U4/U6 small nuclear ribonucleoprotein Pı | PRPF31 | 55.457 | 2 | 53.103413 | yes | 9.722E-08 | 847.39221 | 157.58194 |
| MPACNIMLLGAQI | 13 | Q8WWY3 | U4/U6 small nuclear ribonucleoprotein Pı | PRPF31 | 55.457 | 2 | 53.103413 | yes | 1.422E-07 | 737.87 | 156.88448 |
| ELQGFLDGVK | 10 | Q8WX92 | Negative elongation factor B OS=Homo s | NELFB | 65.698 | 2 | 53.73226 | yes | 9.722E-08 | 553.29803 | 243.8576 |
| MEVELITR | 8 | Q8WX92 | Negative elongation factor B OS=Homo s | NELFB | 65.698 | 2 | 53.73226 | yes | 3.942E-07 | 495.76807 | 212.23328 |
| APVSYPNTLPESFT | 15 | Q8WX92 | Negative elongation factor B OS=Homo s | NELFB | 65.698 | 2 | 53.73226 | yes | 9.722E-08 | 825.9223 | 156.50714 |

| | | | | | | | | | | | |
|---|---|---|---|---|---|---|---|---|---|---|---|
| ALEPTGQSGEAVK | 13 | Q8WX92 | Negative elongation factor B OS=Homo s | NELFB | 65.698 | 2 | 53.73226 | yes | 1.193E-07 | 643.83295 | 148.04161 |
| KLEDLLEK | 8 | Q8WX92 | Negative elongation factor B OS=Homo s | NELFB | 65.698 | 2 | 53.73226 | yes | 4.992E-07 | 494.28967 | 120.43403 |
| FTWCLDACIR | 10 | Q8WX92 | Negative elongation factor B OS=Homo s | NELFB | 65.698 | 2 | 53.73226 | yes | 3.101E-07 | 671.30756 | 117.64853 |
| MPSLQPVVMCVM | 13 | Q8WX92 | Negative elongation factor B OS=Homo s | NELFB | 65.698 | 2 | 53.73226 | yes | 2.151E-07 | 760.37878 | 106.786 |
| LLIHLHPR | 8 | Q8WX92 | Negative elongation factor B OS=Homo s | NELFB | 65.698 | 2 | 53.73226 | yes | 1.164E-07 | 499.81653 | 64.520119 |
| ETLTNCTEPLK | 11 | Q8WX92 | Negative elongation factor B OS=Homo s | NELFB | 65.698 | 2 | 53.73226 | yes | 1.214E-07 | 653.32141 | 57.400654 |
| LEFHQSVFDELR | 12 | Q8WX92 | Negative elongation factor B OS=Homo s | NELFB | 65.698 | 2 | 53.73226 | yes | 9.722E-08 | 760.38062 | 42.559071 |
| HLQELVGQETLPR | 13 | Q8WX92 | Negative elongation factor B OS=Homo s | NELFB | 65.698 | 2 | 53.73226 | yes | 9.722E-08 | 760.41498 | 37.373745 |
| LEQLDHR | 7 | Q8WX92 | Negative elongation factor B OS=Homo s | NELFB | 65.698 | 2 | 53.73226 | yes | 1.4E-06 | 455.74069 | 34.466717 |
| YAALSDQGLDIK | 12 | Q8WX93 | Palladin OS=Homo sapiens OX=9606 GN | PALLD | 150.564 | 2 | 55.866962 | yes | 1.171E-07 | 647.33789 | 227.49368 |
| NEAGIVSCTAR | 11 | Q8WX93 | Palladin OS=Homo sapiens OX=9606 GN | PALLD | 150.564 | 2 | 55.866962 | no | 9.722E-08 | 589.28516 | 204.43613 |
| LMVQAVNQR | 9 | Q8WX93 | Palladin OS=Homo sapiens OX=9606 GN | PALLD | 150.564 | 2 | 55.866962 | yes | 3.241E-07 | 529.79242 | 188.63591 |
| SDHYTIQR | 8 | Q8WX93 | Palladin OS=Homo sapiens OX=9606 GN | PALLD | 150.564 | 2 | 55.866962 | yes | 2.419E-07 | 510.24887 | 177.79915 |
| EDAGWYTVSAK | 11 | Q8WX93 | Palladin OS=Homo sapiens OX=9606 GN | PALLD | 150.564 | 2 | 55.866962 | yes | 9.733E-08 | 613.78802 | 168.6291 |
| GAPPLQVQWFR | 11 | Q8WX93 | Palladin OS=Homo sapiens OX=9606 GN | PALLD | 150.564 | 2 | 55.866962 | yes | 1.611E-07 | 649.85382 | 159.83473 |
| DAGIYTCIATNR | 12 | Q8WX93 | Palladin OS=Homo sapiens OX=9606 GN | PALLD | 150.564 | 2 | 55.866962 | yes | 9.723E-08 | 677.82465 | 147.83493 |
| LISEIEYR | 8 | Q8WX93 | Palladin OS=Homo sapiens OX=9606 GN | PALLD | 150.564 | 2 | 55.866962 | yes | 2.385E-07 | 511.77948 | 146.22865 |
| IFEGMPVTFTCR | 12 | Q8WX93 | Palladin OS=Homo sapiens OX=9606 GN | PALLD | 150.564 | 2 | 55.866962 | yes | 9.722E-08 | 729.34943 | 138.74571 |
| LQQLQNQIR | 9 | Q8WX93 | Palladin OS=Homo sapiens OX=9606 GN | PALLD | 150.564 | 2 | 55.866962 | yes | 4.479E-07 | 570.82782 | 134.83876 |
| SAPAMQSSGSFNY | 15 | Q8WX93 | Palladin OS=Homo sapiens OX=9606 GN | PALLD | 150.564 | 2 | 55.866962 | yes | 2.415E-07 | 787.35681 | 120.78324 |
| ENESLTHSTDR | 11 | Q8WX93 | Palladin OS=Homo sapiens OX=9606 GN | PALLD | 150.564 | 2 | 55.866962 | yes | 9.722E-08 | 644.79181 | 83.523682 |
| IASDEEIQGTK | 11 | Q8WX93 | Palladin OS=Homo sapiens OX=9606 GN | PALLD | 150.564 | 2 | 55.866962 | yes | 3.176E-07 | 595.79858 | 78.67878 |
| LQNTGVADGYPVI | 13 | Q8WX93 | Palladin OS=Homo sapiens OX=9606 GN | PALLD | 150.564 | 2 | 55.866962 | yes | 9.722E-08 | 695.35968 | 78.427063 |
| SLPTPAVLLSPTK | 13 | Q8WX93 | Palladin OS=Homo sapiens OX=9606 GN | PALLD | 150.564 | 2 | 55.866962 | yes | 2.305E-07 | 662.39777 | 62.620579 |
| EAHKPPVFIEK | 11 | Q8WX93 | Palladin OS=Homo sapiens OX=9606 GN | PALLD | 150.564 | 2 | 55.866962 | yes | 1.07E-07 | 647.86133 | 60.11694 |
| QGSEIQDSPDFR | 12 | Q8WX93 | Palladin OS=Homo sapiens OX=9606 GN | PALLD | 150.564 | 2 | 55.866962 | yes | 2.267E-07 | 689.81531 | 36.53294 |
| AGQNSFSLELVVA | 15 | Q8WX93 | Palladin OS=Homo sapiens OX=9606 GN | PALLD | 150.564 | 2 | 55.866962 | yes | 6.386E-07 | 767.41718 | 31.800245 |
| EVDDKDAPR | 9 | Q8WXA9 | Splicing regulatory glutamine/lysine-rich | SREK1 | 59.381 | 2 | 52.843063 | yes | 9.722E-08 | 522.75146 | 147.09404 |
| FAFVEFADQNSVP | 14 | Q8WXA9 | Splicing regulatory glutamine/lysine-rich | SREK1 | 59.381 | 2 | 52.843063 | yes | 2.573E-07 | 813.89917 | 84.193741 |
| INHSNNAIVKPPEM | 21 | Q8WXA9 | Splicing regulatory glutamine/lysine-rich | SREK1 | 59.381 | 3 | 52.843063 | yes | 3.581E-06 | 744.39154 | 34.854118 |
| ALAFNGVMFGDR | 12 | Q8WXA9 | Splicing regulatory glutamine/lysine-rich | SREK1 | 59.381 | 2 | 52.843063 | yes | 3.791E-07 | 649.32153 | 33.133755 |
| VQDLIEGHLTASQ | 13 | Q8WXD5 | Gem-associated protein 6 OS=Homo sapi | GEMIN6 | 18.825 | 2 | 52.51915 | yes | 9.722E-08 | 705.86481 | 1463.364 |
| LMHLFTSGDCK | 11 | Q8WXD5 | Gem-associated protein 6 OS=Homo sapi | GEMIN6 | 18.825 | 2 | 52.51915 | yes | 1.101E-07 | 654.80737 | 1219.2694 |
| AYSPEDLEER | 10 | Q8WXD5 | Gem-associated protein 6 OS=Homo sapi | GEMIN6 | 18.825 | 2 | 52.51915 | yes | 1.01E-07 | 604.77515 | 1092.9137 |
| NHIPITEQGDAPR | 13 | Q8WXD5 | Gem-associated protein 6 OS=Homo sapi | GEMIN6 | 18.825 | 2 | 52.51915 | yes | 4.618E-07 | 724.36804 | 453.20337 |
| NHIPITEQGDAPR | 13 | Q8WXD5 | Gem-associated protein 6 OS=Homo sapi | GEMIN6 | 18.825 | 2 | 52.51915 | yes | 5.089E-07 | 483.2478 | 453.20337 |
| NHIPITEQGDAPR | 13 | Q8WXD5 | Gem-associated protein 6 OS=Homo sapi | GEMIN6 | 18.825 | 3 | 52.51915 | yes | 4.618E-07 | 724.36804 | 453.20337 |
| NHIPITEQGDAPR | 13 | Q8WXD5 | Gem-associated protein 6 OS=Homo sapi | GEMIN6 | 18.825 | 3 | 52.51915 | yes | 5.089E-07 | 483.2478 | 453.20337 |
| GPLEWQDYIYK | 11 | Q8WXD5 | Gem-associated protein 6 OS=Homo sapi | GEMIN6 | 18.825 | 2 | 52.51915 | yes | 9.722E-08 | 706.34827 | 278.64798 |

| | | | | | | | | | | | |
|---|---|---|---|---|---|---|---|---|---|---|---|
| KGPLEWQDYIYK | 12 | Q8WXD5 | Gem-associated protein 6 OS=Homo sapi | GEMIN6 | 18.825 | 2 | 52.51915 | yes | 3.129E-07 | 770.39575 | 87.073135 |
| VILFNPLDACR | 11 | Q8WXE9 | Stonin-2 OS=Homo sapiens OX=9606 Gl | STON2 | 101.166 | 2 | 28.50403 | yes | 2.733E-06 | 659.35284 | 52.319305 |
| LGLNDILVK | 9 | Q8WXE9 | Stonin-2 OS=Homo sapiens OX=9606 Gl | STON2 | 101.166 | 2 | 28.50403 | yes | 3.128E-06 | 492.80804 | 41.998386 |
| YGEPSEVFINR | 11 | Q8WXF1 | Paraspeckle component 1 OS=Homo sapi | PSPC1 | 58.744 | 2 | 57.122116 | yes | 9.722E-08 | 655.82239 | 2465.8193 |
| SFLKPGEK | 8 | Q8WXF1 | Paraspeckle component 1 OS=Homo sapi | PSPC1 | 58.744 | 2 | 57.122116 | yes | 1.107E-07 | 453.25818 | 1291.4668 |
| FAQPGTFEFEYASI | 14 | Q8WXF1 | Paraspeckle component 1 OS=Homo sapi | PSPC1 | 58.744 | 2 | 57.122116 | yes | 1.098E-07 | 825.38336 | 1193.343 |
| CGDGAFLLTTTPR | 13 | Q8WXF1 | Paraspeckle component 1 OS=Homo sapi | PSPC1 | 58.744 | 2 | 57.122116 | yes | 1.132E-07 | 704.84808 | 1184.1766 |
| EKLEAEMEAAR | 11 | Q8WXF1 | Paraspeckle component 1 OS=Homo sapi | PSPC1 | 58.744 | 2 | 57.122116 | yes | 9.722E-08 | 638.81372 | 889.53528 |
| FATHGAALTVK | 11 | Q8WXF1 | Paraspeckle component 1 OS=Homo sapi | PSPC1 | 58.744 | 2 | 57.122116 | yes | 1.075E-07 | 558.31403 | 680.30237 |
| AELDGTILK | 9 | Q8WXF1 | Paraspeckle component 1 OS=Homo sapi | PSPC1 | 58.744 | 2 | 57.122116 | yes | 9.838E-08 | 480.27402 | 551.85815 |
| LEAEMEAAR | 9 | Q8WXF1 | Paraspeckle component 1 OS=Homo sapi | PSPC1 | 58.744 | 2 | 57.122116 | yes | 2.671E-07 | 510.24493 | 304.35721 |
| AVVVVDDR | 8 | Q8WXF1 | Paraspeckle component 1 OS=Homo sapi | PSPC1 | 58.744 | 2 | 57.122116 | yes | 3.284E-07 | 436.74545 | 243.57646 |
| HEHQLMLMR | 9 | Q8WXF1 | Paraspeckle component 1 OS=Homo sapi | PSPC1 | 58.744 | 2 | 57.122116 | yes | 9.722E-08 | 597.79718 | 168.44997 |
| GSQGGNFEGPNKF | 13 | Q8WXF1 | Paraspeckle component 1 OS=Homo sapi | PSPC1 | 58.744 | 2 | 57.122116 | yes | 9.722E-08 | 674.32367 | 126.66981 |
| REEEMIR | 7 | Q8WXF1 | Paraspeckle component 1 OS=Homo sapi | PSPC1 | 58.744 | 2 | 57.122116 | yes | 1.125E-07 | 481.73984 | 108.71135 |
| FPQGPPSQMGSPM | 16 | Q8WXF1 | Paraspeckle component 1 OS=Homo sapi | PSPC1 | 58.744 | 2 | 57.122116 | yes | 8.905E-07 | 830.88214 | 102.68291 |
| AHSLKPSIK | 9 | Q8WXG6 | MAP kinase-activating death domain prot | MADD | 183.304 | 2 | 39.49096 | yes | 4.554E-07 | 490.798 | 28.409859 |
| VIAPNPAQLQGQR | 13 | Q8WXI9 | Transcriptional repressor p66-beta OS=H | GATAD2B | 65.261 | 2 | 50.4107 | yes | 9.809E-08 | 696.3913 | 243.84666 |
| ENINDEPVDMSAR | 13 | Q8WXI9 | Transcriptional repressor p66-beta OS=H | GATAD2B | 65.261 | 2 | 50.4107 | yes | 3.152E-07 | 745.33301 | 127.96509 |
| VEPFVCAQCR | 10 | Q8WXI9 | Transcriptional repressor p66-beta OS=H | GATAD2B | 65.261 | 2 | 50.4107 | yes | 1.022E-06 | 633.29193 | 102.21914 |
| QAPQPQSSLQR | 11 | Q8WXI9 | Transcriptional repressor p66-beta OS=H | GATAD2B | 65.261 | 2 | 50.4107 | yes | 2.695E-07 | 620.32562 | 76.221207 |
| GPPKPGLVR | 9 | Q8WXI9 | Transcriptional repressor p66-beta OS=H | GATAD2B | 65.261 | 2 | 50.4107 | yes | 2.622E-07 | 460.78745 | 50.054474 |
| ILCEQCMTSNQK | 12 | Q8WXI9 | Transcriptional repressor p66-beta OS=H | GATAD2B | 65.261 | 2 | 50.4107 | yes | 2.976E-06 | 756.33618 | 16.383015 |
| EMDNFLAQMEAK | 12 | Q8WXX5 | DnaJ homolog subfamily C member 9 OS | DNAJC9 | 29.91 | 2 | 51.138298 | yes | 1.189E-07 | 713.82068 | 2352.1851 |
| VSLQVHPDR | 9 | Q8WXX5 | DnaJ homolog subfamily C member 9 OS | DNAJC9 | 29.91 | 2 | 51.138298 | yes | 2.041E-07 | 525.78815 | 1581.3546 |
| VYSVLSDR | 8 | Q8WXX5 | DnaJ homolog subfamily C member 9 OS | DNAJC9 | 29.91 | 2 | 51.138298 | yes | 1.423E-07 | 469.75073 | 1045.4767 |
| RFQILGK | 7 | Q8WXX5 | DnaJ homolog subfamily C member 9 OS | DNAJC9 | 29.91 | 2 | 51.138298 | yes | 2.601E-07 | 431.26889 | 569.51105 |
| ISLEDIQAFEK | 11 | Q8WXX5 | DnaJ homolog subfamily C member 9 OS | DNAJC9 | 29.91 | 2 | 51.138298 | yes | 1.118E-07 | 646.84027 | 537.80548 |
| GSEEELADIK | 10 | Q8WXX5 | DnaJ homolog subfamily C member 9 OS | DNAJC9 | 29.91 | 2 | 51.138298 | yes | 2.046E-07 | 545.76678 | 515.87396 |
| KISLEDIQAFEK | 12 | Q8WXX5 | DnaJ homolog subfamily C member 9 OS | DNAJC9 | 29.91 | 2 | 51.138298 | yes | 9.722E-08 | 710.88776 | 180.3035 |
| QAYLDFK | 7 | Q8WXX5 | DnaJ homolog subfamily C member 9 OS | DNAJC9 | 29.91 | 2 | 51.138298 | yes | 2.736E-07 | 442.72925 | 101.4411 |
| KELGLDEGVDSLK | 13 | Q8WXX5 | DnaJ homolog subfamily C member 9 OS | DNAJC9 | 29.91 | 2 | 51.138298 | yes | 1.276E-07 | 701.87482 | 89.52623 |
| IIISDVPSR | 9 | Q8WY36 | HMG box transcription factor BBX OS=I | BBX | 105.131 | 2 | 50.832905 | yes | 3.368E-07 | 500.29529 | 903.46362 |
| TLVSEGMLTSLR | 12 | Q8WY36 | HMG box transcription factor BBX OS=I | BBX | 105.131 | 2 | 50.832905 | yes | 9.892E-08 | 653.85541 | 880.63757 |
| NISGETPEGIK | 11 | Q8WY36 | HMG box transcription factor BBX OS=I | BBX | 105.131 | 2 | 50.832905 | yes | 1.96E-07 | 572.79584 | 650.552 |
| LWAFPSDSSR | 10 | Q8WY36 | HMG box transcription factor BBX OS=I | BBX | 105.131 | 2 | 50.832905 | yes | 1.482E-07 | 583.28546 | 531.52515 |
| STPLTHDGQPK | 11 | Q8WY36 | HMG box transcription factor BBX OS=I | BBX | 105.131 | 2 | 50.832905 | yes | 1.248E-07 | 590.80145 | 340.54581 |
| NFEALQIDDIMAIK | 14 | Q8WY36 | HMG box transcription factor BBX OS=I | BBX | 105.131 | 2 | 50.832905 | yes | 1.225E-07 | 810.9187 | 290.57767 |
| SPTPTVNPR | 9 | Q8WY36 | HMG box transcription factor BBX OS=I | BBX | 105.131 | 2 | 50.832905 | yes | 1.01E-07 | 484.76163 | 271.04895 |

| | | | | | | | | | | |
|---|---|---|---|---|---|---|---|---|---|---|
| EMPQAPVLISCAD | 14 | Q8WY36 | HMG box transcription factor BBX OS=I | BBX | 105.131 | 2 | 50.832905 | yes | 9.903E-08 | 779.86566 | 263.62961 |
| FNSLPQYSPVTFDI | 14 | Q8WY36 | HMG box transcription factor BBX OS=I | BBX | 105.131 | 2 | 50.832905 | yes | 9.722E-08 | 835.91229 | 240.06467 |
| GSILDAKPPK | 10 | Q8WY36 | HMG box transcription factor BBX OS=I | BBX | 105.131 | 2 | 50.832905 | yes | 2.303E-07 | 513.3031 | 220.62682 |
| WCPTTNKPVK | 10 | Q8WY36 | HMG box transcription factor BBX OS=I | BBX | 105.131 | 2 | 50.832905 | yes | 5.436E-07 | 615.8186 | 190.87672 |
| DHMCHPHGIMIIEI | 23 | Q8WY36 | HMG box transcription factor BBX OS=I | BBX | 105.131 | 4 | 50.832905 | yes | 2.793E-07 | 664.07257 | 188.54193 |
| EDCLLGSAK | 9 | Q8WY36 | HMG box transcription factor BBX OS=I | BBX | 105.131 | 2 | 50.832905 | yes | 2.067E-07 | 496.7395 | 113.18774 |
| SPLFQFAEISSSTSI | 21 | Q8WY36 | HMG box transcription factor BBX OS=I | BBX | 105.131 | 3 | 50.832905 | yes | 3.713E-07 | 743.02197 | 75.596741 |
| KDHMCHPHGIMII | 24 | Q8WY36 | HMG box transcription factor BBX OS=I | BBX | 105.131 | 4 | 50.832905 | yes | 1.576E-07 | 696.09631 | 39.772381 |
| VSVPYPSSLLSPSR | 14 | Q8WY91 | Peroxynitrite isomerase THAP4 OS=Hon | THAP4 | 62.89 | 2 | 52.67589 | yes | 3.284E-07 | 744.90649 | 110.74633 |
| SQVCCLR | 7 | Q8WY91 | Peroxynitrite isomerase THAP4 OS=Hon | THAP4 | 62.89 | 2 | 52.67589 | yes | 1.624E-07 | 461.7153 | 102.80887 |
| DVEPSCSGSSLGPI | 15 | Q8WY91 | Peroxynitrite isomerase THAP4 OS=Hon | THAP4 | 62.89 | 2 | 52.67589 | yes | 9.722E-08 | 767.83813 | 94.067581 |
| QAALQGEATPR | 11 | Q8WY91 | Peroxynitrite isomerase THAP4 OS=Hon | THAP4 | 62.89 | 2 | 52.67589 | yes | 9.722E-08 | 571.30164 | 87.854828 |
| YSFLCSEHFTK | 11 | Q8WY91 | Peroxynitrite isomerase THAP4 OS=Hon | THAP4 | 62.89 | 2 | 52.67589 | yes | 2.964E-07 | 709.8241 | 82.440292 |
| TPGDGLATMVAG | 16 | Q8WY91 | Peroxynitrite isomerase THAP4 OS=Hon | THAP4 | 62.89 | 2 | 52.67589 | yes | 1.128E-07 | 745.36938 | 42.180958 |
| LLKPTAVPSIFHLT | 16 | Q8WY91 | Peroxynitrite isomerase THAP4 OS=Hon | THAP4 | 62.89 | 3 | 52.67589 | yes | 3.084E-06 | 598.68896 | 26.658098 |
| RLDADLAR | 8 | Q8WYH8 | Inhibitor of growth protein 5 OS=Homo s | ING5 | 27.751 | 2 | 48.473034 | yes | 1.398E-07 | 465.26178 | 151.80153 |
| VQLAMQTYEMVD | 13 | Q8WYH8 | Inhibitor of growth protein 5 OS=Homo s | ING5 | 27.751 | 2 | 48.473034 | no | 1.143E-07 | 778.37836 | 84.048653 |
| AEIDILAAEYISTVI | 15 | Q8WYH8 | Inhibitor of growth protein 5 OS=Homo s | ING5 | 27.751 | 2 | 48.473034 | yes | 1.21E-07 | 818.44562 | 80.842255 |
| KAEIDILAAEYIST | 16 | Q8WYH8 | Inhibitor of growth protein 5 OS=Homo s | ING5 | 27.751 | 3 | 48.473034 | yes | 7.332E-07 | 588.66449 | 27.501911 |
| SGPSLNELIPDGYN | 15 | Q8WYP5 | Protein ELYS OS=Homo sapiens OX=96 | AHCTF1 | 252.499 | 2 | 55.508766 | yes | 1.348E-07 | 816.40485 | 158.15694 |
| CLVAGLLSPR | 10 | Q8WYP5 | Protein ELYS OS=Homo sapiens OX=96 | AHCTF1 | 252.499 | 2 | 55.508766 | yes | 9.722E-08 | 543.31042 | 118.39233 |
| VDTLPYVPEPIK | 12 | Q8WYP5 | Protein ELYS OS=Homo sapiens OX=96 | AHCTF1 | 252.499 | 2 | 55.508766 | yes | 1.182E-07 | 685.88196 | 117.08633 |
| FSGVNEQPPVVLA | 15 | Q8WYP5 | Protein ELYS OS=Homo sapiens OX=96 | AHCTF1 | 252.499 | 2 | 55.508766 | yes | 1.34E-07 | 792.44324 | 116.271 |
| EISEASENIYSDVR | 14 | Q8WYP5 | Protein ELYS OS=Homo sapiens OX=96 | AHCTF1 | 252.499 | 2 | 55.508766 | yes | 9.722E-08 | 806.37848 | 111.62293 |
| WITEEQPNSATNLI | 14 | Q8WYP5 | Protein ELYS OS=Homo sapiens OX=96 | AHCTF1 | 252.499 | 2 | 55.508766 | yes | 1.21E-07 | 829.91028 | 110.21835 |
| LLDLVVQPVPR | 11 | Q8WYP5 | Protein ELYS OS=Homo sapiens OX=96 | AHCTF1 | 252.499 | 2 | 55.508766 | yes | 9.722E-08 | 624.88733 | 95.767311 |
| KINPSEDVGSK | 11 | Q8WYP5 | Protein ELYS OS=Homo sapiens OX=96 | AHCTF1 | 252.499 | 2 | 55.508766 | yes | 4.647E-07 | 587.30914 | 86.849251 |
| SENVDVGKPALGI | 13 | Q8WYP5 | Protein ELYS OS=Homo sapiens OX=96 | AHCTF1 | 252.499 | 2 | 55.508766 | yes | 1.114E-07 | 657.35663 | 74.41687 |
| LLATASFTK | 9 | Q8WYP5 | Protein ELYS OS=Homo sapiens OX=96 | AHCTF1 | 252.499 | 2 | 55.508766 | yes | 1.908E-07 | 476.27911 | 66.890488 |
| QVVTGTVLTR | 10 | Q8WYP5 | Protein ELYS OS=Homo sapiens OX=96 | AHCTF1 | 252.499 | 2 | 55.508766 | yes | 1.004E-07 | 537.31934 | 66.744995 |
| SSSTALTTNVTEQ | 16 | Q8WYP5 | Protein ELYS OS=Homo sapiens OX=96 | AHCTF1 | 252.499 | 2 | 55.508766 | yes | 2.156E-07 | 848.91541 | 62.628651 |
| TTEVTGTGLGR | 11 | Q8WYP5 | Protein ELYS OS=Homo sapiens OX=96 | AHCTF1 | 252.499 | 2 | 55.508766 | yes | 1.827E-07 | 546.28821 | 59.03455 |
| VAIAENLLDVIK | 12 | Q8WYP5 | Protein ELYS OS=Homo sapiens OX=96 | AHCTF1 | 252.499 | 2 | 55.508766 | yes | 4.682E-07 | 649.38995 | 58.880241 |
| YTLDLTGGMFPLR | 13 | Q8WYP5 | Protein ELYS OS=Homo sapiens OX=96 | AHCTF1 | 252.499 | 2 | 55.508766 | yes | 6.807E-07 | 742.38434 | 50.96524 |
| NSILDQYGK | 9 | Q8WYP5 | Protein ELYS OS=Homo sapiens OX=96 | AHCTF1 | 252.499 | 2 | 55.508766 | yes | 3.751E-06 | 519.26672 | 44.535706 |
| VTAIEPIINHGGAS | 25 | Q8WYP5 | Protein ELYS OS=Homo sapiens OX=96 | AHCTF1 | 252.499 | 4 | 55.508766 | yes | 1.618E-06 | 652.3504 | 43.664333 |
| KIEVPAQLK | 9 | Q8WYP5 | Protein ELYS OS=Homo sapiens OX=96 | AHCTF1 | 252.499 | 2 | 55.508766 | yes | 8.883E-07 | 513.32129 | 36.051662 |
| SAQQEASADVATI | 14 | Q8WYP5 | Protein ELYS OS=Homo sapiens OX=96 | AHCTF1 | 252.499 | 2 | 55.508766 | yes | 2.287E-07 | 701.84406 | 28.170773 |
| DVFASEVTPSDLQ | 14 | Q8WYP5 | Protein ELYS OS=Homo sapiens OX=96 | AHCTF1 | 252.499 | 2 | 55.508766 | yes | 5.336E-06 | 768.38306 | 23.079109 |

| | | | | | | | | | | | |
|---|---|---|---|---|---|---|---|---|---|---|---|
| ASELHLLETPLVVI | 14 | Q8WYP5 | Protein ELYS OS=Homo sapiens OX=96 | AHCTF1 | 252.499 | 2 | 55.508766 | yes | 9.722E-08 | 774.95343 | 14.380718 |
| PSQCSEFIQQSSMK | 14 | Q8WYP5 | Protein ELYS OS=Homo sapiens OX=96 | AHCTF1 | 252.499 | 2 | 55.508766 | yes | 3.282E-06 | 828.87146 | 12.934923 |
| PYFLGTGSIR | 10 | Q8WYQ5 | Microprocessor complex subunit DGCR8 | DGCR8 | 86.046 | 2 | 52.636726 | yes | 9.722E-08 | 555.80072 | 842.06061 |
| LITLSVQDAPTK | 12 | Q8WYQ5 | Microprocessor complex subunit DGCR8 | DGCR8 | 86.046 | 2 | 52.636726 | yes | 1.108E-07 | 643.37177 | 458.70932 |
| SEYVMACGK | 9 | Q8WYQ5 | Microprocessor complex subunit DGCR8 | DGCR8 | 86.046 | 2 | 52.636726 | yes | 9.881E-08 | 522.72809 | 320.28006 |
| SEVCILHEYMQR | 12 | Q8WYQ5 | Microprocessor complex subunit DGCR8 | DGCR8 | 86.046 | 2 | 52.636726 | yes | 1.624E-07 | 782.86597 | 309.91016 |
| ISVSFTESCR | 10 | Q8WYQ5 | Microprocessor complex subunit DGCR8 | DGCR8 | 86.046 | 2 | 52.636726 | yes | 9.722E-08 | 593.28204 | 287.20633 |
| ILQLLHPHVK | 10 | Q8WYQ5 | Microprocessor complex subunit DGCR8 | DGCR8 | 86.046 | 2 | 52.636726 | yes | 9.722E-08 | 599.37695 | 250.85899 |
| GRPPTEPLPDGWI | 27 | Q8WYQ5 | Microprocessor complex subunit DGCR8 | DGCR8 | 86.046 | 4 | 52.636726 | yes | 1.628E-07 | 769.14209 | 249.57597 |
| KFSPDLK | 7 | Q8WYQ5 | Microprocessor complex subunit DGCR8 | DGCR8 | 86.046 | 2 | 52.636726 | yes | 2.713E-07 | 417.73962 | 222.88361 |
| SVIELQQYAK | 10 | Q8WYQ5 | Microprocessor complex subunit DGCR8 | DGCR8 | 86.046 | 2 | 52.636726 | yes | 1.066E-07 | 589.8244 | 219.97742 |
| FDFEQVTVK | 9 | Q8WYQ5 | Microprocessor complex subunit DGCR8 | DGCR8 | 86.046 | 2 | 52.636726 | yes | 2.617E-07 | 556.78479 | 215.09302 |
| RFDFEQVTVK | 10 | Q8WYQ5 | Microprocessor complex subunit DGCR8 | DGCR8 | 86.046 | 2 | 52.636726 | yes | 1.054E-07 | 634.83533 | 203.32373 |
| DESVDLEEFR | 10 | Q8WYQ5 | Microprocessor complex subunit DGCR8 | DGCR8 | 86.046 | 2 | 52.636726 | yes | 1.476E-07 | 619.7804 | 202.67534 |
| NHGMGDTSIK | 10 | Q8WYQ5 | Microprocessor complex subunit DGCR8 | DGCR8 | 86.046 | 2 | 52.636726 | yes | 9.723E-08 | 530.24805 | 185.42395 |
| LLIDPNCSGHSPR | 13 | Q8WYQ5 | Microprocessor complex subunit DGCR8 | DGCR8 | 86.046 | 3 | 52.636726 | yes | 2.676E-07 | 489.24539 | 178.33821 |
| HDPPLSSIPCLHYK | 14 | Q8WYQ5 | Microprocessor complex subunit DGCR8 | DGCR8 | 86.046 | 3 | 52.636726 | yes | 3.209E-06 | 555.28021 | 149.08685 |
| KVLYTGAER | 9 | Q8WYQ5 | Microprocessor complex subunit DGCR8 | DGCR8 | 86.046 | 2 | 52.636726 | yes | 1.1E-07 | 518.79291 | 139.97006 |
| VYELTSK | 7 | Q8WYQ5 | Microprocessor complex subunit DGCR8 | DGCR8 | 86.046 | 2 | 52.636726 | yes | 2.826E-07 | 420.2291 | 92.602669 |
| QAESERPILPANQK | 14 | Q8WYQ5 | Microprocessor complex subunit DGCR8 | DGCR8 | 86.046 | 3 | 52.636726 | yes | 3.172E-07 | 527.61786 | 45.687469 |
| QAESERPILPANQK | 14 | Q8WYQ5 | Microprocessor complex subunit DGCR8 | DGCR8 | 86.046 | 3 | 52.636726 | yes | 6.119E-07 | 790.92316 | 45.687469 |
| QAESERPILPANQK | 14 | Q8WYQ5 | Microprocessor complex subunit DGCR8 | DGCR8 | 86.046 | 2 | 52.636726 | yes | 3.172E-07 | 527.61786 | 45.687469 |
| QAESERPILPANQK | 14 | Q8WYQ5 | Microprocessor complex subunit DGCR8 | DGCR8 | 86.046 | 2 | 52.636726 | yes | 6.119E-07 | 790.92316 | 45.687469 |
| NKPNLHILSK | 10 | Q8WYQ5 | Microprocessor complex subunit DGCR8 | DGCR8 | 86.046 | 2 | 52.636726 | yes | 3.858E-07 | 582.34839 | 53.026703 |
| AGLLSPYQILHECI | 15 | Q8WYQ5 | Microprocessor complex subunit DGCR8 | DGCR8 | 86.046 | 3 | 52.636726 | yes | 4.33E-05 | 581.31506 | 9.1585503 |
| SPALSSWGDPVLL | 14 | Q8WZA1 | Protein O-linked-mannose beta-1,2-N-ace | POMGNT1 | 75.252 | 2 | 49.700832 | yes | 1.993E-06 | 735.40356 | 51.480663 |
| VETMPGLGWVLR | 12 | Q8WZA1 | Protein O-linked-mannose beta-1,2-N-ace | POMGNT1 | 75.252 | 2 | 49.700832 | yes | 1.009E-07 | 679.36847 | 40.872456 |
| CGEMLHIR | 8 | Q92482 | Aquaporin-3 OS=Homo sapiens OX=960 | AQP3 | 31.544 | 2 | 47.985725 | yes | 1.219E-07 | 508.24423 | 22.467327 |
| FLICTDVAAR | 10 | Q92499 | ATP-dependent RNA helicase DDX1 OS | DDX1 | 82.433 | 2 | 53.79254 | yes | 9.819E-08 | 583.30536 | 50161.676 |
| NQALFPACVLK | 11 | Q92499 | ATP-dependent RNA helicase DDX1 OS | DDX1 | 82.433 | 2 | 53.79254 | yes | 1.013E-07 | 630.8421 | 44460.559 |
| GSAFAIGSDGLCC( | 16 | Q92499 | ATP-dependent RNA helicase DDX1 OS | DDX1 | 82.433 | 2 | 53.79254 | yes | 9.722E-08 | 843.37213 | 35157.492 |
| ALIVEPSR | 8 | Q92499 | ATP-dependent RNA helicase DDX1 OS | DDX1 | 82.433 | 2 | 53.79254 | yes | 2.062E-07 | 442.76364 | 28245.656 |
| WQMNPYDR | 8 | Q92499 | ATP-dependent RNA helicase DDX1 OS | DDX1 | 82.433 | 2 | 53.79254 | yes | 1.641E-07 | 555.2453 | 13602.307 |
| WQMNPYDR | 8 | Q92499 | ATP-dependent RNA helicase DDX1 OS | DDX1 | 82.433 | 2 | 53.79254 | yes | 1.064E-07 | 563.24274 | 13602.307 |
| MGLAISLVATEK | 12 | Q92499 | ATP-dependent RNA helicase DDX1 OS | DDX1 | 82.433 | 2 | 53.79254 | yes | 9.722E-08 | 616.84961 | 13380.05 |
| MGLAISLVATEK | 12 | Q92499 | ATP-dependent RNA helicase DDX1 OS | DDX1 | 82.433 | 2 | 53.79254 | yes | 1.133E-07 | 624.84705 | 13380.05 |
| ELAEQTLNNIK | 11 | Q92499 | ATP-dependent RNA helicase DDX1 OS | DDX1 | 82.433 | 2 | 53.79254 | yes | 9.849E-08 | 636.84332 | 25939.516 |
| MDQAIIFCR | 9 | Q92499 | ATP-dependent RNA helicase DDX1 OS | DDX1 | 82.433 | 2 | 53.79254 | yes | 1.238E-07 | 577.27826 | 23808.721 |
| ELLIIGGVAAR | 11 | Q92499 | ATP-dependent RNA helicase DDX1 OS | DDX1 | 82.433 | 2 | 53.79254 | yes | 1.006E-07 | 556.34534 | 23150.012 |

| | | | | | | | | | | | |
|---|---|---|---|---|---|---|---|---|---|---|---|
| FNFGEEEFK | 9 | Q92499 | ATP-dependent RNA helicase DDX1 OS | DDX1 | 82.433 | 2 | 53.79254 | yes | 2.006E-06 | 573.75873 | 21431.307 |
| IMHFPTWVDLK | 11 | Q92499 | ATP-dependent RNA helicase DDX1 OS | DDX1 | 82.433 | 2 | 53.79254 | yes | 6.082E-07 | 693.86554 | 20778.484 |
| VPVDEFDGK | 9 | Q92499 | ATP-dependent RNA helicase DDX1 OS | DDX1 | 82.433 | 2 | 53.79254 | yes | 2.358E-07 | 503.24802 | 20663.617 |
| LDDLVSTGK | 9 | Q92499 | ATP-dependent RNA helicase DDX1 OS | DDX1 | 82.433 | 2 | 53.79254 | yes | 1.534E-07 | 474.25583 | 18039.957 |
| MHNQIPQVTSDGK | 13 | Q92499 | ATP-dependent RNA helicase DDX1 OS | DDX1 | 82.433 | 2 | 53.79254 | yes | 9.722E-08 | 727.85645 | 5928.542 |
| MHNQIPQVTSDGK | 13 | Q92499 | ATP-dependent RNA helicase DDX1 OS | DDX1 | 82.433 | 2 | 53.79254 | yes | 1.061E-06 | 735.85394 | 5928.542 |
| MHNQIPQVTSDGK | 13 | Q92499 | ATP-dependent RNA helicase DDX1 OS | DDX1 | 82.433 | 2 | 53.79254 | yes | 1.971E-07 | 485.57339 | 5928.542 |
| MHNQIPQVTSDGK | 13 | Q92499 | ATP-dependent RNA helicase DDX1 OS | DDX1 | 82.433 | 3 | 53.79254 | yes | 9.722E-08 | 727.85645 | 5928.542 |
| MHNQIPQVTSDGK | 13 | Q92499 | ATP-dependent RNA helicase DDX1 OS | DDX1 | 82.433 | 3 | 53.79254 | yes | 1.061E-06 | 735.85394 | 5928.542 |
| MHNQIPQVTSDGK | 13 | Q92499 | ATP-dependent RNA helicase DDX1 OS | DDX1 | 82.433 | 3 | 53.79254 | yes | 1.971E-07 | 485.57339 | 5928.542 |
| VWYHVCSSR | 9 | Q92499 | ATP-dependent RNA helicase DDX1 OS | DDX1 | 82.433 | 2 | 53.79254 | yes | 2.99E-07 | 597.27966 | 12750.139 |
| KYIDNPK | 7 | Q92499 | ATP-dependent RNA helicase DDX1 OS | DDX1 | 82.433 | 2 | 53.79254 | yes | 1.006E-07 | 439.24252 | 9739.4756 |
| GHVDILAPTVQEL | 18 | Q92499 | ATP-dependent RNA helicase DDX1 OS | DDX1 | 82.433 | 3 | 53.79254 | yes | 9.819E-08 | 635.35443 | 9657.3994 |
| GIDIHGVPYVINVT | 19 | Q92499 | ATP-dependent RNA helicase DDX1 OS | DDX1 | 82.433 | 3 | 53.79254 | yes | 1.138E-07 | 693.70886 | 8011.6182 |
| APDGYIVK | 8 | Q92499 | ATP-dependent RNA helicase DDX1 OS | DDX1 | 82.433 | 2 | 53.79254 | yes | 1.096E-07 | 431.73709 | 5206.1353 |
| GEDSVPDTVHHVV | 19 | Q92499 | ATP-dependent RNA helicase DDX1 OS | DDX1 | 82.433 | 3 | 53.79254 | yes | 1.11E-07 | 675.68481 | 2528.8235 |
| GEDSVPDTVHHVV | 19 | Q92499 | ATP-dependent RNA helicase DDX1 OS | DDX1 | 82.433 | 3 | 53.79254 | yes | 1.542E-07 | 507.01544 | 2528.8235 |
| GEDSVPDTVHHVV | 19 | Q92499 | ATP-dependent RNA helicase DDX1 OS | DDX1 | 82.433 | 4 | 53.79254 | yes | 1.11E-07 | 675.68481 | 2528.8235 |
| GEDSVPDTVHHVV | 19 | Q92499 | ATP-dependent RNA helicase DDX1 OS | DDX1 | 82.433 | 4 | 53.79254 | yes | 1.542E-07 | 507.01544 | 2528.8235 |
| DLGLAFEIPPHMK | 13 | Q92499 | ATP-dependent RNA helicase DDX1 OS | DDX1 | 82.433 | 2 | 53.79254 | yes | 1.875E-06 | 734.38684 | 2865.4756 |
| LQVIVCSATLHSFI | 16 | Q92499 | ATP-dependent RNA helicase DDX1 OS | DDX1 | 82.433 | 3 | 53.79254 | yes | 1.164E-07 | 606.32568 | 2784.6465 |
| PGANSPEMWSEAI | 14 | Q92499 | ATP-dependent RNA helicase DDX1 OS | DDX1 | 82.433 | 2 | 53.79254 | yes | 1.408E-07 | 758.85864 | 1132.0374 |
| HYYEVSCHDQGLC | 14 | Q92499 | ATP-dependent RNA helicase DDX1 OS | DDX1 | 82.433 | 3 | 53.79254 | yes | 1.056E-07 | 608.59113 | 1033.3314 |
| EKVWYHVCSSR | 11 | Q92499 | ATP-dependent RNA helicase DDX1 OS | DDX1 | 82.433 | 3 | 53.79254 | yes | 2.095E-07 | 484.23471 | 495.14642 |
| EKVWYHVCSSR | 11 | Q92499 | ATP-dependent RNA helicase DDX1 OS | DDX1 | 82.433 | 3 | 53.79254 | yes | 9.722E-08 | 725.84845 | 495.14642 |
| EKVWYHVCSSR | 11 | Q92499 | ATP-dependent RNA helicase DDX1 OS | DDX1 | 82.433 | 2 | 53.79254 | yes | 2.095E-07 | 484.23471 | 495.14642 |
| EKVWYHVCSSR | 11 | Q92499 | ATP-dependent RNA helicase DDX1 OS | DDX1 | 82.433 | 2 | 53.79254 | yes | 9.722E-08 | 725.84845 | 495.14642 |
| FNFGEEEFKFPPK | 13 | Q92499 | ATP-dependent RNA helicase DDX1 OS | DDX1 | 82.433 | 2 | 53.79254 | yes | 3.083E-06 | 808.39319 | 340.78986 |
| FNFGEEEFKFPPK | 13 | Q92499 | ATP-dependent RNA helicase DDX1 OS | DDX1 | 82.433 | 2 | 53.79254 | yes | 3.264E-06 | 539.26459 | 340.78986 |
| FNFGEEEFKFPPK | 13 | Q92499 | ATP-dependent RNA helicase DDX1 OS | DDX1 | 82.433 | 3 | 53.79254 | yes | 3.083E-06 | 808.39319 | 340.78986 |
| FNFGEEEFKFPPK | 13 | Q92499 | ATP-dependent RNA helicase DDX1 OS | DDX1 | 82.433 | 3 | 53.79254 | yes | 3.264E-06 | 539.26459 | 340.78986 |
| IDCDNLEQYFIQQC | 20 | Q92499 | ATP-dependent RNA helicase DDX1 OS | DDX1 | 82.433 | 3 | 53.79254 | yes | 1.324E-07 | 775.69873 | 502.11115 |
| FGFGFGGTGK | 10 | Q92499 | ATP-dependent RNA helicase DDX1 OS | DDX1 | 82.433 | 2 | 53.79254 | yes | 2.918E-07 | 487.74017 | 463.59091 |
| RLQVIVCSATLHSI | 17 | Q92499 | ATP-dependent RNA helicase DDX1 OS | DDX1 | 82.433 | 3 | 53.79254 | yes | 1.826E-07 | 658.35944 | 459.23697 |
| MHNQIPQVTSDGK | 14 | Q92499 | ATP-dependent RNA helicase DDX1 OS | DDX1 | 82.433 | 2 | 53.79254 | yes | 1.549E-07 | 805.90704 | 140.97025 |
| SQHSGNAQVTQTK | 13 | Q92499 | ATP-dependent RNA helicase DDX1 OS | DDX1 | 82.433 | 2 | 53.79254 | yes | 9.107E-07 | 693.34204 | 111.35242 |
| FLPNAPK | 7 | Q92499 | ATP-dependent RNA helicase DDX1 OS | DDX1 | 82.433 | 1 | 53.79254 | yes | 3.745E-06 | 786.45087 | 91.985855 |
| GHQFSCVCLHGDF | 13 | Q92499 | ATP-dependent RNA helicase DDX1 OS | DDX1 | 82.433 | 2 | 53.79254 | yes | 9.722E-08 | 786.84338 | 22.39876 |
| DGPVRPQNAEEEK | 13 | Q92504 | Zinc transporter SLC39A7 OS=Homo sap | SLC39A7 | 50.119 | 2 | 51.990887 | yes | 2.239E-07 | 734.85498 | 1193.3026 |

| | | | | | | | | | | | |
|---|---|---|---|---|---|---|---|---|---|---|---|
| GHGHDHEHSHGG | 22 | Q92504 | Zinc transporter SLC39A7 OS=Homo sap | SLC39A7 | 50.119 | 4 | 51.990887 | yes | 9.722E-08 | 556.49823 | 714.97845 |
| DGPVRPQNAEEEK | 14 | Q92504 | Zinc transporter SLC39A7 OS=Homo sap | SLC39A7 | 50.119 | 2 | 51.990887 | yes | 1.684E-07 | 812.90552 | 208.84964 |
| DGPVRPQNAEEEK | 14 | Q92504 | Zinc transporter SLC39A7 OS=Homo sap | SLC39A7 | 50.119 | 2 | 51.990887 | yes | 4.76E-07 | 542.27277 | 208.84964 |
| DGPVRPQNAEEEK | 14 | Q92504 | Zinc transporter SLC39A7 OS=Homo sap | SLC39A7 | 50.119 | 3 | 51.990887 | yes | 1.684E-07 | 812.90552 | 208.84964 |
| DGPVRPQNAEEEK | 14 | Q92504 | Zinc transporter SLC39A7 OS=Homo sap | SLC39A7 | 50.119 | 3 | 51.990887 | yes | 4.76E-07 | 542.27277 | 208.84964 |
| ICLEDNVLMSGVK | 13 | Q92520 | Protein FAM3C OS=Homo sapiens OX=9 | FAM3C | 24.681 | 2 | 48.83065 | yes | 3.163E-07 | 739.37311 | 145.34943 |
| MASGAANVVGPK | 12 | Q92520 | Protein FAM3C OS=Homo sapiens OX=9 | FAM3C | 24.681 | 2 | 48.83065 | yes | 1.099E-07 | 551.28967 | 97.372429 |
| SPFEQHIK | 8 | Q92520 | Protein FAM3C OS=Homo sapiens OX=9 | FAM3C | 24.681 | 2 | 48.83065 | yes | 3.166E-06 | 493.25873 | 59.471931 |
| MDASLGNLFAR | 11 | Q92520 | Protein FAM3C OS=Homo sapiens OX=9 | FAM3C | 24.681 | 2 | 48.83065 | yes | 2.209E-06 | 597.80042 | 25.589359 |
| YSQLVVETIR | 10 | Q92522 | Histone H1.10 OS=Homo sapiens OX=96 | H1-10 | 22.488 | 2 | 50.3837 | yes | 2.172E-07 | 604.33771 | 4787.9805 |
| RGAPAAATAPAPT | 16 | Q92522 | Histone H1.10 OS=Homo sapiens OX=96 | H1-10 | 22.488 | 3 | 50.3837 | yes | 1.783E-07 | 496.60742 | 1209.1936 |
| ALVQNDTLLQVK | 12 | Q92522 | Histone H1.10 OS=Homo sapiens OX=96 | H1-10 | 22.488 | 2 | 50.3837 | yes | 1.525E-07 | 671.39044 | 794.07336 |
| GAPAAATAPAPTA | 15 | Q92522 | Histone H1.10 OS=Homo sapiens OX=96 | H1-10 | 22.488 | 2 | 50.3837 | yes | 2.742E-07 | 666.35693 | 691.16168 |
| KVPWFDQQNGR | 11 | Q92522 | Histone H1.10 OS=Homo sapiens OX=96 | H1-10 | 22.488 | 2 | 50.3837 | yes | 9.81E-08 | 687.8493 | 349.52853 |
| VPWFDQQNGR | 10 | Q92522 | Histone H1.10 OS=Homo sapiens OX=96 | H1-10 | 22.488 | 2 | 50.3837 | yes | 1.537E-07 | 623.80182 | 95.823837 |
| ALIDPSSGLPNR | 12 | Q92530 | Proteasome inhibitor PI31 subunit OS=Ho | PSMF1 | 29.817 | 2 | 48.878525 | yes | 1.091E-07 | 620.33826 | 71.301414 |
| FVGTDPASDEVVL | 15 | Q92538 | Golgi-specific brefeldin A-resistance guai | GBF1 | 206.446 | 2 | 25.764359 | yes | 3.23E-06 | 804.40289 | 357.61292 |
| VFQEAVVDER | 10 | Q92540 | Nonsense-mediated mRNA decay factor S | SMG7 | 127.282 | 2 | 56.539986 | yes | 9.722E-08 | 596.30383 | 456.77252 |
| NQITTLQGQAK | 11 | Q92540 | Nonsense-mediated mRNA decay factor S | SMG7 | 127.282 | 2 | 56.539986 | yes | 1.219E-06 | 601.33038 | 443.86304 |
| LIQCENEVGK | 10 | Q92540 | Nonsense-mediated mRNA decay factor S | SMG7 | 127.282 | 2 | 56.539986 | yes | 3.251E-07 | 595.29773 | 355.20715 |
| SSQLGIISNK | 10 | Q92540 | Nonsense-mediated mRNA decay factor S | SMG7 | 127.282 | 2 | 56.539986 | yes | 1.191E-07 | 523.79565 | 334.23685 |
| WIADNQPR | 8 | Q92540 | Nonsense-mediated mRNA decay factor S | SMG7 | 127.282 | 2 | 56.539986 | yes | 2.398E-07 | 500.25397 | 325.06659 |
| LGPAEVWTSR | 10 | Q92540 | Nonsense-mediated mRNA decay factor S | SMG7 | 127.282 | 2 | 56.539986 | yes | 1.152E-07 | 558.29584 | 264.34995 |
| MLVTDLEYALDK | 12 | Q92540 | Nonsense-mediated mRNA decay factor S | SMG7 | 127.282 | 2 | 56.539986 | yes | 9.722E-08 | 705.86292 | 255.59703 |
| FPFPAASTNLQK | 12 | Q92540 | Nonsense-mediated mRNA decay factor S | SMG7 | 127.282 | 2 | 56.539986 | yes | 3.525E-07 | 660.85095 | 216.59911 |
| QLWNPPQVQGPL( | 14 | Q92540 | Nonsense-mediated mRNA decay factor S | SMG7 | 127.282 | 2 | 56.539986 | yes | 1.199E-06 | 781.42792 | 206.34366 |
| EKLEEQFK | 8 | Q92540 | Nonsense-mediated mRNA decay factor S | SMG7 | 127.282 | 2 | 56.539986 | yes | 2.294E-07 | 525.77692 | 196.12631 |
| WGVSDFIK | 8 | Q92540 | Nonsense-mediated mRNA decay factor S | SMG7 | 127.282 | 2 | 56.539986 | yes | 9.722E-08 | 476.25037 | 149.60539 |
| GDHLTTIFYYCR | 12 | Q92540 | Nonsense-mediated mRNA decay factor S | SMG7 | 127.282 | 2 | 56.539986 | yes | 1.002E-07 | 773.36139 | 140.44202 |
| SPGVFRPEQDPVPI | 14 | Q92540 | Nonsense-mediated mRNA decay factor S | SMG7 | 127.282 | 3 | 56.539986 | yes | 2.711E-07 | 527.61084 | 66.271042 |
| SPGVFRPEQDPVPI | 14 | Q92540 | Nonsense-mediated mRNA decay factor S | SMG7 | 127.282 | 3 | 56.539986 | yes | 2.955E-07 | 790.9126 | 66.271042 |
| SPGVFRPEQDPVPI | 14 | Q92540 | Nonsense-mediated mRNA decay factor S | SMG7 | 127.282 | 2 | 56.539986 | yes | 2.711E-07 | 527.61084 | 66.271042 |
| SPGVFRPEQDPVPI | 14 | Q92540 | Nonsense-mediated mRNA decay factor S | SMG7 | 127.282 | 2 | 56.539986 | yes | 2.955E-07 | 790.9126 | 66.271042 |
| YPNNSMFNEVYGI | 13 | Q92540 | Nonsense-mediated mRNA decay factor S | SMG7 | 127.282 | 2 | 56.539986 | yes | 9.722E-08 | 781.85083 | 90.040085 |
| SVMAQQANIDR | 11 | Q92540 | Nonsense-mediated mRNA decay factor S | SMG7 | 127.282 | 2 | 56.539986 | yes | 9.722E-08 | 616.80621 | 87.23278 |
| NLSNNCDTGEKPV | 17 | Q92540 | Nonsense-mediated mRNA decay factor S | SMG7 | 127.282 | 3 | 56.539986 | yes | 3.571E-07 | 641.6474 | 72.725449 |
| VSMDWLR | 7 | Q92540 | Nonsense-mediated mRNA decay factor S | SMG7 | 127.282 | 2 | 56.539986 | yes | 1.506E-07 | 453.72873 | 66.651726 |
| QPYYLQTQDPIK | 12 | Q92540 | Nonsense-mediated mRNA decay factor S | SMG7 | 127.282 | 2 | 56.539986 | yes | 3.435E-06 | 747.38538 | 32.84721 |
| QNVAVQVK | 8 | Q92540 | Nonsense-mediated mRNA decay factor S | SMG7 | 127.282 | 2 | 56.539986 | yes | 2.283E-07 | 443.26126 | 27.952549 |

| | | | | | | | | | | | |
|---|---|---|---|---|---|---|---|---|---|---|---|
| SQPVSLPEELNR | 12 | Q92541 | RNA polymerase-associated protein RTF | RTF1 | 80.314 | 2 | 43.616028 | yes | 1.459E-06 | 684.8595 | 80.647247 |
| SASDLSEDLFK | 11 | Q92541 | RNA polymerase-associated protein RTF | RTF1 | 80.314 | 2 | 43.616028 | yes | 2.517E-07 | 606.29315 | 29.792305 |
| VILHVDQSR | 9 | Q92547 | DNA topoisomerase 2-binding protein 1 ( | TOPBP1 | 170.679 | 2 | 53.42377 | yes | 2.101E-07 | 533.80383 | 445.60583 |
| NLQLALANSSR | 11 | Q92547 | DNA topoisomerase 2-binding protein 1 ( | TOPBP1 | 170.679 | 2 | 53.42377 | yes | 2.294E-07 | 593.83057 | 365.77124 |
| MSLDISAVQDGR | 12 | Q92547 | DNA topoisomerase 2-binding protein 1 ( | TOPBP1 | 170.679 | 2 | 53.42377 | yes | 9.757E-08 | 646.31915 | 356.30453 |
| QVAASPAVGQPLC | 14 | Q92547 | DNA topoisomerase 2-binding protein 1 ( | TOPBP1 | 170.679 | 2 | 53.42377 | yes | 9.991E-08 | 697.39355 | 255.0983 |
| IYLCGFSGR | 9 | Q92547 | DNA topoisomerase 2-binding protein 1 ( | TOPBP1 | 170.679 | 2 | 53.42377 | yes | 7.878E-07 | 536.76587 | 250.18428 |
| QAVCDPGNIR | 10 | Q92547 | DNA topoisomerase 2-binding protein 1 ( | TOPBP1 | 170.679 | 2 | 53.42377 | yes | 1.328E-07 | 565.2746 | 227.97247 |
| YTDINMEDFK | 10 | Q92547 | DNA topoisomerase 2-binding protein 1 ( | TOPBP1 | 170.679 | 2 | 53.42377 | yes | 9.722E-08 | 638.28174 | 202.85883 |
| GVLTQTLEMR | 10 | Q92547 | DNA topoisomerase 2-binding protein 1 ( | TOPBP1 | 170.679 | 2 | 53.42377 | yes | 9.722E-08 | 574.31061 | 187.58632 |
| WLLECFSK | 8 | Q92547 | DNA topoisomerase 2-binding protein 1 ( | TOPBP1 | 170.679 | 2 | 53.42377 | yes | 1.283E-07 | 541.77063 | 173.80176 |
| EATHLFSDLNK | 11 | Q92547 | DNA topoisomerase 2-binding protein 1 ( | TOPBP1 | 170.679 | 2 | 53.42377 | yes | 2.982E-07 | 637.82239 | 172.03979 |
| HPISEELETPIK | 12 | Q92547 | DNA topoisomerase 2-binding protein 1 ( | TOPBP1 | 170.679 | 2 | 53.42377 | yes | 9.722E-08 | 696.87207 | 166.34859 |
| LFKPSFDVK | 9 | Q92547 | DNA topoisomerase 2-binding protein 1 ( | TOPBP1 | 170.679 | 2 | 53.42377 | yes | 1.596E-07 | 540.80804 | 161.52322 |
| YLVAANLK | 8 | Q92547 | DNA topoisomerase 2-binding protein 1 ( | TOPBP1 | 170.679 | 2 | 53.42377 | yes | 1.245E-07 | 446.26855 | 152.69501 |
| QESGIVEGAFSGW | 14 | Q92547 | DNA topoisomerase 2-binding protein 1 ( | TOPBP1 | 170.679 | 2 | 53.42377 | yes | 9.722E-08 | 747.86481 | 139.17633 |
| DSHLIPTPQAPSIAI | 27 | Q92547 | DNA topoisomerase 2-binding protein 1 ( | TOPBP1 | 170.679 | 4 | 53.42377 | yes | 1.249E-07 | 710.88562 | 134.75055 |
| GFCQDESIYK | 10 | Q92547 | DNA topoisomerase 2-binding protein 1 ( | TOPBP1 | 170.679 | 2 | 53.42377 | yes | 9.722E-08 | 623.77405 | 131.62512 |
| VLPGHSVPLFK | 11 | Q92547 | DNA topoisomerase 2-binding protein 1 ( | TOPBP1 | 170.679 | 2 | 53.42377 | yes | 9.722E-08 | 597.35571 | 129.43466 |
| QCFDPTCTHIVVG | 17 | Q92547 | DNA topoisomerase 2-binding protein 1 ( | TOPBP1 | 170.679 | 3 | 53.42377 | yes | 1.276E-07 | 679.66461 | 62.385273 |
| QCFDPTCTHIVVG | 17 | Q92547 | DNA topoisomerase 2-binding protein 1 ( | TOPBP1 | 170.679 | 3 | 53.42377 | yes | 2.646E-07 | 510.00027 | 62.385273 |
| QCFDPTCTHIVVG | 17 | Q92547 | DNA topoisomerase 2-binding protein 1 ( | TOPBP1 | 170.679 | 4 | 53.42377 | yes | 1.276E-07 | 679.66461 | 62.385273 |
| QCFDPTCTHIVVG | 17 | Q92547 | DNA topoisomerase 2-binding protein 1 ( | TOPBP1 | 170.679 | 4 | 53.42377 | yes | 2.646E-07 | 510.00027 | 62.385273 |
| NAVALSASPQLK | 12 | Q92547 | DNA topoisomerase 2-binding protein 1 ( | TOPBP1 | 170.679 | 2 | 53.42377 | yes | 3.211E-07 | 599.84314 | 113.97321 |
| HGGQYMGQLK | 10 | Q92547 | DNA topoisomerase 2-binding protein 1 ( | TOPBP1 | 170.679 | 2 | 53.42377 | yes | 1.283E-07 | 559.77423 | 108.12859 |
| LLSAVSSTK | 9 | Q92547 | DNA topoisomerase 2-binding protein 1 ( | TOPBP1 | 170.679 | 2 | 53.42377 | yes | 4.052E-06 | 453.26874 | 85.63588 |
| KPILLPSWIK | 10 | Q92547 | DNA topoisomerase 2-binding protein 1 ( | TOPBP1 | 170.679 | 2 | 53.42377 | yes | 4.519E-07 | 597.88409 | 84.382401 |
| IDYCHLIEK | 9 | Q92547 | DNA topoisomerase 2-binding protein 1 ( | TOPBP1 | 170.679 | 2 | 53.42377 | yes | 2.973E-07 | 595.79736 | 79.396133 |
| EPSLHLDTPSK | 11 | Q92547 | DNA topoisomerase 2-binding protein 1 ( | TOPBP1 | 170.679 | 2 | 53.42377 | yes | 1.847E-07 | 612.31696 | 70.442406 |
| QLQEIMSATSIVKI | 18 | Q92547 | DNA topoisomerase 2-binding protein 1 ( | TOPBP1 | 170.679 | 3 | 53.42377 | yes | 2.035E-07 | 672.02985 | 61.031364 |
| IITIEETHEELKK | 13 | Q92547 | DNA topoisomerase 2-binding protein 1 ( | TOPBP1 | 170.679 | 2 | 53.42377 | yes | 3.757E-07 | 791.93799 | 53.424522 |
| KPSTPLSEVIVK | 12 | Q92547 | DNA topoisomerase 2-binding protein 1 ( | TOPBP1 | 170.679 | 2 | 53.42377 | yes | 1.485E-07 | 649.38995 | 52.318016 |
| NDKEPFFVK | 9 | Q92547 | DNA topoisomerase 2-binding protein 1 ( | TOPBP1 | 170.679 | 2 | 53.42377 | yes | 2.209E-07 | 562.29279 | 51.276031 |
| ADESHFLIENSTK | 13 | Q92547 | DNA topoisomerase 2-binding protein 1 ( | TOPBP1 | 170.679 | 2 | 53.42377 | yes | 9.722E-08 | 745.85974 | 35.939274 |
| GMFASTHLILK | 11 | Q92547 | DNA topoisomerase 2-binding protein 1 ( | TOPBP1 | 170.679 | 2 | 53.42377 | yes | 4.77E-07 | 609.33917 | 24.049915 |
| LKPDDSGVNIAEA | 22 | Q92547 | DNA topoisomerase 2-binding protein 1 ( | TOPBP1 | 170.679 | 3 | 53.42377 | yes | 3.57E-06 | 802.06903 | 22.382854 |
| DALAALETPGRPS | 16 | Q92547 | DNA topoisomerase 2-binding protein 1 ( | TOPBP1 | 170.679 | 2 | 53.42377 | yes | 1.379E-06 | 841.44702 | 16.945612 |
| MNECTHLIVQEPK | 13 | Q92547 | DNA topoisomerase 2-binding protein 1 ( | TOPBP1 | 170.679 | 3 | 53.42377 | yes | 1.362E-07 | 533.59369 | 14.841895 |
| VELQQGLR | 8 | Q92552 | Small ribosomal subunit protein mS27 OS | MRPS27 | 47.612 | 2 | 52.583668 | yes | 1.735E-07 | 471.772 | 1824.1846 |

| | | | | | | | | | | | |
|---|---|---|---|---|---|---|---|---|---|---|---|
| KLPVSSLTISR | 11 | Q92552 | Small ribosomal subunit protein mS27 OS | MRPS27 | 47.612 | 2 | 52.583668 | yes | 2.203E-07 | 600.86914 | 1678.9542 |
| LPQYLER | 7 | Q92552 | Small ribosomal subunit protein mS27 OS | MRPS27 | 47.612 | 2 | 52.583668 | yes | 9.722E-08 | 459.7558 | 1283.2826 |
| LVEQLDIEETEQSK | 14 | Q92552 | Small ribosomal subunit protein mS27 OS | MRPS27 | 47.612 | 2 | 52.583668 | yes | 1.388E-07 | 830.91742 | 1036.7706 |
| LIDNISSR | 8 | Q92552 | Small ribosomal subunit protein mS27 OS | MRPS27 | 47.612 | 2 | 52.583668 | yes | 1.378E-07 | 459.25616 | 930.50061 |
| IESEGLLSLTTQLV | 15 | Q92552 | Small ribosomal subunit protein mS27 OS | MRPS27 | 47.612 | 2 | 52.583668 | yes | 9.722E-08 | 815.96674 | 906.3186 |
| EALDVLGAVLK | 11 | Q92552 | Small ribosomal subunit protein mS27 OS | MRPS27 | 47.612 | 2 | 52.583668 | yes | 1.082E-07 | 564.33716 | 797.89386 |
| NFGASLLLPGLK | 12 | Q92552 | Small ribosomal subunit protein mS27 OS | MRPS27 | 47.612 | 2 | 52.583668 | yes | 1.046E-07 | 615.36627 | 787.65948 |
| EEIDHAEYYLYK | 12 | Q92552 | Small ribosomal subunit protein mS27 OS | MRPS27 | 47.612 | 2 | 52.583668 | yes | 1.405E-07 | 786.86444 | 253.54568 |
| EEIDHAEYYLYK | 12 | Q92552 | Small ribosomal subunit protein mS27 OS | MRPS27 | 47.612 | 2 | 52.583668 | yes | 9.722E-08 | 524.91205 | 253.54568 |
| EEIDHAEYYLYK | 12 | Q92552 | Small ribosomal subunit protein mS27 OS | MRPS27 | 47.612 | 3 | 52.583668 | yes | 1.405E-07 | 786.86444 | 253.54568 |
| EEIDHAEYYLYK | 12 | Q92552 | Small ribosomal subunit protein mS27 OS | MRPS27 | 47.612 | 3 | 52.583668 | yes | 9.722E-08 | 524.91205 | 253.54568 |
| VAASPEDIK | 9 | Q92552 | Small ribosomal subunit protein mS27 OS | MRPS27 | 47.612 | 2 | 52.583668 | yes | 1.662E-07 | 465.25055 | 463.47836 |
| AVYHNMPLIWKP( | 17 | Q92552 | Small ribosomal subunit protein mS27 OS | MRPS27 | 47.612 | 3 | 52.583668 | yes | 1.062E-07 | 691.69617 | 229.03706 |
| AVYHNMPLIWKP( | 17 | Q92552 | Small ribosomal subunit protein mS27 OS | MRPS27 | 47.612 | 3 | 52.583668 | yes | 2.389E-06 | 519.02393 | 229.03706 |
| AVYHNMPLIWKP( | 17 | Q92552 | Small ribosomal subunit protein mS27 OS | MRPS27 | 47.612 | 4 | 52.583668 | yes | 1.062E-07 | 691.69617 | 229.03706 |
| AVYHNMPLIWKP( | 17 | Q92552 | Small ribosomal subunit protein mS27 OS | MRPS27 | 47.612 | 4 | 52.583668 | yes | 2.389E-06 | 519.02393 | 229.03706 |
| NWTIHTWIR | 9 | Q92552 | Small ribosomal subunit protein mS27 OS | MRPS27 | 47.612 | 2 | 52.583668 | yes | 4.226E-07 | 613.82507 | 417.63217 |
| HSPNCWYLR | 9 | Q92552 | Small ribosomal subunit protein mS27 OS | MRPS27 | 47.612 | 2 | 52.583668 | yes | 2.521E-07 | 616.7851 | 348.9812 |
| KTDFSWEEER | 10 | Q92552 | Small ribosomal subunit protein mS27 OS | MRPS27 | 47.612 | 2 | 52.583668 | yes | 1.104E-07 | 663.8017 | 328.09821 |
| ALYTLVNK | 8 | Q92552 | Small ribosomal subunit protein mS27 OS | MRPS27 | 47.612 | 2 | 52.583668 | yes | 2.554E-07 | 461.27383 | 226.65881 |
| AVYHNMPLIWK | 11 | Q92552 | Small ribosomal subunit protein mS27 OS | MRPS27 | 47.612 | 2 | 52.583668 | yes | 1.857E-07 | 686.36572 | 196.66272 |
| TDFSWEEER | 9 | Q92552 | Small ribosomal subunit protein mS27 OS | MRPS27 | 47.612 | 2 | 52.583668 | yes | 1.06E-07 | 599.75421 | 194.14455 |
| LPVSSLTISR | 10 | Q92552 | Small ribosomal subunit protein mS27 OS | MRPS27 | 47.612 | 2 | 52.583668 | yes | 2.199E-07 | 536.82166 | 42.372688 |
| EHYCLADLASLMI | 14 | Q92552 | Small ribosomal subunit protein mS27 OS | MRPS27 | 47.612 | 2 | 52.583668 | yes | 9.722E-08 | 833.38422 | 27.267992 |
| LEDLVSPPSIIDK | 13 | Q92570 | Nuclear receptor subfamily 4 group A me | NR4A3 | 68.23 | 2 | 53.184074 | yes | 1.028E-07 | 713.39539 | 1051.6385 |
| VLGALVELR | 9 | Q92570 | Nuclear receptor subfamily 4 group A me | NR4A3 | 68.23 | 2 | 53.184074 | yes | 5.1E-07 | 485.3082 | 874.24078 |
| KICTLGLQR | 9 | Q92570 | Nuclear receptor subfamily 4 group A me | NR4A3 | 68.23 | 2 | 53.184074 | yes | 9.722E-08 | 544.81586 | 265.26935 |
| ICTLGLQR | 8 | Q92570 | Nuclear receptor subfamily 4 group A me | NR4A3 | 68.23 | 2 | 53.184074 | yes | 1.272E-07 | 480.7684 | 157.83833 |
| GFGEWLDSIK | 10 | Q92570 | Nuclear receptor subfamily 4 group A me | NR4A3 | 68.23 | 2 | 53.184074 | yes | 1.159E-06 | 576.29022 | 135.25731 |
| CLSVGMVK | 8 | Q92570 | Nuclear receptor subfamily 4 group A me | NR4A3 | 68.23 | 2 | 53.184074 | yes | 2.594E-07 | 447.23279 | 48.153019 |
| AGGIETIANEFSDR | 14 | Q92598 | Heat shock protein 105 kDa OS=Homo sa | HSPH1 | 96.865 | 2 | 57.76496 | yes | 2.385E-07 | 740.35736 | 557.87158 |
| FVVQNVSAQK | 10 | Q92598 | Heat shock protein 105 kDa OS=Homo sa | HSPH1 | 96.865 | 2 | 57.76496 | yes | 1.269E-07 | 560.31146 | 435.88116 |
| SLDQDPVVR | 9 | Q92598 | Heat shock protein 105 kDa OS=Homo sa | HSPH1 | 96.865 | 2 | 57.76496 | yes | 3.108E-07 | 514.77222 | 425.67535 |
| SQFEELCAELLQK | 13 | Q92598 | Heat shock protein 105 kDa OS=Homo sa | HSPH1 | 96.865 | 2 | 57.76496 | yes | 9.722E-08 | 797.8927 | 263.27515 |
| NAVEEYVYEFR | 11 | Q92598 | Heat shock protein 105 kDa OS=Homo sa | HSPH1 | 96.865 | 2 | 57.76496 | yes | 9.722E-08 | 709.83295 | 261.15399 |
| VLGTAFDPFLGGK | 13 | Q92598 | Heat shock protein 105 kDa OS=Homo sa | HSPH1 | 96.865 | 2 | 57.76496 | yes | 1.102E-07 | 661.36115 | 248.88248 |
| VEDVSAVEIVGGA | 15 | Q92598 | Heat shock protein 105 kDa OS=Homo sa | HSPH1 | 96.865 | 2 | 57.76496 | yes | 9.722E-08 | 751.39648 | 247.77081 |
| DISTTLNADEAVA | 14 | Q92598 | Heat shock protein 105 kDa OS=Homo sa | HSPH1 | 96.865 | 2 | 57.76496 | no | 2.162E-07 | 738.37042 | 225.81216 |
| DLLNMYIETEGK | 12 | Q92598 | Heat shock protein 105 kDa OS=Homo sa | HSPH1 | 96.865 | 2 | 57.76496 | yes | 1.643E-07 | 713.35016 | 206.95964 |

| | | | | | | | | | | | |
|---|---|---|---|---|---|---|---|---|---|---|---|
| MFEELGQR | 8 | Q92598 | Heat shock protein 105 kDa OS=Homo sa | HSPH1 | 96.865 | 2 | 57.76496 | yes | 1.43E-07 | 505.24222 | 174.49042 |
| ELNNTCEPVVTQP | 14 | Q92598 | Heat shock protein 105 kDa OS=Homo sa | HSPH1 | 96.865 | 2 | 57.76496 | yes | 9.722E-08 | 814.90106 | 159.49022 |
| EKENLSYDLVPLK | 13 | Q92598 | Heat shock protein 105 kDa OS=Homo sa | HSPH1 | 96.865 | 2 | 57.76496 | yes | 1.092E-07 | 774.41943 | 102.67577 |
| YNHIDESEMK | 10 | Q92598 | Heat shock protein 105 kDa OS=Homo sa | HSPH1 | 96.865 | 2 | 57.76496 | yes | 2.325E-07 | 633.27698 | 73.111565 |
| LYQECEK | 7 | Q92598 | Heat shock protein 105 kDa OS=Homo sa | HSPH1 | 96.865 | 2 | 57.76496 | no | 2.658E-07 | 485.22095 | 59.002739 |
| LVEHFCAEFK | 10 | Q92598 | Heat shock protein 105 kDa OS=Homo sa | HSPH1 | 96.865 | 2 | 57.76496 | yes | 1.12E-07 | 640.31061 | 52.838757 |
| QDLPSLDEKPR | 11 | Q92598 | Heat shock protein 105 kDa OS=Homo sa | HSPH1 | 96.865 | 2 | 57.76496 | yes | 4.731E-06 | 649.34094 | 41.884583 |
| FICEQDHQNFLR | 12 | Q92598 | Heat shock protein 105 kDa OS=Homo sa | HSPH1 | 96.865 | 2 | 57.76496 | yes | 2.98E-07 | 803.87518 | 29.308895 |
| IEVPLYSLLEQTHL | 15 | Q92598 | Heat shock protein 105 kDa OS=Homo sa | HSPH1 | 96.865 | 3 | 57.76496 | yes | 1.969E-06 | 595.00488 | 9.1927509 |
| NLQEGQVTDPR | 11 | Q92600 | CCR4-NOT transcription complex subuni | CNOT9 | 33.632 | 2 | 51.840717 | yes | 1.161E-07 | 628.81512 | 383.44879 |
| IMESGSELSK | 10 | Q92600 | CCR4-NOT transcription complex subuni | CNOT9 | 33.632 | 2 | 51.840717 | yes | 2.343E-07 | 540.76569 | 240.05672 |
| FSHVAMILGK | 10 | Q92600 | CCR4-NOT transcription complex subuni | CNOT9 | 33.632 | 2 | 51.840717 | yes | 2.702E-07 | 551.8075 | 211.3555 |
| LTSLGVIGALVK | 12 | Q92600 | CCR4-NOT transcription complex subuni | CNOT9 | 33.632 | 2 | 51.840717 | yes | 1.542E-06 | 585.87646 | 165.42065 |
| MHSLATAAPVPTT | 19 | Q92600 | CCR4-NOT transcription complex subuni | CNOT9 | 33.632 | 3 | 51.840717 | yes | 9.722E-08 | 674.35431 | 113.55141 |
| TRPFEYLR | 8 | Q92600 | CCR4-NOT transcription complex subuni | CNOT9 | 33.632 | 2 | 51.840717 | yes | 1.137E-06 | 541.29309 | 100.79092 |
| ENALLELSK | 9 | Q92600 | CCR4-NOT transcription complex subuni | CNOT9 | 33.632 | 2 | 51.840717 | yes | 5.834E-07 | 508.78476 | 98.103386 |
| TVATFILQK | 9 | Q92600 | CCR4-NOT transcription complex subuni | CNOT9 | 33.632 | 2 | 51.840717 | yes | 1.852E-06 | 510.80804 | 84.13414 |
| DTTFAQVLK | 9 | Q92600 | CCR4-NOT transcription complex subuni | CNOT9 | 33.632 | 2 | 51.840717 | yes | 4.168E-07 | 511.77948 | 61.695564 |
| VCNALALLQCVAS | 18 | Q92600 | CCR4-NOT transcription complex subuni | CNOT9 | 33.632 | 3 | 51.840717 | yes | 3.152E-07 | 680.34357 | 38.144981 |
| IILNALVAQQK | 11 | Q92604 | Acyl-CoA:lysophosphatidylglycerol acylt | LPGAT1 | 43.09 | 2 | 46.27062 | yes | 1.605E-07 | 605.87952 | 81.182861 |
| KPTVTHVHYR | 10 | Q92604 | Acyl-CoA:lysophosphatidylglycerol acylt | LPGAT1 | 43.09 | 3 | 46.27062 | yes | 2.355E-07 | 413.23154 | 59.01646 |
| NMQVPQESLEMR | 12 | Q92613 | Protein Jade-3 OS=Homo sapiens OX=96 | JADE3 | 93.809 | 2 | 52.04616 | yes | 1.05E-07 | 731.34485 | 445.97488 |
| ELEEEFYSLVR | 11 | Q92613 | Protein Jade-3 OS=Homo sapiens OX=96 | JADE3 | 93.809 | 2 | 52.04616 | yes | 9.722E-08 | 707.34845 | 237.89955 |
| KPAEVFR | 7 | Q92613 | Protein Jade-3 OS=Homo sapiens OX=96 | JADE3 | 93.809 | 2 | 52.04616 | yes | 1.409E-07 | 423.74524 | 152.32059 |
| VPEGSWLCR | 9 | Q92613 | Protein Jade-3 OS=Homo sapiens OX=96 | JADE3 | 93.809 | 2 | 52.04616 | no | 2.156E-07 | 552.26874 | 131.83865 |
| STVEHFSR | 8 | Q92613 | Protein Jade-3 OS=Homo sapiens OX=96 | JADE3 | 93.809 | 2 | 52.04616 | yes | 9.722E-08 | 481.73816 | 109.45979 |
| NLCYMISR | 8 | Q92613 | Protein Jade-3 OS=Homo sapiens OX=96 | JADE3 | 93.809 | 2 | 52.04616 | yes | 2.745E-07 | 528.75189 | 94.86702 |
| TGACIQCSIK | 10 | Q92613 | Protein Jade-3 OS=Homo sapiens OX=96 | JADE3 | 93.809 | 2 | 52.04616 | yes | 1.137E-07 | 569.27319 | 89.523544 |
| DVLFIRPR | 8 | Q92613 | Protein Jade-3 OS=Homo sapiens OX=96 | JADE3 | 93.809 | 2 | 52.04616 | yes | 2.344E-07 | 508.306 | 84.868324 |
| WALVCNLCK | 9 | Q92613 | Protein Jade-3 OS=Homo sapiens OX=96 | JADE3 | 93.809 | 2 | 52.04616 | yes | 1.899E-07 | 582.28864 | 83.780182 |
| LGEAEYPHHR | 10 | Q92613 | Protein Jade-3 OS=Homo sapiens OX=96 | JADE3 | 93.809 | 2 | 52.04616 | yes | 7.883E-07 | 604.79401 | 58.065113 |
| NTEDLQCYVKPTK | 13 | Q92613 | Protein Jade-3 OS=Homo sapiens OX=96 | JADE3 | 93.809 | 2 | 52.04616 | yes | 1.967E-07 | 798.39032 | 39.928501 |
| VQPIPELSELSR | 12 | Q92614 | Unconventional myosin-XVIIIa OS=Hom | MYO18A | 233.116 | 2 | 55.482334 | yes | 3.326E-07 | 684.38007 | 218.47035 |
| TFVQELER | 8 | Q92614 | Unconventional myosin-XVIIIa OS=Hom | MYO18A | 233.116 | 2 | 55.482334 | yes | 5.735E-07 | 511.26929 | 189.26566 |
| LTAELQDTK | 9 | Q92614 | Unconventional myosin-XVIIIa OS=Hom | MYO18A | 233.116 | 2 | 55.482334 | yes | 2.098E-07 | 509.77438 | 167.07193 |
| ISELTSELTDER | 12 | Q92614 | Unconventional myosin-XVIIIa OS=Hom | MYO18A | 233.116 | 2 | 55.482334 | yes | 2.278E-07 | 696.84625 | 154.09218 |
| LGDLQADSEESQR | 13 | Q92614 | Unconventional myosin-XVIIIa OS=Hom | MYO18A | 233.116 | 2 | 55.482334 | yes | 2.116E-07 | 724.33661 | 104.42127 |
| DMDIAGFTQK | 10 | Q92614 | Unconventional myosin-XVIIIa OS=Hom | MYO18A | 233.116 | 2 | 55.482334 | yes | 2.42E-07 | 563.26587 | 97.432526 |
| LEISNPIPIK | 10 | Q92614 | Unconventional myosin-XVIIIa OS=Hom | MYO18A | 233.116 | 2 | 55.482334 | yes | 9.722E-08 | 562.33972 | 89.000145 |

| | | | | | | | | | | | |
|---|---|---|---|---|---|---|---|---|---|---|---|
| LPPVVPLPPPTLR | 13 | Q92614 | Unconventional myosin-XVIIIa OS=Hon | MYO18A | 233.116 | 2 | 55.482334 | yes | 5.282E-06 | 698.43976 | 79.570038 |
| LHLEGQQVR | 9 | Q92614 | Unconventional myosin-XVIIIa OS=Hon | MYO18A | 233.116 | 2 | 55.482334 | yes | 3.445E-07 | 540.30145 | 78.801231 |
| DLALGLVPGDR | 11 | Q92614 | Unconventional myosin-XVIIIa OS=Hon | MYO18A | 233.116 | 2 | 55.482334 | yes | 1.694E-07 | 563.31677 | 72.162704 |
| QDQSIILLGSSGSG | 15 | Q92614 | Unconventional myosin-XVIIIa OS=Hon | MYO18A | 233.116 | 2 | 55.482334 | yes | 2.763E-07 | 745.39648 | 70.171448 |
| LEMEMER | 7 | Q92614 | Unconventional myosin-XVIIIa OS=Hon | MYO18A | 233.116 | 2 | 55.482334 | yes | 3.597E-07 | 469.20953 | 64.726791 |
| LQVDALIDTIK | 11 | Q92614 | Unconventional myosin-XVIIIa OS=Hon | MYO18A | 233.116 | 2 | 55.482334 | yes | 3.339E-07 | 614.86102 | 60.67038 |
| VKDQEEELDEQA( | 22 | Q92614 | Unconventional myosin-XVIIIa OS=Hon | MYO18A | 233.116 | 3 | 55.482334 | yes | 1.05E-07 | 844.74268 | 58.316505 |
| ELQTQYDALK | 10 | Q92614 | Unconventional myosin-XVIIIa OS=Hon | MYO18A | 233.116 | 2 | 55.482334 | yes | 4.375E-06 | 604.81152 | 54.478813 |
| AVEELLECLDLEK | 13 | Q92614 | Unconventional myosin-XVIIIa OS=Hon | MYO18A | 233.116 | 2 | 55.482334 | yes | 1.949E-07 | 780.8949 | 53.815945 |
| PLIEVQLSEEQIR | 13 | Q92614 | Unconventional myosin-XVIIIa OS=Hon | MYO18A | 233.116 | 2 | 55.482334 | yes | 3.325E-06 | 777.4303 | 50.069103 |
| LEEDQEDMNELMI | 13 | Q92614 | Unconventional myosin-XVIIIa OS=Hon | MYO18A | 233.116 | 2 | 55.482334 | yes | 9.722E-08 | 812.34747 | 48.675503 |
| KIQDLAIR | 8 | Q92614 | Unconventional myosin-XVIIIa OS=Hon | MYO18A | 233.116 | 2 | 55.482334 | yes | 2.826E-06 | 478.798 | 35.266933 |
| QGPEESGLGDGTG | 15 | Q92614 | Unconventional myosin-XVIIIa OS=Hon | MYO18A | 233.116 | 2 | 55.482334 | yes | 1.873E-07 | 714.83368 | 30.073137 |
| NQLEESEFTCAAA | 15 | Q92614 | Unconventional myosin-XVIIIa OS=Hon | MYO18A | 233.116 | 2 | 55.482334 | yes | 1.81E-07 | 848.896 | 29.912458 |
| EKDMLLAEAFSLK | 13 | Q92614 | Unconventional myosin-XVIIIa OS=Hon | MYO18A | 233.116 | 2 | 55.482334 | yes | 1.421E-06 | 747.89728 | 26.636103 |
| NKDEEIQQLR | 10 | Q92614 | Unconventional myosin-XVIIIa OS=Hon | MYO18A | 233.116 | 2 | 55.482334 | yes | 6.342E-07 | 636.83075 | 21.248276 |
| SLPLVQVDEK | 10 | Q92615 | La-related protein 4B OS=Homo sapiens | LARP4B | 80.552 | 2 | 51.80761 | yes | 1.556E-07 | 564.31897 | 6885.6177 |
| LPSALTATACK | 11 | Q92615 | La-related protein 4B OS=Homo sapiens | LARP4B | 80.552 | 2 | 51.80761 | yes | 1.008E-07 | 566.80518 | 5467.8779 |
| TEDLFENR | 8 | Q92615 | La-related protein 4B OS=Homo sapiens | LARP4B | 80.552 | 2 | 51.80761 | yes | 1.094E-07 | 512.24072 | 3837.7102 |
| EPPSSPLQPQK | 11 | Q92615 | La-related protein 4B OS=Homo sapiens | LARP4B | 80.552 | 2 | 51.80761 | yes | 1.562E-07 | 604.31952 | 2514.3015 |
| AIAINTFLPK | 10 | Q92615 | La-related protein 4B OS=Homo sapiens | LARP4B | 80.552 | 2 | 51.80761 | yes | 2.384E-07 | 544.32916 | 2139.9497 |
| KPSYAEICQR | 10 | Q92615 | La-related protein 4B OS=Homo sapiens | LARP4B | 80.552 | 2 | 51.80761 | yes | 9.722E-08 | 626.31116 | 1783.5138 |
| TLEFCLSR | 8 | Q92615 | La-related protein 4B OS=Homo sapiens | LARP4B | 80.552 | 2 | 51.80761 | yes | 6.04E-07 | 513.25787 | 1778.6765 |
| IQNPSAYAK | 9 | Q92615 | La-related protein 4B OS=Homo sapiens | LARP4B | 80.552 | 2 | 51.80761 | yes | 3.305E-07 | 496.26401 | 1719.4075 |
| YREPPALK | 8 | Q92615 | La-related protein 4B OS=Homo sapiens | LARP4B | 80.552 | 2 | 51.80761 | yes | 5.631E-07 | 487.27692 | 1500.9872 |
| LSSLIIGPSK | 10 | Q92615 | La-related protein 4B OS=Homo sapiens | LARP4B | 80.552 | 2 | 51.80761 | yes | 1.073E-07 | 507.81332 | 1250.6613 |
| VVAEPQTQR | 9 | Q92615 | La-related protein 4B OS=Homo sapiens | LARP4B | 80.552 | 2 | 51.80761 | yes | 1.584E-07 | 514.28015 | 1195.047 |
| SPSPAHLPDDPK | 12 | Q92615 | La-related protein 4B OS=Homo sapiens | LARP4B | 80.552 | 2 | 51.80761 | yes | 3.989E-07 | 630.81458 | 1190.152 |
| PLDVSLYAQQR | 11 | Q92615 | La-related protein 4B OS=Homo sapiens | LARP4B | 80.552 | 2 | 51.80761 | yes | 1.612E-07 | 645.34607 | 969.63135 |
| HAIPSAER | 8 | Q92615 | La-related protein 4B OS=Homo sapiens | LARP4B | 80.552 | 2 | 51.80761 | yes | 9.819E-08 | 440.73541 | 618.46478 |
| VEPGSLESSPGLGI | 14 | Q92615 | La-related protein 4B OS=Homo sapiens | LARP4B | 80.552 | 2 | 51.80761 | yes | 1E-07 | 692.85699 | 446.84769 |
| KLAEPAER | 8 | Q92615 | La-related protein 4B OS=Homo sapiens | LARP4B | 80.552 | 2 | 51.80761 | yes | 1.601E-07 | 457.25873 | 319.53043 |
| KNSFGYR | 7 | Q92615 | La-related protein 4B OS=Homo sapiens | LARP4B | 80.552 | 2 | 51.80761 | yes | 9.722E-08 | 436.22467 | 240.79564 |
| RPAGGRPSPSAMC | 14 | Q92615 | La-related protein 4B OS=Homo sapiens | LARP4B | 80.552 | 3 | 51.80761 | yes | 3.29E-07 | 456.91034 | 195.38206 |
| TFQGKPIK | 8 | Q92615 | La-related protein 4B OS=Homo sapiens | LARP4B | 80.552 | 2 | 51.80761 | yes | 3.158E-07 | 459.77402 | 120.10225 |
| PAASPLTSLR | 10 | Q92615 | La-related protein 4B OS=Homo sapiens | LARP4B | 80.552 | 2 | 51.80761 | yes | 1.381E-07 | 506.79294 | 110.05077 |
| KTLEFCLSR | 9 | Q92615 | La-related protein 4B OS=Homo sapiens | LARP4B | 80.552 | 2 | 51.80761 | yes | 1.823E-07 | 577.30536 | 91.624176 |
| KLSTDVDLIVEVLI | 14 | Q92615 | La-related protein 4B OS=Homo sapiens | LARP4B | 80.552 | 2 | 51.80761 | yes | 1.701E-07 | 800.46942 | 10.643242 |
| GIGEYVNIR | 9 | Q92620 | Pre-mRNA-splicing factor ATP-depender | DHX38 | 140.503 | 2 | 48.681126 | yes | 2.453E-07 | 510.77728 | 245.55618 |

| | | | | | | | | | | | |
|---|---|---|---|---|---|---|---|---|---|---|---|
| YMTDGILLR | 9 | Q92620 | Pre-mRNA-splicing factor ATP-depender | DHX38 | 140.503 | 2 | 48.681126 | yes | 1.147E-07 | 541.28918 | 130.38731 |
| EADLDHYSAIIMD | 18 | Q92620 | Pre-mRNA-splicing factor ATP-depender | DHX38 | 140.503 | 3 | 48.681126 | yes | 1.124E-07 | 705.65308 | 120.89366 |
| MLTSGVVHR | 9 | Q92620 | Pre-mRNA-splicing factor ATP-depender | DHX38 | 140.503 | 2 | 48.681126 | yes | 2.985E-07 | 500.27383 | 94.108727 |
| SQWESPSPTPSYR | 13 | Q92620 | Pre-mRNA-splicing factor ATP-depender | DHX38 | 140.503 | 2 | 48.681126 | yes | 1.23E-07 | 761.35205 | 56.689041 |
| LMVEFPLDPALSK | 13 | Q92620 | Pre-mRNA-splicing factor ATP-depender | DHX38 | 140.503 | 2 | 48.681126 | yes | 1.115E-07 | 730.39691 | 48.058865 |
| QPEPVIPVK | 9 | Q92620 | Pre-mRNA-splicing factor ATP-depender | DHX38 | 140.503 | 2 | 48.681126 | yes | 8.986E-07 | 503.80023 | 39.750854 |
| FEDCTSENTLIK | 12 | Q92620 | Pre-mRNA-splicing factor ATP-depender | DHX38 | 140.503 | 2 | 48.681126 | yes | 2.755E-05 | 728.83484 | 35.583809 |
| MSLASCGTDWDIV | 14 | Q92620 | Pre-mRNA-splicing factor ATP-depender | DHX38 | 140.503 | 2 | 48.681126 | yes | 2.529E-07 | 805.86871 | 34.221947 |
| QINEDNER | 8 | Q92620 | Pre-mRNA-splicing factor ATP-depender | DHX38 | 140.503 | 2 | 48.681126 | yes | 6.566E-07 | 509.23343 | 29.909443 |
| DATSDLAIIAR | 11 | Q92620 | Pre-mRNA-splicing factor ATP-depender | DHX38 | 140.503 | 2 | 48.681126 | yes | 3.267E-06 | 573.31171 | 17.679468 |
| LEVDEDFEEDNAA | 14 | Q92620 | Pre-mRNA-splicing factor ATP-depender | DHX38 | 140.503 | 2 | 48.681126 | yes | 2.76E-06 | 812.35468 | 13.970368 |
| ELCQSVMPAGVDI | 13 | Q92621 | Nuclear pore complex protein Nup205 OS | NUP205 | 227.922 | 2 | 51.842937 | yes | 1.767E-06 | 717.3418 | 107.84294 |
| DPPMFIPTPVDR | 12 | Q92621 | Nuclear pore complex protein Nup205 OS | NUP205 | 227.922 | 2 | 51.842937 | yes | 1.893E-07 | 692.8501 | 91.202011 |
| LQDSFASETNLDFI | 14 | Q92621 | Nuclear pore complex protein Nup205 OS | NUP205 | 227.922 | 2 | 51.842937 | yes | 2.05E-07 | 821.88904 | 82.559334 |
| TLQSLLTPQPPLLK | 14 | Q92621 | Nuclear pore complex protein Nup205 OS | NUP205 | 227.922 | 2 | 51.842937 | yes | 7.704E-07 | 774.97162 | 67.821579 |
| DLPSADSVQYR | 11 | Q92621 | Nuclear pore complex protein Nup205 OS | NUP205 | 227.922 | 2 | 51.842937 | yes | 3.142E-07 | 625.8042 | 64.079979 |
| AQIEQVIANCEHK | 13 | Q92621 | Nuclear pore complex protein Nup205 OS | NUP205 | 227.922 | 2 | 51.842937 | yes | 1.058E-07 | 770.38281 | 61.977997 |
| IQQGALELLR | 10 | Q92621 | Nuclear pore complex protein Nup205 OS | NUP205 | 227.922 | 2 | 51.842937 | yes | 9.722E-08 | 570.84039 | 58.73394 |
| ASTEGVAIQGQQG | 15 | Q92621 | Nuclear pore complex protein Nup205 OS | NUP205 | 227.922 | 2 | 51.842937 | yes | 1.006E-07 | 751.88153 | 55.46772 |
| LPGLQATVR | 9 | Q92621 | Nuclear pore complex protein Nup205 OS | NUP205 | 227.922 | 2 | 51.842937 | yes | 4.235E-07 | 477.79019 | 47.197296 |
| SVSGFLHFDTATK | 13 | Q92621 | Nuclear pore complex protein Nup205 OS | NUP205 | 227.922 | 2 | 51.842937 | yes | 1.902E-07 | 705.35663 | 43.267624 |
| IVNEGGIDPLLR | 12 | Q92626 | Peroxidasin homolog OS=Homo sapiens ( | PXDN | 165.275 | 2 | 48.613445 | yes | 1.122E-07 | 648.36951 | 278.53824 |
| HLVLSSGTLR | 10 | Q92626 | Peroxidasin homolog OS=Homo sapiens ( | PXDN | 165.275 | 2 | 48.613445 | yes | 1.373E-07 | 541.81946 | 193.15616 |
| ILGEVGMR | 8 | Q92626 | Peroxidasin homolog OS=Homo sapiens ( | PXDN | 165.275 | 2 | 48.613445 | yes | 2.417E-07 | 437.74438 | 91.632263 |
| VQSDVFR | 7 | Q92626 | Peroxidasin homolog OS=Homo sapiens ( | PXDN | 165.275 | 2 | 48.613445 | yes | 3.048E-07 | 425.72452 | 57.658558 |
| IPSGAFEDLENLK | 13 | Q92626 | Peroxidasin homolog OS=Homo sapiens ( | PXDN | 165.275 | 2 | 48.613445 | yes | 3.622E-07 | 716.86957 | 42.077019 |
| NLNTLLLNNNQIK | 13 | Q92626 | Peroxidasin homolog OS=Homo sapiens ( | PXDN | 165.275 | 2 | 48.613445 | yes | 2.406E-07 | 756.43066 | 39.116779 |
| VWQDCCEDCR | 10 | Q92626 | Peroxidasin homolog OS=Homo sapiens ( | PXDN | 165.275 | 2 | 48.613445 | yes | 2.854E-06 | 714.2605 | 14.44605 |
| VELSTVNVR | 9 | Q92665 | Small ribosomal subunit protein mS31 OS | MRPS31 | 45.319 | 2 | 52.10113 | yes | 9.99E-08 | 508.79037 | 697.91394 |
| ISFSNIISDMK | 11 | Q92665 | Small ribosomal subunit protein mS31 OS | MRPS31 | 45.319 | 2 | 52.10113 | yes | 1.009E-07 | 627.82355 | 368.75967 |
| VEHIEWFR | 8 | Q92665 | Small ribosomal subunit protein mS31 OS | MRPS31 | 45.319 | 2 | 52.10113 | yes | 9.722E-08 | 558.28528 | 242.44012 |
| HFMELVTCGLSK | 12 | Q92665 | Small ribosomal subunit protein mS31 OS | MRPS31 | 45.319 | 2 | 52.10113 | yes | 2.782E-07 | 711.34943 | 80.852341 |
| RLNIFDMMAVTK | 12 | Q92665 | Small ribosomal subunit protein mS31 OS | MRPS31 | 45.319 | 2 | 52.10113 | yes | 2.385E-07 | 719.88092 | 40.610043 |
| DDAAPAPPVADAI | 13 | Q92667 | A-kinase anchor protein 1, mitochondrial | AKAP1 | 97.342 | 2 | 54.74574 | yes | 1.241E-07 | 619.30658 | 1566.0461 |
| VVSTPPSVTEPPEK | 14 | Q92667 | A-kinase anchor protein 1, mitochondrial | AKAP1 | 97.342 | 2 | 54.74574 | yes | 1.387E-07 | 733.8905 | 1387.566 |
| VLEEALLSR | 9 | Q92667 | A-kinase anchor protein 1, mitochondrial | AKAP1 | 97.342 | 2 | 54.74574 | yes | 1.078E-07 | 515.3006 | 1076.8379 |
| LPAEPPALLQTHPI | 16 | Q92667 | A-kinase anchor protein 1, mitochondrial | AKAP1 | 97.342 | 3 | 54.74574 | yes | 1.486E-06 | 599.65405 | 976.60803 |
| SESSGILPNTTDMF | 14 | Q92667 | A-kinase anchor protein 1, mitochondrial | AKAP1 | 97.342 | 2 | 54.74574 | yes | 1.905E-07 | 754.35651 | 907.94092 |
| EPLPVEDVCPK | 11 | Q92667 | A-kinase anchor protein 1, mitochondrial | AKAP1 | 97.342 | 2 | 54.74574 | yes | 9.722E-08 | 641.82104 | 854.94226 |

| | | | | | | | | | | | |
|---|---|---|---|---|---|---|---|---|---|---|---|
| GHVSSHDEQQVE/ | 19 | Q92667 | A-kinase anchor protein 1, mitochondrial | AKAP1 | 97.342 | 3 | 54.74574 | yes | 9.722E-08 | 683.00323 | 363.1636 |
| GHVSSHDEQQVE/ | 19 | Q92667 | A-kinase anchor protein 1, mitochondrial | AKAP1 | 97.342 | 3 | 54.74574 | yes | 2.379E-07 | 512.50427 | 363.1636 |
| GHVSSHDEQQVE/ | 19 | Q92667 | A-kinase anchor protein 1, mitochondrial | AKAP1 | 97.342 | 4 | 54.74574 | yes | 9.722E-08 | 683.00323 | 363.1636 |
| GHVSSHDEQQVE/ | 19 | Q92667 | A-kinase anchor protein 1, mitochondrial | AKAP1 | 97.342 | 4 | 54.74574 | yes | 2.379E-07 | 512.50427 | 363.1636 |
| ETGGAEGTGDAVI | 16 | Q92667 | A-kinase anchor protein 1, mitochondrial | AKAP1 | 97.342 | 2 | 54.74574 | yes | 4.648E-07 | 745.85211 | 721.48199 |
| SIPLECPLSSPK | 12 | Q92667 | A-kinase anchor protein 1, mitochondrial | AKAP1 | 97.342 | 2 | 54.74574 | yes | 9.968E-08 | 664.34998 | 705.62939 |
| VCQASQLQGQK | 11 | Q92667 | A-kinase anchor protein 1, mitochondrial | AKAP1 | 97.342 | 2 | 54.74574 | yes | 1.294E-07 | 623.81409 | 693.85559 |
| FIESAHTELAK | 11 | Q92667 | A-kinase anchor protein 1, mitochondrial | AKAP1 | 97.342 | 2 | 54.74574 | yes | 9.722E-08 | 623.32733 | 615.87512 |
| LELALTGGEAK | 11 | Q92667 | A-kinase anchor protein 1, mitochondrial | AKAP1 | 97.342 | 2 | 54.74574 | yes | 9.722E-08 | 551.31116 | 588.4397 |
| KVQPGYPVVPAEI | 13 | Q92667 | A-kinase anchor protein 1, mitochondrial | AKAP1 | 97.342 | 2 | 54.74574 | yes | 9.722E-08 | 706.40082 | 533.25519 |
| EHVLELENSK | 10 | Q92667 | A-kinase anchor protein 1, mitochondrial | AKAP1 | 97.342 | 2 | 54.74574 | yes | 9.777E-08 | 599.30914 | 397.59027 |
| NSMDSVDSCCSLK | 13 | Q92667 | A-kinase anchor protein 1, mitochondrial | AKAP1 | 97.342 | 2 | 54.74574 | yes | 1.061E-07 | 751.79962 | 330.83453 |
| VQPGYPVVPAEK | 12 | Q92667 | A-kinase anchor protein 1, mitochondrial | AKAP1 | 97.342 | 2 | 54.74574 | yes | 4.513E-07 | 642.35333 | 310.60883 |
| GPSLASLEGEEDK( | 15 | Q92667 | A-kinase anchor protein 1, mitochondrial | AKAP1 | 97.342 | 2 | 54.74574 | yes | 9.809E-08 | 758.87811 | 225.50186 |
| GVEGELGNEESLD | 14 | Q92667 | A-kinase anchor protein 1, mitochondrial | AKAP1 | 97.342 | 2 | 54.74574 | yes | 7.034E-07 | 752.34973 | 116.38345 |
| VKVDVLR | 7 | Q92667 | A-kinase anchor protein 1, mitochondrial | AKAP1 | 97.342 | 2 | 54.74574 | yes | 9.938E-08 | 414.76871 | 105.22874 |
| ADPAIKEPLPVED' | 17 | Q92667 | A-kinase anchor protein 1, mitochondrial | AKAP1 | 97.342 | 3 | 54.74574 | yes | 1.289E-07 | 626.66077 | 97.339119 |
| TESFQNAQAGSNP | 14 | Q92667 | A-kinase anchor protein 1, mitochondrial | AKAP1 | 97.342 | 2 | 54.74574 | yes | 9.722E-08 | 739.84717 | 92.070862 |
| TYVSCLK | 7 | Q92667 | A-kinase anchor protein 1, mitochondrial | AKAP1 | 97.342 | 2 | 54.74574 | yes | 6.84E-07 | 435.72311 | 85.244003 |
| QDSPFSR | 7 | Q92667 | A-kinase anchor protein 1, mitochondrial | AKAP1 | 97.342 | 2 | 54.74574 | yes | 9.887E-08 | 418.69849 | 83.854172 |
| EMSLSDCLNR | 10 | Q92674 | Centromere protein I OS=Homo sapiens ( | CENPI | 86.721 | 2 | 40.286087 | yes | 4.108E-07 | 612.771 | 24.315479 |
| TGESILSLLR | 10 | Q92685 | Dol-P-Man:Man(5)GlcNAc(2)-PP-Dol al | ALG3 | 50.127 | 2 | 39.903057 | yes | 4.318E-07 | 544.81915 | 135.82689 |
| VHSIFVLR | 8 | Q92685 | Dol-P-Man:Man(5)GlcNAc(2)-PP-Dol al | ALG3 | 50.127 | 2 | 39.903057 | yes | 7.644E-07 | 485.79526 | 79.423798 |
| LLPQLTYLDGYDR | 13 | Q92688 | Acidic leucine-rich nuclear phosphoprote | ANP32B | 28.788 | 2 | 45.75946 | no | 1.767E-07 | 783.91174 | 515.7558 |
| IFGGLDMLAEK | 11 | Q92688 | Acidic leucine-rich nuclear phosphoprote | ANP32B | 28.788 | 2 | 45.75946 | yes | 1.749E-07 | 597.31537 | 275.76245 |
| DISTLEPLKK | 10 | Q92688 | Acidic leucine-rich nuclear phosphoprote | ANP32B | 28.788 | 2 | 45.75946 | yes | 3.283E-07 | 572.33459 | 68.906708 |
| FLWECVTSR | 9 | Q92698 | DNA repair and recombination protein R/ | RAD54L | 84.352 | 2 | 45.27809 | yes | 1.899E-07 | 599.28967 | 47.287827 |
| KLPSGPQDSSR | 11 | Q92738 | USP6 N-terminal-like protein OS=Homo | USP6NL | 94.105 | 2 | 43.688465 | yes | 3.195E-07 | 586.30695 | 80.458855 |
| PSPLVLPSSR | 10 | Q92738 | USP6 N-terminal-like protein OS=Homo | USP6NL | 94.105 | 2 | 43.688465 | yes | 6.299E-07 | 526.80859 | 48.443146 |
| AAHPALAVTVPGI | 17 | Q92738 | USP6 N-terminal-like protein OS=Homo | USP6NL | 94.105 | 2 | 43.688465 | yes | 2.487E-07 | 828.46503 | 37.099236 |
| AAHVTLLTSLVEG | 20 | Q92754 | Transcription factor AP-2 gamma OS=Ho | TFAP2C | 49.177 | 3 | 51.92997 | yes | 1.271E-07 | 696.39178 | 1471.0211 |
| NPLNLPCQK | 9 | Q92754 | Transcription factor AP-2 gamma OS=Ho | TFAP2C | 49.177 | 2 | 51.92997 | yes | 2.287E-07 | 542.28442 | 989.63885 |
| EFTELLSQDR | 10 | Q92754 | Transcription factor AP-2 gamma OS=Ho | TFAP2C | 49.177 | 2 | 51.92997 | yes | 9.81E-08 | 619.30658 | 761.64124 |
| PAGLLPHLSGLEA( | 19 | Q92754 | Transcription factor AP-2 gamma OS=Ho | TFAP2C | 49.177 | 3 | 51.92997 | yes | 4.381E-07 | 606.00739 | 682.32611 |
| KNMLLAAQQLCK | 12 | Q92754 | Transcription factor AP-2 gamma OS=Ho | TFAP2C | 49.177 | 2 | 51.92997 | yes | 9.722E-08 | 709.38635 | 80.524956 |
| ELGVLVFENSAK | 12 | Q92759 | General transcription factor IIH subunit 4 | GTF2H4 | 52.187 | 2 | 51.271854 | yes | 1.481E-07 | 653.35608 | 164.4239 |
| FPNMVVAQVTR | 11 | Q92759 | General transcription factor IIH subunit 4 | GTF2H4 | 52.187 | 2 | 51.271854 | yes | 1.155E-07 | 631.33972 | 145.25958 |
| AQEESTGLLSGLR | 13 | Q92759 | General transcription factor IIH subunit 4 | GTF2H4 | 52.187 | 2 | 51.271854 | yes | 9.733E-08 | 680.85699 | 132.49071 |
| QTPVLPPTITDQIR | 14 | Q92759 | General transcription factor IIH subunit 4 | GTF2H4 | 52.187 | 2 | 51.271854 | yes | 1.206E-07 | 789.94611 | 101.03017 |

| | | | | | | | | | | | |
|---|---|---|---|---|---|---|---|---|---|---|---|
| AWSDDTSQLGPDł | 13 | Q92759 | General transcription factor IIH subunit 4 | GTF2H4 | 52.187 | 2 | 51.271854 | yes | 9.722E-08 | 710.323 | 77.976784 |
| EFGLVFQR | 8 | Q92759 | General transcription factor IIH subunit 4 | GTF2H4 | 52.187 | 2 | 51.271854 | yes | 2.374E-07 | 498.26907 | 69.686501 |
| ESVQQAIASGITAC | 21 | Q92759 | General transcription factor IIH subunit 4 | GTF2H4 | 52.187 | 3 | 51.271854 | yes | 7.579E-07 | 770.75702 | 22.405983 |
| VFPWASSLQR | 10 | Q92766 | Ras-responsive element-binding protein 1 | RREB1 | 181.421 | 2 | 39.808502 | yes | 0.0003145 | 595.81946 | 258.16763 |
| APAAPLQAIFK | 11 | Q92766 | Ras-responsive element-binding protein 1 | RREB1 | 181.421 | 2 | 39.808502 | yes | 4.369E-07 | 563.83459 | 51.200409 |
| CDICCVTFR | 9 | Q92766 | Ras-responsive element-binding protein 1 | RREB1 | 181.421 | 2 | 39.808502 | yes | 1.227E-06 | 615.75684 | 25.847895 |
| LGCFNLTVK | 9 | Q92769 | Histone deacetylase 2 OS=Homo sapiens | HDAC2 | 55.365 | 2 | 52.894455 | yes | 1.182E-07 | 526.28387 | 446.73642 |
| YYAVNFPMR | 9 | Q92769 | Histone deacetylase 2 OS=Homo sapiens | HDAC2 | 55.365 | 2 | 52.894455 | yes | 9.722E-08 | 580.78149 | 344.05664 |
| QQTDMAVNWAG( | 17 | Q92769 | Histone deacetylase 2 OS=Homo sapiens | HDAC2 | 55.365 | 3 | 52.894455 | yes | 1.113E-07 | 621.96881 | 172.00661 |
| DGIDDESYGQIFKI | 18 | Q92769 | Histone deacetylase 2 OS=Homo sapiens | HDAC2 | 55.365 | 3 | 52.894455 | yes | 1.93E-07 | 675.67737 | 154.57347 |
| DGIDDESYGQIFK | 13 | Q92769 | Histone deacetylase 2 OS=Homo sapiens | HDAC2 | 55.365 | 2 | 52.894455 | yes | 5.356E-05 | 743.83844 | 55.182991 |
| YHSDEYIK | 8 | Q92769 | Histone deacetylase 2 OS=Homo sapiens | HDAC2 | 55.365 | 2 | 52.894455 | yes | 7.456E-05 | 527.74567 | 22.607971 |
| GPGLASGQLYSYP | 15 | Q92785 | Zinc finger protein ubi-d4 OS=Homo sap | DPF2 | 44.156 | 2 | 49.106724 | yes | 7.115E-07 | 768.89392 | 108.80529 |
| LDASILEDR | 9 | Q92785 | Zinc finger protein ubi-d4 OS=Homo sap | DPF2 | 44.156 | 2 | 49.106724 | yes | 1.201E-07 | 516.27203 | 105.39365 |
| VDDDSLGEFPVTN | 15 | Q92785 | Zinc finger protein ubi-d4 OS=Homo sap | DPF2 | 44.156 | 2 | 49.106724 | yes | 2.749E-07 | 825.88391 | 79.184135 |
| DKPYACDICGK | 11 | Q92785 | Zinc finger protein ubi-d4 OS=Homo sap | DPF2 | 44.156 | 2 | 49.106724 | yes | 1.067E-07 | 663.79449 | 49.985901 |
| AVAAAYTFLQR | 11 | Q92791 | Endoplasmic reticulum protein SC65 OS= | P3H4 | 50.382 | 2 | 45.63461 | yes | 1.79E-07 | 605.83258 | 32.235851 |
| TGKPMINLYTDK | 12 | Q92804 | TATA-binding protein-associated factor 2 | TAF15 | 61.83 | 2 | 52.43798 | yes | 1.66E-07 | 690.86322 | 12588.248 |
| TGKPMINLYTDK | 12 | Q92804 | TATA-binding protein-associated factor 2 | TAF15 | 61.83 | 2 | 52.43798 | yes | 2.064E-07 | 698.86066 | 12588.248 |
| PMINLYTDK | 9 | Q92804 | TATA-binding protein-associated factor 2 | TAF15 | 61.83 | 2 | 52.43798 | yes | 1.767E-07 | 547.78119 | 5655.5347 |
| PMINLYTDK | 9 | Q92804 | TATA-binding protein-associated factor 2 | TAF15 | 61.83 | 2 | 52.43798 | yes | 9.722E-08 | 555.77863 | 5655.5347 |
| SGGGYGGDR | 9 | Q92804 | TATA-binding protein-associated factor 2 | TAF15 | 61.83 | 2 | 52.43798 | yes | 2.53E-07 | 413.17792 | 5146.438 |
| EFHGNIIK | 8 | Q92804 | TATA-binding protein-associated factor 2 | TAF15 | 61.83 | 2 | 52.43798 | yes | 1.528E-07 | 479.26126 | 4242.8652 |
| KTGKPMINLYTDK | 13 | Q92804 | TATA-binding protein-associated factor 2 | TAF15 | 61.83 | 3 | 52.43798 | yes | 1.318E-07 | 503.60956 | 970.42108 |
| KTGKPMINLYTDK | 13 | Q92804 | TATA-binding protein-associated factor 2 | TAF15 | 61.83 | 3 | 52.43798 | yes | 9.722E-08 | 754.91071 | 970.42108 |
| KTGKPMINLYTDK | 13 | Q92804 | TATA-binding protein-associated factor 2 | TAF15 | 61.83 | 3 | 52.43798 | yes | 1.283E-07 | 508.94119 | 970.42108 |
| KTGKPMINLYTDK | 13 | Q92804 | TATA-binding protein-associated factor 2 | TAF15 | 61.83 | 2 | 52.43798 | yes | 1.318E-07 | 503.60956 | 970.42108 |
| KTGKPMINLYTDK | 13 | Q92804 | TATA-binding protein-associated factor 2 | TAF15 | 61.83 | 2 | 52.43798 | yes | 9.722E-08 | 754.91071 | 970.42108 |
| KTGKPMINLYTDK | 13 | Q92804 | TATA-binding protein-associated factor 2 | TAF15 | 61.83 | 2 | 52.43798 | yes | 1.283E-07 | 508.94119 | 970.42108 |
| NSCNQCNEPR | 10 | Q92804 | TATA-binding protein-associated factor 2 | TAF15 | 61.83 | 2 | 52.43798 | yes | 2.83E-07 | 639.75116 | 187.95369 |
| TGKPMINLYTDKD | 16 | Q92804 | TATA-binding protein-associated factor 2 | TAF15 | 61.83 | 3 | 52.43798 | yes | 1.747E-07 | 594.6416 | 183.34006 |
| GYGGSQGGGR | 10 | Q92804 | TATA-binding protein-associated factor 2 | TAF15 | 61.83 | 2 | 52.43798 | yes | 4.574E-07 | 448.20447 | 122.92972 |
| PKGEATVSFDDPP | 16 | Q92804 | TATA-binding protein-associated factor 2 | TAF15 | 61.83 | 2 | 52.43798 | yes | 2.679E-07 | 823.40704 | 38.898849 |
| PKGEATVSFDDPP | 16 | Q92804 | TATA-binding protein-associated factor 2 | TAF15 | 61.83 | 2 | 52.43798 | yes | 8.084E-05 | 549.2738 | 38.898849 |
| PKGEATVSFDDPP | 16 | Q92804 | TATA-binding protein-associated factor 2 | TAF15 | 61.83 | 3 | 52.43798 | yes | 2.679E-07 | 823.40704 | 38.898849 |
| PKGEATVSFDDPP | 16 | Q92804 | TATA-binding protein-associated factor 2 | TAF15 | 61.83 | 3 | 52.43798 | yes | 8.084E-05 | 549.2738 | 38.898849 |
| GPMTGSSGGDR | 11 | Q92804 | TATA-binding protein-associated factor 2 | TAF15 | 61.83 | 2 | 52.43798 | yes | 9.722E-08 | 511.22202 | 55.482094 |
| PMINLYTDKDTGK | 13 | Q92804 | TATA-binding protein-associated factor 2 | TAF15 | 61.83 | 2 | 52.43798 | yes | 9.722E-08 | 748.37671 | 39.779533 |
| PMINLYTDKDTGK | 15 | Q92804 | TATA-binding protein-associated factor 2 | TAF15 | 61.83 | 4 | 52.43798 | yes | 3.589E-07 | 430.97891 | 34.644783 |

| | | | | | | | | | | | |
|---|---|---|---|---|---|---|---|---|---|---|---|
| VIGGICFR | 8 | Q92830 | Histone acetyltransferase KAT2A OS=Hc | KAT2A | 93.926 | 2 | 43.742947 | no | 3.57E-07 | 461.25238 | 170.66248 |
| TLILTHFPK | 9 | Q92830 | Histone acetyltransferase KAT2A OS=Hc | KAT2A | 93.926 | 2 | 43.742947 | no | 3.613E-07 | 535.32385 | 80.29982 |
| KLFVADLQR | 9 | Q92830 | Histone acetyltransferase KAT2A OS=Hc | KAT2A | 93.926 | 2 | 43.742947 | yes | 2.465E-07 | 545.3244 | 59.343361 |
| ELAQQVQQVADD | 15 | Q92841 | Probable ATP-dependent RNA helicase D | DDX17 | 80.273 | 2 | 52.76398 | yes | 1.4E-07 | 846.41541 | 17832.604 |
| NFYVEHPEVAR | 11 | Q92841 | Probable ATP-dependent RNA helicase D | DDX17 | 80.273 | 2 | 52.76398 | yes | 1.736E-07 | 680.83588 | 8466.5137 |
| NFYVEHPEVAR | 11 | Q92841 | Probable ATP-dependent RNA helicase D | DDX17 | 80.273 | 2 | 52.76398 | yes | 1.223E-07 | 454.22632 | 8466.5137 |
| NFYVEHPEVAR | 11 | Q92841 | Probable ATP-dependent RNA helicase D | DDX17 | 80.273 | 3 | 52.76398 | yes | 1.736E-07 | 680.83588 | 8466.5137 |
| NFYVEHPEVAR | 11 | Q92841 | Probable ATP-dependent RNA helicase D | DDX17 | 80.273 | 3 | 52.76398 | yes | 1.223E-07 | 454.22632 | 8466.5137 |
| LMQLVDHR | 8 | Q92841 | Probable ATP-dependent RNA helicase D | DDX17 | 80.273 | 2 | 52.76398 | yes | 2.988E-07 | 506.27383 | 7824.2197 |
| LMQLVDHR | 8 | Q92841 | Probable ATP-dependent RNA helicase D | DDX17 | 80.273 | 2 | 52.76398 | yes | 1.806E-07 | 514.2713 | 7824.2197 |
| CTYLVLDEADR | 11 | Q92841 | Probable ATP-dependent RNA helicase D | DDX17 | 80.273 | 2 | 52.76398 | yes | 1.154E-07 | 677.81903 | 15327.298 |
| GTAYTFFTPGNLK | 13 | Q92841 | Probable ATP-dependent RNA helicase D | DDX17 | 80.273 | 2 | 52.76398 | yes | 1.392E-07 | 708.86151 | 15131.865 |
| VLEEANQAINPK | 12 | Q92841 | Probable ATP-dependent RNA helicase D | DDX17 | 80.273 | 2 | 52.76398 | yes | 1.585E-07 | 663.35663 | 14527.807 |
| DGWPAMCIHGDK | 12 | Q92841 | Probable ATP-dependent RNA helicase D | DDX17 | 80.273 | 2 | 52.76398 | yes | 2.215E-07 | 693.80011 | 14067.482 |
| DWVLNEFR | 8 | Q92841 | Probable ATP-dependent RNA helicase D | DDX17 | 80.273 | 2 | 52.76398 | yes | 1.859E-07 | 539.76947 | 10004.604 |
| DMVGIAQTGSGK | 12 | Q92841 | Probable ATP-dependent RNA helicase D | DDX17 | 80.273 | 2 | 52.76398 | yes | 1.022E-07 | 582.28992 | 9371.3408 |
| SSQSSSQQFSGIGR | 14 | Q92841 | Probable ATP-dependent RNA helicase D | DDX17 | 80.273 | 2 | 52.76398 | yes | 9.722E-08 | 728.34479 | 8242.1885 |
| WDLSELPK | 8 | Q92841 | Probable ATP-dependent RNA helicase D | DDX17 | 80.273 | 2 | 52.76398 | yes | 7.998E-07 | 494.26093 | 6835.8169 |
| APLPDLYPFGTMR | 13 | Q92841 | Probable ATP-dependent RNA helicase D | DDX17 | 80.273 | 2 | 52.76398 | yes | 1.125E-07 | 739.37903 | 2828.7998 |
| APLPDLYPFGTMR | 13 | Q92841 | Probable ATP-dependent RNA helicase D | DDX17 | 80.273 | 2 | 52.76398 | yes | 3.832E-06 | 747.37646 | 2828.7998 |
| LIQLMEEIMAEK | 12 | Q92841 | Probable ATP-dependent RNA helicase D | DDX17 | 80.273 | 2 | 52.76398 | yes | 9.81E-08 | 724.38037 | 2765.3748 |
| LIQLMEEIMAEK | 12 | Q92841 | Probable ATP-dependent RNA helicase D | DDX17 | 80.273 | 2 | 52.76398 | yes | 9.819E-08 | 732.37787 | 2765.3748 |
| KFGNPGER | 8 | Q92841 | Probable ATP-dependent RNA helicase D | DDX17 | 80.273 | 2 | 52.76398 | yes | 3.42E-07 | 452.73541 | 5447.9077 |
| LIDFLESGK | 9 | Q92841 | Probable ATP-dependent RNA helicase D | DDX17 | 80.273 | 2 | 52.76398 | yes | 1.423E-07 | 511.28186 | 4175.436 |
| QLAEDFLR | 8 | Q92841 | Probable ATP-dependent RNA helicase D | DDX17 | 80.273 | 2 | 52.76398 | yes | 2.249E-07 | 496.26401 | 4165.4131 |
| KWDLSELPK | 9 | Q92841 | Probable ATP-dependent RNA helicase D | DDX17 | 80.273 | 2 | 52.76398 | yes | 1.091E-07 | 558.30841 | 3010.1003 |
| TLAYLLPAIVHINH | 20 | Q92841 | Probable ATP-dependent RNA helicase D | DDX17 | 80.273 | 3 | 52.76398 | yes | 1.684E-07 | 787.77411 | 2652.8271 |
| TIIFVETK | 8 | Q92841 | Probable ATP-dependent RNA helicase D | DDX17 | 80.273 | 2 | 52.76398 | yes | 1.912E-07 | 475.78149 | 2352.6895 |
| LTPYEVDELR | 10 | Q92841 | Probable ATP-dependent RNA helicase D | DDX17 | 80.273 | 2 | 52.76398 | yes | 1.283E-07 | 617.81934 | 1886.2062 |
| LTPYEVDELRR | 11 | Q92841 | Probable ATP-dependent RNA helicase D | DDX17 | 80.273 | 2 | 52.76398 | yes | 2.862E-06 | 695.86987 | 755.58014 |
| ESAAPAAAPTAEA | 33 | Q92841 | Probable ATP-dependent RNA helicase D | DDX17 | 80.273 | 4 | 52.76398 | yes | 1.634E-07 | 812.43341 | 347.82446 |
| RCDDLTR | 7 | Q92841 | Probable ATP-dependent RNA helicase D | DDX17 | 80.273 | 2 | 52.76398 | yes | 5.857E-07 | 468.2218 | 264.99625 |
| RCTYLVLDEADR | 12 | Q92841 | Probable ATP-dependent RNA helicase D | DDX17 | 80.273 | 2 | 52.76398 | yes | 1.346E-07 | 755.86957 | 177.46584 |
| RDGWPAMCIHGD | 13 | Q92841 | Probable ATP-dependent RNA helicase D | DDX17 | 80.273 | 3 | 52.76398 | yes | 5.198E-07 | 514.90283 | 79.11573 |
| RDGWPAMCIHGD | 13 | Q92841 | Probable ATP-dependent RNA helicase D | DDX17 | 80.273 | 3 | 52.76398 | yes | 9.403E-07 | 771.85065 | 79.11573 |
| RDGWPAMCIHGD | 13 | Q92841 | Probable ATP-dependent RNA helicase D | DDX17 | 80.273 | 2 | 52.76398 | yes | 5.198E-07 | 514.90283 | 79.11573 |
| RDGWPAMCIHGD | 13 | Q92841 | Probable ATP-dependent RNA helicase D | DDX17 | 80.273 | 2 | 52.76398 | yes | 9.403E-07 | 771.85065 | 79.11573 |
| KKWDLSELPK | 10 | Q92841 | Probable ATP-dependent RNA helicase D | DDX17 | 80.273 | 2 | 52.76398 | yes | 1.073E-07 | 622.3559 | 136.85651 |
| EAATVASATGDSA | 16 | Q92841 | Probable ATP-dependent RNA helicase D | DDX17 | 80.273 | 2 | 52.76398 | yes | 1.021E-07 | 761.8526 | 125.1964 |

| | | | | | | | | | | | |
|---|---|---|---|---|---|---|---|---|---|---|---|
| SLNANTDITSLAR | 13 | Q92845 | Kinesin-associated protein 3 OS=Homo s | KIFAP3 | 91.206 | 2 | 53.57362 | yes | 2.066E-07 | 688.36243 | 56.742767 |
| MLFECSDER | 9 | Q92845 | Kinesin-associated protein 3 OS=Homo s | KIFAP3 | 91.206 | 2 | 53.57362 | yes | 1.955E-07 | 593.74701 | 46.247932 |
| GGNIDVHPSEK | 11 | Q92845 | Kinesin-associated protein 3 OS=Homo s | KIFAP3 | 91.206 | 2 | 53.57362 | yes | 9.722E-08 | 576.78583 | 45.554928 |
| EQVSPLETTLEK | 12 | Q92878 | DNA repair protein RAD50 OS=Homo sa | RAD50 | 153.893 | 2 | 50.7552 | yes | 1.818E-07 | 687.36157 | 171.39439 |
| QKFDEIFSATR | 11 | Q92878 | DNA repair protein RAD50 OS=Homo sa | RAD50 | 153.893 | 2 | 50.7552 | yes | 3.834E-06 | 671.34351 | 166.02618 |
| VIAQLSECEK | 10 | Q92878 | DNA repair protein RAD50 OS=Homo sa | RAD50 | 153.893 | 2 | 50.7552 | yes | 1.682E-07 | 588.80011 | 121.27019 |
| LQGIDLDR | 8 | Q92878 | DNA repair protein RAD50 OS=Homo sa | RAD50 | 153.893 | 2 | 50.7552 | yes | 5.11E-07 | 465.25616 | 115.04433 |
| EMISSLGVSK | 10 | Q92878 | DNA repair protein RAD50 OS=Homo sa | RAD50 | 153.893 | 2 | 50.7552 | yes | 2.617E-07 | 525.77863 | 68.330559 |
| EMGQMQVLQMK | 11 | Q92878 | DNA repair protein RAD50 OS=Homo sa | RAD50 | 153.893 | 2 | 50.7552 | yes | 9.723E-08 | 661.81689 | 64.95977 |
| VLASLIIR | 8 | Q92878 | DNA repair protein RAD50 OS=Homo sa | RAD50 | 153.893 | 2 | 50.7552 | yes | 1.99E-07 | 442.80002 | 64.779259 |
| MEVISLQNEK | 10 | Q92878 | DNA repair protein RAD50 OS=Homo sa | RAD50 | 153.893 | 2 | 50.7552 | yes | 1.693E-07 | 595.80792 | 55.063641 |
| GQDIEYIEIR | 10 | Q92878 | DNA repair protein RAD50 OS=Homo sa | RAD50 | 153.893 | 2 | 50.7552 | yes | 3.992E-07 | 618.31696 | 53.544033 |
| TANQLMNDFAEK | 12 | Q92878 | DNA repair protein RAD50 OS=Homo sa | RAD50 | 153.893 | 2 | 50.7552 | yes | 3.39E-06 | 691.32446 | 31.355156 |
| DIENYIQDGK | 10 | Q92878 | DNA repair protein RAD50 OS=Homo sa | RAD50 | 153.893 | 2 | 50.7552 | yes | 2.844E-05 | 597.78552 | 23.571123 |
| MFVGQVPR | 8 | Q92879 | CUGBP Elav-like family member 1 OS=I | CELF1 | 52.064 | 2 | 51.525978 | yes | 2.105E-07 | 467.25238 | 4530.0078 |
| GCAFVTFTTR | 10 | Q92879 | CUGBP Elav-like family member 1 OS=I | CELF1 | 52.064 | 2 | 51.525978 | yes | 1.048E-07 | 580.28186 | 1794.0042 |
| VMFSSFGQIEECR | 13 | Q92879 | CUGBP Elav-like family member 1 OS=I | CELF1 | 52.064 | 2 | 51.525978 | yes | 9.722E-08 | 795.35797 | 1761.2025 |
| VLPGMHHPIQMK | 12 | Q92879 | CUGBP Elav-like family member 1 OS=I | CELF1 | 52.064 | 3 | 51.525978 | yes | 2.711E-07 | 463.24942 | 517.05328 |
| VLPGMHHPIQMK | 12 | Q92879 | CUGBP Elav-like family member 1 OS=I | CELF1 | 52.064 | 3 | 51.525978 | yes | 2.205E-07 | 694.37048 | 517.05328 |
| VLPGMHHPIQMK | 12 | Q92879 | CUGBP Elav-like family member 1 OS=I | CELF1 | 52.064 | 2 | 51.525978 | yes | 2.711E-07 | 463.24942 | 517.05328 |
| VLPGMHHPIQMK | 12 | Q92879 | CUGBP Elav-like family member 1 OS=I | CELF1 | 52.064 | 2 | 51.525978 | yes | 2.205E-07 | 694.37048 | 517.05328 |
| KLFIGMISK | 9 | Q92879 | CUGBP Elav-like family member 1 OS=I | CELF1 | 52.064 | 2 | 51.525978 | yes | 3.926E-07 | 518.81482 | 772.26758 |
| GCCFVTFYTR | 10 | Q92879 | CUGBP Elav-like family member 1 OS=I | CELF1 | 52.064 | 2 | 51.525978 | no | 1.007E-07 | 655.78644 | 579.21301 |
| VLPGMHHPIQMKI | 18 | Q92879 | CUGBP Elav-like family member 1 OS=I | CELF1 | 52.064 | 3 | 51.525978 | yes | 6.563E-07 | 672.34491 | 74.995155 |
| VLPGMHHPIQMKI | 18 | Q92879 | CUGBP Elav-like family member 1 OS=I | CELF1 | 52.064 | 3 | 51.525978 | yes | 4.517E-07 | 504.5105 | 74.995155 |
| VLPGMHHPIQMKI | 18 | Q92879 | CUGBP Elav-like family member 1 OS=I | CELF1 | 52.064 | 4 | 51.525978 | yes | 6.563E-07 | 672.34491 | 74.995155 |
| VLPGMHHPIQMKI | 18 | Q92879 | CUGBP Elav-like family member 1 OS=I | CELF1 | 52.064 | 4 | 51.525978 | yes | 4.517E-07 | 504.5105 | 74.995155 |
| KAALEAQNALHNI | 14 | Q92879 | CUGBP Elav-like family member 1 OS=I | CELF1 | 52.064 | 3 | 51.525978 | yes | 1.638E-07 | 513.60846 | 70.716354 |
| AMAQTAIK | 8 | Q92879 | CUGBP Elav-like family member 1 OS=I | CELF1 | 52.064 | 2 | 51.525978 | yes | 3.198E-07 | 417.23111 | 24.552326 |
| RQEVISELLVTEA | 17 | Q92888 | Rho guanine nucleotide exchange factor 1 | ARHGEF1 | 102.436 | 3 | 50.05594 | yes | 1.645E-07 | 650.69708 | 155.80368 |
| ADLISEDVQR | 10 | Q92888 | Rho guanine nucleotide exchange factor 1 | ARHGEF1 | 102.436 | 2 | 50.05594 | yes | 5.953E-07 | 573.29346 | 84.322853 |
| GLSSILDAAR | 10 | Q92888 | Rho guanine nucleotide exchange factor 1 | ARHGEF1 | 102.436 | 2 | 50.05594 | yes | 9.919E-08 | 501.78256 | 54.99789 |
| FVQEVVQSQQVA | 15 | Q92888 | Rho guanine nucleotide exchange factor 1 | ARHGEF1 | 102.436 | 2 | 50.05594 | yes | 1.009E-07 | 837.45209 | 42.27652 |
| VYFLIYGGSTEEQI | 14 | Q92889 | DNA repair endonuclease XPF OS=Hom | ERCC4 | 104.486 | 2 | 48.46982 | yes | 4.64E-07 | 831.41211 | 170.34831 |
| YNPGPQDFLLK | 11 | Q92889 | DNA repair endonuclease XPF OS=Hom | ERCC4 | 104.486 | 2 | 48.46982 | yes | 5.932E-07 | 646.33771 | 143.57936 |
| GTASADVSTDTR | 12 | Q92889 | DNA repair endonuclease XPF OS=Hom | ERCC4 | 104.486 | 2 | 48.46982 | yes | 1.856E-07 | 590.77563 | 138.90314 |
| SELPSLIHR | 9 | Q92889 | DNA repair endonuclease XPF OS=Hom | ERCC4 | 104.486 | 2 | 48.46982 | yes | 2.572E-06 | 526.2984 | 107.11758 |
| IEGVEHLPR | 9 | Q92889 | DNA repair endonuclease XPF OS=Hom | ERCC4 | 104.486 | 2 | 48.46982 | yes | 3.296E-06 | 525.29053 | 76.972351 |
| ILVVDFLTDR | 10 | Q92889 | DNA repair endonuclease XPF OS=Hom | ERCC4 | 104.486 | 2 | 48.46982 | yes | 1.144E-07 | 595.84259 | 55.05875 |

| | | | | | | | | | | | |
|---|---|---|---|---|---|---|---|---|---|---|---|
| FVAFSGEGQSLR | 12 | Q92890 | Ubiquitin recognition factor in ER-associ | UFD1 | 34.501 | 2 | 45.047314 | yes | 1.981E-07 | 649.33038 | 57.341042 |
| IIIQESALDYR | 11 | Q92896 | Golgi apparatus protein 1 OS=Homo sapi | GLG1 | 134.552 | 2 | 52.286026 | yes | 2.656E-07 | 660.86151 | 201.59682 |
| GYMVSCLVDHR | 11 | Q92896 | Golgi apparatus protein 1 OS=Homo sapi | GLG1 | 134.552 | 2 | 52.286026 | yes | 1.792E-07 | 668.81049 | 161.44077 |
| MTAIIFSDYR | 10 | Q92896 | Golgi apparatus protein 1 OS=Homo sapi | GLG1 | 134.552 | 2 | 52.286026 | yes | 9.05E-07 | 608.80518 | 155.63298 |
| VAELSSDDFHLDR | 13 | Q92896 | Golgi apparatus protein 1 OS=Homo sapi | GLG1 | 134.552 | 2 | 52.286026 | yes | 9.722E-08 | 752.35736 | 138.95178 |
| DDSELEGQVISCLI | 14 | Q92896 | Golgi apparatus protein 1 OS=Homo sapi | GLG1 | 134.552 | 2 | 52.286026 | yes | 1.331E-07 | 796.87726 | 135.32413 |
| ALNEACESVIQTA | 15 | Q92896 | Golgi apparatus protein 1 OS=Homo sapi | GLG1 | 134.552 | 2 | 52.286026 | yes | 7.996E-07 | 847.39783 | 99.320992 |
| LICGFMDDCK | 10 | Q92896 | Golgi apparatus protein 1 OS=Homo sapi | GLG1 | 134.552 | 2 | 52.286026 | yes | 9.738E-08 | 629.76691 | 78.67646 |
| FCHGILTK | 8 | Q92896 | Golgi apparatus protein 1 OS=Homo sapi | GLG1 | 134.552 | 2 | 52.286026 | yes | 7.821E-07 | 488.25766 | 61.808952 |
| QMSCLMEALEDK | 12 | Q92896 | Golgi apparatus protein 1 OS=Homo sapi | GLG1 | 134.552 | 2 | 52.286026 | yes | 4.938E-07 | 727.81982 | 48.495895 |
| QITQNTDYR | 9 | Q92896 | Golgi apparatus protein 1 OS=Homo sapi | GLG1 | 134.552 | 2 | 52.286026 | yes | 4.82E-07 | 569.77802 | 46.752651 |
| HTWSNNLAVLECI | 17 | Q92896 | Golgi apparatus protein 1 OS=Homo sapi | GLG1 | 134.552 | 3 | 52.286026 | yes | 7.596E-07 | 685.67407 | 40.434151 |
| LLELQYFISR | 10 | Q92896 | Golgi apparatus protein 1 OS=Homo sapi | GLG1 | 134.552 | 2 | 52.286026 | yes | 9.722E-08 | 641.36371 | 40.007488 |
| LSSDCEDQIR | 10 | Q92896 | Golgi apparatus protein 1 OS=Homo sapi | GLG1 | 134.552 | 2 | 52.286026 | yes | 4.739E-07 | 611.77203 | 20.325985 |
| EAIIPGSVYDR | 11 | Q92900 | Regulator of nonsense transcripts 1 OS=H | UPF1 | 124.346 | 2 | 52.936855 | yes | 2.164E-07 | 610.31952 | 23409.709 |
| ADSVVVLLCR | 10 | Q92900 | Regulator of nonsense transcripts 1 OS=H | UPF1 | 124.346 | 2 | 52.936855 | yes | 1.086E-07 | 566.31317 | 16606.039 |
| IPSEQEQLR | 9 | Q92900 | Regulator of nonsense transcripts 1 OS=H | UPF1 | 124.346 | 2 | 52.936855 | yes | 1.536E-07 | 550.29077 | 16167.239 |
| YGVIIVGNPK | 10 | Q92900 | Regulator of nonsense transcripts 1 OS=H | UPF1 | 124.346 | 2 | 52.936855 | yes | 1.68E-07 | 530.31348 | 14807.433 |
| EKDFIILSCVR | 11 | Q92900 | Regulator of nonsense transcripts 1 OS=H | UPF1 | 124.346 | 2 | 52.936855 | yes | 9.722E-08 | 690.37122 | 7341.2354 |
| EKDFIILSCVR | 11 | Q92900 | Regulator of nonsense transcripts 1 OS=H | UPF1 | 124.346 | 2 | 52.936855 | yes | 1.783E-06 | 460.58322 | 7341.2354 |
| EKDFIILSCVR | 11 | Q92900 | Regulator of nonsense transcripts 1 OS=H | UPF1 | 124.346 | 3 | 52.936855 | yes | 9.722E-08 | 690.37122 | 7341.2354 |
| EKDFIILSCVR | 11 | Q92900 | Regulator of nonsense transcripts 1 OS=H | UPF1 | 124.346 | 3 | 52.936855 | yes | 1.783E-06 | 460.58322 | 7341.2354 |
| VLVEGPLNNLR | 11 | Q92900 | Regulator of nonsense transcripts 1 OS=H | UPF1 | 124.346 | 2 | 52.936855 | yes | 9.722E-08 | 612.35895 | 14440.675 |
| LMQGDEICLR | 10 | Q92900 | Regulator of nonsense transcripts 1 OS=H | UPF1 | 124.346 | 2 | 52.936855 | yes | 9.849E-08 | 617.79956 | 6336.293 |
| LMQGDEICLR | 10 | Q92900 | Regulator of nonsense transcripts 1 OS=H | UPF1 | 124.346 | 2 | 52.936855 | yes | 1.455E-07 | 625.797 | 6336.293 |
| AYQHGGVTGLSQ | 13 | Q92900 | Regulator of nonsense transcripts 1 OS=H | UPF1 | 124.346 | 2 | 52.936855 | yes | 2.633E-07 | 690.83075 | 8863.3574 |
| AGLSQSLFER | 10 | Q92900 | Regulator of nonsense transcripts 1 OS=H | UPF1 | 124.346 | 2 | 52.936855 | yes | 1.347E-07 | 554.29327 | 8449.9004 |
| PLSLIQGPPGTGK | 13 | Q92900 | Regulator of nonsense transcripts 1 OS=H | UPF1 | 124.346 | 2 | 52.936855 | yes | 1.151E-07 | 632.86664 | 8289.0029 |
| VPDNYGDEIAIELI | 14 | Q92900 | Regulator of nonsense transcripts 1 OS=H | UPF1 | 124.346 | 2 | 52.936855 | yes | 3.041E-07 | 802.40173 | 8077.21 |
| ESQTQDNITVR | 11 | Q92900 | Regulator of nonsense transcripts 1 OS=H | UPF1 | 124.346 | 2 | 52.936855 | yes | 1.231E-07 | 645.81787 | 7333.3691 |
| YKGDLAPLWK | 10 | Q92900 | Regulator of nonsense transcripts 1 OS=H | UPF1 | 124.346 | 2 | 52.936855 | yes | 1.35E-07 | 595.83203 | 6151.855 |
| LVNTINPGAR | 10 | Q92900 | Regulator of nonsense transcripts 1 OS=H | UPF1 | 124.346 | 2 | 52.936855 | yes | 1.969E-07 | 527.80383 | 6127.8408 |
| DETGELSSADEK | 12 | Q92900 | Regulator of nonsense transcripts 1 OS=H | UPF1 | 124.346 | 2 | 52.936855 | yes | 1.149E-07 | 640.77808 | 5615.354 |
| FMTTAMYDAR | 10 | Q92900 | Regulator of nonsense transcripts 1 OS=H | UPF1 | 124.346 | 2 | 52.936855 | yes | 1.272E-07 | 603.76776 | 1601.2526 |
| FMTTAMYDAR | 10 | Q92900 | Regulator of nonsense transcripts 1 OS=H | UPF1 | 124.346 | 2 | 52.936855 | yes | 1.238E-07 | 611.7652 | 1601.2526 |
| FMTTAMYDAR | 10 | Q92900 | Regulator of nonsense transcripts 1 OS=H | UPF1 | 124.346 | 2 | 52.936855 | yes | 9.974E-08 | 611.7652 | 1601.2526 |
| IAYFTLPK | 8 | Q92900 | Regulator of nonsense transcripts 1 OS=H | UPF1 | 124.346 | 2 | 52.936855 | yes | 3.012E-07 | 476.77875 | 4688.3452 |
| QPCASQSSLK | 10 | Q92900 | Regulator of nonsense transcripts 1 OS=H | UPF1 | 124.346 | 2 | 52.936855 | yes | 8.174E-07 | 553.26898 | 4352.4517 |
| LLGHEVEDVIIK | 12 | Q92900 | Regulator of nonsense transcripts 1 OS=H | UPF1 | 124.346 | 2 | 52.936855 | yes | 2.639E-07 | 682.89282 | 3804.4375 |

| | | | | | | | | | | | |
|---|---|---|---|---|---|---|---|---|---|---|---|
| KLVNTINPGAR | 11 | Q92900 | Regulator of nonsense transcripts 1 OS=H | UPF1 | 124.346 | 2 | 52.936855 | yes | 1.199E-07 | 591.85132 | 3670.3936 |
| LEADYDKK | 8 | Q92900 | Regulator of nonsense transcripts 1 OS=H | UPF1 | 124.346 | 2 | 52.936855 | yes | 1.197E-07 | 491.24802 | 3474.283 |
| ANEHQGIGFLNDP | 14 | Q92900 | Regulator of nonsense transcripts 1 OS=H | UPF1 | 124.346 | 2 | 52.936855 | yes | 1.221E-07 | 784.38422 | 3165.97 |
| EAIDSPVSFLALHN | 17 | Q92900 | Regulator of nonsense transcripts 1 OS=H | UPF1 | 124.346 | 3 | 52.936855 | yes | 1.206E-07 | 637.34253 | 2823.9749 |
| GNTSGSHIVNHLV | 14 | Q92900 | Regulator of nonsense transcripts 1 OS=H | UPF1 | 124.346 | 2 | 52.936855 | yes | 9.722E-08 | 745.89478 | 1756.1573 |
| FTAQGLPDLNHSQ | 18 | Q92900 | Regulator of nonsense transcripts 1 OS=H | UPF1 | 124.346 | 3 | 52.936855 | yes | 1.777E-07 | 663.34607 | 1133.1727 |
| TFAVDETSVSGYI | 16 | Q92900 | Regulator of nonsense transcripts 1 OS=H | UPF1 | 124.346 | 3 | 52.936855 | yes | 9.722E-08 | 606.29657 | 1092.9747 |
| NMDSMPELQK | 10 | Q92900 | Regulator of nonsense transcripts 1 OS=H | UPF1 | 124.346 | 2 | 52.936855 | yes | 2.311E-07 | 596.77045 | 1058.2692 |
| LYQEVEIASVDAF | 16 | Q92900 | Regulator of nonsense transcripts 1 OS=H | UPF1 | 124.346 | 3 | 52.936855 | yes | 9.776E-08 | 608.97571 | 949.09277 |
| DFIILSCVR | 9 | Q92900 | Regulator of nonsense transcripts 1 OS=H | UPF1 | 124.346 | 2 | 52.936855 | yes | 1.71E-07 | 561.80243 | 793.33582 |
| AGAKPDQIGIITPY | 18 | Q92900 | Regulator of nonsense transcripts 1 OS=H | UPF1 | 124.346 | 3 | 52.936855 | yes | 1.345E-07 | 638.67419 | 784.30518 |
| DETGELSSADEKR | 13 | Q92900 | Regulator of nonsense transcripts 1 OS=H | UPF1 | 124.346 | 2 | 52.936855 | yes | 9.722E-08 | 718.82861 | 704.9967 |
| QITAQQINK | 9 | Q92900 | Regulator of nonsense transcripts 1 OS=H | UPF1 | 124.346 | 2 | 52.936855 | yes | 6.587E-07 | 522.29584 | 594.9812 |
| ENPSATLEDLEKP | 25 | Q92900 | Regulator of nonsense transcripts 1 OS=H | UPF1 | 124.346 | 4 | 52.936855 | yes | 2.153E-07 | 704.60815 | 544.51196 |
| CFLSWLVK | 8 | Q92900 | Regulator of nonsense transcripts 1 OS=H | UPF1 | 124.346 | 2 | 52.936855 | yes | 1.674E-07 | 526.78351 | 456.23984 |
| QPLWNHLLNYYK | 12 | Q92900 | Regulator of nonsense transcripts 1 OS=H | UPF1 | 124.346 | 2 | 52.936855 | yes | 4.627E-07 | 794.91718 | 407.31586 |
| RFTAQGLPDLNHS | 19 | Q92900 | Regulator of nonsense transcripts 1 OS=H | UPF1 | 124.346 | 3 | 52.936855 | yes | 1.011E-07 | 715.37976 | 152.46307 |
| RFTAQGLPDLNHS | 19 | Q92900 | Regulator of nonsense transcripts 1 OS=H | UPF1 | 124.346 | 3 | 52.936855 | yes | 1.151E-06 | 536.78662 | 152.46307 |
| RFTAQGLPDLNHS | 19 | Q92900 | Regulator of nonsense transcripts 1 OS=H | UPF1 | 124.346 | 4 | 52.936855 | yes | 1.011E-07 | 715.37976 | 152.46307 |
| RFTAQGLPDLNHS | 19 | Q92900 | Regulator of nonsense transcripts 1 OS=H | UPF1 | 124.346 | 4 | 52.936855 | yes | 1.151E-06 | 536.78662 | 152.46307 |
| TVTSATIVYHLAR | 13 | Q92900 | Regulator of nonsense transcripts 1 OS=H | UPF1 | 124.346 | 3 | 52.936855 | yes | 4.69E-07 | 477.93668 | 147.96533 |
| TVTSATIVYHLAR | 13 | Q92900 | Regulator of nonsense transcripts 1 OS=H | UPF1 | 124.346 | 3 | 52.936855 | yes | 2.845E-06 | 716.40137 | 147.96533 |
| TVTSATIVYHLAR | 13 | Q92900 | Regulator of nonsense transcripts 1 OS=H | UPF1 | 124.346 | 2 | 52.936855 | yes | 4.69E-07 | 477.93668 | 147.96533 |
| TVTSATIVYHLAR | 13 | Q92900 | Regulator of nonsense transcripts 1 OS=H | UPF1 | 124.346 | 2 | 52.936855 | yes | 2.845E-06 | 716.40137 | 147.96533 |
| ESLMQFSK | 8 | Q92900 | Regulator of nonsense transcripts 1 OS=H | UPF1 | 124.346 | 2 | 52.936855 | yes | 2.946E-07 | 485.23914 | 237.31027 |
| LKESQTQDNITVR | 13 | Q92900 | Regulator of nonsense transcripts 1 OS=H | UPF1 | 124.346 | 2 | 52.936855 | yes | 1.034E-07 | 766.40735 | 117.2793 |
| LKESQTQDNITVR | 13 | Q92900 | Regulator of nonsense transcripts 1 OS=H | UPF1 | 124.346 | 2 | 52.936855 | yes | 3.923E-07 | 511.27402 | 117.2793 |
| LKESQTQDNITVR | 13 | Q92900 | Regulator of nonsense transcripts 1 OS=H | UPF1 | 124.346 | 3 | 52.936855 | yes | 1.034E-07 | 766.40735 | 117.2793 |
| LKESQTQDNITVR | 13 | Q92900 | Regulator of nonsense transcripts 1 OS=H | UPF1 | 124.346 | 3 | 52.936855 | yes | 3.923E-07 | 511.27402 | 117.2793 |
| WDLGLNKK | 8 | Q92900 | Regulator of nonsense transcripts 1 OS=H | UPF1 | 124.346 | 2 | 52.936855 | yes | 6.385E-07 | 487.27692 | 196.38245 |
| GFDFQWPQPDK | 11 | Q92900 | Regulator of nonsense transcripts 1 OS=H | UPF1 | 124.346 | 2 | 52.936855 | yes | 9.722E-08 | 682.81714 | 147.96576 |
| SQIDVALSQDSTY | 17 | Q92900 | Regulator of nonsense transcripts 1 OS=H | UPF1 | 124.346 | 3 | 52.936855 | yes | 1.087E-07 | 632.9693 | 113.48936 |
| PDQIGIITPYEGQR | 14 | Q92900 | Regulator of nonsense transcripts 1 OS=H | UPF1 | 124.346 | 2 | 52.936855 | yes | 9.819E-08 | 793.91229 | 112.20072 |
| QLILVGDHCQLGP | 18 | Q92900 | Regulator of nonsense transcripts 1 OS=H | UPF1 | 124.346 | 3 | 52.936855 | yes | 2.45E-07 | 689.68976 | 27.88287 |
| LVVLGIRPIR | 10 | Q92900 | Regulator of nonsense transcripts 1 OS=H | UPF1 | 124.346 | 2 | 52.936855 | yes | 1.227E-06 | 568.38733 | 24.70282 |
| GPSADLLEDLSESI | 14 | Q92908 | Transcription factor GATA-6 OS=Homo s | GATA6 | 60.033 | 2 | 47.476677 | yes | 1.321E-07 | 744.86249 | 501.18307 |
| EPQYSSLSAAR | 11 | Q92908 | Transcription factor GATA-6 OS=Homo s | GATA6 | 60.033 | 2 | 47.476677 | yes | 1.385E-07 | 604.79895 | 387.19019 |
| VMVVLGPQTGR | 11 | Q92917 | G-patch domain and KOW motifs-contair | GPKOW | 52.229 | 2 | 36.08197 | yes | 7.423E-07 | 578.82898 | 56.257416 |
| GCTPSGEGADSEP | 14 | Q92917 | G-patch domain and KOW motifs-contair | GPKOW | 52.229 | 2 | 36.08197 | yes | 2.96E-05 | 710.29388 | 20.113377 |

| | | | | | | | | | | | |
|---|---|---|---|---|---|---|---|---|---|---|---|
| NVEMFMNIEK | 10 | Q92922 | SWI/SNF complex subunit SMARCC1 O | SMARCC1 | 122.868 | 2 | 52.565365 | yes | 1.693E-07 | 627.79651 | 358.05096 |
| LNPQEYLTSTACR | 13 | Q92922 | SWI/SNF complex subunit SMARCC1 O | SMARCC1 | 122.868 | 2 | 52.565365 | no | 3.119E-07 | 776.87482 | 330.25092 |
| HVTNPAFTK | 9 | Q92922 | SWI/SNF complex subunit SMARCC1 O | SMARCC1 | 122.868 | 2 | 52.565365 | yes | 2.69E-07 | 507.772 | 291.59677 |
| NFMIDTYR | 8 | Q92922 | SWI/SNF complex subunit SMARCC1 O | SMARCC1 | 122.868 | 2 | 52.565365 | no | 9.722E-08 | 530.25006 | 149.9978 |
| VREEVPLELVEAH | 15 | Q92922 | SWI/SNF complex subunit SMARCC1 O | SMARCC1 | 122.868 | 3 | 52.565365 | yes | 4.093E-07 | 582.99646 | 137.67708 |
| TLVQNNCLTR | 10 | Q92922 | SWI/SNF complex subunit SMARCC1 O | SMARCC1 | 122.868 | 2 | 52.565365 | yes | 1.443E-07 | 609.81659 | 129.79555 |
| TQDECILHFLR | 11 | Q92922 | SWI/SNF complex subunit SMARCC1 O | SMARCC1 | 122.868 | 2 | 52.565365 | no | 9.722E-08 | 716.35608 | 97.362198 |
| EKPVDLQNFGLR | 12 | Q92922 | SWI/SNF complex subunit SMARCC1 O | SMARCC1 | 122.868 | 2 | 52.565365 | yes | 1.751E-07 | 708.38568 | 85.571968 |
| KVEHEISEGNVAT | 25 | Q92922 | SWI/SNF complex subunit SMARCC1 O | SMARCC1 | 122.868 | 3 | 52.565365 | yes | 1.266E-07 | 804.09058 | 84.903564 |
| YVHADAPTNK | 10 | Q92922 | SWI/SNF complex subunit SMARCC1 O | SMARCC1 | 122.868 | 2 | 52.565365 | yes | 3.53E-07 | 558.27765 | 75.285233 |
| NLTGDVCAVMR | 11 | Q92922 | SWI/SNF complex subunit SMARCC1 O | SMARCC1 | 122.868 | 2 | 52.565365 | yes | 2.176E-07 | 618.29718 | 65.890182 |
| HFEELETIMDR | 11 | Q92922 | SWI/SNF complex subunit SMARCC1 O | SMARCC1 | 122.868 | 2 | 52.565365 | no | 9.722E-08 | 710.33228 | 59.123535 |
| AALEEFSR | 8 | Q92922 | SWI/SNF complex subunit SMARCC1 O | SMARCC1 | 122.868 | 2 | 52.565365 | yes | 1.615E-06 | 461.73508 | 49.786442 |
| MPTTQETDGFQVK | 13 | Q92925 | SWI/SNF-related matrix-associated actin- | SMARCD2 | 58.921 | 2 | 49.955967 | yes | 9.968E-08 | 741.35065 | 41.477142 |
| MMLDDIVSR | 9 | Q92945 | Far upstream element-binding protein 2 C | KHSRP | 73.116 | 2 | 52.56835 | yes | 3.407E-07 | 540.26483 | 4152.7012 |
| VQISPDSGGLPER | 13 | Q92945 | Far upstream element-binding protein 2 C | KHSRP | 73.116 | 2 | 52.56835 | yes | 1.096E-07 | 677.85168 | 4106.4644 |
| IGGGIDVPVPR | 11 | Q92945 | Far upstream element-binding protein 2 C | KHSRP | 73.116 | 2 | 52.56835 | yes | 2.894E-07 | 540.31403 | 3547.8618 |
| IINDLLQSLR | 10 | Q92945 | Far upstream element-binding protein 2 C | KHSRP | 73.116 | 2 | 52.56835 | yes | 9.728E-08 | 592.85352 | 3236.2505 |
| SVSLTGAPESVQK | 13 | Q92945 | Far upstream element-binding protein 2 C | KHSRP | 73.116 | 2 | 52.56835 | yes | 9.722E-08 | 651.84863 | 2775.7493 |
| AINQQTGAFVEISF | 14 | Q92945 | Far upstream element-binding protein 2 C | KHSRP | 73.116 | 2 | 52.56835 | yes | 9.757E-08 | 767.4046 | 2396.8721 |
| IAHIMGPPDR | 10 | Q92945 | Far upstream element-binding protein 2 C | KHSRP | 73.116 | 2 | 52.56835 | yes | 2.457E-07 | 553.79242 | 2255.1846 |
| VPDGMVGLIIGR | 12 | Q92945 | Far upstream element-binding protein 2 C | KHSRP | 73.116 | 2 | 52.56835 | yes | 4.198E-07 | 613.84991 | 2084.7715 |
| MILIQDGSQNTNV | 18 | Q92945 | Far upstream element-binding protein 2 C | KHSRP | 73.116 | 3 | 52.56835 | yes | 1.181E-07 | 681.36151 | 1309.9642 |
| VQQACEMVMDIL | 13 | Q92945 | Far upstream element-binding protein 2 C | KHSRP | 73.116 | 2 | 52.56835 | yes | 9.722E-08 | 796.8833 | 1233.8247 |
| DAFADAVQR | 9 | Q92945 | Far upstream element-binding protein 2 C | KHSRP | 73.116 | 2 | 52.56835 | yes | 1.002E-07 | 496.74344 | 1210.6969 |
| HSVGVVIGR | 9 | Q92945 | Far upstream element-binding protein 2 C | KHSRP | 73.116 | 2 | 52.56835 | yes | 1.823E-07 | 462.27469 | 957.54468 |
| KLASQGDSISSQLC | 21 | Q92945 | Far upstream element-binding protein 2 C | KHSRP | 73.116 | 4 | 52.56835 | yes | 1.624E-07 | 547.04858 | 407.62662 |
| KLASQGDSISSQLC | 21 | Q92945 | Far upstream element-binding protein 2 C | KHSRP | 73.116 | 4 | 52.56835 | yes | 1.845E-07 | 729.06232 | 407.62662 |
| KLASQGDSISSQLC | 21 | Q92945 | Far upstream element-binding protein 2 C | KHSRP | 73.116 | 3 | 52.56835 | yes | 1.624E-07 | 547.04858 | 407.62662 |
| KLASQGDSISSQLC | 21 | Q92945 | Far upstream element-binding protein 2 C | KHSRP | 73.116 | 3 | 52.56835 | yes | 1.845E-07 | 729.06232 | 407.62662 |
| KDAFADAVQR | 10 | Q92945 | Far upstream element-binding protein 2 C | KHSRP | 73.116 | 2 | 52.56835 | yes | 1.313E-07 | 560.79089 | 519.86255 |
| LASQGDSISSQLGF | 20 | Q92945 | Far upstream element-binding protein 2 C | KHSRP | 73.116 | 3 | 52.56835 | yes | 1.048E-07 | 686.36401 | 509.67017 |
| QLEDGDQPESK | 11 | Q92945 | Far upstream element-binding protein 2 C | KHSRP | 73.116 | 2 | 52.56835 | yes | 9.722E-08 | 623.28333 | 493.98123 |
| MILIQDGSQNTNV | 15 | Q92945 | Far upstream element-binding protein 2 C | KHSRP | 73.116 | 2 | 52.56835 | yes | 1.125E-07 | 838.41962 | 482.76898 |
| IIGDPYK | 7 | Q92945 | Far upstream element-binding protein 2 C | KHSRP | 73.116 | 2 | 52.56835 | yes | 1.078E-07 | 403.22635 | 161.1886 |
| TSMTEEYR | 8 | Q92945 | Far upstream element-binding protein 2 C | KHSRP | 73.116 | 2 | 52.56835 | yes | 5.873E-07 | 508.72131 | 74.172585 |
| GSPQQIDHAK | 10 | Q92945 | Far upstream element-binding protein 2 C | KHSRP | 73.116 | 2 | 52.56835 | yes | 5.089E-07 | 540.77527 | 40.98616 |
| ERDQGGFGDR | 10 | Q92945 | Far upstream element-binding protein 2 C | KHSRP | 73.116 | 2 | 52.56835 | yes | 2.309E-06 | 568.75763 | 28.535915 |
| MQFGVPLAR | 9 | Q92947 | Glutaryl-CoA dehydrogenase, mitochondı | GCDH | 48.128 | 2 | 46.105556 | yes | 1.652E-07 | 509.77875 | 198.46127 |

| | | | | | | | | | | |
|---|---|---|---|---|---|---|---|---|---|---|
| AAPEMVSLLK | 10 | Q92947 | Glutaryl-CoA dehydrogenase, mitochondı | GCDH | 48.128 | 2 | 46.105556 | yes | 5.612E-07 | 529.79938 | 106.53748 |
| EPAPTTPK | 8 | Q92954 | Proteoglycan 4 OS=Homo sapiens OX=9 | PRG4 | 151.062 | 2 | 47.57328 | yes | 1.301E-07 | 420.72672 | 250.59146 |
| TIIEWSESHDR | 11 | Q92966 | snRNA-activating protein complex subun | SNAPC3 | 46.753 | 2 | 38.817272 | yes | 4.969E-07 | 686.82825 | 51.059517 |
| TLLENTAITIGR | 12 | Q92973 | Transportin-1 OS=Homo sapiens OX=96 | TNPO1 | 102.356 | 2 | 54.24531 | yes | 9.723E-08 | 651.37482 | 322.96844 |
| QSSFALLGDLTK | 12 | Q92973 | Transportin-1 OS=Homo sapiens OX=96 | TNPO1 | 102.356 | 2 | 54.24531 | no | 9.722E-08 | 640.34827 | 126.97652 |
| ESQSPDTTIQR | 11 | Q92973 | Transportin-1 OS=Homo sapiens OX=96 | TNPO1 | 102.356 | 2 | 54.24531 | yes | 5.317E-06 | 631.30457 | 90.958282 |
| SECLNNIGDSSPLI | 15 | Q92973 | Transportin-1 OS=Homo sapiens OX=96 | TNPO1 | 102.356 | 2 | 54.24531 | yes | 2.759E-07 | 837.90942 | 90.017082 |
| ATVGILITTIASK | 13 | Q92973 | Transportin-1 OS=Homo sapiens OX=96 | TNPO1 | 102.356 | 2 | 54.24531 | yes | 1.119E-06 | 644.39777 | 78.479645 |
| SITCWTLSR | 9 | Q92973 | Transportin-1 OS=Homo sapiens OX=96 | TNPO1 | 102.356 | 2 | 54.24531 | yes | 1.532E-07 | 562.28186 | 71.316429 |
| GELQNWPDLLPK | 12 | Q92973 | Transportin-1 OS=Homo sapiens OX=96 | TNPO1 | 102.356 | 2 | 54.24531 | yes | 1.001E-07 | 705.37482 | 67.33828 |
| FSDQFPLPLK | 10 | Q92973 | Transportin-1 OS=Homo sapiens OX=96 | TNPO1 | 102.356 | 2 | 54.24531 | yes | 1.306E-07 | 596.32404 | 36.02919 |
| YSDIDIILLK | 10 | Q92973 | Transportin-1 OS=Homo sapiens OX=96 | TNPO1 | 102.356 | 2 | 54.24531 | yes | 2.743E-07 | 596.84485 | 22.114578 |
| FLSQLLER | 8 | Q92974 | Rho guanine nucleotide exchange factor 2 | ARHGEF2 | 111.543 | 2 | 53.544666 | yes | 1.625E-07 | 503.29001 | 2086.8943 |
| MQDIPEETESR | 11 | Q92974 | Rho guanine nucleotide exchange factor 2 | ARHGEF2 | 111.543 | 2 | 53.544666 | yes | 1.455E-07 | 667.79828 | 876.84967 |
| MQDIPEETESR | 11 | Q92974 | Rho guanine nucleotide exchange factor 2 | ARHGEF2 | 111.543 | 2 | 53.544666 | yes | 4.588E-07 | 675.79572 | 876.84967 |
| NNTALQSVSLR | 11 | Q92974 | Rho guanine nucleotide exchange factor 2 | ARHGEF2 | 111.543 | 2 | 53.544666 | yes | 1.027E-07 | 601.828 | 1164.0684 |
| ILSQSTDSLNMR | 12 | Q92974 | Rho guanine nucleotide exchange factor 2 | ARHGEF2 | 111.543 | 2 | 53.544666 | yes | 1.4E-07 | 682.84558 | 1114.37 |
| QDLTTALGLVK | 11 | Q92974 | Rho guanine nucleotide exchange factor 2 | ARHGEF2 | 111.543 | 2 | 53.544666 | yes | 1.878E-07 | 579.84009 | 1039.7922 |
| LIHDGCLLWK | 10 | Q92974 | Rho guanine nucleotide exchange factor 2 | ARHGEF2 | 111.543 | 2 | 53.544666 | yes | 9.722E-08 | 627.83679 | 1010.7891 |
| YPLLISR | 7 | Q92974 | Rho guanine nucleotide exchange factor 2 | ARHGEF2 | 111.543 | 2 | 53.544666 | yes | 3.012E-07 | 431.26328 | 742.00494 |
| ATEAGSLEAR | 10 | Q92974 | Rho guanine nucleotide exchange factor 2 | ARHGEF2 | 111.543 | 2 | 53.544666 | yes | 1.021E-07 | 502.754 | 593.66498 |
| LDLPVTTR | 8 | Q92974 | Rho guanine nucleotide exchange factor 2 | ARHGEF2 | 111.543 | 2 | 53.544666 | yes | 1.163E-07 | 457.76892 | 592.57056 |
| QELGSPEER | 9 | Q92974 | Rho guanine nucleotide exchange factor 2 | ARHGEF2 | 111.543 | 2 | 53.544666 | yes | 1.12E-07 | 522.75146 | 383.09473 |
| TYSEFCSR | 8 | Q92974 | Rho guanine nucleotide exchange factor 2 | ARHGEF2 | 111.543 | 2 | 53.544666 | yes | 1.455E-07 | 525.2215 | 343.52759 |
| QATELALLQR | 10 | Q92974 | Rho guanine nucleotide exchange factor 2 | ARHGEF2 | 111.543 | 2 | 53.544666 | yes | 2.184E-07 | 571.83002 | 315.13202 |
| QALCPGSTR | 9 | Q92974 | Rho guanine nucleotide exchange factor 2 | ARHGEF2 | 111.543 | 2 | 53.544666 | yes | 2.902E-07 | 495.24527 | 313.90167 |
| VIQQSVR | 7 | Q92974 | Rho guanine nucleotide exchange factor 2 | ARHGEF2 | 111.543 | 2 | 53.544666 | yes | 1.06E-07 | 415.24814 | 305.68741 |
| QHALLQEELR | 10 | Q92974 | Rho guanine nucleotide exchange factor 2 | ARHGEF2 | 111.543 | 2 | 53.544666 | yes | 9.839E-08 | 618.83838 | 266.9866 |
| ILQHSHGIEEER | 12 | Q92974 | Rho guanine nucleotide exchange factor 2 | ARHGEF2 | 111.543 | 2 | 53.544666 | yes | 9.722E-08 | 724.36804 | 211.79121 |
| TFNGSIELCR | 10 | Q92974 | Rho guanine nucleotide exchange factor 2 | ARHGEF2 | 111.543 | 2 | 53.544666 | yes | 9.722E-08 | 598.79004 | 188.59006 |
| SESLESPR | 8 | Q92974 | Rho guanine nucleotide exchange factor 2 | ARHGEF2 | 111.543 | 2 | 53.544666 | yes | 1.046E-07 | 452.72217 | 128.02547 |
| HGVQECILLVTQR | 13 | Q92974 | Rho guanine nucleotide exchange factor 2 | ARHGEF2 | 111.543 | 2 | 53.544666 | yes | 5.929E-07 | 776.91687 | 79.719772 |
| DFAADSWSLAVDS | 20 | Q92974 | Rho guanine nucleotide exchange factor 2 | ARHGEF2 | 111.543 | 3 | 53.544666 | yes | 2.2E-07 | 751.35883 | 74.701653 |
| KVTRPAVLK | 9 | Q92974 | Rho guanine nucleotide exchange factor 2 | ARHGEF2 | 111.543 | 2 | 53.544666 | yes | 7.177E-07 | 506.33731 | 61.678413 |
| YIFPTLDKPSVVSL | 20 | Q92974 | Rho guanine nucleotide exchange factor 2 | ARHGEF2 | 111.543 | 3 | 53.544666 | yes | 1.584E-07 | 768.11047 | 22.574911 |
| ALVELLR | 7 | Q92974 | Rho guanine nucleotide exchange factor 2 | ARHGEF2 | 111.543 | 2 | 53.544666 | yes | 2.071E-06 | 407.26328 | 15.457449 |
| ERPSSAIYPSDSFR | 14 | Q92974 | Rho guanine nucleotide exchange factor 2 | ARHGEF2 | 111.543 | 3 | 53.544666 | yes | 3.945E-06 | 537.93024 | 8.8083735 |
| NVLIEVNPQTR | 11 | Q92979 | Ribosomal RNA small subunit methyltran | EMG1 | 26.721 | 2 | 52.451366 | yes | 1.248E-07 | 641.85931 | 4225.2275 |
| VGTSFSIPVVSDVI | 14 | Q92979 | Ribosomal RNA small subunit methyltran | EMG1 | 26.721 | 2 | 52.451366 | yes | 1.301E-07 | 731.89862 | 3185.333 |

| | | | | | | | | | | | |
|---|---|---|---|---|---|---|---|---|---|---|---|
| AAPSDGFKPR | 10 | Q92979 | Ribosomal RNA small subunit methyltran | EMG1 | 26.721 | 2 | 52.451366 | yes | 2.426E-07 | 544.28015 | 3038.4299 |
| FCGLMVQLLHK | 11 | Q92979 | Ribosomal RNA small subunit methyltran | EMG1 | 26.721 | 2 | 52.451366 | yes | 1.867E-07 | 673.35962 | 1155.1737 |
| TYELLNCDK | 9 | Q92979 | Ribosomal RNA small subunit methyltran | EMG1 | 26.721 | 2 | 52.451366 | yes | 1.684E-07 | 578.27118 | 1026.793 |
| NPVSDHFPVGCMI | 13 | Q92979 | Ribosomal RNA small subunit methyltran | EMG1 | 26.721 | 2 | 52.451366 | yes | 7.726E-07 | 744.3421 | 404.31201 |
| VSVEYTEK | 8 | Q92979 | Ribosomal RNA small subunit methyltran | EMG1 | 26.721 | 2 | 52.451366 | yes | 1.49E-07 | 477.74255 | 386.27911 |
| SGGEQAQDWDAL | 15 | Q92979 | Ribosomal RNA small subunit methyltran | EMG1 | 26.721 | 2 | 52.451366 | yes | 9.819E-08 | 799.87592 | 189.27798 |
| LIVVLEGASLETVI | 14 | Q92979 | Ribosomal RNA small subunit methyltran | EMG1 | 26.721 | 2 | 52.451366 | yes | 9.722E-08 | 735.9425 | 141.0175 |
| VMVVGPTDVGK | 11 | Q92989 | Polyribonucleotide 5'-hydroxyl-kinase Cl | CLP1 | 47.646 | 2 | 48.041206 | yes | 4.923E-07 | 551.30225 | 325.63095 |
| DLPHFVR | 7 | Q92989 | Polyribonucleotide 5'-hydroxyl-kinase Cl | CLP1 | 47.646 | 2 | 48.041206 | yes | 3.018E-07 | 442.24286 | 220.08513 |
| LLLNYAVR | 8 | Q92989 | Polyribonucleotide 5'-hydroxyl-kinase Cl | CLP1 | 47.646 | 2 | 48.041206 | yes | 1.491E-07 | 481.2951 | 144.61887 |
| ASVSGCVINTCGW | 15 | Q92989 | Polyribonucleotide 5'-hydroxyl-kinase Cl | CLP1 | 47.646 | 2 | 48.041206 | yes | 1.279E-07 | 819.39233 | 101.74371 |
| LADVFNQR | 8 | Q92989 | Polyribonucleotide 5'-hydroxyl-kinase Cl | CLP1 | 47.646 | 2 | 48.041206 | yes | 1.21E-07 | 481.75635 | 55.259464 |
| SHDDIVTR | 8 | Q92993 | Histone acetyltransferase KAT5 OS=Hon | KAT5 | 58.583 | 2 | 38.247513 | yes | 1.361E-06 | 471.7356 | 108.94221 |
| TGTPEKPLSDLGLI | 17 | Q92993 | Histone acetyltransferase KAT5 OS=Hon | KAT5 | 58.583 | 3 | 38.247513 | yes | 5.355E-07 | 616.33514 | 35.351391 |
| FAHLLEFGEK | 10 | Q92994 | Transcription factor IIIB 90 kDa subunit ( | BRF1 | 73.841 | 2 | 25.873682 | yes | 3.212E-06 | 595.81384 | 41.724312 |
| AMASPESGLEVR | 12 | Q92997 | Segment polarity protein dishevelled hom | DVL3 | 78.055 | 2 | 51.10427 | yes | 9.722E-08 | 623.80847 | 146.4377 |
| SHHSLASSLR | 10 | Q92997 | Segment polarity protein dishevelled hom | DVL3 | 78.055 | 2 | 51.10427 | yes | 2.397E-07 | 547.7887 | 68.072929 |
| SMDDDFGVVK | 10 | Q92997 | Segment polarity protein dishevelled hom | DVL3 | 78.055 | 2 | 51.10427 | yes | 2.579E-06 | 556.75006 | 43.882252 |
| EIVHKPGPITLTVA | 15 | Q92997 | Segment polarity protein dishevelled hom | DVL3 | 78.055 | 2 | 51.10427 | yes | 9.81E-08 | 801.98254 | 21.879631 |
| EIVHKPGPITLTVA | 15 | Q92997 | Segment polarity protein dishevelled hom | DVL3 | 78.055 | 2 | 51.10427 | yes | 1.325E-06 | 534.99078 | 21.879631 |
| EIVHKPGPITLTVA | 15 | Q92997 | Segment polarity protein dishevelled hom | DVL3 | 78.055 | 3 | 51.10427 | yes | 9.81E-08 | 801.98254 | 21.879631 |
| EIVHKPGPITLTVA | 15 | Q92997 | Segment polarity protein dishevelled hom | DVL3 | 78.055 | 3 | 51.10427 | yes | 1.325E-06 | 534.99078 | 21.879631 |
| NNFLPNADMETR | 12 | Q93008 | Probable ubiquitin carboxyl-terminal hyd | USP9X | 290.463 | 2 | 51.585155 | yes | 1.431E-07 | 711.32751 | 112.63937 |
| MDTIDQDDELIR | 12 | Q93008 | Probable ubiquitin carboxyl-terminal hyd | USP9X | 290.463 | 2 | 51.585155 | yes | 1.435E-07 | 732.33777 | 102.87915 |
| LAQQISDEASR | 11 | Q93008 | Probable ubiquitin carboxyl-terminal hyd | USP9X | 290.463 | 2 | 51.585155 | yes | 9.722E-08 | 609.30969 | 95.164207 |
| VVIQSNDDIASR | 12 | Q93008 | Probable ubiquitin carboxyl-terminal hyd | USP9X | 290.463 | 2 | 51.585155 | yes | 3.53E-07 | 658.84387 | 90.412025 |
| CLAENAVYLCDR | 12 | Q93008 | Probable ubiquitin carboxyl-terminal hyd | USP9X | 290.463 | 2 | 51.585155 | no | 1.83E-07 | 742.33704 | 88.323723 |
| LWGEPVNLR | 9 | Q93008 | Probable ubiquitin carboxyl-terminal hyd | USP9X | 290.463 | 2 | 51.585155 | no | 1.231E-07 | 542.3009 | 78.526054 |
| LYSVVSQLIR | 10 | Q93008 | Probable ubiquitin carboxyl-terminal hyd | USP9X | 290.463 | 2 | 51.585155 | no | 2.032E-07 | 589.35059 | 76.5606 |
| VISSVSYYTHR | 11 | Q93008 | Probable ubiquitin carboxyl-terminal hyd | USP9X | 290.463 | 2 | 51.585155 | no | 2.246E-07 | 656.33826 | 74.898727 |
| ELLAFQTSEK | 10 | Q93008 | Probable ubiquitin carboxyl-terminal hyd | USP9X | 290.463 | 2 | 51.585155 | no | 3.988E-06 | 583.30859 | 74.742874 |
| FLFTTGFHTK | 10 | Q93008 | Probable ubiquitin carboxyl-terminal hyd | USP9X | 290.463 | 2 | 51.585155 | no | 1.87E-07 | 599.81641 | 66.577698 |
| VLGGSFADQK | 10 | Q93008 | Probable ubiquitin carboxyl-terminal hyd | USP9X | 290.463 | 2 | 51.585155 | no | 4.038E-07 | 511.26929 | 62.566029 |
| ILDYSCSQDR | 10 | Q93008 | Probable ubiquitin carboxyl-terminal hyd | USP9X | 290.463 | 2 | 51.585155 | no | 4.204E-07 | 628.78241 | 50.959122 |
| SPHVFYR | 7 | Q93008 | Probable ubiquitin carboxyl-terminal hyd | USP9X | 290.463 | 2 | 51.585155 | yes | 3.441E-07 | 453.23505 | 49.487469 |
| WVVPVLPK | 8 | Q93008 | Probable ubiquitin carboxyl-terminal hyd | USP9X | 290.463 | 2 | 51.585155 | no | 9.722E-08 | 469.29712 | 46.62178 |
| IIEDCSNSEETVK | 13 | Q93008 | Probable ubiquitin carboxyl-terminal hyd | USP9X | 290.463 | 2 | 51.585155 | yes | 2.142E-07 | 762.34833 | 29.409708 |
| HSGDYFTLLR | 10 | Q93008 | Probable ubiquitin carboxyl-terminal hyd | USP9X | 290.463 | 2 | 51.585155 | yes | 4.203E-07 | 604.80658 | 19.004108 |
| FDDDVVSR | 8 | Q93009 | Ubiquitin carboxyl-terminal hydrolase 7 ( | USP7 | 128.303 | 2 | 55.038944 | yes | 1.12E-07 | 476.72217 | 356.75006 |

| | | | | | | | | | | | |
|---|---|---|---|---|---|---|---|---|---|---|---|
| LSESVLSPPCFVR | 13 | Q93009 | Ubiquitin carboxyl-terminal hydrolase 7 ( | USP7 | 128.303 | 2 | 55.038944 | yes | 3.451E-07 | 745.88721 | 343.4306 |
| FMYDPQTDQNIK | 12 | Q93009 | Ubiquitin carboxyl-terminal hydrolase 7 ( | USP7 | 128.303 | 2 | 55.038944 | yes | 2.044E-07 | 750.3454 | 303.51624 |
| TIPNDPGFVVTLSN | 15 | Q93009 | Ubiquitin carboxyl-terminal hydrolase 7 ( | USP7 | 128.303 | 2 | 55.038944 | yes | 2.386E-06 | 815.43341 | 219.11465 |
| IQDYDVSLDK | 10 | Q93009 | Ubiquitin carboxyl-terminal hydrolase 7 ( | USP7 | 128.303 | 2 | 55.038944 | yes | 1.26E-07 | 598.29572 | 211.81758 |
| VLLDNVENK | 9 | Q93009 | Ubiquitin carboxyl-terminal hydrolase 7 ( | USP7 | 128.303 | 2 | 55.038944 | yes | 2.407E-07 | 522.29022 | 209.44812 |
| HNYEGTLR | 8 | Q93009 | Ubiquitin carboxyl-terminal hydrolase 7 ( | USP7 | 128.303 | 2 | 55.038944 | yes | 4.415E-07 | 495.24359 | 174.61111 |
| GTCVEGTIPK | 10 | Q93009 | Ubiquitin carboxyl-terminal hydrolase 7 ( | USP7 | 128.303 | 2 | 55.038944 | yes | 9.722E-08 | 531.26843 | 162.77563 |
| SEATFQFTVER | 11 | Q93009 | Ubiquitin carboxyl-terminal hydrolase 7 ( | USP7 | 128.303 | 2 | 55.038944 | yes | 9.722E-08 | 657.81989 | 160.95845 |
| ISHLFFHK | 8 | Q93009 | Ubiquitin carboxyl-terminal hydrolase 7 ( | USP7 | 128.303 | 2 | 55.038944 | yes | 1.69E-07 | 514.78741 | 153.46092 |
| LLEIVSYK | 8 | Q93009 | Ubiquitin carboxyl-terminal hydrolase 7 ( | USP7 | 128.303 | 2 | 55.038944 | yes | 1.146E-07 | 482.78931 | 151.30835 |
| IEEIPLDQVDIDKE | 26 | Q93009 | Ubiquitin carboxyl-terminal hydrolase 7 ( | USP7 | 128.303 | 4 | 55.038944 | yes | 1.301E-07 | 766.39301 | 138.35428 |
| CIWLNSQFR | 9 | Q93009 | Ubiquitin carboxyl-terminal hydrolase 7 ( | USP7 | 128.303 | 2 | 55.038944 | yes | 2.585E-07 | 612.30316 | 138.34663 |
| SVPLALQR | 8 | Q93009 | Ubiquitin carboxyl-terminal hydrolase 7 ( | USP7 | 128.303 | 2 | 55.038944 | yes | 9.032E-07 | 442.27164 | 137.45027 |
| MNYFQVAK | 8 | Q93009 | Ubiquitin carboxyl-terminal hydrolase 7 ( | USP7 | 128.303 | 2 | 55.038944 | yes | 1.065E-07 | 500.74966 | 135.89505 |
| DLLPVMCDR | 9 | Q93009 | Ubiquitin carboxyl-terminal hydrolase 7 ( | USP7 | 128.303 | 2 | 55.038944 | yes | 9.722E-08 | 559.77026 | 130.25906 |
| KLYYQQLK | 8 | Q93009 | Ubiquitin carboxyl-terminal hydrolase 7 ( | USP7 | 128.303 | 2 | 55.038944 | yes | 6.996E-07 | 542.31348 | 123.27033 |
| IQSLLDIQEK | 10 | Q93009 | Ubiquitin carboxyl-terminal hydrolase 7 ( | USP7 | 128.303 | 2 | 55.038944 | yes | 9.722E-08 | 593.83752 | 120.17699 |
| EDYYDIQLSIK | 11 | Q93009 | Ubiquitin carboxyl-terminal hydrolase 7 ( | USP7 | 128.303 | 2 | 55.038944 | yes | 2.465E-07 | 693.84302 | 115.4043 |
| AGFIQDTSLILYEE | 22 | Q93009 | Ubiquitin carboxyl-terminal hydrolase 7 ( | USP7 | 128.303 | 3 | 55.038944 | yes | 3.403E-05 | 846.11462 | 110.03304 |
| VDVIFCDK | 8 | Q93009 | Ubiquitin carboxyl-terminal hydrolase 7 ( | USP7 | 128.303 | 2 | 55.038944 | yes | 6.019E-07 | 498.24695 | 108.44382 |
| MVSYIQCK | 8 | Q93009 | Ubiquitin carboxyl-terminal hydrolase 7 ( | USP7 | 128.303 | 2 | 55.038944 | yes | 2.175E-07 | 514.74884 | 101.23027 |
| DLLEECKK | 8 | Q93009 | Ubiquitin carboxyl-terminal hydrolase 7 ( | USP7 | 128.303 | 2 | 55.038944 | yes | 4.962E-07 | 517.763 | 66.986115 |
| FAIVMMGR | 8 | Q93009 | Ubiquitin carboxyl-terminal hydrolase 7 ( | USP7 | 128.303 | 2 | 55.038944 | yes | 9.722E-08 | 462.74335 | 63.586021 |
| FDKDHDVMLFLK | 12 | Q93009 | Ubiquitin carboxyl-terminal hydrolase 7 ( | USP7 | 128.303 | 2 | 55.038944 | yes | 9.722E-08 | 754.38434 | 59.205994 |
| KHTGYVGLK | 9 | Q93009 | Ubiquitin carboxyl-terminal hydrolase 7 ( | USP7 | 128.303 | 2 | 55.038944 | yes | 2.811E-06 | 501.79019 | 53.21069 |
| RPAMLDNEADGN | 13 | Q93009 | Ubiquitin carboxyl-terminal hydrolase 7 ( | USP7 | 128.303 | 2 | 55.038944 | yes | 2.625E-07 | 715.83826 | 14.658098 |
| EDILEFIK | 8 | Q93034 | Cullin-5 OS=Homo sapiens OX=9606 GN | CUL5 | 90.956 | 2 | 43.926197 | yes | 3.169E-07 | 503.7764 | 142.9093 |
| LATELPDAELR | 11 | Q93034 | Cullin-5 OS=Homo sapiens OX=9606 GN | CUL5 | 90.956 | 2 | 43.926197 | yes | 7.638E-07 | 614.33258 | 48.1548 |
| LGEAFDSQLVIGVI | 14 | Q93034 | Cullin-5 OS=Homo sapiens OX=9606 GN | CUL5 | 90.956 | 2 | 43.926197 | yes | 2.391E-07 | 752.41193 | 45.484547 |
| ILNAGAWSR | 9 | Q93034 | Cullin-5 OS=Homo sapiens OX=9606 GN | CUL5 | 90.956 | 2 | 43.926197 | yes | 2.945E-06 | 494.27216 | 42.368938 |
| KLTSEEIEAK | 10 | Q93034 | Cullin-5 OS=Homo sapiens OX=9606 GN | CUL5 | 90.956 | 2 | 43.926197 | yes | 3.271E-06 | 574.3139 | 16.327824 |
| LGFVAGVINR | 10 | Q93050 | V-type proton ATPase 116 kDa subunit a | ATP6V0A1 | 96.413 | 2 | 33.713123 | yes | 4.555E-06 | 523.31128 | 37.356255 |
| LTFLNSFK | 8 | Q93050 | V-type proton ATPase 116 kDa subunit a | ATP6V0A1 | 96.413 | 2 | 33.713123 | no | 1.168E-06 | 485.27383 | 10.236447 |
| MVIPNQPPLTATK | 13 | Q93052 | Lipoma-preferred partner OS=Homo sapi | LPP | 65.747 | 2 | 46.506565 | yes | 1.541E-07 | 705.39471 | 372.55145 |
| EPIMPAPGQEETVI | 14 | Q93052 | Lipoma-preferred partner OS=Homo sapi | LPP | 65.747 | 2 | 46.506565 | yes | 3.768E-07 | 777.38507 | 123.81422 |
| DFHVHCYR | 8 | Q93052 | Lipoma-preferred partner OS=Homo sapi | LPP | 65.747 | 2 | 46.506565 | yes | 1.846E-07 | 567.25092 | 67.326797 |
| FDPEIPQTLR | 10 | Q93062 | RNA-binding protein with multiple splici | RBPMS | 21.802 | 2 | 48.9846 | yes | 1.081E-07 | 608.32202 | 1611.5189 |
| QPVGFVSFDSR | 11 | Q93062 | RNA-binding protein with multiple splici | RBPMS | 21.802 | 2 | 48.9846 | yes | 2.564E-07 | 619.81183 | 379.96353 |
| TLFVSGLPLDIKPR | 14 | Q93062 | RNA-binding protein with multiple splici | RBPMS | 21.802 | 3 | 48.9846 | yes | 1.147E-07 | 519.31171 | 55.154079 |

| | | | | | | | | | | | |
|---|---|---|---|---|---|---|---|---|---|---|---|
| TLFVSGLPLDIKPR | 14 | Q93062 | RNA-binding protein with multiple splici | RBPMS | 21.802 | 3 | 48.9846 | yes | 1.935E-06 | 778.46393 | 55.154079 |
| TLFVSGLPLDIKPR | 14 | Q93062 | RNA-binding protein with multiple splici | RBPMS | 21.802 | 2 | 48.9846 | yes | 1.147E-07 | 519.31171 | 55.154079 |
| TLFVSGLPLDIKPR | 14 | Q93062 | RNA-binding protein with multiple splici | RBPMS | 21.802 | 2 | 48.9846 | yes | 1.935E-06 | 778.46393 | 55.154079 |
| LLVVWNNQNK | 10 | Q93063 | Exostosin-2 OS=Homo sapiens OX=9606 | EXT2 | 82.255 | 2 | 52.215794 | yes | 1.806E-07 | 614.34583 | 269.58029 |
| LPADSPIPER | 10 | Q93063 | Exostosin-2 OS=Homo sapiens OX=9606 | EXT2 | 82.255 | 2 | 52.215794 | yes | 1.737E-07 | 547.79565 | 194.50488 |
| YVDDFGVSVSNTI | 15 | Q93063 | Exostosin-2 OS=Homo sapiens OX=9606 | EXT2 | 82.255 | 2 | 52.215794 | yes | 9.722E-08 | 829.90466 | 164.5338 |
| CTNLSEGVLSVR | 12 | Q93063 | Exostosin-2 OS=Homo sapiens OX=9606 | EXT2 | 82.255 | 2 | 52.215794 | yes | 9.722E-08 | 667.84027 | 114.67308 |
| FASVFGTMPLK | 11 | Q93063 | Exostosin-2 OS=Homo sapiens OX=9606 | EXT2 | 82.255 | 2 | 52.215794 | yes | 3.897E-07 | 599.32043 | 94.627731 |
| AIALATLQIINDR | 13 | Q93063 | Exostosin-2 OS=Homo sapiens OX=9606 | EXT2 | 82.255 | 2 | 52.215794 | yes | 1.533E-07 | 706.41699 | 80.686775 |
| YFNYLYTYK | 9 | Q93063 | Exostosin-2 OS=Homo sapiens OX=9606 | EXT2 | 82.255 | 2 | 52.215794 | yes | 1.71E-07 | 637.80823 | 64.343803 |
| MSDVYSILQSIPQF | 14 | Q93063 | Exostosin-2 OS=Homo sapiens OX=9606 | EXT2 | 82.255 | 2 | 52.215794 | yes | 9.81E-08 | 818.92181 | 59.883556 |
| LNLVGGMFDTVQI | 13 | Q93074 | Mediator of RNA polymerase II transcript | MED12 | 243.082 | 2 | 57.626766 | no | 1.001E-07 | 725.37958 | 90.141151 |
| DVITCEPQGSLIDT | 15 | Q93074 | Mediator of RNA polymerase II transcript | MED12 | 243.082 | 2 | 57.626766 | yes | 7.838E-07 | 838.414 | 90.111107 |
| FGELFSDFCSK | 11 | Q93074 | Mediator of RNA polymerase II transcript | MED12 | 243.082 | 2 | 57.626766 | yes | 9.722E-08 | 668.79755 | 70.410172 |
| LAYFCTR | 7 | Q93074 | Mediator of RNA polymerase II transcript | MED12 | 243.082 | 2 | 57.626766 | yes | 2.173E-07 | 465.72873 | 57.981674 |
| MTCAYYAAISETK | 13 | Q93074 | Mediator of RNA polymerase II transcript | MED12 | 243.082 | 2 | 57.626766 | yes | 3.09E-06 | 754.84161 | 37.611961 |
| QQQQQQILR | 9 | Q93074 | Mediator of RNA polymerase II transcript | MED12 | 243.082 | 2 | 57.626766 | yes | 1.693E-07 | 585.32294 | 37.046261 |
| SGVWLVAPLIAK | 12 | Q93074 | Mediator of RNA polymerase II transcript | MED12 | 243.082 | 2 | 57.626766 | yes | 1.322E-06 | 627.38446 | 21.613958 |
| YEDAVQFIR | 9 | Q93096 | Protein tyrosine phosphatase type IVA 1 ( | PTP4A1 | 19.816 | 2 | 48.841785 | no | 1.334E-07 | 570.78784 | 125.19915 |
| YGVTTIVR | 8 | Q93096 | Protein tyrosine phosphatase type IVA 1 ( | PTP4A1 | 19.816 | 2 | 48.841785 | yes | 1.097E-07 | 454.76364 | 115.1293 |
| FIEELKK | 7 | Q93096 | Protein tyrosine phosphatase type IVA 1 ( | PTP4A1 | 19.816 | 2 | 48.841785 | yes | 1.267E-06 | 453.76837 | 108.83221 |
| FLITHNPTNATLNK | 14 | Q93096 | Protein tyrosine phosphatase type IVA 1 ( | PTP4A1 | 19.816 | 2 | 48.841785 | no | 1.196E-07 | 792.43066 | 30.936893 |
| EEDIVEDGWTIVR | 13 | Q969E8 | Pre-rRNA-processing protein TSR2 homo | TSR2 | 20.895 | 2 | 36.91092 | yes | 6.486E-07 | 780.88068 | 52.100124 |
| IGIEEADSFFK | 11 | Q969F9 | BLOC-2 complex member HPS3 OS=Ho | HPS3 | 113.736 | 2 | 50.224712 | yes | 9.849E-08 | 628.3139 | 66.092926 |
| MSGLSMAEVLAR | 12 | Q969F9 | BLOC-2 complex member HPS3 OS=Ho | HPS3 | 113.736 | 2 | 50.224712 | yes | 5.983E-07 | 632.82306 | 51.072083 |
| EEDPYMDTTLK | 11 | Q969F9 | BLOC-2 complex member HPS3 OS=Ho | HPS3 | 113.736 | 2 | 50.224712 | yes | 2.021E-06 | 671.29755 | 33.809334 |
| SQSSIVPEEEQAAN | 15 | Q969G3 | SWI/SNF-related matrix-associated actin- | SMARCE1 | 46.65 | 2 | 51.245922 | yes | 9.722E-08 | 808.89172 | 49.219536 |
| YLNAYTGIVLLR | 12 | Q969H6 | Ribonuclease P/MRP protein subunit POI | POP5 | 18.82 | 2 | 53.91493 | yes | 9.722E-08 | 698.40338 | 68.437088 |
| YPCFFNTLHVGGT | 15 | Q969H6 | Ribonuclease P/MRP protein subunit POI | POP5 | 18.82 | 3 | 53.91493 | yes | 4.619E-07 | 594.63123 | 53.396515 |
| QENHSSLVSLGGE | 17 | Q969J2 | Zinc finger protein with KRAB and SCAI | ZKSCAN4 | 61.58 | 3 | 47.5431 | yes | 1.692E-07 | 609.6463 | 78.403229 |
| TSYLVQHQR | 9 | Q969J2 | Zinc finger protein with KRAB and SCAI | ZKSCAN4 | 61.58 | 2 | 47.5431 | yes | 1.307E-07 | 566.29889 | 70.52755 |
| VLQVPGLAQGGC( | 14 | Q969J2 | Zinc finger protein with KRAB and SCAI | ZKSCAN4 | 61.58 | 2 | 47.5431 | yes | 1.712E-06 | 757.88196 | 43.566978 |
| KNAALDAQSAED( | 20 | Q969J2 | Zinc finger protein with KRAB and SCAI | ZKSCAN4 | 61.58 | 3 | 47.5431 | yes | 2.651E-07 | 691.69891 | 28.213541 |
| LQLQDEEDDSLCR | 13 | Q969K3 | E3 ubiquitin-protein ligase RNF34 OS=H | RNF34 | 41.641 | 2 | 36.114223 | yes | 1.195E-06 | 810.86212 | 26.309286 |
| NFVNYSGCCEK | 11 | Q969K3 | E3 ubiquitin-protein ligase RNF34 OS=H | RNF34 | 41.641 | 2 | 36.114223 | yes | 7.382E-07 | 689.28174 | 16.375572 |
| IDNVDAFMNIR | 11 | Q969L4 | U7 snRNA-associated Sm-like protein LS | LSM10 | 14.081 | 2 | 47.671177 | yes | 1.282E-07 | 654.32428 | 118.19658 |
| LDDLFVTGR | 9 | Q969L4 | U7 snRNA-associated Sm-like protein LS | LSM10 | 14.081 | 2 | 47.671177 | yes | 2.82E-07 | 518.2771 | 80.312943 |
| TLTVLHPLK | 9 | Q969M3 | Protein YIPF5 OS=Homo sapiens OX=96 | YIPF5 | 27.99 | 2 | 47.754856 | yes | 1.265E-07 | 511.32385 | 579.88959 |
| QYAGYDYSQQGR | 12 | Q969M3 | Protein YIPF5 OS=Homo sapiens OX=96 | YIPF5 | 27.99 | 2 | 47.754856 | yes | 6.171E-07 | 718.31549 | 69.03949 |

| | | | | | | | | | | | |
|---|---|---|---|---|---|---|---|---|---|---|---|
| TLTEPCPLASESR | 13 | Q969N2 | GPI transamidase component PIG-T OS= | PIGT | 65.7 | 2 | 45.091885 | yes | 1.966E-07 | 730.85614 | 63.462635 |
| VVAILCNHQR | 10 | Q969P6 | DNA topoisomerase I, mitochondrial OS= | TOP1MT | 69.872 | 2 | 51.335033 | yes | 2.043E-06 | 605.32971 | 125.00476 |
| VPVEKPVYK | 9 | Q969P6 | DNA topoisomerase I, mitochondrial OS= | TOP1MT | 69.872 | 2 | 51.335033 | yes | 1.321E-07 | 529.81586 | 42.810318 |
| TYNASITLQEQLR | 13 | Q969P6 | DNA topoisomerase I, mitochondrial OS= | TOP1MT | 69.872 | 2 | 51.335033 | yes | 3.072E-07 | 768.90448 | 32.119572 |
| EDGEAADTVGCCS | 15 | Q969P6 | DNA topoisomerase I, mitochondrial OS= | TOP1MT | 69.872 | 2 | 51.335033 | yes | 9.722E-08 | 820.33795 | 23.001781 |
| LPAEDEVLLQK | 11 | Q969Q6 | Serine/threonine-protein phosphatase 2A | PPP2R3C | 53.316 | 2 | 49.678154 | yes | 1.011E-07 | 627.85065 | 51.957493 |
| VEVLNSDAVLPSR | 13 | Q969R5 | Lethal(3)malignant brain tumor-like prote | L3MBTL2 | 79.111 | 2 | 51.46856 | yes | 3.566E-07 | 699.883 | 298.63913 |
| ISSEPVPGEIIAVR | 14 | Q969R5 | Lethal(3)malignant brain tumor-like prote | L3MBTL2 | 79.111 | 2 | 51.46856 | yes | 1.738E-07 | 733.91431 | 142.08087 |
| ASSPELPVSVENIK | 14 | Q969R5 | Lethal(3)malignant brain tumor-like prote | L3MBTL2 | 79.111 | 2 | 51.46856 | yes | 9.418E-07 | 735.39594 | 91.177307 |
| TLPVDFHIK | 9 | Q969R5 | Lethal(3)malignant brain tumor-like prote | L3MBTL2 | 79.111 | 2 | 51.46856 | yes | 4.286E-07 | 535.30566 | 88.04763 |
| FCSVSCSR | 8 | Q969R5 | Lethal(3)malignant brain tumor-like prote | L3MBTL2 | 79.111 | 2 | 51.46856 | no | 5.333E-07 | 501.71021 | 83.311295 |
| AVYTEGGWFEEGI | 14 | Q969R5 | Lethal(3)malignant brain tumor-like prote | L3MBTL2 | 79.111 | 2 | 51.46856 | yes | 1.617E-07 | 802.35852 | 60.359047 |
| KPLLEDDPQGAR | 12 | Q969R5 | Lethal(3)malignant brain tumor-like prote | L3MBTL2 | 79.111 | 2 | 51.46856 | yes | 9.722E-08 | 669.85425 | 51.30315 |
| ALGWTGSTGAALN | 14 | Q969S3 | Cytoplasmic 60S subunit biogenesis facto | ZNF622 | 54.273 | 2 | 54.43017 | yes | 9.722E-08 | 696.35864 | 1393.0851 |
| LQWFEQQAK | 9 | Q969S3 | Cytoplasmic 60S subunit biogenesis facto | ZNF622 | 54.273 | 2 | 54.43017 | yes | 1.025E-07 | 589.30365 | 957.32257 |
| ATYTCITCR | 9 | Q969S3 | Cytoplasmic 60S subunit biogenesis facto | ZNF622 | 54.273 | 2 | 54.43017 | yes | 1.408E-07 | 594.26282 | 312.51495 |
| ATYTCITCR | 9 | Q969S3 | Cytoplasmic 60S subunit biogenesis facto | ZNF622 | 54.273 | 2 | 54.43017 | yes | 1.063E-07 | 573.25751 | 312.51495 |
| VASMAPVTAEGFC | 15 | Q969S3 | Cytoplasmic 60S subunit biogenesis facto | ZNF622 | 54.273 | 2 | 54.43017 | yes | 9.722E-08 | 796.8905 | 299.53113 |
| VASMAPVTAEGFC | 15 | Q969S3 | Cytoplasmic 60S subunit biogenesis facto | ZNF622 | 54.273 | 2 | 54.43017 | yes | 1.988E-07 | 804.88794 | 299.53113 |
| ICLWCNEK | 8 | Q969S3 | Cytoplasmic 60S subunit biogenesis facto | ZNF622 | 54.273 | 2 | 54.43017 | yes | 1.305E-07 | 561.75714 | 452.37921 |
| GSATYCTVCSK | 11 | Q969S3 | Cytoplasmic 60S subunit biogenesis facto | ZNF622 | 54.273 | 2 | 54.43017 | yes | 2.605E-07 | 617.26556 | 413.54468 |
| FASFNAYENHLK | 12 | Q969S3 | Cytoplasmic 60S subunit biogenesis facto | ZNF622 | 54.273 | 2 | 54.43017 | yes | 1.171E-07 | 720.84894 | 380.04868 |
| DAMNAAIQQAIK | 12 | Q969S3 | Cytoplasmic 60S subunit biogenesis facto | ZNF622 | 54.273 | 2 | 54.43017 | yes | 9.722E-08 | 637.33209 | 266.81027 |
| GLGVDSVDKDAM | 21 | Q969S3 | Cytoplasmic 60S subunit biogenesis facto | ZNF622 | 54.273 | 3 | 54.43017 | yes | 9.722E-08 | 715.37207 | 201.3177 |
| KVEMMNEK | 8 | Q969S3 | Cytoplasmic 60S subunit biogenesis facto | ZNF622 | 54.273 | 2 | 54.43017 | yes | 1.264E-07 | 504.74628 | 191.42374 |
| DPSEKPPR | 8 | Q969S3 | Cytoplasmic 60S subunit biogenesis facto | ZNF622 | 54.273 | 2 | 54.43017 | yes | 1.65E-06 | 463.24051 | 55.260071 |
| NLEYDDETMELIL | 18 | Q969S3 | Cytoplasmic 60S subunit biogenesis facto | ZNF622 | 54.273 | 3 | 54.43017 | yes | 6.046E-07 | 689.32971 | 33.908047 |
| MLQVASQASR | 10 | Q969T9 | WW domain-binding protein 2 OS=Homo | WBP2 | 28.087 | 2 | 51.08043 | yes | 1.107E-07 | 545.78729 | 132.3542 |
| KGTVYLTPYR | 10 | Q969T9 | WW domain-binding protein 2 OS=Homo | WBP2 | 28.087 | 2 | 51.08043 | yes | 9.722E-08 | 599.33496 | 49.730602 |
| AMAAVPQDVVR | 11 | Q969V3 | BOS complex subunit NCLN OS=Homo | NCLN | 62.975 | 2 | 47.35794 | yes | 4.176E-07 | 578.81079 | 114.17116 |
| LLDFSYEQYQK | 11 | Q969V3 | BOS complex subunit NCLN OS=Homo | NCLN | 62.975 | 2 | 47.35794 | yes | 3.786E-07 | 717.35101 | 40.219925 |
| MQQYDLQGQPYG | 14 | Q969V3 | BOS complex subunit NCLN OS=Homo | NCLN | 62.975 | 2 | 47.35794 | yes | 1.346E-07 | 842.89105 | 38.212891 |
| RQLALQEEIQVK | 12 | Q969V4 | Tektin-1 OS=Homo sapiens OX=9606 GI | TEKT1 | 48.284 | 2 | 49.774025 | yes | 1.004E-07 | 727.91992 | 212.87248 |
| PEDRESLK | 8 | Q969V5 | Mitochondrial ubiquitin ligase activator o | MUL1 | 39.8 | 2 | 20.593487 | yes | 4.411E-06 | 487.2511 | 193.52121 |
| EAIIVGQVNYPK | 12 | Q969V6 | Myocardin-related transcription factor A ( | MRTFA | 98.919 | 2 | 44.667763 | yes | 2.112E-07 | 665.87189 | 83.659302 |
| AYQDQISPVPGAP | 14 | Q969V6 | Myocardin-related transcription factor A ( | MRTFA | 98.919 | 2 | 44.667763 | yes | 3.265E-06 | 735.883 | 39.004101 |
| SAGEALGSSGTPP | 15 | Q969V6 | Myocardin-related transcription factor A ( | MRTFA | 98.919 | 2 | 44.667763 | yes | 3.548E-06 | 693.35461 | 23.661995 |
| GVAPPTLITDSTGT | 23 | Q969V6 | Myocardin-related transcription factor A ( | MRTFA | 98.919 | 3 | 44.667763 | yes | 1.255E-06 | 779.10046 | 3.9135807 |
| YDLSPITVK | 9 | Q969X5 | Endoplasmic reticulum-Golgi intermediat | ERGIC1 | 32.593 | 2 | 53.39623 | yes | 2.551E-07 | 518.28967 | 147.90868 |

| | | | | | | | | | | | |
|---|---|---|---|---|---|---|---|---|---|---|---|
| EYVAYSHTGR | 10 | Q969X5 | Endoplasmic reticulum-Golgi intermediat | ERGIC1 | 32.593 | 2 | 53.39623 | yes | 4.793E-07 | 591.78052 | 89.913933 |
| LTSNPLASHDYILK | 14 | Q969X5 | Endoplasmic reticulum-Golgi intermediat | ERGIC1 | 32.593 | 2 | 53.39623 | yes | 9.722E-08 | 786.42505 | 37.845833 |
| LTSNPLASHDYILK | 14 | Q969X5 | Endoplasmic reticulum-Golgi intermediat | ERGIC1 | 32.593 | 2 | 53.39623 | yes | 5.435E-06 | 524.61914 | 37.845833 |
| LTSNPLASHDYILK | 14 | Q969X5 | Endoplasmic reticulum-Golgi intermediat | ERGIC1 | 32.593 | 3 | 53.39623 | yes | 9.722E-08 | 786.42505 | 37.845833 |
| LTSNPLASHDYILK | 14 | Q969X5 | Endoplasmic reticulum-Golgi intermediat | ERGIC1 | 32.593 | 3 | 53.39623 | yes | 5.435E-06 | 524.61914 | 37.845833 |
| YSYQYTVANK | 10 | Q969X5 | Endoplasmic reticulum-Golgi intermediat | ERGIC1 | 32.593 | 2 | 53.39623 | yes | 3.347E-07 | 618.7984 | 68.414742 |
| SALQILFSEDSTK | 13 | Q969X6 | U3 small nucleolar RNA-associated prote | UTP4 | 76.891 | 2 | 54.29034 | yes | 1.063E-07 | 719.87482 | 1085.4818 |
| VQFWDSATGTLVI | 13 | Q969X6 | U3 small nucleolar RNA-associated prote | UTP4 | 76.891 | 2 | 54.29034 | yes | 4.537E-07 | 726.38007 | 477.95001 |
| TVAHSPTALISGGT | 21 | Q969X6 | U3 small nucleolar RNA-associated prote | UTP4 | 76.891 | 3 | 54.29034 | yes | 1.002E-07 | 727.38678 | 287.72638 |
| TVAHSPTALISGGT | 26 | Q969X6 | U3 small nucleolar RNA-associated prote | UTP4 | 76.891 | 4 | 54.29034 | yes | 1.113E-07 | 695.37061 | 271.54376 |
| ATEALCWAEGQR | 12 | Q969X6 | U3 small nucleolar RNA-associated prote | UTP4 | 76.891 | 2 | 54.29034 | yes | 9.722E-08 | 696.32227 | 259.41315 |
| LFVASNQGALHIV | 21 | Q969X6 | U3 small nucleolar RNA-associated prote | UTP4 | 76.891 | 3 | 54.29034 | yes | 3.689E-07 | 725.06366 | 247.37477 |
| NADHLLHLK | 9 | Q969X6 | U3 small nucleolar RNA-associated prote | UTP4 | 76.891 | 2 | 54.29034 | yes | 1.337E-07 | 530.79852 | 111.24519 |
| CVAYNNQSNR | 10 | Q969X6 | U3 small nucleolar RNA-associated prote | UTP4 | 76.891 | 2 | 54.29034 | yes | 1.315E-07 | 613.27258 | 107.03127 |
| FFNYVPSGIR | 10 | Q969X6 | U3 small nucleolar RNA-associated prote | UTP4 | 76.891 | 2 | 54.29034 | yes | 1.689E-07 | 600.31403 | 73.238235 |
| TLVAVERPLDDIIA | 21 | Q969X6 | U3 small nucleolar RNA-associated prote | UTP4 | 76.891 | 3 | 54.29034 | yes | 1.285E-07 | 766.78589 | 60.686306 |
| ELQSVEQEVR | 10 | Q969Z0 | FAST kinase domain-containing protein 4 | TBRG4 | 70.739 | 2 | 38.26006 | yes | 5.346E-07 | 608.81201 | 95.642654 |
| LLGSLYALGIPK | 12 | Q969Z0 | FAST kinase domain-containing protein 4 | TBRG4 | 70.739 | 2 | 38.26006 | yes | 9.282E-07 | 622.88428 | 26.267941 |
| GVPVSEAECTAMC | 15 | Q969Z3 | Mitochondrial amidoxime reducing comp | MTARC2 | 38.024 | 2 | 53.79618 | no | 1.105E-07 | 788.87653 | 302.97739 |
| APDMDQLVLPSK | 12 | Q969Z3 | Mitochondrial amidoxime reducing comp | MTARC2 | 38.024 | 2 | 53.79618 | yes | 9.722E-08 | 657.3421 | 117.36332 |
| LSPLFGIYYSVEK | 13 | Q969Z3 | Mitochondrial amidoxime reducing comp | MTARC2 | 38.024 | 2 | 53.79618 | yes | 9.218E-07 | 758.40833 | 54.478142 |
| HETLTSLNLEK | 11 | Q96A26 | Protein FAM162A OS=Homo sapiens OX | FAM162A | 17.343 | 2 | 31.356806 | yes | 1.868E-06 | 642.84332 | 51.171894 |
| VCCEGMLIQLR | 11 | Q96A33 | PAT complex subunit CCDC47 OS=Hom | CCDC47 | 55.874 | 2 | 52.471355 | yes | 9.722E-08 | 689.83563 | 847.55402 |
| LAQAWFNTHR | 10 | Q96A33 | PAT complex subunit CCDC47 OS=Hom | CCDC47 | 55.874 | 2 | 52.471355 | yes | 9.778E-08 | 622.32019 | 417.83441 |
| MVHFLTHYADK | 11 | Q96A33 | PAT complex subunit CCDC47 OS=Hom | CCDC47 | 55.874 | 2 | 52.471355 | yes | 9.722E-08 | 681.33716 | 326.96338 |
| PVSDQVQIK | 9 | Q96A33 | PAT complex subunit CCDC47 OS=Hom | CCDC47 | 55.874 | 2 | 52.471355 | yes | 4.301E-07 | 507.28494 | 221.76279 |
| QDLLNVLAR | 9 | Q96A33 | PAT complex subunit CCDC47 OS=Hom | CCDC47 | 55.874 | 2 | 52.471355 | yes | 2.154E-07 | 521.30621 | 155.58432 |
| RLEEAALR | 8 | Q96A33 | PAT complex subunit CCDC47 OS=Hom | CCDC47 | 55.874 | 2 | 52.471355 | yes | 4.443E-07 | 479.27744 | 128.62541 |
| IMQEEGQPLK | 10 | Q96A33 | PAT complex subunit CCDC47 OS=Hom | CCDC47 | 55.874 | 2 | 52.471355 | yes | 1.823E-07 | 586.80261 | 73.353127 |
| IESVHFSDQFSGPK | 14 | Q96A33 | PAT complex subunit CCDC47 OS=Hom | CCDC47 | 55.874 | 2 | 52.471355 | yes | 3.238E-05 | 789.38336 | 69.825722 |
| RQDLLNVLAR | 10 | Q96A33 | PAT complex subunit CCDC47 OS=Hom | CCDC47 | 55.874 | 2 | 52.471355 | yes | 3.187E-07 | 599.35675 | 61.575176 |
| IMQEEGQPLKLPD | 15 | Q96A33 | PAT complex subunit CCDC47 OS=Hom | CCDC47 | 55.874 | 3 | 52.471355 | yes | 1.786E-06 | 576.30634 | 31.644928 |
| ESEVLIGNLGDK | 12 | Q96A65 | Exocyst complex component 4 OS=Homo | EXOC4 | 110.499 | 2 | 50.24329 | yes | 1.104E-07 | 637.33533 | 80.530663 |
| GENVTVENQPR | 11 | Q96A65 | Exocyst complex component 4 OS=Homo | EXOC4 | 110.499 | 2 | 50.24329 | yes | 2.825E-05 | 621.80731 | 47.0924 |
| LIPPQDILR | 9 | Q96A65 | Exocyst complex component 4 OS=Homo | EXOC4 | 110.499 | 2 | 50.24329 | yes | 1.326E-06 | 532.82678 | 46.296856 |
| SQTGVGELTTQNT | 14 | Q96A65 | Exocyst complex component 4 OS=Homo | EXOC4 | 110.499 | 2 | 50.24329 | yes | 6.038E-07 | 746.37354 | 38.736774 |
| HVLNLLDEIENIK | 13 | Q96A65 | Exocyst complex component 4 OS=Homo | EXOC4 | 110.499 | 2 | 50.24329 | yes | 9.838E-08 | 775.43286 | 22.606199 |
| LYDMADVWVK | 10 | Q96A65 | Exocyst complex component 4 OS=Homo | EXOC4 | 110.499 | 2 | 50.24329 | yes | 2.844E-05 | 620.30756 | 9.2018528 |
| GCEVTPDVNISGQ | 14 | Q96AC1 | Fermitin family homolog 2 OS=Homo sap | FERMT2 | 77.861 | 2 | 50.20359 | yes | 2.279E-07 | 752.35901 | 66.534462 |

| | | | | | | | | | | | |
|---|---|---|---|---|---|---|---|---|---|---|---|
| LSFICTEVDCK | 11 | Q96AC1 | Fermitin family homolog 2 OS=Homo sap | FERMT2 | 77.861 | 2 | 50.20359 | yes | 9.85E-08 | 686.31781 | 42.536137 |
| FAVGIVIGR | 9 | Q96AE4 | Far upstream element-binding protein 1 C | FUBP1 | 67.561 | 2 | 52.598595 | no | 1.412E-07 | 466.28983 | 1570.0347 |
| KIQNDAGVR | 9 | Q96AE4 | Far upstream element-binding protein 1 C | FUBP1 | 67.561 | 2 | 52.598595 | no | 1.479E-07 | 500.78033 | 1233.3298 |
| IQNDAGVR | 8 | Q96AE4 | Far upstream element-binding protein 1 C | FUBP1 | 67.561 | 2 | 52.598595 | no | 2.367E-07 | 436.73288 | 1102.2794 |
| SCMLTGTPESVQS | 15 | Q96AE4 | Far upstream element-binding protein 1 C | FUBP1 | 67.561 | 2 | 52.598595 | yes | 1.562E-07 | 798.37384 | 433.31885 |
| GTPQQIDYAR | 10 | Q96AE4 | Far upstream element-binding protein 1 C | FUBP1 | 67.561 | 2 | 52.598595 | yes | 1.328E-07 | 574.78839 | 295.53522 |
| AWEEYYK | 7 | Q96AE4 | Far upstream element-binding protein 1 C | FUBP1 | 67.561 | 2 | 52.598595 | no | 1.047E-07 | 494.72418 | 265.83191 |
| IGGNEGIDVPIPR | 13 | Q96AE4 | Far upstream element-binding protein 1 C | FUBP1 | 67.561 | 2 | 52.598595 | yes | 9.903E-08 | 668.86462 | 232.1644 |
| IQIAPDSGGLPER | 13 | Q96AE4 | Far upstream element-binding protein 1 C | FUBP1 | 67.561 | 2 | 52.598595 | yes | 3.36E-07 | 676.86206 | 206.98557 |
| VPDGMVGFIIGR | 12 | Q96AE4 | Far upstream element-binding protein 1 C | FUBP1 | 67.561 | 2 | 52.598595 | yes | 9.722E-08 | 630.8421 | 150.33498 |
| MVMIQDGPQNTG. | 18 | Q96AE4 | Far upstream element-binding protein 1 C | FUBP1 | 67.561 | 3 | 52.598595 | yes | 1.075E-07 | 657.66443 | 130.6414 |
| CQHAAEIITDLLR | 13 | Q96AE4 | Far upstream element-binding protein 1 C | FUBP1 | 67.561 | 2 | 52.598595 | yes | 9.722E-08 | 770.40106 | 111.40982 |
| MVMIQDGPQNTG. | 15 | Q96AE4 | Far upstream element-binding protein 1 C | FUBP1 | 67.561 | 2 | 52.598595 | yes | 6.829E-07 | 802.87396 | 103.99583 |
| LLDQIVEK | 8 | Q96AE4 | Far upstream element-binding protein 1 C | FUBP1 | 67.561 | 2 | 52.598595 | yes | 1.558E-05 | 479.28439 | 73.550819 |
| IQQQVDSPMNLK | 12 | Q96AE7 | Tetratricopeptide repeat protein 17 OS=H | TTC17 | 129.559 | 2 | 51.834003 | yes | 1.003E-07 | 700.86377 | 113.9502 |
| ALVWYESTLK | 10 | Q96AE7 | Tetratricopeptide repeat protein 17 OS=H | TTC17 | 129.559 | 2 | 51.834003 | yes | 9.67E-07 | 605.32935 | 101.16479 |
| DQYPQQSLEQIGT | 14 | Q96AE7 | Tetratricopeptide repeat protein 17 OS=H | TTC17 | 129.559 | 2 | 51.834003 | yes | 2.368E-07 | 831.90771 | 100.52129 |
| LIQEEQILR | 9 | Q96AE7 | Tetratricopeptide repeat protein 17 OS=H | TTC17 | 129.559 | 2 | 51.834003 | yes | 9.722E-08 | 571.3324 | 91.281281 |
| PGFEQAIK | 8 | Q96AE7 | Tetratricopeptide repeat protein 17 OS=H | TTC17 | 129.559 | 2 | 51.834003 | yes | 6.579E-07 | 445.24252 | 57.660187 |
| HPHDLVILMR | 10 | Q96AE7 | Tetratricopeptide repeat protein 17 OS=H | TTC17 | 129.559 | 3 | 51.834003 | yes | 6.163E-07 | 410.89737 | 40.213417 |
| ASLHYTGESQLTE | 20 | Q96AE7 | Tetratricopeptide repeat protein 17 OS=H | TTC17 | 129.559 | 3 | 51.834003 | yes | 1.194E-07 | 730.04633 | 23.745422 |
| LQQLPADFGR | 10 | Q96AG4 | Leucine-rich repeat-containing protein 59 | LRRC59 | 34.931 | 2 | 55.65065 | yes | 9.722E-08 | 572.80908 | 8861.7979 |
| ATILDLSCNK | 10 | Q96AG4 | Leucine-rich repeat-containing protein 59 | LRRC59 | 34.931 | 2 | 55.65065 | yes | 1.014E-07 | 567.7948 | 6340.0942 |
| LVTLPVSFAQLK | 12 | Q96AG4 | Leucine-rich repeat-containing protein 59 | LRRC59 | 34.931 | 2 | 55.65065 | yes | 1.701E-07 | 658.40283 | 3133.7407 |
| DNPLDPVLAK | 10 | Q96AG4 | Leucine-rich repeat-containing protein 59 | LRRC59 | 34.931 | 2 | 55.65065 | yes | 1.111E-07 | 541.29803 | 2303.782 |
| LVNLQHLDLLNNK | 13 | Q96AG4 | Leucine-rich repeat-containing protein 59 | LRRC59 | 34.931 | 2 | 55.65065 | yes | 1.531E-07 | 767.44104 | 1573.1597 |
| VAGDCLDEK | 9 | Q96AG4 | Leucine-rich repeat-containing protein 59 | LRRC59 | 34.931 | 2 | 55.65065 | yes | 1.499E-07 | 503.72913 | 1376.547 |
| LTTLPSDFCGLTHI | 16 | Q96AG4 | Leucine-rich repeat-containing protein 59 | LRRC59 | 34.931 | 3 | 55.65065 | yes | 1.172E-07 | 601.32208 | 491.78549 |
| VAGDCLDEKQCK | 12 | Q96AG4 | Leucine-rich repeat-containing protein 59 | LRRC59 | 34.931 | 2 | 55.65065 | yes | 9.722E-08 | 711.82123 | 428.93701 |
| DKLDGNELDLSLS | 21 | Q96AG4 | Leucine-rich repeat-containing protein 59 | LRRC59 | 34.931 | 3 | 55.65065 | yes | 1.092E-07 | 771.73218 | 151.45947 |
| KEYDALK | 7 | Q96AG4 | Leucine-rich repeat-containing protein 59 | LRRC59 | 34.931 | 2 | 55.65065 | yes | 2.349E-07 | 433.73456 | 104.17915 |
| GCLTLYTGR | 9 | Q96AH0 | SOSS complex subunit B2 OS=Homo sap | NABP1 | 22.424 | 2 | 50.147213 | no | 9.889E-08 | 520.76331 | 243.68741 |
| VNDPLIFIR | 9 | Q96AH0 | SOSS complex subunit B2 OS=Homo sap | NABP1 | 22.424 | 2 | 50.147213 | yes | 3.082E-07 | 543.81897 | 115.49264 |
| TSGIIEALHR | 10 | Q96AP4 | Zinc finger-containing ubiquitin peptidase | ZUP1 | 65.959 | 2 | 53.62278 | yes | 9.722E-08 | 548.80908 | 235.46835 |
| EGFDPQGASQLNN | 14 | Q96AP4 | Zinc finger-containing ubiquitin peptidase | ZUP1 | 65.959 | 2 | 53.62278 | yes | 2.035E-07 | 766.85803 | 121.79504 |
| IQSMIEDAWK | 10 | Q96AP4 | Zinc finger-containing ubiquitin peptidase | ZUP1 | 65.959 | 2 | 53.62278 | yes | 3.489E-06 | 610.80261 | 62.741447 |
| QASQVFTAEK | 10 | Q96AP4 | Zinc finger-containing ubiquitin peptidase | ZUP1 | 65.959 | 2 | 53.62278 | yes | 7.008E-07 | 554.78528 | 57.2304 |
| QYGLDNSGGYK | 11 | Q96AP4 | Zinc finger-containing ubiquitin peptidase | ZUP1 | 65.959 | 2 | 53.62278 | yes | 2.861E-07 | 601.27783 | 45.587753 |
| EQLEAEAQALR | 11 | Q96AQ6 | Pre-B-cell leukemia transcription factor-in | PBXIP1 | 80.644 | 2 | 48.311203 | yes | 1.165E-07 | 629.32532 | 65.215164 |

| | | | | | | | | | | | |
|---|---|---|---|---|---|---|---|---|---|---|---|
| ALESELQQLR | 10 | Q96AQ6 | Pre-B-cell leukemia transcription factor-ir | PBXIP1 | 80.644 | 2 | 48.311203 | yes | 4.477E-07 | 593.82495 | 61.106537 |
| QVLPDPEVLEAVC | 15 | Q96AQ6 | Pre-B-cell leukemia transcription factor-ir | PBXIP1 | 80.644 | 2 | 48.311203 | yes | 4.548E-07 | 818.93066 | 38.715412 |
| ETLQGFVTDITAK | 13 | Q96AY2 | Crossover junction endonuclease EME1 ( | EME1 | 63.252 | 2 | 53.886463 | yes | 9.722E-08 | 711.87738 | 189.75247 |
| ELADFTCAFTK | 11 | Q96AY2 | Crossover junction endonuclease EME1 ( | EME1 | 63.252 | 2 | 53.886463 | yes | 9.722E-08 | 651.80536 | 99.846153 |
| QPEASIGSMVSR | 12 | Q96AY2 | Crossover junction endonuclease EME1 ( | EME1 | 63.252 | 2 | 53.886463 | yes | 1.263E-07 | 631.3139 | 89.740936 |
| ELPTVTTNVQNSQ | 15 | Q96B01 | RAD51-associated protein 1 OS=Homo sa | RAD51AP1 | 38.458 | 2 | 50.84935 | yes | 2.299E-07 | 837.42065 | 145.51543 |
| ITVEDDVGGVQGK | 13 | Q96B01 | RAD51-associated protein 1 OS=Homo sa | RAD51AP1 | 38.458 | 2 | 50.84935 | yes | 9.722E-08 | 658.83826 | 72.095566 |
| CNALVTSVDSAPA | 16 | Q96B01 | RAD51-associated protein 1 OS=Homo sa | RAD51AP1 | 38.458 | 2 | 50.84935 | yes | 2.045E-07 | 801.91144 | 57.158356 |
| KVSLSSDTTR | 10 | Q96B01 | RAD51-associated protein 1 OS=Homo sa | RAD51AP1 | 38.458 | 2 | 50.84935 | yes | 3.698E-07 | 547.29602 | 54.675892 |
| TVEPLEYYR | 9 | Q96B26 | Exosome complex component RRP43 OS | EXOSC8 | 30.04 | 2 | 51.2305 | yes | 1.23E-07 | 585.29547 | 511.89618 |
| LGNTTVICGVK | 11 | Q96B26 | Exosome complex component RRP43 OS | EXOSC8 | 30.04 | 2 | 51.2305 | yes | 1.152E-07 | 581.31842 | 435.08224 |
| AEFAAPSTDAPDK | 13 | Q96B26 | Exosome complex component RRP43 OS | EXOSC8 | 30.04 | 2 | 51.2305 | yes | 9.722E-08 | 660.30933 | 350.74866 |
| LQDCMSR | 7 | Q96B26 | Exosome complex component RRP43 OS | EXOSC8 | 30.04 | 2 | 51.2305 | yes | 1.92E-07 | 455.19949 | 131.40962 |
| KLMDEVIK | 8 | Q96B26 | Exosome complex component RRP43 OS | EXOSC8 | 30.04 | 2 | 51.2305 | yes | 1.027E-07 | 488.28079 | 128.24687 |
| LMDEVIK | 7 | Q96B26 | Exosome complex component RRP43 OS | EXOSC8 | 30.04 | 2 | 51.2305 | yes | 9.722E-08 | 424.23331 | 115.38941 |
| LCCLHKPGGSGLT | 16 | Q96B26 | Exosome complex component RRP43 OS | EXOSC8 | 30.04 | 2 | 51.2305 | yes | 4.127E-07 | 828.42145 | 20.91077 |
| EAEDTQVFGDLPR | 13 | Q96B36 | Proline-rich AKT1 substrate 1 OS=Homo | AKT1S1 | 27.384 | 2 | 50.819218 | yes | 9.722E-08 | 738.85187 | 65.843582 |
| AVGVQCPVCR | 10 | Q96BH1 | E3 ubiquitin-protein ligase RNF25 OS=H | RNF25 | 51.219 | 2 | 42.37016 | yes | 6.225E-06 | 573.28137 | 97.838974 |
| GGIIDLEAER | 10 | Q96BH1 | E3 ubiquitin-protein ligase RNF25 OS=H | RNF25 | 51.219 | 2 | 42.37016 | yes | 3.063E-07 | 536.78528 | 72.177017 |
| GLGAQEQGATDHI | 14 | Q96BK5 | PIN2/TERF1-interacting telomerase inhib | PINX1 | 37.035 | 2 | 54.3974 | yes | 9.722E-08 | 712.86005 | 158.44949 |
| LWDIGGQPR | 9 | Q96BM9 | ADP-ribosylation factor-like protein 8A ( | ARL8A | 21.416 | 2 | 31.04211 | yes | 1.974E-06 | 521.27747 | 38.629311 |
| NLSEALGDNVK | 11 | Q96BN2 | Transcriptional adapter 1 OS=Homo sapi | TADA1 | 37.382 | 2 | 47.707355 | yes | 1.275E-07 | 580.30133 | 97.562248 |
| EEFDLEAHR | 9 | Q96BN2 | Transcriptional adapter 1 OS=Homo sapi | TADA1 | 37.382 | 2 | 47.707355 | yes | 2.023E-07 | 573.26471 | 78.197029 |
| ELLIHER | 7 | Q96BN8 | Ubiquitin thioesterase otulin OS=Homo s | OTULIN | 40.263 | 2 | 36.524857 | yes | 6.892E-07 | 455.26126 | 94.844818 |
| LSVAPEMDIMDYC | 14 | Q96BN8 | Ubiquitin thioesterase otulin OS=Homo s | OTULIN | 40.263 | 2 | 36.524857 | yes | 3.656E-06 | 836.375 | 22.622717 |
| VINVETNR | 8 | Q96BP3 | Peptidylprolyl isomerase domain and WD | PPWD1 | 73.575 | 2 | 40.535793 | yes | 3.974E-07 | 472.76163 | 37.985569 |
| AGMVDFQDEEQV | 13 | Q96BR5 | Cytochrome c oxidase assembly factor 7 ( | COA7 | 25.71 | 2 | 47.866093 | yes | 1.243E-07 | 769.34558 | 131.49524 |
| ALTLHYDR | 8 | Q96BY9 | Store-operated calcium entry-associated r | SARAF | 36.976 | 2 | 51.842133 | yes | 2.139E-07 | 494.76416 | 218.04823 |
| RLDPIPQLK | 9 | Q96BY9 | Store-operated calcium entry-associated r | SARAF | 36.976 | 2 | 51.842133 | yes | 1.381E-07 | 540.33221 | 174.70872 |
| GWDGYDVQWECI | 12 | Q96BY9 | Store-operated calcium entry-associated r | SARAF | 36.976 | 2 | 51.842133 | yes | 9.722E-08 | 771.81952 | 50.858414 |
| CVGGTAGCDSYTI | 14 | Q96BY9 | Store-operated calcium entry-associated r | SARAF | 36.976 | 2 | 51.842133 | yes | 1.532E-07 | 736.81085 | 37.431847 |
| FTNSAGPPPPGFK | 13 | Q96BY9 | Store-operated calcium entry-associated r | SARAF | 36.976 | 2 | 51.842133 | yes | 3.059E-06 | 658.83533 | 19.63825 |
| FCDVSLLVQGR | 11 | Q96C00 | Zinc finger and BTB domain-containing p | ZBTB9 | 50.603 | 2 | 43.68801 | yes | 2.487E-07 | 647.33466 | 33.971436 |
| LMAFQPAPK | 9 | Q96C36 | Pyrroline-5-carboxylate reductase 2 OS=I | PYCR2 | 33.638 | 2 | 46.293804 | yes | 1.599E-07 | 501.7757 | 72.957367 |
| VVMNSAQASIK | 11 | Q96C92 | Endosome-associated-trafficking regulato | ENTR1 | 47.962 | 2 | 55.057537 | yes | 1.193E-07 | 574.31061 | 408.85919 |
| QNADVALQNLR | 11 | Q96C92 | Endosome-associated-trafficking regulato | ENTR1 | 47.962 | 2 | 55.057537 | yes | 1.02E-07 | 621.3335 | 201.15094 |
| CGQGASLTVVK | 11 | Q96C92 | Endosome-associated-trafficking regulato | ENTR1 | 47.962 | 2 | 55.057537 | yes | 1.191E-07 | 560.29498 | 115.82145 |
| LKQEISLLQAQVSI | 17 | Q96C92 | Endosome-associated-trafficking regulato | ENTR1 | 47.962 | 3 | 55.057537 | yes | 9.722E-08 | 668.04083 | 70.710991 |
| DLDSPFFGIR | 10 | Q96C92 | Endosome-associated-trafficking regulato | ENTR1 | 47.962 | 2 | 55.057537 | yes | 1.182E-06 | 583.79565 | 25.945469 |

| | | | | | | | | | | | |
|---|---|---|---|---|---|---|---|---|---|---|---|
| TLQISYDALK | 10 | Q96C92 | Endosome-associated-trafficking regulato | ENTR1 | 47.962 | 2 | 55.057537 | yes | 1.85E-07 | 576.31897 | 14.89969 |
| VLVDVPCTTDR | 11 | Q96CB9 | 5-methylcytosine rRNA methyltransferase | NSUN4 | 43.089 | 2 | 50.70318 | yes | 5.152E-07 | 637.8241 | 433.52954 |
| TLALLQTGCCR | 11 | Q96CB9 | 5-methylcytosine rRNA methyltransferase | NSUN4 | 43.089 | 2 | 50.70318 | yes | 3.046E-07 | 646.82611 | 324.14032 |
| NLAANDLSPSR | 11 | Q96CB9 | 5-methylcytosine rRNA methyltransferase | NSUN4 | 43.089 | 2 | 50.70318 | yes | 2.136E-07 | 579.29907 | 270.97968 |
| ILHSYVPEEIR | 11 | Q96CB9 | 5-methylcytosine rRNA methyltransferase | NSUN4 | 43.089 | 2 | 50.70318 | yes | 1.078E-07 | 678.36951 | 238.437 |
| WGELEGDTYDR | 11 | Q96CB9 | 5-methylcytosine rRNA methyltransferase | NSUN4 | 43.089 | 2 | 50.70318 | yes | 9.728E-08 | 670.79132 | 119.75231 |
| HSLHEEENNIFK | 12 | Q96CB9 | 5-methylcytosine rRNA methyltransferase | NSUN4 | 43.089 | 2 | 50.70318 | yes | 3.347E-07 | 748.86005 | 17.773659 |
| LPNVPSSSLPAQPN | 15 | Q96CF2 | Charged multivesicular body protein 4c C | CHMP4C | 26.411 | 2 | 50.24145 | yes | 9.839E-08 | 788.92572 | 78.954765 |
| QLILPALGSGK | 11 | Q96CM3 | Pseudouridylate synthase RPUSD4, mitoc | RPUSD4 | 42.206 | 2 | 52.110546 | yes | 1.465E-07 | 548.83984 | 1341.1466 |
| YIPLHLHAR | 9 | Q96CM3 | Pseudouridylate synthase RPUSD4, mitoc | RPUSD4 | 42.206 | 2 | 52.110546 | yes | 9.907E-08 | 560.32471 | 1239.2152 |
| AEPLHLCHR | 9 | Q96CM3 | Pseudouridylate synthase RPUSD4, mitoc | RPUSD4 | 42.206 | 2 | 52.110546 | yes | 1.113E-07 | 566.78766 | 1088.7921 |
| FFVHSLHR | 8 | Q96CM3 | Pseudouridylate synthase RPUSD4, mitoc | RPUSD4 | 42.206 | 2 | 52.110546 | yes | 2.664E-07 | 521.78271 | 900.20166 |
| DMAHQVQELFR | 11 | Q96CM3 | Pseudouridylate synthase RPUSD4, mitoc | RPUSD4 | 42.206 | 2 | 52.110546 | yes | 9.722E-08 | 687.33514 | 723.69653 |
| ETTGVMVLAWDK | 12 | Q96CM3 | Pseudouridylate synthase RPUSD4, mitoc | RPUSD4 | 42.206 | 2 | 52.110546 | yes | 9.781E-08 | 675.3421 | 592.57574 |
| EELNLVCK | 8 | Q96CM3 | Pseudouridylate synthase RPUSD4, mitoc | RPUSD4 | 42.206 | 2 | 52.110546 | yes | 6.334E-07 | 502.75769 | 553.31763 |
| RVQEIVR | 7 | Q96CM3 | Pseudouridylate synthase RPUSD4, mitoc | RPUSD4 | 42.206 | 2 | 52.110546 | yes | 2.284E-07 | 450.27469 | 522.13904 |
| KEPVSTNAVQR | 11 | Q96CM3 | Pseudouridylate synthase RPUSD4, mitoc | RPUSD4 | 42.206 | 2 | 52.110546 | yes | 1.105E-07 | 614.83588 | 519.15204 |
| GILHQDK | 7 | Q96CM3 | Pseudouridylate synthase RPUSD4, mitoc | RPUSD4 | 42.206 | 2 | 52.110546 | yes | 3.903E-07 | 405.72705 | 515.23993 |
| MTLSPSYR | 8 | Q96CM3 | Pseudouridylate synthase RPUSD4, mitoc | RPUSD4 | 42.206 | 2 | 52.110546 | yes | 1.883E-07 | 477.73929 | 378.03775 |
| KLGLEQSK | 8 | Q96CM3 | Pseudouridylate synthase RPUSD4, mitoc | RPUSD4 | 42.206 | 2 | 52.110546 | yes | 9.722E-08 | 451.76892 | 238.63719 |
| VHPNVLAK | 8 | Q96CM3 | Pseudouridylate synthase RPUSD4, mitoc | RPUSD4 | 42.206 | 2 | 52.110546 | yes | 8.613E-06 | 439.26636 | 116.49648 |
| GGVPGTPSTQSLG | 15 | Q96CP6 | Protein Aster-A OS=Homo sapiens OX=9 | GRAMD1A | 80.681 | 2 | 42.148956 | yes | 3.17E-07 | 700.86005 | 102.04014 |
| LIVDYSCALQR | 11 | Q96CP6 | Protein Aster-A OS=Homo sapiens OX=9 | GRAMD1A | 80.681 | 2 | 42.148956 | no | 3.433E-07 | 669.34772 | 62.122364 |
| YCILGLAR | 8 | Q96CP6 | Protein Aster-A OS=Homo sapiens OX=9 | GRAMD1A | 80.681 | 2 | 42.148956 | yes | 1.237E-06 | 483.26547 | 48.006729 |
| SDFCPEHVPK | 10 | Q96CP7 | TLC domain-containing protein 1 OS=Ho | TLCD1 | 28.548 | 2 | 37.19153 | yes | 6.213E-07 | 608.27679 | 67.470345 |
| QQQDEAYLASLR | 12 | Q96CS3 | FAS-associated factor 2 OS=Homo sapier | FAF2 | 52.624 | 2 | 50.44977 | yes | 9.809E-08 | 711.35461 | 61.795914 |
| SKDCMLQIVCR | 11 | Q96CS7 | Pleckstrin homology domain-containing f | PLEKHB2 | 24.736 | 2 | 53.301647 | yes | 9.722E-08 | 705.33856 | 53.623997 |
| SPDNPMNQR | 9 | Q96CT7 | Coiled-coil domain-containing protein 12 | CCDC124 | 25.836 | 2 | 46.294636 | yes | 2.289E-07 | 529.73779 | 302.67035 |
| VLEEGSVEAR | 10 | Q96CT7 | Coiled-coil domain-containing protein 12 | CCDC124 | 25.836 | 2 | 46.294636 | yes | 2.274E-07 | 544.78278 | 206.07509 |
| AQIEDTLR | 8 | Q96CT7 | Coiled-coil domain-containing protein 12 | CCDC124 | 25.836 | 2 | 46.294636 | yes | 1.599E-07 | 473.25363 | 187.68021 |
| SHLEVPLEENVNR | 13 | Q96CT7 | Coiled-coil domain-containing protein 12 | CCDC124 | 25.836 | 2 | 46.294636 | yes | 2.849E-07 | 768.39429 | 160.08994 |
| AAFTAFEEAQLPR | 13 | Q96CT7 | Coiled-coil domain-containing protein 12 | CCDC124 | 25.836 | 2 | 46.294636 | yes | 4.394E-07 | 725.86987 | 152.06924 |
| VSLMSPDQLR | 10 | Q96CU9 | FAD-dependent oxidoreductase domain-c | FOXRED1 | 53.812 | 2 | 51.158894 | yes | 1.245E-07 | 573.3028 | 392.2478 |
| FNPSGYLLLASEK | 13 | Q96CU9 | FAD-dependent oxidoreductase domain-c | FOXRED1 | 53.812 | 2 | 51.158894 | yes | 1.456E-07 | 719.88245 | 164.17461 |
| VWPHLALR | 8 | Q96CU9 | FAD-dependent oxidoreductase domain-c | FOXRED1 | 53.812 | 2 | 51.158894 | yes | 1.22E-07 | 496.29544 | 160.82393 |
| VPAFETLK | 8 | Q96CU9 | FAD-dependent oxidoreductase domain-c | FOXRED1 | 53.812 | 2 | 51.158894 | yes | 9.722E-08 | 452.76056 | 126.00188 |
| EGLGSNYLGGR | 11 | Q96CU9 | FAD-dependent oxidoreductase domain-c | FOXRED1 | 53.812 | 2 | 51.158894 | yes | 1.232E-07 | 561.78052 | 97.989853 |
| VQSLGVLFCQGEV | 15 | Q96CU9 | FAD-dependent oxidoreductase domain-c | FOXRED1 | 53.812 | 2 | 51.158894 | yes | 1.022E-07 | 846.94055 | 59.579792 |
| DAAAMESNVK | 10 | Q96CU9 | FAD-dependent oxidoreductase domain-c | FOXRED1 | 53.812 | 2 | 51.158894 | yes | 3.777E-06 | 518.24243 | 57.383385 |

| | | | | | | | | | | | |
|---|---|---|---|---|---|---|---|---|---|---|---|
| GGFSLDWDGK | 10 | Q96CU9 | FAD-dependent oxidoreductase domain-c | FOXRED1 | 53.812 | 2 | 51.158894 | yes | 4.409E-07 | 541.2511 | 44.506897 |
| TFITQQGIK | 9 | Q96CW1 | AP-2 complex subunit mu OS=Homo sap | AP2M1 | 49.655 | 2 | 51.710648 | yes | 1.125E-07 | 518.29529 | 2024.031 |
| SYLSGMPECK | 10 | Q96CW1 | AP-2 complex subunit mu OS=Homo sap | AP2M1 | 49.655 | 2 | 51.710648 | yes | 9.722E-08 | 586.25977 | 1466.863 |
| SNIWLAAVTK | 10 | Q96CW1 | AP-2 complex subunit mu OS=Homo sap | AP2M1 | 49.655 | 2 | 51.710648 | yes | 1.457E-07 | 551.81641 | 1260.3302 |
| VNVIHAR | 7 | Q96CW1 | AP-2 complex subunit mu OS=Homo sap | AP2M1 | 49.655 | 2 | 51.710648 | yes | 1.3E-07 | 404.74304 | 1198.3925 |
| LNYSDHDVIK | 10 | Q96CW1 | AP-2 complex subunit mu OS=Homo sap | AP2M1 | 49.655 | 2 | 51.710648 | yes | 2.091E-07 | 602.30383 | 1017.4411 |
| SNFKPSLLAQK | 11 | Q96CW1 | AP-2 complex subunit mu OS=Homo sap | AP2M1 | 49.655 | 2 | 51.710648 | yes | 1.238E-07 | 616.85352 | 768.06848 |
| TSFFHVK | 7 | Q96CW1 | AP-2 complex subunit mu OS=Homo sap | AP2M1 | 49.655 | 2 | 51.710648 | yes | 3.371E-07 | 433.23196 | 648.15228 |
| MCDVMAAYFGK | 11 | Q96CW1 | AP-2 complex subunit mu OS=Homo sap | AP2M1 | 49.655 | 2 | 51.710648 | yes | 9.91E-08 | 646.77722 | 555.52936 |
| ASENAIVWK | 9 | Q96CW1 | AP-2 complex subunit mu OS=Homo sap | AP2M1 | 49.655 | 2 | 51.710648 | yes | 2.212E-07 | 509.27182 | 385.53647 |
| SPVTNIAR | 8 | Q96CW1 | AP-2 complex subunit mu OS=Homo sap | AP2M1 | 49.655 | 2 | 51.710648 | yes | 1.602E-07 | 429.24561 | 289.59036 |
| SGIYETR | 7 | Q96CW1 | AP-2 complex subunit mu OS=Homo sap | AP2M1 | 49.655 | 2 | 51.710648 | yes | 1.487E-07 | 413.20868 | 136.95607 |
| MIGGLFIYNHK | 11 | Q96CW1 | AP-2 complex subunit mu OS=Homo sap | AP2M1 | 49.655 | 2 | 51.710648 | yes | 2.519E-07 | 654.8421 | 101.05271 |
| YLLLGQGDFIR | 11 | Q96CW5 | Gamma-tubulin complex component 3 OS | TUBGCP3 | 103.572 | 2 | 53.72306 | yes | 1.37E-07 | 647.86133 | 2092.8616 |
| ATNAQFDSPEILR | 13 | Q96CW5 | Gamma-tubulin complex component 3 OS | TUBGCP3 | 103.572 | 2 | 53.72306 | yes | 1.161E-07 | 731.37024 | 1746.7926 |
| IDAAYFETSK | 10 | Q96CW5 | Gamma-tubulin complex component 3 OS | TUBGCP3 | 103.572 | 2 | 53.72306 | yes | 1.234E-07 | 572.77966 | 1513.5598 |
| VIGSNFAPTVER | 12 | Q96CW5 | Gamma-tubulin complex component 3 OS | TUBGCP3 | 103.572 | 2 | 53.72306 | yes | 9.722E-08 | 645.34607 | 1466.8773 |
| PQTLPLSYQDR | 11 | Q96CW5 | Gamma-tubulin complex component 3 OS | TUBGCP3 | 103.572 | 2 | 53.72306 | yes | 1.328E-07 | 659.34351 | 1411.9423 |
| MEYILTDIR | 9 | Q96CW5 | Gamma-tubulin complex component 3 OS | TUBGCP3 | 103.572 | 2 | 53.72306 | yes | 3.076E-06 | 577.29974 | 1280.5269 |
| SPNVLLQNLCCR | 12 | Q96CW5 | Gamma-tubulin complex component 3 OS | TUBGCP3 | 103.572 | 2 | 53.72306 | yes | 1.008E-07 | 737.36865 | 1279.7344 |
| LSELGWLHNK | 10 | Q96CW5 | Gamma-tubulin complex component 3 OS | TUBGCP3 | 103.572 | 2 | 53.72306 | yes | 9.893E-08 | 598.82477 | 1277.3719 |
| HLMDLLKPELVR | 12 | Q96CW5 | Gamma-tubulin complex component 3 OS | TUBGCP3 | 103.572 | 3 | 53.72306 | yes | 1.727E-07 | 488.61829 | 386.78244 |
| HLMDLLKPELVR | 12 | Q96CW5 | Gamma-tubulin complex component 3 OS | TUBGCP3 | 103.572 | 3 | 53.72306 | yes | 9.722E-08 | 732.42377 | 386.78244 |
| HLMDLLKPELVR | 12 | Q96CW5 | Gamma-tubulin complex component 3 OS | TUBGCP3 | 103.572 | 3 | 53.72306 | yes | 1.52E-05 | 493.94992 | 386.78244 |
| HLMDLLKPELVR | 12 | Q96CW5 | Gamma-tubulin complex component 3 OS | TUBGCP3 | 103.572 | 2 | 53.72306 | yes | 1.727E-07 | 488.61829 | 386.78244 |
| HLMDLLKPELVR | 12 | Q96CW5 | Gamma-tubulin complex component 3 OS | TUBGCP3 | 103.572 | 2 | 53.72306 | yes | 9.722E-08 | 732.42377 | 386.78244 |
| HLMDLLKPELVR | 12 | Q96CW5 | Gamma-tubulin complex component 3 OS | TUBGCP3 | 103.572 | 2 | 53.72306 | yes | 1.52E-05 | 493.94992 | 386.78244 |
| EADAALFSELHR | 12 | Q96CW5 | Gamma-tubulin complex component 3 OS | TUBGCP3 | 103.572 | 2 | 53.72306 | yes | 1.92E-07 | 679.83856 | 1107.2985 |
| AALEELQR | 8 | Q96CW5 | Gamma-tubulin complex component 3 OS | TUBGCP3 | 103.572 | 2 | 53.72306 | yes | 1.165E-07 | 465.25616 | 802.90894 |
| SMIPSFMTMDQSR | 13 | Q96CW5 | Gamma-tubulin complex component 3 OS | TUBGCP3 | 103.572 | 2 | 53.72306 | yes | 1.363E-07 | 765.84113 | 783.55298 |
| EIEGQWGVTAAEE | 19 | Q96CW5 | Gamma-tubulin complex component 3 OS | TUBGCP3 | 103.572 | 3 | 53.72306 | yes | 1.42E-07 | 735.34155 | 702.57483 |
| SEADVAQQFQYAV | 14 | Q96CW5 | Gamma-tubulin complex component 3 OS | TUBGCP3 | 103.572 | 2 | 53.72306 | yes | 1.214E-07 | 806.39172 | 699.67212 |
| YLLDVLNK | 8 | Q96CW5 | Gamma-tubulin complex component 3 OS | TUBGCP3 | 103.572 | 2 | 53.72306 | yes | 6.496E-07 | 489.28696 | 697.86401 |
| LLVWTYDPK | 9 | Q96CW5 | Gamma-tubulin complex component 3 OS | TUBGCP3 | 103.572 | 2 | 53.72306 | yes | 1.253E-07 | 567.81335 | 691.65112 |
| LDFNEHYK | 8 | Q96CW5 | Gamma-tubulin complex component 3 OS | TUBGCP3 | 103.572 | 2 | 53.72306 | yes | 1.003E-06 | 533.2536 | 591.5871 |
| CLLDSDSR | 8 | Q96CW5 | Gamma-tubulin complex component 3 OS | TUBGCP3 | 103.572 | 2 | 53.72306 | yes | 5.28E-07 | 483.22147 | 493.15039 |
| DAHSTPYYYAR | 11 | Q96CW5 | Gamma-tubulin complex component 3 OS | TUBGCP3 | 103.572 | 2 | 53.72306 | yes | 9.722E-08 | 672.30438 | 461.25092 |
| YSLLDHMQAMR | 11 | Q96CW5 | Gamma-tubulin complex component 3 OS | TUBGCP3 | 103.572 | 2 | 53.72306 | yes | 9.722E-08 | 682.82611 | 206.71817 |
| YSLLDHMQAMR | 11 | Q96CW5 | Gamma-tubulin complex component 3 OS | TUBGCP3 | 103.572 | 2 | 53.72306 | yes | 1.634E-07 | 455.55316 | 206.71817 |

| | | | | | | | | | | | |
|---|---|---|---|---|---|---|---|---|---|---|---|
| YSLLDHMQAMR | 11 | Q96CW5 | Gamma-tubulin complex component 3 OS | TUBGCP3 | 103.572 | 3 | 53.72306 | yes | 9.722E-08 | 682.82611 | 206.71817 |
| YSLLDHMQAMR | 11 | Q96CW5 | Gamma-tubulin complex component 3 OS | TUBGCP3 | 103.572 | 3 | 53.72306 | yes | 1.634E-07 | 455.55316 | 206.71817 |
| DILYVFQGIDGK | 12 | Q96CW5 | Gamma-tubulin complex component 3 OS | TUBGCP3 | 103.572 | 2 | 53.72306 | yes | 9.723E-08 | 684.36389 | 368.43811 |
| ECMSHYLR | 8 | Q96CW5 | Gamma-tubulin complex component 3 OS | TUBGCP3 | 103.572 | 2 | 53.72306 | yes | 1.499E-07 | 548.23914 | 345.04831 |
| LHSQGVLK | 8 | Q96CW5 | Gamma-tubulin complex component 3 OS | TUBGCP3 | 103.572 | 2 | 53.72306 | yes | 3.67E-07 | 441.26379 | 330.99823 |
| ALLNQLR | 7 | Q96CW5 | Gamma-tubulin complex component 3 OS | TUBGCP3 | 103.572 | 2 | 53.72306 | yes | 4.237E-07 | 414.25851 | 236.87074 |
| SFGLVGQSFCAAL | 18 | Q96CW5 | Gamma-tubulin complex component 3 OS | TUBGCP3 | 103.572 | 3 | 53.72306 | yes | 9.722E-08 | 674.00714 | 189.17889 |
| VSSYATLFAQALP | 14 | Q96CW5 | Gamma-tubulin complex component 3 OS | TUBGCP3 | 103.572 | 2 | 53.72306 | yes | 5.816E-07 | 762.41449 | 160.31923 |
| KLHSQGVLK | 9 | Q96CW5 | Gamma-tubulin complex component 3 OS | TUBGCP3 | 103.572 | 2 | 53.72306 | yes | 1.3E-07 | 505.31128 | 118.40484 |
| READAALFSELHR | 13 | Q96CW5 | Gamma-tubulin complex component 3 OS | TUBGCP3 | 103.572 | 3 | 53.72306 | yes | 9.787E-08 | 505.59518 | 64.32486 |
| PATTLYQHNLTGII | 19 | Q96CW5 | Gamma-tubulin complex component 3 OS | TUBGCP3 | 103.572 | 3 | 53.72306 | yes | 9.722E-08 | 700.04791 | 60.462955 |
| MNNTENCYK | 9 | Q96CW5 | Gamma-tubulin complex component 3 OS | TUBGCP3 | 103.572 | 2 | 53.72306 | yes | 2.197E-05 | 587.23682 | 54.976849 |
| KGGELASAVHAY | 14 | Q96CW5 | Gamma-tubulin complex component 3 OS | TUBGCP3 | 103.572 | 2 | 53.72306 | yes | 1.982E-07 | 716.38318 | 39.40464 |
| AGDGDGWVSLAE | 14 | Q96D15 | Reticulocalbin-3 OS=Homo sapiens OX= | RCN3 | 37.493 | 2 | 51.33209 | yes | 1.033E-07 | 723.35461 | 16220.141 |
| AWIAHTQQR | 9 | Q96D15 | Reticulocalbin-3 OS=Homo sapiens OX= | RCN3 | 37.493 | 2 | 51.33209 | yes | 9.734E-08 | 555.79376 | 9406.1553 |
| VGWEELR | 7 | Q96D15 | Reticulocalbin-3 OS=Homo sapiens OX= | RCN3 | 37.493 | 2 | 51.33209 | yes | 1.244E-07 | 444.73233 | 7956.8335 |
| DSVSAAWDTYDTI | 14 | Q96D15 | Reticulocalbin-3 OS=Homo sapiens OX= | RCN3 | 37.493 | 2 | 51.33209 | yes | 1.668E-07 | 801.33936 | 7604.9912 |
| EFDQLTPEESQAR | 13 | Q96D15 | Reticulocalbin-3 OS=Homo sapiens OX= | RCN3 | 37.493 | 2 | 51.33209 | yes | 3.766E-07 | 775.36011 | 5024.2832 |
| DIVIAETLEDLDR | 13 | Q96D15 | Reticulocalbin-3 OS=Homo sapiens OX= | RCN3 | 37.493 | 2 | 51.33209 | yes | 1.104E-07 | 751.39087 | 1906.8167 |
| VHQAAPLSDAPHI | 29 | Q96D15 | Reticulocalbin-3 OS=Homo sapiens OX= | RCN3 | 37.493 | 5 | 51.33209 | yes | 1.087E-07 | 643.89935 | 1831.6714 |
| VADQDGDSMATR | 12 | Q96D15 | Reticulocalbin-3 OS=Homo sapiens OX= | RCN3 | 37.493 | 2 | 51.33209 | yes | 9.722E-08 | 633.27496 | 1549.4899 |
| EELTAFLHPEEFPH | 16 | Q96D15 | Reticulocalbin-3 OS=Homo sapiens OX= | RCN3 | 37.493 | 4 | 51.33209 | yes | 8.63E-07 | 496.49118 | 100.7182 |
| EELTAFLHPEEFPH | 16 | Q96D15 | Reticulocalbin-3 OS=Homo sapiens OX= | RCN3 | 37.493 | 4 | 51.33209 | yes | 0.0001198 | 666.98413 | 100.7182 |
| EELTAFLHPEEFPH | 16 | Q96D15 | Reticulocalbin-3 OS=Homo sapiens OX= | RCN3 | 37.493 | 3 | 51.33209 | yes | 8.63E-07 | 496.49118 | 100.7182 |
| EELTAFLHPEEFPH | 16 | Q96D15 | Reticulocalbin-3 OS=Homo sapiens OX= | RCN3 | 37.493 | 3 | 51.33209 | yes | 0.0001198 | 666.98413 | 100.7182 |
| DGHLDGSEVGHW | 36 | Q96D15 | Reticulocalbin-3 OS=Homo sapiens OX= | RCN3 | 37.493 | 5 | 51.33209 | yes | 2.036E-07 | 791.98273 | 95.999382 |
| STFSVEIVPICK | 12 | Q96D46 | 60S ribosomal export protein NMD3 OS= | NMD3 | 57.604 | 2 | 53.68509 | yes | 9.722E-08 | 690.3656 | 542.46448 |
| MVEFLQCTVPCR | 12 | Q96D46 | 60S ribosomal export protein NMD3 OS= | NMD3 | 57.604 | 2 | 53.68509 | yes | 1.274E-07 | 770.35944 | 228.56703 |
| ELLALCLK | 8 | Q96D46 | 60S ribosomal export protein NMD3 OS= | NMD3 | 57.604 | 2 | 53.68509 | yes | 2.834E-06 | 480.28336 | 109.57558 |
| DNVVCLSPK | 9 | Q96D46 | 60S ribosomal export protein NMD3 OS= | NMD3 | 57.604 | 2 | 53.68509 | yes | 2.212E-07 | 516.26312 | 99.430504 |
| YGMHQNTLR | 9 | Q96D46 | 60S ribosomal export protein NMD3 OS= | NMD3 | 57.604 | 2 | 53.68509 | yes | 3.515E-07 | 560.27185 | 97.171112 |
| EIHDGLDFYYSSK | 13 | Q96D46 | 60S ribosomal export protein NMD3 OS= | NMD3 | 57.604 | 2 | 53.68509 | yes | 2.642E-07 | 787.36212 | 80.427841 |
| VPDVVLIK | 8 | Q96D46 | 60S ribosomal export protein NMD3 OS= | NMD3 | 57.604 | 2 | 53.68509 | yes | 1.47E-07 | 441.78659 | 77.549179 |
| SKVDISQGIPK | 11 | Q96D46 | 60S ribosomal export protein NMD3 OS= | NMD3 | 57.604 | 2 | 53.68509 | yes | 1.002E-07 | 586.33771 | 66.622589 |
| ENMDTDDER | 9 | Q96D46 | 60S ribosomal export protein NMD3 OS= | NMD3 | 57.604 | 2 | 53.68509 | yes | 2.902E-06 | 562.71167 | 51.899406 |
| LISQDIHSNTYNYK | 14 | Q96D46 | 60S ribosomal export protein NMD3 OS= | NMD3 | 57.604 | 2 | 53.68509 | yes | 1.025E-07 | 848.42047 | 24.885132 |
| LISQDIHSNTYNYK | 14 | Q96D46 | 60S ribosomal export protein NMD3 OS= | NMD3 | 57.604 | 2 | 53.68509 | yes | 7.785E-05 | 565.9494 | 24.885132 |
| LISQDIHSNTYNYK | 14 | Q96D46 | 60S ribosomal export protein NMD3 OS= | NMD3 | 57.604 | 3 | 53.68509 | yes | 1.025E-07 | 848.42047 | 24.885132 |
| LISQDIHSNTYNYK | 14 | Q96D46 | 60S ribosomal export protein NMD3 OS= | NMD3 | 57.604 | 3 | 53.68509 | yes | 7.785E-05 | 565.9494 | 24.885132 |

| | | | | | | | | | | | |
|---|---|---|---|---|---|---|---|---|---|---|---|
| HTLGEVWVQK | 10 | Q96D46 | 60S ribosomal export protein NMD3 OS= | NMD3 | 57.604 | 2 | 53.68509 | yes | 4.952E-05 | 598.82477 | 30.918446 |
| KHTLGEVWVQK | 11 | Q96D46 | 60S ribosomal export protein NMD3 OS= | NMD3 | 57.604 | 3 | 53.68509 | yes | 3.665E-06 | 442.25058 | 26.918236 |
| IQDLIPVLLR | 10 | Q96D53 | Atypical kinase COQ8B, mitochondrial O | COQ8B | 60.07 | 2 | 34.44733 | yes | 1.007E-06 | 590.37665 | 39.762909 |
| TAASAPANVSK | 11 | Q96D71 | RalBP1-associated Eps domain-containin | REPS1 | 86.663 | 2 | 43.87175 | yes | 2.411E-07 | 508.77219 | 62.31461 |
| EAALILGVSPTANI | 14 | Q96DA6 | Mitochondrial import inner membrane tra | DNAJC19 | 12.499 | 2 | 49.291725 | yes | 1.048E-07 | 692.39575 | 141.51915 |
| IMLLNHPDK | 9 | Q96DA6 | Mitochondrial import inner membrane tra | DNAJC19 | 12.499 | 2 | 49.291725 | yes | 2.342E-07 | 540.79718 | 114.97623 |
| QVFQSLPK | 8 | Q96DA6 | Mitochondrial import inner membrane tra | DNAJC19 | 12.499 | 2 | 49.291725 | yes | 4.835E-07 | 473.77145 | 73.588226 |
| SAFSGGYYR | 9 | Q96DA6 | Mitochondrial import inner membrane tra | DNAJC19 | 12.499 | 2 | 49.291725 | yes | 6.562E-07 | 504.2327 | 53.367374 |
| GAGEMLEDGSER | 12 | Q96DE5 | Anaphase-promoting complex subunit 16 | ANAPC16 | 11.668 | 2 | 53.277958 | yes | 9.722E-08 | 625.76953 | 102.51726 |
| FKPIEQLLGFTPSS | 15 | Q96DE5 | Anaphase-promoting complex subunit 16 | ANAPC16 | 11.668 | 2 | 53.277958 | yes | 1.232E-07 | 810.93524 | 72.890121 |
| LAGLVEELEADEW | 14 | Q96DE5 | Anaphase-promoting complex subunit 16 | ANAPC16 | 11.668 | 2 | 53.277958 | yes | 1.056E-07 | 815.40961 | 59.690178 |
| GLSPAMSPALQR | 12 | Q96DF8 | Splicing factor ESS-2 homolog OS=Hom | ESS2 | 52.569 | 2 | 47.351185 | yes | 1.751E-07 | 614.32935 | 165.62701 |
| VEGSETPYVDR | 11 | Q96DF8 | Splicing factor ESS-2 homolog OS=Hom | ESS2 | 52.569 | 2 | 47.351185 | yes | 1.348E-07 | 626.2962 | 136.33023 |
| VTENLASLTPK | 11 | Q96DF8 | Splicing factor ESS-2 homolog OS=Hom | ESS2 | 52.569 | 2 | 47.351185 | yes | 1.577E-07 | 586.82971 | 84.433777 |
| METPGASASSLLL | 21 | Q96DF8 | Splicing factor ESS-2 homolog OS=Hom | ESS2 | 52.569 | 3 | 47.351185 | yes | 1.235E-06 | 717.7124 | 70.549278 |
| EYLEAEENGDLER | 13 | Q96DF8 | Splicing factor ESS-2 homolog OS=Hom | ESS2 | 52.569 | 2 | 47.351185 | yes | 8.443E-07 | 783.84955 | 56.134472 |
| CQLQQAAALNAQ | 14 | Q96DF8 | Splicing factor ESS-2 homolog OS=Hom | ESS2 | 52.569 | 2 | 47.351185 | yes | 1.503E-07 | 790.90173 | 32.846558 |
| VLDEEEYIEGLQT | 17 | Q96DF8 | Splicing factor ESS-2 homolog OS=Hom | ESS2 | 52.569 | 3 | 47.351185 | yes | 3.025E-07 | 678.68451 | 32.391685 |
| HAWLYQAEEEFEI | 13 | Q96DF8 | Splicing factor ESS-2 homolog OS=Hom | ESS2 | 52.569 | 2 | 47.351185 | yes | 3.592E-07 | 840.38867 | 10.370853 |
| VEDAMLMFDK | 10 | Q96DH6 | RNA-binding protein Musashi homolog 2 | MSI2 | 35.197 | 2 | 48.412262 | yes | 1.53E-07 | 599.77777 | 3254.7471 |
| GFGFVTFADPASV | 15 | Q96DH6 | RNA-binding protein Musashi homolog 2 | MSI2 | 35.197 | 2 | 48.412262 | yes | 1.152E-07 | 779.38281 | 2774.8445 |
| IFVGGLSANTVVE | 16 | Q96DH6 | RNA-binding protein Musashi homolog 2 | MSI2 | 35.197 | 2 | 48.412262 | yes | 1.743E-07 | 824.45123 | 1878.8396 |
| QYFEQFGK | 8 | Q96DH6 | RNA-binding protein Musashi homolog 2 | MSI2 | 35.197 | 2 | 48.412262 | no | 1.441E-07 | 523.75073 | 1525.3984 |
| VLGQPHHELDSK | 12 | Q96DH6 | RNA-binding protein Musashi homolog 2 | MSI2 | 35.197 | 2 | 48.412262 | yes | 1.877E-07 | 680.35443 | 765.04144 |
| EVMFPPGTR | 9 | Q96DH6 | RNA-binding protein Musashi homolog 2 | MSI2 | 35.197 | 2 | 48.412262 | yes | 1.47E-06 | 517.26038 | 179.87221 |
| VCEIHFHEINNK | 12 | Q96DH6 | RNA-binding protein Musashi homolog 2 | MSI2 | 35.197 | 2 | 48.412262 | no | 2.883E-07 | 770.37225 | 165.67613 |
| FVYVWDTTSR | 10 | Q96DI7 | U5 small nuclear ribonucleoprotein 40 kD | SNRNP40 | 39.311 | 2 | 51.06192 | yes | 1.146E-07 | 637.31421 | 459.52579 |
| TVAVWDSETGER | 12 | Q96DI7 | U5 small nuclear ribonucleoprotein 40 kD | SNRNP40 | 39.311 | 2 | 51.06192 | yes | 1.248E-07 | 675.32025 | 431.46399 |
| VWDVRPFAPK | 10 | Q96DI7 | U5 small nuclear ribonucleoprotein 40 kD | SNRNP40 | 39.311 | 2 | 51.06192 | yes | 9.722E-08 | 607.83765 | 235.64484 |
| KGPELPLVPVK | 11 | Q96DI7 | U5 small nuclear ribonucleoprotein 40 kD | SNRNP40 | 39.311 | 2 | 51.06192 | yes | 1.214E-07 | 588.87115 | 199.812 |
| GPQLVCTGSDDGT | 15 | Q96DI7 | U5 small nuclear ribonucleoprotein 40 kD | SNRNP40 | 39.311 | 2 | 51.06192 | yes | 3.41E-07 | 767.36432 | 56.945503 |
| CSSLQAPIMLLSGI | 22 | Q96DI7 | U5 small nuclear ribonucleoprotein 40 kD | SNRNP40 | 39.311 | 3 | 51.06192 | yes | 4.738E-07 | 847.0531 | 37.496414 |
| VLFFGTDQFAR | 11 | Q96DP5 | Methionyl-tRNA formyltransferase, mitoc | MTFMT | 43.833 | 2 | 53.427277 | yes | 9.722E-08 | 650.83789 | 363.34296 |
| FPYGILNVHPSCLI | 15 | Q96DP5 | Methionyl-tRNA formyltransferase, mitoc | MTFMT | 43.833 | 3 | 53.427277 | yes | 2.87E-06 | 590.64337 | 150.6257 |
| QQPMEGATYAPK | 12 | Q96DP5 | Methionyl-tRNA formyltransferase, mitoc | MTFMT | 43.833 | 2 | 53.427277 | yes | 2.466E-07 | 660.81628 | 128.64011 |
| LLNEALILK | 9 | Q96DP5 | Methionyl-tRNA formyltransferase, mitoc | MTFMT | 43.833 | 2 | 53.427277 | yes | 6.352E-07 | 513.83154 | 97.915482 |
| LTGQALIPGSVIYH | 15 | Q96DP5 | Methionyl-tRNA formyltransferase, mitoc | MTFMT | 43.833 | 2 | 53.427277 | yes | 5.595E-07 | 798.95905 | 52.360565 |
| WEEQTSEQIFR | 11 | Q96DP5 | Methionyl-tRNA formyltransferase, mitoc | MTFMT | 43.833 | 2 | 53.427277 | yes | 4.343E-07 | 726.84131 | 45.544418 |
| LGWEDCR | 7 | Q96DP5 | Methionyl-tRNA formyltransferase, mitoc | MTFMT | 43.833 | 2 | 53.427277 | yes | 2.012E-06 | 468.20563 | 29.591429 |

| | | | | | | | | | | | |
|---|---|---|---|---|---|---|---|---|---|---|---|
| LGANMLISVLK | 11 | Q96DP5 | Methionyl-tRNA formyltransferase, mitoc | MTFMT | 43.833 | 2 | 53.427277 | yes | 9.722E-08 | 579.84937 | 14.748545 |
| LQCSFQTSWLK | 11 | Q96DT7 | Zinc finger and BTB domain-containing p | ZBTB10 | 94.894 | 2 | 51.4895 | yes | 1.902E-07 | 699.34772 | 301.11151 |
| YGVCVDCADK | 10 | Q96DT7 | Zinc finger and BTB domain-containing p | ZBTB10 | 94.894 | 2 | 51.4895 | yes | 2.985E-07 | 593.74701 | 109.56813 |
| NLTNLASNVK | 10 | Q96DT7 | Zinc finger and BTB domain-containing p | ZBTB10 | 94.894 | 2 | 51.4895 | yes | 5.745E-07 | 537.30109 | 70.466789 |
| HLLIHTGVR | 9 | Q96DT7 | Zinc finger and BTB domain-containing p | ZBTB10 | 94.894 | 2 | 51.4895 | yes | 1E-07 | 523.31689 | 69.487106 |
| YGLMPGPSNDFK | 12 | Q96DT7 | Zinc finger and BTB domain-containing p | ZBTB10 | 94.894 | 2 | 51.4895 | yes | 3.086E-06 | 663.31335 | 65.691757 |
| SEAPESVVVDYNN | 14 | Q96DT7 | Zinc finger and BTB domain-containing p | ZBTB10 | 94.894 | 2 | 51.4895 | yes | 9.722E-08 | 789.87335 | 65.626961 |
| IASFWATR | 8 | Q96DT7 | Zinc finger and BTB domain-containing p | ZBTB10 | 94.894 | 2 | 51.4895 | yes | 4.253E-07 | 476.25598 | 51.395378 |
| GPATVDLELDALE | 15 | Q96DT7 | Zinc finger and BTB domain-containing p | ZBTB10 | 94.894 | 2 | 51.4895 | yes | 1.524E-07 | 764.39868 | 48.665405 |
| GSLNEAEAGALPA | 16 | Q96DU7 | Inositol-trisphosphate 3-kinase C OS=Ho | ITPKC | 75.208 | 2 | 51.44139 | yes | 1.082E-07 | 749.38641 | 246.04843 |
| YVACLEELR | 9 | Q96DU7 | Inositol-trisphosphate 3-kinase C OS=Ho | ITPKC | 75.208 | 2 | 51.44139 | yes | 1.075E-07 | 576.78955 | 147.59991 |
| CSPLCPVPR | 9 | Q96DU7 | Inositol-trisphosphate 3-kinase C OS=Ho | ITPKC | 75.208 | 2 | 51.44139 | yes | 1.217E-07 | 543.26514 | 140.32983 |
| TYLEEELVK | 9 | Q96DU7 | Inositol-trisphosphate 3-kinase C OS=Ho | ITPKC | 75.208 | 2 | 51.44139 | yes | 5.183E-07 | 562.29773 | 106.01245 |
| YSPFVVSFR | 9 | Q96DU7 | Inositol-trisphosphate 3-kinase C OS=Ho | ITPKC | 75.208 | 2 | 51.44139 | yes | 1.807E-07 | 551.29004 | 95.41674 |
| FCQCEQR | 7 | Q96DU7 | Inositol-trisphosphate 3-kinase C OS=Ho | ITPKC | 75.208 | 2 | 51.44139 | yes | 9.722E-08 | 514.20782 | 93.639236 |
| EALEISPFFK | 10 | Q96DU7 | Inositol-trisphosphate 3-kinase C OS=Ho | ITPKC | 75.208 | 2 | 51.44139 | yes | 5.796E-07 | 590.81604 | 57.758102 |
| TQALEQVTK | 9 | Q96DU7 | Inositol-trisphosphate 3-kinase C OS=Ho | ITPKC | 75.208 | 2 | 51.44139 | yes | 3.215E-05 | 509.28238 | 42.654823 |
| VWMIDFGK | 8 | Q96DU7 | Inositol-trisphosphate 3-kinase C OS=Ho | ITPKC | 75.208 | 2 | 51.44139 | no | 1.673E-07 | 498.25458 | 39.781658 |
| TEGSSLHSEPER | 12 | Q96DU7 | Inositol-trisphosphate 3-kinase C OS=Ho | ITPKC | 75.208 | 2 | 51.44139 | yes | 3.444E-06 | 664.8075 | 37.307941 |
| MVAVDPGAPTPEI | 23 | Q96DU7 | Inositol-trisphosphate 3-kinase C OS=Ho | ITPKC | 75.208 | 4 | 51.44139 | yes | 5.749E-07 | 590.30225 | 23.164749 |
| PSPCYQLAQR | 10 | Q96DV4 | Large ribosomal subunit protein mL38 OS | MRPL38 | 44.597 | 2 | 54.331726 | yes | 1.656E-07 | 610.29803 | 306.58588 |
| QDQPIDFSEDAR | 12 | Q96DV4 | Large ribosomal subunit protein mL38 OS | MRPL38 | 44.597 | 2 | 54.331726 | yes | 9.722E-08 | 710.82062 | 204.01997 |
| TASVPLDAVR | 10 | Q96DV4 | Large ribosomal subunit protein mL38 OS | MRPL38 | 44.597 | 2 | 54.331726 | yes | 3.867E-07 | 514.79041 | 163.64729 |
| AEQEAQAPHWWR | 12 | Q96DV4 | Large ribosomal subunit protein mL38 OS | MRPL38 | 44.597 | 2 | 54.331726 | yes | 9.722E-08 | 754.8551 | 110.6341 |
| LAEYYGLYR | 9 | Q96DV4 | Large ribosomal subunit protein mL38 OS | MRPL38 | 44.597 | 2 | 54.331726 | yes | 9.722E-08 | 574.29279 | 101.19127 |
| DLFHGATFVPR | 11 | Q96DV4 | Large ribosomal subunit protein mL38 OS | MRPL38 | 44.597 | 2 | 54.331726 | yes | 9.722E-08 | 630.3302 | 90.435951 |
| TQQLLER | 7 | Q96DV4 | Large ribosomal subunit protein mL38 OS | MRPL38 | 44.597 | 2 | 54.331726 | yes | 1.022E-07 | 444.25089 | 73.543015 |
| IDIGLPPPK | 9 | Q96DV4 | Large ribosomal subunit protein mL38 OS | MRPL38 | 44.597 | 2 | 54.331726 | yes | 7.442E-06 | 475.28949 | 24.142727 |
| DSHEPTYGIY | 10 | Q96DV4 | Large ribosomal subunit protein mL38 OS | MRPL38 | 44.597 | 2 | 54.331726 | yes | 2.04E-07 | 591.25909 | 13.121372 |
| LALNQISQISMK | 12 | Q96E11 | Ribosome-recycling factor, mitochondrial | MRRF | 29.278 | 2 | 41.452038 | yes | 3.502E-07 | 673.37903 | 80.815918 |
| AAQDFCQLVAQK | 12 | Q96E22 | Dehydrodolichyl diphosphate synthase co | NUS1 | 33.225 | 2 | 43.87149 | yes | 2.412E-07 | 689.84283 | 127.78793 |
| QQQELLGLDCSK | 12 | Q96E22 | Dehydrodolichyl diphosphate synthase co | NUS1 | 33.225 | 2 | 43.87149 | yes | 4.367E-07 | 709.85083 | 51.373131 |
| FPQVFNTR | 8 | Q96E29 | Transcription termination factor 3, mitoch | MTERF3 | 47.972 | 2 | 52.531826 | yes | 4.626E-07 | 504.76672 | 1741.4369 |
| HPEAANLLLR | 10 | Q96E29 | Transcription termination factor 3, mitoch | MTERF3 | 47.972 | 2 | 52.531826 | yes | 9.991E-08 | 567.32495 | 1612.3236 |
| DVGIEDNQLGAFL | 15 | Q96E29 | Transcription termination factor 3, mitoch | MTERF3 | 47.972 | 2 | 52.531826 | yes | 1.865E-07 | 810.41742 | 1125.1498 |
| LLTGSLEPVK | 10 | Q96E29 | Transcription termination factor 3, mitoch | MTERF3 | 47.972 | 2 | 52.531826 | yes | 1.121E-07 | 528.8186 | 982.41302 |
| HLFLTYLGR | 9 | Q96E29 | Transcription termination factor 3, mitoch | MTERF3 | 47.972 | 2 | 52.531826 | yes | 1.121E-07 | 560.31909 | 973.28693 |
| ASVQDFEK | 8 | Q96E29 | Transcription termination factor 3, mitoch | MTERF3 | 47.972 | 2 | 52.531826 | yes | 3.95E-07 | 462.22708 | 803.91467 |
| LVLLGVDLSK | 10 | Q96E29 | Transcription termination factor 3, mitoch | MTERF3 | 47.972 | 2 | 52.531826 | yes | 9.722E-08 | 528.83679 | 598.50983 |

| | | | | | | | | | | | |
|---|---|---|---|---|---|---|---|---|---|---|---|
| VAYLHSK | 7 | Q96E29 | Transcription termination factor 3, mitoch | MTERF3 | 47.972 | 2 | 52.531826 | yes | 2.078E-07 | 409.23196 | 519.45471 |
| ADVAQMVR | 8 | Q96E29 | Transcription termination factor 3, mitoch | MTERF3 | 47.972 | 2 | 52.531826 | yes | 1.198E-07 | 445.23166 | 496.18613 |
| HNEIQHMITR | 10 | Q96E29 | Transcription termination factor 3, mitoch | MTERF3 | 47.972 | 2 | 52.531826 | yes | 9.722E-08 | 639.8222 | 312.34317 |
| DYVDHSETLQK | 11 | Q96E29 | Transcription termination factor 3, mitoch | MTERF3 | 47.972 | 2 | 52.531826 | yes | 1.391E-07 | 667.81476 | 270.05145 |
| KAPFLLNFSVER | 12 | Q96E29 | Transcription termination factor 3, mitoch | MTERF3 | 47.972 | 2 | 52.531826 | yes | 9.722E-08 | 710.901 | 94.318024 |
| NHAIFSEDLENLK | 13 | Q96E29 | Transcription termination factor 3, mitoch | MTERF3 | 47.972 | 2 | 52.531826 | yes | 1.565E-05 | 765.38336 | 89.0103 |
| QMLLFLK | 7 | Q96E29 | Transcription termination factor 3, mitoch | MTERF3 | 47.972 | 2 | 52.531826 | yes | 3.629E-06 | 446.76987 | 40.106522 |
| APFLLNFSVER | 11 | Q96E29 | Transcription termination factor 3, mitoch | MTERF3 | 47.972 | 2 | 52.531826 | yes | 2.049E-07 | 646.85352 | 28.16983 |
| FMHVLYPR | 8 | Q96EC8 | Protein YIPF6 OS=Homo sapiens OX=96 | YIPF6 | 26.256 | 2 | 53.031776 | yes | 7.315E-07 | 531.78131 | 87.318054 |
| EFDSSTLNESVR | 12 | Q96EC8 | Protein YIPF6 OS=Homo sapiens OX=96 | YIPF6 | 26.256 | 2 | 53.031776 | yes | 9.722E-08 | 692.323 | 75.77491 |
| IFTLKPVR | 8 | Q96EE3 | Nucleoporin SEH1 OS=Homo sapiens OX | SEH1L | 39.649 | 2 | 49.898724 | yes | 1.34E-07 | 487.31329 | 381.88428 |
| VQIFEYNENTR | 11 | Q96EE3 | Nucleoporin SEH1 OS=Homo sapiens OX | SEH1L | 39.649 | 2 | 49.898724 | yes | 1.605E-07 | 706.84387 | 258.20972 |
| TAAVWEEIVGESN | 15 | Q96EE3 | Nucleoporin SEH1 OS=Homo sapiens OX | SEH1L | 39.649 | 2 | 49.898724 | yes | 2.373E-07 | 824.39667 | 221.09854 |
| SFHILAIATK | 10 | Q96EE3 | Nucleoporin SEH1 OS=Homo sapiens OX | SEH1L | 39.649 | 2 | 49.898724 | yes | 9.996E-08 | 550.82678 | 199.52498 |
| LSCSCISWNPSSSR | 14 | Q96EE3 | Nucleoporin SEH1 OS=Homo sapiens OX | SEH1L | 39.649 | 2 | 49.898724 | yes | 4.785E-07 | 820.86139 | 107.58906 |
| DLIHDVSFDFHGR | 13 | Q96EE3 | Nucleoporin SEH1 OS=Homo sapiens OX | SEH1L | 39.649 | 2 | 49.898724 | yes | 3.627E-07 | 779.37585 | 98.064636 |
| HMGLMLATCSAD | 16 | Q96EE3 | Nucleoporin SEH1 OS=Homo sapiens OX | SEH1L | 39.649 | 3 | 49.898724 | yes | 2.019E-07 | 577.94946 | 52.684788 |
| AETLMTVTDPVHI | 22 | Q96EE3 | Nucleoporin SEH1 OS=Homo sapiens OX | SEH1L | 39.649 | 3 | 49.898724 | yes | 2.179E-05 | 790.07037 | 51.310406 |
| MATCSSDQSVK | 11 | Q96EE3 | Nucleoporin SEH1 OS=Homo sapiens OX | SEH1L | 39.649 | 2 | 49.898724 | yes | 4.381E-06 | 607.263 | 46.559425 |
| DICVIQVPPEMR | 12 | Q96EH3 | Mitochondrial assembly of ribosomal larg | MALSU1 | 26.171 | 2 | 54.87801 | yes | 3.49E-07 | 728.86798 | 528.60028 |
| YTDYFVIVSGTSTI | 14 | Q96EH3 | Mitochondrial assembly of ribosomal larg | MALSU1 | 26.171 | 2 | 54.87801 | yes | 9.722E-08 | 804.89886 | 487.6037 |
| LCSVHFQGGR | 10 | Q96EK4 | THAP domain-containing protein 11 OS= | THAP11 | 34.455 | 2 | 50.880695 | yes | 9.722E-08 | 580.7851 | 251.52203 |
| LLAMAVIR | 8 | Q96EK4 | THAP domain-containing protein 11 OS= | THAP11 | 34.455 | 2 | 50.880695 | yes | 1.021E-07 | 443.78076 | 214.36362 |
| VPTIFPLR | 8 | Q96EK4 | THAP domain-containing protein 11 OS= | THAP11 | 34.455 | 2 | 50.880695 | yes | 2.257E-07 | 471.79221 | 169.21198 |
| DILALMEVK | 9 | Q96EK4 | THAP domain-containing protein 11 OS= | THAP11 | 34.455 | 2 | 50.880695 | yes | 9.79E-08 | 516.29388 | 52.899723 |
| VLAFFETDMER | 11 | Q96EK5 | KIF-binding protein OS=Homo sapiens C | KIFBP | 71.814 | 2 | 51.508896 | yes | 9.722E-08 | 679.32648 | 69.026978 |
| FIVDYCEK | 8 | Q96EK5 | KIF-binding protein OS=Homo sapiens C | KIFBP | 71.814 | 2 | 51.508896 | yes | 1.391E-07 | 537.25226 | 67.259315 |
| HDGSEPCVDVLFC | 17 | Q96EL3 | Large ribosomal subunit protein mL53 OS | MRPL53 | 12.107 | 3 | 49.924923 | yes | 1.622E-07 | 632.94794 | 641.22327 |
| TFLQTVSSEK | 10 | Q96EL3 | Large ribosomal subunit protein mL53 OS | MRPL53 | 12.107 | 2 | 49.924923 | yes | 1.874E-07 | 570.30078 | 491.18146 |
| STNLNCSVIADVR | 13 | Q96EL3 | Large ribosomal subunit protein mL53 OS | MRPL53 | 12.107 | 2 | 49.924923 | yes | 9.976E-08 | 724.86176 | 455.67682 |
| VQFCPFEK | 8 | Q96EL3 | Large ribosomal subunit protein mL53 OS | MRPL53 | 12.107 | 2 | 49.924923 | yes | 1.211E-07 | 527.75494 | 186.23438 |
| LTHDTVEQVR | 10 | Q96EN8 | Molybdenum cofactor sulfurase OS=Hom | MOCOS | 98.12 | 2 | 45.86248 | yes | 1.439E-06 | 599.31476 | 148.91638 |
| LAYGYGPGSLR | 11 | Q96EN8 | Molybdenum cofactor sulfurase OS=Hom | MOCOS | 98.12 | 2 | 45.86248 | yes | 3.219E-06 | 577.30365 | 124.31151 |
| FQVLGPCHR | 9 | Q96EN8 | Molybdenum cofactor sulfurase OS=Hom | MOCOS | 98.12 | 2 | 45.86248 | yes | 1.575E-06 | 557.28473 | 103.13662 |
| CQMICIDQQTGQR | 13 | Q96EN8 | Molybdenum cofactor sulfurase OS=Hom | MOCOS | 98.12 | 2 | 45.86248 | yes | 1.721E-07 | 819.36328 | 39.546341 |
| TGCFCNTGACQR | 12 | Q96EN8 | Molybdenum cofactor sulfurase OS=Hom | MOCOS | 98.12 | 2 | 45.86248 | yes | 2.928E-07 | 716.28174 | 11.821265 |
| NIDPKPCTPR | 10 | Q96EP5 | DAZ-associated protein 1 OS=Homo sapi | DAZAP1 | 43.384 | 2 | 49.329697 | yes | 1.043E-07 | 599.30585 | 3051.4119 |
| SQAPGQPGASQWC | 15 | Q96EP5 | DAZ-associated protein 1 OS=Homo sapi | DAZAP1 | 43.384 | 2 | 49.329697 | yes | 2.151E-07 | 757.36078 | 2645.0945 |
| FKDPNCVGTVLAS | 14 | Q96EP5 | DAZ-associated protein 1 OS=Homo sapi | DAZAP1 | 43.384 | 2 | 49.329697 | yes | 1.248E-07 | 782.40106 | 1556.4291 |

| | | | | | | | | | | | |
|---|---|---|---|---|---|---|---|---|---|---|---|
| DPNCVGTVLASR | 12 | Q96EP5 | DAZ-associated protein 1 OS=Homo sapi | DAZAP1 | 43.384 | 2 | 49.329697 | yes | 2.072E-07 | 644.81934 | 1260.8463 |
| KFGVVTEVVMIYI | 16 | Q96EP5 | DAZ-associated protein 1 OS=Homo sapi | DAZAP1 | 43.384 | 3 | 49.329697 | yes | 4.81E-07 | 609.99023 | 194.42419 |
| MNNSGADEIGK | 11 | Q96EP5 | DAZ-associated protein 1 OS=Homo sapi | DAZAP1 | 43.384 | 2 | 49.329697 | yes | 1.345E-07 | 589.26135 | 130.72537 |
| GQNHNVQGFHPY | 13 | Q96EP5 | DAZ-associated protein 1 OS=Homo sapi | DAZAP1 | 43.384 | 2 | 49.329697 | yes | 2.034E-07 | 777.37146 | 102.89333 |
| SEHNLVNIQK | 10 | Q96ES7 | SAGA-associated factor 29 OS=Homo sa | SGF29 | 33.238 | 2 | 52.29492 | yes | 9.722E-08 | 591.31732 | 64.979752 |
| ADAEAECNILR | 11 | Q96ES7 | SAGA-associated factor 29 OS=Homo sa | SGF29 | 33.238 | 2 | 52.29492 | yes | 6.768E-07 | 631.29572 | 31.297712 |
| KLENFLSR | 8 | Q96EU6 | Ribosomal RNA processing protein 36 hc | RRP36 | 29.824 | 2 | 48.78341 | yes | 1.127E-07 | 503.78763 | 477.09229 |
| DREEDGGGLEPAA | 16 | Q96EU6 | Ribosomal RNA processing protein 36 hc | RRP36 | 29.824 | 3 | 48.78341 | yes | 1.104E-07 | 547.93243 | 178.763 |
| DREEDGGGLEPAA | 16 | Q96EU6 | Ribosomal RNA processing protein 36 hc | RRP36 | 29.824 | 3 | 48.78341 | yes | 4.45E-06 | 821.39502 | 178.763 |
| DREEDGGGLEPAA | 16 | Q96EU6 | Ribosomal RNA processing protein 36 hc | RRP36 | 29.824 | 2 | 48.78341 | yes | 1.104E-07 | 547.93243 | 178.763 |
| DREEDGGGLEPAA | 16 | Q96EU6 | Ribosomal RNA processing protein 36 hc | RRP36 | 29.824 | 2 | 48.78341 | yes | 4.45E-06 | 821.39502 | 178.763 |
| TYQFLNDIR | 9 | Q96EU6 | Ribosomal RNA processing protein 36 hc | RRP36 | 29.824 | 2 | 48.78341 | yes | 1.537E-07 | 585.30109 | 336.49835 |
| LQQLLQR | 7 | Q96EU6 | Ribosomal RNA processing protein 36 hc | RRP36 | 29.824 | 2 | 48.78341 | no | 2.4E-07 | 449.77707 | 298.07874 |
| QQQELHLALK | 10 | Q96EU6 | Ribosomal RNA processing protein 36 hc | RRP36 | 29.824 | 2 | 48.78341 | yes | 2.888E-07 | 604.34332 | 87.14521 |
| LENFLSR | 7 | Q96EU6 | Ribosomal RNA processing protein 36 hc | RRP36 | 29.824 | 2 | 48.78341 | yes | 1.181E-07 | 439.74017 | 69.41523 |
| AQAQQGHRPYFLI | 13 | Q96EU6 | Ribosomal RNA processing protein 36 hc | RRP36 | 29.824 | 3 | 48.78341 | yes | 5.043E-07 | 515.27582 | 65.974136 |
| KQQQELHLALK | 11 | Q96EU6 | Ribosomal RNA processing protein 36 hc | RRP36 | 29.824 | 2 | 48.78341 | yes | 6.631E-07 | 668.39081 | 41.18367 |
| LEDPLLWR | 8 | Q96EX3 | Cytoplasmic dynein 2 intermediate chain | DYNC2I2 | 57.801 | 2 | 52.492683 | yes | 1.734E-07 | 521.29004 | 132.7198 |
| CSLAAGEAALTR | 12 | Q96EX3 | Cytoplasmic dynein 2 intermediate chain | DYNC2I2 | 57.801 | 2 | 52.492683 | yes | 1.37E-07 | 610.30859 | 86.030052 |
| GDVQLFDLQK | 10 | Q96EX3 | Cytoplasmic dynein 2 intermediate chain | DYNC2I2 | 57.801 | 2 | 52.492683 | yes | 2.436E-06 | 581.80878 | 76.005112 |
| SCQTASIATASASA | 17 | Q96EX3 | Cytoplasmic dynein 2 intermediate chain | DYNC2I2 | 57.801 | 2 | 52.492683 | yes | 2.405E-07 | 840.9021 | 46.517948 |
| VWQLSTEFTEQGP | 14 | Q96EX3 | Cytoplasmic dynein 2 intermediate chain | DYNC2I2 | 57.801 | 2 | 52.492683 | yes | 4.997E-07 | 839.41522 | 41.966312 |
| SFVCAWNLDR | 10 | Q96EX3 | Cytoplasmic dynein 2 intermediate chain | DYNC2I2 | 57.801 | 2 | 52.492683 | yes | 9.722E-08 | 634.29803 | 38.944714 |
| VMIPVPAGVEDGQ | 16 | Q96EY1 | DnaJ homolog subfamily A member 3, mi | DNAJA3 | 52.489 | 2 | 51.56971 | yes | 2.646E-07 | 834.44287 | 3147.7344 |
| VMIPVPAGVEDGQ | 16 | Q96EY1 | DnaJ homolog subfamily A member 3, mi | DNAJA3 | 52.489 | 2 | 51.56971 | yes | 9.918E-08 | 556.63104 | 3147.7344 |
| VMIPVPAGVEDGQ | 16 | Q96EY1 | DnaJ homolog subfamily A member 3, mi | DNAJA3 | 52.489 | 3 | 51.56971 | yes | 2.646E-07 | 834.44287 | 3147.7344 |
| VMIPVPAGVEDGQ | 16 | Q96EY1 | DnaJ homolog subfamily A member 3, mi | DNAJA3 | 52.489 | 3 | 51.56971 | yes | 9.918E-08 | 556.63104 | 3147.7344 |
| GGPTVDPEELFR | 12 | Q96EY1 | DnaJ homolog subfamily A member 3, mi | DNAJA3 | 52.489 | 2 | 51.56971 | yes | 1.02E-07 | 658.8277 | 4002.0132 |
| GSIIISPCVVCR | 12 | Q96EY1 | DnaJ homolog subfamily A member 3, mi | DNAJA3 | 52.489 | 2 | 51.56971 | yes | 1.109E-07 | 680.85742 | 1142.5725 |
| EFTVNIMDTCER | 12 | Q96EY1 | DnaJ homolog subfamily A member 3, mi | DNAJA3 | 52.489 | 2 | 51.56971 | yes | 1.032E-07 | 757.83435 | 834.93793 |
| EIFITFR | 7 | Q96EY1 | DnaJ homolog subfamily A member 3, mi | DNAJA3 | 52.489 | 2 | 51.56971 | yes | 1.388E-07 | 463.26074 | 479.12277 |
| INSYGYGDHYIHIK | 14 | Q96EY1 | DnaJ homolog subfamily A member 3, mi | DNAJA3 | 52.489 | 2 | 51.56971 | yes | 3.368E-07 | 840.41248 | 347.33298 |
| KAYYQLAK | 8 | Q96EY1 | DnaJ homolog subfamily A member 3, mi | DNAJA3 | 52.489 | 2 | 51.56971 | yes | 9.722E-08 | 492.7793 | 220.5701 |
| EAGEDEEGFLSK | 12 | Q96EY1 | DnaJ homolog subfamily A member 3, mi | DNAJA3 | 52.489 | 2 | 51.56971 | yes | 1.032E-07 | 655.79095 | 184.31619 |
| AYYQLAK | 7 | Q96EY1 | DnaJ homolog subfamily A member 3, mi | DNAJA3 | 52.489 | 2 | 51.56971 | yes | 9.722E-08 | 428.73181 | 102.70271 |
| EDYYQILGVPR | 11 | Q96EY1 | DnaJ homolog subfamily A member 3, mi | DNAJA3 | 52.489 | 2 | 51.56971 | yes | 6.13E-07 | 676.84589 | 101.15215 |
| REIFITFR | 8 | Q96EY1 | DnaJ homolog subfamily A member 3, mi | DNAJA3 | 52.489 | 2 | 51.56971 | yes | 3.888E-07 | 541.31128 | 55.564796 |
| VQHCHYCGGSGM | 23 | Q96EY1 | DnaJ homolog subfamily A member 3, mi | DNAJA3 | 52.489 | 4 | 51.56971 | yes | 3.11E-05 | 660.29089 | 49.434006 |
| REAGEDEEGFLSK | 13 | Q96EY1 | DnaJ homolog subfamily A member 3, mi | DNAJA3 | 52.489 | 2 | 51.56971 | yes | 1.512E-07 | 733.84155 | 38.223488 |

| | | | | | | | | | | | |
|---|---|---|---|---|---|---|---|---|---|---|---|
| KICANDAIPK | 10 | Q96EY4 | Translation machinery-associated protein | TMA16 | 23.865 | 2 | 50.87718 | yes | 8.708E-07 | 565.30536 | 723.20514 |
| ICANDAIPK | 9 | Q96EY4 | Translation machinery-associated protein | TMA16 | 23.865 | 2 | 50.87718 | yes | 3.887E-07 | 501.25787 | 518.15015 |
| EWDFDLK | 7 | Q96EY4 | Translation machinery-associated protein | TMA16 | 23.865 | 2 | 50.87718 | yes | 2.666E-07 | 476.72418 | 467.46152 |
| KDACELIER | 9 | Q96EY4 | Translation machinery-associated protein | TMA16 | 23.865 | 2 | 50.87718 | yes | 9.722E-08 | 567.28461 | 306.65347 |
| DACELIER | 8 | Q96EY4 | Translation machinery-associated protein | TMA16 | 23.865 | 2 | 50.87718 | yes | 1.712E-07 | 503.23712 | 214.51466 |
| DMQGLSLDAASQI | 15 | Q96EY5 | Multivesicular body subunit 12A OS=Hoı | MVB12A | 28.784 | 2 | 52.83027 | yes | 9.722E-08 | 774.37213 | 90.720062 |
| VAVLQALASTVNI | 13 | Q96EY7 | Small ribosomal subunit protein mS39 O | PTCD3 | 78.55 | 2 | 54.431107 | yes | 9.722E-08 | 671.39606 | 811.12781 |
| LFDQPGDPLK | 10 | Q96EY7 | Small ribosomal subunit protein mS39 O | PTCD3 | 78.55 | 2 | 54.431107 | yes | 1.101E-07 | 565.29803 | 366.78229 |
| SPALQVLR | 8 | Q96EY7 | Small ribosomal subunit protein mS39 O | PTCD3 | 78.55 | 2 | 54.431107 | yes | 1.355E-07 | 442.27164 | 269.73495 |
| AGHQFGVTWR | 10 | Q96EY7 | Small ribosomal subunit protein mS39 O | PTCD3 | 78.55 | 2 | 54.431107 | yes | 9.722E-08 | 579.79376 | 265.46527 |
| SELLNELMDSAK | 12 | Q96EY7 | Small ribosomal subunit protein mS39 O | PTCD3 | 78.55 | 2 | 54.431107 | yes | 4.193E-07 | 675.33447 | 185.60196 |
| NEHSYCTMIR | 10 | Q96EY7 | Small ribosomal subunit protein mS39 O | PTCD3 | 78.55 | 2 | 54.431107 | yes | 9.722E-08 | 655.78442 | 177.48636 |
| VEGTDVTGIEEVV | 16 | Q96EY7 | Small ribosomal subunit protein mS39 O | PTCD3 | 78.55 | 2 | 54.431107 | yes | 1.174E-07 | 842.9538 | 177.05508 |
| FIINSYPK | 8 | Q96EY7 | Small ribosomal subunit protein mS39 O | PTCD3 | 78.55 | 2 | 54.431107 | yes | 1.835E-07 | 491.27383 | 158.62984 |
| VMSDFAINQEQK | 12 | Q96EY7 | Small ribosomal subunit protein mS39 O | PTCD3 | 78.55 | 2 | 54.431107 | yes | 4.634E-06 | 705.34009 | 86.469627 |
| SAYESQPIR | 9 | Q96EY7 | Small ribosomal subunit protein mS39 O | PTCD3 | 78.55 | 2 | 54.431107 | yes | 2.857E-06 | 525.76434 | 57.239693 |
| VKPNLQTFNTILK | 13 | Q96EY7 | Small ribosomal subunit protein mS39 O | PTCD3 | 78.55 | 2 | 54.431107 | yes | 1.983E-07 | 758.4483 | 40.795979 |
| DLELAYQVHGLLK | 13 | Q96EY7 | Small ribosomal subunit protein mS39 O | PTCD3 | 78.55 | 2 | 54.431107 | yes | 1.165E-07 | 749.91687 | 29.862608 |
| WYALLYDPVISK | 12 | Q96EZ8 | Microspherule protein 1 OS=Homo sapie | MCRS1 | 51.804 | 2 | 50.202377 | yes | 1.146E-07 | 734.39777 | 78.632156 |
| LSNNSVVEIASLR | 13 | Q96EZ8 | Microspherule protein 1 OS=Homo sapie | MCRS1 | 51.804 | 2 | 50.202377 | yes | 9.85E-08 | 701.38843 | 62.732189 |
| HPDAFYLAR | 9 | Q96EZ8 | Microspherule protein 1 OS=Homo sapie | MCRS1 | 51.804 | 2 | 50.202377 | yes | 3.423E-06 | 545.27747 | 59.434467 |
| ASSQALGTIPK | 11 | Q96EZ8 | Microspherule protein 1 OS=Homo sapie | MCRS1 | 51.804 | 2 | 50.202377 | yes | 4.265E-07 | 536.80347 | 53.766945 |
| FDDELVESSLAK | 12 | Q96EZ8 | Microspherule protein 1 OS=Homo sapie | MCRS1 | 51.804 | 2 | 50.202377 | yes | 1.29E-06 | 676.83264 | 34.128532 |
| QLHPEAIAAIQSK | 13 | Q96EZ8 | Microspherule protein 1 OS=Homo sapie | MCRS1 | 51.804 | 2 | 50.202377 | yes | 4.073E-07 | 703.39355 | 30.729757 |
| APSTPVPPSPAPAP | 18 | Q96EZ8 | Microspherule protein 1 OS=Homo sapie | MCRS1 | 51.804 | 2 | 50.202377 | yes | 2.013E-06 | 842.96704 | 23.867538 |
| VPGLVETLR | 9 | Q96F44 | E3 ubiquitin-protein ligase TRIM11 OS= | TRIM11 | 52.775 | 2 | 53.92483 | yes | 1.02E-07 | 492.29785 | 653.64581 |
| EPLAAFCGDELR | 12 | Q96F44 | E3 ubiquitin-protein ligase TRIM11 OS= | TRIM11 | 52.775 | 2 | 53.92483 | yes | 9.722E-08 | 689.32703 | 488.34045 |
| QNVLGEFER | 9 | Q96F44 | E3 ubiquitin-protein ligase TRIM11 OS= | TRIM11 | 52.775 | 2 | 53.92483 | yes | 2.98E-07 | 546.27765 | 373.16769 |
| LQPPEVVPMELR | 12 | Q96F44 | E3 ubiquitin-protein ligase TRIM11 OS= | TRIM11 | 52.775 | 2 | 53.92483 | yes | 1.451E-07 | 704.38684 | 336.92673 |
| LLCAACER | 8 | Q96F44 | E3 ubiquitin-protein ligase TRIM11 OS= | TRIM11 | 52.775 | 2 | 53.92483 | yes | 1.066E-07 | 496.73624 | 328.19135 |
| FDPGPCVLGQER | 12 | Q96F44 | E3 ubiquitin-protein ligase TRIM11 OS= | TRIM11 | 52.775 | 2 | 53.92483 | yes | 2.809E-07 | 687.82715 | 234.90413 |
| LEEEELEVLPR | 11 | Q96F44 | E3 ubiquitin-protein ligase TRIM11 OS= | TRIM11 | 52.775 | 2 | 53.92483 | yes | 9.849E-08 | 678.35626 | 231.31998 |
| TSWALGVCR | 9 | Q96F44 | E3 ubiquitin-protein ligase TRIM11 OS= | TRIM11 | 52.775 | 2 | 53.92483 | yes | 9.27E-07 | 525.26349 | 216.43193 |
| LLAEEEQQLLQR | 12 | Q96F44 | E3 ubiquitin-protein ligase TRIM11 OS= | TRIM11 | 52.775 | 2 | 53.92483 | yes | 1.101E-07 | 735.40155 | 180.48393 |
| GDVTLDPDTANPE | 21 | Q96F44 | E3 ubiquitin-protein ligase TRIM11 OS= | TRIM11 | 52.775 | 3 | 53.92483 | yes | 1.14E-07 | 776.05554 | 131.1799 |
| QALPDSPER | 9 | Q96F44 | E3 ubiquitin-protein ligase TRIM11 OS= | TRIM11 | 52.775 | 2 | 53.92483 | yes | 3.893E-07 | 506.75653 | 100.61157 |
| HYWEVEVGDR | 10 | Q96F44 | E3 ubiquitin-protein ligase TRIM11 OS= | TRIM11 | 52.775 | 2 | 53.92483 | no | 9.722E-08 | 645.29907 | 69.496918 |
| NLRPNRPLAK | 10 | Q96F44 | E3 ubiquitin-protein ligase TRIM11 OS= | TRIM11 | 52.775 | 2 | 53.92483 | yes | 1.296E-06 | 589.85944 | 15.046697 |
| CLVPEVTFR | 9 | Q96F86 | Enhancer of mRNA-decapping protein 3 ( | EDC3 | 56.078 | 2 | 51.938232 | yes | 1.196E-07 | 560.79462 | 1476.6519 |

| | | | | | | | | | | | |
|---|---|---|---|---|---|---|---|---|---|---|---|
| VSAVDQVSQTISL' | 15 | Q96F86 | Enhancer of mRNA-decapping protein 3 ( | EDC3 | 56.078 | 2 | 51.938232 | yes | 1.346E-07 | 802.4361 | 1279.0249 |
| AAVFEEIDTYER | 12 | Q96F86 | Enhancer of mRNA-decapping protein 3 ( | EDC3 | 56.078 | 2 | 51.938232 | yes | 1.17E-07 | 721.84357 | 1010.3701 |
| IIVPHNVSK | 9 | Q96F86 | Enhancer of mRNA-decapping protein 3 ( | EDC3 | 56.078 | 2 | 51.938232 | yes | 1.006E-07 | 503.80585 | 929.86438 |
| AAVAWANQNR | 10 | Q96F86 | Enhancer of mRNA-decapping protein 3 ( | EDC3 | 56.078 | 2 | 51.938232 | yes | 3.234E-07 | 550.78345 | 889.82037 |
| HMESLSQSK | 9 | Q96F86 | Enhancer of mRNA-decapping protein 3 ( | EDC3 | 56.078 | 2 | 51.938232 | yes | 1.568E-07 | 523.75043 | 699.95679 |
| HGLTLER | 7 | Q96F86 | Enhancer of mRNA-decapping protein 3 ( | EDC3 | 56.078 | 2 | 51.938232 | yes | 3.708E-07 | 413.23251 | 692.25995 |
| SQDVAVSPQQQQ( | 15 | Q96F86 | Enhancer of mRNA-decapping protein 3 ( | EDC3 | 56.078 | 2 | 51.938232 | yes | 1.108E-07 | 845.39667 | 449.62317 |
| TQGQQVSSLK | 10 | Q96F86 | Enhancer of mRNA-decapping protein 3 ( | EDC3 | 56.078 | 2 | 51.938232 | yes | 9.722E-08 | 538.29077 | 441.24304 |
| EFCTDSGLVVPSIS | 19 | Q96F86 | Enhancer of mRNA-decapping protein 3 ( | EDC3 | 56.078 | 3 | 51.938232 | yes | 1.136E-07 | 727.68927 | 329.2894 |
| FVIPLHSA | 8 | Q96F86 | Enhancer of mRNA-decapping protein 3 ( | EDC3 | 56.078 | 2 | 51.938232 | yes | 1.248E-07 | 442.25543 | 239.1561 |
| GIPNERPTR | 9 | Q96F86 | Enhancer of mRNA-decapping protein 3 ( | EDC3 | 56.078 | 2 | 51.938232 | yes | 3.22E-06 | 520.28577 | 228.75607 |
| MLESITNELSLFSK | 14 | Q96F86 | Enhancer of mRNA-decapping protein 3 ( | EDC3 | 56.078 | 2 | 51.938232 | yes | 1.079E-07 | 806.41858 | 227.61298 |
| KLLSVAEK | 8 | Q96F86 | Enhancer of mRNA-decapping protein 3 ( | EDC3 | 56.078 | 2 | 51.938232 | yes | 1.256E-07 | 444.28165 | 152.25388 |
| KPASSSSAPQNIPK | 14 | Q96F86 | Enhancer of mRNA-decapping protein 3 ( | EDC3 | 56.078 | 2 | 51.938232 | yes | 2.89E-07 | 706.38062 | 143.65671 |
| HDENILESEPIVYR | 14 | Q96F86 | Enhancer of mRNA-decapping protein 3 ( | EDC3 | 56.078 | 3 | 51.938232 | yes | 2.801E-07 | 571.95294 | 78.777954 |
| RLEMTGVCASQM. | 22 | Q96F86 | Enhancer of mRNA-decapping protein 3 ( | EDC3 | 56.078 | 3 | 51.938232 | yes | 9.789E-08 | 792.4046 | 58.949097 |
| AGDITELK | 8 | Q96F86 | Enhancer of mRNA-decapping protein 3 ( | EDC3 | 56.078 | 2 | 51.938232 | yes | 2.386E-06 | 423.73199 | 48.203426 |
| HLANHDVQVILFL | 18 | Q96F86 | Enhancer of mRNA-decapping protein 3 ( | EDC3 | 56.078 | 3 | 51.938232 | yes | 1.172E-07 | 702.06158 | 39.721416 |
| IVAHAVEVPAVQS | 15 | Q96FF9 | Sororin OS=Homo sapiens OX=9606 GN | CDCA5 | 27.601 | 3 | 43.54259 | yes | 4.914E-07 | 524.96674 | 90.244415 |
| LETLGSASTSTPGF | 14 | Q96FF9 | Sororin OS=Homo sapiens OX=9606 GN | CDCA5 | 27.601 | 2 | 43.54259 | yes | 2.546E-07 | 688.85443 | 76.401657 |
| LQEDLQK | 7 | Q96FS4 | Signal-induced proliferation-associated pi | SIPA1 | 112.15 | 2 | 28.793007 | yes | 2.664E-06 | 437.23746 | 139.94322 |
| LMDLQLSDSNFR | 12 | Q96FV9 | THO complex subunit 1 OS=Homo sapie | THOC1 | 75.667 | 2 | 54.72845 | yes | 1.181E-07 | 719.85339 | 405.92407 |
| ILAPYLEMK | 9 | Q96FV9 | THO complex subunit 1 OS=Homo sapie | THOC1 | 75.667 | 2 | 54.72845 | yes | 9.893E-08 | 539.30426 | 250.40314 |
| SLPEYLENMVIK | 12 | Q96FV9 | THO complex subunit 1 OS=Homo sapie | THOC1 | 75.667 | 2 | 54.72845 | yes | 1.531E-07 | 718.37872 | 217.96867 |
| LWNLCPDNMEAC | 13 | Q96FV9 | THO complex subunit 1 OS=Homo sapie | THOC1 | 75.667 | 2 | 54.72845 | yes | 9.722E-08 | 825.8573 | 179.74608 |
| MCNDLLR | 7 | Q96FV9 | THO complex subunit 1 OS=Homo sapie | THOC1 | 75.667 | 2 | 54.72845 | yes | 2.058E-07 | 461.21768 | 163.58011 |
| NPVQCYEK | 8 | Q96FV9 | THO complex subunit 1 OS=Homo sapie | THOC1 | 75.667 | 2 | 54.72845 | yes | 2.26E-07 | 519.23969 | 92.55761 |
| DKPVTGEQIEVFA' | 15 | Q96FV9 | THO complex subunit 1 OS=Homo sapie | THOC1 | 75.667 | 2 | 54.72845 | yes | 9.722E-08 | 837.93848 | 88.227112 |
| TGGEHVYFAK | 10 | Q96FV9 | THO complex subunit 1 OS=Homo sapie | THOC1 | 75.667 | 2 | 54.72845 | yes | 1.408E-07 | 554.77472 | 60.39032 |
| SPTPPLFSLPEAR | 13 | Q96FV9 | THO complex subunit 1 OS=Homo sapie | THOC1 | 75.667 | 2 | 54.72845 | yes | 9.722E-08 | 706.38263 | 36.428082 |
| TGEDEDEEDNDAI | 15 | Q96FV9 | THO complex subunit 1 OS=Homo sapie | THOC1 | 75.667 | 2 | 54.72845 | yes | 1.042E-07 | 846.85779 | 29.256678 |
| QIECDSEDMK | 10 | Q96FV9 | THO complex subunit 1 OS=Homo sapie | THOC1 | 75.667 | 2 | 54.72845 | yes | 2.057E-07 | 627.7525 | 28.299276 |
| YSEEVLAVFK | 10 | Q96FV9 | THO complex subunit 1 OS=Homo sapie | THOC1 | 75.667 | 2 | 54.72845 | yes | 3.454E-06 | 592.81354 | 19.257254 |
| LLTSGYLQR | 9 | Q96FW1 | Ubiquitin thioesterase OTUB1 OS=Homc | OTUB1 | 31.285 | 2 | 53.29629 | yes | 2.308E-07 | 525.80072 | 268.5397 |
| FFEHFIEGGR | 10 | Q96FW1 | Ubiquitin thioesterase OTUB1 OS=Homc | OTUB1 | 31.285 | 2 | 53.29629 | yes | 9.722E-08 | 619.80127 | 83.679329 |
| EYAEDDNIYQQK | 12 | Q96FW1 | Ubiquitin thioesterase OTUB1 OS=Homc | OTUB1 | 31.285 | 2 | 53.29629 | yes | 5.149E-07 | 758.33356 | 71.744797 |
| EFCQQEVEPMCK | 12 | Q96FW1 | Ubiquitin thioesterase OTUB1 OS=Homc | OTUB1 | 31.285 | 2 | 53.29629 | yes | 1.301E-07 | 792.82819 | 46.622414 |
| PGHYDILYK | 9 | Q96FW1 | Ubiquitin thioesterase OTUB1 OS=Homc | OTUB1 | 31.285 | 2 | 53.29629 | yes | 3.095E-07 | 553.28748 | 43.475266 |
| HSVDLIGR | 8 | Q96FX7 | tRNA (adenine(58)-N(1))-methyltransfera | TRMT61A | 31.382 | 2 | 54.06126 | yes | 3.272E-07 | 448.75107 | 407.91016 |

| | | | | | | | | | | | |
|---|---|---|---|---|---|---|---|---|---|---|---|
| FCSFSPCIEQVQR | 13 | Q96FX7 | tRNA (adenine(58)-N(1))-methyltransfera | TRMT61A | 31.382 | 2 | 54.06126 | yes | 1.419E-07 | 829.37671 | 407.61493 |
| TIAPTGHLHTVEFI | 17 | Q96FX7 | tRNA (adenine(58)-N(1))-methyltransfera | TRMT61A | 31.382 | 3 | 54.06126 | yes | 1.142E-07 | 658.00977 | 196.93851 |
| TIAPTGHLHTVEFI | 17 | Q96FX7 | tRNA (adenine(58)-N(1))-methyltransfera | TRMT61A | 31.382 | 3 | 54.06126 | yes | 2.407E-07 | 493.75916 | 196.93851 |
| TIAPTGHLHTVEFI | 17 | Q96FX7 | tRNA (adenine(58)-N(1))-methyltransfera | TRMT61A | 31.382 | 4 | 54.06126 | yes | 1.142E-07 | 658.00977 | 196.93851 |
| TIAPTGHLHTVEFI | 17 | Q96FX7 | tRNA (adenine(58)-N(1))-methyltransfera | TRMT61A | 31.382 | 4 | 54.06126 | yes | 2.407E-07 | 493.75916 | 196.93851 |
| TCQALAAR | 8 | Q96FX7 | tRNA (adenine(58)-N(1))-methyltransfera | TRMT61A | 31.382 | 2 | 54.06126 | yes | 2.886E-07 | 445.72928 | 344.27472 |
| EAVGHTGYLTFAT | 14 | Q96FX7 | tRNA (adenine(58)-N(1))-methyltransfera | TRMT61A | 31.382 | 2 | 54.06126 | yes | 9.722E-08 | 747.883 | 156.84346 |
| PGSVVCESGTGSG | 21 | Q96FX7 | tRNA (adenine(58)-N(1))-methyltransfera | TRMT61A | 31.382 | 3 | 54.06126 | yes | 9.722E-08 | 686.34088 | 80.159378 |
| EGDTAILSLGHGAI | 18 | Q96FX7 | tRNA (adenine(58)-N(1))-methyltransfera | TRMT61A | 31.382 | 3 | 54.06126 | yes | 1.208E-07 | 599.64899 | 64.839241 |
| MPAILDGEEAVSK | 13 | Q96FZ2 | Abasic site processing protein HMCES O | HMCES | 40.575 | 2 | 43.04892 | yes | 2.753E-07 | 680.34485 | 48.045654 |
| SPQSNSPVLLSR | 12 | Q96FZ2 | Abasic site processing protein HMCES O | HMCES | 40.575 | 2 | 43.04892 | yes | 3.345E-07 | 642.84894 | 47.39436 |
| WLDFGEVSTQEAI | 14 | Q96FZ2 | Abasic site processing protein HMCES O | HMCES | 40.575 | 2 | 43.04892 | yes | 4.979E-07 | 811.90668 | 39.60305 |
| VITFANQDDYISFR | 14 | Q96G21 | U3 small nucleolar ribonucleoprotein pro | IMP4 | 33.757 | 2 | 54.328266 | yes | 9.722E-08 | 844.91754 | 317.2356 |
| LVFPGAQR | 8 | Q96G21 | U3 small nucleolar ribonucleoprotein pro | IMP4 | 33.757 | 2 | 54.328266 | yes | 2.296E-07 | 444.25851 | 249.58546 |
| NVELTEVGPR | 10 | Q96G21 | U3 small nucleolar ribonucleoprotein pro | IMP4 | 33.757 | 2 | 54.328266 | yes | 9.722E-08 | 557.29858 | 205.74608 |
| WHPYTNTAR | 9 | Q96G21 | U3 small nucleolar ribonucleoprotein pro | IMP4 | 33.757 | 2 | 54.328266 | yes | 4.523E-07 | 573.27795 | 177.4169 |
| WAGVEDPK | 8 | Q96G21 | U3 small nucleolar ribonucleoprotein pro | IMP4 | 33.757 | 2 | 54.328266 | yes | 2.944E-06 | 451.22433 | 78.918854 |
| YLFPVPK | 7 | Q96G21 | U3 small nucleolar ribonucleoprotein pro | IMP4 | 33.757 | 2 | 54.328266 | yes | 1.9E-07 | 432.25491 | 31.091682 |
| ESESGGLRPNK | 11 | Q96G25 | Mediator of RNA polymerase II transcrip | MED8 | 29.08 | 2 | 43.438465 | yes | 2.591E-07 | 587.29657 | 59.21431 |
| TKPDPEVEEQEK | 12 | Q96G25 | Mediator of RNA polymerase II transcrip | MED8 | 29.08 | 2 | 43.438465 | yes | 5.021E-07 | 714.84625 | 29.494921 |
| GMNQVLDVPLTV | 13 | Q96G46 | tRNA-dihydrouridine(47) synthase [NAD | DUS3L | 72.594 | 2 | 50.82049 | yes | 1.093E-07 | 707.39215 | 277.37759 |
| YVPVGLLER | 9 | Q96G46 | tRNA-dihydrouridine(47) synthase [NAD | DUS3L | 72.594 | 2 | 50.82049 | yes | 3.224E-06 | 523.30566 | 232.06093 |
| CVLFETFGR | 9 | Q96G46 | tRNA-dihydrouridine(47) synthase [NAD | DUS3L | 72.594 | 2 | 50.82049 | yes | 1.009E-07 | 564.77893 | 231.18857 |
| HWDISSSER | 9 | Q96G46 | tRNA-dihydrouridine(47) synthase [NAD | DUS3L | 72.594 | 2 | 50.82049 | yes | 2.299E-07 | 558.75708 | 221.17032 |
| TCGPLTDEDVVR | 12 | Q96G46 | tRNA-dihydrouridine(47) synthase [NAD | DUS3L | 72.594 | 2 | 50.82049 | yes | 2.241E-07 | 681.3219 | 207.12636 |
| DWGVALVTLHGR | 12 | Q96G46 | tRNA-dihydrouridine(47) synthase [NAD | DUS3L | 72.594 | 2 | 50.82049 | yes | 9.722E-08 | 662.36206 | 109.18654 |
| YLETKPADLGPR | 12 | Q96G46 | tRNA-dihydrouridine(47) synthase [NAD | DUS3L | 72.594 | 2 | 50.82049 | yes | 7.417E-07 | 680.367 | 75.646255 |
| AMQTGVTGIMIAR | 13 | Q96G46 | tRNA-dihydrouridine(47) synthase [NAD | DUS3L | 72.594 | 2 | 50.82049 | yes | 1.859E-07 | 674.85742 | 64.224991 |
| EQFHQFLEAK | 10 | Q96G46 | tRNA-dihydrouridine(47) synthase [NAD | DUS3L | 72.594 | 2 | 50.82049 | yes | 3.064E-06 | 638.81964 | 35.783051 |
| HQCEDIFGVQLEG | 21 | Q96G46 | tRNA-dihydrouridine(47) synthase [NAD | DUS3L | 72.594 | 3 | 50.82049 | yes | 1.116E-06 | 808.3717 | 15.199093 |
| LYNYFYDR | 8 | Q96GD4 | Aurora kinase B OS=Homo sapiens OX= | AURKB | 39.311 | 2 | 52.123962 | yes | 9.722E-08 | 577.26929 | 281.28687 |
| VMENSSGTPDILTI | 14 | Q96GD4 | Aurora kinase B OS=Homo sapiens OX= | AURKB | 39.311 | 2 | 52.123962 | yes | 1.104E-07 | 760.37469 | 224.50867 |
| HFTIDDFEIGR | 11 | Q96GD4 | Aurora kinase B OS=Homo sapiens OX= | AURKB | 39.311 | 2 | 52.123962 | yes | 9.722E-08 | 675.32788 | 160.70953 |
| LPLAQVSAHPWVI | 13 | Q96GD4 | Aurora kinase B OS=Homo sapiens OX= | AURKB | 39.311 | 2 | 52.123962 | yes | 3.177E-07 | 737.41986 | 73.12735 |
| LPLAQVSAHPWVI | 13 | Q96GD4 | Aurora kinase B OS=Homo sapiens OX= | AURKB | 39.311 | 2 | 52.123962 | yes | 4.868E-07 | 491.94901 | 73.12735 |
| LPLAQVSAHPWVI | 13 | Q96GD4 | Aurora kinase B OS=Homo sapiens OX= | AURKB | 39.311 | 3 | 52.123962 | yes | 3.177E-07 | 737.41986 | 73.12735 |
| LPLAQVSAHPWVI | 13 | Q96GD4 | Aurora kinase B OS=Homo sapiens OX= | AURKB | 39.311 | 3 | 52.123962 | yes | 4.868E-07 | 491.94901 | 73.12735 |
| IYLILEYAPR | 10 | Q96GD4 | Aurora kinase B OS=Homo sapiens OX= | AURKB | 39.311 | 2 | 52.123962 | yes | 2.272E-07 | 625.86078 | 65.456673 |
| WEVYPAQVQGPR | 12 | Q96GE4 | Centrosomal protein of 95 kDa OS=Hom | CEP95 | 95.297 | 2 | 19.456753 | yes | 1.911E-05 | 715.36475 | 35.597374 |

| | | | | | | | | | | | |
|---|---|---|---|---|---|---|---|---|---|---|---|
| VPDLIVIPR | 9 | Q96GE4 | Centrosomal protein of 95 kDa OS=Homo | CEP95 | 95.297 | 2 | 19.456753 | yes | 4.845E-06 | 511.32385 | 32.380772 |
| TVGFYLGTWR | 10 | Q96GJ1 | tRNA (uracil-5-)-methyltransferase homo | TRMT2B | 56.477 | 2 | 54.418217 | yes | 1.53E-06 | 600.31403 | 47.179497 |
| DGNVVCVQSNHL | 13 | Q96GJ1 | tRNA (uracil-5-)-methyltransferase homo | TRMT2B | 56.477 | 2 | 54.418217 | yes | 1.384E-07 | 735.36188 | 40.728111 |
| NVIELCCPPDPAK | 13 | Q96GJ1 | tRNA (uracil-5-)-methyltransferase homo | TRMT2B | 56.477 | 2 | 54.418217 | yes | 2.228E-07 | 756.86292 | 33.27137 |
| LSQEELHVQK | 10 | Q96GJ1 | tRNA (uracil-5-)-methyltransferase homo | TRMT2B | 56.477 | 2 | 54.418217 | yes | 9.722E-08 | 605.82495 | 18.651382 |
| TMTDVVGNPEEEI | 13 | Q96GM5 | SWI/SNF-related matrix-associated actin- | SMARCD1 | 58.233 | 2 | 46.520916 | yes | 1.537E-07 | 738.83539 | 43.576244 |
| ALLNLPGTQTSGE | 15 | Q96GM8 | Target of EGR1 protein 1 OS=Homo sapi | TOE1 | 56.548 | 2 | 51.99673 | yes | 1.364E-07 | 750.40686 | 1209.6787 |
| SLLNQCIEER | 10 | Q96GM8 | Target of EGR1 protein 1 OS=Homo sapi | TOE1 | 56.548 | 2 | 51.99673 | yes | 1.022E-07 | 631.3139 | 962.44501 |
| YAAEFHAR | 8 | Q96GM8 | Target of EGR1 protein 1 OS=Homo sapi | TOE1 | 56.548 | 2 | 51.99673 | yes | 9.722E-08 | 482.73541 | 776.39917 |
| AARPEIADR | 9 | Q96GM8 | Target of EGR1 protein 1 OS=Homo sapi | TOE1 | 56.548 | 2 | 51.99673 | yes | 2.25E-07 | 499.77252 | 509.6839 |
| FVASYLEYAFR | 11 | Q96GM8 | Target of EGR1 protein 1 OS=Homo sapi | TOE1 | 56.548 | 2 | 51.99673 | yes | 9.722E-08 | 683.34552 | 481.79495 |
| TLFLELIR | 8 | Q96GM8 | Target of EGR1 protein 1 OS=Homo sapi | TOE1 | 56.548 | 2 | 51.99673 | yes | 3.768E-07 | 502.81058 | 182.26555 |
| CCLPPATHR | 9 | Q96GM8 | Target of EGR1 protein 1 OS=Homo sapi | TOE1 | 56.548 | 2 | 51.99673 | yes | 2.728E-06 | 556.26044 | 68.57843 |
| SILSLGLACFK | 11 | Q96GM8 | Target of EGR1 protein 1 OS=Homo sapi | TOE1 | 56.548 | 2 | 51.99673 | yes | 1.392E-06 | 604.83899 | 33.361713 |
| SVSLVGEDER | 10 | Q96GQ7 | Probable ATP-dependent RNA helicase D | DDX27 | 89.836 | 2 | 51.481472 | yes | 1.506E-07 | 545.7724 | 349.30472 |
| ACIPVGLLGK | 10 | Q96GQ7 | Probable ATP-dependent RNA helicase D | DDX27 | 89.836 | 2 | 51.481472 | yes | 2.01E-07 | 514.30206 | 346.57968 |
| TAAFALPVLER | 11 | Q96GQ7 | Probable ATP-dependent RNA helicase D | DDX27 | 89.836 | 2 | 51.481472 | yes | 9.722E-08 | 594.34277 | 294.14114 |
| SADFNPDFVFTEK | 13 | Q96GQ7 | Probable ATP-dependent RNA helicase D | DDX27 | 89.836 | 2 | 51.481472 | yes | 2.967E-06 | 758.85138 | 254.24669 |
| MLDEYFEEQMK | 11 | Q96GQ7 | Probable ATP-dependent RNA helicase D | DDX27 | 89.836 | 2 | 51.481472 | yes | 1.154E-07 | 731.81506 | 211.60709 |
| VLVLVPTR | 8 | Q96GQ7 | Probable ATP-dependent RNA helicase D | DDX27 | 89.836 | 2 | 51.481472 | yes | 9.868E-08 | 448.80002 | 210.66586 |
| SVFDEELTNTSK | 12 | Q96GQ7 | Probable ATP-dependent RNA helicase D | DDX27 | 89.836 | 2 | 51.481472 | yes | 9.722E-08 | 685.3277 | 175.13301 |
| DVYAVLQLEAEEK | 13 | Q96GQ7 | Probable ATP-dependent RNA helicase D | DDX27 | 89.836 | 2 | 51.481472 | yes | 1.097E-07 | 753.88794 | 146.56622 |
| EAIVAALLTR | 10 | Q96GQ7 | Probable ATP-dependent RNA helicase D | DDX27 | 89.836 | 2 | 51.481472 | yes | 1.309E-07 | 528.82422 | 124.516 |
| ALQEFDLALR | 10 | Q96GQ7 | Probable ATP-dependent RNA helicase D | DDX27 | 89.836 | 2 | 51.481472 | yes | 2.141E-07 | 588.32458 | 117.61305 |
| QTMLFSATMTDEV | 14 | Q96GQ7 | Probable ATP-dependent RNA helicase D | DDX27 | 89.836 | 2 | 51.481472 | yes | 1.06E-07 | 801.3811 | 101.13518 |
| EMQQSEAQINTAK | 13 | Q96GQ7 | Probable ATP-dependent RNA helicase D | DDX27 | 89.836 | 2 | 51.481472 | yes | 1.122E-07 | 739.3512 | 100.04469 |
| AAPDILIATPGR | 12 | Q96GQ7 | Probable ATP-dependent RNA helicase D | DDX27 | 89.836 | 2 | 51.481472 | yes | 2.177E-07 | 597.8457 | 95.708344 |
| IFVNSNTDVAPFLI | 14 | Q96GQ7 | Probable ATP-dependent RNA helicase D | DDX27 | 89.836 | 2 | 51.481472 | yes | 1.356E-07 | 796.92517 | 80.827507 |
| TVINFTMPNTIK | 12 | Q96GQ7 | Probable ATP-dependent RNA helicase D | DDX27 | 89.836 | 2 | 51.481472 | yes | 2.203E-06 | 689.8736 | 68.552666 |
| SVSLVGEDERK | 11 | Q96GQ7 | Probable ATP-dependent RNA helicase D | DDX27 | 89.836 | 2 | 51.481472 | yes | 1.537E-07 | 609.81989 | 47.971878 |
| VFGTLLQPVTPSL | 16 | Q96GW9 | Methionine--tRNA ligase, mitochondrial | MARS2 | 66.591 | 2 | 40.632225 | yes | 3.924E-07 | 843.47723 | 58.944374 |
| DLKPDNMLISNEG | 16 | Q96GX5 | Serine/threonine-protein kinase greatwall | MASTL | 97.32 | 3 | 57.138676 | yes | 9.722E-08 | 608.64886 | 140.84361 |
| MLGPPLEVLK | 10 | Q96GX5 | Serine/threonine-protein kinase greatwall | MASTL | 97.32 | 2 | 57.138676 | yes | 5.956E-07 | 548.82538 | 101.03113 |
| DIPWPEGEEK | 10 | Q96GX5 | Serine/threonine-protein kinase greatwall | MASTL | 97.32 | 2 | 57.138676 | yes | 2.026E-07 | 600.28259 | 70.747932 |
| SCLETVASNPGMP | 15 | Q96GX5 | Serine/threonine-protein kinase greatwall | MASTL | 97.32 | 2 | 57.138676 | yes | 2.093E-07 | 795.38672 | 57.872524 |
| NLMCELDEDCEK | 12 | Q96GX5 | Serine/threonine-protein kinase greatwall | MASTL | 97.32 | 2 | 57.138676 | yes | 1.191E-07 | 778.30731 | 45.664825 |
| DYLSSSFLCSDDD | 14 | Q96GX5 | Serine/threonine-protein kinase greatwall | MASTL | 97.32 | 2 | 57.138676 | yes | 2.394E-07 | 840.34631 | 45.055855 |
| GVENPAVQESNQI | 13 | Q96GX5 | Serine/threonine-protein kinase greatwall | MASTL | 97.32 | 2 | 57.138676 | yes | 1.529E-06 | 700.34424 | 15.817115 |
| VNPVTLETVLR | 11 | Q96H55 | Unconventional myosin-XIX OS=Homo s | MYO19 | 109.136 | 2 | 38.08482 | yes | 5.471E-07 | 620.86664 | 57.341789 |

| | | | | | | | | | | | |
|---|---|---|---|---|---|---|---|---|---|---|---|
| SLEEDCFEVTR | 11 | Q96H55 | Unconventional myosin-XIX OS=Homo s | MYO19 | 109.136 | 2 | 38.08482 | yes | 6.247E-07 | 692.80609 | 42.046471 |
| MVCSPVTVR | 9 | Q96HA1 | Nuclear envelope pore membrane protein | POM121 | 127.721 | 2 | 48.393764 | no | 2.095E-07 | 524.76752 | 161.15294 |
| ASLQWFNQALEDI | 13 | Q96HA1 | Nuclear envelope pore membrane protein | POM121 | 127.721 | 2 | 48.393764 | no | 5.339E-07 | 775.38593 | 55.947456 |
| VQLLPSR | 7 | Q96HA1 | Nuclear envelope pore membrane protein | POM121 | 127.721 | 2 | 48.393764 | no | 6.2E-07 | 406.75308 | 53.54686 |
| PGPPQPAAAPEGQ | 16 | Q96HA1 | Nuclear envelope pore membrane protein | POM121 | 127.721 | 2 | 48.393764 | yes | 1.154E-07 | 800.90753 | 36.849754 |
| GPFLVALGK | 9 | Q96HC4 | PDZ and LIM domain protein 5 OS=Hom | PDLIM5 | 63.946 | 2 | 53.07125 | yes | 4.24E-07 | 451.27893 | 128.1028 |
| ILAQITGTEHLK | 12 | Q96HC4 | PDZ and LIM domain protein 5 OS=Hom | PDLIM5 | 63.946 | 2 | 53.07125 | yes | 9.722E-08 | 662.38519 | 125.77225 |
| EVVKPVPITSPAVS | 15 | Q96HC4 | PDZ and LIM domain protein 5 OS=Hom | PDLIM5 | 63.946 | 3 | 53.07125 | yes | 7.25E-07 | 517.64325 | 34.482841 |
| EVVKPVPITSPAVS | 15 | Q96HC4 | PDZ and LIM domain protein 5 OS=Hom | PDLIM5 | 63.946 | 3 | 53.07125 | yes | 9.722E-08 | 775.96124 | 34.482841 |
| EVVKPVPITSPAVS | 15 | Q96HC4 | PDZ and LIM domain protein 5 OS=Hom | PDLIM5 | 63.946 | 2 | 53.07125 | yes | 7.25E-07 | 517.64325 | 34.482841 |
| EVVKPVPITSPAVS | 15 | Q96HC4 | PDZ and LIM domain protein 5 OS=Hom | PDLIM5 | 63.946 | 2 | 53.07125 | yes | 9.722E-08 | 775.96124 | 34.482841 |
| FFAPECGR | 8 | Q96HC4 | PDZ and LIM domain protein 5 OS=Hom | PDLIM5 | 63.946 | 2 | 53.07125 | yes | 9.046E-07 | 492.22382 | 55.833229 |
| GCTGSLNMTLQR | 12 | Q96HC4 | PDZ and LIM domain protein 5 OS=Hom | PDLIM5 | 63.946 | 2 | 53.07125 | yes | 9.722E-08 | 669.31866 | 51.108017 |
| DFNMPLTISSLK | 12 | Q96HC4 | PDZ and LIM domain protein 5 OS=Hom | PDLIM5 | 63.946 | 2 | 53.07125 | yes | 9.722E-08 | 683.35779 | 46.376637 |
| GALYCELCYEK | 11 | Q96HC4 | PDZ and LIM domain protein 5 OS=Hom | PDLIM5 | 63.946 | 2 | 53.07125 | yes | 4.37E-07 | 703.30994 | 43.588505 |
| ASAAPKPEPVPVQ | 14 | Q96HC4 | PDZ and LIM domain protein 5 OS=Hom | PDLIM5 | 63.946 | 2 | 53.07125 | yes | 9.722E-08 | 709.90375 | 39.598778 |
| NTMAYIGFVEEK | 12 | Q96HC4 | PDZ and LIM domain protein 5 OS=Hom | PDLIM5 | 63.946 | 2 | 53.07125 | yes | 2.141E-06 | 701.3396 | 33.811905 |
| ALEELNQQLEK | 11 | Q96HJ3 | Coiled-coil domain-containing protein 34 | CCDC34 | 43.226 | 2 | 50.65527 | yes | 1.015E-07 | 657.84863 | 288.19052 |
| SNLCLGTLCR | 10 | Q96HJ3 | Coiled-coil domain-containing protein 34 | CCDC34 | 43.226 | 2 | 50.65527 | yes | 9.729E-08 | 597.29193 | 244.91264 |
| GQGLEVVR | 8 | Q96HJ3 | Coiled-coil domain-containing protein 34 | CCDC34 | 43.226 | 2 | 50.65527 | yes | 1.893E-07 | 429.24561 | 157.82477 |
| KFQEWLENAK | 10 | Q96HJ3 | Coiled-coil domain-containing protein 34 | CCDC34 | 43.226 | 2 | 50.65527 | yes | 2.042E-06 | 646.83533 | 40.549393 |
| IGQETVQDIVYR | 12 | Q96HR3 | Mediator of RNA polymerase II transcript | MED30 | 20.278 | 2 | 48.771633 | yes | 1.106E-07 | 710.87518 | 107.80075 |
| ETMYFAPSMK | 10 | Q96HR8 | H/ACA ribonucleoprotein complex non-c | NAF1 | 53.717 | 2 | 51.1012 | yes | 9.722E-08 | 602.77246 | 465.73685 |
| FNSSDHIESK | 10 | Q96HR8 | H/ACA ribonucleoprotein complex non-c | NAF1 | 53.717 | 2 | 51.1012 | yes | 4.678E-07 | 582.27002 | 248.85439 |
| DFTQYIFTEK | 10 | Q96HR8 | H/ACA ribonucleoprotein complex non-c | NAF1 | 53.717 | 2 | 51.1012 | yes | 2.127E-07 | 646.3139 | 239.73518 |
| AIETTDIISR | 10 | Q96HS1 | Serine/threonine-protein phosphatase PG | PGAM5 | 32.005 | 2 | 52.101807 | yes | 1.045E-07 | 559.80621 | 5057.832 |
| PAEPPAWAGGAR | 12 | Q96HS1 | Serine/threonine-protein phosphatase PG | PGAM5 | 32.005 | 2 | 52.101807 | yes | 1.092E-07 | 590.29889 | 4595.7256 |
| PGPGVWDPNWDR | 12 | Q96HS1 | Serine/threonine-protein phosphatase PG | PGAM5 | 32.005 | 2 | 52.101807 | yes | 1.022E-07 | 698.32562 | 3638.022 |
| EPLSLINVR | 9 | Q96HS1 | Serine/threonine-protein phosphatase PG | PGAM5 | 32.005 | 2 | 52.101807 | yes | 1.112E-07 | 520.80859 | 2936.4351 |
| TLGDTGFMPPDK | 12 | Q96HS1 | Serine/threonine-protein phosphatase PG | PGAM5 | 32.005 | 2 | 52.101807 | yes | 3.649E-07 | 639.80536 | 1982.4047 |
| NVESGEEELASK | 12 | Q96HS1 | Serine/threonine-protein phosphatase PG | PGAM5 | 32.005 | 2 | 52.101807 | yes | 1.074E-07 | 646.30426 | 1870.8848 |
| ALQFPPEGWLR | 11 | Q96HS1 | Serine/threonine-protein phosphatase PG | PGAM5 | 32.005 | 2 | 52.101807 | yes | 6.321E-07 | 657.3537 | 1277.4604 |
| HLPGVCK | 7 | Q96HS1 | Serine/threonine-protein phosphatase PG | PGAM5 | 32.005 | 2 | 52.101807 | yes | 2.953E-07 | 405.71817 | 1251.0176 |
| EQAELTGLR | 9 | Q96HS1 | Serine/threonine-protein phosphatase PG | PGAM5 | 32.005 | 2 | 52.101807 | yes | 3.062E-07 | 508.77219 | 1222.1825 |
| HSQYHVDGSLEK | 12 | Q96HS1 | Serine/threonine-protein phosphatase PG | PGAM5 | 32.005 | 3 | 52.101807 | yes | 1.425E-07 | 467.22488 | 431.51178 |
| HSQYHVDGSLEK | 12 | Q96HS1 | Serine/threonine-protein phosphatase PG | PGAM5 | 32.005 | 3 | 52.101807 | yes | 9.722E-08 | 700.33368 | 431.51178 |
| HSQYHVDGSLEK | 12 | Q96HS1 | Serine/threonine-protein phosphatase PG | PGAM5 | 32.005 | 2 | 52.101807 | yes | 1.425E-07 | 467.22488 | 431.51178 |
| HSQYHVDGSLEK | 12 | Q96HS1 | Serine/threonine-protein phosphatase PG | PGAM5 | 32.005 | 2 | 52.101807 | yes | 9.722E-08 | 700.33368 | 431.51178 |
| IVHSSMTR | 8 | Q96HS1 | Serine/threonine-protein phosphatase PG | PGAM5 | 32.005 | 2 | 52.101807 | yes | 2.085E-07 | 465.74493 | 195.70184 |

| | | | | | | | | | | | |
|---|---|---|---|---|---|---|---|---|---|---|---|
| QEEDSYEIFICHAN | 17 | Q96HS1 | Serine/threonine-protein phosphatase PG. | PGAM5 | 32.005 | 3 | 52.101807 | yes | 9.722E-08 | 708.33362 | 174.10413 |
| REPLSLINVR | 10 | Q96HS1 | Serine/threonine-protein phosphatase PG. | PGAM5 | 32.005 | 2 | 52.101807 | yes | 3.718E-07 | 598.85913 | 144.86069 |
| VSTDLLR | 7 | Q96HS1 | Serine/threonine-protein phosphatase PG. | PGAM5 | 32.005 | 2 | 52.101807 | yes | 9.819E-08 | 402.23471 | 97.731339 |
| RNVESGEEELASK | 13 | Q96HS1 | Serine/threonine-protein phosphatase PG. | PGAM5 | 32.005 | 2 | 52.101807 | yes | 1.045E-06 | 724.3548 | 27.076742 |
| QVVIIHHMRLQAK | 13 | Q96HW7 | Integrator complex subunit 4 OS=Homo s | INTS4 | 108.171 | 2 | 51.95645 | yes | 9.722E-08 | 794.95886 | 45.660522 |
| LVGRPPLPVPAVG | 28 | Q96HY7 | 2-oxoadipate dehydrogenase complex cor | DHTKD1 | 103.077 | 4 | 49.6999 | yes | 1.009E-07 | 741.67902 | 142.44945 |
| LVGRPPLPVPAVG | 28 | Q96HY7 | 2-oxoadipate dehydrogenase complex cor | DHTKD1 | 103.077 | 4 | 49.6999 | yes | 1.602E-07 | 593.54468 | 142.44945 |
| LVGRPPLPVPAVG | 28 | Q96HY7 | 2-oxoadipate dehydrogenase complex cor | DHTKD1 | 103.077 | 5 | 49.6999 | yes | 1.009E-07 | 741.67902 | 142.44945 |
| LVGRPPLPVPAVG | 28 | Q96HY7 | 2-oxoadipate dehydrogenase complex cor | DHTKD1 | 103.077 | 5 | 49.6999 | yes | 1.602E-07 | 593.54468 | 142.44945 |
| KHDFAIIR | 8 | Q96HY7 | 2-oxoadipate dehydrogenase complex cor | DHTKD1 | 103.077 | 2 | 49.6999 | yes | 5.057E-07 | 500.29034 | 230.17253 |
| SSLYCSDIGK | 10 | Q96HY7 | 2-oxoadipate dehydrogenase complex cor | DHTKD1 | 103.077 | 2 | 49.6999 | yes | 2.364E-07 | 565.26337 | 179.0325 |
| LVTVYCEHGHK | 11 | Q96HY7 | 2-oxoadipate dehydrogenase complex cor | DHTKD1 | 103.077 | 2 | 49.6999 | yes | 4.659E-07 | 671.83228 | 72.081871 |
| AEGFVDALHR | 10 | Q96I24 | Far upstream element-binding protein 3 C | FUBP3 | 61.641 | 2 | 51.848835 | yes | 9.722E-08 | 557.78564 | 7761.4917 |
| RPLDDGVGNQLG. | 18 | Q96I24 | Far upstream element-binding protein 3 C | FUBP3 | 61.641 | 3 | 51.848835 | yes | 1.586E-07 | 649.01691 | 5476.4658 |
| MGGGSIEVSVPR | 12 | Q96I24 | Far upstream element-binding protein 3 C | FUBP3 | 61.641 | 2 | 51.848835 | yes | 1.215E-07 | 594.80573 | 2602.2292 |
| MGGGSIEVSVPR | 12 | Q96I24 | Far upstream element-binding protein 3 C | FUBP3 | 61.641 | 2 | 51.848835 | yes | 9.722E-08 | 602.80316 | 2602.2292 |
| LLGQIVDR | 8 | Q96I24 | Far upstream element-binding protein 3 C | FUBP3 | 61.641 | 2 | 51.848835 | yes | 1.232E-07 | 457.27692 | 4470.3208 |
| EMVLEIIR | 8 | Q96I24 | Far upstream element-binding protein 3 C | FUBP3 | 61.641 | 2 | 51.848835 | yes | 1.418E-07 | 501.78625 | 3542.7615 |
| DGFGGLAAAR | 10 | Q96I24 | Far upstream element-binding protein 3 C | FUBP3 | 61.641 | 2 | 51.848835 | yes | 3.283E-07 | 467.74069 | 3087.5645 |
| GVPQQIEVAR | 10 | Q96I24 | Far upstream element-binding protein 3 C | FUBP3 | 61.641 | 2 | 51.848835 | yes | 9.722E-08 | 548.80908 | 2711.3582 |
| MVMIQDGPLPTG. | 15 | Q96I24 | Far upstream element-binding protein 3 C | FUBP3 | 61.641 | 2 | 51.848835 | yes | 1.171E-07 | 786.89166 | 2625.3599 |
| IQFKPDDGISPER | 13 | Q96I24 | Far upstream element-binding protein 3 C | FUBP3 | 61.641 | 3 | 51.848835 | yes | 1.168E-07 | 501.2597 | 1127.4229 |
| IQFKPDDGISPER | 13 | Q96I24 | Far upstream element-binding protein 3 C | FUBP3 | 61.641 | 3 | 51.848835 | yes | 1.174E-07 | 751.38593 | 1127.4229 |
| IQFKPDDGISPER | 13 | Q96I24 | Far upstream element-binding protein 3 C | FUBP3 | 61.641 | 2 | 51.848835 | yes | 1.168E-07 | 501.2597 | 1127.4229 |
| IQFKPDDGISPER | 13 | Q96I24 | Far upstream element-binding protein 3 C | FUBP3 | 61.641 | 2 | 51.848835 | yes | 1.174E-07 | 751.38593 | 1127.4229 |
| AELVQGQSAPVGN | 14 | Q96I24 | Far upstream element-binding protein 3 C | FUBP3 | 61.641 | 2 | 51.848835 | yes | 9.758E-08 | 728.87683 | 1079.0607 |
| AELVQGQSAPVGN | 14 | Q96I24 | Far upstream element-binding protein 3 C | FUBP3 | 61.641 | 2 | 51.848835 | yes | 4.003E-07 | 736.87433 | 1079.0607 |
| TVITEEFK | 8 | Q96I24 | Far upstream element-binding protein 3 C | FUBP3 | 61.641 | 2 | 51.848835 | yes | 2.421E-07 | 483.76077 | 2116.0181 |
| SSGCFPNMAAK | 11 | Q96I24 | Far upstream element-binding protein 3 C | FUBP3 | 61.641 | 2 | 51.848835 | yes | 1.363E-07 | 585.25751 | 1871.011 |
| EKDQADFR | 8 | Q96I24 | Far upstream element-binding protein 3 C | FUBP3 | 61.641 | 2 | 51.848835 | yes | 1.594E-07 | 504.74088 | 1716.5371 |
| SINQQSGAHVELQ | 14 | Q96I24 | Far upstream element-binding protein 3 C | FUBP3 | 61.641 | 2 | 51.848835 | yes | 1.114E-07 | 783.90277 | 759.12976 |
| SINQQSGAHVELQ | 14 | Q96I24 | Far upstream element-binding protein 3 C | FUBP3 | 61.641 | 2 | 51.848835 | yes | 4.952E-07 | 522.93762 | 759.12976 |
| SINQQSGAHVELQ | 14 | Q96I24 | Far upstream element-binding protein 3 C | FUBP3 | 61.641 | 3 | 51.848835 | yes | 1.114E-07 | 783.90277 | 759.12976 |
| SINQQSGAHVELQ | 14 | Q96I24 | Far upstream element-binding protein 3 C | FUBP3 | 61.641 | 3 | 51.848835 | yes | 4.952E-07 | 522.93762 | 759.12976 |
| CQHAAHIISELILT. | 18 | Q96I24 | Far upstream element-binding protein 3 C | FUBP3 | 61.641 | 3 | 51.848835 | yes | 1.128E-07 | 697.36523 | 1255.0675 |
| TVITEEFKVPDK | 12 | Q96I24 | Far upstream element-binding protein 3 C | FUBP3 | 61.641 | 2 | 51.848835 | yes | 1.231E-07 | 703.38232 | 1160.3535 |
| NPPPNSDPNLR | 11 | Q96I24 | Far upstream element-binding protein 3 C | FUBP3 | 61.641 | 2 | 51.848835 | yes | 1.683E-07 | 610.80457 | 886.71204 |
| MVMIQDGPLPTG. | 18 | Q96I24 | Far upstream element-binding protein 3 C | FUBP3 | 61.641 | 3 | 51.848835 | yes | 3.895E-07 | 647.00952 | 638.8396 |
| AAQVMGPPDR | 10 | Q96I24 | Far upstream element-binding protein 3 C | FUBP3 | 61.641 | 2 | 51.848835 | yes | 2.617E-07 | 521.26093 | 422.36353 |

| | | | | | | | | | | | |
|---|---|---|---|---|---|---|---|---|---|---|---|
| MVGFIIGR | 8 | Q96I24 | Far upstream element-binding protein 3 C | FUBP3 | 61.641 | 2 | 51.848835 | yes | 3.843E-07 | 446.75729 | 140.2411 |
| AWEDYYK | 7 | Q96I24 | Far upstream element-binding protein 3 C | FUBP3 | 61.641 | 2 | 51.848835 | yes | 1.295E-07 | 487.71634 | 129.56616 |
| PCVLTGTPESIEQA | 15 | Q96I24 | Far upstream element-binding protein 3 C | FUBP3 | 61.641 | 2 | 51.848835 | yes | 1.572E-07 | 815.41125 | 95.234673 |
| QQVAFYGQTLGQ/ | 20 | Q96I24 | Far upstream element-binding protein 3 C | FUBP3 | 61.641 | 3 | 51.848835 | yes | 2.035E-07 | 740.35406 | 29.62018 |
| SMGGAAIAPPTSL' | 16 | Q96I25 | Splicing factor 45 OS=Homo sapiens OX | RBM17 | 44.962 | 2 | 52.06444 | yes | 1.539E-07 | 764.90564 | 1790.0787 |
| SMGGAAIAPPTSL' | 16 | Q96I25 | Splicing factor 45 OS=Homo sapiens OX | RBM17 | 44.962 | 2 | 52.06444 | yes | 1.058E-07 | 772.90308 | 1790.0787 |
| LLQSQLQVK | 9 | Q96I25 | Splicing factor 45 OS=Homo sapiens OX | RBM17 | 44.962 | 2 | 52.06444 | yes | 2.88E-07 | 528.82422 | 2669.1204 |
| QSTVLAPVIDLK | 12 | Q96I25 | Splicing factor 45 OS=Homo sapiens OX | RBM17 | 44.962 | 2 | 52.06444 | yes | 9.81E-08 | 642.38208 | 2416.0427 |
| IFLEFER | 7 | Q96I25 | Splicing factor 45 OS=Homo sapiens OX | RBM17 | 44.962 | 2 | 52.06444 | yes | 2.613E-07 | 477.25818 | 1461.0563 |
| DFPYEEDSRPR | 11 | Q96I25 | Splicing factor 45 OS=Homo sapiens OX | RBM17 | 44.962 | 2 | 52.06444 | yes | 4.361E-07 | 705.81787 | 1107.9882 |
| ACFYNLDK | 8 | Q96I25 | Splicing factor 45 OS=Homo sapiens OX | RBM17 | 44.962 | 2 | 52.06444 | yes | 1.707E-07 | 515.73676 | 1103.7778 |
| HEQGLSTALSVEK | 13 | Q96I25 | Splicing factor 45 OS=Homo sapiens OX | RBM17 | 44.962 | 2 | 52.06444 | yes | 1.004E-07 | 699.86481 | 909.63989 |
| HEASGFAR | 8 | Q96I25 | Splicing factor 45 OS=Homo sapiens OX | RBM17 | 44.962 | 2 | 52.06444 | yes | 2.109E-07 | 437.71194 | 817.70288 |
| QIVDTPPHVAAGL | 14 | Q96I25 | Splicing factor 45 OS=Homo sapiens OX | RBM17 | 44.962 | 2 | 52.06444 | yes | 1.713E-07 | 723.40918 | 250.93033 |
| QIVDTPPHVAAGL | 14 | Q96I25 | Splicing factor 45 OS=Homo sapiens OX | RBM17 | 44.962 | 2 | 52.06444 | yes | 7.288E-07 | 482.60855 | 250.93033 |
| QIVDTPPHVAAGL | 14 | Q96I25 | Splicing factor 45 OS=Homo sapiens OX | RBM17 | 44.962 | 3 | 52.06444 | yes | 1.713E-07 | 723.40918 | 250.93033 |
| QIVDTPPHVAAGL | 14 | Q96I25 | Splicing factor 45 OS=Homo sapiens OX | RBM17 | 44.962 | 3 | 52.06444 | yes | 7.288E-07 | 482.60855 | 250.93033 |
| SDSNPLTEILK | 11 | Q96I25 | Splicing factor 45 OS=Homo sapiens OX | RBM17 | 44.962 | 2 | 52.06444 | yes | 6.001E-07 | 608.82465 | 476.29041 |
| AAIPPPVYEEQDRI | 15 | Q96I25 | Splicing factor 45 OS=Homo sapiens OX | RBM17 | 44.962 | 3 | 52.06444 | yes | 1.714E-07 | 579.96881 | 463.35947 |
| KSDSNPLTEILK | 12 | Q96I25 | Splicing factor 45 OS=Homo sapiens OX | RBM17 | 44.962 | 2 | 52.06444 | yes | 2.538E-07 | 672.87207 | 396.32016 |
| SLYDDLGVETSDS | 14 | Q96I25 | Splicing factor 45 OS=Homo sapiens OX | RBM17 | 44.962 | 2 | 52.06444 | yes | 1.305E-07 | 785.85956 | 378.42603 |
| SPTGPSNSFLANM( | 20 | Q96I25 | Splicing factor 45 OS=Homo sapiens OX | RBM17 | 44.962 | 3 | 52.06444 | yes | 9.809E-08 | 658.32306 | 148.53474 |
| DRHEASGFAR | 10 | Q96I25 | Splicing factor 45 OS=Homo sapiens OX | RBM17 | 44.962 | 2 | 52.06444 | yes | 9.722E-08 | 573.27594 | 84.053299 |
| AHSLVLTDR | 9 | Q96I51 | RCC1-like G exchanging factor-like prote | RCC1L | 49.898 | 2 | 54.661407 | yes | 4.479E-07 | 506.28271 | 1052.6143 |
| VTMPGEPVDVAC( | 22 | Q96I51 | RCC1-like G exchanging factor-like prote | RCC1L | 49.898 | 3 | 54.661407 | yes | 1.153E-07 | 776.0462 | 1025.0535 |
| DSQLGFHR | 8 | Q96I51 | RCC1-like G exchanging factor-like prote | RCC1L | 49.898 | 2 | 54.661407 | yes | 4.361E-07 | 480.23831 | 913.05963 |
| VLQVSCGR | 8 | Q96I51 | RCC1-like G exchanging factor-like prote | RCC1L | 49.898 | 2 | 54.661407 | yes | 1.096E-07 | 459.74493 | 660.53253 |
| GELFVWGK | 8 | Q96I51 | RCC1-like G exchanging factor-like prote | RCC1L | 49.898 | 2 | 54.661407 | yes | 1.317E-07 | 468.2529 | 641.99438 |
| IQPVPYR | 7 | Q96I51 | RCC1-like G exchanging factor-like prote | RCC1L | 49.898 | 2 | 54.661407 | yes | 3.561E-07 | 436.75308 | 600.59869 |
| VVENEIYSESHR | 12 | Q96I51 | RCC1-like G exchanging factor-like prote | RCC1L | 49.898 | 2 | 54.661407 | yes | 9.722E-08 | 731.35205 | 539.20996 |
| ISSAACGYGFTLLS | 16 | Q96I51 | RCC1-like G exchanging factor-like prote | RCC1L | 49.898 | 2 | 54.661407 | yes | 2.244E-07 | 831.41382 | 442.7818 |
| LEDQYFPWR | 9 | Q96I51 | RCC1-like G exchanging factor-like prote | RCC1L | 49.898 | 2 | 54.661407 | yes | 9.722E-08 | 627.30109 | 266.05591 |
| CLHFSGVGK | 9 | Q96I51 | RCC1-like G exchanging factor-like prote | RCC1L | 49.898 | 2 | 54.661407 | yes | 9.809E-08 | 502.75275 | 247.96466 |
| VWGMGLNK | 8 | Q96I51 | RCC1-like G exchanging factor-like prote | RCC1L | 49.898 | 2 | 54.661407 | yes | 4.521E-06 | 452.73911 | 199.70883 |
| CGLSHFAALTNK | 12 | Q96I51 | RCC1-like G exchanging factor-like prote | RCC1L | 49.898 | 2 | 54.661407 | yes | 9.722E-08 | 659.83228 | 102.16556 |
| KVVENEIYSESHR | 13 | Q96I51 | RCC1-like G exchanging factor-like prote | RCC1L | 49.898 | 2 | 54.661407 | yes | 1.165E-07 | 795.39954 | 26.60498 |
| TNVLGSILR | 9 | Q96I59 | Probable asparagine--tRNA ligase, mitocl | NARS2 | 54.091 | 2 | 42.701942 | yes | 8.195E-07 | 486.79547 | 44.279682 |
| YLQCILGVDNIK | 12 | Q96I59 | Probable asparagine--tRNA ligase, mitocl | NARS2 | 54.091 | 2 | 42.701942 | yes | 2.911E-07 | 718.38434 | 25.123184 |
| LEPTAPALSPR | 11 | Q96IF1 | LIM domain-containing protein ajuba OS | AJUBA | 56.934 | 2 | 52.507294 | yes | 1.431E-07 | 576.32458 | 261.53436 |

| | | | | | | | | | | | |
|---|---|---|---|---|---|---|---|---|---|---|---|
| YQDELTALLR | 10 | Q96IF1 | LIM domain-containing protein ajuba OS | AJUBA | 56.934 | 2 | 52.507294 | yes | 9.843E-08 | 611.32733 | 122.95913 |
| TSGISMGYDQR | 11 | Q96IF1 | LIM domain-containing protein ajuba OS | AJUBA | 56.934 | 2 | 52.507294 | yes | 9.722E-08 | 607.77716 | 94.071136 |
| ALPLPQSLPPDFR | 13 | Q96IF1 | LIM domain-containing protein ajuba OS | AJUBA | 56.934 | 2 | 52.507294 | yes | 4.93E-07 | 725.90625 | 87.568886 |
| GATGGPGDEPLEP | 15 | Q96IF1 | LIM domain-containing protein ajuba OS | AJUBA | 56.934 | 2 | 52.507294 | yes | 9.722E-08 | 712.34424 | 73.582138 |
| GSLLLDGAGAGGA | 17 | Q96IF1 | LIM domain-containing protein ajuba OS | AJUBA | 56.934 | 2 | 52.507294 | yes | 5.224E-07 | 708.36548 | 65.797058 |
| EDYFGTCIK | 9 | Q96IF1 | LIM domain-containing protein ajuba OS | AJUBA | 56.934 | 2 | 52.507294 | yes | 1.273E-07 | 566.75262 | 50.686054 |
| PAGFGCQESR | 10 | Q96IF1 | LIM domain-containing protein ajuba OS | AJUBA | 56.934 | 2 | 52.507294 | yes | 1.544E-07 | 554.74567 | 42.282398 |
| IAPDLPDYDR | 10 | Q96II8 | DISP complex protein LRCH3 OS=Homo | LRCH3 | 86.084 | 2 | 46.130306 | yes | 1.644E-07 | 587.79059 | 113.23685 |
| EAQLAALQYEEEK | 13 | Q96II8 | DISP complex protein LRCH3 OS=Homo | LRCH3 | 86.084 | 2 | 46.130306 | yes | 5.512E-07 | 761.37518 | 94.628418 |
| NVENFLEACR | 10 | Q96II8 | DISP complex protein LRCH3 OS=Homo | LRCH3 | 86.084 | 2 | 46.130306 | no | 2.41E-07 | 626.29297 | 80.589027 |
| NHLVHLPEELAEL | 17 | Q96II8 | DISP complex protein LRCH3 OS=Homo | LRCH3 | 86.084 | 3 | 46.130306 | yes | 3.45E-07 | 665.04584 | 72.646103 |
| ISHEGSPVKPVAIR | 14 | Q96II8 | DISP complex protein LRCH3 OS=Homo | LRCH3 | 86.084 | 3 | 46.130306 | yes | 5.73E-07 | 497.28769 | 60.771732 |
| AVILIGGPQK | 10 | Q96IJ6 | Mannose-1-phosphate guanyltransferase a | GMPPA | 46.292 | 2 | 36.732742 | yes | 6.67E-07 | 498.31604 | 46.303825 |
| LLPAITILGCR | 11 | Q96IJ6 | Mannose-1-phosphate guanyltransferase a | GMPPA | 46.292 | 2 | 36.732742 | yes | 3.697E-06 | 613.8681 | 37.45969 |
| LQYIISDLASAR | 12 | Q96IK5 | Germ cell-less protein-like 1 OS=Homo s | GMCL1 | 58.686 | 2 | 46.993565 | no | 1.424E-07 | 675.37482 | 232.05728 |
| DQEQVVMNLDSR | 12 | Q96IK5 | Germ cell-less protein-like 1 OS=Homo s | GMCL1 | 58.686 | 2 | 46.993565 | no | 2.26E-07 | 717.33807 | 113.21972 |
| DDVLIKPSR | 9 | Q96IK5 | Germ cell-less protein-like 1 OS=Homo s | GMCL1 | 58.686 | 2 | 46.993565 | no | 2.55E-07 | 521.79822 | 109.02869 |
| TTGYQILTLEK | 11 | Q96IK5 | Germ cell-less protein-like 1 OS=Homo s | GMCL1 | 58.686 | 2 | 46.993565 | yes | 2.112E-07 | 633.85065 | 54.073189 |
| QLLTETDVWFSK | 12 | Q96IK5 | Germ cell-less protein-like 1 OS=Homo s | GMCL1 | 58.686 | 2 | 46.993565 | no | 3.938E-07 | 733.87994 | 26.865978 |
| FFIDFSDIGEQQR | 13 | Q96IP4 | Terminal nucleotidyltransferase 5A OS=H | TENT5A | 49.667 | 2 | 53.03193 | yes | 1.165E-07 | 801.38336 | 285.09897 |
| GFRPASDEIK | 10 | Q96IP4 | Terminal nucleotidyltransferase 5A OS=H | TENT5A | 49.667 | 2 | 53.03193 | yes | 1.51E-07 | 560.29327 | 278.5903 |
| LDGILSETIPIHGR | 14 | Q96IP4 | Terminal nucleotidyltransferase 5A OS=H | TENT5A | 49.667 | 2 | 53.03193 | yes | 1.13E-07 | 760.92517 | 144.5981 |
| GNFPTLELQPSLIV | 15 | Q96IP4 | Terminal nucleotidyltransferase 5A OS=H | TENT5A | 49.667 | 2 | 53.03193 | yes | 2.834E-07 | 828.47198 | 106.4641 |
| YCNLLVR | 7 | Q96IP4 | Terminal nucleotidyltransferase 5A OS=H | TENT5A | 49.667 | 2 | 53.03193 | yes | 1.943E-07 | 469.24985 | 93.67791 |
| WSLISLSNNSGK | 12 | Q96IP4 | Terminal nucleotidyltransferase 5A OS=H | TENT5A | 49.667 | 2 | 53.03193 | yes | 9.723E-08 | 653.34351 | 86.053345 |
| QFEFSVDSFQIK | 12 | Q96IP4 | Terminal nucleotidyltransferase 5A OS=H | TENT5A | 49.667 | 2 | 53.03193 | yes | 9.722E-08 | 737.86426 | 30.280632 |
| DLDLIFCADLR | 11 | Q96IP4 | Terminal nucleotidyltransferase 5A OS=H | TENT5A | 49.667 | 2 | 53.03193 | yes | 3.754E-06 | 675.83972 | 11.598235 |
| THCPPTQSLEGK | 12 | Q96IQ9 | Zinc finger protein 414 OS=Homo sapien | ZNF414 | 32.783 | 2 | 47.21342 | yes | 1.376E-07 | 677.82465 | 61.799587 |
| LLEQMGALAVR | 11 | Q96IU2 | Zinc finger BED domain-containing prote | ZBED3 | 25.133 | 2 | 37.39022 | yes | 6.036E-07 | 600.8421 | 114.09644 |
| EGTIQVQGQALFFI | 14 | Q96IU4 | Putative protein-lysine deacylase ABHD1 | ABHD14B | 22.346 | 2 | 42.47527 | yes | 3.012E-07 | 797.42285 | 72.839073 |
| AGPESDAQYQFTC | 15 | Q96IX5 | ATP synthase membrane subunit K, mito | ATP5MK | 6.458 | 2 | 50.04975 | yes | 1.038E-07 | 806.38611 | 649.94434 |
| YFNSYTLTGR | 10 | Q96IX5 | ATP synthase membrane subunit K, mito | ATP5MK | 6.458 | 2 | 50.04975 | yes | 1.1E-07 | 611.29858 | 241.75185 |
| KYFNSYTLTGR | 11 | Q96IX5 | ATP synthase membrane subunit K, mito | ATP5MK | 6.458 | 2 | 50.04975 | yes | 9.926E-08 | 675.34607 | 42.015282 |
| DLDDIEDENEQLK | 13 | Q96IZ0 | PRKC apoptosis WT1 regulator protein C | PAWR | 36.568 | 2 | 52.81688 | yes | 1.361E-07 | 788.35468 | 126.62878 |
| LKEEIDLLNR | 10 | Q96IZ0 | PRKC apoptosis WT1 regulator protein C | PAWR | 36.568 | 2 | 52.81688 | yes | 1.873E-07 | 621.85626 | 120.98503 |
| DANVSGTLVSSST | 16 | Q96IZ0 | PRKC apoptosis WT1 regulator protein C | PAWR | 36.568 | 2 | 52.81688 | yes | 9.722E-08 | 804.40979 | 107.37592 |
| ILCLPQR | 7 | Q96IZ5 | RNA-binding protein 41 OS=Homo sapie | RBM41 | 47.1 | 2 | 54.498672 | yes | 4.329E-07 | 450.26019 | 255.71207 |
| KGPPIQFR | 8 | Q96IZ5 | RNA-binding protein 41 OS=Homo sapie | RBM41 | 47.1 | 2 | 54.498672 | yes | 4.47E-07 | 471.77963 | 244.72542 |
| ILVIEFGK | 8 | Q96IZ5 | RNA-binding protein 41 OS=Homo sapie | RBM41 | 47.1 | 2 | 54.498672 | yes | 9.722E-08 | 459.78659 | 216.21008 |

| | | | | | | | | | | | |
|---|---|---|---|---|---|---|---|---|---|---|---|
| GQAFITFPNK | 10 | Q96IZ5 | RNA-binding protein 41 OS=Homo sapie | RBM41 | 47.1 | 2 | 54.498672 | yes | 2.542E-07 | 561.80072 | 212.98914 |
| ATPEAIQNR | 9 | Q96IZ5 | RNA-binding protein 41 OS=Homo sapie | RBM41 | 47.1 | 2 | 54.498672 | yes | 5.395E-07 | 500.26453 | 173.66315 |
| LQDIEER | 7 | Q96IZ5 | RNA-binding protein 41 OS=Homo sapie | RBM41 | 47.1 | 2 | 54.498672 | yes | 5.762E-07 | 451.73254 | 132.64044 |
| LEEFQLMR | 8 | Q96IZ5 | RNA-binding protein 41 OS=Homo sapie | RBM41 | 47.1 | 2 | 54.498672 | yes | 1.388E-07 | 533.2735 | 105.30399 |
| ELGLNETEILIWK | 13 | Q96IZ5 | RNA-binding protein 41 OS=Homo sapie | RBM41 | 47.1 | 2 | 54.498672 | yes | 9.722E-08 | 779.42981 | 91.079926 |
| ESFAPGTMYKPFG | 14 | Q96IZ5 | RNA-binding protein 41 OS=Homo sapie | RBM41 | 47.1 | 2 | 54.498672 | yes | 1.963E-07 | 780.38177 | 85.843391 |
| ALYYQDEPQK | 10 | Q96IZ5 | RNA-binding protein 41 OS=Homo sapie | RBM41 | 47.1 | 2 | 54.498672 | yes | 1.915E-07 | 627.80371 | 68.970039 |
| SLLQHQLDTSVSIE | 19 | Q96IZ5 | RNA-binding protein 41 OS=Homo sapie | RBM41 | 47.1 | 3 | 54.498672 | yes | 7.937E-07 | 735.68903 | 39.775288 |
| ALYYQDEPQKK | 11 | Q96IZ5 | RNA-binding protein 41 OS=Homo sapie | RBM41 | 47.1 | 2 | 54.498672 | yes | 2.006E-06 | 691.8512 | 33.286243 |
| EIDPTSIPTAIK | 12 | Q96IZ7 | Serine/Arginine-related protein 53 OS=H | RSRC1 | 38.678 | 2 | 39.8561 | yes | 4.343E-07 | 642.8559 | 93.9991 |
| LQLVLEAAAK | 10 | Q96IZ7 | Serine/Arginine-related protein 53 OS=H | RSRC1 | 38.678 | 2 | 39.8561 | yes | 4.841E-07 | 528.3266 | 35.452995 |
| YVLGMQELFR | 10 | Q96J01 | THO complex subunit 3 OS=Homo sapie | THOC3 | 38.772 | 2 | 52.5428 | yes | 9.722E-08 | 628.3288 | 334.43121 |
| TLSFSHDGK | 9 | Q96J01 | THO complex subunit 3 OS=Homo sapie | THOC3 | 38.772 | 2 | 52.5428 | yes | 1.916E-07 | 496.24582 | 218.22972 |
| RPLLAFACDDKDC | 14 | Q96J01 | THO complex subunit 3 OS=Homo sapie | THOC3 | 38.772 | 3 | 52.5428 | yes | 2.649E-07 | 535.93994 | 208.70935 |
| EFLAHSAK | 8 | Q96J01 | THO complex subunit 3 OS=Homo sapie | THOC3 | 38.772 | 2 | 52.5428 | yes | 2.483E-07 | 451.74017 | 184.75742 |
| MLASASEDHFIDIA | 21 | Q96J01 | THO complex subunit 3 OS=Homo sapie | THOC3 | 38.772 | 3 | 52.5428 | yes | 2.196E-07 | 760.02295 | 111.06191 |
| LWEVQCESPTFTV | 18 | Q96J01 | THO complex subunit 3 OS=Homo sapie | THOC3 | 38.772 | 3 | 52.5428 | yes | 1.387E-07 | 739.02618 | 107.53496 |
| TASVFLLEK | 9 | Q96J01 | THO complex subunit 3 OS=Homo sapie | THOC3 | 38.772 | 2 | 52.5428 | yes | 3.628E-07 | 504.29224 | 55.016495 |
| VHSVAWSCDGR | 11 | Q96J01 | THO complex subunit 3 OS=Homo sapie | THOC3 | 38.772 | 2 | 52.5428 | yes | 9.722E-08 | 637.29077 | 46.673031 |
| GHGDSVDQLCWH | 26 | Q96J01 | THO complex subunit 3 OS=Homo sapie | THOC3 | 38.772 | 4 | 52.5428 | yes | 1.163E-07 | 710.57397 | 43.814857 |
| SQLQGAMQQFNQ | 13 | Q96J02 | E3 ubiquitin-protein ligase Itchy homolog | ITCH | 102.803 | 2 | 45.608326 | yes | 1.798E-07 | 768.3728 | 43.215801 |
| SQLQITVISAK | 11 | Q96J02 | E3 ubiquitin-protein ligase Itchy homolog | ITCH | 102.803 | 2 | 45.608326 | yes | 3.327E-07 | 594.35333 | 31.549961 |
| PTLESVR | 7 | Q96J02 | E3 ubiquitin-protein ligase Itchy homolog | ITCH | 102.803 | 2 | 45.608326 | yes | 2.822E-06 | 401.2269 | 30.805218 |
| LGYPQAPPSGLER | 13 | Q96J84 | Kin of IRRE-like protein 1 OS=Homo sap | KIRREL1 | 83.536 | 2 | 51.78163 | yes | 2.895E-07 | 692.86462 | 92.24852 |
| VECFIGSTPPPDR | 13 | Q96J84 | Kin of IRRE-like protein 1 OS=Homo sap | KIRREL1 | 83.536 | 2 | 51.78163 | yes | 3.408E-07 | 737.85339 | 50.992798 |
| ENFLEVGTLER | 11 | Q96J84 | Kin of IRRE-like protein 1 OS=Homo sap | KIRREL1 | 83.536 | 2 | 51.78163 | yes | 4.214E-07 | 653.83551 | 48.777008 |
| EDDTASVSTATR | 12 | Q96J84 | Kin of IRRE-like protein 1 OS=Homo sap | KIRREL1 | 83.536 | 2 | 51.78163 | yes | 9.722E-08 | 626.78619 | 44.295082 |
| PAATIIWFR | 9 | Q96J84 | Kin of IRRE-like protein 1 OS=Homo sap | KIRREL1 | 83.536 | 2 | 51.78163 | yes | 4.045E-07 | 537.80835 | 25.058746 |
| IGDLGLATLK | 10 | Q96J92 | Serine/threonine-protein kinase WNK4 O | WNK4 | 134.74 | 2 | 31.827974 | yes | 1.712E-06 | 500.80551 | 41.261593 |
| ALQVLVDQSK | 10 | Q96JC1 | Vam6/Vps39-like protein OS=Homo sapi | VPS39 | 101.81 | 2 | 52.009506 | yes | 9.722E-08 | 550.81915 | 376.15524 |
| LLVQSIELQR | 10 | Q96JC1 | Vam6/Vps39-like protein OS=Homo sapi | VPS39 | 101.81 | 2 | 52.009506 | yes | 1.001E-07 | 599.86133 | 298.4549 |
| ALNLLPANTQIND | 15 | Q96JC1 | Vam6/Vps39-like protein OS=Homo sapi | VPS39 | 101.81 | 2 | 52.009506 | yes | 5.639E-07 | 833.46777 | 291.18277 |
| IFTEDLPEVESLPR | 14 | Q96JC1 | Vam6/Vps39-like protein OS=Homo sapi | VPS39 | 101.81 | 2 | 52.009506 | yes | 7.007E-07 | 822.92761 | 276.50644 |
| EYLLSFPAGK | 10 | Q96JC1 | Vam6/Vps39-like protein OS=Homo sapi | VPS39 | 101.81 | 2 | 52.009506 | yes | 1.876E-07 | 562.80298 | 225.60132 |
| QLEPLVAPLADGK | 13 | Q96JC1 | Vam6/Vps39-like protein OS=Homo sapi | VPS39 | 101.81 | 2 | 52.009506 | yes | 1.117E-07 | 675.88501 | 220.20366 |
| LLQIIDTTLLK | 11 | Q96JC1 | Vam6/Vps39-like protein OS=Homo sapi | VPS39 | 101.81 | 2 | 52.009506 | yes | 5.323E-07 | 635.90265 | 146.42003 |
| MHDAFEPVPILEK | 13 | Q96JC1 | Vam6/Vps39-like protein OS=Homo sapi | VPS39 | 101.81 | 2 | 52.009506 | yes | 9.722E-08 | 763.38959 | 124.23088 |
| MAEEYCHK | 8 | Q96JC1 | Vam6/Vps39-like protein OS=Homo sapi | VPS39 | 101.81 | 2 | 52.009506 | yes | 1.034E-07 | 534.2179 | 114.12472 |
| QGHLLLYR | 8 | Q96JC1 | Vam6/Vps39-like protein OS=Homo sapi | VPS39 | 101.81 | 2 | 52.009506 | yes | 2.165E-07 | 500.29034 | 98.751091 |

| | | | | | | | | | | | |
|---|---|---|---|---|---|---|---|---|---|---|---|
| QFELALQLAEMK | 12 | Q96JC1 | Vam6/Vps39-like protein OS=Homo sapi | VPS39 | 101.81 | 2 | 52.009506 | yes | 2.121E-07 | 710.87885 | 85.827156 |
| YSELIILYEK | 10 | Q96JC1 | Vam6/Vps39-like protein OS=Homo sapi | VPS39 | 101.81 | 2 | 52.009506 | yes | 3.634E-07 | 635.8501 | 68.995819 |
| SMAWCENSICVGI | 14 | Q96JC1 | Vam6/Vps39-like protein OS=Homo sapi | VPS39 | 101.81 | 2 | 52.009506 | yes | 1.585E-07 | 844.86511 | 63.514336 |
| NLYAFNLFCQK | 11 | Q96JC1 | Vam6/Vps39-like protein OS=Homo sapi | VPS39 | 101.81 | 2 | 52.009506 | yes | 1.121E-07 | 709.35028 | 60.050339 |
| CYLHTNVALVAPI | 15 | Q96JC1 | Vam6/Vps39-like protein OS=Homo sapi | VPS39 | 101.81 | 3 | 52.009506 | yes | 1.995E-07 | 580.659 | 45.297665 |
| VLGFLIENFK | 10 | Q96JC1 | Vam6/Vps39-like protein OS=Homo sapi | VPS39 | 101.81 | 2 | 52.009506 | yes | 4.086E-06 | 590.34222 | 31.548208 |
| LELLEPK | 7 | Q96JC1 | Vam6/Vps39-like protein OS=Homo sapi | VPS39 | 101.81 | 2 | 52.009506 | yes | 1.059E-05 | 421.2551 | 27.44878 |
| KIQQIHVVSQFK | 12 | Q96JC1 | Vam6/Vps39-like protein OS=Homo sapi | VPS39 | 101.81 | 2 | 52.009506 | yes | 1.314E-07 | 727.92755 | 19.267811 |
| EAFISEEEIAK | 11 | Q96JM3 | Chromosome alignment-maintaining phos | CHAMP1 | 89.1 | 2 | 50.808365 | yes | 2.721E-07 | 633.31665 | 358.44702 |
| ATLSNPKPQK | 10 | Q96JM3 | Chromosome alignment-maintaining phos | CHAMP1 | 89.1 | 2 | 50.808365 | yes | 9.722E-08 | 542.31146 | 199.54007 |
| SIPALSMETQK | 11 | Q96JM3 | Chromosome alignment-maintaining phos | CHAMP1 | 89.1 | 2 | 50.808365 | yes | 5.857E-07 | 602.81573 | 186.38582 |
| MYSNVYYHITSK | 12 | Q96JM3 | Chromosome alignment-maintaining phos | CHAMP1 | 89.1 | 2 | 50.808365 | yes | 9.84E-08 | 753.35828 | 162.05286 |
| SPPLPEHQK | 9 | Q96JM3 | Chromosome alignment-maintaining phos | CHAMP1 | 89.1 | 2 | 50.808365 | yes | 6.238E-07 | 516.77728 | 153.8154 |
| CNFESNFPR | 9 | Q96JM3 | Chromosome alignment-maintaining phos | CHAMP1 | 89.1 | 2 | 50.808365 | yes | 1.674E-06 | 585.75348 | 140.92372 |
| CDILVQEELLASPK | 14 | Q96JM3 | Chromosome alignment-maintaining phos | CHAMP1 | 89.1 | 2 | 50.808365 | yes | 1.243E-07 | 807.92401 | 124.6582 |
| KPALFPEPAK | 10 | Q96JM3 | Chromosome alignment-maintaining phos | CHAMP1 | 89.1 | 2 | 50.808365 | yes | 9.939E-08 | 549.32129 | 121.79162 |
| IPCNSAEPK | 9 | Q96JM3 | Chromosome alignment-maintaining phos | CHAMP1 | 89.1 | 2 | 50.808365 | yes | 2.586E-06 | 508.2475 | 116.01946 |
| LLEDTLFPSSK | 11 | Q96JM3 | Chromosome alignment-maintaining phos | CHAMP1 | 89.1 | 2 | 50.808365 | yes | 1.05E-07 | 625.33734 | 111.31717 |
| GAVLHHLVNK | 10 | Q96JM3 | Chromosome alignment-maintaining phos | CHAMP1 | 89.1 | 2 | 50.808365 | yes | 4.294E-05 | 544.3222 | 92.188683 |
| NVLQFTEEK | 9 | Q96JM3 | Chromosome alignment-maintaining phos | CHAMP1 | 89.1 | 2 | 50.808365 | yes | 3.995E-07 | 554.28766 | 89.205627 |
| TAPTLSPEHWK | 11 | Q96JM3 | Chromosome alignment-maintaining phos | CHAMP1 | 89.1 | 2 | 50.808365 | yes | 3.502E-07 | 633.82751 | 82.214684 |
| SVPVCESQK | 9 | Q96JM3 | Chromosome alignment-maintaining phos | CHAMP1 | 89.1 | 2 | 50.808365 | yes | 9.141E-07 | 517.25275 | 65.374214 |
| TSPASLDFPESQK | 13 | Q96JM3 | Chromosome alignment-maintaining phos | CHAMP1 | 89.1 | 2 | 50.808365 | yes | 3.989E-07 | 703.84357 | 43.50042 |
| LECDHCSFR | 9 | Q96JM3 | Chromosome alignment-maintaining phos | CHAMP1 | 89.1 | 2 | 50.808365 | yes | 1.631E-07 | 612.25024 | 24.268234 |
| YVCPHPSCGR | 10 | Q96JP5 | E3 ubiquitin-protein ligase ZFP91 OS=H | ZFP91 | 63.446 | 2 | 51.952522 | yes | 1.165E-07 | 616.76862 | 236.94054 |
| TGSLQLICK | 9 | Q96JP5 | E3 ubiquitin-protein ligase ZFP91 OS=H | ZFP91 | 63.446 | 2 | 51.952522 | yes | 1.464E-07 | 510.28134 | 157.80026 |
| DETYKPHLER | 10 | Q96JP5 | E3 ubiquitin-protein ligase ZFP91 OS=H | ZFP91 | 63.446 | 2 | 51.952522 | yes | 2.45E-07 | 644.32001 | 77.843864 |
| DETYKPHLER | 10 | Q96JP5 | E3 ubiquitin-protein ligase ZFP91 OS=H | ZFP91 | 63.446 | 2 | 51.952522 | yes | 9.722E-08 | 429.88245 | 77.843864 |
| DETYKPHLER | 10 | Q96JP5 | E3 ubiquitin-protein ligase ZFP91 OS=H | ZFP91 | 63.446 | 3 | 51.952522 | yes | 2.45E-07 | 644.32001 | 77.843864 |
| DETYKPHLER | 10 | Q96JP5 | E3 ubiquitin-protein ligase ZFP91 OS=H | ZFP91 | 63.446 | 3 | 51.952522 | yes | 9.722E-08 | 429.88245 | 77.843864 |
| CEMEGCGTVLAHI | 14 | Q96JP5 | E3 ubiquitin-protein ligase ZFP91 OS=H | ZFP91 | 63.446 | 2 | 51.952522 | yes | 1.368E-07 | 808.85254 | 153.16229 |
| DYICEYCAR | 9 | Q96JP5 | E3 ubiquitin-protein ligase ZFP91 OS=H | ZFP91 | 63.446 | 2 | 51.952522 | yes | 1.921E-07 | 625.25244 | 141.91425 |
| AAAAAAAAAVSR | 12 | Q96JP5 | E3 ubiquitin-protein ligase ZFP91 OS=H | ZFP91 | 63.446 | 2 | 51.952522 | yes | 2.397E-07 | 500.78033 | 133.89503 |
| KKPPIQYVR | 9 | Q96JP5 | E3 ubiquitin-protein ligase ZFP91 OS=H | ZFP91 | 63.446 | 2 | 51.952522 | yes | 3.545E-07 | 564.84802 | 127.60247 |
| KYVCPHPSCGR | 11 | Q96JP5 | E3 ubiquitin-protein ligase ZFP91 OS=H | ZFP91 | 63.446 | 3 | 51.952522 | yes | 2.292E-06 | 454.21313 | 62.96883 |
| YLQHHIK | 7 | Q96JP5 | E3 ubiquitin-protein ligase ZFP91 OS=H | ZFP91 | 63.446 | 2 | 51.952522 | yes | 6.072E-07 | 469.76398 | 58.88205 |
| KPPIQYVR | 8 | Q96JP5 | E3 ubiquitin-protein ligase ZFP91 OS=H | ZFP91 | 63.446 | 2 | 51.952522 | yes | 2.626E-07 | 500.80054 | 58.734344 |
| AAPEEPQQRPPEA | 25 | Q96JP5 | E3 ubiquitin-protein ligase ZFP91 OS=H | ZFP91 | 63.446 | 3 | 51.952522 | yes | 4.807E-06 | 830.41748 | 38.489716 |
| LLQEALEAEER | 11 | Q96JY6 | PDZ and LIM domain protein 2 OS=Hon | PDLIM2 | 37.459 | 2 | 48.12285 | yes | 1.197E-07 | 650.84082 | 54.576519 |

| | | | | | | | | | | | |
|---|---|---|---|---|---|---|---|---|---|---|---|
| ETFSTMTLLR | 10 | Q96K58 | Zinc finger protein 668 OS=Homo sapien | ZNF668 | 67.905 | 2 | 55.09244 | yes | 1.352E-07 | 599.81049 | 252.9315 |
| MFSDPSSFR | 9 | Q96K58 | Zinc finger protein 668 OS=Homo sapien | ZNF668 | 67.905 | 2 | 55.09244 | yes | 9.722E-08 | 537.23969 | 222.29999 |
| VHLASHAGELPFR | 13 | Q96K58 | Zinc finger protein 668 OS=Homo sapien | ZNF668 | 67.905 | 3 | 55.09244 | yes | 5.514E-07 | 478.59311 | 75.209961 |
| VHLASHAGELPFR | 13 | Q96K58 | Zinc finger protein 668 OS=Homo sapien | ZNF668 | 67.905 | 3 | 55.09244 | yes | 4.509E-07 | 717.38605 | 75.209961 |
| VHLASHAGELPFR | 13 | Q96K58 | Zinc finger protein 668 OS=Homo sapien | ZNF668 | 67.905 | 2 | 55.09244 | yes | 5.514E-07 | 478.59311 | 75.209961 |
| VHLASHAGELPFR | 13 | Q96K58 | Zinc finger protein 668 OS=Homo sapien | ZNF668 | 67.905 | 2 | 55.09244 | yes | 4.509E-07 | 717.38605 | 75.209961 |
| GFTQLSSYQSHER | 13 | Q96K58 | Zinc finger protein 668 OS=Homo sapien | ZNF668 | 67.905 | 2 | 55.09244 | yes | 9.728E-08 | 770.36298 | 99.397812 |
| AFLSASDLR | 9 | Q96K58 | Zinc finger protein 668 OS=Homo sapien | ZNF668 | 67.905 | 2 | 55.09244 | yes | 5.361E-07 | 490.26401 | 92.041519 |
| PYACPLCPK | 9 | Q96K58 | Zinc finger protein 668 OS=Homo sapien | ZNF668 | 67.905 | 2 | 55.09244 | yes | 1.304E-07 | 553.26208 | 84.928619 |
| THSGEKPFLCPR | 12 | Q96K58 | Zinc finger protein 668 OS=Homo sapien | ZNF668 | 67.905 | 3 | 55.09244 | yes | 5.277E-06 | 476.90659 | 49.043911 |
| PFLCSECGK | 9 | Q96K58 | Zinc finger protein 668 OS=Homo sapien | ZNF668 | 67.905 | 2 | 55.09244 | yes | 2.067E-07 | 549.24133 | 39.343575 |
| LQLSVYTTTR | 10 | Q96KA5 | Lipid scramblase CLPTM1L OS=Homo s | CLPTM1L | 62.23 | 2 | 44.091774 | yes | 2.324E-07 | 591.3299 | 64.51326 |
| LALNVMADNFVFI | 23 | Q96KA5 | Lipid scramblase CLPTM1L OS=Homo s | CLPTM1L | 62.23 | 3 | 44.091774 | yes | 2.797E-07 | 830.08118 | 26.328112 |
| ASQDPFPAAIILK | 13 | Q96KB5 | Lymphokine-activated killer T-cell-origin | PBK | 36.086 | 2 | 46.37323 | yes | 1.577E-07 | 685.88757 | 57.578575 |
| VIELFSVCTNEDPK | 14 | Q96KB5 | Lymphokine-activated killer T-cell-origin | PBK | 36.086 | 2 | 46.37323 | yes | 2.454E-07 | 825.90582 | 54.219719 |
| TATTLPDDMITQR | 13 | Q96KC8 | DnaJ homolog subfamily C member 1 OS | DNAJC1 | 63.883 | 2 | 47.280777 | yes | 3.964E-07 | 731.86395 | 112.17068 |
| LLELALQQYPR | 11 | Q96KC8 | DnaJ homolog subfamily C member 1 OS | DNAJC1 | 63.883 | 2 | 47.280777 | yes | 1.362E-07 | 672.3877 | 51.387917 |
| SLDAFPEDFCR | 11 | Q96KG9 | N-terminal kinase-like protein OS=Homo | SCYL1 | 89.632 | 2 | 53.541187 | yes | 1.002E-07 | 678.7981 | 617.90198 |
| IGSYLSASTR | 10 | Q96KG9 | N-terminal kinase-like protein OS=Homo | SCYL1 | 89.632 | 2 | 53.541187 | yes | 3.554E-07 | 527.78003 | 497.76413 |
| FLSAEEYQQK | 10 | Q96KG9 | N-terminal kinase-like protein OS=Homo | SCYL1 | 89.632 | 2 | 53.541187 | yes | 1.768E-07 | 621.80371 | 487.8598 |
| LNEANLNVELMK | 12 | Q96KG9 | N-terminal kinase-like protein OS=Homo | SCYL1 | 89.632 | 2 | 53.541187 | yes | 1.63E-07 | 694.36615 | 344.96658 |
| ILPVLCGLTVDPEK | 14 | Q96KG9 | N-terminal kinase-like protein OS=Homo | SCYL1 | 89.632 | 2 | 53.541187 | yes | 1.136E-07 | 777.43402 | 194.64922 |
| ATGSPVSIFVYDVI | 14 | Q96KG9 | N-terminal kinase-like protein OS=Homo | SCYL1 | 89.632 | 2 | 53.541187 | yes | 9.722E-08 | 741.89557 | 144.30475 |
| LASEYNWGGPESS | 15 | Q96KG9 | N-terminal kinase-like protein OS=Homo | SCYL1 | 89.632 | 2 | 53.541187 | yes | 9.935E-08 | 820.36536 | 117.53198 |
| FFQELSK | 7 | Q96KG9 | N-terminal kinase-like protein OS=Homo | SCYL1 | 89.632 | 2 | 53.541187 | yes | 2.555E-06 | 449.73709 | 116.45792 |
| APGGFMSNR | 9 | Q96KG9 | N-terminal kinase-like protein OS=Homo | SCYL1 | 89.632 | 2 | 53.541187 | yes | 1.224E-07 | 468.72144 | 94.550774 |
| SMLLLAPK | 8 | Q96KG9 | N-terminal kinase-like protein OS=Homo | SCYL1 | 89.632 | 2 | 53.541187 | yes | 9.722E-08 | 436.76733 | 64.667 |
| TLVPHYCELVGAN | 15 | Q96KG9 | N-terminal kinase-like protein OS=Homo | SCYL1 | 89.632 | 3 | 53.541187 | yes | 1.043E-06 | 566.6275 | 48.871017 |
| PGAEEQTQVAK | 11 | Q96KG9 | N-terminal kinase-like protein OS=Homo | SCYL1 | 89.632 | 2 | 53.541187 | yes | 3.034E-06 | 579.29346 | 45.08852 |
| ELEISWGLHQIVK | 13 | Q96KG9 | N-terminal kinase-like protein OS=Homo | SCYL1 | 89.632 | 2 | 53.541187 | yes | 0.0001403 | 776.43011 | 15.382111 |
| LVTVTKPVPVTKP | 18 | Q96KM6 | Zinc finger protein 512B OS=Homo sapi | ZNF512B | 97.265 | 3 | 52.0097 | yes | 3.848E-06 | 641.06635 | 403.05762 |
| KVPAAPITVSK | 11 | Q96KM6 | Zinc finger protein 512B OS=Homo sapi | ZNF512B | 97.265 | 2 | 52.0097 | yes | 2.911E-07 | 555.84772 | 153.08014 |
| SVPVTKPVPVTKP | 18 | Q96KM6 | Zinc finger protein 512B OS=Homo sapi | ZNF512B | 97.265 | 3 | 52.0097 | yes | 1.934E-07 | 631.05908 | 140.31432 |
| THAENWFR | 8 | Q96KM6 | Zinc finger protein 512B OS=Homo sapi | ZNF512B | 97.265 | 2 | 52.0097 | yes | 6.437E-07 | 530.75159 | 134.32385 |
| AMPVTKPITVTK | 12 | Q96KM6 | Zinc finger protein 512B OS=Homo sapi | ZNF512B | 97.265 | 2 | 52.0097 | yes | 9.722E-08 | 643.38104 | 90.426659 |
| IWNHMDR | 7 | Q96KM6 | Zinc finger protein 512B OS=Homo sapi | ZNF512B | 97.265 | 2 | 52.0097 | yes | 4.671E-06 | 486.22943 | 39.341724 |
| VIEAVSEELSR | 11 | Q96KP1 | Exocyst complex component 2 OS=Homo | EXOC2 | 104.067 | 2 | 46.338802 | yes | 2.065E-07 | 616.33008 | 88.214653 |
| NALNVLQR | 8 | Q96KP1 | Exocyst complex component 2 OS=Homo | EXOC2 | 104.067 | 2 | 46.338802 | yes | 1.586E-07 | 464.27216 | 40.142212 |
| LLIVLSNCCYLER | 13 | Q96KP1 | Exocyst complex component 2 OS=Homo | EXOC2 | 104.067 | 2 | 46.338802 | yes | 1.787E-06 | 826.92859 | 17.851671 |

| | | | | | | | | | | | |
|---|---|---|---|---|---|---|---|---|---|---|---|
| DPSVEGSVK | 9 | Q96KQ4 | Apoptosis-stimulating of p53 protein 1 O: | PPP1R13B | 119.566 | 2 | 25.093235 | yes | 3.339E-06 | 459.23236 | 124.93748 |
| EDDSYLFDLDNK | 12 | Q96KQ7 | Histone-lysine N-methyltransferase EHM | EHMT2 | 132.371 | 2 | 56.595173 | yes | 9.722E-08 | 737.32263 | 113.71449 |
| GFEELPLCSCR | 11 | Q96KQ7 | Histone-lysine N-methyltransferase EHM | EHMT2 | 132.371 | 2 | 56.595173 | yes | 2.806E-07 | 684.30774 | 107.58211 |
| VALMVLCETHR | 11 | Q96KQ7 | Histone-lysine N-methyltransferase EHM | EHMT2 | 132.371 | 2 | 56.595173 | yes | 1.529E-07 | 664.8443 | 101.80097 |
| SPPSVQSLAMR | 11 | Q96KQ7 | Histone-lysine N-methyltransferase EHM | EHMT2 | 132.371 | 2 | 56.595173 | yes | 1.43E-07 | 586.80823 | 92.987038 |
| TGEELGFDYGDR | 12 | Q96KQ7 | Histone-lysine N-methyltransferase EHM | EHMT2 | 132.371 | 2 | 56.595173 | yes | 2.807E-07 | 679.79657 | 58.016701 |
| EGDTAWDLTPER | 12 | Q96KQ7 | Histone-lysine N-methyltransferase EHM | EHMT2 | 132.371 | 2 | 56.595173 | yes | 5.159E-06 | 695.31769 | 28.757788 |
| VFMLHQDLR | 9 | Q96KQ7 | Histone-lysine N-methyltransferase EHM | EHMT2 | 132.371 | 2 | 56.595173 | yes | 3.049E-06 | 579.80804 | 21.158056 |
| FINHLCDPNIIPVR | 14 | Q96KQ7 | Histone-lysine N-methyltransferase EHM | EHMT2 | 132.371 | 3 | 56.595173 | yes | 1.234E-06 | 569.9715 | 12.976208 |
| AENLLLDAEANIK | 13 | Q96L34 | MAP/microtubule affinity-regulating kina | MARK4 | 82.52 | 2 | 49.99072 | yes | 2.223E-05 | 707.38281 | 59.391571 |
| YNEVTATYLLLGR | 13 | Q96L34 | MAP/microtubule affinity-regulating kina | MARK4 | 82.52 | 2 | 49.99072 | yes | 9.938E-08 | 756.90649 | 44.821205 |
| GCNVDTPVSTLTP | 15 | Q96L50 | Leucine-rich repeat protein 1 OS=Homo s | LRR1 | 46.723 | 2 | 41.155922 | yes | 3.649E-07 | 794.40601 | 31.864265 |
| LDALLAELR | 9 | Q96L58 | Beta-1,3-galactosyltransferase 6 OS=Hon | B3GALT6 | 37.138 | 2 | 51.719753 | yes | 2.573E-07 | 507.30313 | 455.02066 |
| GCSNQYLVTHK | 11 | Q96L58 | Beta-1,3-galactosyltransferase 6 OS=Hon | B3GALT6 | 37.138 | 2 | 51.719753 | yes | 9.722E-08 | 653.81409 | 121.40019 |
| FAVGTAGLGAEER | 13 | Q96L58 | Beta-1,3-galactosyltransferase 6 OS=Hon | B3GALT6 | 37.138 | 2 | 51.719753 | yes | 9.722E-08 | 639.32788 | 105.75314 |
| HGDLLLLPALR | 11 | Q96L58 | Beta-1,3-galactosyltransferase 6 OS=Hon | B3GALT6 | 37.138 | 2 | 51.719753 | yes | 1.591E-07 | 609.37189 | 104.27763 |
| LYWGFFSGR | 9 | Q96L58 | Beta-1,3-galactosyltransferase 6 OS=Hon | B3GALT6 | 37.138 | 2 | 51.719753 | yes | 1.247E-07 | 566.78235 | 58.519466 |
| LMTAQNLVSYR | 11 | Q96L73 | Histone-lysine N-methyltransferase, H3 ly | NSD1 | 296.652 | 2 | 51.439617 | yes | 9.722E-08 | 648.34247 | 75.257553 |
| VLSAVVQTLVAK | 12 | Q96L73 | Histone-lysine N-methyltransferase, H3 ly | NSD1 | 296.652 | 2 | 51.439617 | yes | 4.656E-07 | 614.38721 | 72.138649 |
| VGLFALSDIK | 10 | Q96L73 | Histone-lysine N-methyltransferase, H3 ly | NSD1 | 296.652 | 2 | 51.439617 | yes | 2.492E-06 | 531.81335 | 62.521786 |
| QYYVEAFGDPSER | 13 | Q96L73 | Histone-lysine N-methyltransferase, H3 ly | NSD1 | 296.652 | 2 | 51.439617 | yes | 9.927E-08 | 780.85187 | 50.613976 |
| VQIFTADLSEIPR | 13 | Q96L73 | Histone-lysine N-methyltransferase, H3 ly | NSD1 | 296.652 | 2 | 51.439617 | yes | 1.538E-07 | 744.90649 | 49.993881 |
| QMVGGQQLPALA | 14 | Q96L73 | Histone-lysine N-methyltransferase, H3 ly | NSD1 | 296.652 | 2 | 51.439617 | yes | 5.554E-07 | 706.38995 | 31.447468 |
| VLVSGGSTHNSEK | 13 | Q96L73 | Histone-lysine N-methyltransferase, H3 ly | NSD1 | 296.652 | 2 | 51.439617 | yes | 1.952E-06 | 657.83606 | 25.822025 |
| VALPDGTTVTVR | 12 | Q96L92 | Sorting nexin-27 OS=Homo sapiens OX= | SNX27 | 61.266 | 2 | 52.764927 | yes | 3.092E-07 | 614.84845 | 253.91133 |
| EFANFTFPR | 9 | Q96L92 | Sorting nexin-27 OS=Homo sapiens OX= | SNX27 | 61.266 | 2 | 52.764927 | yes | 9.722E-08 | 564.77728 | 229.18292 |
| SESGYGFNVR | 10 | Q96L92 | Sorting nexin-27 OS=Homo sapiens OX= | SNX27 | 61.266 | 2 | 52.764927 | yes | 1.106E-07 | 558.25946 | 144.26172 |
| QAVPISVPR | 9 | Q96L92 | Sorting nexin-27 OS=Homo sapiens OX= | SNX27 | 61.266 | 2 | 52.764927 | yes | 2.503E-06 | 483.79019 | 77.938629 |
| NSTTDQVYQAIAA | 14 | Q96L92 | Sorting nexin-27 OS=Homo sapiens OX= | SNX27 | 61.266 | 2 | 52.764927 | yes | 9.722E-08 | 755.3808 | 65.88781 |
| LAPNEFPHK | 9 | Q96L92 | Sorting nexin-27 OS=Homo sapiens OX= | SNX27 | 61.266 | 2 | 52.764927 | yes | 9.753E-05 | 526.77985 | 35.36618 |
| WPFSLSEQQLDAR | 13 | Q96L92 | Sorting nexin-27 OS=Homo sapiens OX= | SNX27 | 61.266 | 2 | 52.764927 | yes | 2.078E-07 | 788.89136 | 31.518806 |
| GHELVPLEQER | 11 | Q96LD4 | E3 ubiquitin-protein ligase TRIM47 OS= | TRIM47 | 69.533 | 2 | 55.27933 | yes | 1.636E-07 | 653.84113 | 198.00177 |
| MDELGAGIAQSR | 12 | Q96LD4 | E3 ubiquitin-protein ligase TRIM47 OS= | TRIM47 | 69.533 | 2 | 55.27933 | yes | 9.722E-08 | 624.30609 | 124.21679 |
| LALEDGCGPGPGP | 15 | Q96LD4 | E3 ubiquitin-protein ligase TRIM47 OS= | TRIM47 | 69.533 | 2 | 55.27933 | yes | 9.722E-08 | 746.86426 | 71.898544 |
| VLCPINYPLSPTR | 13 | Q96LD4 | E3 ubiquitin-protein ligase TRIM47 OS= | TRIM47 | 69.533 | 2 | 55.27933 | yes | 1.709E-07 | 765.41089 | 58.654224 |
| LDSHFAGLFTHR | 12 | Q96LD4 | E3 ubiquitin-protein ligase TRIM47 OS= | TRIM47 | 69.533 | 2 | 55.27933 | yes | 2.829E-07 | 700.85712 | 50.213837 |
| VCLCEACAAQEHI | 13 | Q96LD4 | E3 ubiquitin-protein ligase TRIM47 OS= | TRIM47 | 69.533 | 2 | 55.27933 | yes | 5.917E-07 | 802.34235 | 25.309158 |
| LFQLQTLGLK | 10 | Q96LI5 | CCR4-NOT transcription complex subuni | CNOT6L | 63.002 | 2 | 51.27696 | no | 1.163E-07 | 580.85553 | 164.38216 |
| LHNLVYLDLSSNK | 13 | Q96LI5 | CCR4-NOT transcription complex subuni | CNOT6L | 63.002 | 2 | 51.27696 | no | 9.722E-08 | 758.41193 | 101.33271 |

| | | | | | | | | | | | |
|---|---|---|---|---|---|---|---|---|---|---|---|
| GYDGFFSPK | 9 | Q96LI5 | CCR4-NOT transcription complex subuni | CNOT6L | 63.002 | 2 | 51.27696 | yes | 1.636E-07 | 509.23746 | 91.198143 |
| SLPAELGNMVSLR | 13 | Q96LI5 | CCR4-NOT transcription complex subuni | CNOT6L | 63.002 | 2 | 51.27696 | no | 2.063E-05 | 693.87415 | 51.889961 |
| GLLSDEDLR | 9 | Q96LL9 | DnaJ homolog subfamily C member 30, n | DNAJC30 | 25.962 | 2 | 41.600403 | yes | 3.429E-07 | 509.26419 | 149.57059 |
| GQAFIGLPNEK | 11 | Q96LT9 | RNA-binding region-containing protein 3 | RNPC3 | 58.576 | 2 | 46.96838 | yes | 3.461E-07 | 587.31677 | 162.51456 |
| EEMETLSVFR | 10 | Q96LT9 | RNA-binding region-containing protein 3 | RNPC3 | 58.576 | 2 | 46.96838 | yes | 2.436E-07 | 620.79755 | 143.01067 |
| YVDFSSETQR | 10 | Q96LT9 | RNA-binding region-containing protein 3 | RNPC3 | 58.576 | 2 | 46.96838 | yes | 1.429E-07 | 616.28314 | 94.971123 |
| YFGAQSVR | 8 | Q96LT9 | RNA-binding region-containing protein 3 | RNPC3 | 58.576 | 2 | 46.96838 | yes | 1.331E-06 | 464.23779 | 69.605637 |
| GCTSSSSLSPPR | 12 | Q96LT9 | RNA-binding region-containing protein 3 | RNPC3 | 58.576 | 2 | 46.96838 | yes | 7.277E-07 | 618.2879 | 33.056999 |
| PMVVQFAR | 8 | Q96LT9 | RNA-binding region-containing protein 3 | RNPC3 | 58.576 | 2 | 46.96838 | yes | 1.707E-06 | 474.26019 | 29.345276 |
| LLDDFGVNIEGVR | 13 | Q96M94 | Kelch-like protein 15 OS=Homo sapiens ( | KLHL15 | 69.775 | 2 | 47.8403 | yes | 1.248E-07 | 723.883 | 72.773575 |
| TQVVTNCWENK | 11 | Q96M94 | Kelch-like protein 15 OS=Homo sapiens ( | KLHL15 | 69.775 | 2 | 47.8403 | yes | 1.246E-06 | 689.82465 | 30.878407 |
| HTDTIIQNIR | 10 | Q96M94 | Kelch-like protein 15 OS=Homo sapiens ( | KLHL15 | 69.775 | 2 | 47.8403 | yes | 1.923E-06 | 605.83057 | 29.570641 |
| AVQSTINHVK | 10 | Q96MF7 | E3 SUMO-protein ligase NSE2 OS=Hom | NSMCE2 | 27.933 | 2 | 49.35602 | yes | 1.041E-07 | 548.80908 | 239.81699 |
| AMVEFATLDR | 10 | Q96MF7 | E3 SUMO-protein ligase NSE2 OS=Hom | NSMCE2 | 27.933 | 2 | 49.35602 | yes | 1.136E-07 | 576.78955 | 148.03835 |
| QLNHYVK | 7 | Q96MF7 | E3 SUMO-protein ligase NSE2 OS=Hom | NSMCE2 | 27.933 | 2 | 49.35602 | yes | 2.273E-07 | 451.24814 | 85.570145 |
| KSDLIQDEALR | 11 | Q96MF7 | E3 SUMO-protein ligase NSE2 OS=Hom | NSMCE2 | 27.933 | 2 | 49.35602 | yes | 1.351E-06 | 644.34882 | 66.614403 |
| NSDADFQNNEK | 11 | Q96MF7 | E3 SUMO-protein ligase NSE2 OS=Hom | NSMCE2 | 27.933 | 2 | 49.35602 | yes | 2.874E-07 | 641.27075 | 46.922108 |
| VCGHTYEEDAIVR | 13 | Q96MF7 | E3 SUMO-protein ligase NSE2 OS=Hom | NSMCE2 | 27.933 | 2 | 49.35602 | yes | 4.162E-07 | 774.85919 | 34.753605 |
| DGFAEEAPSTSR | 12 | Q96MG7 | Non-structural maintenance of chromosor | NSMCE3 | 34.308 | 2 | 44.91428 | yes | 2.026E-07 | 633.78351 | 189.69572 |
| LITEDFVR | 8 | Q96MG7 | Non-structural maintenance of chromosor | NSMCE3 | 34.308 | 2 | 44.91428 | yes | 4.647E-07 | 496.7742 | 185.16225 |
| LQYVFGYK | 8 | Q96MG7 | Non-structural maintenance of chromosor | NSMCE3 | 34.308 | 2 | 44.91428 | yes | 2.562E-07 | 509.27383 | 119.38156 |
| KLITEDFVR | 9 | Q96MG7 | Non-structural maintenance of chromosor | NSMCE3 | 34.308 | 2 | 44.91428 | yes | 2.135E-07 | 560.82166 | 88.840576 |
| HVIGDYK | 7 | Q96MG7 | Non-structural maintenance of chromosor | NSMCE3 | 34.308 | 2 | 44.91428 | yes | 6.147E-07 | 416.22159 | 45.217438 |
| TQSPGGCSAEAVL | 15 | Q96MH2 | Protein HEXIM2 OS=Homo sapiens OX= | HEXIM2 | 32.419 | 2 | 52.629604 | yes | 1.66E-06 | 752.36462 | 321.61029 |
| LSQAEEETR | 9 | Q96MH2 | Protein HEXIM2 OS=Homo sapiens OX= | HEXIM2 | 32.419 | 2 | 52.629604 | yes | 9.722E-08 | 531.75671 | 255.46927 |
| FHTESLQGR | 9 | Q96MH2 | Protein HEXIM2 OS=Homo sapiens OX= | HEXIM2 | 32.419 | 2 | 52.629604 | yes | 1.275E-06 | 537.76996 | 243.58197 |
| QVEELAAEVQR | 11 | Q96MH2 | Protein HEXIM2 OS=Homo sapiens OX= | HEXIM2 | 32.419 | 2 | 52.629604 | yes | 7.745E-07 | 636.33313 | 120.21518 |
| VREEMFAK | 8 | Q96MH2 | Protein HEXIM2 OS=Homo sapiens OX= | HEXIM2 | 32.419 | 2 | 52.629604 | yes | 2.838E-07 | 505.26041 | 92.991432 |
| LQQLQACTGQQSC | 14 | Q96MH2 | Protein HEXIM2 OS=Homo sapiens OX= | HEXIM2 | 32.419 | 2 | 52.629604 | yes | 9.722E-08 | 839.39343 | 75.202301 |
| LIYQQQVSSLEAQI | 14 | Q96MT8 | Centrosomal protein of 63 kDa OS=Homo | CEP63 | 81.344 | 2 | 53.233368 | yes | 1.107E-07 | 831.94409 | 313.93045 |
| FLEEEELR | 8 | Q96MT8 | Centrosomal protein of 63 kDa OS=Homo | CEP63 | 81.344 | 2 | 53.233368 | yes | 1.72E-06 | 532.7666 | 290.98132 |
| EILDQLESLK | 10 | Q96MT8 | Centrosomal protein of 63 kDa OS=Homo | CEP63 | 81.344 | 2 | 53.233368 | yes | 9.722E-08 | 594.32953 | 247.82538 |
| LLEALQEEK | 9 | Q96MT8 | Centrosomal protein of 63 kDa OS=Homo | CEP63 | 81.344 | 2 | 53.233368 | yes | 7.231E-07 | 536.79785 | 247.74202 |
| SHHILER | 7 | Q96MT8 | Centrosomal protein of 63 kDa OS=Homo | CEP63 | 81.344 | 2 | 53.233368 | yes | 4.663E-07 | 446.24341 | 182.14012 |
| ANDTICANELEIER | 14 | Q96MT8 | Centrosomal protein of 63 kDa OS=Homo | CEP63 | 81.344 | 2 | 53.233368 | yes | 1.067E-07 | 824.3858 | 162.59624 |
| THALETCLK | 9 | Q96MT8 | Centrosomal protein of 63 kDa OS=Homo | CEP63 | 81.344 | 2 | 53.233368 | yes | 9.722E-08 | 536.77643 | 151.84828 |
| QIDIMVAHK | 9 | Q96MT8 | Centrosomal protein of 63 kDa OS=Homo | CEP63 | 81.344 | 2 | 53.233368 | yes | 6.309E-07 | 527.78931 | 132.33456 |
| AALQSQENLIHEAI | 14 | Q96MT8 | Centrosomal protein of 63 kDa OS=Homo | CEP63 | 81.344 | 2 | 53.233368 | yes | 9.722E-08 | 790.41296 | 113.28601 |
| LHEELCILK | 9 | Q96MT8 | Centrosomal protein of 63 kDa OS=Homo | CEP63 | 81.344 | 2 | 53.233368 | yes | 1.643E-07 | 577.81555 | 109.22021 |

| | | | | | | | | | | | |
|---|---|---|---|---|---|---|---|---|---|---|---|
| LDAHIEELKR | 10 | Q96MT8 | Centrosomal protein of 63 kDa OS=Hom | CEP63 | 81.344 | 2 | 53.233368 | yes | 1.539E-07 | 612.34076 | 83.532387 |
| IKQEMTMEYK | 10 | Q96MT8 | Centrosomal protein of 63 kDa OS=Hom | CEP63 | 81.344 | 2 | 53.233368 | yes | 1.175E-07 | 650.81744 | 41.768806 |
| GVWSTLPVNEK | 11 | Q96MU7 | YTH domain-containing protein 1 OS=H | YTHDC1 | 84.701 | 2 | 54.808178 | yes | 9.938E-08 | 615.3299 | 523.55164 |
| KLNLAFR | 7 | Q96MU7 | YTH domain-containing protein 1 OS=H | YTHDC1 | 84.701 | 2 | 54.808178 | yes | 1.435E-07 | 431.26889 | 505.70953 |
| VHDYDMR | 7 | Q96MU7 | YTH domain-containing protein 1 OS=H | YTHDC1 | 84.701 | 2 | 54.808178 | yes | 9.722E-08 | 468.20563 | 390.19568 |
| PLSSSVSNNK | 10 | Q96MU7 | YTH domain-containing protein 1 OS=H | YTHDC1 | 84.701 | 2 | 54.808178 | yes | 1.673E-07 | 516.76965 | 362.06165 |
| IDYPPEFHQR | 10 | Q96MU7 | YTH domain-containing protein 1 OS=H | YTHDC1 | 84.701 | 2 | 54.808178 | yes | 3.259E-06 | 651.31732 | 286.57349 |
| SNNHENVSLAK | 11 | Q96MU7 | YTH domain-containing protein 1 OS=H | YTHDC1 | 84.701 | 2 | 54.808178 | yes | 9.722E-08 | 606.802 | 265.63239 |
| RPEDYDIHNSR | 11 | Q96MU7 | YTH domain-containing protein 1 OS=H | YTHDC1 | 84.701 | 3 | 54.808178 | yes | 1.097E-07 | 467.88837 | 78.44751 |
| GISPIVFDR | 9 | Q96MU7 | YTH domain-containing protein 1 OS=H | YTHDC1 | 84.701 | 2 | 54.808178 | yes | 1.308E-07 | 502.2822 | 67.209518 |
| IGLEVDRR | 8 | Q96MU7 | YTH domain-containing protein 1 OS=H | YTHDC1 | 84.701 | 2 | 54.808178 | yes | 3.397E-07 | 479.27744 | 42.717842 |
| QLVSKPLSSSVSNI | 15 | Q96MU7 | YTH domain-containing protein 1 OS=H | YTHDC1 | 84.701 | 2 | 54.808178 | yes | 1.086E-07 | 794.43866 | 32.201744 |
| SGSSASESYAGSEI | 14 | Q96MU7 | YTH domain-containing protein 1 OS=H | YTHDC1 | 84.701 | 2 | 54.808178 | yes | 2.043E-07 | 673.78894 | 15.596422 |
| AASQPTSLAPEK | 12 | Q96N66 | Lysophospholipid acyltransferase 7 OS=I | MBOAT7 | 52.765 | 2 | 52.231754 | yes | 1.366E-06 | 600.31696 | 116.2064 |
| NIDCYSTDFCVR | 12 | Q96N66 | Lysophospholipid acyltransferase 7 OS=I | MBOAT7 | 52.765 | 2 | 52.231754 | yes | 9.722E-08 | 775.32416 | 47.02356 |
| PGSFLLTSTSGR | 12 | Q96N67 | Dedicator of cytokinesis protein 7 OS=H | DOCK7 | 242.561 | 2 | 57.323566 | yes | 9.722E-08 | 611.82495 | 218.41531 |
| LEEAILGSIGAR | 12 | Q96N67 | Dedicator of cytokinesis protein 7 OS=H | DOCK7 | 242.561 | 2 | 57.323566 | yes | 9.722E-08 | 614.84845 | 211.20595 |
| LYSLPNPSVLVSLI | 14 | Q96N67 | Dedicator of cytokinesis protein 7 OS=H | DOCK7 | 242.561 | 2 | 57.323566 | yes | 1.086E-07 | 779.45361 | 150.812 |
| DVYVPNTTYR | 10 | Q96N67 | Dedicator of cytokinesis protein 7 OS=H | DOCK7 | 242.561 | 2 | 57.323566 | yes | 5.239E-07 | 614.30383 | 149.65071 |
| VLIPIHEANR | 10 | Q96N67 | Dedicator of cytokinesis protein 7 OS=H | DOCK7 | 242.561 | 2 | 57.323566 | yes | 1.03E-07 | 581.34058 | 124.77803 |
| EAYTAVVYHNR | 11 | Q96N67 | Dedicator of cytokinesis protein 7 OS=H | DOCK7 | 242.561 | 2 | 57.323566 | yes | 9.722E-08 | 661.828 | 124.49784 |
| SSISALNSSQLEPV | 16 | Q96N67 | Dedicator of cytokinesis protein 7 OS=H | DOCK7 | 242.561 | 2 | 57.323566 | yes | 9.723E-08 | 843.95471 | 115.23959 |
| ESILGGVLK | 9 | Q96N67 | Dedicator of cytokinesis protein 7 OS=H | DOCK7 | 242.561 | 2 | 57.323566 | yes | 3.627E-07 | 458.27911 | 100.66157 |
| LEGFYGER | 8 | Q96N67 | Dedicator of cytokinesis protein 7 OS=H | DOCK7 | 242.561 | 2 | 57.323566 | yes | 1.474E-07 | 485.73508 | 99.375603 |
| AVLPVTCHR | 9 | Q96N67 | Dedicator of cytokinesis protein 7 OS=H | DOCK7 | 242.561 | 2 | 57.323566 | yes | 2.794E-06 | 526.78711 | 98.081841 |
| PATLTVTNFFK | 11 | Q96N67 | Dedicator of cytokinesis protein 7 OS=H | DOCK7 | 242.561 | 2 | 57.323566 | no | 9.722E-08 | 619.84259 | 91.260773 |
| FGDLDEQEFVYK | 12 | Q96N67 | Dedicator of cytokinesis protein 7 OS=H | DOCK7 | 242.561 | 2 | 57.323566 | no | 9.722E-08 | 745.34589 | 87.778748 |
| QNFEIGNNFAR | 11 | Q96N67 | Dedicator of cytokinesis protein 7 OS=H | DOCK7 | 242.561 | 2 | 57.323566 | yes | 2.065E-07 | 655.31781 | 75.238152 |
| GSWACSIFDLK | 11 | Q96N67 | Dedicator of cytokinesis protein 7 OS=H | DOCK7 | 242.561 | 2 | 57.323566 | yes | 1.364E-07 | 642.30811 | 68.714073 |
| NSLPDALLPNLLDI | 14 | Q96N67 | Dedicator of cytokinesis protein 7 OS=H | DOCK7 | 242.561 | 2 | 57.323566 | yes | 3.825E-07 | 775.93048 | 66.907471 |
| SYTEDWAIVIR | 11 | Q96N67 | Dedicator of cytokinesis protein 7 OS=H | DOCK7 | 242.561 | 2 | 57.323566 | yes | 9.722E-08 | 676.84589 | 64.52697 |
| TILTYAEEDLELR | 13 | Q96N67 | Dedicator of cytokinesis protein 7 OS=H | DOCK7 | 242.561 | 2 | 57.323566 | yes | 9.722E-08 | 783.40649 | 60.519867 |
| IANMFELSVPFR | 12 | Q96N67 | Dedicator of cytokinesis protein 7 OS=H | DOCK7 | 242.561 | 2 | 57.323566 | yes | 9.722E-08 | 712.37378 | 59.891312 |
| TTSGDDACNLTSFI | 14 | Q96N67 | Dedicator of cytokinesis protein 7 OS=H | DOCK7 | 242.561 | 2 | 57.323566 | yes | 3.65E-07 | 772.83594 | 49.575649 |
| SQADQFCQR | 9 | Q96N67 | Dedicator of cytokinesis protein 7 OS=H | DOCK7 | 242.561 | 2 | 57.323566 | yes | 6.982E-07 | 570.24854 | 44.443573 |
| LRPITAQLK | 9 | Q96N67 | Dedicator of cytokinesis protein 7 OS=H | DOCK7 | 242.561 | 2 | 57.323566 | yes | 1.741E-06 | 520.33478 | 43.938095 |
| SPSGSAFGSQENLI | 14 | Q96N67 | Dedicator of cytokinesis protein 7 OS=H | DOCK7 | 242.561 | 2 | 57.323566 | yes | 7.809E-05 | 718.84186 | 37.89341 |
| NADETVLQK | 9 | Q96N67 | Dedicator of cytokinesis protein 7 OS=H | DOCK7 | 242.561 | 2 | 57.323566 | yes | 7.371E-07 | 509.26419 | 32.386688 |
| QISGQYSGSPQLLI | 14 | Q96N67 | Dedicator of cytokinesis protein 7 OS=H | DOCK7 | 242.561 | 2 | 57.323566 | yes | 2.811E-06 | 753.40155 | 10.49013 |

| | | | | | | | | | | | |
|---|---|---|---|---|---|---|---|---|---|---|---|
| LGQLSCALCNTPV | 14 | Q96NB3 | Zinc finger protein 830 OS=Homo sapien | ZNF830 | 41.999 | 2 | 40.389744 | yes | 4.05E-07 | 780.89728 | 57.240578 |
| QIGEIDEQIECYR | 13 | Q96NB3 | Zinc finger protein 830 OS=Homo sapien | ZNF830 | 41.999 | 2 | 40.389744 | yes | 2.171E-06 | 826.88287 | 28.207111 |
| EASQGSSASSAPHS | 16 | Q96NB3 | Zinc finger protein 830 OS=Homo sapien | ZNF830 | 41.999 | 2 | 40.389744 | yes | 2.71E-06 | 765.36316 | 2.4552186 |
| DGKPVQPVK | 9 | Q96NC0 | Zinc finger matrin-type protein 2 OS=Hoi | ZMAT2 | 23.612 | 2 | 48.994675 | yes | 2.271E-06 | 484.2822 | 118.03857 |
| QKDYDFEER | 9 | Q96NC0 | Zinc finger matrin-type protein 2 OS=Hoi | ZMAT2 | 23.612 | 2 | 48.994675 | yes | 1.079E-07 | 615.27527 | 66.049446 |
| KWDKDEYEK | 9 | Q96NC0 | Zinc finger matrin-type protein 2 OS=Hoi | ZMAT2 | 23.612 | 2 | 48.994675 | yes | 1.136E-07 | 620.79584 | 48.180943 |
| GIQIFQNLDEEK | 12 | Q96NE9 | FERM domain-containing protein 6 OS=I | FRMD6 | 72.044 | 2 | 47.65517 | yes | 2.795E-07 | 717.36719 | 142.72136 |
| LYMNLQPVLR | 10 | Q96NE9 | FERM domain-containing protein 6 OS=I | FRMD6 | 72.044 | 2 | 47.65517 | yes | 2.604E-07 | 623.85248 | 123.97423 |
| GQSTDSLPQTICR | 13 | Q96NE9 | FERM domain-containing protein 6 OS=I | FRMD6 | 72.044 | 2 | 47.65517 | yes | 1.965E-07 | 731.85138 | 113.24582 |
| LIYYTGCPMR | 10 | Q96NE9 | FERM domain-containing protein 6 OS=I | FRMD6 | 72.044 | 2 | 47.65517 | yes | 5.225E-07 | 637.30701 | 85.663864 |
| LDDVAVHYYR | 10 | Q96NE9 | FERM domain-containing protein 6 OS=I | FRMD6 | 72.044 | 2 | 47.65517 | yes | 3.562E-06 | 625.81183 | 58.680641 |
| DTGPEDSYSSSAIH | 15 | Q96NE9 | FERM domain-containing protein 6 OS=I | FRMD6 | 72.044 | 2 | 47.65517 | yes | 1.284E-07 | 811.35809 | 30.500622 |
| HSTASHSSSHTSGI | 21 | Q96NE9 | FERM domain-containing protein 6 OS=I | FRMD6 | 72.044 | 4 | 47.65517 | yes | 3.769E-07 | 549.01245 | 30.082924 |
| LYDLPDEPFTR | 11 | Q96NW4 | Ankyrin repeat domain-containing proteir | ANKRD27 | 116.985 | 2 | 45.848267 | yes | 1.725E-07 | 683.33789 | 86.255547 |
| YVGTMEASEDAAI | 15 | Q96NW4 | Ankyrin repeat domain-containing proteir | ANKRD27 | 116.985 | 2 | 45.848267 | yes | 4.87E-06 | 816.86414 | 8.7284899 |
| YSLTPEGLELAQK | 13 | Q96NY9 | Crossover junction endonuclease MUS81 | MUS81 | 61.173 | 2 | 51.30485 | yes | 8.654E-07 | 724.88519 | 356.59695 |
| LDDLCSSIIDGR | 12 | Q96NY9 | Crossover junction endonuclease MUS81 | MUS81 | 61.173 | 2 | 51.30485 | yes | 1.087E-07 | 682.32971 | 227.56259 |
| VLLCVDIGETR | 11 | Q96NY9 | Crossover junction endonuclease MUS81 | MUS81 | 61.173 | 2 | 51.30485 | yes | 1.089E-07 | 637.84229 | 213.38556 |
| ESAAYLALLTR | 11 | Q96NY9 | Crossover junction endonuclease MUS81 | MUS81 | 61.173 | 2 | 51.30485 | yes | 9.722E-08 | 604.33771 | 101.40257 |
| ILQHFGDGLCR | 11 | Q96NY9 | Crossover junction endonuclease MUS81 | MUS81 | 61.173 | 2 | 51.30485 | yes | 9.825E-08 | 658.3324 | 93.433586 |
| KRPLPACPNPLFVI | 14 | Q96NY9 | Crossover junction endonuclease MUS81 | MUS81 | 61.173 | 3 | 51.30485 | yes | 5.14E-07 | 555.6521 | 33.509518 |
| EQETLLSTIK | 10 | Q96NY9 | Crossover junction endonuclease MUS81 | MUS81 | 61.173 | 2 | 51.30485 | yes | 3.804E-07 | 581.32172 | 30.261871 |
| VILLVLYR | 8 | Q96NY9 | Crossover junction endonuclease MUS81 | MUS81 | 61.173 | 2 | 51.30485 | yes | 3.279E-06 | 494.83133 | 9.4742146 |
| LAPALPAWPHR | 11 | Q96P11 | 28S rRNA (cytosine-C(5))-methyltransfer | NSUN5 | 46.692 | 2 | 52.088367 | yes | 1.009E-07 | 614.85114 | 1861.0874 |
| QLYALVCETQR | 11 | Q96P11 | 28S rRNA (cytosine-C(5))-methyltransfer | NSUN5 | 46.692 | 2 | 52.088367 | yes | 9.722E-08 | 690.85065 | 1393.1088 |
| TCSDDVVDYFK | 11 | Q96P11 | 28S rRNA (cytosine-C(5))-methyltransfer | NSUN5 | 46.692 | 2 | 52.088367 | yes | 1.568E-07 | 674.78992 | 1247.1084 |
| LHALAGFQQR | 10 | Q96P11 | 28S rRNA (cytosine-C(5))-methyltransfer | NSUN5 | 46.692 | 2 | 52.088367 | no | 1.113E-07 | 570.81726 | 1152.7522 |
| PGPASQLPR | 9 | Q96P11 | 28S rRNA (cytosine-C(5))-methyltransfer | NSUN5 | 46.692 | 2 | 52.088367 | yes | 2.935E-07 | 461.75888 | 1128.4318 |
| AGHLILQDR | 9 | Q96P11 | 28S rRNA (cytosine-C(5))-methyltransfer | NSUN5 | 46.692 | 2 | 52.088367 | no | 4.464E-07 | 511.79071 | 1105.1636 |
| QLEEPGAGTPSPVI | 14 | Q96P11 | 28S rRNA (cytosine-C(5))-methyltransfer | NSUN5 | 46.692 | 2 | 52.088367 | no | 1.187E-07 | 719.37024 | 1020.0997 |
| DALQQNPGAFR | 11 | Q96P11 | 28S rRNA (cytosine-C(5))-methyltransfer | NSUN5 | 46.692 | 2 | 52.088367 | yes | 9.809E-08 | 608.80707 | 987.78571 |
| NEDLLEVGSR | 10 | Q96P11 | 28S rRNA (cytosine-C(5))-methyltransfer | NSUN5 | 46.692 | 2 | 52.088367 | yes | 9.722E-08 | 566.28564 | 943.42426 |
| LASMATLLAR | 10 | Q96P11 | 28S rRNA (cytosine-C(5))-methyltransfer | NSUN5 | 46.692 | 2 | 52.088367 | yes | 2.391E-07 | 523.80499 | 796.19135 |
| GLSTFPGAEHCLR | 13 | Q96P11 | 28S rRNA (cytosine-C(5))-methyltransfer | NSUN5 | 46.692 | 2 | 52.088367 | no | 4.796E-07 | 722.8537 | 757.6864 |
| GLVYSSNFQNVK | 12 | Q96P11 | 28S rRNA (cytosine-C(5))-methyltransfer | NSUN5 | 46.692 | 2 | 52.088367 | yes | 2.425E-07 | 678.35132 | 483.98904 |
| TSHLAALLK | 9 | Q96P11 | 28S rRNA (cytosine-C(5))-methyltransfer | NSUN5 | 46.692 | 2 | 52.088367 | no | 2.723E-07 | 477.29257 | 313.28857 |
| ASSLDDLR | 8 | Q96P11 | 28S rRNA (cytosine-C(5))-methyltransfer | NSUN5 | 46.692 | 2 | 52.088367 | yes | 2.48E-07 | 438.7247 | 298.25839 |
| VLVYELLLGK | 10 | Q96P11 | 28S rRNA (cytosine-C(5))-methyltransfer | NSUN5 | 46.692 | 2 | 52.088367 | yes | 3.183E-07 | 573.86029 | 202.01691 |
| ALCHALTFPSLQR | 13 | Q96P11 | 28S rRNA (cytosine-C(5))-methyltransfer | NSUN5 | 46.692 | 2 | 52.088367 | no | 9.722E-08 | 757.40082 | 170.83917 |

| | | | | | | | | | | | |
|---|---|---|---|---|---|---|---|---|---|---|---|
| LRPHLAK | 7 | Q96P11 | 28S rRNA (cytosine-C(5))-methyltransfer | NSUN5 | 46.692 | 2 | 52.088367 | yes | 1.811E-06 | 417.76904 | 166.65952 |
| GLYAAAAGVLAG | 16 | Q96P11 | 28S rRNA (cytosine-C(5))-methyltransfer | NSUN5 | 46.692 | 2 | 52.088367 | yes | 1.253E-07 | 752.90955 | 85.633629 |
| VGIVGNLSSGK | 11 | Q96P47 | Arf-GAP with GTPase, ANK repeat and I | AGAP3 | 95.045 | 2 | 51.493576 | yes | 4.867E-07 | 515.79822 | 2277.2292 |
| VFQDVAQK | 8 | Q96P47 | Arf-GAP with GTPase, ANK repeat and I | AGAP3 | 95.045 | 2 | 51.493576 | no | 9.722E-08 | 467.75327 | 179.31273 |
| IEDLALQNQIR | 11 | Q96P47 | Arf-GAP with GTPase, ANK repeat and I | AGAP3 | 95.045 | 2 | 51.493576 | yes | 9.722E-08 | 656.86462 | 123.88496 |
| LGNQNAALAVQA | 14 | Q96P47 | Arf-GAP with GTPase, ANK repeat and I | AGAP3 | 95.045 | 2 | 51.493576 | yes | 1.231E-07 | 712.90204 | 120.57407 |
| ATPATAPGTSPR | 12 | Q96P47 | Arf-GAP with GTPase, ANK repeat and I | AGAP3 | 95.045 | 2 | 51.493576 | yes | 1.991E-07 | 563.7962 | 117.35753 |
| IETIAASSTPTPIR | 14 | Q96P47 | Arf-GAP with GTPase, ANK repeat and I | AGAP3 | 95.045 | 2 | 51.493576 | yes | 3.354E-07 | 728.90393 | 79.360641 |
| IFTMAEVYGIR | 11 | Q96P70 | Importin-9 OS=Homo sapiens OX=9606 | IPO9 | 115.963 | 2 | 54.41664 | yes | 9.722E-08 | 650.34192 | 56.79343 |
| NVDILWQAAEICR | 13 | Q96PE3 | Inositol polyphosphate-4-phosphatase typ | INPP4A | 109.956 | 2 | 39.746952 | yes | 8.23E-05 | 794.40106 | 551.03613 |
| EIIAQINTLK | 10 | Q96PE3 | Inositol polyphosphate-4-phosphatase typ | INPP4A | 109.956 | 2 | 39.746952 | yes | 6.218E-06 | 571.84259 | 86.646698 |
| SVFGGAICR | 9 | Q96PE3 | Inositol polyphosphate-4-phosphatase typ | INPP4A | 109.956 | 2 | 39.746952 | yes | 7.03E-07 | 483.74493 | 69.840134 |
| SQTCLPELLR | 10 | Q96PE3 | Inositol polyphosphate-4-phosphatase typ | INPP4A | 109.956 | 2 | 39.746952 | yes | 4.403E-07 | 608.82135 | 51.224636 |
| KLEFVPTNLHIQR | 13 | Q96PE3 | Inositol polyphosphate-4-phosphatase typ | INPP4A | 109.956 | 3 | 39.746952 | yes | 3.192E-06 | 532.30701 | 22.74954 |
| AQPSVSLGAPYR | 12 | Q96PK6 | RNA-binding protein 14 OS=Homo sapie | RBM14 | 69.492 | 2 | 54.498886 | yes | 1.191E-07 | 623.33295 | 14153.535 |
| ASYVAPLTAQPAT | 15 | Q96PK6 | RNA-binding protein 14 OS=Homo sapie | RBM14 | 69.492 | 2 | 54.498886 | yes | 2.102E-07 | 804.92267 | 12380.519 |
| AQPSVSLGAAYR | 12 | Q96PK6 | RNA-binding protein 14 OS=Homo sapie | RBM14 | 69.492 | 2 | 54.498886 | yes | 1.759E-07 | 610.32513 | 12158.438 |
| DYAFVHMEK | 9 | Q96PK6 | RNA-binding protein 14 OS=Homo sapie | RBM14 | 69.492 | 2 | 54.498886 | yes | 1.149E-07 | 570.26312 | 5786.8022 |
| DYAFVHMEK | 9 | Q96PK6 | RNA-binding protein 14 OS=Homo sapie | RBM14 | 69.492 | 2 | 54.498886 | yes | 2.013E-07 | 578.26062 | 5786.8022 |
| RLPDAHSDYAR | 11 | Q96PK6 | RNA-binding protein 14 OS=Homo sapie | RBM14 | 69.492 | 3 | 54.498886 | yes | 1.439E-07 | 434.21796 | 3332.7395 |
| RLPDAHSDYAR | 11 | Q96PK6 | RNA-binding protein 14 OS=Homo sapie | RBM14 | 69.492 | 3 | 54.498886 | yes | 9.722E-08 | 650.8233 | 3332.7395 |
| RLPDAHSDYAR | 11 | Q96PK6 | RNA-binding protein 14 OS=Homo sapie | RBM14 | 69.492 | 2 | 54.498886 | yes | 1.439E-07 | 434.21796 | 3332.7395 |
| RLPDAHSDYAR | 11 | Q96PK6 | RNA-binding protein 14 OS=Homo sapie | RBM14 | 69.492 | 2 | 54.498886 | yes | 9.722E-08 | 650.8233 | 3332.7395 |
| ALVVEMSR | 8 | Q96PK6 | RNA-binding protein 14 OS=Homo sapie | RBM14 | 69.492 | 2 | 54.498886 | yes | 1.146E-07 | 452.74966 | 6116.5688 |
| YSGSYNDYLR | 10 | Q96PK6 | RNA-binding protein 14 OS=Homo sapie | RBM14 | 69.492 | 2 | 54.498886 | yes | 1.362E-07 | 619.27783 | 5736.7109 |
| LSESQLSFR | 9 | Q96PK6 | RNA-binding protein 14 OS=Homo sapie | RBM14 | 69.492 | 2 | 54.498886 | yes | 1.415E-07 | 533.78003 | 5346.71 |
| TQPMTAQAASYR | 12 | Q96PK6 | RNA-binding protein 14 OS=Homo sapie | RBM14 | 69.492 | 2 | 54.498886 | yes | 2.012E-07 | 662.81934 | 2321.416 |
| TQPMTAQAASYR | 12 | Q96PK6 | RNA-binding protein 14 OS=Homo sapie | RBM14 | 69.492 | 2 | 54.498886 | yes | 9.809E-08 | 670.81677 | 2321.416 |
| LAELSDYR | 8 | Q96PK6 | RNA-binding protein 14 OS=Homo sapie | RBM14 | 69.492 | 2 | 54.498886 | yes | 1.642E-07 | 483.74817 | 3848.3286 |
| VIECDVVK | 8 | Q96PK6 | RNA-binding protein 14 OS=Homo sapie | RBM14 | 69.492 | 2 | 54.498886 | yes | 1.283E-07 | 481.25479 | 3525.4441 |
| AAQMHSGYQR | 10 | Q96PK6 | RNA-binding protein 14 OS=Homo sapie | RBM14 | 69.492 | 2 | 54.498886 | yes | 1.183E-07 | 574.76691 | 2859.135 |
| QPTPPFFGR | 9 | Q96PK6 | RNA-binding protein 14 OS=Homo sapie | RBM14 | 69.492 | 2 | 54.498886 | yes | 1.043E-07 | 523.77454 | 2403.0596 |
| KGPGLAVQSGDK | 12 | Q96PK6 | RNA-binding protein 14 OS=Homo sapie | RBM14 | 69.492 | 2 | 54.498886 | yes | 1.074E-07 | 578.81964 | 2306.5923 |
| ASYDDPYKK | 9 | Q96PK6 | RNA-binding protein 14 OS=Homo sapie | RBM14 | 69.492 | 2 | 54.498886 | yes | 1.677E-07 | 543.75873 | 2226.8369 |
| QFAFVHMR | 8 | Q96PK6 | RNA-binding protein 14 OS=Homo sapie | RBM14 | 69.492 | 2 | 54.498886 | yes | 1.915E-07 | 518.26331 | 2097.1101 |
| AQPSASLGVGYR | 12 | Q96PK6 | RNA-binding protein 14 OS=Homo sapie | RBM14 | 69.492 | 2 | 54.498886 | yes | 1.609E-07 | 603.31732 | 2033.1375 |
| LPDAHSDYAR | 10 | Q96PK6 | RNA-binding protein 14 OS=Homo sapie | RBM14 | 69.492 | 2 | 54.498886 | yes | 9.722E-08 | 572.77271 | 1451.9795 |
| PRPLNTWK | 8 | Q96PK6 | RNA-binding protein 14 OS=Homo sapie | RBM14 | 69.492 | 2 | 54.498886 | yes | 3.226E-07 | 506.29034 | 1368.9636 |
| INVELSTK | 8 | Q96PK6 | RNA-binding protein 14 OS=Homo sapie | RBM14 | 69.492 | 2 | 54.498886 | yes | 1.113E-07 | 452.26093 | 1331.7026 |

| | | | | | | | | | | | |
|---|---|---|---|---|---|---|---|---|---|---|---|
| RINVELSTK | 9 | Q96PK6 | RNA-binding protein 14 OS=Homo sapie | RBM14 | 69.492 | 2 | 54.498886 | yes | 3.38E-07 | 530.31146 | 1009.8611 |
| GPGLAVQSGDK | 11 | Q96PK6 | RNA-binding protein 14 OS=Homo sapie | RBM14 | 69.492 | 2 | 54.498886 | yes | 2.126E-07 | 514.77222 | 852.28778 |
| AIEALHGHELRPGI | 14 | Q96PK6 | RNA-binding protein 14 OS=Homo sapie | RBM14 | 69.492 | 3 | 54.498886 | yes | 3.629E-07 | 519.28656 | 412.68365 |
| AIEALHGHELRPGI | 14 | Q96PK6 | RNA-binding protein 14 OS=Homo sapie | RBM14 | 69.492 | 3 | 54.498886 | yes | 9.722E-08 | 778.42621 | 412.68365 |
| AIEALHGHELRPGI | 14 | Q96PK6 | RNA-binding protein 14 OS=Homo sapie | RBM14 | 69.492 | 2 | 54.498886 | yes | 3.629E-07 | 519.28656 | 412.68365 |
| AIEALHGHELRPGI | 14 | Q96PK6 | RNA-binding protein 14 OS=Homo sapie | RBM14 | 69.492 | 2 | 54.498886 | yes | 9.722E-08 | 778.42621 | 412.68365 |
| RLAELSDYR | 9 | Q96PK6 | RNA-binding protein 14 OS=Homo sapie | RBM14 | 69.492 | 2 | 54.498886 | yes | 1.018E-07 | 561.79877 | 331.48877 |
| TQSSASLAASYAA | 24 | Q96PK6 | RNA-binding protein 14 OS=Homo sapie | RBM14 | 69.492 | 3 | 54.498886 | yes | 1.305E-07 | 822.39832 | 156.28183 |
| AIEALHGHELR | 11 | Q96PK6 | RNA-binding protein 14 OS=Homo sapie | RBM14 | 69.492 | 2 | 54.498886 | yes | 1.121E-07 | 623.33856 | 125.27699 |
| ALVVEMSRPR | 10 | Q96PK6 | RNA-binding protein 14 OS=Homo sapie | RBM14 | 69.492 | 2 | 54.498886 | yes | 3.081E-07 | 579.3266 | 59.821892 |
| RLSESQLSFR | 10 | Q96PK6 | RNA-binding protein 14 OS=Homo sapie | RBM14 | 69.492 | 2 | 54.498886 | yes | 1.782E-07 | 611.83057 | 58.93771 |
| LGPPTPGEPEAPA( | 16 | Q96PM9 | Zinc finger protein 385A OS=Homo sapi | ZNF385A | 40.454 | 2 | 53.125607 | yes | 2.655E-07 | 816.40485 | 190.93195 |
| DGVAGKPNPLLSR | 13 | Q96PM9 | Zinc finger protein 385A OS=Homo sapi | ZNF385A | 40.454 | 2 | 53.125607 | yes | 1.021E-07 | 662.37262 | 184.45973 |
| LLYCALCK | 8 | Q96PM9 | Zinc finger protein 385A OS=Homo sapi | ZNF385A | 40.454 | 2 | 53.125607 | yes | 1.166E-07 | 520.76703 | 137.82652 |
| GAGELAGTLTFSK | 13 | Q96PM9 | Zinc finger protein 385A OS=Homo sapi | ZNF385A | 40.454 | 2 | 53.125607 | yes | 1.613E-07 | 626.33258 | 113.40119 |
| TAHGPILFSPY | 11 | Q96PM9 | Zinc finger protein 385A OS=Homo sapi | ZNF385A | 40.454 | 2 | 53.125607 | yes | 3.768E-07 | 601.81384 | 106.5228 |
| FNSQSQAEAHYK | 12 | Q96PM9 | Zinc finger protein 385A OS=Homo sapi | ZNF385A | 40.454 | 2 | 53.125607 | yes | 9.722E-08 | 705.32587 | 75.607193 |
| TFHCEICNVK | 10 | Q96PM9 | Zinc finger protein 385A OS=Homo sapi | ZNF385A | 40.454 | 2 | 53.125607 | yes | 9.722E-08 | 654.29718 | 48.453465 |
| PAPAAPLLQGPPIT | 26 | Q96PM9 | Zinc finger protein 385A OS=Homo sapi | ZNF385A | 40.454 | 4 | 53.125607 | yes | 5.405E-07 | 657.88385 | 41.263824 |
| VAVNSLSQLEAHN | 14 | Q96PM9 | Zinc finger protein 385A OS=Homo sapi | ZNF385A | 40.454 | 2 | 53.125607 | yes | 9.729E-08 | 755.4046 | 37.660553 |
| GEGGTPAPASLPG( | 16 | Q96PM9 | Zinc finger protein 385A OS=Homo sapi | ZNF385A | 40.454 | 2 | 53.125607 | yes | 4.11E-07 | 691.84918 | 36.619179 |
| VNSEVQLK | 8 | Q96PM9 | Zinc finger protein 385A OS=Homo sapi | ZNF385A | 40.454 | 2 | 53.125607 | yes | 4.239E-07 | 458.75854 | 31.694725 |
| AQVFSCPACR | 10 | Q96PU4 | E3 ubiquitin-protein ligase UHRF2 OS=I | UHRF2 | 89.986 | 2 | 51.18682 | no | 9.824E-08 | 598.271 | 1861.4342 |
| LQYPAGYPSDK | 11 | Q96PU4 | E3 ubiquitin-protein ligase UHRF2 OS=I | UHRF2 | 89.986 | 2 | 51.18682 | yes | 2.213E-07 | 619.80621 | 163.33069 |
| IGAPSADQTLTNM | 15 | Q96PU4 | E3 ubiquitin-protein ligase UHRF2 OS=I | UHRF2 | 89.986 | 2 | 51.18682 | yes | 9.722E-08 | 794.89105 | 150.60329 |
| IFLGGSEGTLNDCI | 14 | Q96PU4 | E3 ubiquitin-protein ligase UHRF2 OS=I | UHRF2 | 89.986 | 2 | 51.18682 | yes | 2.877E-07 | 755.86395 | 129.52621 |
| VGPSNQPSTSAR | 12 | Q96PU4 | E3 ubiquitin-protein ligase UHRF2 OS=I | UHRF2 | 89.986 | 2 | 51.18682 | yes | 5.825E-07 | 600.802 | 119.08114 |
| IERPGAHPLSFADC | 15 | Q96PU4 | E3 ubiquitin-protein ligase UHRF2 OS=I | UHRF2 | 89.986 | 3 | 51.18682 | yes | 4.41E-07 | 532.28271 | 107.80255 |
| ALALNCDAPLDDI | 13 | Q96PU4 | E3 ubiquitin-protein ligase UHRF2 OS=I | UHRF2 | 89.986 | 2 | 51.18682 | yes | 3.708E-07 | 708.3454 | 103.10596 |
| LLFEVFDENR | 10 | Q96PU5 | E3 ubiquitin-protein ligase NEDD4-like ( | NEDD4L | 111.932 | 2 | 46.670784 | yes | 1.499E-07 | 641.32733 | 31.571447 |
| LLVPAAEGEDSLK | 13 | Q96PU8 | KH domain-containing RNA-binding pro | QKI | 37.671 | 2 | 49.604954 | yes | 1.062E-07 | 671.36664 | 853.84991 |
| LLDEEISR | 8 | Q96PU8 | KH domain-containing RNA-binding pro | QKI | 37.671 | 2 | 49.604954 | yes | 1.702E-07 | 487.76129 | 387.49405 |
| QLEAETGCK | 9 | Q96PU8 | KH domain-containing RNA-binding pro | QKI | 37.671 | 2 | 49.604954 | yes | 1.018E-07 | 518.24243 | 245.34837 |
| LLVPAAEGEDSLK | 14 | Q96PU8 | KH domain-containing RNA-binding pro | QKI | 37.671 | 2 | 49.604954 | yes | 1.146E-07 | 735.41412 | 114.78354 |
| LLVPAAEGEDSLK | 14 | Q96PU8 | KH domain-containing RNA-binding pro | QKI | 37.671 | 2 | 49.604954 | yes | 1.769E-07 | 490.61185 | 114.78354 |
| LLVPAAEGEDSLK | 14 | Q96PU8 | KH domain-containing RNA-binding pro | QKI | 37.671 | 3 | 49.604954 | yes | 1.146E-07 | 735.41412 | 114.78354 |
| LLVPAAEGEDSLK | 14 | Q96PU8 | KH domain-containing RNA-binding pro | QKI | 37.671 | 3 | 49.604954 | yes | 1.769E-07 | 490.61185 | 114.78354 |
| IITGPAPVLPPAAL | 15 | Q96PU8 | KH domain-containing RNA-binding pro | QKI | 37.671 | 2 | 49.604954 | yes | 5.857E-07 | 743.46124 | 108.05737 |
| EYPDFNFVGR | 10 | Q96PU8 | KH domain-containing RNA-binding pro | QKI | 37.671 | 2 | 49.604954 | yes | 1.274E-07 | 622.29077 | 78.075882 |

| | | | | | | | | | | | |
|---|---|---|---|---|---|---|---|---|---|---|---|
| GKPNWEHLNEDLI | 24 | Q96PU8 | KH domain-containing RNA-binding prot | QKI | 37.671 | 4 | 49.604954 | yes | 3.906E-07 | 707.60852 | 64.838234 |
| TPTPAGPTIMPLIR | 14 | Q96PU8 | KH domain-containing RNA-binding prot | QKI | 37.671 | 2 | 49.604954 | yes | 4.635E-07 | 732.91577 | 37.842361 |
| ADPCVAHALALR | 12 | Q96PV6 | Leukocyte receptor cluster member 8 OS= | LENG8 | 88.158 | 2 | 52.13084 | yes | 1.112E-07 | 647.34027 | 812.90515 |
| TAWALGNYHR | 10 | Q96PV6 | Leukocyte receptor cluster member 8 OS= | LENG8 | 88.158 | 2 | 52.13084 | yes | 9.722E-08 | 594.79907 | 671.14661 |
| QDLTVQGIR | 9 | Q96PV6 | Leukocyte receptor cluster member 8 OS= | LENG8 | 88.158 | 2 | 52.13084 | yes | 5.95E-07 | 515.28802 | 487.82422 |
| NSGDITTELAYLTF | 14 | Q96PV6 | Leukocyte receptor cluster member 8 OS= | LENG8 | 88.158 | 2 | 52.13084 | yes | 1.098E-07 | 777.39392 | 480.58545 |
| MKPAPGTGGLK | 11 | Q96PV6 | Leukocyte receptor cluster member 8 OS= | LENG8 | 88.158 | 2 | 52.13084 | yes | 1.104E-07 | 528.79718 | 417.10846 |
| TEFTVEVYETHAR | 13 | Q96PV6 | Leukocyte receptor cluster member 8 OS= | LENG8 | 88.158 | 2 | 52.13084 | yes | 9.723E-08 | 791.3808 | 372.99664 |
| LTCAPDPSTVRPV. | 17 | Q96PV6 | Leukocyte receptor cluster member 8 OS= | LENG8 | 88.158 | 3 | 52.13084 | yes | 2.184E-07 | 608.6731 | 344.16571 |
| QDYAFACEQMK | 11 | Q96PV6 | Leukocyte receptor cluster member 8 OS= | LENG8 | 88.158 | 2 | 52.13084 | yes | 9.722E-08 | 695.79193 | 185.85066 |
| WEAASSLHPPR | 11 | Q96PV6 | Leukocyte receptor cluster member 8 OS= | LENG8 | 88.158 | 2 | 52.13084 | yes | 2.766E-07 | 625.81744 | 182.92908 |
| LQDGSAYTIDWSR | 13 | Q96PV6 | Leukocyte receptor cluster member 8 OS= | LENG8 | 88.158 | 2 | 52.13084 | yes | 9.722E-08 | 756.35986 | 179.81853 |
| LTCAPDPSTVR | 11 | Q96PV6 | Leukocyte receptor cluster member 8 OS= | LENG8 | 88.158 | 2 | 52.13084 | yes | 6.203E-07 | 608.80316 | 163.47601 |
| KGQQLWNR | 8 | Q96PV6 | Leukocyte receptor cluster member 8 OS= | LENG8 | 88.158 | 2 | 52.13084 | yes | 9.722E-08 | 515.28308 | 125.60827 |
| ILYYIFTK | 8 | Q96PV6 | Leukocyte receptor cluster member 8 OS= | LENG8 | 88.158 | 2 | 52.13084 | yes | 1.179E-07 | 530.8075 | 85.038086 |
| MAALECEDPER | 11 | Q96PV6 | Leukocyte receptor cluster member 8 OS= | LENG8 | 88.158 | 2 | 52.13084 | yes | 1.841E-07 | 660.78156 | 63.092979 |
| RWEAASSLHPPR | 12 | Q96PV6 | Leukocyte receptor cluster member 8 OS= | LENG8 | 88.158 | 3 | 52.13084 | yes | 3.948E-07 | 469.58112 | 45.532349 |
| CFTACESEEDKDR | 13 | Q96PV6 | Leukocyte receptor cluster member 8 OS= | LENG8 | 88.158 | 2 | 52.13084 | yes | 9.819E-08 | 823.82471 | 37.688576 |
| GGTLGGIPGEPAVI | 16 | Q96PV7 | Protein FAM193B OS=Homo sapiens OX | FAM193B | 96.543 | 2 | 52.665657 | yes | 5.134E-07 | 766.89441 | 142.8694 |
| LNTVPNLSR | 9 | Q96PV7 | Protein FAM193B OS=Homo sapiens OX | FAM193B | 96.543 | 2 | 52.665657 | yes | 5.881E-07 | 507.29056 | 116.80153 |
| VAVNWTNFSLK | 11 | Q96PV7 | Protein FAM193B OS=Homo sapiens OX | FAM193B | 96.543 | 2 | 52.665657 | yes | 9.722E-08 | 639.8457 | 107.09639 |
| VNSFLSSR | 8 | Q96PV7 | Protein FAM193B OS=Homo sapiens OX | FAM193B | 96.543 | 2 | 52.665657 | yes | 1.901E-07 | 455.24307 | 98.878868 |
| DMDGVEMDETDR | 12 | Q96PV7 | Protein FAM193B OS=Homo sapiens OX | FAM193B | 96.543 | 2 | 52.665657 | yes | 9.722E-08 | 706.76886 | 93.514862 |
| FCLDSAK | 7 | Q96PV7 | Protein FAM193B OS=Homo sapiens OX | FAM193B | 96.543 | 2 | 52.665657 | yes | 3.657E-07 | 420.69965 | 65.919739 |
| AQLAAEALK | 9 | Q96PV7 | Protein FAM193B OS=Homo sapiens OX | FAM193B | 96.543 | 2 | 52.665657 | yes | 5.58E-06 | 457.76892 | 47.292904 |
| EHAVAISLSHTSCK | 14 | Q96PV7 | Protein FAM193B OS=Homo sapiens OX | FAM193B | 96.543 | 2 | 52.665657 | yes | 1.006E-07 | 770.38281 | 43.503937 |
| TPKPGYPSSEEPSS | 15 | Q96PV7 | Protein FAM193B OS=Homo sapiens OX | FAM193B | 96.543 | 2 | 52.665657 | yes | 3.263E-07 | 795.88593 | 11.552758 |
| LQSLDLLLDTK | 11 | Q96PY5 | Formin-like protein 2 OS=Homo sapiens | FMNL2 | 123.321 | 2 | 50.908413 | no | 9.722E-08 | 629.86627 | 123.23878 |
| LPMPEPGELEER | 12 | Q96PY5 | Formin-like protein 2 OS=Homo sapiens | FMNL2 | 123.321 | 2 | 50.908413 | yes | 4.318E-07 | 698.84247 | 102.32014 |
| DANTQVHTLR | 10 | Q96PY5 | Formin-like protein 2 OS=Homo sapiens | FMNL2 | 123.321 | 2 | 50.908413 | yes | 2.526E-07 | 577.79926 | 99.080627 |
| VTLLEANR | 8 | Q96PY5 | Formin-like protein 2 OS=Homo sapiens | FMNL2 | 123.321 | 2 | 50.908413 | no | 5.047E-07 | 458.26654 | 88.715225 |
| IAQDAFDDVVK | 11 | Q96PY5 | Formin-like protein 2 OS=Homo sapiens | FMNL2 | 123.321 | 2 | 50.908413 | yes | 4.521E-07 | 610.81152 | 67.522278 |
| LGLDEYLDK | 9 | Q96PY5 | Formin-like protein 2 OS=Homo sapiens | FMNL2 | 123.321 | 2 | 50.908413 | yes | 1.053E-06 | 533.27679 | 62.732338 |
| QQQELIAELR | 10 | Q96PY5 | Formin-like protein 2 OS=Homo sapiens | FMNL2 | 123.321 | 2 | 50.908413 | yes | 9.809E-08 | 614.33826 | 60.296238 |
| KDDVHVCIMCLR | 12 | Q96PY5 | Formin-like protein 2 OS=Homo sapiens | FMNL2 | 123.321 | 3 | 50.908413 | yes | 3.921E-07 | 515.91602 | 28.623533 |
| KDDVHVCIMCLR | 12 | Q96PY5 | Formin-like protein 2 OS=Homo sapiens | FMNL2 | 123.321 | 3 | 50.908413 | yes | 8.201E-07 | 773.37036 | 28.623533 |
| KDDVHVCIMCLR | 12 | Q96PY5 | Formin-like protein 2 OS=Homo sapiens | FMNL2 | 123.321 | 2 | 50.908413 | yes | 3.921E-07 | 515.91602 | 28.623533 |
| KDDVHVCIMCLR | 12 | Q96PY5 | Formin-like protein 2 OS=Homo sapiens | FMNL2 | 123.321 | 2 | 50.908413 | yes | 8.201E-07 | 773.37036 | 28.623533 |
| LLEQEALMEQQDI | 14 | Q96PY5 | Formin-like protein 2 OS=Homo sapiens | FMNL2 | 123.321 | 2 | 50.908413 | yes | 1.335E-07 | 836.41656 | 53.519791 |

| | | | | | | | | | | | |
|---|---|---|---|---|---|---|---|---|---|---|---|
| VHLQYEFTK | 9 | Q96PY5 | Formin-like protein 2 OS=Homo sapiens | FMNL2 | 123.321 | 2 | 50.908413 | no | 1.417E-07 | 582.80603 | 48.767349 |
| VQESTQVLR | 9 | Q96PY5 | Formin-like protein 2 OS=Homo sapiens | FMNL2 | 123.321 | 2 | 50.908413 | no | 3.286E-06 | 530.29327 | 39.478378 |
| VEELEENISHLSEK | 14 | Q96PY5 | Formin-like protein 2 OS=Homo sapiens | FMNL2 | 123.321 | 2 | 50.908413 | yes | 1.121E-07 | 828.40979 | 36.333862 |
| NPPHTYIQK | 9 | Q96PY5 | Formin-like protein 2 OS=Homo sapiens | FMNL2 | 123.321 | 2 | 50.908413 | no | 4.696E-06 | 549.29053 | 30.741571 |
| AAAVSLENVLLD\ | 14 | Q96PY5 | Formin-like protein 2 OS=Homo sapiens | FMNL2 | 123.321 | 2 | 50.908413 | no | 3.69E-07 | 721.41669 | 26.929491 |
| LQDTENEAMSK | 11 | Q96PY5 | Formin-like protein 2 OS=Homo sapiens | FMNL2 | 123.321 | 2 | 50.908413 | yes | 4.182E-07 | 633.28754 | 15.273387 |
| HKIQEAYR | 8 | Q96PZ0 | Pseudouridylate synthase 7 homolog OS= | PUS7 | 75.036 | 2 | 56.610306 | yes | 1.599E-06 | 522.7829 | 431.26328 |
| FGTTAVPTYQVGF | 13 | Q96PZ0 | Pseudouridylate synthase 7 homolog OS= | PUS7 | 75.036 | 2 | 56.610306 | yes | 1.151E-07 | 698.86462 | 313.7449 |
| ETSVAIEVIEDTK | 13 | Q96PZ0 | Pseudouridylate synthase 7 homolog OS= | PUS7 | 75.036 | 2 | 56.610306 | yes | 1.084E-07 | 717.37213 | 221.40486 |
| TKDPTAALR | 9 | Q96PZ0 | Pseudouridylate synthase 7 homolog OS= | PUS7 | 75.036 | 2 | 56.610306 | yes | 5.844E-07 | 486.77728 | 191.84622 |
| IPLFNTDVDNLEGI | 14 | Q96PZ0 | Pseudouridylate synthase 7 homolog OS= | PUS7 | 75.036 | 2 | 56.610306 | yes | 1.241E-07 | 787.90668 | 140.02853 |
| AITVQEIAVLK | 11 | Q96PZ0 | Pseudouridylate synthase 7 homolog OS= | PUS7 | 75.036 | 2 | 56.610306 | yes | 9.722E-08 | 592.86609 | 106.90621 |
| TIIHQAIK | 8 | Q96PZ0 | Pseudouridylate synthase 7 homolog OS= | PUS7 | 75.036 | 2 | 56.610306 | yes | 1.915E-07 | 462.28726 | 102.77494 |
| LEELQLFK | 8 | Q96PZ0 | Pseudouridylate synthase 7 homolog OS= | PUS7 | 75.036 | 2 | 56.610306 | yes | 2.111E-07 | 510.29224 | 94.374557 |
| IIIRPQNVSWEVVA | 19 | Q96PZ0 | Pseudouridylate synthase 7 homolog OS= | PUS7 | 75.036 | 3 | 56.610306 | yes | 3.19E-07 | 748.06708 | 92.864723 |
| EMLTADNLDIDNM | 14 | Q96PZ0 | Pseudouridylate synthase 7 homolog OS= | PUS7 | 75.036 | 2 | 56.610306 | yes | 2.511E-07 | 825.87671 | 81.688728 |
| HGLTEADVGITK | 12 | Q96PZ0 | Pseudouridylate synthase 7 homolog OS= | PUS7 | 75.036 | 2 | 56.610306 | yes | 9.722E-08 | 620.83026 | 81.657707 |
| EIGFINYYGMQR | 12 | Q96PZ0 | Pseudouridylate synthase 7 homolog OS= | PUS7 | 75.036 | 2 | 56.610306 | yes | 9.623E-07 | 745.85846 | 47.972042 |
| DTMDAINVLSK | 11 | Q96PZ0 | Pseudouridylate synthase 7 homolog OS= | PUS7 | 75.036 | 2 | 56.610306 | yes | 1.016E-07 | 603.80536 | 44.10944 |
| FVSSHQGFSGILK | 13 | Q96PZ0 | Pseudouridylate synthase 7 homolog OS= | PUS7 | 75.036 | 2 | 56.610306 | yes | 1.138E-07 | 703.875 | 35.59948 |
| GALVVEDNDSGVI | 19 | Q96PZ0 | Pseudouridylate synthase 7 homolog OS= | PUS7 | 75.036 | 3 | 56.610306 | yes | 2.044E-06 | 662.67236 | 26.118628 |
| GSYCHFVLYK | 10 | Q96PZ0 | Pseudouridylate synthase 7 homolog OS= | PUS7 | 75.036 | 2 | 56.610306 | yes | 1.257E-07 | 637.30536 | 25.199301 |
| LGELQGNHFTVVL | 14 | Q96PZ0 | Pseudouridylate synthase 7 homolog OS= | PUS7 | 75.036 | 2 | 56.610306 | yes | 1.047E-07 | 791.93866 | 24.21312 |
| SHLMELGLTAAK | 12 | Q96Q15 | Serine/threonine-protein kinase SMG1 OS | SMG1 | 410.501 | 2 | 48.858925 | yes | 7.102E-07 | 635.84485 | 169.40623 |
| VDTGLEHPIGSSEV | 20 | Q96Q15 | Serine/threonine-protein kinase SMG1 OS | SMG1 | 410.501 | 3 | 48.858925 | yes | 2.662E-07 | 726.03931 | 84.297462 |
| SVLTLANAGR | 10 | Q96Q15 | Serine/threonine-protein kinase SMG1 OS | SMG1 | 410.501 | 2 | 48.858925 | yes | 5.469E-07 | 501.29056 | 62.749218 |
| HGFDLLTEMK | 10 | Q96Q15 | Serine/threonine-protein kinase SMG1 OS | SMG1 | 410.501 | 2 | 48.858925 | yes | 4.243E-07 | 595.79736 | 55.856838 |
| AQDTFQTIEGIIR | 13 | Q96Q15 | Serine/threonine-protein kinase SMG1 OS | SMG1 | 410.501 | 2 | 48.858925 | yes | 1.095E-07 | 746.39374 | 53.879776 |
| NTPASASLEGLAQ | 17 | Q96Q45 | Transmembrane protein 237 OS=Homo sa | TMEM237 | 45.526 | 2 | 42.00774 | yes | 3.238E-07 | 822.42102 | 85.213783 |
| PSYVFSADPIAR | 12 | Q96Q89 | Kinesin-like protein KIF20B OS=Homo s | KIF20B | 210.63 | 2 | 38.405846 | yes | 4.445E-06 | 661.84058 | 53.202011 |
| TLNVLFDSLQER | 12 | Q96Q89 | Kinesin-like protein KIF20B OS=Homo s | KIF20B | 210.63 | 2 | 38.405846 | yes | 5.476E-07 | 717.883 | 44.345242 |
| ETGNINTSLLTLGK | 14 | Q96Q89 | Kinesin-like protein KIF20B OS=Homo s | KIF20B | 210.63 | 2 | 38.405846 | yes | 5.244E-07 | 730.90137 | 17.037855 |
| STGLELETPSLVPV | 15 | Q96QC0 | Serine/threonine-protein phosphatase 1 re | PPP1R10 | 99.058 | 2 | 54.62808 | yes | 1.211E-07 | 785.44037 | 1928.4126 |
| VPWVCPR | 7 | Q96QC0 | Serine/threonine-protein phosphatase 1 re | PPP1R10 | 99.058 | 2 | 54.62808 | yes | 2.65E-07 | 457.23926 | 1492.4348 |
| QSNVAAPGDATPF | 16 | Q96QC0 | Serine/threonine-protein phosphatase 1 re | PPP1R10 | 99.058 | 2 | 54.62808 | yes | 9.722E-08 | 776.88373 | 1126.5372 |
| LLNNWLTYSK | 10 | Q96QC0 | Serine/threonine-protein phosphatase 1 re | PPP1R10 | 99.058 | 2 | 54.62808 | yes | 1.083E-07 | 626.34021 | 825.17163 |
| IKDFGEAAK | 9 | Q96QC0 | Serine/threonine-protein phosphatase 1 re | PPP1R10 | 99.058 | 2 | 54.62808 | yes | 9.722E-08 | 489.76639 | 686.46039 |
| HAFETAR | 7 | Q96QC0 | Serine/threonine-protein phosphatase 1 re | PPP1R10 | 99.058 | 2 | 54.62808 | yes | 1.849E-07 | 416.20901 | 647.32422 |
| YKPLNTTPNATK | 12 | Q96QC0 | Serine/threonine-protein phosphatase 1 re | PPP1R10 | 99.058 | 2 | 54.62808 | yes | 1.517E-07 | 674.367 | 561.20294 |

| | | | | | | | | | | | |
|---|---|---|---|---|---|---|---|---|---|---|---|
| CTYLNILLQTR | 11 | Q96QC0 | Serine/threonine-protein phosphatase 1 re | PPP1R10 | 99.058 | 2 | 54.62808 | yes | 1.071E-07 | 697.87665 | 517.69379 |
| SVTWPEEGK | 9 | Q96QC0 | Serine/threonine-protein phosphatase 1 re | PPP1R10 | 99.058 | 2 | 54.62808 | yes | 1.384E-07 | 516.75348 | 472.5192 |
| GGNEPPPPPPPFR | 13 | Q96QC0 | Serine/threonine-protein phosphatase 1 re | PPP1R10 | 99.058 | 2 | 54.62808 | yes | 1.165E-07 | 679.84619 | 314.92383 |
| GGPGPGPGPYHR | 12 | Q96QC0 | Serine/threonine-protein phosphatase 1 re | PPP1R10 | 99.058 | 2 | 54.62808 | yes | 1.096E-07 | 574.78345 | 308.15964 |
| TTLPERPLTEVK | 12 | Q96QC0 | Serine/threonine-protein phosphatase 1 re | PPP1R10 | 99.058 | 2 | 54.62808 | yes | 1.045E-07 | 692.39575 | 273.37436 |
| GILQELFLNK | 10 | Q96QC0 | Serine/threonine-protein phosphatase 1 re | PPP1R10 | 99.058 | 2 | 54.62808 | yes | 5.495E-07 | 587.84515 | 262.91452 |
| PVESPGDPNQLTR | 13 | Q96QC0 | Serine/threonine-protein phosphatase 1 re | PPP1R10 | 99.058 | 2 | 54.62808 | yes | 9.722E-08 | 705.35461 | 177.12231 |
| EYFYFELDETER | 12 | Q96QC0 | Serine/threonine-protein phosphatase 1 re | PPP1R10 | 99.058 | 2 | 54.62808 | yes | 9.722E-08 | 820.85938 | 139.84869 |
| SSEDEELRK | 9 | Q96QC0 | Serine/threonine-protein phosphatase 1 re | PPP1R10 | 99.058 | 2 | 54.62808 | yes | 3.41E-07 | 546.76202 | 87.884781 |
| KVLSPTAAK | 9 | Q96QC0 | Serine/threonine-protein phosphatase 1 re | PPP1R10 | 99.058 | 2 | 54.62808 | yes | 9.964E-08 | 457.78711 | 66.294426 |
| RLSHDNMEEK | 10 | Q96QC0 | Serine/threonine-protein phosphatase 1 re | PPP1R10 | 99.058 | 3 | 54.62808 | yes | 1.018E-07 | 420.19968 | 30.710846 |
| RLSHDNMEEK | 10 | Q96QC0 | Serine/threonine-protein phosphatase 1 re | PPP1R10 | 99.058 | 3 | 54.62808 | yes | 2.384E-05 | 629.7959 | 30.710846 |
| RLSHDNMEEK | 10 | Q96QC0 | Serine/threonine-protein phosphatase 1 re | PPP1R10 | 99.058 | 2 | 54.62808 | yes | 1.018E-07 | 420.19968 | 30.710846 |
| RLSHDNMEEK | 10 | Q96QC0 | Serine/threonine-protein phosphatase 1 re | PPP1R10 | 99.058 | 2 | 54.62808 | yes | 2.384E-05 | 629.7959 | 30.710846 |
| LSHDNMEEK | 9 | Q96QC0 | Serine/threonine-protein phosphatase 1 re | PPP1R10 | 99.058 | 2 | 54.62808 | yes | 7.606E-07 | 551.7453 | 57.713478 |
| GLDSFLNR | 8 | Q96QC0 | Serine/threonine-protein phosphatase 1 re | PPP1R10 | 99.058 | 2 | 54.62808 | yes | 4.767E-07 | 461.24307 | 47.597256 |
| PLVLPSPLVTPGSN | 18 | Q96QC0 | Serine/threonine-protein phosphatase 1 re | PPP1R10 | 99.058 | 3 | 54.62808 | yes | 1.473E-07 | 631.01428 | 46.552902 |
| LLQQSGAQQFR | 11 | Q96QD9 | UAP56-interacting factor OS=Homo sapi | FYTTD1 | 35.819 | 2 | 45.61095 | yes | 1.797E-07 | 638.34387 | 59.003746 |
| IDMSLDDIIK | 10 | Q96QD9 | UAP56-interacting factor OS=Homo sapi | FYTTD1 | 35.819 | 2 | 45.61095 | yes | 2.254E-06 | 581.80481 | 42.075455 |
| NIEQYFPVLK | 10 | Q96QD9 | UAP56-interacting factor OS=Homo sapi | FYTTD1 | 35.819 | 2 | 45.61095 | yes | 4.829E-05 | 625.84259 | 33.275425 |
| NLLLEEIR | 8 | Q96QE3 | ATPase family AAA domain-containing p | ATAD5 | 207.571 | 2 | 44.080673 | yes | 5.846E-07 | 500.29529 | 111.11433 |
| IMENSGIQMVSK | 12 | Q96QE3 | ATPase family AAA domain-containing p | ATAD5 | 207.571 | 2 | 44.080673 | yes | 3.942E-06 | 668.83362 | 70.997009 |
| SEATDGGFTSQIR | 13 | Q96QE3 | ATPase family AAA domain-containing p | ATAD5 | 207.571 | 2 | 44.080673 | yes | 4.138E-06 | 684.82312 | 55.654541 |
| GIDSDDVQDNSQL | 14 | Q96QE3 | ATPase family AAA domain-containing p | ATAD5 | 207.571 | 2 | 44.080673 | yes | 2.767E-07 | 767.35498 | 43.865208 |
| NFCGPSVTVDASA | 16 | Q96QE3 | ATPase family AAA domain-containing p | ATAD5 | 207.571 | 2 | 44.080673 | yes | 2.328E-07 | 812.88544 | 23.783342 |
| NVQLVCSEHGLDN | 14 | Q96QE3 | ATPase family AAA domain-containing p | ATAD5 | 207.571 | 2 | 44.080673 | yes | 3.308E-07 | 806.89105 | 8.9600983 |
| IAWAHLDR | 8 | Q96QE5 | Transcription elongation factor, mitochon | TEFM | 41.677 | 2 | 56.40195 | yes | 1.298E-07 | 491.26688 | 596.24341 |
| SINIVEHR | 8 | Q96QE5 | Transcription elongation factor, mitochon | TEFM | 41.677 | 2 | 56.40195 | yes | 9.723E-08 | 484.26962 | 543.06165 |
| STVQVCNSILCPK | 13 | Q96QE5 | Transcription elongation factor, mitochon | TEFM | 41.677 | 2 | 56.40195 | yes | 1.279E-07 | 753.37616 | 421.69604 |
| QMFLSTELQR | 10 | Q96QE5 | Transcription elongation factor, mitochon | TEFM | 41.677 | 2 | 56.40195 | yes | 9.722E-08 | 626.82135 | 301.63916 |
| HFELMIGDSR | 10 | Q96QE5 | Transcription elongation factor, mitochon | TEFM | 41.677 | 2 | 56.40195 | yes | 2.347E-07 | 602.7926 | 215.10553 |
| KLTVLDWQQSDR | 12 | Q96QE5 | Transcription elongation factor, mitochon | TEFM | 41.677 | 2 | 56.40195 | yes | 9.722E-08 | 744.89392 | 189.12309 |
| LTVLDWQQSDR | 11 | Q96QE5 | Transcription elongation factor, mitochon | TEFM | 41.677 | 2 | 56.40195 | yes | 9.722E-08 | 680.84644 | 145.1283 |
| TFAQDGQHQVLSN | 15 | Q96QE5 | Transcription elongation factor, mitochon | TEFM | 41.677 | 3 | 56.40195 | yes | 9.722E-08 | 577.94592 | 122.39587 |
| QFLFDSILK | 9 | Q96QE5 | Transcription elongation factor, mitochon | TEFM | 41.677 | 2 | 56.40195 | yes | 2.397E-07 | 555.81335 | 92.440712 |
| ADFYVLEK | 8 | Q96QE5 | Transcription elongation factor, mitochon | TEFM | 41.677 | 2 | 56.40195 | yes | 1.026E-07 | 492.75546 | 60.195599 |
| KLLKPDIER | 9 | Q96QE5 | Transcription elongation factor, mitochon | TEFM | 41.677 | 2 | 56.40195 | yes | 8.579E-07 | 556.34534 | 48.770809 |
| ILVGTNLVR | 9 | Q96QK1 | Vacuolar protein sorting-associated protei | VPS35 | 91.708 | 2 | 46.620235 | yes | 1.512E-07 | 492.81366 | 145.35367 |
| HASNMLGELR | 10 | Q96QK1 | Vacuolar protein sorting-associated protei | VPS35 | 91.708 | 2 | 46.620235 | yes | 3.997E-07 | 564.28491 | 142.9053 |

| | | | | | | | | | | | |
|---|---|---|---|---|---|---|---|---|---|---|---|
| LSQLEGVNVER | 11 | Q96QK1 | Vacuolar protein sorting-associated protei | VPS35 | 91.708 | 2 | 46.620235 | yes | 3.048E-07 | 622.33569 | 129.7196 |
| AELAELPLR | 9 | Q96QK1 | Vacuolar protein sorting-associated protei | VPS35 | 91.708 | 2 | 46.620235 | yes | 3.045E-06 | 506.29529 | 25.311588 |
| VSEVKPSYR | 9 | Q96QR8 | Transcriptional activator protein Pur-beta | PURB | 33.241 | 2 | 53.31246 | yes | 1.436E-07 | 532.79041 | 1099.7714 |
| FFFDVGCNK | 9 | Q96QR8 | Transcriptional activator protein Pur-beta | PURB | 33.241 | 2 | 53.31246 | yes | 1.042E-07 | 567.25787 | 1065.9711 |
| RFYLDVK | 7 | Q96QR8 | Transcriptional activator protein Pur-beta | PURB | 33.241 | 2 | 53.31246 | no | 1.368E-07 | 470.76617 | 944.79871 |
| RLDIQNK | 7 | Q96QR8 | Transcriptional activator protein Pur-beta | PURB | 33.241 | 2 | 53.31246 | yes | 2.798E-07 | 443.75888 | 701.79565 |
| KYYLDLK | 7 | Q96QR8 | Transcriptional activator protein Pur-beta | PURB | 33.241 | 2 | 53.31246 | no | 1.473E-07 | 471.76837 | 473.78842 |
| GGGEQETQELASK | 13 | Q96QR8 | Transcriptional activator protein Pur-beta | PURB | 33.241 | 2 | 53.31246 | yes | 9.722E-08 | 667.31512 | 437.52109 |
| YADEMKEIQER | 11 | Q96QR8 | Transcriptional activator protein Pur-beta | PURB | 33.241 | 2 | 53.31246 | yes | 9.722E-08 | 706.32971 | 104.82371 |
| LTLSMAVAAEFR | 12 | Q96QR8 | Transcriptional activator protein Pur-beta | PURB | 33.241 | 2 | 53.31246 | yes | 3.076E-06 | 654.85266 | 51.067093 |
| NAITVPFK | 8 | Q96QR8 | Transcriptional activator protein Pur-beta | PURB | 33.241 | 2 | 53.31246 | yes | 1.106E-07 | 445.26074 | 41.773712 |
| RFFFDVGCNK | 10 | Q96QR8 | Transcriptional activator protein Pur-beta | PURB | 33.241 | 2 | 53.31246 | yes | 1.446E-07 | 645.30841 | 22.895477 |
| TLLSIPEVVR | 10 | Q96R06 | Sperm-associated antigen 5 OS=Homo sa | SPAG5 | 134.422 | 2 | 52.864365 | yes | 1.239E-07 | 563.84515 | 5624.7139 |
| QSLSLPSMLR | 10 | Q96R06 | Sperm-associated antigen 5 OS=Homo sa | SPAG5 | 134.422 | 2 | 52.864365 | yes | 1.268E-07 | 566.31317 | 5046.5938 |
| LASTIADNQEQDL | 15 | Q96R06 | Sperm-associated antigen 5 OS=Homo sa | SPAG5 | 134.422 | 2 | 52.864365 | yes | 9.722E-08 | 837.91266 | 3638.4055 |
| DAQTQLVGLHAK | 12 | Q96R06 | Sperm-associated antigen 5 OS=Homo sa | SPAG5 | 134.422 | 2 | 52.864365 | yes | 1.001E-07 | 640.8515 | 3576.8391 |
| AETLVCCCFDLLK | 13 | Q96R06 | Sperm-associated antigen 5 OS=Homo sa | SPAG5 | 134.422 | 2 | 52.864365 | yes | 1.548E-07 | 814.87769 | 3567.1926 |
| VWLSQEVDK | 9 | Q96R06 | Sperm-associated antigen 5 OS=Homo sa | SPAG5 | 134.422 | 2 | 52.864365 | yes | 1.241E-07 | 552.29022 | 3429.4658 |
| ELQEVIQQQNEK | 12 | Q96R06 | Sperm-associated antigen 5 OS=Homo sa | SPAG5 | 134.422 | 2 | 52.864365 | yes | 9.722E-08 | 743.3808 | 3241.1238 |
| TSEEAVDPLGNYM | 15 | Q96R06 | Sperm-associated antigen 5 OS=Homo sa | SPAG5 | 134.422 | 2 | 52.864365 | yes | 1.313E-07 | 826.89545 | 3132.532 |
| AQLQILANMDSQL | 14 | Q96R06 | Sperm-associated antigen 5 OS=Homo sa | SPAG5 | 134.422 | 2 | 52.864365 | yes | 1.028E-07 | 786.92432 | 3132.4358 |
| LDDIVQHIYK | 10 | Q96R06 | Sperm-associated antigen 5 OS=Homo sa | SPAG5 | 134.422 | 2 | 52.864365 | yes | 1.641E-07 | 622.33771 | 3038.8101 |
| LNQALCLR | 8 | Q96R06 | Sperm-associated antigen 5 OS=Homo sa | SPAG5 | 134.422 | 2 | 52.864365 | yes | 1.177E-07 | 494.27383 | 2916.3687 |
| LSLSLSPSPQTGK | 13 | Q96R06 | Sperm-associated antigen 5 OS=Homo sa | SPAG5 | 134.422 | 2 | 52.864365 | yes | 1.083E-07 | 657.86682 | 2678.3298 |
| NILEENLR | 8 | Q96R06 | Sperm-associated antigen 5 OS=Homo sa | SPAG5 | 134.422 | 2 | 52.864365 | yes | 2.065E-07 | 500.77472 | 2565.052 |
| ISQLEQDLASMR | 12 | Q96R06 | Sperm-associated antigen 5 OS=Homo sa | SPAG5 | 134.422 | 2 | 52.864365 | yes | 1.022E-07 | 695.85339 | 2456.2905 |
| EEVTHLTR | 8 | Q96R06 | Sperm-associated antigen 5 OS=Homo sa | SPAG5 | 134.422 | 2 | 52.864365 | yes | 9.722E-08 | 492.75909 | 2423.3059 |
| KICELQAR | 8 | Q96R06 | Sperm-associated antigen 5 OS=Homo sa | SPAG5 | 134.422 | 2 | 52.864365 | yes | 9.722E-08 | 509.27911 | 2329.9963 |
| STNTSQTGLVGTK | 13 | Q96R06 | Sperm-associated antigen 5 OS=Homo sa | SPAG5 | 134.422 | 2 | 52.864365 | yes | 9.904E-08 | 647.33588 | 2164.4026 |
| SLQCENLK | 8 | Q96R06 | Sperm-associated antigen 5 OS=Homo sa | SPAG5 | 134.422 | 2 | 52.864365 | yes | 1.136E-07 | 496.2475 | 1724.736 |
| ELTLQPGALTNSG | 14 | Q96R06 | Sperm-associated antigen 5 OS=Homo sa | SPAG5 | 134.422 | 2 | 52.864365 | yes | 1.259E-07 | 714.88831 | 1668.3763 |
| NVMQSWVLISK | 11 | Q96R06 | Sperm-associated antigen 5 OS=Homo sa | SPAG5 | 134.422 | 2 | 52.864365 | yes | 1.119E-07 | 652.85522 | 1187.2186 |
| EEVAPCMGDR | 10 | Q96R06 | Sperm-associated antigen 5 OS=Homo sa | SPAG5 | 134.422 | 2 | 52.864365 | yes | 1.262E-07 | 582.24463 | 1166.4808 |
| SPACSSLTPSLCK | 13 | Q96R06 | Sperm-associated antigen 5 OS=Homo sa | SPAG5 | 134.422 | 2 | 52.864365 | yes | 1.175E-07 | 704.33398 | 1117.3516 |
| VMFLEMK | 7 | Q96R06 | Sperm-associated antigen 5 OS=Homo sa | SPAG5 | 134.422 | 2 | 52.864365 | yes | 2.978E-07 | 449.23227 | 1008.3724 |
| SGELISLR | 8 | Q96R06 | Sperm-associated antigen 5 OS=Homo sa | SPAG5 | 134.422 | 2 | 52.864365 | yes | 1.116E-06 | 437.75327 | 942.48792 |
| ESHEMGQALQQA | 13 | Q96R06 | Sperm-associated antigen 5 OS=Homo sa | SPAG5 | 134.422 | 2 | 52.864365 | yes | 9.722E-08 | 742.84918 | 931.61157 |
| ICELQAR | 7 | Q96R06 | Sperm-associated antigen 5 OS=Homo sa | SPAG5 | 134.422 | 2 | 52.864365 | yes | 3.176E-07 | 445.23166 | 889.53638 |
| SQLAVPHPETQDS | 27 | Q96R06 | Sperm-associated antigen 5 OS=Homo sa | SPAG5 | 134.422 | 4 | 52.864365 | yes | 2.47E-07 | 717.34167 | 722.42169 |

| | | | | | | | | | | | |
|---|---|---|---|---|---|---|---|---|---|---|---|
| TTLEVLR | 7 | Q96R06 | Sperm-associated antigen 5 OS=Homo sa | SPAG5 | 134.422 | 2 | 52.864365 | yes | 9.728E-08 | 416.25037 | 680.8291 |
| HMQAELQQQQAV | 15 | Q96R06 | Sperm-associated antigen 5 OS=Homo sa | SPAG5 | 134.422 | 3 | 52.864365 | yes | 9.722E-08 | 574.97003 | 603.87122 |
| VLEQVSAQLEECK | 13 | Q96R06 | Sperm-associated antigen 5 OS=Homo sa | SPAG5 | 134.422 | 2 | 52.864365 | yes | 9.819E-08 | 766.88489 | 579.79413 |
| SQQALQER | 8 | Q96R06 | Sperm-associated antigen 5 OS=Homo sa | SPAG5 | 134.422 | 2 | 52.864365 | yes | 2.82E-07 | 480.24887 | 567.45624 |
| TDLSSEHFSHSSK | 13 | Q96R06 | Sperm-associated antigen 5 OS=Homo sa | SPAG5 | 134.422 | 2 | 52.864365 | yes | 9.722E-08 | 731.33386 | 487.40955 |
| LQAQEEQHQEVQI | 13 | Q96R06 | Sperm-associated antigen 5 OS=Homo sa | SPAG5 | 134.422 | 2 | 52.864365 | yes | 1.119E-07 | 797.89465 | 439.40912 |
| FSEVAAVSEKPIFQ | 37 | Q96R06 | Sperm-associated antigen 5 OS=Homo sa | SPAG5 | 134.422 | 5 | 52.864365 | yes | 3.207E-07 | 839.80078 | 412.78644 |
| GQTEQLELENSR | 12 | Q96R06 | Sperm-associated antigen 5 OS=Homo sa | SPAG5 | 134.422 | 2 | 52.864365 | yes | 9.722E-08 | 702.34167 | 317.96768 |
| HREEMALR | 8 | Q96R06 | Sperm-associated antigen 5 OS=Homo sa | SPAG5 | 134.422 | 2 | 52.864365 | yes | 7.006E-07 | 521.26654 | 198.48297 |
| FSEVAAVSEK | 10 | Q96R06 | Sperm-associated antigen 5 OS=Homo sa | SPAG5 | 134.422 | 2 | 52.864365 | yes | 3.041E-07 | 533.77441 | 165.23921 |
| RTDLSSEHFSHSSK | 14 | Q96R06 | Sperm-associated antigen 5 OS=Homo sa | SPAG5 | 134.422 | 3 | 52.864365 | yes | 1.33E-07 | 539.92535 | 133.94057 |
| KLSLSLSPSPQTGK | 14 | Q96R06 | Sperm-associated antigen 5 OS=Homo sa | SPAG5 | 134.422 | 2 | 52.864365 | yes | 9.722E-08 | 721.91431 | 106.40834 |
| ILEQIDK | 7 | Q96R06 | Sperm-associated antigen 5 OS=Homo sa | SPAG5 | 134.422 | 2 | 52.864365 | yes | 2.884E-07 | 429.75018 | 96.19101 |
| ELISLLHLSLLHLE | 17 | Q96R06 | Sperm-associated antigen 5 OS=Homo sa | SPAG5 | 134.422 | 3 | 52.864365 | yes | 1.046E-07 | 668.04547 | 82.69381 |
| GKDAAEIVLEAFC | 19 | Q96R06 | Sperm-associated antigen 5 OS=Homo sa | SPAG5 | 134.422 | 3 | 52.864365 | yes | 1.152E-07 | 691.67761 | 64.538017 |
| GVNTSVMLENLR | 12 | Q96R06 | Sperm-associated antigen 5 OS=Homo sa | SPAG5 | 134.422 | 2 | 52.864365 | yes | 3.159E-07 | 666.85065 | 63.335358 |
| RSPACSSLTPSLCK | 14 | Q96R06 | Sperm-associated antigen 5 OS=Homo sa | SPAG5 | 134.422 | 2 | 52.864365 | yes | 2.154E-07 | 782.38452 | 58.028191 |
| ELISLLHLSLLHLE | 25 | Q96R06 | Sperm-associated antigen 5 OS=Homo sa | SPAG5 | 134.422 | 4 | 52.864365 | yes | 9.592E-07 | 723.39349 | 32.475513 |
| YPIVEQYLK | 9 | Q96RD7 | Pannexin-1 OS=Homo sapiens OX=9606 | PANX1 | 48.051 | 2 | 50.72564 | yes | 9.727E-08 | 576.8186 | 139.87729 |
| FAAAPHICSDLK | 12 | Q96RD7 | Pannexin-1 OS=Homo sapiens OX=9606 | PANX1 | 48.051 | 2 | 50.72564 | yes | 7.883E-07 | 665.33466 | 52.652721 |
| VEALPEQVAPESR | 13 | Q96RE7 | Nucleus accumbens-associated protein 1 ( | NACC1 | 57.259 | 2 | 53.02524 | yes | 2.112E-07 | 712.87262 | 224.18069 |
| AVLAASSSYFR | 11 | Q96RE7 | Nucleus accumbens-associated protein 1 ( | NACC1 | 57.259 | 2 | 53.02524 | yes | 1.106E-07 | 586.30896 | 169.41711 |
| LELVTGTNVYITR | 13 | Q96RE7 | Nucleus accumbens-associated protein 1 ( | NACC1 | 57.259 | 2 | 53.02524 | yes | 9.722E-08 | 739.91431 | 151.52341 |
| LQGLYCDVSVVVI | 13 | Q96RE7 | Nucleus accumbens-associated protein 1 ( | NACC1 | 57.259 | 2 | 53.02524 | yes | 1.163E-07 | 740.3974 | 101.33575 |
| AEDDAYTTFISETC | 15 | Q96RE7 | Nucleus accumbens-associated protein 1 ( | NACC1 | 57.259 | 2 | 53.02524 | yes | 9.722E-08 | 824.37286 | 90.619453 |
| NTLANSCGTGIR | 12 | Q96RE7 | Nucleus accumbens-associated protein 1 ( | NACC1 | 57.259 | 2 | 53.02524 | no | 3.355E-06 | 632.30914 | 85.575981 |
| LYDEGDPSEK | 10 | Q96RE7 | Nucleus accumbens-associated protein 1 ( | NACC1 | 57.259 | 2 | 53.02524 | yes | 1.995E-07 | 576.75641 | 40.216442 |
| AQLMNCHVSAGT | 13 | Q96RE7 | Nucleus accumbens-associated protein 1 ( | NACC1 | 57.259 | 3 | 53.02524 | yes | 3.883E-07 | 482.23099 | 33.366234 |
| LLASFFDR | 8 | Q96RE7 | Nucleus accumbens-associated protein 1 ( | NACC1 | 57.259 | 2 | 53.02524 | yes | 3.148E-06 | 484.76364 | 10.235388 |
| GLFPASYVQVIR | 12 | Q96RF0 | Sorting nexin-18 OS=Homo sapiens OX= | SNX18 | 68.895 | 2 | 34.20596 | yes | 1.057E-06 | 675.38245 | 141.23613 |
| LIDFGSAAYLER | 12 | Q96RG2 | PAS domain-containing serine/threonine- | PASK | 142.93 | 2 | 54.44239 | yes | 1.417E-07 | 677.8537 | 96.060623 |
| GSVSCCSLLR | 10 | Q96RG2 | PAS domain-containing serine/threonine- | PASK | 142.93 | 2 | 54.44239 | yes | 5.448E-07 | 569.77081 | 93.530624 |
| VTLEIAILSR | 10 | Q96RG2 | PAS domain-containing serine/threonine- | PASK | 142.93 | 2 | 54.44239 | yes | 4.775E-06 | 557.84515 | 56.778229 |
| LNVQVTSTPVIVM | 14 | Q96RG2 | PAS domain-containing serine/threonine- | PASK | 142.93 | 2 | 54.44239 | yes | 9.722E-08 | 778.94507 | 54.937756 |
| LDEPLASYIFR | 11 | Q96RG2 | PAS domain-containing serine/threonine- | PASK | 142.93 | 2 | 54.44239 | yes | 2.737E-07 | 662.35083 | 49.775948 |
| VLEDCWIEDPK | 11 | Q96RG2 | PAS domain-containing serine/threonine- | PASK | 142.93 | 2 | 54.44239 | yes | 9.362E-07 | 702.32922 | 38.508938 |
| HCLPTLADAK | 10 | Q96RL1 | BRCA1-A complex subunit RAP80 OS=I | UIMC1 | 79.728 | 2 | 52.58691 | yes | 9.85E-08 | 563.28967 | 752.20618 |
| AIAESLNSCR | 10 | Q96RL1 | BRCA1-A complex subunit RAP80 OS=I | UIMC1 | 79.728 | 2 | 52.58691 | yes | 8.169E-07 | 560.77441 | 654.17841 |
| VILCQLEVYQK | 11 | Q96RL1 | BRCA1-A complex subunit RAP80 OS=I | UIMC1 | 79.728 | 2 | 52.58691 | yes | 9.722E-08 | 696.88141 | 585.4624 |

| | | | | | | | | | | | |
|---|---|---|---|---|---|---|---|---|---|---|---|
| MSEQEAR | 7 | Q96RL1 | BRCA1-A complex subunit RAP80 OS=I | UIMC1 | 79.728 | 2 | 52.58691 | yes | 1.092E-06 | 425.68982 | 294.72409 |
| EEPWDHTEK | 9 | Q96RL1 | BRCA1-A complex subunit RAP80 OS=I | UIMC1 | 79.728 | 2 | 52.58691 | yes | 1.161E-07 | 585.75671 | 276.3735 |
| QVTVQPGSR | 9 | Q96RL1 | BRCA1-A complex subunit RAP80 OS=I | UIMC1 | 79.728 | 2 | 52.58691 | yes | 1.065E-07 | 486.26706 | 225.11223 |
| SDSGTAAQTSLDII | 15 | Q96RL1 | BRCA1-A complex subunit RAP80 OS=I | UIMC1 | 79.728 | 2 | 52.58691 | yes | 2.113E-07 | 754.85736 | 175.27899 |
| DVETTSSVSVK | 11 | Q96RL1 | BRCA1-A complex subunit RAP80 OS=I | UIMC1 | 79.728 | 2 | 52.58691 | yes | 2.249E-07 | 576.29315 | 138.51547 |
| EMQSSFTR | 8 | Q96RL1 | BRCA1-A complex subunit RAP80 OS=I | UIMC1 | 79.728 | 2 | 52.58691 | yes | 2.907E-07 | 493.22403 | 86.673065 |
| LLSFLEQSEHK | 11 | Q96RL1 | BRCA1-A complex subunit RAP80 OS=I | UIMC1 | 79.728 | 2 | 52.58691 | yes | 1.204E-07 | 665.8537 | 83.144989 |
| KDVETTSSVSVK | 12 | Q96RL1 | BRCA1-A complex subunit RAP80 OS=I | UIMC1 | 79.728 | 2 | 52.58691 | yes | 1.829E-07 | 640.34064 | 68.217224 |
| VPSPGMEEAGCSR | 13 | Q96RL1 | BRCA1-A complex subunit RAP80 OS=I | UIMC1 | 79.728 | 2 | 52.58691 | yes | 1.211E-07 | 688.80029 | 46.390144 |
| KQVTVQPGSR | 10 | Q96RL1 | BRCA1-A complex subunit RAP80 OS=I | UIMC1 | 79.728 | 2 | 52.58691 | yes | 1.111E-07 | 550.31458 | 33.491726 |
| LLIQVER | 7 | Q96RL7 | Intermembrane lipid transfer protein VPS | VPS13A | 360.277 | 2 | 41.79261 | yes | 3.339E-07 | 435.77402 | 1444.9097 |
| VIQLDTNVPVR | 11 | Q96RL7 | Intermembrane lipid transfer protein VPS | VPS13A | 360.277 | 2 | 41.79261 | yes | 3.359E-06 | 627.36426 | 31.992765 |
| GIVDLIEER | 9 | Q96RP9 | Elongation factor G, mitochondrial OS=H | GFM1 | 83.472 | 2 | 55.107204 | yes | 3.451E-07 | 522.29022 | 158.93201 |
| ETITAPVPFDFTHK | 14 | Q96RP9 | Elongation factor G, mitochondrial OS=H | GFM1 | 83.472 | 2 | 55.107204 | yes | 3.512E-06 | 801.91174 | 156.05428 |
| VIGVLEPLDPEDY | 15 | Q96RP9 | Elongation factor G, mitochondrial OS=H | GFM1 | 83.472 | 2 | 55.107204 | yes | 1.815E-07 | 844.44312 | 142.98022 |
| DGVGAVMDSMEL | 14 | Q96RP9 | Elongation factor G, mitochondrial OS=H | GFM1 | 83.472 | 2 | 55.107204 | yes | 9.722E-08 | 754.8396 | 142.40858 |
| YGEIPAELR | 9 | Q96RP9 | Elongation factor G, mitochondrial OS=H | GFM1 | 83.472 | 2 | 55.107204 | yes | 1.92E-06 | 524.2771 | 136.88258 |
| NIGISAHIDSGK | 12 | Q96RP9 | Elongation factor G, mitochondrial OS=H | GFM1 | 83.472 | 2 | 55.107204 | yes | 2.512E-07 | 606.32257 | 100.05679 |
| YLEATGQLPVK | 11 | Q96RP9 | Elongation factor G, mitochondrial OS=H | GFM1 | 83.472 | 2 | 55.107204 | yes | 2.019E-06 | 609.84009 | 73.380814 |
| DNSHPFVGLAFK | 12 | Q96RP9 | Elongation factor G, mitochondrial OS=H | GFM1 | 83.472 | 2 | 55.107204 | yes | 1.75E-06 | 666.34076 | 68.101257 |
| VYFDTENK | 8 | Q96RP9 | Elongation factor G, mitochondrial OS=H | GFM1 | 83.472 | 2 | 55.107204 | yes | 1.017E-06 | 508.2402 | 61.139851 |
| AIYFDGDFGQIVR | 13 | Q96RP9 | Elongation factor G, mitochondrial OS=H | GFM1 | 83.472 | 2 | 55.107204 | yes | 6.636E-07 | 750.87769 | 42.397659 |
| SFTPVFLGSALK | 12 | Q96RP9 | Elongation factor G, mitochondrial OS=H | GFM1 | 83.472 | 2 | 55.107204 | yes | 9.722E-08 | 633.85828 | 40.91745 |
| KGDTIYNTR | 9 | Q96RP9 | Elongation factor G, mitochondrial OS=H | GFM1 | 83.472 | 2 | 55.107204 | yes | 3.768E-07 | 534.27765 | 40.631573 |
| VMLTQFAEGR | 10 | Q96RR1 | Twinkle mtDNA helicase OS=Homo sapi | TWNK | 77.155 | 2 | 53.518784 | yes | 9.722E-08 | 576.29755 | 706.11896 |
| ASQEADNVLILQD | 14 | Q96RR1 | Twinkle mtDNA helicase OS=Homo sapi | TWNK | 77.155 | 2 | 53.518784 | yes | 1.036E-07 | 786.40485 | 658.63086 |
| IKDDTGPVAK | 10 | Q96RR1 | Twinkle mtDNA helicase OS=Homo sapi | TWNK | 77.155 | 2 | 53.518784 | yes | 1.048E-07 | 522.29022 | 551.52106 |
| ALSPFAESSQLK | 12 | Q96RR1 | Twinkle mtDNA helicase OS=Homo sapi | TWNK | 77.155 | 2 | 53.518784 | yes | 1.373E-07 | 639.34045 | 455.74133 |
| IAAQDYIIGVFR | 12 | Q96RR1 | Twinkle mtDNA helicase OS=Homo sapi | TWNK | 77.155 | 2 | 53.518784 | yes | 9.849E-08 | 683.37988 | 323.44138 |
| FDGDVGVFPLEFN | 14 | Q96RR1 | Twinkle mtDNA helicase OS=Homo sapi | TWNK | 77.155 | 2 | 53.518784 | yes | 9.722E-08 | 792.39069 | 301.50977 |
| APEFEDSEEVR | 11 | Q96RR1 | Twinkle mtDNA helicase OS=Homo sapi | TWNK | 77.155 | 2 | 53.518784 | yes | 9.722E-08 | 654.29114 | 283.51981 |
| PLEALNGGFNLSR | 13 | Q96RR1 | Twinkle mtDNA helicase OS=Homo sapi | TWNK | 77.155 | 2 | 53.518784 | yes | 1.618E-07 | 694.37006 | 136.37553 |
| GQTGVTTSFSLFID | 15 | Q96RR1 | Twinkle mtDNA helicase OS=Homo sapi | TWNK | 77.155 | 2 | 53.518784 | yes | 1.395E-07 | 800.91449 | 123.38372 |
| VTDDTLKR | 8 | Q96RR1 | Twinkle mtDNA helicase OS=Homo sapi | TWNK | 77.155 | 2 | 53.518784 | yes | 3.161E-07 | 474.26144 | 98.255417 |
| KGELTVFTGPTGS | 15 | Q96RR1 | Twinkle mtDNA helicase OS=Homo sapi | TWNK | 77.155 | 2 | 53.518784 | yes | 1.324E-07 | 739.89612 | 61.169178 |
| IVFWLGDDLR | 10 | Q96RR1 | Twinkle mtDNA helicase OS=Homo sapi | TWNK | 77.155 | 2 | 53.518784 | yes | 2.25E-07 | 617.33496 | 55.411701 |
| CFLVRPGDQQPR | 12 | Q96RR1 | Twinkle mtDNA helicase OS=Homo sapi | TWNK | 77.155 | 3 | 53.518784 | yes | 4.646E-06 | 491.58575 | 32.565025 |
| YLDEIVQESWR | 11 | Q96RS0 | Trimethylguanosine synthase OS=Homo s | TGS1 | 96.62 | 2 | 49.388393 | yes | 1.037E-07 | 719.35406 | 135.38799 |
| SMGLPLQFGR | 10 | Q96RS0 | Trimethylguanosine synthase OS=Homo s | TGS1 | 96.62 | 2 | 49.388393 | yes | 3.025E-07 | 553.2948 | 111.80228 |

| | | | | | | | | | | | |
|---|---|---|---|---|---|---|---|---|---|---|---|
| ITNNIVYFLPR | 11 | Q96RS0 | Trimethylguanosine synthase OS=Homo s | TGS1 | 96.62 | 2 | 49.388393 | yes | 3.722E-07 | 675.38245 | 46.862503 |
| KLYNLGLK | 8 | Q96RS0 | Trimethylguanosine synthase OS=Homo s | TGS1 | 96.62 | 2 | 49.388393 | yes | 4.367E-07 | 474.79749 | 36.511612 |
| TITAYFGDLIR | 11 | Q96RS0 | Trimethylguanosine synthase OS=Homo s | TGS1 | 96.62 | 2 | 49.388393 | yes | 7.065E-07 | 635.34552 | 23.811821 |
| ESNSLEISLIK | 11 | Q96RS6 | NudC domain-containing protein 1 OS=H | NUDCD1 | 66.756 | 2 | 43.239464 | yes | 2.672E-07 | 616.84027 | 59.403831 |
| TVQTIVHR | 8 | Q96RT1 | Erbin OS=Homo sapiens OX=9606 GN=I | ERBIN | 158.299 | 2 | 46.64643 | yes | 4.936E-07 | 477.27997 | 58.658974 |
| TPPMMPGSQR | 10 | Q96RT1 | Erbin OS=Homo sapiens OX=9606 GN=I | ERBIN | 158.299 | 2 | 46.64643 | yes | 1.623E-06 | 551.26263 | 58.493778 |
| LQILELR | 7 | Q96RT1 | Erbin OS=Homo sapiens OX=9606 GN=I | ERBIN | 158.299 | 2 | 46.64643 | yes | 4.683E-07 | 442.78183 | 57.047306 |
| MVLTNYMFPQQPI | 13 | Q96RT1 | Erbin OS=Homo sapiens OX=9606 GN=I | ERBIN | 158.299 | 2 | 46.64643 | yes | 1.505E-07 | 812.90234 | 52.102417 |
| PSINEIPER | 9 | Q96RT1 | Erbin OS=Homo sapiens OX=9606 GN=I | ERBIN | 158.299 | 2 | 46.64643 | yes | 1.825E-06 | 527.78003 | 19.69726 |
| VLTIVEASVNPISK | 14 | Q96RT1 | Erbin OS=Homo sapiens OX=9606 GN=I | ERBIN | 158.299 | 2 | 46.64643 | yes | 1.712E-07 | 735.43231 | 16.854969 |
| TALLSHMAGSVQI | 14 | Q96RT7 | Gamma-tubulin complex component 6 OS | TUBGCP6 | 200.499 | 2 | 57.16032 | yes | 2.89E-07 | 759.39832 | 1582.7565 |
| VGENVSDVAPIR | 12 | Q96RT7 | Gamma-tubulin complex component 6 OS | TUBGCP6 | 200.499 | 2 | 57.16032 | yes | 1.593E-07 | 628.33569 | 587.34607 |
| VGENVSDVAPTQI | 14 | Q96RT7 | Gamma-tubulin complex component 6 OS | TUBGCP6 | 200.499 | 2 | 57.16032 | yes | 9.903E-08 | 734.87317 | 462.72882 |
| LATVGDLEEIQR | 12 | Q96RT7 | Gamma-tubulin complex component 6 OS | TUBGCP6 | 200.499 | 2 | 57.16032 | yes | 3.36E-07 | 672.36188 | 375.41119 |
| EVEDCVPVFLK | 11 | Q96RT7 | Gamma-tubulin complex component 6 OS | TUBGCP6 | 200.499 | 2 | 57.16032 | yes | 2.652E-07 | 667.83667 | 292.95078 |
| ETLLPSHPPR | 10 | Q96RT7 | Gamma-tubulin complex component 6 OS | TUBGCP6 | 200.499 | 2 | 57.16032 | yes | 1.028E-07 | 573.81696 | 179.24226 |
| VLSALSDR | 8 | Q96RT7 | Gamma-tubulin complex component 6 OS | TUBGCP6 | 200.499 | 2 | 57.16032 | yes | 3.22E-07 | 430.74545 | 162.40585 |
| WNTHGHVSDASIS | 27 | Q96RT7 | Gamma-tubulin complex component 6 OS | TUBGCP6 | 200.499 | 4 | 57.16032 | yes | 1.863E-07 | 727.60052 | 123.73287 |
| EEPYLTEAGR | 10 | Q96RT7 | Gamma-tubulin complex component 6 OS | TUBGCP6 | 200.499 | 2 | 57.16032 | yes | 3.476E-07 | 582.78021 | 102.45 |
| QEELDDDFSYAR | 12 | Q96RT7 | Gamma-tubulin complex component 6 OS | TUBGCP6 | 200.499 | 2 | 57.16032 | yes | 9.722E-08 | 744.31787 | 101.93241 |
| TSCEPYTR | 8 | Q96RT7 | Gamma-tubulin complex component 6 OS | TUBGCP6 | 200.499 | 2 | 57.16032 | yes | 1.008E-07 | 507.22147 | 97.4328 |
| WNTHGHVSNASIS | 27 | Q96RT7 | Gamma-tubulin complex component 6 OS | TUBGCP6 | 200.499 | 4 | 57.16032 | yes | 4.598E-07 | 719.36151 | 82.771767 |
| WNTHGHVSDASIS | 27 | Q96RT7 | Gamma-tubulin complex component 6 OS | TUBGCP6 | 200.499 | 4 | 57.16032 | yes | 3.187E-07 | 720.0979 | 79.499466 |
| SQLISQAWGPPGG | 15 | Q96RT7 | Gamma-tubulin complex component 6 OS | TUBGCP6 | 200.499 | 2 | 57.16032 | yes | 4.031E-07 | 775.90735 | 72.519714 |
| LKEQFVK | 7 | Q96RT7 | Gamma-tubulin complex component 6 OS | TUBGCP6 | 200.499 | 2 | 57.16032 | yes | 3.18E-07 | 446.26855 | 69.619743 |
| QELIAHAR | 8 | Q96RT7 | Gamma-tubulin complex component 6 OS | TUBGCP6 | 200.499 | 2 | 57.16032 | yes | 2.97E-07 | 469.26434 | 61.380558 |
| HIAHDIYVCGK | 11 | Q96RT7 | Gamma-tubulin complex component 6 OS | TUBGCP6 | 200.499 | 2 | 57.16032 | yes | 2.635E-07 | 656.82697 | 53.808933 |
| VSLFGALVHSR | 11 | Q96RT7 | Gamma-tubulin complex component 6 OS | TUBGCP6 | 200.499 | 2 | 57.16032 | yes | 1.151E-07 | 593.34058 | 52.956718 |
| GYQPHLEDFLLR | 12 | Q96RT7 | Gamma-tubulin complex component 6 OS | TUBGCP6 | 200.499 | 2 | 57.16032 | yes | 9.722E-08 | 744.38568 | 46.557884 |
| WNTHGHVSDTSIR | 13 | Q96RT7 | Gamma-tubulin complex component 6 OS | TUBGCP6 | 200.499 | 3 | 57.16032 | yes | 2.333E-07 | 503.91129 | 21.471338 |
| WNTHGHVSDTSIR | 13 | Q96RT7 | Gamma-tubulin complex component 6 OS | TUBGCP6 | 200.499 | 3 | 57.16032 | yes | 5.323E-07 | 755.36328 | 21.471338 |
| WNTHGHVSDTSIR | 13 | Q96RT7 | Gamma-tubulin complex component 6 OS | TUBGCP6 | 200.499 | 2 | 57.16032 | yes | 2.333E-07 | 503.91129 | 21.471338 |
| WNTHGHVSDTSIR | 13 | Q96RT7 | Gamma-tubulin complex component 6 OS | TUBGCP6 | 200.499 | 2 | 57.16032 | yes | 5.323E-07 | 755.36328 | 21.471338 |
| AALEEGSSQPTER | 13 | Q96RT7 | Gamma-tubulin complex component 6 OS | TUBGCP6 | 200.499 | 2 | 57.16032 | yes | 1.811E-07 | 687.82843 | 40.50742 |
| WNTHGHVSDASIR | 13 | Q96RT7 | Gamma-tubulin complex component 6 OS | TUBGCP6 | 200.499 | 2 | 57.16032 | yes | 6.139E-07 | 740.35803 | 26.178646 |
| LHQGELQLLAGGV | 23 | Q96RT7 | Gamma-tubulin complex component 6 OS | TUBGCP6 | 200.499 | 3 | 57.16032 | yes | 4.776E-07 | 803.47229 | 25.50024 |
| FVIHSDLSK | 9 | Q96RT8 | Gamma-tubulin complex component 5 OS | TUBGCP5 | 118.321 | 2 | 56.15517 | yes | 1.037E-07 | 523.28748 | 287.27142 |
| LTEEFLNAPLPSIK | 14 | Q96RT8 | Gamma-tubulin complex component 5 OS | TUBGCP5 | 118.321 | 2 | 56.15517 | yes | 1.032E-07 | 786.43762 | 280.09219 |
| LDAQQER | 7 | Q96RT8 | Gamma-tubulin complex component 5 OS | TUBGCP5 | 118.321 | 2 | 56.15517 | yes | 3.925E-07 | 430.21704 | 240.96227 |

| | | | | | | | | | | | |
|---|---|---|---|---|---|---|---|---|---|---|---|
| VLVTETQVIR | 10 | Q96RT8 | Gamma-tubulin complex component 5 OS | TUBGCP5 | 118.321 | 2 | 56.15517 | yes | 1.081E-07 | 579.34808 | 184.22263 |
| SCLYPHIDK | 9 | Q96RT8 | Gamma-tubulin complex component 5 OS | TUBGCP5 | 118.321 | 2 | 56.15517 | yes | 9.722E-08 | 566.77643 | 182.06732 |
| FAGGDVCVDR | 10 | Q96RT8 | Gamma-tubulin complex component 5 OS | TUBGCP5 | 118.321 | 2 | 56.15517 | yes | 9.722E-08 | 548.24805 | 169.65198 |
| MYLEQSDFHEK | 11 | Q96RT8 | Gamma-tubulin complex component 5 OS | TUBGCP5 | 118.321 | 2 | 56.15517 | yes | 1.737E-07 | 713.81903 | 168.55913 |
| FLDVNSHK | 8 | Q96RT8 | Gamma-tubulin complex component 5 OS | TUBGCP5 | 118.321 | 2 | 56.15517 | yes | 2.224E-06 | 480.25089 | 144.13086 |
| ASHLLNTLYK | 10 | Q96RT8 | Gamma-tubulin complex component 5 OS | TUBGCP5 | 118.321 | 2 | 56.15517 | yes | 9.81E-08 | 580.32715 | 143.83162 |
| ILHSTGLEFQHQVI | 17 | Q96RT8 | Gamma-tubulin complex component 5 OS | TUBGCP5 | 118.321 | 3 | 56.15517 | yes | 1.692E-07 | 656.0058 | 130.63533 |
| TPLEEQDQNR | 10 | Q96RT8 | Gamma-tubulin complex component 5 OS | TUBGCP5 | 118.321 | 2 | 56.15517 | yes | 9.722E-08 | 615.29144 | 110.66597 |
| LSISFENVDTAK | 12 | Q96RT8 | Gamma-tubulin complex component 5 OS | TUBGCP5 | 118.321 | 2 | 56.15517 | yes | 9.826E-08 | 662.3432 | 81.991913 |
| ETLWLLSGVK | 10 | Q96RT8 | Gamma-tubulin complex component 5 OS | TUBGCP5 | 118.321 | 2 | 56.15517 | yes | 9.722E-08 | 573.33191 | 77.75869 |
| RPFVPAAMFEEHR | 13 | Q96S15 | GATOR2 complex protein WDR24 OS=I | WDR24 | 88.207 | 3 | 44.66025 | yes | 2.114E-07 | 529.60138 | 83.895569 |
| EMHCVQTIASVAR | 13 | Q96S15 | GATOR2 complex protein WDR24 OS=I | WDR24 | 88.207 | 2 | 44.66025 | yes | 2.594E-07 | 751.36615 | 14.991749 |
| SGAPPPSGSAVSTA | 18 | Q96SB4 | SRSF protein kinase 1 OS=Homo sapiens | SRPK1 | 74.325 | 2 | 51.829838 | yes | 2.923E-07 | 818.42047 | 3477.2598 |
| LAAEATEWQR | 10 | Q96SB4 | SRSF protein kinase 1 OS=Homo sapiens | SRPK1 | 74.325 | 2 | 51.829838 | yes | 9.733E-08 | 587.7962 | 3237.8237 |
| SNYQGLPLPCVK | 12 | Q96SB4 | SRSF protein kinase 1 OS=Homo sapiens | SRPK1 | 74.325 | 2 | 51.829838 | yes | 1.017E-07 | 688.35559 | 3118.1118 |
| MQEIEEMEK | 9 | Q96SB4 | SRSF protein kinase 1 OS=Homo sapiens | SRPK1 | 74.325 | 2 | 51.829838 | yes | 1.074E-07 | 583.75702 | 2883.4194 |
| IIHTDIKPENILLSV | 21 | Q96SB4 | SRSF protein kinase 1 OS=Homo sapiens | SRPK1 | 74.325 | 3 | 51.829838 | yes | 1.63E-07 | 836.79889 | 962.21912 |
| IIHTDIKPENILLSV | 21 | Q96SB4 | SRSF protein kinase 1 OS=Homo sapiens | SRPK1 | 74.325 | 3 | 51.829838 | yes | 1.166E-07 | 627.85095 | 962.21912 |
| IIHTDIKPENILLSV | 21 | Q96SB4 | SRSF protein kinase 1 OS=Homo sapiens | SRPK1 | 74.325 | 4 | 51.829838 | yes | 1.63E-07 | 836.79889 | 962.21912 |
| IIHTDIKPENILLSV | 21 | Q96SB4 | SRSF protein kinase 1 OS=Homo sapiens | SRPK1 | 74.325 | 4 | 51.829838 | yes | 1.166E-07 | 627.85095 | 962.21912 |
| EMVVQLLDDFK | 11 | Q96SB4 | SRSF protein kinase 1 OS=Homo sapiens | SRPK1 | 74.325 | 2 | 51.829838 | yes | 1.002E-07 | 668.84448 | 748.38739 |
| EMVVQLLDDFK | 11 | Q96SB4 | SRSF protein kinase 1 OS=Homo sapiens | SRPK1 | 74.325 | 2 | 51.829838 | yes | 2.03E-06 | 676.84198 | 748.38739 |
| KVLALQAR | 8 | Q96SB4 | SRSF protein kinase 1 OS=Homo sapiens | SRPK1 | 74.325 | 2 | 51.829838 | yes | 1.379E-07 | 449.79526 | 887.51068 |
| ATAAECLR | 8 | Q96SB4 | SRSF protein kinase 1 OS=Homo sapiens | SRPK1 | 74.325 | 2 | 51.829838 | yes | 1.877E-07 | 446.22128 | 806.0097 |
| SAEHYTETALDEII | 14 | Q96SB4 | SRSF protein kinase 1 OS=Homo sapiens | SRPK1 | 74.325 | 2 | 51.829838 | yes | 1.125E-07 | 817.88647 | 720.79877 |
| STAGNFLVNPLEPI | 14 | Q96SB4 | SRSF protein kinase 1 OS=Homo sapiens | SRPK1 | 74.325 | 2 | 51.829838 | yes | 2.782E-07 | 743.89862 | 530.00562 |
| QEESESPVER | 10 | Q96SB4 | SRSF protein kinase 1 OS=Homo sapiens | SRPK1 | 74.325 | 2 | 51.829838 | yes | 9.722E-08 | 595.2702 | 241.95181 |
| QEESESPVERPLK | 13 | Q96SB4 | SRSF protein kinase 1 OS=Homo sapiens | SRPK1 | 74.325 | 2 | 51.829838 | yes | 9.722E-08 | 764.38611 | 149.16397 |
| EAFTADGDQVFAC | 14 | Q96SB8 | Structural maintenance of chromosomes p | SMC6 | 126.326 | 2 | 57.675415 | yes | 9.722E-08 | 742.34424 | 241.41461 |
| LLGEIMEHR | 9 | Q96SB8 | Structural maintenance of chromosomes p | SMC6 | 126.326 | 2 | 57.675415 | yes | 1.846E-07 | 549.29224 | 166.0688 |
| LYFDNLLSQR | 10 | Q96SB8 | Structural maintenance of chromosomes p | SMC6 | 126.326 | 2 | 57.675415 | yes | 9.722E-08 | 634.83533 | 110.39407 |
| NETLSISVQPGEGN | 15 | Q96SB8 | Structural maintenance of chromosomes p | SMC6 | 126.326 | 2 | 57.675415 | yes | 9.734E-08 | 786.89685 | 92.859543 |
| SATGSVVSTR | 10 | Q96SB8 | Structural maintenance of chromosomes p | SMC6 | 126.326 | 2 | 57.675415 | yes | 3.741E-07 | 482.75653 | 87.031837 |
| AYNEAEVLYNR | 11 | Q96SB8 | Structural maintenance of chromosomes p | SMC6 | 126.326 | 2 | 57.675415 | yes | 9.722E-08 | 671.32532 | 80.986641 |
| MVEEHMEQQK | 10 | Q96SB8 | Structural maintenance of chromosomes p | SMC6 | 126.326 | 2 | 57.675415 | yes | 9.722E-08 | 644.78668 | 43.159328 |
| DPELALAIESCLK | 13 | Q96SB8 | Structural maintenance of chromosomes p | SMC6 | 126.326 | 2 | 57.675415 | yes | 1.289E-07 | 729.87909 | 14.396563 |
| VILTSPLIR | 9 | Q96SI9 | Spermatid perinuclear RNA-binding prote | STRBP | 73.653 | 2 | 50.639935 | yes | 2.252E-07 | 506.3317 | 1004.0876 |
| LSAFGQIYK | 9 | Q96SI9 | Spermatid perinuclear RNA-binding prote | STRBP | 73.653 | 2 | 50.639935 | yes | 1.611E-07 | 513.78455 | 492.59714 |
| DKPTETLLNTVK | 12 | Q96SI9 | Spermatid perinuclear RNA-binding prote | STRBP | 73.653 | 2 | 50.639935 | yes | 1.112E-07 | 679.87994 | 453.17065 |

| | | | | | | | | | | | |
|---|---|---|---|---|---|---|---|---|---|---|---|
| DNLPIQIQK | 9 | Q96SI9 | Spermatid perinuclear RNA-binding prot‹ | STRBP | 73.653 | 2 | 50.639935 | yes | 1.667E-06 | 534.80603 | 422.23331 |
| TQGPILTASGK | 11 | Q96SI9 | Spermatid perinuclear RNA-binding prot‹ | STRBP | 73.653 | 2 | 50.639935 | yes | 1.642E-07 | 536.80347 | 209.93465 |
| VLEMDPLPSSK | 11 | Q96SI9 | Spermatid perinuclear RNA-binding prot‹ | STRBP | 73.653 | 2 | 50.639935 | yes | 1.175E-07 | 608.31812 | 152.58707 |
| HVSDWLDETNK | 11 | Q96SI9 | Spermatid perinuclear RNA-binding prot‹ | STRBP | 73.653 | 2 | 50.639935 | yes | 1.422E-07 | 672.31494 | 121.53127 |
| ASAALAALEK | 10 | Q96SI9 | Spermatid perinuclear RNA-binding prot‹ | STRBP | 73.653 | 2 | 50.639935 | yes | 2.637E-07 | 472.7742 | 98.438843 |
| MVLLPVMK | 8 | Q96SI9 | Spermatid perinuclear RNA-binding prot‹ | STRBP | 73.653 | 2 | 50.639935 | yes | 3.626E-06 | 465.77939 | 91.9869 |
| CLNALASLR | 9 | Q96SI9 | Spermatid perinuclear RNA-binding prot‹ | STRBP | 73.653 | 2 | 50.639935 | yes | 1.603E-07 | 509.27911 | 77.892174 |
| LNQIRPGLQYK | 11 | Q96SI9 | Spermatid perinuclear RNA-binding prot‹ | STRBP | 73.653 | 2 | 50.639935 | yes | 9.733E-08 | 665.3855 | 67.743698 |
| EDITHSAQHALR | 12 | Q96SI9 | Spermatid perinuclear RNA-binding prot‹ | STRBP | 73.653 | 2 | 50.639935 | yes | 1.555E-07 | 689.34711 | 62.791462 |
| AIDLMNALMR | 10 | Q96SI9 | Spermatid perinuclear RNA-binding prot‹ | STRBP | 73.653 | 2 | 50.639935 | yes | 2.307E-07 | 574.30176 | 45.942886 |
| VPTWAPLK | 8 | Q96SI9 | Spermatid perinuclear RNA-binding prot‹ | STRBP | 73.653 | 2 | 50.639935 | yes | 6.392E-07 | 456.27109 | 41.345886 |
| DPTDALSYMTIQQ | 14 | Q96SI9 | Spermatid perinuclear RNA-binding prot‹ | STRBP | 73.653 | 2 | 50.639935 | yes | 1.079E-06 | 805.89014 | 28.35882 |
| FPTYPVPHYSFF | 12 | Q96SI9 | Spermatid perinuclear RNA-binding prot‹ | STRBP | 73.653 | 2 | 50.639935 | yes | 5.681E-07 | 751.36115 | 21.345673 |
| ILGPLAPEMIDSR | 13 | Q96SN8 | CDK5 regulatory subunit-associated prot‹ | CDK5RAP: | 215.038 | 2 | 54.98115 | yes | 1.607E-07 | 706.38434 | 528.32861 |
| DASVQTVATEGDI | 15 | Q96SN8 | CDK5 regulatory subunit-associated prot‹ | CDK5RAP: | 215.038 | 2 | 54.98115 | yes | 1.058E-07 | 787.90466 | 352.76385 |
| IQALEEDLR | 9 | Q96SN8 | CDK5 regulatory subunit-associated prot‹ | CDK5RAP: | 215.038 | 2 | 54.98115 | yes | 7.681E-07 | 543.79309 | 288.59186 |
| VIFDQLVVTHK | 11 | Q96SN8 | CDK5 regulatory subunit-associated prot‹ | CDK5RAP: | 215.038 | 2 | 54.98115 | yes | 1.622E-07 | 649.87701 | 276.6322 |
| TVSLEHLQR | 9 | Q96SN8 | CDK5 regulatory subunit-associated prot‹ | CDK5RAP: | 215.038 | 2 | 54.98115 | yes | 1.775E-07 | 541.80127 | 255.24623 |
| LPILIKPSR | 9 | Q96SN8 | CDK5 regulatory subunit-associated prot‹ | CDK5RAP: | 215.038 | 2 | 54.98115 | yes | 2.019E-07 | 518.84747 | 234.99841 |
| LSCEAQLVK | 9 | Q96SN8 | CDK5 regulatory subunit-associated prot‹ | CDK5RAP: | 215.038 | 2 | 54.98115 | yes | 1.982E-07 | 524.27881 | 207.69193 |
| LASTEQTELLASK | 13 | Q96SN8 | CDK5 regulatory subunit-associated prot‹ | CDK5RAP: | 215.038 | 2 | 54.98115 | yes | 1.054E-07 | 695.87482 | 197.38507 |
| LTLEEAYR | 8 | Q96SN8 | CDK5 regulatory subunit-associated prot‹ | CDK5RAP: | 215.038 | 2 | 54.98115 | yes | 9.727E-08 | 497.76382 | 196.74866 |
| DVTAAQAELEK | 11 | Q96SN8 | CDK5 regulatory subunit-associated prot‹ | CDK5RAP: | 215.038 | 2 | 54.98115 | yes | 1.147E-07 | 587.80115 | 186.89008 |
| TLEEQISEIR | 10 | Q96SN8 | CDK5 regulatory subunit-associated prot‹ | CDK5RAP: | 215.038 | 2 | 54.98115 | yes | 3.255E-07 | 609.32227 | 147.5219 |
| ITQELSDLQQER | 12 | Q96SN8 | CDK5 regulatory subunit-associated prot‹ | CDK5RAP: | 215.038 | 2 | 54.98115 | yes | 9.722E-08 | 730.37299 | 134.304 |
| EQDIYTHLVK | 10 | Q96SN8 | CDK5 regulatory subunit-associated prot‹ | CDK5RAP: | 215.038 | 2 | 54.98115 | yes | 9.722E-08 | 623.32733 | 127.23312 |
| ATLLLESRPDLLK | 13 | Q96SN8 | CDK5 regulatory subunit-associated prot‹ | CDK5RAP: | 215.038 | 2 | 54.98115 | yes | 9.722E-08 | 734.94031 | 119.27035 |
| GPHPAPLSK | 9 | Q96SN8 | CDK5 regulatory subunit-associated prot‹ | CDK5RAP: | 215.038 | 2 | 54.98115 | yes | 3.001E-07 | 452.25598 | 107.21704 |
| EALIQCLK | 8 | Q96SN8 | CDK5 regulatory subunit-associated prot‹ | CDK5RAP: | 215.038 | 2 | 54.98115 | yes | 1.133E-06 | 487.7706 | 103.94006 |
| DGHGICVISR | 10 | Q96SN8 | CDK5 regulatory subunit-associated prot‹ | CDK5RAP: | 215.038 | 2 | 54.98115 | yes | 1.732E-07 | 557.2771 | 102.53583 |
| LHNQEQVIK | 9 | Q96SN8 | CDK5 regulatory subunit-associated prot‹ | CDK5RAP: | 215.038 | 2 | 54.98115 | yes | 1.008E-07 | 554.80908 | 94.556152 |
| AVESLAEAGGSEI( | 15 | Q96SN8 | CDK5 regulatory subunit-associated prot‹ | CDK5RAP: | 215.038 | 2 | 54.98115 | yes | 1.21E-07 | 758.88373 | 91.835205 |
| HNQQLIQEVR | 10 | Q96SN8 | CDK5 regulatory subunit-associated prot‹ | CDK5RAP: | 215.038 | 2 | 54.98115 | yes | 9.722E-08 | 632.84149 | 91.128571 |
| QLEQDVLSYQNLF | 13 | Q96SN8 | CDK5 regulatory subunit-associated prot‹ | CDK5RAP: | 215.038 | 2 | 54.98115 | yes | 9.733E-08 | 803.41522 | 87.029007 |
| SIETSSTLQSR | 11 | Q96SN8 | CDK5 regulatory subunit-associated prot‹ | CDK5RAP: | 215.038 | 2 | 54.98115 | yes | 1.376E-07 | 604.80951 | 70.576401 |
| QQLLSQNDK | 9 | Q96SN8 | CDK5 regulatory subunit-associated prot‹ | CDK5RAP: | 215.038 | 2 | 54.98115 | yes | 1.982E-07 | 537.2829 | 63.135063 |
| RLEESIK | 7 | Q96SN8 | CDK5 regulatory subunit-associated prot‹ | CDK5RAP: | 215.038 | 2 | 54.98115 | yes | 3.25E-06 | 437.75327 | 60.922176 |
| KVEELNSEIEK | 11 | Q96SN8 | CDK5 regulatory subunit-associated prot‹ | CDK5RAP: | 215.038 | 2 | 54.98115 | yes | 1.876E-07 | 659.34845 | 54.246552 |
| QSSDYEELIQVLK | 13 | Q96SN8 | CDK5 regulatory subunit-associated prot‹ | CDK5RAP: | 215.038 | 2 | 54.98115 | yes | 1.209E-07 | 776.39868 | 47.56395 |

| | | | | | | | | | | | |
|---|---|---|---|---|---|---|---|---|---|---|---|
| TSVMVNSFSQDLL | 21 | Q96SN8 | CDK5 regulatory subunit-associated prot | CDK5RAP | 215.038 | 3 | 54.98115 | yes | 1.38E-06 | 826.41028 | 41.550602 |
| KVSDLIQLVK | 10 | Q96SN8 | CDK5 regulatory subunit-associated prot | CDK5RAP | 215.038 | 2 | 54.98115 | yes | 3.314E-07 | 571.86078 | 25.019604 |
| PIFGLTSNYK | 10 | Q96ST2 | Protein IWS1 homolog OS=Homo sapien | IWS1 | 91.955 | 2 | 50.152782 | yes | 4.722E-07 | 570.30841 | 116.03897 |
| EWLSPLPDR | 9 | Q96ST2 | Protein IWS1 homolog OS=Homo sapien | IWS1 | 91.955 | 2 | 50.152782 | yes | 3.082E-07 | 556.79041 | 67.406921 |
| ILQELPSVSQETLK | 14 | Q96ST2 | Protein IWS1 homolog OS=Homo sapien | IWS1 | 91.955 | 2 | 50.152782 | yes | 9.885E-08 | 792.9458 | 36.421864 |
| WNVEMESSR | 9 | Q96ST2 | Protein IWS1 homolog OS=Homo sapien | IWS1 | 91.955 | 2 | 50.152782 | yes | 3.918E-06 | 569.2533 | 14.428243 |
| LLLEQLEIQQR | 11 | Q96ST8 | Centrosomal protein of 89 kDa OS=Hom | CEP89 | 89.59 | 2 | 28.739477 | yes | 2.677E-06 | 691.90375 | 40.617645 |
| VLEPVRPK | 8 | Q96SZ5 | 2-aminoethanethiol dioxygenase OS=Hor | ADO | 29.751 | 2 | 42.957672 | yes | 3.169E-07 | 469.2951 | 114.67572 |
| AEDLNIAPR | 9 | Q96SZ5 | 2-aminoethanethiol dioxygenase OS=Hor | ADO | 29.751 | 2 | 42.957672 | yes | 4.281E-07 | 499.76691 | 89.923523 |
| EASSSACDLPR | 11 | Q96SZ5 | 2-aminoethanethiol dioxygenase OS=Hor | ADO | 29.751 | 2 | 42.957672 | yes | 2.796E-07 | 596.76678 | 61.587387 |
| QACLTFR | 7 | Q96SZ5 | 2-aminoethanethiol dioxygenase OS=Hor | ADO | 29.751 | 2 | 42.957672 | yes | 4.152E-06 | 448.22635 | 50.806343 |
| MVDILAGPDAYR | 12 | Q96SZ6 | Mitochondrial tRNA methylthiotransfera | CDK5RAP | 67.689 | 2 | 52.002384 | yes | 1.015E-07 | 660.83447 | 260.72598 |
| LKEEILNR | 8 | Q96SZ6 | Mitochondrial tRNA methylthiotransfera | CDK5RAP | 67.689 | 2 | 52.002384 | yes | 4.346E-07 | 507.80075 | 191.80954 |
| FAHLLDQVSR | 10 | Q96SZ6 | Mitochondrial tRNA methylthiotransfera | CDK5RAP | 67.689 | 2 | 52.002384 | yes | 1.277E-07 | 593.32239 | 129.88622 |
| EVTLLGQNVNSFR | 13 | Q96SZ6 | Mitochondrial tRNA methylthiotransfera | CDK5RAP | 67.689 | 2 | 52.002384 | yes | 9.722E-08 | 738.89392 | 98.34256 |
| SRPIASILEEVK | 12 | Q96SZ6 | Mitochondrial tRNA methylthiotransfera | CDK5RAP | 67.689 | 2 | 52.002384 | yes | 9.975E-08 | 671.39044 | 37.367279 |
| SEEPPGTELQR | 11 | Q96T21 | Selenocysteine insertion sequence-bindin | SECISBP2 | 95.462 | 2 | 55.740772 | yes | 1.155E-07 | 621.8017 | 653.08221 |
| TPHNPLDSSAPLM | 14 | Q96T21 | Selenocysteine insertion sequence-bindin | SECISBP2 | 95.462 | 2 | 55.740772 | yes | 7.219E-07 | 754.38232 | 311.17554 |
| TPHNPLDSSAPLM | 14 | Q96T21 | Selenocysteine insertion sequence-bindin | SECISBP2 | 95.462 | 2 | 55.740772 | yes | 1.055E-07 | 503.25729 | 311.17554 |
| TPHNPLDSSAPLM | 14 | Q96T21 | Selenocysteine insertion sequence-bindin | SECISBP2 | 95.462 | 3 | 55.740772 | yes | 7.219E-07 | 754.38232 | 311.17554 |
| TPHNPLDSSAPLM | 14 | Q96T21 | Selenocysteine insertion sequence-bindin | SECISBP2 | 95.462 | 3 | 55.740772 | yes | 1.055E-07 | 503.25729 | 311.17554 |
| EVVKPAAVLSK | 11 | Q96T21 | Selenocysteine insertion sequence-bindin | SECISBP2 | 95.462 | 2 | 55.740772 | yes | 1.496E-07 | 570.85297 | 573.02917 |
| LSADVKPFVPR | 11 | Q96T21 | Selenocysteine insertion sequence-bindin | SECISBP2 | 95.462 | 2 | 55.740772 | yes | 9.903E-08 | 614.85608 | 252.13756 |
| LSADVKPFVPR | 11 | Q96T21 | Selenocysteine insertion sequence-bindin | SECISBP2 | 95.462 | 2 | 55.740772 | yes | 1.733E-07 | 410.23981 | 252.13756 |
| LSADVKPFVPR | 11 | Q96T21 | Selenocysteine insertion sequence-bindin | SECISBP2 | 95.462 | 3 | 55.740772 | yes | 9.903E-08 | 614.85608 | 252.13756 |
| LSADVKPFVPR | 11 | Q96T21 | Selenocysteine insertion sequence-bindin | SECISBP2 | 95.462 | 3 | 55.740772 | yes | 1.733E-07 | 410.23981 | 252.13756 |
| NENTCPLPQEMK | 12 | Q96T21 | Selenocysteine insertion sequence-bindin | SECISBP2 | 95.462 | 2 | 55.740772 | yes | 9.722E-08 | 730.82904 | 471.57684 |
| EVDACVTDLLK | 11 | Q96T21 | Selenocysteine insertion sequence-bindin | SECISBP2 | 95.462 | 2 | 55.740772 | yes | 1.066E-07 | 631.81848 | 450.58911 |
| MVELTVAAR | 9 | Q96T21 | Selenocysteine insertion sequence-bindin | SECISBP2 | 95.462 | 2 | 55.740772 | yes | 4.192E-07 | 495.27606 | 343.75418 |
| YEVLTVQEPPR | 11 | Q96T21 | Selenocysteine insertion sequence-bindin | SECISBP2 | 95.462 | 2 | 55.740772 | yes | 1.18E-07 | 665.8537 | 255.0452 |
| DYCSQMLSK | 9 | Q96T21 | Selenocysteine insertion sequence-bindin | SECISBP2 | 95.462 | 2 | 55.740772 | yes | 1.014E-07 | 566.24408 | 222.50542 |
| GSHHLSIYAENSLI | 14 | Q96T21 | Selenocysteine insertion sequence-bindin | SECISBP2 | 95.462 | 3 | 55.740772 | yes | 9.81E-08 | 519.26697 | 92.108521 |
| GSHHLSIYAENSLI | 14 | Q96T21 | Selenocysteine insertion sequence-bindin | SECISBP2 | 95.462 | 3 | 55.740772 | yes | 9.796E-08 | 778.39679 | 92.108521 |
| GSHHLSIYAENSLI | 14 | Q96T21 | Selenocysteine insertion sequence-bindin | SECISBP2 | 95.462 | 2 | 55.740772 | yes | 9.81E-08 | 519.26697 | 92.108521 |
| GSHHLSIYAENSLI | 14 | Q96T21 | Selenocysteine insertion sequence-bindin | SECISBP2 | 95.462 | 2 | 55.740772 | yes | 9.796E-08 | 778.39679 | 92.108521 |
| ELSWTPMGYVVR | 12 | Q96T21 | Selenocysteine insertion sequence-bindin | SECISBP2 | 95.462 | 2 | 55.740772 | yes | 9.722E-08 | 719.3634 | 178.36768 |
| QSSKPVVVSVGAV | 18 | Q96T21 | Selenocysteine insertion sequence-bindin | SECISBP2 | 95.462 | 3 | 55.740772 | yes | 5.879E-07 | 594.3559 | 129.94966 |
| WGPVHSVSTDISL | 15 | Q96T21 | Selenocysteine insertion sequence-bindin | SECISBP2 | 95.462 | 2 | 55.740772 | yes | 1.021E-07 | 833.94922 | 114.54935 |
| EKEEPHYIEIWK | 12 | Q96T21 | Selenocysteine insertion sequence-bindin | SECISBP2 | 95.462 | 2 | 55.740772 | yes | 6.815E-07 | 800.90393 | 53.62207 |

| | | | | | | | | | | | |
|---|---|---|---|---|---|---|---|---|---|---|---|
| EKEEPHYIEIWK | 12 | Q96T21 | Selenocysteine insertion sequence-bindin; | SECISBP2 | 95.462 | 2 | 55.740772 | yes | 3.721E-07 | 534.27173 | 53.62207 |
| EKEEPHYIEIWK | 12 | Q96T21 | Selenocysteine insertion sequence-bindin; | SECISBP2 | 95.462 | 3 | 55.740772 | yes | 6.815E-07 | 800.90393 | 53.62207 |
| EKEEPHYIEIWK | 12 | Q96T21 | Selenocysteine insertion sequence-bindin; | SECISBP2 | 95.462 | 3 | 55.740772 | yes | 3.721E-07 | 534.27173 | 53.62207 |
| DTEASHLAPNHTT | 16 | Q96T21 | Selenocysteine insertion sequence-bindin; | SECISBP2 | 95.462 | 3 | 55.740772 | yes | 9.722E-08 | 589.28833 | 103.41125 |
| PVVVSVGAVPVLS | 14 | Q96T21 | Selenocysteine insertion sequence-bindin; | SECISBP2 | 95.462 | 2 | 55.740772 | yes | 1.215E-07 | 675.92139 | 78.252007 |
| RLVLGLR | 7 | Q96T21 | Selenocysteine insertion sequence-bindin; | SECISBP2 | 95.462 | 2 | 55.740772 | yes | 1.488E-07 | 413.7847 | 69.946884 |
| QTLSTELSAAPK | 12 | Q96T21 | Selenocysteine insertion sequence-bindin; | SECISBP2 | 95.462 | 2 | 55.740772 | yes | 9.722E-08 | 623.33789 | 65.998306 |
| ASQGSDLEQNEAS | 14 | Q96T21 | Selenocysteine insertion sequence-bindin; | SECISBP2 | 95.462 | 2 | 55.740772 | yes | 9.722E-08 | 746.33716 | 64.751114 |
| NVTSMINLK | 9 | Q96T21 | Selenocysteine insertion sequence-bindin; | SECISBP2 | 95.462 | 2 | 55.740772 | yes | 5.241E-07 | 510.28134 | 62.34227 |
| EEPHYIEIWK | 10 | Q96T21 | Selenocysteine insertion sequence-bindin; | SECISBP2 | 95.462 | 2 | 55.740772 | yes | 1.203E-07 | 672.33514 | 52.50317 |
| AVPVSVVGIFSYD | 21 | Q96T21 | Selenocysteine insertion sequence-bindin; | SECISBP2 | 95.462 | 3 | 55.740772 | yes | 4.255E-06 | 755.38306 | 35.997368 |
| PQAPPSLPTQGPSC | 24 | Q96T21 | Selenocysteine insertion sequence-bindin; | SECISBP2 | 95.462 | 3 | 55.740772 | yes | 0.0001315 | 804.73364 | 29.252041 |
| IEAVYVSR | 8 | Q96T37 | RNA-binding protein 15 OS=Homo sapie | RBM15 | 107.189 | 2 | 53.054188 | yes | 9.722E-08 | 468.76108 | 305.13342 |
| EVDEISPEDDQR | 12 | Q96T37 | RNA-binding protein 15 OS=Homo sapie | RBM15 | 107.189 | 2 | 53.054188 | yes | 1.182E-07 | 716.31537 | 296.4805 |
| LPPPPPPPLPR | 11 | Q96T37 | RNA-binding protein 15 OS=Homo sapie | RBM15 | 107.189 | 2 | 53.054188 | yes | 7.587E-07 | 589.35822 | 264.68396 |
| VDFADTEHR | 9 | Q96T37 | RNA-binding protein 15 OS=Homo sapie | RBM15 | 107.189 | 2 | 53.054188 | yes | 3.586E-07 | 545.25165 | 263.84494 |
| FENLDMSHR | 9 | Q96T37 | RNA-binding protein 15 OS=Homo sapie | RBM15 | 107.189 | 2 | 53.054188 | yes | 9.88E-08 | 574.76129 | 242.97743 |
| VGAGAGAAPFR | 11 | Q96T37 | RNA-binding protein 15 OS=Homo sapie | RBM15 | 107.189 | 2 | 53.054188 | yes | 9.975E-08 | 487.26434 | 240.55017 |
| FGVITEVDIK | 10 | Q96T37 | RNA-binding protein 15 OS=Homo sapie | RBM15 | 107.189 | 2 | 53.054188 | yes | 1.145E-07 | 560.81604 | 223.79651 |
| LHSYSSPSTK | 10 | Q96T37 | RNA-binding protein 15 OS=Homo sapie | RBM15 | 107.189 | 2 | 53.054188 | yes | 1.273E-06 | 553.77747 | 210.92102 |
| LVLYDRPLK | 9 | Q96T37 | RNA-binding protein 15 OS=Homo sapie | RBM15 | 107.189 | 2 | 53.054188 | yes | 1.602E-07 | 558.84241 | 210.36862 |
| ISELGSQLSDEAVE | 22 | Q96T37 | RNA-binding protein 15 OS=Homo sapie | RBM15 | 107.189 | 3 | 53.054188 | yes | 1.735E-07 | 817.39557 | 183.51982 |
| GQTSTYGFLK | 10 | Q96T37 | RNA-binding protein 15 OS=Homo sapie | RBM15 | 107.189 | 2 | 53.054188 | yes | 3.034E-07 | 551.28241 | 172.28076 |
| QDGGTAPVASASP | 14 | Q96T37 | RNA-binding protein 15 OS=Homo sapie | RBM15 | 107.189 | 2 | 53.054188 | yes | 1.711E-07 | 643.32275 | 161.71327 |
| SLSPGGAALGYR | 12 | Q96T37 | RNA-binding protein 15 OS=Homo sapie | RBM15 | 107.189 | 2 | 53.054188 | yes | 1.699E-07 | 574.80658 | 137.29791 |
| QAAGVISLPVGGN | 14 | Q96T37 | RNA-binding protein 15 OS=Homo sapie | RBM15 | 107.189 | 2 | 53.054188 | yes | 5.574E-07 | 655.875 | 117.45157 |
| YQQQYLQPLPLTH | 24 | Q96T37 | RNA-binding protein 15 OS=Homo sapie | RBM15 | 107.189 | 4 | 53.054188 | yes | 1.065E-07 | 730.12122 | 109.68231 |
| LQQLALGR | 8 | Q96T37 | RNA-binding protein 15 OS=Homo sapie | RBM15 | 107.189 | 2 | 53.054188 | yes | 1.596E-06 | 449.77707 | 95.816071 |
| SDGSAPSTSTASSK | 14 | Q96T37 | RNA-binding protein 15 OS=Homo sapie | RBM15 | 107.189 | 2 | 53.054188 | yes | 9.722E-08 | 641.7915 | 33.16507 |
| RLPEESGGR | 9 | Q96T37 | RNA-binding protein 15 OS=Homo sapie | RBM15 | 107.189 | 2 | 53.054188 | yes | 2.69E-06 | 500.76215 | 32.787273 |
| SSSSSAASDTATST | 16 | Q96T37 | RNA-binding protein 15 OS=Homo sapie | RBM15 | 107.189 | 2 | 53.054188 | yes | 1.895E-07 | 772.34515 | 23.427025 |
| GLDDTVSIASR | 11 | Q96T58 | Msx2-interacting protein OS=Homo sapie | SPEN | 402.249 | 2 | 50.193634 | yes | 1.09E-07 | 567.29346 | 127.35044 |
| EQFTLPSVVHR | 11 | Q96T58 | Msx2-interacting protein OS=Homo sapie | SPEN | 402.249 | 2 | 50.193634 | yes | 1.02E-07 | 656.85406 | 124.8118 |
| HLWVGNLPENVR | 12 | Q96T58 | Msx2-interacting protein OS=Homo sapie | SPEN | 402.249 | 2 | 50.193634 | yes | 1.318E-07 | 717.38605 | 117.643 |
| ELQEAAAVPTTPR | 13 | Q96T58 | Msx2-interacting protein OS=Homo sapie | SPEN | 402.249 | 2 | 50.193634 | yes | 9.85E-08 | 691.86737 | 96.903885 |
| QLQVLDDQGPER | 12 | Q96T58 | Msx2-interacting protein OS=Homo sapie | SPEN | 402.249 | 2 | 50.193634 | yes | 1.066E-07 | 699.35461 | 76.795929 |
| SLPLSEGGPPLR | 12 | Q96T58 | Msx2-interacting protein OS=Homo sapie | SPEN | 402.249 | 2 | 50.193634 | yes | 5.114E-07 | 611.84314 | 68.983109 |
| VLPYSNITVR | 10 | Q96T58 | Msx2-interacting protein OS=Homo sapie | SPEN | 402.249 | 2 | 50.193634 | yes | 9.623E-07 | 581.33496 | 64.703094 |
| VTSVQIHGTSEER | 13 | Q96T58 | Msx2-interacting protein OS=Homo sapie | SPEN | 402.249 | 2 | 50.193634 | yes | 3.763E-07 | 721.86536 | 48.127666 |

| | | | | | | | | | | | |
|---|---|---|---|---|---|---|---|---|---|---|---|
| DGLFHEFK | 8 | Q96T58 | Msx2-interacting protein OS=Homo sapie | SPEN | 402.249 | 2 | 50.193634 | yes | 2.835E-06 | 496.74545 | 46.694771 |
| DCQELASISVGSGS | 15 | Q96T58 | Msx2-interacting protein OS=Homo sapie | SPEN | 402.249 | 2 | 50.193634 | yes | 9.215E-07 | 783.36487 | 35.656322 |
| SLVSTPAGPVNVL | 14 | Q96T58 | Msx2-interacting protein OS=Homo sapie | SPEN | 402.249 | 2 | 50.193634 | yes | 1.386E-06 | 691.40613 | 34.161888 |
| EENWSFLDWDSR | 12 | Q96T58 | Msx2-interacting protein OS=Homo sapie | SPEN | 402.249 | 2 | 50.193634 | yes | 4.676E-06 | 792.34167 | 30.325565 |
| FGEIVDIDIK | 10 | Q96T58 | Msx2-interacting protein OS=Homo sapie | SPEN | 402.249 | 2 | 50.193634 | yes | 2.455E-06 | 574.81354 | 29.907375 |
| TQVELVADPETR | 12 | Q96T60 | Bifunctional polynucleotide phosphatase/ | PNKP | 57.077 | 2 | 48.666714 | yes | 1.117E-07 | 679.3515 | 225.60753 |
| VAGFDLDGTLITTI | 14 | Q96T60 | Bifunctional polynucleotide phosphatase/ | PNKP | 57.077 | 2 | 48.666714 | yes | 1.482E-07 | 739.89612 | 204.10138 |
| ILYPEIPR | 8 | Q96T60 | Bifunctional polynucleotide phosphatase/ | PNKP | 57.077 | 2 | 48.666714 | yes | 1.541E-07 | 500.79492 | 149.4032 |
| SNPGWENLEK | 10 | Q96T60 | Bifunctional polynucleotide phosphatase/ | PNKP | 57.077 | 2 | 48.666714 | yes | 1.088E-06 | 587.2804 | 144.48007 |
| CFLFTATLEQAR | 12 | Q96T60 | Bifunctional polynucleotide phosphatase/ | PNKP | 57.077 | 2 | 48.666714 | yes | 2.142E-07 | 728.86627 | 106.57775 |
| SGPLCLPESR | 10 | Q96T60 | Bifunctional polynucleotide phosphatase/ | PNKP | 57.077 | 2 | 48.666714 | yes | 2.891E-07 | 558.2793 | 105.88293 |
| VAIDNTNPDAASR | 13 | Q96T60 | Bifunctional polynucleotide phosphatase/ | PNKP | 57.077 | 2 | 48.666714 | yes | 1.709E-07 | 672.33112 | 54.153236 |
| LVIFTNQMSIGR | 12 | Q96T60 | Bifunctional polynucleotide phosphatase/ | PNKP | 57.077 | 2 | 48.666714 | yes | 2.881E-06 | 689.87921 | 40.100693 |
| LNDTIQLLVR | 10 | Q96T88 | E3 ubiquitin-protein ligase UHRF1 OS=H | UHRF1 | 89.814 | 2 | 55.049015 | yes | 1.462E-07 | 592.85352 | 1720.4203 |
| LWNEVLASLK | 10 | Q96T88 | E3 ubiquitin-protein ligase UHRF1 OS=H | UHRF1 | 89.814 | 2 | 55.049015 | yes | 2.482E-07 | 586.83734 | 1369.2322 |
| VEPYSLTAQQSSLI | 15 | Q96T88 | E3 ubiquitin-protein ligase UHRF1 OS=H | UHRF1 | 89.814 | 2 | 55.049015 | yes | 9.128E-07 | 846.45178 | 1348.6207 |
| IQELFHVEPGLQR | 13 | Q96T88 | E3 ubiquitin-protein ligase UHRF1 OS=H | UHRF1 | 89.814 | 2 | 55.049015 | yes | 2.035E-07 | 783.42535 | 657.03894 |
| IQELFHVEPGLQR | 13 | Q96T88 | E3 ubiquitin-protein ligase UHRF1 OS=H | UHRF1 | 89.814 | 2 | 55.049015 | yes | 3.255E-07 | 522.61932 | 657.03894 |
| IQELFHVEPGLQR | 13 | Q96T88 | E3 ubiquitin-protein ligase UHRF1 OS=H | UHRF1 | 89.814 | 3 | 55.049015 | yes | 2.035E-07 | 783.42535 | 657.03894 |
| IQELFHVEPGLQR | 13 | Q96T88 | E3 ubiquitin-protein ligase UHRF1 OS=H | UHRF1 | 89.814 | 3 | 55.049015 | yes | 3.255E-07 | 522.61932 | 657.03894 |
| DTNMGAWFEAQV | 14 | Q96T88 | E3 ubiquitin-protein ligase UHRF1 OS=H | UHRF1 | 89.814 | 2 | 55.049015 | yes | 2.436E-07 | 812.38281 | 1131.9618 |
| NDASEVVLAGER | 12 | Q96T88 | E3 ubiquitin-protein ligase UHRF1 OS=H | UHRF1 | 89.814 | 2 | 55.049015 | yes | 3.119E-07 | 630.31494 | 1057.2686 |
| GFWYDAEISR | 10 | Q96T88 | E3 ubiquitin-protein ligase UHRF1 OS=H | UHRF1 | 89.814 | 2 | 55.049015 | yes | 9.722E-08 | 622.29077 | 1055.7185 |
| DEPCSSTSRPALEE | 21 | Q96T88 | E3 ubiquitin-protein ligase UHRF1 OS=H | UHRF1 | 89.814 | 3 | 55.049015 | yes | 1.48E-07 | 811.38477 | 997.28986 |
| VCACHLCGGR | 10 | Q96T88 | E3 ubiquitin-protein ligase UHRF1 OS=H | UHRF1 | 89.814 | 2 | 55.049015 | yes | 1.003E-07 | 595.25482 | 917.12927 |
| QTHTVDSLSR | 10 | Q96T88 | E3 ubiquitin-protein ligase UHRF1 OS=H | UHRF1 | 89.814 | 2 | 55.049015 | yes | 9.722E-08 | 572.29126 | 808.4411 |
| QSLVLPHSTK | 10 | Q96T88 | E3 ubiquitin-protein ligase UHRF1 OS=H | UHRF1 | 89.814 | 2 | 55.049015 | yes | 9.722E-08 | 555.31934 | 676.16956 |
| IIFVDEVFK | 9 | Q96T88 | E3 ubiquitin-protein ligase UHRF1 OS=H | UHRF1 | 89.814 | 2 | 55.049015 | yes | 1.348E-07 | 555.31567 | 540.15094 |
| PITTVCQHNVCK | 12 | Q96T88 | E3 ubiquitin-protein ligase UHRF1 OS=H | UHRF1 | 89.814 | 2 | 55.049015 | yes | 9.722E-08 | 728.85541 | 264.34753 |
| KIQELFHVEPGLQI | 14 | Q96T88 | E3 ubiquitin-protein ligase UHRF1 OS=H | UHRF1 | 89.814 | 3 | 55.049015 | yes | 2.708E-07 | 565.31763 | 125.6984 |
| KIQELFHVEPGLQI | 14 | Q96T88 | E3 ubiquitin-protein ligase UHRF1 OS=H | UHRF1 | 89.814 | 3 | 55.049015 | yes | 1.658E-07 | 847.47284 | 125.6984 |
| KIQELFHVEPGLQI | 14 | Q96T88 | E3 ubiquitin-protein ligase UHRF1 OS=H | UHRF1 | 89.814 | 2 | 55.049015 | yes | 2.708E-07 | 565.31763 | 125.6984 |
| KIQELFHVEPGLQI | 14 | Q96T88 | E3 ubiquitin-protein ligase UHRF1 OS=H | UHRF1 | 89.814 | 2 | 55.049015 | yes | 1.658E-07 | 847.47284 | 125.6984 |
| VNEYVDAR | 8 | Q96T88 | E3 ubiquitin-protein ligase UHRF1 OS=H | UHRF1 | 89.814 | 2 | 55.049015 | yes | 9.728E-08 | 483.23798 | 250.76744 |
| YAPAEGNR | 8 | Q96T88 | E3 ubiquitin-protein ligase UHRF1 OS=H | UHRF1 | 89.814 | 2 | 55.049015 | yes | 1.955E-06 | 439.21176 | 235.2103 |
| VQVSESGVHR | 10 | Q96T88 | E3 ubiquitin-protein ligase UHRF1 OS=H | UHRF1 | 89.814 | 2 | 55.049015 | yes | 6.453E-07 | 549.28851 | 192.22194 |
| TKECTIVPSNHYGI | 25 | Q96T88 | E3 ubiquitin-protein ligase UHRF1 OS=H | UHRF1 | 89.814 | 4 | 55.049015 | yes | 9.722E-08 | 703.3584 | 189.14804 |
| RDDDEPGPWTK | 11 | Q96T88 | E3 ubiquitin-protein ligase UHRF1 OS=H | UHRF1 | 89.814 | 2 | 55.049015 | yes | 1.01E-07 | 658.29932 | 169.04428 |
| KLGLTMQYPEGYI | 19 | Q96T88 | E3 ubiquitin-protein ligase UHRF1 OS=H | UHRF1 | 89.814 | 3 | 55.049015 | yes | 1.484E-07 | 723.04541 | 137.51161 |

| | | | | | | | | | | | |
|---|---|---|---|---|---|---|---|---|---|---|---|
| DRPASGSPFQLFLS | 15 | Q96T88 | E3 ubiquitin-protein ligase UHRF1 OS=H | UHRF1 | 89.814 | 2 | 55.049015 | yes | 1.551E-07 | 825.43591 | 44.61961 |
| SSTHGEAAAETDS | 30 | Q96T88 | E3 ubiquitin-protein ligase UHRF1 OS=H | UHRF1 | 89.814 | 4 | 55.049015 | yes | 3.767E-07 | 828.61249 | 43.586643 |
| VQVSESGVHRPHV | 19 | Q96T88 | E3 ubiquitin-protein ligase UHRF1 OS=H | UHRF1 | 89.814 | 4 | 55.049015 | yes | 5.731E-07 | 506.27396 | 28.390966 |
| PGEGSPMVDNPMI | 13 | Q96T88 | E3 ubiquitin-protein ligase UHRF1 OS=H | UHRF1 | 89.814 | 2 | 55.049015 | yes | 0.000111 | 693.81067 | 25.323517 |
| DDDEPGPWTK | 10 | Q96T88 | E3 ubiquitin-protein ligase UHRF1 OS=H | UHRF1 | 89.814 | 2 | 55.049015 | yes | 4.908E-06 | 580.24872 | 20.724401 |
| DEPCSSTSR | 9 | Q96T88 | E3 ubiquitin-protein ligase UHRF1 OS=H | UHRF1 | 89.814 | 2 | 55.049015 | yes | 2.493E-07 | 519.71149 | 18.464176 |
| LAETQNIAPSFVK | 13 | Q96TA2 | ATP-dependent zinc metalloprotease YM | YME1L1 | 86.456 | 2 | 53.05342 | yes | 9.722E-08 | 709.38794 | 882.84686 |
| NIPEAHQDAFK | 11 | Q96TA2 | ATP-dependent zinc metalloprotease YM | YME1L1 | 86.456 | 2 | 53.05342 | yes | 4.702E-07 | 635.31476 | 657.659 |
| FDQSVDPEIIAR | 12 | Q96TA2 | ATP-dependent zinc metalloprotease YM | YME1L1 | 86.456 | 2 | 53.05342 | yes | 1.061E-07 | 695.35406 | 634.48346 |
| LSPETQSAIEQEIR | 14 | Q96TA2 | ATP-dependent zinc metalloprotease YM | YME1L1 | 86.456 | 2 | 53.05342 | yes | 2.429E-07 | 800.91248 | 470.96133 |
| TTTGLDSAVDPVQ | 15 | Q96TA2 | ATP-dependent zinc metalloprotease YM | YME1L1 | 86.456 | 2 | 53.05342 | yes | 9.851E-08 | 781.89014 | 470.87717 |
| LGVMTYSDTGK | 11 | Q96TA2 | ATP-dependent zinc metalloprotease YM | YME1L1 | 86.456 | 2 | 53.05342 | yes | 9.722E-08 | 586.2868 | 438.52637 |
| GILLVGPPGTGK | 12 | Q96TA2 | ATP-dependent zinc metalloprotease YM | YME1L1 | 86.456 | 2 | 53.05342 | yes | 9.823E-08 | 554.83984 | 421.38818 |
| FDMQVTVPRPDVI | 13 | Q96TA2 | ATP-dependent zinc metalloprotease YM | YME1L1 | 86.456 | 2 | 53.05342 | yes | 3.464E-07 | 766.40051 | 330.68176 |
| QELQEVVEFLK | 11 | Q96TA2 | ATP-dependent zinc metalloprotease YM | YME1L1 | 86.456 | 2 | 53.05342 | yes | 9.722E-08 | 681.3692 | 260.01141 |
| EIQIVLEGK | 9 | Q96TA2 | ATP-dependent zinc metalloprotease YM | YME1L1 | 86.456 | 2 | 53.05342 | yes | 1.446E-07 | 514.80298 | 158.55466 |
| AQLLAQMDVSMG | 14 | Q96TA2 | ATP-dependent zinc metalloprotease YM | YME1L1 | 86.456 | 2 | 53.05342 | yes | 9.722E-08 | 738.86853 | 155.82472 |
| NVTFEHVK | 8 | Q96TA2 | ATP-dependent zinc metalloprotease YM | YME1L1 | 86.456 | 2 | 53.05342 | yes | 2.541E-06 | 487.25873 | 150.397 |
| GPTLGHVSLLPENI | 15 | Q96TA2 | ATP-dependent zinc metalloprotease YM | YME1L1 | 86.456 | 2 | 53.05342 | yes | 9.733E-08 | 802.92316 | 101.9106 |
| NLAEALLTYETLD | 15 | Q96TA2 | ATP-dependent zinc metalloprotease YM | YME1L1 | 86.456 | 2 | 53.05342 | yes | 8.274E-07 | 832.94073 | 34.420269 |
| GLAGEIVGEVR | 11 | Q96TC7 | Regulator of microtubule dynamics protei | RMDN3 | 52.119 | 2 | 49.117664 | yes | 1.066E-07 | 550.30896 | 374.91171 |
| AVPGGAGDASVLI | 17 | Q96TC7 | Regulator of microtubule dynamics protei | RMDN3 | 52.119 | 2 | 49.117664 | yes | 1.487E-07 | 782.4281 | 273.29526 |
| EGFQLLLNNK | 10 | Q96TC7 | Regulator of microtubule dynamics protei | RMDN3 | 52.119 | 2 | 49.117664 | yes | 4.44E-07 | 588.32458 | 125.61168 |
| LALELPDVTK | 10 | Q96TC7 | Regulator of microtubule dynamics protei | RMDN3 | 52.119 | 2 | 49.117664 | yes | 1.37E-07 | 549.82391 | 69.68631 |
| LDDAIHVLR | 9 | Q99081 | Transcription factor 12 OS=Homo sapiens | TCF12 | 72.965 | 2 | 46.3097 | no | 9.014E-07 | 526.2984 | 63.538361 |
| GGLQSQSGTVVTT | 16 | Q99081 | Transcription factor 12 OS=Homo sapiens | TCF12 | 72.965 | 2 | 46.3097 | yes | 1.594E-07 | 802.92816 | 36.443897 |
| LAQYEPPQEEK | 11 | Q99417 | c-Myc-binding protein OS=Homo sapiens | MYCBP | 11.967 | 2 | 53.82309 | yes | 1.259E-07 | 666.32751 | 350.39316 |
| LELAEMK | 7 | Q99417 | c-Myc-binding protein OS=Homo sapiens | MYCBP | 11.967 | 2 | 53.82309 | yes | 9.722E-08 | 417.22549 | 169.80954 |
| YEAIVEENKK | 10 | Q99417 | c-Myc-binding protein OS=Homo sapiens | MYCBP | 11.967 | 2 | 53.82309 | yes | 9.722E-08 | 611.81934 | 152.95653 |
| VLVALYEEPEKPN | 20 | Q99417 | c-Myc-binding protein OS=Homo sapiens | MYCBP | 11.967 | 3 | 53.82309 | yes | 2.257E-07 | 759.07886 | 48.889572 |
| EKYEAIVEENKK | 12 | Q99417 | c-Myc-binding protein OS=Homo sapiens | MYCBP | 11.967 | 2 | 53.82309 | yes | 1.722E-06 | 740.38812 | 35.279808 |
| YTISQEAYDQR | 11 | Q99426 | Tubulin-folding cofactor B OS=Homo sap | TBCB | 27.326 | 2 | 51.079414 | yes | 9.722E-08 | 687.32025 | 33.952263 |
| DGTILCTLMNK | 11 | Q99439 | Calponin-2 OS=Homo sapiens OX=9606 | CNN2 | 33.698 | 2 | 53.883564 | yes | 1.513E-07 | 633.31506 | 9810.7129 |
| GLQSGVDIGVK | 11 | Q99439 | Calponin-2 OS=Homo sapiens OX=9606 | CNN2 | 33.698 | 2 | 53.883564 | yes | 2.249E-07 | 536.80347 | 6908.4209 |
| GPSYGLSAEVK | 11 | Q99439 | Calponin-2 OS=Homo sapiens OX=9606 | CNN2 | 33.698 | 2 | 53.883564 | no | 2.314E-07 | 554.28766 | 4734.811 |
| AGQCVIGLQMGTN | 14 | Q99439 | Calponin-2 OS=Homo sapiens OX=9606 | CNN2 | 33.698 | 2 | 53.883564 | yes | 1.284E-07 | 738.86853 | 1132.9949 |
| AGQCVIGLQMGTN | 14 | Q99439 | Calponin-2 OS=Homo sapiens OX=9606 | CNN2 | 33.698 | 2 | 53.883564 | yes | 1.164E-07 | 746.86597 | 1132.9949 |
| CASQSGMTAYGTI | 13 | Q99439 | Calponin-2 OS=Homo sapiens OX=9606 | CNN2 | 33.698 | 2 | 53.883564 | yes | 9.722E-08 | 695.29791 | 533.96094 |
| CASQSGMTAYGTI | 13 | Q99439 | Calponin-2 OS=Homo sapiens OX=9606 | CNN2 | 33.698 | 2 | 53.883564 | yes | 9.722E-08 | 703.29535 | 533.96094 |

| | | | | | | | | | | | |
|---|---|---|---|---|---|---|---|---|---|---|---|
| NFDDATMK | 8 | Q99439 | Calponin-2 OS=Homo sapiens OX=9606 | CNN2 | 33.698 | 2 | 53.883564 | yes | 2.143E-07 | 471.20529 | 577.58185 |
| CASQVGMTAPGTI | 13 | Q99439 | Calponin-2 OS=Homo sapiens OX=9606 | CNN2 | 33.698 | 2 | 53.883564 | yes | 9.722E-08 | 668.31085 | 74.814537 |
| NDFEIVLPENAEK | 13 | Q99459 | Cell division cycle 5-like protein OS=Ho | CDC5L | 92.251 | 2 | 51.81507 | yes | 9.722E-08 | 759.37775 | 1969.0879 |
| ILLGGYQSR | 9 | Q99459 | Cell division cycle 5-like protein OS=Ho | CDC5L | 92.251 | 2 | 51.81507 | yes | 1.33E-07 | 503.78763 | 1856.6708 |
| AQDVLVQEMEVV | 13 | Q99459 | Cell division cycle 5-like protein OS=Ho | CDC5L | 92.251 | 2 | 51.81507 | yes | 1.012E-07 | 744.39233 | 1482.4028 |
| LGLLGLPAPK | 10 | Q99459 | Cell division cycle 5-like protein OS=Ho | CDC5L | 92.251 | 2 | 51.81507 | yes | 2.28E-07 | 489.82095 | 1124.6628 |
| WYEWLDPSIK | 10 | Q99459 | Cell division cycle 5-like protein OS=Ho | CDC5L | 92.251 | 2 | 51.81507 | yes | 9.85E-08 | 668.83228 | 932.16541 |
| IASLLHR | 7 | Q99459 | Cell division cycle 5-like protein OS=Ho | CDC5L | 92.251 | 2 | 51.81507 | yes | 2.636E-07 | 405.25323 | 889.34705 |
| VGQASEIAR | 9 | Q99459 | Cell division cycle 5-like protein OS=Ho | CDC5L | 92.251 | 2 | 51.81507 | yes | 3.224E-07 | 465.75378 | 786.18842 |
| GVDYNAEIPFEK | 12 | Q99459 | Cell division cycle 5-like protein OS=Ho | CDC5L | 92.251 | 2 | 51.81507 | yes | 9.722E-08 | 691.33533 | 708.02209 |
| LKPGEIDPNPETK | 13 | Q99459 | Cell division cycle 5-like protein OS=Ho | CDC5L | 92.251 | 2 | 51.81507 | yes | 9.722E-08 | 719.38281 | 494.06558 |
| DNEEETTDDPR | 11 | Q99459 | Cell division cycle 5-like protein OS=Ho | CDC5L | 92.251 | 2 | 51.81507 | yes | 1.874E-07 | 660.76294 | 490.48254 |
| HEDSAIPR | 8 | Q99459 | Cell division cycle 5-like protein OS=Ho | CDC5L | 92.251 | 2 | 51.81507 | yes | 4.035E-07 | 462.73032 | 451.12616 |
| QQDLDGELR | 9 | Q99459 | Cell division cycle 5-like protein OS=Ho | CDC5L | 92.251 | 2 | 51.81507 | yes | 2.926E-07 | 537.26471 | 410.36652 |
| YADLLLEK | 8 | Q99459 | Cell division cycle 5-like protein OS=Ho | CDC5L | 92.251 | 2 | 51.81507 | yes | 1.018E-07 | 482.77112 | 207.66089 |
| DRIESLEK | 8 | Q99459 | Cell division cycle 5-like protein OS=Ho | CDC5L | 92.251 | 2 | 51.81507 | yes | 9.809E-08 | 495.26672 | 159.31219 |
| LMPTQWR | 7 | Q99459 | Cell division cycle 5-like protein OS=Ho | CDC5L | 92.251 | 2 | 51.81507 | yes | 2.232E-07 | 466.24457 | 124.54816 |
| TPSNGAEGLTPR | 12 | Q99459 | Cell division cycle 5-like protein OS=Ho | CDC5L | 92.251 | 2 | 51.81507 | yes | 5.39E-07 | 600.30438 | 81.789551 |
| QLNDLWDQIEQAI | 17 | Q99459 | Cell division cycle 5-like protein OS=Ho | CDC5L | 92.251 | 3 | 51.81507 | yes | 2.185E-07 | 707.69562 | 73.404114 |
| EMITMLHYDLLHI | 21 | Q99459 | Cell division cycle 5-like protein OS=Ho | CDC5L | 92.251 | 3 | 51.81507 | yes | 9.437E-07 | 842.40326 | 63.040489 |
| SGTTPKPVINSTPG | 15 | Q99459 | Cell division cycle 5-like protein OS=Ho | CDC5L | 92.251 | 2 | 51.81507 | yes | 2.022E-07 | 756.41248 | 61.850731 |
| PLNVEPPLTDLQK | 13 | Q99459 | Cell division cycle 5-like protein OS=Ho | CDC5L | 92.251 | 2 | 51.81507 | yes | 1.389E-07 | 732.40887 | 41.923012 |
| RGVDYNAEIPFEK | 13 | Q99459 | Cell division cycle 5-like protein OS=Ho | CDC5L | 92.251 | 2 | 51.81507 | yes | 6.355E-07 | 769.38593 | 32.395294 |
| DKLNINPEDGMAI | 21 | Q99459 | Cell division cycle 5-like protein OS=Ho | CDC5L | 92.251 | 3 | 51.81507 | yes | 9.722E-08 | 791.03015 | 29.590401 |
| GGLNTPLHESDFS | 19 | Q99459 | Cell division cycle 5-like protein OS=Ho | CDC5L | 92.251 | 3 | 51.81507 | yes | 3.278E-07 | 671.33258 | 11.895288 |
| KIDFLTK | 7 | Q99471 | Prefoldin subunit 5 OS=Homo sapiens O | PFDN5 | 17.329 | 2 | 44.306644 | yes | 4.565E-07 | 432.76309 | 42.851864 |
| IQQLTALGAAQAT | 15 | Q99471 | Prefoldin subunit 5 OS=Homo sapiens O | PFDN5 | 17.329 | 2 | 44.306644 | yes | 3.794E-07 | 742.92517 | 19.801483 |
| QAVMEMMSQK | 10 | Q99471 | Prefoldin subunit 5 OS=Homo sapiens O | PFDN5 | 17.329 | 2 | 44.306644 | yes | 2.244E-07 | 591.76941 | 19.767508 |
| FCADCIITALR | 11 | Q99496 | E3 ubiquitin-protein ligase RING2 OS=H | RNF2 | 37.656 | 2 | 48.102863 | yes | 1.199E-07 | 670.32849 | 306.3566 |
| TWELSLYELQR | 11 | Q99496 | E3 ubiquitin-protein ligase RING2 OS=H | RNF2 | 37.656 | 2 | 48.102863 | yes | 1.551E-07 | 719.37225 | 240.74872 |
| TSGNATVDHLSK | 12 | Q99496 | E3 ubiquitin-protein ligase RING2 OS=H | RNF2 | 37.656 | 2 | 48.102863 | yes | 3.799E-07 | 615.30969 | 111.09302 |
| SLRPDPNFDALISK | 14 | Q99496 | E3 ubiquitin-protein ligase RING2 OS=H | RNF2 | 37.656 | 2 | 48.102863 | no | 1.456E-07 | 786.92267 | 81.846794 |
| GESNQMNLDTASI | 14 | Q99496 | E3 ubiquitin-protein ligase RING2 OS=H | RNF2 | 37.656 | 2 | 48.102863 | yes | 7.23E-07 | 762.33575 | 66.780052 |
| VNKPMELYYAPTI | 13 | Q99496 | E3 ubiquitin-protein ligase RING2 OS=H | RNF2 | 37.656 | 2 | 48.102863 | yes | 1.198E-06 | 777.40527 | 57.059307 |
| HNNQQALSHSIEE | 16 | Q99496 | E3 ubiquitin-protein ligase RING2 OS=H | RNF2 | 37.656 | 3 | 48.102863 | yes | 3.548E-05 | 602.30273 | 36.865337 |
| DVVICPDASLEDA | 14 | Q99497 | Parkinson disease protein 7 OS=Homo sa | PARK7 | 19.892 | 2 | 43.109215 | yes | 2.728E-07 | 766.36908 | 103.25717 |
| GAEEMETVIPVDV | 15 | Q99497 | Parkinson disease protein 7 OS=Homo sa | PARK7 | 19.892 | 2 | 43.109215 | yes | 1.479E-06 | 838.40515 | 36.719646 |
| DNVATFIGWCR | 11 | Q99501 | GAS2-like protein 1 OS=Homo sapiens C | GAS2L1 | 72.717 | 2 | 54.234924 | yes | 3.201E-07 | 669.81659 | 198.7942 |
| VGDSSLLIFVR | 11 | Q99501 | GAS2-like protein 1 OS=Homo sapiens C | GAS2L1 | 72.717 | 2 | 54.234924 | yes | 9.974E-08 | 603.34808 | 142.8347 |

| | | | | | | | | | | | |
|---|---|---|---|---|---|---|---|---|---|---|---|
| ADPVAGIAGSAAK | 13 | Q99501 | GAS2-like protein 1 OS=Homo sapiens C | GAS2L1 | 72.717 | 2 | 54.234924 | yes | 1.979E-07 | 585.31171 | 125.47694 |
| LTTGTPASPR | 10 | Q99501 | GAS2-like protein 1 OS=Homo sapiens C | GAS2L1 | 72.717 | 2 | 54.234924 | yes | 1.584E-07 | 500.77472 | 94.482224 |
| EEQAVLLVR | 9 | Q99501 | GAS2-like protein 1 OS=Homo sapiens C | GAS2L1 | 72.717 | 2 | 54.234924 | yes | 2.014E-07 | 528.80603 | 82.581375 |
| LVQFEQEIER | 10 | Q99501 | GAS2-like protein 1 OS=Homo sapiens C | GAS2L1 | 72.717 | 2 | 54.234924 | yes | 9.722E-08 | 645.83807 | 80.012093 |
| EGPETPPRPR | 10 | Q99501 | GAS2-like protein 1 OS=Homo sapiens C | GAS2L1 | 72.717 | 2 | 54.234924 | yes | 1.195E-07 | 568.29639 | 52.364292 |
| NLDELVR | 7 | Q99501 | GAS2-like protein 1 OS=Homo sapiens C | GAS2L1 | 72.717 | 2 | 54.234924 | yes | 1.21E-06 | 429.73761 | 42.117661 |
| VGGGWDTLEHYL | 14 | Q99501 | GAS2-like protein 1 OS=Homo sapiens C | GAS2L1 | 72.717 | 2 | 54.234924 | yes | 4.146E-06 | 795.38336 | 26.487236 |
| MLLLPSAADGR | 11 | Q99543 | DnaJ homolog subfamily C member 2 OS | DNAJC2 | 71.997 | 2 | 52.956882 | yes | 2.625E-07 | 572.31317 | 498.06104 |
| AAQEQVLNASR | 11 | Q99543 | DnaJ homolog subfamily C member 2 OS | DNAJC2 | 71.997 | 2 | 52.956882 | yes | 2.975E-07 | 593.81238 | 206.6474 |
| TLVDNAYSCDPR | 12 | Q99543 | DnaJ homolog subfamily C member 2 OS | DNAJC2 | 71.997 | 2 | 52.956882 | yes | 9.722E-08 | 705.81952 | 161.58168 |
| EGDNDYFTCITK | 12 | Q99543 | DnaJ homolog subfamily C member 2 OS | DNAJC2 | 71.997 | 2 | 52.956882 | yes | 9.722E-08 | 731.8114 | 158.6076 |
| QIEEINEQIR | 10 | Q99543 | DnaJ homolog subfamily C member 2 OS | DNAJC2 | 71.997 | 2 | 52.956882 | yes | 9.727E-08 | 636.33313 | 123.59868 |
| EFSYLDEEEKEK | 12 | Q99543 | DnaJ homolog subfamily C member 2 OS | DNAJC2 | 71.997 | 2 | 52.956882 | yes | 1.026E-07 | 773.35138 | 85.616318 |
| NWSEDDLQLLIK | 12 | Q99543 | DnaJ homolog subfamily C member 2 OS | DNAJC2 | 71.997 | 2 | 52.956882 | yes | 1.901E-07 | 737.38281 | 77.945633 |
| AYEMLSDPVK | 10 | Q99543 | DnaJ homolog subfamily C member 2 OS | DNAJC2 | 71.997 | 2 | 52.956882 | yes | 3.102E-07 | 576.78394 | 68.315521 |
| NQDHYAVLGLGH | 14 | Q99543 | DnaJ homolog subfamily C member 2 OS | DNAJC2 | 71.997 | 3 | 52.956882 | yes | 2.439E-07 | 526.94269 | 50.619583 |
| TWNHFSDNEAER | 12 | Q99543 | DnaJ homolog subfamily C member 2 OS | DNAJC2 | 71.997 | 2 | 52.956882 | yes | 7.647E-07 | 753.32385 | 34.411518 |
| NASASFQELEDK | 12 | Q99543 | DnaJ homolog subfamily C member 2 OS | DNAJC2 | 71.997 | 2 | 52.956882 | yes | 2.257E-07 | 669.81226 | 13.938761 |
| RYETLVGTIGK | 11 | Q99547 | M-phase phosphoprotein 6 OS=Homo sa | MPHOSPH | 19.024 | 2 | 44.958557 | yes | 2.012E-07 | 618.85095 | 80.406471 |
| LMLQMNAK | 8 | Q99547 | M-phase phosphoprotein 6 OS=Homo sa | MPHOSPH | 19.024 | 2 | 44.958557 | yes | 4.55E-07 | 474.75391 | 50.766609 |
| LMPVSAQTPK | 10 | Q99549 | M-phase phosphoprotein 8 OS=Homo sa | MPHOSPH | 97.183 | 2 | 50.321835 | yes | 2.038E-06 | 536.297 | 190.62404 |
| LVIECGADCNILSK | 14 | Q99549 | M-phase phosphoprotein 8 OS=Homo sa | MPHOSPH | 97.183 | 2 | 50.321835 | yes | 1.066E-07 | 796.39453 | 115.24212 |
| LALLEPVFPIACHR | 14 | Q99549 | M-phase phosphoprotein 8 OS=Homo sa | MPHOSPH | 97.183 | 3 | 50.321835 | yes | 2.082E-07 | 545.97284 | 48.490417 |
| LALLEPVFPIACHR | 14 | Q99549 | M-phase phosphoprotein 8 OS=Homo sa | MPHOSPH | 97.183 | 3 | 50.321835 | yes | 1.376E-07 | 818.45563 | 48.490417 |
| LALLEPVFPIACHR | 14 | Q99549 | M-phase phosphoprotein 8 OS=Homo sa | MPHOSPH | 97.183 | 2 | 50.321835 | yes | 2.082E-07 | 545.97284 | 48.490417 |
| LALLEPVFPIACHR | 14 | Q99549 | M-phase phosphoprotein 8 OS=Homo sa | MPHOSPH | 97.183 | 2 | 50.321835 | yes | 1.376E-07 | 818.45563 | 48.490417 |
| QSNNVLVYDLLK | 12 | Q99549 | M-phase phosphoprotein 8 OS=Homo sa | MPHOSPH | 97.183 | 2 | 50.321835 | yes | 9.819E-08 | 703.38794 | 54.755024 |
| VLDSSISTLK | 10 | Q99551 | Transcription termination factor 1, mitoch | MTERF1 | 45.779 | 2 | 54.234642 | yes | 1.104E-07 | 531.80573 | 183.61594 |
| NPFILIQSTK | 10 | Q99551 | Transcription termination factor 1, mitoch | MTERF1 | 45.779 | 2 | 54.234642 | yes | 9.722E-08 | 580.83734 | 134.34993 |
| LFSLGCTEEEVQK | 13 | Q99551 | Transcription termination factor 1, mitoch | MTERF1 | 45.779 | 2 | 54.234642 | yes | 1.695E-07 | 770.37158 | 73.633194 |
| NLLTMGVDIDMAI | 13 | Q99551 | Transcription termination factor 1, mitoch | MTERF1 | 45.779 | 2 | 54.234642 | yes | 1.521E-07 | 724.86542 | 56.720291 |
| QMVEFLQAAGLSI | 23 | Q99551 | Transcription termination factor 1, mitoch | MTERF1 | 45.779 | 3 | 54.234642 | yes | 2.45E-07 | 839.08478 | 43.260174 |
| SNNNLNLENNIK | 12 | Q99551 | Transcription termination factor 1, mitoch | MTERF1 | 45.779 | 2 | 54.234642 | yes | 1.351E-06 | 693.85223 | 39.356407 |
| IVTSDLEIVNILER | 14 | Q99551 | Transcription termination factor 1, mitoch | MTERF1 | 45.779 | 2 | 54.234642 | yes | 6.558E-06 | 807.45905 | 25.863525 |
| VLAETPDSFEK | 11 | Q99567 | Nuclear pore complex protein Nup88 OS= | NUP88 | 83.542 | 2 | 54.59195 | yes | 9.722E-08 | 618.31134 | 102.22831 |
| ELQLIPDQLR | 10 | Q99567 | Nuclear pore complex protein Nup88 OS= | NUP88 | 83.542 | 2 | 54.59195 | yes | 1.7E-07 | 612.85095 | 90.057182 |
| EDVEVAESPLR | 11 | Q99567 | Nuclear pore complex protein Nup88 OS= | NUP88 | 83.542 | 2 | 54.59195 | yes | 9.722E-08 | 622.31189 | 83.886917 |
| FFTSSTSLTLK | 11 | Q99567 | Nuclear pore complex protein Nup88 OS= | NUP88 | 83.542 | 2 | 54.59195 | yes | 5.876E-07 | 616.33209 | 56.755318 |
| VGSPLTLTDAQTR | 13 | Q99569 | Plakophilin-4 OS=Homo sapiens OX=96 | PKP4 | 131.869 | 2 | 52.9077 | yes | 9.722E-08 | 679.86737 | 494.53146 |

| | | | | | | | | | | |
|---|---|---|---|---|---|---|---|---|---|---|
| EQELQFQR | 8 | Q99569 | Plakophilin-4 OS=Homo sapiens OX=96( | PKP4 | 131.869 | 2 | 52.9077 | yes | 2.709E-06 | 539.26984 | 265.37997 |
| VVSSVATALR | 10 | Q99569 | Plakophilin-4 OS=Homo sapiens OX=96( | PKP4 | 131.869 | 2 | 52.9077 | yes | 3.465E-07 | 501.80075 | 262.89554 |
| LELEVPQAR | 9 | Q99569 | Plakophilin-4 OS=Homo sapiens OX=96( | PKP4 | 131.869 | 2 | 52.9077 | yes | 9.722E-08 | 527.79822 | 243.65335 |
| VSDAVQPNNYLIR | 13 | Q99569 | Plakophilin-4 OS=Homo sapiens OX=96( | PKP4 | 131.869 | 2 | 52.9077 | yes | 6.015E-07 | 744.89392 | 230.58809 |
| TGSVGIGNLQR | 11 | Q99569 | Plakophilin-4 OS=Homo sapiens OX=96( | PKP4 | 131.869 | 2 | 52.9077 | yes | 2.995E-07 | 551.3042 | 224.13297 |
| STTNYVDFYSTK | 12 | Q99569 | Plakophilin-4 OS=Homo sapiens OX=96( | PKP4 | 131.869 | 2 | 52.9077 | yes | 9.722E-08 | 713.33026 | 206.6881 |
| LGAESPSIASTSSTI | 16 | Q99569 | Plakophilin-4 OS=Homo sapiens OX=96( | PKP4 | 131.869 | 2 | 52.9077 | yes | 1.167E-07 | 782.88867 | 179.21536 |
| AQSPSYVISTGVSF | 16 | Q99569 | Plakophilin-4 OS=Homo sapiens OX=96( | PKP4 | 131.869 | 2 | 52.9077 | yes | 4.168E-07 | 818.42047 | 148.16757 |
| TSLGSGFGSPSVTI | 16 | Q99569 | Plakophilin-4 OS=Homo sapiens OX=96( | PKP4 | 131.869 | 2 | 52.9077 | yes | 9.722E-08 | 782.88373 | 146.83342 |
| PSPIYISSYSSPAR | 14 | Q99569 | Plakophilin-4 OS=Homo sapiens OX=96( | PKP4 | 131.869 | 2 | 52.9077 | yes | 1.537E-07 | 762.88831 | 137.55464 |
| AEGQTLVQPSVAN | 14 | Q99569 | Plakophilin-4 OS=Homo sapiens OX=96( | PKP4 | 131.869 | 2 | 52.9077 | yes | 9.722E-08 | 735.38898 | 132.73105 |
| QIVASQLER | 9 | Q99569 | Plakophilin-4 OS=Homo sapiens OX=96( | PKP4 | 131.869 | 2 | 52.9077 | no | 3.892E-07 | 522.29584 | 129.21368 |
| TVENCVCTLR | 10 | Q99569 | Plakophilin-4 OS=Homo sapiens OX=96( | PKP4 | 131.869 | 2 | 52.9077 | yes | 2.462E-07 | 626.29468 | 123.69186 |
| STDVPNTGVSKPR | 13 | Q99569 | Plakophilin-4 OS=Homo sapiens OX=96( | PKP4 | 131.869 | 2 | 52.9077 | yes | 1.084E-07 | 679.35718 | 105.97034 |
| VASPSQGQVGSSS | 15 | Q99569 | Plakophilin-4 OS=Homo sapiens OX=96( | PKP4 | 131.869 | 2 | 52.9077 | yes | 1.266E-07 | 708.35986 | 72.379997 |
| AAAQVLNTLWQY | 13 | Q99569 | Plakophilin-4 OS=Homo sapiens OX=96( | PKP4 | 131.869 | 2 | 52.9077 | yes | 9.722E-08 | 767.41223 | 66.166321 |
| NNYALNTTATYAI | 21 | Q99569 | Plakophilin-4 OS=Homo sapiens OX=96( | PKP4 | 131.869 | 3 | 52.9077 | yes | 4.99E-07 | 840.41528 | 60.23156 |
| PLNPSAYSSTTLPA | 16 | Q99569 | Plakophilin-4 OS=Homo sapiens OX=96( | PKP4 | 131.869 | 2 | 52.9077 | yes | 1.791E-07 | 823.43085 | 55.763256 |
| VVTLLSDPENIVK | 13 | Q99570 | Phosphoinositide 3-kinase regulatory sub | PIK3R4 | 153.104 | 2 | 55.0753 | yes | 9.722E-08 | 713.91119 | 114.16656 |
| VFAIQDPTLPLTSY | 15 | Q99570 | Phosphoinositide 3-kinase regulatory sub | PIK3R4 | 153.104 | 2 | 55.0753 | yes | 1.025E-07 | 846.96399 | 100.05523 |
| LSVEILLDR | 9 | Q99570 | Phosphoinositide 3-kinase regulatory sub | PIK3R4 | 153.104 | 2 | 55.0753 | yes | 1.098E-07 | 529.31622 | 83.130493 |
| ELVTQMIHR | 9 | Q99570 | Phosphoinositide 3-kinase regulatory sub | PIK3R4 | 153.104 | 2 | 55.0753 | yes | 9.722E-08 | 563.80548 | 67.27755 |
| TELQQLIQQK | 10 | Q99570 | Phosphoinositide 3-kinase regulatory sub | PIK3R4 | 153.104 | 2 | 55.0753 | yes | 7.115E-07 | 614.84845 | 30.406033 |
| FWDLAYPER | 9 | Q99570 | Phosphoinositide 3-kinase regulatory sub | PIK3R4 | 153.104 | 2 | 55.0753 | yes | 8.374E-06 | 598.79041 | 28.222153 |
| LCVFFGR | 7 | Q99570 | Phosphoinositide 3-kinase regulatory sub | PIK3R4 | 153.104 | 2 | 55.0753 | yes | 8.068E-07 | 449.73383 | 25.878971 |
| KLVLLSVLK | 9 | Q99570 | Phosphoinositide 3-kinase regulatory sub | PIK3R4 | 153.104 | 2 | 55.0753 | yes | 1.53E-07 | 506.86008 | 23.472536 |
| GSDQEVIQTGKPPI | 14 | Q99570 | Phosphoinositide 3-kinase regulatory sub | PIK3R4 | 153.104 | 2 | 55.0753 | yes | 1.063E-07 | 756.39429 | 10.700191 |
| LLSQGMTEEEEDK | 18 | Q99570 | Phosphoinositide 3-kinase regulatory sub | PIK3R4 | 153.104 | 3 | 55.0753 | yes | 9.938E-07 | 683.02576 | 7.9072585 |
| TDPFCPIFR | 9 | Q99571 | P2X purinoceptor 4 OS=Homo sapiens O | P2RX4 | 43.37 | 2 | 55.31819 | yes | 9.722E-08 | 576.77893 | 207.39194 |
| GYQETDSVVSSVT | 15 | Q99571 | P2X purinoceptor 4 OS=Homo sapiens O | P2RX4 | 43.37 | 2 | 55.31819 | yes | 9.722E-08 | 800.88867 | 107.94547 |
| FDIIVFGK | 8 | Q99571 | P2X purinoceptor 4 OS=Homo sapiens O | P2RX4 | 43.37 | 2 | 55.31819 | yes | 5.095E-07 | 469.77094 | 46.350601 |
| IPILLIQQPGK | 11 | Q99575 | Ribonucleases P/MRP protein subunit PC | POP1 | 114.709 | 2 | 52.823086 | yes | 9.926E-08 | 610.39227 | 646.47784 |
| SSNSLVFQTLPR | 12 | Q99575 | Ribonucleases P/MRP protein subunit PC | POP1 | 114.709 | 2 | 52.823086 | yes | 9.722E-08 | 674.86462 | 571.14142 |
| YITASTFAQAR | 11 | Q99575 | Ribonucleases P/MRP protein subunit PC | POP1 | 114.709 | 2 | 52.823086 | yes | 9.841E-08 | 614.81964 | 427.14307 |
| AGPEGTSQEIPK | 12 | Q99575 | Ribonucleases P/MRP protein subunit PC | POP1 | 114.709 | 2 | 52.823086 | yes | 1.058E-07 | 607.30658 | 414.52875 |
| AACQCVEPIK | 10 | Q99575 | Ribonucleases P/MRP protein subunit PC | POP1 | 114.709 | 2 | 52.823086 | yes | 9.722E-08 | 588.28101 | 348.42542 |
| ALSGMCNIDTGLT | 23 | Q99575 | Ribonucleases P/MRP protein subunit PC | POP1 | 114.709 | 3 | 52.823086 | yes | 8.875E-07 | 808.38873 | 333.52609 |
| EACLSILGHFPR | 12 | Q99575 | Ribonucleases P/MRP protein subunit PC | POP1 | 114.709 | 2 | 52.823086 | yes | 1.401E-07 | 700.36121 | 328.48477 |
| VTLHCSR | 7 | Q99575 | Ribonucleases P/MRP protein subunit PC | POP1 | 114.709 | 2 | 52.823086 | yes | 1.295E-07 | 436.724 | 297.99255 |

| | | | | | | | | | | | |
|---|---|---|---|---|---|---|---|---|---|---|---|
| RLQEIAQK | 8 | Q99575 | Ribonucleases P/MRP protein subunit PC | POP1 | 114.709 | 2 | 52.823086 | yes | 9.722E-08 | 493.29309 | 251.23451 |
| GKEEEILK | 8 | Q99575 | Ribonucleases P/MRP protein subunit PC | POP1 | 114.709 | 2 | 52.823086 | yes | 2.051E-07 | 473.2662 | 190.13243 |
| ITDQEASENHVAA | 22 | Q99575 | Ribonucleases P/MRP protein subunit PC | POP1 | 114.709 | 3 | 52.823086 | yes | 1.431E-07 | 803.39227 | 190.06996 |
| ISDQDLNR | 8 | Q99575 | Ribonucleases P/MRP protein subunit PC | POP1 | 114.709 | 2 | 52.823086 | yes | 2.935E-07 | 480.74088 | 160.98424 |
| NIWLETHIWHAK | 12 | Q99575 | Ribonucleases P/MRP protein subunit PC | POP1 | 114.709 | 2 | 52.823086 | yes | 9.81E-08 | 774.40955 | 147.71741 |
| QGSLVLYR | 8 | Q99575 | Ribonucleases P/MRP protein subunit PC | POP1 | 114.709 | 2 | 52.823086 | yes | 9.821E-08 | 468.26907 | 142.31882 |
| KPDSVSLHCR | 10 | Q99575 | Ribonucleases P/MRP protein subunit PC | POP1 | 114.709 | 2 | 52.823086 | yes | 6.761E-07 | 599.80347 | 124.88008 |
| QLSAWCGPSSEDS | 14 | Q99575 | Ribonucleases P/MRP protein subunit PC | POP1 | 114.709 | 2 | 52.823086 | yes | 1.101E-07 | 790.34393 | 110.9342 |
| SEVPCAPMPK | 10 | Q99575 | Ribonucleases P/MRP protein subunit PC | POP1 | 114.709 | 2 | 52.823086 | yes | 7.466E-07 | 558.26483 | 82.458054 |
| QDILEEIK | 8 | Q99575 | Ribonucleases P/MRP protein subunit PC | POP1 | 114.709 | 2 | 52.823086 | yes | 1.807E-07 | 494.27148 | 81.731155 |
| SELLVPGSQLILGP | 18 | Q99575 | Ribonucleases P/MRP protein subunit PC | POP1 | 114.709 | 3 | 52.823086 | yes | 2.841E-07 | 635.35443 | 76.215797 |
| ESAVHSQYK | 9 | Q99575 | Ribonucleases P/MRP protein subunit PC | POP1 | 114.709 | 2 | 52.823086 | yes | 6.437E-07 | 524.75653 | 72.082092 |
| GSPEPHTMICVPAI | 14 | Q99575 | Ribonucleases P/MRP protein subunit PC | POP1 | 114.709 | 2 | 52.823086 | yes | 6.019E-07 | 762.37091 | 38.592197 |
| HHSGGEKPFQAQK | 13 | Q99575 | Ribonucleases P/MRP protein subunit PC | POP1 | 114.709 | 3 | 52.823086 | yes | 7.428E-07 | 484.24438 | 28.60709 |
| RSEVPCAPMPK | 11 | Q99575 | Ribonucleases P/MRP protein subunit PC | POP1 | 114.709 | 2 | 52.823086 | yes | 9.722E-08 | 636.31537 | 27.188347 |
| NESLTEHPR | 9 | Q99590 | Protein SCAF11 OS=Homo sapiens OX= | SCAF11 | 164.653 | 2 | 58.16271 | yes | 1.096E-07 | 541.76489 | 647.50653 |
| TLPADVQNYYSR | 12 | Q99590 | Protein SCAF11 OS=Homo sapiens OX= | SCAF11 | 164.653 | 2 | 58.16271 | yes | 2.988E-07 | 713.85168 | 638.82581 |
| QDQISGLSQSEVK | 13 | Q99590 | Protein SCAF11 OS=Homo sapiens OX= | SCAF11 | 164.653 | 2 | 58.16271 | yes | 2.034E-07 | 709.85974 | 332.83603 |
| ITESSLVEITEHK | 13 | Q99590 | Protein SCAF11 OS=Homo sapiens OX= | SCAF11 | 164.653 | 2 | 58.16271 | yes | 1.649E-07 | 743.39337 | 305.46915 |
| FVEQQSYK | 8 | Q99590 | Protein SCAF11 OS=Homo sapiens OX= | SCAF11 | 164.653 | 2 | 58.16271 | yes | 1.173E-07 | 514.75598 | 284.58057 |
| TLEEPVSTEK | 10 | Q99590 | Protein SCAF11 OS=Homo sapiens OX= | SCAF11 | 164.653 | 2 | 58.16271 | yes | 9.722E-08 | 566.79022 | 233.17624 |
| SCNEQIEESEK | 11 | Q99590 | Protein SCAF11 OS=Homo sapiens OX= | SCAF11 | 164.653 | 2 | 58.16271 | yes | 9.722E-08 | 676.78534 | 231.68063 |
| VETVSQPSESPK | 12 | Q99590 | Protein SCAF11 OS=Homo sapiens OX= | SCAF11 | 164.653 | 2 | 58.16271 | yes | 1.612E-07 | 644.32501 | 157.17296 |
| LESSEGEIIQTVDR | 14 | Q99590 | Protein SCAF11 OS=Homo sapiens OX= | SCAF11 | 164.653 | 2 | 58.16271 | yes | 9.722E-08 | 788.39667 | 139.41414 |
| SGWTSASSWAVR | 12 | Q99590 | Protein SCAF11 OS=Homo sapiens OX= | SCAF11 | 164.653 | 2 | 58.16271 | yes | 3.285E-07 | 647.81238 | 101.44591 |
| NDCPNWITEK | 10 | Q99590 | Protein SCAF11 OS=Homo sapiens OX= | SCAF11 | 164.653 | 2 | 58.16271 | yes | 9.722E-08 | 638.78497 | 89.175423 |
| KLQIQEK | 7 | Q99590 | Protein SCAF11 OS=Homo sapiens OX= | SCAF11 | 164.653 | 2 | 58.16271 | yes | 9.867E-08 | 443.77145 | 86.62117 |
| ESHWEENR | 8 | Q99590 | Protein SCAF11 OS=Homo sapiens OX= | SCAF11 | 164.653 | 2 | 58.16271 | yes | 2.803E-06 | 543.73358 | 71.360275 |
| VSVAVEASADSSK | 13 | Q99590 | Protein SCAF11 OS=Homo sapiens OX= | SCAF11 | 164.653 | 2 | 58.16271 | yes | 8.506E-07 | 625.31714 | 64.628815 |
| NENSGNSWNK | 10 | Q99590 | Protein SCAF11 OS=Homo sapiens OX= | SCAF11 | 164.653 | 2 | 58.16271 | yes | 1.138E-07 | 575.24963 | 57.061626 |
| SEQEFSFDTPADR | 13 | Q99590 | Protein SCAF11 OS=Homo sapiens OX= | SCAF11 | 164.653 | 2 | 58.16271 | yes | 9.722E-08 | 764.83118 | 50.65744 |
| NDIHLDADDPNSA | 15 | Q99590 | Protein SCAF11 OS=Homo sapiens OX= | SCAF11 | 164.653 | 2 | 58.16271 | yes | 3.59E-07 | 820.36334 | 44.553394 |
| KTLEEPVSTEK | 11 | Q99590 | Protein SCAF11 OS=Homo sapiens OX= | SCAF11 | 164.653 | 2 | 58.16271 | yes | 4.116E-07 | 630.83771 | 24.590635 |
| LAIKPFYQNK | 10 | Q99590 | Protein SCAF11 OS=Homo sapiens OX= | SCAF11 | 164.653 | 3 | 58.16271 | yes | 9.722E-08 | 407.90451 | 12.352505 |
| SFQQELDAR | 9 | Q99598 | Translin-associated protein X OS=Homo | TSNAX | 33.113 | 2 | 55.11568 | yes | 9.508E-07 | 547.26727 | 108.75931 |
| DVNSSSPVMLAFK | 13 | Q99598 | Translin-associated protein X OS=Homo | TSNAX | 33.113 | 2 | 55.11568 | yes | 9.722E-08 | 697.85284 | 99.071152 |
| SLISMDEINK | 10 | Q99598 | Translin-associated protein X OS=Homo | TSNAX | 33.113 | 2 | 55.11568 | yes | 4.929E-07 | 575.29462 | 91.179192 |
| HMLADVFSVK | 10 | Q99598 | Translin-associated protein X OS=Homo | TSNAX | 33.113 | 2 | 55.11568 | yes | 9.881E-08 | 573.80243 | 33.915573 |
| EVGEAFTILSDPK | 13 | Q99615 | DnaJ homolog subfamily C member 7 OS | DNAJC7 | 56.441 | 2 | 55.633038 | yes | 1.401E-07 | 703.36414 | 356.59491 |

| | | | | | | | | | | | |
|---|---|---|---|---|---|---|---|---|---|---|---|
| AECLAMLGR | 9 | Q99615 | DnaJ homolog subfamily C member 7 OS | DNAJC7 | 56.441 | 2 | 55.633038 | yes | 1.16E-07 | 510.75189 | 261.172 |
| MDSTNADALYVR | 12 | Q99615 | DnaJ homolog subfamily C member 7 OS | DNAJC7 | 56.441 | 2 | 55.633038 | yes | 1.668E-07 | 678.31665 | 236.60596 |
| EALGDAQQSVR | 11 | Q99615 | DnaJ homolog subfamily C member 7 OS | DNAJC7 | 56.441 | 2 | 55.633038 | yes | 2.191E-07 | 587.29657 | 173.80525 |
| YPEAQSVASDILR | 13 | Q99615 | DnaJ homolog subfamily C member 7 OS | DNAJC7 | 56.441 | 2 | 55.633038 | yes | 9.722E-08 | 724.87262 | 126.25713 |
| NANAVMEYEK | 10 | Q99615 | DnaJ homolog subfamily C member 7 OS | DNAJC7 | 56.441 | 2 | 55.633038 | yes | 1.102E-07 | 584.7688 | 122.63029 |
| LDDSFVR | 7 | Q99615 | DnaJ homolog subfamily C member 7 OS | DNAJC7 | 56.441 | 2 | 55.633038 | yes | 2.543E-07 | 426.21652 | 88.258438 |
| AVQFFVQALR | 10 | Q99615 | DnaJ homolog subfamily C member 7 OS | DNAJC7 | 56.441 | 2 | 55.633038 | yes | 9.873E-08 | 589.83765 | 86.477135 |
| LDDAIEDCTNAVK | 13 | Q99615 | DnaJ homolog subfamily C member 7 OS | DNAJC7 | 56.441 | 2 | 55.633038 | yes | 9.722E-08 | 732.33777 | 50.00729 |
| GLCLYYEDCIEK | 12 | Q99615 | DnaJ homolog subfamily C member 7 OS | DNAJC7 | 56.441 | 2 | 55.633038 | yes | 1.737E-07 | 781.84692 | 50.001743 |
| VVFCMDR | 7 | Q99615 | DnaJ homolog subfamily C member 7 OS | DNAJC7 | 56.441 | 2 | 55.633038 | yes | 2.769E-07 | 463.71478 | 23.246307 |
| KDYNEAYNYYTK | 12 | Q99615 | DnaJ homolog subfamily C member 7 OS | DNAJC7 | 56.441 | 2 | 55.633038 | yes | 3.38E-06 | 786.35425 | 17.045048 |
| SYEAQDPEIASLSC | 15 | Q99622 | Protein C10 OS=Homo sapiens OX=960( | C12orf57 | 13.179 | 2 | 49.136063 | yes | 1.065E-07 | 797.88336 | 116.80617 |
| ESVFTVEGGHR | 11 | Q99623 | Prohibitin-2 OS=Homo sapiens OX=960( | PHB2 | 33.297 | 2 | 54.850883 | yes | 7.141E-07 | 609.29907 | 203.89462 |
| IGGVQQDTILAEGI | 17 | Q99623 | Prohibitin-2 OS=Homo sapiens OX=960( | PHB2 | 33.297 | 3 | 54.850883 | yes | 9.722E-08 | 618.66711 | 190.28105 |
| IVQAEGEAEAAK | 12 | Q99623 | Prohibitin-2 OS=Homo sapiens OX=960( | PHB2 | 33.297 | 2 | 54.850883 | yes | 9.722E-08 | 608.31439 | 189.12186 |
| LGLDYEER | 8 | Q99623 | Prohibitin-2 OS=Homo sapiens OX=960( | PHB2 | 33.297 | 2 | 54.850883 | yes | 1.408E-07 | 497.74564 | 163.48889 |
| FNASQLITQR | 10 | Q99623 | Prohibitin-2 OS=Homo sapiens OX=960( | PHB2 | 33.297 | 2 | 54.850883 | yes | 1.081E-07 | 589.31982 | 130.98784 |
| VLPSIVNEVLK | 11 | Q99623 | Prohibitin-2 OS=Homo sapiens OX=960( | PHB2 | 33.297 | 2 | 54.850883 | yes | 5.875E-07 | 605.8739 | 105.33043 |
| LLLGAGAVAYGVI | 13 | Q99623 | Prohibitin-2 OS=Homo sapiens OX=960( | PHB2 | 33.297 | 2 | 54.850883 | yes | 1.217E-07 | 630.37714 | 72.418846 |
| AQVSLLIR | 8 | Q99623 | Prohibitin-2 OS=Homo sapiens OX=960( | PHB2 | 33.297 | 2 | 54.850883 | yes | 2.695E-07 | 450.28726 | 66.37439 |
| PNAQELPSMYQR | 12 | Q99623 | Prohibitin-2 OS=Homo sapiens OX=960( | PHB2 | 33.297 | 2 | 54.850883 | yes | 1.887E-07 | 717.3457 | 29.34137 |
| IPELAINPLGDR | 12 | Q99653 | Calcineurin B homologous protein 1 OS= | CHP1 | 22.457 | 2 | 49.624725 | yes | 1.017E-07 | 654.36951 | 249.16905 |
| DELLQVLR | 8 | Q99653 | Calcineurin B homologous protein 1 OS= | CHP1 | 22.457 | 2 | 49.624725 | yes | 7.059E-07 | 493.28748 | 65.379471 |
| DEELEEIKK | 9 | Q99653 | Calcineurin B homologous protein 1 OS= | CHP1 | 22.457 | 2 | 49.624725 | yes | 5.029E-07 | 566.79022 | 44.494465 |
| DNLPLQENVTIQK | 13 | Q99661 | Kinesin-like protein KIF2C OS=Homo sa | KIF2C | 81.313 | 2 | 53.869373 | yes | 1.673E-07 | 756.40686 | 1509.7405 |
| PLVQTIFEGGK | 11 | Q99661 | Kinesin-like protein KIF2C OS=Homo sa | KIF2C | 81.313 | 2 | 53.869373 | yes | 2.817E-07 | 594.83478 | 883.57892 |
| MIDMGSACR | 9 | Q99661 | Kinesin-like protein KIF2C OS=Homo sa | KIF2C | 81.313 | 2 | 53.869373 | yes | 1.977E-07 | 520.71973 | 739.04077 |
| FSLVDLAGNER | 11 | Q99661 | Kinesin-like protein KIF2C OS=Homo sa | KIF2C | 81.313 | 2 | 53.869373 | no | 5.202E-07 | 610.81714 | 699.22668 |
| AESALAQQAK | 10 | Q99661 | Kinesin-like protein KIF2C OS=Homo sa | KIF2C | 81.313 | 2 | 53.869373 | yes | 1.475E-07 | 508.77219 | 661.20361 |
| MSTVSELR | 8 | Q99661 | Kinesin-like protein KIF2C OS=Homo sa | KIF2C | 81.313 | 2 | 53.869373 | yes | 1.083E-07 | 461.73676 | 499.92352 |
| SHACFQIILR | 10 | Q99661 | Kinesin-like protein KIF2C OS=Homo sa | KIF2C | 81.313 | 2 | 53.869373 | yes | 1.274E-07 | 622.83203 | 476.00049 |
| TSGQTFANSNSSR | 13 | Q99661 | Kinesin-like protein KIF2C OS=Homo sa | KIF2C | 81.313 | 2 | 53.869373 | yes | 9.722E-08 | 678.81055 | 365.11639 |
| LAMQLEEQASR | 11 | Q99661 | Kinesin-like protein KIF2C OS=Homo sa | KIF2C | 81.313 | 2 | 53.869373 | yes | 1.14E-07 | 638.32172 | 338.93576 |
| THTMGGDLSGK | 11 | Q99661 | Kinesin-like protein KIF2C OS=Homo sa | KIF2C | 81.313 | 2 | 53.869373 | yes | 1.093E-07 | 552.26111 | 326.8324 |
| ATCFAYGQTGSGK | 13 | Q99661 | Kinesin-like protein KIF2C OS=Homo sa | KIF2C | 81.313 | 2 | 53.869373 | yes | 9.722E-08 | 674.30353 | 276.63168 |
| SCVSVEWAEGGA | 14 | Q99661 | Kinesin-like protein KIF2C OS=Homo sa | KIF2C | 81.313 | 2 | 53.869373 | yes | 1.067E-07 | 740.84045 | 259.75482 |
| EIDVISIPSK | 10 | Q99661 | Kinesin-like protein KIF2C OS=Homo sa | KIF2C | 81.313 | 2 | 53.869373 | yes | 2.082E-07 | 550.81354 | 256.94952 |
| KEIDVISIPSK | 11 | Q99661 | Kinesin-like protein KIF2C OS=Homo sa | KIF2C | 81.313 | 2 | 53.869373 | yes | 9.722E-08 | 614.86102 | 166.93613 |
| MVSEEMEEQVHSI | 14 | Q99661 | Kinesin-like protein KIF2C OS=Homo sa | KIF2C | 81.313 | 3 | 53.869373 | yes | 9.722E-08 | 568.59576 | 131.37405 |

| | | | | | | | | | | | |
|---|---|---|---|---|---|---|---|---|---|---|---|
| PSCPAVAEIPLR | 12 | Q99661 | Kinesin-like protein KIF2C OS=Homo sa | KIF2C | 81.313 | 2 | 53.869373 | yes | 9.722E-08 | 655.35028 | 102.46526 |
| GIYAMASR | 8 | Q99661 | Kinesin-like protein KIF2C OS=Homo sa | KIF2C | 81.313 | 2 | 53.869373 | yes | 4.044E-07 | 434.72092 | 91.039436 |
| SNGLIHSANVR | 11 | Q99661 | Kinesin-like protein KIF2C OS=Homo sa | KIF2C | 81.313 | 2 | 53.869373 | yes | 9.722E-08 | 584.31506 | 73.814438 |
| GSSSANPVNSVR | 12 | Q99661 | Kinesin-like protein KIF2C OS=Homo sa | KIF2C | 81.313 | 2 | 53.869373 | yes | 9.722E-08 | 587.79419 | 44.393131 |
| CLLLVHEPK | 9 | Q99661 | Kinesin-like protein KIF2C OS=Homo sa | KIF2C | 81.313 | 2 | 53.869373 | yes | 3.037E-06 | 554.81281 | 16.7729 |
| MVHILTSVVGSK | 12 | Q99700 | Ataxin-2 OS=Homo sapiens OX=9606 G | ATXN2 | 140.283 | 2 | 51.310287 | yes | 9.819E-08 | 635.86304 | 4979.082 |
| MVHILTSVVGSK | 12 | Q99700 | Ataxin-2 OS=Homo sapiens OX=9606 G | ATXN2 | 140.283 | 2 | 51.310287 | yes | 1.372E-06 | 643.86047 | 4979.082 |
| GISPVVSEHR | 10 | Q99700 | Ataxin-2 OS=Homo sapiens OX=9606 G | ATXN2 | 140.283 | 2 | 51.310287 | yes | 2.354E-07 | 540.79346 | 7479.7173 |
| ANQLAEEIESSAQ | 15 | Q99700 | Ataxin-2 OS=Homo sapiens OX=9606 G | ATXN2 | 140.283 | 2 | 51.310287 | yes | 1.531E-07 | 840.90741 | 6161.2598 |
| TSPSGGTWSSVVS | 17 | Q99700 | Ataxin-2 OS=Homo sapiens OX=9606 G | ATXN2 | 140.283 | 2 | 51.310287 | yes | 1.301E-07 | 830.91809 | 6033.8589 |
| CSDFVVVQFK | 10 | Q99700 | Ataxin-2 OS=Homo sapiens OX=9606 G | ATXN2 | 140.283 | 2 | 51.310287 | yes | 1.183E-07 | 614.80518 | 4721.772 |
| CDLVLDAAHEK | 11 | Q99700 | Ataxin-2 OS=Homo sapiens OX=9606 G | ATXN2 | 140.283 | 2 | 51.310287 | yes | 1.084E-07 | 635.80847 | 3798.7671 |
| STLNPNAK | 8 | Q99700 | Ataxin-2 OS=Homo sapiens OX=9606 G | ATXN2 | 140.283 | 2 | 51.310287 | no | 1.832E-07 | 422.7298 | 3747.7991 |
| VALENDDR | 8 | Q99700 | Ataxin-2 OS=Homo sapiens OX=9606 G | ATXN2 | 140.283 | 2 | 51.310287 | yes | 1.236E-07 | 466.2276 | 2631.7979 |
| DAFTDSAISAK | 11 | Q99700 | Ataxin-2 OS=Homo sapiens OX=9606 G | ATXN2 | 140.283 | 2 | 51.310287 | yes | 9.757E-08 | 563.27478 | 1657.6418 |
| ENIKPNETSPSFSK | 14 | Q99700 | Ataxin-2 OS=Homo sapiens OX=9606 G | ATXN2 | 140.283 | 2 | 51.310287 | yes | 1.041E-07 | 789.39392 | 706.24426 |
| ENIKPNETSPSFSK | 14 | Q99700 | Ataxin-2 OS=Homo sapiens OX=9606 G | ATXN2 | 140.283 | 2 | 51.310287 | yes | 1.13E-07 | 526.59839 | 706.24426 |
| ENIKPNETSPSFSK | 14 | Q99700 | Ataxin-2 OS=Homo sapiens OX=9606 G | ATXN2 | 140.283 | 3 | 51.310287 | yes | 1.041E-07 | 789.39392 | 706.24426 |
| ENIKPNETSPSFSK | 14 | Q99700 | Ataxin-2 OS=Homo sapiens OX=9606 G | ATXN2 | 140.283 | 3 | 51.310287 | yes | 1.13E-07 | 526.59839 | 706.24426 |
| AVPNMPQQR | 9 | Q99700 | Ataxin-2 OS=Homo sapiens OX=9606 G | ATXN2 | 140.283 | 2 | 51.310287 | yes | 3.607E-07 | 520.76892 | 1206.9028 |
| QIDDLKK | 7 | Q99700 | Ataxin-2 OS=Homo sapiens OX=9606 G | ATXN2 | 140.283 | 2 | 51.310287 | yes | 1.909E-07 | 430.2478 | 850.33893 |
| REEIMESILFK | 11 | Q99700 | Ataxin-2 OS=Homo sapiens OX=9606 G | ATXN2 | 140.283 | 2 | 51.310287 | yes | 2.019E-06 | 697.87103 | 834.43384 |
| NGGIYEGVFK | 10 | Q99700 | Ataxin-2 OS=Homo sapiens OX=9606 G | ATXN2 | 140.283 | 2 | 51.310287 | yes | 2.253E-07 | 542.2771 | 602.27734 |
| KDAAEQVR | 8 | Q99700 | Ataxin-2 OS=Homo sapiens OX=9606 G | ATXN2 | 140.283 | 2 | 51.310287 | yes | 9.855E-07 | 458.74597 | 551.27899 |
| PSRPPSHPSAHGSF | 22 | Q99700 | Ataxin-2 OS=Homo sapiens OX=9606 G | ATXN2 | 140.283 | 4 | 51.310287 | yes | 9.919E-08 | 556.28381 | 269.60114 |
| PSRPPSHPSAHGSF | 22 | Q99700 | Ataxin-2 OS=Homo sapiens OX=9606 G | ATXN2 | 140.283 | 4 | 51.310287 | yes | 1.127E-07 | 445.22852 | 269.60114 |
| PSRPPSHPSAHGSF | 22 | Q99700 | Ataxin-2 OS=Homo sapiens OX=9606 G | ATXN2 | 140.283 | 5 | 51.310287 | yes | 9.919E-08 | 556.28381 | 269.60114 |
| PSRPPSHPSAHGSF | 22 | Q99700 | Ataxin-2 OS=Homo sapiens OX=9606 G | ATXN2 | 140.283 | 5 | 51.310287 | yes | 1.127E-07 | 445.22852 | 269.60114 |
| EEIMESILFK | 10 | Q99700 | Ataxin-2 OS=Homo sapiens OX=9606 G | ATXN2 | 140.283 | 2 | 51.310287 | yes | 1.487E-06 | 619.8205 | 407.80841 |
| YQSGPNSLPPR | 11 | Q99700 | Ataxin-2 OS=Homo sapiens OX=9606 G | ATXN2 | 140.283 | 2 | 51.310287 | yes | 1.379E-07 | 608.30945 | 342.18594 |
| AATPTRPPSRPPSR | 14 | Q99700 | Ataxin-2 OS=Homo sapiens OX=9606 G | ATXN2 | 140.283 | 3 | 51.310287 | yes | 9.758E-08 | 497.61105 | 337.62183 |
| DNSEEFLK | 8 | Q99700 | Ataxin-2 OS=Homo sapiens OX=9606 G | ATXN2 | 140.283 | 2 | 51.310287 | yes | 1.123E-06 | 491.22983 | 247.87558 |
| DNSEEFLKR | 9 | Q99700 | Ataxin-2 OS=Homo sapiens OX=9606 G | ATXN2 | 140.283 | 2 | 51.310287 | yes | 1.578E-07 | 569.2804 | 244.66747 |
| DKIEPSAK | 8 | Q99700 | Ataxin-2 OS=Homo sapiens OX=9606 G | ATXN2 | 140.283 | 2 | 51.310287 | yes | 1.392E-07 | 444.24527 | 231.69847 |
| PSTTPTSPR | 9 | Q99700 | Ataxin-2 OS=Homo sapiens OX=9606 G | ATXN2 | 140.283 | 2 | 51.310287 | yes | 4.476E-07 | 472.24582 | 122.78244 |
| RDAFTDSAISAK | 12 | Q99700 | Ataxin-2 OS=Homo sapiens OX=9606 G | ATXN2 | 140.283 | 2 | 51.310287 | yes | 1.131E-07 | 641.32532 | 108.64542 |
| GSISSGLEFVSHNP | 25 | Q99700 | Ataxin-2 OS=Homo sapiens OX=9606 G | ATXN2 | 140.283 | 3 | 51.310287 | yes | 1.045E-07 | 836.42236 | 98.716034 |
| VALENDDRSEEEK | 13 | Q99700 | Ataxin-2 OS=Homo sapiens OX=9606 G | ATXN2 | 140.283 | 2 | 51.310287 | yes | 2.39E-07 | 767.35498 | 72.070137 |
| PNSPSISPSILSNTE | 17 | Q99700 | Ataxin-2 OS=Homo sapiens OX=9606 G | ATXN2 | 140.283 | 3 | 51.310287 | yes | 2.454E-07 | 603.31104 | 63.002472 |

| | | | | | | | | | | | |
|---|---|---|---|---|---|---|---|---|---|---|---|
| RGPEVTSQGVQTS | 21 | Q99700 | Ataxin-2 OS=Homo sapiens OX=9606 Gl | ATXN2 | 140.283 | 3 | 51.310287 | yes | 9.722E-08 | 758.70959 | 53.431679 |
| SFSQPKPSTTPTSPI | 15 | Q99700 | Ataxin-2 OS=Homo sapiens OX=9606 Gl | ATXN2 | 140.283 | 2 | 51.310287 | yes | 4.603E-07 | 809.41522 | 40.263054 |
| IPLLIGGATTSK | 12 | Q99707 | Methionine synthase OS=Homo sapiens ( | MTR | 140.528 | 2 | 48.322544 | yes | 1.164E-07 | 585.85828 | 39.039936 |
| CAVTEEHMR | 9 | Q99708 | DNA endonuclease RBBP8 OS=Homo sa | RBBP8 | 101.942 | 2 | 40.434986 | yes | 4.026E-07 | 566.74731 | 67.206146 |
| ESLETENVLDDIK | 13 | Q99708 | DNA endonuclease RBBP8 OS=Homo sa | RBBP8 | 101.942 | 2 | 40.434986 | yes | 5.437E-07 | 752.8725 | 19.412563 |
| VCNFLASQVPFPSI | 14 | Q99714 | 3-hydroxyacyl-CoA dehydrogenase type-2 | HSD17B10 | 26.924 | 2 | 54.084297 | yes | 4.007E-07 | 811.41138 | 463.80545 |
| VDVAVNCAGIAV/ | 15 | Q99714 | 3-hydroxyacyl-CoA dehydrogenase type-2 | HSD17B10 | 26.924 | 2 | 54.084297 | yes | 9.722E-08 | 737.39014 | 290.71832 |
| VLDVNLMGTFNVI | 14 | Q99714 | 3-hydroxyacyl-CoA dehydrogenase type-2 | HSD17B10 | 26.924 | 2 | 54.084297 | yes | 1.567E-07 | 795.93726 | 74.786133 |
| GGIVGMTLPIAR | 12 | Q99714 | 3-hydroxyacyl-CoA dehydrogenase type-2 | HSD17B10 | 26.924 | 2 | 54.084297 | yes | 6.325E-07 | 592.8446 | 51.955299 |
| ALALGALQNIR | 11 | Q99715 | Collagen alpha-1(XII) chain OS=Homo sa | COL12A1 | 333.147 | 2 | 52.611588 | yes | 2.723E-07 | 570.34839 | 179.86833 |
| VLVVVTDGR | 9 | Q99715 | Collagen alpha-1(XII) chain OS=Homo sa | COL12A1 | 333.147 | 2 | 52.611588 | yes | 1.714E-07 | 479.29001 | 151.15004 |
| VSWDPSPSPVLGY | 14 | Q99715 | Collagen alpha-1(XII) chain OS=Homo sa | COL12A1 | 333.147 | 2 | 52.611588 | yes | 9.722E-08 | 766.39319 | 133.03966 |
| NSDVEIFAVGVK | 12 | Q99715 | Collagen alpha-1(XII) chain OS=Homo sa | COL12A1 | 333.147 | 2 | 52.611588 | yes | 1.14E-06 | 639.34045 | 59.540863 |
| PSESIVVPGNTR | 12 | Q99715 | Collagen alpha-1(XII) chain OS=Homo sa | COL12A1 | 333.147 | 2 | 52.611588 | yes | 4.721E-07 | 628.33569 | 53.475407 |
| IVDDLTINLCNSVK | 14 | Q99715 | Collagen alpha-1(XII) chain OS=Homo sa | COL12A1 | 333.147 | 2 | 52.611588 | yes | 1.619E-07 | 802.42163 | 52.904163 |
| ANFRTVR | 7 | Q99715 | Collagen alpha-1(XII) chain OS=Homo sa | COL12A1 | 333.147 | 2 | 52.611588 | yes | 5.137E-06 | 432.24594 | 45.548847 |
| HVFIVDDFESFEK | 13 | Q99715 | Collagen alpha-1(XII) chain OS=Homo sa | COL12A1 | 333.147 | 2 | 52.611588 | yes | 2.825E-07 | 806.38812 | 39.939606 |
| ITVDPTTDGPTK | 12 | Q99715 | Collagen alpha-1(XII) chain OS=Homo sa | COL12A1 | 333.147 | 2 | 52.611588 | yes | 2.638E-07 | 622.82208 | 38.283478 |
| GGNTMTGDAIDYI | 15 | Q99715 | Collagen alpha-1(XII) chain OS=Homo sa | COL12A1 | 333.147 | 2 | 52.611588 | yes | 2.623E-07 | 777.87708 | 26.632505 |
| LVTAGGGQILSR | 12 | Q99728 | BRCA1-associated RING domain protein | BARD1 | 86.648 | 2 | 55.094906 | yes | 1.175E-07 | 586.34332 | 980.11841 |
| LLLSYGASR | 9 | Q99728 | BRCA1-associated RING domain protein | BARD1 | 86.648 | 2 | 55.094906 | yes | 2.949E-07 | 490.2822 | 488.75964 |
| GETLLHIASIK | 11 | Q99728 | BRCA1-associated RING domain protein | BARD1 | 86.648 | 2 | 55.094906 | yes | 9.722E-08 | 591.34808 | 401.49271 |
| VVELLLQHK | 9 | Q99728 | BRCA1-associated RING domain protein | BARD1 | 86.648 | 2 | 55.094906 | yes | 2.715E-07 | 539.83459 | 304.90308 |
| SLFNDAGNKK | 10 | Q99728 | BRCA1-associated RING domain protein | BARD1 | 86.648 | 2 | 55.094906 | yes | 1.261E-07 | 547.28546 | 135.24437 |
| RDGPLVLIGSGLSS | 18 | Q99728 | BRCA1-associated RING domain protein | BARD1 | 86.648 | 3 | 55.094906 | yes | 4.376E-07 | 628.67767 | 113.14729 |
| QLDSMIQLCSK | 11 | Q99728 | BRCA1-associated RING domain protein | BARD1 | 86.648 | 2 | 55.094906 | yes | 4.724E-07 | 661.82581 | 95.357849 |
| ASVQTQPAIK | 10 | Q99728 | BRCA1-associated RING domain protein | BARD1 | 86.648 | 2 | 55.094906 | yes | 9.722E-08 | 521.79822 | 84.643738 |
| MLSELAVILK | 10 | Q99728 | BRCA1-associated RING domain protein | BARD1 | 86.648 | 2 | 55.094906 | yes | 2.736E-07 | 558.8385 | 64.071457 |
| MFVGGLSWDTSK | 12 | Q99729 | Heterogeneous nuclear ribonucleoprotein | HNRNPAB | 36.225 | 2 | 50.295822 | yes | 4.609E-07 | 664.32117 | 9522.3701 |
| MFVGGLSWDTSK | 12 | Q99729 | Heterogeneous nuclear ribonucleoprotein | HNRNPAB | 36.225 | 2 | 50.295822 | yes | 1.891E-07 | 672.31866 | 9522.3701 |
| RGFVFITFK | 9 | Q99729 | Heterogeneous nuclear ribonucleoprotein | HNRNPAB | 36.225 | 2 | 50.295822 | yes | 9.935E-08 | 557.82404 | 345.64087 |
| DLKDYFTK | 8 | Q99729 | Heterogeneous nuclear ribonucleoprotein | HNRNPAB | 36.225 | 2 | 50.295822 | yes | 2.021E-07 | 515.26624 | 201.91217 |
| EVYQQQQYGSGGI | 13 | Q99729 | Heterogeneous nuclear ribonucleoprotein | HNRNPAB | 36.225 | 2 | 50.295822 | yes | 9.819E-08 | 750.34729 | 193.04718 |
| RGGHQNNYKPY | 11 | Q99729 | Heterogeneous nuclear ribonucleoprotein | HNRNPAB | 36.225 | 2 | 50.295822 | yes | 3.698E-07 | 667.32343 | 123.43304 |
| GFVFITFK | 8 | Q99729 | Heterogeneous nuclear ribonucleoprotein | HNRNPAB | 36.225 | 2 | 50.295822 | yes | 5.795E-06 | 479.77347 | 17.310505 |
| LTDQVMQNPR | 10 | Q99733 | Nucleosome assembly protein 1-like 4 OS | NAP1L4 | 42.824 | 2 | 49.484615 | yes | 2.866E-07 | 601.30334 | 146.83269 |
| VLAALQER | 8 | Q99733 | Nucleosome assembly protein 1-like 4 OS | NAP1L4 | 42.824 | 2 | 49.484615 | yes | 5.375E-07 | 450.26907 | 111.33237 |
| YAALYQPLFDK | 11 | Q99733 | Nucleosome assembly protein 1-like 4 OS | NAP1L4 | 42.824 | 2 | 49.484615 | yes | 1.026E-07 | 664.8479 | 94.581985 |
| NQIVTILQDR | 10 | Q99741 | Cell division control protein 6 homolog C | CDC6 | 62.721 | 2 | 54.349483 | yes | 2.055E-07 | 600.34076 | 764.01727 |

| | | | | | | | | | | | |
|---|---|---|---|---|---|---|---|---|---|---|---|
| LVLNTAVPDR | 10 | Q99741 | Cell division control protein 6 homolog C | CDC6 | 62.721 | 2 | 54.349483 | yes | 9.722E-08 | 549.31934 | 600.86523 |
| LEPTNVQTVTCSPI | 14 | Q99741 | Cell division control protein 6 homolog C | CDC6 | 62.721 | 2 | 54.349483 | yes | 9.735E-08 | 801.40125 | 484.62833 |
| AGSLYLSGAPGTG | 14 | Q99741 | Cell division control protein 6 homolog C | CDC6 | 62.721 | 2 | 54.349483 | yes | 9.722E-08 | 639.83807 | 441.15662 |
| AIEIVESDVK | 10 | Q99741 | Cell division control protein 6 homolog C | CDC6 | 62.721 | 2 | 54.349483 | yes | 2.495E-07 | 551.80316 | 286.84177 |
| SQTILKPLSECK | 12 | Q99741 | Cell division control protein 6 homolog C | CDC6 | 62.721 | 2 | 54.349483 | yes | 9.818E-08 | 702.38177 | 187.45944 |
| DQVLDNAAVQFC. | 14 | Q99741 | Cell division control protein 6 homolog C | CDC6 | 62.721 | 2 | 54.349483 | yes | 1.719E-07 | 803.88574 | 122.6278 |
| TIMLNCMSLR | 10 | Q99741 | Cell division control protein 6 homolog C | CDC6 | 62.721 | 2 | 54.349483 | yes | 4.414E-07 | 619.80634 | 109.81901 |
| LVFDNQLTIK | 10 | Q99741 | Cell division control protein 6 homolog C | CDC6 | 62.721 | 2 | 54.349483 | yes | 2.147E-07 | 595.84259 | 101.06331 |
| EIEHALK | 7 | Q99741 | Cell division control protein 6 homolog C | CDC6 | 62.721 | 2 | 54.349483 | yes | 1.696E-07 | 420.23471 | 85.757744 |
| SQEITTNSEQR | 11 | Q99741 | Cell division control protein 6 homolog C | CDC6 | 62.721 | 2 | 54.349483 | yes | 4.365E-07 | 646.8075 | 77.640671 |
| QEGTCYQQAK | 10 | Q99741 | Cell division control protein 6 homolog C | CDC6 | 62.721 | 2 | 54.349483 | yes | 4.023E-07 | 606.76935 | 38.435467 |
| VGLIHISQVISEVD | 17 | Q99741 | Cell division control protein 6 homolog C | CDC6 | 62.721 | 3 | 54.349483 | yes | 2.879E-07 | 612.67065 | 19.833597 |
| GAIQLGITHTVGSI | 20 | Q99755 | Phosphatidylinositol 4-phosphate 5-kinas | PIP5K1A | 62.634 | 3 | 53.51664 | yes | 1.356E-07 | 689.05151 | 1180.9083 |
| FYGLYCVQAGGK | 12 | Q99755 | Phosphatidylinositol 4-phosphate 5-kinas | PIP5K1A | 62.634 | 2 | 53.51664 | yes | 1.43E-07 | 681.82916 | 880.47058 |
| FMCNTVFK | 8 | Q99755 | Phosphatidylinositol 4-phosphate 5-kinas | PIP5K1A | 62.634 | 2 | 53.51664 | no | 1.847E-07 | 523.74353 | 820.47754 |
| LLPGYYMNLNQNI | 14 | Q99755 | Phosphatidylinositol 4-phosphate 5-kinas | PIP5K1A | 62.634 | 2 | 53.51664 | no | 1.112E-07 | 846.92993 | 764.89453 |
| EKPLPTFK | 8 | Q99755 | Phosphatidylinositol 4-phosphate 5-kinas | PIP5K1A | 62.634 | 2 | 53.51664 | no | 1.561E-07 | 480.28165 | 716.55396 |
| TYAPVAFR | 8 | Q99755 | Phosphatidylinositol 4-phosphate 5-kinas | PIP5K1A | 62.634 | 2 | 53.51664 | no | 3.316E-07 | 462.75052 | 559.07764 |
| EPLSSETQYSVDTI | 14 | Q99755 | Phosphatidylinositol 4-phosphate 5-kinas | PIP5K1A | 62.634 | 2 | 53.51664 | no | 1.417E-07 | 806.37848 | 517.11383 |
| DCLVLQSFK | 9 | Q99755 | Phosphatidylinositol 4-phosphate 5-kinas | PIP5K1A | 62.634 | 2 | 53.51664 | no | 1.883E-07 | 555.28662 | 451.57544 |
| GGTMETDDHMGC | 16 | Q99755 | Phosphatidylinositol 4-phosphate 5-kinas | PIP5K1A | 62.634 | 3 | 53.51664 | no | 2.258E-06 | 548.90833 | 159.4576 |
| GGTMETDDHMGC | 16 | Q99755 | Phosphatidylinositol 4-phosphate 5-kinas | PIP5K1A | 62.634 | 3 | 53.51664 | no | 1.084E-07 | 822.85889 | 159.4576 |
| GGTMETDDHMGC | 16 | Q99755 | Phosphatidylinositol 4-phosphate 5-kinas | PIP5K1A | 62.634 | 2 | 53.51664 | no | 2.258E-06 | 548.90833 | 159.4576 |
| GGTMETDDHMGC | 16 | Q99755 | Phosphatidylinositol 4-phosphate 5-kinas | PIP5K1A | 62.634 | 2 | 53.51664 | no | 1.084E-07 | 822.85889 | 159.4576 |
| ALYSTAMESIQGE. | 15 | Q99755 | Phosphatidylinositol 4-phosphate 5-kinas | PIP5K1A | 62.634 | 2 | 53.51664 | no | 9.722E-08 | 813.89325 | 290.05579 |
| KLEHSWK | 7 | Q99755 | Phosphatidylinositol 4-phosphate 5-kinas | PIP5K1A | 62.634 | 2 | 53.51664 | no | 1.043E-07 | 464.25598 | 283.9595 |
| KIPLKPSPSK | 10 | Q99755 | Phosphatidylinositol 4-phosphate 5-kinas | PIP5K1A | 62.634 | 2 | 53.51664 | no | 2.389E-07 | 547.85022 | 188.77643 |
| IVVMNNLLPR | 10 | Q99755 | Phosphatidylinositol 4-phosphate 5-kinas | PIP5K1A | 62.634 | 2 | 53.51664 | no | 9.804E-08 | 584.84717 | 134.12164 |
| IMDYSLLMSIHNII | 18 | Q99755 | Phosphatidylinositol 4-phosphate 5-kinas | PIP5K1A | 62.634 | 3 | 53.51664 | no | 1.736E-07 | 719.69086 | 117.99019 |
| PGFYAER | 7 | Q99755 | Phosphatidylinositol 4-phosphate 5-kinas | PIP5K1A | 62.634 | 2 | 53.51664 | yes | 1.041E-07 | 420.20596 | 80.037346 |
| ALVHDGDTVSVHI | 13 | Q99755 | Phosphatidylinositol 4-phosphate 5-kinas | PIP5K1A | 62.634 | 2 | 53.51664 | no | 9.819E-08 | 703.36279 | 64.848694 |
| SVDSSGETTYK | 11 | Q99755 | Phosphatidylinositol 4-phosphate 5-kinas | PIP5K1A | 62.634 | 2 | 53.51664 | yes | 9.722E-08 | 587.26715 | 63.171776 |
| IPLKPSPSK | 9 | Q99755 | Phosphatidylinositol 4-phosphate 5-kinas | PIP5K1A | 62.634 | 2 | 53.51664 | no | 2.017E-07 | 483.80276 | 53.187103 |
| ELMNLTGTIPVPYI | 14 | Q99816 | Tumor susceptibility gene 101 protein OS | TSG101 | 43.945 | 2 | 56.22932 | yes | 9.722E-08 | 802.42926 | 201.21411 |
| LEEMVTR | 7 | Q99816 | Tumor susceptibility gene 101 protein OS | TSG101 | 43.945 | 2 | 56.22932 | yes | 4.299E-07 | 439.22601 | 78.845901 |
| ETVNVITLYK | 10 | Q99816 | Tumor susceptibility gene 101 protein OS | TSG101 | 43.945 | 2 | 56.22932 | yes | 3.259E-07 | 590.33459 | 61.425209 |
| AQAELNALK | 9 | Q99816 | Tumor susceptibility gene 101 protein OS | TSG101 | 43.945 | 2 | 56.22932 | yes | 4.51E-07 | 479.27182 | 60.785988 |
| ATISNDGATILK | 12 | Q99832 | T-complex protein 1 subunit eta OS=Hon | CCT7 | 59.367 | 2 | 54.670982 | yes | 1.011E-07 | 602.33258 | 282.59164 |
| SLHDAIMIVR | 10 | Q99832 | T-complex protein 1 subunit eta OS=Hon | CCT7 | 59.367 | 2 | 54.670982 | yes | 9.722E-08 | 577.82117 | 238.33244 |

| | | | | | | | | | | | |
|---|---|---|---|---|---|---|---|---|---|---|---|
| LLDVVHPAAK | 10 | Q99832 | T-complex protein 1 subunit eta OS=Hon | CCT7 | 59.367 | 2 | 54.670982 | yes | 1.885E-07 | 531.81897 | 206.2645 |
| LPIGDVATQYFAD | 14 | Q99832 | T-complex protein 1 subunit eta OS=Hon | CCT7 | 59.367 | 2 | 54.670982 | yes | 1.425E-07 | 783.40155 | 194.14017 |
| YNFFTGCPK | 9 | Q99832 | T-complex protein 1 subunit eta OS=Hon | CCT7 | 59.367 | 2 | 54.670982 | yes | 9.722E-08 | 567.25787 | 157.40794 |
| QVKPYVEEGLHP( | 17 | Q99832 | T-complex protein 1 subunit eta OS=Hon | CCT7 | 59.367 | 3 | 54.670982 | yes | 2.413E-07 | 673.71771 | 89.278641 |
| TFSYAGFEMQPK | 12 | Q99832 | T-complex protein 1 subunit eta OS=Hon | CCT7 | 59.367 | 2 | 54.670982 | yes | 9.722E-08 | 703.32648 | 82.384315 |
| TCTFILR | 7 | Q99832 | T-complex protein 1 subunit eta OS=Hon | CCT7 | 59.367 | 2 | 54.670982 | yes | 2.379E-07 | 455.74438 | 79.450928 |
| QQLLIGAYAK | 10 | Q99832 | T-complex protein 1 subunit eta OS=Hon | CCT7 | 59.367 | 2 | 54.670982 | yes | 5.912E-07 | 552.82422 | 78.15625 |
| AVDPEDDFQR | 10 | Q99848 | Probable rRNA-processing protein EBP2 | EBNA1BP2 | 34.852 | 2 | 48.56982 | yes | 1.295E-07 | 596.26746 | 1198.4418 |
| GLLKPGLNVVLEC | 15 | Q99848 | Probable rRNA-processing protein EBP2 | EBNA1BP2 | 34.852 | 2 | 48.56982 | yes | 4.221E-07 | 767.4718 | 381.28992 |
| GLLKPGLNVVLEC | 15 | Q99848 | Probable rRNA-processing protein EBP2 | EBNA1BP2 | 34.852 | 2 | 48.56982 | yes | 1.315E-07 | 511.98361 | 381.28992 |
| GLLKPGLNVVLEC | 15 | Q99848 | Probable rRNA-processing protein EBP2 | EBNA1BP2 | 34.852 | 3 | 48.56982 | yes | 4.221E-07 | 767.4718 | 381.28992 |
| GLLKPGLNVVLEC | 15 | Q99848 | Probable rRNA-processing protein EBP2 | EBNA1BP2 | 34.852 | 3 | 48.56982 | yes | 1.315E-07 | 511.98361 | 381.28992 |
| ESYDDVSSFR | 10 | Q99848 | Probable rRNA-processing protein EBP2 | EBNA1BP2 | 34.852 | 2 | 48.56982 | yes | 4.156E-07 | 602.75946 | 511.07361 |
| RPTDYFAEMAK | 11 | Q99848 | Probable rRNA-processing protein EBP2 | EBNA1BP2 | 34.852 | 2 | 48.56982 | yes | 1.128E-07 | 664.81879 | 280.44992 |
| KVQTEVLQK | 9 | Q99848 | Probable rRNA-processing protein EBP2 | EBNA1BP2 | 34.852 | 2 | 48.56982 | yes | 3.239E-07 | 536.82166 | 260.80383 |
| QCLAEFK | 7 | Q99848 | Probable rRNA-processing protein EBP2 | EBNA1BP2 | 34.852 | 2 | 48.56982 | yes | 1.584E-07 | 448.22073 | 225.27711 |
| VQTEVLQK | 8 | Q99848 | Probable rRNA-processing protein EBP2 | EBNA1BP2 | 34.852 | 2 | 48.56982 | yes | 4.782E-06 | 472.7742 | 171.17206 |
| PTDYFAEMAK | 10 | Q99848 | Probable rRNA-processing protein EBP2 | EBNA1BP2 | 34.852 | 2 | 48.56982 | yes | 3.691E-06 | 586.76825 | 156.14792 |
| AHMMNAIK | 8 | Q99848 | Probable rRNA-processing protein EBP2 | EBNA1BP2 | 34.852 | 2 | 48.56982 | yes | 1.566E-07 | 458.23059 | 126.40208 |
| ELQDAFSR | 8 | Q99848 | Probable rRNA-processing protein EBP2 | EBNA1BP2 | 34.852 | 2 | 48.56982 | yes | 1.478E-07 | 483.23798 | 121.19684 |
| LDFLEGDQKPLAQ | 14 | Q99848 | Probable rRNA-processing protein EBP2 | EBNA1BP2 | 34.852 | 2 | 48.56982 | yes | 1.223E-07 | 815.43341 | 44.801834 |
| QAQAAVLAVLPR | 12 | Q99848 | Probable rRNA-processing protein EBP2 | EBNA1BP2 | 34.852 | 2 | 48.56982 | yes | 7.851E-06 | 618.87476 | 20.923973 |
| VWPSLQDR | 8 | Q99871 | HAUS augmin-like complex subunit 7 OS | HAUS7 | 39.795 | 2 | 53.1402 | yes | 1.109E-06 | 500.76416 | 213.5657 |
| TIQELLCSPSEYR | 13 | Q99871 | HAUS augmin-like complex subunit 7 OS | HAUS7 | 39.795 | 2 | 53.1402 | yes | 1.155E-07 | 798.39032 | 120.17871 |
| TEYFAQHEQGAA/ | 24 | Q99871 | HAUS augmin-like complex subunit 7 OS | HAUS7 | 39.795 | 3 | 53.1402 | yes | 9.722E-08 | 841.39801 | 74.498589 |
| LEILEWMCTR | 10 | Q99871 | HAUS augmin-like complex subunit 7 OS | HAUS7 | 39.795 | 2 | 53.1402 | yes | 3.809E-07 | 675.83087 | 22.910192 |
| GQLCELSCSTDYR | 13 | Q99873 | Protein arginine N-methyltransferase 1 O: | PRMT1 | 42.462 | 2 | 53.57166 | yes | 9.729E-08 | 794.84015 | 1072.7716 |
| GKVEEVELPVEK | 12 | Q99873 | Protein arginine N-methyltransferase 1 O: | PRMT1 | 42.462 | 2 | 53.57166 | no | 9.722E-08 | 678.37445 | 835.84045 |
| ATLYVTAIEDR | 11 | Q99873 | Protein arginine N-methyltransferase 1 O: | PRMT1 | 42.462 | 2 | 53.57166 | yes | 9.896E-08 | 626.33258 | 777.90393 |
| LDHVVTIIK | 9 | Q99873 | Protein arginine N-methyltransferase 1 O: | PRMT1 | 42.462 | 2 | 53.57166 | yes | 2.471E-07 | 519.32129 | 609.71362 |
| QLVTNACLIK | 10 | Q99873 | Protein arginine N-methyltransferase 1 O: | PRMT1 | 42.462 | 2 | 53.57166 | yes | 9.722E-08 | 580.3288 | 600.2215 |
| WLAPDGLIFPDR | 12 | Q99873 | Protein arginine N-methyltransferase 1 O: | PRMT1 | 42.462 | 2 | 53.57166 | yes | 9.722E-08 | 700.37207 | 510.57269 |
| QTVFYMEDYLTVI | 13 | Q99873 | Protein arginine N-methyltransferase 1 O: | PRMT1 | 42.462 | 2 | 53.57166 | yes | 9.722E-08 | 818.90002 | 388.53693 |
| TGEEIFGTIGMR | 12 | Q99873 | Protein arginine N-methyltransferase 1 O: | PRMT1 | 42.462 | 2 | 53.57166 | yes | 9.819E-08 | 655.8241 | 359.62128 |
| VEEVELPVEK | 10 | Q99873 | Protein arginine N-methyltransferase 1 O: | PRMT1 | 42.462 | 2 | 53.57166 | no | 9.977E-08 | 585.81628 | 302.50455 |
| EPLVDVVDPK | 10 | Q99873 | Protein arginine N-methyltransferase 1 O: | PRMT1 | 42.462 | 2 | 53.57166 | yes | 9.722E-08 | 555.80573 | 232.26546 |
| EVDIYTVK | 8 | Q99873 | Protein arginine N-methyltransferase 1 O: | PRMT1 | 42.462 | 2 | 53.57166 | no | 5.197E-07 | 483.76077 | 181.79039 |
| VIGIECSSISDYAVI | 15 | Q99873 | Protein arginine N-methyltransferase 1 O: | PRMT1 | 42.462 | 2 | 53.57166 | yes | 4.68E-07 | 820.91364 | 154.80658 |
| ANKLDHVVTIIK | 12 | Q99873 | Protein arginine N-methyltransferase 1 O: | PRMT1 | 42.462 | 2 | 53.57166 | yes | 9.809E-08 | 675.90881 | 33.45446 |

| | | | | | | | | | | | |
|---|---|---|---|---|---|---|---|---|---|---|---|
| ANKLDHVVTIIK | 12 | Q99873 | Protein arginine N-methyltransferase 1 O; | PRMT1 | 42.462 | 2 | 53.57166 | yes | 7.638E-07 | 450.94165 | 33.45446 |
| ANKLDHVVTIIK | 12 | Q99873 | Protein arginine N-methyltransferase 1 O; | PRMT1 | 42.462 | 3 | 53.57166 | yes | 9.809E-08 | 675.90881 | 33.45446 |
| ANKLDHVVTIIK | 12 | Q99873 | Protein arginine N-methyltransferase 1 O; | PRMT1 | 42.462 | 3 | 53.57166 | yes | 7.638E-07 | 450.94165 | 33.45446 |
| DVAIKEPLVDVVD | 15 | Q99873 | Protein arginine N-methyltransferase 1 O; | PRMT1 | 42.462 | 2 | 53.57166 | yes | 3.495E-07 | 818.96143 | 61.752499 |
| NSPQEEVELK | 10 | Q99933 | BAG family molecular chaperone regulato | BAG1 | 38.779 | 2 | 53.468643 | yes | 9.722E-08 | 586.79333 | 420.47787 |
| IADQLEELNK | 10 | Q99933 | BAG family molecular chaperone regulato | BAG1 | 38.779 | 2 | 53.468643 | yes | 1.274E-07 | 586.81152 | 184.2243 |
| ELTGIQQGFLPK | 12 | Q99933 | BAG family molecular chaperone regulato | BAG1 | 38.779 | 2 | 53.468643 | yes | 1.664E-07 | 665.87189 | 112.88326 |
| LQAVLADNQQLR | 12 | Q99941 | Cyclic AMP-dependent transcription facto | ATF6B | 76.71 | 2 | 52.078167 | yes | 9.722E-08 | 684.88336 | 104.02621 |
| SQPAFLDAIDR | 11 | Q99941 | Cyclic AMP-dependent transcription facto | ATF6B | 76.71 | 2 | 52.078167 | yes | 2.333E-06 | 616.81714 | 62.595428 |
| AAAEEEDGGPEGP | 15 | Q99942 | E3 ubiquitin-protein ligase RNF5 OS=Ho | RNF5 | 19.882 | 2 | 34.21393 | yes | 1.055E-06 | 770.82916 | 13.853078 |
| SEAASPALPVITK | 13 | Q99958 | Forkhead box protein C2 OS=Homo sapie | FOXC2 | 53.72 | 2 | 51.33148 | yes | 1.086E-07 | 642.36389 | 7256.2295 |
| QGWQNSIR | 8 | Q99958 | Forkhead box protein C2 OS=Homo sapie | FOXC2 | 53.72 | 2 | 51.33148 | no | 2.964E-07 | 494.75159 | 1488.6433 |
| HNLSLNECFVK | 11 | Q99958 | Forkhead box protein C2 OS=Homo sapie | FOXC2 | 53.72 | 2 | 51.33148 | no | 1.204E-07 | 680.83752 | 1367.7589 |
| SYAPYHHHQPAAI | 14 | Q99958 | Forkhead box protein C2 OS=Homo sapie | FOXC2 | 53.72 | 3 | 51.33148 | yes | 2.035E-07 | 535.26367 | 532.54346 |
| SYAPYHHHQPAAI | 14 | Q99958 | Forkhead box protein C2 OS=Homo sapie | FOXC2 | 53.72 | 3 | 51.33148 | yes | 3.551E-07 | 802.39185 | 532.54346 |
| SYAPYHHHQPAAI | 14 | Q99958 | Forkhead box protein C2 OS=Homo sapie | FOXC2 | 53.72 | 2 | 51.33148 | yes | 2.035E-07 | 535.26367 | 532.54346 |
| SYAPYHHHQPAAI | 14 | Q99958 | Forkhead box protein C2 OS=Homo sapie | FOXC2 | 53.72 | 2 | 51.33148 | yes | 3.551E-07 | 802.39185 | 532.54346 |
| VETLSPESALQGSI | 15 | Q99958 | Forkhead box protein C2 OS=Homo sapie | FOXC2 | 53.72 | 2 | 51.33148 | yes | 1.61E-06 | 785.90723 | 1020.0956 |
| EMFNSHR | 7 | Q99958 | Forkhead box protein C2 OS=Homo sapie | FOXC2 | 53.72 | 2 | 51.33148 | yes | 1.003E-07 | 460.70578 | 888.13245 |
| TSPPGGELSPGAGI | 14 | Q99958 | Forkhead box protein C2 OS=Homo sapie | FOXC2 | 53.72 | 2 | 51.33148 | yes | 9.722E-08 | 641.82294 | 883.56287 |
| GAPATPHLADAPK | 13 | Q99958 | Forkhead box protein C2 OS=Homo sapie | FOXC2 | 53.72 | 2 | 51.33148 | yes | 2.65E-07 | 623.33295 | 111.48151 |
| HAAPYSYDCTK | 11 | Q99958 | Forkhead box protein C2 OS=Homo sapie | FOXC2 | 53.72 | 2 | 51.33148 | yes | 9.722E-08 | 656.78497 | 148.89105 |
| AMSLYTGAER | 10 | Q99958 | Forkhead box protein C2 OS=Homo sapie | FOXC2 | 53.72 | 2 | 51.33148 | yes | 5.911E-07 | 549.76605 | 69.941811 |
| HAAPYSYDCTKY | 12 | Q99958 | Forkhead box protein C2 OS=Homo sapie | FOXC2 | 53.72 | 2 | 51.33148 | yes | 6.642E-07 | 738.31665 | 41.240593 |
| PLVPLQPVTQNR | 12 | Q99959 | Plakophilin-2 OS=Homo sapiens OX=960 | PKP2 | 97.415 | 2 | 53.681843 | yes | 1.611E-07 | 681.39862 | 347.71072 |
| IQEQVQQTLAR | 11 | Q99959 | Plakophilin-2 OS=Homo sapiens OX=960 | PKP2 | 97.415 | 2 | 53.681843 | yes | 1.149E-07 | 657.36224 | 255.54419 |
| IMAISAGDAYASN | 14 | Q99959 | Plakophilin-2 OS=Homo sapiens OX=960 | PKP2 | 97.415 | 2 | 53.681843 | yes | 1.007E-07 | 706.34796 | 234.44165 |
| EAGPSVAVDSSGR | 13 | Q99959 | Plakophilin-2 OS=Homo sapiens OX=960 | PKP2 | 97.415 | 2 | 53.681843 | yes | 9.722E-08 | 616.29932 | 203.52429 |
| VQNEDVQR | 8 | Q99959 | Plakophilin-2 OS=Homo sapiens OX=960 | PKP2 | 97.415 | 2 | 53.681843 | yes | 3.173E-07 | 494.24634 | 196.93175 |
| EQYQDVPMPEEK | 12 | Q99959 | Plakophilin-2 OS=Homo sapiens OX=960 | PKP2 | 97.415 | 2 | 53.681843 | yes | 6.872E-07 | 746.83484 | 155.56479 |
| YSQNIYIQNR | 10 | Q99959 | Plakophilin-2 OS=Homo sapiens OX=960 | PKP2 | 97.415 | 2 | 53.681843 | yes | 2.08E-07 | 649.828 | 145.97601 |
| MLHVGDPSVK | 10 | Q99959 | Plakophilin-2 OS=Homo sapiens OX=960 | PKP2 | 97.415 | 2 | 53.681843 | yes | 2.365E-07 | 541.7868 | 137.77451 |
| NLSLQNEIAK | 10 | Q99959 | Plakophilin-2 OS=Homo sapiens OX=960 | PKP2 | 97.415 | 2 | 53.681843 | yes | 2.84E-07 | 565.31421 | 135.71428 |
| ENYLTAGLTVGQV | 14 | Q99959 | Plakophilin-2 OS=Homo sapiens OX=960 | PKP2 | 97.415 | 2 | 53.681843 | yes | 9.722E-08 | 760.90698 | 88.064407 |
| GTIADYQPDDK | 11 | Q99959 | Plakophilin-2 OS=Homo sapiens OX=960 | PKP2 | 97.415 | 2 | 53.681843 | yes | 3.603E-06 | 611.78296 | 76.837448 |
| AHYTHSDYQYSQI | 13 | Q99959 | Plakophilin-2 OS=Homo sapiens OX=960 | PKP2 | 97.415 | 2 | 53.681843 | yes | 1.203E-06 | 828.36346 | 6.6344371 |
| TIEYLQPNPASR | 12 | Q99961 | Endophilin-A2 OS=Homo sapiens OX=96 | SH3GL1 | 41.49 | 2 | 52.954678 | no | 1.665E-07 | 694.86206 | 167.38654 |
| IPDEELR | 7 | Q99961 | Endophilin-A2 OS=Homo sapiens OX=96 | SH3GL1 | 41.49 | 2 | 52.954678 | no | 1.17E-06 | 436.22961 | 149.20724 |
| QAVQILDELAEK | 12 | Q99961 | Endophilin-A2 OS=Homo sapiens OX=96 | SH3GL1 | 41.49 | 2 | 52.954678 | yes | 9.722E-08 | 678.87207 | 106.02228 |

| | | | | | | | | | | | |
|---|---|---|---|---|---|---|---|---|---|---|---|
| QNFIDPLQNLCEK | 13 | Q99961 | Endophilin-A2 OS=Homo sapiens OX=9( | SH3GL1 | 41.49 | 2 | 52.954678 | yes | 3.172E-07 | 809.89832 | 53.384441 |
| ADHMMAMNHGR | 11 | Q99967 | Cbp/p300-interacting transactivator 2 OS: | CITED2 | 28.497 | 2 | 54.808907 | yes | 9.722E-08 | 656.77081 | 186.13527 |
| IIATSQGLLIR | 11 | Q99985 | Semaphorin-3C OS=Homo sapiens OX=9 | SEMA3C | 85.208 | 2 | 53.73861 | yes | 1.146E-07 | 592.8717 | 169.84071 |
| NAAEIVQYGVK | 11 | Q99985 | Semaphorin-3C OS=Homo sapiens OX=9 | SEMA3C | 85.208 | 2 | 53.73861 | yes | 9.722E-08 | 596.32202 | 129.28748 |
| ILLMDEDQDR | 10 | Q99985 | Semaphorin-3C OS=Homo sapiens OX=9 | SEMA3C | 85.208 | 2 | 53.73861 | yes | 9.968E-08 | 624.30048 | 117.11264 |
| DPTHGCGNFVR | 11 | Q99985 | Semaphorin-3C OS=Homo sapiens OX=9 | SEMA3C | 85.208 | 2 | 53.73861 | yes | 1.14E-07 | 630.28296 | 66.875916 |
| VYLTFDELR | 9 | Q99985 | Semaphorin-3C OS=Homo sapiens OX=9 | SEMA3C | 85.208 | 2 | 53.73861 | yes | 3.401E-07 | 578.30585 | 49.123062 |
| YHVLFLGTDR | 10 | Q99985 | Semaphorin-3C OS=Homo sapiens OX=9 | SEMA3C | 85.208 | 2 | 53.73861 | yes | 9.868E-07 | 610.82477 | 45.071407 |
| NHAPITTMK | 9 | Q99985 | Semaphorin-3C OS=Homo sapiens OX=9 | SEMA3C | 85.208 | 2 | 53.73861 | yes | 6.644E-06 | 506.76584 | 41.628262 |
| TSEYFSLSHHPLDY | 15 | Q99985 | Semaphorin-3C OS=Homo sapiens OX=9 | SEMA3C | 85.208 | 3 | 53.73861 | yes | 2.283E-07 | 617.96021 | 34.881245 |
| QQLYVSSNEGVSQ | 18 | Q99985 | Semaphorin-3C OS=Homo sapiens OX=9 | SEMA3C | 85.208 | 3 | 53.73861 | yes | 1.066E-07 | 677.67999 | 23.962969 |
| DSQFETDIVQR | 11 | Q99996 | A-kinase anchor protein 9 OS=Homo sap: | AKAP9 | 452.988 | 2 | 56.658268 | yes | 1.257E-07 | 669.32025 | 443.09253 |
| HVLGILDR | 8 | Q99996 | A-kinase anchor protein 9 OS=Homo sap: | AKAP9 | 452.988 | 2 | 56.658268 | yes | 3.257E-07 | 461.77707 | 403.93622 |
| FAQLEAENSILK | 12 | Q99996 | A-kinase anchor protein 9 OS=Homo sap: | AKAP9 | 452.988 | 2 | 56.658268 | yes | 9.722E-08 | 681.86682 | 370.41406 |
| EDELGSDISALTLR | 14 | Q99996 | A-kinase anchor protein 9 OS=Homo sap: | AKAP9 | 452.988 | 2 | 56.658268 | yes | 9.868E-08 | 759.88593 | 363.88672 |
| LMNVAINELNIK | 12 | Q99996 | A-kinase anchor protein 9 OS=Homo sap: | AKAP9 | 452.988 | 2 | 56.658268 | yes | 2.101E-07 | 686.38684 | 329.09003 |
| FQPISEHQTR | 10 | Q99996 | A-kinase anchor protein 9 OS=Homo sap: | AKAP9 | 452.988 | 2 | 56.658268 | yes | 2.73E-06 | 621.81494 | 327.82053 |
| MLYDAQLSEEQGI | 13 | Q99996 | A-kinase anchor protein 9 OS=Homo sap: | AKAP9 | 452.988 | 2 | 56.658268 | yes | 1.062E-07 | 770.35901 | 324.02399 |
| ETIIEELNTK | 10 | Q99996 | A-kinase anchor protein 9 OS=Homo sap: | AKAP9 | 452.988 | 2 | 56.658268 | yes | 6.517E-07 | 595.31915 | 317.21152 |
| VESTVIPESTIMR | 13 | Q99996 | A-kinase anchor protein 9 OS=Homo sap: | AKAP9 | 452.988 | 2 | 56.658268 | yes | 4.265E-07 | 731.38452 | 304.48273 |
| EFLELTEQSQK | 11 | Q99996 | A-kinase anchor protein 9 OS=Homo sap: | AKAP9 | 452.988 | 2 | 56.658268 | yes | 9.722E-08 | 676.34064 | 302.612 |
| SPLPFQNR | 8 | Q99996 | A-kinase anchor protein 9 OS=Homo sap: | AKAP9 | 452.988 | 2 | 56.658268 | yes | 3.313E-07 | 479.75888 | 274.85925 |
| TGKPTNLLR | 9 | Q99996 | A-kinase anchor protein 9 OS=Homo sap: | AKAP9 | 452.988 | 2 | 56.658268 | yes | 4.999E-07 | 500.3009 | 259.61615 |
| TNIQNLNQLR | 10 | Q99996 | A-kinase anchor protein 9 OS=Homo sap: | AKAP9 | 452.988 | 2 | 56.658268 | yes | 9.722E-08 | 607.336 | 240.86139 |
| GSIINLETR | 9 | Q99996 | A-kinase anchor protein 9 OS=Homo sap: | AKAP9 | 452.988 | 2 | 56.658268 | yes | 6.855E-07 | 501.78256 | 234.82741 |
| LLGELQEQIVQK | 12 | Q99996 | A-kinase anchor protein 9 OS=Homo sap: | AKAP9 | 452.988 | 2 | 56.658268 | yes | 2.006E-07 | 699.40356 | 228.47105 |
| ALNEELHLQR | 10 | Q99996 | A-kinase anchor protein 9 OS=Homo sap: | AKAP9 | 452.988 | 2 | 56.658268 | yes | 3.707E-07 | 611.83057 | 209.92464 |
| LEFGEENLPK | 10 | Q99996 | A-kinase anchor protein 9 OS=Homo sap: | AKAP9 | 452.988 | 2 | 56.658268 | yes | 9.869E-08 | 588.30078 | 205.87349 |
| FLDEQAIDR | 9 | Q99996 | A-kinase anchor protein 9 OS=Homo sap: | AKAP9 | 452.988 | 2 | 56.658268 | yes | 1.356E-07 | 553.77747 | 201.34612 |
| IHQLELQTMK | 10 | Q99996 | A-kinase anchor protein 9 OS=Homo sap: | AKAP9 | 452.988 | 2 | 56.658268 | yes | 9.819E-08 | 620.83954 | 196.23372 |
| EQLAVELSK | 9 | Q99996 | A-kinase anchor protein 9 OS=Homo sap: | AKAP9 | 452.988 | 2 | 56.658268 | yes | 1.846E-07 | 508.78476 | 192.62375 |
| DCVLTISEEMFSK | 13 | Q99996 | A-kinase anchor protein 9 OS=Homo sap: | AKAP9 | 452.988 | 2 | 56.658268 | yes | 9.722E-08 | 779.86005 | 186.80527 |
| SYSNITVNEDQIK | 13 | Q99996 | A-kinase anchor protein 9 OS=Homo sap: | AKAP9 | 452.988 | 2 | 56.658268 | yes | 9.722E-08 | 755.8728 | 180.89044 |
| LSDLSEQLK | 9 | Q99996 | A-kinase anchor protein 9 OS=Homo sap: | AKAP9 | 452.988 | 2 | 56.658268 | yes | 6.235E-07 | 516.78223 | 180.08105 |
| ELEQALLVSADTF( | 15 | Q99996 | A-kinase anchor protein 9 OS=Homo sap: | AKAP9 | 452.988 | 2 | 56.658268 | yes | 1.211E-07 | 846.44617 | 178.40829 |
| EELIFAQEEK | 10 | Q99996 | A-kinase anchor protein 9 OS=Homo sap: | AKAP9 | 452.988 | 2 | 56.658268 | yes | 5.448E-07 | 618.31134 | 174.8434 |
| EVEIDQLNEQVTK | 13 | Q99996 | A-kinase anchor protein 9 OS=Homo sap: | AKAP9 | 452.988 | 2 | 56.658268 | yes | 1.386E-07 | 772.89374 | 172.30269 |
| QLMGTVEELQK | 11 | Q99996 | A-kinase anchor protein 9 OS=Homo sap: | AKAP9 | 452.988 | 2 | 56.658268 | yes | 9.722E-08 | 638.33429 | 170.06517 |
| AEIVSASESR | 10 | Q99996 | A-kinase anchor protein 9 OS=Homo sap: | AKAP9 | 452.988 | 2 | 56.658268 | yes | 1.617E-07 | 524.76709 | 168.30743 |

| | | | | | | | | | | | |
|---|---|---|---|---|---|---|---|---|---|---|---|
| AEGVIDGYADEK | 12 | Q99996 | A-kinase anchor protein 9 OS=Homo sap: | AKAP9 | 452.988 | 2 | 56.658268 | yes | 1.007E-07 | 633.79608 | 166.43822 |
| VSFENMTVGEESK | 13 | Q99996 | A-kinase anchor protein 9 OS=Homo sap: | AKAP9 | 452.988 | 2 | 56.658268 | yes | 9.722E-08 | 728.83484 | 160.877 |
| LLQLESTVSAK | 11 | Q99996 | A-kinase anchor protein 9 OS=Homo sap: | AKAP9 | 452.988 | 2 | 56.658268 | yes | 1.18E-07 | 594.84534 | 156.92316 |
| SEEMTLQINELQK | 13 | Q99996 | A-kinase anchor protein 9 OS=Homo sap: | AKAP9 | 452.988 | 2 | 56.658268 | yes | 2.203E-07 | 781.89014 | 156.47845 |
| MNQLTQELFSLK | 12 | Q99996 | A-kinase anchor protein 9 OS=Homo sap: | AKAP9 | 452.988 | 2 | 56.658268 | yes | 1.324E-07 | 726.38177 | 153.7413 |
| LSSLQDLEK | 9 | Q99996 | A-kinase anchor protein 9 OS=Homo sap: | AKAP9 | 452.988 | 2 | 56.658268 | yes | 1.807E-07 | 516.78223 | 151.54311 |
| DNLGIHYK | 8 | Q99996 | A-kinase anchor protein 9 OS=Homo sap: | AKAP9 | 452.988 | 2 | 56.658268 | yes | 1.297E-07 | 480.25089 | 147.15439 |
| IVELLNETEK | 10 | Q99996 | A-kinase anchor protein 9 OS=Homo sap: | AKAP9 | 452.988 | 2 | 56.658268 | yes | 1.026E-07 | 594.32953 | 144.04973 |
| LQELEQENK | 9 | Q99996 | A-kinase anchor protein 9 OS=Homo sap: | AKAP9 | 452.988 | 2 | 56.658268 | yes | 9.722E-08 | 565.78802 | 141.4747 |
| QTIAEQESK | 9 | Q99996 | A-kinase anchor protein 9 OS=Homo sap: | AKAP9 | 452.988 | 2 | 56.658268 | yes | 2.845E-07 | 517.26166 | 137.20459 |
| VTGSVSININR | 11 | Q99996 | A-kinase anchor protein 9 OS=Homo sap: | AKAP9 | 452.988 | 2 | 56.658268 | yes | 2.993E-07 | 580.32513 | 132.79149 |
| LKEELGLILEEK | 12 | Q99996 | A-kinase anchor protein 9 OS=Homo sap: | AKAP9 | 452.988 | 2 | 56.658268 | yes | 9.926E-08 | 707.41357 | 132.70555 |
| EIEILIEENEDLK | 13 | Q99996 | A-kinase anchor protein 9 OS=Homo sap: | AKAP9 | 452.988 | 2 | 56.658268 | yes | 2.247E-07 | 793.91162 | 128.22607 |
| NLQLNLLLEQQK | 12 | Q99996 | A-kinase anchor protein 9 OS=Homo sap: | AKAP9 | 452.988 | 2 | 56.658268 | yes | 2.26E-07 | 727.4223 | 126.79103 |
| EVEQLANHLK | 10 | Q99996 | A-kinase anchor protein 9 OS=Homo sap: | AKAP9 | 452.988 | 2 | 56.658268 | yes | 9.935E-08 | 590.81964 | 125.48033 |
| ALEIDVEEQVSR | 12 | Q99996 | A-kinase anchor protein 9 OS=Homo sap: | AKAP9 | 452.988 | 2 | 56.658268 | yes | 1.665E-07 | 694.35681 | 120.69546 |
| LQGIMQEFQK | 10 | Q99996 | A-kinase anchor protein 9 OS=Homo sap: | AKAP9 | 452.988 | 2 | 56.658268 | yes | 5.919E-07 | 611.31842 | 116.12712 |
| NLELQVLLESEK | 12 | Q99996 | A-kinase anchor protein 9 OS=Homo sap: | AKAP9 | 452.988 | 2 | 56.658268 | yes | 9.722E-08 | 707.89301 | 115.90842 |
| TTAAVEETIGR | 11 | Q99996 | A-kinase anchor protein 9 OS=Homo sap: | AKAP9 | 452.988 | 2 | 56.658268 | yes | 9.722E-08 | 574.30133 | 115.59873 |
| DGIITQLTANLQQ/ | 15 | Q99996 | A-kinase anchor protein 9 OS=Homo sap: | AKAP9 | 452.988 | 2 | 56.658268 | yes | 1.216E-07 | 821.44958 | 112.76519 |
| GTLEQEVQELQLK | 13 | Q99996 | A-kinase anchor protein 9 OS=Homo sap: | AKAP9 | 452.988 | 2 | 56.658268 | yes | 5.291E-07 | 757.90668 | 111.63836 |
| LLVLQTR | 7 | Q99996 | A-kinase anchor protein 9 OS=Homo sap: | AKAP9 | 452.988 | 2 | 56.658268 | yes | 9.722E-08 | 421.77655 | 108.40533 |
| QQCIQLNEEIEK | 12 | Q99996 | A-kinase anchor protein 9 OS=Homo sap: | AKAP9 | 452.988 | 2 | 56.658268 | yes | 1.263E-07 | 766.37469 | 103.64891 |
| GEESESATDSFPK | 13 | Q99996 | A-kinase anchor protein 9 OS=Homo sap: | AKAP9 | 452.988 | 2 | 56.658268 | yes | 3.579E-07 | 692.29913 | 102.56007 |
| ACMFEPLPIK | 10 | Q99996 | A-kinase anchor protein 9 OS=Homo sap: | AKAP9 | 452.988 | 2 | 56.658268 | yes | 1.202E-07 | 603.30652 | 102.46897 |
| QLMLNEEQLEDMI | 13 | Q99996 | A-kinase anchor protein 9 OS=Homo sap: | AKAP9 | 452.988 | 2 | 56.658268 | yes | 1.683E-07 | 824.88708 | 101.35953 |
| ISSSNQTPQILVK | 13 | Q99996 | A-kinase anchor protein 9 OS=Homo sap: | AKAP9 | 452.988 | 2 | 56.658268 | yes | 1.001E-07 | 707.89862 | 99.871277 |
| DLELTQCYK | 9 | Q99996 | A-kinase anchor protein 9 OS=Homo sap: | AKAP9 | 452.988 | 2 | 56.658268 | yes | 9.722E-08 | 585.27899 | 99.349876 |
| TDSFYHSSGGLEL' | 18 | Q99996 | A-kinase anchor protein 9 OS=Homo sap: | AKAP9 | 452.988 | 3 | 56.658268 | yes | 4.208E-07 | 672.30957 | 98.269562 |
| ALTDYITR | 8 | Q99996 | A-kinase anchor protein 9 OS=Homo sap: | AKAP9 | 452.988 | 2 | 56.658268 | yes | 2.446E-07 | 476.75854 | 94.554955 |
| LQQELANIGQK | 11 | Q99996 | A-kinase anchor protein 9 OS=Homo sap: | AKAP9 | 452.988 | 2 | 56.658268 | yes | 1.866E-07 | 621.34607 | 94.183701 |
| LLEAISETSSQLEH | 16 | Q99996 | A-kinase anchor protein 9 OS=Homo sap: | AKAP9 | 452.988 | 3 | 56.658268 | yes | 9.722E-08 | 585.97565 | 94.091293 |
| FKEEFKPLSK | 10 | Q99996 | A-kinase anchor protein 9 OS=Homo sap: | AKAP9 | 452.988 | 2 | 56.658268 | yes | 9.722E-08 | 626.85046 | 46.784584 |
| FKEEFKPLSK | 10 | Q99996 | A-kinase anchor protein 9 OS=Homo sap: | AKAP9 | 452.988 | 2 | 56.658268 | yes | 3.074E-07 | 418.23605 | 46.784584 |
| FKEEFKPLSK | 10 | Q99996 | A-kinase anchor protein 9 OS=Homo sap: | AKAP9 | 452.988 | 3 | 56.658268 | yes | 9.722E-08 | 626.85046 | 46.784584 |
| FKEEFKPLSK | 10 | Q99996 | A-kinase anchor protein 9 OS=Homo sap: | AKAP9 | 452.988 | 3 | 56.658268 | yes | 3.074E-07 | 418.23605 | 46.784584 |
| LELETTLK | 8 | Q99996 | A-kinase anchor protein 9 OS=Homo sap: | AKAP9 | 452.988 | 2 | 56.658268 | yes | 3.39E-07 | 473.7764 | 92.095917 |
| MAESQEAELER | 11 | Q99996 | A-kinase anchor protein 9 OS=Homo sap: | AKAP9 | 452.988 | 2 | 56.658268 | yes | 5.567E-07 | 646.79297 | 91.920654 |
| VTEEGTELSQR | 11 | Q99996 | A-kinase anchor protein 9 OS=Homo sap: | AKAP9 | 452.988 | 2 | 56.658268 | yes | 9.91E-08 | 624.80695 | 89.956772 |

| | | | | | | | | | | | |
|---|---|---|---|---|---|---|---|---|---|---|---|
| SPQDVEVLK | 9 | Q99996 | A-kinase anchor protein 9 OS=Homo sap | AKAP9 | 452.988 | 2 | 56.658268 | yes | 1.808E-07 | 507.77695 | 87.910736 |
| ETNMTSLQK | 9 | Q99996 | A-kinase anchor protein 9 OS=Homo sap | AKAP9 | 452.988 | 2 | 56.658268 | yes | 1.128E-06 | 526.25806 | 85.595978 |
| LQEELLVK | 8 | Q99996 | A-kinase anchor protein 9 OS=Homo sap | AKAP9 | 452.988 | 2 | 56.658268 | yes | 1.135E-07 | 486.29224 | 84.462929 |
| LIFLDSIK | 8 | Q99996 | A-kinase anchor protein 9 OS=Homo sap | AKAP9 | 452.988 | 2 | 56.658268 | yes | 9.722E-08 | 474.79187 | 81.662926 |
| AYINTISSLK | 10 | Q99996 | A-kinase anchor protein 9 OS=Homo sap | AKAP9 | 452.988 | 2 | 56.658268 | yes | 3.521E-07 | 555.31366 | 79.304893 |
| HGEISFLNEEVK | 12 | Q99996 | A-kinase anchor protein 9 OS=Homo sap | AKAP9 | 452.988 | 2 | 56.658268 | yes | 9.722E-08 | 701.35406 | 74.190453 |
| DLQQSLVNSK | 10 | Q99996 | A-kinase anchor protein 9 OS=Homo sap | AKAP9 | 452.988 | 2 | 56.658268 | yes | 3.778E-07 | 566.30383 | 73.813164 |
| QYQEQLEEEVAK | 12 | Q99996 | A-kinase anchor protein 9 OS=Homo sap | AKAP9 | 452.988 | 2 | 56.658268 | yes | 1.32E-07 | 747.35956 | 72.61692 |
| TFVAETLEMGEVV | 15 | Q99996 | A-kinase anchor protein 9 OS=Homo sap | AKAP9 | 452.988 | 2 | 56.658268 | yes | 1.885E-07 | 841.42133 | 68.558136 |
| QQIDGLQNEMSQK | 13 | Q99996 | A-kinase anchor protein 9 OS=Homo sap | AKAP9 | 452.988 | 2 | 56.658268 | yes | 1.06E-07 | 759.8645 | 67.915276 |
| QLEDLVEELSFSR | 13 | Q99996 | A-kinase anchor protein 9 OS=Homo sap | AKAP9 | 452.988 | 2 | 56.658268 | yes | 3.785E-07 | 782.8963 | 66.832657 |
| VTQTELMR | 8 | Q99996 | A-kinase anchor protein 9 OS=Homo sap | AKAP9 | 452.988 | 2 | 56.658268 | yes | 6.595E-07 | 489.25787 | 59.268162 |
| LKDNLGIHYK | 10 | Q99996 | A-kinase anchor protein 9 OS=Homo sap | AKAP9 | 452.988 | 2 | 56.658268 | yes | 9.935E-08 | 600.84039 | 55.569214 |
| EDFTMQISFLQEK | 13 | Q99996 | A-kinase anchor protein 9 OS=Homo sap | AKAP9 | 452.988 | 2 | 56.658268 | yes | 2.824E-06 | 808.38727 | 50.227745 |
| QNQLILEISK | 10 | Q99996 | A-kinase anchor protein 9 OS=Homo sap | AKAP9 | 452.988 | 2 | 56.658268 | yes | 3.168E-06 | 593.34552 | 49.400921 |
| EITNLEEQLEQFR | 13 | Q99996 | A-kinase anchor protein 9 OS=Homo sap | AKAP9 | 452.988 | 2 | 56.658268 | yes | 5.815E-07 | 824.91248 | 44.099831 |
| EEEIEQLNEVIEK | 13 | Q99996 | A-kinase anchor protein 9 OS=Homo sap | AKAP9 | 452.988 | 2 | 56.658268 | yes | 2.049E-07 | 801.39886 | 41.312443 |
| SVLLAAFR | 8 | Q99996 | A-kinase anchor protein 9 OS=Homo sap | AKAP9 | 452.988 | 2 | 56.658268 | yes | 5.249E-07 | 438.76871 | 40.265072 |
| HQLDVVIAEK | 10 | Q99996 | A-kinase anchor protein 9 OS=Homo sap | AKAP9 | 452.988 | 2 | 56.658268 | yes | 1.157E-07 | 576.32458 | 38.84375 |
| IQSIPENSVNVAID | 18 | Q99996 | A-kinase anchor protein 9 OS=Homo sap | AKAP9 | 452.988 | 3 | 56.658268 | yes | 9.722E-08 | 655.35309 | 37.066231 |
| NSTHSSTAADLLQ | 15 | Q99996 | A-kinase anchor protein 9 OS=Homo sap | AKAP9 | 452.988 | 2 | 56.658268 | yes | 9.722E-08 | 772.38916 | 34.084896 |
| CSELLLSK | 8 | Q99996 | A-kinase anchor protein 9 OS=Homo sap | AKAP9 | 452.988 | 2 | 56.658268 | yes | 2.385E-07 | 475.25479 | 31.42527 |
| EKLEVQCQAEK | 11 | Q99996 | A-kinase anchor protein 9 OS=Homo sap | AKAP9 | 452.988 | 2 | 56.658268 | yes | 2.277E-07 | 681.34009 | 30.438644 |
| TMYPGSCVK | 9 | Q99996 | A-kinase anchor protein 9 OS=Homo sap | AKAP9 | 452.988 | 2 | 56.658268 | yes | 7.23E-07 | 521.73846 | 22.104353 |
| LMFSGLLVSSR | 11 | Q9BPX3 | Condensin complex subunit 3 OS=Homo | NCAPG | 114.335 | 2 | 38.80804 | yes | 4.973E-07 | 605.33667 | 41.435368 |
| TQIVTEIISEIR | 12 | Q9BPX3 | Condensin complex subunit 3 OS=Homo | NCAPG | 114.335 | 2 | 38.80804 | yes | 1.115E-06 | 701.401 | 31.174301 |
| LAYQVLAEK | 9 | Q9BPX3 | Condensin complex subunit 3 OS=Homo | NCAPG | 114.335 | 2 | 38.80804 | yes | 9.11E-06 | 517.79767 | 26.945126 |
| DRPTTGNTLK | 10 | Q9BPX6 | Calcium uptake protein 1, mitochondrial ( | MICU1 | 54.352 | 2 | 55.859894 | yes | 2.456E-07 | 551.7962 | 307.91092 |
| CAQETAWDFALPK | 13 | Q9BPX6 | Calcium uptake protein 1, mitochondrial ( | MICU1 | 54.352 | 2 | 55.859894 | yes | 1.218E-07 | 768.86121 | 205.27856 |
| TADLAPHPEEK | 11 | Q9BPX6 | Calcium uptake protein 1, mitochondrial ( | MICU1 | 54.352 | 2 | 55.859894 | yes | 3.355E-07 | 604.30133 | 204.83249 |
| KVMEYENR | 8 | Q9BPX6 | Calcium uptake protein 1, mitochondrial ( | MICU1 | 54.352 | 2 | 55.859894 | yes | 9.722E-08 | 534.76074 | 113.30527 |
| NFLEFQR | 7 | Q9BPX6 | Calcium uptake protein 1, mitochondrial ( | MICU1 | 54.352 | 2 | 55.859894 | yes | 5.816E-07 | 477.24561 | 104.12447 |
| LQHDVLK | 7 | Q9BPX6 | Calcium uptake protein 1, mitochondrial ( | MICU1 | 54.352 | 2 | 55.859894 | yes | 1.115E-07 | 426.75052 | 63.308353 |
| KLQHDVLK | 8 | Q9BPX6 | Calcium uptake protein 1, mitochondrial ( | MICU1 | 54.352 | 2 | 55.859894 | yes | 1.138E-06 | 490.798 | 52.786057 |
| KTADLAPHPEEK | 12 | Q9BPX6 | Calcium uptake protein 1, mitochondrial ( | MICU1 | 54.352 | 2 | 55.859894 | yes | 6.147E-07 | 668.34882 | 19.964756 |
| CFQEMLEEEEEHE | 19 | Q9BPZ3 | Polyadenylate-binding protein-interacting | PAIP2 | 14.985 | 3 | 50.56013 | yes | 9.766E-08 | 837.03149 | 511.70572 |
| CFQEMLEEEEEHE | 19 | Q9BPZ3 | Polyadenylate-binding protein-interacting | PAIP2 | 14.985 | 3 | 50.56013 | yes | 4.761E-07 | 842.3631 | 511.70572 |
| SNLNPNAK | 8 | Q9BPZ3 | Polyadenylate-binding protein-interacting | PAIP2 | 14.985 | 2 | 50.56013 | yes | 3.136E-07 | 429.22742 | 76.635963 |
| LTYLSNHDYK | 10 | Q9BPZ7 | Target of rapamycin complex 2 subunit M | MAPKAP1 | 59.123 | 2 | 53.655525 | yes | 9.722E-08 | 627.31171 | 167.2531 |

| | | | | | | | | | | | |
|---|---|---|---|---|---|---|---|---|---|---|---|
| KIDVYLPLHSSQDI | 14 | Q9BPZ7 | Target of rapamycin complex 2 subunit M | MAPKAP1 | 59.123 | 3 | 53.655525 | yes | 1.202E-07 | 557.63354 | 81.927971 |
| KIDVYLPLHSSQDI | 14 | Q9BPZ7 | Target of rapamycin complex 2 subunit M | MAPKAP1 | 59.123 | 3 | 53.655525 | yes | 5.206E-07 | 835.94666 | 81.927971 |
| KIDVYLPLHSSQDI | 14 | Q9BPZ7 | Target of rapamycin complex 2 subunit M | MAPKAP1 | 59.123 | 2 | 53.655525 | yes | 1.202E-07 | 557.63354 | 81.927971 |
| KIDVYLPLHSSQDI | 14 | Q9BPZ7 | Target of rapamycin complex 2 subunit M | MAPKAP1 | 59.123 | 2 | 53.655525 | yes | 5.206E-07 | 835.94666 | 81.927971 |
| VNYILESR | 8 | Q9BPZ7 | Target of rapamycin complex 2 subunit M | MAPKAP1 | 59.123 | 2 | 53.655525 | yes | 4.913E-07 | 497.27182 | 141.15718 |
| FGFSTLALVEK | 11 | Q9BPZ7 | Target of rapamycin complex 2 subunit M | MAPKAP1 | 59.123 | 2 | 53.655525 | yes | 9.722E-08 | 606.33716 | 126.46341 |
| FTTDVQLGISGDK' | 23 | Q9BPZ7 | Target of rapamycin complex 2 subunit M | MAPKAP1 | 59.123 | 3 | 53.655525 | yes | 2.638E-07 | 835.43414 | 47.465511 |
| HLYFESDAATVNE | 17 | Q9BPZ7 | Target of rapamycin complex 2 subunit M | MAPKAP1 | 59.123 | 3 | 53.655525 | yes | 1.182E-07 | 650.33868 | 36.190926 |
| GVTYLFPIQVK | 11 | Q9BQ39 | ATP-dependent RNA helicase DDX50 O | DDX50 | 82.566 | 2 | 53.532974 | yes | 9.722E-08 | 632.86865 | 503.5697 |
| VLVATNVAAR | 10 | Q9BQ39 | ATP-dependent RNA helicase DDX50 O | DDX50 | 82.566 | 2 | 53.532974 | yes | 9.722E-08 | 507.30875 | 397.37738 |
| VSSLDTSTHK | 10 | Q9BQ39 | ATP-dependent RNA helicase DDX50 O | DDX50 | 82.566 | 2 | 53.532974 | yes | 9.722E-08 | 537.7749 | 394.15103 |
| VLVLAPTR | 8 | Q9BQ39 | ATP-dependent RNA helicase DDX50 O | DDX50 | 82.566 | 2 | 53.532974 | yes | 1.025E-07 | 434.78436 | 370.37231 |
| LSSNAVSQITR | 11 | Q9BQ39 | ATP-dependent RNA helicase DDX50 O | DDX50 | 82.566 | 2 | 53.532974 | yes | 9.722E-08 | 588.32257 | 331.20557 |
| YEQVDLVGK | 9 | Q9BQ39 | ATP-dependent RNA helicase DDX50 O | DDX50 | 82.566 | 2 | 53.532974 | yes | 4.888E-07 | 525.77692 | 195.81668 |
| LEETLTR | 7 | Q9BQ39 | ATP-dependent RNA helicase DDX50 O | DDX50 | 82.566 | 2 | 53.532974 | yes | 2.272E-07 | 431.23746 | 157.15694 |
| TFGPVYEGK | 9 | Q9BQ39 | ATP-dependent RNA helicase DDX50 O | DDX50 | 82.566 | 2 | 53.532974 | yes | 3.186E-07 | 499.25311 | 124.51807 |
| SSDNKLEETLTR | 12 | Q9BQ39 | ATP-dependent RNA helicase DDX50 O | DDX50 | 82.566 | 2 | 53.532974 | yes | 1.753E-07 | 696.85187 | 106.57502 |
| ENGVTDDLDAPK | 12 | Q9BQ39 | ATP-dependent RNA helicase DDX50 O | DDX50 | 82.566 | 2 | 53.532974 | yes | 1.786E-07 | 637.29895 | 104.69375 |
| AIIFCETK | 8 | Q9BQ39 | ATP-dependent RNA helicase DDX50 O | DDX50 | 82.566 | 2 | 53.532974 | yes | 1.065E-06 | 491.25732 | 76.490028 |
| SRYEQVDLVGK | 11 | Q9BQ39 | ATP-dependent RNA helicase DDX50 O | DDX50 | 82.566 | 2 | 53.532974 | yes | 5.941E-07 | 647.34351 | 73.720901 |
| DLPNGDIDEYEK | 12 | Q9BQ39 | ATP-dependent RNA helicase DDX50 O | DDX50 | 82.566 | 2 | 53.532974 | yes | 4.719E-06 | 704.31738 | 64.86245 |
| LSVACFYGGTSYQ | 20 | Q9BQ39 | ATP-dependent RNA helicase DDX50 O | DDX50 | 82.566 | 3 | 53.532974 | yes | 4.933E-06 | 767.70752 | 44.913918 |
| RVGVPSTMDLVK | 12 | Q9BQ39 | ATP-dependent RNA helicase DDX50 O | DDX50 | 82.566 | 2 | 53.532974 | yes | 1.312E-06 | 651.36591 | 38.058411 |
| ALFAGDIEEMEER | 13 | Q9BQ52 | Zinc phosphodiesterase ELAC protein 2 ( | ELAC2 | 92.219 | 2 | 53.709312 | yes | 9.722E-08 | 755.34814 | 331.9808 |
| VPLFSPNFSEK | 11 | Q9BQ52 | Zinc phosphodiesterase ELAC protein 2 ( | ELAC2 | 92.219 | 2 | 53.709312 | yes | 9.722E-08 | 632.83228 | 259.31473 |
| CVLSGPPQLEK | 11 | Q9BQ52 | Zinc phosphodiesterase ELAC protein 2 ( | ELAC2 | 92.219 | 2 | 53.709312 | yes | 1.446E-07 | 614.32373 | 215.95464 |
| VCFGDFPTMPK | 11 | Q9BQ52 | Zinc phosphodiesterase ELAC protein 2 ( | ELAC2 | 92.219 | 2 | 53.709312 | yes | 1.281E-06 | 649.79901 | 204.97563 |
| CLQEGAEISSPAVI | 15 | Q9BQ52 | Zinc phosphodiesterase ELAC protein 2 ( | ELAC2 | 92.219 | 2 | 53.709312 | yes | 9.722E-08 | 823.39612 | 154.52136 |
| PLHPLLVVAPNQL | 14 | Q9BQ52 | Zinc phosphodiesterase ELAC protein 2 ( | ELAC2 | 92.219 | 3 | 53.709312 | yes | 2.041E-07 | 513.6521 | 77.220665 |
| PLHPLLVVAPNQL | 14 | Q9BQ52 | Zinc phosphodiesterase ELAC protein 2 ( | ELAC2 | 92.219 | 3 | 53.709312 | yes | 1.76E-07 | 769.97449 | 77.220665 |
| PLHPLLVVAPNQL | 14 | Q9BQ52 | Zinc phosphodiesterase ELAC protein 2 ( | ELAC2 | 92.219 | 2 | 53.709312 | yes | 2.041E-07 | 513.6521 | 77.220665 |
| PLHPLLVVAPNQL | 14 | Q9BQ52 | Zinc phosphodiesterase ELAC protein 2 ( | ELAC2 | 92.219 | 2 | 53.709312 | yes | 1.76E-07 | 769.97449 | 77.220665 |
| YLFNCGEGVQR | 11 | Q9BQ52 | Zinc phosphodiesterase ELAC protein 2 ( | ELAC2 | 92.219 | 2 | 53.709312 | yes | 1.717E-07 | 671.81409 | 120.33202 |
| ELAGGLEDGEPQQ | 14 | Q9BQ52 | Zinc phosphodiesterase ELAC protein 2 ( | ELAC2 | 92.219 | 2 | 53.709312 | yes | 1.422E-07 | 735.85718 | 117.56475 |
| DATLLIHEATLED( | 22 | Q9BQ52 | Zinc phosphodiesterase ELAC protein 2 ( | ELAC2 | 92.219 | 3 | 53.709312 | yes | 1.862E-07 | 809.07086 | 100.51778 |
| TCDLEEFQTCLVR | 13 | Q9BQ52 | Zinc phosphodiesterase ELAC protein 2 ( | ELAC2 | 92.219 | 2 | 53.709312 | yes | 9.722E-08 | 835.87927 | 80.193573 |
| LDNIFLTR | 8 | Q9BQ52 | Zinc phosphodiesterase ELAC protein 2 ( | ELAC2 | 92.219 | 2 | 53.709312 | yes | 4.396E-07 | 496.2822 | 43.704769 |
| THSTTSQAISVGMI | 14 | Q9BQ52 | Zinc phosphodiesterase ELAC protein 2 ( | ELAC2 | 92.219 | 2 | 53.709312 | yes | 1.702E-07 | 738.36719 | 42.439404 |
| GGPGSTLSFVGK | 12 | Q9BQ61 | Telomerase RNA component interacting I | TRIR | 18.42 | 2 | 49.98263 | yes | 6.983E-07 | 553.79565 | 516.49512 |

| | | | | | | | | | | | |
|---|---|---|---|---|---|---|---|---|---|---|---|
| TEDEVLTSK | 9 | Q9BQ61 | Telomerase RNA component interacting I | TRIR | 18.42 | 2 | 49.98263 | yes | 2.623E-07 | 511.25604 | 223.88034 |
| EAPGPAGGGGGGS | 14 | Q9BQ61 | Telomerase RNA component interacting I | TRIR | 18.42 | 2 | 49.98263 | yes | 2.218E-07 | 563.76544 | 119.57591 |
| QKTEDEVLTSK | 11 | Q9BQ61 | Telomerase RNA component interacting I | TRIR | 18.42 | 2 | 49.98263 | yes | 9.938E-08 | 639.33282 | 77.498505 |
| KGGPGSTLSFVGK | 13 | Q9BQ61 | Telomerase RNA component interacting I | TRIR | 18.42 | 2 | 49.98263 | yes | 4.624E-07 | 617.84314 | 67.670944 |
| AQTGAPCLSFDIVI | 14 | Q9BQ67 | Glutamate-rich WD repeat-containing pro | GRWD1 | 49.42 | 2 | 51.594925 | yes | 1.38E-07 | 767.88776 | 5160.6313 |
| EPFLLSGGDDGAL | 14 | Q9BQ67 | Glutamate-rich WD repeat-containing pro | GRWD1 | 49.42 | 2 | 51.594925 | yes | 9.722E-08 | 709.86176 | 895.34344 |
| NIHLWTPTDGGSW | 18 | Q9BQ67 | Glutamate-rich WD repeat-containing pro | GRWD1 | 49.42 | 3 | 51.594925 | yes | 1.151E-07 | 707.0083 | 828.11652 |
| PFVGHTR | 7 | Q9BQ67 | Glutamate-rich WD repeat-containing pro | GRWD1 | 49.42 | 2 | 51.594925 | yes | 1.295E-07 | 407.22192 | 726.56732 |
| EGEELVMDEEAY | 17 | Q9BQ67 | Glutamate-rich WD repeat-containing pro | GRWD1 | 49.42 | 3 | 51.594925 | yes | 1.043E-07 | 694.65436 | 709.52386 |
| GQVEVFALR | 9 | Q9BQ67 | Glutamate-rich WD repeat-containing pro | GRWD1 | 49.42 | 2 | 51.594925 | yes | 1.967E-07 | 509.78763 | 408.24658 |
| KPQLELAMVPHY( | 17 | Q9BQ67 | Glutamate-rich WD repeat-containing pro | GRWD1 | 49.42 | 3 | 51.594925 | yes | 9.809E-08 | 641.68048 | 305.11029 |
| QHVAPVTSVEWH | 37 | Q9BQ67 | Glutamate-rich WD repeat-containing pro | GRWD1 | 49.42 | 5 | 51.594925 | yes | 1.951E-07 | 814.59753 | 247.81435 |
| TELPLTLYLCAGT( | 22 | Q9BQ67 | Glutamate-rich WD repeat-containing pro | GRWD1 | 49.42 | 3 | 51.594925 | yes | 9.722E-08 | 808.40057 | 211.69055 |
| REPFLLSGGDDGA | 15 | Q9BQ67 | Glutamate-rich WD repeat-containing pro | GRWD1 | 49.42 | 3 | 51.594925 | yes | 6.783E-07 | 525.6106 | 65.521408 |
| REPFLLSGGDDGA | 15 | Q9BQ67 | Glutamate-rich WD repeat-containing pro | GRWD1 | 49.42 | 3 | 51.594925 | yes | 4.208E-07 | 787.91229 | 65.521408 |
| REPFLLSGGDDGA | 15 | Q9BQ67 | Glutamate-rich WD repeat-containing pro | GRWD1 | 49.42 | 2 | 51.594925 | yes | 6.783E-07 | 525.6106 | 65.521408 |
| REPFLLSGGDDGA | 15 | Q9BQ67 | Glutamate-rich WD repeat-containing pro | GRWD1 | 49.42 | 2 | 51.594925 | yes | 4.208E-07 | 787.91229 | 65.521408 |
| ACMLTTATAHDGI | 21 | Q9BQ67 | Glutamate-rich WD repeat-containing pro | GRWD1 | 49.42 | 3 | 51.594925 | yes | 1.354E-06 | 769.0282 | 8.3739128 |
| HVILSEIK | 8 | Q9BQ70 | Ribosome quality control complex subuni | TCF25 | 76.667 | 2 | 53.114967 | yes | 2.145E-07 | 469.78711 | 310.83499 |
| LFQEWEAHR | 9 | Q9BQ70 | Ribosome quality control complex subuni | TCF25 | 76.667 | 2 | 53.114967 | yes | 9.722E-08 | 608.29889 | 260.25256 |
| GQEPLGPGALHFD | 15 | Q9BQ70 | Ribosome quality control complex subuni | TCF25 | 76.667 | 2 | 53.114967 | yes | 1.751E-06 | 803.92047 | 252.38438 |
| YSKPGLSMR | 9 | Q9BQ70 | Ribosome quality control complex subuni | TCF25 | 76.667 | 2 | 53.114967 | yes | 3.205E-07 | 519.77368 | 251.53366 |
| SGCALTDAVAPGN | 14 | Q9BQ70 | Ribosome quality control complex subuni | TCF25 | 76.667 | 2 | 53.114967 | yes | 1.211E-07 | 680.8299 | 210.30336 |
| CTWLTTPK | 8 | Q9BQ70 | Ribosome quality control complex subuni | TCF25 | 76.667 | 2 | 53.114967 | yes | 1.161E-07 | 503.75494 | 142.62833 |
| QMSFLEK | 7 | Q9BQ70 | Ribosome quality control complex subuni | TCF25 | 76.667 | 2 | 53.114967 | yes | 1.058E-06 | 441.72311 | 80.406723 |
| DDDDAEEEGPK | 11 | Q9BQ70 | Ribosome quality control complex subuni | TCF25 | 76.667 | 2 | 53.114967 | yes | 9.722E-08 | 610.23346 | 66.102295 |
| GLSFFAFEHSEEY( | 19 | Q9BQ70 | Ribosome quality control complex subuni | TCF25 | 76.667 | 3 | 53.114967 | yes | 1.231E-07 | 761.68701 | 60.82021 |
| DDDDAEEEGPKR | 12 | Q9BQ70 | Ribosome quality control complex subuni | TCF25 | 76.667 | 2 | 53.114967 | yes | 1.006E-07 | 688.28406 | 40.772018 |
| SSTGEASENGLEDI | 16 | Q9BQ70 | Ribosome quality control complex subuni | TCF25 | 76.667 | 2 | 53.114967 | yes | 2.893E-06 | 840.3714 | 17.056137 |
| QQECGIAVLEQME | 21 | Q9BQ95 | Evolutionarily conserved signaling interm | ECSIT | 49.149 | 3 | 33.047287 | yes | 1.339E-06 | 804.70831 | 33.154888 |
| FMNVNPFPVPR | 11 | Q9BQ95 | Evolutionarily conserved signaling interm | ECSIT | 49.149 | 2 | 33.047287 | yes | 3.124E-06 | 659.34229 | 20.539295 |
| SDGALLLGASSLS( | 15 | Q9BQA1 | Methylosome protein WDR77 OS=Homo | WDR77 | 36.725 | 2 | 50.599056 | yes | 1.432E-07 | 702.37805 | 1496.2334 |
| DSVFLSCSEDNR | 12 | Q9BQA1 | Methylosome protein WDR77 OS=Homo | WDR77 | 36.725 | 2 | 50.599056 | yes | 1.175E-07 | 714.80664 | 1158.7081 |
| ILLWDTR | 7 | Q9BQA1 | Methylosome protein WDR77 OS=Homo | WDR77 | 36.725 | 2 | 50.599056 | yes | 1.996E-07 | 458.76617 | 1133.0073 |
| ETPPPLVPPAAR | 12 | Q9BQA1 | Methylosome protein WDR77 OS=Homo | WDR77 | 36.725 | 2 | 50.599056 | yes | 2.649E-07 | 622.85352 | 1071.9382 |
| YEHDDIVSTVSVL | 24 | Q9BQA1 | Methylosome protein WDR77 OS=Homo | WDR77 | 36.725 | 3 | 50.599056 | yes | 9.752E-08 | 822.73798 | 910.09937 |
| VWDLAQQVVLSS | 14 | Q9BQA1 | Methylosome protein WDR77 OS=Homo | WDR77 | 36.725 | 2 | 50.599056 | yes | 1.874E-07 | 832.44373 | 664.45801 |
| AHAAQVTCVAASI | 15 | Q9BQA1 | Methylosome protein WDR77 OS=Homo | WDR77 | 36.725 | 3 | 50.599056 | yes | 1.895E-07 | 516.59644 | 232.61763 |
| AHAAQVTCVAASI | 15 | Q9BQA1 | Methylosome protein WDR77 OS=Homo | WDR77 | 36.725 | 3 | 50.599056 | yes | 3.044E-07 | 774.39099 | 232.61763 |

| | | | | | | | | | | | |
|---|---|---|---|---|---|---|---|---|---|---|---|
| AHAAQVTCVAASI | 15 | Q9BQA1 | Methylosome protein WDR77 OS=Homo | WDR77 | 36.725 | 2 | 50.599056 | yes | 1.895E-07 | 516.59644 | 232.61763 |
| AHAAQVTCVAASI | 15 | Q9BQA1 | Methylosome protein WDR77 OS=Homo | WDR77 | 36.725 | 2 | 50.599056 | yes | 3.044E-07 | 774.39099 | 232.61763 |
| EWNLPPNAPACMI | 14 | Q9BQA1 | Methylosome protein WDR77 OS=Homo | WDR77 | 36.725 | 2 | 50.599056 | yes | 2.714E-07 | 842.88214 | 184.52408 |
| ALCDVGTAISCSR | 13 | Q9BQB6 | Vitamin K epoxide reductase complex sul | VKORC1 | 18.235 | 2 | 50.56119 | yes | 9.766E-08 | 705.32922 | 564.42896 |
| GSTWGSPGWVR | 11 | Q9BQB6 | Vitamin K epoxide reductase complex sul | VKORC1 | 18.235 | 2 | 50.56119 | yes | 5.075E-07 | 595.29108 | 92.555283 |
| TIQFVDWCPTGFK | 13 | Q9BQE3 | Tubulin alpha-1C chain OS=Homo sapier | TUBA1C | 49.896 | 2 | 48.94147 | no | 1.084E-07 | 799.88721 | 1334.52 |
| AVCMLSNTTAVAI | 17 | Q9BQE3 | Tubulin alpha-1C chain OS=Homo sapier | TUBA1C | 49.896 | 3 | 48.94147 | yes | 1.612E-07 | 617.63446 | 41.057571 |
| PALETEGLR | 9 | Q9BQE4 | Selenoprotein S OS=Homo sapiens OX=S | SELENOS | 21.163 | 2 | 54.317448 | yes | 1.24E-07 | 493.26929 | 205.51178 |
| IEMWDSMQEGK | 11 | Q9BQE4 | Selenoprotein S OS=Homo sapiens OX=S | SELENOS | 21.163 | 2 | 54.317448 | yes | 1.499E-07 | 677.29431 | 104.57954 |
| AAAAVEPDVVVK | 12 | Q9BQE4 | Selenoprotein S OS=Homo sapiens OX=S | SELENOS | 21.163 | 2 | 54.317448 | yes | 9.722E-08 | 584.83228 | 63.874249 |
| MQEELNAQVEK | 11 | Q9BQE4 | Selenoprotein S OS=Homo sapiens OX=S | SELENOS | 21.163 | 2 | 54.317448 | yes | 2.518E-07 | 659.81903 | 28.291153 |
| ALAEEVEQVHR | 11 | Q9BQE5 | Apolipoprotein L2 OS=Homo sapiens OX | APOL2 | 37.079 | 2 | 47.788548 | yes | 1.259E-07 | 640.83331 | 47.253105 |
| ELEDHIR | 7 | Q9BQE5 | Apolipoprotein L2 OS=Homo sapiens OX | APOL2 | 37.079 | 2 | 47.788548 | yes | 1.367E-06 | 456.2327 | 35.45911 |
| WVTVGDTSLR | 10 | Q9BQE9 | B-cell CLL/lymphoma 7 protein family m | BCL7B | 22.196 | 2 | 48.07122 | no | 1.204E-07 | 567.30109 | 1097.1057 |
| IFKWVPVTDSK | 11 | Q9BQE9 | B-cell CLL/lymphoma 7 protein family m | BCL7B | 22.196 | 2 | 48.07122 | yes | 5.095E-06 | 660.37152 | 118.95781 |
| KWVTVGDTSLR | 11 | Q9BQE9 | B-cell CLL/lymphoma 7 protein family m | BCL7B | 22.196 | 2 | 48.07122 | no | 1.591E-07 | 631.34857 | 29.244072 |
| LVGSVNLFSDENV | 15 | Q9BQG0 | Myb-binding protein 1A OS=Homo sapie | MYBBP1A | 148.855 | 2 | 52.84357 | yes | 9.722E-08 | 823.43085 | 614.59045 |
| EVVEQGLLK | 9 | Q9BQG0 | Myb-binding protein 1A OS=Homo sapie | MYBBP1A | 148.855 | 2 | 52.84357 | yes | 9.722E-08 | 507.79514 | 569.61432 |
| AGALHAQVER | 10 | Q9BQG0 | Myb-binding protein 1A OS=Homo sapie | MYBBP1A | 148.855 | 2 | 52.84357 | yes | 1.065E-07 | 526.28577 | 514.77673 |
| EQLMTVLQAGK | 11 | Q9BQG0 | Myb-binding protein 1A OS=Homo sapie | MYBBP1A | 148.855 | 2 | 52.84357 | yes | 1.464E-07 | 609.33154 | 492.65002 |
| EIPSATQSPISK | 12 | Q9BQG0 | Myb-binding protein 1A OS=Homo sapie | MYBBP1A | 148.855 | 2 | 52.84357 | yes | 5.207E-07 | 629.33789 | 449.56561 |
| ELEAHSAEAR | 10 | Q9BQG0 | Myb-binding protein 1A OS=Homo sapie | MYBBP1A | 148.855 | 2 | 52.84357 | yes | 9.722E-08 | 556.7702 | 422.38681 |
| ALVDILSEVSK | 11 | Q9BQG0 | Myb-binding protein 1A OS=Homo sapie | MYBBP1A | 148.855 | 2 | 52.84357 | yes | 1.103E-07 | 587.3399 | 352.5155 |
| VYSTALSSFLTK | 12 | Q9BQG0 | Myb-binding protein 1A OS=Homo sapie | MYBBP1A | 148.855 | 2 | 52.84357 | yes | 9.919E-08 | 658.85846 | 334.29562 |
| KPTSQIPETK | 10 | Q9BQG0 | Myb-binding protein 1A OS=Homo sapie | MYBBP1A | 148.855 | 2 | 52.84357 | yes | 9.722E-08 | 564.81659 | 332.08044 |
| LLGAALPLLTK | 11 | Q9BQG0 | Myb-binding protein 1A OS=Homo sapie | MYBBP1A | 148.855 | 2 | 52.84357 | yes | 1.497E-07 | 555.36829 | 290.12921 |
| SVFGHICSHLTPR | 13 | Q9BQG0 | Myb-binding protein 1A OS=Homo sapie | MYBBP1A | 148.855 | 3 | 52.84357 | yes | 1.591E-07 | 504.25763 | 122.4443 |
| SVFGHICSHLTPR | 13 | Q9BQG0 | Myb-binding protein 1A OS=Homo sapie | MYBBP1A | 148.855 | 3 | 52.84357 | yes | 3.878E-07 | 755.88281 | 122.4443 |
| SVFGHICSHLTPR | 13 | Q9BQG0 | Myb-binding protein 1A OS=Homo sapie | MYBBP1A | 148.855 | 2 | 52.84357 | yes | 1.591E-07 | 504.25763 | 122.4443 |
| SVFGHICSHLTPR | 13 | Q9BQG0 | Myb-binding protein 1A OS=Homo sapie | MYBBP1A | 148.855 | 2 | 52.84357 | yes | 3.878E-07 | 755.88281 | 122.4443 |
| HQACLLLQK | 9 | Q9BQG0 | Myb-binding protein 1A OS=Homo sapie | MYBBP1A | 148.855 | 2 | 52.84357 | yes | 1.339E-07 | 555.80804 | 236.22935 |
| LHDLYWQAMK | 10 | Q9BQG0 | Myb-binding protein 1A OS=Homo sapie | MYBBP1A | 148.855 | 2 | 52.84357 | yes | 1.7E-07 | 652.82642 | 231.31136 |
| EQLHLVMQGDVIF | 13 | Q9BQG0 | Myb-binding protein 1A OS=Homo sapie | MYBBP1A | 148.855 | 2 | 52.84357 | yes | 1.301E-07 | 769.41138 | 230.05788 |
| HPFSFPLENQAR | 12 | Q9BQG0 | Myb-binding protein 1A OS=Homo sapie | MYBBP1A | 148.855 | 2 | 52.84357 | yes | 9.722E-08 | 721.86243 | 220.84978 |
| KLPAIALDLLR | 11 | Q9BQG0 | Myb-binding protein 1A OS=Homo sapie | MYBBP1A | 148.855 | 2 | 52.84357 | yes | 1.11E-07 | 611.89771 | 214.03665 |
| ATLQEILPEVLK | 12 | Q9BQG0 | Myb-binding protein 1A OS=Homo sapie | MYBBP1A | 148.855 | 2 | 52.84357 | yes | 9.722E-08 | 677.40302 | 208.16228 |
| TLGVQRPK | 8 | Q9BQG0 | Myb-binding protein 1A OS=Homo sapie | MYBBP1A | 148.855 | 2 | 52.84357 | yes | 4.852E-07 | 449.77707 | 192.6532 |
| YCHDLGER | 8 | Q9BQG0 | Myb-binding protein 1A OS=Homo sapie | MYBBP1A | 148.855 | 2 | 52.84357 | yes | 2.897E-07 | 525.22711 | 178.54539 |
| SCFEDPEWK | 9 | Q9BQG0 | Myb-binding protein 1A OS=Homo sapie | MYBBP1A | 148.855 | 2 | 52.84357 | yes | 9.729E-08 | 599.24768 | 166.42157 |

| | | | | | | | | | | | |
|---|---|---|---|---|---|---|---|---|---|---|---|
| VVVTDDSDER | 10 | Q9BQG0 | Myb-binding protein 1A OS=Homo sapie | MYBBP1A | 148.855 | 2 | 52.84357 | yes | 2.5E-07 | 567.76727 | 159.59148 |
| LITGLGVGR | 9 | Q9BQG0 | Myb-binding protein 1A OS=Homo sapie | MYBBP1A | 148.855 | 2 | 52.84357 | yes | 1.041E-06 | 443.27945 | 125.13462 |
| HYGEHVCTAK | 10 | Q9BQG0 | Myb-binding protein 1A OS=Homo sapie | MYBBP1A | 148.855 | 3 | 52.84357 | yes | 2.442E-07 | 401.18549 | 115.76718 |
| QLMGQVLAK | 9 | Q9BQG0 | Myb-binding protein 1A OS=Homo sapie | MYBBP1A | 148.855 | 2 | 52.84357 | yes | 5.369E-07 | 494.28641 | 101.55991 |
| LLQALAQYQNHL( | 17 | Q9BQG0 | Myb-binding protein 1A OS=Homo sapie | MYBBP1A | 148.855 | 3 | 52.84357 | yes | 1.816E-07 | 684.03247 | 70.090591 |
| IFTHHLCR | 8 | Q9BQG0 | Myb-binding protein 1A OS=Homo sapie | MYBBP1A | 148.855 | 2 | 52.84357 | yes | 9.722E-08 | 542.27948 | 62.106594 |
| FCLFHSFFVTK | 11 | Q9BQG0 | Myb-binding protein 1A OS=Homo sapie | MYBBP1A | 148.855 | 2 | 52.84357 | yes | 9.722E-08 | 716.85773 | 53.361164 |
| QPDSPTALYHFNA | 19 | Q9BQG0 | Myb-binding protein 1A OS=Homo sapie | MYBBP1A | 148.855 | 3 | 52.84357 | yes | 3.634E-07 | 736.04791 | 39.211983 |
| EYTSNVFLQGK | 11 | Q9BQP7 | Mitochondrial genome maintenance exon | MGME1 | 39.421 | 2 | 44.59017 | yes | 2.139E-07 | 643.32477 | 111.11296 |
| MILELGEDGFK | 11 | Q9BQP7 | Mitochondrial genome maintenance exon | MGME1 | 39.421 | 2 | 44.59017 | yes | 4.22E-07 | 626.31812 | 38.920715 |
| YSNLVQSVLSSR | 12 | Q9BQP7 | Mitochondrial genome maintenance exon | MGME1 | 39.421 | 2 | 44.59017 | yes | 5.135E-07 | 676.86206 | 26.419006 |
| SGYIESVQHILK | 12 | Q9BQP7 | Mitochondrial genome maintenance exon | MGME1 | 39.421 | 2 | 44.59017 | yes | 6.078E-05 | 687.37482 | 13.020691 |
| VLGLAMVPGHHI\ | 19 | Q9BRA0 | N-alpha-acetyltransferase 38, NatC auxili | NAA38 | 13.515 | 3 | 52.506554 | yes | 1.549E-07 | 685.7226 | 136.65193 |
| TLVGCFLCTDR | 11 | Q9BRA0 | N-alpha-acetyltransferase 38, NatC auxili | NAA38 | 13.515 | 2 | 52.506554 | yes | 9.722E-08 | 671.31812 | 96.821327 |
| TGLVLTDIQR | 10 | Q9BRD0 | BUD13 homolog OS=Homo sapiens OX= | BUD13 | 70.521 | 2 | 51.592598 | yes | 1.732E-07 | 558.32458 | 238.24307 |
| TLGSSDTQQLR | 11 | Q9BRD0 | BUD13 homolog OS=Homo sapiens OX= | BUD13 | 70.521 | 2 | 51.592598 | yes | 9.722E-08 | 603.30969 | 183.53806 |
| EMQKPLAR | 8 | Q9BRD0 | BUD13 homolog OS=Homo sapiens OX= | BUD13 | 70.521 | 2 | 51.592598 | yes | 9.722E-08 | 486.7684 | 75.04216 |
| EGDPMANFIK | 10 | Q9BRD0 | BUD13 homolog OS=Homo sapiens OX= | BUD13 | 70.521 | 2 | 51.592598 | yes | 1.83E-06 | 561.26843 | 70.232315 |
| ESGASHLSFPK | 11 | Q9BRD0 | BUD13 homolog OS=Homo sapiens OX= | BUD13 | 70.521 | 2 | 51.592598 | yes | 1.538E-06 | 580.29077 | 65.896164 |
| QQQQNVEDAMK | 11 | Q9BRD0 | BUD13 homolog OS=Homo sapiens OX= | BUD13 | 70.521 | 2 | 51.592598 | yes | 1.296E-07 | 659.80646 | 55.120628 |
| FNIWPGYR | 8 | Q9BRD0 | BUD13 homolog OS=Homo sapiens OX= | BUD13 | 70.521 | 2 | 51.592598 | yes | 2.481E-07 | 526.76923 | 46.539101 |
| HDSPDLAPNVTYS | 16 | Q9BRD0 | BUD13 homolog OS=Homo sapiens OX= | BUD13 | 70.521 | 3 | 51.592598 | yes | 6.482E-07 | 594.63208 | 36.756374 |
| KAVEELAYK | 9 | Q9BRD0 | BUD13 homolog OS=Homo sapiens OX= | BUD13 | 70.521 | 2 | 51.592598 | yes | 2.233E-06 | 525.7951 | 33.564983 |
| VLGLGLGCLR | 10 | Q9BRJ7 | Tudor-interacting repair regulator protein | NUDT16L1 | 23.339 | 2 | 52.679245 | yes | 1.425E-07 | 529.31299 | 1998.2836 |
| LTEADYLSSHLTE( | 17 | Q9BRJ7 | Tudor-interacting repair regulator protein | NUDT16L1 | 23.339 | 3 | 52.679245 | yes | 1.39E-07 | 642.6499 | 1311.287 |
| LVEALAAATEK | 11 | Q9BRJ7 | Tudor-interacting repair regulator protein | NUDT16L1 | 23.339 | 2 | 52.679245 | yes | 1.02E-07 | 558.31897 | 821.99554 |
| DHGLEVLGLVR | 11 | Q9BRJ7 | Tudor-interacting repair regulator protein | NUDT16L1 | 23.339 | 2 | 52.679245 | no | 1.011E-07 | 604.34332 | 759.09058 |
| FWSLEDGLNR | 10 | Q9BRJ7 | Tudor-interacting repair regulator protein | NUDT16L1 | 23.339 | 2 | 52.679245 | yes | 5.406E-07 | 618.80402 | 756.90527 |
| VVAHLYAR | 8 | Q9BRJ7 | Tudor-interacting repair regulator protein | NUDT16L1 | 23.339 | 2 | 52.679245 | yes | 1.689E-07 | 464.77179 | 580.82928 |
| QLTLEQLHAVEIS/ | 18 | Q9BRJ7 | Tudor-interacting repair regulator protein | NUDT16L1 | 23.339 | 3 | 52.679245 | yes | 9.722E-08 | 677.70422 | 264.23312 |
| QLTLEQLHAVEIS/ | 18 | Q9BRJ7 | Tudor-interacting repair regulator protein | NUDT16L1 | 23.339 | 3 | 52.679245 | yes | 5.011E-07 | 508.53 | 264.23312 |
| QLTLEQLHAVEIS/ | 18 | Q9BRJ7 | Tudor-interacting repair regulator protein | NUDT16L1 | 23.339 | 4 | 52.679245 | yes | 9.722E-08 | 677.70422 | 264.23312 |
| QLTLEQLHAVEIS/ | 18 | Q9BRJ7 | Tudor-interacting repair regulator protein | NUDT16L1 | 23.339 | 4 | 52.679245 | yes | 5.011E-07 | 508.53 | 264.23312 |
| VPLYTQK | 7 | Q9BRJ7 | Tudor-interacting repair regulator protein | NUDT16L1 | 23.339 | 2 | 52.679245 | yes | 1.288E-07 | 424.74747 | 365.48013 |
| FSVLMQMR | 8 | Q9BRJ7 | Tudor-interacting repair regulator protein | NUDT16L1 | 23.339 | 2 | 52.679245 | yes | 1.607E-07 | 506.25934 | 315.55441 |
| VLNMMPEEK | 9 | Q9BRJ7 | Tudor-interacting repair regulator protein | NUDT16L1 | 23.339 | 2 | 52.679245 | yes | 1.09E-07 | 545.76721 | 228.40938 |
| CQLLFALK | 8 | Q9BRJ7 | Tudor-interacting repair regulator protein | NUDT16L1 | 23.339 | 2 | 52.679245 | yes | 1.847E-07 | 496.78351 | 106.23799 |
| VAGGLLGSGTR | 11 | Q9BRK4 | Leucine zipper putative tumor suppressor | LZTS2 | 72.76 | 2 | 49.796696 | yes | 1.902E-07 | 494.28271 | 89.918533 |
| ELELEACSQELQR | 13 | Q9BRK4 | Leucine zipper putative tumor suppressor | LZTS2 | 72.76 | 2 | 49.796696 | yes | 1.003E-07 | 802.88287 | 59.995102 |

| | | | | | | | | | | | |
|---|---|---|---|---|---|---|---|---|---|---|---|
| DLGGFDEDAEPR | 12 | Q9BRK5 | 45 kDa calcium-binding protein OS=Hon | SDF4 | 41.807 | 2 | 57.60111 | yes | 9.722E-08 | 660.78876 | 222.84314 |
| LEMDGHLNR | 9 | Q9BRK5 | 45 kDa calcium-binding protein OS=Hon | SDF4 | 41.807 | 2 | 57.60111 | yes | 4.001E-07 | 542.76385 | 169.6107 |
| VDEETQEVLENLK | 13 | Q9BRK5 | 45 kDa calcium-binding protein OS=Hon | SDF4 | 41.807 | 2 | 57.60111 | yes | 9.722E-08 | 773.38574 | 111.94968 |
| QMIAVADENQNHI | 21 | Q9BRK5 | 45 kDa calcium-binding protein OS=Hon | SDF4 | 41.807 | 3 | 57.60111 | yes | 1.302E-07 | 815.4007 | 80.585526 |
| GFHQEVFLGK | 10 | Q9BRK5 | 45 kDa calcium-binding protein OS=Hon | SDF4 | 41.807 | 2 | 57.60111 | yes | 1.086E-07 | 581.30621 | 40.055672 |
| TAEHFQEAMEESK | 13 | Q9BRK5 | 45 kDa calcium-binding protein OS=Hon | SDF4 | 41.807 | 2 | 57.60111 | yes | 3.452E-07 | 768.83539 | 30.968243 |
| YSEFFTGSK | 9 | Q9BRK5 | 45 kDa calcium-binding protein OS=Hon | SDF4 | 41.807 | 2 | 57.60111 | yes | 2.143E-06 | 533.24799 | 12.837276 |
| IQAGEVSQPSK | 11 | Q9BRP8 | Partner of Y14 and mago OS=Homo sapi | PYM1 | 22.656 | 2 | 49.40474 | yes | 1.665E-07 | 572.30383 | 320.73853 |
| QVEELQQR | 8 | Q9BRP8 | Partner of Y14 and mago OS=Homo sapi | PYM1 | 22.656 | 2 | 49.40474 | yes | 2.442E-07 | 515.26984 | 255.14539 |
| MEAAGSPAATETC | 14 | Q9BRP8 | Partner of Y14 and mago OS=Homo sapi | PYM1 | 22.656 | 2 | 49.40474 | yes | 1.035E-07 | 681.8139 | 177.6098 |
| HSVAVMTVR | 9 | Q9BRS2 | Serine/threonine-protein kinase RIO1 OS | RIOK1 | 65.583 | 2 | 58.20153 | yes | 2.59E-07 | 500.27383 | 625.28503 |
| LNVTDSVINK | 10 | Q9BRS2 | Serine/threonine-protein kinase RIO1 OS | RIOK1 | 65.583 | 2 | 58.20153 | yes | 9.722E-08 | 551.80878 | 493.99493 |
| DKYVSGEFR | 9 | Q9BRS2 | Serine/threonine-protein kinase RIO1 OS | RIOK1 | 65.583 | 2 | 58.20153 | yes | 1.016E-07 | 550.77222 | 290.01849 |
| AMEIASQR | 8 | Q9BRS2 | Serine/threonine-protein kinase RIO1 OS | RIOK1 | 65.583 | 2 | 58.20153 | yes | 1.763E-07 | 453.2291 | 272.13867 |
| VPALLENQVEER | 12 | Q9BRS2 | Serine/threonine-protein kinase RIO1 OS | RIOK1 | 65.583 | 2 | 58.20153 | yes | 9.722E-08 | 698.87518 | 214.48163 |
| ATVEQVLDPR | 10 | Q9BRS2 | Serine/threonine-protein kinase RIO1 OS | RIOK1 | 65.583 | 2 | 58.20153 | yes | 1.998E-06 | 564.3064 | 196.61011 |
| DCANVNDFFMR | 11 | Q9BRS2 | Serine/threonine-protein kinase RIO1 OS | RIOK1 | 65.583 | 2 | 58.20153 | yes | 9.722E-08 | 694.78973 | 175.41681 |
| SSQDHVDEEVFK | 12 | Q9BRS2 | Serine/threonine-protein kinase RIO1 OS | RIOK1 | 65.583 | 2 | 58.20153 | yes | 9.722E-08 | 710.323 | 117.61919 |
| GIITEINGCISTGK | 14 | Q9BRS2 | Serine/threonine-protein kinase RIO1 OS | RIOK1 | 65.583 | 2 | 58.20153 | yes | 1.263E-07 | 731.88214 | 105.88166 |
| SHVLVMSFIGK | 11 | Q9BRS2 | Serine/threonine-protein kinase RIO1 OS | RIOK1 | 65.583 | 2 | 58.20153 | yes | 1.151E-07 | 609.33917 | 80.398232 |
| GYVWNGGSNPQA | 14 | Q9BRS2 | Serine/threonine-protein kinase RIO1 OS | RIOK1 | 65.583 | 2 | 58.20153 | yes | 9.722E-08 | 760.35547 | 71.026413 |
| ISGPNPLSCLK | 11 | Q9BRU9 | rRNA-processing protein UTP23 homolo | UTP23 | 28.403 | 2 | 51.184395 | yes | 9.722E-08 | 593.31842 | 256.17654 |
| CILVASPGFVR | 11 | Q9BRX2 | Protein pelota homolog OS=Homo sapien | PELO | 43.36 | 2 | 58.220314 | yes | 4.663E-07 | 609.83679 | 956.9024 |
| FYEQVVQAIQR | 11 | Q9BRX2 | Protein pelota homolog OS=Homo sapien | PELO | 43.36 | 2 | 58.220314 | yes | 9.809E-08 | 690.86713 | 747.76495 |
| GTNIQENEYVK | 11 | Q9BRX2 | Protein pelota homolog OS=Homo sapien | PELO | 43.36 | 2 | 58.220314 | yes | 4.239E-07 | 647.81732 | 343.5864 |
| EALCDPTVASR | 11 | Q9BRX2 | Protein pelota homolog OS=Homo sapien | PELO | 43.36 | 2 | 58.220314 | yes | 1.812E-07 | 609.79279 | 310.73743 |
| AKVEVNIPR | 9 | Q9BRX2 | Protein pelota homolog OS=Homo sapien | PELO | 43.36 | 2 | 58.220314 | yes | 1.781E-06 | 513.30872 | 232.47791 |
| MLQHEPDR | 8 | Q9BRX2 | Protein pelota homolog OS=Homo sapien | PELO | 43.36 | 2 | 58.220314 | yes | 9.722E-08 | 513.2453 | 225.02342 |
| VQTESSTGSVGSN | 14 | Q9BRX2 | Protein pelota homolog OS=Homo sapien | PELO | 43.36 | 2 | 58.220314 | yes | 1.025E-07 | 704.83679 | 199.12726 |
| KQWDSVVLER | 10 | Q9BRX2 | Protein pelota homolog OS=Homo sapien | PELO | 43.36 | 2 | 58.220314 | yes | 9.722E-08 | 630.34076 | 122.72487 |
| IFSSLHVSGEQLSQ | 23 | Q9BRX2 | Protein pelota homolog OS=Homo sapien | PELO | 43.36 | 3 | 58.220314 | yes | 2.263E-07 | 809.4516 | 100.42256 |
| FLQVHASSGHK | 11 | Q9BRX2 | Protein pelota homolog OS=Homo sapien | PELO | 43.36 | 2 | 58.220314 | yes | 3.617E-07 | 605.82001 | 99.118851 |
| QWDSVVLER | 9 | Q9BRX2 | Protein pelota homolog OS=Homo sapien | PELO | 43.36 | 2 | 58.220314 | yes | 1.254E-07 | 566.29327 | 89.343399 |
| MGAYHTIELEPNR | 13 | Q9BRX2 | Protein pelota homolog OS=Homo sapien | PELO | 43.36 | 2 | 58.220314 | yes | 1.736E-07 | 765.87213 | 69.902489 |
| AHVEAAEEAAR | 11 | Q9BRZ2 | E3 ubiquitin-protein ligase TRIM56 OS= | TRIM56 | 81.488 | 2 | 52.7605 | yes | 9.722E-08 | 577.28345 | 2265.4402 |
| ICLEQLR | 7 | Q9BRZ2 | E3 ubiquitin-protein ligase TRIM56 OS= | TRIM56 | 81.488 | 2 | 52.7605 | yes | 1.863E-07 | 466.25513 | 1792.8264 |
| VGTQVEEAAEGVI | 14 | Q9BRZ2 | E3 ubiquitin-protein ligase TRIM56 OS= | TRIM56 | 81.488 | 2 | 52.7605 | yes | 9.722E-08 | 729.38336 | 1552.5638 |
| VVDLVGYR | 8 | Q9BRZ2 | E3 ubiquitin-protein ligase TRIM56 OS= | TRIM56 | 81.488 | 2 | 52.7605 | yes | 2.284E-07 | 460.76364 | 1154.3459 |
| ITGLCPFGPR | 10 | Q9BRZ2 | E3 ubiquitin-protein ligase TRIM56 OS= | TRIM56 | 81.488 | 2 | 52.7605 | yes | 2.002E-07 | 559.2948 | 1023.6271 |

| | | | | | | | | | | | |
|---|---|---|---|---|---|---|---|---|---|---|---|
| EILVADEQNR | 10 | Q9BRZ2 | E3 ubiquitin-protein ligase TRIM56 OS= | TRIM56 | 81.488 | 2 | 52.7605 | yes | 1.659E-06 | 593.80676 | 1021.1142 |
| QEVLGQLR | 8 | Q9BRZ2 | E3 ubiquitin-protein ligase TRIM56 OS= | TRIM56 | 81.488 | 2 | 52.7605 | yes | 2.234E-07 | 471.772 | 907.02777 |
| ETVPVPPEGVASFI | 14 | Q9BRZ2 | E3 ubiquitin-protein ligase TRIM56 OS= | TRIM56 | 81.488 | 2 | 52.7605 | yes | 1.186E-07 | 728.88776 | 796.15985 |
| LYLINPNGEVQWR | 13 | Q9BRZ2 | E3 ubiquitin-protein ligase TRIM56 OS= | TRIM56 | 81.488 | 2 | 52.7605 | yes | 1.073E-07 | 801.42535 | 777.70984 |
| FLCQPCSQLLCR | 12 | Q9BRZ2 | E3 ubiquitin-protein ligase TRIM56 OS= | TRIM56 | 81.488 | 2 | 52.7605 | yes | 9.722E-08 | 791.37036 | 629.02429 |
| PIFYCSFPTR | 10 | Q9BRZ2 | E3 ubiquitin-protein ligase TRIM56 OS= | TRIM56 | 81.488 | 2 | 52.7605 | yes | 1.025E-07 | 644.31317 | 559.07422 |
| EEGAQTLEEDR | 11 | Q9BRZ2 | E3 ubiquitin-protein ligase TRIM56 OS= | TRIM56 | 81.488 | 2 | 52.7605 | yes | 1.041E-07 | 638.78619 | 495.0206 |
| LSFEEQQPQK | 10 | Q9BRZ2 | E3 ubiquitin-protein ligase TRIM56 OS= | TRIM56 | 81.488 | 2 | 52.7605 | yes | 1.084E-07 | 617.30914 | 441.78726 |
| ALSLSQASHAVAA | 19 | Q9BRZ2 | E3 ubiquitin-protein ligase TRIM56 OS= | TRIM56 | 81.488 | 3 | 52.7605 | yes | 1.3E-07 | 617.66217 | 385.31857 |
| EAEILSLEGAIAQR | 14 | Q9BRZ2 | E3 ubiquitin-protein ligase TRIM56 OS= | TRIM56 | 81.488 | 2 | 52.7605 | yes | 9.722E-08 | 750.40686 | 379.78275 |
| VAVSVAGHVEVY | 21 | Q9BRZ2 | E3 ubiquitin-protein ligase TRIM56 OS= | TRIM56 | 81.488 | 3 | 52.7605 | yes | 9.776E-08 | 730.37213 | 370.18298 |
| GSLLGDFLTAYHG | 18 | Q9BRZ2 | E3 ubiquitin-protein ligase TRIM56 OS= | TRIM56 | 81.488 | 3 | 52.7605 | yes | 1.573E-07 | 658.68628 | 235.07593 |
| AGWYDEEAR | 9 | Q9BRZ2 | E3 ubiquitin-protein ligase TRIM56 OS= | TRIM56 | 81.488 | 2 | 52.7605 | yes | 2.584E-07 | 548.73834 | 195.18835 |
| LDPHLDHPCLPLA | 17 | Q9BRZ2 | E3 ubiquitin-protein ligase TRIM56 OS= | TRIM56 | 81.488 | 4 | 52.7605 | yes | 1.222E-07 | 489.00574 | 182.7457 |
| QGGVQPQAGDGA | 16 | Q9BRZ2 | E3 ubiquitin-protein ligase TRIM56 OS= | TRIM56 | 81.488 | 2 | 52.7605 | yes | 2.621E-06 | 769.88153 | 150.27608 |
| AQTPHEDGGPQPF | 14 | Q9BRZ2 | E3 ubiquitin-protein ligase TRIM56 OS= | TRIM56 | 81.488 | 2 | 52.7605 | yes | 2.245E-07 | 763.8584 | 145.96675 |
| GPGLHGCQPGSVS | 16 | Q9BRZ2 | E3 ubiquitin-protein ligase TRIM56 OS= | TRIM56 | 81.488 | 2 | 52.7605 | yes | 1.028E-07 | 797.88574 | 122.9109 |
| FSLNGDYK | 8 | Q9BRZ2 | E3 ubiquitin-protein ligase TRIM56 OS= | TRIM56 | 81.488 | 2 | 52.7605 | yes | 6.274E-07 | 472.22961 | 107.07204 |
| KGYIFLTLR | 9 | Q9BRZ2 | E3 ubiquitin-protein ligase TRIM56 OS= | TRIM56 | 81.488 | 2 | 52.7605 | yes | 9.722E-08 | 555.83716 | 75.922035 |
| QAAQCPQHPGEAI | 14 | Q9BRZ2 | E3 ubiquitin-protein ligase TRIM56 OS= | TRIM56 | 81.488 | 2 | 52.7605 | yes | 3.473E-07 | 781.87823 | 54.779736 |
| GYIFLTLR | 8 | Q9BRZ2 | E3 ubiquitin-protein ligase TRIM56 OS= | TRIM56 | 81.488 | 2 | 52.7605 | yes | 3.272E-07 | 491.78964 | 43.634907 |
| VTTMVDGR | 8 | Q9BRZ2 | E3 ubiquitin-protein ligase TRIM56 OS= | TRIM56 | 81.488 | 2 | 52.7605 | yes | 4.706E-07 | 439.72366 | 24.04636 |
| NIIGYFEQK | 9 | Q9BS26 | Endoplasmic reticulum resident protein 4 | ERP44 | 46.972 | 2 | 23.40007 | yes | 3.654E-06 | 556.29279 | 40.897308 |
| STYLGIHER | 9 | Q9BS34 | Zinc finger protein 670 OS=Homo sapien | ZNF670 | 44.604 | 2 | 53.117775 | yes | 2.291E-06 | 538.28015 | 140.30313 |
| AFISLTSVDR | 10 | Q9BS34 | Zinc finger protein 670 OS=Homo sapien | ZNF670 | 44.604 | 2 | 53.117775 | yes | 1.826E-07 | 554.80347 | 102.8149 |
| NLSSHVVER | 9 | Q9BS34 | Zinc finger protein 670 OS=Homo sapien | ZNF670 | 44.604 | 2 | 53.117775 | yes | 2.916E-07 | 520.77783 | 73.848686 |
| YSSNLCEHER | 10 | Q9BS34 | Zinc finger protein 670 OS=Homo sapien | ZNF670 | 44.604 | 2 | 53.117775 | yes | 9.722E-08 | 647.77765 | 49.860493 |
| GKPLALVEEIR | 11 | Q9BSD7 | Cancer-related nucleoside-triphosphatase | NTPCR | 20.714 | 2 | 53.680965 | yes | 1.318E-07 | 612.86914 | 956.42743 |
| IGFDVVTLSGTR | 12 | Q9BSD7 | Cancer-related nucleoside-triphosphatase | NTPCR | 20.714 | 2 | 53.680965 | yes | 9.722E-08 | 632.84845 | 742.74231 |
| SSGVPVDGFYTEE | 15 | Q9BSD7 | Cancer-related nucleoside-triphosphatase | NTPCR | 20.714 | 2 | 53.680965 | yes | 2.623E-07 | 821.39136 | 389.5141 |
| VCVIDEIGK | 9 | Q9BSD7 | Cancer-related nucleoside-triphosphatase | NTPCR | 20.714 | 2 | 53.680965 | yes | 6.161E-07 | 516.77332 | 315.02945 |
| HVFLTGPPGVGK | 12 | Q9BSD7 | Cancer-related nucleoside-triphosphatase | NTPCR | 20.714 | 2 | 53.680965 | yes | 9.722E-08 | 604.84296 | 214.29169 |
| SLVPLMGEVR | 10 | Q9BSE5 | Guanidino acid hydrolase, mitochondrial | AGMAT | 37.661 | 2 | 42.48912 | yes | 2.586E-06 | 550.81024 | 66.051117 |
| VVLAEDCWMK | 10 | Q9BSE5 | Guanidino acid hydrolase, mitochondrial | AGMAT | 37.661 | 2 | 42.48912 | yes | 3.005E-07 | 625.79901 | 36.03241 |
| GVIVVEVEDR | 10 | Q9BSH4 | Translational activator of cytochrome c o | TACO1 | 32.478 | 2 | 52.043343 | yes | 3.554E-06 | 557.80878 | 224.65712 |
| FICDASSLHQVR | 12 | Q9BSH4 | Translational activator of cytochrome c o | TACO1 | 32.478 | 2 | 52.043343 | yes | 9.722E-08 | 716.8537 | 97.648666 |
| SKDTYLLYEGR | 11 | Q9BSH4 | Translational activator of cytochrome c o | TACO1 | 32.478 | 2 | 52.043343 | yes | 9.722E-08 | 672.84332 | 63.715988 |
| ALEMAIEAGAEDV | 22 | Q9BSH4 | Translational activator of cytochrome c o | TACO1 | 32.478 | 3 | 52.043343 | yes | 2.363E-07 | 822.03925 | 51.460682 |
| DTYLLYEGR | 9 | Q9BSH4 | Translational activator of cytochrome c o | TACO1 | 32.478 | 2 | 52.043343 | yes | 9.793E-05 | 565.27985 | 23.747303 |

| | | | | | | | | | | | |
|---|---|---|---|---|---|---|---|---|---|---|---|
| AASAPYFEVLEK | 12 | Q9BSJ2 | Gamma-tubulin complex component 2 OS | TUBGCP2 | 102.535 | 2 | 54.0333 | yes | 1.675E-07 | 662.84283 | 1674.339 |
| YTIVQQQIPSFLQK | 14 | Q9BSJ2 | Gamma-tubulin complex component 2 OS | TUBGCP2 | 102.535 | 2 | 54.0333 | yes | 1.006E-07 | 846.9696 | 1609.8944 |
| ISMQELEELR | 10 | Q9BSJ2 | Gamma-tubulin complex component 2 OS | TUBGCP2 | 102.535 | 2 | 54.0333 | yes | 1.004E-07 | 624.31866 | 666.74994 |
| VLLDFLMEEK | 10 | Q9BSJ2 | Gamma-tubulin complex component 2 OS | TUBGCP2 | 102.535 | 2 | 54.0333 | yes | 1.084E-07 | 618.83087 | 319.48468 |
| VLLDFLMEEK | 10 | Q9BSJ2 | Gamma-tubulin complex component 2 OS | TUBGCP2 | 102.535 | 2 | 54.0333 | yes | 7.364E-07 | 626.82831 | 319.48468 |
| YVSAQPLAGR | 10 | Q9BSJ2 | Gamma-tubulin complex component 2 OS | TUBGCP2 | 102.535 | 2 | 54.0333 | yes | 1.107E-07 | 531.29053 | 611.06653 |
| TFLVDPNLDLSIR | 13 | Q9BSJ2 | Gamma-tubulin complex component 2 OS | TUBGCP2 | 102.535 | 2 | 54.0333 | yes | 2.688E-07 | 751.91431 | 459.82214 |
| IHHDVNELLSLLR | 13 | Q9BSJ2 | Gamma-tubulin complex component 2 OS | TUBGCP2 | 102.535 | 2 | 54.0333 | yes | 1.081E-07 | 779.93866 | 427.01892 |
| QGLLSLQK | 8 | Q9BSJ2 | Gamma-tubulin complex component 2 OS | TUBGCP2 | 102.535 | 2 | 54.0333 | yes | 2.986E-06 | 443.77145 | 422.58228 |
| GECLGGSTLSLLHI | 15 | Q9BSJ2 | Gamma-tubulin complex component 2 OS | TUBGCP2 | 102.535 | 2 | 54.0333 | yes | 1.012E-07 | 807.89886 | 369.94324 |
| WPLSLIINR | 9 | Q9BSJ2 | Gamma-tubulin complex component 2 OS | TUBGCP2 | 102.535 | 2 | 54.0333 | yes | 9.722E-08 | 556.33478 | 364.68652 |
| ILPVAASYSAVTR | 13 | Q9BSJ2 | Gamma-tubulin complex component 2 OS | TUBGCP2 | 102.535 | 2 | 54.0333 | yes | 9.722E-08 | 674.38519 | 356.89014 |
| QLCSVWISNK | 10 | Q9BSJ2 | Gamma-tubulin complex component 2 OS | TUBGCP2 | 102.535 | 2 | 54.0333 | yes | 9.722E-08 | 617.8161 | 329.37589 |
| VHGGDGAEVYIDI | 16 | Q9BSJ2 | Gamma-tubulin complex component 2 OS | TUBGCP2 | 102.535 | 3 | 54.0333 | yes | 9.809E-08 | 571.96509 | 323.76263 |
| IQEDYNDK | 8 | Q9BSJ2 | Gamma-tubulin complex component 2 OS | TUBGCP2 | 102.535 | 2 | 54.0333 | yes | 1.275E-07 | 512.73273 | 276.90707 |
| KHLAEHADTVQLV | 22 | Q9BSJ2 | Gamma-tubulin complex component 2 OS | TUBGCP2 | 102.535 | 3 | 54.0333 | yes | 1.425E-07 | 803.42383 | 128.14052 |
| KHLAEHADTVQLV | 22 | Q9BSJ2 | Gamma-tubulin complex component 2 OS | TUBGCP2 | 102.535 | 3 | 54.0333 | yes | 1.608E-07 | 602.81964 | 128.14052 |
| KHLAEHADTVQLV | 22 | Q9BSJ2 | Gamma-tubulin complex component 2 OS | TUBGCP2 | 102.535 | 4 | 54.0333 | yes | 1.425E-07 | 803.42383 | 128.14052 |
| KHLAEHADTVQLV | 22 | Q9BSJ2 | Gamma-tubulin complex component 2 OS | TUBGCP2 | 102.535 | 4 | 54.0333 | yes | 1.608E-07 | 602.81964 | 128.14052 |
| KPVEDITPPR | 10 | Q9BSJ2 | Gamma-tubulin complex component 2 OS | TUBGCP2 | 102.535 | 2 | 54.0333 | yes | 9.823E-08 | 576.32458 | 253.4386 |
| DCMLTNPELLK | 11 | Q9BSJ2 | Gamma-tubulin complex component 2 OS | TUBGCP2 | 102.535 | 2 | 54.0333 | yes | 9.722E-08 | 667.32819 | 213.28755 |
| TMDILASLATSVDI | 14 | Q9BSJ2 | Gamma-tubulin complex component 2 OS | TUBGCP2 | 102.535 | 2 | 54.0333 | yes | 5.675E-07 | 732.88434 | 99.177338 |
| TMDILASLATSVDI | 14 | Q9BSJ2 | Gamma-tubulin complex component 2 OS | TUBGCP2 | 102.535 | 2 | 54.0333 | yes | 3.752E-07 | 740.88184 | 99.177338 |
| MSTANTDPFKDDI | 14 | Q9BSJ2 | Gamma-tubulin complex component 2 OS | TUBGCP2 | 102.535 | 2 | 54.0333 | yes | 1.021E-07 | 791.87451 | 197.07784 |
| ATPQVPVLR | 9 | Q9BSJ2 | Gamma-tubulin complex component 2 OS | TUBGCP2 | 102.535 | 2 | 54.0333 | yes | 7.633E-07 | 490.798 | 190.5843 |
| ETLQYLQQNAK | 11 | Q9BSJ2 | Gamma-tubulin complex component 2 OS | TUBGCP2 | 102.535 | 2 | 54.0333 | yes | 1.37E-07 | 668.34882 | 188.14252 |
| LEALLELALR | 10 | Q9BSJ2 | Gamma-tubulin complex component 2 OS | TUBGCP2 | 102.535 | 2 | 54.0333 | yes | 2.372E-06 | 570.85297 | 182.29326 |
| TPEDFLK | 7 | Q9BSJ2 | Gamma-tubulin complex component 2 OS | TUBGCP2 | 102.535 | 2 | 54.0333 | yes | 1.743E-07 | 425.22128 | 162.9653 |
| GIIHDPYSEFMVEI | 18 | Q9BSJ2 | Gamma-tubulin complex component 2 OS | TUBGCP2 | 102.535 | 3 | 54.0333 | yes | 9.722E-08 | 734.34924 | 156.11838 |
| HLAEHADTVQLVS | 21 | Q9BSJ2 | Gamma-tubulin complex component 2 OS | TUBGCP2 | 102.535 | 3 | 54.0333 | yes | 9.809E-08 | 760.72546 | 75.583527 |
| HLAEHADTVQLVS | 21 | Q9BSJ2 | Gamma-tubulin complex component 2 OS | TUBGCP2 | 102.535 | 3 | 54.0333 | yes | 6.295E-07 | 570.7959 | 75.583527 |
| HLAEHADTVQLVS | 21 | Q9BSJ2 | Gamma-tubulin complex component 2 OS | TUBGCP2 | 102.535 | 4 | 54.0333 | yes | 9.809E-08 | 760.72546 | 75.583527 |
| HLAEHADTVQLVS | 21 | Q9BSJ2 | Gamma-tubulin complex component 2 OS | TUBGCP2 | 102.535 | 4 | 54.0333 | yes | 6.295E-07 | 570.7959 | 75.583527 |
| ECGHDVTCPVAK | 12 | Q9BSJ2 | Gamma-tubulin complex component 2 OS | TUBGCP2 | 102.535 | 3 | 54.0333 | yes | 1.463E-07 | 458.20432 | 66.433823 |
| ECGHDVTCPVAK | 12 | Q9BSJ2 | Gamma-tubulin complex component 2 OS | TUBGCP2 | 102.535 | 3 | 54.0333 | yes | 9.722E-08 | 686.80286 | 66.433823 |
| ECGHDVTCPVAK | 12 | Q9BSJ2 | Gamma-tubulin complex component 2 OS | TUBGCP2 | 102.535 | 2 | 54.0333 | yes | 1.463E-07 | 458.20432 | 66.433823 |
| ECGHDVTCPVAK | 12 | Q9BSJ2 | Gamma-tubulin complex component 2 OS | TUBGCP2 | 102.535 | 2 | 54.0333 | yes | 9.722E-08 | 686.80286 | 66.433823 |
| MSTANTDPFK | 10 | Q9BSJ2 | Gamma-tubulin complex component 2 OS | TUBGCP2 | 102.535 | 2 | 54.0333 | yes | 9.728E-08 | 556.25806 | 112.94442 |
| LDFNGFYTER | 10 | Q9BSJ2 | Gamma-tubulin complex component 2 OS | TUBGCP2 | 102.535 | 2 | 54.0333 | yes | 1.88E-07 | 631.29602 | 91.347404 |

| | | | | | | | | | | | |
|---|---|---|---|---|---|---|---|---|---|---|---|
| TPYVTTTVSAHSA | 14 | Q9BSJ2 | Gamma-tubulin complex component 2 OS | TUBGCP2 | 102.535 | 2 | 54.0333 | yes | 4.809E-07 | 731.88043 | 75.390411 |
| ERAELAAAAVGSS | 25 | Q9BSJ2 | Gamma-tubulin complex component 2 OS | TUBGCP2 | 102.535 | 3 | 54.0333 | yes | 1.712E-06 | 791.4187 | 23.368763 |
| WLVETQVK | 8 | Q9BSJ6 | Protein PIMREG OS=Homo sapiens OX= | PIMREG | 27.48 | 2 | 46.28738 | yes | 3.538E-07 | 501.78458 | 223.74913 |
| ELQPVVSHQETSV | 21 | Q9BSJ6 | Protein PIMREG OS=Homo sapiens OX= | PIMREG | 27.48 | 3 | 46.28738 | yes | 4.061E-07 | 756.38647 | 173.36871 |
| MGSHAHPLR | 9 | Q9BSJ6 | Protein PIMREG OS=Homo sapiens OX= | PIMREG | 27.48 | 2 | 46.28738 | yes | 8.966E-07 | 503.25598 | 159.28482 |
| LETPEPGQQGLQA | 16 | Q9BSJ6 | Protein PIMREG OS=Homo sapiens OX= | PIMREG | 27.48 | 2 | 46.28738 | yes | 3.065E-07 | 833.4314 | 150.39073 |
| LSGAAPAHSAADP | 16 | Q9BSJ6 | Protein PIMREG OS=Homo sapiens OX= | PIMREG | 27.48 | 2 | 46.28738 | yes | 1.89E-07 | 804.39429 | 96.365158 |
| SLQHQEQLEDSK | 12 | Q9BSJ6 | Protein PIMREG OS=Homo sapiens OX= | PIMREG | 27.48 | 2 | 46.28738 | yes | 1.601E-07 | 721.34955 | 82.180092 |
| WQNMGTSVR | 9 | Q9BSJ6 | Protein PIMREG OS=Homo sapiens OX= | PIMREG | 27.48 | 2 | 46.28738 | yes | 5.346E-07 | 539.75854 | 54.669643 |
| FEWELPLDEAQR | 12 | Q9BSJ8 | Extended synaptotagmin-1 OS=Homo sap | ESYT1 | 122.857 | 2 | 53.286903 | yes | 2.713E-07 | 766.87262 | 73.949585 |
| LLAETVAPAVR | 11 | Q9BSJ8 | Extended synaptotagmin-1 OS=Homo sap | ESYT1 | 122.857 | 2 | 53.286903 | yes | 2.026E-07 | 570.34277 | 73.219566 |
| IHVLEAQDLIAK | 12 | Q9BSJ8 | Extended synaptotagmin-1 OS=Homo sap | ESYT1 | 122.857 | 2 | 53.286903 | yes | 9.722E-08 | 675.39301 | 59.931488 |
| GMQLHGVLR | 9 | Q9BSJ8 | Extended synaptotagmin-1 OS=Homo sap | ESYT1 | 122.857 | 2 | 53.286903 | yes | 1.971E-06 | 505.78183 | 40.793129 |
| VELGEKPLR | 9 | Q9BSJ8 | Extended synaptotagmin-1 OS=Homo sap | ESYT1 | 122.857 | 2 | 53.286903 | yes | 1.686E-07 | 520.80859 | 38.084724 |
| GLGPNLVGVAPSR | 13 | Q9BSK2 | Solute carrier family 25 member 33 OS=H | SLC25A33 | 35.376 | 2 | 54.858894 | no | 9.722E-08 | 618.85657 | 95.654167 |
| PTSVTPGLFQVLK | 13 | Q9BSK2 | Solute carrier family 25 member 33 OS=H | SLC25A33 | 35.376 | 2 | 54.858894 | yes | 2.702E-06 | 693.9032 | 61.494328 |
| NQLQSVYK | 8 | Q9BSM1 | Polycomb group RING finger protein 1 O | PCGF1 | 30.347 | 2 | 32.118507 | yes | 1.968E-06 | 490.26401 | 68.798683 |
| PSPLLLQYSVK | 11 | Q9BSM1 | Polycomb group RING finger protein 1 O | PCGF1 | 30.347 | 2 | 32.118507 | yes | 1.618E-06 | 622.86609 | 62.721577 |
| LMFDAFNQR | 9 | Q9BSU1 | Phagosome assembly factor 1 OS=Homo | PHAF1 | 47.525 | 2 | 48.13801 | yes | 1.193E-07 | 571.27661 | 95.32637 |
| HVVAVVLGDVGR | 12 | Q9BT22 | Chitobiosyldiphosphodolichol beta-mann | ALG1 | 52.519 | 2 | 40.31218 | yes | 4.093E-07 | 610.85913 | 42.51009 |
| TFATALDTTR | 10 | Q9BT25 | HAUS augmin-like complex subunit 8 OS | HAUS8 | 44.858 | 2 | 50.99842 | yes | 1.164E-07 | 548.78528 | 410.27908 |
| MENNLAEFER | 10 | Q9BT25 | HAUS augmin-like complex subunit 8 OS | HAUS8 | 44.858 | 2 | 50.99842 | yes | 3.115E-07 | 626.78497 | 173.80089 |
| SFAQVLELSAEASI | 14 | Q9BT25 | HAUS augmin-like complex subunit 8 OS | HAUS8 | 44.858 | 2 | 50.99842 | yes | 1.049E-06 | 740.38812 | 106.85918 |
| WYFNQDSACR | 10 | Q9BT25 | HAUS augmin-like complex subunit 8 OS | HAUS8 | 44.858 | 2 | 50.99842 | yes | 9.722E-08 | 673.78278 | 33.119614 |
| NLLIMCK | 7 | Q9BT25 | HAUS augmin-like complex subunit 8 OS | HAUS8 | 44.858 | 2 | 50.99842 | yes | 2.272E-07 | 446.24316 | 22.041021 |
| KHDPLLR | 7 | Q9BT40 | Inositol polyphosphate 5-phosphatase K ( | INPP5K | 51.091 | 2 | 52.68186 | yes | 1.041E-07 | 439.76398 | 369.75287 |
| STPTGLFGYWGNK | 13 | Q9BT40 | Inositol polyphosphate 5-phosphatase K ( | INPP5K | 51.091 | 2 | 52.68186 | yes | 1.121E-07 | 714.35132 | 303.2146 |
| IEDFGLHFVR | 10 | Q9BT40 | Inositol polyphosphate 5-phosphatase K ( | INPP5K | 51.091 | 2 | 52.68186 | yes | 9.722E-08 | 616.82477 | 109.424 |
| IEDFGLHFVR | 10 | Q9BT40 | Inositol polyphosphate 5-phosphatase K ( | INPP5K | 51.091 | 2 | 52.68186 | yes | 1.632E-07 | 411.55228 | 109.424 |
| IEDFGLHFVR | 10 | Q9BT40 | Inositol polyphosphate 5-phosphatase K ( | INPP5K | 51.091 | 3 | 52.68186 | yes | 9.722E-08 | 616.82477 | 109.424 |
| IEDFGLHFVR | 10 | Q9BT40 | Inositol polyphosphate 5-phosphatase K ( | INPP5K | 51.091 | 3 | 52.68186 | yes | 1.632E-07 | 411.55228 | 109.424 |
| PFQIPPGSLR | 10 | Q9BT40 | Inositol polyphosphate 5-phosphatase K ( | INPP5K | 51.091 | 2 | 52.68186 | yes | 2.767E-06 | 556.31659 | 176.78566 |
| YQHLPYIQILSTK | 13 | Q9BT40 | Inositol polyphosphate 5-phosphatase K ( | INPP5K | 51.091 | 2 | 52.68186 | yes | 1.357E-07 | 802.44574 | 88.568878 |
| CYGGLWEK | 8 | Q9BT40 | Inositol polyphosphate 5-phosphatase K ( | INPP5K | 51.091 | 2 | 52.68186 | yes | 1.028E-06 | 506.73148 | 75.805168 |
| GGVNICLK | 8 | Q9BT40 | Inositol polyphosphate 5-phosphatase K ( | INPP5K | 51.091 | 2 | 52.68186 | yes | 1.676E-07 | 430.73657 | 70.196251 |
| VISFEEQVASIR | 12 | Q9BT78 | COP9 signalosome complex subunit 4 OS | COPS4 | 46.269 | 2 | 53.134598 | yes | 9.722E-08 | 689.37225 | 163.10645 |
| NAAQVLVGIPLET( | 16 | Q9BT78 | COP9 signalosome complex subunit 4 OS | COPS4 | 46.269 | 2 | 53.134598 | yes | 1.588E-07 | 819.46466 | 54.966614 |
| AVIEHNLLSASK | 12 | Q9BT78 | COP9 signalosome complex subunit 4 OS | COPS4 | 46.269 | 2 | 53.134598 | yes | 5.066E-07 | 641.36169 | 43.800888 |
| LQAEALLQQMEEI | 14 | Q9BT92 | Trichoplein keratin filament-binding prot | TCHP | 61.072 | 2 | 50.094128 | yes | 9.903E-08 | 822.43726 | 94.686119 |

| | | | | | | | | | | | |
|---|---|---|---|---|---|---|---|---|---|---|---|
| EMELDLHQK | 9 | Q9BT92 | Trichoplein keratin filament-binding prot | TCHP | 61.072 | 2 | 50.094128 | yes | 1.294E-07 | 571.77917 | 93.59539 |
| LSMNLQER | 8 | Q9BT92 | Trichoplein keratin filament-binding prot | TCHP | 61.072 | 2 | 50.094128 | yes | 3.517E-07 | 495.75549 | 92.86911 |
| EQVMADVAWMK | 11 | Q9BT92 | Trichoplein keratin filament-binding prot | TCHP | 61.072 | 2 | 50.094128 | yes | 7.125E-07 | 654.30975 | 63.496567 |
| QAIEEQLQLER | 11 | Q9BT92 | Trichoplein keratin filament-binding prot | TCHP | 61.072 | 2 | 50.094128 | yes | 3.397E-07 | 678.8595 | 62.085087 |
| HVVNSWEMQK | 10 | Q9BT92 | Trichoplein keratin filament-binding prot | TCHP | 61.072 | 2 | 50.094128 | yes | 2.688E-07 | 629.30585 | 58.916531 |
| KQELEAQVAER | 11 | Q9BT92 | Trichoplein keratin filament-binding prot | TCHP | 61.072 | 2 | 50.094128 | yes | 6.015E-07 | 650.84644 | 35.129589 |
| TAAPSPSLLYK | 11 | Q9BTC0 | Death-inducer obliterator 1 OS=Homo sa | DIDO1 | 243.873 | 2 | 53.7371 | yes | 1.359E-07 | 574.32153 | 598.82367 |
| SGPQSASTAVK | 11 | Q9BTC0 | Death-inducer obliterator 1 OS=Homo sa | DIDO1 | 243.873 | 2 | 53.7371 | yes | 1.422E-07 | 516.76965 | 565.42346 |
| GGLVGQAPMPVPI | 15 | Q9BTC0 | Death-inducer obliterator 1 OS=Homo sa | DIDO1 | 243.873 | 2 | 53.7371 | yes | 1.251E-07 | 754.89252 | 380.87646 |
| LKPEELVSK | 9 | Q9BTC0 | Death-inducer obliterator 1 OS=Homo sa | DIDO1 | 243.873 | 2 | 53.7371 | yes | 1.446E-07 | 521.81079 | 349.35565 |
| IFQPVIEAPGASK | 13 | Q9BTC0 | Death-inducer obliterator 1 OS=Homo sa | DIDO1 | 243.873 | 2 | 53.7371 | yes | 9.926E-08 | 678.8797 | 287.46973 |
| AHLFDLNCK | 9 | Q9BTC0 | Death-inducer obliterator 1 OS=Homo sa | DIDO1 | 243.873 | 2 | 53.7371 | yes | 9.766E-08 | 559.27661 | 234.19188 |
| VATVPQSEK | 9 | Q9BTC0 | Death-inducer obliterator 1 OS=Homo sa | DIDO1 | 243.873 | 2 | 53.7371 | yes | 1.294E-06 | 479.76382 | 211.08722 |
| AFDTQLVER | 9 | Q9BTC0 | Death-inducer obliterator 1 OS=Homo sa | DIDO1 | 243.873 | 2 | 53.7371 | yes | 1.06E-07 | 539.78003 | 209.21339 |
| SSFANVAAATPAII | 14 | Q9BTC0 | Death-inducer obliterator 1 OS=Homo sa | DIDO1 | 243.873 | 2 | 53.7371 | yes | 9.722E-08 | 674.367 | 204.31604 |
| LGAMSAAPSQPNS | 16 | Q9BTC0 | Death-inducer obliterator 1 OS=Homo sa | DIDO1 | 243.873 | 2 | 53.7371 | yes | 1.002E-07 | 814.41467 | 203.68307 |
| GFINMQSVAK | 10 | Q9BTC0 | Death-inducer obliterator 1 OS=Homo sa | DIDO1 | 243.873 | 2 | 53.7371 | yes | 3.445E-07 | 547.7868 | 174.97714 |
| KFPGSAALVGAVF | 13 | Q9BTC0 | Death-inducer obliterator 1 OS=Homo sa | DIDO1 | 243.873 | 2 | 53.7371 | yes | 3.051E-07 | 636.87476 | 168.2245 |
| VEQFLTIAR | 9 | Q9BTC0 | Death-inducer obliterator 1 OS=Homo sa | DIDO1 | 243.873 | 2 | 53.7371 | yes | 1.539E-07 | 538.80859 | 163.20963 |
| FMICCDR | 7 | Q9BTC0 | Death-inducer obliterator 1 OS=Homo sa | DIDO1 | 243.873 | 2 | 53.7371 | yes | 1.143E-07 | 501.2037 | 147.05247 |
| QLEEQEEALR | 10 | Q9BTC0 | Death-inducer obliterator 1 OS=Homo sa | DIDO1 | 243.873 | 2 | 53.7371 | yes | 9.834E-08 | 622.80951 | 136.44199 |
| SPPEGDTTLFLSR | 13 | Q9BTC0 | Death-inducer obliterator 1 OS=Homo sa | DIDO1 | 243.873 | 2 | 53.7371 | yes | 3.214E-06 | 710.35938 | 133.68265 |
| QAGPAPAAATAAS | 14 | Q9BTC0 | Death-inducer obliterator 1 OS=Homo sa | DIDO1 | 243.873 | 2 | 53.7371 | yes | 3.895E-07 | 606.32257 | 130.03549 |
| EPPGSTAGLPQEPI | 14 | Q9BTC0 | Death-inducer obliterator 1 OS=Homo sa | DIDO1 | 243.873 | 2 | 53.7371 | yes | 2.323E-07 | 704.35938 | 126.1304 |
| FGVVANNNR | 9 | Q9BTC0 | Death-inducer obliterator 1 OS=Homo sa | DIDO1 | 243.873 | 2 | 53.7371 | yes | 1.693E-07 | 495.75943 | 117.77634 |
| ALGSAQYEDPR | 11 | Q9BTC0 | Death-inducer obliterator 1 OS=Homo sa | DIDO1 | 243.873 | 2 | 53.7371 | yes | 1.18E-07 | 603.79114 | 106.26098 |
| EMFNLFQVTDNR | 12 | Q9BTC0 | Death-inducer obliterator 1 OS=Homo sa | DIDO1 | 243.873 | 2 | 53.7371 | yes | 9.861E-08 | 757.35883 | 104.26378 |
| LLPFEGPGLESPR | 13 | Q9BTC0 | Death-inducer obliterator 1 OS=Homo sa | DIDO1 | 243.873 | 2 | 53.7371 | yes | 4.115E-07 | 706.38263 | 83.304626 |
| DPFTCPGFASQDK | 13 | Q9BTC0 | Death-inducer obliterator 1 OS=Homo sa | DIDO1 | 243.873 | 2 | 53.7371 | yes | 2.031E-07 | 735.3219 | 77.111702 |
| PVLTSVMVPK | 10 | Q9BTC0 | Death-inducer obliterator 1 OS=Homo sa | DIDO1 | 243.873 | 2 | 53.7371 | yes | 2.609E-06 | 535.81757 | 68.648621 |
| EKPLEEPDAQGR | 12 | Q9BTC0 | Death-inducer obliterator 1 OS=Homo sa | DIDO1 | 243.873 | 2 | 53.7371 | yes | 2.336E-07 | 684.84131 | 60.647495 |
| LQPEEADVPAYPK | 13 | Q9BTC0 | Death-inducer obliterator 1 OS=Homo sa | DIDO1 | 243.873 | 2 | 53.7371 | yes | 2.693E-07 | 728.86957 | 59.592979 |
| KPVVPSVPMASPA | 16 | Q9BTC0 | Death-inducer obliterator 1 OS=Homo sa | DIDO1 | 243.873 | 2 | 53.7371 | yes | 1.535E-07 | 795.44525 | 29.192001 |
| KPVVPSVPMASPA | 16 | Q9BTC0 | Death-inducer obliterator 1 OS=Homo sa | DIDO1 | 243.873 | 2 | 53.7371 | yes | 6.369E-05 | 530.63257 | 29.192001 |
| KPVVPSVPMASPA | 16 | Q9BTC0 | Death-inducer obliterator 1 OS=Homo sa | DIDO1 | 243.873 | 3 | 53.7371 | yes | 1.535E-07 | 795.44525 | 29.192001 |
| KPVVPSVPMASPA | 16 | Q9BTC0 | Death-inducer obliterator 1 OS=Homo sa | DIDO1 | 243.873 | 3 | 53.7371 | yes | 6.369E-05 | 530.63257 | 29.192001 |
| QDVPKPVLTSVM | 15 | Q9BTC0 | Death-inducer obliterator 1 OS=Homo sa | DIDO1 | 243.873 | 3 | 53.7371 | yes | 1.743E-07 | 546.64801 | 29.153496 |
| QDVPKPVLTSVM | 15 | Q9BTC0 | Death-inducer obliterator 1 OS=Homo sa | DIDO1 | 243.873 | 3 | 53.7371 | yes | 2.555E-07 | 819.46838 | 29.153496 |
| QDVPKPVLTSVM | 15 | Q9BTC0 | Death-inducer obliterator 1 OS=Homo sa | DIDO1 | 243.873 | 2 | 53.7371 | yes | 1.743E-07 | 546.64801 | 29.153496 |

| | | | | | | | | | | | |
|---|---|---|---|---|---|---|---|---|---|---|---|
| QDVPKPVLTSVM | 15 | Q9BTC0 | Death-inducer obliterator 1 OS=Homo sa | DIDO1 | 243.873 | 2 | 53.7371 | yes | 2.555E-07 | 819.46838 | 29.153496 |
| AAQDIKDEEPGDL | 15 | Q9BTC0 | Death-inducer obliterator 1 OS=Homo sa | DIDO1 | 243.873 | 2 | 53.7371 | yes | 1.563E-07 | 807.39191 | 51.622978 |
| EQEPTERPLK | 10 | Q9BTC0 | Death-inducer obliterator 1 OS=Homo sa | DIDO1 | 243.873 | 2 | 53.7371 | yes | 1.453E-07 | 613.82239 | 33.625366 |
| SHLDSPEAR | 9 | Q9BTD8 | RNA-binding protein 42 OS=Homo sapie | RBM42 | 50.415 | 2 | 46.87845 | yes | 1.451E-07 | 506.24634 | 700.99933 |
| ELGLGLGLGLK | 11 | Q9BTD8 | RNA-binding protein 42 OS=Homo sapie | RBM42 | 50.415 | 2 | 46.87845 | yes | 2.538E-07 | 535.33441 | 389.19189 |
| IPELLSLR | 8 | Q9BTD8 | RNA-binding protein 42 OS=Homo sapie | RBM42 | 50.415 | 2 | 46.87845 | yes | 1.816E-07 | 470.79492 | 71.078201 |
| IIQHLVPASFR | 11 | Q9BTE3 | Mini-chromosome maintenance complex- | MCMBP | 72.981 | 2 | 53.097397 | yes | 9.722E-08 | 640.87732 | 116.93253 |
| IIQHLVPASFR | 11 | Q9BTE3 | Mini-chromosome maintenance complex- | MCMBP | 72.981 | 2 | 53.097397 | yes | 2.894E-06 | 427.58731 | 116.93253 |
| IIQHLVPASFR | 11 | Q9BTE3 | Mini-chromosome maintenance complex- | MCMBP | 72.981 | 3 | 53.097397 | yes | 9.722E-08 | 640.87732 | 116.93253 |
| IIQHLVPASFR | 11 | Q9BTE3 | Mini-chromosome maintenance complex- | MCMBP | 72.981 | 3 | 53.097397 | yes | 2.894E-06 | 427.58731 | 116.93253 |
| CLSLSAGQTTLSR | 13 | Q9BTE3 | Mini-chromosome maintenance complex- | MCMBP | 72.981 | 2 | 53.097397 | yes | 9.722E-08 | 697.35883 | 165.52101 |
| IHVILAQK | 8 | Q9BTE3 | Mini-chromosome maintenance complex- | MCMBP | 72.981 | 2 | 53.097397 | yes | 9.722E-08 | 461.29764 | 127.8055 |
| VHSPPASLVPR | 11 | Q9BTE3 | Mini-chromosome maintenance complex- | MCMBP | 72.981 | 2 | 53.097397 | yes | 7.745E-07 | 580.33276 | 116.62469 |
| AVEDDFVEMR | 10 | Q9BTE3 | Mini-chromosome maintenance complex- | MCMBP | 72.981 | 2 | 53.097397 | yes | 2.607E-07 | 605.77405 | 110.99961 |
| LQMTIENMNHLK | 12 | Q9BTE3 | Mini-chromosome maintenance complex- | MCMBP | 72.981 | 2 | 53.097397 | yes | 2.095E-07 | 736.37341 | 73.648308 |
| NSTFTEHLYR | 10 | Q9BTE3 | Mini-chromosome maintenance complex- | MCMBP | 72.981 | 2 | 53.097397 | yes | 5.029E-07 | 634.30695 | 62.37851 |
| VYEDWDCFK | 9 | Q9BTE3 | Mini-chromosome maintenance complex- | MCMBP | 72.981 | 2 | 53.097397 | yes | 1.283E-07 | 631.26337 | 61.294975 |
| IYLTLLR | 7 | Q9BTE3 | Mini-chromosome maintenance complex- | MCMBP | 72.981 | 2 | 53.097397 | yes | 8.259E-07 | 446.28674 | 31.047966 |
| EAYVNANQAR | 10 | Q9BTE3 | Mini-chromosome maintenance complex- | MCMBP | 72.981 | 2 | 53.097397 | yes | 8.385E-07 | 568.2782 | 24.461479 |
| SYEDDDDMDLQP | 14 | Q9BTE3 | Mini-chromosome maintenance complex- | MCMBP | 72.981 | 2 | 53.097397 | yes | 1.055E-07 | 842.83575 | 23.305004 |
| TNLIFLSGNR | 10 | Q9BTE6 | Alanyl-tRNA editing protein Aarsd1 OS= | AARSD1 | 45.48 | 2 | 50.885582 | yes | 2.697E-07 | 567.81696 | 119.76308 |
| GTINDISVLR | 10 | Q9BTE6 | Alanyl-tRNA editing protein Aarsd1 OS= | AARSD1 | 45.48 | 2 | 50.885582 | yes | 3.997E-07 | 544.30896 | 106.478 |
| ELSLDDPEVEQVS | 15 | Q9BTE6 | Alanyl-tRNA editing protein Aarsd1 OS= | AARSD1 | 45.48 | 2 | 50.885582 | yes | 9.722E-08 | 836.90485 | 82.601364 |
| SQLNDISSFK | 10 | Q9BTE7 | DCN1-like protein 5 OS=Homo sapiens | DCUN1D5 | 27.509 | 2 | 49.9835 | yes | 2.131E-07 | 569.79059 | 141.0596 |
| LISGEEHFSSK | 11 | Q9BTE7 | DCN1-like protein 5 OS=Homo sapiens | DCUN1D5 | 27.509 | 2 | 49.9835 | yes | 1.694E-07 | 617.30914 | 98.357109 |
| LEAESMGFFTK | 11 | Q9BTE7 | DCN1-like protein 5 OS=Homo sapiens | DCUN1D5 | 27.509 | 2 | 49.9835 | yes | 1.921E-07 | 630.30249 | 88.329964 |
| YAFDFAR | 7 | Q9BTE7 | DCN1-like protein 5 OS=Homo sapiens | DCUN1D5 | 27.509 | 2 | 49.9835 | no | 9.938E-08 | 445.21378 | 52.701942 |
| DQWYNVLEFSR | 11 | Q9BTE7 | DCN1-like protein 5 OS=Homo sapiens | DCUN1D5 | 27.509 | 2 | 49.9835 | yes | 1.88E-07 | 728.84644 | 43.695271 |
| GMTSLQCDCTEK | 12 | Q9BTE7 | DCN1-like protein 5 OS=Homo sapiens | DCUN1D5 | 27.509 | 2 | 49.9835 | yes | 1.11E-07 | 715.29144 | 22.468983 |
| KSPGVAAAVAED | 16 | Q9BTE7 | DCN1-like protein 5 OS=Homo sapiens | DCUN1D5 | 27.509 | 2 | 49.9835 | yes | 4.251E-06 | 735.40155 | 18.3631 |
| SWGNNYPQHR | 10 | Q9BTL3 | RNA guanine-N7 methyltransferase activ | RAMAC | 14.382 | 2 | 52.40284 | no | 9.722E-08 | 629.78925 | 260.35455 |
| FEEMFASR | 8 | Q9BTL3 | RNA guanine-N7 methyltransferase activ | RAMAC | 14.382 | 2 | 52.40284 | yes | 2.035E-07 | 508.72894 | 62.151695 |
| MSEQYFPALQK | 11 | Q9BTP7 | Fanconi anemia core complex-associated | FAAP24 | 23.897 | 2 | 51.209606 | yes | 9.722E-08 | 671.32904 | 58.151119 |
| SLLIQELSK | 9 | Q9BTT4 | Mediator of RNA polymerase II transcrip | MED10 | 15.688 | 2 | 51.692703 | yes | 9.722E-08 | 515.81079 | 176.13095 |
| LVHIIGALR | 9 | Q9BTV4 | Transmembrane protein 43 OS=Homo sa | TMEM43 | 44.876 | 2 | 49.64638 | yes | 1.014E-07 | 496.32419 | 79.539398 |
| LLSDPNYGVHLPA | 15 | Q9BTV4 | Transmembrane protein 43 OS=Homo sa | TMEM43 | 44.876 | 3 | 49.64638 | yes | 1.595E-07 | 541.63489 | 20.323837 |
| LLSDPNYGVHLPA | 15 | Q9BTV4 | Transmembrane protein 43 OS=Homo sa | TMEM43 | 44.876 | 3 | 49.64638 | yes | 3.258E-07 | 811.94867 | 20.323837 |
| LLSDPNYGVHLPA | 15 | Q9BTV4 | Transmembrane protein 43 OS=Homo sa | TMEM43 | 44.876 | 2 | 49.64638 | yes | 1.595E-07 | 541.63489 | 20.323837 |
| LLSDPNYGVHLPA | 15 | Q9BTV4 | Transmembrane protein 43 OS=Homo sa | TMEM43 | 44.876 | 2 | 49.64638 | yes | 3.258E-07 | 811.94867 | 20.323837 |

| | | | | | | | | | | | |
|---|---|---|---|---|---|---|---|---|---|---|---|
| ISGSLVDTSYK | 11 | Q9BTX1 | Nucleoporin NDC1 OS=Homo sapiens O | NDC1 | 76.305 | 2 | 44.75452 | yes | 2.081E-07 | 585.30603 | 161.13867 |
| VSSSYPVEPK | 10 | Q9BTX1 | Nucleoporin NDC1 OS=Homo sapiens O | NDC1 | 76.305 | 2 | 44.75452 | yes | 2.01E-05 | 546.78223 | 49.255589 |
| ITTTFGEHLNAVQ/ | 20 | Q9BTX1 | Nucleoporin NDC1 OS=Homo sapiens O | NDC1 | 76.305 | 3 | 44.75452 | yes | 3.578E-07 | 728.03442 | 23.625168 |
| ELMPCDFPGCGR | 12 | Q9BU19 | Zinc finger protein 692 OS=Homo sapien | ZNF692 | 56.969 | 2 | 48.00319 | yes | 1.215E-07 | 719.79926 | 158.9823 |
| DYICEFCAR | 9 | Q9BU19 | Zinc finger protein 692 OS=Homo sapien | ZNF692 | 56.969 | 2 | 48.00319 | yes | 3.211E-07 | 617.255 | 141.46013 |
| TSSNLVIHR | 9 | Q9BU19 | Zinc finger protein 692 OS=Homo sapien | ZNF692 | 56.969 | 2 | 48.00319 | yes | 1.632E-07 | 513.78815 | 61.302292 |
| LLPSPVTCTPK | 11 | Q9BU19 | Zinc finger protein 692 OS=Homo sapien | ZNF692 | 56.969 | 2 | 48.00319 | yes | 5.442E-05 | 606.83649 | 55.159603 |
| SWCSEATSGQELA | 22 | Q9BU19 | Zinc finger protein 692 OS=Homo sapien | ZNF692 | 56.969 | 3 | 48.00319 | yes | 1.76E-07 | 846.02246 | 46.508835 |
| SFSCPEPACGK | 11 | Q9BU19 | Zinc finger protein 692 OS=Homo sapien | ZNF692 | 56.969 | 2 | 48.00319 | yes | 1.53E-07 | 620.26025 | 29.681383 |
| EAAQAMYIDSYNS | 14 | Q9BU61 | NADH dehydrogenase [ubiquinone] 1 alp | NDUFAF3 | 20.351 | 2 | 46.026104 | yes | 1.674E-07 | 809.86194 | 19.619925 |
| LLGLGSASGSVGR | 13 | Q9BU76 | Multiple myeloma tumor-associated prote | MMTAG2 | 29.412 | 2 | 51.92288 | yes | 2.045E-07 | 587.33295 | 2273.5144 |
| EALLAALGYK | 10 | Q9BU76 | Multiple myeloma tumor-associated prote | MMTAG2 | 29.412 | 2 | 51.92288 | yes | 2.524E-07 | 524.80548 | 1373.0526 |
| ENYLGNSLMAPV( | 14 | Q9BU76 | Multiple myeloma tumor-associated prote | MMTAG2 | 29.412 | 2 | 51.92288 | yes | 9.789E-08 | 760.87994 | 1186.0905 |
| GGQDQFNWEDVK | 12 | Q9BU76 | Multiple myeloma tumor-associated prote | MMTAG2 | 29.412 | 2 | 51.92288 | yes | 1.211E-07 | 711.81787 | 1093.0886 |
| VESGGPGTSAASA | 14 | Q9BU76 | Multiple myeloma tumor-associated prote | MMTAG2 | 29.412 | 2 | 51.92288 | yes | 9.722E-08 | 623.80475 | 818.96906 |
| EEELAAVR | 8 | Q9BU76 | Multiple myeloma tumor-associated prote | MMTAG2 | 29.412 | 2 | 51.92288 | yes | 1.096E-07 | 458.74036 | 175.8154 |
| EDFAEVCKR | 9 | Q9BU76 | Multiple myeloma tumor-associated prote | MMTAG2 | 29.412 | 2 | 51.92288 | yes | 1.259E-07 | 577.26898 | 153.84979 |
| EDFAEVCK | 8 | Q9BU76 | Multiple myeloma tumor-associated prote | MMTAG2 | 29.412 | 2 | 51.92288 | yes | 2.522E-07 | 499.21841 | 79.627548 |
| AFQYADGLEQLR | 12 | Q9BU89 | Deoxyhypusine hydroxylase OS=Homo sa | DOHH | 32.905 | 2 | 51.474236 | yes | 9.722E-08 | 705.85425 | 349.43512 |
| PACLAALQAHADI | 16 | Q9BU89 | Deoxyhypusine hydroxylase OS=Homo sa | DOHH | 32.905 | 3 | 51.474236 | yes | 1.062E-07 | 578.9458 | 41.389076 |
| HECAEALGAIAR | 12 | Q9BU89 | Deoxyhypusine hydroxylase OS=Homo sa | DOHH | 32.905 | 2 | 51.474236 | yes | 6.27E-06 | 649.31952 | 14.974575 |
| SLLVNPVLFPNR | 12 | Q9BUB7 | Transmembrane protein 70, mitochondria | TMEM70 | 28.97 | 2 | 53.790855 | yes | 9.722E-08 | 684.90356 | 86.615509 |
| LSSEFVFNNPNIK | 13 | Q9BUE6 | Iron-sulfur cluster assembly 1 homolog, n | ISCA1 | 14.18 | 2 | 52.244595 | yes | 9.722E-08 | 754.89081 | 136.88545 |
| AALTLTPSAVNK | 12 | Q9BUE6 | Iron-sulfur cluster assembly 1 homolog, n | ISCA1 | 14.18 | 2 | 52.244595 | yes | 5.273E-07 | 593.34552 | 72.676529 |
| AALVDLEPGTMDS | 15 | Q9BUF5 | Tubulin beta-6 chain OS=Homo sapiens ( | TUBB6 | 49.858 | 2 | 52.546646 | yes | 1.204E-07 | 787.39813 | 1802.8418 |
| AALVDLEPGTMDS | 15 | Q9BUF5 | Tubulin beta-6 chain OS=Homo sapiens ( | TUBB6 | 49.858 | 2 | 52.546646 | yes | 6.51E-07 | 795.39563 | 1802.8418 |
| YLTVATVFR | 9 | Q9BUF5 | Tubulin beta-6 chain OS=Homo sapiens ( | TUBB6 | 49.858 | 2 | 52.546646 | no | 2.657E-07 | 535.30566 | 1636.8738 |
| INVYYNESSSQK | 12 | Q9BUF5 | Tubulin beta-6 chain OS=Homo sapiens ( | TUBB6 | 49.858 | 2 | 52.546646 | yes | 9.722E-08 | 716.34119 | 720.63129 |
| ISEQFSAMFR | 10 | Q9BUF5 | Tubulin beta-6 chain OS=Homo sapiens ( | TUBB6 | 49.858 | 2 | 52.546646 | yes | 9.722E-08 | 608.29498 | 219.59692 |
| ISEQFSAMFR | 10 | Q9BUF5 | Tubulin beta-6 chain OS=Homo sapiens ( | TUBB6 | 49.858 | 2 | 52.546646 | yes | 8.697E-06 | 616.29242 | 219.59692 |
| IREEFPDR | 8 | Q9BUF5 | Tubulin beta-6 chain OS=Homo sapiens ( | TUBB6 | 49.858 | 2 | 52.546646 | no | 3.553E-07 | 531.27234 | 361.95709 |
| EVDEQMLAIQSK | 12 | Q9BUF5 | Tubulin beta-6 chain OS=Homo sapiens ( | TUBB6 | 49.858 | 2 | 52.546646 | no | 1.804E-07 | 695.84778 | 284.5484 |
| IMNTFSVMPSPK | 12 | Q9BUF5 | Tubulin beta-6 chain OS=Homo sapiens ( | TUBB6 | 49.858 | 2 | 52.546646 | no | 3.365E-07 | 676.34106 | 137.74496 |
| VAVCDIPPR | 9 | Q9BUF5 | Tubulin beta-6 chain OS=Homo sapiens ( | TUBB6 | 49.858 | 2 | 52.546646 | no | 2.156E-07 | 513.77368 | 124.8241 |
| SGPFGQLFR | 9 | Q9BUF5 | Tubulin beta-6 chain OS=Homo sapiens ( | TUBB6 | 49.858 | 2 | 52.546646 | yes | 2.529E-05 | 504.76672 | 30.777754 |
| VSELPSDFQER | 11 | Q9BUH8 | Brain-enriched guanylate kinase-associate | BEGAIN | 64.804 | 2 | 53.340416 | yes | 1.702E-07 | 653.81732 | 445.84549 |
| TMSPYPAETFR | 11 | Q9BUH8 | Brain-enriched guanylate kinase-associate | BEGAIN | 64.804 | 2 | 53.340416 | yes | 1.612E-07 | 650.30554 | 224.9232 |
| LETEFDSTR | 9 | Q9BUH8 | Brain-enriched guanylate kinase-associate | BEGAIN | 64.804 | 2 | 53.340416 | yes | 2.571E-07 | 549.25909 | 200.87427 |
| HYLEIELR | 8 | Q9BUH8 | Brain-enriched guanylate kinase-associate | BEGAIN | 64.804 | 2 | 53.340416 | yes | 2.785E-07 | 536.79291 | 168.99165 |

| | | | | | | | | | | |
|---|---|---|---|---|---|---|---|---|---|---|
| LSYTTHK | 7 | Q9BUH8 | Brain-enriched guanylate kinase-associate | BEGAIN | 64.804 | 2 | 53.340416 | yes | 2.46E-07 | 425.2269 | 165.15974 |
| VSLHMEK | 7 | Q9BUH8 | Brain-enriched guanylate kinase-associate | BEGAIN | 64.804 | 2 | 53.340416 | yes | 5.333E-07 | 422.2233 | 151.51077 |
| LPGEDMR | 7 | Q9BUH8 | Brain-enriched guanylate kinase-associate | BEGAIN | 64.804 | 2 | 53.340416 | yes | 1.225E-07 | 409.19727 | 119.43164 |
| LSALQEQK | 8 | Q9BUH8 | Brain-enriched guanylate kinase-associate | BEGAIN | 64.804 | 2 | 53.340416 | yes | 5.186E-07 | 458.75854 | 96.363945 |
| IQSNYMALQR | 10 | Q9BUH8 | Brain-enriched guanylate kinase-associate | BEGAIN | 64.804 | 2 | 53.340416 | yes | 2.164E-07 | 612.31372 | 93.480347 |
| ALSHEIVALNSHLI | 17 | Q9BUH8 | Brain-enriched guanylate kinase-associate | BEGAIN | 64.804 | 3 | 53.340416 | yes | 1.455E-07 | 615.67914 | 91.21698 |
| LSEDNELYR | 9 | Q9BUH8 | Brain-enriched guanylate kinase-associate | BEGAIN | 64.804 | 2 | 53.340416 | yes | 9.722E-08 | 569.7724 | 79.57737 |
| KPPATTYEGSPR | 12 | Q9BUH8 | Brain-enriched guanylate kinase-associate | BEGAIN | 64.804 | 2 | 53.340416 | yes | 3.189E-07 | 652.33569 | 70.161743 |
| LSPQQAFPR | 9 | Q9BUH8 | Brain-enriched guanylate kinase-associate | BEGAIN | 64.804 | 2 | 53.340416 | yes | 9.726E-07 | 522.28528 | 43.933578 |
| INQELEDK | 8 | Q9BUH8 | Brain-enriched guanylate kinase-associate | BEGAIN | 64.804 | 2 | 53.340416 | yes | 6.195E-05 | 494.75092 | 40.730869 |
| ASPCSFSER | 9 | Q9BUH8 | Brain-enriched guanylate kinase-associate | BEGAIN | 64.804 | 2 | 53.340416 | yes | 7.719E-07 | 520.72693 | 26.918247 |
| SGDSGGGMYVINI | 15 | Q9BUI4 | DNA-directed RNA polymerase III subun | POLR3C | 60.612 | 2 | 54.6785 | yes | 4.49E-05 | 767.86957 | 2005.6499 |
| TMDYAEVSNTFVI | 13 | Q9BUI4 | DNA-directed RNA polymerase III subun | POLR3C | 60.612 | 2 | 54.6785 | yes | 1.392E-07 | 766.85614 | 100.06004 |
| SIANLIER | 8 | Q9BUI4 | DNA-directed RNA polymerase III subun | POLR3C | 60.612 | 2 | 54.6785 | yes | 1.459E-06 | 458.26654 | 67.553917 |
| GVVEYEAQCSR | 11 | Q9BUI4 | DNA-directed RNA polymerase III subun | POLR3C | 60.612 | 2 | 54.6785 | yes | 9.722E-08 | 649.29572 | 65.388908 |
| MDQTSSEIVR | 10 | Q9BUI4 | DNA-directed RNA polymerase III subun | POLR3C | 60.612 | 2 | 54.6785 | yes | 8.392E-07 | 583.27954 | 48.895115 |
| DQAIVSAVANR | 11 | Q9BUI4 | DNA-directed RNA polymerase III subun | POLR3C | 60.612 | 2 | 54.6785 | yes | 2.97E-07 | 572.30945 | 26.624456 |
| NCAVEFNFGQR | 11 | Q9BUJ2 | Heterogeneous nuclear ribonucleoprotein | HNRNPUL | 95.739 | 2 | 52.22891 | yes | 1.108E-07 | 671.30383 | 13424.334 |
| HLPSTEPDPHVVR | 13 | Q9BUJ2 | Heterogeneous nuclear ribonucleoprotein | HNRNPUL | 95.739 | 3 | 52.22891 | yes | 1.571E-07 | 495.25992 | 5375.6963 |
| HLPSTEPDPHVVR | 13 | Q9BUJ2 | Heterogeneous nuclear ribonucleoprotein | HNRNPUL | 95.739 | 3 | 52.22891 | yes | 1.802E-07 | 742.38623 | 5375.6963 |
| HLPSTEPDPHVVR | 13 | Q9BUJ2 | Heterogeneous nuclear ribonucleoprotein | HNRNPUL | 95.739 | 2 | 52.22891 | yes | 1.571E-07 | 495.25992 | 5375.6963 |
| HLPSTEPDPHVVR | 13 | Q9BUJ2 | Heterogeneous nuclear ribonucleoprotein | HNRNPUL | 95.739 | 2 | 52.22891 | yes | 1.802E-07 | 742.38623 | 5375.6963 |
| AIVICPTDEDLK | 12 | Q9BUJ2 | Heterogeneous nuclear ribonucleoprotein | HNRNPUL | 95.739 | 2 | 52.22891 | yes | 9.722E-08 | 687.35266 | 10238.604 |
| YNILGTNAIMDK | 12 | Q9BUJ2 | Heterogeneous nuclear ribonucleoprotein | HNRNPUL | 95.739 | 2 | 52.22891 | yes | 1E-07 | 676.8476 | 3841.718 |
| YNILGTNAIMDK | 12 | Q9BUJ2 | Heterogeneous nuclear ribonucleoprotein | HNRNPUL | 95.739 | 2 | 52.22891 | yes | 9.722E-08 | 684.84503 | 3841.718 |
| DVPDHAVLEMK | 11 | Q9BUJ2 | Heterogeneous nuclear ribonucleoprotein | HNRNPUL | 95.739 | 2 | 52.22891 | yes | 1.859E-07 | 627.31335 | 5117.0903 |
| QNQFYDTQVIK | 11 | Q9BUJ2 | Heterogeneous nuclear ribonucleoprotein | HNRNPUL | 95.739 | 2 | 52.22891 | yes | 9.722E-08 | 692.34882 | 4786.3599 |
| QYNEEGR | 7 | Q9BUJ2 | Heterogeneous nuclear ribonucleoprotein | HNRNPUL | 95.739 | 2 | 52.22891 | yes | 2.275E-07 | 448.19885 | 3951.5547 |
| EALGGQALYPHVI | 15 | Q9BUJ2 | Heterogeneous nuclear ribonucleoprotein | HNRNPUL | 95.739 | 2 | 52.22891 | yes | 1.499E-07 | 797.95123 | 1649.3916 |
| EALGGQALYPHVI | 15 | Q9BUJ2 | Heterogeneous nuclear ribonucleoprotein | HNRNPUL | 95.739 | 2 | 52.22891 | yes | 2.066E-07 | 532.30322 | 1649.3916 |
| EALGGQALYPHVI | 15 | Q9BUJ2 | Heterogeneous nuclear ribonucleoprotein | HNRNPUL | 95.739 | 3 | 52.22891 | yes | 1.499E-07 | 797.95123 | 1649.3916 |
| EALGGQALYPHVI | 15 | Q9BUJ2 | Heterogeneous nuclear ribonucleoprotein | HNRNPUL | 95.739 | 3 | 52.22891 | yes | 2.066E-07 | 532.30322 | 1649.3916 |
| AIVICPTDEDLKDF | 14 | Q9BUJ2 | Heterogeneous nuclear ribonucleoprotein | HNRNPUL | 95.739 | 2 | 52.22891 | yes | 1.084E-07 | 822.91669 | 2696.6362 |
| AECEILMMVGLPA | 16 | Q9BUJ2 | Heterogeneous nuclear ribonucleoprotein | HNRNPUL | 95.739 | 2 | 52.22891 | yes | 1.086E-07 | 845.42224 | 2228.71 |
| MRPFEGFQR | 9 | Q9BUJ2 | Heterogeneous nuclear ribonucleoprotein | HNRNPUL | 95.739 | 2 | 52.22891 | yes | 1.461E-07 | 584.29004 | 1669.4982 |
| KYNILGTNAIMDK | 13 | Q9BUJ2 | Heterogeneous nuclear ribonucleoprotein | HNRNPUL | 95.739 | 2 | 52.22891 | yes | 1.551E-07 | 740.89508 | 1475.8881 |
| RTDEEGKDVPDHA | 18 | Q9BUJ2 | Heterogeneous nuclear ribonucleoprotein | HNRNPUL | 95.739 | 4 | 52.22891 | yes | 1.862E-07 | 518.00464 | 1353.302 |
| NPPGASTYNK | 10 | Q9BUJ2 | Heterogeneous nuclear ribonucleoprotein | HNRNPUL | 95.739 | 2 | 52.22891 | yes | 2.319E-07 | 524.75653 | 1170.3859 |
| APQQQPPPQQPPPI | 31 | Q9BUJ2 | Heterogeneous nuclear ribonucleoprotein | HNRNPUL | 95.739 | 4 | 52.22891 | yes | 8.942E-07 | 832.927 | 908.61121 |

| | | | | | | | | | | | |
|---|---|---|---|---|---|---|---|---|---|---|---|
| FENYGDK | 7 | Q9BUJ2 | Heterogeneous nuclear ribonucleoprotein | HNRNPUL | 95.739 | 2 | 52.22891 | yes | 2.212E-07 | 436.69287 | 860.33075 |
| LQAALEAEEPDDE | 14 | Q9BUJ2 | Heterogeneous nuclear ribonucleoprotein | HNRNPUL | 95.739 | 2 | 52.22891 | yes | 9.722E-08 | 793.37067 | 622.02301 |
| INEEISVK | 8 | Q9BUJ2 | Heterogeneous nuclear ribonucleoprotein | HNRNPUL | 95.739 | 2 | 52.22891 | yes | 2.212E-07 | 466.25839 | 398.8027 |
| GRVCFEMK | 8 | Q9BUJ2 | Heterogeneous nuclear ribonucleoprotein | HNRNPUL | 95.739 | 2 | 52.22891 | yes | 1.463E-07 | 513.74658 | 320.9826 |
| QENESGYER | 9 | Q9BUJ2 | Heterogeneous nuclear ribonucleoprotein | HNRNPUL | 95.739 | 2 | 52.22891 | yes | 1.084E-07 | 556.23615 | 317.93219 |
| SKAECEILMMVGL | 18 | Q9BUJ2 | Heterogeneous nuclear ribonucleoprotein | HNRNPUL | 95.739 | 3 | 52.22891 | yes | 9.722E-08 | 635.65961 | 161.03815 |
| WMGIAFR | 7 | Q9BUJ2 | Heterogeneous nuclear ribonucleoprotein | HNRNPUL | 95.739 | 2 | 52.22891 | yes | 6.647E-07 | 440.72855 | 120.35279 |
| KAIVICPTDEDLK | 13 | Q9BUJ2 | Heterogeneous nuclear ribonucleoprotein | HNRNPUL | 95.739 | 2 | 52.22891 | yes | 1.695E-06 | 751.40015 | 76.849907 |
| QGAPTSFLPPEAS( | 19 | Q9BUJ2 | Heterogeneous nuclear ribonucleoprotein | HNRNPUL | 95.739 | 3 | 52.22891 | yes | 2.274E-06 | 680.35675 | 70.412567 |
| LKVNELR | 7 | Q9BUJ2 | Heterogeneous nuclear ribonucleoprotein | HNRNPUL | 95.739 | 2 | 52.22891 | yes | 5.001E-07 | 436.27164 | 46.12384 |
| LGPGAGAGFGVLF | 13 | Q9BUL5 | PHD finger protein 23 OS=Homo sapiens | PHF23 | 43.819 | 2 | 30.08767 | yes | 2.29E-06 | 586.33276 | 44.065151 |
| IPDEINDR | 8 | Q9BUL8 | Programmed cell death protein 10 OS=Hc | PDCD10 | 24.702 | 2 | 49.04709 | yes | 1.842E-07 | 486.24326 | 77.818504 |
| VNLSAAQTLR | 10 | Q9BUL8 | Programmed cell death protein 10 OS=Hc | PDCD10 | 24.702 | 2 | 49.04709 | yes | 1.01E-06 | 536.80908 | 73.420105 |
| ENPGLTQDIIMK | 12 | Q9BUL8 | Programmed cell death protein 10 OS=Hc | PDCD10 | 24.702 | 2 | 49.04709 | yes | 2.001E-06 | 679.85284 | 65.824493 |
| SVEVNFTESLLR | 12 | Q9BUL8 | Programmed cell death protein 10 OS=Hc | PDCD10 | 24.702 | 2 | 49.04709 | yes | 1.074E-07 | 697.36975 | 64.029427 |
| LIHQTNLILQTFK | 13 | Q9BUL8 | Programmed cell death protein 10 OS=Hc | PDCD10 | 24.702 | 3 | 49.04709 | yes | 1.057E-05 | 523.64349 | 37.21241 |
| SLGEPAAGEGSAK | 13 | Q9BUL9 | Ribonuclease P protein subunit p25 OS=I | RPP25 | 20.633 | 2 | 50.80551 | yes | 9.722E-08 | 587.29095 | 376.06052 |
| LAGLHQVTR | 9 | Q9BUL9 | Ribonuclease P protein subunit p25 OS=I | RPP25 | 20.633 | 2 | 50.80551 | yes | 1.34E-07 | 497.79324 | 212.96118 |
| AIVFSGCGR | 9 | Q9BUL9 | Ribonuclease P protein subunit p25 OS=I | RPP25 | 20.633 | 2 | 50.80551 | yes | 1.082E-07 | 483.74493 | 161.55035 |
| NVPGLAILLSK | 11 | Q9BUL9 | Ribonuclease P protein subunit p25 OS=I | RPP25 | 20.633 | 2 | 50.80551 | yes | 3.933E-06 | 562.85553 | 38.727787 |
| TVTCAEILK | 9 | Q9BUL9 | Ribonuclease P protein subunit p25 OS=I | RPP25 | 20.633 | 2 | 50.80551 | yes | 5.06E-07 | 517.78119 | 20.078497 |
| NLMAFATASMAQ | 16 | Q9BUL9 | Ribonuclease P protein subunit p25 OS=I | RPP25 | 20.633 | 2 | 50.80551 | yes | 3.856E-06 | 840.91345 | 16.798218 |
| NFLSTPQFLYR | 11 | Q9BUN8 | Derlin-1 OS=Homo sapiens OX=9606 GI | DERL1 | 28.801 | 2 | 46.53258 | yes | 1.535E-07 | 693.36426 | 359.08694 |
| HNWGQGFR | 8 | Q9BUN8 | Derlin-1 OS=Homo sapiens OX=9606 GI | DERL1 | 28.801 | 2 | 46.53258 | yes | 4.909E-07 | 501.23865 | 224.68755 |
| LGAPQTHLGLK | 11 | Q9BUP0 | EF-hand domain-containing protein D1 C | EFHD1 | 26.928 | 2 | 48.07048 | no | 1.489E-07 | 567.83514 | 308.62543 |
| LSEIDVALEGVK | 12 | Q9BUP0 | EF-hand domain-containing protein D1 C | EFHD1 | 26.928 | 2 | 48.07048 | yes | 1.204E-07 | 636.8559 | 67.934814 |
| VFNPYTEFPEFSR | 13 | Q9BUP0 | EF-hand domain-containing protein D1 C | EFHD1 | 26.928 | 2 | 48.07048 | yes | 5.562E-07 | 816.88831 | 27.939404 |
| AAAGELQEDSGLM | 17 | Q9BUP0 | EF-hand domain-containing protein D1 C | EFHD1 | 26.928 | 2 | 48.07048 | yes | 3.636E-05 | 837.922 | 10.985875 |
| DIIGVAETGSGK | 12 | Q9BUQ8 | Probable ATP-dependent RNA helicase D | DDX23 | 95.583 | 2 | 53.691463 | yes | 1.145E-07 | 573.80371 | 849.15747 |
| TVAVIGGISR | 10 | Q9BUQ8 | Probable ATP-dependent RNA helicase D | DDX23 | 95.583 | 2 | 53.691463 | yes | 2.191E-07 | 486.79547 | 700.36932 |
| DILVATDVAGR | 11 | Q9BUQ8 | Probable ATP-dependent RNA helicase D | DDX23 | 95.583 | 2 | 53.691463 | yes | 1.042E-07 | 565.31421 | 664.73651 |
| MIDMGFEPDVQK | 12 | Q9BUQ8 | Probable ATP-dependent RNA helicase D | DDX23 | 95.583 | 2 | 53.691463 | yes | 9.722E-08 | 705.32562 | 647.08917 |
| LIDVLENR | 8 | Q9BUQ8 | Probable ATP-dependent RNA helicase D | DDX23 | 95.583 | 2 | 53.691463 | yes | 5.136E-06 | 486.27966 | 535.23438 |
| FYGDLMEK | 8 | Q9BUQ8 | Probable ATP-dependent RNA helicase D | DDX23 | 95.583 | 2 | 53.691463 | yes | 2.004E-07 | 501.73367 | 508.25381 |
| CTYVVLDEADR | 11 | Q9BUQ8 | Probable ATP-dependent RNA helicase D | DDX23 | 95.583 | 2 | 53.691463 | yes | 9.722E-08 | 670.81116 | 442.57599 |
| HQVQLLGR | 8 | Q9BUQ8 | Probable ATP-dependent RNA helicase D | DDX23 | 95.583 | 2 | 53.691463 | yes | 1.666E-07 | 475.78015 | 393.23013 |
| AQPLSLEELLAK | 12 | Q9BUQ8 | Probable ATP-dependent RNA helicase D | DDX23 | 95.583 | 2 | 53.691463 | yes | 3.619E-07 | 656.37958 | 349.9552 |
| QAIPIGLQNR | 10 | Q9BUQ8 | Probable ATP-dependent RNA helicase D | DDX23 | 95.583 | 2 | 53.691463 | yes | 1.171E-07 | 555.32495 | 344.10272 |
| EDSAVFYELK | 10 | Q9BUQ8 | Probable ATP-dependent RNA helicase D | DDX23 | 95.583 | 2 | 53.691463 | yes | 5.585E-07 | 600.79279 | 300.34213 |

| | | | | | | | | | | | |
|---|---|---|---|---|---|---|---|---|---|---|---|
| MLEDPQER | 8 | Q9BUQ8 | Probable ATP-dependent RNA helicase D | DDX23 | 95.583 | 2 | 53.691463 | yes | 2.393E-07 | 509.23712 | 247.87695 |
| SGVAITFLTK | 10 | Q9BUQ8 | Probable ATP-dependent RNA helicase D | DDX23 | 95.583 | 2 | 53.691463 | yes | 4.195E-07 | 518.80548 | 217.71701 |
| ELAQQIEEETIK | 12 | Q9BUQ8 | Probable ATP-dependent RNA helicase D | DDX23 | 95.583 | 2 | 53.691463 | yes | 9.729E-08 | 715.87231 | 195.37631 |
| MGCEIVIATPGR | 12 | Q9BUQ8 | Probable ATP-dependent RNA helicase D | DDX23 | 95.583 | 2 | 53.691463 | yes | 9.722E-08 | 652.32849 | 181.6348 |
| NIEDYIHR | 8 | Q9BUQ8 | Probable ATP-dependent RNA helicase D | DDX23 | 95.583 | 2 | 53.691463 | yes | 1.318E-07 | 530.26453 | 169.36589 |
| MGYNACTLHGGK | 12 | Q9BUQ8 | Probable ATP-dependent RNA helicase D | DDX23 | 95.583 | 2 | 53.691463 | yes | 9.722E-08 | 654.7948 | 134.80391 |
| FGKPLGIR | 8 | Q9BUQ8 | Probable ATP-dependent RNA helicase D | DDX23 | 95.583 | 2 | 53.691463 | yes | 9.722E-08 | 444.2767 | 88.079254 |
| QTVMFTATMPPAV | 15 | Q9BUQ8 | Probable ATP-dependent RNA helicase D | DDX23 | 95.583 | 2 | 53.691463 | yes | 7.651E-06 | 839.91821 | 66.766083 |
| KGCDVLAK | 8 | Q9BUQ8 | Probable ATP-dependent RNA helicase D | DDX23 | 95.583 | 2 | 53.691463 | yes | 1E-07 | 445.74185 | 44.977104 |
| VFLMSESEK | 9 | Q9BUQ8 | Probable ATP-dependent RNA helicase D | DDX23 | 95.583 | 2 | 53.691463 | yes | 3.255E-07 | 535.26532 | 39.646889 |
| QFQDLGR | 7 | Q9BUQ8 | Probable ATP-dependent RNA helicase D | DDX23 | 95.583 | 2 | 53.691463 | yes | 3.121E-07 | 432.22214 | 39.371761 |
| LSPDPVAGSAVSQ | 16 | Q9BUR4 | Telomerase Cajal body protein 1 OS=Hor | WRAP53 | 59.31 | 2 | 47.86704 | yes | 2.279E-07 | 813.42828 | 183.20393 |
| QSGYPLWSLGR | 11 | Q9BUR4 | Telomerase Cajal body protein 1 OS=Hor | WRAP53 | 59.31 | 2 | 47.86704 | yes | 1.242E-07 | 632.32764 | 110.6684 |
| NFQKPGTWR | 9 | Q9BUZ4 | TNF receptor-associated factor 4 OS=Hor | TRAF4 | 53.543 | 2 | 53.06591 | yes | 3.288E-07 | 567.29614 | 947.2804 |
| LLAQHATSECPK | 12 | Q9BUZ4 | TNF receptor-associated factor 4 OS=Hor | TRAF4 | 53.543 | 2 | 53.06591 | yes | 9.819E-08 | 677.84283 | 420.77301 |
| LLAQHATSECPK | 12 | Q9BUZ4 | TNF receptor-associated factor 4 OS=Hor | TRAF4 | 53.543 | 2 | 53.06591 | yes | 9.723E-08 | 452.23096 | 420.77301 |
| LLAQHATSECPK | 12 | Q9BUZ4 | TNF receptor-associated factor 4 OS=Hor | TRAF4 | 53.543 | 3 | 53.06591 | yes | 9.819E-08 | 677.84283 | 420.77301 |
| LLAQHATSECPK | 12 | Q9BUZ4 | TNF receptor-associated factor 4 OS=Hor | TRAF4 | 53.543 | 3 | 53.06591 | yes | 9.723E-08 | 452.23096 | 420.77301 |
| DSCNTALVLCPFK | 13 | Q9BUZ4 | TNF receptor-associated factor 4 OS=Hor | TRAF4 | 53.543 | 2 | 53.06591 | yes | 1.317E-07 | 762.86292 | 798.85229 |
| CPEDQLPLDYAK | 12 | Q9BUZ4 | TNF receptor-associated factor 4 OS=Hor | TRAF4 | 53.543 | 2 | 53.06591 | yes | 1.165E-07 | 724.83997 | 795.55096 |
| FISHQDIR | 8 | Q9BUZ4 | TNF receptor-associated factor 4 OS=Hor | TRAF4 | 53.543 | 2 | 53.06591 | yes | 2.078E-07 | 508.26962 | 775.21954 |
| GSLDESSLGFGYPI | 14 | Q9BUZ4 | TNF receptor-associated factor 4 OS=Hor | TRAF4 | 53.543 | 2 | 53.06591 | yes | 1.001E-07 | 728.85138 | 760.40613 |
| EDLPGHLK | 8 | Q9BUZ4 | TNF receptor-associated factor 4 OS=Hor | TRAF4 | 53.543 | 2 | 53.06591 | yes | 1.405E-07 | 454.74545 | 647.29785 |
| EPVQVSTCGHR | 11 | Q9BUZ4 | TNF receptor-associated factor 4 OS=Hor | TRAF4 | 53.543 | 2 | 53.06591 | yes | 3.355E-07 | 635.30383 | 560.4928 |
| DLPAHLQHDCPK | 12 | Q9BUZ4 | TNF receptor-associated factor 4 OS=Hor | TRAF4 | 53.543 | 2 | 53.06591 | yes | 9.722E-08 | 715.84589 | 264.88992 |
| DLPAHLQHDCPK | 12 | Q9BUZ4 | TNF receptor-associated factor 4 OS=Hor | TRAF4 | 53.543 | 2 | 53.06591 | yes | 1.004E-07 | 477.56635 | 264.88992 |
| DLPAHLQHDCPK | 12 | Q9BUZ4 | TNF receptor-associated factor 4 OS=Hor | TRAF4 | 53.543 | 3 | 53.06591 | yes | 9.722E-08 | 715.84589 | 264.88992 |
| DLPAHLQHDCPK | 12 | Q9BUZ4 | TNF receptor-associated factor 4 OS=Hor | TRAF4 | 53.543 | 3 | 53.06591 | yes | 1.004E-07 | 477.56635 | 264.88992 |
| LLCPLCGK | 8 | Q9BUZ4 | TNF receptor-associated factor 4 OS=Hor | TRAF4 | 53.543 | 2 | 53.06591 | yes | 9.819E-08 | 480.75391 | 378.33105 |
| TQPCTYCTK | 9 | Q9BUZ4 | TNF receptor-associated factor 4 OS=Hor | TRAF4 | 53.543 | 2 | 53.06591 | yes | 2.559E-07 | 579.74951 | 357.74625 |
| VTFSLLDQSDPGL | 15 | Q9BUZ4 | TNF receptor-associated factor 4 OS=Hor | TRAF4 | 53.543 | 2 | 53.06591 | yes | 9.722E-08 | 795.9223 | 297.36743 |
| HLQGHLNTCSFNV | 19 | Q9BUZ4 | TNF receptor-associated factor 4 OS=Hor | TRAF4 | 53.543 | 4 | 53.06591 | yes | 1.715E-07 | 566.77576 | 271.75143 |
| RLLCPLCGK | 9 | Q9BUZ4 | TNF receptor-associated factor 4 OS=Hor | TRAF4 | 53.543 | 2 | 53.06591 | yes | 1.722E-07 | 558.80444 | 181.13788 |
| HVEESVKPHLAMI | 19 | Q9BUZ4 | TNF receptor-associated factor 4 OS=Hor | TRAF4 | 53.543 | 4 | 53.06591 | yes | 1.278E-07 | 549.27869 | 160.8385 |
| RDLPAHLQHDCPK | 13 | Q9BUZ4 | TNF receptor-associated factor 4 OS=Hor | TRAF4 | 53.543 | 3 | 53.06591 | yes | 1.003E-07 | 529.60004 | 145.49963 |
| PQHVTETFHPDPN | 15 | Q9BUZ4 | TNF receptor-associated factor 4 OS=Hor | TRAF4 | 53.543 | 4 | 53.06591 | yes | 1.41E-06 | 458.97333 | 117.27339 |
| SFHPEQDAGK | 10 | Q9BV38 | WD repeat-containing protein 18 OS=Ho | WDR18 | 47.406 | 2 | 42.947826 | yes | 3.41E-07 | 558.25946 | 79.447014 |
| VATSSLDQTVK | 11 | Q9BV38 | WD repeat-containing protein 18 OS=Ho | WDR18 | 47.406 | 2 | 42.947826 | yes | 5.701E-07 | 574.81152 | 77.852783 |
| TEQLQAVLCSTME | 14 | Q9BV38 | WD repeat-containing protein 18 OS=Ho | WDR18 | 47.406 | 2 | 42.947826 | yes | 2.801E-07 | 819.39728 | 62.134945 |

| | | | | | | | | | | |
|---|---|---|---|---|---|---|---|---|---|---|
| NWNLYPACLR | 10 | Q9BV44 | tRNA (guanine(6)-N2)-methyltransferase | THUMPD3 | 57.003 | 2 | 48.012344 | yes | 1.353E-07 | 653.82172 | 108.85506 |
| AANNIASLLTK | 11 | Q9BV44 | tRNA (guanine(6)-N2)-methyltransferase | THUMPD3 | 57.003 | 2 | 48.012344 | yes | 2.726E-07 | 558.32458 | 63.051903 |
| AVLLTQDTK | 9 | Q9BV44 | tRNA (guanine(6)-N2)-methyltransferase | THUMPD3 | 57.003 | 2 | 48.012344 | yes | 1.139E-06 | 494.78729 | 58.740288 |
| TGSVDIIVTDLPFG | 15 | Q9BV44 | tRNA (guanine(6)-N2)-methyltransferase | THUMPD3 | 57.003 | 2 | 48.012344 | yes | 1.214E-07 | 781.42725 | 43.774021 |
| GDMVTLPAGIYHF | 13 | Q9BV57 | Acireductone dioxygenase OS=Homo sap | ADI1 | 21.499 | 2 | 51.910725 | yes | 9.819E-08 | 715.36646 | 354.58472 |
| VQAWYMDDAPGI | 14 | Q9BV57 | Acireductone dioxygenase OS=Homo sap | ADI1 | 21.499 | 2 | 51.910725 | yes | 5.817E-07 | 810.85919 | 323.25262 |
| DKEDQWIR | 8 | Q9BV57 | Acireductone dioxygenase OS=Homo sap | ADI1 | 21.499 | 2 | 51.910725 | yes | 1.587E-07 | 545.26984 | 267.08319 |
| DKLPNYEEK | 9 | Q9BV57 | Acireductone dioxygenase OS=Homo sap | ADI1 | 21.499 | 2 | 51.910725 | yes | 9.722E-08 | 568.28516 | 179.5979 |
| QPHRPDPGRPVGL | 17 | Q9BV57 | Acireductone dioxygenase OS=Homo sap | ADI1 | 21.499 | 4 | 51.910725 | yes | 1.339E-07 | 488.76978 | 172.44691 |
| LDADKYENDPELE | 14 | Q9BV57 | Acireductone dioxygenase OS=Homo sap | ADI1 | 21.499 | 2 | 51.910725 | yes | 9.722E-08 | 839.89398 | 121.52208 |
| RLGVLYWK | 8 | Q9BV57 | Acireductone dioxygenase OS=Homo sap | ADI1 | 21.499 | 2 | 51.910725 | yes | 7.405E-07 | 517.81091 | 43.701061 |
| NYSWMDIITICK | 12 | Q9BV57 | Acireductone dioxygenase OS=Homo sap | ADI1 | 21.499 | 2 | 51.910725 | yes | 5.955E-07 | 772.3678 | 32.620682 |
| LPDYICPR | 8 | Q9BV68 | E3 ubiquitin-protein ligase RNF126 OS= | RNF126 | 33.862 | 2 | 43.842144 | yes | 2.424E-07 | 517.26038 | 221.59444 |
| YFCHCCSVEIVPR | 13 | Q9BV68 | E3 ubiquitin-protein ligase RNF126 OS= | RNF126 | 33.862 | 3 | 43.842144 | yes | 3.621E-07 | 576.25482 | 25.445368 |
| LDLCTTVLPQIEK | 13 | Q9BVA0 | Katanin p80 WD40 repeat-containing sub | KATNB1 | 72.334 | 2 | 39.944237 | yes | 4.556E-06 | 765.41583 | 64.176056 |
| VYGWEPER | 8 | Q9BVA0 | Katanin p80 WD40 repeat-containing sub | KATNB1 | 72.334 | 2 | 39.944237 | yes | 4.295E-07 | 518.24835 | 37.285179 |
| CFDVVLVNWGK | 11 | Q9BVA0 | Katanin p80 WD40 repeat-containing sub | KATNB1 | 72.334 | 2 | 39.944237 | yes | 4.556E-07 | 668.83954 | 33.834572 |
| ELHLLMASLD | 10 | Q9BVA0 | Katanin p80 WD40 repeat-containing sub | KATNB1 | 72.334 | 2 | 39.944237 | yes | 6.959E-07 | 571.29974 | 21.109314 |
| LLMENPYEGPDSQ | 14 | Q9BVC3 | Sister chromatid cohesion protein DCC1 ( | DSCC1 | 44.826 | 2 | 44.95883 | yes | 3.471E-07 | 810.88232 | 55.503143 |
| FNSLFSLR | 8 | Q9BVC3 | Sister chromatid cohesion protein DCC1 ( | DSCC1 | 44.826 | 2 | 44.95883 | yes | 2.012E-07 | 492.26907 | 46.022625 |
| GDKDEQAVLCSK | 12 | Q9BVC3 | Sister chromatid cohesion protein DCC1 ( | DSCC1 | 44.826 | 2 | 44.95883 | yes | 1.076E-06 | 675.3219 | 39.010075 |
| ILEFDYEMK | 9 | Q9BVC3 | Sister chromatid cohesion protein DCC1 ( | DSCC1 | 44.826 | 2 | 44.95883 | yes | 0.0003456 | 594.28625 | 12.695373 |
| NIASVGFHEDGR | 12 | Q9BVC4 | Target of rapamycin complex subunit LST | MLST8 | 35.877 | 2 | 44.3338 | yes | 2.232E-07 | 651.31531 | 89.158867 |
| TSNFSLMTELSIK | 13 | Q9BVC4 | Target of rapamycin complex subunit LST | MLST8 | 35.877 | 2 | 44.3338 | yes | 2.547E-06 | 735.87909 | 69.598747 |
| LWCVETGEIK | 10 | Q9BVC4 | Target of rapamycin complex subunit LST | MLST8 | 35.877 | 2 | 44.3338 | yes | 4.554E-07 | 617.81049 | 48.005432 |
| WMYTGGEDCTAR | 12 | Q9BVC4 | Target of rapamycin complex subunit LST | MLST8 | 35.877 | 2 | 44.3338 | yes | 4.785E-07 | 723.79248 | 34.520763 |
| PAPAPASPAAR | 11 | Q9BVG8 | Kinesin-like protein KIFC3 OS=Homo sa | KIFC3 | 92.775 | 2 | 52.886776 | yes | 2.828E-07 | 503.27744 | 405.85031 |
| TNLLTLQPALR | 11 | Q9BVG8 | Kinesin-like protein KIFC3 OS=Homo sa | KIFC3 | 92.775 | 2 | 52.886776 | yes | 9.722E-08 | 620.37463 | 246.43137 |
| LISQAQEVSR | 10 | Q9BVG8 | Kinesin-like protein KIFC3 OS=Homo sa | KIFC3 | 92.775 | 2 | 52.886776 | yes | 9.722E-08 | 565.81183 | 234.72165 |
| TLMVVQVSPVEK | 12 | Q9BVG8 | Kinesin-like protein KIFC3 OS=Homo sa | KIFC3 | 92.775 | 2 | 52.886776 | yes | 1.375E-07 | 665.37598 | 223.83234 |
| GMLSELNLEVQQK | 13 | Q9BVG8 | Kinesin-like protein KIFC3 OS=Homo sa | KIFC3 | 92.775 | 2 | 52.886776 | yes | 1.118E-07 | 744.88995 | 214.96225 |
| LTYLLQDSLSGDSI | 14 | Q9BVG8 | Kinesin-like protein KIFC3 OS=Homo sa | KIFC3 | 92.775 | 2 | 52.886776 | yes | 3.354E-07 | 770.39868 | 209.57544 |
| DSHETIASLR | 10 | Q9BVG8 | Kinesin-like protein KIFC3 OS=Homo sa | KIFC3 | 92.775 | 2 | 52.886776 | yes | 1.27E-07 | 564.78583 | 155.54886 |
| AVHENLAGVR | 10 | Q9BVG8 | Kinesin-like protein KIFC3 OS=Homo sa | KIFC3 | 92.775 | 2 | 52.886776 | yes | 2.336E-07 | 533.29364 | 120.29669 |
| VFEFGHTNR | 9 | Q9BVG8 | Kinesin-like protein KIFC3 OS=Homo sa | KIFC3 | 92.775 | 2 | 52.886776 | yes | 9.722E-08 | 553.77252 | 113.42853 |
| AQIAMYESELER | 12 | Q9BVG8 | Kinesin-like protein KIFC3 OS=Homo sa | KIFC3 | 92.775 | 2 | 52.886776 | yes | 9.722E-08 | 720.3454 | 91.053436 |
| LSQLQLEMAESK | 12 | Q9BVG8 | Kinesin-like protein KIFC3 OS=Homo sa | KIFC3 | 92.775 | 2 | 52.886776 | yes | 3.656E-06 | 688.85815 | 73.540886 |
| QGHVPFR | 7 | Q9BVG8 | Kinesin-like protein KIFC3 OS=Homo sa | KIFC3 | 92.775 | 2 | 52.886776 | yes | 3.709E-07 | 420.72739 | 57.471775 |
| ALQLLFSEVQEK | 12 | Q9BVG8 | Kinesin-like protein KIFC3 OS=Homo sa | KIFC3 | 92.775 | 2 | 52.886776 | yes | 9.722E-08 | 702.89032 | 55.519844 |

| | | | | | | | | | | | |
|---|---|---|---|---|---|---|---|---|---|---|---|
| TTEFTNLNEHSSR | 13 | Q9BVG8 | Kinesin-like protein KIFC3 OS=Homo sa | KIFC3 | 92.775 | 2 | 52.886776 | yes | 3.92E-07 | 768.35785 | 53.453762 |
| AHGQMLEEMQSL | 18 | Q9BVG8 | Kinesin-like protein KIFC3 OS=Homo sa | KIFC3 | 92.775 | 3 | 52.886776 | yes | 1.903E-07 | 715.6615 | 23.191948 |
| LFGALLER | 8 | Q9BVI4 | Nucleolar complex protein 4 homolog OS | NOC4L | 58.468 | 2 | 53.533867 | yes | 1.627E-07 | 459.77402 | 1076.2654 |
| ALESSLWELQALQ | 14 | Q9BVI4 | Nucleolar complex protein 4 homolog OS | NOC4L | 58.468 | 2 | 53.533867 | yes | 1.013E-07 | 822.44122 | 743.42139 |
| WEGNYLFPR | 9 | Q9BVI4 | Nucleolar complex protein 4 homolog OS | NOC4L | 58.468 | 2 | 53.533867 | yes | 9.729E-08 | 591.29053 | 720.52271 |
| LGELLGHPSFQVK | 13 | Q9BVI4 | Nucleolar complex protein 4 homolog OS | NOC4L | 58.468 | 2 | 53.533867 | yes | 9.99E-08 | 712.89844 | 339.18723 |
| LGELLGHPSFQVK | 13 | Q9BVI4 | Nucleolar complex protein 4 homolog OS | NOC4L | 58.468 | 2 | 53.533867 | yes | 9.285E-07 | 475.60138 | 339.18723 |
| LGELLGHPSFQVK | 13 | Q9BVI4 | Nucleolar complex protein 4 homolog OS | NOC4L | 58.468 | 3 | 53.533867 | yes | 9.99E-08 | 712.89844 | 339.18723 |
| LGELLGHPSFQVK | 13 | Q9BVI4 | Nucleolar complex protein 4 homolog OS | NOC4L | 58.468 | 3 | 53.533867 | yes | 9.285E-07 | 475.60138 | 339.18723 |
| LYGLLDPSVFHVK | 13 | Q9BVI4 | Nucleolar complex protein 4 homolog OS | NOC4L | 58.468 | 2 | 53.533867 | yes | 9.722E-08 | 744.4165 | 295.63074 |
| LYGLLDPSVFHVK | 13 | Q9BVI4 | Nucleolar complex protein 4 homolog OS | NOC4L | 58.468 | 2 | 53.533867 | yes | 2.189E-07 | 496.6134 | 295.63074 |
| LYGLLDPSVFHVK | 13 | Q9BVI4 | Nucleolar complex protein 4 homolog OS | NOC4L | 58.468 | 3 | 53.533867 | yes | 9.722E-08 | 744.4165 | 295.63074 |
| LYGLLDPSVFHVK | 13 | Q9BVI4 | Nucleolar complex protein 4 homolog OS | NOC4L | 58.468 | 3 | 53.533867 | yes | 2.189E-07 | 496.6134 | 295.63074 |
| YHTMQAAVDAVA | 13 | Q9BVI4 | Nucleolar complex protein 4 homolog OS | NOC4L | 58.468 | 2 | 53.533867 | yes | 1.66E-07 | 716.8537 | 553.30951 |
| FVQLEGAHPLEK | 12 | Q9BVI4 | Nucleolar complex protein 4 homolog OS | NOC4L | 58.468 | 2 | 53.533867 | yes | 1.05E-07 | 684.36951 | 470.00931 |
| HNLEYPDFYR | 10 | Q9BVI4 | Nucleolar complex protein 4 homolog OS | NOC4L | 58.468 | 2 | 53.533867 | yes | 9.722E-08 | 677.31476 | 335.30902 |
| PGELCAQHFTLS | 12 | Q9BVI4 | Nucleolar complex protein 4 homolog OS | NOC4L | 58.468 | 2 | 53.533867 | yes | 1.103E-07 | 680.32172 | 304.89566 |
| EYLDYDDTR | 9 | Q9BVI4 | Nucleolar complex protein 4 homolog OS | NOC4L | 58.468 | 2 | 53.533867 | yes | 9.722E-08 | 595.25403 | 212.30753 |
| REPTVSSFYVK | 11 | Q9BVI4 | Nucleolar complex protein 4 homolog OS | NOC4L | 58.468 | 2 | 53.533867 | yes | 1.751E-07 | 656.84845 | 207.92766 |
| KLYGLLDPSVFHV | 14 | Q9BVI4 | Nucleolar complex protein 4 homolog OS | NOC4L | 58.468 | 3 | 53.533867 | yes | 1.286E-07 | 539.31171 | 96.436165 |
| KLYGLLDPSVFHV | 14 | Q9BVI4 | Nucleolar complex protein 4 homolog OS | NOC4L | 58.468 | 3 | 53.533867 | yes | 9.81E-08 | 808.46393 | 96.436165 |
| KLYGLLDPSVFHV | 14 | Q9BVI4 | Nucleolar complex protein 4 homolog OS | NOC4L | 58.468 | 2 | 53.533867 | yes | 1.286E-07 | 539.31171 | 96.436165 |
| KLYGLLDPSVFHV | 14 | Q9BVI4 | Nucleolar complex protein 4 homolog OS | NOC4L | 58.468 | 2 | 53.533867 | yes | 9.81E-08 | 808.46393 | 96.436165 |
| RLEAVLASR | 9 | Q9BVI4 | Nucleolar complex protein 4 homolog OS | NOC4L | 58.468 | 2 | 53.533867 | yes | 4.065E-07 | 507.80637 | 184.13916 |
| RAELWDTWK | 9 | Q9BVI4 | Nucleolar complex protein 4 homolog OS | NOC4L | 58.468 | 2 | 53.533867 | yes | 9.862E-08 | 602.80908 | 172.97035 |
| GELFVGQLPSEEM | 23 | Q9BVI4 | Nucleolar complex protein 4 homolog OS | NOC4L | 58.468 | 3 | 53.533867 | yes | 3.579E-07 | 808.72266 | 141.26787 |
| SKWEGNYLFPR | 11 | Q9BVI4 | Nucleolar complex protein 4 homolog OS | NOC4L | 58.468 | 2 | 53.533867 | yes | 2.721E-06 | 698.85406 | 89.95475 |
| VLVHRPHGPELDA | 27 | Q9BVI4 | Nucleolar complex protein 4 homolog OS | NOC4L | 58.468 | 4 | 53.533867 | yes | 1.05E-07 | 749.85876 | 89.537025 |
| LEAVLASR | 8 | Q9BVI4 | Nucleolar complex protein 4 homolog OS | NOC4L | 58.468 | 2 | 53.533867 | yes | 1.28E-07 | 429.7558 | 67.069237 |
| AELWDTWK | 8 | Q9BVI4 | Nucleolar complex protein 4 homolog OS | NOC4L | 58.468 | 2 | 53.533867 | yes | 1.018E-07 | 524.75854 | 48.3657 |
| ASSEGTIPQVQR | 12 | Q9BVJ6 | U3 small nucleolar RNA-associated prote | UTP14A | 87.978 | 2 | 52.313892 | yes | 9.723E-08 | 636.83075 | 850.56274 |
| VSEFNVSSEGSGE | 14 | Q9BVJ6 | U3 small nucleolar RNA-associated prote | UTP14A | 87.978 | 2 | 52.313892 | yes | 9.722E-08 | 728.33356 | 546.97089 |
| EAFAGDDVIR | 10 | Q9BVJ6 | U3 small nucleolar RNA-associated prote | UTP14A | 87.978 | 2 | 52.313892 | no | 9.722E-08 | 546.76965 | 532.61249 |
| NLPNVIINEK | 10 | Q9BVJ6 | U3 small nucleolar RNA-associated prote | UTP14A | 87.978 | 2 | 52.313892 | yes | 1.018E-07 | 577.3324 | 351.02707 |
| TIQTPIGSTWNTQR | 14 | Q9BVJ6 | U3 small nucleolar RNA-associated prote | UTP14A | 87.978 | 2 | 52.313892 | no | 9.722E-08 | 801.91534 | 323.44647 |
| DSGSQEVLSELR | 12 | Q9BVJ6 | U3 small nucleolar RNA-associated prote | UTP14A | 87.978 | 2 | 52.313892 | yes | 1.655E-07 | 660.3255 | 276.91537 |
| LLEAISSLDGK | 11 | Q9BVJ6 | U3 small nucleolar RNA-associated prote | UTP14A | 87.978 | 2 | 52.313892 | yes | 1.413E-07 | 573.32428 | 252.04721 |
| QAEQLVFPLEK | 11 | Q9BVJ6 | U3 small nucleolar RNA-associated prote | UTP14A | 87.978 | 2 | 52.313892 | yes | 2.516E-07 | 651.35864 | 186.93228 |
| QPVTDPLLTPVEK | 13 | Q9BVJ6 | U3 small nucleolar RNA-associated prote | UTP14A | 87.978 | 2 | 52.313892 | yes | 9.722E-08 | 718.90338 | 139.36659 |

| | | | | | | | | | | | |
|---|---|---|---|---|---|---|---|---|---|---|---|
| VQTLEELEELGK | 12 | Q9BVJ6 | U3 small nucleolar RNA-associated prote | UTP14A | 87.978 | 2 | 52.313892 | no | 4.656E-07 | 694.36938 | 125.32961 |
| PVLEGQQSER | 10 | Q9BVJ6 | U3 small nucleolar RNA-associated prote | UTP14A | 87.978 | 2 | 52.313892 | no | 1.658E-07 | 571.79364 | 102.52774 |
| NKQPVTDPLLTPV | 15 | Q9BVJ6 | U3 small nucleolar RNA-associated prote | UTP14A | 87.978 | 2 | 52.313892 | yes | 1.675E-07 | 839.97235 | 88.903915 |
| AMSLEEAK | 8 | Q9BVJ6 | U3 small nucleolar RNA-associated prote | UTP14A | 87.978 | 2 | 52.313892 | yes | 1.699E-07 | 439.71805 | 88.62722 |
| SDLSVIQR | 8 | Q9BVJ6 | U3 small nucleolar RNA-associated prote | UTP14A | 87.978 | 2 | 52.313892 | yes | 2.992E-07 | 459.25616 | 88.211891 |
| KVNPAAALEELEK | 13 | Q9BVJ6 | U3 small nucleolar RNA-associated prote | UTP14A | 87.978 | 2 | 52.313892 | yes | 1.149E-06 | 706.39319 | 82.053841 |
| VNPAAALEELEK | 12 | Q9BVJ6 | U3 small nucleolar RNA-associated prote | UTP14A | 87.978 | 2 | 52.313892 | yes | 9.722E-08 | 642.3457 | 75.665512 |
| NIHAAAHQVR | 10 | Q9BVJ6 | U3 small nucleolar RNA-associated prote | UTP14A | 87.978 | 2 | 52.313892 | yes | 3.349E-07 | 558.80469 | 67.823669 |
| TPNNRPDAPK | 10 | Q9BVJ6 | U3 small nucleolar RNA-associated prote | UTP14A | 87.978 | 2 | 52.313892 | no | 9.81E-08 | 555.28851 | 65.329124 |
| ALQSYYEAK | 9 | Q9BVJ6 | U3 small nucleolar RNA-associated prote | UTP14A | 87.978 | 2 | 52.313892 | no | 3.987E-05 | 536.7691 | 63.721272 |
| KTVELPLNK | 9 | Q9BVJ6 | U3 small nucleolar RNA-associated prote | UTP14A | 87.978 | 2 | 52.313892 | yes | 9.427E-07 | 521.31879 | 59.302536 |
| TPLEQEIFNLLHK | 13 | Q9BVJ6 | U3 small nucleolar RNA-associated prote | UTP14A | 87.978 | 2 | 52.313892 | no | 3.569E-07 | 791.43542 | 45.284424 |
| VQTLEELEELGKE | 19 | Q9BVJ6 | U3 small nucleolar RNA-associated prote | UTP14A | 87.978 | 3 | 52.313892 | yes | 2.138E-07 | 775.04224 | 41.67524 |
| WDPVVLK | 7 | Q9BVJ6 | U3 small nucleolar RNA-associated prote | UTP14A | 87.978 | 2 | 52.313892 | yes | 9.722E-08 | 428.75 | 40.335701 |
| DVDLTLPGWGEW | 22 | Q9BVJ6 | U3 small nucleolar RNA-associated prote | UTP14A | 87.978 | 3 | 52.313892 | no | 1.593E-07 | 761.39868 | 36.06908 |
| GLAAGDAIAEIR | 12 | Q9BVJ7 | Dual specificity protein phosphatase 23 O | DUSP23 | 16.589 | 2 | 47.736713 | yes | 2.45E-07 | 578.81964 | 180.12848 |
| TGTMLACYLVK | 11 | Q9BVJ7 | Dual specificity protein phosphatase 23 O | DUSP23 | 16.589 | 2 | 47.736713 | yes | 1.269E-07 | 628.82251 | 166.96217 |
| HLVSLTER | 8 | Q9BVJ7 | Dual specificity protein phosphatase 23 O | DUSP23 | 16.589 | 2 | 47.736713 | yes | 3.768E-07 | 477.772 | 131.65277 |
| GVQPPNFSWVLPC | 14 | Q9BVJ7 | Dual specificity protein phosphatase 23 O | DUSP23 | 16.589 | 2 | 47.736713 | yes | 1.6E-07 | 777.41479 | 119.90614 |
| AVFQFYQR | 8 | Q9BVJ7 | Dual specificity protein phosphatase 23 O | DUSP23 | 16.589 | 2 | 47.736713 | yes | 1.044E-06 | 529.77454 | 71.10614 |
| IPDFCPPAPDQIDR | 14 | Q9BVJ7 | Dual specificity protein phosphatase 23 O | DUSP23 | 16.589 | 2 | 47.736713 | yes | 4.879E-07 | 820.8905 | 55.216469 |
| FSLFAGGMLR | 10 | Q9BVK6 | Transmembrane emp24 domain-containin | TMED9 | 27.278 | 2 | 41.265152 | yes | 3.596E-07 | 549.79187 | 69.686974 |
| CIPLWVGAR | 9 | Q9BVK8 | BOS complex subunit TMEM147 OS=Ho | TMEM147 | 25.261 | 2 | 40.493313 | yes | 3.995E-07 | 536.29205 | 83.430038 |
| VGVIGFPNVGK | 11 | Q9BVP2 | Guanine nucleotide-binding protein-like 3 | GNL3 | 61.994 | 2 | 53.726 | yes | 5.198E-06 | 543.81897 | 4398.7578 |
| LLGGFQETCSK | 11 | Q9BVP2 | Guanine nucleotide-binding protein-like 3 | GNL3 | 61.994 | 2 | 53.726 | yes | 1.067E-07 | 620.30554 | 1842.6084 |
| SPASIEVVKPMEA | 24 | Q9BVP2 | Guanine nucleotide-binding protein-like 3 | GNL3 | 61.994 | 3 | 53.726 | yes | 1.153E-07 | 814.42834 | 1378.2513 |
| GGIPNVEGAAK | 11 | Q9BVP2 | Guanine nucleotide-binding protein-like 3 | GNL3 | 61.994 | 2 | 53.726 | yes | 9.927E-08 | 506.77472 | 1303.1292 |
| CPQVEEAIVQSGQ | 14 | Q9BVP2 | Guanine nucleotide-binding protein-like 3 | GNL3 | 61.994 | 2 | 53.726 | yes | 1.178E-06 | 786.88794 | 1070.717 |
| KDPGVPNSAPFK | 12 | Q9BVP2 | Guanine nucleotide-binding protein-like 3 | GNL3 | 61.994 | 2 | 53.726 | yes | 9.722E-08 | 628.83533 | 856.77191 |
| SMQVVPLDK | 9 | Q9BVP2 | Guanine nucleotide-binding protein-like 3 | GNL3 | 61.994 | 2 | 53.726 | yes | 1.376E-07 | 508.77588 | 703.40607 |
| SGFNLEELEK | 10 | Q9BVP2 | Guanine nucleotide-binding protein-like 3 | GNL3 | 61.994 | 2 | 53.726 | yes | 2.082E-07 | 583.29041 | 646.75323 |
| ENLESWLNYLK | 11 | Q9BVP2 | Guanine nucleotide-binding protein-like 3 | GNL3 | 61.994 | 2 | 53.726 | yes | 9.728E-08 | 704.85901 | 423.91699 |
| KLYCQELK | 8 | Q9BVP2 | Guanine nucleotide-binding protein-like 3 | GNL3 | 61.994 | 2 | 53.726 | yes | 1.709E-07 | 541.28918 | 352.16443 |
| SSIINSLK | 8 | Q9BVP2 | Guanine nucleotide-binding protein-like 3 | GNL3 | 61.994 | 2 | 53.726 | yes | 4.004E-07 | 431.25565 | 250.11215 |
| ELPTVVFR | 8 | Q9BVP2 | Guanine nucleotide-binding protein-like 3 | GNL3 | 61.994 | 2 | 53.726 | yes | 2.01E-07 | 480.7793 | 204.19565 |
| YTVPGYR | 7 | Q9BVP2 | Guanine nucleotide-binding protein-like 3 | GNL3 | 61.994 | 2 | 53.726 | yes | 5.562E-07 | 428.22159 | 151.72354 |
| KLVLILNK | 8 | Q9BVP2 | Guanine nucleotide-binding protein-like 3 | GNL3 | 61.994 | 2 | 53.726 | yes | 1.755E-07 | 470.83133 | 133.92667 |
| KELPTVVFR | 9 | Q9BVP2 | Guanine nucleotide-binding protein-like 3 | GNL3 | 61.994 | 2 | 53.726 | yes | 2.158E-07 | 544.82678 | 97.342957 |
| PSNVEPMEK | 9 | Q9BVP2 | Guanine nucleotide-binding protein-like 3 | GNL3 | 61.994 | 2 | 53.726 | yes | 3.626E-06 | 515.74731 | 47.026775 |

| | | | | | | | | | | | |
|---|---|---|---|---|---|---|---|---|---|---|---|
| GPHLANSILFQSSG | 23 | Q9BVP2 | Guanine nucleotide-binding protein-like 3 | GNL3 | 61.994 | 3 | 53.726 | yes | 2.953E-06 | 809.09558 | 31.187296 |
| LLLVPCPPLR | 10 | Q9BVQ7 | ATPase family gene 2 protein homolog B | AFG2B | 80.711 | 2 | 54.441097 | yes | 1.957E-07 | 589.35992 | 347.34872 |
| GVLLAGPPGVGK | 12 | Q9BVQ7 | ATPase family gene 2 protein homolog B | AFG2B | 80.711 | 2 | 54.441097 | yes | 1.585E-07 | 532.82678 | 275.89606 |
| EVVIGTPTLK | 10 | Q9BVQ7 | ATPase family gene 2 protein homolog B | AFG2B | 80.711 | 2 | 54.441097 | yes | 2.492E-07 | 528.8186 | 265.33835 |
| SSQQEFQEVFNR | 12 | Q9BVQ7 | ATPase family gene 2 protein homolog B | AFG2B | 80.711 | 2 | 54.441097 | yes | 9.722E-08 | 749.84967 | 195.23438 |
| GVLLYGPPGCAK | 12 | Q9BVQ7 | ATPase family gene 2 protein homolog B | AFG2B | 80.711 | 2 | 54.441097 | yes | 4.798E-07 | 616.3288 | 194.94583 |
| TVKPSLSCK | 9 | Q9BVQ7 | ATPase family gene 2 protein homolog B | AFG2B | 80.711 | 2 | 54.441097 | yes | 6.785E-07 | 510.28134 | 136.7178 |
| LGPAALHALGAR | 12 | Q9BVQ7 | ATPase family gene 2 protein homolog B | AFG2B | 80.711 | 2 | 54.441097 | yes | 9.722E-08 | 573.84076 | 94.332207 |
| DLALYENLFK | 10 | Q9BVQ7 | ATPase family gene 2 protein homolog B | AFG2B | 80.711 | 2 | 54.441097 | yes | 2.263E-07 | 613.32678 | 74.803886 |
| EILQVITSK | 9 | Q9BVQ7 | ATPase family gene 2 protein homolog B | AFG2B | 80.711 | 2 | 54.441097 | yes | 2.411E-07 | 515.81079 | 62.721741 |
| VAVWPVLR | 8 | Q9BVQ7 | ATPase family gene 2 protein homolog B | AFG2B | 80.711 | 2 | 54.441097 | yes | 3.345E-07 | 470.29236 | 41.418938 |
| FSYESELFPTFK | 12 | Q9BVS4 | Serine/threonine-protein kinase RIO2 OS | RIOK2 | 63.284 | 2 | 56.713627 | yes | 2.664E-07 | 747.86121 | 1386.7263 |
| VLTAVEMGMK | 10 | Q9BVS4 | Serine/threonine-protein kinase RIO2 OS | RIOK2 | 63.284 | 2 | 56.713627 | yes | 3.801E-07 | 539.7854 | 1103.389 |
| HNVSWLYLSR | 10 | Q9BVS4 | Serine/threonine-protein kinase RIO2 OS | RIOK2 | 63.284 | 2 | 56.713627 | yes | 2.585E-07 | 637.83563 | 793.84277 |
| QVVESVGNQMGV | 14 | Q9BVS4 | Serine/threonine-protein kinase RIO2 OS | RIOK2 | 63.284 | 2 | 56.713627 | yes | 9.722E-08 | 716.36664 | 703.50476 |
| LTNAGYDYLALK | 12 | Q9BVS4 | Serine/threonine-protein kinase RIO2 OS | RIOK2 | 63.284 | 2 | 56.713627 | yes | 1.023E-07 | 671.35608 | 641.48425 |
| DEENVGAMNQYR | 12 | Q9BVS4 | Serine/threonine-protein kinase RIO2 OS | RIOK2 | 63.284 | 2 | 56.713627 | yes | 1.073E-07 | 713.30682 | 497.44968 |
| EMQADDELLHPLC | 17 | Q9BVS4 | Serine/threonine-protein kinase RIO2 OS | RIOK2 | 63.284 | 3 | 56.713627 | yes | 1.018E-07 | 641.63153 | 423.01215 |
| NHEIVPGSLIASIAS | 17 | Q9BVS4 | Serine/threonine-protein kinase RIO2 OS | RIOK2 | 63.284 | 3 | 56.713627 | yes | 9.819E-08 | 583.66833 | 298.91092 |
| REDTLDVEVSASG | 16 | Q9BVS4 | Serine/threonine-protein kinase RIO2 OS | RIOK2 | 63.284 | 3 | 56.713627 | yes | 9.722E-08 | 590.62323 | 218.29126 |
| AEVYGSENESER | 12 | Q9BVS4 | Serine/threonine-protein kinase RIO2 OS | RIOK2 | 63.284 | 2 | 56.713627 | yes | 1.001E-07 | 685.29694 | 149.74231 |
| EDSLSEESADAR | 12 | Q9BVS4 | Serine/threonine-protein kinase RIO2 OS | RIOK2 | 63.284 | 2 | 56.713627 | yes | 1.004E-07 | 654.78113 | 145.54604 |
| EGSEFSFSDGEVAI | 15 | Q9BVS4 | Serine/threonine-protein kinase RIO2 OS | RIOK2 | 63.284 | 2 | 56.713627 | yes | 9.722E-08 | 809.34937 | 125.106 |
| ENMQNIK | 7 | Q9BVS4 | Serine/threonine-protein kinase RIO2 OS | RIOK2 | 63.284 | 2 | 56.713627 | yes | 2.496E-07 | 438.71582 | 63.668175 |
| RPALEDYVVLMK | 12 | Q9BVS5 | tRNA (adenine(58)-N(1))-methyltransfera | TRMT61B | 52.966 | 2 | 51.401196 | yes | 1.728E-07 | 717.39471 | 218.52922 |
| INTDVQLDSQEK | 12 | Q9BVS5 | tRNA (adenine(58)-N(1))-methyltransfera | TRMT61B | 52.966 | 2 | 51.401196 | yes | 9.722E-08 | 695.34644 | 157.66373 |
| ELEDSSGDQGR | 11 | Q9BVS5 | tRNA (adenine(58)-N(1))-methyltransfera | TRMT61B | 52.966 | 2 | 51.401196 | yes | 5.126E-06 | 596.75745 | 145.75082 |
| VISFEVR | 7 | Q9BVS5 | tRNA (adenine(58)-N(1))-methyltransfera | TRMT61B | 52.966 | 2 | 51.401196 | yes | 3.579E-07 | 425.24509 | 111.17265 |
| TCELALSCEK | 10 | Q9BVS5 | tRNA (adenine(58)-N(1))-methyltransfera | TRMT61B | 52.966 | 2 | 51.401196 | yes | 2.434E-07 | 605.77576 | 102.24414 |
| ERPFQAGELILAET | 20 | Q9BVS5 | tRNA (adenine(58)-N(1))-methyltransfera | TRMT61B | 52.966 | 3 | 51.401196 | yes | 1.462E-07 | 725.70245 | 61.516167 |
| ENPGSGEYDFR | 11 | Q9BVV7 | Mitochondrial import inner membrane tra | TIMM21 | 28.203 | 2 | 50.837036 | yes | 9.722E-08 | 635.77039 | 266.452 |
| FYIEGSEPGK | 10 | Q9BVV7 | Mitochondrial import inner membrane tra | TIMM21 | 28.203 | 2 | 50.837036 | yes | 1.994E-07 | 563.77441 | 232.18398 |
| SHPEVIGVFGESVI | 14 | Q9BVV7 | Mitochondrial import inner membrane tra | TIMM21 | 28.203 | 2 | 50.837036 | yes | 4.801E-07 | 742.89081 | 195.19299 |
| QGTVYAQVK | 9 | Q9BVV7 | Mitochondrial import inner membrane tra | TIMM21 | 28.203 | 2 | 50.837036 | yes | 5.497E-07 | 497.27182 | 65.857491 |
| YIFVEIESYPR | 11 | Q9BVV7 | Mitochondrial import inner membrane tra | TIMM21 | 28.203 | 2 | 50.837036 | yes | 7.213E-07 | 708.36389 | 48.587738 |
| GTLSGAPAPPTR | 12 | Q9BW19 | Kinesin-like protein KIFC1 OS=Homo sa | KIFC1 | 73.748 | 2 | 52.792744 | yes | 9.722E-08 | 562.80658 | 732.2356 |
| ACVLELEER | 9 | Q9BW19 | Kinesin-like protein KIFC1 OS=Homo sa | KIFC1 | 73.748 | 2 | 52.792744 | yes | 1.799E-07 | 559.77917 | 569.1311 |
| QVELQEER | 8 | Q9BW19 | Kinesin-like protein KIFC1 OS=Homo sa | KIFC1 | 73.748 | 2 | 52.792744 | yes | 2.471E-07 | 515.76184 | 460.7489 |
| AGPGSEELTVTNA | 14 | Q9BW19 | Kinesin-like protein KIFC1 OS=Homo sa | KIFC1 | 73.748 | 2 | 52.792744 | yes | 9.722E-08 | 701.35205 | 402.44138 |

| | | | | | | | | | | | |
|---|---|---|---|---|---|---|---|---|---|---|---|
| LDPGLALGPGER | 12 | Q9BW19 | Kinesin-like protein KIFC1 OS=Homo sa | KIFC1 | 73.748 | 2 | 52.792744 | yes | 1.128E-07 | 597.82751 | 286.65579 |
| EVDALLHLAR | 10 | Q9BW19 | Kinesin-like protein KIFC1 OS=Homo sa | KIFC1 | 73.748 | 2 | 52.792744 | yes | 1.088E-07 | 568.82477 | 257.55054 |
| CSTAIATGLK | 10 | Q9BW19 | Kinesin-like protein KIFC1 OS=Homo sa | KIFC1 | 73.748 | 2 | 52.792744 | yes | 1.279E-07 | 511.27097 | 254.37447 |
| VNQCVIGTAQANF | 13 | Q9BW19 | Kinesin-like protein KIFC1 OS=Homo sa | KIFC1 | 73.748 | 2 | 52.792744 | yes | 1.041E-07 | 715.86206 | 249.93048 |
| NQKPVPAVPVQK | 12 | Q9BW19 | Kinesin-like protein KIFC1 OS=Homo sa | KIFC1 | 73.748 | 2 | 52.792744 | yes | 1.036E-07 | 652.88788 | 241.4635 |
| HDFSFDR | 7 | Q9BW19 | Kinesin-like protein KIFC1 OS=Homo sa | KIFC1 | 73.748 | 2 | 52.792744 | yes | 1.546E-07 | 462.20395 | 229.78709 |
| LSTQEGLVQELQK | 13 | Q9BW19 | Kinesin-like protein KIFC1 OS=Homo sa | KIFC1 | 73.748 | 2 | 52.792744 | yes | 1.503E-07 | 736.90137 | 228.65892 |
| GQLCDLNAELK | 11 | Q9BW19 | Kinesin-like protein KIFC1 OS=Homo sa | KIFC1 | 73.748 | 2 | 52.792744 | yes | 1.505E-07 | 630.81628 | 216.11781 |
| QETVAQAALLTER | 13 | Q9BW19 | Kinesin-like protein KIFC1 OS=Homo sa | KIFC1 | 73.748 | 2 | 52.792744 | yes | 9.777E-08 | 715.38593 | 208.60255 |
| TTLEGHLAK | 9 | Q9BW19 | Kinesin-like protein KIFC1 OS=Homo sa | KIFC1 | 73.748 | 2 | 52.792744 | yes | 1.284E-06 | 485.27182 | 173.87263 |
| VQAQAEQGQQELI | 13 | Q9BW19 | Kinesin-like protein KIFC1 OS=Homo sa | KIFC1 | 73.748 | 2 | 52.792744 | yes | 4.979E-07 | 728.87317 | 125.68279 |
| LHNQLQELK | 9 | Q9BW19 | Kinesin-like protein KIFC1 OS=Homo sa | KIFC1 | 73.748 | 2 | 52.792744 | yes | 1.982E-07 | 561.81696 | 125.46315 |
| ESHVPYR | 7 | Q9BW19 | Kinesin-like protein KIFC1 OS=Homo sa | KIFC1 | 73.748 | 2 | 52.792744 | yes | 9.722E-08 | 444.22214 | 122.65018 |
| SHSVFQLQISGEHS | 16 | Q9BW19 | Kinesin-like protein KIFC1 OS=Homo sa | KIFC1 | 73.748 | 3 | 52.792744 | yes | 9.722E-08 | 600.29919 | 102.70483 |
| APSQLPLSGSR | 11 | Q9BW19 | Kinesin-like protein KIFC1 OS=Homo sa | KIFC1 | 73.748 | 2 | 52.792744 | yes | 1.13E-07 | 556.80658 | 96.891403 |
| GLMSQLEEK | 9 | Q9BW19 | Kinesin-like protein KIFC1 OS=Homo sa | KIFC1 | 73.748 | 2 | 52.792744 | yes | 5.102E-07 | 517.763 | 75.117432 |
| RPDQMEDGLEPEK | 13 | Q9BW19 | Kinesin-like protein KIFC1 OS=Homo sa | KIFC1 | 73.748 | 2 | 52.792744 | yes | 1.072E-06 | 772.35651 | 32.46196 |
| DVDVYSQILR | 10 | Q9BW27 | Nuclear pore complex protein Nup85 OS= | NUP85 | 75.02 | 2 | 41.044353 | yes | 4.759E-07 | 604.31952 | 77.119034 |
| IMLGDEAALLEQK | 13 | Q9BW27 | Nuclear pore complex protein Nup85 OS= | NUP85 | 75.02 | 2 | 41.044353 | yes | 3.706E-07 | 715.88159 | 55.217522 |
| KPVLSVSAR | 9 | Q9BW66 | Cyclin-dependent kinase 2-interacting prc | CINP | 24.324 | 2 | 48.6108 | yes | 3.431E-07 | 478.798 | 195.42894 |
| LCEELQATLDGLT | 14 | Q9BW66 | Cyclin-dependent kinase 2-interacting prc | CINP | 24.324 | 2 | 48.6108 | yes | 1.123E-07 | 795.90582 | 127.55868 |
| DNAADWHNLILK | 12 | Q9BW66 | Cyclin-dependent kinase 2-interacting prc | CINP | 24.324 | 2 | 48.6108 | yes | 8.041E-07 | 705.36224 | 31.12344 |
| DKIELDSSSPASK | 13 | Q9BW66 | Cyclin-dependent kinase 2-interacting prc | CINP | 24.324 | 2 | 48.6108 | yes | 3.271E-06 | 688.84882 | 21.868088 |
| LEMEVLENLQELK | 13 | Q9BW85 | Splicing factor YJU2 OS=Homo sapiens ( | YJU2 | 37.086 | 2 | 46.608875 | yes | 1.515E-07 | 794.41858 | 35.451515 |
| LGEWDLPEAVQR | 12 | Q9BW92 | Threonine--tRNA ligase, mitochondrial O | TARS2 | 81.037 | 2 | 52.135044 | yes | 1.009E-07 | 706.86206 | 616.88074 |
| VSGISFPTTELLR | 13 | Q9BW92 | Threonine--tRNA ligase, mitochondrial O | TARS2 | 81.037 | 2 | 52.135044 | yes | 1.017E-07 | 710.39575 | 340.51874 |
| LADFGALHR | 9 | Q9BW92 | Threonine--tRNA ligase, mitochondrial O | TARS2 | 81.037 | 2 | 52.135044 | yes | 9.722E-08 | 500.27216 | 265.60864 |
| LLGVLAESCGGK | 12 | Q9BW92 | Threonine--tRNA ligase, mitochondrial O | TARS2 | 81.037 | 2 | 52.135044 | yes | 2.04E-07 | 602.32373 | 257.87863 |
| ICQELTAAAR | 10 | Q9BW92 | Threonine--tRNA ligase, mitochondrial O | TARS2 | 81.037 | 2 | 52.135044 | yes | 1.101E-07 | 566.7926 | 217.63943 |
| LVELQNTR | 8 | Q9BW92 | Threonine--tRNA ligase, mitochondrial O | TARS2 | 81.037 | 2 | 52.135044 | yes | 1.171E-07 | 486.77728 | 214.13773 |
| LLSNSSSLWR | 10 | Q9BW92 | Threonine--tRNA ligase, mitochondrial O | TARS2 | 81.037 | 2 | 52.135044 | yes | 1.751E-07 | 581.81439 | 126.18023 |
| FLTFDSPEGK | 10 | Q9BW92 | Threonine--tRNA ligase, mitochondrial O | TARS2 | 81.037 | 2 | 52.135044 | yes | 2.433E-07 | 570.78223 | 107.55268 |
| LHLIEEK | 7 | Q9BW92 | Threonine--tRNA ligase, mitochondrial O | TARS2 | 81.037 | 2 | 52.135044 | yes | 9.722E-08 | 441.25818 | 68.332306 |
| HTGQIGGLK | 9 | Q9BW92 | Threonine--tRNA ligase, mitochondrial O | TARS2 | 81.037 | 2 | 52.135044 | yes | 1.788E-06 | 455.75888 | 35.403976 |
| GPLAMGLAQPR | 11 | Q9BWE0 | Replication initiator 1 OS=Homo sapiens | REPIN1 | 63.575 | 2 | 52.778934 | yes | 9.809E-08 | 555.80804 | 316.47186 |
| VHVAEALEEAAAI | 13 | Q9BWE0 | Replication initiator 1 OS=Homo sapiens | REPIN1 | 63.575 | 2 | 52.778934 | yes | 9.722E-08 | 669.35663 | 276.57202 |
| LLSGPSQESPQTLC | 15 | Q9BWE0 | Replication initiator 1 OS=Homo sapiens | REPIN1 | 63.575 | 2 | 52.778934 | yes | 1.059E-07 | 771.41211 | 245.79932 |
| HAPFLALHR | 9 | Q9BWE0 | Replication initiator 1 OS=Homo sapiens | REPIN1 | 63.575 | 2 | 52.778934 | yes | 6.681E-07 | 531.30377 | 181.14523 |
| LPLPCPECGR | 10 | Q9BWE0 | Replication initiator 1 OS=Homo sapiens | REPIN1 | 63.575 | 2 | 52.778934 | yes | 3.437E-07 | 599.789 | 176.58151 |

| | | | | | | | | | | | |
|---|---|---|---|---|---|---|---|---|---|---|---|
| PFICGNCGR | 9 | Q9BWE0 | Replication initiator 1 OS=Homo sapiens | REPIN1 | 63.575 | 2 | 52.778934 | yes | 9.728E-08 | 540.73932 | 170.25011 |
| SFAQWDQLVAHK | 12 | Q9BWE0 | Replication initiator 1 OS=Homo sapiens | REPIN1 | 63.575 | 2 | 52.778934 | yes | 2.554E-07 | 715.36475 | 120.68215 |
| SNLITHR | 7 | Q9BWE0 | Replication initiator 1 OS=Homo sapiens | REPIN1 | 63.575 | 2 | 52.778934 | yes | 8.481E-07 | 420.73795 | 52.709709 |
| FTNKPYLTSHR | 11 | Q9BWE0 | Replication initiator 1 OS=Homo sapiens | REPIN1 | 63.575 | 2 | 52.778934 | yes | 1.306E-07 | 682.3595 | 45.613403 |
| PFTCAECGK | 9 | Q9BWE0 | Replication initiator 1 OS=Homo sapiens | REPIN1 | 63.575 | 2 | 52.778934 | yes | 5.578E-05 | 535.22571 | 7.4288688 |
| LVLESPAPVEVNL | 14 | Q9BWF2 | E3 ubiquitin-protein ligase TRAIP OS=H | TRAIP | 53.295 | 2 | 43.74237 | yes | 2.465E-07 | 754.44012 | 122.34043 |
| LQTVYSELDQAK | 12 | Q9BWF2 | E3 ubiquitin-protein ligase TRAIP OS=H | TRAIP | 53.295 | 2 | 43.74237 | yes | 4.167E-07 | 697.86176 | 73.266808 |
| LFIGNLPR | 8 | Q9BWF3 | RNA-binding protein 4 OS=Homo sapien | RBM4 | 40.314 | 2 | 51.026497 | no | 1.965E-07 | 465.28198 | 15542.476 |
| DYAFVHMER | 9 | Q9BWF3 | RNA-binding protein 4 OS=Homo sapien | RBM4 | 40.314 | 2 | 51.026497 | no | 9.926E-08 | 584.26624 | 3970.7612 |
| DYAFVHMER | 9 | Q9BWF3 | RNA-binding protein 4 OS=Homo sapien | RBM4 | 40.314 | 2 | 51.026497 | no | 2.041E-07 | 592.26367 | 3970.7612 |
| MHVQLSTSR | 9 | Q9BWF3 | RNA-binding protein 4 OS=Homo sapien | RBM4 | 40.314 | 2 | 51.026497 | no | 1.415E-07 | 529.77423 | 2638.6975 |
| MHVQLSTSR | 9 | Q9BWF3 | RNA-binding protein 4 OS=Homo sapien | RBM4 | 40.314 | 2 | 51.026497 | no | 1.729E-07 | 537.77167 | 2638.6975 |
| NYGFVHIEDK | 10 | Q9BWF3 | RNA-binding protein 4 OS=Homo sapien | RBM4 | 40.314 | 2 | 51.026497 | no | 6.51E-07 | 611.29858 | 3806.7627 |
| VLECDIIK | 8 | Q9BWF3 | RNA-binding protein 4 OS=Homo sapien | RBM4 | 40.314 | 2 | 51.026497 | no | 1.091E-07 | 495.27045 | 3509.2764 |
| LHGVNINVEASK | 12 | Q9BWF3 | RNA-binding protein 4 OS=Homo sapien | RBM4 | 40.314 | 2 | 51.026497 | no | 9.722E-08 | 640.8515 | 3257.6814 |
| AEDAVEAIR | 9 | Q9BWF3 | RNA-binding protein 4 OS=Homo sapien | RBM4 | 40.314 | 2 | 51.026497 | no | 2.344E-07 | 487.2511 | 3225.4873 |
| TAPGMGDQSGCYI | 13 | Q9BWF3 | RNA-binding protein 4 OS=Homo sapien | RBM4 | 40.314 | 2 | 51.026497 | no | 1.101E-07 | 700.2901 | 3119.1401 |
| GLDNTEFQGK | 10 | Q9BWF3 | RNA-binding protein 4 OS=Homo sapien | RBM4 | 40.314 | 2 | 51.026497 | no | 2.365E-07 | 554.76709 | 2875.6284 |
| LHVGNISPTCTNK | 13 | Q9BWF3 | RNA-binding protein 4 OS=Homo sapien | RBM4 | 40.314 | 2 | 51.026497 | yes | 5.373E-07 | 720.86682 | 2145.6802 |
| TAAEDAIR | 8 | Q9BWF3 | RNA-binding protein 4 OS=Homo sapien | RBM4 | 40.314 | 2 | 51.026497 | no | 1.44E-07 | 423.71942 | 1965.9828 |
| SLFEQYGK | 8 | Q9BWF3 | RNA-binding protein 4 OS=Homo sapien | RBM4 | 40.314 | 2 | 51.026497 | no | 2.215E-07 | 486.24527 | 1528.9957 |
| EATEQEIR | 8 | Q9BWF3 | RNA-binding protein 4 OS=Homo sapien | RBM4 | 40.314 | 2 | 51.026497 | no | 9.894E-08 | 488.24072 | 1267.809 |
| NSLYDMAR | 8 | Q9BWF3 | RNA-binding protein 4 OS=Homo sapien | RBM4 | 40.314 | 2 | 51.026497 | no | 1.441E-07 | 485.22656 | 420.6004 |
| NSLYDMAR | 8 | Q9BWF3 | RNA-binding protein 4 OS=Homo sapien | RBM4 | 40.314 | 2 | 51.026497 | no | 1.35E-07 | 493.22403 | 420.6004 |
| AEAPLLTQTR | 10 | Q9BWG6 | Sodium channel modifier 1 OS=Homo saj | SCNM1 | 25.95 | 2 | 51.30978 | yes | 9.722E-08 | 550.30896 | 551.29907 |
| ALDHYLTLR | 9 | Q9BWG6 | Sodium channel modifier 1 OS=Homo saj | SCNM1 | 25.95 | 2 | 51.30978 | yes | 1.108E-07 | 551.30621 | 422.62878 |
| LITQSALHR | 9 | Q9BWG6 | Sodium channel modifier 1 OS=Homo saj | SCNM1 | 25.95 | 2 | 51.30978 | yes | 2.764E-07 | 519.80634 | 273.75241 |
| EGDDWSQLNVLK | 12 | Q9BWG6 | Sodium channel modifier 1 OS=Homo saj | SCNM1 | 25.95 | 2 | 51.30978 | yes | 1.032E-07 | 702.34369 | 200.54514 |
| HLSSLQLFYGK | 11 | Q9BWG6 | Sodium channel modifier 1 OS=Homo saj | SCNM1 | 25.95 | 2 | 51.30978 | yes | 4.005E-07 | 646.85352 | 135.10196 |
| FACAICPHRPVLDT | 22 | Q9BWG6 | Sodium channel modifier 1 OS=Homo saj | SCNM1 | 25.95 | 4 | 51.30978 | yes | 1.87E-07 | 638.3277 | 38.197098 |
| MLQPCGPPADKPE | 15 | Q9BWJ5 | Splicing factor 3B subunit 5 OS=Homo sa | SF3B5 | 10.136 | 2 | 46.51825 | yes | 1.538E-07 | 841.87927 | 243.28896 |
| YTIHSQLEHLQSK | 13 | Q9BWJ5 | Splicing factor 3B subunit 5 OS=Homo sa | SF3B5 | 10.136 | 2 | 46.51825 | yes | 3.622E-07 | 792.41248 | 15.369978 |
| ELQVGIPVADEAG | 15 | Q9BWM7 | Sideroflexin-3 OS=Homo sapiens OX=96 | SFXN3 | 35.504 | 2 | 51.89524 | yes | 1.789E-07 | 791.41522 | 279.27875 |
| AGVVTPGITEDQL | 15 | Q9BWM7 | Sideroflexin-3 OS=Homo sapiens OX=96 | SFXN3 | 35.504 | 2 | 51.89524 | yes | 9.722E-08 | 821.43341 | 162.39452 |
| NLLLSGAQLEASR | 13 | Q9BWM7 | Sideroflexin-3 OS=Homo sapiens OX=96 | SFXN3 | 35.504 | 2 | 51.89524 | yes | 1.161E-07 | 686.38318 | 140.7482 |
| MGELPLDINIQEPR | 14 | Q9BWM7 | Sideroflexin-3 OS=Homo sapiens OX=96 | SFXN3 | 35.504 | 2 | 51.89524 | yes | 9.722E-08 | 812.92181 | 64.271416 |
| QGLAVTDEILQGK | 13 | Q9BWT3 | Poly(A) polymerase gamma OS=Homo sa | PAPOLG | 82.803 | 2 | 47.949554 | yes | 0.0002089 | 686.37756 | 135.40196 |
| TVSPPTVCTIPTVV | 16 | Q9BWT3 | Poly(A) polymerase gamma OS=Homo sa | PAPOLG | 82.803 | 2 | 47.949554 | yes | 1.974E-06 | 842.45856 | 82.504402 |
| LAIQTISDNLDLR | 13 | Q9BWT3 | Poly(A) polymerase gamma OS=Homo sa | PAPOLG | 82.803 | 2 | 47.949554 | yes | 4.454E-07 | 736.40936 | 76.699425 |

| | | | | | | | | | | | |
|---|---|---|---|---|---|---|---|---|---|---|---|
| LSLDSSCLDSSR | 12 | Q9BWT3 | Poly(A) polymerase gamma OS=Homo sa | PAPOLG | 82.803 | 2 | 47.949554 | yes | 3.26E-07 | 670.31152 | 49.303112 |
| GADIDALCVAPR | 12 | Q9BWT3 | Poly(A) polymerase gamma OS=Homo sa | PAPOLG | 82.803 | 2 | 47.949554 | no | 2.539E-06 | 629.31647 | 41.573673 |
| SDFFQSFFEK | 10 | Q9BWT3 | Poly(A) polymerase gamma OS=Homo sa | PAPOLG | 82.803 | 2 | 47.949554 | yes | 1.226E-07 | 641.29297 | 15.135758 |
| LVNPSQYR | 8 | Q9BX10 | GTP-binding protein 2 OS=Homo sapiens | GTPBP2 | 65.769 | 2 | 53.576225 | yes | 4.13E-07 | 488.76416 | 369.64352 |
| AGQAATLALGDFI | 14 | Q9BX10 | GTP-binding protein 2 OS=Homo sapiens | GTPBP2 | 65.769 | 2 | 53.576225 | yes | 9.722E-08 | 703.35718 | 291.56473 |
| VAVLGNVDSGK | 11 | Q9BX10 | GTP-binding protein 2 OS=Homo sapiens | GTPBP2 | 65.769 | 2 | 53.576225 | yes | 3.976E-06 | 529.79565 | 207.12952 |
| KITEVLVR | 8 | Q9BX10 | GTP-binding protein 2 OS=Homo sapiens | GTPBP2 | 65.769 | 2 | 53.576225 | yes | 5.01E-07 | 479.3082 | 158.32065 |
| FEHLVTQMK | 9 | Q9BX10 | GTP-binding protein 2 OS=Homo sapiens | GTPBP2 | 65.769 | 2 | 53.576225 | yes | 9.722E-08 | 566.79462 | 148.77605 |
| VPFFIVVSK | 9 | Q9BX10 | GTP-binding protein 2 OS=Homo sapiens | GTPBP2 | 65.769 | 2 | 53.576225 | yes | 1.077E-07 | 518.31549 | 128.79796 |
| VFLNILPPLTNSK | 13 | Q9BX10 | GTP-binding protein 2 OS=Homo sapiens | GTPBP2 | 65.769 | 2 | 53.576225 | yes | 6.882E-07 | 728.43213 | 79.123199 |
| EVDYDSDMPR | 10 | Q9BX10 | GTP-binding protein 2 OS=Homo sapiens | GTPBP2 | 65.769 | 2 | 53.576225 | yes | 1.023E-07 | 613.75336 | 67.334801 |
| KVPDNQQFLDLR | 12 | Q9BX10 | GTP-binding protein 2 OS=Homo sapiens | GTPBP2 | 65.769 | 2 | 53.576225 | yes | 8.035E-07 | 736.89642 | 67.169159 |
| EHLGLALALK | 10 | Q9BX10 | GTP-binding protein 2 OS=Homo sapiens | GTPBP2 | 65.769 | 2 | 53.576225 | yes | 3.43E-07 | 532.82678 | 62.255447 |
| SFFDNISSELK | 11 | Q9BX40 | Protein LSM14 homolog B OS=Homo sa | LSM14B | 42.071 | 2 | 55.19487 | yes | 1.006E-07 | 643.81677 | 690.80328 |
| EEIYEYIIFR | 10 | Q9BX40 | Protein LSM14 homolog B OS=Homo sa | LSM14B | 42.071 | 2 | 55.19487 | yes | 9.722E-08 | 687.85065 | 138.5667 |
| KLNTETFGVSGR | 12 | Q9BX40 | Protein LSM14 homolog B OS=Homo sa | LSM14B | 42.071 | 2 | 55.19487 | yes | 1.109E-07 | 654.84894 | 130.96899 |
| EELDKEFK | 8 | Q9BX40 | Protein LSM14 homolog B OS=Homo sa | LSM14B | 42.071 | 2 | 55.19487 | yes | 3.898E-07 | 519.26111 | 107.8801 |
| DITVCEPPK | 9 | Q9BX40 | Protein LSM14 homolog B OS=Homo sa | LSM14B | 42.071 | 2 | 55.19487 | yes | 2.75E-07 | 529.763 | 105.99485 |
| LNTETFGVSGR | 11 | Q9BX40 | Protein LSM14 homolog B OS=Homo sa | LSM14B | 42.071 | 2 | 55.19487 | yes | 2.182E-07 | 590.80145 | 63.452774 |
| ESASYSVTEVQLR | 13 | Q9BX63 | Fanconi anemia group J protein OS=Hom | BRIP1 | 140.868 | 2 | 50.435352 | yes | 2.602E-07 | 734.86755 | 105.5719 |
| EVPVISASTQIMLK | 14 | Q9BX63 | Fanconi anemia group J protein OS=Hom | BRIP1 | 140.868 | 2 | 50.435352 | yes | 3.158E-07 | 758.42621 | 89.971489 |
| VQTIVLTSGTLSPM | 15 | Q9BX63 | Fanconi anemia group J protein OS=Hom | BRIP1 | 140.868 | 2 | 50.435352 | yes | 2.274E-07 | 787.94476 | 88.649811 |
| NSQVWVGTIGSGP | 14 | Q9BX63 | Fanconi anemia group J protein OS=Hom | BRIP1 | 140.868 | 2 | 50.435352 | yes | 3.678E-06 | 715.37537 | 85.75946 |
| DELDSMVNNNIR | 12 | Q9BX63 | Fanconi anemia group J protein OS=Hom | BRIP1 | 140.868 | 2 | 50.435352 | yes | 9.819E-08 | 710.33026 | 84.241966 |
| TVIVEPQGGEK | 11 | Q9BX63 | Fanconi anemia group J protein OS=Hom | BRIP1 | 140.868 | 2 | 50.435352 | yes | 1.319E-07 | 578.81403 | 75.345917 |
| VSEGLDFSDDNAR | 13 | Q9BX63 | Fanconi anemia group J protein OS=Hom | BRIP1 | 140.868 | 2 | 50.435352 | yes | 9.809E-08 | 712.81805 | 55.594379 |
| ICVQELQCPK | 10 | Q9BX63 | Fanconi anemia group J protein OS=Hom | BRIP1 | 140.868 | 2 | 50.435352 | yes | 3.181E-06 | 637.81519 | 40.835945 |
| IWSGNEMLLTLHK | 13 | Q9BX63 | Fanconi anemia group J protein OS=Hom | BRIP1 | 140.868 | 3 | 50.435352 | yes | 1.351E-06 | 514.60968 | 36.410797 |
| HFNYPSTPPSER | 12 | Q9BX63 | Fanconi anemia group J protein OS=Hom | BRIP1 | 140.868 | 2 | 50.435352 | yes | 6.238E-07 | 716.33624 | 20.378546 |
| ADNAFMLLTQAR | 12 | Q9BX70 | BTB/POZ domain-containing protein 2 O | BTBD2 | 55.932 | 2 | 52.19905 | yes | 9.722E-08 | 675.84534 | 87.778702 |
| ALLDLQGHLDAAS | 14 | Q9BXF6 | Rab11 family-interacting protein 5 OS=H | RAB11FIP: | 70.416 | 3 | 44.526394 | yes | 3.665E-06 | 493.93533 | 42.246319 |
| VLAPGASHPGEEE | 16 | Q9BXF6 | Rab11 family-interacting protein 5 OS=H | RAB11FIP: | 70.416 | 2 | 44.526394 | yes | 2.162E-07 | 788.88934 | 25.22574 |
| LKDSAVLDQSAK | 12 | Q9BXF6 | Rab11 family-interacting protein 5 OS=H | RAB11FIP: | 70.416 | 2 | 44.526394 | yes | 2.541E-07 | 637.8512 | 18.596422 |
| YYHLTHDELISLLI | 16 | Q9BXF6 | Rab11 family-interacting protein 5 OS=H | RAB11FIP: | 70.416 | 3 | 44.526394 | yes | 9.977E-07 | 672.03009 | 16.967213 |
| APAAGESLSGTR | 12 | Q9BXI6 | TBC1 domain family member 10A OS=H | TBC1D10A | 57.118 | 2 | 50.137913 | yes | 2.473E-05 | 558.78583 | 138.2617 |
| EHLIQLR | 7 | Q9BXI6 | TBC1 domain family member 10A OS=H | TBC1D10A | 57.118 | 2 | 50.137913 | yes | 9.893E-08 | 454.76926 | 52.677071 |
| VWDMFFCEGVK | 11 | Q9BXI6 | TBC1 domain family member 10A OS=H | TBC1D10A | 57.118 | 2 | 50.137913 | no | 1.912E-06 | 709.31757 | 27.521927 |
| YQLLQLR | 7 | Q9BXJ9 | N-alpha-acetyltransferase 15, NatA auxili | NAA15 | 101.273 | 2 | 41.998363 | no | 4.528E-07 | 467.27945 | 71.870132 |
| GELLLQLCR | 9 | Q9BXJ9 | N-alpha-acetyltransferase 15, NatA auxili | NAA15 | 101.273 | 2 | 41.998363 | yes | 4.354E-07 | 551.30786 | 68.74865 |

| | | | | | | | | | | |
|---|---|---|---|---|---|---|---|---|---|---|
| GCPPVFNTLR | 10 | Q9BXJ9 | N-alpha-acetyltransferase 15, NatA auxili | NAA15 | 101.273 | 2 | 41.998363 | yes | 3.242E-07 | 580.79767 | 64.286942 |
| MVYYLDPSSQK | 11 | Q9BXJ9 | N-alpha-acetyltransferase 15, NatA auxili | NAA15 | 101.273 | 2 | 41.998363 | yes | 3.498E-06 | 665.82104 | 39.584965 |
| WMDEAQALDTAD | 13 | Q9BXJ9 | N-alpha-acetyltransferase 15, NatA auxili | NAA15 | 101.273 | 2 | 41.998363 | yes | 9.529E-07 | 761.33557 | 34.133335 |
| RLPLNFLSGEK | 11 | Q9BXJ9 | N-alpha-acetyltransferase 15, NatA auxili | NAA15 | 101.273 | 2 | 41.998363 | yes | 3.475E-06 | 637.36676 | 7.7680302 |
| VALAGEYGAVTYI | 13 | Q9BXM7 | Serine/threonine-protein kinase PINK1, n | PINK1 | 62.77 | 2 | 51.08937 | yes | 9.809E-08 | 685.35913 | 3929.4622 |
| LHPEGLGHGR | 10 | Q9BXM7 | Serine/threonine-protein kinase PINK1, n | PINK1 | 62.77 | 2 | 51.08937 | yes | 3.897E-07 | 536.78595 | 2997.8325 |
| SKPGPDPLDTR | 11 | Q9BXM7 | Serine/threonine-protein kinase PINK1, n | PINK1 | 62.77 | 2 | 51.08937 | yes | 9.722E-08 | 591.80933 | 1336.8323 |
| LEEYLIGQSIGK | 12 | Q9BXM7 | Serine/threonine-protein kinase PINK1, n | PINK1 | 62.77 | 2 | 51.08937 | yes | 9.722E-08 | 675.3692 | 1296.5167 |
| NYPCTLR | 7 | Q9BXM7 | Serine/threonine-protein kinase PINK1, n | PINK1 | 62.77 | 2 | 51.08937 | yes | 2.664E-07 | 462.22382 | 991.48553 |
| APGAPAFPLAIK | 12 | Q9BXM7 | Serine/threonine-protein kinase PINK1, n | PINK1 | 62.77 | 2 | 51.08937 | yes | 6.628E-07 | 576.84241 | 978.55054 |
| AVSACQEIQAIFTQ | 15 | Q9BXM7 | Serine/threonine-protein kinase PINK1, n | PINK1 | 62.77 | 2 | 51.08937 | yes | 1.367E-07 | 847.43256 | 900.57843 |
| QSVAGLAAR | 9 | Q9BXM7 | Serine/threonine-protein kinase PINK1, n | PINK1 | 62.77 | 2 | 51.08937 | yes | 2.263E-07 | 436.75107 | 790.46814 |
| AWGCAGPCGR | 10 | Q9BXM7 | Serine/threonine-protein kinase PINK1, n | PINK1 | 62.77 | 2 | 51.08937 | yes | 9.893E-08 | 546.22913 | 725.59137 |
| GGNGCLMAPEVST | 15 | Q9BXM7 | Serine/threonine-protein kinase PINK1, n | PINK1 | 62.77 | 2 | 51.08937 | yes | 1.039E-07 | 760.35321 | 622.06732 |
| QYLCVNTPSPR | 11 | Q9BXM7 | Serine/threonine-protein kinase PINK1, n | PINK1 | 62.77 | 2 | 51.08937 | yes | 2.007E-07 | 667.82971 | 589.80872 |
| GPGTSAPGEGQER | 13 | Q9BXM7 | Serine/threonine-protein kinase PINK1, n | PINK1 | 62.77 | 2 | 51.08937 | yes | 1.384E-07 | 621.78912 | 535.95349 |
| PGWAAGPGAEPR | 12 | Q9BXM7 | Serine/threonine-protein kinase PINK1, n | PINK1 | 62.77 | 2 | 51.08937 | yes | 9.919E-08 | 583.29108 | 368.9725 |
| VAANVLHLSLWGI | 19 | Q9BXM7 | Serine/threonine-protein kinase PINK1, n | PINK1 | 62.77 | 3 | 51.08937 | yes | 1.461E-07 | 695.40533 | 299.33408 |
| PGPAAGCVR | 9 | Q9BXM7 | Serine/threonine-protein kinase PINK1, n | PINK1 | 62.77 | 2 | 51.08937 | yes | 1.068E-07 | 442.724 | 250.26436 |
| VGLGLPNR | 8 | Q9BXM7 | Serine/threonine-protein kinase PINK1, n | PINK1 | 62.77 | 2 | 51.08937 | yes | 1.418E-07 | 413.2507 | 111.09072 |
| RVGLGLPNR | 9 | Q9BXM7 | Serine/threonine-protein kinase PINK1, n | PINK1 | 62.77 | 2 | 51.08937 | yes | 2.79E-07 | 491.30124 | 92.719826 |
| QLAPHPNIIR | 10 | Q9BXM7 | Serine/threonine-protein kinase PINK1, n | PINK1 | 62.77 | 2 | 51.08937 | yes | 4.07E-07 | 579.84076 | 53.970112 |
| LTEKCCVETK | 10 | Q9BXM7 | Serine/threonine-protein kinase PINK1, n | PINK1 | 62.77 | 2 | 51.08937 | yes | 1.543E-07 | 634.30469 | 51.861237 |
| VALSEPQPER | 10 | Q9BXP5 | Serrate RNA effector molecule homolog ( | SRRT | 100.667 | 2 | 53.21806 | yes | 1.074E-07 | 563.29858 | 3327.3691 |
| FVTSNTQELGK | 11 | Q9BXP5 | Serrate RNA effector molecule homolog ( | SRRT | 100.667 | 2 | 53.21806 | yes | 1.063E-07 | 612.31696 | 1741.3434 |
| EGNPAEINVER | 11 | Q9BXP5 | Serrate RNA effector molecule homolog ( | SRRT | 100.667 | 2 | 53.21806 | yes | 9.722E-08 | 614.30182 | 1566.0182 |
| EICWNLQNIR | 10 | Q9BXP5 | Serrate RNA effector molecule homolog ( | SRRT | 100.667 | 2 | 53.21806 | yes | 1.281E-07 | 673.33771 | 1334.3088 |
| AGAGLGDGER | 10 | Q9BXP5 | Serrate RNA effector molecule homolog ( | SRRT | 100.667 | 2 | 53.21806 | yes | 3.65E-07 | 451.71997 | 1168.8054 |
| FKGPEFVR | 8 | Q9BXP5 | Serrate RNA effector molecule homolog ( | SRRT | 100.667 | 2 | 53.21806 | yes | 3.707E-07 | 490.27164 | 1165.2936 |
| MEGGTENDLR | 10 | Q9BXP5 | Serrate RNA effector molecule homolog ( | SRRT | 100.667 | 2 | 53.21806 | yes | 1.874E-07 | 561.24823 | 1033.9702 |
| ECELSPGVNR | 10 | Q9BXP5 | Serrate RNA effector molecule homolog ( | SRRT | 100.667 | 2 | 53.21806 | yes | 9.809E-08 | 580.77185 | 999.15326 |
| LIHTLDDR | 8 | Q9BXP5 | Serrate RNA effector molecule homolog ( | SRRT | 100.667 | 2 | 53.21806 | yes | 1.261E-07 | 491.76944 | 959.75403 |
| EVAFFNNFLTDAK | 13 | Q9BXP5 | Serrate RNA effector molecule homolog ( | SRRT | 100.667 | 2 | 53.21806 | yes | 9.722E-08 | 758.37756 | 931.66803 |
| KDPEQEVEK | 9 | Q9BXP5 | Serrate RNA effector molecule homolog ( | SRRT | 100.667 | 2 | 53.21806 | yes | 9.733E-08 | 551.27478 | 929.31982 |
| CGIIHVR | 7 | Q9BXP5 | Serrate RNA effector molecule homolog ( | SRRT | 100.667 | 2 | 53.21806 | yes | 1.009E-07 | 427.73691 | 921.0293 |
| DAAGLECKPR | 10 | Q9BXP5 | Serrate RNA effector molecule homolog ( | SRRT | 100.667 | 2 | 53.21806 | yes | 1.485E-07 | 558.77698 | 825.89087 |
| TCSLFMR | 7 | Q9BXP5 | Serrate RNA effector molecule homolog ( | SRRT | 100.667 | 2 | 53.21806 | yes | 2.724E-07 | 457.71478 | 720.62744 |
| ESLSEEEAQK | 10 | Q9BXP5 | Serrate RNA effector molecule homolog ( | SRRT | 100.667 | 2 | 53.21806 | yes | 1.117E-07 | 575.26715 | 701.5094 |
| ISHGEVLEWQK | 11 | Q9BXP5 | Serrate RNA effector molecule homolog ( | SRRT | 100.667 | 2 | 53.21806 | yes | 9.819E-08 | 663.34607 | 343.08838 |

| | | | | | | | | | | | |
|---|---|---|---|---|---|---|---|---|---|---|---|
| ISHGEVLEWQK | 11 | Q9BXP5 | Serrate RNA effector molecule homolog ( | SRRT | 100.667 | 2 | 53.21806 | yes | 1.136E-07 | 442.56647 | 343.08838 |
| ISHGEVLEWQK | 11 | Q9BXP5 | Serrate RNA effector molecule homolog ( | SRRT | 100.667 | 3 | 53.21806 | yes | 9.819E-08 | 663.34607 | 343.08838 |
| ISHGEVLEWQK | 11 | Q9BXP5 | Serrate RNA effector molecule homolog ( | SRRT | 100.667 | 3 | 53.21806 | yes | 1.136E-07 | 442.56647 | 343.08838 |
| AEIISLCK | 8 | Q9BXP5 | Serrate RNA effector molecule homolog ( | SRRT | 100.667 | 2 | 53.21806 | yes | 9.849E-08 | 467.25732 | 620.95313 |
| RPALPEIK | 8 | Q9BXP5 | Serrate RNA effector molecule homolog ( | SRRT | 100.667 | 2 | 53.21806 | yes | 2.968E-07 | 462.28726 | 441.53619 |
| EFLLSLDDSVDETI | 18 | Q9BXP5 | Serrate RNA effector molecule homolog ( | SRRT | 100.667 | 3 | 53.21806 | yes | 4.705E-07 | 689.34778 | 420.44473 |
| WLCPLSGK | 8 | Q9BXP5 | Serrate RNA effector molecule homolog ( | SRRT | 100.667 | 2 | 53.21806 | yes | 3.452E-07 | 480.75223 | 411.33994 |
| MLDAAVIK | 8 | Q9BXP5 | Serrate RNA effector molecule homolog ( | SRRT | 100.667 | 2 | 53.21806 | yes | 1.166E-07 | 430.74915 | 354.0715 |
| LTPLLSVR | 8 | Q9BXP5 | Serrate RNA effector molecule homolog ( | SRRT | 100.667 | 2 | 53.21806 | yes | 2.025E-07 | 449.78964 | 327.87424 |
| RYPGFMR | 7 | Q9BXP5 | Serrate RNA effector molecule homolog ( | SRRT | 100.667 | 2 | 53.21806 | yes | 1.747E-07 | 463.73691 | 208.71382 |
| RGDDWNDR | 8 | Q9BXP5 | Serrate RNA effector molecule homolog ( | SRRT | 100.667 | 2 | 53.21806 | yes | 9.722E-08 | 517.22595 | 199.58275 |
| QQMQDFFLAHK | 11 | Q9BXP5 | Serrate RNA effector molecule homolog ( | SRRT | 100.667 | 2 | 53.21806 | yes | 1.367E-07 | 696.84009 | 184.98943 |
| RGWVTFDR | 8 | Q9BXP5 | Serrate RNA effector molecule homolog ( | SRRT | 100.667 | 2 | 53.21806 | yes | 2.819E-07 | 518.76978 | 123.43081 |
| DKWLCPLSGK | 10 | Q9BXP5 | Serrate RNA effector molecule homolog ( | SRRT | 100.667 | 2 | 53.21806 | yes | 2.294E-07 | 602.31317 | 118.77109 |
| NINGITQHK | 9 | Q9BXP5 | Serrate RNA effector molecule homolog ( | SRRT | 100.667 | 2 | 53.21806 | yes | 4.093E-07 | 512.78033 | 88.952858 |
| DAAGLECKPRPLH | 14 | Q9BXP5 | Serrate RNA effector molecule homolog ( | SRRT | 100.667 | 3 | 53.21806 | yes | 1.212E-07 | 531.28394 | 43.068954 |
| GDSDDEYDRR | 10 | Q9BXP5 | Serrate RNA effector molecule homolog ( | SRRT | 100.667 | 2 | 53.21806 | yes | 1.006E-07 | 635.25256 | 37.265022 |
| EEEWEKPK | 8 | Q9BXP5 | Serrate RNA effector molecule homolog ( | SRRT | 100.667 | 2 | 53.21806 | yes | 1.422E-07 | 537.75873 | 21.366751 |
| LRECELSPGVNR | 12 | Q9BXP5 | Serrate RNA effector molecule homolog ( | SRRT | 100.667 | 2 | 53.21806 | yes | 4.164E-07 | 715.36444 | 15.696785 |
| EEELYACQR | 9 | Q9BXS4 | Transmembrane protein 59 OS=Homo sa | TMEM59 | 36.224 | 2 | 41.195335 | yes | 3.629E-07 | 599.26385 | 31.068098 |
| SGYQALPWVR | 10 | Q9BXS5 | AP-1 complex subunit mu-1 OS=Homo sa | AP1M1 | 48.587 | 2 | 51.579037 | no | 1.073E-07 | 588.81165 | 190.35223 |
| VLFDNTGR | 8 | Q9BXS5 | AP-1 complex subunit mu-1 OS=Homo sa | AP1M1 | 48.587 | 2 | 51.579037 | yes | 3.498E-07 | 461.24307 | 188.66359 |
| VFLSGMPELR | 10 | Q9BXS5 | AP-1 complex subunit mu-1 OS=Homo sa | AP1M1 | 48.587 | 2 | 51.579037 | no | 9.722E-08 | 574.81024 | 151.3707 |
| AHFGLPSVEAEDK | 13 | Q9BXS5 | AP-1 complex subunit mu-1 OS=Homo sa | AP1M1 | 48.587 | 2 | 51.579037 | yes | 3.057E-07 | 700.34625 | 100.98563 |
| HNNLYLVATSK | 11 | Q9BXS5 | AP-1 complex subunit mu-1 OS=Homo sa | AP1M1 | 48.587 | 2 | 51.579037 | yes | 3.946E-07 | 630.34076 | 82.959007 |
| FEIPYFTTSGIQVR | 14 | Q9BXS5 | AP-1 complex subunit mu-1 OS=Homo sa | AP1M1 | 48.587 | 2 | 51.579037 | yes | 2.305E-06 | 829.43286 | 57.903782 |
| LSVASTPISQR | 11 | Q9BXS6 | Nucleolar and spindle-associated protein | NUSAP1 | 49.452 | 2 | 54.17814 | yes | 1.246E-07 | 579.82751 | 1789.0182 |
| EMESIDQYIER | 11 | Q9BXS6 | Nucleolar and spindle-associated protein | NUSAP1 | 49.452 | 2 | 54.17814 | yes | 9.722E-08 | 706.82178 | 883.30566 |
| LTTEATQTPVSNK | 13 | Q9BXS6 | Nucleolar and spindle-associated protein | NUSAP1 | 49.452 | 2 | 54.17814 | yes | 9.902E-08 | 695.36462 | 543.72546 |
| SAHVTVSGGTPK | 12 | Q9BXS6 | Nucleolar and spindle-associated protein | NUSAP1 | 49.452 | 2 | 54.17814 | yes | 9.722E-08 | 570.80402 | 534.81311 |
| LKPWGQSK | 8 | Q9BXS6 | Nucleolar and spindle-associated protein | NUSAP1 | 49.452 | 2 | 54.17814 | yes | 2.164E-07 | 472.27164 | 469.2659 |
| GEAVLGTHK | 9 | Q9BXS6 | Nucleolar and spindle-associated protein | NUSAP1 | 49.452 | 2 | 54.17814 | yes | 2.007E-06 | 456.25089 | 384.3396 |
| SCGPASQSTLGLK | 13 | Q9BXS6 | Nucleolar and spindle-associated protein | NUSAP1 | 49.452 | 2 | 54.17814 | yes | 2.061E-07 | 653.32703 | 361.75226 |
| KPVFDLK | 7 | Q9BXS6 | Nucleolar and spindle-associated protein | NUSAP1 | 49.452 | 2 | 54.17814 | yes | 4.469E-07 | 423.75781 | 354.36639 |
| YSDLQNLAK | 9 | Q9BXS6 | Nucleolar and spindle-associated protein | NUSAP1 | 49.452 | 2 | 54.17814 | yes | 1.043E-07 | 526.27454 | 334.3598 |
| PLNYEPHK | 8 | Q9BXS6 | Nucleolar and spindle-associated protein | NUSAP1 | 49.452 | 2 | 54.17814 | yes | 9.722E-08 | 499.25873 | 289.76074 |
| TAITTPNFK | 9 | Q9BXS6 | Nucleolar and spindle-associated protein | NUSAP1 | 49.452 | 2 | 54.17814 | yes | 9.795E-08 | 496.7742 | 209.59306 |
| MIIPSLEELDSLK | 13 | Q9BXS6 | Nucleolar and spindle-associated protein | NUSAP1 | 49.452 | 2 | 54.17814 | yes | 1.653E-07 | 744.40491 | 206.47186 |
| SLYTDESSKPGK | 12 | Q9BXS6 | Nucleolar and spindle-associated protein | NUSAP1 | 49.452 | 2 | 54.17814 | yes | 9.728E-08 | 656.32501 | 167.25533 |

| | | | | | | | | | | | |
|---|---|---|---|---|---|---|---|---|---|---|---|
| ISNPTEFQNHEK | 12 | Q9BXS6 | Nucleolar and spindle-associated protein | NUSAP1 | 49.452 | 2 | 54.17814 | yes | 2.021E-07 | 722.3468 | 116.90564 |
| TITGNSAAVITPFK | 14 | Q9BXS6 | Nucleolar and spindle-associated protein | NUSAP1 | 49.452 | 2 | 54.17814 | yes | 9.722E-08 | 710.39575 | 67.214249 |
| QPLGHVTK | 8 | Q9BXS6 | Nucleolar and spindle-associated protein | NUSAP1 | 49.452 | 2 | 54.17814 | yes | 1.612E-07 | 440.25598 | 59.911488 |
| KLHEAHFK | 8 | Q9BXS6 | Nucleolar and spindle-associated protein | NUSAP1 | 49.452 | 2 | 54.17814 | yes | 2.769E-06 | 505.28253 | 59.88599 |
| HFEEHNSMNELK | 12 | Q9BXS6 | Nucleolar and spindle-associated protein | NUSAP1 | 49.452 | 2 | 54.17814 | yes | 1.053E-07 | 757.83826 | 50.562767 |
| ENNYLNQHVNR | 11 | Q9BXS6 | Nucleolar and spindle-associated protein | NUSAP1 | 49.452 | 2 | 54.17814 | yes | 9.722E-08 | 700.83691 | 24.272955 |
| SLNFGGSTGTVPA | 15 | Q9BY12 | S phase cyclin A-associated protein in the | SCAPER | 158.288 | 2 | 55.30979 | yes | 9.727E-08 | 730.87823 | 680.74634 |
| VSLATEATR | 9 | Q9BY12 | S phase cyclin A-associated protein in the | SCAPER | 158.288 | 2 | 55.30979 | yes | 9.918E-08 | 474.26144 | 629.47162 |
| LNELQEER | 8 | Q9BY12 | S phase cyclin A-associated protein in the | SCAPER | 158.288 | 2 | 55.30979 | yes | 9.892E-08 | 515.76184 | 587.40546 |
| HMEQIEQR | 8 | Q9BY12 | S phase cyclin A-associated protein in the | SCAPER | 158.288 | 2 | 55.30979 | yes | 1.43E-07 | 535.75604 | 401.63559 |
| TQFTVSTLDDVK | 12 | Q9BY12 | S phase cyclin A-associated protein in the | SCAPER | 158.288 | 2 | 55.30979 | yes | 9.722E-08 | 677.34845 | 393.72104 |
| EVLMMLDNYVR | 11 | Q9BY12 | S phase cyclin A-associated protein in the | SCAPER | 158.288 | 2 | 55.30979 | yes | 1.006E-07 | 691.84399 | 377.20105 |
| ALIDWIQLQEK | 11 | Q9BY12 | S phase cyclin A-associated protein in the | SCAPER | 158.288 | 2 | 55.30979 | yes | 1.186E-07 | 678.8797 | 306.40451 |
| VSAVVLGCLIANR | 24 | Q9BY12 | S phase cyclin A-associated protein in the | SCAPER | 158.288 | 3 | 55.30979 | yes | 2.107E-07 | 846.43695 | 301.25455 |
| LCQLPFQYFSDPR | 13 | Q9BY12 | S phase cyclin A-associated protein in the | SCAPER | 158.288 | 2 | 55.30979 | yes | 9.935E-08 | 835.90338 | 260.7868 |
| SYSIFDNNR | 9 | Q9BY12 | S phase cyclin A-associated protein in the | SCAPER | 158.288 | 2 | 55.30979 | yes | 9.728E-08 | 558.25946 | 245.29457 |
| NLIAWSVPLESK | 12 | Q9BY12 | S phase cyclin A-associated protein in the | SCAPER | 158.288 | 2 | 55.30979 | yes | 1.212E-07 | 678.8797 | 241.8934 |
| LAPTGVSWADK | 11 | Q9BY12 | S phase cyclin A-associated protein in the | SCAPER | 158.288 | 2 | 55.30979 | yes | 1.258E-07 | 572.80347 | 201.69302 |
| PTSLAWEVK | 9 | Q9BY12 | S phase cyclin A-associated protein in the | SCAPER | 158.288 | 2 | 55.30979 | yes | 2.39E-07 | 515.78204 | 196.03391 |
| QVFEGLTTGLLK | 12 | Q9BY12 | S phase cyclin A-associated protein in the | SCAPER | 158.288 | 2 | 55.30979 | yes | 9.722E-08 | 653.37427 | 166.42551 |
| HVIPSPSTDR | 10 | Q9BY12 | S phase cyclin A-associated protein in the | SCAPER | 158.288 | 2 | 55.30979 | yes | 2.613E-07 | 554.79089 | 161.72395 |
| HPTVLQK | 7 | Q9BY12 | S phase cyclin A-associated protein in the | SCAPER | 158.288 | 2 | 55.30979 | yes | 1.751E-07 | 411.74524 | 143.96365 |
| FPQQAWEEAR | 10 | Q9BY12 | S phase cyclin A-associated protein in the | SCAPER | 158.288 | 2 | 55.30979 | yes | 1.475E-07 | 631.30164 | 136.54811 |
| EVQLQAIVK | 9 | Q9BY12 | S phase cyclin A-associated protein in the | SCAPER | 158.288 | 2 | 55.30979 | yes | 1.363E-07 | 514.31097 | 136.33255 |
| LLHAEFK | 7 | Q9BY12 | S phase cyclin A-associated protein in the | SCAPER | 158.288 | 2 | 55.30979 | yes | 5.263E-07 | 429.24762 | 117.61527 |
| WKEELLDQR | 9 | Q9BY12 | S phase cyclin A-associated protein in the | SCAPER | 158.288 | 2 | 55.30979 | yes | 1.423E-07 | 608.81964 | 110.81438 |
| CLGSQDYLELANR | 13 | Q9BY12 | S phase cyclin A-associated protein in the | SCAPER | 158.288 | 2 | 55.30979 | yes | 1.164E-07 | 769.867 | 107.57645 |
| HANTDYAPK | 9 | Q9BY12 | S phase cyclin A-associated protein in the | SCAPER | 158.288 | 2 | 55.30979 | yes | 3.904E-07 | 508.74344 | 104.56284 |
| INVTSNAR | 8 | Q9BY12 | S phase cyclin A-associated protein in the | SCAPER | 158.288 | 2 | 55.30979 | yes | 2.051E-07 | 437.74069 | 78.271637 |
| QVQVQDSGSWAN | 14 | Q9BY12 | S phase cyclin A-associated protein in the | SCAPER | 158.288 | 2 | 55.30979 | yes | 9.809E-08 | 780.87372 | 75.784424 |
| DAEGWETVQR | 10 | Q9BY12 | S phase cyclin A-associated protein in the | SCAPER | 158.288 | 2 | 55.30979 | yes | 3.85E-06 | 595.77545 | 67.498367 |
| EELLDQR | 7 | Q9BY12 | S phase cyclin A-associated protein in the | SCAPER | 158.288 | 2 | 55.30979 | yes | 1.592E-06 | 451.73254 | 62.903759 |
| AKEYESLMETK | 11 | Q9BY12 | S phase cyclin A-associated protein in the | SCAPER | 158.288 | 2 | 55.30979 | yes | 1.568E-07 | 664.82373 | 62.018181 |
| DSLHSCDHPLAEK | 13 | Q9BY12 | S phase cyclin A-associated protein in the | SCAPER | 158.288 | 3 | 55.30979 | yes | 2.725E-05 | 503.56479 | 30.990097 |
| DSLHSCDHPLAEK | 13 | Q9BY12 | S phase cyclin A-associated protein in the | SCAPER | 158.288 | 3 | 55.30979 | yes | 1.106E-07 | 754.84357 | 30.990097 |
| DSLHSCDHPLAEK | 13 | Q9BY12 | S phase cyclin A-associated protein in the | SCAPER | 158.288 | 2 | 55.30979 | yes | 2.725E-05 | 503.56479 | 30.990097 |
| DSLHSCDHPLAEK | 13 | Q9BY12 | S phase cyclin A-associated protein in the | SCAPER | 158.288 | 2 | 55.30979 | yes | 1.106E-07 | 754.84357 | 30.990097 |
| VLFPSLIAACYNNI | 17 | Q9BY12 | S phase cyclin A-associated protein in the | SCAPER | 158.288 | 3 | 55.30979 | yes | 1.404E-07 | 663.67188 | 56.82151 |
| KDAEGWETVQR | 11 | Q9BY12 | S phase cyclin A-associated protein in the | SCAPER | 158.288 | 2 | 55.30979 | yes | 1.754E-07 | 659.82294 | 56.724155 |

| | | | | | | | | | | | |
|---|---|---|---|---|---|---|---|---|---|---|---|
| CCGCVFSER | 9 | Q9BY42 | Replication termination factor 2 OS=Hon | RTF2 | 33.887 | 2 | 50.800865 | yes | 9.722E-08 | 587.72552 | 65.700546 |
| FICPVVGLEMNGR | 13 | Q9BY42 | Replication termination factor 2 OS=Hon | RTF2 | 33.887 | 2 | 50.800865 | yes | 1.065E-07 | 746.37598 | 51.897636 |
| LSDNPAWEGDK | 11 | Q9BY42 | Replication termination factor 2 OS=Hon | RTF2 | 33.887 | 2 | 50.800865 | yes | 2.84E-07 | 616.28314 | 46.994877 |
| RPIVACELGR | 10 | Q9BY42 | Replication termination factor 2 OS=Hon | RTF2 | 33.887 | 2 | 50.800865 | yes | 6.522E-07 | 585.82422 | 28.507227 |
| EAIENATTNAEVLI | 14 | Q9BY43 | Charged multivesicular body protein 4a C | CHMP4A | 25.099 | 2 | 47.843994 | yes | 1.248E-07 | 765.89154 | 105.68642 |
| QEFLEQK | 7 | Q9BY43 | Charged multivesicular body protein 4a C | CHMP4A | 25.099 | 2 | 47.843994 | yes | 3.351E-06 | 461.23746 | 85.396385 |
| DGTAGIPNLQLYD | 15 | Q9BY44 | Eukaryotic translation initiation factor 2A | EIF2A | 64.991 | 2 | 52.515076 | yes | 1.654E-07 | 802.41992 | 1576.4869 |
| INDFVLSPGPQPYI | 14 | Q9BY44 | Eukaryotic translation initiation factor 2A | EIF2A | 64.991 | 2 | 52.515076 | yes | 1.359E-07 | 787.91431 | 1311.1377 |
| CDPVFDFGTGPR | 12 | Q9BY44 | Eukaryotic translation initiation factor 2A | EIF2A | 64.991 | 2 | 52.515076 | yes | 9.727E-08 | 684.30609 | 630.125 |
| IWHYTGSILHK | 11 | Q9BY44 | Eukaryotic translation initiation factor 2A | EIF2A | 64.991 | 2 | 52.515076 | yes | 1.804E-07 | 677.86694 | 616.86481 |
| SPDLAPTPAPQSTI | 15 | Q9BY44 | Eukaryotic translation initiation factor 2A | EIF2A | 64.991 | 2 | 52.515076 | yes | 1.234E-07 | 767.89661 | 613.64539 |
| VNIISVTNK | 9 | Q9BY44 | Eukaryotic translation initiation factor 2A | EIF2A | 64.991 | 2 | 52.515076 | yes | 9.881E-08 | 494.29529 | 539.34363 |
| GQMEVWDVK | 9 | Q9BY44 | Eukaryotic translation initiation factor 2A | EIF2A | 64.991 | 2 | 52.515076 | yes | 9.722E-08 | 546.26312 | 193.26793 |
| AVCLEFSPK | 9 | Q9BY44 | Eukaryotic translation initiation factor 2A | EIF2A | 64.991 | 2 | 52.515076 | yes | 1.688E-07 | 525.76807 | 177.02097 |
| DGTLFAWGNGEK | 12 | Q9BY44 | Eukaryotic translation initiation factor 2A | EIF2A | 64.991 | 2 | 52.515076 | yes | 9.809E-08 | 647.80676 | 173.92676 |
| KATAVLVIASTDV | 15 | Q9BY44 | Eukaryotic translation initiation factor 2A | EIF2A | 64.991 | 2 | 52.515076 | yes | 1.002E-07 | 765.94049 | 69.742607 |
| KATAVLVIASTDV | 15 | Q9BY44 | Eukaryotic translation initiation factor 2A | EIF2A | 64.991 | 2 | 52.515076 | yes | 2.399E-07 | 510.96277 | 69.742607 |
| KATAVLVIASTDV | 15 | Q9BY44 | Eukaryotic translation initiation factor 2A | EIF2A | 64.991 | 3 | 52.515076 | yes | 1.002E-07 | 765.94049 | 69.742607 |
| KATAVLVIASTDV | 15 | Q9BY44 | Eukaryotic translation initiation factor 2A | EIF2A | 64.991 | 3 | 52.515076 | yes | 2.399E-07 | 510.96277 | 69.742607 |
| LISKPVASDSTYFA | 30 | Q9BY44 | Eukaryotic translation initiation factor 2A | EIF2A | 64.991 | 4 | 52.515076 | yes | 1.682E-07 | 841.65887 | 135.50856 |
| VAVYVPGSK | 9 | Q9BY44 | Eukaryotic translation initiation factor 2A | EIF2A | 64.991 | 2 | 52.515076 | yes | 3.976E-07 | 460.26602 | 97.802887 |
| NTVLATWQPYTTS | 14 | Q9BY44 | Eukaryotic translation initiation factor 2A | EIF2A | 64.991 | 2 | 52.515076 | yes | 2.126E-07 | 805.41467 | 84.458458 |
| NTVSQSISGDPEID | 15 | Q9BY44 | Eukaryotic translation initiation factor 2A | EIF2A | 64.991 | 2 | 52.515076 | yes | 9.722E-08 | 795.38629 | 62.9189 |
| VATAYRPPALR | 11 | Q9BY44 | Eukaryotic translation initiation factor 2A | EIF2A | 64.991 | 2 | 52.515076 | yes | 3.757E-07 | 607.85382 | 62.405556 |
| NCKVCIFSK | 9 | Q9BY44 | Eukaryotic translation initiation factor 2A | EIF2A | 64.991 | 2 | 52.515076 | yes | 2.29E-07 | 578.28607 | 53.251324 |
| GSEGLYMVNGPPI | 22 | Q9BY44 | Eukaryotic translation initiation factor 2A | EIF2A | 64.991 | 3 | 52.515076 | yes | 2.954E-07 | 808.05475 | 28.714418 |
| GLLHSFDLLK | 10 | Q9BY44 | Eukaryotic translation initiation factor 2A | EIF2A | 64.991 | 2 | 52.515076 | yes | 1.728E-07 | 571.83203 | 27.073429 |
| APSTPLLTVR | 10 | Q9BY44 | Eukaryotic translation initiation factor 2A | EIF2A | 64.991 | 2 | 52.515076 | yes | 2.899E-06 | 527.81641 | 20.96759 |
| ELLELGSNVVIASI | 14 | Q9BY49 | Peroxisomal trans-2-enoyl-CoA reductase | PECR | 32.545 | 2 | 41.57083 | yes | 3.444E-07 | 750.42505 | 71.565041 |
| LVHPGVAEVVFVI | 13 | Q9BY77 | Polymerase delta-interacting protein 3 OS | POLDIP3 | 46.09 | 2 | 54.010452 | yes | 1.923E-07 | 697.41376 | 2166.6514 |
| LVHPGVAEVVFVI | 13 | Q9BY77 | Polymerase delta-interacting protein 3 OS | POLDIP3 | 46.09 | 2 | 54.010452 | yes | 8.092E-07 | 465.27826 | 2166.6514 |
| LVHPGVAEVVFVI | 13 | Q9BY77 | Polymerase delta-interacting protein 3 OS | POLDIP3 | 46.09 | 3 | 54.010452 | yes | 1.923E-07 | 697.41376 | 2166.6514 |
| LVHPGVAEVVFVI | 13 | Q9BY77 | Polymerase delta-interacting protein 3 OS | POLDIP3 | 46.09 | 3 | 54.010452 | yes | 8.092E-07 | 465.27826 | 2166.6514 |
| ELPAAEPVLSPLEC | 16 | Q9BY77 | Polymerase delta-interacting protein 3 OS | POLDIP3 | 46.09 | 2 | 54.010452 | yes | 2.115E-07 | 825.95105 | 3905.2715 |
| VGIQQGLLSQSTR | 13 | Q9BY77 | Polymerase delta-interacting protein 3 OS | POLDIP3 | 46.09 | 2 | 54.010452 | yes | 1.473E-07 | 693.88861 | 3361.8108 |
| AMAPLHPHPAGM | 13 | Q9BY77 | Polymerase delta-interacting protein 3 OS | POLDIP3 | 46.09 | 3 | 54.010452 | yes | 2.277E-07 | 462.56915 | 574.45465 |
| AMAPLHPHPAGM | 13 | Q9BY77 | Polymerase delta-interacting protein 3 OS | POLDIP3 | 46.09 | 3 | 54.010452 | yes | 1.001E-07 | 693.3501 | 574.45465 |
| AMAPLHPHPAGM | 13 | Q9BY77 | Polymerase delta-interacting protein 3 OS | POLDIP3 | 46.09 | 3 | 54.010452 | yes | 9.157E-07 | 467.90079 | 574.45465 |
| AMAPLHPHPAGM | 13 | Q9BY77 | Polymerase delta-interacting protein 3 OS | POLDIP3 | 46.09 | 2 | 54.010452 | yes | 2.277E-07 | 462.56915 | 574.45465 |

| | | | | | | | | | | | |
|---|---|---|---|---|---|---|---|---|---|---|---|
| AMAPLHPHPAGM | 13 | Q9BY77 | Polymerase delta-interacting protein 3 OS | POLDIP3 | 46.09 | 2 | 54.010452 | yes | 1.001E-07 | 693.3501 | 574.45465 |
| AMAPLHPHPAGM | 13 | Q9BY77 | Polymerase delta-interacting protein 3 OS | POLDIP3 | 46.09 | 2 | 54.010452 | yes | 9.157E-07 | 467.90079 | 574.45465 |
| FAASGGFLHHMA( | 18 | Q9BY77 | Polymerase delta-interacting protein 3 OS | POLDIP3 | 46.09 | 3 | 54.010452 | yes | 2.216E-07 | 602.29816 | 1640.3589 |
| SSGASVTTQPTEFI | 14 | Q9BY77 | Polymerase delta-interacting protein 3 OS | POLDIP3 | 46.09 | 2 | 54.010452 | yes | 9.722E-08 | 720.35425 | 1627.2012 |
| MTVNNLHPR | 9 | Q9BY77 | Polymerase delta-interacting protein 3 OS | POLDIP3 | 46.09 | 2 | 54.010452 | yes | 1.612E-07 | 541.28223 | 829.11072 |
| INVVNNHQAK | 10 | Q9BY77 | Polymerase delta-interacting protein 3 OS | POLDIP3 | 46.09 | 2 | 54.010452 | yes | 1.894E-07 | 568.81219 | 789.59045 |
| QQTTVPQKPR | 10 | Q9BY77 | Polymerase delta-interacting protein 3 OS | POLDIP3 | 46.09 | 2 | 54.010452 | yes | 9.722E-08 | 591.83313 | 781.45184 |
| CLDGQPMK | 8 | Q9BY77 | Polymerase delta-interacting protein 3 OS | POLDIP3 | 46.09 | 2 | 54.010452 | yes | 2.311E-07 | 474.7175 | 779.98364 |
| KESELPR | 7 | Q9BY77 | Polymerase delta-interacting protein 3 OS | POLDIP3 | 46.09 | 2 | 54.010452 | yes | 2.553E-06 | 429.73761 | 440.414 |
| KDDAITAYK | 9 | Q9BY77 | Polymerase delta-interacting protein 3 OS | POLDIP3 | 46.09 | 2 | 54.010452 | yes | 2.092E-07 | 512.7691 | 422.50967 |
| EEPPKELPAAEPVI | 21 | Q9BY77 | Polymerase delta-interacting protein 3 OS | POLDIP3 | 46.09 | 3 | 54.010452 | yes | 9.722E-08 | 744.39838 | 149.58231 |
| TATFQQR | 7 | Q9BY77 | Polymerase delta-interacting protein 3 OS | POLDIP3 | 46.09 | 2 | 54.010452 | yes | 1.055E-07 | 426.22214 | 125.30068 |
| TIQVPQQK | 8 | Q9BY77 | Polymerase delta-interacting protein 3 OS | POLDIP3 | 46.09 | 2 | 54.010452 | yes | 9.722E-08 | 471.27435 | 69.605896 |
| VVQNDAYTAPALI | 17 | Q9BY77 | Polymerase delta-interacting protein 3 OS | POLDIP3 | 46.09 | 3 | 54.010452 | yes | 2.908E-07 | 601.31964 | 45.071266 |
| LSDSPSMK | 8 | Q9BY77 | Polymerase delta-interacting protein 3 OS | POLDIP3 | 46.09 | 2 | 54.010452 | yes | 1.395E-07 | 432.71021 | 29.497255 |
| EQIENYK | 7 | Q9BYC5 | Alpha-(1,6)-fucosyltransferase OS=Homc | FUT8 | 66.516 | 2 | 36.737156 | yes | 6.666E-07 | 462.22708 | 86.044968 |
| NLEGNELQR | 9 | Q9BYC5 | Alpha-(1,6)-fucosyltransferase OS=Homc | FUT8 | 66.516 | 2 | 36.737156 | yes | 4.252E-06 | 536.77271 | 74.484444 |
| GYAFVEFESEDVA | 14 | Q9BYG3 | MKI67 FHA domain-interacting nucleola | NIFK | 34.223 | 2 | 51.939644 | yes | 1.197E-07 | 795.86975 | 2130.5891 |
| IVAETMNNYLFGE | 14 | Q9BYG3 | MKI67 FHA domain-interacting nucleola | NIFK | 34.223 | 2 | 51.939644 | yes | 9.722E-08 | 828.90613 | 1753.3171 |
| GIDYDFPSLILQK | 13 | Q9BYG3 | MKI67 FHA domain-interacting nucleola | NIFK | 34.223 | 2 | 51.939644 | yes | 1.102E-07 | 754.90338 | 1048.2152 |
| EEIQETQTPTHSR | 13 | Q9BYG3 | MKI67 FHA domain-interacting nucleola | NIFK | 34.223 | 2 | 51.939644 | yes | 9.861E-08 | 778.37097 | 802.06885 |
| KQEQLTPGVVYVI | 13 | Q9BYG3 | MKI67 FHA domain-interacting nucleola | NIFK | 34.223 | 2 | 51.939644 | yes | 1.15E-07 | 758.92773 | 188.64328 |
| KQEQLTPGVVYVI | 13 | Q9BYG3 | MKI67 FHA domain-interacting nucleola | NIFK | 34.223 | 2 | 51.939644 | yes | 1.151E-07 | 506.2876 | 188.64328 |
| KQEQLTPGVVYVI | 13 | Q9BYG3 | MKI67 FHA domain-interacting nucleola | NIFK | 34.223 | 3 | 51.939644 | yes | 1.15E-07 | 758.92773 | 188.64328 |
| KQEQLTPGVVYVI | 13 | Q9BYG3 | MKI67 FHA domain-interacting nucleola | NIFK | 34.223 | 3 | 51.939644 | yes | 1.151E-07 | 506.2876 | 188.64328 |
| LLECHFMPPEK | 11 | Q9BYG3 | MKI67 FHA domain-interacting nucleola | NIFK | 34.223 | 2 | 51.939644 | yes | 4.67E-07 | 700.83868 | 188.29776 |
| LLECHFMPPEK | 11 | Q9BYG3 | MKI67 FHA domain-interacting nucleola | NIFK | 34.223 | 2 | 51.939644 | yes | 5.812E-07 | 467.56155 | 188.29776 |
| LLECHFMPPEK | 11 | Q9BYG3 | MKI67 FHA domain-interacting nucleola | NIFK | 34.223 | 3 | 51.939644 | yes | 4.67E-07 | 700.83868 | 188.29776 |
| LLECHFMPPEK | 11 | Q9BYG3 | MKI67 FHA domain-interacting nucleola | NIFK | 34.223 | 3 | 51.939644 | yes | 5.812E-07 | 467.56155 | 188.29776 |
| QEQLTPGVVYVR | 12 | Q9BYG3 | MKI67 FHA domain-interacting nucleola | NIFK | 34.223 | 2 | 51.939644 | yes | 1.116E-07 | 694.88025 | 327.42795 |
| KVSGTLDTPEK | 11 | Q9BYG3 | MKI67 FHA domain-interacting nucleola | NIFK | 34.223 | 2 | 51.939644 | yes | 9.722E-08 | 587.81934 | 183.31024 |
| VSGTLDTPEK | 10 | Q9BYG3 | MKI67 FHA domain-interacting nucleola | NIFK | 34.223 | 2 | 51.939644 | yes | 1.132E-07 | 523.77185 | 165.40227 |
| DWNIPFK | 7 | Q9BYG3 | MKI67 FHA domain-interacting nucleola | NIFK | 34.223 | 2 | 51.939644 | yes | 5.827E-07 | 460.23724 | 112.46454 |
| QPISCVK | 7 | Q9BYG3 | MKI67 FHA domain-interacting nucleola | NIFK | 34.223 | 2 | 51.939644 | yes | 9.077E-07 | 416.2233 | 76.225388 |
| KGIDYDFPSLILQK | 14 | Q9BYG3 | MKI67 FHA domain-interacting nucleola | NIFK | 34.223 | 2 | 51.939644 | yes | 1.611E-07 | 818.95087 | 52.175339 |
| AVSTANPLLR | 10 | Q9BYG5 | Partitioning defective 6 homolog beta OS | PARD6B | 41.183 | 2 | 53.710793 | yes | 1.768E-07 | 521.30621 | 179.92903 |
| EEADYSAFGTDTL | 15 | Q9BYG5 | Partitioning defective 6 homolog beta OS | PARD6B | 41.183 | 2 | 53.710793 | yes | 9.722E-08 | 830.39105 | 116.35848 |
| VICWLCER | 8 | Q9BYJ4 | E3 ubiquitin-protein ligase TRIM34 OS= | TRIM34 | 56.864 | 2 | 47.550686 | no | 4.15E-07 | 568.27301 | 85.890701 |
| FAEAEDELVQQK | 12 | Q9BYJ4 | E3 ubiquitin-protein ligase TRIM34 OS= | TRIM34 | 56.864 | 2 | 47.550686 | yes | 1.305E-07 | 703.84357 | 57.57486 |

| | | | | | | | | | | | |
|---|---|---|---|---|---|---|---|---|---|---|---|
| HYWEVDVSK | 9 | Q9BYJ4 | E3 ubiquitin-protein ligase TRIM34 OS= | TRIM34 | 56.864 | 2 | 47.550686 | no | 3.962E-07 | 581.78003 | 53.014156 |
| SGPVMGGGLPPPP | 15 | Q9BYJ9 | YTH domain-containing family protein 1 | YTHDF1 | 60.874 | 2 | 50.258553 | yes | 5.526E-07 | 702.3894 | 2212.385 |
| SGPVMGGGLPPPP | 15 | Q9BYJ9 | YTH domain-containing family protein 1 | YTHDF1 | 60.874 | 2 | 50.258553 | yes | 1.486E-07 | 710.38684 | 2212.385 |
| APGMNSLEQGMV | 15 | Q9BYJ9 | YTH domain-containing family protein 1 | YTHDF1 | 60.874 | 2 | 50.258553 | yes | 1.001E-07 | 766.38397 | 1228.1527 |
| APGMNSLEQGMV | 15 | Q9BYJ9 | YTH domain-containing family protein 1 | YTHDF1 | 60.874 | 2 | 50.258553 | yes | 1.662E-06 | 774.38147 | 1228.1527 |
| HNMDIGTWDNK | 11 | Q9BYJ9 | YTH domain-containing family protein 1 | YTHDF1 | 60.874 | 2 | 50.258553 | no | 1.011E-07 | 665.7959 | 1211.6266 |
| HNMDIGTWDNK | 11 | Q9BYJ9 | YTH domain-containing family protein 1 | YTHDF1 | 60.874 | 2 | 50.258553 | no | 1.869E-05 | 673.79333 | 1211.6266 |
| SATSVDTQR | 9 | Q9BYJ9 | YTH domain-containing family protein 1 | YTHDF1 | 60.874 | 2 | 50.258553 | yes | 1.44E-07 | 503.74362 | 1618.8754 |
| SPVDYGTSAGVWS | 16 | Q9BYJ9 | YTH domain-containing family protein 1 | YTHDF1 | 60.874 | 2 | 50.258553 | yes | 1.971E-07 | 848.89429 | 1393.2217 |
| PTSWAAIASK | 10 | Q9BYJ9 | YTH domain-containing family protein 1 | YTHDF1 | 60.874 | 2 | 50.258553 | yes | 1.682E-07 | 516.27966 | 1293.3875 |
| RLDSAFR | 7 | Q9BYJ9 | YTH domain-containing family protein 1 | YTHDF1 | 60.874 | 2 | 50.258553 | yes | 2.084E-07 | 432.73795 | 1031.0245 |
| GKFDVQWIFVK | 11 | Q9BYJ9 | YTH domain-containing family protein 1 | YTHDF1 | 60.874 | 2 | 50.258553 | yes | 1.222E-07 | 683.87952 | 847.76831 |
| QEEEEVVR | 8 | Q9BYJ9 | YTH domain-containing family protein 1 | YTHDF1 | 60.874 | 2 | 50.258553 | yes | 1.707E-07 | 509.246 | 607.85388 |
| RQEEEEVVR | 9 | Q9BYJ9 | YTH domain-containing family protein 1 | YTHDF1 | 60.874 | 2 | 50.258553 | yes | 9.825E-08 | 587.29657 | 488.62723 |
| EFEWNLK | 7 | Q9BYJ9 | YTH domain-containing family protein 1 | YTHDF1 | 60.874 | 2 | 50.258553 | yes | 1.385E-07 | 483.23999 | 252.79727 |
| DPSQVPIYSLR | 11 | Q9BYK8 | Helicase with zinc finger domain 2 OS=H | HELZ2 | 294.651 | 2 | 53.958668 | yes | 2.394E-07 | 637.84058 | 715.15051 |
| LEGLPAPTASPR | 12 | Q9BYK8 | Helicase with zinc finger domain 2 OS=H | HELZ2 | 294.651 | 2 | 53.958668 | yes | 9.809E-08 | 604.83533 | 469.94644 |
| GPALALEFNR | 10 | Q9BYK8 | Helicase with zinc finger domain 2 OS=H | HELZ2 | 294.651 | 2 | 53.958668 | yes | 1.142E-07 | 544.2984 | 453.84964 |
| LQNLGLDR | 8 | Q9BYK8 | Helicase with zinc finger domain 2 OS=H | HELZ2 | 294.651 | 2 | 53.958668 | yes | 2.715E-07 | 464.76416 | 450.14978 |
| ANLEEVAEVVR | 11 | Q9BYK8 | Helicase with zinc finger domain 2 OS=H | HELZ2 | 294.651 | 2 | 53.958668 | yes | 9.722E-08 | 614.83026 | 425.10049 |
| SSVASGPISPTNYR | 14 | Q9BYK8 | Helicase with zinc finger domain 2 OS=H | HELZ2 | 294.651 | 2 | 53.958668 | yes | 9.722E-08 | 718.36243 | 379.13324 |
| HELAFVCR | 8 | Q9BYK8 | Helicase with zinc finger domain 2 OS=H | HELZ2 | 294.651 | 2 | 53.958668 | yes | 2.337E-06 | 516.25818 | 377.0694 |
| EGIAGVAVSSITK | 13 | Q9BYK8 | Helicase with zinc finger domain 2 OS=H | HELZ2 | 294.651 | 2 | 53.958668 | yes | 9.809E-08 | 616.34827 | 357.90546 |
| DLGQVSVGSFEILI | 16 | Q9BYK8 | Helicase with zinc finger domain 2 OS=H | HELZ2 | 294.651 | 2 | 53.958668 | yes | 9.722E-08 | 837.44647 | 356.07312 |
| QGFYYPLGIVR | 11 | Q9BYK8 | Helicase with zinc finger domain 2 OS=H | HELZ2 | 294.651 | 2 | 53.958668 | yes | 9.722E-08 | 656.85608 | 351.23654 |
| YVLVSTVR | 8 | Q9BYK8 | Helicase with zinc finger domain 2 OS=H | HELZ2 | 294.651 | 2 | 53.958668 | yes | 1.678E-07 | 468.7793 | 347.37836 |
| LVFHENYR | 8 | Q9BYK8 | Helicase with zinc finger domain 2 OS=H | HELZ2 | 294.651 | 2 | 53.958668 | yes | 9.722E-08 | 539.27747 | 346.90796 |
| ILGLEYSLR | 9 | Q9BYK8 | Helicase with zinc finger domain 2 OS=H | HELZ2 | 294.651 | 2 | 53.958668 | yes | 4.142E-07 | 532.31097 | 336.62402 |
| LVAEFLVGSECTR | 13 | Q9BYK8 | Helicase with zinc finger domain 2 OS=H | HELZ2 | 294.651 | 2 | 53.958668 | yes | 9.722E-08 | 740.87683 | 330.94528 |
| LLAEYQR | 7 | Q9BYK8 | Helicase with zinc finger domain 2 OS=H | HELZ2 | 294.651 | 2 | 53.958668 | yes | 9.722E-08 | 446.74799 | 329.08435 |
| QLLELVELQR | 10 | Q9BYK8 | Helicase with zinc finger domain 2 OS=H | HELZ2 | 294.651 | 2 | 53.958668 | yes | 9.722E-08 | 620.86664 | 292.73703 |
| VVLLGDHK | 8 | Q9BYK8 | Helicase with zinc finger domain 2 OS=H | HELZ2 | 294.651 | 2 | 53.958668 | yes | 1.063E-07 | 440.76617 | 290.5199 |
| LGGPCILYCGPSNI | 14 | Q9BYK8 | Helicase with zinc finger domain 2 OS=H | HELZ2 | 294.651 | 2 | 53.958668 | yes | 9.809E-08 | 768.37091 | 282.05374 |
| EVLPEASTWEQGL | 14 | Q9BYK8 | Helicase with zinc finger domain 2 OS=H | HELZ2 | 294.651 | 2 | 53.958668 | yes | 1.251E-07 | 807.90973 | 273.29379 |
| SDLDQRPTK | 9 | Q9BYK8 | Helicase with zinc finger domain 2 OS=H | HELZ2 | 294.651 | 2 | 53.958668 | yes | 9.722E-08 | 530.27509 | 271.41504 |
| VPSSTADFQVGVR | 13 | Q9BYK8 | Helicase with zinc finger domain 2 OS=H | HELZ2 | 294.651 | 2 | 53.958668 | yes | 7.098E-07 | 681.85425 | 269.32486 |
| YLDVVLQR | 8 | Q9BYK8 | Helicase with zinc finger domain 2 OS=H | HELZ2 | 294.651 | 2 | 53.958668 | yes | 9.974E-08 | 503.29001 | 266.28394 |
| CCLALGLQPEAR | 12 | Q9BYK8 | Helicase with zinc finger domain 2 OS=H | HELZ2 | 294.651 | 2 | 53.958668 | yes | 9.722E-08 | 694.34467 | 258.50168 |
| DGVLDVEAR | 9 | Q9BYK8 | Helicase with zinc finger domain 2 OS=H | HELZ2 | 294.651 | 2 | 53.958668 | yes | 4.403E-07 | 487.2511 | 254.65988 |

| | | | | | | | | | | | |
|---|---|---|---|---|---|---|---|---|---|---|---|
| FAPSVVQSDR | 10 | Q9BYK8 | Helicase with zinc finger domain 2 OS=H | HELZ2 | 294.651 | 2 | 53.958668 | yes | 1.633E-06 | 553.28546 | 250.40334 |
| LNPSQNVAVR | 10 | Q9BYK8 | Helicase with zinc finger domain 2 OS=H | HELZ2 | 294.651 | 2 | 53.958668 | yes | 1.172E-07 | 549.3067 | 239.24963 |
| LVLAGDHMQVTPI | 13 | Q9BYK8 | Helicase with zinc finger domain 2 OS=H | HELZ2 | 294.651 | 3 | 53.958668 | yes | 1.727E-07 | 479.59415 | 101.64482 |
| LVLAGDHMQVTPI | 13 | Q9BYK8 | Helicase with zinc finger domain 2 OS=H | HELZ2 | 294.651 | 3 | 53.958668 | yes | 5.641E-07 | 718.88757 | 101.64482 |
| LVLAGDHMQVTPI | 13 | Q9BYK8 | Helicase with zinc finger domain 2 OS=H | HELZ2 | 294.651 | 2 | 53.958668 | yes | 1.727E-07 | 479.59415 | 101.64482 |
| LVLAGDHMQVTPI | 13 | Q9BYK8 | Helicase with zinc finger domain 2 OS=H | HELZ2 | 294.651 | 2 | 53.958668 | yes | 5.641E-07 | 718.88757 | 101.64482 |
| SLLVSTDEGNENSI | 14 | Q9BYK8 | Helicase with zinc finger domain 2 OS=H | HELZ2 | 294.651 | 2 | 53.958668 | yes | 3.284E-07 | 746.85992 | 202.01881 |
| HCSFVPETFER | 11 | Q9BYK8 | Helicase with zinc finger domain 2 OS=H | HELZ2 | 294.651 | 2 | 53.958668 | yes | 9.722E-08 | 704.81934 | 193.29776 |
| VVVTTTSQAR | 10 | Q9BYK8 | Helicase with zinc finger domain 2 OS=H | HELZ2 | 294.651 | 2 | 53.958668 | yes | 2.667E-07 | 531.30109 | 190.15869 |
| TPQGQLQGAFR | 11 | Q9BYK8 | Helicase with zinc finger domain 2 OS=H | HELZ2 | 294.651 | 2 | 53.958668 | yes | 9.722E-08 | 601.81744 | 186.75552 |
| SHCGHFLEVAR | 11 | Q9BYK8 | Helicase with zinc finger domain 2 OS=H | HELZ2 | 294.651 | 3 | 53.958668 | yes | 9.722E-08 | 438.21204 | 184.02391 |
| CTDAIVSFISR | 11 | Q9BYK8 | Helicase with zinc finger domain 2 OS=H | HELZ2 | 294.651 | 2 | 53.958668 | yes | 1.727E-07 | 634.81879 | 181.27948 |
| MELKPLR | 7 | Q9BYK8 | Helicase with zinc finger domain 2 OS=H | HELZ2 | 294.651 | 2 | 53.958668 | yes | 9.722E-08 | 443.76257 | 164.61256 |
| ELLDESQK | 8 | Q9BYK8 | Helicase with zinc finger domain 2 OS=H | HELZ2 | 294.651 | 2 | 53.958668 | yes | 3.767E-06 | 481.24548 | 162.95483 |
| VLNTVLTR | 8 | Q9BYK8 | Helicase with zinc finger domain 2 OS=H | HELZ2 | 294.651 | 2 | 53.958668 | yes | 2.024E-07 | 458.28473 | 155.16573 |
| QTYNIPGGR | 9 | Q9BYK8 | Helicase with zinc finger domain 2 OS=H | HELZ2 | 294.651 | 2 | 53.958668 | yes | 9.723E-08 | 503.25925 | 148.62712 |
| GLEEASPLVTSIAL | 16 | Q9BYK8 | Helicase with zinc finger domain 2 OS=H | HELZ2 | 294.651 | 2 | 53.958668 | yes | 1.164E-06 | 806.94885 | 126.74406 |
| LHLLASLWK | 9 | Q9BYK8 | Helicase with zinc finger domain 2 OS=H | HELZ2 | 294.651 | 2 | 53.958668 | yes | 1.564E-07 | 540.83185 | 114.79062 |
| RPVLLQK | 7 | Q9BYK8 | Helicase with zinc finger domain 2 OS=H | HELZ2 | 294.651 | 2 | 53.958668 | yes | 1.973E-07 | 427.28455 | 111.37595 |
| QLSYEEAEEVIR | 12 | Q9BYK8 | Helicase with zinc finger domain 2 OS=H | HELZ2 | 294.651 | 2 | 53.958668 | yes | 9.722E-08 | 733.36212 | 110.13913 |
| PVPQPLCR | 8 | Q9BYK8 | Helicase with zinc finger domain 2 OS=H | HELZ2 | 294.651 | 2 | 53.958668 | yes | 6.531E-07 | 483.76312 | 108.83397 |
| YHTELGR | 7 | Q9BYK8 | Helicase with zinc finger domain 2 OS=H | HELZ2 | 294.651 | 2 | 53.958668 | yes | 9.722E-08 | 438.22214 | 107.75515 |
| AAEHTLLHR | 9 | Q9BYK8 | Helicase with zinc finger domain 2 OS=H | HELZ2 | 294.651 | 2 | 53.958668 | yes | 1.061E-07 | 524.28833 | 106.64924 |
| ETLPDPCPVPYGSI | 26 | Q9BYK8 | Helicase with zinc finger domain 2 OS=H | HELZ2 | 294.651 | 4 | 53.958668 | yes | 1.003E-07 | 707.10645 | 102.82917 |
| AHSAQELQEWVR | 12 | Q9BYK8 | Helicase with zinc finger domain 2 OS=H | HELZ2 | 294.651 | 2 | 53.958668 | yes | 9.722E-08 | 727.36279 | 101.44949 |
| LWESFIR | 7 | Q9BYK8 | Helicase with zinc finger domain 2 OS=H | HELZ2 | 294.651 | 2 | 53.958668 | yes | 5.037E-07 | 475.75836 | 94.133011 |
| EGRPNQSLR | 9 | Q9BYK8 | Helicase with zinc finger domain 2 OS=H | HELZ2 | 294.651 | 2 | 53.958668 | yes | 2.078E-07 | 528.78088 | 87.278564 |
| QLRPVVK | 7 | Q9BYK8 | Helicase with zinc finger domain 2 OS=H | HELZ2 | 294.651 | 2 | 53.958668 | yes | 2.751E-07 | 420.2767 | 83.015053 |
| WPEAAALIQEK | 11 | Q9BYK8 | Helicase with zinc finger domain 2 OS=H | HELZ2 | 294.651 | 2 | 53.958668 | yes | 1.703E-06 | 628.33771 | 78.611382 |
| EYMIQFNR | 8 | Q9BYK8 | Helicase with zinc finger domain 2 OS=H | HELZ2 | 294.651 | 2 | 53.958668 | yes | 4.013E-07 | 550.76331 | 77.634445 |
| VPPLLIYGPFGTGK | 14 | Q9BYK8 | Helicase with zinc finger domain 2 OS=H | HELZ2 | 294.651 | 2 | 53.958668 | yes | 5.995E-07 | 729.92139 | 71.405777 |
| LGFVVDVEAGSR | 12 | Q9BYK8 | Helicase with zinc finger domain 2 OS=H | HELZ2 | 294.651 | 2 | 53.958668 | yes | 1.738E-07 | 624.83276 | 71.176857 |
| SLHLAVQLK | 9 | Q9BYK8 | Helicase with zinc finger domain 2 OS=H | HELZ2 | 294.651 | 2 | 53.958668 | yes | 3.676E-07 | 504.81366 | 69.941597 |
| SNQEQVQPGGPPR | 13 | Q9BYK8 | Helicase with zinc finger domain 2 OS=H | HELZ2 | 294.651 | 2 | 53.958668 | yes | 1.182E-07 | 697.34454 | 62.521141 |
| QILLALGHGGSAY | 16 | Q9BYK8 | Helicase with zinc finger domain 2 OS=H | HELZ2 | 294.651 | 3 | 53.958668 | yes | 6.967E-07 | 538.63013 | 50.936089 |
| NLALGHPEEMER | 12 | Q9BYK8 | Helicase with zinc finger domain 2 OS=H | HELZ2 | 294.651 | 2 | 53.958668 | yes | 9.722E-08 | 698.33789 | 62.141403 |
| EYFHSHVSGGHPE | 18 | Q9BYK8 | Helicase with zinc finger domain 2 OS=H | HELZ2 | 294.651 | 4 | 53.958668 | yes | 3.156E-07 | 505.99606 | 53.786621 |
| RPPSVLGHAGK | 11 | Q9BYK8 | Helicase with zinc finger domain 2 OS=H | HELZ2 | 294.651 | 2 | 53.958668 | yes | 3.634E-06 | 559.82507 | 37.808983 |
| KQELAVALIAGWC | 18 | Q9BYK8 | Helicase with zinc finger domain 2 OS=H | HELZ2 | 294.651 | 3 | 53.958668 | yes | 1.435E-07 | 613.33551 | 36.170563 |

| | | | | | | | | | | | |
|---|---|---|---|---|---|---|---|---|---|---|---|
| CICVVSHGAQVSA | 15 | Q9BYK8 | Helicase with zinc finger domain 2 OS=H | HELZ2 | 294.651 | 3 | 53.958668 | yes | 5.364E-06 | 552.94849 | 32.550476 |
| AQEGLCLIGDHLLI | 15 | Q9BYK8 | Helicase with zinc finger domain 2 OS=H | HELZ2 | 294.651 | 3 | 53.958668 | yes | 2.445E-06 | 569.97852 | 14.46359 |
| HSVCPEGLSMEQV | 20 | Q9BYK8 | Helicase with zinc finger domain 2 OS=H | HELZ2 | 294.651 | 3 | 53.958668 | yes | 1.286E-07 | 747.68439 | 5.3079715 |
| ETIVEVGSDLPDSC | 15 | Q9BYW2 | Histone-lysine N-methyltransferase SETD | SETD2 | 287.597 | 2 | 56.017868 | yes | 1.374E-07 | 773.38574 | 198.97881 |
| LMSMPVMTVDYS | 13 | Q9BYW2 | Histone-lysine N-methyltransferase SETD | SETD2 | 287.597 | 2 | 56.017868 | yes | 1.864E-07 | 751.35858 | 165.87643 |
| SEDLGKPSR | 9 | Q9BYW2 | Histone-lysine N-methyltransferase SETD | SETD2 | 287.597 | 2 | 56.017868 | yes | 4.301E-07 | 494.75653 | 159.55878 |
| IYYYHVITR | 9 | Q9BYW2 | Histone-lysine N-methyltransferase SETD | SETD2 | 287.597 | 2 | 56.017868 | yes | 9.722E-08 | 614.32965 | 145.79539 |
| TETSYLEMER | 10 | Q9BYW2 | Histone-lysine N-methyltransferase SETD | SETD2 | 287.597 | 2 | 56.017868 | yes | 9.838E-08 | 629.78461 | 134.08063 |
| NDEIIDATQK | 10 | Q9BYW2 | Histone-lysine N-methyltransferase SETD | SETD2 | 287.597 | 2 | 56.017868 | yes | 3.766E-07 | 573.78552 | 132.48344 |
| TLEHLPIPTK | 10 | Q9BYW2 | Histone-lysine N-methyltransferase SETD | SETD2 | 287.597 | 2 | 56.017868 | yes | 2.162E-07 | 574.83734 | 124.81917 |
| ETEPLVSPHQDK | 12 | Q9BYW2 | Histone-lysine N-methyltransferase SETD | SETD2 | 287.597 | 2 | 56.017868 | yes | 9.789E-08 | 690.34369 | 112.38173 |
| FGAVYKPK | 8 | Q9BYW2 | Histone-lysine N-methyltransferase SETD | SETD2 | 287.597 | 2 | 56.017868 | yes | 1.415E-07 | 455.26328 | 111.81499 |
| LLMIECSSR | 9 | Q9BYW2 | Histone-lysine N-methyltransferase SETD | SETD2 | 287.597 | 2 | 56.017868 | yes | 1.545E-07 | 554.77808 | 110.56596 |
| GRPEISLDER | 10 | Q9BYW2 | Histone-lysine N-methyltransferase SETD | SETD2 | 287.597 | 2 | 56.017868 | yes | 8.476E-07 | 586.30695 | 101.42812 |
| MQCECTPLSK | 10 | Q9BYW2 | Histone-lysine N-methyltransferase SETD | SETD2 | 287.597 | 2 | 56.017868 | yes | 5.108E-07 | 627.26978 | 100.46625 |
| NPEISFTQSSR | 11 | Q9BYW2 | Histone-lysine N-methyltransferase SETD | SETD2 | 287.597 | 2 | 56.017868 | yes | 1.114E-07 | 633.30969 | 99.539001 |
| TVDISDLATK | 10 | Q9BYW2 | Histone-lysine N-methyltransferase SETD | SETD2 | 287.597 | 2 | 56.017868 | yes | 6.477E-07 | 531.78748 | 98.48101 |
| NSFCCTELNEEIK | 13 | Q9BYW2 | Histone-lysine N-methyltransferase SETD | SETD2 | 287.597 | 2 | 56.017868 | yes | 9.722E-08 | 822.35565 | 96.642288 |
| LSTEADTDTPK | 11 | Q9BYW2 | Histone-lysine N-methyltransferase SETD | SETD2 | 287.597 | 2 | 56.017868 | yes | 1.52E-07 | 589.28278 | 94.207611 |
| LNDSPTLK | 8 | Q9BYW2 | Histone-lysine N-methyltransferase SETD | SETD2 | 287.597 | 2 | 56.017868 | yes | 1.455E-07 | 444.24527 | 73.263275 |
| NPEDLECNENVK | 12 | Q9BYW2 | Histone-lysine N-methyltransferase SETD | SETD2 | 287.597 | 2 | 56.017868 | yes | 9.722E-08 | 730.81976 | 67.614548 |
| LPELSHPQSDGVD | 21 | Q9BYW2 | Histone-lysine N-methyltransferase SETD | SETD2 | 287.597 | 3 | 56.017868 | yes | 3.7E-07 | 750.36084 | 65.683296 |
| AHTPLNTPDPSTK | 13 | Q9BYW2 | Histone-lysine N-methyltransferase SETD | SETD2 | 287.597 | 2 | 56.017868 | yes | 9.849E-08 | 689.85168 | 56.901981 |
| YQQNAEQYGGTR | 12 | Q9BYW2 | Histone-lysine N-methyltransferase SETD | SETD2 | 287.597 | 2 | 56.017868 | yes | 2.194E-07 | 707.82092 | 53.032951 |
| LPTSEPEADAEIEP | 15 | Q9BYW2 | Histone-lysine N-methyltransferase SETD | SETD2 | 287.597 | 2 | 56.017868 | yes | 1.523E-07 | 813.39886 | 41.427814 |
| KLFEQEVAQR | 10 | Q9BYW2 | Histone-lysine N-methyltransferase SETD | SETD2 | 287.597 | 2 | 56.017868 | yes | 1.624E-07 | 624.34076 | 38.705818 |
| SDPLGHPNSEETVI | 14 | Q9BYW2 | Histone-lysine N-methyltransferase SETD | SETD2 | 287.597 | 2 | 56.017868 | yes | 4.96E-07 | 755.36261 | 38.542313 |
| TAEADTSSELAK | 12 | Q9BYW2 | Histone-lysine N-methyltransferase SETD | SETD2 | 287.597 | 2 | 56.017868 | yes | 6.147E-07 | 611.79352 | 36.916702 |
| AQGEIACGEDCLN | 14 | Q9BYW2 | Histone-lysine N-methyltransferase SETD | SETD2 | 287.597 | 2 | 56.017868 | yes | 1.784E-06 | 796.8432 | 23.3482 |
| SEFITHDSHDSIK | 13 | Q9BYW2 | Histone-lysine N-methyltransferase SETD | SETD2 | 287.597 | 2 | 56.017868 | yes | 2.688E-07 | 758.35736 | 19.261639 |
| LQSVVVVPK | 9 | Q9BYW2 | Histone-lysine N-methyltransferase SETD | SETD2 | 287.597 | 2 | 56.017868 | yes | 3.094E-05 | 484.81058 | 9.818018 |
| LIEPLDYENVIVQK | 14 | Q9BZ29 | Dedicator of cytokinesis protein 9 OS=Hc | DOCK9 | 236.446 | 2 | 35.353886 | yes | 3.799E-05 | 836.96143 | 39.642273 |
| GSLISTDSGNSLPE | 15 | Q9BZ29 | Dedicator of cytokinesis protein 9 OS=Hc | DOCK9 | 236.446 | 2 | 35.353886 | yes | 8.454E-07 | 766.88116 | 30.750118 |
| EQVETVPQATVK | 12 | Q9BZ95 | Histone-lysine N-methyltransferase NSD3 | NSD3 | 161.613 | 2 | 53.767124 | yes | 9.722E-08 | 664.85645 | 230.76656 |
| SLPASITMHK | 10 | Q9BZ95 | Histone-lysine N-methyltransferase NSD3 | NSD3 | 161.613 | 2 | 53.767124 | yes | 2.527E-07 | 542.79462 | 146.07153 |
| AQWDIGIAHAEK | 12 | Q9BZ95 | Histone-lysine N-methyltransferase NSD3 | NSD3 | 161.613 | 2 | 53.767124 | yes | 1.523E-07 | 669.84369 | 104.09449 |
| QYEELLAEATK | 11 | Q9BZ95 | Histone-lysine N-methyltransferase NSD3 | NSD3 | 161.613 | 2 | 53.767124 | yes | 9.722E-08 | 647.8299 | 66.827461 |
| ELFESSLCGDLLNI | 22 | Q9BZ95 | Histone-lysine N-methyltransferase NSD3 | NSD3 | 161.613 | 3 | 53.767124 | yes | 1.466E-07 | 836.06372 | 66.311966 |
| IEQYTFIYIDK | 11 | Q9BZ95 | Histone-lysine N-methyltransferase NSD3 | NSD3 | 161.613 | 2 | 53.767124 | yes | 2.024E-06 | 716.87158 | 54.291412 |

| | | | | | | | | | | | |
|---|---|---|---|---|---|---|---|---|---|---|---|
| FIDQFVYSTK | 10 | Q9BZ95 | Histone-lysine N-methyltransferase NSD3 | NSD3 | 161.613 | 2 | 53.767124 | yes | 5.848E-06 | 624.31897 | 42.543175 |
| VQIQVADLSEIPR | 13 | Q9BZ95 | Histone-lysine N-methyltransferase NSD3 | NSD3 | 161.613 | 2 | 53.767124 | yes | 5.312E-06 | 734.41193 | 22.104769 |
| LHLAIPTR | 8 | Q9BZE4 | GTP-binding protein 4 OS=Homo sapiens | GTPBP4 | 73.965 | 2 | 53.346222 | yes | 1.3E-07 | 460.78745 | 3574.4348 |
| TLLLCGYPNVGK | 12 | Q9BZE4 | GTP-binding protein 4 OS=Homo sapiens | GTPBP4 | 73.965 | 2 | 53.346222 | yes | 1.034E-07 | 667.86047 | 3115.6326 |
| LPTIDPNTR | 9 | Q9BZE4 | GTP-binding protein 4 OS=Homo sapiens | GTPBP4 | 73.965 | 2 | 53.346222 | yes | 3.715E-07 | 513.78253 | 2695.2209 |
| LSQILTDFPK | 10 | Q9BZE4 | GTP-binding protein 4 OS=Homo sapiens | GTPBP4 | 73.965 | 2 | 53.346222 | yes | 9.901E-08 | 581.32935 | 1913.3611 |
| YWDLMNLSEK | 10 | Q9BZE4 | GTP-binding protein 4 OS=Homo sapiens | GTPBP4 | 73.965 | 2 | 53.346222 | yes | 3.782E-07 | 649.80792 | 1911.7067 |
| SLFVGHMDYK | 10 | Q9BZE4 | GTP-binding protein 4 OS=Homo sapiens | GTPBP4 | 73.965 | 2 | 53.346222 | yes | 1.4E-07 | 598.79205 | 914.48163 |
| SLFVGHMDYK | 10 | Q9BZE4 | GTP-binding protein 4 OS=Homo sapiens | GTPBP4 | 73.965 | 2 | 53.346222 | yes | 1.611E-07 | 606.78955 | 914.48163 |
| LALGQINIAK | 10 | Q9BZE4 | GTP-binding protein 4 OS=Homo sapiens | GTPBP4 | 73.965 | 2 | 53.346222 | yes | 2.391E-07 | 520.82678 | 1769.63 |
| IAELSEDDQK | 10 | Q9BZE4 | GTP-binding protein 4 OS=Homo sapiens | GTPBP4 | 73.965 | 2 | 53.346222 | yes | 1.005E-07 | 574.27753 | 1754.3488 |
| ADVDVQPYAFTTK | 13 | Q9BZE4 | GTP-binding protein 4 OS=Homo sapiens | GTPBP4 | 73.965 | 2 | 53.346222 | yes | 9.809E-08 | 727.86176 | 1462.1038 |
| EQLELFQNIR | 10 | Q9BZE4 | GTP-binding protein 4 OS=Homo sapiens | GTPBP4 | 73.965 | 2 | 53.346222 | yes | 9.732E-08 | 645.34607 | 1447.3528 |
| QSLEYLEQVR | 10 | Q9BZE4 | GTP-binding protein 4 OS=Homo sapiens | GTPBP4 | 73.965 | 2 | 53.346222 | yes | 1.232E-07 | 632.83026 | 1350.9596 |
| PLFINKPLIVVANK | 14 | Q9BZE4 | GTP-binding protein 4 OS=Homo sapiens | GTPBP4 | 73.965 | 3 | 53.346222 | yes | 6.92E-07 | 522.66412 | 435.04691 |
| PLFINKPLIVVANK | 14 | Q9BZE4 | GTP-binding protein 4 OS=Homo sapiens | GTPBP4 | 73.965 | 3 | 53.346222 | yes | 3.676E-07 | 783.49249 | 435.04691 |
| PLFINKPLIVVANK | 14 | Q9BZE4 | GTP-binding protein 4 OS=Homo sapiens | GTPBP4 | 73.965 | 2 | 53.346222 | yes | 6.92E-07 | 522.66412 | 435.04691 |
| PLFINKPLIVVANK | 14 | Q9BZE4 | GTP-binding protein 4 OS=Homo sapiens | GTPBP4 | 73.965 | 2 | 53.346222 | yes | 3.676E-07 | 783.49249 | 435.04691 |
| SLGVDMDDKDDA | 19 | Q9BZE4 | GTP-binding protein 4 OS=Homo sapiens | GTPBP4 | 73.965 | 3 | 53.346222 | yes | 1.049E-07 | 702.65674 | 838.38672 |
| KITVVPSAK | 9 | Q9BZE4 | GTP-binding protein 4 OS=Homo sapiens | GTPBP4 | 73.965 | 2 | 53.346222 | yes | 1.206E-06 | 471.80276 | 683.38281 |
| EDSAPPSSVAR | 11 | Q9BZE4 | GTP-binding protein 4 OS=Homo sapiens | GTPBP4 | 73.965 | 2 | 53.346222 | yes | 7.147E-07 | 558.27002 | 569.89233 |
| WQVVDTPGILDHI | 17 | Q9BZE4 | GTP-binding protein 4 OS=Homo sapiens | GTPBP4 | 73.965 | 3 | 53.346222 | yes | 2.823E-07 | 664.0058 | 431.68671 |
| DFIDLTLSK | 9 | Q9BZE4 | GTP-binding protein 4 OS=Homo sapiens | GTPBP4 | 73.965 | 2 | 53.346222 | yes | 1.682E-07 | 526.28711 | 417.8551 |
| YGDSLYR | 7 | Q9BZE4 | GTP-binding protein 4 OS=Homo sapiens | GTPBP4 | 73.965 | 2 | 53.346222 | yes | 9.722E-08 | 437.20868 | 398.71683 |
| HVFDMKPK | 8 | Q9BZE4 | GTP-binding protein 4 OS=Homo sapiens | GTPBP4 | 73.965 | 2 | 53.346222 | yes | 2.877E-07 | 501.26547 | 386.21051 |
| ITVVPSAK | 8 | Q9BZE4 | GTP-binding protein 4 OS=Homo sapiens | GTPBP4 | 73.965 | 2 | 53.346222 | yes | 2.344E-07 | 407.75528 | 271.98505 |
| NLVDNVAK | 8 | Q9BZE4 | GTP-binding protein 4 OS=Homo sapiens | GTPBP4 | 73.965 | 2 | 53.346222 | yes | 1.996E-07 | 436.74545 | 268.76202 |
| METEESR | 7 | Q9BZE4 | GTP-binding protein 4 OS=Homo sapiens | GTPBP4 | 73.965 | 2 | 53.346222 | yes | 1.585E-06 | 441.1871 | 192.51167 |
| HDKIPEIWEGHNIA | 23 | Q9BZE4 | GTP-binding protein 4 OS=Homo sapiens | GTPBP4 | 73.965 | 4 | 53.346222 | yes | 9.722E-08 | 677.09027 | 174.48695 |
| KLEELEKEEELR | 12 | Q9BZE4 | GTP-binding protein 4 OS=Homo sapiens | GTPBP4 | 73.965 | 2 | 53.346222 | yes | 1.232E-07 | 772.91193 | 153.2977 |
| ERPPFIPEGVVAR | 13 | Q9BZE4 | GTP-binding protein 4 OS=Homo sapiens | GTPBP4 | 73.965 | 2 | 53.346222 | yes | 3.498E-06 | 733.90936 | 140.5696 |
| PLIVVANK | 8 | Q9BZE4 | GTP-binding protein 4 OS=Homo sapiens | GTPBP4 | 73.965 | 2 | 53.346222 | yes | 3.972E-07 | 427.27893 | 102.07539 |
| ERDLELEMGDDYI | 18 | Q9BZE4 | GTP-binding protein 4 OS=Homo sapiens | GTPBP4 | 73.965 | 3 | 53.346222 | yes | 3.104E-06 | 732.35608 | 76.333633 |
| LEELEKEEELR | 11 | Q9BZE4 | GTP-binding protein 4 OS=Homo sapiens | GTPBP4 | 73.965 | 2 | 53.346222 | yes | 3.531E-06 | 708.86444 | 13.588562 |
| LLQNLEILQR | 10 | Q9BZF3 | Oxysterol-binding protein-related protein | OSBPL6 | 106.306 | 2 | 52.649967 | yes | 9.722E-08 | 620.37463 | 147.98746 |
| FFVLDNGMLK | 10 | Q9BZF3 | Oxysterol-binding protein-related protein | OSBPL6 | 106.306 | 2 | 52.649967 | yes | 3.326E-07 | 592.31262 | 34.771435 |
| ASETDDPYER | 10 | Q9BZF3 | Oxysterol-binding protein-related protein | OSBPL6 | 106.306 | 2 | 52.649967 | yes | 7.416E-06 | 591.74908 | 19.22966 |
| SLTTITELQR | 10 | Q9BZF9 | Uveal autoantigen with coiled-coil domai | UACA | 162.506 | 2 | 44.330166 | yes | 2.234E-07 | 581.32733 | 95.580429 |
| SDLETQISSLNEK | 13 | Q9BZF9 | Uveal autoantigen with coiled-coil domai | UACA | 162.506 | 2 | 44.330166 | yes | 6.109E-07 | 732.36481 | 62.638744 |

| | | | | | | | | | | | |
|---|---|---|---|---|---|---|---|---|---|---|---|
| SHDAIIDDLNR | 11 | Q9BZF9 | Uveal autoantigen with coiled-coil domai | UACA | 162.506 | 2 | 44.330166 | yes | 1.416E-06 | 634.81512 | 41.868977 |
| ELSMCEMER | 9 | Q9BZF9 | Uveal autoantigen with coiled-coil domai | UACA | 162.506 | 2 | 44.330166 | yes | 1.263E-06 | 592.74084 | 12.372897 |
| VLAELPQCLR | 10 | Q9BZG1 | Ras-related protein Rab-34 OS=Homo sa | RAB34 | 29.045 | 2 | 52.399525 | yes | 1.087E-07 | 599.83429 | 537.04248 |
| VIVVGDLSVGK | 11 | Q9BZG1 | Ras-related protein Rab-34 OS=Homo sa | RAB34 | 29.045 | 2 | 52.399525 | yes | 8.396E-07 | 543.33191 | 219.8817 |
| DALQVAQEMK | 10 | Q9BZG1 | Ras-related protein Rab-34 OS=Homo sa | RAB34 | 29.045 | 2 | 52.399525 | yes | 9.722E-08 | 566.78699 | 211.58328 |
| AEYWAVSSLTGEN | 15 | Q9BZG1 | Ras-related protein Rab-34 OS=Homo sa | RAB34 | 29.045 | 2 | 52.399525 | yes | 1.298E-07 | 841.41266 | 126.2478 |
| QWLADALK | 8 | Q9BZG1 | Ras-related protein Rab-34 OS=Homo sa | RAB34 | 29.045 | 2 | 52.399525 | yes | 1.088E-07 | 472.76364 | 114.48016 |
| INSDDSNLYLTASI | 14 | Q9BZG1 | Ras-related protein Rab-34 OS=Homo sa | RAB34 | 29.045 | 2 | 52.399525 | yes | 9.722E-08 | 770.87811 | 73.891556 |
| ATIGVDFEMER | 11 | Q9BZG1 | Ras-related protein Rab-34 OS=Homo sa | RAB34 | 29.045 | 2 | 52.399525 | yes | 9.722E-08 | 634.30298 | 72.271797 |
| VILASDDGCIR | 11 | Q9BZH6 | WD repeat-containing protein 11 OS=Ho | WDR11 | 136.685 | 2 | 51.42248 | yes | 9.927E-08 | 609.81097 | 3250.3142 |
| DKPLELWDVR | 10 | Q9BZH6 | WD repeat-containing protein 11 OS=Ho | WDR11 | 136.685 | 2 | 51.42248 | yes | 1.276E-07 | 635.84314 | 2833.2214 |
| NELQLVDLPTGR | 12 | Q9BZH6 | WD repeat-containing protein 11 OS=Ho | WDR11 | 136.685 | 2 | 51.42248 | yes | 1.367E-07 | 677.86987 | 2446.6416 |
| LITAIYADYAR | 11 | Q9BZH6 | WD repeat-containing protein 11 OS=Ho | WDR11 | 136.685 | 2 | 51.42248 | yes | 1.003E-07 | 635.34552 | 2444.2891 |
| MCPPLTTK | 8 | Q9BZH6 | WD repeat-containing protein 11 OS=Ho | WDR11 | 136.685 | 2 | 51.42248 | yes | 1.605E-07 | 474.23807 | 2093.3489 |
| CTDQLLLLGQTDR | 13 | Q9BZH6 | WD repeat-containing protein 11 OS=Ho | WDR11 | 136.685 | 2 | 51.42248 | yes | 1.063E-07 | 766.8905 | 2074.8892 |
| LPDLSLDNMIGQS. | 22 | Q9BZH6 | WD repeat-containing protein 11 OS=Ho | WDR11 | 136.685 | 3 | 51.42248 | yes | 1.127E-07 | 788.39349 | 1954.6584 |
| SELSQNISAR | 10 | Q9BZH6 | WD repeat-containing protein 11 OS=Ho | WDR11 | 136.685 | 2 | 51.42248 | yes | 1.084E-07 | 552.78583 | 1793.7109 |
| ILDVDWCTSDK | 11 | Q9BZH6 | WD repeat-containing protein 11 OS=Ho | WDR11 | 136.685 | 2 | 51.42248 | yes | 1.014E-07 | 676.31354 | 1713.394 |
| EVQMVSSLR | 9 | Q9BZH6 | WD repeat-containing protein 11 OS=Ho | WDR11 | 136.685 | 2 | 51.42248 | yes | 9.809E-08 | 524.77643 | 1586.6932 |
| AVQLLLETSADNQ | 21 | Q9BZH6 | WD repeat-containing protein 11 OS=Ho | WDR11 | 136.685 | 3 | 51.42248 | yes | 1.594E-07 | 823.39691 | 1203.1202 |
| NHMLLLYPR | 9 | Q9BZH6 | WD repeat-containing protein 11 OS=Ho | WDR11 | 136.685 | 2 | 51.42248 | yes | 1.075E-07 | 578.81842 | 1192.8372 |
| GNDESAIEMIK | 11 | Q9BZH6 | WD repeat-containing protein 11 OS=Ho | WDR11 | 136.685 | 2 | 51.42248 | yes | 3.371E-07 | 603.78717 | 1076.3296 |
| QYLAVVFR | 8 | Q9BZH6 | WD repeat-containing protein 11 OS=Ho | WDR11 | 136.685 | 2 | 51.42248 | yes | 2.322E-07 | 498.28726 | 1068.3993 |
| DLLNELESPK | 10 | Q9BZH6 | WD repeat-containing protein 11 OS=Ho | WDR11 | 136.685 | 2 | 51.42248 | yes | 1.747E-07 | 579.30603 | 863.09509 |
| TLTGALNAHNK | 11 | Q9BZH6 | WD repeat-containing protein 11 OS=Ho | WDR11 | 136.685 | 2 | 51.42248 | yes | 9.722E-08 | 570.31201 | 816.18982 |
| YGAFEVTEDTEK | 12 | Q9BZH6 | WD repeat-containing protein 11 OS=Ho | WDR11 | 136.685 | 2 | 51.42248 | yes | 1.009E-07 | 694.81445 | 709.19122 |
| LNPEECADVLR | 11 | Q9BZH6 | WD repeat-containing protein 11 OS=Ho | WDR11 | 136.685 | 2 | 51.42248 | yes | 3.011E-07 | 658.31915 | 678.1496 |
| LAEGVQLLCLIDK | 13 | Q9BZH6 | WD repeat-containing protein 11 OS=Ho | WDR11 | 136.685 | 2 | 51.42248 | yes | 2.69E-07 | 736.41309 | 598.5036 |
| ELSIHSCEVK | 10 | Q9BZH6 | WD repeat-containing protein 11 OS=Ho | WDR11 | 136.685 | 2 | 51.42248 | yes | 9.722E-08 | 601.29773 | 452.58179 |
| AALFVEACLK | 10 | Q9BZH6 | WD repeat-containing protein 11 OS=Ho | WDR11 | 136.685 | 2 | 51.42248 | yes | 1.022E-07 | 561.30481 | 428.32416 |
| SQCDAIR | 7 | Q9BZH6 | WD repeat-containing protein 11 OS=Ho | WDR11 | 136.685 | 2 | 51.42248 | yes | 2.066E-07 | 425.19781 | 373.74326 |
| IIVWDVAAGVAQC | 20 | Q9BZH6 | WD repeat-containing protein 11 OS=Ho | WDR11 | 136.685 | 3 | 51.42248 | yes | 1.21E-07 | 746.38428 | 287.53888 |
| YLQTYGEWNR | 10 | Q9BZH6 | WD repeat-containing protein 11 OS=Ho | WDR11 | 136.685 | 2 | 51.42248 | yes | 1.069E-07 | 665.31476 | 241.608 |
| VYISSPHSSPAHNI | 14 | Q9BZH6 | WD repeat-containing protein 11 OS=Ho | WDR11 | 136.685 | 2 | 51.42248 | yes | 1.381E-07 | 762.38367 | 116.42345 |
| VYISSPHSSPAHNI | 14 | Q9BZH6 | WD repeat-containing protein 11 OS=Ho | WDR11 | 136.685 | 2 | 51.42248 | yes | 4.621E-07 | 508.59155 | 116.42345 |
| VYISSPHSSPAHNI | 14 | Q9BZH6 | WD repeat-containing protein 11 OS=Ho | WDR11 | 136.685 | 3 | 51.42248 | yes | 1.381E-07 | 762.38367 | 116.42345 |
| VYISSPHSSPAHNI | 14 | Q9BZH6 | WD repeat-containing protein 11 OS=Ho | WDR11 | 136.685 | 3 | 51.42248 | yes | 4.621E-07 | 508.59155 | 116.42345 |
| LASADVNGK | 9 | Q9BZH6 | WD repeat-containing protein 11 OS=Ho | WDR11 | 136.685 | 2 | 51.42248 | yes | 2.2E-07 | 437.73508 | 200.69832 |
| ILITQEK | 7 | Q9BZH6 | WD repeat-containing protein 11 OS=Ho | WDR11 | 136.685 | 2 | 51.42248 | yes | 4.322E-07 | 422.76056 | 180.27046 |

| | | | | | | | | | | | |
|---|---|---|---|---|---|---|---|---|---|---|---|
| KCTDQLLLLGQTD | 14 | Q9BZH6 | WD repeat-containing protein 11 OS=Ho | WDR11 | 136.685 | 3 | 51.42248 | yes | 2.025E-07 | 554.29443 | 85.438759 |
| KCTDQLLLLGQTD | 14 | Q9BZH6 | WD repeat-containing protein 11 OS=Ho | WDR11 | 136.685 | 3 | 51.42248 | yes | 1.012E-07 | 830.93799 | 85.438759 |
| KCTDQLLLLGQTD | 14 | Q9BZH6 | WD repeat-containing protein 11 OS=Ho | WDR11 | 136.685 | 2 | 51.42248 | yes | 2.025E-07 | 554.29443 | 85.438759 |
| KCTDQLLLLGQTD | 14 | Q9BZH6 | WD repeat-containing protein 11 OS=Ho | WDR11 | 136.685 | 2 | 51.42248 | yes | 1.012E-07 | 830.93799 | 85.438759 |
| NFPTITALEWSPSH | 17 | Q9BZH6 | WD repeat-containing protein 11 OS=Ho | WDR11 | 136.685 | 3 | 51.42248 | yes | 1.589E-07 | 652.33887 | 115.75089 |
| WVDHLCSPQVNQ | 13 | Q9BZH6 | WD repeat-containing protein 11 OS=Ho | WDR11 | 136.685 | 2 | 51.42248 | yes | 2.014E-07 | 805.89081 | 76.422119 |
| LVATNMIANGK | 11 | Q9BZH6 | WD repeat-containing protein 11 OS=Ho | WDR11 | 136.685 | 2 | 51.42248 | yes | 9.722E-08 | 566.31317 | 49.578907 |
| IPPDGSMGSITCIA | 16 | Q9BZH6 | WD repeat-containing protein 11 OS=Ho | WDR11 | 136.685 | 3 | 51.42248 | yes | 1.323E-05 | 578.28473 | 46.358898 |
| VLEMSMK | 7 | Q9BZH6 | WD repeat-containing protein 11 OS=Ho | WDR11 | 136.685 | 2 | 51.42248 | yes | 8.682E-07 | 419.21408 | 23.145832 |
| NQEDIILFR | 9 | Q9BZI7 | Regulator of nonsense transcripts 3B OS= | UPF3B | 57.763 | 2 | 54.082207 | yes | 1.076E-07 | 574.30896 | 4765.4214 |
| VGTIDDDPEYR | 11 | Q9BZI7 | Regulator of nonsense transcripts 3B OS= | UPF3B | 57.763 | 2 | 54.082207 | yes | 9.722E-08 | 640.2937 | 2653.4941 |
| LCPPDDSTK | 9 | Q9BZI7 | Regulator of nonsense transcripts 3B OS= | UPF3B | 57.763 | 2 | 54.082207 | yes | 3.286E-06 | 516.73694 | 2422.8684 |
| MTSTPETLLEEIEA | 15 | Q9BZI7 | Regulator of nonsense transcripts 3B OS= | UPF3B | 57.763 | 2 | 54.082207 | yes | 1.004E-07 | 846.42401 | 1518.261 |
| ASGQSCTLPK | 10 | Q9BZI7 | Regulator of nonsense transcripts 3B OS= | UPF3B | 57.763 | 2 | 54.082207 | yes | 9.722E-08 | 524.75824 | 1080.5081 |
| FDGYVFLDNK | 10 | Q9BZI7 | Regulator of nonsense transcripts 3B OS= | UPF3B | 57.763 | 2 | 54.082207 | yes | 1.296E-07 | 609.29547 | 1009.9706 |
| DRFDGYVFLDNK | 12 | Q9BZI7 | Regulator of nonsense transcripts 3B OS= | UPF3B | 57.763 | 2 | 54.082207 | yes | 5.418E-07 | 744.8595 | 821.02679 |
| DRPAMQLYQPGA | 13 | Q9BZI7 | Regulator of nonsense transcripts 3B OS= | UPF3B | 57.763 | 2 | 54.082207 | yes | 2.174E-07 | 751.88025 | 289.04584 |
| FLESYATDNEK | 11 | Q9BZI7 | Regulator of nonsense transcripts 3B OS= | UPF3B | 57.763 | 2 | 54.082207 | yes | 1.048E-07 | 658.80389 | 275.36951 |
| KTTPLLSFLK | 10 | Q9BZI7 | Regulator of nonsense transcripts 3B OS= | UPF3B | 57.763 | 2 | 54.082207 | yes | 1.566E-07 | 574.35791 | 241.49156 |
| EQLQEHLQPMPEH | 32 | Q9BZI7 | Regulator of nonsense transcripts 3B OS= | UPF3B | 57.763 | 5 | 54.082207 | yes | 9.722E-08 | 800.76062 | 223.25516 |
| KAESTESIGSSEK | 13 | Q9BZI7 | Regulator of nonsense transcripts 3B OS= | UPF3B | 57.763 | 2 | 54.082207 | yes | 1.602E-07 | 676.83063 | 196.54581 |
| KFLESYATDNEK | 12 | Q9BZI7 | Regulator of nonsense transcripts 3B OS= | UPF3B | 57.763 | 2 | 54.082207 | yes | 9.809E-08 | 722.85138 | 192.4892 |
| RLPPTLTK | 8 | Q9BZI7 | Regulator of nonsense transcripts 3B OS= | UPF3B | 57.763 | 2 | 54.082207 | yes | 3.025E-06 | 463.2951 | 166.74355 |
| TTPLLSFLK | 9 | Q9BZI7 | Regulator of nonsense transcripts 3B OS= | UPF3B | 57.763 | 2 | 54.082207 | yes | 1.183E-07 | 510.31042 | 159.07693 |
| PAMQLYQPGAR | 11 | Q9BZI7 | Regulator of nonsense transcripts 3B OS= | UPF3B | 57.763 | 2 | 54.082207 | yes | 2.744E-06 | 616.31622 | 108.31809 |
| RVTLLTPAGATGS | 23 | Q9BZI7 | Regulator of nonsense transcripts 3B OS= | UPF3B | 57.763 | 3 | 54.082207 | yes | 4.376E-07 | 693.02216 | 87.228409 |
| LDKENLSDER | 10 | Q9BZI7 | Regulator of nonsense transcripts 3B OS= | UPF3B | 57.763 | 2 | 54.082207 | yes | 9.722E-08 | 609.8017 | 76.101517 |
| IDRIPER | 7 | Q9BZI7 | Regulator of nonsense transcripts 3B OS= | UPF3B | 57.763 | 2 | 54.082207 | yes | 3.753E-07 | 449.75888 | 59.959129 |
| NRLCPPDDSTK | 11 | Q9BZI7 | Regulator of nonsense transcripts 3B OS= | UPF3B | 57.763 | 2 | 54.082207 | yes | 4.913E-07 | 651.80896 | 40.988941 |
| LMLLESWR | 8 | Q9BZJ0 | Crooked neck-like protein 1 OS=Homo sa | CRNKL1 | 100.452 | 2 | 53.113033 | yes | 4.4E-07 | 524.28644 | 178.47757 |
| LVESDAEAEAVR | 12 | Q9BZJ0 | Crooked neck-like protein 1 OS=Homo sa | CRNKL1 | 100.452 | 2 | 53.113033 | yes | 9.722E-08 | 644.82263 | 148.2019 |
| FVLVHPDVK | 9 | Q9BZJ0 | Crooked neck-like protein 1 OS=Homo sa | CRNKL1 | 100.452 | 2 | 53.113033 | yes | 2.144E-07 | 527.30823 | 83.538048 |
| APAEVQITAEQLLI | 14 | Q9BZJ0 | Crooked neck-like protein 1 OS=Homo sa | CRNKL1 | 100.452 | 2 | 53.113033 | yes | 2.607E-07 | 769.93048 | 73.066055 |
| ELELLPPPPQQK | 12 | Q9BZJ0 | Crooked neck-like protein 1 OS=Homo sa | CRNKL1 | 100.452 | 2 | 53.113033 | yes | 4.853E-07 | 694.89282 | 55.783459 |
| SFEEEFGTASDK | 12 | Q9BZJ0 | Crooked neck-like protein 1 OS=Homo sa | CRNKL1 | 100.452 | 2 | 53.113033 | yes | 1.984E-07 | 673.79095 | 51.670795 |
| TTIIWDAHTGEAK | 13 | Q9BZK7 | F-box-like/WD repeat-containing protein | TBL1XR1 | 55.595 | 2 | 47.432358 | no | 8.933E-07 | 721.86737 | 167.42409 |
| YLASGSFDK | 9 | Q9BZK7 | F-box-like/WD repeat-containing protein | TBL1XR1 | 55.595 | 2 | 47.432358 | no | 4.268E-07 | 494.24274 | 93.520851 |
| GICIHTLTK | 9 | Q9BZK7 | F-box-like/WD repeat-containing protein | TBL1XR1 | 55.595 | 2 | 47.432358 | yes | 1.688E-07 | 521.78931 | 84.036385 |
| LAQQQAAAAAAA | 23 | Q9BZK7 | F-box-like/WD repeat-containing protein | TBL1XR1 | 55.595 | 3 | 47.432358 | yes | 1.331E-07 | 685.6908 | 69.785606 |

| | | | | | | | | | | | |
|---|---|---|---|---|---|---|---|---|---|---|---|
| GTGGIFEVCWNAA | 16 | Q9BZK7 | F-box-like/WD repeat-containing protein | TBL1XR1 | 55.595 | 2 | 47.432358 | yes | 4.595E-07 | 841.38556 | 32.307583 |
| NEVAILQSLR | 10 | Q9BZL6 | Serine/threonine-protein kinase D2 OS=H | PRKD2 | 96.723 | 2 | 45.90796 | yes | 1.707E-07 | 571.83002 | 174.04076 |
| LFLNEVAK | 8 | Q9BZR9 | E3 ubiquitin-protein ligase TRIM8 OS=H | TRIM8 | 61.489 | 2 | 52.259785 | yes | 4.039E-07 | 467.27383 | 454.65207 |
| EQDIEDQLYK | 10 | Q9BZR9 | E3 ubiquitin-protein ligase TRIM8 OS=H | TRIM8 | 61.489 | 2 | 52.259785 | yes | 9.722E-08 | 640.80389 | 437.92603 |
| GCIGEAWAK | 9 | Q9BZR9 | E3 ubiquitin-protein ligase TRIM8 OS=H | TRIM8 | 61.489 | 2 | 52.259785 | yes | 2.497E-06 | 496.23694 | 393.25986 |
| LHQLLDEDLR | 10 | Q9BZR9 | E3 ubiquitin-protein ligase TRIM8 OS=H | TRIM8 | 61.489 | 2 | 52.259785 | yes | 1.073E-07 | 626.33826 | 344.36182 |
| GHLLVEADDVR | 11 | Q9BZR9 | E3 ubiquitin-protein ligase TRIM8 OS=H | TRIM8 | 61.489 | 2 | 52.259785 | yes | 1.602E-07 | 612.32257 | 257.82767 |
| TQTCTSSSLSPTK | 13 | Q9BZR9 | E3 ubiquitin-protein ligase TRIM8 OS=H | TRIM8 | 61.489 | 2 | 52.259785 | yes | 1.786E-07 | 699.33246 | 172.66943 |
| LTNIVEK | 7 | Q9BZR9 | E3 ubiquitin-protein ligase TRIM8 OS=H | TRIM8 | 61.489 | 2 | 52.259785 | yes | 3.241E-07 | 408.7449 | 86.500214 |
| RGPPLPAQK | 9 | Q9BZR9 | E3 ubiquitin-protein ligase TRIM8 OS=H | TRIM8 | 61.489 | 2 | 52.259785 | yes | 1.651E-07 | 482.29034 | 84.365097 |
| FNALHVEKPPAAL | 19 | Q9BZR9 | E3 ubiquitin-protein ligase TRIM8 OS=H | TRIM8 | 61.489 | 3 | 52.259785 | yes | 1.915E-06 | 756.0506 | 77.130005 |
| TEDVSFMK | 8 | Q9BZR9 | E3 ubiquitin-protein ligase TRIM8 OS=H | TRIM8 | 61.489 | 2 | 52.259785 | yes | 4.718E-07 | 478.72333 | 67.777039 |
| FCSENAAQALHLG | 15 | Q9BZR9 | E3 ubiquitin-protein ligase TRIM8 OS=H | TRIM8 | 61.489 | 3 | 52.259785 | yes | 6.668E-07 | 568.27039 | 62.635208 |
| ILVCSVDNCYCSS | 25 | Q9BZR9 | E3 ubiquitin-protein ligase TRIM8 OS=H | TRIM8 | 61.489 | 4 | 52.259785 | yes | 4E-07 | 723.32593 | 52.22876 |
| AWSCPQHNAYR | 11 | Q9BZR9 | E3 ubiquitin-protein ligase TRIM8 OS=H | TRIM8 | 61.489 | 2 | 52.259785 | yes | 9.722E-08 | 695.30945 | 41.32119 |
| IVQLLGQNEVDYR | 13 | Q9BZX2 | Uridine-cytidine kinase 2 OS=Homo sapi | UCK2 | 29.3 | 2 | 53.99298 | yes | 1.655E-07 | 773.91486 | 651.36615 |
| LFVDTDADTR | 10 | Q9BZX2 | Uridine-cytidine kinase 2 OS=Homo sapi | UCK2 | 29.3 | 2 | 53.99298 | yes | 3.184E-07 | 576.78021 | 628.05292 |
| QVVILSQDSFYR | 12 | Q9BZX2 | Uridine-cytidine kinase 2 OS=Homo sapi | UCK2 | 29.3 | 2 | 53.99298 | yes | 9.722E-08 | 727.88556 | 213.28267 |
| YADVIIPR | 8 | Q9BZX2 | Uridine-cytidine kinase 2 OS=Homo sapi | UCK2 | 29.3 | 2 | 53.99298 | no | 1.478E-07 | 473.77145 | 125.77023 |
| ELPQFATGENLPR | 13 | Q9BZZ5 | Apoptosis inhibitor 5 OS=Homo sapiens ( | API5 | 59.005 | 2 | 51.319393 | yes | 1.148E-06 | 736.38062 | 99.670471 |
| PTVEELYR | 8 | Q9BZZ5 | Apoptosis inhibitor 5 OS=Homo sapiens ( | API5 | 59.005 | 2 | 51.319393 | yes | 2.152E-07 | 503.76382 | 87.216942 |
| LLAEMSSFCGDME | 14 | Q9BZZ5 | Apoptosis inhibitor 5 OS=Homo sapiens ( | API5 | 59.005 | 2 | 51.319393 | yes | 9.722E-08 | 809.3515 | 61.77404 |
| ITNNINVLIK | 10 | Q9BZZ5 | Apoptosis inhibitor 5 OS=Homo sapiens ( | API5 | 59.005 | 2 | 51.319393 | yes | 3.927E-07 | 571.35059 | 46.971195 |
| EVEELILTESK | 11 | Q9BZZ5 | Apoptosis inhibitor 5 OS=Homo sapiens ( | API5 | 59.005 | 2 | 51.319393 | yes | 1.329E-07 | 645.3454 | 37.213226 |
| QLAAVATLLR | 10 | Q9C029 | E3 ubiquitin-protein ligase TRIM7 OS=H | TRIM7 | 56.631 | 2 | 53.08419 | yes | 2.689E-07 | 528.33221 | 173.41673 |
| EHAVLPLDEAVQE | 15 | Q9C029 | E3 ubiquitin-protein ligase TRIM7 OS=H | TRIM7 | 56.631 | 2 | 53.08419 | yes | 1.12E-07 | 824.93066 | 169.21889 |
| AICVVCDR | 8 | Q9C029 | E3 ubiquitin-protein ligase TRIM7 OS=H | TRIM7 | 56.631 | 2 | 53.08419 | yes | 2.007E-07 | 496.73624 | 165.44395 |
| AFLVEQEGR | 9 | Q9C029 | E3 ubiquitin-protein ligase TRIM7 OS=H | TRIM7 | 56.631 | 2 | 53.08419 | yes | 1.199E-07 | 524.77472 | 155.52905 |
| ELEDCEVFR | 9 | Q9C029 | E3 ubiquitin-protein ligase TRIM7 OS=H | TRIM7 | 56.631 | 2 | 53.08419 | yes | 9.722E-08 | 598.76624 | 123.05965 |
| VELTLDPDTANPR | 13 | Q9C029 | E3 ubiquitin-protein ligase TRIM7 OS=H | TRIM7 | 56.631 | 2 | 53.08419 | yes | 9.722E-08 | 720.87012 | 93.067398 |
| SPLSCGHLSR | 10 | Q9C029 | E3 ubiquitin-protein ligase TRIM7 OS=H | TRIM7 | 56.631 | 2 | 53.08419 | yes | 1.383E-07 | 557.2771 | 87.719429 |
| LYCQDDGR | 8 | Q9C029 | E3 ubiquitin-protein ligase TRIM7 OS=H | TRIM7 | 56.631 | 2 | 53.08419 | yes | 9.825E-08 | 513.71912 | 76.172241 |
| VLASCGFSSGR | 11 | Q9C029 | E3 ubiquitin-protein ligase TRIM7 OS=H | TRIM7 | 56.631 | 2 | 53.08419 | yes | 2.947E-07 | 570.77698 | 58.80632 |
| SVPTFTVIPEGPR | 13 | Q9C086 | INO80 complex subunit B OS=Homo sap | INO80B | 38.638 | 2 | 49.19765 | yes | 1.06E-07 | 700.38263 | 235.99207 |
| TGQALCSLQCYR | 12 | Q9C086 | INO80 complex subunit B OS=Homo sap | INO80B | 38.638 | 2 | 49.19765 | yes | 6.409E-07 | 728.83722 | 205.02576 |
| CSVPGCPHPR | 10 | Q9C086 | INO80 complex subunit B OS=Homo sap | INO80B | 38.638 | 2 | 49.19765 | yes | 1.531E-06 | 583.76331 | 152.56161 |
| GELDDNGDLKK | 11 | Q9C086 | INO80 complex subunit B OS=Homo sap | INO80B | 38.638 | 2 | 49.19765 | yes | 1.506E-07 | 602.2962 | 74.538879 |
| ASAAGAECELAR | 12 | Q9C0B0 | RING finger protein unkempt homolog O | UNK | 88.085 | 2 | 53.51529 | yes | 1.048E-07 | 603.28259 | 915.53381 |
| SQEQPLLQPK | 10 | Q9C0B0 | RING finger protein unkempt homolog O | UNK | 88.085 | 2 | 53.51529 | yes | 1.36E-07 | 584.32202 | 710.02606 |

| | | | | | | | | | | | |
|---|---|---|---|---|---|---|---|---|---|---|---|
| TEQQFHPEIYK | 11 | Q9C0B0 | RING finger protein unkempt homolog O | UNK | 88.085 | 2 | 53.51529 | yes | 9.722E-08 | 710.34882 | 527.22479 |
| TEQCPLFVQHK | 11 | Q9C0B0 | RING finger protein unkempt homolog O | UNK | 88.085 | 2 | 53.51529 | yes | 1.022E-07 | 693.84534 | 423.05923 |
| AVFHMQSVK | 9 | Q9C0B0 | RING finger protein unkempt homolog O | UNK | 88.085 | 2 | 53.51529 | yes | 1.642E-07 | 523.77625 | 333.83084 |
| WQETAYVLGNYK | 12 | Q9C0B0 | RING finger protein unkempt homolog O | UNK | 88.085 | 2 | 53.51529 | yes | 1.246E-07 | 736.36444 | 312.95404 |
| ILSEEPR | 7 | Q9C0B0 | RING finger protein unkempt homolog O | UNK | 88.085 | 2 | 53.51529 | yes | 3.776E-06 | 422.23218 | 270.04898 |
| KLQEELER | 8 | Q9C0B0 | RING finger protein unkempt homolog O | UNK | 88.085 | 2 | 53.51529 | yes | 1.082E-07 | 522.78784 | 262.96237 |
| TGICIHETDSK | 11 | Q9C0B0 | RING finger protein unkempt homolog O | UNK | 88.085 | 2 | 53.51529 | yes | 9.823E-08 | 630.7981 | 234.90314 |
| EDQVGAEYLK | 10 | Q9C0B0 | RING finger protein unkempt homolog O | UNK | 88.085 | 2 | 53.51529 | yes | 4.429E-07 | 576.28259 | 172.40729 |
| DGTFNYSPDVYCT | 14 | Q9C0B0 | RING finger protein unkempt homolog O | UNK | 88.085 | 2 | 53.51529 | yes | 9.722E-08 | 833.85632 | 166.58638 |
| QELDEANSTIK | 11 | Q9C0B0 | RING finger protein unkempt homolog O | UNK | 88.085 | 2 | 53.51529 | yes | 9.722E-08 | 624.30933 | 138.86818 |
| LRQELDEANSTIK | 13 | Q9C0B0 | RING finger protein unkempt homolog O | UNK | 88.085 | 2 | 53.51529 | yes | 1.013E-07 | 758.90192 | 109.29553 |
| DALEVQVK | 8 | Q9C0B0 | RING finger protein unkempt homolog O | UNK | 88.085 | 2 | 53.51529 | yes | 3.145E-06 | 451.25311 | 55.070999 |
| LQEELER | 7 | Q9C0B0 | RING finger protein unkempt homolog O | UNK | 88.085 | 2 | 53.51529 | no | 1.967E-06 | 458.74036 | 52.211884 |
| GVGSPEPGPTAPY | 16 | Q9C0B5 | Palmitoyltransferase ZDHHC5 OS=Homo | ZDHHC5 | 77.545 | 2 | 42.074017 | yes | 3.205E-07 | 777.89917 | 41.788837 |
| VSAPGVLTAQDR | 12 | Q9C0C2 | 182 kDa tankyrase-1-binding protein OS= | TNKS1BP1 | 181.796 | 2 | 56.49554 | yes | 9.722E-08 | 607.33038 | 1300.6316 |
| VNLFPGLSPSALK | 13 | Q9C0C2 | 182 kDa tankyrase-1-binding protein OS= | TNKS1BP1 | 181.796 | 2 | 56.49554 | yes | 1.29E-07 | 671.89008 | 496.01532 |
| DFCIEASER | 9 | Q9C0C2 | 182 kDa tankyrase-1-binding protein OS= | TNKS1BP1 | 181.796 | 2 | 56.49554 | yes | 1.014E-07 | 563.7453 | 442.19641 |
| DLAEVGEGGGHS( | 15 | Q9C0C2 | 182 kDa tankyrase-1-binding protein OS= | TNKS1BP1 | 181.796 | 2 | 56.49554 | yes | 9.722E-08 | 741.85022 | 403.1676 |
| FAATTVEEILAK | 12 | Q9C0C2 | 182 kDa tankyrase-1-binding protein OS= | TNKS1BP1 | 181.796 | 2 | 56.49554 | yes | 9.722E-08 | 646.85846 | 349.90317 |
| EAAFSPGQQDWSF | 13 | Q9C0C2 | 182 kDa tankyrase-1-binding protein OS= | TNKS1BP1 | 181.796 | 2 | 56.49554 | yes | 1.254E-07 | 739.83661 | 345.33487 |
| ESAVGQMGWSGG | 16 | Q9C0C2 | 182 kDa tankyrase-1-binding protein OS= | TNKS1BP1 | 181.796 | 2 | 56.49554 | yes | 1.074E-07 | 817.90137 | 342.75824 |
| GVGQADWTPDLG | 14 | Q9C0C2 | 182 kDa tankyrase-1-binding protein OS= | TNKS1BP1 | 181.796 | 2 | 56.49554 | yes | 1.047E-07 | 742.87823 | 334.07431 |
| SSGSEGSSPNWLQ | 16 | Q9C0C2 | 182 kDa tankyrase-1-binding protein OS= | TNKS1BP1 | 181.796 | 2 | 56.49554 | yes | 9.79E-08 | 824.40228 | 329.36472 |
| SQEADVQDWEFR | 12 | Q9C0C2 | 182 kDa tankyrase-1-binding protein OS= | TNKS1BP1 | 181.796 | 2 | 56.49554 | yes | 9.722E-08 | 755.33386 | 314.55713 |
| AAEPQEQEFGK | 11 | Q9C0C2 | 182 kDa tankyrase-1-binding protein OS= | TNKS1BP1 | 181.796 | 2 | 56.49554 | yes | 1.722E-07 | 617.29095 | 309.54633 |
| KIPSVEDSLGEGSF | 14 | Q9C0C2 | 182 kDa tankyrase-1-binding protein OS= | TNKS1BP1 | 181.796 | 2 | 56.49554 | yes | 9.722E-08 | 737.3808 | 251.74271 |
| DVSLGDWEFGK | 11 | Q9C0C2 | 182 kDa tankyrase-1-binding protein OS= | TNKS1BP1 | 181.796 | 2 | 56.49554 | yes | 1.795E-07 | 626.79584 | 242.47382 |
| TTAGSVDWTDQL( | 15 | Q9C0C2 | 182 kDa tankyrase-1-binding protein OS= | TNKS1BP1 | 181.796 | 2 | 56.49554 | yes | 1.031E-07 | 810.40485 | 223.9279 |
| GWVGEFSLSVGPQ | 14 | Q9C0C2 | 182 kDa tankyrase-1-binding protein OS= | TNKS1BP1 | 181.796 | 2 | 56.49554 | yes | 9.722E-08 | 759.88861 | 222.33702 |
| DWTSDVNVK | 9 | Q9C0C2 | 182 kDa tankyrase-1-binding protein OS= | TNKS1BP1 | 181.796 | 2 | 56.49554 | yes | 3.001E-07 | 532.25635 | 208.80531 |
| SYQFGIIGNDR | 11 | Q9C0C2 | 182 kDa tankyrase-1-binding protein OS= | TNKS1BP1 | 181.796 | 2 | 56.49554 | yes | 1.006E-07 | 635.31476 | 180.94743 |
| FSEGVLQSPSQDQ | 15 | Q9C0C2 | 182 kDa tankyrase-1-binding protein OS= | TNKS1BP1 | 181.796 | 2 | 56.49554 | yes | 9.722E-08 | 839.8996 | 171.48006 |
| APVIRPGGTLGLSE | 30 | Q9C0C2 | 182 kDa tankyrase-1-binding protein OS= | TNKS1BP1 | 181.796 | 4 | 56.49554 | yes | 2.085E-07 | 777.64026 | 146.99559 |
| DVSLGTYGSR | 10 | Q9C0C2 | 182 kDa tankyrase-1-binding protein OS= | TNKS1BP1 | 181.796 | 2 | 56.49554 | yes | 9.722E-08 | 527.76184 | 140.733 |
| MNMLAGPQPYGG | 14 | Q9C0C2 | 182 kDa tankyrase-1-binding protein OS= | TNKS1BP1 | 181.796 | 2 | 56.49554 | yes | 9.722E-08 | 725.84448 | 137.49409 |
| VVGKPAQLGTQR | 12 | Q9C0C2 | 182 kDa tankyrase-1-binding protein OS= | TNKS1BP1 | 181.796 | 2 | 56.49554 | yes | 1.154E-07 | 627.36987 | 129.99051 |
| MEGGHFVPPGK | 11 | Q9C0C2 | 182 kDa tankyrase-1-binding protein OS= | TNKS1BP1 | 181.796 | 2 | 56.49554 | yes | 1.214E-07 | 578.28442 | 121.64574 |
| SAEEGELAESK | 11 | Q9C0C2 | 182 kDa tankyrase-1-binding protein OS= | TNKS1BP1 | 181.796 | 2 | 56.49554 | yes | 9.722E-08 | 575.26715 | 116.45551 |
| KAPAPFRPASER | 12 | Q9C0C2 | 182 kDa tankyrase-1-binding protein OS= | TNKS1BP1 | 181.796 | 3 | 56.49554 | yes | 2.889E-07 | 442.91409 | 90.262611 |

| | | | | | | | | | | | |
|---|---|---|---|---|---|---|---|---|---|---|---|
| LTFNHDGSSR | 10 | Q9C0C2 | 182 kDa tankyrase-1-binding protein OS= | TNKS1BP1 | 181.796 | 2 | 56.49554 | yes | 1.417E-07 | 567.27032 | 89.845932 |
| DAELQDQEFGK | 11 | Q9C0C2 | 182 kDa tankyrase-1-binding protein OS= | TNKS1BP1 | 181.796 | 2 | 56.49554 | yes | 9.722E-08 | 640.2937 | 84.542328 |
| VSGAGFSPSSK | 11 | Q9C0C2 | 182 kDa tankyrase-1-binding protein OS= | TNKS1BP1 | 181.796 | 2 | 56.49554 | yes | 4.559E-07 | 512.25891 | 76.929123 |
| RPLPFAPR | 8 | Q9C0C2 | 182 kDa tankyrase-1-binding protein OS= | TNKS1BP1 | 181.796 | 2 | 56.49554 | yes | 9.81E-08 | 477.2876 | 71.555405 |
| DSLGTYSSR | 9 | Q9C0C2 | 182 kDa tankyrase-1-binding protein OS= | TNKS1BP1 | 181.796 | 2 | 56.49554 | yes | 1.685E-07 | 493.23291 | 68.742493 |
| PSLLVPVGPRPPR | 13 | Q9C0C2 | 182 kDa tankyrase-1-binding protein OS= | TNKS1BP1 | 181.796 | 3 | 56.49554 | yes | 3.142E-06 | 462.28561 | 31.871357 |
| NLEVSSCVGSGGS | 17 | Q9C0C2 | 182 kDa tankyrase-1-binding protein OS= | TNKS1BP1 | 181.796 | 2 | 56.49554 | yes | 1.094E-07 | 848.38379 | 26.434259 |
| ALPAKPALPAK | 11 | Q9C0C2 | 182 kDa tankyrase-1-binding protein OS= | TNKS1BP1 | 181.796 | 2 | 56.49554 | yes | 8.651E-07 | 538.84497 | 23.593134 |
| DEDGSTLFR | 9 | Q9C0C2 | 182 kDa tankyrase-1-binding protein OS= | TNKS1BP1 | 181.796 | 2 | 56.49554 | yes | 4.461E-07 | 520.23816 | 16.525232 |
| LIGTNCIIYPVNSK | 14 | Q9C0C9 | (E3-independent) E2 ubiquitin-conjugatir | UBE2O | 141.294 | 2 | 39.964527 | yes | 4.284E-07 | 796.42926 | 47.121376 |
| VEVVWADNSK | 10 | Q9C0C9 | (E3-independent) E2 ubiquitin-conjugatir | UBE2O | 141.294 | 2 | 39.964527 | yes | 3.21E-06 | 573.79309 | 31.623386 |
| LEDLSYLDGQR | 11 | Q9C0D5 | Protein TANC1 OS=Homo sapiens OX=9 | TANC1 | 202.22 | 2 | 51.15563 | yes | 1.259E-07 | 654.82513 | 171.83633 |
| GAVVVGNVGFGK | 12 | Q9C0D5 | Protein TANC1 OS=Homo sapiens OX=9 | TANC1 | 202.22 | 2 | 51.15563 | yes | 2.932E-07 | 552.31403 | 147.28827 |
| TNDFGMAEEFASK | 13 | Q9C0D5 | Protein TANC1 OS=Homo sapiens OX=9 | TANC1 | 202.22 | 2 | 51.15563 | yes | 4.49E-06 | 723.8139 | 140.61848 |
| GVLEPLTNLR | 10 | Q9C0D5 | Protein TANC1 OS=Homo sapiens OX=9 | TANC1 | 202.22 | 2 | 51.15563 | yes | 4.5E-07 | 556.32715 | 124.28052 |
| FMTQSAFER | 9 | Q9C0D5 | Protein TANC1 OS=Homo sapiens OX=9 | TANC1 | 202.22 | 2 | 51.15563 | yes | 1.009E-07 | 558.76074 | 121.76829 |
| GVPPLFCAAR | 10 | Q9C0D5 | Protein TANC1 OS=Homo sapiens OX=9 | TANC1 | 202.22 | 2 | 51.15563 | yes | 1.747E-06 | 544.28949 | 98.947884 |
| LMEEGNVMYK | 10 | Q9C0D5 | Protein TANC1 OS=Homo sapiens OX=9 | TANC1 | 202.22 | 2 | 51.15563 | yes | 1.349E-07 | 607.2832 | 86.160637 |
| VSLYLNLSR | 9 | Q9C0D5 | Protein TANC1 OS=Homo sapiens OX=9 | TANC1 | 202.22 | 2 | 51.15563 | yes | 3.664E-07 | 532.80859 | 85.055107 |
| GVSMSLPSSPLLPF | 14 | Q9C0D5 | Protein TANC1 OS=Homo sapiens OX=9 | TANC1 | 202.22 | 2 | 51.15563 | yes | 3.166E-07 | 720.89758 | 83.20784 |
| LTLDLFQR | 8 | Q9C0D5 | Protein TANC1 OS=Homo sapiens OX=9 | TANC1 | 202.22 | 2 | 51.15563 | yes | 2.681E-07 | 503.29001 | 73.587296 |
| EGLSALSWACLK | 12 | Q9C0D5 | Protein TANC1 OS=Homo sapiens OX=9 | TANC1 | 202.22 | 2 | 51.15563 | yes | 2.916E-07 | 667.84229 | 69.197266 |
| VPGDAVIMPFR | 11 | Q9C0D5 | Protein TANC1 OS=Homo sapiens OX=9 | TANC1 | 202.22 | 2 | 51.15563 | yes | 2.093E-07 | 601.32355 | 57.428017 |
| EPQLQSMLSLR | 11 | Q9C0D5 | Protein TANC1 OS=Homo sapiens OX=9 | TANC1 | 202.22 | 2 | 51.15563 | yes | 2.751E-06 | 651.34772 | 55.853973 |
| GKLEVCELLLGHG | 18 | Q9C0D5 | Protein TANC1 OS=Homo sapiens OX=9 | TANC1 | 202.22 | 3 | 51.15563 | yes | 5.465E-07 | 637.01563 | 55.122837 |
| SCVQDPVAAFK | 11 | Q9C0D5 | Protein TANC1 OS=Homo sapiens OX=9 | TANC1 | 202.22 | 2 | 51.15563 | yes | 3.309E-07 | 611.30023 | 54.536285 |
| EPVAQPGLLLQPSI | 14 | Q9C0D5 | Protein TANC1 OS=Homo sapiens OX=9 | TANC1 | 202.22 | 2 | 51.15563 | yes | 3.452E-07 | 738.92468 | 51.128086 |
| LSLDDFPDNK | 10 | Q9C0D5 | Protein TANC1 OS=Homo sapiens OX=9 | TANC1 | 202.22 | 2 | 51.15563 | yes | 7.053E-06 | 582.28259 | 44.810043 |
| QFVAALADLQEAV | 14 | Q9C0D5 | Protein TANC1 OS=Homo sapiens OX=9 | TANC1 | 202.22 | 2 | 51.15563 | yes | 9.722E-08 | 751.91431 | 40.114727 |
| NTSVVVALLR | 10 | Q9C0D5 | Protein TANC1 OS=Homo sapiens OX=9 | TANC1 | 202.22 | 2 | 51.15563 | yes | 2.108E-06 | 536.32965 | 28.761406 |
| SQPSSSVPSSYIR | 13 | Q9C0D5 | Protein TANC1 OS=Homo sapiens OX=9 | TANC1 | 202.22 | 2 | 51.15563 | yes | 1.915E-07 | 697.84918 | 26.675716 |
| SIVEDIHQGK | 10 | Q9C0D6 | FH2 domain-containing protein 1 OS=Ho | FHDC1 | 124.762 | 2 | 46.831467 | yes | 1.461E-07 | 563.29858 | 72.957558 |
| GPQSLSASSSSLTP | 17 | Q9C0D6 | FH2 domain-containing protein 1 OS=Ho | FHDC1 | 124.762 | 2 | 46.831467 | yes | 3.471E-06 | 831.90942 | 35.255104 |
| PISPSSPSYRPPNTI | 15 | Q9C0D6 | FH2 domain-containing protein 1 OS=Ho | FHDC1 | 124.762 | 3 | 46.831467 | yes | 1.763E-06 | 552.62152 | 31.363113 |
| ISAEAVGVDISGR | 13 | Q9C0F1 | Centrosomal protein of 44 kDa OS=Homo | CEP44 | 44.14 | 2 | 55.28945 | yes | 1.021E-07 | 637.34094 | 383.47263 |
| VTLLETEMLLSK | 12 | Q9C0F1 | Centrosomal protein of 44 kDa OS=Homo | CEP44 | 44.14 | 2 | 55.28945 | yes | 1.32E-07 | 688.88892 | 190.6143 |
| DQFNYKPILTK | 11 | Q9C0F1 | Centrosomal protein of 44 kDa OS=Homo | CEP44 | 44.14 | 2 | 55.28945 | yes | 9.722E-08 | 683.87189 | 170.04475 |
| MFEETAELLK | 10 | Q9C0F1 | Centrosomal protein of 44 kDa OS=Homo | CEP44 | 44.14 | 2 | 55.28945 | yes | 9.722E-08 | 605.80481 | 127.0851 |
| VMVDENTWTNLL | 14 | Q9C0F1 | Centrosomal protein of 44 kDa OS=Homo | CEP44 | 44.14 | 2 | 55.28945 | yes | 9.722E-08 | 839.41687 | 60.867611 |

| | | | | | | | | | | | |
|---|---|---|---|---|---|---|---|---|---|---|---|
| QFIQCGFAEWK | 11 | Q9C0F1 | Centrosomal protein of 44 kDa OS=Hom | CEP44 | 44.14 | 2 | 55.28945 | yes | 2.926E-07 | 707.33466 | 47.960075 |
| SSHLIQHQR | 9 | Q9C0F3 | Zinc finger protein 436 OS=Homo sapien | ZNF436 | 54.277 | 2 | 52.817394 | yes | 7.225E-05 | 553.29669 | 41.989983 |
| GFTQSSNLITHQR | 13 | Q9C0F3 | Zinc finger protein 436 OS=Homo sapien | ZNF436 | 54.277 | 2 | 52.817394 | yes | 9.722E-08 | 744.88135 | 27.192783 |
| VPIPAPNEVLNDR | 13 | Q9C0J8 | pre-mRNA 3' end processing protein WD | WDR33 | 145.892 | 2 | 53.255478 | yes | 2.731E-07 | 717.39099 | 510.99197 |
| TIDYNPSVIK | 10 | Q9C0J8 | pre-mRNA 3' end processing protein WD | WDR33 | 145.892 | 2 | 53.255478 | yes | 2.324E-07 | 575.31116 | 291.18811 |
| IPPLNPGQGPGPNI | 14 | Q9C0J8 | pre-mRNA 3' end processing protein WD | WDR33 | 145.892 | 2 | 53.255478 | yes | 1.608E-06 | 693.38043 | 251.57034 |
| GPHPPGGLLGHGP | 17 | Q9C0J8 | pre-mRNA 3' end processing protein WD | WDR33 | 145.892 | 3 | 53.255478 | yes | 1.874E-07 | 579.62653 | 247.34674 |
| TPLLGDGPR | 9 | Q9C0J8 | pre-mRNA 3' end processing protein WD | WDR33 | 145.892 | 2 | 53.255478 | yes | 4.976E-07 | 463.25873 | 223.12268 |
| PIPAQFQQAWMQ | 14 | Q9C0J8 | pre-mRNA 3' end processing protein WD | WDR33 | 145.892 | 2 | 53.255478 | yes | 1.002E-07 | 843.92468 | 221.51282 |
| EASFSPTDNK | 10 | Q9C0J8 | pre-mRNA 3' end processing protein WD | WDR33 | 145.892 | 2 | 53.255478 | yes | 1.516E-06 | 548.25128 | 212.21971 |
| EELQVFR | 7 | Q9C0J8 | pre-mRNA 3' end processing protein WD | WDR33 | 145.892 | 2 | 53.255478 | yes | 2.995E-06 | 460.74545 | 208.67363 |
| YWQSNMNNVK | 10 | Q9C0J8 | pre-mRNA 3' end processing protein WD | WDR33 | 145.892 | 2 | 53.255478 | yes | 9.722E-08 | 642.29553 | 146.74402 |
| GLLPTPDEFPR | 11 | Q9C0J8 | pre-mRNA 3' end processing protein WD | WDR33 | 145.892 | 2 | 53.255478 | yes | 9.722E-08 | 621.3299 | 133.90611 |
| NLKEELQVFR | 10 | Q9C0J8 | pre-mRNA 3' end processing protein WD | WDR33 | 145.892 | 2 | 53.255478 | yes | 9.722E-08 | 638.35645 | 56.424438 |
| NLKEELQVFR | 10 | Q9C0J8 | pre-mRNA 3' end processing protein WD | WDR33 | 145.892 | 2 | 53.255478 | yes | 3.344E-06 | 425.90671 | 56.424438 |
| NLKEELQVFR | 10 | Q9C0J8 | pre-mRNA 3' end processing protein WD | WDR33 | 145.892 | 3 | 53.255478 | yes | 9.722E-08 | 638.35645 | 56.424438 |
| NLKEELQVFR | 10 | Q9C0J8 | pre-mRNA 3' end processing protein WD | WDR33 | 145.892 | 3 | 53.255478 | yes | 3.344E-06 | 425.90671 | 56.424438 |
| CVDWHPTK | 8 | Q9C0J8 | pre-mRNA 3' end processing protein WD | WDR33 | 145.892 | 2 | 53.255478 | yes | 2.748E-07 | 521.74237 | 88.974983 |
| LAMEQEQMGK | 10 | Q9C0J8 | pre-mRNA 3' end processing protein WD | WDR33 | 145.892 | 2 | 53.255478 | yes | 1.603E-07 | 582.77301 | 85.900871 |
| CPVFVVR | 7 | Q9C0J8 | pre-mRNA 3' end processing protein WD | WDR33 | 145.892 | 2 | 53.255478 | yes | 1.061E-06 | 438.74164 | 73.757767 |
| FQHQAPR | 7 | Q9C0J8 | pre-mRNA 3' end processing protein WD | WDR33 | 145.892 | 2 | 53.255478 | yes | 1.474E-07 | 442.23029 | 64.255112 |
| TGQSLATLHAHK | 12 | Q9C0J8 | pre-mRNA 3' end processing protein WD | WDR33 | 145.892 | 2 | 53.255478 | yes | 2.549E-07 | 632.34387 | 34.457733 |
| AMTWSHNDMWM | 21 | Q9C0J8 | pre-mRNA 3' end processing protein WD | WDR33 | 145.892 | 3 | 53.255478 | yes | 2.972E-07 | 817.69415 | 22.558001 |
| VFNCISYSPLCK | 12 | Q9GZL7 | Ribosome biogenesis protein WDR12 OS | WDR12 | 47.708 | 2 | 52.489407 | yes | 9.722E-08 | 744.35468 | 292.94385 |
| APLYDLAAHEDK | 12 | Q9GZL7 | Ribosome biogenesis protein WDR12 OS | WDR12 | 47.708 | 2 | 52.489407 | yes | 6.439E-07 | 671.83551 | 280.77765 |
| GAEEWILTGSYDK | 13 | Q9GZL7 | Ribosome biogenesis protein WDR12 OS | WDR12 | 47.708 | 2 | 52.489407 | yes | 2.73E-07 | 734.85138 | 263.71155 |
| HVEFDFLIK | 9 | Q9GZL7 | Ribosome biogenesis protein WDR12 OS | WDR12 | 47.708 | 2 | 52.489407 | yes | 9.722E-08 | 574.31097 | 213.23056 |
| VWDVESGSLK | 10 | Q9GZL7 | Ribosome biogenesis protein WDR12 OS | WDR12 | 47.708 | 2 | 52.489407 | yes | 4.199E-07 | 560.28766 | 179.74983 |
| FCSGSWDK | 8 | Q9GZL7 | Ribosome biogenesis protein WDR12 OS | WDR12 | 47.708 | 2 | 52.489407 | yes | 3.249E-06 | 493.70547 | 97.738243 |
| GHAGSVDSIAVDC | 17 | Q9GZL7 | Ribosome biogenesis protein WDR12 OS | WDR12 | 47.708 | 2 | 52.489407 | yes | 2.565E-07 | 779.37878 | 19.400858 |
| ELEQANDDLER | 11 | Q9GZM8 | Nuclear distribution protein nudE-like 1 ( | NDEL1 | 38.375 | 2 | 52.873856 | yes | 9.722E-08 | 666.30731 | 42.324947 |
| LEYLQIPPVSR | 11 | Q9GZP9 | Derlin-2 OS=Homo sapiens OX=9606 GI | DERL2 | 27.568 | 2 | 42.496998 | yes | 3.001E-07 | 657.87445 | 46.185692 |
| EQAFGGLTR | 9 | Q9GZR2 | RNA exonuclease 4 OS=Homo sapiens O | REXO4 | 46.672 | 2 | 52.38029 | yes | 2.131E-07 | 489.75378 | 267.79205 |
| VLFLDHPK | 8 | Q9GZR2 | RNA exonuclease 4 OS=Homo sapiens O | REXO4 | 46.672 | 2 | 52.38029 | yes | 3.245E-07 | 484.78183 | 151.98357 |
| ALALDCEMVGVG | 14 | Q9GZR2 | RNA exonuclease 4 OS=Homo sapiens O | REXO4 | 46.672 | 2 | 52.38029 | yes | 2.76E-07 | 730.3678 | 145.26917 |
| QGEELEVVQK | 10 | Q9GZR2 | RNA exonuclease 4 OS=Homo sapiens O | REXO4 | 46.672 | 2 | 52.38029 | yes | 1.337E-05 | 579.80371 | 117.49758 |
| ILGLQVQQAEHCS | 22 | Q9GZR2 | RNA exonuclease 4 OS=Homo sapiens O | REXO4 | 46.672 | 3 | 52.38029 | yes | 3.083E-06 | 823.07751 | 91.399109 |
| QLGQSEGSVSLSL | 15 | Q9GZR2 | RNA exonuclease 4 OS=Homo sapiens O | REXO4 | 46.672 | 2 | 52.38029 | yes | 9.722E-08 | 766.41992 | 70.517853 |
| VSIVNQYGK | 9 | Q9GZR2 | RNA exonuclease 4 OS=Homo sapiens O | REXO4 | 46.672 | 2 | 52.38029 | yes | 5.275E-05 | 504.27966 | 58.378807 |

| | | | | | | | | | | | |
|---|---|---|---|---|---|---|---|---|---|---|---|
| APEDFSQNWK | 10 | Q9GZR2 | RNA exonuclease 4 OS=Homo sapiens O | REXO4 | 46.672 | 2 | 52.38029 | yes | 9.722E-08 | 611.2804 | 50.153309 |
| VSVAAGSEEDHK | 12 | Q9GZS1 | DNA-directed RNA polymerase I subunit | POLR1E | 47.261 | 2 | 54.179707 | yes | 2.715E-06 | 614.79382 | 342.13403 |
| SALGPGVPHIINTK | 14 | Q9GZS1 | DNA-directed RNA polymerase I subunit | POLR1E | 47.261 | 2 | 54.179707 | yes | 1.021E-07 | 702.40393 | 250.47087 |
| LGTLSLPLPPAQTS | 16 | Q9GZS1 | DNA-directed RNA polymerase I subunit | POLR1E | 47.261 | 2 | 54.179707 | yes | 3.014E-06 | 833.46216 | 196.3161 |
| LSYVGNNFGTGAL | 14 | Q9GZS1 | DNA-directed RNA polymerase I subunit | POLR1E | 47.261 | 2 | 54.179707 | yes | 9.722E-08 | 720.87769 | 159.45506 |
| WQYCGAPDGSQR | 12 | Q9GZS1 | DNA-directed RNA polymerase I subunit | POLR1E | 47.261 | 2 | 54.179707 | yes | 9.722E-08 | 712.8042 | 157.17595 |
| MDSCIEAFGTTK | 12 | Q9GZS1 | DNA-directed RNA polymerase I subunit | POLR1E | 47.261 | 2 | 54.179707 | yes | 9.81E-08 | 680.29962 | 155.27193 |
| HFVGILNK | 8 | Q9GZS1 | DNA-directed RNA polymerase I subunit | POLR1E | 47.261 | 2 | 54.179707 | yes | 3.162E-07 | 464.27417 | 132.84775 |
| NVTSEEILK | 9 | Q9GZS1 | DNA-directed RNA polymerase I subunit | POLR1E | 47.261 | 2 | 54.179707 | yes | 4.358E-07 | 516.78223 | 110.14517 |
| AAETIIDTK | 9 | Q9GZS1 | DNA-directed RNA polymerase I subunit | POLR1E | 47.261 | 2 | 54.179707 | yes | 1.774E-07 | 481.26367 | 71.102875 |
| FILSIAYSPDGK | 12 | Q9GZS3 | Superkiller complex protein 8 OS=Homo | SKIC8 | 33.581 | 2 | 54.375107 | yes | 9.722E-08 | 655.85315 | 392.10641 |
| VNIFGVESGK | 10 | Q9GZS3 | Superkiller complex protein 8 OS=Homo | SKIC8 | 33.581 | 2 | 54.375107 | yes | 6.119E-07 | 525.28491 | 309.612 |
| LLHTLEGHAMPIR | 13 | Q9GZS3 | Superkiller complex protein 8 OS=Homo | SKIC8 | 33.581 | 3 | 54.375107 | yes | 9.81E-08 | 496.60989 | 51.619022 |
| LLHTLEGHAMPIR | 13 | Q9GZS3 | Superkiller complex protein 8 OS=Homo | SKIC8 | 33.581 | 3 | 54.375107 | yes | 9.722E-08 | 744.41119 | 51.619022 |
| LLHTLEGHAMPIR | 13 | Q9GZS3 | Superkiller complex protein 8 OS=Homo | SKIC8 | 33.581 | 2 | 54.375107 | yes | 9.81E-08 | 496.60989 | 51.619022 |
| LLHTLEGHAMPIR | 13 | Q9GZS3 | Superkiller complex protein 8 OS=Homo | SKIC8 | 33.581 | 2 | 54.375107 | yes | 9.722E-08 | 744.41119 | 51.619022 |
| QEQAHDDAIWSVA | 18 | Q9GZS3 | Superkiller complex protein 8 OS=Homo | SKIC8 | 33.581 | 3 | 54.375107 | yes | 1.01E-07 | 685.98883 | 65.843651 |
| LPQTSDDEK | 9 | Q9GZT3 | SRA stem-loop-interacting RNA-binding | SLIRP | 12.35 | 2 | 50.93255 | yes | 1.468E-07 | 516.74585 | 1334.6898 |
| IPWTAASSQLK | 11 | Q9GZT3 | SRA stem-loop-interacting RNA-binding | SLIRP | 12.35 | 2 | 50.93255 | yes | 1.164E-07 | 601.3324 | 1212.4863 |
| EHFAQFGHVR | 10 | Q9GZT3 | SRA stem-loop-interacting RNA-binding | SLIRP | 12.35 | 2 | 50.93255 | yes | 9.722E-08 | 614.3045 | 1140.0201 |
| GLGWVQFSSEEGL | 14 | Q9GZT3 | SRA stem-loop-interacting RNA-binding | SLIRP | 12.35 | 2 | 50.93255 | yes | 1.074E-07 | 782.89136 | 1039.7166 |
| SINQPVAFVR | 10 | Q9GZT3 | SRA stem-loop-interacting RNA-binding | SLIRP | 12.35 | 2 | 50.93255 | yes | 1.683E-07 | 565.81946 | 595.20111 |
| CILPFDK | 7 | Q9GZT3 | SRA stem-loop-interacting RNA-binding | SLIRP | 12.35 | 2 | 50.93255 | yes | 1.802E-07 | 446.73349 | 305.76181 |
| RIPWTAASSQLK | 12 | Q9GZT3 | SRA stem-loop-interacting RNA-binding | SLIRP | 12.35 | 2 | 50.93255 | yes | 1.022E-07 | 679.383 | 49.710495 |
| RIPWTAASSQLK | 12 | Q9GZT3 | SRA stem-loop-interacting RNA-binding | SLIRP | 12.35 | 2 | 50.93255 | yes | 2.136E-05 | 453.25775 | 49.710495 |
| RIPWTAASSQLK | 12 | Q9GZT3 | SRA stem-loop-interacting RNA-binding | SLIRP | 12.35 | 3 | 50.93255 | yes | 1.022E-07 | 679.383 | 49.710495 |
| RIPWTAASSQLK | 12 | Q9GZT3 | SRA stem-loop-interacting RNA-binding | SLIRP | 12.35 | 3 | 50.93255 | yes | 2.136E-05 | 453.25775 | 49.710495 |
| NALQQENHIIDGV | 14 | Q9GZT3 | SRA stem-loop-interacting RNA-binding | SLIRP | 12.35 | 2 | 50.93255 | yes | 4.971E-07 | 789.91534 | 43.263584 |
| LALEYIVPCMNK | 12 | Q9GZT9 | Egl nine homolog 1 OS=Homo sapiens O | EGLN1 | 46.021 | 2 | 51.238697 | yes | 9.722E-08 | 725.87524 | 357.93619 |
| ETGQQIGDEVR | 11 | Q9GZT9 | Egl nine homolog 1 OS=Homo sapiens O | EGLN1 | 46.021 | 2 | 51.238697 | yes | 9.722E-08 | 616.29932 | 267.36133 |
| FTDGQLVSQK | 10 | Q9GZT9 | Egl nine homolog 1 OS=Homo sapiens O | EGLN1 | 46.021 | 2 | 51.238697 | yes | 2.369E-07 | 561.79309 | 118.38284 |
| CVTCIYYLNK | 10 | Q9GZT9 | Egl nine homolog 1 OS=Homo sapiens O | EGLN1 | 46.021 | 2 | 51.238697 | yes | 5.743E-05 | 667.31757 | 79.294281 |
| VELNKPSDSVGK | 12 | Q9GZT9 | Egl nine homolog 1 OS=Homo sapiens O | EGLN1 | 46.021 | 2 | 51.238697 | yes | 9.976E-08 | 636.84332 | 66.203011 |
| AAAGGQGSAVAA | 24 | Q9GZT9 | Egl nine homolog 1 OS=Homo sapiens O | EGLN1 | 46.021 | 3 | 51.238697 | yes | 2.212E-07 | 741.03296 | 39.980877 |
| HGICVVDDFLGK | 12 | Q9GZT9 | Egl nine homolog 1 OS=Homo sapiens O | EGLN1 | 46.021 | 2 | 51.238697 | yes | 1.098E-07 | 680.3399 | 29.293232 |
| YFLNHIEK | 8 | Q9GZV5 | WW domain-containing transcription regu | WWTR1 | 44.102 | 2 | 26.970488 | yes | 3.036E-06 | 532.28217 | 59.705921 |
| ALLLGHLAR | 9 | Q9GZZ6 | Neuronal acetylcholine receptor subunit a | CHRNA10 | 49.705 | 2 | 44.086437 | yes | 2.326E-07 | 482.30853 | 45.039288 |
| MSSEVVDSNPYSR | 13 | Q9GZZ9 | Ubiquitin-like modifier-activating enzyme | UBA5 | 44.864 | 2 | 48.338478 | yes | 1.163E-07 | 735.83008 | 76.914215 |
| LQVLHPFDPEQR | 12 | Q9H000 | E3 ubiquitin-protein ligase makorin-2 OS | MKRN2 | 46.941 | 2 | 48.73082 | yes | 1.837E-07 | 739.89117 | 811.06262 |

| | | | | | | | | | | | |
|---|---|---|---|---|---|---|---|---|---|---|---|
| LQVLHPFDPEQR | 12 | Q9H000 | E3 ubiquitin-protein ligase makorin-2 OS | MKRN2 | 46.941 | 2 | 48.73082 | yes | 1.517E-07 | 493.59653 | 811.06262 |
| LQVLHPFDPEQR | 12 | Q9H000 | E3 ubiquitin-protein ligase makorin-2 OS | MKRN2 | 46.941 | 3 | 48.73082 | yes | 1.837E-07 | 739.89117 | 811.06262 |
| LQVLHPFDPEQR | 12 | Q9H000 | E3 ubiquitin-protein ligase makorin-2 OS | MKRN2 | 46.941 | 3 | 48.73082 | yes | 1.517E-07 | 493.59653 | 811.06262 |
| LWDFIENR | 8 | Q9H000 | E3 ubiquitin-protein ligase makorin-2 OS | MKRN2 | 46.941 | 2 | 48.73082 | yes | 1.477E-07 | 546.77728 | 1319.9484 |
| AFAFQASQDK | 10 | Q9H000 | E3 ubiquitin-protein ligase makorin-2 OS | MKRN2 | 46.941 | 2 | 48.73082 | yes | 1.11E-07 | 556.77222 | 1084.9067 |
| ICMLTFEHEMEK | 12 | Q9H000 | E3 ubiquitin-protein ligase makorin-2 OS | MKRN2 | 46.941 | 2 | 48.73082 | yes | 1.164E-07 | 784.35132 | 502.8389 |
| VCSICMEVILEK | 12 | Q9H000 | E3 ubiquitin-protein ligase makorin-2 OS | MKRN2 | 46.941 | 2 | 48.73082 | yes | 1.603E-07 | 740.86407 | 481.17758 |
| QFENPIIK | 8 | Q9H000 | E3 ubiquitin-protein ligase makorin-2 OS | MKRN2 | 46.941 | 2 | 48.73082 | yes | 3.107E-07 | 494.77673 | 427.86307 |
| YFMHGVCR | 8 | Q9H000 | E3 ubiquitin-protein ligase makorin-2 OS | MKRN2 | 46.941 | 2 | 48.73082 | yes | 3.184E-07 | 535.23895 | 344.09641 |
| QLSSQGTVR | 9 | Q9H000 | E3 ubiquitin-protein ligase makorin-2 OS | MKRN2 | 46.941 | 2 | 48.73082 | yes | 2.343E-07 | 488.26453 | 267.87997 |
| NELIEAFK | 8 | Q9H000 | E3 ubiquitin-protein ligase makorin-2 OS | MKRN2 | 46.941 | 2 | 48.73082 | yes | 9.765E-07 | 482.26093 | 159.05672 |
| KNELIEAFK | 9 | Q9H000 | E3 ubiquitin-protein ligase makorin-2 OS | MKRN2 | 46.941 | 2 | 48.73082 | yes | 2.133E-07 | 546.30841 | 157.78246 |
| PHSYLDAIR | 9 | Q9H000 | E3 ubiquitin-protein ligase makorin-2 OS | MKRN2 | 46.941 | 2 | 48.73082 | yes | 4.075E-07 | 536.28271 | 78.053009 |
| VISEFVIPSVYWVI | 19 | Q9H000 | E3 ubiquitin-protein ligase makorin-2 OS | MKRN2 | 46.941 | 3 | 48.73082 | yes | 7.079E-07 | 760.73523 | 42.070854 |
| GYVLGETSNLPSP | 15 | Q9H040 | DNA-dependent metalloprotease SPRTN | SPRTN | 55.134 | 2 | 46.798195 | yes | 1.467E-07 | 759.89355 | 68.155708 |
| IENVVLDANCSR | 12 | Q9H074 | Polyadenylate-binding protein-interacting | PAIP1 | 53.525 | 2 | 52.311222 | yes | 9.722E-08 | 695.3432 | 4584.2603 |
| LTGSVLEDAWK | 11 | Q9H074 | Polyadenylate-binding protein-interacting | PAIP1 | 53.525 | 2 | 52.311222 | yes | 9.919E-08 | 609.8219 | 3519.0664 |
| MDMEEIIQR | 9 | Q9H074 | Polyadenylate-binding protein-interacting | PAIP1 | 53.525 | 2 | 52.311222 | yes | 5.235E-07 | 582.77301 | 2190.7334 |
| PQVVVAPVLMSK | 12 | Q9H074 | Polyadenylate-binding protein-interacting | PAIP1 | 53.525 | 2 | 52.311222 | yes | 9.722E-08 | 634.37579 | 1529.9507 |
| YQELLER | 7 | Q9H074 | Polyadenylate-binding protein-interacting | PAIP1 | 53.525 | 2 | 52.311222 | yes | 1.426E-07 | 475.75073 | 1372.8369 |
| ADILQVGLR | 9 | Q9H074 | Polyadenylate-binding protein-interacting | PAIP1 | 53.525 | 2 | 52.311222 | yes | 1.111E-07 | 492.79547 | 956.05499 |
| FCLESER | 7 | Q9H074 | Polyadenylate-binding protein-interacting | PAIP1 | 53.525 | 2 | 52.311222 | yes | 1.821E-07 | 470.71329 | 941.10065 |
| APGFLQPPPLR | 11 | Q9H074 | Polyadenylate-binding protein-interacting | PAIP1 | 53.525 | 2 | 52.311222 | yes | 1.186E-07 | 596.84552 | 859.40234 |
| GKMDMEEIIQR | 11 | Q9H074 | Polyadenylate-binding protein-interacting | PAIP1 | 53.525 | 2 | 52.311222 | yes | 9.722E-08 | 675.33124 | 368.18808 |
| IPQQNSESAMAK | 12 | Q9H074 | Polyadenylate-binding protein-interacting | PAIP1 | 53.525 | 2 | 52.311222 | yes | 1.157E-06 | 652.31915 | 168.62013 |
| LCNYLSHHLTISPQ | 19 | Q9H074 | Polyadenylate-binding protein-interacting | PAIP1 | 53.525 | 4 | 52.311222 | yes | 2.535E-07 | 561.77991 | 110.40432 |
| CRTEYEVK | 8 | Q9H074 | Polyadenylate-binding protein-interacting | PAIP1 | 53.525 | 2 | 52.311222 | yes | 2.236E-06 | 542.75824 | 96.177017 |
| ELLDGGANPLQR | 12 | Q9H078 | Mitochondrial disaggregase OS=Homo sa | CLPB | 78.729 | 2 | 54.40058 | yes | 1.649E-07 | 641.84113 | 536.16473 |
| LDMSEFQER | 9 | Q9H078 | Mitochondrial disaggregase OS=Homo sa | CLPB | 78.729 | 2 | 54.40058 | yes | 1.736E-07 | 577.76099 | 452.2059 |
| QEALEMSR | 8 | Q9H078 | Mitochondrial disaggregase OS=Homo sa | CLPB | 78.729 | 2 | 54.40058 | yes | 3.431E-07 | 482.23184 | 236.62816 |
| SPELPSPQAEK | 11 | Q9H078 | Mitochondrial disaggregase OS=Homo sa | CLPB | 78.729 | 2 | 54.40058 | yes | 2.562E-07 | 591.80371 | 222.56439 |
| IAENLGDVQISDK | 13 | Q9H078 | Mitochondrial disaggregase OS=Homo sa | CLPB | 78.729 | 2 | 54.40058 | yes | 1.067E-07 | 701.36462 | 213.24815 |
| ENVIRPILK | 9 | Q9H078 | Mitochondrial disaggregase OS=Homo sa | CLPB | 78.729 | 2 | 54.40058 | yes | 1.574E-07 | 541.34003 | 150.53307 |
| RDEFLGR | 7 | Q9H078 | Mitochondrial disaggregase OS=Homo sa | CLPB | 78.729 | 2 | 54.40058 | yes | 1.115E-07 | 446.73541 | 129.3376 |
| ITVEDSDK | 8 | Q9H078 | Mitochondrial disaggregase OS=Homo sa | CLPB | 78.729 | 2 | 54.40058 | yes | 2.583E-07 | 453.72437 | 106.00571 |
| ANNMQEVSR | 9 | Q9H078 | Mitochondrial disaggregase OS=Homo sa | CLPB | 78.729 | 2 | 54.40058 | yes | 6.041E-07 | 524.74567 | 100.21358 |
| QCPNAVVLFDEVI | 14 | Q9H078 | Mitochondrial disaggregase OS=Homo sa | CLPB | 78.729 | 2 | 54.40058 | yes | 1.444E-07 | 817.39813 | 83.127686 |
| LKEHIIGQESAIAT | 20 | Q9H078 | Mitochondrial disaggregase OS=Homo sa | CLPB | 78.729 | 3 | 54.40058 | yes | 2.624E-07 | 693.06366 | 51.916534 |
| NEMGHTPLDYAR | 12 | Q9H078 | Mitochondrial disaggregase OS=Homo sa | CLPB | 78.729 | 2 | 54.40058 | yes | 9.722E-08 | 702.32227 | 40.417431 |

| | | | | | | | | | | | |
|---|---|---|---|---|---|---|---|---|---|---|---|
| EHIIGQESAIATVG | 18 | Q9H078 | Mitochondrial disaggregase OS=Homo sa | CLPB | 78.729 | 3 | 54.40058 | yes | 3.27E-06 | 612.67065 | 32.169022 |
| TEIINDGNIQILK | 13 | Q9H079 | KATNB1-like protein 1 OS=Homo sapie | KATNBL1 | 34.767 | 2 | 52.10684 | yes | 9.722E-08 | 735.91174 | 29.446489 |
| SDIVVQIVDAR | 11 | Q9H089 | Large subunit GTPase 1 homolog OS=Ho | LSG1 | 75.226 | 2 | 53.06352 | yes | 3.535E-07 | 607.84058 | 5027.7241 |
| TGLLSFEESQR | 11 | Q9H089 | Large subunit GTPase 1 homolog OS=Ho | LSG1 | 75.226 | 2 | 53.06352 | yes | 9.818E-08 | 633.81989 | 4217.437 |
| VSVSATPGHTK | 11 | Q9H089 | Large subunit GTPase 1 homolog OS=Ho | LSG1 | 75.226 | 2 | 53.06352 | yes | 1.11E-07 | 542.29327 | 2093.2075 |
| LILTPFER | 8 | Q9H089 | Large subunit GTPase 1 homolog OS=Ho | LSG1 | 75.226 | 2 | 53.06352 | yes | 3.933E-07 | 494.79492 | 2045.2272 |
| CEDLECYVK | 9 | Q9H089 | Large subunit GTPase 1 homolog OS=Ho | LSG1 | 75.226 | 2 | 53.06352 | yes | 9.722E-08 | 608.25464 | 1577.3788 |
| TFFHQENVR | 9 | Q9H089 | Large subunit GTPase 1 homolog OS=Ho | LSG1 | 75.226 | 2 | 53.06352 | yes | 1.705E-07 | 589.29108 | 1356.4949 |
| HVLEATYGINIITP | 15 | Q9H089 | Large subunit GTPase 1 homolog OS=Ho | LSG1 | 75.226 | 3 | 53.06352 | yes | 1.492E-07 | 566.31757 | 641.67944 |
| HVLEATYGINIITP | 15 | Q9H089 | Large subunit GTPase 1 homolog OS=Ho | LSG1 | 75.226 | 3 | 53.06352 | yes | 2.338E-07 | 848.97266 | 641.67944 |
| HVLEATYGINIITP | 15 | Q9H089 | Large subunit GTPase 1 homolog OS=Ho | LSG1 | 75.226 | 2 | 53.06352 | yes | 1.492E-07 | 566.31757 | 641.67944 |
| HVLEATYGINIITP | 15 | Q9H089 | Large subunit GTPase 1 homolog OS=Ho | LSG1 | 75.226 | 2 | 53.06352 | yes | 2.338E-07 | 848.97266 | 641.67944 |
| LLYCHPPPGR | 10 | Q9H089 | Large subunit GTPase 1 homolog OS=Ho | LSG1 | 75.226 | 2 | 53.06352 | yes | 5.561E-07 | 605.31348 | 547.77216 |
| LLYCHPPPGR | 10 | Q9H089 | Large subunit GTPase 1 homolog OS=Ho | LSG1 | 75.226 | 2 | 53.06352 | yes | 1.566E-07 | 403.87808 | 547.77216 |
| LLYCHPPPGR | 10 | Q9H089 | Large subunit GTPase 1 homolog OS=Ho | LSG1 | 75.226 | 3 | 53.06352 | yes | 5.561E-07 | 605.31348 | 547.77216 |
| LLYCHPPPGR | 10 | Q9H089 | Large subunit GTPase 1 homolog OS=Ho | LSG1 | 75.226 | 3 | 53.06352 | yes | 1.566E-07 | 403.87808 | 547.77216 |
| SSTINTIMGNK | 11 | Q9H089 | Large subunit GTPase 1 homolog OS=Ho | LSG1 | 75.226 | 2 | 53.06352 | yes | 1.145E-07 | 583.29773 | 1065.728 |
| ADLLTAEQR | 9 | Q9H089 | Large subunit GTPase 1 homolog OS=Ho | LSG1 | 75.226 | 2 | 53.06352 | yes | 1.436E-07 | 508.77219 | 989.41083 |
| ENVILINK | 8 | Q9H089 | Large subunit GTPase 1 homolog OS=Ho | LSG1 | 75.226 | 2 | 53.06352 | yes | 1.35E-07 | 471.78458 | 950.5675 |
| GFMTAHGQPDQP | 13 | Q9H089 | Large subunit GTPase 1 homolog OS=Ho | LSG1 | 75.226 | 2 | 53.06352 | yes | 9.722E-08 | 721.33569 | 423.84113 |
| GFMTAHGQPDQP | 13 | Q9H089 | Large subunit GTPase 1 homolog OS=Ho | LSG1 | 75.226 | 2 | 53.06352 | yes | 1.037E-07 | 481.22623 | 423.84113 |
| GFMTAHGQPDQP | 13 | Q9H089 | Large subunit GTPase 1 homolog OS=Ho | LSG1 | 75.226 | 3 | 53.06352 | yes | 9.722E-08 | 721.33569 | 423.84113 |
| GFMTAHGQPDQP | 13 | Q9H089 | Large subunit GTPase 1 homolog OS=Ho | LSG1 | 75.226 | 3 | 53.06352 | yes | 1.037E-07 | 481.22623 | 423.84113 |
| DGQLTVGLVGYP | 16 | Q9H089 | Large subunit GTPase 1 homolog OS=Ho | LSG1 | 75.226 | 2 | 53.06352 | yes | 1.156E-07 | 808.93579 | 579.55487 |
| QFLCIPR | 7 | Q9H089 | Large subunit GTPase 1 homolog OS=Ho | LSG1 | 75.226 | 2 | 53.06352 | yes | 3.207E-07 | 467.25238 | 569.66833 |
| DPVTFQHQHQR | 11 | Q9H089 | Large subunit GTPase 1 homolog OS=Ho | LSG1 | 75.226 | 2 | 53.06352 | yes | 9.722E-08 | 696.84198 | 490.07449 |
| SAWAMYFEK | 9 | Q9H089 | Large subunit GTPase 1 homolog OS=Ho | LSG1 | 75.226 | 2 | 53.06352 | yes | 9.903E-08 | 566.76025 | 375.66254 |
| QIENIVDK | 8 | Q9H089 | Large subunit GTPase 1 homolog OS=Ho | LSG1 | 75.226 | 2 | 53.06352 | yes | 2.88E-07 | 479.76382 | 289.33585 |
| QIHNFSHLVSK | 11 | Q9H089 | Large subunit GTPase 1 homolog OS=Ho | LSG1 | 75.226 | 2 | 53.06352 | yes | 3.515E-07 | 655.35419 | 256.89124 |
| RAPAGGSLGR | 10 | Q9H089 | Large subunit GTPase 1 homolog OS=Ho | LSG1 | 75.226 | 2 | 53.06352 | yes | 9.577E-07 | 471.2674 | 246.87889 |
| VKDGQLTVGLVG | 18 | Q9H089 | Large subunit GTPase 1 homolog OS=Ho | LSG1 | 75.226 | 3 | 53.06352 | yes | 9.722E-08 | 615.34741 | 228.1234 |
| DHVPPVSLVCQNI | 15 | Q9H089 | Large subunit GTPase 1 homolog OS=Ho | LSG1 | 75.226 | 3 | 53.06352 | yes | 1.178E-07 | 577.63843 | 174.3793 |
| GVQAVMGYK | 9 | Q9H089 | Large subunit GTPase 1 homolog OS=Ho | LSG1 | 75.226 | 2 | 53.06352 | yes | 3.7E-07 | 476.74966 | 120.70802 |
| SGPTPSAAELAR | 12 | Q9H093 | NUAK family SNF1-like kinase 2 OS=H | NUAK2 | 69.613 | 2 | 41.949715 | yes | 6.558E-07 | 578.80145 | 82.452568 |
| EQKPPQASGLLLH | 14 | Q9H093 | NUAK family SNF1-like kinase 2 OS=H | NUAK2 | 69.613 | 3 | 41.949715 | yes | 3.265E-07 | 525.29846 | 43.924156 |
| IVIVMEYASR | 10 | Q9H093 | NUAK family SNF1-like kinase 2 OS=H | NUAK2 | 69.613 | 2 | 41.949715 | yes | 2.274E-06 | 590.82336 | 32.705364 |
| SLFVVVGDR | 9 | Q9H0A0 | RNA cytidine acetyltransferase OS=Hom | NAT10 | 115.73 | 2 | 54.029274 | yes | 1.133E-07 | 496.2822 | 739.50928 |
| AGPNASIISLK | 11 | Q9H0A0 | RNA cytidine acetyltransferase OS=Hom | NAT10 | 115.73 | 2 | 54.029274 | yes | 1.132E-07 | 535.81384 | 708.49115 |
| SMDLSEYIIR | 10 | Q9H0A0 | RNA cytidine acetyltransferase OS=Hom | NAT10 | 115.73 | 2 | 54.029274 | yes | 1.096E-07 | 613.80792 | 706.58813 |

| | | | | | | | | | | | |
|---|---|---|---|---|---|---|---|---|---|---|---|
| AGFVPVYLR | 9 | Q9H0A0 | RNA cytidine acetyltransferase OS=Hom | NAT10 | 115.73 | 2 | 54.029274 | yes | 4.774E-07 | 511.2951 | 504.6767 |
| FILSLASCK | 9 | Q9H0A0 | RNA cytidine acetyltransferase OS=Hom | NAT10 | 115.73 | 2 | 54.029274 | yes | 2.277E-07 | 519.78625 | 375.53616 |
| ASEVFLQR | 8 | Q9H0A0 | RNA cytidine acetyltransferase OS=Hom | NAT10 | 115.73 | 2 | 54.029274 | yes | 7.241E-07 | 475.25873 | 350.08771 |
| FIEGISEK | 8 | Q9H0A0 | RNA cytidine acetyltransferase OS=Hom | NAT10 | 115.73 | 2 | 54.029274 | yes | 1.474E-07 | 461.74765 | 347.57217 |
| LFNEVQEK | 8 | Q9H0A0 | RNA cytidine acetyltransferase OS=Hom | NAT10 | 115.73 | 2 | 54.029274 | yes | 1.544E-07 | 503.76382 | 329.02243 |
| KDLPPLLLK | 9 | Q9H0A0 | RNA cytidine acetyltransferase OS=Hom | NAT10 | 115.73 | 2 | 54.029274 | yes | 2.136E-06 | 518.84186 | 323.05984 |
| TLSDDLDEAAK | 11 | Q9H0A0 | RNA cytidine acetyltransferase OS=Hom | NAT10 | 115.73 | 2 | 54.029274 | yes | 4.927E-07 | 589.28278 | 299.00854 |
| AIEEQMVAAK | 10 | Q9H0A0 | RNA cytidine acetyltransferase OS=Hom | NAT10 | 115.73 | 2 | 54.029274 | yes | 8.105E-07 | 545.28406 | 266.93756 |
| ILIENGVAER | 10 | Q9H0A0 | RNA cytidine acetyltransferase OS=Hom | NAT10 | 115.73 | 2 | 54.029274 | yes | 9.722E-08 | 557.31677 | 242.00464 |
| LDYLGVSYGLTPR | 13 | Q9H0A0 | RNA cytidine acetyltransferase OS=Hom | NAT10 | 115.73 | 2 | 54.029274 | yes | 9.722E-08 | 727.38794 | 174.11967 |
| TEAHQDVVGR | 10 | Q9H0A0 | RNA cytidine acetyltransferase OS=Hom | NAT10 | 115.73 | 2 | 54.029274 | yes | 4.004E-07 | 556.2782 | 168.96246 |
| QTIQYIHPADAVK | 13 | Q9H0A0 | RNA cytidine acetyltransferase OS=Hom | NAT10 | 115.73 | 2 | 54.029274 | yes | 9.789E-08 | 742.3988 | 160.46715 |
| DVVMEPTMK | 9 | Q9H0A0 | RNA cytidine acetyltransferase OS=Hom | NAT10 | 115.73 | 2 | 54.029274 | yes | 8.858E-07 | 525.25391 | 159.47415 |
| ALQLLQMYYEGR | 12 | Q9H0A0 | RNA cytidine acetyltransferase OS=Hom | NAT10 | 115.73 | 2 | 54.029274 | yes | 1.093E-07 | 742.88196 | 145.69411 |
| QLYTVTMDVHSR | 12 | Q9H0A0 | RNA cytidine acetyltransferase OS=Hom | NAT10 | 115.73 | 2 | 54.029274 | yes | 2.808E-07 | 725.36139 | 120.02171 |
| YCYYNETHK | 9 | Q9H0A0 | RNA cytidine acetyltransferase OS=Hom | NAT10 | 115.73 | 2 | 54.029274 | yes | 9.722E-08 | 639.26642 | 118.08941 |
| ARPSVLWCYK | 10 | Q9H0A0 | RNA cytidine acetyltransferase OS=Hom | NAT10 | 115.73 | 2 | 54.029274 | yes | 1.002E-07 | 640.33441 | 110.24432 |
| DQVVILHHMLSK | 12 | Q9H0A0 | RNA cytidine acetyltransferase OS=Hom | NAT10 | 115.73 | 2 | 54.029274 | yes | 9.722E-08 | 710.39246 | 94.227562 |
| YAPGDAVEK | 9 | Q9H0A0 | RNA cytidine acetyltransferase OS=Hom | NAT10 | 115.73 | 2 | 54.029274 | yes | 5.636E-07 | 475.23489 | 90.126053 |
| QQSAQSQVSTTAE | 15 | Q9H0A0 | RNA cytidine acetyltransferase OS=Hom | NAT10 | 115.73 | 2 | 54.029274 | yes | 9.809E-08 | 803.88702 | 79.44944 |
| DTLFCYHK | 8 | Q9H0A0 | RNA cytidine acetyltransferase OS=Hom | NAT10 | 115.73 | 2 | 54.029274 | yes | 1.841E-07 | 542.25006 | 55.450874 |
| QSILNSLSR | 9 | Q9H0A0 | RNA cytidine acetyltransferase OS=Hom | NAT10 | 115.73 | 2 | 54.029274 | yes | 1.637E-06 | 509.28799 | 53.248291 |
| PSVLWCYK | 8 | Q9H0A0 | RNA cytidine acetyltransferase OS=Hom | NAT10 | 115.73 | 2 | 54.029274 | yes | 5.282E-06 | 526.76532 | 19.377048 |
| YEVAVPLCK | 9 | Q9H0B6 | Kinesin light chain 2 OS=Homo sapiens ( | KLC2 | 68.935 | 2 | 52.18665 | no | 6.162E-07 | 539.78369 | 134.75473 |
| AVIQGLETLR | 10 | Q9H0B6 | Kinesin light chain 2 OS=Homo sapiens ( | KLC2 | 68.935 | 2 | 52.18665 | yes | 2.652E-07 | 550.32715 | 94.251991 |
| LGPDDPNVAK | 10 | Q9H0B6 | Kinesin light chain 2 OS=Homo sapiens ( | KLC2 | 68.935 | 2 | 52.18665 | no | 3.016E-07 | 513.26672 | 71.918312 |
| EAAHLLNDALAIR | 13 | Q9H0B6 | Kinesin light chain 2 OS=Homo sapiens ( | KLC2 | 68.935 | 2 | 52.18665 | yes | 1.05E-07 | 703.89117 | 69.188278 |
| SEQAVAQLEEEK | 12 | Q9H0B6 | Kinesin light chain 2 OS=Homo sapiens ( | KLC2 | 68.935 | 2 | 52.18665 | no | 9.722E-08 | 680.83319 | 49.280972 |
| QLSNLALLCQNQC | 14 | Q9H0B6 | Kinesin light chain 2 OS=Homo sapiens ( | KLC2 | 68.935 | 2 | 52.18665 | yes | 5.689E-07 | 793.91962 | 28.836617 |
| ALEIYATR | 8 | Q9H0B6 | Kinesin light chain 2 OS=Homo sapiens ( | KLC2 | 68.935 | 2 | 52.18665 | yes | 1.682E-05 | 468.76108 | 26.230186 |
| DSAPYGEYGSWYI | 13 | Q9H0B6 | Kinesin light chain 2 OS=Homo sapiens ( | KLC2 | 68.935 | 2 | 52.18665 | yes | 1.987E-07 | 761.82788 | 20.728914 |
| NSPGSQVASNPR | 12 | Q9H0D6 | 5'-3' exoribonuclease 2 OS=Homo sapien: | XRN2 | 108.583 | 2 | 53.17401 | yes | 1.163E-07 | 607.29962 | 1599.3079 |
| GVGAEPLLPWNR | 12 | Q9H0D6 | 5'-3' exoribonuclease 2 OS=Homo sapien: | XRN2 | 108.583 | 2 | 53.17401 | yes | 4.382E-07 | 654.85657 | 1054.2343 |
| VREEILAK | 8 | Q9H0D6 | 5'-3' exoribonuclease 2 OS=Homo sapien: | XRN2 | 108.583 | 2 | 53.17401 | yes | 1.234E-07 | 479.29001 | 965.81763 |
| NSLGGDVLFVGK | 12 | Q9H0D6 | 5'-3' exoribonuclease 2 OS=Homo sapien: | XRN2 | 108.583 | 2 | 53.17401 | yes | 1.521E-07 | 603.3299 | 938.31775 |
| DLTQNTVVSINFK | 13 | Q9H0D6 | 5'-3' exoribonuclease 2 OS=Homo sapien: | XRN2 | 108.583 | 2 | 53.17401 | yes | 1.043E-07 | 739.89612 | 857.53009 |
| RPVHLDQAAFR | 11 | Q9H0D6 | 5'-3' exoribonuclease 2 OS=Homo sapien: | XRN2 | 108.583 | 3 | 53.17401 | yes | 1.117E-07 | 437.24231 | 343.61325 |
| RPVHLDQAAFR | 11 | Q9H0D6 | 5'-3' exoribonuclease 2 OS=Homo sapien: | XRN2 | 108.583 | 3 | 53.17401 | yes | 9.722E-08 | 655.3598 | 343.61325 |
| RPVHLDQAAFR | 11 | Q9H0D6 | 5'-3' exoribonuclease 2 OS=Homo sapien: | XRN2 | 108.583 | 2 | 53.17401 | yes | 1.117E-07 | 437.24231 | 343.61325 |

| | | | | | | | | | | | |
|---|---|---|---|---|---|---|---|---|---|---|---|
| RPVHLDQAAFR | 11 | Q9H0D6 | 5'-3' exoribonuclease 2 OS=Homo sapien: | XRN2 | 108.583 | 2 | 53.17401 | yes | 9.722E-08 | 655.3598 | 343.61325 |
| DPQFAEDYIFK | 11 | Q9H0D6 | 5'-3' exoribonuclease 2 OS=Homo sapien: | XRN2 | 108.583 | 2 | 53.17401 | yes | 1.598E-07 | 686.82465 | 677.56079 |
| KPAAVLKPSDWEI | 13 | Q9H0D6 | 5'-3' exoribonuclease 2 OS=Homo sapien: | XRN2 | 108.583 | 3 | 53.17401 | yes | 9.722E-08 | 490.27679 | 278.1936 |
| KPAAVLKPSDWEI | 13 | Q9H0D6 | 5'-3' exoribonuclease 2 OS=Homo sapien: | XRN2 | 108.583 | 3 | 53.17401 | yes | 9.722E-08 | 734.91156 | 278.1936 |
| KPAAVLKPSDWEI | 13 | Q9H0D6 | 5'-3' exoribonuclease 2 OS=Homo sapien: | XRN2 | 108.583 | 2 | 53.17401 | yes | 9.722E-08 | 490.27679 | 278.1936 |
| KPAAVLKPSDWEI | 13 | Q9H0D6 | 5'-3' exoribonuclease 2 OS=Homo sapien: | XRN2 | 108.583 | 2 | 53.17401 | yes | 9.722E-08 | 734.91156 | 278.1936 |
| TGGYLTESGYVNL | 15 | Q9H0D6 | 5'-3' exoribonuclease 2 OS=Homo sapien: | XRN2 | 108.583 | 2 | 53.17401 | yes | 1.31E-07 | 829.41266 | 541.65033 |
| LFSIVRPR | 8 | Q9H0D6 | 5'-3' exoribonuclease 2 OS=Homo sapien: | XRN2 | 108.583 | 2 | 53.17401 | yes | 2.925E-07 | 494.30853 | 500.7099 |
| QWKPQLGFNR | 10 | Q9H0D6 | 5'-3' exoribonuclease 2 OS=Homo sapien: | XRN2 | 108.583 | 2 | 53.17401 | yes | 1.026E-07 | 637.34363 | 250.22702 |
| QWKPQLGFNR | 10 | Q9H0D6 | 5'-3' exoribonuclease 2 OS=Homo sapien: | XRN2 | 108.583 | 2 | 53.17401 | yes | 9.722E-08 | 425.23154 | 250.22702 |
| QWKPQLGFNR | 10 | Q9H0D6 | 5'-3' exoribonuclease 2 OS=Homo sapien: | XRN2 | 108.583 | 3 | 53.17401 | yes | 1.026E-07 | 637.34363 | 250.22702 |
| QWKPQLGFNR | 10 | Q9H0D6 | 5'-3' exoribonuclease 2 OS=Homo sapien: | XRN2 | 108.583 | 3 | 53.17401 | yes | 9.722E-08 | 425.23154 | 250.22702 |
| KDDEDSFR | 8 | Q9H0D6 | 5'-3' exoribonuclease 2 OS=Homo sapien: | XRN2 | 108.583 | 2 | 53.17401 | yes | 3.729E-07 | 506.22253 | 324.90521 |
| KYPSIIVNCVEEKI | 15 | Q9H0D6 | 5'-3' exoribonuclease 2 OS=Homo sapien: | XRN2 | 108.583 | 3 | 53.17401 | yes | 1.584E-07 | 601.99396 | 210.1035 |
| NKFDVDAADEK | 11 | Q9H0D6 | 5'-3' exoribonuclease 2 OS=Homo sapien: | XRN2 | 108.583 | 2 | 53.17401 | yes | 9.732E-08 | 626.2962 | 205.0773 |
| FDVDAADEK | 9 | Q9H0D6 | 5'-3' exoribonuclease 2 OS=Homo sapien: | XRN2 | 108.583 | 2 | 53.17401 | yes | 1.106E-07 | 505.22726 | 195.32607 |
| EEFKPNKPK | 9 | Q9H0D6 | 5'-3' exoribonuclease 2 OS=Homo sapien: | XRN2 | 108.583 | 2 | 53.17401 | yes | 1.101E-07 | 558.80603 | 186.44174 |
| QAAYEMR | 7 | Q9H0D6 | 5'-3' exoribonuclease 2 OS=Homo sapien: | XRN2 | 108.583 | 2 | 53.17401 | yes | 2.442E-07 | 434.70273 | 176.89891 |
| TLGHVMPR | 8 | Q9H0D6 | 5'-3' exoribonuclease 2 OS=Homo sapien: | XRN2 | 108.583 | 2 | 53.17401 | yes | 1.027E-07 | 455.75 | 176.76031 |
| NLTVILSDASAPGI | 18 | Q9H0D6 | 5'-3' exoribonuclease 2 OS=Homo sapien: | XRN2 | 108.583 | 3 | 53.17401 | yes | 1.92E-07 | 613.31451 | 131.64053 |
| YYYQGCASWK | 10 | Q9H0D6 | 5'-3' exoribonuclease 2 OS=Homo sapien: | XRN2 | 108.583 | 2 | 53.17401 | yes | 9.722E-08 | 663.28461 | 94.383507 |
| LLYMAIDGVAPR | 12 | Q9H0D6 | 5'-3' exoribonuclease 2 OS=Homo sapien: | XRN2 | 108.583 | 2 | 53.17401 | yes | 9.722E-08 | 659.86304 | 75.068161 |
| PVHLDQAAFR | 10 | Q9H0D6 | 5'-3' exoribonuclease 2 OS=Homo sapien: | XRN2 | 108.583 | 2 | 53.17401 | yes | 1.717E-07 | 577.30927 | 75.011932 |
| PCGLCNQFGHEVK | 13 | Q9H0D6 | 5'-3' exoribonuclease 2 OS=Homo sapien: | XRN2 | 108.583 | 2 | 53.17401 | yes | 1.113E-07 | 773.35046 | 35.676632 |
| PCGLCNQFGHEVK | 13 | Q9H0D6 | 5'-3' exoribonuclease 2 OS=Homo sapien: | XRN2 | 108.583 | 2 | 53.17401 | yes | 1.96E-06 | 515.90277 | 35.676632 |
| PCGLCNQFGHEVK | 13 | Q9H0D6 | 5'-3' exoribonuclease 2 OS=Homo sapien: | XRN2 | 108.583 | 3 | 53.17401 | yes | 1.113E-07 | 773.35046 | 35.676632 |
| PCGLCNQFGHEVK | 13 | Q9H0D6 | 5'-3' exoribonuclease 2 OS=Homo sapien: | XRN2 | 108.583 | 3 | 53.17401 | yes | 1.96E-06 | 515.90277 | 35.676632 |
| GKHDELADSLPCA | 21 | Q9H0D6 | 5'-3' exoribonuclease 2 OS=Homo sapien: | XRN2 | 108.583 | 4 | 53.17401 | yes | 9.722E-08 | 601.79547 | 25.667253 |
| EVGVQSSER | 9 | Q9H0G5 | Nuclear speckle splicing regulatory protei | NSRP1 | 66.391 | 2 | 50.021336 | yes | 1.181E-07 | 495.74619 | 828.80945 |
| AAALEACLDVTK | 12 | Q9H0G5 | Nuclear speckle splicing regulatory protei | NSRP1 | 66.391 | 2 | 50.021336 | yes | 9.932E-08 | 631.32648 | 617.77783 |
| VIETPENDFK | 10 | Q9H0G5 | Nuclear speckle splicing regulatory protei | NSRP1 | 66.391 | 2 | 50.021336 | yes | 3.011E-07 | 596.29822 | 525.02112 |
| YIHNLLK | 7 | Q9H0G5 | Nuclear speckle splicing regulatory protei | NSRP1 | 66.391 | 2 | 50.021336 | yes | 1.132E-07 | 450.76871 | 294.87204 |
| VEENPDADSDFDA | 14 | Q9H0G5 | Nuclear speckle splicing regulatory protei | NSRP1 | 66.391 | 2 | 50.021336 | yes | 9.977E-08 | 776.32593 | 232.36761 |
| CILQTDVK | 8 | Q9H0G5 | Nuclear speckle splicing regulatory protei | NSRP1 | 66.391 | 2 | 50.021336 | yes | 2.124E-07 | 488.76022 | 212.74371 |
| TQQLHPVLQK | 10 | Q9H0G5 | Nuclear speckle splicing regulatory protei | NSRP1 | 66.391 | 2 | 50.021336 | yes | 1.455E-07 | 596.34583 | 62.03001 |
| ALAEDATVYEYDS | 20 | Q9H0G5 | Nuclear speckle splicing regulatory protei | NSRP1 | 66.391 | 3 | 50.021336 | yes | 1.182E-07 | 785.35089 | 50.015068 |
| EQERPPEAVSK | 11 | Q9H0G5 | Nuclear speckle splicing regulatory protei | NSRP1 | 66.391 | 2 | 50.021336 | yes | 4.116E-07 | 635.32532 | 44.308933 |
| KTQQLHPVLQK | 11 | Q9H0G5 | Nuclear speckle splicing regulatory protei | NSRP1 | 66.391 | 2 | 50.021336 | yes | 2.354E-07 | 660.39331 | 21.443649 |
| KTQQLHPVLQK | 11 | Q9H0G5 | Nuclear speckle splicing regulatory protei | NSRP1 | 66.391 | 2 | 50.021336 | yes | 1.307E-06 | 440.59799 | 21.443649 |

| | | | | | | | | | | | |
|---|---|---|---|---|---|---|---|---|---|---|---|
| KTQQLHPVLQK | 11 | Q9H0G5 | Nuclear speckle splicing regulatory protei | NSRP1 | 66.391 | 3 | 50.021336 | yes | 2.354E-07 | 660.39331 | 21.443649 |
| KTQQLHPVLQK | 11 | Q9H0G5 | Nuclear speckle splicing regulatory protei | NSRP1 | 66.391 | 3 | 50.021336 | yes | 1.307E-06 | 440.59799 | 21.443649 |
| HLLNQAVGEEEVI | 14 | Q9H0G5 | Nuclear speckle splicing regulatory protei | NSRP1 | 66.391 | 2 | 50.021336 | yes | 2.519E-07 | 781.91229 | 35.694225 |
| TDESLDWDSSLVK | 13 | Q9H0H5 | Rac GTPase-activating protein 1 OS=Hor | RACGAP1 | 71.027 | 2 | 52.801586 | yes | 1.285E-07 | 747.85156 | 4156.7451 |
| SIGSAVDQGNESIV | 16 | Q9H0H5 | Rac GTPase-activating protein 1 OS=Hor | RACGAP1 | 71.027 | 2 | 52.801586 | yes | 1.108E-07 | 787.90466 | 3769.7537 |
| GLTETGLYR | 9 | Q9H0H5 | Rac GTPase-activating protein 1 OS=Hor | RACGAP1 | 71.027 | 2 | 52.801586 | yes | 1.496E-07 | 505.26929 | 3092.7551 |
| NLFEQLVR | 8 | Q9H0H5 | Rac GTPase-activating protein 1 OS=Hor | RACGAP1 | 71.027 | 2 | 52.801586 | yes | 2.261E-07 | 509.78763 | 2234.853 |
| NLKEPLLTFR | 10 | Q9H0H5 | Rac GTPase-activating protein 1 OS=Hor | RACGAP1 | 71.027 | 2 | 52.801586 | yes | 9.722E-08 | 615.86389 | 1003.0504 |
| NLKEPLLTFR | 10 | Q9H0H5 | Rac GTPase-activating protein 1 OS=Hor | RACGAP1 | 71.027 | 2 | 52.801586 | yes | 9.723E-08 | 410.91168 | 1003.0504 |
| NLKEPLLTFR | 10 | Q9H0H5 | Rac GTPase-activating protein 1 OS=Hor | RACGAP1 | 71.027 | 3 | 52.801586 | yes | 9.722E-08 | 615.86389 | 1003.0504 |
| NLKEPLLTFR | 10 | Q9H0H5 | Rac GTPase-activating protein 1 OS=Hor | RACGAP1 | 71.027 | 3 | 52.801586 | yes | 9.723E-08 | 410.91168 | 1003.0504 |
| TVIKPESCVPCGK | 13 | Q9H0H5 | Rac GTPase-activating protein 1 OS=Hor | RACGAP1 | 71.027 | 2 | 52.801586 | yes | 1.069E-07 | 737.87329 | 785.91254 |
| TVIKPESCVPCGK | 13 | Q9H0H5 | Rac GTPase-activating protein 1 OS=Hor | RACGAP1 | 71.027 | 2 | 52.801586 | yes | 1.643E-06 | 492.25128 | 785.91254 |
| TVIKPESCVPCGK | 13 | Q9H0H5 | Rac GTPase-activating protein 1 OS=Hor | RACGAP1 | 71.027 | 3 | 52.801586 | yes | 1.069E-07 | 737.87329 | 785.91254 |
| TVIKPESCVPCGK | 13 | Q9H0H5 | Rac GTPase-activating protein 1 OS=Hor | RACGAP1 | 71.027 | 3 | 52.801586 | yes | 1.643E-06 | 492.25128 | 785.91254 |
| QFVDGPPGPVK | 11 | Q9H0H5 | Rac GTPase-activating protein 1 OS=Hor | RACGAP1 | 71.027 | 2 | 52.801586 | yes | 1.016E-07 | 570.80603 | 1510.7738 |
| TPSSSSLSQR | 10 | Q9H0H5 | Rac GTPase-activating protein 1 OS=Hor | RACGAP1 | 71.027 | 2 | 52.801586 | yes | 2.318E-07 | 525.26471 | 1412.6056 |
| NQVDVEIK | 8 | Q9H0H5 | Rac GTPase-activating protein 1 OS=Hor | RACGAP1 | 71.027 | 2 | 52.801586 | yes | 1.584E-07 | 472.75601 | 1358.8353 |
| EPLLTFR | 7 | Q9H0H5 | Rac GTPase-activating protein 1 OS=Hor | RACGAP1 | 71.027 | 2 | 52.801586 | yes | 2.721E-07 | 438.2529 | 1302.4302 |
| MDVANLAK | 8 | Q9H0H5 | Rac GTPase-activating protein 1 OS=Hor | RACGAP1 | 71.027 | 2 | 52.801586 | yes | 2.656E-07 | 431.22858 | 844.0835 |
| QGNFFASPMLK | 11 | Q9H0H5 | Rac GTPase-activating protein 1 OS=Hor | RACGAP1 | 71.027 | 2 | 52.801586 | yes | 6.873E-07 | 620.31317 | 735.99329 |
| CPLPCIPTLIGTPVI | 15 | Q9H0H5 | Rac GTPase-activating protein 1 OS=Hor | RACGAP1 | 71.027 | 2 | 52.801586 | yes | 9.722E-08 | 833.45697 | 472.79272 |
| RVEILSEGNEVQFI | 18 | Q9H0H5 | Rac GTPase-activating protein 1 OS=Hor | RACGAP1 | 71.027 | 3 | 52.801586 | yes | 9.845E-08 | 691.71613 | 453.97934 |
| DFEDFRK | 7 | Q9H0H5 | Rac GTPase-activating protein 1 OS=Hor | RACGAP1 | 71.027 | 2 | 52.801586 | yes | 1.616E-07 | 478.72726 | 337.93982 |
| YKDLLMK | 7 | Q9H0H5 | Rac GTPase-activating protein 1 OS=Hor | RACGAP1 | 71.027 | 2 | 52.801586 | yes | 1.29E-07 | 455.75696 | 172.6723 |
| AEADCEKLER | 10 | Q9H0H5 | Rac GTPase-activating protein 1 OS=Hor | RACGAP1 | 71.027 | 2 | 52.801586 | yes | 9.722E-08 | 610.78241 | 144.75037 |
| SALAFLNR | 8 | Q9H0H5 | Rac GTPase-activating protein 1 OS=Hor | RACGAP1 | 71.027 | 2 | 52.801586 | yes | 2.601E-07 | 446.25598 | 80.222832 |
| LHDFVSK | 7 | Q9H0H5 | Rac GTPase-activating protein 1 OS=Hor | RACGAP1 | 71.027 | 2 | 52.801586 | yes | 2.054E-07 | 423.22943 | 62.896427 |
| MDTMMLNVR | 9 | Q9H0H5 | Rac GTPase-activating protein 1 OS=Hor | RACGAP1 | 71.027 | 2 | 52.801586 | yes | 4.26E-07 | 576.76416 | 61.884808 |
| TETDSVGTPQSNG | 16 | Q9H0H5 | Rac GTPase-activating protein 1 OS=Hor | RACGAP1 | 71.027 | 2 | 52.801586 | yes | 9.722E-08 | 818.86523 | 40.845123 |
| VSLLGPVTTPEHQ | 16 | Q9H0H5 | Rac GTPase-activating protein 1 OS=Hor | RACGAP1 | 71.027 | 3 | 52.801586 | yes | 2.845E-06 | 578.00494 | 25.364283 |
| ELVLLPAGADSLR | 13 | Q9H0I2 | Enkurin domain-containing protein 1 OS= | ENKD1 | 38.76 | 2 | 43.405025 | yes | 2.606E-07 | 677.39044 | 133.89391 |
| LDLLTSDR | 8 | Q9H0I2 | Enkurin domain-containing protein 1 OS= | ENKD1 | 38.76 | 2 | 43.405025 | yes | 6.534E-07 | 466.75601 | 88.07663 |
| VLVGEFVR | 8 | Q9H0J9 | Protein mono-ADP-ribosyltransferase PA | PARP12 | 79.064 | 2 | 56.512794 | yes | 1.494E-07 | 459.77402 | 1900.094 |
| VVLAASPLR | 9 | Q9H0J9 | Protein mono-ADP-ribosyltransferase PA | PARP12 | 79.064 | 2 | 56.512794 | yes | 2.067E-07 | 463.2951 | 1687.8445 |
| LGMSSDLVSR | 10 | Q9H0J9 | Protein mono-ADP-ribosyltransferase PA | PARP12 | 79.064 | 2 | 56.512794 | yes | 1.149E-07 | 532.77386 | 1686.1332 |
| MGLSADALER | 10 | Q9H0J9 | Protein mono-ADP-ribosyltransferase PA | PARP12 | 79.064 | 2 | 56.512794 | yes | 1.229E-07 | 531.76605 | 998.18207 |
| GNASFVRPPAK | 11 | Q9H0J9 | Protein mono-ADP-ribosyltransferase PA | PARP12 | 79.064 | 2 | 56.512794 | yes | 1.139E-07 | 572.31708 | 927.36157 |
| ITLSSSSEEYQK | 12 | Q9H0J9 | Protein mono-ADP-ribosyltransferase PA | PARP12 | 79.064 | 2 | 56.512794 | yes | 1.343E-07 | 686.33557 | 856.1543 |

| | | | | | | | | | | | |
|---|---|---|---|---|---|---|---|---|---|---|---|
| HNYELDFK | 8 | Q9H0J9 | Protein mono-ADP-ribosyltransferase PA | PARP12 | 79.064 | 2 | 56.512794 | yes | 3.346E-07 | 533.2536 | 816.09802 |
| TLPFYFVQK | 9 | Q9H0J9 | Protein mono-ADP-ribosyltransferase PA | PARP12 | 79.064 | 2 | 56.512794 | yes | 9.722E-08 | 571.81586 | 541.56696 |
| VCGVHGTSYGK | 11 | Q9H0J9 | Protein mono-ADP-ribosyltransferase PA | PARP12 | 79.064 | 2 | 56.512794 | yes | 9.732E-08 | 582.77698 | 480.78583 |
| DAAYSHHYSK | 10 | Q9H0J9 | Protein mono-ADP-ribosyltransferase PA | PARP12 | 79.064 | 2 | 56.512794 | yes | 1.111E-07 | 589.76489 | 240.41805 |
| SDTQTHTMFLAR | 12 | Q9H0J9 | Protein mono-ADP-ribosyltransferase PA | PARP12 | 79.064 | 2 | 56.512794 | yes | 2.554E-07 | 704.33789 | 187.97482 |
| VPPLFVPQGTSER | 13 | Q9H0J9 | Protein mono-ADP-ribosyltransferase PA | PARP12 | 79.064 | 2 | 56.512794 | yes | 1.086E-06 | 713.88806 | 108.84885 |
| GKWEDLDNMELII | 20 | Q9H0J9 | Protein mono-ADP-ribosyltransferase PA | PARP12 | 79.064 | 3 | 56.512794 | yes | 9.722E-08 | 818.70325 | 99.472893 |
| PGCVGLCAQLHLC | 14 | Q9H0J9 | Protein mono-ADP-ribosyltransferase PA | PARP12 | 79.064 | 2 | 56.512794 | yes | 1.033E-07 | 820.89453 | 77.600952 |
| NLVYGTTK | 8 | Q9H0J9 | Protein mono-ADP-ribosyltransferase PA | PARP12 | 79.064 | 2 | 56.512794 | yes | 1.134E-06 | 448.2478 | 71.694717 |
| SHDFSNSENLEK | 12 | Q9H0J9 | Protein mono-ADP-ribosyltransferase PA | PARP12 | 79.064 | 2 | 56.512794 | yes | 9.722E-08 | 703.81274 | 52.780144 |
| DLNFLEDNPSLK | 12 | Q9H0K1 | Serine/threonine-protein kinase SIK2 OS= | SIK2 | 103.916 | 2 | 52.988834 | yes | 9.722E-08 | 702.85388 | 84.004929 |
| WMLIEVPVQR | 10 | Q9H0K1 | Serine/threonine-protein kinase SIK2 OS= | SIK2 | 103.916 | 2 | 52.988834 | yes | 3.834E-07 | 635.85248 | 38.548622 |
| VGFYDIEGTLGK | 12 | Q9H0K1 | Serine/threonine-protein kinase SIK2 OS= | SIK2 | 103.916 | 2 | 52.988834 | yes | 2.671E-07 | 649.83496 | 27.270086 |
| AQNTCQLYCK | 10 | Q9H0K1 | Serine/threonine-protein kinase SIK2 OS= | SIK2 | 103.916 | 2 | 52.988834 | yes | 3.288E-06 | 643.2868 | 21.321323 |
| IMEAIENVK | 9 | Q9H0K6 | Pseudouridylate synthase PUS7L OS=Ho | PUS7L | 80.701 | 2 | 48.161274 | yes | 1.189E-07 | 523.78119 | 56.089165 |
| IHVTPLIPGK | 10 | Q9H0L4 | Cleavage stimulation factor subunit 2 tau | CSTF2T | 64.437 | 2 | 38.127384 | yes | 5.44E-07 | 537.83716 | 49.485298 |
| NQLQGAMQQFNQ | 13 | Q9H0M0 | NEDD4-like E3 ubiquitin-protein ligase \ | WWP1 | 105.202 | 2 | 46.878258 | yes | 7.498E-07 | 781.87823 | 73.408127 |
| LQLQVTVSSAK | 11 | Q9H0M0 | NEDD4-like E3 ubiquitin-protein ligase \ | WWP1 | 105.202 | 2 | 46.878258 | yes | 1.451E-07 | 587.34552 | 30.138086 |
| IQIEAIPLALQGR | 13 | Q9H0S4 | Probable ATP-dependent RNA helicase D | DDX47 | 50.647 | 2 | 52.29773 | yes | 1.572E-07 | 711.42737 | 2876.5071 |
| YLVMDEADR | 9 | Q9H0S4 | Probable ATP-dependent RNA helicase D | DDX47 | 50.647 | 2 | 52.29773 | yes | 1.122E-07 | 556.25806 | 2383.9624 |
| DIIGLAETGSGK | 12 | Q9H0S4 | Probable ATP-dependent RNA helicase D | DDX47 | 50.647 | 2 | 52.29773 | yes | 1.104E-07 | 580.81152 | 1879.5441 |
| ILNMDFETEVDK | 12 | Q9H0S4 | Probable ATP-dependent RNA helicase D | DDX47 | 50.647 | 2 | 52.29773 | yes | 1.001E-07 | 727.3476 | 925.86365 |
| ILNMDFETEVDK | 12 | Q9H0S4 | Probable ATP-dependent RNA helicase D | DDX47 | 50.647 | 2 | 52.29773 | yes | 3.006E-06 | 735.34503 | 925.86365 |
| LFALVLTPTR | 10 | Q9H0S4 | Probable ATP-dependent RNA helicase D | DDX47 | 50.647 | 2 | 52.29773 | yes | 1.916E-07 | 565.85022 | 1264.616 |
| TFLFSATMTK | 10 | Q9H0S4 | Probable ATP-dependent RNA helicase D | DDX47 | 50.647 | 2 | 52.29773 | yes | 9.893E-08 | 573.79681 | 1047.4127 |
| GLDIPHVDVVVNF | 20 | Q9H0S4 | Probable ATP-dependent RNA helicase D | DDX47 | 50.647 | 3 | 52.29773 | yes | 1.164E-07 | 734.72333 | 992.16016 |
| LIDHLENTK | 9 | Q9H0S4 | Probable ATP-dependent RNA helicase D | DDX47 | 50.647 | 2 | 52.29773 | yes | 1.376E-07 | 541.79565 | 887.7793 |
| LQQYYIFIPSK | 11 | Q9H0S4 | Probable ATP-dependent RNA helicase D | DDX47 | 50.647 | 2 | 52.29773 | yes | 1.21E-07 | 700.38464 | 858.68542 |
| SILLATDVASR | 11 | Q9H0S4 | Probable ATP-dependent RNA helicase D | DDX47 | 50.647 | 2 | 52.29773 | yes | 9.722E-08 | 573.3299 | 460.73004 |
| KPHIIIATPGR | 11 | Q9H0S4 | Probable ATP-dependent RNA helicase D | DDX47 | 50.647 | 2 | 52.29773 | yes | 1.119E-07 | 601.87201 | 379.5842 |
| IEHLIGK | 7 | Q9H0S4 | Probable ATP-dependent RNA helicase D | DDX47 | 50.647 | 2 | 52.29773 | yes | 9.722E-08 | 405.24762 | 95.626259 |
| KTFLFSATMTK | 11 | Q9H0S4 | Probable ATP-dependent RNA helicase D | DDX47 | 50.647 | 2 | 52.29773 | yes | 4.414E-07 | 637.8443 | 84.133598 |
| NLGFTAIPLHGQM | 17 | Q9H0S4 | Probable ATP-dependent RNA helicase D | DDX47 | 50.647 | 3 | 52.29773 | yes | 1.573E-07 | 610.31732 | 59.424667 |
| EDAGDNDDTEGAI | 16 | Q9H0S4 | Probable ATP-dependent RNA helicase D | DDX47 | 50.647 | 2 | 52.29773 | yes | 1.81E-07 | 817.35046 | 58.598743 |
| VSQLMEWTNK | 10 | Q9H0U3 | Magnesium transporter protein 1 OS=Hoı | MAGT1 | 38.037 | 2 | 50.120567 | yes | 9.897E-08 | 618.30811 | 930.8064 |
| GFSAEQIAR | 9 | Q9H0U3 | Magnesium transporter protein 1 OS=Hoı | MAGT1 | 38.037 | 2 | 50.120567 | yes | 1.74E-07 | 489.75378 | 380.41412 |
| RGDTYELQVR | 10 | Q9H0U3 | Magnesium transporter protein 1 OS=Hoı | MAGT1 | 38.037 | 2 | 50.120567 | yes | 1.22E-07 | 618.82019 | 251.38577 |
| SNMEFLFNK | 9 | Q9H0U3 | Magnesium transporter protein 1 OS=Hoı | MAGT1 | 38.037 | 2 | 50.120567 | yes | 2.611E-07 | 565.271 | 94.02018 |
| SNMEFLFNK | 9 | Q9H0U3 | Magnesium transporter protein 1 OS=Hoı | MAGT1 | 38.037 | 2 | 50.120567 | yes | 3.157E-07 | 573.26843 | 94.02018 |

| | | | | | | | | | | | |
|---|---|---|---|---|---|---|---|---|---|---|---|
| KEMVLSEK | 8 | Q9H0U3 | Magnesium transporter protein 1 OS=Hoı | MAGT1 | 38.037 | 2 | 50.120567 | yes | 1.373E-07 | 482.2626 | 154.92033 |
| YHGYPYSFLMS | 11 | Q9H0U3 | Magnesium transporter protein 1 OS=Hoı | MAGT1 | 38.037 | 2 | 50.120567 | yes | 1.166E-07 | 682.80261 | 65.36293 |
| YHGYPYSFLMS | 11 | Q9H0U3 | Magnesium transporter protein 1 OS=Hoı | MAGT1 | 38.037 | 2 | 50.120567 | yes | 3.601E-06 | 690.80011 | 65.36293 |
| NVVACESIGR | 10 | Q9H0U6 | Large ribosomal subunit protein uL18m C | MRPL18 | 20.577 | 2 | 47.251328 | yes | 1.369E-07 | 552.77698 | 175.59828 |
| LQSAMTEGGVVLI | 13 | Q9H0U6 | Large ribosomal subunit protein uL18m C | MRPL18 | 20.577 | 2 | 47.251328 | yes | 2.73E-07 | 680.86627 | 128.30251 |
| NLELLSVAR | 9 | Q9H0U6 | Large ribosomal subunit protein uL18m C | MRPL18 | 20.577 | 2 | 47.251328 | yes | 1.917E-07 | 507.80075 | 107.40489 |
| GMGTAAPAAMDP | 18 | Q9H0W8 | Nonsense-mediated mRNA decay factor S | SMG9 | 57.651 | 2 | 39.12236 | yes | 4.776E-07 | 836.41327 | 104.26354 |
| ANVNLAYLGAK | 11 | Q9H0Z9 | RNA-binding protein 38 OS=Homo sapie | RBM38 | 25.498 | 2 | 53.669662 | no | 9.722E-08 | 567.31934 | 122.73544 |
| ACKDPNPIIDGR | 12 | Q9H0Z9 | RNA-binding protein 38 OS=Homo sapie | RBM38 | 25.498 | 2 | 53.669662 | no | 1.17E-06 | 678.34045 | 98.976425 |
| GYGFVTMADR | 10 | Q9H0Z9 | RNA-binding protein 38 OS=Homo sapie | RBM38 | 25.498 | 2 | 53.669662 | no | 4.229E-07 | 558.76074 | 66.112373 |
| PLAAPGAMHGSQI | 13 | Q9H0Z9 | RNA-binding protein 38 OS=Homo sapie | RBM38 | 25.498 | 2 | 53.669662 | yes | 9.81E-08 | 632.82697 | 56.461384 |
| KYFEGFGDIEEAV | 18 | Q9H0Z9 | RNA-binding protein 38 OS=Homo sapie | RBM38 | 25.498 | 3 | 53.669662 | yes | 3.895E-07 | 696.68097 | 36.027847 |
| SLQTGFAIGVQQLI | 20 | Q9H0Z9 | RNA-binding protein 38 OS=Homo sapie | RBM38 | 25.498 | 3 | 53.669662 | yes | 5.069E-07 | 736.41467 | 24.645781 |
| KANVNLAYLGAK | 12 | Q9H0Z9 | RNA-binding protein 38 OS=Homo sapie | RBM38 | 25.498 | 2 | 53.669662 | no | 9.722E-08 | 631.36676 | 21.992157 |
| SFVLPSWMVEK | 11 | Q9H147 | Deoxynucleotidyltransferase terminal-inte | DNTTIP1 | 37.014 | 2 | 54.07453 | yes | 2.128E-07 | 661.8443 | 393.46796 |
| LNESTTFVLGSR | 12 | Q9H147 | Deoxynucleotidyltransferase terminal-inte | DNTTIP1 | 37.014 | 2 | 54.07453 | yes | 1.062E-07 | 662.34882 | 341.37308 |
| MAYLLIEEDIR | 11 | Q9H147 | Deoxynucleotidyltransferase terminal-inte | DNTTIP1 | 37.014 | 2 | 54.07453 | yes | 9.722E-08 | 683.35779 | 311.90018 |
| AAAGMVWKPK | 10 | Q9H147 | Deoxynucleotidyltransferase terminal-inte | DNTTIP1 | 37.014 | 2 | 54.07453 | yes | 1.3E-07 | 529.79443 | 183.15228 |
| LTHELPGIK | 9 | Q9H147 | Deoxynucleotidyltransferase terminal-inte | DNTTIP1 | 37.014 | 2 | 54.07453 | yes | 1.56E-07 | 504.29785 | 166.69476 |
| DLAASDDYR | 9 | Q9H147 | Deoxynucleotidyltransferase terminal-inte | DNTTIP1 | 37.014 | 2 | 54.07453 | yes | 3.347E-07 | 513.23035 | 102.58675 |
| LLFSDGEK | 8 | Q9H147 | Deoxynucleotidyltransferase terminal-inte | DNTTIP1 | 37.014 | 2 | 54.07453 | yes | 2.893E-06 | 454.73984 | 100.49448 |
| KYMETLR | 7 | Q9H147 | Deoxynucleotidyltransferase terminal-inte | DNTTIP1 | 37.014 | 2 | 54.07453 | yes | 3.473E-07 | 470.74966 | 59.182888 |
| GRPPGHILSSDR | 12 | Q9H147 | Deoxynucleotidyltransferase terminal-inte | DNTTIP1 | 37.014 | 3 | 54.07453 | yes | 1.797E-06 | 431.2337 | 33.820755 |
| WVDSNVPQIIR | 11 | Q9H1A4 | Anaphase-promoting complex subunit 1 ( | ANAPC1 | 216.5 | 2 | 51.41995 | yes | 9.722E-08 | 663.86188 | 226.82054 |
| VQYVVDHAMK | 10 | Q9H1A4 | Anaphase-promoting complex subunit 1 ( | ANAPC1 | 216.5 | 2 | 51.41995 | yes | 4.159E-07 | 595.30536 | 112.44371 |
| EMSETSELWQIK | 12 | Q9H1A4 | Anaphase-promoting complex subunit 1 ( | ANAPC1 | 216.5 | 2 | 51.41995 | yes | 1.238E-07 | 740.85303 | 110.59074 |
| TQLIFGSVTNIPAK | 14 | Q9H1A4 | Anaphase-promoting complex subunit 1 ( | ANAPC1 | 216.5 | 2 | 51.41995 | yes | 2.274E-07 | 744.92468 | 110.38604 |
| LGTMDMSITR | 10 | Q9H1A4 | Anaphase-promoting complex subunit 1 ( | ANAPC1 | 216.5 | 2 | 51.41995 | yes | 2.061E-07 | 562.77557 | 109.869 |
| YHLQALR | 7 | Q9H1A4 | Anaphase-promoting complex subunit 1 ( | ANAPC1 | 216.5 | 2 | 51.41995 | yes | 1.385E-07 | 450.75613 | 106.05833 |
| LLQSAHPVR | 9 | Q9H1A4 | Anaphase-promoting complex subunit 1 ( | ANAPC1 | 216.5 | 2 | 51.41995 | yes | 2.89E-07 | 510.80109 | 101.38112 |
| LLQLCQR | 7 | Q9H1A4 | Anaphase-promoting complex subunit 1 ( | ANAPC1 | 216.5 | 2 | 51.41995 | yes | 1.064E-07 | 465.76312 | 99.103363 |
| VFITSDLCGQK | 11 | Q9H1A4 | Anaphase-promoting complex subunit 1 ( | ANAPC1 | 216.5 | 2 | 51.41995 | yes | 1.022E-07 | 634.32117 | 96.733315 |
| NFDFEGSLSPVIAP | 15 | Q9H1A4 | Anaphase-promoting complex subunit 1 ( | ANAPC1 | 216.5 | 2 | 51.41995 | yes | 1.193E-07 | 810.91705 | 85.197319 |
| LQVEQEENR | 9 | Q9H1A4 | Anaphase-promoting complex subunit 1 ( | ANAPC1 | 216.5 | 2 | 51.41995 | yes | 2.417E-06 | 572.78326 | 84.836365 |
| YGLLFER | 7 | Q9H1A4 | Anaphase-promoting complex subunit 1 ( | ANAPC1 | 216.5 | 2 | 51.41995 | yes | 1.975E-07 | 449.24509 | 54.082138 |
| LVLEFFSSR | 9 | Q9H1A4 | Anaphase-promoting complex subunit 1 ( | ANAPC1 | 216.5 | 2 | 51.41995 | yes | 1.306E-07 | 549.3031 | 51.159878 |
| AHSPALGVHSFSG | 16 | Q9H1A4 | Anaphase-promoting complex subunit 1 ( | ANAPC1 | 216.5 | 3 | 51.41995 | yes | 4.286E-07 | 550.62177 | 47.083778 |
| FSEQGGTPQNVAT | 22 | Q9H1A4 | Anaphase-promoting complex subunit 1 ( | ANAPC1 | 216.5 | 3 | 51.41995 | yes | 1.295E-07 | 763.38037 | 42.947563 |
| FNISSHNQSPK | 11 | Q9H1A4 | Anaphase-promoting complex subunit 1 ( | ANAPC1 | 216.5 | 2 | 51.41995 | yes | 3.776E-07 | 629.81238 | 41.158218 |

| | | | | | | | | | | | |
|---|---|---|---|---|---|---|---|---|---|---|---|
| PSTPLDGVSTPK | 12 | Q9H1A4 | Anaphase-promoting complex subunit 1 ( | ANAPC1 | 216.5 | 2 | 51.41995 | yes | 3.678E-06 | 599.81934 | 35.737259 |
| HSTSVSSLAER | 11 | Q9H1A4 | Anaphase-promoting complex subunit 1 ( | ANAPC1 | 216.5 | 2 | 51.41995 | yes | 7.43E-07 | 587.29657 | 26.098709 |
| IAPASQIDSAWIVY | 18 | Q9H1A4 | Anaphase-promoting complex subunit 1 ( | ANAPC1 | 216.5 | 3 | 51.41995 | yes | 5.009E-06 | 667.6983 | 18.842081 |
| IAYMLVVHGR | 10 | Q9H1B5 | Xylosyltransferase 2 OS=Homo sapiens C | XYLT2 | 96.768 | 2 | 46.106632 | yes | 6.073E-07 | 579.82623 | 64.540276 |
| LLQFWEPLGETR | 12 | Q9H1B5 | Xylosyltransferase 2 OS=Homo sapiens C | XYLT2 | 96.768 | 2 | 46.106632 | yes | 1.651E-07 | 744.89594 | 49.589291 |
| LVYQIIEDAGNK | 12 | Q9H1D9 | DNA-directed RNA polymerase III subun | POLR3F | 35.684 | 2 | 48.693474 | yes | 1.31E-07 | 681.86682 | 94.747261 |
| VQPPDADPVEIEN | 14 | Q9H1D9 | DNA-directed RNA polymerase III subun | POLR3F | 35.684 | 2 | 48.693474 | yes | 1.624E-07 | 789.89154 | 61.56847 |
| LLSMGQLDLLR | 11 | Q9H1D9 | DNA-directed RNA polymerase III subun | POLR3F | 35.684 | 2 | 48.693474 | yes | 1.113E-07 | 629.86304 | 43.582729 |
| SNTGLLYR | 8 | Q9H1D9 | DNA-directed RNA polymerase III subun | POLR3F | 35.684 | 2 | 48.693474 | yes | 0.0001298 | 462.25089 | 24.16861 |
| TSPALHPEQK | 10 | Q9H1I8 | Activating signal cointegrator 1 complex : | ASCC2 | 86.361 | 2 | 49.841908 | yes | 2.138E-07 | 554.29327 | 3705.0361 |
| YFVLYKPPPK | 10 | Q9H1I8 | Activating signal cointegrator 1 complex : | ASCC2 | 86.361 | 2 | 49.841908 | yes | 1.002E-07 | 626.36041 | 1783.3181 |
| YFVLYKPPPK | 10 | Q9H1I8 | Activating signal cointegrator 1 complex : | ASCC2 | 86.361 | 2 | 49.841908 | yes | 5.33E-07 | 417.90939 | 1783.3181 |
| YFVLYKPPPK | 10 | Q9H1I8 | Activating signal cointegrator 1 complex : | ASCC2 | 86.361 | 3 | 49.841908 | yes | 1.002E-07 | 626.36041 | 1783.3181 |
| YFVLYKPPPK | 10 | Q9H1I8 | Activating signal cointegrator 1 complex : | ASCC2 | 86.361 | 3 | 49.841908 | yes | 5.33E-07 | 417.90939 | 1783.3181 |
| LAPTLSQLDR | 10 | Q9H1I8 | Activating signal cointegrator 1 complex : | ASCC2 | 86.361 | 2 | 49.841908 | yes | 1.457E-07 | 557.31677 | 3122.8333 |
| LTPSDMPLLELK | 12 | Q9H1I8 | Activating signal cointegrator 1 complex : | ASCC2 | 86.361 | 2 | 49.841908 | yes | 1.148E-07 | 678.87579 | 1332.2388 |
| LTPSDMPLLELK | 12 | Q9H1I8 | Activating signal cointegrator 1 complex : | ASCC2 | 86.361 | 2 | 49.841908 | yes | 3.347E-07 | 686.87323 | 1332.2388 |
| EMKPDPTPLLTSR | 13 | Q9H1I8 | Activating signal cointegrator 1 complex : | ASCC2 | 86.361 | 2 | 49.841908 | yes | 1.185E-06 | 742.89252 | 1516.7424 |
| LLGDLWQR | 8 | Q9H1I8 | Activating signal cointegrator 1 complex : | ASCC2 | 86.361 | 2 | 49.841908 | yes | 2.479E-07 | 500.78235 | 1469.3102 |
| PALPLDQLQITHK | 13 | Q9H1I8 | Activating signal cointegrator 1 complex : | ASCC2 | 86.361 | 2 | 49.841908 | yes | 2.848E-06 | 737.4248 | 502.31595 |
| PALPLDQLQITHK | 13 | Q9H1I8 | Activating signal cointegrator 1 complex : | ASCC2 | 86.361 | 2 | 49.841908 | yes | 4.396E-06 | 491.9523 | 502.31595 |
| PALPLDQLQITHK | 13 | Q9H1I8 | Activating signal cointegrator 1 complex : | ASCC2 | 86.361 | 3 | 49.841908 | yes | 2.848E-06 | 737.4248 | 502.31595 |
| PALPLDQLQITHK | 13 | Q9H1I8 | Activating signal cointegrator 1 complex : | ASCC2 | 86.361 | 3 | 49.841908 | yes | 4.396E-06 | 491.9523 | 502.31595 |
| KFDEGVASAPEVV | 17 | Q9H1I8 | Activating signal cointegrator 1 complex : | ASCC2 | 86.361 | 3 | 49.841908 | yes | 1.106E-07 | 617.30487 | 608.17017 |
| CLDSYLR | 7 | Q9H1I8 | Activating signal cointegrator 1 complex : | ASCC2 | 86.361 | 2 | 49.841908 | yes | 2.633E-07 | 463.72366 | 447.02228 |
| RPFTIPQVLR | 10 | Q9H1I8 | Activating signal cointegrator 1 complex : | ASCC2 | 86.361 | 2 | 49.841908 | yes | 2.565E-07 | 613.87201 | 391.66565 |
| GNSPLLQK | 8 | Q9H1I8 | Activating signal cointegrator 1 complex : | ASCC2 | 86.361 | 2 | 49.841908 | yes | 4.161E-07 | 428.74799 | 286.34717 |
| SLLNDKR | 7 | Q9H1I8 | Activating signal cointegrator 1 complex : | ASCC2 | 86.361 | 2 | 49.841908 | yes | 4.805E-07 | 423.24561 | 149.59412 |
| PALPLDQLQITHKI | 16 | Q9H1I8 | Activating signal cointegrator 1 complex : | ASCC2 | 86.361 | 3 | 49.841908 | yes | 3.768E-07 | 605.34387 | 122.05116 |
| SVFLTFLR | 8 | Q9H1I8 | Activating signal cointegrator 1 complex : | ASCC2 | 86.361 | 2 | 49.841908 | yes | 1.681E-06 | 491.78964 | 45.790173 |
| LSALEVQFHR | 10 | Q9H1J1 | Regulator of nonsense transcripts 3A OS= | UPF3A | 54.697 | 2 | 54.309845 | yes | 9.722E-08 | 600.3302 | 342.08218 |
| TTPLLEYIK | 9 | Q9H1J1 | Regulator of nonsense transcripts 3A OS= | UPF3A | 54.697 | 2 | 54.309845 | yes | 2.06E-07 | 539.31317 | 314.54315 |
| TSANPETLLGEME | 15 | Q9H1J1 | Regulator of nonsense transcripts 3A OS= | UPF3A | 54.697 | 2 | 54.309845 | yes | 1.053E-07 | 795.88763 | 164.31091 |
| FLETYCVEEEK | 11 | Q9H1J1 | Regulator of nonsense transcripts 3A OS= | UPF3A | 54.697 | 2 | 54.309845 | yes | 9.819E-08 | 723.82648 | 145.51285 |
| QESCAPGAVVK | 11 | Q9H1J1 | Regulator of nonsense transcripts 3A OS= | UPF3A | 54.697 | 2 | 54.309845 | yes | 1.339E-07 | 573.28461 | 96.74015 |
| NPDDILLFR | 9 | Q9H1J1 | Regulator of nonsense transcripts 3A OS= | UPF3A | 54.697 | 2 | 54.309845 | yes | 1.073E-07 | 551.79822 | 66.091675 |
| GSQDSGAPGEAMI | 14 | Q9H1J1 | Regulator of nonsense transcripts 3A OS= | UPF3A | 54.697 | 2 | 54.309845 | yes | 7.932E-07 | 696.29645 | 14.31607 |
| NVGTGLVGAPAC( | 17 | Q9H1K1 | Iron-sulfur cluster assembly enzyme ISCU | ISCU | 17.999 | 2 | 44.93114 | yes | 2.021E-07 | 823.40546 | 68.946777 |
| EYQISAGDFPEVK | 13 | Q9H223 | EH domain-containing protein 4 OS=Hon | EHD4 | 61.176 | 2 | 46.02116 | yes | 3.028E-07 | 741.85919 | 90.505745 |

| | | | | | | | | | | | |
|---|---|---|---|---|---|---|---|---|---|---|---|
| SISVIDSPGILSGEK | 15 | Q9H223 | EH domain-containing protein 4 OS=Hor | EHD4 | 61.176 | 2 | 46.02116 | yes | 3.046E-07 | 751.40906 | 84.120224 |
| AGGADAVQTVTG | 15 | Q9H223 | EH domain-containing protein 4 OS=Hor | EHD4 | 61.176 | 2 | 46.02116 | yes | 1.675E-07 | 686.86261 | 47.036179 |
| FMCSQLPNQVLK | 12 | Q9H223 | EH domain-containing protein 4 OS=Hor | EHD4 | 61.176 | 2 | 46.02116 | yes | 5.732E-07 | 732.87054 | 41.101532 |
| LPEIYIQLQR | 10 | Q9H223 | EH domain-containing protein 4 OS=Hor | EHD4 | 61.176 | 2 | 46.02116 | yes | 4.203E-07 | 636.86914 | 31.199896 |
| ATSEVASDYK | 10 | Q9H254 | Spectrin beta chain, non-erythrocytic 4 O | SPTBN4 | 288.986 | 2 | 36.660736 | yes | 1.666E-05 | 535.75366 | 437.15366 |
| ALLQQALR | 8 | Q9H254 | Spectrin beta chain, non-erythrocytic 4 O | SPTBN4 | 288.986 | 2 | 36.660736 | yes | 6.746E-07 | 456.78491 | 90.696289 |
| LGDTPGVSYSDIA | 15 | Q9H269 | Vacuolar protein sorting-associated protei | VPS16 | 94.694 | 2 | 48.805065 | yes | 3.013E-06 | 761.3808 | 106.18187 |
| VAALQTAADAFYI | 13 | Q9H269 | Vacuolar protein sorting-associated protei | VPS16 | 94.694 | 2 | 48.805065 | yes | 4.359E-07 | 684.86151 | 56.917557 |
| ALLLVGDVAQAAI | 19 | Q9H269 | Vacuolar protein sorting-associated protei | VPS16 | 94.694 | 3 | 48.805065 | yes | 1.382E-06 | 654.36536 | 41.160564 |
| IFHTEFGSGVAILT | 18 | Q9H269 | Vacuolar protein sorting-associated protei | VPS16 | 94.694 | 3 | 48.805065 | yes | 1.102E-07 | 638.33911 | 36.697449 |
| LCEFNCNIR | 9 | Q9H269 | Vacuolar protein sorting-associated protei | VPS16 | 94.694 | 2 | 48.805065 | yes | 2.902E-07 | 613.27625 | 34.01712 |
| LEPSYVIR | 8 | Q9H270 | Vacuolar protein sorting-associated protei | VPS11 | 107.837 | 2 | 51.409428 | yes | 5.59E-07 | 488.77673 | 143.09579 |
| LHAEAISLLK | 10 | Q9H270 | Vacuolar protein sorting-associated protei | VPS11 | 107.837 | 2 | 51.409428 | yes | 4.262E-07 | 547.83203 | 135.48969 |
| LTSSLEAGLQR | 11 | Q9H270 | Vacuolar protein sorting-associated protei | VPS11 | 107.837 | 2 | 51.409428 | yes | 7.294E-07 | 587.82495 | 112.52521 |
| GPCFAFEGHK | 10 | Q9H270 | Vacuolar protein sorting-associated protei | VPS11 | 107.837 | 2 | 51.409428 | yes | 2.295E-07 | 575.26093 | 68.397041 |
| LPFEQAESNMK | 11 | Q9H270 | Vacuolar protein sorting-associated protei | VPS11 | 107.837 | 2 | 51.409428 | yes | 1.912E-06 | 647.31079 | 62.430687 |
| GNYPVTGLAFR | 11 | Q9H270 | Vacuolar protein sorting-associated protei | VPS11 | 107.837 | 2 | 51.409428 | yes | 4.331E-07 | 597.81696 | 49.790688 |
| QILNIYDLCNK | 11 | Q9H270 | Vacuolar protein sorting-associated protei | VPS11 | 107.837 | 2 | 51.409428 | yes | 9.722E-08 | 697.36084 | 49.267784 |
| LTLLTDPPTAR | 11 | Q9H270 | Vacuolar protein sorting-associated protei | VPS11 | 107.837 | 2 | 51.409428 | yes | 9.722E-08 | 599.34552 | 32.990715 |
| LMLQYDAVVNSPI | 14 | Q9H2D1 | Solute carrier family 25 member 32 OS=I | SLC25A32 | 35.408 | 2 | 47.132076 | yes | 1.648E-07 | 821.92212 | 92.511948 |
| GMFDTLVK | 8 | Q9H2D1 | Solute carrier family 25 member 32 OS=I | SLC25A32 | 35.408 | 2 | 47.132076 | yes | 1.393E-07 | 455.73877 | 69.780045 |
| IFAVAATYPYQVV | 14 | Q9H2D1 | Solute carrier family 25 member 32 OS=I | SLC25A32 | 35.408 | 2 | 47.132076 | yes | 2.797E-06 | 799.44049 | 32.823898 |
| ELQVLSEQYSQK | 12 | Q9H2D6 | TRIO and F-actin-binding protein OS=H | TRIOBP | 261.376 | 2 | 34.896484 | yes | 2.155E-06 | 726.37244 | 77.939781 |
| SCTDVTEYAVQR | 12 | Q9H2D6 | TRIO and F-actin-binding protein OS=H | TRIOBP | 261.376 | 2 | 34.896484 | yes | 9.217E-07 | 714.82483 | 18.967279 |
| NDLWVNPLR | 9 | Q9H2G4 | Testis-specific Y-encoded-like protein 2 ( | TSPYL2 | 79.436 | 2 | 51.33737 | yes | 3.503E-07 | 563.80383 | 125.10261 |
| YLTNLQVQDLR | 11 | Q9H2G4 | Testis-specific Y-encoded-like protein 2 ( | TSPYL2 | 79.436 | 2 | 51.33737 | yes | 9.722E-08 | 681.87244 | 110.80659 |
| SVTLHTDVGDIK | 12 | Q9H2H8 | Peptidyl-prolyl cis-trans isomerase-like 3 | PPIL3 | 18.155 | 2 | 51.000538 | yes | 1.022E-07 | 663.84863 | 604.26013 |
| IEVFCER | 7 | Q9H2H8 | Peptidyl-prolyl cis-trans isomerase-like 3 | PPIL3 | 18.155 | 2 | 51.000538 | yes | 1.143E-07 | 476.73148 | 456.28427 |
| VIDGLETLDELEK | 13 | Q9H2H8 | Peptidyl-prolyl cis-trans isomerase-like 3 | PPIL3 | 18.155 | 2 | 51.000538 | yes | 1.331E-07 | 737.38776 | 215.08257 |
| TYRPLNDVHIK | 11 | Q9H2H8 | Peptidyl-prolyl cis-trans isomerase-like 3 | PPIL3 | 18.155 | 2 | 51.000538 | yes | 1.276E-07 | 678.37512 | 186.5957 |
| KFEDEYSEYLK | 11 | Q9H2H8 | Peptidyl-prolyl cis-trans isomerase-like 3 | PPIL3 | 18.155 | 2 | 51.000538 | yes | 9.722E-08 | 725.84045 | 150.03888 |
| VIDGLETLDELEKI | 19 | Q9H2H8 | Peptidyl-prolyl cis-trans isomerase-like 3 | PPIL3 | 18.155 | 3 | 51.000538 | yes | 1.161E-07 | 718.72284 | 105.28619 |
| GFMVQTGDPTGT | 14 | Q9H2H8 | Peptidyl-prolyl cis-trans isomerase-like 3 | PPIL3 | 18.155 | 2 | 51.000538 | yes | 2.445E-07 | 712.33539 | 73.296974 |
| LGEQVFGTTGK | 11 | Q9H2H9 | Sodium-coupled neutral amino acid symp | SLC38A1 | 54.048 | 2 | 30.902111 | yes | 2.022E-06 | 568.80096 | 57.400486 |
| TALDLADPSAK | 11 | Q9H2K2 | Poly [ADP-ribose] polymerase tankyrase- | TNKS2 | 126.918 | 2 | 44.269035 | yes | 4.673E-07 | 551.29297 | 96.934578 |
| SAIDLAPTPQLK | 12 | Q9H2K2 | Poly [ADP-ribose] polymerase tankyrase- | TNKS2 | 126.918 | 2 | 44.269035 | yes | 1.301E-05 | 627.35864 | 78.824997 |
| DGDTDIQDLLR | 11 | Q9H2K2 | Poly [ADP-ribose] polymerase tankyrase- | TNKS2 | 126.918 | 2 | 44.269035 | yes | 3.132E-06 | 630.80695 | 45.175152 |
| AAYCGHLQTCR | 11 | Q9H2K2 | Poly [ADP-ribose] polymerase tankyrase- | TNKS2 | 126.918 | 2 | 44.269035 | yes | 2.257E-07 | 668.79791 | 33.979759 |
| VLIDLIQR | 8 | Q9H2K8 | Serine/threonine-protein kinase TAO3 O | TAOK3 | 105.406 | 2 | 46.908245 | yes | 1.443E-07 | 485.3082 | 70.943237 |

| | | | | | | | | | | | |
|---|---|---|---|---|---|---|---|---|---|---|---|
| SLPSQQMVNR | 10 | Q9H2P0 | Activity-dependent neuroprotector homec | ADNP | 123.563 | 2 | 56.571587 | yes | 5.065E-07 | 580.29803 | 180.36447 |
| ILSDIGLEYCK | 11 | Q9H2P0 | Activity-dependent neuroprotector homec | ADNP | 123.563 | 2 | 56.571587 | yes | 1.022E-06 | 655.83667 | 161.52347 |
| VPAVANYIMK | 10 | Q9H2P0 | Activity-dependent neuroprotector homec | ADNP | 123.563 | 2 | 56.571587 | yes | 2.613E-07 | 553.30737 | 158.92761 |
| MVHIDEEMGPK | 11 | Q9H2P0 | Activity-dependent neuroprotector homec | ADNP | 123.563 | 2 | 56.571587 | yes | 9.722E-08 | 643.29938 | 114.28016 |
| LNQSPSLAPVK | 11 | Q9H2P0 | Activity-dependent neuroprotector homec | ADNP | 123.563 | 2 | 56.571587 | yes | 4.24E-07 | 577.3324 | 87.286407 |
| TYEQMEFPLLK | 11 | Q9H2P0 | Activity-dependent neuroprotector homec | ADNP | 123.563 | 2 | 56.571587 | yes | 1.258E-07 | 699.85229 | 82.966499 |
| SVGQGYSVGQSMI | 13 | Q9H2P0 | Activity-dependent neuroprotector homec | ADNP | 123.563 | 2 | 56.571587 | yes | 1.006E-07 | 678.32227 | 80.475502 |
| SPSLSQSQASR | 11 | Q9H2P0 | Activity-dependent neuroprotector homec | ADNP | 123.563 | 2 | 56.571587 | yes | 4.161E-07 | 574.28876 | 55.383942 |
| SKPLMLIAPK | 10 | Q9H2P0 | Activity-dependent neuroprotector homec | ADNP | 123.563 | 2 | 56.571587 | yes | 3.993E-07 | 549.34119 | 51.443886 |
| QADSVEQAVYYCI | 13 | Q9H2P0 | Activity-dependent neuroprotector homec | ADNP | 123.563 | 2 | 56.571587 | yes | 1.122E-07 | 780.85358 | 51.024429 |
| SKPLMLIAPKPQDI | 14 | Q9H2P0 | Activity-dependent neuroprotector homec | ADNP | 123.563 | 3 | 56.571587 | yes | 9.722E-08 | 522.64099 | 41.584648 |
| NVHSEDFENR | 10 | Q9H2P0 | Activity-dependent neuroprotector homec | ADNP | 123.563 | 2 | 56.571587 | yes | 3.757E-05 | 623.776 | 31.6702 |
| HEMDFDAEWLFEI | 17 | Q9H2P0 | Activity-dependent neuroprotector homec | ADNP | 123.563 | 3 | 56.571587 | yes | 2.375E-05 | 731.30573 | 3.8516304 |
| ELDILLQEK | 9 | Q9H2U1 | ATP-dependent DNA/RNA helicase DH> | DHX36 | 114.761 | 2 | 55.30749 | yes | 1.31E-07 | 550.81354 | 6676.1182 |
| TPLEELCLQIK | 11 | Q9H2U1 | ATP-dependent DNA/RNA helicase DH> | DHX36 | 114.761 | 2 | 55.30749 | yes | 9.722E-08 | 672.3656 | 4858.6172 |
| LMDPPSNEAVLLS | 15 | Q9H2U1 | ATP-dependent DNA/RNA helicase DH> | DHX36 | 114.761 | 2 | 55.30749 | yes | 1.849E-07 | 827.94525 | 2316.9182 |
| LMDPPSNEAVLLS | 15 | Q9H2U1 | ATP-dependent DNA/RNA helicase DH> | DHX36 | 114.761 | 2 | 55.30749 | yes | 9.722E-08 | 835.94275 | 2316.9182 |
| VILMSATLNAEK | 12 | Q9H2U1 | ATP-dependent DNA/RNA helicase DH> | DHX36 | 114.761 | 2 | 55.30749 | yes | 9.722E-08 | 645.36029 | 4613.4785 |
| EKLPSYGMQK | 10 | Q9H2U1 | ATP-dependent DNA/RNA helicase DH> | DHX36 | 114.761 | 2 | 55.30749 | yes | 1.128E-07 | 590.80518 | 4471.4741 |
| TDGLVAVHPK | 10 | Q9H2U1 | ATP-dependent DNA/RNA helicase DH> | DHX36 | 114.761 | 2 | 55.30749 | yes | 2.135E-07 | 518.79291 | 4191.4365 |
| LPVEPHIGK | 9 | Q9H2U1 | ATP-dependent DNA/RNA helicase DH> | DHX36 | 114.761 | 2 | 55.30749 | yes | 2.183E-07 | 495.29257 | 3375.0154 |
| AVVHMDER | 8 | Q9H2U1 | ATP-dependent DNA/RNA helicase DH> | DHX36 | 114.761 | 2 | 55.30749 | yes | 1.801E-07 | 478.73456 | 3248.6152 |
| YIEMQHFR | 8 | Q9H2U1 | ATP-dependent DNA/RNA helicase DH> | DHX36 | 114.761 | 2 | 55.30749 | yes | 1.499E-07 | 562.2713 | 1601.7179 |
| YIEMQHFR | 8 | Q9H2U1 | ATP-dependent DNA/RNA helicase DH> | DHX36 | 114.761 | 2 | 55.30749 | yes | 1.074E-07 | 570.26874 | 1601.7179 |
| GFMQGHVNR | 9 | Q9H2U1 | ATP-dependent DNA/RNA helicase DH> | DHX36 | 114.761 | 2 | 55.30749 | yes | 1.086E-07 | 523.25342 | 3055.1833 |
| HLMELNALDK | 10 | Q9H2U1 | ATP-dependent DNA/RNA helicase DH> | DHX36 | 114.761 | 2 | 55.30749 | yes | 1.064E-07 | 592.31061 | 2885.0549 |
| IVCTQPR | 7 | Q9H2U1 | ATP-dependent DNA/RNA helicase DH> | DHX36 | 114.761 | 2 | 55.30749 | yes | 1.18E-07 | 437.23419 | 2862.0083 |
| ASLLDDYQLPEILF | 14 | Q9H2U1 | ATP-dependent DNA/RNA helicase DH> | DHX36 | 114.761 | 2 | 55.30749 | yes | 9.722E-08 | 823.44342 | 2790.656 |
| QEELTPLGVHLAR | 13 | Q9H2U1 | ATP-dependent DNA/RNA helicase DH> | DHX36 | 114.761 | 2 | 55.30749 | yes | 1.699E-07 | 731.90424 | 1154.4086 |
| QEELTPLGVHLAR | 13 | Q9H2U1 | ATP-dependent DNA/RNA helicase DH> | DHX36 | 114.761 | 2 | 55.30749 | yes | 4.58E-07 | 488.27194 | 1154.4086 |
| QEELTPLGVHLAR | 13 | Q9H2U1 | ATP-dependent DNA/RNA helicase DH> | DHX36 | 114.761 | 3 | 55.30749 | yes | 1.699E-07 | 731.90424 | 1154.4086 |
| QEELTPLGVHLAR | 13 | Q9H2U1 | ATP-dependent DNA/RNA helicase DH> | DHX36 | 114.761 | 3 | 55.30749 | yes | 4.58E-07 | 488.27194 | 1154.4086 |
| AVICAGLYPK | 10 | Q9H2U1 | ATP-dependent DNA/RNA helicase DH> | DHX36 | 114.761 | 2 | 55.30749 | yes | 1.097E-07 | 546.2995 | 2238.707 |
| LGGIAYFLSR | 10 | Q9H2U1 | ATP-dependent DNA/RNA helicase DH> | DHX36 | 114.761 | 2 | 55.30749 | yes | 1.099E-07 | 548.8111 | 2025.8859 |
| DPFVIPLGK | 9 | Q9H2U1 | ATP-dependent DNA/RNA helicase DH> | DHX36 | 114.761 | 2 | 55.30749 | yes | 2.722E-07 | 493.28949 | 1718.7271 |
| EEQIVQLLNSVQA | 14 | Q9H2U1 | ATP-dependent DNA/RNA helicase DH> | DHX36 | 114.761 | 2 | 55.30749 | yes | 1.258E-07 | 799.94104 | 1672.9624 |
| SYDYHQNWGR | 10 | Q9H2U1 | ATP-dependent DNA/RNA helicase DH> | DHX36 | 114.761 | 2 | 55.30749 | yes | 1.076E-07 | 684.29181 | 1480.7113 |
| NLQSDVLMTVVK | 12 | Q9H2U1 | ATP-dependent DNA/RNA helicase DH> | DHX36 | 114.761 | 2 | 55.30749 | yes | 1.456E-07 | 673.87103 | 703.58032 |
| NLQSDVLMTVVK | 12 | Q9H2U1 | ATP-dependent DNA/RNA helicase DH> | DHX36 | 114.761 | 2 | 55.30749 | yes | 2.486E-07 | 681.86853 | 703.58032 |

| | | | | | | | | | | | |
|---|---|---|---|---|---|---|---|---|---|---|---|
| IESPHPVDWNDTK | 13 | Q9H2U1 | ATP-dependent DNA/RNA helicase DH | DHX36 | 114.761 | 2 | 55.30749 | yes | 9.988E-08 | 769.36774 | 665.59222 |
| IESPHPVDWNDTK | 13 | Q9H2U1 | ATP-dependent DNA/RNA helicase DH | DHX36 | 114.761 | 2 | 55.30749 | yes | 1.578E-07 | 513.24756 | 665.59222 |
| IESPHPVDWNDTK | 13 | Q9H2U1 | ATP-dependent DNA/RNA helicase DH | DHX36 | 114.761 | 3 | 55.30749 | yes | 9.988E-08 | 769.36774 | 665.59222 |
| IESPHPVDWNDTK | 13 | Q9H2U1 | ATP-dependent DNA/RNA helicase DH | DHX36 | 114.761 | 3 | 55.30749 | yes | 1.578E-07 | 513.24756 | 665.59222 |
| SVNVEQTDFHYNV | 19 | Q9H2U1 | ATP-dependent DNA/RNA helicase DH | DHX36 | 114.761 | 3 | 55.30749 | yes | 1.69E-07 | 802.73407 | 1188.8394 |
| EIGMWYAK | 8 | Q9H2U1 | ATP-dependent DNA/RNA helicase DH | DHX36 | 114.761 | 2 | 55.30749 | yes | 2.356E-07 | 499.24423 | 538.00385 |
| EIGMWYAK | 8 | Q9H2U1 | ATP-dependent DNA/RNA helicase DH | DHX36 | 114.761 | 2 | 55.30749 | yes | 1.897E-07 | 507.24167 | 538.00385 |
| GQFAEHLLGAGFV | 16 | Q9H2U1 | ATP-dependent DNA/RNA helicase DH | DHX36 | 114.761 | 2 | 55.30749 | yes | 1.237E-07 | 838.43115 | 534.02319 |
| GQFAEHLLGAGFV | 16 | Q9H2U1 | ATP-dependent DNA/RNA helicase DH | DHX36 | 114.761 | 2 | 55.30749 | yes | 5.962E-07 | 559.28986 | 534.02319 |
| GQFAEHLLGAGFV | 16 | Q9H2U1 | ATP-dependent DNA/RNA helicase DH | DHX36 | 114.761 | 3 | 55.30749 | yes | 1.237E-07 | 838.43115 | 534.02319 |
| GQFAEHLLGAGFV | 16 | Q9H2U1 | ATP-dependent DNA/RNA helicase DH | DHX36 | 114.761 | 3 | 55.30749 | yes | 5.962E-07 | 559.28986 | 534.02319 |
| VQPGHCYHLYNGI | 14 | Q9H2U1 | ATP-dependent DNA/RNA helicase DH | DHX36 | 114.761 | 3 | 55.30749 | yes | 1.113E-07 | 571.94745 | 523.68439 |
| VQPGHCYHLYNGI | 14 | Q9H2U1 | ATP-dependent DNA/RNA helicase DH | DHX36 | 114.761 | 3 | 55.30749 | yes | 2.101E-06 | 429.2124 | 523.68439 |
| VQPGHCYHLYNGI | 14 | Q9H2U1 | ATP-dependent DNA/RNA helicase DH | DHX36 | 114.761 | 4 | 55.30749 | yes | 1.113E-07 | 571.94745 | 523.68439 |
| VQPGHCYHLYNGI | 14 | Q9H2U1 | ATP-dependent DNA/RNA helicase DH | DHX36 | 114.761 | 4 | 55.30749 | yes | 2.101E-06 | 429.2124 | 523.68439 |
| LPSYGMQK | 8 | Q9H2U1 | ATP-dependent DNA/RNA helicase DH | DHX36 | 114.761 | 2 | 55.30749 | yes | 1.046E-07 | 462.23639 | 538.04614 |
| DPESNINSDNEK | 12 | Q9H2U1 | ATP-dependent DNA/RNA helicase DH | DHX36 | 114.761 | 2 | 55.30749 | yes | 7.145E-07 | 681.29443 | 523.20105 |
| KELDILLQEK | 10 | Q9H2U1 | ATP-dependent DNA/RNA helicase DH | DHX36 | 114.761 | 2 | 55.30749 | yes | 9.722E-08 | 614.86102 | 259.36264 |
| HLMELNALDKQEI | 23 | Q9H2U1 | ATP-dependent DNA/RNA helicase DH | DHX36 | 114.761 | 4 | 55.30749 | yes | 1.092E-07 | 657.6048 | 161.2663 |
| ELVNLIDNHQVTV | 22 | Q9H2U1 | ATP-dependent DNA/RNA helicase DH | DHX36 | 114.761 | 3 | 55.30749 | yes | 9.722E-08 | 795.06891 | 146.40506 |
| REEQIVQLLNSVQ | 15 | Q9H2U1 | ATP-dependent DNA/RNA helicase DH | DHX36 | 114.761 | 3 | 55.30749 | yes | 9.722E-08 | 585.66351 | 76.083122 |
| NDKESEAQISWFA | 26 | Q9H2U1 | ATP-dependent DNA/RNA helicase DH | DHX36 | 114.761 | 4 | 55.30749 | yes | 1.47E-06 | 732.08478 | 49.373547 |
| NTPCSENKLDIQEI | 14 | Q9H2U1 | ATP-dependent DNA/RNA helicase DH | DHX36 | 114.761 | 2 | 55.30749 | yes | 1.06E-07 | 838.40143 | 5.3099389 |
| YVANIFPYK | 9 | Q9H2U2 | Inorganic pyrophosphatase 2, mitochondr | PPA2 | 37.921 | 2 | 53.256737 | yes | 1.001E-07 | 557.80023 | 519.70068 |
| LIAINANDPEASK | 13 | Q9H2U2 | Inorganic pyrophosphatase 2, mitochondr | PPA2 | 37.921 | 2 | 53.256737 | yes | 9.722E-08 | 678.36188 | 153.42648 |
| SDISQPSGPLLPELS | 16 | Q9H2Y7 | Zinc finger protein 106 OS=Homo sapien | ZNF106 | 208.883 | 2 | 53.801422 | yes | 4.862E-07 | 834.44617 | 259.49719 |
| AVSCLATAQEGAF | 13 | Q9H2Y7 | Zinc finger protein 106 OS=Homo sapien | ZNF106 | 208.883 | 2 | 53.801422 | yes | 9.722E-08 | 667.33008 | 257.63959 |
| VNCLLVTQTSGK | 12 | Q9H2Y7 | Zinc finger protein 106 OS=Homo sapien | ZNF106 | 208.883 | 2 | 53.801422 | yes | 1E-07 | 660.35303 | 201.06647 |
| NAALYTGSSDHTII | 14 | Q9H2Y7 | Zinc finger protein 106 OS=Homo sapien | ZNF106 | 208.883 | 2 | 53.801422 | yes | 9.722E-08 | 753.37079 | 129.0934 |
| LGTVATYR | 8 | Q9H2Y7 | Zinc finger protein 106 OS=Homo sapien | ZNF106 | 208.883 | 2 | 53.801422 | yes | 2.829E-07 | 440.74799 | 99.205307 |
| SLSESSVIMDR | 11 | Q9H2Y7 | Zinc finger protein 106 OS=Homo sapien | ZNF106 | 208.883 | 2 | 53.801422 | yes | 6.58E-07 | 612.30048 | 98.097809 |
| IGTLGSATTELLSG | 17 | Q9H2Y7 | Zinc finger protein 106 OS=Homo sapien | ZNF106 | 208.883 | 2 | 53.801422 | yes | 9.722E-08 | 832.44672 | 82.284515 |
| ECVEQLQLEDR | 11 | Q9H2Y7 | Zinc finger protein 106 OS=Homo sapien | ZNF106 | 208.883 | 2 | 53.801422 | yes | 3.604E-07 | 709.83264 | 80.265305 |
| NCDAFFTAR | 9 | Q9H2Y7 | Zinc finger protein 106 OS=Homo sapien | ZNF106 | 208.883 | 2 | 53.801422 | yes | 1.399E-07 | 551.24274 | 72.993233 |
| SLQCMDNNLLQAI | 13 | Q9H2Y7 | Zinc finger protein 106 OS=Homo sapien | ZNF106 | 208.883 | 2 | 53.801422 | yes | 2.438E-07 | 781.87433 | 71.126274 |
| SQVDQLLNISLR | 12 | Q9H2Y7 | Zinc finger protein 106 OS=Homo sapien | ZNF106 | 208.883 | 2 | 53.801422 | yes | 1.083E-07 | 693.39099 | 62.520519 |
| VCGVTEVGLSAYA | 14 | Q9H2Y7 | Zinc finger protein 106 OS=Homo sapien | ZNF106 | 208.883 | 2 | 53.801422 | yes | 1.64E-07 | 727.3714 | 55.336605 |
| ASSDSAASFEVVR | 13 | Q9H2Y7 | Zinc finger protein 106 OS=Homo sapien | ZNF106 | 208.883 | 2 | 53.801422 | yes | 9.863E-08 | 663.32025 | 54.377129 |
| ESELQMTSAASPH | 21 | Q9H2Y7 | Zinc finger protein 106 OS=Homo sapien | ZNF106 | 208.883 | 3 | 53.801422 | yes | 4.636E-06 | 746.38757 | 46.881668 |

| | | | | | | | | | | | |
|---|---|---|---|---|---|---|---|---|---|---|---|
| LSFASETPLEK | 11 | Q9H2Y7 | Zinc finger protein 106 OS=Homo sapien | ZNF106 | 208.883 | 2 | 53.801422 | yes | 1.443E-07 | 611.32172 | 38.137268 |
| VTTSTWEDSR | 10 | Q9H2Y7 | Zinc finger protein 106 OS=Homo sapien | ZNF106 | 208.883 | 2 | 53.801422 | yes | 1.991E-06 | 591.27527 | 32.891571 |
| NVNWEQVIQQVTI | 13 | Q9H2Y7 | Zinc finger protein 106 OS=Homo sapien | ZNF106 | 208.883 | 2 | 53.801422 | yes | 1.838E-07 | 793.42029 | 28.135448 |
| TNFFSPGEHSNPSN | 28 | Q9H2Y7 | Zinc finger protein 106 OS=Homo sapien | ZNF106 | 208.883 | 4 | 53.801422 | yes | 2.771E-06 | 775.86536 | 21.760962 |
| YFTSNPASK | 9 | Q9H300 | Presenilin-associated rhomboid-like prote | PARL | 42.191 | 2 | 49.73661 | yes | 4.297E-07 | 507.74817 | 134.95454 |
| ADWLDSIRPQK | 11 | Q9H300 | Presenilin-associated rhomboid-like prote | PARL | 42.191 | 2 | 49.73661 | yes | 1.006E-07 | 664.8515 | 112.75854 |
| SDPGTSGEAYK | 11 | Q9H300 | Presenilin-associated rhomboid-like prote | PARL | 42.191 | 2 | 49.73661 | yes | 1.449E-06 | 556.24872 | 108.79869 |
| LEVQAEEER | 9 | Q9H307 | Pinin OS=Homo sapiens OX=9606 GN=I | PNN | 81.628 | 2 | 53.810345 | yes | 1.041E-07 | 551.7724 | 638.54016 |
| LTEVPVEPVLTVH | 17 | Q9H307 | Pinin OS=Homo sapiens OX=9606 GN=I | PNN | 81.628 | 3 | 53.810345 | yes | 1.122E-07 | 625.34717 | 541.2464 |
| RIEFAEQINK | 10 | Q9H307 | Pinin OS=Homo sapiens OX=9606 GN=I | PNN | 81.628 | 2 | 53.810345 | yes | 2.011E-07 | 624.34076 | 173.17891 |
| DLIQDQNMDEK | 11 | Q9H307 | Pinin OS=Homo sapiens OX=9606 GN=I | PNN | 81.628 | 2 | 53.810345 | yes | 2.972E-07 | 674.80609 | 164.10443 |
| MEEETEVR | 8 | Q9H307 | Pinin OS=Homo sapiens OX=9606 GN=I | PNN | 81.628 | 2 | 53.810345 | yes | 5.939E-07 | 511.72659 | 162.28973 |
| HVIADQEVMETNI | 13 | Q9H307 | Pinin OS=Homo sapiens OX=9606 GN=I | PNN | 81.628 | 2 | 53.810345 | yes | 9.722E-08 | 771.3725 | 124.75285 |
| IFGLLMGTLQK | 11 | Q9H307 | Pinin OS=Homo sapiens OX=9606 GN=I | PNN | 81.628 | 2 | 53.810345 | yes | 2.695E-07 | 610.85724 | 74.82431 |
| KPALQSSVVATSK | 13 | Q9H307 | Pinin OS=Homo sapiens OX=9606 GN=I | PNN | 81.628 | 2 | 53.810345 | yes | 3.783E-07 | 658.38263 | 36.28075 |
| LLALSGPGGGR | 11 | Q9H307 | Pinin OS=Homo sapiens OX=9606 GN=I | PNN | 81.628 | 2 | 53.810345 | yes | 1.064E-07 | 499.29309 | 27.149317 |
| FKQESTVATER | 11 | Q9H307 | Pinin OS=Homo sapiens OX=9606 GN=I | PNN | 81.628 | 2 | 53.810345 | yes | 6.632E-07 | 648.33313 | 23.571325 |
| ASNPVIAAQLCSK | 13 | Q9H329 | Band 4.1-like protein 4B OS=Homo sapie | EPB41L4B | 99.713 | 2 | 54.10515 | yes | 1.274E-07 | 679.85846 | 342.37811 |
| YYSSEPNNLR | 10 | Q9H329 | Band 4.1-like protein 4B OS=Homo sapie | EPB41L4B | 99.713 | 2 | 54.10515 | yes | 9.722E-08 | 621.79114 | 268.38733 |
| WLEMYGVDMHV | 13 | Q9H329 | Band 4.1-like protein 4B OS=Homo sapie | EPB41L4B | 99.713 | 2 | 54.10515 | yes | 9.722E-08 | 817.89453 | 191.34845 |
| SPAQAELSYLNK | 12 | Q9H329 | Band 4.1-like protein 4B OS=Homo sapie | EPB41L4B | 99.713 | 2 | 54.10515 | yes | 1.428E-07 | 660.84332 | 158.35739 |
| LTLVVVEDDDQGI | 13 | Q9H329 | Band 4.1-like protein 4B OS=Homo sapie | EPB41L4B | 99.713 | 2 | 54.10515 | yes | 2.029E-07 | 729.87537 | 133.34515 |
| YLFVLQLR | 8 | Q9H329 | Band 4.1-like protein 4B OS=Homo sapie | EPB41L4B | 99.713 | 2 | 54.10515 | yes | 9.733E-08 | 526.3186 | 112.89555 |
| GPDNALLEATGGA | 16 | Q9H3F6 | BTB/POZ domain-containing adapter for | KCTD10 | 35.433 | 2 | 50.30097 | yes | 9.819E-08 | 735.37079 | 162.18356 |
| VEFPEAR | 7 | Q9H3F6 | BTB/POZ domain-containing adapter for | KCTD10 | 35.433 | 2 | 50.30097 | no | 7.127E-07 | 424.21906 | 49.758701 |
| REIEELLAEAK | 11 | Q9H3F6 | BTB/POZ domain-containing adapter for | KCTD10 | 35.433 | 2 | 50.30097 | yes | 1.287E-07 | 650.85901 | 43.095989 |
| SHHLDEDEER | 10 | Q9H3F6 | BTB/POZ domain-containing adapter for | KCTD10 | 35.433 | 2 | 50.30097 | yes | 4.118E-07 | 633.77094 | 24.587166 |
| LIATSNKPAVK | 11 | Q9H3F6 | BTB/POZ domain-containing adapter for | KCTD10 | 35.433 | 2 | 50.30097 | yes | 1.543E-07 | 571.35059 | 22.031305 |
| PITPGTLR | 8 | Q9H3H3 | UPF0696 protein C11orf68 OS=Homo sa | C11orf68 | 31.431 | 2 | 35.699802 | yes | 7.942E-07 | 427.75836 | 32.201023 |
| TLTPDQWAR | 9 | Q9H3K6 | BolA-like protein 2 OS=Homo sapiens O | BOLA2 | 10.117 | 2 | 48.88249 | yes | 1.091E-07 | 544.28015 | 796.90424 |
| FEGKPLLQR | 9 | Q9H3K6 | BolA-like protein 2 OS=Homo sapiens O | BOLA2 | 10.117 | 2 | 48.88249 | yes | 1.785E-06 | 544.31659 | 296.69733 |
| LVNACLAEELPHII | 19 | Q9H3K6 | BolA-like protein 2 OS=Homo sapiens O | BOLA2 | 10.117 | 3 | 48.88249 | yes | 3.389E-07 | 740.38074 | 70.856308 |
| LMILTVTQK | 9 | Q9H3L0 | Cobalamin trafficking protein CblD OS=I | MMADHC | 32.941 | 2 | 49.99422 | yes | 9.937E-08 | 523.81757 | 78.246964 |
| ENPCQEQGDVIQII | 14 | Q9H3P2 | Negative elongation factor A OS=Homo s | NELFA | 57.277 | 2 | 53.540802 | yes | 1.708E-07 | 829.39612 | 145.12547 |
| SPTAPSVFSPTGNI | 14 | Q9H3P2 | Negative elongation factor A OS=Homo s | NELFA | 57.277 | 2 | 53.540802 | yes | 1.2E-07 | 709.35718 | 119.19286 |
| STETAQQLK | 9 | Q9H3P2 | Negative elongation factor A OS=Homo s | NELFA | 57.277 | 2 | 53.540802 | yes | 4.004E-07 | 503.26419 | 82.752037 |
| NALTTLAGPLTPPV | 15 | Q9H3P2 | Negative elongation factor A OS=Homo s | NELFA | 57.277 | 2 | 53.540802 | yes | 2.14E-06 | 746.94031 | 66.539703 |
| EQMFAAQEMFK | 11 | Q9H3P2 | Negative elongation factor A OS=Homo s | NELFA | 57.277 | 2 | 53.540802 | yes | 9.722E-08 | 680.30725 | 64.590469 |
| LLDISELDMVGAG | 14 | Q9H3P2 | Negative elongation factor A OS=Homo s | NELFA | 57.277 | 2 | 53.540802 | yes | 3.437E-07 | 744.88995 | 50.418488 |

| | | | | | | | | | | | |
|---|---|---|---|---|---|---|---|---|---|---|---|
| LSEHTEDLPK | 10 | Q9H3P2 | Negative elongation factor A OS=Homo s | NELFA | 57.277 | 2 | 53.540802 | yes | 2.779E-07 | 584.79584 | 41.198303 |
| LLLGTLHLPR | 10 | Q9H3P2 | Negative elongation factor A OS=Homo s | NELFA | 57.277 | 2 | 53.540802 | yes | 1.152E-07 | 566.86371 | 18.522161 |
| QQEVVVAGSSLPT | 16 | Q9H3P7 | Golgi resident protein GCP60 OS=Homo | ACBD3 | 60.594 | 2 | 46.766758 | yes | 1.475E-07 | 808.92816 | 43.783699 |
| ESLPVIAAPSMWT | 14 | Q9H3P7 | Golgi resident protein GCP60 OS=Homo | ACBD3 | 60.594 | 2 | 46.766758 | yes | 2.673E-07 | 779.40833 | 40.291931 |
| QLVSSSVHSK | 10 | Q9H3Q1 | Cdc42 effector protein 4 OS=Homo sapie | CDC42EP4 | 37.98 | 2 | 49.41254 | yes | 1.034E-07 | 536.29327 | 115.64229 |
| NAMSLPQLNEK | 11 | Q9H3Q1 | Cdc42 effector protein 4 OS=Homo sapie | CDC42EP4 | 37.98 | 2 | 49.41254 | yes | 3.138E-07 | 622.81879 | 75.235687 |
| IDLDSMENSER | 11 | Q9H3R5 | Centromere protein H OS=Homo sapiens | CENPH | 28.481 | 2 | 37.31936 | yes | 6.1E-07 | 654.79047 | 23.838936 |
| ISAQVVDYYK | 10 | Q9H3S7 | Tyrosine-protein phosphatase non-recepto | PTPN23 | 178.974 | 2 | 51.7457 | yes | 3.323E-07 | 593.31116 | 100.6968 |
| GQPDTVQDALR | 11 | Q9H3S7 | Tyrosine-protein phosphatase non-recepto | PTPN23 | 178.974 | 2 | 51.7457 | yes | 3.408E-07 | 600.30438 | 96.796227 |
| ALENPDTASLLGR | 13 | Q9H3S7 | Tyrosine-protein phosphatase non-recepto | PTPN23 | 178.974 | 2 | 51.7457 | yes | 9.728E-08 | 678.8595 | 71.959602 |
| FCYEAVVR | 8 | Q9H3S7 | Tyrosine-protein phosphatase non-recepto | PTPN23 | 178.974 | 2 | 51.7457 | yes | 2.997E-07 | 522.25256 | 65.037178 |
| ASFTNSELHR | 10 | Q9H3S7 | Tyrosine-protein phosphatase non-recepto | PTPN23 | 178.974 | 2 | 51.7457 | yes | 8.506E-07 | 581.28601 | 59.120586 |
| VAYFQSALDK | 10 | Q9H3S7 | Tyrosine-protein phosphatase non-recepto | PTPN23 | 178.974 | 2 | 51.7457 | yes | 7.944E-07 | 571.29803 | 58.185848 |
| DLLEEDELLEQK | 12 | Q9H3S7 | Tyrosine-protein phosphatase non-recepto | PTPN23 | 178.974 | 2 | 51.7457 | yes | 5.409E-07 | 737.36957 | 51.547665 |
| HVEQVLQR | 8 | Q9H3S7 | Tyrosine-protein phosphatase non-recepto | PTPN23 | 178.974 | 2 | 51.7457 | yes | 4.026E-07 | 504.7829 | 50.761761 |
| GQPMVHGALSLAl | 17 | Q9H3S7 | Tyrosine-protein phosphatase non-recepto | PTPN23 | 178.974 | 3 | 51.7457 | yes | 9.722E-08 | 574.98218 | 48.42749 |
| NYGENPEAYNEEL | 14 | Q9H3S7 | Tyrosine-protein phosphatase non-recepto | PTPN23 | 178.974 | 2 | 51.7457 | yes | 1.846E-07 | 835.37067 | 22.171768 |
| SMQQLDPSPAR | 11 | Q9H410 | Kinetochore-associated protein DSN1 hoi | DSN1 | 40.068 | 2 | 40.048725 | yes | 4.239E-07 | 615.30078 | 123.2628 |
| EALENANTNTEVL | 14 | Q9H444 | Charged multivesicular body protein 4b C | CHMP4B | 24.951 | 2 | 51.838055 | yes | 9.722E-08 | 773.39136 | 407.2049 |
| GGPTPQEAIQR | 11 | Q9H444 | Charged multivesicular body protein 4b C | CHMP4B | 24.951 | 2 | 51.838055 | yes | 9.723E-08 | 577.30164 | 256.13651 |
| KQEFLEK | 7 | Q9H444 | Charged multivesicular body protein 4b C | CHMP4B | 24.951 | 2 | 51.838055 | no | 9.728E-08 | 461.25565 | 153.26569 |
| KIEQELTAAK | 10 | Q9H444 | Charged multivesicular body protein 4b C | CHMP4B | 24.951 | 2 | 51.838055 | yes | 8.97E-07 | 565.8244 | 149.07167 |
| AAHDNMDIDK | 10 | Q9H444 | Charged multivesicular body protein 4b C | CHMP4B | 24.951 | 2 | 51.838055 | no | 3.676E-07 | 565.25073 | 53.302917 |
| AKDLVELDPEVEC | 31 | Q9H4A5 | Golgi phosphoprotein 3-like OS=Homo s: | GOLPH3L | 32.768 | 4 | 24.391312 | yes | 3.462E-06 | 843.94037 | 21.106665 |
| SDAPTGDVLLDEA | 15 | Q9H4A6 | Golgi phosphoprotein 3 OS=Homo sapier | GOLPH3 | 33.811 | 2 | 43.62268 | yes | 2.514E-07 | 772.39612 | 59.34243 |
| LQLEACGMR | 9 | Q9H4A6 | Golgi phosphoprotein 3 OS=Homo sapier | GOLPH3 | 33.811 | 2 | 43.62268 | yes | 2.663E-07 | 539.26263 | 30.154375 |
| LSAPSYLAR | 9 | Q9H4B6 | Protein salvador homolog 1 OS=Homo sa | SAV1 | 44.634 | 2 | 45.799747 | yes | 5.307E-07 | 489.27435 | 217.63275 |
| YYYSDNFFDGQR | 12 | Q9H4B6 | Protein salvador homolog 1 OS=Homo sa | SAV1 | 44.634 | 2 | 45.799747 | yes | 2.383E-07 | 787.83093 | 145.56242 |
| QALLTELENR | 10 | Q9H4B6 | Protein salvador homolog 1 OS=Homo sa | SAV1 | 44.634 | 2 | 45.799747 | yes | 1.737E-07 | 593.82495 | 99.496872 |
| NEVSKPAEVQGK | 12 | Q9H4B6 | Protein salvador homolog 1 OS=Homo sa | SAV1 | 44.634 | 2 | 45.799747 | yes | 3.921E-07 | 643.34094 | 28.644213 |
| NSSCFVEWIPNNV | 14 | Q9H4B7 | Tubulin beta-1 chain OS=Homo sapiens ( | TUBB1 | 50.327 | 2 | 41.677486 | yes | 3.394E-07 | 847.40375 | 65.556389 |
| EGIVIPECAPGGLY | 15 | Q9H4F8 | SPARC-related modular calcium-binding | SMOC1 | 48.163 | 2 | 54.156925 | yes | 7.691E-07 | 801.91345 | 112.74181 |
| QSALEEAQQNPR | 12 | Q9H4F8 | SPARC-related modular calcium-binding | SMOC1 | 48.163 | 2 | 54.156925 | yes | 1.666E-07 | 685.83661 | 72.221588 |
| DPTLGVVHR | 9 | Q9H4F8 | SPARC-related modular calcium-binding | SMOC1 | 48.163 | 2 | 54.156925 | yes | 4.075E-07 | 497.27744 | 69.13678 |
| FTDYCDLNK | 9 | Q9H4F8 | SPARC-related modular calcium-binding | SMOC1 | 48.163 | 2 | 54.156925 | yes | 3.822E-07 | 588.25549 | 62.466297 |
| SYESMCEYQR | 10 | Q9H4F8 | SPARC-related modular calcium-binding | SMOC1 | 48.163 | 2 | 54.156925 | yes | 9.722E-08 | 676.76593 | 58.562233 |
| TTEADDPFKDR | 11 | Q9H4F8 | SPARC-related modular calcium-binding | SMOC1 | 48.163 | 2 | 54.156925 | yes | 2.062E-07 | 647.79913 | 23.2472 |
| ELTLGNLLR | 9 | Q9H4H8 | Protein FAM83D OS=Homo sapiens OX= | FAM83D | 64.425 | 2 | 50.951534 | yes | 1.12E-07 | 514.80859 | 1976.9574 |
| FLNPDEVHAILR | 12 | Q9H4H8 | Protein FAM83D OS=Homo sapiens OX= | FAM83D | 64.425 | 2 | 50.951534 | yes | 1.28E-07 | 712.38824 | 1927.4972 |

| | | | | | | | | | | | |
|---|---|---|---|---|---|---|---|---|---|---|---|
| FTLIDGIR | 8 | Q9H4H8 | Protein FAM83D OS=Homo sapiens OX= | FAM83D | 64.425 | 2 | 50.951534 | yes | 2.552E-07 | 467.77145 | 1564.6866 |
| TITGNIYYAR | 10 | Q9H4H8 | Protein FAM83D OS=Homo sapiens OX= | FAM83D | 64.425 | 2 | 50.951534 | yes | 9.728E-08 | 586.30896 | 1091.0825 |
| VETHFQPR | 8 | Q9H4H8 | Protein FAM83D OS=Homo sapiens OX= | FAM83D | 64.425 | 2 | 50.951534 | yes | 4.442E-07 | 507.26178 | 859.49628 |
| IASSQTTIWSR | 11 | Q9H4H8 | Protein FAM83D OS=Homo sapiens OX= | FAM83D | 64.425 | 2 | 50.951534 | yes | 1.121E-07 | 625.33038 | 778.61865 |
| KADLDPEMPAEGI | 13 | Q9H4H8 | Protein FAM83D OS=Homo sapiens OX= | FAM83D | 64.425 | 2 | 50.951534 | yes | 1.713E-07 | 700.83997 | 374.19589 |
| KADLDPEMPAEGI | 13 | Q9H4H8 | Protein FAM83D OS=Homo sapiens OX= | FAM83D | 64.425 | 2 | 50.951534 | yes | 4.68E-07 | 467.56238 | 374.19589 |
| KADLDPEMPAEGI | 13 | Q9H4H8 | Protein FAM83D OS=Homo sapiens OX= | FAM83D | 64.425 | 3 | 50.951534 | yes | 1.713E-07 | 700.83997 | 374.19589 |
| KADLDPEMPAEGI | 13 | Q9H4H8 | Protein FAM83D OS=Homo sapiens OX= | FAM83D | 64.425 | 3 | 50.951534 | yes | 4.68E-07 | 467.56238 | 374.19589 |
| VATGSYSFTWTDC | 14 | Q9H4H8 | Protein FAM83D OS=Homo sapiens OX= | FAM83D | 64.425 | 2 | 50.951534 | yes | 1.396E-07 | 760.35681 | 711.72546 |
| LLSHFQSSNK | 10 | Q9H4H8 | Protein FAM83D OS=Homo sapiens OX= | FAM83D | 64.425 | 2 | 50.951534 | yes | 1.241E-07 | 580.80658 | 683.85974 |
| FDHLTNR | 7 | Q9H4H8 | Protein FAM83D OS=Homo sapiens OX= | FAM83D | 64.425 | 2 | 50.951534 | yes | 1.055E-07 | 451.72757 | 512.33142 |
| LNHMLAMLSR | 10 | Q9H4H8 | Protein FAM83D OS=Homo sapiens OX= | FAM83D | 64.425 | 2 | 50.951534 | yes | 1.001E-07 | 593.31519 | 492.92126 |
| GAGEGGPYGCK | 11 | Q9H4H8 | Protein FAM83D OS=Homo sapiens OX= | FAM83D | 64.425 | 2 | 50.951534 | yes | 9.831E-07 | 526.72693 | 257.38248 |
| TTDFHNPGYPK | 11 | Q9H4H8 | Protein FAM83D OS=Homo sapiens OX= | FAM83D | 64.425 | 2 | 50.951534 | yes | 9.722E-08 | 638.80145 | 199.67303 |
| QFHFAGIR | 8 | Q9H4H8 | Protein FAM83D OS=Homo sapiens OX= | FAM83D | 64.425 | 2 | 50.951534 | yes | 1.337E-07 | 488.2616 | 139.62465 |
| GTQSTEGSPVSK | 12 | Q9H4H8 | Protein FAM83D OS=Homo sapiens OX= | FAM83D | 64.425 | 2 | 50.951534 | yes | 1.618E-07 | 589.28839 | 132.27782 |
| ADLDPEMPAEGK | 12 | Q9H4H8 | Protein FAM83D OS=Homo sapiens OX= | FAM83D | 64.425 | 2 | 50.951534 | yes | 2.745E-07 | 636.79248 | 104.51386 |
| KPHDCESSTVSEEI | 20 | Q9H4H8 | Protein FAM83D OS=Homo sapiens OX= | FAM83D | 64.425 | 3 | 50.951534 | yes | 1.734E-07 | 799.6731 | 32.988358 |
| HQELPGK | 7 | Q9H4K7 | Mitochondrial ribosome-associated GTPa | MTG2 | 43.956 | 2 | 49.77903 | yes | 7.71E-07 | 404.71924 | 88.733521 |
| VLHLELK | 7 | Q9H4K7 | Mitochondrial ribosome-associated GTPa | MTG2 | 43.956 | 2 | 49.77903 | yes | 4.074E-07 | 426.27109 | 72.836632 |
| IDLPEAQANLSQLI | 14 | Q9H4K7 | Mitochondrial ribosome-associated GTPa | MTG2 | 43.956 | 2 | 49.77903 | yes | 3.303E-05 | 784.42554 | 58.111423 |
| YELEMYEK | 8 | Q9H4K7 | Mitochondrial ribosome-associated GTPa | MTG2 | 43.956 | 2 | 49.77903 | yes | 3.598E-07 | 552.74951 | 55.098785 |
| VLYDAYAEAELGC | 15 | Q9H4K7 | Mitochondrial ribosome-associated GTPa | MTG2 | 43.956 | 2 | 49.77903 | yes | 1.475E-07 | 827.90723 | 49.737896 |
| SGAVLYIR | 8 | Q9H4K7 | Mitochondrial ribosome-associated GTPa | MTG2 | 43.956 | 2 | 49.77903 | yes | 2.043E-06 | 439.75836 | 40.019176 |
| APVTCTPGQPGQC | 14 | Q9H4K7 | Mitochondrial ribosome-associated GTPa | MTG2 | 43.956 | 2 | 49.77903 | yes | 1.004E-07 | 748.86737 | 27.1621 |
| TITYFDSQR | 9 | Q9H4L4 | Sentrin-specific protease 3 OS=Homo sap | SENP3 | 65.01 | 2 | 45.060654 | yes | 3.242E-07 | 565.77747 | 271.42719 |
| STSLTFHWK | 9 | Q9H4L4 | Sentrin-specific protease 3 OS=Homo sap | SENP3 | 65.01 | 2 | 45.060654 | yes | 3.945E-07 | 553.7851 | 124.86019 |
| GLVLQLIQSYQR | 12 | Q9H4L4 | Sentrin-specific protease 3 OS=Homo sap | SENP3 | 65.01 | 2 | 45.060654 | yes | 1.976E-07 | 709.41174 | 56.723667 |
| VHFFNSFFYDK | 11 | Q9H4L4 | Sentrin-specific protease 3 OS=Homo sap | SENP3 | 65.01 | 2 | 45.060654 | yes | 5.636E-05 | 725.84314 | 43.268696 |
| SGGGFGPDPGSGT | 18 | Q9H4L4 | Sentrin-specific protease 3 OS=Homo sap | SENP3 | 65.01 | 2 | 45.060654 | yes | 2.994E-07 | 808.88678 | 36.364906 |
| NFVFWQDVR | 9 | Q9H4L5 | Oxysterol-binding protein-related protein | OSBPL3 | 101.224 | 2 | 51.67378 | yes | 1.286E-07 | 605.80383 | 2156.4238 |
| FLEEGNLEEAEIQK | 14 | Q9H4L5 | Oxysterol-binding protein-related protein | OSBPL3 | 101.224 | 2 | 51.67378 | yes | 1.001E-07 | 824.90686 | 1492.8181 |
| AAQIPSPLER | 10 | Q9H4L5 | Oxysterol-binding protein-related protein | OSBPL3 | 101.224 | 2 | 51.67378 | yes | 4.549E-07 | 541.30365 | 1464.9076 |
| QTLGPVLDSGR | 11 | Q9H4L5 | Oxysterol-binding protein-related protein | OSBPL3 | 101.224 | 2 | 51.67378 | yes | 1.823E-07 | 571.81183 | 1418.1074 |
| NALSSALAQNTDL | 14 | Q9H4L5 | Oxysterol-binding protein-related protein | OSBPL3 | 101.224 | 2 | 51.67378 | yes | 1.286E-07 | 723.38336 | 1234.2142 |
| QDSWEVVEGLR | 11 | Q9H4L5 | Oxysterol-binding protein-related protein | OSBPL3 | 101.224 | 2 | 51.67378 | yes | 1.084E-07 | 659.32532 | 1230.3787 |
| LCEELEYSELLDK | 13 | Q9H4L5 | Oxysterol-binding protein-related protein | OSBPL3 | 101.224 | 2 | 51.67378 | yes | 1.105E-07 | 820.88983 | 1025.3862 |
| GEMNYTQEPPVQI | 13 | Q9H4L5 | Oxysterol-binding protein-related protein | OSBPL3 | 101.224 | 2 | 51.67378 | yes | 9.722E-08 | 760.85614 | 964.81482 |
| VTSCIHNILSGQR | 13 | Q9H4L5 | Oxysterol-binding protein-related protein | OSBPL3 | 101.224 | 2 | 51.67378 | yes | 3.265E-07 | 742.88556 | 629.18359 |

| | | | | | | | | | | | |
|---|---|---|---|---|---|---|---|---|---|---|---|
| VPLWLQSSEDMEI | 13 | Q9H4L5 | Oxysterol-binding protein-related protein | OSBPL3 | 101.224 | 2 | 51.67378 | yes | 1.18E-07 | 781.38196 | 624.57074 |
| CIDLDTEEHIYHLK | 14 | Q9H4L5 | Oxysterol-binding protein-related protein | OSBPL3 | 101.224 | 3 | 51.67378 | yes | 9.809E-08 | 595.95404 | 617.6626 |
| VLEENHVEHQPR | 12 | Q9H4L5 | Oxysterol-binding protein-related protein | OSBPL3 | 101.224 | 3 | 51.67378 | yes | 1.7E-07 | 496.25143 | 301.98328 |
| VLEENHVEHQPR | 12 | Q9H4L5 | Oxysterol-binding protein-related protein | OSBPL3 | 101.224 | 3 | 51.67378 | yes | 2.964E-07 | 743.87347 | 301.98328 |
| VLEENHVEHQPR | 12 | Q9H4L5 | Oxysterol-binding protein-related protein | OSBPL3 | 101.224 | 2 | 51.67378 | yes | 1.7E-07 | 496.25143 | 301.98328 |
| VLEENHVEHQPR | 12 | Q9H4L5 | Oxysterol-binding protein-related protein | OSBPL3 | 101.224 | 2 | 51.67378 | yes | 2.964E-07 | 743.87347 | 301.98328 |
| IHAESLLLDSPAVA | 15 | Q9H4L5 | Oxysterol-binding protein-related protein | OSBPL3 | 101.224 | 2 | 51.67378 | yes | 3.127E-07 | 782.44067 | 544.82959 |
| SEEVFDEWVSK | 11 | Q9H4L5 | Oxysterol-binding protein-related protein | OSBPL3 | 101.224 | 2 | 51.67378 | yes | 9.819E-08 | 677.81171 | 516.10895 |
| NLHDDSCYCK | 10 | Q9H4L5 | Oxysterol-binding protein-related protein | OSBPL3 | 101.224 | 2 | 51.67378 | yes | 9.722E-08 | 656.25824 | 473.81848 |
| ALVHQLSNESR | 11 | Q9H4L5 | Oxysterol-binding protein-related protein | OSBPL3 | 101.224 | 2 | 51.67378 | yes | 1.892E-07 | 627.3335 | 301.84714 |
| PFNPVLGETYECIF | 14 | Q9H4L5 | Oxysterol-binding protein-related protein | OSBPL3 | 101.224 | 2 | 51.67378 | no | 9.911E-08 | 847.914 | 292.10678 |
| SLLPPTDTR | 9 | Q9H4L5 | Oxysterol-binding protein-related protein | OSBPL3 | 101.224 | 2 | 51.67378 | yes | 1.044E-06 | 500.2771 | 291.21216 |
| AGSKPFNPVLGET | 18 | Q9H4L5 | Oxysterol-binding protein-related protein | OSBPL3 | 101.224 | 3 | 51.67378 | no | 1.199E-07 | 680.00696 | 186.13046 |
| MQEDLCHIAHK | 11 | Q9H4L5 | Oxysterol-binding protein-related protein | OSBPL3 | 101.224 | 2 | 51.67378 | yes | 1.533E-07 | 691.32117 | 140.0578 |
| SGDNLAEENSR | 11 | Q9H4L5 | Oxysterol-binding protein-related protein | OSBPL3 | 101.224 | 2 | 51.67378 | yes | 9.491E-07 | 596.26544 | 133.3217 |
| LHGCIDVGLSVMS | 15 | Q9H4L5 | Oxysterol-binding protein-related protein | OSBPL3 | 101.224 | 3 | 51.67378 | yes | 2.892E-07 | 538.94958 | 38.252579 |
| LHGCIDVGLSVMS | 15 | Q9H4L5 | Oxysterol-binding protein-related protein | OSBPL3 | 101.224 | 3 | 51.67378 | yes | 4.075E-06 | 807.92072 | 38.252579 |
| LHGCIDVGLSVMS | 15 | Q9H4L5 | Oxysterol-binding protein-related protein | OSBPL3 | 101.224 | 2 | 51.67378 | yes | 2.892E-07 | 538.94958 | 38.252579 |
| LHGCIDVGLSVMS | 15 | Q9H4L5 | Oxysterol-binding protein-related protein | OSBPL3 | 101.224 | 2 | 51.67378 | yes | 4.075E-06 | 807.92072 | 38.252579 |
| DLAHCHAYLVEM | 24 | Q9H4L5 | Oxysterol-binding protein-related protein | OSBPL3 | 101.224 | 4 | 51.67378 | yes | 2.307E-07 | 717.3504 | 66.555984 |
| SMEIVPIGTTHVTL | 25 | Q9H4L5 | Oxysterol-binding protein-related protein | OSBPL3 | 101.224 | 4 | 51.67378 | yes | 3.465E-07 | 714.36127 | 30.825333 |
| RGPPSEAGGGKPP | 14 | Q9H4M7 | Pleckstrin homology domain-containing f | PLEKHA4 | 85.401 | 3 | 46.800823 | yes | 1.467E-07 | 454.91275 | 69.622849 |
| LLPLEEHYR | 9 | Q9H4M9 | EH domain-containing protein 1 OS=Hon | EHD1 | 60.627 | 2 | 47.25686 | no | 1.367E-07 | 585.31934 | 283.41272 |
| IILLFDAHK | 9 | Q9H4M9 | EH domain-containing protein 1 OS=Hon | EHD1 | 60.627 | 2 | 47.25686 | no | 1.451E-07 | 535.32385 | 207.64043 |
| VYGALMWSLGK | 11 | Q9H4M9 | EH domain-containing protein 1 OS=Hon | EHD1 | 60.627 | 2 | 47.25686 | no | 2.451E-07 | 612.82593 | 99.975052 |
| LDISDEFSEVIK | 12 | Q9H4M9 | EH domain-containing protein 1 OS=Hon | EHD1 | 60.627 | 2 | 47.25686 | yes | 1.382E-07 | 697.85614 | 57.426399 |
| SWPQGLATLETR | 12 | Q9H4P4 | E3 ubiquitin-protein ligase NRDP1 OS=I | RNF41 | 35.906 | 2 | 49.791508 | yes | 1.408E-07 | 679.85681 | 671.62195 |
| SVVQQQQTR | 9 | Q9H4P4 | E3 ubiquitin-protein ligase NRDP1 OS=I | RNF41 | 35.906 | 2 | 49.791508 | yes | 1.652E-07 | 537.28851 | 102.87773 |
| DELPNHNCIK | 10 | Q9H4P4 | E3 ubiquitin-protein ligase NRDP1 OS=I | RNF41 | 35.906 | 2 | 49.791508 | yes | 1.414E-07 | 620.29297 | 95.948563 |
| RYYENYVAK | 9 | Q9H4P4 | E3 ubiquitin-protein ligase NRDP1 OS=I | RNF41 | 35.906 | 2 | 49.791508 | yes | 1.003E-07 | 603.30109 | 77.994301 |
| LQIACDNAVFGCS | 17 | Q9H4P4 | E3 ubiquitin-protein ligase NRDP1 OS=I | RNF41 | 35.906 | 3 | 49.791508 | yes | 5.035E-07 | 627.30994 | 10.522335 |
| LSSFPCPVCGR | 11 | Q9H4Z2 | Zinc finger protein 335 OS=Homo sapien | ZNF335 | 144.893 | 2 | 49.5488 | yes | 1.022E-07 | 640.29974 | 130.28664 |
| EDFLLSHVAVK | 11 | Q9H4Z2 | Zinc finger protein 335 OS=Homo sapien | ZNF335 | 144.893 | 2 | 49.5488 | yes | 1.464E-07 | 629.34552 | 103.46733 |
| CPDCPFSAR | 9 | Q9H4Z2 | Zinc finger protein 335 OS=Homo sapien | ZNF335 | 144.893 | 2 | 49.5488 | yes | 1.562E-06 | 555.22876 | 67.993164 |
| CDECSYTSVYR | 11 | Q9H4Z2 | Zinc finger protein 335 OS=Homo sapien | ZNF335 | 144.893 | 2 | 49.5488 | yes | 2.58E-07 | 720.28192 | 46.418419 |
| WPLLQCGGLPR | 11 | Q9H4Z2 | Zinc finger protein 335 OS=Homo sapien | ZNF335 | 144.893 | 2 | 49.5488 | yes | 1.278E-07 | 648.84772 | 36.965317 |
| HASSFEEWGR | 10 | Q9H4Z2 | Zinc finger protein 335 OS=Homo sapien | ZNF335 | 144.893 | 2 | 49.5488 | yes | 4.895E-07 | 603.27032 | 22.173166 |
| LALPQSDASLLSR | 13 | Q9H583 | HEAT repeat-containing protein 1 OS=H | HEATR1 | 242.371 | 2 | 54.958416 | yes | 2.058E-07 | 685.88556 | 423.12408 |
| VFAEYPGSSAQLR | 13 | Q9H583 | HEAT repeat-containing protein 1 OS=H | HEATR1 | 242.371 | 2 | 54.958416 | yes | 1.88E-07 | 712.86206 | 232.03699 |

| | | | | | | | | | | | |
|---|---|---|---|---|---|---|---|---|---|---|---|
| GISVNAEQVR | 10 | Q9H583 | HEAT repeat-containing protein 1 OS=H | HEATR1 | 242.371 | 2 | 54.958416 | yes | 1.108E-07 | 536.79089 | 214.73839 |
| AKPLGTVQQK | 10 | Q9H583 | HEAT repeat-containing protein 1 OS=H | HEATR1 | 242.371 | 2 | 54.958416 | yes | 1.422E-07 | 535.32184 | 141.84804 |
| SGICSLHPLLR | 11 | Q9H583 | HEAT repeat-containing protein 1 OS=H | HEATR1 | 242.371 | 2 | 54.958416 | yes | 1.064E-07 | 626.84515 | 121.53577 |
| VVESGGPEILK | 11 | Q9H583 | HEAT repeat-containing protein 1 OS=H | HEATR1 | 242.371 | 2 | 54.958416 | yes | 9.722E-08 | 564.31897 | 119.70087 |
| DEAMVLHLTLGK | 12 | Q9H583 | HEAT repeat-containing protein 1 OS=H | HEATR1 | 242.371 | 2 | 54.958416 | yes | 9.722E-08 | 663.85791 | 93.727501 |
| YNEFSVSLLNEDP | 14 | Q9H583 | HEAT repeat-containing protein 1 OS=H | HEATR1 | 242.371 | 2 | 54.958416 | yes | 1.076E-07 | 827.90161 | 88.217644 |
| LDDTYSFQVINK | 12 | Q9H583 | HEAT repeat-containing protein 1 OS=H | HEATR1 | 242.371 | 2 | 54.958416 | yes | 9.722E-08 | 721.86176 | 82.61615 |
| ITSEMGSASQANI | 14 | Q9H583 | HEAT repeat-containing protein 1 OS=H | HEATR1 | 242.371 | 2 | 54.958416 | yes | 9.722E-08 | 732.85919 | 71.452484 |
| GLVGNPLPSVR | 11 | Q9H583 | HEAT repeat-containing protein 1 OS=H | HEATR1 | 242.371 | 2 | 54.958416 | yes | 9.776E-07 | 554.82727 | 55.084412 |
| CLEWLIHR | 8 | Q9H583 | HEAT repeat-containing protein 1 OS=H | HEATR1 | 242.371 | 2 | 54.958416 | yes | 9.722E-08 | 563.79492 | 54.875847 |
| LLTFYNLADCIAE | 14 | Q9H583 | HEAT repeat-containing protein 1 OS=H | HEATR1 | 242.371 | 2 | 54.958416 | yes | 4.796E-07 | 835.92657 | 35.325771 |
| IISVFVDALPHVPE | 16 | Q9H583 | HEAT repeat-containing protein 1 OS=H | HEATR1 | 242.371 | 3 | 54.958416 | yes | 3.824E-07 | 610.34045 | 33.101364 |
| AQHSENDLEEVG | 13 | Q9H583 | HEAT repeat-containing protein 1 OS=H | HEATR1 | 242.371 | 2 | 54.958416 | yes | 4.624E-06 | 728.33917 | 20.926796 |
| AFADSSYLLR | 10 | Q9H5H4 | Zinc finger protein 768 OS=Homo sapien | ZNF768 | 60.229 | 2 | 51.896088 | yes | 9.722E-08 | 571.79565 | 259.49438 |
| AFGDSSYLLR | 10 | Q9H5H4 | Zinc finger protein 768 OS=Homo sapien | ZNF768 | 60.229 | 2 | 51.896088 | yes | 1.23E-07 | 564.78784 | 136.92502 |
| CYSQNSSLR | 9 | Q9H5H4 | Zinc finger protein 768 OS=Homo sapien | ZNF768 | 60.229 | 2 | 51.896088 | yes | 1.109E-07 | 557.75092 | 133.85193 |
| AFSQSSDLIR | 10 | Q9H5H4 | Zinc finger protein 768 OS=Homo sapien | ZNF768 | 60.229 | 2 | 51.896088 | yes | 2.582E-07 | 562.29077 | 129.41759 |
| AFSQSSDLIK | 10 | Q9H5H4 | Zinc finger protein 768 OS=Homo sapien | ZNF768 | 60.229 | 2 | 51.896088 | yes | 2.921E-06 | 548.28766 | 54.356667 |
| SPGYESESSR | 10 | Q9H5H4 | Zinc finger protein 768 OS=Homo sapien | ZNF768 | 60.229 | 2 | 51.896088 | yes | 4.835E-07 | 549.73853 | 53.136536 |
| TQSPEFEAQSSK | 12 | Q9H5H4 | Zinc finger protein 768 OS=Homo sapien | ZNF768 | 60.229 | 2 | 51.896088 | yes | 9.992E-08 | 669.81226 | 37.924206 |
| ARPPLPR | 7 | Q9H5P4 | PDZ domain-containing protein 7 OS=H | PDZD7 | 111.752 | 2 | 45.842636 | yes | 1.727E-07 | 403.75342 | 130.79745 |
| LAETLAQIYLGK | 12 | Q9H5Q4 | Dimethyladenosine transferase 2, mitoch | TFB2M | 45.349 | 2 | 52.306133 | yes | 9.722E-08 | 660.38208 | 656.46008 |
| TFIPHLESLGK | 11 | Q9H5Q4 | Dimethyladenosine transferase 2, mitoch | TFB2M | 45.349 | 2 | 52.306133 | yes | 1.536E-07 | 621.34808 | 633.85706 |
| SATVIDHLR | 9 | Q9H5Q4 | Dimethyladenosine transferase 2, mitoch | TFB2M | 45.349 | 2 | 52.306133 | yes | 1.653E-07 | 506.28271 | 523.1192 |
| ELLDQLQQK | 9 | Q9H5Q4 | Dimethyladenosine transferase 2, mitoch | TFB2M | 45.349 | 2 | 52.306133 | yes | 9.722E-08 | 557.80878 | 461.32339 |
| WLYDETLEDR | 10 | Q9H5Q4 | Dimethyladenosine transferase 2, mitoch | TFB2M | 45.349 | 2 | 52.306133 | yes | 9.722E-08 | 670.31189 | 366.42615 |
| LYLIQMIPR | 9 | Q9H5Q4 | Dimethyladenosine transferase 2, mitoch | TFB2M | 45.349 | 2 | 52.306133 | yes | 4.194E-07 | 573.83881 | 316.26703 |
| LAYDLYSCTSIYK | 13 | Q9H5Q4 | Dimethyladenosine transferase 2, mitoch | TFB2M | 45.349 | 2 | 52.306133 | yes | 9.759E-08 | 798.88434 | 264.16104 |
| VVNMHPQDFK | 10 | Q9H5Q4 | Dimethyladenosine transferase 2, mitoch | TFB2M | 45.349 | 2 | 52.306133 | yes | 5.151E-07 | 607.80298 | 217.59232 |
| SGGVIKPPAMSSR | 13 | Q9H5Q4 | Dimethyladenosine transferase 2, mitoch | TFB2M | 45.349 | 2 | 52.306133 | yes | 1.143E-07 | 643.8479 | 183.63351 |
| KGPLENPK | 8 | Q9H5Q4 | Dimethyladenosine transferase 2, mitoch | TFB2M | 45.349 | 2 | 52.306133 | yes | 9.779E-08 | 441.7558 | 154.40808 |
| IEVNMFIGEK | 10 | Q9H5Q4 | Dimethyladenosine transferase 2, mitoch | TFB2M | 45.349 | 2 | 52.306133 | yes | 1.424E-07 | 590.30756 | 149.83783 |
| VIHCDFFK | 8 | Q9H5Q4 | Dimethyladenosine transferase 2, mitoch | TFB2M | 45.349 | 2 | 52.306133 | yes | 5.47E-07 | 533.26294 | 146.40189 |
| VVGMFPSR | 8 | Q9H5Q4 | Dimethyladenosine transferase 2, mitoch | TFB2M | 45.349 | 2 | 52.306133 | yes | 4.523E-07 | 446.73911 | 139.412 |
| VVALESDK | 8 | Q9H5Q4 | Dimethyladenosine transferase 2, mitoch | TFB2M | 45.349 | 2 | 52.306133 | yes | 2.466E-06 | 430.73984 | 76.215508 |
| PSRPPHLLLECNP | 28 | Q9H5Q4 | Dimethyladenosine transferase 2, mitoch | TFB2M | 45.349 | 4 | 52.306133 | yes | 1.481E-07 | 738.15491 | 57.815887 |
| FIELPLTQR | 9 | Q9H5U6 | rRNA N6-adenosine-methyltransferase Z | ZCCHC4 | 59.01 | 2 | 52.392643 | yes | 3.662E-07 | 558.82422 | 757.76398 |
| FCSPCQR | 7 | Q9H5U6 | rRNA N6-adenosine-methyltransferase Z | ZCCHC4 | 59.01 | 2 | 52.392643 | yes | 1.431E-06 | 477.69965 | 215.93199 |
| VLCVGTPR | 8 | Q9H5U6 | rRNA N6-adenosine-methyltransferase Z | ZCCHC4 | 59.01 | 2 | 52.392643 | yes | 9.792E-08 | 451.24985 | 156.06117 |

| | | | | | | | | | | | |
|---|---|---|---|---|---|---|---|---|---|---|---|
| FYACSACR | 8 | Q9H5U6 | rRNA N6-adenosine-methyltransferase Z( | ZCCHC4 | 59.01 | 2 | 52.392643 | yes | 9.809E-08 | 517.71277 | 136.97845 |
| DCNFFQWEDEK | 11 | Q9H5U6 | rRNA N6-adenosine-methyltransferase Z( | ZCCHC4 | 59.01 | 2 | 52.392643 | yes | 9.722E-08 | 759.30371 | 119.12778 |
| TNAQYLFADR | 10 | Q9H5U6 | rRNA N6-adenosine-methyltransferase Z( | ZCCHC4 | 59.01 | 2 | 52.392643 | yes | 9.722E-08 | 599.7962 | 112.76162 |
| CQPPLSR | 7 | Q9H5U6 | rRNA N6-adenosine-methyltransferase Z( | ZCCHC4 | 59.01 | 2 | 52.392643 | yes | 5.296E-07 | 429.21854 | 81.078217 |
| STCPNIATSK | 10 | Q9H5U6 | rRNA N6-adenosine-methyltransferase Z( | ZCCHC4 | 59.01 | 2 | 52.392643 | yes | 2.517E-07 | 539.76349 | 70.4617 |
| RPSQLLYPLENK | 12 | Q9H5U6 | rRNA N6-adenosine-methyltransferase Z( | ZCCHC4 | 59.01 | 2 | 52.392643 | yes | 1.916E-07 | 729.40918 | 67.601387 |
| NGFEAVEAEGSAC | 15 | Q9H5U6 | rRNA N6-adenosine-methyltransferase Z( | ZCCHC4 | 59.01 | 2 | 52.392643 | yes | 1.841E-07 | 777.33606 | 28.644112 |
| NAPVTFIVDGAVV | 14 | Q9H5V9 | STING ER exit protein OS=Homo sapien | STEEP1 | 25.625 | 2 | 48.294956 | yes | 1.986E-07 | 715.40613 | 449.24084 |
| LQELAELEAK | 10 | Q9H5V9 | STING ER exit protein OS=Homo sapien | STEEP1 | 25.625 | 2 | 48.294956 | yes | 1.166E-07 | 572.31641 | 207.37474 |
| CGLPLFYQSQPK | 12 | Q9H5V9 | STING ER exit protein OS=Homo sapien | STEEP1 | 25.625 | 2 | 48.294956 | yes | 4.079E-07 | 719.3634 | 153.85071 |
| EVADSYAQNAK | 11 | Q9H5V9 | STING ER exit protein OS=Homo sapien | STEEP1 | 25.625 | 2 | 48.294956 | yes | 2.563E-07 | 598.28314 | 78.841736 |
| ALGIDNVLR | 9 | Q9H5Z1 | Probable ATP-dependent RNA helicase D | DHX35 | 78.911 | 2 | 37.992107 | yes | 5.539E-07 | 485.78763 | 52.99572 |
| EMMVDPLLTK | 10 | Q9H5Z1 | Probable ATP-dependent RNA helicase D | DHX35 | 78.911 | 2 | 37.992107 | yes | 2.782E-06 | 588.80377 | 39.163048 |
| GCSSLLVPGLTCT( | 15 | Q9H633 | Ribonuclease P protein subunit p21 OS=H | RPP21 | 17.571 | 2 | 53.37089 | yes | 3.166E-07 | 824.91089 | 274.54279 |
| WTVQTCLTCQR | 11 | Q9H633 | Ribonuclease P protein subunit p21 OS=H | RPP21 | 17.571 | 2 | 53.37089 | yes | 9.722E-08 | 726.83978 | 183.33049 |
| FYCYTER | 7 | Q9H633 | Ribonuclease P protein subunit p21 OS=H | RPP21 | 17.571 | 2 | 53.37089 | yes | 4.867E-07 | 519.72113 | 70.014961 |
| MLELPTIYR | 9 | Q9H668 | CST complex subunit STN1 OS=Homo sa | STN1 | 42.12 | 2 | 42.907204 | yes | 1.709E-06 | 568.31262 | 52.836166 |
| DDGFDNLYYVTR | 12 | Q9H668 | CST complex subunit STN1 OS=Homo sa | STN1 | 42.12 | 2 | 42.907204 | yes | 2.818E-07 | 739.33331 | 32.797813 |
| IMEETNTQIAWPSI | 14 | Q9H694 | Protein bicaudal C homolog 1 OS=Homo | BICC1 | 104.844 | 2 | 52.478897 | yes | 9.722E-08 | 824.40601 | 279.52728 |
| MYGATVIVR | 9 | Q9H694 | Protein bicaudal C homolog 1 OS=Homo | BICC1 | 104.844 | 2 | 52.478897 | yes | 1.21E-06 | 505.27859 | 216.75923 |
| YGAISTSSLGEK | 12 | Q9H694 | Protein bicaudal C homolog 1 OS=Homo | BICC1 | 104.844 | 2 | 52.478897 | yes | 4.097E-07 | 606.80896 | 200.36737 |
| EMIMSVLDTK | 10 | Q9H694 | Protein bicaudal C homolog 1 OS=Homo | BICC1 | 104.844 | 2 | 52.478897 | yes | 1.012E-07 | 583.7934 | 154.01314 |
| MLLAISELNK | 10 | Q9H694 | Protein bicaudal C homolog 1 OS=Homo | BICC1 | 104.844 | 2 | 52.478897 | yes | 5.99E-07 | 566.32574 | 151.51822 |
| QTVELLQGTK | 10 | Q9H694 | Protein bicaudal C homolog 1 OS=Homo | BICC1 | 104.844 | 2 | 52.478897 | yes | 1.181E-07 | 558.81659 | 128.29242 |
| ELGITTFGAR | 10 | Q9H694 | Protein bicaudal C homolog 1 OS=Homo | BICC1 | 104.844 | 2 | 52.478897 | yes | 3.302E-07 | 532.79041 | 99.375984 |
| KPVVTEVR | 8 | Q9H694 | Protein bicaudal C homolog 1 OS=Homo | BICC1 | 104.844 | 2 | 52.478897 | yes | 1.843E-07 | 464.28473 | 95.912949 |
| MDVSHTEHSHVIC | 14 | Q9H694 | Protein bicaudal C homolog 1 OS=Homo | BICC1 | 104.844 | 2 | 52.478897 | yes | 9.819E-08 | 788.88049 | 33.233978 |
| ANHVSYKPTMTT | 23 | Q9H694 | Protein bicaudal C homolog 1 OS=Homo | BICC1 | 104.844 | 4 | 52.478897 | yes | 2.341E-07 | 637.8006 | 29.393208 |
| SAAYSQVLLR | 10 | Q9H6D7 | HAUS augmin-like complex subunit 4 OS | HAUS4 | 42.4 | 2 | 49.969067 | yes | 1.758E-06 | 554.31146 | 119.19205 |
| LQQEVEEQLK | 10 | Q9H6D7 | HAUS augmin-like complex subunit 4 OS | HAUS4 | 42.4 | 2 | 49.969067 | yes | 9.963E-08 | 622.33008 | 89.584633 |
| VIQELLVDYYVK | 12 | Q9H6D7 | HAUS augmin-like complex subunit 4 OS | HAUS4 | 42.4 | 2 | 49.969067 | yes | 5.37E-07 | 741.41614 | 32.853016 |
| LLLNLSQHVDESG | 20 | Q9H6D7 | HAUS augmin-like complex subunit 4 OS | HAUS4 | 42.4 | 3 | 49.969067 | yes | 5.534E-07 | 717.73883 | 24.263622 |
| FVHPSLSR | 8 | Q9H6H4 | Receptor expression-enhancing protein 4 | REEP4 | 29.396 | 2 | 51.825813 | yes | 1.803E-07 | 471.76141 | 955.95477 |
| SYETVLSFGK | 10 | Q9H6H4 | Receptor expression-enhancing protein 4 | REEP4 | 29.396 | 2 | 51.825813 | yes | 1.473E-07 | 565.79004 | 781.51007 |
| SFSMQDLR | 8 | Q9H6H4 | Receptor expression-enhancing protein 4 | REEP4 | 29.396 | 2 | 51.825813 | yes | 1.595E-07 | 492.23438 | 467.54834 |
| RPPIGYR | 7 | Q9H6H4 | Receptor expression-enhancing protein 4 | REEP4 | 29.396 | 2 | 51.825813 | yes | 1.653E-07 | 429.75085 | 347.87222 |
| KFVHPSLSR | 9 | Q9H6H4 | Receptor expression-enhancing protein 4 | REEP4 | 29.396 | 2 | 51.825813 | yes | 2.303E-07 | 535.8089 | 237.66223 |
| EIDAYIVQAK | 10 | Q9H6H4 | Receptor expression-enhancing protein 4 | REEP4 | 29.396 | 2 | 51.825813 | yes | 9.722E-08 | 575.31116 | 108.72659 |
| SISDAPAPAYHDPI | 23 | Q9H6H4 | Receptor expression-enhancing protein 4 | REEP4 | 29.396 | 4 | 51.825813 | yes | 2.82E-06 | 646.06342 | 21.430546 |

| | | | | | | | | | | | |
|---|---|---|---|---|---|---|---|---|---|---|---|
| SPQQVMGSLVK | 11 | Q9H6Q4 | Cytosolic iron-sulfur assembly componen | CIAO3 | 53.021 | 2 | 33.035095 | yes | 1.343E-06 | 587.31842 | 43.359951 |
| MTVPEIQR | 8 | Q9H6R0 | ATP-dependent RNA helicase DHX33 O | DHX33 | 78.875 | 2 | 50.881546 | yes | 8.454E-07 | 487.26041 | 247.24225 |
| QGIIAVTQPR | 10 | Q9H6R0 | ATP-dependent RNA helicase DHX33 O | DHX33 | 78.875 | 2 | 50.881546 | yes | 9.722E-08 | 541.81946 | 167.73499 |
| FDDVTSEDTR | 10 | Q9H6R0 | ATP-dependent RNA helicase DHX33 O | DHX33 | 78.875 | 2 | 50.881546 | yes | 1.217E-07 | 592.75696 | 164.75229 |
| TIHTDVLFGVVK | 12 | Q9H6R0 | ATP-dependent RNA helicase DHX33 O | DHX33 | 78.875 | 2 | 50.881546 | yes | 9.881E-08 | 664.88226 | 144.54395 |
| FLTDGMLLR | 9 | Q9H6R0 | ATP-dependent RNA helicase DHX33 O | DHX33 | 78.875 | 2 | 50.881546 | yes | 5.921E-07 | 533.29169 | 140.88602 |
| NMTLVAEVR | 9 | Q9H6R0 | ATP-dependent RNA helicase DHX33 O | DHX33 | 78.875 | 2 | 50.881546 | yes | 1.108E-07 | 516.77893 | 97.424942 |
| VAAISLATR | 9 | Q9H6R0 | ATP-dependent RNA helicase DHX33 O | DHX33 | 78.875 | 2 | 50.881546 | yes | 2.98E-07 | 451.27692 | 80.578232 |
| FISSEGDHMTLLNI | 16 | Q9H6R0 | ATP-dependent RNA helicase DHX33 O | DHX33 | 78.875 | 3 | 50.881546 | yes | 2.406E-06 | 632.64886 | 78.962219 |
| EAISDSLLR | 9 | Q9H6R0 | ATP-dependent RNA helicase DHX33 O | DHX33 | 78.875 | 2 | 50.881546 | yes | 1.484E-07 | 502.27457 | 75.095093 |
| SLPIFQAR | 8 | Q9H6R0 | ATP-dependent RNA helicase DHX33 O | DHX33 | 78.875 | 2 | 50.881546 | yes | 5.319E-07 | 466.27164 | 73.631897 |
| NLDNAVLIGETGS | 15 | Q9H6R0 | ATP-dependent RNA helicase DHX33 O | DHX33 | 78.875 | 2 | 50.881546 | yes | 1.766E-07 | 744.38867 | 69.190689 |
| YNPDSGLEVLAV | 14 | Q9H6R0 | ATP-dependent RNA helicase DHX33 O | DHX33 | 78.875 | 2 | 50.881546 | yes | 0.0003559 | 780.90448 | 66.603706 |
| YVVDTGMVK | 9 | Q9H6R0 | ATP-dependent RNA helicase DHX33 O | DHX33 | 78.875 | 2 | 50.881546 | yes | 2.841E-07 | 506.2626 | 66.554077 |
| YSCVILDEAHER | 12 | Q9H6R0 | ATP-dependent RNA helicase DHX33 O | DHX33 | 78.875 | 2 | 50.881546 | yes | 1.886E-07 | 746.34845 | 49.838573 |
| EDSGICYR | 8 | Q9H6R0 | ATP-dependent RNA helicase DHX33 O | DHX33 | 78.875 | 2 | 50.881546 | yes | 6.761E-07 | 500.21365 | 45.445385 |
| LQVEELLK | 8 | Q9H6R4 | Nucleolar protein 6 OS=Homo sapiens O | NOL6 | 127.594 | 2 | 55.37085 | yes | 3.176E-07 | 486.29224 | 388.55975 |
| EAVVWEAASMSQ | 13 | Q9H6R4 | Nucleolar protein 6 OS=Homo sapiens O | NOL6 | 127.594 | 2 | 55.37085 | yes | 1.084E-07 | 718.34796 | 378.18063 |
| ATATHTDVLK | 10 | Q9H6R4 | Nucleolar protein 6 OS=Homo sapiens O | NOL6 | 127.594 | 2 | 55.37085 | yes | 1.375E-07 | 528.78784 | 202.317 |
| AQLPVMVIVTPQD | 14 | Q9H6R4 | Nucleolar protein 6 OS=Homo sapiens O | NOL6 | 127.594 | 2 | 55.37085 | yes | 2.293E-07 | 783.93726 | 192.5325 |
| ETEILFHSSLLR | 12 | Q9H6R4 | Nucleolar protein 6 OS=Homo sapiens O | NOL6 | 127.594 | 2 | 55.37085 | yes | 3.005E-07 | 722.89337 | 178.30054 |
| IHTTMSGYQVLR | 12 | Q9H6R4 | Nucleolar protein 6 OS=Homo sapiens O | NOL6 | 127.594 | 2 | 55.37085 | yes | 9.722E-08 | 703.36646 | 157.85745 |
| PAFSFYETLR | 10 | Q9H6R4 | Nucleolar protein 6 OS=Homo sapiens O | NOL6 | 127.594 | 2 | 55.37085 | yes | 9.722E-08 | 615.81134 | 123.97586 |
| PTSFQPQPFK | 10 | Q9H6R4 | Nucleolar protein 6 OS=Homo sapiens O | NOL6 | 127.594 | 2 | 55.37085 | yes | 3.652E-07 | 588.80603 | 117.98061 |
| LAELLTQQHGLQC | 14 | Q9H6R4 | Nucleolar protein 6 OS=Homo sapiens O | NOL6 | 127.594 | 2 | 55.37085 | yes | 9.722E-08 | 833.93829 | 53.306862 |
| LAELLTQQHGLQC | 14 | Q9H6R4 | Nucleolar protein 6 OS=Homo sapiens O | NOL6 | 127.594 | 2 | 55.37085 | yes | 1.112E-07 | 556.29462 | 53.306862 |
| LAELLTQQHGLQC | 14 | Q9H6R4 | Nucleolar protein 6 OS=Homo sapiens O | NOL6 | 127.594 | 3 | 55.37085 | yes | 9.722E-08 | 833.93829 | 53.306862 |
| LAELLTQQHGLQC | 14 | Q9H6R4 | Nucleolar protein 6 OS=Homo sapiens O | NOL6 | 127.594 | 3 | 55.37085 | yes | 1.112E-07 | 556.29462 | 53.306862 |
| EVQSPEGMISLR | 12 | Q9H6R4 | Nucleolar protein 6 OS=Homo sapiens O | NOL6 | 127.594 | 2 | 55.37085 | yes | 9.728E-08 | 673.34265 | 86.683998 |
| GPAPAGEQLR | 10 | Q9H6R4 | Nucleolar protein 6 OS=Homo sapiens O | NOL6 | 127.594 | 2 | 55.37085 | yes | 1.417E-07 | 498.26706 | 86.073036 |
| ADDGFHLLLMTPK | 13 | Q9H6R4 | Nucleolar protein 6 OS=Homo sapiens O | NOL6 | 127.594 | 2 | 55.37085 | yes | 9.728E-08 | 729.37646 | 74.677734 |
| LSMMLLDSR | 9 | Q9H6R4 | Nucleolar protein 6 OS=Homo sapiens O | NOL6 | 127.594 | 2 | 55.37085 | yes | 2.58E-07 | 533.27521 | 63.555389 |
| YTEVFPPTPVR | 11 | Q9H6R4 | Nucleolar protein 6 OS=Homo sapiens O | NOL6 | 127.594 | 2 | 55.37085 | yes | 1.252E-06 | 653.34552 | 41.891159 |
| ETSSTGEEALVAA | 15 | Q9H6R4 | Nucleolar protein 6 OS=Homo sapiens O | NOL6 | 127.594 | 2 | 55.37085 | yes | 1.093E-07 | 760.38354 | 36.678417 |
| QAVDSPAASFCR | 12 | Q9H6R4 | Nucleolar protein 6 OS=Homo sapiens O | NOL6 | 127.594 | 2 | 55.37085 | yes | 3.242E-07 | 654.80371 | 32.753834 |
| AFDHVLHLR | 9 | Q9H6R4 | Nucleolar protein 6 OS=Homo sapiens O | NOL6 | 127.594 | 2 | 55.37085 | yes | 1.673E-06 | 554.30652 | 26.658361 |
| LTDTTFLPSSK | 11 | Q9H6R7 | WD repeat and coiled-coil-containing pro | WDCP | 79.136 | 2 | 27.93571 | yes | 2.857E-06 | 605.32172 | 70.095245 |
| VVLIVGETGSGK | 12 | Q9H6S0 | 3'-5' RNA helicase YTHDC2 OS=Homo s | YTHDC2 | 160.249 | 2 | 54.085217 | yes | 1.249E-07 | 579.84009 | 2993.0044 |
| LASNALQEPSSFR | 13 | Q9H6S0 | 3'-5' RNA helicase YTHDC2 OS=Homo s | YTHDC2 | 160.249 | 2 | 54.085217 | yes | 9.722E-08 | 710.36499 | 2368.1309 |

| | | | | | | | | | | | |
|---|---|---|---|---|---|---|---|---|---|---|---|
| DPFVLPTQASQK | 12 | Q9H6S0 | 3'-5' RNA helicase YTHDC2 OS=Homo s | YTHDC2 | 160.249 | 2 | 54.085217 | yes | 9.722E-08 | 665.8537 | 1947.1093 |
| GNVFAVEAENR | 11 | Q9H6S0 | 3'-5' RNA helicase YTHDC2 OS=Homo s | YTHDC2 | 160.249 | 2 | 54.085217 | yes | 3.117E-07 | 603.29907 | 1926.1923 |
| TQLLGQLR | 8 | Q9H6S0 | 3'-5' RNA helicase YTHDC2 OS=Homo s | YTHDC2 | 160.249 | 2 | 54.085217 | yes | 2.464E-07 | 464.78235 | 1835.736 |
| LAAIAVAER | 9 | Q9H6S0 | 3'-5' RNA helicase YTHDC2 OS=Homo s | YTHDC2 | 160.249 | 2 | 54.085217 | yes | 2.247E-07 | 457.27692 | 1822.1295 |
| YFGSCPVIYIQGR | 13 | Q9H6S0 | 3'-5' RNA helicase YTHDC2 OS=Homo s | YTHDC2 | 160.249 | 2 | 54.085217 | yes | 1.102E-07 | 780.38739 | 1477.1436 |
| SQDWGSAGLGGV | 14 | Q9H6S0 | 3'-5' RNA helicase YTHDC2 OS=Homo s | YTHDC2 | 160.249 | 2 | 54.085217 | yes | 9.722E-08 | 704.84644 | 1466.4955 |
| MVLCAVVLK | 9 | Q9H6S0 | 3'-5' RNA helicase YTHDC2 OS=Homo s | YTHDC2 | 160.249 | 2 | 54.085217 | yes | 2.627E-07 | 516.80084 | 1462.9764 |
| IFCTQPR | 7 | Q9H6S0 | 3'-5' RNA helicase YTHDC2 OS=Homo s | YTHDC2 | 160.249 | 2 | 54.085217 | yes | 1.897E-07 | 461.23419 | 1415.4951 |
| WHSLFLR | 7 | Q9H6S0 | 3'-5' RNA helicase YTHDC2 OS=Homo s | YTHDC2 | 160.249 | 2 | 54.085217 | yes | 9.722E-08 | 479.76651 | 1375.5463 |
| GIWSTTPSNER | 11 | Q9H6S0 | 3'-5' RNA helicase YTHDC2 OS=Homo s | YTHDC2 | 160.249 | 2 | 54.085217 | yes | 1.287E-07 | 624.30438 | 1251.446 |
| IGQTIGYQIR | 10 | Q9H6S0 | 3'-5' RNA helicase YTHDC2 OS=Homo s | YTHDC2 | 160.249 | 2 | 54.085217 | yes | 1.214E-07 | 574.82477 | 1185.2856 |
| ENLVLTGPK | 9 | Q9H6S0 | 3'-5' RNA helicase YTHDC2 OS=Homo s | YTHDC2 | 160.249 | 2 | 54.085217 | yes | 9.491E-07 | 485.78201 | 1078.9954 |
| DVNCLEPWLIK | 11 | Q9H6S0 | 3'-5' RNA helicase YTHDC2 OS=Homo s | YTHDC2 | 160.249 | 2 | 54.085217 | yes | 1.153E-07 | 693.85791 | 1028.7162 |
| TLMAGDSTLSTVT | 23 | Q9H6S0 | 3'-5' RNA helicase YTHDC2 OS=Homo s | YTHDC2 | 160.249 | 3 | 54.085217 | yes | 1.286E-07 | 837.42334 | 1024.5018 |
| APEPPPALIVR | 11 | Q9H6S0 | 3'-5' RNA helicase YTHDC2 OS=Homo s | YTHDC2 | 160.249 | 2 | 54.085217 | yes | 1.761E-07 | 580.34534 | 929.98108 |
| ILFDDKR | 7 | Q9H6S0 | 3'-5' RNA helicase YTHDC2 OS=Homo s | YTHDC2 | 160.249 | 2 | 54.085217 | yes | 1.399E-07 | 453.7558 | 913.67493 |
| LSQSLGLVSK | 10 | Q9H6S0 | 3'-5' RNA helicase YTHDC2 OS=Homo s | YTHDC2 | 160.249 | 2 | 54.085217 | yes | 2.18E-07 | 516.30841 | 895.18445 |
| AIIAVLSTEEQSAG | 24 | Q9H6S0 | 3'-5' RNA helicase YTHDC2 OS=Homo s | YTHDC2 | 160.249 | 3 | 54.085217 | yes | 1.297E-07 | 818.43799 | 792.422 |
| DVNTNSENWAVV | 13 | Q9H6S0 | 3'-5' RNA helicase YTHDC2 OS=Homo s | YTHDC2 | 160.249 | 2 | 54.085217 | yes | 9.722E-08 | 738.35986 | 763.52759 |
| PMSSEELPLASSW | 14 | Q9H6S0 | 3'-5' RNA helicase YTHDC2 OS=Homo s | YTHDC2 | 160.249 | 2 | 54.085217 | yes | 9.722E-08 | 795.38507 | 750.09863 |
| ASNGWMALDWAI | 12 | Q9H6S0 | 3'-5' RNA helicase YTHDC2 OS=Homo s | YTHDC2 | 160.249 | 2 | 54.085217 | yes | 3.674E-06 | 675.31897 | 702.57813 |
| EMDACLSDIWLHK | 13 | Q9H6S0 | 3'-5' RNA helicase YTHDC2 OS=Homo s | YTHDC2 | 160.249 | 2 | 54.085217 | yes | 1.562E-07 | 809.3736 | 673.65076 |
| QSLPVFEK | 8 | Q9H6S0 | 3'-5' RNA helicase YTHDC2 OS=Homo s | YTHDC2 | 160.249 | 2 | 54.085217 | yes | 2.177E-07 | 474.26346 | 618.77509 |
| GESEFDSFR | 9 | Q9H6S0 | 3'-5' RNA helicase YTHDC2 OS=Homo s | YTHDC2 | 160.249 | 2 | 54.085217 | yes | 4.873E-07 | 537.23035 | 615.23151 |
| EMEFPSSLTSTER | 13 | Q9H6S0 | 3'-5' RNA helicase YTHDC2 OS=Homo s | YTHDC2 | 160.249 | 2 | 54.085217 | yes | 2.336E-07 | 757.34558 | 538.86957 |
| SPSPALHPPQK | 11 | Q9H6S0 | 3'-5' RNA helicase YTHDC2 OS=Homo s | YTHDC2 | 160.249 | 2 | 54.085217 | yes | 1.173E-07 | 579.81696 | 493.60123 |
| HSETSATALMVAA | 15 | Q9H6S0 | 3'-5' RNA helicase YTHDC2 OS=Homo s | YTHDC2 | 160.249 | 3 | 54.085217 | yes | 1.691E-07 | 501.25244 | 202.20796 |
| HSETSATALMVAA | 15 | Q9H6S0 | 3'-5' RNA helicase YTHDC2 OS=Homo s | YTHDC2 | 160.249 | 3 | 54.085217 | yes | 9.722E-08 | 751.375 | 202.20796 |
| HSETSATALMVAA | 15 | Q9H6S0 | 3'-5' RNA helicase YTHDC2 OS=Homo s | YTHDC2 | 160.249 | 2 | 54.085217 | yes | 1.691E-07 | 501.25244 | 202.20796 |
| HSETSATALMVAA | 15 | Q9H6S0 | 3'-5' RNA helicase YTHDC2 OS=Homo s | YTHDC2 | 160.249 | 2 | 54.085217 | yes | 9.722E-08 | 751.375 | 202.20796 |
| MPLQELCLHTK | 11 | Q9H6S0 | 3'-5' RNA helicase YTHDC2 OS=Homo s | YTHDC2 | 160.249 | 2 | 54.085217 | yes | 1.362E-07 | 685.35199 | 386.57416 |
| IAVNIALER | 9 | Q9H6S0 | 3'-5' RNA helicase YTHDC2 OS=Homo s | YTHDC2 | 160.249 | 2 | 54.085217 | yes | 2.123E-07 | 499.80328 | 361.45566 |
| FTAGAFSDHMALI | 14 | Q9H6S0 | 3'-5' RNA helicase YTHDC2 OS=Homo s | YTHDC2 | 160.249 | 3 | 54.085217 | yes | 1.214E-07 | 512.92578 | 162.99855 |
| FTAGAFSDHMALI | 14 | Q9H6S0 | 3'-5' RNA helicase YTHDC2 OS=Homo s | YTHDC2 | 160.249 | 3 | 54.085217 | yes | 9.722E-08 | 768.88501 | 162.99855 |
| FTAGAFSDHMALI | 14 | Q9H6S0 | 3'-5' RNA helicase YTHDC2 OS=Homo s | YTHDC2 | 160.249 | 2 | 54.085217 | yes | 1.214E-07 | 512.92578 | 162.99855 |
| FTAGAFSDHMALI | 14 | Q9H6S0 | 3'-5' RNA helicase YTHDC2 OS=Homo s | YTHDC2 | 160.249 | 2 | 54.085217 | yes | 9.722E-08 | 768.88501 | 162.99855 |
| CRPGICFR | 8 | Q9H6S0 | 3'-5' RNA helicase YTHDC2 OS=Homo s | YTHDC2 | 160.249 | 2 | 54.085217 | yes | 2.059E-07 | 533.25769 | 316.04108 |
| LNNGIPQIPVK | 11 | Q9H6S0 | 3'-5' RNA helicase YTHDC2 OS=Homo s | YTHDC2 | 160.249 | 2 | 54.085217 | yes | 1.623E-07 | 596.85608 | 289.35608 |
| FHPASVLSQPQYK | 13 | Q9H6S0 | 3'-5' RNA helicase YTHDC2 OS=Homo s | YTHDC2 | 160.249 | 2 | 54.085217 | yes | 6.003E-05 | 751.39355 | 284.49496 |

| | | | | | | | | | | | |
|---|---|---|---|---|---|---|---|---|---|---|---|
| TLLTFCTNGVLLR | 13 | Q9H6S0 | 3'-5' RNA helicase YTHDC2 OS=Homo s | YTHDC2 | 160.249 | 2 | 54.085217 | yes | 9.722E-08 | 754.4187 | 259.66217 |
| TTANLAALK | 9 | Q9H6S0 | 3'-5' RNA helicase YTHDC2 OS=Homo s | YTHDC2 | 160.249 | 2 | 54.085217 | yes | 2.119E-07 | 451.76892 | 197.17317 |
| AALVAGMYPNLV | 16 | Q9H6S0 | 3'-5' RNA helicase YTHDC2 OS=Homo s | YTHDC2 | 160.249 | 3 | 54.085217 | yes | 1.087E-07 | 575.97504 | 182.88132 |
| APSKPWSQVDEAT | 15 | Q9H6S0 | 3'-5' RNA helicase YTHDC2 OS=Homo s | YTHDC2 | 160.249 | 2 | 54.085217 | yes | 2.372E-07 | 842.93628 | 167.47147 |
| GFASQVEQLISMG | 19 | Q9H6S0 | 3'-5' RNA helicase YTHDC2 OS=Homo s | YTHDC2 | 160.249 | 3 | 54.085217 | yes | 9.935E-08 | 668.33868 | 165.13869 |
| DGSETAHAMMTC | 19 | Q9H6S0 | 3'-5' RNA helicase YTHDC2 OS=Homo s | YTHDC2 | 160.249 | 3 | 54.085217 | yes | 1.139E-07 | 706.97156 | 155.37595 |
| EMFLEDILR | 9 | Q9H6S0 | 3'-5' RNA helicase YTHDC2 OS=Homo s | YTHDC2 | 160.249 | 2 | 54.085217 | yes | 1.355E-07 | 583.29974 | 116.27814 |
| SRPSSVSPR | 9 | Q9H6S0 | 3'-5' RNA helicase YTHDC2 OS=Homo s | YTHDC2 | 160.249 | 2 | 54.085217 | yes | 1.362E-07 | 507.76999 | 88.102707 |
| AYHHSFDDEK | 10 | Q9H6S0 | 3'-5' RNA helicase YTHDC2 OS=Homo s | YTHDC2 | 160.249 | 2 | 54.085217 | yes | 1.389E-07 | 624.76764 | 67.893661 |
| NLEISQQK | 8 | Q9H6S0 | 3'-5' RNA helicase YTHDC2 OS=Homo s | YTHDC2 | 160.249 | 2 | 54.085217 | yes | 3.15E-06 | 480.26144 | 62.653336 |
| RGESEFDSFR | 10 | Q9H6S0 | 3'-5' RNA helicase YTHDC2 OS=Homo s | YTHDC2 | 160.249 | 2 | 54.085217 | yes | 3.734E-07 | 615.28088 | 62.344654 |
| FSDFLLTK | 8 | Q9H6S0 | 3'-5' RNA helicase YTHDC2 OS=Homo s | YTHDC2 | 160.249 | 2 | 54.085217 | yes | 1.734E-07 | 485.76584 | 56.503654 |
| NLNPPSSNWEVEK | 13 | Q9H6S1 | 5-azacytidine-induced protein 2 OS=Hom | AZI2 | 44.935 | 2 | 57.00761 | yes | 3.072E-07 | 757.36774 | 170.38045 |
| EVELLQLR | 8 | Q9H6S1 | 5-azacytidine-induced protein 2 OS=Hom | AZI2 | 44.935 | 2 | 57.00761 | yes | 9.722E-08 | 500.29529 | 142.33159 |
| VLNEQLQSK | 9 | Q9H6S1 | 5-azacytidine-induced protein 2 OS=Hom | AZI2 | 44.935 | 2 | 57.00761 | yes | 4.717E-07 | 529.79565 | 94.373238 |
| ACAPVGCSEDLGR | 13 | Q9H6S1 | 5-azacytidine-induced protein 2 OS=Hom | AZI2 | 44.935 | 2 | 57.00761 | yes | 9.893E-08 | 696.30573 | 70.912216 |
| TEVETQQVMR | 10 | Q9H6S1 | 5-azacytidine-induced protein 2 OS=Hom | AZI2 | 44.935 | 2 | 57.00761 | yes | 9.722E-08 | 610.8006 | 38.969818 |
| QTDPYQEDNLK | 11 | Q9H6S1 | 5-azacytidine-induced protein 2 OS=Hom | AZI2 | 44.935 | 2 | 57.00761 | yes | 1.35E-06 | 675.81226 | 15.172853 |
| YVEVVPCSTEEMS | 14 | Q9H6T0 | Epithelial splicing regulatory protein 2 OS | ESRP2 | 78.402 | 2 | 52.095734 | yes | 1.003E-07 | 843.37909 | 973.99414 |
| STAAEVQQVLNR | 12 | Q9H6T0 | Epithelial splicing regulatory protein 2 OS | ESRP2 | 78.402 | 2 | 52.095734 | no | 1.088E-07 | 658.35187 | 791.29877 |
| IAGGTSLEVAR | 11 | Q9H6T0 | Epithelial splicing regulatory protein 2 OS | ESRP2 | 78.402 | 2 | 52.095734 | yes | 2.761E-07 | 537.30109 | 785.26282 |
| VLMGGTLGR | 9 | Q9H6T0 | Epithelial splicing regulatory protein 2 OS | ESRP2 | 78.402 | 2 | 52.095734 | yes | 2.267E-07 | 452.25766 | 696.02039 |
| ADVVDSETVVR | 11 | Q9H6T0 | Epithelial splicing regulatory protein 2 OS | ESRP2 | 78.402 | 2 | 52.095734 | yes | 1.05E-07 | 595.30658 | 691.23303 |
| GGVALCLNAQGR | 12 | Q9H6T0 | Epithelial splicing regulatory protein 2 OS | ESRP2 | 78.402 | 2 | 52.095734 | yes | 1.562E-07 | 608.31677 | 672.16882 |
| GLPWQSSDQDVAI | 13 | Q9H6T0 | Epithelial splicing regulatory protein 2 OS | ESRP2 | 78.402 | 2 | 52.095734 | yes | 2.02E-07 | 729.85223 | 538.03729 |
| EASGLSADSLAR | 12 | Q9H6T0 | Epithelial splicing regulatory protein 2 OS | ESRP2 | 78.402 | 2 | 52.095734 | yes | 9.722E-08 | 588.79639 | 442.57263 |
| QVLHPEASR | 9 | Q9H6T0 | Epithelial splicing regulatory protein 2 OS | ESRP2 | 78.402 | 2 | 52.095734 | yes | 3.365E-07 | 518.78033 | 426.29977 |
| MQGVPYTAGMK | 11 | Q9H6T0 | Epithelial splicing regulatory protein 2 OS | ESRP2 | 78.402 | 2 | 52.095734 | yes | 1.159E-07 | 591.78595 | 361.23599 |
| FVDSEQR | 7 | Q9H6T0 | Epithelial splicing regulatory protein 2 OS | ESRP2 | 78.402 | 2 | 52.095734 | yes | 1.162E-06 | 440.71161 | 216.72369 |
| EPSSQLFSKPEVIK | 14 | Q9H6T0 | Epithelial splicing regulatory protein 2 OS | ESRP2 | 78.402 | 2 | 52.095734 | yes | 9.722E-08 | 794.93268 | 92.442963 |
| EFHMQHPSTCPAR | 13 | Q9H6T0 | Epithelial splicing regulatory protein 2 OS | ESRP2 | 78.402 | 2 | 52.095734 | yes | 2.349E-07 | 799.35358 | 34.833103 |
| AEAAALSTQCR | 11 | Q9H6T0 | Epithelial splicing regulatory protein 2 OS | ESRP2 | 78.402 | 2 | 52.095734 | yes | 3.367E-07 | 589.28516 | 27.881086 |
| GMDADPYNPVLPT | 15 | Q9H6T3 | RNA polymerase II-associated protein 3 ( | RPAP3 | 75.719 | 2 | 50.119892 | yes | 1.066E-07 | 830.39343 | 139.43225 |
| SSPDMLYQYLK | 11 | Q9H6T3 | RNA polymerase II-associated protein 3 ( | RPAP3 | 75.719 | 2 | 50.119892 | yes | 9.897E-08 | 672.82886 | 121.68083 |
| YDEAIDCYTK | 10 | Q9H6T3 | RNA polymerase II-associated protein 3 ( | RPAP3 | 75.719 | 2 | 50.119892 | yes | 1.234E-07 | 639.27136 | 95.298317 |
| QNAEELQDFMR | 11 | Q9H6T3 | RNA polymerase II-associated protein 3 ( | RPAP3 | 75.719 | 2 | 50.119892 | yes | 2.715E-07 | 690.81427 | 78.40242 |
| ILHDFYIEK | 9 | Q9H6T3 | RNA polymerase II-associated protein 3 ( | RPAP3 | 75.719 | 2 | 50.119892 | yes | 1.036E-07 | 589.31622 | 53.76305 |
| NSSQDDLFPTSDTI | 15 | Q9H6T3 | RNA polymerase II-associated protein 3 ( | RPAP3 | 75.719 | 2 | 50.119892 | yes | 1.715E-07 | 840.37903 | 49.480816 |
| QDFETVLLLEPGN | 14 | Q9H6T3 | RNA polymerase II-associated protein 3 ( | RPAP3 | 75.719 | 2 | 50.119892 | yes | 1.063E-07 | 801.9223 | 49.022427 |

| | | | | | | | | | | | |
|---|---|---|---|---|---|---|---|---|---|---|---|
| ALFNHIDK | 8 | Q9H6T3 | RNA polymerase II-associated protein 3 ( | RPAP3 | 75.719 | 2 | 50.119892 | yes | 2.971E-07 | 479.26126 | 39.0821 |
| ILPAPQFGAQK | 11 | Q9H6U6 | BCAS3 microtubule associated cell migra | BCAS3 | 101.237 | 2 | 43.18129 | yes | 2.774E-06 | 585.33752 | 107.90611 |
| ILVVVLQEK | 9 | Q9H6U6 | BCAS3 microtubule associated cell migra | BCAS3 | 101.237 | 2 | 43.18129 | yes | 2.696E-07 | 520.83936 | 63.797855 |
| LWMGPQFQFK | 10 | Q9H6U6 | BCAS3 microtubule associated cell migra | BCAS3 | 101.237 | 2 | 43.18129 | yes | 7.973E-07 | 641.32605 | 26.56867 |
| ALSVYGLPIR | 10 | Q9H6W3 | Ribosomal oxygenase 1 OS=Homo sapier | RIOX1 | 71.086 | 2 | 49.451096 | yes | 1.184E-06 | 544.82678 | 263.54565 |
| LGHFAPVDAVAD( | 14 | Q9H6W3 | Ribosomal oxygenase 1 OS=Homo sapier | RIOX1 | 71.086 | 2 | 49.451096 | yes | 2.181E-07 | 748.38623 | 98.331055 |
| LVGEGGHLFLYYT | 18 | Q9H6W3 | Ribosomal oxygenase 1 OS=Homo sapier | RIOX1 | 71.086 | 3 | 49.451096 | yes | 3.945E-07 | 685.34955 | 72.770401 |
| LVEVPAAPVR | 10 | Q9H6W3 | Ribosomal oxygenase 1 OS=Homo sapier | RIOX1 | 71.086 | 2 | 49.451096 | yes | 2.638E-07 | 525.81897 | 68.701813 |
| DFIHDSLPPVLTDF | 14 | Q9H6W3 | Ribosomal oxygenase 1 OS=Homo sapier | RIOX1 | 71.086 | 2 | 49.451096 | yes | 1.486E-07 | 812.9201 | 65.346718 |
| DFMDYMGAQHSD | 14 | Q9H6W3 | Ribosomal oxygenase 1 OS=Homo sapier | RIOX1 | 71.086 | 2 | 49.451096 | yes | 1.029E-07 | 816.32684 | 54.249275 |
| MDGLQASAGPLR | 12 | Q9H6W3 | Ribosomal oxygenase 1 OS=Homo sapier | RIOX1 | 71.086 | 2 | 49.451096 | yes | 1.952E-06 | 629.31647 | 37.579823 |
| QSIYIQR | 7 | Q9H707 | Zinc finger protein 552 OS=Homo sapien | ZNF552 | 46.199 | 2 | 53.140217 | no | 2.606E-07 | 454.25345 | 518.84351 |
| DILSQHER | 8 | Q9H707 | Zinc finger protein 552 OS=Homo sapien | ZNF552 | 46.199 | 2 | 53.140217 | yes | 3.469E-07 | 499.25671 | 241.98465 |
| SGLLQQEATHTGK | 13 | Q9H707 | Zinc finger protein 552 OS=Homo sapien | ZNF552 | 46.199 | 2 | 53.140217 | yes | 9.722E-08 | 685.35718 | 182.49902 |
| SFTHSSTFR | 9 | Q9H707 | Zinc finger protein 552 OS=Homo sapien | ZNF552 | 46.199 | 2 | 53.140217 | yes | 1.941E-07 | 535.25671 | 150.25244 |
| YHLIAHQR | 8 | Q9H707 | Zinc finger protein 552 OS=Homo sapien | ZNF552 | 46.199 | 2 | 53.140217 | yes | 2.224E-07 | 519.28558 | 138.95456 |
| GSVEEALFAK | 10 | Q9H707 | Zinc finger protein 552 OS=Homo sapien | ZNF552 | 46.199 | 2 | 53.140217 | no | 1.585E-07 | 525.77692 | 84.166237 |
| TECVSLFHGGK | 11 | Q9H707 | Zinc finger protein 552 OS=Homo sapien | ZNF552 | 46.199 | 2 | 53.140217 | yes | 1.846E-07 | 617.79785 | 80.696938 |
| LHVSGESSVFSES( | 15 | Q9H707 | Zinc finger protein 552 OS=Homo sapien | ZNF552 | 46.199 | 2 | 53.140217 | no | 9.722E-08 | 775.3783 | 74.778603 |
| FPVQGTVTFEDVA | 15 | Q9H707 | Zinc finger protein 552 OS=Homo sapien | ZNF552 | 46.199 | 2 | 53.140217 | yes | 9.722E-08 | 818.93268 | 34.413166 |
| YDSFSNHQGVHTF | 13 | Q9H707 | Zinc finger protein 552 OS=Homo sapien | ZNF552 | 46.199 | 2 | 53.140217 | yes | 2.805E-07 | 774.35291 | 7.6347275 |
| FSFSPEPTLEDIR | 13 | Q9H773 | dCTP pyrophosphatase 1 OS=Homo sapi( | DCTPP1 | 18.681 | 2 | 46.515335 | yes | 1.122E-06 | 769.38031 | 146.77907 |
| TDGEPGPQGWSPF | 13 | Q9H773 | dCTP pyrophosphatase 1 OS=Homo sapi( | DCTPP1 | 18.681 | 2 | 46.515335 | yes | 3.874E-06 | 692.31799 | 105.46531 |
| VDLPLAVLSK | 10 | Q9H773 | dCTP pyrophosphatase 1 OS=Homo sapi( | DCTPP1 | 18.681 | 2 | 46.515335 | yes | 1.539E-07 | 527.82898 | 35.649918 |
| THLASDDLYK | 10 | Q9H7B2 | Ribosome production factor 2 homolog O | RPF2 | 35.583 | 2 | 49.645683 | yes | 1.388E-07 | 581.79059 | 337.90826 |
| SDCSLFMFGSHNK | 13 | Q9H7B2 | Ribosome production factor 2 homolog O | RPF2 | 35.583 | 2 | 49.645683 | yes | 3.049E-07 | 765.32922 | 210.37761 |
| KPYGVLYK | 8 | Q9H7B2 | Ribosome production factor 2 homolog O | RPF2 | 35.583 | 2 | 49.645683 | yes | 1.122E-07 | 484.28421 | 104.55779 |
| NISHDTFGTTYGR | 13 | Q9H7B2 | Ribosome production factor 2 homolog O | RPF2 | 35.583 | 2 | 49.645683 | yes | 1.014E-07 | 734.84442 | 101.95609 |
| GPTVSNIR | 8 | Q9H7B2 | Ribosome production factor 2 homolog O | RPF2 | 35.583 | 2 | 49.645683 | yes | 1.666E-07 | 422.23779 | 50.349297 |
| PFEDQTSLEFFSK | 13 | Q9H7B2 | Ribosome production factor 2 homolog O | RPF2 | 35.583 | 2 | 49.645683 | yes | 1.118E-07 | 787.87231 | 47.389469 |
| RTHLASDDLYK | 11 | Q9H7B2 | Ribosome production factor 2 homolog O | RPF2 | 35.583 | 2 | 49.645683 | yes | 2.329E-05 | 659.84113 | 11.552453 |
| LYSFYDLESNINK | 13 | Q9H7B4 | Histone-lysine N-methyltransferase SMY | SMYD3 | 49.098 | 2 | 50.713173 | yes | 9.728E-08 | 803.39337 | 49.093445 |
| LPDINIYQLK | 10 | Q9H7B4 | Histone-lysine N-methyltransferase SMY | SMYD3 | 49.098 | 2 | 50.713173 | yes | 1.197E-07 | 608.85046 | 43.487679 |
| DADMLTGDEQVW | 13 | Q9H7B4 | Histone-lysine N-methyltransferase SMY | SMYD3 | 49.098 | 2 | 50.713173 | yes | 1.627E-07 | 754.34033 | 24.914816 |
| TVLASDTHQR | 10 | Q9H7D7 | WD repeat-containing protein 26 OS=Ho | WDR26 | 72.124 | 2 | 49.51373 | yes | 1.023E-07 | 564.29382 | 89.258385 |
| VLEALQVLR | 9 | Q9H7D7 | WD repeat-containing protein 26 OS=Ho | WDR26 | 72.124 | 2 | 49.51373 | yes | 1.727E-07 | 520.82678 | 79.962479 |
| FLLLQQK | 7 | Q9H7D7 | WD repeat-containing protein 26 OS=Ho | WDR26 | 72.124 | 2 | 49.51373 | yes | 4.076E-07 | 445.27893 | 33.438351 |
| SVLEGSGLPR | 10 | Q9H7E2 | Tudor domain-containing protein 3 OS=F | TDRD3 | 73.185 | 2 | 53.01694 | yes | 3.14E-07 | 507.78256 | 5258.0811 |
| SQTINNEAFSGIK | 13 | Q9H7E2 | Tudor domain-containing protein 3 OS=F | TDRD3 | 73.185 | 2 | 53.01694 | yes | 9.722E-08 | 704.85699 | 2869.7002 |

| | | | | | | | | | | | |
|---|---|---|---|---|---|---|---|---|---|---|---|
| MGTLNVEEPK | 10 | Q9H7E2 | Tudor domain-containing protein 3 OS=H | TDRD3 | 73.185 | 2 | 53.01694 | yes | 3.06E-07 | 559.28156 | 2451.2104 |
| SQPQQLHQGQYR | 12 | Q9H7E2 | Tudor domain-containing protein 3 OS=H | TDRD3 | 73.185 | 2 | 53.01694 | yes | 1.081E-07 | 735.36584 | 1090.9011 |
| HITEMGFSK | 9 | Q9H7E2 | Tudor domain-containing protein 3 OS=H | TDRD3 | 73.185 | 2 | 53.01694 | yes | 1.603E-07 | 525.25787 | 1003.8367 |
| SNLNMNAAGNR | 11 | Q9H7E2 | Tudor domain-containing protein 3 OS=H | TDRD3 | 73.185 | 2 | 53.01694 | yes | 9.81E-08 | 581.27509 | 982.38593 |
| AEVEALHSSGMTA | 16 | Q9H7E2 | Tudor domain-containing protein 3 OS=H | TDRD3 | 73.185 | 2 | 53.01694 | yes | 1.006E-07 | 814.91925 | 438.58173 |
| AEVEALHSSGMTA | 16 | Q9H7E2 | Tudor domain-containing protein 3 OS=H | TDRD3 | 73.185 | 2 | 53.01694 | yes | 9.722E-08 | 822.91669 | 438.58173 |
| LQMTDGHISCTAV | 20 | Q9H7E2 | Tudor domain-containing protein 3 OS=H | TDRD3 | 73.185 | 3 | 53.01694 | yes | 4.749E-07 | 769.01477 | 820.58008 |
| CVSHVQVDSR | 10 | Q9H7E2 | Tudor domain-containing protein 3 OS=H | TDRD3 | 73.185 | 2 | 53.01694 | yes | 4.355E-07 | 593.78528 | 777.57556 |
| IFYNSGPK | 8 | Q9H7E2 | Tudor domain-containing protein 3 OS=H | TDRD3 | 73.185 | 2 | 53.01694 | yes | 1.862E-07 | 463.24252 | 638.88104 |
| QKPVMGPPLR | 10 | Q9H7E2 | Tudor domain-containing protein 3 OS=H | TDRD3 | 73.185 | 2 | 53.01694 | yes | 1.929E-07 | 561.82623 | 575.0191 |
| SEDEEDLGNAR | 11 | Q9H7E2 | Tudor domain-containing protein 3 OS=H | TDRD3 | 73.185 | 2 | 53.01694 | yes | 1.063E-07 | 617.76276 | 498.04498 |
| ISLNTPPGTK | 10 | Q9H7E2 | Tudor domain-containing protein 3 OS=H | TDRD3 | 73.185 | 2 | 53.01694 | yes | 3.133E-07 | 514.29279 | 495.87936 |
| IGPIKPAGPVTAVP | 26 | Q9H7E2 | Tudor domain-containing protein 3 OS=H | TDRD3 | 73.185 | 4 | 53.01694 | yes | 3.015E-07 | 686.11371 | 477.86014 |
| HFNVNTDYQNPVI | 13 | Q9H7E2 | Tudor domain-containing protein 3 OS=H | TDRD3 | 73.185 | 2 | 53.01694 | yes | 1.101E-07 | 802.38422 | 383.71613 |
| ESQTSIPDYFYDR | 13 | Q9H7E2 | Tudor domain-containing protein 3 OS=H | TDRD3 | 73.185 | 2 | 53.01694 | yes | 1.02E-07 | 810.86249 | 236.63257 |
| PSTSSVSEVWAED | 14 | Q9H7E2 | Tudor domain-containing protein 3 OS=H | TDRD3 | 73.185 | 2 | 53.01694 | yes | 9.975E-08 | 775.36011 | 219.98763 |
| ILESSIPMEYAK | 12 | Q9H7E2 | Tudor domain-containing protein 3 OS=H | TDRD3 | 73.185 | 2 | 53.01694 | yes | 3.383E-07 | 690.8576 | 218.21051 |
| PTQQFYQPPR | 10 | Q9H7E2 | Tudor domain-containing protein 3 OS=H | TDRD3 | 73.185 | 2 | 53.01694 | yes | 3.121E-07 | 631.31982 | 216.00211 |
| PAGPVTAVPCDDK | 13 | Q9H7E2 | Tudor domain-containing protein 3 OS=H | TDRD3 | 73.185 | 2 | 53.01694 | yes | 3.401E-07 | 663.82153 | 141.47478 |
| TAAIAEVAK | 9 | Q9H7E2 | Tudor domain-containing protein 3 OS=H | TDRD3 | 73.185 | 2 | 53.01694 | yes | 2.021E-07 | 437.25565 | 80.497635 |
| SGPIKPEK | 8 | Q9H7E2 | Tudor domain-containing protein 3 OS=H | TDRD3 | 73.185 | 2 | 53.01694 | yes | 5.416E-07 | 428.25037 | 50.98299 |
| DTSYPLGSQHSDG | 16 | Q9H7E2 | Tudor domain-containing protein 3 OS=H | TDRD3 | 73.185 | 3 | 53.01694 | yes | 1.031E-06 | 570.59705 | 32.458092 |
| STRPTQQFYQPPR | 13 | Q9H7E2 | Tudor domain-containing protein 3 OS=H | TDRD3 | 73.185 | 2 | 53.01694 | yes | 2.408E-06 | 803.41022 | 20.935947 |
| ATLDMPDEEFR | 11 | Q9H7E9 | UPF0488 protein C8orf33 OS=Homo sap | C8orf33 | 24.993 | 2 | 51.05559 | yes | 1.084E-07 | 662.29791 | 2233.8713 |
| AAAYSAQVQPVD | 16 | Q9H7E9 | UPF0488 protein C8orf33 OS=Homo sap | C8orf33 | 24.993 | 2 | 51.05559 | yes | 9.722E-08 | 802.90497 | 1582.6678 |
| LPGPVSSAR | 9 | Q9H7E9 | UPF0488 protein C8orf33 OS=Homo sap | C8orf33 | 24.993 | 2 | 51.05559 | yes | 9.722E-08 | 442.25345 | 1086.4838 |
| AHPLGDEGGTASK | 13 | Q9H7E9 | UPF0488 protein C8orf33 OS=Homo sap | C8orf33 | 24.993 | 2 | 51.05559 | yes | 1.483E-07 | 620.30182 | 414.77444 |
| EQAIGAIR | 8 | Q9H7E9 | UPF0488 protein C8orf33 OS=Homo sap | C8orf33 | 24.993 | 2 | 51.05559 | yes | 3.082E-07 | 429.24561 | 340.5603 |
| AALGHLAGEAAA | 23 | Q9H7E9 | UPF0488 protein C8orf33 OS=Homo sap | C8orf33 | 24.993 | 3 | 51.05559 | yes | 1.543E-07 | 715.68835 | 235.49625 |
| QKEQAIGAIR | 10 | Q9H7E9 | UPF0488 protein C8orf33 OS=Homo sap | C8orf33 | 24.993 | 2 | 51.05559 | yes | 9.722E-08 | 557.32239 | 58.608986 |
| FLENPDLSQTEEK | 13 | Q9H7H0 | Methyltransferase-like protein 17, mitoch | METTL17 | 50.734 | 2 | 50.690586 | yes | 3.53E-07 | 775.37268 | 177.22418 |
| QFLPVSPK | 8 | Q9H7H0 | Methyltransferase-like protein 17, mitoch | METTL17 | 50.734 | 2 | 50.690586 | yes | 1.853E-07 | 458.26855 | 129.0972 |
| FSMVILAR | 8 | Q9H7H0 | Methyltransferase-like protein 17, mitoch | METTL17 | 50.734 | 2 | 50.690586 | yes | 1.133E-07 | 468.77042 | 106.37437 |
| AGHSLLMDAR | 10 | Q9H7H0 | Methyltransferase-like protein 17, mitoch | METTL17 | 50.734 | 2 | 50.690586 | yes | 4.192E-07 | 535.77423 | 93.063927 |
| TEVVQTLWR | 9 | Q9H7H0 | Methyltransferase-like protein 17, mitoch | METTL17 | 50.734 | 2 | 50.690586 | yes | 9.728E-08 | 566.31146 | 71.435112 |
| GAVLHALR | 8 | Q9H7H0 | Methyltransferase-like protein 17, mitoch | METTL17 | 50.734 | 2 | 50.690586 | yes | 1.407E-07 | 418.7587 | 29.571775 |
| DAASAGLGSIGVK | 13 | Q9H7N4 | Splicing factor, arginine/serine-rich 19 OS | SCAF1 | 139.27 | 2 | 48.526714 | yes | 2.398E-06 | 573.31171 | 50.957798 |
| SGGGSGGSGGQVS | 15 | Q9H7N4 | Splicing factor, arginine/serine-rich 19 OS | SCAF1 | 139.27 | 2 | 48.526714 | yes | 2.808E-07 | 617.80475 | 45.665489 |
| EVLYDSEGLSGEE | 14 | Q9H7N4 | Splicing factor, arginine/serine-rich 19 OS | SCAF1 | 139.27 | 2 | 48.526714 | yes | 3.215E-06 | 791.86517 | 45.25161 |

| | | | | | | | | | | | |
|---|---|---|---|---|---|---|---|---|---|---|---|
| SPFLKPDER | 9 | Q9H7N4 | Splicing factor, arginine/serine-rich 19 OS | SCAF1 | 139.27 | 2 | 48.526714 | yes | 1.136E-07 | 544.79041 | 40.53381 |
| GAEETSWSGEER | 12 | Q9H7N4 | Splicing factor, arginine/serine-rich 19 OS | SCAF1 | 139.27 | 2 | 48.526714 | yes | 3.27E-07 | 669.28387 | 33.471848 |
| APAPAPPPSGSTSC | 17 | Q9H7N4 | Splicing factor, arginine/serine-rich 19 OS | SCAF1 | 139.27 | 2 | 48.526714 | yes | 6.466E-05 | 812.87286 | 8.8649731 |
| MLTLLPGER | 9 | Q9H7T9 | Aurora kinase A- and ninein-interacting p | AUNIP | 40.253 | 2 | 50.128147 | yes | 1.296E-07 | 515.29169 | 267.17108 |
| SPVSVFSWDNEK | 12 | Q9H7T9 | Aurora kinase A- and ninein-interacting p | AUNIP | 40.253 | 2 | 50.128147 | yes | 1.231E-07 | 697.83295 | 235.8038 |
| ANIYFTQR | 8 | Q9H7T9 | Aurora kinase A- and ninein-interacting p | AUNIP | 40.253 | 2 | 50.128147 | yes | 2.021E-07 | 506.76416 | 223.70488 |
| QAPVLLQTYR | 10 | Q9H7T9 | Aurora kinase A- and ninein-interacting p | AUNIP | 40.253 | 2 | 50.128147 | yes | 1.165E-07 | 594.84039 | 137.46727 |
| SIASFFTLQPGK | 12 | Q9H7T9 | Aurora kinase A- and ninein-interacting p | AUNIP | 40.253 | 2 | 50.128147 | yes | 9.893E-08 | 648.35333 | 80.102959 |
| ESWNGENIESVK | 12 | Q9H7T9 | Aurora kinase A- and ninein-interacting p | AUNIP | 40.253 | 2 | 50.128147 | yes | 4.231E-07 | 696.3255 | 52.161339 |
| SVSSHTESQINK | 12 | Q9H7T9 | Aurora kinase A- and ninein-interacting p | AUNIP | 40.253 | 2 | 50.128147 | yes | 2.696E-07 | 658.82568 | 49.023342 |
| VQTHLIKPGTK | 11 | Q9H7T9 | Aurora kinase A- and ninein-interacting p | AUNIP | 40.253 | 2 | 50.128147 | yes | 3.934E-06 | 611.36932 | 6.7737303 |
| TEPPSSEPISFIPVK | 15 | Q9H7Z3 | Nuclear exosome regulator NRDE2 OS=I | NRDE2 | 132.673 | 2 | 40.76281 | yes | 5.795E-06 | 814.4325 | 94.085945 |
| EIHATIPETGLMHF | 14 | Q9H7Z3 | Nuclear exosome regulator NRDE2 OS=I | NRDE2 | 132.673 | 3 | 40.76281 | yes | 4.868E-07 | 535.61194 | 74.157501 |
| LTESSPYGPYTGQ' | 21 | Q9H7Z3 | Nuclear exosome regulator NRDE2 OS=I | NRDE2 | 132.673 | 3 | 40.76281 | yes | 3.854E-07 | 758.41071 | 70.625328 |
| YLSELAEQPER | 11 | Q9H7Z6 | Histone acetyltransferase KAT8 OS=Hom | KAT8 | 52.403 | 2 | 42.961956 | yes | 2.794E-07 | 667.83295 | 75.060997 |
| LWLCEYCLK | 9 | Q9H7Z6 | Histone acetyltransferase KAT8 OS=Hom | KAT8 | 52.403 | 2 | 42.961956 | yes | 4.689E-07 | 642.80939 | 48.16832 |
| NTPSISEEQIK | 11 | Q9H814 | Phosphorylated adapter RNA export prote | PHAX | 44.403 | 2 | 52.84681 | yes | 2.239E-07 | 623.3197 | 1944.8243 |
| DIFYIENQK | 9 | Q9H814 | Phosphorylated adapter RNA export prote | PHAX | 44.403 | 2 | 52.84681 | yes | 1.084E-07 | 585.29547 | 1492.2461 |
| VLGGDSAMR | 9 | Q9H814 | Phosphorylated adapter RNA export prote | PHAX | 44.403 | 2 | 52.84681 | yes | 2.598E-07 | 453.2291 | 1221.8083 |
| YEITAEDSQEK | 11 | Q9H814 | Phosphorylated adapter RNA export prote | PHAX | 44.403 | 2 | 52.84681 | yes | 9.893E-08 | 656.79877 | 1143.9131 |
| AFQNTATACAPVS | 16 | Q9H814 | Phosphorylated adapter RNA export prote | PHAX | 44.403 | 3 | 52.84681 | yes | 9.722E-08 | 598.6178 | 831.38422 |
| TPGGVFLNLLK | 11 | Q9H814 | Phosphorylated adapter RNA export prote | PHAX | 44.403 | 2 | 52.84681 | yes | 9.899E-08 | 579.84772 | 796.7428 |
| QSETYNYLLAK | 11 | Q9H814 | Phosphorylated adapter RNA export prote | PHAX | 44.403 | 2 | 52.84681 | yes | 1.042E-07 | 665.33789 | 765.60571 |
| SLNFQEDDDTSR | 12 | Q9H814 | Phosphorylated adapter RNA export prote | PHAX | 44.403 | 2 | 52.84681 | yes | 9.722E-08 | 713.80768 | 694.39429 |
| ELDEYMHGGK | 10 | Q9H814 | Phosphorylated adapter RNA export prote | PHAX | 44.403 | 2 | 52.84681 | yes | 2.207E-07 | 589.76099 | 646.77582 |
| LGNRPEMNYK | 10 | Q9H814 | Phosphorylated adapter RNA export prote | PHAX | 44.403 | 2 | 52.84681 | yes | 9.809E-08 | 611.30585 | 286.31653 |
| MALEVGDMEDGQ | 27 | Q9H814 | Phosphorylated adapter RNA export prote | PHAX | 44.403 | 4 | 52.84681 | yes | 2.817E-07 | 718.55457 | 142.44685 |
| CFNPPPKPEPFQFC | 26 | Q9H814 | Phosphorylated adapter RNA export prote | PHAX | 44.403 | 4 | 52.84681 | yes | 3.579E-06 | 707.35626 | 60.426353 |
| VADEISFR | 8 | Q9H814 | Phosphorylated adapter RNA export prote | PHAX | 44.403 | 2 | 52.84681 | yes | 1.214E-06 | 468.74289 | 30.007545 |
| GRYEITAEDSQEK | 13 | Q9H814 | Phosphorylated adapter RNA export prote | PHAX | 44.403 | 2 | 52.84681 | yes | 1.787E-07 | 763.36011 | 26.761391 |
| DLDKELDEYMHG | 14 | Q9H814 | Phosphorylated adapter RNA export prote | PHAX | 44.403 | 2 | 52.84681 | yes | 2.62E-05 | 825.37744 | 10.617301 |
| TESDFSNLDSEK | 12 | Q9H825 | tRNA N(3)-methylcytidine methyltransfer | METTL8 | 33.387 | 2 | 49.940903 | yes | 4.499E-07 | 686.29913 | 301.60724 |
| VLLEEQVK | 8 | Q9H825 | tRNA N(3)-methylcytidine methyltransfer | METTL8 | 33.387 | 2 | 49.940903 | yes | 3.39E-07 | 479.28439 | 294.68518 |
| GPMETGLFPGSNA | 16 | Q9H825 | tRNA N(3)-methylcytidine methyltransfer | METTL8 | 33.387 | 2 | 49.940903 | yes | 1.959E-07 | 841.40375 | 130.18269 |
| MHCPTVPDEK | 10 | Q9H825 | tRNA N(3)-methylcytidine methyltransfer | METTL8 | 33.387 | 2 | 49.940903 | yes | 9.974E-08 | 607.27063 | 19.895056 |
| CSDIISYTFKP | 11 | Q9H840 | Gem-associated protein 7 OS=Homo sapi | GEMIN7 | 14.537 | 2 | 51.76806 | yes | 1.398E-07 | 665.82104 | 620.33069 |
| SLLAMVGHQVSF1 | 19 | Q9H840 | Gem-associated protein 7 OS=Homo sapi | GEMIN7 | 14.537 | 3 | 51.76806 | yes | 1.584E-07 | 694.37018 | 102.66496 |
| RAPLRPEVPEIQEC | 27 | Q9H840 | Gem-associated protein 7 OS=Homo sapi | GEMIN7 | 14.537 | 4 | 51.76806 | yes | 3.133E-07 | 798.15436 | 86.362297 |
| MQTPVNIPVPVLR | 13 | Q9H840 | Gem-associated protein 7 OS=Homo sapi | GEMIN7 | 14.537 | 2 | 51.76806 | yes | 9.722E-08 | 732.42377 | 82.150299 |

| | | | | | | | | | | | |
|---|---|---|---|---|---|---|---|---|---|---|---|
| TEVVDSDGSVK | 11 | Q9H845 | Complex I assembly factor ACAD9, mito | ACAD9 | 68.761 | 2 | 46.203876 | yes | 3.514E-07 | 568.27753 | 87.286957 |
| VAMNILNSGR | 10 | Q9H845 | Complex I assembly factor ACAD9, mito | ACAD9 | 68.761 | 2 | 46.203876 | yes | 1.622E-07 | 537.78986 | 64.374893 |
| IPVENILGEVGDGI | 15 | Q9H845 | Complex I assembly factor ACAD9, mito | ACAD9 | 68.761 | 2 | 46.203876 | yes | 3.412E-07 | 793.92487 | 50.731571 |
| LIEMTAEYACTR | 12 | Q9H845 | Complex I assembly factor ACAD9, mito | ACAD9 | 68.761 | 2 | 46.203876 | yes | 3.147E-07 | 729.3418 | 30.716602 |
| FSMGSVVAGLLK | 12 | Q9H845 | Complex I assembly factor ACAD9, mito | ACAD9 | 68.761 | 2 | 46.203876 | yes | 6.859E-05 | 604.83899 | 4.7388945 |
| VMMPSPASSMFR | 12 | Q9H869 | YY1-associated protein 1 OS=Homo sapi | YY1AP1 | 87.945 | 2 | 47.727486 | yes | 1.271E-07 | 670.81165 | 109.55231 |
| QSIPVLLEGR | 10 | Q9H8H2 | Probable ATP-dependent RNA helicase D | DDX31 | 94.088 | 2 | 50.970444 | yes | 9.722E-08 | 556.32715 | 199.47652 |
| GVLLCTDVAAR | 11 | Q9H8H2 | Probable ATP-dependent RNA helicase D | DDX31 | 94.088 | 2 | 50.970444 | yes | 2.864E-06 | 587.8161 | 177.29987 |
| SDGPYALVLVPTR | 13 | Q9H8H2 | Probable ATP-dependent RNA helicase D | DDX31 | 94.088 | 2 | 50.970444 | yes | 1.348E-07 | 694.38263 | 162.95152 |
| TAVFQEFSHSR | 11 | Q9H8H2 | Probable ATP-dependent RNA helicase D | DDX31 | 94.088 | 2 | 50.970444 | yes | 1.276E-07 | 654.82019 | 124.7584 |
| QNVLLSATLTEGV | 15 | Q9H8H2 | Probable ATP-dependent RNA helicase D | DDX31 | 94.088 | 2 | 50.970444 | yes | 1.065E-07 | 801.44647 | 110.15312 |
| WLVFDEADR | 9 | Q9H8H2 | Probable ATP-dependent RNA helicase D | DDX31 | 94.088 | 2 | 50.970444 | yes | 1.512E-07 | 575.78003 | 88.099594 |
| DITVILNAVNAEC( | 15 | Q9H8H2 | Probable ATP-dependent RNA helicase D | DDX31 | 94.088 | 2 | 50.970444 | yes | 1.02E-07 | 844.43781 | 84.987831 |
| MEDILCVLTR | 10 | Q9H8H2 | Probable ATP-dependent RNA helicase D | DDX31 | 94.088 | 2 | 50.970444 | yes | 3.946E-07 | 625.31757 | 63.915489 |
| ILDLGFEK | 8 | Q9H8H2 | Probable ATP-dependent RNA helicase D | DDX31 | 94.088 | 2 | 50.970444 | yes | 3.912E-07 | 467.76584 | 62.791245 |
| SHAVGPQEIR | 10 | Q9H8H2 | Probable ATP-dependent RNA helicase D | DDX31 | 94.088 | 2 | 50.970444 | yes | 4.57E-06 | 547.29108 | 60.397873 |
| ELALQSFDTVQK | 12 | Q9H8H2 | Probable ATP-dependent RNA helicase D | DDX31 | 94.088 | 2 | 50.970444 | yes | 1.039E-07 | 689.86426 | 56.228878 |
| SLHLGHVAK | 9 | Q9H8H2 | Probable ATP-dependent RNA helicase D | DDX31 | 94.088 | 2 | 50.970444 | yes | 9.722E-08 | 481.28253 | 54.050148 |
| IGCHGSSLLILAPSI | 26 | Q9H8H2 | Probable ATP-dependent RNA helicase D | DDX31 | 94.088 | 4 | 50.970444 | yes | 2.923E-06 | 689.10364 | 27.341471 |
| KVEVEPPPDRPVR | 13 | Q9H8M2 | Bromodomain-containing protein 9 OS=I | BRD9 | 67.001 | 3 | 22.666197 | yes | 3.817E-06 | 506.61932 | 41.73431 |
| YLPLGDER | 8 | Q9H8V3 | Protein ECT2 OS=Homo sapiens OX=96 | ECT2 | 103.505 | 2 | 53.87372 | yes | 1.05E-07 | 481.75073 | 814.72784 |
| TSLADSSIFDSK | 12 | Q9H8V3 | Protein ECT2 OS=Homo sapiens OX=96 | ECT2 | 103.505 | 2 | 53.87372 | yes | 2.176E-07 | 635.81171 | 656.76532 |
| VIGPPVVLNCSQK | 13 | Q9H8V3 | Protein ECT2 OS=Homo sapiens OX=96 | ECT2 | 103.505 | 2 | 53.87372 | yes | 1.043E-07 | 705.89233 | 632.09174 |
| GGPILAPEEIK | 11 | Q9H8V3 | Protein ECT2 OS=Homo sapiens OX=96 | ECT2 | 103.505 | 2 | 53.87372 | yes | 9.722E-08 | 562.32153 | 438.57224 |
| SVSMLSLNTPNSN | 14 | Q9H8V3 | Protein ECT2 OS=Homo sapiens OX=96 | ECT2 | 103.505 | 2 | 53.87372 | yes | 1.02E-07 | 760.38031 | 399.35138 |
| TIFGSIPDIFDVHTI | 15 | Q9H8V3 | Protein ECT2 OS=Homo sapiens OX=96 | ECT2 | 103.505 | 2 | 53.87372 | yes | 1.468E-07 | 845.44598 | 322.09088 |
| VILVQEAGK | 9 | Q9H8V3 | Protein ECT2 OS=Homo sapiens OX=96 | ECT2 | 103.505 | 2 | 53.87372 | yes | 2.594E-07 | 478.79239 | 313.99042 |
| AGETMYLYEK | 10 | Q9H8V3 | Protein ECT2 OS=Homo sapiens OX=96 | ECT2 | 103.505 | 2 | 53.87372 | yes | 1.482E-07 | 602.78137 | 290.95688 |
| ETLAQLSR | 8 | Q9H8V3 | Protein ECT2 OS=Homo sapiens OX=96 | ECT2 | 103.505 | 2 | 53.87372 | yes | 9.819E-08 | 459.25616 | 278.43808 |
| SPHGQTRPPASLK | 13 | Q9H8V3 | Protein ECT2 OS=Homo sapiens OX=96 | ECT2 | 103.505 | 3 | 53.87372 | yes | 2.626E-07 | 459.25287 | 138.91501 |
| SPHGQTRPPASLK | 13 | Q9H8V3 | Protein ECT2 OS=Homo sapiens OX=96 | ECT2 | 103.505 | 3 | 53.87372 | yes | 6.766E-07 | 688.37567 | 138.91501 |
| SPHGQTRPPASLK | 13 | Q9H8V3 | Protein ECT2 OS=Homo sapiens OX=96 | ECT2 | 103.505 | 2 | 53.87372 | yes | 2.626E-07 | 459.25287 | 138.91501 |
| SPHGQTRPPASLK | 13 | Q9H8V3 | Protein ECT2 OS=Homo sapiens OX=96 | ECT2 | 103.505 | 2 | 53.87372 | yes | 6.766E-07 | 688.37567 | 138.91501 |
| ALMTSHGSVEGR | 12 | Q9H8V3 | Protein ECT2 OS=Homo sapiens OX=96 | ECT2 | 103.505 | 2 | 53.87372 | yes | 9.85E-08 | 622.80621 | 252.60791 |
| VETISLGEHPCDR | 13 | Q9H8V3 | Protein ECT2 OS=Homo sapiens OX=96 | ECT2 | 103.505 | 2 | 53.87372 | yes | 9.809E-08 | 756.85919 | 123.91193 |
| VETISLGEHPCDR | 13 | Q9H8V3 | Protein ECT2 OS=Homo sapiens OX=96 | ECT2 | 103.505 | 2 | 53.87372 | yes | 2.756E-07 | 504.90857 | 123.91193 |
| VETISLGEHPCDR | 13 | Q9H8V3 | Protein ECT2 OS=Homo sapiens OX=96 | ECT2 | 103.505 | 3 | 53.87372 | yes | 9.809E-08 | 756.85919 | 123.91193 |
| VETISLGEHPCDR | 13 | Q9H8V3 | Protein ECT2 OS=Homo sapiens OX=96 | ECT2 | 103.505 | 3 | 53.87372 | yes | 2.756E-07 | 504.90857 | 123.91193 |
| CTHLVVEENIVK | 12 | Q9H8V3 | Protein ECT2 OS=Homo sapiens OX=96 | ECT2 | 103.505 | 2 | 53.87372 | yes | 1.058E-07 | 720.87939 | 119.53162 |

| | | | | | | | | | | | |
|---|---|---|---|---|---|---|---|---|---|---|---|
| CTHLVVEENIVK | 12 | Q9H8V3 | Protein ECT2 OS=Homo sapiens OX=96 | ECT2 | 103.505 | 2 | 53.87372 | yes | 3.093E-06 | 480.92203 | 119.53162 |
| CTHLVVEENIVK | 12 | Q9H8V3 | Protein ECT2 OS=Homo sapiens OX=96 | ECT2 | 103.505 | 3 | 53.87372 | yes | 1.058E-07 | 720.87939 | 119.53162 |
| CTHLVVEENIVK | 12 | Q9H8V3 | Protein ECT2 OS=Homo sapiens OX=96 | ECT2 | 103.505 | 3 | 53.87372 | yes | 3.093E-06 | 480.92203 | 119.53162 |
| HIHLMPLSQIK | 11 | Q9H8V3 | Protein ECT2 OS=Homo sapiens OX=96 | ECT2 | 103.505 | 3 | 53.87372 | yes | 3.989E-07 | 439.58844 | 112.93175 |
| HIHLMPLSQIK | 11 | Q9H8V3 | Protein ECT2 OS=Homo sapiens OX=96 | ECT2 | 103.505 | 3 | 53.87372 | yes | 1.865E-07 | 658.87903 | 112.93175 |
| HIHLMPLSQIK | 11 | Q9H8V3 | Protein ECT2 OS=Homo sapiens OX=96 | ECT2 | 103.505 | 2 | 53.87372 | yes | 3.989E-07 | 439.58844 | 112.93175 |
| HIHLMPLSQIK | 11 | Q9H8V3 | Protein ECT2 OS=Homo sapiens OX=96 | ECT2 | 103.505 | 2 | 53.87372 | yes | 1.865E-07 | 658.87903 | 112.93175 |
| LVTLVHHMGGVII | 13 | Q9H8V3 | Protein ECT2 OS=Homo sapiens OX=96 | ECT2 | 103.505 | 3 | 53.87372 | yes | 1.018E-07 | 477.94662 | 106.63457 |
| LVTLVHHMGGVII | 13 | Q9H8V3 | Protein ECT2 OS=Homo sapiens OX=96 | ECT2 | 103.505 | 3 | 53.87372 | yes | 9.722E-08 | 716.41626 | 106.63457 |
| LVTLVHHMGGVII | 13 | Q9H8V3 | Protein ECT2 OS=Homo sapiens OX=96 | ECT2 | 103.505 | 2 | 53.87372 | yes | 1.018E-07 | 477.94662 | 106.63457 |
| LVTLVHHMGGVII | 13 | Q9H8V3 | Protein ECT2 OS=Homo sapiens OX=96 | ECT2 | 103.505 | 2 | 53.87372 | yes | 9.722E-08 | 716.41626 | 106.63457 |
| VTHLVANCTQGEI | 13 | Q9H8V3 | Protein ECT2 OS=Homo sapiens OX=96 | ECT2 | 103.505 | 2 | 53.87372 | yes | 1.137E-07 | 728.86426 | 191.40854 |
| SIGDIFLK | 8 | Q9H8V3 | Protein ECT2 OS=Homo sapiens OX=96 | ECT2 | 103.505 | 2 | 53.87372 | yes | 3.813E-07 | 446.76056 | 180.50821 |
| TNMEEMTEMQGG | 13 | Q9H8V3 | Protein ECT2 OS=Homo sapiens OX=96 | ECT2 | 103.505 | 2 | 53.87372 | yes | 9.826E-08 | 743.30457 | 176.33777 |
| VAVSLGTPIMKPE | 17 | Q9H8V3 | Protein ECT2 OS=Homo sapiens OX=96 | ECT2 | 103.505 | 3 | 53.87372 | yes | 1.805E-07 | 644.6936 | 140.83743 |
| AENSVLTSTTGR | 12 | Q9H8V3 | Protein ECT2 OS=Homo sapiens OX=96 | ECT2 | 103.505 | 2 | 53.87372 | yes | 9.722E-08 | 639.32025 | 123.04972 |
| QSLVELLIR | 9 | Q9H8V3 | Protein ECT2 OS=Homo sapiens OX=96 | ECT2 | 103.505 | 2 | 53.87372 | yes | 6.393E-07 | 535.83203 | 118.55191 |
| EVMTHINEDK | 10 | Q9H8V3 | Protein ECT2 OS=Homo sapiens OX=96 | ECT2 | 103.505 | 2 | 53.87372 | yes | 1.082E-07 | 608.28735 | 118.31894 |
| LKETLAQLSR | 10 | Q9H8V3 | Protein ECT2 OS=Homo sapiens OX=96 | ECT2 | 103.505 | 2 | 53.87372 | yes | 1.938E-07 | 579.8457 | 63.005714 |
| INQAKPECGR | 10 | Q9H8V3 | Protein ECT2 OS=Homo sapiens OX=96 | ECT2 | 103.505 | 2 | 53.87372 | yes | 2.475E-07 | 586.79565 | 62.41465 |
| NEQDFYAAVDDFI | 13 | Q9H8V3 | Protein ECT2 OS=Homo sapiens OX=96 | ECT2 | 103.505 | 2 | 53.87372 | yes | 9.722E-08 | 795.34698 | 60.004051 |
| EVMTHINEDKR | 11 | Q9H8V3 | Protein ECT2 OS=Homo sapiens OX=96 | ECT2 | 103.505 | 2 | 53.87372 | yes | 9.901E-08 | 686.33789 | 37.011017 |
| AGTLSPGLGPQGP | 14 | Q9H8X2 | Inositol-pentakisphosphate 2-kinase OS=I | IPPK | 56.017 | 2 | 52.73507 | yes | 1.532E-07 | 654.35693 | 315.625 |
| YCPLDLYSGNK | 11 | Q9H8X2 | Inositol-pentakisphosphate 2-kinase OS=I | IPPK | 56.017 | 2 | 52.73507 | yes | 2.82E-07 | 665.31079 | 142.23241 |
| SLLQEAQNNLK | 11 | Q9H8X2 | Inositol-pentakisphosphate 2-kinase OS=I | IPPK | 56.017 | 2 | 52.73507 | yes | 9.722E-08 | 629.34351 | 130.0054 |
| ELVHVITR | 8 | Q9H8X2 | Inositol-pentakisphosphate 2-kinase OS=I | IPPK | 56.017 | 2 | 52.73507 | yes | 6.538E-07 | 483.79019 | 124.94682 |
| YLEEFPEER | 9 | Q9H8X2 | Inositol-pentakisphosphate 2-kinase OS=I | IPPK | 56.017 | 2 | 52.73507 | yes | 5.46E-07 | 606.28259 | 121.55815 |
| SLVVAHAQR | 9 | Q9H8X2 | Inositol-pentakisphosphate 2-kinase OS=I | IPPK | 56.017 | 2 | 52.73507 | yes | 4.769E-07 | 490.78543 | 91.659523 |
| AQVPGPLTPEMEA | 14 | Q9H8Y5 | tRNA endonuclease ANKZF1 OS=Homo | ANKZF1 | 80.927 | 2 | 52.861305 | yes | 5.462E-07 | 748.38232 | 1415.8318 |
| DLLAGPSWAK | 10 | Q9H8Y5 | tRNA endonuclease ANKZF1 OS=Homo | ANKZF1 | 80.927 | 2 | 52.861305 | yes | 1.6E-07 | 529.28748 | 905.40692 |
| LWDIPLATR | 9 | Q9H8Y5 | tRNA endonuclease ANKZF1 OS=Homo | ANKZF1 | 80.927 | 2 | 52.861305 | yes | 2.023E-07 | 542.8111 | 549.49731 |
| DKPLLSALDFEK | 12 | Q9H8Y5 | tRNA endonuclease ANKZF1 OS=Homo | ANKZF1 | 80.927 | 2 | 52.861305 | yes | 1.022E-07 | 688.37701 | 532.01587 |
| EALGQNEESPK | 11 | Q9H8Y5 | tRNA endonuclease ANKZF1 OS=Homo | ANKZF1 | 80.927 | 2 | 52.861305 | yes | 9.722E-08 | 601.28839 | 492.31845 |
| LLLEAGADPTVQD | 15 | Q9H8Y5 | tRNA endonuclease ANKZF1 OS=Homo | ANKZF1 | 80.927 | 2 | 52.861305 | yes | 9.722E-08 | 792.91504 | 432.09027 |
| GGPSHSAGANLR | 12 | Q9H8Y5 | tRNA endonuclease ANKZF1 OS=Homo | ANKZF1 | 80.927 | 2 | 52.861305 | yes | 1.209E-07 | 562.28381 | 418.65222 |
| LLQGPMDISEK | 11 | Q9H8Y5 | tRNA endonuclease ANKZF1 OS=Homo | ANKZF1 | 80.927 | 2 | 52.861305 | yes | 1.093E-06 | 615.82355 | 378.78964 |
| ALEEAGTILLR | 11 | Q9H8Y5 | tRNA endonuclease ANKZF1 OS=Homo | ANKZF1 | 80.927 | 2 | 52.861305 | yes | 2.311E-07 | 593.34552 | 348.66763 |
| ARPPYTVAADK | 11 | Q9H8Y5 | tRNA endonuclease ANKZF1 OS=Homo | ANKZF1 | 80.927 | 2 | 52.861305 | yes | 4.221E-07 | 594.8222 | 282.53607 |
| LQLAPSPADPR | 11 | Q9H8Y5 | tRNA endonuclease ANKZF1 OS=Homo | ANKZF1 | 80.927 | 2 | 52.861305 | yes | 1.113E-07 | 582.8222 | 278.7558 |

| | | | | | | | | | | | |
|---|---|---|---|---|---|---|---|---|---|---|---|
| RYNEATLYK | 9 | Q9H8Y5 | tRNA endonuclease ANKZF1 OS=Homo | ANKZF1 | 80.927 | 2 | 52.861305 | yes | 9.722E-08 | 579.30109 | 251.04106 |
| LTTLHVYEEDPR | 12 | Q9H8Y5 | tRNA endonuclease ANKZF1 OS=Homo | ANKZF1 | 80.927 | 2 | 52.861305 | yes | 1.704E-07 | 736.87262 | 165.14235 |
| RPTFQELQR | 9 | Q9H8Y5 | tRNA endonuclease ANKZF1 OS=Homo | ANKZF1 | 80.927 | 2 | 52.861305 | yes | 1.391E-07 | 587.82001 | 145.51997 |
| LAAQLGAPTSPIPD | 21 | Q9H8Y5 | tRNA endonuclease ANKZF1 OS=Homo | ANKZF1 | 80.927 | 3 | 52.861305 | yes | 1.304E-07 | 698.05139 | 98.092926 |
| YNEATLYK | 8 | Q9H8Y5 | tRNA endonuclease ANKZF1 OS=Homo | ANKZF1 | 80.927 | 2 | 52.861305 | yes | 3.677E-07 | 501.25055 | 82.046738 |
| NPDAYDYNK | 9 | Q9H8Y5 | tRNA endonuclease ANKZF1 OS=Homo | ANKZF1 | 80.927 | 2 | 52.861305 | yes | 7.32E-07 | 550.23816 | 76.206703 |
| LSRPPGFYPHR | 11 | Q9H8Y5 | tRNA endonuclease ANKZF1 OS=Homo | ANKZF1 | 80.927 | 3 | 52.861305 | yes | 1.093E-07 | 442.90704 | 58.274052 |
| GSGIFDESTPVQTR | 14 | Q9H910 | Jupiter microtubule associated homolog 2 | JPT2 | 20.064 | 2 | 50.231762 | yes | 9.847E-08 | 747.36517 | 68.976044 |
| TSDIFGSPVTATSR | 14 | Q9H910 | Jupiter microtubule associated homolog 2 | JPT2 | 20.064 | 2 | 50.231762 | yes | 3.6E-07 | 719.86224 | 39.51482 |
| GVNEDTYSGILDC | 15 | Q9H936 | Mitochondrial glutamate carrier 1 OS=Hc | SLC25A22 | 34.471 | 2 | 52.3437 | yes | 9.722E-08 | 835.37793 | 338.8978 |
| LAANDFFR | 8 | Q9H936 | Mitochondrial glutamate carrier 1 OS=Hc | SLC25A22 | 34.471 | 2 | 52.3437 | no | 9.722E-08 | 477.24561 | 289.64465 |
| VYTSMSDCLIK | 11 | Q9H936 | Mitochondrial glutamate carrier 1 OS=Hc | SLC25A22 | 34.471 | 2 | 52.3437 | yes | 9.722E-08 | 658.81488 | 178.05313 |
| SEGYFGMYR | 9 | Q9H936 | Mitochondrial glutamate carrier 1 OS=Hc | SLC25A22 | 34.471 | 2 | 52.3437 | yes | 9.722E-08 | 555.23969 | 162.25304 |
| IQLQDAGR | 8 | Q9H936 | Mitochondrial glutamate carrier 1 OS=Hc | SLC25A22 | 34.471 | 2 | 52.3437 | no | 2.019E-06 | 450.7485 | 119.60371 |
| GAAVNLTLVTPEK | 13 | Q9H936 | Mitochondrial glutamate carrier 1 OS=Hc | SLC25A22 | 34.471 | 2 | 52.3437 | no | 0.0001718 | 656.8772 | 11.232589 |
| GVTCVSQMPVAE( | 14 | Q9H944 | Mediator of RNA polymerase II transcript | MED20 | 23.222 | 2 | 48.47804 | yes | 1.142E-07 | 731.85504 | 79.297089 |
| SVQQTVELLTR | 11 | Q9H944 | Mediator of RNA polymerase II transcript | MED20 | 23.222 | 2 | 48.47804 | yes | 2.518E-07 | 637.35913 | 44.982143 |
| KQQQVPVAGIR | 11 | Q9H944 | Mediator of RNA polymerase II transcript | MED20 | 23.222 | 2 | 48.47804 | yes | 1.329E-06 | 612.36456 | 38.796165 |
| SLNSENSYVSPR | 12 | Q9H992 | E3 ubiquitin-protein ligase MARCHF7 O | MARCHF7 | 78.051 | 2 | 44.16694 | yes | 3.617E-07 | 676.82568 | 41.237572 |
| SGDFTTSSYVQDR | 13 | Q9H992 | E3 ubiquitin-protein ligase MARCHF7 O | MARCHF7 | 78.051 | 2 | 44.16694 | yes | 4.676E-07 | 731.82587 | 40.052944 |
| GSSLNDTYHSR | 11 | Q9H992 | E3 ubiquitin-protein ligase MARCHF7 O | MARCHF7 | 78.051 | 2 | 44.16694 | yes | 2.295E-07 | 618.78381 | 31.277512 |
| NALLLLPEEQQDP | 14 | Q9H9A5 | CCR4-NOT transcription complex subuni | CNOT10 | 82.31 | 2 | 51.787777 | yes | 9.722E-08 | 804.43561 | 879.1991 |
| QYTEAISVGEK | 11 | Q9H9A5 | CCR4-NOT transcription complex subuni | CNOT10 | 82.31 | 2 | 51.787777 | yes | 9.881E-08 | 612.80896 | 491.78998 |
| FIPAPPSSPLR | 11 | Q9H9A5 | CCR4-NOT transcription complex subuni | CNOT10 | 82.31 | 2 | 51.787777 | yes | 4.171E-07 | 591.33752 | 444.05258 |
| SNQTTTDNLR | 10 | Q9H9A5 | CCR4-NOT transcription complex subuni | CNOT10 | 82.31 | 2 | 51.787777 | yes | 2.134E-07 | 575.27838 | 344.58557 |
| ALHLLAVLEK | 10 | Q9H9A5 | CCR4-NOT transcription complex subuni | CNOT10 | 82.31 | 2 | 51.787777 | yes | 1.349E-07 | 553.85022 | 311.22693 |
| HNLGIFYFK | 9 | Q9H9A5 | CCR4-NOT transcription complex subuni | CNOT10 | 82.31 | 2 | 51.787777 | yes | 2.149E-07 | 569.80585 | 271.16266 |
| LAECCIAANK | 10 | Q9H9A5 | CCR4-NOT transcription complex subuni | CNOT10 | 82.31 | 2 | 51.787777 | yes | 1.352E-07 | 575.27319 | 267.64987 |
| SVMNTAGNSAPSL | 16 | Q9H9A5 | CCR4-NOT transcription complex subuni | CNOT10 | 82.31 | 2 | 51.787777 | yes | 1.052E-07 | 818.92181 | 249.66266 |
| AESGALIEAAK | 11 | Q9H9A5 | CCR4-NOT transcription complex subuni | CNOT10 | 82.31 | 2 | 51.787777 | yes | 1.02E-07 | 530.28766 | 222.16423 |
| LYQFIEPFEEK | 11 | Q9H9A5 | CCR4-NOT transcription complex subuni | CNOT10 | 82.31 | 2 | 51.787777 | yes | 9.978E-08 | 721.86377 | 161.35437 |
| KKPVFQPVHPIQPI | 24 | Q9H9A5 | CCR4-NOT transcription complex subuni | CNOT10 | 82.31 | 4 | 51.787777 | yes | 1.85E-07 | 697.6427 | 121.67329 |
| PMCTLLTNK | 9 | Q9H9A5 | CCR4-NOT transcription complex subuni | CNOT10 | 82.31 | 2 | 51.787777 | yes | 2.465E-07 | 539.27521 | 118.97407 |
| KQELENLK | 8 | Q9H9A5 | CCR4-NOT transcription complex subuni | CNOT10 | 82.31 | 2 | 51.787777 | yes | 9.85E-08 | 501.28494 | 96.592606 |
| TVMLFNLGSAYCI | 14 | Q9H9A5 | CCR4-NOT transcription complex subuni | CNOT10 | 82.31 | 2 | 51.787777 | yes | 1.18E-07 | 822.91547 | 96.004356 |
| LLNSSNIAEHPGFN | 15 | Q9H9A5 | CCR4-NOT transcription complex subuni | CNOT10 | 82.31 | 2 | 51.787777 | yes | 9.722E-08 | 829.42194 | 84.153816 |
| SNFEYLR | 7 | Q9H9A5 | CCR4-NOT transcription complex subuni | CNOT10 | 82.31 | 2 | 51.787777 | yes | 1.996E-07 | 464.7298 | 56.232292 |
| NQLLPAVK | 8 | Q9H9A5 | CCR4-NOT transcription complex subuni | CNOT10 | 82.31 | 2 | 51.787777 | yes | 1.709E-07 | 441.77402 | 18.803364 |
| ELQPLTFNR | 9 | Q9H9A7 | RecQ-mediated genome instability protei | RMI1 | 70.145 | 2 | 55.23072 | yes | 9.722E-08 | 559.30365 | 140.60118 |

| | | | | | | | | | | | |
|---|---|---|---|---|---|---|---|---|---|---|---|
| QSSFEPEFVISPR | 13 | Q9H9A7 | RecQ-mediated genome instability protei | RMI1 | 70.145 | 2 | 55.23072 | yes | 9.722E-08 | 761.88043 | 130.01328 |
| KDPLQYQK | 8 | Q9H9A7 | RecQ-mediated genome instability protei | RMI1 | 70.145 | 2 | 55.23072 | yes | 6.927E-07 | 510.27966 | 86.771835 |
| ILIYGNISFR | 10 | Q9H9A7 | RecQ-mediated genome instability protei | RMI1 | 70.145 | 2 | 55.23072 | yes | 3.191E-07 | 598.34534 | 59.735615 |
| NVSEQMTNEDK | 11 | Q9H9A7 | RecQ-mediated genome instability protei | RMI1 | 70.145 | 2 | 55.23072 | yes | 9.722E-08 | 647.78265 | 48.273521 |
| SIFSVHCNVPLAHI | 18 | Q9H9A7 | RecQ-mediated genome instability protei | RMI1 | 70.145 | 3 | 55.23072 | yes | 2.462E-06 | 696.01068 | 37.505272 |
| DGEVYCIDAR | 10 | Q9H9B1 | Histone-lysine N-methyltransferase EHM | EHMT1 | 141.467 | 2 | 48.901024 | no | 3.426E-07 | 599.26385 | 128.37575 |
| VFMAHQDLR | 9 | Q9H9B1 | Histone-lysine N-methyltransferase EHM | EHMT1 | 141.467 | 2 | 48.901024 | yes | 4.604E-05 | 558.78455 | 107.01019 |
| VNPQDGTNTLTR | 12 | Q9H9B1 | Histone-lysine N-methyltransferase EHM | EHMT1 | 141.467 | 2 | 48.901024 | yes | 1.298E-07 | 658.33368 | 95.747879 |
| ADTTSTVTPVPGQ | 15 | Q9H9B1 | Histone-lysine N-methyltransferase EHM | EHMT1 | 141.467 | 2 | 48.901024 | yes | 1.929E-07 | 765.88593 | 93.119759 |
| SVVGLHAASK | 10 | Q9H9B1 | Histone-lysine N-methyltransferase EHM | EHMT1 | 141.467 | 2 | 48.901024 | yes | 1.052E-06 | 484.77982 | 70.336334 |
| DAEGSTCLHLAAK | 13 | Q9H9B1 | Histone-lysine N-methyltransferase EHM | EHMT1 | 141.467 | 2 | 48.901024 | yes | 1.088E-07 | 686.8299 | 40.248173 |
| FINHHCEPNLVPVI | 14 | Q9H9B1 | Histone-lysine N-methyltransferase EHM | EHMT1 | 141.467 | 3 | 48.901024 | yes | 1.848E-07 | 577.96313 | 23.529844 |
| EITTLANNQCMAT | 22 | Q9H9B1 | Histone-lysine N-methyltransferase EHM | EHMT1 | 141.467 | 3 | 48.901024 | yes | 2.48E-07 | 830.3847 | 23.373556 |
| HVSPLIGR | 8 | Q9H9B4 | Sideroflexin-1 OS=Homo sapiens OX=96 | SFXN1 | 35.62 | 2 | 52.216663 | yes | 4.701E-07 | 439.76398 | 159.41869 |
| QAITQVVVSR | 10 | Q9H9B4 | Sideroflexin-1 OS=Homo sapiens OX=96 | SFXN1 | 35.62 | 2 | 52.216663 | yes | 9.722E-08 | 550.82477 | 104.41054 |
| YIYDSAFHPDTGEI | 14 | Q9H9B4 | Sideroflexin-1 OS=Homo sapiens OX=96 | SFXN1 | 35.62 | 2 | 52.216663 | yes | 9.723E-08 | 821.8728 | 66.950813 |
| NILLTNEQLESAR | 13 | Q9H9B4 | Sideroflexin-1 OS=Homo sapiens OX=96 | SFXN1 | 35.62 | 2 | 52.216663 | yes | 1.631E-07 | 750.90448 | 43.142578 |
| VGIPVTDENGNR | 12 | Q9H9B4 | Sideroflexin-1 OS=Homo sapiens OX=96 | SFXN1 | 35.62 | 2 | 52.216663 | yes | 5.932E-07 | 635.82294 | 42.831123 |
| VLPAPMLQYGGR | 12 | Q9H9G7 | Protein argonaute-3 OS=Homo sapiens O | AGO3 | 97.361 | 2 | 53.593517 | no | 1.394E-07 | 651.35535 | 1513.9623 |
| AVQIHQDTLR | 10 | Q9H9G7 | Protein argonaute-3 OS=Homo sapiens O | AGO3 | 97.361 | 2 | 53.593517 | yes | 1.031E-07 | 590.82526 | 727.60358 |
| VSWHLLHEVLTGI | 13 | Q9H9G7 | Protein argonaute-3 OS=Homo sapiens O | AGO3 | 97.361 | 2 | 53.593517 | yes | 1.066E-07 | 773.9281 | 346.00488 |
| VSWHLLHEVLTGI | 13 | Q9H9G7 | Protein argonaute-3 OS=Homo sapiens O | AGO3 | 97.361 | 2 | 53.593517 | yes | 1.771E-07 | 516.28778 | 346.00488 |
| VSWHLLHEVLTGI | 13 | Q9H9G7 | Protein argonaute-3 OS=Homo sapiens O | AGO3 | 97.361 | 3 | 53.593517 | yes | 1.066E-07 | 773.9281 | 346.00488 |
| VSWHLLHEVLTGI | 13 | Q9H9G7 | Protein argonaute-3 OS=Homo sapiens O | AGO3 | 97.361 | 3 | 53.593517 | yes | 1.771E-07 | 516.28778 | 346.00488 |
| MWAIACFATQR | 11 | Q9H9G7 | Protein argonaute-3 OS=Homo sapiens O | AGO3 | 97.361 | 2 | 53.593517 | yes | 9.722E-08 | 677.82336 | 676.72784 |
| KPSIAAVVGSMDA | 17 | Q9H9G7 | Protein argonaute-3 OS=Homo sapiens O | AGO3 | 97.361 | 3 | 53.593517 | yes | 1.007E-07 | 574.97003 | 635.4364 |
| TVATPSHGVWDM | 13 | Q9H9G7 | Protein argonaute-3 OS=Homo sapiens O | AGO3 | 97.361 | 2 | 53.593517 | yes | 1.014E-07 | 728.8537 | 312.77924 |
| TVATPSHGVWDM | 13 | Q9H9G7 | Protein argonaute-3 OS=Homo sapiens O | AGO3 | 97.361 | 2 | 53.593517 | yes | 1.21E-07 | 486.23825 | 312.77924 |
| TVATPSHGVWDM | 13 | Q9H9G7 | Protein argonaute-3 OS=Homo sapiens O | AGO3 | 97.361 | 3 | 53.593517 | yes | 1.014E-07 | 728.8537 | 312.77924 |
| TVATPSHGVWDM | 13 | Q9H9G7 | Protein argonaute-3 OS=Homo sapiens O | AGO3 | 97.361 | 3 | 53.593517 | yes | 1.21E-07 | 486.23825 | 312.77924 |
| VRDEMAHVTGR | 11 | Q9H9G7 | Protein argonaute-3 OS=Homo sapiens O | AGO3 | 97.361 | 3 | 53.593517 | yes | 2.419E-07 | 424.21555 | 288.80698 |
| VRDEMAHVTGR | 11 | Q9H9G7 | Protein argonaute-3 OS=Homo sapiens O | AGO3 | 97.361 | 3 | 53.593517 | yes | 1.63E-07 | 635.8197 | 288.80698 |
| VRDEMAHVTGR | 11 | Q9H9G7 | Protein argonaute-3 OS=Homo sapiens O | AGO3 | 97.361 | 2 | 53.593517 | yes | 2.419E-07 | 424.21555 | 288.80698 |
| VRDEMAHVTGR | 11 | Q9H9G7 | Protein argonaute-3 OS=Homo sapiens O | AGO3 | 97.361 | 2 | 53.593517 | yes | 1.63E-07 | 635.8197 | 288.80698 |
| VEVTHCGTMR | 10 | Q9H9G7 | Protein argonaute-3 OS=Homo sapiens O | AGO3 | 97.361 | 2 | 53.593517 | yes | 9.819E-08 | 595.27625 | 548.4679 |
| EACISLEK | 8 | Q9H9G7 | Protein argonaute-3 OS=Homo sapiens O | AGO3 | 97.361 | 2 | 53.593517 | yes | 1.464E-07 | 475.2366 | 343.75342 |
| RPVYDGK | 7 | Q9H9G7 | Protein argonaute-3 OS=Homo sapiens O | AGO3 | 97.361 | 2 | 53.593517 | yes | 1.106E-07 | 417.72705 | 330.0929 |
| QFHTGVEIK | 9 | Q9H9G7 | Protein argonaute-3 OS=Homo sapiens O | AGO3 | 97.361 | 2 | 53.593517 | yes | 2.845E-07 | 529.7851 | 292.15051 |
| PSVFQQPVIFLGAI | 24 | Q9H9G7 | Protein argonaute-3 OS=Homo sapiens O | AGO3 | 97.361 | 3 | 53.593517 | no | 1.283E-07 | 826.43231 | 234.01253 |

| | | | | | | | | | | | |
|---|---|---|---|---|---|---|---|---|---|---|---|
| LLANCFQVEIPK | 12 | Q9H9G7 | Protein argonaute-3 OS=Homo sapiens O | AGO3 | 97.361 | 2 | 53.593517 | yes | 9.722E-08 | 716.38684 | 152.77777 |
| SVSIPAPAYYAHL' | 17 | Q9H9G7 | Protein argonaute-3 OS=Homo sapiens O | AGO3 | 97.361 | 3 | 53.593517 | no | 2.506E-06 | 621.33685 | 141.92915 |
| RPGYGTMGK | 9 | Q9H9G7 | Protein argonaute-3 OS=Homo sapiens O | AGO3 | 97.361 | 2 | 53.593517 | yes | 5.413E-07 | 483.74493 | 71.960258 |
| QVLYYELLAIR | 11 | Q9H9G7 | Protein argonaute-3 OS=Homo sapiens O | AGO3 | 97.361 | 2 | 53.593517 | yes | 9.81E-08 | 690.89795 | 71.169388 |
| TLPEPLELDKPISTI | 25 | Q9H9G7 | Protein argonaute-3 OS=Homo sapiens O | AGO3 | 97.361 | 4 | 53.593517 | yes | 6.639E-07 | 688.88654 | 68.057732 |
| EVVDSMVQHFK | 11 | Q9H9G7 | Protein argonaute-3 OS=Homo sapiens O | AGO3 | 97.361 | 2 | 53.593517 | yes | 1.107E-06 | 659.82666 | 23.93606 |
| IEEDTENMAEK | 11 | Q9H9H4 | Vacuolar protein sorting-associated protei | VPS37B | 31.308 | 2 | 41.21864 | yes | 3.618E-07 | 654.78485 | 25.893494 |
| DFLITQMTGK | 10 | Q9H9J2 | Large ribosomal subunit protein mL44 O! | MRPL44 | 37.536 | 2 | 47.79829 | yes | 1.257E-07 | 577.29974 | 909.18866 |
| LYGFTENR | 8 | Q9H9J2 | Large ribosomal subunit protein mL44 O! | MRPL44 | 37.536 | 2 | 47.79829 | yes | 2.115E-07 | 500.24835 | 773.72791 |
| LQENFSLDLLK | 11 | Q9H9J2 | Large ribosomal subunit protein mL44 O! | MRPL44 | 37.536 | 2 | 47.79829 | yes | 1.434E-07 | 660.36389 | 653.21204 |
| CPPPPVR | 7 | Q9H9J2 | Large ribosomal subunit protein mL44 O! | MRPL44 | 37.536 | 2 | 47.79829 | yes | 1.044E-06 | 411.71817 | 309.22983 |
| TAFVNSCYIK | 10 | Q9H9J2 | Large ribosomal subunit protein mL44 O! | MRPL44 | 37.536 | 2 | 47.79829 | yes | 2.459E-07 | 601.79736 | 180.49028 |
| RPWNYSK | 7 | Q9H9J2 | Large ribosomal subunit protein mL44 O! | MRPL44 | 37.536 | 2 | 47.79829 | yes | 4.943E-07 | 475.74579 | 52.26503 |
| KLIAEGPGETVLV/ | 20 | Q9H9J2 | Large ribosomal subunit protein mL44 O! | MRPL44 | 37.536 | 3 | 47.79829 | yes | 1.268E-07 | 694.7074 | 39.38353 |
| IHVLPTNR | 8 | Q9H9L4 | KAT8 regulatory NSL complex subunit 2 | KANSL2 | 55.043 | 2 | 46.877792 | yes | 1.451E-07 | 475.28253 | 114.38291 |
| YGVEALLHR | 9 | Q9H9L4 | KAT8 regulatory NSL complex subunit 2 | KANSL2 | 55.043 | 2 | 46.877792 | yes | 1.809E-07 | 529.29309 | 66.507568 |
| LQSLYIDQFK | 10 | Q9H9L4 | KAT8 regulatory NSL complex subunit 2 | KANSL2 | 55.043 | 2 | 46.877792 | yes | 4.176E-07 | 627.84009 | 20.896309 |
| QVCDAWQR | 8 | Q9H9P5 | Putative E3 ubiquitin-protein ligase UNK | UNKL | 73.828 | 2 | 46.121964 | yes | 2.775E-07 | 531.74292 | 70.051193 |
| SLAGSAPVAIPGSI | 16 | Q9H9P5 | Putative E3 ubiquitin-protein ligase UNK | UNKL | 73.828 | 2 | 46.121964 | yes | 3.417E-07 | 746.92773 | 68.146057 |
| QWEESWQQVK | 10 | Q9H9P5 | Putative E3 ubiquitin-protein ligase UNK | UNKL | 73.828 | 2 | 46.121964 | yes | 1.647E-07 | 674.32001 | 60.167706 |
| WQDANFVLGSYK | 12 | Q9H9P5 | Putative E3 ubiquitin-protein ligase UNK | UNKL | 73.828 | 2 | 46.121964 | yes | 1.156E-06 | 714.35132 | 53.516304 |
| DGTFNYSPDVYCS | 14 | Q9H9P5 | Putative E3 ubiquitin-protein ligase UNK | UNKL | 73.828 | 2 | 46.121964 | yes | 2.72E-06 | 826.84851 | 25.567102 |
| SIDGMQYPIVIK | 12 | Q9H9P8 | L-2-hydroxyglutarate dehydrogenase, mit | L2HGDH | 50.317 | 2 | 47.976063 | yes | 1.221E-07 | 682.36816 | 46.893871 |
| ELATLLHMK | 9 | Q9H9Q4 | Non-homologous end-joining factor 1 OS | NHEJ1 | 33.338 | 2 | 38.462753 | yes | 5.203E-07 | 528.2995 | 89.262764 |
| FELGGGVSIVR | 11 | Q9H9T3 | Elongator complex protein 3 OS=Homo s | ELP3 | 62.259 | 2 | 53.239773 | yes | 1.332E-07 | 567.31934 | 730.39191 |
| IAVISGVGTR | 10 | Q9H9T3 | Elongator complex protein 3 OS=Homo s | ELP3 | 62.259 | 2 | 53.239773 | yes | 1.465E-07 | 486.79547 | 516.13019 |
| LYPTLVIR | 8 | Q9H9T3 | Elongator complex protein 3 OS=Homo s | ELP3 | 62.259 | 2 | 53.239773 | yes | 2.6E-07 | 487.8053 | 491.96643 |
| SYSPSDLVELVAR | 13 | Q9H9T3 | Elongator complex protein 3 OS=Homo s | ELP3 | 62.259 | 2 | 53.239773 | yes | 9.722E-08 | 718.375 | 422.59683 |
| YDPFLQTR | 8 | Q9H9T3 | Elongator complex protein 3 OS=Homo s | ELP3 | 62.259 | 2 | 53.239773 | yes | 1.067E-07 | 520.26398 | 280.48822 |
| AVCESFHLAK | 10 | Q9H9T3 | Elongator complex protein 3 OS=Homo s | ELP3 | 62.259 | 2 | 53.239773 | yes | 1.73E-07 | 581.28967 | 270.07803 |
| LEIGVQSVYEDVA | 14 | Q9H9T3 | Elongator complex protein 3 OS=Homo s | ELP3 | 62.259 | 2 | 53.239773 | yes | 1.002E-07 | 789.41211 | 248.43138 |
| ELHVYGSVVPVSS | 14 | Q9H9T3 | Elongator complex protein 3 OS=Homo s | ELP3 | 62.259 | 3 | 53.239773 | yes | 1.188E-07 | 510.27545 | 114.27234 |
| ELHVYGSVVPVSS | 14 | Q9H9T3 | Elongator complex protein 3 OS=Homo s | ELP3 | 62.259 | 3 | 53.239773 | yes | 4.454E-07 | 764.90955 | 114.27234 |
| ELHVYGSVVPVSS | 14 | Q9H9T3 | Elongator complex protein 3 OS=Homo s | ELP3 | 62.259 | 2 | 53.239773 | yes | 1.188E-07 | 510.27545 | 114.27234 |
| ELHVYGSVVPVSS | 14 | Q9H9T3 | Elongator complex protein 3 OS=Homo s | ELP3 | 62.259 | 2 | 53.239773 | yes | 4.454E-07 | 764.90955 | 114.27234 |
| YGLSAQPR | 8 | Q9H9T3 | Elongator complex protein 3 OS=Homo s | ELP3 | 62.259 | 2 | 53.239773 | yes | 9.881E-08 | 446.23779 | 223.16209 |
| LVDIIAAVPPQYR | 13 | Q9H9T3 | Elongator complex protein 3 OS=Homo s | ELP3 | 62.259 | 2 | 53.239773 | yes | 1.032E-06 | 727.92194 | 109.69003 |
| ILALVPPWTR | 10 | Q9H9T3 | Elongator complex protein 3 OS=Homo s | ELP3 | 62.259 | 2 | 53.239773 | yes | 1.988E-06 | 583.35822 | 105.41104 |
| GTGLYELWK | 9 | Q9H9T3 | Elongator complex protein 3 OS=Homo s | ELP3 | 62.259 | 2 | 53.239773 | yes | 9.722E-08 | 533.78204 | 96.654694 |

| | | | | | | | | | | | |
|---|---|---|---|---|---|---|---|---|---|---|---|
| HLSDMLTYGCTR | 12 | Q9H9T3 | Elongator complex protein 3 OS=Homo s | ELP3 | 62.259 | 2 | 53.239773 | yes | 1.21E-07 | 727.33179 | 91.603783 |
| LQGPYMVK | 8 | Q9H9T3 | Elongator complex protein 3 OS=Homo s | ELP3 | 62.259 | 2 | 53.239773 | yes | 2.221E-07 | 468.25458 | 79.067566 |
| VRPYQVELVR | 10 | Q9H9T3 | Elongator complex protein 3 OS=Homo s | ELP3 | 62.259 | 2 | 53.239773 | yes | 7.093E-07 | 629.86694 | 49.019337 |
| QLIEAHEQGK | 10 | Q9H9T3 | Elongator complex protein 3 OS=Homo s | ELP3 | 62.259 | 2 | 53.239773 | yes | 1.236E-06 | 576.80402 | 27.439392 |
| HLTDPSYGIPR | 11 | Q9H9Y6 | DNA-directed RNA polymerase I subunit | POLR1B | 128.23 | 2 | 54.99176 | yes | 1.36E-07 | 628.32513 | 294.08588 |
| LQTPQSPLVR | 10 | Q9H9Y6 | DNA-directed RNA polymerase I subunit | POLR1B | 128.23 | 2 | 54.99176 | yes | 9.728E-08 | 569.83258 | 257.69357 |
| AALQELTR | 8 | Q9H9Y6 | DNA-directed RNA polymerase I subunit | POLR1B | 128.23 | 2 | 54.99176 | yes | 3.233E-07 | 451.25873 | 186.35568 |
| DLAPGIADSLR | 11 | Q9H9Y6 | DNA-directed RNA polymerase I subunit | POLR1B | 128.23 | 2 | 54.99176 | yes | 5.246E-07 | 564.3064 | 183.00981 |
| IVMEEGCSTQK | 11 | Q9H9Y6 | DNA-directed RNA polymerase I subunit | POLR1B | 128.23 | 2 | 54.99176 | yes | 9.722E-08 | 641.29431 | 175.22113 |
| QVLNYLGECFR | 11 | Q9H9Y6 | DNA-directed RNA polymerase I subunit | POLR1B | 128.23 | 2 | 54.99176 | yes | 9.722E-08 | 699.84534 | 168.80243 |
| TSVSMNTDNLMR | 12 | Q9H9Y6 | DNA-directed RNA polymerase I subunit | POLR1B | 128.23 | 2 | 54.99176 | yes | 9.722E-08 | 684.81592 | 162.44104 |
| EANVYPAECR | 10 | Q9H9Y6 | DNA-directed RNA polymerase I subunit | POLR1B | 128.23 | 2 | 54.99176 | yes | 1.732E-06 | 604.77185 | 102.31435 |
| ENCVVDNIK | 9 | Q9H9Y6 | DNA-directed RNA polymerase I subunit | POLR1B | 128.23 | 2 | 54.99176 | yes | 1.436E-06 | 545.76349 | 95.139183 |
| SEFIDLSEK | 9 | Q9H9Y6 | DNA-directed RNA polymerase I subunit | POLR1B | 128.23 | 2 | 54.99176 | yes | 2.424E-07 | 534.26642 | 92.950317 |
| AAGYNFYGTER | 11 | Q9H9Y6 | DNA-directed RNA polymerase I subunit | POLR1B | 128.23 | 2 | 54.99176 | yes | 2.796E-07 | 624.78583 | 90.960106 |
| EKLEGWLVSIK | 11 | Q9H9Y6 | DNA-directed RNA polymerase I subunit | POLR1B | 128.23 | 2 | 54.99176 | yes | 1.28E-07 | 651.37683 | 88.085312 |
| LFNCSDR | 7 | Q9H9Y6 | DNA-directed RNA polymerase I subunit | POLR1B | 128.23 | 2 | 54.99176 | yes | 1.113E-07 | 456.20563 | 87.161469 |
| QTMGFPLLTYQDF | 13 | Q9H9Y6 | DNA-directed RNA polymerase I subunit | POLR1B | 128.23 | 2 | 54.99176 | yes | 1.022E-06 | 785.39014 | 84.390274 |
| QFLGYVPIMVK | 11 | Q9H9Y6 | DNA-directed RNA polymerase I subunit | POLR1B | 128.23 | 2 | 54.99176 | yes | 4.398E-07 | 647.86505 | 81.281921 |
| GPGYTQYGVSMH | 15 | Q9H9Y6 | DNA-directed RNA polymerase I subunit | POLR1B | 128.23 | 3 | 54.99176 | yes | 3.439E-05 | 571.2608 | 62.640453 |
| GFAHGSVYK | 9 | Q9H9Y6 | DNA-directed RNA polymerase I subunit | POLR1B | 128.23 | 2 | 54.99176 | yes | 2.174E-07 | 483.24561 | 60.807034 |
| LVRPVQNLALGK | 12 | Q9H9Y6 | DNA-directed RNA polymerase I subunit | POLR1B | 128.23 | 2 | 54.99176 | yes | 3.115E-07 | 654.4115 | 47.009972 |
| LDDDGLPFIGAK | 12 | Q9H9Y6 | DNA-directed RNA polymerase I subunit | POLR1B | 128.23 | 2 | 54.99176 | yes | 1.284E-06 | 630.82715 | 33.79298 |
| QGDSSLVFGIK | 11 | Q9H9Y6 | DNA-directed RNA polymerase I subunit | POLR1B | 128.23 | 2 | 54.99176 | yes | 7.27E-07 | 575.80878 | 23.889292 |
| LEGWLVSIK | 9 | Q9H9Y6 | DNA-directed RNA polymerase I subunit | POLR1B | 128.23 | 2 | 54.99176 | yes | 2.256E-05 | 522.80804 | 21.965029 |
| NMYQCQMGK | 9 | Q9H9Y6 | DNA-directed RNA polymerase I subunit | POLR1B | 128.23 | 2 | 54.99176 | yes | 1.324E-07 | 580.23828 | 5.8882418 |
| VILATENDYCK | 11 | Q9HA38 | Zinc finger matrin-type protein 3 OS=Hoı | ZMAT3 | 32.06 | 2 | 44.612133 | yes | 2.131E-07 | 663.32391 | 63.382492 |
| LGAQLADLHLDNI | 13 | Q9HA64 | Ketosamine-3-kinase OS=Homo sapiens ( | FN3KRP | 34.413 | 2 | 53.586414 | yes | 9.722E-08 | 704.38318 | 61.530937 |
| VLDAPGGGSVLVN | 19 | Q9HA64 | Ketosamine-3-kinase OS=Homo sapiens ( | FN3KRP | 34.413 | 3 | 53.586414 | yes | 1.067E-07 | 671.98999 | 31.226641 |
| AAPPQIPDTR | 10 | Q9HAF1 | Chromatin modification-related protein M | MEAF6 | 21.635 | 2 | 52.957554 | yes | 2.809E-07 | 533.28802 | 722.90253 |
| KQELAETLANLER | 13 | Q9HAF1 | Chromatin modification-related protein M | MEAF6 | 21.635 | 3 | 52.957554 | yes | 1.088E-07 | 505.61063 | 79.639938 |
| KQELAETLANLER | 13 | Q9HAF1 | Chromatin modification-related protein M | MEAF6 | 21.635 | 3 | 52.957554 | yes | 2.441E-06 | 757.91229 | 79.639938 |
| KQELAETLANLER | 13 | Q9HAF1 | Chromatin modification-related protein M | MEAF6 | 21.635 | 2 | 52.957554 | yes | 1.088E-07 | 505.61063 | 79.639938 |
| KQELAETLANLER | 13 | Q9HAF1 | Chromatin modification-related protein M | MEAF6 | 21.635 | 2 | 52.957554 | yes | 2.441E-06 | 757.91229 | 79.639938 |
| QELAETLANLER | 12 | Q9HAF1 | Chromatin modification-related protein M | MEAF6 | 21.635 | 2 | 52.957554 | yes | 9.722E-08 | 693.86481 | 129.47964 |
| WVEVDTWESLQK | 12 | Q9HAN9 | Nicotinamide/nicotinic acid mononucleot | NMNAT1 | 31.933 | 2 | 52.99356 | yes | 9.722E-08 | 760.375 | 1338.9504 |
| VIMAELATK | 9 | Q9HAN9 | Nicotinamide/nicotinic acid mononucleot | NMNAT1 | 31.933 | 2 | 52.99356 | yes | 2.689E-07 | 488.28079 | 1226.4479 |
| GLIPAYHR | 8 | Q9HAN9 | Nicotinamide/nicotinic acid mononucleot | NMNAT1 | 31.933 | 2 | 52.99356 | yes | 9.855E-08 | 463.76398 | 644.24652 |
| NAGVILAPLQR | 11 | Q9HAN9 | Nicotinamide/nicotinic acid mononucleot | NMNAT1 | 31.933 | 2 | 52.99356 | yes | 9.722E-08 | 576.34839 | 638.66046 |

| | | | | | | | | | | | |
|---|---|---|---|---|---|---|---|---|---|---|---|
| GIISPVGDAYK | 11 | Q9HAN9 | Nicotinamide/nicotinic acid mononucleot | NMNAT1 | 31.933 | 2 | 52.99356 | yes | 1.69E-07 | 560.30585 | 572.6087 |
| HNLYSSESEDR | 11 | Q9HAN9 | Nicotinamide/nicotinic acid mononucleot | NMNAT1 | 31.933 | 2 | 52.99356 | yes | 9.722E-08 | 668.79181 | 406.98764 |
| YLVPDLVQEYIEK | 13 | Q9HAN9 | Nicotinamide/nicotinic acid mononucleot | NMNAT1 | 31.933 | 2 | 52.99356 | yes | 1.924E-07 | 804.92963 | 187.44319 |
| TEVVLLACGSFNP | 20 | Q9HAN9 | Nicotinamide/nicotinic acid mononucleot | NMNAT1 | 31.933 | 3 | 52.99356 | yes | 9.722E-08 | 758.05743 | 136.60229 |
| WTETQDSSQK | 10 | Q9HAN9 | Nicotinamide/nicotinic acid mononucleot | NMNAT1 | 31.933 | 2 | 52.99356 | yes | 9.722E-08 | 605.27277 | 121.0358 |
| KWTETQDSSQK | 11 | Q9HAN9 | Nicotinamide/nicotinic acid mononucleot | NMNAT1 | 31.933 | 2 | 52.99356 | yes | 1.786E-07 | 669.32025 | 108.84504 |
| FIYESDVLWK | 10 | Q9HAN9 | Nicotinamide/nicotinic acid mononucleot | NMNAT1 | 31.933 | 2 | 52.99356 | yes | 3.281E-06 | 650.33459 | 74.456879 |
| INLVDLAGSER | 11 | Q9HAQ2 | Kinesin-like protein KIF9 OS=Homo sapi | KIF9 | 89.987 | 2 | 49.397343 | yes | 1.036E-07 | 593.82495 | 541.35675 |
| LLPETVTIDSER | 12 | Q9HAS0 | Protein Njmu-R1 OS=Homo sapiens OX= | C17orf75 | 44.622 | 2 | 55.261734 | yes | 9.819E-08 | 686.86957 | 1212.1948 |
| VVLLFQEK | 8 | Q9HAS0 | Protein Njmu-R1 OS=Homo sapiens OX= | C17orf75 | 44.622 | 2 | 55.261734 | yes | 2.488E-07 | 488.2973 | 735.48181 |
| GAVFEGLGNVASV | 16 | Q9HAS0 | Protein Njmu-R1 OS=Homo sapiens OX= | C17orf75 | 44.622 | 2 | 55.261734 | yes | 9.722E-08 | 795.4303 | 526.32715 |
| NCGSGDILLK | 10 | Q9HAS0 | Protein Njmu-R1 OS=Homo sapiens OX= | C17orf75 | 44.622 | 2 | 55.261734 | yes | 3.201E-06 | 538.77386 | 292.34061 |
| AIQDTNNLK | 9 | Q9HAS0 | Protein Njmu-R1 OS=Homo sapiens OX= | C17orf75 | 44.622 | 2 | 55.261734 | yes | 9.722E-08 | 508.77219 | 286.91724 |
| MLPSLQESMDGDI | 14 | Q9HAS0 | Protein Njmu-R1 OS=Homo sapiens OX= | C17orf75 | 44.622 | 2 | 55.261734 | yes | 2.033E-07 | 790.35254 | 119.37716 |
| MLPSLQESMDGDI | 14 | Q9HAS0 | Protein Njmu-R1 OS=Homo sapiens OX= | C17orf75 | 44.622 | 2 | 55.261734 | yes | 1.772E-07 | 811.35785 | 119.37716 |
| GGNPFLFR | 8 | Q9HAS0 | Protein Njmu-R1 OS=Homo sapiens OX= | C17orf75 | 44.622 | 2 | 55.261734 | yes | 6.94E-07 | 454.24286 | 225.41042 |
| VGCYYCLFQNEK | 12 | Q9HAS0 | Protein Njmu-R1 OS=Homo sapiens OX= | C17orf75 | 44.622 | 2 | 55.261734 | yes | 1.729E-07 | 790.84723 | 211.3006 |
| QVLENFK | 7 | Q9HAS0 | Protein Njmu-R1 OS=Homo sapiens OX= | C17orf75 | 44.622 | 2 | 55.261734 | yes | 9.81E-08 | 439.24252 | 182.52516 |
| VEHEEMPEAK | 10 | Q9HAS0 | Protein Njmu-R1 OS=Homo sapiens OX= | C17orf75 | 44.622 | 2 | 55.261734 | yes | 1.532E-07 | 599.77405 | 175.24265 |
| NPSEYVVCFLGGS | 15 | Q9HAS0 | Protein Njmu-R1 OS=Homo sapiens OX= | C17orf75 | 44.622 | 2 | 55.261734 | yes | 9.722E-08 | 843.39563 | 121.59333 |
| QAEMNHYALFK | 11 | Q9HAS0 | Protein Njmu-R1 OS=Homo sapiens OX= | C17orf75 | 44.622 | 2 | 55.261734 | yes | 9.722E-08 | 676.32678 | 64.446358 |
| MTVEEQMER | 9 | Q9HAU0 | Pleckstrin homology domain-containing f | PLEKHA5 | 127.464 | 2 | 56.708927 | yes | 3.941E-07 | 576.75482 | 300.58597 |
| IVNVSLADLR | 10 | Q9HAU0 | Pleckstrin homology domain-containing f | PLEKHA5 | 127.464 | 2 | 56.708927 | yes | 1.091E-06 | 550.32715 | 142.35504 |
| SEPVSEIETSVVK | 13 | Q9HAU0 | Pleckstrin homology domain-containing f | PLEKHA5 | 127.464 | 2 | 56.708927 | yes | 1.066E-07 | 702.36682 | 128.54358 |
| TNSMQQLEQWIK | 12 | Q9HAU0 | Pleckstrin homology domain-containing f | PLEKHA5 | 127.464 | 2 | 56.708927 | yes | 9.722E-08 | 753.37451 | 116.91802 |
| GVISYQTLPR | 10 | Q9HAU0 | Pleckstrin homology domain-containing f | PLEKHA5 | 127.464 | 2 | 56.708927 | yes | 3.834E-07 | 567.31934 | 116.3576 |
| STTWLHPVTGEAV | 18 | Q9HAU0 | Pleckstrin homology domain-containing f | PLEKHA5 | 127.464 | 3 | 56.708927 | yes | 1.465E-07 | 650.00604 | 108.55148 |
| VFFINEEAK | 9 | Q9HAU0 | Pleckstrin homology domain-containing f | PLEKHA5 | 127.464 | 2 | 56.708927 | yes | 4.544E-07 | 548.78729 | 101.58333 |
| AMLDAALVQTEP | 14 | Q9HAU0 | Pleckstrin homology domain-containing f | PLEKHA5 | 127.464 | 2 | 56.708927 | yes | 5.35E-07 | 743.40271 | 77.123047 |
| DDLQNGLLSTCR | 12 | Q9HAU0 | Pleckstrin homology domain-containing f | PLEKHA5 | 127.464 | 2 | 56.708927 | yes | 3.963E-07 | 696.33282 | 63.447453 |
| ETEPQNVDFSK | 11 | Q9HAU0 | Pleckstrin homology domain-containing f | PLEKHA5 | 127.464 | 2 | 56.708927 | yes | 4.228E-05 | 647.30151 | 50.500957 |
| TYYFCTDTGK | 10 | Q9HAU0 | Pleckstrin homology domain-containing f | PLEKHA5 | 127.464 | 2 | 56.708927 | yes | 3.832E-06 | 628.26862 | 46.547703 |
| PINLSSSENK | 10 | Q9HAU0 | Pleckstrin homology domain-containing f | PLEKHA5 | 127.464 | 2 | 56.708927 | yes | 2.69E-07 | 544.78278 | 40.302128 |
| DGQDRPLTK | 9 | Q9HAU0 | Pleckstrin homology domain-containing f | PLEKHA5 | 127.464 | 2 | 56.708927 | yes | 2.43E-06 | 515.26984 | 34.392719 |
| RNPNAPVVR | 9 | Q9HAU0 | Pleckstrin homology domain-containing f | PLEKHA5 | 127.464 | 2 | 56.708927 | yes | 2.719E-06 | 511.79633 | 29.507957 |
| EEIFEESYR | 9 | Q9HAU4 | E3 ubiquitin-protein ligase SMURF2 OS= | SMURF2 | 86.196 | 2 | 51.344727 | no | 1.022E-07 | 601.27222 | 137.07115 |
| VYFVDHNNR | 9 | Q9HAU4 | E3 ubiquitin-protein ligase SMURF2 OS= | SMURF2 | 86.196 | 2 | 51.344727 | yes | 4.118E-07 | 582.28326 | 113.0957 |
| AVEFFDEER | 9 | Q9HAU4 | E3 ubiquitin-protein ligase SMURF2 OS= | SMURF2 | 86.196 | 2 | 51.344727 | yes | 2.02E-07 | 571.26166 | 95.54837 |
| QGAGFLGCVR | 10 | Q9HAU4 | E3 ubiquitin-protein ligase SMURF2 OS= | SMURF2 | 86.196 | 2 | 51.344727 | no | 1.763E-07 | 532.76892 | 92.101784 |

| | | | | | | | | | | | |
|---|---|---|---|---|---|---|---|---|---|---|---|
| GQIVVSLQSR | 10 | Q9HAU4 | E3 ubiquitin-protein ligase SMURF2 OS= | SMURF2 | 86.196 | 2 | 51.344727 | yes | 9.722E-08 | 543.81696 | 69.858376 |
| GIEAQFLALQK | 11 | Q9HAU4 | E3 ubiquitin-protein ligase SMURF2 OS= | SMURF2 | 86.196 | 2 | 51.344727 | no | 1.89E-06 | 609.34808 | 45.355534 |
| TTQWERPTRPASE | 19 | Q9HAU4 | E3 ubiquitin-protein ligase SMURF2 OS= | SMURF2 | 86.196 | 3 | 51.344727 | yes | 0.0002332 | 736.02606 | 30.521049 |
| EGLSLIYEQLK | 11 | Q9HAU5 | Regulator of nonsense transcripts 2 OS=H | UPF2 | 147.811 | 2 | 55.202396 | yes | 1.201E-07 | 646.85846 | 3833.5293 |
| MMLENLQQR | 9 | Q9HAU5 | Regulator of nonsense transcripts 2 OS=H | UPF2 | 147.811 | 2 | 55.202396 | yes | 1.566E-07 | 581.789 | 3309.4185 |
| LCNSLEESIR | 10 | Q9HAU5 | Regulator of nonsense transcripts 2 OS=H | UPF2 | 147.811 | 2 | 55.202396 | yes | 9.722E-08 | 610.8006 | 2367.0564 |
| LTLDINER | 8 | Q9HAU5 | Regulator of nonsense transcripts 2 OS=H | UPF2 | 147.811 | 2 | 55.202396 | yes | 9.722E-08 | 487.26929 | 2056.6084 |
| FLGELYNYR | 9 | Q9HAU5 | Regulator of nonsense transcripts 2 OS=H | UPF2 | 147.811 | 2 | 55.202396 | yes | 1.283E-07 | 587.79822 | 2055.4551 |
| QQPFQNLLK | 9 | Q9HAU5 | Regulator of nonsense transcripts 2 OS=H | UPF2 | 147.811 | 2 | 55.202396 | yes | 2.348E-07 | 558.31403 | 1896.3052 |
| MLLSDFSHHHIEM | 23 | Q9HAU5 | Regulator of nonsense transcripts 2 OS=H | UPF2 | 147.811 | 4 | 55.202396 | yes | 9.793E-08 | 690.31964 | 1762.0579 |
| KLPWQDQEVK | 10 | Q9HAU5 | Regulator of nonsense transcripts 2 OS=H | UPF2 | 147.811 | 2 | 55.202396 | yes | 3.158E-07 | 635.84314 | 1699.9426 |
| LGMEVNQPK | 9 | Q9HAU5 | Regulator of nonsense transcripts 2 OS=H | UPF2 | 147.811 | 2 | 55.202396 | yes | 3.036E-07 | 508.26569 | 1590.0571 |
| MVESAVIFR | 9 | Q9HAU5 | Regulator of nonsense transcripts 2 OS=H | UPF2 | 147.811 | 2 | 55.202396 | yes | 2.25E-07 | 526.28387 | 1409.9667 |
| VHQLDVAIPLHLK | 13 | Q9HAU5 | Regulator of nonsense transcripts 2 OS=H | UPF2 | 147.811 | 3 | 55.202396 | yes | 2.081E-07 | 494.96457 | 679.25964 |
| VHQLDVAIPLHLK | 13 | Q9HAU5 | Regulator of nonsense transcripts 2 OS=H | UPF2 | 147.811 | 3 | 55.202396 | yes | 1.006E-07 | 741.94318 | 679.25964 |
| VHQLDVAIPLHLK | 13 | Q9HAU5 | Regulator of nonsense transcripts 2 OS=H | UPF2 | 147.811 | 2 | 55.202396 | yes | 2.081E-07 | 494.96457 | 679.25964 |
| VHQLDVAIPLHLK | 13 | Q9HAU5 | Regulator of nonsense transcripts 2 OS=H | UPF2 | 147.811 | 2 | 55.202396 | yes | 1.006E-07 | 741.94318 | 679.25964 |
| HCGDDIAGLVPR | 12 | Q9HAU5 | Regulator of nonsense transcripts 2 OS=H | UPF2 | 147.811 | 2 | 55.202396 | yes | 1.173E-07 | 655.31952 | 1125.2028 |
| LLDEQEQEDEEAS | 19 | Q9HAU5 | Regulator of nonsense transcripts 2 OS=H | UPF2 | 147.811 | 3 | 55.202396 | yes | 1.198E-07 | 719.99762 | 1118.3439 |
| QYEEFAMSYQK | 11 | Q9HAU5 | Regulator of nonsense transcripts 2 OS=H | UPF2 | 147.811 | 2 | 55.202396 | yes | 1.192E-07 | 712.31354 | 1087.9773 |
| NDTLHCLK | 8 | Q9HAU5 | Regulator of nonsense transcripts 2 OS=H | UPF2 | 147.811 | 2 | 55.202396 | yes | 1.083E-07 | 500.74765 | 1076.08 |
| LDLLPFYAR | 9 | Q9HAU5 | Regulator of nonsense transcripts 2 OS=H | UPF2 | 147.811 | 2 | 55.202396 | yes | 1.692E-07 | 554.31348 | 1049.5088 |
| DQINIETK | 8 | Q9HAU5 | Regulator of nonsense transcripts 2 OS=H | UPF2 | 147.811 | 2 | 55.202396 | yes | 2.585E-07 | 480.75348 | 822.92688 |
| AAMDFCMNMNTK | 12 | Q9HAU5 | Regulator of nonsense transcripts 2 OS=H | UPF2 | 147.811 | 2 | 55.202396 | yes | 9.722E-08 | 717.28766 | 402.85486 |
| AAMDFCMNMNTK | 12 | Q9HAU5 | Regulator of nonsense transcripts 2 OS=H | UPF2 | 147.811 | 2 | 55.202396 | yes | 2.366E-06 | 725.2851 | 402.85486 |
| QVQDLER | 7 | Q9HAU5 | Regulator of nonsense transcripts 2 OS=H | UPF2 | 147.811 | 2 | 55.202396 | yes | 1.531E-07 | 444.2327 | 714.24182 |
| YIAEAVASIVEAK | 13 | Q9HAU5 | Regulator of nonsense transcripts 2 OS=H | UPF2 | 147.811 | 2 | 55.202396 | yes | 1.911E-07 | 682.37701 | 645.80371 |
| NFYENLIDLK | 10 | Q9HAU5 | Regulator of nonsense transcripts 2 OS=H | UPF2 | 147.811 | 2 | 55.202396 | yes | 1.613E-07 | 634.82971 | 561.73364 |
| KLDCFLVYFQR | 11 | Q9HAU5 | Regulator of nonsense transcripts 2 OS=H | UPF2 | 147.811 | 2 | 55.202396 | yes | 1.048E-07 | 744.88702 | 560.8421 |
| QAMHLDAR | 8 | Q9HAU5 | Regulator of nonsense transcripts 2 OS=H | UPF2 | 147.811 | 2 | 55.202396 | yes | 1.453E-07 | 471.23471 | 487.94644 |
| QQEEEAAAQMK | 11 | Q9HAU5 | Regulator of nonsense transcripts 2 OS=H | UPF2 | 147.811 | 2 | 55.202396 | yes | 3.081E-07 | 631.78772 | 473.47867 |
| TSVLLEQMMR | 10 | Q9HAU5 | Regulator of nonsense transcripts 2 OS=H | UPF2 | 147.811 | 2 | 55.202396 | yes | 7.097E-07 | 604.31232 | 473.11685 |
| EYFTSLTK | 8 | Q9HAU5 | Regulator of nonsense transcripts 2 OS=H | UPF2 | 147.811 | 2 | 55.202396 | yes | 1.483E-07 | 494.75293 | 377.49707 |
| LPWQDQEVK | 9 | Q9HAU5 | Regulator of nonsense transcripts 2 OS=H | UPF2 | 147.811 | 2 | 55.202396 | yes | 9.849E-08 | 571.79565 | 350.78311 |
| LGLVNDKDSK | 10 | Q9HAU5 | Regulator of nonsense transcripts 2 OS=H | UPF2 | 147.811 | 2 | 55.202396 | yes | 1.09E-07 | 544.80096 | 349.4502 |
| RPPLQEYVR | 9 | Q9HAU5 | Regulator of nonsense transcripts 2 OS=H | UPF2 | 147.811 | 2 | 55.202396 | yes | 9.722E-08 | 579.32495 | 303.04691 |
| KLLDEQEQEDEEA | 20 | Q9HAU5 | Regulator of nonsense transcripts 2 OS=H | UPF2 | 147.811 | 3 | 55.202396 | yes | 9.722E-08 | 762.69592 | 290.09479 |
| YADFAPSLLQVWK | 13 | Q9HAU5 | Regulator of nonsense transcripts 2 OS=H | UPF2 | 147.811 | 2 | 55.202396 | yes | 1.514E-06 | 769.40613 | 264.55731 |
| TITEQQR | 7 | Q9HAU5 | Regulator of nonsense transcripts 2 OS=H | UPF2 | 147.811 | 2 | 55.202396 | yes | 9.539E-05 | 438.2327 | 209.78467 |

| | | | | | | | | | | | |
|---|---|---|---|---|---|---|---|---|---|---|---|
| DSLSHDFNGLNLS | 14 | Q9HAU5 | Regulator of nonsense transcripts 2 OS=H | UPF2 | 147.811 | 2 | 55.202396 | yes | 1.387E-07 | 773.87842 | 194.25815 |
| PEENFFSR | 8 | Q9HAU5 | Regulator of nonsense transcripts 2 OS=H | UPF2 | 147.811 | 2 | 55.202396 | yes | 9.722E-08 | 513.23798 | 168.32697 |
| KLTLDINER | 9 | Q9HAU5 | Regulator of nonsense transcripts 2 OS=H | UPF2 | 147.811 | 2 | 55.202396 | yes | 1.584E-07 | 551.31677 | 164.33481 |
| HVPCVEDEDFIQA | 16 | Q9HAU5 | Regulator of nonsense transcripts 2 OS=H | UPF2 | 147.811 | 3 | 55.202396 | yes | 9.722E-08 | 638.96826 | 139.59674 |
| GAPNADLIFK | 10 | Q9HAU5 | Regulator of nonsense transcripts 2 OS=H | UPF2 | 147.811 | 2 | 55.202396 | yes | 2.394E-07 | 523.28748 | 123.70502 |
| KDQINIETK | 9 | Q9HAU5 | Regulator of nonsense transcripts 2 OS=H | UPF2 | 147.811 | 2 | 55.202396 | yes | 2.176E-07 | 544.80096 | 92.773758 |
| LDCFLVYFQR | 10 | Q9HAU5 | Regulator of nonsense transcripts 2 OS=H | UPF2 | 147.811 | 2 | 55.202396 | yes | 2.4E-07 | 680.83954 | 66.836899 |
| AFVPAILFK | 9 | Q9HAU5 | Regulator of nonsense transcripts 2 OS=H | UPF2 | 147.811 | 2 | 55.202396 | yes | 6.247E-07 | 503.31021 | 56.02718 |
| KKDQINIETK | 10 | Q9HAU5 | Regulator of nonsense transcripts 2 OS=H | UPF2 | 147.811 | 2 | 55.202396 | yes | 9.722E-08 | 608.84845 | 54.669754 |
| SLEVWTK | 7 | Q9HAU5 | Regulator of nonsense transcripts 2 OS=H | UPF2 | 147.811 | 2 | 55.202396 | yes | 1.929E-07 | 431.73709 | 48.114002 |
| RQQEEEAAAQMK | 12 | Q9HAU5 | Regulator of nonsense transcripts 2 OS=H | UPF2 | 147.811 | 2 | 55.202396 | yes | 2.987E-07 | 709.83826 | 34.622162 |
| ENETDEENTEVMI | 14 | Q9HAU5 | Regulator of nonsense transcripts 2 OS=H | UPF2 | 147.811 | 2 | 55.202396 | yes | 9.722E-08 | 840.86707 | 23.808384 |
| MHAIPLR | 7 | Q9HAV0 | Guanine nucleotide-binding protein subu | GNB4 | 37.568 | 2 | 50.217983 | yes | 9.459E-07 | 419.24182 | 129.87204 |
| TFVSGACDASSK | 12 | Q9HAV0 | Guanine nucleotide-binding protein subu | GNB4 | 37.568 | 2 | 50.217983 | yes | 9.849E-08 | 615.27698 | 34.720276 |
| WQVINQR | 7 | Q9HAV4 | Exportin-5 OS=Homo sapiens OX=9606 | XPO5 | 136.312 | 2 | 47.747448 | yes | 2.706E-06 | 472.25906 | 48.372086 |
| SCDPGLEDPCGLN | 14 | Q9HAV4 | Exportin-5 OS=Homo sapiens OX=9606 | XPO5 | 136.312 | 2 | 47.747448 | yes | 1.266E-07 | 795.33777 | 28.265224 |
| ETQIITGSDESCR | 13 | Q9HAW4 | Claspin OS=Homo sapiens OX=9606 GN | CLSPN | 151.094 | 2 | 54.84239 | yes | 9.722E-08 | 748.33832 | 120.98853 |
| MGYFPTEEK | 9 | Q9HAW4 | Claspin OS=Homo sapiens OX=9606 GN | CLSPN | 151.094 | 2 | 54.84239 | yes | 2.807E-07 | 551.24969 | 108.28116 |
| TMLDDDKR | 8 | Q9HAW4 | Claspin OS=Homo sapiens OX=9606 GN | CLSPN | 151.094 | 2 | 54.84239 | yes | 4.351E-06 | 497.23712 | 103.38435 |
| LDEDDSCSLLTK | 12 | Q9HAW4 | Claspin OS=Homo sapiens OX=9606 GN | CLSPN | 151.094 | 2 | 54.84239 | yes | 4.038E-07 | 698.31903 | 89.379608 |
| TIHDFFK | 7 | Q9HAW4 | Claspin OS=Homo sapiens OX=9606 GN | CLSPN | 151.094 | 2 | 54.84239 | yes | 7.225E-07 | 454.23724 | 89.266975 |
| SLSSDSTLLLFK | 12 | Q9HAW4 | Claspin OS=Homo sapiens OX=9606 GN | CLSPN | 151.094 | 2 | 54.84239 | yes | 1.365E-07 | 655.86377 | 70.918472 |
| NFVFHTLSPVK | 11 | Q9HAW4 | Claspin OS=Homo sapiens OX=9606 GN | CLSPN | 151.094 | 2 | 54.84239 | yes | 1.622E-07 | 644.85608 | 60.477623 |
| ENMEELLNLCSGK | 13 | Q9HAW4 | Claspin OS=Homo sapiens OX=9606 GN | CLSPN | 151.094 | 2 | 54.84239 | yes | 9.722E-08 | 768.85529 | 45.843719 |
| TVADSDESYMEK | 12 | Q9HAW4 | Claspin OS=Homo sapiens OX=9606 GN | CLSPN | 151.094 | 2 | 54.84239 | yes | 6.215E-07 | 687.7901 | 40.494713 |
| AGQTVNVNVIVK | 12 | Q9HAW4 | Claspin OS=Homo sapiens OX=9606 GN | CLSPN | 151.094 | 2 | 54.84239 | yes | 4.104E-07 | 621.36426 | 16.764494 |
| FQVDPSGEIVELAI | 14 | Q9HB07 | MYG1 exonuclease OS=Homo sapiens O | MYG1 | 42.478 | 2 | 50.696175 | yes | 1.494E-07 | 766.40375 | 410.67584 |
| AMDLVQEEFLQR | 12 | Q9HB07 | MYG1 exonuclease OS=Homo sapiens O | MYG1 | 42.478 | 2 | 50.696175 | yes | 1.369E-07 | 739.86902 | 311.24435 |
| ALVEEALAQR | 10 | Q9HB07 | MYG1 exonuclease OS=Homo sapiens O | MYG1 | 42.478 | 2 | 50.696175 | yes | 1.727E-07 | 550.30896 | 308.17322 |
| LPLPEPWR | 8 | Q9HB07 | MYG1 exonuclease OS=Homo sapiens O | MYG1 | 42.478 | 2 | 50.696175 | yes | 9.728E-08 | 504.28726 | 210.39427 |
| LDFYQHSWLPAR | 12 | Q9HB07 | MYG1 exonuclease OS=Homo sapiens O | MYG1 | 42.478 | 3 | 50.696175 | yes | 3.456E-07 | 511.59302 | 83.249283 |
| LDFYQHSWLPAR | 12 | Q9HB07 | MYG1 exonuclease OS=Homo sapiens O | MYG1 | 42.478 | 3 | 50.696175 | yes | 4.863E-07 | 766.88586 | 83.249283 |
| LDFYQHSWLPAR | 12 | Q9HB07 | MYG1 exonuclease OS=Homo sapiens O | MYG1 | 42.478 | 2 | 50.696175 | yes | 3.456E-07 | 511.59302 | 83.249283 |
| LDFYQHSWLPAR | 12 | Q9HB07 | MYG1 exonuclease OS=Homo sapiens O | MYG1 | 42.478 | 2 | 50.696175 | yes | 4.863E-07 | 766.88586 | 83.249283 |
| YALTTTLSAR | 10 | Q9HB07 | MYG1 exonuclease OS=Homo sapiens O | MYG1 | 42.478 | 2 | 50.696175 | yes | 1.248E-07 | 548.80347 | 163.70004 |
| THECLVK | 7 | Q9HB19 | Pleckstrin homology domain-containing f | PLEKHA2 | 47.256 | 2 | 46.340134 | yes | 1.673E-07 | 443.7262 | 87.399734 |
| SQSYIPTSGCR | 11 | Q9HB19 | Pleckstrin homology domain-containing f | PLEKHA2 | 47.256 | 2 | 46.340134 | yes | 5.023E-06 | 628.29041 | 83.590851 |
| DNLFEIITSSR | 11 | Q9HB19 | Pleckstrin homology domain-containing f | PLEKHA2 | 47.256 | 2 | 46.340134 | yes | 1.585E-07 | 647.83551 | 58.446453 |
| EIGAAVQALK | 10 | Q9HB19 | Pleckstrin homology domain-containing f | PLEKHA2 | 47.256 | 2 | 46.340134 | yes | 2.963E-07 | 500.29529 | 41.50161 |

| | | | | | | | | | | | |
|---|---|---|---|---|---|---|---|---|---|---|---|
| DNLFEIVTTSR | 11 | Q9HB21 | Pleckstrin homology domain-containing f | PLEKHA1 | 45.554 | 2 | 44.1754 | yes | 2.293E-07 | 647.83551 | 28.723467 |
| EYPNLVTIYR | 10 | Q9HB58 | Sp110 nuclear body protein OS=Homo sa | SP110 | 78.396 | 2 | 50.126015 | yes | 1.697E-07 | 634.33771 | 155.37592 |
| VVHNILTQLER | 11 | Q9HB58 | Sp110 nuclear body protein OS=Homo sa | SP110 | 78.396 | 2 | 50.126015 | yes | 1.339E-07 | 661.383 | 119.43609 |
| CEGMTLGELLK | 11 | Q9HB58 | Sp110 nuclear body protein OS=Homo sa | SP110 | 78.396 | 2 | 50.126015 | yes | 9.894E-08 | 625.80957 | 72.76413 |
| SDTVDFHCSK | 10 | Q9HB58 | Sp110 nuclear body protein OS=Homo sa | SP110 | 78.396 | 2 | 50.126015 | yes | 4.634E-07 | 598.25604 | 41.64669 |
| MYMESLEACR | 10 | Q9HB58 | Sp110 nuclear body protein OS=Homo sa | SP110 | 78.396 | 2 | 50.126015 | yes | 1.012E-07 | 645.26978 | 36.25808 |
| SEEIIDGTSEMNEG | 15 | Q9HB58 | Sp110 nuclear body protein OS=Homo sa | SP110 | 78.396 | 2 | 50.126015 | yes | 3.521E-07 | 819.86182 | 35.654007 |
| VTQGAASPGHGIQ | 15 | Q9HB58 | Sp110 nuclear body protein OS=Homo sa | SP110 | 78.396 | 2 | 50.126015 | yes | 1.117E-07 | 740.38116 | 26.298218 |
| TDTVLILCR | 9 | Q9HB71 | Calcyclin-binding protein OS=Homo sapi | CACYBP | 26.21 | 2 | 52.837257 | yes | 1.071E-07 | 545.79987 | 573.90515 |
| WDYLTQVEK | 9 | Q9HB71 | Calcyclin-binding protein OS=Homo sapi | CACYBP | 26.21 | 2 | 52.837257 | yes | 9.989E-08 | 591.29547 | 186.14929 |
| ISNYGWDQSDK | 11 | Q9HB71 | Calcyclin-binding protein OS=Homo sapi | CACYBP | 26.21 | 2 | 52.837257 | yes | 1.018E-07 | 656.79382 | 89.211166 |
| IYEDGDDDMKR | 11 | Q9HB71 | Calcyclin-binding protein OS=Homo sapi | CACYBP | 26.21 | 2 | 52.837257 | yes | 9.722E-08 | 678.79047 | 51.423016 |
| IYEDGDDDMK | 10 | Q9HB71 | Calcyclin-binding protein OS=Homo sapi | CACYBP | 26.21 | 2 | 52.837257 | yes | 5.521E-07 | 600.73987 | 40.915382 |
| AELLDNEKPAAVV | 23 | Q9HB71 | Calcyclin-binding protein OS=Homo sapi | CACYBP | 26.21 | 3 | 52.837257 | yes | 3.563E-06 | 800.77222 | 31.876278 |
| GCQFLGPAMQEEA | 15 | Q9HBD1 | Roquin-2 OS=Homo sapiens OX=9606 G | RC3H2 | 131.669 | 2 | 55.683857 | no | 2.67E-07 | 839.90002 | 1764.0597 |
| FIVTDLSGHR | 10 | Q9HBD1 | Roquin-2 OS=Homo sapiens OX=9606 G | RC3H2 | 131.669 | 2 | 55.683857 | yes | 9.723E-08 | 572.80908 | 1620.699 |
| SVQELTIVLQR | 11 | Q9HBD1 | Roquin-2 OS=Homo sapiens OX=9606 G | RC3H2 | 131.669 | 2 | 55.683857 | yes | 9.722E-08 | 643.37738 | 1188.8048 |
| LVLLALEDGSALSI | 14 | Q9HBD1 | Roquin-2 OS=Homo sapiens OX=9606 G | RC3H2 | 131.669 | 2 | 55.683857 | no | 1.352E-07 | 728.92212 | 945.5918 |
| VLVLFVVQR | 9 | Q9HBD1 | Roquin-2 OS=Homo sapiens OX=9606 G | RC3H2 | 131.669 | 2 | 55.683857 | no | 2.141E-07 | 536.84747 | 701.33435 |
| VHLFETQR | 8 | Q9HBD1 | Roquin-2 OS=Homo sapiens OX=9606 G | RC3H2 | 131.669 | 2 | 55.683857 | yes | 3.249E-07 | 515.27747 | 669.11041 |
| SHMQSIIDK | 9 | Q9HBD1 | Roquin-2 OS=Homo sapiens OX=9606 G | RC3H2 | 131.669 | 2 | 55.683857 | no | 3.63E-07 | 529.76862 | 632.02356 |
| RDEDSSLMQLK | 11 | Q9HBD1 | Roquin-2 OS=Homo sapiens OX=9606 G | RC3H2 | 131.669 | 2 | 55.683857 | no | 2.246E-07 | 661.32446 | 526.97131 |
| SNSLLLQR | 8 | Q9HBD1 | Roquin-2 OS=Homo sapiens OX=9606 G | RC3H2 | 131.669 | 2 | 55.683857 | yes | 7.039E-07 | 465.772 | 510.11728 |
| WNSLDEGR | 8 | Q9HBD1 | Roquin-2 OS=Homo sapiens OX=9606 G | RC3H2 | 131.669 | 2 | 55.683857 | yes | 3.627E-07 | 488.72778 | 318.26312 |
| TSCEEQIR | 8 | Q9HBD1 | Roquin-2 OS=Homo sapiens OX=9606 G | RC3H2 | 131.669 | 2 | 55.683857 | yes | 3.046E-06 | 511.73221 | 256.44629 |
| GVASLNQSALSR | 12 | Q9HBD1 | Roquin-2 OS=Homo sapiens OX=9606 G | RC3H2 | 131.669 | 2 | 55.683857 | yes | 2.495E-07 | 601.828 | 200.19273 |
| DEDSSLMQLK | 10 | Q9HBD1 | Roquin-2 OS=Homo sapiens OX=9606 G | RC3H2 | 131.669 | 2 | 55.683857 | no | 9.722E-08 | 583.27393 | 188.68147 |
| ADFSESVSGTK | 11 | Q9HBD1 | Roquin-2 OS=Homo sapiens OX=9606 G | RC3H2 | 131.669 | 2 | 55.683857 | yes | 9.809E-08 | 564.2644 | 185.33582 |
| WSSYGNEATSSAH | 17 | Q9HBD1 | Roquin-2 OS=Homo sapiens OX=9606 G | RC3H2 | 131.669 | 3 | 55.683857 | yes | 3.751E-07 | 648.62213 | 148.82748 |
| HLTLNLLSK | 9 | Q9HBD1 | Roquin-2 OS=Homo sapiens OX=9606 G | RC3H2 | 131.669 | 2 | 55.683857 | yes | 9.991E-07 | 519.81897 | 79.726822 |
| VGVNNTVTTTAGN | 26 | Q9HBD1 | Roquin-2 OS=Homo sapiens OX=9606 G | RC3H2 | 131.669 | 3 | 55.683857 | yes | 9.722E-08 | 840.77655 | 78.504898 |
| KVLVLFVVQR | 10 | Q9HBD1 | Roquin-2 OS=Homo sapiens OX=9606 G | RC3H2 | 131.669 | 2 | 55.683857 | no | 9.722E-08 | 600.89496 | 71.926323 |
| HSSTGDLLSLELQC | 16 | Q9HBD1 | Roquin-2 OS=Homo sapiens OX=9606 G | RC3H2 | 131.669 | 3 | 55.683857 | yes | 5.19E-07 | 576.30383 | 62.568665 |
| IWRPPMYQR | 9 | Q9HBD1 | Roquin-2 OS=Homo sapiens OX=9606 G | RC3H2 | 131.669 | 3 | 55.683857 | yes | 1.017E-06 | 416.22198 | 14.188071 |
| DVYHLNR | 7 | Q9HBE1 | POZ-, AT hook-, and zinc finger-containi | PATZ1 | 74.06 | 2 | 48.78827 | no | 2.45E-07 | 458.73541 | 1460.0844 |
| EAGILPCGLCGK | 12 | Q9HBE1 | POZ-, AT hook-, and zinc finger-containi | PATZ1 | 74.06 | 2 | 48.78827 | yes | 3.008E-07 | 637.81519 | 116.80234 |
| FCDVLLR | 7 | Q9HBE1 | POZ-, AT hook-, and zinc finger-containi | PATZ1 | 74.06 | 2 | 48.78827 | yes | 1.606E-07 | 461.74438 | 70.721497 |
| VFGDILDFAYTSR | 13 | Q9HBE1 | POZ-, AT hook-, and zinc finger-containi | PATZ1 | 74.06 | 2 | 48.78827 | yes | 1.104E-07 | 752.37756 | 43.017574 |
| ANLLDSMFGSPGC | 15 | Q9HBE1 | POZ-, AT hook-, and zinc finger-containi | PATZ1 | 74.06 | 2 | 48.78827 | yes | 2.279E-07 | 767.88776 | 31.612514 |

| | | | | | | | | | | | |
|---|---|---|---|---|---|---|---|---|---|---|---|
| LGENGLPISEDPDC | 16 | Q9HBE1 | POZ-, AT hook-, and zinc finger-containi | PATZ1 | 74.06 | 2 | 48.78827 | yes | 2.934E-06 | 833.40759 | 25.6394 |
| QSNVQILVPPAR | 12 | Q9HBE1 | POZ-, AT hook-, and zinc finger-containi | PATZ1 | 74.06 | 2 | 48.78827 | yes | 2.089E-06 | 661.383 | 22.495453 |
| GIPMVLIGCK | 10 | Q9HBH0 | Rho-related GTP-binding protein RhoF C | RHOF | 23.626 | 2 | 46.52066 | yes | 1.291E-06 | 544.30377 | 220.40222 |
| AALYLECSAK | 10 | Q9HBH0 | Rho-related GTP-binding protein RhoF C | RHOF | 23.626 | 2 | 46.52066 | yes | 2.27E-07 | 563.28406 | 145.97742 |
| ENVEDVFR | 8 | Q9HBH0 | Rho-related GTP-binding protein RhoF C | RHOF | 23.626 | 2 | 46.52066 | yes | 1.537E-07 | 504.24326 | 123.65639 |
| IVIVGDGGCGK | 11 | Q9HBH0 | Rho-related GTP-binding protein RhoF C | RHOF | 23.626 | 2 | 46.52066 | yes | 4.241E-07 | 537.78424 | 91.121056 |
| AAQLEPITYMQGL | 20 | Q9HBH0 | Rho-related GTP-binding protein RhoF C | RHOF | 23.626 | 3 | 46.52066 | yes | 2.356E-07 | 760.37683 | 49.96278 |
| VFVNPSLR | 8 | Q9HBH1 | Peptide deformylase, mitochondrial OS=I | PDF | 27.014 | 2 | 50.34518 | yes | 2.612E-07 | 466.27164 | 374.54553 |
| AQLGGPELQR | 10 | Q9HBH1 | Peptide deformylase, mitochondrial OS=I | PDF | 27.014 | 2 | 50.34518 | yes | 1.282E-07 | 534.79346 | 345.66632 |
| CVGLSAPQLGVPR | 13 | Q9HBH1 | Peptide deformylase, mitochondrial OS=I | PDF | 27.014 | 2 | 50.34518 | yes | 3.42E-07 | 677.36902 | 191.21036 |
| QMEPFPLR | 8 | Q9HBH1 | Peptide deformylase, mitochondrial OS=I | PDF | 27.014 | 2 | 50.34518 | yes | 9.818E-08 | 509.26294 | 144.93317 |
| TFTNVYWMK | 9 | Q9HBH1 | Peptide deformylase, mitochondrial OS=I | PDF | 27.014 | 2 | 50.34518 | yes | 2.941E-07 | 595.28918 | 20.52953 |
| VATTPGSPSLGR | 12 | Q9HBL0 | Tensin-1 OS=Homo sapiens OX=9606 G | TNS1 | 197.023 | 2 | 47.005505 | yes | 1.422E-07 | 571.81183 | 173.85828 |
| VKPCGICR | 8 | Q9HBL0 | Tensin-1 OS=Homo sapiens OX=9606 G | TNS1 | 197.023 | 2 | 47.005505 | yes | 3.53E-07 | 495.25461 | 131.67261 |
| AHLESLVASR | 10 | Q9HBL0 | Tensin-1 OS=Homo sapiens OX=9606 G | TNS1 | 197.023 | 2 | 47.005505 | yes | 4.867E-07 | 541.80127 | 76.877876 |
| VCSFSCHR | 8 | Q9HBL0 | Tensin-1 OS=Homo sapiens OX=9606 G | TNS1 | 197.023 | 2 | 47.005505 | yes | 2.038E-07 | 526.72363 | 70.977814 |
| TPLSALGLKPHNPA | 25 | Q9HBL0 | Tensin-1 OS=Homo sapiens OX=9606 G | TNS1 | 197.023 | 4 | 47.005505 | yes | 8.933E-06 | 661.86969 | 66.976898 |
| VSAQGITLTDNQR | 13 | Q9HBL0 | Tensin-1 OS=Homo sapiens OX=9606 G | TNS1 | 197.023 | 2 | 47.005505 | no | 1.498E-07 | 701.86786 | 58.146461 |
| LLSGFGLER | 9 | Q9HBL0 | Tensin-1 OS=Homo sapiens OX=9606 G | TNS1 | 197.023 | 2 | 47.005505 | yes | 2.203E-07 | 496.2822 | 49.615505 |
| VEFVFSYGPEK | 11 | Q9HBL0 | Tensin-1 OS=Homo sapiens OX=9606 G | TNS1 | 197.023 | 2 | 47.005505 | yes | 7.755E-07 | 651.32428 | 42.37077 |
| AGPAHAGHTAPMI | 27 | Q9HBL0 | Tensin-1 OS=Homo sapiens OX=9606 G | TNS1 | 197.023 | 4 | 47.005505 | yes | 2.758E-06 | 696.08728 | 40.458176 |
| QVITQEGCTCK | 11 | Q9HBL0 | Tensin-1 OS=Homo sapiens OX=9606 G | TNS1 | 197.023 | 2 | 47.005505 | yes | 3.2E-07 | 662.30524 | 24.082588 |
| SQLTTIVLAAR | 11 | Q9HC07 | Putative divalent cation/proton antiporter | TMEM165 | 34.906 | 2 | 39.264896 | yes | 4.688E-07 | 586.85352 | 79.222252 |
| LQVGGFEHAR | 10 | Q9HC21 | Mitochondrial thiamine pyrophosphate ca | SLC25A19 | 35.512 | 2 | 53.837776 | yes | 1.136E-07 | 557.29364 | 388.36301 |
| YHGILQASR | 9 | Q9HC21 | Mitochondrial thiamine pyrophosphate ca | SLC25A19 | 35.512 | 2 | 53.837776 | yes | 6.689E-07 | 522.7829 | 253.72034 |
| HAVGTMYR | 8 | Q9HC21 | Mitochondrial thiamine pyrophosphate ca | SLC25A19 | 35.512 | 2 | 53.837776 | yes | 3.218E-07 | 467.73181 | 192.24997 |
| ALISPFDVIK | 10 | Q9HC21 | Mitochondrial thiamine pyrophosphate ca | SLC25A19 | 35.512 | 2 | 53.837776 | yes | 1.216E-07 | 551.82898 | 173.09818 |
| FQLQHER | 7 | Q9HC21 | Mitochondrial thiamine pyrophosphate ca | SLC25A19 | 35.512 | 2 | 53.837776 | yes | 4.355E-07 | 479.24869 | 144.08038 |
| SEGPQVFYK | 9 | Q9HC21 | Mitochondrial thiamine pyrophosphate ca | SLC25A19 | 35.512 | 2 | 53.837776 | yes | 3.774E-07 | 527.76385 | 123.4592 |
| TLTYPLDLFK | 10 | Q9HC21 | Mitochondrial thiamine pyrophosphate ca | SLC25A19 | 35.512 | 2 | 53.837776 | yes | 3.085E-06 | 605.83954 | 87.765411 |
| QILQEEGPTAFWK | 13 | Q9HC21 | Mitochondrial thiamine pyrophosphate ca | SLC25A19 | 35.512 | 2 | 53.837776 | yes | 9.722E-08 | 773.89862 | 65.472816 |
| VGYDPKPDGR | 10 | Q9HC21 | Mitochondrial thiamine pyrophosphate ca | SLC25A19 | 35.512 | 2 | 53.837776 | yes | 3.113E-07 | 552.27765 | 64.643982 |
| DLLLTCAQDR | 10 | Q9HC35 | Echinoderm microtubule-associated prote | EML4 | 108.917 | 2 | 48.811165 | yes | 1.102E-07 | 602.80316 | 71.20723 |
| DVIINQEGEYIK | 12 | Q9HC35 | Echinoderm microtubule-associated prote | EML4 | 108.917 | 2 | 48.811165 | yes | 2.127E-07 | 710.86957 | 47.359077 |
| AALADVLR | 8 | Q9HC35 | Echinoderm microtubule-associated prote | EML4 | 108.917 | 2 | 48.811165 | yes | 1.154E-07 | 414.75052 | 46.340027 |
| VIAVADDFCK | 10 | Q9HC35 | Echinoderm microtubule-associated prote | EML4 | 108.917 | 2 | 48.811165 | yes | 2.372E-06 | 569.28406 | 43.630375 |
| LVDEPGHCADFHF | 26 | Q9HC35 | Echinoderm microtubule-associated prote | EML4 | 108.917 | 4 | 48.811165 | yes | 6.825E-07 | 679.82788 | 36.121693 |
| DPGNLGTILR | 10 | Q9HC36 | rRNA methyltransferase 3, mitochondrial | MRM3 | 47.021 | 2 | 55.076202 | yes | 1.964E-07 | 528.29584 | 1251.2063 |
| APSTWEESGLR | 11 | Q9HC36 | rRNA methyltransferase 3, mitochondrial | MRM3 | 47.021 | 2 | 55.076202 | yes | 1.001E-07 | 616.79895 | 832.573 |

| | | | | | | | | | | | |
|---|---|---|---|---|---|---|---|---|---|---|---|
| VVFPSGEVVEQK | 12 | Q9HC36 | rRNA methyltransferase 3, mitochondrial | MRM3 | 47.021 | 2 | 55.076202 | yes | 1.086E-07 | 659.35608 | 761.6748 |
| AGMGAHFR | 8 | Q9HC36 | rRNA methyltransferase 3, mitochondrial | MRM3 | 47.021 | 2 | 55.076202 | yes | 2.098E-07 | 423.7056 | 406.7377 |
| TQLQHSLPLLLICD | 17 | Q9HC36 | rRNA methyltransferase 3, mitochondrial | MRM3 | 47.021 | 3 | 55.076202 | yes | 9.796E-08 | 678.71033 | 290.6142 |
| GCVDAWEPK | 9 | Q9HC36 | rRNA methyltransferase 3, mitochondrial | MRM3 | 47.021 | 2 | 55.076202 | yes | 1.034E-07 | 531.23969 | 253.09076 |
| EKQPLEESASR | 11 | Q9HC36 | rRNA methyltransferase 3, mitochondrial | MRM3 | 47.021 | 2 | 55.076202 | yes | 4.207E-07 | 637.32275 | 235.92496 |
| QPLEESASR | 9 | Q9HC36 | rRNA methyltransferase 3, mitochondrial | MRM3 | 47.021 | 2 | 55.076202 | yes | 1.737E-07 | 508.754 | 181.21581 |
| VKFEDIK | 7 | Q9HC36 | rRNA methyltransferase 3, mitochondrial | MRM3 | 47.021 | 2 | 55.076202 | yes | 9.722E-08 | 439.75275 | 169.76772 |
| ASDHGWVCDQR | 11 | Q9HC36 | rRNA methyltransferase 3, mitochondrial | MRM3 | 47.021 | 2 | 55.076202 | yes | 9.722E-08 | 665.78326 | 157.68164 |
| LSSVMTIVK | 9 | Q9HC36 | rRNA methyltransferase 3, mitochondrial | MRM3 | 47.021 | 2 | 55.076202 | yes | 1.564E-07 | 489.28864 | 103.06587 |
| VFAAEALLK | 9 | Q9HC52 | Chromobox protein homolog 8 OS=Homo | CBX8 | 43.396 | 2 | 52.27784 | yes | 1.298E-07 | 481.28949 | 1124.9955 |
| LLAAFEER | 8 | Q9HC52 | Chromobox protein homolog 8 OS=Homo | CBX8 | 43.396 | 2 | 52.27784 | yes | 1.079E-07 | 474.76108 | 805.66974 |
| KLDDTPSGAGK | 11 | Q9HC52 | Chromobox protein homolog 8 OS=Homo | CBX8 | 43.396 | 2 | 52.27784 | yes | 9.722E-08 | 544.78278 | 548.82941 |
| ESNTDQGFFK | 10 | Q9HC52 | Chromobox protein homolog 8 OS=Homo | CBX8 | 43.396 | 2 | 52.27784 | yes | 1.067E-07 | 586.76459 | 273.5853 |
| LAVDTFPAR | 9 | Q9HC52 | Chromobox protein homolog 8 OS=Homo | CBX8 | 43.396 | 2 | 52.27784 | yes | 1.264E-07 | 495.27435 | 243.92747 |
| EMELYGPK | 8 | Q9HC52 | Chromobox protein homolog 8 OS=Homo | CBX8 | 43.396 | 2 | 52.27784 | yes | 9.809E-08 | 483.73367 | 207.36322 |
| SPQDLASTSR | 10 | Q9HC52 | Chromobox protein homolog 8 OS=Homo | CBX8 | 43.396 | 2 | 52.27784 | yes | 1.15E-07 | 531.26471 | 159.29292 |
| VDDKPSSPGDSSK | 13 | Q9HC52 | Chromobox protein homolog 8 OS=Homo | CBX8 | 43.396 | 2 | 52.27784 | yes | 1.55E-07 | 659.80969 | 140.0903 |
| FPAGHSVIQLAR | 12 | Q9HC52 | Chromobox protein homolog 8 OS=Homo | CBX8 | 43.396 | 2 | 52.27784 | yes | 2.068E-07 | 648.36456 | 94.369713 |
| GYQLEPDLSEEVS | 15 | Q9HC62 | Sentrin-specific protease 2 OS=Homo sap | SENP2 | 67.855 | 2 | 41.087856 | yes | 3.683E-07 | 846.90741 | 57.147911 |
| TDDLLELTEDMEK | 13 | Q9HC62 | Sentrin-specific protease 2 OS=Homo sap | SENP2 | 67.855 | 2 | 41.087856 | yes | 3.75E-07 | 776.35834 | 39.49929 |
| SQIQMLVR | 8 | Q9HC77 | Centromere protein J OS=Homo sapiens ( | CENPJ | 153.001 | 2 | 54.576187 | yes | 2.042E-07 | 487.77621 | 93.935783 |
| LNISQDQPPGDNA | 14 | Q9HC77 | Centromere protein J OS=Homo sapiens ( | CENPJ | 153.001 | 2 | 54.576187 | yes | 2.415E-07 | 762.87372 | 84.316109 |
| SGSVTLSQKPK | 11 | Q9HC77 | Centromere protein J OS=Homo sapiens ( | CENPJ | 153.001 | 2 | 54.576187 | yes | 9.722E-08 | 566.32202 | 79.102531 |
| AAVDISTSFPIK | 12 | Q9HC77 | Centromere protein J OS=Homo sapiens ( | CENPJ | 153.001 | 2 | 54.576187 | yes | 9.722E-08 | 624.84534 | 65.855438 |
| FQNSQISSGTQVEK | 14 | Q9HC77 | Centromere protein J OS=Homo sapiens ( | CENPJ | 153.001 | 2 | 54.576187 | yes | 2.342E-07 | 776.88373 | 36.43018 |
| VTEQIQEANDK | 11 | Q9HC77 | Centromere protein J OS=Homo sapiens ( | CENPJ | 153.001 | 2 | 54.576187 | yes | 2.04E-06 | 637.81476 | 29.680239 |
| AGVILNR | 7 | Q9HC77 | Centromere protein J OS=Homo sapiens ( | CENPJ | 153.001 | 1 | 54.576187 | yes | 5.151E-05 | 742.45697 | 0.7702624 |
| MNLFSLCQK | 9 | Q9HC98 | Serine/threonine-protein kinase Nek6 OS= | NEK6 | 35.715 | 2 | 45.822727 | yes | 1.732E-07 | 570.78064 | 80.82695 |
| HPNTLSFR | 8 | Q9HC98 | Serine/threonine-protein kinase Nek6 OS= | NEK6 | 35.715 | 2 | 45.822727 | yes | 2.356E-06 | 486.2565 | 52.160423 |
| VQIFEMMDAK | 10 | Q9HC98 | Serine/threonine-protein kinase Nek6 OS= | NEK6 | 35.715 | 2 | 45.822727 | yes | 3.134E-07 | 606.29358 | 47.14373 |
| ALVNQLHER | 9 | Q9HCC0 | Methylcrotonoyl-CoA carboxylase beta cl | MCCC2 | 61.334 | 2 | 46.075012 | yes | 4.309E-07 | 540.30145 | 77.405861 |
| AFYGDTLVTGFAR | 13 | Q9HCC0 | Methylcrotonoyl-CoA carboxylase beta cl | MCCC2 | 61.334 | 2 | 46.075012 | yes | 1.66E-07 | 709.35913 | 59.205189 |
| GGAYYPVTVK | 10 | Q9HCC0 | Methylcrotonoyl-CoA carboxylase beta cl | MCCC2 | 61.334 | 2 | 46.075012 | yes | 2.56E-07 | 527.78204 | 39.43111 |
| QGTIFLAGPPLVK | 13 | Q9HCC0 | Methylcrotonoyl-CoA carboxylase beta cl | MCCC2 | 61.334 | 2 | 46.075012 | yes | 2.687E-06 | 670.90045 | 27.634689 |
| AATGEEVSAEDLG | 20 | Q9HCC0 | Methylcrotonoyl-CoA carboxylase beta cl | MCCC2 | 61.334 | 3 | 46.075012 | yes | 1.4E-06 | 686.64459 | 21.730776 |
| MVAAVACAQVPK | 12 | Q9HCC0 | Methylcrotonoyl-CoA carboxylase beta cl | MCCC2 | 61.334 | 2 | 46.075012 | yes | 3.417E-06 | 622.82813 | 10.634861 |
| MADEAILQER | 10 | Q9HCD5 | Nuclear receptor coactivator 5 OS=Homo | NCOA5 | 65.537 | 2 | 51.90752 | yes | 9.935E-08 | 588.28992 | 591.29205 |
| YLTAEETDK | 9 | Q9HCD5 | Nuclear receptor coactivator 5 OS=Homo | NCOA5 | 65.537 | 2 | 51.90752 | yes | 3.368E-07 | 535.25604 | 368.11975 |
| QYFEEIQR | 8 | Q9HCD5 | Nuclear receptor coactivator 5 OS=Homo | NCOA5 | 65.537 | 2 | 51.90752 | yes | 3.408E-07 | 556.77222 | 320.66675 |

| | | | | | | | | | | | |
|---|---|---|---|---|---|---|---|---|---|---|---|
| GPPGPESQSR | 10 | Q9HCD5 | Nuclear receptor coactivator 5 OS=Homo | NCOA5 | 65.537 | 2 | 51.90752 | yes | 2.718E-07 | 506.24634 | 287.02649 |
| SSTDSLPGPISR | 12 | Q9HCD5 | Nuclear receptor coactivator 5 OS=Homo | NCOA5 | 65.537 | 2 | 51.90752 | yes | 1.71E-07 | 608.81201 | 279.79395 |
| NMPQADAMVLVA | 13 | Q9HCD5 | Nuclear receptor coactivator 5 OS=Homo | NCOA5 | 65.537 | 2 | 51.90752 | yes | 1.707E-07 | 708.36035 | 126.18681 |
| PGAPSQGLFGQPS: | 15 | Q9HCD5 | Nuclear receptor coactivator 5 OS=Homo | NCOA5 | 65.537 | 2 | 51.90752 | yes | 9.722E-08 | 743.37585 | 98.25251 |
| LAPASNMTSQR | 11 | Q9HCD5 | Nuclear receptor coactivator 5 OS=Homo | NCOA5 | 65.537 | 2 | 51.90752 | yes | 4.799E-07 | 588.29553 | 91.495094 |
| QPLGATSGASLK | 12 | Q9HCD5 | Nuclear receptor coactivator 5 OS=Homo | NCOA5 | 65.537 | 2 | 51.90752 | yes | 3.101E-06 | 565.31421 | 86.751884 |
| SQASGNQPPSILGC | 24 | Q9HCD5 | Nuclear receptor coactivator 5 OS=Homo | NCOA5 | 65.537 | 3 | 51.90752 | yes | 1.75E-07 | 780.37677 | 68.644279 |
| PVSSTGINFDNPSV | 16 | Q9HCD5 | Nuclear receptor coactivator 5 OS=Homo | NCOA5 | 65.537 | 2 | 51.90752 | yes | 1.037E-07 | 845.42578 | 31.309517 |
| MFVHPSFR | 8 | Q9HCD6 | Protein TANC2 OS=Homo sapiens OX=9 | TANC2 | 219.65 | 2 | 53.936672 | yes | 4.002E-07 | 510.75784 | 221.90054 |
| LVALSCHGTR | 10 | Q9HCD6 | Protein TANC2 OS=Homo sapiens OX=9 | TANC2 | 219.65 | 2 | 53.936672 | yes | 2.918E-06 | 557.29529 | 130.24385 |
| VSLLLNLSR | 9 | Q9HCD6 | Protein TANC2 OS=Homo sapiens OX=9 | TANC2 | 219.65 | 2 | 53.936672 | yes | 1.043E-07 | 507.81894 | 110.26114 |
| LLLEQGAAVAQPN | 14 | Q9HCD6 | Protein TANC2 OS=Homo sapiens OX=9 | TANC2 | 219.65 | 2 | 53.936672 | yes | 2.64E-07 | 740.41754 | 84.532356 |
| TLPVAQAYQDNLY | 14 | Q9HCD6 | Protein TANC2 OS=Homo sapiens OX=9 | TANC2 | 219.65 | 2 | 53.936672 | yes | 1.46E-06 | 826.42554 | 81.493362 |
| VSQASSYPDVK | 11 | Q9HCD6 | Protein TANC2 OS=Homo sapiens OX=9 | TANC2 | 219.65 | 2 | 53.936672 | yes | 4.907E-07 | 590.79584 | 75.449829 |
| SSMDSCLYR | 9 | Q9HCD6 | Protein TANC2 OS=Homo sapiens OX=9 | TANC2 | 219.65 | 2 | 53.936672 | yes | 2.087E-06 | 559.73389 | 65.968208 |
| TLSQGSYLYLK | 11 | Q9HCD6 | Protein TANC2 OS=Homo sapiens OX=9 | TANC2 | 219.65 | 2 | 53.936672 | yes | 9.722E-08 | 636.84534 | 60.401455 |
| EGLTALSWACLK | 12 | Q9HCD6 | Protein TANC2 OS=Homo sapiens OX=9 | TANC2 | 219.65 | 2 | 53.936672 | yes | 1.099E-06 | 674.8501 | 54.373283 |
| LMEEGDMFYK | 10 | Q9HCD6 | Protein TANC2 OS=Homo sapiens OX=9 | TANC2 | 219.65 | 2 | 53.936672 | yes | 1.164E-07 | 631.77521 | 51.920509 |
| SSSQLGSPDVSHLI | 15 | Q9HCD6 | Protein TANC2 OS=Homo sapiens OX=9 | TANC2 | 219.65 | 2 | 53.936672 | yes | 9.722E-08 | 791.91284 | 49.905613 |
| LVYQGSIGGIVGD( | 15 | Q9HCD6 | Protein TANC2 OS=Homo sapiens OX=9 | TANC2 | 219.65 | 2 | 53.936672 | yes | 8.766E-07 | 745.90173 | 49.696327 |
| LGFLLGEK | 8 | Q9HCD6 | Protein TANC2 OS=Homo sapiens OX=9 | TANC2 | 219.65 | 2 | 53.936672 | yes | 1.381E-07 | 438.76309 | 48.784615 |
| MNDFGMAEEFATI | 13 | Q9HCD6 | Protein TANC2 OS=Homo sapiens OX=9 | TANC2 | 219.65 | 2 | 53.936672 | yes | 9.722E-08 | 745.81818 | 39.447533 |
| PNSVAATSSAHLE | 22 | Q9HCD6 | Protein TANC2 OS=Homo sapiens OX=9 | TANC2 | 219.65 | 3 | 53.936672 | yes | 4.878E-07 | 796.38733 | 33.986721 |
| LTFDLIEK | 8 | Q9HCD6 | Protein TANC2 OS=Homo sapiens OX=9 | TANC2 | 219.65 | 2 | 53.936672 | yes | 2.217E-06 | 489.77896 | 33.871132 |
| VDENMTASTYSLN | 14 | Q9HCD6 | Protein TANC2 OS=Homo sapiens OX=9 | TANC2 | 219.65 | 2 | 53.936672 | yes | 2.835E-06 | 786.86414 | 26.120926 |
| NTSVVVTLLK | 10 | Q9HCD6 | Protein TANC2 OS=Homo sapiens OX=9 | TANC2 | 219.65 | 2 | 53.936672 | yes | 4.041E-07 | 537.33191 | 25.562759 |
| QFAAALEDLNEAI | 14 | Q9HCD6 | Protein TANC2 OS=Homo sapiens OX=9 | TANC2 | 219.65 | 2 | 53.936672 | yes | 3.015E-06 | 766.90137 | 24.875395 |
| VHLPSSIYR | 9 | Q9HCE1 | Helicase MOV-10 OS=Homo sapiens OX | MOV10 | 113.672 | 2 | 50.557953 | yes | 1.196E-07 | 536.3009 | 12543.589 |
| EGEGGLSLQVEPE | 15 | Q9HCE1 | Helicase MOV-10 OS=Homo sapiens OX | MOV10 | 113.672 | 2 | 50.557953 | yes | 1.739E-07 | 843.4101 | 9384.0176 |
| SVGVISPYR | 9 | Q9HCE1 | Helicase MOV-10 OS=Homo sapiens OX | MOV10 | 113.672 | 2 | 50.557953 | yes | 5.723E-07 | 489.27435 | 9014.2656 |
| LLTLEVPGVTESR | 13 | Q9HCE1 | Helicase MOV-10 OS=Homo sapiens OX | MOV10 | 113.672 | 2 | 50.557953 | yes | 2.789E-07 | 707.401 | 8410.9932 |
| ITGNPVVTNR | 10 | Q9HCE1 | Helicase MOV-10 OS=Homo sapiens OX | MOV10 | 113.672 | 2 | 50.557953 | yes | 1.721E-07 | 535.80127 | 7512.2964 |
| VGSVEEFQGQER | 12 | Q9HCE1 | Helicase MOV-10 OS=Homo sapiens OX | MOV10 | 113.672 | 2 | 50.557953 | yes | 1.584E-07 | 682.82568 | 7254.7871 |
| ETGDPGGQLVLAC | 16 | Q9HCE1 | Helicase MOV-10 OS=Homo sapiens OX | MOV10 | 113.672 | 2 | 50.557953 | yes | 1.503E-07 | 791.39697 | 6927.7974 |
| LLLHLEELQMEHD | 15 | Q9HCE1 | Helicase MOV-10 OS=Homo sapiens OX | MOV10 | 113.672 | 3 | 50.557953 | yes | 1.375E-07 | 630.33649 | 3122.8401 |
| LLLHLEELQMEHD | 15 | Q9HCE1 | Helicase MOV-10 OS=Homo sapiens OX | MOV10 | 113.672 | 3 | 50.557953 | yes | 2.679E-07 | 473.00418 | 3122.8401 |
| LLLHLEELQMEHD | 15 | Q9HCE1 | Helicase MOV-10 OS=Homo sapiens OX | MOV10 | 113.672 | 4 | 50.557953 | yes | 1.375E-07 | 630.33649 | 3122.8401 |
| LLLHLEELQMEHD | 15 | Q9HCE1 | Helicase MOV-10 OS=Homo sapiens OX | MOV10 | 113.672 | 4 | 50.557953 | yes | 2.679E-07 | 473.00418 | 3122.8401 |
| SVILISTVR | 9 | Q9HCE1 | Helicase MOV-10 OS=Homo sapiens OX | MOV10 | 113.672 | 2 | 50.557953 | yes | 1.319E-07 | 494.31351 | 6222.8794 |

| | | | | | | | | | | | |
|---|---|---|---|---|---|---|---|---|---|---|---|
| VLITTLITAGR | 11 | Q9HCE1 | Helicase MOV-10 OS=Homo sapiens OX | MOV10 | 113.672 | 2 | 50.557953 | yes | 1.532E-07 | 579.36627 | 6030.0474 |
| EAGQCFESFLVVR | 13 | Q9HCE1 | Helicase MOV-10 OS=Homo sapiens OX | MOV10 | 113.672 | 2 | 50.557953 | yes | 2.671E-07 | 771.37451 | 5257.9038 |
| LLTYNSLYK | 9 | Q9HCE1 | Helicase MOV-10 OS=Homo sapiens OX | MOV10 | 113.672 | 2 | 50.557953 | yes | 1.166E-07 | 557.81079 | 4370.9375 |
| GLDMETDR | 8 | Q9HCE1 | Helicase MOV-10 OS=Homo sapiens OX | MOV10 | 113.672 | 2 | 50.557953 | yes | 1.909E-07 | 468.70819 | 4361.0938 |
| PAPYIIFGPPGTGK | 14 | Q9HCE1 | Helicase MOV-10 OS=Homo sapiens OX | MOV10 | 113.672 | 2 | 50.557953 | yes | 9.144E-07 | 707.89008 | 3986.6377 |
| LSFSMSLLSR | 10 | Q9HCE1 | Helicase MOV-10 OS=Homo sapiens OX | MOV10 | 113.672 | 2 | 50.557953 | yes | 1.286E-07 | 570.80774 | 1990.4994 |
| LSFSMSLLSR | 10 | Q9HCE1 | Helicase MOV-10 OS=Homo sapiens OX | MOV10 | 113.672 | 2 | 50.557953 | yes | 1.284E-07 | 578.80518 | 1990.4994 |
| IANLAYVTK | 9 | Q9HCE1 | Helicase MOV-10 OS=Homo sapiens OX | MOV10 | 113.672 | 2 | 50.557953 | yes | 1.054E-07 | 496.79239 | 3433.5188 |
| DVPLLPSDVK | 10 | Q9HCE1 | Helicase MOV-10 OS=Homo sapiens OX | MOV10 | 113.672 | 2 | 50.557953 | yes | 2.905E-07 | 541.80823 | 3280.877 |
| LDLQQGQNLLQGI | 15 | Q9HCE1 | Helicase MOV-10 OS=Homo sapiens OX | MOV10 | 113.672 | 2 | 50.557953 | yes | 6.753E-07 | 827.95978 | 2791.1536 |
| HGLGYSLLER | 10 | Q9HCE1 | Helicase MOV-10 OS=Homo sapiens OX | MOV10 | 113.672 | 2 | 50.557953 | yes | 1.541E-07 | 572.80908 | 2523.8967 |
| AQLETALK | 8 | Q9HCE1 | Helicase MOV-10 OS=Homo sapiens OX | MOV10 | 113.672 | 2 | 50.557953 | yes | 1.356E-07 | 437.25565 | 2394.772 |
| HGVDVEVQGPHE. | 14 | Q9HCE1 | Helicase MOV-10 OS=Homo sapiens OX | MOV10 | 113.672 | 3 | 50.557953 | yes | 9.809E-08 | 510.5867 | 981.00812 |
| HGVDVEVQGPHE. | 14 | Q9HCE1 | Helicase MOV-10 OS=Homo sapiens OX | MOV10 | 113.672 | 3 | 50.557953 | yes | 9.893E-08 | 765.3764 | 981.00812 |
| HGVDVEVQGPHE. | 14 | Q9HCE1 | Helicase MOV-10 OS=Homo sapiens OX | MOV10 | 113.672 | 2 | 50.557953 | yes | 9.809E-08 | 510.5867 | 981.00812 |
| HGVDVEVQGPHE. | 14 | Q9HCE1 | Helicase MOV-10 OS=Homo sapiens OX | MOV10 | 113.672 | 2 | 50.557953 | yes | 9.893E-08 | 765.3764 | 981.00812 |
| KGPDGYDPQFITK | 13 | Q9HCE1 | Helicase MOV-10 OS=Homo sapiens OX | MOV10 | 113.672 | 2 | 50.557953 | yes | 1.549E-07 | 733.36975 | 1852.621 |
| GPDGYDPQFITK | 12 | Q9HCE1 | Helicase MOV-10 OS=Homo sapiens OX | MOV10 | 113.672 | 2 | 50.557953 | yes | 9.888E-08 | 669.32227 | 1789.9395 |
| KGEYVFPAK | 9 | Q9HCE1 | Helicase MOV-10 OS=Homo sapiens OX | MOV10 | 113.672 | 2 | 50.557953 | yes | 1.404E-07 | 519.78455 | 1255.0151 |
| RFNVAVTR | 8 | Q9HCE1 | Helicase MOV-10 OS=Homo sapiens OX | MOV10 | 113.672 | 2 | 50.557953 | yes | 2.059E-07 | 481.78015 | 1171.9259 |
| LLTLEVPGVTESRI | 18 | Q9HCE1 | Helicase MOV-10 OS=Homo sapiens OX | MOV10 | 113.672 | 3 | 50.557953 | yes | 1.864E-07 | 656.04926 | 1035.9202 |
| VNFTFNR | 7 | Q9HCE1 | Helicase MOV-10 OS=Homo sapiens OX | MOV10 | 113.672 | 2 | 50.557953 | yes | 1.62E-07 | 449.23251 | 877.55115 |
| AEYLHGK | 7 | Q9HCE1 | Helicase MOV-10 OS=Homo sapiens OX | MOV10 | 113.672 | 2 | 50.557953 | yes | 1.37E-07 | 409.21378 | 817.88751 |
| KEVLTLR | 7 | Q9HCE1 | Helicase MOV-10 OS=Homo sapiens OX | MOV10 | 113.672 | 2 | 50.557953 | yes | 1.108E-07 | 429.77402 | 766.20782 |
| SLESNPEQLQAMR | 13 | Q9HCE1 | Helicase MOV-10 OS=Homo sapiens OX | MOV10 | 113.672 | 2 | 50.557953 | yes | 9.766E-08 | 751.867 | 742.54156 |
| QGFPIIFHGVMGK | 13 | Q9HCE1 | Helicase MOV-10 OS=Homo sapiens OX | MOV10 | 113.672 | 2 | 50.557953 | yes | 1.063E-07 | 715.88428 | 661.01294 |
| LLLAPSSK | 8 | Q9HCE1 | Helicase MOV-10 OS=Homo sapiens OX | MOV10 | 113.672 | 2 | 50.557953 | yes | 3.569E-07 | 414.76309 | 522.20251 |
| FLAAVAHSPLAAQ | 21 | Q9HCE1 | Helicase MOV-10 OS=Homo sapiens OX | MOV10 | 113.672 | 3 | 50.557953 | yes | 9.977E-08 | 746.75378 | 200.59906 |
| FLAAVAHSPLAAQ | 21 | Q9HCE1 | Helicase MOV-10 OS=Homo sapiens OX | MOV10 | 113.672 | 3 | 50.557953 | yes | 3.838E-07 | 752.08539 | 200.59906 |
| TVTLVEAIK | 9 | Q9HCE1 | Helicase MOV-10 OS=Homo sapiens OX | MOV10 | 113.672 | 2 | 50.557953 | yes | 1.021E-07 | 487.30005 | 325.42871 |
| IEEGERPDR | 9 | Q9HCE1 | Helicase MOV-10 OS=Homo sapiens OX | MOV10 | 113.672 | 2 | 50.557953 | yes | 1.031E-07 | 550.7702 | 303.16379 |
| ENGGYTGCPFPAK | 13 | Q9HCE1 | Helicase MOV-10 OS=Homo sapiens OX | MOV10 | 113.672 | 2 | 50.557953 | yes | 1.059E-07 | 699.31134 | 232.5045 |
| AHILACAPSNSGAI | 19 | Q9HCE1 | Helicase MOV-10 OS=Homo sapiens OX | MOV10 | 113.672 | 3 | 50.557953 | yes | 1.156E-07 | 685.33502 | 195.39847 |
| VFLEFCK | 7 | Q9HCE1 | Helicase MOV-10 OS=Homo sapiens OX | MOV10 | 113.672 | 2 | 50.557953 | yes | 2.169E-07 | 471.7413 | 171.97339 |
| FLAAVAHSPLAAQ | 15 | Q9HCE1 | Helicase MOV-10 OS=Homo sapiens OX | MOV10 | 113.672 | 3 | 50.557953 | yes | 3.492E-07 | 512.96808 | 85.339401 |
| HYDLESVPMTWD. | 19 | Q9HCE1 | Helicase MOV-10 OS=Homo sapiens OX | MOV10 | 113.672 | 3 | 50.557953 | yes | 1.674E-07 | 767.01978 | 77.104103 |
| LNPSDTDAVR | 10 | Q9HCE7 | E3 ubiquitin-protein ligase SMURF1 OS= | SMURF1 | 86.114 | 2 | 46.94822 | yes | 5.386E-07 | 544.27252 | 96.28611 |
| IGTGGSVVDCR | 11 | Q9HCE7 | E3 ubiquitin-protein ligase SMURF1 OS= | SMURF1 | 86.114 | 2 | 46.94822 | yes | 1.434E-07 | 560.77441 | 43.27906 |
| TEINLVSFR | 9 | Q9HCG8 | Pre-mRNA-splicing factor CWC22 homo | CWC22 | 105.467 | 2 | 53.255062 | yes | 3.316E-07 | 539.79822 | 72.758209 |

| | | | | | | | | | | | |
|---|---|---|---|---|---|---|---|---|---|---|---|
| EYMESFEGIFK | 11 | Q9HCG8 | Pre-mRNA-splicing factor CWC22 homo | CWC22 | 105.467 | 2 | 53.255062 | yes | 9.722E-08 | 690.31305 | 60.797424 |
| FPQIGELILK | 10 | Q9HCG8 | Pre-mRNA-splicing factor CWC22 homo | CWC22 | 105.467 | 2 | 53.255062 | yes | 6.938E-06 | 579.3501 | 23.865614 |
| AQILEVLR | 8 | Q9HCH0 | Nck-associated protein 5-like OS=Homo : | NCKAP5L | 139.462 | 2 | 44.196075 | yes | 2.285E-07 | 471.29257 | 54.072746 |
| LTLGINVTSIAVR | 13 | Q9HCI7 | E3 ubiquitin-protein ligase MSL2 OS=Ho | MSL2 | 62.541 | 2 | 25.784876 | yes | 3.227E-06 | 678.91412 | 20.37252 |
| QTGTGWIGGPVPV | 14 | Q9HCJ0 | Trinucleotide repeat-containing gene 6C p | TNRC6C | 175.965 | 2 | 55.62024 | yes | 4.264E-07 | 698.88281 | 105.66592 |
| VLSNTGWGQTPVI | 13 | Q9HCJ0 | Trinucleotide repeat-containing gene 6C p | TNRC6C | 175.965 | 2 | 55.62024 | yes | 9.722E-08 | 693.87244 | 100.52238 |
| LQIQQQMLQAQR | 12 | Q9HCJ0 | Trinucleotide repeat-containing gene 6C p | TNRC6C | 175.965 | 2 | 55.62024 | yes | 3.876E-07 | 742.90375 | 56.71516 |
| HGAIPGGLSIGPPG | 15 | Q9HCJ0 | Trinucleotide repeat-containing gene 6C p | TNRC6C | 175.965 | 2 | 55.62024 | yes | 3.089E-06 | 679.383 | 48.548813 |
| QLAQALLVK | 9 | Q9HCJ0 | Trinucleotide repeat-containing gene 6C p | TNRC6C | 175.965 | 2 | 55.62024 | yes | 5.186E-07 | 492.31604 | 41.416161 |
| QNTAWEFEESPR | 12 | Q9HCJ0 | Trinucleotide repeat-containing gene 6C p | TNRC6C | 175.965 | 2 | 55.62024 | yes | 2.246E-07 | 747.33643 | 37.1185 |
| MDTMIGDGR | 9 | Q9HCJ0 | Trinucleotide repeat-containing gene 6C p | TNRC6C | 175.965 | 2 | 55.62024 | yes | 3.678E-07 | 498.21786 | 25.155182 |
| SNPAWSAGGGDW | 26 | Q9HCJ0 | Trinucleotide repeat-containing gene 6C p | TNRC6C | 175.965 | 3 | 55.62024 | yes | 1.618E-06 | 843.38605 | 24.086033 |
| ATGSAQGNFTGHT | 14 | Q9HCJ0 | Trinucleotide repeat-containing gene 6C p | TNRC6C | 175.965 | 2 | 55.62024 | yes | 0.0002107 | 709.83661 | 21.958811 |
| SPTWGEPPKPK | 11 | Q9HCJ0 | Trinucleotide repeat-containing gene 6C p | TNRC6C | 175.965 | 2 | 55.62024 | yes | 2.661E-07 | 612.32458 | 20.142776 |
| TVATPNQGVWDM | 13 | Q9HCK5 | Protein argonaute-4 OS=Homo sapiens O | AGO4 | 97.097 | 2 | 41.107666 | yes | 3.673E-07 | 737.85901 | 25.587517 |
| IVLQGNQLAALTQ | 15 | Q9HCK8 | Chromodomain-helicase-DNA-binding pr | CHD8 | 290.52 | 2 | 50.126503 | yes | 9.893E-08 | 784.46198 | 46.219379 |
| IDLVCQAVLSGK | 12 | Q9HCK8 | Chromodomain-helicase-DNA-binding pr | CHD8 | 290.52 | 2 | 50.126503 | yes | 5.409E-06 | 651.85791 | 38.37466 |
| LNTITPVVGK | 10 | Q9HCK8 | Chromodomain-helicase-DNA-binding pr | CHD8 | 290.52 | 2 | 50.126503 | yes | 7.448E-07 | 521.31879 | 24.320236 |
| VVTIQLQVQQPQQ | 14 | Q9HCK8 | Chromodomain-helicase-DNA-binding pr | CHD8 | 290.52 | 2 | 50.126503 | yes | 1.263E-07 | 818.97266 | 16.120632 |
| FFNPSIPSLGLR | 12 | Q9HCL2 | Glycerol-3-phosphate acyltransferase 1, m | GPAM | 93.795 | 2 | 52.80372 | yes | 1.074E-07 | 674.37463 | 206.3344 |
| KLPEPLSWR | 9 | Q9HCL2 | Glycerol-3-phosphate acyltransferase 1, m | GPAM | 93.795 | 2 | 52.80372 | yes | 1.068E-06 | 563.3244 | 155.54913 |
| ALLHGHIVELLR | 12 | Q9HCL2 | Glycerol-3-phosphate acyltransferase 1, m | GPAM | 93.795 | 3 | 52.80372 | yes | 3.256E-07 | 457.61374 | 116.81485 |
| LSYVLFIQER | 10 | Q9HCL2 | Glycerol-3-phosphate acyltransferase 1, m | GPAM | 93.795 | 2 | 52.80372 | yes | 1.733E-07 | 634.3559 | 69.35228 |
| KNESLWSVAR | 10 | Q9HCL2 | Glycerol-3-phosphate acyltransferase 1, m | GPAM | 93.795 | 2 | 52.80372 | yes | 9.722E-08 | 595.31982 | 67.882225 |
| QQQFLEIFLEGTR | 13 | Q9HCL2 | Glycerol-3-phosphate acyltransferase 1, m | GPAM | 93.795 | 2 | 52.80372 | yes | 0.0003145 | 804.92267 | 59.522503 |
| NVIYINETHTR | 11 | Q9HCL2 | Glycerol-3-phosphate acyltransferase 1, m | GPAM | 93.795 | 2 | 52.80372 | yes | 5.526E-07 | 680.35443 | 54.491295 |
| RLDETPDGR | 9 | Q9HCL2 | Glycerol-3-phosphate acyltransferase 1, m | GPAM | 93.795 | 2 | 52.80372 | yes | 1.345E-07 | 529.76489 | 52.04723 |
| NVAVYAESATYCI | 15 | Q9HCL2 | Glycerol-3-phosphate acyltransferase 1, m | GPAM | 93.795 | 2 | 52.80372 | yes | 6.452E-07 | 844.42163 | 25.910587 |
| VSLLDGTDVSVDL | 15 | Q9HCM4 | Band 4.1-like protein 5 OS=Homo sapien | EPB41L5 | 81.857 | 2 | 56.292038 | yes | 1.124E-07 | 779.42218 | 511.04123 |
| FYSSEPNNLR | 10 | Q9HCM4 | Band 4.1-like protein 5 OS=Homo sapien | EPB41L5 | 81.857 | 2 | 56.292038 | yes | 1.32E-07 | 613.79364 | 449.18219 |
| AATHIPAAGDSK | 12 | Q9HCM4 | Band 4.1-like protein 5 OS=Homo sapien | EPB41L5 | 81.857 | 2 | 56.292038 | yes | 9.722E-08 | 569.7962 | 358.90665 |
| GQTPAQAETNYLN | 14 | Q9HCM4 | Band 4.1-like protein 5 OS=Homo sapien | EPB41L5 | 81.857 | 2 | 56.292038 | yes | 1.456E-07 | 767.87842 | 314.57709 |
| EQEHTFVFR | 9 | Q9HCM4 | Band 4.1-like protein 5 OS=Homo sapien | EPB41L5 | 81.857 | 2 | 56.292038 | no | 9.722E-08 | 596.79089 | 205.44473 |
| IGSPYCLHLR | 10 | Q9HCM4 | Band 4.1-like protein 5 OS=Homo sapien | EPB41L5 | 81.857 | 2 | 56.292038 | yes | 1.05E-07 | 608.31879 | 172.59061 |
| LTLVVVEDDDQGI | 13 | Q9HCM4 | Band 4.1-like protein 5 OS=Homo sapien | EPB41L5 | 81.857 | 2 | 56.292038 | yes | 1.691E-07 | 715.87231 | 93.441879 |
| CAVEHHAFFR | 10 | Q9HCM4 | Band 4.1-like protein 5 OS=Homo sapien | EPB41L5 | 81.857 | 2 | 56.292038 | no | 9.722E-08 | 637.2984 | 73.903824 |
| YLFVLQLK | 8 | Q9HCM4 | Band 4.1-like protein 5 OS=Homo sapien | EPB41L5 | 81.857 | 2 | 56.292038 | yes | 5.376E-07 | 512.31549 | 65.129013 |
| WLEMYGVDMHV | 13 | Q9HCM4 | Band 4.1-like protein 5 OS=Homo sapien | EPB41L5 | 81.857 | 3 | 56.292038 | yes | 1.001E-07 | 536.26337 | 24.532146 |
| WLEMYGVDMHV | 13 | Q9HCM4 | Band 4.1-like protein 5 OS=Homo sapien | EPB41L5 | 81.857 | 3 | 56.292038 | yes | 1.191E-06 | 803.89148 | 24.532146 |

| | | | | | | | | | | | |
|---|---|---|---|---|---|---|---|---|---|---|---|
| WLEMYGVDMHV' | 13 | Q9HCM4 | Band 4.1-like protein 5 OS=Homo sapien | EPB41L5 | 81.857 | 2 | 56.292038 | yes | 1.001E-07 | 536.26337 | 24.532146 |
| WLEMYGVDMHV' | 13 | Q9HCM4 | Band 4.1-like protein 5 OS=Homo sapien | EPB41L5 | 81.857 | 2 | 56.292038 | yes | 1.191E-06 | 803.89148 | 24.532146 |
| EHVIHIIDFGLAK | 13 | Q9HCP0 | Casein kinase I isoform gamma-1 OS=Ho | CSNK1G1 | 48.512 | 2 | 38.098373 | yes | 5.462E-07 | 746.41956 | 23.544964 |
| EHVIHIIDFGLAK | 13 | Q9HCP0 | Casein kinase I isoform gamma-1 OS=Ho | CSNK1G1 | 48.512 | 2 | 38.098373 | yes | 1.379E-06 | 497.94879 | 23.544964 |
| EHVIHIIDFGLAK | 13 | Q9HCP0 | Casein kinase I isoform gamma-1 OS=Ho | CSNK1G1 | 48.512 | 3 | 38.098373 | yes | 5.462E-07 | 746.41956 | 23.544964 |
| EHVIHIIDFGLAK | 13 | Q9HCP0 | Casein kinase I isoform gamma-1 OS=Ho | CSNK1G1 | 48.512 | 3 | 38.098373 | yes | 1.379E-06 | 497.94879 | 23.544964 |
| TLFTDLFEK | 9 | Q9HCP0 | Casein kinase I isoform gamma-1 OS=Ho | CSNK1G1 | 48.512 | 2 | 38.098373 | yes | 8.618E-07 | 557.29498 | 34.327515 |
| VQSLNVDAIIR | 11 | Q9HCS7 | Pre-mRNA-splicing factor SYF1 OS=Ho | XAB2 | 100.01 | 2 | 52.883392 | yes | 9.807E-08 | 614.35645 | 368.13943 |
| AIEVLSDEHAR | 11 | Q9HCS7 | Pre-mRNA-splicing factor SYF1 OS=Ho | XAB2 | 100.01 | 2 | 52.883392 | yes | 9.722E-08 | 620.32001 | 291.40451 |
| LEQQSVPAAVFGS | 15 | Q9HCS7 | Pre-mRNA-splicing factor SYF1 OS=Ho | XAB2 | 100.01 | 2 | 52.883392 | yes | 1.499E-07 | 787.43286 | 225.79921 |
| AAEIYGVTHTR | 11 | Q9HCS7 | Pre-mRNA-splicing factor SYF1 OS=Ho | XAB2 | 100.01 | 2 | 52.883392 | yes | 1.591E-07 | 609.31732 | 216.07077 |
| LSPESAEEYIEYLK | 14 | Q9HCS7 | Pre-mRNA-splicing factor SYF1 OS=Ho | XAB2 | 100.01 | 2 | 52.883392 | yes | 7.036E-07 | 835.91162 | 185.25356 |
| LNQLYER | 7 | Q9HCS7 | Pre-mRNA-splicing factor SYF1 OS=Ho | XAB2 | 100.01 | 2 | 52.883392 | yes | 3.716E-07 | 468.25089 | 174.9061 |
| LATVVNDER | 9 | Q9HCS7 | Pre-mRNA-splicing factor SYF1 OS=Ho | XAB2 | 100.01 | 2 | 52.883392 | yes | 8.185E-07 | 508.77219 | 151.00568 |
| SHPLPETAVR | 10 | Q9HCS7 | Pre-mRNA-splicing factor SYF1 OS=Ho | XAB2 | 100.01 | 2 | 52.883392 | yes | 2.202E-07 | 553.80127 | 140.2963 |
| HENYDEALR | 9 | Q9HCS7 | Pre-mRNA-splicing factor SYF1 OS=Ho | XAB2 | 100.01 | 2 | 52.883392 | yes | 2.209E-07 | 573.76233 | 107.02679 |
| AIYSFCSQICDPR | 13 | Q9HCS7 | Pre-mRNA-splicing factor SYF1 OS=Ho | XAB2 | 100.01 | 2 | 52.883392 | yes | 1.727E-07 | 808.8634 | 78.773071 |
| DLFEQALDGCPPK | 13 | Q9HCS7 | Pre-mRNA-splicing factor SYF1 OS=Ho | XAB2 | 100.01 | 2 | 52.883392 | yes | 2.229E-07 | 745.35321 | 74.853935 |
| AEYFDGSEPVQNR | 13 | Q9HCS7 | Pre-mRNA-splicing factor SYF1 OS=Ho | XAB2 | 100.01 | 2 | 52.883392 | yes | 2.073E-07 | 756.34167 | 57.753002 |
| AEQLAAEAER | 10 | Q9HCS7 | Pre-mRNA-splicing factor SYF1 OS=Ho | XAB2 | 100.01 | 2 | 52.883392 | yes | 6.984E-07 | 544.27252 | 39.925911 |
| FYEDNGQLDDAR | 12 | Q9HCS7 | Pre-mRNA-splicing factor SYF1 OS=Ho | XAB2 | 100.01 | 2 | 52.883392 | yes | 3.619E-07 | 721.81274 | 35.1749 |
| CVTDPAYEDVNN( | 16 | Q9HCS7 | Pre-mRNA-splicing factor SYF1 OS=Ho | XAB2 | 100.01 | 3 | 52.883392 | yes | 3.525E-07 | 660.27209 | 10.356167 |
| FGQVPDQPAGLR | 12 | Q9HCU5 | Prolactin regulatory element-binding prot | PREB | 45.469 | 2 | 50.688232 | yes | 9.81E-08 | 642.83838 | 615.30243 |
| LFTVQIPHK | 9 | Q9HCU5 | Prolactin regulatory element-binding prot | PREB | 45.469 | 2 | 50.688232 | yes | 7.991E-07 | 541.82147 | 140.63972 |
| CQLHLLPSR | 9 | Q9HCU5 | Prolactin regulatory element-binding prot | PREB | 45.469 | 2 | 50.688232 | yes | 1.791E-07 | 562.30566 | 128.20294 |
| RAPELYR | 7 | Q9HCU5 | Prolactin regulatory element-binding prot | PREB | 45.469 | 2 | 50.688232 | yes | 9.728E-08 | 452.7536 | 105.30899 |
| GPELLGSHETALFS | 20 | Q9HCU5 | Prolactin regulatory element-binding prot | PREB | 45.469 | 3 | 50.688232 | yes | 1.193E-07 | 695.69189 | 92.13916 |
| EAHGIVVTDVAFL | 16 | Q9HCU5 | Prolactin regulatory element-binding prot | PREB | 45.469 | 3 | 50.688232 | yes | 2.075E-07 | 575.64545 | 49.562931 |
| VYAENAIR | 8 | Q9HD42 | Charged multivesicular body protein 1a C | CHMP1A | 21.703 | 2 | 51.36214 | yes | 9.849E-08 | 468.25089 | 757.96814 |
| ALSTMDLQK | 9 | Q9HD42 | Charged multivesicular body protein 1a C | CHMP1A | 21.703 | 2 | 51.36214 | yes | 9.722E-08 | 503.7655 | 446.79099 |
| NEGVNWLR | 8 | Q9HD42 | Charged multivesicular body protein 1a C | CHMP1A | 21.703 | 2 | 51.36214 | yes | 1.31E-07 | 494.25397 | 328.31903 |
| SQEDQLSR | 8 | Q9HD42 | Charged multivesicular body protein 1a C | CHMP1A | 21.703 | 2 | 51.36214 | yes | 2.011E-07 | 481.73053 | 272.52802 |
| KNEGVNWLR | 9 | Q9HD42 | Charged multivesicular body protein 1a C | CHMP1A | 21.703 | 2 | 51.36214 | yes | 1.172E-07 | 558.30145 | 153.09772 |
| VQTAVTMK | 8 | Q9HD42 | Charged multivesicular body protein 1a C | CHMP1A | 21.703 | 2 | 51.36214 | yes | 9.722E-08 | 439.24423 | 118.27578 |
| LAATSEVGDLPWK | 13 | Q9HD67 | Unconventional myosin-X OS=Homo sap | MYO10 | 237.347 | 2 | 54.33081 | yes | 1.175E-07 | 693.86682 | 165.66907 |
| YSGMLETVR | 9 | Q9HD67 | Unconventional myosin-X OS=Homo sap | MYO10 | 237.347 | 2 | 54.33081 | yes | 3.103E-07 | 528.26312 | 144.55061 |
| VEGQEFIVR | 9 | Q9HD67 | Unconventional myosin-X OS=Homo sap | MYO10 | 237.347 | 2 | 54.33081 | yes | 3.059E-06 | 538.79041 | 119.18626 |
| RPFQDFYK | 8 | Q9HD67 | Unconventional myosin-X OS=Homo sap | MYO10 | 237.347 | 2 | 54.33081 | yes | 4.82E-07 | 550.77985 | 85.05349 |
| REEEVSHAAMVIR | 13 | Q9HD67 | Unconventional myosin-X OS=Homo sap | MYO10 | 237.347 | 3 | 54.33081 | yes | 9.722E-08 | 509.59631 | 79.229866 |

| | | | | | | | | | | | |
|---|---|---|---|---|---|---|---|---|---|---|---|
| FVQLNICQK | 9 | Q9HD67 | Unconventional myosin-X OS=Homo sap | MYO10 | 237.347 | 2 | 54.33081 | yes | 1.452E-07 | 575.30786 | 78.724785 |
| WSSAIQNVTDTK | 12 | Q9HD67 | Unconventional myosin-X OS=Homo sap | MYO10 | 237.347 | 2 | 54.33081 | yes | 9.722E-08 | 675.33844 | 77.573471 |
| LMYFENDSEEK | 11 | Q9HD67 | Unconventional myosin-X OS=Homo sap | MYO10 | 237.347 | 2 | 54.33081 | yes | 1.222E-07 | 702.80304 | 75.929573 |
| LYCFLDTDNVPK | 12 | Q9HD67 | Unconventional myosin-X OS=Homo sap | MYO10 | 237.347 | 2 | 54.33081 | yes | 2.154E-07 | 742.85815 | 71.760338 |
| ELLFETSEVVDVA | 14 | Q9HD67 | Unconventional myosin-X OS=Homo sap | MYO10 | 237.347 | 2 | 54.33081 | yes | 1.73E-06 | 789.91669 | 68.25367 |
| DEIEALIHR | 9 | Q9HD67 | Unconventional myosin-X OS=Homo sap | MYO10 | 237.347 | 2 | 54.33081 | yes | 2.526E-07 | 548.29327 | 66.806404 |
| APIDTPTQQLIQDI | 15 | Q9HD67 | Unconventional myosin-X OS=Homo sap | MYO10 | 237.347 | 2 | 54.33081 | yes | 1.836E-07 | 840.96198 | 64.649719 |
| HYAGEVQYDVR | 11 | Q9HD67 | Unconventional myosin-X OS=Homo sap | MYO10 | 237.347 | 2 | 54.33081 | yes | 4.325E-07 | 668.81769 | 59.64106 |
| AHVLGFLAR | 9 | Q9HD67 | Unconventional myosin-X OS=Homo sap | MYO10 | 237.347 | 2 | 54.33081 | yes | 3.081E-06 | 492.29291 | 39.690048 |
| AILESSPIMEAFGN | 16 | Q9HD67 | Unconventional myosin-X OS=Homo sap | MYO10 | 237.347 | 2 | 54.33081 | yes | 1.671E-06 | 839.42944 | 31.450649 |
| QSVPYGMATVIR | 12 | Q9HDC5 | Junctophilin-1 OS=Homo sapiens OX=96 | JPH1 | 71.686 | 2 | 50.43383 | yes | 1.509E-07 | 661.35028 | 228.71028 |
| ADAADQAALAAR | 12 | Q9HDC5 | Junctophilin-1 OS=Homo sapiens OX=96 | JPH1 | 71.686 | 2 | 50.43383 | yes | 2.101E-07 | 572.29126 | 152.67799 |
| HGLGVETK | 8 | Q9HDC5 | Junctophilin-1 OS=Homo sapiens OX=96 | JPH1 | 71.686 | 2 | 50.43383 | yes | 9.809E-08 | 420.73233 | 149.57347 |
| GEWSHGFK | 8 | Q9HDC5 | Junctophilin-1 OS=Homo sapiens OX=96 | JPH1 | 71.686 | 2 | 50.43383 | yes | 3.505E-07 | 474.22214 | 146.77539 |
| SVADEQVTAIVNK | 18 | Q9HDC5 | Junctophilin-1 OS=Homo sapiens OX=96 | JPH1 | 71.686 | 3 | 50.43383 | yes | 1.745E-07 | 644.01532 | 127.00848 |
| QSLCTPAR | 8 | Q9HDC5 | Junctophilin-1 OS=Homo sapiens OX=96 | JPH1 | 71.686 | 2 | 50.43383 | yes | 1.087E-07 | 466.73456 | 93.229546 |
| HGYGCTVFPDGSK | 13 | Q9HDC5 | Junctophilin-1 OS=Homo sapiens OX=96 | JPH1 | 71.686 | 2 | 50.43383 | yes | 1.313E-07 | 712.81677 | 75.826073 |
| ESPHFYR | 7 | Q9HDC5 | Junctophilin-1 OS=Homo sapiens OX=96 | JPH1 | 71.686 | 2 | 50.43383 | yes | 1.206E-07 | 468.22214 | 65.373055 |
| NPASNDSCPALEK | 13 | Q9HDC5 | Junctophilin-1 OS=Homo sapiens OX=96 | JPH1 | 71.686 | 2 | 50.43383 | yes | 3.868E-07 | 701.81702 | 32.316654 |
| AKADAADQAALA | 14 | Q9HDC5 | Junctophilin-1 OS=Homo sapiens OX=96 | JPH1 | 71.686 | 2 | 50.43383 | yes | 1.655E-05 | 671.8573 | 17.018711 |
| LFLLQDSGR | 9 | Q9NNW5 | tRNA (34-2'-O)-methyltransferase regulat | WDR6 | 121.725 | 2 | 53.234436 | yes | 3.309E-06 | 524.79291 | 2290.5107 |
| ILSPSIMVSASIDQI | 15 | Q9NNW5 | tRNA (34-2'-O)-methyltransferase regulat | WDR6 | 121.725 | 2 | 53.234436 | yes | 1.174E-07 | 808.93744 | 1439.3423 |
| VHSLSWALR | 9 | Q9NNW5 | tRNA (34-2'-O)-methyltransferase regulat | WDR6 | 121.725 | 2 | 53.234436 | yes | 9.722E-08 | 534.80109 | 1435.8605 |
| AVTLAGSWR | 9 | Q9NNW5 | tRNA (34-2'-O)-methyltransferase regulat | WDR6 | 121.725 | 2 | 53.234436 | yes | 4.993E-07 | 480.76672 | 1399.0093 |
| QAWVITGGDDSGI | 14 | Q9NNW5 | tRNA (34-2'-O)-methyltransferase regulat | WDR6 | 121.725 | 2 | 53.234436 | yes | 1.435E-07 | 737.86786 | 1362.1682 |
| GLGVSALCFK | 10 | Q9NNW5 | tRNA (34-2'-O)-methyltransferase regulat | WDR6 | 121.725 | 2 | 53.234436 | yes | 2.452E-07 | 526.28387 | 1322.9468 |
| LWHLVGR | 7 | Q9NNW5 | tRNA (34-2'-O)-methyltransferase regulat | WDR6 | 121.725 | 2 | 53.234436 | yes | 1.847E-07 | 440.76123 | 941.94122 |
| CTLSSACLIGDAW | 14 | Q9NNW5 | tRNA (34-2'-O)-methyltransferase regulat | WDR6 | 121.725 | 2 | 53.234436 | yes | 1.006E-07 | 791.3736 | 897.1568 |
| DGQLQPVLR | 9 | Q9NNW5 | tRNA (34-2'-O)-methyltransferase regulat | WDR6 | 121.725 | 2 | 53.234436 | yes | 1.122E-07 | 513.29053 | 769.46564 |
| GSVLLFPSRPGLLK | 14 | Q9NNW5 | tRNA (34-2'-O)-methyltransferase regulat | WDR6 | 121.725 | 2 | 53.234436 | yes | 1.362E-07 | 742.45337 | 701.67792 |
| GLLATASEDR | 10 | Q9NNW5 | tRNA (34-2'-O)-methyltransferase regulat | WDR6 | 121.725 | 2 | 53.234436 | yes | 1.01E-06 | 516.76965 | 639.64355 |
| LHIVNCGGGHR | 11 | Q9NNW5 | tRNA (34-2'-O)-methyltransferase regulat | WDR6 | 121.725 | 2 | 53.234436 | yes | 9.722E-08 | 610.30927 | 356.70886 |
| ILQLLAETFHHK | 12 | Q9NNW5 | tRNA (34-2'-O)-methyltransferase regulat | WDR6 | 121.725 | 2 | 53.234436 | yes | 9.722E-08 | 725.41425 | 329.97858 |
| GMNWLAGLR | 9 | Q9NNW5 | tRNA (34-2'-O)-methyltransferase regulat | WDR6 | 121.725 | 2 | 53.234436 | yes | 1.061E-07 | 509.26855 | 161.09045 |
| GMNWLAGLR | 9 | Q9NNW5 | tRNA (34-2'-O)-methyltransferase regulat | WDR6 | 121.725 | 2 | 53.234436 | yes | 2.451E-06 | 517.26599 | 161.09045 |
| ALGGCTRPHVILR | 13 | Q9NNW5 | tRNA (34-2'-O)-methyltransferase regulat | WDR6 | 121.725 | 3 | 53.234436 | yes | 2.174E-07 | 483.94174 | 285.30237 |
| VHSFTHEAPNQR | 12 | Q9NNW5 | tRNA (34-2'-O)-methyltransferase regulat | WDR6 | 121.725 | 3 | 53.234436 | yes | 9.722E-08 | 474.9006 | 94.942825 |
| VHSFTHEAPNQR | 12 | Q9NNW5 | tRNA (34-2'-O)-methyltransferase regulat | WDR6 | 121.725 | 3 | 53.234436 | yes | 9.722E-08 | 711.84729 | 94.942825 |
| VHSFTHEAPNQR | 12 | Q9NNW5 | tRNA (34-2'-O)-methyltransferase regulat | WDR6 | 121.725 | 2 | 53.234436 | yes | 9.722E-08 | 474.9006 | 94.942825 |

| | | | | | | | | | | | |
|---|---|---|---|---|---|---|---|---|---|---|---|
| VHSFTHEAPNQR | 12 | Q9NNW5 | tRNA (34-2'-O)-methyltransferase regulat | WDR6 | 121.725 | 2 | 53.234436 | yes | 9.722E-08 | 711.84729 | 94.942825 |
| LLAGEGPDVLVYS | 21 | Q9NNW5 | tRNA (34-2'-O)-methyltransferase regulat | WDR6 | 121.725 | 3 | 53.234436 | yes | 2.643E-07 | 743.3952 | 180.18268 |
| VVPINTPTAAVDQ | 19 | Q9NNW5 | tRNA (34-2'-O)-methyltransferase regulat | WDR6 | 121.725 | 3 | 53.234436 | yes | 1.554E-07 | 656.69818 | 175.51428 |
| GSVLLFPSR | 9 | Q9NNW5 | tRNA (34-2'-O)-methyltransferase regulat | WDR6 | 121.725 | 2 | 53.234436 | yes | 9.804E-08 | 488.28473 | 158.69844 |
| LDEYWDR | 7 | Q9NNW5 | tRNA (34-2'-O)-methyltransferase regulat | WDR6 | 121.725 | 2 | 53.234436 | yes | 1.465E-07 | 498.7247 | 106.6862 |
| CWEQLLEDK | 9 | Q9NNW5 | tRNA (34-2'-O)-methyltransferase regulat | WDR6 | 121.725 | 2 | 53.234436 | yes | 9.722E-08 | 610.78442 | 104.21134 |
| DGDVMLYR | 8 | Q9NNW5 | tRNA (34-2'-O)-methyltransferase regulat | WDR6 | 121.725 | 2 | 53.234436 | yes | 4.849E-07 | 484.72894 | 99.878891 |
| RGSVLLFPSRPGLI | 15 | Q9NNW5 | tRNA (34-2'-O)-methyltransferase regulat | WDR6 | 121.725 | 3 | 53.234436 | yes | 5.067E-07 | 547.33838 | 63.169834 |
| DGAYYQLFVR | 10 | Q9NNW5 | tRNA (34-2'-O)-methyltransferase regulat | WDR6 | 121.725 | 2 | 53.234436 | yes | 1.041E-07 | 616.30896 | 37.521858 |
| VQNIGHCFGHSAR | 13 | Q9NNW5 | tRNA (34-2'-O)-methyltransferase regulat | WDR6 | 121.725 | 2 | 53.234436 | yes | 9.723E-08 | 741.85455 | 31.638248 |
| NWNHVVDAR | 9 | Q9NP50 | SIN3-HDAC complex-associated factor C | SINHCAF | 24.853 | 2 | 53.371628 | yes | 5.344E-07 | 555.77557 | 129.35907 |
| ICCGIIYK | 8 | Q9NP50 | SIN3-HDAC complex-associated factor C | SINHCAF | 24.853 | 2 | 53.371628 | yes | 3.561E-07 | 513.75916 | 91.409103 |
| SIEGCCICR | 9 | Q9NP50 | SIN3-HDAC complex-associated factor C | SINHCAF | 24.853 | 2 | 53.371628 | yes | 9.722E-08 | 577.74121 | 60.468498 |
| AISSDMYFGR | 10 | Q9NP61 | ADP-ribosylation factor GTPase-activatir | ARFGAP3 | 56.929 | 2 | 41.740284 | yes | 1.495E-05 | 573.76605 | 64.720299 |
| ATLEVSSIIK | 10 | Q9NP61 | ADP-ribosylation factor GTPase-activatir | ARFGAP3 | 56.929 | 2 | 41.740284 | yes | 3.363E-07 | 530.81604 | 62.386295 |
| LSVFANGVVTSIQI | 15 | Q9NP61 | ADP-ribosylation factor GTPase-activatir | ARFGAP3 | 56.929 | 2 | 41.740284 | yes | 5.459E-05 | 803.43341 | 30.408409 |
| DLDPNNVIIEQEEF | 14 | Q9NP64 | Zinc finger CCHC domain-containing prc | ZCCHC17 | 27.57 | 2 | 53.62248 | yes | 9.81E-08 | 842.41284 | 1006.395 |
| ITLEAVLNTTCK | 12 | Q9NP64 | Zinc finger CCHC domain-containing prc | ZCCHC17 | 27.57 | 2 | 53.62248 | yes | 9.722E-08 | 681.86853 | 742.20282 |
| DCFMQPGGTK | 10 | Q9NP64 | Zinc finger CCHC domain-containing prc | ZCCHC17 | 27.57 | 2 | 53.62248 | yes | 9.722E-08 | 570.74426 | 562.10016 |
| VDKPSEIVDVGDK | 13 | Q9NP64 | Zinc finger CCHC domain-containing prc | ZCCHC17 | 27.57 | 2 | 53.62248 | yes | 9.722E-08 | 700.867 | 297.89957 |
| SFQDYTGQK | 9 | Q9NP64 | Zinc finger CCHC domain-containing prc | ZCCHC17 | 27.57 | 2 | 53.62248 | yes | 9.722E-08 | 537.24854 | 85.76989 |
| SAEFEKPDPTR | 11 | Q9NP64 | Zinc finger CCHC domain-containing prc | ZCCHC17 | 27.57 | 2 | 53.62248 | yes | 9.722E-08 | 638.81201 | 84.426262 |
| LEVDVIQER | 9 | Q9NP66 | High mobility group protein 20A OS=Ho | HMG20A | 40.144 | 2 | 43.091652 | yes | 2.736E-07 | 550.80096 | 109.97334 |
| ILIIGESGVGK | 11 | Q9NP72 | Ras-related protein Rab-18 OS=Homo saj | RAB18 | 22.978 | 2 | 52.316757 | yes | 9.722E-08 | 543.33191 | 554.20612 |
| LAIWDTAGQER | 11 | Q9NP72 | Ras-related protein Rab-18 OS=Homo saj | RAB18 | 22.978 | 2 | 52.316757 | yes | 1.003E-07 | 630.32257 | 269.89859 |
| NDIVNMLVGNK | 11 | Q9NP72 | Ras-related protein Rab-18 OS=Homo saj | RAB18 | 22.978 | 2 | 52.316757 | yes | 9.722E-08 | 608.82135 | 151.95158 |
| TLTPSYYR | 8 | Q9NP72 | Ras-related protein Rab-18 OS=Homo saj | RAB18 | 22.978 | 2 | 52.316757 | yes | 1.959E-06 | 500.75854 | 99.594666 |
| GAQGVILVYDVTF | 13 | Q9NP72 | Ras-related protein Rab-18 OS=Homo saj | RAB18 | 22.978 | 2 | 52.316757 | yes | 4.427E-07 | 695.88806 | 40.031002 |
| ALDPEIYR | 8 | Q9NP73 | Putative bifunctional UDP-N-acetylgluco: | ALG13 | 126.057 | 2 | 50.869358 | yes | 5.227E-07 | 488.75854 | 109.81458 |
| YHEEYLYR | 8 | Q9NP73 | Putative bifunctional UDP-N-acetylgluco: | ALG13 | 126.057 | 2 | 50.869358 | yes | 9.722E-08 | 586.77222 | 77.448753 |
| NVEFDVWLDSR | 11 | Q9NP73 | Putative bifunctional UDP-N-acetylgluco: | ALG13 | 126.057 | 2 | 50.869358 | yes | 2.52E-06 | 690.33313 | 64.13707 |
| GKPLVVVINEK | 11 | Q9NP73 | Putative bifunctional UDP-N-acetylgluco: | ALG13 | 126.057 | 2 | 50.869358 | yes | 5.53E-07 | 598.37408 | 49.513618 |
| SDYMEYAGR | 9 | Q9NP73 | Putative bifunctional UDP-N-acetylgluco: | ALG13 | 126.057 | 2 | 50.869358 | yes | 1.143E-06 | 546.22675 | 45.71032 |
| CCQSYDNFSYR | 11 | Q9NP73 | Putative bifunctional UDP-N-acetylgluco: | ALG13 | 126.057 | 2 | 50.869358 | yes | 9.052E-06 | 750.28754 | 41.162743 |
| NNAVTGSEDAHTI | 15 | Q9NP73 | Putative bifunctional UDP-N-acetylgluco: | ALG13 | 126.057 | 2 | 50.869358 | yes | 1.303E-07 | 811.35809 | 37.097389 |
| VYDQVVEDLNSR | 12 | Q9NP77 | RNA polymerase II subunit A C-terminal | SSU72 | 22.575 | 2 | 46.06773 | yes | 1.662E-07 | 718.85443 | 63.723423 |
| LYAMQTGMK | 9 | Q9NP79 | Vacuolar protein sorting-associated protei | VTA1 | 33.88 | 2 | 50.385063 | yes | 2.228E-07 | 521.75665 | 166.35667 |
| LTPEDFAR | 8 | Q9NP79 | Vacuolar protein sorting-associated protei | VTA1 | 33.88 | 2 | 50.385063 | yes | 9.81E-08 | 474.74289 | 133.37024 |
| DPVVAYYCR | 9 | Q9NP79 | Vacuolar protein sorting-associated protei | VTA1 | 33.88 | 2 | 50.385063 | yes | 1.055E-06 | 571.76862 | 123.00076 |

| | | | | | | | | | | | |
|---|---|---|---|---|---|---|---|---|---|---|---|
| LMDQLEALK | 9 | Q9NP79 | Vacuolar protein sorting-associated protei | VTA1 | 33.88 | 2 | 50.385063 | yes | 4.609E-07 | 530.789 | 110.81902 |
| MFLYADNEDR | 10 | Q9NP79 | Vacuolar protein sorting-associated protei | VTA1 | 33.88 | 2 | 50.385063 | yes | 1.545E-07 | 637.27954 | 109.62766 |
| LMDQLEALKK | 10 | Q9NP79 | Vacuolar protein sorting-associated protei | VTA1 | 33.88 | 2 | 50.385063 | yes | 4.79E-07 | 594.83649 | 26.380989 |
| EGYSALPQLDIER | 13 | Q9NP81 | Serine--tRNA ligase, mitochondrial OS=I | SARS2 | 58.283 | 2 | 52.47616 | yes | 1.142E-07 | 745.87793 | 659.87268 |
| NLLYEYAR | 8 | Q9NP81 | Serine--tRNA ligase, mitochondrial OS=I | SARS2 | 58.283 | 2 | 52.47616 | yes | 1.528E-07 | 521.27185 | 393.75436 |
| FCACPEEAAHALE | 15 | Q9NP81 | Serine--tRNA ligase, mitochondrial OS=I | SARS2 | 58.283 | 3 | 52.47616 | yes | 9.722E-08 | 591.93976 | 302.5777 |
| KPGLPGQPAVS | 11 | Q9NP81 | Serine--tRNA ligase, mitochondrial OS=I | SARS2 | 58.283 | 2 | 52.47616 | yes | 3.082E-07 | 525.80072 | 278.13 |
| KFDIEAWMPGR | 11 | Q9NP81 | Serine--tRNA ligase, mitochondrial OS=I | SARS2 | 58.283 | 2 | 52.47616 | yes | 1.086E-07 | 675.33716 | 217.55124 |
| MVCSSTCYR | 9 | Q9NP81 | Serine--tRNA ligase, mitochondrial OS=I | SARS2 | 58.283 | 2 | 52.47616 | yes | 1.327E-07 | 582.23572 | 165.15599 |
| LLIALLESNQQK | 12 | Q9NP81 | Serine--tRNA ligase, mitochondrial OS=I | SARS2 | 58.283 | 2 | 52.47616 | yes | 9.722E-08 | 685.40613 | 120.88895 |
| QLQEQIR | 7 | Q9NP81 | Serine--tRNA ligase, mitochondrial OS=I | SARS2 | 58.283 | 2 | 52.47616 | yes | 9.729E-08 | 457.75635 | 115.97497 |
| GHLEIGEK | 8 | Q9NP81 | Serine--tRNA ligase, mitochondrial OS=I | SARS2 | 58.283 | 2 | 52.47616 | yes | 1.294E-07 | 441.73761 | 91.603928 |
| VLHMVGDKPVFSI | 16 | Q9NP81 | Serine--tRNA ligase, mitochondrial OS=I | SARS2 | 58.283 | 3 | 52.47616 | yes | 1.727E-07 | 619.6662 | 72.03656 |
| ELVHLYPR | 8 | Q9NP81 | Serine--tRNA ligase, mitochondrial OS=I | SARS2 | 58.283 | 2 | 52.47616 | yes | 1.477E-07 | 513.79016 | 39.468445 |
| TFFQAASEEDIFR | 13 | Q9NP87 | DNA-directed DNA/RNA polymerase mu | POLM | 54.816 | 2 | 51.798443 | yes | 1.499E-07 | 780.87012 | 117.64188 |
| AGLQHHQDLSTPV | 15 | Q9NP87 | DNA-directed DNA/RNA polymerase mu | POLM | 54.816 | 3 | 51.798443 | yes | 9.962E-08 | 557.96906 | 64.872086 |
| LFTQIFGVGVK | 11 | Q9NP87 | DNA-directed DNA/RNA polymerase mu | POLM | 54.816 | 2 | 51.798443 | yes | 9.722E-08 | 604.85553 | 56.771732 |
| DIDPQNDLTFLR | 12 | Q9NP97 | Dynein light chain roadblock-type 1 OS=I | DYNLRB1 | 10.922 | 2 | 51.482006 | no | 9.898E-08 | 723.86481 | 116.73554 |
| GVQGIIVVNTEGIP | 16 | Q9NP97 | Dynein light chain roadblock-type 1 OS=I | DYNLRB1 | 10.922 | 2 | 51.482006 | yes | 9.722E-08 | 818.98523 | 112.04374 |
| TFLAQHASL | 9 | Q9NPA8 | Transcription and mRNA export factor EI | ENY2 | 11.529 | 2 | 42.52821 | yes | 2.985E-07 | 494.26654 | 233.82394 |
| EWYAHNPYPSPR | 12 | Q9NPC8 | Homeobox protein SIX2 OS=Homo sapie | SIX2 | 32.287 | 3 | 50.97041 | no | 1.34E-06 | 506.23712 | 62.09008 |
| EWYAHNPYPSPR | 12 | Q9NPC8 | Homeobox protein SIX2 OS=Homo sapie | SIX2 | 32.287 | 3 | 50.97041 | no | 9.722E-08 | 758.85205 | 62.09008 |
| EWYAHNPYPSPR | 12 | Q9NPC8 | Homeobox protein SIX2 OS=Homo sapie | SIX2 | 32.287 | 2 | 50.97041 | no | 1.34E-06 | 506.23712 | 62.09008 |
| EWYAHNPYPSPR | 12 | Q9NPC8 | Homeobox protein SIX2 OS=Homo sapie | SIX2 | 32.287 | 2 | 50.97041 | no | 9.722E-08 | 758.85205 | 62.09008 |
| GRPLGAVGK | 9 | Q9NPC8 | Homeobox protein SIX2 OS=Homo sapie | SIX2 | 32.287 | 2 | 50.97041 | no | 2.01E-07 | 427.76398 | 121.44767 |
| AHYIEAEK | 8 | Q9NPC8 | Homeobox protein SIX2 OS=Homo sapie | SIX2 | 32.287 | 2 | 50.97041 | yes | 2.283E-07 | 480.74289 | 115.06615 |
| SIWDGEETSYCFK | 13 | Q9NPC8 | Homeobox protein SIX2 OS=Homo sapie | SIX2 | 32.287 | 2 | 50.97041 | yes | 2.841E-07 | 811.34558 | 56.925117 |
| SCEMGLQLR | 9 | Q9NPD3 | Exosome complex component RRP41 OS | EXOSC4 | 26.383 | 2 | 47.274372 | yes | 1.408E-07 | 547.26007 | 569.05255 |
| VLEAAAQAAR | 10 | Q9NPD3 | Exosome complex component RRP41 OS | EXOSC4 | 26.383 | 2 | 47.274372 | yes | 2.271E-07 | 500.28271 | 520.6521 |
| DVHTLLDR | 8 | Q9NPD3 | Exosome complex component RRP41 OS | EXOSC4 | 26.383 | 2 | 47.274372 | yes | 1.364E-07 | 484.76163 | 299.80997 |
| ALAVVYGPHEIR | 12 | Q9NPD3 | Exosome complex component RRP41 OS | EXOSC4 | 26.383 | 3 | 47.274372 | yes | 1.854E-07 | 442.25058 | 44.181393 |
| ALPDRALVNCQYS | 22 | Q9NPD3 | Exosome complex component RRP41 OS | EXOSC4 | 26.383 | 3 | 47.274372 | yes | 2.603E-06 | 815.06055 | 28.054098 |
| TLTWEAMEQIR | 11 | Q9NPE2 | Neugrin OS=Homo sapiens OX=9606 GN | NGRN | 32.408 | 2 | 52.286617 | yes | 9.968E-08 | 689.34521 | 1768.5286 |
| LAEGFDVSTDVIR | 13 | Q9NPE2 | Neugrin OS=Homo sapiens OX=9606 GN | NGRN | 32.408 | 2 | 52.286617 | yes | 1.079E-07 | 711.36719 | 1371.9265 |
| EIQDLEESFVPVAA | 20 | Q9NPE2 | Neugrin OS=Homo sapiens OX=9606 GN | NGRN | 32.408 | 3 | 52.286617 | yes | 1.552E-07 | 735.38306 | 1080.157 |
| FLPTLEQK | 8 | Q9NPE2 | Neugrin OS=Homo sapiens OX=9606 GN | NGRN | 32.408 | 2 | 52.286617 | yes | 3.234E-07 | 488.27911 | 1046.498 |
| ELQEVESTLK | 10 | Q9NPE2 | Neugrin OS=Homo sapiens OX=9606 GN | NGRN | 32.408 | 2 | 52.286617 | yes | 1.058E-07 | 588.31134 | 1023.946 |
| GTGSGALPSGQK | 12 | Q9NPE2 | Neugrin OS=Homo sapiens OX=9606 GN | NGRN | 32.408 | 2 | 52.286617 | yes | 9.733E-08 | 530.27509 | 581.46613 |
| LLPAGHSVSGSLLI | 22 | Q9NPE2 | Neugrin OS=Homo sapiens OX=9606 GN | NGRN | 32.408 | 3 | 52.286617 | yes | 1.708E-06 | 725.7124 | 563.96887 |

| | | | | | | | | | | | |
|---|---|---|---|---|---|---|---|---|---|---|---|
| YSSDSESPR | 9 | Q9NPE2 | Neugrin OS=Homo sapiens OX=9606 GN | NGRN | 32.408 | 2 | 52.286617 | yes | 1.251E-07 | 514.21997 | 389.19189 |
| EFFDSNGNFLYR | 12 | Q9NPE2 | Neugrin OS=Homo sapiens OX=9606 GN | NGRN | 32.408 | 2 | 52.286617 | yes | 9.722E-08 | 754.84387 | 333.08182 |
| QMEAPGAPPR | 10 | Q9NPE2 | Neugrin OS=Homo sapiens OX=9606 GN | NGRN | 32.408 | 2 | 52.286617 | yes | 1.133E-07 | 527.26093 | 198.85144 |
| EPDPDSDWEPEER | 13 | Q9NPE2 | Neugrin OS=Homo sapiens OX=9606 GN | NGRN | 32.408 | 2 | 52.286617 | yes | 9.819E-08 | 800.82355 | 188.10745 |
| MFLQYYLNEQGDI | 13 | Q9NPE3 | H/ACA ribonucleoprotein complex subun | NOP10 | 7.706 | 2 | 49.82625 | yes | 1.002E-07 | 838.8905 | 1136.4362 |
| VLMTQQPR | 8 | Q9NPE3 | H/ACA ribonucleoprotein complex subun | NOP10 | 7.706 | 2 | 49.82625 | yes | 1.809E-07 | 486.7684 | 108.27686 |
| KFDPMGQQTCSAI | 16 | Q9NPE3 | H/ACA ribonucleoprotein complex subun | NOP10 | 7.706 | 3 | 49.82625 | yes | 2.215E-07 | 610.95062 | 61.116634 |
| LRPLGLLVAMK | 11 | Q9NPF0 | CD320 antigen OS=Homo sapiens OX=9 | CD320 | 28.992 | 2 | 20.401146 | yes | 4.478E-06 | 605.88885 | 16.636059 |
| DAPPLLPSDTGQG | 15 | Q9NPF5 | DNA methyltransferase 1-associated prot | DMAP1 | 52.993 | 2 | 51.927494 | yes | 1.699E-07 | 793.8941 | 657.04413 |
| LITAADTTAEQR | 12 | Q9NPF5 | DNA methyltransferase 1-associated prot | DMAP1 | 52.993 | 2 | 51.927494 | yes | 1.079E-07 | 645.33844 | 500.13773 |
| IPVFDAGHER | 10 | Q9NPF5 | DNA methyltransferase 1-associated prot | DMAP1 | 52.993 | 2 | 51.927494 | yes | 9.722E-08 | 570.79346 | 490.13467 |
| FVVIHDR | 7 | Q9NPF5 | DNA methyltransferase 1-associated prot | DMAP1 | 52.993 | 2 | 51.927494 | yes | 1.079E-07 | 443.2507 | 258.04343 |
| EAEKPAVPETAGII | 14 | Q9NPF5 | DNA methyltransferase 1-associated prot | DMAP1 | 52.993 | 2 | 51.927494 | yes | 1.354E-07 | 720.39069 | 143.41454 |
| AETDHLFDLSR | 11 | Q9NPF5 | DNA methyltransferase 1-associated prot | DMAP1 | 52.993 | 2 | 51.927494 | yes | 5.796E-07 | 652.3175 | 138.19307 |
| MKLPSSVGQK | 10 | Q9NPF5 | DNA methyltransferase 1-associated prot | DMAP1 | 52.993 | 2 | 51.927494 | yes | 2.367E-06 | 537.80243 | 77.854843 |
| EVYALLYSDK | 10 | Q9NPF5 | DNA methyltransferase 1-associated prot | DMAP1 | 52.993 | 2 | 51.927494 | yes | 1.13E-07 | 600.81097 | 66.011337 |
| DGAMFFHWR | 9 | Q9NPF5 | DNA methyltransferase 1-associated prot | DMAP1 | 52.993 | 2 | 51.927494 | yes | 2.111E-07 | 583.76367 | 47.840729 |
| EQEQLDRIVK | 10 | Q9NPI1 | Bromodomain-containing protein 7 OS=I | BRD7 | 74.139 | 2 | 25.690676 | yes | 3.242E-06 | 629.34351 | 111.59661 |
| EQPAVVGLDSEEM | 15 | Q9NPI6 | mRNA-decapping enzyme 1A OS=Homo | DCP1A | 63.279 | 2 | 53.61563 | yes | 1.736E-07 | 844.89343 | 842.10553 |
| TDIEGTLFVYR | 11 | Q9NPI6 | mRNA-decapping enzyme 1A OS=Homo | DCP1A | 63.279 | 2 | 53.61563 | yes | 9.804E-08 | 657.34045 | 798.98792 |
| ASSPSPLTIGTPESC | 16 | Q9NPI6 | mRNA-decapping enzyme 1A OS=Homo | DCP1A | 63.279 | 2 | 53.61563 | yes | 1.006E-07 | 814.41791 | 582.21326 |
| AGQEMSLAALK | 11 | Q9NPI6 | mRNA-decapping enzyme 1A OS=Homo | DCP1A | 63.279 | 2 | 53.61563 | yes | 9.915E-08 | 559.79736 | 438.10941 |
| LMADVVEEETR | 11 | Q9NPI6 | mRNA-decapping enzyme 1A OS=Homo | DCP1A | 63.279 | 2 | 53.61563 | yes | 2.388E-07 | 646.31354 | 261.07089 |
| HLTVEELFGTSLPK | 14 | Q9NPI6 | mRNA-decapping enzyme 1A OS=Homo | DCP1A | 63.279 | 3 | 53.61563 | yes | 1.373E-07 | 524.28735 | 126.82397 |
| HLTVEELFGTSLPK | 14 | Q9NPI6 | mRNA-decapping enzyme 1A OS=Homo | DCP1A | 63.279 | 3 | 53.61563 | yes | 4.006E-07 | 785.92743 | 126.82397 |
| HLTVEELFGTSLPK | 14 | Q9NPI6 | mRNA-decapping enzyme 1A OS=Homo | DCP1A | 63.279 | 2 | 53.61563 | yes | 1.373E-07 | 524.28735 | 126.82397 |
| HLTVEELFGTSLPK | 14 | Q9NPI6 | mRNA-decapping enzyme 1A OS=Homo | DCP1A | 63.279 | 2 | 53.61563 | yes | 4.006E-07 | 785.92743 | 126.82397 |
| LNMHNLVEPVNK | 12 | Q9NPI6 | mRNA-decapping enzyme 1A OS=Homo | DCP1A | 63.279 | 2 | 53.61563 | yes | 9.722E-08 | 704.37427 | 252.34326 |
| KPSIILSK | 8 | Q9NPI6 | mRNA-decapping enzyme 1A OS=Homo | DCP1A | 63.279 | 2 | 53.61563 | yes | 1.391E-07 | 443.29202 | 104.71747 |
| SQLQDTLIHLIK | 12 | Q9NPI6 | mRNA-decapping enzyme 1A OS=Homo | DCP1A | 63.279 | 2 | 53.61563 | yes | 1.375E-07 | 704.91156 | 19.10692 |
| EAEQILATAVYQA | 14 | Q9NPJ6 | Mediator of RNA polymerase II transcript | MED4 | 29.746 | 2 | 56.927185 | yes | 9.722E-08 | 767.90924 | 139.60812 |
| ELIEMLAISR | 10 | Q9NPJ6 | Mediator of RNA polymerase II transcript | MED4 | 29.746 | 2 | 56.927185 | yes | 1.512E-07 | 587.82867 | 84.330246 |
| RPYPTDLEMR | 10 | Q9NPJ6 | Mediator of RNA polymerase II transcript | MED4 | 29.746 | 2 | 56.927185 | yes | 1.016E-06 | 639.31897 | 46.308369 |
| IHHEMQVLEK | 10 | Q9NPJ6 | Mediator of RNA polymerase II transcript | MED4 | 29.746 | 2 | 56.927185 | yes | 1.418E-07 | 632.32935 | 45.938023 |
| KIEALLNLPR | 10 | Q9NPL8 | Complex I assembly factor TIMMDC1, m | TIMMDC1 | 32.178 | 2 | 50.529728 | yes | 9.788E-08 | 583.86639 | 122.52251 |
| LQVTEHLPEK | 10 | Q9NPL8 | Complex I assembly factor TIMMDC1, m | TIMMDC1 | 32.178 | 2 | 50.529728 | yes | 4.341E-07 | 597.3299 | 78.181915 |
| VHELNEEIGK | 10 | Q9NQ29 | Putative RNA-binding protein Luc7-like | LUC7L | 43.728 | 2 | 53.203705 | no | 1.962E-07 | 584.30383 | 5336.1782 |
| NSMPASSFQQQK | 12 | Q9NQ29 | Putative RNA-binding protein Luc7-like | LUC7L | 43.728 | 2 | 53.203705 | no | 1.498E-07 | 676.81677 | 2375.8835 |
| NSMPASSFQQQK | 12 | Q9NQ29 | Putative RNA-binding protein Luc7-like | LUC7L | 43.728 | 2 | 53.203705 | no | 1.665E-07 | 684.81427 | 2375.8835 |

| | | | | | | | | | | | |
|---|---|---|---|---|---|---|---|---|---|---|---|
| LADHFGGK | 8 | Q9NQ29 | Putative RNA-binding protein Luc7-like | LUC7L | 43.728 | 2 | 53.203705 | no | 9.722E-08 | 422.71924 | 2540.5464 |
| ALLDQLMGTAR | 11 | Q9NQ29 | Putative RNA-binding protein Luc7-like | LUC7L | 43.728 | 2 | 53.203705 | yes | 1.555E-07 | 594.82391 | 1810.0496 |
| IHDLALR | 7 | Q9NQ29 | Putative RNA-binding protein Luc7-like | LUC7L | 43.728 | 2 | 53.203705 | yes | 9.722E-08 | 419.2507 | 1770.3427 |
| LHLGFIQIR | 9 | Q9NQ29 | Putative RNA-binding protein Luc7-like | LUC7L | 43.728 | 2 | 53.203705 | yes | 1.107E-07 | 548.8349 | 1550.6432 |
| VCEVCSAYLGLHI | 16 | Q9NQ29 | Putative RNA-binding protein Luc7-like | LUC7L | 43.728 | 3 | 53.203705 | no | 1.23E-07 | 636.61731 | 1461.9503 |
| EKLDQLR | 7 | Q9NQ29 | Putative RNA-binding protein Luc7-like | LUC7L | 43.728 | 2 | 53.203705 | yes | 1.297E-07 | 451.25873 | 697.01184 |
| AEQLGAEGNVDES | 15 | Q9NQ29 | Putative RNA-binding protein Luc7-like | LUC7L | 43.728 | 2 | 53.203705 | yes | 1.105E-07 | 787.86829 | 563.14972 |
| VCEVCSAYLGLHI | 17 | Q9NQ29 | Putative RNA-binding protein Luc7-like | LUC7L | 43.728 | 3 | 53.203705 | no | 1.663E-07 | 688.65106 | 480.68277 |
| MDLGECTK | 8 | Q9NQ29 | Putative RNA-binding protein Luc7-like | LUC7L | 43.728 | 2 | 53.203705 | yes | 1.373E-07 | 477.20697 | 422.32532 |
| LAETQEEISAEVSA | 15 | Q9NQ29 | Putative RNA-binding protein Luc7-like | LUC7L | 43.728 | 2 | 53.203705 | yes | 9.722E-08 | 802.9043 | 422.0798 |
| SAQAQMR | 7 | Q9NQ29 | Putative RNA-binding protein Luc7-like | LUC7L | 43.728 | 2 | 53.203705 | no | 9.722E-08 | 417.20035 | 71.150841 |
| LPGGPTLTFQVK | 12 | Q9NQ55 | Suppressor of SWI4 1 homolog OS=Hom | PPAN | 53.194 | 2 | 48.155056 | yes | 1.283E-07 | 629.36371 | 565.258 |
| TEEELQAILEAK | 12 | Q9NQ55 | Suppressor of SWI4 1 homolog OS=Hom | PPAN | 53.194 | 2 | 48.155056 | yes | 1.191E-07 | 687.36157 | 256.95441 |
| VMFHSFVSK | 9 | Q9NQ55 | Suppressor of SWI4 1 homolog OS=Hom | PPAN | 53.194 | 2 | 48.155056 | yes | 1.402E-07 | 541.27856 | 191.08316 |
| VMEPLTASR | 9 | Q9NQ55 | Suppressor of SWI4 1 homolog OS=Hom | PPAN | 53.194 | 2 | 48.155056 | yes | 2.218E-07 | 502.26569 | 127.0629 |
| TETNVYFK | 8 | Q9NQ55 | Suppressor of SWI4 1 homolog OS=Hom | PPAN | 53.194 | 2 | 48.155056 | yes | 5.806E-07 | 501.25055 | 115.10151 |
| NLEAYAANPHSFV | 16 | Q9NQ55 | Suppressor of SWI4 1 homolog OS=Hom | PPAN | 53.194 | 3 | 48.155056 | yes | 1.483E-07 | 612.97241 | 51.16153 |
| VGGSDEEASGIPSI | 14 | Q9NQ55 | Suppressor of SWI4 1 homolog OS=Hom | PPAN | 53.194 | 2 | 48.155056 | yes | 1.715E-07 | 680.82062 | 48.523727 |
| VNLNTIK | 7 | Q9NQ55 | Suppressor of SWI4 1 homolog OS=Hom | PPAN | 53.194 | 2 | 48.155056 | yes | 2.2E-07 | 401.24509 | 18.145164 |
| SVYGGEFIQQLK | 12 | Q9NQG5 | Regulation of nuclear pre-mRNA domain | RPRD1B | 36.9 | 2 | 46.80878 | yes | 1.465E-07 | 684.86151 | 47.607819 |
| KGPEFTR | 7 | Q9NQG5 | Regulation of nuclear pre-mRNA domain | RPRD1B | 36.9 | 2 | 46.80878 | yes | 3.028E-06 | 417.72705 | 28.930952 |
| ALQDLENAASGDA | 16 | Q9NQG5 | Regulation of nuclear pre-mRNA domain | RPRD1B | 36.9 | 2 | 46.80878 | yes | 2.608E-07 | 815.90521 | 22.186272 |
| AAAEDVNVTFEDO | 15 | Q9NQP4 | Prefoldin subunit 4 OS=Homo sapiens O | PFDN4 | 15.315 | 2 | 48.03604 | yes | 1.21E-07 | 832.89172 | 20.151911 |
| TFLHTVQR | 8 | Q9NQS7 | Inner centromere protein OS=Homo sapie | INCENP | 105.429 | 2 | 52.352665 | yes | 2.402E-07 | 501.27997 | 207.60573 |
| KPIPTWAR | 8 | Q9NQS7 | Inner centromere protein OS=Homo sapie | INCENP | 105.429 | 2 | 52.352665 | yes | 9.988E-07 | 484.78745 | 130.87756 |
| IAQVSPGPR | 9 | Q9NQS7 | Inner centromere protein OS=Homo sapie | INCENP | 105.429 | 2 | 52.352665 | yes | 1.042E-07 | 462.76672 | 108.63815 |
| NQMLMTPTSAPR | 12 | Q9NQS7 | Inner centromere protein OS=Homo sapie | INCENP | 105.429 | 2 | 52.352665 | yes | 1.199E-07 | 673.83142 | 75.82119 |
| LMEFLCNMDNK | 11 | Q9NQS7 | Inner centromere protein OS=Homo sapie | INCENP | 105.429 | 2 | 52.352665 | yes | 9.722E-08 | 707.81183 | 74.568619 |
| TSSAVWNSPPLQG | 15 | Q9NQS7 | Inner centromere protein OS=Homo sapie | INCENP | 105.429 | 2 | 52.352665 | yes | 1.812E-07 | 785.90228 | 64.788124 |
| WLQQEEEER | 9 | Q9NQS7 | Inner centromere protein OS=Homo sapie | INCENP | 105.429 | 2 | 52.352665 | yes | 4.815E-07 | 623.78857 | 51.480354 |
| LLNVEVPQK | 9 | Q9NQS7 | Inner centromere protein OS=Homo sapie | INCENP | 105.429 | 2 | 52.352665 | yes | 5.187E-05 | 520.31097 | 35.48679 |
| QNAEQHVTQLMS | 18 | Q9NQS7 | Inner centromere protein OS=Homo sapie | INCENP | 105.429 | 3 | 52.352665 | yes | 3.096E-06 | 693.68109 | 22.138077 |
| ATLEVILRPK | 10 | Q9NQT4 | Exosome complex component RRP46 OS | EXOSC5 | 25.249 | 2 | 52.228832 | yes | 1.167E-07 | 570.36096 | 857.62347 |
| AVLTFALDSVER | 12 | Q9NQT4 | Exosome complex component RRP46 OS | EXOSC5 | 25.249 | 2 | 52.228832 | yes | 9.722E-08 | 660.86151 | 355.48505 |
| IGLPGVAEK | 9 | Q9NQT4 | Exosome complex component RRP46 OS | EXOSC5 | 25.249 | 2 | 52.228832 | yes | 1.327E-07 | 442.26602 | 203.08337 |
| NTCEAVVLGTLHP | 14 | Q9NQT4 | Exosome complex component RRP46 OS | EXOSC5 | 25.249 | 2 | 52.228832 | yes | 9.862E-08 | 783.90649 | 71.497665 |
| NTCEAVVLGTLHP | 14 | Q9NQT4 | Exosome complex component RRP46 OS | EXOSC5 | 25.249 | 2 | 52.228832 | yes | 8.759E-05 | 522.94006 | 71.497665 |
| NTCEAVVLGTLHP | 14 | Q9NQT4 | Exosome complex component RRP46 OS | EXOSC5 | 25.249 | 3 | 52.228832 | yes | 9.862E-08 | 783.90649 | 71.497665 |
| NTCEAVVLGTLHP | 14 | Q9NQT4 | Exosome complex component RRP46 OS | EXOSC5 | 25.249 | 3 | 52.228832 | yes | 8.759E-05 | 522.94006 | 71.497665 |

| | | | | | | | | | | | |
|---|---|---|---|---|---|---|---|---|---|---|---|
| AEPASVAAESLAG | 15 | Q9NQT5 | Exosome complex component RRP40 OS | EXOSC3 | 29.573 | 2 | 48.99757 | yes | 1.159E-07 | 729.36517 | 770.72302 |
| NRPNVQVGDLIYC | 20 | Q9NQT5 | Exosome complex component RRP40 OS | EXOSC3 | 29.573 | 3 | 48.99757 | yes | 1.662E-07 | 744.40259 | 192.59512 |
| LLAPDCEIIQEVGK | 14 | Q9NQT5 | Exosome complex component RRP40 OS | EXOSC3 | 29.573 | 2 | 48.99757 | yes | 2.229E-07 | 792.9187 | 144.4373 |
| GDHVIGIVTAK | 11 | Q9NQT5 | Exosome complex component RRP40 OS | EXOSC3 | 29.573 | 2 | 48.99757 | yes | 1.079E-07 | 555.31934 | 119.00835 |
| ANGMGVIGQDGLI | 15 | Q9NQT5 | Exosome complex component RRP40 OS | EXOSC3 | 29.573 | 2 | 48.99757 | yes | 5.736E-06 | 760.40051 | 30.911779 |
| VILNPVNTNLSK | 12 | Q9NQT8 | Kinesin-like protein KIF13B OS=Homo s | KIF13B | 202.789 | 2 | 49.929443 | yes | 1.527E-07 | 656.38519 | 54.407661 |
| QGFAQSLLK | 9 | Q9NQT8 | Kinesin-like protein KIF13B OS=Homo s | KIF13B | 202.789 | 2 | 49.929443 | yes | 2.037E-06 | 496.2822 | 50.244213 |
| LIQEMTVTWEEK | 12 | Q9NQT8 | Kinesin-like protein KIF13B OS=Homo s | KIF13B | 202.789 | 2 | 49.929443 | yes | 4.327E-07 | 753.87909 | 45.048996 |
| CNPGYGLLVR | 10 | Q9NQT8 | Kinesin-like protein KIF13B OS=Homo s | KIF13B | 202.789 | 2 | 49.929443 | yes | 5.34E-07 | 574.79767 | 42.148376 |
| ECEEDNPVIR | 10 | Q9NQT8 | Kinesin-like protein KIF13B OS=Homo s | KIF13B | 202.789 | 2 | 49.929443 | yes | 1.207E-06 | 630.77991 | 38.843384 |
| DVPTGGIFQLR | 11 | Q9NQT8 | Kinesin-like protein KIF13B OS=Homo s | KIF13B | 202.789 | 2 | 49.929443 | yes | 9.54E-07 | 601.83002 | 34.451305 |
| LQYAVPIINQK | 11 | Q9NQT8 | Kinesin-like protein KIF13B OS=Homo s | KIF13B | 202.789 | 2 | 49.929443 | yes | 2.269E-06 | 643.87701 | 30.159494 |
| GSLLSEPAIQVR | 12 | Q9NQT8 | Kinesin-like protein KIF13B OS=Homo s | KIF13B | 202.789 | 2 | 49.929443 | yes | 9.975E-08 | 635.36169 | 26.678394 |
| QLESLGISLQSSGII | 15 | Q9NQT8 | Kinesin-like protein KIF13B OS=Homo s | KIF13B | 202.789 | 2 | 49.929443 | yes | 4.805E-07 | 780.43561 | 15.150396 |
| ITLFLEEDK | 9 | Q9NQW6 | Anillin OS=Homo sapiens OX=9606 GN | ANLN | 124.199 | 2 | 51.92133 | yes | 1.043E-07 | 554.30023 | 229.36896 |
| EICLQSQSK | 9 | Q9NQW6 | Anillin OS=Homo sapiens OX=9606 GN | ANLN | 124.199 | 2 | 51.92133 | yes | 1.611E-07 | 546.7713 | 227.44478 |
| INLANCTSR | 9 | Q9NQW6 | Anillin OS=Homo sapiens OX=9606 GN | ANLN | 124.199 | 2 | 51.92133 | yes | 5.691E-07 | 524.76385 | 223.51625 |
| SCEGQNPELLPK | 12 | Q9NQW6 | Anillin OS=Homo sapiens OX=9606 GN | ANLN | 124.199 | 2 | 51.92133 | yes | 5.063E-07 | 686.33228 | 213.11615 |
| CQVNSSVEER | 10 | Q9NQW6 | Anillin OS=Homo sapiens OX=9606 GN | ANLN | 124.199 | 2 | 51.92133 | yes | 1.02E-07 | 604.27222 | 212.68398 |
| LEATAASSVK | 10 | Q9NQW6 | Anillin OS=Homo sapiens OX=9606 GN | ANLN | 124.199 | 2 | 51.92133 | yes | 9.722E-08 | 488.7691 | 169.54553 |
| TLLIDELNK | 9 | Q9NQW6 | Anillin OS=Homo sapiens OX=9606 GN | ANLN | 124.199 | 2 | 51.92133 | yes | 2.437E-07 | 529.80823 | 142.7533 |
| STTPGGTGIKPFLE | 15 | Q9NQW6 | Anillin OS=Homo sapiens OX=9606 GN | ANLN | 124.199 | 2 | 51.92133 | yes | 9.793E-08 | 780.92267 | 113.50011 |
| TGVSKPIVK | 9 | Q9NQW6 | Anillin OS=Homo sapiens OX=9606 GN | ANLN | 124.199 | 2 | 51.92133 | yes | 7.268E-07 | 464.79492 | 98.590309 |
| GSVTLSEIR | 9 | Q9NQW6 | Anillin OS=Homo sapiens OX=9606 GN | ANLN | 124.199 | 2 | 51.92133 | yes | 1.828E-07 | 481.26929 | 94.203636 |
| STLSQTVPSK | 10 | Q9NQW6 | Anillin OS=Homo sapiens OX=9606 GN | ANLN | 124.199 | 2 | 51.92133 | yes | 5.567E-07 | 524.28766 | 92.350143 |
| LWQPDACYKPIGK | 14 | Q9NQW6 | Anillin OS=Homo sapiens OX=9606 GN | ANLN | 124.199 | 2 | 51.92133 | yes | 1.007E-07 | 836.92944 | 87.943291 |
| SNIHSSVMASPGG | 18 | Q9NQW6 | Anillin OS=Homo sapiens OX=9606 GN | ANLN | 124.199 | 3 | 51.92133 | yes | 2.517E-07 | 590.63696 | 86.846733 |
| QNSVQEQPGTACL | 15 | Q9NQW6 | Anillin OS=Homo sapiens OX=9606 GN | ANLN | 124.199 | 2 | 51.92133 | yes | 1.668E-07 | 823.8938 | 73.222885 |
| ASPQSEFMPSK | 11 | Q9NQW6 | Anillin OS=Homo sapiens OX=9606 GN | ANLN | 124.199 | 2 | 51.92133 | yes | 3.609E-07 | 604.78442 | 57.181976 |
| VTENQIPAK | 9 | Q9NQW6 | Anillin OS=Homo sapiens OX=9606 GN | ANLN | 124.199 | 2 | 51.92133 | yes | 7.968E-07 | 500.2771 | 46.040352 |
| EIEMSVDDDDINS | 15 | Q9NQW6 | Anillin OS=Homo sapiens OX=9606 GN | ANLN | 124.199 | 2 | 51.92133 | yes | 3.298E-07 | 848.86456 | 42.762291 |
| NTFELITVR | 9 | Q9NQW6 | Anillin OS=Homo sapiens OX=9606 GN | ANLN | 124.199 | 2 | 51.92133 | yes | 2.521E-07 | 546.80603 | 36.244015 |
| QEATFCSQR | 9 | Q9NQW6 | Anillin OS=Homo sapiens OX=9606 GN | ANLN | 124.199 | 2 | 51.92133 | yes | 2.125E-07 | 563.75092 | 28.494045 |
| VPFLSSLEGHIYLK | 14 | Q9NQW6 | Anillin OS=Homo sapiens OX=9606 GN | ANLN | 124.199 | 2 | 51.92133 | yes | 2.325E-07 | 801.94812 | 19.561255 |
| PVPQVDEAETR | 11 | Q9NQZ2 | Something about silencing protein 10 OS | UTP3 | 54.558 | 2 | 51.18912 | yes | 2.72E-07 | 620.81201 | 368.51279 |
| YSGELSGIR | 9 | Q9NQZ2 | Something about silencing protein 10 OS | UTP3 | 54.558 | 2 | 51.18912 | yes | 4.474E-07 | 491.25363 | 114.08785 |
| LYYDTDYGSK | 10 | Q9NQZ2 | Something about silencing protein 10 OS | UTP3 | 54.558 | 2 | 51.18912 | yes | 8.495E-07 | 612.7746 | 97.917885 |
| VPAHGHPVIER | 11 | Q9NQZ2 | Something about silencing protein 10 OS | UTP3 | 54.558 | 3 | 51.18912 | yes | 9.722E-08 | 404.55963 | 38.789394 |
| KLYYDTDYGSK | 11 | Q9NQZ2 | Something about silencing protein 10 OS | UTP3 | 54.558 | 2 | 51.18912 | yes | 5.448E-07 | 676.82208 | 31.01387 |

| | | | | | | | | | | | |
|---|---|---|---|---|---|---|---|---|---|---|---|
| VVLEGPAPWGFR | 12 | Q9NR12 | PDZ and LIM domain protein 7 OS=Hom | PDLIM7 | 49.845 | 2 | 50.63397 | yes | 2.286E-07 | 664.36151 | 1572.2777 |
| DFNVPLSISR | 10 | Q9NR12 | PDZ and LIM domain protein 7 OS=Hom | PDLIM7 | 49.845 | 2 | 50.63397 | yes | 1.345E-07 | 574.30896 | 461.20459 |
| YTFAPSVSLNK | 11 | Q9NR12 | PDZ and LIM domain protein 7 OS=Hom | PDLIM7 | 49.845 | 2 | 50.63397 | yes | 9.733E-08 | 613.8244 | 450.50583 |
| ITGEIMHALK | 10 | Q9NR12 | PDZ and LIM domain protein 7 OS=Hom | PDLIM7 | 49.845 | 2 | 50.63397 | yes | 1.182E-07 | 556.81024 | 440.24722 |
| AFYMEEGVPYCEI | 13 | Q9NR12 | PDZ and LIM domain protein 7 OS=Hom | PDLIM7 | 49.845 | 2 | 50.63397 | yes | 1.145E-07 | 825.84998 | 409.92648 |
| VLEEGGFFEEK | 11 | Q9NR12 | PDZ and LIM domain protein 7 OS=Hom | PDLIM7 | 49.845 | 2 | 50.63397 | yes | 1.163E-07 | 642.31134 | 362.04449 |
| GAIFCPPCYDVR | 12 | Q9NR12 | PDZ and LIM domain protein 7 OS=Hom | PDLIM7 | 49.845 | 2 | 50.63397 | yes | 1.647E-07 | 727.83142 | 248.70732 |
| ASAPAADPPR | 10 | Q9NR12 | PDZ and LIM domain protein 7 OS=Hom | PDLIM7 | 49.845 | 2 | 50.63397 | yes | 2.722E-07 | 476.74597 | 192.13136 |
| HSQPATPTPLQSR | 13 | Q9NR12 | PDZ and LIM domain protein 7 OS=Hom | PDLIM7 | 49.845 | 2 | 50.63397 | yes | 1.072E-06 | 710.37061 | 130.1274 |
| LMENTEDWRPR | 11 | Q9NR12 | PDZ and LIM domain protein 7 OS=Hom | PDLIM7 | 49.845 | 2 | 50.63397 | yes | 3.057E-06 | 723.84332 | 53.222542 |
| ILAHLTGTEFMQD | 20 | Q9NR12 | PDZ and LIM domain protein 7 OS=Hom | PDLIM7 | 49.845 | 3 | 50.63397 | yes | 9.171E-07 | 775.37909 | 34.638573 |
| IGVPSATEIIK | 11 | Q9NR30 | Nucleolar RNA helicase 2 OS=Homo sap | DDX21 | 87.345 | 2 | 56.469852 | yes | 1.05E-07 | 564.33716 | 4048.0984 |
| EEYQLVQVEQK | 11 | Q9NR30 | Nucleolar RNA helicase 2 OS=Homo sap | DDX21 | 87.345 | 2 | 56.469852 | yes | 9.722E-08 | 696.85388 | 3341.1799 |
| TAITVEHLAIK | 11 | Q9NR30 | Nucleolar RNA helicase 2 OS=Homo sap | DDX21 | 87.345 | 2 | 56.469852 | yes | 9.722E-08 | 598.3559 | 2556.2734 |
| EGAFSNFPISEETII | 15 | Q9NR30 | Nucleolar RNA helicase 2 OS=Homo sap | DDX21 | 87.345 | 2 | 56.469852 | no | 9.722E-08 | 834.90942 | 2507.3804 |
| NGIDILVGTPGR | 12 | Q9NR30 | Nucleolar RNA helicase 2 OS=Homo sap | DDX21 | 87.345 | 2 | 56.469852 | no | 1.004E-07 | 606.34076 | 2332.583 |
| DVESYIHR | 8 | Q9NR30 | Nucleolar RNA helicase 2 OS=Homo sap | DDX21 | 87.345 | 2 | 56.469852 | yes | 2.834E-07 | 509.75125 | 2149.7097 |
| EQLGEEIDSK | 10 | Q9NR30 | Nucleolar RNA helicase 2 OS=Homo sap | DDX21 | 87.345 | 2 | 56.469852 | yes | 1.357E-07 | 574.27753 | 2108.8701 |
| SDAGLESDTAMK | 12 | Q9NR30 | Nucleolar RNA helicase 2 OS=Homo sap | DDX21 | 87.345 | 2 | 56.469852 | yes | 9.722E-08 | 612.77429 | 2000.2444 |
| APQVLVLAPTR | 11 | Q9NR30 | Nucleolar RNA helicase 2 OS=Homo sap | DDX21 | 87.345 | 2 | 56.469852 | yes | 9.723E-08 | 582.85858 | 1903.5715 |
| STYEQVDLIGK | 11 | Q9NR30 | Nucleolar RNA helicase 2 OS=Homo sap | DDX21 | 87.345 | 2 | 56.469852 | yes | 9.977E-08 | 626.82465 | 1444.5045 |
| LHGELQDR | 8 | Q9NR30 | Nucleolar RNA helicase 2 OS=Homo sap | DDX21 | 87.345 | 2 | 56.469852 | yes | 1.396E-07 | 484.25143 | 1414.1263 |
| NGSFGVLVATNV/ | 15 | Q9NR30 | Nucleolar RNA helicase 2 OS=Homo sap | DDX21 | 87.345 | 2 | 56.469852 | yes | 9.915E-08 | 738.40192 | 1323.6854 |
| EAQELSQNSAIK | 12 | Q9NR30 | Nucleolar RNA helicase 2 OS=Homo sap | DDX21 | 87.345 | 2 | 56.469852 | yes | 9.722E-08 | 659.33588 | 1159.8678 |
| GVTFLFPIQAK | 11 | Q9NR30 | Nucleolar RNA helicase 2 OS=Homo sap | DDX21 | 87.345 | 2 | 56.469852 | yes | 9.935E-08 | 610.85553 | 851.10706 |
| QDAQSLHGDIPQK | 13 | Q9NR30 | Nucleolar RNA helicase 2 OS=Homo sap | DDX21 | 87.345 | 2 | 56.469852 | yes | 9.809E-08 | 718.86005 | 351.07721 |
| QDAQSLHGDIPQK | 13 | Q9NR30 | Nucleolar RNA helicase 2 OS=Homo sap | DDX21 | 87.345 | 2 | 56.469852 | yes | 9.722E-08 | 479.57581 | 351.07721 |
| QDAQSLHGDIPQK | 13 | Q9NR30 | Nucleolar RNA helicase 2 OS=Homo sap | DDX21 | 87.345 | 3 | 56.469852 | yes | 9.809E-08 | 718.86005 | 351.07721 |
| QDAQSLHGDIPQK | 13 | Q9NR30 | Nucleolar RNA helicase 2 OS=Homo sap | DDX21 | 87.345 | 3 | 56.469852 | yes | 9.722E-08 | 479.57581 | 351.07721 |
| AEPSEVDMNSPK | 12 | Q9NR30 | Nucleolar RNA helicase 2 OS=Homo sap | DDX21 | 87.345 | 2 | 56.469852 | yes | 2.07E-07 | 652.29535 | 702.11108 |
| TGVCICFYQHK | 11 | Q9NR30 | Nucleolar RNA helicase 2 OS=Homo sap | DDX21 | 87.345 | 2 | 56.469852 | yes | 9.722E-08 | 706.82611 | 572.7157 |
| SDKTEEIAEEEETV | 17 | Q9NR30 | Nucleolar RNA helicase 2 OS=Homo sap | DDX21 | 87.345 | 3 | 56.469852 | yes | 1.025E-07 | 660.98102 | 329.25226 |
| TEEIAEEEETVFPK | 14 | Q9NR30 | Nucleolar RNA helicase 2 OS=Homo sap | DDX21 | 87.345 | 2 | 56.469852 | yes | 3.065E-07 | 825.89087 | 272.22537 |
| TFHHVYSGK | 9 | Q9NR30 | Nucleolar RNA helicase 2 OS=Homo sap | DDX21 | 87.345 | 2 | 56.469852 | yes | 9.722E-08 | 538.26959 | 215.99124 |
| NEEPSEEEIDAPKP | 15 | Q9NR30 | Nucleolar RNA helicase 2 OS=Homo sap | DDX21 | 87.345 | 3 | 56.469852 | yes | 9.722E-08 | 571.27222 | 179.10464 |
| LLDSVPPTAISHFK | 14 | Q9NR30 | Nucleolar RNA helicase 2 OS=Homo sap | DDX21 | 87.345 | 3 | 56.469852 | yes | 3.236E-07 | 508.95221 | 66.966782 |
| LLDSVPPTAISHFK | 14 | Q9NR30 | Nucleolar RNA helicase 2 OS=Homo sap | DDX21 | 87.345 | 3 | 56.469852 | yes | 2.255E-07 | 762.92468 | 66.966782 |
| LLDSVPPTAISHFK | 14 | Q9NR30 | Nucleolar RNA helicase 2 OS=Homo sap | DDX21 | 87.345 | 2 | 56.469852 | yes | 3.236E-07 | 508.95221 | 66.966782 |
| LLDSVPPTAISHFK | 14 | Q9NR30 | Nucleolar RNA helicase 2 OS=Homo sap | DDX21 | 87.345 | 2 | 56.469852 | yes | 2.255E-07 | 762.92468 | 66.966782 |

| | | | | | | | | | | | |
|---|---|---|---|---|---|---|---|---|---|---|---|
| KAEPSEVDMNSPK | 13 | Q9NR30 | Nucleolar RNA helicase 2 OS=Homo sap | DDX21 | 87.345 | 2 | 56.469852 | yes | 2.263E-07 | 716.34283 | 97.669983 |
| ELANQVSK | 8 | Q9NR30 | Nucleolar RNA helicase 2 OS=Homo sap | DDX21 | 87.345 | 2 | 56.469852 | yes | 4.062E-07 | 444.74289 | 91.722641 |
| NEEPSEEEIDAPK | 13 | Q9NR30 | Nucleolar RNA helicase 2 OS=Homo sap | DDX21 | 87.345 | 2 | 56.469852 | yes | 3.976E-07 | 743.83081 | 84.057518 |
| KEAQELSQNSAIK | 13 | Q9NR30 | Nucleolar RNA helicase 2 OS=Homo sap | DDX21 | 87.345 | 2 | 56.469852 | yes | 2.217E-07 | 723.38336 | 52.039837 |
| RIGVPSATEIIK | 12 | Q9NR30 | Nucleolar RNA helicase 2 OS=Homo sap | DDX21 | 87.345 | 2 | 56.469852 | yes | 7.293E-07 | 642.3877 | 40.945099 |
| ELKEQLGEEIDSK | 13 | Q9NR30 | Nucleolar RNA helicase 2 OS=Homo sap | DDX21 | 87.345 | 2 | 56.469852 | yes | 1.152E-07 | 759.38831 | 30.542149 |
| TFSFAIPLIEK | 11 | Q9NR30 | Nucleolar RNA helicase 2 OS=Homo sap | DDX21 | 87.345 | 2 | 56.469852 | yes | 0.0001076 | 633.36066 | 0.40448 |
| LVFLGLDNAGK | 11 | Q9NR31 | GTP-binding protein SAR1a OS=Homo s | SAR1A | 22.367 | 2 | 43.6599 | no | 2.498E-07 | 573.82953 | 108.05819 |
| PMEVFMCSVLK | 11 | Q9NR31 | GTP-binding protein SAR1a OS=Homo s | SAR1A | 22.367 | 2 | 43.6599 | yes | 2.566E-07 | 670.82422 | 36.213299 |
| VGFEEVIVVTTR | 12 | Q9NR50 | Translation initiation factor eIF-2B subun | EIF2B3 | 50.241 | 2 | 45.581184 | yes | 1.806E-07 | 674.8772 | 141.00664 |
| HLVGVDSLIGPET( | 18 | Q9NR50 | Translation initiation factor eIF-2B subun | EIF2B3 | 50.241 | 3 | 45.581184 | yes | 5.868E-06 | 631.34229 | 38.74469 |
| MTDLTSSIPK | 10 | Q9NR50 | Translation initiation factor eIF-2B subun | EIF2B3 | 50.241 | 2 | 45.581184 | yes | 6.029E-07 | 546.78394 | 20.976145 |
| KGQDSIEPVPGQK | 13 | Q9NR50 | Translation initiation factor eIF-2B subun | EIF2B3 | 50.241 | 2 | 45.581184 | yes | 4.346E-06 | 691.86737 | 17.638577 |
| YFHPPAHLQAK | 11 | Q9NR56 | Muscleblind-like protein 1 OS=Homo sap | MBNL1 | 41.818 | 2 | 52.911827 | no | 9.722E-08 | 654.84601 | 1256.5283 |
| YFHPPAHLQAK | 11 | Q9NR56 | Muscleblind-like protein 1 OS=Homo sap | MBNL1 | 41.818 | 2 | 52.911827 | no | 1.133E-07 | 436.89978 | 1256.5283 |
| YFHPPAHLQAK | 11 | Q9NR56 | Muscleblind-like protein 1 OS=Homo sap | MBNL1 | 41.818 | 3 | 52.911827 | no | 9.722E-08 | 654.84601 | 1256.5283 |
| YFHPPAHLQAK | 11 | Q9NR56 | Muscleblind-like protein 1 OS=Homo sap | MBNL1 | 41.818 | 3 | 52.911827 | no | 1.133E-07 | 436.89978 | 1256.5283 |
| YLHPPPHLK | 9 | Q9NR56 | Muscleblind-like protein 1 OS=Homo sap | MBNL1 | 41.818 | 2 | 52.911827 | no | 7.085E-07 | 551.31384 | 1807.5518 |
| VIACFDSLK | 9 | Q9NR56 | Muscleblind-like protein 1 OS=Homo sap | MBNL1 | 41.818 | 2 | 52.911827 | no | 3.938E-07 | 526.77588 | 1093.3256 |
| WLTLEVCR | 8 | Q9NR56 | Muscleblind-like protein 1 OS=Homo sap | MBNL1 | 41.818 | 2 | 52.911827 | no | 1.643E-07 | 538.78149 | 306.50076 |
| RPLEATFDLGIPQA | 21 | Q9NR56 | Muscleblind-like protein 1 OS=Homo sap | MBNL1 | 41.818 | 3 | 52.911827 | yes | 9.722E-08 | 758.10699 | 131.37585 |
| TDRLEVCR | 8 | Q9NR56 | Muscleblind-like protein 1 OS=Homo sap | MBNL1 | 41.818 | 2 | 52.911827 | yes | 3.383E-07 | 524.76385 | 117.9023 |
| NNLIQQK | 7 | Q9NR56 | Muscleblind-like protein 1 OS=Homo sap | MBNL1 | 41.818 | 2 | 52.911827 | no | 4.747E-07 | 429.24561 | 28.879406 |
| QQWLNCIR | 8 | Q9NR81 | Rho guanine nucleotide exchange factor 3 | ARHGEF3 | 59.783 | 2 | 41.325077 | yes | 6.103E-07 | 559.28223 | 90.689728 |
| LLYLEEGQK | 9 | Q9NR81 | Rho guanine nucleotide exchange factor 3 | ARHGEF3 | 59.783 | 2 | 41.325077 | yes | 3.62E-06 | 546.80042 | 52.583916 |
| DLLLEDLQDGEVR | 13 | Q9NR81 | Rho guanine nucleotide exchange factor 3 | ARHGEF3 | 59.783 | 2 | 41.325077 | yes | 4.687E-07 | 757.88849 | 44.418385 |
| VLCCHGELK | 9 | Q9NR81 | Rho guanine nucleotide exchange factor 3 | ARHGEF3 | 59.783 | 2 | 41.325077 | yes | 3.566E-07 | 558.27045 | 41.375965 |
| LDLSLNR | 7 | Q9NR97 | Toll-like receptor 8 OS=Homo sapiens O: | TLR8 | 119.829 | 2 | 34.51699 | yes | 9.929E-07 | 415.74017 | 788.54034 |
| ELDTDRPSLVR | 11 | Q9NRA8 | Eukaryotic translation initiation factor 4E | EIF4ENIF1 | 108.201 | 2 | 53.553295 | yes | 1.602E-07 | 650.84644 | 188.59439 |
| ELDTDRPSLVR | 11 | Q9NRA8 | Eukaryotic translation initiation factor 4E | EIF4ENIF1 | 108.201 | 2 | 53.553295 | yes | 7.316E-07 | 434.23337 | 188.59439 |
| ELDTDRPSLVR | 11 | Q9NRA8 | Eukaryotic translation initiation factor 4E | EIF4ENIF1 | 108.201 | 3 | 53.553295 | yes | 1.602E-07 | 650.84644 | 188.59439 |
| ELDTDRPSLVR | 11 | Q9NRA8 | Eukaryotic translation initiation factor 4E | EIF4ENIF1 | 108.201 | 3 | 53.553295 | yes | 7.316E-07 | 434.23337 | 188.59439 |
| VISVDELEYR | 10 | Q9NRA8 | Eukaryotic translation initiation factor 4E | EIF4ENIF1 | 108.201 | 2 | 53.553295 | yes | 1.229E-07 | 611.81934 | 377.14993 |
| WHASLYPASGR | 11 | Q9NRA8 | Eukaryotic translation initiation factor 4E | EIF4ENIF1 | 108.201 | 2 | 53.553295 | yes | 9.997E-08 | 622.81219 | 255.00769 |
| VDLKPLLSSLSANI | 14 | Q9NRA8 | Eukaryotic translation initiation factor 4E | EIF4ENIF1 | 108.201 | 2 | 53.553295 | yes | 2.11E-07 | 742.93774 | 119.80482 |
| VDLKPLLSSLSANI | 14 | Q9NRA8 | Eukaryotic translation initiation factor 4E | EIF4ENIF1 | 108.201 | 2 | 53.553295 | yes | 5.824E-07 | 495.62759 | 119.80482 |
| VDLKPLLSSLSANI | 14 | Q9NRA8 | Eukaryotic translation initiation factor 4E | EIF4ENIF1 | 108.201 | 3 | 53.553295 | yes | 2.11E-07 | 742.93774 | 119.80482 |
| VDLKPLLSSLSANI | 14 | Q9NRA8 | Eukaryotic translation initiation factor 4E | EIF4ENIF1 | 108.201 | 3 | 53.553295 | yes | 5.824E-07 | 495.62759 | 119.80482 |
| ASGTLPSQPK | 10 | Q9NRA8 | Eukaryotic translation initiation factor 4E | EIF4ENIF1 | 108.201 | 2 | 53.553295 | yes | 3.543E-06 | 493.26929 | 221.53293 |

| | | | | | | | | | | | |
|---|---|---|---|---|---|---|---|---|---|---|---|
| YDSDGVWDPEK | 11 | Q9NRA8 | Eukaryotic translation initiation factor 4E | EIF4ENIF1 | 108.201 | 2 | 53.553295 | yes | 1.142E-07 | 655.7804 | 156.25739 |
| SMGETESGDAFLD | 15 | Q9NRA8 | Eukaryotic translation initiation factor 4E | EIF4ENIF1 | 108.201 | 2 | 53.553295 | yes | 9.722E-08 | 800.36395 | 118.56075 |
| SVLHPPGSGSHAA | 27 | Q9NRA8 | Eukaryotic translation initiation factor 4E | EIF4ENIF1 | 108.201 | 4 | 53.553295 | yes | 6.124E-07 | 671.0993 | 96.306381 |
| ESSHSGVVLSVEE | 19 | Q9NRA8 | Eukaryotic translation initiation factor 4E | EIF4ENIF1 | 108.201 | 3 | 53.553295 | yes | 1.25E-07 | 652.66888 | 92.682053 |
| SSSLGSTPHEELER | 14 | Q9NRA8 | Eukaryotic translation initiation factor 4E | EIF4ENIF1 | 108.201 | 2 | 53.553295 | yes | 1.655E-07 | 764.86554 | 87.783325 |
| EELLDIK | 7 | Q9NRA8 | Eukaryotic translation initiation factor 4E | EIF4ENIF1 | 108.201 | 2 | 53.553295 | yes | 3.04E-06 | 430.24219 | 54.356339 |
| SFGGGCHVTAAVS | 15 | Q9NRA8 | Eukaryotic translation initiation factor 4E | EIF4ENIF1 | 108.201 | 2 | 53.553295 | yes | 1.823E-07 | 746.85168 | 54.063011 |
| NSTPFMAEHLEET | 21 | Q9NRA8 | Eukaryotic translation initiation factor 4E | EIF4ENIF1 | 108.201 | 3 | 53.553295 | yes | 5.164E-07 | 787.70947 | 29.646545 |
| ELLPSFHEVSVYPI | 14 | Q9NRC1 | Suppressor of tumorigenicity 7 protein OS | ST7 | 67.167 | 2 | 51.114567 | yes | 1.081E-07 | 822.93524 | 147.19336 |
| ELLPSFHEVSVYPI | 14 | Q9NRC1 | Suppressor of tumorigenicity 7 protein OS | ST7 | 67.167 | 2 | 51.114567 | yes | 9.722E-08 | 548.95923 | 147.19336 |
| ELLPSFHEVSVYPI | 14 | Q9NRC1 | Suppressor of tumorigenicity 7 protein OS | ST7 | 67.167 | 3 | 51.114567 | yes | 1.081E-07 | 822.93524 | 147.19336 |
| ELLPSFHEVSVYPI | 14 | Q9NRC1 | Suppressor of tumorigenicity 7 protein OS | ST7 | 67.167 | 3 | 51.114567 | yes | 9.722E-08 | 548.95923 | 147.19336 |
| SATICYTAALLK | 12 | Q9NRC1 | Suppressor of tumorigenicity 7 protein OS | ST7 | 67.167 | 2 | 51.114567 | yes | 2.093E-07 | 656.35248 | 148.19809 |
| YDDISLPK | 8 | Q9NRC1 | Suppressor of tumorigenicity 7 protein OS | ST7 | 67.167 | 2 | 51.114567 | no | 3.274E-07 | 475.74512 | 110.80247 |
| NPLNLFR | 7 | Q9NRC1 | Suppressor of tumorigenicity 7 protein OS | ST7 | 67.167 | 2 | 51.114567 | no | 2.066E-07 | 437.2507 | 99.522408 |
| AVEFNPHVPK | 10 | Q9NRC1 | Suppressor of tumorigenicity 7 protein OS | ST7 | 67.167 | 2 | 51.114567 | no | 3.495E-07 | 569.30621 | 80.054008 |
| YTWVTGR | 7 | Q9NRC1 | Suppressor of tumorigenicity 7 protein OS | ST7 | 67.167 | 2 | 51.114567 | yes | 3.639E-06 | 441.72705 | 42.600647 |
| DTNVLVYIK | 9 | Q9NRC1 | Suppressor of tumorigenicity 7 protein OS | ST7 | 67.167 | 2 | 51.114567 | no | 1.007E-06 | 532.80298 | 23.378798 |
| DTIVHFGER | 9 | Q9NRC8 | NAD-dependent protein deacetylase sirtu | SIRT7 | 44.899 | 2 | 50.887753 | yes | 1.381E-07 | 537.27234 | 357.62912 |
| LLAESADLVTELQ | 15 | Q9NRC8 | NAD-dependent protein deacetylase sirtu | SIRT7 | 44.899 | 2 | 50.887753 | yes | 9.722E-08 | 807.93848 | 298.17886 |
| ADTILCLGSSLK | 12 | Q9NRC8 | NAD-dependent protein deacetylase sirtu | SIRT7 | 44.899 | 2 | 50.887753 | yes | 7.928E-07 | 639.3421 | 96.893288 |
| LLMAELGLEIPAY | 15 | Q9NRC8 | NAD-dependent protein deacetylase sirtu | SIRT7 | 44.899 | 2 | 50.887753 | yes | 3.98E-07 | 838.45801 | 69.266281 |
| GPNGVWTLLQK | 11 | Q9NRC8 | NAD-dependent protein deacetylase sirtu | SIRT7 | 44.899 | 2 | 50.887753 | yes | 1.645E-06 | 606.84039 | 37.590504 |
| LISLPLSR | 8 | Q9NRG0 | Chromatin accessibility complex protein 1 | CHRAC1 | 14.711 | 2 | 42.65784 | yes | 2.93E-07 | 449.78964 | 122.99395 |
| LLSASPVDAAIR | 12 | Q9NRG9 | Aladin OS=Homo sapiens OX=9606 GN= | AAAS | 59.575 | 2 | 50.94048 | yes | 1.166E-07 | 606.85095 | 308.90063 |
| ILATTPSAVFR | 11 | Q9NRG9 | Aladin OS=Homo sapiens OX=9606 GN= | AAAS | 59.575 | 2 | 50.94048 | yes | 9.722E-08 | 588.34277 | 214.53769 |
| FAVALLDDSVR | 11 | Q9NRG9 | Aladin OS=Homo sapiens OX=9606 GN= | AAAS | 59.575 | 2 | 50.94048 | yes | 2.64E-07 | 603.3299 | 78.647491 |
| TASGWALALCR | 11 | Q9NRG9 | Aladin OS=Homo sapiens OX=9606 GN= | AAAS | 59.575 | 2 | 50.94048 | yes | 3.948E-07 | 603.30841 | 69.213173 |
| IQVPDYYDIIK | 11 | Q9NRL2 | Bromodomain adjacent to zinc finger dom | BAZ1A | 178.703 | 2 | 52.239956 | yes | 1.894E-07 | 683.86627 | 275.46478 |
| YFVEETVEVIR | 11 | Q9NRL2 | Bromodomain adjacent to zinc finger dom | BAZ1A | 178.703 | 2 | 52.239956 | yes | 9.722E-08 | 692.36139 | 192.38255 |
| HYDDFFER | 8 | Q9NRL2 | Bromodomain adjacent to zinc finger dom | BAZ1A | 178.703 | 2 | 52.239956 | yes | 2.049E-07 | 564.74091 | 185.61765 |
| IQSAIACTNIFPLGI | 15 | Q9NRL2 | Bromodomain adjacent to zinc finger dom | BAZ1A | 178.703 | 2 | 52.239956 | yes | 1.329E-07 | 830.94562 | 162.95276 |
| LLTLVSTR | 8 | Q9NRL2 | Bromodomain adjacent to zinc finger dom | BAZ1A | 178.703 | 2 | 52.239956 | yes | 1.582E-07 | 451.78711 | 150.62309 |
| APLDASDSGR | 10 | Q9NRL2 | Bromodomain adjacent to zinc finger dom | BAZ1A | 178.703 | 2 | 52.239956 | yes | 9.722E-08 | 494.73834 | 141.9496 |
| IYQGTLGAIK | 10 | Q9NRL2 | Bromodomain adjacent to zinc finger dom | BAZ1A | 178.703 | 2 | 52.239956 | yes | 2.231E-07 | 532.31097 | 100.40115 |
| PGLTYQEALESEK | 13 | Q9NRL2 | Bromodomain adjacent to zinc finger dom | BAZ1A | 178.703 | 2 | 52.239956 | yes | 9.849E-08 | 732.86444 | 98.966431 |
| SLDLDSCTLSEILR | 14 | Q9NRL2 | Bromodomain adjacent to zinc finger dom | BAZ1A | 178.703 | 2 | 52.239956 | yes | 1.819E-07 | 811.40869 | 97.98645 |
| KPIALNIIR | 9 | Q9NRL2 | Bromodomain adjacent to zinc finger dom | BAZ1A | 178.703 | 2 | 52.239956 | yes | 9.722E-08 | 519.34515 | 82.721153 |
| DLTEALDEDADPT | 14 | Q9NRL2 | Bromodomain adjacent to zinc finger dom | BAZ1A | 178.703 | 2 | 52.239956 | yes | 1.543E-07 | 766.85175 | 76.320259 |

| | | | | | | | | | | | |
|---|---|---|---|---|---|---|---|---|---|---|---|
| IHISQEDNVANK | 12 | Q9NRL2 | Bromodomain adjacent to zinc finger don | BAZ1A | 178.703 | 2 | 52.239956 | yes | 1.043E-07 | 684.3493 | 73.074661 |
| SSNAYDPSQMCAI | 14 | Q9NRL2 | Bromodomain adjacent to zinc finger don | BAZ1A | 178.703 | 2 | 52.239956 | yes | 3.558E-06 | 794.32434 | 57.573502 |
| LHILASGADVTSAI | 16 | Q9NRL2 | Bromodomain adjacent to zinc finger don | BAZ1A | 178.703 | 2 | 52.239956 | yes | 1.21E-07 | 784.42554 | 48.211147 |
| HDDSWPFLK | 9 | Q9NRL2 | Bromodomain adjacent to zinc finger don | BAZ1A | 178.703 | 2 | 52.239956 | yes | 3.154E-07 | 572.77472 | 47.342064 |
| ELHESAIVK | 9 | Q9NRL2 | Bromodomain adjacent to zinc finger don | BAZ1A | 178.703 | 2 | 52.239956 | yes | 8.918E-07 | 513.28491 | 46.723755 |
| GGFDATDDACME | 14 | Q9NRL2 | Bromodomain adjacent to zinc finger don | BAZ1A | 178.703 | 2 | 52.239956 | yes | 1.662E-07 | 779.31903 | 46.611031 |
| QSPEPSPVTLGR | 12 | Q9NRL2 | Bromodomain adjacent to zinc finger don | BAZ1A | 178.703 | 2 | 52.239956 | yes | 4.222E-07 | 634.33569 | 44.747032 |
| ILHALCGK | 8 | Q9NRL2 | Bromodomain adjacent to zinc finger don | BAZ1A | 178.703 | 2 | 52.239956 | yes | 3.635E-07 | 456.26019 | 43.814396 |
| LHEICDDIFAYVK | 13 | Q9NRL2 | Bromodomain adjacent to zinc finger don | BAZ1A | 178.703 | 2 | 52.239956 | yes | 1.064E-07 | 811.89777 | 39.657471 |
| EPLTADEEEALKQ | 17 | Q9NRL2 | Bromodomain adjacent to zinc finger don | BAZ1A | 178.703 | 3 | 52.239956 | yes | 4.41E-07 | 674.99573 | 28.342312 |
| GDAENMVLCDGC | 14 | Q9NRL2 | Bromodomain adjacent to zinc finger don | BAZ1A | 178.703 | 2 | 52.239956 | yes | 2.67E-06 | 806.31342 | 5.8172588 |
| DIKPDNILIDLDGH | 16 | Q9NRM7 | Serine/threonine-protein kinase LATS2 O | LATS2 | 120.136 | 3 | 53.259766 | yes | 9.722E-08 | 607.00354 | 135.77306 |
| LTDFGLCTGFR | 11 | Q9NRM7 | Serine/threonine-protein kinase LATS2 O | LATS2 | 120.136 | 2 | 53.259766 | no | 1.071E-07 | 643.81354 | 96.543915 |
| AGLCEAEQEQMR | 12 | Q9NRM7 | Serine/threonine-protein kinase LATS2 O | LATS2 | 120.136 | 2 | 53.259766 | yes | 9.722E-08 | 711.31104 | 81.69603 |
| TPPETGGYASLPTI | 14 | Q9NRM7 | Serine/threonine-protein kinase LATS2 O | LATS2 | 120.136 | 2 | 53.259766 | yes | 2.213E-06 | 709.86176 | 29.277496 |
| VPFLVLECPNLK | 12 | Q9NRP0 | Oligosaccharyltransferase complex subun | OSTC | 16.83 | 2 | 55.39528 | yes | 9.722E-08 | 714.8996 | 90.693085 |
| LLFNDPDLR | 9 | Q9NRR4 | Ribonuclease 3 OS=Homo sapiens OX=9 | DROSHA | 159.316 | 2 | 53.706665 | yes | 2.267E-07 | 551.79822 | 1132.5757 |
| LIGYTFQDR | 9 | Q9NRR4 | Ribonuclease 3 OS=Homo sapiens OX=9 | DROSHA | 159.316 | 2 | 53.706665 | yes | 1.893E-07 | 556.79041 | 529.98499 |
| TLQTVGPSHAR | 11 | Q9NRR4 | Ribonuclease 3 OS=Homo sapiens OX=9 | DROSHA | 159.316 | 2 | 53.706665 | yes | 9.722E-08 | 583.81744 | 499.16904 |
| PAQLSYAGDPQYC | 14 | Q9NRR4 | Ribonuclease 3 OS=Homo sapiens OX=9 | DROSHA | 159.316 | 2 | 53.706665 | yes | 1.037E-07 | 783.38336 | 491.28369 |
| QLIETSPVLQK | 11 | Q9NRR4 | Ribonuclease 3 OS=Homo sapiens OX=9 | DROSHA | 159.316 | 2 | 53.706665 | yes | 9.722E-08 | 628.36646 | 451.14627 |
| SQLQQCCLTLR | 11 | Q9NRR4 | Ribonuclease 3 OS=Homo sapiens OX=9 | DROSHA | 159.316 | 2 | 53.706665 | yes | 9.723E-08 | 703.8476 | 444.28705 |
| GPLFEDSPPCCPR | 13 | Q9NRR4 | Ribonuclease 3 OS=Homo sapiens OX=9 | DROSHA | 159.316 | 2 | 53.706665 | yes | 9.728E-08 | 766.33704 | 428.62982 |
| TVGFNHLTLGHNC | 14 | Q9NRR4 | Ribonuclease 3 OS=Homo sapiens OX=9 | DROSHA | 159.316 | 3 | 53.706665 | yes | 1.827E-07 | 531.94635 | 186.49466 |
| TVGFNHLTLGHNC | 14 | Q9NRR4 | Ribonuclease 3 OS=Homo sapiens OX=9 | DROSHA | 159.316 | 3 | 53.706665 | yes | 9.809E-08 | 797.41583 | 186.49466 |
| TVGFNHLTLGHNC | 14 | Q9NRR4 | Ribonuclease 3 OS=Homo sapiens OX=9 | DROSHA | 159.316 | 2 | 53.706665 | yes | 1.827E-07 | 531.94635 | 186.49466 |
| TVGFNHLTLGHNC | 14 | Q9NRR4 | Ribonuclease 3 OS=Homo sapiens OX=9 | DROSHA | 159.316 | 2 | 53.706665 | yes | 9.809E-08 | 797.41583 | 186.49466 |
| EFILNQDWNDPK | 12 | Q9NRR4 | Ribonuclease 3 OS=Homo sapiens OX=9 | DROSHA | 159.316 | 2 | 53.706665 | yes | 1.584E-07 | 759.86481 | 363.00137 |
| EVTVELSSQGFWK | 13 | Q9NRR4 | Ribonuclease 3 OS=Homo sapiens OX=9 | DROSHA | 159.316 | 2 | 53.706665 | yes | 3.219E-07 | 755.38281 | 307.2019 |
| HLLANSPK | 8 | Q9NRR4 | Ribonuclease 3 OS=Homo sapiens OX=9 | DROSHA | 159.316 | 2 | 53.706665 | yes | 1.629E-07 | 440.25598 | 234.47598 |
| FMLYAHGPDLCR | 12 | Q9NRR4 | Ribonuclease 3 OS=Homo sapiens OX=9 | DROSHA | 159.316 | 2 | 53.706665 | yes | 9.722E-08 | 740.34723 | 231.5878 |
| GMIVTNPGTKPSS | 15 | Q9NRR4 | Ribonuclease 3 OS=Homo sapiens OX=9 | DROSHA | 159.316 | 2 | 53.706665 | yes | 1.284E-07 | 772.41669 | 171.74739 |
| HSIYPGEEAIKPCR | 14 | Q9NRR4 | Ribonuclease 3 OS=Homo sapiens OX=9 | DROSHA | 159.316 | 3 | 53.706665 | yes | 1.037E-07 | 552.9436 | 159.32356 |
| NSLSNCGIR | 9 | Q9NRR4 | Ribonuclease 3 OS=Homo sapiens OX=9 | DROSHA | 159.316 | 2 | 53.706665 | yes | 1.814E-06 | 510.7482 | 154.8136 |
| TEGKEPDIPLYK | 12 | Q9NRR4 | Ribonuclease 3 OS=Homo sapiens OX=9 | DROSHA | 159.316 | 2 | 53.706665 | yes | 9.722E-08 | 695.36664 | 127.01373 |
| YNFPQMAHQK | 10 | Q9NRR4 | Ribonuclease 3 OS=Homo sapiens OX=9 | DROSHA | 159.316 | 2 | 53.706665 | yes | 3.317E-07 | 632.3006 | 124.97567 |
| YHQCLMHLDK | 10 | Q9NRR4 | Ribonuclease 3 OS=Homo sapiens OX=9 | DROSHA | 159.316 | 2 | 53.706665 | yes | 9.85E-08 | 672.81299 | 118.38188 |
| HSIYPGEEAIKPCR | 22 | Q9NRR4 | Ribonuclease 3 OS=Homo sapiens OX=9 | DROSHA | 159.316 | 4 | 53.706665 | yes | 4.448E-07 | 625.30615 | 81.388885 |
| FEEELGSR | 8 | Q9NRR4 | Ribonuclease 3 OS=Homo sapiens OX=9 | DROSHA | 159.316 | 2 | 53.706665 | yes | 2.016E-07 | 483.72998 | 80.415031 |

| | | | | | | | | | | | |
|---|---|---|---|---|---|---|---|---|---|---|---|
| EPDETEDIKK | 10 | Q9NRR4 | Ribonuclease 3 OS=Homo sapiens OX=9 | DROSHA | 159.316 | 2 | 53.706665 | yes | 1.583E-06 | 602.29059 | 69.385414 |
| DLYDKFEEELGSR | 13 | Q9NRR4 | Ribonuclease 3 OS=Homo sapiens OX=9 | DROSHA | 159.316 | 2 | 53.706665 | yes | 4.271E-07 | 800.87811 | 54.538284 |
| GINTLINIMSR | 11 | Q9NRR4 | Ribonuclease 3 OS=Homo sapiens OX=9 | DROSHA | 159.316 | 2 | 53.706665 | yes | 1.344E-05 | 616.34497 | 36.120201 |
| LGCCVVEKPQPK | 12 | Q9NRR8 | CDC42 small effector protein 1 OS=Hom | CDC42SE1 | 8.926 | 2 | 37.79594 | yes | 5.693E-07 | 707.86267 | 38.355103 |
| AMYPGTFYFQFK | 12 | Q9NRW3 | DNA dC->dU-editing enzyme APOBEC- | APOBEC3( | 22.826 | 2 | 50.08252 | yes | 1.02E-07 | 750.35504 | 722.41113 |
| AMYPGTFYFQFK | 12 | Q9NRW3 | DNA dC->dU-editing enzyme APOBEC- | APOBEC3( | 22.826 | 2 | 50.08252 | yes | 4.005E-07 | 758.35248 | 722.41113 |
| NLWEANDR | 8 | Q9NRW3 | DNA dC->dU-editing enzyme APOBEC- | APOBEC3( | 22.826 | 2 | 50.08252 | yes | 2.566E-07 | 509.24106 | 1099.558 |
| NETWLCFTVEGIK | 13 | Q9NRW3 | DNA dC->dU-editing enzyme APOBEC- | APOBEC3( | 22.826 | 2 | 50.08252 | yes | 9.91E-08 | 798.88995 | 457.16354 |
| TAICNLILGNPPSK | 14 | Q9NRX1 | RNA-binding protein PNO1 OS=Homo s | PNO1 | 27.925 | 2 | 52.548378 | yes | 9.728E-08 | 749.40833 | 3054.4609 |
| FTIENVTR | 8 | Q9NRX1 | RNA-binding protein PNO1 OS=Homo s | PNO1 | 27.925 | 2 | 52.548378 | yes | 9.992E-07 | 490.26401 | 1553.3165 |
| VHILGSFQNIK | 11 | Q9NRX1 | RNA-binding protein PNO1 OS=Homo s | PNO1 | 27.925 | 2 | 52.548378 | yes | 1.12E-07 | 628.36151 | 1435.417 |
| KIPVPANR | 8 | Q9NRX1 | RNA-binding protein PNO1 OS=Homo s | PNO1 | 27.925 | 2 | 52.548378 | yes | 4.861E-07 | 447.77963 | 642.84686 |
| IFTPIVEHLGLQIR | 14 | Q9NRX1 | RNA-binding protein PNO1 OS=Homo s | PNO1 | 27.925 | 3 | 52.548378 | yes | 1.641E-07 | 545.99091 | 222.06912 |
| IFTPIVEHLGLQIR | 14 | Q9NRX1 | RNA-binding protein PNO1 OS=Homo s | PNO1 | 27.925 | 3 | 52.548378 | yes | 9.732E-08 | 818.48267 | 222.06912 |
| IFTPIVEHLGLQIR | 14 | Q9NRX1 | RNA-binding protein PNO1 OS=Homo s | PNO1 | 27.925 | 2 | 52.548378 | yes | 1.641E-07 | 545.99091 | 222.06912 |
| IFTPIVEHLGLQIR | 14 | Q9NRX1 | RNA-binding protein PNO1 OS=Homo s | PNO1 | 27.925 | 2 | 52.548378 | yes | 9.732E-08 | 818.48267 | 222.06912 |
| MDTEEARPAK | 10 | Q9NRX1 | RNA-binding protein PNO1 OS=Homo s | PNO1 | 27.925 | 2 | 52.548378 | yes | 1.133E-07 | 574.27423 | 341.21591 |
| MDTEEAR | 7 | Q9NRX1 | RNA-binding protein PNO1 OS=Homo s | PNO1 | 27.925 | 2 | 52.548378 | yes | 9.722E-08 | 426.18182 | 186.2419 |
| LISQIQPEVDR | 11 | Q9NRZ9 | Lymphoid-specific helicase OS=Homo sa | HELLS | 97.074 | 2 | 53.178707 | yes | 2.12E-07 | 649.35913 | 347.15317 |
| IDDFPNELEK | 10 | Q9NRZ9 | Lymphoid-specific helicase OS=Homo sa | HELLS | 97.074 | 2 | 53.178707 | yes | 1.426E-07 | 610.29572 | 345.23138 |
| TIANMFGSSEK | 11 | Q9NRZ9 | Lymphoid-specific helicase OS=Homo sa | HELLS | 97.074 | 2 | 53.178707 | yes | 1.818E-07 | 592.78442 | 247.68164 |
| ETIELSPTGR | 10 | Q9NRZ9 | Lymphoid-specific helicase OS=Homo sa | HELLS | 97.074 | 2 | 53.178707 | yes | 3.569E-07 | 551.79059 | 227.31311 |
| IDEELVTNSGK | 11 | Q9NRZ9 | Lymphoid-specific helicase OS=Homo sa | HELLS | 97.074 | 2 | 53.178707 | yes | 9.722E-08 | 602.8064 | 191.38165 |
| YLIVDEGHR | 9 | Q9NRZ9 | Lymphoid-specific helicase OS=Homo sa | HELLS | 97.074 | 2 | 53.178707 | yes | 9.722E-08 | 551.28802 | 147.83746 |
| GGQSGLNLSK | 10 | Q9NRZ9 | Lymphoid-specific helicase OS=Homo sa | HELLS | 97.074 | 2 | 53.178707 | yes | 4.036E-07 | 480.75909 | 146.69716 |
| EVVVYAPLSK | 10 | Q9NRZ9 | Lymphoid-specific helicase OS=Homo sa | HELLS | 97.074 | 2 | 53.178707 | yes | 1.078E-07 | 552.8186 | 140.82935 |
| SDLIDQMNASGPII | 14 | Q9NRZ9 | Lymphoid-specific helicase OS=Homo sa | HELLS | 97.074 | 2 | 53.178707 | yes | 4.994E-07 | 744.87177 | 123.7709 |
| IGQTKPVVVYR | 11 | Q9NRZ9 | Lymphoid-specific helicase OS=Homo sa | HELLS | 97.074 | 2 | 53.178707 | yes | 2.223E-07 | 630.37714 | 79.265701 |
| LDGSMSYSER | 10 | Q9NRZ9 | Lymphoid-specific helicase OS=Homo sa | HELLS | 97.074 | 2 | 53.178707 | yes | 9.722E-08 | 572.75061 | 76.493591 |
| EDESYNISEVMSK | 13 | Q9NRZ9 | Lymphoid-specific helicase OS=Homo sa | HELLS | 97.074 | 2 | 53.178707 | yes | 1.605E-06 | 765.83508 | 62.439159 |
| WYQVEGMEWLR | 11 | Q9NRZ9 | Lymphoid-specific helicase OS=Homo sa | HELLS | 97.074 | 2 | 53.178707 | yes | 9.722E-08 | 748.85321 | 57.921421 |
| NSIDASEEKPVMR | 13 | Q9NRZ9 | Lymphoid-specific helicase OS=Homo sa | HELLS | 97.074 | 2 | 53.178707 | yes | 1.081E-07 | 738.36157 | 57.016884 |
| MEQQQLEEQK | 10 | Q9NRZ9 | Lymphoid-specific helicase OS=Homo sa | HELLS | 97.074 | 2 | 53.178707 | yes | 2.652E-05 | 645.80334 | 52.063053 |
| NALQHCYWK | 9 | Q9NRZ9 | Lymphoid-specific helicase OS=Homo sa | HELLS | 97.074 | 2 | 53.178707 | yes | 1.475E-07 | 610.28748 | 32.292099 |
| RLQHLLEK | 8 | Q9NRZ9 | Lymphoid-specific helicase OS=Homo sa | HELLS | 97.074 | 2 | 53.178707 | yes | 2.236E-07 | 518.81671 | 18.324123 |
| CNGQPVPFQQPK | 12 | Q9NRZ9 | Lymphoid-specific helicase OS=Homo sa | HELLS | 97.074 | 2 | 53.178707 | yes | 4.126E-06 | 700.34296 | 15.727356 |
| SRPHGACVPQR | 11 | Q9NS15 | Latent-transforming growth factor beta-bi | LTBP3 | 139.36 | 3 | 49.884293 | yes | 1E-07 | 422.21576 | 89.908417 |
| HGQVYLR | 7 | Q9NS23 | Ras association domain-containing protei | RASSF1 | 39.22 | 2 | 39.57008 | yes | 4.507E-07 | 436.74051 | 57.598526 |
| VSVEFCSACAR | 11 | Q9NS37 | CREB/ATF bZIP transcription factor OS= | CREBZF | 37.134 | 2 | 49.176586 | yes | 1.062E-07 | 643.2868 | 11.915234 |

| | | | | | | | | | | | |
|---|---|---|---|---|---|---|---|---|---|---|---|
| LWGLTEMFPER | 11 | Q9NS69 | Mitochondrial import receptor subunit TC | TOMM22 | 15.522 | 2 | 50.80743 | yes | 2.38E-07 | 689.84485 | 155.64218 |
| LQMEQQQQLQQR | 12 | Q9NS69 | Mitochondrial import receptor subunit TC | TOMM22 | 15.522 | 2 | 50.80743 | yes | 9.722E-08 | 779.39374 | 136.06734 |
| SGNYPSSLSNETDI | 14 | Q9NS86 | LanC-like protein 2 OS=Homo sapiens O | LANCL2 | 50.855 | 2 | 52.731304 | yes | 2.386E-07 | 763.83954 | 336.54193 |
| EVVNAIIESGK | 11 | Q9NS86 | LanC-like protein 2 OS=Homo sapiens O | LANCL2 | 50.855 | 2 | 52.731304 | yes | 1.345E-07 | 579.8219 | 295.37457 |
| VTCDQTYLLR | 10 | Q9NS86 | LanC-like protein 2 OS=Homo sapiens O | LANCL2 | 50.855 | 2 | 52.731304 | yes | 1.517E-07 | 634.81879 | 213.09081 |
| VTFLCGDAGPLAV | 20 | Q9NS86 | LanC-like protein 2 OS=Homo sapiens O | LANCL2 | 50.855 | 3 | 52.731304 | yes | 2.628E-07 | 696.70306 | 179.76396 |
| EAMECSDVIWQR | 12 | Q9NS86 | LanC-like protein 2 OS=Homo sapiens O | LANCL2 | 50.855 | 2 | 52.731304 | yes | 9.722E-08 | 762.33447 | 124.00152 |
| FPAFELDSSK | 10 | Q9NS86 | LanC-like protein 2 OS=Homo sapiens O | LANCL2 | 50.855 | 2 | 52.731304 | yes | 2.58E-07 | 570.78223 | 74.522881 |
| IKDLLQQMEEGLK | 13 | Q9NS86 | LanC-like protein 2 OS=Homo sapiens O | LANCL2 | 50.855 | 2 | 52.731304 | yes | 9.722E-08 | 772.92126 | 66.049667 |
| DLLQQMEEGLK | 11 | Q9NS86 | LanC-like protein 2 OS=Homo sapiens O | LANCL2 | 50.855 | 2 | 52.731304 | yes | 9.819E-08 | 652.33173 | 62.282845 |
| PSIDYVR | 7 | Q9NS86 | LanC-like protein 2 OS=Homo sapiens O | LANCL2 | 50.855 | 2 | 52.731304 | yes | 6.872E-07 | 425.2269 | 53.971893 |
| SDCESQECVTK | 11 | Q9NS86 | LanC-like protein 2 OS=Homo sapiens O | LANCL2 | 50.855 | 2 | 52.731304 | yes | 1.05E-07 | 671.76611 | 24.082579 |
| LMDNFLIR | 8 | Q9NS91 | E3 ubiquitin-protein ligase RAD18 OS=F | RAD18 | 56.223 | 2 | 43.35956 | yes | 2.626E-07 | 511.27859 | 59.760109 |
| SEFQLLVDQAR | 11 | Q9NS91 | E3 ubiquitin-protein ligase RAD18 OS=F | RAD18 | 56.223 | 2 | 43.35956 | yes | 4.546E-05 | 653.34351 | 54.328609 |
| EMSGSTSELLIK | 12 | Q9NS91 | E3 ubiquitin-protein ligase RAD18 OS=F | RAD18 | 56.223 | 2 | 43.35956 | yes | 2.625E-07 | 647.8316 | 52.108559 |
| NIFIECTGTDFTK | 13 | Q9NSD9 | Phenylalanine--tRNA ligase beta subunit | FARSB | 66.116 | 2 | 54.97003 | yes | 9.722E-08 | 773.36633 | 69.574486 |
| LFEISDIVIK | 10 | Q9NSD9 | Phenylalanine--tRNA ligase beta subunit | FARSB | 66.116 | 2 | 54.97003 | yes | 2.467E-05 | 588.84735 | 33.716274 |
| SHTFPELAYR | 10 | Q9NSD9 | Phenylalanine--tRNA ligase beta subunit | FARSB | 66.116 | 2 | 54.97003 | yes | 5.605E-07 | 610.80658 | 14.979856 |
| LGVLHPDVITK | 11 | Q9NSD9 | Phenylalanine--tRNA ligase beta subunit | FARSB | 66.116 | 2 | 54.97003 | yes | 2.607E-06 | 596.35846 | 14.166908 |
| FPLLKPSPK | 9 | Q9NSE4 | Isoleucine--tRNA ligase, mitochondrial O | IARS2 | 113.792 | 2 | 49.68651 | yes | 5.898E-07 | 513.82092 | 89.051666 |
| AVLEEGTDVVIK | 12 | Q9NSE4 | Isoleucine--tRNA ligase, mitochondrial O | IARS2 | 113.792 | 2 | 49.68651 | yes | 1.011E-07 | 636.8559 | 55.354343 |
| YTAESSDTLCPR | 12 | Q9NSE4 | Isoleucine--tRNA ligase, mitochondrial O | IARS2 | 113.792 | 2 | 49.68651 | yes | 1.567E-06 | 700.31152 | 36.590672 |
| VCFLATGAGR | 10 | Q9NSP4 | Centromere protein M OS=Homo sapiens | CENPM | 19.737 | 2 | 45.311817 | yes | 1.888E-07 | 526.2713 | 50.136795 |
| NESVPEVEALLAR | 13 | Q9NSV4 | Protein diaphanous homolog 3 OS=Homo | DIAPH3 | 136.926 | 2 | 53.008736 | yes | 1.108E-07 | 713.88043 | 95.404404 |
| FFCIVEGLR | 9 | Q9NSV4 | Protein diaphanous homolog 3 OS=Homo | DIAPH3 | 136.926 | 2 | 53.008736 | yes | 2.647E-07 | 570.79718 | 88.848335 |
| INELQAELQAFK | 12 | Q9NSV4 | Protein diaphanous homolog 3 OS=Homo | DIAPH3 | 136.926 | 2 | 53.008736 | yes | 1.451E-07 | 702.38007 | 79.451736 |
| ALMNTQYGLER | 11 | Q9NSV4 | Protein diaphanous homolog 3 OS=Homo | DIAPH3 | 136.926 | 2 | 53.008736 | yes | 9.722E-08 | 648.32428 | 75.167679 |
| LAESMIQNLIK | 11 | Q9NSV4 | Protein diaphanous homolog 3 OS=Homo | DIAPH3 | 136.926 | 2 | 53.008736 | yes | 1.675E-07 | 630.35504 | 58.400105 |
| NDGLDIQLK | 9 | Q9NSV4 | Protein diaphanous homolog 3 OS=Homo | DIAPH3 | 136.926 | 2 | 53.008736 | yes | 4.888E-07 | 508.27457 | 57.44838 |
| EMVMQYINTASK | 12 | Q9NSV4 | Protein diaphanous homolog 3 OS=Homo | DIAPH3 | 136.926 | 2 | 53.008736 | yes | 1.648E-07 | 707.83887 | 44.799149 |
| VSVETLEK | 8 | Q9NSV4 | Protein diaphanous homolog 3 OS=Homo | DIAPH3 | 136.926 | 2 | 53.008736 | yes | 6.708E-06 | 452.75293 | 42.961941 |
| HPNMMTDVVK | 10 | Q9NSV4 | Protein diaphanous homolog 3 OS=Homo | DIAPH3 | 136.926 | 2 | 53.008736 | yes | 1.488E-07 | 586.28357 | 35.455833 |
| LENTFCCQQK | 10 | Q9NSV4 | Protein diaphanous homolog 3 OS=Homo | DIAPH3 | 136.926 | 2 | 53.008736 | yes | 5.567E-07 | 664.29211 | 34.073975 |
| ELNYNLDTHTSTG | 14 | Q9NSV4 | Protein diaphanous homolog 3 OS=Homo | DIAPH3 | 136.926 | 2 | 53.008736 | yes | 9.722E-08 | 810.88428 | 25.388891 |
| ITDTIGPTETSIAPR | 15 | Q9NSY1 | BMP-2-inducible protein kinase OS=Hon | BMP2K | 129.172 | 2 | 45.97465 | yes | 3.548E-07 | 786.41742 | 92.489037 |
| VLQQLQQGDWR | 11 | Q9NSY1 | BMP-2-inducible protein kinase OS=Hon | BMP2K | 129.172 | 2 | 45.97465 | yes | 1.688E-07 | 685.86243 | 64.994659 |
| LWLFGYDEQR | 10 | Q9NT62 | Ubiquitin-like-conjugating enzyme ATG3 | ATG3 | 35.865 | 2 | 49.346294 | yes | 1.042E-07 | 663.82751 | 52.500324 |
| TDAGGEDAILQTR | 13 | Q9NT62 | Ubiquitin-like-conjugating enzyme ATG3 | ATG3 | 35.865 | 2 | 49.346294 | yes | 1.515E-07 | 673.83093 | 45.420067 |
| VAEAALQIFK | 10 | Q9NTI5 | Sister chromatid cohesion protein PDS5 h | PDS5B | 164.668 | 2 | 51.48313 | yes | 9.722E-08 | 545.31879 | 127.36638 |

| | | | | | | | | | | | |
|---|---|---|---|---|---|---|---|---|---|---|---|
| FNDIHVPIR | 9 | Q9NTI5 | Sister chromatid cohesion protein PDS5 h | PDS5B | 164.668 | 2 | 51.48313 | yes | 1.022E-07 | 555.80634 | 114.8398 |
| LGMDDLTK | 8 | Q9NTI5 | Sister chromatid cohesion protein PDS5 h | PDS5B | 164.668 | 2 | 51.48313 | yes | 9.722E-08 | 446.72586 | 106.78565 |
| IEEDFPHIR | 9 | Q9NTI5 | Sister chromatid cohesion protein PDS5 h | PDS5B | 164.668 | 2 | 51.48313 | yes | 5.773E-07 | 578.29327 | 86.1959 |
| VQDIEQLK | 8 | Q9NTI5 | Sister chromatid cohesion protein PDS5 h | PDS5B | 164.668 | 2 | 51.48313 | yes | 2.912E-07 | 486.77167 | 63.823635 |
| HDVIVSIVTAAK | 12 | Q9NTI5 | Sister chromatid cohesion protein PDS5 h | PDS5B | 164.668 | 2 | 51.48313 | yes | 1.89E-06 | 626.86664 | 50.820702 |
| SGPPAPEEEEEEEF | 14 | Q9NTI5 | Sister chromatid cohesion protein PDS5 h | PDS5B | 164.668 | 2 | 51.48313 | yes | 5.331E-07 | 792.83661 | 19.995369 |
| TTNVLGAVNKPLS | 17 | Q9NTI5 | Sister chromatid cohesion protein PDS5 h | PDS5B | 164.668 | 2 | 51.48313 | yes | 1.036E-07 | 828.96759 | 15.492989 |
| FTQLDLEDVQVR | 12 | Q9NTJ3 | Structural maintenance of chromosomes p | SMC4 | 147.182 | 2 | 52.875813 | yes | 2.467E-07 | 731.88043 | 541.91266 |
| LLEENVSAFK | 10 | Q9NTJ3 | Structural maintenance of chromosomes p | SMC4 | 147.182 | 2 | 52.875813 | yes | 1.305E-07 | 575.31116 | 373.25552 |
| EALIAASETLK | 11 | Q9NTJ3 | Structural maintenance of chromosomes p | SMC4 | 147.182 | 2 | 52.875813 | yes | 9.722E-08 | 573.32428 | 368.9794 |
| LPQTEQELK | 9 | Q9NTJ3 | Structural maintenance of chromosomes p | SMC4 | 147.182 | 2 | 52.875813 | yes | 4.392E-07 | 543.29547 | 294.93826 |
| DTLVADNLDQATI | 13 | Q9NTJ3 | Structural maintenance of chromosomes p | SMC4 | 147.182 | 2 | 52.875813 | yes | 2.023E-07 | 716.35736 | 259.905 |
| ELEANVLATAPDK | 13 | Q9NTJ3 | Structural maintenance of chromosomes p | SMC4 | 147.182 | 2 | 52.875813 | yes | 1.417E-07 | 685.86176 | 195.70593 |
| VLDAIIQEK | 9 | Q9NTJ3 | Structural maintenance of chromosomes p | SMC4 | 147.182 | 2 | 52.875813 | yes | 9.766E-08 | 514.80298 | 167.54936 |
| EVDDLTAELK | 10 | Q9NTJ3 | Structural maintenance of chromosomes p | SMC4 | 147.182 | 2 | 52.875813 | yes | 1.061E-07 | 566.79022 | 162.3429 |
| DIQSCTVEVHFQK | 13 | Q9NTJ3 | Structural maintenance of chromosomes p | SMC4 | 147.182 | 2 | 52.875813 | yes | 9.809E-08 | 795.88269 | 155.09912 |
| SNILSNEMK | 9 | Q9NTJ3 | Structural maintenance of chromosomes p | SMC4 | 147.182 | 2 | 52.875813 | yes | 5.417E-07 | 518.26062 | 149.78157 |
| MDVAQSELDIYLS | 14 | Q9NTJ3 | Structural maintenance of chromosomes p | SMC4 | 147.182 | 2 | 52.875813 | yes | 9.927E-08 | 820.40344 | 147.21848 |
| VIQENEHALQK | 11 | Q9NTJ3 | Structural maintenance of chromosomes p | SMC4 | 147.182 | 2 | 52.875813 | yes | 9.722E-08 | 654.84894 | 144.49651 |
| NTNAAEESLPEIQK | 14 | Q9NTJ3 | Structural maintenance of chromosomes p | SMC4 | 147.182 | 2 | 52.875813 | yes | 9.809E-08 | 772.38354 | 118.33273 |
| SNNIINETTTR | 11 | Q9NTJ3 | Structural maintenance of chromosomes p | SMC4 | 147.182 | 2 | 52.875813 | yes | 2.455E-07 | 631.82037 | 105.57816 |
| LIEQEEYLNVQVK | 13 | Q9NTJ3 | Structural maintenance of chromosomes p | SMC4 | 147.182 | 2 | 52.875813 | yes | 9.722E-08 | 802.93018 | 97.984573 |
| QNIGVATFIGLDK | 13 | Q9NTJ3 | Structural maintenance of chromosomes p | SMC4 | 147.182 | 2 | 52.875813 | yes | 9.722E-08 | 688.38263 | 95.740173 |
| NNMFEISDR | 9 | Q9NTJ3 | Structural maintenance of chromosomes p | SMC4 | 147.182 | 2 | 52.875813 | yes | 4.609E-07 | 563.2533 | 91.258682 |
| ILGPFHK | 7 | Q9NTJ3 | Structural maintenance of chromosomes p | SMC4 | 147.182 | 2 | 52.875813 | yes | 4.248E-06 | 406.24487 | 90.289955 |
| SLVHDLFQK | 9 | Q9NTJ3 | Structural maintenance of chromosomes p | SMC4 | 147.182 | 2 | 52.875813 | yes | 2.761E-07 | 543.80072 | 84.596153 |
| DNTSVYHISGK | 11 | Q9NTJ3 | Structural maintenance of chromosomes p | SMC4 | 147.182 | 2 | 52.875813 | yes | 3.9E-07 | 610.79895 | 83.399185 |
| SHGIDLDHNR | 10 | Q9NTJ3 | Structural maintenance of chromosomes p | SMC4 | 147.182 | 2 | 52.875813 | yes | 1.675E-07 | 582.28125 | 67.822449 |
| QAFYFALR | 8 | Q9NTJ3 | Structural maintenance of chromosomes p | SMC4 | 147.182 | 2 | 52.875813 | yes | 4.751E-07 | 508.27164 | 66.540695 |
| IFNLSGGEK | 9 | Q9NTJ3 | Structural maintenance of chromosomes p | SMC4 | 147.182 | 2 | 52.875813 | yes | 1.701E-06 | 482.75854 | 55.765617 |
| LHNTIVEINNHK | 12 | Q9NTJ3 | Structural maintenance of chromosomes p | SMC4 | 147.182 | 2 | 52.875813 | yes | 9.722E-08 | 716.38879 | 55.192818 |
| MTEIQTPENTPR | 12 | Q9NTJ3 | Structural maintenance of chromosomes p | SMC4 | 147.182 | 2 | 52.875813 | yes | 1.92E-07 | 708.84302 | 54.446266 |
| KKEELYLQR | 9 | Q9NTJ3 | Structural maintenance of chromosomes p | SMC4 | 147.182 | 2 | 52.875813 | yes | 1.249E-06 | 603.8457 | 31.699324 |
| EKELMGFSK | 9 | Q9NTJ3 | Structural maintenance of chromosomes p | SMC4 | 147.182 | 2 | 52.875813 | yes | 2.337E-06 | 534.77332 | 29.526327 |
| LEEQDEFEK | 9 | Q9NTJ5 | Phosphatidylinositol-3-phosphatase SAC | SACM1L | 66.967 | 2 | 50.918255 | yes | 9.722E-08 | 583.76422 | 48.324165 |
| QVIINLINQK | 10 | Q9NTJ5 | Phosphatidylinositol-3-phosphatase SAC | SACM1L | 66.967 | 2 | 50.918255 | yes | 8.69E-07 | 591.86389 | 22.384813 |
| VSTEVTLAVK | 10 | Q9NTJ5 | Phosphatidylinositol-3-phosphatase SAC | SACM1L | 66.967 | 2 | 50.918255 | yes | 3.073E-06 | 523.80823 | 10.71031 |
| IGIVGLPNVGK | 11 | Q9NTK5 | Obg-like ATPase 1 OS=Homo sapiens OX | OLA1 | 44.744 | 2 | 41.55997 | yes | 3.449E-07 | 533.83459 | 124.63142 |
| DEEMIGPIIDK | 11 | Q9NTK5 | Obg-like ATPase 1 OS=Homo sapiens OX | OLA1 | 44.744 | 2 | 41.55997 | yes | 3.751E-07 | 630.31305 | 34.877121 |

| | | | | | | | | | | | |
|---|---|---|---|---|---|---|---|---|---|---|---|
| LNCCYPGCQFK | 11 | Q9NTW7 | Zinc finger protein 64 OS=Homo sapiens | ZFP64 | 72.217 | 2 | 44.106163 | yes | 1.155E-06 | 723.80176 | 51.991207 |
| NSSQLTVHLR | 10 | Q9NTW7 | Zinc finger protein 64 OS=Homo sapiens | ZFP64 | 72.217 | 2 | 44.106163 | yes | 2.319E-07 | 577.81744 | 38.509869 |
| HCDVLAVPVK | 10 | Q9NUD5 | Zinc finger CCHC domain-containing prc | ZCCHC3 | 43.548 | 2 | 52.136505 | yes | 9.995E-08 | 569.30786 | 1127.0851 |
| ICFQGDEGACPTR | 13 | Q9NUD5 | Zinc finger CCHC domain-containing prc | ZCCHC3 | 43.548 | 2 | 52.136505 | yes | 1.284E-07 | 755.82428 | 842.3924 |
| HLPGAFFLGAER | 12 | Q9NUD5 | Zinc finger CCHC domain-containing prc | ZCCHC3 | 43.548 | 2 | 52.136505 | yes | 4.534E-07 | 657.85132 | 682.34021 |
| FGIWTGEYK | 9 | Q9NUD5 | Zinc finger CCHC domain-containing prc | ZCCHC3 | 43.548 | 2 | 52.136505 | yes | 1.015E-07 | 550.77417 | 600.61029 |
| AVHNSVAAQLTG' | 16 | Q9NUD5 | Zinc finger CCHC domain-containing prc | ZCCHC3 | 43.548 | 2 | 52.136505 | yes | 9.796E-08 | 766.4024 | 221.96567 |
| AVHNSVAAQLTG' | 16 | Q9NUD5 | Zinc finger CCHC domain-containing prc | ZCCHC3 | 43.548 | 2 | 52.136505 | yes | 3.671E-07 | 511.27069 | 221.96567 |
| AVHNSVAAQLTG' | 16 | Q9NUD5 | Zinc finger CCHC domain-containing prc | ZCCHC3 | 43.548 | 3 | 52.136505 | yes | 9.796E-08 | 766.4024 | 221.96567 |
| AVHNSVAAQLTG' | 16 | Q9NUD5 | Zinc finger CCHC domain-containing prc | ZCCHC3 | 43.548 | 3 | 52.136505 | yes | 3.671E-07 | 511.27069 | 221.96567 |
| DPAGEAVDPR | 10 | Q9NUD5 | Zinc finger CCHC domain-containing prc | ZCCHC3 | 43.548 | 2 | 52.136505 | yes | 1.692E-07 | 513.74615 | 371.3381 |
| GRPQLLPPAR | 10 | Q9NUD5 | Zinc finger CCHC domain-containing prc | ZCCHC3 | 43.548 | 2 | 52.136505 | yes | 9.722E-08 | 552.83545 | 347.56433 |
| KGIVCNLCGK | 10 | Q9NUD5 | Zinc finger CCHC domain-containing prc | ZCCHC3 | 43.548 | 2 | 52.136505 | yes | 9.727E-08 | 574.79938 | 274.47345 |
| GHAFAQCPK | 9 | Q9NUD5 | Zinc finger CCHC domain-containing prc | ZCCHC3 | 43.548 | 2 | 52.136505 | yes | 1.443E-07 | 508.24255 | 273.71921 |
| GIVCNLCGK | 9 | Q9NUD5 | Zinc finger CCHC domain-containing prc | ZCCHC3 | 43.548 | 2 | 52.136505 | yes | 1.009E-07 | 510.75189 | 222.15942 |
| CGEEGHLSPYCR | 12 | Q9NUD5 | Zinc finger CCHC domain-containing prc | ZCCHC3 | 43.548 | 2 | 52.136505 | yes | 2.041E-07 | 732.80334 | 193.7103 |
| RHCDVLAVPVK | 11 | Q9NUD5 | Zinc finger CCHC domain-containing prc | ZCCHC3 | 43.548 | 3 | 52.136505 | yes | 9.722E-08 | 431.90805 | 64.416512 |
| RHCDVLAVPVK | 11 | Q9NUD5 | Zinc finger CCHC domain-containing prc | ZCCHC3 | 43.548 | 3 | 52.136505 | yes | 9.722E-08 | 647.35846 | 64.416512 |
| RHCDVLAVPVK | 11 | Q9NUD5 | Zinc finger CCHC domain-containing prc | ZCCHC3 | 43.548 | 2 | 52.136505 | yes | 9.722E-08 | 431.90805 | 64.416512 |
| RHCDVLAVPVK | 11 | Q9NUD5 | Zinc finger CCHC domain-containing prc | ZCCHC3 | 43.548 | 2 | 52.136505 | yes | 9.722E-08 | 647.35846 | 64.416512 |
| HLFCPDLLR | 9 | Q9NUI1 | Peroxisomal 2,4-dienoyl-CoA reductase [ | DECR2 | 30.778 | 2 | 53.34881 | yes | 9.722E-08 | 585.80804 | 339.97269 |
| VNSLAPGPISGTEC | 16 | Q9NUI1 | Peroxisomal 2,4-dienoyl-CoA reductase [ | DECR2 | 30.778 | 2 | 53.34881 | yes | 6.229E-07 | 784.42554 | 260.05685 |
| GLPDFASFSAK | 11 | Q9NUI1 | Peroxisomal 2,4-dienoyl-CoA reductase [ | DECR2 | 30.778 | 2 | 53.34881 | yes | 9.722E-08 | 570.29022 | 202.33389 |
| LGGPQASLSTK | 11 | Q9NUI1 | Peroxisomal 2,4-dienoyl-CoA reductase [ | DECR2 | 30.778 | 2 | 53.34881 | yes | 9.81E-08 | 529.79565 | 167.37717 |
| TVMDIDTSGTFNV | 15 | Q9NUI1 | Peroxisomal 2,4-dienoyl-CoA reductase [ | DECR2 | 30.778 | 2 | 53.34881 | yes | 1.102E-07 | 821.89069 | 149.03519 |
| HLAVEWGPQNIR | 12 | Q9NUI1 | Peroxisomal 2,4-dienoyl-CoA reductase [ | DECR2 | 30.778 | 2 | 53.34881 | yes | 9.722E-08 | 710.37823 | 114.33068 |
| VAFITGGGSGIGFR | 14 | Q9NUI1 | Peroxisomal 2,4-dienoyl-CoA reductase [ | DECR2 | 30.778 | 2 | 53.34881 | yes | 5.148E-07 | 669.86188 | 98.810524 |
| GQALQVHAGSAK | 12 | Q9NUI1 | Peroxisomal 2,4-dienoyl-CoA reductase [ | DECR2 | 30.778 | 2 | 53.34881 | yes | 4.659E-06 | 583.81744 | 52.156883 |
| APPAVMAAVDQA | 14 | Q9NUI1 | Peroxisomal 2,4-dienoyl-CoA reductase [ | DECR2 | 30.778 | 2 | 53.34881 | yes | 5.871E-07 | 691.37903 | 41.022999 |
| YFHPPAHLQAR | 11 | Q9NUK0 | Muscleblind-like protein 3 OS=Homo sap | MBNL3 | 38.532 | 3 | 48.550377 | yes | 1.211E-07 | 446.23517 | 429.14169 |
| YFHPPAHLQAR | 11 | Q9NUK0 | Muscleblind-like protein 3 OS=Homo sap | MBNL3 | 38.532 | 3 | 48.550377 | yes | 2.849E-07 | 668.84912 | 429.14169 |
| YFHPPAHLQAR | 11 | Q9NUK0 | Muscleblind-like protein 3 OS=Homo sap | MBNL3 | 38.532 | 2 | 48.550377 | yes | 1.211E-07 | 446.23517 | 429.14169 |
| YFHPPAHLQAR | 11 | Q9NUK0 | Muscleblind-like protein 3 OS=Homo sap | MBNL3 | 38.532 | 2 | 48.550377 | yes | 2.849E-07 | 668.84912 | 429.14169 |
| VVACFDSLK | 9 | Q9NUK0 | Muscleblind-like protein 3 OS=Homo sap | MBNL3 | 38.532 | 2 | 48.550377 | yes | 1.196E-07 | 519.76807 | 771.0614 |
| SDKLEVCR | 8 | Q9NUK0 | Muscleblind-like protein 3 OS=Homo sap | MBNL3 | 38.532 | 2 | 48.550377 | yes | 1.132E-07 | 503.75293 | 580.8064 |
| TAVNVALIR | 9 | Q9NUK0 | Muscleblind-like protein 3 OS=Homo sap | MBNL3 | 38.532 | 2 | 48.550377 | yes | 2.737E-07 | 499.80328 | 163.97267 |
| ALQALQNEPIPER | 13 | Q9NUL3 | Double-stranded RNA-binding protein St | STAU2 | 62.609 | 2 | 52.993885 | yes | 1.066E-07 | 739.90173 | 4072.7991 |
| NMPVSFEVIK | 10 | Q9NUL3 | Double-stranded RNA-binding protein St | STAU2 | 62.609 | 2 | 52.993885 | yes | 1.738E-07 | 582.31012 | 2998.7495 |
| KLPPLPVVEKPK | 12 | Q9NUL3 | Double-stranded RNA-binding protein St | STAU2 | 62.609 | 3 | 52.993885 | yes | 1.23E-07 | 448.95862 | 1308.9281 |

| | | | | | | | | | | | |
|---|---|---|---|---|---|---|---|---|---|---|---|
| KLPPLPVVEKPK | 12 | Q9NUL3 | Double-stranded RNA-binding protein St | STAU2 | 62.609 | 3 | 52.993885 | yes | 4.421E-07 | 672.93433 | 1308.9281 |
| KLPPLPVVEKPK | 12 | Q9NUL3 | Double-stranded RNA-binding protein St | STAU2 | 62.609 | 2 | 52.993885 | yes | 1.23E-07 | 448.95862 | 1308.9281 |
| KLPPLPVVEKPK | 12 | Q9NUL3 | Double-stranded RNA-binding protein St | STAU2 | 62.609 | 2 | 52.993885 | yes | 4.421E-07 | 672.93433 | 1308.9281 |
| NAAEAMLLQLGYI | 13 | Q9NUL3 | Double-stranded RNA-binding protein St | STAU2 | 62.609 | 2 | 52.993885 | yes | 9.849E-08 | 711.37646 | 1307.296 |
| NAAEAMLLQLGYI | 13 | Q9NUL3 | Double-stranded RNA-binding protein St | STAU2 | 62.609 | 2 | 52.993885 | yes | 4.422E-06 | 719.37396 | 1307.296 |
| ASTNLQDQLEK | 11 | Q9NUL3 | Double-stranded RNA-binding protein St | STAU2 | 62.609 | 2 | 52.993885 | yes | 1.002E-07 | 623.81732 | 2509.4114 |
| AATTVLQELK | 10 | Q9NUL3 | Double-stranded RNA-binding protein St | STAU2 | 62.609 | 2 | 52.993885 | yes | 1.714E-07 | 537.31366 | 2268.4697 |
| VSVGEFSAEGEGN | 15 | Q9NUL3 | Double-stranded RNA-binding protein St | STAU2 | 62.609 | 2 | 52.993885 | yes | 9.722E-08 | 748.8468 | 2114.9465 |
| RGEPAIYRPLDPK | 13 | Q9NUL3 | Double-stranded RNA-binding protein St | STAU2 | 62.609 | 3 | 52.993885 | yes | 1.902E-07 | 504.61581 | 973.46796 |
| RGEPAIYRPLDPK | 13 | Q9NUL3 | Double-stranded RNA-binding protein St | STAU2 | 62.609 | 3 | 52.993885 | yes | 1.7E-07 | 756.4201 | 973.46796 |
| RGEPAIYRPLDPK | 13 | Q9NUL3 | Double-stranded RNA-binding protein St | STAU2 | 62.609 | 2 | 52.993885 | yes | 1.902E-07 | 504.61581 | 973.46796 |
| RGEPAIYRPLDPK | 13 | Q9NUL3 | Double-stranded RNA-binding protein St | STAU2 | 62.609 | 2 | 52.993885 | yes | 1.7E-07 | 756.4201 | 973.46796 |
| PGFPEPTNNTPK | 12 | Q9NUL3 | Double-stranded RNA-binding protein St | STAU2 | 62.609 | 2 | 52.993885 | yes | 9.722E-08 | 649.82239 | 1683.2247 |
| EKEPDYVLLSER | 12 | Q9NUL3 | Double-stranded RNA-binding protein St | STAU2 | 62.609 | 2 | 52.993885 | yes | 9.722E-08 | 739.38031 | 1561.8174 |
| EPDYVLLSER | 10 | Q9NUL3 | Double-stranded RNA-binding protein St | STAU2 | 62.609 | 2 | 52.993885 | yes | 9.722E-08 | 610.81152 | 1533.1731 |
| GSSPTPPCSPVQPS | 15 | Q9NUL3 | Double-stranded RNA-binding protein St | STAU2 | 62.609 | 2 | 52.993885 | yes | 1.375E-07 | 763.36938 | 1413.072 |
| AGPEYGQGMNPIS | 14 | Q9NUL3 | Double-stranded RNA-binding protein St | STAU2 | 62.609 | 2 | 52.993885 | yes | 1.002E-07 | 738.84863 | 626.04083 |
| AGPEYGQGMNPIS | 14 | Q9NUL3 | Double-stranded RNA-binding protein St | STAU2 | 62.609 | 2 | 52.993885 | yes | 9.819E-08 | 746.84607 | 626.04083 |
| EFVMQVK | 7 | Q9NUL3 | Double-stranded RNA-binding protein St | STAU2 | 62.609 | 2 | 52.993885 | no | 2.259E-07 | 440.73349 | 1144.2212 |
| VGNEVATGTGPNI | 13 | Q9NUL3 | Double-stranded RNA-binding protein St | STAU2 | 62.609 | 2 | 52.993885 | yes | 4.092E-07 | 622.3175 | 1095.8621 |
| QLEYLAR | 7 | Q9NUL3 | Double-stranded RNA-binding protein St | STAU2 | 62.609 | 2 | 52.993885 | yes | 1.905E-07 | 446.74799 | 960.96002 |
| LAQIQQAK | 8 | Q9NUL3 | Double-stranded RNA-binding protein St | STAU2 | 62.609 | 2 | 52.993885 | no | 2.702E-07 | 450.26907 | 943.56116 |
| YHCPVPK | 7 | Q9NUL3 | Double-stranded RNA-binding protein St | STAU2 | 62.609 | 2 | 52.993885 | yes | 1.641E-07 | 450.72345 | 689.72791 |
| IQGFQAALSALK | 12 | Q9NUL3 | Double-stranded RNA-binding protein St | STAU2 | 62.609 | 2 | 52.993885 | yes | 1.532E-07 | 623.86133 | 686.37317 |
| GEPAIYRPLDPK | 12 | Q9NUL3 | Double-stranded RNA-binding protein St | STAU2 | 62.609 | 2 | 52.993885 | yes | 1.037E-07 | 678.36951 | 661.27582 |
| KNAAEAMLLQLG | 14 | Q9NUL3 | Double-stranded RNA-binding protein St | STAU2 | 62.609 | 3 | 52.993885 | yes | 2.697E-07 | 517.2851 | 298.39282 |
| KNAAEAMLLQLG | 14 | Q9NUL3 | Double-stranded RNA-binding protein St | STAU2 | 62.609 | 3 | 52.993885 | yes | 9.722E-08 | 775.42395 | 298.39282 |
| KNAAEAMLLQLG | 14 | Q9NUL3 | Double-stranded RNA-binding protein St | STAU2 | 62.609 | 2 | 52.993885 | yes | 2.697E-07 | 517.2851 | 298.39282 |
| KNAAEAMLLQLG | 14 | Q9NUL3 | Double-stranded RNA-binding protein St | STAU2 | 62.609 | 2 | 52.993885 | yes | 9.722E-08 | 775.42395 | 298.39282 |
| TAMCLVNELAR | 11 | Q9NUL3 | Double-stranded RNA-binding protein St | STAU2 | 62.609 | 2 | 52.993885 | yes | 9.722E-08 | 639.32068 | 565.21729 |
| ALTESTLPK | 9 | Q9NUL3 | Double-stranded RNA-binding protein St | STAU2 | 62.609 | 2 | 52.993885 | yes | 1.465E-07 | 480.27402 | 298.79492 |
| VGNEVATGTGPNI | 14 | Q9NUL3 | Double-stranded RNA-binding protein St | STAU2 | 62.609 | 2 | 52.993885 | yes | 1.705E-07 | 686.36499 | 257.46332 |
| ADNNQAPPGSIAQ | 16 | Q9NUL3 | Double-stranded RNA-binding protein St | STAU2 | 62.609 | 2 | 52.993885 | yes | 1.215E-07 | 843.38104 | 119.19969 |
| KLPPLPVVEK | 10 | Q9NUL3 | Double-stranded RNA-binding protein St | STAU2 | 62.609 | 2 | 52.993885 | yes | 1.335E-06 | 560.36041 | 97.646362 |
| RGEPAIYR | 8 | Q9NUL3 | Double-stranded RNA-binding protein St | STAU2 | 62.609 | 2 | 52.993885 | yes | 3.747E-07 | 481.26434 | 63.491367 |
| VISGTTLGYLSPK | 13 | Q9NUL3 | Double-stranded RNA-binding protein St | STAU2 | 62.609 | 2 | 52.993885 | yes | 4.235E-07 | 668.37958 | 51.781471 |
| AQDDLIPAVDR | 11 | Q9NUL5 | Shiftless antiviral inhibitor of ribosomal f | SHFL | 33.111 | 2 | 50.381927 | yes | 1.204E-07 | 606.81458 | 952.66992 |
| REPHVPGTSCAHP | 14 | Q9NUL5 | Shiftless antiviral inhibitor of ribosomal f | SHFL | 33.111 | 3 | 50.381927 | yes | 5.383E-07 | 524.92816 | 315.50467 |
| WDRDPDR | 7 | Q9NUL5 | Shiftless antiviral inhibitor of ribosomal f | SHFL | 33.111 | 2 | 50.381927 | yes | 1.247E-07 | 480.22012 | 193.99698 |

| | | | | | | | | | | | |
|---|---|---|---|---|---|---|---|---|---|---|---|
| RYEPVPADK | 9 | Q9NUL5 | Shiftless antiviral inhibitor of ribosomal f | SHFL | 33.111 | 2 | 50.381927 | yes | 2.222E-07 | 537.78253 | 152.95506 |
| EPHVPGTSCAHPK | 13 | Q9NUL5 | Shiftless antiviral inhibitor of ribosomal f | SHFL | 33.111 | 2 | 50.381927 | yes | 9.81E-08 | 708.83807 | 142.33856 |
| VASPDAVTTITSSK | 14 | Q9NUL7 | Probable ATP-dependent RNA helicase D | DDX28 | 59.582 | 2 | 50.50684 | yes | 1.063E-07 | 688.867 | 2199.085 |
| PGPLLVSAR | 9 | Q9NUL7 | Probable ATP-dependent RNA helicase D | DDX28 | 59.582 | 2 | 50.50684 | yes | 9.79E-08 | 455.27945 | 1566.1423 |
| LQGQMPALMR | 10 | Q9NUL7 | Probable ATP-dependent RNA helicase D | DDX28 | 59.582 | 2 | 50.50684 | yes | 5.855E-07 | 572.80194 | 1304.2502 |
| GSFADLGLEPR | 11 | Q9NUL7 | Probable ATP-dependent RNA helicase D | DDX28 | 59.582 | 2 | 50.50684 | yes | 7.721E-07 | 581.29858 | 1271.9398 |
| APLASQGWK | 9 | Q9NUL7 | Probable ATP-dependent RNA helicase D | DDX28 | 59.582 | 2 | 50.50684 | yes | 1.261E-07 | 479.26126 | 924.71576 |
| DLEGGHGMR | 9 | Q9NUL7 | Probable ATP-dependent RNA helicase D | DDX28 | 59.582 | 2 | 50.50684 | yes | 1.987E-07 | 486.2218 | 855.67273 |
| HVVCAAETGSGK | 12 | Q9NUL7 | Probable ATP-dependent RNA helicase D | DDX28 | 59.582 | 2 | 50.50684 | yes | 9.819E-08 | 608.29297 | 843.32861 |
| QPSADVLVATPGA | 16 | Q9NUL7 | Probable ATP-dependent RNA helicase D | DDX28 | 59.582 | 2 | 50.50684 | yes | 1.437E-07 | 826.95398 | 813.84961 |
| LLGQPSLDSLPIPA | 16 | Q9NUL7 | Probable ATP-dependent RNA helicase D | DDX28 | 59.582 | 2 | 50.50684 | yes | 1.686E-07 | 837.48285 | 756.41827 |
| DILLCTDIASR | 11 | Q9NUL7 | Probable ATP-dependent RNA helicase D | DDX28 | 59.582 | 2 | 50.50684 | yes | 1.02E-07 | 638.83191 | 374.11307 |
| AQQEAPAVR | 9 | Q9NUL7 | Probable ATP-dependent RNA helicase D | DDX28 | 59.582 | 2 | 50.50684 | yes | 1.204E-07 | 485.25925 | 261.64584 |
| VAELVHILK | 9 | Q9NUL7 | Probable ATP-dependent RNA helicase D | DDX28 | 59.582 | 2 | 50.50684 | yes | 1.12E-07 | 511.32385 | 218.07732 |
| VGIFQSFQK | 9 | Q9NUL7 | Probable ATP-dependent RNA helicase D | DDX28 | 59.582 | 2 | 50.50684 | yes | 1.638E-07 | 527.29004 | 200.49825 |
| SHIAEGPADLEDPI | 17 | Q9NUL7 | Probable ATP-dependent RNA helicase D | DDX28 | 59.582 | 3 | 50.50684 | yes | 5.382E-07 | 612.96362 | 155.05731 |
| GLVLVPSR | 8 | Q9NUL7 | Probable ATP-dependent RNA helicase D | DDX28 | 59.582 | 2 | 50.50684 | yes | 6.004E-07 | 420.76871 | 133.80217 |
| RPELNQPAR | 9 | Q9NUL7 | Probable ATP-dependent RNA helicase D | DDX28 | 59.582 | 2 | 50.50684 | yes | 2.753E-07 | 540.79907 | 99.84507 |
| ELAQQVR | 7 | Q9NUL7 | Probable ATP-dependent RNA helicase D | DDX28 | 59.582 | 2 | 50.50684 | yes | 5.667E-07 | 422.23779 | 81.207474 |
| SAYVSYDVQK | 10 | Q9NUM4 | Transmembrane protein 106B OS=Homo | TMEM106B | 31.128 | 2 | 48.54762 | yes | 1.132E-07 | 580.28516 | 275.91812 |
| EDAYDGVTSENMI | 13 | Q9NUM4 | Transmembrane protein 106B OS=Homo | TMEM106B | 31.128 | 2 | 48.54762 | yes | 1.325E-07 | 743.80939 | 43.954731 |
| VLQSEFCNAVR | 11 | Q9NUP9 | Protein lin-7 homolog C OS=Homo sapie | LIN7C | 21.834 | 2 | 50.0681 | yes | 1.507E-07 | 661.82971 | 410.53937 |
| VLEEMESR | 8 | Q9NUP9 | Protein lin-7 homolog C OS=Homo sapie | LIN7C | 21.834 | 2 | 50.0681 | yes | 1.255E-07 | 496.7395 | 403.03656 |
| TEEGLGFNIMGGK | 13 | Q9NUP9 | Protein lin-7 homolog C OS=Homo sapie | LIN7C | 21.834 | 2 | 50.0681 | yes | 1.022E-07 | 676.82941 | 327.79578 |
| EQNSPIYISR | 10 | Q9NUP9 | Protein lin-7 homolog C OS=Homo sapie | LIN7C | 21.834 | 2 | 50.0681 | no | 2.461E-07 | 603.80933 | 263.39005 |
| EVVYEHVYETVDIS | 19 | Q9NUP9 | Protein lin-7 homolog C OS=Homo sapie | LIN7C | 21.834 | 3 | 50.0681 | yes | 9.918E-08 | 746.69019 | 241.48677 |
| ATVAAFAASEGHS | 16 | Q9NUP9 | Protein lin-7 homolog C OS=Homo sapie | LIN7C | 21.834 | 3 | 50.0681 | no | 1.973E-07 | 536.93408 | 22.737421 |
| ATVAAFAASEGHS | 16 | Q9NUP9 | Protein lin-7 homolog C OS=Homo sapie | LIN7C | 21.834 | 3 | 50.0681 | no | 4.54E-06 | 804.89752 | 22.737421 |
| ATVAAFAASEGHS | 16 | Q9NUP9 | Protein lin-7 homolog C OS=Homo sapie | LIN7C | 21.834 | 2 | 50.0681 | no | 1.973E-07 | 536.93408 | 22.737421 |
| ATVAAFAASEGHS | 16 | Q9NUP9 | Protein lin-7 homolog C OS=Homo sapie | LIN7C | 21.834 | 2 | 50.0681 | no | 4.54E-06 | 804.89752 | 22.737421 |
| VLSASQAFAAQR | 12 | Q9NUQ2 | 1-acyl-sn-glycerol-3-phosphate acyltransf | AGPAT5 | 42.073 | 2 | 55.076824 | yes | 9.844E-07 | 624.83838 | 38.096481 |
| MLIEFYESPDPER | 13 | Q9NUQ2 | 1-acyl-sn-glycerol-3-phosphate acyltransf | AGPAT5 | 42.073 | 2 | 55.076824 | yes | 9.722E-08 | 813.37946 | 32.110195 |
| ALLQMAEEK | 9 | Q9NUQ3 | Gamma-taxilin OS=Homo sapiens OX=9 | TXLNG | 60.586 | 2 | 51.547333 | yes | 1.3E-07 | 516.77332 | 308.6521 |
| SNELFTTFR | 9 | Q9NUQ3 | Gamma-taxilin OS=Homo sapiens OX=9 | TXLNG | 60.586 | 2 | 51.547333 | yes | 9.758E-08 | 557.78003 | 245.57321 |
| YADLLEESR | 9 | Q9NUQ3 | Gamma-taxilin OS=Homo sapiens OX=9 | TXLNG | 60.586 | 2 | 51.547333 | yes | 9.722E-08 | 548.26947 | 245.36613 |
| FEEFQTTMAK | 10 | Q9NUQ3 | Gamma-taxilin OS=Homo sapiens OX=9 | TXLNG | 60.586 | 2 | 51.547333 | yes | 9.819E-08 | 616.2868 | 228.77434 |
| NLVSPAYCTQESR | 13 | Q9NUQ3 | Gamma-taxilin OS=Homo sapiens OX=9 | TXLNG | 60.586 | 2 | 51.547333 | yes | 9.722E-08 | 762.85919 | 194.3027 |
| LQQTTQLIK | 9 | Q9NUQ3 | Gamma-taxilin OS=Homo sapiens OX=9 | TXLNG | 60.586 | 2 | 51.547333 | yes | 9.81E-08 | 536.82166 | 127.20853 |
| LIEQYALR | 8 | Q9NUQ3 | Gamma-taxilin OS=Homo sapiens OX=9 | TXLNG | 60.586 | 2 | 51.547333 | yes | 9.729E-07 | 503.29001 | 111.87379 |

| | | | | | | | | | | | |
|---|---|---|---|---|---|---|---|---|---|---|---|
| QENIELGEK | 9 | Q9NUQ3 | Gamma-taxilin OS=Homo sapiens OX=9( | TXLNG | 60.586 | 2 | 51.547333 | yes | 4.973E-07 | 530.26947 | 95.820648 |
| ELQQQLVDAK | 10 | Q9NUQ3 | Gamma-taxilin OS=Homo sapiens OX=9( | TXLNG | 60.586 | 2 | 51.547333 | yes | 1.136E-07 | 586.31952 | 74.741211 |
| QQEVQLK | 7 | Q9NUQ3 | Gamma-taxilin OS=Homo sapiens OX=9( | TXLNG | 60.586 | 2 | 51.547333 | yes | 6.9E-07 | 436.74545 | 59.399658 |
| KLIEQYALR | 9 | Q9NUQ3 | Gamma-taxilin OS=Homo sapiens OX=9( | TXLNG | 60.586 | 2 | 51.547333 | yes | 7.462E-07 | 567.33752 | 33.821678 |
| LRQENIELGEK | 11 | Q9NUQ3 | Gamma-taxilin OS=Homo sapiens OX=9( | TXLNG | 60.586 | 2 | 51.547333 | yes | 4.419E-07 | 664.86206 | 27.701439 |
| GGGAEEATEAGR | 12 | Q9NUQ3 | Gamma-taxilin OS=Homo sapiens OX=9( | TXLNG | 60.586 | 2 | 51.547333 | yes | 2.942E-06 | 552.74945 | 2.2139883 |
| TASWLAAELSK | 11 | Q9NUU7 | ATP-dependent RNA helicase DDX19A ( | DDX19A | 53.976 | 2 | 49.10931 | yes | 2.072E-06 | 588.81659 | 63.230942 |
| VFVLDEADVMIAT | 22 | Q9NUU7 | ATP-dependent RNA helicase DDX19A ( | DDX19A | 53.976 | 3 | 49.10931 | yes | 1.067E-07 | 824.74445 | 49.402336 |
| DPNSPLYSVK | 10 | Q9NUU7 | ATP-dependent RNA helicase DDX19A ( | DDX19A | 53.976 | 2 | 49.10931 | yes | 5.176E-07 | 560.28766 | 48.088425 |
| GNPFQFYLTR | 10 | Q9NUW8 | Tyrosyl-DNA phosphodiesterase 1 OS=H | TDP1 | 68.421 | 2 | 47.186794 | yes | 1.381E-07 | 621.81696 | 115.47479 |
| EWIDVIHK | 8 | Q9NUW8 | Tyrosyl-DNA phosphodiesterase 1 OS=H | TDP1 | 68.421 | 2 | 47.186794 | yes | 1.707E-07 | 520.28217 | 88.712051 |
| HDLSETNVYLIGST | 17 | Q9NUW8 | Tyrosyl-DNA phosphodiesterase 1 OS=H | TDP1 | 68.421 | 3 | 47.186794 | yes | 2.393E-07 | 620.31464 | 40.234749 |
| APDTHGNMWVPS | 12 | Q9NUW8 | Tyrosyl-DNA phosphodiesterase 1 OS=H | TDP1 | 68.421 | 2 | 47.186794 | yes | 1.315E-06 | 656.29297 | 38.98233 |
| LATVLSGACDGEV | 14 | Q9NV06 | DDB1- and CUL4-associated factor 13 O | DCAF13 | 51.403 | 2 | 54.969334 | yes | 1.702E-07 | 724.36414 | 521.1499 |
| SIYSQIQEQR | 10 | Q9NV06 | DDB1- and CUL4-associated factor 13 O | DCAF13 | 51.403 | 2 | 54.969334 | yes | 9.722E-08 | 626.32001 | 462.53152 |
| YIMCGSDEMNIR | 12 | Q9NV06 | DDB1- and CUL4-associated factor 13 O | DCAF13 | 51.403 | 2 | 54.969334 | yes | 3.705E-07 | 744.81763 | 427.08768 |
| EFVSASFDK | 9 | Q9NV06 | DDB1- and CUL4-associated factor 13 O | DCAF13 | 51.403 | 2 | 54.969334 | yes | 1.835E-07 | 515.24799 | 399.32849 |
| NYDPALHPFEVPR | 13 | Q9NV06 | DDB1- and CUL4-associated factor 13 O | DCAF13 | 51.403 | 2 | 54.969334 | yes | 5.131E-07 | 777.88861 | 196.63251 |
| NYDPALHPFEVPR | 13 | Q9NV06 | DDB1- and CUL4-associated factor 13 O | DCAF13 | 51.403 | 2 | 54.969334 | yes | 1.724E-07 | 518.92816 | 196.63251 |
| NYDPALHPFEVPR | 13 | Q9NV06 | DDB1- and CUL4-associated factor 13 O | DCAF13 | 51.403 | 3 | 54.969334 | yes | 5.131E-07 | 777.88861 | 196.63251 |
| NYDPALHPFEVPR | 13 | Q9NV06 | DDB1- and CUL4-associated factor 13 O | DCAF13 | 51.403 | 3 | 54.969334 | yes | 1.724E-07 | 518.92816 | 196.63251 |
| TIQAHEGFVR | 10 | Q9NV06 | DDB1- and CUL4-associated factor 13 O | DCAF13 | 51.403 | 2 | 54.969334 | yes | 9.722E-08 | 579.3067 | 299.91748 |
| MQHVICVK | 8 | Q9NV06 | DDB1- and CUL4-associated factor 13 O | DCAF13 | 51.403 | 2 | 54.969334 | yes | 4.013E-07 | 507.7648 | 298.88953 |
| IWNLTQR | 7 | Q9NV06 | DDB1- and CUL4-associated factor 13 O | DCAF13 | 51.403 | 2 | 54.969334 | yes | 3.333E-06 | 465.76141 | 273.96851 |
| PGSVPLVSEK | 10 | Q9NV06 | DDB1- and CUL4-associated factor 13 O | DCAF13 | 51.403 | 2 | 54.969334 | yes | 2.136E-07 | 506.78729 | 245.79132 |
| FCGTSFFTVGDDK | 13 | Q9NV06 | DDB1- and CUL4-associated factor 13 O | DCAF13 | 51.403 | 2 | 54.969334 | yes | 1.555E-07 | 740.82428 | 222.45963 |
| HSKPGSVPLVSEK | 13 | Q9NV06 | DDB1- and CUL4-associated factor 13 O | DCAF13 | 51.403 | 3 | 54.969334 | yes | 2.965E-07 | 455.58926 | 96.213791 |
| HSKPGSVPLVSEK | 13 | Q9NV06 | DDB1- and CUL4-associated factor 13 O | DCAF13 | 51.403 | 3 | 54.969334 | yes | 1.839E-07 | 682.88025 | 96.213791 |
| HSKPGSVPLVSEK | 13 | Q9NV06 | DDB1- and CUL4-associated factor 13 O | DCAF13 | 51.403 | 2 | 54.969334 | yes | 2.965E-07 | 455.58926 | 96.213791 |
| HSKPGSVPLVSEK | 13 | Q9NV06 | DDB1- and CUL4-associated factor 13 O | DCAF13 | 51.403 | 2 | 54.969334 | yes | 1.839E-07 | 682.88025 | 96.213791 |
| NIVLYDMR | 8 | Q9NV06 | DDB1- and CUL4-associated factor 13 O | DCAF13 | 51.403 | 2 | 54.969334 | yes | 3.21E-06 | 512.26825 | 187.0513 |
| TVYTGIDHHWK | 11 | Q9NV06 | DDB1- and CUL4-associated factor 13 O | DCAF13 | 51.403 | 2 | 54.969334 | yes | 9.722E-08 | 678.83838 | 181.7382 |
| PFLASLDGHR | 10 | Q9NV06 | DDB1- and CUL4-associated factor 13 O | DCAF13 | 51.403 | 2 | 54.969334 | yes | 1.492E-07 | 556.79602 | 137.84486 |
| FQHYPHIK | 8 | Q9NV06 | DDB1- and CUL4-associated factor 13 O | DCAF13 | 51.403 | 2 | 54.969334 | yes | 4.225E-07 | 535.28253 | 73.766212 |
| KVILDMR | 7 | Q9NV06 | DDB1- and CUL4-associated factor 13 O | DCAF13 | 51.403 | 2 | 54.969334 | yes | 2.865E-06 | 437.76257 | 29.486977 |
| LYALGLVPTR | 10 | Q9NV31 | U3 small nucleolar ribonucleoprotein pro | IMP3 | 21.851 | 2 | 51.635242 | yes | 4.921E-07 | 551.83459 | 592.80762 |
| VGPDVVTDPAFLV | 15 | Q9NV31 | U3 small nucleolar ribonucleoprotein pro | IMP3 | 21.851 | 2 | 51.635242 | yes | 9.722E-08 | 793.43286 | 570.33063 |
| MAQHLQAAVAFV | 18 | Q9NV31 | U3 small nucleolar ribonucleoprotein pro | IMP3 | 21.851 | 3 | 51.635242 | yes | 2.082E-07 | 664.67926 | 202.79257 |
| SMEDFVTWVDSSI | 13 | Q9NV31 | U3 small nucleolar ribonucleoprotein pro | IMP3 | 21.851 | 2 | 51.635242 | yes | 2.008E-07 | 765.84265 | 135.44933 |

| | | | | | | | | | | | |
|---|---|---|---|---|---|---|---|---|---|---|---|
| VLTANSNPSSPSA/ | 15 | Q9NV56 | MRG/MORF4L-binding protein OS=Hon | MRGBP | 22.418 | 2 | 54.061718 | yes | 2.012E-07 | 722.37555 | 75.229065 |
| NFVLPEEIIQEVR | 13 | Q9NV56 | MRG/MORF4L-binding protein OS=Hon | MRGBP | 22.418 | 2 | 54.061718 | yes | 9.722E-08 | 793.43286 | 62.421604 |
| ILGDLSYR | 8 | Q9NVC6 | Mediator of RNA polymerase II transcrip | MED17 | 72.891 | 2 | 50.934746 | yes | 1.266E-07 | 468.76108 | 122.65983 |
| VLITSQGYEQICK | 13 | Q9NVC6 | Mediator of RNA polymerase II transcrip | MED17 | 72.891 | 2 | 50.934746 | yes | 4.716E-07 | 769.89777 | 94.455383 |
| LEAAQNVLLCK | 11 | Q9NVC6 | Mediator of RNA polymerase II transcrip | MED17 | 72.891 | 2 | 50.934746 | yes | 3.25E-06 | 629.84485 | 65.336174 |
| AAATIDSLASR | 11 | Q9NVC6 | Mediator of RNA polymerase II transcrip | MED17 | 72.891 | 2 | 50.934746 | yes | 3.551E-07 | 538.29077 | 64.663666 |
| QAPDIGDLGTVNL | 15 | Q9NVC6 | Mediator of RNA polymerase II transcrip | MED17 | 72.891 | 2 | 50.934746 | yes | 1.65E-07 | 794.42249 | 62.706676 |
| QNPQTLQLISK | 11 | Q9NVC6 | Mediator of RNA polymerase II transcrip | MED17 | 72.891 | 2 | 50.934746 | yes | 2.464E-07 | 635.36169 | 59.941788 |
| NEINSLQSSEGLLE | 15 | Q9NVC6 | Mediator of RNA polymerase II transcrip | MED17 | 72.891 | 2 | 50.934746 | yes | 2.177E-07 | 830.92303 | 46.729839 |
| EIFAQLSR | 8 | Q9NVC6 | Mediator of RNA polymerase II transcrip | MED17 | 72.891 | 2 | 50.934746 | yes | 9.722E-08 | 482.26654 | 28.538029 |
| MDLVCQR | 7 | Q9NVF7 | F-box only protein 28 OS=Homo sapiens | FBXO28 | 41.15 | 2 | 52.632523 | yes | 9.722E-08 | 461.21768 | 142.46503 |
| HADILAAVETR | 11 | Q9NVF7 | F-box only protein 28 OS=Homo sapiens | FBXO28 | 41.15 | 2 | 52.632523 | yes | 9.722E-08 | 598.32513 | 101.3281 |
| LLEQTQIIGEQNAR | 14 | Q9NVF7 | F-box only protein 28 OS=Homo sapiens | FBXO28 | 41.15 | 2 | 52.632523 | yes | 9.722E-08 | 806.93628 | 71.097595 |
| AHEVLQELR | 9 | Q9NVF7 | F-box only protein 28 OS=Homo sapiens | FBXO28 | 41.15 | 2 | 52.632523 | yes | 3.035E-07 | 547.80127 | 52.026318 |
| DISSMAMEYFDEK | 13 | Q9NVF7 | F-box only protein 28 OS=Homo sapiens | FBXO28 | 41.15 | 2 | 52.632523 | yes | 9.722E-08 | 783.32855 | 46.259628 |
| EFQAPIGSEEGLR | 13 | Q9NVH0 | Exonuclease 3'-5' domain-containing prot | EXD2 | 70.353 | 2 | 53.408688 | yes | 1.193E-07 | 716.85699 | 500.76685 |
| LLQDYGLVVR | 10 | Q9NVH0 | Exonuclease 3'-5' domain-containing prot | EXD2 | 70.353 | 2 | 53.408688 | yes | 9.722E-08 | 588.34277 | 461.59076 |
| ENLCVVCGK | 9 | Q9NVH0 | Exonuclease 3'-5' domain-containing prot | EXD2 | 70.353 | 2 | 53.408688 | yes | 2.475E-07 | 539.75464 | 268.15723 |
| GIGELVSEEPFVVK | 14 | Q9NVH0 | Exonuclease 3'-5' domain-containing prot | EXD2 | 70.353 | 2 | 53.408688 | yes | 9.722E-08 | 751.90869 | 250.89067 |
| KEELLQALR | 9 | Q9NVH0 | Exonuclease 3'-5' domain-containing prot | EXD2 | 70.353 | 2 | 53.408688 | yes | 9.758E-08 | 550.32715 | 249.16681 |
| NVIPHEYR | 8 | Q9NVH0 | Exonuclease 3'-5' domain-containing prot | EXD2 | 70.353 | 2 | 53.408688 | yes | 1.002E-07 | 514.26959 | 243.90862 |
| TSPVTQQPQQK | 11 | Q9NVH0 | Exonuclease 3'-5' domain-containing prot | EXD2 | 70.353 | 2 | 53.408688 | yes | 1.06E-07 | 621.32788 | 190.40659 |
| VGVGCSEDASK | 11 | Q9NVH0 | Exonuclease 3'-5' domain-containing prot | EXD2 | 70.353 | 2 | 53.408688 | yes | 1.254E-07 | 554.75061 | 176.6745 |
| SLMQLESR | 8 | Q9NVH0 | Exonuclease 3'-5' domain-containing prot | EXD2 | 70.353 | 2 | 53.408688 | yes | 1.624E-07 | 482.25003 | 176.1368 |
| MDGMVPGNHQGF | 12 | Q9NVH0 | Exonuclease 3'-5' domain-containing prot | EXD2 | 70.353 | 2 | 53.408688 | yes | 2.734E-07 | 649.79004 | 150.75 |
| ISNENYVPHGLK | 12 | Q9NVH0 | Exonuclease 3'-5' domain-containing prot | EXD2 | 70.353 | 2 | 53.408688 | yes | 2.139E-07 | 685.85681 | 118.28823 |
| VVQCHSQGGLR | 11 | Q9NVH0 | Exonuclease 3'-5' domain-containing prot | EXD2 | 70.353 | 2 | 53.408688 | yes | 9.722E-08 | 620.81439 | 104.96896 |
| ALLNAESLPTQR | 12 | Q9NVH0 | Exonuclease 3'-5' domain-containing prot | EXD2 | 70.353 | 2 | 53.408688 | yes | 1.577E-07 | 656.86462 | 101.91318 |
| NDDHSSWR | 8 | Q9NVH0 | Exonuclease 3'-5' domain-containing prot | EXD2 | 70.353 | 2 | 53.408688 | yes | 1.42E-07 | 508.71268 | 81.198776 |
| SLAETVLNFPLDK | 13 | Q9NVH0 | Exonuclease 3'-5' domain-containing prot | EXD2 | 70.353 | 2 | 53.408688 | yes | 1.823E-07 | 723.89557 | 67.990524 |
| KFGEDLPIQLS | 11 | Q9NVH0 | Exonuclease 3'-5' domain-containing prot | EXD2 | 70.353 | 2 | 53.408688 | yes | 2.381E-06 | 623.83752 | 53.171532 |
| ELPPPEDDQLHSS/ | 16 | Q9NVH0 | Exonuclease 3'-5' domain-containing prot | EXD2 | 70.353 | 3 | 53.408688 | yes | 0.0001022 | 596.62347 | 31.643978 |
| NNLLCNGLSLK | 11 | Q9NVH0 | Exonuclease 3'-5' domain-containing prot | EXD2 | 70.353 | 2 | 53.408688 | yes | 3.846E-06 | 623.33466 | 27.607338 |
| CQGVVDIPFR | 10 | Q9NVH0 | Exonuclease 3'-5' domain-containing prot | EXD2 | 70.353 | 2 | 53.408688 | yes | 9.722E-08 | 595.80298 | 22.469341 |
| AGFFGTVVEYGAE | 14 | Q9NVH1 | DnaJ homolog subfamily C member 11 O | DNAJC11 | 63.279 | 2 | 51.063076 | yes | 9.722E-08 | 751.86737 | 122.70277 |
| WGIQSAMNTSIVR | 13 | Q9NVH1 | DnaJ homolog subfamily C member 11 O | DNAJC11 | 63.279 | 2 | 51.063076 | yes | 2.83E-07 | 731.8772 | 91.035637 |
| AGLPGFYDPCVGE | 15 | Q9NVH1 | DnaJ homolog subfamily C member 11 O | DNAJC11 | 63.279 | 2 | 51.063076 | yes | 2.869E-06 | 819.87708 | 68.527603 |
| LILTEASK | 8 | Q9NVH1 | DnaJ homolog subfamily C member 11 O | DNAJC11 | 63.279 | 2 | 51.063076 | yes | 4.359E-07 | 437.76584 | 67.024864 |
| GLEMEGWEVVER | 12 | Q9NVH1 | DnaJ homolog subfamily C member 11 O | DNAJC11 | 63.279 | 2 | 51.063076 | yes | 1.056E-07 | 717.34009 | 46.756802 |

| | | | | | | | | | | | |
|---|---|---|---|---|---|---|---|---|---|---|---|
| VIDVTVPLQCLVK | 13 | Q9NVH1 | DnaJ homolog subfamily C member 11 O | DNAJC11 | 63.279 | 2 | 51.063076 | yes | 8.138E-07 | 742.43127 | 46.491096 |
| NILMYGPPGTGK | 12 | Q9NVI7 | ATPase family AAA domain-containing p | ATAD3A | 71.37 | 2 | 53.948174 | yes | 1.74E-07 | 624.32629 | 1933.073 |
| QTVLESIR | 8 | Q9NVI7 | ATPase family AAA domain-containing p | ATAD3A | 71.37 | 2 | 53.948174 | no | 1.041E-07 | 473.27182 | 1878.0669 |
| ITVLEALR | 8 | Q9NVI7 | ATPase family AAA domain-containing p | ATAD3A | 71.37 | 2 | 53.948174 | no | 3.394E-07 | 457.78711 | 1862.4012 |
| LAQFDYGR | 8 | Q9NVI7 | ATPase family AAA domain-containing p | ATAD3A | 71.37 | 2 | 53.948174 | no | 6.846E-07 | 485.24307 | 1709.3463 |
| GLGDRPAPK | 9 | Q9NVI7 | ATPase family AAA domain-containing p | ATAD3A | 71.37 | 2 | 53.948174 | no | 2.687E-07 | 455.75888 | 1543.9818 |
| LGKPSLVR | 8 | Q9NVI7 | ATPase family AAA domain-containing p | ATAD3A | 71.37 | 2 | 53.948174 | no | 3.557E-07 | 435.28198 | 1505.2555 |
| QQQLLNEENLR | 11 | Q9NVI7 | ATPase family AAA domain-containing p | ATAD3A | 71.37 | 2 | 53.948174 | no | 9.722E-08 | 692.86261 | 1499.7701 |
| HPIQVSR | 7 | Q9NVI7 | ATPase family AAA domain-containing p | ATAD3A | 71.37 | 2 | 53.948174 | no | 1.318E-07 | 418.74051 | 1166.0707 |
| TAGTLFGEGFR | 11 | Q9NVI7 | ATPase family AAA domain-containing p | ATAD3A | 71.37 | 2 | 53.948174 | no | 6.145E-07 | 578.29327 | 1111.5645 |
| ATLNAFLYR | 9 | Q9NVI7 | ATPase family AAA domain-containing p | ATAD3A | 71.37 | 2 | 53.948174 | no | 1.003E-07 | 534.79547 | 976.76624 |
| YVLKPATEGK | 10 | Q9NVI7 | ATPase family AAA domain-containing p | ATAD3A | 71.37 | 2 | 53.948174 | yes | 9.809E-08 | 553.31622 | 849.31714 |
| LLSRPQDALEGVV | 21 | Q9NVI7 | ATPase family AAA domain-containing p | ATAD3A | 71.37 | 3 | 53.948174 | yes | 2.02E-07 | 750.75323 | 741.59747 |
| DKWSNFDPTGLEF | 13 | Q9NVI7 | ATPase family AAA domain-containing p | ATAD3A | 71.37 | 2 | 53.948174 | no | 1.66E-07 | 782.87317 | 446.34894 |
| LALHSGMDYAIMT | 22 | Q9NVI7 | ATPase family AAA domain-containing p | ATAD3A | 71.37 | 3 | 53.948174 | no | 9.722E-08 | 755.0271 | 173.68507 |
| LALHSGMDYAIMT | 22 | Q9NVI7 | ATPase family AAA domain-containing p | ATAD3A | 71.37 | 3 | 53.948174 | no | 9.903E-08 | 760.35876 | 173.68507 |
| ENADIIR | 7 | Q9NVI7 | ATPase family AAA domain-containing p | ATAD3A | 71.37 | 2 | 53.948174 | no | 4.229E-07 | 415.72195 | 132.29465 |
| QRYEDQLK | 8 | Q9NVI7 | ATPase family AAA domain-containing p | ATAD3A | 71.37 | 2 | 53.948174 | no | 1.134E-06 | 540.27765 | 96.663284 |
| LFDWANTSR | 9 | Q9NVI7 | ATPase family AAA domain-containing p | ATAD3A | 71.37 | 2 | 53.948174 | no | 9.722E-08 | 555.27234 | 81.534546 |
| PQDALEGVVLSPS | 17 | Q9NVI7 | ATPase family AAA domain-containing p | ATAD3A | 71.37 | 3 | 53.948174 | yes | 1.274E-06 | 594.31952 | 41.648151 |
| MNLSAIQDR | 9 | Q9NVJ2 | ADP-ribosylation factor-like protein 8B ( | ARL8B | 21.54 | 2 | 49.955734 | no | 1.284E-07 | 524.26624 | 464.11594 |
| DLPNALDEK | 9 | Q9NVJ2 | ADP-ribosylation factor-like protein 8B ( | ARL8B | 21.54 | 2 | 49.955734 | yes | 1.604E-07 | 507.75876 | 186.43071 |
| EICCYSISCK | 10 | Q9NVJ2 | ADP-ribosylation factor-like protein 8B ( | ARL8B | 21.54 | 2 | 49.955734 | no | 9.968E-08 | 660.27509 | 50.046124 |
| EEMELTLVGLQYS | 15 | Q9NVJ2 | ADP-ribosylation factor-like protein 8B ( | ARL8B | 21.54 | 2 | 49.955734 | no | 1.664E-06 | 848.92676 | 45.679142 |
| EKDNIDITLQWLIQ | 17 | Q9NVJ2 | ADP-ribosylation factor-like protein 8B ( | ARL8B | 21.54 | 3 | 49.955734 | no | 3.737E-07 | 694.37238 | 26.421968 |
| SILEDPPSISEGCGC | 16 | Q9NVM9 | Integrator complex subunit 13 OS=Homo | INTS13 | 80.226 | 2 | 50.21811 | yes | 2.893E-07 | 837.39362 | 101.66048 |
| ITDFGEFMR | 9 | Q9NVM9 | Integrator complex subunit 13 OS=Homo | INTS13 | 80.226 | 2 | 50.21811 | yes | 9.849E-08 | 558.26312 | 78.555283 |
| IMWNELETLVR | 11 | Q9NVM9 | Integrator complex subunit 13 OS=Homo | INTS13 | 80.226 | 2 | 50.21811 | yes | 4.124E-07 | 702.37122 | 43.655537 |
| TLLMENAER | 9 | Q9NVM9 | Integrator complex subunit 13 OS=Homo | INTS13 | 80.226 | 2 | 50.21811 | yes | 2.056E-06 | 538.77386 | 42.154945 |
| VLECLMACR | 9 | Q9NVM9 | Integrator complex subunit 13 OS=Homo | INTS13 | 80.226 | 2 | 50.21811 | yes | 2.45E-07 | 576.27216 | 37.99395 |
| ESLTEEDVLNCQK | 13 | Q9NVM9 | Integrator complex subunit 13 OS=Homo | INTS13 | 80.226 | 2 | 50.21811 | yes | 3.708E-07 | 782.86157 | 30.125797 |
| GKEELAEAEIIK | 12 | Q9NVM9 | Integrator complex subunit 13 OS=Homo | INTS13 | 80.226 | 2 | 50.21811 | yes | 1.177E-06 | 665.36664 | 14.91214 |
| CSVPVDQASESLL | 14 | Q9NVN8 | Guanine nucleotide-binding protein-like 3 | GNL3L | 65.574 | 2 | 51.13598 | yes | 1.699E-07 | 766.88489 | 2069.7612 |
| LADPVTPVETILQF | 14 | Q9NVN8 | Guanine nucleotide-binding protein-like 3 | GNL3L | 65.574 | 2 | 51.13598 | yes | 1.171E-07 | 776.44067 | 746.39545 |
| LADPVTPVETILQF | 14 | Q9NVN8 | Guanine nucleotide-binding protein-like 3 | GNL3L | 65.574 | 2 | 51.13598 | yes | 3.085E-06 | 517.96289 | 746.39545 |
| LADPVTPVETILQF | 14 | Q9NVN8 | Guanine nucleotide-binding protein-like 3 | GNL3L | 65.574 | 3 | 51.13598 | yes | 1.171E-07 | 776.44067 | 746.39545 |
| LADPVTPVETILQF | 14 | Q9NVN8 | Guanine nucleotide-binding protein-like 3 | GNL3L | 65.574 | 3 | 51.13598 | yes | 3.085E-06 | 517.96289 | 746.39545 |
| ACFGAENLMR | 10 | Q9NVN8 | Guanine nucleotide-binding protein-like 3 | GNL3L | 65.574 | 2 | 51.13598 | yes | 9.722E-08 | 584.7655 | 1460.177 |
| ACSVGAVPGITK | 12 | Q9NVN8 | Guanine nucleotide-binding protein-like 3 | GNL3L | 65.574 | 2 | 51.13598 | yes | 1.675E-07 | 580.31061 | 1242.6401 |

| | | | | | | | | | | | |
|---|---|---|---|---|---|---|---|---|---|---|---|
| IGDLTGYCTNPNR | 13 | Q9NVN8 | Guanine nucleotide-binding protein-like 3 | GNL3L | 65.574 | 2 | 51.13598 | yes | 2.276E-07 | 740.84607 | 1080.6359 |
| CFQMEEAVLR | 10 | Q9NVN8 | Guanine nucleotide-binding protein-like 3 | GNL3L | 65.574 | 2 | 51.13598 | yes | 1.02E-07 | 641.79956 | 1061.8651 |
| TIESYCQDVLR | 11 | Q9NVN8 | Guanine nucleotide-binding protein-like 3 | GNL3L | 65.574 | 2 | 51.13598 | yes | 1.185E-07 | 692.33228 | 929.96014 |
| FMQEVYLDK | 9 | Q9NVN8 | Guanine nucleotide-binding protein-like 3 | GNL3L | 65.574 | 2 | 51.13598 | yes | 9.722E-08 | 586.78644 | 875.6261 |
| SNSMVDVCSVDR | 12 | Q9NVN8 | Guanine nucleotide-binding protein-like 3 | GNL3L | 65.574 | 2 | 51.13598 | yes | 9.79E-08 | 684.79773 | 843.75714 |
| VGVVGLPNVGK | 11 | Q9NVN8 | Guanine nucleotide-binding protein-like 3 | GNL3L | 65.574 | 2 | 51.13598 | yes | 2.204E-07 | 519.81897 | 609.34955 |
| NELPTVAFK | 9 | Q9NVN8 | Guanine nucleotide-binding protein-like 3 | GNL3L | 65.574 | 2 | 51.13598 | yes | 1.999E-07 | 509.78201 | 460.59821 |
| ISWPYPQPAK | 10 | Q9NVN8 | Guanine nucleotide-binding protein-like 3 | GNL3L | 65.574 | 2 | 51.13598 | yes | 2.887E-07 | 593.81641 | 460.37595 |
| AVLADWVSGK | 10 | Q9NVN8 | Guanine nucleotide-binding protein-like 3 | GNL3L | 65.574 | 2 | 51.13598 | yes | 9.978E-08 | 523.28748 | 446.15756 |
| VLGNYCR | 7 | Q9NVN8 | Guanine nucleotide-binding protein-like 3 | GNL3L | 65.574 | 2 | 51.13598 | yes | 2.26E-07 | 441.21854 | 378.34613 |
| GGLYSQEQAAK | 11 | Q9NVN8 | Guanine nucleotide-binding protein-like 3 | GNL3L | 65.574 | 2 | 51.13598 | yes | 2.021E-07 | 576.28821 | 175.58952 |
| KLVLVLNK | 8 | Q9NVN8 | Guanine nucleotide-binding protein-like 3 | GNL3L | 65.574 | 2 | 51.13598 | yes | 6.622E-07 | 463.82349 | 156.22966 |
| HQMGWAK | 7 | Q9NVN8 | Guanine nucleotide-binding protein-like 3 | GNL3L | 65.574 | 2 | 51.13598 | yes | 2.643E-06 | 429.20798 | 98.681091 |
| KGGLYSQEQAAK | 12 | Q9NVN8 | Guanine nucleotide-binding protein-like 3 | GNL3L | 65.574 | 2 | 51.13598 | yes | 1.413E-07 | 640.33569 | 93.389938 |
| LLHSPMTK | 8 | Q9NVN8 | Guanine nucleotide-binding protein-like 3 | GNL3L | 65.574 | 2 | 51.13598 | yes | 1.768E-07 | 463.76004 | 91.676926 |
| CNLEEISNYYGVS | 28 | Q9NVN8 | Guanine nucleotide-binding protein-like 3 | GNL3L | 65.574 | 4 | 51.13598 | yes | 5.605E-07 | 811.63458 | 67.211929 |
| NLQCLVIDEADR | 12 | Q9NVP1 | ATP-dependent RNA helicase DDX18 O | DDX18 | 75.407 | 2 | 52.96278 | yes | 9.722E-08 | 723.35626 | 940.63025 |
| ILDVGFEEELK | 11 | Q9NVP1 | ATP-dependent RNA helicase DDX18 O | DDX18 | 75.407 | 2 | 52.96278 | yes | 9.794E-08 | 646.34265 | 850.59686 |
| KVEDLAR | 7 | Q9NVP1 | ATP-dependent RNA helicase DDX18 O | DDX18 | 75.407 | 2 | 52.96278 | yes | 3.929E-07 | 415.74017 | 684.63159 |
| ELAMQTFGVLK | 11 | Q9NVP1 | ATP-dependent RNA helicase DDX18 O | DDX18 | 75.407 | 2 | 52.96278 | yes | 9.722E-08 | 618.83649 | 525.88507 |
| ISDIQSQLEK | 10 | Q9NVP1 | ATP-dependent RNA helicase DDX18 O | DDX18 | 75.407 | 2 | 52.96278 | yes | 9.722E-08 | 580.81152 | 462.46988 |
| QTMLFSATQTR | 11 | Q9NVP1 | ATP-dependent RNA helicase DDX18 O | DDX18 | 75.407 | 2 | 52.96278 | yes | 9.893E-08 | 642.32428 | 454.68933 |
| EPLYVGVDDDK | 11 | Q9NVP1 | ATP-dependent RNA helicase DDX18 O | DDX18 | 75.407 | 2 | 52.96278 | yes | 2.286E-07 | 625.30096 | 403.29532 |
| GHALLILRPEELGF | 16 | Q9NVP1 | ATP-dependent RNA helicase DDX18 O | DDX18 | 75.407 | 3 | 52.96278 | yes | 2.816E-07 | 612.02808 | 378.07309 |
| VPPFVDLNVNSNE | 15 | Q9NVP1 | ATP-dependent RNA helicase DDX18 O | DDX18 | 75.407 | 2 | 52.96278 | yes | 9.722E-08 | 814.91754 | 371.64703 |
| VPLSEFDFSWSK | 12 | Q9NVP1 | ATP-dependent RNA helicase DDX18 O | DDX18 | 75.407 | 2 | 52.96278 | yes | 1.144E-07 | 721.35352 | 358.53873 |
| EMGFTNMTEIQHK | 13 | Q9NVP1 | ATP-dependent RNA helicase DDX18 O | DDX18 | 75.407 | 2 | 52.96278 | yes | 9.722E-08 | 783.35797 | 228.53633 |
| SIRPLLEGR | 9 | Q9NVP1 | ATP-dependent RNA helicase DDX18 O | DDX18 | 75.407 | 2 | 52.96278 | yes | 9.722E-08 | 520.81421 | 224.4003 |
| NGTGVLILSPTR | 12 | Q9NVP1 | ATP-dependent RNA helicase DDX18 O | DDX18 | 75.407 | 2 | 52.96278 | yes | 1.212E-07 | 614.35645 | 170.87402 |
| LGNGINIIVATPGR | 14 | Q9NVP1 | ATP-dependent RNA helicase DDX18 O | DDX18 | 75.407 | 2 | 52.96278 | yes | 9.722E-08 | 697.90936 | 159.85327 |
| KEPLYVGVDDDK | 12 | Q9NVP1 | ATP-dependent RNA helicase DDX18 O | DDX18 | 75.407 | 2 | 52.96278 | yes | 4.359E-07 | 689.34845 | 110.51463 |
| SEEESAETTK | 10 | Q9NVP1 | ATP-dependent RNA helicase DDX18 O | DDX18 | 75.407 | 2 | 52.96278 | yes | 4.487E-07 | 555.74347 | 94.450554 |
| LLDHMQNTPGFM | 14 | Q9NVP1 | ATP-dependent RNA helicase DDX18 O | DDX18 | 75.407 | 2 | 52.96278 | yes | 1.112E-07 | 847.90509 | 80.096313 |
| AYDSHSLK | 8 | Q9NVP1 | ATP-dependent RNA helicase DDX18 O | DDX18 | 75.407 | 2 | 52.96278 | yes | 9.722E-08 | 460.72726 | 76.380508 |
| YHYELLNYIDLPV | 19 | Q9NVP1 | ATP-dependent RNA helicase DDX18 O | DDX18 | 75.407 | 3 | 52.96278 | yes | 2.162E-07 | 757.7439 | 74.00013 |
| MVNDAEPDTK | 10 | Q9NVP1 | ATP-dependent RNA helicase DDX18 O | DDX18 | 75.407 | 2 | 52.96278 | yes | 1.603E-05 | 560.25299 | 30.391691 |
| LMVFFSSCMSVK | 12 | Q9NVP1 | ATP-dependent RNA helicase DDX18 O | DDX18 | 75.407 | 2 | 52.96278 | yes | 1.577E-06 | 718.34277 | 18.435074 |
| ANATVDGLEQGY | 19 | Q9NVP1 | ATP-dependent RNA helicase DDX18 O | DDX18 | 75.407 | 3 | 52.96278 | yes | 4.989E-07 | 679.65796 | 8.9437609 |
| AFVEATFQR | 9 | Q9NVU0 | DNA-directed RNA polymerase III subun | POLR3E | 79.899 | 2 | 52.991486 | yes | 1.327E-07 | 534.77728 | 120.63837 |

| | | | | | | | | | | | |
|---|---|---|---|---|---|---|---|---|---|---|---|
| LTQECGEDLSK | 11 | Q9NVU0 | DNA-directed RNA polymerase III subun | POLR3E | 79.899 | 2 | 52.991486 | yes | 9.722E-08 | 640.29535 | 81.871338 |
| TLPLADQIK | 9 | Q9NVU0 | DNA-directed RNA polymerase III subun | POLR3E | 79.899 | 2 | 52.991486 | yes | 2.779E-07 | 499.79767 | 67.642738 |
| DFLEHMAVVR | 10 | Q9NVU0 | DNA-directed RNA polymerase III subun | POLR3E | 79.899 | 2 | 52.991486 | yes | 1.339E-07 | 608.81079 | 42.504631 |
| DCCVSYGGMWYL | 13 | Q9NVU0 | DNA-directed RNA polymerase III subun | POLR3E | 79.899 | 2 | 52.991486 | yes | 3.24E-06 | 819.84113 | 26.625723 |
| GKPTEASIEAR | 11 | Q9NVU7 | Protein SDA1 homolog OS=Homo sapien | SDAD1 | 79.872 | 2 | 49.00214 | yes | 2.972E-07 | 579.80933 | 237.54631 |
| TVNVITTACFSK | 12 | Q9NVU7 | Protein SDA1 homolog OS=Homo sapien | SDAD1 | 79.872 | 2 | 49.00214 | yes | 1.078E-07 | 670.8476 | 104.99936 |
| LPSNLPQLQNLIK | 13 | Q9NVU7 | Protein SDA1 homolog OS=Homo sapien | SDAD1 | 79.872 | 2 | 49.00214 | yes | 1.702E-07 | 739.44049 | 94.012238 |
| TLIHLFR | 7 | Q9NVU7 | Protein SDA1 homolog OS=Homo sapien | SDAD1 | 79.872 | 2 | 49.00214 | yes | 5.79E-07 | 450.2767 | 90.195488 |
| NSGEVMTVGINAI | 14 | Q9NVU7 | Protein SDA1 homolog OS=Homo sapien | SDAD1 | 79.872 | 2 | 49.00214 | yes | 1.369E-06 | 716.87683 | 45.596176 |
| LNSMPMEER | 9 | Q9NVU7 | Protein SDA1 homolog OS=Homo sapien | SDAD1 | 79.872 | 2 | 49.00214 | yes | 4.029E-06 | 553.75208 | 38.258884 |
| EFQLTEENTK | 10 | Q9NVV4 | Poly(A) RNA polymerase, mitochondrial | MTPAP | 66.172 | 2 | 54.386658 | yes | 1.031E-07 | 619.79858 | 2230.0652 |
| TVLIHCPEK | 9 | Q9NVV4 | Poly(A) RNA polymerase, mitochondrial | MTPAP | 66.172 | 2 | 54.386658 | yes | 1.128E-07 | 548.79462 | 2008.8331 |
| SPPILPTLDSLK | 12 | Q9NVV4 | Poly(A) RNA polymerase, mitochondrial | MTPAP | 66.172 | 2 | 54.386658 | yes | 1.316E-07 | 640.87665 | 1975.7168 |
| NVSQSQLQK | 9 | Q9NVV4 | Poly(A) RNA polymerase, mitochondrial | MTPAP | 66.172 | 2 | 54.386658 | yes | 3.275E-07 | 516.27765 | 1275.7959 |
| FSEMQNER | 8 | Q9NVV4 | Poly(A) RNA polymerase, mitochondrial | MTPAP | 66.172 | 2 | 54.386658 | yes | 1.865E-07 | 520.72693 | 1033.3739 |
| PFGSSVNTFGK | 11 | Q9NVV4 | Poly(A) RNA polymerase, mitochondrial | MTPAP | 66.172 | 2 | 54.386658 | yes | 2.671E-07 | 570.78784 | 838.89801 |
| ISGNFLMEFQVK | 12 | Q9NVV4 | Poly(A) RNA polymerase, mitochondrial | MTPAP | 66.172 | 2 | 54.386658 | yes | 9.722E-08 | 706.86578 | 743.48853 |
| CVIEGNNCTFVR | 12 | Q9NVV4 | Poly(A) RNA polymerase, mitochondrial | MTPAP | 66.172 | 2 | 54.386658 | yes | 9.722E-08 | 734.83722 | 714.41345 |
| RSPPILPTLDSLK | 13 | Q9NVV4 | Poly(A) RNA polymerase, mitochondrial | MTPAP | 66.172 | 2 | 54.386658 | yes | 5.075E-07 | 718.92719 | 299.65839 |
| ILSVLGECLDHFGI | 21 | Q9NVV4 | Poly(A) RNA polymerase, mitochondrial | MTPAP | 66.172 | 3 | 54.386658 | yes | 2.187E-06 | 762.38544 | 146.05336 |
| DEQPSGSVETGFEI | 15 | Q9NVV4 | Poly(A) RNA polymerase, mitochondrial | MTPAP | 66.172 | 2 | 54.386658 | yes | 9.819E-08 | 812.85229 | 125.49052 |
| NLLESLK | 7 | Q9NVV4 | Poly(A) RNA polymerase, mitochondrial | MTPAP | 66.172 | 2 | 54.386658 | yes | 2.294E-07 | 408.7449 | 112.05322 |
| FAIETVK | 7 | Q9NVV4 | Poly(A) RNA polymerase, mitochondrial | MTPAP | 66.172 | 2 | 54.386658 | yes | 9.849E-08 | 404.23419 | 84.456848 |
| NIPHLAANLK | 10 | Q9NVX0 | HAUS augmin-like complex subunit 2 OS | HAUS2 | 26.934 | 2 | 51.86504 | yes | 9.722E-08 | 545.82202 | 175.99068 |
| LQQITNIQAEIYQK | 14 | Q9NVX0 | HAUS augmin-like complex subunit 2 OS | HAUS2 | 26.934 | 2 | 51.86504 | yes | 9.722E-08 | 845.46216 | 142.47002 |
| LETHLETIR | 9 | Q9NVX0 | HAUS augmin-like complex subunit 2 OS | HAUS2 | 26.934 | 2 | 51.86504 | yes | 1.073E-07 | 556.30896 | 112.77023 |
| ILAEESYLYK | 10 | Q9NVX0 | HAUS augmin-like complex subunit 2 OS | HAUS2 | 26.934 | 2 | 51.86504 | yes | 1.16E-07 | 614.8266 | 107.97098 |
| TVSCFVNFTR | 10 | Q9NVX0 | HAUS augmin-like complex subunit 2 OS | HAUS2 | 26.934 | 2 | 51.86504 | yes | 1.438E-07 | 615.80042 | 55.780151 |
| WGGDGLLYSASQI | 14 | Q9NVX2 | Notchless protein homolog 1 OS=Homo s | NLE1 | 53.321 | 2 | 51.95377 | yes | 4.561E-07 | 762.85748 | 169.2411 |
| VLDIIYQPQAIFR | 13 | Q9NVX2 | Notchless protein homolog 1 OS=Homo s | NLE1 | 53.321 | 2 | 51.95377 | yes | 9.722E-08 | 788.4483 | 160.5632 |
| LLVSGSSDSTLK | 12 | Q9NVX2 | Notchless protein homolog 1 OS=Homo s | NLE1 | 53.321 | 2 | 51.95377 | yes | 2.081E-07 | 603.83246 | 150.37956 |
| TLESQAVETEK | 11 | Q9NVX2 | Notchless protein homolog 1 OS=Homo s | NLE1 | 53.321 | 2 | 51.95377 | yes | 9.722E-08 | 617.81171 | 131.19075 |
| LAMDLPGHADEV | 23 | Q9NVX2 | Notchless protein homolog 1 OS=Homo s | NLE1 | 53.321 | 3 | 51.95377 | yes | 5.096E-06 | 848.06042 | 58.649792 |
| VMGPMLDAATR | 11 | Q9NW08 | DNA-directed RNA polymerase III subun | POLR3B | 127.785 | 2 | 53.784378 | yes | 1.12E-07 | 581.29138 | 211.38921 |
| VILIQEQLSK | 10 | Q9NW08 | DNA-directed RNA polymerase III subun | POLR3B | 127.785 | 2 | 53.784378 | yes | 9.722E-08 | 585.85828 | 166.2998 |
| SSCHVSSLR | 9 | Q9NW08 | DNA-directed RNA polymerase III subun | POLR3B | 127.785 | 2 | 53.784378 | yes | 1.865E-07 | 516.74817 | 153.15201 |
| VTSDADPMWYLK | 12 | Q9NW08 | DNA-directed RNA polymerase III subun | POLR3B | 127.785 | 2 | 53.784378 | yes | 2.032E-07 | 713.3396 | 106.19813 |
| IIVEADRK | 8 | Q9NW08 | DNA-directed RNA polymerase III subun | POLR3B | 127.785 | 2 | 53.784378 | yes | 1.402E-07 | 472.2822 | 78.576317 |
| QAMGTIGYNQR | 11 | Q9NW08 | DNA-directed RNA polymerase III subun | POLR3B | 127.785 | 2 | 53.784378 | yes | 1.483E-07 | 619.80096 | 45.665863 |

| | | | | | | | | | | | |
|---|---|---|---|---|---|---|---|---|---|---|---|
| HGYNYLGK | 8 | Q9NW08 | DNA-directed RNA polymerase III subun | POLR3B | 127.785 | 2 | 53.784378 | yes | 1.248E-07 | 476.23779 | 40.991219 |
| CIYTAVMVR | 9 | Q9NW08 | DNA-directed RNA polymerase III subun | POLR3B | 127.785 | 2 | 53.784378 | yes | 3.602E-07 | 556.7832 | 32.2136 |
| AQIFTQMQALK | 11 | Q9NW08 | DNA-directed RNA polymerase III subun | POLR3B | 127.785 | 2 | 53.784378 | yes | 1.066E-07 | 639.84735 | 27.421282 |
| VLALPSHR | 8 | Q9NW13 | RNA-binding protein 28 OS=Homo sapie | RBM28 | 85.738 | 2 | 55.048088 | yes | 3.108E-07 | 446.77179 | 482.29303 |
| SEQLEELFSQVGP\ | 15 | Q9NW13 | RNA-binding protein 28 OS=Homo sapie | RBM28 | 85.738 | 2 | 55.048088 | yes | 9.722E-08 | 845.43835 | 439.30997 |
| GCAFAQFMTQEA/ | 15 | Q9NW13 | RNA-binding protein 28 OS=Homo sapie | RBM28 | 85.738 | 2 | 55.048088 | yes | 2.167E-07 | 844.38196 | 426.81036 |
| CLLAASPENEAGG | 15 | Q9NW13 | RNA-binding protein 28 OS=Homo sapie | RBM28 | 85.738 | 2 | 55.048088 | yes | 9.722E-08 | 765.38507 | 426.71838 |
| LINNNPEIFGPLK | 13 | Q9NW13 | RNA-binding protein 28 OS=Homo sapie | RBM28 | 85.738 | 2 | 55.048088 | yes | 1.022E-07 | 734.91156 | 385.2644 |
| KLPSDVNEGK | 10 | Q9NW13 | RNA-binding protein 28 OS=Homo sapie | RBM28 | 85.738 | 2 | 55.048088 | yes | 1.624E-06 | 543.79309 | 362.1954 |
| LLLSATSGEK | 10 | Q9NW13 | RNA-binding protein 28 OS=Homo sapie | RBM28 | 85.738 | 2 | 55.048088 | yes | 1.905E-07 | 509.7926 | 284.06995 |
| AAEGVSAADMAK | 12 | Q9NW13 | RNA-binding protein 28 OS=Homo sapie | RBM28 | 85.738 | 2 | 55.048088 | yes | 2.772E-07 | 560.7688 | 250.26376 |
| IVLHPDTEHSK | 11 | Q9NW13 | RNA-binding protein 28 OS=Homo sapie | RBM28 | 85.738 | 2 | 55.048088 | yes | 1.068E-07 | 638.33826 | 230.13026 |
| VTKPVQIQK | 9 | Q9NW13 | RNA-binding protein 28 OS=Homo sapie | RBM28 | 85.738 | 2 | 55.048088 | yes | 2.39E-07 | 520.82678 | 228.19864 |
| AGSTSWTGFQTK | 12 | Q9NW13 | RNA-binding protein 28 OS=Homo sapie | RBM28 | 85.738 | 2 | 55.048088 | yes | 1.83E-07 | 635.80676 | 175.56046 |
| LCLHNLPK | 8 | Q9NW13 | RNA-binding protein 28 OS=Homo sapie | RBM28 | 85.738 | 2 | 55.048088 | yes | 1.073E-06 | 497.77875 | 170.42543 |
| FNQLVEQYK | 9 | Q9NW13 | RNA-binding protein 28 OS=Homo sapie | RBM28 | 85.738 | 2 | 55.048088 | yes | 5.883E-07 | 584.80347 | 167.27499 |
| ERFELLK | 7 | Q9NW13 | RNA-binding protein 28 OS=Homo sapie | RBM28 | 85.738 | 2 | 55.048088 | yes | 2.39E-07 | 467.77145 | 149.50682 |
| LKDQNIFVSR | 10 | Q9NW13 | RNA-binding protein 28 OS=Homo sapie | RBM28 | 85.738 | 2 | 55.048088 | yes | 9.722E-08 | 610.34332 | 138.87268 |
| AEVEQVELPDGK | 12 | Q9NW13 | RNA-binding protein 28 OS=Homo sapie | RBM28 | 85.738 | 2 | 55.048088 | yes | 9.722E-08 | 657.33282 | 120.21657 |
| QQLSSEQVSR | 10 | Q9NW13 | RNA-binding protein 28 OS=Homo sapie | RBM28 | 85.738 | 2 | 55.048088 | yes | 2.602E-07 | 581.29657 | 115.604 |
| KLLLSATSGEK | 11 | Q9NW13 | RNA-binding protein 28 OS=Homo sapie | RBM28 | 85.738 | 2 | 55.048088 | yes | 1.65E-07 | 573.84009 | 97.585991 |
| QCFVVTEK | 8 | Q9NW13 | RNA-binding protein 28 OS=Homo sapie | RBM28 | 85.738 | 2 | 55.048088 | yes | 1.981E-07 | 505.75241 | 80.505692 |
| DTQSVSAIGEEK | 12 | Q9NW13 | RNA-binding protein 28 OS=Homo sapie | RBM28 | 85.738 | 2 | 55.048088 | yes | 1.429E-07 | 632.30676 | 69.639618 |
| EITTFEGCK | 9 | Q9NW13 | RNA-binding protein 28 OS=Homo sapie | RBM28 | 85.738 | 2 | 55.048088 | yes | 4.705E-07 | 542.75262 | 55.288677 |
| YYGINDPVADK | 11 | Q9NW64 | Pre-mRNA-splicing factor RBM22 OS=H | RBM22 | 46.896 | 2 | 48.01741 | yes | 3.362E-07 | 627.80371 | 1313.9613 |
| NHFYQFGEIR | 10 | Q9NW64 | Pre-mRNA-splicing factor RBM22 OS=H | RBM22 | 46.896 | 2 | 48.01741 | yes | 2.454E-07 | 655.81744 | 806.87396 |
| QQCAFIQFATR | 11 | Q9NW64 | Pre-mRNA-splicing factor RBM22 OS=H | RBM22 | 46.896 | 2 | 48.01741 | yes | 1.86E-07 | 685.33771 | 536.54724 |
| ATSTSDMLLK | 10 | Q9NW64 | Pre-mRNA-splicing factor RBM22 OS=H | RBM22 | 46.896 | 2 | 48.01741 | yes | 1.213E-07 | 533.77606 | 353.78113 |
| EYYTQNMER | 9 | Q9NW64 | Pre-mRNA-splicing factor RBM22 OS=H | RBM22 | 46.896 | 2 | 48.01741 | yes | 1.673E-07 | 617.26385 | 330.42194 |
| NRPHICSFWVK | 11 | Q9NW64 | Pre-mRNA-splicing factor RBM22 OS=H | RBM22 | 46.896 | 3 | 48.01741 | yes | 2.04E-07 | 481.91531 | 239.74603 |
| RGEECPYR | 8 | Q9NW64 | Pre-mRNA-splicing factor RBM22 OS=H | RBM22 | 46.896 | 2 | 48.01741 | yes | 2.32E-07 | 533.74036 | 64.903656 |
| MAVDLPQDISNK | 12 | Q9NW75 | G patch domain-containing protein 2 OS= | GPATCH2 | 58.944 | 2 | 51.986378 | yes | 9.733E-08 | 665.83722 | 1547.8069 |
| GGFAETGDHSR | 11 | Q9NW75 | G patch domain-containing protein 2 OS= | GPATCH2 | 58.944 | 2 | 51.986378 | yes | 9.722E-08 | 567.25214 | 1374.597 |
| QGDDEQSDWFYEI | 13 | Q9NW75 | G patch domain-containing protein 2 OS= | GPATCH2 | 58.944 | 2 | 51.986378 | yes | 1.108E-07 | 823.83392 | 285.4429 |
| AYQYQEFTK | 9 | Q9NW75 | G patch domain-containing protein 2 OS= | GPATCH2 | 58.944 | 2 | 51.986378 | yes | 1.063E-07 | 589.27985 | 255.13264 |
| SGGTPTSMVPIPGI | 18 | Q9NW75 | G patch domain-containing protein 2 OS= | GPATCH2 | 58.944 | 2 | 51.986378 | yes | 1.834E-06 | 848.44037 | 153.82004 |
| RPLWHESDFAVDI | 17 | Q9NW75 | G patch domain-containing protein 2 OS= | GPATCH2 | 58.944 | 4 | 51.986378 | yes | 9.722E-08 | 503.74878 | 134.23012 |
| GLGLGFPLPK | 10 | Q9NW75 | G patch domain-containing protein 2 OS= | GPATCH2 | 58.944 | 2 | 51.986378 | yes | 1.949E-06 | 499.8053 | 91.141037 |
| TMTQPPEGCR | 10 | Q9NW75 | G patch domain-containing protein 2 OS= | GPATCH2 | 58.944 | 2 | 51.986378 | yes | 3.473E-07 | 588.76044 | 90.283569 |

| | | | | | | | | | | | |
|---|---|---|---|---|---|---|---|---|---|---|---|
| MLQNMGWTPGSC | 15 | Q9NW75 | G patch domain-containing protein 2 OS= | GPATCH2 | 58.944 | 2 | 51.986378 | yes | 5.035E-07 | 802.88721 | 56.613834 |
| QPIGAPAAGNSWH | 16 | Q9NW75 | G patch domain-containing protein 2 OS= | GPATCH2 | 58.944 | 3 | 51.986378 | yes | 2.579E-06 | 565.94983 | 43.565655 |
| IQDEGVVLESEETI | 18 | Q9NW75 | G patch domain-containing protein 2 OS= | GPATCH2 | 58.944 | 3 | 51.986378 | yes | 2.965E-07 | 678.32715 | 40.283058 |
| DKMECEEQK | 9 | Q9NW75 | G patch domain-containing protein 2 OS= | GPATCH2 | 58.944 | 2 | 51.986378 | yes | 2.434E-07 | 598.74976 | 38.618855 |
| DHSDSDDQMLVA | 13 | Q9NW75 | G patch domain-containing protein 2 OS= | GPATCH2 | 58.944 | 2 | 51.986378 | yes | 5.833E-07 | 730.81976 | 37.365417 |
| RPSSNLNNNVR | 11 | Q9NW75 | G patch domain-containing protein 2 OS= | GPATCH2 | 58.944 | 2 | 51.986378 | yes | 6.158E-05 | 635.83417 | 32.593979 |
| ELQSLSLQR | 9 | Q9NW81 | Distal membrane-arm assembly complex p | DMAC2 | 29.268 | 2 | 52.72549 | yes | 2.314E-07 | 537.30109 | 386.42429 |
| LGAAVAPEGNQK | 12 | Q9NW81 | Distal membrane-arm assembly complex p | DMAC2 | 29.268 | 2 | 52.72549 | yes | 9.722E-08 | 577.81183 | 231.34048 |
| GLACLHHLQNLR | 12 | Q9NW81 | Distal membrane-arm assembly complex p | DMAC2 | 29.268 | 3 | 52.72549 | yes | 1.06E-07 | 477.92612 | 95.738167 |
| GLACLHHLQNLR | 12 | Q9NW81 | Distal membrane-arm assembly complex p | DMAC2 | 29.268 | 3 | 52.72549 | yes | 1.305E-07 | 716.3855 | 95.738167 |
| GLACLHHLQNLR | 12 | Q9NW81 | Distal membrane-arm assembly complex p | DMAC2 | 29.268 | 2 | 52.72549 | yes | 1.06E-07 | 477.92612 | 95.738167 |
| GLACLHHLQNLR | 12 | Q9NW81 | Distal membrane-arm assembly complex p | DMAC2 | 29.268 | 2 | 52.72549 | yes | 1.305E-07 | 716.3855 | 95.738167 |
| DKEWIRPDK | 9 | Q9NW81 | Distal membrane-arm assembly complex p | DMAC2 | 29.268 | 2 | 52.72549 | yes | 2.523E-07 | 593.81439 | 60.293056 |
| QHGPYGAGAFFILI | 14 | Q9NW81 | Distal membrane-arm assembly complex p | DMAC2 | 29.268 | 2 | 52.72549 | yes | 2.13E-07 | 753.39862 | 20.654381 |
| QHGPYGAGAFFILI | 14 | Q9NW81 | Distal membrane-arm assembly complex p | DMAC2 | 29.268 | 2 | 52.72549 | yes | 5.348E-07 | 502.6015 | 20.654381 |
| QHGPYGAGAFFILI | 14 | Q9NW81 | Distal membrane-arm assembly complex p | DMAC2 | 29.268 | 3 | 52.72549 | yes | 2.13E-07 | 753.39862 | 20.654381 |
| QHGPYGAGAFFILI | 14 | Q9NW81 | Distal membrane-arm assembly complex p | DMAC2 | 29.268 | 3 | 52.72549 | yes | 5.348E-07 | 502.6015 | 20.654381 |
| DGQLIECR | 8 | Q9NWF9 | E3 ubiquitin-protein ligase RNF216 OS=I | RNF216 | 99.406 | 2 | 55.744152 | yes | 1.616E-07 | 495.7373 | 192.24915 |
| GHYAITR | 7 | Q9NWF9 | E3 ubiquitin-protein ligase RNF216 OS=I | RNF216 | 99.406 | 2 | 55.744152 | yes | 1.036E-07 | 409.21939 | 171.32129 |
| EITNQVVPQER | 11 | Q9NWF9 | E3 ubiquitin-protein ligase RNF216 OS=I | RNF216 | 99.406 | 2 | 55.744152 | yes | 9.722E-08 | 656.84644 | 160.75668 |
| CSLWTDPTEDDEK | 13 | Q9NWF9 | E3 ubiquitin-protein ligase RNF216 OS=I | RNF216 | 99.406 | 2 | 55.744152 | yes | 9.722E-08 | 798.33014 | 158.79283 |
| IDFFDYSK | 8 | Q9NWF9 | E3 ubiquitin-protein ligase RNF216 OS=I | RNF216 | 99.406 | 2 | 55.744152 | yes | 2.247E-07 | 517.74512 | 130.72009 |
| FSCPNPHCR | 9 | Q9NWF9 | E3 ubiquitin-protein ligase RNF216 OS=I | RNF216 | 99.406 | 2 | 55.744152 | yes | 1.179E-07 | 587.74768 | 129.93773 |
| YAQEAVFGSGK | 11 | Q9NWF9 | E3 ubiquitin-protein ligase RNF216 OS=I | RNF216 | 99.406 | 2 | 55.744152 | yes | 1.575E-07 | 578.78528 | 112.49272 |
| MFFLENK | 7 | Q9NWF9 | E3 ubiquitin-protein ligase RNF216 OS=I | RNF216 | 99.406 | 2 | 55.744152 | yes | 1.582E-07 | 464.73349 | 99.913582 |
| WQELSPETSGK | 11 | Q9NWF9 | E3 ubiquitin-protein ligase RNF216 OS=I | RNF216 | 99.406 | 2 | 55.744152 | yes | 7.292E-07 | 631.30658 | 98.533859 |
| GQEWINLR | 8 | Q9NWF9 | E3 ubiquitin-protein ligase RNF216 OS=I | RNF216 | 99.406 | 2 | 55.744152 | yes | 4.208E-07 | 508.26962 | 85.10936 |
| KAEEEVAAAYADI | 16 | Q9NWF9 | E3 ubiquitin-protein ligase RNF216 OS=I | RNF216 | 99.406 | 3 | 55.744152 | yes | 1.355E-07 | 588.63184 | 84.979042 |
| IGPPLEKPVEK | 11 | Q9NWF9 | E3 ubiquitin-protein ligase RNF216 OS=I | RNF216 | 99.406 | 2 | 55.744152 | yes | 5.123E-07 | 603.85828 | 63.363789 |
| SPGAPCQECSR | 11 | Q9NWF9 | E3 ubiquitin-protein ligase RNF216 OS=I | RNF216 | 99.406 | 2 | 55.744152 | yes | 1.283E-07 | 624.75842 | 50.678593 |
| DGPITISDSSDEER | 14 | Q9NWF9 | E3 ubiquitin-protein ligase RNF216 OS=I | RNF216 | 99.406 | 2 | 55.744152 | yes | 1.978E-07 | 760.83917 | 24.544575 |
| CGAQMCYLCR | 10 | Q9NWF9 | E3 ubiquitin-protein ligase RNF216 OS=I | RNF216 | 99.406 | 2 | 55.744152 | yes | 7.402E-07 | 659.7616 | 12.548758 |
| ELLSAEENKR | 10 | Q9NWH9 | SAFB-like transcription modulator OS=H | SLTM | 117.149 | 2 | 54.503418 | yes | 9.722E-08 | 594.81458 | 211.92299 |
| FGHGSDYSR | 9 | Q9NWH9 | SAFB-like transcription modulator OS=H | SLTM | 117.149 | 2 | 54.503418 | yes | 4.543E-07 | 513.2254 | 150.12227 |
| REDPSFER | 8 | Q9NWH9 | SAFB-like transcription modulator OS=H | SLTM | 117.149 | 2 | 54.503418 | yes | 1.265E-06 | 518.24634 | 127.24332 |
| KLSLDTDAR | 9 | Q9NWH9 | SAFB-like transcription modulator OS=H | SLTM | 117.149 | 2 | 54.503418 | yes | 5.303E-07 | 509.78 | 88.573448 |
| NIWVSGLSSNTK | 12 | Q9NWH9 | SAFB-like transcription modulator OS=H | SLTM | 117.149 | 2 | 54.503418 | yes | 4.367E-07 | 653.34351 | 87.572998 |
| DYEMNANHK | 9 | Q9NWH9 | SAFB-like transcription modulator OS=H | SLTM | 117.149 | 2 | 54.503418 | yes | 3.72E-06 | 561.23767 | 68.261528 |
| GGGSQHYPEER | 11 | Q9NWH9 | SAFB-like transcription modulator OS=H | SLTM | 117.149 | 2 | 54.503418 | yes | 0.0001164 | 608.77069 | 57.024586 |

| | | | | | | | | | | | |
|---|---|---|---|---|---|---|---|---|---|---|---|
| CYGIVTMSSSTEV! | 15 | Q9NWH9 | SAFB-like transcription modulator OS=H | SLTM | 117.149 | 2 | 54.503418 | yes | 2.616E-07 | 838.88458 | 49.092522 |
| SPGHMVILDQTK | 12 | Q9NWH9 | SAFB-like transcription modulator OS=H | SLTM | 117.149 | 2 | 54.503418 | yes | 9.722E-08 | 663.34772 | 44.958035 |
| DDPYWSENKK | 10 | Q9NWH9 | SAFB-like transcription modulator OS=H | SLTM | 117.149 | 2 | 54.503418 | yes | 1.211E-06 | 641.29095 | 44.386887 |
| IEYMQQNLVR | 10 | Q9NWK9 | Box C/D snoRNA protein 1 OS=Homo sa | ZNHIT6 | 53.919 | 2 | 54.553837 | yes | 1.726E-07 | 647.33466 | 248.03915 |
| SQTGVQILMK | 10 | Q9NWK9 | Box C/D snoRNA protein 1 OS=Homo sa | ZNHIT6 | 53.919 | 2 | 54.553837 | yes | 1.127E-07 | 552.80774 | 209.33405 |
| VIIEYPTLHVVLK | 13 | Q9NWK9 | Box C/D snoRNA protein 1 OS=Homo sa | ZNHIT6 | 53.919 | 2 | 54.553837 | yes | 1.214E-07 | 762.46344 | 96.149689 |
| VIIEYPTLHVVLK | 13 | Q9NWK9 | Box C/D snoRNA protein 1 OS=Homo sa | ZNHIT6 | 53.919 | 2 | 54.553837 | yes | 3.343E-06 | 508.64471 | 96.149689 |
| VIIEYPTLHVVLK | 13 | Q9NWK9 | Box C/D snoRNA protein 1 OS=Homo sa | ZNHIT6 | 53.919 | 3 | 54.553837 | yes | 1.214E-07 | 762.46344 | 96.149689 |
| VIIEYPTLHVVLK | 13 | Q9NWK9 | Box C/D snoRNA protein 1 OS=Homo sa | ZNHIT6 | 53.919 | 3 | 54.553837 | yes | 3.343E-06 | 508.64471 | 96.149689 |
| VKEEPPINHPVGC! | 14 | Q9NWK9 | Box C/D snoRNA protein 1 OS=Homo sa | ZNHIT6 | 53.919 | 3 | 54.553837 | yes | 1.019E-07 | 535.28021 | 69.886909 |
| VKEEPPINHPVGC! | 14 | Q9NWK9 | Box C/D snoRNA protein 1 OS=Homo sa | ZNHIT6 | 53.919 | 3 | 54.553837 | yes | 4.665E-07 | 802.41669 | 69.886909 |
| VKEEPPINHPVGC! | 14 | Q9NWK9 | Box C/D snoRNA protein 1 OS=Homo sa | ZNHIT6 | 53.919 | 2 | 54.553837 | yes | 1.019E-07 | 535.28021 | 69.886909 |
| VKEEPPINHPVGC! | 14 | Q9NWK9 | Box C/D snoRNA protein 1 OS=Homo sa | ZNHIT6 | 53.919 | 2 | 54.553837 | yes | 4.665E-07 | 802.41669 | 69.886909 |
| QEIIDWPGTEGR | 12 | Q9NWK9 | Box C/D snoRNA protein 1 OS=Homo sa | ZNHIT6 | 53.919 | 2 | 54.553837 | yes | 1.438E-06 | 700.84387 | 113.26993 |
| YSCSLPCVK | 9 | Q9NWK9 | Box C/D snoRNA protein 1 OS=Homo sa | ZNHIT6 | 53.919 | 2 | 54.553837 | yes | 6.426E-07 | 557.25702 | 105.03082 |
| TINEILKPYIDPEK | 14 | Q9NWK9 | Box C/D snoRNA protein 1 OS=Homo sa | ZNHIT6 | 53.919 | 2 | 54.553837 | yes | 2.006E-07 | 836.96143 | 97.807831 |
| EELMHGECVK | 10 | Q9NWK9 | Box C/D snoRNA protein 1 OS=Homo sa | ZNHIT6 | 53.919 | 2 | 54.553837 | yes | 9.722E-08 | 616.27594 | 94.47551 |
| DENAGVLEVK | 10 | Q9NWK9 | Box C/D snoRNA protein 1 OS=Homo sa | ZNHIT6 | 53.919 | 2 | 54.553837 | yes | 1.871E-07 | 537.27728 | 85.140366 |
| FLEDVAR | 7 | Q9NWK9 | Box C/D snoRNA protein 1 OS=Homo sa | ZNHIT6 | 53.919 | 2 | 54.553837 | yes | 4.354E-07 | 425.2269 | 77.910995 |
| AELTCNGVR | 9 | Q9NWK9 | Box C/D snoRNA protein 1 OS=Homo sa | ZNHIT6 | 53.919 | 2 | 54.553837 | yes | 4.905E-07 | 510.25058 | 74.250694 |
| SGGGLHSVAEGVF | 13 | Q9NWK9 | Box C/D snoRNA protein 1 OS=Homo sa | ZNHIT6 | 53.919 | 2 | 54.553837 | yes | 1.985E-07 | 613.31781 | 42.256615 |
| LQFPQSQAEYIEK | 13 | Q9NWK9 | Box C/D snoRNA protein 1 OS=Homo sa | ZNHIT6 | 53.919 | 2 | 54.553837 | yes | 1.403E-05 | 790.90137 | 34.107964 |
| ETSINTLGTER | 11 | Q9NWQ4 | G patch domain-containing protein 2-like | GPATCH2I | 54.26 | 2 | 54.611187 | yes | 1.468E-07 | 610.80951 | 222.26595 |
| LGELWEEMALSPF | 13 | Q9NWQ4 | G patch domain-containing protein 2-like | GPATCH2I | 54.26 | 2 | 54.611187 | yes | 9.722E-08 | 765.8847 | 181.80536 |
| WAPDHCSEVER | 11 | Q9NWQ4 | G patch domain-containing protein 2-like | GPATCH2I | 54.26 | 2 | 54.611187 | yes | 1.288E-07 | 693.29877 | 153.81227 |
| ATDATTATFFK | 11 | Q9NWQ4 | G patch domain-containing protein 2-like | GPATCH2I | 54.26 | 2 | 54.611187 | yes | 2.167E-07 | 587.29297 | 145.23056 |
| VTSEVAASLQQK | 12 | Q9NWQ4 | G patch domain-containing protein 2-like | GPATCH2I | 54.26 | 2 | 54.611187 | yes | 2.138E-07 | 630.84332 | 123.87699 |
| VSDWSYER | 8 | Q9NWQ4 | G patch domain-containing protein 2-like | GPATCH2I | 54.26 | 2 | 54.611187 | yes | 2.192E-07 | 521.23541 | 90.184204 |
| FSDSTFLLPSR | 11 | Q9NWQ4 | G patch domain-containing protein 2-like | GPATCH2I | 54.26 | 2 | 54.611187 | yes | 1.911E-07 | 635.32733 | 63.684158 |
| QHSWHESDSFTEN | 20 | Q9NWQ4 | G patch domain-containing protein 2-like | GPATCH2I | 54.26 | 4 | 54.611187 | yes | 3.496E-07 | 614.28058 | 49.400589 |
| SPSSEWLVR | 9 | Q9NWQ4 | G patch domain-containing protein 2-like | GPATCH2I | 54.26 | 2 | 54.611187 | yes | 5.279E-06 | 530.77472 | 43.756809 |
| KPVATASLSSPSA\ | 30 | Q9NWQ4 | G patch domain-containing protein 2-like | GPATCH2I | 54.26 | 4 | 54.611187 | yes | 1.483E-07 | 744.63403 | 42.659775 |
| QANVHWGPPCSR | 12 | Q9NWQ4 | G patch domain-containing protein 2-like | GPATCH2I | 54.26 | 2 | 54.611187 | yes | 2.315E-05 | 704.83057 | 14.92839 |
| HGLPLVTLTLPSR | 13 | Q9NWR8 | Calcium uniporter regulatory subunit MC | MCUB | 39.083 | 2 | 41.44231 | yes | 3.507E-07 | 702.42212 | 18.963221 |
| FEELLLQASK | 10 | Q9NWS0 | PIH1 domain-containing protein 1 OS=H | PIH1D1 | 32.363 | 2 | 39.701912 | yes | 4.429E-07 | 589.32678 | 104.3045 |
| LTCLGVWLDK | 10 | Q9NWT1 | p21-activated protein kinase-interacting p | PAK1IP1 | 43.964 | 2 | 51.357014 | yes | 1.058E-07 | 602.82336 | 931.51166 |
| GQVTFLSIHPSGK | 13 | Q9NWT1 | p21-activated protein kinase-interacting p | PAK1IP1 | 43.964 | 2 | 51.357014 | yes | 9.722E-08 | 685.875 | 771.3396 |
| DETIHIYDMK | 10 | Q9NWT1 | p21-activated protein kinase-interacting p | PAK1IP1 | 43.964 | 2 | 51.357014 | yes | 9.722E-08 | 632.79755 | 737.27069 |
| GEQYVVIIQNK | 11 | Q9NWT1 | p21-activated protein kinase-interacting p | PAK1IP1 | 43.964 | 2 | 51.357014 | yes | 1.122E-07 | 645.85626 | 370.45905 |

| | | | | | | | | | | | |
|---|---|---|---|---|---|---|---|---|---|---|---|
| ESLPPAAEPSPVSK | 14 | Q9NWT1 | p21-activated protein kinase-interacting p | PAK1IP1 | 43.964 | 2 | 51.357014 | yes | 3.316E-07 | 704.86957 | 353.59479 |
| TWNLVEGR | 8 | Q9NWT1 | p21-activated protein kinase-interacting p | PAK1IP1 | 43.964 | 2 | 51.357014 | yes | 1.538E-07 | 487.75635 | 325.19916 |
| HLISGAEDGLICIW | 17 | Q9NWT1 | p21-activated protein kinase-interacting p | PAK1IP1 | 43.964 | 3 | 51.357014 | yes | 9.722E-08 | 633.32074 | 285.87622 |
| LALSVGTDK | 9 | Q9NWT1 | p21-activated protein kinase-interacting p | PAK1IP1 | 43.964 | 2 | 51.357014 | yes | 6.777E-07 | 452.26093 | 256.8848 |
| QNAHIVEWSPR | 11 | Q9NWT1 | p21-activated protein kinase-interacting p | PAK1IP1 | 43.964 | 2 | 51.357014 | yes | 3.164E-06 | 668.84149 | 160.68547 |
| VPPSLLCEINTNAF | 14 | Q9NWT1 | p21-activated protein kinase-interacting p | PAK1IP1 | 43.964 | 2 | 51.357014 | yes | 9.722E-08 | 792.41412 | 124.32793 |
| FLSESVLAVAGDE | 17 | Q9NWT1 | p21-activated protein kinase-interacting p | PAK1IP1 | 43.964 | 3 | 51.357014 | yes | 3.206E-06 | 611.99127 | 26.127991 |
| EAPEGWQTPK | 10 | Q9NWT8 | Small ribosomal subunit protein mS38 OS | AURKAIP1 | 22.355 | 2 | 50.462982 | yes | 9.809E-08 | 571.77747 | 1063.5774 |
| MSVSPLESWLTAR | 13 | Q9NWT8 | Small ribosomal subunit protein mS38 OS | AURKAIP1 | 22.355 | 2 | 50.462982 | yes | 3.445E-06 | 738.87939 | 34.479305 |
| EVLLEAQDMAVR | 12 | Q9NWU5 | Large ribosomal subunit protein uL22m C | MRPL22 | 23.641 | 2 | 50.09048 | yes | 1.269E-07 | 687.35828 | 1083.3682 |
| EVLLEAQDMAVR | 12 | Q9NWU5 | Large ribosomal subunit protein uL22m C | MRPL22 | 23.641 | 2 | 50.09048 | yes | 3.989E-07 | 695.35577 | 1083.3682 |
| LVEGPPPPPEPPK | 13 | Q9NWU5 | Large ribosomal subunit protein uL22m C | MRPL22 | 23.641 | 2 | 50.09048 | yes | 1.979E-06 | 677.37427 | 1274.7324 |
| IVYPPQLPGEPR | 12 | Q9NWU5 | Large ribosomal subunit protein uL22m C | MRPL22 | 23.641 | 2 | 50.09048 | yes | 2.983E-06 | 683.37988 | 948.24396 |
| DHNVEFR | 7 | Q9NWU5 | Large ribosomal subunit protein uL22m C | MRPL22 | 23.641 | 2 | 50.09048 | yes | 2.232E-07 | 458.71722 | 532.75336 |
| SNLYIAESTSGR | 12 | Q9NWU5 | Large ribosomal subunit protein uL22m C | MRPL22 | 23.641 | 2 | 50.09048 | yes | 9.967E-08 | 649.32275 | 503.02081 |
| EYIQQLR | 7 | Q9NWU5 | Large ribosomal subunit protein uL22m C | MRPL22 | 23.641 | 2 | 50.09048 | yes | 2.307E-07 | 475.25873 | 447.24268 |
| VYCHYFVK | 8 | Q9NWU5 | Large ribosomal subunit protein uL22m C | MRPL22 | 23.641 | 2 | 50.09048 | yes | 9.904E-08 | 558.27075 | 256.04324 |
| LLAHPLQR | 8 | Q9NWV8 | BRISC and BRCA1-A complex member | BABAM1 | 36.561 | 2 | 51.81968 | yes | 1.398E-07 | 474.29291 | 295.39789 |
| SWQVPPPAPEVQI | 14 | Q9NWV8 | BRISC and BRCA1-A complex member | BABAM1 | 36.561 | 2 | 51.81968 | yes | 4.04E-06 | 802.43317 | 157.21254 |
| DMFAFMGSLDTK | 12 | Q9NWV8 | BRISC and BRCA1-A complex member | BABAM1 | 36.561 | 2 | 51.81968 | yes | 9.722E-08 | 681.80707 | 154.88541 |
| TNALNVSQK | 9 | Q9NWV8 | BRISC and BRCA1-A complex member | BABAM1 | 36.561 | 2 | 51.81968 | yes | 3.363E-07 | 487.76691 | 104.40881 |
| MIEMFVR | 7 | Q9NWV8 | BRISC and BRCA1-A complex member | BABAM1 | 36.561 | 2 | 51.81968 | yes | 1.86E-07 | 463.23535 | 88.630287 |
| IIPGIGNFGDR | 11 | Q9NWZ5 | Uridine-cytidine kinase-like 1 OS=Homo | UCKL1 | 61.142 | 2 | 36.953968 | yes | 6.443E-07 | 579.81696 | 73.975525 |
| QYFAETER | 8 | Q9NWZ8 | Gem-associated protein 8 OS=Homo sapi | GEMIN8 | 28.637 | 2 | 53.61301 | yes | 9.722E-08 | 522.24329 | 1255.6616 |
| ATRPWYSHPVYAI | 13 | Q9NWZ8 | Gem-associated protein 8 OS=Homo sapi | GEMIN8 | 28.637 | 3 | 53.61301 | yes | 9.851E-08 | 535.27582 | 1202.4436 |
| IQAMEAAVQLSFD | 14 | Q9NWZ8 | Gem-associated protein 8 OS=Homo sapi | GEMIN8 | 28.637 | 2 | 53.61301 | yes | 9.722E-08 | 775.89777 | 1036.4596 |
| YWPVIPLK | 8 | Q9NWZ8 | Gem-associated protein 8 OS=Homo sapi | GEMIN8 | 28.637 | 2 | 53.61301 | yes | 6.982E-07 | 508.3024 | 576.90656 |
| QQQLDAER | 8 | Q9NWZ8 | Gem-associated protein 8 OS=Homo sapi | GEMIN8 | 28.637 | 2 | 53.61301 | yes | 9.56E-07 | 494.24634 | 271.71545 |
| SVEAPTER | 8 | Q9NWZ8 | Gem-associated protein 8 OS=Homo sapi | GEMIN8 | 28.637 | 2 | 53.61301 | yes | 8.629E-07 | 444.7247 | 119.12342 |
| LYGDSAAK | 8 | Q9NWZ8 | Gem-associated protein 8 OS=Homo sapi | GEMIN8 | 28.637 | 2 | 53.61301 | yes | 9.722E-08 | 412.71106 | 99.003342 |
| SVEAPTERPGER | 12 | Q9NWZ8 | Gem-associated protein 8 OS=Homo sapi | GEMIN8 | 28.637 | 2 | 53.61301 | yes | 9.722E-08 | 664.33368 | 89.625824 |
| PLLAVQEALK | 10 | Q9NX04 | AFG2-interacting ribosome maturation fa | AIRIM | 23.373 | 2 | 49.30515 | yes | 2.269E-07 | 541.33441 | 296.12314 |
| QLVAGDIVLDK | 11 | Q9NX04 | AFG2-interacting ribosome maturation fa | AIRIM | 23.373 | 2 | 49.30515 | yes | 1.047E-07 | 585.84009 | 94.710724 |
| FEDVPALR | 8 | Q9NX04 | AFG2-interacting ribosome maturation fa | AIRIM | 23.373 | 2 | 49.30515 | yes | 1.029E-05 | 473.75327 | 10.382149 |
| ALTECQGAVGLGS | 14 | Q9NX07 | tRNA selenocysteine 1-associated protein | TRNAU1A | 32.499 | 2 | 52.71938 | yes | 9.722E-08 | 695.85339 | 60.481647 |
| GALLDVCVEQGK | 12 | Q9NX09 | DNA damage-inducible transcript 4 prote | DDIT4 | 25.371 | 2 | 40.392838 | yes | 4.048E-07 | 644.83191 | 75.055748 |
| LAYSEPCGLR | 10 | Q9NX09 | DNA damage-inducible transcript 4 prote | DDIT4 | 25.371 | 2 | 40.392838 | yes | 9.917E-07 | 583.28717 | 56.211109 |
| FSSSSTSSSPSSLPR | 15 | Q9NX09 | DNA damage-inducible transcript 4 prote | DDIT4 | 25.371 | 2 | 40.392838 | yes | 9.218E-07 | 757.36011 | 17.937458 |
| RPTCVIMVK | 9 | Q9NX24 | H/ACA ribonucleoprotein complex subun | NHP2 | 17.202 | 2 | 46.875217 | yes | 1.452E-07 | 552.30682 | 503.64636 |

| | | | | | | | | | | |
|---|---|---|---|---|---|---|---|---|---|---|
| TEALYYTDDTAM/ | 14 | Q9NX46 | ADP-ribosylhydrolase ARH3 OS=Homo | ADPRS | 38.947 | 2 | 40.025646 | yes | 4.249E-07 | 810.86414 | 37.690998 |
| SYNISIWDTAGR | 12 | Q9NX57 | Ras-related protein Rab-20 OS=Homo sa| RAB20 | 26.278 | 2 | 49.81002 | yes | 1.783E-07 | 691.83856 | 249.10551 |
| QSLVELEDR | 9 | Q9NX57 | Ras-related protein Rab-20 OS=Homo sa| RAB20 | 26.278 | 2 | 49.81002 | yes | 1.481E-07 | 544.78278 | 125.75685 |
| VDLTEEGALAGQE | 14 | Q9NX57 | Ras-related protein Rab-20 OS=Homo sa| RAB20 | 26.278 | 2 | 49.81002 | yes | 8.728E-07 | 730.36737 | 85.170822 |
| QVQLEDAVALYK | 12 | Q9NX57 | Ras-related protein Rab-20 OS=Homo sa| RAB20 | 26.278 | 2 | 49.81002 | yes | 1.002E-07 | 688.87463 | 73.478996 |
| IVLLGDMNVGK | 11 | Q9NX57 | Ras-related protein Rab-20 OS=Homo sa| RAB20 | 26.278 | 2 | 49.81002 | yes | 2.91E-07 | 579.83118 | 72.156738 |
| ELLEQISAFDNVPF | 14 | Q9NX58 | Cell growth-regulating nucleolar protein ( | LYAR | 43.634 | 2 | 54.579662 | yes | 1.108E-07 | 815.92542 | 1160.9641 |
| VFFTCNACGESVK | 13 | Q9NX58 | Cell growth-regulating nucleolar protein ( | LYAR | 43.634 | 2 | 54.579662 | yes | 9.722E-08 | 759.83942 | 724.83435 |
| SEEELLVIFNK | 11 | Q9NX58 | Cell growth-regulating nucleolar protein ( | LYAR | 43.634 | 2 | 54.579662 | yes | 1.398E-07 | 660.8559 | 409.58923 |
| NCECLSCIDCGK | 12 | Q9NX58 | Cell growth-regulating nucleolar protein ( | LYAR | 43.634 | 2 | 54.579662 | yes | 9.722E-08 | 758.28839 | 390.16443 |
| VKDAVEQQGEVK | 12 | Q9NX58 | Cell growth-regulating nucleolar protein ( | LYAR | 43.634 | 2 | 54.579662 | yes | 9.722E-08 | 665.35406 | 332.19757 |
| QAPDNEITIK | 10 | Q9NX58 | Cell growth-regulating nucleolar protein ( | LYAR | 43.634 | 2 | 54.579662 | yes | 1.074E-07 | 564.7984 | 268.68768 |
| DAVEQQGEVK | 10 | Q9NX58 | Cell growth-regulating nucleolar protein ( | LYAR | 43.634 | 2 | 54.579662 | yes | 8.626E-07 | 551.7724 | 87.812698 |
| PLHPVANPHAEIST | 15 | Q9NX58 | Cell growth-regulating nucleolar protein ( | LYAR | 43.634 | 3 | 54.579662 | yes | 1.41E-07 | 537.62646 | 22.398094 |
| PLHPVANPHAEIST | 15 | Q9NX58 | Cell growth-regulating nucleolar protein ( | LYAR | 43.634 | 3 | 54.579662 | yes | 1.5E-07 | 805.9361 | 22.398094 |
| PLHPVANPHAEIST | 15 | Q9NX58 | Cell growth-regulating nucleolar protein ( | LYAR | 43.634 | 2 | 54.579662 | yes | 1.41E-07 | 537.62646 | 22.398094 |
| PLHPVANPHAEIST | 15 | Q9NX58 | Cell growth-regulating nucleolar protein ( | LYAR | 43.634 | 2 | 54.579662 | yes | 1.5E-07 | 805.9361 | 22.398094 |
| RVAEELALEQAK | 12 | Q9NX63 | MICOS complex subunit MIC19 OS=Hor | CHCHD3 | 26.153 | 2 | 55.10231 | yes | 3.472E-06 | 678.87769 | 190.25745 |
| VAEELALEQAK | 11 | Q9NX63 | MICOS complex subunit MIC19 OS=Hor | CHCHD3 | 26.153 | 2 | 55.10231 | yes | 9.722E-08 | 600.82715 | 164.20071 |
| LSENVIDR | 8 | Q9NX63 | MICOS complex subunit MIC19 OS=Hor | CHCHD3 | 26.153 | 2 | 55.10231 | yes | 1.667E-06 | 473.25363 | 101.67004 |
| YSGAYGASVSDEE | 15 | Q9NX63 | MICOS complex subunit MIC19 OS=Hor | CHCHD3 | 26.153 | 2 | 55.10231 | yes | 9.722E-08 | 788.3623 | 79.369537 |
| LGEESPSLHK | 10 | Q9NX74 | tRNA-dihydrouridine(20) synthase [NAD | DUS2 | 55.051 | 2 | 53.974216 | yes | 2.726E-06 | 548.78528 | 425.51471 |
| LFSSIVTVAEQK | 12 | Q9NX74 | tRNA-dihydrouridine(20) synthase [NAD | DUS2 | 55.051 | 2 | 53.974216 | yes | 9.722E-08 | 661.37177 | 295.5589 |
| VVFQMGTSDAER | 12 | Q9NX74 | tRNA-dihydrouridine(20) synthase [NAD | DUS2 | 55.051 | 2 | 53.974216 | yes | 9.722E-08 | 670.31915 | 154.18771 |
| MCLLEWCR | 8 | Q9NX74 | tRNA-dihydrouridine(20) synthase [NAD | DUS2 | 55.051 | 2 | 53.974216 | yes | 9.722E-08 | 584.25903 | 105.63316 |
| EAPDQDPGGPR | 11 | Q9NX74 | tRNA-dihydrouridine(20) synthase [NAD | DUS2 | 55.051 | 2 | 53.974216 | yes | 9.722E-08 | 569.75983 | 102.21931 |
| AQELAQPGDLCK | 12 | Q9NX74 | tRNA-dihydrouridine(20) synthase [NAD | DUS2 | 55.051 | 2 | 53.974216 | yes | 2.179E-07 | 665.32703 | 77.753288 |
| LAQPVYETVQR | 11 | Q9NX74 | tRNA-dihydrouridine(20) synthase [NAD | DUS2 | 55.051 | 2 | 53.974216 | yes | 8.644E-06 | 652.35388 | 68.390053 |
| YCLCQMLR | 8 | Q9NX74 | tRNA-dihydrouridine(20) synthase [NAD | DUS2 | 55.051 | 2 | 53.974216 | yes | 1.012E-07 | 572.25903 | 66.24408 |
| LILAPMVR | 8 | Q9NX74 | tRNA-dihydrouridine(20) synthase [NAD | DUS2 | 55.051 | 2 | 53.974216 | yes | 3.783E-07 | 456.7886 | 31.632839 |
| ILPSLEDTLSLVK | 13 | Q9NX74 | tRNA-dihydrouridine(20) synthase [NAD | DUS2 | 55.051 | 2 | 53.974216 | yes | 3.333E-07 | 714.42145 | 30.787746 |
| YAVQYDNHYTNT | 13 | Q9NX74 | tRNA-dihydrouridine(20) synthase [NAD | DUS2 | 55.051 | 2 | 53.974216 | yes | 2.107E-07 | 808.87061 | 20.632433 |
| AAMWNPSIFLK | 11 | Q9NX74 | tRNA-dihydrouridine(20) synthase [NAD | DUS2 | 55.051 | 2 | 53.974216 | yes | 5.014E-07 | 639.33917 | 8.2857924 |
| GGTDPVYNIGALL | 14 | Q9NX95 | Syntabulin OS=Homo sapiens OX=9606 | SYBU | 72.389 | 2 | 48.745564 | yes | 1.108E-07 | 723.39099 | 82.939476 |
| VSPASESPFSEEES | 15 | Q9NX95 | Syntabulin OS=Homo sapiens OX=9606 | SYBU | 72.389 | 2 | 48.745564 | yes | 4.135E-07 | 819.3681 | 76.294647 |
| ELDFAACVEER | 11 | Q9NX95 | Syntabulin OS=Homo sapiens OX=9606 | SYBU | 72.389 | 2 | 48.745564 | yes | 3.411E-07 | 669.80334 | 48.573509 |
| VEAQLALK | 8 | Q9NX95 | Syntabulin OS=Homo sapiens OX=9606 | SYBU | 72.389 | 2 | 48.745564 | no | 2.343E-05 | 436.26602 | 24.617023 |
| YGLDTEQVWR | 10 | Q9NXC5 | GATOR2 complex protein MIOS OS=Ho | MIOS | 98.585 | 2 | 52.592735 | yes | 1.018E-07 | 633.80933 | 104.6039 |
| DGVDLMESYVDR | 12 | Q9NXC5 | GATOR2 complex protein MIOS OS=Ho | MIOS | 98.585 | 2 | 52.592735 | yes | 1.606E-07 | 699.8139 | 103.75007 |

| | | | | | | | | | | | |
|---|---|---|---|---|---|---|---|---|---|---|---|
| AIQILNEGASSEK | 13 | Q9NXC5 | GATOR2 complex protein MIOS OS=Ho | MIOS | 98.585 | 2 | 52.592735 | yes | 9.722E-08 | 680.35938 | 88.62059 |
| AVQGVTVDPYFHI | 14 | Q9NXC5 | GATOR2 complex protein MIOS OS=Ho | MIOS | 98.585 | 2 | 52.592735 | yes | 1.03E-07 | 802.39679 | 65.744682 |
| GFSQYGVSGSPTK | 13 | Q9NXC5 | GATOR2 complex protein MIOS OS=Ho | MIOS | 98.585 | 2 | 52.592735 | yes | 9.722E-08 | 657.81989 | 51.826172 |
| TMSDFTVFER | 10 | Q9NXC5 | GATOR2 complex protein MIOS OS=Ho | MIOS | 98.585 | 2 | 52.592735 | yes | 9.722E-08 | 616.78442 | 43.710304 |
| NHILAGNEDPQLK | 13 | Q9NXC5 | GATOR2 complex protein MIOS OS=Ho | MIOS | 98.585 | 2 | 52.592735 | yes | 1.457E-07 | 724.87823 | 42.825108 |
| DDFLGSFCR | 9 | Q9NXE4 | Sphingomyelin phosphodiesterase 4 OS=I | SMPD4 | 97.81 | 2 | 46.516014 | yes | 1.539E-07 | 558.74255 | 62.61726 |
| VPGYGLQVR | 9 | Q9NXE8 | Pre-mRNA-splicing factor CWC25 homo | CWC25 | 49.648 | 2 | 52.675014 | yes | 8.494E-07 | 494.78235 | 1444.5675 |
| SWHPQTLR | 8 | Q9NXE8 | Pre-mRNA-splicing factor CWC25 homo | CWC25 | 49.648 | 2 | 52.675014 | yes | 2.831E-07 | 512.76978 | 959.32104 |
| LDWMYQGPGGM' | 14 | Q9NXE8 | Pre-mRNA-splicing factor CWC25 homo | CWC25 | 49.648 | 2 | 52.675014 | yes | 9.722E-08 | 812.37396 | 922.34436 |
| NQGLQGPLTAEQK | 13 | Q9NXE8 | Pre-mRNA-splicing factor CWC25 homo | CWC25 | 49.648 | 2 | 52.675014 | yes | 9.938E-08 | 692.36499 | 861.86853 |
| LESASTSSLEDR | 12 | Q9NXE8 | Pre-mRNA-splicing factor CWC25 homo | CWC25 | 49.648 | 2 | 52.675014 | yes | 9.722E-08 | 647.80969 | 847.76459 |
| KLSAEELER | 9 | Q9NXE8 | Pre-mRNA-splicing factor CWC25 homo | CWC25 | 49.648 | 2 | 52.675014 | yes | 9.722E-08 | 537.79309 | 596.75165 |
| KIEELQR | 7 | Q9NXE8 | Pre-mRNA-splicing factor CWC25 homo | CWC25 | 49.648 | 2 | 52.675014 | yes | 3.941E-07 | 458.26654 | 341.60858 |
| MANSSPVLSK | 10 | Q9NXE8 | Pre-mRNA-splicing factor CWC25 homo | CWC25 | 49.648 | 2 | 52.675014 | yes | 1.231E-07 | 517.271 | 302.13083 |
| IREDPLFIIR | 10 | Q9NXE8 | Pre-mRNA-splicing factor CWC25 homo | CWC25 | 49.648 | 2 | 52.675014 | yes | 9.722E-08 | 636.37714 | 229.29884 |
| LSAEELER | 8 | Q9NXE8 | Pre-mRNA-splicing factor CWC25 homo | CWC25 | 49.648 | 2 | 52.675014 | yes | 9.722E-08 | 473.74564 | 215.44113 |
| ELLQMSLEK | 9 | Q9NXE8 | Pre-mRNA-splicing factor CWC25 homo | CWC25 | 49.648 | 2 | 52.675014 | yes | 3.224E-07 | 545.79425 | 196.00012 |
| EDPLFIIR | 8 | Q9NXE8 | Pre-mRNA-splicing factor CWC25 homo | CWC25 | 49.648 | 2 | 52.675014 | yes | 2.589E-07 | 501.78458 | 177.45567 |
| YAEDVGAVK | 9 | Q9NXE8 | Pre-mRNA-splicing factor CWC25 homo | CWC25 | 49.648 | 2 | 52.675014 | yes | 1.768E-07 | 476.24274 | 152.7865 |
| DEYLLGRPIDK | 11 | Q9NXE8 | Pre-mRNA-splicing factor CWC25 homo | CWC25 | 49.648 | 2 | 52.675014 | yes | 1.835E-07 | 659.8537 | 151.0585 |
| RQEMMENAK | 9 | Q9NXE8 | Pre-mRNA-splicing factor CWC25 homo | CWC25 | 49.648 | 2 | 52.675014 | yes | 1.082E-07 | 568.763 | 78.617798 |
| EVLNNPVK | 8 | Q9NXE8 | Pre-mRNA-splicing factor CWC25 homo | CWC25 | 49.648 | 2 | 52.675014 | yes | 4.245E-07 | 456.76108 | 58.208721 |
| MKLESASTSSLEDI | 14 | Q9NXE8 | Pre-mRNA-splicing factor CWC25 homo | CWC25 | 49.648 | 2 | 52.675014 | yes | 1.141E-07 | 777.37744 | 19.526142 |
| VLEAVSANPGR | 11 | Q9NXH9 | tRNA (guanine(26)-N(2))-dimethyltransf | TRMT1 | 72.235 | 2 | 53.869083 | yes | 2.417E-07 | 556.80658 | 408.5466 |
| SALLHADFR | 9 | Q9NXH9 | tRNA (guanine(26)-N(2))-dimethyltransf | TRMT1 | 72.235 | 2 | 53.869083 | yes | 1.833E-07 | 515.27747 | 360.37814 |
| LQANFTIR | 8 | Q9NXH9 | tRNA (guanine(26)-N(2))-dimethyltransf | TRMT1 | 72.235 | 2 | 53.869083 | yes | 9.722E-08 | 481.77454 | 280.83719 |
| DLTCAVITEFAR | 12 | Q9NXH9 | tRNA (guanine(26)-N(2))-dimethyltransf | TRMT1 | 72.235 | 2 | 53.869083 | yes | 1.009E-07 | 698.35046 | 279.6254 |
| TDAPASALWDIMF | 13 | Q9NXH9 | tRNA (guanine(26)-N(2))-dimethyltransf | TRMT1 | 72.235 | 2 | 53.869083 | yes | 9.722E-08 | 723.8559 | 279.01999 |
| LSETSPAFR | 9 | Q9NXH9 | tRNA (guanine(26)-N(2))-dimethyltransf | TRMT1 | 72.235 | 2 | 53.869083 | yes | 1.335E-07 | 504.26144 | 220.98354 |
| FQANPEANWGPR | 12 | Q9NXH9 | tRNA (guanine(26)-N(2))-dimethyltransf | TRMT1 | 72.235 | 2 | 53.869083 | yes | 1.423E-07 | 693.83112 | 215.6306 |
| EVQETTVTEGAAK | 13 | Q9NXH9 | tRNA (guanine(26)-N(2))-dimethyltransf | TRMT1 | 72.235 | 2 | 53.869083 | yes | 9.722E-08 | 681.841 | 167.71402 |
| MLMYQHQR | 8 | Q9NXH9 | tRNA (guanine(26)-N(2))-dimethyltransf | TRMT1 | 72.235 | 2 | 53.869083 | yes | 1.71E-07 | 553.76532 | 136.3493 |
| ACHEMALR | 8 | Q9NXH9 | tRNA (guanine(26)-N(2))-dimethyltransf | TRMT1 | 72.235 | 2 | 53.869083 | yes | 2.506E-07 | 494.22858 | 134.35832 |
| ESENLASGDQPR | 12 | Q9NXH9 | tRNA (guanine(26)-N(2))-dimethyltransf | TRMT1 | 72.235 | 2 | 53.869083 | yes | 2.379E-07 | 651.79968 | 125.73036 |
| SVVANDASTR | 10 | Q9NXH9 | tRNA (guanine(26)-N(2))-dimethyltransf | TRMT1 | 72.235 | 2 | 53.869083 | yes | 1.188E-07 | 510.25943 | 99.615837 |
| IVLHSLDLR | 9 | Q9NXH9 | tRNA (guanine(26)-N(2))-dimethyltransf | TRMT1 | 72.235 | 2 | 53.869083 | yes | 1.115E-07 | 533.3244 | 95.488846 |
| FALEVPGLR | 9 | Q9NXH9 | tRNA (guanine(26)-N(2))-dimethyltransf | TRMT1 | 72.235 | 2 | 53.869083 | yes | 5.498E-07 | 501.29257 | 83.984131 |
| GDQCCYSHSPPTPI | 14 | Q9NXH9 | tRNA (guanine(26)-N(2))-dimethyltransf | TRMT1 | 72.235 | 2 | 53.869083 | yes | 3.452E-06 | 831.34338 | 13.301085 |
| VLEVETTPGSLR | 12 | Q9NXL9 | DNA helicase MCM9 OS=Homo sapiens | MCM9 | 127.313 | 2 | 49.16975 | yes | 1.552E-07 | 650.85901 | 66.400757 |

| | | | | | | | | | | | |
|---|---|---|---|---|---|---|---|---|---|---|---|
| NTVVVSPHPK | 10 | Q9NXL9 | DNA helicase MCM9 OS=Homo sapiens | MCM9 | 127.313 | 2 | 49.16975 | yes | 1.062E-07 | 539.30621 | 65.021751 |
| VILEDDLVDSCK | 12 | Q9NXL9 | DNA helicase MCM9 OS=Homo sapiens | MCM9 | 127.313 | 2 | 49.16975 | yes | 5.168E-06 | 703.3476 | 58.939297 |
| SGDDLTIYGIVMQ | 14 | Q9NXL9 | DNA helicase MCM9 OS=Homo sapiens | MCM9 | 127.313 | 2 | 49.16975 | yes | 1.756E-07 | 784.39288 | 40.116753 |
| LECLVQLVR | 9 | Q9NXR5 | Ankyrin repeat domain-containing proteir | ANKRD10 | 44.768 | 2 | 45.619286 | yes | 1.795E-07 | 565.32355 | 87.769814 |
| CLEDSEDFGVK | 11 | Q9NXR5 | Ankyrin repeat domain-containing proteir | ANKRD10 | 44.768 | 2 | 45.619286 | yes | 7.694E-07 | 649.7821 | 40.598694 |
| AGATLNVSTTR | 11 | Q9NXR5 | Ankyrin repeat domain-containing proteir | ANKRD10 | 44.768 | 2 | 45.619286 | yes | 2.889E-06 | 545.7962 | 30.819118 |
| VGLDATNCLR | 10 | Q9NXR7 | BRISC and BRCA1-A complex member 2 | BABAM2 | 43.552 | 2 | 50.73118 | yes | 9.727E-08 | 559.78479 | 956.04883 |
| LHIPYAGETLK | 11 | Q9NXR7 | BRISC and BRCA1-A complex member 2 | BABAM2 | 43.552 | 2 | 50.73118 | yes | 1.827E-07 | 621.34808 | 545.81879 |
| SGCTSLTPGPNCDI | 14 | Q9NXR7 | BRISC and BRCA1-A complex member 2 | BABAM2 | 43.552 | 2 | 50.73118 | yes | 1.318E-07 | 761.32465 | 495.00488 |
| NNWTGEFSAR | 10 | Q9NXR7 | BRISC and BRCA1-A complex member 2 | BABAM2 | 43.552 | 2 | 50.73118 | yes | 1.02E-07 | 591.27032 | 312.38089 |
| MSPEVALNR | 9 | Q9NXR7 | BRISC and BRCA1-A complex member 2 | BABAM2 | 43.552 | 2 | 50.73118 | yes | 2.401E-07 | 529.76862 | 102.78836 |
| MSPEVALNR | 9 | Q9NXR7 | BRISC and BRCA1-A complex member 2 | BABAM2 | 43.552 | 2 | 50.73118 | yes | 2.785E-07 | 508.76331 | 102.78836 |
| KNNWTGEFSAR | 11 | Q9NXR7 | BRISC and BRCA1-A complex member 2 | BABAM2 | 43.552 | 2 | 50.73118 | yes | 3.644E-07 | 655.31781 | 54.360916 |
| VPLIGSLPEAR | 11 | Q9NXS2 | Glutaminyl-peptide cyclotransferase-like p | QPCTL | 42.925 | 2 | 51.132866 | yes | 1.067E-07 | 576.34277 | 795.25238 |
| VVGQLDPQR | 9 | Q9NXS2 | Glutaminyl-peptide cyclotransferase-like p | QPCTL | 42.925 | 2 | 51.132866 | yes | 2.679E-07 | 506.28271 | 539.44073 |
| KFLEATLR | 8 | Q9NXS2 | Glutaminyl-peptide cyclotransferase-like p | QPCTL | 42.925 | 2 | 51.132866 | yes | 9.722E-08 | 489.29257 | 438.93347 |
| HLTLACHYDSK | 11 | Q9NXS2 | Glutaminyl-peptide cyclotransferase-like p | QPCTL | 42.925 | 2 | 51.132866 | yes | 3.483E-07 | 672.8219 | 106.38136 |
| HLTLACHYDSK | 11 | Q9NXS2 | Glutaminyl-peptide cyclotransferase-like p | QPCTL | 42.925 | 2 | 51.132866 | yes | 9.577E-07 | 448.8837 | 106.38136 |
| HLTLACHYDSK | 11 | Q9NXS2 | Glutaminyl-peptide cyclotransferase-like p | QPCTL | 42.925 | 3 | 51.132866 | yes | 3.483E-07 | 672.8219 | 106.38136 |
| HLTLACHYDSK | 11 | Q9NXS2 | Glutaminyl-peptide cyclotransferase-like p | QPCTL | 42.925 | 3 | 51.132866 | yes | 9.577E-07 | 448.8837 | 106.38136 |
| TEELPLGR | 8 | Q9NXS2 | Glutaminyl-peptide cyclotransferase-like p | QPCTL | 42.925 | 2 | 51.132866 | yes | 3.624E-07 | 457.75073 | 96.481064 |
| GLMEPLLPPK | 10 | Q9NXS2 | Glutaminyl-peptide cyclotransferase-like p | QPCTL | 42.925 | 2 | 51.132866 | yes | 3.541E-07 | 547.81757 | 91.676086 |
| TPGSPGNLQVR | 11 | Q9NXS2 | Glutaminyl-peptide cyclotransferase-like p | QPCTL | 42.925 | 2 | 51.132866 | yes | 1.692E-07 | 563.3042 | 82.052826 |
| LWSTYLR | 7 | Q9NXS2 | Glutaminyl-peptide cyclotransferase-like p | QPCTL | 42.925 | 2 | 51.132866 | yes | 3.241E-07 | 469.75836 | 71.314949 |
| RTEELPLGR | 9 | Q9NXS2 | Glutaminyl-peptide cyclotransferase-like p | QPCTL | 42.925 | 2 | 51.132866 | yes | 1.414E-07 | 535.80127 | 66.74881 |
| GGIGAGLGGGLCR | 13 | Q9NXV2 | BTB/POZ domain-containing protein KC | KCTD5 | 26.093 | 2 | 47.448112 | yes | 1.333E-07 | 572.79822 | 160.347 |
| CSAGLGALAQR | 11 | Q9NXV2 | BTB/POZ domain-containing protein KC | KCTD5 | 26.093 | 2 | 47.448112 | yes | 1.585E-07 | 552.28491 | 109.32483 |
| LNVGGTYFLTTR | 12 | Q9NXV2 | BTB/POZ domain-containing protein KC | KCTD5 | 26.093 | 2 | 47.448112 | yes | 3.427E-07 | 671.36169 | 100.74618 |
| AENHCELLSPAR | 12 | Q9NXV2 | BTB/POZ domain-containing protein KC | KCTD5 | 26.093 | 2 | 47.448112 | yes | 1.328E-07 | 719.84082 | 34.613514 |
| VAAWVEALR | 9 | Q9NXV6 | CDKN2A-interacting protein OS=Homo s | CDKN2AIP | 61.125 | 2 | 52.34422 | yes | 3.174E-07 | 507.79019 | 948.02979 |
| SSQESIVCELR | 11 | Q9NXV6 | CDKN2A-interacting protein OS=Homo s | CDKN2AIP | 61.125 | 2 | 52.34422 | yes | 9.722E-08 | 654.31665 | 713.88275 |
| ILSMAEGIK | 9 | Q9NXV6 | CDKN2A-interacting protein OS=Homo s | CDKN2AIP | 61.125 | 2 | 52.34422 | yes | 6.584E-07 | 481.27298 | 561.61127 |
| ELADLPQNK | 9 | Q9NXV6 | CDKN2A-interacting protein OS=Homo s | CDKN2AIP | 61.125 | 2 | 52.34422 | yes | 1.585E-07 | 514.27454 | 458.7171 |
| STSLASVSQLASK | 13 | Q9NXV6 | CDKN2A-interacting protein OS=Homo s | CDKN2AIP | 61.125 | 2 | 52.34422 | yes | 1.115E-07 | 639.84863 | 413.46436 |
| AQEVSEYLSQNPR | 13 | Q9NXV6 | CDKN2A-interacting protein OS=Homo s | CDKN2AIP | 61.125 | 2 | 52.34422 | yes | 2.718E-07 | 781.87592 | 363.06006 |
| STSQVAASLLASK | 13 | Q9NXV6 | CDKN2A-interacting protein OS=Homo s | CDKN2AIP | 61.125 | 2 | 52.34422 | yes | 9.722E-08 | 631.8512 | 312.10895 |
| SVYLGTGCGK | 10 | Q9NXV6 | CDKN2A-interacting protein OS=Homo s | CDKN2AIP | 61.125 | 2 | 52.34422 | yes | 9.722E-08 | 521.25531 | 301.0976 |
| VTDAPTYTTR | 10 | Q9NXV6 | CDKN2A-interacting protein OS=Homo s | CDKN2AIP | 61.125 | 2 | 52.34422 | yes | 1.851E-07 | 562.78278 | 300.20993 |
| SSSQTSGSLVSK | 12 | Q9NXV6 | CDKN2A-interacting protein OS=Homo s | CDKN2AIP | 61.125 | 2 | 52.34422 | yes | 2.219E-07 | 584.2962 | 279.01428 |

| | | | | | | | | | | | |
|---|---|---|---|---|---|---|---|---|---|---|---|
| TSSEASVSSSVAK | 13 | Q9NXV6 | CDKN2A-interacting protein OS=Homo : | CDKN2AII | 61.125 | 2 | 52.34422 | yes | 9.722E-08 | 620.30676 | 243.19954 |
| PSSETASSGLTSK | 13 | Q9NXV6 | CDKN2A-interacting protein OS=Homo : | CDKN2AII | 61.125 | 2 | 52.34422 | yes | 3.297E-07 | 626.30676 | 202.86226 |
| SSSSTNTSLLTSK | 13 | Q9NXV6 | CDKN2A-interacting protein OS=Homo : | CDKN2AII | 61.125 | 2 | 52.34422 | yes | 1.075E-07 | 656.83319 | 200.80869 |
| SSSQTSTSQLPSK | 13 | Q9NXV6 | CDKN2A-interacting protein OS=Homo : | CDKN2AII | 61.125 | 2 | 52.34422 | yes | 9.722E-08 | 669.33081 | 152.23297 |
| HGSASFVSLLK | 11 | Q9NXV6 | CDKN2A-interacting protein OS=Homo : | CDKN2AII | 61.125 | 2 | 52.34422 | yes | 9.935E-08 | 573.31934 | 110.12948 |
| GISSSNEGVEEPSK | 15 | Q9NXV6 | CDKN2A-interacting protein OS=Homo : | CDKN2AII | 61.125 | 2 | 52.34422 | yes | 6.69E-07 | 774.38104 | 9.4748821 |
| NVEDDYIANLR | 11 | Q9NXW2 | DnaJ homolog subfamily B member 12 O | DNAJB12 | 41.86 | 2 | 53.45896 | yes | 9.914E-08 | 661.32275 | 404.17389 |
| LSEVQASLHG | 10 | Q9NXW2 | DnaJ homolog subfamily B member 12 O | DNAJB12 | 41.86 | 2 | 53.45896 | yes | 9.722E-08 | 520.77222 | 304.15894 |
| DYYEILGVSR | 10 | Q9NXW2 | DnaJ homolog subfamily B member 12 O | DNAJB12 | 41.86 | 2 | 53.45896 | yes | 9.846E-08 | 607.80621 | 289.29837 |
| NHAPGATEAFK | 11 | Q9NXW2 | DnaJ homolog subfamily B member 12 O | DNAJB12 | 41.86 | 2 | 53.45896 | yes | 1.462E-07 | 571.78308 | 288.74829 |
| AIGTAYAVLSNPEI | 14 | Q9NXW2 | DnaJ homolog subfamily B member 12 O | DNAJB12 | 41.86 | 2 | 53.45896 | yes | 9.819E-08 | 717.38538 | 215.45772 |
| GYTAEQVAAVK | 11 | Q9NXW2 | DnaJ homolog subfamily B member 12 O | DNAJB12 | 41.86 | 2 | 53.45896 | yes | 1.001E-07 | 568.80096 | 94.537567 |
| MESNKDEAER | 10 | Q9NXW2 | DnaJ homolog subfamily B member 12 O | DNAJB12 | 41.86 | 2 | 53.45896 | yes | 1.804E-07 | 625.76953 | 68.981133 |
| YFGDTDMYHR | 10 | Q9NXW2 | DnaJ homolog subfamily B member 12 O | DNAJB12 | 41.86 | 2 | 53.45896 | yes | 1.274E-07 | 652.77185 | 61.042477 |
| KQYDQFGDDK | 10 | Q9NXW2 | DnaJ homolog subfamily B member 12 O | DNAJB12 | 41.86 | 2 | 53.45896 | yes | 1.473E-07 | 622.28314 | 16.659622 |
| TFEISEPVITPSQR | 14 | Q9NXX6 | Non-structural maintenance of chromosor | NSMCE4A | 44.302 | 2 | 48.901173 | yes | 5.126E-07 | 802.41992 | 239.9584 |
| KVPVIQEER | 9 | Q9NXX6 | Non-structural maintenance of chromosor | NSMCE4A | 44.302 | 2 | 48.901173 | yes | 1.44E-07 | 549.31934 | 216.68886 |
| ALINSVQQNR | 10 | Q9NXX6 | Non-structural maintenance of chromosor | NSMCE4A | 44.302 | 2 | 48.901173 | yes | 3.4E-07 | 571.81744 | 202.99458 |
| SDLSSFDMLR | 10 | Q9NXX6 | Non-structural maintenance of chromosor | NSMCE4A | 44.302 | 2 | 48.901173 | yes | 1.088E-07 | 585.77661 | 142.65959 |
| NQGIIALSYR | 10 | Q9NXX6 | Non-structural maintenance of chromosor | NSMCE4A | 44.302 | 2 | 48.901173 | yes | 4.511E-07 | 567.81696 | 111.95918 |
| ILGLLQTYFR | 10 | Q9NXX6 | Non-structural maintenance of chromosor | NSMCE4A | 44.302 | 2 | 48.901173 | no | 3.997E-07 | 612.36096 | 42.770676 |
| VDEIFGQLR | 9 | Q9NY12 | H/ACA ribonucleoprotein complex subun | GAR1 | 22.348 | 2 | 51.003822 | yes | 2.545E-07 | 538.79041 | 3137.7847 |
| FLPRPPGEK | 9 | Q9NY12 | H/ACA ribonucleoprotein complex subun | GAR1 | 22.348 | 2 | 51.003822 | yes | 1.423E-06 | 520.79803 | 2015.8595 |
| FYIDPYK | 7 | Q9NY12 | H/ACA ribonucleoprotein complex subun | GAR1 | 22.348 | 2 | 51.003822 | yes | 9.722E-08 | 473.23947 | 664.05475 |
| VPYFNAPVYLENK | 13 | Q9NY12 | H/ACA ribonucleoprotein complex subun | GAR1 | 22.348 | 2 | 51.003822 | yes | 9.819E-08 | 777.40356 | 209.57008 |
| FADFTVYR | 8 | Q9NY61 | Protein AATF OS=Homo sapiens OX=96 | AATF | 63.133 | 2 | 49.682076 | yes | 1.988E-07 | 509.75327 | 89.971573 |
| TSSLDPNDQVAM( | 14 | Q9NY61 | Protein AATF OS=Homo sapiens OX=96 | AATF | 63.133 | 2 | 49.682076 | yes | 1.809E-07 | 745.84882 | 87.718742 |
| TPGFSVQSISDFEK | 14 | Q9NY61 | Protein AATF OS=Homo sapiens OX=96 | AATF | 63.133 | 2 | 49.682076 | yes | 1.011E-07 | 771.37775 | 66.121658 |
| DKGGPEFSSALK | 12 | Q9NY61 | Protein AATF OS=Homo sapiens OX=96 | AATF | 63.133 | 2 | 49.682076 | yes | 1.379E-07 | 618.31696 | 58.507725 |
| ATGPVVEQAVR | 11 | Q9NY93 | Probable ATP-dependent RNA helicase I | DDX56 | 61.59 | 2 | 52.660732 | yes | 1.163E-07 | 563.81439 | 1711.5438 |
| IEELLSGENR | 10 | Q9NY93 | Probable ATP-dependent RNA helicase I | DDX56 | 61.59 | 2 | 52.660732 | yes | 9.722E-08 | 580.30133 | 1073.7979 |
| VANVSAAEDSVS( | 14 | Q9NY93 | Probable ATP-dependent RNA helicase I | DDX56 | 61.59 | 2 | 52.660732 | yes | 9.809E-08 | 716.85498 | 841.03296 |
| LLQAVTDLGWSR | 12 | Q9NY93 | Probable ATP-dependent RNA helicase I | DDX56 | 61.59 | 2 | 52.660732 | yes | 1.022E-07 | 679.875 | 785.54993 |
| GPILLPYQFR | 10 | Q9NY93 | Probable ATP-dependent RNA helicase I | DDX56 | 61.59 | 2 | 52.660732 | yes | 1.327E-07 | 602.34784 | 742.0528 |
| SLLFVNTLER | 10 | Q9NY93 | Probable ATP-dependent RNA helicase I | DDX56 | 61.59 | 2 | 52.660732 | yes | 1.04E-07 | 596.34021 | 595.82837 |
| MEEIEGFR | 8 | Q9NY93 | Probable ATP-dependent RNA helicase I | DDX56 | 61.59 | 2 | 52.660732 | yes | 1.762E-07 | 505.73422 | 412.45486 |
| SLLCHLPR | 8 | Q9NY93 | Probable ATP-dependent RNA helicase I | DDX56 | 61.59 | 2 | 52.660732 | yes | 1.283E-07 | 498.2764 | 398.80118 |
| EELLHSEK | 8 | Q9NY93 | Probable ATP-dependent RNA helicase I | DDX56 | 61.59 | 2 | 52.660732 | yes | 9.722E-08 | 492.75348 | 382.56921 |
| PTLIQEK | 7 | Q9NY93 | Probable ATP-dependent RNA helicase I | DDX56 | 61.59 | 2 | 52.660732 | yes | 3.3E-07 | 414.7449 | 170.14827 |

| | | | | | | | | | | | |
|---|---|---|---|---|---|---|---|---|---|---|---|
| HDLPLHPAVVK | 11 | Q9NY93 | Probable ATP-dependent RNA helicase D | DDX56 | 61.59 | 2 | 52.660732 | yes | 1.083E-07 | 613.3562 | 75.05748 |
| HDLPLHPAVVK | 11 | Q9NY93 | Probable ATP-dependent RNA helicase D | DDX56 | 61.59 | 2 | 52.660732 | yes | 6.344E-07 | 409.2399 | 75.05748 |
| HDLPLHPAVVK | 11 | Q9NY93 | Probable ATP-dependent RNA helicase D | DDX56 | 61.59 | 3 | 52.660732 | yes | 1.083E-07 | 613.3562 | 75.05748 |
| HDLPLHPAVVK | 11 | Q9NY93 | Probable ATP-dependent RNA helicase D | DDX56 | 61.59 | 3 | 52.660732 | yes | 6.344E-07 | 409.2399 | 75.05748 |
| ELILHNPVTLK | 11 | Q9NY93 | Probable ATP-dependent RNA helicase D | DDX56 | 61.59 | 2 | 52.660732 | yes | 9.758E-08 | 638.88483 | 144.39445 |
| AIPLALEGK | 9 | Q9NY93 | Probable ATP-dependent RNA helicase D | DDX56 | 61.59 | 2 | 52.660732 | yes | 2.493E-07 | 456.28165 | 106.35117 |
| PHLGHVPDYLVPP | 16 | Q9NY93 | Probable ATP-dependent RNA helicase D | DDX56 | 61.59 | 3 | 52.660732 | yes | 2.283E-07 | 594.33344 | 91.963219 |
| KATGPVVEQAVR | 12 | Q9NY93 | Probable ATP-dependent RNA helicase D | DDX56 | 61.59 | 2 | 52.660732 | yes | 6.281E-07 | 627.86188 | 85.097534 |
| GLVLVPTK | 8 | Q9NY93 | Probable ATP-dependent RNA helicase D | DDX56 | 61.59 | 2 | 52.660732 | yes | 2.141E-07 | 413.77347 | 61.741138 |
| AVLMEKPDVVVG | 16 | Q9NY93 | Probable ATP-dependent RNA helicase D | DDX56 | 61.59 | 3 | 52.660732 | yes | 2.681E-07 | 566.64667 | 16.314556 |
| LAYLPVGR | 8 | Q9NYA1 | Sphingosine kinase 1 OS=Homo sapiens ( | SPHK1 | 42.518 | 2 | 48.3094 | yes | 1.165E-07 | 444.76871 | 40.338108 |
| FGAQNVAR | 8 | Q9NYB0 | Telomeric repeat-binding factor 2-interact | TERF2IP | 44.26 | 2 | 47.81722 | yes | 2.02E-07 | 431.73013 | 120.31078 |
| YLLGDAPVSPSSQ | 14 | Q9NYB0 | Telomeric repeat-binding factor 2-interact | TERF2IP | 44.26 | 2 | 47.81722 | yes | 2.016E-07 | 731.38281 | 96.556183 |
| LELEAYR | 7 | Q9NYB0 | Telomeric repeat-binding factor 2-interact | TERF2IP | 44.26 | 2 | 47.81722 | yes | 5.542E-07 | 447.23999 | 48.652622 |
| LGPASAADTGSEA | 14 | Q9NYB0 | Telomeric repeat-binding factor 2-interact | TERF2IP | 44.26 | 2 | 47.81722 | yes | 9.574E-07 | 637.81476 | 25.321545 |
| SSLTQHSWQSLK | 12 | Q9NYB0 | Telomeric repeat-binding factor 2-interact | TERF2IP | 44.26 | 2 | 47.81722 | yes | 1.252E-07 | 701.35968 | 19.171795 |
| VVAIYDYTK | 9 | Q9NYB9 | Abl interactor 2 OS=Homo sapiens OX=9 | ABI2 | 55.664 | 2 | 50.26077 | no | 2.953E-07 | 536.28967 | 203.15642 |
| EIGILTTNK | 9 | Q9NYB9 | Abl interactor 2 OS=Homo sapiens OX=9 | ABI2 | 55.664 | 2 | 50.26077 | no | 3.096E-07 | 494.78729 | 188.99583 |
| KPIDYTILDDIGHG | 16 | Q9NYB9 | Abl interactor 2 OS=Homo sapiens OX=9 | ABI2 | 55.664 | 3 | 50.26077 | yes | 9.824E-08 | 595.32446 | 106.49296 |
| HTPPTIGGSLPYR | 13 | Q9NYB9 | Abl interactor 2 OS=Homo sapiens OX=9 | ABI2 | 55.664 | 2 | 50.26077 | yes | 7.999E-07 | 698.37262 | 93.584488 |
| ALFDSYTNLER | 11 | Q9NYB9 | Abl interactor 2 OS=Homo sapiens OX=9 | ABI2 | 55.664 | 2 | 50.26077 | yes | 1.23E-06 | 664.8277 | 91.848122 |
| REIGILTTNK | 10 | Q9NYB9 | Abl interactor 2 OS=Homo sapiens OX=9 | ABI2 | 55.664 | 2 | 50.26077 | no | 3.12E-05 | 572.83789 | 33.283035 |
| IIAPANLER | 9 | Q9NYB9 | Abl interactor 2 OS=Homo sapiens OX=9 | ABI2 | 55.664 | 2 | 50.26077 | yes | 8.678E-07 | 498.79547 | 31.275227 |
| AEHLESPQLGGK | 12 | Q9NYD6 | Homeobox protein Hox-C10 OS=Homo s | HOXC10 | 38.073 | 2 | 51.724045 | yes | 5.99E-07 | 633.32788 | 1505.5419 |
| AENTTGNWLTAK | 12 | Q9NYD6 | Homeobox protein Hox-C10 OS=Homo s | HOXC10 | 38.073 | 2 | 51.724045 | yes | 1.876E-07 | 653.32532 | 1478.2684 |
| HQTLELEK | 8 | Q9NYD6 | Homeobox protein Hox-C10 OS=Homo s | HOXC10 | 38.073 | 2 | 51.724045 | no | 4.245E-07 | 499.26929 | 1133.6176 |
| VSFPETPK | 8 | Q9NYD6 | Homeobox protein Hox-C10 OS=Homo s | HOXC10 | 38.073 | 2 | 51.724045 | yes | 1.005E-06 | 452.74237 | 1003.8934 |
| SDSQTPSPNEIK | 12 | Q9NYD6 | Homeobox protein Hox-C10 OS=Homo s | HOXC10 | 38.073 | 2 | 51.724045 | yes | 1.433E-06 | 651.81226 | 913.71332 |
| TEQSLAGPK | 9 | Q9NYD6 | Homeobox protein Hox-C10 OS=Homo s | HOXC10 | 38.073 | 2 | 51.724045 | yes | 1.001E-07 | 465.74817 | 856.88147 |
| EFLFNMYLTR | 10 | Q9NYD6 | Homeobox protein Hox-C10 OS=Homo s | HOXC10 | 38.073 | 2 | 51.724045 | no | 1.163E-07 | 667.33411 | 395.67444 |
| EFLFNMYLTR | 10 | Q9NYD6 | Homeobox protein Hox-C10 OS=Homo s | HOXC10 | 38.073 | 2 | 51.724045 | no | 1.727E-07 | 675.33154 | 395.67444 |
| ASPSYSALDK | 10 | Q9NYD6 | Homeobox protein Hox-C10 OS=Homo s | HOXC10 | 38.073 | 2 | 51.724045 | yes | 3.873E-07 | 519.75873 | 782.58301 |
| GCGLAPSLSK | 10 | Q9NYD6 | Homeobox protein Hox-C10 OS=Homo s | HOXC10 | 38.073 | 2 | 51.724045 | yes | 9.722E-08 | 495.25787 | 699.26343 |
| EENVCCMYSAEK | 12 | Q9NYD6 | Homeobox protein Hox-C10 OS=Homo s | HOXC10 | 38.073 | 2 | 51.724045 | yes | 1.001E-07 | 760.29675 | 289.43689 |
| PLSSCSYPPSVK | 12 | Q9NYD6 | Homeobox protein Hox-C10 OS=Homo s | HOXC10 | 38.073 | 2 | 51.724045 | yes | 4.758E-06 | 661.32648 | 232.50441 |
| MTCPRNVTPNSYA | 24 | Q9NYD6 | Homeobox protein Hox-C10 OS=Homo s | HOXC10 | 38.073 | 4 | 51.724045 | yes | 8.061E-06 | 637.05353 | 164.46681 |
| SGPEAALYSHPLPI | 29 | Q9NYD6 | Homeobox protein Hox-C10 OS=Homo s | HOXC10 | 38.073 | 4 | 51.724045 | yes | 9.671E-07 | 811.1369 | 74.544189 |
| TINLTDR | 7 | Q9NYD6 | Homeobox protein Hox-C10 OS=Homo s | HOXC10 | 38.073 | 2 | 51.724045 | yes | 4.533E-06 | 416.7298 | 40.558998 |
| SAAMTLNER | 9 | Q9NYF8 | Bcl-2-associated transcription factor 1 OS | BCLAF1 | 106.123 | 2 | 52.428097 | yes | 1.37E-07 | 496.74512 | 1747.4998 |

| | | | | | | | | | | | |
|---|---|---|---|---|---|---|---|---|---|---|---|
| SSFYPDGGDQETA | 14 | Q9NYF8 | Bcl-2-associated transcription factor 1 OS | BCLAF1 | 106.123 | 2 | 52.428097 | yes | 2.014E-07 | 751.32568 | 1682.5885 |
| TIAPQNAPR | 9 | Q9NYF8 | Bcl-2-associated transcription factor 1 OS | BCLAF1 | 106.123 | 2 | 52.428097 | yes | 1.156E-07 | 484.26962 | 1145.5844 |
| SFATASHR | 8 | Q9NYF8 | Bcl-2-associated transcription factor 1 OS | BCLAF1 | 106.123 | 2 | 52.428097 | yes | 1.627E-07 | 438.71976 | 740.39825 |
| EYSGFAGVSR | 10 | Q9NYF8 | Bcl-2-associated transcription factor 1 OS | BCLAF1 | 106.123 | 2 | 52.428097 | yes | 3.716E-07 | 536.75653 | 674.62952 |
| CDSADLR | 7 | Q9NYF8 | Bcl-2-associated transcription factor 1 OS | BCLAF1 | 106.123 | 2 | 52.428097 | yes | 9.722E-08 | 418.68198 | 535.60541 |
| ETGYVVERPSTTK | 13 | Q9NYF8 | Bcl-2-associated transcription factor 1 OS | BCLAF1 | 106.123 | 2 | 52.428097 | yes | 9.722E-08 | 733.87793 | 325.55356 |
| ETGYVVER | 8 | Q9NYF8 | Bcl-2-associated transcription factor 1 OS | BCLAF1 | 106.123 | 2 | 52.428097 | yes | 1.287E-06 | 476.74036 | 162.33351 |
| MIASDSHRPEVK | 12 | Q9NYF8 | Bcl-2-associated transcription factor 1 OS | BCLAF1 | 106.123 | 2 | 52.428097 | yes | 9.722E-08 | 685.34827 | 155.95959 |
| FTDEESR | 7 | Q9NYF8 | Bcl-2-associated transcription factor 1 OS | BCLAF1 | 106.123 | 2 | 52.428097 | yes | 1.264E-07 | 442.19324 | 153.70662 |
| IDISPSTLR | 9 | Q9NYF8 | Bcl-2-associated transcription factor 1 OS | BCLAF1 | 106.123 | 2 | 52.428097 | yes | 2.301E-07 | 501.28494 | 134.2646 |
| KSVLADQGK | 9 | Q9NYF8 | Bcl-2-associated transcription factor 1 OS | BCLAF1 | 106.123 | 2 | 52.428097 | yes | 7.7E-06 | 473.27182 | 129.78207 |
| DLFDYSPPLHK | 11 | Q9NYF8 | Bcl-2-associated transcription factor 1 OS | BCLAF1 | 106.123 | 2 | 52.428097 | yes | 1.022E-07 | 666.33514 | 127.73118 |
| LKDLFDYSPPLHK | 13 | Q9NYF8 | Bcl-2-associated transcription factor 1 OS | BCLAF1 | 106.123 | 2 | 52.428097 | yes | 9.819E-08 | 786.92468 | 58.778648 |
| KAEGEPQEESPLK | 13 | Q9NYF8 | Bcl-2-associated transcription factor 1 OS | BCLAF1 | 106.123 | 2 | 52.428097 | yes | 8.325E-06 | 721.36212 | 26.573147 |
| LPELEQLER | 9 | Q9NYH9 | U3 small nucleolar RNA-associated prote | UTP6 | 70.194 | 2 | 54.69439 | yes | 9.722E-08 | 563.80878 | 317.60184 |
| IGLFSHAEIK | 10 | Q9NYH9 | U3 small nucleolar RNA-associated prote | UTP6 | 70.194 | 2 | 54.69439 | yes | 1.496E-07 | 557.81641 | 230.28192 |
| PFSVDFFR | 8 | Q9NYH9 | U3 small nucleolar RNA-associated prote | UTP6 | 70.194 | 2 | 54.69439 | yes | 1.212E-07 | 507.7558 | 206.48123 |
| LLSECQYK | 8 | Q9NYH9 | U3 small nucleolar RNA-associated prote | UTP6 | 70.194 | 2 | 54.69439 | yes | 2.24E-06 | 520.75769 | 112.9325 |
| SQEDTEAVFK | 10 | Q9NYH9 | U3 small nucleolar RNA-associated prote | UTP6 | 70.194 | 2 | 54.69439 | yes | 1.188E-07 | 577.27222 | 111.45296 |
| CYITFCLER | 9 | Q9NYH9 | U3 small nucleolar RNA-associated prote | UTP6 | 70.194 | 2 | 54.69439 | yes | 9.722E-08 | 631.28882 | 104.1714 |
| CCAVYEEAVK | 10 | Q9NYH9 | U3 small nucleolar RNA-associated prote | UTP6 | 70.194 | 2 | 54.69439 | yes | 5.758E-07 | 614.77045 | 60.372143 |
| LQVLIESK | 8 | Q9NYH9 | U3 small nucleolar RNA-associated prote | UTP6 | 70.194 | 2 | 54.69439 | yes | 1.554E-07 | 465.28696 | 50.576687 |
| EALEVAVAGTELF | 14 | Q9NYH9 | U3 small nucleolar RNA-associated prote | UTP6 | 70.194 | 2 | 54.69439 | yes | 1.789E-07 | 752.90393 | 18.881147 |
| EQESCNMANIR | 11 | Q9NYH9 | U3 small nucleolar RNA-associated prote | UTP6 | 70.194 | 2 | 54.69439 | yes | 4.706E-06 | 676.2901 | 12.175312 |
| DLPFETLEVEAK | 12 | Q9NYK5 | Large ribosomal subunit protein mL39 OS | MRPL39 | 38.712 | 2 | 52.737206 | yes | 1.463E-07 | 695.85864 | 1648.2979 |
| IGDFIDVSEGPLIPF | 15 | Q9NYK5 | Large ribosomal subunit protein mL39 OS | MRPL39 | 38.712 | 2 | 52.737206 | yes | 9.722E-08 | 814.43811 | 1621.1362 |
| FQGVSLPVHLR | 11 | Q9NYK5 | Large ribosomal subunit protein mL39 OS | MRPL39 | 38.712 | 2 | 52.737206 | yes | 9.722E-08 | 626.86169 | 1540.457 |
| YKVDFIEEK | 9 | Q9NYK5 | Large ribosomal subunit protein mL39 OS | MRPL39 | 38.712 | 2 | 52.737206 | yes | 1.201E-07 | 585.80573 | 945.59564 |
| VALEIFQHSK | 10 | Q9NYK5 | Large ribosomal subunit protein mL39 OS | MRPL39 | 38.712 | 2 | 52.737206 | yes | 9.819E-08 | 586.32715 | 738.95313 |
| DAHALIYK | 8 | Q9NYK5 | Large ribosomal subunit protein mL39 OS | MRPL39 | 38.712 | 2 | 52.737206 | yes | 9.81E-08 | 465.7558 | 732.50696 |
| AFKDEYMVNLVR | 12 | Q9NYK5 | Large ribosomal subunit protein mL39 OS | MRPL39 | 38.712 | 2 | 52.737206 | yes | 9.722E-08 | 742.88196 | 567.39038 |
| MVTEDQSK | 8 | Q9NYK5 | Large ribosomal subunit protein mL39 OS | MRPL39 | 38.712 | 2 | 52.737206 | yes | 1.734E-07 | 469.21841 | 535.74292 |
| SCAMMMGCVIER | 12 | Q9NYK5 | Large ribosomal subunit protein mL39 OS | MRPL39 | 38.712 | 2 | 52.737206 | yes | 1.703E-07 | 722.7973 | 364.79587 |
| AHFTIWDK | 8 | Q9NYK5 | Large ribosomal subunit protein mL39 OS | MRPL39 | 38.712 | 2 | 52.737206 | yes | 1.977E-07 | 509.26126 | 325.48093 |
| TDPGTVFVMNK | 11 | Q9NYK5 | Large ribosomal subunit protein mL39 OS | MRPL39 | 38.712 | 2 | 52.737206 | yes | 9.722E-08 | 604.80261 | 263.25308 |
| DEYMVNLVR | 9 | Q9NYK5 | Large ribosomal subunit protein mL39 OS | MRPL39 | 38.712 | 2 | 52.737206 | yes | 9.722E-08 | 569.78168 | 255.82901 |
| DCDPGEVNK | 9 | Q9NYK5 | Large ribosomal subunit protein mL39 OS | MRPL39 | 38.712 | 2 | 52.737206 | yes | 3.765E-07 | 517.21637 | 58.283169 |
| VDFIEEK | 7 | Q9NYK5 | Large ribosomal subunit protein mL39 OS | MRPL39 | 38.712 | 2 | 52.737206 | yes | 2.638E-07 | 440.22656 | 37.933891 |
| LDEWMPTK | 8 | Q9NYK5 | Large ribosomal subunit protein mL39 OS | MRPL39 | 38.712 | 2 | 52.737206 | yes | 3.813E-07 | 510.24695 | 28.524132 |

| | | | | | | | | | | | |
|---|---|---|---|---|---|---|---|---|---|---|---|
| SILALVDGQPWDM | 19 | Q9NYK5 | Large ribosomal subunit protein mL39 OS | MRPL39 | 38.712 | 3 | 52.737206 | yes | 4.804E-07 | 725.72211 | 13.758419 |
| EAEILSVLSHR | 11 | Q9NYL2 | Mitogen-activated protein kinase kinase k | MAP3K20 | 91.156 | 2 | 54.60203 | yes | 9.809E-08 | 627.34607 | 752.32397 |
| YQQITPVNQSR | 11 | Q9NYL2 | Mitogen-activated protein kinase kinase k | MAP3K20 | 91.156 | 2 | 54.60203 | yes | 9.722E-08 | 667.34656 | 599.83032 |
| LLLLEEEDLK | 10 | Q9NYL2 | Mitogen-activated protein kinase kinase k | MAP3K20 | 91.156 | 2 | 54.60203 | yes | 2.405E-07 | 607.84753 | 481.78647 |
| DVTFNTNLPDAEII | 15 | Q9NYL2 | Mitogen-activated protein kinase kinase k | MAP3K20 | 91.156 | 2 | 54.60203 | yes | 9.722E-08 | 845.43835 | 335.30356 |
| SSSPTQYGLTK | 11 | Q9NYL2 | Mitogen-activated protein kinase kinase k | MAP3K20 | 91.156 | 2 | 54.60203 | yes | 1.425E-07 | 584.79584 | 318.21024 |
| LTIPSSCPR | 9 | Q9NYL2 | Mitogen-activated protein kinase kinase k | MAP3K20 | 91.156 | 2 | 54.60203 | yes | 1.92E-06 | 515.77112 | 269.96497 |
| MTKPPFVMEK | 10 | Q9NYL2 | Mitogen-activated protein kinase kinase k | MAP3K20 | 91.156 | 2 | 54.60203 | yes | 2.664E-07 | 604.31433 | 230.49156 |
| CEIEATLER | 9 | Q9NYL2 | Mitogen-activated protein kinase kinase k | MAP3K20 | 91.156 | 2 | 54.60203 | yes | 2.757E-07 | 560.7688 | 215.44196 |
| SDSSADCQWLDTI | 14 | Q9NYL2 | Mitogen-activated protein kinase kinase k | MAP3K20 | 91.156 | 2 | 54.60203 | yes | 9.722E-08 | 827.3623 | 179.65601 |
| GMHYLHMEAPVK | 12 | Q9NYL2 | Mitogen-activated protein kinase kinase k | MAP3K20 | 91.156 | 2 | 54.60203 | yes | 1.765E-07 | 706.8443 | 73.52224 |
| GMHYLHMEAPVK | 12 | Q9NYL2 | Mitogen-activated protein kinase kinase k | MAP3K20 | 91.156 | 2 | 54.60203 | yes | 4.253E-06 | 471.56531 | 73.52224 |
| GMHYLHMEAPVK | 12 | Q9NYL2 | Mitogen-activated protein kinase kinase k | MAP3K20 | 91.156 | 3 | 54.60203 | yes | 1.765E-07 | 706.8443 | 73.52224 |
| GMHYLHMEAPVK | 12 | Q9NYL2 | Mitogen-activated protein kinase kinase k | MAP3K20 | 91.156 | 3 | 54.60203 | yes | 4.253E-06 | 471.56531 | 73.52224 |
| NVVIAADGVLK | 11 | Q9NYL2 | Mitogen-activated protein kinase kinase k | MAP3K20 | 91.156 | 2 | 54.60203 | yes | 1.002E-07 | 549.82953 | 145.08688 |
| QIASNTSLQR | 10 | Q9NYL2 | Mitogen-activated protein kinase kinase k | MAP3K20 | 91.156 | 2 | 54.60203 | yes | 2.097E-07 | 559.30164 | 139.02068 |
| GHIIHFK | 7 | Q9NYL2 | Mitogen-activated protein kinase kinase k | MAP3K20 | 91.156 | 2 | 54.60203 | yes | 9.722E-08 | 426.24796 | 136.67603 |
| RPQVPIK | 7 | Q9NYL2 | Mitogen-activated protein kinase kinase k | MAP3K20 | 91.156 | 2 | 54.60203 | yes | 4.594E-07 | 419.26889 | 110.75334 |
| HVHSIQWSR | 9 | Q9NYL2 | Mitogen-activated protein kinase kinase k | MAP3K20 | 91.156 | 2 | 54.60203 | yes | 1.881E-06 | 575.29926 | 82.78199 |
| KGDSSAEMSVYAS | 16 | Q9NYL2 | Mitogen-activated protein kinase kinase k | MAP3K20 | 91.156 | 3 | 54.60203 | yes | 1.82E-07 | 573.94678 | 46.333855 |
| VSQSALNPHQSPD | 15 | Q9NYL2 | Mitogen-activated protein kinase kinase k | MAP3K20 | 91.156 | 2 | 54.60203 | yes | 9.974E-07 | 827.91284 | 12.667 |
| MEHFDASLSTYFK | 13 | Q9NYP7 | Elongation of very long chain fatty acids p | ELOVL5 | 35.294 | 2 | 50.94514 | yes | 9.722E-08 | 809.36633 | 122.57705 |
| VVVCGLLSVGK | 11 | Q9NYS0 | NF-kappa-B inhibitor-interacting Ras-like | NKIRAS1 | 21.643 | 2 | 49.32222 | yes | 1.045E-07 | 565.83374 | 76.02095 |
| LWEVTVTDR | 9 | Q9NYS0 | NF-kappa-B inhibitor-interacting Ras-like | NKIRAS1 | 21.643 | 2 | 49.32222 | yes | 3.497E-07 | 559.79565 | 75.044075 |
| EGFPITAIR | 9 | Q9NYV4 | Cyclin-dependent kinase 12 OS=Homo sa | CDK12 | 164.155 | 2 | 51.4893 | no | 1.422E-07 | 502.2822 | 272.33405 |
| CSNILLNNSGQIK | 13 | Q9NYV4 | Cyclin-dependent kinase 12 OS=Homo sa | CDK12 | 164.155 | 2 | 51.4893 | yes | 1.164E-07 | 730.87994 | 174.38927 |
| CTAEQTLQSDFLK | 13 | Q9NYV4 | Cyclin-dependent kinase 12 OS=Homo sa | CDK12 | 164.155 | 2 | 51.4893 | yes | 9.722E-08 | 770.8692 | 99.133347 |
| VPLALHPVVGQPF | 15 | Q9NYV4 | Cyclin-dependent kinase 12 OS=Homo sa | CDK12 | 164.155 | 3 | 51.4893 | yes | 1.532E-07 | 538.99585 | 43.398941 |
| VPLALHPVVGQPF | 15 | Q9NYV4 | Cyclin-dependent kinase 12 OS=Homo sa | CDK12 | 164.155 | 3 | 51.4893 | yes | 7.871E-07 | 807.99017 | 43.398941 |
| VPLALHPVVGQPF | 15 | Q9NYV4 | Cyclin-dependent kinase 12 OS=Homo sa | CDK12 | 164.155 | 2 | 51.4893 | yes | 1.532E-07 | 538.99585 | 43.398941 |
| VPLALHPVVGQPF | 15 | Q9NYV4 | Cyclin-dependent kinase 12 OS=Homo sa | CDK12 | 164.155 | 2 | 51.4893 | yes | 7.871E-07 | 807.99017 | 43.398941 |
| LPLNSSLGAELSR | 13 | Q9NYV4 | Cyclin-dependent kinase 12 OS=Homo sa | CDK12 | 164.155 | 2 | 51.4893 | yes | 1.544E-07 | 678.87769 | 83.835602 |
| NSGPALTESLVQTI | 16 | Q9NYV4 | Cyclin-dependent kinase 12 OS=Homo sa | CDK12 | 164.155 | 2 | 51.4893 | yes | 9.722E-08 | 828.96198 | 56.52668 |
| DKDTGELVALK | 11 | Q9NYV4 | Cyclin-dependent kinase 12 OS=Homo sa | CDK12 | 164.155 | 2 | 51.4893 | yes | 8.546E-07 | 594.82715 | 37.776207 |
| AEGSSNSVVHAET | 14 | Q9NYV4 | Cyclin-dependent kinase 12 OS=Homo sa | CDK12 | 164.155 | 2 | 51.4893 | yes | 6.919E-05 | 708.34167 | 37.64259 |
| SNEETDDYGK | 10 | Q9NYV4 | Cyclin-dependent kinase 12 OS=Homo sa | CDK12 | 164.155 | 2 | 51.4893 | yes | 3.003E-06 | 579.23328 | 29.989523 |
| CVDKFDIIGIIGEG | 21 | Q9NYV4 | Cyclin-dependent kinase 12 OS=Homo sa | CDK12 | 164.155 | 3 | 51.4893 | no | 2.152E-06 | 792.40326 | 21.833567 |
| QSGVVVEEPPPSK | 13 | Q9NYV4 | Cyclin-dependent kinase 12 OS=Homo sa | CDK12 | 164.155 | 2 | 51.4893 | yes | 3.387E-05 | 676.85645 | 15.766448 |
| MSNLTVHYR | 9 | Q9NYW8 | RB-associated KRAB zinc finger protein | RBAK | 82.995 | 2 | 49.00074 | yes | 1.078E-07 | 560.78204 | 91.632034 |

| | | | | | | | | | | | |
|---|---|---|---|---|---|---|---|---|---|---|---|
| VSYLTIHYR | 9 | Q9NYW8 | RB-associated KRAB zinc finger protein | RBAK | 82.995 | 2 | 49.00074 | yes | 2.874E-07 | 576.31403 | 49.166691 |
| GCLENMPEADCIP | 14 | Q9NYY3 | Serine/threonine-protein kinase PLK2 OS | PLK2 | 78.237 | 2 | 44.900517 | yes | 2.411E-07 | 817.35455 | 111.43776 |
| VGDFGLAAR | 9 | Q9NYY3 | Serine/threonine-protein kinase PLK2 OS | PLK2 | 78.237 | 2 | 44.900517 | no | 2.551E-07 | 453.24561 | 101.98087 |
| TITYQPAASTK | 11 | Q9NYY3 | Serine/threonine-protein kinase PLK2 OS | PLK2 | 78.237 | 2 | 44.900517 | yes | 2.031E-07 | 590.81403 | 96.921936 |
| EQLSTSFQWVTK | 12 | Q9NYY3 | Serine/threonine-protein kinase PLK2 OS | PLK2 | 78.237 | 2 | 44.900517 | yes | 2.854E-07 | 727.36975 | 66.135933 |
| LGNFFINEAMELK | 13 | Q9NYY3 | Serine/threonine-protein kinase PLK2 OS | PLK2 | 78.237 | 2 | 44.900517 | yes | 8.195E-07 | 763.38959 | 58.170647 |
| HDFFLQGFTPDR | 12 | Q9NYY3 | Serine/threonine-protein kinase PLK2 OS | PLK2 | 78.237 | 2 | 44.900517 | yes | 2.114E-07 | 740.35443 | 39.861885 |
| TDEELQPPTTTVAI | 14 | Q9NYY3 | Serine/threonine-protein kinase PLK2 OS | PLK2 | 78.237 | 2 | 44.900517 | yes | 7.605E-07 | 779.39136 | 20.693653 |
| ILVDQQVWK | 9 | Q9NYY8 | FAST kinase domain-containing protein 2 | FASTKD2 | 81.463 | 2 | 52.2481 | yes | 1.215E-07 | 564.82422 | 1508.0875 |
| IVEDPESLNMK | 11 | Q9NYY8 | FAST kinase domain-containing protein 2 | FASTKD2 | 81.463 | 2 | 52.2481 | yes | 9.722E-08 | 637.81848 | 1154.7825 |
| LHTLDTCLK | 9 | Q9NYY8 | FAST kinase domain-containing protein 2 | FASTKD2 | 81.463 | 2 | 52.2481 | yes | 1.253E-07 | 550.79205 | 1109.3784 |
| SLILLDECSK | 10 | Q9NYY8 | FAST kinase domain-containing protein 2 | FASTKD2 | 81.463 | 2 | 52.2481 | yes | 5.426E-07 | 589.3103 | 1015.6068 |
| VVLDNIHGCPLR | 12 | Q9NYY8 | FAST kinase domain-containing protein 2 | FASTKD2 | 81.463 | 2 | 52.2481 | yes | 9.722E-08 | 696.87445 | 495.72485 |
| VVLDNIHGCPLR | 12 | Q9NYY8 | FAST kinase domain-containing protein 2 | FASTKD2 | 81.463 | 2 | 52.2481 | yes | 1.794E-07 | 464.91873 | 495.72485 |
| VVLDNIHGCPLR | 12 | Q9NYY8 | FAST kinase domain-containing protein 2 | FASTKD2 | 81.463 | 3 | 52.2481 | yes | 9.722E-08 | 696.87445 | 495.72485 |
| VVLDNIHGCPLR | 12 | Q9NYY8 | FAST kinase domain-containing protein 2 | FASTKD2 | 81.463 | 3 | 52.2481 | yes | 1.794E-07 | 464.91873 | 495.72485 |
| DIALSLPQLPR | 11 | Q9NYY8 | FAST kinase domain-containing protein 2 | FASTKD2 | 81.463 | 2 | 52.2481 | yes | 3.74E-07 | 611.86133 | 458.55121 |
| DLQYHNLDLFK | 11 | Q9NYY8 | FAST kinase domain-containing protein 2 | FASTKD2 | 81.463 | 2 | 52.2481 | yes | 9.722E-08 | 703.35913 | 437.34949 |
| IEDVFTLQVVMK | 12 | Q9NYY8 | FAST kinase domain-containing protein 2 | FASTKD2 | 81.463 | 2 | 52.2481 | yes | 1.004E-07 | 711.3891 | 431.9946 |
| SAYCLGSSHPR | 11 | Q9NYY8 | FAST kinase domain-containing protein 2 | FASTKD2 | 81.463 | 2 | 52.2481 | yes | 4.76E-07 | 617.78528 | 406.55579 |
| IMINILQSCK | 10 | Q9NYY8 | FAST kinase domain-containing protein 2 | FASTKD2 | 81.463 | 2 | 52.2481 | yes | 1.048E-07 | 610.33051 | 268.38028 |
| LDDTVYLR | 8 | Q9NYY8 | FAST kinase domain-containing protein 2 | FASTKD2 | 81.463 | 2 | 52.2481 | yes | 1.684E-07 | 497.76382 | 225.15457 |
| VAEVLSSLLGGEG | 17 | Q9NYY8 | FAST kinase domain-containing protein 2 | FASTKD2 | 81.463 | 3 | 52.2481 | yes | 1.06E-07 | 577.30884 | 184.83447 |
| DVHLPHNYHIDFE | 15 | Q9NYY8 | FAST kinase domain-containing protein 2 | FASTKD2 | 81.463 | 4 | 52.2481 | yes | 5.405E-07 | 476.99051 | 158.3414 |
| LEMEDAVTFLK | 11 | Q9NYY8 | FAST kinase domain-containing protein 2 | FASTKD2 | 81.463 | 2 | 52.2481 | yes | 9.809E-08 | 648.33124 | 120.93626 |
| PALPAGPVGASSW | 16 | Q9NYZ3 | G2 and S phase-expressed protein 1 OS=l | GTSE1 | 76.645 | 2 | 54.57898 | yes | 1.112E-07 | 768.91211 | 217.0442 |
| AVGSPLCVPAR | 11 | Q9NYZ3 | G2 and S phase-expressed protein 1 OS=l | GTSE1 | 76.645 | 2 | 54.57898 | yes | 2.495E-07 | 563.80548 | 178.40727 |
| LLASSPALPSSGAQ | 16 | Q9NYZ3 | G2 and S phase-expressed protein 1 OS=l | GTSE1 | 76.645 | 2 | 54.57898 | yes | 1.846E-07 | 763.42029 | 129.04967 |
| VTVHSTPVR | 9 | Q9NYZ3 | G2 and S phase-expressed protein 1 OS=l | GTSE1 | 76.645 | 2 | 54.57898 | yes | 3.82E-07 | 498.28528 | 118.84254 |
| VMPTPTNQFK | 10 | Q9NYZ3 | G2 and S phase-expressed protein 1 OS=l | GTSE1 | 76.645 | 2 | 54.57898 | yes | 3.432E-07 | 581.79987 | 77.745621 |
| SIPGAAEKPK | 10 | Q9NYZ3 | G2 and S phase-expressed protein 1 OS=l | GTSE1 | 76.645 | 2 | 54.57898 | yes | 1.048E-07 | 499.28748 | 76.657967 |
| EAHLLALHIESSSR | 14 | Q9NYZ3 | G2 and S phase-expressed protein 1 OS=l | GTSE1 | 76.645 | 2 | 54.57898 | yes | 9.722E-08 | 781.91791 | 47.646759 |
| PLIDLMTNTPDMN | 14 | Q9NYZ3 | G2 and S phase-expressed protein 1 OS=l | GTSE1 | 76.645 | 2 | 54.57898 | yes | 5.135E-07 | 801.89691 | 26.977245 |
| FSHGTMPLR | 9 | Q9NZ01 | Very-long-chain enoyl-CoA reductase OS | TECR | 36.035 | 2 | 54.77429 | yes | 9.85E-08 | 523.26599 | 1242.6864 |
| FSHGTMPLR | 9 | Q9NZ01 | Very-long-chain enoyl-CoA reductase OS | TECR | 36.035 | 2 | 54.77429 | yes | 5.45E-07 | 531.26349 | 1242.6864 |
| LPVGTTATLYFR | 12 | Q9NZ01 | Very-long-chain enoyl-CoA reductase OS | TECR | 36.035 | 2 | 54.77429 | yes | 1.262E-07 | 669.87445 | 2455.9873 |
| THPQWYPAR | 9 | Q9NZ01 | Very-long-chain enoyl-CoA reductase OS | TECR | 36.035 | 2 | 54.77429 | yes | 1.8E-07 | 578.28833 | 2367.3049 |
| LLETLFVHR | 9 | Q9NZ01 | Very-long-chain enoyl-CoA reductase OS | TECR | 36.035 | 2 | 54.77429 | yes | 1.384E-07 | 564.33221 | 2059.2937 |
| SLKDEDVLQK | 10 | Q9NZ01 | Very-long-chain enoyl-CoA reductase OS | TECR | 36.035 | 2 | 54.77429 | yes | 2.252E-07 | 587.81934 | 1303.4264 |

| | | | | | | | | | | | |
|---|---|---|---|---|---|---|---|---|---|---|---|
| VEPHATIAEIK | 11 | Q9NZ01 | Very-long-chain enoyl-CoA reductase OS | TECR | 36.035 | 2 | 54.77429 | yes | 1.073E-07 | 604.33771 | 1157.0079 |
| YDFTSSR | 7 | Q9NZ01 | Very-long-chain enoyl-CoA reductase OS | TECR | 36.035 | 2 | 54.77429 | yes | 3.133E-07 | 438.19833 | 664.17328 |
| KIPYPTK | 7 | Q9NZ01 | Very-long-chain enoyl-CoA reductase OS | TECR | 36.035 | 2 | 54.77429 | yes | 1.651E-07 | 423.75781 | 322.70547 |
| DLRPAGSK | 8 | Q9NZ01 | Very-long-chain enoyl-CoA reductase OS | TECR | 36.035 | 2 | 54.77429 | yes | 9.722E-08 | 422.23779 | 102.0732 |
| VPFIYGHK | 8 | Q9NZ01 | Very-long-chain enoyl-CoA reductase OS | TECR | 36.035 | 2 | 54.77429 | yes | 1.542E-07 | 480.76871 | 66.887848 |
| HYEVEILDAK | 10 | Q9NZ01 | Very-long-chain enoyl-CoA reductase OS | TECR | 36.035 | 2 | 54.77429 | yes | 3.001E-07 | 608.81403 | 36.799431 |
| IVHAFDMEDLGDK | 13 | Q9NZ45 | CDGSH iron-sulfur domain-containing pı | CISD1 | 12.2 | 2 | 48.617695 | yes | 1.662E-07 | 745.35321 | 189.48065 |
| IVHAFDMEDLGDK | 13 | Q9NZ45 | CDGSH iron-sulfur domain-containing pı | CISD1 | 12.2 | 2 | 48.617695 | yes | 4.849E-06 | 753.35065 | 189.48065 |
| AMINLHIQK | 9 | Q9NZ45 | CDGSH iron-sulfur domain-containing pı | CISD1 | 12.2 | 2 | 48.617695 | yes | 1.122E-07 | 534.30511 | 128.20837 |
| HNEETGDNVGPLI | 15 | Q9NZ45 | CDGSH iron-sulfur domain-containing pı | CISD1 | 12.2 | 2 | 48.617695 | yes | 1.92E-06 | 818.42047 | 16.3524 |
| LYEPDQLQELK | 11 | Q9NZB2 | Constitutive coactivator of PPAR-gamma | FAM120A | 121.888 | 2 | 53.57181 | yes | 1.329E-07 | 688.35883 | 9861.0459 |
| LPPEFSPVIIK | 11 | Q9NZB2 | Constitutive coactivator of PPAR-gamma | FAM120A | 121.888 | 2 | 53.57181 | yes | 2.993E-07 | 620.37103 | 9568.709 |
| QSVLEGLSFSR | 11 | Q9NZB2 | Constitutive coactivator of PPAR-gamma | FAM120A | 121.888 | 2 | 53.57181 | yes | 9.722E-08 | 611.82495 | 7542.9111 |
| HCPSAVVPVELQK | 13 | Q9NZB2 | Constitutive coactivator of PPAR-gamma | FAM120A | 121.888 | 2 | 53.57181 | yes | 3.271E-07 | 732.38739 | 3362.1648 |
| HCPSAVVPVELQK | 13 | Q9NZB2 | Constitutive coactivator of PPAR-gamma | FAM120A | 121.888 | 2 | 53.57181 | yes | 1.083E-07 | 488.59402 | 3362.1648 |
| HCPSAVVPVELQK | 13 | Q9NZB2 | Constitutive coactivator of PPAR-gamma | FAM120A | 121.888 | 3 | 53.57181 | yes | 3.271E-07 | 732.38739 | 3362.1648 |
| HCPSAVVPVELQK | 13 | Q9NZB2 | Constitutive coactivator of PPAR-gamma | FAM120A | 121.888 | 3 | 53.57181 | yes | 1.083E-07 | 488.59402 | 3362.1648 |
| HTPLYER | 7 | Q9NZB2 | Constitutive coactivator of PPAR-gamma | FAM120A | 121.888 | 2 | 53.57181 | no | 1.971E-07 | 458.23779 | 6219.917 |
| DLPPAALLYR | 10 | Q9NZB2 | Constitutive coactivator of PPAR-gamma | FAM120A | 121.888 | 2 | 53.57181 | yes | 5.515E-07 | 564.82422 | 5599.2383 |
| NIQDTSDLDAIAK | 13 | Q9NZB2 | Constitutive coactivator of PPAR-gamma | FAM120A | 121.888 | 2 | 53.57181 | yes | 9.914E-08 | 702.35425 | 5569.0161 |
| SLTTSQYLMHEVA | 14 | Q9NZB2 | Constitutive coactivator of PPAR-gamma | FAM120A | 121.888 | 2 | 53.57181 | yes | 1.526E-07 | 804.40851 | 2505.6909 |
| SLTTSQYLMHEVA | 14 | Q9NZB2 | Constitutive coactivator of PPAR-gamma | FAM120A | 121.888 | 2 | 53.57181 | yes | 1.353E-07 | 812.40601 | 2505.6909 |
| AHQLVLPPCDVVI | 14 | Q9NZB2 | Constitutive coactivator of PPAR-gamma | FAM120A | 121.888 | 2 | 53.57181 | no | 1.468E-07 | 794.94763 | 4777.6045 |
| SPQTPELVEALAFI | 14 | Q9NZB2 | Constitutive coactivator of PPAR-gamma | FAM120A | 121.888 | 2 | 53.57181 | yes | 4.539E-07 | 779.41718 | 4349.1973 |
| YMVQWPGAR | 9 | Q9NZB2 | Constitutive coactivator of PPAR-gamma | FAM120A | 121.888 | 2 | 53.57181 | yes | 6.863E-07 | 554.27386 | 4330.9917 |
| IAVSIEDEANK | 11 | Q9NZB2 | Constitutive coactivator of PPAR-gamma | FAM120A | 121.888 | 2 | 53.57181 | yes | 3.729E-07 | 594.80896 | 4160.8594 |
| EAALEAAVLNK | 11 | Q9NZB2 | Constitutive coactivator of PPAR-gamma | FAM120A | 121.888 | 2 | 53.57181 | yes | 1.945E-07 | 564.81659 | 3925.9553 |
| DVFQHSQSR | 9 | Q9NZB2 | Constitutive coactivator of PPAR-gamma | FAM120A | 121.888 | 2 | 53.57181 | yes | 9.893E-08 | 552.26508 | 3841.2319 |
| QLDLNPNR | 8 | Q9NZB2 | Constitutive coactivator of PPAR-gamma | FAM120A | 121.888 | 2 | 53.57181 | yes | 8.653E-07 | 485.25925 | 3518.1191 |
| AEGSSTASSGSQLA | 17 | Q9NZB2 | Constitutive coactivator of PPAR-gamma | FAM120A | 121.888 | 2 | 53.57181 | yes | 1.241E-07 | 783.86572 | 3377.272 |
| AIGYYSATSK | 10 | Q9NZB2 | Constitutive coactivator of PPAR-gamma | FAM120A | 121.888 | 2 | 53.57181 | yes | 3.683E-07 | 530.7691 | 3222.876 |
| GVQGFQDYIEK | 11 | Q9NZB2 | Constitutive coactivator of PPAR-gamma | FAM120A | 121.888 | 2 | 53.57181 | yes | 1.635E-07 | 642.31696 | 3127.0657 |
| EWTCPNLK | 8 | Q9NZB2 | Constitutive coactivator of PPAR-gamma | FAM120A | 121.888 | 2 | 53.57181 | no | 1.313E-07 | 524.25006 | 2863.0486 |
| IENLDPR | 7 | Q9NZB2 | Constitutive coactivator of PPAR-gamma | FAM120A | 121.888 | 2 | 53.57181 | yes | 3.366E-06 | 428.7298 | 2215.3379 |
| PMSFHPPHYLAAR | 13 | Q9NZB2 | Constitutive coactivator of PPAR-gamma | FAM120A | 121.888 | 3 | 53.57181 | yes | 1.034E-07 | 508.59073 | 1047.3402 |
| PMSFHPPHYLAAR | 13 | Q9NZB2 | Constitutive coactivator of PPAR-gamma | FAM120A | 121.888 | 3 | 53.57181 | yes | 1.002E-07 | 762.38245 | 1047.3402 |
| PMSFHPPHYLAAR | 13 | Q9NZB2 | Constitutive coactivator of PPAR-gamma | FAM120A | 121.888 | 2 | 53.57181 | yes | 1.034E-07 | 508.59073 | 1047.3402 |
| PMSFHPPHYLAAR | 13 | Q9NZB2 | Constitutive coactivator of PPAR-gamma | FAM120A | 121.888 | 2 | 53.57181 | yes | 1.002E-07 | 762.38245 | 1047.3402 |
| GPFPLQVVSVGGP | 15 | Q9NZB2 | Constitutive coactivator of PPAR-gamma | FAM120A | 121.888 | 2 | 53.57181 | yes | 1.121E-07 | 740.91718 | 1679.9047 |

| | | | | | | | | | | | |
|---|---|---|---|---|---|---|---|---|---|---|---|
| AFLACMR | 7 | Q9NZB2 | Constitutive coactivator of PPAR-gamma | FAM120A | 121.888 | 2 | 53.57181 | yes | 2.015E-07 | 434.71204 | 1367.3138 |
| QTAQQIVSHVQNK | 13 | Q9NZB2 | Constitutive coactivator of PPAR-gamma | FAM120A | 121.888 | 2 | 53.57181 | yes | 2.416E-07 | 740.89697 | 1301.5188 |
| APSHSESALNNDS | 14 | Q9NZB2 | Constitutive coactivator of PPAR-gamma | FAM120A | 121.888 | 2 | 53.57181 | yes | 1.022E-07 | 728.83679 | 1048.0198 |
| QRPPQTPLR | 9 | Q9NZB2 | Constitutive coactivator of PPAR-gamma | FAM120A | 121.888 | 2 | 53.57181 | yes | 2.138E-07 | 546.81726 | 860.81622 |
| QYVYGVLFSLAES | 14 | Q9NZB2 | Constitutive coactivator of PPAR-gamma | FAM120A | 121.888 | 2 | 53.57181 | yes | 1.109E-07 | 816.42505 | 839.06189 |
| NLTEQNSYSNIPHI | 16 | Q9NZB2 | Constitutive coactivator of PPAR-gamma | FAM120A | 121.888 | 3 | 53.57181 | no | 5.054E-07 | 610.95874 | 781.55487 |
| VAAASGHCGAFSC | 18 | Q9NZB2 | Constitutive coactivator of PPAR-gamma | FAM120A | 121.888 | 3 | 53.57181 | yes | 1.246E-07 | 575.2533 | 731.97687 |
| RQELDAFLAQALS | 15 | Q9NZB2 | Constitutive coactivator of PPAR-gamma | FAM120A | 121.888 | 2 | 53.57181 | yes | 1.128E-07 | 843.96234 | 349.40417 |
| RQELDAFLAQALS | 15 | Q9NZB2 | Constitutive coactivator of PPAR-gamma | FAM120A | 121.888 | 2 | 53.57181 | yes | 1.278E-07 | 562.97729 | 349.40417 |
| RQELDAFLAQALS | 15 | Q9NZB2 | Constitutive coactivator of PPAR-gamma | FAM120A | 121.888 | 3 | 53.57181 | yes | 1.128E-07 | 843.96234 | 349.40417 |
| RQELDAFLAQALS | 15 | Q9NZB2 | Constitutive coactivator of PPAR-gamma | FAM120A | 121.888 | 3 | 53.57181 | yes | 1.278E-07 | 562.97729 | 349.40417 |
| SQGGVQPIPSQGG | 14 | Q9NZB2 | Constitutive coactivator of PPAR-gamma | FAM120A | 121.888 | 2 | 53.57181 | yes | 9.722E-08 | 670.35187 | 522.5141 |
| GVISTPVIR | 9 | Q9NZB2 | Constitutive coactivator of PPAR-gamma | FAM120A | 121.888 | 2 | 53.57181 | yes | 1.552E-07 | 471.29257 | 480.20065 |
| VAQSIEDHHQEVI | 17 | Q9NZB2 | Constitutive coactivator of PPAR-gamma | FAM120A | 121.888 | 4 | 53.57181 | yes | 1.075E-07 | 506.99564 | 410.30618 |
| QELDAFLAQALSP | 14 | Q9NZB2 | Constitutive coactivator of PPAR-gamma | FAM120A | 121.888 | 2 | 53.57181 | yes | 1.306E-07 | 765.91174 | 279.84766 |
| VWFLPPVCMAHC | 14 | Q9NZB2 | Constitutive coactivator of PPAR-gamma | FAM120A | 121.888 | 3 | 53.57181 | yes | 4.616E-07 | 595.9632 | 232.75063 |
| PGPFGMPGMVPPI | 29 | Q9NZB2 | Constitutive coactivator of PPAR-gamma | FAM120A | 121.888 | 4 | 53.57181 | yes | 3.249E-07 | 767.39862 | 182.44016 |
| LEIAGTVVGHWAC | 15 | Q9NZB2 | Constitutive coactivator of PPAR-gamma | FAM120A | 121.888 | 3 | 53.57181 | yes | 1.564E-07 | 518.27917 | 153.44632 |
| IAVSIEDEANKDLI | 21 | Q9NZB2 | Constitutive coactivator of PPAR-gamma | FAM120A | 121.888 | 3 | 53.57181 | yes | 4.199E-07 | 766.74951 | 117.56257 |
| NQAAIQGRPPYAA | 20 | Q9NZB2 | Constitutive coactivator of PPAR-gamma | FAM120A | 121.888 | 3 | 53.57181 | yes | 3.685E-07 | 691.02374 | 116.46581 |
| GLMYPYIFHVLTK | 13 | Q9NZB2 | Constitutive coactivator of PPAR-gamma | FAM120A | 121.888 | 2 | 53.57181 | yes | 1.831E-07 | 791.42853 | 104.25804 |
| REAALEAAVLNK | 12 | Q9NZB2 | Constitutive coactivator of PPAR-gamma | FAM120A | 121.888 | 2 | 53.57181 | yes | 1.562E-07 | 642.86713 | 99.45443 |
| NHMDITTPPLPPV | 19 | Q9NZB2 | Constitutive coactivator of PPAR-gamma | FAM120A | 121.888 | 3 | 53.57181 | yes | 4.608E-07 | 699.71021 | 27.169195 |
| VILLSGTPAMSR | 12 | Q9NZC9 | SWI/SNF-related matrix-associated actin- | SMARCAL | 105.938 | 2 | 52.343067 | yes | 5.054E-07 | 622.85522 | 310.31693 |
| VVLDAITQELER | 12 | Q9NZC9 | SWI/SNF-related matrix-associated actin- | SMARCAL | 105.938 | 2 | 52.343067 | yes | 9.722E-08 | 693.38538 | 295.59525 |
| LLLEEAVMLR | 10 | Q9NZC9 | SWI/SNF-related matrix-associated actin- | SMARCAL | 105.938 | 2 | 52.343067 | yes | 1.209E-07 | 593.84686 | 158.35944 |
| WSFLLEEHSK | 10 | Q9NZC9 | SWI/SNF-related matrix-associated actin- | SMARCAL | 105.938 | 2 | 52.343067 | yes | 9.754E-08 | 638.32202 | 100.00744 |
| KIVVIAPGR | 9 | Q9NZC9 | SWI/SNF-related matrix-associated actin- | SMARCAL | 105.938 | 2 | 52.343067 | yes | 1.16E-07 | 476.81876 | 95.935265 |
| LLLADDMGLGK | 11 | Q9NZC9 | SWI/SNF-related matrix-associated actin- | SMARCAL | 105.938 | 2 | 52.343067 | yes | 1.064E-06 | 573.31537 | 93.929794 |
| KPEEMPTACPGHS | 15 | Q9NZC9 | SWI/SNF-related matrix-associated actin- | SMARCAL | 105.938 | 3 | 52.343067 | yes | 2.355E-07 | 565.26434 | 82.600868 |
| FTWEQAFLR | 9 | Q9NZC9 | SWI/SNF-related matrix-associated actin- | SMARCAL | 105.938 | 2 | 52.343067 | yes | 1.079E-07 | 599.30621 | 73.633377 |
| EDLCQQFQLSER | 12 | Q9NZC9 | SWI/SNF-related matrix-associated actin- | SMARCAL | 105.938 | 2 | 52.343067 | yes | 9.927E-08 | 776.85663 | 70.946342 |
| GTADDYLWPLIQE | 14 | Q9NZC9 | SWI/SNF-related matrix-associated actin- | SMARCAL | 105.938 | 2 | 52.343067 | yes | 1.022E-07 | 824.91449 | 44.698772 |
| AGVNFAIAK | 9 | Q9NZC9 | SWI/SNF-related matrix-associated actin- | SMARCAL | 105.938 | 2 | 52.343067 | yes | 1.088E-07 | 445.75836 | 38.881317 |
| TSSGTSIAGNPFQA | 15 | Q9NZC9 | SWI/SNF-related matrix-associated actin- | SMARCAL | 105.938 | 2 | 52.343067 | yes | 1.086E-07 | 733.36774 | 36.010788 |
| VVIIDESHFLK | 11 | Q9NZC9 | SWI/SNF-related matrix-associated actin- | SMARCAL | 105.938 | 2 | 52.343067 | yes | 1.087E-07 | 650.36902 | 35.375023 |
| TIQAICIAAFYR | 12 | Q9NZC9 | SWI/SNF-related matrix-associated actin- | SMARCAL | 105.938 | 2 | 52.343067 | yes | 5.92E-06 | 713.87921 | 3.5749702 |
| LHCGLWVR | 8 | Q9NZE8 | Large ribosomal subunit protein bL35m C | MRPL35 | 21.515 | 2 | 52.57135 | yes | 2.54E-07 | 520.77655 | 261.88446 |
| NWYVDDPYQK | 10 | Q9NZE8 | Large ribosomal subunit protein bL35m C | MRPL35 | 21.515 | 2 | 52.57135 | yes | 9.722E-08 | 664.30133 | 153.19345 |

| | | | | | | | | | | | |
|---|---|---|---|---|---|---|---|---|---|---|---|
| EDLVQICGAADGI | 14 | Q9NZI7 | Upstream-binding protein 1 OS=Homo sa | UBP1 | 60.492 | 2 | 53.33027 | yes | 9.938E-08 | 758.87482 | 458.70856 |
| LFSNFSGADLLK | 12 | Q9NZI7 | Upstream-binding protein 1 OS=Homo sa | UBP1 | 60.492 | 2 | 53.33027 | yes | 1.387E-07 | 656.35083 | 440.03958 |
| LTIYVCR | 7 | Q9NZI7 | Upstream-binding protein 1 OS=Homo sa | UBP1 | 60.492 | 2 | 53.33027 | yes | 2.509E-07 | 462.75223 | 140.31851 |
| AESSDGIHIILK | 12 | Q9NZI7 | Upstream-binding protein 1 OS=Homo sa | UBP1 | 60.492 | 2 | 53.33027 | yes | 9.722E-08 | 641.8537 | 109.96053 |
| LQYTEHQQLEGWI | 13 | Q9NZI7 | Upstream-binding protein 1 OS=Homo sa | UBP1 | 60.492 | 2 | 53.33027 | yes | 3.183E-06 | 830.40991 | 41.85294 |
| TLPADYGDSLAK | 12 | Q9NZI7 | Upstream-binding protein 1 OS=Homo sa | UBP1 | 60.492 | 2 | 53.33027 | yes | 2.843E-07 | 625.81677 | 28.427933 |
| MILEIMHK | 8 | Q9NZI8 | Insulin-like growth factor 2 mRNA-bindi | IGF2BP1 | 63.481 | 2 | 52.71008 | yes | 2.41E-07 | 507.77737 | 9070.4736 |
| MILEIMHK | 8 | Q9NZI8 | Insulin-like growth factor 2 mRNA-bindi | IGF2BP1 | 63.481 | 2 | 52.71008 | yes | 1.799E-07 | 515.77484 | 9070.4736 |
| MILEIMHK | 8 | Q9NZI8 | Insulin-like growth factor 2 mRNA-bindi | IGF2BP1 | 63.481 | 2 | 52.71008 | yes | 4.403E-07 | 515.77484 | 9070.4736 |
| QQQVDIPLR | 9 | Q9NZI8 | Insulin-like growth factor 2 mRNA-bindi | IGF2BP1 | 63.481 | 2 | 52.71008 | yes | 1.885E-07 | 548.80908 | 21104.613 |
| LKEENFFGPK | 10 | Q9NZI8 | Insulin-like growth factor 2 mRNA-bindi | IGF2BP1 | 63.481 | 2 | 52.71008 | yes | 1.543E-07 | 604.81915 | 8180.5161 |
| LKEENFFGPK | 10 | Q9NZI8 | Insulin-like growth factor 2 mRNA-bindi | IGF2BP1 | 63.481 | 2 | 52.71008 | yes | 2.101E-07 | 403.54852 | 8180.5161 |
| LKEENFFGPK | 10 | Q9NZI8 | Insulin-like growth factor 2 mRNA-bindi | IGF2BP1 | 63.481 | 3 | 52.71008 | yes | 1.543E-07 | 604.81915 | 8180.5161 |
| LKEENFFGPK | 10 | Q9NZI8 | Insulin-like growth factor 2 mRNA-bindi | IGF2BP1 | 63.481 | 3 | 52.71008 | yes | 2.101E-07 | 403.54852 | 8180.5161 |
| ISYSGQFLVK | 10 | Q9NZI8 | Insulin-like growth factor 2 mRNA-bindi | IGF2BP1 | 63.481 | 2 | 52.71008 | yes | 1.21E-07 | 571.31622 | 14638.041 |
| DQTPDENDQVIVK | 13 | Q9NZI8 | Insulin-like growth factor 2 mRNA-bindi | IGF2BP1 | 63.481 | 2 | 52.71008 | yes | 2.038E-07 | 750.86249 | 8725.9844 |
| IAPPETPDSK | 10 | Q9NZI8 | Insulin-like growth factor 2 mRNA-bindi | IGF2BP1 | 63.481 | 2 | 52.71008 | yes | 6.996E-07 | 527.77441 | 7761.8857 |
| IIGHFYASQMAQR | 13 | Q9NZI8 | Insulin-like growth factor 2 mRNA-bindi | IGF2BP1 | 63.481 | 2 | 52.71008 | yes | 9.938E-08 | 761.38519 | 6764.5083 |
| LLVPTQYVGAIIGK | 14 | Q9NZI8 | Insulin-like growth factor 2 mRNA-bindi | IGF2BP1 | 63.481 | 2 | 52.71008 | yes | 1.561E-07 | 736.44775 | 5833.104 |
| RLEIEHSVPK | 10 | Q9NZI8 | Insulin-like growth factor 2 mRNA-bindi | IGF2BP1 | 63.481 | 2 | 52.71008 | yes | 1.011E-07 | 604.34332 | 5720.8506 |
| TADEVPLK | 8 | Q9NZI8 | Insulin-like growth factor 2 mRNA-bindi | IGF2BP1 | 63.481 | 2 | 52.71008 | yes | 1.409E-07 | 436.73984 | 2530.5413 |
| AEQEIMK | 7 | Q9NZI8 | Insulin-like growth factor 2 mRNA-bindi | IGF2BP1 | 63.481 | 2 | 52.71008 | yes | 7.16E-07 | 424.71277 | 1250.0774 |
| AISVHSTPEGCSSA | 16 | Q9NZI8 | Insulin-like growth factor 2 mRNA-bindi | IGF2BP1 | 63.481 | 2 | 52.71008 | yes | 9.722E-08 | 845.87982 | 1226.7677 |
| NIPPQLR | 7 | Q9NZI8 | Insulin-like growth factor 2 mRNA-bindi | IGF2BP1 | 63.481 | 2 | 52.71008 | yes | 2.924E-06 | 419.2507 | 969.26141 |
| GAIENCCR | 8 | Q9NZI8 | Insulin-like growth factor 2 mRNA-bindi | IGF2BP1 | 63.481 | 2 | 52.71008 | yes | 9.732E-08 | 490.20786 | 383.52298 |
| LNGHQLENHALK | 12 | Q9NZI8 | Insulin-like growth factor 2 mRNA-bindi | IGF2BP1 | 63.481 | 2 | 52.71008 | yes | 1.018E-07 | 687.36786 | 336.49762 |
| EENFFGPK | 8 | Q9NZI8 | Insulin-like growth factor 2 mRNA-bindi | IGF2BP1 | 63.481 | 2 | 52.71008 | yes | 4.802E-07 | 484.22961 | 289.9769 |
| IRDILAQVK | 9 | Q9NZI8 | Insulin-like growth factor 2 mRNA-bindi | IGF2BP1 | 63.481 | 2 | 52.71008 | yes | 4.995E-07 | 528.33221 | 286.77075 |
| QGSPVAAGAPAK | 12 | Q9NZI8 | Insulin-like growth factor 2 mRNA-bindi | IGF2BP1 | 63.481 | 2 | 52.71008 | yes | 9.968E-08 | 527.28802 | 252.15248 |
| LEIEHSVPK | 9 | Q9NZI8 | Insulin-like growth factor 2 mRNA-bindi | IGF2BP1 | 63.481 | 2 | 52.71008 | yes | 1.138E-07 | 526.29279 | 124.53001 |
| VSYIPDEQIAQGPE | 17 | Q9NZI8 | Insulin-like growth factor 2 mRNA-bindi | IGF2BP1 | 63.481 | 3 | 52.71008 | yes | 9.722E-08 | 624.97437 | 15.868596 |
| MLFNSVLR | 8 | Q9NZJ0 | Denticleless protein homolog OS=Homo | DTL | 79.468 | 2 | 51.813374 | yes | 1.26E-07 | 490.27332 | 179.33412 |
| LVTAAGDQTAK | 11 | Q9NZJ0 | Denticleless protein homolog OS=Homo | DTL | 79.468 | 2 | 51.813374 | yes | 4.299E-07 | 537.79309 | 175.31987 |
| LYNTESQSFR | 10 | Q9NZJ0 | Denticleless protein homolog OS=Homo | DTL | 79.468 | 2 | 51.813374 | yes | 1.79E-07 | 622.79895 | 89.336678 |
| QPQLGVLR | 8 | Q9NZJ0 | Denticleless protein homolog OS=Homo | DTL | 79.468 | 2 | 51.813374 | yes | 2.447E-06 | 455.77707 | 67.039024 |
| GSVSSVSPKPPSSF | 15 | Q9NZJ0 | Denticleless protein homolog OS=Homo | DTL | 79.468 | 2 | 51.813374 | yes | 9.722E-08 | 745.89612 | 57.979347 |
| TPSSSPPITPPASET | 16 | Q9NZJ0 | Denticleless protein homolog OS=Homo | DTL | 79.468 | 2 | 51.813374 | yes | 2.5E-07 | 798.90942 | 46.012062 |
| PGLVTVTSSQSTPA | 15 | Q9NZJ0 | Denticleless protein homolog OS=Homo | DTL | 79.468 | 2 | 51.813374 | yes | 2.795E-07 | 736.90137 | 22.943119 |
| YLSVQGQLFR | 10 | Q9NZJ7 | Mitochondrial carrier homolog 1 OS=Ho | MTCH1 | 41.545 | 2 | 51.743656 | yes | 1.231E-07 | 605.83258 | 462.05655 |

| | | | | | | | | | | | |
|---|---|---|---|---|---|---|---|---|---|---|---|
| LMSNALSTVTR | 11 | Q9NZJ7 | Mitochondrial carrier homolog 1 OS=Hor | MTCH1 | 41.545 | 2 | 51.743656 | yes | 1.231E-07 | 596.82135 | 220.58455 |
| MLAHPLHVISMR | 12 | Q9NZJ7 | Mitochondrial carrier homolog 1 OS=Hor | MTCH1 | 41.545 | 2 | 51.743656 | yes | 1.262E-07 | 702.88379 | 89.417458 |
| MLAHPLHVISMR | 12 | Q9NZJ7 | Mitochondrial carrier homolog 1 OS=Hor | MTCH1 | 41.545 | 2 | 51.743656 | yes | 1.159E-07 | 468.92493 | 89.417458 |
| MLAHPLHVISMR | 12 | Q9NZJ7 | Mitochondrial carrier homolog 1 OS=Hor | MTCH1 | 41.545 | 3 | 51.743656 | yes | 1.262E-07 | 702.88379 | 89.417458 |
| MLAHPLHVISMR | 12 | Q9NZJ7 | Mitochondrial carrier homolog 1 OS=Hor | MTCH1 | 41.545 | 3 | 51.743656 | yes | 1.159E-07 | 468.92493 | 89.417458 |
| ETSYEMMMQCVS | 13 | Q9NZJ7 | Mitochondrial carrier homolog 1 OS=Hor | MTCH1 | 41.545 | 2 | 51.743656 | yes | 9.722E-08 | 826.33276 | 146.39557 |
| VFPPDEIEQVSNK | 13 | Q9NZJ7 | Mitochondrial carrier homolog 1 OS=Hor | MTCH1 | 41.545 | 2 | 51.743656 | yes | 1.63E-06 | 751.38031 | 109.19541 |
| SWIHCWK | 7 | Q9NZJ7 | Mitochondrial carrier homolog 1 OS=Hor | MTCH1 | 41.545 | 2 | 51.743656 | yes | 1.165E-07 | 508.74219 | 85.928459 |
| YSGVLSSIGK | 10 | Q9NZJ7 | Mitochondrial carrier homolog 1 OS=Hor | MTCH1 | 41.545 | 2 | 51.743656 | yes | 1.665E-07 | 505.77948 | 28.483351 |
| YLEAGAAGLR | 10 | Q9NZL4 | Hsp70-binding protein 1 OS=Homo sapie | HSPBP1 | 39.303 | 2 | 50.198242 | yes | 1.789E-06 | 510.77728 | 173.98868 |
| LDGFSVLMR | 9 | Q9NZL4 | Hsp70-binding protein 1 OS=Homo sapie | HSPBP1 | 39.303 | 2 | 50.198242 | yes | 1.216E-07 | 519.27606 | 61.515831 |
| EQEAGLLQFLR | 11 | Q9NZL4 | Hsp70-binding protein 1 OS=Homo sapie | HSPBP1 | 39.303 | 2 | 50.198242 | yes | 9.85E-08 | 652.35388 | 54.102856 |
| EPELGLEELLR | 11 | Q9NZL4 | Hsp70-binding protein 1 OS=Homo sapie | HSPBP1 | 39.303 | 2 | 50.198242 | yes | 1.857E-07 | 649.35352 | 46.460835 |
| YTLPLTEGK | 9 | Q9NZM1 | Myoferlin OS=Homo sapiens OX=9606 ( | MYOF | 234.709 | 2 | 52.64804 | yes | 2.092E-07 | 511.28186 | 59.580688 |
| GPVGTVSEAQLAR | 13 | Q9NZM1 | Myoferlin OS=Homo sapiens OX=9606 ( | MYOF | 234.709 | 2 | 52.64804 | yes | 9.722E-08 | 642.84894 | 50.404068 |
| MYELSCYLPQEK | 12 | Q9NZM1 | Myoferlin OS=Homo sapiens OX=9606 ( | MYOF | 234.709 | 2 | 52.64804 | yes | 3.699E-06 | 780.8573 | 36.1553 |
| ELPDSVPQECTVR | 13 | Q9NZM1 | Myoferlin OS=Homo sapiens OX=9606 ( | MYOF | 234.709 | 2 | 52.64804 | yes | 2.42E-05 | 765.36688 | 6.5313702 |
| VDLILENTSK | 10 | Q9NZM5 | Ribosome biogenesis protein NOP53 OS= | NOP53 | 54.39 | 2 | 51.75115 | yes | 9.968E-08 | 566.31641 | 625.15283 |
| LFFVDTGSK | 9 | Q9NZM5 | Ribosome biogenesis protein NOP53 OS= | NOP53 | 54.39 | 2 | 51.75115 | yes | 9.722E-08 | 507.26874 | 342.9881 |
| LLNPSATR | 8 | Q9NZM5 | Ribosome biogenesis protein NOP53 OS= | NOP53 | 54.39 | 2 | 51.75115 | yes | 1.464E-07 | 436.25345 | 154.55685 |
| TSGGLLSEAPNEK | 13 | Q9NZM5 | Ribosome biogenesis protein NOP53 OS= | NOP53 | 54.39 | 2 | 51.75115 | yes | 1.317E-07 | 651.83044 | 124.74689 |
| DVLAHQVPNAK | 11 | Q9NZM5 | Ribosome biogenesis protein NOP53 OS= | NOP53 | 54.39 | 2 | 51.75115 | yes | 1.113E-07 | 596.32764 | 109.49763 |
| AKPGPQDTVER | 11 | Q9NZM5 | Ribosome biogenesis protein NOP53 OS= | NOP53 | 54.39 | 2 | 51.75115 | yes | 3.637E-07 | 599.31476 | 80.636848 |
| VVGTPEVLR | 9 | Q9NZN4 | EH domain-containing protein 2 OS=Hor | EHD2 | 61.162 | 2 | 52.587868 | yes | 2.236E-06 | 485.29001 | 89.489204 |
| LFELEEQDLFR | 11 | Q9NZN4 | EH domain-containing protein 2 OS=Hor | EHD2 | 61.162 | 2 | 52.587868 | yes | 9.722E-08 | 719.86426 | 62.368362 |
| GYDFPAVLR | 9 | Q9NZN4 | EH domain-containing protein 2 OS=Hor | EHD2 | 61.162 | 2 | 52.587868 | yes | 9.169E-07 | 519.27435 | 45.84594 |
| FGAFHSPALEDAD | 30 | Q9NZN4 | EH domain-containing protein 2 OS=Hor | EHD2 | 61.162 | 4 | 52.587868 | yes | 2.487E-07 | 789.63403 | 41.744572 |
| TVWQDLICR | 9 | Q9NZN5 | Rho guanine nucleotide exchange factor 1 | ARHGEF1: | 173.233 | 2 | 50.31292 | yes | 9.819E-08 | 595.80298 | 51.211185 |
| EGSGNPTPLINPLA | 16 | Q9NZN8 | CCR4-NOT transcription complex subuni | CNOT2 | 59.738 | 2 | 54.135956 | yes | 9.81E-08 | 796.92316 | 404.82852 |
| APYVGMVTK | 9 | Q9NZN8 | CCR4-NOT transcription complex subuni | CNOT2 | 59.738 | 2 | 54.135956 | yes | 1.68E-07 | 483.25986 | 343.84848 |
| SSPSIICMPK | 10 | Q9NZN8 | CCR4-NOT transcription complex subuni | CNOT2 | 59.738 | 2 | 54.135956 | yes | 9.722E-08 | 560.28046 | 215.93712 |
| GMSNNTPQLNR | 11 | Q9NZN8 | CCR4-NOT transcription complex subuni | CNOT2 | 59.738 | 2 | 54.135956 | yes | 1.314E-07 | 616.29602 | 195.06064 |
| APGMEPTMK | 9 | Q9NZN8 | CCR4-NOT transcription complex subuni | CNOT2 | 59.738 | 2 | 54.135956 | yes | 2.543E-07 | 481.22772 | 171.72282 |
| TNSMSSSGLGSPN | 14 | Q9NZN8 | CCR4-NOT transcription complex subuni | CNOT2 | 59.738 | 2 | 54.135956 | yes | 9.722E-08 | 697.82007 | 147.53906 |
| GILPMNPR | 8 | Q9NZN8 | CCR4-NOT transcription complex subuni | CNOT2 | 59.738 | 2 | 54.135956 | yes | 2.617E-07 | 449.25238 | 108.93159 |
| GIQVLPDGR | 9 | Q9NZN8 | CCR4-NOT transcription complex subuni | CNOT2 | 59.738 | 2 | 54.135956 | yes | 3.271E-06 | 477.772 | 85.726669 |
| KGIQVLPDGR | 10 | Q9NZN8 | CCR4-NOT transcription complex subuni | CNOT2 | 59.738 | 2 | 54.135956 | yes | 9.722E-08 | 541.81946 | 48.238899 |
| GTYYFFDCLNWR | 12 | Q9NZN8 | CCR4-NOT transcription complex subuni | CNOT2 | 59.738 | 2 | 54.135956 | yes | 9.722E-08 | 821.36139 | 30.315689 |
| FVEGVDSDYHDEN | 25 | Q9NZN8 | CCR4-NOT transcription complex subuni | CNOT2 | 59.738 | 4 | 54.135956 | yes | 0.0001305 | 764.81702 | 13.042014 |

| | | | | | | | | | | | |
|---|---|---|---|---|---|---|---|---|---|---|---|
| LGTVITPDTWK | 11 | Q9P021 | Cysteine-rich PDZ-binding protein OS=H | CRIPT | 11.216 | 2 | 45.96857 | yes | 1.69E-07 | 615.84009 | 762.56915 |
| KLGTVITPDTWK | 12 | Q9P021 | Cysteine-rich PDZ-binding protein OS=H | CRIPT | 11.216 | 2 | 45.96857 | yes | 3.57E-07 | 679.88757 | 90.211868 |
| KGICAMCGK | 9 | Q9P021 | Cysteine-rich PDZ-binding protein OS=H | CRIPT | 11.216 | 2 | 45.96857 | yes | 4.283E-06 | 512.74048 | 23.142422 |
| LESEGSPETLTNLR | 14 | Q9P035 | Very-long-chain (3R)-3-hydroxyacyl-CoA | HACD3 | 43.16 | 2 | 49.570328 | yes | 1.021E-07 | 773.39136 | 1775.6957 |
| FSFTLPYPVK | 10 | Q9P035 | Very-long-chain (3R)-3-hydroxyacyl-CoA | HACD3 | 43.16 | 2 | 49.570328 | yes | 1.136E-07 | 599.82898 | 643.47003 |
| KVSQWWER | 8 | Q9P035 | Very-long-chain (3R)-3-hydroxyacyl-CoA | HACD3 | 43.16 | 2 | 49.570328 | yes | 1.518E-07 | 559.79071 | 368.32108 |
| QVNITVQK | 8 | Q9P035 | Very-long-chain (3R)-3-hydroxyacyl-CoA | HACD3 | 43.16 | 2 | 49.570328 | yes | 1.778E-05 | 465.27435 | 150.05554 |
| RPLFLAPDFDR | 11 | Q9P035 | Very-long-chain (3R)-3-hydroxyacyl-CoA | HACD3 | 43.16 | 2 | 49.570328 | yes | 1.432E-07 | 673.86444 | 108.74897 |
| VELSDVQNPAISIT | 21 | Q9P035 | Very-long-chain (3R)-3-hydroxyacyl-CoA | HACD3 | 43.16 | 3 | 49.570328 | yes | 6.244E-07 | 785.08478 | 93.853287 |
| WLDESDAEMELR | 12 | Q9P035 | Very-long-chain (3R)-3-hydroxyacyl-CoA | HACD3 | 43.16 | 2 | 49.570328 | yes | 2.839E-07 | 747.33246 | 84.087357 |
| VSQWWER | 7 | Q9P035 | Very-long-chain (3R)-3-hydroxyacyl-CoA | HACD3 | 43.16 | 2 | 49.570328 | yes | 1.127E-07 | 495.74323 | 29.825569 |
| PLFLAPDFDR | 10 | Q9P035 | Very-long-chain (3R)-3-hydroxyacyl-CoA | HACD3 | 43.16 | 2 | 49.570328 | yes | 2.933E-07 | 595.81384 | 26.688372 |
| ENLGVALEIDGLEI | 15 | Q9P0B6 | Coiled-coil domain-containing protein 16 | CCDC167 | 11.46 | 2 | 39.435535 | yes | 4.587E-07 | 814.92255 | 33.324852 |
| LQIEDFEAR | 9 | Q9P0J0 | NADH dehydrogenase [ubiquinone] 1 alp | NDUFA13 | 16.699 | 2 | 50.93962 | yes | 1.076E-07 | 560.78528 | 248.21822 |
| VGESVFHTTR | 10 | Q9P0J0 | NADH dehydrogenase [ubiquinone] 1 alp | NDUFA13 | 16.699 | 2 | 50.93962 | yes | 9.722E-08 | 566.79089 | 91.839661 |
| ENLEEEAIIMK | 11 | Q9P0J0 | NADH dehydrogenase [ubiquinone] 1 alp | NDUFA13 | 16.699 | 2 | 50.93962 | yes | 1.216E-06 | 659.8316 | 35.555073 |
| QDMPPPGGYGPID | 15 | Q9P0J0 | NADH dehydrogenase [ubiquinone] 1 alp | NDUFA13 | 16.699 | 2 | 50.93962 | yes | 1.732E-06 | 817.87958 | 18.484222 |
| HTTDHPMQCILTR | 13 | Q9P0J7 | E3 ubiquitin-protein ligase KCMF1 OS=I | KCMF1 | 41.946 | 3 | 37.148605 | yes | 2.906E-06 | 537.25732 | 37.91011 |
| HEGVSCDACLK | 11 | Q9P0J7 | E3 ubiquitin-protein ligase KCMF1 OS=I | KCMF1 | 41.946 | 2 | 37.148605 | yes | 6.254E-07 | 638.27643 | 8.1481781 |
| HEQILVLDPPTDLI | 14 | Q9P0L0 | Vesicle-associated membrane protein-ass | VAPA | 27.894 | 2 | 55.75121 | yes | 1.082E-07 | 809.44598 | 662.85431 |
| HEQILVLDPPTDLI | 14 | Q9P0L0 | Vesicle-associated membrane protein-ass | VAPA | 27.894 | 2 | 55.75121 | yes | 1.138E-07 | 539.96643 | 662.85431 |
| HEQILVLDPPTDLI | 14 | Q9P0L0 | Vesicle-associated membrane protein-ass | VAPA | 27.894 | 3 | 55.75121 | yes | 1.082E-07 | 809.44598 | 662.85431 |
| HEQILVLDPPTDLI | 14 | Q9P0L0 | Vesicle-associated membrane protein-ass | VAPA | 27.894 | 3 | 55.75121 | yes | 1.138E-07 | 539.96643 | 662.85431 |
| CVFEMPNENDK | 11 | Q9P0L0 | Vesicle-associated membrane protein-ass | VAPA | 27.894 | 2 | 55.75121 | yes | 9.722E-08 | 691.78937 | 599.15594 |
| EAKPDELMDSK | 11 | Q9P0L0 | Vesicle-associated membrane protein-ass | VAPA | 27.894 | 2 | 55.75121 | yes | 9.858E-08 | 631.80029 | 394.23795 |
| GPFTDVVTTNLK | 12 | Q9P0L0 | Vesicle-associated membrane protein-ass | VAPA | 27.894 | 2 | 55.75121 | no | 9.722E-08 | 646.34827 | 330.03677 |
| VAHSDKPGSTSTA | 16 | Q9P0L0 | Vesicle-associated membrane protein-ass | VAPA | 27.894 | 2 | 55.75121 | yes | 9.722E-08 | 824.4079 | 111.04733 |
| VAHSDKPGSTSTA | 16 | Q9P0L0 | Vesicle-associated membrane protein-ass | VAPA | 27.894 | 2 | 55.75121 | yes | 1.775E-07 | 549.94104 | 111.04733 |
| VAHSDKPGSTSTA | 16 | Q9P0L0 | Vesicle-associated membrane protein-ass | VAPA | 27.894 | 3 | 55.75121 | yes | 9.722E-08 | 824.4079 | 111.04733 |
| VAHSDKPGSTSTA | 16 | Q9P0L0 | Vesicle-associated membrane protein-ass | VAPA | 27.894 | 3 | 55.75121 | yes | 1.775E-07 | 549.94104 | 111.04733 |
| KLMEECK | 7 | Q9P0L0 | Vesicle-associated membrane protein-ass | VAPA | 27.894 | 2 | 55.75121 | yes | 1.061E-07 | 469.22772 | 168.78845 |
| LMEECKR | 7 | Q9P0L0 | Vesicle-associated membrane protein-ass | VAPA | 27.894 | 2 | 55.75121 | yes | 5.381E-07 | 483.23077 | 120.77372 |
| FKGPFTDVVTTNL | 14 | Q9P0L0 | Vesicle-associated membrane protein-ass | VAPA | 27.894 | 2 | 55.75121 | yes | 1.622E-07 | 783.92993 | 61.205654 |
| LNDMEPSK | 8 | Q9P0L0 | Vesicle-associated membrane protein-ass | VAPA | 27.894 | 2 | 55.75121 | yes | 3.759E-07 | 467.22095 | 52.570221 |
| PHSVSLNDTETR | 12 | Q9P0L0 | Vesicle-associated membrane protein-ass | VAPA | 27.894 | 2 | 55.75121 | yes | 4.568E-07 | 678.33112 | 10.540121 |
| SMSTSGHPIK | 10 | Q9P0L2 | Serine/threonine-protein kinase MARK1 ( | MARK1 | 89.003 | 2 | 52.7724 | yes | 1.979E-07 | 522.76074 | 603.95813 |
| DEINDALINQK | 11 | Q9P0L2 | Serine/threonine-protein kinase MARK1 ( | MARK1 | 89.003 | 2 | 52.7724 | yes | 1.045E-07 | 636.82513 | 568.72888 |
| SEMTASPLVGPER | 13 | Q9P0L2 | Serine/threonine-protein kinase MARK1 ( | MARK1 | 89.003 | 2 | 52.7724 | yes | 9.722E-08 | 687.34009 | 489.82446 |
| IADFGFSNEFTVGN | 15 | Q9P0L2 | Serine/threonine-protein kinase MARK1 ( | MARK1 | 89.003 | 2 | 52.7724 | yes | 9.729E-08 | 823.39648 | 416.34244 |

| | | | | | | | | | | | |
|---|---|---|---|---|---|---|---|---|---|---|---|
| LLVLNPIK | 8 | Q9P0L2 | Serine/threonine-protein kinase MARK1 ( | MARK1 | 89.003 | 2 | 52.7724 | yes | 1.22E-07 | 455.31021 | 414.51572 |
| STFHGEQLR | 9 | Q9P0L2 | Serine/threonine-protein kinase MARK1 ( | MARK1 | 89.003 | 2 | 52.7724 | yes | 2.089E-07 | 537.76996 | 363.11996 |
| QDSLVQWEMEVC | 13 | Q9P0L2 | Serine/threonine-protein kinase MARK1 ( | MARK1 | 89.003 | 2 | 52.7724 | yes | 1.186E-07 | 826.37634 | 295.62775 |
| TTSSMDPNDMMR | 12 | Q9P0L2 | Serine/threonine-protein kinase MARK1 ( | MARK1 | 89.003 | 2 | 52.7724 | yes | 2.385E-07 | 693.27832 | 266.18948 |
| AENLLLDGDMNIK | 13 | Q9P0L2 | Serine/threonine-protein kinase MARK1 ( | MARK1 | 89.003 | 2 | 52.7724 | yes | 9.722E-08 | 723.3689 | 266.00873 |
| RGTSTGIISK | 10 | Q9P0L2 | Serine/threonine-protein kinase MARK1 ( | MARK1 | 89.003 | 2 | 52.7724 | yes | 3.166E-07 | 510.29584 | 199.1147 |
| VLDANNCDYEQK | 12 | Q9P0L2 | Serine/threonine-protein kinase MARK1 ( | MARK1 | 89.003 | 2 | 52.7724 | yes | 1.876E-07 | 734.82227 | 162.31799 |
| FLLFCVHGDAR | 11 | Q9P0L2 | Serine/threonine-protein kinase MARK1 ( | MARK1 | 89.003 | 2 | 52.7724 | yes | 1.762E-07 | 667.83734 | 72.748611 |
| FLLFCVHGDAR | 11 | Q9P0L2 | Serine/threonine-protein kinase MARK1 ( | MARK1 | 89.003 | 2 | 52.7724 | yes | 1.926E-07 | 445.56064 | 72.748611 |
| FLLFCVHGDAR | 11 | Q9P0L2 | Serine/threonine-protein kinase MARK1 ( | MARK1 | 89.003 | 3 | 52.7724 | yes | 1.762E-07 | 667.83734 | 72.748611 |
| FLLFCVHGDAR | 11 | Q9P0L2 | Serine/threonine-protein kinase MARK1 ( | MARK1 | 89.003 | 3 | 52.7724 | yes | 1.926E-07 | 445.56064 | 72.748611 |
| YDEVMATYILLGR | 13 | Q9P0L2 | Serine/threonine-protein kinase MARK1 ( | MARK1 | 89.003 | 2 | 52.7724 | yes | 9.722E-08 | 772.3949 | 100.82195 |
| IDIMVTMGFAR | 11 | Q9P0L2 | Serine/threonine-protein kinase MARK1 ( | MARK1 | 89.003 | 2 | 52.7724 | yes | 1.415E-07 | 627.32269 | 49.157352 |
| DTENHTSVDGYTE | 20 | Q9P0L2 | Serine/threonine-protein kinase MARK1 ( | MARK1 | 89.003 | 4 | 52.7724 | yes | 1.007E-07 | 568.01465 | 48.083149 |
| SVAYNGPPASPSH | 22 | Q9P0L2 | Serine/threonine-protein kinase MARK1 ( | MARK1 | 89.003 | 3 | 52.7724 | yes | 7.68E-07 | 742.02252 | 35.160263 |
| VGVSMFNRPH | 10 | Q9P0S9 | Transmembrane protein 14C OS=Homo s | TMEM14C | 11.565 | 2 | 38.41255 | yes | 5.239E-07 | 572.29004 | 46.069748 |
| DAGELAQHSR | 10 | Q9P0T4 | Zinc finger protein 581 OS=Homo sapien | ZNF581 | 22.011 | 2 | 52.257145 | yes | 9.722E-08 | 542.26251 | 209.38206 |
| VFEYMSYLQR | 10 | Q9P0T4 | Zinc finger protein 581 OS=Homo sapien | ZNF581 | 22.011 | 2 | 52.257145 | yes | 2.832E-06 | 668.32373 | 105.06083 |
| FMEQNTLQK | 9 | Q9P0T4 | Zinc finger protein 581 OS=Homo sapien | ZNF581 | 22.011 | 2 | 52.257145 | yes | 5.972E-07 | 569.78168 | 78.20298 |
| CYSCPVCSR | 9 | Q9P0T4 | Zinc finger protein 581 OS=Homo sapien | ZNF581 | 22.011 | 2 | 52.257145 | yes | 3.015E-06 | 594.7334 | 24.504095 |
| ELTSVYDSR | 9 | Q9P0U3 | Sentrin-specific protease 1 OS=Homo sap | SENP1 | 73.481 | 2 | 39.96184 | yes | 4.285E-07 | 535.26166 | 92.426819 |
| MVWEILHR | 8 | Q9P0U3 | Sentrin-specific protease 1 OS=Homo sap | SENP1 | 73.481 | 2 | 39.96184 | yes | 8.164E-07 | 542.29205 | 31.102179 |
| DVIQELHLGLDK | 12 | Q9P0V3 | SH3 domain-binding protein 4 OS=Homo | SH3BP4 | 107.496 | 2 | 53.796944 | yes | 9.823E-08 | 690.38007 | 194.52234 |
| TLLVSEVTR | 9 | Q9P0V3 | SH3 domain-binding protein 4 OS=Homo | SH3BP4 | 107.496 | 2 | 53.796944 | yes | 3.071E-06 | 509.30057 | 167.71042 |
| MDCQGLVVR | 9 | Q9P0V3 | SH3 domain-binding protein 4 OS=Homo | SH3BP4 | 107.496 | 2 | 53.796944 | yes | 2.167E-07 | 539.26263 | 139.59297 |
| ELELLGGWTDDK | 12 | Q9P0V3 | SH3 domain-binding protein 4 OS=Homo | SH3BP4 | 107.496 | 2 | 53.796944 | yes | 1.699E-07 | 688.34064 | 115.17849 |
| TLLMENISSWR | 11 | Q9P0V3 | SH3 domain-binding protein 4 OS=Homo | SH3BP4 | 107.496 | 2 | 53.796944 | yes | 6.628E-07 | 675.34772 | 112.4039 |
| SCSISPVLEVK | 11 | Q9P0V3 | SH3 domain-binding protein 4 OS=Homo | SH3BP4 | 107.496 | 2 | 53.796944 | yes | 1.36E-07 | 609.82355 | 112.11622 |
| STVGLQCLR | 9 | Q9P0V3 | SH3 domain-binding protein 4 OS=Homo | SH3BP4 | 107.496 | 2 | 53.796944 | yes | 1.268E-07 | 517.27661 | 108.17384 |
| ALLDPPLELNSDR | 13 | Q9P0V3 | SH3 domain-binding protein 4 OS=Homo | SH3BP4 | 107.496 | 2 | 53.796944 | yes | 9.722E-08 | 726.88831 | 103.21015 |
| SDAPTSSSFFTGLK | 14 | Q9P0V3 | SH3 domain-binding protein 4 OS=Homo | SH3BP4 | 107.496 | 2 | 53.796944 | yes | 1.75E-07 | 722.85138 | 76.487312 |
| IILSPFATTTK | 11 | Q9P0V3 | SH3 domain-binding protein 4 OS=Homo | SH3BP4 | 107.496 | 2 | 53.796944 | yes | 5.791E-07 | 596.35284 | 76.452026 |
| SYSLSELSVLQAK | 13 | Q9P0V3 | SH3 domain-binding protein 4 OS=Homo | SH3BP4 | 107.496 | 2 | 53.796944 | yes | 9.722E-08 | 712.88519 | 69.030525 |
| DYCPTNFTTLK | 11 | Q9P0V3 | SH3 domain-binding protein 4 OS=Homo | SH3BP4 | 107.496 | 2 | 53.796944 | yes | 5.133E-07 | 680.3161 | 34.706902 |
| TVVTIFGHDCAPK | 13 | Q9P0V3 | SH3 domain-binding protein 4 OS=Homo | SH3BP4 | 107.496 | 2 | 53.796944 | yes | 9.927E-08 | 722.86627 | 28.740772 |
| QAPNPAPVALQLV | 15 | Q9P0V3 | SH3 domain-binding protein 4 OS=Homo | SH3BP4 | 107.496 | 2 | 53.796944 | yes | 3.46E-06 | 795.44354 | 23.846094 |
| VCMFSNMTNYEV | 13 | Q9P0V3 | SH3 domain-binding protein 4 OS=Homo | SH3BP4 | 107.496 | 2 | 53.796944 | yes | 9.722E-08 | 811.85419 | 15.757796 |
| EMLICTNMEDLR | 12 | Q9P0V9 | Septin-10 OS=Homo sapiens OX=9606 G | SEPTIN10 | 52.593 | 2 | 46.608894 | yes | 1.515E-07 | 762.84637 | 50.804134 |
| AQTYELQESNVQL | 14 | Q9P0V9 | Septin-10 OS=Homo sapiens OX=9606 G | SEPTIN10 | 52.593 | 2 | 46.608894 | yes | 4.357E-07 | 825.92029 | 24.667948 |

| | | | | | | | | | | | |
|---|---|---|---|---|---|---|---|---|---|---|---|
| LLLSAELYCR | 10 | Q9P1Y5 | Calmodulin-regulated spectrin-associated | CAMSAP3 | 134.75 | 2 | 51.055573 | yes | 9.722E-08 | 619.33411 | 315.47916 |
| LSAALSSLQR | 10 | Q9P1Y5 | Calmodulin-regulated spectrin-associated | CAMSAP3 | 134.75 | 2 | 51.055573 | yes | 9.722E-08 | 523.30365 | 215.4127 |
| APSPSGLMSPSR | 12 | Q9P1Y5 | Calmodulin-regulated spectrin-associated | CAMSAP3 | 134.75 | 2 | 51.055573 | yes | 4.092E-07 | 593.79785 | 115.82207 |
| RIEAIFAK | 8 | Q9P1Y5 | Calmodulin-regulated spectrin-associated | CAMSAP3 | 134.75 | 2 | 51.055573 | yes | 1.426E-06 | 474.28726 | 97.448303 |
| ASPAAPADGAAPA | 18 | Q9P1Y5 | Calmodulin-regulated spectrin-associated | CAMSAP3 | 134.75 | 2 | 51.055573 | yes | 1.9E-07 | 824.42609 | 91.762276 |
| RGTVPALPERPVR | 13 | Q9P1Y5 | Calmodulin-regulated spectrin-associated | CAMSAP3 | 134.75 | 3 | 51.055573 | yes | 2.2E-06 | 483.28793 | 89.558914 |
| LLFWVDTTVR | 10 | Q9P1Y5 | Calmodulin-regulated spectrin-associated | CAMSAP3 | 134.75 | 2 | 51.055573 | yes | 1.555E-07 | 625.35059 | 55.860703 |
| TVTPAMVEGIYK | 12 | Q9P1Y5 | Calmodulin-regulated spectrin-associated | CAMSAP3 | 134.75 | 2 | 51.055573 | yes | 3.137E-06 | 654.84705 | 52.722046 |
| AVASSPAATNSEV | 14 | Q9P1Y5 | Calmodulin-regulated spectrin-associated | CAMSAP3 | 134.75 | 2 | 51.055573 | yes | 1.009E-07 | 666.34369 | 43.952362 |
| AAASLAWVLR | 10 | Q9P1Y5 | Calmodulin-regulated spectrin-associated | CAMSAP3 | 134.75 | 2 | 51.055573 | yes | 1.077E-06 | 529.31128 | 37.069633 |
| IYSQSTLSLSTVAN | 23 | Q9P1Y5 | Calmodulin-regulated spectrin-associated | CAMSAP3 | 134.75 | 3 | 51.055573 | yes | 6.439E-06 | 816.09058 | 36.898262 |
| SLDQYDFSR | 9 | Q9P1Y5 | Calmodulin-regulated spectrin-associated | CAMSAP3 | 134.75 | 2 | 51.055573 | yes | 3.514E-06 | 565.75928 | 30.963591 |
| SGCCDDSALAR | 11 | Q9P1Y5 | Calmodulin-regulated spectrin-associated | CAMSAP3 | 134.75 | 2 | 51.055573 | yes | 1.172E-07 | 606.24261 | 28.957348 |
| HPLLSSGGPQSPLF | 14 | Q9P1Y5 | Calmodulin-regulated spectrin-associated | CAMSAP3 | 134.75 | 2 | 51.055573 | yes | 7.652E-07 | 723.39661 | 25.833187 |
| ILELEDDIQTISEK | 14 | Q9P1Z2 | Calcium-binding and coiled-coil domain-c | CALCOCO | 77.337 | 2 | 47.437294 | yes | 3.742E-07 | 823.43018 | 48.348587 |
| LAELGLHLK | 9 | Q9P1Z2 | Calcium-binding and coiled-coil domain-c | CALCOCO | 77.337 | 2 | 47.437294 | yes | 1.33E-07 | 497.3082 | 38.970745 |
| LVLSEEAVR | 9 | Q9P209 | Centrosomal protein of 72 kDa OS=Homo | CEP72 | 71.718 | 2 | 51.95488 | yes | 9.722E-08 | 508.29276 | 182.99486 |
| LDLSGEMVPGPLP | 17 | Q9P209 | Centrosomal protein of 72 kDa OS=Homo | CEP72 | 71.718 | 2 | 51.95488 | yes | 9.722E-08 | 839.44763 | 174.06572 |
| LHFASEDSLDSK | 12 | Q9P209 | Centrosomal protein of 72 kDa OS=Homo | CEP72 | 71.718 | 2 | 51.95488 | yes | 5.97E-07 | 674.82263 | 71.737976 |
| HLLSPQLVQYQCC | 17 | Q9P209 | Centrosomal protein of 72 kDa OS=Homo | CEP72 | 71.718 | 3 | 51.95488 | yes | 2.851E-07 | 643.98773 | 61.963913 |
| LHALTELVDVDFR | 13 | Q9P209 | Centrosomal protein of 72 kDa OS=Homo | CEP72 | 71.718 | 2 | 51.95488 | yes | 4.12E-07 | 764.41193 | 50.53875 |
| SLHSNEAFLAQAR | 13 | Q9P209 | Centrosomal protein of 72 kDa OS=Homo | CEP72 | 71.718 | 3 | 51.95488 | yes | 3.72E-07 | 481.91614 | 23.127747 |
| TLSQPEASETEEQF | 14 | Q9P209 | Centrosomal protein of 72 kDa OS=Homo | CEP72 | 71.718 | 2 | 51.95488 | yes | 6.097E-06 | 802.87354 | 19.940355 |
| LGALQGDPEALTL | 15 | Q9P253 | Vacuolar protein sorting-associated protei | VPS18 | 110.186 | 2 | 49.139717 | yes | 1.065E-07 | 808.93579 | 68.132004 |
| AGGAAVVITEPEH | 15 | Q9P258 | Protein RCC2 OS=Homo sapiens OX=96 | RCC2 | 56.085 | 2 | 53.82776 | yes | 1.001E-07 | 740.39374 | 3182.6741 |
| AGGAAVVITEPEH | 15 | Q9P258 | Protein RCC2 OS=Homo sapiens OX=96 | RCC2 | 56.085 | 2 | 53.82776 | yes | 1.38E-07 | 493.93158 | 3182.6741 |
| AGGAAVVITEPEH | 15 | Q9P258 | Protein RCC2 OS=Homo sapiens OX=96 | RCC2 | 56.085 | 3 | 53.82776 | yes | 1.001E-07 | 740.39374 | 3182.6741 |
| AGGAAVVITEPEH | 15 | Q9P258 | Protein RCC2 OS=Homo sapiens OX=96 | RCC2 | 56.085 | 3 | 53.82776 | yes | 1.38E-07 | 493.93158 | 3182.6741 |
| IEYDCELVPR | 10 | Q9P258 | Protein RCC2 OS=Homo sapiens OX=96 | RCC2 | 56.085 | 2 | 53.82776 | yes | 4.239E-07 | 647.31079 | 4558.2334 |
| KLPEYNPR | 8 | Q9P258 | Protein RCC2 OS=Homo sapiens OX=96 | RCC2 | 56.085 | 2 | 53.82776 | yes | 1.116E-07 | 508.77982 | 3281.4209 |
| YGCLAGVR | 8 | Q9P258 | Protein RCC2 OS=Homo sapiens OX=96 | RCC2 | 56.085 | 2 | 53.82776 | yes | 1.313E-07 | 448.22635 | 2955.3352 |
| DGQILPVPNVVVR | 13 | Q9P258 | Protein RCC2 OS=Homo sapiens OX=96 | RCC2 | 56.085 | 2 | 53.82776 | yes | 1.289E-07 | 703.41174 | 2853.5093 |
| LFDFPGR | 7 | Q9P258 | Protein RCC2 OS=Homo sapiens OX=96 | RCC2 | 56.085 | 2 | 53.82776 | yes | 2.009E-07 | 426.22415 | 2537.6851 |
| AVQDLCGWR | 9 | Q9P258 | Protein RCC2 OS=Homo sapiens OX=96 | RCC2 | 56.085 | 2 | 53.82776 | yes | 1.197E-07 | 552.76636 | 1567.4497 |
| LIEGLSHEVIVSAA | 17 | Q9P258 | Protein RCC2 OS=Homo sapiens OX=96 | RCC2 | 56.085 | 3 | 53.82776 | yes | 2.034E-07 | 604.32086 | 1482.5903 |
| NHTLALTETGSVF. | 19 | Q9P258 | Protein RCC2 OS=Homo sapiens OX=96 | RCC2 | 56.085 | 3 | 53.82776 | yes | 9.722E-08 | 679.34094 | 1413.1741 |
| MACGAEFSMIMD( | 14 | Q9P258 | Protein RCC2 OS=Homo sapiens OX=96 | RCC2 | 56.085 | 2 | 53.82776 | yes | 9.722E-08 | 825.82629 | 1189.353 |
| TVVSGSCAAHSLL | 19 | Q9P258 | Protein RCC2 OS=Homo sapiens OX=96 | RCC2 | 56.085 | 3 | 53.82776 | yes | 1.767E-07 | 644.33496 | 1187.542 |
| VFSWGFGGYGR | 11 | Q9P258 | Protein RCC2 OS=Homo sapiens OX=96 | RCC2 | 56.085 | 2 | 53.82776 | yes | 2.348E-07 | 616.79602 | 894.40912 |

| | | | | | | | | | | | |
|---|---|---|---|---|---|---|---|---|---|---|---|
| RVAIFIEK | 8 | Q9P258 | Protein RCC2 OS=Homo sapiens OX=96 | RCC2 | 56.085 | 2 | 53.82776 | yes | 3.172E-07 | 488.30292 | 722.12872 |
| TLDGIFSEQVAMG | 22 | Q9P258 | Protein RCC2 OS=Homo sapiens OX=96 | RCC2 | 56.085 | 3 | 53.82776 | yes | 1.147E-07 | 798.74225 | 647.69989 |
| NLGQNLWGPHR | 11 | Q9P258 | Protein RCC2 OS=Homo sapiens OX=96 | RCC2 | 56.085 | 2 | 53.82776 | yes | 1.584E-07 | 646.33636 | 487.41553 |
| GNLYSFGCPEYGQ | 21 | Q9P258 | Protein RCC2 OS=Homo sapiens OX=96 | RCC2 | 56.085 | 3 | 53.82776 | yes | 2.078E-07 | 767.33942 | 375.8537 |
| LPEYNPR | 7 | Q9P258 | Protein RCC2 OS=Homo sapiens OX=96 | RCC2 | 56.085 | 2 | 53.82776 | yes | 1.535E-07 | 444.73233 | 314.46802 |
| TKDGQILPVPNVV | 15 | Q9P258 | Protein RCC2 OS=Homo sapiens OX=96 | RCC2 | 56.085 | 2 | 53.82776 | yes | 9.722E-08 | 817.98303 | 54.610497 |
| SPAAPSPLALR | 11 | Q9P270 | SLAIN motif-containing protein 2 OS=H | SLAIN2 | 62.544 | 2 | 52.062916 | yes | 4.743E-07 | 540.31403 | 873.53442 |
| SVSPLVWCR | 9 | Q9P270 | SLAIN motif-containing protein 2 OS=H | SLAIN2 | 62.544 | 2 | 52.062916 | yes | 2.213E-07 | 552.28693 | 611.86633 |
| SGAVQGAGSLGPC | 17 | Q9P270 | SLAIN motif-containing protein 2 OS=H | SLAIN2 | 62.544 | 2 | 52.062916 | yes | 9.722E-08 | 748.89441 | 602.8584 |
| TSNTQVDSVK | 10 | Q9P270 | SLAIN motif-containing protein 2 OS=H | SLAIN2 | 62.544 | 2 | 52.062916 | yes | 2.41E-07 | 539.7724 | 466.44565 |
| LNDVTDVQILAR | 12 | Q9P270 | SLAIN motif-containing protein 2 OS=H | SLAIN2 | 62.544 | 2 | 52.062916 | yes | 9.808E-08 | 678.87769 | 395.98822 |
| QLILPGNSGNLK | 12 | Q9P270 | SLAIN motif-containing protein 2 OS=H | SLAIN2 | 62.544 | 2 | 52.062916 | yes | 1.016E-07 | 627.36426 | 357.44333 |
| QVLDYPSPDVECA | 14 | Q9P270 | SLAIN motif-containing protein 2 OS=H | SLAIN2 | 62.544 | 2 | 52.062916 | yes | 1.535E-07 | 810.88232 | 336.05884 |
| MQPQASAIPSPGK | 13 | Q9P270 | SLAIN motif-containing protein 2 OS=H | SLAIN2 | 62.544 | 2 | 52.062916 | yes | 6.171E-07 | 656.3399 | 239.44621 |
| PSAPSAGGIPVPR | 13 | Q9P270 | SLAIN motif-containing protein 2 OS=H | SLAIN2 | 62.544 | 2 | 52.062916 | yes | 5.909E-07 | 603.33551 | 100.13455 |
| SDSNFQVPNGGIPI | 14 | Q9P270 | SLAIN motif-containing protein 2 OS=H | SLAIN2 | 62.544 | 2 | 52.062916 | yes | 9.722E-08 | 744.36548 | 53.664139 |
| KLQELVK | 7 | Q9P270 | SLAIN motif-containing protein 2 OS=H | SLAIN2 | 62.544 | 2 | 52.062916 | yes | 1.897E-07 | 429.27637 | 45.265644 |
| VLFPTER | 7 | Q9P275 | Ubiquitin carboxyl-terminal hydrolase 36 | USP36 | 122.908 | 2 | 51.835995 | yes | 6.767E-07 | 431.24509 | 143.10616 |
| IEFEPASK | 8 | Q9P275 | Ubiquitin carboxyl-terminal hydrolase 36 | USP36 | 122.908 | 2 | 51.835995 | yes | 7.735E-06 | 460.73984 | 141.25522 |
| NFWSVTHPAK | 10 | Q9P275 | Ubiquitin carboxyl-terminal hydrolase 36 | USP36 | 122.908 | 2 | 51.835995 | yes | 3.454E-07 | 593.80383 | 105.51954 |
| MSAVSQDAIEDSR | 13 | Q9P275 | Ubiquitin carboxyl-terminal hydrolase 36 | USP36 | 122.908 | 2 | 51.835995 | yes | 3.199E-06 | 704.82227 | 103.55672 |
| TSNVLTLSLK | 10 | Q9P275 | Ubiquitin carboxyl-terminal hydrolase 36 | USP36 | 122.908 | 2 | 51.835995 | yes | 9.722E-08 | 538.32153 | 63.269176 |
| AVSPAPQSSSR | 11 | Q9P275 | Ubiquitin carboxyl-terminal hydrolase 36 | USP36 | 122.908 | 2 | 51.835995 | yes | 2.455E-06 | 543.78052 | 62.771042 |
| KSPEGLISR | 9 | Q9P275 | Ubiquitin carboxyl-terminal hydrolase 36 | USP36 | 122.908 | 2 | 51.835995 | yes | 2.921E-06 | 493.7851 | 61.787853 |
| ADVLSGENAYMC | 14 | Q9P275 | Ubiquitin carboxyl-terminal hydrolase 36 | USP36 | 122.908 | 2 | 51.835995 | yes | 5.599E-07 | 764.84216 | 52.75996 |
| TSHPVVASTWPVH | 14 | Q9P275 | Ubiquitin carboxyl-terminal hydrolase 36 | USP36 | 122.908 | 2 | 51.835995 | yes | 9.873E-08 | 787.41534 | 38.291294 |
| MGLSQAPPVSWN | 15 | Q9P275 | Ubiquitin carboxyl-terminal hydrolase 36 | USP36 | 122.908 | 2 | 51.835995 | yes | 4.633E-05 | 814.89612 | 30.079275 |
| FGNQEDAHEFLR | 12 | Q9P275 | Ubiquitin carboxyl-terminal hydrolase 36 | USP36 | 122.908 | 2 | 51.835995 | yes | 1.905E-07 | 731.83911 | 28.969585 |
| SFSYQLEALK | 10 | Q9P275 | Ubiquitin carboxyl-terminal hydrolase 36 | USP36 | 122.908 | 2 | 51.835995 | yes | 4.264E-06 | 593.31116 | 26.852152 |
| WSFDDVPMTPLR | 12 | Q9P287 | BRCA2 and CDKN1A-interacting protein | BCCIP | 35.98 | 2 | 51.215015 | yes | 1.204E-06 | 732.35303 | 108.77912 |
| GTQCVEQIQELVL | 14 | Q9P287 | BRCA2 and CDKN1A-interacting protein | BCCIP | 35.98 | 2 | 51.215015 | yes | 9.722E-08 | 836.93799 | 53.232544 |
| LLQQLFLK | 8 | Q9P287 | BRCA2 and CDKN1A-interacting protein | BCCIP | 35.98 | 2 | 51.215015 | yes | 4.114E-06 | 501.82095 | 33.884014 |
| AVESGVPQPPDPP | 16 | Q9P287 | BRCA2 and CDKN1A-interacting protein | BCCIP | 35.98 | 2 | 51.215015 | yes | 3.288E-07 | 836.93628 | 32.939034 |
| SIAVAEAACPGITD | 15 | Q9P289 | Serine/threonine-protein kinase 26 OS=H | STK26 | 46.529 | 2 | 43.28358 | yes | 4.377E-07 | 751.87958 | 77.378212 |
| NQAIEELEK | 9 | Q9P289 | Serine/threonine-protein kinase 26 OS=H | STK26 | 46.529 | 2 | 43.28358 | yes | 2.655E-07 | 537.27728 | 66.805794 |
| TIMEIEESR | 9 | Q9P2D0 | Inhibitor of Bruton tyrosine kinase OS=H | IBTK | 150.528 | 2 | 54.17133 | yes | 6.078E-07 | 554.27118 | 344.67719 |
| VLVSGGHMEYK | 11 | Q9P2D0 | Inhibitor of Bruton tyrosine kinase OS=H | IBTK | 150.528 | 2 | 54.17133 | yes | 1.164E-07 | 610.31061 | 295.96582 |
| ISAPQPIPSNR | 11 | Q9P2D0 | Inhibitor of Bruton tyrosine kinase OS=H | IBTK | 150.528 | 2 | 54.17133 | yes | 2.018E-07 | 590.32764 | 250.28705 |
| QVFISDIALNR | 11 | Q9P2D0 | Inhibitor of Bruton tyrosine kinase OS=H | IBTK | 150.528 | 2 | 54.17133 | yes | 9.722E-08 | 638.35645 | 239.81796 |

| | | | | | | | | | | |
|---|---|---|---|---|---|---|---|---|---|---|
| ICILAMDGAGR | 11 | Q9P2D0 | Inhibitor of Bruton tyrosine kinase OS=H | IBTK | 150.528 | 2 | 54.17133 | yes | 3.472E-07 | 588.79681 | 213.99742 |
| EGLSALDLVMK | 11 | Q9P2D0 | Inhibitor of Bruton tyrosine kinase OS=H | IBTK | 150.528 | 2 | 54.17133 | yes | 1.52E-07 | 588.32068 | 194.72618 |
| KFDFSNLSSR | 10 | Q9P2D0 | Inhibitor of Bruton tyrosine kinase OS=H | IBTK | 150.528 | 2 | 54.17133 | yes | 9.926E-08 | 600.80402 | 192.17264 |
| SNQAQTVSER | 10 | Q9P2D0 | Inhibitor of Bruton tyrosine kinase OS=H | IBTK | 150.528 | 2 | 54.17133 | yes | 1.615E-07 | 560.27307 | 189.64708 |
| YILAVHSDFFQK | 12 | Q9P2D0 | Inhibitor of Bruton tyrosine kinase OS=H | IBTK | 150.528 | 2 | 54.17133 | yes | 9.722E-08 | 734.38519 | 187.724 |
| NEILFVTQDGEGFI | 14 | Q9P2D0 | Inhibitor of Bruton tyrosine kinase OS=H | IBTK | 150.528 | 2 | 54.17133 | yes | 9.722E-08 | 812.90192 | 186.49814 |
| SLDVLSDGVLK | 11 | Q9P2D0 | Inhibitor of Bruton tyrosine kinase OS=H | IBTK | 150.528 | 2 | 54.17133 | yes | 9.722E-08 | 573.32428 | 178.06396 |
| HGVSLYIQDK | 10 | Q9P2D0 | Inhibitor of Bruton tyrosine kinase OS=H | IBTK | 150.528 | 2 | 54.17133 | yes | 1.06E-07 | 580.30896 | 167.5871 |
| HALDVLSVVTK | 11 | Q9P2D0 | Inhibitor of Bruton tyrosine kinase OS=H | IBTK | 150.528 | 2 | 54.17133 | yes | 9.722E-08 | 591.34808 | 165.76982 |
| NALHLVSSCGK | 11 | Q9P2D0 | Inhibitor of Bruton tyrosine kinase OS=H | IBTK | 150.528 | 2 | 54.17133 | yes | 1.378E-07 | 593.30585 | 151.96144 |
| TPQGPLAVPMWN | 13 | Q9P2D0 | Inhibitor of Bruton tyrosine kinase OS=H | IBTK | 150.528 | 2 | 54.17133 | yes | 9.722E-08 | 719.87921 | 151.29773 |
| GDIYLLADYQCK | 12 | Q9P2D0 | Inhibitor of Bruton tyrosine kinase OS=H | IBTK | 150.528 | 2 | 54.17133 | yes | 1.795E-07 | 729.85028 | 130.10355 |
| EICEVALTEK | 10 | Q9P2D0 | Inhibitor of Bruton tyrosine kinase OS=H | IBTK | 150.528 | 2 | 54.17133 | yes | 5.312E-07 | 596.29993 | 96.905266 |
| VNFHEDDNQK | 10 | Q9P2D0 | Inhibitor of Bruton tyrosine kinase OS=H | IBTK | 150.528 | 2 | 54.17133 | yes | 1.217E-07 | 623.27838 | 75.888031 |
| DLQSPDFTTGFHSI | 15 | Q9P2D0 | Inhibitor of Bruton tyrosine kinase OS=H | IBTK | 150.528 | 2 | 54.17133 | yes | 1.982E-07 | 847.88647 | 65.316071 |
| LGHGDEQTCLVPR | 13 | Q9P2D0 | Inhibitor of Bruton tyrosine kinase OS=H | IBTK | 150.528 | 2 | 54.17133 | yes | 1.262E-07 | 741.36188 | 51.433014 |
| DEDSAGCHLFVVE | 14 | Q9P2D0 | Inhibitor of Bruton tyrosine kinase OS=H | IBTK | 150.528 | 2 | 54.17133 | yes | 1.876E-07 | 803.36432 | 40.20631 |
| VDPEHLK | 7 | Q9P2D0 | Inhibitor of Bruton tyrosine kinase OS=H | IBTK | 150.528 | 2 | 54.17133 | yes | 7.992E-07 | 419.2269 | 40.132179 |
| DFLLEEK | 7 | Q9P2D0 | Inhibitor of Bruton tyrosine kinase OS=H | IBTK | 150.528 | 2 | 54.17133 | yes | 6.314E-06 | 447.23438 | 28.555098 |
| KGVLDWLIQK | 10 | Q9P2D0 | Inhibitor of Bruton tyrosine kinase OS=H | IBTK | 150.528 | 2 | 54.17133 | yes | 1.021E-07 | 600.36096 | 24.200399 |
| AFLSSHCYNAATIF | 14 | Q9P2D0 | Inhibitor of Bruton tyrosine kinase OS=H | IBTK | 150.528 | 2 | 54.17133 | yes | 1.276E-07 | 791.88776 | 18.189713 |
| PFIGMSSAPR | 10 | Q9P2D1 | Chromodomain-helicase-DNA-binding pr | CHD7 | 335.928 | 2 | 54.791916 | yes | 4.031E-06 | 531.77368 | 1188.2598 |
| EQVLHHPQLGER | 12 | Q9P2D1 | Chromodomain-helicase-DNA-binding pr | CHD7 | 335.928 | 3 | 54.791916 | yes | 6.989E-07 | 481.58813 | 111.31586 |
| EQVLHHPQLGER | 12 | Q9P2D1 | Chromodomain-helicase-DNA-binding pr | CHD7 | 335.928 | 3 | 54.791916 | yes | 3.097E-05 | 721.8786 | 111.31586 |
| EQVLHHPQLGER | 12 | Q9P2D1 | Chromodomain-helicase-DNA-binding pr | CHD7 | 335.928 | 2 | 54.791916 | yes | 6.989E-07 | 481.58813 | 111.31586 |
| EQVLHHPQLGER | 12 | Q9P2D1 | Chromodomain-helicase-DNA-binding pr | CHD7 | 335.928 | 2 | 54.791916 | yes | 3.097E-05 | 721.8786 | 111.31586 |
| AMSNPAGTPPPQV | 14 | Q9P2D1 | Chromodomain-helicase-DNA-binding pr | CHD7 | 335.928 | 2 | 54.791916 | yes | 2.37E-07 | 711.86157 | 205.83026 |
| DDFPGGVDNQELN | 14 | Q9P2D1 | Chromodomain-helicase-DNA-binding pr | CHD7 | 335.928 | 2 | 54.791916 | yes | 5.56E-07 | 788.35535 | 171.04372 |
| NNLVIDTPR | 9 | Q9P2D1 | Chromodomain-helicase-DNA-binding pr | CHD7 | 335.928 | 2 | 54.791916 | no | 9.722E-08 | 521.28802 | 150.33752 |
| KIEIEAER | 8 | Q9P2D1 | Chromodomain-helicase-DNA-binding pr | CHD7 | 335.928 | 2 | 54.791916 | yes | 2.594E-07 | 494.2771 | 138.48457 |
| ESGEEVEIEEFYVK | 14 | Q9P2D1 | Chromodomain-helicase-DNA-binding pr | CHD7 | 335.928 | 2 | 54.791916 | yes | 9.722E-08 | 843.89087 | 130.70262 |
| LDNICEAVLK | 10 | Q9P2D1 | Chromodomain-helicase-DNA-binding pr | CHD7 | 335.928 | 2 | 54.791916 | yes | 9.722E-08 | 587.81049 | 129.772 |
| CVVIDEAHR | 9 | Q9P2D1 | Chromodomain-helicase-DNA-binding pr | CHD7 | 335.928 | 2 | 54.791916 | yes | 3.514E-07 | 549.77167 | 125.37864 |
| WTDILSHGR | 9 | Q9P2D1 | Chromodomain-helicase-DNA-binding pr | CHD7 | 335.928 | 2 | 54.791916 | yes | 9.722E-08 | 542.78033 | 120.82374 |
| IPVINLEDGTR | 11 | Q9P2D1 | Chromodomain-helicase-DNA-binding pr | CHD7 | 335.928 | 2 | 54.791916 | yes | 9.722E-08 | 613.84058 | 113.44872 |
| TDYHILNDPELSFL | 18 | Q9P2D1 | Chromodomain-helicase-DNA-binding pr | CHD7 | 335.928 | 3 | 54.791916 | yes | 9.722E-08 | 710.01636 | 105.98949 |
| EDALNLSVPR | 10 | Q9P2D1 | Chromodomain-helicase-DNA-binding pr | CHD7 | 335.928 | 2 | 54.791916 | yes | 9.722E-08 | 557.29858 | 85.86927 |
| ALVNHSGLSAPVP | 14 | Q9P2D1 | Chromodomain-helicase-DNA-binding pr | CHD7 | 335.928 | 3 | 54.791916 | yes | 6.405E-06 | 473.26852 | 80.184189 |
| QLTEQDVETICR | 12 | Q9P2D1 | Chromodomain-helicase-DNA-binding pr | CHD7 | 335.928 | 2 | 54.791916 | yes | 9.823E-08 | 746.35901 | 80.022247 |

| | | | | | | | | | | | |
|---|---|---|---|---|---|---|---|---|---|---|---|
| QECEAEASSVK | 11 | Q9P2D1 | Chromodomain-helicase-DNA-binding pr | CHD7 | 335.928 | 2 | 54.791916 | yes | 1.62E-07 | 619.27191 | 78.311081 |
| LLTGPVVR | 8 | Q9P2D1 | Chromodomain-helicase-DNA-binding pr | CHD7 | 335.928 | 2 | 54.791916 | yes | 6.639E-07 | 427.77655 | 71.084656 |
| AFEEDIETPPTR | 12 | Q9P2D1 | Chromodomain-helicase-DNA-binding pr | CHD7 | 335.928 | 2 | 54.791916 | yes | 5.434E-07 | 702.83569 | 66.361053 |
| GEPVTHYLVK | 10 | Q9P2D1 | Chromodomain-helicase-DNA-binding pr | CHD7 | 335.928 | 2 | 54.791916 | yes | 5.47E-07 | 571.81384 | 52.941273 |
| HGQSFVDSSSMW( | 15 | Q9P2D1 | Chromodomain-helicase-DNA-binding pr | CHD7 | 335.928 | 3 | 54.791916 | yes | 9.99E-08 | 559.91943 | 49.541882 |
| VVSTFGVIFDPVK | 13 | Q9P2D1 | Chromodomain-helicase-DNA-binding pr | CHD7 | 335.928 | 2 | 54.791916 | yes | 9.722E-08 | 704.39777 | 41.293499 |
| THTITIESEGK | 11 | Q9P2D1 | Chromodomain-helicase-DNA-binding pr | CHD7 | 335.928 | 2 | 54.791916 | yes | 7.928E-07 | 608.31439 | 35.99688 |
| EATSSTSNFSSLSSI | 15 | Q9P2D1 | Chromodomain-helicase-DNA-binding pr | CHD7 | 335.928 | 2 | 54.791916 | yes | 1.217E-07 | 766.85736 | 33.853664 |
| DGFYMEDGDPSV/ | 19 | Q9P2D1 | Chromodomain-helicase-DNA-binding pr | CHD7 | 335.928 | 3 | 54.791916 | yes | 3.351E-06 | 726.99689 | 28.724689 |
| NVEGLDLLFMSHK | 13 | Q9P2D1 | Chromodomain-helicase-DNA-binding pr | CHD7 | 335.928 | 2 | 54.791916 | yes | 1.759E-07 | 751.88721 | 24.115791 |
| AVEAVVNDTSGEI | 14 | Q9P2D3 | HEAT repeat-containing protein 5B OS=I | HEATR5B | 224.303 | 2 | 41.738003 | yes | 4.421E-07 | 716.84937 | 58.678513 |
| STSVNLNQASGAV | 17 | Q9P2D3 | HEAT repeat-containing protein 5B OS=I | HEATR5B | 224.303 | 2 | 41.738003 | yes | 3.363E-07 | 795.90771 | 53.057552 |
| VFAADCLCR | 9 | Q9P2D3 | HEAT repeat-containing protein 5B OS=I | HEATR5B | 224.303 | 2 | 41.738003 | yes | 1.057E-06 | 556.25482 | 52.523212 |
| VHNLLVSSLDK | 11 | Q9P2D3 | HEAT repeat-containing protein 5B OS=I | HEATR5B | 224.303 | 2 | 41.738003 | yes | 3.623E-06 | 612.85095 | 46.501637 |
| SKPFVAPR | 8 | Q9P2D3 | HEAT repeat-containing protein 5B OS=I | HEATR5B | 224.303 | 2 | 41.738003 | yes | 4.783E-06 | 451.26636 | 35.736488 |
| INVLDPGQWPSK | 12 | Q9P2E3 | NFX1-type zinc finger-containing protein | ZNFX1 | 220.227 | 2 | 55.920544 | yes | 1.113E-07 | 677.36169 | 855.56616 |
| ELAIIQGPPGTGK | 13 | Q9P2E3 | NFX1-type zinc finger-containing protein | ZNFX1 | 220.227 | 2 | 55.920544 | yes | 1.049E-07 | 640.86407 | 770.01294 |
| LSFTSQELSDLR | 12 | Q9P2E3 | NFX1-type zinc finger-containing protein | ZNFX1 | 220.227 | 2 | 55.920544 | yes | 9.722E-08 | 698.35938 | 637.56549 |
| VGFLQISNR | 9 | Q9P2E3 | NFX1-type zinc finger-containing protein | ZNFX1 | 220.227 | 2 | 55.920544 | yes | 3.939E-07 | 517.29309 | 464.71387 |
| ATLPCSGLGISEEE | 15 | Q9P2E3 | NFX1-type zinc finger-containing protein | ZNFX1 | 220.227 | 2 | 55.920544 | yes | 1.372E-07 | 809.89069 | 404.63452 |
| ASGVDIEEETEK | 12 | Q9P2E3 | NFX1-type zinc finger-containing protein | ZNFX1 | 220.227 | 2 | 55.920544 | yes | 1.002E-07 | 653.80408 | 379.61514 |
| NLPMHLR | 7 | Q9P2E3 | NFX1-type zinc finger-containing protein | ZNFX1 | 220.227 | 2 | 55.920544 | yes | 2.407E-07 | 440.74472 | 353.08719 |
| VQTIIEHLQEK | 11 | Q9P2E3 | NFX1-type zinc finger-containing protein | ZNFX1 | 220.227 | 2 | 55.920544 | yes | 9.722E-08 | 669.37482 | 337.96906 |
| NIVECNSHVK | 10 | Q9P2E3 | NFX1-type zinc finger-containing protein | ZNFX1 | 220.227 | 2 | 55.920544 | yes | 9.722E-08 | 600.29553 | 329.89542 |
| LLYGSLVCMSK | 11 | Q9P2E3 | NFX1-type zinc finger-containing protein | ZNFX1 | 220.227 | 2 | 55.920544 | yes | 1.084E-07 | 635.83032 | 322.51221 |
| VQIVSAIGYPR | 11 | Q9P2E3 | NFX1-type zinc finger-containing protein | ZNFX1 | 220.227 | 2 | 55.920544 | yes | 9.729E-08 | 601.84821 | 316.91687 |
| LWLQLYQADTR | 11 | Q9P2E3 | NFX1-type zinc finger-containing protein | ZNFX1 | 220.227 | 2 | 55.920544 | yes | 1.017E-07 | 703.875 | 296.49475 |
| FLESLLQK | 8 | Q9P2E3 | NFX1-type zinc finger-containing protein | ZNFX1 | 220.227 | 2 | 55.920544 | yes | 3.978E-07 | 489.28696 | 295.05115 |
| MCPEIAR | 7 | Q9P2E3 | NFX1-type zinc finger-containing protein | ZNFX1 | 220.227 | 2 | 55.920544 | yes | 9.722E-08 | 438.70694 | 272.66708 |
| FTQEDEQLVQEK | 12 | Q9P2E3 | NFX1-type zinc finger-containing protein | ZNFX1 | 220.227 | 2 | 55.920544 | yes | 1.416E-07 | 747.35956 | 259.6033 |
| ELLSHSSMK | 9 | Q9P2E3 | NFX1-type zinc finger-containing protein | ZNFX1 | 220.227 | 2 | 55.920544 | yes | 9.722E-08 | 516.26312 | 227.87607 |
| GMYCIGNMQMLA | 13 | Q9P2E3 | NFX1-type zinc finger-containing protein | ZNFX1 | 220.227 | 2 | 55.920544 | yes | 9.722E-08 | 758.84259 | 187.01329 |
| VQFDTKPLK | 9 | Q9P2E3 | NFX1-type zinc finger-containing protein | ZNFX1 | 220.227 | 2 | 55.920544 | yes | 9.722E-08 | 538.31097 | 165.09756 |
| LEEIEIIK | 8 | Q9P2E3 | NFX1-type zinc finger-containing protein | ZNFX1 | 220.227 | 2 | 55.920544 | yes | 2.218E-07 | 493.79205 | 144.46515 |
| LLICSHK | 7 | Q9P2E3 | NFX1-type zinc finger-containing protein | ZNFX1 | 220.227 | 2 | 55.920544 | yes | 1.138E-07 | 435.74692 | 138.80194 |
| LTYLVNLLTR | 10 | Q9P2E3 | NFX1-type zinc finger-containing protein | ZNFX1 | 220.227 | 2 | 55.920544 | yes | 9.722E-08 | 603.36627 | 131.58998 |
| AYMSIMTQMK | 10 | Q9P2E3 | NFX1-type zinc finger-containing protein | ZNFX1 | 220.227 | 2 | 55.920544 | yes | 1.28E-07 | 602.28217 | 112.17431 |
| LEQVHEWLAK | 10 | Q9P2E3 | NFX1-type zinc finger-containing protein | ZNFX1 | 220.227 | 2 | 55.920544 | yes | 9.722E-08 | 626.83789 | 111.13335 |
| EQEDLCR | 7 | Q9P2E3 | NFX1-type zinc finger-containing protein | ZNFX1 | 220.227 | 2 | 55.920544 | yes | 2.284E-07 | 475.20581 | 109.91547 |

| | | | | | | | | | | | |
|---|---|---|---|---|---|---|---|---|---|---|---|
| EFQCMKPCQK | 10 | Q9P2E3 | NFX1-type zinc finger-containing protein | ZNFX1 | 220.227 | 2 | 55.920544 | yes | 1.197E-07 | 678.29889 | 103.219 |
| PSANVYDLAK | 10 | Q9P2E3 | NFX1-type zinc finger-containing protein | ZNFX1 | 220.227 | 2 | 55.920544 | yes | 2.809E-07 | 539.28241 | 101.76883 |
| CGHVEGLLYGGLI | 15 | Q9P2E3 | NFX1-type zinc finger-containing protein | ZNFX1 | 220.227 | 3 | 55.920544 | yes | 2.384E-07 | 538.96057 | 49.418602 |
| CGHVEGLLYGGLI | 15 | Q9P2E3 | NFX1-type zinc finger-containing protein | ZNFX1 | 220.227 | 3 | 55.920544 | yes | 2.18E-06 | 807.93726 | 49.418602 |
| CGHVEGLLYGGLI | 15 | Q9P2E3 | NFX1-type zinc finger-containing protein | ZNFX1 | 220.227 | 2 | 55.920544 | yes | 2.384E-07 | 538.96057 | 49.418602 |
| CGHVEGLLYGGLI | 15 | Q9P2E3 | NFX1-type zinc finger-containing protein | ZNFX1 | 220.227 | 2 | 55.920544 | yes | 2.18E-06 | 807.93726 | 49.418602 |
| ENNQIGPMLR | 10 | Q9P2E3 | NFX1-type zinc finger-containing protein | ZNFX1 | 220.227 | 2 | 55.920544 | yes | 1.002E-07 | 586.29803 | 93.401909 |
| LQEDLHILK | 9 | Q9P2E3 | NFX1-type zinc finger-containing protein | ZNFX1 | 220.227 | 2 | 55.920544 | yes | 1.317E-07 | 554.82166 | 88.522224 |
| FVQLEDCSHIFEV( | 18 | Q9P2E3 | NFX1-type zinc finger-containing protein | ZNFX1 | 220.227 | 3 | 55.920544 | yes | 1.284E-07 | 736.02478 | 88.14016 |
| ICVALSR | 7 | Q9P2E3 | NFX1-type zinc finger-containing protein | ZNFX1 | 220.227 | 2 | 55.920544 | yes | 1.11E-07 | 409.73129 | 85.67028 |
| EDFVRPLR | 8 | Q9P2E3 | NFX1-type zinc finger-containing protein | ZNFX1 | 220.227 | 2 | 55.920544 | yes | 9.809E-08 | 516.28528 | 78.024872 |
| IQGSAGEIATSQER | 14 | Q9P2E3 | NFX1-type zinc finger-containing protein | ZNFX1 | 220.227 | 2 | 55.920544 | yes | 1.051E-07 | 723.86279 | 69.838768 |
| EVIGGTNHTLER | 12 | Q9P2E3 | NFX1-type zinc finger-containing protein | ZNFX1 | 220.227 | 2 | 55.920544 | yes | 1.009E-07 | 663.34406 | 65.075821 |
| NFNLEVSLFER | 11 | Q9P2E3 | NFX1-type zinc finger-containing protein | ZNFX1 | 220.227 | 2 | 55.920544 | yes | 2.251E-07 | 684.35132 | 62.111084 |
| QSVLHVLGILK | 11 | Q9P2E3 | NFX1-type zinc finger-containing protein | ZNFX1 | 220.227 | 2 | 55.920544 | yes | 9.722E-08 | 603.88208 | 51.532257 |
| NQANNPPANALR | 12 | Q9P2E3 | NFX1-type zinc finger-containing protein | ZNFX1 | 220.227 | 2 | 55.920544 | yes | 1.486E-07 | 640.32874 | 43.237709 |
| DAQVVGMTTTGA | 14 | Q9P2E3 | NFX1-type zinc finger-containing protein | ZNFX1 | 220.227 | 2 | 55.920544 | yes | 3.622E-06 | 675.34009 | 42.797752 |
| VDTYTLVQPEAEI | 19 | Q9P2E3 | NFX1-type zinc finger-containing protein | ZNFX1 | 220.227 | 3 | 55.920544 | yes | 1.157E-07 | 751.02197 | 33.658241 |
| CGTILDCGHPCPG! | 22 | Q9P2E3 | NFX1-type zinc finger-containing protein | ZNFX1 | 220.227 | 4 | 55.920544 | yes | 3.747E-07 | 641.75525 | 14.333686 |
| HVLEGLQEVQEEI | 18 | Q9P2E3 | NFX1-type zinc finger-containing protein | ZNFX1 | 220.227 | 3 | 55.920544 | yes | 7.649E-06 | 718.02716 | 4.1586962 |
| LLYFTGQR | 8 | Q9P2E5 | Chondroitin sulfate glucuronyltransferase | CHPF2 | 85.949 | 2 | 54.166813 | yes | 4.037E-07 | 499.27692 | 503.11871 |
| LSEELYHR | 8 | Q9P2E5 | Chondroitin sulfate glucuronyltransferase | CHPF2 | 85.949 | 2 | 54.166813 | yes | 1.679E-07 | 523.76691 | 496.52231 |
| LLVAVLTSR | 9 | Q9P2E5 | Chondroitin sulfate glucuronyltransferase | CHPF2 | 85.949 | 2 | 54.166813 | yes | 1.163E-07 | 486.31604 | 475.20551 |
| FSALELER | 8 | Q9P2E5 | Chondroitin sulfate glucuronyltransferase | CHPF2 | 85.949 | 2 | 54.166813 | yes | 2.153E-07 | 482.75854 | 363.38593 |
| AEEFIGAGEQAR | 12 | Q9P2E5 | Chondroitin sulfate glucuronyltransferase | CHPF2 | 85.949 | 2 | 54.166813 | yes | 2.153E-07 | 639.30969 | 361.59119 |
| AYSEIEQLQAQIR | 13 | Q9P2E5 | Chondroitin sulfate glucuronyltransferase | CHPF2 | 85.949 | 2 | 54.166813 | yes | 1.403E-07 | 774.90448 | 341.77954 |
| YIQTELGSR | 9 | Q9P2E5 | Chondroitin sulfate glucuronyltransferase | CHPF2 | 85.949 | 2 | 54.166813 | yes | 1.402E-07 | 533.78003 | 317.48257 |
| FSGLHLFR | 8 | Q9P2E5 | Chondroitin sulfate glucuronyltransferase | CHPF2 | 85.949 | 2 | 54.166813 | yes | 1.526E-07 | 488.77179 | 299.16577 |
| GAPDPFLGVK | 10 | Q9P2E5 | Chondroitin sulfate glucuronyltransferase | CHPF2 | 85.949 | 2 | 54.166813 | yes | 3.655E-07 | 500.77673 | 244.99486 |
| LDQSDEDFKPR | 11 | Q9P2E5 | Chondroitin sulfate glucuronyltransferase | CHPF2 | 85.949 | 2 | 54.166813 | yes | 9.722E-08 | 675.32025 | 163.04007 |
| AVEPGLVQK | 9 | Q9P2E5 | Chondroitin sulfate glucuronyltransferase | CHPF2 | 85.949 | 2 | 54.166813 | yes | 5.693E-07 | 470.77673 | 136.28236 |
| VEILPMPYVTEATI | 14 | Q9P2E5 | Chondroitin sulfate glucuronyltransferase | CHPF2 | 85.949 | 2 | 54.166813 | yes | 2.611E-07 | 809.92908 | 128.49582 |
| YCHGGFGYLLSR | 12 | Q9P2E5 | Chondroitin sulfate glucuronyltransferase | CHPF2 | 85.949 | 2 | 54.166813 | yes | 2.048E-06 | 715.33771 | 100.79842 |
| LAALAGHLSINQD | 18 | Q9P2E5 | Chondroitin sulfate glucuronyltransferase | CHPF2 | 85.949 | 3 | 54.166813 | yes | 1.306E-06 | 642.35828 | 65.614304 |
| PAWLMSETLR | 10 | Q9P2E5 | Chondroitin sulfate glucuronyltransferase | CHPF2 | 85.949 | 2 | 54.166813 | yes | 1.985E-07 | 602.31317 | 60.930862 |
| APAGMQVVSHGD | 14 | Q9P2E5 | Chondroitin sulfate glucuronyltransferase | CHPF2 | 85.949 | 2 | 54.166813 | yes | 9.722E-08 | 727.34625 | 44.639473 |
| AAAAELER | 8 | Q9P2E5 | Chondroitin sulfate glucuronyltransferase | CHPF2 | 85.949 | 2 | 54.166813 | yes | 4.552E-06 | 415.72195 | 17.719189 |
| SIEALLEAGQAR | 12 | Q9P2E9 | Ribosome-binding protein 1 OS=Homo sa | RRBP1 | 152.457 | 2 | 53.456432 | yes | 4.173E-07 | 629.34351 | 4618.2715 |
| EVPMVVVPPVGAI | 13 | Q9P2E9 | Ribosome-binding protein 1 OS=Homo sa | RRBP1 | 152.457 | 2 | 53.456432 | yes | 2.482E-07 | 661.38104 | 2057.9048 |

| | | | | | | | | | | | |
|---|---|---|---|---|---|---|---|---|---|---|---|
| EVPMVVVPPVGAI | 13 | Q9P2E9 | Ribosome-binding protein 1 OS=Homo sa | RRBP1 | 152.457 | 2 | 53.456432 | yes | 2.182E-06 | 669.37848 | 2057.9048 |
| EQEITAVQAR | 10 | Q9P2E9 | Ribosome-binding protein 1 OS=Homo sa | RRBP1 | 152.457 | 2 | 53.456432 | yes | 1.211E-07 | 572.80145 | 2565.0815 |
| DALNQATSQVESK | 13 | Q9P2E9 | Ribosome-binding protein 1 OS=Homo sa | RRBP1 | 152.457 | 2 | 53.456432 | yes | 9.722E-08 | 695.84406 | 2466.45 |
| AGIIQDTWHK | 10 | Q9P2E9 | Ribosome-binding protein 1 OS=Homo sa | RRBP1 | 152.457 | 2 | 53.456432 | yes | 1.4E-07 | 584.80908 | 2427.9514 |
| LQQENSILR | 9 | Q9P2E9 | Ribosome-binding protein 1 OS=Homo sa | RRBP1 | 152.457 | 2 | 53.456432 | no | 2.703E-07 | 550.80658 | 2319.2573 |
| LIEILSEK | 8 | Q9P2E9 | Ribosome-binding protein 1 OS=Homo sa | RRBP1 | 152.457 | 2 | 53.456432 | yes | 1.576E-07 | 472.78677 | 2212.6697 |
| QLLLESQSQLDAA | 14 | Q9P2E9 | Ribosome-binding protein 1 OS=Homo sa | RRBP1 | 152.457 | 2 | 53.456432 | yes | 1.031E-07 | 772.41992 | 2092.1685 |
| AENSQLTER | 9 | Q9P2E9 | Ribosome-binding protein 1 OS=Homo sa | RRBP1 | 152.457 | 2 | 53.456432 | yes | 1.008E-07 | 524.2569 | 1495.7813 |
| LQSSEAEVR | 9 | Q9P2E9 | Ribosome-binding protein 1 OS=Homo sa | RRBP1 | 152.457 | 2 | 53.456432 | yes | 5.143E-07 | 509.76181 | 1380.5291 |
| NTDVAQSPEAPK | 12 | Q9P2E9 | Ribosome-binding protein 1 OS=Homo sa | RRBP1 | 152.457 | 2 | 53.456432 | yes | 1.185E-07 | 628.80951 | 1349.5092 |
| HPPAPAEPSSDLAS | 15 | Q9P2E9 | Ribosome-binding protein 1 OS=Homo sa | RRBP1 | 152.457 | 2 | 53.456432 | yes | 1.441E-07 | 752.37555 | 1250.2792 |
| QSDELALVR | 9 | Q9P2E9 | Ribosome-binding protein 1 OS=Homo sa | RRBP1 | 152.457 | 2 | 53.456432 | yes | 9.722E-08 | 515.78003 | 1223.5109 |
| KLTSDLGR | 8 | Q9P2E9 | Ribosome-binding protein 1 OS=Homo sa | RRBP1 | 152.457 | 2 | 53.456432 | yes | 9.722E-08 | 445.25873 | 1169.9567 |
| ETSYEEALANQR | 12 | Q9P2E9 | Ribosome-binding protein 1 OS=Homo sa | RRBP1 | 152.457 | 2 | 53.456432 | yes | 9.722E-08 | 705.82843 | 1101.0616 |
| QVLQLQASHR | 10 | Q9P2E9 | Ribosome-binding protein 1 OS=Homo sa | RRBP1 | 152.457 | 2 | 53.456432 | yes | 1.745E-07 | 590.33325 | 1004.8961 |
| ELESQVSGLEK | 11 | Q9P2E9 | Ribosome-binding protein 1 OS=Homo sa | RRBP1 | 152.457 | 2 | 53.456432 | yes | 9.734E-08 | 609.81427 | 979.58496 |
| SILAETEGMLR | 11 | Q9P2E9 | Ribosome-binding protein 1 OS=Homo sa | RRBP1 | 152.457 | 2 | 53.456432 | yes | 1.189E-07 | 610.32117 | 922.05731 |
| HLEEIVEK | 8 | Q9P2E9 | Ribosome-binding protein 1 OS=Homo sa | RRBP1 | 152.457 | 2 | 53.456432 | yes | 1.691E-07 | 498.77167 | 856.59631 |
| VGAAEEELQK | 10 | Q9P2E9 | Ribosome-binding protein 1 OS=Homo sa | RRBP1 | 152.457 | 2 | 53.456432 | yes | 1.827E-07 | 537.27728 | 836.56403 |
| IPDHDPAPNVTVLI | 15 | Q9P2E9 | Ribosome-binding protein 1 OS=Homo sa | RRBP1 | 152.457 | 2 | 53.456432 | yes | 3.654E-07 | 828.95703 | 746.2417 |
| RLDEVSR | 7 | Q9P2E9 | Ribosome-binding protein 1 OS=Homo sa | RRBP1 | 152.457 | 2 | 53.456432 | yes | 9.722E-08 | 437.74069 | 743.01508 |
| LLATEQEDAAVAK | 13 | Q9P2E9 | Ribosome-binding protein 1 OS=Homo sa | RRBP1 | 152.457 | 2 | 53.456432 | yes | 3.249E-07 | 679.86176 | 730.33179 |
| KLQEQLEK | 8 | Q9P2E9 | Ribosome-binding protein 1 OS=Homo sa | RRBP1 | 152.457 | 2 | 53.456432 | yes | 1.448E-07 | 508.29276 | 600.21045 |
| LTAEFEEAQTSACI | 14 | Q9P2E9 | Ribosome-binding protein 1 OS=Homo sa | RRBP1 | 152.457 | 2 | 53.456432 | yes | 1.276E-07 | 806.86725 | 586.49554 |
| HMAAASAECQNY | 14 | Q9P2E9 | Ribosome-binding protein 1 OS=Homo sa | RRBP1 | 152.457 | 2 | 53.456432 | yes | 9.722E-08 | 776.33759 | 252.28804 |
| HMAAASAECQNY | 14 | Q9P2E9 | Ribosome-binding protein 1 OS=Homo sa | RRBP1 | 152.457 | 2 | 53.456432 | yes | 1.022E-07 | 517.89417 | 252.28804 |
| HMAAASAECQNY | 14 | Q9P2E9 | Ribosome-binding protein 1 OS=Homo sa | RRBP1 | 152.457 | 3 | 53.456432 | yes | 9.722E-08 | 776.33759 | 252.28804 |
| HMAAASAECQNY | 14 | Q9P2E9 | Ribosome-binding protein 1 OS=Homo sa | RRBP1 | 152.457 | 3 | 53.456432 | yes | 1.022E-07 | 517.89417 | 252.28804 |
| GNTPATGTTQGK | 12 | Q9P2E9 | Ribosome-binding protein 1 OS=Homo sa | RRBP1 | 152.457 | 2 | 53.456432 | yes | 2.126E-07 | 566.78326 | 477.18799 |
| GELESSDQVR | 10 | Q9P2E9 | Ribosome-binding protein 1 OS=Homo sa | RRBP1 | 152.457 | 2 | 53.456432 | yes | 1.324E-07 | 560.26746 | 475.25519 |
| SKCEELSGLHGQL | 17 | Q9P2E9 | Ribosome-binding protein 1 OS=Homo sa | RRBP1 | 152.457 | 3 | 53.456432 | yes | 9.989E-08 | 647.98639 | 433.82925 |
| TTQEQLAR | 8 | Q9P2E9 | Ribosome-binding protein 1 OS=Homo sa | RRBP1 | 152.457 | 2 | 53.456432 | yes | 2.505E-07 | 473.75125 | 427.62219 |
| ESEEALQK | 8 | Q9P2E9 | Ribosome-binding protein 1 OS=Homo sa | RRBP1 | 152.457 | 2 | 53.456432 | yes | 3.38E-07 | 467.22983 | 381.27731 |
| EVQQLQGK | 8 | Q9P2E9 | Ribosome-binding protein 1 OS=Homo sa | RRBP1 | 152.457 | 2 | 53.456432 | yes | 1.803E-07 | 465.25616 | 380.93451 |
| QQLSEMK | 7 | Q9P2E9 | Ribosome-binding protein 1 OS=Homo sa | RRBP1 | 152.457 | 2 | 53.456432 | yes | 1.457E-07 | 432.2182 | 370.35828 |
| LKELESQVSGLEK | 13 | Q9P2E9 | Ribosome-binding protein 1 OS=Homo sa | RRBP1 | 152.457 | 2 | 53.456432 | yes | 9.722E-08 | 730.40375 | 366.94598 |
| LQEELEK | 7 | Q9P2E9 | Ribosome-binding protein 1 OS=Homo sa | RRBP1 | 152.457 | 2 | 53.456432 | yes | 9.902E-07 | 444.73727 | 352.91003 |
| EKGPTLLK | 8 | Q9P2E9 | Ribosome-binding protein 1 OS=Homo sa | RRBP1 | 152.457 | 2 | 53.456432 | yes | 1.355E-07 | 443.27383 | 348.83878 |
| CEELSGLHGQLQE | 15 | Q9P2E9 | Ribosome-binding protein 1 OS=Homo sa | RRBP1 | 152.457 | 3 | 53.456432 | yes | 1.256E-07 | 576.2774 | 336.87518 |

| | | | | | | | | | | | |
|---|---|---|---|---|---|---|---|---|---|---|---|
| LKGELESSDQVR | 12 | Q9P2E9 | Ribosome-binding protein 1 OS=Homo sa | RRBP1 | 152.457 | 2 | 53.456432 | yes | 1.188E-07 | 680.85699 | 334.92926 |
| KVEGAPTQGR | 10 | Q9P2E9 | Ribosome-binding protein 1 OS=Homo sa | RRBP1 | 152.457 | 2 | 53.456432 | yes | 6.002E-07 | 521.78564 | 321.70587 |
| AAILETAPK | 9 | Q9P2E9 | Ribosome-binding protein 1 OS=Homo sa | RRBP1 | 152.457 | 2 | 53.456432 | yes | 3.148E-07 | 457.2713 | 303.62042 |
| AMEALATAEQACl | 13 | Q9P2E9 | Ribosome-binding protein 1 OS=Homo sa | RRBP1 | 152.457 | 2 | 53.456432 | yes | 1.004E-07 | 697.32617 | 289.03046 |
| AQEQQQQMAELH | 14 | Q9P2E9 | Ribosome-binding protein 1 OS=Homo sa | RRBP1 | 152.457 | 2 | 53.456432 | yes | 9.722E-08 | 828.39392 | 220.51956 |
| LQEQLEK | 7 | Q9P2E9 | Ribosome-binding protein 1 OS=Homo sa | RRBP1 | 152.457 | 2 | 53.456432 | yes | 1.112E-07 | 444.24527 | 218.83153 |
| VEGAPTQGR | 9 | Q9P2E9 | Ribosome-binding protein 1 OS=Homo sa | RRBP1 | 152.457 | 2 | 53.456432 | yes | 3.231E-07 | 457.73816 | 170.52502 |
| LRQELSK | 7 | Q9P2E9 | Ribosome-binding protein 1 OS=Homo sa | RRBP1 | 152.457 | 2 | 53.456432 | yes | 4.298E-07 | 437.26126 | 127.4342 |
| KGEGAPIQGK | 10 | Q9P2E9 | Ribosome-binding protein 1 OS=Homo sa | RRBP1 | 152.457 | 2 | 53.456432 | yes | 3.272E-07 | 492.77728 | 124.78526 |
| ELCHTQSSHASLR | 13 | Q9P2E9 | Ribosome-binding protein 1 OS=Homo sa | RRBP1 | 152.457 | 2 | 53.456432 | yes | 1.344E-07 | 763.36243 | 72.21524 |
| SVEEEEQVWR | 10 | Q9P2E9 | Ribosome-binding protein 1 OS=Homo sa | RRBP1 | 152.457 | 2 | 53.456432 | yes | 1.318E-07 | 645.8017 | 60.836037 |
| YLDSDASFTENSD' | 16 | Q9P2G4 | Microtubule-associated protein 10 OS=H | MAP10 | 100.345 | 3 | 24.95942 | yes | 3.362E-06 | 603.25873 | 228.13339 |
| AMFLSEMAEAK | 11 | Q9P2G9 | Kelch-like protein 8 OS=Homo sapiens O | KLHL8 | 68.803 | 2 | 36.336452 | yes | 7.109E-07 | 614.29102 | 61.689079 |
| VVLASQPDLECGF | 15 | Q9P2I0 | Cleavage and polyadenylation specificity | CPSF2 | 88.487 | 2 | 49.699524 | yes | 1.443E-07 | 839.41687 | 304.34531 |
| GDGNVLIAVDTAC | 14 | Q9P2I0 | Cleavage and polyadenylation specificity | CPSF2 | 88.487 | 2 | 49.699524 | yes | 1.009E-07 | 679.35718 | 197.86604 |
| FLIDNPSEK | 9 | Q9P2I0 | Cleavage and polyadenylation specificity | CPSF2 | 88.487 | 2 | 49.699524 | yes | 2.757E-07 | 531.77692 | 164.18187 |
| NSIILTYR | 8 | Q9P2I0 | Cleavage and polyadenylation specificity | CPSF2 | 88.487 | 2 | 49.699524 | yes | 4.22E-06 | 490.2822 | 145.21698 |
| DLFIQWCQDPK | 11 | Q9P2I0 | Cleavage and polyadenylation specificity | CPSF2 | 88.487 | 2 | 49.699524 | yes | 1.352E-07 | 725.34521 | 124.42214 |
| VTYIDYEGR | 9 | Q9P2I0 | Cleavage and polyadenylation specificity | CPSF2 | 88.487 | 2 | 49.699524 | yes | 2.895E-07 | 558.27203 | 118.60658 |
| SYPMFPAPEER | 11 | Q9P2I0 | Cleavage and polyadenylation specificity | CPSF2 | 88.487 | 2 | 49.699524 | yes | 1.443E-07 | 662.30554 | 82.584511 |
| HLSLCHGLSDLAR | 13 | Q9P2I0 | Cleavage and polyadenylation specificity | CPSF2 | 88.487 | 3 | 49.699524 | yes | 1.783E-07 | 493.58926 | 74.856842 |
| MGQMFMYDLYQS | 13 | Q9P2I0 | Cleavage and polyadenylation specificity | CPSF2 | 88.487 | 2 | 49.699524 | yes | 3.275E-07 | 835.36218 | 70.533226 |
| IGLEGCLCQDFYR | 13 | Q9P2I0 | Cleavage and polyadenylation specificity | CPSF2 | 88.487 | 2 | 49.699524 | yes | 1.461E-07 | 815.87128 | 67.759727 |
| FSQIVNLK | 8 | Q9P2I0 | Cleavage and polyadenylation specificity | CPSF2 | 88.487 | 2 | 49.699524 | yes | 2.656E-07 | 474.7793 | 47.477047 |
| LHETVDATSETHI' | 17 | Q9P2I0 | Cleavage and polyadenylation specificity | CPSF2 | 88.487 | 3 | 49.699524 | yes | 2.264E-07 | 667.00085 | 40.092506 |
| VFEVNASNLEK | 11 | Q9P2J5 | Leucine--tRNA ligase, cytoplasmic OS=H | LARS1 | 134.467 | 2 | 53.78463 | yes | 1.045E-07 | 625.32477 | 131.36945 |
| STGNFLTLTQAIDK | 14 | Q9P2J5 | Leucine--tRNA ligase, cytoplasmic OS=H | LARS1 | 134.467 | 2 | 53.78463 | yes | 1.611E-07 | 754.90137 | 93.248741 |
| TGFFEFQAAK | 10 | Q9P2J5 | Leucine--tRNA ligase, cytoplasmic OS=H | LARS1 | 134.467 | 2 | 53.78463 | yes | 2.023E-07 | 573.28491 | 74.071442 |
| VFASELNAGIIK | 12 | Q9P2J5 | Leucine--tRNA ligase, cytoplasmic OS=H | LARS1 | 134.467 | 2 | 53.78463 | yes | 9.722E-08 | 631.36115 | 67.356667 |
| ILDLQLEFDEK | 11 | Q9P2J5 | Leucine--tRNA ligase, cytoplasmic OS=H | LARS1 | 134.467 | 2 | 53.78463 | yes | 1.806E-06 | 681.86121 | 46.843491 |
| WQFLTLR | 7 | Q9P2J5 | Leucine--tRNA ligase, cytoplasmic OS=H | LARS1 | 134.467 | 2 | 53.78463 | yes | 4.431E-07 | 482.27417 | 37.087433 |
| VIASELGSMPELK | 13 | Q9P2J5 | Leucine--tRNA ligase, cytoplasmic OS=H | LARS1 | 134.467 | 2 | 53.78463 | yes | 9.722E-08 | 687.37085 | 34.343979 |
| NLETFCEETR | 10 | Q9P2J5 | Leucine--tRNA ligase, cytoplasmic OS=H | LARS1 | 134.467 | 2 | 53.78463 | yes | 1.204E-07 | 649.78772 | 28.587399 |
| VTDSTGQQVVGEI | 15 | Q9P2L0 | WD repeat-containing protein 35 OS=Ho | WDR35 | 133.548 | 2 | 31.338625 | yes | 1.875E-06 | 787.42523 | 9.2653837 |
| VFDLPESLGGIR | 12 | Q9P2N7 | Kelch-like protein 13 OS=Homo sapiens | KLHL13 | 73.868 | 2 | 48.93566 | no | 2.97E-07 | 651.85626 | 136.86528 |
| GQSDVGVAVFENI | 13 | Q9P2N7 | Kelch-like protein 13 OS=Homo sapiens | KLHL13 | 73.868 | 2 | 48.93566 | no | 2.703E-07 | 675.33844 | 67.991203 |
| NFPALLSTGEFLK | 13 | Q9P2N7 | Kelch-like protein 13 OS=Homo sapiens | KLHL13 | 73.868 | 2 | 48.93566 | no | 6.444E-07 | 718.89282 | 63.346321 |
| ELMCFDPDTDK | 11 | Q9P2N7 | Kelch-like protein 13 OS=Homo sapiens | KLHL13 | 73.868 | 2 | 48.93566 | yes | 1.084E-07 | 685.78375 | 37.731514 |
| IANTYNLTEVDK | 12 | Q9P2N7 | Kelch-like protein 13 OS=Homo sapiens | KLHL13 | 73.868 | 2 | 48.93566 | yes | 4.809E-05 | 690.85388 | 11.779054 |

| | | | | | | | | | | | |
|---|---|---|---|---|---|---|---|---|---|---|---|
| TDLYFMPLAGSK | 12 | Q9P2R6 | Arginine-glutamic acid dipeptide repeats ] | RERE | 172.424 | 2 | 52.935333 | yes | 1.847E-07 | 671.83923 | 124.04201 |
| ELVITDPVIK | 10 | Q9P2R6 | Arginine-glutamic acid dipeptide repeats ] | RERE | 172.424 | 2 | 52.935333 | yes | 1.207E-07 | 563.83954 | 115.51785 |
| ENILLCTDCR | 10 | Q9P2R6 | Arginine-glutamic acid dipeptide repeats ] | RERE | 172.424 | 2 | 52.935333 | yes | 9.722E-08 | 647.29993 | 70.18045 |
| AFQWDPQQPR | 10 | Q9P2S5 | WD repeat-containing protein WRAP73 ( | WRAP73 | 51.589 | 2 | 48.565487 | yes | 1.193E-07 | 636.80963 | 597.35645 |
| ILLYSLDGR | 9 | Q9P2S5 | WD repeat-containing protein WRAP73 ( | WRAP73 | 51.589 | 2 | 48.565487 | yes | 2.921E-07 | 525.3031 | 384.70074 |
| ACLQGITFTR | 10 | Q9P2S5 | WD repeat-containing protein WRAP73 ( | WRAP73 | 51.589 | 2 | 48.565487 | yes | 1.369E-07 | 583.80298 | 322.47845 |
| HILNTTEFHLR | 11 | Q9P2S5 | WD repeat-containing protein WRAP73 ( | WRAP73 | 51.589 | 3 | 48.565487 | yes | 1.862E-07 | 460.9176 | 245.62102 |
| YLASCVQYR | 9 | Q9P2S5 | WD repeat-containing protein WRAP73 ( | WRAP73 | 51.589 | 2 | 48.565487 | yes | 4.385E-07 | 580.28186 | 147.08578 |
| ITVWSLCTK | 9 | Q9P2S5 | WD repeat-containing protein WRAP73 ( | WRAP73 | 51.589 | 2 | 48.565487 | yes | 1.129E-07 | 554.297 | 142.99933 |
| LFAVLEQLSPVR | 12 | Q9P2S5 | WD repeat-containing protein WRAP73 ( | WRAP73 | 51.589 | 2 | 48.565487 | yes | 3.088E-07 | 686.40338 | 130.22574 |
| YMALAER | 7 | Q9P2S5 | WD repeat-containing protein WRAP73 ( | WRAP73 | 51.589 | 2 | 48.565487 | yes | 2.987E-07 | 427.21545 | 127.8354 |
| ILNHVTWK | 8 | Q9P2S5 | WD repeat-containing protein WRAP73 ( | WRAP73 | 51.589 | 2 | 48.565487 | yes | 4.723E-07 | 505.79272 | 87.049629 |
| MITEFGHPAAINDI | 15 | Q9P2S5 | WD repeat-containing protein WRAP73 ( | WRAP73 | 51.589 | 2 | 48.565487 | yes | 1.166E-07 | 820.90869 | 36.862301 |
| MITEFGHPAAINDI | 15 | Q9P2S5 | WD repeat-containing protein WRAP73 ( | WRAP73 | 51.589 | 2 | 48.565487 | yes | 5.413E-07 | 547.60822 | 36.862301 |
| MITEFGHPAAINDI | 15 | Q9P2S5 | WD repeat-containing protein WRAP73 ( | WRAP73 | 51.589 | 3 | 48.565487 | yes | 1.166E-07 | 820.90869 | 36.862301 |
| MITEFGHPAAINDI | 15 | Q9P2S5 | WD repeat-containing protein WRAP73 ( | WRAP73 | 51.589 | 3 | 48.565487 | yes | 5.413E-07 | 547.60822 | 36.862301 |
| DHFCLCFLETEAV | 19 | Q9P2S5 | WD repeat-containing protein WRAP73 ( | WRAP73 | 51.589 | 3 | 48.565487 | yes | 2.972E-07 | 762.34308 | 67.595558 |
| SPQLGLGCLSFPPP | 15 | Q9P2S5 | WD repeat-containing protein WRAP73 ( | WRAP73 | 51.589 | 2 | 48.565487 | yes | 3.011E-06 | 813.42706 | 10.711043 |
| YEIASVPVSLQTLK | 19 | Q9P2S5 | WD repeat-containing protein WRAP73 ( | WRAP73 | 51.589 | 3 | 48.565487 | yes | 1.596E-05 | 706.05981 | 10.299953 |
| ELSSALTTPEMQK | 13 | Q9P2W1 | Homologous-pairing protein 2 homolog ( | PSMC3IP | 24.907 | 2 | 37.387592 | yes | 6.038E-07 | 717.8609 | 67.801033 |
| ELQSQIQEAR | 10 | Q9P2X0 | Dolichol-phosphate mannosyltransferase : | DPM3 | 10.094 | 2 | 47.230724 | yes | 1.373E-07 | 601.31219 | 161.56328 |
| VEQESLLTAFR | 11 | Q9UBB4 | Ataxin-10 OS=Homo sapiens OX=9606 ( | ATXN10 | 53.489 | 2 | 53.74372 | yes | 1.08E-07 | 646.84589 | 2617.749 |
| MEEQGLADASLLK | 13 | Q9UBB4 | Ataxin-10 OS=Homo sapiens OX=9606 ( | ATXN10 | 53.489 | 2 | 53.74372 | yes | 9.849E-08 | 702.85559 | 1869.9783 |
| LIGNLCYK | 8 | Q9UBB4 | Ataxin-10 OS=Homo sapiens OX=9606 ( | ATXN10 | 53.489 | 2 | 53.74372 | yes | 1.156E-07 | 490.76532 | 1684.8274 |
| ETTNIFSNCGCVR | 13 | Q9UBB4 | Ataxin-10 OS=Homo sapiens OX=9606 ( | ATXN10 | 53.489 | 2 | 53.74372 | yes | 9.819E-08 | 779.34283 | 1009.3914 |
| HAELIASTFVDQCI | 14 | Q9UBB4 | Ataxin-10 OS=Homo sapiens OX=9606 ( | ATXN10 | 53.489 | 2 | 53.74372 | yes | 9.722E-08 | 809.89832 | 332.89191 |
| HAELIASTFVDQCI | 14 | Q9UBB4 | Ataxin-10 OS=Homo sapiens OX=9606 ( | ATXN10 | 53.489 | 2 | 53.74372 | yes | 1.404E-07 | 540.26794 | 332.89191 |
| HAELIASTFVDQCI | 14 | Q9UBB4 | Ataxin-10 OS=Homo sapiens OX=9606 ( | ATXN10 | 53.489 | 3 | 53.74372 | yes | 9.722E-08 | 809.89832 | 332.89191 |
| HAELIASTFVDQCI | 14 | Q9UBB4 | Ataxin-10 OS=Homo sapiens OX=9606 ( | ATXN10 | 53.489 | 3 | 53.74372 | yes | 1.404E-07 | 540.26794 | 332.89191 |
| CGLQFLGNIASR | 12 | Q9UBB4 | Ataxin-10 OS=Homo sapiens OX=9606 ( | ATXN10 | 53.489 | 2 | 53.74372 | yes | 1.064E-07 | 668.34552 | 650.88306 |
| SSHAVELACR | 10 | Q9UBB4 | Ataxin-10 OS=Homo sapiens OX=9606 ( | ATXN10 | 53.489 | 2 | 53.74372 | yes | 9.722E-08 | 565.2746 | 617.29144 |
| KVGFEVEK | 8 | Q9UBB4 | Ataxin-10 OS=Homo sapiens OX=9606 ( | ATXN10 | 53.489 | 2 | 53.74372 | yes | 1.22E-07 | 468.26346 | 520.27106 |
| SPELVQAMFPK | 11 | Q9UBB4 | Ataxin-10 OS=Homo sapiens OX=9606 ( | ATXN10 | 53.489 | 2 | 53.74372 | yes | 6.474E-07 | 623.82867 | 482.61902 |
| ITSDEPLTK | 9 | Q9UBB4 | Ataxin-10 OS=Homo sapiens OX=9606 ( | ATXN10 | 53.489 | 2 | 53.74372 | yes | 1.522E-07 | 502.26895 | 410.16168 |
| AEGDISNVANGFK | 13 | Q9UBB4 | Ataxin-10 OS=Homo sapiens OX=9606 ( | ATXN10 | 53.489 | 2 | 53.74372 | yes | 9.722E-08 | 661.32275 | 333.39465 |
| AAPRPPPAR | 9 | Q9UBB4 | Ataxin-10 OS=Homo sapiens OX=9606 ( | ATXN10 | 53.489 | 2 | 53.74372 | yes | 7.176E-07 | 487.78015 | 295.02997 |
| NACIECSVNQNSIF | 14 | Q9UBB4 | Ataxin-10 OS=Homo sapiens OX=9606 ( | ATXN10 | 53.489 | 2 | 53.74372 | yes | 9.722E-08 | 832.87762 | 185.41727 |
| DDIPVFLR | 8 | Q9UBB4 | Ataxin-10 OS=Homo sapiens OX=9606 ( | ATXN10 | 53.489 | 2 | 53.74372 | yes | 2.297E-07 | 487.76892 | 119.25053 |
| NPAVWLNTSQPLC | 14 | Q9UBB5 | Methyl-CpG-binding domain protein 2 OS | MBD2 | 43.255 | 2 | 51.943146 | yes | 9.734E-08 | 814.41669 | 81.60675 |

| | | | | | | | | | | | |
|---|---|---|---|---|---|---|---|---|---|---|---|
| GKPDLNTTLPIR | 12 | Q9UBB5 | Methyl-CpG-binding domain protein 2 O: | MBD2 | 43.255 | 2 | 51.943146 | yes | 1.515E-06 | 662.88281 | 72.555649 |
| LQGLSASDVTEQII | 15 | Q9UBB5 | Methyl-CpG-binding domain protein 2 O: | MBD2 | 43.255 | 2 | 51.943146 | yes | 9.722E-08 | 801.44086 | 49.193691 |
| YLGNTVDLSSFDF | 14 | Q9UBB5 | Methyl-CpG-binding domain protein 2 O: | MBD2 | 43.255 | 2 | 51.943146 | yes | 6.634E-07 | 817.39648 | 38.539368 |
| LLSTSPALQGTPAS | 15 | Q9UBB6 | Neurochondrin OS=Homo sapiens OX=9 | NCDN | 78.865 | 2 | 41.987804 | yes | 3.247E-07 | 749.91486 | 64.610916 |
| IFDAVGFTFPNR | 12 | Q9UBB6 | Neurochondrin OS=Homo sapiens OX=9 | NCDN | 78.865 | 2 | 41.987804 | yes | 7.78E-07 | 692.35645 | 36.571056 |
| IVIYIDR | 7 | Q9UBB9 | Tuftelin-interacting protein 11 OS=Homo | TFIP11 | 96.82 | 2 | 53.998543 | yes | 9.722E-08 | 446.26855 | 352.26126 |
| GIGVAASSVPMNF | 14 | Q9UBB9 | Tuftelin-interacting protein 11 OS=Homo | TFIP11 | 96.82 | 2 | 53.998543 | yes | 9.722E-08 | 689.3634 | 314.67294 |
| DYSAPVNFISAGLI | 14 | Q9UBB9 | Tuftelin-interacting protein 11 OS=Homo | TFIP11 | 96.82 | 2 | 53.998543 | yes | 2.01E-07 | 741.38538 | 309.17841 |
| SLSHLYR | 7 | Q9UBB9 | Tuftelin-interacting protein 11 OS=Homo | TFIP11 | 96.82 | 2 | 53.998543 | yes | 1.128E-07 | 459.24561 | 291.45572 |
| LEPLYSPIR | 9 | Q9UBB9 | Tuftelin-interacting protein 11 OS=Homo | TFIP11 | 96.82 | 2 | 53.998543 | yes | 9.722E-08 | 544.31097 | 284.17261 |
| NAQGIINPIEAK | 12 | Q9UBB9 | Tuftelin-interacting protein 11 OS=Homo | TFIP11 | 96.82 | 2 | 53.998543 | yes | 1.026E-07 | 634.35388 | 188.99654 |
| NIVTQWQPR | 9 | Q9UBB9 | Tuftelin-interacting protein 11 OS=Homo | TFIP11 | 96.82 | 2 | 53.998543 | yes | 8.374E-07 | 571.30927 | 172.14323 |
| SFMDFGSWER | 10 | Q9UBB9 | Tuftelin-interacting protein 11 OS=Homo | TFIP11 | 96.82 | 2 | 53.998543 | yes | 9.722E-08 | 631.26898 | 150.65517 |
| AVSSNVGAYMQP | 15 | Q9UBB9 | Tuftelin-interacting protein 11 OS=Homo | TFIP11 | 96.82 | 2 | 53.998543 | yes | 9.796E-08 | 754.36975 | 143.11404 |
| SMFSDQVLAHPSV | 14 | Q9UBB9 | Tuftelin-interacting protein 11 OS=Homo | TFIP11 | 96.82 | 2 | 53.998543 | yes | 1.575E-07 | 773.39014 | 135.92754 |
| IFETLQDK | 8 | Q9UBB9 | Tuftelin-interacting protein 11 OS=Homo | TFIP11 | 96.82 | 2 | 53.998543 | yes | 3.926E-06 | 497.2662 | 108.50383 |
| MTEVLDHEER | 10 | Q9UBB9 | Tuftelin-interacting protein 11 OS=Homo | TFIP11 | 96.82 | 2 | 53.998543 | yes | 5.015E-07 | 629.79022 | 92.171539 |
| AEEHNIVFMPVIGI | 14 | Q9UBB9 | Tuftelin-interacting protein 11 OS=Homo | TFIP11 | 96.82 | 3 | 53.998543 | yes | 1.543E-07 | 528.61322 | 42.433258 |
| AEEHNIVFMPVIGI | 14 | Q9UBB9 | Tuftelin-interacting protein 11 OS=Homo | TFIP11 | 96.82 | 3 | 53.998543 | yes | 3.674E-07 | 792.41614 | 42.433258 |
| AEEHNIVFMPVIGI | 14 | Q9UBB9 | Tuftelin-interacting protein 11 OS=Homo | TFIP11 | 96.82 | 2 | 53.998543 | yes | 1.543E-07 | 528.61322 | 42.433258 |
| AEEHNIVFMPVIGI | 14 | Q9UBB9 | Tuftelin-interacting protein 11 OS=Homo | TFIP11 | 96.82 | 2 | 53.998543 | yes | 3.674E-07 | 792.41614 | 42.433258 |
| DKFNEALDIMNR | 12 | Q9UBB9 | Tuftelin-interacting protein 11 OS=Homo | TFIP11 | 96.82 | 2 | 53.998543 | yes | 9.722E-08 | 733.35883 | 84.008369 |
| DFQYEAMQER | 10 | Q9UBB9 | Tuftelin-interacting protein 11 OS=Homo | TFIP11 | 96.82 | 2 | 53.998543 | yes | 9.81E-08 | 658.78241 | 65.766197 |
| SLLENDQLLSHGG | 23 | Q9UBB9 | Tuftelin-interacting protein 11 OS=Homo | TFIP11 | 96.82 | 3 | 53.998543 | yes | 1.56E-07 | 841.74561 | 58.621857 |
| VLEMVEECER | 10 | Q9UBB9 | Tuftelin-interacting protein 11 OS=Homo | TFIP11 | 96.82 | 2 | 53.998543 | yes | 2.331E-07 | 647.29431 | 56.146721 |
| QLYTFGR | 7 | Q9UBB9 | Tuftelin-interacting protein 11 OS=Homo | TFIP11 | 96.82 | 2 | 53.998543 | yes | 2.995E-07 | 442.73486 | 51.901886 |
| VYYSYSQISHK | 11 | Q9UBB9 | Tuftelin-interacting protein 11 OS=Homo | TFIP11 | 96.82 | 2 | 53.998543 | yes | 2.558E-05 | 687.83807 | 47.968376 |
| DMVVNLFHELEK | 12 | Q9UBB9 | Tuftelin-interacting protein 11 OS=Homo | TFIP11 | 96.82 | 2 | 53.998543 | yes | 0.0001134 | 737.37396 | 33.08107 |
| ETSSLQELEAQK | 12 | Q9UBC2 | Epidermal growth factor receptor substrat | EPS15L1 | 94.256 | 2 | 50.06337 | yes | 6.529E-07 | 681.841 | 74.018036 |
| NDPFTSDPFTK | 11 | Q9UBC2 | Epidermal growth factor receptor substrat | EPS15L1 | 94.256 | 2 | 50.06337 | yes | 1.385E-07 | 634.79333 | 53.00779 |
| VGASEAALFLK | 11 | Q9UBC2 | Epidermal growth factor receptor substrat | EPS15L1 | 94.256 | 2 | 50.06337 | yes | 9.919E-08 | 553.31622 | 48.271755 |
| ELDDISQEIAQLQR | 14 | Q9UBC2 | Epidermal growth factor receptor substrat | EPS15L1 | 94.256 | 2 | 50.06337 | yes | 4.172E-07 | 829.42322 | 42.454857 |
| LISQLMDCCVDIK | 13 | Q9UBD5 | Origin recognition complex subunit 3 OS | ORC3 | 82.255 | 2 | 48.285744 | yes | 1.167E-07 | 797.8855 | 90.615868 |
| EHLTTVLDK | 9 | Q9UBD5 | Origin recognition complex subunit 3 OS | ORC3 | 82.255 | 2 | 48.285744 | yes | 5.897E-06 | 528.29022 | 64.941994 |
| TDLYHLQK | 8 | Q9UBD5 | Origin recognition complex subunit 3 OS | ORC3 | 82.255 | 2 | 48.285744 | yes | 4.865E-06 | 509.27182 | 43.102112 |
| SYCENHLGSTAK | 12 | Q9UBD5 | Origin recognition complex subunit 3 OS | ORC3 | 82.255 | 2 | 48.285744 | yes | 1.379E-07 | 683.80646 | 31.548317 |
| INFLSNNQCENIR | 13 | Q9UBD5 | Origin recognition complex subunit 3 OS | ORC3 | 82.255 | 2 | 48.285744 | yes | 3.799E-07 | 811.39117 | 29.743795 |
| AVSELELLGFIKPT | 15 | Q9UBD5 | Origin recognition complex subunit 3 OS | ORC3 | 82.255 | 2 | 48.285744 | yes | 2.252E-07 | 822.98218 | 15.123428 |
| ATFYLNVLQQR | 11 | Q9UBF2 | Coatomer subunit gamma-2 OS=Homo sa | COPG2 | 97.623 | 2 | 39.10982 | yes | 4.784E-07 | 676.86969 | 60.058216 |

| | | | | | | | | | | | |
|---|---|---|---|---|---|---|---|---|---|---|---|
| CPLCQQDWVVQR | 12 | Q9UBF6 | RING-box protein 2 OS=Homo sapiens C | RNF7 | 12.684 | 2 | 49.743744 | yes | 1.006E-07 | 794.87158 | 46.58849 |
| ASADLMSYCEEHA | 14 | Q9UBI6 | Guanine nucleotide-binding protein G(I)/( | GNG12 | 8.007 | 2 | 47.20794 | yes | 1.377E-07 | 820.34558 | 29.222366 |
| YETFISDVLQR | 11 | Q9UBK9 | Protein UXT OS=Homo sapiens OX=960 | UXT | 18.247 | 2 | 54.574223 | yes | 9.722E-08 | 685.8512 | 89.547577 |
| VLLALHDR | 8 | Q9UBL3 | Set1/Ash2 histone methyltransferase com | ASH2L | 68.723 | 2 | 50.420815 | yes | 9.809E-08 | 468.78491 | 70.916626 |
| YTAAVPYR | 8 | Q9UBM7 | 7-dehydrocholesterol reductase OS=Hom | DHCR7 | 54.49 | 2 | 42.221756 | yes | 1.021E-06 | 470.74799 | 45.611004 |
| VIECSYTSADGQR | 13 | Q9UBM7 | 7-dehydrocholesterol reductase OS=Hom | DHCR7 | 54.49 | 2 | 42.221756 | yes | 3.135E-07 | 743.33557 | 35.288277 |
| VYVSLPNEETR | 11 | Q9UBP0 | Spastin OS=Homo sapiens OX=9606 GN | SPAST | 67.198 | 2 | 52.60295 | yes | 3.296E-07 | 653.83551 | 209.95149 |
| FDDIAGQDLAK | 11 | Q9UBP0 | Spastin OS=Homo sapiens OX=9606 GN | SPAST | 67.198 | 2 | 52.60295 | yes | 2.143E-07 | 596.79584 | 193.05086 |
| SVSPQTLEAYIR | 12 | Q9UBP0 | Spastin OS=Homo sapiens OX=9606 GN | SPAST | 67.198 | 2 | 52.60295 | yes | 1.647E-07 | 682.36444 | 183.15002 |
| TNKPSTPTTATR | 12 | Q9UBP0 | Spastin OS=Homo sapiens OX=9606 GN | SPAST | 67.198 | 2 | 52.60295 | yes | 1.527E-07 | 637.83856 | 153.09418 |
| GIAVIVTGQGEQCI | 15 | Q9UBP0 | Spastin OS=Homo sapiens OX=9606 GN | SPAST | 67.198 | 2 | 52.60295 | yes | 1.018E-07 | 808.90668 | 141.84181 |
| MTDGYSGSDLTAI | 15 | Q9UBP0 | Spastin OS=Homo sapiens OX=9606 GN | SPAST | 67.198 | 2 | 52.60295 | yes | 9.903E-08 | 765.36127 | 139.39319 |
| ELKPEQVK | 8 | Q9UBP0 | Spastin OS=Homo sapiens OX=9606 GN | SPAST | 67.198 | 2 | 52.60295 | yes | 9.722E-08 | 485.78201 | 130.68152 |
| APSYSGLSMVSGV | 14 | Q9UBP0 | Spastin OS=Homo sapiens OX=9606 GN | SPAST | 67.198 | 2 | 52.60295 | yes | 2.669E-07 | 691.85284 | 117.1879 |
| QAFEYISIALR | 11 | Q9UBP0 | Spastin OS=Homo sapiens OX=9606 GN | SPAST | 67.198 | 2 | 52.60295 | yes | 2.518E-07 | 655.85876 | 115.26969 |
| TLAGDVHIVR | 10 | Q9UBQ0 | Vacuolar protein sorting-associated protei | VPS29 | 20.506 | 2 | 24.487799 | yes | 3.445E-06 | 540.81165 | 26.577213 |
| YNPENLATLER | 11 | Q9UBQ5 | Eukaryotic translation initiation factor 3 s | EIF3K | 25.06 | 2 | 51.06208 | yes | 1.065E-07 | 660.33313 | 3464.7893 |
| ALTNLPHTDFTLCI | 14 | Q9UBQ5 | Eukaryotic translation initiation factor 3 s | EIF3K | 25.06 | 2 | 51.06208 | yes | 1.512E-07 | 815.9165 | 1204.0881 |
| ENAYDLEANLAVI | 14 | Q9UBQ5 | Eukaryotic translation initiation factor 3 s | EIF3K | 25.06 | 2 | 51.06208 | yes | 9.722E-08 | 781.90668 | 1139.1702 |
| FICHVVGITYQHID | 15 | Q9UBQ5 | Eukaryotic translation initiation factor 3 s | EIF3K | 25.06 | 3 | 51.06208 | yes | 1.022E-07 | 619.98578 | 225.99069 |
| FICHVVGITYQHID | 15 | Q9UBQ5 | Eukaryotic translation initiation factor 3 s | EIF3K | 25.06 | 3 | 51.06208 | yes | 2.964E-07 | 465.24118 | 225.99069 |
| FICHVVGITYQHID | 15 | Q9UBQ5 | Eukaryotic translation initiation factor 3 s | EIF3K | 25.06 | 4 | 51.06208 | yes | 1.022E-07 | 619.98578 | 225.99069 |
| FICHVVGITYQHID | 15 | Q9UBQ5 | Eukaryotic translation initiation factor 3 s | EIF3K | 25.06 | 4 | 51.06208 | yes | 2.964E-07 | 465.24118 | 225.99069 |
| VIVVWNNIGEK | 11 | Q9UBQ6 | Exostosin-like 2 OS=Homo sapiens OX=( | EXTL2 | 37.466 | 2 | 43.841732 | yes | 2.424E-07 | 635.86133 | 44.182411 |
| NVDGVNYASITR | 12 | Q9UBR2 | Cathepsin Z OS=Homo sapiens OX=960( | CTSZ | 33.868 | 2 | 45.18782 | yes | 1.931E-07 | 654.83075 | 703.12885 |
| VGDYGSLSGR | 10 | Q9UBR2 | Cathepsin Z OS=Homo sapiens OX=960( | CTSZ | 33.868 | 2 | 45.18782 | yes | 2.087E-07 | 505.74872 | 473.93585 |
| STYPRPHEYLSPAI | 17 | Q9UBR2 | Cathepsin Z OS=Homo sapiens OX=960( | CTSZ | 33.868 | 3 | 45.18782 | yes | 1.508E-06 | 657.67053 | 58.652485 |
| NSWGEPWGER | 10 | Q9UBR2 | Cathepsin Z OS=Homo sapiens OX=960( | CTSZ | 33.868 | 2 | 45.18782 | yes | 7.755E-07 | 609.27032 | 20.374153 |
| VVQVWFQNR | 9 | Q9UBR4 | LIM/homeobox protein Lhx3 OS=Homo | LHX3 | 43.358 | 2 | 50.243633 | no | 9.838E-08 | 588.31964 | 143.00603 |
| QRWGQYFR | 8 | Q9UBR4 | LIM/homeobox protein Lhx3 OS=Homo | LHX3 | 43.358 | 2 | 50.243633 | yes | 2.202E-06 | 570.78851 | 53.114075 |
| DLKPENIMLSSQGI | 16 | Q9UBS0 | Ribosomal protein S6 kinase beta-2 OS=I | RPS6KB2 | 53.456 | 3 | 46.175797 | yes | 1.63E-07 | 603.98907 | 65.715431 |
| TLEVEIEPGVR | 11 | Q9UBS4 | DnaJ homolog subfamily B member 11 O | DNAJB11 | 40.514 | 2 | 49.55759 | yes | 1.021E-07 | 621.34045 | 962.23718 |
| LALQLHPDR | 9 | Q9UBS4 | DnaJ homolog subfamily B member 11 O | DNAJB11 | 40.514 | 2 | 49.55759 | yes | 2.047E-07 | 531.80634 | 195.50172 |
| QYDTYGEEGLK | 11 | Q9UBS4 | DnaJ homolog subfamily B member 11 O | DNAJB11 | 40.514 | 2 | 49.55759 | yes | 1.033E-07 | 651.79608 | 106.79372 |
| FAMELEQSR | 9 | Q9UBT6 | DNA polymerase kappa OS=Homo sapie | POLK | 98.809 | 2 | 52.625458 | yes | 1.362E-07 | 555.76605 | 163.31117 |
| TTLTASAGIAPNTM | 17 | Q9UBT6 | DNA polymerase kappa OS=Homo sapie | POLK | 98.809 | 2 | 52.625458 | yes | 9.722E-08 | 830.95056 | 149.43355 |
| ALGIITCTELYQQR | 14 | Q9UBT6 | DNA polymerase kappa OS=Homo sapie | POLK | 98.809 | 2 | 52.625458 | yes | 9.722E-08 | 833.43506 | 124.01656 |
| AQGCISLEALNK | 12 | Q9UBT6 | DNA polymerase kappa OS=Homo sapie | POLK | 98.809 | 2 | 52.625458 | yes | 2.737E-07 | 652.33734 | 99.500366 |
| HTLDIFFK | 8 | Q9UBT6 | DNA polymerase kappa OS=Homo sapie | POLK | 98.809 | 2 | 52.625458 | yes | 5.744E-07 | 510.7793 | 94.919044 |

| | | | | | | | | | | | |
|---|---|---|---|---|---|---|---|---|---|---|---|
| QSFQTSQPFQVLK | 13 | Q9UBT6 | DNA polymerase kappa OS=Homo sapie | POLK | 98.809 | 2 | 52.625458 | yes | 5.464E-07 | 769.40411 | 54.411072 |
| TEFITQEENR | 10 | Q9UBU7 | Protein DBF4 homolog A OS=Homo sapi | DBF4 | 76.858 | 2 | 52.002117 | yes | 9.722E-08 | 633.8017 | 154.54524 |
| CSMLSTAEDDIR | 12 | Q9UBU7 | Protein DBF4 homolog A OS=Homo sapi | DBF4 | 76.858 | 2 | 52.002117 | yes | 9.722E-08 | 699.30542 | 137.9771 |
| VEDMSQLYR | 9 | Q9UBU7 | Protein DBF4 homolog A OS=Homo sapi | DBF4 | 76.858 | 2 | 52.002117 | yes | 3.206E-07 | 570.7713 | 130.61121 |
| ENLEPNAEFDKR | 12 | Q9UBU7 | Protein DBF4 homolog A OS=Homo sapi | DBF4 | 76.858 | 2 | 52.002117 | yes | 9.722E-08 | 731.35205 | 93.351677 |
| YSVGSLSPVSASVI | 15 | Q9UBU7 | Protein DBF4 homolog A OS=Homo sapi | DBF4 | 76.858 | 2 | 52.002117 | yes | 9.722E-08 | 747.41412 | 84.264374 |
| GHFQGGIQVK | 10 | Q9UBU7 | Protein DBF4 homolog A OS=Homo sapi | DBF4 | 76.858 | 2 | 52.002117 | yes | 5.095E-07 | 535.79071 | 46.837128 |
| LVFDFVEYEK | 10 | Q9UBU7 | Protein DBF4 homolog A OS=Homo sapi | DBF4 | 76.858 | 2 | 52.002117 | yes | 3.456E-07 | 644.8266 | 39.192928 |
| VFYLDLPSVTISEK | 14 | Q9UBU7 | Protein DBF4 homolog A OS=Homo sapi | DBF4 | 76.858 | 2 | 52.002117 | yes | 4.086E-07 | 805.93744 | 28.890621 |
| GYCECCLQK | 9 | Q9UBU7 | Protein DBF4 homolog A OS=Homo sapi | DBF4 | 76.858 | 2 | 52.002117 | yes | 8.528E-07 | 609.24103 | 18.465853 |
| VDPTVENEETFMN | 14 | Q9UBU8 | Mortality factor 4-like protein 1 OS=Hon | MORF4L1 | 41.474 | 2 | 50.281174 | yes | 1.517E-07 | 840.88031 | 475.45245 |
| NVDSILEDYANYK | 13 | Q9UBU8 | Mortality factor 4-like protein 1 OS=Hon | MORF4L1 | 41.474 | 2 | 50.281174 | yes | 1.06E-07 | 772.36737 | 223.29962 |
| VLCFHGPLLYEAK | 13 | Q9UBU8 | Mortality factor 4-like protein 1 OS=Hon | MORF4L1 | 41.474 | 2 | 50.281174 | yes | 9.819E-08 | 773.90796 | 176.97015 |
| QLFYLPAK | 8 | Q9UBU8 | Mortality factor 4-like protein 1 OS=Hon | MORF4L1 | 41.474 | 2 | 50.281174 | yes | 1.736E-07 | 490.28421 | 156.0657 |
| ANQEQYAEGK | 10 | Q9UBU8 | Mortality factor 4-like protein 1 OS=Hon | MORF4L1 | 41.474 | 2 | 50.281174 | yes | 8.529E-07 | 569.26221 | 99.129868 |
| FERPQYAEILADHI | 29 | Q9UBU8 | Mortality factor 4-like protein 1 OS=Hon | MORF4L1 | 41.474 | 4 | 50.281174 | yes | 3.61E-07 | 831.1673 | 72.467155 |
| NSATLFSASDYEV. | 20 | Q9UBU8 | Mortality factor 4-like protein 1 OS=Hon | MORF4L1 | 41.474 | 3 | 50.281174 | yes | 2.619E-06 | 752.01855 | 10.415089 |
| GSYFGTENLK | 10 | Q9UBU9 | Nuclear RNA export factor 1 OS=Homo s | NXF1 | 70.183 | 2 | 50.435905 | yes | 1.285E-07 | 558.27203 | 1054.4926 |
| LDDMSSIVQK | 10 | Q9UBU9 | Nuclear RNA export factor 1 OS=Homo s | NXF1 | 70.183 | 2 | 50.435905 | yes | 1.426E-07 | 568.2868 | 1020.653 |
| AQFFVEDASTASA | 15 | Q9UBU9 | Nuclear RNA export factor 1 OS=Homo s | NXF1 | 70.183 | 2 | 50.435905 | yes | 1.078E-07 | 792.89886 | 693.45563 |
| SAQAFTHLK | 9 | Q9UBU9 | Nuclear RNA export factor 1 OS=Homo s | NXF1 | 70.183 | 2 | 50.435905 | yes | 1.861E-07 | 501.772 | 651.94519 |
| NASSEEIQR | 9 | Q9UBU9 | Nuclear RNA export factor 1 OS=Homo s | NXF1 | 70.183 | 2 | 50.435905 | yes | 1.113E-07 | 517.24908 | 637.95612 |
| DQSTYISAIR | 10 | Q9UBU9 | Nuclear RNA export factor 1 OS=Homo s | NXF1 | 70.183 | 2 | 50.435905 | yes | 1.354E-07 | 577.29602 | 610.68848 |
| ITIPYGR | 7 | Q9UBU9 | Nuclear RNA export factor 1 OS=Homo s | NXF1 | 70.183 | 2 | 50.435905 | yes | 1.004E-07 | 410.23981 | 591.86938 |
| GEIPEVAFMK | 10 | Q9UBU9 | Nuclear RNA export factor 1 OS=Homo s | NXF1 | 70.183 | 2 | 50.435905 | yes | 3.959E-07 | 560.789 | 306.14694 |
| ILNLSGNELK | 10 | Q9UBU9 | Nuclear RNA export factor 1 OS=Homo s | NXF1 | 70.183 | 2 | 50.435905 | yes | 5.35E-07 | 550.81915 | 290.29834 |
| AKGEIPEVAFMK | 12 | Q9UBU9 | Nuclear RNA export factor 1 OS=Homo s | NXF1 | 70.183 | 2 | 50.435905 | yes | 1.44E-07 | 660.35504 | 288.54364 |
| AWLLSMIQSK | 10 | Q9UBU9 | Nuclear RNA export factor 1 OS=Homo s | NXF1 | 70.183 | 2 | 50.435905 | yes | 1.004E-07 | 588.82593 | 221.70042 |
| CSVPFTPIEFHYEN | 16 | Q9UBU9 | Nuclear RNA export factor 1 OS=Homo s | NXF1 | 70.183 | 3 | 50.435905 | yes | 9.809E-08 | 666.31226 | 188.85651 |
| RYDGSQQALDLK | 12 | Q9UBU9 | Nuclear RNA export factor 1 OS=Homo s | NXF1 | 70.183 | 2 | 50.435905 | yes | 9.963E-08 | 697.35718 | 181.6282 |
| LKDPTLR | 7 | Q9UBU9 | Nuclear RNA export factor 1 OS=Homo s | NXF1 | 70.183 | 2 | 50.435905 | yes | 2.124E-06 | 421.75836 | 157.10399 |
| CLQDNNWDYTR | 11 | Q9UBU9 | Nuclear RNA export factor 1 OS=Homo s | NXF1 | 70.183 | 2 | 50.435905 | yes | 1.734E-07 | 742.81482 | 151.21419 |
| SCMAATLR | 8 | Q9UBU9 | Nuclear RNA export factor 1 OS=Homo s | NXF1 | 70.183 | 2 | 50.435905 | yes | 4.297E-07 | 455.21768 | 146.34827 |
| YDGSQQALDLK | 11 | Q9UBU9 | Nuclear RNA export factor 1 OS=Homo s | NXF1 | 70.183 | 2 | 50.435905 | yes | 2.077E-05 | 619.30658 | 115.23338 |
| SSLAEYFK | 8 | Q9UBU9 | Nuclear RNA export factor 1 OS=Homo s | NXF1 | 70.183 | 2 | 50.435905 | yes | 2.017E-06 | 472.73984 | 54.914494 |
| AAQLPWEDGR | 10 | Q9UBV7 | Beta-1,4-galactosyltransferase 7 OS=Hon | B4GALT7 | 37.406 | 2 | 46.76587 | yes | 1.475E-07 | 571.78308 | 84.950935 |
| PSGITTGYK | 9 | Q9UBV7 | Beta-1,4-galactosyltransferase 7 OS=Hon | B4GALT7 | 37.406 | 2 | 46.76587 | yes | 3.683E-07 | 462.24527 | 37.251297 |
| LSFEDFVTMTASR | 13 | Q9UBV8 | Peflin OS=Homo sapiens OX=9606 GN= | PEF1 | 30.381 | 2 | 50.81941 | yes | 9.722E-08 | 752.36102 | 237.78462 |
| SANPAMQLDR | 10 | Q9UBV8 | Peflin OS=Homo sapiens OX=9606 GN= | PEF1 | 30.381 | 2 | 50.81941 | yes | 2.314E-07 | 551.7691 | 233.71616 |

| | | | | | | | | | | | |
|---|---|---|---|---|---|---|---|---|---|---|---|
| NLFQQYDR | 8 | Q9UBV8 | Peflin OS=Homo sapiens OX=9606 GN= | PEF1 | 30.381 | 2 | 50.81941 | yes | 1.812E-06 | 542.26453 | 217.84534 |
| IDVYGFSALWK | 11 | Q9UBV8 | Peflin OS=Homo sapiens OX=9606 GN= | PEF1 | 30.381 | 2 | 50.81941 | yes | 1.986E-07 | 649.84259 | 48.025154 |
| EKDTAVQGNIR | 11 | Q9UBV8 | Peflin OS=Homo sapiens OX=9606 GN= | PEF1 | 30.381 | 2 | 50.81941 | yes | 3.233E-07 | 615.8255 | 43.368416 |
| VTTAAAAAATSQI | 22 | Q9UBW8 | COP9 signalosome complex subunit 7a O | COPS7A | 30.277 | 3 | 46.174614 | yes | 1.63E-07 | 761.05212 | 31.576151 |
| LFSGATMASSR | 11 | Q9UBX3 | Mitochondrial dicarboxylate carrier OS=I | SLC25A10 | 31.283 | 2 | 50.45693 | yes | 1.24E-07 | 564.2793 | 1183.5164 |
| GSQGPLPFHEK | 11 | Q9UBX3 | Mitochondrial dicarboxylate carrier OS=I | SLC25A10 | 31.283 | 2 | 50.45693 | yes | 9.809E-08 | 598.80658 | 1162.3179 |
| FAIYETVR | 8 | Q9UBX3 | Mitochondrial dicarboxylate carrier OS=I | SLC25A10 | 31.283 | 2 | 50.45693 | yes | 1.05E-07 | 499.76892 | 1146.4967 |
| VHLQTQQEVK | 10 | Q9UBX3 | Mitochondrial dicarboxylate carrier OS=I | SLC25A10 | 31.283 | 2 | 50.45693 | yes | 9.823E-08 | 605.33295 | 745.31567 |
| WYFGGLASCGAA | 22 | Q9UBX3 | Mitochondrial dicarboxylate carrier OS=I | SLC25A10 | 31.283 | 3 | 50.45693 | yes | 1.568E-06 | 833.05353 | 55.207066 |
| QMTYSLTR | 8 | Q9UBX3 | Mitochondrial dicarboxylate carrier OS=I | SLC25A10 | 31.283 | 2 | 50.45693 | yes | 4.213E-06 | 500.25003 | 50.297035 |
| DQPFTILYR | 9 | Q9UBX5 | Fibulin-5 OS=Homo sapiens OX=9606 G | FBLN5 | 50.181 | 2 | 48.607796 | yes | 1.123E-07 | 576.80603 | 67.029999 |
| QSGQCLDIDECR | 12 | Q9UBX5 | Fibulin-5 OS=Homo sapiens OX=9606 G | FBLN5 | 50.181 | 2 | 48.607796 | yes | 8.698E-07 | 740.8114 | 23.434198 |
| FLPEFAGTQLK | 11 | Q9UBZ4 | DNA-(apurinic or apyrimidinic site) endo | APEX2 | 57.401 | 2 | 47.33926 | yes | 1.414E-07 | 625.84259 | 433.35831 |
| LDYVLGDR | 8 | Q9UBZ4 | DNA-(apurinic or apyrimidinic site) endo | APEX2 | 57.401 | 2 | 47.33926 | yes | 1.351E-07 | 475.75073 | 193.24521 |
| AEALLAAGSHVIIL | 22 | Q9UBZ4 | DNA-(apurinic or apyrimidinic site) endo | APEX2 | 57.401 | 3 | 47.33926 | yes | 7.724E-07 | 748.0816 | 87.785126 |
| SVLAGPLR | 8 | Q9UBZ4 | DNA-(apurinic or apyrimidinic site) endo | APEX2 | 57.401 | 2 | 47.33926 | yes | 7.376E-07 | 406.75308 | 68.752182 |
| CNFFLWSRPS | 10 | Q9UBZ4 | DNA-(apurinic or apyrimidinic site) endo | APEX2 | 57.401 | 2 | 47.33926 | yes | 2.447E-06 | 657.30841 | 40.15419 |
| QEGAFTCWSAVT | 15 | Q9UBZ4 | DNA-(apurinic or apyrimidinic site) endo | APEX2 | 57.401 | 2 | 47.33926 | yes | 3.41E-07 | 820.87793 | 23.691891 |
| DLQELSIQVEEER | 13 | Q9UDW3 | Zinc finger matrin-type protein 5 OS=Ho | ZMAT5 | 19.971 | 2 | 42.241623 | yes | 3.126E-07 | 794.39667 | 64.000694 |
| LANELPDWFQTAI | 13 | Q9UDY2 | Tight junction protein ZO-2 OS=Homo sa | TJP2 | 133.959 | 2 | 54.166264 | yes | 1.885E-07 | 766.89081 | 2531.5383 |
| AFEVMDEFDGR | 11 | Q9UDY2 | Tight junction protein ZO-2 OS=Homo sa | TJP2 | 133.959 | 2 | 54.166264 | yes | 9.85E-08 | 658.28479 | 1633.561 |
| GEMVTILAQSR | 11 | Q9UDY2 | Tight junction protein ZO-2 OS=Homo sa | TJP2 | 133.959 | 2 | 54.166264 | yes | 9.722E-08 | 602.82135 | 1129.386 |
| MQELQEAQNAR | 11 | Q9UDY2 | Tight junction protein ZO-2 OS=Homo sa | TJP2 | 133.959 | 2 | 54.166264 | yes | 9.85E-08 | 659.31445 | 1083.6132 |
| ETPQSLAFTR | 10 | Q9UDY2 | Tight junction protein ZO-2 OS=Homo sa | TJP2 | 133.959 | 2 | 54.166264 | yes | 1.606E-07 | 575.29858 | 1022.1829 |
| GLDHDFGPSR | 10 | Q9UDY2 | Tight junction protein ZO-2 OS=Homo sa | TJP2 | 133.959 | 2 | 54.166264 | yes | 9.728E-08 | 550.75958 | 1013.9658 |
| MEGMDDDPEDR | 11 | Q9UDY2 | Tight junction protein ZO-2 OS=Homo sa | TJP2 | 133.959 | 2 | 54.166264 | yes | 1.113E-07 | 655.23718 | 906.83514 |
| DGNLHEGDIILK | 12 | Q9UDY2 | Tight junction protein ZO-2 OS=Homo sa | TJP2 | 133.959 | 2 | 54.166264 | yes | 9.722E-08 | 662.34882 | 822.71387 |
| HPDIYAVPIK | 10 | Q9UDY2 | Tight junction protein ZO-2 OS=Homo sa | TJP2 | 133.959 | 2 | 54.166264 | yes | 1.332E-07 | 576.82422 | 779.64551 |
| LQLVVLR | 7 | Q9UDY2 | Tight junction protein ZO-2 OS=Homo sa | TJP2 | 133.959 | 2 | 54.166264 | yes | 1.011E-07 | 420.78693 | 777.43311 |
| GFGIAVSGGR | 10 | Q9UDY2 | Tight junction protein ZO-2 OS=Homo sa | TJP2 | 133.959 | 2 | 54.166264 | yes | 3.307E-07 | 460.75107 | 717.59644 |
| SIDQDYER | 8 | Q9UDY2 | Tight junction protein ZO-2 OS=Homo sa | TJP2 | 133.959 | 2 | 54.166264 | yes | 1.079E-07 | 513.23035 | 574.93347 |
| STGDIAGTVVPETI | 15 | Q9UDY2 | Tight junction protein ZO-2 OS=Homo sa | TJP2 | 133.959 | 2 | 54.166264 | yes | 1.116E-07 | 744.88068 | 549.05774 |
| LGSQIFVK | 8 | Q9UDY2 | Tight junction protein ZO-2 OS=Homo sa | TJP2 | 133.959 | 2 | 54.166264 | yes | 1.388E-07 | 446.26855 | 502.70258 |
| AYDPDYER | 8 | Q9UDY2 | Tight junction protein ZO-2 OS=Homo sa | TJP2 | 133.959 | 2 | 54.166264 | yes | 1.638E-07 | 514.7196 | 418.7511 |
| GRPGPIGVLLMK | 12 | Q9UDY2 | Tight junction protein ZO-2 OS=Homo sa | TJP2 | 133.959 | 2 | 54.166264 | yes | 1.252E-07 | 619.3761 | 179.90245 |
| GRPGPIGVLLMK | 12 | Q9UDY2 | Tight junction protein ZO-2 OS=Homo sa | TJP2 | 133.959 | 2 | 54.166264 | yes | 2.224E-07 | 413.25317 | 179.90245 |
| GRPGPIGVLLMK | 12 | Q9UDY2 | Tight junction protein ZO-2 OS=Homo sa | TJP2 | 133.959 | 3 | 54.166264 | yes | 1.252E-07 | 619.3761 | 179.90245 |
| GRPGPIGVLLMK | 12 | Q9UDY2 | Tight junction protein ZO-2 OS=Homo sa | TJP2 | 133.959 | 3 | 54.166264 | yes | 2.224E-07 | 413.25317 | 179.90245 |
| VVDTLYDGK | 9 | Q9UDY2 | Tight junction protein ZO-2 OS=Homo sa | TJP2 | 133.959 | 2 | 54.166264 | yes | 5.703E-07 | 505.26367 | 350.39655 |

| | | | | | | | | | | | |
|---|---|---|---|---|---|---|---|---|---|---|---|
| THKPDPGTPQHTS | 21 | Q9UDY2 | Tight junction protein ZO-2 OS=Homo sa | TJP2 | 133.959 | 4 | 54.166264 | yes | 9.722E-08 | 581.2951 | 140.41135 |
| THKPDPGTPQHTS | 21 | Q9UDY2 | Tight junction protein ZO-2 OS=Homo sa | TJP2 | 133.959 | 4 | 54.166264 | yes | 4.854E-07 | 774.72437 | 140.41135 |
| THKPDPGTPQHTS | 21 | Q9UDY2 | Tight junction protein ZO-2 OS=Homo sa | TJP2 | 133.959 | 3 | 54.166264 | yes | 9.722E-08 | 581.2951 | 140.41135 |
| THKPDPGTPQHTS | 21 | Q9UDY2 | Tight junction protein ZO-2 OS=Homo sa | TJP2 | 133.959 | 3 | 54.166264 | yes | 4.854E-07 | 774.72437 | 140.41135 |
| KLFDQANK | 8 | Q9UDY2 | Tight junction protein ZO-2 OS=Homo sa | TJP2 | 133.959 | 2 | 54.166264 | yes | 5.833E-07 | 482.26654 | 280.50122 |
| GSYGSDAEEEEYR | 13 | Q9UDY2 | Tight junction protein ZO-2 OS=Homo sa | TJP2 | 133.959 | 2 | 54.166264 | yes | 1.023E-07 | 746.29712 | 275.67554 |
| HALLDVTPK | 9 | Q9UDY2 | Tight junction protein ZO-2 OS=Homo sa | TJP2 | 133.959 | 2 | 54.166264 | yes | 1.781E-07 | 497.29001 | 273.27335 |
| STGDIAGTVVPETI | 18 | Q9UDY2 | Tight junction protein ZO-2 OS=Homo sa | TJP2 | 133.959 | 3 | 54.166264 | yes | 1.006E-07 | 624.32166 | 247.68864 |
| GLDQDHAR | 8 | Q9UDY2 | Tight junction protein ZO-2 OS=Homo sa | TJP2 | 133.959 | 2 | 54.166264 | yes | 1.118E-07 | 456.22012 | 216.94754 |
| QIIEQDK | 7 | Q9UDY2 | Tight junction protein ZO-2 OS=Homo sa | TJP2 | 133.959 | 2 | 54.166264 | yes | 1.498E-07 | 437.23746 | 175.72609 |
| YQEDPPAPQPK | 11 | Q9UDY2 | Tight junction protein ZO-2 OS=Homo sa | TJP2 | 133.959 | 2 | 54.166264 | yes | 1.141E-07 | 635.30914 | 162.91379 |
| INGTVTENMSLTD | 15 | Q9UDY2 | Tight junction protein ZO-2 OS=Homo sa | TJP2 | 133.959 | 2 | 54.166264 | yes | 1.151E-07 | 811.39612 | 152.593 |
| AEQMASVQNAQR | 12 | Q9UDY2 | Tight junction protein ZO-2 OS=Homo sa | TJP2 | 133.959 | 2 | 54.166264 | yes | 3.118E-07 | 666.81989 | 127.01573 |
| SSEPVQHEESIR | 12 | Q9UDY2 | Tight junction protein ZO-2 OS=Homo sa | TJP2 | 133.959 | 2 | 54.166264 | yes | 1.764E-07 | 699.33643 | 111.44907 |
| RPVVLFGPIADIAN | 16 | Q9UDY2 | Tight junction protein ZO-2 OS=Homo sa | TJP2 | 133.959 | 3 | 54.166264 | yes | 9.722E-08 | 585.99902 | 98.079887 |
| SNPSAVAGNETPG | 17 | Q9UDY2 | Tight junction protein ZO-2 OS=Homo sa | TJP2 | 133.959 | 2 | 54.166264 | yes | 2.049E-07 | 794.38409 | 94.773445 |
| RGFGIAVSGGR | 11 | Q9UDY2 | Tight junction protein ZO-2 OS=Homo sa | TJP2 | 133.959 | 2 | 54.166264 | yes | 9.864E-07 | 538.80164 | 43.03001 |
| SREDLTAVVSVST | 14 | Q9UDY2 | Tight junction protein ZO-2 OS=Homo sa | TJP2 | 133.959 | 2 | 54.166264 | yes | 3.629E-07 | 746.4043 | 29.795292 |
| TQNKEESYDFSK | 12 | Q9UDY2 | Tight junction protein ZO-2 OS=Homo sa | TJP2 | 133.959 | 2 | 54.166264 | yes | 1.104E-07 | 738.33606 | 15.988995 |
| EIYDQFGEEGLK | 12 | Q9UDY4 | DnaJ homolog subfamily B member 4 OS | DNAJB4 | 37.807 | 2 | 47.753067 | yes | 1.265E-07 | 714.33807 | 321.61188 |
| EVAEAYEVLSDPK | 13 | Q9UDY4 | DnaJ homolog subfamily B member 4 OS | DNAJB4 | 37.807 | 2 | 47.753067 | yes | 7.664E-07 | 725.35901 | 293.75897 |
| LKQDPPVIHELR | 12 | Q9UDY4 | DnaJ homolog subfamily B member 4 OS | DNAJB4 | 37.807 | 3 | 47.753067 | yes | 2.04E-07 | 482.28055 | 142.36494 |
| LKQDPPVIHELR | 12 | Q9UDY4 | DnaJ homolog subfamily B member 4 OS | DNAJB4 | 37.807 | 3 | 47.753067 | yes | 2.47E-07 | 722.91718 | 142.36494 |
| LKQDPPVIHELR | 12 | Q9UDY4 | DnaJ homolog subfamily B member 4 OS | DNAJB4 | 37.807 | 2 | 47.753067 | yes | 2.04E-07 | 482.28055 | 142.36494 |
| LKQDPPVIHELR | 12 | Q9UDY4 | DnaJ homolog subfamily B member 4 OS | DNAJB4 | 37.807 | 2 | 47.753067 | yes | 2.47E-07 | 722.91718 | 142.36494 |
| IIGYGLPFPK | 10 | Q9UDY4 | DnaJ homolog subfamily B member 4 OS | DNAJB4 | 37.807 | 2 | 47.753067 | yes | 2.086E-07 | 552.82623 | 188.27675 |
| ILTIEIK | 7 | Q9UDY4 | DnaJ homolog subfamily B member 4 OS | DNAJB4 | 37.807 | 2 | 47.753067 | yes | 7.924E-05 | 415.27332 | 10.303391 |
| GTPEETGSYLVSK | 13 | Q9UDY8 | Mucosa-associated lymphoid tissue lympl | MALT1 | 92.273 | 2 | 50.411304 | yes | 1.402E-07 | 684.33807 | 229.60272 |
| LSCLDLEQCSLK | 12 | Q9UDY8 | Mucosa-associated lymphoid tissue lympl | MALT1 | 92.273 | 2 | 50.411304 | yes | 1.276E-07 | 733.35492 | 198.12863 |
| LSELLDQAPEGR | 12 | Q9UDY8 | Mucosa-associated lymphoid tissue lympl | MALT1 | 92.273 | 2 | 50.411304 | yes | 1.288E-07 | 664.34625 | 195.44188 |
| NELPLTHETK | 10 | Q9UDY8 | Mucosa-associated lymphoid tissue lympl | MALT1 | 92.273 | 2 | 50.411304 | yes | 1.657E-06 | 591.31171 | 187.07245 |
| SENCLCVQNILK | 12 | Q9UDY8 | Mucosa-associated lymphoid tissue lympl | MALT1 | 92.273 | 2 | 50.411304 | yes | 9.809E-08 | 739.36053 | 169.77396 |
| LQICVEPTSQK | 11 | Q9UDY8 | Mucosa-associated lymphoid tissue lympl | MALT1 | 92.273 | 2 | 50.411304 | yes | 1.143E-07 | 651.83972 | 159.72902 |
| VLEPEGSPSLCLLK | 14 | Q9UDY8 | Mucosa-associated lymphoid tissue lympl | MALT1 | 92.273 | 2 | 50.411304 | yes | 1.791E-07 | 771.41583 | 151.19566 |
| KVEIIIGR | 8 | Q9UDY8 | Mucosa-associated lymphoid tissue lympl | MALT1 | 92.273 | 2 | 50.411304 | yes | 1.197E-07 | 464.30292 | 118.96451 |
| ITVLLDEVAEDMG | 14 | Q9UDY8 | Mucosa-associated lymphoid tissue lympl | MALT1 | 92.273 | 2 | 50.411304 | yes | 1.41E-07 | 766.89746 | 58.304283 |
| ATGHPFVQYQWFI | 13 | Q9UDY8 | Mucosa-associated lymphoid tissue lympl | MALT1 | 92.273 | 2 | 50.411304 | yes | 1.823E-07 | 804.90155 | 55.237915 |
| AHELPESMCLK | 11 | Q9UDY8 | Mucosa-associated lymphoid tissue lympl | MALT1 | 92.273 | 2 | 50.411304 | yes | 3.029E-06 | 657.81268 | 52.606667 |
| LREPLLR | 7 | Q9UDY8 | Mucosa-associated lymphoid tissue lympl | MALT1 | 92.273 | 2 | 50.411304 | yes | 1.2E-05 | 448.78745 | 40.889236 |

| | | | | | | | | | | | |
|---|---|---|---|---|---|---|---|---|---|---|---|
| LAELAGSR | 8 | Q9UDY8 | Mucosa-associated lymphoid tissue lympl | MALT1 | 92.273 | 2 | 50.411304 | yes | 2.336E-06 | 408.73233 | 32.805729 |
| VVSLLDLTEYEMR | 13 | Q9UDY8 | Mucosa-associated lymphoid tissue lympl | MALT1 | 92.273 | 2 | 50.411304 | yes | 1.467E-07 | 784.40546 | 30.403433 |
| QEVNVGKPLIAK | 12 | Q9UDY8 | Mucosa-associated lymphoid tissue lympl | MALT1 | 92.273 | 2 | 50.411304 | yes | 1.731E-07 | 648.3877 | 18.75473 |
| SSSVLLEHLR | 10 | Q9UEG4 | Zinc finger protein 629 OS=Homo sapien | ZNF629 | 96.621 | 2 | 53.174603 | yes | 1.795E-07 | 570.8222 | 168.05769 |
| SFIQSSELTQHQR | 13 | Q9UEG4 | Zinc finger protein 629 OS=Homo sapien | ZNF629 | 96.621 | 2 | 53.174603 | yes | 1.722E-07 | 780.89191 | 85.945084 |
| SFIMSSTLIR | 10 | Q9UEG4 | Zinc finger protein 629 OS=Homo sapien | ZNF629 | 96.621 | 2 | 53.174603 | yes | 1.173E-07 | 577.81555 | 72.52317 |
| TFTLSATLLR | 10 | Q9UEG4 | Zinc finger protein 629 OS=Homo sapien | ZNF629 | 96.621 | 2 | 53.174603 | yes | 9.722E-08 | 561.82953 | 64.013519 |
| SFSQSSNLITHVR | 13 | Q9UEG4 | Zinc finger protein 629 OS=Homo sapien | ZNF629 | 96.621 | 2 | 53.174603 | yes | 1.496E-07 | 738.38367 | 56.584236 |
| AFTQSTNLIK | 10 | Q9UEG4 | Zinc finger protein 629 OS=Homo sapien | ZNF629 | 96.621 | 2 | 53.174603 | yes | 3.006E-06 | 561.81134 | 45.14875 |
| FGQNHNLLK | 9 | Q9UEG4 | Zinc finger protein 629 OS=Homo sapien | ZNF629 | 96.621 | 2 | 53.174603 | yes | 7.063E-06 | 535.79071 | 41.076859 |
| LFEEFLELCK | 10 | Q9UER7 | Death domain-associated protein 6 OS=H | DAXX | 81.373 | 2 | 53.710293 | yes | 9.722E-08 | 664.33374 | 166.04584 |
| QQLQLMAQDAFR | 12 | Q9UER7 | Death domain-associated protein 6 OS=H | DAXX | 81.373 | 2 | 53.710293 | yes | 5.24E-07 | 724.86938 | 102.90328 |
| LYVYINELCTVLK | 13 | Q9UER7 | Death domain-associated protein 6 OS=H | DAXX | 81.373 | 2 | 53.710293 | yes | 9.722E-08 | 814.44183 | 102.15452 |
| SPMSSLQISNEK | 12 | Q9UER7 | Death domain-associated protein 6 OS=H | DAXX | 81.373 | 2 | 53.710293 | yes | 1.802E-07 | 660.82684 | 69.103539 |
| QTGSGPLGNSYVE | 14 | Q9UER7 | Death domain-associated protein 6 OS=H | DAXX | 81.373 | 2 | 53.710293 | yes | 5.983E-07 | 732.85748 | 34.557907 |
| LSQTPHSQPPRPGT | 16 | Q9UER7 | Death domain-associated protein 6 OS=H | DAXX | 81.373 | 3 | 53.710293 | yes | 4.56E-07 | 597.63708 | 32.054783 |
| EFQLFQGVTR | 10 | Q9UET6 | tRNA (cytidine(32)/guanosine(34)-2'-O)- | FTSJ1 | 36.08 | 2 | 48.79611 | yes | 1.402E-07 | 612.8222 | 1198.8057 |
| SYPLDLEGGSEYK | 13 | Q9UET6 | tRNA (cytidine(32)/guanosine(34)-2'-O)- | FTSJ1 | 36.08 | 2 | 48.79611 | yes | 1.733E-07 | 729.34338 | 1171.1354 |
| EIIQHFK | 7 | Q9UET6 | tRNA (cytidine(32)/guanosine(34)-2'-O)- | FTSJ1 | 36.08 | 2 | 48.79611 | yes | 2.917E-07 | 457.75836 | 775.88281 |
| EIRPQDCPISR | 11 | Q9UET6 | tRNA (cytidine(32)/guanosine(34)-2'-O)- | FTSJ1 | 36.08 | 2 | 48.79611 | yes | 1.104E-07 | 685.84589 | 100.31663 |
| EIRPQDCPISR | 11 | Q9UET6 | tRNA (cytidine(32)/guanosine(34)-2'-O)- | FTSJ1 | 36.08 | 2 | 48.79611 | yes | 1.884E-06 | 457.56635 | 100.31663 |
| EIRPQDCPISR | 11 | Q9UET6 | tRNA (cytidine(32)/guanosine(34)-2'-O)- | FTSJ1 | 36.08 | 3 | 48.79611 | yes | 1.104E-07 | 685.84589 | 100.31663 |
| EIRPQDCPISR | 11 | Q9UET6 | tRNA (cytidine(32)/guanosine(34)-2'-O)- | FTSJ1 | 36.08 | 3 | 48.79611 | yes | 1.884E-06 | 457.56635 | 100.31663 |
| ASSAASSEHFEK | 12 | Q9UEU0 | Vesicle transport through interaction with | VTI1B | 26.689 | 2 | 41.081482 | yes | 3.686E-07 | 646.79132 | 34.654243 |
| VGEIEFEGLMR | 11 | Q9UEY8 | Gamma-adducin OS=Homo sapiens OX= | ADD3 | 79.155 | 2 | 55.01233 | yes | 9.722E-08 | 640.32117 | 268.07007 |
| EDLECLIQEQMK | 12 | Q9UEY8 | Gamma-adducin OS=Homo sapiens OX= | ADD3 | 79.155 | 2 | 55.01233 | yes | 9.722E-08 | 768.35767 | 202.82883 |
| GLSFSEATASNLVI | 14 | Q9UEY8 | Gamma-adducin OS=Homo sapiens OX= | ADD3 | 79.155 | 2 | 55.01233 | yes | 2.411E-07 | 712.375 | 192.54645 |
| VTQILQSPAFR | 11 | Q9UEY8 | Gamma-adducin OS=Homo sapiens OX= | ADD3 | 79.155 | 2 | 55.01233 | yes | 9.809E-08 | 630.35895 | 182.63396 |
| LSTSTTIENIEITIK | 15 | Q9UEY8 | Gamma-adducin OS=Homo sapiens OX= | ADD3 | 79.155 | 2 | 55.01233 | yes | 9.722E-08 | 831.96167 | 110.07757 |
| TLDNLGYR | 8 | Q9UEY8 | Gamma-adducin OS=Homo sapiens OX= | ADD3 | 79.155 | 2 | 55.01233 | yes | 5.401E-07 | 476.24835 | 101.73895 |
| VNVPEESR | 8 | Q9UEY8 | Gamma-adducin OS=Homo sapiens OX= | ADD3 | 79.155 | 2 | 55.01233 | yes | 5.707E-07 | 465.23798 | 86.685333 |
| VNIIGEVVDQGSTI | 16 | Q9UEY8 | Gamma-adducin OS=Homo sapiens OX= | ADD3 | 79.155 | 2 | 55.01233 | yes | 9.722E-08 | 843.45703 | 62.769611 |
| IEEVLSPEGSPSK | 13 | Q9UEY8 | Gamma-adducin OS=Homo sapiens OX= | ADD3 | 79.155 | 2 | 55.01233 | yes | 5.424E-07 | 686.35376 | 62.629738 |
| QDFNMMEQR | 9 | Q9UEY8 | Gamma-adducin OS=Homo sapiens OX= | ADD3 | 79.155 | 2 | 55.01233 | yes | 9.722E-08 | 599.75262 | 59.659893 |
| EQDHIIIIPR | 10 | Q9UEY8 | Gamma-adducin OS=Homo sapiens OX= | ADD3 | 79.155 | 3 | 55.01233 | yes | 1.467E-07 | 411.9032 | 48.135002 |
| FMATLGPEEEAEK | 13 | Q9UFC0 | Leucine-rich repeat and WD repeat-conta | LRWD1 | 70.862 | 2 | 41.339798 | yes | 3.559E-07 | 726.33978 | 103.02769 |
| DASSTYSQVENLN | 14 | Q9UFC0 | Leucine-rich repeat and WD repeat-conta | LRWD1 | 70.862 | 2 | 41.339798 | yes | 3.283E-06 | 792.36847 | 23.194422 |
| ELFFEDSIDDAK | 12 | Q9UFF9 | CCR4-NOT transcription complex subuni | CNOT8 | 33.541 | 2 | 54.167587 | yes | 1.224E-06 | 714.83008 | 176.40468 |
| GGLQEVADQLDL( | 14 | Q9UFF9 | CCR4-NOT transcription complex subuni | CNOT8 | 33.541 | 2 | 54.167587 | yes | 9.722E-08 | 771.39954 | 129.29733 |

| | | | | | | | | | | | |
|---|---|---|---|---|---|---|---|---|---|---|---|
| SSIDYQYQLLR | 11 | Q9UFF9 | CCR4-NOT transcription complex subuni | CNOT8 | 33.541 | 2 | 54.167587 | yes | 3.478E-07 | 693.35663 | 126.73489 |
| IIQLGLTFTNEK | 12 | Q9UFF9 | CCR4-NOT transcription complex subuni | CNOT8 | 33.541 | 2 | 54.167587 | yes | 9.722E-08 | 688.89282 | 113.35727 |
| CNVDLLK | 7 | Q9UFF9 | CCR4-NOT transcription complex subuni | CNOT8 | 33.541 | 2 | 54.167587 | no | 1.754E-06 | 431.22858 | 68.537613 |
| TALYVTPLDR | 10 | Q9UFW8 | CGG triplet repeat-binding protein 1 OS= | CGGBP1 | 18.821 | 2 | 52.610317 | yes | 1.562E-06 | 574.81915 | 169.5088 |
| VTEFGGELHEDGC | 14 | Q9UFW8 | CGG triplet repeat-binding protein 1 OS= | CGGBP1 | 18.821 | 2 | 52.610317 | yes | 9.795E-08 | 737.84406 | 150.92226 |
| FVVTAPPAR | 9 | Q9UFW8 | CGG triplet repeat-binding protein 1 OS= | CGGBP1 | 18.821 | 2 | 52.610317 | yes | 1.301E-06 | 479.27945 | 143.18109 |
| SAISDHLK | 8 | Q9UFW8 | CGG triplet repeat-binding protein 1 OS= | CGGBP1 | 18.821 | 2 | 52.610317 | yes | 1.018E-07 | 435.73761 | 132.25723 |
| MCLEANIPLEK | 11 | Q9UFW8 | CGG triplet repeat-binding protein 1 OS= | CGGBP1 | 18.821 | 2 | 52.610317 | yes | 9.722E-08 | 659.33069 | 70.34539 |
| VSVIQDFVK | 9 | Q9UFW8 | CGG triplet repeat-binding protein 1 OS= | CGGBP1 | 18.821 | 2 | 52.610317 | yes | 1.93E-07 | 517.79767 | 54.488907 |
| AEFEEQNVR | 9 | Q9UFW8 | CGG triplet repeat-binding protein 1 OS= | CGGBP1 | 18.821 | 2 | 52.610317 | yes | 2.404E-07 | 561.26471 | 43.166611 |
| KAEFEEQNVR | 10 | Q9UFW8 | CGG triplet repeat-binding protein 1 OS= | CGGBP1 | 18.821 | 2 | 52.610317 | yes | 1.294E-07 | 625.31219 | 40.415901 |
| DLHTNSPR | 8 | Q9UG56 | Phosphatidylserine decarboxylase proenzy | PISD | 46.672 | 2 | 43.496033 | yes | 2.565E-07 | 470.23578 | 114.69034 |
| ELFCHNER | 8 | Q9UG56 | Phosphatidylserine decarboxylase proenzy | PISD | 46.672 | 2 | 43.496033 | yes | 3.134E-06 | 552.74817 | 67.506538 |
| HGFFSLTAVGATN | 18 | Q9UG56 | Phosphatidylserine decarboxylase proenzy | PISD | 46.672 | 3 | 43.496033 | yes | 3.689E-07 | 611.9917 | 59.225471 |
| VEILPSYIPVR | 11 | Q9UGJ1 | Gamma-tubulin complex component 4 OS | TUBGCP4 | 76.09 | 2 | 56.560577 | yes | 1.576E-07 | 643.37933 | 855.15649 |
| LMVEESDLLGQLK | 13 | Q9UGJ1 | Gamma-tubulin complex component 4 OS | TUBGCP4 | 76.09 | 2 | 56.560577 | yes | 1.617E-07 | 737.89471 | 393.95126 |
| LMVEESDLLGQLK | 13 | Q9UGJ1 | Gamma-tubulin complex component 4 OS | TUBGCP4 | 76.09 | 2 | 56.560577 | yes | 9.894E-08 | 745.89215 | 393.95126 |
| LIEEENMLAPSLK | 13 | Q9UGJ1 | Gamma-tubulin complex component 4 OS | TUBGCP4 | 76.09 | 2 | 56.560577 | yes | 1.067E-07 | 743.89471 | 171.82671 |
| LIEEENMLAPSLK | 13 | Q9UGJ1 | Gamma-tubulin complex component 4 OS | TUBGCP4 | 76.09 | 2 | 56.560577 | yes | 6.878E-07 | 751.89215 | 171.82671 |
| AFCTGLDSVLQPY | 14 | Q9UGJ1 | Gamma-tubulin complex component 4 OS | TUBGCP4 | 76.09 | 2 | 56.560577 | yes | 9.722E-08 | 813.90088 | 214.33571 |
| STVAEHLWK | 9 | Q9UGJ1 | Gamma-tubulin complex component 4 OS | TUBGCP4 | 76.09 | 2 | 56.560577 | yes | 1.666E-07 | 535.7851 | 213.98172 |
| EAPASGWAALGLS | 15 | Q9UGJ1 | Gamma-tubulin complex component 4 OS | TUBGCP4 | 76.09 | 2 | 56.560577 | yes | 9.722E-08 | 760.89081 | 182.57295 |
| ILAVCHGVMYK | 11 | Q9UGJ1 | Gamma-tubulin complex component 4 OS | TUBGCP4 | 76.09 | 2 | 56.560577 | yes | 9.722E-08 | 645.8385 | 157.90004 |
| NHQINSDLAQLLL | 14 | Q9UGJ1 | Gamma-tubulin complex component 4 OS | TUBGCP4 | 76.09 | 2 | 56.560577 | yes | 9.722E-08 | 817.95227 | 141.76801 |
| TPPTAVTEHDVNV | 21 | Q9UGJ1 | Gamma-tubulin complex component 4 OS | TUBGCP4 | 76.09 | 3 | 56.560577 | yes | 2.054E-07 | 759.71344 | 79.386848 |
| LGTDYIR | 7 | Q9UGJ1 | Gamma-tubulin complex component 4 OS | TUBGCP4 | 76.09 | 2 | 56.560577 | yes | 1.695E-07 | 419.2269 | 79.148598 |
| FTEFIEQYTGHVQ( | 33 | Q9UGJ1 | Gamma-tubulin complex component 4 OS | TUBGCP4 | 76.09 | 5 | 56.560577 | yes | 1.084E-07 | 767.97229 | 74.885384 |
| HSCGGLPPVR | 10 | Q9UGJ1 | Gamma-tubulin complex component 4 OS | TUBGCP4 | 76.09 | 2 | 56.560577 | yes | 1.564E-07 | 540.27435 | 54.60527 |
| GAAQLSILVK | 10 | Q9UGJ1 | Gamma-tubulin complex component 4 OS | TUBGCP4 | 76.09 | 2 | 56.560577 | yes | 4.876E-07 | 500.31351 | 31.734676 |
| LLSPLQDVDIK | 11 | Q9UGL1 | Lysine-specific demethylase 5B OS=Hom | KDM5B | 175.658 | 2 | 38.15634 | yes | 5.419E-07 | 620.86102 | 46.787979 |
| MLLWHGSR | 8 | Q9UGN5 | Poly [ADP-ribose] polymerase 2 OS=Hor | PARP2 | 66.206 | 2 | 53.917957 | yes | 1.383E-07 | 500.26328 | 766.7738 |
| SPLKPESQLDLR | 12 | Q9UGN5 | Poly [ADP-ribose] polymerase 2 OS=Hor | PARP2 | 66.206 | 2 | 53.917957 | yes | 1.809E-07 | 691.88556 | 285.86954 |
| SPLKPESQLDLR | 12 | Q9UGN5 | Poly [ADP-ribose] polymerase 2 OS=Hor | PARP2 | 66.206 | 2 | 53.917957 | yes | 2.093E-07 | 461.59277 | 285.86954 |
| SPLKPESQLDLR | 12 | Q9UGN5 | Poly [ADP-ribose] polymerase 2 OS=Hor | PARP2 | 66.206 | 3 | 53.917957 | yes | 1.809E-07 | 691.88556 | 285.86954 |
| SPLKPESQLDLR | 12 | Q9UGN5 | Poly [ADP-ribose] polymerase 2 OS=Hor | PARP2 | 66.206 | 3 | 53.917957 | yes | 2.093E-07 | 461.59277 | 285.86954 |
| ALMEACNEFYTR | 12 | Q9UGN5 | Poly [ADP-ribose] polymerase 2 OS=Hor | PARP2 | 66.206 | 2 | 53.917957 | yes | 1.041E-07 | 752.8316 | 511.2041 |
| IPHDFGLR | 8 | Q9UGN5 | Poly [ADP-ribose] polymerase 2 OS=Hor | PARP2 | 66.206 | 2 | 53.917957 | yes | 1.118E-07 | 477.76141 | 432.9949 |
| APVDPECTAK | 10 | Q9UGN5 | Poly [ADP-ribose] polymerase 2 OS=Hor | PARP2 | 66.206 | 2 | 53.917957 | yes | 9.722E-08 | 544.25806 | 389.07837 |
| MGQHSLVACSGNI | 15 | Q9UGN5 | Poly [ADP-ribose] polymerase 2 OS=Hor | PARP2 | 66.206 | 3 | 53.917957 | yes | 9.722E-08 | 539.2608 | 153.78824 |

| | | | | | | | | | | | |
|---|---|---|---|---|---|---|---|---|---|---|---|
| MGQHSLVACSGN | 15 | Q9UGN5 | Poly [ADP-ribose] polymerase 2 OS=Hor | PARP2 | 66.206 | 3 | 53.917957 | yes | 1.456E-07 | 808.38757 | 153.78824 |
| MGQHSLVACSGN | 15 | Q9UGN5 | Poly [ADP-ribose] polymerase 2 OS=Hor | PARP2 | 66.206 | 2 | 53.917957 | yes | 9.722E-08 | 539.2608 | 153.78824 |
| MGQHSLVACSGN | 15 | Q9UGN5 | Poly [ADP-ribose] polymerase 2 OS=Hor | PARP2 | 66.206 | 2 | 53.917957 | yes | 1.456E-07 | 808.38757 | 153.78824 |
| KIEDCIR | 7 | Q9UGN5 | Poly [ADP-ribose] polymerase 2 OS=Hor | PARP2 | 66.206 | 2 | 53.917957 | yes | 4.437E-07 | 467.24475 | 281.75821 |
| GIYFADMSSK | 10 | Q9UGN5 | Poly [ADP-ribose] polymerase 2 OS=Hor | PARP2 | 66.206 | 2 | 53.917957 | yes | 1.008E-07 | 559.763 | 211.36787 |
| SANYCFASR | 9 | Q9UGN5 | Poly [ADP-ribose] polymerase 2 OS=Hor | PARP2 | 66.206 | 2 | 53.917957 | yes | 1.011E-07 | 538.23492 | 160.15318 |
| YYLIQLLEDDAQR | 13 | Q9UGN5 | Poly [ADP-ribose] polymerase 2 OS=Hor | PARP2 | 66.206 | 2 | 53.917957 | yes | 1.385E-07 | 820.41992 | 148.73636 |
| GKAPVDPECTAK | 12 | Q9UGN5 | Poly [ADP-ribose] polymerase 2 OS=Hor | PARP2 | 66.206 | 2 | 53.917957 | yes | 2.944E-07 | 636.81628 | 71.377419 |
| NLHCALRPLDHES | 17 | Q9UGN5 | Poly [ADP-ribose] polymerase 2 OS=Hor | PARP2 | 66.206 | 4 | 53.917957 | yes | 4.13E-07 | 533.01129 | 34.228897 |
| FIEDLNMYEASK | 12 | Q9UGP4 | LIM domain-containing protein 1 OS=Hc | LIMD1 | 72.191 | 2 | 53.674393 | yes | 9.722E-08 | 730.34235 | 427.2525 |
| TPSVSAPLALSCPF | 14 | Q9UGP4 | LIM domain-containing protein 1 OS=Hc | LIMD1 | 72.191 | 2 | 53.674393 | yes | 1.06E-07 | 728.38483 | 405.68951 |
| LPCQPLVPGPELRI | 20 | Q9UGP4 | LIM domain-containing protein 1 OS=Hc | LIMD1 | 72.191 | 3 | 53.674393 | yes | 7.273E-07 | 724.73303 | 318.0499 |
| QLSLSSSR | 8 | Q9UGP4 | LIM domain-containing protein 1 OS=Hc | LIMD1 | 72.191 | 2 | 53.674393 | yes | 2.102E-06 | 439.24051 | 294.26141 |
| LEALTQR | 7 | Q9UGP4 | LIM domain-containing protein 1 OS=Hc | LIMD1 | 72.191 | 2 | 53.674393 | yes | 1.501E-07 | 415.74017 | 288.89667 |
| YDDLGLEASK | 10 | Q9UGP4 | LIM domain-containing protein 1 OS=Hc | LIMD1 | 72.191 | 2 | 53.674393 | yes | 2.134E-07 | 555.76929 | 199.8429 |
| RPSSTALHQHHF | 12 | Q9UGP4 | LIM domain-containing protein 1 OS=Hc | LIMD1 | 72.191 | 3 | 53.674393 | yes | 2.733E-07 | 473.24097 | 182.32172 |
| PGCTDLGTGPK | 11 | Q9UGP4 | LIM domain-containing protein 1 OS=Hc | LIMD1 | 72.191 | 2 | 53.674393 | yes | 1.002E-07 | 551.76349 | 178.22775 |
| GAGNNPEFEETR | 12 | Q9UGP4 | LIM domain-containing protein 1 OS=Hc | LIMD1 | 72.191 | 2 | 53.674393 | yes | 9.722E-08 | 660.79437 | 170.70259 |
| ADYFGACVK | 9 | Q9UGP4 | LIM domain-containing protein 1 OS=Hc | LIMD1 | 72.191 | 2 | 53.674393 | yes | 9.722E-08 | 515.73676 | 161.69998 |
| PNVDPQPWFQDGI | 14 | Q9UGP4 | LIM domain-containing protein 1 OS=Hc | LIMD1 | 72.191 | 2 | 53.674393 | yes | 9.79E-08 | 812.89136 | 95.815056 |
| LPCQPLVPGPELR | 13 | Q9UGP4 | LIM domain-containing protein 1 OS=Hc | LIMD1 | 72.191 | 2 | 53.674393 | yes | 2.994E-06 | 738.40558 | 76.066475 |
| EGPLGWSSDGSLG | 24 | Q9UGP4 | LIM domain-containing protein 1 OS=Hc | LIMD1 | 72.191 | 3 | 53.674393 | yes | 2.731E-06 | 800.72742 | 60.193268 |
| SSEGSLGGQNSGIC | 16 | Q9UGP4 | LIM domain-containing protein 1 OS=Hc | LIMD1 | 72.191 | 2 | 53.674393 | yes | 1.078E-07 | 731.84766 | 55.702312 |
| SNSGLGGEVSGVN | 15 | Q9UGP4 | LIM domain-containing protein 1 OS=Hc | LIMD1 | 72.191 | 2 | 53.674393 | yes | 3.917E-06 | 704.84045 | 44.73027 |
| SSEKPTGLWSTASS | 16 | Q9UGP4 | LIM domain-containing protein 1 OS=Hc | LIMD1 | 72.191 | 3 | 53.674393 | yes | 9.722E-08 | 574.61993 | 39.429543 |
| LSQMAVQGLQQFI | 13 | Q9UGP8 | Translocation protein SEC63 homolog OS | SEC63 | 87.997 | 2 | 51.3156 | yes | 3.621E-07 | 739.3952 | 152.70212 |
| APTLASLENCMK | 12 | Q9UGP8 | Translocation protein SEC63 homolog OS | SEC63 | 87.997 | 2 | 51.3156 | yes | 9.722E-08 | 667.82581 | 114.8176 |
| QTMAEVFEK | 9 | Q9UGP8 | Translocation protein SEC63 homolog OS | SEC63 | 87.997 | 2 | 51.3156 | yes | 1.042E-07 | 541.763 | 80.798523 |
| PTDNILIPQLIR | 12 | Q9UGP8 | Translocation protein SEC63 homolog OS | SEC63 | 87.997 | 2 | 51.3156 | yes | 8.609E-07 | 696.91412 | 70.19989 |
| YSGDQILIR | 9 | Q9UGP8 | Translocation protein SEC63 homolog OS | SEC63 | 87.997 | 2 | 51.3156 | yes | 2.04E-07 | 532.79041 | 46.200916 |
| SDSYMGLDQIKPL | 14 | Q9UGP8 | Translocation protein SEC63 homolog OS | SEC63 | 87.997 | 2 | 51.3156 | yes | 6.829E-07 | 797.91089 | 35.980789 |
| SPLLQLPHIEEDNL | 15 | Q9UGP8 | Translocation protein SEC63 homolog OS | SEC63 | 87.997 | 3 | 51.3156 | yes | 4.138E-07 | 591.98798 | 28.749697 |
| LDALDSFGSTR | 11 | Q9UGR2 | Zinc finger CCCH domain-containing prc | ZC3H7B | 109.858 | 2 | 52.79982 | yes | 1.584E-07 | 591.29346 | 1531.2865 |
| GLQFIQSTLPLK | 12 | Q9UGR2 | Zinc finger CCCH domain-containing prc | ZC3H7B | 109.858 | 2 | 52.79982 | yes | 1.221E-07 | 672.89795 | 891.69403 |
| DSPSVCSNLAAK | 12 | Q9UGR2 | Zinc finger CCCH domain-containing prc | ZC3H7B | 109.858 | 2 | 52.79982 | yes | 1.073E-07 | 624.7981 | 854.11774 |
| LVQNLFAEGNDLF | 14 | Q9UGR2 | Zinc finger CCCH domain-containing prc | ZC3H7B | 109.858 | 2 | 52.79982 | yes | 9.722E-08 | 818.4281 | 759.61243 |
| ALGLDSESIR | 10 | Q9UGR2 | Zinc finger CCCH domain-containing prc | ZC3H7B | 109.858 | 2 | 52.79982 | yes | 2.099E-07 | 530.78528 | 734.41833 |
| AACYFTMGLYEK | 12 | Q9UGR2 | Zinc finger CCCH domain-containing prc | ZC3H7B | 109.858 | 2 | 52.79982 | yes | 1.455E-07 | 727.32819 | 648.38281 |
| NPLAATHEFK | 10 | Q9UGR2 | Zinc finger CCCH domain-containing prc | ZC3H7B | 109.858 | 2 | 52.79982 | yes | 1.021E-07 | 564.29584 | 631.87671 |

| | | | | | | | | | | |
|---|---|---|---|---|---|---|---|---|---|---|
| CLVHIVR | 7 | Q9UGR2 | Zinc finger CCCH domain-containing prc | ZC3H7B | 109.858 | 2 | 52.79982 | yes | 1.724E-07 | 448.76038 | 485.49707 |
| ILDMQQTYDMWL | 13 | Q9UGR2 | Zinc finger CCCH domain-containing prc | ZC3H7B | 109.858 | 2 | 52.79982 | yes | 9.728E-08 | 842.90729 | 472.03931 |
| VFTSDSDASGWAI | 14 | Q9UGR2 | Zinc finger CCCH domain-containing prc | ZC3H7B | 109.858 | 2 | 52.79982 | yes | 4.025E-07 | 773.35205 | 471.54633 |
| TSFVGSYYLCK | 11 | Q9UGR2 | Zinc finger CCCH domain-containing prc | ZC3H7B | 109.858 | 2 | 52.79982 | yes | 9.722E-08 | 662.81573 | 396.82037 |
| FVCGQCWR | 8 | Q9UGR2 | Zinc finger CCCH domain-containing prc | ZC3H7B | 109.858 | 2 | 52.79982 | yes | 9.745E-08 | 556.74188 | 387.61481 |
| QEEYEAFLLK | 10 | Q9UGR2 | Zinc finger CCCH domain-containing prc | ZC3H7B | 109.858 | 2 | 52.79982 | yes | 9.722E-08 | 635.32172 | 384.46008 |
| CSLALPHDESVTQ | 21 | Q9UGR2 | Zinc finger CCCH domain-containing prc | ZC3H7B | 109.858 | 3 | 52.79982 | yes | 9.722E-08 | 775.38989 | 321.64716 |
| KDMLLCPR | 8 | Q9UGR2 | Zinc finger CCCH domain-containing prc | ZC3H7B | 109.858 | 2 | 52.79982 | yes | 1.379E-07 | 516.77008 | 288.50842 |
| DMLLCPR | 7 | Q9UGR2 | Zinc finger CCCH domain-containing prc | ZC3H7B | 109.858 | 2 | 52.79982 | yes | 1.854E-07 | 452.7226 | 262.09003 |
| THGPSTFDLQMK | 12 | Q9UGR2 | Zinc finger CCCH domain-containing prc | ZC3H7B | 109.858 | 2 | 52.79982 | yes | 1.301E-07 | 681.32953 | 248.1331 |
| RGSLTIAK | 8 | Q9UGR2 | Zinc finger CCCH domain-containing prc | ZC3H7B | 109.858 | 2 | 52.79982 | yes | 1.033E-07 | 423.26379 | 191.85861 |
| QACQLCYPK | 9 | Q9UGR2 | Zinc finger CCCH domain-containing prc | ZC3H7B | 109.858 | 2 | 52.79982 | yes | 9.722E-08 | 584.26788 | 184.851 |
| NGQVVEPDKDLK | 12 | Q9UGR2 | Zinc finger CCCH domain-containing prc | ZC3H7B | 109.858 | 2 | 52.79982 | yes | 1.164E-07 | 671.35406 | 170.01205 |
| YWQQMEAHAGK | 11 | Q9UGR2 | Zinc finger CCCH domain-containing prc | ZC3H7B | 109.858 | 2 | 52.79982 | yes | 1.187E-07 | 674.80878 | 150.95493 |
| HNPGKPGEGTPISS | 15 | Q9UGR2 | Zinc finger CCCH domain-containing prc | ZC3H7B | 109.858 | 3 | 52.79982 | yes | 1.409E-07 | 511.93045 | 142.47078 |
| CQYVGNCSFAHSI | 16 | Q9UGR2 | Zinc finger CCCH domain-containing prc | ZC3H7B | 109.858 | 3 | 52.79982 | no | 9.927E-08 | 647.60535 | 80.804596 |
| KWVSVRPLPSIR | 12 | Q9UGR2 | Zinc finger CCCH domain-containing prc | ZC3H7B | 109.858 | 3 | 52.79982 | yes | 3.587E-07 | 479.96115 | 38.4604 |
| QWQQHIQSEK | 10 | Q9UGR2 | Zinc finger CCCH domain-containing prc | ZC3H7B | 109.858 | 2 | 52.79982 | yes | 1.767E-07 | 656.32562 | 38.172821 |
| KYWQQMEAHAGI | 12 | Q9UGR2 | Zinc finger CCCH domain-containing prc | ZC3H7B | 109.858 | 2 | 52.79982 | yes | 2.039E-07 | 738.85626 | 34.048325 |
| WVSVRPLPSIR | 11 | Q9UGR2 | Zinc finger CCCH domain-containing prc | ZC3H7B | 109.858 | 2 | 52.79982 | yes | 1.508E-07 | 655.39063 | 27.115194 |
| EHQGIFTFLCEICFI | 19 | Q9UGR2 | Zinc finger CCCH domain-containing prc | ZC3H7B | 109.858 | 3 | 52.79982 | yes | 2.047E-07 | 795.37689 | 22.915506 |
| GSLDKPDSFMEET | 26 | Q9UGR2 | Zinc finger CCCH domain-containing prc | ZC3H7B | 109.858 | 4 | 52.79982 | yes | 4.105E-07 | 715.33069 | 11.573093 |
| KQWQQHIQSEK | 11 | Q9UGR2 | Zinc finger CCCH domain-containing prc | ZC3H7B | 109.858 | 2 | 52.79982 | yes | 4.202E-07 | 720.37311 | 5.7308078 |
| DLLFDPLGGVK | 11 | Q9UGR2 | Zinc finger CCCH domain-containing prc | ZC3H7B | 109.858 | 2 | 52.79982 | yes | 7.591E-05 | 587.32935 | 3.7683821 |
| SVICDISPLR | 10 | Q9UGU0 | Transcription factor 20 OS=Homo sapien: | TCF20 | 211.772 | 2 | 49.544456 | yes | 1.153E-07 | 580.31061 | 129.53963 |
| QAVPIVEPQEPEIK | 14 | Q9UGU0 | Transcription factor 20 OS=Homo sapien: | TCF20 | 211.772 | 2 | 49.544456 | yes | 1.74E-07 | 788.93268 | 92.217712 |
| NVSGFPQYPTGQE | 14 | Q9UGU0 | Transcription factor 20 OS=Homo sapien: | TCF20 | 211.772 | 2 | 49.544456 | yes | 4.163E-07 | 776.37555 | 72.586136 |
| LYGLQEALEIAR | 12 | Q9UGU0 | Transcription factor 20 OS=Homo sapien: | TCF20 | 211.772 | 2 | 49.544456 | yes | 2.888E-07 | 688.38263 | 54.957157 |
| AHAYGDPNAGLN: | 18 | Q9UGU0 | Transcription factor 20 OS=Homo sapien: | TCF20 | 211.772 | 3 | 49.544456 | yes | 1.022E-07 | 652.65015 | 42.738342 |
| QINLTDYPIPR | 11 | Q9UGU0 | Transcription factor 20 OS=Homo sapien: | TCF20 | 211.772 | 2 | 49.544456 | yes | 2.599E-06 | 665.36169 | 25.006771 |
| LLGLTPPEGGAGD | 14 | Q9UGY1 | Nucleolar protein 12 OS=Homo sapiens ( | NOL12 | 24.663 | 2 | 50.07438 | yes | 3.051E-07 | 676.86206 | 1026.6903 |
| EEALEEADELDR | 12 | Q9UGY1 | Nucleolar protein 12 OS=Homo sapiens ( | NOL12 | 24.663 | 2 | 50.07438 | yes | 9.915E-08 | 709.81769 | 883.00446 |
| LVLSFDEEK | 9 | Q9UGY1 | Nucleolar protein 12 OS=Homo sapiens ( | NOL12 | 24.663 | 2 | 50.07438 | yes | 1.229E-07 | 540.28461 | 675.93512 |
| SRDPLLSQR | 9 | Q9UGY1 | Nucleolar protein 12 OS=Homo sapiens ( | NOL12 | 24.663 | 2 | 50.07438 | yes | 1.503E-05 | 536.29889 | 47.786392 |
| FDENYAFLHR | 10 | Q9UH17 | DNA dC->dU-editing enzyme APOBEC- | APOBEC3I | 45.925 | 2 | 52.77704 | yes | 9.722E-08 | 656.30945 | 975.73627 |
| SNLLWDTGVFR | 11 | Q9UH17 | DNA dC->dU-editing enzyme APOBEC- | APOBEC3I | 45.925 | 2 | 52.77704 | no | 1E-07 | 654.34076 | 868.54413 |
| AFLQENTHVR | 10 | Q9UH17 | DNA dC->dU-editing enzyme APOBEC- | APOBEC3I | 45.925 | 2 | 52.77704 | no | 9.722E-08 | 607.81744 | 642.69702 |
| NLLCGFYGR | 9 | Q9UH17 | DNA dC->dU-editing enzyme APOBEC- | APOBEC3I | 45.925 | 2 | 52.77704 | no | 1.137E-07 | 550.2713 | 559.87549 |
| LAEFLSEHPNVTL7 | 19 | Q9UH17 | DNA dC->dU-editing enzyme APOBEC- | APOBEC3I | 45.925 | 3 | 52.77704 | yes | 9.722E-08 | 690.37238 | 536.85327 |

| | | | | | | | | | | | |
|---|---|---|---|---|---|---|---|---|---|---|---|
| SYTWLCYEVK | 10 | Q9UH17 | DNA dC->dU-editing enzyme APOBEC- | APOBEC3I | 45.925 | 2 | 52.77704 | no | 9.722E-08 | 674.81573 | 479.75049 |
| QTYLCYEVER | 10 | Q9UH17 | DNA dC->dU-editing enzyme APOBEC- | APOBEC3I | 45.925 | 2 | 52.77704 | yes | 1.415E-07 | 680.81372 | 240.78519 |
| IYDYDPLYK | 9 | Q9UH17 | DNA dC->dU-editing enzyme APOBEC- | APOBEC3I | 45.925 | 2 | 52.77704 | no | 1.812E-07 | 595.29242 | 204.18092 |
| LYYYWER | 7 | Q9UH17 | DNA dC->dU-editing enzyme APOBEC- | APOBEC3I | 45.925 | 2 | 52.77704 | no | 1.28E-07 | 546.76111 | 136.68655 |
| IQEELSALR | 9 | Q9UH99 | SUN domain-containing protein 2 OS=H | SUN2 | 80.311 | 2 | 52.33096 | yes | 9.819E-08 | 529.79565 | 316.51291 |
| IRPTAVTLEHVPK | 13 | Q9UH99 | SUN domain-containing protein 2 OS=H | SUN2 | 80.311 | 3 | 52.33096 | yes | 9.722E-08 | 487.6243 | 145.8521 |
| ILTHVAEMQGK | 11 | Q9UH99 | SUN domain-containing protein 2 OS=H | SUN2 | 80.311 | 2 | 52.33096 | yes | 9.722E-08 | 613.83173 | 107.04046 |
| EEMQAQLR | 8 | Q9UH99 | SUN domain-containing protein 2 OS=H | SUN2 | 80.311 | 2 | 52.33096 | yes | 4.298E-07 | 502.74512 | 104.05067 |
| EAAASLSLTLQK | 12 | Q9UH99 | SUN domain-containing protein 2 OS=H | SUN2 | 80.311 | 2 | 52.33096 | yes | 1.204E-07 | 616.34827 | 65.388451 |
| RLEALAAEFSSNW | 15 | Q9UH99 | SUN domain-containing protein 2 OS=H | SUN2 | 80.311 | 3 | 52.33096 | yes | 1.591E-07 | 583.96881 | 51.386574 |
| ILTNWGHPEYTCI | 15 | Q9UH99 | SUN domain-containing protein 2 OS=H | SUN2 | 80.311 | 3 | 52.33096 | yes | 6.87E-07 | 641.64545 | 48.921898 |
| AEHQQDSEDLFK | 12 | Q9UH99 | SUN domain-containing protein 2 OS=H | SUN2 | 80.311 | 2 | 52.33096 | yes | 6.194E-07 | 723.82843 | 36.206947 |
| DSSPHFQAEQR | 11 | Q9UH99 | SUN domain-containing protein 2 OS=H | SUN2 | 80.311 | 2 | 52.33096 | yes | 2.775E-06 | 651.29712 | 10.393569 |
| ELSVEEQIK | 9 | Q9UHB6 | LIM domain and actin-binding protein 1 ( | LIMA1 | 85.226 | 2 | 52.974236 | yes | 1.172E-07 | 537.78748 | 1446.3445 |
| VGVLAASMEAK | 11 | Q9UHB6 | LIM domain and actin-binding protein 1 ( | LIMA1 | 85.226 | 2 | 52.974236 | yes | 1.146E-07 | 538.29443 | 1000.9696 |
| SQDVELWEGEVVI | 13 | Q9UHB6 | LIM domain and actin-binding protein 1 ( | LIMA1 | 85.226 | 2 | 52.974236 | yes | 1.443E-07 | 759.37775 | 964.49652 |
| YNVPLNR | 7 | Q9UHB6 | LIM domain and actin-binding protein 1 ( | LIMA1 | 85.226 | 2 | 52.974236 | yes | 1.739E-07 | 438.24033 | 725.58893 |
| ISANENSLAVR | 11 | Q9UHB6 | LIM domain and actin-binding protein 1 ( | LIMA1 | 85.226 | 2 | 52.974236 | yes | 9.722E-08 | 587.31476 | 416.7142 |
| SEVQQPVHPK | 10 | Q9UHB6 | LIM domain and actin-binding protein 1 ( | LIMA1 | 85.226 | 2 | 52.974236 | yes | 1.639E-07 | 574.80658 | 413.56052 |
| TVYPMER | 7 | Q9UHB6 | LIM domain and actin-binding protein 1 ( | LIMA1 | 85.226 | 2 | 52.974236 | no | 9.722E-08 | 448.22073 | 366.83371 |
| NENEEILER | 9 | Q9UHB6 | LIM domain and actin-binding protein 1 ( | LIMA1 | 85.226 | 2 | 52.974236 | yes | 9.804E-08 | 573.27527 | 264.76208 |
| QSSSTNYTNELK | 12 | Q9UHB6 | LIM domain and actin-binding protein 1 ( | LIMA1 | 85.226 | 2 | 52.974236 | yes | 9.722E-08 | 686.323 | 244.46124 |
| PHFNQLFK | 8 | Q9UHB6 | LIM domain and actin-binding protein 1 ( | LIMA1 | 85.226 | 2 | 52.974236 | yes | 4.327E-07 | 515.7771 | 191.68839 |
| IYCKPHFNQLFK | 12 | Q9UHB6 | LIM domain and actin-binding protein 1 ( | LIMA1 | 85.226 | 2 | 52.974236 | yes | 3.682E-06 | 797.91357 | 72.260117 |
| IYCKPHFNQLFK | 12 | Q9UHB6 | LIM domain and actin-binding protein 1 ( | LIMA1 | 85.226 | 2 | 52.974236 | yes | 4.76E-07 | 532.27814 | 72.260117 |
| IYCKPHFNQLFK | 12 | Q9UHB6 | LIM domain and actin-binding protein 1 ( | LIMA1 | 85.226 | 3 | 52.974236 | yes | 3.682E-06 | 797.91357 | 72.260117 |
| IYCKPHFNQLFK | 12 | Q9UHB6 | LIM domain and actin-binding protein 1 ( | LIMA1 | 85.226 | 3 | 52.974236 | yes | 4.76E-07 | 532.27814 | 72.260117 |
| ASSLSESSPPK | 11 | Q9UHB6 | LIM domain and actin-binding protein 1 ( | LIMA1 | 85.226 | 2 | 52.974236 | yes | 1.043E-07 | 545.27478 | 144.45367 |
| ETPHSPGVEDAPIA | 15 | Q9UHB6 | LIM domain and actin-binding protein 1 ( | LIMA1 | 85.226 | 2 | 52.974236 | yes | 2.526E-07 | 774.38867 | 116.74498 |
| SRPFTVAASFQSTS | 16 | Q9UHB6 | LIM domain and actin-binding protein 1 ( | LIMA1 | 85.226 | 3 | 52.974236 | yes | 9.722E-08 | 571.63702 | 83.993042 |
| QQSPQEPK | 8 | Q9UHB6 | LIM domain and actin-binding protein 1 ( | LIMA1 | 85.226 | 2 | 52.974236 | yes | 5.776E-07 | 471.23798 | 68.044922 |
| SEISENTDASGK | 12 | Q9UHB6 | LIM domain and actin-binding protein 1 ( | LIMA1 | 85.226 | 2 | 52.974236 | yes | 9.722E-08 | 619.28076 | 51.574371 |
| GNYDEGFGHRPHI | 13 | Q9UHB6 | LIM domain and actin-binding protein 1 ( | LIMA1 | 85.226 | 2 | 52.974236 | yes | 1.468E-07 | 757.35016 | 36.234592 |
| SSAIVEIFSK | 10 | Q9UHB6 | LIM domain and actin-binding protein 1 ( | LIMA1 | 85.226 | 2 | 52.974236 | yes | 1.364E-06 | 540.80042 | 35.199669 |
| SNTENLSQHFR | 11 | Q9UHB6 | LIM domain and actin-binding protein 1 ( | LIMA1 | 85.226 | 2 | 52.974236 | yes | 2.59E-07 | 666.81818 | 16.111092 |
| IDLNLLTR | 8 | Q9UHB7 | AF4/FMR2 family member 4 OS=Homo : | AFF4 | 127.46 | 2 | 53.886547 | yes | 2.195E-07 | 479.29001 | 420.2999 |
| WQLDNWLNK | 9 | Q9UHB7 | AF4/FMR2 family member 4 OS=Homo : | AFF4 | 127.46 | 2 | 53.886547 | yes | 1.09E-07 | 608.80908 | 144.90869 |
| WTPVGPAPSTSQS | 15 | Q9UHB7 | AF4/FMR2 family member 4 OS=Homo : | AFF4 | 127.46 | 2 | 53.886547 | yes | 7.673E-07 | 785.89661 | 140.29175 |
| MFSPMEEK | 8 | Q9UHB7 | AF4/FMR2 family member 4 OS=Homo : | AFF4 | 127.46 | 2 | 53.886547 | yes | 3.245E-07 | 499.71973 | 128.37704 |

| | | | | | | | | | | | |
|---|---|---|---|---|---|---|---|---|---|---|---|
| NYLAPDATAADKI | 13 | Q9UHB7 | AF4/FMR2 family member 4 OS=Homo : | AFF4 | 127.46 | 2 | 53.886547 | yes | 9.786E-08 | 703.35718 | 109.15402 |
| SNPNFFEQR | 9 | Q9UHB7 | AF4/FMR2 family member 4 OS=Homo : | AFF4 | 127.46 | 2 | 53.886547 | yes | 9.722E-08 | 569.76746 | 108.36907 |
| NSYNNSQAPSPGL | 16 | Q9UHB7 | AF4/FMR2 family member 4 OS=Homo : | AFF4 | 127.46 | 2 | 53.886547 | yes | 3.553E-07 | 810.88428 | 49.359165 |
| QGLHWLR | 7 | Q9UHB7 | AF4/FMR2 family member 4 OS=Homo : | AFF4 | 127.46 | 2 | 53.886547 | yes | 2.041E-07 | 455.25632 | 49.190182 |
| NYSADHYLQEAK | 12 | Q9UHB7 | AF4/FMR2 family member 4 OS=Homo : | AFF4 | 127.46 | 2 | 53.886547 | yes | 5.844E-07 | 719.8335 | 31.845039 |
| SPFPMYSETVDLIK | 14 | Q9UHB7 | AF4/FMR2 family member 4 OS=Homo : | AFF4 | 127.46 | 2 | 53.886547 | yes | 6.87E-07 | 813.90784 | 23.429735 |
| LTVLCLR | 7 | Q9UHB7 | AF4/FMR2 family member 4 OS=Homo : | AFF4 | 127.46 | 2 | 53.886547 | yes | 2.98E-06 | 437.76257 | 23.29619 |
| LEALITQTR | 9 | Q9UHB9 | Signal recognition particle subunit SRP68 | SRP68 | 70.73 | 2 | 51.94969 | yes | 1.609E-07 | 522.80603 | 1184.9099 |
| LFESMLSECR | 10 | Q9UHB9 | Signal recognition particle subunit SRP68 | SRP68 | 70.73 | 2 | 51.94969 | yes | 1.607E-07 | 636.29156 | 594.9942 |
| DNKPLVER | 8 | Q9UHB9 | Signal recognition particle subunit SRP68 | SRP68 | 70.73 | 2 | 51.94969 | yes | 9.926E-08 | 485.76944 | 450.89844 |
| YANEVNSDAGAFI | 13 | Q9UHB9 | Signal recognition particle subunit SRP68 | SRP68 | 70.73 | 2 | 51.94969 | yes | 1.127E-07 | 693.32025 | 411.60904 |
| WSEALVLYDR | 10 | Q9UHB9 | Signal recognition particle subunit SRP68 | SRP68 | 70.73 | 2 | 51.94969 | yes | 1.32E-07 | 626.32202 | 361.4097 |
| ALLQQQPEDDSK | 12 | Q9UHB9 | Signal recognition particle subunit SRP68 | SRP68 | 70.73 | 2 | 51.94969 | yes | 9.722E-08 | 686.34119 | 332.3891 |
| QAATMSEVEWR | 11 | Q9UHB9 | Signal recognition particle subunit SRP68 | SRP68 | 70.73 | 2 | 51.94969 | yes | 1.349E-07 | 654.30609 | 260.74445 |
| VTEELLTDNR | 10 | Q9UHB9 | Signal recognition particle subunit SRP68 | SRP68 | 70.73 | 2 | 51.94969 | yes | 5.231E-06 | 595.30658 | 199.74365 |
| FETFCLDPSLVTK | 13 | Q9UHB9 | Signal recognition particle subunit SRP68 | SRP68 | 70.73 | 2 | 51.94969 | yes | 1.209E-07 | 778.8869 | 189.66589 |
| CFFIAQSYVLVK | 12 | Q9UHB9 | Signal recognition particle subunit SRP68 | SRP68 | 70.73 | 2 | 51.94969 | yes | 1.066E-07 | 737.89178 | 109.21547 |
| DYILEGEPGK | 10 | Q9UHB9 | Signal recognition particle subunit SRP68 | SRP68 | 70.73 | 2 | 51.94969 | yes | 3.4E-06 | 560.77966 | 101.7196 |
| KWSEALVLYDR | 11 | Q9UHB9 | Signal recognition particle subunit SRP68 | SRP68 | 70.73 | 2 | 51.94969 | yes | 9.722E-08 | 690.36951 | 90.468063 |
| KVTEELLTDNR | 11 | Q9UHB9 | Signal recognition particle subunit SRP68 | SRP68 | 70.73 | 2 | 51.94969 | yes | 9.759E-08 | 659.35406 | 88.756142 |
| DLPDVQELITQVR | 13 | Q9UHB9 | Signal recognition particle subunit SRP68 | SRP68 | 70.73 | 2 | 51.94969 | yes | 1.012E-07 | 763.41467 | 82.750336 |
| SPRPQDLIR | 9 | Q9UHB9 | Signal recognition particle subunit SRP68 | SRP68 | 70.73 | 2 | 51.94969 | yes | 1.059E-07 | 541.30927 | 67.819893 |
| YCAYNIGDQSAIN | 19 | Q9UHB9 | Signal recognition particle subunit SRP68 | SRP68 | 70.73 | 3 | 51.94969 | yes | 5.384E-05 | 759.67474 | 40.060139 |
| VSNLQYLHSYLTY | 15 | Q9UHB9 | Signal recognition particle subunit SRP68 | SRP68 | 70.73 | 3 | 51.94969 | yes | 9.809E-08 | 614.66473 | 30.27146 |
| TAPSCTEAPLQGSV | 16 | Q9UHC7 | E3 ubiquitin-protein ligase makorin-1 OS | MKRN1 | 53.35 | 2 | 51.518448 | yes | 1.899E-07 | 823.90637 | 3319.304 |
| YFMHGVCK | 8 | Q9UHC7 | E3 ubiquitin-protein ligase makorin-1 OS | MKRN1 | 53.35 | 2 | 51.518448 | yes | 1.092E-07 | 521.2359 | 965.06104 |
| QEEATATELTTK | 12 | Q9UHC7 | E3 ubiquitin-protein ligase makorin-1 OS | MKRN1 | 53.35 | 2 | 51.518448 | no | 1.003E-07 | 661.3277 | 944.31848 |
| DMELSFAVQR | 10 | Q9UHC7 | E3 ubiquitin-protein ligase makorin-1 OS | MKRN1 | 53.35 | 2 | 51.518448 | no | 9.947E-08 | 598.29242 | 816.78271 |
| GYCIYGDR | 8 | Q9UHC7 | E3 ubiquitin-protein ligase makorin-1 OS | MKRN1 | 53.35 | 2 | 51.518448 | yes | 2.056E-07 | 502.21872 | 606.95996 |
| NHFWELIEER | 10 | Q9UHC7 | E3 ubiquitin-protein ligase makorin-1 OS | MKRN1 | 53.35 | 2 | 51.518448 | yes | 9.722E-08 | 686.83588 | 524.84186 |
| HAYPDGR | 7 | Q9UHC7 | E3 ubiquitin-protein ligase makorin-1 OS | MKRN1 | 53.35 | 2 | 51.518448 | no | 9.728E-08 | 408.19339 | 524.66089 |
| GSCPFGGNCFYK | 12 | Q9UHC7 | E3 ubiquitin-protein ligase makorin-1 OS | MKRN1 | 53.35 | 2 | 51.518448 | yes | 9.85E-08 | 697.2868 | 381.21082 |
| EESEKEQTAVETK | 13 | Q9UHC7 | E3 ubiquitin-protein ligase makorin-1 OS | MKRN1 | 53.35 | 2 | 51.518448 | yes | 1.076E-07 | 754.35974 | 116.16402 |
| LFAIEEETTTR | 11 | Q9UHD2 | Serine/threonine-protein kinase TBK1 OS | TBK1 | 83.643 | 2 | 52.26141 | yes | 2.113E-07 | 655.33533 | 451.66797 |
| TTEENPIFVVSR | 12 | Q9UHD2 | Serine/threonine-protein kinase TBK1 OS | TBK1 | 83.643 | 2 | 52.26141 | yes | 1.48E-07 | 696.36188 | 310.83228 |
| AITGVVCYACR | 11 | Q9UHD2 | Serine/threonine-protein kinase TBK1 OS | TBK1 | 83.643 | 2 | 52.26141 | yes | 1.414E-07 | 635.30756 | 282.61664 |
| VIGEDGQSVYK | 11 | Q9UHD2 | Serine/threonine-protein kinase TBK1 OS | TBK1 | 83.643 | 2 | 52.26141 | yes | 9.722E-08 | 597.80371 | 212.07552 |
| DVVGGMNHLR | 10 | Q9UHD2 | Serine/threonine-protein kinase TBK1 OS | TBK1 | 83.643 | 2 | 52.26141 | yes | 3.238E-07 | 549.27966 | 208.02602 |
| LSPGGSLADAWAI | 20 | Q9UHD2 | Serine/threonine-protein kinase TBK1 OS | TBK1 | 83.643 | 3 | 52.26141 | yes | 2.083E-07 | 687.00458 | 203.69942 |

| | | | | | | | | | | | |
|---|---|---|---|---|---|---|---|---|---|---|---|
| EPLNTIGLIYEK | 12 | Q9UHD2 | Serine/threonine-protein kinase TBK1 OS | TBK1 | 83.643 | 2 | 52.26141 | yes | 9.722E-08 | 695.38483 | 179.2469 |
| IISSNQELIYEGR | 13 | Q9UHD2 | Serine/threonine-protein kinase TBK1 OS | TBK1 | 83.643 | 2 | 52.26141 | yes | 3.698E-07 | 761.39899 | 166.58481 |
| YDLDGDASMAK | 11 | Q9UHD2 | Serine/threonine-protein kinase TBK1 OS | TBK1 | 83.643 | 2 | 52.26141 | yes | 1.108E-07 | 593.25824 | 136.47946 |
| IASTLLLYQELMR | 13 | Q9UHD2 | Serine/threonine-protein kinase TBK1 OS | TBK1 | 83.643 | 2 | 52.26141 | yes | 1.41E-07 | 775.9342 | 129.17821 |
| LYYHATK | 7 | Q9UHD2 | Serine/threonine-protein kinase TBK1 OS | TBK1 | 83.643 | 2 | 52.26141 | yes | 1.214E-07 | 448.23724 | 111.90825 |
| LAYNEEQIHK | 10 | Q9UHD2 | Serine/threonine-protein kinase TBK1 OS | TBK1 | 83.643 | 2 | 52.26141 | yes | 1.163E-07 | 622.81714 | 100.83798 |
| KYEAFLNK | 8 | Q9UHD2 | Serine/threonine-protein kinase TBK1 OS | TBK1 | 83.643 | 2 | 52.26141 | yes | 9.925E-08 | 506.77673 | 94.054268 |
| LKEEMEGVVK | 10 | Q9UHD2 | Serine/threonine-protein kinase TBK1 OS | TBK1 | 83.643 | 2 | 52.26141 | yes | 1.249E-06 | 581.31281 | 67.098999 |
| RLVLEPGR | 8 | Q9UHD2 | Serine/threonine-protein kinase TBK1 OS | TBK1 | 83.643 | 2 | 52.26141 | yes | 4.749E-06 | 470.29034 | 20.010254 |
| VPEVPTAPATDAA | 15 | Q9UHD8 | Septin-9 OS=Homo sapiens OX=9606 GI | SEPTIN9 | 65.402 | 2 | 51.303165 | yes | 1.744E-07 | 732.39069 | 211.91278 |
| VVNIVPVIAK | 10 | Q9UHD8 | Septin-9 OS=Homo sapiens OX=9606 GI | SEPTIN9 | 65.402 | 2 | 51.303165 | yes | 9.722E-08 | 526.34735 | 206.05753 |
| DITSSIHFEAYR | 12 | Q9UHD8 | Septin-9 OS=Homo sapiens OX=9606 GI | SEPTIN9 | 65.402 | 2 | 51.303165 | yes | 1.248E-07 | 719.85168 | 148.10782 |
| YLQEEVNINR | 10 | Q9UHD8 | Septin-9 OS=Homo sapiens OX=9606 GI | SEPTIN9 | 65.402 | 2 | 51.303165 | yes | 1.022E-07 | 639.32788 | 133.52501 |
| PQPPVAEATPR | 11 | Q9UHD8 | Septin-9 OS=Homo sapiens OX=9606 GI | SEPTIN9 | 65.402 | 2 | 51.303165 | yes | 1.251E-07 | 581.81439 | 123.08881 |
| LGDSSGPALK | 10 | Q9UHD8 | Septin-9 OS=Homo sapiens OX=9606 GI | SEPTIN9 | 65.402 | 2 | 51.303165 | yes | 1.288E-07 | 472.75601 | 95.727493 |
| EFDEDSEDR | 9 | Q9UHD8 | Septin-9 OS=Homo sapiens OX=9606 GI | SEPTIN9 | 65.402 | 2 | 51.303165 | yes | 9.849E-08 | 571.21765 | 94.521362 |
| QVENAGAIGPSR | 12 | Q9UHD8 | Septin-9 OS=Homo sapiens OX=9606 GI | SEPTIN9 | 65.402 | 2 | 51.303165 | yes | 1.167E-07 | 599.81238 | 86.689285 |
| SITHDIEEK | 9 | Q9UHD8 | Septin-9 OS=Homo sapiens OX=9606 GI | SEPTIN9 | 65.402 | 2 | 51.303165 | yes | 1.986E-07 | 536.26947 | 71.903015 |
| RTELSIDISSK | 11 | Q9UHD8 | Septin-9 OS=Homo sapiens OX=9606 GI | SEPTIN9 | 65.402 | 2 | 51.303165 | yes | 9.819E-08 | 624.84332 | 69.086319 |
| VQTPLLR | 7 | Q9UHD8 | Septin-9 OS=Homo sapiens OX=9606 GI | SEPTIN9 | 65.402 | 2 | 51.303165 | yes | 8.021E-07 | 413.76089 | 41.432354 |
| STLINTLFK | 9 | Q9UHD8 | Septin-9 OS=Homo sapiens OX=9606 GI | SEPTIN9 | 65.402 | 2 | 51.303165 | yes | 1.37E-07 | 518.80548 | 36.454197 |
| RVQTPLLR | 8 | Q9UHD8 | Septin-9 OS=Homo sapiens OX=9606 GI | SEPTIN9 | 65.402 | 2 | 51.303165 | yes | 2.865E-06 | 491.81146 | 19.652584 |
| SQTDQLVLIFAGK | 13 | Q9UHD9 | Ubiquilin-2 OS=Homo sapiens OX=9606 | UBQLN2 | 65.697 | 2 | 51.200703 | yes | 9.722E-08 | 710.39575 | 55.244232 |
| GPAAAQGSAAAPA | 16 | Q9UHD9 | Ubiquilin-2 OS=Homo sapiens OX=9606 | UBQLN2 | 65.697 | 2 | 51.200703 | yes | 2.808E-07 | 697.35718 | 24.550541 |
| NPAMMQEMMR | 10 | Q9UHD9 | Ubiquilin-2 OS=Homo sapiens OX=9606 | UBQLN2 | 65.697 | 2 | 51.200703 | no | 3.074E-06 | 619.76105 | 14.836178 |
| VLISTDLTSR | 10 | Q9UHI6 | Probable ATP-dependent RNA helicase D | DDX20 | 92.241 | 2 | 52.92738 | yes | 1.254E-07 | 552.81659 | 5009.8311 |
| LFILDEADK | 9 | Q9UHI6 | Probable ATP-dependent RNA helicase D | DDX20 | 92.241 | 2 | 52.92738 | yes | 1.745E-07 | 532.28711 | 3625.2888 |
| LNSSDPSLIGLK | 12 | Q9UHI6 | Probable ATP-dependent RNA helicase D | DDX20 | 92.241 | 2 | 52.92738 | yes | 1.301E-07 | 622.34827 | 3148.3567 |
| VVNSYPLAHK | 10 | Q9UHI6 | Probable ATP-dependent RNA helicase D | DDX20 | 92.241 | 2 | 52.92738 | yes | 9.733E-08 | 564.31403 | 2336.4478 |
| YQESPGIQMK | 10 | Q9UHI6 | Probable ATP-dependent RNA helicase D | DDX20 | 92.241 | 2 | 52.92738 | yes | 5.358E-07 | 590.78699 | 2290.2507 |
| TQHLQELFSR | 10 | Q9UHI6 | Probable ATP-dependent RNA helicase D | DDX20 | 92.241 | 2 | 52.92738 | yes | 9.722E-08 | 629.83057 | 1096.2992 |
| TQHLQELFSR | 10 | Q9UHI6 | Probable ATP-dependent RNA helicase D | DDX20 | 92.241 | 2 | 52.92738 | yes | 1.111E-07 | 420.22281 | 1096.2992 |
| TQHLQELFSR | 10 | Q9UHI6 | Probable ATP-dependent RNA helicase D | DDX20 | 92.241 | 3 | 52.92738 | yes | 9.722E-08 | 629.83057 | 1096.2992 |
| TQHLQELFSR | 10 | Q9UHI6 | Probable ATP-dependent RNA helicase D | DDX20 | 92.241 | 3 | 52.92738 | yes | 1.111E-07 | 420.22281 | 1096.2992 |
| FGTLGLTVTYCCR | 13 | Q9UHI6 | Probable ATP-dependent RNA helicase D | DDX20 | 92.241 | 2 | 52.92738 | yes | 9.849E-08 | 774.37091 | 1709.3013 |
| CGLDLIVQAK | 10 | Q9UHI6 | Probable ATP-dependent RNA helicase D | DDX20 | 92.241 | 2 | 52.92738 | yes | 1.748E-07 | 558.80774 | 1647.7615 |
| AQHLADILSSK | 11 | Q9UHI6 | Probable ATP-dependent RNA helicase D | DDX20 | 92.241 | 2 | 52.92738 | yes | 9.819E-08 | 591.82751 | 1637.3003 |
| QLIELDYLNPGSIR | 14 | Q9UHI6 | Probable ATP-dependent RNA helicase D | DDX20 | 92.241 | 2 | 52.92738 | yes | 9.722E-08 | 815.9436 | 1521.1335 |
| EGLEKPVEIIR | 11 | Q9UHI6 | Probable ATP-dependent RNA helicase D | DDX20 | 92.241 | 2 | 52.92738 | yes | 9.733E-08 | 641.87189 | 1448.3302 |

| | | | | | | | | | | | |
|---|---|---|---|---|---|---|---|---|---|---|---|
| SHSECGIIEK | 10 | Q9UHI6 | Probable ATP-dependent RNA helicase D | DDX20 | 92.241 | 2 | 52.92738 | yes | 1.022E-07 | 580.27423 | 1187.0782 |
| SYLEGSSDNQLK | 12 | Q9UHI6 | Probable ATP-dependent RNA helicase D | DDX20 | 92.241 | 2 | 52.92738 | yes | 9.722E-08 | 670.82007 | 1150.1207 |
| AAVHTYGIASVPN | 17 | Q9UHI6 | Probable ATP-dependent RNA helicase D | DDX20 | 92.241 | 3 | 52.92738 | yes | 9.722E-08 | 589.32471 | 996.49646 |
| AAVHTYGIASVPN | 18 | Q9UHI6 | Probable ATP-dependent RNA helicase D | DDX20 | 92.241 | 3 | 52.92738 | yes | 1.508E-07 | 632.02301 | 910.9848 |
| NNSVSGLSVK | 10 | Q9UHI6 | Probable ATP-dependent RNA helicase D | DDX20 | 92.241 | 2 | 52.92738 | yes | 3.098E-07 | 502.77219 | 685.01166 |
| GEEENMMMR | 9 | Q9UHI6 | Probable ATP-dependent RNA helicase D | DDX20 | 92.241 | 2 | 52.92738 | yes | 1.006E-07 | 563.71991 | 594.88336 |
| IPFNQALVFSNLHS | 15 | Q9UHI6 | Probable ATP-dependent RNA helicase D | DDX20 | 92.241 | 3 | 52.92738 | yes | 1.543E-07 | 581.64941 | 462.87256 |
| AAGFERPSPVQLK | 13 | Q9UHI6 | Probable ATP-dependent RNA helicase D | DDX20 | 92.241 | 2 | 52.92738 | yes | 1.173E-07 | 700.38824 | 235.12546 |
| TAQDLSSPR | 9 | Q9UHI6 | Probable ATP-dependent RNA helicase D | DDX20 | 92.241 | 2 | 52.92738 | yes | 1.091E-07 | 487.74872 | 228.3573 |
| EIAVQIHSVITAIGI | 16 | Q9UHI6 | Probable ATP-dependent RNA helicase D | DDX20 | 92.241 | 2 | 52.92738 | yes | 9.722E-08 | 846.50635 | 151.80038 |
| NSVQTPVENSTNS | 18 | Q9UHI6 | Probable ATP-dependent RNA helicase D | DDX20 | 92.241 | 3 | 52.92738 | yes | 1.022E-07 | 666.32764 | 150.16217 |
| CHIAVGSPGR | 10 | Q9UHI6 | Probable ATP-dependent RNA helicase D | DDX20 | 92.241 | 2 | 52.92738 | yes | 3.391E-07 | 527.26654 | 128.93184 |
| NVILQCVR | 8 | Q9UHK0 | FMR1-interacting protein NUFIP1 OS=H | NUFIP1 | 56.3 | 2 | 48.69791 | yes | 5.799E-07 | 501.28168 | 532.85748 |
| NYPTLANIER | 10 | Q9UHK0 | FMR1-interacting protein NUFIP1 OS=H | NUFIP1 | 56.3 | 2 | 48.69791 | yes | 2.286E-07 | 595.81183 | 338.89514 |
| GAVLTTTQYGK | 11 | Q9UHK0 | FMR1-interacting protein NUFIP1 OS=H | NUFIP1 | 56.3 | 2 | 48.69791 | yes | 1.743E-07 | 569.80878 | 205.96387 |
| DFFGLDTNSAK | 11 | Q9UHK0 | FMR1-interacting protein NUFIP1 OS=H | NUFIP1 | 56.3 | 2 | 48.69791 | yes | 1.365E-07 | 607.78802 | 153.58853 |
| DYHNYQTLFEPR | 12 | Q9UHK0 | FMR1-interacting protein NUFIP1 OS=H | NUFIP1 | 56.3 | 2 | 48.69791 | yes | 1.267E-07 | 791.86786 | 65.05468 |
| CPELDCSFTAHEK | 13 | Q9UHK0 | FMR1-interacting protein NUFIP1 OS=H | NUFIP1 | 56.3 | 2 | 48.69791 | yes | 1.349E-06 | 797.33722 | 61.453426 |
| NMHAPGMK | 8 | Q9UHK0 | FMR1-interacting protein NUFIP1 OS=H | NUFIP1 | 56.3 | 2 | 48.69791 | yes | 2.736E-07 | 443.20712 | 50.083328 |
| AVTGSGSHLCDLK | 13 | Q9UHK0 | FMR1-interacting protein NUFIP1 OS=H | NUFIP1 | 56.3 | 2 | 48.69791 | yes | 1.113E-07 | 672.83246 | 32.053497 |
| VDAGQPELLTER | 12 | Q9UHN1 | DNA polymerase subunit gamma-2, mito | POLG2 | 54.912 | 2 | 45.476055 | yes | 1.839E-07 | 664.34625 | 106.30674 |
| EIEYVETVTSR | 11 | Q9UHR4 | Brain-specific angiogenesis inhibitor 1-as | BAIAP2L1 | 56.883 | 2 | 53.52728 | yes | 9.927E-08 | 663.33282 | 709.65417 |
| DGWLYGEHDVSK | 12 | Q9UHR4 | Brain-specific angiogenesis inhibitor 1-as | BAIAP2L1 | 56.883 | 2 | 53.52728 | yes | 9.722E-08 | 703.32275 | 550.91565 |
| IMNMIEEIK | 9 | Q9UHR4 | Brain-specific angiogenesis inhibitor 1-as | BAIAP2L1 | 56.883 | 2 | 53.52728 | yes | 9.722E-08 | 560.79065 | 209.98605 |
| TIFPHTAGSNK | 11 | Q9UHR4 | Brain-specific angiogenesis inhibitor 1-as | BAIAP2L1 | 56.883 | 2 | 53.52728 | yes | 1.058E-07 | 586.80658 | 168.81268 |
| KIELDVK | 7 | Q9UHR4 | Brain-specific angiogenesis inhibitor 1-as | BAIAP2L1 | 56.883 | 2 | 53.52728 | yes | 1.535E-07 | 422.76056 | 141.14705 |
| SVSVATGLNMMK | 12 | Q9UHR4 | Brain-specific angiogenesis inhibitor 1-as | BAIAP2L1 | 56.883 | 2 | 53.52728 | yes | 3.429E-07 | 619.31757 | 133.22368 |
| VNNSTGTSEDPSL( | 15 | Q9UHR4 | Brain-specific angiogenesis inhibitor 1-as | BAIAP2L1 | 56.883 | 2 | 53.52728 | yes | 9.722E-08 | 802.87915 | 131.98679 |
| GWFPSSYTK | 9 | Q9UHR4 | Brain-specific angiogenesis inhibitor 1-as | BAIAP2L1 | 56.883 | 2 | 53.52728 | yes | 3.403E-07 | 536.75854 | 125.95393 |
| EIIHELEK | 8 | Q9UHR4 | Brain-specific angiogenesis inhibitor 1-as | BAIAP2L1 | 56.883 | 2 | 53.52728 | yes | 1.224E-07 | 505.77948 | 93.166664 |
| NVMEQFNPGLR | 11 | Q9UHR4 | Brain-specific angiogenesis inhibitor 1-as | BAIAP2L1 | 56.883 | 2 | 53.52728 | yes | 3.081E-07 | 652.8244 | 86.463791 |
| WQETCVDAIK | 10 | Q9UHR4 | Brain-specific angiogenesis inhibitor 1-as | BAIAP2L1 | 56.883 | 2 | 53.52728 | yes | 1.174E-07 | 625.29773 | 64.251556 |
| LNESLDENFK | 10 | Q9UHR4 | Brain-specific angiogenesis inhibitor 1-as | BAIAP2L1 | 56.883 | 2 | 53.52728 | yes | 1.453E-07 | 604.79333 | 48.524357 |
| TTVISAVGTIVK | 12 | Q9UHR5 | SAP30-binding protein OS=Homo sapien | SAP30BP | 33.871 | 2 | 54.571697 | yes | 1.112E-07 | 594.86353 | 314.53354 |
| GGLVSDAYGEDDI | 15 | Q9UHR5 | SAP30-binding protein OS=Homo sapien | SAP30BP | 33.871 | 2 | 54.571697 | yes | 1.482E-07 | 794.34973 | 196.18376 |
| EGMDMNYIIQR | 11 | Q9UHR5 | SAP30-binding protein OS=Homo sapien | SAP30BP | 33.871 | 2 | 54.571697 | yes | 9.722E-08 | 685.31555 | 34.280594 |
| MVGGVLVER | 9 | Q9UHV9 | Prefoldin subunit 2 OS=Homo sapiens O: | PFDN2 | 16.648 | 2 | 49.031677 | yes | 3.668E-07 | 480.27078 | 317.64035 |
| LMGEDEKPAAK | 11 | Q9UHV9 | Prefoldin subunit 2 OS=Homo sapiens O: | PFDN2 | 16.648 | 2 | 49.031677 | yes | 2.084E-07 | 594.80011 | 106.13352 |
| GAVSAEQVIAGFN | 14 | Q9UHV9 | Prefoldin subunit 2 OS=Homo sapiens O: | PFDN2 | 16.648 | 2 | 49.031677 | yes | 2.748E-07 | 709.87299 | 87.753174 |

| | | | | | | | | | | | |
|---|---|---|---|---|---|---|---|---|---|---|---|
| IIETLTQQLQAK | 12 | Q9UHV9 | Prefoldin subunit 2 OS=Homo sapiens O | PFDN2 | 16.648 | 2 | 49.031677 | yes | 1.075E-07 | 693.40356 | 72.171478 |
| SIDMSWDSVTMK | 12 | Q9UHX1 | Poly(U)-binding-splicing factor PUF60 O | PUF60 | 59.876 | 2 | 53.070168 | yes | 1.402E-07 | 700.31525 | 1140.5856 |
| SIDMSWDSVTMK | 12 | Q9UHX1 | Poly(U)-binding-splicing factor PUF60 O | PUF60 | 59.876 | 2 | 53.070168 | yes | 2.292E-07 | 708.31268 | 1140.5856 |
| IFVEFSIASETHK | 13 | Q9UHX1 | Poly(U)-binding-splicing factor PUF60 O | PUF60 | 59.876 | 2 | 53.070168 | yes | 9.722E-08 | 754.39319 | 2162.7397 |
| GYGFIEYEK | 9 | Q9UHX1 | Poly(U)-binding-splicing factor PUF60 O | PUF60 | 59.876 | 2 | 53.070168 | yes | 1.1E-07 | 553.26367 | 1729.8313 |
| QAFAPFGPIK | 10 | Q9UHX1 | Poly(U)-binding-splicing factor PUF60 O | PUF60 | 59.876 | 2 | 53.070168 | yes | 4.91E-07 | 538.30035 | 1562.7896 |
| WKPPQGTDSIK | 11 | Q9UHX1 | Poly(U)-binding-splicing factor PUF60 O | PUF60 | 59.876 | 2 | 53.070168 | yes | 6.069E-07 | 628.83533 | 1555.276 |
| VIIYQEK | 7 | Q9UHX1 | Poly(U)-binding-splicing factor PUF60 O | PUF60 | 59.876 | 2 | 53.070168 | yes | 2.004E-07 | 446.76056 | 1323.8221 |
| VVAEVYDQER | 10 | Q9UHX1 | Poly(U)-binding-splicing factor PUF60 O | PUF60 | 59.876 | 2 | 53.070168 | yes | 1.338E-07 | 604.30133 | 1021.587 |
| KQESTVMVLR | 10 | Q9UHX1 | Poly(U)-binding-splicing factor PUF60 O | PUF60 | 59.876 | 2 | 53.070168 | yes | 9.722E-08 | 595.83173 | 898.85748 |
| QGEEEDAEIIVK | 12 | Q9UHX1 | Poly(U)-binding-splicing factor PUF60 O | PUF60 | 59.876 | 2 | 53.070168 | yes | 1.347E-07 | 680.33557 | 638.67108 |
| KVVAEVYDQER | 11 | Q9UHX1 | Poly(U)-binding-splicing factor PUF60 O | PUF60 | 59.876 | 2 | 53.070168 | yes | 1.024E-07 | 668.34882 | 443.02887 |
| ALAIMCR | 7 | Q9UHX1 | Poly(U)-binding-splicing factor PUF60 O | PUF60 | 59.876 | 2 | 53.070168 | yes | 1.405E-07 | 417.71985 | 278.23639 |
| KYAMEQSIK | 9 | Q9UHX1 | Poly(U)-binding-splicing factor PUF60 O | PUF60 | 59.876 | 2 | 53.070168 | yes | 2.031E-07 | 549.28662 | 156.99898 |
| YAMEQSIK | 8 | Q9UHX1 | Poly(U)-binding-splicing factor PUF60 O | PUF60 | 59.876 | 2 | 53.070168 | yes | 2.17E-07 | 485.23914 | 108.25857 |
| EEEELFPESERPEM | 29 | Q9UHX1 | Poly(U)-binding-splicing factor PUF60 O | PUF60 | 59.876 | 4 | 53.070168 | yes | 3.284E-07 | 838.37512 | 47.280743 |
| GLQQITGHGGLR | 12 | Q9UI09 | NADH dehydrogenase [ubiquinone] 1 alp | NDUFA12 | 17.115 | 2 | 50.025623 | yes | 9.927E-08 | 618.84399 | 40.60202 |
| VGTAQLALVAR | 11 | Q9UI10 | Translation initiation factor eIF-2B subun | EIF2B4 | 57.558 | 2 | 55.421684 | yes | 9.823E-08 | 549.83514 | 274.64075 |
| IVLAAQAISR | 10 | Q9UI10 | Translation initiation factor eIF-2B subun | EIF2B4 | 57.558 | 2 | 55.421684 | yes | 1.381E-07 | 521.3244 | 210.40042 |
| VSLFSHLPQYSR | 12 | Q9UI10 | Translation initiation factor eIF-2B subun | EIF2B4 | 57.558 | 2 | 55.421684 | yes | 9.722E-08 | 717.38043 | 183.72928 |
| ELPESGIQLGTPR | 13 | Q9UI10 | Translation initiation factor eIF-2B subun | EIF2B4 | 57.558 | 2 | 55.421684 | yes | 3.265E-07 | 698.87518 | 179.89037 |
| ASPSTAGETPSGVI | 14 | Q9UI10 | Translation initiation factor eIF-2B subun | EIF2B4 | 57.558 | 2 | 55.421684 | yes | 4.899E-07 | 644.82263 | 165.41112 |
| LGLQYSQGLVSGS | 16 | Q9UI10 | Translation initiation factor eIF-2B subun | EIF2B4 | 57.558 | 2 | 55.421684 | yes | 9.722E-08 | 825.4339 | 155.01373 |
| ILQEAWTEGR | 10 | Q9UI10 | Translation initiation factor eIF-2B subun | EIF2B4 | 57.558 | 2 | 55.421684 | yes | 2.041E-07 | 601.81183 | 126.70592 |
| LPEYPQVDDLLLR | 13 | Q9UI10 | Translation initiation factor eIF-2B subun | EIF2B4 | 57.558 | 2 | 55.421684 | yes | 4.059E-07 | 785.92743 | 32.289383 |
| ICPVEFNPNFVAR | 13 | Q9UI30 | Multifunctional methyltransferase subunit | TRMT112 | 14.2 | 2 | 50.049095 | yes | 9.926E-08 | 781.89282 | 2004.4564 |
| GPVEGYEENEEFL | 14 | Q9UI30 | Multifunctional methyltransferase subunit | TRMT112 | 14.2 | 2 | 50.049095 | yes | 1.326E-07 | 834.38104 | 489.53656 |
| VEWSAFLEAADNI | 14 | Q9UI30 | Multifunctional methyltransferase subunit | TRMT112 | 14.2 | 2 | 50.049095 | yes | 5.106E-07 | 810.90448 | 334.72601 |
| LQATEVR | 7 | Q9UI30 | Multifunctional methyltransferase subunit | TRMT112 | 14.2 | 2 | 50.049095 | yes | 2.841E-07 | 408.73233 | 236.71539 |
| GIPNMLLSEEETES | 14 | Q9UI30 | Multifunctional methyltransferase subunit | TRMT112 | 14.2 | 2 | 50.049095 | yes | 1.876E-07 | 774.85852 | 81.012939 |
| ADVILSDMAPNAT | 16 | Q9UI43 | rRNA methyltransferase 2, mitochondrial | MRM2 | 27.424 | 2 | 50.358963 | yes | 2.007E-06 | 839.41687 | 215.82591 |
| IIKPEASR | 8 | Q9UI43 | rRNA methyltransferase 2, mitochondrial | MRM2 | 27.424 | 2 | 50.358963 | yes | 1.046E-07 | 457.27692 | 203.24817 |
| LTEEFQNVR | 9 | Q9UI43 | rRNA methyltransferase 2, mitochondrial | MRM2 | 27.424 | 2 | 50.358963 | yes | 1.025E-07 | 568.29077 | 114.48781 |
| TGAEHLWLTR | 10 | Q9UI43 | rRNA methyltransferase 2, mitochondrial | MRM2 | 27.424 | 2 | 50.358963 | yes | 9.81E-08 | 592.31458 | 62.578033 |
| NQNYLVPSPVLR | 12 | Q9UID6 | Zinc finger protein 639 OS=Homo sapien | ZNF639 | 56.055 | 2 | 51.936344 | yes | 9.722E-08 | 700.38824 | 242.2014 |
| SQALNVTAQQK | 11 | Q9UID6 | Zinc finger protein 639 OS=Homo sapien | ZNF639 | 56.055 | 2 | 51.936344 | yes | 9.722E-08 | 594.32257 | 145.70064 |
| NFFVCQVCGFR | 11 | Q9UID6 | Zinc finger protein 639 OS=Homo sapien | ZNF639 | 56.055 | 2 | 51.936344 | yes | 1.262E-07 | 717.32629 | 82.777443 |
| ESFSTNMLLIEHAI | 14 | Q9UID6 | Zinc finger protein 639 OS=Homo sapien | ZNF639 | 56.055 | 2 | 51.936344 | yes | 1.553E-07 | 810.40851 | 56.21336 |
| CELCEFNSK | 9 | Q9UID6 | Zinc finger protein 639 OS=Homo sapien | ZNF639 | 56.055 | 2 | 51.936344 | yes | 9.39E-07 | 593.74701 | 50.239021 |

| | | | | | | | | | | | |
|---|---|---|---|---|---|---|---|---|---|---|---|
| LHEEDPYICK | 10 | Q9UID6 | Zinc finger protein 639 OS=Homo sapien | ZNF639 | 56.055 | 2 | 51.936344 | yes | 1.17E-06 | 652.30298 | 28.114599 |
| MPVGDFFEER | 10 | Q9UIF9 | Bromodomain adjacent to zinc finger don | BAZ2A | 211.198 | 2 | 47.81842 | yes | 1.252E-07 | 613.77917 | 40.930687 |
| IISNVPADSLIR | 12 | Q9UIG0 | Tyrosine-protein kinase BAZ1B OS=Hon | BAZ1B | 170.903 | 2 | 54.773224 | yes | 1.004E-07 | 649.37738 | 337.83823 |
| GGLGYVEETSEFE | 15 | Q9UIG0 | Tyrosine-protein kinase BAZ1B OS=Hon | BAZ1B | 170.903 | 2 | 54.773224 | yes | 9.733E-08 | 822.38104 | 110.81194 |
| LVLEMVHHNTASI | 15 | Q9UIG0 | Tyrosine-protein kinase BAZ1B OS=Hon | BAZ1B | 170.903 | 3 | 54.773224 | yes | 2.386E-06 | 574.30652 | 92.917053 |
| SCDGNQELLNFLR | 13 | Q9UIG0 | Tyrosine-protein kinase BAZ1B OS=Hon | BAZ1B | 170.903 | 2 | 54.773224 | yes | 9.722E-08 | 783.3725 | 82.340652 |
| LETSEFFELTSEEK | 14 | Q9UIG0 | Tyrosine-protein kinase BAZ1B OS=Hon | BAZ1B | 170.903 | 2 | 54.773224 | yes | 6.039E-07 | 844.89868 | 76.767159 |
| WASMSEEQR | 9 | Q9UIG0 | Tyrosine-protein kinase BAZ1B OS=Hon | BAZ1B | 170.903 | 2 | 54.773224 | yes | 1.527E-07 | 562.24548 | 70.694786 |
| YAVGEECDFEVGK | 13 | Q9UIG0 | Tyrosine-protein kinase BAZ1B OS=Hon | BAZ1B | 170.903 | 2 | 54.773224 | yes | 5.345E-07 | 751.82703 | 65.284576 |
| FSDFLLDPYK | 10 | Q9UIG0 | Tyrosine-protein kinase BAZ1B OS=Hon | BAZ1B | 170.903 | 2 | 54.773224 | yes | 2.736E-07 | 622.81354 | 60.445358 |
| QGQNLWFLCDSQI | 13 | Q9UIG0 | Tyrosine-protein kinase BAZ1B OS=Hon | BAZ1B | 170.903 | 2 | 54.773224 | yes | 2.044E-07 | 812.38281 | 56.200764 |
| YEDQELTGK | 9 | Q9UIG0 | Tyrosine-protein kinase BAZ1B OS=Hon | BAZ1B | 170.903 | 2 | 54.773224 | yes | 9.734E-08 | 541.75366 | 54.204872 |
| YQDIIHSIHLAR | 12 | Q9UIG0 | Tyrosine-protein kinase BAZ1B OS=Hon | BAZ1B | 170.903 | 3 | 54.773224 | yes | 1.027E-07 | 489.26852 | 40.634438 |
| SPEEHLEEMMK | 11 | Q9UIG0 | Tyrosine-protein kinase BAZ1B OS=Hon | BAZ1B | 170.903 | 2 | 54.773224 | yes | 1.146E-06 | 680.29962 | 30.113939 |
| ENSSQIAQDHQK | 12 | Q9UIG0 | Tyrosine-protein kinase BAZ1B OS=Hon | BAZ1B | 170.903 | 2 | 54.773224 | yes | 1.207E-06 | 692.82623 | 24.740532 |
| LVDTAWLEIMTK | 12 | Q9UIG0 | Tyrosine-protein kinase BAZ1B OS=Hon | BAZ1B | 170.903 | 2 | 54.773224 | yes | 9.809E-08 | 710.38123 | 18.247025 |
| QVFTNAEVYNCR | 12 | Q9UIG0 | Tyrosine-protein kinase BAZ1B OS=Hon | BAZ1B | 170.903 | 2 | 54.773224 | yes | 2.798E-07 | 750.84863 | 16.613535 |
| NASMNTQHGTATI | 22 | Q9UIG0 | Tyrosine-protein kinase BAZ1B OS=Hon | BAZ1B | 170.903 | 3 | 54.773224 | yes | 4.098E-07 | 763.3656 | 15.28936 |
| TIPTLNEGTVK | 11 | Q9UII4 | E3 ISG15--protein ligase HERC5 OS=Ho | HERC5 | 116.853 | 2 | 52.197334 | yes | 1.375E-07 | 586.82971 | 1016.067 |
| YINYIFNDSVK | 11 | Q9UII4 | E3 ISG15--protein ligase HERC5 OS=Ho | HERC5 | 116.853 | 2 | 52.197334 | yes | 9.722E-08 | 688.34827 | 718.28644 |
| DQLMPLPVK | 9 | Q9UII4 | E3 ISG15--protein ligase HERC5 OS=Ho | HERC5 | 116.853 | 2 | 52.197334 | yes | 2.656E-07 | 520.79407 | 664.33026 |
| LLDQMPSLEDLK | 12 | Q9UII4 | E3 ISG15--protein ligase HERC5 OS=Ho | HERC5 | 116.853 | 2 | 52.197334 | yes | 6.71E-07 | 701.36835 | 651.87036 |
| FLVFLTGTDR | 10 | Q9UII4 | E3 ISG15--protein ligase HERC5 OS=Ho | HERC5 | 116.853 | 2 | 52.197334 | yes | 9.722E-08 | 584.82166 | 615.77838 |
| LNFFVEVCR | 9 | Q9UII4 | E3 ISG15--protein ligase HERC5 OS=Ho | HERC5 | 116.853 | 2 | 52.197334 | yes | 4.844E-07 | 592.30005 | 591.65729 |
| DDPSLIEGLDNQK | 13 | Q9UII4 | E3 ISG15--protein ligase HERC5 OS=Ho | HERC5 | 116.853 | 2 | 52.197334 | yes | 1.041E-07 | 722.35175 | 491.99054 |
| LLHTDTLLK | 9 | Q9UII4 | E3 ISG15--protein ligase HERC5 OS=Ho | HERC5 | 116.853 | 2 | 52.197334 | yes | 2.481E-07 | 527.31879 | 407.01617 |
| FESILQEK | 8 | Q9UII4 | E3 ISG15--protein ligase HERC5 OS=Ho | HERC5 | 116.853 | 2 | 52.197334 | yes | 2.202E-07 | 497.2662 | 378.18573 |
| QLCCSPGR | 8 | Q9UII4 | E3 ISG15--protein ligase HERC5 OS=Ho | HERC5 | 116.853 | 2 | 52.197334 | yes | 3.295E-07 | 489.2182 | 310.24365 |
| VTQIACGR | 8 | Q9UII4 | E3 ISG15--protein ligase HERC5 OS=Ho | HERC5 | 116.853 | 2 | 52.197334 | yes | 9.792E-08 | 452.73709 | 282.72922 |
| TTEMMPVYLDLNI | 13 | Q9UII4 | E3 ISG15--protein ligase HERC5 OS=Ho | HERC5 | 116.853 | 2 | 52.197334 | yes | 9.722E-08 | 777.88074 | 271.16992 |
| SPGAQLWLFPSAA | 17 | Q9UII4 | E3 ISG15--protein ligase HERC5 OS=Ho | HERC5 | 116.853 | 3 | 52.197334 | yes | 4.42E-07 | 603.32495 | 241.63527 |
| GGAGVQVHQLLA | 18 | Q9UII4 | E3 ISG15--protein ligase HERC5 OS=Ho | HERC5 | 116.853 | 3 | 52.197334 | yes | 2.553E-07 | 545.62891 | 119.34105 |
| GGAGVQVHQLLA | 18 | Q9UII4 | E3 ISG15--protein ligase HERC5 OS=Ho | HERC5 | 116.853 | 3 | 52.197334 | yes | 1.042E-07 | 817.9397 | 119.34105 |
| GGAGVQVHQLLA | 18 | Q9UII4 | E3 ISG15--protein ligase HERC5 OS=Ho | HERC5 | 116.853 | 2 | 52.197334 | yes | 2.553E-07 | 545.62891 | 119.34105 |
| GGAGVQVHQLLA | 18 | Q9UII4 | E3 ISG15--protein ligase HERC5 OS=Ho | HERC5 | 116.853 | 2 | 52.197334 | yes | 1.042E-07 | 817.9397 | 119.34105 |
| SAAIEEER | 8 | Q9UII4 | E3 ISG15--protein ligase HERC5 OS=Ho | HERC5 | 116.853 | 2 | 52.197334 | yes | 3.197E-06 | 452.72217 | 171.37032 |
| ITFCCPESWNER | 12 | Q9UII4 | E3 ISG15--protein ligase HERC5 OS=Ho | HERC5 | 116.853 | 2 | 52.197334 | yes | 1.032E-07 | 799.83997 | 157.62685 |
| NECGQLGLGHTES | 14 | Q9UII4 | E3 ISG15--protein ligase HERC5 OS=Ho | HERC5 | 116.853 | 2 | 52.197334 | yes | 1.375E-07 | 765.35431 | 154.06148 |
| DVIVGNTDYDWK | 12 | Q9UII4 | E3 ISG15--protein ligase HERC5 OS=Ho | HERC5 | 116.853 | 2 | 52.197334 | yes | 9.722E-08 | 712.83826 | 142.31894 |

| | | | | | | | | | | | |
|---|---|---|---|---|---|---|---|---|---|---|---|
| MCDEDIIK | 8 | Q9UII4 | E3 ISG15--protein ligase HERC5 OS=Ho | HERC5 | 116.853 | 2 | 52.197334 | yes | 1.28E-07 | 512.22791 | 137.12292 |
| WHTLAYVSDLGK | 12 | Q9UII4 | E3 ISG15--protein ligase HERC5 OS=Ho | HERC5 | 116.853 | 2 | 52.197334 | yes | 9.722E-08 | 695.36169 | 128.03035 |
| ELSPDLGK | 8 | Q9UII4 | E3 ISG15--protein ligase HERC5 OS=Ho | HERC5 | 116.853 | 2 | 52.197334 | yes | 3.286E-07 | 429.73199 | 120.90276 |
| KENSYVNLK | 9 | Q9UII4 | E3 ISG15--protein ligase HERC5 OS=Ho | HERC5 | 116.853 | 2 | 52.197334 | yes | 7.718E-07 | 547.79565 | 80.05368 |
| KLLDQMPSLEDLK | 13 | Q9UII4 | E3 ISG15--protein ligase HERC5 OS=Ho | HERC5 | 116.853 | 2 | 52.197334 | yes | 4.41E-07 | 765.41583 | 44.545547 |
| LALHELK | 7 | Q9UIJ7 | GTP:AMP phosphotransferase AK3, mito | AK3 | 25.566 | 2 | 51.441986 | yes | 9.722E-08 | 412.25543 | 150.30086 |
| EMFFEDHIDDAK | 12 | Q9UIV1 | CCR4-NOT transcription complex subuni | CNOT7 | 32.746 | 2 | 49.92002 | yes | 9.977E-08 | 748.82178 | 244.0101 |
| SNADYQYQLLR | 11 | Q9UIV1 | CCR4-NOT transcription complex subuni | CNOT7 | 32.746 | 2 | 49.92002 | yes | 1.074E-07 | 685.83856 | 173.0152 |
| GGLQEVAEQLELE | 14 | Q9UIV1 | CCR4-NOT transcription complex subuni | CNOT7 | 32.746 | 2 | 49.92002 | yes | 1.083E-07 | 785.90723 | 130.66508 |
| IAEGAQQGDPLSR | 13 | Q9UJ70 | N-acetyl-D-glucosamine kinase OS=Hom | NAGK | 37.376 | 2 | 42.28575 | yes | 3.104E-07 | 671.34149 | 56.134605 |
| EGFLLALTQGR | 11 | Q9UJ70 | N-acetyl-D-glucosamine kinase OS=Hom | NAGK | 37.376 | 2 | 42.28575 | yes | 2.094E-06 | 602.83789 | 50.848698 |
| FISALNNVAER | 11 | Q9UJA3 | DNA helicase MCM8 OS=Homo sapiens | MCM8 | 93.697 | 2 | 53.277683 | yes | 1.764E-07 | 617.33295 | 153.34891 |
| VYNYEPLTQLK | 11 | Q9UJA3 | DNA helicase MCM8 OS=Homo sapiens | MCM8 | 93.697 | 2 | 53.277683 | yes | 3.659E-07 | 684.36389 | 122.01347 |
| IQELMSDDQR | 10 | Q9UJA3 | DNA helicase MCM8 OS=Homo sapiens | MCM8 | 93.697 | 2 | 53.277683 | yes | 3.187E-07 | 617.79022 | 108.85025 |
| GDPHILVVGDPGL | 15 | Q9UJA3 | DNA helicase MCM8 OS=Homo sapiens | MCM8 | 93.697 | 2 | 53.277683 | yes | 1.118E-07 | 737.40662 | 101.27482 |
| VLQDFYLELR | 10 | Q9UJA3 | DNA helicase MCM8 OS=Homo sapiens | MCM8 | 93.697 | 2 | 53.277683 | yes | 9.722E-08 | 648.35333 | 95.332893 |
| TSEQTPQFLLSTK | 13 | Q9UJA3 | DNA helicase MCM8 OS=Homo sapiens | MCM8 | 93.697 | 2 | 53.277683 | yes | 3.156E-07 | 740.38812 | 72.959457 |
| VSNIKPLCTK | 10 | Q9UJA3 | DNA helicase MCM8 OS=Homo sapiens | MCM8 | 93.697 | 2 | 53.277683 | yes | 2.246E-07 | 580.3288 | 68.947357 |
| LNSSPITTR | 9 | Q9UJA3 | DNA helicase MCM8 OS=Homo sapiens | MCM8 | 93.697 | 2 | 53.277683 | yes | 1.394E-06 | 494.77472 | 68.029449 |
| MNSQDSNTSVLEV | 22 | Q9UJA3 | DNA helicase MCM8 OS=Homo sapiens | MCM8 | 93.697 | 3 | 53.277683 | yes | 2.17E-07 | 817.06738 | 59.537277 |
| SQMLQAACNVAPI | 13 | Q9UJA3 | DNA helicase MCM8 OS=Homo sapiens | MCM8 | 93.697 | 2 | 53.277683 | yes | 4.652E-07 | 723.35303 | 58.184109 |
| LIVNSLCPVIFGHE | 17 | Q9UJA3 | DNA helicase MCM8 OS=Homo sapiens | MCM8 | 93.697 | 3 | 53.277683 | yes | 2.545E-07 | 646.70087 | 38.569656 |
| HIDLYDKDEIER | 12 | Q9UJA3 | DNA helicase MCM8 OS=Homo sapiens | MCM8 | 93.697 | 3 | 53.277683 | yes | 8.104E-07 | 515.92297 | 35.463886 |
| TSIIAAANPVGGHY | 16 | Q9UJA3 | DNA helicase MCM8 OS=Homo sapiens | MCM8 | 93.697 | 2 | 53.277683 | yes | 5.442E-07 | 806.92572 | 30.430201 |
| HLEAAALLSER | 11 | Q9UJA5 | tRNA (adenine(58)-N(1))-methyltransfera | TRMT6 | 55.8 | 2 | 51.667984 | yes | 1.008E-07 | 605.33295 | 958.1593 |
| NADGLIVASR | 10 | Q9UJA5 | tRNA (adenine(58)-N(1))-methyltransfera | TRMT6 | 55.8 | 2 | 51.667984 | yes | 5.413E-07 | 508.28018 | 377.17163 |
| KYEAIITVVKPSTR | 14 | Q9UJA5 | tRNA (adenine(58)-N(1))-methyltransfera | TRMT6 | 55.8 | 3 | 51.667984 | yes | 9.722E-08 | 535.65051 | 157.92111 |
| KYEAIITVVKPSTR | 14 | Q9UJA5 | tRNA (adenine(58)-N(1))-methyltransfera | TRMT6 | 55.8 | 3 | 51.667984 | yes | 9.722E-08 | 802.97217 | 157.92111 |
| KYEAIITVVKPSTR | 14 | Q9UJA5 | tRNA (adenine(58)-N(1))-methyltransfera | TRMT6 | 55.8 | 2 | 51.667984 | yes | 9.722E-08 | 535.65051 | 157.92111 |
| KYEAIITVVKPSTR | 14 | Q9UJA5 | tRNA (adenine(58)-N(1))-methyltransfera | TRMT6 | 55.8 | 2 | 51.667984 | yes | 9.722E-08 | 802.97217 | 157.92111 |
| AATACFGFPK | 10 | Q9UJA5 | tRNA (adenine(58)-N(1))-methyltransfera | TRMT6 | 55.8 | 2 | 51.667984 | yes | 9.919E-08 | 535.26038 | 291.39777 |
| EPLLECYTK | 9 | Q9UJA5 | tRNA (adenine(58)-N(1))-methyltransfera | TRMT6 | 55.8 | 2 | 51.667984 | yes | 9.723E-08 | 576.78394 | 270.40799 |
| IRDGDFVVLK | 10 | Q9UJA5 | tRNA (adenine(58)-N(1))-methyltransfera | TRMT6 | 55.8 | 2 | 51.667984 | yes | 1.021E-07 | 581.33496 | 180.09438 |
| ILSIMYYAR | 9 | Q9UJA5 | tRNA (adenine(58)-N(1))-methyltransfera | TRMT6 | 55.8 | 2 | 51.667984 | yes | 2.149E-06 | 565.30737 | 170.90324 |
| NYQVLPDR | 8 | Q9UJA5 | tRNA (adenine(58)-N(1))-methyltransfera | TRMT6 | 55.8 | 2 | 51.667984 | yes | 1.39E-07 | 502.76163 | 158.27133 |
| SFLSGLYEFPLNK | 13 | Q9UJA5 | tRNA (adenine(58)-N(1))-methyltransfera | TRMT6 | 55.8 | 2 | 51.667984 | yes | 1.414E-07 | 757.89813 | 147.44275 |
| YEAIITVVKPSTR | 13 | Q9UJA5 | tRNA (adenine(58)-N(1))-methyltransfera | TRMT6 | 55.8 | 2 | 51.667984 | yes | 1.029E-07 | 738.92468 | 134.22621 |
| DSALVEESNGTLE | 15 | Q9UJA5 | tRNA (adenine(58)-N(1))-methyltransfera | TRMT6 | 55.8 | 2 | 51.667984 | yes | 1.228E-05 | 810.88361 | 130.82217 |
| DKTEFAQDK | 9 | Q9UJA5 | tRNA (adenine(58)-N(1))-methyltransfera | TRMT6 | 55.8 | 2 | 51.667984 | yes | 9.722E-08 | 541.26166 | 78.464531 |

| | | | | | | | | | | | |
|---|---|---|---|---|---|---|---|---|---|---|---|
| PFVVYCQYK | 9 | Q9UJA5 | tRNA (adenine(58)-N(1))-methyltransfera | TRMT6 | 55.8 | 2 | 51.667984 | yes | 3.835E-07 | 602.297 | 75.91642 |
| KYEAIITVVK | 10 | Q9UJA5 | tRNA (adenine(58)-N(1))-methyltransfera | TRMT6 | 55.8 | 2 | 51.667984 | yes | 1.026E-07 | 582.35535 | 67.326851 |
| LSETWLR | 7 | Q9UJA5 | tRNA (adenine(58)-N(1))-methyltransfera | TRMT6 | 55.8 | 2 | 51.667984 | yes | 7.12E-07 | 452.74799 | 50.536617 |
| ALKDEIDVLR | 10 | Q9UJC3 | Protein Hook homolog 1 OS=Homo sapie | HOOK1 | 84.648 | 2 | 47.467472 | yes | 1.324E-07 | 586.33771 | 54.422089 |
| HIPLALPR | 8 | Q9UJF2 | Ras GTPase-activating protein nGAP OS= | RASAL2 | 128.558 | 2 | 56.126568 | yes | 3.501E-06 | 458.78998 | 427.32462 |
| SLTNPTPIQQQLR | 13 | Q9UJF2 | Ras GTPase-activating protein nGAP OS= | RASAL2 | 128.558 | 2 | 56.126568 | yes | 1.043E-07 | 748.41498 | 229.38722 |
| LMAVEEELKK | 10 | Q9UJF2 | Ras GTPase-activating protein nGAP OS= | RASAL2 | 128.558 | 2 | 56.126568 | yes | 2.452E-06 | 595.32849 | 197.20082 |
| IINSYCVFPR | 10 | Q9UJF2 | Ras GTPase-activating protein nGAP OS= | RASAL2 | 128.558 | 2 | 56.126568 | no | 2.156E-07 | 634.82642 | 170.20615 |
| LWIIEAK | 7 | Q9UJF2 | Ras GTPase-activating protein nGAP OS= | RASAL2 | 128.558 | 2 | 56.126568 | yes | 3.268E-07 | 436.76566 | 129.07901 |
| ASIDSSLENLSTAS | 16 | Q9UJF2 | Ras GTPase-activating protein nGAP OS= | RASAL2 | 128.558 | 2 | 56.126568 | yes | 9.733E-08 | 819.40247 | 118.53313 |
| FQTITILPMEQYK | 13 | Q9UJF2 | Ras GTPase-activating protein nGAP OS= | RASAL2 | 128.558 | 2 | 56.126568 | yes | 3.655E-07 | 806.42621 | 116.08557 |
| LLVQEQQMQK | 10 | Q9UJF2 | Ras GTPase-activating protein nGAP OS= | RASAL2 | 128.558 | 2 | 56.126568 | yes | 5.593E-07 | 622.83698 | 98.633011 |
| LSGPSNSSMEDFTI | 14 | Q9UJF2 | Ras GTPase-activating protein nGAP OS= | RASAL2 | 128.558 | 2 | 56.126568 | yes | 3.954E-07 | 750.33777 | 97.568481 |
| APPSLPHSASLR | 12 | Q9UJF2 | Ras GTPase-activating protein nGAP OS= | RASAL2 | 128.558 | 2 | 56.126568 | yes | 2.895E-07 | 616.84094 | 97.037689 |
| LMSALTQVK | 9 | Q9UJF2 | Ras GTPase-activating protein nGAP OS= | RASAL2 | 128.558 | 2 | 56.126568 | yes | 4.066E-07 | 495.78625 | 92.849846 |
| LTGSQLSITQVASI | 15 | Q9UJF2 | Ras GTPase-activating protein nGAP OS= | RASAL2 | 128.558 | 2 | 56.126568 | yes | 9.722E-08 | 773.44598 | 82.727943 |
| WYPVSTPTPNK | 11 | Q9UJF2 | Ras GTPase-activating protein nGAP OS= | RASAL2 | 128.558 | 2 | 56.126568 | yes | 3.335E-06 | 645.3299 | 66.093872 |
| CGEHDVLIFR | 10 | Q9UJF2 | Ras GTPase-activating protein nGAP OS= | RASAL2 | 128.558 | 2 | 56.126568 | yes | 1.855E-07 | 623.30585 | 64.27874 |
| ETQSTPQSAPQVR | 13 | Q9UJF2 | Ras GTPase-activating protein nGAP OS= | RASAL2 | 128.558 | 2 | 56.126568 | yes | 2.545E-07 | 714.85748 | 62.769459 |
| GENSFLQATVAK | 12 | Q9UJF2 | Ras GTPase-activating protein nGAP OS= | RASAL2 | 128.558 | 2 | 56.126568 | yes | 3.423E-07 | 632.83026 | 60.897938 |
| IFEDPTDSDLHK | 12 | Q9UJF2 | Ras GTPase-activating protein nGAP OS= | RASAL2 | 128.558 | 2 | 56.126568 | yes | 9.722E-08 | 708.83569 | 58.408005 |
| MCCELAFCK | 9 | Q9UJF2 | Ras GTPase-activating protein nGAP OS= | RASAL2 | 128.558 | 2 | 56.126568 | no | 2.491E-07 | 609.74237 | 44.080284 |
| RLEEYER | 7 | Q9UJF2 | Ras GTPase-activating protein nGAP OS= | RASAL2 | 128.558 | 2 | 56.126568 | yes | 2.736E-05 | 497.75125 | 37.149986 |
| NKEELACALVHIL | 18 | Q9UJF2 | Ras GTPase-activating protein nGAP OS= | RASAL2 | 128.558 | 3 | 56.126568 | yes | 2.344E-07 | 680.36157 | 35.048576 |
| ALYESDENCEVDP | 29 | Q9UJF2 | Ras GTPase-activating protein nGAP OS= | RASAL2 | 128.558 | 4 | 56.126568 | yes | 1.543E-06 | 839.3739 | 23.870449 |
| SPSQDNTDSYFR | 12 | Q9UJF2 | Ras GTPase-activating protein nGAP OS= | RASAL2 | 128.558 | 2 | 56.126568 | yes | 0.0002759 | 708.80493 | 8.2890873 |
| YLNIFGESQPNPK | 13 | Q9UJS0 | Electrogenic aspartate/glutamate antiporte | SLC25A13 | 74.176 | 2 | 51.32813 | yes | 1.608E-07 | 753.883 | 1006.1332 |
| GLLPQLLGVAPEK | 13 | Q9UJS0 | Electrogenic aspartate/glutamate antiporte | SLC25A13 | 74.176 | 2 | 51.32813 | yes | 1.242E-07 | 667.90576 | 403.83331 |
| VSALSVVR | 8 | Q9UJS0 | Electrogenic aspartate/glutamate antiporte | SLC25A13 | 74.176 | 2 | 51.32813 | yes | 1.161E-06 | 415.75836 | 331.11996 |
| LAVATFAGIENK | 12 | Q9UJS0 | Electrogenic aspartate/glutamate antiporte | SLC25A13 | 74.176 | 2 | 51.32813 | yes | 9.722E-08 | 617.34552 | 323.77344 |
| STGSFVGELMYK | 12 | Q9UJS0 | Electrogenic aspartate/glutamate antiporte | SLC25A13 | 74.176 | 2 | 51.32813 | yes | 1.296E-07 | 659.82104 | 286.48618 |
| PVLLQVAESAYR | 12 | Q9UJS0 | Electrogenic aspartate/glutamate antiporte | SLC25A13 | 74.176 | 2 | 51.32813 | yes | 1.731E-07 | 673.37738 | 259.71915 |
| MTLADIER | 8 | Q9UJS0 | Electrogenic aspartate/glutamate antiporte | SLC25A13 | 74.176 | 2 | 51.32813 | yes | 5.129E-07 | 474.7446 | 252.94614 |
| INLPAPNPDHVGG | 15 | Q9UJS0 | Electrogenic aspartate/glutamate antiporte | SLC25A13 | 74.176 | 2 | 51.32813 | yes | 1.673E-07 | 796.41498 | 186.70323 |
| EEFVLAAQK | 9 | Q9UJS0 | Electrogenic aspartate/glutamate antiporte | SLC25A13 | 74.176 | 2 | 51.32813 | yes | 2.976E-07 | 517.77948 | 175.51056 |
| TVELLSGVVDQTK | 13 | Q9UJS0 | Electrogenic aspartate/glutamate antiporte | SLC25A13 | 74.176 | 2 | 51.32813 | yes | 9.722E-08 | 694.88519 | 175.25003 |
| RADPAELR | 8 | Q9UJS0 | Electrogenic aspartate/glutamate antiporte | SLC25A13 | 74.176 | 2 | 51.32813 | yes | 1.365E-07 | 464.25397 | 147.76343 |
| VTAIDFR | 7 | Q9UJS0 | Electrogenic aspartate/glutamate antiporte | SLC25A13 | 74.176 | 2 | 51.32813 | yes | 9.722E-08 | 411.22943 | 104.93044 |
| DLGFFGIYK | 9 | Q9UJS0 | Electrogenic aspartate/glutamate antiporte | SLC25A13 | 74.176 | 2 | 51.32813 | yes | 4.522E-06 | 530.27911 | 44.365513 |

| | | | | | | | | | | | |
|---|---|---|---|---|---|---|---|---|---|---|---|
| HLTYAEFTQFLLEI | 20 | Q9UJS0 | Electrogenic aspartate/glutamate antiporte | SLC25A13 | 74.176 | 3 | 51.32813 | yes | 1.151E-07 | 811.09338 | 31.771992 |
| QVFGQTTIHQHIPF | 27 | Q9UJS0 | Electrogenic aspartate/glutamate antiporte | SLC25A13 | 74.176 | 4 | 51.32813 | yes | 2.045E-07 | 810.90582 | 31.429293 |
| HEESIMNIIR | 10 | Q9UJT1 | Tubulin delta chain OS=Homo sapiens O. | TUBD1 | 51.035 | 2 | 48.090656 | yes | 3.456E-07 | 621.31897 | 38.271782 |
| HVWPPLSGLPPLSI | 14 | Q9UJT1 | Tubulin delta chain OS=Homo sapiens O. | TUBD1 | 51.035 | 2 | 48.090656 | yes | 1.201E-07 | 764.43774 | 36.506279 |
| FQDVGPQAPVGSV | 16 | Q9UJU6 | Drebrin-like protein OS=Homo sapiens O | DBNL | 48.208 | 3 | 49.433723 | yes | 5.528E-06 | 573.96155 | 65.185661 |
| AEEDVEPECIMEK | 13 | Q9UJU6 | Drebrin-like protein OS=Homo sapiens O | DBNL | 48.208 | 2 | 49.433723 | yes | 1.031E-07 | 789.83673 | 58.770451 |
| LLQEDSSPLLR | 11 | Q9UJV9 | Probable ATP-dependent RNA helicase D | DDX41 | 69.838 | 2 | 55.03034 | yes | 9.722E-08 | 635.8537 | 616.89563 |
| MIDMGFEGDIR | 11 | Q9UJV9 | Probable ATP-dependent RNA helicase D | DDX41 | 69.838 | 2 | 55.03034 | yes | 9.722E-08 | 642.29156 | 573.21057 |
| TPPPVLIFAEK | 11 | Q9UJV9 | Probable ATP-dependent RNA helicase D | DDX41 | 69.838 | 2 | 55.03034 | yes | 1.314E-07 | 606.35535 | 491.03046 |
| DVLVATDVASK | 11 | Q9UJV9 | Probable ATP-dependent RNA helicase D | DDX41 | 69.838 | 2 | 55.03034 | yes | 2.43E-07 | 559.30859 | 468.92441 |
| YLALDEADR | 9 | Q9UJV9 | Probable ATP-dependent RNA helicase D | DDX41 | 69.838 | 2 | 55.03034 | yes | 1.932E-07 | 533.26422 | 452.29318 |
| MVSLDICR | 8 | Q9UJV9 | Probable ATP-dependent RNA helicase D | DDX41 | 69.838 | 2 | 55.03034 | yes | 5.878E-07 | 497.24643 | 408.29559 |
| MVYLLECLQK | 10 | Q9UJV9 | Probable ATP-dependent RNA helicase D | DDX41 | 69.838 | 2 | 55.03034 | yes | 1.032E-07 | 648.83813 | 404.95306 |
| YVLSMSEER | 9 | Q9UJV9 | Probable ATP-dependent RNA helicase D | DDX41 | 69.838 | 2 | 55.03034 | yes | 1.09E-07 | 557.26587 | 399.85074 |
| EGPYGLIICPSR | 12 | Q9UJV9 | Probable ATP-dependent RNA helicase D | DDX41 | 69.838 | 2 | 55.03034 | yes | 9.722E-08 | 681.34772 | 354.64755 |
| AGAASLDVIQEVE | 16 | Q9UJV9 | Probable ATP-dependent RNA helicase D | DDX41 | 69.838 | 2 | 55.03034 | yes | 9.722E-08 | 846.44617 | 282.18527 |
| DMIGIAFTGSGK | 12 | Q9UJV9 | Probable ATP-dependent RNA helicase D | DDX41 | 69.838 | 2 | 55.03034 | yes | 9.722E-08 | 598.80261 | 271.94833 |
| ILESVAEGR | 9 | Q9UJV9 | Probable ATP-dependent RNA helicase D | DDX41 | 69.838 | 2 | 55.03034 | yes | 1.105E-07 | 487.26929 | 242.97705 |
| GVEAVAIHGGK | 11 | Q9UJV9 | Probable ATP-dependent RNA helicase D | DDX41 | 69.838 | 2 | 55.03034 | yes | 3.123E-07 | 519.29053 | 209.20798 |
| CALCIGGMSVK | 11 | Q9UJV9 | Probable ATP-dependent RNA helicase D | DDX41 | 69.838 | 2 | 55.03034 | yes | 9.722E-08 | 598.28522 | 184.23076 |
| KADVDAIHEYLLL | 14 | Q9UJV9 | Probable ATP-dependent RNA helicase D | DDX41 | 69.838 | 3 | 55.03034 | yes | 1.265E-07 | 543.30664 | 91.335449 |
| KADVDAIHEYLLL | 14 | Q9UJV9 | Probable ATP-dependent RNA helicase D | DDX41 | 69.838 | 3 | 55.03034 | yes | 3.989E-07 | 814.45636 | 91.335449 |
| KADVDAIHEYLLL | 14 | Q9UJV9 | Probable ATP-dependent RNA helicase D | DDX41 | 69.838 | 2 | 55.03034 | yes | 1.265E-07 | 543.30664 | 91.335449 |
| KADVDAIHEYLLL | 14 | Q9UJV9 | Probable ATP-dependent RNA helicase D | DDX41 | 69.838 | 2 | 55.03034 | yes | 3.989E-07 | 814.45636 | 91.335449 |
| SGNTGIATTFINK | 13 | Q9UJV9 | Probable ATP-dependent RNA helicase D | DDX41 | 69.838 | 2 | 55.03034 | yes | 9.722E-08 | 662.34882 | 176.38853 |
| QTLLFSATMPK | 11 | Q9UJV9 | Probable ATP-dependent RNA helicase D | DDX41 | 69.838 | 2 | 55.03034 | yes | 9.722E-08 | 618.83649 | 144.87814 |
| ACDESVLMDLK | 11 | Q9UJV9 | Probable ATP-dependent RNA helicase D | DDX41 | 69.838 | 2 | 55.03034 | yes | 9.722E-08 | 640.79669 | 129.35054 |
| GITYDDPIK | 9 | Q9UJV9 | Probable ATP-dependent RNA helicase D | DDX41 | 69.838 | 2 | 55.03034 | yes | 1.122E-07 | 511.26367 | 107.75797 |
| GCAFCGGLGHR | 11 | Q9UJV9 | Probable ATP-dependent RNA helicase D | DDX41 | 69.838 | 2 | 55.03034 | yes | 7.011E-07 | 596.26093 | 104.90054 |
| SALVKPVTINVGR | 13 | Q9UJV9 | Probable ATP-dependent RNA helicase D | DDX41 | 69.838 | 2 | 55.03034 | yes | 1.965E-07 | 677.41425 | 104.44449 |
| YHILVEGDGIPPPII | 15 | Q9UJV9 | Probable ATP-dependent RNA helicase D | DDX41 | 69.838 | 2 | 55.03034 | yes | 7.023E-06 | 824.45886 | 53.045338 |
| HGVHMMVATPGR | 12 | Q9UJV9 | Probable ATP-dependent RNA helicase D | DDX41 | 69.838 | 3 | 55.03034 | yes | 1.322E-07 | 431.54987 | 43.246101 |
| KDVLVATDVASK | 12 | Q9UJV9 | Probable ATP-dependent RNA helicase D | DDX41 | 69.838 | 2 | 55.03034 | yes | 4.684E-07 | 623.35608 | 31.070972 |
| VPPVLQVLHCGDE | 22 | Q9UJV9 | Probable ATP-dependent RNA helicase D | DDX41 | 69.838 | 3 | 55.03034 | yes | 0.0003307 | 807.73486 | 0.40448 |
| IMSIPNLR | 8 | Q9UJW0 | Dynactin subunit 4 OS=Homo sapiens O> | DCTN4 | 52.338 | 2 | 51.134514 | yes | 7.63E-07 | 472.27332 | 121.9245 |
| AYYLACGFCR | 10 | Q9UJW0 | Dynactin subunit 4 OS=Homo sapiens O> | DCTN4 | 52.338 | 2 | 51.134514 | yes | 2.041E-07 | 640.78119 | 97.166595 |
| ATSISTQLPDDPAK | 14 | Q9UJW0 | Dynactin subunit 4 OS=Homo sapiens O> | DCTN4 | 52.338 | 2 | 51.134514 | yes | 1.779E-07 | 722.36993 | 90.63887 |
| IQLVAVNYIPEVR | 13 | Q9UJW0 | Dynactin subunit 4 OS=Homo sapiens O> | DCTN4 | 52.338 | 2 | 51.134514 | yes | 9.722E-08 | 757.44049 | 75.604004 |
| LIEYYQQLAQK | 11 | Q9UJW0 | Dynactin subunit 4 OS=Homo sapiens O> | DCTN4 | 52.338 | 2 | 51.134514 | yes | 2.605E-07 | 698.8772 | 73.344925 |

| | | | | | | | | | | | |
|---|---|---|---|---|---|---|---|---|---|---|---|
| APLSQLYFCR | 10 | Q9UJW0 | Dynactin subunit 4 OS=Homo sapiens OX | DCTN4 | 52.338 | 2 | 51.134514 | yes | 5.598E-07 | 627.8186 | 52.24905 |
| EEGEVTVCFK | 10 | Q9UJW0 | Dynactin subunit 4 OS=Homo sapiens OX | DCTN4 | 52.338 | 2 | 51.134514 | yes | 3.564E-06 | 599.27643 | 14.814902 |
| AGASISTLAGLSLK | 14 | Q9UJW0 | Dynactin subunit 4 OS=Homo sapiens OX | DCTN4 | 52.338 | 2 | 51.134514 | yes | 7.086E-05 | 644.8772 | 13.746393 |
| FLSLPDTWMK | 10 | Q9UJX2 | Cell division cycle protein 23 homolog O | CDC23 | 68.835 | 2 | 54.372303 | yes | 9.722E-08 | 619.31793 | 114.38406 |
| YQNLIDVGFSK | 11 | Q9UJX2 | Cell division cycle protein 23 homolog O | CDC23 | 68.835 | 2 | 54.372303 | yes | 1.49E-07 | 642.33514 | 97.908043 |
| MLVALGECYEK | 11 | Q9UJX2 | Cell division cycle protein 23 homolog O | CDC23 | 68.835 | 2 | 54.372303 | yes | 1.059E-07 | 656.81744 | 60.046738 |
| QLLLIAGLTR | 10 | Q9UJX2 | Cell division cycle protein 23 homolog O | CDC23 | 68.835 | 2 | 54.372303 | yes | 7.887E-07 | 549.35571 | 55.110786 |
| AAHFLHGCNSK | 11 | Q9UJX2 | Cell division cycle protein 23 homolog O | CDC23 | 68.835 | 2 | 54.372303 | yes | 3.265E-07 | 621.29584 | 37.825356 |
| LWDEASTCAQK | 11 | Q9UJX2 | Cell division cycle protein 23 homolog O | CDC23 | 68.835 | 2 | 54.372303 | yes | 1.345E-07 | 654.7981 | 15.644584 |
| MNVIDHVR | 8 | Q9UJX3 | Anaphase-promoting complex subunit 7 ( | ANAPC7 | 63.134 | 2 | 54.77702 | yes | 9.906E-07 | 513.26349 | 272.79129 |
| VQEAIIHFR | 9 | Q9UJX3 | Anaphase-promoting complex subunit 7 ( | ANAPC7 | 63.134 | 2 | 54.77702 | yes | 1.446E-07 | 556.81421 | 234.68155 |
| YEDGIALLR | 9 | Q9UJX3 | Anaphase-promoting complex subunit 7 ( | ANAPC7 | 63.134 | 2 | 54.77702 | yes | 4.225E-07 | 525.28491 | 175.87946 |
| LEDVENLGCR | 10 | Q9UJX3 | Anaphase-promoting complex subunit 7 ( | ANAPC7 | 63.134 | 2 | 54.77702 | yes | 1.001E-07 | 602.78497 | 175.29434 |
| EAMVMANNVYK | 11 | Q9UJX3 | Anaphase-promoting complex subunit 7 ( | ANAPC7 | 63.134 | 2 | 54.77702 | yes | 9.722E-08 | 635.30194 | 145.20091 |
| FEQAQMLDPYLIK | 13 | Q9UJX3 | Anaphase-promoting complex subunit 7 ( | ANAPC7 | 63.134 | 2 | 54.77702 | yes | 1.836E-07 | 798.41052 | 114.4918 |
| MAECYTMLK | 9 | Q9UJX3 | Anaphase-promoting complex subunit 7 ( | ANAPC7 | 63.134 | 2 | 54.77702 | yes | 9.722E-08 | 573.75323 | 97.492462 |
| INMMLANLYK | 10 | Q9UJX3 | Anaphase-promoting complex subunit 7 ( | ANAPC7 | 63.134 | 2 | 54.77702 | yes | 1.272E-07 | 605.81976 | 92.485336 |
| ALTQRPDYIK | 10 | Q9UJX3 | Anaphase-promoting complex subunit 7 ( | ANAPC7 | 63.134 | 2 | 54.77702 | yes | 3.617E-07 | 602.83789 | 90.566849 |
| AISTICSLEK | 10 | Q9UJX3 | Anaphase-promoting complex subunit 7 ( | ANAPC7 | 63.134 | 2 | 54.77702 | yes | 3.294E-07 | 561.29718 | 88.881805 |
| DMAAAGLHSNVR | 12 | Q9UJX3 | Anaphase-promoting complex subunit 7 ( | ANAPC7 | 63.134 | 2 | 54.77702 | yes | 1.018E-07 | 621.3064 | 77.443726 |
| GMDVYGYLLAR | 11 | Q9UJX3 | Anaphase-promoting complex subunit 7 ( | ANAPC7 | 63.134 | 2 | 54.77702 | yes | 2.343E-06 | 629.31842 | 49.428417 |
| NALANQSDCVLHI | 13 | Q9UJX3 | Anaphase-promoting complex subunit 7 ( | ANAPC7 | 63.134 | 2 | 54.77702 | yes | 2.751E-06 | 749.36499 | 48.889671 |
| TGEGAVSLMER | 11 | Q9UJX4 | Anaphase-promoting complex subunit 5 ( | ANAPC5 | 85.077 | 2 | 49.920322 | yes | 2.723E-07 | 575.28204 | 174.85605 |
| DSDLLHWK | 8 | Q9UJX4 | Anaphase-promoting complex subunit 5 ( | ANAPC5 | 85.077 | 2 | 49.920322 | yes | 9.977E-08 | 507.25616 | 107.61902 |
| CQVASAASYDQPK | 13 | Q9UJX4 | Anaphase-promoting complex subunit 5 ( | ANAPC5 | 85.077 | 2 | 49.920322 | yes | 4.415E-07 | 712.82739 | 69.206772 |
| TVEDADMELTSR | 12 | Q9UJX4 | Anaphase-promoting complex subunit 5 ( | ANAPC5 | 85.077 | 2 | 49.920322 | yes | 1.054E-07 | 683.8114 | 55.325203 |
| QAEFFLSQQASLLI | 14 | Q9UJX4 | Anaphase-promoting complex subunit 5 ( | ANAPC5 | 85.077 | 2 | 49.920322 | yes | 1.613E-07 | 805.43286 | 46.432312 |
| AVVLQAQNQMSE | 15 | Q9UJX4 | Anaphase-promoting complex subunit 5 ( | ANAPC5 | 85.077 | 2 | 49.920322 | yes | 3.888E-07 | 827.42249 | 15.416682 |
| ANELLQVIDSSMK | 13 | Q9UJX5 | Anaphase-promoting complex subunit 4 ( | ANAPC4 | 92.117 | 2 | 50.02956 | yes | 2.4E-07 | 724.37671 | 96.223915 |
| SMNQAICIPLYR | 12 | Q9UJX5 | Anaphase-promoting complex subunit 4 ( | ANAPC4 | 92.117 | 2 | 50.02956 | yes | 9.927E-08 | 733.36816 | 93.325691 |
| FGSFTYATTEK | 11 | Q9UJX5 | Anaphase-promoting complex subunit 4 ( | ANAPC4 | 92.117 | 2 | 50.02956 | yes | 3.654E-07 | 626.29822 | 72.393776 |
| LGQSIESSYSSIQK | 14 | Q9UJX5 | Anaphase-promoting complex subunit 4 ( | ANAPC4 | 92.117 | 2 | 50.02956 | yes | 1.543E-07 | 763.88849 | 43.511082 |
| LDEQCSAIPTR | 11 | Q9UJX5 | Anaphase-promoting complex subunit 4 ( | ANAPC4 | 92.117 | 2 | 50.02956 | yes | 2.277E-07 | 645.31134 | 43.164089 |
| VSCVLSSNLR | 10 | Q9UJX5 | Anaphase-promoting complex subunit 4 ( | ANAPC4 | 92.117 | 2 | 50.02956 | yes | 4.718E-07 | 567.80042 | 39.453125 |
| LVISHLQSGSESLL | 21 | Q9UJX5 | Anaphase-promoting complex subunit 4 ( | ANAPC4 | 92.117 | 3 | 50.02956 | yes | 1.356E-07 | 785.09692 | 34.872269 |
| FPTCFPSFR | 9 | Q9UJX5 | Anaphase-promoting complex subunit 4 ( | ANAPC4 | 92.117 | 2 | 50.02956 | yes | 3.371E-07 | 579.77368 | 30.31926 |
| LSSAWPGTLR | 10 | Q9UJY1 | Heat shock protein beta-8 OS=Homo sapi | HSPB8 | 21.605 | 2 | 49.154087 | yes | 1.063E-07 | 544.2984 | 487.52689 |
| TPPPFPGEPWK | 11 | Q9UJY1 | Heat shock protein beta-8 OS=Homo sapi | HSPB8 | 21.605 | 2 | 49.154087 | yes | 1.758E-07 | 626.82166 | 138.42468 |
| VCVNVHSFKPEEL | 16 | Q9UJY1 | Heat shock protein beta-8 OS=Homo sapi | HSPB8 | 21.605 | 3 | 49.154087 | yes | 2.324E-07 | 639.33044 | 89.957779 |

| | | | | | | | | | | | |
|---|---|---|---|---|---|---|---|---|---|---|---|
| ADGQMPFSCHYPS | 14 | Q9UJY1 | Heat shock protein beta-8 OS=Homo sapi | HSPB8 | 21.605 | 2 | 49.154087 | yes | 2.482E-06 | 847.85614 | 34.654877 |
| QQEGGIVSK | 9 | Q9UJY1 | Heat shock protein beta-8 OS=Homo sapi | HSPB8 | 21.605 | 2 | 49.154087 | yes | 5.077E-06 | 473.25363 | 29.378609 |
| AEQINQAAGEASA | 17 | Q9UJZ1 | Stomatin-like protein 2, mitochondrial OS | STOML2 | 38.535 | 2 | 53.143574 | yes | 9.722E-08 | 835.93903 | 247.14362 |
| ESMQMQVEAER | 11 | Q9UJZ1 | Stomatin-like protein 2, mitochondrial OS | STOML2 | 38.535 | 2 | 53.143574 | yes | 9.722E-08 | 669.29486 | 161.83835 |
| ESAINVAEGK | 10 | Q9UJZ1 | Stomatin-like protein 2, mitochondrial OS | STOML2 | 38.535 | 2 | 53.143574 | yes | 5.584E-07 | 509.26419 | 128.08527 |
| DVQGTDASLDEEL | 15 | Q9UJZ1 | Stomatin-like protein 2, mitochondrial OS | STOML2 | 38.535 | 2 | 53.143574 | yes | 2.101E-07 | 831.87628 | 85.812019 |
| APVPGTPDSLSSGS | 16 | Q9UJZ1 | Stomatin-like protein 2, mitochondrial OS | STOML2 | 38.535 | 2 | 53.143574 | yes | 5.58E-07 | 757.87592 | 75.966476 |
| LDCPLAMER | 9 | Q9UK41 | Vacuolar protein sorting-associated protei | VPS28 | 25.426 | 2 | 48.923134 | yes | 1.086E-07 | 552.76245 | 144.25479 |
| QLGLVVCR | 8 | Q9UK45 | U6 snRNA-associated Sm-like protein LS | LSM7 | 11.603 | 2 | 48.222668 | yes | 1.178E-07 | 472.77094 | 470.45026 |
| ESILDLSK | 8 | Q9UK45 | U6 snRNA-associated Sm-like protein LS | LSM7 | 11.603 | 2 | 48.222668 | yes | 2.18E-07 | 452.75293 | 84.459587 |
| KESILDLSK | 9 | Q9UK45 | U6 snRNA-associated Sm-like protein LS | LSM7 | 11.603 | 2 | 48.222668 | yes | 5.237E-07 | 516.80042 | 39.580219 |
| ALQIPLPTR | 9 | Q9UK58 | Cyclin-L1 OS=Homo sapiens OX=9606 ( | CCNL1 | 59.634 | 2 | 38.23911 | yes | 5.361E-07 | 504.81366 | 16.004593 |
| ATDDLINVTGR | 11 | Q9UK59 | Lariat debranching enzyme OS=Homo sa | DBR1 | 61.555 | 2 | 49.246628 | yes | 1.053E-07 | 587.80676 | 586.86676 |
| IYETLALAER | 10 | Q9UK59 | Lariat debranching enzyme OS=Homo sa | DBR1 | 61.555 | 2 | 49.246628 | yes | 1.393E-07 | 589.8244 | 249.37236 |
| WDYSATEEGMK | 11 | Q9UK59 | Lariat debranching enzyme OS=Homo sa | DBR1 | 61.555 | 2 | 49.246628 | yes | 1.279E-07 | 658.77679 | 225.58505 |
| FAALMQHQAK | 10 | Q9UK59 | Lariat debranching enzyme OS=Homo sa | DBR1 | 61.555 | 2 | 49.246628 | yes | 1.327E-07 | 572.80023 | 213.31854 |
| HMQTFYR | 7 | Q9UK59 | Lariat debranching enzyme OS=Homo sa | DBR1 | 61.555 | 2 | 49.246628 | yes | 2.423E-07 | 491.73181 | 205.34351 |
| QPIDIFLSHDWPR | 13 | Q9UK59 | Lariat debranching enzyme OS=Homo sa | DBR1 | 61.555 | 2 | 49.246628 | yes | 4.046E-07 | 812.41754 | 45.76091 |
| QPIDIFLSHDWPR | 13 | Q9UK59 | Lariat debranching enzyme OS=Homo sa | DBR1 | 61.555 | 2 | 49.246628 | yes | 2.513E-07 | 541.94745 | 45.76091 |
| QPIDIFLSHDWPR | 13 | Q9UK59 | Lariat debranching enzyme OS=Homo sa | DBR1 | 61.555 | 3 | 49.246628 | yes | 4.046E-07 | 812.41754 | 45.76091 |
| QPIDIFLSHDWPR | 13 | Q9UK59 | Lariat debranching enzyme OS=Homo sa | DBR1 | 61.555 | 3 | 49.246628 | yes | 2.513E-07 | 541.94745 | 45.76091 |
| ELINLIQCR | 9 | Q9UK61 | Protein TASOR OS=Homo sapiens OX=9 | TASOR | 189.032 | 2 | 55.596725 | yes | 1.709E-07 | 579.8186 | 254.02512 |
| YQVQTICEK | 9 | Q9UK61 | Protein TASOR OS=Homo sapiens OX=9 | TASOR | 189.032 | 2 | 55.596725 | yes | 1.783E-07 | 584.78699 | 202.08211 |
| ALGLSTDDAYEEL | 14 | Q9UK61 | Protein TASOR OS=Homo sapiens OX=9 | TASOR | 189.032 | 2 | 55.596725 | yes | 1.004E-07 | 776.87811 | 179.41495 |
| NAPELDCLIR | 10 | Q9UK61 | Protein TASOR OS=Homo sapiens OX=9 | TASOR | 189.032 | 2 | 55.596725 | yes | 1.438E-07 | 600.80573 | 143.07576 |
| EFIMFPYDSR | 10 | Q9UK61 | Protein TASOR OS=Homo sapiens OX=9 | TASOR | 189.032 | 2 | 55.596725 | yes | 2.931E-07 | 652.80261 | 139.71457 |
| ELSESYAFLMVDR | 13 | Q9UK61 | Protein TASOR OS=Homo sapiens OX=9 | TASOR | 189.032 | 2 | 55.596725 | yes | 1.772E-07 | 780.37415 | 137.312 |
| GTQSLLLEVDATSI | 14 | Q9UK61 | Protein TASOR OS=Homo sapiens OX=9 | TASOR | 189.032 | 2 | 55.596725 | yes | 1.61E-07 | 731.39337 | 129.80077 |
| IASNPLER | 8 | Q9UK61 | Protein TASOR OS=Homo sapiens OX=9 | TASOR | 189.032 | 2 | 55.596725 | yes | 5.555E-07 | 450.25089 | 128.10205 |
| LIPITGGNAR | 10 | Q9UK61 | Protein TASOR OS=Homo sapiens OX=9 | TASOR | 189.032 | 2 | 55.596725 | yes | 1.263E-06 | 506.3009 | 102.69449 |
| ETYLGPNEVLK | 11 | Q9UK61 | Protein TASOR OS=Homo sapiens OX=9 | TASOR | 189.032 | 2 | 55.596725 | yes | 4.992E-07 | 631.83496 | 95.887878 |
| LQAQNIQQR | 9 | Q9UK61 | Protein TASOR OS=Homo sapiens OX=9 | TASOR | 189.032 | 2 | 55.596725 | yes | 1.113E-07 | 549.80438 | 95.411652 |
| KLEDLPENMR | 10 | Q9UK61 | Protein TASOR OS=Homo sapiens OX=9 | TASOR | 189.032 | 2 | 55.596725 | yes | 9.722E-08 | 622.81879 | 88.164246 |
| IGSNMESLLQK | 11 | Q9UK61 | Protein TASOR OS=Homo sapiens OX=9 | TASOR | 189.032 | 2 | 55.596725 | yes | 1.011E-07 | 610.32117 | 79.094704 |
| LVLVKPLGDR | 10 | Q9UK61 | Protein TASOR OS=Homo sapiens OX=9 | TASOR | 189.032 | 2 | 55.596725 | yes | 1.02E-07 | 555.35571 | 69.644768 |
| YSVAISTSEVGTDI | 15 | Q9UK61 | Protein TASOR OS=Homo sapiens OX=9 | TASOR | 189.032 | 2 | 55.596725 | yes | 1.025E-07 | 797.39136 | 63.281761 |
| SIYDPMGVK | 9 | Q9UK61 | Protein TASOR OS=Homo sapiens OX=9 | TASOR | 189.032 | 2 | 55.596725 | yes | 1.959E-06 | 505.25479 | 56.276649 |
| ITILGSPSMGVYLS | 15 | Q9UK61 | Protein TASOR OS=Homo sapiens OX=9 | TASOR | 189.032 | 2 | 55.596725 | yes | 4.314E-07 | 797.43707 | 52.799923 |
| LLCYISLR | 8 | Q9UK61 | Protein TASOR OS=Homo sapiens OX=9 | TASOR | 189.032 | 2 | 55.596725 | yes | 3.756E-07 | 519.29425 | 46.366238 |

| | | | | | | | | | | | |
|---|---|---|---|---|---|---|---|---|---|---|---|
| HVIPVSSSDFNNK | 13 | Q9UK61 | Protein TASOR OS=Homo sapiens OX=9 | TASOR | 189.032 | 2 | 55.596725 | yes | 1.157E-06 | 722.36499 | 45.510017 |
| SNNVEETEIHWK | 12 | Q9UK61 | Protein TASOR OS=Homo sapiens OX=9 | TASOR | 189.032 | 2 | 55.596725 | yes | 2.032E-07 | 743.35205 | 41.497047 |
| TVLGEYNLFSR | 11 | Q9UK61 | Protein TASOR OS=Homo sapiens OX=9 | TASOR | 189.032 | 2 | 55.596725 | yes | 1.088E-07 | 649.84058 | 40.392941 |
| FYIHEEEESVLCK | 13 | Q9UK61 | Protein TASOR OS=Homo sapiens OX=9 | TASOR | 189.032 | 2 | 55.596725 | yes | 3.919E-07 | 841.89014 | 26.560816 |
| STTGVSEILLK | 11 | Q9UK73 | Protein fem-1 homolog B OS=Homo sapi | FEM1B | 70.265 | 2 | 42.963665 | yes | 2.793E-07 | 574.33209 | 20.844074 |
| ILVLHLNR | 8 | Q9UK80 | Ubiquitin carboxyl-terminal hydrolase 21 | USP21 | 62.656 | 2 | 43.114044 | yes | 2.725E-07 | 489.31638 | 27.346609 |
| SLFIDTYSDVGR | 12 | Q9UK99 | F-box only protein 3 OS=Homo sapiens ( | FBXO3 | 54.561 | 2 | 49.033516 | yes | 1.075E-07 | 686.84082 | 91.62838 |
| NVVSGGFPIIR | 11 | Q9UK99 | F-box only protein 3 OS=Homo sapiens ( | FBXO3 | 54.561 | 2 | 49.033516 | yes | 9.946E-07 | 579.83514 | 89.780136 |
| YWLISEEEK | 9 | Q9UK99 | F-box only protein 3 OS=Homo sapiens ( | FBXO3 | 54.561 | 2 | 49.033516 | yes | 1.407E-06 | 598.79535 | 19.523956 |
| VTNILMLK | 8 | Q9UKA9 | Polypyrimidine tract-binding protein 2 OS | PTBP2 | 57.491 | 2 | 45.961605 | yes | 1.692E-07 | 466.28589 | 15920.932 |
| EGLDDQGLTK | 10 | Q9UKA9 | Polypyrimidine tract-binding protein 2 OS | PTBP2 | 57.491 | 2 | 45.961605 | yes | 2.431E-07 | 538.26691 | 116.47195 |
| LWDIECGACLR | 11 | Q9UKB1 | F-box/WD repeat-containing protein 11 C | FBXW11 | 62.091 | 2 | 52.874065 | no | 9.722E-08 | 696.82355 | 319.82669 |
| LVVSGSSDNTIR | 12 | Q9UKB1 | F-box/WD repeat-containing protein 11 C | FBXW11 | 62.091 | 2 | 52.874065 | no | 3.059E-07 | 624.33313 | 286.14511 |
| APASTLCLR | 9 | Q9UKB1 | F-box/WD repeat-containing protein 11 C | FBXW11 | 62.091 | 2 | 52.874065 | yes | 1.559E-07 | 494.76584 | 273.99484 |
| GIACLQYR | 8 | Q9UKB1 | F-box/WD repeat-containing protein 11 C | FBXW11 | 62.091 | 2 | 52.874065 | no | 5.786E-07 | 490.75275 | 192.80353 |
| VWSTSTCEFVR | 11 | Q9UKB1 | F-box/WD repeat-containing protein 11 C | FBXW11 | 62.091 | 2 | 52.874065 | yes | 9.722E-08 | 686.32172 | 140.95297 |
| SLCAAELVCK | 10 | Q9UKB1 | F-box/WD repeat-containing protein 11 C | FBXW11 | 62.091 | 2 | 52.874065 | no | 1.121E-07 | 575.78339 | 133.51225 |
| IIQDIETIESNWR | 13 | Q9UKB1 | F-box/WD repeat-containing protein 11 C | FBXW11 | 62.091 | 2 | 52.874065 | no | 7.708E-07 | 808.91754 | 95.747292 |
| AAVNVVDFDDK | 11 | Q9UKB1 | F-box/WD repeat-containing protein 11 C | FBXW11 | 62.091 | 2 | 52.874065 | no | 4.673E-07 | 596.79584 | 91.933174 |
| GVYCLQYDDEK | 11 | Q9UKB1 | F-box/WD repeat-containing protein 11 C | FBXW11 | 62.091 | 2 | 52.874065 | yes | 2.626E-07 | 695.30322 | 65.462547 |
| RGWDQYLFK | 9 | Q9UKB1 | F-box/WD repeat-containing protein 11 C | FBXW11 | 62.091 | 2 | 52.874065 | yes | 1E-06 | 606.81165 | 40.580902 |
| LLYTGDFSR | 9 | Q9UKF6 | Cleavage and polyadenylation specificity | CPSF3 | 77.486 | 2 | 47.133945 | yes | 1.554E-07 | 536.2771 | 224.73146 |
| TANLNLETR | 9 | Q9UKF6 | Cleavage and polyadenylation specificity | CPSF3 | 77.486 | 2 | 47.133945 | yes | 2.65E-07 | 516.27765 | 145.76274 |
| EMVELAAQR | 9 | Q9UKF6 | Cleavage and polyadenylation specificity | CPSF3 | 77.486 | 2 | 47.133945 | yes | 1.475E-07 | 523.76862 | 131.84471 |
| LYEALTPVH | 9 | Q9UKF6 | Cleavage and polyadenylation specificity | CPSF3 | 77.486 | 2 | 47.133945 | yes | 3.356E-07 | 521.78204 | 108.68706 |
| QININNPFVFK | 11 | Q9UKF6 | Cleavage and polyadenylation specificity | CPSF3 | 77.486 | 2 | 47.133945 | yes | 3.995E-07 | 667.36676 | 56.07251 |
| WLLSDYVK | 8 | Q9UKF6 | Cleavage and polyadenylation specificity | CPSF3 | 77.486 | 2 | 47.133945 | yes | 1.846E-06 | 512.27911 | 51.085674 |
| NTEAVTLNFR | 10 | Q9UKF6 | Cleavage and polyadenylation specificity | CPSF3 | 77.486 | 2 | 47.133945 | yes | 1.393E-07 | 582.80402 | 51.02494 |
| FCNTVHDIVNR | 11 | Q9UKF6 | Cleavage and polyadenylation specificity | CPSF3 | 77.486 | 3 | 47.133945 | yes | 3.883E-07 | 458.89096 | 39.293709 |
| SAIPAEESDQLLIR | 14 | Q9UKF6 | Cleavage and polyadenylation specificity | CPSF3 | 77.486 | 2 | 47.133945 | yes | 0.0001914 | 792.41742 | 25.478981 |
| LSCEPVMEEK | 10 | Q9UKI2 | Cdc42 effector protein 3 OS=Homo sapie | CDC42EP3 | 27.679 | 2 | 37.550503 | yes | 5.895E-07 | 611.27814 | 46.509827 |
| LFACGIGSTANR | 12 | Q9UKK3 | Protein mono-ADP-ribosyltransferase PA | PARP4 | 192.595 | 2 | 53.092358 | yes | 9.722E-08 | 633.81659 | 312.14993 |
| VNIIQFGTGYK | 11 | Q9UKK3 | Protein mono-ADP-ribosyltransferase PA | PARP4 | 192.595 | 2 | 53.092358 | yes | 1.392E-07 | 620.34021 | 308.92496 |
| LELGNDWDSATK | 12 | Q9UKK3 | Protein mono-ADP-ribosyltransferase PA | PARP4 | 192.595 | 2 | 53.092358 | yes | 1.167E-07 | 674.82263 | 146.36246 |
| YLSLLYPAR | 9 | Q9UKK3 | Protein mono-ADP-ribosyltransferase PA | PARP4 | 192.595 | 2 | 53.092358 | yes | 6.054E-07 | 548.31348 | 130.39009 |
| SHVPIEAK | 8 | Q9UKK3 | Protein mono-ADP-ribosyltransferase PA | PARP4 | 192.595 | 2 | 53.092358 | yes | 1.284E-07 | 440.74799 | 113.95226 |
| FSMPGDQIK | 9 | Q9UKK3 | Protein mono-ADP-ribosyltransferase PA | PARP4 | 192.595 | 2 | 53.092358 | yes | 7.563E-07 | 511.75241 | 106.967 |
| KADLCQLIR | 9 | Q9UKK3 | Protein mono-ADP-ribosyltransferase PA | PARP4 | 192.595 | 2 | 53.092358 | yes | 1.975E-07 | 558.81335 | 105.9928 |
| LLLICDVALGK | 11 | Q9UKK3 | Protein mono-ADP-ribosyltransferase PA | PARP4 | 192.595 | 2 | 53.092358 | yes | 9.722E-08 | 607.86249 | 93.953529 |

| | | | | | | | | | | | |
|---|---|---|---|---|---|---|---|---|---|---|---|
| TEGQYPSICPR | 11 | Q9UKK3 | Protein mono-ADP-ribosyltransferase PA | PARP4 | 192.595 | 2 | 53.092358 | yes | 9.722E-08 | 654.30609 | 88.4916 |
| NQSLLASSEWPEL | 14 | Q9UKK3 | Protein mono-ADP-ribosyltransferase PA | PARP4 | 192.595 | 2 | 53.092358 | yes | 9.722E-08 | 815.41522 | 80.921028 |
| VNETTEFLSK | 10 | Q9UKK3 | Protein mono-ADP-ribosyltransferase PA | PARP4 | 192.595 | 2 | 53.092358 | yes | 2.94E-07 | 584.29822 | 69.110199 |
| MMTEFYR | 7 | Q9UKK3 | Protein mono-ADP-ribosyltransferase PA | PARP4 | 192.595 | 2 | 53.092358 | yes | 4.187E-07 | 489.2146 | 64.967766 |
| EVNLGLLAK | 9 | Q9UKK3 | Protein mono-ADP-ribosyltransferase PA | PARP4 | 192.595 | 2 | 53.092358 | yes | 1.5E-06 | 478.79239 | 49.923439 |
| NIPWAFEAIK | 10 | Q9UKK3 | Protein mono-ADP-ribosyltransferase PA | PARP4 | 192.595 | 2 | 53.092358 | yes | 6.579E-07 | 594.82422 | 40.44154 |
| LCSPSCHSVSVK | 12 | Q9UKK3 | Protein mono-ADP-ribosyltransferase PA | PARP4 | 192.595 | 2 | 53.092358 | yes | 2.463E-07 | 680.82104 | 37.744595 |
| SPVDVLQIFR | 10 | Q9UKK3 | Protein mono-ADP-ribosyltransferase PA | PARP4 | 192.595 | 2 | 53.092358 | yes | 1.786E-07 | 587.33496 | 35.532688 |
| QLLGLQPISTVSPL | 16 | Q9UKK3 | Protein mono-ADP-ribosyltransferase PA | PARP4 | 192.595 | 3 | 53.092358 | yes | 5.543E-07 | 587.01233 | 35.355091 |
| AEGILLLVK | 9 | Q9UKK3 | Protein mono-ADP-ribosyltransferase PA | PARP4 | 192.595 | 2 | 53.092358 | yes | 3.99E-07 | 478.31296 | 21.334988 |
| DFHPSDHTELEEY | 22 | Q9UKK3 | Protein mono-ADP-ribosyltransferase PA | PARP4 | 192.595 | 4 | 53.092358 | yes | 3.863E-07 | 678.55566 | 15.640368 |
| QYIISEELISEGK | 13 | Q9UKK9 | ADP-sugar pyrophosphatase OS=Homo s | NUDT5 | 24.328 | 2 | 47.898754 | yes | 2.69E-06 | 754.89575 | 127.44643 |
| EQTADGVAVIPVL | 15 | Q9UKK9 | ADP-sugar pyrophosphatase OS=Homo s | NUDT5 | 24.328 | 2 | 47.898754 | yes | 1.236E-07 | 798.44122 | 99.649994 |
| PKPGDGEFVEVISI | 16 | Q9UKK9 | ADP-sugar pyrophosphatase OS=Homo s | NUDT5 | 24.328 | 3 | 47.898754 | yes | 5.345E-07 | 571.31366 | 21.452747 |
| PFEVPFLK | 8 | Q9UKK9 | ADP-sugar pyrophosphatase OS=Homo s | NUDT5 | 24.328 | 2 | 47.898754 | yes | 1.185E-05 | 488.77875 | 14.202264 |
| DFQAISDVIGNK | 12 | Q9UKL0 | REST corepressor 1 OS=Homo sapiens O | RCOR1 | 53.328 | 2 | 48.506187 | yes | 1.138E-07 | 653.83551 | 64.925438 |
| WTTEEQLLAVQAI | 14 | Q9UKL0 | REST corepressor 1 OS=Homo sapiens O | RCOR1 | 53.328 | 2 | 48.506187 | yes | 4.507E-07 | 829.44904 | 23.37779 |
| DVDVNLFESTIR | 12 | Q9UKM7 | Endoplasmic reticulum mannosyl-oligosa | MAN1B1 | 79.58 | 2 | 44.100758 | yes | 2.321E-07 | 704.35938 | 60.642593 |
| VTQHIHGLSGK | 11 | Q9UKM7 | Endoplasmic reticulum mannosyl-oligosa | MAN1B1 | 79.58 | 2 | 44.100758 | yes | 3.084E-05 | 588.82782 | 12.532706 |
| GYAFVQYSNER | 11 | Q9UKM9 | RNA-binding protein Raly OS=Homo sap | RALY | 32.464 | 2 | 55.109722 | yes | 9.722E-08 | 667.31219 | 841.83124 |
| SNIDALLSR | 9 | Q9UKM9 | RNA-binding protein Raly OS=Homo sap | RALY | 32.464 | 2 | 55.109722 | yes | 1.002E-07 | 494.77472 | 832.98462 |
| LEQIAAEQK | 9 | Q9UKM9 | RNA-binding protein Raly OS=Homo sap | RALY | 32.464 | 2 | 55.109722 | yes | 2.284E-07 | 515.28241 | 659.81342 |
| LQASNVTNK | 9 | Q9UKM9 | RNA-binding protein Raly OS=Homo sap | RALY | 32.464 | 2 | 55.109722 | yes | 1.063E-07 | 487.76691 | 625.86359 |
| VFIGNLNTALVK | 12 | Q9UKM9 | RNA-binding protein Raly OS=Homo sap | RALY | 32.464 | 2 | 55.109722 | yes | 1.212E-07 | 644.88483 | 528.1344 |
| KSDVETIFSK | 10 | Q9UKM9 | RNA-binding protein Raly OS=Homo sap | RALY | 32.464 | 2 | 55.109722 | yes | 1.082E-07 | 577.30859 | 302.76923 |
| SSELQAIK | 8 | Q9UKM9 | RNA-binding protein Raly OS=Homo sap | RALY | 32.464 | 2 | 55.109722 | yes | 1.197E-07 | 438.24527 | 178.78383 |
| LSPVPVPR | 8 | Q9UKM9 | RNA-binding protein Raly OS=Homo sap | RALY | 32.464 | 2 | 55.109722 | yes | 1.236E-07 | 432.76871 | 124.99525 |
| LTIQANLNR | 9 | Q9UKN8 | General transcription factor 3C polypepti | GTF3C4 | 91.983 | 2 | 52.94299 | yes | 9.722E-08 | 521.80383 | 423.33337 |
| IEAVEMHLTR | 10 | Q9UKN8 | General transcription factor 3C polypepti | GTF3C4 | 91.983 | 2 | 52.94299 | yes | 1.284E-07 | 599.8161 | 383.16559 |
| VLIGHISK | 8 | Q9UKN8 | General transcription factor 3C polypepti | GTF3C4 | 91.983 | 2 | 52.94299 | yes | 9.722E-08 | 433.77655 | 280.70456 |
| MSGLIVGSAFGPIK | 14 | Q9UKN8 | General transcription factor 3C polypepti | GTF3C4 | 91.983 | 2 | 52.94299 | yes | 9.722E-08 | 688.88397 | 254.62154 |
| MEWSGICTTQQVK | 13 | Q9UKN8 | General transcription factor 3C polypepti | GTF3C4 | 91.983 | 2 | 52.94299 | yes | 1.668E-07 | 784.36578 | 240.36917 |
| EMDQLPVHSIK | 11 | Q9UKN8 | General transcription factor 3C polypepti | GTF3C4 | 91.983 | 2 | 52.94299 | yes | 9.722E-08 | 648.83447 | 204.23837 |
| QVDLIDLVR | 9 | Q9UKN8 | General transcription factor 3C polypepti | GTF3C4 | 91.983 | 2 | 52.94299 | yes | 1.132E-07 | 535.81384 | 195.33916 |
| CLLHDSIAR | 9 | Q9UKN8 | General transcription factor 3C polypepti | GTF3C4 | 91.983 | 2 | 52.94299 | yes | 7.06E-07 | 542.78204 | 171.75356 |
| CLLAALTMDNR | 11 | Q9UKN8 | General transcription factor 3C polypepti | GTF3C4 | 91.983 | 2 | 52.94299 | yes | 1.133E-07 | 639.32068 | 171.61356 |
| LQYAVSGLEPLAW | 18 | Q9UKN8 | General transcription factor 3C polypepti | GTF3C4 | 91.983 | 3 | 52.94299 | yes | 9.722E-08 | 691.01294 | 157.42412 |
| FEHQLIK | 7 | Q9UKN8 | General transcription factor 3C polypepti | GTF3C4 | 91.983 | 2 | 52.94299 | yes | 4.586E-06 | 457.75836 | 156.21042 |
| TSVPAPLNSCLLK | 13 | Q9UKN8 | General transcription factor 3C polypepti | GTF3C4 | 91.983 | 2 | 52.94299 | yes | 4.79E-07 | 700.38434 | 130.69669 |

| | | | | | | | | | | | |
|---|---|---|---|---|---|---|---|---|---|---|---|
| NYQVQFVTLK | 10 | Q9UKN8 | General transcription factor 3C polypeptic | GTF3C4 | 91.983 | 2 | 52.94299 | yes | 3.496E-07 | 620.34021 | 128.90327 |
| EPAADAAPGPSAA | 15 | Q9UKN8 | General transcription factor 3C polypeptic | GTF3C4 | 91.983 | 2 | 52.94299 | yes | 2.508E-06 | 714.3493 | 122.36702 |
| HSMQTPVR | 8 | Q9UKN8 | General transcription factor 3C polypeptic | GTF3C4 | 91.983 | 2 | 52.94299 | yes | 7.901E-07 | 478.24255 | 115.20577 |
| ILYQSMQK | 8 | Q9UKN8 | General transcription factor 3C polypeptic | GTF3C4 | 91.983 | 2 | 52.94299 | yes | 1.326E-06 | 505.7706 | 107.80892 |
| HPAPEDPDWIK | 11 | Q9UKN8 | General transcription factor 3C polypeptic | GTF3C4 | 91.983 | 2 | 52.94299 | yes | 9.722E-08 | 652.81714 | 98.877121 |
| LSDVFGSVR | 9 | Q9UKN8 | General transcription factor 3C polypeptic | GTF3C4 | 91.983 | 2 | 52.94299 | yes | 2.846E-06 | 490.26401 | 83.316856 |
| CSCSLVVAAR | 10 | Q9UKN8 | General transcription factor 3C polypeptic | GTF3C4 | 91.983 | 2 | 52.94299 | yes | 8.561E-07 | 561.77338 | 60.326462 |
| TPSEALWKPTHED | 15 | Q9UKN8 | General transcription factor 3C polypeptic | GTF3C4 | 91.983 | 3 | 52.94299 | yes | 1.876E-07 | 575.95294 | 23.076511 |
| ADSAVSQEQLR | 11 | Q9UKS6 | Protein kinase C and casein kinase substra | PACSIN3 | 48.487 | 2 | 53.561653 | yes | 9.722E-08 | 602.30182 | 955.82129 |
| GPQYGTLEK | 9 | Q9UKS6 | Protein kinase C and casein kinase substra | PACSIN3 | 48.487 | 2 | 53.561653 | yes | 4.208E-07 | 496.75601 | 785.20374 |
| AYAQQLADWAR | 11 | Q9UKS6 | Protein kinase C and casein kinase substra | PACSIN3 | 48.487 | 2 | 53.561653 | yes | 9.722E-08 | 646.82275 | 713.37708 |
| SPDEVTLTSIVPTR | 14 | Q9UKS6 | Protein kinase C and casein kinase substra | PACSIN3 | 48.487 | 2 | 53.561653 | yes | 8.324E-07 | 757.90668 | 691.50995 |
| AWHAFFTAAER | 11 | Q9UKS6 | Protein kinase C and casein kinase substra | PACSIN3 | 48.487 | 2 | 53.561653 | yes | 9.722E-08 | 653.82001 | 642.65259 |
| LSALHLEVR | 9 | Q9UKS6 | Protein kinase C and casein kinase substra | PACSIN3 | 48.487 | 2 | 53.561653 | yes | 5.87E-07 | 519.30872 | 637.29712 |
| AQYEQTLAELHR | 12 | Q9UKS6 | Protein kinase C and casein kinase substra | PACSIN3 | 48.487 | 2 | 53.561653 | yes | 1.102E-07 | 729.87042 | 468.03015 |
| LCGDLVSCFQER | 12 | Q9UKS6 | Protein kinase C and casein kinase substra | PACSIN3 | 48.487 | 2 | 53.561653 | yes | 9.722E-08 | 742.33704 | 353.38385 |
| AQKPWLK | 7 | Q9UKS6 | Protein kinase C and casein kinase substra | PACSIN3 | 48.487 | 2 | 53.561653 | yes | 9.722E-08 | 435.76343 | 135.44951 |
| KAQKPWLK | 8 | Q9UKS6 | Protein kinase C and casein kinase substra | PACSIN3 | 48.487 | 2 | 53.561653 | yes | 3.406E-07 | 499.81091 | 116.53217 |
| IGLYPANYVECVG | 14 | Q9UKS6 | Protein kinase C and casein kinase substra | PACSIN3 | 48.487 | 2 | 53.561653 | yes | 9.722E-08 | 763.3714 | 109.31746 |
| LQGQDSER | 8 | Q9UKS6 | Protein kinase C and casein kinase substra | PACSIN3 | 48.487 | 2 | 53.561653 | yes | 6.425E-07 | 466.72525 | 106.57135 |
| LSCIGSPIVSPR | 12 | Q9UKT4 | F-box only protein 5 OS=Homo sapiens ( | FBXO5 | 50.147 | 2 | 45.08882 | yes | 1.967E-07 | 643.35028 | 100.93228 |
| HNEFSEVAK | 9 | Q9UKT4 | F-box only protein 5 OS=Homo sapiens ( | FBXO5 | 50.147 | 2 | 45.08882 | yes | 4.726E-07 | 530.75653 | 60.677467 |
| LVSYTPAYLEGSC | 14 | Q9UKT4 | F-box only protein 5 OS=Homo sapiens ( | FBXO5 | 50.147 | 2 | 45.08882 | yes | 2.32E-07 | 794.38977 | 45.229076 |
| KVTLGDTLTR | 10 | Q9UKV3 | Apoptotic chromatin condensation induce | ACIN1 | 151.862 | 2 | 53.182808 | yes | 9.879E-08 | 552.32458 | 333.16968 |
| VTLGDTLTR | 9 | Q9UKV3 | Apoptotic chromatin condensation induce | ACIN1 | 151.862 | 2 | 53.182808 | yes | 2.658E-07 | 488.2771 | 285.58649 |
| SQEQEVLER | 9 | Q9UKV3 | Apoptotic chromatin condensation induce | ACIN1 | 151.862 | 2 | 53.182808 | yes | 2.664E-07 | 559.27783 | 217.64432 |
| TGTLVEEAFWIDK | 13 | Q9UKV3 | Apoptotic chromatin condensation induce | ACIN1 | 151.862 | 2 | 53.182808 | yes | 1.902E-07 | 754.88519 | 217.48553 |
| GLLVDRPSETK | 11 | Q9UKV3 | Apoptotic chromatin condensation induce | ACIN1 | 151.862 | 2 | 53.182808 | yes | 9.722E-08 | 607.84058 | 168.82491 |
| VPEESVLPLVQK | 12 | Q9UKV3 | Apoptotic chromatin condensation induce | ACIN1 | 151.862 | 2 | 53.182808 | yes | 9.722E-08 | 669.38739 | 136.95891 |
| ASHTLLPSHR | 10 | Q9UKV3 | Apoptotic chromatin condensation induce | ACIN1 | 151.862 | 2 | 53.182808 | yes | 4.346E-07 | 559.80688 | 102.1535 |
| SGVSITIDDPVR | 12 | Q9UKV3 | Apoptotic chromatin condensation induce | ACIN1 | 151.862 | 2 | 53.182808 | yes | 4.726E-07 | 629.83551 | 99.326553 |
| TTSPLEEEER | 10 | Q9UKV3 | Apoptotic chromatin condensation induce | ACIN1 | 151.862 | 2 | 53.182808 | yes | 1.375E-07 | 595.7804 | 99.1884 |
| STLADYSAQK | 10 | Q9UKV3 | Apoptotic chromatin condensation induce | ACIN1 | 151.862 | 2 | 53.182808 | yes | 9.722E-08 | 542.26947 | 88.890404 |
| IEELALAK | 8 | Q9UKV3 | Apoptotic chromatin condensation induce | ACIN1 | 151.862 | 2 | 53.182808 | yes | 2.258E-07 | 443.76584 | 85.558372 |
| GALMLENLQK | 10 | Q9UKV3 | Apoptotic chromatin condensation induce | ACIN1 | 151.862 | 2 | 53.182808 | yes | 1.201E-07 | 558.80774 | 85.110199 |
| KPSISITTESLK | 12 | Q9UKV3 | Apoptotic chromatin condensation induce | ACIN1 | 151.862 | 2 | 53.182808 | yes | 1.745E-07 | 652.37701 | 77.175125 |
| PFTLGQLK | 8 | Q9UKV3 | Apoptotic chromatin condensation induce | ACIN1 | 151.862 | 2 | 53.182808 | yes | 2.196E-07 | 452.26855 | 57.251652 |
| DLEPESDR | 8 | Q9UKV3 | Apoptotic chromatin condensation induce | ACIN1 | 151.862 | 2 | 53.182808 | yes | 5.579E-07 | 480.71707 | 41.338581 |
| VLQPPSILYGGR | 12 | Q9UKV8 | Protein argonaute-2 OS=Homo sapiens O | AGO2 | 97.209 | 2 | 54.45324 | yes | 2.44E-06 | 650.37463 | 4756.5522 |

| | | | | | | | | | | | |
|---|---|---|---|---|---|---|---|---|---|---|---|
| AIATPVQGVWDM | 13 | Q9UKV8 | Protein argonaute-2 OS=Homo sapiens O | AGO2 | 97.209 | 2 | 54.45324 | yes | 2.43E-06 | 722.37427 | 2049.3462 |
| AIATPVQGVWDM | 13 | Q9UKV8 | Protein argonaute-2 OS=Homo sapiens O | AGO2 | 97.209 | 2 | 54.45324 | yes | 1.423E-07 | 730.37177 | 2049.3462 |
| NLYTAMPLPIGR | 12 | Q9UKV8 | Protein argonaute-2 OS=Homo sapiens O | AGO2 | 97.209 | 2 | 54.45324 | yes | 1.078E-07 | 673.36847 | 3961.0959 |
| SGNIPAGTTVDTK | 13 | Q9UKV8 | Protein argonaute-2 OS=Homo sapiens O | AGO2 | 97.209 | 2 | 54.45324 | yes | 1.566E-07 | 630.82513 | 3905.7449 |
| SASFNTDPYVR | 11 | Q9UKV8 | Protein argonaute-2 OS=Homo sapiens O | AGO2 | 97.209 | 2 | 54.45324 | yes | 1.874E-07 | 628.79895 | 3384.8254 |
| TTPQTLSNLCLK | 12 | Q9UKV8 | Protein argonaute-2 OS=Homo sapiens O | AGO2 | 97.209 | 2 | 54.45324 | yes | 1.09E-07 | 688.36615 | 2887.948 |
| DYQPGITFIVVQK | 13 | Q9UKV8 | Protein argonaute-2 OS=Homo sapiens O | AGO2 | 97.209 | 2 | 54.45324 | yes | 1.77E-07 | 754.41138 | 2634.7026 |
| VWAIACFAPQR | 11 | Q9UKV8 | Protein argonaute-2 OS=Homo sapiens O | AGO2 | 97.209 | 2 | 54.45324 | yes | 1.937E-07 | 659.8399 | 2431.02 |
| KPSIAAVVGSMDA | 17 | Q9UKV8 | Protein argonaute-2 OS=Homo sapiens O | AGO2 | 97.209 | 3 | 54.45324 | yes | 2.012E-07 | 583.97369 | 2374.7495 |
| LTDNQTSTMIR | 11 | Q9UKV8 | Protein argonaute-2 OS=Homo sapiens O | AGO2 | 97.209 | 2 | 54.45324 | yes | 9.722E-08 | 640.31915 | 2360.5061 |
| VEITHCGQMK | 10 | Q9UKV8 | Protein argonaute-2 OS=Homo sapiens O | AGO2 | 97.209 | 2 | 54.45324 | yes | 1.007E-07 | 601.78644 | 1836.7242 |
| SFTEQLR | 7 | Q9UKV8 | Protein argonaute-2 OS=Homo sapiens O | AGO2 | 97.209 | 2 | 54.45324 | yes | 9.728E-08 | 440.7298 | 1772.5828 |
| LQANFFEMDIPK | 12 | Q9UKV8 | Protein argonaute-2 OS=Homo sapiens O | AGO2 | 97.209 | 2 | 54.45324 | yes | 2.112E-07 | 726.86322 | 1746.1033 |
| DKVELEVTLPGEG | 14 | Q9UKV8 | Protein argonaute-2 OS=Homo sapiens O | AGO2 | 97.209 | 2 | 54.45324 | yes | 1.341E-07 | 757.40906 | 1177.9744 |
| LFCTDKNER | 9 | Q9UKV8 | Protein argonaute-2 OS=Homo sapiens O | AGO2 | 97.209 | 2 | 54.45324 | yes | 1.235E-07 | 591.78223 | 1049.9756 |
| QCTEVHLK | 8 | Q9UKV8 | Protein argonaute-2 OS=Homo sapiens O | AGO2 | 97.209 | 2 | 54.45324 | yes | 2.314E-07 | 507.75549 | 965.45911 |
| EIVEHMVQHFK | 11 | Q9UKV8 | Protein argonaute-2 OS=Homo sapiens O | AGO2 | 97.209 | 2 | 54.45324 | yes | 1.133E-07 | 698.85571 | 919.39154 |
| DEMTDVTGR | 9 | Q9UKV8 | Protein argonaute-2 OS=Homo sapiens O | AGO2 | 97.209 | 2 | 54.45324 | yes | 7.047E-07 | 512.22424 | 882.62439 |
| QEIIQDLAAMVR | 12 | Q9UKV8 | Protein argonaute-2 OS=Homo sapiens O | AGO2 | 97.209 | 2 | 54.45324 | yes | 1.073E-07 | 693.87415 | 360.55469 |
| QEIIQDLAAMVR | 12 | Q9UKV8 | Protein argonaute-2 OS=Homo sapiens O | AGO2 | 97.209 | 2 | 54.45324 | yes | 9.809E-08 | 701.87158 | 360.55469 |
| QFHTGIEIK | 9 | Q9UKV8 | Protein argonaute-2 OS=Homo sapiens O | AGO2 | 97.209 | 2 | 54.45324 | yes | 9.722E-08 | 536.79291 | 567.70465 |
| DGVSEGQFQQVLF | 20 | Q9UKV8 | Protein argonaute-2 OS=Homo sapiens O | AGO2 | 97.209 | 3 | 54.45324 | yes | 3.502E-07 | 759.39758 | 260.02832 |
| DGVSEGQFQQVLF | 20 | Q9UKV8 | Protein argonaute-2 OS=Homo sapiens O | AGO2 | 97.209 | 3 | 54.45324 | yes | 1.129E-06 | 569.79999 | 260.02832 |
| DGVSEGQFQQVLF | 20 | Q9UKV8 | Protein argonaute-2 OS=Homo sapiens O | AGO2 | 97.209 | 4 | 54.45324 | yes | 3.502E-07 | 759.39758 | 260.02832 |
| DGVSEGQFQQVLF | 20 | Q9UKV8 | Protein argonaute-2 OS=Homo sapiens O | AGO2 | 97.209 | 4 | 54.45324 | yes | 1.129E-06 | 569.79999 | 260.02832 |
| SIEEQQKPLTDSQF | 14 | Q9UKV8 | Protein argonaute-2 OS=Homo sapiens O | AGO2 | 97.209 | 2 | 54.45324 | yes | 9.904E-08 | 829.92084 | 497.64151 |
| WVSCVSLQALHD | 17 | Q9UKV8 | Protein argonaute-2 OS=Homo sapiens O | AGO2 | 97.209 | 3 | 54.45324 | yes | 9.777E-08 | 633.65625 | 388.24133 |
| PSHYHVLWDDNR | 12 | Q9UKV8 | Protein argonaute-2 OS=Homo sapiens O | AGO2 | 97.209 | 2 | 54.45324 | yes | 1.012E-07 | 769.86041 | 196.53056 |
| KLTDNQTSTMIR | 12 | Q9UKV8 | Protein argonaute-2 OS=Homo sapiens O | AGO2 | 97.209 | 2 | 54.45324 | yes | 9.722E-08 | 704.36664 | 136.87752 |
| FSSDELQILTYQLC | 19 | Q9UKV8 | Protein argonaute-2 OS=Homo sapiens O | AGO2 | 97.209 | 3 | 54.45324 | yes | 1.248E-07 | 791.72296 | 130.46902 |
| PSIAAVVGSMDAF | 16 | Q9UKV8 | Protein argonaute-2 OS=Homo sapiens O | AGO2 | 97.209 | 3 | 54.45324 | yes | 4.055E-07 | 541.27533 | 87.672958 |
| PPVFQQPVIFLGAI | 24 | Q9UKV8 | Protein argonaute-2 OS=Homo sapiens O | AGO2 | 97.209 | 3 | 54.45324 | yes | 1.823E-07 | 829.77252 | 62.90657 |
| KNLYTAMPLPIGR | 13 | Q9UKV8 | Protein argonaute-2 OS=Homo sapiens O | AGO2 | 97.209 | 2 | 54.45324 | yes | 1.187E-07 | 737.41595 | 46.263779 |
| ITHPTEFDFYLCSH | 22 | Q9UKV8 | Protein argonaute-2 OS=Homo sapiens O | AGO2 | 97.209 | 3 | 54.45324 | yes | 2.504E-07 | 846.40076 | 45.905933 |
| LQTESGEFSR | 10 | Q9UKX2 | Myosin-2 OS=Homo sapiens OX=9606 C | MYH2 | 223.045 | 2 | 34.93067 | yes | 9.157E-07 | 577.27783 | 142.01222 |
| VAAETQSPSLFGST | 15 | Q9UKX7 | Nuclear pore complex protein Nup50 OS= | NUP50 | 50.145 | 2 | 51.42047 | yes | 2.338E-07 | 761.89099 | 110.30357 |
| QLAALNCSVR | 10 | Q9UKX7 | Nuclear pore complex protein Nup50 OS= | NUP50 | 50.145 | 2 | 51.42047 | yes | 1.558E-07 | 566.3006 | 76.431152 |
| NVGFESDTGGAFK | 13 | Q9UKX7 | Nuclear pore complex protein Nup50 OS= | NUP50 | 50.145 | 2 | 51.42047 | yes | 1.606E-07 | 664.80951 | 67.209099 |
| TDPSSLGATSASFN | 17 | Q9UKX7 | Nuclear pore complex protein Nup50 OS= | NUP50 | 50.145 | 2 | 51.42047 | yes | 1.173E-06 | 843.9021 | 30.909811 |

| | | | | | | | | | | | |
|---|---|---|---|---|---|---|---|---|---|---|---|
| YLANIEQQHGNSG | 14 | Q9UKX7 | Nuclear pore complex protein Nup50 OS= | NUP50 | 50.145 | 2 | 51.42047 | yes | 9.722E-08 | 793.88715 | 18.585052 |
| FIDLESDNLCR | 11 | Q9UKZ9 | Procollagen C-endopeptidase enhancer 2 | PCOLCE2 | 45.717 | 2 | 47.888397 | yes | 1.238E-07 | 691.32446 | 83.392258 |
| EGNLAIQQAGK | 11 | Q9UKZ9 | Procollagen C-endopeptidase enhancer 2 | PCOLCE2 | 45.717 | 2 | 47.888397 | yes | 3.401E-07 | 564.80402 | 76.463509 |
| GLNYIIMGQVGED | 15 | Q9UKZ9 | Procollagen C-endopeptidase enhancer 2 | PCOLCE2 | 45.717 | 2 | 47.888397 | yes | 2.724E-07 | 811.40375 | 55.418003 |
| VVVLNFR | 7 | Q9UKZ9 | Procollagen C-endopeptidase enhancer 2 | PCOLCE2 | 45.717 | 2 | 47.888397 | yes | 3.796E-06 | 423.76343 | 48.367157 |
| YCGDSPPAPIVSEF | 14 | Q9UKZ9 | Procollagen C-endopeptidase enhancer 2 | PCOLCE2 | 45.717 | 2 | 47.888397 | yes | 1.489E-07 | 774.36157 | 14.534818 |
| YLDLEEEADTTK | 12 | Q9UL15 | BAG family molecular chaperone regulato | BAG5 | 51.2 | 2 | 53.599857 | yes | 7.369E-07 | 713.83282 | 122.09959 |
| ICAVQEIIEDCMK | 13 | Q9UL15 | BAG family molecular chaperone regulato | BAG5 | 51.2 | 2 | 53.599857 | yes | 9.722E-08 | 804.87512 | 79.34111 |
| QLLALDAVDPQGI | 15 | Q9UL15 | BAG family molecular chaperone regulato | BAG5 | 51.2 | 2 | 53.599857 | yes | 6.378E-06 | 813.42267 | 33.562069 |
| LAQNILSYLDLK | 12 | Q9UL15 | BAG family molecular chaperone regulato | BAG5 | 51.2 | 2 | 53.599857 | yes | 9.722E-08 | 695.90063 | 29.554604 |
| KQPSLPLSEDAHPS | 17 | Q9UL15 | BAG family molecular chaperone regulato | BAG5 | 51.2 | 3 | 53.599857 | yes | 2.838E-06 | 601.99152 | 16.513111 |
| AVQVHQDTLR | 10 | Q9UL18 | Protein argonaute-1 OS=Homo sapiens O | AGO1 | 97.215 | 2 | 56.79019 | no | 9.722E-08 | 583.81744 | 2689.9653 |
| YAQGADSVEPMFI | 13 | Q9UL18 | Protein argonaute-1 OS=Homo sapiens O | AGO1 | 97.215 | 2 | 56.79019 | no | 1.399E-07 | 735.83771 | 2665.9827 |
| EVWFGFHQSVR | 11 | Q9UL18 | Protein argonaute-1 OS=Homo sapiens O | AGO1 | 97.215 | 2 | 56.79019 | no | 9.722E-08 | 696.34637 | 2402.0801 |
| VLPAPILQYGGR | 12 | Q9UL18 | Protein argonaute-1 OS=Homo sapiens O | AGO1 | 97.215 | 2 | 56.79019 | yes | 9.722E-08 | 642.37714 | 1778.6691 |
| TSPQTLSNLCLK | 12 | Q9UL18 | Protein argonaute-1 OS=Homo sapiens O | AGO1 | 97.215 | 2 | 56.79019 | no | 2.204E-07 | 681.35828 | 1614.5692 |
| DAGMPIQGQPCFC | 14 | Q9UL18 | Protein argonaute-1 OS=Homo sapiens O | AGO1 | 97.215 | 2 | 56.79019 | no | 9.809E-08 | 804.85199 | 1454.1283 |
| NIYTVTALPIGNER | 14 | Q9UL18 | Protein argonaute-1 OS=Homo sapiens O | AGO1 | 97.215 | 2 | 56.79019 | yes | 1.063E-07 | 780.92267 | 1226.5635 |
| ELLIQFYK | 8 | Q9UL18 | Protein argonaute-1 OS=Homo sapiens O | AGO1 | 97.215 | 2 | 56.79019 | no | 1.425E-07 | 527.30261 | 1218.911 |
| TPVYAEVK | 8 | Q9UL18 | Protein argonaute-1 OS=Homo sapiens O | AGO1 | 97.215 | 2 | 56.79019 | no | 1.011E-07 | 453.75018 | 1000.8025 |
| HTYLPLEVCNIVA | 16 | Q9UL18 | Protein argonaute-1 OS=Homo sapiens O | AGO1 | 97.215 | 3 | 56.79019 | no | 9.728E-08 | 623.99286 | 966.89636 |
| LTDNQTSTMIK | 11 | Q9UL18 | Protein argonaute-1 OS=Homo sapiens O | AGO1 | 97.215 | 2 | 56.79019 | no | 9.771E-08 | 626.3161 | 964.03906 |
| DYQPGITYIVVQK | 13 | Q9UL18 | Protein argonaute-1 OS=Homo sapiens O | AGO1 | 97.215 | 2 | 56.79019 | no | 1.557E-07 | 762.40887 | 799.92017 |
| SVSIPAPAYYAR | 12 | Q9UL18 | Protein argonaute-1 OS=Homo sapiens O | AGO1 | 97.215 | 2 | 56.79019 | no | 2.945E-07 | 647.84314 | 793.81366 |
| MMLNIDVSATAFY | 14 | Q9UL18 | Protein argonaute-1 OS=Homo sapiens O | AGO1 | 97.215 | 2 | 56.79019 | no | 1.27E-07 | 802.39655 | 346.09277 |
| MMLNIDVSATAFY | 14 | Q9UL18 | Protein argonaute-1 OS=Homo sapiens O | AGO1 | 97.215 | 2 | 56.79019 | no | 2.918E-06 | 810.39404 | 346.09277 |
| VDFEVTIPGEGK | 12 | Q9UL18 | Protein argonaute-1 OS=Homo sapiens O | AGO1 | 97.215 | 2 | 56.79019 | yes | 9.722E-08 | 645.83246 | 682.98047 |
| KPSITAVVGSMDA | 17 | Q9UL18 | Protein argonaute-1 OS=Homo sapiens O | AGO1 | 97.215 | 3 | 56.79019 | yes | 1.08E-07 | 584.97357 | 660.80981 |
| VKDDMTEVTGR | 11 | Q9UL18 | Protein argonaute-1 OS=Homo sapiens O | AGO1 | 97.215 | 2 | 56.79019 | yes | 9.722E-08 | 625.80591 | 557.4693 |
| LLANYFEVDIPK | 12 | Q9UL18 | Protein argonaute-1 OS=Homo sapiens O | AGO1 | 97.215 | 2 | 56.79019 | yes | 9.722E-08 | 711.38739 | 391.24374 |
| VWAIACFAPQK | 11 | Q9UL18 | Protein argonaute-1 OS=Homo sapiens O | AGO1 | 97.215 | 2 | 56.79019 | yes | 9.873E-08 | 645.83679 | 354.94485 |
| SAVFQQPVIFLGAI | 24 | Q9UL18 | Protein argonaute-1 OS=Homo sapiens O | AGO1 | 97.215 | 3 | 56.79019 | yes | 1.591E-07 | 817.76044 | 330.93857 |
| QEIIEDLSYMVR | 12 | Q9UL18 | Protein argonaute-1 OS=Homo sapiens O | AGO1 | 97.215 | 2 | 56.79019 | yes | 2.575E-07 | 748.37671 | 171.43872 |
| NFTDQLR | 7 | Q9UL18 | Protein argonaute-1 OS=Homo sapiens O | AGO1 | 97.215 | 2 | 56.79019 | yes | 7.049E-07 | 447.22742 | 169.86586 |
| WLAIVSWR | 8 | Q9UL18 | Protein argonaute-1 OS=Homo sapiens O | AGO1 | 97.215 | 2 | 56.79019 | yes | 9.722E-08 | 515.79529 | 155.4101 |
| NIDEQPKPLTDSQI | 14 | Q9UL18 | Protein argonaute-1 OS=Homo sapiens O | AGO1 | 97.215 | 3 | 56.79019 | yes | 2.689E-07 | 547.61279 | 64.069038 |
| NIDEQPKPLTDSQI | 14 | Q9UL18 | Protein argonaute-1 OS=Homo sapiens O | AGO1 | 97.215 | 3 | 56.79019 | yes | 4.093E-07 | 820.91559 | 64.069038 |
| NIDEQPKPLTDSQI | 14 | Q9UL18 | Protein argonaute-1 OS=Homo sapiens O | AGO1 | 97.215 | 2 | 56.79019 | yes | 2.689E-07 | 547.61279 | 64.069038 |
| NIDEQPKPLTDSQI | 14 | Q9UL18 | Protein argonaute-1 OS=Homo sapiens O | AGO1 | 97.215 | 2 | 56.79019 | yes | 4.093E-07 | 820.91559 | 64.069038 |

| | | | | | | | | | | | |
|---|---|---|---|---|---|---|---|---|---|---|---|
| PSHYYVLWDDNR | 12 | Q9UL18 | Protein argonaute-1 OS=Homo sapiens O | AGO1 | 97.215 | 2 | 56.79019 | yes | 9.722E-08 | 782.86261 | 118.16142 |
| VEVTHCGQMK | 10 | Q9UL18 | Protein argonaute-1 OS=Homo sapiens O | AGO1 | 97.215 | 2 | 56.79019 | no | 1.175E-06 | 594.77863 | 115.46658 |
| AIATPNQGVWDM | 13 | Q9UL18 | Protein argonaute-1 OS=Homo sapiens O | AGO1 | 97.215 | 2 | 56.79019 | yes | 4.249E-07 | 729.86157 | 103.66946 |
| FTADELQILTYQL( | 19 | Q9UL18 | Protein argonaute-1 OS=Homo sapiens O | AGO1 | 97.215 | 3 | 56.79019 | yes | 1.178E-07 | 791.06323 | 84.659668 |
| RPASHQTFPLQLES | 27 | Q9UL18 | Protein argonaute-1 OS=Homo sapiens O | AGO1 | 97.215 | 4 | 56.79019 | yes | 9.722E-08 | 781.38953 | 80.353966 |
| KLTDNQTSTMIK | 12 | Q9UL18 | Protein argonaute-1 OS=Homo sapiens O | AGO1 | 97.215 | 2 | 56.79019 | no | 2.088E-07 | 690.36359 | 50.326458 |
| RVGDTLLGMATQ( | 17 | Q9UL18 | Protein argonaute-1 OS=Homo sapiens O | AGO1 | 97.215 | 3 | 56.79019 | no | 1.065E-07 | 625.9975 | 49.989471 |
| LGGINNILVPHQR | 13 | Q9UL18 | Protein argonaute-1 OS=Homo sapiens O | AGO1 | 97.215 | 2 | 56.79019 | no | 3.97E-06 | 715.91498 | 41.466667 |
| RPGIGTVGK | 9 | Q9UL18 | Protein argonaute-1 OS=Homo sapiens O | AGO1 | 97.215 | 2 | 56.79019 | yes | 1.236E-07 | 442.76926 | 36.180561 |
| VVLLGEGCVGK | 11 | Q9UL25 | Ras-related protein Rab-21 OS=Homo sa | RAB21 | 24.348 | 2 | 41.94355 | yes | 9.965E-07 | 565.81555 | 88.783928 |
| MLGNEICLCIVGNI | 14 | Q9UL25 | Ras-related protein Rab-21 OS=Homo sa | RAB21 | 24.348 | 2 | 41.94355 | yes | 3.267E-07 | 810.89893 | 41.578468 |
| VNLAIWDTAGQEI | 13 | Q9UL25 | Ras-related protein Rab-21 OS=Homo sa | RAB21 | 24.348 | 2 | 41.94355 | yes | 2.8E-06 | 736.87823 | 31.953152 |
| STPTENELK | 9 | Q9UL33 | Trafficking protein particle complex subu | TRAPPC2I | 16.146 | 2 | 48.209015 | yes | 1.22E-07 | 509.7562 | 101.23388 |
| FVMVVDSSNTALI | 13 | Q9UL33 | Trafficking protein particle complex subu | TRAPPC2I | 16.146 | 2 | 48.209015 | yes | 1.181E-07 | 719.87158 | 93.023224 |
| VYGYVTNSK | 9 | Q9UL33 | Trafficking protein particle complex subu | TRAPPC2I | 16.146 | 2 | 48.209015 | yes | 2.496E-07 | 515.76385 | 85.929047 |
| LWEAVSGAQPVGI | 13 | Q9UL40 | Zinc finger protein 346 OS=Homo sapien | ZNF346 | 32.933 | 2 | 51.9479 | yes | 2.74E-07 | 685.36475 | 501.33328 |
| YLAIHGMETLK | 11 | Q9UL40 | Zinc finger protein 346 OS=Homo sapien | ZNF346 | 32.933 | 2 | 51.9479 | yes | 1.113E-07 | 638.34192 | 310.50046 |
| VCCALLISESQK | 12 | Q9UL40 | Zinc finger protein 346 OS=Homo sapien | ZNF346 | 32.933 | 2 | 51.9479 | yes | 9.819E-08 | 704.35217 | 288.90668 |
| LADPAVTDFPAGK | 13 | Q9UL40 | Zinc finger protein 346 OS=Homo sapien | ZNF346 | 32.933 | 2 | 51.9479 | yes | 1.006E-07 | 651.34045 | 275.66934 |
| NQCLFTNTQCK | 11 | Q9UL40 | Zinc finger protein 346 OS=Homo sapien | ZNF346 | 32.933 | 2 | 51.9479 | yes | 9.722E-08 | 707.3161 | 241.88837 |
| EEVEHMIQK | 9 | Q9UL40 | Zinc finger protein 346 OS=Homo sapien | ZNF346 | 32.933 | 2 | 51.9479 | yes | 2.372E-07 | 571.77917 | 210.07062 |
| AGNILLSEPGLVK | 13 | Q9UL54 | Serine/threonine-protein kinase TAO2 OS | TAOK2 | 138.252 | 2 | 53.509396 | yes | 9.722E-08 | 655.88757 | 322.55038 |
| ELAALLEAQK | 10 | Q9UL54 | Serine/threonine-protein kinase TAO2 OS | TAOK2 | 138.252 | 2 | 53.509396 | yes | 1.315E-06 | 543.31366 | 243.0576 |
| DLECALLLR | 9 | Q9UL54 | Serine/threonine-protein kinase TAO2 OS | TAOK2 | 138.252 | 2 | 53.509396 | yes | 9.722E-08 | 551.79987 | 139.56992 |
| QYFELQCR | 8 | Q9UL54 | Serine/threonine-protein kinase TAO2 OS | TAOK2 | 138.252 | 2 | 53.509396 | yes | 1.075E-07 | 572.26624 | 108.63229 |
| ILGEESGAPSPSPQ | 15 | Q9UL54 | Serine/threonine-protein kinase TAO2 OS | TAOK2 | 138.252 | 2 | 53.509396 | yes | 9.722E-08 | 748.88318 | 69.424431 |
| HPNTIQYR | 8 | Q9UL54 | Serine/threonine-protein kinase TAO2 OS | TAOK2 | 138.252 | 2 | 53.509396 | yes | 4.443E-07 | 514.76721 | 59.427963 |
| EELQENPSTPK | 11 | Q9UL54 | Serine/threonine-protein kinase TAO2 OS | TAOK2 | 138.252 | 2 | 53.509396 | yes | 4.924E-05 | 636.30933 | 28.334068 |
| VPLQDLMVR | 9 | Q9ULA0 | Aspartyl aminopeptidase OS=Homo sapie | DNPEP | 53.411 | 2 | 39.177097 | yes | 7.993E-07 | 535.80499 | 51.969955 |
| IPHLAIHLQR | 10 | Q9ULA0 | Aspartyl aminopeptidase OS=Homo sapie | DNPEP | 53.411 | 2 | 39.177097 | yes | 4.742E-07 | 599.36438 | 6.7658811 |
| EGPFYPTLR | 9 | Q9ULC4 | Malignant T-cell-amplified sequence 1 OS | MCTS1 | 20.556 | 2 | 41.48077 | yes | 3.772E-07 | 540.27966 | 137.77391 |
| MSAEDIEK | 8 | Q9ULC4 | Malignant T-cell-amplified sequence 1 OS | MCTS1 | 20.556 | 2 | 41.48077 | no | 3.489E-07 | 461.71295 | 74.262955 |
| GIGIENIHYLNDGL | 18 | Q9ULC4 | Malignant T-cell-amplified sequence 1 OS | MCTS1 | 20.556 | 3 | 41.48077 | no | 3.544E-07 | 704.02283 | 56.628979 |
| KAEILINK | 8 | Q9ULD2 | Microtubule-associated tumor suppressor | MTUS1 | 141.397 | 2 | 55.464092 | yes | 5.311E-07 | 464.79492 | 2176.4758 |
| TLSQELVNLR | 10 | Q9ULD2 | Microtubule-associated tumor suppressor | MTUS1 | 141.397 | 2 | 55.464092 | yes | 1.223E-07 | 586.83533 | 434.55035 |
| GELVTASTTCEK | 12 | Q9ULD2 | Microtubule-associated tumor suppressor | MTUS1 | 141.397 | 2 | 55.464092 | yes | 1.283E-07 | 648.31104 | 357.19034 |
| TMCMSTPVLEPTK | 13 | Q9ULD2 | Microtubule-associated tumor suppressor | MTUS1 | 141.397 | 2 | 55.464092 | yes | 9.722E-08 | 747.8537 | 349.18042 |
| LVDNNTALVDK | 11 | Q9ULD2 | Microtubule-associated tumor suppressor | MTUS1 | 141.397 | 2 | 55.464092 | yes | 1.224E-07 | 601.32477 | 291.00024 |
| VGPPVSCLR | 9 | Q9ULD2 | Microtubule-associated tumor suppressor | MTUS1 | 141.397 | 2 | 55.464092 | yes | 6.64E-07 | 492.7684 | 271.37607 |

| | | | | | | | | | | | |
|---|---|---|---|---|---|---|---|---|---|---|---|
| SEAQVLNPEHK | 11 | Q9ULD2 | Microtubule-associated tumor suppressor | MTUS1 | 141.397 | 2 | 55.464092 | yes | 1.815E-07 | 626.32001 | 247.38551 |
| TAQSSWVNLPR | 11 | Q9ULD2 | Microtubule-associated tumor suppressor | MTUS1 | 141.397 | 2 | 55.464092 | yes | 9.722E-08 | 629.83057 | 234.39093 |
| SPTSSAIPLQSPR | 13 | Q9ULD2 | Microtubule-associated tumor suppressor | MTUS1 | 141.397 | 2 | 55.464092 | yes | 9.722E-08 | 670.86206 | 207.4664 |
| FQQENEELK | 9 | Q9ULD2 | Microtubule-associated tumor suppressor | MTUS1 | 141.397 | 2 | 55.464092 | yes | 1.464E-06 | 582.78021 | 197.1618 |
| LSMENEELLWK | 11 | Q9ULD2 | Microtubule-associated tumor suppressor | MTUS1 | 141.397 | 2 | 55.464092 | yes | 1.002E-07 | 696.34741 | 180.94016 |
| VTFSVSPIEATEK | 13 | Q9ULD2 | Microtubule-associated tumor suppressor | MTUS1 | 141.397 | 2 | 55.464092 | yes | 1.327E-07 | 704.37195 | 180.77216 |
| AYEASLSEIK | 10 | Q9ULD2 | Microtubule-associated tumor suppressor | MTUS1 | 141.397 | 2 | 55.464092 | yes | 2.823E-07 | 555.78748 | 180.01648 |
| NLFTALNAVEK | 11 | Q9ULD2 | Microtubule-associated tumor suppressor | MTUS1 | 141.397 | 2 | 55.464092 | yes | 9.838E-08 | 610.33771 | 164.02121 |
| NISKPDSCGLR | 11 | Q9ULD2 | Microtubule-associated tumor suppressor | MTUS1 | 141.397 | 2 | 55.464092 | yes | 4.269E-07 | 623.81409 | 151.85089 |
| KTEIISYPR | 9 | Q9ULD2 | Microtubule-associated tumor suppressor | MTUS1 | 141.397 | 2 | 55.464092 | yes | 3.103E-06 | 553.81384 | 133.90263 |
| VTETEDTQMVSK | 12 | Q9ULD2 | Microtubule-associated tumor suppressor | MTUS1 | 141.397 | 2 | 55.464092 | yes | 9.722E-08 | 684.32159 | 131.14943 |
| SENDALNEK | 9 | Q9ULD2 | Microtubule-associated tumor suppressor | MTUS1 | 141.397 | 2 | 55.464092 | yes | 3.537E-06 | 510.23563 | 120.07345 |
| LEIEASHSEK | 10 | Q9ULD2 | Microtubule-associated tumor suppressor | MTUS1 | 141.397 | 2 | 55.464092 | yes | 1.004E-07 | 571.78802 | 101.79858 |
| AHVHLMK | 7 | Q9ULD2 | Microtubule-associated tumor suppressor | MTUS1 | 141.397 | 2 | 55.464092 | yes | 3.232E-07 | 418.23398 | 99.246628 |
| CENQSGFILQLK | 12 | Q9ULD2 | Microtubule-associated tumor suppressor | MTUS1 | 141.397 | 2 | 55.464092 | yes | 9.722E-08 | 718.86377 | 81.191933 |
| QQTVLSR | 7 | Q9ULD2 | Microtubule-associated tumor suppressor | MTUS1 | 141.397 | 2 | 55.464092 | yes | 1.067E-07 | 416.23779 | 59.430592 |
| DSICQCPALVGTEI | 16 | Q9ULD2 | Microtubule-associated tumor suppressor | MTUS1 | 141.397 | 3 | 55.464092 | yes | 5.664E-07 | 601.63068 | 58.869446 |
| SLEDLLSEK | 9 | Q9ULD2 | Microtubule-associated tumor suppressor | MTUS1 | 141.397 | 2 | 55.464092 | yes | 2.385E-07 | 517.27423 | 58.65448 |
| MQLQEQFDNLNA | 18 | Q9ULD2 | Microtubule-associated tumor suppressor | MTUS1 | 141.397 | 3 | 55.464092 | yes | 1.546E-07 | 701.99701 | 43.478714 |
| DLGTQNHTSELILS | 20 | Q9ULD2 | Microtubule-associated tumor suppressor | MTUS1 | 141.397 | 3 | 55.464092 | yes | 1.304E-07 | 708.0307 | 41.099483 |
| PQQTSASSPSSVNS | 15 | Q9ULD2 | Microtubule-associated tumor suppressor | MTUS1 | 141.397 | 2 | 55.464092 | yes | 1.624E-07 | 766.86859 | 35.009621 |
| LRDTYIEEAEK | 11 | Q9ULD2 | Microtubule-associated tumor suppressor | MTUS1 | 141.397 | 2 | 55.464092 | yes | 1.781E-07 | 683.84607 | 18.241953 |
| IVEVDIDGR | 9 | Q9ULD4 | Bromodomain and PHD finger-containing | BRPF3 | 135.745 | 2 | 49.983097 | yes | 2.363E-07 | 508.27457 | 353.42273 |
| SPLQLGNEPLQR | 12 | Q9ULD4 | Bromodomain and PHD finger-containing | BRPF3 | 135.745 | 2 | 49.983097 | yes | 1.355E-07 | 676.37006 | 346.58041 |
| ETLTYAQAQR | 10 | Q9ULD4 | Bromodomain and PHD finger-containing | BRPF3 | 135.745 | 2 | 49.983097 | no | 2.107E-07 | 590.80145 | 266.90363 |
| LAQPPPPQPPSLNK | 14 | Q9ULD4 | Bromodomain and PHD finger-containing | BRPF3 | 135.745 | 2 | 49.983097 | yes | 1.21E-06 | 742.41699 | 240.57155 |
| SLLLPFEDR | 9 | Q9ULD4 | Bromodomain and PHD finger-containing | BRPF3 | 135.745 | 2 | 49.983097 | yes | 1.584E-06 | 545.3006 | 152.34943 |
| GLGAAIQCHK | 10 | Q9ULD4 | Bromodomain and PHD finger-containing | BRPF3 | 135.745 | 2 | 49.983097 | yes | 1.127E-07 | 527.77673 | 148.63838 |
| ICSGLSFQR | 9 | Q9ULD4 | Bromodomain and PHD finger-containing | BRPF3 | 135.745 | 2 | 49.983097 | yes | 1.765E-07 | 534.26874 | 146.35606 |
| AHLSPEVQLK | 10 | Q9ULD4 | Bromodomain and PHD finger-containing | BRPF3 | 135.745 | 2 | 49.983097 | yes | 1.772E-07 | 561.31934 | 124.86668 |
| GYPSYPALIIDPK | 13 | Q9ULD4 | Bromodomain and PHD finger-containing | BRPF3 | 135.745 | 2 | 49.983097 | no | 2.891E-07 | 717.38739 | 124.8407 |
| LHNYWLLK | 8 | Q9ULD4 | Bromodomain and PHD finger-containing | BRPF3 | 135.745 | 2 | 49.983097 | yes | 9.938E-08 | 543.80835 | 102.28872 |
| LDLVSAMR | 8 | Q9ULD4 | Bromodomain and PHD finger-containing | BRPF3 | 135.745 | 2 | 49.983097 | yes | 1.043E-07 | 452.74966 | 95.850548 |
| GPHSFVTSSYL | 11 | Q9ULD4 | Bromodomain and PHD finger-containing | BRPF3 | 135.745 | 2 | 49.983097 | yes | 1.825E-07 | 597.79309 | 73.907883 |
| KVEEDELLEK | 10 | Q9ULD4 | Bromodomain and PHD finger-containing | BRPF3 | 135.745 | 2 | 49.983097 | yes | 2.947E-07 | 616.32446 | 70.678986 |
| LTCYICK | 7 | Q9ULD4 | Bromodomain and PHD finger-containing | BRPF3 | 135.745 | 2 | 49.983097 | no | 2.29E-07 | 479.23026 | 68.678185 |
| QAENIGYDPER | 11 | Q9ULD4 | Bromodomain and PHD finger-containing | BRPF3 | 135.745 | 2 | 49.983097 | yes | 9.67E-07 | 646.29932 | 59.470116 |
| TTLDLLQEK | 9 | Q9ULD4 | Bromodomain and PHD finger-containing | BRPF3 | 135.745 | 2 | 49.983097 | yes | 1.076E-07 | 530.79785 | 44.586044 |
| ENSEQPQFPGK | 11 | Q9ULD4 | Bromodomain and PHD finger-containing | BRPF3 | 135.745 | 2 | 49.983097 | yes | 5.081E-07 | 630.79639 | 40.829369 |

| | | | | | | | | | | | |
|---|---|---|---|---|---|---|---|---|---|---|---|
| TAYCEAHSPPGAA | 16 | Q9ULD4 | Bromodomain and PHD finger-containing | BRPF3 | 135.745 | 2 | 49.983097 | yes | 7.277E-07 | 830.3808 | 37.07423 |
| PMDFSTMR | 8 | Q9ULD4 | Bromodomain and PHD finger-containing | BRPF3 | 135.745 | 2 | 49.983097 | yes | 1.559E-07 | 492.7175 | 33.308571 |
| LLIELIR | 7 | Q9ULD4 | Bromodomain and PHD finger-containing | BRPF3 | 135.745 | 2 | 49.983097 | yes | 4.245E-07 | 435.29456 | 28.658745 |
| ISIYDPLK | 8 | Q9ULD4 | Bromodomain and PHD finger-containing | BRPF3 | 135.745 | 2 | 49.983097 | yes | 4.84E-07 | 474.77368 | 23.48786 |
| TNLATENQYLR | 11 | Q9ULG6 | Cell cycle progression protein 1 OS=Hom | CCPG1 | 87.341 | 2 | 53.802666 | yes | 2.743E-07 | 661.83856 | 280.60873 |
| GELQQLSGSQLHG | 14 | Q9ULG6 | Cell cycle progression protein 1 OS=Hom | CCPG1 | 87.341 | 2 | 53.802666 | yes | 9.722E-08 | 741.38898 | 238.27255 |
| ALSSLQEELNK | 11 | Q9ULG6 | Cell cycle progression protein 1 OS=Hom | CCPG1 | 87.341 | 2 | 53.802666 | yes | 9.722E-08 | 616.33008 | 181.01462 |
| LVAEQETEPSK | 11 | Q9ULG6 | Cell cycle progression protein 1 OS=Hom | CCPG1 | 87.341 | 2 | 53.802666 | yes | 9.722E-08 | 615.81427 | 172.98828 |
| CWTLTEAEK | 9 | Q9ULG6 | Cell cycle progression protein 1 OS=Hom | CCPG1 | 87.341 | 2 | 53.802666 | yes | 1.165E-07 | 569.26587 | 138.56595 |
| EYEVDNDGVFEK | 12 | Q9ULG6 | Cell cycle progression protein 1 OS=Hom | CCPG1 | 87.341 | 2 | 53.802666 | yes | 9.722E-08 | 722.31738 | 128.1613 |
| IHEDELNDMK | 10 | Q9ULG6 | Cell cycle progression protein 1 OS=Hom | CCPG1 | 87.341 | 2 | 53.802666 | yes | 9.722E-08 | 622.28479 | 114.28527 |
| KLTFEQQR | 8 | Q9ULG6 | Cell cycle progression protein 1 OS=Hom | CCPG1 | 87.341 | 2 | 53.802666 | yes | 2.004E-07 | 525.29053 | 110.10048 |
| LFTDFVNDVK | 10 | Q9ULG6 | Cell cycle progression protein 1 OS=Hom | CCPG1 | 87.341 | 2 | 53.802666 | yes | 1.133E-05 | 599.31116 | 47.084583 |
| LDEYIYR | 7 | Q9ULG6 | Cell cycle progression protein 1 OS=Hom | CCPG1 | 87.341 | 2 | 53.802666 | yes | 4.656E-07 | 486.24527 | 35.51078 |
| LTCLECMR | 8 | Q9ULJ3 | Zinc finger and BTB domain-containing p | ZBTB21 | 118.871 | 2 | 51.280865 | yes | 2.04E-07 | 541.74323 | 138.76364 |
| EASLWPCEK | 9 | Q9ULJ3 | Zinc finger and BTB domain-containing p | ZBTB21 | 118.871 | 2 | 51.280865 | yes | 5.653E-07 | 560.26062 | 136.93364 |
| TEPSSPLSDPSDIIR | 15 | Q9ULJ3 | Zinc finger and BTB domain-containing p | ZBTB21 | 118.871 | 2 | 51.280865 | yes | 1.122E-07 | 807.40448 | 129.27609 |
| NAAVCPYCSLR | 11 | Q9ULJ3 | Zinc finger and BTB domain-containing p | ZBTB21 | 118.871 | 2 | 51.280865 | yes | 1.043E-07 | 655.80261 | 116.16779 |
| NAVLPSKPLQDR | 12 | Q9ULJ3 | Zinc finger and BTB domain-containing p | ZBTB21 | 118.871 | 2 | 51.280865 | yes | 1.385E-07 | 669.38043 | 110.53223 |
| VFVEDDENSSQK | 12 | Q9ULJ3 | Zinc finger and BTB domain-containing p | ZBTB21 | 118.871 | 2 | 51.280865 | yes | 9.722E-08 | 698.815 | 95.790443 |
| EHAPLASPVENK | 12 | Q9ULJ3 | Zinc finger and BTB domain-containing p | ZBTB21 | 118.871 | 2 | 51.280865 | yes | 5.148E-07 | 646.33569 | 84.571159 |
| FFSPELK | 7 | Q9ULJ3 | Zinc finger and BTB domain-containing p | ZBTB21 | 118.871 | 2 | 51.280865 | yes | 3.656E-07 | 434.23419 | 62.055473 |
| SFSASQSTDR | 10 | Q9ULJ3 | Zinc finger and BTB domain-containing p | ZBTB21 | 118.871 | 2 | 51.280865 | yes | 2.329E-07 | 543.24652 | 61.514751 |
| KLIDIVR | 7 | Q9ULJ3 | Zinc finger and BTB domain-containing p | ZBTB21 | 118.871 | 2 | 51.280865 | yes | 7.422E-07 | 428.78436 | 55.982723 |
| NPSPASSSHAVLD | 15 | Q9ULJ3 | Zinc finger and BTB domain-containing p | ZBTB21 | 118.871 | 2 | 51.280865 | yes | 2.829E-07 | 769.87592 | 26.695749 |
| SLEHSGSLDDPNR | 13 | Q9ULJ3 | Zinc finger and BTB domain-containing p | ZBTB21 | 118.871 | 2 | 51.280865 | yes | 2.272E-07 | 713.83148 | 24.543228 |
| TAFSLWSHEQTHN | 13 | Q9ULJ3 | Zinc finger and BTB domain-containing p | ZBTB21 | 118.871 | 2 | 51.280865 | yes | 3.296E-06 | 779.35767 | 21.309114 |
| EAMDDKPGVSGQ | 15 | Q9ULJ3 | Zinc finger and BTB domain-containing p | ZBTB21 | 118.871 | 2 | 51.280865 | yes | 2.989E-06 | 786.39032 | 12.436813 |
| DMVCNMLGLNK | 11 | Q9ULK4 | Mediator of RNA polymerase II transcrip | MED23 | 156.474 | 2 | 40.447636 | yes | 5.721E-06 | 647.80127 | 38.167278 |
| EVGNALLNVVLK | 12 | Q9ULK4 | Mediator of RNA polymerase II transcrip | MED23 | 156.474 | 2 | 40.447636 | yes | 4.019E-07 | 634.88226 | 23.689774 |
| FLELLPVSK | 9 | Q9ULK4 | Mediator of RNA polymerase II transcrip | MED23 | 156.474 | 2 | 40.447636 | yes | 8.359E-06 | 523.31824 | 18.74029 |
| SWIMLQEPDR | 10 | Q9ULM6 | CCR4-NOT transcription complex subuni | CNOT6 | 63.307 | 2 | 51.48384 | yes | 9.722E-08 | 637.81354 | 46.703693 |
| TVTPDASVYSDIV | 15 | Q9ULP9 | TBC1 domain family member 24 OS=Ho | TBC1D24 | 62.92 | 2 | 34.014275 | yes | 1.098E-06 | 776.39868 | 45.378708 |
| VAQIQNAGLGEFR | 13 | Q9ULR0 | Pre-mRNA-splicing factor ISY1 homolog | ISY1 | 32.993 | 2 | 51.929367 | yes | 9.722E-08 | 701.87549 | 159.95453 |
| ILYIVASDPYSR | 12 | Q9ULT8 | E3 ubiquitin-protein ligase HECTD1 OS= | HECTD1 | 289.369 | 2 | 53.293453 | yes | 9.722E-08 | 698.8772 | 78.382065 |
| LVVVELNNR | 9 | Q9ULT8 | E3 ubiquitin-protein ligase HECTD1 OS= | HECTD1 | 289.369 | 2 | 53.293453 | yes | 1.905E-07 | 528.31403 | 74.526276 |
| KLGPDSSVR | 9 | Q9ULT8 | E3 ubiquitin-protein ligase HECTD1 OS= | HECTD1 | 289.369 | 2 | 53.293453 | yes | 3.144E-06 | 479.76944 | 70.698372 |
| EDLPGFVFESNR | 12 | Q9ULT8 | E3 ubiquitin-protein ligase HECTD1 OS= | HECTD1 | 289.369 | 2 | 53.293453 | yes | 9.722E-08 | 705.33844 | 56.191784 |
| GSPVVTHDLLR | 11 | Q9ULT8 | E3 ubiquitin-protein ligase HECTD1 OS= | HECTD1 | 289.369 | 2 | 53.293453 | yes | 2.313E-06 | 597.33551 | 22.358536 |

| | | | | | | | | | | | |
|---|---|---|---|---|---|---|---|---|---|---|---|
| IPSTDTQVQPK | 11 | Q9ULV3 | Cip1-interacting zinc finger protein OS=I | CIZ1 | 100.046 | 2 | 44.8657 | yes | 2.041E-07 | 607.32477 | 91.88884 |
| EMPGVWGAGGSL | 13 | Q9ULV3 | Cip1-interacting zinc finger protein OS=I | CIZ1 | 100.046 | 2 | 44.8657 | yes | 3.739E-07 | 644.82135 | 69.419853 |
| KFVEHVK | 7 | Q9ULV3 | Cip1-interacting zinc finger protein OS=I | CIZ1 | 100.046 | 2 | 44.8657 | yes | 8.149E-07 | 443.76089 | 56.909176 |
| QVQLQQEAEPLK | 12 | Q9ULV3 | Cip1-interacting zinc finger protein OS=I | CIZ1 | 100.046 | 2 | 44.8657 | yes | 4.106E-07 | 705.883 | 32.536655 |
| VTWDSSFCAVNPF | 13 | Q9ULV4 | Coronin-1C OS=Homo sapiens OX=9606 | CORO1C | 53.25 | 2 | 49.20943 | yes | 1.154E-07 | 769.85645 | 293.41006 |
| NADPILISLK | 10 | Q9ULV4 | Coronin-1C OS=Homo sapiens OX=9606 | CORO1C | 53.25 | 2 | 49.20943 | yes | 4.42E-07 | 542.32404 | 247.70935 |
| VGIVAWHPTAR | 11 | Q9ULV4 | Coronin-1C OS=Homo sapiens OX=9606 | CORO1C | 53.25 | 2 | 49.20943 | yes | 1.204E-07 | 603.84076 | 143.27942 |
| NILDSKPTANK | 11 | Q9ULV4 | Coronin-1C OS=Homo sapiens OX=9606 | CORO1C | 53.25 | 2 | 49.20943 | yes | 4.34E-07 | 600.83276 | 134.24655 |
| KCEPIIMTVPR | 11 | Q9ULV4 | Coronin-1C OS=Homo sapiens OX=9606 | CORO1C | 53.25 | 2 | 49.20943 | no | 1.581E-07 | 672.3623 | 123.43348 |
| RVGIVAWHPTAR | 12 | Q9ULV4 | Coronin-1C OS=Homo sapiens OX=9606 | CORO1C | 53.25 | 3 | 49.20943 | yes | 1.276E-06 | 454.92996 | 109.67349 |
| NDQCYDDIR | 9 | Q9ULV4 | Coronin-1C OS=Homo sapiens OX=9606 | CORO1C | 53.25 | 2 | 49.20943 | yes | 1.059E-07 | 599.74329 | 89.087753 |
| SSDQPLTVPVSPK | 13 | Q9ULW0 | Targeting protein for Xklp2 OS=Homo sa | TPX2 | 85.654 | 2 | 52.905148 | yes | 9.722E-08 | 677.86426 | 1664.0885 |
| ILEDVVGVPEK | 11 | Q9ULW0 | Targeting protein for Xklp2 OS=Homo sa | TPX2 | 85.654 | 2 | 52.905148 | yes | 1.253E-07 | 599.3399 | 1590.2515 |
| SVDFHFR | 7 | Q9ULW0 | Targeting protein for Xklp2 OS=Homo sa | TPX2 | 85.654 | 2 | 52.905148 | yes | 1.686E-07 | 454.22467 | 1308.9098 |
| STAELEAEELEK | 12 | Q9ULW0 | Targeting protein for Xklp2 OS=Homo sa | TPX2 | 85.654 | 2 | 52.905148 | yes | 9.722E-08 | 674.82758 | 1150.0206 |
| ILEGGPILPK | 10 | Q9ULW0 | Targeting protein for Xklp2 OS=Homo sa | TPX2 | 85.654 | 2 | 52.905148 | yes | 2.89E-07 | 518.82367 | 1091.1283 |
| MQQEVVEMR | 9 | Q9ULW0 | Targeting protein for Xklp2 OS=Homo sa | TPX2 | 85.654 | 2 | 52.905148 | yes | 9.729E-08 | 575.27319 | 948.27759 |
| LALAGIGQPVK | 11 | Q9ULW0 | Targeting protein for Xklp2 OS=Homo sa | TPX2 | 85.654 | 2 | 52.905148 | yes | 1.223E-07 | 533.83459 | 726.7287 |
| EVNFTSELR | 9 | Q9ULW0 | Targeting protein for Xklp2 OS=Homo sa | TPX2 | 85.654 | 2 | 52.905148 | yes | 9.766E-08 | 547.77747 | 723.19391 |
| AQQLEEAR | 8 | Q9ULW0 | Targeting protein for Xklp2 OS=Homo sa | TPX2 | 85.654 | 2 | 52.905148 | yes | 4.792E-07 | 472.74344 | 640.02386 |
| DPQTPVLQTK | 10 | Q9ULW0 | Targeting protein for Xklp2 OS=Homo sa | TPX2 | 85.654 | 2 | 52.905148 | yes | 1.188E-07 | 563.80878 | 522.88287 |
| HQLEEELR | 8 | Q9ULW0 | Targeting protein for Xklp2 OS=Homo sa | TPX2 | 85.654 | 2 | 52.905148 | yes | 4.16E-07 | 527.26984 | 482.41904 |
| MAEVEAQK | 8 | Q9ULW0 | Targeting protein for Xklp2 OS=Homo sa | TPX2 | 85.654 | 2 | 52.905148 | yes | 1.45E-07 | 453.22348 | 457.44193 |
| CATPVIIDEILPSK | 14 | Q9ULW0 | Targeting protein for Xklp2 OS=Homo sa | TPX2 | 85.654 | 2 | 52.905148 | yes | 1.963E-07 | 778.42365 | 405.42014 |
| STEEQELEK | 9 | Q9ULW0 | Targeting protein for Xklp2 OS=Homo sa | TPX2 | 85.654 | 2 | 52.905148 | yes | 1.025E-07 | 546.75641 | 394.66632 |
| KYQGLEIK | 8 | Q9ULW0 | Targeting protein for Xklp2 OS=Homo sa | TPX2 | 85.654 | 2 | 52.905148 | yes | 1.344E-07 | 489.78458 | 376.38675 |
| ARPNTVISQEPFVI | 15 | Q9ULW0 | Targeting protein for Xklp2 OS=Homo sa | TPX2 | 85.654 | 3 | 52.905148 | yes | 1.05E-07 | 561.64569 | 120.7038 |
| ARPNTVISQEPFVI | 15 | Q9ULW0 | Targeting protein for Xklp2 OS=Homo sa | TPX2 | 85.654 | 3 | 52.905148 | yes | 1.396E-07 | 841.96484 | 120.7038 |
| ARPNTVISQEPFVI | 15 | Q9ULW0 | Targeting protein for Xklp2 OS=Homo sa | TPX2 | 85.654 | 2 | 52.905148 | yes | 1.05E-07 | 561.64569 | 120.7038 |
| ARPNTVISQEPFVI | 15 | Q9ULW0 | Targeting protein for Xklp2 OS=Homo sa | TPX2 | 85.654 | 2 | 52.905148 | yes | 1.396E-07 | 841.96484 | 120.7038 |
| TVEICPFSFDSR | 12 | Q9ULW0 | Targeting protein for Xklp2 OS=Homo sa | TPX2 | 85.654 | 2 | 52.905148 | yes | 1.029E-07 | 729.34009 | 198.80922 |
| GCTIVKPFNLSQGI | 14 | Q9ULW0 | Targeting protein for Xklp2 OS=Homo sa | TPX2 | 85.654 | 2 | 52.905148 | yes | 1.021E-07 | 774.91376 | 186.10982 |
| AQPVPHYGVPFKF | 19 | Q9ULW0 | Targeting protein for Xklp2 OS=Homo sa | TPX2 | 85.654 | 4 | 52.905148 | yes | 2.033E-07 | 533.54163 | 82.569717 |
| AQPVPHYGVPFKF | 19 | Q9ULW0 | Targeting protein for Xklp2 OS=Homo sa | TPX2 | 85.654 | 4 | 52.905148 | yes | 1.819E-07 | 711.0531 | 82.569717 |
| AQPVPHYGVPFKF | 19 | Q9ULW0 | Targeting protein for Xklp2 OS=Homo sa | TPX2 | 85.654 | 3 | 52.905148 | yes | 2.033E-07 | 533.54163 | 82.569717 |
| AQPVPHYGVPFKF | 19 | Q9ULW0 | Targeting protein for Xklp2 OS=Homo sa | TPX2 | 85.654 | 3 | 52.905148 | yes | 1.819E-07 | 711.0531 | 82.569717 |
| MPTKEDEEEDEPV | 16 | Q9ULW0 | Targeting protein for Xklp2 OS=Homo sa | TPX2 | 85.654 | 3 | 52.905148 | yes | 1.341E-07 | 629.9679 | 145.42804 |
| KDDINLLPSK | 10 | Q9ULW0 | Targeting protein for Xklp2 OS=Homo sa | TPX2 | 85.654 | 2 | 52.905148 | yes | 9.722E-08 | 571.8244 | 136.69513 |
| YQGLEIK | 7 | Q9ULW0 | Targeting protein for Xklp2 OS=Homo sa | TPX2 | 85.654 | 2 | 52.905148 | yes | 1.026E-06 | 425.73709 | 114.10925 |

| | | | | | | | | | | | |
|---|---|---|---|---|---|---|---|---|---|---|---|
| HTVPCMPPAK | 10 | Q9ULW0 | Targeting protein for Xklp2 OS=Homo sa | TPX2 | 85.654 | 2 | 52.905148 | yes | 3.437E-07 | 569.28082 | 105.60501 |
| EDEEEDEPVVIK | 12 | Q9ULW0 | Targeting protein for Xklp2 OS=Homo sa | TPX2 | 85.654 | 2 | 52.905148 | yes | 1.436E-07 | 715.83032 | 68.474777 |
| KPPVKPPTEPIGFD | 19 | Q9ULW0 | Targeting protein for Xklp2 OS=Homo sa | TPX2 | 85.654 | 3 | 52.905148 | yes | 2.615E-07 | 712.06464 | 59.391174 |
| TEDEHFEFHSR | 11 | Q9ULW0 | Targeting protein for Xklp2 OS=Homo sa | TPX2 | 85.654 | 2 | 52.905148 | yes | 4.273E-07 | 717.30768 | 50.24675 |
| KANLQQAIVTPLK | 21 | Q9ULW0 | Targeting protein for Xklp2 OS=Homo sa | TPX2 | 85.654 | 4 | 52.905148 | yes | 3.323E-07 | 601.8363 | 42.773224 |
| ICRDPQTPVLQTK | 13 | Q9ULW0 | Targeting protein for Xklp2 OS=Homo sa | TPX2 | 85.654 | 2 | 52.905148 | yes | 1.649E-07 | 778.41669 | 36.65992 |
| KLALAGIGQPVK | 12 | Q9ULW0 | Targeting protein for Xklp2 OS=Homo sa | TPX2 | 85.654 | 2 | 52.905148 | yes | 9.722E-08 | 597.88208 | 27.739677 |
| ILEDVVGVPEKK | 12 | Q9ULW0 | Targeting protein for Xklp2 OS=Homo sa | TPX2 | 85.654 | 2 | 52.905148 | yes | 1.19E-06 | 663.38739 | 25.328415 |
| ALPLPHFDTINLPE | 15 | Q9ULW0 | Targeting protein for Xklp2 OS=Homo sa | TPX2 | 85.654 | 3 | 52.905148 | yes | 2.822E-05 | 568.98218 | 18.687311 |
| ETDFYLQSVER | 11 | Q9ULW3 | Activator of basal transcription 1 OS=Ho | ABT1 | 31.079 | 2 | 53.41374 | yes | 1.039E-07 | 693.83044 | 300.41901 |
| NLLSAYGEVGR | 11 | Q9ULW3 | Activator of basal transcription 1 OS=Ho | ABT1 | 31.079 | 2 | 53.41374 | yes | 2.404E-06 | 589.81183 | 142.40674 |
| VAASLHNTPMGAI | 13 | Q9ULW3 | Activator of basal transcription 1 OS=Ho | ABT1 | 31.079 | 2 | 53.41374 | yes | 9.722E-08 | 662.84314 | 108.68336 |
| DYTEGWVEFR | 10 | Q9ULW3 | Activator of basal transcription 1 OS=Ho | ABT1 | 31.079 | 2 | 53.41374 | yes | 9.862E-07 | 651.29346 | 107.62124 |
| VFFQAEDR | 8 | Q9ULW3 | Activator of basal transcription 1 OS=Ho | ABT1 | 31.079 | 2 | 53.41374 | yes | 1.49E-07 | 506.24835 | 77.008591 |
| VMLVGDSGVGK | 11 | Q9ULW5 | Ras-related protein Rab-26 OS=Homo sa | RAB26 | 27.9 | 2 | 46.460556 | yes | 1.553E-07 | 531.28662 | 73.529106 |
| LQMWDTAGQER | 11 | Q9ULW5 | Ras-related protein Rab-26 OS=Homo sa | RAB26 | 27.9 | 2 | 46.460556 | yes | 2.649E-06 | 667.81152 | 34.467796 |
| LAVLPYELR | 9 | Q9ULX3 | RNA-binding protein NOB1 OS=Homo s | NOB1 | 46.675 | 2 | 53.841064 | yes | 1.738E-07 | 537.32129 | 5267.9526 |
| FKEPLPEYVR | 10 | Q9ULX3 | RNA-binding protein NOB1 OS=Homo s | NOB1 | 46.675 | 2 | 53.841064 | yes | 3.859E-07 | 639.34808 | 2006.6365 |
| FKEPLPEYVR | 10 | Q9ULX3 | RNA-binding protein NOB1 OS=Homo s | NOB1 | 46.675 | 2 | 53.841064 | yes | 1.976E-07 | 426.56781 | 2006.6365 |
| FKEPLPEYVR | 10 | Q9ULX3 | RNA-binding protein NOB1 OS=Homo s | NOB1 | 46.675 | 3 | 53.841064 | yes | 3.859E-07 | 639.34808 | 2006.6365 |
| FKEPLPEYVR | 10 | Q9ULX3 | RNA-binding protein NOB1 OS=Homo s | NOB1 | 46.675 | 3 | 53.841064 | yes | 1.976E-07 | 426.56781 | 2006.6365 |
| HAALQDIGK | 9 | Q9ULX3 | RNA-binding protein NOB1 OS=Homo s | NOB1 | 46.675 | 2 | 53.841064 | yes | 1.163E-07 | 476.76416 | 1106.6005 |
| APVEHVVADAGA | 15 | Q9ULX3 | RNA-binding protein NOB1 OS=Homo s | NOB1 | 46.675 | 2 | 53.841064 | yes | 9.722E-08 | 776.41754 | 1023.6275 |
| YAINPHLTEDQR | 12 | Q9ULX3 | RNA-binding protein NOB1 OS=Homo s | NOB1 | 46.675 | 2 | 53.841064 | yes | 9.722E-08 | 728.86261 | 327.32468 |
| YAINPHLTEDQR | 12 | Q9ULX3 | RNA-binding protein NOB1 OS=Homo s | NOB1 | 46.675 | 2 | 53.841064 | yes | 1.206E-07 | 486.24417 | 327.32468 |
| YAINPHLTEDQR | 12 | Q9ULX3 | RNA-binding protein NOB1 OS=Homo s | NOB1 | 46.675 | 3 | 53.841064 | yes | 9.722E-08 | 728.86261 | 327.32468 |
| YAINPHLTEDQR | 12 | Q9ULX3 | RNA-binding protein NOB1 OS=Homo s | NOB1 | 46.675 | 3 | 53.841064 | yes | 1.206E-07 | 486.24417 | 327.32468 |
| EVVTEIR | 7 | Q9ULX3 | RNA-binding protein NOB1 OS=Homo s | NOB1 | 46.675 | 2 | 53.841064 | yes | 1.063E-07 | 423.23999 | 494.96927 |
| EPLPEYVR | 8 | Q9ULX3 | RNA-binding protein NOB1 OS=Homo s | NOB1 | 46.675 | 2 | 53.841064 | yes | 5.653E-07 | 501.76639 | 343.73862 |
| VFCSHCGNK | 9 | Q9ULX3 | RNA-binding protein NOB1 OS=Homo s | NOB1 | 46.675 | 2 | 53.841064 | yes | 1.001E-07 | 554.73676 | 299.19189 |
| VSSSIQHPETPLHIS | 22 | Q9ULX3 | RNA-binding protein NOB1 OS=Homo s | NOB1 | 46.675 | 4 | 53.841064 | yes | 1.132E-07 | 619.32611 | 149.75429 |
| VSSSIQHPETPLHIS | 31 | Q9ULX3 | RNA-binding protein NOB1 OS=Homo s | NOB1 | 46.675 | 5 | 53.841064 | yes | 1.998E-07 | 702.57031 | 104.40477 |
| VSVTVSDDGTLHN | 17 | Q9ULX3 | RNA-binding protein NOB1 OS=Homo s | NOB1 | 46.675 | 3 | 53.841064 | yes | 5.333E-07 | 629.97211 | 52.998299 |
| IQFVCSLCK | 9 | Q9ULX6 | A-kinase anchor protein 8-like OS=Homo | AKAP8L | 71.64 | 2 | 52.81045 | yes | 1.163E-07 | 577.78845 | 2062.7854 |
| TVEDLDGLIQQIYI | 14 | Q9ULX6 | A-kinase anchor protein 8-like OS=Homo | AKAP8L | 71.64 | 2 | 52.81045 | yes | 1E-07 | 831.93848 | 1642.9283 |
| TFYEDEMASHLDS | 14 | Q9ULX6 | A-kinase anchor protein 8-like OS=Homo | AKAP8L | 71.64 | 2 | 52.81045 | yes | 1.062E-07 | 836.86157 | 1509.5881 |
| MWEDPMGAR | 9 | Q9ULX6 | A-kinase anchor protein 8-like OS=Homo | AKAP8L | 71.64 | 2 | 52.81045 | yes | 1.152E-07 | 546.7337 | 645.96307 |
| QTADFLQEYVTNK | 13 | Q9ULX6 | A-kinase anchor protein 8-like OS=Homo | AKAP8L | 71.64 | 2 | 52.81045 | yes | 1.238E-07 | 778.88318 | 494.00705 |
| GALTTQDENGQTK | 13 | Q9ULX6 | A-kinase anchor protein 8-like OS=Homo | AKAP8L | 71.64 | 2 | 52.81045 | yes | 9.722E-08 | 681.82843 | 253.38449 |

| | | | | | | | | | | | |
|---|---|---|---|---|---|---|---|---|---|---|---|
| GIPGLDVEDDEEG | 18 | Q9ULX6 | A-kinase anchor protein 8-like OS=Homo | AKAP8L | 71.64 | 2 | 52.81045 | yes | 5.894E-07 | 842.37085 | 139.17636 |
| GQCMSGASR | 9 | Q9ULX6 | A-kinase anchor protein 8-like OS=Homo | AKAP8L | 71.64 | 2 | 52.81045 | yes | 3.567E-07 | 477.20001 | 90.205788 |
| YDSYESCDSR | 10 | Q9ULX6 | A-kinase anchor protein 8-like OS=Homo | AKAP8L | 71.64 | 2 | 52.81045 | yes | 1.249E-06 | 641.23804 | 44.537994 |
| PMASGYGR | 8 | Q9ULX6 | A-kinase anchor protein 8-like OS=Homo | AKAP8L | 71.64 | 2 | 52.81045 | yes | 7.307E-07 | 419.69745 | 21.621656 |
| FGFGFGNGMK | 10 | Q9ULX6 | A-kinase anchor protein 8-like OS=Homo | AKAP8L | 71.64 | 2 | 52.81045 | yes | 3.166E-06 | 531.24731 | 9.2160549 |
| GPATLVAPASVITI | 16 | Q9ULX9 | Transcription factor MafF OS=Homo sap | MAFF | 17.761 | 2 | 42.688892 | yes | 2.916E-07 | 768.97162 | 27.560879 |
| ILGLAPSR | 8 | Q9UM00 | Calcium load-activated calcium channel C | TMCO1 | 27.08 | 2 | 50.58627 | yes | 3.649E-07 | 413.76089 | 780.03979 |
| QAGGFLGPPPPSGI | 14 | Q9UM00 | Calcium load-activated calcium channel C | TMCO1 | 27.08 | 2 | 50.58627 | yes | 9.758E-08 | 655.34857 | 502.02124 |
| KETITESAGR | 10 | Q9UM00 | Calcium load-activated calcium channel C | TMCO1 | 27.08 | 2 | 50.58627 | yes | 2.247E-07 | 546.28821 | 306.97913 |
| VGALAWNAEQLS | 16 | Q9UM11 | Fizzy-related protein homolog OS=Homo | FZR1 | 55.18 | 2 | 47.09161 | yes | 1.402E-07 | 823.41827 | 106.2442 |
| LQQFFNER | 8 | Q9UM54 | Unconventional myosin-VI OS=Homo sa | MYO6 | 149.692 | 2 | 52.74649 | yes | 2.657E-07 | 541.2749 | 114.61563 |
| SSEELLSALQK | 11 | Q9UM54 | Unconventional myosin-VI OS=Homo sa | MYO6 | 149.692 | 2 | 52.74649 | yes | 1.171E-07 | 602.82465 | 111.51136 |
| STMMTQEQIQK | 11 | Q9UM54 | Unconventional myosin-VI OS=Homo sa | MYO6 | 149.692 | 2 | 52.74649 | yes | 1.164E-07 | 662.81543 | 102.25145 |
| FVEIHFNEK | 9 | Q9UM54 | Unconventional myosin-VI OS=Homo sa | MYO6 | 149.692 | 2 | 52.74649 | yes | 9.722E-08 | 581.79822 | 99.05719 |
| IQAEVEAQLAR | 11 | Q9UM54 | Unconventional myosin-VI OS=Homo sa | MYO6 | 149.692 | 2 | 52.74649 | yes | 1.191E-07 | 614.33826 | 98.945526 |
| VMLTTAGGTK | 10 | Q9UM54 | Unconventional myosin-VI OS=Homo sa | MYO6 | 149.692 | 2 | 52.74649 | yes | 1.967E-07 | 489.76807 | 88.491669 |
| SDPDHLAELVK | 11 | Q9UM54 | Unconventional myosin-VI OS=Homo sa | MYO6 | 149.692 | 2 | 52.74649 | yes | 1.564E-07 | 612.31696 | 86.049004 |
| NYHIFYR | 7 | Q9UM54 | Unconventional myosin-VI OS=Homo sa | MYO6 | 149.692 | 2 | 52.74649 | yes | 4.51E-07 | 506.7536 | 84.747787 |
| IVEANPLLEAFGN | 15 | Q9UM54 | Unconventional myosin-VI OS=Homo sa | MYO6 | 149.692 | 2 | 52.74649 | yes | 9.722E-08 | 793.43286 | 80.541199 |
| TQLNLLLDK | 9 | Q9UM54 | Unconventional myosin-VI OS=Homo sa | MYO6 | 149.692 | 2 | 52.74649 | yes | 1.34E-07 | 529.31622 | 79.879471 |
| MSQSIIVSGESGAC | 15 | Q9UM54 | Unconventional myosin-VI OS=Homo sa | MYO6 | 149.692 | 2 | 52.74649 | yes | 2.873E-07 | 725.86395 | 54.714287 |
| TVYSHLFDHVVNI | 13 | Q9UM54 | Unconventional myosin-VI OS=Homo sa | MYO6 | 149.692 | 3 | 52.74649 | yes | 6.3E-07 | 529.6073 | 36.058563 |
| TVPEELVKPEELSI | 14 | Q9UMS4 | Pre-mRNA-processing factor 19 OS=Hon | PRPF19 | 55.181 | 2 | 55.03553 | yes | 2.25E-07 | 799.43781 | 1755.8691 |
| TVPEELVKPEELSI | 14 | Q9UMS4 | Pre-mRNA-processing factor 19 OS=Hon | PRPF19 | 55.181 | 2 | 55.03553 | yes | 6.038E-07 | 533.29431 | 1755.8691 |
| TVPEELVKPEELSI | 14 | Q9UMS4 | Pre-mRNA-processing factor 19 OS=Hon | PRPF19 | 55.181 | 3 | 55.03553 | yes | 2.25E-07 | 799.43781 | 1755.8691 |
| TVPEELVKPEELSI | 14 | Q9UMS4 | Pre-mRNA-processing factor 19 OS=Hon | PRPF19 | 55.181 | 3 | 55.03553 | yes | 6.038E-07 | 533.29431 | 1755.8691 |
| IWSVPNASCVQVV | 14 | Q9UMS4 | Pre-mRNA-processing factor 19 OS=Hon | PRPF19 | 55.181 | 2 | 55.03553 | yes | 2.89E-07 | 807.92468 | 2902.9985 |
| TLQLDNNFEVK | 11 | Q9UMS4 | Pre-mRNA-processing factor 19 OS=Hon | PRPF19 | 55.181 | 2 | 55.03553 | yes | 9.722E-08 | 660.84332 | 2058.6919 |
| ATVLTTER | 8 | Q9UMS4 | Pre-mRNA-processing factor 19 OS=Hon | PRPF19 | 55.181 | 2 | 55.03553 | yes | 1.8E-07 | 445.75073 | 1042.9882 |
| SSEQILATLK | 10 | Q9UMS4 | Pre-mRNA-processing factor 19 OS=Hon | PRPF19 | 55.181 | 2 | 55.03553 | yes | 9.722E-08 | 545.31116 | 1005.9076 |
| PKPPSATSIPAILK | 14 | Q9UMS4 | Pre-mRNA-processing factor 19 OS=Hon | PRPF19 | 55.181 | 2 | 55.03553 | yes | 3.8E-07 | 710.43213 | 450.78485 |
| PKPPSATSIPAILK | 14 | Q9UMS4 | Pre-mRNA-processing factor 19 OS=Hon | PRPF19 | 55.181 | 2 | 55.03553 | yes | 1.215E-07 | 473.95718 | 450.78485 |
| PKPPSATSIPAILK | 14 | Q9UMS4 | Pre-mRNA-processing factor 19 OS=Hon | PRPF19 | 55.181 | 3 | 55.03553 | yes | 3.8E-07 | 710.43213 | 450.78485 |
| PKPPSATSIPAILK | 14 | Q9UMS4 | Pre-mRNA-processing factor 19 OS=Hon | PRPF19 | 55.181 | 3 | 55.03553 | yes | 1.215E-07 | 473.95718 | 450.78485 |
| QVASHVGLHSASI | 28 | Q9UMS4 | Pre-mRNA-processing factor 19 OS=Hon | PRPF19 | 55.181 | 4 | 55.03553 | yes | 1.916E-07 | 725.87885 | 877.82343 |
| KVTSVVFHPSQDL | 24 | Q9UMS4 | Pre-mRNA-processing factor 19 OS=Hon | PRPF19 | 55.181 | 4 | 55.03553 | yes | 1.061E-07 | 651.09723 | 520.09222 |
| VAHPIRPKPPSATS | 20 | Q9UMS4 | Pre-mRNA-processing factor 19 OS=Hon | PRPF19 | 55.181 | 3 | 55.03553 | yes | 2.106E-07 | 698.42462 | 441.70313 |
| QQLQTTR | 7 | Q9UMS4 | Pre-mRNA-processing factor 19 OS=Hon | PRPF19 | 55.181 | 2 | 55.03553 | yes | 2.538E-07 | 437.74069 | 297.95374 |
| QWTEILHFTEHSGI | 25 | Q9UMS4 | Pre-mRNA-processing factor 19 OS=Hon | PRPF19 | 55.181 | 4 | 55.03553 | yes | 1.911E-07 | 701.85303 | 103.44046 |

| | | | | | | | | | | | |
|---|---|---|---|---|---|---|---|---|---|---|---|
| QWTEILHFTEHSGI | 25 | Q9UMS4 | Pre-mRNA-processing factor 19 OS=Hon | PRPF19 | 55.181 | 4 | 55.03553 | yes | 9.722E-08 | 561.6839 | 103.44046 |
| QWTEILHFTEHSGI | 25 | Q9UMS4 | Pre-mRNA-processing factor 19 OS=Hon | PRPF19 | 55.181 | 5 | 55.03553 | yes | 1.911E-07 | 701.85303 | 103.44046 |
| QWTEILHFTEHSGI | 25 | Q9UMS4 | Pre-mRNA-processing factor 19 OS=Hon | PRPF19 | 55.181 | 5 | 55.03553 | yes | 9.722E-08 | 561.6839 | 103.44046 |
| VTSVVFHPSQDLV | 23 | Q9UMS4 | Pre-mRNA-processing factor 19 OS=Hon | PRPF19 | 55.181 | 3 | 55.03553 | yes | 9.722E-08 | 825.09558 | 178.43903 |
| FIASTGMDR | 9 | Q9UMS4 | Pre-mRNA-processing factor 19 OS=Hon | PRPF19 | 55.181 | 2 | 55.03553 | yes | 2.332E-07 | 499.24222 | 146.05408 |
| ALQDEWDAVMLH | 17 | Q9UMS4 | Pre-mRNA-processing factor 19 OS=Hon | PRPF19 | 55.181 | 3 | 55.03553 | yes | 1.64E-06 | 678.00342 | 11.309099 |
| AAVQFLNNAWGI( | 14 | Q9UMY1 | Nucleolar protein 7 OS=Homo sapiens O. | NOL7 | 29.427 | 2 | 53.904392 | yes | 5.56E-07 | 780.4201 | 840.27679 |
| LLPDTILEK | 9 | Q9UMY1 | Nucleolar protein 7 OS=Homo sapiens O. | NOL7 | 29.427 | 2 | 53.904392 | yes | 1.884E-07 | 521.31317 | 608.72992 |
| LTTASQTNIK | 10 | Q9UMY1 | Nucleolar protein 7 OS=Homo sapiens O. | NOL7 | 29.427 | 2 | 53.904392 | yes | 2.842E-07 | 538.80096 | 508.57892 |
| EELFIEQK | 8 | Q9UMY1 | Nucleolar protein 7 OS=Homo sapiens O. | NOL7 | 29.427 | 2 | 53.904392 | yes | 4.574E-05 | 518.27148 | 396.65439 |
| KLLPDTILEK | 10 | Q9UMY1 | Nucleolar protein 7 OS=Homo sapiens O. | NOL7 | 29.427 | 2 | 53.904392 | yes | 9.722E-08 | 585.36066 | 266.60727 |
| REELFIEQK | 9 | Q9UMY1 | Nucleolar protein 7 OS=Homo sapiens O. | NOL7 | 29.427 | 2 | 53.904392 | yes | 1.93E-07 | 596.32202 | 143.00035 |
| QQAAQAFIHNSLY | 19 | Q9UMY1 | Nucleolar protein 7 OS=Homo sapiens O. | NOL7 | 29.427 | 3 | 53.904392 | yes | 2.523E-07 | 691.68018 | 55.664227 |
| ETLGQIWALANR | 12 | Q9UMZ2 | Synergin gamma OS=Homo sapiens OX= | SYNRG | 140.655 | 2 | 49.146717 | yes | 5.496E-07 | 686.37262 | 56.870426 |
| ILETTMTPTGIDTA | 15 | Q9UMZ2 | Synergin gamma OS=Homo sapiens OX= | SYNRG | 140.655 | 2 | 49.146717 | yes | 1.399E-07 | 796.41602 | 55.247887 |
| SAGTDDGFTDFK | 12 | Q9UMZ2 | Synergin gamma OS=Homo sapiens OX= | SYNRG | 140.655 | 2 | 49.146717 | yes | 1.064E-07 | 630.77258 | 39.594501 |
| IYIPLPEEAAR | 11 | Q9UN37 | Vacuolar protein sorting-associated protei | VPS4A | 48.898 | 2 | 53.104008 | yes | 7.738E-07 | 636.35333 | 601.8952 |
| GILLFGPPGTGK | 12 | Q9UN37 | Vacuolar protein sorting-associated protei | VPS4A | 48.898 | 2 | 53.104008 | no | 9.722E-08 | 578.83984 | 467.50565 |
| LLEPVVCMSDMLI | 13 | Q9UN37 | Vacuolar protein sorting-associated protei | VPS4A | 48.898 | 2 | 53.104008 | yes | 1.167E-07 | 781.89063 | 234.89952 |
| WNDVAGLEGAK | 11 | Q9UN37 | Vacuolar protein sorting-associated protei | VPS4A | 48.898 | 2 | 53.104008 | yes | 1.301E-07 | 580.29077 | 190.01202 |
| CVQYLDR | 7 | Q9UN37 | Vacuolar protein sorting-associated protei | VPS4A | 48.898 | 2 | 53.104008 | yes | 1.248E-07 | 477.2291 | 124.96351 |
| DSLMQPVR | 8 | Q9UN37 | Vacuolar protein sorting-associated protei | VPS4A | 48.898 | 2 | 53.104008 | yes | 8.211E-07 | 473.24475 | 105.35098 |
| NLFELAR | 7 | Q9UN37 | Vacuolar protein sorting-associated protei | VPS4A | 48.898 | 2 | 53.104008 | yes | 3.261E-07 | 431.74271 | 88.641876 |
| TEGYSGADISIIVR | 14 | Q9UN37 | Vacuolar protein sorting-associated protei | VPS4A | 48.898 | 2 | 53.104008 | yes | 2.824E-07 | 740.88574 | 88.405716 |
| KTEGYSGADISIIV | 15 | Q9UN37 | Vacuolar protein sorting-associated protei | VPS4A | 48.898 | 3 | 53.104008 | yes | 1.467E-07 | 536.95789 | 60.020741 |
| PTVNADDLLK | 10 | Q9UN37 | Vacuolar protein sorting-associated protei | VPS4A | 48.898 | 2 | 53.104008 | yes | 5.06E-07 | 543.29547 | 49.736958 |
| LHLGSTPHNLTDA | 20 | Q9UN37 | Vacuolar protein sorting-associated protei | VPS4A | 48.898 | 3 | 53.104008 | yes | 9.722E-08 | 737.05396 | 47.446632 |
| LTADLSAETLQAR | 13 | Q9UN81 | LINE-1 retrotransposable element ORF1 | L1RE1 | 40.056 | 2 | 53.478413 | yes | 1.5E-07 | 694.87262 | 271.60999 |
| NLEECITR | 8 | Q9UN81 | LINE-1 retrotransposable element ORF1 | L1RE1 | 40.056 | 2 | 53.478413 | yes | 9.722E-08 | 517.75043 | 224.98204 |
| LSFISEGEIK | 10 | Q9UN81 | LINE-1 retrotransposable element ORF1 | L1RE1 | 40.056 | 2 | 53.478413 | yes | 9.722E-08 | 561.80573 | 214.50763 |
| VSAMEDEMNEMK | 12 | Q9UN81 | LINE-1 retrotransposable element ORF1 | L1RE1 | 40.056 | 2 | 53.478413 | yes | 9.722E-08 | 707.28839 | 164.20784 |
| NEQSLQEIWDYVK | 13 | Q9UN81 | LINE-1 retrotransposable element ORF1 | L1RE1 | 40.056 | 2 | 53.478413 | yes | 9.722E-08 | 826.40173 | 121.52999 |
| SRCDQLEER | 9 | Q9UN81 | LINE-1 retrotransposable element ORF1 | L1RE1 | 40.056 | 2 | 53.478413 | yes | 4E-07 | 596.7724 | 66.718719 |
| REWGPIFNILK | 11 | Q9UN81 | LINE-1 retrotransposable element ORF1 | L1RE1 | 40.056 | 2 | 53.478413 | yes | 9.938E-08 | 686.89044 | 53.797066 |
| LIGVPESDVENGTI | 14 | Q9UN81 | LINE-1 retrotransposable element ORF1 | L1RE1 | 40.056 | 2 | 53.478413 | yes | 3.081E-07 | 729.37775 | 30.984993 |
| VMEKPSPLLVGR | 12 | Q9UN86 | Ras GTPase-activating protein-binding pr | G3BP2 | 54.122 | 2 | 51.86807 | no | 2.312E-07 | 663.38409 | 17301.799 |
| VMEKPSPLLVGR | 12 | Q9UN86 | Ras GTPase-activating protein-binding pr | G3BP2 | 54.122 | 2 | 51.86807 | no | 2.097E-07 | 442.59183 | 17301.799 |
| VMEKPSPLLVGR | 12 | Q9UN86 | Ras GTPase-activating protein-binding pr | G3BP2 | 54.122 | 2 | 51.86807 | no | 1.46E-07 | 671.38159 | 17301.799 |
| VMEKPSPLLVGR | 12 | Q9UN86 | Ras GTPase-activating protein-binding pr | G3BP2 | 54.122 | 2 | 51.86807 | no | 1.23E-07 | 447.92349 | 17301.799 |

| | | | | | | | | | | |
|---|---|---|---|---|---|---|---|---|---|---|
| VMEKPSPLLVGR | 12 | Q9UN86 | Ras GTPase-activating protein-binding pr | G3BP2 | 54.122 | 2 | 51.86807 | no | 1.17E-07 | 684.3894 | 17301.799 |
| VMEKPSPLLVGR | 12 | Q9UN86 | Ras GTPase-activating protein-binding pr | G3BP2 | 54.122 | 3 | 51.86807 | no | 2.312E-07 | 663.38409 | 17301.799 |
| VMEKPSPLLVGR | 12 | Q9UN86 | Ras GTPase-activating protein-binding pr | G3BP2 | 54.122 | 3 | 51.86807 | no | 2.097E-07 | 442.59183 | 17301.799 |
| VMEKPSPLLVGR | 12 | Q9UN86 | Ras GTPase-activating protein-binding pr | G3BP2 | 54.122 | 3 | 51.86807 | no | 1.46E-07 | 671.38159 | 17301.799 |
| VMEKPSPLLVGR | 12 | Q9UN86 | Ras GTPase-activating protein-binding pr | G3BP2 | 54.122 | 3 | 51.86807 | no | 1.23E-07 | 447.92349 | 17301.799 |
| VMEKPSPLLVGR | 12 | Q9UN86 | Ras GTPase-activating protein-binding pr | G3BP2 | 54.122 | 3 | 51.86807 | no | 1.17E-07 | 684.3894 | 17301.799 |
| FYVHNDMFR | 9 | Q9UN86 | Ras GTPase-activating protein-binding pr | G3BP2 | 54.122 | 2 | 51.86807 | yes | 1.238E-07 | 614.78204 | 11922.228 |
| FYVHNDMFR | 9 | Q9UN86 | Ras GTPase-activating protein-binding pr | G3BP2 | 54.122 | 2 | 51.86807 | yes | 9.838E-08 | 410.19046 | 11922.228 |
| FYVHNDMFR | 9 | Q9UN86 | Ras GTPase-activating protein-binding pr | G3BP2 | 54.122 | 2 | 51.86807 | yes | 1.29E-07 | 622.77948 | 11922.228 |
| FYVHNDMFR | 9 | Q9UN86 | Ras GTPase-activating protein-binding pr | G3BP2 | 54.122 | 3 | 51.86807 | yes | 1.238E-07 | 614.78204 | 11922.228 |
| FYVHNDMFR | 9 | Q9UN86 | Ras GTPase-activating protein-binding pr | G3BP2 | 54.122 | 3 | 51.86807 | yes | 9.838E-08 | 410.19046 | 11922.228 |
| FYVHNDMFR | 9 | Q9UN86 | Ras GTPase-activating protein-binding pr | G3BP2 | 54.122 | 3 | 51.86807 | yes | 1.29E-07 | 622.77948 | 11922.228 |
| VLSLNFSECHTK | 12 | Q9UN86 | Ras GTPase-activating protein-binding pr | G3BP2 | 54.122 | 2 | 51.86807 | yes | 9.722E-08 | 717.8559 | 35757.695 |
| TEELKPQVEEK | 11 | Q9UN86 | Ras GTPase-activating protein-binding pr | G3BP2 | 54.122 | 2 | 51.86807 | yes | 9.722E-08 | 665.34845 | 9578.0703 |
| TEELKPQVEEK | 11 | Q9UN86 | Ras GTPase-activating protein-binding pr | G3BP2 | 54.122 | 2 | 51.86807 | yes | 1.009E-07 | 443.9014 | 9578.0703 |
| TEELKPQVEEK | 11 | Q9UN86 | Ras GTPase-activating protein-binding pr | G3BP2 | 54.122 | 3 | 51.86807 | yes | 9.722E-08 | 665.34845 | 9578.0703 |
| TEELKPQVEEK | 11 | Q9UN86 | Ras GTPase-activating protein-binding pr | G3BP2 | 54.122 | 3 | 51.86807 | yes | 1.009E-07 | 443.9014 | 9578.0703 |
| VEAKPEVQSQPPR | 13 | Q9UN86 | Ras GTPase-activating protein-binding pr | G3BP2 | 54.122 | 3 | 51.86807 | yes | 9.722E-08 | 488.9317 | 9274.2324 |
| VEAKPEVQSQPPR | 13 | Q9UN86 | Ras GTPase-activating protein-binding pr | G3BP2 | 54.122 | 3 | 51.86807 | yes | 1.244E-07 | 732.89392 | 9274.2324 |
| VEAKPEVQSQPPR | 13 | Q9UN86 | Ras GTPase-activating protein-binding pr | G3BP2 | 54.122 | 2 | 51.86807 | yes | 9.722E-08 | 488.9317 | 9274.2324 |
| VEAKPEVQSQPPR | 13 | Q9UN86 | Ras GTPase-activating protein-binding pr | G3BP2 | 54.122 | 2 | 51.86807 | yes | 1.244E-07 | 732.89392 | 9274.2324 |
| GIVGGGMMR | 9 | Q9UN86 | Ras GTPase-activating protein-binding pr | G3BP2 | 54.122 | 2 | 51.86807 | yes | 1.821E-07 | 439.22278 | 5922.6514 |
| GIVGGGMMR | 9 | Q9UN86 | Ras GTPase-activating protein-binding pr | G3BP2 | 54.122 | 2 | 51.86807 | yes | 3.158E-06 | 447.22021 | 5922.6514 |
| GIVGGGMMR | 9 | Q9UN86 | Ras GTPase-activating protein-binding pr | G3BP2 | 54.122 | 2 | 51.86807 | yes | 1.146E-07 | 447.22021 | 5922.6514 |
| PSPLLVGR | 8 | Q9UN86 | Ras GTPase-activating protein-binding pr | G3BP2 | 54.122 | 2 | 51.86807 | no | 1.686E-07 | 419.76089 | 13309.732 |
| NLEELEEK | 8 | Q9UN86 | Ras GTPase-activating protein-binding pr | G3BP2 | 54.122 | 2 | 51.86807 | yes | 2.38E-07 | 502.25076 | 12991.465 |
| APEYLHR | 7 | Q9UN86 | Ras GTPase-activating protein-binding pr | G3BP2 | 54.122 | 2 | 51.86807 | yes | 2.25E-07 | 443.23251 | 9643.8359 |
| QYYTLLNK | 8 | Q9UN86 | Ras GTPase-activating protein-binding pr | G3BP2 | 54.122 | 2 | 51.86807 | yes | 1.23E-07 | 521.78204 | 8910.7803 |
| NSSYVHGGVDAS( | 28 | Q9UN86 | Ras GTPase-activating protein-binding pr | G3BP2 | 54.122 | 4 | 51.86807 | yes | 3.234E-07 | 749.35767 | 2976.9563 |
| NSSYVHGGVDAS( | 28 | Q9UN86 | Ras GTPase-activating protein-binding pr | G3BP2 | 54.122 | 4 | 51.86807 | yes | 1.01E-07 | 599.68762 | 2976.9563 |
| NSSYVHGGVDAS( | 28 | Q9UN86 | Ras GTPase-activating protein-binding pr | G3BP2 | 54.122 | 5 | 51.86807 | yes | 3.234E-07 | 749.35767 | 2976.9563 |
| NSSYVHGGVDAS( | 28 | Q9UN86 | Ras GTPase-activating protein-binding pr | G3BP2 | 54.122 | 5 | 51.86807 | yes | 1.01E-07 | 599.68762 | 2976.9563 |
| YPDSHQLFVGNLP | 22 | Q9UN86 | Ras GTPase-activating protein-binding pr | G3BP2 | 54.122 | 4 | 51.86807 | yes | 2.577E-07 | 645.81458 | 4732.521 |
| ERPGFPPR | 8 | Q9UN86 | Ras GTPase-activating protein-binding pr | G3BP2 | 54.122 | 2 | 51.86807 | yes | 2.873E-06 | 478.25906 | 2659.6746 |
| AFSWASVTSK | 10 | Q9UN86 | Ras GTPase-activating protein-binding pr | G3BP2 | 54.122 | 2 | 51.86807 | yes | 1.368E-07 | 542.2771 | 1841.4967 |
| EFFMSFGNVVELR | 13 | Q9UN86 | Ras GTPase-activating protein-binding pr | G3BP2 | 54.122 | 2 | 51.86807 | yes | 9.722E-08 | 795.8847 | 1181.6204 |
| HVDAHATLSDGV' | 28 | Q9UN86 | Ras GTPase-activating protein-binding pr | G3BP2 | 54.122 | 4 | 51.86807 | yes | 1.852E-07 | 729.87128 | 671.65936 |
| PEVQSQPPR | 9 | Q9UN86 | Ras GTPase-activating protein-binding pr | G3BP2 | 54.122 | 2 | 51.86807 | yes | 2.59E-07 | 519.27234 | 503.63611 |
| GDMEQNDSDNR | 11 | Q9UN86 | Ras GTPase-activating protein-binding pr | G3BP2 | 54.122 | 2 | 51.86807 | yes | 9.722E-08 | 640.74402 | 309.33762 |

| | | | | | | | | | | | |
|---|---|---|---|---|---|---|---|---|---|---|---|
| PQEAVYGQNDIHH | 14 | Q9UN86 | Ras GTPase-activating protein-binding pr | G3BP2 | 54.122 | 2 | 51.86807 | yes | 3.513E-07 | 818.39734 | 283.78305 |
| MVMEKPSPLLVGI | 13 | Q9UN86 | Ras GTPase-activating protein-binding pr | G3BP2 | 54.122 | 2 | 51.86807 | no | 1.465E-07 | 749.90967 | 259.80377 |
| KFMQTFVLAPEGS | 17 | Q9UN86 | Ras GTPase-activating protein-binding pr | G3BP2 | 54.122 | 3 | 51.86807 | yes | 9.722E-08 | 631.66949 | 109.98982 |
| KFMQTFVLAPEGS | 17 | Q9UN86 | Ras GTPase-activating protein-binding pr | G3BP2 | 54.122 | 3 | 51.86807 | yes | 1.434E-07 | 637.0011 | 109.98982 |
| NSSYVHGGVDAS( | 14 | Q9UN86 | Ras GTPase-activating protein-binding pr | G3BP2 | 54.122 | 2 | 51.86807 | yes | 1.084E-07 | 689.3233 | 68.530678 |
| NSSYVHGGVDAS( | 14 | Q9UN86 | Ras GTPase-activating protein-binding pr | G3BP2 | 54.122 | 2 | 51.86807 | yes | 1.149E-07 | 459.88464 | 68.530678 |
| NSSYVHGGVDAS( | 14 | Q9UN86 | Ras GTPase-activating protein-binding pr | G3BP2 | 54.122 | 3 | 51.86807 | yes | 1.084E-07 | 689.3233 | 68.530678 |
| NSSYVHGGVDAS( | 14 | Q9UN86 | Ras GTPase-activating protein-binding pr | G3BP2 | 54.122 | 3 | 51.86807 | yes | 1.149E-07 | 459.88464 | 68.530678 |
| LPNFGFVVFDDSE | 17 | Q9UN86 | Ras GTPase-activating protein-binding pr | G3BP2 | 54.122 | 3 | 51.86807 | yes | 3.431E-05 | 655.995 | 15.632596 |
| NPLVAVYYTNR | 11 | Q9UNE7 | E3 ubiquitin-protein ligase CHIP OS=Ho | STUB1 | 34.857 | 2 | 52.022694 | yes | 9.722E-08 | 655.34857 | 160.78786 |
| LNFGDDIPSALR | 12 | Q9UNE7 | E3 ubiquitin-protein ligase CHIP OS=Ho | STUB1 | 34.857 | 2 | 52.022694 | yes | 1.869E-07 | 659.34351 | 139.84746 |
| VGHFDPVTR | 9 | Q9UNE7 | E3 ubiquitin-protein ligase CHIP OS=Ho | STUB1 | 34.857 | 2 | 52.022694 | yes | 1.117E-07 | 514.26959 | 133.29584 |
| IHQESELHSYLSR | 13 | Q9UNE7 | E3 ubiquitin-protein ligase CHIP OS=Ho | STUB1 | 34.857 | 3 | 52.022694 | yes | 2.559E-07 | 533.60223 | 43.659241 |
| IHQESELHSYLSR | 13 | Q9UNE7 | E3 ubiquitin-protein ligase CHIP OS=Ho | STUB1 | 34.857 | 3 | 52.022694 | yes | 7.084E-05 | 799.89972 | 43.659241 |
| IHQESELHSYLSR | 13 | Q9UNE7 | E3 ubiquitin-protein ligase CHIP OS=Ho | STUB1 | 34.857 | 2 | 52.022694 | yes | 2.559E-07 | 533.60223 | 43.659241 |
| IHQESELHSYLSR | 13 | Q9UNE7 | E3 ubiquitin-protein ligase CHIP OS=Ho | STUB1 | 34.857 | 2 | 52.022694 | yes | 7.084E-05 | 799.89972 | 43.659241 |
| DIEEHLQR | 8 | Q9UNE7 | E3 ubiquitin-protein ligase CHIP OS=Ho | STUB1 | 34.857 | 2 | 52.022694 | yes | 4.752E-06 | 520.26196 | 61.869556 |
| KYPEAAACYGR | 11 | Q9UNE7 | E3 ubiquitin-protein ligase CHIP OS=Ho | STUB1 | 34.857 | 2 | 52.022694 | yes | 9.722E-08 | 643.30334 | 37.957661 |
| LCSDLMNCLHER | 12 | Q9UNF0 | Protein kinase C and casein kinase substr | PACSIN2 | 55.739 | 2 | 53.206623 | yes | 9.722E-08 | 774.3418 | 24.24172 |
| EYTDVYPEIIER | 12 | Q9UNF1 | Melanoma-associated antigen D2 OS=Ho | MAGED2 | 64.955 | 2 | 54.336502 | no | 4.285E-07 | 763.87231 | 246.26935 |
| ALEVSEDVK | 9 | Q9UNF1 | Melanoma-associated antigen D2 OS=Ho | MAGED2 | 64.955 | 2 | 54.336502 | yes | 8.783E-07 | 495.26111 | 122.69189 |
| SDTSESGAGLTR | 12 | Q9UNF1 | Melanoma-associated antigen D2 OS=Ho | MAGED2 | 64.955 | 2 | 54.336502 | yes | 1.186E-07 | 611.78094 | 97.424385 |
| GPIAFWAR | 8 | Q9UNF1 | Melanoma-associated antigen D2 OS=Ho | MAGED2 | 64.955 | 2 | 54.336502 | yes | 9.722E-08 | 459.25323 | 94.433502 |
| SSEAVIWEVLR | 11 | Q9UNF1 | Melanoma-associated antigen D2 OS=Ho | MAGED2 | 64.955 | 2 | 54.336502 | yes | 9.823E-08 | 644.84845 | 73.32766 |
| GSWCEQQLCACDI | 13 | Q9UNK4 | Group IID secretory phospholipase A2 OS | PLA2G2D | 16.547 | 2 | 37.98052 | yes | 5.548E-07 | 821.32635 | 27.191916 |
| RLDTDLAR | 8 | Q9UNL4 | Inhibitor of growth protein 4 OS=Homo s | ING4 | 28.531 | 2 | 48.761234 | yes | 1.107E-07 | 480.26706 | 75.220818 |
| LATEYMSSAR | 10 | Q9UNL4 | Inhibitor of growth protein 4 OS=Homo s | ING4 | 28.531 | 2 | 48.761234 | yes | 2.385E-07 | 564.7713 | 39.445633 |
| HFGSVVAQTIR | 11 | Q9UNN5 | FAS-associated factor 1 OS=Homo sapier | FAF1 | 73.955 | 2 | 45.991238 | yes | 3.818E-07 | 607.83563 | 27.228596 |
| QILENELQIPVSK | 13 | Q9UNN5 | FAS-associated factor 1 OS=Homo sapier | FAF1 | 73.955 | 2 | 45.991238 | yes | 1.684E-07 | 755.92743 | 16.47298 |
| VLYVGGLAEEVDI | 14 | Q9UNP9 | Peptidyl-prolyl cis-trans isomerase E OS= | PPIE | 33.431 | 2 | 52.511295 | yes | 9.728E-08 | 753.88794 | 178.52795 |
| KFDDENFILK | 10 | Q9UNP9 | Peptidyl-prolyl cis-trans isomerase E OS= | PPIE | 33.431 | 2 | 52.511295 | yes | 1.012E-07 | 634.82971 | 104.44595 |
| PVWSDDDWLK | 10 | Q9UNP9 | Peptidyl-prolyl cis-trans isomerase E OS= | PPIE | 33.431 | 2 | 52.511295 | yes | 9.722E-08 | 630.7984 | 89.780594 |
| SDVVPMTAENFR | 12 | Q9UNP9 | Peptidyl-prolyl cis-trans isomerase E OS= | PPIE | 33.431 | 2 | 52.511295 | yes | 7.705E-07 | 683.32703 | 83.933853 |
| HVVFGEVTEGLD\ | 15 | Q9UNP9 | Peptidyl-prolyl cis-trans isomerase E OS= | PPIE | 33.431 | 3 | 52.511295 | yes | 9.722E-08 | 557.30176 | 39.858574 |
| HVVFGEVTEGLD\ | 15 | Q9UNP9 | Peptidyl-prolyl cis-trans isomerase E OS= | PPIE | 33.431 | 3 | 52.511295 | yes | 2.466E-07 | 835.44904 | 39.858574 |
| HVVFGEVTEGLD\ | 15 | Q9UNP9 | Peptidyl-prolyl cis-trans isomerase E OS= | PPIE | 33.431 | 2 | 52.511295 | yes | 9.722E-08 | 557.30176 | 39.858574 |
| HVVFGEVTEGLD\ | 15 | Q9UNP9 | Peptidyl-prolyl cis-trans isomerase E OS= | PPIE | 33.431 | 2 | 52.511295 | yes | 2.466E-07 | 835.44904 | 39.858574 |
| LSINTQLLAR | 10 | Q9UNQ2 | Probable dimethyladenosine transferase C | DIMT1 | 35.237 | 2 | 52.37304 | yes | 1.346E-07 | 564.84039 | 3463.8433 |
| IQQILTSTGFSDK | 13 | Q9UNQ2 | Probable dimethyladenosine transferase C | DIMT1 | 35.237 | 2 | 52.37304 | yes | 1.152E-07 | 719.38281 | 3123.0806 |

| | | | | | | | | | | | |
|---|---|---|---|---|---|---|---|---|---|---|---|
| SSAVQQLLEK | 10 | Q9UNQ2 | Probable dimethyladenosine transferase C | DIMT1 | 35.237 | 2 | 52.37304 | yes | 1.197E-07 | 551.80878 | 2769.8896 |
| AALRPTDVVLEVC | 22 | Q9UNQ2 | Probable dimethyladenosine transferase C | DIMT1 | 35.237 | 3 | 52.37304 | yes | 2.603E-07 | 742.40344 | 1283.1418 |
| AALRPTDVVLEVC | 22 | Q9UNQ2 | Probable dimethyladenosine transferase C | DIMT1 | 35.237 | 3 | 52.37304 | yes | 4.328E-07 | 747.73511 | 1283.1418 |
| SMDIDDFIR | 9 | Q9UNQ2 | Probable dimethyladenosine transferase C | DIMT1 | 35.237 | 2 | 52.37304 | yes | 1.021E-07 | 556.25806 | 2533.053 |
| LQVLVGDVLK | 10 | Q9UNQ2 | Probable dimethyladenosine transferase C | DIMT1 | 35.237 | 2 | 52.37304 | yes | 1.398E-07 | 542.34222 | 2369.824 |
| LVAELHK | 7 | Q9UNQ2 | Probable dimethyladenosine transferase C | DIMT1 | 35.237 | 2 | 52.37304 | yes | 9.81E-08 | 405.24762 | 1459.1847 |
| NPLIINSIIDK | 11 | Q9UNQ2 | Probable dimethyladenosine transferase C | DIMT1 | 35.237 | 2 | 52.37304 | yes | 9.722E-08 | 620.36902 | 1373.1863 |
| CAILMFQR | 8 | Q9UNQ2 | Probable dimethyladenosine transferase C | DIMT1 | 35.237 | 2 | 52.37304 | yes | 1.852E-07 | 519.76477 | 1295.4821 |
| LLHGFNAEGIHFS | 13 | Q9UNQ2 | Probable dimethyladenosine transferase C | DIMT1 | 35.237 | 2 | 52.37304 | yes | 2.219E-07 | 721.36475 | 787.70062 |
| VVACELDPR | 9 | Q9UNQ2 | Probable dimethyladenosine transferase C | DIMT1 | 35.237 | 2 | 52.37304 | yes | 1.158E-06 | 529.76862 | 649.67975 |
| KVVACELDPR | 10 | Q9UNQ2 | Probable dimethyladenosine transferase C | DIMT1 | 35.237 | 2 | 52.37304 | yes | 3.034E-07 | 593.8161 | 478.25192 |
| NNFRPPPK | 8 | Q9UNQ2 | Probable dimethyladenosine transferase C | DIMT1 | 35.237 | 2 | 52.37304 | yes | 4.416E-07 | 485.26688 | 380.64996 |
| IHCSVHNIIIPEDFS | 19 | Q9UNQ2 | Probable dimethyladenosine transferase C | DIMT1 | 35.237 | 4 | 52.37304 | yes | 1.073E-07 | 552.78339 | 62.245335 |
| IQQILTSTGFSDKR | 14 | Q9UNQ2 | Probable dimethyladenosine transferase C | DIMT1 | 35.237 | 2 | 52.37304 | yes | 9.722E-08 | 797.43341 | 55.875584 |
| IMEAIELYR | 9 | Q9UNX4 | WD repeat-containing protein 3 OS=Hom | WDR3 | 106.099 | 2 | 54.262436 | yes | 9.722E-08 | 569.30225 | 374.86847 |
| LITGASDSELR | 11 | Q9UNX4 | WD repeat-containing protein 3 OS=Hom | WDR3 | 106.099 | 2 | 54.262436 | yes | 1.348E-07 | 581.30914 | 360.82855 |
| GNIVFVTLR | 9 | Q9UNX4 | WD repeat-containing protein 3 OS=Hom | WDR3 | 106.099 | 2 | 54.262436 | yes | 1.423E-07 | 509.80585 | 275.44183 |
| SFDLIHSPHGELK | 13 | Q9UNX4 | WD repeat-containing protein 3 OS=Hom | WDR3 | 106.099 | 2 | 54.262436 | yes | 1.122E-07 | 740.38318 | 112.06606 |
| SFDLIHSPHGELK | 13 | Q9UNX4 | WD repeat-containing protein 3 OS=Hom | WDR3 | 106.099 | 2 | 54.262436 | yes | 3.455E-07 | 493.92453 | 112.06606 |
| SFDLIHSPHGELK | 13 | Q9UNX4 | WD repeat-containing protein 3 OS=Hom | WDR3 | 106.099 | 3 | 54.262436 | yes | 1.122E-07 | 740.38318 | 112.06606 |
| SFDLIHSPHGELK | 13 | Q9UNX4 | WD repeat-containing protein 3 OS=Hom | WDR3 | 106.099 | 3 | 54.262436 | yes | 3.455E-07 | 493.92453 | 112.06606 |
| SHLFFTAGK | 9 | Q9UNX4 | WD repeat-containing protein 3 OS=Hom | WDR3 | 106.099 | 2 | 54.262436 | yes | 4.092E-07 | 504.26907 | 162.80988 |
| IWGLDFGDCHK | 11 | Q9UNX4 | WD repeat-containing protein 3 OS=Hom | WDR3 | 106.099 | 2 | 54.262436 | yes | 9.722E-08 | 674.31116 | 139.55679 |
| DAITQALFLR | 10 | Q9UNX4 | WD repeat-containing protein 3 OS=Hom | WDR3 | 106.099 | 2 | 54.262436 | yes | 9.794E-08 | 574.32715 | 139.31157 |
| FWDFELVK | 8 | Q9UNX4 | WD repeat-containing protein 3 OS=Hom | WDR3 | 106.099 | 2 | 54.262436 | yes | 3.634E-07 | 542.27911 | 109.44279 |
| SLFAHDDSVMYLC | 17 | Q9UNX4 | WD repeat-containing protein 3 OS=Hom | WDR3 | 106.099 | 3 | 54.262436 | yes | 2.793E-07 | 666.33276 | 64.565346 |
| IFYVDTLK | 8 | Q9UNX4 | WD repeat-containing protein 3 OS=Hom | WDR3 | 106.099 | 2 | 54.262436 | yes | 3.256E-07 | 499.78149 | 54.53735 |
| LLAVSLLDCTVK | 12 | Q9UNX4 | WD repeat-containing protein 3 OS=Hom | WDR3 | 106.099 | 2 | 54.262436 | yes | 1.002E-07 | 666.38379 | 47.822849 |
| ILILQGLK | 8 | Q9UNX4 | WD repeat-containing protein 3 OS=Hom | WDR3 | 106.099 | 2 | 54.262436 | yes | 1.607E-07 | 449.31021 | 40.393379 |
| YVASAVFGVIGSQ | 14 | Q9UNX4 | WD repeat-containing protein 3 OS=Hom | WDR3 | 106.099 | 2 | 54.262436 | yes | 5.991E-05 | 713.39044 | 14.100563 |
| TEVWGLVLLSEEK | 13 | Q9UNX4 | WD repeat-containing protein 3 OS=Hom | WDR3 | 106.099 | 2 | 54.262436 | yes | 2.305E-05 | 751.90869 | 14.086901 |
| VALEFGSEEPR | 11 | Q9UNY4 | Transcription termination factor 2 OS=Hc | TTF2 | 129.589 | 2 | 39.870213 | yes | 1.593E-05 | 617.30914 | 71.743134 |
| VYLYHGPNR | 9 | Q9UNY4 | Transcription termination factor 2 OS=Hc | TTF2 | 129.589 | 2 | 39.870213 | yes | 4.336E-07 | 559.79071 | 45.710358 |
| LTVEEAVR | 8 | Q9UPN3 | Microtubule-actin cross-linking factor 1, i | MACF1 | 838.308 | 2 | 56.122227 | no | 1.921E-07 | 458.75854 | 938.15198 |
| HWITIIR | 7 | Q9UPN3 | Microtubule-actin cross-linking factor 1, i | MACF1 | 838.308 | 2 | 56.122227 | no | 1.555E-07 | 469.78217 | 537.71332 |
| QQVEAAETIK | 10 | Q9UPN3 | Microtubule-actin cross-linking factor 1, i | MACF1 | 838.308 | 2 | 56.122227 | no | 6.743E-07 | 558.7984 | 325.97513 |
| NDDITDGNPK | 10 | Q9UPN3 | Microtubule-actin cross-linking factor 1, i | MACF1 | 838.308 | 2 | 56.122227 | no | 1.204E-07 | 544.74634 | 130.48622 |
| MPPLIPAEVDK | 11 | Q9UPN3 | Microtubule-actin cross-linking factor 1, i | MACF1 | 838.308 | 2 | 56.122227 | no | 7.652E-07 | 605.33105 | 115.78957 |
| IEDEVTR | 7 | Q9UPN3 | Microtubule-actin cross-linking factor 1, i | MACF1 | 838.308 | 2 | 56.122227 | no | 3.135E-07 | 431.21927 | 108.55624 |

| | | | | | | | | | | |
|---|---|---|---|---|---|---|---|---|---|---|
| LVSDTIGQR | 9 | Q9UPN3 | Microtubule-actin cross-linking factor 1, i | MACF1 | 838.308 | 2 | 56.122227 | no | 2.662E-07 | 494.77472 | 107.65201 |
| VDEIDAAIQR | 10 | Q9UPN3 | Microtubule-actin cross-linking factor 1, i | MACF1 | 838.308 | 2 | 56.122227 | no | 2.439E-07 | 565.29602 | 104.78197 |
| ENLEQAFEVAER | 12 | Q9UPN3 | Microtubule-actin cross-linking factor 1, i | MACF1 | 838.308 | 2 | 56.122227 | yes | 1.198E-07 | 717.84662 | 96.080933 |
| LCTMPPVGTDLNT | 15 | Q9UPN3 | Microtubule-actin cross-linking factor 1, i | MACF1 | 838.308 | 2 | 56.122227 | no | 5.693E-07 | 823.41803 | 95.245949 |
| VEVYQQQIEMEK | 12 | Q9UPN3 | Microtubule-actin cross-linking factor 1, i | MACF1 | 838.308 | 2 | 56.122227 | no | 1.219E-07 | 762.37415 | 83.391838 |
| FEQLCLQQQEK | 11 | Q9UPN3 | Microtubule-actin cross-linking factor 1, i | MACF1 | 838.308 | 2 | 56.122227 | yes | 2.397E-07 | 725.85339 | 67.575539 |
| LEQDQTTAQLQVC | 14 | Q9UPN3 | Microtubule-actin cross-linking factor 1, i | MACF1 | 838.308 | 2 | 56.122227 | no | 9.722E-08 | 815.42578 | 63.854923 |
| ELNPEEGEMVEEK | 13 | Q9UPN3 | Microtubule-actin cross-linking factor 1, i | MACF1 | 838.308 | 2 | 56.122227 | no | 9.722E-08 | 766.8429 | 61.774174 |
| LEDLLQDSMDEK | 12 | Q9UPN3 | Microtubule-actin cross-linking factor 1, i | MACF1 | 838.308 | 2 | 56.122227 | yes | 2.916E-07 | 718.33472 | 61.210457 |
| LNHQGELMLK | 10 | Q9UPN3 | Microtubule-actin cross-linking factor 1, i | MACF1 | 838.308 | 2 | 56.122227 | no | 5.444E-07 | 591.8186 | 61.11462 |
| ITGQLESLESR | 11 | Q9UPN3 | Microtubule-actin cross-linking factor 1, i | MACF1 | 838.308 | 2 | 56.122227 | no | 7.12E-07 | 616.8277 | 60.790688 |
| LECTNLYR | 8 | Q9UPN3 | Microtubule-actin cross-linking factor 1, i | MACF1 | 838.308 | 2 | 56.122227 | yes | 8.538E-07 | 534.76074 | 59.242214 |
| FLDVLELAEK | 10 | Q9UPN3 | Microtubule-actin cross-linking factor 1, i | MACF1 | 838.308 | 2 | 56.122227 | no | 2.274E-06 | 588.82916 | 57.024582 |
| LDIEASEAECR | 11 | Q9UPN3 | Microtubule-actin cross-linking factor 1, i | MACF1 | 838.308 | 2 | 56.122227 | yes | 9.722E-08 | 646.79297 | 57.01342 |
| VFANEVNAHR | 10 | Q9UPN3 | Microtubule-actin cross-linking factor 1, i | MACF1 | 838.308 | 2 | 56.122227 | no | 9.722E-08 | 578.79651 | 55.003048 |
| LSVQSAISTQPEAV | 15 | Q9UPN3 | Microtubule-actin cross-linking factor 1, i | MACF1 | 838.308 | 2 | 56.122227 | yes | 9.22E-07 | 779.4278 | 52.754509 |
| FQSTYEELTGWLR | 13 | Q9UPN3 | Microtubule-actin cross-linking factor 1, i | MACF1 | 838.308 | 2 | 56.122227 | no | 2.711E-07 | 815.39899 | 50.896667 |
| DLDLVQTWNLEK | 12 | Q9UPN3 | Microtubule-actin cross-linking factor 1, i | MACF1 | 838.308 | 2 | 56.122227 | yes | 9.722E-08 | 737.38281 | 50.013714 |
| LVSDANEQYK | 10 | Q9UPN3 | Microtubule-actin cross-linking factor 1, i | MACF1 | 838.308 | 2 | 56.122227 | no | 8.165E-06 | 583.78802 | 46.628494 |
| TEQSVALLEQK | 11 | Q9UPN3 | Microtubule-actin cross-linking factor 1, i | MACF1 | 838.308 | 2 | 56.122227 | no | 6.308E-07 | 623.33789 | 42.725086 |
| ALLELVPWR | 9 | Q9UPN3 | Microtubule-actin cross-linking factor 1, i | MACF1 | 838.308 | 2 | 56.122227 | no | 6.719E-07 | 548.82928 | 41.768768 |
| DENYYQLEELAFR | 13 | Q9UPN3 | Microtubule-actin cross-linking factor 1, i | MACF1 | 838.308 | 2 | 56.122227 | yes | 3.42E-07 | 845.39136 | 40.112354 |
| IAQLQEALLHCGK | 13 | Q9UPN3 | Microtubule-actin cross-linking factor 1, i | MACF1 | 838.308 | 2 | 56.122227 | no | 9.722E-08 | 740.9007 | 39.959873 |
| GGSSPETLPSLLK | 13 | Q9UPN3 | Microtubule-actin cross-linking factor 1, i | MACF1 | 838.308 | 2 | 56.122227 | yes | 5.139E-06 | 643.35352 | 39.673637 |
| LEMTAVADIFDR | 12 | Q9UPN3 | Microtubule-actin cross-linking factor 1, i | MACF1 | 838.308 | 2 | 56.122227 | no | 1.25E-07 | 690.84503 | 37.765823 |
| LELLLQIANK | 10 | Q9UPN3 | Microtubule-actin cross-linking factor 1, i | MACF1 | 838.308 | 2 | 56.122227 | yes | 3.219E-06 | 577.86078 | 35.031582 |
| HQLEIFDALGSQAC | 17 | Q9UPN3 | Microtubule-actin cross-linking factor 1, i | MACF1 | 838.308 | 3 | 56.122227 | yes | 2.847E-06 | 639.97565 | 34.706142 |
| LLLWTQK | 7 | Q9UPN3 | Microtubule-actin cross-linking factor 1, i | MACF1 | 838.308 | 2 | 56.122227 | yes | 5.712E-07 | 451.27893 | 34.608498 |
| SIDEMNNAWENLN | 14 | Q9UPN3 | Microtubule-actin cross-linking factor 1, i | MACF1 | 838.308 | 2 | 56.122227 | no | 5.894E-07 | 839.38049 | 33.918819 |
| LLDAEDVDVPSPD | 15 | Q9UPN3 | Microtubule-actin cross-linking factor 1, i | MACF1 | 838.308 | 2 | 56.122227 | yes | 1.036E-06 | 821.39636 | 33.527771 |
| ATVDMLQAEGGR | 12 | Q9UPN3 | Microtubule-actin cross-linking factor 1, i | MACF1 | 838.308 | 2 | 56.122227 | no | 4.817E-06 | 624.30609 | 33.278343 |
| MFNALIHR | 8 | Q9UPN3 | Microtubule-actin cross-linking factor 1, i | MACF1 | 838.308 | 2 | 56.122227 | yes | 5.335E-07 | 501.27112 | 27.528645 |
| VNSGCVMMENK | 11 | Q9UPN3 | Microtubule-actin cross-linking factor 1, i | MACF1 | 838.308 | 2 | 56.122227 | yes | 5.475E-07 | 634.77521 | 26.106449 |
| EQYSTSLAQSEAEI | 15 | Q9UPN3 | Microtubule-actin cross-linking factor 1, i | MACF1 | 838.308 | 2 | 56.122227 | yes | 9.988E-08 | 842.40723 | 23.781672 |
| LQSQLQENEEFQK | 13 | Q9UPN3 | Microtubule-actin cross-linking factor 1, i | MACF1 | 838.308 | 2 | 56.122227 | yes | 2.966E-07 | 810.89685 | 22.131714 |
| GQLGEAEGENLR | 12 | Q9UPN4 | Centrosomal protein of 131 kDa OS=Hon | CEP131 | 122.149 | 2 | 56.9228 | yes | 1.075E-07 | 636.81256 | 195.8317 |
| AAIQELQQK | 9 | Q9UPN4 | Centrosomal protein of 131 kDa OS=Hon | CEP131 | 122.149 | 2 | 56.9228 | yes | 1.188E-06 | 514.79041 | 155.40208 |
| RPGSAATTKPIVR | 13 | Q9UPN4 | Centrosomal protein of 131 kDa OS=Hon | CEP131 | 122.149 | 3 | 56.9228 | yes | 9.722E-08 | 451.93689 | 145.85223 |
| LAASEEETR | 9 | Q9UPN4 | Centrosomal protein of 131 kDa OS=Hon | CEP131 | 122.149 | 2 | 56.9228 | yes | 1.67E-06 | 503.246 | 134.94727 |

| | | | | | | | | | | | |
|---|---|---|---|---|---|---|---|---|---|---|---|
| EVTEEEAER | 9 | Q9UPN4 | Centrosomal protein of 131 kDa OS=Hon | CEP131 | 122.149 | 2 | 56.9228 | yes | 9.892E-08 | 546.24622 | 129.32877 |
| SSSALDSPAGPR | 12 | Q9UPN4 | Centrosomal protein of 131 kDa OS=Hon | CEP131 | 122.149 | 2 | 56.9228 | yes | 1.109E-07 | 572.78326 | 107.79614 |
| EHYEATIQR | 9 | Q9UPN4 | Centrosomal protein of 131 kDa OS=Hon | CEP131 | 122.149 | 2 | 56.9228 | yes | 3.221E-07 | 573.78052 | 106.6794 |
| SGEGTLLDLHQQK | 13 | Q9UPN4 | Centrosomal protein of 131 kDa OS=Hon | CEP131 | 122.149 | 2 | 56.9228 | yes | 1.009E-07 | 713.37024 | 83.338181 |
| GFTLPSNAR | 9 | Q9UPN4 | Centrosomal protein of 131 kDa OS=Hon | CEP131 | 122.149 | 2 | 56.9228 | yes | 2.832E-06 | 481.75635 | 78.475288 |
| HLAFIDQLIEDK | 12 | Q9UPN4 | Centrosomal protein of 131 kDa OS=Hon | CEP131 | 122.149 | 2 | 56.9228 | yes | 1.551E-07 | 721.38794 | 74.260582 |
| QPAQELSPTPGGT. | 19 | Q9UPN4 | Centrosomal protein of 131 kDa OS=Hon | CEP131 | 122.149 | 3 | 56.9228 | yes | 9.728E-08 | 644.33759 | 62.738609 |
| ECTVALAPNFTAN | 15 | Q9UPN4 | Centrosomal protein of 131 kDa OS=Hon | CEP131 | 122.149 | 2 | 56.9228 | yes | 1.021E-07 | 839.4043 | 59.968742 |
| SNLAQVIR | 8 | Q9UPN4 | Centrosomal protein of 131 kDa OS=Hon | CEP131 | 122.149 | 2 | 56.9228 | yes | 2.876E-07 | 450.76672 | 53.21994 |
| QQLELEEVHR | 10 | Q9UPN4 | Centrosomal protein of 131 kDa OS=Hon | CEP131 | 122.149 | 2 | 56.9228 | yes | 9.722E-08 | 640.83331 | 51.657967 |
| SSNQTAPSLNNIIK | 14 | Q9UPN4 | Centrosomal protein of 131 kDa OS=Hon | CEP131 | 122.149 | 2 | 56.9228 | yes | 3.607E-07 | 743.89661 | 50.035114 |
| HQMELNTLK | 9 | Q9UPN4 | Centrosomal protein of 131 kDa OS=Hon | CEP131 | 122.149 | 2 | 56.9228 | yes | 1.497E-06 | 557.28967 | 48.912628 |
| KEEAWLLNR | 9 | Q9UPN4 | Centrosomal protein of 131 kDa OS=Hon | CEP131 | 122.149 | 2 | 56.9228 | yes | 5.525E-07 | 579.81696 | 42.070961 |
| SNSTTQVSQPR | 11 | Q9UPN4 | Centrosomal protein of 131 kDa OS=Hon | CEP131 | 122.149 | 2 | 56.9228 | yes | 9.722E-08 | 602.79944 | 35.905731 |
| SVSVVTGSEQK | 11 | Q9UPN4 | Centrosomal protein of 131 kDa OS=Hon | CEP131 | 122.149 | 2 | 56.9228 | yes | 2.767E-07 | 560.79584 | 26.80113 |
| VFDYFEGATSQR | 12 | Q9UPN6 | SR-related and CTD-associated factor 8 C | SCAF8 | 140.52 | 2 | 57.247448 | yes | 9.722E-08 | 710.33063 | 464.72272 |
| GCAYVCMVHR | 10 | Q9UPN6 | SR-related and CTD-associated factor 8 C | SCAF8 | 140.52 | 2 | 57.247448 | yes | 5.478E-07 | 626.77283 | 375.5592 |
| GFGQEVHR | 8 | Q9UPN6 | SR-related and CTD-associated factor 8 C | SCAF8 | 140.52 | 2 | 57.247448 | yes | 1.39E-07 | 465.23303 | 341.26761 |
| VPGLYVIDSIVR | 12 | Q9UPN6 | SR-related and CTD-associated factor 8 C | SCAF8 | 140.52 | 2 | 57.247448 | no | 9.722E-08 | 665.89008 | 166.88951 |
| VLPVYGGPK | 9 | Q9UPN6 | SR-related and CTD-associated factor 8 C | SCAF8 | 140.52 | 2 | 57.247448 | yes | 4.112E-07 | 465.27637 | 121.81355 |
| DHFGRPPVDIR | 11 | Q9UPN6 | SR-related and CTD-associated factor 8 C | SCAF8 | 140.52 | 3 | 57.247448 | yes | 3.176E-07 | 436.89844 | 70.43251 |
| QSVDNVTNPEK | 11 | Q9UPN6 | SR-related and CTD-associated factor 8 C | SCAF8 | 140.52 | 2 | 57.247448 | yes | 1.012E-07 | 615.8017 | 64.061157 |
| IPLGNDNIQQEGDI | 14 | Q9UPN6 | SR-related and CTD-associated factor 8 C | SCAF8 | 140.52 | 2 | 57.247448 | yes | 1.981E-06 | 784.88678 | 44.968624 |
| QQLLEQQQPQK | 11 | Q9UPN6 | SR-related and CTD-associated factor 8 C | SCAF8 | 140.52 | 2 | 57.247448 | yes | 2.026E-07 | 684.36749 | 40.203194 |
| ETVQTTQSPTPVEI | 14 | Q9UPN6 | SR-related and CTD-associated factor 8 C | SCAF8 | 140.52 | 2 | 57.247448 | yes | 1.637E-07 | 772.89374 | 31.679075 |
| VAGALVQNTEK | 11 | Q9UPN7 | Serine/threonine-protein phosphatase 6 re | PPP6R1 | 96.724 | 2 | 49.64568 | yes | 1.014E-07 | 565.31421 | 835.49719 |
| VMGILINR | 8 | Q9UPN7 | Serine/threonine-protein phosphatase 6 re | PPP6R1 | 96.724 | 2 | 49.64568 | yes | 1.915E-07 | 458.27585 | 657.85919 |
| ILTSWEENDR | 10 | Q9UPN7 | Serine/threonine-protein phosphatase 6 re | PPP6R1 | 96.724 | 2 | 49.64568 | yes | 1.367E-07 | 631.8042 | 462.91675 |
| KTDQLVSFLR | 10 | Q9UPN7 | Serine/threonine-protein phosphatase 6 re | PPP6R1 | 96.724 | 2 | 49.64568 | yes | 2.874E-07 | 603.8457 | 217.03375 |
| GYMGHLTR | 8 | Q9UPN7 | Serine/threonine-protein phosphatase 6 re | PPP6R1 | 96.724 | 2 | 49.64568 | yes | 4.756E-07 | 467.73181 | 187.70172 |
| KGYMGHLTR | 9 | Q9UPN7 | Serine/threonine-protein phosphatase 6 re | PPP6R1 | 96.724 | 2 | 49.64568 | yes | 1.889E-06 | 531.7793 | 113.6097 |
| QDVVNWLNEEK | 11 | Q9UPN7 | Serine/threonine-protein phosphatase 6 re | PPP6R1 | 96.724 | 2 | 49.64568 | yes | 4.411E-07 | 687.33844 | 76.934341 |
| LIEQIHPSK | 9 | Q9UPN7 | Serine/threonine-protein phosphatase 6 re | PPP6R1 | 96.724 | 2 | 49.64568 | yes | 1.763E-07 | 532.80859 | 71.480751 |
| GPNAEQLR | 8 | Q9UPN7 | Serine/threonine-protein phosphatase 6 re | PPP6R1 | 96.724 | 2 | 49.64568 | yes | 3.179E-07 | 442.73288 | 51.473583 |
| LLTCVERPQLR | 11 | Q9UPN7 | Serine/threonine-protein phosphatase 6 re | PPP6R1 | 96.724 | 2 | 49.64568 | yes | 1.104E-07 | 692.89008 | 48.726574 |
| STTPMAPGVFLTQ | 14 | Q9UPP1 | Histone lysine demethylase PHF8 OS=Hc | PHF8 | 117.865 | 2 | 53.041077 | yes | 1.434E-07 | 753.3927 | 736.50153 |
| DGLGMTLPSPSFT | 15 | Q9UPP1 | Histone lysine demethylase PHF8 OS=Hc | PHF8 | 117.865 | 2 | 53.041077 | yes | 1.415E-07 | 789.40326 | 547.41815 |
| LPYDVTR | 7 | Q9UPP1 | Histone lysine demethylase PHF8 OS=Hc | PHF8 | 117.865 | 2 | 53.041077 | no | 5.539E-07 | 432.23471 | 390.47681 |
| LSTADLFR | 8 | Q9UPP1 | Histone lysine demethylase PHF8 OS=Hc | PHF8 | 117.865 | 2 | 53.041077 | yes | 9.722E-08 | 461.75327 | 274.27811 |

| | | | | | | | | | | | |
|---|---|---|---|---|---|---|---|---|---|---|---|
| LVEDIFQQNVGK | 12 | Q9UPP1 | Histone lysine demethylase PHF8 OS=Hc | PHF8 | 117.865 | 2 | 53.041077 | yes | 9.838E-08 | 695.37225 | 230.7529 |
| VLNVISLEFSDTR | 13 | Q9UPP1 | Histone lysine demethylase PHF8 OS=Hc | PHF8 | 117.865 | 2 | 53.041077 | yes | 9.722E-08 | 746.90393 | 229.89893 |
| YCLMSVR | 7 | Q9UPP1 | Histone lysine demethylase PHF8 OS=Hc | PHF8 | 117.865 | 2 | 53.041077 | no | 1.974E-07 | 464.7226 | 205.54636 |
| PNAFGMAQANR | 11 | Q9UPP1 | Histone lysine demethylase PHF8 OS=Hc | PHF8 | 117.865 | 2 | 53.041077 | yes | 1.509E-07 | 588.78259 | 184.76077 |
| ATLIIRPK | 8 | Q9UPP1 | Histone lysine demethylase PHF8 OS=Hc | PHF8 | 117.865 | 2 | 53.041077 | yes | 2.272E-07 | 456.30548 | 179.66458 |
| IFPAGSIPLTR | 11 | Q9UPP1 | Histone lysine demethylase PHF8 OS=Hc | PHF8 | 117.865 | 2 | 53.041077 | yes | 4.013E-07 | 586.34534 | 173.41394 |
| VASIETGLAAAAA | 14 | Q9UPP1 | Histone lysine demethylase PHF8 OS=Hc | PHF8 | 117.865 | 2 | 53.041077 | no | 1.372E-07 | 636.86151 | 132.98141 |
| LAQQELQK | 8 | Q9UPP1 | Histone lysine demethylase PHF8 OS=Hc | PHF8 | 117.865 | 2 | 53.041077 | yes | 1.385E-07 | 479.27182 | 107.02341 |
| LSNLVETPK | 9 | Q9UPP1 | Histone lysine demethylase PHF8 OS=Hc | PHF8 | 117.865 | 2 | 53.041077 | yes | 2.637E-07 | 500.78729 | 106.37715 |
| EALPDHEDEIPETV | 15 | Q9UPP1 | Histone lysine demethylase PHF8 OS=Hc | PHF8 | 117.865 | 3 | 53.041077 | yes | 1.004E-07 | 583.94788 | 94.420952 |
| NSDDAPWSPK | 10 | Q9UPP1 | Histone lysine demethylase PHF8 OS=Hc | PHF8 | 117.865 | 2 | 53.041077 | yes | 9.722E-08 | 558.75146 | 89.595169 |
| AADIDLYHCPNCE | 22 | Q9UPP1 | Histone lysine demethylase PHF8 OS=Hc | PHF8 | 117.865 | 4 | 53.041077 | yes | 3.155E-06 | 635.79871 | 46.918526 |
| KEALPDHEDEIPET | 16 | Q9UPP1 | Histone lysine demethylase PHF8 OS=Hc | PHF8 | 117.865 | 3 | 53.041077 | yes | 9.722E-08 | 626.64618 | 42.136112 |
| LLAGEDGTSER | 11 | Q9UPQ0 | LIM and calponin homology domains-cor | LIMCH1 | 121.868 | 2 | 52.824436 | yes | 2.518E-07 | 574.28314 | 286.15936 |
| GQLGDAVSGTDVI | 13 | Q9UPQ0 | LIM and calponin homology domains-cor | LIMCH1 | 121.868 | 2 | 52.824436 | yes | 3.056E-07 | 637.82037 | 228.7 |
| QTPSPDVVLR | 10 | Q9UPQ0 | LIM and calponin homology domains-cor | LIMCH1 | 121.868 | 2 | 52.824436 | yes | 1.127E-07 | 556.30896 | 228.45126 |
| SAVPFNQYLPNK | 12 | Q9UPQ0 | LIM and calponin homology domains-cor | LIMCH1 | 121.868 | 2 | 52.824436 | yes | 3.848E-07 | 689.36169 | 222.82161 |
| SNQTAYVPAPLR | 12 | Q9UPQ0 | LIM and calponin homology domains-cor | LIMCH1 | 121.868 | 2 | 52.824436 | yes | 9.733E-08 | 658.8515 | 215.02713 |
| SFQGDDSDLLLK | 12 | Q9UPQ0 | LIM and calponin homology domains-cor | LIMCH1 | 121.868 | 2 | 52.824436 | yes | 1.674E-07 | 669.33282 | 190.17767 |
| WQQEQER | 7 | Q9UPQ0 | LIM and calponin homology domains-cor | LIMCH1 | 121.868 | 2 | 52.824436 | yes | 1.231E-06 | 502.23322 | 179.72614 |
| FTATVETTIAR | 11 | Q9UPQ0 | LIM and calponin homology domains-cor | LIMCH1 | 121.868 | 2 | 52.824436 | yes | 1.131E-07 | 605.32733 | 174.07399 |
| DSGYIDCWDSER | 12 | Q9UPQ0 | LIM and calponin homology domains-cor | LIMCH1 | 121.868 | 2 | 52.824436 | yes | 9.722E-08 | 751.79626 | 165.84428 |
| VHGSPLELK | 9 | Q9UPQ0 | LIM and calponin homology domains-cor | LIMCH1 | 121.868 | 2 | 52.824436 | yes | 3.34E-07 | 490.2822 | 161.01346 |
| IDLGNCQDEK | 10 | Q9UPQ0 | LIM and calponin homology domains-cor | LIMCH1 | 121.868 | 2 | 52.824436 | yes | 9.722E-08 | 596.26917 | 152.92221 |
| DDSFDSLDSFGSR | 13 | Q9UPQ0 | LIM and calponin homology domains-cor | LIMCH1 | 121.868 | 2 | 52.824436 | yes | 9.722E-08 | 724.30225 | 120.25562 |
| EFEGLLAQMR | 10 | Q9UPQ0 | LIM and calponin homology domains-cor | LIMCH1 | 121.868 | 2 | 52.824436 | yes | 1.025E-07 | 597.3028 | 107.19586 |
| GSLTEGALAHSGN | 17 | Q9UPQ0 | LIM and calponin homology domains-cor | LIMCH1 | 121.868 | 2 | 52.824436 | yes | 9.727E-08 | 812.91809 | 104.17108 |
| VELVLSQK | 8 | Q9UPQ0 | LIM and calponin homology domains-cor | LIMCH1 | 121.868 | 2 | 52.824436 | yes | 9.722E-08 | 458.27911 | 103.45528 |
| ASVLDTSMSAGSC | 17 | Q9UPQ0 | LIM and calponin homology domains-cor | LIMCH1 | 121.868 | 2 | 52.824436 | yes | 2.207E-07 | 791.37488 | 102.83183 |
| QEQLQLINNQLR | 12 | Q9UPQ0 | LIM and calponin homology domains-cor | LIMCH1 | 121.868 | 2 | 52.824436 | yes | 9.722E-08 | 748.9126 | 80.224991 |
| KDTDDIESPK | 10 | Q9UPQ0 | LIM and calponin homology domains-cor | LIMCH1 | 121.868 | 2 | 52.824436 | yes | 9.722E-08 | 574.27753 | 78.390213 |
| VSVNGETVHR | 10 | Q9UPQ0 | LIM and calponin homology domains-cor | LIMCH1 | 121.868 | 2 | 52.824436 | yes | 5.344E-07 | 549.28851 | 63.591133 |
| AANSCTSYSGTTL | 16 | Q9UPQ0 | LIM and calponin homology domains-cor | LIMCH1 | 121.868 | 2 | 52.824436 | yes | 9.722E-08 | 844.40143 | 58.972176 |
| SPEPEATLTFPFLD | 15 | Q9UPQ0 | LIM and calponin homology domains-cor | LIMCH1 | 121.868 | 2 | 52.824436 | yes | 2.739E-07 | 846.42999 | 49.047913 |
| GSDSESDLPHR | 11 | Q9UPQ0 | LIM and calponin homology domains-cor | LIMCH1 | 121.868 | 2 | 52.824436 | yes | 1.867E-07 | 600.26801 | 44.863255 |
| SINHQIESPSER | 12 | Q9UPQ0 | LIM and calponin homology domains-cor | LIMCH1 | 121.868 | 2 | 52.824436 | yes | 8.003E-07 | 698.84442 | 41.286907 |
| GVHEDHQLDTEAC | 46 | Q9UPQ0 | LIM and calponin homology domains-cor | LIMCH1 | 121.868 | 6 | 52.824436 | yes | 9.722E-08 | 832.04828 | 41.282585 |
| FTISEAVLER | 10 | Q9UPQ0 | LIM and calponin homology domains-cor | LIMCH1 | 121.868 | 2 | 52.824436 | yes | 3.284E-07 | 582.81659 | 38.715324 |
| EEDDKWQDDLAR | 12 | Q9UPQ0 | LIM and calponin homology domains-cor | LIMCH1 | 121.868 | 2 | 52.824436 | yes | 1.074E-07 | 760.33661 | 33.647274 |

| | | | | | | | | | | | |
|---|---|---|---|---|---|---|---|---|---|---|---|
| VEEPVLQNQIR | 11 | Q9UPQ3 | Arf-GAP with GTPase, ANK repeat and I | AGAP1 | 94.471 | 2 | 50.769135 | yes | 1.873E-07 | 662.86462 | 242.50137 |
| TALHLACR | 8 | Q9UPQ3 | Arf-GAP with GTPase, ANK repeat and I | AGAP1 | 94.471 | 2 | 50.769135 | yes | 9.722E-08 | 471.2529 | 144.94693 |
| ATSACAPISSPK | 12 | Q9UPQ3 | Arf-GAP with GTPase, ANK repeat and I | AGAP1 | 94.471 | 2 | 50.769135 | yes | 1.008E-07 | 595.29773 | 139.81091 |
| LTSQSEAMALQSII | 14 | Q9UPQ3 | Arf-GAP with GTPase, ANK repeat and I | AGAP1 | 94.471 | 2 | 50.769135 | yes | 1.114E-07 | 767.89832 | 91.172546 |
| GMPFGMGLGNTSI | 13 | Q9UPQ9 | Trinucleotide repeat-containing gene 6B p | TNRC6B | 194.003 | 2 | 53.75079 | yes | 1.208E-07 | 662.81049 | 1020.0328 |
| GMPFGMGLGNTSI | 13 | Q9UPQ9 | Trinucleotide repeat-containing gene 6B p | TNRC6B | 194.003 | 2 | 53.75079 | yes | 7.042E-07 | 670.80792 | 1020.0328 |
| MDLSVGSLSDK | 11 | Q9UPQ9 | Trinucleotide repeat-containing gene 6B p | TNRC6B | 194.003 | 2 | 53.75079 | yes | 9.793E-08 | 576.28424 | 1700.9501 |
| AMNLGDFNDIMR | 12 | Q9UPQ9 | Trinucleotide repeat-containing gene 6B p | TNRC6B | 194.003 | 2 | 53.75079 | yes | 1.338E-07 | 698.82104 | 1495.8075 |
| VLSNTGWGQTQIK | 13 | Q9UPQ9 | Trinucleotide repeat-containing gene 6B p | TNRC6B | 194.003 | 2 | 53.75079 | yes | 1.214E-07 | 716.38318 | 1456.532 |
| MENAGVNFVVSG | 13 | Q9UPQ9 | Trinucleotide repeat-containing gene 6B p | TNRC6B | 194.003 | 2 | 53.75079 | yes | 9.968E-08 | 690.34045 | 1237.6135 |
| DMGTTDSGPYFEK | 13 | Q9UPQ9 | Trinucleotide repeat-containing gene 6B p | TNRC6B | 194.003 | 2 | 53.75079 | yes | 9.722E-08 | 724.30591 | 1236.5234 |
| SGWGELSASTEWK | 13 | Q9UPQ9 | Trinucleotide repeat-containing gene 6B p | TNRC6B | 194.003 | 2 | 53.75079 | yes | 9.819E-08 | 719.33588 | 1051.0575 |
| LASASTWSDGGSV | 14 | Q9UPQ9 | Trinucleotide repeat-containing gene 6B p | TNRC6B | 194.003 | 2 | 53.75079 | yes | 9.722E-08 | 697.33893 | 988.56982 |
| MTSGVSQGEWK | 11 | Q9UPQ9 | Trinucleotide repeat-containing gene 6B p | TNRC6B | 194.003 | 2 | 53.75079 | yes | 9.722E-08 | 605.28204 | 969.66125 |
| GTEGWESAATQTK | 13 | Q9UPQ9 | Trinucleotide repeat-containing gene 6B p | TNRC6B | 194.003 | 2 | 53.75079 | yes | 9.722E-08 | 683.31769 | 935.31342 |
| EQAQIHNTDGPK | 12 | Q9UPQ9 | Trinucleotide repeat-containing gene 6B p | TNRC6B | 194.003 | 2 | 53.75079 | yes | 1.002E-07 | 669.32587 | 914.27319 |
| STLPGSTTSNK | 11 | Q9UPQ9 | Trinucleotide repeat-containing gene 6B p | TNRC6B | 194.003 | 2 | 53.75079 | yes | 1.844E-07 | 546.78021 | 881.48254 |
| GGNNDSWMNPLA | 13 | Q9UPQ9 | Trinucleotide repeat-containing gene 6B p | TNRC6B | 194.003 | 2 | 53.75079 | yes | 9.722E-08 | 702.32227 | 813.79718 |
| QQQEQQLAR | 9 | Q9UPQ9 | Trinucleotide repeat-containing gene 6B p | TNRC6B | 194.003 | 2 | 53.75079 | yes | 9.722E-08 | 564.79144 | 662.23816 |
| PSSPWSSTAPR | 11 | Q9UPQ9 | Trinucleotide repeat-containing gene 6B p | TNRC6B | 194.003 | 2 | 53.75079 | yes | 9.926E-08 | 586.78839 | 654.4137 |
| TPSSWNENPSK | 11 | Q9UPQ9 | Trinucleotide repeat-containing gene 6B p | TNRC6B | 194.003 | 2 | 53.75079 | yes | 1.023E-07 | 623.78857 | 378.48837 |
| EPNLPTPMTSK | 11 | Q9UPQ9 | Trinucleotide repeat-containing gene 6B p | TNRC6B | 194.003 | 2 | 53.75079 | yes | 6.081E-07 | 607.80792 | 331.77747 |
| GWGTQDSR | 8 | Q9UPQ9 | Trinucleotide repeat-containing gene 6B p | TNRC6B | 194.003 | 2 | 53.75079 | yes | 2.763E-07 | 453.70685 | 270.37415 |
| TGSVGSWGAAR | 11 | Q9UPQ9 | Trinucleotide repeat-containing gene 6B p | TNRC6B | 194.003 | 2 | 53.75079 | yes | 2.429E-06 | 524.76215 | 258.97775 |
| SGFRPPNSK | 9 | Q9UPQ9 | Trinucleotide repeat-containing gene 6B p | TNRC6B | 194.003 | 2 | 53.75079 | yes | 2.304E-06 | 495.26178 | 205.10564 |
| GGWEDCKR | 8 | Q9UPQ9 | Trinucleotide repeat-containing gene 6B p | TNRC6B | 194.003 | 2 | 53.75079 | yes | 9.722E-08 | 504.2218 | 142.58722 |
| QDTVWDIEEVPRP | 16 | Q9UPQ9 | Trinucleotide repeat-containing gene 6B p | TNRC6B | 194.003 | 3 | 53.75079 | yes | 5.25E-07 | 633.31451 | 99.575653 |
| NTTPLPRPPPGLTN | 16 | Q9UPQ9 | Trinucleotide repeat-containing gene 6B p | TNRC6B | 194.003 | 3 | 53.75079 | yes | 1.378E-07 | 567.32117 | 95.69635 |
| PSNSSWSSGPQPA | 16 | Q9UPQ9 | Trinucleotide repeat-containing gene 6B p | TNRC6B | 194.003 | 2 | 53.75079 | yes | 1.547E-07 | 814.38916 | 89.991539 |
| NSGGWGDAPSQSI | 16 | Q9UPQ9 | Trinucleotide repeat-containing gene 6B p | TNRC6B | 194.003 | 2 | 53.75079 | yes | 1.119E-07 | 832.36011 | 87.44899 |
| EEQLMEDK | 8 | Q9UPQ9 | Trinucleotide repeat-containing gene 6B p | TNRC6B | 194.003 | 2 | 53.75079 | yes | 2.938E-06 | 511.22897 | 87.445633 |
| NGWGEEVDQTK | 11 | Q9UPQ9 | Trinucleotide repeat-containing gene 6B p | TNRC6B | 194.003 | 2 | 53.75079 | yes | 1.043E-07 | 631.78601 | 84.455666 |
| KTGSVGSWGAAR | 12 | Q9UPQ9 | Trinucleotide repeat-containing gene 6B p | TNRC6B | 194.003 | 2 | 53.75079 | yes | 3.321E-07 | 588.80963 | 83.911827 |
| VAVPNGQPPSAAR | 13 | Q9UPQ9 | Trinucleotide repeat-containing gene 6B p | TNRC6B | 194.003 | 2 | 53.75079 | yes | 4.721E-07 | 632.34387 | 72.571999 |
| EREDSWK | 7 | Q9UPQ9 | Trinucleotide repeat-containing gene 6B p | TNRC6B | 194.003 | 2 | 53.75079 | yes | 1.536E-07 | 475.22232 | 67.956749 |
| QWTSMMEGLPSV | 21 | Q9UPQ9 | Trinucleotide repeat-containing gene 6B p | TNRC6B | 194.003 | 3 | 53.75079 | yes | 3.212E-05 | 792.69757 | 52.790276 |
| EQEREEQLMEDK | 12 | Q9UPQ9 | Trinucleotide repeat-containing gene 6B p | TNRC6B | 194.003 | 2 | 53.75079 | yes | 1.971E-07 | 782.35138 | 31.058403 |
| NTGGWNDYK | 9 | Q9UPQ9 | Trinucleotide repeat-containing gene 6B p | TNRC6B | 194.003 | 2 | 53.75079 | yes | 3.914E-05 | 527.73309 | 24.12985 |
| KAEELLWR | 8 | Q9UPR3 | Nonsense-mediated mRNA decay factor S | SMG5 | 113.928 | 2 | 53.54245 | yes | 2.133E-07 | 522.79547 | 395.87024 |

| | | | | | | | | | | | |
|---|---|---|---|---|---|---|---|---|---|---|---|
| LMFLHPVDYGR | 11 | Q9UPR3 | Nonsense-mediated mRNA decay factor S | SMG5 | 113.928 | 2 | 53.54245 | yes | 1.136E-07 | 674.34753 | 388.93369 |
| SFGHFIAR | 8 | Q9UPR3 | Nonsense-mediated mRNA decay factor S | SMG5 | 113.928 | 2 | 53.54245 | yes | 1.363E-07 | 467.74832 | 326.70367 |
| AVVEAVHR | 8 | Q9UPR3 | Nonsense-mediated mRNA decay factor S | SMG5 | 113.928 | 2 | 53.54245 | yes | 4.176E-07 | 440.7536 | 310.82535 |
| EMDWAQMACHR | 11 | Q9UPR3 | Nonsense-mediated mRNA decay factor S | SMG5 | 113.928 | 2 | 53.54245 | yes | 9.722E-08 | 717.78918 | 156.07756 |
| KVYYEVIQLIK | 11 | Q9UPR3 | Nonsense-mediated mRNA decay factor S | SMG5 | 113.928 | 2 | 53.54245 | yes | 9.902E-08 | 698.41595 | 149.37167 |
| LDLILCNK | 8 | Q9UPR3 | Nonsense-mediated mRNA decay factor S | SMG5 | 113.928 | 2 | 53.54245 | yes | 5.652E-07 | 494.77844 | 88.279976 |
| VYYEVIQLIK | 10 | Q9UPR3 | Nonsense-mediated mRNA decay factor S | SMG5 | 113.928 | 2 | 53.54245 | yes | 9.722E-08 | 634.36847 | 54.4286 |
| YYNVEAMYCYLR | 12 | Q9UPR3 | Nonsense-mediated mRNA decay factor S | SMG5 | 113.928 | 2 | 53.54245 | yes | 2.995E-07 | 822.86292 | 54.256172 |
| RQDADAWTLYK | 11 | Q9UPR3 | Nonsense-mediated mRNA decay factor S | SMG5 | 113.928 | 2 | 53.54245 | yes | 4.157E-07 | 683.84113 | 40.595863 |
| LQVLMAEGLLPAV | 14 | Q9UPR3 | Nonsense-mediated mRNA decay factor S | SMG5 | 113.928 | 2 | 53.54245 | yes | 3.236E-06 | 741.44165 | 30.680128 |
| NMVELGQVK | 9 | Q9UPS8 | Ankyrin repeat domain-containing proteir | ANKRD26 | 196.411 | 2 | 52.112495 | yes | 0.0003106 | 509.2735 | 112.44959 |
| DNNEILSQQLFK | 12 | Q9UPS8 | Ankyrin repeat domain-containing proteir | ANKRD26 | 196.411 | 2 | 52.112495 | yes | 9.722E-08 | 724.87262 | 89.273079 |
| MNFDVSNLK | 9 | Q9UPS8 | Ankyrin repeat domain-containing proteir | ANKRD26 | 196.411 | 2 | 52.112495 | yes | 5.167E-07 | 534.26312 | 56.187424 |
| NSMLQEEIAMLR | 12 | Q9UPS8 | Ankyrin repeat domain-containing proteir | ANKRD26 | 196.411 | 2 | 52.112495 | yes | 1.371E-07 | 717.8576 | 55.566463 |
| LMTASQQSR | 9 | Q9UPS8 | Ankyrin repeat domain-containing proteir | ANKRD26 | 196.411 | 2 | 52.112495 | yes | 1.325E-06 | 511.25839 | 48.691154 |
| VQQILLLR | 8 | Q9UPS8 | Ankyrin repeat domain-containing proteir | ANKRD26 | 196.411 | 2 | 52.112495 | yes | 4.426E-07 | 491.82404 | 44.531181 |
| LSQLQSENMLLR | 12 | Q9UPS8 | Ankyrin repeat domain-containing proteir | ANKRD26 | 196.411 | 2 | 52.112495 | no | 6.726E-06 | 716.38483 | 41.05188 |
| EEPTKPGIAK | 10 | Q9UPS8 | Ankyrin repeat domain-containing proteir | ANKRD26 | 196.411 | 2 | 52.112495 | yes | 1.121E-06 | 535.29803 | 37.998463 |
| IKDLESELSK | 10 | Q9UPS8 | Ankyrin repeat domain-containing proteir | ANKRD26 | 196.411 | 2 | 52.112495 | yes | 4.752E-07 | 581.32172 | 33.09222 |
| KLEQEVINLR | 10 | Q9UPS8 | Ankyrin repeat domain-containing proteir | ANKRD26 | 196.411 | 2 | 52.112495 | yes | 1.173E-07 | 621.36426 | 25.90097 |
| MLADDAEAGAED | 21 | Q9UPT8 | Zinc finger CCCH domain-containing prc | ZC3H4 | 140.257 | 3 | 54.26317 | yes | 1.683E-07 | 774.03351 | 1197.8066 |
| ATEPAADTGAQPK | 13 | Q9UPT8 | Zinc finger CCCH domain-containing prc | ZC3H4 | 140.257 | 2 | 54.26317 | yes | 9.722E-08 | 628.80951 | 1161.3378 |
| AVNIPLDPLPGHPI | 15 | Q9UPT8 | Zinc finger CCCH domain-containing prc | ZC3H4 | 140.257 | 2 | 54.26317 | yes | 1.325E-07 | 804.96466 | 570.56995 |
| AVNIPLDPLPGHPI | 15 | Q9UPT8 | Zinc finger CCCH domain-containing prc | ZC3H4 | 140.257 | 2 | 54.26317 | yes | 1.341E-07 | 536.97888 | 570.56995 |
| AVNIPLDPLPGHPI | 15 | Q9UPT8 | Zinc finger CCCH domain-containing prc | ZC3H4 | 140.257 | 3 | 54.26317 | yes | 1.325E-07 | 804.96466 | 570.56995 |
| AVNIPLDPLPGHPI | 15 | Q9UPT8 | Zinc finger CCCH domain-containing prc | ZC3H4 | 140.257 | 3 | 54.26317 | yes | 1.341E-07 | 536.97888 | 570.56995 |
| FYITGFCAR | 9 | Q9UPT8 | Zinc finger CCCH domain-containing prc | ZC3H4 | 140.257 | 2 | 54.26317 | yes | 1.113E-07 | 567.77368 | 931.27374 |
| PGAGVPDFLPSAQ | 14 | Q9UPT8 | Zinc finger CCCH domain-containing prc | ZC3H4 | 140.257 | 2 | 54.26317 | yes | 1.65E-07 | 706.37006 | 697.802 |
| AKEPPFVR | 8 | Q9UPT8 | Zinc finger CCCH domain-containing prc | ZC3H4 | 140.257 | 2 | 54.26317 | yes | 1.947E-06 | 472.27164 | 477.68701 |
| LQKPTDSTASSR | 12 | Q9UPT8 | Zinc finger CCCH domain-containing prc | ZC3H4 | 140.257 | 2 | 54.26317 | yes | 4.626E-07 | 645.83606 | 204.93831 |
| LQKPTDSTASSR | 12 | Q9UPT8 | Zinc finger CCCH domain-containing prc | ZC3H4 | 140.257 | 2 | 54.26317 | yes | 6.209E-07 | 430.89313 | 204.93831 |
| LQKPTDSTASSR | 12 | Q9UPT8 | Zinc finger CCCH domain-containing prc | ZC3H4 | 140.257 | 3 | 54.26317 | yes | 4.626E-07 | 645.83606 | 204.93831 |
| LQKPTDSTASSR | 12 | Q9UPT8 | Zinc finger CCCH domain-containing prc | ZC3H4 | 140.257 | 3 | 54.26317 | yes | 6.209E-07 | 430.89313 | 204.93831 |
| PKPGAGVPDFLPS. | 16 | Q9UPT8 | Zinc finger CCCH domain-containing prc | ZC3H4 | 140.257 | 2 | 54.26317 | yes | 1.464E-07 | 818.94391 | 146.87126 |
| PKPGAGVPDFLPS. | 16 | Q9UPT8 | Zinc finger CCCH domain-containing prc | ZC3H4 | 140.257 | 2 | 54.26317 | yes | 1.248E-07 | 546.2984 | 146.87126 |
| PKPGAGVPDFLPS. | 16 | Q9UPT8 | Zinc finger CCCH domain-containing prc | ZC3H4 | 140.257 | 3 | 54.26317 | yes | 1.464E-07 | 818.94391 | 146.87126 |
| PKPGAGVPDFLPS. | 16 | Q9UPT8 | Zinc finger CCCH domain-containing prc | ZC3H4 | 140.257 | 3 | 54.26317 | yes | 1.248E-07 | 546.2984 | 146.87126 |
| SQLQQFSHIK | 10 | Q9UPT8 | Zinc finger CCCH domain-containing prc | ZC3H4 | 140.257 | 2 | 54.26317 | yes | 9.722E-08 | 608.32764 | 274.48334 |
| KIPSLFEIVVR | 11 | Q9UPT8 | Zinc finger CCCH domain-containing prc | ZC3H4 | 140.257 | 2 | 54.26317 | yes | 9.722E-08 | 650.90302 | 219.33241 |

| | | | | | | | | | | | |
|---|---|---|---|---|---|---|---|---|---|---|---|
| EGEQDAASLK | 10 | Q9UPT8 | Zinc finger CCCH domain-containing prc | ZC3H4 | 140.257 | 2 | 54.26317 | yes | 1.805E-06 | 524.25128 | 204.37073 |
| YNSYNRPR | 8 | Q9UPT8 | Zinc finger CCCH domain-containing prc | ZC3H4 | 140.257 | 2 | 54.26317 | yes | 6.7E-07 | 535.26233 | 199.58698 |
| MLADDAEAGAED | 20 | Q9UPT8 | Zinc finger CCCH domain-containing prc | ZC3H4 | 140.257 | 3 | 54.26317 | yes | 3.52E-07 | 731.33521 | 193.77373 |
| MLADDAEAGAED | 14 | Q9UPT8 | Zinc finger CCCH domain-containing prc | ZC3H4 | 140.257 | 2 | 54.26317 | yes | 9.722E-08 | 732.81158 | 144.07962 |
| KIPSLFEIVVRPTG( | 19 | Q9UPT8 | Zinc finger CCCH domain-containing prc | ZC3H4 | 140.257 | 3 | 54.26317 | yes | 9.722E-08 | 709.08417 | 118.40087 |
| KSALEQPETGK | 11 | Q9UPT8 | Zinc finger CCCH domain-containing prc | ZC3H4 | 140.257 | 2 | 54.26317 | yes | 1.272E-07 | 594.31696 | 83.134415 |
| AENCPYMHGDFP( | 14 | Q9UPT8 | Zinc finger CCCH domain-containing prc | ZC3H4 | 140.257 | 3 | 54.26317 | yes | 1.399E-07 | 575.8988 | 53.152824 |
| GFDPTASPFCQ | 11 | Q9UPT8 | Zinc finger CCCH domain-containing prc | ZC3H4 | 140.257 | 2 | 54.26317 | yes | 1.127E-07 | 613.76099 | 34.490673 |
| TGSGSPFAGNSPAI | 14 | Q9UPT8 | Zinc finger CCCH domain-containing prc | ZC3H4 | 140.257 | 2 | 54.26317 | yes | 9.918E-08 | 653.31274 | 24.492939 |
| ITSNCTIGLR | 10 | Q9UPT9 | Ubiquitin carboxyl-terminal hydrolase 22 | USP22 | 59.961 | 2 | 31.290644 | yes | 1.89E-06 | 567.80042 | 131.67276 |
| TLLDISGYR | 9 | Q9UPU9 | Protein Smaug homolog 1 OS=Homo sap | SAMD4A | 79.415 | 2 | 56.095898 | yes | 5.409E-07 | 519.28491 | 224.75943 |
| NTTATPTIMK | 10 | Q9UPU9 | Protein Smaug homolog 1 OS=Homo sap | SAMD4A | 79.415 | 2 | 56.095898 | yes | 3.98E-07 | 539.28406 | 223.21109 |
| DSGICINASNWQD | 14 | Q9UPU9 | Protein Smaug homolog 1 OS=Homo sap | SAMD4A | 79.415 | 2 | 56.095898 | yes | 1.061E-07 | 804.35956 | 186.15744 |
| IPLQELHQMILTPII | 15 | Q9UPU9 | Protein Smaug homolog 1 OS=Homo sap | SAMD4A | 79.415 | 3 | 56.095898 | yes | 9.722E-08 | 592.01471 | 111.6433 |
| CLIHEAFTETQK | 12 | Q9UPU9 | Protein Smaug homolog 1 OS=Homo sap | SAMD4A | 79.415 | 2 | 56.095898 | yes | 9.722E-08 | 738.86121 | 97.973862 |
| AYSSPSTTPEAR | 12 | Q9UPU9 | Protein Smaug homolog 1 OS=Homo sap | SAMD4A | 79.415 | 2 | 56.095898 | yes | 1.172E-07 | 633.8017 | 87.019676 |
| NTFQEEGSGMK | 11 | Q9UPU9 | Protein Smaug homolog 1 OS=Homo sap | SAMD4A | 79.415 | 2 | 56.095898 | yes | 9.722E-08 | 614.26917 | 73.698418 |
| QYQIPSR | 7 | Q9UPU9 | Protein Smaug homolog 1 OS=Homo sap | SAMD4A | 79.415 | 2 | 56.095898 | yes | 3.764E-07 | 446.23779 | 72.883179 |
| DQVGVLAGWFK | 11 | Q9UPU9 | Protein Smaug homolog 1 OS=Homo sap | SAMD4A | 79.415 | 2 | 56.095898 | yes | 7.737E-07 | 610.32715 | 70.261864 |
| SALAMWLNHLEDI | 13 | Q9UPU9 | Protein Smaug homolog 1 OS=Homo sap | SAMD4A | 79.415 | 2 | 56.095898 | yes | 2.484E-07 | 778.38794 | 52.727661 |
| ILAHSIEHNQHIEES | 16 | Q9UPU9 | Protein Smaug homolog 1 OS=Homo sap | SAMD4A | 79.415 | 4 | 56.095898 | yes | 1.279E-06 | 478.99606 | 21.312927 |
| QPSLMGPESQSPD( | 15 | Q9UPU9 | Protein Smaug homolog 1 OS=Homo sap | SAMD4A | 79.415 | 2 | 56.095898 | yes | 5.953E-06 | 830.8689 | 18.840736 |
| LALLEVQEETAR | 12 | Q9UPV0 | Centrosomal protein of 164 kDa OS=Hon | CEP164 | 164.314 | 2 | 47.374893 | yes | 1.343E-07 | 686.37756 | 70.000328 |
| LGYMSASEQLR | 11 | Q9UPV0 | Centrosomal protein of 164 kDa OS=Hon | CEP164 | 164.314 | 2 | 47.374893 | yes | 5.532E-06 | 627.81097 | 51.624489 |
| SFHGLDFGFR | 10 | Q9UPV0 | Centrosomal protein of 164 kDa OS=Hon | CEP164 | 164.314 | 2 | 47.374893 | yes | 2.165E-06 | 591.78815 | 35.154701 |
| FCVGLLR | 7 | Q9UPW5 | Cytosolic carboxypeptidase 1 OS=Homo | AGTPBP1 | 138.449 | 2 | 41.36569 | yes | 3.978E-07 | 432.74164 | 60.426083 |
| TLQQQPGDQNR | 11 | Q9UPW5 | Cytosolic carboxypeptidase 1 OS=Homo | AGTPBP1 | 138.449 | 2 | 41.36569 | yes | 3.546E-07 | 642.81818 | 59.117088 |
| FDLPTEYR | 8 | Q9UPY3 | Endoribonuclease Dicer OS=Homo sapiei | DICER1 | 218.682 | 2 | 55.030483 | yes | 1.184E-07 | 520.75598 | 297.46393 |
| ELLEMEPETAK | 11 | Q9UPY3 | Endoribonuclease Dicer OS=Homo sapiei | DICER1 | 218.682 | 2 | 55.030483 | yes | 9.722E-08 | 645.3183 | 233.64349 |
| VVALICCEK | 9 | Q9UPY3 | Endoribonuclease Dicer OS=Homo sapiei | DICER1 | 218.682 | 2 | 55.030483 | yes | 2.344E-07 | 546.28302 | 195.39859 |
| SNAETATDLVVLD | 14 | Q9UPY3 | Endoribonuclease Dicer OS=Homo sapiei | DICER1 | 218.682 | 2 | 55.030483 | yes | 1.106E-07 | 752.38611 | 170.85909 |
| YPNLDFGWK | 9 | Q9UPY3 | Endoribonuclease Dicer OS=Homo sapiei | DICER1 | 218.682 | 2 | 55.030483 | yes | 9.809E-08 | 570.27966 | 164.93196 |
| NLSVSCAAASVAS | 15 | Q9UPY3 | Endoribonuclease Dicer OS=Homo sapiei | DICER1 | 218.682 | 2 | 55.030483 | yes | 1.013E-07 | 740.36462 | 155.27065 |
| LHCLLTAEELR | 11 | Q9UPY3 | Endoribonuclease Dicer OS=Homo sapiei | DICER1 | 218.682 | 2 | 55.030483 | yes | 9.722E-08 | 677.86102 | 136.19051 |
| LEDYQDAVIIPR | 12 | Q9UPY3 | Endoribonuclease Dicer OS=Homo sapiei | DICER1 | 218.682 | 2 | 55.030483 | yes | 2.089E-07 | 716.37756 | 109.42979 |
| LEMLGDSFLK | 10 | Q9UPY3 | Endoribonuclease Dicer OS=Homo sapiei | DICER1 | 218.682 | 2 | 55.030483 | yes | 9.975E-08 | 576.80212 | 108.79823 |
| YTAVVLNR | 8 | Q9UPY3 | Endoribonuclease Dicer OS=Homo sapiei | DICER1 | 218.682 | 2 | 55.030483 | yes | 2.39E-07 | 468.26907 | 98.573837 |
| VTINTAIGHINR | 12 | Q9UPY3 | Endoribonuclease Dicer OS=Homo sapiei | DICER1 | 218.682 | 2 | 55.030483 | yes | 1.613E-07 | 654.87274 | 90.642166 |
| AVLVVLGPWCAD | 13 | Q9UPY3 | Endoribonuclease Dicer OS=Homo sapiei | DICER1 | 218.682 | 2 | 55.030483 | yes | 2.511E-07 | 714.3894 | 86.41935 |

| | | | | | | | | | | | |
|---|---|---|---|---|---|---|---|---|---|---|---|
| AQTASDAGVGVR | 12 | Q9UPY3 | Endoribonuclease Dicer OS=Homo sapiei | DICER1 | 218.682 | 2 | 55.030483 | yes | 2.918E-07 | 566.29126 | 85.945427 |
| DFCQGNQLNYYK | 12 | Q9UPY3 | Endoribonuclease Dicer OS=Homo sapiei | DICER1 | 218.682 | 2 | 55.030483 | yes | 1.14E-07 | 775.34064 | 82.876266 |
| FIDNMLMGSGAFV | 14 | Q9UPY3 | Endoribonuclease Dicer OS=Homo sapiei | DICER1 | 218.682 | 2 | 55.030483 | yes | 1.022E-07 | 765.37817 | 80.895866 |
| MDSEQSPSIGYSSF | 14 | Q9UPY3 | Endoribonuclease Dicer OS=Homo sapiei | DICER1 | 218.682 | 2 | 55.030483 | yes | 2.05E-07 | 772.33832 | 77.779953 |
| APISNYIMLADTDI | 14 | Q9UPY3 | Endoribonuclease Dicer OS=Homo sapiei | DICER1 | 218.682 | 2 | 55.030483 | yes | 1.023E-07 | 776.38977 | 75.885056 |
| VSNCNLYR | 8 | Q9UPY3 | Endoribonuclease Dicer OS=Homo sapiei | DICER1 | 218.682 | 2 | 55.030483 | yes | 2.194E-07 | 513.2453 | 65.598579 |
| SALVNNTIFASLAV | 15 | Q9UPY3 | Endoribonuclease Dicer OS=Homo sapiei | DICER1 | 218.682 | 2 | 55.030483 | yes | 4.722E-07 | 774.44324 | 61.48613 |
| AAQLFLCSLGLK | 12 | Q9UPY3 | Endoribonuclease Dicer OS=Homo sapiei | DICER1 | 218.682 | 2 | 55.030483 | yes | 9.722E-08 | 660.87085 | 61.352814 |
| NEMQGMDSELR | 11 | Q9UPY3 | Endoribonuclease Dicer OS=Homo sapiei | DICER1 | 218.682 | 2 | 55.030483 | yes | 9.722E-08 | 655.27917 | 42.399719 |
| IGELDDHLMPVGK | 13 | Q9UPY3 | Endoribonuclease Dicer OS=Homo sapiei | DICER1 | 218.682 | 2 | 55.030483 | yes | 9.935E-08 | 712.36615 | 40.338188 |
| TFIAVLLTK | 9 | Q9UPY3 | Endoribonuclease Dicer OS=Homo sapiei | DICER1 | 218.682 | 2 | 55.030483 | yes | 3.452E-07 | 503.3208 | 38.784649 |
| PIPQIPHFPVYTR | 13 | Q9UPY3 | Endoribonuclease Dicer OS=Homo sapiei | DICER1 | 218.682 | 3 | 55.030483 | yes | 6.108E-07 | 522.29266 | 34.060825 |
| LNSLTLAR | 8 | Q9UQ13 | Leucine-rich repeat protein SHOC-2 OS= | SHOC2 | 64.889 | 2 | 37.226948 | yes | 6.182E-07 | 444.26907 | 66.723526 |
| TPAIPTAVNLADSI | 14 | Q9UQ35 | Serine/arginine repetitive matrix protein 2 | SRRM2 | 299.616 | 2 | 52.463924 | yes | 5.62E-06 | 713.38843 | 1107.1851 |
| TAVAPSAVNLADF | 14 | Q9UQ35 | Serine/arginine repetitive matrix protein 2 | SRRM2 | 299.616 | 2 | 52.463924 | yes | 4.463E-07 | 691.37537 | 880.44153 |
| TPQAPASANLVGP | 14 | Q9UQ35 | Serine/arginine repetitive matrix protein 2 | SRRM2 | 299.616 | 2 | 52.463924 | yes | 1.363E-07 | 689.87549 | 858.0545 |
| ENSFGSPLEFR | 11 | Q9UQ35 | Serine/arginine repetitive matrix protein 2 | SRRM2 | 299.616 | 2 | 52.463924 | yes | 9.722E-08 | 641.80676 | 837.01154 |
| TPTAPAVNLAGAR | 13 | Q9UQ35 | Serine/arginine repetitive matrix protein 2 | SRRM2 | 299.616 | 2 | 52.463924 | yes | 1.418E-07 | 619.84619 | 784.26819 |
| SAHATAPVNIAGS | 14 | Q9UQ35 | Serine/arginine repetitive matrix protein 2 | SRRM2 | 299.616 | 2 | 52.463924 | yes | 9.722E-08 | 676.35748 | 361.09744 |
| SAHATAPVNIAGS | 14 | Q9UQ35 | Serine/arginine repetitive matrix protein 2 | SRRM2 | 299.616 | 2 | 52.463924 | yes | 3.626E-05 | 451.24075 | 361.09744 |
| SAHATAPVNIAGS | 14 | Q9UQ35 | Serine/arginine repetitive matrix protein 2 | SRRM2 | 299.616 | 3 | 52.463924 | yes | 9.722E-08 | 676.35748 | 361.09744 |
| SAHATAPVNIAGS | 14 | Q9UQ35 | Serine/arginine repetitive matrix protein 2 | SRRM2 | 299.616 | 3 | 52.463924 | yes | 3.626E-05 | 451.24075 | 361.09744 |
| VPLSAYER | 8 | Q9UQ35 | Serine/arginine repetitive matrix protein 2 | SRRM2 | 299.616 | 2 | 52.463924 | yes | 1.602E-07 | 467.75327 | 644.60016 |
| PSMSPTPLDR | 10 | Q9UQ35 | Serine/arginine repetitive matrix protein 2 | SRRM2 | 299.616 | 2 | 52.463924 | yes | 1.542E-07 | 550.77386 | 639.43213 |
| SPVPSAFSDQSR | 12 | Q9UQ35 | Serine/arginine repetitive matrix protein 2 | SRRM2 | 299.616 | 2 | 52.463924 | yes | 9.722E-08 | 639.30969 | 528.08588 |
| IPAASAAAMNLAS | 15 | Q9UQ35 | Serine/arginine repetitive matrix protein 2 | SRRM2 | 299.616 | 2 | 52.463924 | yes | 1.382E-07 | 707.8772 | 457.13651 |
| TAAALAPASLTSA | 14 | Q9UQ35 | Serine/arginine repetitive matrix protein 2 | SRRM2 | 299.616 | 2 | 52.463924 | yes | 1.062E-07 | 650.86462 | 442.52679 |
| SPGMLEPLGSSR | 12 | Q9UQ35 | Serine/arginine repetitive matrix protein 2 | SRRM2 | 299.616 | 2 | 52.463924 | yes | 9.734E-08 | 615.81097 | 340.34039 |
| KPIDSLR | 7 | Q9UQ35 | Serine/arginine repetitive matrix protein 2 | SRRM2 | 299.616 | 2 | 52.463924 | yes | 9.9E-08 | 414.75052 | 303.98199 |
| HSLSGSSPGMK | 11 | Q9UQ35 | Serine/arginine repetitive matrix protein 2 | SRRM2 | 299.616 | 2 | 52.463924 | yes | 3.657E-06 | 544.26367 | 263.23151 |
| TPPVALNSSR | 10 | Q9UQ35 | Serine/arginine repetitive matrix protein 2 | SRRM2 | 299.616 | 2 | 52.463924 | yes | 3.73E-07 | 521.28802 | 260.6897 |
| SCFESSPDPELK | 12 | Q9UQ35 | Serine/arginine repetitive matrix protein 2 | SRRM2 | 299.616 | 2 | 52.463924 | yes | 9.823E-08 | 698.30847 | 224.36008 |
| TSVPENHAQSR | 11 | Q9UQ35 | Serine/arginine repetitive matrix protein 2 | SRRM2 | 299.616 | 2 | 52.463924 | yes | 1.903E-07 | 613.29962 | 213.48326 |
| SLSYSPVER | 9 | Q9UQ35 | Serine/arginine repetitive matrix protein 2 | SRRM2 | 299.616 | 2 | 52.463924 | yes | 5.196E-07 | 519.26672 | 206.5451 |
| RLEAALVK | 8 | Q9UQ35 | Serine/arginine repetitive matrix protein 2 | SRRM2 | 299.616 | 2 | 52.463924 | yes | 1.188E-07 | 450.28726 | 204.71597 |
| GEFSASPMLK | 10 | Q9UQ35 | Serine/arginine repetitive matrix protein 2 | SRRM2 | 299.616 | 2 | 52.463924 | yes | 2.142E-07 | 533.7655 | 193.82968 |
| TPPSAPSQSR | 10 | Q9UQ35 | Serine/arginine repetitive matrix protein 2 | SRRM2 | 299.616 | 2 | 52.463924 | yes | 3.539E-07 | 514.26196 | 165.18129 |
| TPAAAAAMNLASI | 14 | Q9UQ35 | Serine/arginine repetitive matrix protein 2 | SRRM2 | 299.616 | 2 | 52.463924 | yes | 1.479E-07 | 671.35083 | 115.67521 |
| EGRPPEPTPAK | 11 | Q9UQ35 | Serine/arginine repetitive matrix protein 2 | SRRM2 | 299.616 | 2 | 52.463924 | yes | 1.155E-07 | 589.81183 | 104.92766 |

| | | | | | | | | | | | |
|---|---|---|---|---|---|---|---|---|---|---|---|
| VKPETPPR | 8 | Q9UQ35 | Serine/arginine repetitive matrix protein 2 | SRRM2 | 299.616 | 2 | 52.463924 | yes | 3.601E-07 | 462.26907 | 88.077316 |
| RVPSPTPAPK | 10 | Q9UQ35 | Serine/arginine repetitive matrix protein 2 | SRRM2 | 299.616 | 2 | 52.463924 | yes | 7.237E-07 | 525.30872 | 66.278267 |
| RPNPDILDHER | 11 | Q9UQ35 | Serine/arginine repetitive matrix protein 2 | SRRM2 | 299.616 | 3 | 52.463924 | yes | 1.245E-07 | 454.56888 | 31.010185 |
| RPNPDILDHER | 11 | Q9UQ35 | Serine/arginine repetitive matrix protein 2 | SRRM2 | 299.616 | 3 | 52.463924 | yes | 9.893E-08 | 681.34967 | 31.010185 |
| RPNPDILDHER | 11 | Q9UQ35 | Serine/arginine repetitive matrix protein 2 | SRRM2 | 299.616 | 2 | 52.463924 | yes | 1.245E-07 | 454.56888 | 31.010185 |
| RPNPDILDHER | 11 | Q9UQ35 | Serine/arginine repetitive matrix protein 2 | SRRM2 | 299.616 | 2 | 52.463924 | yes | 9.893E-08 | 681.34967 | 31.010185 |
| QSHSGSISPYPK | 12 | Q9UQ35 | Serine/arginine repetitive matrix protein 2 | SRRM2 | 299.616 | 2 | 52.463924 | yes | 1.115E-07 | 644.32001 | 53.896156 |
| SVSPCSNVESR | 11 | Q9UQ35 | Serine/arginine repetitive matrix protein 2 | SRRM2 | 299.616 | 2 | 52.463924 | yes | 1.509E-06 | 611.28003 | 45.325745 |
| PAVTETHQLAELN | 15 | Q9UQ35 | Serine/arginine repetitive matrix protein 2 | SRRM2 | 299.616 | 2 | 52.463924 | yes | 9.722E-08 | 840.43359 | 39.260574 |
| SSGHSSSELSPDA\ | 16 | Q9UQ35 | Serine/arginine repetitive matrix protein 2 | SRRM2 | 299.616 | 2 | 52.463924 | yes | 1.51E-07 | 808.87354 | 17.992527 |
| SSGHSSSELSPDA\ | 16 | Q9UQ35 | Serine/arginine repetitive matrix protein 2 | SRRM2 | 299.616 | 2 | 52.463924 | yes | 6.788E-07 | 539.58478 | 17.992527 |
| SSGHSSSELSPDA\ | 16 | Q9UQ35 | Serine/arginine repetitive matrix protein 2 | SRRM2 | 299.616 | 3 | 52.463924 | yes | 1.51E-07 | 808.87354 | 17.992527 |
| SSGHSSSELSPDA\ | 16 | Q9UQ35 | Serine/arginine repetitive matrix protein 2 | SRRM2 | 299.616 | 3 | 52.463924 | yes | 6.788E-07 | 539.58478 | 17.992527 |
| SEMEVQDAELK | 11 | Q9UQ80 | Proliferation-associated protein 2G4 OS= | PA2G4 | 43.787 | 2 | 53.469395 | yes | 9.722E-08 | 639.79773 | 338.841 |
| TIIQNPTDQQK | 11 | Q9UQ80 | Proliferation-associated protein 2G4 OS= | PA2G4 | 43.787 | 2 | 53.469395 | yes | 9.722E-08 | 643.34094 | 218.05316 |
| ITSGPFEPDLYK | 12 | Q9UQ80 | Proliferation-associated protein 2G4 OS= | PA2G4 | 43.787 | 2 | 53.469395 | yes | 9.728E-08 | 683.84808 | 201.62338 |
| AAHLCAEAALR | 11 | Q9UQ80 | Proliferation-associated protein 2G4 OS= | PA2G4 | 43.787 | 2 | 53.469395 | yes | 9.85E-08 | 591.80603 | 161.17543 |
| EGEFVAQFK | 9 | Q9UQ80 | Proliferation-associated protein 2G4 OS= | PA2G4 | 43.787 | 2 | 53.469395 | yes | 1.113E-07 | 527.76385 | 115.73641 |
| MGVVECAK | 8 | Q9UQ80 | Proliferation-associated protein 2G4 OS= | PA2G4 | 43.787 | 2 | 53.469395 | yes | 3.247E-07 | 447.2146 | 107.35319 |
| HELLQPFNVLYEK | 13 | Q9UQ80 | Proliferation-associated protein 2G4 OS= | PA2G4 | 43.787 | 2 | 53.469395 | yes | 9.722E-08 | 815.43542 | 35.952309 |
| TISPPTLGTLR | 11 | Q9UQ84 | Exonuclease 1 OS=Homo sapiens OX=96 | EXO1 | 94.104 | 2 | 52.271523 | yes | 1.844E-07 | 578.34021 | 280.38885 |
| SANVSSIWHR | 10 | Q9UQ84 | Exonuclease 1 OS=Homo sapiens OX=96 | EXO1 | 94.104 | 2 | 52.271523 | yes | 1.276E-07 | 578.79651 | 246.45782 |
| EASEPIHVR | 9 | Q9UQ84 | Exonuclease 1 OS=Homo sapiens OX=96 | EXO1 | 94.104 | 2 | 52.271523 | yes | 1.443E-07 | 519.27234 | 232.13373 |
| SCFSWSGGLGDFS | 14 | Q9UQ84 | Exonuclease 1 OS=Homo sapiens OX=96 | EXO1 | 94.104 | 2 | 52.271523 | yes | 9.722E-08 | 781.83826 | 229.40218 |
| VSIQPLDETAVTDI | 14 | Q9UQ84 | Exonuclease 1 OS=Homo sapiens OX=96 | EXO1 | 94.104 | 2 | 52.271523 | yes | 1.566E-07 | 758.39868 | 210.93123 |
| PILVFDGCTLPSK | 13 | Q9UQ84 | Exonuclease 1 OS=Homo sapiens OX=96 | EXO1 | 94.104 | 2 | 52.271523 | yes | 1.473E-07 | 723.88672 | 201.71584 |
| SSSADSLSTTK | 11 | Q9UQ84 | Exonuclease 1 OS=Homo sapiens OX=96 | EXO1 | 94.104 | 2 | 52.271523 | yes | 1.295E-07 | 542.26184 | 191.2637 |
| QLGDVFTEEK | 10 | Q9UQ84 | Exonuclease 1 OS=Homo sapiens OX=96 | EXO1 | 94.104 | 2 | 52.271523 | yes | 1.228E-07 | 583.29041 | 136.96063 |
| SINITHAMAHK | 11 | Q9UQ84 | Exonuclease 1 OS=Homo sapiens OX=96 | EXO1 | 94.104 | 2 | 52.271523 | yes | 1.197E-07 | 611.82172 | 83.134163 |
| LANNPDIVK | 9 | Q9UQ84 | Exonuclease 1 OS=Homo sapiens OX=96 | EXO1 | 94.104 | 2 | 52.271523 | yes | 9.722E-08 | 492.27966 | 81.310982 |
| PESGTVSDAPQLK | 13 | Q9UQ84 | Exonuclease 1 OS=Homo sapiens OX=96 | EXO1 | 94.104 | 2 | 52.271523 | yes | 1.166E-07 | 664.83826 | 79.677628 |
| MDQFGNGLEIDQA | 14 | Q9UQ84 | Exonuclease 1 OS=Homo sapiens OX=96 | EXO1 | 94.104 | 2 | 52.271523 | yes | 1.018E-07 | 797.36993 | 67.604881 |
| GIQGLLQFIK | 10 | Q9UQ84 | Exonuclease 1 OS=Homo sapiens OX=96 | EXO1 | 94.104 | 2 | 52.271523 | yes | 1.313E-07 | 558.84241 | 39.153664 |
| LLDSQSDQTSK | 11 | Q9UQ84 | Exonuclease 1 OS=Homo sapiens OX=96 | EXO1 | 94.104 | 2 | 52.271523 | yes | 6.752E-07 | 611.30151 | 35.448704 |
| FFCSSDSTDCVSNI | 14 | Q9UQ84 | Exonuclease 1 OS=Homo sapiens OX=96 | EXO1 | 94.104 | 2 | 52.271523 | yes | 2.437E-07 | 827.32959 | 27.62023 |
| LSDSYSNTLPVR | 12 | Q9UQB8 | Brain-specific angiogenesis inhibitor 1-as | BAIAP2 | 60.868 | 2 | 53.39932 | yes | 9.722E-08 | 676.34625 | 500.06137 |
| VLDSDGSDR | 9 | Q9UQB8 | Brain-specific angiogenesis inhibitor 1-as | BAIAP2 | 60.868 | 2 | 53.39932 | yes | 1.05E-07 | 482.22253 | 425.04642 |
| LHMSLQQGK | 9 | Q9UQB8 | Brain-specific angiogenesis inhibitor 1-as | BAIAP2 | 60.868 | 2 | 53.39932 | yes | 1.446E-07 | 521.27911 | 358.15335 |
| TIMEQFNPSLR | 11 | Q9UQB8 | Brain-specific angiogenesis inhibitor 1-as | BAIAP2 | 60.868 | 2 | 53.39932 | yes | 9.722E-08 | 668.3399 | 342.57764 |

| | | | | | | | | | | | |
|---|---|---|---|---|---|---|---|---|---|---|---|
| ELGDVLFQMAEVI | 14 | Q9UQB8 | Brain-specific angiogenesis inhibitor 1-as | BAIAP2 | 60.868 | 2 | 53.39932 | yes | 9.722E-08 | 822.41412 | 277.46713 |
| LPLWQQACADPSI | 13 | Q9UQB8 | Brain-specific angiogenesis inhibitor 1-as | BAIAP2 | 60.868 | 2 | 53.39932 | yes | 9.733E-06 | 757.37701 | 201.35799 |
| ELQYIDAISNK | 11 | Q9UQB8 | Brain-specific angiogenesis inhibitor 1-as | BAIAP2 | 60.868 | 2 | 53.39932 | yes | 9.722E-08 | 647.33789 | 198.72736 |
| MGELASESQGSK | 12 | Q9UQB8 | Brain-specific angiogenesis inhibitor 1-as | BAIAP2 | 60.868 | 2 | 53.39932 | yes | 9.789E-07 | 612.28229 | 170.63759 |
| SNLVISDPIPGAK | 13 | Q9UQB8 | Brain-specific angiogenesis inhibitor 1-as | BAIAP2 | 60.868 | 2 | 53.39932 | yes | 1.527E-07 | 655.86938 | 145.40265 |
| QGELENYVSDGYI | 13 | Q9UQB8 | Brain-specific angiogenesis inhibitor 1-as | BAIAP2 | 60.868 | 2 | 53.39932 | yes | 3.916E-07 | 751.34393 | 143.96126 |
| AFPAQTASGFK | 11 | Q9UQB8 | Brain-specific angiogenesis inhibitor 1-as | BAIAP2 | 60.868 | 2 | 53.39932 | yes | 2.738E-06 | 562.79041 | 137.04778 |
| SLSPPQSQSK | 10 | Q9UQB8 | Brain-specific angiogenesis inhibitor 1-as | BAIAP2 | 60.868 | 2 | 53.39932 | yes | 6.998E-07 | 529.77747 | 117.20795 |
| SFHNELLTQLEQK | 13 | Q9UQB8 | Brain-specific angiogenesis inhibitor 1-as | BAIAP2 | 60.868 | 2 | 53.39932 | yes | 3.721E-07 | 793.91229 | 114.70216 |
| ALAGVTYAAK | 10 | Q9UQB8 | Brain-specific angiogenesis inhibitor 1-as | BAIAP2 | 60.868 | 2 | 53.39932 | yes | 2.303E-06 | 482.77673 | 75.661644 |
| PLPVPPELAPFVGI | 14 | Q9UQB8 | Brain-specific angiogenesis inhibitor 1-as | BAIAP2 | 60.868 | 2 | 53.39932 | yes | 8.467E-05 | 744.93231 | 52.480358 |
| FCFLVEK | 7 | Q9UQB8 | Brain-specific angiogenesis inhibitor 1-as | BAIAP2 | 60.868 | 2 | 53.39932 | yes | 3.661E-07 | 471.7413 | 27.913082 |
| VETYLNENLR | 10 | Q9UQE7 | Structural maintenance of chromosomes p | SMC3 | 141.542 | 2 | 52.41912 | yes | 1.001E-07 | 625.82239 | 281.73007 |
| QVIIQGFR | 8 | Q9UQE7 | Structural maintenance of chromosomes p | SMC3 | 141.542 | 2 | 52.41912 | yes | 9.809E-08 | 480.78491 | 184.37454 |
| DQTIVDPFSSK | 11 | Q9UQE7 | Structural maintenance of chromosomes p | SMC3 | 141.542 | 2 | 52.41912 | yes | 9.722E-08 | 618.80896 | 165.71159 |
| ALDQFVNFSEQK | 12 | Q9UQE7 | Structural maintenance of chromosomes p | SMC3 | 141.542 | 2 | 52.41912 | yes | 9.722E-08 | 713.35406 | 154.22202 |
| IKLEGIITR | 9 | Q9UQE7 | Structural maintenance of chromosomes p | SMC3 | 141.542 | 2 | 52.41912 | yes | 9.722E-08 | 521.83459 | 118.1387 |
| ELGSLPQEAFEK | 12 | Q9UQE7 | Structural maintenance of chromosomes p | SMC3 | 141.542 | 2 | 52.41912 | yes | 3.685E-07 | 674.3432 | 114.74144 |
| EHMDAINHDTK | 11 | Q9UQE7 | Structural maintenance of chromosomes p | SMC3 | 141.542 | 2 | 52.41912 | yes | 9.722E-08 | 655.79333 | 113.8112 |
| MNLPGEVTFLPLN | 14 | Q9UQE7 | Structural maintenance of chromosomes p | SMC3 | 141.542 | 2 | 52.41912 | yes | 1.511E-07 | 786.92633 | 109.4847 |
| EMQQLSGGQK | 10 | Q9UQE7 | Structural maintenance of chromosomes p | SMC3 | 141.542 | 2 | 52.41912 | yes | 5.045E-07 | 553.26898 | 106.29352 |
| NDVMNLLESAGFS | 14 | Q9UQE7 | Structural maintenance of chromosomes p | SMC3 | 141.542 | 2 | 52.41912 | yes | 9.809E-08 | 776.87482 | 100.27374 |
| VSHIDVITAEMAK | 13 | Q9UQE7 | Structural maintenance of chromosomes p | SMC3 | 141.542 | 2 | 52.41912 | yes | 4.038E-07 | 707.37396 | 87.548637 |
| LPIDKEEVSLR | 11 | Q9UQE7 | Structural maintenance of chromosomes p | SMC3 | 141.542 | 2 | 52.41912 | yes | 9.974E-07 | 649.86938 | 33.344379 |
| LPIDKEEVSLR | 11 | Q9UQE7 | Structural maintenance of chromosomes p | SMC3 | 141.542 | 2 | 52.41912 | yes | 2.054E-07 | 433.582 | 33.344379 |
| LPIDKEEVSLR | 11 | Q9UQE7 | Structural maintenance of chromosomes p | SMC3 | 141.542 | 3 | 52.41912 | yes | 9.974E-07 | 649.86938 | 33.344379 |
| LPIDKEEVSLR | 11 | Q9UQE7 | Structural maintenance of chromosomes p | SMC3 | 141.542 | 3 | 52.41912 | yes | 2.054E-07 | 433.582 | 33.344379 |
| DLQDELAGNSEQI | 13 | Q9UQE7 | Structural maintenance of chromosomes p | SMC3 | 141.542 | 2 | 52.41912 | yes | 1.147E-07 | 737.84204 | 63.880138 |
| KYEAIQLTFK | 10 | Q9UQE7 | Structural maintenance of chromosomes p | SMC3 | 141.542 | 2 | 52.41912 | yes | 1.871E-07 | 620.85046 | 53.350391 |
| KAEEELGELEAK | 12 | Q9UQE7 | Structural maintenance of chromosomes p | SMC3 | 141.542 | 2 | 52.41912 | yes | 9.722E-08 | 673.34589 | 45.89286 |
| EENAEQQALAAK | 12 | Q9UQE7 | Structural maintenance of chromosomes p | SMC3 | 141.542 | 2 | 52.41912 | yes | 6.535E-07 | 651.32025 | 43.674404 |
| LFYHIVDSDEVSTI | 14 | Q9UQE7 | Structural maintenance of chromosomes p | SMC3 | 141.542 | 2 | 52.41912 | yes | 1.284E-07 | 826.91193 | 43.386482 |
| SMEVSTQLAR | 10 | Q9UQE7 | Structural maintenance of chromosomes p | SMC3 | 141.542 | 2 | 52.41912 | yes | 3.663E-07 | 561.28461 | 37.856323 |
| AFTMDCITLEGDQ | 17 | Q9UQE7 | Structural maintenance of chromosomes p | SMC3 | 141.542 | 3 | 52.41912 | yes | 1.041E-06 | 660.63654 | 34.688908 |
| INQMATAPDSQR | 12 | Q9UQE7 | Structural maintenance of chromosomes p | SMC3 | 141.542 | 2 | 52.41912 | yes | 4.902E-07 | 666.32227 | 24.18725 |
| TLQTMQNFQK | 10 | Q9UQN3 | Charged multivesicular body protein 2b C | CHMP2B | 23.907 | 2 | 47.96233 | yes | 1.223E-07 | 619.81354 | 118.85506 |
| ATISDEEIER | 10 | Q9UQN3 | Charged multivesicular body protein 2b C | CHMP2B | 23.907 | 2 | 47.96233 | yes | 2.483E-07 | 581.78296 | 118.44011 |
| VTSMSTQTK | 9 | Q9UQN3 | Charged multivesicular body protein 2b C | CHMP2B | 23.907 | 2 | 47.96233 | yes | 4.981E-07 | 491.74731 | 108.64153 |
| SVPGTTSSPLVGDI | 17 | Q9UQR0 | Sex comb on midleg-like protein 2 OS=H | SCML2 | 77.257 | 2 | 48.081367 | yes | 3.559E-07 | 821.43835 | 175.51297 |

| | | | | | | | | | | | |
|---|---|---|---|---|---|---|---|---|---|---|---|
| LVDSPDIQPVGTCI | 15 | Q9UQR0 | Sex comb on midleg-like protein 2 OS=H | SCML2 | 77.257 | 2 | 48.081367 | yes | 3.307E-07 | 829.40869 | 107.28944 |
| IVQACVDCALETK | 13 | Q9UQR0 | Sex comb on midleg-like protein 2 OS=H | SCML2 | 77.257 | 2 | 48.081367 | yes | 2.534E-07 | 753.86816 | 73.467499 |
| IKPPGPTAVPK | 11 | Q9UQR0 | Sex comb on midleg-like protein 2 OS=H | SCML2 | 77.257 | 2 | 48.081367 | yes | 1.532E-07 | 552.84241 | 45.688347 |
| ETGSISAPSECFR | 13 | Q9UQR0 | Sex comb on midleg-like protein 2 OS=H | SCML2 | 77.257 | 2 | 48.081367 | yes | 1.202E-07 | 720.82483 | 32.913147 |
| SSSLNSGNYLNPA( | 15 | Q9UQR0 | Sex comb on midleg-like protein 2 OS=H | SCML2 | 77.257 | 2 | 48.081367 | yes | 4.377E-07 | 820.3783 | 27.19833 |
| FLQQALDR | 8 | Q9UQR1 | Zinc finger protein 148 OS=Homo sapien | ZNF148 | 88.977 | 2 | 50.86614 | yes | 2.076E-07 | 495.772 | 413.78192 |
| TNYHLQR | 7 | Q9UQR1 | Zinc finger protein 148 OS=Homo sapien | ZNF148 | 88.977 | 2 | 50.86614 | yes | 2.303E-07 | 466.24084 | 184.96358 |
| YLQAASNNSR | 10 | Q9UQR1 | Zinc finger protein 148 OS=Homo sapien | ZNF148 | 88.977 | 2 | 50.86614 | yes | 1.281E-07 | 562.2782 | 110.2442 |
| ANMLQEYSK | 9 | Q9UQR1 | Zinc finger protein 148 OS=Homo sapien | ZNF148 | 88.977 | 2 | 50.86614 | yes | 2.53E-07 | 542.26062 | 99.232567 |
| ILTINEDGSLGLK | 13 | Q9UQR1 | Zinc finger protein 148 OS=Homo sapien | ZNF148 | 88.977 | 2 | 50.86614 | yes | 1.108E-07 | 686.88776 | 93.241074 |
| YAFELVDK | 8 | Q9UQR1 | Zinc finger protein 148 OS=Homo sapien | ZNF148 | 88.977 | 2 | 50.86614 | yes | 3.629E-07 | 492.75546 | 79.621933 |
| DKNDYLPLYSSST | 14 | Q9UQR1 | Zinc finger protein 148 OS=Homo sapien | ZNF148 | 88.977 | 2 | 50.86614 | yes | 9.819E-08 | 815.90161 | 59.661247 |
| QEITFTDVSEQLMI | 14 | Q9UQR1 | Zinc finger protein 148 OS=Homo sapien | ZNF148 | 88.977 | 2 | 50.86614 | yes | 9.722E-08 | 848.91418 | 58.024246 |
| QVYATMPINSFR | 12 | Q9UQR1 | Zinc finger protein 148 OS=Homo sapien | ZNF148 | 88.977 | 2 | 50.86614 | yes | 6.616E-07 | 713.86102 | 57.21656 |
| HVFIHTGEKPFQC: | 19 | Q9UQR1 | Zinc finger protein 148 OS=Homo sapien | ZNF148 | 88.977 | 4 | 50.86614 | yes | 5.548E-07 | 595.0213 | 50.02256 |
| LEGLFLK | 7 | Q9UQR1 | Zinc finger protein 148 OS=Homo sapien | ZNF148 | 88.977 | 2 | 50.86614 | yes | 1.657E-07 | 410.25238 | 16.434992 |
| LHVTALDYLAPYA | 14 | Q9Y221 | 60S ribosome subunit biogenesis protein | NIP7 | 20.463 | 2 | 50.852535 | yes | 1.028E-07 | 787.9325 | 188.84933 |
| LVSLGTCFGK | 10 | Q9Y221 | 60S ribosome subunit biogenesis protein | NIP7 | 20.463 | 2 | 50.852535 | yes | 8.411E-07 | 541.28918 | 125.0125 |
| VWIKPGAEQSFLY | 19 | Q9Y221 | 60S ribosome subunit biogenesis protein | NIP7 | 20.463 | 4 | 50.852535 | yes | 9.722E-08 | 547.29938 | 20.928835 |
| ILGVGISTGGR | 11 | Q9Y223 | Bifunctional UDP-N-acetylglucosamine 2 | GNE | 79.275 | 2 | 52.336323 | yes | 2.293E-07 | 515.30621 | 242.8273 |
| ILLAGCPSYDK | 11 | Q9Y223 | Bifunctional UDP-N-acetylglucosamine 2 | GNE | 79.275 | 2 | 52.336323 | yes | 2.042E-07 | 618.8183 | 210.66682 |
| EVGAFGTPVINLG' | 15 | Q9Y223 | Bifunctional UDP-N-acetylglucosamine 2 | GNE | 79.275 | 2 | 52.336323 | yes | 1.09E-07 | 765.91736 | 185.38797 |
| SIDLQEPLQK | 10 | Q9Y223 | Bifunctional UDP-N-acetylglucosamine 2 | GNE | 79.275 | 2 | 52.336323 | yes | 1.083E-07 | 585.8219 | 117.66305 |
| KLHDEDLLLVEGM | 17 | Q9Y223 | Bifunctional UDP-N-acetylglucosamine 2 | GNE | 79.275 | 3 | 52.336323 | yes | 9.722E-08 | 641.68005 | 104.33339 |
| ETGENVLHVR | 10 | Q9Y223 | Bifunctional UDP-N-acetylglucosamine 2 | GNE | 79.275 | 2 | 52.336323 | yes | 9.722E-08 | 577.30164 | 98.307854 |
| MIEQDDFDINTR | 12 | Q9Y223 | Bifunctional UDP-N-acetylglucosamine 2 | GNE | 79.275 | 2 | 52.336323 | yes | 7.873E-07 | 748.83795 | 88.156174 |
| EGIVLHSTK | 9 | Q9Y223 | Bifunctional UDP-N-acetylglucosamine 2 | GNE | 79.275 | 2 | 52.336323 | yes | 1.952E-07 | 492.27966 | 84.906288 |
| LIQEWNSVDLR | 11 | Q9Y223 | Bifunctional UDP-N-acetylglucosamine 2 | GNE | 79.275 | 2 | 52.336323 | yes | 1.539E-07 | 686.86462 | 72.157005 |
| SKDYIVALQHPVT | 17 | Q9Y223 | Bifunctional UDP-N-acetylglucosamine 2 | GNE | 79.275 | 3 | 52.336323 | yes | 1.771E-07 | 643.35443 | 62.427986 |
| LAHYHVCCTR | 10 | Q9Y223 | Bifunctional UDP-N-acetylglucosamine 2 | GNE | 79.275 | 2 | 52.336323 | yes | 9.722E-08 | 658.80298 | 28.849121 |
| LAHYHVCCTR | 10 | Q9Y223 | Bifunctional UDP-N-acetylglucosamine 2 | GNE | 79.275 | 2 | 52.336323 | yes | 5.145E-07 | 439.53775 | 28.849121 |
| LAHYHVCCTR | 10 | Q9Y223 | Bifunctional UDP-N-acetylglucosamine 2 | GNE | 79.275 | 3 | 52.336323 | yes | 9.722E-08 | 658.80298 | 28.849121 |
| LAHYHVCCTR | 10 | Q9Y223 | Bifunctional UDP-N-acetylglucosamine 2 | GNE | 79.275 | 3 | 52.336323 | yes | 5.145E-07 | 439.53775 | 28.849121 |
| MWLGDDVK | 8 | Q9Y223 | Bifunctional UDP-N-acetylglucosamine 2 | GNE | 79.275 | 2 | 52.336323 | yes | 3.584E-07 | 482.23386 | 54.445137 |
| LKPDIMIVHGDR | 12 | Q9Y223 | Bifunctional UDP-N-acetylglucosamine 2 | GNE | 79.275 | 3 | 52.336323 | yes | 9.255E-07 | 465.25885 | 45.899082 |
| TLVLFPNIDAGSK | 13 | Q9Y223 | Bifunctional UDP-N-acetylglucosamine 2 | GNE | 79.275 | 2 | 52.336323 | yes | 3.158E-05 | 687.88501 | 12.340944 |
| NIHSSDWPK | 9 | Q9Y224 | RNA transcription, translation and transp | RTRAF | 28.069 | 2 | 52.650043 | yes | 1.12E-06 | 542.26453 | 21971.129 |
| NFIVWLEDQK | 10 | Q9Y224 | RNA transcription, translation and transp | RTRAF | 28.069 | 2 | 52.650043 | yes | 1.15E-07 | 646.33771 | 18144.377 |
| LTQDAVAK | 8 | Q9Y224 | RNA transcription, translation and transp | RTRAF | 28.069 | 2 | 52.650043 | yes | 9.976E-08 | 423.23999 | 18046.158 |

| | | | | | | | | | | | |
|---|---|---|---|---|---|---|---|---|---|---|---|
| LEYGDNAEK | 9 | Q9Y224 | RNA transcription, translation and transp‹ | RTRAF | 28.069 | 2 | 52.650043 | yes | 2.248E-07 | 519.74054 | 14459.907 |
| EGLPVALDK | 9 | Q9Y224 | RNA transcription, translation and transp‹ | RTRAF | 28.069 | 2 | 52.650043 | yes | 1.102E-07 | 471.26874 | 13070.705 |
| YKDLVPDNSK | 10 | Q9Y224 | RNA transcription, translation and transp‹ | RTRAF | 28.069 | 2 | 52.650043 | yes | 1.66E-07 | 589.80621 | 12845.647 |
| LLHIEELR | 8 | Q9Y224 | RNA transcription, translation and transp‹ | RTRAF | 28.069 | 2 | 52.650043 | yes | 1.94E-07 | 511.80328 | 12676.749 |
| INEAIVAVQAIIAD | 16 | Q9Y224 | RNA transcription, translation and transp‹ | RTRAF | 28.069 | 2 | 52.650043 | yes | 1.096E-07 | 832.98273 | 4355.062 |
| INEAIVAVQAIIAD | 16 | Q9Y224 | RNA transcription, translation and transp‹ | RTRAF | 28.069 | 2 | 52.650043 | yes | 9.722E-08 | 555.65759 | 4355.062 |
| INEAIVAVQAIIAD | 16 | Q9Y224 | RNA transcription, translation and transp‹ | RTRAF | 28.069 | 3 | 52.650043 | yes | 1.096E-07 | 832.98273 | 4355.062 |
| INEAIVAVQAIIAD | 16 | Q9Y224 | RNA transcription, translation and transp‹ | RTRAF | 28.069 | 3 | 52.650043 | yes | 9.722E-08 | 555.65759 | 4355.062 |
| LTALDYHNPAGFN | 21 | Q9Y224 | RNA transcription, translation and transp‹ | RTRAF | 28.069 | 4 | 52.650043 | yes | 1.022E-07 | 625.29059 | 1446.3102 |
| HDDYLVMLK | 9 | Q9Y224 | RNA transcription, translation and transp‹ | RTRAF | 28.069 | 2 | 52.650043 | yes | 3.548E-07 | 567.28662 | 1196.3071 |
| KLTALDYHNPAGI | 22 | Q9Y224 | RNA transcription, translation and transp‹ | RTRAF | 28.069 | 4 | 52.650043 | yes | 9.722E-08 | 657.31433 | 978.26611 |
| AGVMALANLLQIC | 14 | Q9Y224 | RNA transcription, translation and transp‹ | RTRAF | 28.069 | 2 | 52.650043 | yes | 4.561E-07 | 757.42957 | 94.49675 |
| DLVPDNSK | 8 | Q9Y224 | RNA transcription, translation and transp‹ | RTRAF | 28.069 | 2 | 52.650043 | yes | 2.689E-05 | 444.22708 | 17.020546 |
| LLIVSTTPYSEK | 12 | Q9Y230 | RuvB-like 2 OS=Homo sapiens OX=960‹ | RUVBL2 | 51.157 | 2 | 56.650497 | yes | 1.036E-07 | 675.87939 | 989.85248 |
| AAGVVLEMIR | 10 | Q9Y230 | RuvB-like 2 OS=Homo sapiens OX=960‹ | RUVBL2 | 51.157 | 2 | 56.650497 | yes | 1.204E-07 | 529.80499 | 666.38239 |
| GLGLDDALEPR | 11 | Q9Y230 | RuvB-like 2 OS=Homo sapiens OX=960‹ | RUVBL2 | 51.157 | 2 | 56.650497 | yes | 9.968E-08 | 578.30383 | 649.06726 |
| FVQCPDGELQK | 11 | Q9Y230 | RuvB-like 2 OS=Homo sapiens OX=960‹ | RUVBL2 | 51.157 | 2 | 56.650497 | yes | 1.585E-07 | 660.81628 | 555.79907 |
| IGLETSLR | 8 | Q9Y230 | RuvB-like 2 OS=Homo sapiens OX=960‹ | RUVBL2 | 51.157 | 2 | 56.650497 | yes | 1.202E-07 | 444.76108 | 431.18454 |
| GTEVQVDDIK | 10 | Q9Y230 | RuvB-like 2 OS=Homo sapiens OX=960‹ | RUVBL2 | 51.157 | 2 | 56.650497 | yes | 1.162E-05 | 552.28259 | 379.01898 |
| QASQGMVGQLAA | 13 | Q9Y230 | RuvB-like 2 OS=Homo sapiens OX=960‹ | RUVBL2 | 51.157 | 2 | 56.650497 | yes | 9.722E-08 | 658.84064 | 349.89459 |
| TEALTQAFR | 9 | Q9Y230 | RuvB-like 2 OS=Homo sapiens OX=960‹ | RUVBL2 | 51.157 | 2 | 56.650497 | yes | 2.642E-07 | 518.77472 | 236.11508 |
| TTEMETIYDLGTK | 13 | Q9Y230 | RuvB-like 2 OS=Homo sapiens OX=960‹ | RUVBL2 | 51.157 | 2 | 56.650497 | yes | 9.722E-08 | 751.35815 | 221.86642 |
| EYQDAFLFNELK | 12 | Q9Y230 | RuvB-like 2 OS=Homo sapiens OX=960‹ | RUVBL2 | 51.157 | 2 | 56.650497 | yes | 9.722E-08 | 758.86957 | 197.11977 |
| AVLIAGQPGTGK | 12 | Q9Y230 | RuvB-like 2 OS=Homo sapiens OX=960‹ | RUVBL2 | 51.157 | 2 | 56.650497 | yes | 9.722E-08 | 556.32715 | 170.74376 |
| VYSLFLDESR | 10 | Q9Y230 | RuvB-like 2 OS=Homo sapiens OX=960‹ | RUVBL2 | 51.157 | 2 | 56.650497 | yes | 9.722E-08 | 614.81403 | 157.1125 |
| GTEVQVDDIKR | 11 | Q9Y230 | RuvB-like 2 OS=Homo sapiens OX=960‹ | RUVBL2 | 51.157 | 2 | 56.650497 | yes | 1.002E-07 | 630.33313 | 129.10123 |
| VQAGDVITIDK | 11 | Q9Y230 | RuvB-like 2 OS=Homo sapiens OX=960‹ | RUVBL2 | 51.157 | 2 | 56.650497 | yes | 6.514E-07 | 579.8219 | 112.63625 |
| DYDAMGSQTK | 10 | Q9Y230 | RuvB-like 2 OS=Homo sapiens OX=960‹ | RUVBL2 | 51.157 | 2 | 56.650497 | yes | 9.722E-08 | 558.2373 | 87.260002 |
| DKVQAGDVITIDK | 13 | Q9Y230 | RuvB-like 2 OS=Homo sapiens OX=960‹ | RUVBL2 | 51.157 | 2 | 56.650497 | yes | 1.136E-07 | 701.38281 | 69.311287 |
| TQGFLALFSGDTG | 16 | Q9Y230 | RuvB-like 2 OS=Homo sapiens OX=960‹ | RUVBL2 | 51.157 | 2 | 56.650497 | yes | 7.66E-07 | 842.43304 | 32.566593 |
| EVQSALSTAAADD | 15 | Q9Y232 | Chromodomain Y-like protein OS=Homo | CDYL | 66.482 | 2 | 48.058395 | yes | 1.206E-07 | 746.85992 | 63.707306 |
| ALNAVQPQLDVQ | 15 | Q9Y234 | Lipoyltransferase 1, mitochondrial OS=H | LIPT1 | 42.48 | 2 | 51.463158 | yes | 9.722E-08 | 798.44122 | 70.794174 |
| VLMASVQGSK | 10 | Q9Y251 | Heparanase OS=Homo sapiens OX=9606 | HPSE | 61.149 | 2 | 53.353386 | yes | 9.722E-08 | 510.28134 | 561.44788 |
| LYGPDVGQPR | 10 | Q9Y251 | Heparanase OS=Homo sapiens OX=9606 | HPSE | 61.149 | 2 | 53.353386 | yes | 9.722E-08 | 551.28802 | 470.21808 |
| MVDDQTLPPLMEI | 13 | Q9Y251 | Heparanase OS=Homo sapiens OX=9606 | HPSE | 61.149 | 2 | 53.353386 | yes | 9.722E-08 | 758.87292 | 453.96423 |
| PLGPHGLLSK | 10 | Q9Y251 | Heparanase OS=Homo sapiens OX=9606 | HPSE | 61.149 | 2 | 53.353386 | yes | 5.384E-07 | 509.80585 | 175.49704 |
| VFQVVESTRPGK | 12 | Q9Y251 | Heparanase OS=Homo sapiens OX=9606 | HPSE | 61.149 | 2 | 53.353386 | yes | 3.072E-07 | 673.875 | 175.16002 |
| GLSPAYLR | 8 | Q9Y251 | Heparanase OS=Homo sapiens OX=9606 | HPSE | 61.149 | 2 | 53.353386 | yes | 1.122E-07 | 438.75052 | 124.58368 |
| FLILLGSPK | 9 | Q9Y251 | Heparanase OS=Homo sapiens OX=9606 | HPSE | 61.149 | 2 | 53.353386 | yes | 9.107E-07 | 494.31549 | 96.743042 |

| | | | | | | | | | | | |
|---|---|---|---|---|---|---|---|---|---|---|---|
| VYLHCTNTDNPR | 12 | Q9Y251 | Heparanase OS=Homo sapiens OX=9606 | HPSE | 61.149 | 2 | 53.353386 | yes | 9.722E-08 | 745.34625 | 96.373604 |
| YLLRPLGPHGLLSI | 14 | Q9Y251 | Heparanase OS=Homo sapiens OX=9606 | HPSE | 61.149 | 3 | 53.353386 | yes | 1.498E-07 | 521.98383 | 43.54512 |
| YLLRPLGPHGLLSI | 14 | Q9Y251 | Heparanase OS=Homo sapiens OX=9606 | HPSE | 61.149 | 3 | 53.353386 | yes | 4.12E-06 | 782.47211 | 43.54512 |
| YLLRPLGPHGLLSI | 14 | Q9Y251 | Heparanase OS=Homo sapiens OX=9606 | HPSE | 61.149 | 2 | 53.353386 | yes | 1.498E-07 | 521.98383 | 43.54512 |
| YLLRPLGPHGLLSI | 14 | Q9Y251 | Heparanase OS=Homo sapiens OX=9606 | HPSE | 61.149 | 2 | 53.353386 | yes | 4.12E-06 | 782.47211 | 43.54512 |
| TDFLIFDPK | 9 | Q9Y251 | Heparanase OS=Homo sapiens OX=9606 | HPSE | 61.149 | 2 | 53.353386 | yes | 1.38E-07 | 548.28967 | 84.806885 |
| LEWPYQEQLLLR | 12 | Q9Y251 | Heparanase OS=Homo sapiens OX=9606 | HPSE | 61.149 | 2 | 53.353386 | yes | 3.632E-06 | 794.43011 | 57.799255 |
| TQGSGPAVTATK | 12 | Q9Y253 | DNA polymerase eta OS=Homo sapiens ( | POLH | 78.414 | 2 | 35.322266 | yes | 8.503E-07 | 559.29602 | 76.749672 |
| EASVEVMEIMSR | 12 | Q9Y253 | DNA polymerase eta OS=Homo sapiens ( | POLH | 78.414 | 2 | 35.322266 | yes | 2.495E-06 | 690.82849 | 73.892311 |
| ATTSLESFFQK | 11 | Q9Y253 | DNA polymerase eta OS=Homo sapiens ( | POLH | 78.414 | 2 | 35.322266 | yes | 3.099E-06 | 629.81934 | 38.693481 |
| MSHDAFTVIK | 10 | Q9Y253 | DNA polymerase eta OS=Homo sapiens ( | POLH | 78.414 | 2 | 35.322266 | yes | 4.315E-06 | 574.79205 | 34.213825 |
| VLENIELNK | 9 | Q9Y262 | Eukaryotic translation initiation factor 3 s | EIF3L | 66.727 | 2 | 53.51623 | yes | 1.429E-07 | 536.30585 | 1944.1034 |
| VYELQASR | 8 | Q9Y262 | Eukaryotic translation initiation factor 3 s | EIF3L | 66.727 | 2 | 53.51623 | yes | 9.722E-08 | 483.25616 | 1797.6362 |
| DMIHIADTK | 9 | Q9Y262 | Eukaryotic translation initiation factor 3 s | EIF3L | 66.727 | 2 | 53.51623 | yes | 1.15E-07 | 522.26312 | 1412.7491 |
| IDESIHLQLR | 10 | Q9Y262 | Eukaryotic translation initiation factor 3 s | EIF3L | 66.727 | 2 | 53.51623 | yes | 9.722E-08 | 612.34076 | 1347.1361 |
| NFIQYFHK | 8 | Q9Y262 | Eukaryotic translation initiation factor 3 s | EIF3L | 66.727 | 2 | 53.51623 | yes | 2.388E-07 | 548.78235 | 1252.9834 |
| LYTTMPVAK | 9 | Q9Y262 | Eukaryotic translation initiation factor 3 s | EIF3L | 66.727 | 2 | 53.51623 | yes | 1.736E-07 | 512.28082 | 1105.9282 |
| TVSDLIDQK | 9 | Q9Y262 | Eukaryotic translation initiation factor 3 s | EIF3L | 66.727 | 2 | 53.51623 | yes | 3.977E-06 | 509.77438 | 1032.1658 |
| VSSDVIDQK | 9 | Q9Y262 | Eukaryotic translation initiation factor 3 s | EIF3L | 66.727 | 2 | 53.51623 | yes | 2.918E-07 | 495.75876 | 764.65717 |
| LAGFLDLTEQEFR | 13 | Q9Y262 | Eukaryotic translation initiation factor 3 s | EIF3L | 66.727 | 2 | 53.51623 | yes | 1.343E-07 | 769.89612 | 735.375 |
| KSEEEIDFLR | 10 | Q9Y262 | Eukaryotic translation initiation factor 3 s | EIF3L | 66.727 | 2 | 53.51623 | yes | 1.728E-07 | 633.32227 | 555.52222 |
| VLENIELNKK | 10 | Q9Y262 | Eukaryotic translation initiation factor 3 s | EIF3L | 66.727 | 2 | 53.51623 | yes | 9.722E-08 | 600.35333 | 420.85739 |
| RYQDAIR | 7 | Q9Y262 | Eukaryotic translation initiation factor 3 s | EIF3L | 66.727 | 2 | 53.51623 | yes | 1.455E-07 | 461.24869 | 418.20642 |
| LHSLLGDYYQAIK | 13 | Q9Y262 | Eukaryotic translation initiation factor 3 s | EIF3L | 66.727 | 2 | 53.51623 | yes | 3.675E-07 | 760.909 | 254.52284 |
| EPFLQQLK | 8 | Q9Y262 | Eukaryotic translation initiation factor 3 s | EIF3L | 66.727 | 2 | 53.51623 | yes | 1.178E-07 | 501.78458 | 192.47212 |
| RYGDFFIR | 8 | Q9Y262 | Eukaryotic translation initiation factor 3 s | EIF3L | 66.727 | 2 | 53.51623 | yes | 1.621E-07 | 537.27997 | 143.15042 |
| SEEEIDFLR | 9 | Q9Y262 | Eukaryotic translation initiation factor 3 s | EIF3L | 66.727 | 2 | 53.51623 | yes | 9.722E-08 | 569.27478 | 106.70314 |
| QDLAYER | 7 | Q9Y262 | Eukaryotic translation initiation factor 3 s | EIF3L | 66.727 | 2 | 53.51623 | yes | 5.656E-07 | 447.71942 | 86.896011 |
| LDPSIFESLQK | 11 | Q9Y265 | RuvB-like 1 OS=Homo sapiens OX=960( | RUVBL1 | 50.228 | 2 | 52.924717 | yes | 9.722E-08 | 638.84283 | 578.35162 |
| GLGLDESGLAK | 11 | Q9Y265 | RuvB-like 1 OS=Homo sapiens OX=960( | RUVBL1 | 50.228 | 2 | 52.924717 | yes | 3.251E-07 | 530.28766 | 489.51303 |
| AVLLAGPPGTGK | 12 | Q9Y265 | RuvB-like 1 OS=Homo sapiens OX=960( | RUVBL1 | 50.228 | 2 | 52.924717 | yes | 1.419E-07 | 540.82422 | 402.40161 |
| YSVQLLTPANLLA | 14 | Q9Y265 | RuvB-like 1 OS=Homo sapiens OX=960( | RUVBL1 | 50.228 | 2 | 52.924717 | yes | 9.722E-08 | 765.94812 | 352.38763 |
| KTEVLMENFR | 10 | Q9Y265 | RuvB-like 1 OS=Homo sapiens OX=960( | RUVBL1 | 50.228 | 2 | 52.924717 | yes | 9.975E-08 | 633.82916 | 293.84067 |
| EACGVIVELIK | 11 | Q9Y265 | RuvB-like 1 OS=Homo sapiens OX=960( | RUVBL1 | 50.228 | 2 | 52.924717 | yes | 1.241E-07 | 615.84174 | 279.06119 |
| TMLYTPQEMK | 10 | Q9Y265 | RuvB-like 1 OS=Homo sapiens OX=960( | RUVBL1 | 50.228 | 2 | 52.924717 | yes | 9.722E-08 | 621.29889 | 249.42105 |
| QAASGLVGQENAI | 13 | Q9Y265 | RuvB-like 1 OS=Homo sapiens OX=960( | RUVBL1 | 50.228 | 2 | 52.924717 | yes | 1.039E-07 | 650.83386 | 217.64636 |
| TISHVIIGLK | 10 | Q9Y265 | RuvB-like 1 OS=Homo sapiens OX=960( | RUVBL1 | 50.228 | 2 | 52.924717 | yes | 1.257E-07 | 540.84241 | 90.451912 |
| EIIQDVTLHDLDV/ | 17 | Q9Y265 | RuvB-like 1 OS=Homo sapiens OX=960( | RUVBL1 | 50.228 | 3 | 52.924717 | yes | 1.274E-07 | 641.33746 | 35.960094 |
| AQTEGINISEEALN | 22 | Q9Y265 | RuvB-like 1 OS=Homo sapiens OX=960( | RUVBL1 | 50.228 | 3 | 52.924717 | yes | 2.409E-07 | 775.39581 | 32.866508 |

| | | | | | | | | | | | |
|---|---|---|---|---|---|---|---|---|---|---|---|
| LSDLDSETR | 9 | Q9Y266 | Nuclear migration protein nudC OS=Hom | NUDC | 38.243 | 2 | 50.087673 | yes | 3.353E-07 | 518.25128 | 196.10443 |
| LVSSDPEINTK | 11 | Q9Y266 | Nuclear migration protein nudC OS=Hom | NUDC | 38.243 | 2 | 50.087673 | yes | 6.145E-07 | 601.81677 | 107.3871 |
| VEESSWLIEDGK | 12 | Q9Y266 | Nuclear migration protein nudC OS=Hom | NUDC | 38.243 | 2 | 50.087673 | yes | 1.916E-07 | 696.33807 | 102.17181 |
| TDFFIGGEEGMAE | 14 | Q9Y266 | Nuclear migration protein nudC OS=Hom | NUDC | 38.243 | 2 | 50.087673 | yes | 1.171E-07 | 765.84265 | 95.827301 |
| ELTDEEAER | 9 | Q9Y266 | Nuclear migration protein nudC OS=Hom | NUDC | 38.243 | 2 | 50.087673 | yes | 9.905E-08 | 546.24622 | 93.354584 |
| GQPAIIDGELYNEV | 15 | Q9Y266 | Nuclear migration protein nudC OS=Hom | NUDC | 38.243 | 2 | 50.087673 | yes | 5.67E-07 | 823.42523 | 85.170189 |
| FMDQHPEMDFSK | 12 | Q9Y266 | Nuclear migration protein nudC OS=Hom | NUDC | 38.243 | 2 | 50.087673 | yes | 1.13E-07 | 756.3183 | 57.074856 |
| LTLDTIFVPNTGK | 13 | Q9Y277 | Voltage-dependent anion-selective chann | VDAC3 | 30.659 | 2 | 43.88254 | yes | 2.408E-07 | 709.89813 | 67.860046 |
| DPSAVHTVR | 9 | Q9Y283 | Inversin OS=Homo sapiens OX=9606 GN | INVS | 117.827 | 2 | 45.985485 | yes | 1.685E-07 | 491.25925 | 90.74501 |
| YTPLDYALLGER | 12 | Q9Y283 | Inversin OS=Homo sapiens OX=9606 GN | INVS | 117.827 | 2 | 45.985485 | yes | 3.422E-06 | 705.86682 | 86.772408 |
| TPLMYCVLADR | 11 | Q9Y283 | Inversin OS=Homo sapiens OX=9606 GN | INVS | 117.827 | 2 | 45.985485 | yes | 7.567E-07 | 669.83087 | 77.624191 |
| VVDSMEDEVQR | 11 | Q9Y285 | Phenylalanine--tRNA ligase alpha subuni | FARSA | 57.564 | 2 | 51.078705 | yes | 9.779E-08 | 653.80084 | 533.86725 |
| YGINNIR | 7 | Q9Y285 | Phenylalanine--tRNA ligase alpha subuni | FARSA | 57.564 | 2 | 51.078705 | yes | 1.681E-07 | 425.23251 | 311.63095 |
| DQHDTFFLR | 9 | Q9Y285 | Phenylalanine--tRNA ligase alpha subuni | FARSA | 57.564 | 2 | 51.078705 | yes | 9.722E-08 | 589.78308 | 290.28384 |
| VNLQMVYDSPLCI | 13 | Q9Y285 | Phenylalanine--tRNA ligase alpha subuni | FARSA | 57.564 | 2 | 51.078705 | yes | 1.72E-07 | 797.88947 | 216.94095 |
| RLEASDGGLDSAE | 30 | Q9Y285 | Phenylalanine--tRNA ligase alpha subuni | FARSA | 57.564 | 4 | 51.078705 | yes | 2.265E-07 | 756.633 | 148.59418 |
| LDAEPRPPPTQEA | 14 | Q9Y285 | Phenylalanine--tRNA ligase alpha subuni | FARSA | 57.564 | 2 | 51.078705 | yes | 9.903E-07 | 746.37555 | 145.32753 |
| HWELTAEGEEIAR | 13 | Q9Y285 | Phenylalanine--tRNA ligase alpha subuni | FARSA | 57.564 | 2 | 51.078705 | yes | 2.017E-07 | 770.87317 | 117.43396 |
| THSQGGYGSQGYI | 13 | Q9Y285 | Phenylalanine--tRNA ligase alpha subuni | FARSA | 57.564 | 2 | 51.078705 | yes | 9.722E-08 | 685.31018 | 108.7702 |
| KLLAEVTLK | 9 | Q9Y285 | Phenylalanine--tRNA ligase alpha subuni | FARSA | 57.564 | 2 | 51.078705 | yes | 1.001E-07 | 507.83151 | 77.782669 |
| LLAEVTLK | 8 | Q9Y285 | Phenylalanine--tRNA ligase alpha subuni | FARSA | 57.564 | 2 | 51.078705 | yes | 3.171E-07 | 443.78403 | 54.040329 |
| ENPPVKPDFSK | 11 | Q9Y294 | Histone chaperone ASF1A OS=Homo sap | ASF1A | 22.969 | 2 | 47.15927 | yes | 1.602E-07 | 629.32733 | 878.71808 |
| FHINWEDNTEK | 11 | Q9Y294 | Histone chaperone ASF1A OS=Homo sap | ASF1A | 22.969 | 2 | 47.15927 | yes | 1.388E-07 | 716.82825 | 518.21692 |
| IAEIEAEMAR | 10 | Q9Y295 | Developmentally-regulated GTP-binding | DRG1 | 40.543 | 2 | 50.572533 | yes | 9.759E-08 | 566.78699 | 1449.452 |
| IIENELEGFGIR | 12 | Q9Y295 | Developmentally-regulated GTP-binding | DRG1 | 40.543 | 2 | 50.572533 | yes | 1.131E-07 | 695.37225 | 1406.3639 |
| SDATADDLIDVVE | 16 | Q9Y295 | Developmentally-regulated GTP-binding | DRG1 | 40.543 | 2 | 50.572533 | yes | 1.641E-07 | 845.39996 | 1338.1284 |
| VYIPCIYVLNK | 11 | Q9Y295 | Developmentally-regulated GTP-binding | DRG1 | 40.543 | 2 | 50.572533 | yes | 1.024E-07 | 691.38104 | 1013.3501 |
| IHNADVTLR | 9 | Q9Y295 | Developmentally-regulated GTP-binding | DRG1 | 40.543 | 2 | 50.572533 | yes | 1.808E-07 | 519.78815 | 730.61285 |
| LNSKPPNIGFK | 11 | Q9Y295 | Developmentally-regulated GTP-binding | DRG1 | 40.543 | 2 | 50.572533 | yes | 1.49E-07 | 607.84821 | 708.78223 |
| IGFVGFPSVGK | 11 | Q9Y295 | Developmentally-regulated GTP-binding | DRG1 | 40.543 | 2 | 50.572533 | yes | 1.087E-07 | 554.31348 | 696.12463 |
| TTVEDFCMK | 9 | Q9Y295 | Developmentally-regulated GTP-binding | DRG1 | 40.543 | 2 | 50.572533 | yes | 1.136E-07 | 565.74646 | 455.66589 |
| GGGGGGPGEGFDV | 15 | Q9Y295 | Developmentally-regulated GTP-binding | DRG1 | 40.543 | 2 | 50.572533 | yes | 4.426E-07 | 631.29401 | 361.173 |
| YALVWGLSVK | 10 | Q9Y295 | Developmentally-regulated GTP-binding | DRG1 | 40.543 | 2 | 50.572533 | yes | 1.457E-07 | 568.32916 | 334.80258 |
| WNFDDLLEK | 9 | Q9Y295 | Developmentally-regulated GTP-binding | DRG1 | 40.543 | 2 | 50.572533 | yes | 1.93E-07 | 590.28766 | 299.27808 |
| ATAHHLGLLK | 10 | Q9Y295 | Developmentally-regulated GTP-binding | DRG1 | 40.543 | 2 | 50.572533 | yes | 1.043E-07 | 530.81671 | 296.66495 |
| IDQISIEELDIIYK | 14 | Q9Y295 | Developmentally-regulated GTP-binding | DRG1 | 40.543 | 2 | 50.572533 | yes | 9.819E-08 | 846.45874 | 197.75813 |
| DHTLEDEDVIQIVI | 14 | Q9Y295 | Developmentally-regulated GTP-binding | DRG1 | 40.543 | 2 | 50.572533 | yes | 1.283E-07 | 827.42017 | 165.77246 |
| VPHCVPISAHHR | 12 | Q9Y295 | Developmentally-regulated GTP-binding | DRG1 | 40.543 | 3 | 50.572533 | yes | 3.506E-07 | 470.57883 | 62.565685 |
| VPHCVPISAHHR | 12 | Q9Y295 | Developmentally-regulated GTP-binding | DRG1 | 40.543 | 3 | 50.572533 | yes | 4.722E-07 | 705.36456 | 62.565685 |

| | | | | | | | | | | | |
|---|---|---|---|---|---|---|---|---|---|---|---|
| VPHCVPISAHHR | 12 | Q9Y295 | Developmentally-regulated GTP-binding | DRG1 | 40.543 | 2 | 50.572533 | yes | 3.506E-07 | 470.57883 | 62.565685 |
| VPHCVPISAHHR | 12 | Q9Y295 | Developmentally-regulated GTP-binding | DRG1 | 40.543 | 2 | 50.572533 | yes | 4.722E-07 | 705.36456 | 62.565685 |
| APAGTLCLR | 9 | Q9Y297 | F-box/WD repeat-containing protein 1A ( | BTRC | 68.867 | 2 | 40.9121 | yes | 3.774E-07 | 479.76056 | 50.150997 |
| VWNTSTCEFVR | 11 | Q9Y297 | F-box/WD repeat-containing protein 1A ( | BTRC | 68.867 | 2 | 40.9121 | yes | 1.491E-06 | 699.82715 | 37.135223 |
| ELATVLSDQPGLL( | 16 | Q9Y2A7 | Nck-associated protein 1 OS=Homo sapie | NCKAP1 | 128.79 | 2 | 51.73884 | yes | 1.722E-07 | 819.45905 | 514.25104 |
| LVVENVDVLTQM | 13 | Q9Y2A7 | Nck-associated protein 1 OS=Homo sapie | NCKAP1 | 128.79 | 2 | 51.73884 | yes | 1.505E-07 | 758.41364 | 492.22192 |
| SLSELLGPYGMK | 12 | Q9Y2A7 | Nck-associated protein 1 OS=Homo sapie | NCKAP1 | 128.79 | 2 | 51.73884 | yes | 9.722E-08 | 647.83923 | 447.39359 |
| AAEDLFVNIR | 10 | Q9Y2A7 | Nck-associated protein 1 OS=Homo sapie | NCKAP1 | 128.79 | 2 | 51.73884 | yes | 1.082E-07 | 574.30896 | 435.82892 |
| KFPAVETR | 8 | Q9Y2A7 | Nck-associated protein 1 OS=Homo sapie | NCKAP1 | 128.79 | 2 | 51.73884 | yes | 2.374E-06 | 474.26907 | 427.51425 |
| AKPSYLIDK | 9 | Q9Y2A7 | Nck-associated protein 1 OS=Homo sapie | NCKAP1 | 128.79 | 2 | 51.73884 | yes | 2.934E-07 | 517.79767 | 309.87552 |
| MFQQCLELPSQSR | 13 | Q9Y2A7 | Nck-associated protein 1 OS=Homo sapie | NCKAP1 | 128.79 | 2 | 51.73884 | yes | 1.795E-07 | 812.38452 | 276.78006 |
| ASLGLADHR | 9 | Q9Y2A7 | Nck-associated protein 1 OS=Homo sapie | NCKAP1 | 128.79 | 2 | 51.73884 | yes | 2.295E-07 | 470.25397 | 256.12442 |
| EKPGVESMR | 9 | Q9Y2A7 | Nck-associated protein 1 OS=Homo sapie | NCKAP1 | 128.79 | 2 | 51.73884 | yes | 5.505E-07 | 516.76074 | 223.93977 |
| EYLTSHLEIR | 10 | Q9Y2A7 | Nck-associated protein 1 OS=Homo sapie | NCKAP1 | 128.79 | 2 | 51.73884 | yes | 2.451E-07 | 630.83276 | 197.59033 |
| QVEDGEVFDFR | 11 | Q9Y2A7 | Nck-associated protein 1 OS=Homo sapie | NCKAP1 | 128.79 | 2 | 51.73884 | yes | 1.091E-07 | 670.80951 | 197.46101 |
| NNNQQLAQLQK | 11 | Q9Y2A7 | Nck-associated protein 1 OS=Homo sapie | NCKAP1 | 128.79 | 2 | 51.73884 | yes | 9.722E-08 | 649.84418 | 148.34677 |
| SLAQEALR | 8 | Q9Y2A7 | Nck-associated protein 1 OS=Homo sapie | NCKAP1 | 128.79 | 2 | 51.73884 | yes | 3.282E-07 | 444.25089 | 145.10916 |
| PSELLTSVR | 9 | Q9Y2A7 | Nck-associated protein 1 OS=Homo sapie | NCKAP1 | 128.79 | 2 | 51.73884 | yes | 2.32E-07 | 501.28494 | 137.08185 |
| LQAYTSVSK | 9 | Q9Y2A7 | Nck-associated protein 1 OS=Homo sapie | NCKAP1 | 128.79 | 2 | 51.73884 | yes | 1.854E-06 | 498.77167 | 129.22563 |
| LTILNDR | 7 | Q9Y2A7 | Nck-associated protein 1 OS=Homo sapie | NCKAP1 | 128.79 | 2 | 51.73884 | no | 1.466E-07 | 422.74799 | 125.19031 |
| SLSLCNMFLDEMA | 14 | Q9Y2A7 | Nck-associated protein 1 OS=Homo sapie | NCKAP1 | 128.79 | 2 | 51.73884 | yes | 9.722E-08 | 829.883 | 124.69498 |
| EFLALASSSLLK | 12 | Q9Y2A7 | Nck-associated protein 1 OS=Homo sapie | NCKAP1 | 128.79 | 2 | 51.73884 | yes | 3.483E-07 | 639.86884 | 124.5485 |
| KYGPVMQR | 8 | Q9Y2A7 | Nck-associated protein 1 OS=Homo sapie | NCKAP1 | 128.79 | 2 | 51.73884 | yes | 1.16E-07 | 489.76312 | 109.9444 |
| LSSVDSVLK | 9 | Q9Y2A7 | Nck-associated protein 1 OS=Homo sapie | NCKAP1 | 128.79 | 2 | 51.73884 | yes | 1.491E-07 | 474.27402 | 95.152771 |
| MMEEFVPHSK | 10 | Q9Y2A7 | Nck-associated protein 1 OS=Homo sapie | NCKAP1 | 128.79 | 2 | 51.73884 | yes | 1.511E-07 | 617.78339 | 92.630905 |
| TSFDKPDQMAALI | 14 | Q9Y2A7 | Nck-associated protein 1 OS=Homo sapie | NCKAP1 | 128.79 | 2 | 51.73884 | yes | 2.102E-07 | 799.89777 | 72.158524 |
| WVLEHISK | 8 | Q9Y2C4 | Nuclease EXOG, mitochondrial OS=Hon | EXOG | 41.085 | 2 | 52.012054 | yes | 9.722E-08 | 506.28473 | 68.245934 |
| NAEIEPDDYFMSR | 13 | Q9Y2C4 | Nuclease EXOG, mitochondrial OS=Hon | EXOG | 41.085 | 2 | 52.012054 | yes | 5.858E-07 | 793.8432 | 29.188557 |
| CYTNHALSYDQAI | 13 | Q9Y2C4 | Nuclease EXOG, mitochondrial OS=Hon | EXOG | 41.085 | 2 | 52.012054 | yes | 5.285E-06 | 785.85138 | 2.8946614 |
| VLHDEILNKPMET | 15 | Q9Y2D2 | UDP-N-acetylglucosamine transporter OS | SLC35A3 | 35.985 | 3 | 39.785828 | yes | 4.382E-07 | 593.99396 | 50.574108 |
| LQNIIASR | 8 | Q9Y2D8 | Afadin- and alpha-actinin-binding protein | SSX2IP | 71.237 | 2 | 52.967133 | yes | 9.722E-08 | 457.77454 | 1225.6334 |
| ESMWDLSCETVR | 12 | Q9Y2D8 | Afadin- and alpha-actinin-binding protein | SSX2IP | 71.237 | 2 | 52.967133 | yes | 1.214E-07 | 756.82654 | 977.99597 |
| EMIGLQER | 8 | Q9Y2D8 | Afadin- and alpha-actinin-binding protein | SSX2IP | 71.237 | 2 | 52.967133 | yes | 2.801E-07 | 488.25003 | 501.8667 |
| NLLAQENVETQNI | 14 | Q9Y2D8 | Afadin- and alpha-actinin-binding protein | SSX2IP | 71.237 | 2 | 52.967133 | yes | 9.81E-08 | 807.42828 | 494.47 |
| LKEEWSLFK | 9 | Q9Y2D8 | Afadin- and alpha-actinin-binding protein | SSX2IP | 71.237 | 2 | 52.967133 | yes | 1.383E-07 | 590.32404 | 396.37955 |
| EMISLLSPQK | 10 | Q9Y2D8 | Afadin- and alpha-actinin-binding protein | SSX2IP | 71.237 | 2 | 52.967133 | yes | 1.042E-07 | 573.31537 | 303.85449 |
| QILMENAELK | 10 | Q9Y2D8 | Afadin- and alpha-actinin-binding protein | SSX2IP | 71.237 | 2 | 52.967133 | yes | 1.324E-07 | 594.8183 | 292.85876 |
| TISQYTSETK | 10 | Q9Y2D8 | Afadin- and alpha-actinin-binding protein | SSX2IP | 71.237 | 2 | 52.967133 | yes | 1.163E-07 | 579.28784 | 209.89767 |
| LKEQLETSR | 9 | Q9Y2D8 | Afadin- and alpha-actinin-binding protein | SSX2IP | 71.237 | 2 | 52.967133 | yes | 1.51E-07 | 552.3064 | 208.78864 |

| | | | | | | | | | | |
|---|---|---|---|---|---|---|---|---|---|---|
| SLPASPSTSDFCQT | 15 | Q9Y2D8 | Afadin- and alpha-actinin-binding protein | SSX2IP | 71.237 | 2 | 52.967133 | yes | 1.577E-07 | 827.38049 | 183.49426 |
| ILLNDYEYR | 9 | Q9Y2D8 | Afadin- and alpha-actinin-binding protein | SSX2IP | 71.237 | 2 | 52.967133 | yes | 3.119E-07 | 599.80878 | 183.46976 |
| VHLEGFNDEDVIS | 14 | Q9Y2D8 | Afadin- and alpha-actinin-binding protein | SSX2IP | 71.237 | 2 | 52.967133 | yes | 1.054E-07 | 815.39697 | 164.23093 |
| LELEIQQCK | 9 | Q9Y2D8 | Afadin- and alpha-actinin-binding protein | SSX2IP | 71.237 | 2 | 52.967133 | yes | 1.02E-06 | 580.80261 | 159.19652 |
| IAMDILNYVGR | 11 | Q9Y2D8 | Afadin- and alpha-actinin-binding protein | SSX2IP | 71.237 | 2 | 52.967133 | yes | 9.722E-08 | 632.83954 | 154.14313 |
| LHQLVMNK | 8 | Q9Y2D8 | Afadin- and alpha-actinin-binding protein | SSX2IP | 71.237 | 2 | 52.967133 | yes | 1.792E-07 | 491.77875 | 120.54416 |
| NLHQLLK | 7 | Q9Y2D8 | Afadin- and alpha-actinin-binding protein | SSX2IP | 71.237 | 2 | 52.967133 | yes | 3.128E-07 | 433.26636 | 95.156479 |
| KNLLAQENVETQN | 15 | Q9Y2D8 | Afadin- and alpha-actinin-binding protein | SSX2IP | 71.237 | 3 | 52.967133 | yes | 2.815E-07 | 581.31964 | 14.606962 |
| NLPGPASAMIGFK | 13 | Q9Y2F5 | Little elongation complex subunit 1 OS=H | ICE1 | 247.891 | 2 | 49.815067 | yes | 1.023E-07 | 651.84735 | 64.032349 |
| SHVYVGNISK | 10 | Q9Y2F5 | Little elongation complex subunit 1 OS=H | ICE1 | 247.891 | 2 | 49.815067 | yes | 4.567E-07 | 552.29584 | 58.318802 |
| ILDTMYPELSAR | 12 | Q9Y2F5 | Little elongation complex subunit 1 OS=H | ICE1 | 247.891 | 2 | 49.815067 | yes | 1.002E-07 | 704.86066 | 57.868549 |
| IINTDNLLTEYQK | 13 | Q9Y2F5 | Little elongation complex subunit 1 OS=H | ICE1 | 247.891 | 2 | 49.815067 | yes | 3.579E-07 | 782.91449 | 52.377769 |
| EDFLCQNVEK | 10 | Q9Y2F5 | Little elongation complex subunit 1 OS=H | ICE1 | 247.891 | 2 | 49.815067 | yes | 5.609E-07 | 641.2926 | 49.105091 |
| VTTSGLETFTATLF | 14 | Q9Y2F5 | Little elongation complex subunit 1 OS=H | ICE1 | 247.891 | 2 | 49.815067 | yes | 1.147E-07 | 748.90137 | 28.326653 |
| ACVNFLETSLSAK | 13 | Q9Y2F9 | BTB/POZ domain-containing protein 3 O | BTBD3 | 58.42 | 2 | 43.342583 | yes | 2.631E-07 | 720.36359 | 81.048065 |
| FVHVYSNR | 8 | Q9Y2G8 | DnaJ homolog subfamily C member 16 O | DNAJC16 | 90.592 | 2 | 52.986366 | yes | 6.425E-07 | 511.26434 | 671.16718 |
| QFVESLLPGNLVE | 14 | Q9Y2G8 | DnaJ homolog subfamily C member 16 O | DNAJC16 | 90.592 | 2 | 52.986366 | yes | 5.757E-07 | 786.93524 | 295.91882 |
| LLEGSLQR | 8 | Q9Y2G8 | DnaJ homolog subfamily C member 16 O | DNAJC16 | 90.592 | 2 | 52.986366 | yes | 3.662E-07 | 458.26654 | 291.22699 |
| ISFFHNAVVR | 10 | Q9Y2G8 | DnaJ homolog subfamily C member 16 O | DNAJC16 | 90.592 | 2 | 52.986366 | yes | 9.722E-08 | 595.32745 | 246.30563 |
| QIIDDFITR | 9 | Q9Y2G8 | DnaJ homolog subfamily C member 16 O | DNAJC16 | 90.592 | 2 | 52.986366 | yes | 2.164E-07 | 560.80347 | 214.27136 |
| AYEILSNEEK | 10 | Q9Y2G8 | DnaJ homolog subfamily C member 16 O | DNAJC16 | 90.592 | 2 | 52.986366 | yes | 3.255E-07 | 598.29572 | 211.95462 |
| TLEDPWIGSESDK | 13 | Q9Y2G8 | DnaJ homolog subfamily C member 16 O | DNAJC16 | 90.592 | 2 | 52.986366 | yes | 1.177E-07 | 738.84625 | 180.66647 |
| LFHELCPVK | 9 | Q9Y2G8 | DnaJ homolog subfamily C member 16 O | DNAJC16 | 90.592 | 2 | 52.986366 | yes | 2.193E-07 | 571.80499 | 159.51651 |
| EHINRPADVIQAR | 13 | Q9Y2G8 | DnaJ homolog subfamily C member 16 O | DNAJC16 | 90.592 | 3 | 52.986366 | yes | 3.254E-06 | 506.94269 | 89.637558 |
| YCVVLLTAETTK | 12 | Q9Y2G8 | DnaJ homolog subfamily C member 16 O | DNAJC16 | 90.592 | 2 | 52.986366 | yes | 2.165E-07 | 699.37085 | 77.828209 |
| LSLWMER | 7 | Q9Y2G8 | DnaJ homolog subfamily C member 16 O | DNAJC16 | 90.592 | 2 | 52.986366 | yes | 2.825E-07 | 467.74438 | 47.282532 |
| ITSDWCFSCIHIEPV | 17 | Q9Y2G8 | DnaJ homolog subfamily C member 16 O | DNAJC16 | 90.592 | 3 | 52.986366 | yes | 5.313E-07 | 726.67639 | 45.396217 |
| LACETAGVIR | 10 | Q9Y2G9 | Protein strawberry notch homolog 2 OS=I | SBNO2 | 150.276 | 2 | 46.152637 | yes | 2.045E-06 | 545.28967 | 33.872314 |
| ALWFSVSNDLK | 11 | Q9Y2G9 | Protein strawberry notch homolog 2 OS=I | SBNO2 | 150.276 | 2 | 46.152637 | no | 1.637E-07 | 640.33771 | 31.678429 |
| TQPVVLGTASADH | 24 | Q9Y2I8 | WD repeat-containing protein 37 OS=Ho | WDR37 | 54.666 | 3 | 31.727364 | yes | 1.745E-06 | 832.44287 | 22.719753 |
| LYQVMETER | 9 | Q9Y2K2 | Serine/threonine-protein kinase SIK3 OS= | SIK3 | 144.852 | 2 | 49.257538 | yes | 1.165E-07 | 584.78699 | 226.10809 |
| IGYYEIDR | 8 | Q9Y2K2 | Serine/threonine-protein kinase SIK3 OS= | SIK3 | 144.852 | 2 | 49.257538 | yes | 1.325E-07 | 514.75598 | 181.11122 |
| LPPTEFAQLIK | 11 | Q9Y2K2 | Serine/threonine-protein kinase SIK3 OS= | SIK3 | 144.852 | 2 | 49.257538 | yes | 8.75E-07 | 628.86609 | 164.91982 |
| LIAECQQLK | 9 | Q9Y2K2 | Serine/threonine-protein kinase SIK3 OS= | SIK3 | 144.852 | 2 | 49.257538 | yes | 4.297E-07 | 551.79987 | 145.2244 |
| TQLDEENLK | 9 | Q9Y2K2 | Serine/threonine-protein kinase SIK3 OS= | SIK3 | 144.852 | 2 | 49.257538 | yes | 1.179E-07 | 545.27478 | 139.83777 |
| AENLLLDANLNIK | 13 | Q9Y2K2 | Serine/threonine-protein kinase SIK3 OS= | SIK3 | 144.852 | 2 | 49.257538 | yes | 1.209E-07 | 720.90649 | 122.27238 |
| FSDGAASIQAFK | 12 | Q9Y2K2 | Serine/threonine-protein kinase SIK3 OS= | SIK3 | 144.852 | 2 | 49.257538 | yes | 1.488E-07 | 621.31171 | 122.11268 |
| MYGGQIDER | 9 | Q9Y2K2 | Serine/threonine-protein kinase SIK3 OS= | SIK3 | 144.852 | 2 | 49.257538 | yes | 1.625E-07 | 534.74255 | 111.07653 |
| EPVIGNCMDR | 10 | Q9Y2K2 | Serine/threonine-protein kinase SIK3 OS= | SIK3 | 144.852 | 2 | 49.257538 | yes | 1.051E-07 | 595.76825 | 98.178337 |

| | | | | | | | | | | | |
|---|---|---|---|---|---|---|---|---|---|---|---|
| DSNTLHLPTER | 11 | Q9Y2K2 | Serine/threonine-protein kinase SIK3 OS= | SIK3 | 144.852 | 2 | 49.257538 | yes | 3.356E-07 | 641.82294 | 82.07724 |
| LSMEQICK | 8 | Q9Y2K2 | Serine/threonine-protein kinase SIK3 OS= | SIK3 | 144.852 | 2 | 49.257538 | yes | 5.277E-07 | 504.74628 | 80.457329 |
| QIVTAVYFCHCR | 12 | Q9Y2K2 | Serine/threonine-protein kinase SIK3 OS= | SIK3 | 144.852 | 2 | 49.257538 | yes | 8.054E-07 | 777.37122 | 61.374023 |
| LLPPPAPGSPAAPA | 31 | Q9Y2K2 | Serine/threonine-protein kinase SIK3 OS= | SIK3 | 144.852 | 4 | 49.257538 | yes | 3.618E-06 | 717.39758 | 44.206768 |
| HMNQGDAGSLAPS | 22 | Q9Y2K2 | Serine/threonine-protein kinase SIK3 OS= | SIK3 | 144.852 | 3 | 49.257538 | yes | 2.93E-07 | 748.6759 | 35.235775 |
| QLQQECEQLQK | 11 | Q9Y2K2 | Serine/threonine-protein kinase SIK3 OS= | SIK3 | 144.852 | 2 | 49.257538 | yes | 3.786E-06 | 716.34845 | 8.9112911 |
| VLEVTDLPEGITR | 13 | Q9Y2K5 | R3H domain-containing protein 2 OS=Hc | R3HDM2 | 106.999 | 2 | 50.91504 | yes | 9.722E-08 | 721.3985 | 554.1535 |
| ASSFSGISILTR | 12 | Q9Y2K5 | R3H domain-containing protein 2 OS=Hc | R3HDM2 | 106.999 | 2 | 50.91504 | yes | 3.288E-07 | 619.84058 | 359.73404 |
| SASTDLGTADVVL | 15 | Q9Y2K5 | R3H domain-containing protein 2 OS=Hc | R3HDM2 | 106.999 | 2 | 50.91504 | yes | 9.766E-08 | 731.3808 | 307.79324 |
| KLVEESVNK | 9 | Q9Y2K5 | R3H domain-containing protein 2 OS=Hc | R3HDM2 | 106.999 | 2 | 50.91504 | yes | 1.232E-07 | 523.29803 | 57.715199 |
| NTEFQQR | 7 | Q9Y2K5 | R3H domain-containing protein 2 OS=Hc | R3HDM2 | 106.999 | 2 | 50.91504 | yes | 9.944E-07 | 461.7225 | 42.07618 |
| YYSMDQR | 7 | Q9Y2K5 | R3H domain-containing protein 2 OS=Hc | R3HDM2 | 106.999 | 2 | 50.91504 | yes | 4.633E-07 | 481.70547 | 30.544891 |
| LQAITASSANLR | 12 | Q9Y2K7 | Lysine-specific demethylase 2A OS=Hon | KDM2A | 132.793 | 2 | 56.33355 | yes | 1.526E-07 | 622.8515 | 155.21669 |
| LAGLDITDATLR | 12 | Q9Y2K7 | Lysine-specific demethylase 2A OS=Hon | KDM2A | 132.793 | 2 | 56.33355 | yes | 1.861E-07 | 629.8537 | 149.42239 |
| LTDQTLIYLR | 10 | Q9Y2K7 | Lysine-specific demethylase 2A OS=Hon | KDM2A | 132.793 | 2 | 56.33355 | yes | 9.722E-08 | 618.35333 | 146.61365 |
| VLVQHCPAR | 9 | Q9Y2K7 | Lysine-specific demethylase 2A OS=Hon | KDM2A | 132.793 | 2 | 56.33355 | yes | 5.987E-07 | 540.29254 | 101.83506 |
| GSYLTVTLQR | 10 | Q9Y2K7 | Lysine-specific demethylase 2A OS=Hon | KDM2A | 132.793 | 2 | 56.33355 | yes | 1.354E-07 | 569.31677 | 63.329369 |
| ILLEELANSDPK | 12 | Q9Y2K7 | Lysine-specific demethylase 2A OS=Hon | KDM2A | 132.793 | 2 | 56.33355 | yes | 3.952E-06 | 671.36664 | 62.32663 |
| CYQEDSSEK | 9 | Q9Y2K7 | Lysine-specific demethylase 2A OS=Hon | KDM2A | 132.793 | 2 | 56.33355 | yes | 1.292E-06 | 573.22443 | 59.542351 |
| AIVPQALSGIIK | 12 | Q9Y2K7 | Lysine-specific demethylase 2A OS=Hon | KDM2A | 132.793 | 2 | 56.33355 | yes | 1.859E-06 | 605.3819 | 59.396229 |
| ELHGTSIVPK | 10 | Q9Y2K7 | Lysine-specific demethylase 2A OS=Hon | KDM2A | 132.793 | 2 | 56.33355 | yes | 5.88E-07 | 540.80603 | 58.724171 |
| YSLTELNMAGCNK | 13 | Q9Y2K7 | Lysine-specific demethylase 2A OS=Hon | KDM2A | 132.793 | 2 | 56.33355 | yes | 3.778E-07 | 750.84473 | 32.186321 |
| GSWAQDGDESWM | 14 | Q9Y2K7 | Lysine-specific demethylase 2A OS=Hon | KDM2A | 132.793 | 2 | 56.33355 | yes | 7.315E-07 | 826.84155 | 18.312416 |
| SGTYLQGTFR | 10 | Q9Y2L1 | Exosome complex exonuclease RRP44 O | DIS3 | 109.003 | 2 | 50.474068 | yes | 1.03E-07 | 565.28546 | 139.34535 |
| IIVAIDGWPR | 10 | Q9Y2L1 | Exosome complex exonuclease RRP44 O | DIS3 | 109.003 | 2 | 50.474068 | yes | 1.193E-07 | 570.33221 | 121.89571 |
| IEDTVFHVFDK | 11 | Q9Y2L1 | Exosome complex exonuclease RRP44 O | DIS3 | 109.003 | 2 | 50.474068 | yes | 1.439E-06 | 675.34045 | 73.42585 |
| YADVIVHR | 8 | Q9Y2L1 | Exosome complex exonuclease RRP44 O | DIS3 | 109.003 | 2 | 50.474068 | yes | 2.742E-07 | 486.76672 | 73.269997 |
| SLTANPELIDR | 11 | Q9Y2L1 | Exosome complex exonuclease RRP44 O | DIS3 | 109.003 | 2 | 50.474068 | yes | 1.02E-07 | 614.83026 | 70.274139 |
| LACLSEEGNEIESC | 15 | Q9Y2L1 | Exosome complex exonuclease RRP44 O | DIS3 | 109.003 | 2 | 50.474068 | yes | 5.082E-07 | 818.38019 | 52.902252 |
| IDSANMNDDITTSI | 15 | Q9Y2L1 | Exosome complex exonuclease RRP44 O | DIS3 | 109.003 | 2 | 50.474068 | yes | 9.809E-08 | 833.39105 | 45.413441 |
| GALTLSSPEVR | 11 | Q9Y2L1 | Exosome complex exonuclease RRP44 O | DIS3 | 109.003 | 2 | 50.474068 | yes | 2.959E-07 | 565.31421 | 38.738731 |
| NVIVLQTVLQEVR | 13 | Q9Y2L1 | Exosome complex exonuclease RRP44 O | DIS3 | 109.003 | 2 | 50.474068 | yes | 1.739E-07 | 755.95123 | 35.938416 |
| YGLEGTVFFEEK | 12 | Q9Y2L1 | Exosome complex exonuclease RRP44 O | DIS3 | 109.003 | 2 | 50.474068 | yes | 1.215E-07 | 709.84552 | 35.535561 |
| GVTNDQVDPSVDY | 15 | Q9Y2P8 | RNA 3'-terminal phosphate cyclase-like p | RCL1 | 40.843 | 2 | 52.469498 | yes | 1.215E-07 | 793.40704 | 1181.6312 |
| QFGIDGESFELK | 12 | Q9Y2P8 | RNA 3'-terminal phosphate cyclase-like p | RCL1 | 40.843 | 2 | 52.469498 | yes | 9.722E-08 | 685.33533 | 782.2243 |
| LLLEEIYR | 8 | Q9Y2P8 | RNA 3'-terminal phosphate cyclase-like p | RCL1 | 40.843 | 2 | 52.469498 | yes | 1.464E-07 | 524.80548 | 726.25012 |
| VLKPIQLTDPGK | 12 | Q9Y2P8 | RNA 3'-terminal phosphate cyclase-like p | RCL1 | 40.843 | 2 | 52.469498 | yes | 9.809E-08 | 654.89795 | 353.79541 |
| VLKPIQLTDPGK | 12 | Q9Y2P8 | RNA 3'-terminal phosphate cyclase-like p | RCL1 | 40.843 | 2 | 52.469498 | yes | 3.108E-07 | 436.93439 | 353.79541 |
| VLKPIQLTDPGK | 12 | Q9Y2P8 | RNA 3'-terminal phosphate cyclase-like p | RCL1 | 40.843 | 3 | 52.469498 | yes | 9.809E-08 | 654.89795 | 353.79541 |

| | | | | | | | | | | |
|---|---|---|---|---|---|---|---|---|---|---|
| VLKPIQLTDPGK | 12 | Q9Y2P8 | RNA 3'-terminal phosphate cyclase-like p | RCL1 | 40.843 | 3 | 52.469498 | yes | 3.108E-07 | 436.93439 | 353.79541 |
| LVLSTLSGR | 9 | Q9Y2P8 | RNA 3'-terminal phosphate cyclase-like p | RCL1 | 40.843 | 2 | 52.469498 | yes | 2.042E-07 | 473.29001 | 494.44437 |
| VLMTCVGIGFSNL | 15 | Q9Y2P8 | RNA 3'-terminal phosphate cyclase-like p | RCL1 | 40.843 | 2 | 52.469498 | yes | 9.722E-08 | 813.4231 | 338.44824 |
| IETKPCGEELK | 11 | Q9Y2P8 | RNA 3'-terminal phosphate cyclase-like p | RCL1 | 40.843 | 2 | 52.469498 | yes | 1.084E-07 | 652.33173 | 285.63477 |
| DFEASFIR | 8 | Q9Y2P8 | RNA 3'-terminal phosphate cyclase-like p | RCL1 | 40.843 | 2 | 52.469498 | yes | 4.539E-07 | 492.74289 | 223.67978 |
| FIPDIYIYTDHMK | 13 | Q9Y2P8 | RNA 3'-terminal phosphate cyclase-like p | RCL1 | 40.843 | 2 | 52.469498 | yes | 2.227E-07 | 828.41052 | 169.83134 |
| ATALPLLK | 8 | Q9Y2P8 | RNA 3'-terminal phosphate cyclase-like p | RCL1 | 40.843 | 2 | 52.469498 | yes | 1.083E-07 | 413.77347 | 52.330177 |
| KVLKPIQLTDPGK | 13 | Q9Y2P8 | RNA 3'-terminal phosphate cyclase-like p | RCL1 | 40.843 | 2 | 52.469498 | yes | 9.722E-08 | 718.94537 | 26.723585 |
| AGMSAEQAQGLLI | 14 | Q9Y2Q3 | Glutathione S-transferase kappa 1 OS=Hc | GSTK1 | 25.497 | 2 | 41.736404 | yes | 3.364E-07 | 716.85864 | 84.45639 |
| FILMDCMEGR | 10 | Q9Y2Q5 | Ragulator complex protein LAMTOR2 O | LAMTOR2 | 13.508 | 2 | 46.20241 | yes | 1.623E-07 | 636.28271 | 65.160797 |
| ELLASAPTGSGK | 12 | Q9Y2R4 | Probable ATP-dependent RNA helicase D | DDX52 | 67.467 | 2 | 50.61796 | yes | 1.223E-07 | 565.80621 | 480.09192 |
| MDVHDLFR | 8 | Q9Y2R4 | Probable ATP-dependent RNA helicase D | DDX52 | 67.467 | 2 | 50.61796 | yes | 2.641E-07 | 537.75549 | 455.75696 |
| DNTVHSFR | 8 | Q9Y2R4 | Probable ATP-dependent RNA helicase D | DDX52 | 67.467 | 2 | 50.61796 | yes | 2.28E-07 | 488.23578 | 279.25287 |
| ELASQIHR | 8 | Q9Y2R4 | Probable ATP-dependent RNA helicase D | DDX52 | 67.467 | 2 | 50.61796 | yes | 4.848E-07 | 477.26178 | 235.29797 |
| AITFFTEDDKPLLR | 14 | Q9Y2R4 | Probable ATP-dependent RNA helicase D | DDX52 | 67.467 | 2 | 50.61796 | yes | 2.511E-07 | 833.44598 | 76.982056 |
| AITFFTEDDKPLLR | 14 | Q9Y2R4 | Probable ATP-dependent RNA helicase D | DDX52 | 67.467 | 2 | 50.61796 | yes | 1.069E-07 | 555.96643 | 76.982056 |
| AITFFTEDDKPLLR | 14 | Q9Y2R4 | Probable ATP-dependent RNA helicase D | DDX52 | 67.467 | 3 | 50.61796 | yes | 2.511E-07 | 833.44598 | 76.982056 |
| AITFFTEDDKPLLR | 14 | Q9Y2R4 | Probable ATP-dependent RNA helicase D | DDX52 | 67.467 | 3 | 50.61796 | yes | 1.069E-07 | 555.96643 | 76.982056 |
| ALIISPTR | 8 | Q9Y2R4 | Probable ATP-dependent RNA helicase D | DDX52 | 67.467 | 2 | 50.61796 | yes | 3.408E-07 | 435.77402 | 119.70798 |
| KFDILVTTPNR | 11 | Q9Y2R4 | Probable ATP-dependent RNA helicase D | DDX52 | 67.467 | 2 | 50.61796 | yes | 1.084E-07 | 652.37207 | 114.58865 |
| KEESLTER | 8 | Q9Y2R4 | Probable ATP-dependent RNA helicase D | DDX52 | 67.467 | 2 | 50.61796 | yes | 1.771E-07 | 496.25638 | 108.46162 |
| LNLDNVISVSIGAF | 14 | Q9Y2R4 | Probable ATP-dependent RNA helicase D | DDX52 | 67.467 | 2 | 50.61796 | yes | 9.734E-08 | 735.91736 | 95.612389 |
| ESISTTPK | 8 | Q9Y2R4 | Probable ATP-dependent RNA helicase D | DDX52 | 67.467 | 2 | 50.61796 | yes | 2.379E-07 | 431.72946 | 54.687214 |
| GVNLVINYDFPTS | 20 | Q9Y2R4 | Probable ATP-dependent RNA helicase D | DDX52 | 67.467 | 3 | 50.61796 | yes | 1.121E-07 | 775.06207 | 50.626251 |
| ELFHELIYEGINVD | 20 | Q9Y2R4 | Probable ATP-dependent RNA helicase D | DDX52 | 67.467 | 3 | 50.61796 | yes | 1.464E-07 | 799.41302 | 24.654402 |
| DQLASIFLACTSHK | 14 | Q9Y2R4 | Probable ATP-dependent RNA helicase D | DDX52 | 67.467 | 2 | 50.61796 | yes | 0.0001337 | 795.90088 | 15.930445 |
| GVVLFPWQAR | 10 | Q9Y2S7 | Polymerase delta-interacting protein 2 OS | POLDIP2 | 42.034 | 2 | 49.55003 | yes | 1.154E-06 | 586.8324 | 120.2567 |
| IFSLSGTLETVR | 12 | Q9Y2S7 | Polymerase delta-interacting protein 2 OS | POLDIP2 | 42.034 | 2 | 49.55003 | yes | 1.022E-07 | 661.86938 | 87.299568 |
| FERPDGSHFDVR | 12 | Q9Y2S7 | Polymerase delta-interacting protein 2 OS | POLDIP2 | 42.034 | 3 | 49.55003 | yes | 4.502E-07 | 487.90051 | 78.025276 |
| THTYYQVLIDAR | 12 | Q9Y2S7 | Polymerase delta-interacting protein 2 OS | POLDIP2 | 42.034 | 2 | 49.55003 | yes | 1.369E-07 | 740.38318 | 66.411316 |
| LENLDSDVVQLR | 12 | Q9Y2S7 | Polymerase delta-interacting protein 2 OS | POLDIP2 | 42.034 | 2 | 49.55003 | yes | 4.566E-07 | 700.87262 | 54.992157 |
| VLETVGVFEVPK | 12 | Q9Y2S7 | Polymerase delta-interacting protein 2 OS | POLDIP2 | 42.034 | 2 | 49.55003 | yes | 8.862E-07 | 658.87665 | 12.285917 |
| IQQLAISGLK | 10 | Q9Y2T2 | AP-3 complex subunit mu-1 OS=Homo sa | AP3M1 | 46.94 | 2 | 54.24669 | no | 1.26E-07 | 535.83203 | 430.09286 |
| VSSQNLVAIPVYV | 14 | Q9Y2T2 | AP-3 complex subunit mu-1 OS=Homo sa | AP3M1 | 46.94 | 2 | 54.24669 | yes | 9.722E-08 | 758.94031 | 200.62737 |
| LLDDVSFHPCIR | 12 | Q9Y2T2 | AP-3 complex subunit mu-1 OS=Homo sa | AP3M1 | 46.94 | 3 | 54.24669 | yes | 3.23E-07 | 491.25024 | 71.759308 |
| ELIKPPTILR | 10 | Q9Y2T2 | AP-3 complex subunit mu-1 OS=Homo sa | AP3M1 | 46.94 | 2 | 54.24669 | no | 1.846E-06 | 590.37665 | 25.624058 |
| FQNIDFAEEVYTR | 13 | Q9Y2T3 | Guanine deaminase OS=Homo sapiens O | GDA | 51.004 | 2 | 45.23377 | yes | 8.086E-07 | 816.38867 | 72.900581 |
| IVFLEEASQQEK | 12 | Q9Y2T3 | Guanine deaminase OS=Homo sapiens O | GDA | 51.004 | 2 | 45.23377 | yes | 1.915E-07 | 710.86957 | 49.678944 |
| PVSGDLSYLR | 10 | Q9Y2U8 | Inner nuclear membrane protein Man1 OS | LEMD3 | 99.998 | 2 | 42.718018 | yes | 3.429E-07 | 553.79565 | 74.767815 |

| | | | | | | | | | | | |
|---|---|---|---|---|---|---|---|---|---|---|---|
| TLSVQEAAAYLK | 12 | Q9Y2U8 | Inner nuclear membrane protein Man1 OS | LEMD3 | 99.998 | 2 | 42.718018 | yes | 3.574E-07 | 647.35608 | 47.945251 |
| WIQPSASCDK | 10 | Q9Y2U8 | Inner nuclear membrane protein Man1 OS | LEMD3 | 99.998 | 2 | 42.718018 | yes | 3.721E-05 | 596.27679 | 26.729361 |
| AVDFLAANESR | 11 | Q9Y2U8 | Inner nuclear membrane protein Man1 OS | LEMD3 | 99.998 | 2 | 42.718018 | yes | 4.622E-07 | 596.80145 | 25.735718 |
| PLMSFWCR | 8 | Q9Y2U8 | Inner nuclear membrane protein Man1 OS | LEMD3 | 99.998 | 2 | 42.718018 | yes | 2.903E-07 | 548.75696 | 17.174212 |
| YGLSPGPVTESTRI | 18 | Q9Y2U8 | Inner nuclear membrane protein Man1 OS | LEMD3 | 99.998 | 3 | 42.718018 | yes | 4.326E-07 | 655.35443 | 13.726499 |
| IDISPSTFR | 9 | Q9Y2W1 | Thyroid hormone receptor-associated prot | THRAP3 | 108.666 | 2 | 53.634094 | yes | 0.0001085 | 518.2771 | 12709.533 |
| PSGLLAQER | 9 | Q9Y2W1 | Thyroid hormone receptor-associated prot | THRAP3 | 108.666 | 2 | 53.634094 | yes | 1.128E-07 | 485.76944 | 2748.499 |
| WEGLVYAPPGK | 11 | Q9Y2W1 | Thyroid hormone receptor-associated prot | THRAP3 | 108.666 | 2 | 53.634094 | yes | 1.127E-07 | 608.82166 | 2435.0874 |
| SPLQSVVVR | 9 | Q9Y2W1 | Thyroid hormone receptor-associated prot | THRAP3 | 108.666 | 2 | 53.634094 | yes | 2.16E-07 | 492.79547 | 2387.7393 |
| MDSFDEDLAR | 10 | Q9Y2W1 | Thyroid hormone receptor-associated prot | THRAP3 | 108.666 | 2 | 53.634094 | yes | 4.47E-07 | 599.75586 | 1127.8799 |
| MDSFDEDLAR | 10 | Q9Y2W1 | Thyroid hormone receptor-associated prot | THRAP3 | 108.666 | 2 | 53.634094 | yes | 9.722E-08 | 607.75336 | 1127.8799 |
| YKDDPVDLR | 9 | Q9Y2W1 | Thyroid hormone receptor-associated prot | THRAP3 | 108.666 | 2 | 53.634094 | yes | 1.951E-07 | 560.78528 | 1976.8533 |
| GSFSDTGLGDGK | 12 | Q9Y2W1 | Thyroid hormone receptor-associated prot | THRAP3 | 108.666 | 2 | 53.634094 | yes | 9.722E-08 | 570.76202 | 1491.0372 |
| EESAASGGAAYTK | 13 | Q9Y2W1 | Thyroid hormone receptor-associated prot | THRAP3 | 108.666 | 2 | 53.634094 | yes | 9.722E-08 | 621.28583 | 1302.9047 |
| LRDDFEK | 7 | Q9Y2W1 | Thyroid hormone receptor-associated prot | THRAP3 | 108.666 | 2 | 53.634094 | yes | 1.308E-07 | 461.73508 | 898.8056 |
| TDSEKPFR | 8 | Q9Y2W1 | Thyroid hormone receptor-associated prot | THRAP3 | 108.666 | 2 | 53.634094 | yes | 8.188E-07 | 490.24582 | 850.77045 |
| SIFQHIQSAQSQR | 13 | Q9Y2W1 | Thyroid hormone receptor-associated prot | THRAP3 | 108.666 | 2 | 53.634094 | yes | 1.225E-07 | 765.39459 | 358.75488 |
| SIFQHIQSAQSQR | 13 | Q9Y2W1 | Thyroid hormone receptor-associated prot | THRAP3 | 108.666 | 2 | 53.634094 | yes | 2.092E-06 | 510.59882 | 358.75488 |
| SIFQHIQSAQSQR | 13 | Q9Y2W1 | Thyroid hormone receptor-associated prot | THRAP3 | 108.666 | 3 | 53.634094 | yes | 1.225E-07 | 765.39459 | 358.75488 |
| SIFQHIQSAQSQR | 13 | Q9Y2W1 | Thyroid hormone receptor-associated prot | THRAP3 | 108.666 | 3 | 53.634094 | yes | 2.092E-06 | 510.59882 | 358.75488 |
| EEEWDPEYTPK | 11 | Q9Y2W1 | Thyroid hormone receptor-associated prot | THRAP3 | 108.666 | 2 | 53.634094 | no | 9.722E-08 | 711.80664 | 538.02869 |
| EHHFGSSGMTLHE | 14 | Q9Y2W1 | Thyroid hormone receptor-associated prot | THRAP3 | 108.666 | 3 | 53.634094 | yes | 1.05E-07 | 542.24768 | 230.94739 |
| EHHFGSSGMTLHE | 14 | Q9Y2W1 | Thyroid hormone receptor-associated prot | THRAP3 | 108.666 | 3 | 53.634094 | yes | 1.146E-07 | 812.86786 | 230.94739 |
| EHHFGSSGMTLHE | 14 | Q9Y2W1 | Thyroid hormone receptor-associated prot | THRAP3 | 108.666 | 2 | 53.634094 | yes | 1.05E-07 | 542.24768 | 230.94739 |
| EHHFGSSGMTLHE | 14 | Q9Y2W1 | Thyroid hormone receptor-associated prot | THRAP3 | 108.666 | 2 | 53.634094 | yes | 1.146E-07 | 812.86786 | 230.94739 |
| YYLHDDR | 7 | Q9Y2W1 | Thyroid hormone receptor-associated prot | THRAP3 | 108.666 | 2 | 53.634094 | yes | 2E-06 | 491.22488 | 412.41241 |
| ASAVSELSPR | 10 | Q9Y2W1 | Thyroid hormone receptor-associated prot | THRAP3 | 108.666 | 2 | 53.634094 | yes | 1.672E-07 | 508.77219 | 369.58167 |
| SPSELFAQHIVTIV | 18 | Q9Y2W1 | Thyroid hormone receptor-associated prot | THRAP3 | 108.666 | 3 | 53.634094 | yes | 3.278E-07 | 681.37756 | 286.33316 |
| ESEFDDEPK | 9 | Q9Y2W1 | Thyroid hormone receptor-associated prot | THRAP3 | 108.666 | 2 | 53.634094 | yes | 1.153E-07 | 548.22748 | 263.13464 |
| EAQVNVR | 7 | Q9Y2W1 | Thyroid hormone receptor-associated prot | THRAP3 | 108.666 | 2 | 53.634094 | yes | 1.154E-07 | 408.22214 | 238.54056 |
| NREEEWDPEYTPK | 13 | Q9Y2W1 | Thyroid hormone receptor-associated prot | THRAP3 | 108.666 | 2 | 53.634094 | yes | 1.185E-07 | 846.87866 | 237.11874 |
| SPPSTGSTYGSSQK | 14 | Q9Y2W1 | Thyroid hormone receptor-associated prot | THRAP3 | 108.666 | 2 | 53.634094 | yes | 9.722E-08 | 692.323 | 212.3819 |
| KYYLHDDR | 8 | Q9Y2W1 | Thyroid hormone receptor-associated prot | THRAP3 | 108.666 | 2 | 53.634094 | yes | 9.722E-08 | 555.27234 | 203.96414 |
| EESTTGFDK | 9 | Q9Y2W1 | Thyroid hormone receptor-associated prot | THRAP3 | 108.666 | 2 | 53.634094 | yes | 3.372E-07 | 507.22473 | 202.53333 |
| KESEFDDEPK | 10 | Q9Y2W1 | Thyroid hormone receptor-associated prot | THRAP3 | 108.666 | 2 | 53.634094 | yes | 9.722E-08 | 612.27496 | 125.66485 |
| TEELEEESFPER | 12 | Q9Y2W1 | Thyroid hormone receptor-associated prot | THRAP3 | 108.666 | 2 | 53.634094 | yes | 9.722E-08 | 747.83337 | 106.44916 |
| EQTFSGGTSQDTK | 13 | Q9Y2W1 | Thyroid hormone receptor-associated prot | THRAP3 | 108.666 | 2 | 53.634094 | yes | 1.002E-07 | 693.31262 | 82.21022 |
| EEKDNIQPTTE | 11 | Q9Y2W1 | Thyroid hormone receptor-associated prot | THRAP3 | 108.666 | 2 | 53.634094 | yes | 2.537E-07 | 652.30426 | 76.446426 |
| KTEELEEESFPER | 13 | Q9Y2W1 | Thyroid hormone receptor-associated prot | THRAP3 | 108.666 | 2 | 53.634094 | yes | 9.722E-08 | 811.88086 | 27.705055 |

| | | | | | | | | | | | |
|---|---|---|---|---|---|---|---|---|---|---|---|
| KTEELEEESFPER | 13 | Q9Y2W1 | Thyroid hormone receptor-associated prot | THRAP3 | 108.666 | 2 | 53.634094 | yes | 3.265E-07 | 541.58966 | 27.705055 |
| KTEELEEESFPER | 13 | Q9Y2W1 | Thyroid hormone receptor-associated prot | THRAP3 | 108.666 | 3 | 53.634094 | yes | 9.722E-08 | 811.88086 | 27.705055 |
| KTEELEEESFPER | 13 | Q9Y2W1 | Thyroid hormone receptor-associated prot | THRAP3 | 108.666 | 3 | 53.634094 | yes | 3.265E-07 | 541.58966 | 27.705055 |
| AQLSQYFDAVK | 11 | Q9Y2W2 | WW domain-binding protein 11 OS=Hon | WBP11 | 69.998 | 2 | 44.651886 | yes | 3.249E-07 | 635.32733 | 151.52251 |
| SGPSVPVSVQTK | 12 | Q9Y2W2 | WW domain-binding protein 11 OS=Hon | WBP11 | 69.998 | 2 | 44.651886 | yes | 2.116E-07 | 593.32733 | 113.76804 |
| KLEVEYEQK | 9 | Q9Y2W2 | WW domain-binding protein 11 OS=Hon | WBP11 | 69.998 | 2 | 44.651886 | yes | 4.845E-05 | 583.30859 | 54.502201 |
| TSAYGPPTR | 9 | Q9Y2W2 | WW domain-binding protein 11 OS=Hon | WBP11 | 69.998 | 2 | 44.651886 | yes | 5.898E-07 | 475.24051 | 54.489182 |
| DEDMLYSPELAQF | 13 | Q9Y2W2 | WW domain-binding protein 11 OS=Hon | WBP11 | 69.998 | 2 | 44.651886 | yes | 2.466E-07 | 783.85889 | 47.569008 |
| TAGPAVHLK | 9 | Q9Y2X0 | Mediator of RNA polymerase II transcript | MED16 | 96.794 | 2 | 42.793526 | yes | 3.637E-07 | 447.26379 | 57.323154 |
| DKFPAITHLK | 10 | Q9Y2X0 | Mediator of RNA polymerase II transcript | MED16 | 96.794 | 2 | 42.793526 | yes | 3.948E-07 | 585.33752 | 45.598392 |
| LPISLTNTDLK | 11 | Q9Y2X0 | Mediator of RNA polymerase II transcript | MED16 | 96.794 | 2 | 42.793526 | yes | 2.872E-07 | 607.85315 | 36.374916 |
| SQMDGLIPGVEPR | 13 | Q9Y2X3 | Nucleolar protein 58 OS=Homo sapiens ( | NOP58 | 59.579 | 2 | 53.39258 | yes | 9.734E-08 | 699.8559 | 2035.5385 |
| LSELLPEEVEAEVI | 14 | Q9Y2X3 | Nucleolar protein 58 OS=Homo sapiens ( | NOP58 | 59.579 | 2 | 53.39258 | yes | 1.9E-07 | 792.922 | 1542.7631 |
| TQLYEYLQNR | 10 | Q9Y2X3 | Nucleolar protein 58 OS=Homo sapiens ( | NOP58 | 59.579 | 2 | 53.39258 | yes | 1.025E-07 | 664.33569 | 1104.7136 |
| LQEVDSLWK | 9 | Q9Y2X3 | Nucleolar protein 58 OS=Homo sapiens ( | NOP58 | 59.579 | 2 | 53.39258 | yes | 1.026E-07 | 559.29803 | 922.65356 |
| IISDNLTYCK | 10 | Q9Y2X3 | Nucleolar protein 58 OS=Homo sapiens ( | NOP58 | 59.579 | 2 | 53.39258 | yes | 1.423E-07 | 613.80792 | 862.86853 |
| HAASTVQILGAEK | 13 | Q9Y2X3 | Nucleolar protein 58 OS=Homo sapiens ( | NOP58 | 59.579 | 2 | 53.39258 | yes | 9.722E-08 | 662.86462 | 713.71008 |
| EAHEPLAVADAK | 12 | Q9Y2X3 | Nucleolar protein 58 OS=Homo sapiens ( | NOP58 | 59.579 | 2 | 53.39258 | yes | 1.122E-06 | 625.82239 | 629.07892 |
| TYDPSGDSTLPTCS | 15 | Q9Y2X3 | Nucleolar protein 58 OS=Homo sapiens ( | NOP58 | 59.579 | 2 | 53.39258 | yes | 1.014E-07 | 814.85907 | 262.95581 |
| VKVEEEEEEKVAE | 20 | Q9Y2X3 | Nucleolar protein 58 OS=Homo sapiens ( | NOP58 | 59.579 | 3 | 53.39258 | yes | 1.214E-07 | 783.71124 | 188.18871 |
| ELNNYIMR | 8 | Q9Y2X3 | Nucleolar protein 58 OS=Homo sapiens ( | NOP58 | 59.579 | 2 | 53.39258 | yes | 2.482E-07 | 526.76331 | 185.55177 |
| EWYGWHFPELGK | 12 | Q9Y2X3 | Nucleolar protein 58 OS=Homo sapiens ( | NOP58 | 59.579 | 2 | 53.39258 | yes | 9.739E-08 | 774.86713 | 156.73541 |
| VAEEEETSVK | 10 | Q9Y2X3 | Nucleolar protein 58 OS=Homo sapiens ( | NOP58 | 59.579 | 2 | 53.39258 | yes | 1.168E-07 | 560.77203 | 145.15382 |
| KLQEVDSLWK | 10 | Q9Y2X3 | Nucleolar protein 58 OS=Homo sapiens ( | NOP58 | 59.579 | 2 | 53.39258 | yes | 1.864E-06 | 623.34552 | 97.020447 |
| YGLIYHASLVGQT | 16 | Q9Y2X3 | Nucleolar protein 58 OS=Homo sapiens ( | NOP58 | 59.579 | 3 | 53.39258 | yes | 2.079E-07 | 578.64557 | 85.841934 |
| VEEEEEEKVAEEE | 18 | Q9Y2X3 | Nucleolar protein 58 OS=Homo sapiens ( | NOP58 | 59.579 | 3 | 53.39258 | yes | 1.545E-07 | 707.99011 | 85.816322 |
| LIAHAGSLLNLAK | 13 | Q9Y2X3 | Nucleolar protein 58 OS=Homo sapiens ( | NOP58 | 59.579 | 3 | 53.39258 | yes | 1.101E-07 | 440.93811 | 73.513153 |
| VKVEEEEEEK | 10 | Q9Y2X3 | Nucleolar protein 58 OS=Homo sapiens ( | NOP58 | 59.579 | 2 | 53.39258 | yes | 1.301E-07 | 624.30371 | 53.711452 |
| LPPVTSFTR | 9 | Q9Y2X9 | Zinc finger protein 281 OS=Homo sapien | ZNF281 | 96.915 | 2 | 52.610962 | yes | 1.219E-07 | 509.29001 | 1784.8754 |
| EDPFNIAEPR | 10 | Q9Y2X9 | Zinc finger protein 281 OS=Homo sapien | ZNF281 | 96.915 | 2 | 52.610962 | yes | 1.992E-07 | 594.28821 | 1704.3657 |
| HVLIHTGER | 9 | Q9Y2X9 | Zinc finger protein 281 OS=Homo sapien | ZNF281 | 96.915 | 2 | 52.610962 | yes | 9.722E-08 | 531.29614 | 1195.8726 |
| YLQQAFEK | 8 | Q9Y2X9 | Zinc finger protein 281 OS=Homo sapien | ZNF281 | 96.915 | 2 | 52.610962 | yes | 1.241E-07 | 513.76636 | 1190.8218 |
| EPAASAAAFPSQR | 13 | Q9Y2X9 | Zinc finger protein 281 OS=Homo sapien | ZNF281 | 96.915 | 2 | 52.610962 | yes | 9.722E-08 | 651.8255 | 1059.6687 |
| ASMLQEYSK | 9 | Q9Y2X9 | Zinc finger protein 281 OS=Homo sapien | ZNF281 | 96.915 | 2 | 52.610962 | yes | 2.329E-07 | 528.75513 | 677.88892 |
| CDTCQQYFSR | 10 | Q9Y2X9 | Zinc finger protein 281 OS=Homo sapien | ZNF281 | 96.915 | 2 | 52.610962 | yes | 1.003E-07 | 682.77155 | 590.89447 |
| KEPAASAAAFPSQ | 14 | Q9Y2X9 | Zinc finger protein 281 OS=Homo sapien | ZNF281 | 96.915 | 3 | 52.610962 | yes | 1.287E-07 | 477.58441 | 284.64987 |
| KEPAASAAAFPSQ | 14 | Q9Y2X9 | Zinc finger protein 281 OS=Homo sapien | ZNF281 | 96.915 | 3 | 52.610962 | yes | 3.187E-07 | 715.87299 | 284.64987 |
| KEPAASAAAFPSQ | 14 | Q9Y2X9 | Zinc finger protein 281 OS=Homo sapien | ZNF281 | 96.915 | 2 | 52.610962 | yes | 1.287E-07 | 477.58441 | 284.64987 |
| KEPAASAAAFPSQ | 14 | Q9Y2X9 | Zinc finger protein 281 OS=Homo sapien | ZNF281 | 96.915 | 2 | 52.610962 | yes | 3.187E-07 | 715.87299 | 284.64987 |

| | | | | | | | | | | | |
|---|---|---|---|---|---|---|---|---|---|---|---|
| NLETSSAFQSSSQK | 14 | Q9Y2X9 | Zinc finger protein 281 OS=Homo sapien | ZNF281 | 96.915 | 2 | 52.610962 | yes | 5.785E-07 | 757.36011 | 357.07413 |
| TCGEVIVK | 8 | Q9Y2X9 | Zinc finger protein 281 OS=Homo sapien | ZNF281 | 96.915 | 2 | 52.610962 | yes | 9.722E-08 | 453.24167 | 303.85193 |
| EKPFGCDQCSMK | 12 | Q9Y2X9 | Zinc finger protein 281 OS=Homo sapien | ZNF281 | 96.915 | 2 | 52.610962 | yes | 1.968E-07 | 743.80981 | 284.16348 |
| DVLLSSSSR | 9 | Q9Y2X9 | Zinc finger protein 281 OS=Homo sapien | ZNF281 | 96.915 | 2 | 52.610962 | yes | 5.643E-07 | 482.25891 | 279.72229 |
| RTCGEVIVK | 9 | Q9Y2X9 | Zinc finger protein 281 OS=Homo sapien | ZNF281 | 96.915 | 2 | 52.610962 | yes | 6.201E-07 | 531.29224 | 205.88914 |
| PSGSLGIVSNNSVE | 25 | Q9Y2X9 | Zinc finger protein 281 OS=Homo sapien | ZNF281 | 96.915 | 3 | 52.610962 | yes | 9.798E-08 | 815.76587 | 190.45872 |
| SPGLGGGEGGSHG | 34 | Q9Y2X9 | Zinc finger protein 281 OS=Homo sapien | ZNF281 | 96.915 | 5 | 52.610962 | yes | 1.478E-07 | 713.16193 | 133.07462 |
| PSLVGDGEGAILSI | 29 | Q9Y2X9 | Zinc finger protein 281 OS=Homo sapien | ZNF281 | 96.915 | 5 | 52.610962 | yes | 1.936E-07 | 622.10248 | 58.969368 |
| PSLVGDGEGAILSI | 29 | Q9Y2X9 | Zinc finger protein 281 OS=Homo sapien | ZNF281 | 96.915 | 5 | 52.610962 | yes | 2.209E-06 | 777.37628 | 58.969368 |
| PSLVGDGEGAILSI | 29 | Q9Y2X9 | Zinc finger protein 281 OS=Homo sapien | ZNF281 | 96.915 | 4 | 52.610962 | yes | 1.936E-07 | 622.10248 | 58.969368 |
| PSLVGDGEGAILSI | 29 | Q9Y2X9 | Zinc finger protein 281 OS=Homo sapien | ZNF281 | 96.915 | 4 | 52.610962 | yes | 2.209E-06 | 777.37628 | 58.969368 |
| PHICDHCSAAFR | 12 | Q9Y2X9 | Zinc finger protein 281 OS=Homo sapien | ZNF281 | 96.915 | 3 | 52.610962 | yes | 8.302E-07 | 490.8837 | 45.926456 |
| AYFMQLQTR | 9 | Q9Y2Y1 | DNA-directed RNA polymerase III subun | POLR3K | 12.321 | 2 | 50.324883 | yes | 9.819E-08 | 579.29224 | 153.50121 |
| SADEPMTTFYK | 11 | Q9Y2Y1 | DNA-directed RNA polymerase III subun | POLR3K | 12.321 | 2 | 50.324883 | yes | 3.02E-07 | 645.28955 | 138.72177 |
| TGSLSDYSR | 9 | Q9Y2Y9 | Krueppel-like factor 13 OS=Homo sapien | KLF13 | 31.18 | 2 | 43.960773 | yes | 3.451E-07 | 493.23291 | 154.44765 |
| DSASLFVVAR | 10 | Q9Y2Y9 | Krueppel-like factor 13 OS=Homo sapien | KLF13 | 31.18 | 2 | 43.960773 | yes | 2.393E-07 | 532.79041 | 135.57564 |
| KFSCPICEK | 9 | Q9Y2Y9 | Krueppel-like factor 13 OS=Homo sapien | KLF13 | 31.18 | 2 | 43.960773 | yes | 1.915E-06 | 584.77808 | 54.257744 |
| EGPESRPEGAAVA | 20 | Q9Y2Y9 | Krueppel-like factor 13 OS=Homo sapien | KLF13 | 31.18 | 3 | 43.960773 | yes | 2.378E-07 | 669.34821 | 43.013 |
| LGDQPSIGLAQTLI | 15 | Q9Y2Z2 | Protein MTO1 homolog, mitochondrial O | MTO1 | 79.964 | 2 | 48.277317 | yes | 1.169E-07 | 785.4278 | 42.537895 |
| AGHNVIALVGGAT | 15 | Q9Y2Z4 | Tyrosine--tRNA ligase, mitochondrial OS | YARS2 | 53.2 | 2 | 55.085022 | yes | 1.099E-07 | 703.89679 | 1209.5217 |
| AGHNVIALVGGAT | 15 | Q9Y2Z4 | Tyrosine--tRNA ligase, mitochondrial OS | YARS2 | 53.2 | 2 | 55.085022 | yes | 1.308E-07 | 469.60028 | 1209.5217 |
| AGHNVIALVGGAT | 15 | Q9Y2Z4 | Tyrosine--tRNA ligase, mitochondrial OS | YARS2 | 53.2 | 3 | 55.085022 | yes | 1.099E-07 | 703.89679 | 1209.5217 |
| AGHNVIALVGGAT | 15 | Q9Y2Z4 | Tyrosine--tRNA ligase, mitochondrial OS | YARS2 | 53.2 | 3 | 55.085022 | yes | 1.308E-07 | 469.60028 | 1209.5217 |
| IELPELFDR | 9 | Q9Y2Z4 | Tyrosine--tRNA ligase, mitochondrial OS | YARS2 | 53.2 | 2 | 55.085022 | yes | 1.464E-07 | 566.30585 | 758.9364 |
| AHSGAQGLLAAQI | 13 | Q9Y2Z4 | Tyrosine--tRNA ligase, mitochondrial OS | YARS2 | 53.2 | 2 | 55.085022 | yes | 1.002E-07 | 626.34387 | 692.7771 |
| SAGNAVWLNR | 10 | Q9Y2Z4 | Tyrosine--tRNA ligase, mitochondrial OS | YARS2 | 53.2 | 2 | 55.085022 | yes | 1.861E-07 | 544.28577 | 602.08069 |
| DFFPETGTK | 9 | Q9Y2Z4 | Tyrosine--tRNA ligase, mitochondrial OS | YARS2 | 53.2 | 2 | 55.085022 | yes | 3.069E-07 | 521.24799 | 583.92938 |
| CTQALYHSSIDALI | 22 | Q9Y2Z4 | Tyrosine--tRNA ligase, mitochondrial OS | YARS2 | 53.2 | 3 | 55.085022 | yes | 9.722E-08 | 846.73322 | 381.33575 |
| EALETER | 7 | Q9Y2Z4 | Tyrosine--tRNA ligase, mitochondrial OS | YARS2 | 53.2 | 2 | 55.085022 | yes | 3.299E-07 | 424.21143 | 276.28748 |
| QHLVDFLAAVGGI | 15 | Q9Y2Z4 | Tyrosine--tRNA ligase, mitochondrial OS | YARS2 | 53.2 | 3 | 55.085022 | yes | 1.206E-07 | 556.29858 | 216.06798 |
| KANAIPDGPR | 10 | Q9Y2Z4 | Tyrosine--tRNA ligase, mitochondrial OS | YARS2 | 53.2 | 2 | 55.085022 | yes | 1.009E-07 | 519.78815 | 163.24409 |
| QPDDSVER | 8 | Q9Y2Z4 | Tyrosine--tRNA ligase, mitochondrial OS | YARS2 | 53.2 | 2 | 55.085022 | yes | 9.722E-08 | 473.21725 | 98.592033 |
| MITEGGVSINHQQ | 29 | Q9Y2Z4 | Tyrosine--tRNA ligase, mitochondrial OS | YARS2 | 53.2 | 4 | 55.085022 | yes | 9.722E-08 | 786.17358 | 70.671196 |
| EAILEYILHQK | 11 | Q9Y314 | Nitric oxide synthase-interacting protein ( | NOSIP | 33.172 | 2 | 52.203766 | yes | 9.809E-08 | 678.8797 | 644.22632 |
| PSGAVVTLECVEK | 13 | Q9Y314 | Nitric oxide synthase-interacting protein ( | NOSIP | 33.172 | 2 | 52.203766 | yes | 1.127E-07 | 694.85815 | 285.62506 |
| DTAASGYGTQNIR | 13 | Q9Y314 | Nitric oxide synthase-interacting protein ( | NOSIP | 33.172 | 2 | 52.203766 | yes | 9.722E-08 | 677.3233 | 277.4054 |
| DSLSNATPCAVLR | 13 | Q9Y314 | Nitric oxide synthase-interacting protein ( | NOSIP | 33.172 | 2 | 52.203766 | yes | 2.881E-07 | 702.35101 | 216.37801 |
| NCTAGAVYTYHEI | 13 | Q9Y314 | Nitric oxide synthase-interacting protein ( | NOSIP | 33.172 | 2 | 52.203766 | yes | 1.062E-07 | 757.34064 | 130.14534 |
| GGTGFAGSGVK | 11 | Q9Y314 | Nitric oxide synthase-interacting protein ( | NOSIP | 33.172 | 2 | 52.203766 | yes | 1.12E-07 | 469.24051 | 114.03804 |

| | | | | | | | | | | | |
|---|---|---|---|---|---|---|---|---|---|---|---|
| TVTCPMSGKPLR | 12 | Q9Y314 | Nitric oxide synthase-interacting protein ( | NOSIP | 33.172 | 2 | 52.203766 | yes | 9.562E-07 | 673.84961 | 79.621307 |
| KDMVDPVTGDK | 11 | Q9Y314 | Nitric oxide synthase-interacting protein ( | NOSIP | 33.172 | 2 | 52.203766 | yes | 1.013E-06 | 602.79755 | 25.678183 |
| IFILGPSHHVPLSR | 14 | Q9Y316 | Protein MEMO1 OS=Homo sapiens OX= | MEMO1 | 33.734 | 3 | 52.861366 | yes | 9.722E-08 | 524.9718 | 138.45058 |
| EFNLNELYQR | 10 | Q9Y320 | Thioredoxin-related transmembrane prote | TMX2 | 34.038 | 2 | 38.31265 | yes | 5.311E-07 | 663.32788 | 37.025116 |
| CYIVATVDR | 9 | Q9Y324 | rRNA-processing protein FCF1 homolog | FCF1 | 23.37 | 2 | 53.764736 | yes | 9.722E-08 | 548.77643 | 262.51724 |
| GTYADDCLVQR | 11 | Q9Y324 | rRNA-processing protein FCF1 homolog | FCF1 | 23.37 | 2 | 53.764736 | yes | 9.722E-08 | 649.29572 | 230.6682 |
| MPDDYGAPR | 9 | Q9Y324 | rRNA-processing protein FCF1 homolog | FCF1 | 23.37 | 2 | 53.764736 | yes | 4.978E-07 | 511.22403 | 104.30621 |
| LTDISVTDPEK | 11 | Q9Y333 | U6 snRNA-associated Sm-like protein LS | LSM2 | 10.835 | 2 | 41.96354 | yes | 3.258E-07 | 609.31665 | 250.47223 |
| YPHMLSVK | 8 | Q9Y333 | U6 snRNA-associated Sm-like protein LS | LSM2 | 10.835 | 2 | 41.96354 | yes | 6.775E-07 | 487.76004 | 234.31158 |
| MLFYSFFK | 8 | Q9Y333 | U6 snRNA-associated Sm-like protein LS | LSM2 | 10.835 | 2 | 41.96354 | yes | 6.476E-05 | 541.77264 | 33.263657 |
| NDLSICGTLHSVD( | 19 | Q9Y333 | U6 snRNA-associated Sm-like protein LS | LSM2 | 10.835 | 3 | 41.96354 | yes | 4.646E-07 | 730.7002 | 28.265633 |
| LHLGFIEIR | 9 | Q9Y383 | Putative RNA-binding protein Luc7-like : | LUC7L2 | 46.514 | 2 | 52.16696 | yes | 3.036E-06 | 549.3269 | 9336.499 |
| AMLDQLMGTSR | 11 | Q9Y383 | Putative RNA-binding protein Luc7-like : | LUC7L2 | 46.514 | 2 | 52.16696 | yes | 9.722E-08 | 611.79956 | 2170.853 |
| AMLDQLMGTSR | 11 | Q9Y383 | Putative RNA-binding protein Luc7-like : | LUC7L2 | 46.514 | 2 | 52.16696 | yes | 1.603E-07 | 619.797 | 2170.853 |
| AMLDQLMGTSR | 11 | Q9Y383 | Putative RNA-binding protein Luc7-like : | LUC7L2 | 46.514 | 2 | 52.16696 | yes | 1.444E-07 | 619.797 | 2170.853 |
| VHDLALR | 7 | Q9Y383 | Putative RNA-binding protein Luc7-like : | LUC7L2 | 46.514 | 2 | 52.16696 | yes | 2.122E-07 | 412.24286 | 4462.4165 |
| MDLGECLK | 8 | Q9Y383 | Putative RNA-binding protein Luc7-like : | LUC7L2 | 46.514 | 2 | 52.16696 | yes | 2.963E-07 | 483.22516 | 3965.8054 |
| LAETQEEISAEVA/ | 15 | Q9Y383 | Putative RNA-binding protein Luc7-like : | LUC7L2 | 46.514 | 2 | 52.16696 | yes | 2.353E-07 | 794.90686 | 1709.0676 |
| EKLEELKR | 8 | Q9Y383 | Putative RNA-binding protein Luc7-like : | LUC7L2 | 46.514 | 2 | 52.16696 | yes | 1.544E-07 | 522.80603 | 1381.7207 |
| VEQLGAEGNVEES | 15 | Q9Y383 | Putative RNA-binding protein Luc7-like : | LUC7L2 | 46.514 | 2 | 52.16696 | yes | 1.2E-07 | 808.89172 | 1096.5847 |
| EKLEELK | 7 | Q9Y383 | Putative RNA-binding protein Luc7-like : | LUC7L2 | 46.514 | 2 | 52.16696 | yes | 2.247E-07 | 444.75546 | 558.8446 |
| VMDEVEK | 7 | Q9Y383 | Putative RNA-binding protein Luc7-like : | LUC7L2 | 46.514 | 2 | 52.16696 | yes | 2.157E-07 | 425.20477 | 151.94405 |
| FRDQDLASCDR | 11 | Q9Y383 | Putative RNA-binding protein Luc7-like : | LUC7L2 | 46.514 | 2 | 52.16696 | yes | 9.722E-08 | 691.80951 | 74.580994 |
| EAEEVYR | 7 | Q9Y383 | Putative RNA-binding protein Luc7-like : | LUC7L2 | 46.514 | 2 | 52.16696 | yes | 1.272E-07 | 448.21143 | 69.652802 |
| DQDLASCDRDR | 11 | Q9Y383 | Putative RNA-binding protein Luc7-like : | LUC7L2 | 46.514 | 2 | 52.16696 | yes | 9.722E-08 | 675.78876 | 24.466513 |
| GFCFLCYEDQR | 11 | Q9Y388 | RNA-binding motif protein, X-linked 2 O | RBMX2 | 37.336 | 2 | 52.6417 | yes | 1.125E-07 | 747.81067 | 505.56976 |
| LINELNER | 8 | Q9Y388 | RNA-binding motif protein, X-linked 2 O | RBMX2 | 37.336 | 2 | 52.6417 | yes | 2.287E-07 | 500.77472 | 460.45505 |
| VDHVSNYR | 8 | Q9Y388 | RNA-binding motif protein, X-linked 2 O | RBMX2 | 37.336 | 2 | 52.6417 | yes | 1.871E-07 | 495.24359 | 255.46164 |
| DSEEIDDVTR | 10 | Q9Y388 | RNA-binding motif protein, X-linked 2 O | RBMX2 | 37.336 | 2 | 52.6417 | yes | 9.722E-08 | 589.76221 | 169.30719 |
| TAYSGGAEDLER | 12 | Q9Y388 | RNA-binding motif protein, X-linked 2 O | RBMX2 | 37.336 | 2 | 52.6417 | yes | 9.722E-08 | 634.79132 | 155.34824 |
| VSWHSEYK | 8 | Q9Y388 | RNA-binding motif protein, X-linked 2 O | RBMX2 | 37.336 | 2 | 52.6417 | yes | 1.113E-07 | 518.24835 | 152.38683 |
| STILAVDNFNGIK | 13 | Q9Y388 | RNA-binding motif protein, X-linked 2 O | RBMX2 | 37.336 | 2 | 52.6417 | yes | 2.423E-07 | 696.38007 | 97.873642 |
| QFSYLIENMAR | 11 | Q9Y399 | Small ribosomal subunit protein uS2m OS | MRPS2 | 33.25 | 2 | 50.44566 | yes | 9.809E-08 | 686.3399 | 990.61694 |
| LLFGPTVR | 8 | Q9Y399 | Small ribosomal subunit protein uS2m OS | MRPS2 | 33.25 | 2 | 50.44566 | yes | 2.393E-07 | 451.77655 | 962.60352 |
| LFQTAITR | 8 | Q9Y399 | Small ribosomal subunit protein uS2m OS | MRPS2 | 33.25 | 2 | 50.44566 | yes | 2.031E-07 | 475.27692 | 689.83435 |
| QQVEALYR | 8 | Q9Y399 | Small ribosomal subunit protein uS2m OS | MRPS2 | 33.25 | 2 | 50.44566 | yes | 1.257E-07 | 503.76944 | 618.37494 |
| HSDFFNVK | 8 | Q9Y399 | Small ribosomal subunit protein uS2m OS | MRPS2 | 33.25 | 2 | 50.44566 | yes | 1.111E-07 | 497.24307 | 515.664 |
| DCGEYAHTR | 9 | Q9Y399 | Small ribosomal subunit protein uS2m OS | MRPS2 | 33.25 | 2 | 50.44566 | yes | 1.859E-07 | 554.72748 | 410.11667 |
| FMEPYIFGSR | 10 | Q9Y399 | Small ribosomal subunit protein uS2m OS | MRPS2 | 33.25 | 2 | 50.44566 | yes | 1.156E-07 | 623.79987 | 287.00165 |

| | | | | | | | | | | | |
|---|---|---|---|---|---|---|---|---|---|---|---|
| KGIILFISR | 9 | Q9Y399 | Small ribosomal subunit protein uS2m OS | MRPS2 | 33.25 | 2 | 50.44566 | yes | 4.299E-07 | 523.83966 | 238.62405 |
| ILNEPLK | 7 | Q9Y399 | Small ribosomal subunit protein uS2m OS | MRPS2 | 33.25 | 2 | 50.44566 | yes | 1.624E-07 | 413.75528 | 66.737602 |
| ESEDSTDFNDK | 11 | Q9Y399 | Small ribosomal subunit protein uS2m OS | MRPS2 | 33.25 | 2 | 50.44566 | yes | 5.01E-07 | 643.75458 | 19.304152 |
| LQDGVHIIK | 9 | Q9Y3A2 | Probable U3 small nucleolar RNA-associa | UTP11 | 30.447 | 2 | 47.413773 | yes | 4.609E-07 | 511.80328 | 115.09081 |
| HVFFFDTK | 8 | Q9Y3A2 | Probable U3 small nucleolar RNA-associa | UTP11 | 30.447 | 2 | 47.413773 | yes | 9.17E-07 | 520.76361 | 114.71696 |
| SELHLLDFQGK | 11 | Q9Y3A2 | Probable U3 small nucleolar RNA-associa | UTP11 | 30.447 | 2 | 47.413773 | yes | 1.335E-07 | 643.84058 | 90.523811 |
| EEVTPEQLK | 9 | Q9Y3A2 | Probable U3 small nucleolar RNA-associa | UTP11 | 30.447 | 2 | 47.413773 | yes | 2.695E-06 | 536.77966 | 81.00795 |
| HLGLLEK | 7 | Q9Y3A2 | Probable U3 small nucleolar RNA-associa | UTP11 | 30.447 | 2 | 47.413773 | yes | 3.863E-07 | 405.24762 | 73.060715 |
| ETVNSPAIYK | 10 | Q9Y3A2 | Probable U3 small nucleolar RNA-associa | UTP11 | 30.447 | 2 | 47.413773 | yes | 7.68E-07 | 561.29547 | 59.207138 |
| GPLLVSTESHPVK | 13 | Q9Y3A4 | Ribosomal RNA-processing protein 7 hor | RRP7A | 32.335 | 2 | 51.875698 | yes | 1.229E-07 | 682.38263 | 1354.4148 |
| VEVDTFMEAYDQ | 13 | Q9Y3A4 | Ribosomal RNA-processing protein 7 hor | RRP7A | 32.335 | 2 | 51.875698 | yes | 1.037E-07 | 787.85577 | 664.00171 |
| IPSPLGYAAIPIK | 13 | Q9Y3A4 | Ribosomal RNA-processing protein 7 hor | RRP7A | 32.335 | 2 | 51.875698 | yes | 9.722E-08 | 670.40283 | 350.68893 |
| QQASHYLYVR | 10 | Q9Y3A4 | Ribosomal RNA-processing protein 7 hor | RRP7A | 32.335 | 2 | 51.875698 | yes | 1.084E-07 | 632.82526 | 106.47384 |
| QQASHYLYVR | 10 | Q9Y3A4 | Ribosomal RNA-processing protein 7 hor | RRP7A | 32.335 | 2 | 51.875698 | yes | 9.722E-08 | 422.21927 | 106.47384 |
| QQASHYLYVR | 10 | Q9Y3A4 | Ribosomal RNA-processing protein 7 hor | RRP7A | 32.335 | 3 | 51.875698 | yes | 1.084E-07 | 632.82526 | 106.47384 |
| QQASHYLYVR | 10 | Q9Y3A4 | Ribosomal RNA-processing protein 7 hor | RRP7A | 32.335 | 3 | 51.875698 | yes | 9.722E-08 | 422.21927 | 106.47384 |
| PSGVSAALALK | 11 | Q9Y3A4 | Ribosomal RNA-processing protein 7 hor | RRP7A | 32.335 | 2 | 51.875698 | yes | 6.63E-07 | 507.30313 | 95.456253 |
| SFRPGDIVLAK | 11 | Q9Y3B2 | Exosome complex component CSL4 OS= | EXOSC1 | 21.452 | 2 | 51.045097 | yes | 1.3E-07 | 601.84821 | 420.61563 |
| SSENGALPVVSVV | 14 | Q9Y3B2 | Exosome complex component CSL4 OS= | EXOSC1 | 21.452 | 2 | 51.045097 | yes | 2.977E-06 | 707.38843 | 340.12131 |
| YCIPGER | 7 | Q9Y3B2 | Exosome complex component CSL4 OS= | EXOSC1 | 21.452 | 2 | 51.045097 | yes | 9.722E-08 | 447.71054 | 271.05484 |
| VHILYVGSMPLK | 12 | Q9Y3B2 | Exosome complex component CSL4 OS= | EXOSC1 | 21.452 | 2 | 51.045097 | yes | 3.637E-07 | 678.88904 | 244.40665 |
| DKVEIYK | 7 | Q9Y3B2 | Exosome complex component CSL4 OS= | EXOSC1 | 21.452 | 2 | 51.045097 | yes | 1.197E-07 | 447.75018 | 143.89011 |
| GTAYVVYEDIFDA | 14 | Q9Y3B4 | Splicing factor 3B subunit 6 OS=Homo sa | SF3B6 | 14.585 | 2 | 53.385822 | yes | 1.045E-07 | 795.88794 | 1948.1649 |
| ITAEEMYDIFGK | 12 | Q9Y3B4 | Splicing factor 3B subunit 6 OS=Homo sa | SF3B6 | 14.585 | 2 | 53.385822 | yes | 1.5E-07 | 708.83942 | 252.72397 |
| YGINTDPPK | 9 | Q9Y3B4 | Splicing factor 3B subunit 6 OS=Homo sa | SF3B6 | 14.585 | 2 | 53.385822 | yes | 5.815E-07 | 502.75601 | 238.43919 |
| YLVVLYYNANR | 11 | Q9Y3B4 | Splicing factor 3B subunit 6 OS=Homo sa | SF3B6 | 14.585 | 2 | 53.385822 | yes | 1.651E-07 | 694.37207 | 74.594101 |
| NACDHLSGFNVCN | 14 | Q9Y3B4 | Splicing factor 3B subunit 6 OS=Homo sa | SF3B6 | 14.585 | 2 | 53.385822 | yes | 9.722E-08 | 832.35681 | 59.116936 |
| IGQPTVSYFLK | 11 | Q9Y3B7 | Large ribosomal subunit protein uL11m C | MRPL11 | 20.684 | 2 | 53.58354 | yes | 1.36E-07 | 626.85046 | 1957.3921 |
| AGLAMPGPPLGPV | 17 | Q9Y3B7 | Large ribosomal subunit protein uL11m C | MRPL11 | 20.684 | 2 | 53.58354 | yes | 2.486E-07 | 815.9585 | 1656.6476 |
| EADLAAQEEAAK | 12 | Q9Y3B7 | Large ribosomal subunit protein uL11m C | MRPL11 | 20.684 | 2 | 53.58354 | yes | 9.819E-08 | 623.30151 | 1192.7212 |
| DLSSEELAAFQK | 12 | Q9Y3B7 | Large ribosomal subunit protein uL11m C | MRPL11 | 20.684 | 2 | 53.58354 | yes | 1.113E-07 | 669.33282 | 1060.3593 |
| HVYEIAR | 7 | Q9Y3B7 | Large ribosomal subunit protein uL11m C | MRPL11 | 20.684 | 2 | 53.58354 | yes | 1.02E-07 | 444.24033 | 696.51141 |
| EVAGLVTLK | 9 | Q9Y3B7 | Large ribosomal subunit protein uL11m C | MRPL11 | 20.684 | 2 | 53.58354 | yes | 9.722E-08 | 465.28696 | 619.12561 |
| KPEVGGVIR | 9 | Q9Y3B7 | Large ribosomal subunit protein uL11m C | MRPL11 | 20.684 | 2 | 53.58354 | yes | 1.517E-07 | 477.79019 | 498.83032 |
| GVSINQFCK | 9 | Q9Y3B7 | Large ribosomal subunit protein uL11m C | MRPL11 | 20.684 | 2 | 53.58354 | yes | 3.729E-07 | 526.76331 | 390.03079 |
| AIFLAAQK | 8 | Q9Y3B7 | Large ribosomal subunit protein uL11m C | MRPL11 | 20.684 | 2 | 53.58354 | yes | 9.81E-08 | 431.26328 | 134.62228 |
| EADLAAQEEAAKI | 13 | Q9Y3B7 | Large ribosomal subunit protein uL11m C | MRPL11 | 20.684 | 2 | 53.58354 | yes | 9.722E-08 | 687.349 | 39.01247 |
| DFMIQGGDPTGTG | 14 | Q9Y3C6 | Peptidyl-prolyl cis-trans isomerase-like 1 | PPIL1 | 18.237 | 2 | 50.36326 | yes | 1.659E-07 | 726.33282 | 2451.3359 |
| VCQGIGMVNR | 10 | Q9Y3C6 | Peptidyl-prolyl cis-trans isomerase-like 1 | PPIL1 | 18.237 | 2 | 50.36326 | yes | 1.034E-07 | 567.28137 | 1303.8964 |

| | | | | | | | | | | | |
|---|---|---|---|---|---|---|---|---|---|---|---|
| QFEDELHPDLK | 11 | Q9Y3C6 | Peptidyl-prolyl cis-trans isomerase-like 1 | PPIL1 | 18.237 | 2 | 50.36326 | yes | 9.81E-08 | 685.83295 | 698.88232 |
| NFAELAR | 7 | Q9Y3C6 | Peptidyl-prolyl cis-trans isomerase-like 1 | PPIL1 | 18.237 | 2 | 50.36326 | yes | 1.585E-07 | 410.71924 | 182.9743 |
| AAAVAMETDDAG | 14 | Q9Y3C7 | Mediator of RNA polymerase II transcript | MED31 | 15.806 | 2 | 50.53987 | yes | 9.781E-08 | 717.31989 | 217.56667 |
| LKEEYQSLIR | 10 | Q9Y3C8 | Ubiquitin-fold modifier-conjugating enzy | UFC1 | 19.459 | 2 | 49.468246 | yes | 1.027E-07 | 639.85626 | 53.00975 |
| LVALNLDR | 8 | Q9Y3D3 | Small ribosomal subunit protein bS16m C | MRPS16 | 15.345 | 2 | 52.55853 | yes | 6.128E-07 | 457.27692 | 1297.0454 |
| EVLLASQK | 8 | Q9Y3D3 | Small ribosomal subunit protein bS16m C | MRPS16 | 15.345 | 2 | 52.55853 | yes | 1.102E-07 | 444.26346 | 530.56915 |
| HWIGCGAHLSK | 11 | Q9Y3D3 | Small ribosomal subunit protein bS16m C | MRPS16 | 15.345 | 2 | 52.55853 | yes | 9.722E-08 | 633.31403 | 153.20929 |
| HWIGCGAHLSKPM | 15 | Q9Y3D3 | Small ribosomal subunit protein bS16m C | MRPS16 | 15.345 | 3 | 52.55853 | yes | 1.113E-07 | 584.2887 | 61.781963 |
| STQFEYAWCLVR | 12 | Q9Y3D6 | Mitochondrial fission 1 protein OS=Homo | FIS1 | 16.938 | 2 | 30.801414 | yes | 2.056E-06 | 780.3692 | 46.833195 |
| SVGGSFYLQSK | 11 | Q9Y3D7 | Mitochondrial import inner membrane tra | PAM16 | 13.825 | 2 | 49.193855 | yes | 1.06E-07 | 586.80096 | 367.86011 |
| NYEHLFK | 7 | Q9Y3D7 | Mitochondrial import inner membrane tra | PAM16 | 13.825 | 2 | 49.193855 | yes | 1.971E-06 | 475.74017 | 66.541962 |
| YINVGDLAR | 9 | Q9Y3D8 | Adenylate kinase isoenzyme 6 OS=Homo | AK6 | 20.062 | 2 | 47.913742 | yes | 1.233E-07 | 510.77728 | 236.84041 |
| TDTNVLYER | 9 | Q9Y3D8 | Adenylate kinase isoenzyme 6 OS=Homo | AK6 | 20.062 | 2 | 47.913742 | yes | 1.678E-07 | 555.7749 | 108.65279 |
| LETVGSIFSR | 10 | Q9Y3D9 | Small ribosomal subunit protein mS23 OS | MRPS23 | 21.771 | 2 | 50.634083 | yes | 1.841E-07 | 554.80347 | 1664.5144 |
| AFDLFNPNFK | 10 | Q9Y3D9 | Small ribosomal subunit protein mS23 OS | MRPS23 | 21.771 | 2 | 50.634083 | yes | 7.241E-07 | 606.80603 | 1537.89 |
| ALLAEGVILR | 10 | Q9Y3D9 | Small ribosomal subunit protein mS23 OS | MRPS23 | 21.771 | 2 | 50.634083 | yes | 1.143E-07 | 527.83459 | 978.87628 |
| FYSVYGSGQR | 10 | Q9Y3D9 | Small ribosomal subunit protein mS23 OS | MRPS23 | 21.771 | 2 | 50.634083 | yes | 2.843E-07 | 582.27765 | 756.39996 |
| APIQDIWYHEDR | 12 | Q9Y3D9 | Small ribosomal subunit protein mS23 OS | MRPS23 | 21.771 | 2 | 50.634083 | yes | 1.397E-07 | 771.87042 | 481.59628 |
| EPVFQRPR | 8 | Q9Y3D9 | Small ribosomal subunit protein mS23 OS | MRPS23 | 21.771 | 2 | 50.634083 | yes | 9.733E-08 | 514.78546 | 398.64069 |
| LGETDEEK | 8 | Q9Y3D9 | Small ribosomal subunit protein mS23 OS | MRPS23 | 21.771 | 2 | 50.634083 | yes | 1.643E-07 | 460.71402 | 326.14594 |
| KSEHLSVRPQTAL | 20 | Q9Y3D9 | Small ribosomal subunit protein mS23 OS | MRPS23 | 21.771 | 4 | 50.634083 | yes | 1.362E-07 | 581.80145 | 72.77021 |
| PQTALEENETQK | 12 | Q9Y3D9 | Small ribosomal subunit protein mS23 OS | MRPS23 | 21.771 | 2 | 50.634083 | yes | 9.809E-08 | 694.33862 | 38.553364 |
| EVPQDQHLEAPAI | 16 | Q9Y3D9 | Small ribosomal subunit protein mS23 OS | MRPS23 | 21.771 | 3 | 50.634083 | yes | 2.921E-06 | 597.95514 | 15.606886 |
| RNPEMLK | 7 | Q9Y3E5 | Peptidyl-tRNA hydrolase 2, mitochondria | PTRH2 | 19.194 | 2 | 50.08531 | yes | 1.089E-07 | 444.242 | 103.73461 |
| MLGLTVSLIQDAG | 14 | Q9Y3E5 | Peptidyl-tRNA hydrolase 2, mitochondria | PTRH2 | 19.194 | 2 | 50.08531 | yes | 3.81E-07 | 737.40833 | 50.088596 |
| THTDTESEASILGI | 19 | Q9Y3E5 | Peptidyl-tRNA hydrolase 2, mitochondria | PTRH2 | 19.194 | 3 | 50.08531 | yes | 3.427E-07 | 680.63953 | 47.652702 |
| VAAQCSHAAVSA | 14 | Q9Y3E5 | Peptidyl-tRNA hydrolase 2, mitochondria | PTRH2 | 19.194 | 2 | 50.08531 | yes | 9.908E-08 | 731.85901 | 15.425114 |
| IPEIQATMR | 9 | Q9Y3E7 | Charged multivesicular body protein 3 OS | CHMP3 | 25.074 | 2 | 47.400547 | yes | 2.693E-07 | 529.7868 | 340.04163 |
| ILFEITAGALGK | 12 | Q9Y3E7 | Charged multivesicular body protein 3 OS | CHMP3 | 25.074 | 2 | 47.400547 | yes | 1.339E-07 | 616.86609 | 147.23573 |
| AHMNSVLMGMK | 11 | Q9Y3E7 | Charged multivesicular body protein 3 OS | CHMP3 | 25.074 | 2 | 47.400547 | yes | 1.488E-06 | 609.79321 | 106.89636 |
| CVLPEEDSGELAK | 15 | Q9Y3F4 | Serine-threonine kinase receptor-associate | STRAP | 38.439 | 2 | 53.507862 | yes | 1.721E-07 | 836.41656 | 2860.2427 |
| CVLPEEDSGELAK | 15 | Q9Y3F4 | Serine-threonine kinase receptor-associate | STRAP | 38.439 | 2 | 53.507862 | yes | 1.146E-07 | 557.94678 | 2860.2427 |
| CVLPEEDSGELAK | 15 | Q9Y3F4 | Serine-threonine kinase receptor-associate | STRAP | 38.439 | 3 | 53.507862 | yes | 1.721E-07 | 836.41656 | 2860.2427 |
| CVLPEEDSGELAK | 15 | Q9Y3F4 | Serine-threonine kinase receptor-associate | STRAP | 38.439 | 3 | 53.507862 | yes | 1.146E-07 | 557.94678 | 2860.2427 |
| GAVWGATLNK | 10 | Q9Y3F4 | Serine-threonine kinase receptor-associate | STRAP | 38.439 | 2 | 53.507862 | yes | 1.553E-07 | 508.77982 | 3982.365 |
| EFLVAGGEDFK | 11 | Q9Y3F4 | Serine-threonine kinase receptor-associate | STRAP | 38.439 | 2 | 53.507862 | yes | 1.489E-07 | 606.30078 | 3662.4009 |
| DGKPMLR | 7 | Q9Y3F4 | Serine-threonine kinase receptor-associate | STRAP | 38.439 | 2 | 53.507862 | yes | 1.259E-07 | 408.72345 | 3519.134 |
| SIAFHSAVSLDPIK | 14 | Q9Y3F4 | Serine-threonine kinase receptor-associate | STRAP | 38.439 | 2 | 53.507862 | yes | 1.699E-07 | 742.909 | 1734.8162 |
| SIAFHSAVSLDPIK | 14 | Q9Y3F4 | Serine-threonine kinase receptor-associate | STRAP | 38.439 | 2 | 53.507862 | yes | 2.174E-07 | 495.60843 | 1734.8162 |

| | | | | | | | | | | | |
|---|---|---|---|---|---|---|---|---|---|---|---|
| SIAFHSAVSLDPIK | 14 | Q9Y3F4 | Serine-threonine kinase receptor-associat | STRAP | 38.439 | 3 | 53.507862 | yes | 1.699E-07 | 742.909 | 1734.8162 |
| SIAFHSAVSLDPIK | 14 | Q9Y3F4 | Serine-threonine kinase receptor-associat | STRAP | 38.439 | 3 | 53.507862 | yes | 2.174E-07 | 495.60843 | 1734.8162 |
| YDYNSGEELESYK | 13 | Q9Y3F4 | Serine-threonine kinase receptor-associat | STRAP | 38.439 | 2 | 53.507862 | yes | 1.166E-07 | 798.83862 | 3405.2104 |
| EISGHTSGIK | 10 | Q9Y3F4 | Serine-threonine kinase receptor-associat | STRAP | 38.439 | 2 | 53.507862 | yes | 1.121E-07 | 514.77222 | 2669.9575 |
| LWDHATMTEVK | 11 | Q9Y3F4 | Serine-threonine kinase receptor-associat | STRAP | 38.439 | 2 | 53.507862 | yes | 9.766E-08 | 665.82666 | 787.15649 |
| LWDHATMTEVK | 11 | Q9Y3F4 | Serine-threonine kinase receptor-associat | STRAP | 38.439 | 2 | 53.507862 | yes | 9.722E-08 | 444.22018 | 787.15649 |
| LWDHATMTEVK | 11 | Q9Y3F4 | Serine-threonine kinase receptor-associat | STRAP | 38.439 | 2 | 53.507862 | yes | 9.722E-08 | 673.8241 | 787.15649 |
| LWDHATMTEVK | 11 | Q9Y3F4 | Serine-threonine kinase receptor-associat | STRAP | 38.439 | 3 | 53.507862 | yes | 9.766E-08 | 665.82666 | 787.15649 |
| LWDHATMTEVK | 11 | Q9Y3F4 | Serine-threonine kinase receptor-associat | STRAP | 38.439 | 3 | 53.507862 | yes | 9.722E-08 | 444.22018 | 787.15649 |
| LWDHATMTEVK | 11 | Q9Y3F4 | Serine-threonine kinase receptor-associat | STRAP | 38.439 | 3 | 53.507862 | yes | 9.722E-08 | 673.8241 | 787.15649 |
| VWDAVSGDELMT | 16 | Q9Y3F4 | Serine-threonine kinase receptor-associat | STRAP | 38.439 | 3 | 53.507862 | yes | 9.819E-08 | 591.29431 | 1010.4387 |
| VWDAVSGDELMT | 16 | Q9Y3F4 | Serine-threonine kinase receptor-associat | STRAP | 38.439 | 3 | 53.507862 | yes | 3.424E-05 | 596.62592 | 1010.4387 |
| QGDTGDWIGTFLC | 15 | Q9Y3F4 | Serine-threonine kinase receptor-associat | STRAP | 38.439 | 2 | 53.507862 | yes | 9.722E-08 | 816.39429 | 1752.4139 |
| GHFGPIHCVR | 10 | Q9Y3F4 | Serine-threonine kinase receptor-associat | STRAP | 38.439 | 2 | 53.507862 | yes | 1.104E-07 | 590.29565 | 1276.452 |
| IYDLNKPEAEPK | 12 | Q9Y3F4 | Serine-threonine kinase receptor-associat | STRAP | 38.439 | 3 | 53.507862 | yes | 1.025E-07 | 472.91714 | 633.79309 |
| IYDLNKPEAEPK | 12 | Q9Y3F4 | Serine-threonine kinase receptor-associat | STRAP | 38.439 | 3 | 53.507862 | yes | 9.722E-08 | 708.87207 | 633.79309 |
| IYDLNKPEAEPK | 12 | Q9Y3F4 | Serine-threonine kinase receptor-associat | STRAP | 38.439 | 2 | 53.507862 | yes | 1.025E-07 | 472.91714 | 633.79309 |
| IYDLNKPEAEPK | 12 | Q9Y3F4 | Serine-threonine kinase receptor-associat | STRAP | 38.439 | 2 | 53.507862 | yes | 9.722E-08 | 708.87207 | 633.79309 |
| QTPLTCSGHTR | 11 | Q9Y3F4 | Serine-threonine kinase receptor-associat | STRAP | 38.439 | 2 | 53.507862 | yes | 3.372E-07 | 629.30383 | 1230.2445 |
| CVLPEEDSGELAK | 13 | Q9Y3F4 | Serine-threonine kinase receptor-associat | STRAP | 38.439 | 2 | 53.507862 | yes | 4.798E-07 | 723.84265 | 1145.6259 |
| SFEAPATINSASLH | 17 | Q9Y3F4 | Serine-threonine kinase receptor-associat | STRAP | 38.439 | 3 | 53.507862 | yes | 1.712E-07 | 600.30383 | 568.42328 |
| ALWCSEDK | 8 | Q9Y3F4 | Serine-threonine kinase receptor-associat | STRAP | 38.439 | 2 | 53.507862 | yes | 1.002E-07 | 504.72638 | 378.3801 |
| AATAAADFTAK | 11 | Q9Y3F4 | Serine-threonine kinase receptor-associat | STRAP | 38.439 | 2 | 53.507862 | yes | 5.719E-07 | 519.26672 | 328.13181 |
| LWQTVVGK | 8 | Q9Y3F4 | Serine-threonine kinase receptor-associat | STRAP | 38.439 | 2 | 53.507862 | yes | 1.097E-07 | 465.77402 | 178.70001 |
| GGGVGGFLPAMK | 12 | Q9Y3I0 | RNA-splicing ligase RtcB homolog OS=I | RTCB | 55.211 | 2 | 52.86701 | yes | 1.265E-07 | 545.78931 | 28258.82 |
| GGGVGGFLPAMK | 12 | Q9Y3I0 | RNA-splicing ligase RtcB homolog OS=I | RTCB | 55.211 | 2 | 52.86701 | yes | 1.355E-07 | 553.7868 | 28258.82 |
| LVMEEAPESYK | 11 | Q9Y3I0 | RNA-splicing ligase RtcB homolog OS=I | RTCB | 55.211 | 2 | 52.86701 | yes | 1.527E-07 | 648.31305 | 26219.23 |
| LVMEEAPESYK | 11 | Q9Y3I0 | RNA-splicing ligase RtcB homolog OS=I | RTCB | 55.211 | 2 | 52.86701 | yes | 2.354E-07 | 656.31049 | 26219.23 |
| TNLDESDVQPVK | 12 | Q9Y3I0 | RNA-splicing ligase RtcB homolog OS=I | RTCB | 55.211 | 2 | 52.86701 | yes | 9.722E-08 | 672.83569 | 36795.137 |
| GMAAAGNYAWVI | 13 | Q9Y3I0 | RNA-splicing ligase RtcB homolog OS=I | RTCB | 55.211 | 2 | 52.86701 | yes | 9.823E-08 | 690.82751 | 17927.369 |
| GMAAAGNYAWVI | 13 | Q9Y3I0 | RNA-splicing ligase RtcB homolog OS=I | RTCB | 55.211 | 2 | 52.86701 | yes | 9.722E-08 | 698.82495 | 17927.369 |
| SYNDELQFLEK | 11 | Q9Y3I0 | RNA-splicing ligase RtcB homolog OS=I | RTCB | 55.211 | 2 | 52.86701 | yes | 9.722E-08 | 693.33282 | 28425.105 |
| NLDFQDVLDK | 10 | Q9Y3I0 | RNA-splicing ligase RtcB homolog OS=I | RTCB | 55.211 | 2 | 52.86701 | yes | 1.267E-07 | 603.80371 | 28014.744 |
| QIGNVAALPGIVHI | 14 | Q9Y3I0 | RNA-splicing ligase RtcB homolog OS=I | RTCB | 55.211 | 3 | 52.86701 | yes | 1.501E-07 | 482.2843 | 13543.595 |
| QIGNVAALPGIVHI | 14 | Q9Y3I0 | RNA-splicing ligase RtcB homolog OS=I | RTCB | 55.211 | 3 | 52.86701 | yes | 5.982E-07 | 722.92279 | 13543.595 |
| QIGNVAALPGIVHI | 14 | Q9Y3I0 | RNA-splicing ligase RtcB homolog OS=I | RTCB | 55.211 | 2 | 52.86701 | yes | 1.501E-07 | 482.2843 | 13543.595 |
| QIGNVAALPGIVHI | 14 | Q9Y3I0 | RNA-splicing ligase RtcB homolog OS=I | RTCB | 55.211 | 2 | 52.86701 | yes | 5.982E-07 | 722.92279 | 13543.595 |
| IASPEGQDYLK | 11 | Q9Y3I0 | RNA-splicing ligase RtcB homolog OS=I | RTCB | 55.211 | 2 | 52.86701 | yes | 9.722E-08 | 610.81152 | 23425.596 |
| LMFEELR | 7 | Q9Y3I0 | RNA-splicing ligase RtcB homolog OS=I | RTCB | 55.211 | 2 | 52.86701 | yes | 1.962E-07 | 469.24423 | 9157.877 |

| | | | | | | | | | | | |
|---|---|---|---|---|---|---|---|---|---|---|---|
| LMFEELR | 7 | Q9Y3I0 | RNA-splicing ligase RtcB homolog OS=H | RTCB | 55.211 | 2 | 52.86701 | yes | 4.153E-06 | 477.24167 | 9157.877 |
| MLQADPNK | 8 | Q9Y3I0 | RNA-splicing ligase RtcB homolog OS=H | RTCB | 55.211 | 2 | 52.86701 | yes | 2.552E-07 | 458.73148 | 11868.287 |
| GLGHQVATDALVA | 16 | Q9Y3I0 | RNA-splicing ligase RtcB homolog OS=H | RTCB | 55.211 | 2 | 52.86701 | yes | 1.337E-07 | 820.42725 | 3819.2261 |
| GLGHQVATDALVA | 16 | Q9Y3I0 | RNA-splicing ligase RtcB homolog OS=H | RTCB | 55.211 | 2 | 52.86701 | yes | 3.914E-07 | 828.42468 | 3819.2261 |
| GLGHQVATDALVA | 16 | Q9Y3I0 | RNA-splicing ligase RtcB homolog OS=H | RTCB | 55.211 | 2 | 52.86701 | yes | 9.722E-08 | 552.6189 | 3819.2261 |
| GLGHQVATDALVA | 16 | Q9Y3I0 | RNA-splicing ligase RtcB homolog OS=H | RTCB | 55.211 | 3 | 52.86701 | yes | 1.337E-07 | 820.42725 | 3819.2261 |
| GLGHQVATDALVA | 16 | Q9Y3I0 | RNA-splicing ligase RtcB homolog OS=H | RTCB | 55.211 | 3 | 52.86701 | yes | 3.914E-07 | 828.42468 | 3819.2261 |
| GLGHQVATDALVA | 16 | Q9Y3I0 | RNA-splicing ligase RtcB homolog OS=H | RTCB | 55.211 | 3 | 52.86701 | yes | 9.722E-08 | 552.6189 | 3819.2261 |
| LADMGIAIR | 9 | Q9Y3I0 | RNA-splicing ligase RtcB homolog OS=H | RTCB | 55.211 | 2 | 52.86701 | yes | 2E-07 | 480.27078 | 5588.7632 |
| LADMGIAIR | 9 | Q9Y3I0 | RNA-splicing ligase RtcB homolog OS=H | RTCB | 55.211 | 2 | 52.86701 | yes | 1.737E-07 | 488.26822 | 5588.7632 |
| GQVCVMIHSGSR | 12 | Q9Y3I0 | RNA-splicing ligase RtcB homolog OS=H | RTCB | 55.211 | 2 | 52.86701 | yes | 1.59E-06 | 665.82135 | 5252.542 |
| GQVCVMIHSGSR | 12 | Q9Y3I0 | RNA-splicing ligase RtcB homolog OS=H | RTCB | 55.211 | 2 | 52.86701 | yes | 9.722E-08 | 673.81885 | 5252.542 |
| LRPIAVIK | 8 | Q9Y3I0 | RNA-splicing ligase RtcB homolog OS=H | RTCB | 55.211 | 2 | 52.86701 | yes | 9.058E-06 | 455.31583 | 3174.885 |
| EGYAWAEDK | 9 | Q9Y3I0 | RNA-splicing ligase RtcB homolog OS=H | RTCB | 55.211 | 2 | 52.86701 | yes | 1.297E-07 | 534.73529 | 2750.7993 |
| VEQHVVDGK | 9 | Q9Y3I0 | RNA-splicing ligase RtcB homolog OS=H | RTCB | 55.211 | 2 | 52.86701 | yes | 1.995E-07 | 505.76691 | 2658.0676 |
| DKIIVNDR | 8 | Q9Y3I0 | RNA-splicing ligase RtcB homolog OS=H | RTCB | 55.211 | 2 | 52.86701 | yes | 4.117E-07 | 486.77728 | 2496.0625 |
| SSMTFLTR | 8 | Q9Y3I0 | RNA-splicing ligase RtcB homolog OS=H | RTCB | 55.211 | 2 | 52.86701 | yes | 1.202E-06 | 471.73929 | 1228.1686 |
| SSMTFLTR | 8 | Q9Y3I0 | RNA-splicing ligase RtcB homolog OS=H | RTCB | 55.211 | 2 | 52.86701 | yes | 1.437E-07 | 479.73676 | 1228.1686 |
| RNLDFQDVLDK | 11 | Q9Y3I0 | RNA-splicing ligase RtcB homolog OS=H | RTCB | 55.211 | 2 | 52.86701 | yes | 1.218E-07 | 681.85425 | 1365.314 |
| KGFVPNMQVEGV | 21 | Q9Y3I0 | RNA-splicing ligase RtcB homolog OS=H | RTCB | 55.211 | 3 | 52.86701 | yes | 3.154E-07 | 795.40332 | 791.35468 |
| EHCEEYGR | 8 | Q9Y3I0 | RNA-splicing ligase RtcB homolog OS=H | RTCB | 55.211 | 2 | 52.86701 | yes | 2.358E-07 | 540.21417 | 650.97253 |
| EGYAWAEDKEHC | 17 | Q9Y3I0 | RNA-splicing ligase RtcB homolog OS=H | RTCB | 55.211 | 4 | 52.86701 | yes | 1.169E-07 | 532.97211 | 136.43515 |
| EGYAWAEDKEHC | 17 | Q9Y3I0 | RNA-splicing ligase RtcB homolog OS=H | RTCB | 55.211 | 4 | 52.86701 | yes | 2.455E-05 | 710.2937 | 136.43515 |
| EGYAWAEDKEHC | 17 | Q9Y3I0 | RNA-splicing ligase RtcB homolog OS=H | RTCB | 55.211 | 3 | 52.86701 | yes | 1.169E-07 | 532.97211 | 136.43515 |
| EGYAWAEDKEHC | 17 | Q9Y3I0 | RNA-splicing ligase RtcB homolog OS=H | RTCB | 55.211 | 3 | 52.86701 | yes | 2.455E-05 | 710.2937 | 136.43515 |
| GFVPNMQVEGVFY | 20 | Q9Y3I0 | RNA-splicing ligase RtcB homolog OS=H | RTCB | 55.211 | 3 | 52.86701 | yes | 1.39E-07 | 752.70502 | 99.825706 |
| RGLPQLGTLGAGN | 31 | Q9Y3I0 | RNA-splicing ligase RtcB homolog OS=H | RTCB | 55.211 | 4 | 52.86701 | yes | 1.283E-07 | 844.18481 | 81.302094 |
| INKNCWR | 7 | Q9Y3I0 | RNA-splicing ligase RtcB homolog OS=H | RTCB | 55.211 | 2 | 52.86701 | yes | 7.408E-07 | 495.75055 | 44.987686 |
| LQLLPESFICK | 11 | Q9Y3I1 | F-box only protein 7 OS=Homo sapiens ( | FBXO7 | 58.503 | 2 | 47.326687 | yes | 1.354E-07 | 674.37067 | 61.159725 |
| TGQPLPQSIQQALI | 14 | Q9Y3M8 | StAR-related lipid transfer protein 13 OS= | STARD13 | 124.968 | 2 | 34.835217 | yes | 9.325E-07 | 768.92828 | 42.633255 |
| VDDLYTLLPR | 10 | Q9Y3M8 | StAR-related lipid transfer protein 13 OS= | STARD13 | 124.968 | 2 | 34.835217 | yes | 1.687E-05 | 602.83228 | 24.56485 |
| LPICAQCGK | 9 | Q9Y3S2 | Zinc finger protein 330 OS=Homo sapien | ZNF330 | 36.202 | 2 | 53.28455 | yes | 4.336E-07 | 523.7597 | 421.10138 |
| ACFCDDHTR | 9 | Q9Y3S2 | Zinc finger protein 330 OS=Homo sapien | ZNF330 | 36.202 | 2 | 53.28455 | yes | 9.722E-08 | 591.22675 | 142.91675 |
| AFCYFCNSVQK | 11 | Q9Y3S2 | Zinc finger protein 330 OS=Homo sapien | ZNF330 | 36.202 | 2 | 53.28455 | yes | 9.722E-08 | 712.3103 | 81.882339 |
| TYASGYAHYEEQI | 14 | Q9Y3S2 | Zinc finger protein 330 OS=Homo sapien | ZNF330 | 36.202 | 2 | 53.28455 | yes | 2.614E-07 | 831.33936 | 12.991979 |
| LSNVNLQEK | 9 | Q9Y3T9 | Nucleolar complex protein 2 homolog OS | NOC2L | 84.92 | 2 | 46.668243 | yes | 2.551E-07 | 522.78784 | 219.20874 |
| LEDLNFPEIK | 10 | Q9Y3T9 | Nucleolar complex protein 2 homolog OS | NOC2L | 84.92 | 2 | 46.668243 | yes | 3.437E-07 | 609.32428 | 175.09428 |
| LFHEVVQAFR | 10 | Q9Y3T9 | Nucleolar complex protein 2 homolog OS | NOC2L | 84.92 | 2 | 46.668243 | yes | 1.841E-07 | 623.34058 | 87.202171 |
| LFHEVVQAFR | 10 | Q9Y3T9 | Nucleolar complex protein 2 homolog OS | NOC2L | 84.92 | 2 | 46.668243 | yes | 2.069E-07 | 415.89615 | 87.202171 |

| | | | | | | | | | | | |
|---|---|---|---|---|---|---|---|---|---|---|---|
| LFHEVVQAFR | 10 | Q9Y3T9 | Nucleolar complex protein 2 homolog OS | NOC2L | 84.92 | 3 | 46.668243 | yes | 1.841E-07 | 623.34058 | 87.202171 |
| LFHEVVQAFR | 10 | Q9Y3T9 | Nucleolar complex protein 2 homolog OS | NOC2L | 84.92 | 3 | 46.668243 | yes | 2.069E-07 | 415.89615 | 87.202171 |
| NSVPVTVAMVER | 12 | Q9Y3T9 | Nucleolar complex protein 2 homolog OS | NOC2L | 84.92 | 2 | 46.668243 | yes | 9.085E-07 | 651.34772 | 159.89403 |
| PINFSVILK | 9 | Q9Y3T9 | Nucleolar complex protein 2 homolog OS | NOC2L | 84.92 | 2 | 46.668243 | yes | 1.688E-07 | 515.81842 | 125.7933 |
| VQENSAYICSR | 11 | Q9Y3T9 | Nucleolar complex protein 2 homolog OS | NOC2L | 84.92 | 2 | 46.668243 | yes | 1.178E-06 | 663.80896 | 120.74797 |
| DTFLGPVLK | 9 | Q9Y3T9 | Nucleolar complex protein 2 homolog OS | NOC2L | 84.92 | 2 | 46.668243 | yes | 3.301E-07 | 495.28696 | 94.527748 |
| DRDPEFYK | 8 | Q9Y3T9 | Nucleolar complex protein 2 homolog OS | NOC2L | 84.92 | 2 | 46.668243 | yes | 1.5E-07 | 535.2511 | 68.274529 |
| GILRPLSTR | 9 | Q9Y3T9 | Nucleolar complex protein 2 homolog OS | NOC2L | 84.92 | 2 | 46.668243 | yes | 3.945E-07 | 506.81674 | 39.570316 |
| EELSNVLAAMR | 11 | Q9Y3U8 | Large ribosomal subunit protein eL36 OS | RPL36 | 12.254 | 2 | 51.95422 | yes | 4.854E-07 | 616.81879 | 14460.578 |
| EELSNVLAAMR | 11 | Q9Y3U8 | Large ribosomal subunit protein eL36 OS | RPL36 | 12.254 | 2 | 51.95422 | yes | 9.722E-08 | 624.81628 | 14460.578 |
| EVCGFAPYER | 10 | Q9Y3U8 | Large ribosomal subunit protein eL36 OS | RPL36 | 12.254 | 2 | 51.95422 | yes | 2.113E-07 | 614.27679 | 23265.172 |
| YPMAVGLNK | 9 | Q9Y3U8 | Large ribosomal subunit protein eL36 OS | RPL36 | 12.254 | 2 | 51.95422 | yes | 1.198E-07 | 496.76532 | 7435.9873 |
| YPMAVGLNK | 9 | Q9Y3U8 | Large ribosomal subunit protein eL36 OS | RPL36 | 12.254 | 2 | 51.95422 | yes | 2.485E-07 | 504.76279 | 7435.9873 |
| KREELSNVLAAM | 13 | Q9Y3U8 | Large ribosomal subunit protein eL36 OS | RPL36 | 12.254 | 3 | 51.95422 | yes | 1.664E-07 | 506.28033 | 95.545395 |
| KREELSNVLAAM | 13 | Q9Y3U8 | Large ribosomal subunit protein eL36 OS | RPL36 | 12.254 | 3 | 51.95422 | yes | 1.999E-07 | 758.91687 | 95.545395 |
| KREELSNVLAAM | 13 | Q9Y3U8 | Large ribosomal subunit protein eL36 OS | RPL36 | 12.254 | 2 | 51.95422 | yes | 1.664E-07 | 506.28033 | 95.545395 |
| KREELSNVLAAM | 13 | Q9Y3U8 | Large ribosomal subunit protein eL36 OS | RPL36 | 12.254 | 2 | 51.95422 | yes | 1.999E-07 | 758.91687 | 95.545395 |
| REELSNVLAAMR | 12 | Q9Y3U8 | Large ribosomal subunit protein eL36 OS | RPL36 | 12.254 | 2 | 51.95422 | yes | 1.48E-07 | 694.86938 | 39.047352 |
| NKQPTPVNIR | 10 | Q9Y3Y2 | Chromatin target of PRMT1 protein OS= | CHTOP | 26.397 | 2 | 52.999947 | yes | 1.122E-07 | 583.83563 | 945.77747 |
| ASMQQQQQLASA | 13 | Q9Y3Y2 | Chromatin target of PRMT1 protein OS= | CHTOP | 26.397 | 2 | 52.999947 | yes | 9.722E-08 | 723.85956 | 722.34973 |
| EQLDNQLDAYMS | 13 | Q9Y3Y2 | Chromatin target of PRMT1 protein OS= | CHTOP | 26.397 | 2 | 52.999947 | yes | 1.1E-07 | 777.85889 | 630.13458 |
| QPTPVNIR | 8 | Q9Y3Y2 | Chromatin target of PRMT1 protein OS= | CHTOP | 26.397 | 2 | 52.999947 | yes | 2.317E-07 | 462.76672 | 256.18451 |
| LAQQMENR | 8 | Q9Y3Y2 | Chromatin target of PRMT1 protein OS= | CHTOP | 26.397 | 2 | 52.999947 | yes | 3.322E-07 | 495.24527 | 127.43659 |
| IIDTPQFQR | 9 | Q9Y3Z3 | Deoxynucleoside triphosphate triphospho | SAMHD1 | 72.201 | 2 | 47.56853 | yes | 3.969E-06 | 559.30365 | 105.70168 |
| SFLYEIVSNK | 10 | Q9Y3Z3 | Deoxynucleoside triphosphate triphospho | SAMHD1 | 72.201 | 2 | 47.56853 | yes | 5.349E-07 | 600.31897 | 82.975601 |
| FENLGVSSLGER | 12 | Q9Y3Z3 | Deoxynucleoside triphosphate triphospho | SAMHD1 | 72.201 | 2 | 47.56853 | yes | 3.596E-07 | 654.33313 | 79.713692 |
| YVGETQPTGQIK | 12 | Q9Y3Z3 | Deoxynucleoside triphosphate triphospho | SAMHD1 | 72.201 | 2 | 47.56853 | yes | 1.301E-07 | 660.84332 | 26.735308 |
| LSSGFDDIDLPSAV | 15 | Q9Y446 | Plakophilin-3 OS=Homo sapiens OX=96 | PKP3 | 87.083 | 2 | 52.581142 | yes | 1.156E-07 | 782.39868 | 2066.0903 |
| LLQLGQQPR | 9 | Q9Y446 | Plakophilin-3 OS=Homo sapiens OX=96 | PKP3 | 87.083 | 2 | 52.581142 | yes | 3.282E-07 | 526.81421 | 1633.4697 |
| GVGGAVPGAVLE | 16 | Q9Y446 | Plakophilin-3 OS=Homo sapiens OX=96 | PKP3 | 87.083 | 2 | 52.581142 | yes | 1.464E-07 | 724.91461 | 1548.3427 |
| AGGLDWPEATEV | 16 | Q9Y446 | Plakophilin-3 OS=Homo sapiens OX=96 | PKP3 | 87.083 | 2 | 52.581142 | yes | 9.809E-08 | 836.40228 | 1545.2428 |
| PAYSPASWSSR | 11 | Q9Y446 | Plakophilin-3 OS=Homo sapiens OX=96 | PKP3 | 87.083 | 2 | 52.581142 | yes | 9.85E-08 | 604.78839 | 1343.0262 |
| ILNPLLDR | 8 | Q9Y446 | Plakophilin-3 OS=Homo sapiens OX=96 | PKP3 | 87.083 | 2 | 52.581142 | yes | 1.786E-07 | 477.29257 | 1307.5511 |
| YLMASDPNLQVL | 20 | Q9Y446 | Plakophilin-3 OS=Homo sapiens OX=96 | PKP3 | 87.083 | 3 | 52.581142 | yes | 2.658E-07 | 744.72089 | 1285.0757 |
| LALEQER | 7 | Q9Y446 | Plakophilin-3 OS=Homo sapiens OX=96 | PKP3 | 87.083 | 2 | 52.581142 | yes | 3.717E-07 | 429.73761 | 1062.3984 |
| ELPLAADALTFAE | 16 | Q9Y446 | Plakophilin-3 OS=Homo sapiens OX=96 | PKP3 | 87.083 | 2 | 52.581142 | yes | 1.23E-07 | 837.95105 | 763.53625 |
| LYDEMPPSALQR | 12 | Q9Y446 | Plakophilin-3 OS=Homo sapiens OX=96 | PKP3 | 87.083 | 2 | 52.581142 | yes | 1.471E-07 | 710.35046 | 726.78363 |
| HTTEAAAGALQN | 18 | Q9Y446 | Plakophilin-3 OS=Homo sapiens OX=96 | PKP3 | 87.083 | 3 | 52.581142 | yes | 9.848E-08 | 599.6344 | 707.56781 |
| AASSLLANLWQY | 14 | Q9Y446 | Plakophilin-3 OS=Homo sapiens OX=96 | PKP3 | 87.083 | 2 | 52.581142 | yes | 9.722E-08 | 789.91736 | 660.08643 |

| | | | | | | | | | | | |
|---|---|---|---|---|---|---|---|---|---|---|---|
| ADYDTLSLR | 9 | Q9Y446 | Plakophilin-3 OS=Homo sapiens OX=96( | PKP3 | 87.083 | 2 | 52.581142 | yes | 1.735E-07 | 527.26422 | 612.99158 |
| LGPGGLDDR | 9 | Q9Y446 | Plakophilin-3 OS=Homo sapiens OX=96( | PKP3 | 87.083 | 2 | 52.581142 | yes | 1.477E-07 | 450.2327 | 607.40399 |
| VVSHLIEK | 8 | Q9Y446 | Plakophilin-3 OS=Homo sapiens OX=96( | PKP3 | 87.083 | 2 | 52.581142 | yes | 1.446E-07 | 462.7793 | 502.78802 |
| LFNHANQEVQR | 11 | Q9Y446 | Plakophilin-3 OS=Homo sapiens OX=96( | PKP3 | 87.083 | 2 | 52.581142 | yes | 9.722E-08 | 678.34436 | 475.49542 |
| DLLYFDGLR | 9 | Q9Y446 | Plakophilin-3 OS=Homo sapiens OX=96( | PKP3 | 87.083 | 2 | 52.581142 | yes | 9.722E-08 | 556.29279 | 430.21109 |
| GQYHTLQAGFSSR | 13 | Q9Y446 | Plakophilin-3 OS=Homo sapiens OX=96( | PKP3 | 87.083 | 2 | 52.581142 | yes | 3.017E-07 | 726.35492 | 183.69321 |
| GQYHTLQAGFSSR | 13 | Q9Y446 | Plakophilin-3 OS=Homo sapiens OX=96( | PKP3 | 87.083 | 2 | 52.581142 | yes | 2.128E-07 | 484.57239 | 183.69321 |
| GQYHTLQAGFSSR | 13 | Q9Y446 | Plakophilin-3 OS=Homo sapiens OX=96( | PKP3 | 87.083 | 3 | 52.581142 | yes | 3.017E-07 | 726.35492 | 183.69321 |
| GQYHTLQAGFSSR | 13 | Q9Y446 | Plakophilin-3 OS=Homo sapiens OX=96( | PKP3 | 87.083 | 3 | 52.581142 | yes | 2.128E-07 | 484.57239 | 183.69321 |
| NLIYDNADNK | 10 | Q9Y446 | Plakophilin-3 OS=Homo sapiens OX=96( | PKP3 | 87.083 | 2 | 52.581142 | yes | 9.819E-08 | 590.28564 | 357.12088 |
| NLSSASQATR | 10 | Q9Y446 | Plakophilin-3 OS=Homo sapiens OX=96( | PKP3 | 87.083 | 2 | 52.581142 | yes | 9.722E-08 | 517.76489 | 323.40369 |
| SVENAVCVLR | 10 | Q9Y446 | Plakophilin-3 OS=Homo sapiens OX=96( | PKP3 | 87.083 | 2 | 52.581142 | yes | 1.152E-07 | 573.80042 | 273.68707 |
| TSGFRPIAK | 9 | Q9Y446 | Plakophilin-3 OS=Homo sapiens OX=96( | PKP3 | 87.083 | 2 | 52.581142 | yes | 1.018E-07 | 488.78235 | 156.48816 |
| PVSFHER | 7 | Q9Y446 | Plakophilin-3 OS=Homo sapiens OX=96( | PKP3 | 87.083 | 2 | 52.581142 | yes | 4.199E-07 | 436.22467 | 121.2272 |
| SAVDLSCSR | 9 | Q9Y446 | Plakophilin-3 OS=Homo sapiens OX=96( | PKP3 | 87.083 | 2 | 52.581142 | yes | 2.224E-07 | 497.73474 | 118.87016 |
| HTTEAAAGALQNI | 19 | Q9Y446 | Plakophilin-3 OS=Homo sapiens OX=96( | PKP3 | 87.083 | 3 | 52.581142 | yes | 2.628E-06 | 651.66809 | 44.656963 |
| GLDPALGSETLASI | 14 | Q9Y448 | Small kinetochore-associated protein OS= | KNSTRN | 35.439 | 2 | 50.710754 | yes | 1.244E-07 | 693.86481 | 9783.626 |
| NQLLEAVNK | 9 | Q9Y448 | Small kinetochore-associated protein OS= | KNSTRN | 35.439 | 2 | 50.710754 | yes | 9.728E-08 | 514.79041 | 3249.5728 |
| DNCLAILESK | 10 | Q9Y448 | Small kinetochore-associated protein OS= | KNSTRN | 35.439 | 2 | 50.710754 | yes | 1.364E-07 | 581.79224 | 1989.2863 |
| QMEELQALK | 9 | Q9Y448 | Small kinetochore-associated protein OS= | KNSTRN | 35.439 | 2 | 50.710754 | yes | 1.002E-07 | 545.28406 | 1970.1716 |
| LTETQGELK | 9 | Q9Y448 | Small kinetochore-associated protein OS= | KNSTRN | 35.439 | 2 | 50.710754 | yes | 2.323E-07 | 509.77438 | 1899.3541 |
| KQMEELQALK | 10 | Q9Y448 | Small kinetochore-associated protein OS= | KNSTRN | 35.439 | 2 | 50.710754 | yes | 1.046E-07 | 609.33154 | 770.56494 |
| FRDNCLAILESK | 12 | Q9Y448 | Small kinetochore-associated protein OS= | KNSTRN | 35.439 | 2 | 50.710754 | yes | 1.37E-07 | 733.37701 | 445.58936 |
| DKNQLLEAVNK | 11 | Q9Y448 | Small kinetochore-associated protein OS= | KNSTRN | 35.439 | 2 | 50.710754 | yes | 4.163E-07 | 636.35132 | 311.29764 |
| QLHSGGPENDVTK | 13 | Q9Y448 | Small kinetochore-associated protein OS= | KNSTRN | 35.439 | 2 | 50.710754 | yes | 1.076E-07 | 691.33893 | 238.39894 |
| SEEELKDK | 8 | Q9Y448 | Small kinetochore-associated protein OS= | KNSTRN | 35.439 | 2 | 50.710754 | yes | 1.213E-07 | 489.24292 | 133.98709 |
| KGYKPLSK | 8 | Q9Y448 | Small kinetochore-associated protein OS= | KNSTRN | 35.439 | 2 | 50.710754 | yes | 1.857E-07 | 460.78183 | 121.70551 |
| LFNETAK | 7 | Q9Y448 | Small kinetochore-associated protein OS= | KNSTRN | 35.439 | 2 | 50.710754 | yes | 1.083E-07 | 411.72144 | 77.612259 |
| VITLMDAPGHK | 11 | Q9Y450 | HBS1-like protein OS=Homo sapiens OX | HBS1L | 75.473 | 2 | 53.36759 | yes | 9.722E-08 | 591.32098 | 3205.2175 |
| INVGCIFCGPK | 11 | Q9Y450 | HBS1-like protein OS=Homo sapiens OX | HBS1L | 75.473 | 2 | 53.36759 | yes | 1.26E-07 | 632.8125 | 3043.2402 |
| SLGVTQLAVAVNI | 13 | Q9Y450 | HBS1-like protein OS=Homo sapiens OX | HBS1L | 75.473 | 2 | 53.36759 | yes | 3.234E-07 | 650.38519 | 2891.7302 |
| EHGLLVR | 7 | Q9Y450 | HBS1-like protein OS=Homo sapiens OX | HBS1L | 75.473 | 2 | 53.36759 | yes | 2.442E-07 | 412.24286 | 1980.9138 |
| ALSGVLEQDR | 10 | Q9Y450 | HBS1-like protein OS=Homo sapiens OX | HBS1L | 75.473 | 2 | 53.36759 | yes | 1.022E-07 | 544.29077 | 1865.0612 |
| SIDKPFR | 7 | Q9Y450 | HBS1-like protein OS=Homo sapiens OX | HBS1L | 75.473 | 2 | 53.36759 | yes | 1.3E-07 | 431.74271 | 1807.4596 |
| IEAGYIQTGDR | 11 | Q9Y450 | HBS1-like protein OS=Homo sapiens OX | HBS1L | 75.473 | 2 | 53.36759 | yes | 1.083E-07 | 611.80676 | 1776.649 |
| QLLNLVVIGHVDA | 15 | Q9Y450 | HBS1-like protein OS=Homo sapiens OX | HBS1L | 75.473 | 2 | 53.36759 | yes | 1.465E-07 | 788.46448 | 876.56067 |
| QLLNLVVIGHVDA | 15 | Q9Y450 | HBS1-like protein OS=Homo sapiens OX | HBS1L | 75.473 | 2 | 53.36759 | yes | 1.281E-07 | 525.97876 | 876.56067 |
| QLLNLVVIGHVDA | 15 | Q9Y450 | HBS1-like protein OS=Homo sapiens OX | HBS1L | 75.473 | 3 | 53.36759 | yes | 1.465E-07 | 788.46448 | 876.56067 |
| QLLNLVVIGHVDA | 15 | Q9Y450 | HBS1-like protein OS=Homo sapiens OX | HBS1L | 75.473 | 3 | 53.36759 | yes | 1.281E-07 | 525.97876 | 876.56067 |

| | | | | | | | | | | | |
|---|---|---|---|---|---|---|---|---|---|---|---|
| GQNALVELQTQR | 12 | Q9Y450 | HBS1-like protein OS=Homo sapiens OX | HBS1L | 75.473 | 2 | 53.36759 | yes | 9.722E-08 | 678.86511 | 1355.2816 |
| MDQVNWQQER | 10 | Q9Y450 | HBS1-like protein OS=Homo sapiens OX | HBS1L | 75.473 | 2 | 53.36759 | yes | 1.011E-07 | 667.30133 | 1316.6001 |
| PIALELYK | 8 | Q9Y450 | HBS1-like protein OS=Homo sapiens OX | HBS1L | 75.473 | 2 | 53.36759 | yes | 2.522E-07 | 473.78403 | 1216.793 |
| LYSCLDHMR | 9 | Q9Y450 | HBS1-like protein OS=Homo sapiens OX | HBS1L | 75.473 | 2 | 53.36759 | yes | 1.277E-07 | 597.77338 | 1097.6714 |
| YGGSTIAAGVVTE | 16 | Q9Y450 | HBS1-like protein OS=Homo sapiens OX | HBS1L | 75.473 | 2 | 53.36759 | yes | 9.926E-08 | 797.9198 | 1089.4958 |
| LCVSDVFK | 8 | Q9Y450 | HBS1-like protein OS=Homo sapiens OX | HBS1L | 75.473 | 2 | 53.36759 | yes | 2.716E-07 | 484.24951 | 1069.3135 |
| DQGSGFCITGK | 11 | Q9Y450 | HBS1-like protein OS=Homo sapiens OX | HBS1L | 75.473 | 2 | 53.36759 | yes | 9.722E-08 | 585.26642 | 1018.0627 |
| LLAMPPNETCTVK | 13 | Q9Y450 | HBS1-like protein OS=Homo sapiens OX | HBS1L | 75.473 | 2 | 53.36759 | yes | 1.37E-07 | 737.37561 | 966.91724 |
| GVTMDVGMTK | 10 | Q9Y450 | HBS1-like protein OS=Homo sapiens OX | HBS1L | 75.473 | 2 | 53.36759 | yes | 1.642E-06 | 519.75153 | 412.25421 |
| YGGSTIAAGVVTE | 15 | Q9Y450 | HBS1-like protein OS=Homo sapiens OX | HBS1L | 75.473 | 2 | 53.36759 | yes | 9.722E-08 | 733.3985 | 386.37396 |
| GFPVLLHYQTVSE | 18 | Q9Y450 | HBS1-like protein OS=Homo sapiens OX | HBS1L | 75.473 | 3 | 53.36759 | yes | 1.05E-07 | 666.70679 | 220.40987 |
| GLCLLEQIDSFKPP | 16 | Q9Y450 | HBS1-like protein OS=Homo sapiens OX | HBS1L | 75.473 | 3 | 53.36759 | yes | 4.422E-07 | 634.33649 | 143.08592 |
| GITLHDEPVDWAA | 29 | Q9Y450 | HBS1-like protein OS=Homo sapiens OX | HBS1L | 75.473 | 4 | 53.36759 | yes | 2.409E-07 | 769.14703 | 114.71631 |
| GPPIEDAIASSDVL | 19 | Q9Y450 | HBS1-like protein OS=Homo sapiens OX | HBS1L | 75.473 | 3 | 53.36759 | yes | 2.988E-07 | 633.98969 | 73.334419 |
| SANPPHTIQASEEQ | 22 | Q9Y450 | HBS1-like protein OS=Homo sapiens OX | HBS1L | 75.473 | 3 | 53.36759 | yes | 9.722E-08 | 759.37665 | 71.139923 |
| RLISVLNK | 8 | Q9Y450 | HBS1-like protein OS=Homo sapiens OX | HBS1L | 75.473 | 2 | 53.36759 | yes | 5.46E-07 | 471.80838 | 61.769806 |
| SWPAAIPHLR | 10 | Q9Y483 | Metal-response element-binding transcrip | MTF2 | 67.09 | 2 | 53.549007 | yes | 9.722E-08 | 574.3222 | 108.36107 |
| ALQTQNSEIVK | 11 | Q9Y483 | Metal-response element-binding transcrip | MTF2 | 67.09 | 2 | 53.549007 | yes | 3.962E-07 | 615.83807 | 56.552086 |
| YEHVLEALNDYK | 12 | Q9Y483 | Metal-response element-binding transcrip | MTF2 | 67.09 | 2 | 53.549007 | yes | 3.392E-07 | 747.36719 | 49.213226 |
| AQEACGPLEMDSA | 22 | Q9Y490 | Talin-1 OS=Homo sapiens OX=9606 GN | TLN1 | 269.768 | 3 | 53.0562 | yes | 8.495E-07 | 802.38226 | 4137.0845 |
| AVAEQIPLLVQGV | 14 | Q9Y490 | Talin-1 OS=Homo sapiens OX=9606 GN | TLN1 | 269.768 | 2 | 53.0562 | yes | 5.244E-07 | 746.94592 | 135.79602 |
| ALEATTEHIR | 10 | Q9Y490 | Talin-1 OS=Homo sapiens OX=9606 GN | TLN1 | 269.768 | 2 | 53.0562 | yes | 9.722E-08 | 570.80402 | 100.16332 |
| NLGTALAELR | 10 | Q9Y490 | Talin-1 OS=Homo sapiens OX=9606 GN | TLN1 | 269.768 | 2 | 53.0562 | yes | 7.659E-07 | 529.30365 | 95.560059 |
| TMLESAGGLIQTA | 14 | Q9Y490 | Talin-1 OS=Homo sapiens OX=9606 GN | TLN1 | 269.768 | 2 | 53.0562 | yes | 1.229E-07 | 724.38232 | 80.8573 |
| VMVTNVTSLLK | 11 | Q9Y490 | Talin-1 OS=Homo sapiens OX=9606 GN | TLN1 | 269.768 | 2 | 53.0562 | no | 1.345E-07 | 602.85211 | 64.926811 |
| LAQAAQSSVATITI | 14 | Q9Y490 | Talin-1 OS=Homo sapiens OX=9606 GN | TLN1 | 269.768 | 2 | 53.0562 | yes | 2.745E-07 | 708.89392 | 35.122662 |
| TVTDMLMTICAR | 12 | Q9Y490 | Talin-1 OS=Homo sapiens OX=9606 GN | TLN1 | 269.768 | 2 | 53.0562 | yes | 6.084E-07 | 706.34076 | 34.815254 |
| LLAALLEDEGGSG | 21 | Q9Y490 | Talin-1 OS=Homo sapiens OX=9606 GN | TLN1 | 269.768 | 3 | 53.0562 | yes | 3.026E-06 | 708.06708 | 24.59621 |
| EIEGLLENIR | 10 | Q9Y496 | Kinesin-like protein KIF3A OS=Homo sa | KIF3A | 80.041 | 2 | 50.81642 | yes | 1.003E-07 | 593.32733 | 113.80534 |
| VIVGGVDLLAK | 11 | Q9Y496 | Kinesin-like protein KIF3A OS=Homo sa | KIF3A | 80.041 | 2 | 50.81642 | yes | 6.449E-07 | 542.34222 | 93.079781 |
| TFTFDTVFGPESK | 13 | Q9Y496 | Kinesin-like protein KIF3A OS=Homo sa | KIF3A | 80.041 | 2 | 50.81642 | yes | 9.894E-08 | 738.35626 | 72.02047 |
| LLEESNMELEER | 12 | Q9Y496 | Kinesin-like protein KIF3A OS=Homo sa | KIF3A | 80.041 | 2 | 50.81642 | yes | 9.722E-08 | 746.35339 | 67.630775 |
| VSYLEIYNEEVR | 12 | Q9Y496 | Kinesin-like protein KIF3A OS=Homo sa | KIF3A | 80.041 | 2 | 50.81642 | yes | 1.75E-06 | 757.38031 | 38.741516 |
| AMLIEAGSPFR | 11 | Q9Y4A5 | Transformation/transcription domain-assc | TRRAP | 437.601 | 2 | 52.94587 | yes | 2.687E-07 | 596.31317 | 145.24672 |
| LFDESILIGSGYTAI | 15 | Q9Y4A5 | Transformation/transcription domain-assc | TRRAP | 437.601 | 2 | 52.94587 | yes | 5.158E-07 | 821.4278 | 143.72885 |
| SVTVIMEPHK | 10 | Q9Y4A5 | Transformation/transcription domain-assc | TRRAP | 437.601 | 2 | 52.94587 | yes | 2.387E-07 | 570.80774 | 133.21133 |
| VIYEGLTNYEK | 11 | Q9Y4A5 | Transformation/transcription domain-assc | TRRAP | 437.601 | 2 | 52.94587 | yes | 1.166E-07 | 664.84027 | 105.25376 |
| VYPQAVYFPIR | 11 | Q9Y4A5 | Transformation/transcription domain-assc | TRRAP | 437.601 | 2 | 52.94587 | yes | 1.249E-07 | 676.8717 | 95.357742 |
| YQLSAIFK | 8 | Q9Y4A5 | Transformation/transcription domain-assc | TRRAP | 437.601 | 2 | 52.94587 | yes | 4.626E-06 | 485.27383 | 87.029076 |

| | | | | | | | | | | | |
|---|---|---|---|---|---|---|---|---|---|---|---|
| TATGAISAVFGR | 12 | Q9Y4A5 | Transformation/transcription domain-assc | TRRAP | 437.601 | 2 | 52.94587 | yes | 2.164E-07 | 575.81439 | 85.863228 |
| QAMAILTPAVPAR | 13 | Q9Y4A5 | Transformation/transcription domain-assc | TRRAP | 437.601 | 2 | 52.94587 | yes | 2.144E-07 | 669.88177 | 85.551651 |
| TIPNVIISHR | 10 | Q9Y4A5 | Transformation/transcription domain-assc | TRRAP | 437.601 | 2 | 52.94587 | yes | 1.423E-07 | 575.34058 | 81.866676 |
| IIIELHK | 7 | Q9Y4A5 | Transformation/transcription domain-assc | TRRAP | 437.601 | 2 | 52.94587 | yes | 6.663E-07 | 433.27893 | 81.847961 |
| APGEAQFIPNK | 11 | Q9Y4A5 | Transformation/transcription domain-assc | TRRAP | 437.601 | 2 | 52.94587 | yes | 2.105E-07 | 586.30896 | 80.71434 |
| LDLQFQAIK | 9 | Q9Y4A5 | Transformation/transcription domain-assc | TRRAP | 437.601 | 2 | 52.94587 | yes | 1.718E-07 | 538.31097 | 72.598488 |
| LGFTPSVTIEQR | 12 | Q9Y4A5 | Transformation/transcription domain-assc | TRRAP | 437.601 | 2 | 52.94587 | yes | 3.556E-07 | 674.367 | 65.423904 |
| LEPAFLSGLR | 10 | Q9Y4A5 | Transformation/transcription domain-assc | TRRAP | 437.601 | 2 | 52.94587 | yes | 9.722E-08 | 551.81641 | 64.023438 |
| EALVQVEVSCPK | 12 | Q9Y4A5 | Transformation/transcription domain-assc | TRRAP | 437.601 | 2 | 52.94587 | yes | 1.219E-06 | 679.85284 | 59.961197 |
| TITWGITSCK | 10 | Q9Y4A5 | Transformation/transcription domain-assc | TRRAP | 437.601 | 2 | 52.94587 | yes | 1.127E-06 | 583.79736 | 59.417828 |
| LVLEIIHR | 8 | Q9Y4A5 | Transformation/transcription domain-assc | TRRAP | 437.601 | 2 | 52.94587 | yes | 9.722E-08 | 496.81619 | 57.336147 |
| AELMQALWR | 9 | Q9Y4A5 | Transformation/transcription domain-assc | TRRAP | 437.601 | 2 | 52.94587 | yes | 3.553E-07 | 559.2948 | 54.001888 |
| YEELECLYAAVGK | 13 | Q9Y4A5 | Transformation/transcription domain-assc | TRRAP | 437.601 | 2 | 52.94587 | yes | 1.521E-07 | 772.86871 | 50.257286 |
| SACSNNPSYIDR | 12 | Q9Y4A5 | Transformation/transcription domain-assc | TRRAP | 437.601 | 2 | 52.94587 | yes | 2.571E-07 | 692.30151 | 39.024632 |
| LAVDLSEVVIK | 11 | Q9Y4A5 | Transformation/transcription domain-assc | TRRAP | 437.601 | 2 | 52.94587 | yes | 9.722E-08 | 593.35809 | 34.75198 |
| SLPSLVPLR | 9 | Q9Y4A5 | Transformation/transcription domain-assc | TRRAP | 437.601 | 2 | 52.94587 | yes | 3.291E-07 | 491.3082 | 32.277241 |
| GTQASHQVLR | 10 | Q9Y4A5 | Transformation/transcription domain-assc | TRRAP | 437.601 | 2 | 52.94587 | yes | 5.693E-07 | 548.79651 | 30.028469 |
| QHTDTVVNFLIR | 12 | Q9Y4A5 | Transformation/transcription domain-assc | TRRAP | 437.601 | 2 | 52.94587 | yes | 6.967E-07 | 721.89117 | 29.306833 |
| GIAACMTCGNTK | 12 | Q9Y4A5 | Transformation/transcription domain-assc | TRRAP | 437.601 | 2 | 52.94587 | yes | 3.515E-07 | 642.2807 | 28.641691 |
| FVDFNDPNFGDEL | 14 | Q9Y4A5 | Transformation/transcription domain-assc | TRRAP | 437.601 | 2 | 52.94587 | yes | 8.549E-07 | 828.88068 | 27.466173 |
| ELYADFIAAQIK | 12 | Q9Y4A5 | Transformation/transcription domain-assc | TRRAP | 437.601 | 2 | 52.94587 | yes | 4.581E-06 | 691.37177 | 26.84931 |
| KLVLEIIHR | 9 | Q9Y4A5 | Transformation/transcription domain-assc | TRRAP | 437.601 | 2 | 52.94587 | yes | 1.41E-07 | 560.86371 | 21.090239 |
| LVPAAEEEELLR | 12 | Q9Y4B5 | Microtubule cross-linking factor 1 OS=H | MTCL1 | 209.526 | 2 | 54.34099 | yes | 1.921E-07 | 684.87207 | 12640.422 |
| LVEEEANILGR | 11 | Q9Y4B5 | Microtubule cross-linking factor 1 OS=H | MTCL1 | 209.526 | 2 | 54.34099 | no | 9.722E-08 | 621.83807 | 7731.0854 |
| PAAPGPAVPSSGR | 13 | Q9Y4B5 | Microtubule cross-linking factor 1 OS=H | MTCL1 | 209.526 | 2 | 54.34099 | yes | 9.722E-08 | 582.31201 | 7223.1782 |
| VAETGQVDGELIR | 13 | Q9Y4B5 | Microtubule cross-linking factor 1 OS=H | MTCL1 | 209.526 | 2 | 54.34099 | yes | 9.968E-08 | 693.86481 | 5143.4341 |
| AGKPPGAEPPSAA | 15 | Q9Y4B5 | Microtubule cross-linking factor 1 OS=H | MTCL1 | 209.526 | 2 | 54.34099 | yes | 1.845E-07 | 674.86462 | 2235.2551 |
| AGKPPGAEPPSAA | 15 | Q9Y4B5 | Microtubule cross-linking factor 1 OS=H | MTCL1 | 209.526 | 2 | 54.34099 | yes | 9.759E-08 | 450.24548 | 2235.2551 |
| AGKPPGAEPPSAA | 15 | Q9Y4B5 | Microtubule cross-linking factor 1 OS=H | MTCL1 | 209.526 | 3 | 54.34099 | yes | 1.845E-07 | 674.86462 | 2235.2551 |
| AGKPPGAEPPSAA | 15 | Q9Y4B5 | Microtubule cross-linking factor 1 OS=H | MTCL1 | 209.526 | 3 | 54.34099 | yes | 9.759E-08 | 450.24548 | 2235.2551 |
| EQLEWQLGPAR | 11 | Q9Y4B5 | Microtubule cross-linking factor 1 OS=H | MTCL1 | 209.526 | 2 | 54.34099 | yes | 1.17E-07 | 663.84369 | 4445.917 |
| APPSPGSLAAPGR | 13 | Q9Y4B5 | Microtubule cross-linking factor 1 OS=H | MTCL1 | 209.526 | 2 | 54.34099 | yes | 1.901E-07 | 589.31982 | 3967.6006 |
| IVELEVENR | 9 | Q9Y4B5 | Microtubule cross-linking factor 1 OS=H | MTCL1 | 209.526 | 2 | 54.34099 | no | 1.054E-07 | 550.80096 | 3948.0347 |
| LGELGSSAESK | 11 | Q9Y4B5 | Microtubule cross-linking factor 1 OS=H | MTCL1 | 209.526 | 2 | 54.34099 | yes | 1.091E-07 | 539.27478 | 3144.2263 |
| EKVELLDR | 8 | Q9Y4B5 | Microtubule cross-linking factor 1 OS=H | MTCL1 | 209.526 | 2 | 54.34099 | yes | 1.557E-07 | 501.28494 | 3110.2437 |
| SISEIEDHNR | 10 | Q9Y4B5 | Microtubule cross-linking factor 1 OS=H | MTCL1 | 209.526 | 2 | 54.34099 | yes | 9.734E-08 | 600.28619 | 3053.3413 |
| AVSVSSMSEFQR | 12 | Q9Y4B5 | Microtubule cross-linking factor 1 OS=H | MTCL1 | 209.526 | 2 | 54.34099 | yes | 3.431E-07 | 664.31915 | 1447.4709 |
| AVSVSSMSEFQR | 12 | Q9Y4B5 | Microtubule cross-linking factor 1 OS=H | MTCL1 | 209.526 | 2 | 54.34099 | yes | 2.364E-07 | 672.31665 | 1447.4709 |
| LQISELSGK | 9 | Q9Y4B5 | Microtubule cross-linking factor 1 OS=H | MTCL1 | 209.526 | 2 | 54.34099 | yes | 2.184E-07 | 487.77948 | 2828.9568 |

| | | | | | | | | | | | |
|---|---|---|---|---|---|---|---|---|---|---|---|
| CDLAAHLGLR | 10 | Q9Y4B5 | Microtubule cross-linking factor 1 OS=H | MTCL1 | 209.526 | 2 | 54.34099 | yes | 1.036E-07 | 563.29529 | 2733.1355 |
| NAICSGPGELQVK | 13 | Q9Y4B5 | Microtubule cross-linking factor 1 OS=H | MTCL1 | 209.526 | 2 | 54.34099 | yes | 1.286E-07 | 686.84808 | 2728.251 |
| SHVLTEQSGLR | 11 | Q9Y4B5 | Microtubule cross-linking factor 1 OS=H | MTCL1 | 209.526 | 2 | 54.34099 | yes | 9.819E-08 | 613.828 | 2545.4954 |
| QALQNELER | 9 | Q9Y4B5 | Microtubule cross-linking factor 1 OS=H | MTCL1 | 209.526 | 2 | 54.34099 | yes | 9.722E-08 | 550.78839 | 2504.3848 |
| LMDISPFLPEK | 11 | Q9Y4B5 | Microtubule cross-linking factor 1 OS=H | MTCL1 | 209.526 | 2 | 54.34099 | yes | 9.722E-08 | 645.34412 | 1251.9802 |
| LMDISPFLPEK | 11 | Q9Y4B5 | Microtubule cross-linking factor 1 OS=H | MTCL1 | 209.526 | 2 | 54.34099 | yes | 5.733E-07 | 653.34161 | 1251.9802 |
| NLMQQELR | 8 | Q9Y4B5 | Microtubule cross-linking factor 1 OS=H | MTCL1 | 209.526 | 2 | 54.34099 | yes | 1.813E-07 | 516.26874 | 2308.1658 |
| VQIGDHSLR | 9 | Q9Y4B5 | Microtubule cross-linking factor 1 OS=H | MTCL1 | 209.526 | 2 | 54.34099 | yes | 6.12E-07 | 512.78033 | 2306.3242 |
| LLADSHSLVMDLR | 13 | Q9Y4B5 | Microtubule cross-linking factor 1 OS=H | MTCL1 | 209.526 | 2 | 54.34099 | yes | 1.115E-07 | 735.3927 | 2123.9949 |
| QQMIEVEISK | 10 | Q9Y4B5 | Microtubule cross-linking factor 1 OS=H | MTCL1 | 209.526 | 2 | 54.34099 | yes | 9.722E-08 | 602.81573 | 1973.7491 |
| QMEEEGEEFTEGE | 20 | Q9Y4B5 | Microtubule cross-linking factor 1 OS=H | MTCL1 | 209.526 | 3 | 54.34099 | yes | 1.559E-07 | 788.66925 | 982.66016 |
| QMEEEGEEFTEGE | 20 | Q9Y4B5 | Microtubule cross-linking factor 1 OS=H | MTCL1 | 209.526 | 3 | 54.34099 | yes | 1.189E-07 | 794.00092 | 982.66016 |
| TFNQDDSADLR | 11 | Q9Y4B5 | Microtubule cross-linking factor 1 OS=H | MTCL1 | 209.526 | 2 | 54.34099 | yes | 9.803E-08 | 641.28894 | 1808.5436 |
| WQIHHSEK | 8 | Q9Y4B5 | Microtubule cross-linking factor 1 OS=H | MTCL1 | 209.526 | 2 | 54.34099 | yes | 1.423E-07 | 532.76721 | 1767.8311 |
| QVVENQQLFSAFK | 13 | Q9Y4B5 | Microtubule cross-linking factor 1 OS=H | MTCL1 | 209.526 | 2 | 54.34099 | yes | 9.849E-08 | 769.40411 | 1673.3379 |
| EPSLGEQSR | 9 | Q9Y4B5 | Microtubule cross-linking factor 1 OS=H | MTCL1 | 209.526 | 2 | 54.34099 | yes | 1.886E-07 | 501.74619 | 1585.1356 |
| WPCTSPR | 7 | Q9Y4B5 | Microtubule cross-linking factor 1 OS=H | MTCL1 | 209.526 | 2 | 54.34099 | yes | 1.781E-07 | 452.21072 | 1540.9965 |
| EEAFLMR | 7 | Q9Y4B5 | Microtubule cross-linking factor 1 OS=H | MTCL1 | 209.526 | 2 | 54.34099 | yes | 2.214E-07 | 448.22073 | 1521.4487 |
| LITDTDSFLHDAGI | 15 | Q9Y4B5 | Microtubule cross-linking factor 1 OS=H | MTCL1 | 209.526 | 2 | 54.34099 | yes | 1.662E-07 | 837.42828 | 1494.4351 |
| EDVTPPLSPDDLK | 13 | Q9Y4B5 | Microtubule cross-linking factor 1 OS=H | MTCL1 | 209.526 | 2 | 54.34099 | yes | 8.701E-07 | 713.35901 | 1470.8951 |
| KIVELEVENR | 10 | Q9Y4B5 | Microtubule cross-linking factor 1 OS=H | MTCL1 | 209.526 | 2 | 54.34099 | no | 9.917E-08 | 614.84845 | 1455.6024 |
| GYSESAWAR | 9 | Q9Y4B5 | Microtubule cross-linking factor 1 OS=H | MTCL1 | 209.526 | 2 | 54.34099 | yes | 1.682E-07 | 513.7356 | 1405.541 |
| LLEHALK | 7 | Q9Y4B5 | Microtubule cross-linking factor 1 OS=H | MTCL1 | 209.526 | 2 | 54.34099 | yes | 2.453E-07 | 412.25543 | 1345.4276 |
| AGGGATPVSSPSR | 13 | Q9Y4B5 | Microtubule cross-linking factor 1 OS=H | MTCL1 | 209.526 | 2 | 54.34099 | yes | 1.735E-07 | 572.29126 | 1328.4738 |
| ELEMHSLALQNTL | 16 | Q9Y4B5 | Microtubule cross-linking factor 1 OS=H | MTCL1 | 209.526 | 3 | 54.34099 | yes | 1.101E-07 | 640.99139 | 1264.5653 |
| ELQAFLEQVNR | 11 | Q9Y4B5 | Microtubule cross-linking factor 1 OS=H | MTCL1 | 209.526 | 2 | 54.34099 | yes | 1.304E-07 | 673.85681 | 1258.5055 |
| AEDENETLR | 9 | Q9Y4B5 | Microtubule cross-linking factor 1 OS=H | MTCL1 | 209.526 | 2 | 54.34099 | yes | 2.144E-07 | 538.74634 | 1256.222 |
| IPAVTLAVTSVAG | 17 | Q9Y4B5 | Microtubule cross-linking factor 1 OS=H | MTCL1 | 209.526 | 2 | 54.34099 | yes | 1.339E-07 | 805.46722 | 616.63367 |
| IPAVTLAVTSVAG | 17 | Q9Y4B5 | Microtubule cross-linking factor 1 OS=H | MTCL1 | 209.526 | 2 | 54.34099 | yes | 1.792E-06 | 537.3139 | 616.63367 |
| IPAVTLAVTSVAG | 17 | Q9Y4B5 | Microtubule cross-linking factor 1 OS=H | MTCL1 | 209.526 | 3 | 54.34099 | yes | 1.339E-07 | 805.46722 | 616.63367 |
| IPAVTLAVTSVAG | 17 | Q9Y4B5 | Microtubule cross-linking factor 1 OS=H | MTCL1 | 209.526 | 3 | 54.34099 | yes | 1.792E-06 | 537.3139 | 616.63367 |
| METLNGPAGGGAI | 16 | Q9Y4B5 | Microtubule cross-linking factor 1 OS=H | MTCL1 | 209.526 | 2 | 54.34099 | yes | 1.53E-07 | 764.35901 | 1184.6394 |
| VAPAEPLSR | 9 | Q9Y4B5 | Microtubule cross-linking factor 1 OS=H | MTCL1 | 209.526 | 2 | 54.34099 | yes | 2.581E-07 | 470.26654 | 1122.074 |
| EKDELEQELQK | 11 | Q9Y4B5 | Microtubule cross-linking factor 1 OS=H | MTCL1 | 209.526 | 2 | 54.34099 | yes | 1.066E-07 | 694.84882 | 1095.2698 |
| VLHSPPAVR | 9 | Q9Y4B5 | Microtubule cross-linking factor 1 OS=H | MTCL1 | 209.526 | 2 | 54.34099 | yes | 1.313E-07 | 488.29034 | 1061.0399 |
| TVGPPTPAAR | 10 | Q9Y4B5 | Microtubule cross-linking factor 1 OS=H | MTCL1 | 209.526 | 2 | 54.34099 | yes | 1.378E-07 | 483.772 | 1019.1481 |
| HLQFVEEEAELLR | 13 | Q9Y4B5 | Microtubule cross-linking factor 1 OS=H | MTCL1 | 209.526 | 2 | 54.34099 | no | 1.002E-07 | 806.9201 | 450.64368 |
| HLQFVEEEAELLR | 13 | Q9Y4B5 | Microtubule cross-linking factor 1 OS=H | MTCL1 | 209.526 | 2 | 54.34099 | no | 4.006E-07 | 538.28247 | 450.64368 |
| HLQFVEEEAELLR | 13 | Q9Y4B5 | Microtubule cross-linking factor 1 OS=H | MTCL1 | 209.526 | 3 | 54.34099 | no | 1.002E-07 | 806.9201 | 450.64368 |

| | | | | | | | | | | | |
|---|---|---|---|---|---|---|---|---|---|---|---|
| HLQFVEEEAELLR | 13 | Q9Y4B5 | Microtubule cross-linking factor 1 OS=H( | MTCL1 | 209.526 | 3 | 54.34099 | no | 4.006E-07 | 538.28247 | 450.64368 |
| QLTHELSK | 8 | Q9Y4B5 | Microtubule cross-linking factor 1 OS=H( | MTCL1 | 209.526 | 2 | 54.34099 | yes | 9.729E-08 | 478.26401 | 894.07153 |
| AAWDVEWAVLK | 11 | Q9Y4B5 | Microtubule cross-linking factor 1 OS=H( | MTCL1 | 209.526 | 2 | 54.34099 | yes | 9.722E-08 | 644.34021 | 888.74805 |
| LPEEEENHK | 9 | Q9Y4B5 | Microtubule cross-linking factor 1 OS=H( | MTCL1 | 209.526 | 2 | 54.34099 | yes | 1.417E-07 | 562.76459 | 858.53876 |
| LLPAASSDAESGT( | 22 | Q9Y4B5 | Microtubule cross-linking factor 1 OS=H( | MTCL1 | 209.526 | 3 | 54.34099 | yes | 3.91E-07 | 734.02075 | 831.22797 |
| HHHLHPVAER | 10 | Q9Y4B5 | Microtubule cross-linking factor 1 OS=H( | MTCL1 | 209.526 | 3 | 54.34099 | yes | 3.123E-07 | 411.55136 | 828.63239 |
| FKFEPPR | 7 | Q9Y4B5 | Microtubule cross-linking factor 1 OS=H( | MTCL1 | 209.526 | 2 | 54.34099 | yes | 1.095E-06 | 460.75308 | 798.51257 |
| QNIFLFYVK | 9 | Q9Y4B5 | Microtubule cross-linking factor 1 OS=H( | MTCL1 | 209.526 | 2 | 54.34099 | yes | 1.182E-07 | 586.32916 | 764.05206 |
| GGSFLCDQK | 9 | Q9Y4B5 | Microtubule cross-linking factor 1 OS=H( | MTCL1 | 209.526 | 2 | 54.34099 | yes | 1.006E-07 | 506.23184 | 749.16815 |
| EDSEYLVTLK | 10 | Q9Y4B5 | Microtubule cross-linking factor 1 OS=H( | MTCL1 | 209.526 | 2 | 54.34099 | yes | 9.722E-08 | 598.80591 | 675.13226 |
| RGGSFLCDQK | 10 | Q9Y4B5 | Microtubule cross-linking factor 1 OS=H( | MTCL1 | 209.526 | 2 | 54.34099 | yes | 1.943E-07 | 584.28241 | 570.98413 |
| VDSITAAGGEGPFI | 17 | Q9Y4B5 | Microtubule cross-linking factor 1 OS=H( | MTCL1 | 209.526 | 2 | 54.34099 | yes | 1.658E-07 | 831.4101 | 559.84064 |
| ALLEDFR | 7 | Q9Y4B5 | Microtubule cross-linking factor 1 OS=H( | MTCL1 | 209.526 | 2 | 54.34099 | yes | 1.256E-07 | 432.23471 | 535.15424 |
| AKAEDENETLR | 11 | Q9Y4B5 | Microtubule cross-linking factor 1 OS=H( | MTCL1 | 209.526 | 2 | 54.34099 | yes | 1.009E-07 | 638.31238 | 512.60352 |
| AREDSEYLVTLK | 12 | Q9Y4B5 | Microtubule cross-linking factor 1 OS=H( | MTCL1 | 209.526 | 2 | 54.34099 | yes | 1.31E-07 | 712.375 | 508.18915 |
| LQQQYASDK | 9 | Q9Y4B5 | Microtubule cross-linking factor 1 OS=H( | MTCL1 | 209.526 | 2 | 54.34099 | yes | 2.13E-07 | 540.76965 | 428.31052 |
| PSEASEPCPTELLK | 14 | Q9Y4B5 | Microtubule cross-linking factor 1 OS=H( | MTCL1 | 209.526 | 2 | 54.34099 | yes | 1.931E-07 | 779.37689 | 386.90198 |
| CQLQFAK | 7 | Q9Y4B5 | Microtubule cross-linking factor 1 OS=H( | MTCL1 | 209.526 | 2 | 54.34099 | yes | 1.803E-07 | 447.72873 | 373.63672 |
| LKEQLEWQLGPAI | 13 | Q9Y4B5 | Microtubule cross-linking factor 1 OS=H( | MTCL1 | 209.526 | 2 | 54.34099 | yes | 1.328E-07 | 784.43317 | 175.32761 |
| LKEQLEWQLGPAI | 13 | Q9Y4B5 | Microtubule cross-linking factor 1 OS=H( | MTCL1 | 209.526 | 2 | 54.34099 | yes | 1.047E-06 | 523.2912 | 175.32761 |
| LKEQLEWQLGPAI | 13 | Q9Y4B5 | Microtubule cross-linking factor 1 OS=H( | MTCL1 | 209.526 | 3 | 54.34099 | yes | 1.328E-07 | 784.43317 | 175.32761 |
| LKEQLEWQLGPAI | 13 | Q9Y4B5 | Microtubule cross-linking factor 1 OS=H( | MTCL1 | 209.526 | 3 | 54.34099 | yes | 1.047E-06 | 523.2912 | 175.32761 |
| DGNVRPFPHQGSL | 14 | Q9Y4B5 | Microtubule cross-linking factor 1 OS=H( | MTCL1 | 209.526 | 3 | 54.34099 | yes | 1.443E-07 | 527.27448 | 168.05801 |
| DGNVRPFPHQGSL | 14 | Q9Y4B5 | Microtubule cross-linking factor 1 OS=H( | MTCL1 | 209.526 | 3 | 54.34099 | yes | 3.932E-07 | 790.40802 | 168.05801 |
| DGNVRPFPHQGSL | 14 | Q9Y4B5 | Microtubule cross-linking factor 1 OS=H( | MTCL1 | 209.526 | 2 | 54.34099 | yes | 1.443E-07 | 527.27448 | 168.05801 |
| DGNVRPFPHQGSL | 14 | Q9Y4B5 | Microtubule cross-linking factor 1 OS=H( | MTCL1 | 209.526 | 2 | 54.34099 | yes | 3.932E-07 | 790.40802 | 168.05801 |
| APSPARPFLK | 10 | Q9Y4B5 | Microtubule cross-linking factor 1 OS=H( | MTCL1 | 209.526 | 2 | 54.34099 | yes | 9.722E-08 | 542.31909 | 291.00726 |
| GGPPEPMLSR | 10 | Q9Y4B5 | Microtubule cross-linking factor 1 OS=H( | MTCL1 | 209.526 | 2 | 54.34099 | yes | 2.042E-07 | 520.76331 | 261.62354 |
| SLEQDLK | 7 | Q9Y4B5 | Microtubule cross-linking factor 1 OS=H( | MTCL1 | 209.526 | 2 | 54.34099 | no | 1.418E-07 | 416.72418 | 239.5683 |
| LQHENHALLSNIQ | 14 | Q9Y4B5 | Microtubule cross-linking factor 1 OS=H( | MTCL1 | 209.526 | 2 | 54.34099 | yes | 9.722E-08 | 836.94751 | 218.96594 |
| TMGTQTVQTISVG | 20 | Q9Y4B5 | Microtubule cross-linking factor 1 OS=H( | MTCL1 | 209.526 | 3 | 54.34099 | yes | 9.722E-08 | 712.04395 | 206.66646 |
| EEGGEGTPVK | 10 | Q9Y4B5 | Microtubule cross-linking factor 1 OS=H( | MTCL1 | 209.526 | 2 | 54.34099 | yes | 1.194E-07 | 501.74054 | 191.65614 |
| SPPERPVPGTPK | 12 | Q9Y4B5 | Microtubule cross-linking factor 1 OS=H( | MTCL1 | 209.526 | 2 | 54.34099 | yes | 9.838E-08 | 631.34857 | 87.036873 |
| SPPERPVPGTPK | 12 | Q9Y4B5 | Microtubule cross-linking factor 1 OS=H( | MTCL1 | 209.526 | 2 | 54.34099 | yes | 4.123E-07 | 421.23483 | 87.036873 |
| SPPERPVPGTPK | 12 | Q9Y4B5 | Microtubule cross-linking factor 1 OS=H( | MTCL1 | 209.526 | 3 | 54.34099 | yes | 9.838E-08 | 631.34857 | 87.036873 |
| SPPERPVPGTPK | 12 | Q9Y4B5 | Microtubule cross-linking factor 1 OS=H( | MTCL1 | 209.526 | 3 | 54.34099 | yes | 4.123E-07 | 421.23483 | 87.036873 |
| RPLDSPLCTSLGFA | 24 | Q9Y4B5 | Microtubule cross-linking factor 1 OS=H( | MTCL1 | 209.526 | 4 | 54.34099 | yes | 1.078E-07 | 661.58716 | 170.93323 |
| LQPPGQHHR | 9 | Q9Y4B5 | Microtubule cross-linking factor 1 OS=H( | MTCL1 | 209.526 | 2 | 54.34099 | yes | 2.184E-07 | 535.28613 | 159.90901 |
| KLLEHALK | 8 | Q9Y4B5 | Microtubule cross-linking factor 1 OS=H( | MTCL1 | 209.526 | 2 | 54.34099 | yes | 9.722E-08 | 476.30292 | 147.19409 |

| | | | | | | | | | | | |
|---|---|---|---|---|---|---|---|---|---|---|---|
| NRLPEEEENHK | 11 | Q9Y4B5 | Microtubule cross-linking factor 1 OS=H | MTCL1 | 209.526 | 2 | 54.34099 | yes | 9.722E-08 | 697.83661 | 122.66335 |
| KTFNQDDSADLR | 12 | Q9Y4B5 | Microtubule cross-linking factor 1 OS=H | MTCL1 | 209.526 | 2 | 54.34099 | yes | 9.822E-08 | 705.33643 | 104.86983 |
| SPSPIGVGSEMCR | 13 | Q9Y4B5 | Microtubule cross-linking factor 1 OS=H | MTCL1 | 209.526 | 2 | 54.34099 | yes | 1.791E-07 | 688.81848 | 103.24364 |
| GCGFPVGEHSPHS | 14 | Q9Y4B5 | Microtubule cross-linking factor 1 OS=H | MTCL1 | 209.526 | 2 | 54.34099 | yes | 1.024E-07 | 762.34406 | 102.71305 |
| RVDSITAAGGEGP | 18 | Q9Y4B5 | Microtubule cross-linking factor 1 OS=H | MTCL1 | 209.526 | 3 | 54.34099 | yes | 1.002E-07 | 606.64288 | 99.150497 |
| ELGPDLQSR | 9 | Q9Y4B5 | Microtubule cross-linking factor 1 OS=H | MTCL1 | 209.526 | 2 | 54.34099 | yes | 1.371E-07 | 507.76437 | 96.003242 |
| PFPHQGSLR | 9 | Q9Y4B5 | Microtubule cross-linking factor 1 OS=H | MTCL1 | 209.526 | 2 | 54.34099 | yes | 1.159E-06 | 519.77759 | 89.590279 |
| DRQEWER | 7 | Q9Y4B5 | Microtubule cross-linking factor 1 OS=H | MTCL1 | 209.526 | 2 | 54.34099 | yes | 5.098E-07 | 509.73868 | 86.763344 |
| DKPPPGAGAR | 10 | Q9Y4B5 | Microtubule cross-linking factor 1 OS=H | MTCL1 | 209.526 | 2 | 54.34099 | yes | 2.601E-07 | 483.26178 | 69.318916 |
| CRLEQLEEK | 9 | Q9Y4B5 | Microtubule cross-linking factor 1 OS=H | MTCL1 | 209.526 | 2 | 54.34099 | yes | 9.722E-08 | 602.80316 | 57.193348 |
| PVAMWPCADADS | 22 | Q9Y4B5 | Microtubule cross-linking factor 1 OS=H | MTCL1 | 209.526 | 3 | 54.34099 | yes | 1.711E-07 | 834.73126 | 54.572781 |
| SWDYTPNR | 8 | Q9Y4B5 | Microtubule cross-linking factor 1 OS=H | MTCL1 | 209.526 | 2 | 54.34099 | yes | 2.778E-07 | 519.7356 | 36.889664 |
| LGELGVQGGHQA | 21 | Q9Y4B5 | Microtubule cross-linking factor 1 OS=H | MTCL1 | 209.526 | 4 | 54.34099 | yes | 8.512E-07 | 540.74799 | 23.246231 |
| EVATSLAYLPEWK | 13 | Q9Y4C2 | TRPM8 channel-associated factor 1 OS=I | TCAF1 | 102.126 | 2 | 55.764843 | yes | 9.84E-08 | 753.89557 | 455.24542 |
| TLENPEPLLR | 10 | Q9Y4C2 | TRPM8 channel-associated factor 1 OS=I | TCAF1 | 102.126 | 2 | 55.764843 | yes | 1.009E-07 | 591.3299 | 401.76236 |
| NQTNLPTENVDK | 12 | Q9Y4C2 | TRPM8 channel-associated factor 1 OS=I | TCAF1 | 102.126 | 2 | 55.764843 | yes | 9.935E-08 | 686.83881 | 332.24243 |
| STLAEFQVIMGR | 12 | Q9Y4C2 | TRPM8 channel-associated factor 1 OS=I | TCAF1 | 102.126 | 2 | 55.764843 | yes | 9.722E-08 | 676.35559 | 301.18536 |
| LWDEVMQAVAR | 11 | Q9Y4C2 | TRPM8 channel-associated factor 1 OS=I | TCAF1 | 102.126 | 2 | 55.764843 | yes | 1.269E-07 | 659.33466 | 298.84552 |
| LGPFLLNAVR | 10 | Q9Y4C2 | TRPM8 channel-associated factor 1 OS=I | TCAF1 | 102.126 | 2 | 55.764843 | yes | 1.196E-07 | 550.33478 | 277.80353 |
| MFSHQVQK | 8 | Q9Y4C2 | TRPM8 channel-associated factor 1 OS=I | TCAF1 | 102.126 | 2 | 55.764843 | yes | 1.735E-07 | 502.75275 | 230.26239 |
| LGETTLEEWK | 10 | Q9Y4C2 | TRPM8 channel-associated factor 1 OS=I | TCAF1 | 102.126 | 2 | 55.764843 | yes | 9.722E-08 | 603.30603 | 188.55096 |
| ENPVINDCCR | 10 | Q9Y4C2 | TRPM8 channel-associated factor 1 OS=I | TCAF1 | 102.126 | 2 | 55.764843 | yes | 9.722E-08 | 638.77411 | 181.92097 |
| VVVQTELR | 8 | Q9Y4C2 | TRPM8 channel-associated factor 1 OS=I | TCAF1 | 102.126 | 2 | 55.764843 | yes | 1.02E-07 | 472.2822 | 168.18535 |
| CCLDKPTK | 8 | Q9Y4C2 | TRPM8 channel-associated factor 1 OS=I | TCAF1 | 102.126 | 2 | 55.764843 | yes | 2.911E-07 | 511.2439 | 163.33536 |
| NPGVSPLAR | 9 | Q9Y4C2 | TRPM8 channel-associated factor 1 OS=I | TCAF1 | 102.126 | 2 | 55.764843 | yes | 2.611E-07 | 455.75888 | 162.71124 |
| YCWMSTGLYIPGR | 13 | Q9Y4C2 | TRPM8 channel-associated factor 1 OS=I | TCAF1 | 102.126 | 2 | 55.764843 | yes | 9.728E-08 | 802.37341 | 161.0251 |
| GLWGPVHELGR | 11 | Q9Y4C2 | TRPM8 channel-associated factor 1 OS=I | TCAF1 | 102.126 | 2 | 55.764843 | yes | 9.809E-08 | 610.83038 | 157.16467 |
| ANIALWPPVR | 10 | Q9Y4C2 | TRPM8 channel-associated factor 1 OS=I | TCAF1 | 102.126 | 2 | 55.764843 | yes | 2.297E-07 | 568.8324 | 156.02988 |
| LYLLTQMPH | 9 | Q9Y4C2 | TRPM8 channel-associated factor 1 OS=I | TCAF1 | 102.126 | 2 | 55.764843 | yes | 1.802E-07 | 558.2995 | 133.89674 |
| IQIGCHTDDLTR | 12 | Q9Y4C2 | TRPM8 channel-associated factor 1 OS=I | TCAF1 | 102.126 | 2 | 55.764843 | yes | 9.722E-08 | 714.84863 | 57.289082 |
| NQLTSVPSLISGLG | 15 | Q9Y4C4 | Malignant fibrous histiocytoma-amplified | MFHAS1 | 116.95 | 2 | 40.60407 | yes | 3.938E-07 | 771.43591 | 15.740252 |
| VALGETQLVQEVI | 13 | Q9Y4C8 | Probable RNA-binding protein 19 OS=H | RBM19 | 107.332 | 2 | 53.349167 | yes | 1.175E-07 | 721.40411 | 298.2771 |
| AYSEVDGQVFQGI | 13 | Q9Y4C8 | Probable RNA-binding protein 19 OS=H | RBM19 | 107.332 | 2 | 53.349167 | yes | 3.99E-07 | 728.3468 | 258.44891 |
| NLPYTSTEEDLEK | 13 | Q9Y4C8 | Probable RNA-binding protein 19 OS=H | RBM19 | 107.332 | 2 | 53.349167 | yes | 1.059E-07 | 769.86469 | 240.97952 |
| ATKPAVTLAR | 10 | Q9Y4C8 | Probable RNA-binding protein 19 OS=H | RBM19 | 107.332 | 2 | 53.349167 | yes | 2.614E-07 | 514.31659 | 170.53082 |
| MLHVLPSTIK | 10 | Q9Y4C8 | Probable RNA-binding protein 19 OS=H | RBM19 | 107.332 | 2 | 53.349167 | yes | 9.819E-08 | 569.8363 | 159.9474 |
| ELFSTFGELK | 10 | Q9Y4C8 | Probable RNA-binding protein 19 OS=H | RBM19 | 107.332 | 2 | 53.349167 | yes | 1.042E-07 | 585.80573 | 154.14626 |
| AFNALCHSTHLYG | 14 | Q9Y4C8 | Probable RNA-binding protein 19 OS=H | RBM19 | 107.332 | 3 | 53.349167 | yes | 1.71E-07 | 549.60004 | 49.000282 |
| AFNALCHSTHLYG | 14 | Q9Y4C8 | Probable RNA-binding protein 19 OS=H | RBM19 | 107.332 | 3 | 53.349167 | yes | 3.008E-07 | 823.89642 | 49.000282 |

| | | | | | | | | | | | |
|---|---|---|---|---|---|---|---|---|---|---|---|
| AFNALCHSTHLYG | 14 | Q9Y4C8 | Probable RNA-binding protein 19 OS=H | RBM19 | 107.332 | 2 | 53.349167 | yes | 1.71E-07 | 549.60004 | 49.000282 |
| AFNALCHSTHLYG | 14 | Q9Y4C8 | Probable RNA-binding protein 19 OS=H | RBM19 | 107.332 | 2 | 53.349167 | yes | 3.008E-07 | 823.89642 | 49.000282 |
| QLFAAFGTLTDCSI | 15 | Q9Y4C8 | Probable RNA-binding protein 19 OS=H | RBM19 | 107.332 | 2 | 53.349167 | yes | 9.722E-08 | 836.42419 | 85.469597 |
| GAGQEQGMPAGK | 12 | Q9Y4C8 | Probable RNA-binding protein 19 OS=H | RBM19 | 107.332 | 2 | 53.349167 | yes | 1.965E-07 | 565.7666 | 77.806702 |
| ELSDMDYLK | 9 | Q9Y4C8 | Probable RNA-binding protein 19 OS=H | RBM19 | 107.332 | 2 | 53.349167 | yes | 3.524E-07 | 557.26025 | 51.800236 |
| EASEDASALGSSSY | 15 | Q9Y4C8 | Probable RNA-binding protein 19 OS=H | RBM19 | 107.332 | 2 | 53.349167 | yes | 1.069E-06 | 751.3363 | 50.79438 |
| VLVYGLGHK | 9 | Q9Y4D8 | Probable E3 ubiquitin-protein ligase HEC | HECTD4 | 439.345 | 2 | 46.150475 | yes | 1.638E-07 | 493.2951 | 45.779114 |
| TSQASDTLTIPLSR | 14 | Q9Y4D8 | Probable E3 ubiquitin-protein ligase HEC | HECTD4 | 439.345 | 2 | 46.150475 | yes | 1.963E-07 | 745.39648 | 36.92263 |
| SDEEEQQYVIK | 11 | Q9Y4E5 | E3 SUMO-protein ligase ZNF451 OS=H | ZNF451 | 121.485 | 2 | 51.384384 | yes | 9.722E-08 | 684.31989 | 142.22517 |
| FVTQQQYR | 8 | Q9Y4E5 | E3 SUMO-protein ligase ZNF451 OS=H | ZNF451 | 121.485 | 2 | 51.384384 | yes | 5.252E-07 | 535.2749 | 85.973907 |
| SHEGVACVQK | 10 | Q9Y4E5 | E3 SUMO-protein ligase ZNF451 OS=H | ZNF451 | 121.485 | 2 | 51.384384 | yes | 3.745E-05 | 557.7691 | 52.983158 |
| PILCPIMHCNK | 11 | Q9Y4E5 | E3 SUMO-protein ligase ZNF451 OS=H | ZNF451 | 121.485 | 3 | 51.384384 | yes | 2.926E-06 | 461.5629 | 27.85125 |
| AAYVLFYQR | 9 | Q9Y4E8 | Ubiquitin carboxyl-terminal hydrolase 15 | USP15 | 112.42 | 2 | 48.460262 | no | 1.752E-07 | 565.80328 | 106.12315 |
| MIVTDIYNHR | 10 | Q9Y4E8 | Ubiquitin carboxyl-terminal hydrolase 15 | USP15 | 112.42 | 2 | 48.460262 | yes | 2.672E-06 | 631.32153 | 45.855808 |
| VEVYLTELK | 9 | Q9Y4E8 | Ubiquitin carboxyl-terminal hydrolase 15 | USP15 | 112.42 | 2 | 48.460262 | yes | 1.145E-07 | 547.31061 | 36.901974 |
| LYNLLLLR | 8 | Q9Y4E8 | Ubiquitin carboxyl-terminal hydrolase 15 | USP15 | 112.42 | 2 | 48.460262 | yes | 2.193E-07 | 509.32642 | 33.150375 |
| TEDTEHVIIPVCLR | 14 | Q9Y4E8 | Ubiquitin carboxyl-terminal hydrolase 15 | USP15 | 112.42 | 2 | 48.460262 | yes | 2.157E-06 | 841.43256 | 31.92037 |
| TLEVYLVR | 8 | Q9Y4E8 | Ubiquitin carboxyl-terminal hydrolase 15 | USP15 | 112.42 | 2 | 48.460262 | yes | 1.693E-06 | 496.79239 | 25.921137 |
| YFDENAAEDFEK | 12 | Q9Y4E8 | Ubiquitin carboxyl-terminal hydrolase 15 | USP15 | 112.42 | 2 | 48.460262 | yes | 1.257E-07 | 739.30951 | 21.429298 |
| LCENGNMNNVVT | 13 | Q9Y4E8 | Ubiquitin carboxyl-terminal hydrolase 15 | USP15 | 112.42 | 2 | 48.460262 | yes | 5.409E-06 | 760.85083 | 13.204464 |
| VSAGEPGSHPSPAI | 15 | Q9Y4F1 | FERM, ARHGEF and pleckstrin domain- | FARP1 | 118.633 | 3 | 52.5709 | yes | 1.167E-07 | 482.5759 | 263.57513 |
| VSAGEPGSHPSPAI | 15 | Q9Y4F1 | FERM, ARHGEF and pleckstrin domain- | FARP1 | 118.633 | 3 | 52.5709 | yes | 9.722E-08 | 723.36023 | 263.57513 |
| VSAGEPGSHPSPAI | 15 | Q9Y4F1 | FERM, ARHGEF and pleckstrin domain- | FARP1 | 118.633 | 2 | 52.5709 | yes | 1.167E-07 | 482.5759 | 263.57513 |
| VSAGEPGSHPSPAI | 15 | Q9Y4F1 | FERM, ARHGEF and pleckstrin domain- | FARP1 | 118.633 | 2 | 52.5709 | yes | 9.722E-08 | 723.36023 | 263.57513 |
| YIPQQDALEDK | 11 | Q9Y4F1 | FERM, ARHGEF and pleckstrin domain- | FARP1 | 118.633 | 2 | 52.5709 | yes | 4.485E-07 | 660.32751 | 442.0925 |
| LFEEPKPK | 8 | Q9Y4F1 | FERM, ARHGEF and pleckstrin domain- | FARP1 | 118.633 | 2 | 52.5709 | yes | 1.543E-07 | 494.27911 | 428.45602 |
| LGAPENSGISTLER | 14 | Q9Y4F1 | FERM, ARHGEF and pleckstrin domain- | FARP1 | 118.633 | 2 | 52.5709 | yes | 1.147E-07 | 722.37555 | 413.47937 |
| SLIFPNFEPLHK | 12 | Q9Y4F1 | FERM, ARHGEF and pleckstrin domain- | FARP1 | 118.633 | 2 | 52.5709 | yes | 1.503E-07 | 721.39557 | 378.01529 |
| AESEYTFER | 9 | Q9Y4F1 | FERM, ARHGEF and pleckstrin domain- | FARP1 | 118.633 | 2 | 52.5709 | yes | 1.131E-07 | 566.25128 | 318.94418 |
| IVEFHHNHIGQTPA | 25 | Q9Y4F1 | FERM, ARHGEF and pleckstrin domain- | FARP1 | 118.633 | 4 | 52.5709 | yes | 9.975E-08 | 726.12152 | 258.43839 |
| EDAMPEALK | 9 | Q9Y4F1 | FERM, ARHGEF and pleckstrin domain- | FARP1 | 118.633 | 2 | 52.5709 | yes | 1.313E-07 | 502.24188 | 245.92293 |
| DLIGIDNLVVPGR | 13 | Q9Y4F1 | FERM, ARHGEF and pleckstrin domain- | FARP1 | 118.633 | 2 | 52.5709 | yes | 9.758E-08 | 690.89594 | 219.19063 |
| YLFALQVK | 8 | Q9Y4F1 | FERM, ARHGEF and pleckstrin domain- | FARP1 | 118.633 | 2 | 52.5709 | yes | 1.536E-07 | 491.29202 | 179.00891 |
| FHTNFLK | 7 | Q9Y4F1 | FERM, ARHGEF and pleckstrin domain- | FARP1 | 118.633 | 2 | 52.5709 | yes | 5.465E-05 | 453.74524 | 178.21883 |
| SHVYYFR | 7 | Q9Y4F1 | FERM, ARHGEF and pleckstrin domain- | FARP1 | 118.633 | 2 | 52.5709 | yes | 2.559E-06 | 486.24033 | 153.2235 |
| HLAAHLWK | 8 | Q9Y4F1 | FERM, ARHGEF and pleckstrin domain- | FARP1 | 118.633 | 2 | 52.5709 | yes | 9.766E-08 | 488.27979 | 103.45656 |
| WVEDIQMAIDLAE | 14 | Q9Y4F1 | FERM, ARHGEF and pleckstrin domain- | FARP1 | 118.633 | 2 | 52.5709 | yes | 2.393E-07 | 830.9162 | 99.776733 |
| LALWEGR | 7 | Q9Y4F1 | FERM, ARHGEF and pleckstrin domain- | FARP1 | 118.633 | 2 | 52.5709 | yes | 1.041E-07 | 422.73743 | 80.511841 |
| QSIIVAASSR | 10 | Q9Y4F1 | FERM, ARHGEF and pleckstrin domain- | FARP1 | 118.633 | 2 | 52.5709 | yes | 1.47E-07 | 516.29584 | 65.691277 |

| | | | | | | | | | | | | |
|---|---|---|---|---|---|---|---|---|---|---|---|---|
| ICVEHHAFFR | 10 | Q9Y4F1 | FERM, ARHGEF and pleckstrin domain- | FARP1 | 118.633 | 3 | 52.5709 | yes | 9.733E-08 | 439.21701 | 32.798134 |
| ICVEHHAFFR | 10 | Q9Y4F1 | FERM, ARHGEF and pleckstrin domain- | FARP1 | 118.633 | 3 | 52.5709 | yes | 1.248E-06 | 658.32184 | 32.798134 |
| ICVEHHAFFR | 10 | Q9Y4F1 | FERM, ARHGEF and pleckstrin domain- | FARP1 | 118.633 | 2 | 52.5709 | yes | 9.733E-08 | 439.21701 | 32.798134 |
| ICVEHHAFFR | 10 | Q9Y4F1 | FERM, ARHGEF and pleckstrin domain- | FARP1 | 118.633 | 2 | 52.5709 | yes | 1.248E-06 | 658.32184 | 32.798134 |
| INAFNWAK | 8 | Q9Y4F1 | FERM, ARHGEF and pleckstrin domain- | FARP1 | 118.633 | 2 | 52.5709 | yes | 2.212E-07 | 482.25598 | 57.044998 |
| RLEMYGIR | 8 | Q9Y4F1 | FERM, ARHGEF and pleckstrin domain- | FARP1 | 118.633 | 2 | 52.5709 | yes | 4.736E-07 | 519.28168 | 23.006479 |
| ILVSLATGAASK | 12 | Q9Y4F3 | Meiosis regulator and mRNA stability fac | MARF1 | 192.86 | 2 | 53.61044 | yes | 9.722E-08 | 565.84259 | 345.59937 |
| ILEVPESHTASELK | 14 | Q9Y4F3 | Meiosis regulator and mRNA stability fac | MARF1 | 192.86 | 2 | 53.61044 | yes | 1.304E-07 | 776.91449 | 101.1524 |
| ILEVPESHTASELK | 14 | Q9Y4F3 | Meiosis regulator and mRNA stability fac | MARF1 | 192.86 | 2 | 53.61044 | yes | 6.082E-07 | 518.27875 | 101.1524 |
| ILEVPESHTASELK | 14 | Q9Y4F3 | Meiosis regulator and mRNA stability fac | MARF1 | 192.86 | 3 | 53.61044 | yes | 1.304E-07 | 776.91449 | 101.1524 |
| ILEVPESHTASELK | 14 | Q9Y4F3 | Meiosis regulator and mRNA stability fac | MARF1 | 192.86 | 3 | 53.61044 | yes | 6.082E-07 | 518.27875 | 101.1524 |
| LLTLTHR | 7 | Q9Y4F3 | Meiosis regulator and mRNA stability fac | MARF1 | 192.86 | 2 | 53.61044 | yes | 9.722E-08 | 427.26636 | 198.383 |
| DVPSPLHAGSK | 11 | Q9Y4F3 | Meiosis regulator and mRNA stability fac | MARF1 | 192.86 | 2 | 53.61044 | yes | 2.968E-07 | 554.29327 | 166.95529 |
| VLSITGCSAILR | 12 | Q9Y4F3 | Meiosis regulator and mRNA stability fac | MARF1 | 192.86 | 2 | 53.61044 | yes | 1.234E-07 | 645.36591 | 130.77257 |
| LAANFSLAPITK | 12 | Q9Y4F3 | Meiosis regulator and mRNA stability fac | MARF1 | 192.86 | 2 | 53.61044 | yes | 1.259E-07 | 623.36371 | 104.00825 |
| NSGVAEPVYK | 10 | Q9Y4F3 | Meiosis regulator and mRNA stability fac | MARF1 | 192.86 | 2 | 53.61044 | yes | 3.752E-06 | 532.27454 | 83.373398 |
| LQDYDVSSISALT( | 15 | Q9Y4F3 | Meiosis regulator and mRNA stability fac | MARF1 | 192.86 | 2 | 53.61044 | yes | 7.429E-07 | 834.42798 | 80.739952 |
| QVSPLLASQSWSS | 14 | Q9Y4F3 | Meiosis regulator and mRNA stability fac | MARF1 | 192.86 | 2 | 53.61044 | yes | 9.722E-08 | 773.4046 | 73.082458 |
| MENEDVFGNR | 10 | Q9Y4F3 | Meiosis regulator and mRNA stability fac | MARF1 | 192.86 | 2 | 53.61044 | yes | 1.312E-07 | 605.76147 | 71.418175 |
| LVVPTHGNSSAAV | 17 | Q9Y4F3 | Meiosis regulator and mRNA stability fac | MARF1 | 192.86 | 4 | 53.61044 | yes | 8.543E-07 | 416.97961 | 57.014374 |
| LTSLYLFAK | 9 | Q9Y4F3 | Meiosis regulator and mRNA stability fac | MARF1 | 192.86 | 2 | 53.61044 | yes | 3.166E-06 | 528.31042 | 55.623013 |
| RFTQDLLK | 8 | Q9Y4F3 | Meiosis regulator and mRNA stability fac | MARF1 | 192.86 | 2 | 53.61044 | yes | 1.41E-05 | 510.79547 | 47.972031 |
| ELQQLLQEAFAR | 12 | Q9Y4F3 | Meiosis regulator and mRNA stability fac | MARF1 | 192.86 | 2 | 53.61044 | yes | 2.261E-06 | 723.39099 | 47.159882 |
| EAEFICVCDISK | 12 | Q9Y4F3 | Meiosis regulator and mRNA stability fac | MARF1 | 192.86 | 2 | 53.61044 | yes | 2.925E-07 | 735.8338 | 42.951248 |
| DFSEHEFDPDSYK | 13 | Q9Y4F3 | Meiosis regulator and mRNA stability fac | MARF1 | 192.86 | 2 | 53.61044 | yes | 8.458E-07 | 808.33099 | 27.212601 |
| ALAAQFVK | 8 | Q9Y4F3 | Meiosis regulator and mRNA stability fac | MARF1 | 192.86 | 2 | 53.61044 | yes | 5.815E-07 | 424.25543 | 18.204702 |
| VDGLVNFEK | 9 | Q9Y4G8 | Rap guanine nucleotide exchange factor 2 | RAPGEF2 | 167.417 | 2 | 49.021877 | no | 3.22E-07 | 510.77167 | 111.27914 |
| LLNIACAAK | 9 | Q9Y4G8 | Rap guanine nucleotide exchange factor 2 | RAPGEF2 | 167.417 | 2 | 49.021877 | no | 1.076E-07 | 487.27859 | 74.975044 |
| MASVNMDPALMF | 13 | Q9Y4G8 | Rap guanine nucleotide exchange factor 2 | RAPGEF2 | 167.417 | 2 | 49.021877 | yes | 2.439E-07 | 741.84875 | 67.950684 |
| GFGIFVDSVDSGSI | 14 | Q9Y4G8 | Rap guanine nucleotide exchange factor 2 | RAPGEF2 | 167.417 | 2 | 49.021877 | yes | 3.263E-07 | 707.84607 | 58.74041 |
| KLYEDAQMAR | 10 | Q9Y4G8 | Rap guanine nucleotide exchange factor 2 | RAPGEF2 | 167.417 | 2 | 49.021877 | no | 1.116E-07 | 612.80573 | 57.772881 |
| NIEPTEYIDDLFK | 13 | Q9Y4G8 | Rap guanine nucleotide exchange factor 2 | RAPGEF2 | 167.417 | 2 | 49.021877 | no | 4.872E-07 | 798.89325 | 45.221134 |
| KPPDYNVALQR | 11 | Q9Y4G8 | Rap guanine nucleotide exchange factor 2 | RAPGEF2 | 167.417 | 2 | 49.021877 | yes | 3.89E-06 | 650.85406 | 30.724321 |
| DLTFLHEGNDSK | 12 | Q9Y4G8 | Rap guanine nucleotide exchange factor 2 | RAPGEF2 | 167.417 | 2 | 49.021877 | yes | 3.141E-06 | 688.32806 | 28.289747 |
| TFLSSPMEVGK | 11 | Q9Y4G8 | Rap guanine nucleotide exchange factor 2 | RAPGEF2 | 167.417 | 2 | 49.021877 | yes | 3.138E-07 | 598.30499 | 15.247623 |
| PVSVAGSPLSPGP\ | 15 | Q9Y4H2 | Insulin receptor substrate 2 OS=Homo sa| | IRS2 | 137.335 | 2 | 52.158783 | yes | 9.809E-08 | 710.40137 | 620.04114 |
| GVPGCCYSSLPR | 12 | Q9Y4H2 | Insulin receptor substrate 2 OS=Homo sa| | IRS2 | 137.335 | 2 | 52.158783 | yes | 4.798E-07 | 676.80792 | 451.51312 |
| ALTDLVSEGR | 10 | Q9Y4H2 | Insulin receptor substrate 2 OS=Homo sa| | IRS2 | 137.335 | 2 | 52.158783 | yes | 1.631E-07 | 530.78528 | 374.08057 |
| QLQPAPPLAPQGR | 13 | Q9Y4H2 | Insulin receptor substrate 2 OS=Homo sa| | IRS2 | 137.335 | 2 | 52.158783 | yes | 3.957E-07 | 686.88843 | 331.17783 |

| | | | | | | | | | | | |
|---|---|---|---|---|---|---|---|---|---|---|---|
| SNTPESIAETPPAR | 14 | Q9Y4H2 | Insulin receptor substrate 2 OS=Homo saj | IRS2 | 137.335 | 2 | 52.158783 | yes | 1.073E-07 | 735.36517 | 305.4216 |
| VAYHPYPEDYGDI | 19 | Q9Y4H2 | Insulin receptor substrate 2 OS=Homo saj | IRS2 | 137.335 | 3 | 52.158783 | yes | 9.789E-08 | 740.01154 | 282.6312 |
| TDSLAATPPAAK | 12 | Q9Y4H2 | Insulin receptor substrate 2 OS=Homo saj | IRS2 | 137.335 | 2 | 52.158783 | yes | 9.722E-08 | 571.80621 | 194.39726 |
| LSMEHADGK | 9 | Q9Y4H2 | Insulin receptor substrate 2 OS=Homo saj | IRS2 | 137.335 | 2 | 52.158783 | yes | 1.166E-07 | 494.23184 | 186.01315 |
| EVWQVNLKPK | 10 | Q9Y4H2 | Insulin receptor substrate 2 OS=Homo saj | IRS2 | 137.335 | 2 | 52.158783 | yes | 9.722E-08 | 620.85608 | 178.1203 |
| YLIALYTK | 8 | Q9Y4H2 | Insulin receptor substrate 2 OS=Homo saj | IRS2 | 137.335 | 2 | 52.158783 | yes | 1.382E-07 | 492.79187 | 170.46077 |
| SDDYMPMSPASVS | 16 | Q9Y4H2 | Insulin receptor substrate 2 OS=Homo saj | IRS2 | 137.335 | 2 | 52.158783 | yes | 3.166E-07 | 841.87366 | 144.67374 |
| LEYYESEK | 8 | Q9Y4H2 | Insulin receptor substrate 2 OS=Homo saj | IRS2 | 137.335 | 2 | 52.158783 | yes | 1.741E-07 | 530.7453 | 143.30759 |
| VIALDCCLNINK | 12 | Q9Y4H2 | Insulin receptor substrate 2 OS=Homo saj | IRS2 | 137.335 | 2 | 52.158783 | yes | 7.341E-07 | 716.86798 | 115.51109 |
| VSGDAAQDLDR | 11 | Q9Y4H2 | Insulin receptor substrate 2 OS=Homo saj | IRS2 | 137.335 | 2 | 52.158783 | yes | 1.485E-07 | 573.77289 | 81.169746 |
| HNSASVENVSLR | 12 | Q9Y4H2 | Insulin receptor substrate 2 OS=Homo saj | IRS2 | 137.335 | 2 | 52.158783 | yes | 9.722E-08 | 656.83386 | 80.996361 |
| CGHSDSFFFIEVGF | 14 | Q9Y4H2 | Insulin receptor substrate 2 OS=Homo saj | IRS2 | 137.335 | 2 | 52.158783 | yes | 2.594E-06 | 829.375 | 29.141947 |
| MLPELFQDDEK | 11 | Q9Y4I1 | Unconventional myosin-Va OS=Homo sa | MYO5A | 215.405 | 2 | 34.70601 | yes | 9.563E-07 | 682.82373 | 63.395981 |
| CSGTELSLAHCR | 12 | Q9Y4K0 | Lysyl oxidase homolog 2 OS=Homo sapi | LOXL2 | 86.725 | 2 | 49.47893 | yes | 2.78E-07 | 695.81372 | 489.6304 |
| AYKPEQPLVR | 10 | Q9Y4K0 | Lysyl oxidase homolog 2 OS=Homo sapi | LOXL2 | 86.725 | 2 | 49.47893 | yes | 1.152E-07 | 600.84039 | 206.88901 |
| LGPQVSLDPMK | 11 | Q9Y4K0 | Lysyl oxidase homolog 2 OS=Homo sapi | LOXL2 | 86.725 | 2 | 49.47893 | yes | 1.026E-07 | 592.8208 | 180.3046 |
| TPVMEGYVEVK | 11 | Q9Y4K0 | Lysyl oxidase homolog 2 OS=Homo sapi | LOXL2 | 86.725 | 2 | 49.47893 | yes | 1.638E-07 | 626.31812 | 82.354988 |
| VEVLVER | 7 | Q9Y4K0 | Lysyl oxidase homolog 2 OS=Homo sapi | LOXL2 | 86.725 | 2 | 49.47893 | yes | 1.732E-07 | 422.25037 | 64.057663 |
| HTEDVGVVCSDK | 12 | Q9Y4K0 | Lysyl oxidase homolog 2 OS=Homo sapi | LOXL2 | 86.725 | 2 | 49.47893 | yes | 6.19E-07 | 673.30627 | 47.825485 |
| GANILLTDHGDVK | 13 | Q9Y4K4 | Mitogen-activated protein kinase kinase k | MAP4K5 | 95.024 | 2 | 53.543434 | yes | 9.722E-08 | 676.86206 | 85.87471 |
| YLGEMEPPALPR | 12 | Q9Y4R8 | Telomere length regulation protein TEL2 | TELO2 | 91.748 | 2 | 18.85523 | yes | 5.121E-06 | 686.8501 | 49.962288 |
| ETWELEEFR | 9 | Q9Y4W2 | Ribosomal biogenesis protein LAS1L OS | LAS1L | 83.065 | 2 | 55.312973 | yes | 9.722E-08 | 619.78802 | 118.058 |
| FVNLISER | 8 | Q9Y4W2 | Ribosomal biogenesis protein LAS1L OS | LAS1L | 83.065 | 2 | 55.312973 | yes | 1.924E-07 | 489.27435 | 90.960579 |
| LLYGMALVR | 9 | Q9Y4W2 | Ribosomal biogenesis protein LAS1L OS | LAS1L | 83.065 | 2 | 55.312973 | yes | 2.021E-07 | 518.30463 | 65.750702 |
| AMGQGLPDEEQEI | 13 | Q9Y4W2 | Ribosomal biogenesis protein LAS1L OS | LAS1L | 83.065 | 2 | 55.312973 | yes | 4.245E-07 | 716.32465 | 54.86919 |
| LLDVTGGLGTDEL | 14 | Q9Y4W2 | Ribosomal biogenesis protein LAS1L OS | LAS1L | 83.065 | 2 | 55.312973 | yes | 4.661E-07 | 729.89355 | 52.394341 |
| WTVELIVANTK | 11 | Q9Y4W2 | Ribosomal biogenesis protein LAS1L OS | LAS1L | 83.065 | 2 | 55.312973 | yes | 1.709E-07 | 637.36115 | 49.247086 |
| FSAGQWEAR | 9 | Q9Y4W2 | Ribosomal biogenesis protein LAS1L OS | LAS1L | 83.065 | 2 | 55.312973 | yes | 3.748E-06 | 526.2514 | 38.453484 |
| LFNCSASLDWPR | 12 | Q9Y4W2 | Ribosomal biogenesis protein LAS1L OS | LAS1L | 83.065 | 2 | 55.312973 | yes | 4.352E-07 | 733.34827 | 28.158045 |
| VGQISFDLPR | 10 | Q9Y4W6 | AFG3-like protein 2 OS=Homo sapiens C | AFG3L2 | 88.584 | 2 | 50.682903 | yes | 4.184E-07 | 566.31146 | 352.15073 |
| QIFIGPPDIK | 10 | Q9Y4W6 | AFG3-like protein 2 OS=Homo sapiens C | AFG3L2 | 88.584 | 2 | 50.682903 | yes | 9.819E-08 | 564.3266 | 257.8815 |
| GMGGLFSVGETTA | 14 | Q9Y4W6 | AFG3-like protein 2 OS=Homo sapiens C | AFG3L2 | 88.584 | 2 | 50.682903 | yes | 9.927E-08 | 677.83722 | 222.10382 |
| GAILTGPPGTGK | 12 | Q9Y4W6 | AFG3-like protein 2 OS=Homo sapiens C | AFG3L2 | 88.584 | 2 | 50.682903 | yes | 1.69E-07 | 534.80603 | 219.91251 |
| ILINDAYK | 8 | Q9Y4W6 | AFG3-like protein 2 OS=Homo sapiens C | AFG3L2 | 88.584 | 2 | 50.682903 | yes | 1.055E-07 | 475.2713 | 206.29074 |
| VLKDEIDVK | 9 | Q9Y4W6 | AFG3-like protein 2 OS=Homo sapiens C | AFG3L2 | 88.584 | 2 | 50.682903 | yes | 9.728E-08 | 529.80823 | 189.93472 |
| TQVLQPEEK | 9 | Q9Y4W6 | AFG3-like protein 2 OS=Homo sapiens C | AFG3L2 | 88.584 | 2 | 50.682903 | yes | 1.626E-07 | 536.28766 | 167.22038 |
| HFEQAIER | 8 | Q9Y4W6 | AFG3-like protein 2 OS=Homo sapiens C | AFG3L2 | 88.584 | 2 | 50.682903 | yes | 5.784E-07 | 515.25928 | 155.55626 |
| VHLRPLK | 7 | Q9Y4W6 | AFG3-like protein 2 OS=Homo sapiens C | AFG3L2 | 88.584 | 2 | 50.682903 | yes | 4.126E-07 | 431.7847 | 80.869843 |
| TVALLTEK | 8 | Q9Y4W6 | AFG3-like protein 2 OS=Homo sapiens C | AFG3L2 | 88.584 | 2 | 50.682903 | yes | 1.316E-06 | 437.76584 | 29.459082 |

| | | | | | | | | | | | |
|---|---|---|---|---|---|---|---|---|---|---|---|
| KTQVLQPEEK | 10 | Q9Y4W6 | AFG3-like protein 2 OS=Homo sapiens C | AFG3L2 | 88.584 | 2 | 50.682903 | yes | 1.7E-07 | 600.33514 | 28.979815 |
| EVNEVIQNPATITF | 14 | Q9Y4X5 | E3 ubiquitin-protein ligase ARIH1 OS=H | ARIH1 | 64.118 | 2 | 47.800507 | yes | 1.256E-07 | 792.42303 | 195.02301 |
| DISQDSLQDIK | 11 | Q9Y4X5 | E3 ubiquitin-protein ligase ARIH1 OS=H | ARIH1 | 64.118 | 2 | 47.800507 | yes | 2.54E-07 | 631.31714 | 154.71326 |
| LFAECHVINPSK | 12 | Q9Y4X5 | E3 ubiquitin-protein ligase ARIH1 OS=H | ARIH1 | 64.118 | 2 | 47.800507 | yes | 3.697E-06 | 707.86102 | 62.340057 |
| LFAECHVINPSK | 12 | Q9Y4X5 | E3 ubiquitin-protein ligase ARIH1 OS=H | ARIH1 | 64.118 | 2 | 47.800507 | yes | 3.623E-07 | 472.2431 | 62.340057 |
| LFAECHVINPSK | 12 | Q9Y4X5 | E3 ubiquitin-protein ligase ARIH1 OS=H | ARIH1 | 64.118 | 3 | 47.800507 | yes | 3.697E-06 | 707.86102 | 62.340057 |
| LFAECHVINPSK | 12 | Q9Y4X5 | E3 ubiquitin-protein ligase ARIH1 OS=H | ARIH1 | 64.118 | 3 | 47.800507 | yes | 3.623E-07 | 472.2431 | 62.340057 |
| YLFYCNR | 7 | Q9Y4X5 | E3 ubiquitin-protein ligase ARIH1 OS=H | ARIH1 | 64.118 | 2 | 47.800507 | yes | 6.372E-07 | 518.23944 | 77.302879 |
| YFDGNLEK | 8 | Q9Y4X5 | E3 ubiquitin-protein ligase ARIH1 OS=H | ARIH1 | 64.118 | 2 | 47.800507 | yes | 2.272E-07 | 493.23489 | 61.187096 |
| AVDVLCQCR | 9 | Q9Y4X5 | E3 ubiquitin-protein ligase ARIH1 OS=H | ARIH1 | 64.118 | 2 | 47.800507 | yes | 5.097E-07 | 560.7655 | 61.179863 |
| IPDEIIDMVK | 10 | Q9Y4Z0 | U6 snRNA-associated Sm-like protein LS | LSM4 | 15.35 | 2 | 50.27311 | yes | 9.85E-08 | 586.81519 | 1248.3046 |
| TAQNHPMLVELK | 12 | Q9Y4Z0 | U6 snRNA-associated Sm-like protein LS | LSM4 | 15.35 | 2 | 50.27311 | yes | 1.104E-07 | 690.86884 | 275.77823 |
| TAQNHPMLVELK | 12 | Q9Y4Z0 | U6 snRNA-associated Sm-like protein LS | LSM4 | 15.35 | 2 | 50.27311 | yes | 9.823E-08 | 460.91498 | 275.77823 |
| TAQNHPMLVELK | 12 | Q9Y4Z0 | U6 snRNA-associated Sm-like protein LS | LSM4 | 15.35 | 3 | 50.27311 | yes | 1.104E-07 | 690.86884 | 275.77823 |
| TAQNHPMLVELK | 12 | Q9Y4Z0 | U6 snRNA-associated Sm-like protein LS | LSM4 | 15.35 | 3 | 50.27311 | yes | 9.823E-08 | 460.91498 | 275.77823 |
| MPECYIR | 7 | Q9Y4Z0 | U6 snRNA-associated Sm-like protein LS | LSM4 | 15.35 | 2 | 50.27311 | yes | 1.114E-07 | 484.72006 | 492.13559 |
| YQNYIMEGVK | 10 | Q9Y508 | E3 ubiquitin-protein ligase RNF114 OS=I | RNF114 | 25.695 | 2 | 47.111435 | yes | 1.398E-07 | 622.80261 | 537.45337 |
| NFDQEGLVEHCK | 12 | Q9Y508 | E3 ubiquitin-protein ligase RNF114 OS=I | RNF114 | 25.695 | 2 | 47.111435 | yes | 2.677E-07 | 738.33282 | 359.46475 |
| YTFPCPYCPEK | 11 | Q9Y508 | E3 ubiquitin-protein ligase RNF114 OS=I | RNF114 | 25.695 | 2 | 47.111435 | yes | 1.135E-06 | 731.3125 | 359.24228 |
| FTCPVCLEVYEK | 12 | Q9Y508 | E3 ubiquitin-protein ligase RNF114 OS=I | RNF114 | 25.695 | 2 | 47.111435 | yes | 3.793E-07 | 772.8598 | 209.3569 |
| LFHSTDTK | 8 | Q9Y508 | E3 ubiquitin-protein ligase RNF114 OS=I | RNF114 | 25.695 | 2 | 47.111435 | yes | 1.816E-07 | 474.74289 | 200.20947 |
| KPVCGVCR | 8 | Q9Y508 | E3 ubiquitin-protein ligase RNF114 OS=I | RNF114 | 25.695 | 2 | 47.111435 | yes | 4.305E-07 | 488.24677 | 123.01302 |
| ICDGVQFGAGIR | 12 | Q9Y512 | Sorting and assembly machinery compone | SAMM50 | 51.977 | 2 | 43.68924 | yes | 2.486E-07 | 646.8244 | 82.107605 |
| WSYGAGIVLR | 10 | Q9Y512 | Sorting and assembly machinery compone | SAMM50 | 51.977 | 2 | 43.68924 | yes | 2.815E-07 | 561.30872 | 77.273857 |
| VTGQFPWSSLR | 11 | Q9Y512 | Sorting and assembly machinery compone | SAMM50 | 51.977 | 2 | 43.68924 | yes | 2.829E-07 | 639.33551 | 25.21489 |
| GMSAEYSFPIWK | 12 | Q9Y512 | Sorting and assembly machinery compone | SAMM50 | 51.977 | 2 | 43.68924 | yes | 3.678E-07 | 708.33685 | 20.663305 |
| PGVAAPPEVAPAP | 14 | Q9Y520 | Protein PRRC2C OS=Homo sapiens OX= | PRRC2C | 316.912 | 2 | 53.288067 | yes | 2.921E-06 | 650.86664 | 24587.709 |
| SSSQIPAQPSVAK | 13 | Q9Y520 | Protein PRRC2C OS=Homo sapiens OX= | PRRC2C | 316.912 | 2 | 53.288067 | yes | 1.512E-07 | 650.34882 | 18776.598 |
| AEHIPSGPLR | 10 | Q9Y520 | Protein PRRC2C OS=Homo sapiens OX= | PRRC2C | 316.912 | 2 | 53.288067 | yes | 1.013E-06 | 538.79602 | 12618.605 |
| LPDLSPVENK | 10 | Q9Y520 | Protein PRRC2C OS=Homo sapiens OX= | PRRC2C | 316.912 | 2 | 53.288067 | yes | 1.004E-07 | 556.30334 | 12125.824 |
| DHAISLSEPR | 10 | Q9Y520 | Protein PRRC2C OS=Homo sapiens OX= | PRRC2C | 316.912 | 2 | 53.288067 | yes | 2.199E-07 | 562.78839 | 9341.1982 |
| VAPPAVLNDISK | 12 | Q9Y520 | Protein PRRC2C OS=Homo sapiens OX= | PRRC2C | 316.912 | 2 | 53.288067 | yes | 4.925E-07 | 612.35333 | 8810.2705 |
| RHEQFIPIAADK | 12 | Q9Y520 | Protein PRRC2C OS=Homo sapiens OX= | PRRC2C | 316.912 | 3 | 53.288067 | yes | 3.773E-07 | 475.59302 | 2830.3813 |
| RHEQFIPIAADK | 12 | Q9Y520 | Protein PRRC2C OS=Homo sapiens OX= | PRRC2C | 316.912 | 3 | 53.288067 | yes | 3.057E-07 | 712.88586 | 2830.3813 |
| RHEQFIPIAADK | 12 | Q9Y520 | Protein PRRC2C OS=Homo sapiens OX= | PRRC2C | 316.912 | 2 | 53.288067 | yes | 3.773E-07 | 475.59302 | 2830.3813 |
| RHEQFIPIAADK | 12 | Q9Y520 | Protein PRRC2C OS=Homo sapiens OX= | PRRC2C | 316.912 | 2 | 53.288067 | yes | 3.057E-07 | 712.88586 | 2830.3813 |
| QQSEISAAVER | 11 | Q9Y520 | Protein PRRC2C OS=Homo sapiens OX= | PRRC2C | 316.912 | 2 | 53.288067 | yes | 1.001E-07 | 609.30969 | 5513.1431 |
| ATEEPEDVR | 9 | Q9Y520 | Protein PRRC2C OS=Homo sapiens OX= | PRRC2C | 316.912 | 2 | 53.288067 | yes | 1.889E-07 | 523.24347 | 5249.0132 |
| NADLNAQTVVK | 11 | Q9Y520 | Protein PRRC2C OS=Homo sapiens OX= | PRRC2C | 316.912 | 2 | 53.288067 | yes | 9.722E-08 | 586.81714 | 4868.2192 |

| | | | | | | | | | | | |
|---|---|---|---|---|---|---|---|---|---|---|---|
| GNDPNVNIVPK | 11 | Q9Y520 | Protein PRRC2C OS=Homo sapiens OX= | PRRC2C | 316.912 | 2 | 53.288067 | yes | 2.161E-06 | 583.81183 | 4735.915 |
| PAVLSGYFK | 9 | Q9Y520 | Protein PRRC2C OS=Homo sapiens OX= | PRRC2C | 316.912 | 2 | 53.288067 | yes | 2.514E-07 | 491.27383 | 4334.7534 |
| YATLSLFNTYK | 11 | Q9Y520 | Protein PRRC2C OS=Homo sapiens OX= | PRRC2C | 316.912 | 2 | 53.288067 | yes | 1.063E-07 | 660.84534 | 4287.1987 |
| MLWGSDPYPHAEI | 20 | Q9Y520 | Protein PRRC2C OS=Homo sapiens OX= | PRRC2C | 316.912 | 3 | 53.288067 | yes | 5.194E-07 | 752.02478 | 2003.8118 |
| MLWGSDPYPHAEI | 20 | Q9Y520 | Protein PRRC2C OS=Homo sapiens OX= | PRRC2C | 316.912 | 3 | 53.288067 | yes | 3.533E-07 | 757.35638 | 2003.8118 |
| TVNQQTMAAPVV | 13 | Q9Y520 | Protein PRRC2C OS=Homo sapiens OX= | PRRC2C | 316.912 | 2 | 53.288067 | yes | 9.881E-08 | 693.87415 | 1867.1692 |
| TVNQQTMAAPVV | 13 | Q9Y520 | Protein PRRC2C OS=Homo sapiens OX= | PRRC2C | 316.912 | 2 | 53.288067 | yes | 1.166E-07 | 701.87158 | 1867.1692 |
| LLAQQHPPPDR | 11 | Q9Y520 | Protein PRRC2C OS=Homo sapiens OX= | PRRC2C | 316.912 | 2 | 53.288067 | yes | 5.383E-07 | 636.34637 | 1644.8489 |
| LLAQQHPPPDR | 11 | Q9Y520 | Protein PRRC2C OS=Homo sapiens OX= | PRRC2C | 316.912 | 2 | 53.288067 | yes | 1.354E-07 | 424.56668 | 1644.8489 |
| LLAQQHPPPDR | 11 | Q9Y520 | Protein PRRC2C OS=Homo sapiens OX= | PRRC2C | 316.912 | 3 | 53.288067 | yes | 5.383E-07 | 636.34637 | 1644.8489 |
| LLAQQHPPPDR | 11 | Q9Y520 | Protein PRRC2C OS=Homo sapiens OX= | PRRC2C | 316.912 | 3 | 53.288067 | yes | 1.354E-07 | 424.56668 | 1644.8489 |
| AEVLQSTQR | 9 | Q9Y520 | Protein PRRC2C OS=Homo sapiens OX= | PRRC2C | 316.912 | 2 | 53.288067 | yes | 1.542E-07 | 516.27765 | 3106.9768 |
| EPNLEPMVEK | 10 | Q9Y520 | Protein PRRC2C OS=Homo sapiens OX= | PRRC2C | 316.912 | 2 | 53.288067 | yes | 1.995E-07 | 593.29462 | 1551.52 |
| EPNLEPMVEK | 10 | Q9Y520 | Protein PRRC2C OS=Homo sapiens OX= | PRRC2C | 316.912 | 2 | 53.288067 | yes | 8.608E-07 | 601.29211 | 1551.52 |
| AWENSPNVR | 9 | Q9Y520 | Protein PRRC2C OS=Homo sapiens OX= | PRRC2C | 316.912 | 2 | 53.288067 | yes | 1.335E-07 | 536.76215 | 2774.1345 |
| GTSSHLPPPPK | 11 | Q9Y520 | Protein PRRC2C OS=Homo sapiens OX= | PRRC2C | 316.912 | 2 | 53.288067 | yes | 9.81E-08 | 559.30365 | 2585.8093 |
| SADKIPEPK | 9 | Q9Y520 | Protein PRRC2C OS=Homo sapiens OX= | PRRC2C | 316.912 | 2 | 53.288067 | yes | 1.584E-07 | 492.77167 | 2515.4336 |
| EPIERPEEKPK | 11 | Q9Y520 | Protein PRRC2C OS=Homo sapiens OX= | PRRC2C | 316.912 | 3 | 53.288067 | yes | 1.002E-07 | 451.24539 | 1229.7285 |
| EPIERPEEKPK | 11 | Q9Y520 | Protein PRRC2C OS=Homo sapiens OX= | PRRC2C | 316.912 | 3 | 53.288067 | yes | 1.018E-07 | 676.36444 | 1229.7285 |
| EPIERPEEKPK | 11 | Q9Y520 | Protein PRRC2C OS=Homo sapiens OX= | PRRC2C | 316.912 | 2 | 53.288067 | yes | 1.002E-07 | 451.24539 | 1229.7285 |
| EPIERPEEKPK | 11 | Q9Y520 | Protein PRRC2C OS=Homo sapiens OX= | PRRC2C | 316.912 | 2 | 53.288067 | yes | 1.018E-07 | 676.36444 | 1229.7285 |
| PAQVSQPFR | 9 | Q9Y520 | Protein PRRC2C OS=Homo sapiens OX= | PRRC2C | 316.912 | 2 | 53.288067 | yes | 9.722E-08 | 515.27747 | 2395.7908 |
| GPSFNQER | 8 | Q9Y520 | Protein PRRC2C OS=Homo sapiens OX= | PRRC2C | 316.912 | 2 | 53.288067 | yes | 2.693E-07 | 467.7225 | 2340.9541 |
| MTYPPLHGPMR | 11 | Q9Y520 | Protein PRRC2C OS=Homo sapiens OX= | PRRC2C | 316.912 | 2 | 53.288067 | yes | 1.073E-07 | 650.3205 | 1095.3295 |
| MTYPPLHGPMR | 11 | Q9Y520 | Protein PRRC2C OS=Homo sapiens OX= | PRRC2C | 316.912 | 2 | 53.288067 | yes | 3.241E-07 | 433.88275 | 1095.3295 |
| MTYPPLHGPMR | 11 | Q9Y520 | Protein PRRC2C OS=Homo sapiens OX= | PRRC2C | 316.912 | 3 | 53.288067 | yes | 1.073E-07 | 650.3205 | 1095.3295 |
| MTYPPLHGPMR | 11 | Q9Y520 | Protein PRRC2C OS=Homo sapiens OX= | PRRC2C | 316.912 | 3 | 53.288067 | yes | 3.241E-07 | 433.88275 | 1095.3295 |
| STTPTSSPFR | 10 | Q9Y520 | Protein PRRC2C OS=Homo sapiens OX= | PRRC2C | 316.912 | 2 | 53.288067 | yes | 2.45E-07 | 540.76965 | 2056.9883 |
| QQVADEDEIWK | 11 | Q9Y520 | Protein PRRC2C OS=Homo sapiens OX= | PRRC2C | 316.912 | 2 | 53.288067 | yes | 1.512E-07 | 680.82263 | 1922.9121 |
| TLSAPQEER | 9 | Q9Y520 | Protein PRRC2C OS=Homo sapiens OX= | PRRC2C | 316.912 | 2 | 53.288067 | yes | 9.722E-08 | 515.76184 | 1807.9199 |
| AMMPPYMFQQYP | 13 | Q9Y520 | Protein PRRC2C OS=Homo sapiens OX= | PRRC2C | 316.912 | 2 | 53.288067 | yes | 5.214E-07 | 830.37769 | 1702.3813 |
| SESSDFEVVPK | 11 | Q9Y520 | Protein PRRC2C OS=Homo sapiens OX= | PRRC2C | 316.912 | 2 | 53.288067 | yes | 9.722E-08 | 612.29315 | 1617.3871 |
| QPSPEEIR | 8 | Q9Y520 | Protein PRRC2C OS=Homo sapiens OX= | PRRC2C | 316.912 | 2 | 53.288067 | yes | 3.275E-07 | 478.24582 | 1542.0587 |
| PLTSFGSAPSSEGA | 15 | Q9Y520 | Protein PRRC2C OS=Homo sapiens OX= | PRRC2C | 316.912 | 2 | 53.288067 | yes | 3.991E-07 | 718.35681 | 1533.3885 |
| VEEKPPPAPSIATK | 14 | Q9Y520 | Protein PRRC2C OS=Homo sapiens OX= | PRRC2C | 316.912 | 2 | 53.288067 | yes | 3.859E-07 | 732.40887 | 696.40594 |
| VEEKPPPAPSIATK | 14 | Q9Y520 | Protein PRRC2C OS=Homo sapiens OX= | PRRC2C | 316.912 | 2 | 53.288067 | yes | 4.237E-07 | 488.60834 | 696.40594 |
| VEEKPPPAPSIATK | 14 | Q9Y520 | Protein PRRC2C OS=Homo sapiens OX= | PRRC2C | 316.912 | 3 | 53.288067 | yes | 3.859E-07 | 732.40887 | 696.40594 |
| VEEKPPPAPSIATK | 14 | Q9Y520 | Protein PRRC2C OS=Homo sapiens OX= | PRRC2C | 316.912 | 3 | 53.288067 | yes | 4.237E-07 | 488.60834 | 696.40594 |
| FFSEQQQSK | 9 | Q9Y520 | Protein PRRC2C OS=Homo sapiens OX= | PRRC2C | 316.912 | 2 | 53.288067 | yes | 1.87E-07 | 564.76965 | 1370.0536 |

| | | | | | | | | | | | |
|---|---|---|---|---|---|---|---|---|---|---|---|
| MNSIVYQK | 8 | Q9Y520 | Protein PRRC2C OS=Homo sapiens OX= | PRRC2C | 316.912 | 2 | 53.288067 | yes | 3.445E-07 | 491.75494 | 1257.7778 |
| QFQSAPATVR | 10 | Q9Y520 | Protein PRRC2C OS=Homo sapiens OX= | PRRC2C | 316.912 | 2 | 53.288067 | yes | 9.722E-08 | 552.79346 | 1257.2828 |
| QAVPGRPGPFPSK | 13 | Q9Y520 | Protein PRRC2C OS=Homo sapiens OX= | PRRC2C | 316.912 | 2 | 53.288067 | yes | 1.073E-07 | 669.36987 | 585.28784 |
| QAVPGRPGPFPSK | 13 | Q9Y520 | Protein PRRC2C OS=Homo sapiens OX= | PRRC2C | 316.912 | 2 | 53.288067 | yes | 1.043E-07 | 446.58234 | 585.28784 |
| QAVPGRPGPFPSK | 13 | Q9Y520 | Protein PRRC2C OS=Homo sapiens OX= | PRRC2C | 316.912 | 3 | 53.288067 | yes | 1.073E-07 | 669.36987 | 585.28784 |
| QAVPGRPGPFPSK | 13 | Q9Y520 | Protein PRRC2C OS=Homo sapiens OX= | PRRC2C | 316.912 | 3 | 53.288067 | yes | 1.043E-07 | 446.58234 | 585.28784 |
| MSEMELK | 7 | Q9Y520 | Protein PRRC2C OS=Homo sapiens OX= | PRRC2C | 316.912 | 2 | 53.288067 | yes | 2.487E-07 | 434.20117 | 882.76453 |
| SIQTPQSHGTLTAE | 19 | Q9Y520 | Protein PRRC2C OS=Homo sapiens OX= | PRRC2C | 316.912 | 3 | 53.288067 | yes | 1.799E-07 | 709.35529 | 878.90063 |
| DNQWNPR | 7 | Q9Y520 | Protein PRRC2C OS=Homo sapiens OX= | PRRC2C | 316.912 | 2 | 53.288067 | yes | 2.763E-07 | 465.21484 | 788.66626 |
| STTTSDPPNICK | 12 | Q9Y520 | Protein PRRC2C OS=Homo sapiens OX= | PRRC2C | 316.912 | 2 | 53.288067 | yes | 1.327E-07 | 660.80865 | 768.91626 |
| DLVIERPR | 8 | Q9Y520 | Protein PRRC2C OS=Homo sapiens OX= | PRRC2C | 316.912 | 2 | 53.288067 | yes | 1.777E-07 | 499.29309 | 616.09796 |
| GLIPAGTQHSMIAT | 17 | Q9Y520 | Protein PRRC2C OS=Homo sapiens OX= | PRRC2C | 316.912 | 2 | 53.288067 | yes | 2.656E-07 | 841.94836 | 589.39624 |
| GPCGSLSGVR | 10 | Q9Y520 | Protein PRRC2C OS=Homo sapiens OX= | PRRC2C | 316.912 | 2 | 53.288067 | yes | 9.913E-08 | 495.24527 | 568.36707 |
| HEQFIPIAADK | 11 | Q9Y520 | Protein PRRC2C OS=Homo sapiens OX= | PRRC2C | 316.912 | 2 | 53.288067 | yes | 5.777E-07 | 634.83533 | 560.78406 |
| EEQVIQVWNK | 10 | Q9Y520 | Protein PRRC2C OS=Homo sapiens OX= | PRRC2C | 316.912 | 2 | 53.288067 | yes | 1.119E-07 | 636.83276 | 508.74725 |
| SFSSQRPVDR | 10 | Q9Y520 | Protein PRRC2C OS=Homo sapiens OX= | PRRC2C | 316.912 | 2 | 53.288067 | yes | 1.334E-07 | 589.79926 | 404.00092 |
| TPEVPPAQPK | 10 | Q9Y520 | Protein PRRC2C OS=Homo sapiens OX= | PRRC2C | 316.912 | 2 | 53.288067 | yes | 3.882E-06 | 532.29279 | 386.48914 |
| KAWENSPNVR | 10 | Q9Y520 | Protein PRRC2C OS=Homo sapiens OX= | PRRC2C | 316.912 | 2 | 53.288067 | yes | 3.46E-07 | 600.80963 | 370.6481 |
| QSEVPKPEDGEPPI | 14 | Q9Y520 | Protein PRRC2C OS=Homo sapiens OX= | PRRC2C | 316.912 | 3 | 53.288067 | yes | 3.012E-07 | 522.25824 | 176.69102 |
| QSEVPKPEDGEPPI | 14 | Q9Y520 | Protein PRRC2C OS=Homo sapiens OX= | PRRC2C | 316.912 | 3 | 53.288067 | yes | 3.367E-06 | 782.88373 | 176.69102 |
| QSEVPKPEDGEPPI | 14 | Q9Y520 | Protein PRRC2C OS=Homo sapiens OX= | PRRC2C | 316.912 | 2 | 53.288067 | yes | 3.012E-07 | 522.25824 | 176.69102 |
| QSEVPKPEDGEPPI | 14 | Q9Y520 | Protein PRRC2C OS=Homo sapiens OX= | PRRC2C | 316.912 | 2 | 53.288067 | yes | 3.367E-06 | 782.88373 | 176.69102 |
| KNADLNAQTVVK | 12 | Q9Y520 | Protein PRRC2C OS=Homo sapiens OX= | PRRC2C | 316.912 | 2 | 53.288067 | yes | 9.809E-08 | 650.86462 | 344.45935 |
| LNGQQAALASQYI | 13 | Q9Y520 | Protein PRRC2C OS=Homo sapiens OX= | PRRC2C | 316.912 | 2 | 53.288067 | yes | 1.076E-07 | 710.37061 | 329.68466 |
| GNQAQAALK | 9 | Q9Y520 | Protein PRRC2C OS=Homo sapiens OX= | PRRC2C | 316.912 | 2 | 53.288067 | yes | 2.57E-07 | 450.7485 | 327.89374 |
| PLYQPMQPHPQHI | 21 | Q9Y520 | Protein PRRC2C OS=Homo sapiens OX= | PRRC2C | 316.912 | 4 | 53.288067 | yes | 3.347E-07 | 612.54913 | 296.07043 |
| VGENVLPPK | 9 | Q9Y520 | Protein PRRC2C OS=Homo sapiens OX= | PRRC2C | 316.912 | 2 | 53.288067 | yes | 2.298E-06 | 476.77673 | 292.62799 |
| EEEPVFTR | 8 | Q9Y520 | Protein PRRC2C OS=Homo sapiens OX= | PRRC2C | 316.912 | 2 | 53.288067 | yes | 3.612E-07 | 503.74564 | 289.50208 |
| KKEEQVIQVWNK | 12 | Q9Y520 | Protein PRRC2C OS=Homo sapiens OX= | PRRC2C | 316.912 | 2 | 53.288067 | yes | 9.81E-08 | 764.92773 | 245.57259 |
| PSILSASELK | 10 | Q9Y520 | Protein PRRC2C OS=Homo sapiens OX= | PRRC2C | 316.912 | 2 | 53.288067 | yes | 1.384E-07 | 522.80042 | 232.7368 |
| AFGSGIDIK | 9 | Q9Y520 | Protein PRRC2C OS=Homo sapiens OX= | PRRC2C | 316.912 | 2 | 53.288067 | yes | 1.071E-07 | 454.2478 | 225.20975 |
| FPSTETATLAQK | 12 | Q9Y520 | Protein PRRC2C OS=Homo sapiens OX= | PRRC2C | 316.912 | 2 | 53.288067 | yes | 1.354E-07 | 647.33789 | 212.50479 |
| EEVNDRPVR | 9 | Q9Y520 | Protein PRRC2C OS=Homo sapiens OX= | PRRC2C | 316.912 | 2 | 53.288067 | yes | 1.128E-07 | 557.28601 | 208.91 |
| ISAVESQPSR | 10 | Q9Y520 | Protein PRRC2C OS=Homo sapiens OX= | PRRC2C | 316.912 | 2 | 53.288067 | yes | 1.076E-07 | 537.2829 | 200.05772 |
| REEVNDRPVR | 10 | Q9Y520 | Protein PRRC2C OS=Homo sapiens OX= | PRRC2C | 316.912 | 2 | 53.288067 | yes | 1.238E-07 | 635.33655 | 92.747215 |
| REEVNDRPVR | 10 | Q9Y520 | Protein PRRC2C OS=Homo sapiens OX= | PRRC2C | 316.912 | 2 | 53.288067 | yes | 8.211E-07 | 423.89346 | 92.747215 |
| REEVNDRPVR | 10 | Q9Y520 | Protein PRRC2C OS=Homo sapiens OX= | PRRC2C | 316.912 | 3 | 53.288067 | yes | 1.238E-07 | 635.33655 | 92.747215 |
| REEVNDRPVR | 10 | Q9Y520 | Protein PRRC2C OS=Homo sapiens OX= | PRRC2C | 316.912 | 3 | 53.288067 | yes | 8.211E-07 | 423.89346 | 92.747215 |
| EQEKECELEK | 10 | Q9Y520 | Protein PRRC2C OS=Homo sapiens OX= | PRRC2C | 316.912 | 2 | 53.288067 | yes | 1.206E-07 | 661.30066 | 169.87804 |

| | | | | | | | | | | | |
|---|---|---|---|---|---|---|---|---|---|---|---|
| PSRPSTLR | 8 | Q9Y520 | Protein PRRC2C OS=Homo sapiens OX= | PRRC2C | 316.912 | 2 | 53.288067 | yes | 2.004E-07 | 457.26434 | 154.7429 |
| LLEKPYVR | 8 | Q9Y520 | Protein PRRC2C OS=Homo sapiens OX= | PRRC2C | 316.912 | 2 | 53.288067 | yes | 5.483E-07 | 509.3082 | 151.79373 |
| EPIERPEEK | 9 | Q9Y520 | Protein PRRC2C OS=Homo sapiens OX= | PRRC2C | 316.912 | 2 | 53.288067 | yes | 9.722E-08 | 563.79059 | 149.25433 |
| KYATLSLFNTYK | 12 | Q9Y520 | Protein PRRC2C OS=Homo sapiens OX= | PRRC2C | 316.912 | 2 | 53.288067 | yes | 1.705E-07 | 724.89282 | 137.09299 |
| PGTPPIAGR | 9 | Q9Y520 | Protein PRRC2C OS=Homo sapiens OX= | PRRC2C | 316.912 | 2 | 53.288067 | yes | 1.402E-07 | 433.24814 | 122.63764 |
| RLQDEER | 7 | Q9Y520 | Protein PRRC2C OS=Homo sapiens OX= | PRRC2C | 316.912 | 2 | 53.288067 | yes | 5.868E-07 | 473.24106 | 109.16499 |
| VGENVLPPKR | 10 | Q9Y520 | Protein PRRC2C OS=Homo sapiens OX= | PRRC2C | 316.912 | 2 | 53.288067 | yes | 4.453E-06 | 554.82727 | 103.47614 |
| KEEQVIQVWNK | 11 | Q9Y520 | Protein PRRC2C OS=Homo sapiens OX= | PRRC2C | 316.912 | 2 | 53.288067 | yes | 1.055E-07 | 700.88025 | 100.09343 |
| GPPPSWASEPER | 12 | Q9Y520 | Protein PRRC2C OS=Homo sapiens OX= | PRRC2C | 316.912 | 2 | 53.288067 | yes | 1.784E-06 | 655.31219 | 94.786469 |
| DLVIERPRPDSR | 12 | Q9Y520 | Protein PRRC2C OS=Homo sapiens OX= | PRRC2C | 316.912 | 3 | 53.288067 | yes | 1.759E-07 | 484.93542 | 77.703903 |
| WLMMQSYMDPR | 11 | Q9Y520 | Protein PRRC2C OS=Homo sapiens OX= | PRRC2C | 316.912 | 2 | 53.288067 | yes | 9.722E-08 | 729.32233 | 73.680618 |
| MKEQEKECELEK | 12 | Q9Y520 | Protein PRRC2C OS=Homo sapiens OX= | PRRC2C | 316.912 | 2 | 53.288067 | yes | 9.722E-08 | 790.86835 | 63.730938 |
| ERDWFPDQGYR | 11 | Q9Y520 | Protein PRRC2C OS=Homo sapiens OX= | PRRC2C | 316.912 | 2 | 53.288067 | yes | 1.102E-07 | 734.83386 | 46.013554 |
| PVQNPLQTTSQSSI | 14 | Q9Y520 | Protein PRRC2C OS=Homo sapiens OX= | PRRC2C | 316.912 | 2 | 53.288067 | yes | 2.499E-07 | 757.8941 | 45.849678 |
| VEEKPPPAPSIATK | 17 | Q9Y520 | Protein PRRC2C OS=Homo sapiens OX= | PRRC2C | 316.912 | 3 | 53.288067 | yes | 8.07E-07 | 606.01575 | 21.133804 |
| VEEKPPPAPSIATK | 17 | Q9Y520 | Protein PRRC2C OS=Homo sapiens OX= | PRRC2C | 316.912 | 3 | 53.288067 | yes | 2.251E-06 | 454.76364 | 21.133804 |
| VEEKPPPAPSIATK | 17 | Q9Y520 | Protein PRRC2C OS=Homo sapiens OX= | PRRC2C | 316.912 | 4 | 53.288067 | yes | 8.07E-07 | 606.01575 | 21.133804 |
| VEEKPPPAPSIATK | 17 | Q9Y520 | Protein PRRC2C OS=Homo sapiens OX= | PRRC2C | 316.912 | 4 | 53.288067 | yes | 2.251E-06 | 454.76364 | 21.133804 |
| EKETNDDNYGPGI | 24 | Q9Y520 | Protein PRRC2C OS=Homo sapiens OX= | PRRC2C | 316.912 | 4 | 53.288067 | yes | 7.112E-07 | 693.82532 | 40.826488 |
| TGPIKPQAIK | 10 | Q9Y520 | Protein PRRC2C OS=Homo sapiens OX= | PRRC2C | 316.912 | 2 | 53.288067 | yes | 5.629E-07 | 526.82678 | 37.519642 |
| RDQMEGSPNSSES | 19 | Q9Y520 | Protein PRRC2C OS=Homo sapiens OX= | PRRC2C | 316.912 | 4 | 53.288067 | yes | 1.018E-07 | 544.99738 | 29.991362 |
| VKDAQQVEPEGQI | 21 | Q9Y520 | Protein PRRC2C OS=Homo sapiens OX= | PRRC2C | 316.912 | 4 | 53.288067 | yes | 3.376E-05 | 574.05005 | 14.256578 |
| VKDAQQVEPEGQI | 21 | Q9Y520 | Protein PRRC2C OS=Homo sapiens OX= | PRRC2C | 316.912 | 4 | 53.288067 | yes | 1.341E-06 | 765.06427 | 14.256578 |
| VKDAQQVEPEGQI | 21 | Q9Y520 | Protein PRRC2C OS=Homo sapiens OX= | PRRC2C | 316.912 | 3 | 53.288067 | yes | 3.376E-05 | 574.05005 | 14.256578 |
| VKDAQQVEPEGQI | 21 | Q9Y520 | Protein PRRC2C OS=Homo sapiens OX= | PRRC2C | 316.912 | 3 | 53.288067 | yes | 1.341E-06 | 765.06427 | 14.256578 |
| AFGSGIDIKPGTPP | 18 | Q9Y520 | Protein PRRC2C OS=Homo sapiens OX= | PRRC2C | 316.912 | 3 | 53.288067 | yes | 1.258E-07 | 585.32471 | 27.022516 |
| DDDKAKPGFLPK | 12 | Q9Y520 | Protein PRRC2C OS=Homo sapiens OX= | PRRC2C | 316.912 | 2 | 53.288067 | yes | 6.26E-07 | 665.8537 | 26.523634 |
| PLEPVSTVQVEPA | 15 | Q9Y520 | Protein PRRC2C OS=Homo sapiens OX= | PRRC2C | 316.912 | 2 | 53.288067 | yes | 1.548E-06 | 796.94836 | 19.397484 |
| SVEDVRPHHTDAN | 25 | Q9Y520 | Protein PRRC2C OS=Homo sapiens OX= | PRRC2C | 316.912 | 4 | 53.288067 | yes | 1.078E-06 | 713.82178 | 4.0899553 |
| GLSNLFLSCPIPK | 13 | Q9Y570 | Protein phosphatase methylesterase 1 OS= | PPME1 | 42.316 | 2 | 30.945292 | yes | 2.007E-06 | 723.39471 | 41.892006 |
| TMDNMTSSAQIIQ | 14 | Q9Y580 | RNA-binding protein 7 OS=Homo sapien | RBM7 | 30.504 | 2 | 50.855408 | yes | 1.401E-07 | 798.37946 | 642.69861 |
| TLFVGNLETK | 10 | Q9Y580 | RNA-binding protein 7 OS=Homo sapien | RBM7 | 30.504 | 2 | 50.855408 | yes | 3.619E-07 | 561.31366 | 620.30322 |
| SFSSPENFQR | 10 | Q9Y580 | RNA-binding protein 7 OS=Homo sapien | RBM7 | 30.504 | 2 | 50.855408 | yes | 3.214E-07 | 599.77802 | 292.47098 |
| MNSYPYLADR | 10 | Q9Y580 | RNA-binding protein 7 OS=Homo sapien | RBM7 | 30.504 | 2 | 50.855408 | yes | 3.371E-07 | 615.28461 | 287.45938 |
| QFAFVNFK | 8 | Q9Y580 | RNA-binding protein 7 OS=Homo sapien | RBM7 | 30.504 | 2 | 50.855408 | yes | 9.722E-08 | 500.76617 | 106.79402 |
| LGSTEVASNVPK | 12 | Q9Y5A9 | YTH domain-containing family protein 2 | YTHDF2 | 62.335 | 2 | 50.937172 | yes | 9.728E-08 | 601.32477 | 5097.4209 |
| SASSLLEQRPK | 11 | Q9Y5A9 | YTH domain-containing family protein 2 | YTHDF2 | 62.335 | 2 | 50.937172 | yes | 1.005E-07 | 629.34351 | 3104.9124 |
| HTTSIFDDFSHYEK | 14 | Q9Y5A9 | YTH domain-containing family protein 2 | YTHDF2 | 62.335 | 3 | 50.937172 | yes | 1.182E-07 | 576.26178 | 2744.5515 |
| APGMNTIDQGMA | 15 | Q9Y5A9 | YTH domain-containing family protein 2 | YTHDF2 | 62.335 | 2 | 50.937172 | yes | 1.173E-07 | 759.37616 | 1183.4843 |

| | | | | | | | | | | | |
|---|---|---|---|---|---|---|---|---|---|---|---|
| APGMNTIDQGMA. | 15 | Q9Y5A9 | YTH domain-containing family protein 2 | YTHDF2 | 62.335 | 2 | 50.937172 | yes | 2.062E-07 | 767.3736 | 1183.4843 |
| YNIWCSTEHGNK | 12 | Q9Y5A9 | YTH domain-containing family protein 2 | YTHDF2 | 62.335 | 2 | 50.937172 | yes | 1.121E-07 | 754.83301 | 2211.396 |
| SASSLLEQR | 9 | Q9Y5A9 | YTH domain-containing family protein 2 | YTHDF2 | 62.335 | 2 | 50.937172 | yes | 9.722E-08 | 516.76965 | 1833.9012 |
| SINNYNPK | 8 | Q9Y5A9 | YTH domain-containing family protein 2 | YTHDF2 | 62.335 | 2 | 50.937172 | yes | 3.67E-07 | 475.24051 | 1036.0404 |
| NGIAGSSLPPPPIK | 14 | Q9Y5A9 | YTH domain-containing family protein 2 | YTHDF2 | 62.335 | 2 | 50.937172 | yes | 1.846E-07 | 674.38519 | 726.60028 |
| DVPNSQLR | 8 | Q9Y5A9 | YTH domain-containing family protein 2 | YTHDF2 | 62.335 | 2 | 50.937172 | yes | 1.729E-07 | 464.74597 | 239.66986 |
| LENNENKPVTNSR | 13 | Q9Y5A9 | YTH domain-containing family protein 2 | YTHDF2 | 62.335 | 2 | 50.937172 | yes | 1.281E-07 | 757.88153 | 120.003 |
| PASWADIASK | 10 | Q9Y5A9 | YTH domain-containing family protein 2 | YTHDF2 | 62.335 | 2 | 50.937172 | yes | 3.312E-07 | 523.26929 | 73.375595 |
| GSGFGHNGVDGN( | 34 | Q9Y5A9 | YTH domain-containing family protein 2 | YTHDF2 | 62.335 | 4 | 50.937172 | yes | 6.913E-07 | 812.63043 | 60.679695 |
| ALMAPNLDSFGR | 12 | Q9Y5B6 | PAX3- and PAX7-binding protein 1 OS= | PAXBP1 | 104.804 | 2 | 53.462307 | yes | 1.408E-07 | 646.32678 | 230.18204 |
| AQNVINCFPK | 10 | Q9Y5B6 | PAX3- and PAX7-binding protein 1 OS= | PAXBP1 | 104.804 | 2 | 53.462307 | yes | 3.103E-07 | 595.80298 | 174.59601 |
| TISQLENFCR | 10 | Q9Y5B6 | PAX3- and PAX7-binding protein 1 OS= | PAXBP1 | 104.804 | 2 | 53.462307 | yes | 2.756E-07 | 634.30859 | 168.59558 |
| ALDHAMSVASDH | 15 | Q9Y5B6 | PAX3- and PAX7-binding protein 1 OS= | PAXBP1 | 104.804 | 3 | 53.462307 | yes | 1.025E-07 | 532.25958 | 150.60855 |
| TPSNEMTPVTIDL\ | 15 | Q9Y5B6 | PAX3- and PAX7-binding protein 1 OS= | PAXBP1 | 104.804 | 2 | 53.462307 | yes | 1.371E-07 | 822.92926 | 127.23964 |
| DAYIGLCLPK | 10 | Q9Y5B6 | PAX3- and PAX7-binding protein 1 OS= | PAXBP1 | 104.804 | 2 | 53.462307 | yes | 9.758E-08 | 575.30225 | 126.53987 |
| WEQEQIR | 7 | Q9Y5B6 | PAX3- and PAX7-binding protein 1 OS= | PAXBP1 | 104.804 | 2 | 53.462307 | yes | 3.146E-07 | 494.74597 | 85.972549 |
| NSGPYLFFQR | 10 | Q9Y5B6 | PAX3- and PAX7-binding protein 1 OS= | PAXBP1 | 104.804 | 2 | 53.462307 | yes | 1.287E-07 | 614.80908 | 79.585953 |
| TLQELSIDGLLNR | 13 | Q9Y5B6 | PAX3- and PAX7-binding protein 1 OS= | PAXBP1 | 104.804 | 2 | 53.462307 | yes | 1.06E-07 | 736.40936 | 67.465424 |
| TLDDDVFMPLYPK | 13 | Q9Y5B6 | PAX3- and PAX7-binding protein 1 OS= | PAXBP1 | 104.804 | 2 | 53.462307 | yes | 3.161E-07 | 777.38147 | 61.81934 |
| NSIGCSDVEK | 10 | Q9Y5B6 | PAX3- and PAX7-binding protein 1 OS= | PAXBP1 | 104.804 | 2 | 53.462307 | yes | 2.738E-07 | 554.75061 | 59.87545 |
| EQEKDDVDVALLI | 18 | Q9Y5B6 | PAX3- and PAX7-binding protein 1 OS= | PAXBP1 | 104.804 | 3 | 53.462307 | yes | 9.722E-08 | 681.02814 | 56.811459 |
| RYELLFYPGDGSV | 19 | Q9Y5B8 | Nucleoside diphosphate kinase 7 OS=Hoı | NME7 | 42.492 | 3 | 53.39745 | yes | 9.722E-08 | 752.36487 | 332.38306 |
| VNVEEFYEVYK | 11 | Q9Y5B8 | Nucleoside diphosphate kinase 7 OS=Hoı | NME7 | 42.492 | 2 | 53.39745 | yes | 9.722E-08 | 709.84552 | 129.2023 |
| QLVLIDYGDQYTA | 14 | Q9Y5B8 | Nucleoside diphosphate kinase 7 OS=Hoı | NME7 | 42.492 | 2 | 53.39745 | yes | 1.877E-07 | 827.92542 | 112.07627 |
| EFCGPADPEIAR | 12 | Q9Y5B8 | Nucleoside diphosphate kinase 7 OS=Hoı | NME7 | 42.492 | 2 | 53.39745 | yes | 9.722E-08 | 681.31134 | 57.407818 |
| MAVTLDKDAYYR | 12 | Q9Y5B9 | FACT complex subunit SPT16 OS=Hom | SUPT16H | 119.914 | 2 | 53.061817 | yes | 0.0003225 | 731.35577 | 992.75867 |
| EELEFEVPFR | 10 | Q9Y5B9 | FACT complex subunit SPT16 OS=Hom | SUPT16H | 119.914 | 2 | 53.061817 | yes | 9.722E-08 | 647.81934 | 341.39899 |
| NEGNIFPNPEATF\ | 15 | Q9Y5B9 | FACT complex subunit SPT16 OS=Hom | SUPT16H | 119.914 | 2 | 53.061817 | yes | 1.106E-07 | 838.91754 | 193.49461 |
| ELAAQLNEEAK | 11 | Q9Y5B9 | FACT complex subunit SPT16 OS=Hom | SUPT16H | 119.914 | 2 | 53.061817 | yes | 1.06E-07 | 608.31439 | 177.69234 |
| INFYCPGSALGR | 12 | Q9Y5B9 | FACT complex subunit SPT16 OS=Hom | SUPT16H | 119.914 | 2 | 53.061817 | yes | 1.236E-07 | 677.83228 | 173.21574 |
| DKFPGEFMK | 9 | Q9Y5B9 | FACT complex subunit SPT16 OS=Hom | SUPT16H | 119.914 | 2 | 53.061817 | yes | 0.0001315 | 549.76807 | 115.70371 |
| ICDVYNAVMDVV | 13 | Q9Y5B9 | FACT complex subunit SPT16 OS=Hom | SUPT16H | 119.914 | 2 | 53.061817 | yes | 1.905E-06 | 763.37311 | 83.096588 |
| YTEGVQSLNWTK | 12 | Q9Y5B9 | FACT complex subunit SPT16 OS=Hom | SUPT16H | 119.914 | 2 | 53.061817 | yes | 3.5E-07 | 713.35406 | 60.717014 |
| NLGFGMGIEFR | 11 | Q9Y5B9 | FACT complex subunit SPT16 OS=Hom | SUPT16H | 119.914 | 2 | 53.061817 | yes | 2.769E-07 | 620.81079 | 59.609592 |
| GDKVDILYNNIK | 12 | Q9Y5B9 | FACT complex subunit SPT16 OS=Hom | SUPT16H | 119.914 | 2 | 53.061817 | yes | 5.316E-07 | 696.38007 | 40.256653 |
| NISMSVEGDYTYL | 14 | Q9Y5B9 | FACT complex subunit SPT16 OS=Hom | SUPT16H | 119.914 | 2 | 53.061817 | yes | 0.0002281 | 824.38782 | 6.1778712 |
| TGNFISTSTSLPR | 13 | Q9Y5J1 | U3 small nucleolar RNA-associated prote | UTP18 | 62.004 | 2 | 51.232655 | yes | 1E-07 | 690.8595 | 982.76318 |
| IQSIYLER | 8 | Q9Y5J1 | U3 small nucleolar RNA-associated prote | UTP18 | 62.004 | 2 | 51.232655 | yes | 3.503E-07 | 511.28748 | 874.29626 |
| LIPVHQVR | 8 | Q9Y5J1 | U3 small nucleolar RNA-associated prote | UTP18 | 62.004 | 2 | 51.232655 | yes | 1.248E-07 | 481.30072 | 740.3598 |

| | | | | | | | | | | | |
|---|---|---|---|---|---|---|---|---|---|---|---|
| VLYVYDMLAGK | 11 | Q9Y5J1 | U3 small nucleolar RNA-associated prote | UTP18 | 62.004 | 2 | 51.232655 | yes | 2.149E-07 | 636.33887 | 532.55707 |
| SGYFALGNEK | 10 | Q9Y5J1 | U3 small nucleolar RNA-associated prote | UTP18 | 62.004 | 2 | 51.232655 | yes | 1.226E-07 | 543.26672 | 530.90833 |
| NISHVHTMDFSPR | 13 | Q9Y5J1 | U3 small nucleolar RNA-associated prote | UTP18 | 62.004 | 3 | 51.232655 | yes | 1.069E-07 | 514.24902 | 176.87854 |
| NISHVHTMDFSPR | 13 | Q9Y5J1 | U3 small nucleolar RNA-associated prote | UTP18 | 62.004 | 3 | 51.232655 | yes | 4.719E-06 | 770.86987 | 176.87854 |
| NISHVHTMDFSPR | 13 | Q9Y5J1 | U3 small nucleolar RNA-associated prote | UTP18 | 62.004 | 2 | 51.232655 | yes | 1.069E-07 | 514.24902 | 176.87854 |
| NISHVHTMDFSPR | 13 | Q9Y5J1 | U3 small nucleolar RNA-associated prote | UTP18 | 62.004 | 2 | 51.232655 | yes | 4.719E-06 | 770.86987 | 176.87854 |
| LEEDKPAVER | 10 | Q9Y5J1 | U3 small nucleolar RNA-associated prote | UTP18 | 62.004 | 2 | 51.232655 | yes | 9.722E-08 | 593.30914 | 291.3255 |
| LRLEEDKPAVER | 12 | Q9Y5J1 | U3 small nucleolar RNA-associated prote | UTP18 | 62.004 | 3 | 51.232655 | yes | 1.21E-07 | 485.60358 | 198.40913 |
| EEFQHAMGGVPA' | 18 | Q9Y5J1 | U3 small nucleolar RNA-associated prote | UTP18 | 62.004 | 3 | 51.232655 | yes | 1.34E-06 | 663.64386 | 127.64143 |
| AGAGPGGPPQKPA | 18 | Q9Y5J1 | U3 small nucleolar RNA-associated prote | UTP18 | 62.004 | 3 | 51.232655 | yes | 1.152E-07 | 553.95691 | 41.711617 |
| AGAGPGGPPQKPA | 18 | Q9Y5J1 | U3 small nucleolar RNA-associated prote | UTP18 | 62.004 | 3 | 51.232655 | yes | 1.465E-07 | 830.4317 | 41.711617 |
| AGAGPGGPPQKPA | 18 | Q9Y5J1 | U3 small nucleolar RNA-associated prote | UTP18 | 62.004 | 2 | 51.232655 | yes | 1.152E-07 | 553.95691 | 41.711617 |
| AGAGPGGPPQKPA | 18 | Q9Y5J1 | U3 small nucleolar RNA-associated prote | UTP18 | 62.004 | 2 | 51.232655 | yes | 1.465E-07 | 830.4317 | 41.711617 |
| LHHYSDF | 7 | Q9Y5J1 | U3 small nucleolar RNA-associated prote | UTP18 | 62.004 | 2 | 51.232655 | yes | 2.071E-06 | 459.70886 | 82.188278 |
| NCQHANAERPTV/ | 14 | Q9Y5J1 | U3 small nucleolar RNA-associated prote | UTP18 | 62.004 | 4 | 51.232655 | yes | 1.614E-07 | 406.69968 | 60.407211 |
| KPPARPSAAAAAL | 22 | Q9Y5J1 | U3 small nucleolar RNA-associated prote | UTP18 | 62.004 | 4 | 51.232655 | yes | 1.402E-07 | 537.54462 | 60.07399 |
| PSAAAAAIAVAAA | 17 | Q9Y5J1 | U3 small nucleolar RNA-associated prote | UTP18 | 62.004 | 3 | 51.232655 | yes | 5.76E-07 | 533.27753 | 29.922529 |
| PSAAAAAIAVAAA | 17 | Q9Y5J1 | U3 small nucleolar RNA-associated prote | UTP18 | 62.004 | 3 | 51.232655 | yes | 0.0002141 | 799.41266 | 29.922529 |
| PSAAAAAIAVAAA | 17 | Q9Y5J1 | U3 small nucleolar RNA-associated prote | UTP18 | 62.004 | 2 | 51.232655 | yes | 5.76E-07 | 533.27753 | 29.922529 |
| PSAAAAAIAVAAA | 17 | Q9Y5J1 | U3 small nucleolar RNA-associated prote | UTP18 | 62.004 | 2 | 51.232655 | yes | 0.0002141 | 799.41266 | 29.922529 |
| ACFSANGEEVLAT | 18 | Q9Y5J1 | U3 small nucleolar RNA-associated prote | UTP18 | 62.004 | 3 | 51.232655 | yes | 1.331E-07 | 636.96338 | 59.312908 |
| GLALSNSDVIR | 11 | Q9Y5K5 | Ubiquitin carboxyl-terminal hydrolase iso | UCHL5 | 37.607 | 2 | 41.247177 | yes | 3.604E-07 | 572.81964 | 74.994255 |
| LKPVHGLIFLFK | 12 | Q9Y5K5 | Ubiquitin carboxyl-terminal hydrolase iso | UCHL5 | 37.607 | 3 | 41.247177 | yes | 4.811E-06 | 471.29898 | 5.5730839 |
| SVDFDSLTVR | 10 | Q9Y5K6 | CD2-associated protein OS=Homo sapien | CD2AP | 71.452 | 2 | 48.57321 | yes | 3.454E-06 | 569.79059 | 40.024254 |
| ANTTAFLTPLEIK | 13 | Q9Y5K6 | CD2-associated protein OS=Homo sapien | CD2AP | 71.452 | 2 | 48.57321 | yes | 1.128E-07 | 709.89813 | 36.292801 |
| EGEIIHLISK | 10 | Q9Y5K6 | CD2-associated protein OS=Homo sapien | CD2AP | 71.452 | 2 | 48.57321 | yes | 1.168E-07 | 569.82697 | 21.793987 |
| LDPGLIMEQVK | 11 | Q9Y5L4 | Mitochondrial import inner membrane tra | TIMM13 | 10.501 | 2 | 47.785175 | yes | 1.495E-07 | 621.84174 | 413.93561 |
| VQIAVANAQELLC | 14 | Q9Y5L4 | Mitochondrial import inner membrane tra | TIMM13 | 10.501 | 2 | 47.785175 | yes | 1.259E-07 | 776.94391 | 156.16806 |
| YMDAWNTVSR | 10 | Q9Y5L4 | Mitochondrial import inner membrane tra | TIMM13 | 10.501 | 2 | 47.785175 | yes | 4.182E-06 | 621.78223 | 66.695595 |
| ACGCPLYWK | 9 | Q9Y5P8 | Serine/threonine-protein phosphatase 2A | PPP2R3B | 65.061 | 2 | 45.15592 | yes | 3.585E-06 | 577.7597 | 45.48542 |
| TPTSIEYWFR | 10 | Q9Y5P8 | Serine/threonine-protein phosphatase 2A | PPP2R3B | 65.061 | 2 | 45.15592 | yes | 1.942E-07 | 650.32202 | 38.283176 |
| MVCVEYPGVVR | 11 | Q9Y5Q8 | General transcription factor 3C polypeptic | GTF3C5 | 59.571 | 2 | 54.310764 | yes | 9.733E-08 | 654.82556 | 593.19055 |
| MLPTLGGEEGVSR | 13 | Q9Y5Q8 | General transcription factor 3C polypeptic | GTF3C5 | 59.571 | 2 | 54.310764 | yes | 9.989E-08 | 673.34265 | 362.38995 |
| QGLGPSGTSGAR | 12 | Q9Y5Q8 | General transcription factor 3C polypeptic | GTF3C5 | 59.571 | 2 | 54.310764 | yes | 6.615E-07 | 544.2782 | 237.19528 |
| VLMLRPEK | 8 | Q9Y5Q8 | General transcription factor 3C polypeptic | GTF3C5 | 59.571 | 2 | 54.310764 | yes | 9.722E-08 | 493.29678 | 228.02901 |
| IYQVLDFR | 8 | Q9Y5Q8 | General transcription factor 3C polypeptic | GTF3C5 | 59.571 | 2 | 54.310764 | yes | 2.093E-07 | 527.29004 | 169.43527 |
| STYNYSLPITVK | 12 | Q9Y5Q8 | General transcription factor 3C polypeptic | GTF3C5 | 59.571 | 2 | 54.310764 | yes | 1.912E-07 | 693.3692 | 153.68562 |
| QPLEAAAQTWR | 11 | Q9Y5Q8 | General transcription factor 3C polypeptic | GTF3C5 | 59.571 | 2 | 54.310764 | yes | 9.722E-08 | 635.83057 | 126.25677 |
| HGYAPSDLPVK | 11 | Q9Y5Q8 | General transcription factor 3C polypeptic | GTF3C5 | 59.571 | 2 | 54.310764 | yes | 9.917E-08 | 592.30896 | 114.26928 |

| | | | | | | | | | | | |
|---|---|---|---|---|---|---|---|---|---|---|---|
| VCTNPVDR | 8 | Q9Y5Q8 | General transcription factor 3C polypepti | GTF3C5 | 59.571 | 2 | 54.310764 | yes | 3.025E-07 | 480.73199 | 106.54372 |
| LKDSVYIFR | 9 | Q9Y5Q8 | General transcription factor 3C polypepti | GTF3C5 | 59.571 | 2 | 54.310764 | yes | 5.375E-07 | 570.82422 | 90.942101 |
| DPYCHPVCANR | 11 | Q9Y5Q8 | General transcription factor 3C polypepti | GTF3C5 | 59.571 | 2 | 54.310764 | yes | 2.834E-07 | 694.79535 | 53.68681 |
| ILNLLSPSDGER | 12 | Q9Y5Q9 | General transcription factor 3C polypepti | GTF3C3 | 101.273 | 2 | 52.96355 | yes | 9.722E-08 | 657.35663 | 506.06747 |
| ISFEEFER | 8 | Q9Y5Q9 | General transcription factor 3C polypepti | GTF3C3 | 101.273 | 2 | 52.96355 | yes | 2.029E-07 | 528.75348 | 362.30701 |
| SSLYEQMGDHK | 11 | Q9Y5Q9 | General transcription factor 3C polypepti | GTF3C3 | 101.273 | 2 | 52.96355 | yes | 3.358E-07 | 647.79022 | 215.59247 |
| ISLSTLQQQLGQPE | 15 | Q9Y5Q9 | General transcription factor 3C polypepti | GTF3C3 | 101.273 | 2 | 52.96355 | yes | 1.165E-07 | 835.45959 | 195.39835 |
| GLMGEANIR | 9 | Q9Y5Q9 | General transcription factor 3C polypepti | GTF3C3 | 101.273 | 2 | 52.96355 | yes | 6.028E-07 | 480.75021 | 178.0136 |
| STLLFSQGK | 9 | Q9Y5Q9 | General transcription factor 3C polypepti | GTF3C3 | 101.273 | 2 | 52.96355 | yes | 1.2E-07 | 490.7742 | 126.42598 |
| GPCQESFYNLGR | 12 | Q9Y5Q9 | General transcription factor 3C polypepti | GTF3C3 | 101.273 | 2 | 52.96355 | yes | 9.722E-08 | 714.32227 | 106.48651 |
| AQVCLISSSK | 10 | Q9Y5Q9 | General transcription factor 3C polypepti | GTF3C3 | 101.273 | 2 | 52.96355 | yes | 8.955E-07 | 546.78955 | 98.974747 |
| LAEMSLEQDNIK | 12 | Q9Y5Q9 | General transcription factor 3C polypepti | GTF3C3 | 101.273 | 2 | 52.96355 | yes | 7.402E-07 | 695.84778 | 76.967682 |
| QAIFCYTK | 8 | Q9Y5Q9 | General transcription factor 3C polypepti | GTF3C3 | 101.273 | 2 | 52.96355 | yes | 3.816E-07 | 515.75494 | 75.439903 |
| AIYSLCDLSR | 10 | Q9Y5Q9 | General transcription factor 3C polypepti | GTF3C3 | 101.273 | 2 | 52.96355 | yes | 9.722E-08 | 599.30023 | 67.485626 |
| VVDLAPLHLDAR | 12 | Q9Y5Q9 | General transcription factor 3C polypepti | GTF3C3 | 101.273 | 3 | 52.96355 | yes | 9.722E-08 | 440.25412 | 36.001354 |
| ELEYFGLSAAILDK | 14 | Q9Y5Q9 | General transcription factor 3C polypepti | GTF3C3 | 101.273 | 2 | 52.96355 | yes | 4.735E-07 | 784.91394 | 28.487473 |
| LFLYDLPEGK | 10 | Q9Y5S2 | Serine/threonine-protein kinase MRCK b | CDC42BPI | 194.316 | 2 | 51.04889 | yes | 1.089E-06 | 597.82391 | 111.63295 |
| MTEELEALR | 9 | Q9Y5S2 | Serine/threonine-protein kinase MRCK b | CDC42BPI | 194.316 | 2 | 51.04889 | no | 1.401E-07 | 546.27368 | 103.53721 |
| VTASLLGAPSK | 11 | Q9Y5S2 | Serine/threonine-protein kinase MRCK b | CDC42BPI | 194.316 | 2 | 51.04889 | yes | 1.874E-07 | 522.30841 | 96.0364 |
| TMEWVQTIGLR | 11 | Q9Y5S2 | Serine/threonine-protein kinase MRCK b | CDC42BPI | 194.316 | 2 | 51.04889 | yes | 1.67E-07 | 667.35028 | 85.572899 |
| TSSLLILTENENEK | 14 | Q9Y5S2 | Serine/threonine-protein kinase MRCK b | CDC42BPI | 194.316 | 2 | 51.04889 | yes | 1.389E-06 | 795.91467 | 83.270386 |
| VLFYEEELVR | 10 | Q9Y5S2 | Serine/threonine-protein kinase MRCK b | CDC42BPI | 194.316 | 2 | 51.04889 | yes | 9.775E-08 | 648.84534 | 75.52269 |
| IVILLCGR | 8 | Q9Y5S2 | Serine/threonine-protein kinase MRCK b | CDC42BPI | 194.316 | 2 | 51.04889 | yes | 1.604E-07 | 472.2915 | 71.190887 |
| LALQEFSELNER | 12 | Q9Y5S2 | Serine/threonine-protein kinase MRCK b | CDC42BPI | 194.316 | 2 | 51.04889 | yes | 1.749E-06 | 724.87262 | 63.115326 |
| LELQSALEAEIR | 12 | Q9Y5S2 | Serine/threonine-protein kinase MRCK b | CDC42BPI | 194.316 | 2 | 51.04889 | no | 2.789E-07 | 686.37756 | 62.270798 |
| KLNEEIER | 8 | Q9Y5S2 | Serine/threonine-protein kinase MRCK b | CDC42BPI | 194.316 | 2 | 51.04889 | yes | 7.886E-07 | 515.78003 | 46.500511 |
| LILCYEIQR | 9 | Q9Y5S2 | Serine/threonine-protein kinase MRCK b | CDC42BPI | 194.316 | 2 | 51.04889 | yes | 9.722E-08 | 604.3288 | 46.233688 |
| FSGAVLNVPDTSD | 16 | Q9Y5S2 | Serine/threonine-protein kinase MRCK b | CDC42BPI | 194.316 | 2 | 51.04889 | yes | 1.429E-07 | 825.9021 | 45.807682 |
| AYAVVCDCK | 9 | Q9Y5S2 | Serine/threonine-protein kinase MRCK b | CDC42BPI | 194.316 | 2 | 51.04889 | yes | 1.756E-07 | 543.24133 | 44.797501 |
| PFTQLVK | 7 | Q9Y5S2 | Serine/threonine-protein kinase MRCK b | CDC42BPI | 194.316 | 2 | 51.04889 | yes | 2.451E-07 | 416.75 | 44.76849 |
| ELEAQLDDAVAEA | 15 | Q9Y5S2 | Serine/threonine-protein kinase MRCK b | CDC42BPI | 194.316 | 2 | 51.04889 | yes | 3.483E-06 | 794.88867 | 43.586788 |
| GYLQALASK | 9 | Q9Y5S2 | Serine/threonine-protein kinase MRCK b | CDC42BPI | 194.316 | 2 | 51.04889 | no | 1.977E-07 | 475.76892 | 42.947422 |
| GCQLMATATLK | 11 | Q9Y5S2 | Serine/threonine-protein kinase MRCK b | CDC42BPI | 194.316 | 2 | 51.04889 | yes | 1.737E-07 | 597.3045 | 37.204178 |
| STQPGVIASQVLDI | 15 | Q9Y5S2 | Serine/threonine-protein kinase MRCK b | CDC42BPI | 194.316 | 2 | 51.04889 | yes | 4.705E-06 | 792.44122 | 32.017986 |
| PGPAPTNLAR | 10 | Q9Y5S2 | Serine/threonine-protein kinase MRCK b | CDC42BPI | 194.316 | 2 | 51.04889 | yes | 1.549E-06 | 497.27744 | 25.594105 |
| HAFFEGLNWENIR | 13 | Q9Y5S2 | Serine/threonine-protein kinase MRCK b | CDC42BPI | 194.316 | 3 | 51.04889 | yes | 1.414E-06 | 544.93542 | 13.541736 |
| LCSFVDKLTAQNR | 13 | Q9Y5S2 | Serine/threonine-protein kinase MRCK b | CDC42BPI | 194.316 | 3 | 51.04889 | yes | 5.155E-05 | 517.93646 | 3.5432162 |
| NIHLNLDR | 8 | Q9Y5S9 | RNA-binding protein 8A OS=Homo sapi | RBM8A | 19.889 | 2 | 51.29379 | yes | 1.499E-07 | 497.77505 | 8660.0938 |
| GYTLVEYETYK | 11 | Q9Y5S9 | RNA-binding protein 8A OS=Homo sapi | RBM8A | 19.889 | 2 | 51.29379 | yes | 9.722E-08 | 683.33228 | 2444.603 |

| | | | | | | | | | | | |
|---|---|---|---|---|---|---|---|---|---|---|---|
| FAEYGEIK | 8 | Q9Y5S9 | RNA-binding protein 8A OS=Homo sapi | RBM8A | 19.889 | 2 | 51.29379 | yes | 1.16E-07 | 478.73984 | 1434.0988 |
| VMILNHPDK | 9 | Q9Y5T4 | DnaJ homolog subfamily C member 15 O | DNAJC15 | 16.383 | 2 | 51.139668 | yes | 2.45E-07 | 533.78931 | 141.66298 |
| EAGLILGVSPSAGI | 14 | Q9Y5T4 | DnaJ homolog subfamily C member 15 O | DNAJC15 | 16.383 | 2 | 51.139668 | yes | 9.722E-08 | 649.86938 | 116.00584 |
| ISTPSFSSYYK | 11 | Q9Y5T4 | DnaJ homolog subfamily C member 15 O | DNAJC15 | 16.383 | 2 | 51.139668 | yes | 5.211E-07 | 640.3139 | 114.32287 |
| IWKPLEQVITETAI | 14 | Q9Y5T4 | DnaJ homolog subfamily C member 15 O | DNAJC15 | 16.383 | 2 | 51.139668 | yes | 2.04E-07 | 828.47198 | 50.529037 |
| IWKPLEQVITETAI | 14 | Q9Y5T4 | DnaJ homolog subfamily C member 15 O | DNAJC15 | 16.383 | 2 | 51.139668 | yes | 4.182E-06 | 552.65039 | 50.529037 |
| IWKPLEQVITETAI | 14 | Q9Y5T4 | DnaJ homolog subfamily C member 15 O | DNAJC15 | 16.383 | 3 | 51.139668 | yes | 2.04E-07 | 828.47198 | 50.529037 |
| IWKPLEQVITETAI | 14 | Q9Y5T4 | DnaJ homolog subfamily C member 15 O | DNAJC15 | 16.383 | 3 | 51.139668 | yes | 4.182E-06 | 552.65039 | 50.529037 |
| YAEYLQPSAK | 10 | Q9Y5T4 | DnaJ homolog subfamily C member 15 O | DNAJC15 | 16.383 | 2 | 51.139668 | yes | 2.689E-07 | 585.29547 | 56.434502 |
| LLCEVCTR | 8 | Q9Y5T5 | Ubiquitin carboxyl-terminal hydrolase 16 | USP16 | 93.571 | 2 | 48.61421 | yes | 1.122E-07 | 525.75714 | 382.17731 |
| ENPPMNSPCQITVI | 14 | Q9Y5T5 | Ubiquitin carboxyl-terminal hydrolase 16 | USP16 | 93.571 | 2 | 48.61421 | yes | 1.67E-07 | 807.88434 | 294.3082 |
| MIESVTDNQK | 10 | Q9Y5T5 | Ubiquitin carboxyl-terminal hydrolase 16 | USP16 | 93.571 | 2 | 48.61421 | yes | 2.776E-07 | 582.78192 | 194.98076 |
| GYQQQDSQELLR | 12 | Q9Y5T5 | Ubiquitin carboxyl-terminal hydrolase 16 | USP16 | 93.571 | 2 | 48.61421 | yes | 1.866E-07 | 732.85748 | 171.65614 |
| FQQAGFNLR | 9 | Q9Y5T5 | Ubiquitin carboxyl-terminal hydrolase 16 | USP16 | 93.571 | 2 | 48.61421 | yes | 4.508E-07 | 540.7829 | 161.14114 |
| ELFSQVCK | 8 | Q9Y5T5 | Ubiquitin carboxyl-terminal hydrolase 16 | USP16 | 93.571 | 2 | 48.61421 | yes | 2.201E-06 | 505.75241 | 95.692635 |
| KGLEQGNLK | 9 | Q9Y5T5 | Ubiquitin carboxyl-terminal hydrolase 16 | USP16 | 93.571 | 2 | 48.61421 | yes | 1.368E-07 | 493.7851 | 51.656128 |
| VLNSQAYLLFYER | 13 | Q9Y5T5 | Ubiquitin carboxyl-terminal hydrolase 16 | USP16 | 93.571 | 2 | 48.61421 | yes | 5.284E-07 | 808.42755 | 49.042377 |
| GMSSTFSQR | 9 | Q9Y5U2 | U5 small nuclear ribonucleoprotein TSSC | TSSC4 | 34.326 | 2 | 44.498047 | yes | 5.382E-07 | 500.72946 | 34.858723 |
| DIFDCLEGAAR | 11 | Q9Y5U2 | U5 small nuclear ribonucleoprotein TSSC | TSSC4 | 34.326 | 2 | 44.498047 | yes | 2.172E-07 | 633.79279 | 30.677702 |
| VPPVPDYVAHPER | 13 | Q9Y5U2 | U5 small nuclear ribonucleoprotein TSSC | TSSC4 | 34.326 | 3 | 44.498047 | yes | 2.96E-06 | 492.5929 | 23.866119 |
| STFFGTSDEFIEK | 13 | Q9Y5W9 | Sorting nexin-11 OS=Homo sapiens OX= | SNX11 | 30.434 | 2 | 45.6798 | yes | 1.776E-07 | 754.3512 | 53.264091 |
| FGSAIPIPSLPDK | 13 | Q9Y5X1 | Sorting nexin-9 OS=Homo sapiens OX=9 | SNX9 | 66.592 | 2 | 51.50734 | yes | 6.339E-07 | 671.37427 | 7464.6323 |
| IYDYNSVIR | 9 | Q9Y5X1 | Sorting nexin-9 OS=Homo sapiens OX=9 | SNX9 | 66.592 | 2 | 51.50734 | yes | 1.153E-07 | 571.79565 | 2487.3132 |
| GFLGCFPDIIGTHK | 14 | Q9Y5X1 | Sorting nexin-9 OS=Homo sapiens OX=9 | SNX9 | 66.592 | 2 | 51.50734 | yes | 1.145E-07 | 781.3952 | 1674.0319 |
| PGTEQYLLAK | 10 | Q9Y5X1 | Sorting nexin-9 OS=Homo sapiens OX=9 | SNX9 | 66.592 | 2 | 51.50734 | yes | 2.584E-07 | 560.30585 | 1265.9868 |
| HFDWLYER | 8 | Q9Y5X1 | Sorting nexin-9 OS=Homo sapiens OX=9 | SNX9 | 66.592 | 2 | 51.50734 | yes | 2.092E-07 | 583.2749 | 1009.7626 |
| LQAWMTR | 7 | Q9Y5X1 | Sorting nexin-9 OS=Homo sapiens OX=9 | SNX9 | 66.592 | 2 | 51.50734 | yes | 1.128E-07 | 453.23672 | 999.39508 |
| TYEEIASLVAEQPI | 14 | Q9Y5X1 | Sorting nexin-9 OS=Homo sapiens OX=9 | SNX9 | 66.592 | 2 | 51.50734 | yes | 2.627E-07 | 789.40649 | 857.95074 |
| HPVISESEVFQQFL | 17 | Q9Y5X1 | Sorting nexin-9 OS=Homo sapiens OX=9 | SNX9 | 66.592 | 3 | 51.50734 | yes | 1.696E-07 | 693.02155 | 510.38547 |
| FEEEFIK | 7 | Q9Y5X1 | Sorting nexin-9 OS=Homo sapiens OX=9 | SNX9 | 66.592 | 2 | 51.50734 | yes | 4.184E-06 | 471.23438 | 478.26346 |
| VSIMSYALQAEMI | 19 | Q9Y5X1 | Sorting nexin-9 OS=Homo sapiens OX=9 | SNX9 | 66.592 | 3 | 51.50734 | yes | 1.063E-07 | 745.68597 | 241.32729 |
| RCTGPLPK | 8 | Q9Y5X1 | Sorting nexin-9 OS=Homo sapiens OX=9 | SNX9 | 66.592 | 2 | 51.50734 | yes | 3.6E-07 | 464.75528 | 152.01201 |
| KDLHFLMECNHE | 14 | Q9Y5X1 | Sorting nexin-9 OS=Homo sapiens OX=9 | SNX9 | 66.592 | 3 | 51.50734 | yes | 9.722E-08 | 621.95538 | 79.658455 |
| ESAQCVGDEFLNC | 14 | Q9Y5X2 | Sorting nexin-8 OS=Homo sapiens OX=9 | SNX8 | 52.57 | 2 | 39.613186 | yes | 4.993E-07 | 828.85327 | 59.991241 |
| FVNLVAR | 7 | Q9Y5X2 | Sorting nexin-8 OS=Homo sapiens OX=9 | SNX8 | 52.57 | 2 | 39.613186 | yes | 4.481E-07 | 409.7478 | 52.630699 |
| IVEQENAIQTMELI | 14 | Q9Y5X2 | Sorting nexin-8 OS=Homo sapiens OX=9 | SNX8 | 52.57 | 2 | 39.613186 | yes | 4.913E-07 | 837.42999 | 28.039442 |
| MEAAAEPGNLAG | 14 | Q9Y5Y2 | Cytosolic Fe-S cluster assembly factor NU | NUBP2 | 28.826 | 2 | 49.68128 | yes | 1.362E-07 | 714.35101 | 330.43683 |
| GWAPVFLDR | 9 | Q9Y5Y2 | Cytosolic Fe-S cluster assembly factor NU | NUBP2 | 28.826 | 2 | 49.68128 | yes | 3.762E-07 | 530.78235 | 218.32452 |
| VGILDVDLCGPSIP | 15 | Q9Y5Y2 | Cytosolic Fe-S cluster assembly factor NU | NUBP2 | 28.826 | 2 | 49.68128 | yes | 1.011E-07 | 805.93219 | 65.77607 |

| | | | | | | | | | | | |
|---|---|---|---|---|---|---|---|---|---|---|---|
| AGLQTSPPIVPLDF | 14 | Q9Y5Y5 | Peroxisomal membrane protein PEX16 O | PEX16 | 38.629 | 2 | 54.738743 | yes | 9.722E-08 | 732.41449 | 311.79428 |
| HWGAPQQR | 8 | Q9Y5Y5 | Peroxisomal membrane protein PEX16 O | PEX16 | 38.629 | 2 | 54.738743 | yes | 1.63E-07 | 490.24646 | 178.06671 |
| HPAATAQLETAVF | 13 | Q9Y5Y5 | Peroxisomal membrane protein PEX16 O | PEX16 | 38.629 | 2 | 54.738743 | yes | 1.055E-07 | 682.86768 | 151.19164 |
| KLPVSLSQQK | 10 | Q9Y5Y5 | Peroxisomal membrane protein PEX16 O | PEX16 | 38.629 | 2 | 54.738743 | yes | 3.553E-07 | 564.34277 | 113.47021 |
| YQEYVTR | 7 | Q9Y5Y5 | Peroxisomal membrane protein PEX16 O | PEX16 | 38.629 | 2 | 54.738743 | yes | 1.97E-07 | 479.73508 | 59.053589 |
| GFSYLLAGR | 9 | Q9Y5Y5 | Peroxisomal membrane protein PEX16 O | PEX16 | 38.629 | 2 | 54.738743 | yes | 2.894E-07 | 492.26907 | 51.315655 |
| LWEMDNMLIQIK | 12 | Q9Y605 | MORF4 family-associated protein 1 OS=l | MRFAP1 | 14.65 | 2 | 46.799294 | no | 3.085E-07 | 767.39386 | 333.44083 |
| MAEMLVELVR | 10 | Q9Y605 | MORF4 family-associated protein 1 OS=l | MRFAP1 | 14.65 | 2 | 46.799294 | yes | 1.467E-07 | 595.8172 | 325.70508 |
| EDIASLTR | 8 | Q9Y605 | MORF4 family-associated protein 1 OS=l | MRFAP1 | 14.65 | 2 | 46.799294 | yes | 7.33E-07 | 452.74036 | 226.07133 |
| GYAPESVLER | 10 | Q9Y606 | Pseudouridylate synthase 1 homolog OS= | PUS1 | 47.471 | 2 | 51.083477 | yes | 9.722E-08 | 560.78528 | 596.22772 |
| EGLEFAVIR | 9 | Q9Y606 | Pseudouridylate synthase 1 homolog OS= | PUS1 | 47.471 | 2 | 51.083477 | yes | 2.53E-07 | 517.28748 | 581.69177 |
| GVSAAGQVVSLK | 12 | Q9Y606 | Pseudouridylate synthase 1 homolog OS= | PUS1 | 47.471 | 2 | 51.083477 | yes | 9.809E-08 | 558.32458 | 549.33807 |
| LSAETLQQVNR | 11 | Q9Y606 | Pseudouridylate synthase 1 homolog OS= | PUS1 | 47.471 | 2 | 51.083477 | yes | 1.117E-07 | 629.84113 | 502.76312 |
| EEHIYPTIIGTER | 13 | Q9Y606 | Pseudouridylate synthase 1 homolog OS= | PUS1 | 47.471 | 2 | 51.083477 | yes | 5.454E-07 | 779.39899 | 274.27634 |
| VWEDGEHPAK | 10 | Q9Y606 | Pseudouridylate synthase 1 homolog OS= | PUS1 | 47.471 | 2 | 51.083477 | yes | 1.37E-07 | 584.27509 | 217.61447 |
| INSHLPSHIR | 10 | Q9Y606 | Pseudouridylate synthase 1 homolog OS= | PUS1 | 47.471 | 2 | 51.083477 | yes | 9.727E-08 | 587.328 | 193.12108 |
| APGLGLVLER | 10 | Q9Y606 | Pseudouridylate synthase 1 homolog OS= | PUS1 | 47.471 | 2 | 51.083477 | yes | 1.084E-07 | 512.8111 | 166.67278 |
| TIEDDLVSALVR | 12 | Q9Y606 | Pseudouridylate synthase 1 homolog OS= | PUS1 | 47.471 | 2 | 51.083477 | yes | 9.722E-08 | 665.86426 | 156.83606 |
| MVGLVVAIVK | 10 | Q9Y606 | Pseudouridylate synthase 1 homolog OS= | PUS1 | 47.471 | 2 | 51.083477 | yes | 3.706E-07 | 514.83044 | 51.437794 |
| IVLLMAYSGK | 10 | Q9Y606 | Pseudouridylate synthase 1 homolog OS= | PUS1 | 47.471 | 2 | 51.083477 | yes | 1.885E-06 | 547.81757 | 47.224586 |
| SGCIPENHGEDMR | 13 | Q9Y606 | Pseudouridylate synthase 1 homolog OS= | PUS1 | 47.471 | 2 | 51.083477 | yes | 2.864E-07 | 751.31158 | 11.643147 |
| FSAEDEALSNIAR | 13 | Q9Y608 | Leucine-rich repeat flightless-interacting ] | LRRFIP2 | 82.172 | 2 | 53.11575 | yes | 9.937E-08 | 711.84662 | 411.43384 |
| LQLEEER | 7 | Q9Y608 | Leucine-rich repeat flightless-interacting ] | LRRFIP2 | 82.172 | 2 | 53.11575 | yes | 4.593E-07 | 458.74036 | 271.31348 |
| AEQDITTLEQSISR | 14 | Q9Y608 | Leucine-rich repeat flightless-interacting ] | LRRFIP2 | 82.172 | 2 | 53.11575 | yes | 1.594E-07 | 795.9021 | 242.84167 |
| AMVSNAQLDNEK | 12 | Q9Y608 | Leucine-rich repeat flightless-interacting ] | LRRFIP2 | 82.172 | 2 | 53.11575 | no | 9.722E-08 | 660.31665 | 165.95062 |
| EELLSQIR | 8 | Q9Y608 | Leucine-rich repeat flightless-interacting ] | LRRFIP2 | 82.172 | 2 | 53.11575 | yes | 2.011E-07 | 494.2771 | 120.79225 |
| NNLIYQVDTLK | 11 | Q9Y608 | Leucine-rich repeat flightless-interacting ] | LRRFIP2 | 82.172 | 2 | 53.11575 | yes | 1.278E-07 | 660.86151 | 114.64671 |
| IEEMEMTNSHLAK | 13 | Q9Y608 | Leucine-rich repeat flightless-interacting ] | LRRFIP2 | 82.172 | 2 | 53.11575 | yes | 9.722E-08 | 766.85779 | 105.89408 |
| DQIQDVEGR | 9 | Q9Y608 | Leucine-rich repeat flightless-interacting ] | LRRFIP2 | 82.172 | 2 | 53.11575 | yes | 9.077E-07 | 530.2569 | 93.304214 |
| TALDKIEEMEMTN | 18 | Q9Y608 | Leucine-rich repeat flightless-interacting ] | LRRFIP2 | 82.172 | 3 | 53.11575 | yes | 3.251E-07 | 687.6712 | 37.61068 |
| FLENVAAAETEK | 12 | Q9Y613 | FH1/FH2 domain-containing protein 1 O! | FHOD1 | 126.552 | 2 | 51.55582 | yes | 1.235E-07 | 661.33533 | 217.45236 |
| GDGEPVSVVTVR | 12 | Q9Y613 | FH1/FH2 domain-containing protein 1 O! | FHOD1 | 126.552 | 2 | 51.55582 | yes | 9.722E-08 | 607.82239 | 189.03468 |
| TPQSPAPCVLLR | 12 | Q9Y613 | FH1/FH2 domain-containing protein 1 O! | FHOD1 | 126.552 | 2 | 51.55582 | yes | 9.85E-08 | 669.86353 | 131.35411 |
| KPTLILR | 7 | Q9Y613 | FH1/FH2 domain-containing protein 1 O! | FHOD1 | 126.552 | 2 | 51.55582 | yes | 5.901E-07 | 420.78693 | 127.99045 |
| VGMEQLVQNATFI | 13 | Q9Y613 | FH1/FH2 domain-containing protein 1 O! | FHOD1 | 126.552 | 2 | 51.55582 | yes | 4.435E-07 | 746.88245 | 101.60544 |
| LDYDSMER | 8 | Q9Y613 | FH1/FH2 domain-containing protein 1 O! | FHOD1 | 126.552 | 2 | 51.55582 | yes | 9.927E-08 | 514.72131 | 86.011688 |
| AALLNFDEFAVSK | 13 | Q9Y613 | FH1/FH2 domain-containing protein 1 O! | FHOD1 | 126.552 | 2 | 51.55582 | yes | 1.134E-06 | 712.87463 | 72.393768 |
| APTCSLDGALPLG. | 21 | Q9Y613 | FH1/FH2 domain-containing protein 1 O! | FHOD1 | 126.552 | 3 | 51.55582 | yes | 9.722E-08 | 715.38086 | 67.567688 |
| LLTMMPTEEER | 11 | Q9Y613 | FH1/FH2 domain-containing protein 1 O! | FHOD1 | 126.552 | 2 | 51.55582 | yes | 1.003E-07 | 675.32562 | 64.443329 |

| | | | | | | | | | | | |
|---|---|---|---|---|---|---|---|---|---|---|---|
| QLWDSPETAPAAF | 13 | Q9Y613 | FH1/FH2 domain-containing protein 1 OS | FHOD1 | 126.552 | 2 | 51.55582 | yes | 9.981E-08 | 721.35718 | 54.096485 |
| TQLVLYENALK | 11 | Q9Y613 | FH1/FH2 domain-containing protein 1 OS | FHOD1 | 126.552 | 2 | 51.55582 | yes | 7.258E-07 | 646.36646 | 38.086697 |
| LQTMYEMETSGK | 12 | Q9Y619 | Mitochondrial ornithine transporter 1 OS= | SLC25A15 | 32.737 | 2 | 45.025536 | yes | 1.988E-07 | 709.32056 | 24.191576 |
| GLTDCCLK | 8 | Q9Y619 | Mitochondrial ornithine transporter 1 OS= | SLC25A15 | 32.737 | 2 | 45.025536 | yes | 5.765E-07 | 483.72281 | 23.242025 |
| LDGQTPISQR | 10 | Q9Y620 | DNA repair and recombination protein RA | RAD54B | 102.967 | 2 | 50.756428 | yes | 2.683E-07 | 557.7962 | 95.63588 |
| PGALQIELYR | 10 | Q9Y620 | DNA repair and recombination protein RA | RAD54B | 102.967 | 2 | 50.756428 | yes | 9.723E-08 | 580.32715 | 54.702988 |
| TLIVTPGSLVNNW | 14 | Q9Y620 | DNA repair and recombination protein RA | RAD54B | 102.967 | 2 | 50.756428 | yes | 2.671E-06 | 771.43793 | 42.717178 |
| FDLLICDEGHR | 11 | Q9Y620 | DNA repair and recombination protein RA | RAD54B | 102.967 | 2 | 50.756428 | yes | 1.334E-07 | 687.82715 | 10.627686 |
| EPGEVVDSLVGK | 12 | Q9Y657 | Spindlin-1 OS=Homo sapiens OX=9606 ( | SPIN1 | 29.601 | 2 | 45.65486 | yes | 1.783E-07 | 614.82465 | 77.379158 |
| DHGLLIYHIPQVEF | 15 | Q9Y676 | Small ribosomal subunit protein mS40 OS | MRPS18B | 29.396 | 3 | 51.4221 | yes | 2.45E-07 | 596.32483 | 1852.4252 |
| YLESEEYQER | 10 | Q9Y676 | Small ribosomal subunit protein mS40 OS | MRPS18B | 29.396 | 2 | 51.4221 | yes | 9.927E-08 | 673.29895 | 911.94403 |
| PVWADYR | 7 | Q9Y676 | Small ribosomal subunit protein mS40 OS | MRPS18B | 29.396 | 2 | 51.4221 | yes | 1.266E-07 | 453.72705 | 828.44702 |
| VVGNPCPICR | 10 | Q9Y676 | Small ribosomal subunit protein mS40 OS | MRPS18B | 29.396 | 2 | 51.4221 | yes | 1.075E-07 | 586.28918 | 549.67932 |
| RLTQAIQK | 8 | Q9Y676 | Small ribosomal subunit protein mS40 OS | MRPS18B | 29.396 | 2 | 51.4221 | yes | 9.722E-08 | 479.29562 | 211.69968 |
| TPAEASSTGQTGP( | 17 | Q9Y676 | Small ribosomal subunit protein mS40 OS | MRPS18B | 29.396 | 2 | 51.4221 | yes | 1.215E-07 | 801.88391 | 128.16225 |
| VFNETPINPR | 10 | Q9Y678 | Coatomer subunit gamma-1 OS=Homo sa | COPG1 | 97.719 | 2 | 51.00954 | yes | 5.73E-07 | 593.81439 | 245.42184 |
| SVPLATAPMAEQF | 13 | Q9Y678 | Coatomer subunit gamma-1 OS=Homo sa | COPG1 | 97.719 | 2 | 51.00954 | yes | 9.81E-08 | 685.85846 | 237.69008 |
| LFQSNDPTLR | 10 | Q9Y678 | Coatomer subunit gamma-1 OS=Homo sa | COPG1 | 97.719 | 2 | 51.00954 | yes | 9.722E-08 | 595.81183 | 173.9588 |
| ATFYLNVLEQK | 11 | Q9Y678 | Coatomer subunit gamma-1 OS=Homo sa | COPG1 | 97.719 | 2 | 51.00954 | yes | 1.883E-07 | 663.35864 | 138.29602 |
| VVLEHEEVR | 9 | Q9Y678 | Coatomer subunit gamma-1 OS=Homo sa | COPG1 | 97.719 | 2 | 51.00954 | yes | 5.711E-07 | 555.30109 | 125.64257 |
| NTHTLLLAGVFR | 12 | Q9Y678 | Coatomer subunit gamma-1 OS=Homo sa | COPG1 | 97.719 | 2 | 51.00954 | yes | 1.831E-07 | 671.3855 | 26.388634 |
| MCYLTIK | 7 | Q9Y678 | Coatomer subunit gamma-1 OS=Homo sa | COPG1 | 97.719 | 2 | 51.00954 | no | 7.361E-05 | 464.73517 | 16.193373 |
| EVLPHVPLGVIQR | 13 | Q9Y679 | Lipid droplet-regulating VLDL assembly | AUP1 | 45.787 | 2 | 50.551834 | yes | 4.476E-07 | 728.93536 | 118.49368 |
| TMCAVLGLVAR | 11 | Q9Y679 | Lipid droplet-regulating VLDL assembly | AUP1 | 45.787 | 2 | 50.551834 | yes | 9.773E-08 | 595.82281 | 72.632607 |
| GTQSLPTASASK | 12 | Q9Y679 | Lipid droplet-regulating VLDL assembly | AUP1 | 45.787 | 2 | 50.551834 | yes | 2.414E-07 | 574.30133 | 68.568855 |
| IPLDVNDMK | 9 | Q9Y697 | Cysteine desulfurase OS=Homo sapiens ( | NFS1 | 50.196 | 2 | 45.197834 | yes | 5.856E-06 | 522.77332 | 151.91719 |
| AIGTDEDLAHSSIR | 14 | Q9Y697 | Cysteine desulfurase OS=Homo sapiens ( | NFS1 | 50.196 | 2 | 45.197834 | yes | 1.928E-07 | 742.87061 | 69.137344 |
| FTTEEEVDYTVEK | 13 | Q9Y697 | Cysteine desulfurase OS=Homo sapiens ( | NFS1 | 50.196 | 2 | 45.197834 | yes | 2.653E-07 | 795.3645 | 42.223324 |
| THAYGWESEAAM | 14 | Q9Y697 | Cysteine desulfurase OS=Homo sapiens ( | NFS1 | 50.196 | 2 | 45.197834 | yes | 2.344E-07 | 819.35431 | 38.314392 |
| LYLPVQNK | 8 | Q9Y6A4 | Cilia- and flagella-associated protein 20 ( | CFAP20 | 22.775 | 2 | 54.165176 | yes | 1.253E-06 | 487.78711 | 699.9303 |
| LYSEDELPAEFK | 12 | Q9Y6A4 | Cilia- and flagella-associated protein 20 ( | CFAP20 | 22.775 | 2 | 54.165176 | yes | 1.426E-07 | 720.84827 | 500.13605 |
| AYGTNYIETLR | 11 | Q9Y6A4 | Cilia- and flagella-associated protein 20 ( | CFAP20 | 22.775 | 2 | 54.165176 | yes | 1.143E-07 | 650.83026 | 419.67209 |
| VKPFICTMPMR | 11 | Q9Y6A4 | Cilia- and flagella-associated protein 20 ( | CFAP20 | 22.775 | 2 | 54.165176 | yes | 1.211E-07 | 690.35345 | 246.60999 |
| VQIHANCR | 8 | Q9Y6A4 | Cilia- and flagella-associated protein 20 ( | CFAP20 | 22.775 | 2 | 54.165176 | yes | 1.388E-07 | 499.25345 | 204.36765 |
| YFTFEVQVLDDK | 12 | Q9Y6A4 | Cilia- and flagella-associated protein 20 ( | CFAP20 | 22.775 | 2 | 54.165176 | yes | 9.722E-08 | 752.37195 | 103.54692 |
| ASNYQSTTR | 9 | Q9Y6A4 | Cilia- and flagella-associated protein 20 ( | CFAP20 | 22.775 | 2 | 54.165176 | yes | 2.508E-07 | 514.24377 | 72.435219 |
| GLFTGLTPR | 9 | Q9Y6C9 | Mitochondrial carrier homolog 2 OS=Hon | MTCH2 | 33.331 | 2 | 53.220448 | yes | 1.186E-07 | 481.27692 | 188.63759 |
| EVSSSFDHVIK | 11 | Q9Y6C9 | Mitochondrial carrier homolog 2 OS=Hon | MTCH2 | 33.331 | 2 | 53.220448 | yes | 2.022E-07 | 624.31696 | 79.342789 |
| SAATLITHPFHVIT | 16 | Q9Y6C9 | Mitochondrial carrier homolog 2 OS=Hon | MTCH2 | 33.331 | 3 | 53.220448 | yes | 9.722E-08 | 593.00885 | 54.928013 |

| | | | | | | | | | | | |
|---|---|---|---|---|---|---|---|---|---|---|---|
| LADFGVAGQLTDT | 16 | Q9Y6E0 | Serine/threonine-protein kinase 24 OS=H | STK24 | 49.308 | 2 | 50.056915 | no | 9.919E-08 | 838.94635 | 327.37872 |
| GLDYLHSEK | 9 | Q9Y6E0 | Serine/threonine-protein kinase 24 OS=H | STK24 | 49.308 | 2 | 50.056915 | no | 4.417E-07 | 531.26672 | 122.89872 |
| AANVLLSEHGEVK | 13 | Q9Y6E0 | Serine/threonine-protein kinase 24 OS=H | STK24 | 49.308 | 2 | 50.056915 | yes | 1.256E-07 | 683.86987 | 28.006943 |
| EELVAEQALK | 10 | Q9Y6E2 | eIF5-mimic protein 1 OS=Homo sapiens ( | BZW2 | 48.163 | 2 | 45.735558 | yes | 4.248E-06 | 565.30859 | 81.792419 |
| LLELFPVNR | 9 | Q9Y6E2 | eIF5-mimic protein 1 OS=Homo sapiens ( | BZW2 | 48.163 | 2 | 45.735558 | yes | 1.758E-07 | 550.82678 | 64.495491 |
| TIVCYHPSVDIPYE | 21 | Q9Y6G3 | Large ribosomal subunit protein mL42 OS | MRPL42 | 16.662 | 4 | 41.816093 | yes | 3.327E-07 | 631.32697 | 87.163681 |
| SVSSNVASVSPIPA | 17 | Q9Y6G9 | Cytoplasmic dynein 1 light intermediate c | DYNC1LI1 | 56.58 | 2 | 36.183537 | yes | 2.657E-06 | 793.92285 | 43.178398 |
| FSLDAVSLK | 9 | Q9Y6G9 | Cytoplasmic dynein 1 light intermediate c | DYNC1LI1 | 56.58 | 2 | 36.183537 | yes | 7.297E-07 | 490.27658 | 24.865093 |
| AVLGDDPQLMK | 11 | Q9Y6J9 | TAF6-like RNA polymerase II p300/CBP | TAF6L | 67.815 | 2 | 51.042934 | yes | 3.591E-07 | 593.81049 | 163.29454 |
| EGELYFPEDR | 10 | Q9Y6J9 | TAF6-like RNA polymerase II p300/CBP | TAF6L | 67.815 | 2 | 51.042934 | yes | 1.171E-07 | 627.78552 | 96.816765 |
| ELYAFFGDSLATR | 13 | Q9Y6J9 | TAF6-like RNA polymerase II p300/CBP | TAF6L | 67.815 | 2 | 51.042934 | yes | 9.722E-08 | 745.36975 | 82.346542 |
| VHVSYLDGK | 9 | Q9Y6J9 | TAF6-like RNA polymerase II p300/CBP | TAF6L | 67.815 | 2 | 51.042934 | yes | 4.353E-07 | 509.27182 | 77.000267 |
| LTVEDFNR | 8 | Q9Y6J9 | TAF6-like RNA polymerase II p300/CBP | TAF6L | 67.815 | 2 | 51.042934 | yes | 2.018E-07 | 497.25363 | 61.084316 |
| KMPQLTASAIVSPI | 20 | Q9Y6J9 | TAF6-like RNA polymerase II p300/CBP | TAF6L | 67.815 | 3 | 51.042934 | yes | 3.059E-07 | 707.69678 | 45.755585 |
| SVSHDLEQLHR | 11 | Q9Y6J9 | TAF6-like RNA polymerase II p300/CBP | TAF6L | 67.815 | 2 | 51.042934 | yes | 1.041E-07 | 660.83636 | 35.247425 |
| EATQNSSQFMK | 11 | Q9Y6J9 | TAF6-like RNA polymerase II p300/CBP | TAF6L | 67.815 | 2 | 51.042934 | yes | 2.103E-07 | 635.79022 | 31.516298 |
| GSGGGGPASASGP | 23 | Q9Y6J9 | TAF6-like RNA polymerase II p300/CBP | TAF6L | 67.815 | 3 | 51.042934 | yes | 1.844E-06 | 675.00323 | 17.543005 |
| LFQLPTPPLSR | 11 | Q9Y6K0 | Choline/ethanolaminephosphotransferase | CEPT1 | 46.554 | 2 | 38.9756 | yes | 4.868E-07 | 634.8717 | 563.38055 |
| TVLELVTQYR | 10 | Q9Y6K5 | 2'-5'-oligoadenylate synthase 3 OS=Homo | OAS3 | 121.171 | 2 | 51.497597 | yes | 2.222E-07 | 611.34552 | 1449.2117 |
| FISEFLQPNR | 10 | Q9Y6K5 | 2'-5'-oligoadenylate synthase 3 OS=Homo | OAS3 | 121.171 | 2 | 51.497597 | yes | 9.722E-08 | 625.83002 | 1241.8325 |
| LTFPEQSVPGALQI | 15 | Q9Y6K5 | 2'-5'-oligoadenylate synthase 3 OS=Homo | OAS3 | 121.171 | 2 | 51.497597 | yes | 2.83E-07 | 845.4516 | 866.18799 |
| MHLLGQLR | 8 | Q9Y6K5 | 2'-5'-oligoadenylate synthase 3 OS=Homo | OAS3 | 121.171 | 2 | 51.497597 | yes | 4.881E-07 | 484.27893 | 818.45331 |
| GMGDPVQSWK | 10 | Q9Y6K5 | 2'-5'-oligoadenylate synthase 3 OS=Homo | OAS3 | 121.171 | 2 | 51.497597 | yes | 9.862E-08 | 552.76074 | 476.68063 |
| AQLEACQQER | 10 | Q9Y6K5 | 2'-5'-oligoadenylate synthase 3 OS=Homo | OAS3 | 121.171 | 2 | 51.497597 | yes | 6.688E-07 | 616.78802 | 465.40924 |
| EAAACTSALCCMC | 14 | Q9Y6K5 | 2'-5'-oligoadenylate synthase 3 OS=Homo | OAS3 | 121.171 | 2 | 51.497597 | yes | 9.781E-08 | 779.31744 | 389.46521 |
| QFLAQVNK | 8 | Q9Y6K5 | 2'-5'-oligoadenylate synthase 3 OS=Homo | OAS3 | 121.171 | 2 | 51.497597 | yes | 1.966E-07 | 474.26907 | 321.4407 |
| RAEILDEMR | 9 | Q9Y6K5 | 2'-5'-oligoadenylate synthase 3 OS=Homo | OAS3 | 121.171 | 2 | 51.497597 | yes | 9.722E-08 | 566.7926 | 320.23618 |
| LLTSGCQEGEHK | 12 | Q9Y6K5 | 2'-5'-oligoadenylate synthase 3 OS=Homo | OAS3 | 121.171 | 2 | 51.497597 | yes | 9.814E-08 | 679.82208 | 314.51413 |
| RAEILSEMR | 9 | Q9Y6K5 | 2'-5'-oligoadenylate synthase 3 OS=Homo | OAS3 | 121.171 | 2 | 51.497597 | yes | 2.802E-06 | 552.79517 | 289.70297 |
| DSQFNMAEGFR | 11 | Q9Y6K5 | 2'-5'-oligoadenylate synthase 3 OS=Homo | OAS3 | 121.171 | 2 | 51.497597 | yes | 9.722E-08 | 651.28259 | 288.76849 |
| AVDTICSFLK | 10 | Q9Y6K5 | 2'-5'-oligoadenylate synthase 3 OS=Homo | OAS3 | 121.171 | 2 | 51.497597 | yes | 9.732E-08 | 577.29974 | 267.66052 |
| NFVNIRPAK | 9 | Q9Y6K5 | 2'-5'-oligoadenylate synthase 3 OS=Homo | OAS3 | 121.171 | 2 | 51.497597 | yes | 1.373E-07 | 529.8089 | 248.07036 |
| ALGALAAALR | 10 | Q9Y6K5 | 2'-5'-oligoadenylate synthase 3 OS=Homo | OAS3 | 121.171 | 2 | 51.497597 | yes | 1.151E-07 | 463.79272 | 207.93092 |
| SLNAVYPR | 8 | Q9Y6K5 | 2'-5'-oligoadenylate synthase 3 OS=Homo | OAS3 | 121.171 | 2 | 51.497597 | yes | 2.703E-07 | 460.25345 | 190.49358 |
| PIILDPADPTGNLG | 18 | Q9Y6K5 | 2'-5'-oligoadenylate synthase 3 OS=Homo | OAS3 | 121.171 | 3 | 51.497597 | yes | 2.851E-07 | 624.33051 | 187.81567 |
| GSLPPQHGLELLTV | 22 | Q9Y6K5 | 2'-5'-oligoadenylate synthase 3 OS=Homo | OAS3 | 121.171 | 3 | 51.497597 | yes | 8.63E-07 | 794.08136 | 172.29195 |
| QDCFNMAQGFR | 11 | Q9Y6K5 | 2'-5'-oligoadenylate synthase 3 OS=Homo | OAS3 | 121.171 | 2 | 51.497597 | yes | 1.611E-07 | 687.28992 | 165.16556 |
| RAEIISEIR | 9 | Q9Y6K5 | 2'-5'-oligoadenylate synthase 3 OS=Homo | OAS3 | 121.171 | 2 | 51.497597 | yes | 1.238E-06 | 543.81696 | 136.48946 |
| DFIISRPTK | 9 | Q9Y6K5 | 2'-5'-oligoadenylate synthase 3 OS=Homo | OAS3 | 121.171 | 2 | 51.497597 | yes | 1.025E-07 | 538.80859 | 135.55925 |

| | | | | | | | | | | | |
|---|---|---|---|---|---|---|---|---|---|---|---|
| AGCSGLGHPIQLD | 17 | Q9Y6K5 | 2'-5'-oligoadenylate synthase 3 OS=Homo | OAS3 | 121.171 | 3 | 51.497597 | yes | 3.298E-07 | 597.96509 | 128.18956 |
| NFMNIRPVK | 9 | Q9Y6K5 | 2'-5'-oligoadenylate synthase 3 OS=Homo | OAS3 | 121.171 | 2 | 51.497597 | yes | 2.947E-07 | 559.81061 | 107.52198 |
| AEILSEMR | 8 | Q9Y6K5 | 2'-5'-oligoadenylate synthase 3 OS=Homo | OAS3 | 121.171 | 2 | 51.497597 | yes | 3.525E-07 | 474.7446 | 79.898804 |
| HWYQQCTK | 8 | Q9Y6K5 | 2'-5'-oligoadenylate synthase 3 OS=Homo | OAS3 | 121.171 | 2 | 51.497597 | yes | 4.504E-06 | 575.75854 | 68.076225 |
| AEILDEMR | 8 | Q9Y6K5 | 2'-5'-oligoadenylate synthase 3 OS=Homo | OAS3 | 121.171 | 2 | 51.497597 | yes | 8.657E-07 | 488.74203 | 46.452824 |
| DKTVGDFLK | 9 | Q9Y6K5 | 2'-5'-oligoadenylate synthase 3 OS=Homo | OAS3 | 121.171 | 2 | 51.497597 | yes | 3.286E-06 | 511.77948 | 45.104004 |
| MDLYSTPAAALDI | 13 | Q9Y6K5 | 2'-5'-oligoadenylate synthase 3 OS=Homo | OAS3 | 121.171 | 2 | 51.497597 | yes | 2.735E-07 | 733.35321 | 41.651741 |
| PKPQVYSTLLNSG | 27 | Q9Y6K5 | 2'-5'-oligoadenylate synthase 3 OS=Homo | OAS3 | 121.171 | 4 | 51.497597 | yes | 3.346E-07 | 755.86426 | 14.827857 |
| QLCIYWTINYNAK | 13 | Q9Y6K5 | 2'-5'-oligoadenylate synthase 3 OS=Homo | OAS3 | 121.171 | 2 | 51.497597 | yes | 2.645E-06 | 843.91907 | 9.1165323 |
| IVGGEDAELGR | 11 | Q9Y6M0 | Testisin OS=Homo sapiens OX=9606 GN | PRSS21 | 34.884 | 2 | 45.201874 | yes | 1.926E-07 | 558.28821 | 167.59529 |
| YLGNSPYDIALVK | 13 | Q9Y6M0 | Testisin OS=Homo sapiens OX=9606 GN | PRSS21 | 34.884 | 2 | 45.201874 | yes | 2.75E-07 | 726.89032 | 78.877968 |
| DACFGDSGGPLAC | 15 | Q9Y6M0 | Testisin OS=Homo sapiens OX=9606 GN | PRSS21 | 34.884 | 2 | 45.201874 | yes | 4.156E-06 | 784.82703 | 19.705496 |
| TDCWVTGWGYIK | 12 | Q9Y6M0 | Testisin OS=Homo sapiens OX=9606 GN | PRSS21 | 34.884 | 2 | 45.201874 | yes | 8.35E-06 | 743.34521 | 6.938674 |
| KLPLAGQVLLK | 11 | Q9Y6M1 | Insulin-like growth factor 2 mRNA-bindi | IGF2BP2 | 66.122 | 2 | 52.919773 | yes | 2.242E-06 | 590.39484 | 3976.6348 |
| IMEVDYSVSK | 10 | Q9Y6M1 | Insulin-like growth factor 2 mRNA-bindi | IGF2BP2 | 66.122 | 2 | 52.919773 | yes | 9.81E-08 | 585.78918 | 3833.7212 |
| LKEENFFNPK | 10 | Q9Y6M1 | Insulin-like growth factor 2 mRNA-bindi | IGF2BP2 | 66.122 | 2 | 52.919773 | yes | 9.722E-08 | 633.3299 | 1617.1219 |
| LKEENFFNPK | 10 | Q9Y6M1 | Insulin-like growth factor 2 mRNA-bindi | IGF2BP2 | 66.122 | 2 | 52.919773 | yes | 4.163E-07 | 422.55566 | 1617.1219 |
| LKEENFFNPK | 10 | Q9Y6M1 | Insulin-like growth factor 2 mRNA-bindi | IGF2BP2 | 66.122 | 3 | 52.919773 | yes | 9.722E-08 | 633.3299 | 1617.1219 |
| LKEENFFNPK | 10 | Q9Y6M1 | Insulin-like growth factor 2 mRNA-bindi | IGF2BP2 | 66.122 | 3 | 52.919773 | yes | 4.163E-07 | 422.55566 | 1617.1219 |
| DQTPDENEEVIVR | 13 | Q9Y6M1 | Insulin-like growth factor 2 mRNA-bindi | IGF2BP2 | 66.122 | 2 | 52.919773 | yes | 1.127E-07 | 772.36536 | 3052.0098 |
| MILEIMQK | 8 | Q9Y6M1 | Insulin-like growth factor 2 mRNA-bindi | IGF2BP2 | 66.122 | 2 | 52.919773 | yes | 2.203E-07 | 503.27722 | 2843.7607 |
| IIGHFFASQTAQR | 13 | Q9Y6M1 | Insulin-like growth factor 2 mRNA-bindi | IGF2BP2 | 66.122 | 2 | 52.919773 | yes | 1.248E-07 | 738.3913 | 2587.8938 |
| IAPAEGPDVSER | 12 | Q9Y6M1 | Insulin-like growth factor 2 mRNA-bindi | IGF2BP2 | 66.122 | 2 | 52.919773 | yes | 1.799E-07 | 620.81201 | 2552.1208 |
| YPQGVASQR | 9 | Q9Y6M1 | Insulin-like growth factor 2 mRNA-bindi | IGF2BP2 | 66.122 | 2 | 52.919773 | yes | 1.003E-07 | 503.25925 | 1770.4083 |
| QIDFPLR | 7 | Q9Y6M1 | Insulin-like growth factor 2 mRNA-bindi | IGF2BP2 | 66.122 | 2 | 52.919773 | yes | 2.393E-07 | 444.75052 | 1114.4453 |
| LSGHQFENYSFK | 12 | Q9Y6M1 | Insulin-like growth factor 2 mRNA-bindi | IGF2BP2 | 66.122 | 2 | 52.919773 | yes | 1.051E-07 | 728.84644 | 725.99585 |
| LAEEIPLK | 8 | Q9Y6M1 | Insulin-like growth factor 2 mRNA-bindi | IGF2BP2 | 66.122 | 2 | 52.919773 | yes | 1.377E-07 | 456.77368 | 615.9032 |
| GTVEACASAEIEIM | 15 | Q9Y6M1 | Insulin-like growth factor 2 mRNA-bindi | IGF2BP2 | 66.122 | 2 | 52.919773 | yes | 1.795E-07 | 804.88403 | 425.35245 |
| EIVQQVK | 7 | Q9Y6M1 | Insulin-like growth factor 2 mRNA-bindi | IGF2BP2 | 66.122 | 2 | 52.919773 | yes | 1.012E-07 | 422.25037 | 355.33371 |
| ILAHNGLVGR | 10 | Q9Y6M1 | Insulin-like growth factor 2 mRNA-bindi | IGF2BP2 | 66.122 | 2 | 52.919773 | yes | 9.885E-08 | 525.31433 | 219.65308 |
| LPLAGQVLLK | 10 | Q9Y6M1 | Insulin-like growth factor 2 mRNA-bindi | IGF2BP2 | 66.122 | 2 | 52.919773 | yes | 2.953E-07 | 526.34735 | 146.88031 |
| ENSGAAEKPVTIH | 24 | Q9Y6M1 | Insulin-like growth factor 2 mRNA-bindi | IGF2BP2 | 66.122 | 4 | 52.919773 | yes | 9.722E-08 | 628.79877 | 134.56381 |
| TVNELQNLTSAEV | 17 | Q9Y6M1 | Insulin-like growth factor 2 mRNA-bindi | IGF2BP2 | 66.122 | 3 | 52.919773 | yes | 9.722E-08 | 628.34595 | 38.498585 |
| KENSGAAEKPVTI | 25 | Q9Y6M1 | Insulin-like growth factor 2 mRNA-bindi | IGF2BP2 | 66.122 | 4 | 52.919773 | yes | 2.945E-06 | 660.82251 | 25.897524 |
| KLFTDLFDR | 9 | Q9Y6M4 | Casein kinase I isoform gamma-3 OS=Ho | CSNK1G3 | 51.389 | 2 | 54.504513 | no | 9.722E-08 | 577.81384 | 514.74432 |
| APQLHLEYR | 9 | Q9Y6M4 | Casein kinase I isoform gamma-3 OS=Ho | CSNK1G3 | 51.389 | 2 | 54.504513 | no | 9.722E-08 | 563.80383 | 353.89438 |
| IGCGNFGELR | 10 | Q9Y6M4 | Casein kinase I isoform gamma-3 OS=Ho | CSNK1G3 | 51.389 | 2 | 54.504513 | no | 1.048E-07 | 561.77167 | 282.15811 |
| DVKPENFLIGR | 11 | Q9Y6M4 | Casein kinase I isoform gamma-3 OS=Ho | CSNK1G3 | 51.389 | 2 | 54.504513 | no | 1.309E-06 | 644.35645 | 266.28864 |
| MEYVHSK | 7 | Q9Y6M4 | Casein kinase I isoform gamma-3 OS=Ho | CSNK1G3 | 51.389 | 2 | 54.504513 | no | 2.609E-07 | 447.21292 | 204.5258 |

| | | | | | | | | | | | |
|---|---|---|---|---|---|---|---|---|---|---|---|
| RLDFFEKPDYDYL | 14 | Q9Y6M4 | Casein kinase I isoform gamma-3 OS=Ho | CSNK1G3 | 51.389 | 3 | 54.504513 | no | 9.722E-08 | 626.31244 | 201.14848 |
| YMSINTHLGK | 10 | Q9Y6M4 | Casein kinase I isoform gamma-3 OS=Ho | CSNK1G3 | 51.389 | 2 | 54.504513 | no | 2.398E-07 | 582.29755 | 158.09456 |
| EYIDPETK | 8 | Q9Y6M4 | Casein kinase I isoform gamma-3 OS=Ho | CSNK1G3 | 51.389 | 2 | 54.504513 | no | 1.976E-07 | 497.74002 | 131.14851 |
| LFTDLFDR | 8 | Q9Y6M4 | Casein kinase I isoform gamma-3 OS=Ho | CSNK1G3 | 51.389 | 2 | 54.504513 | no | 9.722E-08 | 513.76636 | 129.2025 |
| GTGSSSSGVLMVC | 17 | Q9Y6M4 | Casein kinase I isoform gamma-3 OS=Ho | CSNK1G3 | 51.389 | 2 | 54.504513 | yes | 9.722E-08 | 826.90668 | 112.43285 |
| TQQVIHIIDFGLAK | 14 | Q9Y6M4 | Casein kinase I isoform gamma-3 OS=Ho | CSNK1G3 | 51.389 | 2 | 54.504513 | yes | 0.000304 | 791.95123 | 31.778908 |
| AMYPDYFAK | 9 | Q9Y6M9 | NADH dehydrogenase [ubiquinone] 1 bet | NDUFB9 | 21.831 | 2 | 53.387558 | yes | 9.85E-06 | 553.25476 | 192.48193 |
| EAEEEFWYR | 9 | Q9Y6M9 | NADH dehydrogenase [ubiquinone] 1 bet | NDUFB9 | 21.831 | 2 | 53.387558 | yes | 9.722E-08 | 629.7724 | 138.55087 |
| YFACLMR | 7 | Q9Y6M9 | NADH dehydrogenase [ubiquinone] 1 bet | NDUFB9 | 21.831 | 2 | 53.387558 | yes | 1.695E-07 | 480.72513 | 51.245956 |
| EGNAIFTFPNTPVK | 14 | Q9Y6N5 | Sulfide:quinone oxidoreductase, mitochoi | SQOR | 49.961 | 2 | 53.26379 | yes | 1.842E-07 | 767.89862 | 462.90045 |
| TAAAVAAQSGILD | 14 | Q9Y6N5 | Sulfide:quinone oxidoreductase, mitochoi | SQOR | 49.961 | 2 | 53.26379 | yes | 9.722E-08 | 672.36749 | 412.20438 |
| AEPLETFPFDQSK | 13 | Q9Y6N5 | Sulfide:quinone oxidoreductase, mitochoi | SQOR | 49.961 | 2 | 53.26379 | yes | 1.046E-06 | 754.867 | 318.21857 |
| IMYLSEAYFR | 10 | Q9Y6N5 | Sulfide:quinone oxidoreductase, mitochoi | SQOR | 49.961 | 2 | 53.26379 | yes | 1E-07 | 646.8208 | 232.85092 |
| VILAEFDYK | 9 | Q9Y6N5 | Sulfide:quinone oxidoreductase, mitochoi | SQOR | 49.961 | 2 | 53.26379 | yes | 1.716E-07 | 549.29749 | 165.83795 |
| HFYQPIWTLVGAC | 15 | Q9Y6N5 | Sulfide:quinone oxidoreductase, mitochoi | SQOR | 49.961 | 2 | 53.26379 | yes | 1.24E-07 | 844.45135 | 123.94614 |
| GYWGGPAFLR | 10 | Q9Y6N5 | Sulfide:quinone oxidoreductase, mitochoi | SQOR | 49.961 | 2 | 53.26379 | yes | 1.507E-06 | 562.28778 | 115.47556 |
| YADALQEIIQER | 12 | Q9Y6N5 | Sulfide:quinone oxidoreductase, mitochoi | SQOR | 49.961 | 2 | 53.26379 | yes | 1.096E-07 | 724.87262 | 111.4572 |
| GLPEGFAHPK | 10 | Q9Y6N5 | Sulfide:quinone oxidoreductase, mitochoi | SQOR | 49.961 | 2 | 53.26379 | yes | 2.579E-06 | 526.77985 | 109.98422 |
| PTASVIPSGVEWIK | 14 | Q9Y6N5 | Sulfide:quinone oxidoreductase, mitochoi | SQOR | 49.961 | 2 | 53.26379 | yes | 8.846E-07 | 742.41138 | 81.883408 |
| NCIHTDDDEK | 10 | Q9Y6N5 | Sulfide:quinone oxidoreductase, mitochoi | SQOR | 49.961 | 2 | 53.26379 | yes | 9.826E-07 | 623.75385 | 14.752645 |
| LNELPSTLQR | 10 | Q9Y6R0 | Numb-like protein OS=Homo sapiens OX | NUMBL | 64.892 | 2 | 51.75344 | yes | 1.037E-07 | 585.82751 | 450.11871 |
| HAPLEQLVR | 9 | Q9Y6R0 | Numb-like protein OS=Homo sapiens OX | NUMBL | 64.892 | 2 | 51.75344 | yes | 8.543E-07 | 531.80634 | 394.4787 |
| WLEEVSQVAK | 10 | Q9Y6R0 | Numb-like protein OS=Homo sapiens OX | NUMBL | 64.892 | 2 | 51.75344 | yes | 9.809E-08 | 594.81659 | 392.16058 |
| AGAFPPPAIPSAPG | 18 | Q9Y6R0 | Numb-like protein OS=Homo sapiens OX | NUMBL | 64.892 | 2 | 51.75344 | yes | 4.445E-07 | 846.44684 | 167.69159 |
| ECGVTAAFDASR | 12 | Q9Y6R0 | Numb-like protein OS=Homo sapiens OX | NUMBL | 64.892 | 2 | 51.75344 | yes | 9.722E-08 | 642.2879 | 125.3904 |
| RTDFQVK | 7 | Q9Y6R0 | Numb-like protein OS=Homo sapiens OX | NUMBL | 64.892 | 2 | 51.75344 | yes | 3.266E-07 | 447.24561 | 110.61385 |
| GEAGTPVAAGTTA | 18 | Q9Y6R0 | Numb-like protein OS=Homo sapiens OX | NUMBL | 64.892 | 2 | 51.75344 | yes | 1.088E-07 | 805.92847 | 106.52974 |
| SVLWVSADGLR | 11 | Q9Y6R0 | Numb-like protein OS=Homo sapiens OX | NUMBL | 64.892 | 2 | 51.75344 | yes | 9.722E-08 | 601.83002 | 80.929352 |
| INSPHYLTK | 9 | Q9Y6W3 | Calpain-7 OS=Homo sapiens OX=9606 C | CAPN7 | 92.653 | 2 | 46.127724 | yes | 1.645E-07 | 536.79291 | 630.32184 |
| TGPLLIELR | 9 | Q9Y6W3 | Calpain-7 OS=Homo sapiens OX=9606 C | CAPN7 | 92.653 | 2 | 46.127724 | yes | 3.873E-07 | 506.31351 | 63.144276 |
| ISSTSVKPKPPPVR | 14 | Q9Y6W3 | Calpain-7 OS=Homo sapiens OX=9606 C | CAPN7 | 92.653 | 3 | 46.127724 | yes | 1.697E-06 | 498.29971 | 37.956039 |
| LTADPDSEVATTSI | 15 | Q9Y6X2 | E3 SUMO-protein ligase PIAS3 OS=Hon | PIAS3 | 68.018 | 2 | 37.854507 | yes | 5.646E-07 | 788.39667 | 64.378082 |
| VSLMCPLGK | 9 | Q9Y6X2 | E3 SUMO-protein ligase PIAS3 OS=Hon | PIAS3 | 68.018 | 2 | 37.854507 | no | 1.645E-06 | 502.767 | 46.989014 |
| CDYTIQVQLR | 10 | Q9Y6X2 | E3 SUMO-protein ligase PIAS3 OS=Hon | PIAS3 | 68.018 | 2 | 37.854507 | yes | 8.802E-07 | 648.32428 | 38.415325 |
| FDYVPTDTTPR | 11 | Q9Y6X9 | ATPase MORC2 OS=Homo sapiens OX= | MORC2 | 117.824 | 2 | 56.244835 | yes | 2.439E-07 | 656.31439 | 444.07065 |
| TLPFQLSSVEK | 11 | Q9Y6X9 | ATPase MORC2 OS=Homo sapiens OX= | MORC2 | 117.824 | 2 | 56.244835 | yes | 9.722E-08 | 624.84534 | 260.5293 |
| DLDGMFIYNCSR | 12 | Q9Y6X9 | ATPase MORC2 OS=Homo sapiens OX= | MORC2 | 117.824 | 2 | 56.244835 | yes | 9.722E-08 | 745.82379 | 228.82094 |
| ELNFVFGVNIEHR | 13 | Q9Y6X9 | ATPase MORC2 OS=Homo sapiens OX= | MORC2 | 117.824 | 2 | 56.244835 | yes | 1.122E-07 | 787.40973 | 226.73517 |
| TNIVALLQK | 9 | Q9Y6X9 | ATPase MORC2 OS=Homo sapiens OX= | MORC2 | 117.824 | 2 | 56.244835 | yes | 2.02E-07 | 500.31351 | 225.29326 |

| | | | | | | | | | | | |
|---|---|---|---|---|---|---|---|---|---|---|---|
| IFIHGHK | 7 | Q9Y6X9 | ATPase MORC2 OS=Homo sapiens OX= | MORC2 | 117.824 | 2 | 56.244835 | yes | 1.113E-07 | 426.24796 | 171.19725 |
| KLPLEVTTR | 9 | Q9Y6X9 | ATPase MORC2 OS=Homo sapiens OX= | MORC2 | 117.824 | 2 | 56.244835 | yes | 1.166E-07 | 528.82422 | 163.59656 |
| KAPVISSTPK | 10 | Q9Y6X9 | ATPase MORC2 OS=Homo sapiens OX= | MORC2 | 117.824 | 2 | 56.244835 | yes | 2.096E-07 | 514.31097 | 153.88353 |
| LSCCLYKPR | 9 | Q9Y6X9 | ATPase MORC2 OS=Homo sapiens OX= | MORC2 | 117.824 | 2 | 56.244835 | yes | 3.785E-06 | 598.79938 | 153.48227 |
| FAIETELIYK | 10 | Q9Y6X9 | ATPase MORC2 OS=Homo sapiens OX= | MORC2 | 117.824 | 2 | 56.244835 | yes | 9.722E-08 | 613.83698 | 118.80463 |
| NAPSRPPSLPTPR | 13 | Q9Y6X9 | ATPase MORC2 OS=Homo sapiens OX= | MORC2 | 117.824 | 3 | 56.244835 | yes | 2.143E-07 | 463.92474 | 109.48268 |
| AFTNYSSLNR | 10 | Q9Y6X9 | ATPase MORC2 OS=Homo sapiens OX= | MORC2 | 117.824 | 2 | 56.244835 | yes | 9.722E-08 | 607.79364 | 102.58351 |
| YFLPPSFPISK | 11 | Q9Y6X9 | ATPase MORC2 OS=Homo sapiens OX= | MORC2 | 117.824 | 2 | 56.244835 | yes | 1.968E-06 | 648.35535 | 80.506271 |
| PSTEEPVR | 8 | Q9Y6X9 | ATPase MORC2 OS=Homo sapiens OX= | MORC2 | 117.824 | 2 | 56.244835 | yes | 1.272E-07 | 457.73254 | 77.946754 |
| AMGEHLAQYWK | 11 | Q9Y6X9 | ATPase MORC2 OS=Homo sapiens OX= | MORC2 | 117.824 | 2 | 56.244835 | yes | 1.44E-07 | 667.32153 | 68.816566 |
| VKFDYVPTDTTPR | 13 | Q9Y6X9 | ATPase MORC2 OS=Homo sapiens OX= | MORC2 | 117.824 | 2 | 56.244835 | yes | 9.722E-08 | 769.89612 | 65.714088 |
| TEEEVMTQFMK | 11 | Q9Y6X9 | ATPase MORC2 OS=Homo sapiens OX= | MORC2 | 117.824 | 2 | 56.244835 | yes | 2.294E-07 | 686.80981 | 41.059578 |
| TREPVTDNVEK | 11 | Q9Y6X9 | ATPase MORC2 OS=Homo sapiens OX= | MORC2 | 117.824 | 2 | 56.244835 | yes | 6.282E-07 | 644.33063 | 32.858166 |
| LNPEYFQTR | 9 | Q9Y6Y1 | Calmodulin-binding transcription activatc | CAMTA1 | 183.672 | 2 | 37.747227 | yes | 5.733E-07 | 584.29327 | 222.99118 |
| LRSSLLTK | 8 | Q9Y6Y1 | Calmodulin-binding transcription activatc | CAMTA1 | 183.672 | 2 | 37.747227 | yes | 2.757E-06 | 459.29257 | 49.370285 |
| SIIECVDDFR | 10 | Q9Y6Y8 | SEC23-interacting protein OS=Homo sap | SEC23IP | 111.077 | 2 | 20.224403 | yes | 4.542E-06 | 627.29517 | 21.832525 |
| SEQEDEVLLVSSSI | 14 | A0A024RBG1 | Diphosphoinositol polyphosphate phosph | NUDT4B | 20.435 | 2 | 44.229782 | yes | 2.272E-07 | 789.38629 | 135.26849 |
| YVDIAIPCNNK | 11 | A0A8I5KQE6 | Small ribosomal subunit protein uS2B OS | RPSA2 | 32.91 | 2 | 51.5342 | yes | 7.05E-07 | 653.82666 | 13309.147 |
| FAAATGATPIAGR | 13 | A0A8I5KQE6 | Small ribosomal subunit protein uS2B OS | RPSA2 | 32.91 | 2 | 51.5342 | yes | 9.722E-08 | 602.32764 | 5523.5137 |
| SDGIYIINLK | 10 | A0A8I5KQE6 | Small ribosomal subunit protein uS2B OS | RPSA2 | 32.91 | 2 | 51.5342 | yes | 5.025E-07 | 568.32153 | 5112.1191 |
| KSDGIYIINLK | 11 | A0A8I5KQE6 | Small ribosomal subunit protein uS2B OS | RPSA2 | 32.91 | 2 | 51.5342 | yes | 9.722E-08 | 632.36902 | 839.0791 |
| LLVVTDPR | 8 | A0A8I5KQE6 | Small ribosomal subunit protein uS2B OS | RPSA2 | 32.91 | 2 | 51.5342 | yes | 4.202E-07 | 456.7793 | 720.66522 |
| SGALDVLQMK | 10 | A0A8I5KQE6 | Small ribosomal subunit protein uS2B OS | RPSA2 | 32.91 | 2 | 51.5342 | yes | 1.77E-07 | 552.29187 | 257.90112 |
| SGALDVLQMK | 10 | A0A8I5KQE6 | Small ribosomal subunit protein uS2B OS | RPSA2 | 32.91 | 2 | 51.5342 | yes | 4.88E-06 | 560.28937 | 257.90112 |
| EHPWEVMPDLYF' | 14 | A0A8I5KQE6 | Small ribosomal subunit protein uS2B OS | RPSA2 | 32.91 | 3 | 51.5342 | yes | 6.488E-07 | 627.95917 | 57.758888 |
| DVLLFEHDR | 9 | A0PJW6 | Transmembrane protein 223 OS=Homo sa | TMEM223 | 22.049 | 2 | 48.558205 | yes | 1.13E-07 | 572.29327 | 79.735062 |
| GEVPAMLPLK | 10 | A0PJW6 | Transmembrane protein 223 OS=Homo sa | TMEM223 | 22.049 | 2 | 48.558205 | yes | 4.08E-07 | 527.80188 | 58.63765 |
| PSLVVSGIMER | 11 | A1L020 | RNA-binding protein MEX3A OS=Homo | MEX3A | 54.174 | 2 | 46.836845 | yes | 4.555E-06 | 594.32629 | 109.483 |
| IQQQTNTYIITPSR | 14 | A1L020 | RNA-binding protein MEX3A OS=Homo | MEX3A | 54.174 | 2 | 46.836845 | yes | 1.46E-07 | 831.94409 | 105.94456 |
| SPATSAGPELAGLI | 15 | A1L020 | RNA-binding protein MEX3A OS=Homo | MEX3A | 54.174 | 2 | 46.836845 | yes | 1.615E-07 | 712.38062 | 83.447258 |
| GTQEFSPLCLR | 11 | A6NHQ4 | Elongin BC and Polycomb repressive con | EPOP | 39.322 | 2 | 50.396805 | yes | 9.809E-08 | 654.32428 | 94.179222 |
| ASSLGPGPGELAA | 14 | A6NHQ4 | Elongin BC and Polycomb repressive con | EPOP | 39.322 | 2 | 50.396805 | yes | 2.307E-07 | 641.84113 | 86.956413 |
| AEVASFVADR | 10 | A6NKF1 | SAC3 domain-containing protein 1 OS=F | SAC3D1 | 43.554 | 2 | 44.496178 | yes | 1.06E-06 | 532.77222 | 68.254402 |
| YLAGEVAESADIA | 14 | A6NKF1 | SAC3 domain-containing protein 1 OS=F | SAC3D1 | 43.554 | 2 | 44.496178 | yes | 2.172E-07 | 732.87012 | 59.424473 |
| ESPLLNDYR | 9 | A6ZKI3 | Retrotransposon Gag-like protein 8C OS= | RTL8C | 13.172 | 2 | 37.710545 | no | 5.764E-07 | 553.77747 | 159.48238 |
| ALLALPIRPATR | 12 | A6ZKI3 | Retrotransposon Gag-like protein 8C OS= | RTL8C | 13.172 | 3 | 37.710545 | yes | 1.729E-06 | 431.27847 | 29.464787 |
| ALLALPIRPATR | 12 | A6ZKI3 | Retrotransposon Gag-like protein 8C OS= | RTL8C | 13.172 | 3 | 37.710545 | yes | 6.202E-07 | 646.41406 | 29.464787 |
| ALLALPIRPATR | 12 | A6ZKI3 | Retrotransposon Gag-like protein 8C OS= | RTL8C | 13.172 | 2 | 37.710545 | yes | 1.729E-06 | 431.27847 | 29.464787 |
| ALLALPIRPATR | 12 | A6ZKI3 | Retrotransposon Gag-like protein 8C OS= | RTL8C | 13.172 | 2 | 37.710545 | yes | 6.202E-07 | 646.41406 | 29.464787 |

| | | | | | | | | | | | |
|---|---|---|---|---|---|---|---|---|---|---|---|
| IKPVTDEAQLVYR | 13 | A8K855 | EF-hand calcium-binding domain-contain | EFCAB7 | 71.982 | 2 | 49.142746 | yes | 1.064E-07 | 766.42755 | 94.959793 |
| DQLYLALQHAGR | 12 | A8K855 | EF-hand calcium-binding domain-contain | EFCAB7 | 71.982 | 2 | 49.142746 | yes | 1.165E-07 | 692.87024 | 54.052292 |
| SSLENIISK | 9 | A8K855 | EF-hand calcium-binding domain-contain | EFCAB7 | 71.982 | 2 | 49.142746 | yes | 2.9E-07 | 495.77695 | 24.371765 |
| ATFGPAIAAVQK | 12 | A8MPP1 | Putative ATP-dependent RNA helicase D | DDX11L8 | 101.811 | 2 | 31.414774 | yes | 1.849E-06 | 587.33496 | 40.308891 |
| PAGEPAWVTQFV( | 14 | A8MPP1 | Putative ATP-dependent RNA helicase D | DDX11L8 | 101.811 | 2 | 31.414774 | no | 5.132E-05 | 779.40662 | 11.106343 |
| MSIHLIHMR | 9 | A8MV81 | HIG1 domain family member 1C OS=Hoı | HIGD1C | 11.079 | 2 | 50.96698 | yes | 9.722E-08 | 569.30463 | 190.32448 |
| NILFVITKPDVYK | 13 | E9PAV3 | Nascent polypeptide-associated complex s | NACA | 205.422 | 2 | 49.491035 | no | 1.068E-07 | 775.45306 | 1052.8137 |
| IEDLSQQAQLAAA | 15 | E9PAV3 | Nascent polypeptide-associated complex s | NACA | 205.422 | 2 | 49.491035 | yes | 1.31E-07 | 807.92029 | 789.66064 |
| DIELVMSQANVSR | 13 | E9PAV3 | Nascent polypeptide-associated complex s | NACA | 205.422 | 2 | 49.491035 | yes | 1.036E-07 | 731.37195 | 731.10699 |
| SPASDTYIVFGEAI | 14 | E9PAV3 | Nascent polypeptide-associated complex s | NACA | 205.422 | 2 | 49.491035 | yes | 1.025E-07 | 742.867 | 205.68204 |
| ENPTGVVLLLK | 11 | G9CGD6 | CNK3/IPCEF1 fusion protein OS=Homo | CNK3/IPCI | 100.365 | 2 | 41.038845 | yes | 3.708E-07 | 591.85828 | 38.698467 |
| APFTGITDFSVTK | 13 | G9CGD6 | CNK3/IPCEF1 fusion protein OS=Homo | CNK3/IPCI | 100.365 | 2 | 41.038845 | yes | 3.984E-07 | 692.36139 | 34.412769 |
| DDVIVTASNFSSEI | 14 | O00178 | GTP-binding protein 1 OS=Homo sapien | GTPBP1 | 72.454 | 2 | 54.23002 | yes | 1.021E-07 | 770.36792 | 1297.1182 |
| KIPVLVQSK | 9 | O00178 | GTP-binding protein 1 OS=Homo sapien | GTPBP1 | 72.454 | 2 | 54.23002 | yes | 1.571E-07 | 506.3317 | 1065.3231 |
| VGDNDFLEVR | 10 | O00178 | GTP-binding protein 1 OS=Homo sapien | GTPBP1 | 72.454 | 2 | 54.23002 | yes | 1.084E-07 | 582.28821 | 721.52478 |
| VAVVGNVDAGK | 11 | O00178 | GTP-binding protein 1 OS=Homo sapien | GTPBP1 | 72.454 | 2 | 54.23002 | yes | 4.79E-07 | 514.79041 | 584.54468 |
| TPEYLHIDQR | 10 | O00178 | GTP-binding protein 1 OS=Homo sapien | GTPBP1 | 72.454 | 2 | 54.23002 | yes | 2.941E-07 | 636.32257 | 450.71826 |
| SMAEQIEADVILLI | 14 | O00178 | GTP-binding protein 1 OS=Homo sapien | GTPBP1 | 72.454 | 2 | 54.23002 | yes | 1.021E-07 | 794.42419 | 435.71652 |
| MFLNLLSPR | 9 | O00178 | GTP-binding protein 1 OS=Homo sapien | GTPBP1 | 72.454 | 2 | 54.23002 | yes | 1.107E-07 | 545.8075 | 395.64307 |
| IDMCPANILQETLI | 14 | O00178 | GTP-binding protein 1 OS=Homo sapien | GTPBP1 | 72.454 | 2 | 54.23002 | yes | 1.208E-07 | 823.41803 | 315.16013 |
| YQAMVHCGSIR | 11 | O00178 | GTP-binding protein 1 OS=Homo sapien | GTPBP1 | 72.454 | 2 | 54.23002 | yes | 9.722E-08 | 661.31061 | 290.18259 |
| QTATILSMDK | 10 | O00178 | GTP-binding protein 1 OS=Homo sapien | GTPBP1 | 72.454 | 2 | 54.23002 | yes | 1.417E-07 | 554.28937 | 247.13777 |
| GGQTASFALK | 10 | O00178 | GTP-binding protein 1 OS=Homo sapien | GTPBP1 | 72.454 | 2 | 54.23002 | yes | 1.379E-07 | 490.26401 | 177.42467 |
| GMVMVSPR | 8 | O00178 | GTP-binding protein 1 OS=Homo sapien | GTPBP1 | 72.454 | 2 | 54.23002 | yes | 1.202E-07 | 438.72513 | 153.73418 |
| LNPQASWEFEAEII | 25 | O00178 | GTP-binding protein 1 OS=Homo sapien | GTPBP1 | 72.454 | 4 | 54.23002 | yes | 1.348E-07 | 722.12909 | 137.83289 |
| LLQTTNNSPMNSK | 13 | O00178 | GTP-binding protein 1 OS=Homo sapien | GTPBP1 | 72.454 | 2 | 54.23002 | yes | 9.722E-08 | 724.36414 | 107.3286 |
| VITFIDLAGHEK | 12 | O00178 | GTP-binding protein 1 OS=Homo sapien | GTPBP1 | 72.454 | 2 | 54.23002 | yes | 9.823E-08 | 671.87189 | 65.404442 |
| IPVLVQSK | 8 | O00178 | GTP-binding protein 1 OS=Homo sapien | GTPBP1 | 72.454 | 2 | 54.23002 | yes | 3.52E-07 | 442.28421 | 64.72039 |
| KGMVMVSPR | 9 | O00178 | GTP-binding protein 1 OS=Homo sapien | GTPBP1 | 72.454 | 2 | 54.23002 | yes | 6.383E-07 | 502.77261 | 47.829967 |
| MTLTDSIER | 9 | O00458 | Interferon-related developmental regulato | IFRD1 | 50.269 | 2 | 53.880886 | yes | 2.829E-07 | 533.26587 | 456.44324 |
| IICDGSASMQAR | 12 | O00458 | Interferon-related developmental regulato | IFRD1 | 50.269 | 2 | 53.880886 | yes | 9.722E-08 | 654.80536 | 293.31281 |
| LPSLLSCDDVNMR | 13 | O00458 | Interferon-related developmental regulato | IFRD1 | 50.269 | 2 | 53.880886 | yes | 1.273E-07 | 760.36578 | 292.23627 |
| MYIDCWVK | 8 | O00458 | Interferon-related developmental regulato | IFRD1 | 50.269 | 2 | 53.880886 | yes | 1.805E-07 | 557.75665 | 258.52191 |
| MLYEFILER | 9 | O00458 | Interferon-related developmental regulato | IFRD1 | 50.269 | 2 | 53.880886 | yes | 9.722E-08 | 607.31793 | 213.10352 |
| HLYNSAAFK | 9 | O00458 | Interferon-related developmental regulato | IFRD1 | 50.269 | 2 | 53.880886 | yes | 1.445E-06 | 525.77197 | 192.55731 |
| QAALEGIK | 8 | O00458 | Interferon-related developmental regulato | IFRD1 | 50.269 | 2 | 53.880886 | yes | 2.06E-07 | 415.24252 | 100.25495 |
| VTPAAANYR | 9 | O14545 | TRAF-type zinc finger domain-containing | TRAFD1 | 64.842 | 2 | 55.090748 | yes | 1.18E-07 | 481.75635 | 237.02713 |
| QIEALDPPMR | 10 | O14545 | TRAF-type zinc finger domain-containing | TRAFD1 | 64.842 | 2 | 55.090748 | yes | 2.711E-07 | 585.3028 | 172.11067 |
| PGCQPSSPCVPK | 12 | O14545 | TRAF-type zinc finger domain-containing | TRAFD1 | 64.842 | 2 | 55.090748 | yes | 1.602E-07 | 657.30249 | 150.38574 |

| | | | | | | | | | | | |
|---|---|---|---|---|---|---|---|---|---|---|---|
| AEFLDDQETR | 10 | O14545 | TRAF-type zinc finger domain-containing | TRAFD1 | 64.842 | 2 | 55.090748 | yes | 9.722E-08 | 633.28583 | 144.90244 |
| AFESDVFHNR | 10 | O14545 | TRAF-type zinc finger domain-containing | TRAFD1 | 64.842 | 2 | 55.090748 | yes | 1.857E-07 | 611.28601 | 133.30829 |
| NIGMCPTCK | 9 | O14545 | TRAF-type zinc finger domain-containing | TRAFD1 | 64.842 | 2 | 55.090748 | yes | 2.138E-07 | 540.73535 | 104.85565 |
| LSNSDSQDIQGR | 12 | O14545 | TRAF-type zinc finger domain-containing | TRAFD1 | 64.842 | 2 | 55.090748 | yes | 1.34E-07 | 660.31293 | 80.317726 |
| TELCGNCGR | 9 | O14545 | TRAF-type zinc finger domain-containing | TRAFD1 | 64.842 | 2 | 55.090748 | yes | 1.305E-07 | 533.72388 | 65.583542 |
| AYGELPEHAK | 10 | O14602 | Eukaryotic translation initiation factor 1A | EIF1AY | 16.443 | 2 | 50.91926 | yes | 9.722E-08 | 557.78003 | 292.63483 |
| GVSFNESAADNLK | 13 | O15063 | Granule associated Rac and RHOG effect | GARRE1 | 116.02 | 2 | 52.23008 | yes | 3.292E-06 | 676.32806 | 87.553383 |
| YVAGVSQAMQQK | 12 | O15063 | Granule associated Rac and RHOG effect | GARRE1 | 116.02 | 2 | 52.23008 | yes | 1.164E-07 | 655.33209 | 77.626038 |
| YLDALNVFCR | 10 | O15063 | Granule associated Rac and RHOG effect | GARRE1 | 116.02 | 2 | 52.23008 | yes | 9.722E-08 | 635.8161 | 76.041855 |
| LQSILNIGNFPR | 12 | O15063 | Granule associated Rac and RHOG effect | GARRE1 | 116.02 | 2 | 52.23008 | yes | 3.293E-06 | 686.39081 | 61.953087 |
| IVVQVPSTWCLK | 12 | O15063 | Granule associated Rac and RHOG effect | GARRE1 | 116.02 | 2 | 52.23008 | yes | 1.953E-07 | 715.39722 | 54.351002 |
| NIFIAGCSEEK | 11 | O15063 | Granule associated Rac and RHOG effect | GARRE1 | 116.02 | 2 | 52.23008 | yes | 3.617E-07 | 634.30298 | 43.840385 |
| LGPTEGRPQLK | 11 | O15235 | Small ribosomal subunit protein uS12m C | MRPS12 | 15.173 | 2 | 51.11513 | yes | 9.722E-08 | 598.34332 | 943.65259 |
| GVVLCTFTR | 9 | O15235 | Small ribosomal subunit protein uS12m C | MRPS12 | 15.173 | 2 | 51.11513 | yes | 1.647E-07 | 526.78149 | 708.89606 |
| YDCGHVQK | 8 | O15235 | Small ribosomal subunit protein uS12m C | MRPS12 | 15.173 | 2 | 51.11513 | yes | 3.227E-07 | 503.72418 | 364.82733 |
| EAVCFIPGEGHTL( | 25 | O15235 | Small ribosomal subunit protein uS12m C | MRPS12 | 15.173 | 4 | 51.11513 | yes | 2.625E-07 | 694.60254 | 259.21368 |
| KLGPTEGRPQLK | 12 | O15235 | Small ribosomal subunit protein uS12m C | MRPS12 | 15.173 | 3 | 51.11513 | yes | 1.154E-07 | 441.92963 | 50.670929 |
| KLGPTEGRPQLK | 12 | O15235 | Small ribosomal subunit protein uS12m C | MRPS12 | 15.173 | 3 | 51.11513 | yes | 2.405E-07 | 662.39081 | 50.670929 |
| KLGPTEGRPQLK | 12 | O15235 | Small ribosomal subunit protein uS12m C | MRPS12 | 15.173 | 2 | 51.11513 | yes | 1.154E-07 | 441.92963 | 50.670929 |
| KLGPTEGRPQLK | 12 | O15235 | Small ribosomal subunit protein uS12m C | MRPS12 | 15.173 | 2 | 51.11513 | yes | 2.405E-07 | 662.39081 | 50.670929 |
| TQDLPGVK | 8 | O15235 | Small ribosomal subunit protein uS12m C | MRPS12 | 15.173 | 2 | 51.11513 | yes | 9.722E-08 | 429.23999 | 61.678791 |
| LKPGPQLPGR | 10 | O15370 | Transcription factor SOX-12 OS=Homo s | SOX12 | 34.122 | 2 | 41.3906 | yes | 4.303E-07 | 531.82458 | 155.41641 |
| HMADYPDYK | 9 | O15370 | Transcription factor SOX-12 OS=Homo s | SOX12 | 34.122 | 2 | 41.3906 | no | 3.533E-07 | 570.24493 | 60.25885 |
| WQLLQDSEK | 9 | O15370 | Transcription factor SOX-12 OS=Homo s | SOX12 | 34.122 | 2 | 41.3906 | yes | 3.7E-05 | 573.79309 | 54.579929 |
| EGLLDPNVK | 9 | O43414 | ERI1 exoribonuclease 3 OS=Homo sapie | ERI3 | 37.239 | 2 | 51.79671 | yes | 3.198E-06 | 492.77167 | 515.58093 |
| SIFVTCGDWDLK | 12 | O43414 | ERI1 exoribonuclease 3 OS=Homo sapie | ERI3 | 37.239 | 2 | 51.79671 | yes | 9.722E-08 | 720.84503 | 435.46744 |
| GLSLQHIGR | 9 | O43414 | ERI1 exoribonuclease 3 OS=Homo sapie | ERI3 | 37.239 | 2 | 51.79671 | yes | 1.496E-07 | 490.78543 | 327.32758 |
| AYSFAMGCWPK | 11 | O43414 | ERI1 exoribonuclease 3 OS=Homo sapie | ERI3 | 37.239 | 2 | 51.79671 | yes | 9.634E-07 | 659.29138 | 56.817448 |
| IGIGELITR | 9 | O75153 | Clustered mitochondria protein homolog | CLUH | 146.67 | 2 | 53.776443 | yes | 1.541E-07 | 486.29785 | 3184.894 |
| VTAQSIIPGILER | 13 | O75153 | Clustered mitochondria protein homolog | CLUH | 146.67 | 2 | 53.776443 | yes | 6.05E-07 | 698.91156 | 2055.9106 |
| VVEEPYTVR | 9 | O75153 | Clustered mitochondria protein homolog | CLUH | 146.67 | 2 | 53.776443 | yes | 1.249E-07 | 546.29022 | 2020.3341 |
| AVGSISSTAFDIR | 13 | O75153 | Clustered mitochondria protein homolog | CLUH | 146.67 | 2 | 53.776443 | yes | 5.907E-07 | 662.34882 | 1957.5271 |
| HYILDLLR | 8 | O75153 | Clustered mitochondria protein homolog | CLUH | 146.67 | 2 | 53.776443 | yes | 1.201E-06 | 521.80585 | 1786.051 |
| DWKPLQCLK | 9 | O75153 | Clustered mitochondria protein homolog | CLUH | 146.67 | 2 | 53.776443 | yes | 3.314E-07 | 594.31573 | 1329.0433 |
| DQEQSVIYGSIDF( | 15 | O75153 | Clustered mitochondria protein homolog | CLUH | 146.67 | 2 | 53.776443 | yes | 6.102E-07 | 843.40448 | 1309.0126 |
| CLTQQAVALQR | 11 | O75153 | Clustered mitochondria protein homolog | CLUH | 146.67 | 2 | 53.776443 | yes | 9.766E-08 | 644.34552 | 1257.3888 |
| QELVDAFVEHR | 11 | O75153 | Clustered mitochondria protein homolog | CLUH | 146.67 | 2 | 53.776443 | yes | 9.722E-08 | 671.84113 | 1239.3114 |
| VHSDFTAAATR | 11 | O75153 | Clustered mitochondria protein homolog | CLUH | 146.67 | 2 | 53.776443 | yes | 1.049E-07 | 588.29382 | 1219.5648 |
| QVSITASTR | 9 | O75153 | Clustered mitochondria protein homolog | CLUH | 146.67 | 2 | 53.776443 | yes | 9.789E-08 | 481.76691 | 714.83423 |

| | | | | | | | | | | | |
|---|---|---|---|---|---|---|---|---|---|---|---|
| LGYEEHIPGQTR | 12 | O75153 | Clustered mitochondria protein homolog | CLUH | 146.67 | 2 | 53.776443 | yes | 9.722E-08 | 700.35187 | 704.90491 |
| ELAETIAADDGTD | 15 | O75153 | Clustered mitochondria protein homolog | CLUH | 146.67 | 2 | 53.776443 | yes | 9.722E-08 | 787.37067 | 687.20569 |
| DCMEHAVLPVDG | 20 | O75153 | Clustered mitochondria protein homolog | CLUH | 146.67 | 3 | 53.776443 | yes | 1.42E-07 | 738.68323 | 671.03424 |
| VLTMSGWNPPPGI | 14 | O75153 | Clustered mitochondria protein homolog | CLUH | 146.67 | 2 | 53.776443 | yes | 1.311E-07 | 763.38263 | 628.0495 |
| SVEGLQEGSVLR | 12 | O75153 | Clustered mitochondria protein homolog | CLUH | 146.67 | 2 | 53.776443 | yes | 2.436E-07 | 637.34094 | 575.84979 |
| VALSHHLVAR | 10 | O75153 | Clustered mitochondria protein homolog | CLUH | 146.67 | 2 | 53.776443 | yes | 9.904E-08 | 551.82764 | 451.92264 |
| ASDAFHFFQSGQA | 14 | O75153 | Clustered mitochondria protein homolog | CLUH | 146.67 | 2 | 53.776443 | yes | 2.734E-07 | 770.86261 | 444.54919 |
| YLELLER | 7 | O75153 | Clustered mitochondria protein homolog | CLUH | 146.67 | 2 | 53.776443 | yes | 9.722E-08 | 468.26346 | 356.73468 |
| VLELVLR | 7 | O75153 | Clustered mitochondria protein homolog | CLUH | 146.67 | 2 | 53.776443 | no | 1.408E-07 | 421.27893 | 334.45334 |
| HKPAFTEEDVLNII | 18 | O75153 | Clustered mitochondria protein homolog | CLUH | 146.67 | 3 | 53.776443 | yes | 9.722E-08 | 695.04559 | 287.92654 |
| TMNEIYR | 7 | O75153 | Clustered mitochondria protein homolog | CLUH | 146.67 | 2 | 53.776443 | yes | 1.603E-07 | 463.72366 | 253.02914 |
| LHYIMGDYAEALS | 17 | O75153 | Clustered mitochondria protein homolog | CLUH | 146.67 | 3 | 53.776443 | yes | 1.839E-07 | 660.98767 | 203.48816 |
| FPESCQDEVR | 10 | O75153 | Clustered mitochondria protein homolog | CLUH | 146.67 | 2 | 53.776443 | yes | 2.916E-07 | 633.7746 | 185.92041 |
| TCFSLHLDGNVLD | 19 | O75153 | Clustered mitochondria protein homolog | CLUH | 146.67 | 3 | 53.776443 | yes | 9.722E-08 | 754.03204 | 166.48093 |
| EAAAAEPPR | 9 | O75153 | Clustered mitochondria protein homolog | CLUH | 146.67 | 2 | 53.776443 | yes | 4.531E-07 | 456.2327 | 162.41666 |
| FLENALAVSTK | 11 | O75153 | Clustered mitochondria protein homolog | CLUH | 146.67 | 2 | 53.776443 | yes | 5.401E-07 | 596.83228 | 157.84958 |
| AEDAYTSR | 8 | O75153 | Clustered mitochondria protein homolog | CLUH | 146.67 | 2 | 53.776443 | yes | 3.588E-07 | 456.70651 | 129.80147 |
| DWNEELQTTR | 10 | O75153 | Clustered mitochondria protein homolog | CLUH | 146.67 | 2 | 53.776443 | yes | 1.214E-07 | 646.29932 | 69.113541 |
| KNFAVLQK | 8 | O75153 | Clustered mitochondria protein homolog | CLUH | 146.67 | 2 | 53.776443 | yes | 4.09E-07 | 474.28726 | 56.873779 |
| AVLMSER | 7 | O75153 | Clustered mitochondria protein homolog | CLUH | 146.67 | 2 | 53.776443 | yes | 2.894E-06 | 403.21545 | 53.912773 |
| DEEVELCSSVECK | 13 | O75153 | Clustered mitochondria protein homolog | CLUH | 146.67 | 2 | 53.776443 | yes | 2.398E-07 | 792.33185 | 34.793163 |
| AEEPMATEPAPAG | 26 | O75153 | Clustered mitochondria protein homolog | CLUH | 146.67 | 3 | 53.776443 | yes | 1.074E-07 | 820.72858 | 24.512896 |
| MHGDLMYLFVITA | 16 | O75153 | Clustered mitochondria protein homolog | CLUH | 146.67 | 3 | 53.776443 | yes | 4.03E-06 | 637.64282 | 13.401663 |
| AAEADGPLKR | 10 | O75352 | Mannose-P-dolichol utilization defect 1 p | MPDU1 | 26.638 | 2 | 43.962 | yes | 2.378E-07 | 535.28546 | 109.8343 |
| TGTCGYCGLQFR | 12 | O75380 | NADH dehydrogenase [ubiquinone] iron- | NDUFS6 | 13.712 | 2 | 50.22754 | yes | 9.849E-08 | 710.31085 | 358.82971 |
| VTHTGQVYDDK | 11 | O75380 | NADH dehydrogenase [ubiquinone] iron- | NDUFS6 | 13.712 | 2 | 50.22754 | yes | 1.004E-07 | 631.8042 | 90.757591 |
| MVSEAGTGFCFNT | 14 | O75394 | Large ribosomal subunit protein bL33m C | MRPL33 | 7.62 | 2 | 50.947174 | yes | 9.722E-08 | 774.84473 | 154.57419 |
| SSFLIEHQR | 9 | O75820 | Zinc finger protein 189 OS=Homo sapien | ZNF189 | 72.976 | 2 | 46.020203 | yes | 1.675E-07 | 558.79346 | 118.48071 |
| TFSVSSTLIR | 10 | O75820 | Zinc finger protein 189 OS=Homo sapien | ZNF189 | 72.976 | 2 | 46.020203 | yes | 3.97E-07 | 555.81134 | 71.708611 |
| AGLVDDFEK | 9 | O75947 | ATP synthase subunit d, mitochondrial O | ATP5PD | 18.492 | 2 | 49.482285 | yes | 1.026E-07 | 497.24802 | 480.65866 |
| SWNETLTSR | 9 | O75947 | ATP synthase subunit d, mitochondrial O | ATP5PD | 18.492 | 2 | 49.482285 | yes | 3.661E-07 | 547.26727 | 250.97935 |
| YTAQVDAEEK | 10 | O75947 | ATP synthase subunit d, mitochondrial O | ATP5PD | 18.492 | 2 | 49.482285 | yes | 1.689E-07 | 577.27222 | 91.297073 |
| YTAQVDAEEKEDV | 14 | O75947 | ATP synthase subunit d, mitochondrial O | ATP5PD | 18.492 | 2 | 49.482285 | yes | 1.503E-07 | 812.88867 | 27.41573 |
| VELVPPTPAEIPR | 13 | O75964 | ATP synthase subunit g, mitochondrial O | ATP5MG | 11.429 | 2 | 53.57656 | yes | 1.001E-07 | 709.40613 | 645.841 |
| IVNSAQTGSFK | 11 | O75964 | ATP synthase subunit g, mitochondrial O | ATP5MG | 11.429 | 2 | 53.57656 | yes | 1.065E-07 | 576.3064 | 174.44144 |
| LATFWYYAK | 9 | O75964 | ATP synthase subunit g, mitochondrial O | ATP5MG | 11.429 | 2 | 53.57656 | yes | 1.343E-07 | 581.80023 | 95.493103 |
| TPALVNAAVTYSK | 13 | O75964 | ATP synthase subunit g, mitochondrial O | ATP5MG | 11.429 | 2 | 53.57656 | yes | 9.722E-08 | 667.86938 | 84.572899 |
| DFLLNYPFSTSVK | 13 | O95084 | Serine protease 23 OS=Homo sapiens OX | PRSS23 | 43.002 | 2 | 51.9777 | yes | 9.722E-08 | 765.89557 | 397.28561 |
| LSTGCTGTLVAEK | 13 | O95084 | Serine protease 23 OS=Homo sapiens OX | PRSS23 | 43.002 | 2 | 51.9777 | yes | 1.001E-07 | 668.84247 | 351.83322 |

| | | | | | | | | | | | |
|---|---|---|---|---|---|---|---|---|---|---|---|
| YAQICYWIK | 9 | O95084 | Serine protease 23 OS=Homo sapiens OX | PRSS23 | 43.002 | 2 | 51.9777 | yes | 3.046E-06 | 622.81024 | 222.91035 |
| LPVVLPQSTLNLA | 14 | O95084 | Serine protease 23 OS=Homo sapiens OX | PRSS23 | 43.002 | 2 | 51.9777 | yes | 1.006E-07 | 746.9585 | 219.41437 |
| HVLTAAHCIHDGK | 13 | O95084 | Serine protease 23 OS=Homo sapiens OX | PRSS23 | 43.002 | 3 | 51.9777 | yes | 1.084E-07 | 486.91388 | 47.473988 |
| HVLTAAHCIHDGK | 13 | O95084 | Serine protease 23 OS=Homo sapiens OX | PRSS23 | 43.002 | 3 | 51.9777 | yes | 1.316E-06 | 729.86719 | 47.473988 |
| HVLTAAHCIHDGK | 13 | O95084 | Serine protease 23 OS=Homo sapiens OX | PRSS23 | 43.002 | 2 | 51.9777 | yes | 1.084E-07 | 486.91388 | 47.473988 |
| HVLTAAHCIHDGK | 13 | O95084 | Serine protease 23 OS=Homo sapiens OX | PRSS23 | 43.002 | 2 | 51.9777 | yes | 1.316E-06 | 729.86719 | 47.473988 |
| LPVVLPQSTLNLA | 22 | O95084 | Serine protease 23 OS=Homo sapiens OX | PRSS23 | 43.002 | 3 | 51.9777 | yes | 1.761E-07 | 770.10187 | 51.873264 |
| LEVSSSCGPQCHK | 13 | O95084 | Serine protease 23 OS=Homo sapiens OX | PRSS23 | 43.002 | 2 | 51.9777 | yes | 9.722E-08 | 744.83215 | 49.024368 |
| SVNEGAYIR | 9 | O95625 | Zinc finger and BTB domain-containing p | ZBTB11 | 119.384 | 2 | 51.336872 | yes | 1.936E-07 | 504.75909 | 297.32498 |
| LSEVAEAIQTVK | 12 | O95625 | Zinc finger and BTB domain-containing p | ZBTB11 | 119.384 | 2 | 51.336872 | yes | 2.516E-07 | 644.36139 | 92.992661 |
| SVLSANSEYFR | 11 | O95625 | Zinc finger and BTB domain-containing p | ZBTB11 | 119.384 | 2 | 51.336872 | yes | 9.728E-08 | 636.81458 | 72.143639 |
| QFQCELCVK | 9 | O95625 | Zinc finger and BTB domain-containing p | ZBTB11 | 119.384 | 2 | 51.336872 | yes | 1.183E-07 | 606.28101 | 64.78688 |
| TDFGPDDDTYR | 11 | O95625 | Zinc finger and BTB domain-containing p | ZBTB11 | 119.384 | 2 | 51.336872 | yes | 9.722E-08 | 651.26764 | 51.082813 |
| CGECGMVFQR | 10 | O95625 | Zinc finger and BTB domain-containing p | ZBTB11 | 119.384 | 2 | 51.336872 | yes | 9.722E-08 | 622.25446 | 36.197247 |
| GPSSAVLPSALGSI | 14 | O95685 | Protein phosphatase 1 regulatory subunit | PPP1R3D | 32.56 | 2 | 44.76895 | yes | 4.009E-07 | 649.85681 | 63.187275 |
| VTCSDLGISGTVR | 13 | O95685 | Protein phosphatase 1 regulatory subunit | PPP1R3D | 32.56 | 2 | 44.76895 | yes | 2.076E-07 | 682.84558 | 41.478989 |
| LKETSENALK | 10 | O95757 | Heat shock 70 kDa protein 4L OS=Homo | HSPA4L | 94.513 | 2 | 47.029728 | yes | 1.599E-07 | 566.81403 | 101.37962 |
| SGGIETIANEYSDR | 14 | O95757 | Heat shock 70 kDa protein 4L OS=Homo | HSPA4L | 94.513 | 2 | 47.029728 | yes | 1.417E-07 | 756.35223 | 36.809532 |
| NAVEEYVYDFR | 11 | O95757 | Heat shock 70 kDa protein 4L OS=Homo | HSPA4L | 94.513 | 2 | 47.029728 | yes | 3.506E-06 | 702.82513 | 34.391602 |
| ATSILEFYR | 9 | P09001 | Large ribosomal subunit protein uL3m OS | MRPL3 | 38.633 | 2 | 50.042866 | yes | 1.083E-07 | 550.29279 | 1043.3026 |
| HVVTLLQVQDCH | 15 | P09001 | Large ribosomal subunit protein uL3m OS | MRPL3 | 38.633 | 3 | 50.042866 | yes | 1.001E-07 | 596.9978 | 881.15491 |
| LGMMPLWTK | 9 | P09001 | Large ribosomal subunit protein uL3m OS | MRPL3 | 38.633 | 2 | 50.042866 | yes | 1.849E-07 | 538.78522 | 683.2019 |
| PGQYVDVTAK | 10 | P09001 | Large ribosomal subunit protein uL3m OS | MRPL3 | 38.633 | 2 | 50.042866 | yes | 2.152E-07 | 539.28241 | 657.21527 |
| IFNITDNAAIKPGT | 22 | P09001 | Large ribosomal subunit protein uL3m OS | MRPL3 | 38.633 | 3 | 50.042866 | yes | 3.001E-07 | 810.7691 | 208.01367 |
| IFNITDNAAIKPGT | 22 | P09001 | Large ribosomal subunit protein uL3m OS | MRPL3 | 38.633 | 3 | 50.042866 | yes | 1.078E-07 | 608.32867 | 208.01367 |
| IFNITDNAAIKPGT | 22 | P09001 | Large ribosomal subunit protein uL3m OS | MRPL3 | 38.633 | 4 | 50.042866 | yes | 3.001E-07 | 810.7691 | 208.01367 |
| IFNITDNAAIKPGT | 22 | P09001 | Large ribosomal subunit protein uL3m OS | MRPL3 | 38.633 | 4 | 50.042866 | yes | 1.078E-07 | 608.32867 | 208.01367 |
| RPGAVATGDIGR | 12 | P09001 | Large ribosomal subunit protein uL3m OS | MRPL3 | 38.633 | 2 | 50.042866 | yes | 1.845E-07 | 585.32294 | 398.07587 |
| KATSILEFYR | 10 | P09001 | Large ribosomal subunit protein uL3m OS | MRPL3 | 38.633 | 2 | 50.042866 | yes | 1.075E-07 | 614.34021 | 318.5452 |
| IFNITDNAAIK | 11 | P09001 | Large ribosomal subunit protein uL3m OS | MRPL3 | 38.633 | 2 | 50.042866 | yes | 1.001E-07 | 610.33771 | 279.76999 |
| LCPLKDEPWPIHP | 20 | P09001 | Large ribosomal subunit protein uL3m OS | MRPL3 | 38.633 | 4 | 50.042866 | yes | 9.926E-08 | 616.05847 | 146.16856 |
| DEPWPIHPWEPGS | 15 | P09001 | Large ribosomal subunit protein uL3m OS | MRPL3 | 38.633 | 3 | 50.042866 | yes | 2.678E-07 | 617.2934 | 62.386723 |
| QLVSDEDKAQLAS | 14 | P09001 | Large ribosomal subunit protein uL3m OS | MRPL3 | 38.633 | 2 | 50.042866 | yes | 4.488E-06 | 766.40173 | 12.616667 |
| LSCAASGFTFSR | 12 | P0DOX5 | Immunoglobulin gamma-1 heavy chain O | -- | 49.329 | 2 | 45.490356 | yes | 1.834E-07 | 652.30859 | 218.8822 |
| PSSDNPNVLVLK | 12 | P16260 | Solute carrier family 25 member 16 OS=H | SLC25A16 | 36.224 | 2 | 35.557312 | yes | 8.144E-07 | 641.8537 | 18.395315 |
| EGGFFGFYR | 9 | P16260 | Solute carrier family 25 member 16 OS=H | SLC25A16 | 36.224 | 2 | 35.557312 | yes | 0.0002209 | 540.25092 | 11.285881 |
| NSYLILHR | 8 | P17029 | Zinc finger protein with KRAB and SCA | ZKSCAN1 | 63.631 | 2 | 48.86295 | yes | 1.094E-07 | 508.28781 | 118.18077 |
| EATGLSPQAAQEK | 13 | P17029 | Zinc finger protein with KRAB and SCA | ZKSCAN1 | 63.631 | 2 | 48.86295 | yes | 9.036E-07 | 665.33588 | 107.58032 |
| DQAMASALFTADS | 18 | P17029 | Zinc finger protein with KRAB and SCA | ZKSCAN1 | 63.631 | 3 | 48.86295 | yes | 6.786E-07 | 634.29749 | 93.357185 |

| | | | | | | | | | | | |
|---|---|---|---|---|---|---|---|---|---|---|---|
| AFSQQSQLVR | 10 | P17097 | Zinc finger protein 7 OS=Homo sapiens ( | ZNF7 | 77.887 | 2 | 34.756424 | yes | 9.47E-07 | 582.31201 | 60.031261 |
| TTCLFLHLR | 9 | P17098 | Zinc finger protein 8 OS=Homo sapiens ( | ZNF8 | 64.971 | 2 | 27.623072 | yes | 2.918E-06 | 580.81586 | 90.355019 |
| EGFPTDAPYPTTL( | 15 | P17098 | Zinc finger protein 8 OS=Homo sapiens ( | ZNF8 | 64.971 | 2 | 27.623072 | yes | 4.104E-06 | 797.39337 | 87.490669 |
| RPDVLFK | 7 | P31277 | Homeobox protein Hox-D11 OS=Homo s | HOXD11 | 35.198 | 2 | 48.69886 | yes | 1.113E-07 | 437.76089 | 190.54832 |
| THLMSEEEWR | 10 | P33552 | Cyclin-dependent kinases regulatory subu | CKS2 | 9.861 | 2 | 38.118275 | yes | 5.447E-07 | 659.29822 | 176.03932 |
| GNYDAAQR | 8 | P35542 | Serum amyloid A-4 protein OS=Homo sa | SAA4 | 14.747 | 2 | 40.02654 | yes | 4.249E-07 | 447.70685 | 115.3739 |
| ALGPEQLLR | 9 | P36915 | Guanine nucleotide-binding protein-like 1 | GNL1 | 68.661 | 2 | 46.05047 | yes | 1.783E-07 | 498.79547 | 184.15723 |
| VDLAPPALVVAW | 13 | P36915 | Guanine nucleotide-binding protein-like 1 | GNL1 | 68.661 | 2 | 46.05047 | yes | 5.081E-07 | 689.90826 | 61.40519 |
| IPVQALLHLR | 10 | P36915 | Guanine nucleotide-binding protein-like 1 | GNL1 | 68.661 | 2 | 46.05047 | yes | 2.946E-06 | 580.36914 | 45.061985 |
| YFQTYFLTPSVK | 12 | P36915 | Guanine nucleotide-binding protein-like 1 | GNL1 | 68.661 | 2 | 46.05047 | yes | 1.667E-07 | 747.38739 | 39.274086 |
| LNQQPSQGLGPR | 12 | P36915 | Guanine nucleotide-binding protein-like 1 | GNL1 | 68.661 | 2 | 46.05047 | yes | 3.975E-06 | 647.84674 | 19.533895 |
| GTWESHPETTELV | 18 | P36915 | Guanine nucleotide-binding protein-like 1 | GNL1 | 68.661 | 3 | 46.05047 | yes | 1.667E-06 | 680.3446 | 18.926405 |
| TVFVGNLPVTCNK | 13 | P42696 | RNA-binding protein 34 OS=Homo sapie | RBM34 | 48.565 | 2 | 49.926247 | yes | 9.976E-08 | 724.88196 | 246.87065 |
| SLIPAEGTLSK | 11 | P42696 | RNA-binding protein 34 OS=Homo sapie | RBM34 | 48.565 | 2 | 49.926247 | yes | 5.555E-07 | 558.31897 | 215.66002 |
| ILDDTEDTVVSQR | 13 | P42696 | RNA-binding protein 34 OS=Homo sapie | RBM34 | 48.565 | 2 | 49.926247 | yes | 2.583E-07 | 745.8703 | 156.66028 |
| VDLASETSSR | 10 | P42696 | RNA-binding protein 34 OS=Homo sapie | RBM34 | 48.565 | 2 | 49.926247 | yes | 6.72E-07 | 532.76459 | 127.97942 |
| LGQVASSLFR | 10 | P42696 | RNA-binding protein 34 OS=Homo sapie | RBM34 | 48.565 | 2 | 49.926247 | yes | 1.123E-07 | 539.30621 | 124.08421 |
| SVFVGNLPYK | 10 | P42696 | RNA-binding protein 34 OS=Homo sapie | RBM34 | 48.565 | 2 | 49.926247 | yes | 3.982E-07 | 562.31097 | 57.096771 |
| GLGATFILR | 9 | P49406 | Large ribosomal subunit protein bL19m ( | MRPL19 | 33.535 | 2 | 49.692272 | yes | 2.549E-07 | 474.28726 | 2096.1665 |
| ISQFLGICIQR | 11 | P49406 | Large ribosomal subunit protein bL19m ( | MRPL19 | 33.535 | 2 | 49.692272 | yes | 1.01E-07 | 667.86609 | 2069.2573 |
| FDLCLTEQQMK | 11 | P49406 | Large ribosomal subunit protein bL19m ( | MRPL19 | 33.535 | 2 | 49.692272 | yes | 1.166E-07 | 706.83105 | 1904.2151 |
| FLSPEFIPR | 9 | P49406 | Large ribosomal subunit protein bL19m ( | MRPL19 | 33.535 | 2 | 49.692272 | yes | 2.94E-07 | 553.30566 | 1717.4781 |
| VLHIPEFYVGSILR | 14 | P49406 | Large ribosomal subunit protein bL19m ( | MRPL19 | 33.535 | 2 | 49.692272 | yes | 1.05E-07 | 821.96942 | 908.44519 |
| LDDSLLYLR | 9 | P49406 | Large ribosomal subunit protein bL19m ( | MRPL19 | 33.535 | 2 | 49.692272 | yes | 5.68E-07 | 554.30585 | 357.72614 |
| RLDDSLLYLR | 10 | P49406 | Large ribosomal subunit protein bL19m ( | MRPL19 | 33.535 | 2 | 49.692272 | yes | 3.368E-07 | 632.35645 | 210.79974 |
| VTTADPYASGK | 11 | P49406 | Large ribosomal subunit protein bL19m ( | MRPL19 | 33.535 | 2 | 49.692272 | yes | 1.605E-07 | 555.27728 | 206.77908 |
| DALPEYSTFDVNM | 14 | P49406 | Large ribosomal subunit protein bL19m ( | MRPL19 | 33.535 | 2 | 49.692272 | yes | 1.851E-07 | 815.37689 | 158.18541 |
| PVVQEPNQK | 9 | P49406 | Large ribosomal subunit protein bL19m ( | MRPL19 | 33.535 | 2 | 49.692272 | yes | 7.927E-06 | 519.78253 | 148.54155 |
| AIGFVVGQTDWEK | 13 | P49750 | YLP motif-containing protein 1 OS=Hom | YLPM1 | 241.646 | 2 | 53.781284 | yes | 9.722E-08 | 725.37225 | 2256.4856 |
| MPLPAPSLSHQPPI | 17 | P49750 | YLP motif-containing protein 1 OS=Hom | YLPM1 | 241.646 | 3 | 53.781284 | yes | 1.311E-07 | 598.32239 | 1968.6071 |
| VEQIPYGER | 9 | P49750 | YLP motif-containing protein 1 OS=Hom | YLPM1 | 241.646 | 2 | 53.781284 | yes | 1.117E-07 | 545.78003 | 1441.7476 |
| ITDESGHLAEK | 11 | P49750 | YLP motif-containing protein 1 OS=Hom | YLPM1 | 241.646 | 2 | 53.781284 | yes | 9.722E-08 | 600.29877 | 1105.4183 |
| VGFQYQGIMQK | 11 | P49750 | YLP motif-containing protein 1 OS=Hom | YLPM1 | 241.646 | 2 | 53.781284 | yes | 1.022E-07 | 649.83173 | 981.61774 |
| GHEEFPLDGR | 10 | P49750 | YLP motif-containing protein 1 OS=Hom | YLPM1 | 241.646 | 2 | 53.781284 | yes | 9.722E-08 | 578.77271 | 915.9884 |
| DYQDDTLELYNR | 12 | P49750 | YLP motif-containing protein 1 OS=Hom | YLPM1 | 241.646 | 2 | 53.781284 | yes | 1.061E-07 | 772.8468 | 820.27759 |
| DKEVEFGGPAPR | 12 | P49750 | YLP motif-containing protein 1 OS=Hom | YLPM1 | 241.646 | 2 | 53.781284 | yes | 1.062E-07 | 651.32788 | 756.17603 |
| ITLRPDPLPER | 11 | P49750 | YLP motif-containing protein 1 OS=Hom | YLPM1 | 241.646 | 2 | 53.781284 | yes | 1.822E-07 | 653.8775 | 368.828 |
| ITLRPDPLPER | 11 | P49750 | YLP motif-containing protein 1 OS=Hom | YLPM1 | 241.646 | 2 | 53.781284 | yes | 9.722E-08 | 436.25412 | 368.828 |
| ITLRPDPLPER | 11 | P49750 | YLP motif-containing protein 1 OS=Hom | YLPM1 | 241.646 | 3 | 53.781284 | yes | 1.822E-07 | 653.8775 | 368.828 |

| | | | | | | | | | | | |
|---|---|---|---|---|---|---|---|---|---|---|---|
| ITLRPDPLPER | 11 | P49750 | YLP motif-containing protein 1 OS=Hom | YLPM1 | 241.646 | 3 | 53.781284 | yes | 9.722E-08 | 436.25412 | 368.828 |
| TTVQQEPLESGAK | 13 | P49750 | YLP motif-containing protein 1 OS=Hom | YLPM1 | 241.646 | 2 | 53.781284 | yes | 1.041E-07 | 694.35681 | 688.36835 |
| MADHWETAPR | 10 | P49750 | YLP motif-containing protein 1 OS=Hom | YLPM1 | 241.646 | 2 | 53.781284 | yes | 1.276E-07 | 607.2746 | 327.76633 |
| MADHWETAPR | 10 | P49750 | YLP motif-containing protein 1 OS=Hom | YLPM1 | 241.646 | 2 | 53.781284 | yes | 1.356E-07 | 405.18549 | 327.76633 |
| MADHWETAPR | 10 | P49750 | YLP motif-containing protein 1 OS=Hom | YLPM1 | 241.646 | 3 | 53.781284 | yes | 1.276E-07 | 607.2746 | 327.76633 |
| MADHWETAPR | 10 | P49750 | YLP motif-containing protein 1 OS=Hom | YLPM1 | 241.646 | 3 | 53.781284 | yes | 1.356E-07 | 405.18549 | 327.76633 |
| SQLLAPPPPSAPPC | 16 | P49750 | YLP motif-containing protein 1 OS=Hom | YLPM1 | 241.646 | 2 | 53.781284 | yes | 4.506E-07 | 785.93304 | 654.75397 |
| NVDDILKPPGR | 11 | P49750 | YLP motif-containing protein 1 OS=Hom | YLPM1 | 241.646 | 2 | 53.781284 | yes | 9.722E-08 | 612.34076 | 462.18973 |
| MQSAAFSIAADVK | 13 | P49750 | YLP motif-containing protein 1 OS=Hom | YLPM1 | 241.646 | 2 | 53.781284 | yes | 1.128E-07 | 669.83972 | 446.42972 |
| QHLLSLQQR | 9 | P49750 | YLP motif-containing protein 1 OS=Hom | YLPM1 | 241.646 | 2 | 53.781284 | yes | 6.481E-07 | 561.82257 | 436.70032 |
| APWNHGEER | 9 | P49750 | YLP motif-containing protein 1 OS=Hom | YLPM1 | 241.646 | 2 | 53.781284 | yes | 1.714E-07 | 548.25195 | 366.35944 |
| SSYLESPR | 8 | P49750 | YLP motif-containing protein 1 OS=Hom | YLPM1 | 241.646 | 2 | 53.781284 | yes | 5.405E-07 | 469.73254 | 360.55392 |
| HFDQFWSAAK | 10 | P49750 | YLP motif-containing protein 1 OS=Hom | YLPM1 | 241.646 | 2 | 53.781284 | yes | 6.544E-07 | 618.79346 | 338.27457 |
| RFEDLGSR | 8 | P49750 | YLP motif-containing protein 1 OS=Hom | YLPM1 | 241.646 | 2 | 53.781284 | yes | 2.58E-07 | 490.25143 | 337.15887 |
| SLDNEWDR | 8 | P49750 | YLP motif-containing protein 1 OS=Hom | YLPM1 | 241.646 | 2 | 53.781284 | yes | 1.806E-07 | 517.73053 | 337.09012 |
| STFETEHAGQR | 11 | P49750 | YLP motif-containing protein 1 OS=Hom | YLPM1 | 241.646 | 2 | 53.781284 | yes | 9.722E-08 | 631.79163 | 332.70746 |
| DREPYFDR | 8 | P49750 | YLP motif-containing protein 1 OS=Hom | YLPM1 | 241.646 | 2 | 53.781284 | yes | 4.952E-07 | 549.25415 | 324.10773 |
| AQAVTQPVPLANI | 24 | P49750 | YLP motif-containing protein 1 OS=Hom | YLPM1 | 241.646 | 3 | 53.781284 | yes | 2.279E-06 | 826.12347 | 315.67999 |
| WYPSDVDR | 8 | P49750 | YLP motif-containing protein 1 OS=Hom | YLPM1 | 241.646 | 2 | 53.781284 | yes | 1.001E-07 | 519.23798 | 293.2066 |
| QSNVIADHR | 9 | P49750 | YLP motif-containing protein 1 OS=Hom | YLPM1 | 241.646 | 2 | 53.781284 | yes | 1.452E-07 | 520.26758 | 249.21376 |
| EPLADTSSNQQK | 12 | P49750 | YLP motif-containing protein 1 OS=Hom | YLPM1 | 241.646 | 2 | 53.781284 | yes | 9.722E-08 | 659.31769 | 244.23622 |
| EQHLAQLQQLQQI | 16 | P49750 | YLP motif-containing protein 1 OS=Hom | YLPM1 | 241.646 | 3 | 53.781284 | yes | 9.722E-08 | 663.34253 | 240.60254 |
| PMDMYDR | 7 | P49750 | YLP motif-containing protein 1 OS=Hom | YLPM1 | 241.646 | 2 | 53.781284 | yes | 9.722E-08 | 464.1886 | 237.48918 |
| DWEAIASR | 8 | P49750 | YLP motif-containing protein 1 OS=Hom | YLPM1 | 241.646 | 2 | 53.781284 | yes | 1.252E-07 | 474.2327 | 226.61935 |
| PAFGQQHQQQPK | 12 | P49750 | YLP motif-containing protein 1 OS=Hom | YLPM1 | 241.646 | 3 | 53.781284 | yes | 1.113E-07 | 465.23721 | 111.28529 |
| PAFGQQHQQQPK | 12 | P49750 | YLP motif-containing protein 1 OS=Hom | YLPM1 | 241.646 | 3 | 53.781284 | yes | 2.423E-07 | 697.35217 | 111.28529 |
| PAFGQQHQQQPK | 12 | P49750 | YLP motif-containing protein 1 OS=Hom | YLPM1 | 241.646 | 2 | 53.781284 | yes | 1.113E-07 | 465.23721 | 111.28529 |
| PAFGQQHQQQPK | 12 | P49750 | YLP motif-containing protein 1 OS=Hom | YLPM1 | 241.646 | 2 | 53.781284 | yes | 2.423E-07 | 697.35217 | 111.28529 |
| DQLQEYEK | 8 | P49750 | YLP motif-containing protein 1 OS=Hom | YLPM1 | 241.646 | 2 | 53.781284 | yes | 2.056E-07 | 526.74835 | 219.30099 |
| WADLEEK | 7 | P49750 | YLP motif-containing protein 1 OS=Hom | YLPM1 | 241.646 | 2 | 53.781284 | yes | 1.456E-07 | 445.71634 | 214.66414 |
| SKWEMDTSEAK | 11 | P49750 | YLP motif-containing protein 1 OS=Hom | YLPM1 | 241.646 | 2 | 53.781284 | yes | 9.722E-08 | 656.29791 | 212.99251 |
| VNSFQNMK | 8 | P49750 | YLP motif-containing protein 1 OS=Hom | YLPM1 | 241.646 | 2 | 53.781284 | yes | 1.017E-07 | 484.23694 | 211.58104 |
| EKVPGGLQGSQDI | 13 | P49750 | YLP motif-containing protein 1 OS=Hom | YLPM1 | 241.646 | 2 | 53.781284 | yes | 1.765E-07 | 685.8548 | 208.29393 |
| GLVRPGSSR | 9 | P49750 | YLP motif-containing protein 1 OS=Hom | YLPM1 | 241.646 | 2 | 53.781284 | yes | 9.722E-08 | 464.76978 | 196.16478 |
| ATQSYLQEK | 9 | P49750 | YLP motif-containing protein 1 OS=Hom | YLPM1 | 241.646 | 2 | 53.781284 | yes | 8.436E-07 | 534.27203 | 184.65894 |
| ERLDDWDR | 8 | P49750 | YLP motif-containing protein 1 OS=Hom | YLPM1 | 241.646 | 2 | 53.781284 | yes | 2.261E-07 | 552.75708 | 158.14401 |
| DQDMDEDYNR | 10 | P49750 | YLP motif-containing protein 1 OS=Hom | YLPM1 | 241.646 | 2 | 53.781284 | yes | 9.752E-08 | 650.74097 | 134.30095 |
| GPWWDDWER | 9 | P49750 | YLP motif-containing protein 1 OS=Hom | YLPM1 | 241.646 | 2 | 53.781284 | yes | 9.722E-08 | 623.76746 | 117.12276 |
| WEMDTSEAK | 9 | P49750 | YLP motif-containing protein 1 OS=Hom | YLPM1 | 241.646 | 2 | 53.781284 | yes | 1.311E-07 | 548.73444 | 112.90322 |

| | | | | | | | | | | | |
|---|---|---|---|---|---|---|---|---|---|---|---|
| ISRPMDMYDR | 10 | P49750 | YLP motif-containing protein 1 OS=Hom | YLPM1 | 241.646 | 2 | 53.781284 | yes | 9.722E-08 | 642.29718 | 108.27055 |
| QHEIILK | 7 | P49750 | YLP motif-containing protein 1 OS=Hom | YLPM1 | 241.646 | 2 | 53.781284 | yes | 1.383E-07 | 440.76617 | 107.23045 |
| VLSLDDYFITEVEI | 14 | P49750 | YLP motif-containing protein 1 OS=Hom | YLPM1 | 241.646 | 2 | 53.781284 | yes | 1.724E-07 | 835.92981 | 104.97554 |
| LDKLDGLR | 8 | P49750 | YLP motif-containing protein 1 OS=Hom | YLPM1 | 241.646 | 2 | 53.781284 | yes | 3.404E-07 | 465.27435 | 80.347443 |
| VPGGLQGSQDR | 11 | P49750 | YLP motif-containing protein 1 OS=Hom | YLPM1 | 241.646 | 2 | 53.781284 | yes | 3.719E-07 | 557.28601 | 58.192894 |
| PLDEQESQFR | 10 | P49750 | YLP motif-containing protein 1 OS=Hom | YLPM1 | 241.646 | 2 | 53.781284 | yes | 2.613E-07 | 624.79639 | 56.549583 |
| DRGPFRPEPGDGG | 15 | P49750 | YLP motif-containing protein 1 OS=Hom | YLPM1 | 241.646 | 3 | 53.781284 | yes | 1.08E-06 | 538.59375 | 19.437691 |
| DRGPFRPEPGDGG | 15 | P49750 | YLP motif-containing protein 1 OS=Hom | YLPM1 | 241.646 | 3 | 53.781284 | yes | 3.611E-07 | 807.38696 | 19.437691 |
| DRGPFRPEPGDGG | 15 | P49750 | YLP motif-containing protein 1 OS=Hom | YLPM1 | 241.646 | 2 | 53.781284 | yes | 1.08E-06 | 538.59375 | 19.437691 |
| DRGPFRPEPGDGG | 15 | P49750 | YLP motif-containing protein 1 OS=Hom | YLPM1 | 241.646 | 2 | 53.781284 | yes | 3.611E-07 | 807.38696 | 19.437691 |
| VLSLDDYFITEVEI | 17 | P49750 | YLP motif-containing protein 1 OS=Hom | YLPM1 | 241.646 | 3 | 53.781284 | yes | 1.196E-06 | 686.349 | 34.282288 |
| DAEEEESELGYIPK | 14 | P49750 | YLP motif-containing protein 1 OS=Hom | YLPM1 | 241.646 | 2 | 53.781284 | yes | 9.997E-08 | 804.86743 | 34.030811 |
| ELAEDDSILK | 10 | P56385 | ATP synthase subunit e, mitochondrial O: | ATP5ME | 7.934 | 2 | 50.442577 | yes | 9.809E-08 | 566.79022 | 269.65817 |
| YNYLKPR | 7 | P56385 | ATP synthase subunit e, mitochondrial O: | ATP5ME | 7.934 | 2 | 50.442577 | yes | 7.975E-07 | 477.26379 | 113.22848 |
| VPPVQVSPLIK | 11 | P56385 | ATP synthase subunit e, mitochondrial O: | ATP5ME | 7.934 | 2 | 50.442577 | yes | 6.213E-07 | 588.87115 | 80.349503 |
| YSALFLGVAYGAT | 14 | P56385 | ATP synthase subunit e, mitochondrial O: | ATP5ME | 7.934 | 2 | 50.442577 | yes | 1.865E-07 | 744.89594 | 79.494339 |
| FTSCDLLWR | 9 | P57775 | F-box/WD repeat-containing protein 4 OS | FBXW4 | 46.338 | 2 | 48.56058 | yes | 1.13E-07 | 599.28967 | 95.362541 |
| LGTDLMTSVPVK | 12 | P57775 | F-box/WD repeat-containing protein 4 OS | FBXW4 | 46.338 | 2 | 48.56058 | yes | 4.285E-07 | 630.84705 | 52.887585 |
| FYLLTEHFK | 9 | P58317 | Zinc finger protein 121 OS=Homo sapien | ZNF121 | 44.695 | 2 | 54.778767 | yes | 1.05E-07 | 599.31879 | 1005.7311 |
| IHTGEKPYECK | 11 | P58317 | Zinc finger protein 121 OS=Homo sapien | ZNF121 | 44.695 | 3 | 54.778767 | no | 1.1E-07 | 454.55545 | 373.01508 |
| THTGEKPYICK | 11 | P58317 | Zinc finger protein 121 OS=Homo sapien | ZNF121 | 44.695 | 3 | 54.778767 | no | 1.814E-07 | 445.22382 | 161.70979 |
| THTGEKPYICK | 11 | P58317 | Zinc finger protein 121 OS=Homo sapien | ZNF121 | 44.695 | 3 | 54.778767 | no | 2.487E-07 | 667.33209 | 161.70979 |
| THTGEKPYICK | 11 | P58317 | Zinc finger protein 121 OS=Homo sapien | ZNF121 | 44.695 | 2 | 54.778767 | no | 1.814E-07 | 445.22382 | 161.70979 |
| THTGEKPYICK | 11 | P58317 | Zinc finger protein 121 OS=Homo sapien | ZNF121 | 44.695 | 2 | 54.778767 | no | 2.487E-07 | 667.33209 | 161.70979 |
| SFEYSDCEEAFVD | 21 | P58317 | Zinc finger protein 121 OS=Homo sapien | ZNF121 | 44.695 | 3 | 54.778767 | yes | 1.693E-07 | 844.69592 | 314.08264 |
| AFTYSTSHAVSVK | 13 | P58317 | Zinc finger protein 121 OS=Homo sapien | ZNF121 | 44.695 | 2 | 54.778767 | yes | 9.722E-08 | 699.35663 | 309.31552 |
| YSSYLNSHMR | 10 | P58317 | Zinc finger protein 121 OS=Homo sapien | ZNF121 | 44.695 | 2 | 54.778767 | yes | 9.722E-08 | 629.28766 | 220.05919 |
| IHTGIKPYK | 9 | P58317 | Zinc finger protein 121 OS=Homo sapien | ZNF121 | 44.695 | 2 | 54.778767 | no | 1.004E-07 | 528.81366 | 197.29263 |
| ITHNGETLYEQK | 12 | P58317 | Zinc finger protein 121 OS=Homo sapien | ZNF121 | 44.695 | 2 | 54.778767 | yes | 1.28E-07 | 716.85699 | 189.67134 |
| AFTVSSSLHNHVK | 13 | P58317 | Zinc finger protein 121 OS=Homo sapien | ZNF121 | 44.695 | 2 | 54.778767 | yes | 1.084E-07 | 713.87549 | 169.38754 |
| MHTVEKPYECK | 11 | P58317 | Zinc finger protein 121 OS=Homo sapien | ZNF121 | 44.695 | 2 | 54.778767 | yes | 9.722E-08 | 711.33124 | 83.915291 |
| MHTVEKPYECK | 11 | P58317 | Zinc finger protein 121 OS=Homo sapien | ZNF121 | 44.695 | 2 | 54.778767 | yes | 9.722E-08 | 474.55658 | 83.915291 |
| MHTVEKPYECK | 11 | P58317 | Zinc finger protein 121 OS=Homo sapien | ZNF121 | 44.695 | 3 | 54.778767 | yes | 9.722E-08 | 711.33124 | 83.915291 |
| MHTVEKPYECK | 11 | P58317 | Zinc finger protein 121 OS=Homo sapien | ZNF121 | 44.695 | 3 | 54.778767 | yes | 9.722E-08 | 474.55658 | 83.915291 |
| THTEEKPFECK | 11 | P58317 | Zinc finger protein 121 OS=Homo sapien | ZNF121 | 44.695 | 2 | 54.778767 | yes | 9.722E-08 | 703.32446 | 104.23055 |
| PYICNECGK | 9 | P58317 | Zinc finger protein 121 OS=Homo sapien | ZNF121 | 44.695 | 2 | 54.778767 | no | 1.447E-07 | 570.74426 | 58.254593 |
| THTGEKPYECK | 11 | P58317 | Zinc finger protein 121 OS=Homo sapien | ZNF121 | 44.695 | 3 | 54.778767 | no | 3.619E-07 | 450.54333 | 35.043198 |
| NIQHINR | 7 | P58557 | Endoribonuclease YbeY OS=Homo sapie | YBEY | 19.299 | 2 | 54.305447 | yes | 4.844E-07 | 447.74884 | 218.91628 |
| FDLGIICVDNK | 11 | P58557 | Endoribonuclease YbeY OS=Homo sapie | YBEY | 19.299 | 2 | 54.305447 | yes | 9.722E-08 | 647.32904 | 122.60685 |

| | | | | | | | | | | | |
|---|---|---|---|---|---|---|---|---|---|---|---|
| AVLDELGR | 8 | P58557 | Endoribonuclease YbeY OS=Homo sapie | YBEY | 19.299 | 2 | 54.305447 | yes | 2.294E-07 | 436.74545 | 101.87605 |
| TLQYLSQGNVVFK | 13 | P82663 | Small ribosomal subunit protein mS25 OS | MRPS25 | 20.117 | 2 | 53.215355 | yes | 9.722E-08 | 748.909 | 1515.9812 |
| EIMEHIR | 7 | P82663 | Small ribosomal subunit protein mS25 OS | MRPS25 | 20.117 | 2 | 53.215355 | yes | 1.871E-07 | 464.23947 | 522.57123 |
| QLSHPANFGPR | 11 | P82663 | Small ribosomal subunit protein mS25 OS | MRPS25 | 20.117 | 2 | 53.215355 | yes | 9.722E-08 | 612.31763 | 145.75339 |
| QLSHPANFGPR | 11 | P82663 | Small ribosomal subunit protein mS25 OS | MRPS25 | 20.117 | 2 | 53.215355 | yes | 2.318E-07 | 408.54752 | 145.75339 |
| QLSHPANFGPR | 11 | P82663 | Small ribosomal subunit protein mS25 OS | MRPS25 | 20.117 | 3 | 53.215355 | yes | 9.722E-08 | 612.31763 | 145.75339 |
| QLSHPANFGPR | 11 | P82663 | Small ribosomal subunit protein mS25 OS | MRPS25 | 20.117 | 3 | 53.215355 | yes | 2.318E-07 | 408.54752 | 145.75339 |
| VMTVNYNTHGEL | 17 | P82663 | Small ribosomal subunit protein mS25 OS | MRPS25 | 20.117 | 3 | 53.215355 | yes | 1.388E-07 | 616.62836 | 230.59563 |
| KFVFFNIPQIQYK | 13 | P82663 | Small ribosomal subunit protein mS25 OS | MRPS25 | 20.117 | 2 | 53.215355 | yes | 9.722E-08 | 836.46649 | 84.787628 |
| HFPIEIDSTDYVSS | 19 | P82673 | Small ribosomal subunit protein mS35 OS | MRPS35 | 36.845 | 3 | 50.938015 | yes | 9.722E-08 | 702.67615 | 647.60553 |
| LSSLNLDDHAK | 11 | P82673 | Small ribosomal subunit protein mS35 OS | MRPS35 | 36.845 | 2 | 50.938015 | yes | 1.319E-07 | 606.81458 | 598.80194 |
| IPNFLHLTPVAIK | 13 | P82673 | Small ribosomal subunit protein mS35 OS | MRPS35 | 36.845 | 2 | 50.938015 | yes | 9.734E-08 | 731.94269 | 142.36673 |
| IPNFLHLTPVAIK | 13 | P82673 | Small ribosomal subunit protein mS35 OS | MRPS35 | 36.845 | 2 | 50.938015 | yes | 5.388E-07 | 488.29755 | 142.36673 |
| IPNFLHLTPVAIK | 13 | P82673 | Small ribosomal subunit protein mS35 OS | MRPS35 | 36.845 | 3 | 50.938015 | yes | 9.734E-08 | 731.94269 | 142.36673 |
| IPNFLHLTPVAIK | 13 | P82673 | Small ribosomal subunit protein mS35 OS | MRPS35 | 36.845 | 3 | 50.938015 | yes | 5.388E-07 | 488.29755 | 142.36673 |
| PSAVPLPVR | 9 | P82673 | Small ribosomal subunit protein mS35 OS | MRPS35 | 36.845 | 2 | 50.938015 | yes | 1.312E-07 | 468.28726 | 45.081322 |
| EGNLELLK | 8 | P82673 | Small ribosomal subunit protein mS35 OS | MRPS35 | 36.845 | 2 | 50.938015 | yes | 2.411E-06 | 458.26093 | 44.718689 |
| TTDVLTIK | 8 | P82673 | Small ribosomal subunit protein mS35 OS | MRPS35 | 36.845 | 2 | 50.938015 | yes | 1.248E-05 | 445.76331 | 35.354752 |
| NEEENENSISQYK | 13 | P82673 | Small ribosomal subunit protein mS35 OS | MRPS35 | 36.845 | 2 | 50.938015 | yes | 1.942E-07 | 792.3446 | 23.399227 |
| EECGPLPIVVASPR | 14 | P82675 | Small ribosomal subunit protein uS5m OS | MRPS5 | 48.007 | 2 | 55.906605 | yes | 9.79E-08 | 762.39795 | 1554.2139 |
| GLHVVEIR | 8 | P82675 | Small ribosomal subunit protein uS5m OS | MRPS5 | 48.007 | 2 | 55.906605 | yes | 1.269E-07 | 461.77707 | 487.66754 |
| YEDHTIFHDISLR | 13 | P82675 | Small ribosomal subunit protein uS5m OS | MRPS5 | 48.007 | 3 | 55.906605 | yes | 9.758E-08 | 549.27051 | 234.12836 |
| YEDHTIFHDISLR | 13 | P82675 | Small ribosomal subunit protein uS5m OS | MRPS5 | 48.007 | 3 | 55.906605 | yes | 9.722E-08 | 823.4021 | 234.12836 |
| YEDHTIFHDISLR | 13 | P82675 | Small ribosomal subunit protein uS5m OS | MRPS5 | 48.007 | 2 | 55.906605 | yes | 9.758E-08 | 549.27051 | 234.12836 |
| YEDHTIFHDISLR | 13 | P82675 | Small ribosomal subunit protein uS5m OS | MRPS5 | 48.007 | 2 | 55.906605 | yes | 9.722E-08 | 823.4021 | 234.12836 |
| LTADELWK | 8 | P82675 | Small ribosomal subunit protein uS5m OS | MRPS5 | 48.007 | 2 | 55.906605 | yes | 1.261E-07 | 488.26093 | 366.76376 |
| AVHHLHYIER | 10 | P82675 | Small ribosomal subunit protein uS5m OS | MRPS5 | 48.007 | 2 | 55.906605 | yes | 9.809E-08 | 637.84125 | 316.1593 |
| KDPEPEDEVPDVK | 13 | P82675 | Small ribosomal subunit protein uS5m OS | MRPS5 | 48.007 | 2 | 55.906605 | yes | 4.779E-07 | 748.85938 | 208.65282 |
| GALAETGAGAK | 11 | P82675 | Small ribosomal subunit protein uS5m OS | MRPS5 | 48.007 | 2 | 55.906605 | yes | 3.846E-06 | 473.25363 | 120.45689 |
| VSGSINMLSLTQGI | 16 | P82675 | Small ribosomal subunit protein uS5m OS | MRPS5 | 48.007 | 3 | 55.906605 | yes | 2.072E-06 | 580.31006 | 51.861404 |
| DPEPEDEVPDVK | 12 | P82675 | Small ribosomal subunit protein uS5m OS | MRPS5 | 48.007 | 2 | 55.906605 | yes | 4.357E-07 | 684.81189 | 25.878494 |
| FSIYPPIPGEESSLR | 15 | P82912 | Small ribosomal subunit protein uS11m C | MRPS11 | 20.616 | 2 | 55.307507 | yes | 6.822E-07 | 846.43561 | 966.40039 |
| GTGIAAQTAGIAA | 16 | P82912 | Small ribosomal subunit protein uS11m C | MRPS11 | 20.616 | 2 | 55.307507 | yes | 1.4E-07 | 700.38623 | 397.33936 |
| KGTGIAAQTAGIA | 17 | P82912 | Small ribosomal subunit protein uS11m C | MRPS11 | 20.616 | 2 | 55.307507 | yes | 9.722E-08 | 764.43372 | 75.20211 |
| KGTGIAAQTAGIA | 17 | P82912 | Small ribosomal subunit protein uS11m C | MRPS11 | 20.616 | 2 | 55.307507 | yes | 4.646E-07 | 509.95825 | 75.20211 |
| KGTGIAAQTAGIA | 17 | P82912 | Small ribosomal subunit protein uS11m C | MRPS11 | 20.616 | 3 | 55.307507 | yes | 9.722E-08 | 764.43372 | 75.20211 |
| KGTGIAAQTAGIA | 17 | P82912 | Small ribosomal subunit protein uS11m C | MRPS11 | 20.616 | 3 | 55.307507 | yes | 4.646E-07 | 509.95825 | 75.20211 |
| KFEEIPIAHIK | 11 | P82912 | Small ribosomal subunit protein uS11m C | MRPS11 | 20.616 | 2 | 55.307507 | yes | 2.765E-06 | 662.88483 | 148.08417 |
| FEEIPIAHIK | 10 | P82912 | Small ribosomal subunit protein uS11m C | MRPS11 | 20.616 | 2 | 55.307507 | yes | 4.027E-06 | 598.83734 | 64.943977 |

| | | | | | | | | | | | |
|---|---|---|---|---|---|---|---|---|---|---|---|
| TVMVQEGNVESA | 14 | P82921 | Small ribosomal subunit protein bS21m C | MRPS21 | 10.689 | 2 | 53.760036 | yes | 9.722E-08 | 791.88013 | 323.45233 |
| KINFLMR | 7 | P82921 | Small ribosomal subunit protein bS21m C | MRPS21 | 10.689 | 2 | 53.760036 | yes | 8.169E-07 | 461.27057 | 122.93343 |
| IYNMEMAR | 8 | P82921 | Small ribosomal subunit protein bS21m C | MRPS21 | 10.689 | 2 | 53.760036 | yes | 2.378E-07 | 514.23859 | 75.573784 |
| HPEQIFPR | 8 | P82933 | Small ribosomal subunit protein uS9m OS | MRPS9 | 45.835 | 2 | 52.97128 | yes | 2.463E-07 | 512.27216 | 1239.0441 |
| HDVTCTVSGGGR | 12 | P82933 | Small ribosomal subunit protein uS9m OS | MRPS9 | 45.835 | 2 | 52.97128 | yes | 9.722E-08 | 623.28564 | 843.11707 |
| QAGLLTTDPR | 10 | P82933 | Small ribosomal subunit protein uS9m OS | MRPS9 | 45.835 | 2 | 52.97128 | yes | 1.794E-07 | 536.29327 | 836.17554 |
| LSDLDYMQFIR | 11 | P82933 | Small ribosomal subunit protein uS9m OS | MRPS9 | 45.835 | 2 | 52.97128 | yes | 9.722E-08 | 700.8476 | 556.27136 |
| RETYTEDFIK | 10 | P82933 | Small ribosomal subunit protein uS9m OS | MRPS9 | 45.835 | 2 | 52.97128 | yes | 1.121E-07 | 651.32227 | 301.62384 |
| EQLMFPFHFVDR | 12 | P82933 | Small ribosomal subunit protein uS9m OS | MRPS9 | 45.835 | 2 | 52.97128 | yes | 1.215E-07 | 783.38214 | 231.55199 |
| QIEEFNIGK | 9 | P82933 | Small ribosomal subunit protein uS9m OS | MRPS9 | 45.835 | 2 | 52.97128 | yes | 1.466E-07 | 539.28241 | 225.70856 |
| EELEEMLVEK | 10 | P82933 | Small ribosomal subunit protein uS9m OS | MRPS9 | 45.835 | 2 | 52.97128 | yes | 1.121E-07 | 624.80505 | 170.72585 |
| ETYTEDFIK | 9 | P82933 | Small ribosomal subunit protein uS9m OS | MRPS9 | 45.835 | 2 | 52.97128 | yes | 8.315E-07 | 573.27167 | 87.617134 |
| LGCLLER | 7 | Q02509 | Otoconin-90 OS=Homo sapiens OX=960 | OC90 | 51.728 | 2 | 52.113342 | yes | 9.722E-08 | 430.73657 | 148.39952 |
| MLSALLTGVNR | 11 | Q03701 | CCAAT/enhancer-binding protein zeta OS | CEBPZ | 120.974 | 2 | 54.474964 | yes | 1.335E-07 | 587.83429 | 382.41779 |
| AIVSSGTLGDR | 11 | Q03701 | CCAAT/enhancer-binding protein zeta OS | CEBPZ | 120.974 | 2 | 54.474964 | yes | 1.059E-07 | 538.29077 | 301.94104 |
| ALLVQVVNK | 9 | Q03701 | CCAAT/enhancer-binding protein zeta OS | CEBPZ | 120.974 | 2 | 54.474964 | yes | 2.127E-07 | 492.31604 | 122.74243 |
| ASHLLETLLCK | 11 | Q03701 | CCAAT/enhancer-binding protein zeta OS | CEBPZ | 120.974 | 2 | 54.474964 | yes | 4.865E-07 | 642.85266 | 117.24738 |
| KPEVASWVHFDN | 14 | Q03701 | CCAAT/enhancer-binding protein zeta OS | CEBPZ | 120.974 | 3 | 54.474964 | yes | 1.974E-07 | 557.29474 | 112.52937 |
| ELLITDLLPDNR | 12 | Q03701 | CCAAT/enhancer-binding protein zeta OS | CEBPZ | 120.974 | 2 | 54.474964 | yes | 9.722E-08 | 706.39319 | 106.93423 |
| MLDPGLMTCSK | 11 | Q03701 | CCAAT/enhancer-binding protein zeta OS | CEBPZ | 120.974 | 2 | 54.474964 | yes | 9.722E-08 | 626.79034 | 99.826736 |
| IFSQRPFDK | 9 | Q03701 | CCAAT/enhancer-binding protein zeta OS | CEBPZ | 120.974 | 2 | 54.474964 | yes | 3.74E-07 | 569.30621 | 84.287956 |
| QAMFLNLVYK | 10 | Q03701 | CCAAT/enhancer-binding protein zeta OS | CEBPZ | 120.974 | 2 | 54.474964 | yes | 9.722E-08 | 613.83374 | 78.443588 |
| ENTDSVVMQPK | 11 | Q03701 | CCAAT/enhancer-binding protein zeta OS | CEBPZ | 120.974 | 2 | 54.474964 | yes | 1.901E-07 | 624.30048 | 63.571342 |
| QTLLLRPGGK | 10 | Q03701 | CCAAT/enhancer-binding protein zeta OS | CEBPZ | 120.974 | 2 | 54.474964 | yes | 3.064E-07 | 541.83765 | 60.989677 |
| QQCLMALDTFK | 11 | Q03701 | CCAAT/enhancer-binding protein zeta OS | CEBPZ | 120.974 | 2 | 54.474964 | yes | 4.368E-06 | 677.82831 | 56.26424 |
| QNIFEFFER | 9 | Q03701 | CCAAT/enhancer-binding protein zeta OS | CEBPZ | 120.974 | 2 | 54.474964 | yes | 9.722E-08 | 615.30109 | 52.427971 |
| GTDDFDFAGSFQG | 15 | Q03701 | CCAAT/enhancer-binding protein zeta OS | CEBPZ | 120.974 | 2 | 54.474964 | yes | 1.642E-07 | 808.85242 | 40.611641 |
| EPLEFHAK | 8 | Q03701 | CCAAT/enhancer-binding protein zeta OS | CEBPZ | 120.974 | 2 | 54.474964 | yes | 4.507E-07 | 485.75327 | 39.704727 |
| AYPYSQTGDDK | 11 | Q03701 | CCAAT/enhancer-binding protein zeta OS | CEBPZ | 120.974 | 2 | 54.474964 | yes | 9.735E-07 | 622.77515 | 39.017738 |
| YIQEAEMLR | 9 | Q05048 | Cleavage stimulation factor subunit 1 OS= | CSTF1 | 48.358 | 2 | 44.124714 | yes | 3.135E-07 | 576.78955 | 458.5954 |
| LWEISTGR | 8 | Q05048 | Cleavage stimulation factor subunit 1 OS= | CSTF1 | 48.358 | 2 | 44.124714 | yes | 3.458E-07 | 481.25873 | 118.36916 |
| AHDGAEVCSAIFS | 14 | Q05048 | Cleavage stimulation factor subunit 1 OS= | CSTF1 | 48.358 | 2 | 44.124714 | yes | 4.986E-07 | 746.34845 | 90.374596 |
| SISFHPSGDFILVG | 21 | Q05048 | Cleavage stimulation factor subunit 1 OS= | CSTF1 | 48.358 | 3 | 44.124714 | yes | 2.312E-07 | 770.40607 | 73.554413 |
| TISLCCWDSR | 10 | Q05048 | Cleavage stimulation factor subunit 1 OS= | CSTF1 | 48.358 | 2 | 44.124714 | yes | 2.54E-07 | 649.2868 | 70.566788 |
| FTVLFPSGTPLPAR | 14 | Q0VDF9 | Heat shock 70 kDa protein 14 OS=Homo | HSPA14 | 54.795 | 2 | 51.829086 | yes | 1.13E-07 | 751.92194 | 472.59683 |
| FELLCSPLFNK | 11 | Q0VDF9 | Heat shock 70 kDa protein 14 OS=Homo | HSPA14 | 54.795 | 2 | 51.829086 | yes | 9.722E-08 | 684.35504 | 208.99829 |
| YEIDTGEETK | 10 | Q0VDF9 | Heat shock 70 kDa protein 14 OS=Homo | HSPA14 | 54.795 | 2 | 51.829086 | yes | 1.799E-07 | 592.76953 | 194.38321 |
| AAGFNVLR | 8 | Q0VDF9 | Heat shock 70 kDa protein 14 OS=Homo | HSPA14 | 54.795 | 2 | 51.829086 | yes | 4.075E-07 | 424.24286 | 166.69051 |
| FAQVVLQDLDK | 11 | Q0VDF9 | Heat shock 70 kDa protein 14 OS=Homo | HSPA14 | 54.795 | 2 | 51.829086 | yes | 2.049E-07 | 638.35083 | 86.274368 |

| | | | | | | | | | | | |
|---|---|---|---|---|---|---|---|---|---|---|---|
| FVNPEDVAR | 9 | Q0VDF9 | Heat shock 70 kDa protein 14 OS=Homo | HSPA14 | 54.795 | 2 | 51.829086 | yes | 1.227E-06 | 523.76691 | 69.881897 |
| LIHEPSAALLAYGI | 22 | Q0VDF9 | Heat shock 70 kDa protein 14 OS=Homo | HSPA14 | 54.795 | 3 | 51.829086 | yes | 0.0001919 | 746.73035 | 64.701523 |
| GLLDQNGFTADDI | 15 | Q0VDF9 | Heat shock 70 kDa protein 14 OS=Homo | HSPA14 | 54.795 | 2 | 51.829086 | yes | 3.127E-07 | 810.89685 | 44.450554 |
| IEALCDEER | 9 | Q12872 | Splicing factor, suppressor of white-aprico | SFSWAP | 104.822 | 2 | 48.84126 | yes | 1.362E-07 | 567.75842 | 166.7365 |
| HSLPSAYR | 8 | Q12872 | Splicing factor, suppressor of white-aprico | SFSWAP | 104.822 | 2 | 48.84126 | yes | 3.484E-06 | 465.74323 | 142.85312 |
| DYQLSEEEAR | 10 | Q12872 | Splicing factor, suppressor of white-aprico | SFSWAP | 104.822 | 2 | 48.84126 | yes | 2.043E-07 | 620.27802 | 112.29815 |
| VVDPDHPLAALVI | 13 | Q12872 | Splicing factor, suppressor of white-aprico | SFSWAP | 104.822 | 2 | 48.84126 | yes | 3.31E-07 | 701.39606 | 50.855225 |
| VVDPDHPLAALVI | 13 | Q12872 | Splicing factor, suppressor of white-aprico | SFSWAP | 104.822 | 2 | 48.84126 | yes | 2.867E-05 | 467.93314 | 50.855225 |
| VVDPDHPLAALVI | 13 | Q12872 | Splicing factor, suppressor of white-aprico | SFSWAP | 104.822 | 3 | 48.84126 | yes | 3.31E-07 | 701.39606 | 50.855225 |
| VVDPDHPLAALVI | 13 | Q12872 | Splicing factor, suppressor of white-aprico | SFSWAP | 104.822 | 3 | 48.84126 | yes | 2.867E-05 | 467.93314 | 50.855225 |
| EAQISSAIVSSVQS | 15 | Q12872 | Splicing factor, suppressor of white-aprico | SFSWAP | 104.822 | 2 | 48.84126 | yes | 1.097E-07 | 767.40961 | 98.304657 |
| LEELMKPLK | 9 | Q12872 | Splicing factor, suppressor of white-aprico | SFSWAP | 104.822 | 2 | 48.84126 | yes | 1.768E-07 | 550.82281 | 95.841179 |
| NSQFDFLR | 8 | Q12872 | Splicing factor, suppressor of white-aprico | SFSWAP | 104.822 | 2 | 48.84126 | yes | 4.796E-07 | 513.75378 | 60.696911 |
| VELLVFGYACK | 11 | Q12872 | Splicing factor, suppressor of white-aprico | SFSWAP | 104.822 | 2 | 48.84126 | yes | 1.952E-07 | 649.8443 | 39.785248 |
| KPQLTQEELEAK | 12 | Q12872 | Splicing factor, suppressor of white-aprico | SFSWAP | 104.822 | 2 | 48.84126 | yes | 1.73E-07 | 707.38281 | 9.0698185 |
| LLPDFLLER | 9 | Q12894 | Interferon-related developmental regulato | IFRD2 | 48.048 | 2 | 50.651367 | yes | 1.067E-07 | 558.3266 | 314.15921 |
| LPQLLSSESVNLR | 13 | Q12894 | Interferon-related developmental regulato | IFRD2 | 48.048 | 2 | 50.651367 | yes | 1.263E-07 | 728.41193 | 297.3609 |
| LTLADALEK | 9 | Q12894 | Interferon-related developmental regulato | IFRD2 | 48.048 | 2 | 50.651367 | yes | 4.861E-07 | 487.28186 | 187.07675 |
| RLTLADALEK | 10 | Q12894 | Interferon-related developmental regulato | IFRD2 | 48.048 | 2 | 50.651367 | yes | 9.732E-08 | 565.3324 | 149.48509 |
| QGALESLR | 8 | Q12894 | Interferon-related developmental regulato | IFRD2 | 48.048 | 2 | 50.651367 | yes | 4.08E-07 | 437.24307 | 146.29662 |
| EYVDCLTDK | 9 | Q12894 | Interferon-related developmental regulato | IFRD2 | 48.048 | 2 | 50.651367 | yes | 4.972E-07 | 571.75537 | 82.943588 |
| LKEYVDCLTDK | 11 | Q12894 | Interferon-related developmental regulato | IFRD2 | 48.048 | 2 | 50.651367 | yes | 2.468E-07 | 692.34485 | 37.176514 |
| ILVDQVTGISR | 11 | Q12926 | ELAV-like protein 2 OS=Homo sapiens ( | ELAVL2 | 39.504 | 2 | 51.313774 | yes | 9.722E-08 | 600.85095 | 3654.7861 |
| ELEQLFSQYGR | 11 | Q12926 | ELAV-like protein 2 OS=Homo sapiens ( | ELAVL2 | 39.504 | 2 | 51.313774 | no | 1.031E-07 | 685.34094 | 2403.6211 |
| SLFGSIGEIESCK | 13 | Q12926 | ELAV-like protein 2 OS=Homo sapiens ( | ELAVL2 | 39.504 | 2 | 51.313774 | no | 9.722E-08 | 713.84778 | 2254.9128 |
| AINTLNGLR | 9 | Q12926 | ELAV-like protein 2 OS=Homo sapiens ( | ELAVL2 | 39.504 | 2 | 51.313774 | no | 1.444E-07 | 486.28528 | 2054.5283 |
| DANLYVSGLPK | 11 | Q12926 | ELAV-like protein 2 OS=Homo sapiens ( | ELAVL2 | 39.504 | 2 | 51.313774 | no | 2.468E-07 | 588.81659 | 828.52625 |
| YPGPLAQQAQR | 11 | Q12926 | ELAV-like protein 2 OS=Homo sapiens ( | ELAVL2 | 39.504 | 2 | 51.313774 | yes | 1.22E-07 | 614.82526 | 699.2193 |
| LDNLLNMAYGVK | 12 | Q12926 | ELAV-like protein 2 OS=Homo sapiens ( | ELAVL2 | 39.504 | 2 | 51.313774 | no | 1.003E-07 | 675.85791 | 369.93277 |
| RIEAEEAIK | 9 | Q12926 | ELAV-like protein 2 OS=Homo sapiens ( | ELAVL2 | 39.504 | 2 | 51.313774 | no | 1.256E-07 | 529.79565 | 220.08379 |
| IEAEEAIK | 8 | Q12926 | ELAV-like protein 2 OS=Homo sapiens ( | ELAVL2 | 39.504 | 2 | 51.313774 | no | 3.19E-07 | 451.74512 | 181.6944 |
| FANNPSQK | 8 | Q12926 | ELAV-like protein 2 OS=Homo sapiens ( | ELAVL2 | 39.504 | 2 | 51.313774 | no | 1.737E-07 | 453.22742 | 122.77609 |
| GVEGLIDIENPNR | 13 | Q13442 | 28 kDa heat- and acid-stable phosphoprot | PDAP1 | 20.631 | 2 | 41.637302 | yes | 3.411E-07 | 713.37024 | 103.15976 |
| DKDFAIDIIK | 10 | Q15181 | Inorganic pyrophosphatase OS=Homo sa | PPA1 | 32.661 | 2 | 52.28351 | yes | 1.775E-07 | 589.32678 | 485.98001 |
| DVFHMVVEVPR | 11 | Q15181 | Inorganic pyrophosphatase OS=Homo sa | PPA1 | 32.661 | 2 | 52.28351 | yes | 9.722E-08 | 664.34497 | 270.53668 |
| GISCMNTTLSESPF | 15 | Q15181 | Inorganic pyrophosphatase OS=Homo sa | PPA1 | 32.661 | 2 | 52.28351 | yes | 1.647E-07 | 836.38947 | 240.72333 |
| AAPFSLEYR | 9 | Q15181 | Inorganic pyrophosphatase OS=Homo sa | PPA1 | 32.661 | 2 | 52.28351 | yes | 2.555E-07 | 527.27185 | 46.571541 |
| IGTNLSLIQCPR | 12 | Q15390 | Mitochondrial fission regulator 1 OS=Ho | MTFR1 | 37.001 | 2 | 49.947914 | yes | 1.189E-07 | 686.37427 | 171.12103 |
| ICALENELAALR | 12 | Q15390 | Mitochondrial fission regulator 1 OS=Ho | MTFR1 | 37.001 | 2 | 49.947914 | yes | 9.968E-08 | 686.86627 | 105.5257 |

| | | | | | | | | | | | |
|---|---|---|---|---|---|---|---|---|---|---|---|
| LQILSLR | 7 | Q15404 | Ras suppressor protein 1 OS=Homo sapie | RSU1 | 31.541 | 2 | 52.80288 | yes | 9.722E-08 | 421.77655 | 53.434765 |
| LHEEWLLR | 8 | Q15695 | Putative U2 small nuclear ribonucleoprot | ZRSR2P1 | 57.644 | 2 | 49.79486 | yes | 1.041E-06 | 548.3009 | 129.7637 |
| SMFTTFGMEQCR | 12 | Q15695 | Putative U2 small nuclear ribonucleoprot | ZRSR2P1 | 57.644 | 2 | 49.79486 | yes | 1.003E-07 | 747.81232 | 80.259956 |
| QLQCEFCPVTR | 11 | Q15695 | Putative U2 small nuclear ribonucleoprot | ZRSR2P1 | 57.644 | 2 | 49.79486 | yes | 1.136E-07 | 719.33429 | 59.786366 |
| KHNFPTSSPTLLIK | 14 | Q15695 | Putative U2 small nuclear ribonucleoprot | ZRSR2P1 | 57.644 | 3 | 49.79486 | yes | 3.235E-07 | 528.30322 | 32.5779 |
| HCNFLHVFR | 9 | Q15695 | Putative U2 small nuclear ribonucleoprot | ZRSR2P1 | 57.644 | 3 | 49.79486 | yes | 2.075E-06 | 410.53806 | 16.199364 |
| VYQETSEMR | 9 | Q15773 | Myeloid leukemia factor 2 OS=Homo sap | MLF2 | 28.148 | 2 | 53.770164 | yes | 9.722E-08 | 571.76099 | 136.12576 |
| LAIQGPEDSPSR | 12 | Q15773 | Myeloid leukemia factor 2 OS=Homo sap | MLF2 | 28.148 | 2 | 53.770164 | yes | 2.592E-07 | 635.32532 | 96.224854 |
| QQRPLEFR | 8 | Q15773 | Myeloid leukemia factor 2 OS=Homo sap | MLF2 | 28.148 | 2 | 53.770164 | yes | 4.771E-06 | 537.29614 | 52.309044 |
| WTEPYCVIAAVK | 12 | Q17RY6 | Lymphocyte antigen 6K OS=Homo sapie | LY6K | 18.673 | 2 | 50.25957 | yes | 1.471E-07 | 718.86578 | 274.36548 |
| ENTFECQNPR | 10 | Q17RY6 | Lymphocyte antigen 6K OS=Homo sapie | LY6K | 18.673 | 2 | 50.25957 | yes | 9.824E-08 | 647.77765 | 177.35757 |
| VWCHVCER | 8 | Q17RY6 | Lymphocyte antigen 6K OS=Homo sapie | LY6K | 18.673 | 2 | 50.25957 | yes | 9.849E-08 | 573.25256 | 90.430428 |
| QCSAGCAAMER | 11 | Q17RY6 | Lymphocyte antigen 6K OS=Homo sapie | LY6K | 18.673 | 2 | 50.25957 | yes | 2.092E-07 | 620.74701 | 14.403969 |
| MEAGPHPR | 8 | Q2NKX9 | UPF0561 protein C2orf68 OS=Homo sap | C2orf68 | 18.751 | 2 | 51.391068 | yes | 9.722E-08 | 468.72144 | 352.64789 |
| VSAHTPLDPPMR | 12 | Q2NKX9 | UPF0561 protein C2orf68 OS=Homo sap | C2orf68 | 18.751 | 2 | 51.391068 | yes | 2.838E-07 | 660.84009 | 155.88707 |
| VSAHTPLDPPMR | 12 | Q2NKX9 | UPF0561 protein C2orf68 OS=Homo sap | C2orf68 | 18.751 | 2 | 51.391068 | yes | 2.972E-07 | 440.89581 | 155.88707 |
| VSAHTPLDPPMR | 12 | Q2NKX9 | UPF0561 protein C2orf68 OS=Homo sap | C2orf68 | 18.751 | 3 | 51.391068 | yes | 2.838E-07 | 660.84009 | 155.88707 |
| VSAHTPLDPPMR | 12 | Q2NKX9 | UPF0561 protein C2orf68 OS=Homo sap | C2orf68 | 18.751 | 3 | 51.391068 | yes | 2.972E-07 | 440.89581 | 155.88707 |
| KPDLQVYLPR | 10 | Q2NKX9 | UPF0561 protein C2orf68 OS=Homo sap | C2orf68 | 18.751 | 2 | 51.391068 | yes | 1.15E-07 | 614.85608 | 197.46664 |
| LEEMFPDEVDTPR | 13 | Q2NL82 | Pre-rRNA-processing protein TSR1 homo | TSR1 | 91.811 | 2 | 52.45582 | yes | 9.722E-08 | 789.36127 | 5653.6138 |
| QIDAPGDPFPLNPF | 14 | Q2NL82 | Pre-rRNA-processing protein TSR1 homo | TSR1 | 91.811 | 2 | 52.45582 | yes | 1.059E-07 | 768.89392 | 4010.7051 |
| DGPPHQVLVVPLF | 15 | Q2NL82 | Pre-rRNA-processing protein TSR1 homo | TSR1 | 91.811 | 3 | 52.45582 | yes | 9.722E-08 | 550.974 | 1238.3102 |
| DGPPHQVLVVPLF | 15 | Q2NL82 | Pre-rRNA-processing protein TSR1 homo | TSR1 | 91.811 | 3 | 52.45582 | yes | 9.889E-08 | 825.95734 | 1238.3102 |
| DGPPHQVLVVPLF | 15 | Q2NL82 | Pre-rRNA-processing protein TSR1 homo | TSR1 | 91.811 | 3 | 52.45582 | yes | 4.772E-07 | 413.48233 | 1238.3102 |
| DGPPHQVLVVPLF | 15 | Q2NL82 | Pre-rRNA-processing protein TSR1 homo | TSR1 | 91.811 | 2 | 52.45582 | yes | 9.722E-08 | 550.974 | 1238.3102 |
| DGPPHQVLVVPLF | 15 | Q2NL82 | Pre-rRNA-processing protein TSR1 homo | TSR1 | 91.811 | 2 | 52.45582 | yes | 9.889E-08 | 825.95734 | 1238.3102 |
| DGPPHQVLVVPLF | 15 | Q2NL82 | Pre-rRNA-processing protein TSR1 homo | TSR1 | 91.811 | 2 | 52.45582 | yes | 4.772E-07 | 413.48233 | 1238.3102 |
| DGPPHQVLVVPLF | 15 | Q2NL82 | Pre-rRNA-processing protein TSR1 homo | TSR1 | 91.811 | 4 | 52.45582 | yes | 9.722E-08 | 550.974 | 1238.3102 |
| DGPPHQVLVVPLF | 15 | Q2NL82 | Pre-rRNA-processing protein TSR1 homo | TSR1 | 91.811 | 4 | 52.45582 | yes | 9.889E-08 | 825.95734 | 1238.3102 |
| DGPPHQVLVVPLF | 15 | Q2NL82 | Pre-rRNA-processing protein TSR1 homo | TSR1 | 91.811 | 4 | 52.45582 | yes | 4.772E-07 | 413.48233 | 1238.3102 |
| MSVLNMVVR | 9 | Q2NL82 | Pre-rRNA-processing protein TSR1 homo | TSR1 | 91.811 | 2 | 52.45582 | yes | 1.135E-07 | 524.78571 | 3245.7114 |
| EDVLWFKPVELR | 12 | Q2NL82 | Pre-rRNA-processing protein TSR1 homo | TSR1 | 91.811 | 2 | 52.45582 | yes | 1.297E-07 | 765.91937 | 1615.9072 |
| EDVLWFKPVELR | 12 | Q2NL82 | Pre-rRNA-processing protein TSR1 homo | TSR1 | 91.811 | 2 | 52.45582 | yes | 7.483E-07 | 510.94867 | 1615.9072 |
| EDVLWFKPVELR | 12 | Q2NL82 | Pre-rRNA-processing protein TSR1 homo | TSR1 | 91.811 | 3 | 52.45582 | yes | 1.297E-07 | 765.91937 | 1615.9072 |
| EDVLWFKPVELR | 12 | Q2NL82 | Pre-rRNA-processing protein TSR1 homo | TSR1 | 91.811 | 3 | 52.45582 | yes | 7.483E-07 | 510.94867 | 1615.9072 |
| SQDTVLMNLYK | 11 | Q2NL82 | Pre-rRNA-processing protein TSR1 homo | TSR1 | 91.811 | 2 | 52.45582 | yes | 9.722E-08 | 656.33429 | 3160.2942 |
| PGDLHVVLDMAK | 12 | Q2NL82 | Pre-rRNA-processing protein TSR1 homo | TSR1 | 91.811 | 2 | 52.45582 | yes | 1.181E-07 | 647.84485 | 1558.25 |
| PGDLHVVLDMAK | 12 | Q2NL82 | Pre-rRNA-processing protein TSR1 homo | TSR1 | 91.811 | 2 | 52.45582 | yes | 1.661E-07 | 432.2323 | 1558.25 |
| PGDLHVVLDMAK | 12 | Q2NL82 | Pre-rRNA-processing protein TSR1 homo | TSR1 | 91.811 | 3 | 52.45582 | yes | 1.181E-07 | 647.84485 | 1558.25 |

| | | | | | | | | | | | |
|---|---|---|---|---|---|---|---|---|---|---|---|
| PGDLHVVLDMAK | 12 | Q2NL82 | Pre-rRNA-processing protein TSR1 homc | TSR1 | 91.811 | 3 | 52.45582 | yes | 1.661E-07 | 432.2323 | 1558.25 |
| ASPLFSQHTAADK | 13 | Q2NL82 | Pre-rRNA-processing protein TSR1 homc | TSR1 | 91.811 | 2 | 52.45582 | yes | 3.502E-07 | 686.84644 | 3031.01 |
| LLHIVGYGDFQMK | 13 | Q2NL82 | Pre-rRNA-processing protein TSR1 homc | TSR1 | 91.811 | 2 | 52.45582 | yes | 1.012E-07 | 760.90015 | 1320.4825 |
| LLHIVGYGDFQMK | 13 | Q2NL82 | Pre-rRNA-processing protein TSR1 homc | TSR1 | 91.811 | 2 | 52.45582 | yes | 2.205E-07 | 507.60251 | 1320.4825 |
| LLHIVGYGDFQMK | 13 | Q2NL82 | Pre-rRNA-processing protein TSR1 homc | TSR1 | 91.811 | 3 | 52.45582 | yes | 1.012E-07 | 760.90015 | 1320.4825 |
| LLHIVGYGDFQMK | 13 | Q2NL82 | Pre-rRNA-processing protein TSR1 homc | TSR1 | 91.811 | 3 | 52.45582 | yes | 2.205E-07 | 507.60251 | 1320.4825 |
| IFQFQNFTNTR | 11 | Q2NL82 | Pre-rRNA-processing protein TSR1 homc | TSR1 | 91.811 | 2 | 52.45582 | yes | 9.722E-08 | 708.35693 | 2272.9548 |
| ENLPQDYAR | 9 | Q2NL82 | Pre-rRNA-processing protein TSR1 homc | TSR1 | 91.811 | 2 | 52.45582 | yes | 1.25E-06 | 553.26727 | 2185.292 |
| EELIFHCGFR | 10 | Q2NL82 | Pre-rRNA-processing protein TSR1 homc | TSR1 | 91.811 | 2 | 52.45582 | yes | 1.128E-07 | 654.31372 | 2091.2312 |
| VVLSGHPFK | 9 | Q2NL82 | Pre-rRNA-processing protein TSR1 homc | TSR1 | 91.811 | 2 | 52.45582 | yes | 1.403E-06 | 492.28726 | 1892.2855 |
| ISLPEAMQLLQDR | 13 | Q2NL82 | Pre-rRNA-processing protein TSR1 homc | TSR1 | 91.811 | 2 | 52.45582 | yes | 2.327E-07 | 757.40576 | 907.13678 |
| WFFTSAR | 7 | Q2NL82 | Pre-rRNA-processing protein TSR1 homc | TSR1 | 91.811 | 2 | 52.45582 | yes | 1.386E-07 | 457.72958 | 738.23932 |
| AKEELIFHCGFR | 12 | Q2NL82 | Pre-rRNA-processing protein TSR1 homc | TSR1 | 91.811 | 2 | 52.45582 | yes | 1.117E-07 | 753.87976 | 458.62796 |
| QQHLAFR | 7 | Q2NL82 | Pre-rRNA-processing protein TSR1 homc | TSR1 | 91.811 | 2 | 52.45582 | yes | 9.823E-08 | 450.24594 | 387.85779 |
| EPLGTHGHMK | 10 | Q2NL82 | Pre-rRNA-processing protein TSR1 homc | TSR1 | 91.811 | 2 | 52.45582 | yes | 1.02E-07 | 553.77423 | 353.94006 |
| RDPGNTEPVK | 10 | Q2NL82 | Pre-rRNA-processing protein TSR1 homc | TSR1 | 91.811 | 2 | 52.45582 | yes | 1.73E-05 | 556.78839 | 256.3573 |
| SEISSTVPQGGME | 13 | Q2NL82 | Pre-rRNA-processing protein TSR1 homc | TSR1 | 91.811 | 2 | 52.45582 | yes | 3.274E-07 | 661.30066 | 198.8298 |
| DPGNTEPVK | 9 | Q2NL82 | Pre-rRNA-processing protein TSR1 homc | TSR1 | 91.811 | 2 | 52.45582 | yes | 1.814E-07 | 478.73782 | 136.88329 |
| AYLFAHAVDFVPS | 23 | Q2NL82 | Pre-rRNA-processing protein TSR1 homc | TSR1 | 91.811 | 3 | 52.45582 | yes | 1.317E-06 | 845.43573 | 116.59869 |
| RVVLSGHPFK | 10 | Q2NL82 | Pre-rRNA-processing protein TSR1 homc | TSR1 | 91.811 | 2 | 52.45582 | yes | 3.027E-07 | 570.33783 | 114.10337 |
| QGTPLIAFSLLPHE | 16 | Q2NL82 | Pre-rRNA-processing protein TSR1 homc | TSR1 | 91.811 | 3 | 52.45582 | yes | 2.88E-07 | 593.66486 | 51.806244 |
| DTGTVHLNELGNT | 22 | Q2NL82 | Pre-rRNA-processing protein TSR1 homc | TSR1 | 91.811 | 3 | 52.45582 | yes | 1.471E-07 | 844.07739 | 47.82682 |
| RSPLTSR | 7 | Q2T9L4 | Inhibitory synaptic factor 1 OS=Homo sa| | INSYN1 | 31.929 | 2 | 49.171486 | yes | 1.062E-07 | 408.73795 | 918.21442 |
| LDSDAAGVFCIR | 12 | Q3B726 | DNA-directed RNA polymerase I subunit | POLR1F | 37.433 | 2 | 51.812855 | yes | 9.722E-08 | 662.32172 | 229.6629 |
| RLEEIMK | 7 | Q3KQU3 | MAP7 domain-containing protein 1 OS=I | MAP7D1 | 92.82 | 2 | 53.515995 | no | 9.445E-07 | 459.75751 | 502.67368 |
| AELGAGAPPAVVA | 14 | Q3KQU3 | MAP7 domain-containing protein 1 OS=I | MAP7D1 | 92.82 | 2 | 53.515995 | yes | 1.603E-07 | 639.86188 | 424.42484 |
| SASASPLTPCSVTR | 14 | Q3KQU3 | MAP7 domain-containing protein 1 OS=I | MAP7D1 | 92.82 | 2 | 53.515995 | yes | 1.018E-07 | 717.35626 | 259.65912 |
| SLQLSAWESSIVDI | 14 | Q3KQU3 | MAP7 domain-containing protein 1 OS=I | MAP7D1 | 92.82 | 2 | 53.515995 | yes | 9.722E-08 | 795.90973 | 236.26346 |
| LSASTASELSPK | 12 | Q3KQU3 | MAP7 domain-containing protein 1 OS=I | MAP7D1 | 92.82 | 2 | 53.515995 | yes | 3.171E-07 | 595.81677 | 228.1395 |
| HVDSIINK | 8 | Q3KQU3 | MAP7 domain-containing protein 1 OS=I | MAP7D1 | 92.82 | 2 | 53.515995 | yes | 8.046E-07 | 463.25873 | 217.41649 |
| CSVSAVNLPK | 10 | Q3KQU3 | MAP7 domain-containing protein 1 OS=I | MAP7D1 | 92.82 | 2 | 53.515995 | yes | 4.49E-07 | 537.78424 | 144.93282 |
| LMTPTLSFLAR | 11 | Q3KQU3 | MAP7 domain-containing protein 1 OS=I | MAP7D1 | 92.82 | 2 | 53.515995 | yes | 9.722E-08 | 625.35229 | 110.91313 |
| QLPLEPESPSGQVG | 16 | Q3KQU3 | MAP7 domain-containing protein 1 OS=I | MAP7D1 | 92.82 | 2 | 53.515995 | yes | 9.926E-08 | 845.94159 | 97.571732 |
| AVVQSPQVTEVL | 12 | Q3KQU3 | MAP7 domain-containing protein 1 OS=I | MAP7D1 | 92.82 | 2 | 53.515995 | yes | 9.722E-08 | 635.35608 | 89.408119 |
| QSLPASPR | 8 | Q3KQU3 | MAP7 domain-containing protein 1 OS=I | MAP7D1 | 92.82 | 2 | 53.515995 | yes | 1.883E-05 | 428.23779 | 83.292732 |
| EANANGSSPEPVK | 13 | Q3KQU3 | MAP7 domain-containing protein 1 OS=I | MAP7D1 | 92.82 | 2 | 53.515995 | yes | 1.194E-05 | 650.31238 | 37.805984 |
| KPNAGGSPAPVR | 12 | Q3KQU3 | MAP7 domain-containing protein 1 OS=I | MAP7D1 | 92.82 | 2 | 53.515995 | yes | 1.039E-07 | 575.82001 | 35.653198 |
| RKPNAGGSPAPVR | 13 | Q3KQU3 | MAP7 domain-containing protein 1 OS=I | MAP7D1 | 92.82 | 3 | 53.515995 | yes | 3.286E-06 | 436.24945 | 17.627426 |
| AEDNLLALGLK | 11 | Q3T8J9 | GON-4-like protein OS=Homo sapiens O | GON4L | 248.621 | 2 | 45.76316 | yes | 1.748E-07 | 578.83228 | 61.472637 |

| | | | | | | | | | | | |
|---|---|---|---|---|---|---|---|---|---|---|---|
| ELGTFAQSSIALHI | 19 | Q3T8J9 | GON-4-like protein OS=Homo sapiens O | GON4L | 248.621 | 3 | 45.76316 | no | 3.763E-07 | 714.36401 | 41.648544 |
| VSIQMPTIIR | 10 | Q3ZCT1 | Zinc finger protein 260 OS=Homo sapien | ZNF260 | 47.223 | 2 | 42.103687 | yes | 1.405E-06 | 579.33917 | 33.704498 |
| ITSLIVHVR | 9 | Q3ZCT1 | Zinc finger protein 260 OS=Homo sapien | ZNF260 | 47.223 | 2 | 42.103687 | yes | 3.191E-07 | 519.3269 | 26.806217 |
| MFADDLHNLNK | 11 | Q4U2R6 | Large ribosomal subunit protein mL51 OS | MRPL51 | 15.095 | 2 | 43.64523 | yes | 2.504E-07 | 659.31647 | 654.01514 |
| AGQSEAGCCLLER | 13 | Q53HC5 | Kelch-like protein 26 OS=Homo sapiens | KLHL26 | 68.14 | 2 | 51.4022 | yes | 2.825E-07 | 725.82428 | 155.8615 |
| IYIVGGYNWR | 10 | Q53HC5 | Kelch-like protein 26 OS=Homo sapiens | KLHL26 | 68.14 | 2 | 51.4022 | yes | 9.722E-08 | 620.82727 | 58.058159 |
| TWGHAGAASGGR | 12 | Q53HC5 | Kelch-like protein 26 OS=Homo sapiens | KLHL26 | 68.14 | 2 | 51.4022 | yes | 3.178E-06 | 564.27069 | 37.240082 |
| SPVPLTPPGCVALI | 16 | Q567V2 | Mpv17-like protein 2 OS=Homo sapiens | MPV17L2 | 23.181 | 2 | 46.518227 | yes | 1.683E-07 | 840.44287 | 121.70299 |
| LLSAGQLLFQGR | 12 | Q567V2 | Mpv17-like protein 2 OS=Homo sapiens | MPV17L2 | 23.181 | 2 | 46.518227 | yes | 1.538E-07 | 651.88007 | 102.83285 |
| SLIINTFYSNK | 11 | Q58FF6 | Putative heat shock protein HSP 90-beta | HSP90AB4 | 58.265 | 2 | 50.068333 | yes | 9.918E-08 | 650.35083 | 209.31064 |
| HSQFLGYPITLYLI | 15 | Q58FF8 | Putative heat shock protein HSP 90-beta | HSP90AB2 | 44.349 | 3 | 49.490353 | yes | 1.025E-07 | 603.65759 | 53.884033 |
| LLASSVPELR | 10 | Q5H9U9 | Probable ATP-dependent RNA helicase I | DDX60L | 197.674 | 2 | 55.557217 | yes | 9.722E-08 | 542.82166 | 404.74539 |
| VLCVTSCSLSLR | 12 | Q5H9U9 | Probable ATP-dependent RNA helicase I | DDX60L | 197.674 | 2 | 55.557217 | yes | 9.722E-08 | 697.86017 | 335.67242 |
| DPHVLDFLK | 9 | Q5H9U9 | Probable ATP-dependent RNA helicase I | DDX60L | 197.674 | 2 | 55.557217 | yes | 1.066E-07 | 542.29529 | 210.03226 |
| ILEISEDCTYADVK | 14 | Q5H9U9 | Probable ATP-dependent RNA helicase I | DDX60L | 197.674 | 2 | 55.557217 | yes | 9.722E-08 | 828.39526 | 209.29895 |
| AALYEYNLAVMK | 12 | Q5H9U9 | Probable ATP-dependent RNA helicase I | DDX60L | 197.674 | 2 | 55.557217 | yes | 9.722E-08 | 693.36029 | 186.27968 |
| FYAYTMESTDR | 11 | Q5H9U9 | Probable ATP-dependent RNA helicase I | DDX60L | 197.674 | 2 | 55.557217 | yes | 1.037E-07 | 692.29791 | 185.75423 |
| NLSPDSLSSSK | 11 | Q5H9U9 | Probable ATP-dependent RNA helicase I | DDX60L | 197.674 | 2 | 55.557217 | yes | 3.048E-06 | 567.78552 | 182.57701 |
| DHAVFFR | 7 | Q5H9U9 | Probable ATP-dependent RNA helicase I | DDX60L | 197.674 | 2 | 55.557217 | yes | 1.357E-07 | 446.22723 | 175.57864 |
| VILAELPEDFK | 11 | Q5H9U9 | Probable ATP-dependent RNA helicase I | DDX60L | 197.674 | 2 | 55.557217 | yes | 1.388E-07 | 637.35553 | 149.60474 |
| EMTQLILNEMPK | 12 | Q5H9U9 | Probable ATP-dependent RNA helicase I | DDX60L | 197.674 | 2 | 55.557217 | yes | 9.722E-08 | 723.87018 | 149.01297 |
| IHSLLER | 7 | Q5H9U9 | Probable ATP-dependent RNA helicase I | DDX60L | 197.674 | 2 | 55.557217 | yes | 9.722E-08 | 434.25598 | 132.53519 |
| GSQQFSQDVMEK | 12 | Q5H9U9 | Probable ATP-dependent RNA helicase I | DDX60L | 197.674 | 2 | 55.557217 | yes | 9.722E-08 | 692.31409 | 131.64877 |
| AEYINFLENLK | 11 | Q5H9U9 | Probable ATP-dependent RNA helicase I | DDX60L | 197.674 | 2 | 55.557217 | yes | 9.722E-08 | 677.35608 | 113.14149 |
| HSLLSFK | 7 | Q5H9U9 | Probable ATP-dependent RNA helicase I | DDX60L | 197.674 | 2 | 55.557217 | no | 9.722E-08 | 416.23981 | 104.9669 |
| VIVTQTTRPK | 10 | Q5H9U9 | Probable ATP-dependent RNA helicase I | DDX60L | 197.674 | 2 | 55.557217 | yes | 1.098E-07 | 571.84821 | 102.63268 |
| ENETVIQSAYK | 11 | Q5H9U9 | Probable ATP-dependent RNA helicase I | DDX60L | 197.674 | 2 | 55.557217 | yes | 1.641E-07 | 641.3197 | 101.58956 |
| YVIFDEVHYLGR | 12 | Q5H9U9 | Probable ATP-dependent RNA helicase I | DDX60L | 197.674 | 2 | 55.557217 | yes | 9.722E-08 | 755.88806 | 100.98448 |
| LLSDDYDR | 8 | Q5H9U9 | Probable ATP-dependent RNA helicase I | DDX60L | 197.674 | 2 | 55.557217 | yes | 1.163E-07 | 498.73526 | 92.117012 |
| SLIQHLEEIR | 10 | Q5H9U9 | Probable ATP-dependent RNA helicase I | DDX60L | 197.674 | 2 | 55.557217 | yes | 6.959E-07 | 619.34857 | 84.22213 |
| VITCSWIR | 8 | Q5H9U9 | Probable ATP-dependent RNA helicase I | DDX60L | 197.674 | 2 | 55.557217 | yes | 3.932E-07 | 517.77625 | 73.251114 |
| AQELCPEEFILFK | 13 | Q5H9U9 | Probable ATP-dependent RNA helicase I | DDX60L | 197.674 | 2 | 55.557217 | yes | 1.441E-07 | 812.40802 | 70.433563 |
| GIGYHHSSMYFK | 12 | Q5H9U9 | Probable ATP-dependent RNA helicase I | DDX60L | 197.674 | 3 | 55.557217 | yes | 8.466E-07 | 476.22394 | 31.50742 |
| GIGYHHSSMYFK | 12 | Q5H9U9 | Probable ATP-dependent RNA helicase I | DDX60L | 197.674 | 3 | 55.557217 | yes | 5.504E-07 | 713.83228 | 31.50742 |
| GIGYHHSSMYFK | 12 | Q5H9U9 | Probable ATP-dependent RNA helicase I | DDX60L | 197.674 | 2 | 55.557217 | yes | 8.466E-07 | 476.22394 | 31.50742 |
| GIGYHHSSMYFK | 12 | Q5H9U9 | Probable ATP-dependent RNA helicase I | DDX60L | 197.674 | 2 | 55.557217 | yes | 5.504E-07 | 713.83228 | 31.50742 |
| NESAVIVAPTSSGI | 14 | Q5H9U9 | Probable ATP-dependent RNA helicase I | DDX60L | 197.674 | 2 | 55.557217 | no | 2.197E-07 | 680.35938 | 55.719334 |
| KYSIDIGPAR | 10 | Q5H9U9 | Probable ATP-dependent RNA helicase I | DDX60L | 197.674 | 2 | 55.557217 | yes | 1.019E-06 | 560.31146 | 55.636303 |
| VNVVLSSGHESDT | 15 | Q5H9U9 | Probable ATP-dependent RNA helicase I | DDX60L | 197.674 | 2 | 55.557217 | yes | 9.722E-08 | 806.91809 | 54.748779 |

| | | | | | | | | | | | |
|---|---|---|---|---|---|---|---|---|---|---|---|
| VVTATETLALGIHI | 17 | Q5H9U9 | Probable ATP-dependent RNA helicase D | DDX60L | 197.674 | 3 | 55.557217 | yes | 1.443E-07 | 614.32684 | 19.208544 |
| MPPPANLPSLK | 11 | Q5JSZ5 | Protein PRRC2B OS=Homo sapiens OX= | PRRC2B | 242.967 | 2 | 52.842957 | no | 6.355E-07 | 582.82593 | 2053.3284 |
| MPPPANLPSLK | 11 | Q5JSZ5 | Protein PRRC2B OS=Homo sapiens OX= | PRRC2B | 242.967 | 2 | 52.842957 | no | 4.551E-06 | 590.82336 | 2053.3284 |
| HGLQSLGK | 8 | Q5JSZ5 | Protein PRRC2B OS=Homo sapiens OX= | PRRC2B | 242.967 | 2 | 52.842957 | no | 1.062E-07 | 420.24033 | 2844.6348 |
| RMPPPANLPSLK | 12 | Q5JSZ5 | Protein PRRC2B OS=Homo sapiens OX= | PRRC2B | 242.967 | 2 | 52.842957 | no | 2.297E-07 | 668.8739 | 1265.4304 |
| RMPPPANLPSLK | 12 | Q5JSZ5 | Protein PRRC2B OS=Homo sapiens OX= | PRRC2B | 242.967 | 2 | 52.842957 | no | 2.487E-06 | 660.87646 | 1265.4304 |
| FIQSEMSEAVER | 12 | Q5JSZ5 | Protein PRRC2B OS=Homo sapiens OX= | PRRC2B | 242.967 | 2 | 52.842957 | yes | 9.722E-08 | 713.33759 | 841.4472 |
| FIQSEMSEAVER | 12 | Q5JSZ5 | Protein PRRC2B OS=Homo sapiens OX= | PRRC2B | 242.967 | 2 | 52.842957 | yes | 3.215E-06 | 721.33502 | 841.4472 |
| GSFPLPQLR | 9 | Q5JSZ5 | Protein PRRC2B OS=Homo sapiens OX= | PRRC2B | 242.967 | 2 | 52.842957 | yes | 1.871E-06 | 507.79019 | 1630.9904 |
| SSDTLAMDMR | 10 | Q5JSZ5 | Protein PRRC2B OS=Homo sapiens OX= | PRRC2B | 242.967 | 2 | 52.842957 | yes | 1.158E-07 | 563.74701 | 1156.1489 |
| CSPLEPDFVPDEK | 13 | Q5JSZ5 | Protein PRRC2B OS=Homo sapiens OX= | PRRC2B | 242.967 | 2 | 52.842957 | yes | 1.242E-07 | 766.85052 | 1148.9031 |
| YSTLSLFDK | 9 | Q5JSZ5 | Protein PRRC2B OS=Homo sapiens OX= | PRRC2B | 242.967 | 2 | 52.842957 | yes | 2.644E-07 | 537.2793 | 1084.6433 |
| EGTAFNISSWDK | 12 | Q5JSZ5 | Protein PRRC2B OS=Homo sapiens OX= | PRRC2B | 242.967 | 2 | 52.842957 | yes | 1.017E-07 | 677.81732 | 990.75983 |
| LLSFSPEEFPTLK | 13 | Q5JSZ5 | Protein PRRC2B OS=Homo sapiens OX= | PRRC2B | 242.967 | 2 | 52.842957 | yes | 9.722E-08 | 754.40576 | 749.63977 |
| GNDPNIVIVPK | 11 | Q5JSZ5 | Protein PRRC2B OS=Homo sapiens OX= | PRRC2B | 242.967 | 2 | 52.842957 | yes | 9.722E-08 | 583.3324 | 723.2442 |
| PTIINAENLK | 10 | Q5JSZ5 | Protein PRRC2B OS=Homo sapiens OX= | PRRC2B | 242.967 | 2 | 52.842957 | yes | 1.025E-07 | 556.81915 | 720.89948 |
| DYVPDSYR | 8 | Q5JSZ5 | Protein PRRC2B OS=Homo sapiens OX= | PRRC2B | 242.967 | 2 | 52.842957 | yes | 9.722E-08 | 507.72998 | 658.01752 |
| GEEYLSAFDK | 10 | Q5JSZ5 | Protein PRRC2B OS=Homo sapiens OX= | PRRC2B | 242.967 | 2 | 52.842957 | yes | 9.722E-08 | 579.76929 | 657.09924 |
| EVPWSPSAEK | 10 | Q5JSZ5 | Protein PRRC2B OS=Homo sapiens OX= | PRRC2B | 242.967 | 2 | 52.842957 | yes | 1.286E-07 | 565.27985 | 642.29895 |
| AQADFDSCISSQR | 13 | Q5JSZ5 | Protein PRRC2B OS=Homo sapiens OX= | PRRC2B | 242.967 | 2 | 52.842957 | yes | 2.058E-07 | 742.82538 | 618.06873 |
| EQAVQVPVK | 9 | Q5JSZ5 | Protein PRRC2B OS=Homo sapiens OX= | PRRC2B | 242.967 | 2 | 52.842957 | yes | 9.819E-08 | 499.28748 | 574.77979 |
| AAHASADLPEASS | 14 | Q5JSZ5 | Protein PRRC2B OS=Homo sapiens OX= | PRRC2B | 242.967 | 2 | 52.842957 | yes | 9.722E-08 | 677.8335 | 452.38815 |
| LQEAPSAASQMK | 12 | Q5JSZ5 | Protein PRRC2B OS=Homo sapiens OX= | PRRC2B | 242.967 | 2 | 52.842957 | yes | 1.074E-07 | 630.81628 | 447.44632 |
| GYPLPHPK | 8 | Q5JSZ5 | Protein PRRC2B OS=Homo sapiens OX= | PRRC2B | 242.967 | 2 | 52.842957 | yes | 4.742E-07 | 454.75308 | 425.32678 |
| GPGHATFGR | 9 | Q5JSZ5 | Protein PRRC2B OS=Homo sapiens OX= | PRRC2B | 242.967 | 2 | 52.842957 | yes | 1.414E-07 | 450.22775 | 421.47379 |
| KAPDPQPPPR | 10 | Q5JSZ5 | Protein PRRC2B OS=Homo sapiens OX= | PRRC2B | 242.967 | 2 | 52.842957 | yes | 1.024E-07 | 551.80383 | 407.66446 |
| SIYCSSQR | 8 | Q5JSZ5 | Protein PRRC2B OS=Homo sapiens OX= | PRRC2B | 242.967 | 2 | 52.842957 | yes | 2.451E-07 | 500.72946 | 383.71329 |
| LHGWAPGPDYQK | 12 | Q5JSZ5 | Protein PRRC2B OS=Homo sapiens OX= | PRRC2B | 242.967 | 2 | 52.842957 | yes | 9.722E-08 | 684.83838 | 361.98651 |
| HPDAFGGR | 8 | Q5JSZ5 | Protein PRRC2B OS=Homo sapiens OX= | PRRC2B | 242.967 | 2 | 52.842957 | yes | 9.759E-08 | 428.70667 | 323.44611 |
| QLSMSSADSADAK | 13 | Q5JSZ5 | Protein PRRC2B OS=Homo sapiens OX= | PRRC2B | 242.967 | 2 | 52.842957 | yes | 1.017E-07 | 655.79828 | 300.95224 |
| GCSEDHSGLDAK | 12 | Q5JSZ5 | Protein PRRC2B OS=Homo sapiens OX= | PRRC2B | 242.967 | 2 | 52.842957 | yes | 9.819E-08 | 638.26715 | 296.91034 |
| SSQGDSGVDLSAE | 15 | Q5JSZ5 | Protein PRRC2B OS=Homo sapiens OX= | PRRC2B | 242.967 | 2 | 52.842957 | yes | 9.722E-08 | 747.83698 | 292.71082 |
| QELGEESTR | 9 | Q5JSZ5 | Protein PRRC2B OS=Homo sapiens OX= | PRRC2B | 242.967 | 2 | 52.842957 | yes | 2.914E-07 | 524.7489 | 270.02789 |
| NESSFSASLGR | 11 | Q5JSZ5 | Protein PRRC2B OS=Homo sapiens OX= | PRRC2B | 242.967 | 2 | 52.842957 | yes | 9.722E-08 | 577.77545 | 254.44835 |
| IKQELGEESTR | 11 | Q5JSZ5 | Protein PRRC2B OS=Homo sapiens OX= | PRRC2B | 242.967 | 2 | 52.842957 | yes | 9.722E-08 | 645.33844 | 247.47722 |
| TPVDFYPSALHPS( | 17 | Q5JSZ5 | Protein PRRC2B OS=Homo sapiens OX= | PRRC2B | 242.967 | 3 | 52.842957 | yes | 9.722E-08 | 620.65021 | 240.10417 |
| ESPHWQSK | 8 | Q5JSZ5 | Protein PRRC2B OS=Homo sapiens OX= | PRRC2B | 242.967 | 2 | 52.842957 | yes | 2.916E-07 | 499.73816 | 232.53735 |
| SEDQNCVPPLQER | 13 | Q5JSZ5 | Protein PRRC2B OS=Homo sapiens OX= | PRRC2B | 242.967 | 2 | 52.842957 | yes | 1.947E-06 | 786.35956 | 216.91772 |
| VDEKPSLGAVK | 11 | Q5JSZ5 | Protein PRRC2B OS=Homo sapiens OX= | PRRC2B | 242.967 | 2 | 52.842957 | yes | 1.039E-07 | 571.8244 | 208.09875 |

| | | | | | | | | | | |
|---|---|---|---|---|---|---|---|---|---|---|
| QNVPSGGPVPSPQ | 16 | Q5JSZ5 | Protein PRRC2B OS=Homo sapiens OX= | PRRC2B | 242.967 | 2 | 52.842957 | yes | 3.735E-07 | 842.42609 | 180.53419 |
| KLEPGGFGEK | 10 | Q5JSZ5 | Protein PRRC2B OS=Homo sapiens OX= | PRRC2B | 242.967 | 2 | 52.842957 | yes | 7.357E-07 | 531.28491 | 149.64209 |
| QFQMNDQDGK | 10 | Q5JSZ5 | Protein PRRC2B OS=Homo sapiens OX= | PRRC2B | 242.967 | 2 | 52.842957 | yes | 5.924E-07 | 605.76147 | 141.16231 |
| GFHFADSK | 8 | Q5JSZ5 | Protein PRRC2B OS=Homo sapiens OX= | PRRC2B | 242.967 | 2 | 52.842957 | yes | 9.722E-08 | 454.71667 | 138.01933 |
| IPPPGSQPPVLNTS | 15 | Q5JSZ5 | Protein PRRC2B OS=Homo sapiens OX= | PRRC2B | 242.967 | 2 | 52.842957 | yes | 1.119E-07 | 780.43066 | 134.59477 |
| VASETHSEGSEYE | 17 | Q5JSZ5 | Protein PRRC2B OS=Homo sapiens OX= | PRRC2B | 242.967 | 3 | 52.842957 | yes | 2.944E-07 | 631.29004 | 131.95604 |
| PVGHEGGLR | 9 | Q5JSZ5 | Protein PRRC2B OS=Homo sapiens OX= | PRRC2B | 242.967 | 2 | 52.842957 | yes | 1.025E-07 | 461.24869 | 128.76489 |
| PMMPQESLNGTG | 14 | Q5JSZ5 | Protein PRRC2B OS=Homo sapiens OX= | PRRC2B | 242.967 | 2 | 52.842957 | yes | 1.924E-07 | 789.34711 | 127.40488 |
| HIISATSLSTSPTEL | 18 | Q5JSZ5 | Protein PRRC2B OS=Homo sapiens OX= | PRRC2B | 242.967 | 3 | 52.842957 | yes | 9.722E-08 | 619.66193 | 123.60518 |
| GEEYLSAFDKK | 11 | Q5JSZ5 | Protein PRRC2B OS=Homo sapiens OX= | PRRC2B | 242.967 | 2 | 52.842957 | yes | 1.859E-07 | 643.81677 | 120.83037 |
| KPECGSWDVSHQ | 24 | Q5JSZ5 | Protein PRRC2B OS=Homo sapiens OX= | PRRC2B | 242.967 | 4 | 52.842957 | yes | 1.97E-07 | 674.05566 | 119.83323 |
| GSENGNEGSLLER | 13 | Q5JSZ5 | Protein PRRC2B OS=Homo sapiens OX= | PRRC2B | 242.967 | 2 | 52.842957 | yes | 9.722E-08 | 681.31824 | 104.07346 |
| DPSLRPAQPVR | 11 | Q5JSZ5 | Protein PRRC2B OS=Homo sapiens OX= | PRRC2B | 242.967 | 3 | 52.842957 | yes | 3.738E-07 | 412.56668 | 101.75635 |
| TFYPHHPQMLGFD | 15 | Q5JSZ5 | Protein PRRC2B OS=Homo sapiens OX= | PRRC2B | 242.967 | 3 | 52.842957 | yes | 2.377E-07 | 614.96301 | 101.74459 |
| LSNCGYGR | 8 | Q5JSZ5 | Protein PRRC2B OS=Homo sapiens OX= | PRRC2B | 242.967 | 2 | 52.842957 | yes | 1.785E-07 | 463.71109 | 98.828773 |
| FSDDEEEEEVVK | 12 | Q5JSZ5 | Protein PRRC2B OS=Homo sapiens OX= | PRRC2B | 242.967 | 2 | 52.842957 | yes | 9.722E-08 | 727.81207 | 90.281906 |
| EEGKDWAEAVGA | 14 | Q5JSZ5 | Protein PRRC2B OS=Homo sapiens OX= | PRRC2B | 242.967 | 2 | 52.842957 | yes | 9.722E-08 | 752.85498 | 80.888214 |
| SSPYGTLKPEEMS | 21 | Q5JSZ5 | Protein PRRC2B OS=Homo sapiens OX= | PRRC2B | 242.967 | 3 | 52.842957 | yes | 2.087E-07 | 725.69281 | 80.699539 |
| KVTPIDSPPVWSP | 22 | Q5JSZ5 | Protein PRRC2B OS=Homo sapiens OX= | PRRC2B | 242.967 | 3 | 52.842957 | yes | 2.947E-07 | 810.75061 | 80.085312 |
| WMMMPSYMDPR | 11 | Q5JSZ5 | Protein PRRC2B OS=Homo sapiens OX= | PRRC2B | 242.967 | 2 | 52.842957 | yes | 1.903E-07 | 722.79767 | 72.1231 |
| EFARPEDCPR | 10 | Q5JSZ5 | Protein PRRC2B OS=Homo sapiens OX= | PRRC2B | 242.967 | 2 | 52.842957 | yes | 9.819E-08 | 638.79059 | 67.934341 |
| PSGSAVNMGSVQ | 20 | Q5JSZ5 | Protein PRRC2B OS=Homo sapiens OX= | PRRC2B | 242.967 | 3 | 52.842957 | yes | 1.163E-07 | 682.3338 | 57.839306 |
| GVLDLSYGPGPSL | 14 | Q5JSZ5 | Protein PRRC2B OS=Homo sapiens OX= | PRRC2B | 242.967 | 2 | 52.842957 | yes | 1.932E-07 | 715.88556 | 55.191097 |
| KLHGWAPGPDYQ | 13 | Q5JSZ5 | Protein PRRC2B OS=Homo sapiens OX= | PRRC2B | 242.967 | 2 | 52.842957 | yes | 1.841E-07 | 748.88586 | 51.981308 |
| RLLEEER | 7 | Q5JSZ5 | Protein PRRC2B OS=Homo sapiens OX= | PRRC2B | 242.967 | 2 | 52.842957 | yes | 4.595E-07 | 472.76163 | 51.921898 |
| SFFQDEHVADSEN | 20 | Q5JSZ5 | Protein PRRC2B OS=Homo sapiens OX= | PRRC2B | 242.967 | 3 | 52.842957 | yes | 1.595E-07 | 799.02814 | 36.214478 |
| APRPTIINAENLK | 13 | Q5JSZ5 | Protein PRRC2B OS=Homo sapiens OX= | PRRC2B | 242.967 | 2 | 52.842957 | yes | 4.371E-07 | 718.91461 | 31.970955 |
| QQSIEDKEDKPPP | 14 | Q5JSZ5 | Protein PRRC2B OS=Homo sapiens OX= | PRRC2B | 242.967 | 3 | 52.842957 | yes | 5.051E-07 | 556.28467 | 26.819855 |
| AAGTCGVLLR | 10 | Q5T653 | Large ribosomal subunit protein uL2m OS | MRPL2 | 33.301 | 2 | 45.499863 | yes | 1.831E-07 | 509.27911 | 165.20543 |
| QMQVLETCVATV | 14 | Q5T653 | Large ribosomal subunit protein uL2m OS | MRPL2 | 33.301 | 2 | 45.499863 | yes | 9.583E-07 | 796.40015 | 151.21861 |
| WIIATENMQAGDT | 22 | Q5T653 | Large ribosomal subunit protein uL2m OS | MRPL2 | 33.301 | 3 | 45.499863 | yes | 2.236E-07 | 818.74463 | 106.96418 |
| IRPLPPMK | 8 | Q5T653 | Large ribosomal subunit protein uL2m OS | MRPL2 | 33.301 | 2 | 45.499863 | yes | 4.773E-06 | 476.29404 | 47.254139 |
| AISSTEAVLNNR | 12 | Q5T8P6 | RNA-binding protein 26 OS=Homo sapie | RBM26 | 113.598 | 2 | 57.19036 | no | 6.505E-07 | 637.83856 | 677.10352 |
| TLEPICDADPSALA | 15 | Q5T8P6 | RNA-binding protein 26 OS=Homo sapie | RBM26 | 113.598 | 2 | 57.19036 | yes | 1.517E-07 | 800.89801 | 510.71213 |
| VEFFPHQEK | 9 | Q5T8P6 | RNA-binding protein 26 OS=Homo sapie | RBM26 | 113.598 | 2 | 57.19036 | yes | 9.723E-08 | 580.79041 | 412.69672 |
| ELLDTELDLYK | 11 | Q5T8P6 | RNA-binding protein 26 OS=Homo sapie | RBM26 | 113.598 | 2 | 57.19036 | yes | 9.991E-08 | 676.35321 | 305.93546 |
| VMQPLVQQPILPV | 15 | Q5T8P6 | RNA-binding protein 26 OS=Homo sapie | RBM26 | 113.598 | 2 | 57.19036 | yes | 1.001E-07 | 845.01019 | 301.93262 |
| MQAGEEVTELR | 11 | Q5T8P6 | RNA-binding protein 26 OS=Homo sapie | RBM26 | 113.598 | 2 | 57.19036 | yes | 9.722E-08 | 631.80591 | 298.08813 |
| TLLVSTSAVDNNE | 16 | Q5T8P6 | RNA-binding protein 26 OS=Homo sapie | RBM26 | 113.598 | 2 | 57.19036 | yes | 1.081E-07 | 845.43634 | 267.56204 |

| | | | | | | | | | | | |
|---|---|---|---|---|---|---|---|---|---|---|---|
| EGSTQQLQTTSPK | 13 | Q5T8P6 | RNA-binding protein 26 OS=Homo sapie | RBM26 | 113.598 | 2 | 57.19036 | yes | 1.003E-07 | 702.85187 | 218.71562 |
| TIGSGEPGVPTK | 12 | Q5T8P6 | RNA-binding protein 26 OS=Homo sapie | RBM26 | 113.598 | 2 | 57.19036 | yes | 3.657E-07 | 571.80621 | 216.98964 |
| SYLPPPEQPSSGSL | 15 | Q5T8P6 | RNA-binding protein 26 OS=Homo sapie | RBM26 | 113.598 | 2 | 57.19036 | yes | 1.217E-06 | 793.90668 | 196.14467 |
| TVYNPAALK | 9 | Q5T8P6 | RNA-binding protein 26 OS=Homo sapie | RBM26 | 113.598 | 2 | 57.19036 | yes | 2.359E-07 | 488.77673 | 186.95534 |
| LNEHFSR | 7 | Q5T8P6 | RNA-binding protein 26 OS=Homo sapie | RBM26 | 113.598 | 2 | 57.19036 | yes | 1.656E-07 | 451.72757 | 176.34393 |
| LFDAVNTK | 8 | Q5T8P6 | RNA-binding protein 26 OS=Homo sapie | RBM26 | 113.598 | 2 | 57.19036 | yes | 9.849E-08 | 454.2478 | 142.57253 |
| YTELQLEAAK | 10 | Q5T8P6 | RNA-binding protein 26 OS=Homo sapie | RBM26 | 113.598 | 2 | 57.19036 | yes | 1.475E-07 | 583.30859 | 119.40993 |
| ALEISAFTESDR | 12 | Q5T8P6 | RNA-binding protein 26 OS=Homo sapie | RBM26 | 113.598 | 2 | 57.19036 | yes | 1.352E-07 | 669.83044 | 113.44627 |
| TWFDKPNFNR | 10 | Q5T8P6 | RNA-binding protein 26 OS=Homo sapie | RBM26 | 113.598 | 2 | 57.19036 | yes | 9.722E-08 | 662.82526 | 97.465897 |
| KQEILEK | 7 | Q5T8P6 | RNA-binding protein 26 OS=Homo sapie | RBM26 | 113.598 | 2 | 57.19036 | yes | 1.043E-07 | 444.26346 | 83.483948 |
| IVVDSESR | 8 | Q5T8P6 | RNA-binding protein 26 OS=Homo sapie | RBM26 | 113.598 | 2 | 57.19036 | yes | 1.44E-07 | 452.74036 | 76.341255 |
| AEAEAAAVHGAR | 12 | Q5T8P6 | RNA-binding protein 26 OS=Homo sapie | RBM26 | 113.598 | 2 | 57.19036 | yes | 1.517E-07 | 576.79144 | 61.036457 |
| KYTELQLEAAK | 11 | Q5T8P6 | RNA-binding protein 26 OS=Homo sapie | RBM26 | 113.598 | 2 | 57.19036 | yes | 2.636E-07 | 647.35608 | 58.441715 |
| ALCIDQLDVFLQK | 13 | Q5T8P6 | RNA-binding protein 26 OS=Homo sapie | RBM26 | 113.598 | 2 | 57.19036 | yes | 5.401E-07 | 781.91595 | 57.861664 |
| SEDKAEIMK | 9 | Q5T8P6 | RNA-binding protein 26 OS=Homo sapie | RBM26 | 113.598 | 2 | 57.19036 | yes | 4.098E-06 | 525.76044 | 27.462523 |
| RVTIMPK | 7 | Q5TEC6 | Histone H3-7 OS=Homo sapiens OX=96( | H3-7 | 15.431 | 2 | 46.98932 | no | 2.765E-07 | 422.75729 | 6243.1392 |
| RVTIMPK | 7 | Q5TEC6 | Histone H3-7 OS=Homo sapiens OX=96( | H3-7 | 15.431 | 2 | 46.98932 | no | 1.812E-07 | 430.75476 | 6243.1392 |
| YRPGTVALR | 9 | Q5TEC6 | Histone H3-7 OS=Homo sapiens OX=96( | H3-7 | 15.431 | 2 | 46.98932 | no | 1.425E-07 | 516.80109 | 11359.66 |
| EAYLVGLFEDTNL | 19 | Q5TEC6 | Histone H3-7 OS=Homo sapiens OX=96( | H3-7 | 15.431 | 3 | 46.98932 | yes | 1.637E-07 | 722.0296 | 80.482216 |
| STELLIR | 7 | Q5TEC6 | Histone H3-7 OS=Homo sapiens OX=96( | H3-7 | 15.431 | 2 | 46.98932 | no | 8.835E-07 | 416.25037 | 71.869713 |
| YLDVVNR | 7 | Q5VU97 | VWFA and cache domain-containing pro | CACHD1 | 142.29 | 2 | 44.85661 | yes | 2.044E-07 | 439.74017 | 396.91205 |
| TFSDISNFGAHQR | 13 | Q5VV52 | Zinc finger protein 691 OS=Homo sapien | ZNF691 | 35.81 | 2 | 49.502224 | yes | 2.991E-07 | 740.35242 | 178.93152 |
| SDLTTHQQDHLGK | 13 | Q5VV52 | Zinc finger protein 691 OS=Homo sapien | ZNF691 | 35.81 | 2 | 49.502224 | yes | 1.025E-07 | 740.36298 | 67.516121 |
| MLLSWDGGESR | 11 | Q658Y4 | Protein FAM91A1 OS=Homo sapiens OX | FAM91A1 | 93.909 | 2 | 55.644054 | yes | 4.351E-07 | 625.79535 | 1536.0626 |
| MLLSWDGGESR | 11 | Q658Y4 | Protein FAM91A1 OS=Homo sapiens OX | FAM91A1 | 93.909 | 2 | 55.644054 | yes | 5.205E-07 | 633.79279 | 1536.0626 |
| CESLLGLDPATCSI | 14 | Q658Y4 | Protein FAM91A1 OS=Homo sapiens OX | FAM91A1 | 93.909 | 2 | 55.644054 | yes | 2.2E-07 | 789.86615 | 2724.1411 |
| NQYIDLMNQCR | 11 | Q658Y4 | Protein FAM91A1 OS=Homo sapiens OX | FAM91A1 | 93.909 | 2 | 55.644054 | yes | 9.809E-08 | 727.82941 | 1754.8137 |
| YFDHALTLR | 9 | Q658Y4 | Protein FAM91A1 OS=Homo sapiens OX | FAM91A1 | 93.909 | 2 | 55.644054 | yes | 1.074E-07 | 568.2984 | 1744.2295 |
| VQGDYFETLLYK | 12 | Q658Y4 | Protein FAM91A1 OS=Homo sapiens OX | FAM91A1 | 93.909 | 2 | 55.644054 | yes | 1.602E-07 | 738.37445 | 1494.5742 |
| DGVLSEWSGR | 10 | Q658Y4 | Protein FAM91A1 OS=Homo sapiens OX | FAM91A1 | 93.909 | 2 | 55.644054 | yes | 1.113E-07 | 553.26727 | 1369.4221 |
| IIDSGPQLSGSLDY | 23 | Q658Y4 | Protein FAM91A1 OS=Homo sapiens OX | FAM91A1 | 93.909 | 3 | 55.644054 | yes | 1.408E-07 | 840.43073 | 1321.9358 |
| ESLQNLLHSSR | 11 | Q658Y4 | Protein FAM91A1 OS=Homo sapiens OX | FAM91A1 | 93.909 | 2 | 55.644054 | yes | 1.138E-07 | 642.33875 | 1160.9412 |
| KLPDIFQSYDR | 11 | Q658Y4 | Protein FAM91A1 OS=Homo sapiens OX | FAM91A1 | 93.909 | 2 | 55.644054 | no | 1E-07 | 691.35913 | 1045.5271 |
| SHAVTMFEVGK | 11 | Q658Y4 | Protein FAM91A1 OS=Homo sapiens OX | FAM91A1 | 93.909 | 2 | 55.644054 | yes | 1.001E-07 | 603.3028 | 972.94214 |
| ICTLPEK | 7 | Q658Y4 | Protein FAM91A1 OS=Homo sapiens OX | FAM91A1 | 93.909 | 2 | 55.644054 | yes | 1.051E-07 | 430.73096 | 905.61151 |
| IAAHGLCR | 8 | Q658Y4 | Protein FAM91A1 OS=Homo sapiens OX | FAM91A1 | 93.909 | 2 | 55.644054 | yes | 1.467E-07 | 449.23981 | 801.65625 |
| MNIDVEFHIR | 10 | Q658Y4 | Protein FAM91A1 OS=Homo sapiens OX | FAM91A1 | 93.909 | 2 | 55.644054 | yes | 2.562E-07 | 658.32678 | 327.86838 |
| MNIDVEFHIR | 10 | Q658Y4 | Protein FAM91A1 OS=Homo sapiens OX | FAM91A1 | 93.909 | 2 | 55.644054 | yes | 1.03E-05 | 666.32428 | 327.86838 |
| SPSSLLIANLHLQ | 13 | Q658Y4 | Protein FAM91A1 OS=Homo sapiens OX | FAM91A1 | 93.909 | 2 | 55.644054 | yes | 1.15E-07 | 696.89594 | 620.69653 |

| | | | | | | | | | | | |
|---|---|---|---|---|---|---|---|---|---|---|---|
| VDLQHLCGYVTM | 22 | Q658Y4 | Protein FAM91A1 OS=Homo sapiens OX | FAM91A1 | 93.909 | 3 | 55.644054 | yes | 9.722E-08 | 831.07379 | 565.49091 |
| NTILFLR | 7 | Q658Y4 | Protein FAM91A1 OS=Homo sapiens OX | FAM91A1 | 93.909 | 2 | 55.644054 | yes | 1.84E-07 | 438.76871 | 546.26221 |
| RYYEELLK | 8 | Q658Y4 | Protein FAM91A1 OS=Homo sapiens OX | FAM91A1 | 93.909 | 2 | 55.644054 | yes | 1.764E-07 | 557.3006 | 499.37311 |
| GQVINLDQLHSSW | 14 | Q658Y4 | Protein FAM91A1 OS=Homo sapiens OX | FAM91A1 | 93.909 | 2 | 55.644054 | yes | 9.722E-08 | 812.92572 | 410.35168 |
| YYEELLK | 7 | Q658Y4 | Protein FAM91A1 OS=Homo sapiens OX | FAM91A1 | 93.909 | 2 | 55.644054 | yes | 2.595E-07 | 479.25003 | 249.79845 |
| NAVSMYCR | 8 | Q658Y4 | Protein FAM91A1 OS=Homo sapiens OX | FAM91A1 | 93.909 | 2 | 55.644054 | yes | 1.259E-07 | 500.72058 | 183.15977 |
| KLSDASDER | 9 | Q658Y4 | Protein FAM91A1 OS=Homo sapiens OX | FAM91A1 | 93.909 | 2 | 55.644054 | yes | 6.615E-07 | 510.75146 | 148.06592 |
| QVVLYSIR | 8 | Q658Y4 | Protein FAM91A1 OS=Homo sapiens OX | FAM91A1 | 93.909 | 2 | 55.644054 | yes | 1.948E-07 | 489.29257 | 142.76295 |
| KESLQNLLHSSR | 12 | Q658Y4 | Protein FAM91A1 OS=Homo sapiens OX | FAM91A1 | 93.909 | 2 | 55.644054 | yes | 9.722E-08 | 706.38623 | 98.895889 |
| YHFAALEK | 8 | Q69YN2 | CWF19-like protein 1 OS=Homo sapiens | CWF19L1 | 60.62 | 2 | 51.689636 | yes | 4.602E-07 | 489.7558 | 128.49252 |
| EVLASEAILNVPDI | 14 | Q69YN2 | CWF19-like protein 1 OS=Homo sapiens | CWF19L1 | 60.62 | 2 | 51.689636 | yes | 9.722E-08 | 749.41156 | 88.658684 |
| WCVVFER | 7 | Q69YN2 | CWF19-like protein 1 OS=Homo sapiens | CWF19L1 | 60.62 | 2 | 51.689636 | yes | 2.425E-07 | 498.242 | 23.603254 |
| CVGNFGNSSGEVD | 15 | Q69YN2 | CWF19-like protein 1 OS=Homo sapiens | CWF19L1 | 60.62 | 2 | 51.689636 | yes | 2.819E-06 | 785.84375 | 18.64489 |
| ETMTNLHPK | 9 | Q6IPR3 | tRNA wybutosine-synthesizing protein 3 | TYW3 | 29.795 | 2 | 42.82771 | yes | 7.641E-07 | 535.76862 | 113.37981 |
| DQFFTTSSCAGR | 12 | Q6IPR3 | tRNA wybutosine-synthesizing protein 3 | TYW3 | 29.795 | 2 | 42.82771 | yes | 2.854E-07 | 688.79858 | 108.53966 |
| FYNCLQHALER | 11 | Q6IPR3 | tRNA wybutosine-synthesizing protein 3 | TYW3 | 29.795 | 2 | 42.82771 | yes | 1.246E-06 | 725.84845 | 41.622879 |
| SQGAITER | 8 | Q6NUK4 | Receptor expression-enhancing protein 3 | REEP3 | 29.264 | 2 | 48.307873 | yes | 4.973E-07 | 431.22488 | 598.74011 |
| PYQPLPEAK | 9 | Q6NUK4 | Receptor expression-enhancing protein 3 | REEP3 | 29.264 | 2 | 48.307873 | yes | 6.992E-07 | 521.78204 | 365.13135 |
| SKPAPSESAGYGIF | 16 | Q6NUK4 | Receptor expression-enhancing protein 3 | REEP3 | 29.264 | 2 | 48.307873 | yes | 1.166E-07 | 801.4303 | 215.50705 |
| GYETMVNFGR | 10 | Q6NUK4 | Receptor expression-enhancing protein 3 | REEP3 | 29.264 | 2 | 48.307873 | yes | 2.479E-07 | 587.27148 | 146.86606 |
| KFLHPLLSSK | 10 | Q6NUK4 | Receptor expression-enhancing protein 3 | REEP3 | 29.264 | 2 | 48.307873 | yes | 1.165E-07 | 585.35571 | 140.30183 |
| QGLNLAATAAVTA | 16 | Q6NUK4 | Receptor expression-enhancing protein 3 | REEP3 | 29.264 | 2 | 48.307873 | yes | 7.318E-07 | 749.93304 | 77.489235 |
| EIDDYIVQAK | 10 | Q6NUK4 | Receptor expression-enhancing protein 3 | REEP3 | 29.264 | 2 | 48.307873 | yes | 2.256E-07 | 597.30603 | 71.997124 |
| SFSMHDLTTIQGDI | 28 | Q6NUK4 | Receptor expression-enhancing protein 3 | REEP3 | 29.264 | 4 | 48.307873 | yes | 2.69E-07 | 786.13818 | 66.464684 |
| KRPQVYF | 7 | Q6NUK4 | Receptor expression-enhancing protein 3 | REEP3 | 29.264 | 2 | 48.307873 | yes | 2.054E-07 | 469.26636 | 52.234253 |
| EREIDDYIVQAK | 12 | Q6NUK4 | Receptor expression-enhancing protein 3 | REEP3 | 29.264 | 2 | 48.307873 | yes | 3.515E-07 | 739.87793 | 44.344532 |
| LVCVGPECSLFYSI | 14 | Q6NVV7 | Cysteine-rich DPF motif domain-containi | CDPF1 | 13.878 | 2 | 48.761158 | yes | 1.107E-07 | 829.89948 | 61.923687 |
| LFLINEQSPR | 10 | Q6NW34 | Nucleolus and neural progenitor protein ( | NEPRO | 64.552 | 2 | 53.391243 | yes | 3.061E-07 | 608.83789 | 169.69258 |
| QLLNSGVSMPVIQ | 15 | Q6NW34 | Nucleolus and neural progenitor protein ( | NEPRO | 64.552 | 2 | 53.391243 | yes | 1.065E-07 | 807.94781 | 76.669701 |
| ATQETSFDFK | 10 | Q6NW34 | Nucleolus and neural progenitor protein ( | NEPRO | 64.552 | 2 | 53.391243 | yes | 6.818E-07 | 587.27478 | 59.959808 |
| ASEETLLGISK | 11 | Q6NW34 | Nucleolus and neural progenitor protein ( | NEPRO | 64.552 | 2 | 53.391243 | yes | 3.518E-07 | 574.3139 | 59.532879 |
| INVQNNVDLGQPV | 14 | Q6NW34 | Nucleolus and neural progenitor protein ( | NEPRO | 64.552 | 2 | 53.391243 | yes | 1.326E-07 | 769.42029 | 52.9865 |
| HLEAQGTSLPK | 11 | Q6NW34 | Nucleolus and neural progenitor protein ( | NEPRO | 64.552 | 2 | 53.391243 | yes | 9.722E-08 | 590.81964 | 46.909096 |
| VMTFDDDIR | 9 | Q6NXE6 | Armadillo repeat-containing protein 6 OS | ARMC6 | 54.142 | 2 | 31.566822 | yes | 1.798E-06 | 556.25806 | 73.652786 |
| DQSGVQELVK | 10 | Q6NXE6 | Armadillo repeat-containing protein 6 OS | ARMC6 | 54.142 | 2 | 31.566822 | yes | 2.324E-06 | 551.79059 | 59.720837 |
| TPIPTSLR | 8 | Q6P1L8 | Large ribosomal subunit protein uL14m ( | MRPL14 | 15.948 | 2 | 49.767204 | yes | 9.946E-07 | 442.76364 | 884.57996 |
| VGDQILLAIK | 10 | Q6P1L8 | Large ribosomal subunit protein uL14m ( | MRPL14 | 15.948 | 2 | 49.767204 | yes | 1.899E-07 | 535.33441 | 706.15955 |
| VVDNSALGNSPYH | 14 | Q6P1L8 | Large ribosomal subunit protein uL14m ( | MRPL14 | 15.948 | 2 | 49.767204 | yes | 1.004E-07 | 764.87878 | 193.18825 |
| LQGFEFWSR | 9 | Q6P1R4 | tRNA-dihydrouridine(16/17) synthase [N | DUS1L | 53.231 | 2 | 47.14606 | yes | 1.39E-07 | 585.29053 | 77.178322 |

| | | | | | | | | | | | |
|---|---|---|---|---|---|---|---|---|---|---|---|
| ALEEEEGGTEVLSl | 14 | Q6P1R4 | tRNA-dihydrouridine(16/17) synthase [N | DUS1L | 53.231 | 2 | 47.14606 | yes | 4.119E-07 | 745.86469 | 47.465607 |
| TFDPSLKPK | 9 | Q6P1R4 | tRNA-dihydrouridine(16/17) synthase [N | DUS1L | 53.231 | 2 | 47.14606 | yes | 2.817E-06 | 516.78986 | 40.777534 |
| DKGSAALGGVLAI | 16 | Q6P582 | Mitotic-spindle organizing protein 2A OS | MZT2A | 16.221 | 2 | 25.676088 | yes | 3.244E-06 | 764.4281 | 19.785173 |
| FQAEIPLMR | 9 | Q6PJG2 | Mitotic deacetylase-associated SANT dom | MIDEAS | 114.989 | 2 | 53.101124 | yes | 1.066E-07 | 552.79718 | 419.64941 |
| TSAAMLSQQVAS\ | 14 | Q6PJG2 | Mitotic deacetylase-associated SANT dom | MIDEAS | 114.989 | 2 | 53.101124 | yes | 9.722E-08 | 710.87683 | 247.87703 |
| ALAAADPHK | 9 | Q6PJG2 | Mitotic deacetylase-associated SANT dom | MIDEAS | 114.989 | 2 | 53.101124 | yes | 1.716E-06 | 447.24561 | 195.10075 |
| CGVEFSEPSLATK | 13 | Q6PJG2 | Mitotic deacetylase-associated SANT dom | MIDEAS | 114.989 | 2 | 53.101124 | yes | 9.722E-08 | 712.83997 | 187.53241 |
| AVSTGDCGQVLR | 12 | Q6PJG2 | Mitotic deacetylase-associated SANT dom | MIDEAS | 114.989 | 2 | 53.101124 | yes | 3.316E-07 | 631.81152 | 159.26294 |
| AVELASLQNAK | 11 | Q6PJG2 | Mitotic deacetylase-associated SANT dom | MIDEAS | 114.989 | 2 | 53.101124 | yes | 1.467E-07 | 572.32202 | 151.91563 |
| ASQEANLLTLAQK | 13 | Q6PJG2 | Mitotic deacetylase-associated SANT dom | MIDEAS | 114.989 | 2 | 53.101124 | yes | 4.228E-07 | 693.883 | 139.80157 |
| LEGEPSVR | 8 | Q6PJG2 | Mitotic deacetylase-associated SANT dom | MIDEAS | 114.989 | 2 | 53.101124 | yes | 7.446E-07 | 443.73508 | 102.99335 |
| TVAQCVEFYYTYk | 13 | Q6PJG2 | Mitotic deacetylase-associated SANT dom | MIDEAS | 114.989 | 2 | 53.101124 | yes | 9.722E-08 | 836.38977 | 100.90579 |
| EAPAMGSEEGMR | 12 | Q6PJG2 | Mitotic deacetylase-associated SANT dom | MIDEAS | 114.989 | 2 | 53.101124 | yes | 1.418E-07 | 632.76843 | 89.453163 |
| EQPPPLQPPQQSIR | 14 | Q6PJG2 | Mitotic deacetylase-associated SANT dom | MIDEAS | 114.989 | 2 | 53.101124 | yes | 2.807E-07 | 807.93353 | 78.767136 |
| WPNSVMAPGR | 10 | Q6PJG2 | Mitotic deacetylase-associated SANT dom | MIDEAS | 114.989 | 2 | 53.101124 | yes | 4.153E-06 | 557.77673 | 72.888634 |
| KEGEEEVPEIQEK | 13 | Q6PJG2 | Mitotic deacetylase-associated SANT dom | MIDEAS | 114.989 | 2 | 53.101124 | yes | 5.062E-07 | 772.37793 | 39.329674 |
| NGTLTFGDVDTSD | 15 | Q6PJG2 | Mitotic deacetylase-associated SANT dom | MIDEAS | 114.989 | 2 | 53.101124 | yes | 1.635E-06 | 799.86267 | 35.899349 |
| TQNQENTFPCK | 11 | Q6PJG2 | Mitotic deacetylase-associated SANT dom | MIDEAS | 114.989 | 2 | 53.101124 | yes | 3.116E-06 | 683.80646 | 35.438751 |
| QRPRPEPLIIPTK | 13 | Q6PJG2 | Mitotic deacetylase-associated SANT dom | MIDEAS | 114.989 | 3 | 53.101124 | yes | 2.813E-05 | 515.64722 | 25.902353 |
| IPGTDAQAQAEDM | 16 | Q6PJG2 | Mitotic deacetylase-associated SANT dom | MIDEAS | 114.989 | 2 | 53.101124 | yes | 2.943E-07 | 844.40143 | 13.811671 |
| IVEILPEHLK | 10 | Q6UXM1 | Leucine-rich repeats and immunoglobulin | LRIG3 | 123.434 | 2 | 50.41008 | yes | 9.809E-08 | 595.86078 | 47.133415 |
| CLGDLLDCSR | 10 | Q6UXM1 | Leucine-rich repeats and immunoglobulin | LRIG3 | 123.434 | 2 | 50.41008 | yes | 4.457E-06 | 604.77356 | 19.286568 |
| AQGGEVMEYTTIL | 14 | Q6UXM1 | Leucine-rich repeats and immunoglobulin | LRIG3 | 123.434 | 2 | 50.41008 | yes | 2.243E-06 | 784.39288 | 3.8220077 |
| ITSLYVECR | 9 | Q6ZNB6 | NF-X1-type zinc finger protein NFXL1 C | NFXL1 | 101.34 | 2 | 54.83756 | yes | 9.728E-08 | 570.78955 | 731.81262 |
| GSCYPCPETVDVK | 13 | Q6ZNB6 | NF-X1-type zinc finger protein NFXL1 C | NFXL1 | 101.34 | 2 | 54.83756 | yes | 1.183E-07 | 756.32886 | 509.26837 |
| SRPLGCLHPCILR | 13 | Q6ZNB6 | NF-X1-type zinc finger protein NFXL1 C | NFXL1 | 101.34 | 3 | 54.83756 | yes | 2.65E-07 | 526.95001 | 361.04538 |
| VLECGIHR | 8 | Q6ZNB6 | NF-X1-type zinc finger protein NFXL1 C | NFXL1 | 101.34 | 2 | 54.83756 | yes | 2.213E-07 | 492.25818 | 300.0802 |
| QQAELEAFENR | 11 | Q6ZNB6 | NF-X1-type zinc finger protein NFXL1 C | NFXL1 | 101.34 | 2 | 54.83756 | yes | 1.616E-07 | 667.82037 | 273.40085 |
| HSPAGSQALQTTA | 21 | Q6ZNB6 | NF-X1-type zinc finger protein NFXL1 C | NFXL1 | 101.34 | 3 | 54.83756 | yes | 1.568E-07 | 715.01978 | 234.62032 |
| MVTTTCYCK | 9 | Q6ZNB6 | NF-X1-type zinc finger protein NFXL1 C | NFXL1 | 101.34 | 2 | 54.83756 | yes | 1.121E-07 | 582.24829 | 229.06841 |
| ITTADVNEK | 9 | Q6ZNB6 | NF-X1-type zinc finger protein NFXL1 C | NFXL1 | 101.34 | 2 | 54.83756 | yes | 2.541E-07 | 495.75876 | 184.95213 |
| DCPWPCPK | 8 | Q6ZNB6 | NF-X1-type zinc finger protein NFXL1 C | NFXL1 | 101.34 | 2 | 54.83756 | yes | 1.166E-07 | 530.22296 | 180.04654 |
| ILDCQNHTCMK | 11 | Q6ZNB6 | NF-X1-type zinc finger protein NFXL1 C | NFXL1 | 101.34 | 2 | 54.83756 | yes | 9.722E-08 | 710.3125 | 166.62892 |
| CGHLCPAPCHDQA | 16 | Q6ZNB6 | NF-X1-type zinc finger protein NFXL1 C | NFXL1 | 101.34 | 3 | 54.83756 | yes | 9.722E-08 | 626.29199 | 154.50449 |
| EWSCQLPCGQK | 11 | Q6ZNB6 | NF-X1-type zinc finger protein NFXL1 C | NFXL1 | 101.34 | 2 | 54.83756 | yes | 9.722E-08 | 696.80536 | 135.56384 |
| CLLLCHPGPCPPCI | 15 | Q6ZNB6 | NF-X1-type zinc finger protein NFXL1 C | NFXL1 | 101.34 | 3 | 54.83756 | yes | 1.101E-07 | 602.61542 | 121.52428 |
| KITTADVNEK | 10 | Q6ZNB6 | NF-X1-type zinc finger protein NFXL1 C | NFXL1 | 101.34 | 2 | 54.83756 | yes | 1.186E-07 | 559.80621 | 70.053871 |
| QYVNEAFQAGAM | 22 | Q6ZNB6 | NF-X1-type zinc finger protein NFXL1 C | NFXL1 | 101.34 | 3 | 54.83756 | yes | 9.722E-08 | 825.40076 | 54.697151 |
| GPCETCR | 7 | Q6ZNB6 | NF-X1-type zinc finger protein NFXL1 C | NFXL1 | 101.34 | 2 | 54.83756 | yes | 9.219E-07 | 440.17603 | 43.440872 |

| | | | | | | | | | | | |
|---|---|---|---|---|---|---|---|---|---|---|---|
| NLLSCCK | 7 | Q6ZNB6 | NF-X1-type zinc finger protein NFXL1 C | NFXL1 | 101.34 | 2 | 54.83756 | yes | 2.318E-06 | 447.71222 | 38.445377 |
| ENQVSIECDTTCK | 13 | Q6ZNB6 | NF-X1-type zinc finger protein NFXL1 C | NFXL1 | 101.34 | 2 | 54.83756 | yes | 2.144E-07 | 792.33746 | 26.53055 |
| MPCHKPYLCETK | 12 | Q6ZNB6 | NF-X1-type zinc finger protein NFXL1 C | NFXL1 | 101.34 | 2 | 54.83756 | yes | 2.281E-07 | 782.35944 | 25.844212 |
| YYCYCGK | 7 | Q6ZNB6 | NF-X1-type zinc finger protein NFXL1 C | NFXL1 | 101.34 | 2 | 54.83756 | yes | 1.751E-06 | 507.19641 | 17.263908 |
| SSREFIER | 8 | Q6ZWK6 | Transmembrane protease serine 11F OS=I | TMPRSS11 | 49.41 | 2 | 41.30832 | yes | 3.574E-07 | 512.26453 | 216.41034 |
| LLSVGDGTR | 9 | Q7L2K0 | Tubulin epsilon and delta complex protei | TEDC2 | 46.403 | 2 | 43.51919 | yes | 2.556E-07 | 459.25616 | 95.691612 |
| QLQLEQSLR | 9 | Q7L2K0 | Tubulin epsilon and delta complex protei | TEDC2 | 46.403 | 2 | 43.51919 | yes | 4.934E-07 | 557.81439 | 77.427811 |
| VFQPQNLR | 8 | Q7L3T8 | Probable proline--tRNA ligase, mitochon | PARS2 | 53.263 | 2 | 51.26312 | yes | 4.621E-06 | 501.27997 | 207.6196 |
| GIEVGHTFYLGTK | 13 | Q7L3T8 | Probable proline--tRNA ligase, mitochon | PARS2 | 53.263 | 2 | 51.26312 | yes | 1.024E-07 | 711.37482 | 142.57033 |
| FGYPFVIIAGK | 11 | Q7L3T8 | Probable proline--tRNA ligase, mitochon | PARS2 | 53.263 | 2 | 51.26312 | yes | 3.176E-07 | 606.34479 | 114.02277 |
| VIDQEMQAIGGQK | 13 | Q7L3T8 | Probable proline--tRNA ligase, mitochon | PARS2 | 53.263 | 2 | 51.26312 | yes | 9.722E-08 | 708.86121 | 93.362686 |
| THLTIGNR | 8 | Q7L3T8 | Probable proline--tRNA ligase, mitochon | PARS2 | 53.263 | 2 | 51.26312 | yes | 1.068E-07 | 456.2565 | 91.215706 |
| PTLAEMGCYGLGV | 15 | Q7L3T8 | Probable proline--tRNA ligase, mitochon | PARS2 | 53.263 | 2 | 51.26312 | yes | 2.158E-07 | 812.89471 | 48.791519 |
| QLPFLLYQVTR | 11 | Q7L3T8 | Probable proline--tRNA ligase, mitochon | PARS2 | 53.263 | 2 | 51.26312 | yes | 5.635E-06 | 689.39807 | 18.257122 |
| EEVEVAQVQVPTI | 15 | Q7Z2T5 | TRMT1-like protein OS=Homo sapiens C | TRMT1L | 81.748 | 2 | 52.863205 | yes | 9.722E-08 | 826.4361 | 970.30804 |
| VSVEFEGYR | 9 | Q7Z2T5 | TRMT1-like protein OS=Homo sapiens C | TRMT1L | 81.748 | 2 | 52.863205 | yes | 1.128E-07 | 543.26672 | 803.32965 |
| THFDPMGVR | 9 | Q7Z2T5 | TRMT1-like protein OS=Homo sapiens C | TRMT1L | 81.748 | 2 | 52.863205 | yes | 9.722E-08 | 530.25568 | 706.1524 |
| QACPLCPK | 8 | Q7Z2T5 | TRMT1-like protein OS=Homo sapiens C | TRMT1L | 81.748 | 2 | 52.863205 | yes | 3.152E-07 | 487.23334 | 679.61395 |
| MDANVLMHLR | 10 | Q7Z2T5 | TRMT1-like protein OS=Homo sapiens C | TRMT1L | 81.748 | 2 | 52.863205 | yes | 9.722E-08 | 600.30481 | 548.97888 |
| TDMLGHVR | 8 | Q7Z2T5 | TRMT1-like protein OS=Homo sapiens C | TRMT1L | 81.748 | 2 | 52.863205 | yes | 1.325E-07 | 464.73709 | 539.68298 |
| SSYIAASTAKPPK | 13 | Q7Z2T5 | TRMT1-like protein OS=Homo sapiens C | TRMT1L | 81.748 | 2 | 52.863205 | yes | 1.371E-07 | 660.86151 | 410.74591 |
| FLCYLSQAGFR | 11 | Q7Z2T5 | TRMT1-like protein OS=Homo sapiens C | TRMT1L | 81.748 | 2 | 52.863205 | yes | 1.339E-07 | 681.33716 | 376.73633 |
| SDDILEEGEK | 10 | Q7Z2T5 | TRMT1-like protein OS=Homo sapiens C | TRMT1L | 81.748 | 2 | 52.863205 | yes | 3.091E-07 | 567.76166 | 308.17123 |
| TDAPLMQFK | 9 | Q7Z2T5 | TRMT1-like protein OS=Homo sapiens C | TRMT1L | 81.748 | 2 | 52.863205 | yes | 1.421E-06 | 525.76807 | 262.60391 |
| HYGCNIVR | 8 | Q7Z2T5 | TRMT1-like protein OS=Homo sapiens C | TRMT1L | 81.748 | 2 | 52.863205 | yes | 5.278E-07 | 509.74799 | 228.26178 |
| SNEMITNLGK | 10 | Q7Z2T5 | TRMT1-like protein OS=Homo sapiens C | TRMT1L | 81.748 | 2 | 52.863205 | yes | 1.784E-06 | 553.77917 | 218.87978 |
| HLQNLHWK | 8 | Q7Z2T5 | TRMT1-like protein OS=Homo sapiens C | TRMT1L | 81.748 | 2 | 52.863205 | yes | 1.14E-07 | 538.2934 | 178.02272 |
| KPLECLDAFGATG | 21 | Q7Z2T5 | TRMT1-like protein OS=Homo sapiens C | TRMT1L | 81.748 | 3 | 52.863205 | yes | 9.722E-08 | 769.39392 | 133.65366 |
| MCICHLPCR | 9 | Q7Z2T5 | TRMT1-like protein OS=Homo sapiens C | TRMT1L | 81.748 | 2 | 52.863205 | yes | 1.214E-07 | 623.76923 | 132.98495 |
| TTDDTTTDNYIAQ | 15 | Q7Z2T5 | TRMT1-like protein OS=Homo sapiens C | TRMT1L | 81.748 | 2 | 52.863205 | yes | 9.722E-08 | 822.37341 | 131.82767 |
| TDSYFNPK | 8 | Q7Z2T5 | TRMT1-like protein OS=Homo sapiens C | TRMT1L | 81.748 | 2 | 52.863205 | yes | 4.486E-07 | 486.22708 | 112.51698 |
| DGNMVEENPYR | 11 | Q7Z2T5 | TRMT1-like protein OS=Homo sapiens C | TRMT1L | 81.748 | 2 | 52.863205 | yes | 2.004E-07 | 662.28534 | 102.0274 |
| EKSDDILEEGEK | 12 | Q7Z2T5 | TRMT1-like protein OS=Homo sapiens C | TRMT1L | 81.748 | 2 | 52.863205 | yes | 9.722E-08 | 696.33044 | 70.969421 |
| QLPCNCHGSMPGI | 13 | Q7Z2T5 | TRMT1-like protein OS=Homo sapiens C | TRMT1L | 81.748 | 3 | 52.863205 | yes | 1.106E-07 | 495.8847 | 60.481571 |
| KSNEMITNLGK | 11 | Q7Z2T5 | TRMT1-like protein OS=Homo sapiens C | TRMT1L | 81.748 | 2 | 52.863205 | yes | 2.559E-07 | 617.82666 | 59.669044 |
| IVTTPQTVLR | 10 | Q7Z2W9 | Large ribosomal subunit protein bL21m C | MRPL21 | 22.815 | 2 | 52.689335 | yes | 1.643E-07 | 564.34277 | 256.93723 |
| LFAVVHFASR | 10 | Q7Z2W9 | Large ribosomal subunit protein bL21m C | MRPL21 | 22.815 | 2 | 52.689335 | yes | 1.02E-07 | 573.82458 | 169.80498 |
| VLLVGADNFTLLC | 14 | Q7Z2W9 | Large ribosomal subunit protein bL21m C | MRPL21 | 22.815 | 2 | 52.689335 | yes | 1.066E-07 | 730.42957 | 154.74908 |
| VNEMIVTGQYGR | 12 | Q7Z2W9 | Large ribosomal subunit protein bL21m C | MRPL21 | 22.815 | 2 | 52.689335 | yes | 9.722E-08 | 683.84283 | 113.93525 |

| | | | | | | | | | | | |
|---|---|---|---|---|---|---|---|---|---|---|---|
| KVNEMIVTGQYGI | 13 | Q7Z2W9 | Large ribosomal subunit protein bL21m C | MRPL21 | 22.815 | 2 | 52.689335 | yes | 1.248E-07 | 747.89032 | 32.878979 |
| PLPTCVAECGGTV | 14 | Q7Z408 | CUB and sushi domain-containing protei | CSMD2 | 380.04 | 2 | 28.715206 | yes | 2.683E-06 | 758.86597 | 36.24794 |
| FIAYQETVEK | 10 | Q7Z4Q2 | HEAT repeat-containing protein 3 OS=H | HEATR3 | 74.583 | 2 | 52.260426 | yes | 1.018E-07 | 614.31641 | 168.59642 |
| SQAEIINALLK | 11 | Q7Z4Q2 | HEAT repeat-containing protein 3 OS=H | HEATR3 | 74.583 | 2 | 52.260426 | yes | 9.722E-08 | 600.35333 | 130.96806 |
| LGPLLLDPSLAVR | 13 | Q7Z4Q2 | HEAT repeat-containing protein 3 OS=H | HEATR3 | 74.583 | 2 | 52.260426 | yes | 1.916E-07 | 682.41901 | 128.01999 |
| TLVAGTIWNLK | 11 | Q7Z4Q2 | HEAT repeat-containing protein 3 OS=H | HEATR3 | 74.583 | 2 | 52.260426 | yes | 9.722E-08 | 608.35846 | 127.57116 |
| ECACAGLAR | 9 | Q7Z4Q2 | HEAT repeat-containing protein 3 OS=H | HEATR3 | 74.583 | 2 | 52.260426 | yes | 2.781E-07 | 504.22348 | 107.67665 |
| AGIHSSNVGVR | 11 | Q7Z4Q2 | HEAT repeat-containing protein 3 OS=H | HEATR3 | 74.583 | 2 | 52.260426 | yes | 2.031E-07 | 548.79651 | 91.596512 |
| TFVSDLLPPTDK | 12 | Q7Z4Q2 | HEAT repeat-containing protein 3 OS=H | HEATR3 | 74.583 | 2 | 52.260426 | yes | 7.851E-07 | 666.8559 | 89.851135 |
| TAFPNSIAVDLCSF | 14 | Q7Z4Q2 | HEAT repeat-containing protein 3 OS=H | HEATR3 | 74.583 | 2 | 52.260426 | yes | 1.92E-06 | 775.88519 | 47.379772 |
| NIGCFLLEVTTK | 12 | Q7Z4Q2 | HEAT repeat-containing protein 3 OS=H | HEATR3 | 74.583 | 2 | 52.260426 | yes | 4.338E-07 | 697.87103 | 24.633549 |
| GGGLGSPGEGGAL | 15 | Q7Z7A3 | Cytoplasmic tRNA 2-thiolation protein 1 | CTU1 | 36.45 | 2 | 52.19417 | yes | 1.022E-07 | 641.33093 | 179.48613 |
| ALCQACALLDGLN | 14 | Q7Z7A3 | Cytoplasmic tRNA 2-thiolation protein 1 | CTU1 | 36.45 | 2 | 52.19417 | yes | 1.841E-07 | 787.89252 | 156.83936 |
| SCCTFCGVLR | 10 | Q7Z7A3 | Cytoplasmic tRNA 2-thiolation protein 1 | CTU1 | 36.45 | 2 | 52.19417 | yes | 9.722E-08 | 630.27014 | 114.12788 |
| DSTVLAHVLR | 10 | Q7Z7A3 | Cytoplasmic tRNA 2-thiolation protein 1 | CTU1 | 36.45 | 2 | 52.19417 | yes | 1.604E-07 | 555.81696 | 73.102997 |
| EVVLYAHFR | 9 | Q7Z7A3 | Cytoplasmic tRNA 2-thiolation protein 1 | CTU1 | 36.45 | 2 | 52.19417 | yes | 2.985E-07 | 567.30872 | 53.14159 |
| QDGSTIHIR | 9 | Q7Z7F7 | Large ribosomal subunit protein mL55 OS | MRPL55 | 15.129 | 2 | 47.606228 | yes | 1.295E-07 | 513.76996 | 96.019264 |
| PGLIWIPPPGR | 11 | Q86SX3 | Tubulin epsilon and delta complex protei | TEDC1 | 54.232 | 2 | 55.60649 | yes | 9.722E-08 | 601.85583 | 221.56779 |
| LALAQLPEDGSQG | 15 | Q86SX3 | Tubulin epsilon and delta complex protei | TEDC1 | 54.232 | 2 | 55.60649 | yes | 2.474E-07 | 771.39954 | 172.02104 |
| GPVPEQMLAQAR | 12 | Q86SX3 | Tubulin epsilon and delta complex protei | TEDC1 | 54.232 | 2 | 55.60649 | yes | 2.661E-07 | 648.84009 | 166.21915 |
| ELQALEEELR | 10 | Q86SX3 | Tubulin epsilon and delta complex protei | TEDC1 | 54.232 | 2 | 55.60649 | yes | 9.722E-08 | 615.32227 | 128.65247 |
| SLPSGPSPEIFR | 12 | Q86SX3 | Tubulin epsilon and delta complex protei | TEDC1 | 54.232 | 2 | 55.60649 | yes | 4.395E-07 | 643.84058 | 117.45624 |
| SQEACLEAVLR | 11 | Q86SX3 | Tubulin epsilon and delta complex protei | TEDC1 | 54.232 | 2 | 55.60649 | yes | 9.722E-08 | 638.32172 | 89.118896 |
| ELGALQQCWER | 11 | Q86SX3 | Tubulin epsilon and delta complex protei | TEDC1 | 54.232 | 2 | 55.60649 | yes | 9.722E-08 | 695.33264 | 76.082588 |
| HVQWLMGK | 8 | Q86SX3 | Tubulin epsilon and delta complex protei | TEDC1 | 54.232 | 2 | 55.60649 | yes | 2.612E-05 | 499.76566 | 44.196968 |
| GFSAEDVQR | 9 | Q86TN4 | tRNA 2'-phosphotransferase 1 OS=Homo | TRPT1 | 27.742 | 2 | 51.506973 | yes | 1.392E-07 | 504.74088 | 132.13864 |
| FALQLGDPSTGLLI | 15 | Q86TN4 | tRNA 2'-phosphotransferase 1 OS=Homo | TRPT1 | 27.742 | 2 | 51.506973 | yes | 9.722E-08 | 800.95648 | 57.268547 |
| VLDAIWSGAALR | 12 | Q86U38 | Nucleolar protein 9 OS=Homo sapiens O | NOP9 | 69.438 | 2 | 51.088127 | yes | 1.13E-07 | 636.35895 | 328.23187 |
| EVETQALALSTNR | 13 | Q86U38 | Nucleolar protein 9 OS=Homo sapiens O | NOP9 | 69.438 | 2 | 51.088127 | yes | 9.722E-08 | 716.37555 | 299.21283 |
| DLMVHNIMK | 9 | Q86U38 | Nucleolar protein 9 OS=Homo sapiens O | NOP9 | 69.438 | 2 | 51.088127 | yes | 9.809E-08 | 550.7832 | 236.70518 |
| GQYVALACSR | 10 | Q86U38 | Nucleolar protein 9 OS=Homo sapiens O | NOP9 | 69.438 | 2 | 51.088127 | yes | 1.453E-07 | 562.77948 | 181.74422 |
| DPFGHHVAR | 9 | Q86U38 | Nucleolar protein 9 OS=Homo sapiens O | NOP9 | 69.438 | 2 | 51.088127 | yes | 1.313E-07 | 518.25958 | 144.93663 |
| EAWEQQQGAVAK | 12 | Q86U38 | Nucleolar protein 9 OS=Homo sapiens O | NOP9 | 69.438 | 2 | 51.088127 | yes | 1.028E-07 | 672.83075 | 58.121639 |
| CGVHVLQSALLQI | 15 | Q86U38 | Nucleolar protein 9 OS=Homo sapiens O | NOP9 | 69.438 | 2 | 51.088127 | yes | 3.496E-06 | 845.97467 | 25.638355 |
| CGVHVLQSALLQI | 15 | Q86U38 | Nucleolar protein 9 OS=Homo sapiens O | NOP9 | 69.438 | 2 | 51.088127 | yes | 5.437E-07 | 564.31891 | 25.638355 |
| CGVHVLQSALLQI | 15 | Q86U38 | Nucleolar protein 9 OS=Homo sapiens O | NOP9 | 69.438 | 3 | 51.088127 | yes | 3.496E-06 | 845.97467 | 25.638355 |
| CGVHVLQSALLQI | 15 | Q86U38 | Nucleolar protein 9 OS=Homo sapiens O | NOP9 | 69.438 | 3 | 51.088127 | yes | 5.437E-07 | 564.31891 | 25.638355 |
| ENLQDQIR | 8 | Q86XH1 | Dynein regulatory complex protein 11 OS | IQCA1 | 95.342 | 2 | 34.310005 | yes | 1.035E-06 | 508.26199 | 138.09633 |
| ATDLILDHIR | 10 | Q86Y79 | Peptidyl-tRNA hydrolase OS=Homo sapi | PTRH1 | 22.937 | 2 | 56.956417 | yes | 4.856E-07 | 583.83002 | 1371.1274 |

| | | | | | | | | | | | |
|---|---|---|---|---|---|---|---|---|---|---|---|
| LGVAESWTR | 9 | Q86Y79 | Peptidyl-tRNA hydrolase OS=Homo sapi | PTRH1 | 22.937 | 2 | 56.956417 | yes | 6.102E-07 | 509.76944 | 629.02704 |
| CVLEPRPPGK | 10 | Q86Y79 | Peptidyl-tRNA hydrolase OS=Homo sapi | PTRH1 | 22.937 | 2 | 56.956417 | yes | 1.264E-07 | 576.81335 | 436.81696 |
| WMVAGLGNPGLP | 15 | Q86Y79 | Peptidyl-tRNA hydrolase OS=Homo sapi | PTRH1 | 22.937 | 2 | 56.956417 | yes | 1.406E-07 | 763.40082 | 360.4278 |
| HSVGMAVLGQLA | 13 | Q86Y79 | Peptidyl-tRNA hydrolase OS=Homo sapi | PTRH1 | 22.937 | 2 | 56.956417 | yes | 1.203E-07 | 669.86914 | 206.19562 |
| SCISCLNSNAMPR | 13 | Q86Y79 | Peptidyl-tRNA hydrolase OS=Homo sapi | PTRH1 | 22.937 | 2 | 56.956417 | yes | 9.722E-08 | 755.33398 | 106.77209 |
| HCAADLALAPLGI | 22 | Q86Y79 | Peptidyl-tRNA hydrolase OS=Homo sapi | PTRH1 | 22.937 | 4 | 56.956417 | yes | 6.001E-07 | 593.33234 | 45.859333 |
| HCAADLALAPLGI | 22 | Q86Y79 | Peptidyl-tRNA hydrolase OS=Homo sapi | PTRH1 | 22.937 | 4 | 56.956417 | yes | 5.066E-07 | 790.77399 | 45.859333 |
| HCAADLALAPLGI | 22 | Q86Y79 | Peptidyl-tRNA hydrolase OS=Homo sapi | PTRH1 | 22.937 | 3 | 56.956417 | yes | 6.001E-07 | 593.33234 | 45.859333 |
| HCAADLALAPLGI | 22 | Q86Y79 | Peptidyl-tRNA hydrolase OS=Homo sapi | PTRH1 | 22.937 | 3 | 56.956417 | yes | 5.066E-07 | 790.77399 | 45.859333 |
| DRHCAADLALAPI | 24 | Q86Y79 | Peptidyl-tRNA hydrolase OS=Homo sapi | PTRH1 | 22.937 | 4 | 56.956417 | yes | 3.293E-07 | 661.11432 | 64.407974 |
| VLDALVAR | 8 | Q8IUR0 | Trafficking protein particle complex subu | TRAPPC5 | 20.784 | 2 | 37.426125 | yes | 1.42E-06 | 428.76617 | 53.607468 |
| EPLINTYISVPK | 12 | Q8IUR0 | Trafficking protein particle complex subu | TRAPPC5 | 20.784 | 2 | 37.426125 | yes | 6.003E-07 | 687.38739 | 43.825108 |
| FEEAVIAR | 8 | Q8IUR0 | Trafficking protein particle complex subu | TRAPPC5 | 20.784 | 2 | 37.426125 | yes | 3.265E-06 | 467.75327 | 24.756283 |
| CSLAVPQDEHVIK | 13 | Q8IWR0 | Zinc finger CCCH domain-containing prc | ZC3H7A | 110.538 | 2 | 52.713158 | yes | 2.388E-07 | 748.38232 | 1290.2736 |
| CSLAVPQDEHVIK | 13 | Q8IWR0 | Zinc finger CCCH domain-containing prc | ZC3H7A | 110.538 | 2 | 52.713158 | yes | 1.36E-07 | 499.25729 | 1290.2736 |
| CSLAVPQDEHVIK | 13 | Q8IWR0 | Zinc finger CCCH domain-containing prc | ZC3H7A | 110.538 | 3 | 52.713158 | yes | 2.388E-07 | 748.38232 | 1290.2736 |
| CSLAVPQDEHVIK | 13 | Q8IWR0 | Zinc finger CCCH domain-containing prc | ZC3H7A | 110.538 | 3 | 52.713158 | yes | 1.36E-07 | 499.25729 | 1290.2736 |
| FVCAQCLR | 8 | Q8IWR0 | Zinc finger CCCH domain-containing prc | ZC3H7A | 110.538 | 2 | 52.713158 | yes | 2.301E-07 | 527.25208 | 1732.8514 |
| FPTGYFSICDR | 11 | Q8IWR0 | Zinc finger CCCH domain-containing prc | ZC3H7A | 110.538 | 2 | 52.713158 | yes | 9.935E-08 | 681.81097 | 1549.4255 |
| TNYEGPYYICK | 11 | Q8IWR0 | Zinc finger CCCH domain-containing prc | ZC3H7A | 110.538 | 2 | 52.713158 | yes | 9.722E-08 | 704.3161 | 1279.0464 |
| CLVHILR | 7 | Q8IWR0 | Zinc finger CCCH domain-containing prc | ZC3H7A | 110.538 | 2 | 52.713158 | yes | 2.393E-07 | 455.76819 | 1119.4515 |
| ALNHSVEDIEPDLI | 17 | Q8IWR0 | Zinc finger CCCH domain-containing prc | ZC3H7A | 110.538 | 3 | 52.713158 | yes | 1.156E-07 | 640.3338 | 1071.0824 |
| QACQICFVK | 9 | Q8IWR0 | Zinc finger CCCH domain-containing prc | ZC3H7A | 110.538 | 2 | 52.713158 | yes | 1.613E-07 | 577.27826 | 911.60486 |
| HPLEGTHELR | 10 | Q8IWR0 | Zinc finger CCCH domain-containing prc | ZC3H7A | 110.538 | 2 | 52.713158 | yes | 1.083E-07 | 594.80963 | 899.02301 |
| NLFNEGNDVYR | 11 | Q8IWR0 | Zinc finger CCCH domain-containing prc | ZC3H7A | 110.538 | 2 | 52.713158 | yes | 1.202E-07 | 670.81512 | 734.28644 |
| IACYSNMGFHDK | 12 | Q8IWR0 | Zinc finger CCCH domain-containing prc | ZC3H7A | 110.538 | 2 | 52.713158 | yes | 1.089E-07 | 721.81323 | 727.26123 |
| SEDIASQSNK | 10 | Q8IWR0 | Zinc finger CCCH domain-containing prc | ZC3H7A | 110.538 | 2 | 52.713158 | yes | 1.814E-07 | 539.75421 | 657.12646 |
| QWQGHISSEK | 10 | Q8IWR0 | Zinc finger CCCH domain-containing prc | ZC3H7A | 110.538 | 2 | 52.713158 | yes | 9.722E-08 | 600.29382 | 516.52936 |
| FAHGNAELHEWEI | 14 | Q8IWR0 | Zinc finger CCCH domain-containing prc | ZC3H7A | 110.538 | 3 | 52.713158 | yes | 9.79E-08 | 575.59735 | 514.57147 |
| LTQELAQK | 8 | Q8IWR0 | Zinc finger CCCH domain-containing prc | ZC3H7A | 110.538 | 2 | 52.713158 | yes | 9.722E-08 | 465.76639 | 484.54773 |
| SFHGQCQLDLCR | 12 | Q8IWR0 | Zinc finger CCCH domain-containing prc | ZC3H7A | 110.538 | 2 | 52.713158 | yes | 1.418E-07 | 760.84027 | 454.74243 |
| NDFGNFFGSAVTK | 13 | Q8IWR0 | Zinc finger CCCH domain-containing prc | ZC3H7A | 110.538 | 2 | 52.713158 | yes | 1.046E-07 | 702.33313 | 414.55258 |
| WMNIRPLPTK | 10 | Q8IWR0 | Zinc finger CCCH domain-containing prc | ZC3H7A | 110.538 | 2 | 52.713158 | yes | 1.343E-07 | 628.3526 | 328.11078 |
| DHLIGPNDNDFGK | 13 | Q8IWR0 | Zinc finger CCCH domain-containing prc | ZC3H7A | 110.538 | 2 | 52.713158 | yes | 1.025E-07 | 721.33893 | 229.03815 |
| KDILIGR | 7 | Q8IWR0 | Zinc finger CCCH domain-containing prc | ZC3H7A | 110.538 | 2 | 52.713158 | yes | 1.05E-07 | 407.76089 | 166.06824 |
| KDHLIGPNDNDFG | 14 | Q8IWR0 | Zinc finger CCCH domain-containing prc | ZC3H7A | 110.538 | 2 | 52.713158 | yes | 9.722E-08 | 785.38641 | 165.2384 |
| NGQVIEPDKNR | 11 | Q8IWR0 | Zinc finger CCCH domain-containing prc | ZC3H7A | 110.538 | 2 | 52.713158 | yes | 9.722E-08 | 635.33093 | 143.9704 |
| DVAAEEECR | 9 | Q8IWR0 | Zinc finger CCCH domain-containing prc | ZC3H7A | 110.538 | 2 | 52.713158 | yes | 3.61E-07 | 539.72711 | 105.56863 |
| SEEILIPK | 8 | Q8IWR0 | Zinc finger CCCH domain-containing prc | ZC3H7A | 110.538 | 2 | 52.713158 | yes | 2.188E-07 | 464.77112 | 91.080437 |

| | | | | | | | | | | | |
|---|---|---|---|---|---|---|---|---|---|---|---|
| DLSTSTSR | 8 | Q8IWR0 | Zinc finger CCCH domain-containing prc | ZC3H7A | 110.538 | 2 | 52.713158 | yes | 2.004E-06 | 433.71432 | 27.55411 |
| YGCLREDECFYAF | 19 | Q8IWR0 | Zinc finger CCCH domain-containing prc | ZC3H7A | 110.538 | 3 | 52.713158 | yes | 1.313E-06 | 797.03656 | 10.19038 |
| PYQCGECGK | 9 | Q8IWY8 | Zinc finger and SCAN domain-containing | ZSCAN29 | 96.72 | 2 | 47.63726 | no | 1.287E-07 | 549.72076 | 67.318878 |
| DISEEVQLHR | 10 | Q8IWY8 | Zinc finger and SCAN domain-containing | ZSCAN29 | 96.72 | 2 | 47.63726 | yes | 2.028E-07 | 613.31219 | 36.289955 |
| ELPSGSHIGDR | 11 | Q8IWY8 | Zinc finger and SCAN domain-containing | ZSCAN29 | 96.72 | 2 | 47.63726 | yes | 1.897E-07 | 584.29126 | 31.941954 |
| CLAIHDVIR | 9 | Q8IX18 | Probable ATP-dependent RNA helicase D | DHX40 | 88.561 | 2 | 55.038673 | yes | 9.722E-08 | 548.80023 | 214.18019 |
| FSVIILDEAHER | 12 | Q8IX18 | Probable ATP-dependent RNA helicase D | DHX40 | 88.561 | 2 | 55.038673 | yes | 9.722E-08 | 714.87769 | 182.19815 |
| HILGDPNLTK | 10 | Q8IX18 | Probable ATP-dependent RNA helicase D | DHX40 | 88.561 | 2 | 55.038673 | yes | 6.189E-07 | 554.31146 | 153.36696 |
| LSAVCIADR | 9 | Q8IX18 | Probable ATP-dependent RNA helicase D | DHX40 | 88.561 | 2 | 55.038673 | yes | 2.035E-07 | 502.76331 | 139.40237 |
| FPYLDPPNER | 10 | Q8IX18 | Probable ATP-dependent RNA helicase D | DHX40 | 88.561 | 2 | 55.038673 | yes | 8.955E-07 | 624.3064 | 111.34554 |
| YMTDGCLLK | 9 | Q8IX18 | Probable ATP-dependent RNA helicase D | DHX40 | 88.561 | 2 | 55.038673 | yes | 5.183E-07 | 550.7594 | 103.25706 |
| KVAAISVAQR | 10 | Q8IX18 | Probable ATP-dependent RNA helicase D | DHX40 | 88.561 | 2 | 55.038673 | yes | 3.684E-07 | 521.82202 | 80.172806 |
| DNSFLIVTGNTGSC | 15 | Q8IX18 | Probable ATP-dependent RNA helicase D | DHX40 | 88.561 | 2 | 55.038673 | yes | 9.722E-08 | 755.3808 | 76.275398 |
| QLYQCDAIDR | 10 | Q8IX18 | Probable ATP-dependent RNA helicase D | DHX40 | 88.561 | 2 | 55.038673 | yes | 2.112E-06 | 641.29822 | 67.090385 |
| FCNLIGPR | 8 | Q8IX18 | Probable ATP-dependent RNA helicase D | DHX40 | 88.561 | 2 | 55.038673 | yes | 2.994E-06 | 488.75528 | 49.773705 |
| VVVMSATMELAK | 12 | Q8IX18 | Probable ATP-dependent RNA helicase D | DHX40 | 88.561 | 2 | 55.038673 | yes | 2.324E-06 | 647.84088 | 47.836781 |
| TSLTSVVLTLK | 11 | Q8IX18 | Probable ATP-dependent RNA helicase D | DHX40 | 88.561 | 2 | 55.038673 | yes | 9.819E-08 | 581.35809 | 39.526455 |
| YLYEAGFSQHGMI | 19 | Q8IX18 | Probable ATP-dependent RNA helicase D | DHX40 | 88.561 | 3 | 55.038673 | yes | 1.221E-06 | 718.6861 | 27.205915 |
| TLYVHWTDAGQLT | 20 | Q8IY67 | Ribonucleoprotein PTB-binding 1 OS=Ho | RAVER1 | 63.878 | 3 | 51.08569 | yes | 2.786E-07 | 760.40259 | 516.37347 |
| TLYVHWTDAGQLT | 20 | Q8IY67 | Ribonucleoprotein PTB-binding 1 OS=Ho | RAVER1 | 63.878 | 3 | 51.08569 | yes | 2.063E-07 | 570.55371 | 516.37347 |
| TLYVHWTDAGQLT | 20 | Q8IY67 | Ribonucleoprotein PTB-binding 1 OS=Ho | RAVER1 | 63.878 | 4 | 51.08569 | yes | 2.786E-07 | 760.40259 | 516.37347 |
| TLYVHWTDAGQLT | 20 | Q8IY67 | Ribonucleoprotein PTB-binding 1 OS=Ho | RAVER1 | 63.878 | 4 | 51.08569 | yes | 2.063E-07 | 570.55371 | 516.37347 |
| LPPGFNDVDALCR | 13 | Q8IY67 | Ribonucleoprotein PTB-binding 1 OS=Ho | RAVER1 | 63.878 | 2 | 51.08569 | yes | 1.325E-07 | 737.36139 | 876.33801 |
| SDLLGKPLGPR | 11 | Q8IY67 | Ribonucleoprotein PTB-binding 1 OS=Ho | RAVER1 | 63.878 | 2 | 51.08569 | yes | 9.935E-08 | 576.84039 | 761.30444 |
| CFLVYSER | 8 | Q8IY67 | Ribonucleoprotein PTB-binding 1 OS=Ho | RAVER1 | 63.878 | 2 | 51.08569 | yes | 2.096E-07 | 537.25787 | 692.40881 |
| SGAEVEAGDAAEF | 13 | Q8IY67 | Ribonucleoprotein PTB-binding 1 OS=Ho | RAVER1 | 63.878 | 2 | 51.08569 | yes | 1.722E-07 | 631.28638 | 494.78 |
| GYGFAEYMK | 9 | Q8IY67 | Ribonucleoprotein PTB-binding 1 OS=Ho | RAVER1 | 63.878 | 2 | 51.08569 | yes | 2.254E-07 | 533.23914 | 428.81311 |
| GLPGDVTNQEVHI | 21 | Q8IY67 | Ribonucleoprotein PTB-binding 1 OS=Ho | RAVER1 | 63.878 | 3 | 51.08569 | yes | 1.08E-06 | 781.38855 | 392.2883 |
| AADVSVTHRPPLS | 15 | Q8IY67 | Ribonucleoprotein PTB-binding 1 OS=Ho | RAVER1 | 63.878 | 3 | 51.08569 | yes | 1.041E-07 | 539.63 | 102.69647 |
| AADVSVTHRPPLS | 15 | Q8IY67 | Ribonucleoprotein PTB-binding 1 OS=Ho | RAVER1 | 63.878 | 3 | 51.08569 | yes | 9.722E-08 | 808.94141 | 102.69647 |
| AADVSVTHRPPLS | 15 | Q8IY67 | Ribonucleoprotein PTB-binding 1 OS=Ho | RAVER1 | 63.878 | 2 | 51.08569 | yes | 1.041E-07 | 539.63 | 102.69647 |
| AADVSVTHRPPLS | 15 | Q8IY67 | Ribonucleoprotein PTB-binding 1 OS=Ho | RAVER1 | 63.878 | 2 | 51.08569 | yes | 9.722E-08 | 808.94141 | 102.69647 |
| VSFCAPGPPGR | 11 | Q8IY67 | Ribonucleoprotein PTB-binding 1 OS=Ho | RAVER1 | 63.878 | 2 | 51.08569 | yes | 9.722E-08 | 572.78204 | 82.345604 |
| PFGSLER | 7 | Q8IY67 | Ribonucleoprotein PTB-binding 1 OS=Ho | RAVER1 | 63.878 | 2 | 51.08569 | yes | 5.682E-06 | 403.21378 | 67.982841 |
| EALGLGPPAAQLT | 22 | Q8IY67 | Ribonucleoprotein PTB-binding 1 OS=Ho | RAVER1 | 63.878 | 3 | 51.08569 | yes | 7.574E-06 | 708.07214 | 23.660437 |
| ECGGLQTLLR | 10 | Q8IYL2 | Probable tRNA (uracil-O(2)-)-methyltran: | TRMT44 | 84.629 | 2 | 48.794964 | yes | 1.104E-07 | 573.80042 | 37.856255 |
| WSAALPCAEAR | 11 | Q8IYL2 | Probable tRNA (uracil-O(2)-)-methyltran: | TRMT44 | 84.629 | 2 | 48.794964 | yes | 2.877E-06 | 616.29803 | 32.468967 |
| SISLEPLQK | 9 | Q8N0T1 | Ribosomal biogenesis factor OS=Homo sa | RBIS | 11.457 | 2 | 50.189457 | yes | 3.07E-07 | 507.79514 | 298.49655 |
| AKPVTTNLK | 9 | Q8N0T1 | Ribosomal biogenesis factor OS=Homo sa | RBIS | 11.457 | 2 | 50.189457 | yes | 1.118E-07 | 486.29785 | 261.66254 |

| | | | | | | | | | | | |
|---|---|---|---|---|---|---|---|---|---|---|---|
| NVFHIASQK | 9 | Q8N0T1 | Ribosomal biogenesis factor OS=Homo sa | RBIS | 11.457 | 2 | 50.189457 | yes | 8.288E-07 | 522.28528 | 99.223457 |
| HESKPVNVDEATF | 13 | Q8N0T1 | Ribosomal biogenesis factor OS=Homo sa | RBIS | 11.457 | 3 | 50.189457 | yes | 9.854E-08 | 494.58295 | 95.480331 |
| INIMNEEK | 8 | Q8N0T1 | Ribosomal biogenesis factor OS=Homo sa | RBIS | 11.457 | 2 | 50.189457 | yes | 9.212E-07 | 495.74988 | 72.245712 |
| ELIPQQR | 7 | Q8N0T1 | Ribosomal biogenesis factor OS=Homo sa | RBIS | 11.457 | 2 | 50.189457 | yes | 9.574E-07 | 442.25345 | 63.224491 |
| SGRPPFPPEVLAEA | 20 | Q8N0Z8 | tRNA pseudouridine synthase-like 1 OS= | PUSL1 | 33.233 | 3 | 47.40171 | yes | 9.585E-07 | 734.39905 | 29.992443 |
| VAPAHGLFLK | 10 | Q8N0Z8 | tRNA pseudouridine synthase-like 1 OS= | PUSL1 | 33.233 | 2 | 47.40171 | yes | 1.339E-07 | 526.81622 | 16.203251 |
| LSPTAATVQFK | 11 | Q8N143 | B-cell CLL/lymphoma 6 member B protei | BCL6B | 51.532 | 2 | 39.71404 | yes | 4.422E-07 | 581.82697 | 57.560783 |
| ELLLTGPGLEER | 12 | Q8N1G4 | Leucine-rich repeat-containing protein 47 | LRRC47 | 63.473 | 2 | 52.81899 | yes | 9.722E-08 | 663.86682 | 1089.3381 |
| DGPSLLVVEQVR | 12 | Q8N1G4 | Leucine-rich repeat-containing protein 47 | LRRC47 | 63.473 | 2 | 52.81899 | yes | 9.722E-08 | 656.367 | 826.86285 |
| ADLATAPPHVTVV | 14 | Q8N1G4 | Leucine-rich repeat-containing protein 47 | LRRC47 | 63.473 | 2 | 52.81899 | yes | 9.722E-08 | 723.9068 | 449.77313 |
| GMDLQPGNALK | 11 | Q8N1G4 | Leucine-rich repeat-containing protein 47 | LRRC47 | 63.473 | 2 | 52.81899 | yes | 1.111E-07 | 572.29498 | 387.12753 |
| ELSPDIAHLASLK | 13 | Q8N1G4 | Leucine-rich repeat-containing protein 47 | LRRC47 | 63.473 | 2 | 52.81899 | yes | 1.448E-07 | 697.38794 | 382.47757 |
| VLHVSENPVPLTV | 14 | Q8N1G4 | Leucine-rich repeat-containing protein 47 | LRRC47 | 63.473 | 2 | 52.81899 | yes | 9.722E-08 | 780.44885 | 292.20313 |
| QLQLEAEEQR | 10 | Q8N1G4 | Leucine-rich repeat-containing protein 47 | LRRC47 | 63.473 | 2 | 52.81899 | yes | 1.817E-07 | 622.3175 | 236.92937 |
| QSVSGLHR | 8 | Q8N1G4 | Leucine-rich repeat-containing protein 47 | LRRC47 | 63.473 | 2 | 52.81899 | yes | 2.349E-07 | 442.24084 | 132.30945 |
| GPLLYCARPPQDL | 14 | Q8N1G4 | Leucine-rich repeat-containing protein 47 | LRRC47 | 63.473 | 2 | 52.81899 | yes | 7.531E-07 | 814.43488 | 26.550488 |
| AFSQSTSLCIHQR | 13 | Q8N3J9 | Zinc finger protein 664 OS=Homo sapien | ZNF664 | 30.284 | 2 | 50.18167 | yes | 9.861E-08 | 767.87518 | 33.493053 |
| FEFLGINR | 8 | Q8N3X1 | Formin-binding protein 4 OS=Homo sapi | FNBP4 | 110.266 | 2 | 51.8527 | yes | 4.376E-07 | 498.26907 | 321.27106 |
| QQQLVSGMAER | 11 | Q8N3X1 | Formin-binding protein 4 OS=Homo sapi | FNBP4 | 110.266 | 2 | 51.8527 | yes | 2.149E-07 | 623.81409 | 236.39613 |
| ATEISTAVVQR | 11 | Q8N3X1 | Formin-binding protein 4 OS=Homo sapi | FNBP4 | 110.266 | 2 | 51.8527 | yes | 1.245E-07 | 587.82495 | 218.90601 |
| FQIGELANTLTSK | 13 | Q8N3X1 | Formin-binding protein 4 OS=Homo sapi | FNBP4 | 110.266 | 2 | 51.8527 | yes | 9.81E-08 | 711.38538 | 198.92191 |
| LQDAAEQLK | 9 | Q8N3X1 | Formin-binding protein 4 OS=Homo sapi | FNBP4 | 110.266 | 2 | 51.8527 | yes | 5.296E-05 | 508.27457 | 100.75558 |
| KLQDAAEQLK | 10 | Q8N3X1 | Formin-binding protein 4 OS=Homo sapi | FNBP4 | 110.266 | 2 | 51.8527 | yes | 1.823E-07 | 572.32202 | 68.497101 |
| NANFEALPEDWR | 12 | Q8N3X1 | Formin-binding protein 4 OS=Homo sapi | FNBP4 | 110.266 | 2 | 51.8527 | yes | 1.102E-07 | 731.34149 | 57.761635 |
| SDISQPASQDGMR | 13 | Q8N3X1 | Formin-binding protein 4 OS=Homo sapi | FNBP4 | 110.266 | 2 | 51.8527 | yes | 3.571E-07 | 696.31464 | 53.496681 |
| KATEISTAVVQR | 12 | Q8N3X1 | Formin-binding protein 4 OS=Homo sapi | FNBP4 | 110.266 | 2 | 51.8527 | yes | 1.466E-07 | 651.87244 | 53.407269 |
| QYEINATPK | 9 | Q8N3X1 | Formin-binding protein 4 OS=Homo sapi | FNBP4 | 110.266 | 2 | 51.8527 | yes | 9.722E-08 | 532.27454 | 52.980652 |
| VVQNPPKPVMTTF | 13 | Q8N3X1 | Formin-binding protein 4 OS=Homo sapi | FNBP4 | 110.266 | 2 | 51.8527 | yes | 1.16E-07 | 733.91101 | 29.287895 |
| GVAASLLAPLLPE( | 21 | Q8N3X1 | Formin-binding protein 4 OS=Homo sapi | FNBP4 | 110.266 | 3 | 51.8527 | yes | 1.179E-06 | 741.07397 | 28.119465 |
| LLELTGTIDK | 10 | Q8N442 | Translation factor GUF1, mitochondrial ( | GUF1 | 74.328 | 2 | 43.87957 | yes | 2.875E-07 | 551.82135 | 111.63365 |
| ALVFDSTFDQYR | 12 | Q8N442 | Translation factor GUF1, mitochondrial ( | GUF1 | 74.328 | 2 | 43.87957 | yes | 2.409E-07 | 731.35406 | 83.163132 |
| IMMLCEAR | 8 | Q8N442 | Translation factor GUF1, mitochondrial ( | GUF1 | 74.328 | 2 | 43.87957 | yes | 3.639E-07 | 512.24286 | 53.251759 |
| GVIANVALFDGVV | 15 | Q8N442 | Translation factor GUF1, mitochondrial ( | GUF1 | 74.328 | 2 | 43.87957 | yes | 2.501E-06 | 744.92468 | 24.70686 |
| LLLMSTDQEELSR | 13 | Q8N4T4 | Rho guanine nucleotide exchange factor 3 | ARHGEF39 | 38.296 | 2 | 51.86312 | yes | 9.732E-08 | 767.8927 | 305.05585 |
| SQTTLQEQLK | 10 | Q8N4T4 | Rho guanine nucleotide exchange factor 3 | ARHGEF39 | 38.296 | 2 | 51.86312 | yes | 9.722E-08 | 588.31696 | 166.37851 |
| ELLETER | 7 | Q8N4T4 | Rho guanine nucleotide exchange factor 3 | ARHGEF39 | 38.296 | 2 | 51.86312 | yes | 4.547E-07 | 445.23489 | 149.84303 |
| PRPPLHLLR | 9 | Q8N4T4 | Rho guanine nucleotide exchange factor 3 | ARHGEF39 | 38.296 | 2 | 51.86312 | yes | 2.531E-06 | 549.84839 | 56.854031 |
| HVVFSGSGR | 9 | Q8N5L8 | Ribonuclease P protein subunit p25-like p | RPP25L | 17.632 | 2 | 46.470566 | yes | 2.496E-07 | 473.24869 | 589.52277 |
| AVSCAEIVK | 9 | Q8N5L8 | Ribonuclease P protein subunit p25-like p | RPP25L | 17.632 | 2 | 46.470566 | yes | 1.551E-07 | 488.76022 | 495.00833 |

| | | | | | | | | | | | |
|---|---|---|---|---|---|---|---|---|---|---|---|
| HVPAVWVLLSR | 11 | Q8N5L8 | Ribonuclease P protein subunit p25-like p | RPP25L | 17.632 | 2 | 46.470566 | yes | 3.147E-07 | 638.87988 | 293.17484 |
| VPGLHQLTK | 9 | Q8N5L8 | Ribonuclease P protein subunit p25-like p | RPP25L | 17.632 | 2 | 46.470566 | yes | 2.256E-06 | 496.798 | 148.22417 |
| RVPGLHQLTK | 10 | Q8N5L8 | Ribonuclease P protein subunit p25-like p | RPP25L | 17.632 | 2 | 46.470566 | yes | 2.615E-07 | 574.84857 | 136.15152 |
| NLLGLALGR | 9 | Q8N5L8 | Ribonuclease P protein subunit p25-like p | RPP25L | 17.632 | 2 | 46.470566 | yes | 6.393E-07 | 463.79272 | 44.110535 |
| DVLDFYNVPIQDR | 13 | Q8N5N7 | Large ribosomal subunit protein mL50 OS | MRPL50 | 18.325 | 2 | 51.247196 | yes | 9.722E-08 | 797.39899 | 2671.0508 |
| AYTPPEDLQSR | 11 | Q8N5N7 | Large ribosomal subunit protein mL50 OS | MRPL50 | 18.325 | 2 | 51.247196 | yes | 2.411E-07 | 638.81201 | 1524.5872 |
| KEPILVCPPLR | 11 | Q8N5N7 | Large ribosomal subunit protein mL50 OS | MRPL50 | 18.325 | 2 | 51.247196 | yes | 9.722E-08 | 661.38666 | 292.40875 |
| KEPILVCPPLR | 11 | Q8N5N7 | Large ribosomal subunit protein mL50 OS | MRPL50 | 18.325 | 2 | 51.247196 | yes | 3.907E-07 | 441.26019 | 292.40875 |
| KEPILVCPPLR | 11 | Q8N5N7 | Large ribosomal subunit protein mL50 OS | MRPL50 | 18.325 | 3 | 51.247196 | yes | 9.722E-08 | 661.38666 | 292.40875 |
| KEPILVCPPLR | 11 | Q8N5N7 | Large ribosomal subunit protein mL50 OS | MRPL50 | 18.325 | 3 | 51.247196 | yes | 3.907E-07 | 441.26019 | 292.40875 |
| EKEPVVVETVEEK | 13 | Q8N5N7 | Large ribosomal subunit protein mL50 OS | MRPL50 | 18.325 | 2 | 51.247196 | yes | 2.175E-07 | 757.90106 | 528.95154 |
| EPVVVETVEEK | 11 | Q8N5N7 | Large ribosomal subunit protein mL50 OS | MRPL50 | 18.325 | 2 | 51.247196 | yes | 2.025E-07 | 629.33228 | 330.52429 |
| FDELSASNLPPNLK | 14 | Q8N5N7 | Large ribosomal subunit protein mL50 OS | MRPL50 | 18.325 | 2 | 51.247196 | yes | 2.464E-07 | 772.90137 | 287.25974 |
| VFMWTVSGTPCR | 12 | Q8N5N7 | Large ribosomal subunit protein mL50 OS | MRPL50 | 18.325 | 2 | 51.247196 | yes | 1.202E-07 | 720.84174 | 92.59668 |
| LSCWVYK | 7 | Q8N7A1 | Kelch domain-containing protein 1 OS=H | KLHDC1 | 46.72 | 2 | 31.806744 | yes | 1.719E-06 | 478.23895 | 249.73116 |
| LLQSSGADFHK | 11 | Q8N8A2 | Serine/threonine-protein phosphatase 6 re | ANKRD44 | 107.604 | 2 | 48.787136 | yes | 1.429E-07 | 601.81183 | 74.243889 |
| QCLELLLER | 9 | Q8N8A2 | Serine/threonine-protein phosphatase 6 re | ANKRD44 | 107.604 | 2 | 48.787136 | yes | 1.104E-07 | 587.31842 | 58.153957 |
| LLLEIADNPEAVD | 15 | Q8N8A2 | Serine/threonine-protein phosphatase 6 re | ANKRD44 | 107.604 | 2 | 48.787136 | yes | 1.286E-07 | 819.95105 | 47.733047 |
| TEDVNTLDSEK | 11 | Q8N8A2 | Serine/threonine-protein phosphatase 6 re | ANKRD44 | 107.604 | 2 | 48.787136 | yes | 1.995E-07 | 625.79095 | 43.929779 |
| GHTECVEALINQG | 19 | Q8N8A2 | Serine/threonine-protein phosphatase 6 re | ANKRD44 | 107.604 | 3 | 48.787136 | yes | 1.347E-06 | 691.68597 | 23.817572 |
| TALDLAAFK | 9 | Q8N8A2 | Serine/threonine-protein phosphatase 6 re | ANKRD44 | 107.604 | 2 | 48.787136 | yes | 1.458E-05 | 475.2713 | 20.13422 |
| VFNIYTDATPLR | 12 | Q8N8A6 | ATP-dependent RNA helicase DDX51 OS | DDX51 | 72.458 | 2 | 46.71279 | yes | 1.894E-07 | 705.37482 | 77.386307 |
| LLFSATLTQNPEK | 13 | Q8N8A6 | ATP-dependent RNA helicase DDX51 OS | DDX51 | 72.458 | 2 | 46.71279 | yes | 8.043E-07 | 731.401 | 70.53437 |
| MLTEAGAPELQR | 12 | Q8N8A6 | ATP-dependent RNA helicase DDX51 OS | DDX51 | 72.458 | 2 | 46.71279 | yes | 1.489E-07 | 658.33734 | 55.1064 |
| SGGGIVEPIPLNIK | 14 | Q8N954 | G patch domain-containing protein 11 OS | GPATCH11 | 33.278 | 2 | 50.23634 | yes | 9.843E-08 | 697.40613 | 462.26764 |
| LQILTSYLR | 9 | Q8N954 | G patch domain-containing protein 11 OS | GPATCH11 | 33.278 | 2 | 50.23634 | yes | 1.594E-07 | 553.83203 | 236.84117 |
| ACQQLDVQK | 9 | Q8N954 | G patch domain-containing protein 11 OS | GPATCH11 | 33.278 | 2 | 50.23634 | yes | 1.423E-07 | 545.27148 | 132.28717 |
| SLKEEEQER | 9 | Q8N954 | G patch domain-containing protein 11 OS | GPATCH11 | 33.278 | 2 | 50.23634 | yes | 2.419E-07 | 574.28314 | 116.21469 |
| SGIGHEASLK | 10 | Q8N954 | G patch domain-containing protein 11 OS | GPATCH11 | 33.278 | 2 | 50.23634 | yes | 2.518E-07 | 499.76691 | 81.559395 |
| NKQDEMK | 7 | Q8N954 | G patch domain-containing protein 11 OS | GPATCH11 | 33.278 | 2 | 50.23634 | yes | 2.581E-05 | 446.71329 | 33.870621 |
| ESESEEAEAGAAE | 15 | Q8NAF0 | Zinc finger protein 579 OS=Homo sapien | ZNF579 | 60.51 | 2 | 46.994556 | yes | 1.424E-07 | 789.34991 | 42.363445 |
| YLVVPPLETGLK | 12 | Q8NB91 | Fanconi anemia group B protein OS=Hon | FANCB | 97.727 | 2 | 53.30505 | yes | 9.722E-08 | 664.89484 | 83.699684 |
| VFLSLEDLSTGK | 12 | Q8NB91 | Fanconi anemia group B protein OS=Hon | FANCB | 97.727 | 2 | 53.30505 | yes | 1.155E-07 | 654.8559 | 75.841728 |
| TPILHVR | 7 | Q8NB91 | Fanconi anemia group B protein OS=Hon | FANCB | 97.727 | 2 | 53.30505 | yes | 2.818E-07 | 418.26105 | 66.098839 |
| ISLIALTR | 8 | Q8NB91 | Fanconi anemia group B protein OS=Hon | FANCB | 97.727 | 2 | 53.30505 | yes | 3.251E-06 | 443.78964 | 43.14859 |
| LSDNFQDSEQLVE | 14 | Q8NB91 | Fanconi anemia group B protein OS=Hon | FANCB | 97.727 | 2 | 53.30505 | yes | 2.85E-07 | 826.3941 | 33.622368 |
| IMCCNCVSDFR | 11 | Q8NB91 | Fanconi anemia group B protein OS=Hon | FANCB | 97.727 | 2 | 53.30505 | yes | 9.722E-08 | 731.29071 | 19.532919 |
| TSISSQYVIR | 10 | Q8NBA8 | tRNA-uridine aminocarboxypropyltransfe | DTWD2 | 33.416 | 2 | 43.368347 | yes | 2.621E-07 | 577.31421 | 219.08859 |
| TVPLLAACLPQDK | 13 | Q8NBA8 | tRNA-uridine aminocarboxypropyltransfe | DTWD2 | 33.416 | 2 | 43.368347 | yes | 4.149E-07 | 713.39215 | 139.59171 |

| | | | | | | | | | | | |
|---|---|---|---|---|---|---|---|---|---|---|---|
| GGGLSLLPPPPGGI | 14 | Q8NC96 | Adaptin ear-binding coat-associated prote | NECAP1 | 29.738 | 2 | 49.9932 | yes | 1.137E-07 | 623.86133 | 244.60338 |
| SAFIGIGFTDR | 11 | Q8NC96 | Adaptin ear-binding coat-associated prote | NECAP1 | 29.738 | 2 | 49.9932 | yes | 9.937E-08 | 592.30896 | 176.2113 |
| GTNTSSPYDAGAD | 16 | Q8ND82 | Zinc finger protein 280C OS=Homo sapie | ZNF280C | 83.096 | 2 | 38.709934 | yes | 5.038E-07 | 844.38153 | 32.07246 |
| GITLVCLK | 8 | Q8ND82 | Zinc finger protein 280C OS=Homo sapie | ZNF280C | 83.096 | 2 | 38.709934 | yes | 2.141E-06 | 452.27023 | 25.101526 |
| MMLSQIASK | 9 | Q8NDX6 | Zinc finger protein 740 OS=Homo sapien | ZNF740 | 21.857 | 2 | 46.696842 | yes | 3.313E-07 | 504.76447 | 141.1814 |
| HILIHTGEKPFECD | 19 | Q8NDX6 | Zinc finger protein 740 OS=Homo sapien | ZNF740 | 21.857 | 4 | 46.696842 | yes | 2.548E-07 | 593.53027 | 38.786446 |
| VHSGEKPYQCER | 12 | Q8NDX6 | Zinc finger protein 740 OS=Homo sapien | ZNF740 | 21.857 | 3 | 46.696842 | yes | 3.259E-07 | 497.23328 | 35.213612 |
| DDDSLSEASHSK | 12 | Q8NDX6 | Zinc finger protein 740 OS=Homo sapien | ZNF740 | 21.857 | 2 | 46.696842 | yes | 1.493E-07 | 645.77588 | 29.096174 |
| NFVCEHCFGAFR | 12 | Q8NDX6 | Zinc finger protein 740 OS=Homo sapien | ZNF740 | 21.857 | 2 | 46.696842 | yes | 1.758E-06 | 772.33209 | 25.408211 |
| LLEEIHAMK | 9 | Q8NEF9 | Serum response factor-binding protein 1 ( | SRFBP1 | 48.634 | 2 | 52.857273 | yes | 1.376E-07 | 542.297 | 376.63498 |
| SALGDDINFEK | 11 | Q8NEF9 | Serum response factor-binding protein 1 ( | SRFBP1 | 48.634 | 2 | 52.857273 | yes | 9.935E-08 | 604.79333 | 369.02679 |
| EQQSNIAVFQGK | 12 | Q8NEF9 | Serum response factor-binding protein 1 ( | SRFBP1 | 48.634 | 2 | 52.857273 | yes | 9.722E-08 | 674.84644 | 235.11833 |
| LAVHPLLK | 8 | Q8NEF9 | Serum response factor-binding protein 1 ( | SRFBP1 | 48.634 | 2 | 52.857273 | yes | 9.789E-08 | 445.79474 | 234.41133 |
| DSVVSLESQK | 10 | Q8NEF9 | Serum response factor-binding protein 1 ( | SRFBP1 | 48.634 | 2 | 52.857273 | yes | 1.196E-07 | 546.28259 | 201.53064 |
| SLDFPQNEPQIK | 12 | Q8NEF9 | Serum response factor-binding protein 1 ( | SRFBP1 | 48.634 | 2 | 52.857273 | yes | 3.512E-07 | 708.36188 | 146.77589 |
| ELKPDIVTK | 9 | Q8NEF9 | Serum response factor-binding protein 1 ( | SRFBP1 | 48.634 | 2 | 52.857273 | yes | 2.555E-07 | 521.81079 | 142.02274 |
| EATVISEQK | 9 | Q8NEF9 | Serum response factor-binding protein 1 ( | SRFBP1 | 48.634 | 2 | 52.857273 | yes | 2.635E-06 | 502.76657 | 124.22356 |
| QNVAEVESSK | 10 | Q8NEF9 | Serum response factor-binding protein 1 ( | SRFBP1 | 48.634 | 2 | 52.857273 | yes | 5.813E-07 | 545.7724 | 82.950104 |
| KLESVFFHSLSGSK | 14 | Q8NEF9 | Serum response factor-binding protein 1 ( | SRFBP1 | 48.634 | 2 | 52.857273 | yes | 9.723E-08 | 783.41974 | 63.885952 |
| AQPGTLNLNNEVV | 14 | Q8NEF9 | Serum response factor-binding protein 1 ( | SRFBP1 | 48.634 | 2 | 52.857273 | yes | 4.882E-07 | 769.91229 | 60.973835 |
| LCEAFLAGDER | 11 | Q8NFI3 | Cytosolic endo-beta-N-acetylglucosamini | ENGASE | 83.987 | 2 | 53.79833 | yes | 9.722E-08 | 640.8006 | 224.39114 |
| YLTTQLHR | 8 | Q8NFI3 | Cytosolic endo-beta-N-acetylglucosamini | ENGASE | 83.987 | 2 | 53.79833 | yes | 2.902E-07 | 516.28528 | 179.74402 |
| EVVSFSPDPLPVR | 13 | Q8NFI3 | Cytosolic endo-beta-N-acetylglucosamini | ENGASE | 83.987 | 2 | 53.79833 | yes | 9.733E-08 | 721.38794 | 170.63803 |
| IVDLLVEAAGPGQ | 15 | Q8NFI3 | Cytosolic endo-beta-N-acetylglucosamini | ENGASE | 83.987 | 2 | 53.79833 | yes | 1.214E-07 | 776.9201 | 80.714592 |
| VPQAEWGR | 8 | Q8NFI3 | Cytosolic endo-beta-N-acetylglucosamini | ENGASE | 83.987 | 2 | 53.79833 | yes | 2.188E-07 | 471.74323 | 77.364563 |
| RMEFLVEPVPK | 11 | Q8NFI3 | Cytosolic endo-beta-N-acetylglucosamini | ENGASE | 83.987 | 2 | 53.79833 | yes | 3.91E-06 | 672.87085 | 71.070564 |
| SIKDEEEETVFR | 12 | Q8NFI3 | Cytosolic endo-beta-N-acetylglucosamini | ENGASE | 83.987 | 2 | 53.79833 | yes | 4.338E-07 | 741.35956 | 67.643211 |
| LVQITQFFR | 9 | Q8NFI3 | Cytosolic endo-beta-N-acetylglucosamini | ENGASE | 83.987 | 2 | 53.79833 | yes | 5.044E-07 | 576.33221 | 65.152374 |
| HSDMFLLKPEDGR | 13 | Q8TBB5 | Kelch domain-containing protein 4 OS=H | KLHDC4 | 57.892 | 3 | 53.686832 | yes | 1.37E-07 | 515.58905 | 293.60199 |
| QVLTAPGSAGQPR | 13 | Q8TBB5 | Kelch domain-containing protein 4 OS=H | KLHDC4 | 57.892 | 2 | 53.686832 | yes | 2.167E-07 | 641.34912 | 271.37143 |
| TQTVELPCPPPSPR | 14 | Q8TBB5 | Kelch domain-containing protein 4 OS=H | KLHDC4 | 57.892 | 2 | 53.686832 | yes | 9.249E-07 | 789.90088 | 191.97104 |
| LSPSGTGPTPR | 11 | Q8TBB5 | Kelch domain-containing protein 4 OS=H | KLHDC4 | 57.892 | 2 | 53.686832 | yes | 1.643E-07 | 535.28546 | 186.63326 |
| DLWVLHLATK | 10 | Q8TBB5 | Kelch domain-containing protein 4 OS=H | KLHDC4 | 57.892 | 2 | 53.686832 | yes | 9.722E-08 | 598.34534 | 151.43764 |
| TFLYNELYVYNTR | 13 | Q8TBB5 | Kelch domain-containing protein 4 OS=H | KLHDC4 | 57.892 | 2 | 53.686832 | yes | 1.02E-07 | 848.42249 | 122.67912 |
| MNPSGVKPTPR | 11 | Q8TBB5 | Kelch domain-containing protein 4 OS=H | KLHDC4 | 57.892 | 2 | 53.686832 | yes | 9.722E-08 | 592.31622 | 120.82639 |
| HGVLYVYGGMFE | 16 | Q8TBB5 | Kelch domain-containing protein 4 OS=H | KLHDC4 | 57.892 | 3 | 53.686832 | yes | 1.165E-07 | 590.94714 | 70.443314 |
| KEEEDLEALIAHF( | 19 | Q8TBB5 | Kelch domain-containing protein 4 OS=H | KLHDC4 | 57.892 | 3 | 53.686832 | yes | 9.722E-08 | 734.0426 | 66.349213 |
| WFEGQLK | 7 | Q8TBB5 | Kelch domain-containing protein 4 OS=H | KLHDC4 | 57.892 | 2 | 53.686832 | yes | 3.171E-06 | 454.23724 | 57.741798 |
| EVVAEDGTVVTIK | 13 | Q8TBB5 | Kelch domain-containing protein 4 OS=H | KLHDC4 | 57.892 | 2 | 53.686832 | yes | 5.822E-05 | 680.37195 | 40.775547 |

| | | | | | | | | | | | |
|---|---|---|---|---|---|---|---|---|---|---|---|
| CYECGESGHLSYA | 16 | Q8TBF4 | Zinc finger CCHC-type and RNA-binding | ZCRB1 | 24.592 | 3 | 32.754288 | yes | 2.039E-06 | 639.92365 | 30.426159 |
| WKPSSGVPSTSDD | 15 | Q8TBF4 | Zinc finger CCHC-type and RNA-binding | ZCRB1 | 24.592 | 2 | 32.754288 | yes | 1.423E-06 | 803.37878 | 12.617508 |
| VFQASPEDHEK | 11 | Q8TCC3 | Large ribosomal subunit protein uL30m C | MRPL30 | 18.546 | 2 | 50.424335 | yes | 9.809E-08 | 643.8042 | 198.98955 |
| LFVINEVCEMK | 11 | Q8TDN6 | Ribosome biogenesis protein BRX1 homo | BRIX1 | 41.402 | 2 | 50.83431 | yes | 9.722E-08 | 691.34637 | 1639.9784 |
| RNEIDAEPPAK | 11 | Q8TDN6 | Ribosome biogenesis protein BRX1 homo | BRIX1 | 41.402 | 2 | 50.83431 | yes | 1.069E-06 | 620.32001 | 201.23395 |
| CIYFEAK | 7 | Q8TDN6 | Ribosome biogenesis protein BRX1 homo | BRIX1 | 41.402 | 2 | 50.83431 | yes | 5.882E-07 | 465.72311 | 117.398 |
| FLVQNIHTLAELK | 13 | Q8TDN6 | Ribosome biogenesis protein BRX1 homo | BRIX1 | 41.402 | 3 | 50.83431 | yes | 9.838E-08 | 509.29608 | 86.702965 |
| HATAEEVEEEER | 12 | Q8TDN6 | Ribosome biogenesis protein BRX1 homo | BRIX1 | 41.402 | 2 | 50.83431 | yes | 1.665E-07 | 714.81549 | 28.029373 |
| FVLNLIK | 7 | Q8TDN6 | Ribosome biogenesis protein BRX1 homo | BRIX1 | 41.402 | 2 | 50.83431 | yes | 3.12E-06 | 423.776 | 13.492077 |
| ILIFSSR | 7 | Q8TDN6 | Ribosome biogenesis protein BRX1 homo | BRIX1 | 41.402 | 2 | 50.83431 | yes | 3.867E-06 | 418.25543 | 8.7985106 |
| PLLSFDPAFDELPH | 19 | Q8TDN6 | Ribosome biogenesis protein BRX1 homo | BRIX1 | 41.402 | 3 | 50.83431 | yes | 0.000102 | 729.38928 | 0.30336 |
| MQAVVPLNK | 9 | Q8TF39 | Zinc finger protein 483 OS=Homo sapien | ZNF483 | 85.099 | 2 | 36.257748 | yes | 7.206E-07 | 521.29169 | 59.405617 |
| AFTQLSNLQSHR | 12 | Q8TF68 | Zinc finger protein 384 OS=Homo sapien | ZNF384 | 63.22 | 2 | 50.77502 | yes | 9.91E-08 | 701.3653 | 260.48843 |
| VYTCTICSR | 9 | Q8TF68 | Zinc finger protein 384 OS=Homo sapien | ZNF384 | 63.22 | 2 | 50.77502 | yes | 1.076E-07 | 580.26538 | 246.61708 |
| AYTSETYLMK | 10 | Q8TF68 | Zinc finger protein 384 OS=Homo sapien | ZNF384 | 63.22 | 2 | 50.77502 | no | 6.071E-07 | 603.78918 | 149.36964 |
| PYSCNFCEK | 9 | Q8TF68 | Zinc finger protein 384 OS=Homo sapien | ZNF384 | 63.22 | 2 | 50.77502 | yes | 4.714E-07 | 602.7417 | 103.41586 |
| MKDQLLPEK | 9 | Q8TF68 | Zinc finger protein 384 OS=Homo sapien | ZNF384 | 63.22 | 2 | 50.77502 | yes | 9.722E-08 | 551.30225 | 102.04976 |
| VASTLTEEGGGGG | 25 | Q8TF68 | Zinc finger protein 384 OS=Homo sapien | ZNF384 | 63.22 | 3 | 50.77502 | yes | 1.5E-07 | 732.37323 | 41.359451 |
| TFANSSYLAQHIR | 13 | Q8TF68 | Zinc finger protein 384 OS=Homo sapien | ZNF384 | 63.22 | 2 | 50.77502 | yes | 3.289E-07 | 754.38623 | 25.377415 |
| DFIIQTGDPTGTGR | 14 | Q8WUA2 | Peptidyl-prolyl cis-trans isomerase-like 4 | PPIL4 | 57.226 | 2 | 51.59497 | yes | 1.021E-07 | 739.36774 | 590.6286 |
| IGADEEIDDFK | 11 | Q8WUA2 | Peptidyl-prolyl cis-trans isomerase-like 4 | PPIL4 | 57.226 | 2 | 51.59497 | yes | 1.528E-07 | 626.29059 | 566.16003 |
| MDNVLIDDR | 9 | Q8WUA2 | Peptidyl-prolyl cis-trans isomerase-like 4 | PPIL4 | 57.226 | 2 | 51.59497 | yes | 9.722E-08 | 545.76349 | 467.24631 |
| IHVDFSQSVAK | 11 | Q8WUA2 | Peptidyl-prolyl cis-trans isomerase-like 4 | PPIL4 | 57.226 | 2 | 51.59497 | yes | 9.722E-08 | 615.82751 | 403.46301 |
| YQTDLYER | 8 | Q8WUA2 | Peptidyl-prolyl cis-trans isomerase-like 4 | PPIL4 | 57.226 | 2 | 51.59497 | yes | 9.722E-08 | 544.25635 | 375.84689 |
| YDLILDEQAEDSK | 13 | Q8WUA2 | Peptidyl-prolyl cis-trans isomerase-like 4 | PPIL4 | 57.226 | 2 | 51.59497 | yes | 2.101E-07 | 769.86469 | 290.98343 |
| EQDKPPNLVLK | 11 | Q8WUA2 | Peptidyl-prolyl cis-trans isomerase-like 4 | PPIL4 | 57.226 | 2 | 51.59497 | yes | 1.058E-07 | 640.86407 | 226.86707 |
| YYNYCLIHNVQR | 12 | Q8WUA2 | Peptidyl-prolyl cis-trans isomerase-like 4 | PPIL4 | 57.226 | 2 | 51.59497 | yes | 1.055E-07 | 821.89337 | 117.11366 |
| ACLNFLK | 7 | Q8WUA2 | Peptidyl-prolyl cis-trans isomerase-like 4 | PPIL4 | 57.226 | 2 | 51.59497 | yes | 2.236E-07 | 433.23364 | 101.92451 |
| TGEPFVFNYR | 10 | Q8WUF8 | Cotranscriptional regulator FAM172A OS | FAM172A | 47.973 | 2 | 30.693893 | yes | 2.092E-06 | 615.30109 | 35.028233 |
| SLKEEDGEEIVTYI | 18 | Q8WUH1 | Protein Churchill OS=Homo sapiens OX= | CHURC1 | 12.887 | 3 | 46.036587 | yes | 1.672E-07 | 722.34039 | 89.320572 |
| WVPVVDPQEEER | 12 | Q8WUZ0 | B-cell CLL/lymphoma 7 protein family m | BCL7C | 23.469 | 2 | 46.77642 | yes | 1.473E-07 | 741.86481 | 196.77083 |
| HVFSEEVLAK | 10 | Q8WV60 | Pentatricopeptide repeat-containing prote | PTCD2 | 43.968 | 2 | 50.806114 | yes | 9.722E-08 | 579.81134 | 1036.0397 |
| GELITLLHLCESR | 13 | Q8WV60 | Pentatricopeptide repeat-containing prote | PTCD2 | 43.968 | 2 | 50.806114 | yes | 1.568E-07 | 770.91125 | 624.37915 |
| SALQVLIEMK | 10 | Q8WV60 | Pentatricopeptide repeat-containing prote | PTCD2 | 43.968 | 2 | 50.806114 | yes | 1.256E-07 | 566.32574 | 569.0437 |
| DHVELAK | 7 | Q8WV60 | Pentatricopeptide repeat-containing prote | PTCD2 | 43.968 | 2 | 50.806114 | yes | 1.063E-07 | 406.21906 | 284.03842 |
| SHTLLLNK | 8 | Q8WV60 | Pentatricopeptide repeat-containing prote | PTCD2 | 43.968 | 2 | 50.806114 | yes | 3.182E-07 | 463.27692 | 114.82468 |
| VAVACNLSGTK | 11 | Q8WV60 | Pentatricopeptide repeat-containing prote | PTCD2 | 43.968 | 2 | 50.806114 | yes | 1.514E-07 | 560.29498 | 100.85997 |
| VKDVPALVAK | 10 | Q8WV60 | Pentatricopeptide repeat-containing prote | PTCD2 | 43.968 | 2 | 50.806114 | yes | 2.746E-06 | 520.32916 | 68.105492 |
| KVAVACNLSGTK | 12 | Q8WV60 | Pentatricopeptide repeat-containing prote | PTCD2 | 43.968 | 2 | 50.806114 | yes | 2.082E-07 | 624.34247 | 43.956161 |

| | | | | | | | | | | | |
|---|---|---|---|---|---|---|---|---|---|---|---|
| VGQETFYDVYR | 11 | Q8WVI0 | Ubiquinol-cytochrome c reductase compl | UQCC5 | 8.697 | 2 | 52.70676 | yes | 9.722E-08 | 688.8277 | 498.0004 |
| LCQLYPEENLEK | 12 | Q8WYQ9 | Zinc finger CCHC domain-containing pro | ZCCHC14 | 100.043 | 2 | 50.857307 | yes | 9.722E-08 | 768.37415 | 288.63019 |
| SSSEVTEFISK | 11 | Q8WYQ9 | Zinc finger CCHC domain-containing pro | ZCCHC14 | 100.043 | 2 | 50.857307 | yes | 1.878E-06 | 607.30096 | 37.521442 |
| VPTEPPATSVAAR | 13 | Q92610 | Zinc finger protein 592 OS=Homo sapien | ZNF592 | 137.529 | 2 | 39.299618 | yes | 4.667E-07 | 648.35132 | 90.247765 |
| LPFDTPPIGK | 10 | Q93073 | Selenocysteine insertion sequence-bindin | SECISBP2L | 121.776 | 2 | 49.86062 | yes | 1.489E-07 | 542.80548 | 1601.3524 |
| LSAEVEPFIPQK | 12 | Q93073 | Selenocysteine insertion sequence-bindin | SECISBP2L | 121.776 | 2 | 49.86062 | yes | 1.062E-07 | 679.37177 | 1148.796 |
| LVELTEEAR | 9 | Q93073 | Selenocysteine insertion sequence-bindin | SECISBP2L | 121.776 | 2 | 49.86062 | yes | 1.446E-07 | 530.28766 | 845.68701 |
| CVIISPNCEK | 10 | Q93073 | Selenocysteine insertion sequence-bindin | SECISBP2L | 121.776 | 2 | 49.86062 | no | 1.001E-07 | 610.29413 | 775.12622 |
| IESWVSETQR | 10 | Q93073 | Selenocysteine insertion sequence-bindin | SECISBP2L | 121.776 | 2 | 49.86062 | yes | 1.478E-07 | 617.80676 | 735.38318 |
| SLSETTATMLWK | 12 | Q93073 | Selenocysteine insertion sequence-bindin | SECISBP2L | 121.776 | 2 | 49.86062 | yes | 1.125E-07 | 684.34741 | 591.68158 |
| TMETLQLGK | 9 | Q93073 | Selenocysteine insertion sequence-bindin | SECISBP2L | 121.776 | 2 | 49.86062 | yes | 1.831E-07 | 510.77335 | 421.88727 |
| EYCNQVLCK | 9 | Q93073 | Selenocysteine insertion sequence-bindin | SECISBP2L | 121.776 | 2 | 49.86062 | yes | 1.168E-07 | 607.27063 | 379.52719 |
| QPSLVATGSTTSAT | 18 | Q93073 | Selenocysteine insertion sequence-bindin | SECISBP2L | 121.776 | 2 | 49.86062 | yes | 1.727E-07 | 832.42853 | 368.40979 |
| QFPLYNNDIR | 10 | Q93073 | Selenocysteine insertion sequence-bindin | SECISBP2L | 121.776 | 2 | 49.86062 | yes | 1.136E-07 | 640.32513 | 287.47028 |
| QITNTRPLSYTVVT | 21 | Q93073 | Selenocysteine insertion sequence-bindin | SECISBP2L | 121.776 | 3 | 49.86062 | yes | 1.238E-07 | 779.08502 | 130.9039 |
| HSNNQPAAGALR | 12 | Q93073 | Selenocysteine insertion sequence-bindin | SECISBP2L | 121.776 | 2 | 49.86062 | yes | 1.026E-07 | 618.31561 | 97.206322 |
| RLVMGLR | 7 | Q93073 | Selenocysteine insertion sequence-bindin | SECISBP2L | 121.776 | 2 | 49.86062 | yes | 3.398E-07 | 422.76291 | 73.526558 |
| KLQEALSK | 8 | Q93073 | Selenocysteine insertion sequence-bindin | SECISBP2L | 121.776 | 2 | 49.86062 | yes | 1.242E-07 | 458.77673 | 53.285549 |
| ALAAALATAQEYS | 20 | Q93073 | Selenocysteine insertion sequence-bindin | SECISBP2L | 121.776 | 3 | 49.86062 | yes | 1.967E-07 | 709.02094 | 31.351006 |
| LKRPTALK | 8 | Q93073 | Selenocysteine insertion sequence-bindin | SECISBP2L | 121.776 | 2 | 49.86062 | yes | 1.063E-06 | 463.81091 | 24.038883 |
| DSLYAQGR | 8 | Q969Q0 | Ribosomal protein eL42-like OS=Homo s | RPL36AL | 12.469 | 2 | 54.06536 | yes | 9.722E-08 | 455.22488 | 1012.3507 |
| GKDSLYAQGR | 10 | Q969Q0 | Ribosomal protein eL42-like OS=Homo s | RPL36AL | 12.469 | 2 | 54.06536 | yes | 1.659E-07 | 547.78308 | 949.87897 |
| AFSSGSNFTR | 10 | Q969W8 | Zinc finger protein 566 OS=Homo sapien | ZNF566 | 49.22 | 2 | 39.36893 | yes | 4.626E-07 | 537.25415 | 124.24113 |
| GQWPVLESR | 9 | Q969W8 | Zinc finger protein 566 OS=Homo sapien | ZNF566 | 49.22 | 2 | 39.36893 | yes | 3.972E-06 | 536.28271 | 98.767357 |
| TFICGSDLTR | 10 | Q969W8 | Zinc finger protein 566 OS=Homo sapien | ZNF566 | 49.22 | 2 | 39.36893 | yes | 5.143E-07 | 585.28461 | 98.060707 |
| GTMIPSEAPLLHR | 13 | Q96A35 | Large ribosomal subunit protein uL24m C | MRPL24 | 24.915 | 2 | 50.96317 | yes | 1.434E-07 | 711.38214 | 1852.813 |
| ADGIVPETWIDGPK | 14 | Q96A35 | Large ribosomal subunit protein uL24m C | MRPL24 | 24.915 | 2 | 50.96317 | yes | 1.565E-07 | 749.38281 | 1273.6506 |
| KPTEIEWR | 8 | Q96A35 | Large ribosomal subunit protein uL24m C | MRPL24 | 24.915 | 2 | 50.96317 | yes | 1.952E-07 | 529.7851 | 859.95087 |
| YGMSPPGSVADK | 12 | Q96A35 | Large ribosomal subunit protein uL24m C | MRPL24 | 24.915 | 2 | 50.96317 | yes | 1.968E-07 | 604.78442 | 489.49036 |
| KNPPWIR | 7 | Q96A35 | Large ribosomal subunit protein uL24m C | MRPL24 | 24.915 | 2 | 50.96317 | yes | 1.294E-07 | 455.76651 | 423.44061 |
| TYVPCLK | 7 | Q96A35 | Large ribosomal subunit protein uL24m C | MRPL24 | 24.915 | 2 | 50.96317 | yes | 9.722E-08 | 440.73349 | 400.30664 |
| TLQEEVMEAMGIK | 13 | Q96A35 | Large ribosomal subunit protein uL24m C | MRPL24 | 24.915 | 2 | 50.96317 | yes | 1.635E-07 | 739.86511 | 383.41672 |
| DTSVEDALER | 10 | Q96A35 | Large ribosomal subunit protein uL24m C | MRPL24 | 24.915 | 2 | 50.96317 | yes | 9.809E-08 | 567.76727 | 301.31189 |
| YGMSPPGSVADKI | 13 | Q96A35 | Large ribosomal subunit protein uL24m C | MRPL24 | 24.915 | 2 | 50.96317 | yes | 6.695E-07 | 682.83502 | 92.290947 |
| NWVVVGGLNTHY | 13 | Q96A35 | Large ribosomal subunit protein uL24m C | MRPL24 | 24.915 | 2 | 50.96317 | yes | 2.032E-07 | 757.89679 | 26.110668 |
| QLLPGDEFSLR | 11 | Q96A47 | Insulin gene enhancer protein ISL-2 OS=I | ISL2 | 39.769 | 2 | 48.619965 | yes | 1.361E-07 | 637.84058 | 693.33905 |
| GLHLPDAGSGR | 11 | Q96A47 | Insulin gene enhancer protein ISL-2 OS=I | ISL2 | 39.769 | 2 | 48.619965 | yes | 2.41E-07 | 540.28326 | 582.1806 |
| EQLVEMTGLSPR | 12 | Q96A47 | Insulin gene enhancer protein ISL-2 OS=I | ISL2 | 39.769 | 2 | 48.619965 | no | 1.262E-07 | 680.35046 | 508.87543 |
| VSPDLEWHAACLK | 13 | Q96A47 | Insulin gene enhancer protein ISL-2 OS=I | ISL2 | 39.769 | 2 | 48.619965 | no | 2.442E-07 | 763.37701 | 498.40268 |

| | | | | | | | | | | | |
|---|---|---|---|---|---|---|---|---|---|---|---|
| ADHGLLLER | 9 | Q96A47 | Insulin gene enhancer protein ISL-2 OS=I | ISL2 | 39.769 | 2 | 48.619965 | yes | 3.873E-07 | 512.28271 | 488.3183 |
| CQVGFSSSDLVMF | 13 | Q96A47 | Insulin gene enhancer protein ISL-2 OS=I | ISL2 | 39.769 | 2 | 48.619965 | yes | 1.276E-07 | 743.34485 | 268.29651 |
| DSVYHIECFR | 10 | Q96A47 | Insulin gene enhancer protein ISL-2 OS=I | ISL2 | 39.769 | 2 | 48.619965 | yes | 3.042E-07 | 663.30078 | 228.82217 |
| TCYAANPRPDALN | 14 | Q96A47 | Insulin gene enhancer protein ISL-2 OS=I | ISL2 | 39.769 | 3 | 48.619965 | no | 5.134E-07 | 536.59381 | 63.585625 |
| TCYAANPRPDALN | 14 | Q96A47 | Insulin gene enhancer protein ISL-2 OS=I | ISL2 | 39.769 | 3 | 48.619965 | no | 1.467E-07 | 804.38708 | 63.585625 |
| TCYAANPRPDALN | 14 | Q96A47 | Insulin gene enhancer protein ISL-2 OS=I | ISL2 | 39.769 | 2 | 48.619965 | no | 5.134E-07 | 536.59381 | 63.585625 |
| TCYAANPRPDALN | 14 | Q96A47 | Insulin gene enhancer protein ISL-2 OS=I | ISL2 | 39.769 | 2 | 48.619965 | no | 1.467E-07 | 804.38708 | 63.585625 |
| SPGPLPGAR | 9 | Q96A47 | Insulin gene enhancer protein ISL-2 OS=I | ISL2 | 39.769 | 2 | 48.619965 | yes | 3.274E-06 | 426.24033 | 72.805435 |
| HENAVQGSAVEV( | 20 | Q96A47 | Insulin gene enhancer protein ISL-2 OS=I | ISL2 | 39.769 | 3 | 48.619965 | yes | 1.122E-07 | 756.7063 | 71.923538 |
| EHELLCR | 7 | Q96A47 | Insulin gene enhancer protein ISL-2 OS=I | ISL2 | 39.769 | 2 | 48.619965 | yes | 6.248E-07 | 478.73456 | 51.125351 |
| NMPDSENPTSVFS | 14 | Q96BU1 | S100P-binding protein OS=Homo sapiens | S100PBP | 45.582 | 2 | 45.874214 | yes | 2.938E-06 | 790.85413 | 163.67094 |
| YSLTQWVDR | 9 | Q96BU1 | S100P-binding protein OS=Homo sapiens | S100PBP | 45.582 | 2 | 45.874214 | yes | 1.717E-07 | 584.29327 | 104.54694 |
| GSQILLDTPR | 10 | Q96BU1 | S100P-binding protein OS=Homo sapiens | S100PBP | 45.582 | 2 | 45.874214 | yes | 1.079E-06 | 550.30896 | 70.584511 |
| ADLNQGIGEPQSPS | 15 | Q96C19 | EF-hand domain-containing protein D2 O | EFHD2 | 26.698 | 2 | 52.775383 | yes | 1.138E-07 | 784.88678 | 130.8273 |
| LSEIDVSSEGVK | 12 | Q96C19 | EF-hand domain-containing protein D2 O | EFHD2 | 26.698 | 2 | 52.775383 | yes | 9.722E-08 | 631.82733 | 110.96692 |
| LEQWEEPWVEDR | 12 | Q96C28 | Zinc finger protein 707 OS=Homo sapien | ZNF707 | 43.088 | 2 | 46.92924 | yes | 3.087E-06 | 808.37299 | 55.402328 |
| AVELSFICGTCGK | 13 | Q96C28 | Zinc finger protein 707 OS=Homo sapien | ZNF707 | 43.088 | 2 | 46.92924 | yes | 1.439E-07 | 721.34436 | 34.243649 |
| LTVSEDGPGVR | 11 | Q96CW6 | Probable RNA polymerase II nuclear loca | SLC7A6OS | 35.028 | 2 | 51.73769 | yes | 1.243E-07 | 565.29602 | 338.28955 |
| SAEPAEALVLACK | 13 | Q96CW6 | Probable RNA polymerase II nuclear loca | SLC7A6OS | 35.028 | 2 | 51.73769 | yes | 9.722E-08 | 679.85284 | 167.67905 |
| YPLDVQK | 7 | Q96CW6 | Probable RNA polymerase II nuclear loca | SLC7A6OS | 35.028 | 2 | 51.73769 | yes | 3.776E-06 | 431.73709 | 25.513634 |
| NFSTSNLNTDFLAI | 14 | Q96DY7 | Mdm2-binding protein OS=Homo sapiens | MTBP | 102.194 | 2 | 47.188 | yes | 1.381E-07 | 786.38867 | 112.68201 |
| HVLQNLETFEK | 11 | Q96DY7 | Mdm2-binding protein OS=Homo sapiens | MTBP | 102.194 | 2 | 47.188 | yes | 3.234E-06 | 679.35913 | 20.833897 |
| QEVPMCTGPEAR | 12 | Q96EK7 | Constitutive coactivator of peroxisome pr | FAM120B | 103.784 | 2 | 51.10198 | yes | 2.051E-07 | 687.81067 | 282.57166 |
| EWFVYPGNPLR | 11 | Q96EK7 | Constitutive coactivator of peroxisome pr | FAM120B | 103.784 | 2 | 51.10198 | yes | 1.453E-07 | 689.35114 | 194.38347 |
| LEALMCTNPEIK | 12 | Q96EK7 | Constitutive coactivator of peroxisome pr | FAM120B | 103.784 | 2 | 51.10198 | yes | 9.722E-08 | 709.85455 | 193.82785 |
| ILWLNQEPEIQVR | 13 | Q96EK7 | Constitutive coactivator of peroxisome pr | FAM120B | 103.784 | 2 | 51.10198 | yes | 9.81E-08 | 819.4541 | 192.91875 |
| LPVATDFEFK | 10 | Q96EK7 | Constitutive coactivator of peroxisome pr | FAM120B | 103.784 | 2 | 51.10198 | yes | 1.459E-07 | 583.80823 | 176.38618 |
| GYAVEVLLEQNR | 12 | Q96EK7 | Constitutive coactivator of peroxisome pr | FAM120B | 103.784 | 2 | 51.10198 | yes | 1.031E-07 | 695.86987 | 136.1658 |
| VLYLYQGEK | 9 | Q96EK7 | Constitutive coactivator of peroxisome pr | FAM120B | 103.784 | 2 | 51.10198 | yes | 1.547E-07 | 556.80298 | 135.25034 |
| GNIILAVSDHISK | 13 | Q96EK7 | Constitutive coactivator of peroxisome pr | FAM120B | 103.784 | 2 | 51.10198 | yes | 1.175E-07 | 683.88806 | 115.90675 |
| EEVPMCSDAESR | 12 | Q96EK7 | Constitutive coactivator of peroxisome pr | FAM120B | 103.784 | 2 | 51.10198 | yes | 8.539E-07 | 705.28723 | 108.78469 |
| QQVTMVSDTEILK | 13 | Q96EK7 | Constitutive coactivator of peroxisome pr | FAM120B | 103.784 | 2 | 51.10198 | yes | 9.818E-08 | 746.38977 | 92.891251 |
| KLEEILPLGPNK | 12 | Q96EK7 | Constitutive coactivator of peroxisome pr | FAM120B | 103.784 | 2 | 51.10198 | yes | 1.041E-07 | 675.9032 | 83.09523 |
| EVPVYTDSEPR | 11 | Q96EK7 | Constitutive coactivator of peroxisome pr | FAM120B | 103.784 | 2 | 51.10198 | yes | 2.669E-07 | 646.31189 | 82.771729 |
| EVPMCSDPEPR | 11 | Q96EK7 | Constitutive coactivator of peroxisome pr | FAM120B | 103.784 | 2 | 51.10198 | yes | 1.874E-07 | 658.78412 | 60.909004 |
| TSDFMPWNVFDGI | 13 | Q96EK7 | Constitutive coactivator of peroxisome pr | FAM120B | 103.784 | 2 | 51.10198 | yes | 2.823E-07 | 772.34796 | 60.898968 |
| QVISTSSDAESR | 12 | Q96EK7 | Constitutive coactivator of peroxisome pr | FAM120B | 103.784 | 2 | 51.10198 | yes | 3.565E-07 | 640.30988 | 55.043041 |
| LIFFFDGMVEQDK | 13 | Q96EK7 | Constitutive coactivator of peroxisome pr | FAM120B | 103.784 | 2 | 51.10198 | yes | 2.103E-06 | 794.8894 | 47.88343 |
| QEVPMYTGSEPR | 12 | Q96EK7 | Constitutive coactivator of peroxisome pr | FAM120B | 103.784 | 2 | 51.10198 | yes | 9.968E-08 | 697.32446 | 34.80447 |

| | | | | | | | | | | | |
|---|---|---|---|---|---|---|---|---|---|---|---|
| GLQGFVGSTCPHI( | 20 | Q96EK7 | Constitutive coactivator of peroxisome pr | FAM120B | 103.784 | 3 | 51.10198 | yes | 3.033E-06 | 741.03461 | 31.111738 |
| QEDPTNVGPEVK | 12 | Q96EK7 | Constitutive coactivator of peroxisome pr | FAM120B | 103.784 | 2 | 51.10198 | yes | 8.111E-07 | 656.82263 | 29.186508 |
| TGIPLNVLPK | 10 | Q96GA3 | Protein LTV1 homolog OS=Homo sapien | LTV1 | 54.855 | 2 | 51.771713 | yes | 2.811E-06 | 526.32916 | 2023.2953 |
| DPLAADESAPQR | 12 | Q96GA3 | Protein LTV1 homolog OS=Homo sapien | LTV1 | 54.855 | 2 | 51.771713 | yes | 2.773E-07 | 635.30713 | 1527.1926 |
| LQEVLNDYYK | 10 | Q96GA3 | Protein LTV1 homolog OS=Homo sapien | LTV1 | 54.855 | 2 | 51.771713 | yes | 9.722E-08 | 642.82715 | 1236.535 |
| FTEYSMTSSVMR | 12 | Q96GA3 | Protein LTV1 homolog OS=Homo sapien | LTV1 | 54.855 | 2 | 51.771713 | yes | 9.974E-08 | 719.82068 | 580.34607 |
| FTEYSMTSSVMR | 12 | Q96GA3 | Protein LTV1 homolog OS=Homo sapien | LTV1 | 54.855 | 2 | 51.771713 | yes | 3.553E-06 | 727.81818 | 580.34607 |
| ATGEEEGMDIQK | 12 | Q96GA3 | Protein LTV1 homolog OS=Homo sapien | LTV1 | 54.855 | 2 | 51.771713 | yes | 9.722E-08 | 654.29285 | 1108.4833 |
| AVSFHLVHR | 9 | Q96GA3 | Protein LTV1 homolog OS=Homo sapien | LTV1 | 54.855 | 2 | 51.771713 | yes | 3.006E-07 | 533.30127 | 1056.9679 |
| AIADHLFWSEETK | 13 | Q96GA3 | Protein LTV1 homolog OS=Homo sapien | LTV1 | 54.855 | 2 | 51.771713 | yes | 1.011E-07 | 773.88043 | 962.27429 |
| EETLVIPSTGIK | 12 | Q96GA3 | Protein LTV1 homolog OS=Homo sapien | LTV1 | 54.855 | 2 | 51.771713 | yes | 1.501E-07 | 643.86377 | 500.50281 |
| NEQLTLHDER | 10 | Q96GA3 | Protein LTV1 homolog OS=Homo sapien | LTV1 | 54.855 | 2 | 51.771713 | yes | 4.04E-07 | 627.80731 | 464.00317 |
| IQMINGSDLPK | 11 | Q96GA3 | Protein LTV1 homolog OS=Homo sapien | LTV1 | 54.855 | 2 | 51.771713 | yes | 9.722E-08 | 608.32373 | 298.85349 |
| EPSGPSELIPSSTFS | 19 | Q96GA3 | Protein LTV1 homolog OS=Homo sapien | LTV1 | 54.855 | 3 | 51.771713 | yes | 1.846E-07 | 671.66058 | 286.39944 |
| EEKEETLVIPSTGII | 15 | Q96GA3 | Protein LTV1 homolog OS=Homo sapien | LTV1 | 54.855 | 2 | 51.771713 | yes | 1.002E-07 | 836.9538 | 253.47673 |
| GDSNDDYDSAGLI | 24 | Q96GA3 | Protein LTV1 homolog OS=Homo sapien | LTV1 | 54.855 | 4 | 51.771713 | yes | 1.308E-06 | 637.50879 | 78.051102 |
| ARPELEELLAK | 11 | Q96GC5 | Large ribosomal subunit protein mL48 OS | MRPL48 | 23.935 | 2 | 43.224724 | yes | 2.678E-07 | 634.86407 | 201.82947 |
| FCCECGIR | 8 | Q96GY0 | Zinc finger C2HC domain-containing pro | ZC2HC1A | 35.093 | 2 | 51.726704 | yes | 9.722E-08 | 551.21735 | 720.04028 |
| VSSSSSSLGNK | 11 | Q96GY0 | Zinc finger C2HC domain-containing pro | ZC2HC1A | 35.093 | 2 | 51.726704 | yes | 1.307E-07 | 526.76459 | 521.32147 |
| FNENAADR | 8 | Q96GY0 | Zinc finger C2HC domain-containing pro | ZC2HC1A | 35.093 | 2 | 51.726704 | yes | 1.26E-07 | 468.71213 | 469.36243 |
| LQTLSPSHK | 9 | Q96GY0 | Zinc finger C2HC domain-containing pro | ZC2HC1A | 35.093 | 2 | 51.726704 | yes | 2.048E-07 | 505.7851 | 391.94788 |
| NPAPGVLTNK | 10 | Q96GY0 | Zinc finger C2HC domain-containing pro | ZC2HC1A | 35.093 | 2 | 51.726704 | yes | 5.005E-06 | 505.7851 | 385.44614 |
| KHEEFIATIR | 10 | Q96GY0 | Zinc finger C2HC domain-containing pro | ZC2HC1A | 35.093 | 3 | 51.726704 | yes | 1.81E-07 | 415.23129 | 188.8625 |
| KHEEFIATIR | 10 | Q96GY0 | Zinc finger C2HC domain-containing pro | ZC2HC1A | 35.093 | 3 | 51.726704 | yes | 3.094E-07 | 622.34332 | 188.8625 |
| KHEEFIATIR | 10 | Q96GY0 | Zinc finger C2HC domain-containing pro | ZC2HC1A | 35.093 | 2 | 51.726704 | yes | 1.81E-07 | 415.23129 | 188.8625 |
| KHEEFIATIR | 10 | Q96GY0 | Zinc finger C2HC domain-containing pro | ZC2HC1A | 35.093 | 2 | 51.726704 | yes | 3.094E-07 | 622.34332 | 188.8625 |
| HEEFIATIR | 9 | Q96GY0 | Zinc finger C2HC domain-containing pro | ZC2HC1A | 35.093 | 2 | 51.726704 | yes | 2.263E-07 | 558.29584 | 211.57841 |
| GIEGHSPGNLPK | 12 | Q96GY0 | Zinc finger C2HC domain-containing pro | ZC2HC1A | 35.093 | 2 | 51.726704 | yes | 3.27E-07 | 603.31732 | 202.76387 |
| GIAAPHAGANVKF | 14 | Q96GY0 | Zinc finger C2HC domain-containing pro | ZC2HC1A | 35.093 | 3 | 51.726704 | yes | 2.943E-07 | 453.59323 | 201.75693 |
| TFFPVALK | 8 | Q96GY0 | Zinc finger C2HC domain-containing pro | ZC2HC1A | 35.093 | 2 | 51.726704 | yes | 2.336E-07 | 461.77347 | 154.27188 |
| AEGTDIPTVK | 10 | Q96GY0 | Zinc finger C2HC domain-containing pro | ZC2HC1A | 35.093 | 2 | 51.726704 | yes | 1.835E-07 | 515.77441 | 122.12452 |
| TQVYKPPALK | 10 | Q96GY0 | Zinc finger C2HC domain-containing pro | ZC2HC1A | 35.093 | 2 | 51.726704 | yes | 9.722E-08 | 572.83984 | 85.79998 |
| YPVEWAK | 7 | Q96GY0 | Zinc finger C2HC domain-containing pro | ZC2HC1A | 35.093 | 2 | 51.726704 | yes | 5.797E-07 | 446.73181 | 72.803093 |
| LTNVPACVLITPGI | 16 | Q96HE9 | Proline-rich protein 11 OS=Homo sapien: | PRR11 | 40.086 | 2 | 50.965687 | yes | 9.722E-08 | 842.95056 | 380.90875 |
| LITPPPPPPSPER | 13 | Q96HE9 | Proline-rich protein 11 OS=Homo sapien: | PRR11 | 40.086 | 2 | 50.965687 | yes | 6.975E-07 | 699.39301 | 239.23401 |
| VGISSIDISQSR | 12 | Q96HE9 | Proline-rich protein 11 OS=Homo sapien: | PRR11 | 40.086 | 2 | 50.965687 | yes | 1.128E-07 | 631.34094 | 234.57724 |
| VTNVLITPGK | 10 | Q96HE9 | Proline-rich protein 11 OS=Homo sapien: | PRR11 | 40.086 | 2 | 50.965687 | yes | 1.553E-07 | 521.31879 | 215.40146 |
| QQFCILESK | 9 | Q96HE9 | Proline-rich protein 11 OS=Homo sapien: | PRR11 | 40.086 | 2 | 50.965687 | yes | 1.204E-07 | 576.78955 | 66.878807 |
| AAPAAQAEPER | 11 | Q96JN8 | Neuralized-like protein 4 OS=Homo sapi( | NEURL4 | 166.907 | 2 | 51.17319 | yes | 1.06E-07 | 555.78052 | 87.210335 |

| | | | | | | | | | | | |
|---|---|---|---|---|---|---|---|---|---|---|---|
| TTWMVSSCEVR | 11 | Q96JN8 | Neuralized-like protein 4 OS=Homo sapi | NEURL4 | 166.907 | 2 | 51.17319 | yes | 1.083E-07 | 678.30774 | 80.355141 |
| SEFNDAIVISNR | 12 | Q96JN8 | Neuralized-like protein 4 OS=Homo sapi | NEURL4 | 166.907 | 2 | 51.17319 | yes | 9.722E-08 | 682.84387 | 52.206627 |
| VWAVVDLYGK | 10 | Q96JN8 | Neuralized-like protein 4 OS=Homo sapi | NEURL4 | 166.907 | 2 | 51.17319 | yes | 2.193E-07 | 575.31879 | 30.653826 |
| ESVCWGPPPAASP | 15 | Q96JN8 | Neuralized-like protein 4 OS=Homo sapi | NEURL4 | 166.907 | 2 | 51.17319 | yes | 9.413E-07 | 798.39795 | 29.556229 |
| EDITSSAQFALR | 12 | Q96KR1 | Zinc finger RNA-binding protein OS=Ho | ZFR | 117.013 | 2 | 53.235844 | yes | 9.722E-08 | 669.33844 | 1070.8147 |
| STPVTSAVQIPEVK | 14 | Q96KR1 | Zinc finger RNA-binding protein OS=Ho | ZFR | 117.013 | 2 | 53.235844 | yes | 2.077E-07 | 728.40631 | 947.59589 |
| AISSASSPQSPGDA | 16 | Q96KR1 | Zinc finger RNA-binding protein OS=Ho | ZFR | 117.013 | 2 | 53.235844 | yes | 1.022E-07 | 772.38916 | 943.69592 |
| VLGMDPLPQMSQI | 13 | Q96KR1 | Zinc finger RNA-binding protein OS=Ho | ZFR | 117.013 | 2 | 53.235844 | yes | 9.919E-08 | 736.37341 | 887.21991 |
| NVNLVLLCSEKPS | 14 | Q96KR1 | Zinc finger RNA-binding protein OS=Ho | ZFR | 117.013 | 2 | 53.235844 | yes | 9.96E-08 | 800.94 | 538.7442 |
| LVSDSLSEHEK | 11 | Q96KR1 | Zinc finger RNA-binding protein OS=Ho | ZFR | 117.013 | 2 | 53.235844 | yes | 4.951E-07 | 622.31189 | 528.17047 |
| IVSITER | 7 | Q96KR1 | Zinc finger RNA-binding protein OS=Ho | ZFR | 117.013 | 2 | 53.235844 | yes | 1.112E-07 | 409.24252 | 428.50476 |
| LCECSFNDPNAK | 12 | Q96KR1 | Zinc finger RNA-binding protein OS=Ho | ZFR | 117.013 | 2 | 53.235844 | yes | 9.722E-08 | 727.8056 | 420.90631 |
| CLDALAALR | 9 | Q96KR1 | Zinc finger RNA-binding protein OS=Ho | ZFR | 117.013 | 2 | 53.235844 | yes | 2.093E-07 | 501.77368 | 412.21252 |
| STAPAVAYDSK | 11 | Q96KR1 | Zinc finger RNA-binding protein OS=Ho | ZFR | 117.013 | 2 | 53.235844 | yes | 1.014E-07 | 555.27728 | 376.76703 |
| VSAVPTNMAAK | 11 | Q96KR1 | Zinc finger RNA-binding protein OS=Ho | ZFR | 117.013 | 2 | 53.235844 | yes | 1.554E-07 | 544.79205 | 366.69553 |
| VNPDLQVEVKPSI | 14 | Q96KR1 | Zinc finger RNA-binding protein OS=Ho | ZFR | 117.013 | 3 | 53.235844 | yes | 1.96E-07 | 531.97015 | 180.10043 |
| VNPDLQVEVKPSI | 14 | Q96KR1 | Zinc finger RNA-binding protein OS=Ho | ZFR | 117.013 | 3 | 53.235844 | yes | 9.722E-08 | 797.4516 | 180.10043 |
| VNPDLQVEVKPSI | 14 | Q96KR1 | Zinc finger RNA-binding protein OS=Ho | ZFR | 117.013 | 2 | 53.235844 | yes | 1.96E-07 | 531.97015 | 180.10043 |
| VNPDLQVEVKPSI | 14 | Q96KR1 | Zinc finger RNA-binding protein OS=Ho | ZFR | 117.013 | 2 | 53.235844 | yes | 9.722E-08 | 797.4516 | 180.10043 |
| VFECISSGIILK | 12 | Q96KR1 | Zinc finger RNA-binding protein OS=Ho | ZFR | 117.013 | 2 | 53.235844 | yes | 9.722E-08 | 683.37598 | 348.70496 |
| ISCAGPQTYK | 10 | Q96KR1 | Zinc finger RNA-binding protein OS=Ho | ZFR | 117.013 | 2 | 53.235844 | yes | 1.785E-07 | 562.77386 | 281.2843 |
| QLAVISPEK | 9 | Q96KR1 | Zinc finger RNA-binding protein OS=Ho | ZFR | 117.013 | 2 | 53.235844 | yes | 2.679E-07 | 492.78986 | 279.51004 |
| MQVTITLTSPIIR | 13 | Q96KR1 | Zinc finger RNA-binding protein OS=Ho | ZFR | 117.013 | 2 | 53.235844 | yes | 1.23E-07 | 736.92889 | 274.71176 |
| CELCDVSCTGADA | 19 | Q96KR1 | Zinc finger RNA-binding protein OS=Ho | ZFR | 117.013 | 3 | 53.235844 | yes | 9.722E-08 | 723.64343 | 250.33475 |
| FNIHNNR | 7 | Q96KR1 | Zinc finger RNA-binding protein OS=Ho | ZFR | 117.013 | 2 | 53.235844 | yes | 2.189E-07 | 457.73322 | 187.62805 |
| KVNPDLQVEVKPS | 15 | Q96KR1 | Zinc finger RNA-binding protein OS=Ho | ZFR | 117.013 | 3 | 53.235844 | yes | 9.722E-08 | 574.66846 | 181.30673 |
| GSPGLLDPCEK | 11 | Q96KR1 | Zinc finger RNA-binding protein OS=Ho | ZFR | 117.013 | 2 | 53.235844 | yes | 1.66E-07 | 586.78442 | 111.24333 |
| ANGLQSCVIIIR | 12 | Q96KR1 | Zinc finger RNA-binding protein OS=Ho | ZFR | 117.013 | 2 | 53.235844 | yes | 1.62E-07 | 672.37683 | 107.14906 |
| YEEDMYWR | 8 | Q96KR1 | Zinc finger RNA-binding protein OS=Ho | ZFR | 117.013 | 2 | 53.235844 | yes | 1.771E-07 | 596.24243 | 103.28859 |
| GLTTTGNSSLNSTS | 17 | Q96KR1 | Zinc finger RNA-binding protein OS=Ho | ZFR | 117.013 | 2 | 53.235844 | yes | 1.035E-07 | 841.91321 | 72.22744 |
| CAVSEAAIILNSCV | 17 | Q96KR1 | Zinc finger RNA-binding protein OS=Ho | ZFR | 117.013 | 3 | 53.235844 | yes | 1.061E-07 | 620.97797 | 51.250057 |
| DSDGVDGFEAEGK | 13 | Q96KR1 | Zinc finger RNA-binding protein OS=Ho | ZFR | 117.013 | 2 | 53.235844 | yes | 2.152E-07 | 663.2782 | 48.726074 |
| DPPDVLDR | 8 | Q96KR1 | Zinc finger RNA-binding protein OS=Ho | ZFR | 117.013 | 2 | 53.235844 | yes | 1.548E-06 | 463.73254 | 39.684086 |
| HATIYPTEEELQAV | 16 | Q96KR1 | Zinc finger RNA-binding protein OS=Ho | ZFR | 117.013 | 3 | 53.235844 | yes | 1.577E-07 | 619.65112 | 29.948183 |
| PAPVAVAAAATA | 32 | Q96KR1 | Zinc finger RNA-binding protein OS=Ho | ZFR | 117.013 | 4 | 53.235844 | yes | 4.685E-07 | 789.38617 | 13.39164 |
| LQCETAAQQK | 10 | Q96M89 | Coiled-coil domain-containing protein 13 | CCDC138 | 76.219 | 2 | 34.032593 | yes | 1.094E-06 | 588.78754 | 84.303452 |
| GFNYTVFDCK | 10 | Q96MX6 | Dynein axonemal assembly factor 10 OS= | DNAAF10 | 39.741 | 2 | 43.86705 | yes | 2.414E-07 | 625.77917 | 40.25872 |
| VSFPSYLASPAGSI | 14 | Q96PE2 | Rho guanine nucleotide exchange factor 1 | ARHGEF1 | 221.673 | 2 | 38.552425 | yes | 5.142E-07 | 719.86987 | 70.458488 |
| ILCIGAVPGLQPR | 13 | Q96PE2 | Rho guanine nucleotide exchange factor 1 | ARHGEF1 | 221.673 | 2 | 38.552425 | yes | 2.535E-06 | 697.40283 | 57.008434 |

| | | | | | | | | | | | |
|---|---|---|---|---|---|---|---|---|---|---|---|
| DFNAPALLER | 10 | Q96SL8 | Flt3-interacting zinc finger protein 1 OS= | FIZ1 | 51.996 | 2 | 51.669083 | yes | 1.354E-07 | 573.30109 | 229.8436 |
| LLLGPAGGGVPK | 12 | Q96SL8 | Flt3-interacting zinc finger protein 1 OS= | FIZ1 | 51.996 | 2 | 51.669083 | yes | 9.722E-08 | 539.83459 | 185.77202 |
| PFPCHICGK | 9 | Q96SL8 | Flt3-interacting zinc finger protein 1 OS= | FIZ1 | 51.996 | 2 | 51.669083 | yes | 3.167E-07 | 558.25989 | 130.46613 |
| HVLVHTGEKPFPC | 18 | Q96SL8 | Flt3-interacting zinc finger protein 1 OS= | FIZ1 | 51.996 | 4 | 51.669083 | yes | 6.145E-07 | 527.76587 | 100.38351 |
| IFGCSECEK | 9 | Q96SL8 | Flt3-interacting zinc finger protein 1 OS= | FIZ1 | 51.996 | 2 | 51.669083 | yes | 1.364E-07 | 565.23627 | 76.487961 |
| ESVVDYCNR | 9 | Q99442 | Translocation protein SEC62 OS=Homo s | SEC62 | 45.862 | 2 | 57.5447 | yes | 1.983E-07 | 571.25073 | 115.3653 |
| IQEVGEPSKEEK | 12 | Q99442 | Translocation protein SEC62 OS=Homo s | SEC62 | 45.862 | 2 | 57.5447 | yes | 9.722E-08 | 686.85138 | 68.878761 |
| DACVISSDFHER | 12 | Q99805 | Transmembrane 9 superfamily member 2 | TM9SF2 | 75.776 | 2 | 47.742027 | yes | 1.267E-07 | 718.31714 | 118.45199 |
| VQVWFQNR | 8 | Q99811 | Paired mesoderm homeobox protein 2 OS | PRRX2 | 27.08 | 2 | 55.70088 | no | 1.096E-07 | 538.78546 | 316.29266 |
| TTFNSSQLQALER | 13 | Q99811 | Paired mesoderm homeobox protein 2 OS | PRRX2 | 27.08 | 2 | 55.70088 | no | 3.079E-07 | 747.88098 | 185.57523 |
| EFSLHHSQVPTVN | 13 | Q99811 | Paired mesoderm homeobox protein 2 OS | PRRX2 | 27.08 | 2 | 55.70088 | yes | 9.722E-08 | 747.87042 | 94.079216 |
| SYSQEAAIEQPVAl | 15 | Q99811 | Paired mesoderm homeobox protein 2 OS | PRRX2 | 27.08 | 2 | 55.70088 | yes | 9.728E-08 | 823.41266 | 68.361465 |
| RVNLSEAR | 8 | Q99811 | Paired mesoderm homeobox protein 2 OS | PRRX2 | 27.08 | 2 | 55.70088 | yes | 3.877E-06 | 472.76724 | 53.845284 |
| IQESIER | 7 | Q9BPW4 | Apolipoprotein L4 OS=Homo sapiens OX | APOL4 | 39.165 | 2 | 45.496525 | yes | 1.832E-07 | 437.73508 | 106.37386 |
| RFTEEVIEYFQKK | 13 | Q9BPW4 | Apolipoprotein L4 OS=Homo sapiens OX | APOL4 | 39.165 | 3 | 45.496525 | yes | 4.151E-06 | 572.97003 | 15.080924 |
| FIADQLDHLNVTK | 13 | Q9BQC6 | Large ribosomal subunit protein mL63 OS | MRPL57 | 12.267 | 2 | 45.851894 | yes | 1.724E-07 | 757.40411 | 48.989429 |
| FTPPIYQPK | 9 | Q9BRJ2 | Large ribosomal subunit protein mL45 OS | MRPL45 | 35.351 | 2 | 50.251568 | yes | 1.592E-07 | 545.80023 | 1200.0745 |
| DVLEYVVFEK | 10 | Q9BRJ2 | Large ribosomal subunit protein mL45 OS | MRPL45 | 35.351 | 2 | 50.251568 | yes | 9.885E-08 | 620.8266 | 476.3555 |
| QLTNPYGSWR | 10 | Q9BRJ2 | Large ribosomal subunit protein mL45 OS | MRPL45 | 35.351 | 2 | 50.251568 | yes | 1.083E-07 | 611.3042 | 389.81238 |
| QTLAIYDR | 8 | Q9BRJ2 | Large ribosomal subunit protein mL45 OS | MRPL45 | 35.351 | 2 | 50.251568 | yes | 1.617E-07 | 490.26401 | 304.54855 |
| IKDYDANFK | 9 | Q9BRJ2 | Large ribosomal subunit protein mL45 OS | MRPL45 | 35.351 | 2 | 50.251568 | yes | 1.376E-07 | 557.28241 | 281.9657 |
| IVPPWAPPK | 9 | Q9BRJ2 | Large ribosomal subunit protein mL45 OS | MRPL45 | 35.351 | 2 | 50.251568 | yes | 1.192E-07 | 502.80002 | 187.39702 |
| TMASQVSIR | 9 | Q9BRJ2 | Large ribosomal subunit protein mL45 OS | MRPL45 | 35.351 | 2 | 50.251568 | yes | 9.834E-08 | 496.76331 | 73.02597 |
| IKDFPEK | 7 | Q9BRJ2 | Large ribosomal subunit protein mL45 OS | MRPL45 | 35.351 | 2 | 50.251568 | yes | 3.127E-07 | 438.7449 | 55.149574 |
| KAGLVIPPEK | 10 | Q9BRJ2 | Large ribosomal subunit protein mL45 OS | MRPL45 | 35.351 | 2 | 50.251568 | yes | 3.41E-06 | 526.32916 | 37.347794 |
| AIVTTDDFASK | 11 | Q9BRR8 | G patch domain-containing protein 1 OS= | GPATCH1 | 103.345 | 2 | 38.813835 | yes | 4.971E-07 | 584.29822 | 28.112911 |
| LSVGFELLR | 9 | Q9BRR8 | G patch domain-containing protein 1 OS= | GPATCH1 | 103.345 | 2 | 38.813835 | yes | 6.738E-07 | 517.30566 | 19.157297 |
| LDGDVLMK | 8 | Q9BRT6 | Protein LLP homolog OS=Homo sapiens | LLPH | 15.226 | 2 | 51.927563 | yes | 9.722E-08 | 445.73624 | 596.48804 |
| DVQEIATVVVPK | 12 | Q9BRT6 | Protein LLP homolog OS=Homo sapiens | LLPH | 15.226 | 2 | 51.927563 | yes | 1.882E-07 | 649.37177 | 37.077442 |
| ILDLIDDAWR | 10 | Q9BS18 | Anaphase-promoting complex subunit 13 | ANAPC13 | 8.522 | 2 | 41.371258 | yes | 3.543E-07 | 615.3299 | 63.529984 |
| VLDVGFVGR | 9 | Q9BSF4 | Mitochondrial import inner membrane tra | TIMM29 | 29.234 | 2 | 51.586643 | yes | 9.722E-08 | 481.27692 | 146.37192 |
| LALSQAHSLVQAE | 16 | Q9BSF4 | Mitochondrial import inner membrane tra | TIMM29 | 29.234 | 3 | 51.586643 | yes | 4.485E-07 | 564.31268 | 89.556526 |
| LGIEGYPTYK | 10 | Q9BSF8 | BTB/POZ domain-containing protein 10 ( | BTBD10 | 53.78 | 2 | 31.515915 | yes | 1.815E-06 | 570.80042 | 82.783112 |
| LSQVSLLER | 9 | Q9BSJ5 | Uncharacterized protein C17orf80 OS=H | C17orf80 | 67.316 | 2 | 51.795986 | yes | 6.426E-07 | 522.80603 | 72.33268 |
| TFPLPAVGLER | 11 | Q9BSJ5 | Uncharacterized protein C17orf80 OS=H | C17orf80 | 67.316 | 2 | 51.795986 | yes | 4.14E-06 | 600.34277 | 56.151318 |
| ALMESPEGQLSLE | 15 | Q9BSJ5 | Uncharacterized protein C17orf80 OS=H | C17orf80 | 67.316 | 2 | 51.795986 | yes | 1.064E-07 | 814.91364 | 55.925335 |
| NTESLILSLK | 10 | Q9BSJ5 | Uncharacterized protein C17orf80 OS=H | C17orf80 | 67.316 | 2 | 51.795986 | yes | 5.327E-07 | 559.32678 | 43.15255 |
| LLDVPTGDCHISPI | 14 | Q9BSJ5 | Uncharacterized protein C17orf80 OS=H | C17orf80 | 67.316 | 2 | 51.795986 | yes | 3.458E-06 | 776.39539 | 41.029804 |
| TVMSHGCENFNTI | 13 | Q9BSJ5 | Uncharacterized protein C17orf80 OS=H | C17orf80 | 67.316 | 2 | 51.795986 | yes | 9.722E-08 | 776.83521 | 15.693996 |

| | | | | | | | | | | |
|---|---|---|---|---|---|---|---|---|---|---|
| RAPSTSPSFEGTQE | 25 | Q9BUT9 | MAPK regulated corepressor interacting p | MCRIP2 | 17.828 | 4 | 51.276634 | yes | 1.108E-07 | 699.08234 | 920.27289 |
| PVQYVER | 7 | Q9BUT9 | MAPK regulated corepressor interacting p | MCRIP2 | 17.828 | 2 | 51.276634 | yes | 1.781E-07 | 445.74017 | 857.23901 |
| TGPTQQQVEGR | 11 | Q9BUT9 | MAPK regulated corepressor interacting p | MCRIP2 | 17.828 | 2 | 51.276634 | yes | 9.722E-08 | 600.802 | 707.22913 |
| QPAPPTSQPPR | 11 | Q9BUT9 | MAPK regulated corepressor interacting p | MCRIP2 | 17.828 | 2 | 51.276634 | yes | 1.279E-07 | 588.31201 | 694.90454 |
| RTGPTQQQVEGR | 12 | Q9BUT9 | MAPK regulated corepressor interacting p | MCRIP2 | 17.828 | 2 | 51.276634 | yes | 3.26E-07 | 678.85254 | 15.38221 |
| SPPLSPVGTTPVK | 13 | Q9BVC5 | Ashwin OS=Homo sapiens OX=9606 GN | C2orf49 | 25.859 | 2 | 49.396927 | yes | 1.036E-07 | 640.36646 | 145.98071 |
| ILDTLGLLR | 9 | Q9BWU0 | Kanadaptin OS=Homo sapiens OX=9606 | SLC4A1AP | 88.815 | 2 | 50.320198 | yes | 4.265E-07 | 507.32132 | 167.42097 |
| IVKPAEIPELK | 11 | Q9BWU0 | Kanadaptin OS=Homo sapiens OX=9606 | SLC4A1AP | 88.815 | 2 | 50.320198 | yes | 9.819E-08 | 618.88171 | 163.37193 |
| EAMIQCSLEACR | 12 | Q9BWU0 | Kanadaptin OS=Homo sapiens OX=9606 | SLC4A1AP | 88.815 | 2 | 50.320198 | yes | 9.863E-08 | 734.32306 | 138.37096 |
| ELSEISER | 8 | Q9BWU0 | Kanadaptin OS=Homo sapiens OX=9606 | SLC4A1AP | 88.815 | 2 | 50.320198 | yes | 1.878E-06 | 481.7431 | 138.19543 |
| LFILQGPEEDR | 11 | Q9BWU0 | Kanadaptin OS=Homo sapiens OX=9606 | SLC4A1AP | 88.815 | 2 | 50.320198 | yes | 1.591E-07 | 658.84589 | 105.21403 |
| IDEKPETFESLVAK | 14 | Q9BWU0 | Kanadaptin OS=Homo sapiens OX=9606 | SLC4A1AP | 88.815 | 2 | 50.320198 | yes | 1.06E-07 | 803.42218 | 97.144409 |
| APASSSSNPEEVQI | 14 | Q9BWU0 | Kanadaptin OS=Homo sapiens OX=9606 | SLC4A1AP | 88.815 | 2 | 50.320198 | yes | 4.119E-07 | 715.84155 | 25.297163 |
| KQQQILLEK | 9 | Q9BWU0 | Kanadaptin OS=Homo sapiens OX=9606 | SLC4A1AP | 88.815 | 2 | 50.320198 | yes | 1.005E-06 | 564.34277 | 21.145748 |
| CPFPDCAK | 8 | Q9BXK1 | Krueppel-like factor 16 OS=Homo sapien | KLF16 | 25.431 | 2 | 52.28868 | yes | 1.684E-07 | 497.70969 | 347.14868 |
| HPGFHPDLLR | 10 | Q9BXK1 | Krueppel-like factor 16 OS=Homo sapien | KLF16 | 25.431 | 2 | 52.28868 | yes | 1.325E-07 | 594.81726 | 198.85706 |
| RHPGFHPDLLR | 11 | Q9BXK1 | Krueppel-like factor 16 OS=Homo sapien | KLF16 | 25.431 | 3 | 52.28868 | yes | 9.722E-08 | 448.91431 | 139.17519 |
| FSCPLCSK | 8 | Q9BXK1 | Krueppel-like factor 16 OS=Homo sapien | KLF16 | 25.431 | 2 | 52.28868 | yes | 1.376E-07 | 499.72534 | 72.187363 |
| RFSCPLCSK | 9 | Q9BXK1 | Krueppel-like factor 16 OS=Homo sapien | KLF16 | 25.431 | 2 | 52.28868 | yes | 4.715E-07 | 577.77588 | 37.584644 |
| APGAAPSAAAK | 11 | Q9BXK1 | Krueppel-like factor 16 OS=Homo sapien | KLF16 | 25.431 | 2 | 52.28868 | yes | 3.909E-06 | 456.25089 | 28.332867 |
| PFACDWQGCDK | 11 | Q9BXK1 | Krueppel-like factor 16 OS=Homo sapien | KLF16 | 25.431 | 2 | 52.28868 | yes | 7.276E-07 | 692.27643 | 7.7293496 |
| NNIDVCPECGHLK | 13 | Q9BYC8 | Large ribosomal subunit protein bL32m C | MRPL32 | 21.405 | 2 | 51.367275 | yes | 1.284E-07 | 778.35321 | 398.32455 |
| HVLCAYCYEK | 10 | Q9BYC8 | Large ribosomal subunit protein bL32m C | MRPL32 | 21.405 | 2 | 51.367275 | yes | 9.722E-08 | 671.79956 | 281.59464 |
| KRPSWFTQN | 9 | Q9BYC8 | Large ribosomal subunit protein bL32m C | MRPL32 | 21.405 | 2 | 51.367275 | yes | 2.694E-07 | 582.30145 | 125.79647 |
| YPALIGNLVK | 10 | Q9BYC9 | Large ribosomal subunit protein bL20m C | MRPL20 | 17.443 | 2 | 44.031673 | yes | 2.391E-06 | 544.32916 | 85.416847 |
| VLADLAIYEPK | 11 | Q9BYC9 | Large ribosomal subunit protein bL20m C | MRPL20 | 17.443 | 2 | 44.031673 | yes | 2.348E-07 | 616.35028 | 52.283241 |
| LAIYGMLPK | 9 | Q9BYD1 | Large ribosomal subunit protein uL13m C | MRPL13 | 20.692 | 2 | 53.27874 | yes | 1.025E-07 | 503.2937 | 119.684 |
| APQQWATFAR | 10 | Q9BYD1 | Large ribosomal subunit protein uL13m C | MRPL13 | 20.692 | 2 | 53.27874 | yes | 8.092E-07 | 588.30145 | 92.146835 |
| IWYLLDGK | 8 | Q9BYD1 | Large ribosomal subunit protein uL13m C | MRPL13 | 20.692 | 2 | 53.27874 | yes | 9.722E-08 | 504.28165 | 72.883118 |
| LWTPPEDYR | 9 | Q9BYD1 | Large ribosomal subunit protein uL13m C | MRPL13 | 20.692 | 2 | 53.27874 | yes | 8.34E-07 | 588.78784 | 55.097279 |
| NLVEELPQPR | 10 | Q9BYD1 | Large ribosomal subunit protein uL13m C | MRPL13 | 20.692 | 2 | 53.27874 | yes | 7.432E-07 | 597.82751 | 44.792564 |
| QVTAAQLHLR | 10 | Q9BYD1 | Large ribosomal subunit protein uL13m C | MRPL13 | 20.692 | 2 | 53.27874 | yes | 3.662E-07 | 568.83038 | 33.595295 |
| VYSSHTGYPGGFR | 13 | Q9BYD1 | Large ribosomal subunit protein uL13m C | MRPL13 | 20.692 | 2 | 53.27874 | yes | 9.722E-08 | 714.33875 | 17.57461 |
| WELNPEIVAR | 10 | Q9BYD2 | Large ribosomal subunit protein bL9m OS | MRPL9 | 30.244 | 2 | 52.1498 | yes | 9.722E-08 | 613.83002 | 1086.0433 |
| LPEEPITR | 8 | Q9BYD2 | Large ribosomal subunit protein bL9m OS | MRPL9 | 30.244 | 2 | 52.1498 | yes | 1.695E-07 | 477.76639 | 677.51685 |
| YWLAQQAAK | 9 | Q9BYD2 | Large ribosomal subunit protein bL9m OS | MRPL9 | 30.244 | 2 | 52.1498 | yes | 3.16E-07 | 539.78766 | 529.15955 |
| LLPQGLAVYASPE | 15 | Q9BYD2 | Large ribosomal subunit protein bL9m OS | MRPL9 | 30.244 | 2 | 52.1498 | yes | 5.597E-07 | 800.44067 | 301.88586 |
| KLFEEEK | 7 | Q9BYD2 | Large ribosomal subunit protein bL9m OS | MRPL9 | 30.244 | 2 | 52.1498 | yes | 3.038E-07 | 461.74765 | 191.69084 |
| VGLADLHPDVFAT | 16 | Q9BYD3 | Large ribosomal subunit protein uL4m OS | MRPL4 | 34.92 | 3 | 51.16567 | yes | 1.083E-07 | 560.30188 | 1465.4692 |

| | | | | | | | | | | | |
|---|---|---|---|---|---|---|---|---|---|---|---|
| VGLADLHPDVFAT | 16 | Q9BYD3 | Large ribosomal subunit protein uL4m OS | MRPL4 | 34.92 | 3 | 51.16567 | yes | 3.204E-07 | 839.94922 | 1465.4692 |
| VGLADLHPDVFAT | 16 | Q9BYD3 | Large ribosomal subunit protein uL4m OS | MRPL4 | 34.92 | 2 | 51.16567 | yes | 1.083E-07 | 560.30188 | 1465.4692 |
| VGLADLHPDVFAT | 16 | Q9BYD3 | Large ribosomal subunit protein uL4m OS | MRPL4 | 34.92 | 2 | 51.16567 | yes | 3.204E-07 | 839.94922 | 1465.4692 |
| LLWQDSR | 7 | Q9BYD3 | Large ribosomal subunit protein uL4m OS | MRPL4 | 34.92 | 2 | 51.16567 | yes | 2.009E-07 | 459.24561 | 1138.9645 |
| GPTSYYYMLPMK | 12 | Q9BYD3 | Large ribosomal subunit protein uL4m OS | MRPL4 | 34.92 | 2 | 51.16567 | yes | 1.068E-07 | 725.84088 | 348.27658 |
| GPTSYYYMLPMK | 12 | Q9BYD3 | Large ribosomal subunit protein uL4m OS | MRPL4 | 34.92 | 2 | 51.16567 | yes | 2.992E-07 | 733.83838 | 348.27658 |
| GPTSYYYMLPMK | 12 | Q9BYD3 | Large ribosomal subunit protein uL4m OS | MRPL4 | 34.92 | 2 | 51.16567 | yes | 1.454E-07 | 733.83838 | 348.27658 |
| KVELPVPTHR | 10 | Q9BYD3 | Large ribosomal subunit protein uL4m OS | MRPL4 | 34.92 | 2 | 51.16567 | yes | 9.722E-08 | 588.34839 | 703.83447 |
| LDILHQVAMWQK | 12 | Q9BYD3 | Large ribosomal subunit protein uL4m OS | MRPL4 | 34.92 | 2 | 51.16567 | yes | 4.493E-07 | 741.40033 | 518.10828 |
| RPVQAWVESLR | 11 | Q9BYD3 | Large ribosomal subunit protein uL4m OS | MRPL4 | 34.92 | 2 | 51.16567 | yes | 9.722E-08 | 670.87531 | 126.58033 |
| VELPVPTHR | 9 | Q9BYD3 | Large ribosomal subunit protein uL4m OS | MRPL4 | 34.92 | 2 | 51.16567 | yes | 1.763E-06 | 524.3009 | 88.546234 |
| LAQDDLHIMDSLE | 29 | Q9BYD3 | Large ribosomal subunit protein uL4m OS | MRPL4 | 34.92 | 4 | 51.16567 | yes | 3.245E-06 | 839.41321 | 13.90206 |
| MEFVSEVQR | 9 | Q9BYN8 | Small ribosomal subunit protein mS26 OS | MRPS26 | 24.212 | 2 | 50.851677 | yes | 1.025E-07 | 562.77386 | 2443.5779 |
| EVLQLQEEVK | 10 | Q9BYN8 | Small ribosomal subunit protein mS26 OS | MRPS26 | 24.212 | 2 | 50.851677 | yes | 1.189E-07 | 607.83496 | 1588.3011 |
| ELMAWNQAENR | 11 | Q9BYN8 | Small ribosomal subunit protein mS26 OS | MRPS26 | 24.212 | 2 | 50.851677 | yes | 1.41E-07 | 681.31696 | 1200.5291 |
| KAEEVQAWAQR | 11 | Q9BYN8 | Small ribosomal subunit protein mS26 OS | MRPS26 | 24.212 | 2 | 50.851677 | yes | 9.722E-08 | 658.34131 | 391.1792 |
| NYNWAITR | 8 | Q9BYN8 | Small ribosomal subunit protein mS26 OS | MRPS26 | 24.212 | 2 | 50.851677 | yes | 1.671E-07 | 519.26178 | 338.72455 |
| KNYNWAITR | 9 | Q9BYN8 | Small ribosomal subunit protein mS26 OS | MRPS26 | 24.212 | 2 | 50.851677 | yes | 1.612E-07 | 583.30927 | 263.60645 |
| EGLVVRPQR | 9 | Q9BYN8 | Small ribosomal subunit protein mS26 OS | MRPS26 | 24.212 | 2 | 50.851677 | yes | 1.062E-07 | 527.31183 | 215.23325 |
| QALEQAR | 7 | Q9BYN8 | Small ribosomal subunit protein mS26 OS | MRPS26 | 24.212 | 2 | 50.851677 | yes | 5.332E-07 | 408.22214 | 71.836319 |
| VYEIPGLEPITFAG | 15 | Q9BZE1 | Large ribosomal subunit protein mL37 OS | MRPL37 | 48.118 | 2 | 48.881767 | yes | 7.323E-07 | 817.44543 | 116.70965 |
| VLEQPVVVQSVGT | 16 | Q9BZE1 | Large ribosomal subunit protein mL37 OS | MRPL37 | 48.118 | 2 | 48.881767 | yes | 1.091E-07 | 841.95721 | 95.178268 |
| FLALYLHGAA | 10 | Q9BZE1 | Large ribosomal subunit protein mL37 OS | MRPL37 | 48.118 | 2 | 48.881767 | yes | 1.891E-06 | 538.30035 | 64.397186 |
| VDTGPREDLIK | 11 | Q9C075 | Keratin, type I cytoskeletal 23 OS=Homo | KRT23 | 48.132 | 2 | 23.877525 | yes | 3.558E-06 | 621.83807 | 177.39774 |
| FMPPDDPLGR | 10 | Q9C0D7 | Probable ribonuclease ZC3H12C OS=Ho | ZC3H12C | 99.341 | 2 | 54.746452 | no | 6.739E-07 | 572.77643 | 321.99014 |
| ILVFTPSR | 8 | Q9C0D7 | Probable ribonuclease ZC3H12C OS=Ho | ZC3H12C | 99.341 | 2 | 54.746452 | no | 1.032E-07 | 466.78183 | 243.67017 |
| HGPSLDNFLR | 10 | Q9C0D7 | Probable ribonuclease ZC3H12C OS=Ho | ZC3H12C | 99.341 | 2 | 54.746452 | no | 9.722E-08 | 578.29889 | 210.18484 |
| LGYSEEQVQLVLN | 14 | Q9C0D7 | Probable ribonuclease ZC3H12C OS=Ho | ZC3H12C | 99.341 | 2 | 54.746452 | yes | 9.722E-08 | 810.43561 | 205.06656 |
| LLMYSFVNDK | 10 | Q9C0D7 | Probable ribonuclease ZC3H12C OS=Ho | ZC3H12C | 99.341 | 2 | 54.746452 | no | 9.722E-08 | 615.31537 | 179.47873 |
| TQVYQDLEEK | 10 | Q9C0D7 | Probable ribonuclease ZC3H12C OS=Ho | ZC3H12C | 99.341 | 2 | 54.746452 | yes | 1.721E-07 | 626.8064 | 130.64012 |
| FSLDTDYR | 8 | Q9C0D7 | Probable ribonuclease ZC3H12C OS=Ho | ZC3H12C | 99.341 | 2 | 54.746452 | yes | 2.287E-06 | 508.73782 | 122.22891 |
| SEADQTVSTINTIT | 15 | Q9C0D7 | Probable ribonuclease ZC3H12C OS=Ho | ZC3H12C | 99.341 | 2 | 54.746452 | yes | 7.048E-07 | 818.41284 | 116.2867 |
| LAVDWFLER | 9 | Q9C0D7 | Probable ribonuclease ZC3H12C OS=Ho | ZC3H12C | 99.341 | 2 | 54.746452 | yes | 9.722E-08 | 574.80859 | 115.92838 |
| KPPDVVR | 7 | Q9C0D7 | Probable ribonuclease ZC3H12C OS=Ho | ZC3H12C | 99.341 | 2 | 54.746452 | yes | 1.294E-06 | 405.74524 | 100.84589 |
| GQSYSHEEPK | 10 | Q9C0D7 | Probable ribonuclease ZC3H12C OS=Ho | ZC3H12C | 99.341 | 2 | 54.746452 | yes | 9.722E-08 | 581.26221 | 92.941063 |
| NPHMTDAQQLAA | 18 | Q9C0D7 | Probable ribonuclease ZC3H12C OS=Ho | ZC3H12C | 99.341 | 3 | 54.746452 | yes | 5.84E-07 | 650.67529 | 48.62579 |
| RVVCYDDR | 8 | Q9C0D7 | Probable ribonuclease ZC3H12C OS=Ho | ZC3H12C | 99.341 | 2 | 54.746452 | no | 2.281E-06 | 541.75604 | 35.62941 |
| DITVFVPAWR | 10 | Q9C0D7 | Probable ribonuclease ZC3H12C OS=Ho | ZC3H12C | 99.341 | 2 | 54.746452 | no | 1.796E-07 | 602.32965 | 20.420918 |
| NHGTPMPYEQYPF | 13 | Q9C0D7 | Probable ribonuclease ZC3H12C OS=Ho | ZC3H12C | 99.341 | 2 | 54.746452 | yes | 6.049E-07 | 781.35883 | 17.945509 |

| | | | | | | | | | | | |
|---|---|---|---|---|---|---|---|---|---|---|---|
| FLSLEYSPAGK | 11 | Q9C0H5 | Rho GTPase-activating protein 39 OS=H | ARHGAP3 | 121.286 | 2 | 44.556515 | yes | 5.424E-07 | 606.31897 | 87.830994 |
| QGPPSPCQQLVLT | 14 | Q9C0H5 | Rho GTPase-activating protein 39 OS=H | ARHGAP3 | 121.286 | 2 | 44.556515 | yes | 1.116E-06 | 776.91125 | 80.694954 |
| QLVYVEQAGSSPK | 13 | Q9C0H5 | Rho GTPase-activating protein 39 OS=H | ARHGAP3 | 121.286 | 2 | 44.556515 | yes | 2.151E-07 | 703.36975 | 68.388878 |
| DELYIQLCR | 9 | Q9C0H5 | Rho GTPase-activating protein 39 OS=H | ARHGAP3 | 121.286 | 2 | 44.556515 | yes | 4.338E-06 | 605.30023 | 58.284565 |
| LGFPQILLEK | 10 | Q9C0H5 | Rho GTPase-activating protein 39 OS=H | ARHGAP3 | 121.286 | 2 | 44.556515 | yes | 3.265E-07 | 579.3501 | 42.81514 |
| FLQVFVQPANVAV | 15 | Q9C0H5 | Rho GTPase-activating protein 39 OS=H | ARHGAP3 | 121.286 | 2 | 44.556515 | yes | 2.664E-07 | 830.97467 | 31.734987 |
| EGSTSSSLEPEPDT | 16 | Q9C0H5 | Rho GTPase-activating protein 39 OS=H | ARHGAP3 | 121.286 | 2 | 44.556515 | yes | 5.696E-07 | 846.87598 | 8.020813 |
| VHQVTPQTHFIS | 12 | Q9GZP4 | PITH domain-containing protein 1 OS=H | PITHD1 | 24.178 | 2 | 52.649235 | yes | 2.456E-07 | 697.36475 | 60.873177 |
| HEVTICNYEASAN | 18 | Q9GZP4 | PITH domain-containing protein 1 OS=H | PITHD1 | 24.178 | 3 | 52.649235 | yes | 9.722E-08 | 695.31317 | 40.87846 |
| LEDCVLLATDVAA | 14 | Q9GZR7 | ATP-dependent RNA helicase DDX24 OS | DDX24 | 96.332 | 2 | 50.952724 | yes | 1.117E-07 | 773.4007 | 471.04083 |
| VQHVIHYQVPR | 11 | Q9GZR7 | ATP-dependent RNA helicase DDX24 OS | DDX24 | 96.332 | 3 | 50.952724 | yes | 1.425E-07 | 459.25797 | 402.35587 |
| CLVVDEADR | 9 | Q9GZR7 | ATP-dependent RNA helicase DDX24 OS | DDX24 | 96.332 | 2 | 50.952724 | yes | 1.595E-07 | 538.75568 | 375.634 |
| DKLDILGAAETGS | 15 | Q9GZR7 | ATP-dependent RNA helicase DDX24 OS | DDX24 | 96.332 | 2 | 50.952724 | yes | 9.85E-08 | 737.89099 | 254.57233 |
| GLEPSQSTAAK | 11 | Q9GZR7 | ATP-dependent RNA helicase DDX24 OS | DDX24 | 96.332 | 2 | 50.952724 | yes | 3.051E-07 | 544.78278 | 177.53767 |
| KDEDIPLFPVQTK | 13 | Q9GZR7 | ATP-dependent RNA helicase DDX24 OS | DDX24 | 96.332 | 2 | 50.952724 | yes | 2.56E-07 | 765.41412 | 172.38887 |
| SLVFANSISCIK | 12 | Q9GZR7 | ATP-dependent RNA helicase DDX24 OS | DDX24 | 96.332 | 2 | 50.952724 | yes | 1.612E-07 | 669.85791 | 149.89661 |
| NEATVETLTETK | 12 | Q9GZR7 | ATP-dependent RNA helicase DDX24 OS | DDX24 | 96.332 | 2 | 50.952724 | yes | 9.85E-08 | 668.33557 | 143.20149 |
| DEDIPLFPVQTK | 12 | Q9GZR7 | ATP-dependent RNA helicase DDX24 OS | DDX24 | 96.332 | 2 | 50.952724 | yes | 1.905E-07 | 701.36664 | 101.4247 |
| HLLSQPLFTESQK | 13 | Q9GZR7 | ATP-dependent RNA helicase DDX24 OS | DDX24 | 96.332 | 2 | 50.952724 | yes | 9.722E-08 | 764.41193 | 96.135231 |
| RPEIVVATPGR | 11 | Q9GZR7 | ATP-dependent RNA helicase DDX24 OS | DDX24 | 96.332 | 2 | 50.952724 | yes | 3.189E-06 | 597.85132 | 21.657681 |
| RPLLGLVLTPTR | 12 | Q9GZR7 | ATP-dependent RNA helicase DDX24 OS | DDX24 | 96.332 | 2 | 50.952724 | yes | 1.704E-07 | 668.42719 | 20.981667 |
| AQAVSEEEEEEEG | 14 | Q9GZR7 | ATP-dependent RNA helicase DDX24 OS | DDX24 | 96.332 | 2 | 50.952724 | yes | 5.401E-07 | 782.33649 | 17.886877 |
| KINQLPEAER | 10 | Q9H061 | Transmembrane protein 126A OS=Homo | TMEM126 | 21.528 | 2 | 44.880234 | yes | 2.037E-07 | 599.33295 | 295.9201 |
| VVSLSTPIIWR | 11 | Q9H0U9 | Testis-specific Y-encoded-like protein 1 ( | TSPYL1 | 49.193 | 2 | 48.301685 | yes | 2.071E-07 | 635.87952 | 252.18318 |
| SASELTAGAEAEA | 17 | Q9H0U9 | Testis-specific Y-encoded-like protein 1 ( | TSPYL1 | 49.193 | 2 | 48.301685 | yes | 1.166E-07 | 846.40216 | 153.0322 |
| YITNLEVK | 8 | Q9H0U9 | Testis-specific Y-encoded-like protein 1 ( | TSPYL1 | 49.193 | 2 | 48.301685 | yes | 2.272E-07 | 490.27658 | 140.76097 |
| AFQQLEHK | 8 | Q9H0U9 | Testis-specific Y-encoded-like protein 1 ( | TSPYL1 | 49.193 | 2 | 48.301685 | yes | 1.982E-07 | 500.76416 | 72.089706 |
| CATVSAAVAER | 11 | Q9H0U9 | Testis-specific Y-encoded-like protein 1 ( | TSPYL1 | 49.193 | 2 | 48.301685 | yes | 1.244E-07 | 567.78223 | 48.134037 |
| DLEAEIHPLK | 10 | Q9H0X4 | Protein FAM234A OS=Homo sapiens OX | FAM234A | 59.66 | 2 | 38.73928 | yes | 5.018E-07 | 582.81659 | 36.459484 |
| LLDLGAGDGEVTK | 13 | Q9H1A3 | Protein-L-histidine N-pros-methyltransfer | METTL9 | 36.537 | 2 | 51.48904 | yes | 1.065E-07 | 644.3432 | 981.09082 |
| GSMFVFSPDQFQR | 13 | Q9H1A3 | Protein-L-histidine N-pros-methyltransfer | METTL9 | 36.537 | 2 | 51.48904 | yes | 9.722E-08 | 773.36139 | 318.16824 |
| GSMFVFSPDQFQR | 13 | Q9H1A3 | Protein-L-histidine N-pros-methyltransfer | METTL9 | 36.537 | 2 | 51.48904 | yes | 2.664E-07 | 781.35883 | 318.16824 |
| WEKPSEILEIK | 11 | Q9H1A3 | Protein-L-histidine N-pros-methyltransfer | METTL9 | 36.537 | 2 | 51.48904 | yes | 1.064E-07 | 686.37958 | 314.91669 |
| CDQPLTLLK | 9 | Q9H1A3 | Protein-L-histidine N-pros-methyltransfer | METTL9 | 36.537 | 2 | 51.48904 | yes | 1.145E-07 | 544.29443 | 185.07536 |
| AGFVIEAFTR | 10 | Q9H1A3 | Protein-L-histidine N-pros-methyltransfer | METTL9 | 36.537 | 2 | 51.48904 | yes | 9.722E-08 | 555.80072 | 137.30742 |
| ALLLTGDFSQAGN | 14 | Q9H2W6 | Large ribosomal subunit protein mL46 OS | MRPL46 | 31.706 | 2 | 52.817043 | yes | 1.79E-07 | 717.883 | 491.34161 |
| DELGDYLKPK | 10 | Q9H2W6 | Large ribosomal subunit protein mL46 OS | MRPL46 | 31.706 | 2 | 52.817043 | yes | 1.104E-07 | 589.30859 | 289.237 |
| TLATLSENNMEAK | 13 | Q9H2W6 | Large ribosomal subunit protein mL46 OS | MRPL46 | 31.706 | 2 | 52.817043 | yes | 9.722E-08 | 711.35065 | 217.03667 |
| GHHVWVTK | 8 | Q9H2W6 | Large ribosomal subunit protein mL46 OS | MRPL46 | 31.706 | 2 | 52.817043 | yes | 1.46E-07 | 482.2616 | 117.22214 |

| | | | | | | | | | | | |
|---|---|---|---|---|---|---|---|---|---|---|---|
| LLGALCLQRPPVV | 15 | Q9H2W6 | Large ribosomal subunit protein mL46 OS | MRPL46 | 31.706 | 3 | 52.817043 | yes | 1.857E-07 | 550.99567 | 37.30209 |
| LLGALCLQRPPVV | 15 | Q9H2W6 | Large ribosomal subunit protein mL46 OS | MRPL46 | 31.706 | 3 | 52.817043 | yes | 1.803E-05 | 825.98981 | 37.30209 |
| LLGALCLQRPPVV | 15 | Q9H2W6 | Large ribosomal subunit protein mL46 OS | MRPL46 | 31.706 | 2 | 52.817043 | yes | 1.857E-07 | 550.99567 | 37.30209 |
| LLGALCLQRPPVV | 15 | Q9H2W6 | Large ribosomal subunit protein mL46 OS | MRPL46 | 31.706 | 2 | 52.817043 | yes | 1.803E-05 | 825.98981 | 37.30209 |
| SLYSDHELR | 9 | Q9H2W6 | Large ribosomal subunit protein mL46 OS | MRPL46 | 31.706 | 2 | 52.817043 | yes | 3.635E-06 | 560.27509 | 61.44136 |
| NLVLLVR | 7 | Q9H2W6 | Large ribosomal subunit protein mL46 OS | MRPL46 | 31.706 | 2 | 52.817043 | yes | 1.38E-06 | 413.77908 | 43.723724 |
| AYNILIGELDCSK | 13 | Q9H3C7 | Gametogenetin-binding protein 2 OS=Ho | GGNBP2 | 79.087 | 2 | 53.181305 | yes | 6.686E-07 | 748.37671 | 3017.9727 |
| GGQPLPWFEHR | 11 | Q9H3C7 | Gametogenetin-binding protein 2 OS=Ho | GGNBP2 | 79.087 | 2 | 53.181305 | yes | 9.722E-08 | 662.33325 | 2441.2957 |
| LGSCITDPGNR | 11 | Q9H3C7 | Gametogenetin-binding protein 2 OS=Ho | GGNBP2 | 79.087 | 2 | 53.181305 | yes | 1.963E-07 | 595.28516 | 1443.9299 |
| GYCAALYEGLR | 11 | Q9H3C7 | Gametogenetin-binding protein 2 OS=Ho | GGNBP2 | 79.087 | 2 | 53.181305 | yes | 1.11E-07 | 636.80573 | 1316.8003 |
| QQDLSIAMVVTSR | 13 | Q9H3C7 | Gametogenetin-binding protein 2 OS=Ho | GGNBP2 | 79.087 | 2 | 53.181305 | yes | 9.722E-08 | 724.38232 | 1242.7129 |
| RPICSGWLTTAGA | 14 | Q9H3C7 | Gametogenetin-binding protein 2 OS=Ho | GGNBP2 | 79.087 | 2 | 53.181305 | yes | 1.108E-07 | 752.37225 | 858.88916 |
| ETDFIENSSCK | 11 | Q9H3C7 | Gametogenetin-binding protein 2 OS=Ho | GGNBP2 | 79.087 | 2 | 53.181305 | yes | 1.122E-07 | 665.28497 | 821.24042 |
| LNDMIDAIPK | 10 | Q9H3C7 | Gametogenetin-binding protein 2 OS=Ho | GGNBP2 | 79.087 | 2 | 53.181305 | yes | 1.325E-07 | 565.29974 | 716.44775 |
| SFEMTVEK | 8 | Q9H3C7 | Gametogenetin-binding protein 2 OS=Ho | GGNBP2 | 79.087 | 2 | 53.181305 | yes | 7.037E-07 | 485.73114 | 181.49503 |
| HIHVCCETDFIAHI | 17 | Q9H3C7 | Gametogenetin-binding protein 2 OS=Ho | GGNBP2 | 79.087 | 3 | 53.181305 | yes | 9.722E-08 | 693.34015 | 139.35948 |
| LYTLFYVHGSK | 11 | Q9H3C7 | Gametogenetin-binding protein 2 OS=Ho | GGNBP2 | 79.087 | 2 | 53.181305 | yes | 9.722E-08 | 664.3559 | 124.42424 |
| ETSGNTMHTVFHF | 13 | Q9H3C7 | Gametogenetin-binding protein 2 OS=Ho | GGNBP2 | 79.087 | 2 | 53.181305 | yes | 1.305E-07 | 758.85168 | 118.13817 |
| KSFEMTVEK | 9 | Q9H3C7 | Gametogenetin-binding protein 2 OS=Ho | GGNBP2 | 79.087 | 2 | 53.181305 | yes | 2.656E-06 | 549.77863 | 114.2597 |
| LFSQLVESGNPALI | 21 | Q9H3C7 | Gametogenetin-binding protein 2 OS=Ho | GGNBP2 | 79.087 | 3 | 53.181305 | yes | 1.361E-07 | 732.73523 | 92.329048 |
| TIDIAQEEVLTCLG | 20 | Q9H3C7 | Gametogenetin-binding protein 2 OS=Ho | GGNBP2 | 79.087 | 3 | 53.181305 | yes | 1.429E-07 | 791.74213 | 81.343918 |
| CQLHSLDTHKPK | 12 | Q9H3C7 | Gametogenetin-binding protein 2 OS=Ho | GGNBP2 | 79.087 | 2 | 53.181305 | yes | 5.362E-06 | 732.37482 | 12.212641 |
| FIPDDITFDDEPK | 13 | Q9H501 | ESF1 homolog OS=Homo sapiens OX=9 | ESF1 | 98.797 | 2 | 53.3222 | yes | 5.455E-07 | 776.36432 | 296.32834 |
| TDLDNSIGIK | 10 | Q9H501 | ESF1 homolog OS=Homo sapiens OX=9 | ESF1 | 98.797 | 2 | 53.3222 | yes | 2.328E-07 | 538.28516 | 206.22919 |
| TLDSGTSEIVK | 11 | Q9H501 | ESF1 homolog OS=Homo sapiens OX=9 | ESF1 | 98.797 | 2 | 53.3222 | yes | 3.877E-07 | 575.30353 | 198.09378 |
| QLLQVIQEK | 9 | Q9H501 | ESF1 homolog OS=Homo sapiens OX=9 | ESF1 | 98.797 | 2 | 53.3222 | yes | 1.21E-07 | 549.82953 | 129.34039 |
| LAVCNMDWDR | 10 | Q9H501 | ESF1 homolog OS=Homo sapiens OX=9 | ESF1 | 98.797 | 2 | 53.3222 | yes | 9.722E-08 | 640.28156 | 116.36153 |
| DVASEVNLTAYKI | 14 | Q9H501 | ESF1 homolog OS=Homo sapiens OX=9 | ESF1 | 98.797 | 2 | 53.3222 | yes | 1.46E-07 | 767.90924 | 107.82594 |
| DGTSPEEEIEIER | 13 | Q9H501 | ESF1 homolog OS=Homo sapiens OX=9 | ESF1 | 98.797 | 2 | 53.3222 | yes | 1.018E-06 | 752.34412 | 89.772293 |
| SIDPALSMLIK | 11 | Q9H501 | ESF1 homolog OS=Homo sapiens OX=9 | ESF1 | 98.797 | 2 | 53.3222 | yes | 9.723E-08 | 594.33887 | 87.736694 |
| GRPISHSTTEDLKF | 14 | Q9H501 | ESF1 homolog OS=Homo sapiens OX=9 | ESF1 | 98.797 | 3 | 53.3222 | yes | 9.728E-08 | 532.95325 | 72.991943 |
| DKLTPWEQFLEK | 12 | Q9H501 | ESF1 homolog OS=Homo sapiens OX=9 | ESF1 | 98.797 | 2 | 53.3222 | yes | 9.723E-08 | 767.401 | 71.828026 |
| SQKDDEEQIAK | 11 | Q9H501 | ESF1 homolog OS=Homo sapiens OX=9 | ESF1 | 98.797 | 2 | 53.3222 | yes | 7.042E-07 | 645.81226 | 31.841618 |
| NIVQHTTDSSLEEI | 14 | Q9H501 | ESF1 homolog OS=Homo sapiens OX=9 | ESF1 | 98.797 | 2 | 53.3222 | yes | 9.722E-08 | 800.89429 | 21.798733 |
| FWEMPEK | 7 | Q9H501 | ESF1 homolog OS=Homo sapiens OX=9 | ESF1 | 98.797 | 2 | 53.3222 | yes | 8.083E-07 | 483.72311 | 17.312138 |
| MAIFELVSK | 9 | Q9H5L6 | DNA transposase THAP9 OS=Homo sapi | THAP9 | 103.411 | 2 | 37.70097 | yes | 5.772E-07 | 519.28864 | 20.930315 |
| KAEVVQVIR | 9 | Q9H6F5 | Coiled-coil domain-containing protein 86 | CCDC86 | 40.236 | 2 | 51.35812 | yes | 1.07E-07 | 521.3244 | 765.37347 |
| ALVEFESNPEETR | 13 | Q9H6F5 | Coiled-coil domain-containing protein 86 | CCDC86 | 40.236 | 2 | 51.35812 | yes | 1.011E-07 | 760.86499 | 510.55319 |
| FSQMLQDKPLR | 11 | Q9H6F5 | Coiled-coil domain-containing protein 86 | CCDC86 | 40.236 | 2 | 51.35812 | yes | 9.935E-08 | 681.86353 | 301.85037 |

| | | | | | | | | | | | |
|---|---|---|---|---|---|---|---|---|---|---|---|
| AGLGSPERPPK | 11 | Q9H6F5 | Coiled-coil domain-containing protein 86 | CCDC86 | 40.236 | 2 | 51.35812 | yes | 1.28E-07 | 554.80908 | 294.54697 |
| CQPKPSEEAPK | 11 | Q9H6F5 | Coiled-coil domain-containing protein 86 | CCDC86 | 40.236 | 2 | 51.35812 | yes | 1.06E-07 | 635.80847 | 123.41018 |
| QPPQQPAAK | 9 | Q9H6F5 | Coiled-coil domain-containing protein 86 | CCDC86 | 40.236 | 2 | 51.35812 | yes | 1.381E-07 | 482.76416 | 122.47498 |
| FSQMLQDK | 8 | Q9H6F5 | Coiled-coil domain-containing protein 86 | CCDC86 | 40.236 | 2 | 51.35812 | yes | 9.722E-08 | 498.7446 | 115.55767 |
| QQDLHLESPQR | 11 | Q9H6F5 | Coiled-coil domain-containing protein 86 | CCDC86 | 40.236 | 2 | 51.35812 | yes | 2.704E-07 | 675.84167 | 92.438866 |
| EELPVIPK | 8 | Q9H6F5 | Coiled-coil domain-containing protein 86 | CCDC86 | 40.236 | 2 | 51.35812 | yes | 2.828E-06 | 462.77368 | 35.143768 |
| YTLLLGQDENSVII | 14 | Q9H8H0 | Nucleolar protein 11 OS=Homo sapiens ( | NOL11 | 81.124 | 2 | 53.54432 | yes | 1.165E-07 | 796.93018 | 310.8266 |
| TDQFLVTDSGR | 11 | Q9H8H0 | Nucleolar protein 11 OS=Homo sapiens ( | NOL11 | 81.124 | 2 | 53.54432 | yes | 9.722E-08 | 619.8042 | 258.02606 |
| ILSVQGTEPLVLFK | 14 | Q9H8H0 | Nucleolar protein 11 OS=Homo sapiens ( | NOL11 | 81.124 | 2 | 53.54432 | yes | 9.722E-08 | 772.45831 | 196.40549 |
| IWNNEDVNLDK | 11 | Q9H8H0 | Nucleolar protein 11 OS=Homo sapiens ( | NOL11 | 81.124 | 2 | 53.54432 | yes | 1.076E-07 | 680.33063 | 172.86533 |
| SQISVYSELNK | 11 | Q9H8H0 | Nucleolar protein 11 OS=Homo sapiens ( | NOL11 | 81.124 | 2 | 53.54432 | yes | 1.149E-07 | 634.33008 | 162.99478 |
| GLEALLADPQQK | 12 | Q9H8H0 | Nucleolar protein 11 OS=Homo sapiens ( | NOL11 | 81.124 | 2 | 53.54432 | yes | 1.002E-07 | 641.8537 | 159.32898 |
| AEPSFYPR | 8 | Q9H8H0 | Nucleolar protein 11 OS=Homo sapiens ( | NOL11 | 81.124 | 2 | 53.54432 | yes | 2.702E-06 | 483.73761 | 136.24281 |
| CEVSSLAGALGK | 12 | Q9H8H0 | Nucleolar protein 11 OS=Homo sapiens ( | NOL11 | 81.124 | 2 | 53.54432 | yes | 1.011E-06 | 596.30554 | 126.70534 |
| ECLSVWNIK | 9 | Q9H8H0 | Nucleolar protein 11 OS=Homo sapiens ( | NOL11 | 81.124 | 2 | 53.54432 | yes | 1.639E-07 | 574.79205 | 119.38428 |
| ESTSCPVVQK | 10 | Q9H8H0 | Nucleolar protein 11 OS=Homo sapiens ( | NOL11 | 81.124 | 2 | 53.54432 | yes | 3.256E-07 | 567.77661 | 118.31306 |
| ATLSAEVYR | 9 | Q9H8H0 | Nucleolar protein 11 OS=Homo sapiens ( | NOL11 | 81.124 | 2 | 53.54432 | yes | 1.667E-06 | 505.26929 | 96.376488 |
| VSDQKPLGSWSVI | 13 | Q9H8H0 | Nucleolar protein 11 OS=Homo sapiens ( | NOL11 | 81.124 | 2 | 53.54432 | yes | 4.147E-07 | 715.88556 | 81.007874 |
| HPVLIFITEK | 10 | Q9H8H0 | Nucleolar protein 11 OS=Homo sapiens ( | NOL11 | 81.124 | 2 | 53.54432 | yes | 9.722E-08 | 598.85553 | 59.705471 |
| CNNCDQELNK | 10 | Q9H8H0 | Nucleolar protein 11 OS=Homo sapiens ( | NOL11 | 81.124 | 2 | 53.54432 | yes | 1.22E-07 | 647.76117 | 29.296005 |
| DTHIYDAR | 8 | Q9H967 | WD repeat-containing protein 76 OS=Ho | WDR76 | 69.77 | 2 | 53.16288 | yes | 1.746E-07 | 495.7356 | 101.54086 |
| IPLLTTIR | 8 | Q9H967 | WD repeat-containing protein 76 OS=Ho | WDR76 | 69.77 | 2 | 53.16288 | yes | 1.539E-07 | 463.8053 | 85.878975 |
| VVTTCADCNLR | 11 | Q9H967 | WD repeat-containing protein 76 OS=Ho | WDR76 | 69.77 | 2 | 53.16288 | yes | 9.722E-08 | 654.80536 | 77.260277 |
| FGQVGLCDLTQQF | 14 | Q9H967 | WD repeat-containing protein 76 OS=Ho | WDR76 | 69.77 | 2 | 53.16288 | yes | 4.36E-07 | 795.90088 | 76.943283 |
| QYFITAGLR | 9 | Q9H967 | WD repeat-containing protein 76 OS=Ho | WDR76 | 69.77 | 2 | 53.16288 | yes | 1.097E-07 | 534.79547 | 69.868378 |
| AIFEEVYR | 8 | Q9H967 | WD repeat-containing protein 76 OS=Ho | WDR76 | 69.77 | 2 | 53.16288 | yes | 3.737E-07 | 513.76636 | 68.684639 |
| SIASAYFSPLTGNR | 14 | Q9H967 | WD repeat-containing protein 76 OS=Ho | WDR76 | 69.77 | 2 | 53.16288 | yes | 1.197E-07 | 742.38062 | 67.809677 |
| ENQNIAYVSLR | 11 | Q9H967 | WD repeat-containing protein 76 OS=Ho | WDR76 | 69.77 | 2 | 53.16288 | yes | 2.805E-07 | 653.84113 | 28.12594 |
| ILITTSDRPHGR | 12 | Q9H9Y2 | Ribosome production factor 1 OS=Homo | RPF1 | 40.111 | 3 | 46.56198 | yes | 1.528E-07 | 455.92474 | 114.12131 |
| DFTDLIVINEDR | 12 | Q9H9Y2 | Ribosome production factor 1 OS=Homo | RPF1 | 40.111 | 2 | 46.56198 | yes | 3.285E-07 | 725.36462 | 113.76002 |
| AAAFPPGFSISEIK | 14 | Q9H9Y2 | Ribosome production factor 1 OS=Homo | RPF1 | 40.111 | 2 | 46.56198 | yes | 3.042E-06 | 717.88501 | 72.544365 |
| LAEFGAAGVNR | 11 | Q9HA92 | Radical S-adenosyl methionine domain-co | RSAD1 | 48.714 | 2 | 33.172947 | yes | 1.305E-06 | 552.79346 | 100.45983 |
| IQTLEPDNWMK | 11 | Q9HA92 | Radical S-adenosyl methionine domain-co | RSAD1 | 48.714 | 2 | 33.172947 | yes | 1.593E-05 | 687.83972 | 51.629147 |
| TGAELVTCGSVLK | 13 | Q9HCN8 | Stromal cell-derived factor 2-like protein | SDF2L1 | 23.599 | 2 | 54.513847 | yes | 9.722E-08 | 667.85284 | 186.7784 |
| GLEEFFDDPK | 10 | Q9HD33 | Large ribosomal subunit protein uL29m ( | MRPL47 | 29.451 | 2 | 51.63483 | yes | 9.838E-08 | 598.77716 | 851.56036 |
| SGAAWTCQQLR | 11 | Q9HD33 | Large ribosomal subunit protein uL29m ( | MRPL47 | 29.451 | 2 | 51.63483 | yes | 1.194E-07 | 639.3064 | 636.36005 |
| VVDSMDALDK | 10 | Q9HD33 | Large ribosomal subunit protein uL29m ( | MRPL47 | 29.451 | 2 | 51.63483 | yes | 2.909E-07 | 546.76569 | 601.60748 |
| LPMPSPER | 8 | Q9HD33 | Large ribosomal subunit protein uL29m ( | MRPL47 | 29.451 | 2 | 51.63483 | yes | 1.101E-07 | 463.74185 | 474.4462 |
| QWVIPWHLNK | 10 | Q9HD33 | Large ribosomal subunit protein uL29m ( | MRPL47 | 29.451 | 2 | 51.63483 | yes | 9.722E-08 | 660.8642 | 462.42743 |

| | | | | | | | | | | | |
|---|---|---|---|---|---|---|---|---|---|---|---|
| NMLLTLEQEAK | 11 | Q9HD33 | Large ribosomal subunit protein uL29m C | MRPL47 | 29.451 | 2 | 51.63483 | yes | 1.272E-07 | 645.3421 | 460.95135 |
| KFPHLAEAQK | 10 | Q9HD33 | Large ribosomal subunit protein uL29m C | MRPL47 | 29.451 | 2 | 51.63483 | yes | 1.229E-07 | 584.82727 | 209.33075 |
| ARPGAWR | 7 | Q9HD33 | Large ribosomal subunit protein uL29m C | MRPL47 | 29.451 | 2 | 51.63483 | yes | 1.23E-07 | 407.22754 | 150.37466 |
| LLQTGQER | 8 | Q9HD33 | Large ribosomal subunit protein uL29m C | MRPL47 | 29.451 | 2 | 51.63483 | yes | 1.186E-07 | 472.76163 | 95.563828 |
| FFALPYVDHFLR | 12 | Q9HD33 | Large ribosomal subunit protein uL29m C | MRPL47 | 29.451 | 3 | 51.63483 | yes | 3.758E-06 | 508.93811 | 12.71777 |
| PLYETIEDNDVK | 12 | Q9NP92 | Large ribosomal subunit protein mL65 OS | MRPS30 | 50.365 | 2 | 55.5751 | yes | 3.055E-07 | 718.3512 | 111.33957 |
| QNCADQIEVVFR | 12 | Q9NP92 | Large ribosomal subunit protein mL65 OS | MRPS30 | 50.365 | 2 | 55.5751 | yes | 9.722E-08 | 739.85645 | 95.650299 |
| WYQYFTK | 7 | Q9NP92 | Large ribosomal subunit protein mL65 OS | MRPS30 | 50.365 | 2 | 55.5751 | yes | 2.837E-07 | 518.25037 | 60.930614 |
| YQIDDKPNNQIR | 12 | Q9NP92 | Large ribosomal subunit protein mL65 OS | MRPS30 | 50.365 | 2 | 55.5751 | yes | 1.85E-07 | 752.38116 | 38.437683 |
| HSLFNAMNR | 9 | Q9NPA3 | Mid1-interacting protein 1 OS=Homo sap | MID1IP1 | 20.202 | 2 | 38.310764 | yes | 5.312E-07 | 545.26654 | 127.86935 |
| DMYSHYVLLK | 10 | Q9NPA3 | Mid1-interacting protein 1 OS=Homo sap | MID1IP1 | 20.202 | 2 | 38.310764 | yes | 3.238E-06 | 634.8208 | 34.192245 |
| DILVDLGHLEGAD | 28 | Q9NPA3 | Mid1-interacting protein 1 OS=Homo sap | MID1IP1 | 20.202 | 4 | 38.310764 | yes | 6.774E-06 | 805.63898 | 4.085248 |
| ALQELPLCLR | 10 | Q9NQS1 | Cell death regulator Aven OS=Homo sapi | AVEN | 38.507 | 2 | 53.87244 | yes | 9.722E-08 | 606.8421 | 463.87302 |
| EGDNILPDQTSQDI | 15 | Q9NQS1 | Cell death regulator Aven OS=Homo sapi | AVEN | 38.507 | 2 | 53.87244 | yes | 9.722E-08 | 836.90485 | 236.03319 |
| SVAAGCPVLLGK | 12 | Q9NQS1 | Cell death regulator Aven OS=Homo sapi | AVEN | 38.507 | 2 | 53.87244 | yes | 2.532E-07 | 586.3288 | 145.1142 |
| EWDSEASCPK | 10 | Q9NQS1 | Cell death regulator Aven OS=Homo sapi | AVEN | 38.507 | 2 | 53.87244 | yes | 1.196E-07 | 604.74805 | 91.347504 |
| QNSAFYVDSELLV | 14 | Q9NQS1 | Cell death regulator Aven OS=Homo sapi | AVEN | 38.507 | 2 | 53.87244 | yes | 9.722E-08 | 820.91754 | 56.71944 |
| LNVAAELVQGTVI | 22 | Q9NQS1 | Cell death regulator Aven OS=Homo sapi | AVEN | 38.507 | 3 | 53.87244 | yes | 9.722E-08 | 777.12115 | 46.979866 |
| SELGLNFYQQVK | 12 | Q9NRM2 | Zinc finger protein 277 OS=Homo sapien | ZNF277 | 52.788 | 2 | 54.253853 | yes | 4.137E-07 | 713.37225 | 383.83185 |
| EILEQQQQER | 10 | Q9NRM2 | Zinc finger protein 277 OS=Homo sapien | ZNF277 | 52.788 | 2 | 54.253853 | yes | 1.508E-07 | 650.82825 | 291.15591 |
| SVILNHMAR | 9 | Q9NRM2 | Zinc finger protein 277 OS=Homo sapien | ZNF277 | 52.788 | 2 | 54.253853 | yes | 6.514E-07 | 520.78711 | 243.86325 |
| FYVINYLELGK | 11 | Q9NRM2 | Zinc finger protein 277 OS=Homo sapien | ZNF277 | 52.788 | 2 | 54.253853 | yes | 9.722E-08 | 679.8714 | 190.43379 |
| LYVHMEDAHEFDI | 15 | Q9NRM2 | Zinc finger protein 277 OS=Homo sapien | ZNF277 | 52.788 | 3 | 54.253853 | yes | 9.903E-08 | 620.63806 | 167.56323 |
| HTSLLPDR | 8 | Q9NRM2 | Zinc finger protein 277 OS=Homo sapien | ZNF277 | 52.788 | 2 | 54.253853 | yes | 4.036E-06 | 469.75635 | 70.331604 |
| SWEEVQLEDDR | 11 | Q9NRM2 | Zinc finger protein 277 OS=Homo sapien | ZNF277 | 52.788 | 2 | 54.253853 | yes | 9.722E-08 | 703.31512 | 67.45018 |
| MADFWLTEK | 9 | Q9NRX2 | Large ribosomal subunit protein bL17m C | MRPL17 | 20.051 | 2 | 50.611767 | yes | 1.245E-07 | 570.77332 | 1002.8862 |
| GNCLPPLPLPR | 11 | Q9NRX2 | Large ribosomal subunit protein bL17m C | MRPL17 | 20.051 | 2 | 50.611767 | yes | 7.319E-07 | 617.34229 | 787.78937 |
| LFQVLAPR | 8 | Q9NRX2 | Large ribosomal subunit protein bL17m C | MRPL17 | 20.051 | 2 | 50.611767 | yes | 5.094E-07 | 472.28983 | 555.75446 |
| IEAPWAR | 7 | Q9NRX2 | Large ribosomal subunit protein bL17m C | MRPL17 | 20.051 | 2 | 50.611767 | yes | 2.87E-07 | 421.72958 | 481.32886 |
| MAVIEYK | 7 | Q9NRX2 | Large ribosomal subunit protein bL17m C | MRPL17 | 20.051 | 2 | 50.611767 | yes | 9.741E-08 | 427.22803 | 304.6008 |
| MGLGPESR | 8 | Q9NRX2 | Large ribosomal subunit protein bL17m C | MRPL17 | 20.051 | 2 | 50.611767 | yes | 2.699E-07 | 423.71054 | 242.55298 |
| MLQIPNR | 7 | Q9NRX2 | Large ribosomal subunit protein bL17m C | MRPL17 | 20.051 | 2 | 50.611767 | yes | 1.865E-07 | 436.24457 | 125.63484 |
| DALPELLGLEAGSI | 14 | Q9NSI2 | Ribosome biogenesis protein SLX9 homo | SLX9 | 25.457 | 2 | 51.00664 | yes | 1.004E-07 | 720.88831 | 1473.7263 |
| QQLLEEER | 8 | Q9NSI2 | Ribosome biogenesis protein SLX9 homo | SLX9 | 25.457 | 2 | 51.00664 | yes | 2.176E-07 | 522.76965 | 1270.8019 |
| ASPLVAIGQTLAR | 13 | Q9NSI2 | Ribosome biogenesis protein SLX9 homo | SLX9 | 25.457 | 2 | 51.00664 | yes | 1.593E-07 | 648.88531 | 880.10809 |
| FQELLASPAYR | 11 | Q9NSI2 | Ribosome biogenesis protein SLX9 homo | SLX9 | 25.457 | 2 | 51.00664 | yes | 9.722E-08 | 647.84314 | 852.88251 |
| IDPSALVQK | 9 | Q9NSI2 | Ribosome biogenesis protein SLX9 homo | SLX9 | 25.457 | 2 | 51.00664 | yes | 1.157E-07 | 485.78201 | 733.98914 |
| RATVVVGDLHPLI | 13 | Q9NSI2 | Ribosome biogenesis protein SLX9 homo | SLX9 | 25.457 | 3 | 51.00664 | yes | 1.48E-07 | 478.2843 | 139.42494 |
| ATVVVGDLHPLR | 12 | Q9NSI2 | Ribosome biogenesis protein SLX9 homo | SLX9 | 25.457 | 2 | 51.00664 | yes | 2.754E-07 | 638.87225 | 67.704903 |

| | | | | | | | | | | | |
|---|---|---|---|---|---|---|---|---|---|---|---|
| FLSPQDTFEALK | 12 | Q9NTZ6 | RNA-binding protein 12 OS=Homo sapie | RBM12 | 97.396 | 2 | 46.087955 | yes | 2.173E-07 | 698.36139 | 103.71272 |
| FIQVHPITK | 9 | Q9NTZ6 | RNA-binding protein 12 OS=Homo sapie | RBM12 | 97.396 | 2 | 46.087955 | yes | 3.58E-07 | 541.82147 | 80.112328 |
| DFFHGLR | 7 | Q9NTZ6 | RNA-binding protein 12 OS=Homo sapie | RBM12 | 97.396 | 2 | 46.087955 | yes | 2.618E-07 | 446.22723 | 71.58168 |
| VDAVHLLK | 8 | Q9NTZ6 | RNA-binding protein 12 OS=Homo sapie | RBM12 | 97.396 | 2 | 46.087955 | yes | 1.657E-07 | 447.77402 | 42.701347 |
| FSCDIEQLK | 9 | Q9NUQ6 | SPATS2-like protein OS=Homo sapiens ( | SPATS2L | 61.729 | 2 | 51.62974 | yes | 9.722E-08 | 570.27368 | 3948.2573 |
| AELNTHVNVK | 10 | Q9NUQ6 | SPATS2-like protein OS=Homo sapiens ( | SPATS2L | 61.729 | 2 | 51.62974 | yes | 1.03E-07 | 583.81183 | 2640.5713 |
| FNPQYHNNR | 9 | Q9NUQ6 | SPATS2-like protein OS=Homo sapiens ( | SPATS2L | 61.729 | 2 | 51.62974 | yes | 1.115E-07 | 595.2785 | 2589.4138 |
| EVSLMAEMDK | 10 | Q9NUQ6 | SPATS2-like protein OS=Homo sapiens ( | SPATS2L | 61.729 | 2 | 51.62974 | yes | 1.277E-07 | 576.7674 | 2195.9021 |
| TPEAPAHSEKPR | 12 | Q9NUQ6 | SPATS2-like protein OS=Homo sapiens ( | SPATS2L | 61.729 | 3 | 51.62974 | yes | 1.937E-07 | 440.56158 | 910.92096 |
| TPEAPAHSEKPR | 12 | Q9NUQ6 | SPATS2-like protein OS=Homo sapiens ( | SPATS2L | 61.729 | 3 | 51.62974 | yes | 1.851E-07 | 660.33875 | 910.92096 |
| TPEAPAHSEKPR | 12 | Q9NUQ6 | SPATS2-like protein OS=Homo sapiens ( | SPATS2L | 61.729 | 2 | 51.62974 | yes | 1.937E-07 | 440.56158 | 910.92096 |
| TPEAPAHSEKPR | 12 | Q9NUQ6 | SPATS2-like protein OS=Homo sapiens ( | SPATS2L | 61.729 | 2 | 51.62974 | yes | 1.851E-07 | 660.33875 | 910.92096 |
| VKEEAMEILTAR | 12 | Q9NUQ6 | SPATS2-like protein OS=Homo sapiens ( | SPATS2L | 61.729 | 2 | 51.62974 | yes | 9.854E-08 | 695.37396 | 1598.5604 |
| KYDEELGK | 8 | Q9NUQ6 | SPATS2-like protein OS=Homo sapiens ( | SPATS2L | 61.729 | 2 | 51.62974 | yes | 1.564E-07 | 491.24802 | 1529.6016 |
| ISILEEPSK | 9 | Q9NUQ6 | SPATS2-like protein OS=Homo sapiens ( | SPATS2L | 61.729 | 2 | 51.62974 | yes | 4.208E-07 | 508.28714 | 1498.178 |
| AAFAELHNCIIDK | 13 | Q9NUQ6 | SPATS2-like protein OS=Homo sapiens ( | SPATS2L | 61.729 | 2 | 51.62974 | yes | 4.292E-07 | 751.37701 | 1346.8744 |
| AQIMLCGEITHPK | 13 | Q9NUQ6 | SPATS2-like protein OS=Homo sapiens ( | SPATS2L | 61.729 | 2 | 51.62974 | yes | 1.13E-07 | 749.38123 | 1144.0352 |
| TPCSSLLPLLNAH/ | 19 | Q9NUQ6 | SPATS2-like protein OS=Homo sapiens ( | SPATS2L | 61.729 | 3 | 51.62974 | yes | 1.73E-07 | 646.67529 | 1114.4358 |
| KISILEEPSK | 10 | Q9NUQ6 | SPATS2-like protein OS=Homo sapiens ( | SPATS2L | 61.729 | 2 | 51.62974 | yes | 1.241E-07 | 572.33459 | 899.11493 |
| EEAMEILTAR | 10 | Q9NUQ6 | SPATS2-like protein OS=Homo sapiens ( | SPATS2L | 61.729 | 2 | 51.62974 | yes | 1.132E-07 | 581.79224 | 698.18359 |
| NQEASLGMK | 9 | Q9NUQ6 | SPATS2-like protein OS=Homo sapiens ( | SPATS2L | 61.729 | 2 | 51.62974 | yes | 9.819E-08 | 489.23965 | 692.48712 |
| CTVSLTR | 7 | Q9NUQ6 | SPATS2-like protein OS=Homo sapiens ( | SPATS2L | 61.729 | 2 | 51.62974 | yes | 1.083E-07 | 418.71838 | 527.47461 |
| SQGSGNEAEPLGK | 13 | Q9NUQ6 | SPATS2-like protein OS=Homo sapiens ( | SPATS2L | 61.729 | 2 | 51.62974 | yes | 9.722E-08 | 637.30457 | 363.16818 |
| AVQAFVDGSAIQV | 15 | Q9NUQ6 | SPATS2-like protein OS=Homo sapiens ( | SPATS2L | 61.729 | 2 | 51.62974 | yes | 9.722E-08 | 773.43542 | 189.74606 |
| LSLDGNPK | 8 | Q9NUQ6 | SPATS2-like protein OS=Homo sapiens ( | SPATS2L | 61.729 | 2 | 51.62974 | yes | 1.022E-07 | 422.23218 | 146.88515 |
| VEGGFDQYR | 9 | Q9NUQ8 | ATP-binding cassette sub-family F memb | ABCF3 | 79.745 | 2 | 49.448853 | yes | 2.004E-07 | 535.74872 | 122.16148 |
| LLLGDLAPVR | 10 | Q9NUQ8 | ATP-binding cassette sub-family F memb | ABCF3 | 79.745 | 2 | 49.448853 | yes | 6.658E-07 | 533.83459 | 109.83198 |
| ALLQEQFR | 8 | Q9NUQ8 | ATP-binding cassette sub-family F memb | ABCF3 | 79.745 | 2 | 49.448853 | yes | 6.857E-07 | 502.77982 | 98.448616 |
| IENFDVSFGDR | 11 | Q9NUQ8 | ATP-binding cassette sub-family F memb | ABCF3 | 79.745 | 2 | 49.448853 | yes | 1.098E-07 | 649.8042 | 91.27562 |
| PLASLSGGQK | 10 | Q9NUQ8 | ATP-binding cassette sub-family F memb | ABCF3 | 79.745 | 2 | 49.448853 | yes | 2.695E-06 | 479.27182 | 88.150673 |
| AEGSEAAELAEIY/ | 15 | Q9NUQ8 | ATP-binding cassette sub-family F memb | ABCF3 | 79.745 | 2 | 49.448853 | yes | 1.364E-07 | 776.38049 | 82.462036 |
| VLLAGADVNLAW | 14 | Q9NUQ8 | ATP-binding cassette sub-family F memb | ABCF3 | 79.745 | 2 | 49.448853 | yes | 1.03E-07 | 727.90936 | 73.511017 |
| GGVILVSHDER | 11 | Q9NUQ8 | ATP-binding cassette sub-family F memb | ABCF3 | 79.745 | 2 | 49.448853 | yes | 1.072E-07 | 591.31732 | 50.378136 |
| LPELKPVDK | 9 | Q9NUQ8 | ATP-binding cassette sub-family F memb | ABCF3 | 79.745 | 2 | 49.448853 | yes | 5.635E-07 | 519.81335 | 42.734993 |
| ELWVCEGGGVTR | 12 | Q9NUQ8 | ATP-binding cassette sub-family F memb | ABCF3 | 79.745 | 2 | 49.448853 | yes | 6.124E-07 | 681.82715 | 30.16914 |
| EYEAQQQYR | 9 | Q9NUQ8 | ATP-binding cassette sub-family F memb | ABCF3 | 79.745 | 2 | 49.448853 | yes | 2.627E-07 | 607.77545 | 21.503975 |
| LLCPVPDSYTLK | 12 | Q9NVE5 | Ubiquitin carboxyl-terminal hydrolase 40 | USP40 | 140.13 | 2 | 51.20736 | yes | 2.752E-06 | 703.37341 | 171.55128 |
| GENLGPQDVLLR | 12 | Q9NVE5 | Ubiquitin carboxyl-terminal hydrolase 40 | USP40 | 140.13 | 2 | 51.20736 | yes | 1.006E-07 | 655.85681 | 145.22369 |
| NSSLQAESDFQR | 12 | Q9NVE5 | Ubiquitin carboxyl-terminal hydrolase 40 | USP40 | 140.13 | 2 | 51.20736 | yes | 1.01E-07 | 691.32074 | 67.890648 |

| | | | | | | | | | | | |
|---|---|---|---|---|---|---|---|---|---|---|---|
| IEICLEPLQK | 10 | Q9NVE5 | Ubiquitin carboxyl-terminal hydrolase 40 | USP40 | 140.13 | 2 | 51.20736 | yes | 1.557E-06 | 621.84174 | 53.422722 |
| KLPPFLTVSLLR | 12 | Q9NVE5 | Ubiquitin carboxyl-terminal hydrolase 40 | USP40 | 140.13 | 2 | 51.20736 | yes | 9.722E-08 | 692.43976 | 43.847363 |
| IIPLQLQR | 8 | Q9NVE5 | Ubiquitin carboxyl-terminal hydrolase 40 | USP40 | 140.13 | 2 | 51.20736 | yes | 1.879E-06 | 490.81619 | 42.259342 |
| NLLIDDDDDFSTIR | 14 | Q9NVE5 | Ubiquitin carboxyl-terminal hydrolase 40 | USP40 | 140.13 | 2 | 51.20736 | yes | 3.196E-07 | 826.3941 | 21.88529 |
| TSLIGEEWPSPR | 12 | Q9NVR7 | TBCC domain-containing protein 1 OS=I | TBCCD1 | 63.531 | 2 | 46.53415 | yes | 1.534E-07 | 686.34882 | 102.05582 |
| SSDTLAGAHVK | 11 | Q9NVR7 | TBCC domain-containing protein 1 OS=I | TBCCD1 | 63.531 | 2 | 46.53415 | yes | 1.823E-06 | 543.2829 | 42.589424 |
| ITGLCQEEHR | 10 | Q9NVS2 | Large ribosomal subunit protein mL66 OS | MRPS18A | 22.184 | 2 | 47.15738 | yes | 6.552E-07 | 621.7984 | 110.74146 |
| LPEGVVPK | 8 | Q9NVS2 | Large ribosomal subunit protein mL66 OS | MRPS18A | 22.184 | 2 | 47.15738 | yes | 1.388E-07 | 419.75528 | 48.98988 |
| LDPATSAVTVTR | 12 | Q9NVX7 | Kelch repeat and BTB domain-containing | KBTBD4 | 58.144 | 2 | 48.564674 | yes | 3.22E-06 | 615.83807 | 91.916954 |
| LVLSAQSCFFR | 11 | Q9NVX7 | Kelch repeat and BTB domain-containing | KBTBD4 | 58.144 | 2 | 48.564674 | yes | 1.129E-07 | 664.34497 | 63.117657 |
| AREEENYSFLPLVI | 18 | Q9NWA0 | Mediator of RNA polymerase II transcript | MED9 | 16.403 | 3 | 47.0438 | yes | 1.414E-07 | 724.71979 | 81.968719 |
| RVEELVAK | 8 | Q9NWB6 | Arginine and glutamate-rich protein 1 OS | ARGLU1 | 33.217 | 2 | 53.56432 | yes | 1.329E-07 | 472.2822 | 811.44043 |
| QLLEELER | 8 | Q9NWB6 | Arginine and glutamate-rich protein 1 OS | ARGLU1 | 33.217 | 2 | 53.56432 | yes | 1.142E-07 | 515.28241 | 595.52637 |
| RVEEELEK | 8 | Q9NWB6 | Arginine and glutamate-rich protein 1 OS | ARGLU1 | 33.217 | 2 | 53.56432 | yes | 3.561E-07 | 516.27203 | 363.91922 |
| ILEENNR | 7 | Q9NWB6 | Arginine and glutamate-rich protein 1 OS | ARGLU1 | 33.217 | 2 | 53.56432 | yes | 1.743E-07 | 444.2327 | 257.99817 |
| LAEEQLR | 7 | Q9NWB6 | Arginine and glutamate-rich protein 1 OS | ARGLU1 | 33.217 | 2 | 53.56432 | yes | 9.722E-08 | 429.73761 | 249.7023 |
| LIEEETAR | 8 | Q9NWB6 | Arginine and glutamate-rich protein 1 OS | ARGLU1 | 33.217 | 2 | 53.56432 | yes | 4.384E-07 | 480.75348 | 85.288727 |
| VPAPFKPITR | 10 | Q9NX20 | Large ribosomal subunit protein uL16m C | MRPL16 | 28.45 | 2 | 50.672646 | yes | 9.728E-08 | 563.34259 | 617.14429 |
| TLLPVPSFEDVSIP | 18 | Q9NX20 | Large ribosomal subunit protein uL16m C | MRPL16 | 28.45 | 3 | 50.672646 | yes | 1.944E-07 | 666.03821 | 358.4649 |
| VLSPYDLTHK | 10 | Q9NX20 | Large ribosomal subunit protein uL16m C | MRPL16 | 28.45 | 2 | 50.672646 | yes | 2.024E-07 | 586.81915 | 310.05484 |
| GAIDHYVTPVK | 11 | Q9NX20 | Large ribosomal subunit protein uL16m C | MRPL16 | 28.45 | 2 | 50.672646 | yes | 2.067E-07 | 600.32458 | 210.60512 |
| NNQNPWTFER | 10 | Q9NX20 | Large ribosomal subunit protein uL16m C | MRPL16 | 28.45 | 2 | 50.672646 | yes | 2.889E-07 | 653.30219 | 64.564674 |
| LLLQHAPLR | 9 | Q9NXF1 | Testis-expressed protein 10 OS=Homo sa | TEX10 | 105.675 | 2 | 46.289204 | yes | 1.6E-07 | 530.8349 | 94.910233 |
| SQSWILSVNPNR | 12 | Q9NXF1 | Testis-expressed protein 10 OS=Homo sa | TEX10 | 105.675 | 2 | 46.289204 | yes | 2.945E-07 | 700.86768 | 86.778397 |
| SQNFDILQSAISK | 13 | Q9NXF1 | Testis-expressed protein 10 OS=Homo sa | TEX10 | 105.675 | 2 | 46.289204 | yes | 4.517E-07 | 725.88043 | 86.632416 |
| EELTWLQSLR | 10 | Q9NXF1 | Testis-expressed protein 10 OS=Homo sa | TEX10 | 105.675 | 2 | 46.289204 | yes | 4.921E-07 | 637.84058 | 59.919098 |
| VGPEELPVVGQLL | 14 | Q9NXF1 | Testis-expressed protein 10 OS=Homo sa | TEX10 | 105.675 | 2 | 46.289204 | yes | 1.661E-07 | 753.43793 | 48.455593 |
| DCSWIEMIR | 9 | Q9NXF1 | Testis-expressed protein 10 OS=Homo sa | TEX10 | 105.675 | 2 | 46.289204 | yes | 2.491E-07 | 605.27319 | 41.526756 |
| LMLQSLR | 7 | Q9NXF1 | Testis-expressed protein 10 OS=Homo sa | TEX10 | 105.675 | 2 | 46.289204 | yes | 1.126E-06 | 430.75476 | 24.327702 |
| FGYQFTQQGPR | 11 | Q9NYL9 | Tropomodulin-3 OS=Homo sapiens OX= | TMOD3 | 39.595 | 2 | 51.756527 | yes | 1.015E-07 | 664.82275 | 512.84998 |
| NIPIPTLK | 8 | Q9NYL9 | Tropomodulin-3 OS=Homo sapiens OX= | TMOD3 | 39.595 | 2 | 51.756527 | no | 1.41E-07 | 448.28421 | 340.11481 |
| MLEENTNILK | 10 | Q9NYL9 | Tropomodulin-3 OS=Homo sapiens OX= | TMOD3 | 39.595 | 2 | 51.756527 | yes | 1.276E-07 | 602.81573 | 338.12885 |
| EHLLSYLEK | 9 | Q9NYL9 | Tropomodulin-3 OS=Homo sapiens OX= | TMOD3 | 39.595 | 2 | 51.756527 | yes | 9.722E-08 | 566.30585 | 172.2756 |
| QKPVQTFTEEK | 11 | Q9NYL9 | Tropomodulin-3 OS=Homo sapiens OX= | TMOD3 | 39.595 | 2 | 51.756527 | yes | 1.499E-07 | 667.8512 | 164.31689 |
| SNDPVATAFAEMI | 14 | Q9NYL9 | Tropomodulin-3 OS=Homo sapiens OX= | TMOD3 | 39.595 | 2 | 51.756527 | yes | 2.043E-06 | 747.3689 | 126.49951 |
| CFSLAATR | 8 | Q9NYL9 | Tropomodulin-3 OS=Homo sapiens OX= | TMOD3 | 39.595 | 2 | 51.756527 | yes | 3.424E-07 | 463.23166 | 123.44201 |
| DREDYVPYTGEK | 12 | Q9NYL9 | Tropomodulin-3 OS=Homo sapiens OX= | TMOD3 | 39.595 | 2 | 51.756527 | yes | 1.324E-07 | 736.33862 | 42.711048 |
| DREDYVPYTGEK | 12 | Q9NYL9 | Tropomodulin-3 OS=Homo sapiens OX= | TMOD3 | 39.595 | 2 | 51.756527 | yes | 3.705E-05 | 491.22818 | 42.711048 |
| DREDYVPYTGEK | 12 | Q9NYL9 | Tropomodulin-3 OS=Homo sapiens OX= | TMOD3 | 39.595 | 3 | 51.756527 | yes | 1.324E-07 | 736.33862 | 42.711048 |

| | | | | | | | | | | | |
|---|---|---|---|---|---|---|---|---|---|---|---|
| DREDYVPYTGEK | 12 | Q9NYL9 | Tropomodulin-3 OS=Homo sapiens OX= | TMOD3 | 39.595 | 3 | 51.756527 | yes | 3.705E-05 | 491.22818 | 42.711048 |
| ENDAHLVEVNLNI | 15 | Q9NYL9 | Tropomodulin-3 OS=Homo sapiens OX= | TMOD3 | 39.595 | 3 | 51.756527 | yes | 1.337E-07 | 574.63208 | 44.922539 |
| TKENDAHLVEVNI | 17 | Q9NYL9 | Tropomodulin-3 OS=Homo sapiens OX= | TMOD3 | 39.595 | 3 | 51.756527 | yes | 8.657E-05 | 651.01294 | 9.7316513 |
| TLVLIGAQGVGR | 12 | Q9NZW5 | Protein PALS2 OS=Homo sapiens OX=9 | PALS2 | 61.117 | 2 | 40.838573 | no | 1.928E-06 | 592.36151 | 43.933029 |
| VENNDLVIAR | 10 | Q9NZW5 | Protein PALS2 OS=Homo sapiens OX=9 | PALS2 | 61.117 | 2 | 40.838573 | yes | 3.813E-07 | 571.81183 | 31.347605 |
| LQYLIDLGR | 9 | Q9P015 | Large ribosomal subunit protein uL15m ( | MRPL15 | 33.42 | 2 | 56.829033 | yes | 1.127E-07 | 545.81641 | 1456.6945 |
| YGYILPDITK | 10 | Q9P015 | Large ribosomal subunit protein uL15m ( | MRPL15 | 33.42 | 2 | 56.829033 | yes | 9.899E-08 | 591.82391 | 1081.176 |
| ILKPTDENLLK | 11 | Q9P015 | Large ribosomal subunit protein uL15m ( | MRPL15 | 33.42 | 2 | 56.829033 | yes | 2.396E-07 | 642.38208 | 1054.1683 |
| QYKPLSLNR | 9 | Q9P015 | Large ribosomal subunit protein uL15m ( | MRPL15 | 33.42 | 2 | 56.829033 | yes | 1.322E-07 | 559.81946 | 650.23792 |
| YGFNEGHSFR | 10 | Q9P015 | Large ribosomal subunit protein uL15m ( | MRPL15 | 33.42 | 2 | 56.829033 | yes | 1.481E-07 | 607.27289 | 317.62256 |
| YGFNEGHSFR | 10 | Q9P015 | Large ribosomal subunit protein uL15m ( | MRPL15 | 33.42 | 2 | 56.829033 | yes | 6.929E-07 | 405.18436 | 317.62256 |
| YGFNEGHSFR | 10 | Q9P015 | Large ribosomal subunit protein uL15m ( | MRPL15 | 33.42 | 3 | 56.829033 | yes | 1.481E-07 | 607.27289 | 317.62256 |
| YGFNEGHSFR | 10 | Q9P015 | Large ribosomal subunit protein uL15m ( | MRPL15 | 33.42 | 3 | 56.829033 | yes | 6.929E-07 | 405.18436 | 317.62256 |
| NGGVVTTAFYDPI | 13 | Q9P015 | Large ribosomal subunit protein uL15m ( | MRPL15 | 33.42 | 2 | 56.829033 | yes | 9.819E-08 | 698.84644 | 492.86734 |
| VSLANLKPNPGSK | 13 | Q9P015 | Large ribosomal subunit protein uL15m ( | MRPL15 | 33.42 | 2 | 56.829033 | yes | 9.722E-08 | 662.88281 | 389.16391 |
| KYGYILPDITK | 11 | Q9P015 | Large ribosomal subunit protein uL15m ( | MRPL15 | 33.42 | 2 | 56.829033 | yes | 2.867E-07 | 655.8714 | 240.93036 |
| GVTIQPLKR | 9 | Q9P015 | Large ribosomal subunit protein uL15m ( | MRPL15 | 33.42 | 2 | 56.829033 | yes | 9.722E-08 | 506.31912 | 118.34516 |
| GVTIQPLK | 8 | Q9P015 | Large ribosomal subunit protein uL15m ( | MRPL15 | 33.42 | 2 | 56.829033 | yes | 1.14E-06 | 428.26855 | 40.571907 |
| AQTSLESQFTDR | 12 | Q9P031 | Thyroid transcription factor 1-associated | CCDC59 | 28.67 | 2 | 52.72967 | yes | 9.722E-08 | 691.83093 | 494.62256 |
| AQEEYEQIQAK | 11 | Q9P031 | Thyroid transcription factor 1-associated | CCDC59 | 28.67 | 2 | 52.72967 | yes | 9.722E-08 | 668.82263 | 264.89499 |
| WRPGGIEAR | 9 | Q9P031 | Thyroid transcription factor 1-associated | CCDC59 | 28.67 | 2 | 52.72967 | yes | 2.641E-07 | 521.28308 | 180.18979 |
| GEGVSTVGYR | 10 | Q9P031 | Thyroid transcription factor 1-associated | CCDC59 | 28.67 | 2 | 52.72967 | yes | 1.084E-07 | 512.75653 | 179.03114 |
| EGQGFAFR | 8 | Q9P031 | Thyroid transcription factor 1-associated | CCDC59 | 28.67 | 2 | 52.72967 | yes | 9.927E-08 | 456.22214 | 51.965645 |
| CKDCYLVK | 8 | Q9P0J6 | Large ribosomal subunit protein bL36m ( | MRPL36 | 11.785 | 2 | 49.319736 | yes | 1.045E-07 | 543.25952 | 675.13409 |
| TFVHVVPAKPEGT | 15 | Q9P0M9 | Large ribosomal subunit protein bL27m ( | MRPL27 | 16.073 | 3 | 51.132423 | yes | 1.083E-07 | 552.9751 | 114.91122 |
| TFVHVVPAKPEGT | 15 | Q9P0M9 | Large ribosomal subunit protein bL27m ( | MRPL27 | 16.073 | 3 | 51.132423 | yes | 9.722E-08 | 828.95905 | 114.91122 |
| TFVHVVPAKPEGT | 15 | Q9P0M9 | Large ribosomal subunit protein bL27m ( | MRPL27 | 16.073 | 2 | 51.132423 | yes | 1.083E-07 | 552.9751 | 114.91122 |
| TFVHVVPAKPEGT | 15 | Q9P0M9 | Large ribosomal subunit protein bL27m ( | MRPL27 | 16.073 | 2 | 51.132423 | yes | 9.722E-08 | 828.95905 | 114.91122 |
| CLYALEEGIVR | 11 | Q9P0M9 | Large ribosomal subunit protein bL27m ( | MRPL27 | 16.073 | 2 | 51.132423 | yes | 1.107E-07 | 661.84229 | 200.13953 |
| NTEAVDLITR | 10 | Q9P0M9 | Large ribosomal subunit protein bL27m ( | MRPL27 | 16.073 | 2 | 51.132423 | yes | 1.181E-07 | 566.30383 | 147.20886 |
| EVYVPHPR | 8 | Q9P0M9 | Large ribosomal subunit protein bL27m ( | MRPL27 | 16.073 | 2 | 51.132423 | yes | 2.532E-07 | 498.76672 | 135.58618 |
| SGPGLLPSEITR | 12 | Q9P1Y6 | PHD and RING finger domain-containing | PHRF1 | 178.667 | 2 | 44.36267 | yes | 2.222E-07 | 613.84058 | 165.20113 |
| EGAWDMEDVAPT | 15 | Q9P1Y6 | PHD and RING finger domain-containing | PHRF1 | 178.667 | 2 | 44.36267 | yes | 3.883E-07 | 816.86975 | 63.675575 |
| GVGSTFESFR | 10 | Q9P1Y6 | PHD and RING finger domain-containing | PHRF1 | 178.667 | 2 | 44.36267 | yes | 2.466E-07 | 543.76434 | 38.659153 |
| SGPQILTPLGDR | 12 | Q9P206 | Uncharacterized protein KIAA1522 OS=I | KIAA1522 | 107.095 | 2 | 51.945606 | yes | 1.047E-07 | 627.34607 | 1357.3954 |
| SVGAPGGAPTPAL | 23 | Q9P206 | Uncharacterized protein KIAA1522 OS=I | KIAA1522 | 107.095 | 3 | 51.945606 | yes | 2.88E-07 | 709.39539 | 861.31091 |
| STVLGLPQHVQK | 12 | Q9P206 | Uncharacterized protein KIAA1522 OS=I | KIAA1522 | 107.095 | 2 | 51.945606 | yes | 9.934E-08 | 653.8775 | 667.38019 |
| ASPVPAPSSGLHA | 16 | Q9P206 | Uncharacterized protein KIAA1522 OS=I | KIAA1522 | 107.095 | 2 | 51.945606 | yes | 1.173E-07 | 758.91516 | 515.85956 |
| PVSPETQADLQR | 12 | Q9P206 | Uncharacterized protein KIAA1522 OS=I | KIAA1522 | 107.095 | 2 | 51.945606 | yes | 1.513E-07 | 670.84387 | 506.71075 |

| | | | | | | | | | | | |
|---|---|---|---|---|---|---|---|---|---|---|---|
| VPAPFSPPPSKPR | 13 | Q9P206 | Uncharacterized protein KIAA1522 OS=I | KIAA1522 | 107.095 | 2 | 51.945606 | yes | 9.722E-08 | 688.88788 | 474.28967 |
| CSLHSASPASVR | 12 | Q9P206 | Uncharacterized protein KIAA1522 OS=I | KIAA1522 | 107.095 | 2 | 51.945606 | yes | 9.722E-08 | 636.31171 | 107.20412 |
| CSLHSASPASVR | 12 | Q9P206 | Uncharacterized protein KIAA1522 OS=I | KIAA1522 | 107.095 | 2 | 51.945606 | yes | 1.013E-07 | 424.54355 | 107.20412 |
| CSLHSASPASVR | 12 | Q9P206 | Uncharacterized protein KIAA1522 OS=I | KIAA1522 | 107.095 | 3 | 51.945606 | yes | 9.722E-08 | 636.31171 | 107.20412 |
| CSLHSASPASVR | 12 | Q9P206 | Uncharacterized protein KIAA1522 OS=I | KIAA1522 | 107.095 | 3 | 51.945606 | yes | 1.013E-07 | 424.54355 | 107.20412 |
| KPSVGVPPPASPSY | 16 | Q9P206 | Uncharacterized protein KIAA1522 OS=I | KIAA1522 | 107.095 | 3 | 51.945606 | yes | 6.863E-07 | 545.96661 | 103.44804 |
| KPSVGVPPPASPSY | 16 | Q9P206 | Uncharacterized protein KIAA1522 OS=I | KIAA1522 | 107.095 | 3 | 51.945606 | yes | 7.155E-07 | 818.44629 | 103.44804 |
| KPSVGVPPPASPSY | 16 | Q9P206 | Uncharacterized protein KIAA1522 OS=I | KIAA1522 | 107.095 | 2 | 51.945606 | yes | 6.863E-07 | 545.96661 | 103.44804 |
| KPSVGVPPPASPSY | 16 | Q9P206 | Uncharacterized protein KIAA1522 OS=I | KIAA1522 | 107.095 | 2 | 51.945606 | yes | 7.155E-07 | 818.44629 | 103.44804 |
| MGLPGSDSQK | 10 | Q9P206 | Uncharacterized protein KIAA1522 OS=I | KIAA1522 | 107.095 | 2 | 51.945606 | yes | 3.631E-06 | 510.24493 | 176.69353 |
| FSSVSSPQPR | 10 | Q9P206 | Uncharacterized protein KIAA1522 OS=I | KIAA1522 | 107.095 | 2 | 51.945606 | yes | 1.041E-07 | 546.27765 | 137.53925 |
| FVIPPHPK | 8 | Q9P206 | Uncharacterized protein KIAA1522 OS=I | KIAA1522 | 107.095 | 2 | 51.945606 | yes | 6.604E-07 | 467.77908 | 128.66139 |
| VSLQALEAEAEAG | 22 | Q9P206 | Uncharacterized protein KIAA1522 OS=I | KIAA1522 | 107.095 | 3 | 51.945606 | yes | 1.151E-07 | 773.0495 | 121.12659 |
| IPTVDGRPR | 9 | Q9P206 | Uncharacterized protein KIAA1522 OS=I | KIAA1522 | 107.095 | 2 | 51.945606 | yes | 3.403E-07 | 505.79071 | 46.215717 |
| GSPSGGSTAEASDT | 18 | Q9P206 | Uncharacterized protein KIAA1522 OS=I | KIAA1522 | 107.095 | 2 | 51.945606 | yes | 6.281E-07 | 846.9054 | 43.512642 |
| SISEQRPPQAPK | 12 | Q9P206 | Uncharacterized protein KIAA1522 OS=I | KIAA1522 | 107.095 | 2 | 51.945606 | yes | 4.976E-07 | 669.36224 | 29.971188 |
| MVNAAVNTYGAA | 17 | Q9P242 | Neuronal tyrosine-phosphorylated phosph | NYAP2 | 70.548 | 2 | 45.179527 | yes | 2.327E-07 | 818.39905 | 41.116486 |
| LSTSSETVSSTAAS | 15 | Q9P242 | Neuronal tyrosine-phosphorylated phosph | NYAP2 | 70.548 | 2 | 45.179527 | yes | 1.934E-07 | 728.3623 | 36.789627 |
| SSLMDTADGVPVS | 15 | Q9P265 | Disco-interacting protein 2 homolog B OS | DIP2B | 171.492 | 2 | 55.729652 | yes | 9.722E-08 | 761.36432 | 256.17203 |
| AACILTSQTLMR | 12 | Q9P265 | Disco-interacting protein 2 homolog B OS | DIP2B | 171.492 | 2 | 55.729652 | yes | 9.722E-08 | 682.85486 | 217.49229 |
| DLGQIEENDLVR | 12 | Q9P265 | Disco-interacting protein 2 homolog B OS | DIP2B | 171.492 | 2 | 55.729652 | yes | 1.626E-07 | 700.85443 | 151.53828 |
| LQCELYSSR | 9 | Q9P265 | Disco-interacting protein 2 homolog B OS | DIP2B | 171.492 | 2 | 55.729652 | yes | 8.679E-07 | 578.27679 | 150.06398 |
| EGSVMGVTVSR | 11 | Q9P265 | Disco-interacting protein 2 homolog B OS | DIP2B | 171.492 | 2 | 55.729652 | yes | 2.045E-07 | 561.28461 | 119.13455 |
| GLGNQVEVLK | 10 | Q9P265 | Disco-interacting protein 2 homolog B OS | DIP2B | 171.492 | 2 | 55.729652 | yes | 1.209E-07 | 528.80603 | 113.26943 |
| LAYTLLNK | 8 | Q9P265 | Disco-interacting protein 2 homolog B OS | DIP2B | 171.492 | 2 | 55.729652 | no | 1.573E-07 | 468.28165 | 104.66769 |
| LSFGDAAQTLWAI | 13 | Q9P265 | Disco-interacting protein 2 homolog B OS | DIP2B | 171.492 | 2 | 55.729652 | yes | 2.219E-07 | 718.37006 | 102.53809 |
| GTTVCTASCLQLH | 14 | Q9P265 | Disco-interacting protein 2 homolog B OS | DIP2B | 171.492 | 2 | 55.729652 | yes | 9.722E-08 | 788.38452 | 83.820251 |
| RHNADDIVATGLA | 18 | Q9P265 | Disco-interacting protein 2 homolog B OS | DIP2B | 171.492 | 3 | 55.729652 | yes | 9.722E-08 | 637.00946 | 80.296677 |
| NQTPAPSAAQTSA | 16 | Q9P265 | Disco-interacting protein 2 homolog B OS | DIP2B | 171.492 | 2 | 55.729652 | yes | 1.525E-07 | 778.38916 | 78.022011 |
| TDEIGEICVSSR | 12 | Q9P265 | Disco-interacting protein 2 homolog B OS | DIP2B | 171.492 | 2 | 55.729652 | yes | 1.306E-07 | 683.3194 | 77.755424 |
| TGYLGFVR | 8 | Q9P265 | Disco-interacting protein 2 homolog B OS | DIP2B | 171.492 | 2 | 55.729652 | yes | 2.323E-07 | 456.75052 | 76.760735 |
| YHPIDIETSVSR | 12 | Q9P265 | Disco-interacting protein 2 homolog B OS | DIP2B | 171.492 | 2 | 55.729652 | yes | 1.21E-06 | 708.8595 | 70.084145 |
| SDIHTEAVQAALA | 14 | Q9P265 | Disco-interacting protein 2 homolog B OS | DIP2B | 171.492 | 2 | 55.729652 | yes | 9.722E-08 | 727.38593 | 68.039192 |
| TGGMMYFGLAGV | 14 | Q9P265 | Disco-interacting protein 2 homolog B OS | DIP2B | 171.492 | 2 | 55.729652 | yes | 5.837E-07 | 716.85181 | 53.282574 |
| VVIVNPETK | 9 | Q9P265 | Disco-interacting protein 2 homolog B OS | DIP2B | 171.492 | 2 | 55.729652 | yes | 6.259E-07 | 499.79767 | 45.433773 |
| TWPTIIDTDDLPR | 13 | Q9P265 | Disco-interacting protein 2 homolog B OS | DIP2B | 171.492 | 2 | 55.729652 | yes | 9.44E-07 | 771.89355 | 42.893944 |
| NEPVLKPGDR | 10 | Q9P265 | Disco-interacting protein 2 homolog B OS | DIP2B | 171.492 | 2 | 55.729652 | yes | 1.557E-07 | 562.80658 | 34.366146 |
| SIQGSSTSSSASSTI | 21 | Q9P265 | Disco-interacting protein 2 homolog B OS | DIP2B | 171.492 | 3 | 55.729652 | yes | 4.78E-07 | 679.66254 | 19.563946 |
| LMLPDSPLVEEGR | 13 | Q9P2F8 | Signal-induced proliferation-associated 1- | SIPA1L2 | 190.438 | 2 | 54.36365 | yes | 1.477E-06 | 728.37921 | 98.982956 |

| | | | | | | | | | | | |
|---|---|---|---|---|---|---|---|---|---|---|---|
| SNDLVLSCPYFR | 12 | Q9P2F8 | Signal-induced proliferation-associated 1· | SIPA1L2 | 190.438 | 2 | 54.36365 | yes | 1.19E-07 | 735.8559 | 64.363945 |
| SAVFRDFLLAK | 11 | Q9P2F8 | Signal-induced proliferation-associated 1· | SIPA1L2 | 190.438 | 2 | 54.36365 | yes | 1.125E-05 | 633.86389 | 49.265938 |
| NEFWFSDGSLSDK | 13 | Q9P2F8 | Signal-induced proliferation-associated 1· | SIPA1L2 | 190.438 | 2 | 54.36365 | yes | 9.114E-07 | 766.33862 | 31.921658 |
| AVLQAEVQHLR | 11 | Q9P2F8 | Signal-induced proliferation-associated 1· | SIPA1L2 | 190.438 | 2 | 54.36365 | yes | 9.722E-08 | 632.36206 | 21.241434 |
| HSEAAPGPVSCTR | 15 | Q9P2Z0 | THAP domain-containing protein 10 OS= | THAP10 | 28.351 | 3 | 52.0491 | yes | 9.722E-08 | 541.26697 | 44.788364 |
| SEQSDLSYMAVQV | 14 | Q9P2Z0 | THAP domain-containing protein 10 OS= | THAP10 | 28.351 | 2 | 52.0491 | yes | 4.261E-07 | 792.88232 | 6.7881308 |
| ELTVDNSFEFEK | 12 | Q9UHA3 | Probable ribosome biogenesis protein RL | RSL24D1 | 19.622 | 2 | 49.974964 | yes | 1.108E-07 | 729.34338 | 265.71774 |
| QNIHLIR | 7 | Q9UHA3 | Probable ribosome biogenesis protein RL | RSL24D1 | 19.622 | 2 | 49.974964 | yes | 9.957E-08 | 447.26941 | 236.37187 |
| CNAPYCSLR | 9 | Q9UHR6 | Zinc finger HIT domain-containing protei | ZNHIT2 | 42.884 | 2 | 47.043674 | yes | 4.813E-07 | 570.74988 | 96.962151 |
| VLGDVPGACTPVV | 16 | Q9UHR6 | Zinc finger HIT domain-containing protei | ZNHIT2 | 42.884 | 2 | 47.043674 | yes | 1.414E-07 | 819.43762 | 32.257851 |
| YAQLVMGPAGSG | 13 | Q9UHW5 | GPN-loop GTPase 3 OS=Homo sapiens ( | GPN3 | 32.762 | 2 | 55.354862 | yes | 9.722E-08 | 639.82916 | 146.61217 |
| AICGLIDDYSMVR | 13 | Q9UHW5 | GPN-loop GTPase 3 OS=Homo sapiens ( | GPN3 | 32.762 | 2 | 55.354862 | yes | 9.722E-08 | 756.86292 | 62.499626 |
| FGGLVLSPVGK | 11 | Q9UJK0 | 18S rRNA aminocarboxypropyltransferas | TSR3 | 33.596 | 2 | 50.27695 | yes | 3.398E-07 | 537.32129 | 877.46051 |
| QYASPADR | 8 | Q9UJK0 | 18S rRNA aminocarboxypropyltransferas | TSR3 | 33.596 | 2 | 50.27695 | yes | 9.82E-08 | 454.21704 | 203.41216 |
| LLPYLVAANPVNY | 15 | Q9UJK0 | 18S rRNA aminocarboxypropyltransferas | TSR3 | 33.596 | 2 | 50.27695 | yes | 4.66E-06 | 830.46448 | 94.971275 |
| LDETPFGK | 8 | Q9UJK0 | 18S rRNA aminocarboxypropyltransferas | TSR3 | 33.596 | 2 | 50.27695 | yes | 1.518E-07 | 453.73199 | 80.581474 |
| GFLDLNR | 7 | Q9UJK0 | 18S rRNA aminocarboxypropyltransferas | TSR3 | 33.596 | 2 | 50.27695 | yes | 1.887E-07 | 417.72705 | 76.715195 |
| YAACGSPEEVLQA | 22 | Q9UJK0 | 18S rRNA aminocarboxypropyltransferas | TSR3 | 33.596 | 3 | 50.27695 | yes | 1.236E-07 | 809.0495 | 74.739601 |
| GEVGLLFTNR | 10 | Q9UKD2 | mRNA turnover protein 4 homolog OS=I | MRTO4 | 27.561 | 2 | 53.40101 | yes | 1.188E-07 | 553.30365 | 2678.4944 |
| EGDVLTPEQAR | 11 | Q9UKD2 | mRNA turnover protein 4 homolog OS=I | MRTO4 | 27.561 | 2 | 53.40101 | yes | 1.084E-07 | 607.8042 | 2073.4834 |
| GVVTLLSDYEVCK | 13 | Q9UKD2 | mRNA turnover protein 4 homolog OS=I | MRTO4 | 27.561 | 2 | 53.40101 | yes | 9.722E-08 | 741.87909 | 1252.6877 |
| YMWDSQSGR | 9 | Q9UKD2 | mRNA turnover protein 4 homolog OS=I | MRTO4 | 27.561 | 2 | 53.40101 | yes | 9.722E-08 | 565.24017 | 1083.095 |
| VMMVALGR | 8 | Q9UKD2 | mRNA turnover protein 4 homolog OS=I | MRTO4 | 27.561 | 2 | 53.40101 | yes | 2.953E-07 | 438.74335 | 915.76227 |
| LFGYEMAEFK | 10 | Q9UKD2 | mRNA turnover protein 4 homolog OS=I | MRTO4 | 27.561 | 2 | 53.40101 | yes | 1.056E-07 | 617.79425 | 879.04529 |
| QNLIEELR | 8 | Q9UKD2 | mRNA turnover protein 4 homolog OS=I | MRTO4 | 27.561 | 2 | 53.40101 | yes | 2.264E-07 | 507.78256 | 807.5426 |
| TKEEVNEWFTK | 11 | Q9UKD2 | mRNA turnover protein 4 homolog OS=I | MRTO4 | 27.561 | 2 | 53.40101 | yes | 1.062E-07 | 705.84863 | 664.72559 |
| YTEMDYAR | 8 | Q9UKD2 | mRNA turnover protein 4 homolog OS=I | MRTO4 | 27.561 | 2 | 53.40101 | yes | 1.865E-07 | 524.72388 | 574.2146 |
| QLGLPTALK | 9 | Q9UKD2 | mRNA turnover protein 4 homolog OS=I | MRTO4 | 27.561 | 2 | 53.40101 | yes | 1.75E-07 | 470.79492 | 527.16028 |
| QNLIEELRK | 9 | Q9UKD2 | mRNA turnover protein 4 homolog OS=I | MRTO4 | 27.561 | 2 | 53.40101 | yes | 1.882E-07 | 571.83002 | 60.921051 |
| EEVNEWFTK | 9 | Q9UKD2 | mRNA turnover protein 4 homolog OS=I | MRTO4 | 27.561 | 2 | 53.40101 | yes | 1.401E-06 | 591.27728 | 25.1511 |
| LHELAEQR | 8 | Q9UKJ3 | G patch domain-containing protein 8 OS= | GPATCH8 | 164.198 | 2 | 54.666218 | yes | 1.81E-07 | 498.26706 | 132.15825 |
| DHLQGLDPGEPNK | 13 | Q9UKJ3 | G patch domain-containing protein 8 OS= | GPATCH8 | 164.198 | 2 | 54.666218 | yes | 9.366E-07 | 710.3468 | 105.68814 |
| ANFYCELCDK | 10 | Q9UKJ3 | G patch domain-containing protein 8 OS= | GPATCH8 | 164.198 | 2 | 54.666218 | no | 9.722E-08 | 660.27338 | 85.181038 |
| ASEQMDGDNTTHI | 14 | Q9UKJ3 | G patch domain-containing protein 8 OS= | GPATCH8 | 164.198 | 2 | 54.666218 | yes | 1.19E-07 | 765.82806 | 76.224998 |
| MDAPASGSACSGL | 15 | Q9UKJ3 | G patch domain-containing protein 8 OS= | GPATCH8 | 164.198 | 2 | 54.666218 | yes | 9.722E-08 | 733.32416 | 69.183464 |
| ETSLATPAGK | 10 | Q9UKJ3 | G patch domain-containing protein 8 OS= | GPATCH8 | 164.198 | 2 | 54.666218 | yes | 9.365E-07 | 487.76129 | 68.901855 |
| HPTGPFFPVLSK | 12 | Q9UKJ3 | G patch domain-containing protein 8 OS= | GPATCH8 | 164.198 | 2 | 54.666218 | yes | 5.895E-07 | 663.86389 | 51.144047 |
| TVSEVSEQPK | 10 | Q9UKJ3 | G patch domain-containing protein 8 OS= | GPATCH8 | 164.198 | 2 | 54.666218 | yes | 4.186E-06 | 552.28259 | 34.734486 |
| VLEVEKEDTEELR | 13 | Q9UKJ3 | G patch domain-containing protein 8 OS= | GPATCH8 | 164.198 | 2 | 54.666218 | yes | 3.445E-07 | 794.90686 | 29.256927 |

| | | | | | | | | | | | |
|---|---|---|---|---|---|---|---|---|---|---|---|
| LVYHIGLTPAK | 11 | Q9UKZ1 | CCR4-NOT transcription complex subuni | CNOT11 | 55.216 | 2 | 51.20519 | yes | 9.722E-08 | 606.36096 | 115.18231 |
| TLDTGETPSETK | 12 | Q9UKZ1 | CCR4-NOT transcription complex subuni | CNOT11 | 55.216 | 2 | 51.20519 | yes | 1.191E-07 | 639.80664 | 89.821533 |
| SPLSSPQQTQLLGI | 20 | Q9UKZ1 | CCR4-NOT transcription complex subuni | CNOT11 | 55.216 | 3 | 51.20519 | yes | 3.839E-07 | 732.39001 | 79.293678 |
| GGASGPGSGSGGP | 19 | Q9UKZ1 | CCR4-NOT transcription complex subuni | CNOT11 | 55.216 | 2 | 51.20519 | yes | 2.821E-06 | 720.33472 | 11.55683 |
| LEPLKPLK | 8 | Q9ULL5 | Proline-rich protein 12 OS=Homo sapien | PRR12 | 211.045 | 2 | 49.11249 | yes | 1.066E-07 | 469.30768 | 107.8406 |
| SLATGQPATSR | 11 | Q9ULL5 | Proline-rich protein 12 OS=Homo sapien | PRR12 | 211.045 | 2 | 49.11249 | yes | 3.563E-07 | 544.78839 | 102.74358 |
| YHLQSVIR | 8 | Q9ULL5 | Proline-rich protein 12 OS=Homo sapien | PRR12 | 211.045 | 2 | 49.11249 | yes | 3.737E-06 | 508.28781 | 85.818764 |
| VGVHLLEPATR | 11 | Q9ULL5 | Proline-rich protein 12 OS=Homo sapien | PRR12 | 211.045 | 2 | 49.11249 | yes | 1.684E-07 | 596.34583 | 68.722969 |
| SHGLEPAAPSPR | 12 | Q9ULL5 | Proline-rich protein 12 OS=Homo sapien | PRR12 | 211.045 | 2 | 49.11249 | yes | 1.166E-07 | 609.81494 | 21.184406 |
| LCYYIDR | 7 | Q9UN30 | Sex comb on midleg-like protein 1 OS=H | SCML1 | 37.448 | 2 | 46.63465 | yes | 1.508E-07 | 501.73929 | 98.736298 |
| LLCASPVEYFR | 11 | Q9UNK9 | Protein angel homolog 1 OS=Homo sapie | ANGEL1 | 75.276 | 2 | 49.225468 | yes | 1.174E-07 | 677.84485 | 147.22226 |
| FCSIACQR | 8 | Q9UNK9 | Protein angel homolog 1 OS=Homo sapie | ANGEL1 | 75.276 | 2 | 49.225468 | yes | 5.527E-07 | 521.23389 | 133.83365 |
| ALSDAFFTCR | 10 | Q9UNK9 | Protein angel homolog 1 OS=Homo sapie | ANGEL1 | 75.276 | 2 | 49.225468 | yes | 1.056E-07 | 594.2793 | 78.356941 |
| ESGSPYSVLVDTK | 13 | Q9Y2D9 | Zinc finger protein 652 OS=Homo sapien | ZNF652 | 69.745 | 2 | 49.637463 | yes | 1.966E-07 | 691.34589 | 96.083664 |
| FSCEICEK | 8 | Q9Y2D9 | Zinc finger protein 652 OS=Homo sapien | ZNF652 | 69.745 | 2 | 49.637463 | no | 1.076E-07 | 536.72552 | 89.876495 |
| DMPFTCETCGK | 11 | Q9Y2D9 | Zinc finger protein 652 OS=Homo sapien | ZNF652 | 69.745 | 2 | 49.637463 | no | 1.015E-07 | 673.26471 | 64.325188 |
| VHSLQHSGEKPFR | 13 | Q9Y2D9 | Zinc finger protein 652 OS=Homo sapien | ZNF652 | 69.745 | 3 | 49.637463 | no | 2.571E-06 | 507.93555 | 39.674583 |
| ALVAEHLR | 8 | Q9Y2L6 | FERM domain-containing protein 4B OS | FRMD4B | 118.047 | 2 | 37.43911 | yes | 5.992E-07 | 454.76926 | 85.395363 |
| LLDLELTSR | 9 | Q9Y2Q9 | Small ribosomal subunit protein bS1m O | MRPS28 | 20.843 | 2 | 51.657204 | yes | 1.117E-07 | 530.30585 | 2806.4688 |
| AGGFASALER | 10 | Q9Y2Q9 | Small ribosomal subunit protein bS1m O | MRPS28 | 20.843 | 2 | 51.657204 | yes | 9.722E-08 | 489.75378 | 1399.6406 |
| HSPLTQMGPAK | 11 | Q9Y2Q9 | Small ribosomal subunit protein bS1m O | MRPS28 | 20.843 | 2 | 51.657204 | yes | 4.752E-07 | 583.80298 | 595.12927 |
| HSELLQK | 7 | Q9Y2Q9 | Small ribosomal subunit protein bS1m O | MRPS28 | 20.843 | 2 | 51.657204 | yes | 1.542E-07 | 427.74017 | 316.01282 |
| HELAEIVFK | 9 | Q9Y2R5 | Small ribosomal subunit protein uS17m C | MRPS17 | 14.503 | 2 | 48.4702 | yes | 1.158E-06 | 543.3031 | 1379.0723 |
| TYFAHDALQQCTV | 20 | Q9Y2R5 | Small ribosomal subunit protein uS17m C | MRPS17 | 14.503 | 3 | 48.4702 | yes | 1.144E-07 | 774.06335 | 700.03101 |
| FYQVPVPLPDR | 11 | Q9Y2R9 | Small ribosomal subunit protein uS7m O | MRPS7 | 28.135 | 2 | 52.898552 | yes | 1.106E-07 | 665.86133 | 3960.2405 |
| TSSVFEDPVISK | 12 | Q9Y2R9 | Small ribosomal subunit protein uS7m O | MRPS7 | 28.135 | 2 | 52.898552 | yes | 1.038E-07 | 654.83771 | 3268.7327 |
| SLMIQTLEAVK | 11 | Q9Y2R9 | Small ribosomal subunit protein uS7m O | MRPS7 | 28.135 | 2 | 52.898552 | yes | 9.882E-08 | 616.84961 | 1521.7765 |
| NPYTIFHQALK | 11 | Q9Y2R9 | Small ribosomal subunit protein uS7m O | MRPS7 | 28.135 | 2 | 52.898552 | yes | 3.976E-07 | 666.35895 | 1298.7031 |
| LLEAFHNQGPVIK | 13 | Q9Y2R9 | Small ribosomal subunit protein uS7m O | MRPS7 | 28.135 | 2 | 52.898552 | yes | 1.182E-07 | 733.41174 | 517.0755 |
| LLEAFHNQGPVIK | 13 | Q9Y2R9 | Small ribosomal subunit protein uS7m O | MRPS7 | 28.135 | 2 | 52.898552 | yes | 2.233E-07 | 489.27692 | 517.0755 |
| LLEAFHNQGPVIK | 13 | Q9Y2R9 | Small ribosomal subunit protein uS7m O | MRPS7 | 28.135 | 3 | 52.898552 | yes | 1.182E-07 | 733.41174 | 517.0755 |
| LLEAFHNQGPVIK | 13 | Q9Y2R9 | Small ribosomal subunit protein uS7m O | MRPS7 | 28.135 | 3 | 52.898552 | yes | 2.233E-07 | 489.27692 | 517.0755 |
| NCEPMIGLVPILK | 13 | Q9Y2R9 | Small ribosomal subunit protein uS7m O | MRPS7 | 28.135 | 2 | 52.898552 | yes | 1.318E-07 | 742.40417 | 887.52081 |
| WMITECR | 7 | Q9Y2R9 | Small ribosomal subunit protein uS7m O | MRPS7 | 28.135 | 2 | 52.898552 | yes | 1.307E-06 | 498.22549 | 879.74542 |
| YSPEFKDPLIDK | 12 | Q9Y2R9 | Small ribosomal subunit protein uS7m O | MRPS7 | 28.135 | 2 | 52.898552 | yes | 1.361E-07 | 726.37445 | 758.96912 |
| KPVEELTEEEK | 11 | Q9Y2R9 | Small ribosomal subunit protein uS7m O | MRPS7 | 28.135 | 2 | 52.898552 | yes | 1.477E-07 | 665.84045 | 387.09875 |
| YHAASAEEQATIE | 14 | Q9Y2R9 | Small ribosomal subunit protein uS7m O | MRPS7 | 28.135 | 2 | 52.898552 | yes | 1.221E-07 | 788.37354 | 211.9088 |
| FTNMMMIGGNK | 11 | Q9Y2R9 | Small ribosomal subunit protein uS7m O | MRPS7 | 28.135 | 2 | 52.898552 | yes | 9.722E-08 | 622.28522 | 131.1539 |
| EIVGASTNWR | 10 | Q9Y305 | Acyl-coenzyme A thioesterase 9, mitocho | ACOT9 | 49.902 | 2 | 52.81505 | yes | 1.171E-07 | 566.79089 | 571.04645 |

| | | | | | | | | | | | |
|---|---|---|---|---|---|---|---|---|---|---|---|
| TYGESMLYLDGQI | 13 | Q9Y305 | Acyl-coenzyme A thioesterase 9, mitocho | ACOT9 | 49.902 | 2 | 52.81505 | yes | 1.032E-07 | 766.85614 | 563.97601 |
| EVPLVFPK | 8 | Q9Y305 | Acyl-coenzyme A thioesterase 9, mitocho | ACOT9 | 49.902 | 2 | 52.81505 | yes | 1.011E-07 | 464.77875 | 441.54181 |
| SLEICHPQER | 10 | Q9Y305 | Acyl-coenzyme A thioesterase 9, mitocho | ACOT9 | 49.902 | 2 | 52.81505 | yes | 9.722E-08 | 634.80621 | 393.75992 |
| YLTVQNTVR | 9 | Q9Y305 | Acyl-coenzyme A thioesterase 9, mitocho | ACOT9 | 49.902 | 2 | 52.81505 | yes | 2.858E-07 | 547.30365 | 376.23059 |
| VLPSNAVWMENS | 13 | Q9Y305 | Acyl-coenzyme A thioesterase 9, mitocho | ACOT9 | 49.902 | 2 | 52.81505 | yes | 8.492E-07 | 737.87158 | 287.61447 |
| IAFSSTSLLK | 10 | Q9Y305 | Acyl-coenzyme A thioesterase 9, mitocho | ACOT9 | 49.902 | 2 | 52.81505 | yes | 1.041E-07 | 533.81079 | 253.412 |
| FSGHVSWVGK | 10 | Q9Y305 | Acyl-coenzyme A thioesterase 9, mitocho | ACOT9 | 49.902 | 2 | 52.81505 | yes | 1.732E-07 | 552.28528 | 242.70622 |
| VHSEVASLQEK | 11 | Q9Y305 | Acyl-coenzyme A thioesterase 9, mitocho | ACOT9 | 49.902 | 2 | 52.81505 | yes | 1.281E-07 | 613.82239 | 223.72943 |
| HFNSMSGPATLR | 12 | Q9Y305 | Acyl-coenzyme A thioesterase 9, mitocho | ACOT9 | 49.902 | 2 | 52.81505 | yes | 9.901E-08 | 659.32208 | 141.32103 |
| TTIHEMFLSTLDPK | 14 | Q9Y305 | Acyl-coenzyme A thioesterase 9, mitocho | ACOT9 | 49.902 | 2 | 52.81505 | yes | 1.448E-07 | 816.9187 | 135.44716 |
| LLHSFLAK | 8 | Q9Y305 | Acyl-coenzyme A thioesterase 9, mitocho | ACOT9 | 49.902 | 2 | 52.81505 | yes | 9.935E-08 | 464.78436 | 125.10234 |
| KLLHSFLAK | 9 | Q9Y305 | Acyl-coenzyme A thioesterase 9, mitocho | ACOT9 | 49.902 | 2 | 52.81505 | yes | 1.136E-07 | 528.83185 | 32.916656 |
| AMEVDIEERPK | 11 | Q9Y3C1 | Nucleolar protein 16 OS=Homo sapiens ( | NOP16 | 21.189 | 2 | 51.17566 | yes | 9.893E-08 | 658.82941 | 2115.2432 |
| QNLAEMGLAVDPI | 14 | Q9Y3C1 | Nucleolar protein 16 OS=Homo sapiens ( | NOP16 | 21.189 | 2 | 51.17566 | yes | 2.008E-07 | 764.38281 | 1684.1691 |
| YMVENHGEDYK | 11 | Q9Y3C1 | Nucleolar protein 16 OS=Homo sapiens ( | NOP16 | 21.189 | 2 | 51.17566 | yes | 1.791E-07 | 692.79553 | 670.24542 |
| NYYQDTPK | 8 | Q9Y3C1 | Nucleolar protein 16 OS=Homo sapiens ( | NOP16 | 21.189 | 2 | 51.17566 | yes | 1.732E-07 | 514.73779 | 668.16644 |
| KPYVLNDLEAEAS | 18 | Q9Y3C1 | Nucleolar protein 16 OS=Homo sapiens ( | NOP16 | 21.189 | 3 | 51.17566 | yes | 9.722E-08 | 682.03687 | 311.16617 |
| KPYVLNDLEAEAS | 17 | Q9Y3C1 | Nucleolar protein 16 OS=Homo sapiens ( | NOP16 | 21.189 | 3 | 51.17566 | yes | 9.795E-08 | 639.33856 | 264.84018 |
| HAWDHAK | 7 | Q9Y3C1 | Nucleolar protein 16 OS=Homo sapiens ( | NOP16 | 21.189 | 2 | 51.17566 | yes | 6.233E-07 | 432.7092 | 139.3853 |
| AMEVDIEER | 9 | Q9Y3C1 | Nucleolar protein 16 OS=Homo sapiens ( | NOP16 | 21.189 | 2 | 51.17566 | yes | 9.722E-08 | 546.25549 | 121.29166 |
| EGVLEPNPVR | 10 | Q9Y3X0 | Coiled-coil domain-containing protein 9 ( | CCDC9 | 59.703 | 2 | 49.478703 | yes | 1.05E-07 | 555.30109 | 1208.1913 |
| NWGGPDFER | 9 | Q9Y3X0 | Coiled-coil domain-containing protein 9 ( | CCDC9 | 59.703 | 2 | 49.478703 | yes | 1.032E-07 | 539.24103 | 1100.0881 |
| AGLGSAGDMTLSN | 16 | Q9Y3X0 | Coiled-coil domain-containing protein 9 ( | CCDC9 | 59.703 | 2 | 49.478703 | yes | 2.184E-07 | 762.8609 | 683.85718 |
| KAELEGVAVTAPR | 13 | Q9Y3X0 | Coiled-coil domain-containing protein 9 ( | CCDC9 | 59.703 | 2 | 49.478703 | yes | 1.083E-07 | 670.88025 | 235.21317 |
| KAELEGVAVTAPR | 13 | Q9Y3X0 | Coiled-coil domain-containing protein 9 ( | CCDC9 | 59.703 | 2 | 49.478703 | yes | 1.113E-07 | 447.58926 | 235.21317 |
| KAELEGVAVTAPR | 13 | Q9Y3X0 | Coiled-coil domain-containing protein 9 ( | CCDC9 | 59.703 | 3 | 49.478703 | yes | 1.083E-07 | 670.88025 | 235.21317 |
| KAELEGVAVTAPR | 13 | Q9Y3X0 | Coiled-coil domain-containing protein 9 ( | CCDC9 | 59.703 | 3 | 49.478703 | yes | 1.113E-07 | 447.58926 | 235.21317 |
| DGPVPAHEPSHR | 12 | Q9Y3X0 | Coiled-coil domain-containing protein 9 ( | CCDC9 | 59.703 | 2 | 49.478703 | yes | 1.026E-07 | 649.81543 | 421.05582 |
| ENVAVESEK | 9 | Q9Y3X0 | Coiled-coil domain-containing protein 9 ( | CCDC9 | 59.703 | 2 | 49.478703 | yes | 1.78E-07 | 502.74838 | 376.6199 |
| YQEIEEDR | 8 | Q9Y3X0 | Coiled-coil domain-containing protein 9 ( | CCDC9 | 59.703 | 2 | 49.478703 | yes | 2.226E-07 | 541.24347 | 308.99786 |
| RSPGTPRPPGASK | 13 | Q9Y3X0 | Coiled-coil domain-containing protein 9 ( | CCDC9 | 59.703 | 3 | 49.478703 | yes | 0.0001736 | 436.57748 | 291.03427 |
| REWDAEK | 7 | Q9Y3X0 | Coiled-coil domain-containing protein 9 ( | CCDC9 | 59.703 | 2 | 49.478703 | yes | 1.809E-07 | 467.22488 | 245.18021 |
| MNEEMEK | 7 | Q9Y3X0 | Coiled-coil domain-containing protein 9 ( | CCDC9 | 59.703 | 2 | 49.478703 | yes | 2.033E-07 | 455.68588 | 222.92355 |
| SPGTPRPPGASK | 12 | Q9Y3X0 | Coiled-coil domain-containing protein 9 ( | CCDC9 | 59.703 | 2 | 49.478703 | yes | 7.281E-07 | 576.31201 | 203.19441 |
| AELEGVAVTAPR | 12 | Q9Y3X0 | Coiled-coil domain-containing protein 9 ( | CCDC9 | 59.703 | 2 | 49.478703 | yes | 1.903E-06 | 606.83276 | 121.36801 |
| RYQEIEEDR | 9 | Q9Y3X0 | Coiled-coil domain-containing protein 9 ( | CCDC9 | 59.703 | 2 | 49.478703 | yes | 4.303E-07 | 619.29401 | 11.955926 |
| AYSDHDDRWETK | 12 | Q9Y3X0 | Coiled-coil domain-containing protein 9 ( | CCDC9 | 59.703 | 2 | 49.478703 | yes | 4.905E-07 | 761.83148 | 4.7186408 |
| HLDTLTEHYDIPK | 13 | Q9Y421 | Protein FAM32A OS=Homo sapiens OX= | FAM32A | 13.179 | 3 | 50.605637 | yes | 4.101E-07 | 527.93512 | 124.52685 |
| HLDTLTEHYDIPK | 13 | Q9Y421 | Protein FAM32A OS=Homo sapiens OX= | FAM32A | 13.179 | 3 | 50.605637 | yes | 3.221E-07 | 791.39899 | 124.52685 |

| | | | | | | | | | | | |
|---|---|---|---|---|---|---|---|---|---|---|---|
| HLDTLTEHYDIPK | 13 | Q9Y421 | Protein FAM32A OS=Homo sapiens OX= | FAM32A | 13.179 | 2 | 50.605637 | yes | 4.101E-07 | 527.93512 | 124.52685 |
| HLDTLTEHYDIPK | 13 | Q9Y421 | Protein FAM32A OS=Homo sapiens OX= | FAM32A | 13.179 | 2 | 50.605637 | yes | 3.221E-07 | 791.39899 | 124.52685 |
| TPAQAAFEK | 9 | Q9Y421 | Protein FAM32A OS=Homo sapiens OX= | FAM32A | 13.179 | 2 | 50.605637 | yes | 1.369E-07 | 481.75073 | 237.2038 |
| LLEAMGTSK | 9 | Q9Y421 | Protein FAM32A OS=Homo sapiens OX= | FAM32A | 13.179 | 2 | 50.605637 | yes | 9.743E-08 | 475.25479 | 198.80188 |
| MEAYEQVQK | 9 | Q9Y421 | Protein FAM32A OS=Homo sapiens OX= | FAM32A | 13.179 | 2 | 50.605637 | yes | 1.447E-07 | 584.27118 | 42.621628 |
| SLASSAQPGLGK | 12 | Q9Y4F5 | Centrosomal protein of 170 kDa protein E | CEP170B | 171.689 | 2 | 51.59131 | yes | 3.649E-07 | 558.3064 | 108.66869 |
| LAQDFMAQCLR | 11 | Q9Y4F5 | Centrosomal protein of 170 kDa protein E | CEP170B | 171.689 | 2 | 51.59131 | yes | 1.014E-07 | 676.82611 | 108.26624 |
| GSLDWPSEER | 10 | Q9Y4F5 | Centrosomal protein of 170 kDa protein E | CEP170B | 171.689 | 2 | 51.59131 | yes | 9.722E-08 | 588.27002 | 91.436119 |
| FGYDSNMYVLER | 12 | Q9Y4F5 | Centrosomal protein of 170 kDa protein E | CEP170B | 171.689 | 2 | 51.59131 | yes | 1.144E-07 | 747.34009 | 88.03949 |
| EATPGEMVSAETK | 13 | Q9Y4F5 | Centrosomal protein of 170 kDa protein E | CEP170B | 171.689 | 2 | 51.59131 | yes | 1.061E-06 | 675.31628 | 86.118813 |
| GSLGNPEPVGR | 11 | Q9Y4F5 | Centrosomal protein of 170 kDa protein E | CEP170B | 171.689 | 2 | 51.59131 | yes | 2.898E-07 | 541.78308 | 81.845917 |
| AASAAGVPLEASG | 17 | Q9Y4F5 | Centrosomal protein of 170 kDa protein E | CEP170B | 171.689 | 2 | 51.59131 | yes | 7.039E-07 | 806.91809 | 80.389275 |
| EECELMLQSR | 10 | Q9Y4F5 | Centrosomal protein of 170 kDa protein E | CEP170B | 171.689 | 2 | 51.59131 | yes | 9.722E-08 | 647.79193 | 75.42733 |
| MALLQEFASR | 10 | Q9Y4F5 | Centrosomal protein of 170 kDa protein E | CEP170B | 171.689 | 2 | 51.59131 | yes | 7.095E-07 | 583.30536 | 62.842857 |
| APAEPQGCSFR | 11 | Q9Y4F5 | Centrosomal protein of 170 kDa protein E | CEP170B | 171.689 | 2 | 51.59131 | yes | 3.411E-07 | 610.27985 | 54.777523 |
| INAENEVPILK | 11 | Q9Y4F5 | Centrosomal protein of 170 kDa protein E | CEP170B | 171.689 | 2 | 51.59131 | yes | 4.951E-06 | 620.35083 | 53.905743 |
| AGSFTGTSDPEAA | 16 | Q9Y4F5 | Centrosomal protein of 170 kDa protein E | CEP170B | 171.689 | 2 | 51.59131 | yes | 3.929E-07 | 767.86023 | 46.178646 |
| REPSYFEIPTK | 11 | Q9Y4F5 | Centrosomal protein of 170 kDa protein E | CEP170B | 171.689 | 2 | 51.59131 | yes | 3.591E-07 | 683.8537 | 35.736351 |
| SQSFTHSPSGDPK | 13 | Q9Y4F5 | Centrosomal protein of 170 kDa protein E | CEP170B | 171.689 | 2 | 51.59131 | yes | 0.0002732 | 687.81787 | 20.726379 |
| FVASCGFTPDVK | 12 | Q9Y4P3 | Transducin beta-like protein 2 OS=Homo | TBL2 | 49.798 | 2 | 53.35533 | yes | 9.802E-08 | 664.32117 | 2066.449 |
| AMVEEMQGHLK | 11 | Q9Y4P3 | Transducin beta-like protein 2 OS=Homo | TBL2 | 49.798 | 2 | 53.35533 | yes | 9.722E-08 | 636.80737 | 1731.0753 |
| ANVELDHATLVR | 12 | Q9Y4P3 | Transducin beta-like protein 2 OS=Homo | TBL2 | 49.798 | 2 | 53.35533 | yes | 9.722E-08 | 669.36224 | 851.76935 |
| ANVELDHATLVR | 12 | Q9Y4P3 | Transducin beta-like protein 2 OS=Homo | TBL2 | 49.798 | 2 | 53.35533 | yes | 9.722E-08 | 446.57724 | 851.76935 |
| ANVELDHATLVR | 12 | Q9Y4P3 | Transducin beta-like protein 2 OS=Homo | TBL2 | 49.798 | 3 | 53.35533 | yes | 9.722E-08 | 669.36224 | 851.76935 |
| ANVELDHATLVR | 12 | Q9Y4P3 | Transducin beta-like protein 2 OS=Homo | TBL2 | 49.798 | 3 | 53.35533 | yes | 9.722E-08 | 446.57724 | 851.76935 |
| APVIDIGIANTGK | 13 | Q9Y4P3 | Transducin beta-like protein 2 OS=Homo | TBL2 | 49.798 | 2 | 53.35533 | yes | 9.964E-08 | 634.86407 | 1655.5266 |
| LFHNTPGHR | 9 | Q9Y4P3 | Transducin beta-like protein 2 OS=Homo | TBL2 | 49.798 | 2 | 53.35533 | yes | 1.204E-07 | 539.7807 | 1416.9404 |
| LWDTDVEYK | 9 | Q9Y4P3 | Transducin beta-like protein 2 OS=Homo | TBL2 | 49.798 | 2 | 53.35533 | yes | 9.964E-08 | 584.77966 | 1166.7546 |
| LQQQLTQAQETLK | 13 | Q9Y4P3 | Transducin beta-like protein 2 OS=Homo | TBL2 | 49.798 | 2 | 53.35533 | yes | 1.191E-07 | 764.9201 | 570.27417 |
| FLASCGDR | 8 | Q9Y4P3 | Transducin beta-like protein 2 OS=Homo | TBL2 | 49.798 | 2 | 53.35533 | yes | 9.809E-08 | 463.21344 | 454.62109 |
| KGEFQEVVR | 9 | Q9Y4P3 | Transducin beta-like protein 2 OS=Homo | TBL2 | 49.798 | 2 | 53.35533 | yes | 4.383E-07 | 546.29584 | 448.58054 |
| KQDPYLLK | 8 | Q9Y4P3 | Transducin beta-like protein 2 OS=Homo | TBL2 | 49.798 | 2 | 53.35533 | yes | 1.01E-07 | 502.79239 | 395.57465 |
| LALSPNAQVLALA | 24 | Q9Y4P3 | Transducin beta-like protein 2 OS=Homo | TBL2 | 49.798 | 3 | 53.35533 | yes | 1.387E-07 | 832.79047 | 378.90405 |
| GEFQEVVR | 8 | Q9Y4P3 | Transducin beta-like protein 2 OS=Homo | TBL2 | 49.798 | 2 | 53.35533 | yes | 9.741E-08 | 482.24835 | 300.5275 |
| REDGGYTFTATPE | 17 | Q9Y4P3 | Transducin beta-like protein 2 OS=Homo | TBL2 | 49.798 | 3 | 53.35533 | yes | 9.722E-08 | 644.29877 | 300.30164 |
| YLATCADDR | 9 | Q9Y4P3 | Transducin beta-like protein 2 OS=Homo | TBL2 | 49.798 | 2 | 53.35533 | yes | 1.691E-07 | 542.74005 | 124.85842 |
| AFIVWLANGDTLR | 13 | Q9Y4P3 | Transducin beta-like protein 2 OS=Homo | TBL2 | 49.798 | 2 | 53.35533 | yes | 9.722E-08 | 738.40393 | 99.438469 |
| QDPYLLK | 7 | Q9Y4P3 | Transducin beta-like protein 2 OS=Homo | TBL2 | 49.798 | 2 | 53.35533 | yes | 1.086E-07 | 438.7449 | 86.224899 |
| HKAPVIDIGIANTC | 15 | Q9Y4P3 | Transducin beta-like protein 2 OS=Homo | TBL2 | 49.798 | 2 | 53.35533 | yes | 1.3E-07 | 767.44104 | 56.098675 |

| | | | | | | | | | | | |
|---|---|---|---|---|---|---|---|---|---|---|---|
| GEKEECFER | 9 | Q9Y4P3 | Transducin beta-like protein 2 OS=Homo | TBL2 | 49.798 | 2 | 53.35533 | yes | 1.771E-06 | 592.25604 | 13.649877 |
| GHSAAVHSFAFSN | 16 | Q9Y4P3 | Transducin beta-like protein 2 OS=Homo | TBL2 | 49.798 | 2 | 53.35533 | yes | 1.189E-06 | 845.39001 | 12.720895 |
| FIDTTLAITSR | 11 | Q9Y5J9 | Mitochondrial import inner membrane tra | TIMM8B | 9.344 | 2 | 49.372612 | yes | 1.039E-07 | 619.34296 | 86.022408 |
| TENCLSSCVDR | 11 | Q9Y5J9 | Mitochondrial import inner membrane tra | TIMM8B | 9.344 | 2 | 49.372612 | yes | 3.043E-07 | 670.7821 | 35.784592 |
| AVLLVGLCDSGK | 12 | Q9Y5M8 | Signal recognition particle receptor subun | SRPRB | 29.703 | 2 | 54.604458 | yes | 1.025E-07 | 616.33936 | 640.9552 |
| EFEFSQLPLK | 10 | Q9Y5M8 | Signal recognition particle receptor subun | SRPRB | 29.703 | 2 | 54.604458 | yes | 1.366E-07 | 619.32678 | 574.24817 |
| DTQTSITDSCAVYI | 14 | Q9Y5M8 | Signal recognition particle receptor subun | SRPRB | 29.703 | 2 | 54.604458 | yes | 1.164E-07 | 808.86469 | 558.03125 |
| LIQQQLEK | 8 | Q9Y5M8 | Signal recognition particle receptor subun | SRPRB | 29.703 | 2 | 54.604458 | yes | 9.722E-08 | 500.29529 | 479.9498 |
| NTPSFLIACNK | 11 | Q9Y5M8 | Signal recognition particle receptor subun | SRPRB | 29.703 | 2 | 54.604458 | yes | 9.722E-08 | 632.82135 | 404.50381 |
| GDVGSADIQDLEK | 13 | Q9Y5M8 | Signal recognition particle receptor subun | SRPRB | 29.703 | 2 | 54.604458 | yes | 3.156E-07 | 673.82532 | 377.19348 |
| AIVFVVDSAAFQR | 13 | Q9Y5M8 | Signal recognition particle receptor subun | SRPRB | 29.703 | 2 | 54.604458 | yes | 9.722E-08 | 711.89063 | 305.05524 |
| VEFLECSAK | 9 | Q9Y5M8 | Signal recognition particle receptor subun | SRPRB | 29.703 | 2 | 54.604458 | yes | 9.722E-08 | 541.763 | 235.03456 |
| GKEFEFSQLPLK | 12 | Q9Y5M8 | Signal recognition particle receptor subun | SRPRB | 29.703 | 2 | 54.604458 | yes | 1.266E-07 | 711.88501 | 56.674431 |
| LLTGLYR | 7 | Q9Y5M8 | Signal recognition particle receptor subun | SRPRB | 29.703 | 2 | 54.604458 | yes | 1.977E-06 | 418.25543 | 48.871513 |
| ILIATDVASR | 10 | Q9Y6V7 | Probable ATP-dependent RNA helicase D | DDX49 | 54.227 | 2 | 56.137844 | yes | 1.061E-07 | 529.81384 | 14112.104 |
| ELAYQIAEQFR | 11 | Q9Y6V7 | Probable ATP-dependent RNA helicase D | DDX49 | 54.227 | 2 | 56.137844 | yes | 9.722E-08 | 684.35132 | 8818.1094 |
| DAYLVHLIQR | 10 | Q9Y6V7 | Probable ATP-dependent RNA helicase D | DDX49 | 54.227 | 2 | 56.137844 | yes | 9.722E-08 | 614.34583 | 7901.625 |
| TPSGSHSGPVPSQ( | 16 | Q9Y6V7 | Probable ATP-dependent RNA helicase D | DDX49 | 54.227 | 2 | 56.137844 | yes | 1.43E-07 | 753.88098 | 5645.0313 |
| FLVMDEADR | 9 | Q9Y6V7 | Probable ATP-dependent RNA helicase D | DDX49 | 54.227 | 2 | 56.137844 | yes | 1.498E-07 | 548.26062 | 4375.4829 |
| LEAAHFDEK | 9 | Q9Y6V7 | Probable ATP-dependent RNA helicase D | DDX49 | 54.227 | 2 | 56.137844 | yes | 1.286E-07 | 530.25891 | 3034.3079 |
| KPHVVIATPGR | 11 | Q9Y6V7 | Probable ATP-dependent RNA helicase D | DDX49 | 54.227 | 2 | 56.137844 | yes | 1.242E-07 | 587.85638 | 2091.4121 |
| QTLLFSATLTDTLF | 14 | Q9Y6V7 | Probable ATP-dependent RNA helicase D | DDX49 | 54.227 | 2 | 56.137844 | yes | 1.079E-07 | 790.43811 | 1947.7405 |
| TAAFVLPILQK | 11 | Q9Y6V7 | Probable ATP-dependent RNA helicase D | DDX49 | 54.227 | 2 | 56.137844 | yes | 5.244E-07 | 600.87115 | 1620.5029 |
| YLLVPEK | 7 | Q9Y6V7 | Probable ATP-dependent RNA helicase D | DDX49 | 54.227 | 2 | 56.137844 | yes | 1.442E-07 | 431.25766 | 1308.9515 |
| QLILEGKDPDLEAI | 14 | Q9Y6V7 | Probable ATP-dependent RNA helicase D | DDX49 | 54.227 | 2 | 56.137844 | yes | 1.084E-07 | 784.93018 | 422.78174 |
| QLILEGKDPDLEAI | 14 | Q9Y6V7 | Probable ATP-dependent RNA helicase D | DDX49 | 54.227 | 2 | 56.137844 | yes | 9.934E-08 | 523.6225 | 422.78174 |
| QLILEGKDPDLEAI | 14 | Q9Y6V7 | Probable ATP-dependent RNA helicase D | DDX49 | 54.227 | 3 | 56.137844 | yes | 1.084E-07 | 784.93018 | 422.78174 |
| QLILEGKDPDLEAI | 14 | Q9Y6V7 | Probable ATP-dependent RNA helicase D | DDX49 | 54.227 | 3 | 56.137844 | yes | 9.934E-08 | 523.6225 | 422.78174 |
| FQDEHEDWSIIIFT | 18 | Q9Y6V7 | Probable ATP-dependent RNA helicase D | DDX49 | 54.227 | 3 | 56.137844 | yes | 3.906E-07 | 761.68439 | 627.81256 |
| TCQILCMMLR | 10 | Q9Y6V7 | Probable ATP-dependent RNA helicase D | DDX49 | 54.227 | 2 | 56.137844 | yes | 9.722E-08 | 663.31348 | 512.70471 |
| EKVEETLK | 8 | Q9Y6V7 | Probable ATP-dependent RNA helicase D | DDX49 | 54.227 | 2 | 56.137844 | yes | 1.872E-07 | 488.27148 | 415.05679 |
| RQTLLFSATLTDTI | 15 | Q9Y6V7 | Probable ATP-dependent RNA helicase D | DDX49 | 54.227 | 3 | 56.137844 | yes | 9.722E-08 | 579.32825 | 99.785782 |
| VEETLKR | 7 | Q9Y6V7 | Probable ATP-dependent RNA helicase D | DDX49 | 54.227 | 2 | 56.137844 | yes | 3.561E-07 | 437.75327 | 39.649155 |
| VKDAYLVHLIQR | 12 | Q9Y6V7 | Probable ATP-dependent RNA helicase D | DDX49 | 54.227 | 2 | 56.137844 | yes | 9.722E-08 | 727.92755 | 20.966005 |
| LLIYGASTR | 9 | A0A075B6H7 | Probable non-functional immunoglobulin | IGKV3-7 | 12.784 | 2 | 48.510216 | yes | 1.138E-07 | 497.29001 | 5571.3477 |
| NTLYLQMGSLR | 11 | A0A075B6Q5 | Immunoglobulin heavy variable 3-64 OS= | IGHV3-64 | 12.891 | 2 | 44.09693 | yes | 2.322E-07 | 648.34247 | 47.864841 |
| LLIYKVSNR | 9 | A0A075B6R9 | Probable non-functional immunoglobulin | IGKV2D-2 | 13.079 | 2 | 49.11704 | yes | 1.066E-07 | 553.34003 | 2345.4038 |
| FSGSGAGTDFTLK | 13 | A0A075B6R9 | Probable non-functional immunoglobulin | IGKV2D-2 | 13.079 | 2 | 49.11704 | no | 4.893E-07 | 644.31439 | 722.64117 |
| FSGSGSGTDFTLK | 13 | A0A075B6S2 | Immunoglobulin kappa variable 2D-29 O | IGKV2D-2 | 13.144 | 2 | 54.11979 | no | 3.345E-07 | 652.31189 | 34094.293 |

| | | | | | | | | | | | |
|---|---|---|---|---|---|---|---|---|---|---|---|
| SSQSLLHSDGK | 11 | A0A075B6S2 | Immunoglobulin kappa variable 2D-29 O | IGKV2D-2 | 13.144 | 2 | 54.11979 | yes | 9.722E-08 | 579.79114 | 226.99821 |
| LLIYGASIR | 9 | A0A087WSY6 | Immunoglobulin kappa variable 3D-15 O | IGKV3D-1 | 12.535 | 2 | 50.200718 | yes | 9.85E-08 | 503.3082 | 237.53035 |
| SEDTAMYYCAR | 11 | A0A0A0MS14 | Immunoglobulin heavy variable 1-45 OS= | IGHV1-45 | 13.509 | 2 | 52.193905 | yes | 9.722E-08 | 683.77374 | 699.29346 |
| SEDTAMYYCAR | 11 | A0A0A0MS14 | Immunoglobulin heavy variable 1-45 OS= | IGHV1-45 | 13.509 | 2 | 52.193905 | yes | 3.328E-07 | 691.77118 | 699.29346 |
| TEDTAVYYCTR | 11 | A0A0A0MS15 | Immunoglobulin heavy variable 3-49 OS= | IGHV3-49 | 13.056 | 2 | 48.63744 | yes | 1.121E-07 | 689.80084 | 180.91304 |
| FTISRDDSK | 9 | A0A0A0MS15 | Immunoglobulin heavy variable 3-49 OS= | IGHV3-49 | 13.056 | 2 | 48.63744 | no | 1.464E-07 | 534.76965 | 121.6334 |
| NTLYLQMNSLR | 11 | A0A0B4J1X5 | Immunoglobulin heavy variable 3-74 OS= | IGHV3-74 | 12.84 | 2 | 50.19753 | no | 4.402E-07 | 676.85321 | 190.77589 |
| AEDTAVYYCAR | 11 | A0A0B4J1X5 | Immunoglobulin heavy variable 3-74 OS= | IGHV3-74 | 12.84 | 2 | 50.19753 | no | 9.85E-08 | 659.79022 | 103.06982 |
| GRFTISR | 7 | A0A0B4J1X5 | Immunoglobulin heavy variable 3-74 OS= | IGHV3-74 | 12.84 | 2 | 50.19753 | no | 2.53E-07 | 418.74051 | 98.870743 |
| SEDTAVYYCAR | 11 | A0A0B4J2H0 | Immunoglobulin heavy variable 1-69D O | IGHV1-69I | 12.661 | 2 | 50.88182 | yes | 9.722E-08 | 667.78772 | 126.08485 |
| FSGSGSGTDFTLTI | 16 | A0A0C4DH25 | Immunoglobulin kappa variable 3D-20 O | IGKV3D-2 | 12.516 | 2 | 49.635296 | yes | 1.015E-07 | 816.89685 | 74.455872 |
| SDDTAVYYCAR | 11 | A0A0C4DH31 | Immunoglobulin heavy variable 1-18 OS= | IGHV1-18 | 12.821 | 2 | 56.43467 | yes | 9.722E-08 | 660.77991 | 29.451866 |
| NTLYLQMSSLR | 11 | A0A0J9YX35 | Immunoglobulin heavy variable 3-64D O | IGHV3-64I | 12.823 | 2 | 50.768456 | yes | 9.722E-08 | 663.34772 | 1238.1682 |
| HVQGILR | 7 | A8MWD9 | Putative small nuclear ribonucleoprotein | SNRPGP15 | 8.545 | 2 | 48.586056 | yes | 1.823E-07 | 411.75085 | 5268.1304 |
| GNSIIMLEALER | 12 | A8MWD9 | Putative small nuclear ribonucleoprotein | SNRPGP15 | 8.545 | 2 | 48.586056 | yes | 1.127E-07 | 673.36084 | 4901.2573 |
| GCILTLVER | 9 | B5ME19 | Eukaryotic translation initiation factor 3 s | EIF3CL | 105.473 | 2 | 52.22095 | yes | 1.157E-07 | 530.79462 | 4347.3135 |
| LGSLVENNER | 10 | B5ME19 | Eukaryotic translation initiation factor 3 s | EIF3CL | 105.473 | 2 | 52.22095 | yes | 9.741E-08 | 565.79364 | 2941.3491 |
| ELLGQGLLLR | 10 | B5ME19 | Eukaryotic translation initiation factor 3 s | EIF3CL | 105.473 | 2 | 52.22095 | yes | 1.108E-07 | 556.34534 | 2365.4573 |
| TMVQLGICAFR | 11 | B5ME19 | Eukaryotic translation initiation factor 3 s | EIF3CL | 105.473 | 2 | 52.22095 | yes | 1.186E-07 | 648.33356 | 927.18573 |
| TMVQLGICAFR | 11 | B5ME19 | Eukaryotic translation initiation factor 3 s | EIF3CL | 105.473 | 2 | 52.22095 | yes | 1.065E-07 | 656.33105 | 927.18573 |
| QPLLGPPESMR | 11 | B5ME19 | Eukaryotic translation initiation factor 3 s | EIF3CL | 105.473 | 2 | 52.22095 | yes | 1.108E-06 | 612.82391 | 1812.8468 |
| VWDLFPEADK | 10 | B5ME19 | Eukaryotic translation initiation factor 3 s | EIF3CL | 105.473 | 2 | 52.22095 | yes | 1.236E-07 | 610.30334 | 1559.7797 |
| GTEITHAVVIK | 11 | B5ME19 | Eukaryotic translation initiation factor 3 s | EIF3CL | 105.473 | 2 | 52.22095 | yes | 6.203E-07 | 584.34021 | 1508.5962 |
| FEELTNLIR | 9 | B5ME19 | Eukaryotic translation initiation factor 3 s | EIF3CL | 105.473 | 2 | 52.22095 | yes | 1.101E-07 | 567.81134 | 1272.9637 |
| KIQEESLR | 8 | B5ME19 | Eukaryotic translation initiation factor 3 s | EIF3CL | 105.473 | 2 | 52.22095 | yes | 1.341E-07 | 501.78256 | 1264.3501 |
| QGTYGGYFR | 9 | B5ME19 | Eukaryotic translation initiation factor 3 s | EIF3CL | 105.473 | 2 | 52.22095 | yes | 3.584E-07 | 524.74597 | 1168.084 |
| DFESHITSYK | 10 | B5ME19 | Eukaryotic translation initiation factor 3 s | EIF3CL | 105.473 | 2 | 52.22095 | yes | 1.367E-07 | 613.78802 | 1078.3083 |
| LNEILQAR | 8 | B5ME19 | Eukaryotic translation initiation factor 3 s | EIF3CL | 105.473 | 2 | 52.22095 | yes | 9.722E-08 | 478.77982 | 993.11578 |
| DAHNALLDIQSSG | 14 | B5ME19 | Eukaryotic translation initiation factor 3 s | EIF3CL | 105.473 | 2 | 52.22095 | yes | 1.104E-07 | 748.87622 | 926.73846 |
| KLNEILQAR | 9 | B5ME19 | Eukaryotic translation initiation factor 3 s | EIF3CL | 105.473 | 2 | 52.22095 | yes | 1.076E-07 | 542.82727 | 883.91516 |
| DEAQVCAIIER | 11 | B5ME19 | Eukaryotic translation initiation factor 3 s | EIF3CL | 105.473 | 2 | 52.22095 | yes | 9.79E-08 | 652.31915 | 699.47968 |
| CLEEFELLGK | 10 | B5ME19 | Eukaryotic translation initiation factor 3 s | EIF3CL | 105.473 | 2 | 52.22095 | yes | 1.15E-07 | 619.3103 | 501.89966 |
| GTTEEVCR | 8 | B5ME19 | Eukaryotic translation initiation factor 3 s | EIF3CL | 105.473 | 2 | 52.22095 | yes | 3.409E-06 | 476.21365 | 410.34024 |
| QLTPPEGSSK | 10 | B5ME19 | Eukaryotic translation initiation factor 3 s | EIF3CL | 105.473 | 2 | 52.22095 | yes | 9.861E-08 | 522.27203 | 244.53767 |
| ILHTYYK | 7 | B5ME19 | Eukaryotic translation initiation factor 3 s | EIF3CL | 105.473 | 2 | 52.22095 | yes | 9.733E-08 | 469.26074 | 228.65356 |
| RFEELTNLIR | 10 | B5ME19 | Eukaryotic translation initiation factor 3 s | EIF3CL | 105.473 | 2 | 52.22095 | yes | 8.334E-07 | 645.86188 | 216.40335 |
| TCHSFIINEK | 10 | B5ME19 | Eukaryotic translation initiation factor 3 s | EIF3CL | 105.473 | 2 | 52.22095 | yes | 2.115E-07 | 624.80573 | 184.3092 |
| QPLLLSEDEEDTKI | 14 | B5ME19 | Eukaryotic translation initiation factor 3 s | EIF3CL | 105.473 | 2 | 52.22095 | yes | 1.152E-07 | 836.92303 | 121.31699 |
| KNEGYMR | 7 | B5ME19 | Eukaryotic translation initiation factor 3 s | EIF3CL | 105.473 | 2 | 52.22095 | yes | 7.273E-07 | 449.216 | 50.144501 |

| | | | | | | | | | | | |
|---|---|---|---|---|---|---|---|---|---|---|---|
| TCAILCHIYHHALI | 16 | B5ME19 | Eukaryotic translation initiation factor 3 s | EIF3CL | 105.473 | 4 | 52.22095 | yes | 1.957E-06 | 497.9978 | 6.4702148 |
| TCAILCHIYHHALI | 16 | B5ME19 | Eukaryotic translation initiation factor 3 s | EIF3CL | 105.473 | 4 | 52.22095 | yes | 4.014E-06 | 663.66132 | 6.4702148 |
| TCAILCHIYHHALI | 16 | B5ME19 | Eukaryotic translation initiation factor 3 s | EIF3CL | 105.473 | 3 | 52.22095 | yes | 1.957E-06 | 497.9978 | 6.4702148 |
| TCAILCHIYHHALI | 16 | B5ME19 | Eukaryotic translation initiation factor 3 s | EIF3CL | 105.473 | 3 | 52.22095 | yes | 4.014E-06 | 663.66132 | 6.4702148 |
| VTVLGQPK | 8 | B9A064 | Immunoglobulin lambda-like polypeptide | IGLL5 | 23.064 | 2 | 45.285374 | yes | 1.897E-07 | 421.26074 | 317.33917 |
| YFSLEGYLR | 9 | E7ETH6 | Zinc finger protein 587B OS=Homo sapie | ZNF587B | 45.542 | 2 | 48.615997 | yes | 1.122E-07 | 574.29279 | 41.275337 |
| SFSSNVNLK | 9 | E7ETH6 | Zinc finger protein 587B OS=Homo sapie | ZNF587B | 45.542 | 2 | 48.615997 | yes | 1.352E-06 | 498.26144 | 23.124826 |
| IVVDELSNLK | 10 | L0R819 | ASNSD1 upstream open reading frame pr | ASDURF | 11.25 | 2 | 45.896873 | yes | 3.492E-07 | 565.32678 | 132.5159 |
| QQQNSNIFFLADR | 13 | L0R819 | ASNSD1 upstream open reading frame pr | ASDURF | 11.25 | 2 | 45.896873 | yes | 1.71E-07 | 790.89441 | 44.36319 |
| EEIALGGGGFCVH | 14 | O15037 | Protein KHNYN OS=Homo sapiens OX= | KHNYN | 74.534 | 2 | 48.9052 | yes | 1.478E-07 | 751.36444 | 131.03828 |
| FAVSAEAENK | 10 | O15037 | Protein KHNYN OS=Homo sapiens OX= | KHNYN | 74.534 | 2 | 48.9052 | yes | 1.088E-07 | 533.26422 | 128.01862 |
| ELPGEEAWER | 10 | O15037 | Protein KHNYN OS=Homo sapiens OX= | KHNYN | 74.534 | 2 | 48.9052 | yes | 5.064E-07 | 608.28564 | 127.48295 |
| SSALLGAQCQGVI | 13 | O15037 | Protein KHNYN OS=Homo sapiens OX= | KHNYN | 74.534 | 2 | 48.9052 | yes | 4.688E-07 | 673.84589 | 59.750179 |
| ESLDTGSMGPGDC | 14 | O15037 | Protein KHNYN OS=Homo sapiens OX= | KHNYN | 74.534 | 2 | 48.9052 | yes | 2.436E-07 | 741.30341 | 57.466473 |
| LYSLSLLSLTPSR | 13 | O15037 | Protein KHNYN OS=Homo sapiens OX= | KHNYN | 74.534 | 2 | 48.9052 | yes | 5.639E-07 | 725.41919 | 45.962257 |
| GIAIAVQYFWDR | 12 | O15037 | Protein KHNYN OS=Homo sapiens OX= | KHNYN | 74.534 | 2 | 48.9052 | yes | 3.432E-07 | 719.8775 | 43.75293 |
| IFSVGVSVLPK | 11 | O15037 | Protein KHNYN OS=Homo sapiens OX= | KHNYN | 74.534 | 2 | 48.9052 | yes | 4.225E-07 | 573.3501 | 38.23769 |
| LAEAADLLNR | 10 | O15050 | TPR and ankyrin repeat-containing protei | TRANK1 | 336.222 | 2 | 54.82216 | yes | 9.722E-08 | 543.30109 | 105.9453 |
| IIATEQNTCAMEK | 13 | O15050 | TPR and ankyrin repeat-containing protei | TRANK1 | 336.222 | 2 | 54.82216 | yes | 2.54E-07 | 754.85779 | 83.80162 |
| VSDMVEDLYR | 10 | O15050 | TPR and ankyrin repeat-containing protei | TRANK1 | 336.222 | 2 | 54.82216 | yes | 2.921E-07 | 613.78973 | 82.11171 |
| PLEPIILIGR | 10 | O15050 | TPR and ankyrin repeat-containing protei | TRANK1 | 336.222 | 2 | 54.82216 | yes | 9.926E-08 | 560.85809 | 81.286919 |
| YCQIAQNDPGPILI | 14 | O15050 | TPR and ankyrin repeat-containing protei | TRANK1 | 336.222 | 2 | 54.82216 | yes | 1.664E-07 | 822.91174 | 75.84166 |
| DENFPLFVTSK | 11 | O15050 | TPR and ankyrin repeat-containing protei | TRANK1 | 336.222 | 2 | 54.82216 | yes | 5.712E-07 | 648.82715 | 60.026043 |
| KEPGALRPCSLR | 12 | O15050 | TPR and ankyrin repeat-containing protei | TRANK1 | 336.222 | 3 | 54.82216 | yes | 4.465E-07 | 461.92236 | 54.226234 |
| TAYNPALIWK | 10 | O15050 | TPR and ankyrin repeat-containing protei | TRANK1 | 336.222 | 2 | 54.82216 | yes | 2.517E-07 | 588.82422 | 51.049992 |
| VGELEYAVIDLNP | 14 | O15050 | TPR and ankyrin repeat-containing protei | TRANK1 | 336.222 | 2 | 54.82216 | yes | 1.154E-05 | 794.42249 | 38.759121 |
| CYVEDTEAEK | 10 | O15050 | TPR and ankyrin repeat-containing protei | TRANK1 | 336.222 | 2 | 54.82216 | yes | 1.181E-06 | 622.26099 | 37.978951 |
| ALESEKDER | 9 | O15050 | TPR and ankyrin repeat-containing protei | TRANK1 | 336.222 | 2 | 54.82216 | yes | 1.715E-06 | 538.76459 | 34.920795 |
| LSYSASQFYLEAA | 15 | O15050 | TPR and ankyrin repeat-containing protei | TRANK1 | 336.222 | 2 | 54.82216 | yes | 3.826E-07 | 824.91449 | 33.415051 |
| SCFEFFGISQVDAI | 14 | O15050 | TPR and ankyrin repeat-containing protei | TRANK1 | 336.222 | 2 | 54.82216 | yes | 3.368E-06 | 817.87958 | 26.475721 |
| ALTGDAAIQPR | 11 | O43824 | Putative GTP-binding protein 6 OS=Hom | GTPBP6 | 56.898 | 2 | 51.515373 | yes | 9.967E-08 | 556.80658 | 408.87372 |
| GEPLLPAGTQR | 11 | O43824 | Putative GTP-binding protein 6 OS=Hom | GTPBP6 | 56.898 | 2 | 51.515373 | yes | 2.54E-07 | 569.81439 | 227.3121 |
| LAGAQLSWLYK | 11 | O43824 | Putative GTP-binding protein 6 OS=Hom | GTPBP6 | 56.898 | 2 | 51.515373 | yes | 9.723E-08 | 625.35059 | 178.08656 |
| EFPVISVVGYTNC | 15 | O43824 | Putative GTP-binding protein 6 OS=Hom | GTPBP6 | 56.898 | 2 | 51.515373 | yes | 9.722E-08 | 835.41632 | 170.40089 |
| DVSHPEAELQK | 11 | O43824 | Putative GTP-binding protein 6 OS=Hom | GTPBP6 | 56.898 | 2 | 51.515373 | yes | 3.077E-07 | 626.81201 | 157.77266 |
| GNFEHLTEK | 9 | O43824 | Putative GTP-binding protein 6 OS=Hom | GTPBP6 | 56.898 | 2 | 51.515373 | yes | 2.709E-07 | 537.76434 | 117.67975 |
| GSPDITCVFLNVEI | 14 | O43824 | Putative GTP-binding protein 6 OS=Hom | GTPBP6 | 56.898 | 2 | 51.515373 | yes | 2.555E-07 | 803.89832 | 89.938812 |
| ELEAAWGVEVFDI | 13 | O43824 | Putative GTP-binding protein 6 OS=Hom | GTPBP6 | 56.898 | 2 | 51.515373 | yes | 3.081E-07 | 760.87262 | 78.419823 |
| FTVVLHIFR | 9 | O43824 | Putative GTP-binding protein 6 OS=Hom | GTPBP6 | 56.898 | 2 | 51.515373 | yes | 1.121E-07 | 566.33728 | 69.435852 |

| | | | | | | | | | | | |
|---|---|---|---|---|---|---|---|---|---|---|---|
| CSVLSTLR | 8 | O43824 | Putative GTP-binding protein 6 OS=Hom | GTPBP6 | 56.898 | 2 | 51.515373 | yes | 1.362E-07 | 468.25256 | 68.92482 |
| VCLVHPDVK | 9 | O43824 | Putative GTP-binding protein 6 OS=Hom | GTPBP6 | 56.898 | 2 | 51.515373 | yes | 1.161E-07 | 533.78931 | 67.153221 |
| AELDAAVLK | 9 | O43824 | Putative GTP-binding protein 6 OS=Hom | GTPBP6 | 56.898 | 2 | 51.515373 | yes | 6.172E-07 | 465.26874 | 47.685455 |
| PAGVSSAVIPVLR | 13 | O60287 | Nucleolar pre-ribosomal-associated protei | URB1 | 254.39 | 2 | 49.061596 | yes | 3.618E-05 | 633.38245 | 114.46228 |
| YSEAITQSVLK | 11 | O60287 | Nucleolar pre-ribosomal-associated protei | URB1 | 254.39 | 2 | 49.061596 | yes | 1.073E-07 | 619.83496 | 56.286194 |
| LICESLYASGYR | 12 | O60287 | Nucleolar pre-ribosomal-associated protei | URB1 | 254.39 | 2 | 49.061596 | yes | 1.181E-07 | 716.35046 | 49.347618 |
| SHILPHPVVVADLI | 15 | O60287 | Nucleolar pre-ribosomal-associated protei | URB1 | 254.39 | 3 | 49.061596 | yes | 3.216E-06 | 546.66278 | 34.26915 |
| DFVDLSTGEDAHK | 13 | O75794 | Cell division cycle protein 123 homolog ( | CDC123 | 39.135 | 2 | 55.423866 | yes | 1.272E-07 | 717.33081 | 289.47513 |
| DYLLDDGTLVVSC | 14 | O75794 | Cell division cycle protein 123 homolog ( | CDC123 | 39.135 | 2 | 55.423866 | yes | 9.81E-08 | 761.89099 | 235.16638 |
| WCELIPGAEFR | 11 | O75794 | Cell division cycle protein 123 homolog ( | CDC123 | 39.135 | 2 | 55.423866 | yes | 9.722E-08 | 689.33466 | 190.18785 |
| VQEAINSLGGSVFI | 15 | O75794 | Cell division cycle protein 123 homolog ( | CDC123 | 39.135 | 2 | 55.423866 | yes | 9.722E-08 | 773.41718 | 171.07495 |
| SVILPLPQNVK | 11 | O75794 | Cell division cycle protein 123 homolog ( | CDC123 | 39.135 | 2 | 55.423866 | yes | 1.382E-07 | 604.37408 | 148.15807 |
| DAYWIAMNSSLK | 12 | O75794 | Cell division cycle protein 123 homolog ( | CDC123 | 39.135 | 2 | 55.423866 | yes | 9.722E-08 | 699.83972 | 110.51158 |
| DYTQYYDHISK | 11 | O75794 | Cell division cycle protein 123 homolog ( | CDC123 | 39.135 | 2 | 55.423866 | yes | 3.775E-07 | 716.82263 | 55.354393 |
| VCFVGDGFTR | 10 | O95478 | Ribosome biogenesis protein NSA2 homo | NSA2 | 30.066 | 2 | 50.736164 | yes | 1.163E-07 | 579.27405 | 915.02832 |
| TPQGAVPAYLLDR | 13 | O95478 | Ribosome biogenesis protein NSA2 homo | NSA2 | 30.066 | 2 | 50.736164 | yes | 1.068E-07 | 700.88025 | 356.93286 |
| NPSSPLYTTLGVIT | 15 | O95478 | Ribosome biogenesis protein NSA2 homo | NSA2 | 30.066 | 2 | 50.736164 | yes | 9.81E-08 | 795.94049 | 117.93756 |
| ATFCLPILGVK | 11 | O95478 | Ribosome biogenesis protein NSA2 homo | NSA2 | 30.066 | 2 | 50.736164 | yes | 4.277E-07 | 609.84937 | 84.521149 |
| PQNEYIELHR | 10 | O95478 | Ribosome biogenesis protein NSA2 homo | NSA2 | 30.066 | 2 | 50.736164 | yes | 9.723E-08 | 649.828 | 81.462509 |
| AHVTHPELK | 9 | O95478 | Ribosome biogenesis protein NSA2 homo | NSA2 | 30.066 | 2 | 50.736164 | yes | 1.489E-07 | 516.28528 | 77.811203 |
| WEVPLPK | 7 | O95478 | Ribosome biogenesis protein NSA2 homo | NSA2 | 30.066 | 2 | 50.736164 | yes | 1.46E-06 | 434.75 | 40.523865 |
| SGFLGTNPAPR | 11 | P0CG12 | Decreased expression in renal and prostat | DERPC | 51.392 | 2 | 36.60483 | yes | 7.154E-07 | 558.79346 | 59.094666 |
| GMNPTGTGAVSFF | 14 | P0CG12 | Decreased expression in renal and prostat | DERPC | 51.392 | 2 | 36.60483 | yes | 6.806E-07 | 696.34045 | 58.737862 |
| MEEDQELER | 9 | P0DPB5 | Protein POLR1D, isoform 2 OS=Homo sa | POLR1D | 14.333 | 2 | 45.82229 | no | 2.045E-07 | 610.75861 | 108.09528 |
| AETMGPMGWMK | 11 | P0DPB5 | Protein POLR1D, isoform 2 OS=Homo sa | POLR1D | 14.333 | 2 | 45.82229 | yes | 1.732E-07 | 619.77197 | 41.988758 |
| TFSSLGNLR | 9 | P16415 | Zinc finger protein 823 OS=Homo sapien | ZNF823 | 70.271 | 2 | 40.47115 | yes | 4.006E-07 | 497.76944 | 105.39409 |
| AFSCPTYFR | 9 | P17017 | Zinc finger protein 14 OS=Homo sapiens | ZNF14 | 75.354 | 2 | 46.464916 | yes | 1.552E-07 | 574.76331 | 50.705368 |
| TFSFSSSLQR | 10 | P17017 | Zinc finger protein 14 OS=Homo sapiens | ZNF14 | 75.354 | 2 | 46.464916 | yes | 3.291E-06 | 580.29077 | 26.539139 |
| CGETTSQMPNVNI | 15 | P17017 | Zinc finger protein 14 OS=Homo sapiens | ZNF14 | 75.354 | 2 | 46.464916 | yes | 4.555E-06 | 846.88763 | 7.9501591 |
| GVNTGAVGSYIYD | 14 | P52943 | Cysteine-rich protein 2 OS=Homo sapien | CRIP2 | 22.493 | 2 | 48.7052 | yes | 1.113E-07 | 736.36243 | 112.53496 |
| TLTPGGHAEHDGK | 28 | P52943 | Cysteine-rich protein 2 OS=Homo sapien | CRIP2 | 22.493 | 4 | 48.7052 | yes | 1.207E-06 | 781.62866 | 7.5347023 |
| SISTFLGTLENEHL | 15 | P78312 | Protein FAM193A OS=Homo sapiens OX | FAM193A | 139.988 | 2 | 54.285778 | yes | 1.055E-07 | 844.94635 | 346.91965 |
| DIDLDSVDMDETE | 14 | P78312 | Protein FAM193A OS=Homo sapiens OX | FAM193A | 139.988 | 2 | 54.285778 | yes | 1.03E-07 | 826.85144 | 303.29303 |
| VAEEWLECQK | 10 | P78312 | Protein FAM193A OS=Homo sapiens OX | FAM193A | 139.988 | 2 | 54.285778 | yes | 9.722E-08 | 646.30298 | 292.89697 |
| VVDLMSITEQK | 11 | P78312 | Protein FAM193A OS=Homo sapiens OX | FAM193A | 139.988 | 2 | 54.285778 | yes | 2.182E-07 | 631.83667 | 252.55252 |
| AVTGENNFTDTMI | 13 | P78312 | Protein FAM193A OS=Homo sapiens OX | FAM193A | 139.988 | 2 | 54.285778 | yes | 9.722E-08 | 728.33026 | 228.81673 |
| FIEEQLTNK | 9 | P78312 | Protein FAM193A OS=Homo sapiens OX | FAM193A | 139.988 | 2 | 54.285778 | yes | 3.339E-06 | 561.29547 | 226.92668 |
| MLTEDWELFK | 10 | P78312 | Protein FAM193A OS=Homo sapiens OX | FAM193A | 139.988 | 2 | 54.285778 | yes | 9.722E-08 | 656.31812 | 210.61156 |
| FISLWGSEVMNDK | 13 | P78312 | Protein FAM193A OS=Homo sapiens OX | FAM193A | 139.988 | 2 | 54.285778 | yes | 9.722E-08 | 763.3714 | 196.48921 |

| | | | | | | | | | | | |
|---|---|---|---|---|---|---|---|---|---|---|---|
| LSMPDCPNCNYR | 12 | P78312 | Protein FAM193A OS=Homo sapiens OX | FAM193A | 139.988 | 2 | 54.285778 | yes | 2.844E-07 | 763.81287 | 185.23322 |
| FCLDSAR | 7 | P78312 | Protein FAM193A OS=Homo sapiens OX | FAM193A | 139.988 | 2 | 54.285778 | yes | 1.523E-07 | 434.70273 | 140.63368 |
| PGLGADGDAADPV | 16 | P78312 | Protein FAM193A OS=Homo sapiens OX | FAM193A | 139.988 | 2 | 54.285778 | yes | 1.202E-07 | 763.85773 | 130.60289 |
| LSINWSNFSLK | 11 | P78312 | Protein FAM193A OS=Homo sapiens OX | FAM193A | 139.988 | 2 | 54.285778 | yes | 9.722E-08 | 654.85095 | 113.0479 |
| EHLHLQEEQR | 10 | P78312 | Protein FAM193A OS=Homo sapiens OX | FAM193A | 139.988 | 2 | 54.285778 | yes | 2.154E-07 | 659.82855 | 51.591503 |
| LVLAESPQPK | 10 | P78312 | Protein FAM193A OS=Homo sapiens OX | FAM193A | 139.988 | 2 | 54.285778 | yes | 1.701E-07 | 541.31622 | 36.697243 |
| IDAYVDEQMTMK | 12 | P78312 | Protein FAM193A OS=Homo sapiens OX | FAM193A | 139.988 | 2 | 54.285778 | yes | 2.286E-06 | 722.32837 | 27.959827 |
| RPPSVGDVFHGISI | 14 | P78312 | Protein FAM193A OS=Homo sapiens OX | FAM193A | 139.988 | 3 | 54.285778 | yes | 7.591E-07 | 499.27203 | 18.879883 |
| NLPEGVAMEVTK | 12 | P82664 | Small ribosomal subunit protein uS10m C | MRPS10 | 23 | 2 | 54.09316 | yes | 1.951E-07 | 644.33429 | 631.13239 |
| TQLEQLPEHIK | 11 | P82664 | Small ribosomal subunit protein uS10m C | MRPS10 | 23 | 2 | 54.09316 | yes | 9.722E-08 | 668.367 | 517.73468 |
| CLELEHLTGSTADY | 21 | P82664 | Small ribosomal subunit protein uS10m C | MRPS10 | 23 | 3 | 54.09316 | yes | 9.849E-08 | 837.4126 | 123.11264 |
| RLSVLVK | 7 | P82664 | Small ribosomal subunit protein uS10m C | MRPS10 | 23 | 2 | 54.09316 | yes | 2.087E-07 | 407.77908 | 17.915951 |
| ILGDINSDFTR | 11 | P86790 | Vacuolar fusion protein CCZ1 homolog B | CCZ1B | 55.867 | 2 | 41.325233 | yes | 4.435E-07 | 625.82239 | 65.918282 |
| ELYVILNQK | 9 | P86790 | Vacuolar fusion protein CCZ1 homolog B | CCZ1B | 55.867 | 2 | 41.325233 | yes | 3.566E-07 | 560.32404 | 59.143806 |
| NVGLCEAIVQFTR | 13 | P86790 | Vacuolar fusion protein CCZ1 homolog B | CCZ1B | 55.867 | 2 | 41.325233 | yes | 2.479E-05 | 753.89032 | 24.315382 |
| VVYTPESLYK | 10 | Q08AF3 | Schlafen family member 5 OS=Homo sap | SLFN5 | 101.055 | 2 | 47.242325 | yes | 3.999E-07 | 599.82135 | 78.810425 |
| ENHFLIFVK | 9 | Q08AF3 | Schlafen family member 5 OS=Homo sap | SLFN5 | 101.055 | 2 | 47.242325 | yes | 1.37E-07 | 573.81897 | 41.983341 |
| YFPVFSDR | 8 | Q08AF3 | Schlafen family member 5 OS=Homo sap | SLFN5 | 101.055 | 2 | 47.242325 | yes | 2.108E-06 | 515.7533 | 26.259285 |
| VLEITELPDGITR | 13 | Q15032 | R3H domain-containing protein 1 OS=Ho | R3HDM1 | 120.697 | 2 | 52.443806 | yes | 1.258E-07 | 728.40631 | 5242.7563 |
| ASSFSGISVLTR | 12 | Q15032 | R3H domain-containing protein 1 OS=Ho | R3HDM1 | 120.697 | 2 | 52.443806 | yes | 1.093E-07 | 612.83276 | 2884.6345 |
| LQDEDASSTQQR | 12 | Q15032 | R3H domain-containing protein 1 OS=Ho | R3HDM1 | 120.697 | 2 | 52.443806 | yes | 9.81E-08 | 689.31567 | 1546.1765 |
| AASTDLGAGETVV | 15 | Q15032 | R3H domain-containing protein 1 OS=Ho | R3HDM1 | 120.697 | 2 | 52.443806 | yes | 9.893E-08 | 688.35681 | 1501.605 |
| IQIQLTQSFEK | 11 | Q15032 | R3H domain-containing protein 1 OS=Ho | R3HDM1 | 120.697 | 2 | 52.443806 | yes | 1.173E-07 | 667.86938 | 1311.9308 |
| PWSSTDSDSSLR | 12 | Q15032 | R3H domain-containing protein 1 OS=Ho | R3HDM1 | 120.697 | 2 | 52.443806 | yes | 1.096E-07 | 669.30206 | 890.76208 |
| PFVPGQGDSR | 10 | Q15032 | R3H domain-containing protein 1 OS=Ho | R3HDM1 | 120.697 | 2 | 52.443806 | yes | 4.506E-07 | 530.26453 | 854.05872 |
| DDKGEDFQK | 9 | Q15032 | R3H domain-containing protein 1 OS=Ho | R3HDM1 | 120.697 | 2 | 52.443806 | yes | 9.722E-08 | 541.24347 | 603.12756 |
| YYCDHQR | 7 | Q15032 | R3H domain-containing protein 1 OS=Ho | R3HDM1 | 120.697 | 2 | 52.443806 | yes | 9.722E-08 | 521.21399 | 507.20688 |
| HYDFHILER | 9 | Q15032 | R3H domain-containing protein 1 OS=Ho | R3HDM1 | 120.697 | 2 | 52.443806 | yes | 9.904E-08 | 615.3067 | 425.94052 |
| STNSHQSSTENELI | 14 | Q15032 | R3H domain-containing protein 1 OS=Ho | R3HDM1 | 120.697 | 2 | 52.443806 | yes | 9.722E-08 | 781.35809 | 178.72112 |
| KAASTDLGAGETV | 16 | Q15032 | R3H domain-containing protein 1 OS=Ho | R3HDM1 | 120.697 | 2 | 52.443806 | yes | 3.427E-07 | 752.4043 | 54.314529 |
| KAASTDLGAGETV | 16 | Q15032 | R3H domain-containing protein 1 OS=Ho | R3HDM1 | 120.697 | 2 | 52.443806 | yes | 3.589E-06 | 501.93863 | 54.314529 |
| KAASTDLGAGETV | 16 | Q15032 | R3H domain-containing protein 1 OS=Ho | R3HDM1 | 120.697 | 3 | 52.443806 | yes | 3.427E-07 | 752.4043 | 54.314529 |
| KAASTDLGAGETV | 16 | Q15032 | R3H domain-containing protein 1 OS=Ho | R3HDM1 | 120.697 | 3 | 52.443806 | yes | 3.589E-06 | 501.93863 | 54.314529 |
| LEQEILDFIGNNES | 16 | Q15032 | R3H domain-containing protein 1 OS=Ho | R3HDM1 | 120.697 | 3 | 52.443806 | yes | 1.848E-07 | 625.31451 | 48.144428 |
| TPPGVPAPSDPLLC | 15 | Q15434 | RNA-binding motif, single-stranded-inter | RBMS2 | 43.959 | 2 | 46.877335 | yes | 1.522E-07 | 774.90814 | 3634.3113 |
| GYGFVDFDSPSAA | 15 | Q15434 | RNA-binding motif, single-stranded-inter | RBMS2 | 43.959 | 2 | 46.877335 | yes | 1.577E-07 | 794.86755 | 2163.9277 |
| PGISTFGYNR | 10 | Q15434 | RNA-binding motif, single-stranded-inter | RBMS2 | 43.959 | 2 | 46.877335 | yes | 1.451E-07 | 556.28015 | 1594.3397 |
| CEAIITHFNGK | 11 | Q15434 | RNA-binding motif, single-stranded-inter | RBMS2 | 43.959 | 2 | 46.877335 | yes | 2.102E-07 | 645.31897 | 209.96289 |
| CILTGHELPCR | 11 | Q15527 | Surfeit locus protein 2 OS=Homo sapiens | SURF2 | 29.618 | 2 | 50.910465 | yes | 9.722E-08 | 678.33154 | 102.55136 |

| | | | | | | | | | | | |
|---|---|---|---|---|---|---|---|---|---|---|---|
| NPHQLFCK | 8 | Q15527 | Surfeit locus protein 2 OS=Homo sapiens | SURF2 | 29.618 | 2 | 50.910465 | yes | 4.434E-07 | 522.25818 | 84.386795 |
| LPELQVYTR | 9 | Q15527 | Surfeit locus protein 2 OS=Homo sapiens | SURF2 | 29.618 | 2 | 50.910465 | yes | 9.819E-08 | 559.81384 | 83.168076 |
| CPEHVLR | 7 | Q15527 | Surfeit locus protein 2 OS=Homo sapiens | SURF2 | 29.618 | 2 | 50.910465 | yes | 6.049E-07 | 455.73181 | 47.099155 |
| QGVEYVPACLVHI | 13 | Q15527 | Surfeit locus protein 2 OS=Homo sapiens | SURF2 | 29.618 | 2 | 50.910465 | yes | 2.935E-05 | 764.39044 | 5.8623676 |
| ALQEEIDR | 8 | Q1ED39 | Lysine-rich nucleolar protein 1 OS=Homo | KNOP1 | 51.59 | 2 | 51.003124 | yes | 1.01E-07 | 487.2511 | 229.95074 |
| AADSLQQNLQR | 11 | Q1ED39 | Lysine-rich nucleolar protein 1 OS=Homo | KNOP1 | 51.59 | 2 | 51.003124 | yes | 9.722E-08 | 622.32312 | 194.55409 |
| VEQPVIEEPALK | 12 | Q1ED39 | Lysine-rich nucleolar protein 1 OS=Homo | KNOP1 | 51.59 | 2 | 51.003124 | yes | 4.657E-07 | 676.37701 | 109.674 |
| VEAPEYIPISDDPK | 14 | Q1ED39 | Lysine-rich nucleolar protein 1 OS=Homo | KNOP1 | 51.59 | 2 | 51.003124 | yes | 1.183E-06 | 786.89325 | 93.464912 |
| DVGDTCSVGK | 10 | Q1ED39 | Lysine-rich nucleolar protein 1 OS=Homo | KNOP1 | 51.59 | 2 | 51.003124 | yes | 9.612E-06 | 519.23206 | 86.57428 |
| KVEQPVIEEPALK | 13 | Q1ED39 | Lysine-rich nucleolar protein 1 OS=Homo | KNOP1 | 51.59 | 2 | 51.003124 | yes | 1.063E-07 | 740.4245 | 61.23296 |
| GAGLGFSTAPNK | 12 | Q1ED39 | Lysine-rich nucleolar protein 1 OS=Homo | KNOP1 | 51.59 | 2 | 51.003124 | yes | 1.294E-06 | 560.29327 | 52.733227 |
| GNMDEAHIDQVR | 12 | Q1ED39 | Lysine-rich nucleolar protein 1 OS=Homo | KNOP1 | 51.59 | 2 | 51.003124 | yes | 2.093E-07 | 692.81732 | 30.525307 |
| KDEEQAALGQK | 11 | Q1ED39 | Lysine-rich nucleolar protein 1 OS=Homo | KNOP1 | 51.59 | 2 | 51.003124 | yes | 2.352E-06 | 608.81201 | 28.575094 |
| YNVLGAETVLNQI | 14 | Q1KMD3 | Heterogeneous nuclear ribonucleoprotein | HNRNPUL | 85.105 | 2 | 51.910126 | yes | 9.722E-08 | 804.41412 | 911.44781 |
| EGCTEVSLLR | 10 | Q1KMD3 | Heterogeneous nuclear ribonucleoprotein | HNRNPUL | 85.105 | 2 | 51.910126 | yes | 9.819E-08 | 582.28992 | 752.22437 |
| DLLVQQASQCLSK | 13 | Q1KMD3 | Heterogeneous nuclear ribonucleoprotein | HNRNPUL | 85.105 | 2 | 51.910126 | yes | 4.126E-07 | 745.38757 | 596.21442 |
| YGGQPLFSEK | 10 | Q1KMD3 | Heterogeneous nuclear ribonucleoprotein | HNRNPUL | 85.105 | 2 | 51.910126 | yes | 9.722E-08 | 563.28241 | 463.616 |
| ALLPHVLCK | 9 | Q1KMD3 | Heterogeneous nuclear ribonucleoprotein | HNRNPUL | 85.105 | 2 | 51.910126 | yes | 1.022E-07 | 525.81006 | 356.41998 |
| GQGYVGGQR | 9 | Q1KMD3 | Heterogeneous nuclear ribonucleoprotein | HNRNPUL | 85.105 | 2 | 51.910126 | yes | 6.248E-07 | 461.2305 | 342.84274 |
| GLEEPEMDPK | 10 | Q1KMD3 | Heterogeneous nuclear ribonucleoprotein | HNRNPUL | 85.105 | 2 | 51.910126 | yes | 6.554E-07 | 572.76318 | 323.1308 |
| VVVVVPNEEDWK | 12 | Q1KMD3 | Heterogeneous nuclear ribonucleoprotein | HNRNPUL | 85.105 | 2 | 51.910126 | yes | 2.609E-07 | 706.87463 | 313.98932 |
| KVVVVVPNEEDW | 13 | Q1KMD3 | Heterogeneous nuclear ribonucleoprotein | HNRNPUL | 85.105 | 2 | 51.910126 | yes | 9.722E-08 | 770.92212 | 138.76755 |
| KVVVVVPNEEDW | 13 | Q1KMD3 | Heterogeneous nuclear ribonucleoprotein | HNRNPUL | 85.105 | 2 | 51.910126 | yes | 2.286E-06 | 514.28381 | 138.76755 |
| KVVVVVPNEEDW | 13 | Q1KMD3 | Heterogeneous nuclear ribonucleoprotein | HNRNPUL | 85.105 | 3 | 51.910126 | yes | 9.722E-08 | 770.92212 | 138.76755 |
| KVVVVVPNEEDW | 13 | Q1KMD3 | Heterogeneous nuclear ribonucleoprotein | HNRNPUL | 85.105 | 3 | 51.910126 | yes | 2.286E-06 | 514.28381 | 138.76755 |
| NCVVELNFGQK | 11 | Q1KMD3 | Heterogeneous nuclear ribonucleoprotein | HNRNPUL | 85.105 | 2 | 51.910126 | yes | 1.062E-07 | 654.32428 | 254.95731 |
| SGDETPGSEVPGD | 14 | Q1KMD3 | Heterogeneous nuclear ribonucleoprotein | HNRNPUL | 85.105 | 2 | 51.910126 | yes | 1.312E-07 | 687.80463 | 112.49091 |
| VGWSVDFSR | 9 | Q1KMD3 | Heterogeneous nuclear ribonucleoprotein | HNRNPUL | 85.105 | 2 | 51.910126 | yes | 1.066E-07 | 526.7616 | 93.204849 |
| EEAQPIVTK | 9 | Q1KMD3 | Heterogeneous nuclear ribonucleoprotein | HNRNPUL | 85.105 | 2 | 51.910126 | yes | 2.048E-06 | 507.77695 | 76.974464 |
| NGEDLGVAFWISK | 13 | Q1KMD3 | Heterogeneous nuclear ribonucleoprotein | HNRNPUL | 85.105 | 2 | 51.910126 | yes | 2.384E-07 | 718.36444 | 64.025436 |
| FPTLWSGAR | 9 | Q1KMD3 | Heterogeneous nuclear ribonucleoprotein | HNRNPUL | 85.105 | 2 | 51.910126 | yes | 1.847E-07 | 517.77454 | 58.002583 |
| KEVEGDDVPESIM | 17 | Q1KMD3 | Heterogeneous nuclear ribonucleoprotein | HNRNPUL | 85.105 | 3 | 51.910126 | yes | 7.458E-07 | 650.31293 | 53.565044 |
| LEDQVIGVALQAS | 14 | Q2KHR3 | Glutamine and serine-rich protein 1 OS=I | QSER1 | 189.973 | 2 | 49.076942 | yes | 1.071E-07 | 735.91174 | 88.378395 |
| ELGSVPLR | 8 | Q2M296 | Methenyltetrahydrofolate synthase domai | MTHFSD | 42.173 | 2 | 48.1769 | yes | 2.311E-07 | 435.7558 | 297.68076 |
| AFLHYPDSAAAQC | 22 | Q2M296 | Methenyltetrahydrofolate synthase domai | MTHFSD | 42.173 | 3 | 48.1769 | yes | 1.187E-07 | 801.73401 | 265.79944 |
| DVTLQGEHQHLPE | 26 | Q2M296 | Methenyltetrahydrofolate synthase domai | MTHFSD | 42.173 | 4 | 48.1769 | yes | 1.204E-07 | 721.36078 | 207.19353 |
| ISLEMMEK | 8 | Q2M296 | Methenyltetrahydrofolate synthase domai | MTHFSD | 42.173 | 2 | 48.1769 | yes | 3.041E-07 | 490.74319 | 192.27142 |
| TLLVPTPR | 8 | Q2M296 | Methenyltetrahydrofolate synthase domai | MTHFSD | 42.173 | 2 | 48.1769 | yes | 1.652E-07 | 448.78183 | 176.87367 |
| GSYLACQNIK | 10 | Q2M296 | Methenyltetrahydrofolate synthase domai | MTHFSD | 42.173 | 2 | 48.1769 | yes | 7.486E-07 | 577.28717 | 115.58315 |

| | | | | | | | | | | | |
|---|---|---|---|---|---|---|---|---|---|---|---|
| NYSVPIGLDSR | 11 | Q2M296 | Methenyltetrahydrofolate synthase domai | MTHFSD | 42.173 | 2 | 48.1769 | yes | 1.926E-06 | 610.81714 | 82.237152 |
| KTLLVPTPR | 9 | Q2M296 | Methenyltetrahydrofolate synthase domai | MTHFSD | 42.173 | 2 | 48.1769 | yes | 3.046E-06 | 512.82928 | 8.6561499 |
| CPLCTYETR | 9 | Q2TB10 | Zinc finger protein 800 OS=Homo sapien | ZNF800 | 75.236 | 2 | 46.22293 | yes | 3.355E-07 | 600.26282 | 67.280968 |
| KPIDFVLNK | 9 | Q2TB10 | Zinc finger protein 800 OS=Homo sapien | ZNF800 | 75.236 | 2 | 46.22293 | yes | 1.617E-07 | 537.32129 | 53.596764 |
| SGIQQIIECFR | 11 | Q2TB10 | Zinc finger protein 800 OS=Homo sapien | ZNF800 | 75.236 | 2 | 46.22293 | yes | 1.741E-07 | 675.84534 | 49.02702 |
| SFCYSSVLLR | 10 | Q499Z4 | Zinc finger protein 672 OS=Homo sapien | ZNF672 | 50.225 | 2 | 51.11853 | yes | 9.964E-08 | 616.31061 | 126.94241 |
| QSALLFHQAR | 10 | Q499Z4 | Zinc finger protein 672 OS=Homo sapien | ZNF672 | 50.225 | 2 | 51.11853 | yes | 1.387E-07 | 585.82257 | 113.62566 |
| SDLVVHQR | 8 | Q499Z4 | Zinc finger protein 672 OS=Homo sapien | ZNF672 | 50.225 | 2 | 51.11853 | yes | 2.052E-07 | 477.26178 | 102.75458 |
| GFLESATLVR | 10 | Q499Z4 | Zinc finger protein 672 OS=Homo sapien | ZNF672 | 50.225 | 2 | 51.11853 | yes | 1.317E-07 | 546.80603 | 102.07895 |
| VSDAHQCGVCGK | 12 | Q499Z4 | Zinc finger protein 672 OS=Homo sapien | ZNF672 | 50.225 | 2 | 51.11853 | yes | 9.722E-08 | 659.28735 | 27.553984 |
| AHPLGTTSDPAAP | 16 | Q499Z4 | Zinc finger protein 672 OS=Homo sapien | ZNF672 | 50.225 | 2 | 51.11853 | yes | 3.916E-07 | 812.91315 | 15.334335 |
| CVDVCSDASR | 10 | Q49AG3 | Zinc finger BED domain-containing prote | ZBED5 | 78.911 | 2 | 45.465664 | yes | 5.089E-07 | 584.73969 | 61.514858 |
| LSDCLTNSSWLLR | 13 | Q49AG3 | Zinc finger BED domain-containing prote | ZBED5 | 78.911 | 2 | 45.465664 | yes | 5.057E-07 | 782.89307 | 59.080135 |
| IAEAVTLIK | 9 | Q49AG3 | Zinc finger BED domain-containing prote | ZBED5 | 78.911 | 2 | 45.465664 | yes | 3.201E-06 | 479.30258 | 56.36668 |
| LNEVNLSMQGK | 11 | Q49AG3 | Zinc finger BED domain-containing prote | ZBED5 | 78.911 | 2 | 45.465664 | yes | 6.699E-07 | 616.81879 | 54.083199 |
| DKDISFFK | 8 | Q49AG3 | Zinc finger BED domain-containing prote | ZBED5 | 78.911 | 2 | 45.465664 | yes | 1.841E-07 | 500.26093 | 51.169758 |
| LAYLADIFTK | 10 | Q49AG3 | Zinc finger BED domain-containing prote | ZBED5 | 78.911 | 2 | 45.465664 | yes | 1.926E-06 | 577.82642 | 47.239426 |
| VGILQSEDK | 9 | Q49AG3 | Zinc finger BED domain-containing prote | ZBED5 | 78.911 | 2 | 45.465664 | yes | 7.66E-06 | 494.7691 | 40.083019 |
| ELLVFMDSAFR | 11 | Q49AG3 | Zinc finger BED domain-containing prote | ZBED5 | 78.911 | 2 | 45.465664 | yes | 0.0001478 | 664.33936 | 11.710246 |
| ALEFTNIYVK | 10 | Q4VXU2 | Polyadenylate-binding protein 1-like OS= | PABPC1L | 68.393 | 2 | 41.454086 | yes | 3.502E-07 | 599.32935 | 39.50544 |
| HCLLTCEECK | 10 | Q56VL3 | OCIA domain-containing protein 2 OS=H | OCIAD2 | 16.954 | 2 | 25.745628 | yes | 3.233E-06 | 675.28595 | 12.699386 |
| LWNFTEEEVSER | 12 | Q5SWX8 | Protein odr-4 homolog OS=Homo sapiens | ODR4 | 51.104 | 2 | 38.378887 | yes | 5.262E-07 | 769.85974 | 39.636982 |
| STATVQICSGSVNI | 15 | Q5SWX8 | Protein odr-4 homolog OS=Homo sapiens | ODR4 | 51.104 | 2 | 38.378887 | yes | 8.097E-07 | 782.90363 | 28.726622 |
| DFAVEPAENVSEA | 15 | Q5SYE7 | NHS-like protein 1 OS=Homo sapiens OX | NHSL1 | 170.669 | 2 | 47.377438 | yes | 1.343E-07 | 823.90466 | 169.12663 |
| PAPVAQPPAAAPP | 24 | Q5T1J5 | Putative coiled-coil-helix-coiled-coil-heli | CHCHD2P | 15.49 | 3 | 19.475168 | yes | 4.837E-06 | 713.3833 | 70.143463 |
| ILTDEMLLQACEG | 14 | Q5T3I0 | G patch domain-containing protein 4 OS= | GPATCH4 | 50.382 | 2 | 52.759636 | yes | 1.065E-07 | 824.90527 | 1870.5289 |
| LEAQEQAFLAR | 11 | Q5T3I0 | G patch domain-containing protein 4 OS= | GPATCH4 | 50.382 | 2 | 52.759636 | yes | 1.02E-07 | 638.33826 | 1013.5472 |
| EDAAGTSGLGELN | 15 | Q5T3I0 | G patch domain-containing protein 4 OS= | GPATCH4 | 50.382 | 2 | 52.759636 | yes | 9.722E-08 | 738.84985 | 560.00574 |
| AYADPCSR | 8 | Q5T3I0 | G patch domain-containing protein 4 OS= | GPATCH4 | 50.382 | 2 | 52.759636 | yes | 9.722E-08 | 470.20309 | 284.5249 |
| MATLTSGGEKPNK | 13 | Q5T3I0 | G patch domain-containing protein 4 OS= | GPATCH4 | 50.382 | 2 | 52.759636 | yes | 1.937E-07 | 667.34265 | 197.10699 |
| GEETVLGGGTR | 11 | Q5T3I0 | G patch domain-containing protein 4 OS= | GPATCH4 | 50.382 | 2 | 52.759636 | yes | 1.343E-07 | 538.27252 | 154.29422 |
| QQQEEEDLNLEDR | 13 | Q5T3I0 | G patch domain-containing protein 4 OS= | GPATCH4 | 50.382 | 2 | 52.759636 | yes | 9.904E-08 | 823.36865 | 140.36661 |
| PNLLYQK | 7 | Q5T3I0 | G patch domain-containing protein 4 OS= | GPATCH4 | 50.382 | 2 | 52.759636 | yes | 2.941E-05 | 438.2529 | 70.936859 |
| QQHQEEEDILDVR | 13 | Q5T3I0 | G patch domain-containing protein 4 OS= | GPATCH4 | 50.382 | 2 | 52.759636 | yes | 1.618E-07 | 819.88953 | 38.361553 |
| MGVLEEGGK | 9 | Q5T3I0 | G patch domain-containing protein 4 OS= | GPATCH4 | 50.382 | 2 | 52.759636 | yes | 3.924E-07 | 460.23132 | 17.62215 |
| LNYEGLPIGK | 10 | Q5VTE6 | Protein angel homolog 2 OS=Homo sapie | ANGEL2 | 62.339 | 2 | 50.467663 | yes | 9.809E-08 | 552.30841 | 61.534191 |
| TDSDLTQTQLK | 11 | Q5VTE6 | Protein angel homolog 2 OS=Homo sapie | ANGEL2 | 62.339 | 2 | 50.467663 | yes | 3.075E-06 | 625.31714 | 28.341707 |
| AGGGCVMTIGEMI | 14 | Q5VTL8 | Pre-mRNA-splicing factor 38B OS=Hom | PRPF38B | 64.468 | 2 | 51.857147 | yes | 9.722E-08 | 726.34381 | 2441.9949 |
| VTHVEPWEK | 9 | Q5VTL8 | Pre-mRNA-splicing factor 38B OS=Hom | PRPF38B | 64.468 | 2 | 51.857147 | yes | 1.074E-07 | 562.79041 | 1749.6548 |

| | | | | | | | | | | | |
|---|---|---|---|---|---|---|---|---|---|---|---|
| QGNVLPLWGNEK | 12 | Q5VTL8 | Pre-mRNA-splicing factor 38B OS=Hom | PRPF38B | 64.468 | 2 | 51.857147 | yes | 1.431E-07 | 677.85931 | 1741.8397 |
| VQLYELK | 7 | Q5VTL8 | Pre-mRNA-splicing factor 38B OS=Hom | PRPF38B | 64.468 | 2 | 51.857147 | yes | 3.125E-07 | 446.76056 | 748.85614 |
| QVMGLITHTDSPY | 15 | Q5VTL8 | Pre-mRNA-splicing factor 38B OS=Hom | PRPF38B | 64.468 | 3 | 51.857147 | yes | 9.722E-08 | 577.63464 | 335.43988 |
| EGHGSSSFDR | 10 | Q5VTL8 | Pre-mRNA-splicing factor 38B OS=Hom | PRPF38B | 64.468 | 2 | 51.857147 | yes | 2.47E-07 | 539.73108 | 322.96545 |
| ALGFMYIR | 8 | Q5VTL8 | Pre-mRNA-splicing factor 38B OS=Hom | PRPF38B | 64.468 | 2 | 51.857147 | yes | 1.327E-07 | 485.76257 | 296.06879 |
| TYHEVVDEIYFK | 12 | Q5VTL8 | Pre-mRNA-splicing factor 38B OS=Hom | PRPF38B | 64.468 | 2 | 51.857147 | yes | 6.716E-07 | 771.87738 | 264.00049 |
| DGKEGAEEIDR | 11 | Q5VTL8 | Pre-mRNA-splicing factor 38B OS=Hom | PRPF38B | 64.468 | 2 | 51.857147 | yes | 3.954E-06 | 609.78351 | 86.910812 |
| TAGQTGMCGGVR | 12 | Q5VTL8 | Pre-mRNA-splicing factor 38B OS=Hom | PRPF38B | 64.468 | 2 | 51.857147 | yes | 1.23E-07 | 597.77136 | 63.250275 |
| ELEVDSQCVR | 10 | Q5VWQ0 | Lysine-specific demethylase 9 OS=Homo | RSBN1 | 90.073 | 2 | 53.28448 | yes | 9.974E-08 | 617.79022 | 200.004 |
| TVSAVCSLAWHIR | 13 | Q5VWQ0 | Lysine-specific demethylase 9 OS=Homo | RSBN1 | 90.073 | 2 | 53.28448 | no | 9.722E-08 | 750.39301 | 158.6338 |
| MTLPWGTLSSLR | 12 | Q5VWQ0 | Lysine-specific demethylase 9 OS=Homo | RSBN1 | 90.073 | 2 | 53.28448 | yes | 9.722E-08 | 681.36591 | 138.0052 |
| EVQTVCAGLTR | 11 | Q5VWQ0 | Lysine-specific demethylase 9 OS=Homo | RSBN1 | 90.073 | 2 | 53.28448 | yes | 2.041E-07 | 617.31647 | 130.57547 |
| NLQYLPR | 7 | Q5VWQ0 | Lysine-specific demethylase 9 OS=Homo | RSBN1 | 90.073 | 2 | 53.28448 | no | 3.25E-06 | 452.25598 | 130.41269 |
| IQLCDNDIYFIPR | 13 | Q5VWQ0 | Lysine-specific demethylase 9 OS=Homo | RSBN1 | 90.073 | 2 | 53.28448 | yes | 1.422E-07 | 833.9165 | 112.99388 |
| PGEQMIPTADMPK | 13 | Q5VWQ0 | Lysine-specific demethylase 9 OS=Homo | RSBN1 | 90.073 | 2 | 53.28448 | yes | 1.305E-07 | 707.83887 | 97.556656 |
| AVQFGEWSDQPR | 12 | Q5VWQ0 | Lysine-specific demethylase 9 OS=Homo | RSBN1 | 90.073 | 2 | 53.28448 | yes | 1.69E-07 | 710.33624 | 85.31208 |
| SQSDDGPIMWVR | 12 | Q5VWQ0 | Lysine-specific demethylase 9 OS=Homo | RSBN1 | 90.073 | 2 | 53.28448 | yes | 9.819E-08 | 695.82465 | 73.104446 |
| TYCCGTYR | 8 | Q5VWQ0 | Lysine-specific demethylase 9 OS=Homo | RSBN1 | 90.073 | 2 | 53.28448 | yes | 4.494E-07 | 540.71552 | 38.411121 |
| AHADHVGQGFDW | 22 | Q5VWQ0 | Lysine-specific demethylase 9 OS=Homo | RSBN1 | 90.073 | 3 | 53.28448 | yes | 9.294E-07 | 765.38196 | 25.344648 |
| VLPASELSDPGLLI | 15 | Q5VYV7 | Protein SLX4IP OS=Homo sapiens OX=9 | SLX4IP | 45.552 | 2 | 32.358868 | yes | 3.383E-06 | 776.45325 | 69.229362 |
| STQNAELCVFPDR | 13 | Q5VYV7 | Protein SLX4IP OS=Homo sapiens OX=9 | SLX4IP | 45.552 | 2 | 32.358868 | yes | 1.542E-06 | 768.85919 | 52.575035 |
| AAVQVVGSWPSV | 15 | Q6AI08 | HEAT repeat-containing protein 6 OS=H | HEATR6 | 128.782 | 2 | 48.40802 | yes | 1.152E-07 | 811.9361 | 120.73894 |
| LSDLAQSDPEVR | 12 | Q6AI08 | HEAT repeat-containing protein 6 OS=H | HEATR6 | 128.782 | 2 | 48.40802 | yes | 3.91E-07 | 665.33588 | 92.346153 |
| YLPELLGNTGLLM | 14 | Q6AI08 | HEAT repeat-containing protein 6 OS=H | HEATR6 | 128.782 | 2 | 48.40802 | yes | 4.132E-07 | 781.43652 | 32.631565 |
| PYVCLECGK | 9 | Q6DD87 | Zinc finger protein 787 OS=Homo sapien | ZNF787 | 40.429 | 2 | 50.912605 | yes | 1.029E-07 | 563.25702 | 1075.2982 |
| LHPELSGPGVAAK | 13 | Q6DD87 | Zinc finger protein 787 OS=Homo sapien | ZNF787 | 40.429 | 2 | 50.912605 | yes | 1.745E-07 | 638.35645 | 1065.0719 |
| AQHGDGLGAAGG | 33 | Q6DD87 | Zinc finger protein 787 OS=Homo sapien | ZNF787 | 40.429 | 4 | 50.912605 | yes | 1.612E-07 | 830.88477 | 906.63354 |
| SHSGLKPFVCPR | 12 | Q6DD87 | Zinc finger protein 787 OS=Homo sapien | ZNF787 | 40.429 | 3 | 50.912605 | yes | 1.136E-07 | 462.24332 | 390.13809 |
| SHSGLKPFVCPR | 12 | Q6DD87 | Zinc finger protein 787 OS=Homo sapien | ZNF787 | 40.429 | 3 | 50.912605 | yes | 2.105E-07 | 692.86133 | 390.13809 |
| SHSGLKPFVCPR | 12 | Q6DD87 | Zinc finger protein 787 OS=Homo sapien | ZNF787 | 40.429 | 2 | 50.912605 | yes | 1.136E-07 | 462.24332 | 390.13809 |
| SHSGLKPFVCPR | 12 | Q6DD87 | Zinc finger protein 787 OS=Homo sapien | ZNF787 | 40.429 | 2 | 50.912605 | yes | 2.105E-07 | 692.86133 | 390.13809 |
| LSPPQSAPPAGPPP | 15 | Q6DD87 | Zinc finger protein 787 OS=Homo sapien | ZNF787 | 40.429 | 2 | 50.912605 | yes | 2.611E-07 | 734.89899 | 455.50208 |
| SFSHWSK | 7 | Q6DD87 | Zinc finger protein 787 OS=Homo sapien | ZNF787 | 40.429 | 2 | 50.912605 | yes | 3.393E-07 | 439.7114 | 235.60239 |
| GFGHGAGLLAHQI | 13 | Q6DD87 | Zinc finger protein 787 OS=Homo sapien | ZNF787 | 40.429 | 3 | 50.912605 | yes | 6.779E-07 | 440.90219 | 192.78992 |
| TFSQSSHLVQHR | 12 | Q6DD87 | Zinc finger protein 787 OS=Homo sapien | ZNF787 | 40.429 | 2 | 50.912605 | yes | 9.722E-08 | 713.86292 | 136.79068 |
| PYACLECGK | 9 | Q6DD87 | Zinc finger protein 787 OS=Homo sapien | ZNF787 | 40.429 | 2 | 50.912605 | yes | 9.445E-07 | 549.24133 | 134.27794 |
| FSWSSNLMQHQR | 12 | Q6DD87 | Zinc finger protein 787 OS=Homo sapien | ZNF787 | 40.429 | 2 | 50.912605 | yes | 4.02E-06 | 760.85681 | 124.09687 |
| IDGVSLLVQR | 10 | Q6DKI1 | Ribosomal protein uL30-like OS=Homo s | RPL7L1 | 29.67 | 2 | 53.929253 | yes | 1.283E-07 | 550.32715 | 2039.4714 |
| HSLAFVVR | 8 | Q6DKI1 | Ribosomal protein uL30-like OS=Homo s | RPL7L1 | 29.67 | 2 | 53.929253 | yes | 2.17E-07 | 464.77179 | 726.49573 |

| | | | | | | | | | | | |
|---|---|---|---|---|---|---|---|---|---|---|---|
| LESFLHDSWR | 10 | Q6DKI1 | Ribosomal protein uL30-like OS=Homo s | RPL7L1 | 29.67 | 2 | 53.929253 | yes | 9.722E-08 | 645.31732 | 455.70532 |
| KIPLVPENLLK | 11 | Q6DKI1 | Ribosomal protein uL30-like OS=Homo s | RPL7L1 | 29.67 | 2 | 53.929253 | yes | 1.166E-07 | 632.4054 | 286.6189 |
| IVEPYVTWGFPNL | 14 | Q6DKI1 | Ribosomal protein uL30-like OS=Homo s | RPL7L1 | 29.67 | 2 | 53.929253 | yes | 9.722E-08 | 831.94812 | 233.73599 |
| IFSGVFVK | 8 | Q6DKI1 | Ribosomal protein uL30-like OS=Homo s | RPL7L1 | 29.67 | 2 | 53.929253 | yes | 4.431E-07 | 448.76566 | 51.09647 |
| LEVKPHALELPDK | 13 | Q6DKI1 | Ribosomal protein uL30-like OS=Homo s | RPL7L1 | 29.67 | 3 | 53.929253 | yes | 2.967E-06 | 496.95221 | 27.385786 |
| KIFSGVFVK | 9 | Q6DKI1 | Ribosomal protein uL30-like OS=Homo s | RPL7L1 | 29.67 | 2 | 53.929253 | yes | 3.01E-06 | 512.81311 | 22.27261 |
| NNYQGFLAELR | 11 | Q6ICC9 | Retrotransposon Gag-like protein 6 OS=H | RTL6 | 26.155 | 2 | 47.488743 | yes | 1.544E-07 | 662.83588 | 67.814697 |
| LAGFLMQMDR | 10 | Q6ICC9 | Retrotransposon Gag-like protein 6 OS=H | RTL6 | 26.155 | 2 | 47.488743 | yes | 1.318E-07 | 591.29395 | 56.572411 |
| LDYMVTCAVCTR | 12 | Q6ICG6 | Uncharacterized protein KIAA0930 OS=I | KIAA0930 | 45.795 | 2 | 49.045208 | yes | 1.074E-07 | 744.83582 | 98.76664 |
| ILTDILEVR | 9 | Q6ICG6 | Uncharacterized protein KIAA0930 OS=I | KIAA0930 | 45.795 | 2 | 49.045208 | yes | 2.683E-07 | 536.32404 | 95.846329 |
| QDDMLFYVR | 9 | Q6ICG6 | Uncharacterized protein KIAA0930 OS=I | KIAA0930 | 45.795 | 2 | 49.045208 | yes | 1.475E-06 | 593.78168 | 77.572594 |
| YSNMEFVR | 8 | Q6ICG6 | Uncharacterized protein KIAA0930 OS=I | KIAA0930 | 45.795 | 2 | 49.045208 | yes | 2.832E-07 | 523.24219 | 73.258232 |
| TNTFQGVIFQGSIR | 14 | Q6ICG6 | Uncharacterized protein KIAA0930 OS=I | KIAA0930 | 45.795 | 2 | 49.045208 | yes | 1.252E-06 | 784.41498 | 65.064667 |
| EVCGLGCFK | 9 | Q6ICG6 | Uncharacterized protein KIAA0930 OS=I | KIAA0930 | 45.795 | 2 | 49.045208 | yes | 1.187E-06 | 535.2439 | 36.505089 |
| LAYSGSESGADGR | 13 | Q6ICG6 | Uncharacterized protein KIAA0930 OS=I | KIAA0930 | 45.795 | 2 | 49.045208 | yes | 2.768E-07 | 635.28894 | 31.352156 |
| KLDNFQSEK | 9 | Q6IQ21 | Zinc finger protein 770 OS=Homo sapien | ZNF770 | 80.008 | 2 | 46.17367 | yes | 2.515E-05 | 554.78528 | 103.71159 |
| NIHACTICGK | 10 | Q6IQ21 | Zinc finger protein 770 OS=Homo sapien | ZNF770 | 80.008 | 2 | 46.17367 | yes | 1.631E-07 | 587.27881 | 31.876303 |
| ELSPVSFQYR | 10 | Q6NW29 | RWD domain-containing protein 4 OS=H | RWDD4 | 21.251 | 2 | 50.107197 | yes | 1.013E-07 | 613.31421 | 141.84196 |
| SIYEGDESFR | 10 | Q6NW29 | RWD domain-containing protein 4 OS=H | RWDD4 | 21.251 | 2 | 50.107197 | yes | 9.901E-08 | 601.76984 | 114.15574 |
| SGPEEPPSLVLGK | 13 | Q6NX49 | Zinc finger protein 544 OS=Homo sapien | ZNF544 | 81.742 | 2 | 21.615509 | yes | 4.089E-06 | 655.35352 | 49.873787 |
| TLEELDPESR | 10 | Q6P161 | Large ribosomal subunit protein mL54 OS | MRPL54 | 15.82 | 2 | 49.85748 | yes | 1.001E-07 | 594.79077 | 1300.0587 |
| HEPIASVR | 8 | Q6PCB5 | Lysine-specific demethylase RSBN1L OS | RSBN1L | 94.87 | 2 | 53.434483 | yes | 9.722E-08 | 454.75107 | 464.52298 |
| PGEQMIPVADMPK | 13 | Q6PCB5 | Lysine-specific demethylase RSBN1L OS | RSBN1L | 94.87 | 2 | 53.434483 | yes | 1.029E-07 | 706.84924 | 310.13974 |
| CTLPWGTLSSLK | 12 | Q6PCB5 | Lysine-specific demethylase RSBN1L OS | RSBN1L | 94.87 | 2 | 53.434483 | yes | 1.849E-07 | 681.85791 | 294.44269 |
| YELQQIK | 7 | Q6PCB5 | Lysine-specific demethylase RSBN1L OS | RSBN1L | 94.87 | 2 | 53.434483 | yes | 4.674E-07 | 461.25565 | 200.71815 |
| IPENSEFPFVSLK | 13 | Q6PCB5 | Lysine-specific demethylase RSBN1L OS | RSBN1L | 94.87 | 2 | 53.434483 | yes | 4.358E-07 | 753.89557 | 192.49068 |
| IQLYDNDIYFIPR | 13 | Q6PCB5 | Lysine-specific demethylase RSBN1L OS | RSBN1L | 94.87 | 2 | 53.434483 | yes | 9.722E-08 | 835.43286 | 190.61394 |
| IKEEPVNVNIPEK | 13 | Q6PCB5 | Lysine-specific demethylase RSBN1L OS | RSBN1L | 94.87 | 2 | 53.434483 | yes | 9.727E-08 | 754.91956 | 84.613525 |
| IKEEPVNVNIPEK | 13 | Q6PCB5 | Lysine-specific demethylase RSBN1L OS | RSBN1L | 94.87 | 2 | 53.434483 | yes | 4.934E-06 | 503.61548 | 84.613525 |
| IKEEPVNVNIPEK | 13 | Q6PCB5 | Lysine-specific demethylase RSBN1L OS | RSBN1L | 94.87 | 3 | 53.434483 | yes | 9.727E-08 | 754.91956 | 84.613525 |
| IKEEPVNVNIPEK | 13 | Q6PCB5 | Lysine-specific demethylase RSBN1L OS | RSBN1L | 94.87 | 3 | 53.434483 | yes | 4.934E-06 | 503.61548 | 84.613525 |
| RLDTLEFK | 8 | Q6PCB5 | Lysine-specific demethylase RSBN1L OS | RSBN1L | 94.87 | 2 | 53.434483 | yes | 2.548E-07 | 511.28748 | 161.97151 |
| ASLDSVFSDK | 10 | Q6PCB5 | Lysine-specific demethylase RSBN1L OS | RSBN1L | 94.87 | 2 | 53.434483 | yes | 9.722E-08 | 534.76404 | 147.06619 |
| LDHVQFAEFK | 10 | Q6PCB5 | Lysine-specific demethylase RSBN1L OS | RSBN1L | 94.87 | 2 | 53.434483 | yes | 3.316E-07 | 617.31677 | 142.75429 |
| AVHCGEWPDQPR | 12 | Q6PCB5 | Lysine-specific demethylase RSBN1L OS | RSBN1L | 94.87 | 2 | 53.434483 | yes | 9.729E-08 | 726.32788 | 118.35881 |
| LLVPPTLLHAQPH | 29 | Q6PCB5 | Lysine-specific demethylase RSBN1L OS | RSBN1L | 94.87 | 4 | 53.434483 | yes | 6.875E-07 | 727.42365 | 46.73822 |
| NVVHQFK | 7 | Q6PCB5 | Lysine-specific demethylase RSBN1L OS | RSBN1L | 94.87 | 2 | 53.434483 | yes | 3.551E-07 | 436.24286 | 45.531956 |
| QTTAAVGVLK | 10 | Q6PCB5 | Lysine-specific demethylase RSBN1L OS | RSBN1L | 94.87 | 2 | 53.434483 | yes | 1.841E-07 | 494.29529 | 33.409542 |
| KDSDDGPIMWVR | 12 | Q6PCB5 | Lysine-specific demethylase RSBN1L OS | RSBN1L | 94.87 | 2 | 53.434483 | yes | 5.315E-07 | 709.84027 | 22.198196 |

| | | | | | | | | | | | |
|---|---|---|---|---|---|---|---|---|---|---|---|
| FGEVVLQPPELTA | 14 | Q6PK04 | Coiled-coil domain-containing protein 13 | CCDC137 | 33.232 | 2 | 52.853317 | yes | 1.53E-06 | 778.42755 | 1388.7325 |
| LEQELLR | 7 | Q6PK04 | Coiled-coil domain-containing protein 13 | CCDC137 | 33.232 | 2 | 52.853317 | yes | 1.526E-07 | 450.76108 | 801.57465 |
| NQDEQEIPFR | 10 | Q6PK04 | Coiled-coil domain-containing protein 13 | CCDC137 | 33.232 | 2 | 52.853317 | yes | 1.429E-07 | 638.30182 | 396.68405 |
| GEEPDIAVPK | 10 | Q6PK04 | Coiled-coil domain-containing protein 13 | CCDC137 | 33.232 | 2 | 52.853317 | yes | 1.231E-07 | 527.77441 | 362.81143 |
| MQQEAQHVLFLSI | 13 | Q6PK04 | Coiled-coil domain-containing protein 13 | CCDC137 | 33.232 | 2 | 52.853317 | yes | 9.722E-08 | 779.90594 | 214.43076 |
| KGESDGAYIHR | 11 | Q6PK04 | Coiled-coil domain-containing protein 13 | CCDC137 | 33.232 | 2 | 52.853317 | yes | 9.893E-08 | 616.80457 | 180.97929 |
| QQVQPLGK | 8 | Q6PK04 | Coiled-coil domain-containing protein 13 | CCDC137 | 33.232 | 2 | 52.853317 | yes | 1.211E-07 | 449.26126 | 162.88365 |
| QPEVQAAPK | 9 | Q6PK04 | Coiled-coil domain-containing protein 13 | CCDC137 | 33.232 | 2 | 52.853317 | yes | 4.347E-07 | 484.26401 | 133.97467 |
| GESDGAYIHR | 10 | Q6PK04 | Coiled-coil domain-containing protein 13 | CCDC137 | 33.232 | 2 | 52.853317 | yes | 2.188E-07 | 552.75708 | 94.093575 |
| QQQLHGERPHLTS | 14 | Q6PK04 | Coiled-coil domain-containing protein 13 | CCDC137 | 33.232 | 4 | 52.853317 | yes | 8.784E-07 | 422.47504 | 27.57815 |
| YCQPEEEVAR | 10 | Q6Y2X3 | DnaJ homolog subfamily C member 14 O | DNAJC14 | 78.569 | 2 | 52.199226 | yes | 9.722E-08 | 640.78241 | 131.12883 |
| AAWDIVSNAEK | 11 | Q6Y2X3 | DnaJ homolog subfamily C member 14 O | DNAJC14 | 78.569 | 2 | 52.199226 | yes | 9.722E-08 | 602.30383 | 104.12492 |
| VWMGVWTGR | 9 | Q6Y2X3 | DnaJ homolog subfamily C member 14 O | DNAJC14 | 78.569 | 2 | 52.199226 | yes | 3.254E-07 | 546.27637 | 101.97117 |
| VGISPDTHR | 9 | Q6Y2X3 | DnaJ homolog subfamily C member 14 O | DNAJC14 | 78.569 | 2 | 52.199226 | yes | 4.394E-07 | 491.25925 | 73.150597 |
| VYDITEWAGCQR | 12 | Q6Y2X3 | DnaJ homolog subfamily C member 14 O | DNAJC14 | 78.569 | 2 | 52.199226 | yes | 1.056E-07 | 749.3432 | 59.680618 |
| ITYFALMDGK | 10 | Q6Y2X3 | DnaJ homolog subfamily C member 14 O | DNAJC14 | 78.569 | 2 | 52.199226 | yes | 2.226E-06 | 579.79681 | 28.531919 |
| EAMNTMMCSR | 10 | Q6Y2X3 | DnaJ homolog subfamily C member 14 O | DNAJC14 | 78.569 | 2 | 52.199226 | yes | 6.959E-07 | 615.74036 | 26.259651 |
| LWDTDFKPITK | 11 | Q6ZMW3 | Echinoderm microtubule-associated prote | EML6 | 217.9 | 2 | 39.98435 | yes | 4.271E-07 | 682.36646 | 57.675968 |
| LHTGEKPFK | 9 | Q6ZN06 | Zinc finger protein 813 OS=Homo sapien | ZNF813 | 71.721 | 2 | 47.73566 | no | 1.269E-07 | 528.79547 | 238.07899 |
| STLQDCSQLR | 10 | Q6ZUT6 | Coiled-coil domain-containing protein 9B | CCDC9B | 57.325 | 2 | 38.496407 | yes | 5.181E-07 | 604.29041 | 115.89191 |
| SGAQQGLAPR | 10 | Q6ZUT6 | Coiled-coil domain-containing protein 9B | CCDC9B | 57.325 | 2 | 38.496407 | yes | 1.69E-06 | 492.76471 | 51.134605 |
| SPPTQVAISSDSAR | 14 | Q6ZUT6 | Coiled-coil domain-containing protein 9B | CCDC9B | 57.325 | 2 | 38.496407 | yes | 3.159E-06 | 708.35986 | 38.256123 |
| KGEMETESQPK | 11 | Q7Z309 | PABIR family member 2 OS=Homo sapie | PABIR2 | 26.929 | 2 | 49.655434 | yes | 1.013E-07 | 632.29791 | 35.20293 |
| ILGSASPEEEQEKP | 21 | Q7Z422 | SUZ domain-containing protein 1 OS=Hc | SZRD1 | 16.998 | 3 | 47.478325 | yes | 1.32E-07 | 789.08344 | 143.58191 |
| ILGSASPEEEQEKP | 21 | Q7Z422 | SUZ domain-containing protein 1 OS=Hc | SZRD1 | 16.998 | 3 | 47.478325 | yes | 3.316E-07 | 592.06439 | 143.58191 |
| ILGSASPEEEQEKP | 21 | Q7Z422 | SUZ domain-containing protein 1 OS=Hc | SZRD1 | 16.998 | 4 | 47.478325 | yes | 1.32E-07 | 789.08344 | 143.58191 |
| ILGSASPEEEQEKP | 21 | Q7Z422 | SUZ domain-containing protein 1 OS=Hc | SZRD1 | 16.998 | 4 | 47.478325 | yes | 3.316E-07 | 592.06439 | 143.58191 |
| ISQPEDSR | 8 | Q7Z422 | SUZ domain-containing protein 1 OS=Hc | SZRD1 | 16.998 | 2 | 47.478325 | yes | 1.483E-07 | 466.2276 | 138.23071 |
| QPLGPDGSQGFK | 12 | Q7Z422 | SUZ domain-containing protein 1 OS=Hc | SZRD1 | 16.998 | 2 | 47.478325 | yes | 1.464E-07 | 615.80933 | 113.88829 |
| EAEYAEAR | 8 | Q7Z422 | SUZ domain-containing protein 1 OS=Hc | SZRD1 | 16.998 | 2 | 47.478325 | yes | 3.937E-07 | 469.71432 | 94.426247 |
| TEQALPDSELAEY | 15 | Q7Z736 | Pleckstrin homology domain-containing f | PLEKHH3 | 85.318 | 2 | 50.97379 | yes | 7.924E-07 | 846.90741 | 102.64989 |
| GLPEDDPDIVVK | 12 | Q7Z736 | Pleckstrin homology domain-containing f | PLEKHH3 | 85.318 | 2 | 50.97379 | yes | 2.352E-06 | 648.83771 | 93.109268 |
| ALAGGTLVADVLT | 14 | Q7Z736 | Pleckstrin homology domain-containing f | PLEKHH3 | 85.318 | 2 | 50.97379 | yes | 9.722E-08 | 678.89594 | 75.204491 |
| DSLDQFSSSGK | 11 | Q7Z736 | Pleckstrin homology domain-containing f | PLEKHH3 | 85.318 | 2 | 50.97379 | yes | 1.654E-07 | 585.76727 | 70.804832 |
| GPLEVTLTQPVR | 12 | Q7Z736 | Pleckstrin homology domain-containing f | PLEKHH3 | 85.318 | 2 | 50.97379 | yes | 2.008E-07 | 655.37738 | 56.728989 |
| APLETPTQLLLR | 12 | Q7Z736 | Pleckstrin homology domain-containing f | PLEKHH3 | 85.318 | 2 | 50.97379 | yes | 1.779E-06 | 676.40082 | 53.131023 |
| GHLLGHLER | 9 | Q7Z736 | Pleckstrin homology domain-containing f | PLEKHH3 | 85.318 | 2 | 50.97379 | yes | 1.974E-07 | 516.29089 | 46.952362 |
| YDVLELSTEPGR | 12 | Q7Z736 | Pleckstrin homology domain-containing f | PLEKHH3 | 85.318 | 2 | 50.97379 | yes | 6.981E-06 | 689.84607 | 45.126698 |
| ALDPADQHLR | 10 | Q7Z7K0 | COX assembly mitochondrial protein hon | CMC1 | 12.49 | 2 | 36.15318 | yes | 7.333E-07 | 589.30164 | 116.1627 |

| | | | | | | | | | | | |
|---|---|---|---|---|---|---|---|---|---|---|---|
| VVTCEVDAQPPEL | 15 | Q86VU5 | Catechol O-methyltransferase domain-cor | COMTD1 | 28.809 | 2 | 41.77405 | yes | 3.347E-07 | 835.41431 | 114.93988 |
| MAGNEALSPTSPF | 14 | Q86WR7 | Proline and serine-rich protein 2 OS=Hon | PROSER2 | 45.802 | 2 | 52.672024 | yes | 9.722E-08 | 739.35883 | 199.49983 |
| SVPTPLVMAQK | 11 | Q86WR7 | Proline and serine-rich protein 2 OS=Hon | PROSER2 | 45.802 | 2 | 52.672024 | yes | 9.734E-08 | 585.83118 | 191.70055 |
| SGDPGPGPSHPAQ | 15 | Q86WR7 | Proline and serine-rich protein 2 OS=Hon | PROSER2 | 45.802 | 2 | 52.672024 | yes | 9.722E-08 | 714.84692 | 98.137611 |
| PGPALPSTR | 9 | Q86WR7 | Proline and serine-rich protein 2 OS=Hon | PROSER2 | 45.802 | 2 | 52.672024 | yes | 4.381E-07 | 448.25345 | 40.361477 |
| SAPSSFAPAGK | 11 | Q86WR7 | Proline and serine-rich protein 2 OS=Hon | PROSER2 | 45.802 | 2 | 52.672024 | yes | 3.215E-07 | 510.26144 | 39.907005 |
| PQGITVQFAGR | 11 | Q86WR7 | Proline and serine-rich protein 2 OS=Hon | PROSER2 | 45.802 | 2 | 52.672024 | yes | 5.313E-07 | 587.32239 | 25.863724 |
| IHTGEKPYK | 9 | Q86XN6 | Zinc finger protein 761 OS=Homo sapien | ZNF761 | 87.717 | 2 | 55.284946 | no | 1.069E-07 | 536.79291 | 420.92896 |
| DVMLENYR | 8 | Q86XN6 | Zinc finger protein 761 OS=Homo sapien | ZNF761 | 87.717 | 2 | 55.284946 | no | 9.766E-08 | 520.2475 | 136.72478 |
| DVAIEFSQEEWK | 12 | Q86XN6 | Zinc finger protein 761 OS=Homo sapien | ZNF761 | 87.717 | 2 | 55.284946 | no | 9.722E-08 | 740.85138 | 121.87663 |
| TFNQQLTLK | 9 | Q86XN6 | Zinc finger protein 761 OS=Homo sapien | ZNF761 | 87.717 | 2 | 55.284946 | yes | 7.546E-07 | 546.80603 | 71.321487 |
| AFNYSSLLR | 9 | Q86XN6 | Zinc finger protein 761 OS=Homo sapien | ZNF761 | 87.717 | 2 | 55.284946 | no | 2.701E-07 | 535.7851 | 41.138226 |
| AEAMEILLSR | 10 | Q86XZ4 | Spermatogenesis-associated serine-rich pr | SPATS2 | 59.545 | 2 | 51.83579 | yes | 1.376E-07 | 566.80518 | 4486.8589 |
| EVALLAEMDK | 10 | Q86XZ4 | Spermatogenesis-associated serine-rich pr | SPATS2 | 59.545 | 2 | 51.83579 | yes | 9.723E-08 | 559.79175 | 3505.8792 |
| YQSAPSQAPGNTII | 15 | Q86XZ4 | Spermatogenesis-associated serine-rich pr | SPATS2 | 59.545 | 2 | 51.83579 | yes | 1.065E-07 | 809.89465 | 2789.2947 |
| FTCDVETLK | 9 | Q86XZ4 | Spermatogenesis-associated serine-rich pr | SPATS2 | 59.545 | 2 | 51.83579 | yes | 9.9E-08 | 556.76825 | 2505.0715 |
| TVQAFMEGSASEV | 15 | Q86XZ4 | Spermatogenesis-associated serine-rich pr | SPATS2 | 59.545 | 2 | 51.83579 | yes | 1.211E-07 | 798.90051 | 2400.6682 |
| KLSSNIEK | 8 | Q86XZ4 | Spermatogenesis-associated serine-rich pr | SPATS2 | 59.545 | 2 | 51.83579 | yes | 1.495E-07 | 459.76639 | 2129.4731 |
| KYDEDLGR | 8 | Q86XZ4 | Spermatogenesis-associated serine-rich pr | SPATS2 | 59.545 | 2 | 51.83579 | yes | 3.294E-07 | 498.24326 | 1411.0834 |
| QAFAELESCLMDR | 13 | Q86XZ4 | Spermatogenesis-associated serine-rich pr | SPATS2 | 59.545 | 2 | 51.83579 | yes | 1.047E-07 | 785.35541 | 1274.7842 |
| SIDSFGQVSHPK | 12 | Q86XZ4 | Spermatogenesis-associated serine-rich pr | SPATS2 | 59.545 | 2 | 51.83579 | yes | 5.346E-07 | 651.32788 | 699.20331 |
| CTVSLAR | 7 | Q86XZ4 | Spermatogenesis-associated serine-rich pr | SPATS2 | 59.545 | 2 | 51.83579 | yes | 9.722E-08 | 403.71307 | 481.4942 |
| TGSMLQNGVSDFI | 15 | Q86XZ4 | Spermatogenesis-associated serine-rich pr | SPATS2 | 59.545 | 2 | 51.83579 | yes | 1.702E-07 | 807.37744 | 337.35559 |
| SNNEIILVLQHFDN | 18 | Q86XZ4 | Spermatogenesis-associated serine-rich pr | SPATS2 | 59.545 | 3 | 51.83579 | yes | 1.53E-07 | 720.02478 | 207.05554 |
| EWTVTGK | 7 | Q86XZ4 | Spermatogenesis-associated serine-rich pr | SPATS2 | 59.545 | 2 | 51.83579 | yes | 9.81E-08 | 410.71362 | 174.75052 |
| PAAEPSNGIPDSSK | 14 | Q86XZ4 | Spermatogenesis-associated serine-rich pr | SPATS2 | 59.545 | 2 | 51.83579 | yes | 1.127E-07 | 685.33331 | 96.098427 |
| SVSIQEEQSAPSSE | 15 | Q86XZ4 | Spermatogenesis-associated serine-rich pr | SPATS2 | 59.545 | 2 | 51.83579 | yes | 9.051E-07 | 803.38373 | 40.283195 |
| AVGEIVQDYDSDK | 13 | Q86YQ8 | Copine-8 OS=Homo sapiens OX=9606 G | CPNE8 | 63.108 | 2 | 44.520897 | yes | 2.164E-07 | 719.83844 | 86.005714 |
| GTDLTDEQIR | 10 | Q8IXT5 | RNA-binding protein 12B OS=Homo sap | RBM12B | 118.103 | 2 | 53.015568 | yes | 1.083E-07 | 574.28314 | 1332.871 |
| GLPYLVNEDDVR | 12 | Q8IXT5 | RNA-binding protein 12B OS=Homo sap | RBM12B | 118.103 | 2 | 53.015568 | yes | 1.086E-07 | 695.35406 | 1329.6858 |
| FLGTEVLLR | 9 | Q8IXT5 | RNA-binding protein 12B OS=Homo sap | RBM12B | 118.103 | 2 | 53.015568 | yes | 2.005E-06 | 524.31348 | 1223.5133 |
| WLPEEDFR | 8 | Q8IXT5 | RNA-binding protein 12B OS=Homo sap | RBM12B | 118.103 | 2 | 53.015568 | yes | 2.978E-07 | 546.26147 | 1159.8628 |
| NLSLSIDER | 9 | Q8IXT5 | RNA-binding protein 12B OS=Homo sap | RBM12B | 118.103 | 2 | 53.015568 | yes | 3.296E-07 | 523.77747 | 1032.0471 |
| DYNTALSLHK | 10 | Q8IXT5 | RNA-binding protein 12B OS=Homo sap | RBM12B | 118.103 | 2 | 53.015568 | yes | 6.471E-07 | 581.29858 | 894.30389 |
| HPPDEDFR | 8 | Q8IXT5 | RNA-binding protein 12B OS=Homo sap | RBM12B | 118.103 | 2 | 53.015568 | yes | 1.034E-06 | 506.72778 | 824.1521 |
| RPLEEDFR | 8 | Q8IXT5 | RNA-binding protein 12B OS=Homo sap | RBM12B | 118.103 | 2 | 53.015568 | yes | 1.604E-07 | 531.27234 | 758.23657 |
| QLPEEDFR | 8 | Q8IXT5 | RNA-binding protein 12B OS=Homo sap | RBM12B | 118.103 | 2 | 53.015568 | yes | 3.53E-07 | 517.2511 | 688.54883 |
| RPLEEDWR | 8 | Q8IXT5 | RNA-binding protein 12B OS=Homo sap | RBM12B | 118.103 | 2 | 53.015568 | yes | 1.866E-07 | 550.77783 | 588.48315 |
| DSSVELFLSSK | 11 | Q8IXT5 | RNA-binding protein 12B OS=Homo sap | RBM12B | 118.103 | 2 | 53.015568 | yes | 1.104E-07 | 606.31134 | 568.17828 |

| | | | | | | | | | | | |
|---|---|---|---|---|---|---|---|---|---|---|---|
| AENPYLFLR | 9 | Q8IXT5 | RNA-binding protein 12B OS=Homo sap | RBM12B | 118.103 | 2 | 53.015568 | yes | 1.465E-07 | 561.80072 | 515.38721 |
| RPPEDDFR | 8 | Q8IXT5 | RNA-binding protein 12B OS=Homo sap | RBM12B | 118.103 | 2 | 53.015568 | yes | 8.992E-07 | 516.2489 | 515.34912 |
| FASCVDASGGLK | 12 | Q8IXT5 | RNA-binding protein 12B OS=Homo sap | RBM12B | 118.103 | 2 | 53.015568 | yes | 9.722E-08 | 606.28992 | 449.36639 |
| DPPIYSVGAFENFF | 14 | Q8IXT5 | RNA-binding protein 12B OS=Homo sap | RBM12B | 118.103 | 2 | 53.015568 | yes | 9.722E-08 | 806.39374 | 447.31183 |
| RPSEEDFR | 8 | Q8IXT5 | RNA-binding protein 12B OS=Homo sap | RBM12B | 118.103 | 2 | 53.015568 | yes | 1.055E-07 | 518.24634 | 439.34021 |
| RPPEEDFR | 8 | Q8IXT5 | RNA-binding protein 12B OS=Homo sap | RBM12B | 118.103 | 2 | 53.015568 | yes | 8.715E-07 | 523.25671 | 437.81265 |
| RPPPEHFR | 8 | Q8IXT5 | RNA-binding protein 12B OS=Homo sap | RBM12B | 118.103 | 2 | 53.015568 | yes | 9.722E-08 | 518.27777 | 432.37695 |
| GVGLGEALVK | 10 | Q8IXT5 | RNA-binding protein 12B OS=Homo sap | RBM12B | 118.103 | 2 | 53.015568 | yes | 9.722E-08 | 471.78458 | 416.83511 |
| PVHIDPISR | 9 | Q8IXT5 | RNA-binding protein 12B OS=Homo sap | RBM12B | 118.103 | 2 | 53.015568 | yes | 9.89E-08 | 517.29309 | 389.85553 |
| RPWEEDFR | 8 | Q8IXT5 | RNA-binding protein 12B OS=Homo sap | RBM12B | 118.103 | 2 | 53.015568 | yes | 1.06E-07 | 567.76996 | 332.71924 |
| RPLQGEWR | 8 | Q8IXT5 | RNA-binding protein 12B OS=Homo sap | RBM12B | 118.103 | 2 | 53.015568 | yes | 9.722E-08 | 521.28308 | 271.30695 |
| ANVNEILDFFHGY | 14 | Q8IXT5 | RNA-binding protein 12B OS=Homo sap | RBM12B | 118.103 | 2 | 53.015568 | yes | 1.695E-07 | 847.91791 | 250.74725 |
| RPPEEDWR | 8 | Q8IXT5 | RNA-binding protein 12B OS=Homo sap | RBM12B | 118.103 | 2 | 53.015568 | yes | 1.081E-07 | 542.76215 | 248.62225 |
| HSSEDFR | 7 | Q8IXT5 | RNA-binding protein 12B OS=Homo sap | RBM12B | 118.103 | 2 | 53.015568 | yes | 3.411E-07 | 439.19357 | 233.33611 |
| HSPEDFR | 7 | Q8IXT5 | RNA-binding protein 12B OS=Homo sap | RBM12B | 118.103 | 2 | 53.015568 | yes | 9.79E-08 | 444.20395 | 198.36739 |
| HNMGSFPEGR | 10 | Q8IXT5 | RNA-binding protein 12B OS=Homo sap | RBM12B | 118.103 | 2 | 53.015568 | yes | 1.334E-07 | 566.25366 | 193.06479 |
| QPPEEDLR | 8 | Q8IXT5 | RNA-binding protein 12B OS=Homo sap | RBM12B | 118.103 | 2 | 53.015568 | yes | 5.571E-07 | 492.24326 | 179.713 |
| LPDNFRPPGEDFR | 13 | Q8IXT5 | RNA-binding protein 12B OS=Homo sap | RBM12B | 118.103 | 3 | 53.015568 | yes | 4.772E-07 | 520.59155 | 88.780388 |
| LPDNFRPPGEDFR | 13 | Q8IXT5 | RNA-binding protein 12B OS=Homo sap | RBM12B | 118.103 | 3 | 53.015568 | yes | 3.441E-07 | 780.38367 | 88.780388 |
| LPDNFRPPGEDFR | 13 | Q8IXT5 | RNA-binding protein 12B OS=Homo sap | RBM12B | 118.103 | 2 | 53.015568 | yes | 4.772E-07 | 520.59155 | 88.780388 |
| LPDNFRPPGEDFR | 13 | Q8IXT5 | RNA-binding protein 12B OS=Homo sap | RBM12B | 118.103 | 2 | 53.015568 | yes | 3.441E-07 | 780.38367 | 88.780388 |
| DLNDRPVGPR | 10 | Q8IXT5 | RNA-binding protein 12B OS=Homo sap | RBM12B | 118.103 | 2 | 53.015568 | yes | 2.049E-07 | 569.80182 | 167.0475 |
| GDHSHLFDSK | 10 | Q8IXT5 | RNA-binding protein 12B OS=Homo sap | RBM12B | 118.103 | 2 | 53.015568 | yes | 9.722E-08 | 571.76489 | 159.22183 |
| EAPEEDPRLPDNFI | 21 | Q8IXT5 | RNA-binding protein 12B OS=Homo sap | RBM12B | 118.103 | 4 | 53.015568 | yes | 2.151E-07 | 621.5451 | 128.88995 |
| SREEDFR | 7 | Q8IXT5 | RNA-binding protein 12B OS=Homo sap | RBM12B | 118.103 | 2 | 53.015568 | yes | 2.467E-06 | 469.71997 | 111.59142 |
| HSPEEDFR | 8 | Q8IXT5 | RNA-binding protein 12B OS=Homo sap | RBM12B | 118.103 | 2 | 53.015568 | yes | 1.857E-07 | 508.72525 | 108.33849 |
| RPPQEHFR | 8 | Q8IXT5 | RNA-binding protein 12B OS=Homo sap | RBM12B | 118.103 | 2 | 53.015568 | yes | 3.513E-07 | 533.7807 | 79.25827 |
| CPSDEDFR | 8 | Q8IXT5 | RNA-binding protein 12B OS=Homo sap | RBM12B | 118.103 | 2 | 53.015568 | yes | 8.608E-07 | 513.20331 | 24.601278 |
| AVANAVQQEVK | 11 | Q8IXX5 | Transmembrane protein 183A OS=Homo | TMEM183. | 42.827 | 2 | 50.860016 | yes | 1.823E-07 | 578.81964 | 102.15225 |
| VTVADYANSDPA' | 15 | Q8IXX5 | Transmembrane protein 183A OS=Homo | TMEM183. | 42.827 | 2 | 50.860016 | yes | 9.722E-08 | 788.90192 | 87.849594 |
| HYTLDASLPLR | 11 | Q8IXX5 | Transmembrane protein 183A OS=Homo | TMEM183. | 42.827 | 2 | 50.860016 | no | 1.156E-06 | 643.34857 | 69.305122 |
| CLLFWCR | 7 | Q8IXX5 | Transmembrane protein 183A OS=Homo | TMEM183. | 42.827 | 2 | 50.860016 | no | 1.592E-07 | 527.75171 | 52.302238 |
| LASAQEVAGSTSA | 14 | Q8IYL3 | UPF0688 protein C1orf174 OS=Homo sa | C1orf174 | 25.977 | 2 | 50.959755 | yes | 9.722E-08 | 660.34369 | 62.94471 |
| AFNESSNLTTHK | 12 | Q8IYN0 | Zinc finger protein 100 OS=Homo sapien | ZNF100 | 62.745 | 2 | 49.291805 | no | 1.048E-07 | 674.82825 | 20.154261 |
| ITPAGLQDLPCR | 12 | Q8N531 | F-box/LRR-repeat protein 6 OS=Homo sa | FBXL6 | 58.559 | 2 | 53.77962 | yes | 1.177E-07 | 670.85321 | 185.04701 |
| NSIPLQLPVEALQK | 14 | Q8N531 | F-box/LRR-repeat protein 6 OS=Homo sa | FBXL6 | 58.559 | 2 | 53.77962 | yes | 9.722E-08 | 775.45105 | 99.497681 |
| ELDLSGQGFSEK | 12 | Q8N531 | F-box/LRR-repeat protein 6 OS=Homo sa | FBXL6 | 58.559 | 2 | 53.77962 | yes | 9.722E-08 | 655.31714 | 99.365921 |
| GCPQLQVLR | 9 | Q8N531 | F-box/LRR-repeat protein 6 OS=Homo sa | FBXL6 | 58.559 | 2 | 53.77962 | yes | 3.419E-07 | 535.79242 | 95.073914 |
| LTLIHWK | 7 | Q8N531 | F-box/LRR-repeat protein 6 OS=Homo sa | FBXL6 | 58.559 | 2 | 53.77962 | yes | 1.371E-07 | 455.77908 | 52.070465 |

| | | | | | | | | | | | |
|---|---|---|---|---|---|---|---|---|---|---|---|
| GSGYHLLQSDSMI | 23 | Q8N531 | F-box/LRR-repeat protein 6 OS=Homo sa | FBXL6 | 58.559 | 3 | 53.77962 | yes | 2.446E-07 | 809.74939 | 45.600521 |
| VDPALGEFPFAK | 12 | Q8N567 | Zinc finger CCHC domain-containing pro | ZCCHC9 | 30.477 | 2 | 50.4978 | yes | 1.183E-07 | 645.84009 | 537.21136 |
| LCGSVEHLK | 9 | Q8N567 | Zinc finger CCHC domain-containing pro | ZCCHC9 | 30.477 | 2 | 50.4978 | yes | 4.963E-07 | 521.77112 | 416.27966 |
| CFVCGEMGHLSR | 12 | Q8N567 | Zinc finger CCHC domain-containing pro | ZCCHC9 | 30.477 | 2 | 50.4978 | yes | 9.795E-08 | 726.81268 | 372.14133 |
| NAMVCFHCR | 9 | Q8N567 | Zinc finger CCHC domain-containing pro | ZCCHC9 | 30.477 | 2 | 50.4978 | yes | 9.862E-08 | 597.75189 | 261.33609 |
| GLYADGGGCKLC( | 19 | Q8N567 | Zinc finger CCHC domain-containing pro | ZCCHC9 | 30.477 | 3 | 50.4978 | yes | 1.429E-07 | 674.32416 | 66.854813 |
| RPLPATSWEDMKI | 13 | Q8N567 | Zinc finger CCHC domain-containing pro | ZCCHC9 | 30.477 | 3 | 50.4978 | yes | 1.093E-06 | 520.27307 | 27.721157 |
| LIYPLLCR | 8 | Q8N5C6 | S1 RNA-binding domain-containing prot | SRBD1 | 111.776 | 2 | 55.31473 | yes | 2.895E-07 | 524.30463 | 836.22449 |
| LKVEEETSK | 9 | Q8N5C6 | S1 RNA-binding domain-containing prot | SRBD1 | 111.776 | 2 | 55.31473 | yes | 2.428E-07 | 531.78748 | 545.13666 |
| NAPQISDGSEVVV | 15 | Q8N5C6 | S1 RNA-binding domain-containing prot | SRBD1 | 111.776 | 2 | 55.31473 | yes | 1.796E-07 | 771.41211 | 474.72705 |
| FLSSIGGTLYEVGK | 20 | Q8N5C6 | S1 RNA-binding domain-containing prot | SRBD1 | 111.776 | 3 | 55.31473 | yes | 1.154E-07 | 738.04877 | 472.7525 |
| ELINNLDADSLR | 12 | Q8N5C6 | S1 RNA-binding domain-containing prot | SRBD1 | 111.776 | 2 | 55.31473 | yes | 1.191E-07 | 686.85699 | 459.85074 |
| EVQQTLEELR | 10 | Q8N5C6 | S1 RNA-binding domain-containing prot | SRBD1 | 111.776 | 2 | 55.31473 | yes | 9.979E-08 | 622.8277 | 451.32162 |
| VEVQVLNIDIPR | 12 | Q8N5C6 | S1 RNA-binding domain-containing prot | SRBD1 | 111.776 | 2 | 55.31473 | yes | 9.722E-08 | 697.90375 | 437.86545 |
| VQVQDVVLK | 9 | Q8N5C6 | S1 RNA-binding domain-containing prot | SRBD1 | 111.776 | 2 | 55.31473 | yes | 9.728E-08 | 514.31097 | 409.32098 |
| VQDPLAELVK | 10 | Q8N5C6 | S1 RNA-binding domain-containing prot | SRBD1 | 111.776 | 2 | 55.31473 | yes | 9.722E-08 | 556.32153 | 369.29947 |
| SFQQCAGFIR | 10 | Q8N5C6 | S1 RNA-binding domain-containing prot | SRBD1 | 111.776 | 2 | 55.31473 | yes | 9.722E-08 | 607.29279 | 362.17508 |
| DKDTLDFIR | 9 | Q8N5C6 | S1 RNA-binding domain-containing prot | SRBD1 | 111.776 | 2 | 55.31473 | yes | 3.543E-07 | 561.79309 | 348.83951 |
| TDFDKPDFK | 9 | Q8N5C6 | S1 RNA-binding domain-containing prot | SRBD1 | 111.776 | 2 | 55.31473 | yes | 3.445E-07 | 556.7666 | 320.12781 |
| QLGLEGAAR | 9 | Q8N5C6 | S1 RNA-binding domain-containing prot | SRBD1 | 111.776 | 2 | 55.31473 | yes | 1.922E-07 | 457.75635 | 318.34259 |
| RVQDPLAELVK | 11 | Q8N5C6 | S1 RNA-binding domain-containing prot | SRBD1 | 111.776 | 2 | 55.31473 | yes | 1.236E-07 | 634.37207 | 314.44272 |
| ILYNSLNDSFK | 11 | Q8N5C6 | S1 RNA-binding domain-containing prot | SRBD1 | 111.776 | 2 | 55.31473 | yes | 9.722E-08 | 657.34045 | 304.78351 |
| NIHHHQILAINR | 12 | Q8N5C6 | S1 RNA-binding domain-containing prot | SRBD1 | 111.776 | 3 | 55.31473 | yes | 9.926E-08 | 489.276 | 142.59409 |
| NIHHHQILAINR | 12 | Q8N5C6 | S1 RNA-binding domain-containing prot | SRBD1 | 111.776 | 3 | 55.31473 | yes | 9.722E-08 | 733.4104 | 142.59409 |
| NIHHHQILAINR | 12 | Q8N5C6 | S1 RNA-binding domain-containing prot | SRBD1 | 111.776 | 2 | 55.31473 | yes | 9.926E-08 | 489.276 | 142.59409 |
| NIHHHQILAINR | 12 | Q8N5C6 | S1 RNA-binding domain-containing prot | SRBD1 | 111.776 | 2 | 55.31473 | yes | 9.722E-08 | 733.4104 | 142.59409 |
| LDTVQTLK | 8 | Q8N5C6 | S1 RNA-binding domain-containing prot | SRBD1 | 111.776 | 2 | 55.31473 | yes | 9.722E-08 | 459.26874 | 276.40109 |
| TFEELEHVSAPYK | 13 | Q8N5C6 | S1 RNA-binding domain-containing prot | SRBD1 | 111.776 | 2 | 55.31473 | yes | 9.722E-08 | 775.38031 | 247.36258 |
| EMPGLDPNLR | 10 | Q8N5C6 | S1 RNA-binding domain-containing prot | SRBD1 | 111.776 | 2 | 55.31473 | yes | 9.809E-08 | 571.28717 | 230.82072 |
| HVCIQSSLAK | 10 | Q8N5C6 | S1 RNA-binding domain-containing prot | SRBD1 | 111.776 | 2 | 55.31473 | yes | 9.722E-08 | 571.80298 | 204.31276 |
| TLMGVDPGYK | 10 | Q8N5C6 | S1 RNA-binding domain-containing prot | SRBD1 | 111.776 | 2 | 55.31473 | yes | 2.5E-07 | 540.77332 | 204.17168 |
| ETEAYFADLIMK | 12 | Q8N5C6 | S1 RNA-binding domain-containing prot | SRBD1 | 111.776 | 2 | 55.31473 | yes | 9.722E-08 | 715.84723 | 172.50658 |
| FLLYQHFSCNIR | 12 | Q8N5C6 | S1 RNA-binding domain-containing prot | SRBD1 | 111.776 | 3 | 55.31473 | yes | 7.296E-07 | 533.26965 | 62.227371 |
| FLLYQHFSCNIR | 12 | Q8N5C6 | S1 RNA-binding domain-containing prot | SRBD1 | 111.776 | 3 | 55.31473 | yes | 3.687E-07 | 799.40082 | 62.227371 |
| FLLYQHFSCNIR | 12 | Q8N5C6 | S1 RNA-binding domain-containing prot | SRBD1 | 111.776 | 2 | 55.31473 | yes | 7.296E-07 | 533.26965 | 62.227371 |
| FLLYQHFSCNIR | 12 | Q8N5C6 | S1 RNA-binding domain-containing prot | SRBD1 | 111.776 | 2 | 55.31473 | yes | 3.687E-07 | 799.40082 | 62.227371 |
| QLLLTSPVPGR | 11 | Q8N5C6 | S1 RNA-binding domain-containing prot | SRBD1 | 111.776 | 2 | 55.31473 | yes | 1.92E-07 | 590.85608 | 117.54205 |
| LFNDDNTIPFIIR | 13 | Q8N5C6 | S1 RNA-binding domain-containing prot | SRBD1 | 111.776 | 2 | 55.31473 | yes | 9.722E-08 | 789.41974 | 114.39597 |
| NKLDTVQTLK | 10 | Q8N5C6 | S1 RNA-binding domain-containing prot | SRBD1 | 111.776 | 2 | 55.31473 | yes | 9.722E-08 | 580.33771 | 106.51885 |

| | | | | | | | | | | | |
|---|---|---|---|---|---|---|---|---|---|---|---|
| TETYPQGQPVK | 11 | Q8N5C6 | S1 RNA-binding domain-containing prot | SRBD1 | 111.776 | 2 | 55.31473 | yes | 3.948E-07 | 624.31696 | 82.277573 |
| TAVNVLLKPNPLD | 27 | Q8N5C6 | S1 RNA-binding domain-containing prot | SRBD1 | 111.776 | 4 | 55.31473 | yes | 4.006E-07 | 774.64679 | 37.443142 |
| ITLDLIR | 7 | Q8N5C6 | S1 RNA-binding domain-containing prot | SRBD1 | 111.776 | 2 | 55.31473 | yes | 5.532E-07 | 422.26855 | 34.859966 |
| LAIISPTSQILHTDV | 25 | Q8N5C6 | S1 RNA-binding domain-containing prot | SRBD1 | 111.776 | 4 | 55.31473 | yes | 4.239E-07 | 707.12463 | 29.631258 |
| ELLTAGSDGR | 10 | Q8N5I9 | NOP protein chaperone 1 OS=Homo sapi | NOPCHAP | 20.124 | 2 | 48.81326 | yes | 4.704E-07 | 509.76181 | 150.98537 |
| FNIENIDGPHSK | 12 | Q8N5I9 | NOP protein chaperone 1 OS=Homo sapi | NOPCHAP | 20.124 | 2 | 48.81326 | yes | 1.102E-07 | 685.83856 | 147.43076 |
| IEVLDSPASK | 10 | Q8N5I9 | NOP protein chaperone 1 OS=Homo sapi | NOPCHAP | 20.124 | 2 | 48.81326 | yes | 4.598E-07 | 529.79004 | 121.32195 |
| TSTLQTVR | 8 | Q8N5I9 | NOP protein chaperone 1 OS=Homo sapi | NOPCHAP | 20.124 | 2 | 48.81326 | yes | 2.875E-07 | 453.25616 | 64.627411 |
| SNYVPPPETYTTEI | 14 | Q8N6N3 | UPF0690 protein C1orf52 OS=Homo sap | C1orf52 | 20.599 | 2 | 37.273956 | yes | 6.139E-07 | 813.38831 | 60.462841 |
| LPGPDELFR | 9 | Q8N6N3 | UPF0690 protein C1orf52 OS=Homo sap | C1orf52 | 20.599 | 2 | 37.273956 | yes | 2.46E-06 | 522.27966 | 45.314873 |
| KPPPPELDMAIK | 12 | Q8N6N3 | UPF0690 protein C1orf52 OS=Homo sap | C1orf52 | 20.599 | 2 | 37.273956 | yes | 4.052E-06 | 668.37067 | 23.508343 |
| LQPLTQAAYVGEA | 14 | Q8N8D1 | Programmed cell death protein 7 OS=Hor | PDCD7 | 54.7 | 2 | 32.507107 | yes | 1.496E-06 | 758.90955 | 48.277287 |
| AQEEADYIEWLK | 12 | Q8N9T8 | Protein KRI1 homolog OS=Homo sapien | KRI1 | 82.599 | 2 | 52.927856 | yes | 1.512E-07 | 747.85919 | 281.07797 |
| EYWNDPELDEGEI | 13 | Q8N9T8 | Protein KRI1 homolog OS=Homo sapien | KRI1 | 82.599 | 2 | 52.927856 | yes | 9.722E-08 | 826.34717 | 145.23015 |
| LLGPTVMLGGCEF | 15 | Q8N9T8 | Protein KRI1 homolog OS=Homo sapien | KRI1 | 82.599 | 2 | 52.927856 | yes | 1.642E-07 | 818.9129 | 139.22498 |
| LDYEDIIDDLPCR | 13 | Q8N9T8 | Protein KRI1 homolog OS=Homo sapien | KRI1 | 82.599 | 2 | 52.927856 | yes | 9.722E-08 | 818.87982 | 123.69888 |
| VRPMYLK | 7 | Q8N9T8 | Protein KRI1 homolog OS=Homo sapien | KRI1 | 82.599 | 2 | 52.927856 | yes | 2.121E-07 | 453.76514 | 92.935982 |
| LQETSSQSYVEEQ | 14 | Q8N9T8 | Protein KRI1 homolog OS=Homo sapien | KRI1 | 82.599 | 2 | 52.927856 | yes | 1.006E-07 | 828.3916 | 83.372704 |
| EEAETPAEATGKP | 15 | Q8N9T8 | Protein KRI1 homolog OS=Homo sapien | KRI1 | 82.599 | 3 | 52.927856 | yes | 5.674E-06 | 538.59705 | 38.354809 |
| EEAETPAEATGKP | 15 | Q8N9T8 | Protein KRI1 homolog OS=Homo sapien | KRI1 | 82.599 | 3 | 52.927856 | yes | 6.28E-07 | 807.39191 | 38.354809 |
| EEAETPAEATGKP | 15 | Q8N9T8 | Protein KRI1 homolog OS=Homo sapien | KRI1 | 82.599 | 2 | 52.927856 | yes | 5.674E-06 | 538.59705 | 38.354809 |
| EEAETPAEATGKP | 15 | Q8N9T8 | Protein KRI1 homolog OS=Homo sapien | KRI1 | 82.599 | 2 | 52.927856 | yes | 6.28E-07 | 807.39191 | 38.354809 |
| LQAFGLNPK | 9 | Q8N9T8 | Protein KRI1 homolog OS=Homo sapien | KRI1 | 82.599 | 2 | 52.927856 | yes | 2.513E-07 | 494.28473 | 50.667652 |
| TLAETLVNLDVR | 12 | Q8NB46 | Serine/threonine-protein phosphatase 6 re | ANKRD52 | 115.077 | 2 | 52.39243 | yes | 1.236E-07 | 672.38007 | 413.41901 |
| ENINVLDQER | 10 | Q8NB46 | Serine/threonine-protein phosphatase 6 re | ANKRD52 | 115.077 | 2 | 52.39243 | yes | 2.054E-07 | 615.30969 | 264.88898 |
| TALFLATER | 9 | Q8NB46 | Serine/threonine-protein phosphatase 6 re | ANKRD52 | 115.077 | 2 | 52.39243 | yes | 1.368E-07 | 511.28748 | 176.63083 |
| FGNTPLHVAAR | 11 | Q8NB46 | Serine/threonine-protein phosphatase 6 re | ANKRD52 | 115.077 | 2 | 52.39243 | yes | 1.306E-07 | 591.82257 | 164.54173 |
| GSTADAADLR | 10 | Q8NB46 | Serine/threonine-protein phosphatase 6 re | ANKRD52 | 115.077 | 2 | 52.39243 | yes | 1.343E-07 | 488.73834 | 158.21233 |
| ADLTVLDENK | 10 | Q8NB46 | Serine/threonine-protein phosphatase 6 re | ANKRD52 | 115.077 | 2 | 52.39243 | yes | 2.023E-07 | 559.29041 | 145.27562 |
| SPLHMAAIHGR | 11 | Q8NB46 | Serine/threonine-protein phosphatase 6 re | ANKRD52 | 115.077 | 2 | 52.39243 | yes | 9.722E-08 | 595.31659 | 52.771416 |
| TPIHLASACGHTA | 16 | Q8NB46 | Serine/threonine-protein phosphatase 6 re | ANKRD52 | 115.077 | 3 | 52.39243 | yes | 3.145E-07 | 568.63849 | 51.239418 |
| PILSVQR | 7 | Q8NE00 | Transmembrane protein 104 OS=Homo s | TMEM104 | 55.679 | 2 | 38.83608 | yes | 4.956E-07 | 406.75308 | 219.38 |
| LHYDILEDR | 9 | Q8NEM2 | SHC SH2 domain-binding protein 1 OS= | SHCBP1 | 75.691 | 2 | 52.425503 | yes | 9.722E-08 | 587.29858 | 2075.2288 |
| LIENPLLR | 8 | Q8NEM2 | SHC SH2 domain-binding protein 1 OS= | SHCBP1 | 75.691 | 2 | 52.425503 | yes | 1.5E-07 | 484.30038 | 1850.1647 |
| AVWHTNVFK | 9 | Q8NEM2 | SHC SH2 domain-binding protein 1 OS= | SHCBP1 | 75.691 | 2 | 52.425503 | yes | 1.285E-07 | 551.29565 | 860.65094 |
| ASEVQEFTAEFLEI | 14 | Q8NEM2 | SHC SH2 domain-binding protein 1 OS= | SHCBP1 | 75.691 | 2 | 52.425503 | yes | 9.722E-08 | 814.39612 | 761.14484 |
| GDTFVDCTGADIK | 13 | Q8NEM2 | SHC SH2 domain-binding protein 1 OS= | SHCBP1 | 75.691 | 2 | 52.425503 | yes | 9.865E-08 | 699.8139 | 710.41797 |
| DFLDEHYDIPK | 11 | Q8NEM2 | SHC SH2 domain-binding protein 1 OS= | SHCBP1 | 75.691 | 2 | 52.425503 | yes | 1E-07 | 696.32751 | 629.09375 |
| AYQDYILADCK | 11 | Q8NEM2 | SHC SH2 domain-binding protein 1 OS= | SHCBP1 | 75.691 | 2 | 52.425503 | yes | 1.341E-07 | 680.3161 | 605.09869 |

| | | | | | | | | | | | |
|---|---|---|---|---|---|---|---|---|---|---|---|
| VLVEITDVDFAAL | 14 | Q8NEM2 | SHC SH2 domain-binding protein 1 OS= | SHCBP1 | 75.691 | 2 | 52.425503 | yes | 1.186E-07 | 766.93213 | 505.435 |
| FFYQNIWR | 8 | Q8NEM2 | SHC SH2 domain-binding protein 1 OS= | SHCBP1 | 75.691 | 2 | 52.425503 | yes | 2.234E-06 | 587.29565 | 389.17673 |
| VLEPSGWR | 8 | Q8NEM2 | SHC SH2 domain-binding protein 1 OS= | SHCBP1 | 75.691 | 2 | 52.425503 | yes | 1.625E-07 | 472.25345 | 371.69299 |
| ITHVVSSTMMAGL | 15 | Q8NEM2 | SHC SH2 domain-binding protein 1 OS= | SHCBP1 | 75.691 | 2 | 52.425503 | yes | 9.722E-08 | 808.43658 | 156.14655 |
| ITHVVSSTMMAGL | 15 | Q8NEM2 | SHC SH2 domain-binding protein 1 OS= | SHCBP1 | 75.691 | 2 | 52.425503 | yes | 1.26E-07 | 539.29346 | 156.14655 |
| ITHVVSSTMMAGL | 15 | Q8NEM2 | SHC SH2 domain-binding protein 1 OS= | SHCBP1 | 75.691 | 3 | 52.425503 | yes | 9.722E-08 | 808.43658 | 156.14655 |
| ITHVVSSTMMAGL | 15 | Q8NEM2 | SHC SH2 domain-binding protein 1 OS= | SHCBP1 | 75.691 | 3 | 52.425503 | yes | 1.26E-07 | 539.29346 | 156.14655 |
| IQTSGEPDVAER | 12 | Q8NEM2 | SHC SH2 domain-binding protein 1 OS= | SHCBP1 | 75.691 | 2 | 52.425503 | yes | 1.108E-07 | 651.32025 | 269.64853 |
| DKPGSSLQSFMPE | 15 | Q8NEM2 | SHC SH2 domain-binding protein 1 OS= | SHCBP1 | 75.691 | 2 | 52.425503 | yes | 9.722E-08 | 804.39032 | 243.0844 |
| LCQEPGEEER | 10 | Q8NEM2 | SHC SH2 domain-binding protein 1 OS= | SHCBP1 | 75.691 | 2 | 52.425503 | yes | 2.297E-07 | 623.77203 | 165.28993 |
| GKGDTFVDCTGAI | 15 | Q8NEM2 | SHC SH2 domain-binding protein 1 OS= | SHCBP1 | 75.691 | 2 | 52.425503 | yes | 1.076E-07 | 792.37213 | 113.28469 |
| ISMVNNIIHNNEG | 20 | Q8NEM2 | SHC SH2 domain-binding protein 1 OS= | SHCBP1 | 75.691 | 3 | 52.425503 | yes | 3.807E-07 | 738.39642 | 85.884163 |
| LAEPYLCDSQVST | 20 | Q8NEM2 | SHC SH2 domain-binding protein 1 OS= | SHCBP1 | 75.691 | 3 | 52.425503 | yes | 1.176E-07 | 804.01929 | 34.163208 |
| ADFQYQESQMR | 11 | Q8TAV0 | Protein FAM76A OS=Homo sapiens OX= | FAM76A | 35.05 | 2 | 47.78197 | yes | 1.26E-07 | 701.80646 | 50.660046 |
| TDCVMISTR | 9 | Q8TBR7 | TLC domain-containing protein 3A OS=I | TLCD3A | 29.383 | 2 | 41.63239 | yes | 3.414E-07 | 541.75208 | 86.146004 |
| LVSSVHAVLATGS | 19 | Q8TBR7 | TLC domain-containing protein 3A OS=I | TLCD3A | 29.383 | 3 | 41.63239 | yes | 7.83E-07 | 631.38245 | 38.023724 |
| TAVTTVPSMGIGL | 15 | Q8WUH6 | Transmembrane protein 263 OS=Homo sa | TMEM263 | 11.749 | 2 | 44.942158 | yes | 2.018E-07 | 737.4209 | 93.660889 |
| LWNETVELFR | 10 | Q8WUY9 | DEP domain-containing protein 1B OS=I | DEPDC1B | 61.772 | 2 | 50.658188 | yes | 2.12E-07 | 653.84314 | 120.83385 |
| SMTLSYLQK | 9 | Q8WUY9 | DEP domain-containing protein 1B OS=I | DEPDC1B | 61.772 | 2 | 50.658188 | yes | 6.88E-07 | 535.78119 | 100.1938 |
| VPLALQTSIEER | 12 | Q8WUY9 | DEP domain-containing protein 1B OS=I | DEPDC1B | 61.772 | 2 | 50.658188 | yes | 1.559E-07 | 678.38007 | 89.403168 |
| KLQLLMR | 7 | Q8WUY9 | DEP domain-containing protein 1B OS=I | DEPDC1B | 61.772 | 2 | 50.658188 | no | 1.588E-07 | 451.28622 | 78.898666 |
| SYPEVYQER | 9 | Q8WUY9 | DEP domain-containing protein 1B OS=I | DEPDC1B | 61.772 | 2 | 50.658188 | yes | 3.191E-07 | 585.7749 | 69.887344 |
| DEVDLDELLAAR | 12 | Q8WUY9 | DEP domain-containing protein 1B OS=I | DEPDC1B | 61.772 | 2 | 50.658188 | yes | 9.729E-08 | 679.84357 | 43.96653 |
| CSQNFGPEVTR | 11 | Q8WUY9 | DEP domain-containing protein 1B OS=I | DEPDC1B | 61.772 | 2 | 50.658188 | yes | 2.469E-06 | 647.7959 | 42.949959 |
| TPLDLACEFGR | 11 | Q8WXE0 | Caskin-2 OS=Homo sapiens OX=9606 G | CASKIN2 | 126.784 | 2 | 47.127293 | no | 1.395E-07 | 639.81097 | 70.701599 |
| AGDVITVLEQHPD | 15 | Q8WXE0 | Caskin-2 OS=Homo sapiens OX=9606 G | CASKIN2 | 126.784 | 2 | 47.127293 | yes | 2.635E-07 | 803.91284 | 62.9039 |
| VGLSPDSPAGDR | 12 | Q8WXE0 | Caskin-2 OS=Homo sapiens OX=9606 G | CASKIN2 | 126.784 | 2 | 47.127293 | yes | 3.111E-05 | 585.79114 | 25.920767 |
| TTIISSLPEFEK | 12 | Q92545 | Transmembrane protein 131 OS=Homo sa | TMEM131 | 205.138 | 2 | 42.736694 | yes | 2.895E-07 | 682.86902 | 220.49966 |
| IVGISSEGNLNTLS | 21 | Q92545 | Transmembrane protein 131 OS=Homo sa | TMEM131 | 205.138 | 3 | 42.736694 | yes | 4.463E-07 | 738.3587 | 127.48272 |
| IWSPTIGR | 8 | Q92545 | Transmembrane protein 131 OS=Homo sa | TMEM131 | 205.138 | 2 | 42.736694 | yes | 1.868E-06 | 465.26379 | 106.2011 |
| EAFVQSESIIEVLR | 14 | Q92545 | Transmembrane protein 131 OS=Homo sa | TMEM131 | 205.138 | 2 | 42.736694 | yes | 3.204E-07 | 810.43561 | 80.939644 |
| LVDNRPPALAK | 11 | Q92545 | Transmembrane protein 131 OS=Homo sa | TMEM131 | 205.138 | 2 | 42.736694 | yes | 1.49E-06 | 597.3537 | 46.903549 |
| TLEFQVFR | 8 | Q92545 | Transmembrane protein 131 OS=Homo sa | TMEM131 | 205.138 | 2 | 42.736694 | yes | 4.024E-07 | 520.28217 | 40.71418 |
| LSFEASNPPFDVGI | 19 | Q92545 | Transmembrane protein 131 OS=Homo sa | TMEM131 | 205.138 | 3 | 42.736694 | yes | 8.272E-07 | 722.03223 | 32.262608 |
| GSYSSIVNSSSSSD | 16 | Q92545 | Transmembrane protein 131 OS=Homo sa | TMEM131 | 205.138 | 2 | 42.736694 | yes | 3.788E-07 | 801.3681 | 30.20118 |
| LSPAPLAHPSHPEF | 14 | Q92545 | Transmembrane protein 131 OS=Homo sa | TMEM131 | 205.138 | 2 | 42.736694 | yes | 8.327E-07 | 754.90204 | 13.753619 |
| DMYLIPLGATDK | 12 | Q92576 | PHD finger protein 3 OS=Homo sapiens ( | PHF3 | 229.482 | 2 | 52.80293 | yes | 1.041E-07 | 668.84448 | 252.41176 |
| TGETVVEEMIATR | 13 | Q92576 | PHD finger protein 3 OS=Homo sapiens ( | PHF3 | 229.482 | 2 | 52.80293 | yes | 9.722E-08 | 718.35852 | 208.27789 |
| LMAQEQVR | 8 | Q92576 | PHD finger protein 3 OS=Homo sapiens ( | PHF3 | 229.482 | 2 | 52.80293 | yes | 4.597E-07 | 487.75803 | 198.37167 |

| | | | | | | | | | | | |
|---|---|---|---|---|---|---|---|---|---|---|---|
| ISPQTVWDYVEK | 12 | Q92576 | PHD finger protein 3 OS=Homo sapiens ( | PHF3 | 229.482 | 2 | 52.80293 | yes | 4.479E-07 | 732.87207 | 136.42291 |
| HSDPWGR | 7 | Q92576 | PHD finger protein 3 OS=Homo sapiens ( | PHF3 | 229.482 | 2 | 52.80293 | yes | 6.409E-07 | 427.69882 | 109.75436 |
| IEPLGYCEDAESNI | 14 | Q92576 | PHD finger protein 3 OS=Homo sapiens ( | PHF3 | 229.482 | 2 | 52.80293 | yes | 2.251E-07 | 826.86469 | 109.16747 |
| MSPEELASK | 9 | Q92576 | PHD finger protein 3 OS=Homo sapiens ( | PHF3 | 229.482 | 2 | 52.80293 | yes | 0.0001412 | 496.24188 | 107.20612 |
| NSQSSSVSYLESK | 13 | Q92576 | PHD finger protein 3 OS=Homo sapiens ( | PHF3 | 229.482 | 2 | 52.80293 | yes | 1.176E-07 | 708.33606 | 98.856674 |
| SLEKPEGSEK | 10 | Q92576 | PHD finger protein 3 OS=Homo sapiens ( | PHF3 | 229.482 | 2 | 52.80293 | yes | 2.761E-06 | 552.28259 | 83.66156 |
| EQEAAMEIQEPAA | 15 | Q92576 | PHD finger protein 3 OS=Homo sapiens ( | PHF3 | 229.482 | 2 | 52.80293 | yes | 9.722E-08 | 829.88812 | 75.880455 |
| NSCVQQSDNLK | 11 | Q92576 | PHD finger protein 3 OS=Homo sapiens ( | PHF3 | 229.482 | 2 | 52.80293 | yes | 1.12E-07 | 646.79858 | 73.738365 |
| ESTTVTCTGEK | 11 | Q92576 | PHD finger protein 3 OS=Homo sapiens ( | PHF3 | 229.482 | 2 | 52.80293 | yes | 1.982E-06 | 606.77429 | 54.075752 |
| EWEQESER | 8 | Q92576 | PHD finger protein 3 OS=Homo sapiens ( | PHF3 | 229.482 | 2 | 52.80293 | yes | 1.438E-07 | 546.73328 | 45.84948 |
| GVLNVHPAASASK | 20 | Q92576 | PHD finger protein 3 OS=Homo sapiens ( | PHF3 | 229.482 | 3 | 52.80293 | yes | 1.345E-07 | 673.36407 | 39.202744 |
| WQLAPLR | 7 | Q92576 | PHD finger protein 3 OS=Homo sapiens ( | PHF3 | 229.482 | 2 | 52.80293 | yes | 5.003E-06 | 442.26105 | 29.596613 |
| EHLTEQINVEEK | 12 | Q92576 | PHD finger protein 3 OS=Homo sapiens ( | PHF3 | 229.482 | 2 | 52.80293 | yes | 6.884E-06 | 734.86755 | 28.138918 |
| SPQFINLK | 8 | Q92576 | PHD finger protein 3 OS=Homo sapiens ( | PHF3 | 229.482 | 2 | 52.80293 | yes | 2.576E-07 | 473.77145 | 21.488443 |
| KNEALMECK | 9 | Q92576 | PHD finger protein 3 OS=Homo sapiens ( | PHF3 | 229.482 | 2 | 52.80293 | yes | 4.059E-07 | 561.76776 | 16.821442 |
| EFQGCISDFCDPR | 13 | Q93075 | Putative deoxyribonuclease TATDN2 OS | TATDN2 | 85.024 | 2 | 53.84413 | yes | 9.722E-08 | 815.83484 | 87.950035 |
| IIVETDAPYFLPR | 13 | Q93075 | Putative deoxyribonuclease TATDN2 OS | TATDN2 | 85.024 | 2 | 53.84413 | yes | 2.282E-07 | 767.41919 | 84.214188 |
| QLQLAVSLK | 9 | Q93075 | Putative deoxyribonuclease TATDN2 OS | TATDN2 | 85.024 | 2 | 53.84413 | yes | 3.86E-07 | 500.31351 | 76.558945 |
| LSFQGTFTK | 9 | Q93075 | Putative deoxyribonuclease TATDN2 OS | TATDN2 | 85.024 | 2 | 53.84413 | yes | 1.649E-07 | 514.77417 | 71.327866 |
| CTTPVPEQHK | 10 | Q93075 | Putative deoxyribonuclease TATDN2 OS | TATDN2 | 85.024 | 2 | 53.84413 | yes | 5.993E-07 | 598.79004 | 64.028419 |
| AVAFGEMGLDYS | 14 | Q93075 | Putative deoxyribonuclease TATDN2 OS | TATDN2 | 85.024 | 2 | 53.84413 | yes | 2.57E-07 | 775.8634 | 48.217243 |
| KPLVIHCR | 8 | Q93075 | Putative deoxyribonuclease TATDN2 OS | TATDN2 | 85.024 | 2 | 53.84413 | yes | 9.687E-07 | 511.80002 | 30.494465 |
| VSVSLVHPSEGIR | 13 | Q969Y0 | NXPE family member 3 OS=Homo sapie | NXPE3 | 63.8 | 2 | 41.616173 | yes | 3.422E-07 | 690.38568 | 50.537308 |
| KVVHLFGDSTIR | 12 | Q969Y0 | NXPE family member 3 OS=Homo sapie | NXPE3 | 63.8 | 3 | 41.616173 | yes | 4.376E-07 | 457.92963 | 37.875648 |
| VALEDDGFR | 9 | Q96C57 | Protein CUSTOS OS=Homo sapiens OX= | CUSTOS | 28.171 | 2 | 49.20064 | yes | 1.284E-07 | 511.2511 | 151.79944 |
| AGAANSQLSTSQP | 16 | Q96C57 | Protein CUSTOS OS=Homo sapiens OX= | CUSTOS | 28.171 | 2 | 49.20064 | yes | 1.183E-07 | 794.4079 | 117.94666 |
| LFFTSVPGGR | 10 | Q96C57 | Protein CUSTOS OS=Homo sapiens OX= | CUSTOS | 28.171 | 2 | 49.20064 | yes | 1.06E-07 | 540.79547 | 111.40918 |
| EAAVSASDILQESA | 23 | Q96C57 | Protein CUSTOS OS=Homo sapiens OX= | CUSTOS | 28.171 | 3 | 49.20064 | yes | 1.454E-07 | 780.39569 | 63.899086 |
| VEQATKPSFER | 11 | Q96E39 | RNA binding motif protein, X-linked-like | RBMXL1 | 42.142 | 2 | 53.999287 | yes | 9.722E-08 | 646.33569 | 106.14186 |
| IVEVLLIK | 8 | Q96E39 | RNA binding motif protein, X-linked-like | RBMXL1 | 42.142 | 2 | 53.999287 | yes | 1.341E-07 | 463.81787 | 104.19044 |
| SDLYSSCDR | 9 | Q96E39 | RNA binding motif protein, X-linked-like | RBMXL1 | 42.142 | 2 | 53.999287 | yes | 1.001E-07 | 551.72711 | 72.773148 |
| RPAEDDVPR | 9 | Q96EI5 | Transcription elongation factor A protein- | TCEAL4 | 24.648 | 2 | 51.57485 | yes | 4.95E-07 | 527.76746 | 125.1114 |
| GLAHYLK | 7 | Q96EI5 | Transcription elongation factor A protein- | TCEAL4 | 24.648 | 2 | 51.57485 | yes | 1.108E-07 | 401.23453 | 44.421101 |
| KPEVTCTLEDK | 11 | Q96EI5 | Transcription elongation factor A protein- | TCEAL4 | 24.648 | 2 | 51.57485 | yes | 9.722E-08 | 660.32922 | 36.722946 |
| LGAFLWMQR | 9 | Q96EI5 | Transcription elongation factor A protein- | TCEAL4 | 24.648 | 2 | 51.57485 | no | 2.749E-07 | 561.29987 | 33.803833 |
| LYSENEGMASNQ | 14 | Q96EI5 | Transcription elongation factor A protein- | TCEAL4 | 24.648 | 2 | 51.57485 | yes | 9.848E-08 | 764.34082 | 28.414242 |
| GNQLEICAVVLR | 12 | Q96EL2 | Small ribosomal subunit protein uS3m OS | MRPS24 | 19.016 | 2 | 52.343216 | yes | 9.722E-08 | 686.37427 | 840.28833 |
| LHLQTVPSK | 9 | Q96EL2 | Small ribosomal subunit protein uS3m OS | MRPS24 | 19.016 | 2 | 52.343216 | yes | 7.985E-07 | 511.80328 | 433.50305 |
| TVEDVFLR | 8 | Q96EL2 | Small ribosomal subunit protein uS3m OS | MRPS24 | 19.016 | 2 | 52.343216 | yes | 5.289E-07 | 489.76639 | 210.44284 |

| | | | | | | | | | | | |
|---|---|---|---|---|---|---|---|---|---|---|---|
| GWLSLHTGNLDGI | 19 | Q96EL2 | Small ribosomal subunit protein uS3m OS | MRPS24 | 19.016 | 3 | 52.343216 | yes | 1.128E-06 | 693.3277 | 29.247831 |
| SHWVFESR | 8 | Q96ET8 | Golgi apparatus membrane protein TVP2. | TVP23C | 31.105 | 2 | 48.239716 | yes | 1.175E-07 | 524.25397 | 71.848282 |
| WWNHIDEDGK | 10 | Q96ET8 | Golgi apparatus membrane protein TVP2. | TVP23C | 31.105 | 2 | 48.239716 | yes | 2.162E-06 | 650.29126 | 51.092503 |
| AALLEFEER | 9 | Q96EV2 | RNA-binding protein 33 OS=Homo sapie | RBM33 | 129.986 | 2 | 52.623398 | yes | 1.33E-07 | 539.28241 | 590.42249 |
| NQDVSISNVQPK | 12 | Q96EV2 | RNA-binding protein 33 OS=Homo sapie | RBM33 | 129.986 | 2 | 52.623398 | yes | 9.722E-08 | 664.84387 | 572.37146 |
| TVPQSQTQPLHK | 12 | Q96EV2 | RNA-binding protein 33 OS=Homo sapie | RBM33 | 129.986 | 2 | 52.623398 | yes | 1.263E-07 | 682.37006 | 349.45459 |
| SIQGIHPAK | 9 | Q96EV2 | RNA-binding protein 33 OS=Homo sapie | RBM33 | 129.986 | 2 | 52.623398 | yes | 1.013E-07 | 475.77454 | 218.91211 |
| VLPIKPADVEEPA' | 19 | Q96EV2 | RNA-binding protein 33 OS=Homo sapie | RBM33 | 129.986 | 3 | 52.623398 | yes | 2.206E-07 | 686.05273 | 200.85387 |
| TNSGGGDGPHISSI | 14 | Q96EV2 | RNA-binding protein 33 OS=Homo sapie | RBM33 | 129.986 | 2 | 52.623398 | yes | 9.722E-08 | 657.30768 | 192.46509 |
| VKPASPVAQPK | 11 | Q96EV2 | RNA-binding protein 33 OS=Homo sapie | RBM33 | 129.986 | 2 | 52.623398 | yes | 1.252E-07 | 561.33752 | 186.94722 |
| NIPETLELSAEAK | 13 | Q96EV2 | RNA-binding protein 33 OS=Homo sapie | RBM33 | 129.986 | 2 | 52.623398 | yes | 1.03E-07 | 707.87482 | 125.05231 |
| FKEPAHALAFQQK | 13 | Q96EV2 | RNA-binding protein 33 OS=Homo sapie | RBM33 | 129.986 | 3 | 52.623398 | yes | 6.36E-07 | 505.60867 | 84.442703 |
| LPQPPEVGPQPAR | 13 | Q96EV2 | RNA-binding protein 33 OS=Homo sapie | RBM33 | 129.986 | 2 | 52.623398 | yes | 4.243E-06 | 693.38043 | 64.621117 |
| NIHINPHFK | 9 | Q96EV2 | RNA-binding protein 33 OS=Homo sapie | RBM33 | 129.986 | 2 | 52.623398 | yes | 0.0003011 | 560.30652 | 62.321346 |
| TETEFPDEDEETR | 13 | Q96EV2 | RNA-binding protein 33 OS=Homo sapie | RBM33 | 129.986 | 2 | 52.623398 | yes | 1.063E-06 | 799.32867 | 23.377913 |
| ADFYCAEVAR | 10 | Q96F63 | Coiled-coil domain-containing protein 97 | CCDC97 | 38.947 | 2 | 44.03296 | yes | 2.348E-07 | 601.26898 | 72.521042 |
| SVSPLLSTHVLGK | 13 | Q96H12 | Myb/SANT-like DNA-binding domain-cc | MSANTD3 | 32.364 | 2 | 52.546085 | yes | 3.621E-06 | 669.39301 | 44.257248 |
| ECEMAEEEHR | 10 | Q96H12 | Myb/SANT-like DNA-binding domain-cc | MSANTD3 | 32.364 | 2 | 52.546085 | yes | 9.722E-08 | 660.25317 | 34.078472 |
| SILLALVEK | 9 | Q96H12 | Myb/SANT-like DNA-binding domain-cc | MSANTD3 | 32.364 | 2 | 52.546085 | yes | 1.509E-05 | 493.31824 | 10.945455 |
| LLVADQDFSFK | 11 | Q96HP4 | Oxidoreductase NAD-binding domain-co | OXNAD1 | 34.855 | 2 | 41.59582 | yes | 3.431E-07 | 641.83752 | 124.21304 |
| SSNLIIHQR | 9 | Q96JF6 | Zinc finger protein 594 OS=Homo sapien | ZNF594 | 93.907 | 2 | 50.773567 | yes | 9.722E-08 | 534.30145 | 282.70392 |
| IKPAATSHVEGSG | 18 | Q96ME7 | Zinc finger protein 512 OS=Homo sapien | ZNF512 | 64.683 | 3 | 49.725044 | yes | 1.007E-07 | 565.97235 | 149.53342 |
| VLQDLVPDDR | 10 | Q96ME7 | Zinc finger protein 512 OS=Homo sapien | ZNF512 | 64.683 | 2 | 49.725044 | yes | 1.604E-06 | 585.31171 | 147.09335 |
| AGLAYHLR | 8 | Q96ME7 | Zinc finger protein 512 OS=Homo sapien | ZNF512 | 64.683 | 2 | 49.725044 | yes | 3.925E-06 | 450.75613 | 68.889282 |
| EPEQEPVPAQFQK | 13 | Q96ME7 | Zinc finger protein 512 OS=Homo sapien | ZNF512 | 64.683 | 2 | 49.725044 | yes | 4.14E-07 | 763.87793 | 58.778721 |
| NNEELVVSASCK | 12 | Q96ME7 | Zinc finger protein 512 OS=Homo sapien | ZNF512 | 64.683 | 2 | 49.725044 | yes | 4.135E-07 | 675.3219 | 49.459015 |
| MDHLLVENIER | 11 | Q96MN5 | Transcription elongation factor A N-termi | TCEANC2 | 24.151 | 2 | 48.399323 | yes | 4.006E-07 | 684.85065 | 84.522247 |
| LLSEALELK | 9 | Q96MN5 | Transcription elongation factor A N-termi | TCEANC2 | 24.151 | 2 | 48.399323 | yes | 4.623E-07 | 508.30533 | 52.733997 |
| ENLVEALQELK | 11 | Q96MN5 | Transcription elongation factor A N-termi | TCEANC2 | 24.151 | 2 | 48.399323 | yes | 1.153E-07 | 643.35352 | 44.998474 |
| KDECYPVPANTSA | 19 | Q96N46 | Tetratricopeptide repeat protein 14 OS=H | TTC14 | 88.319 | 3 | 46.41054 | yes | 5.221E-07 | 726.68097 | 83.067169 |
| LGVISSEELPLYYR | 14 | Q96N46 | Tetratricopeptide repeat protein 14 OS=H | TTC14 | 88.319 | 2 | 46.41054 | yes | 3.196E-07 | 819.94049 | 28.623201 |
| SLLGSAAEPAR | 11 | Q96N46 | Tetratricopeptide repeat protein 14 OS=H | TTC14 | 88.319 | 2 | 46.41054 | yes | 2.916E-07 | 536.29327 | 23.315987 |
| LNTNQGEYER | 10 | Q96N46 | Tetratricopeptide repeat protein 14 OS=H | TTC14 | 88.319 | 2 | 46.41054 | yes | 1.566E-07 | 612.28619 | 21.684464 |
| SLQSLGTPSDTQEI | 15 | Q96NA8 | t-SNARE domain-containing protein 1 O: | TSNARE1 | 55.949 | 2 | 47.400223 | yes | 1.339E-07 | 816.41541 | 59.825275 |
| DGEGDLGPAGTPI | 16 | Q96NA8 | t-SNARE domain-containing protein 1 O: | TSNARE1 | 55.949 | 2 | 47.400223 | yes | 1.759E-06 | 775.8941 | 23.095037 |
| ISPQSNVDFDLMLI | 14 | Q96S19 | Methyltransferase-like 26 OS=Homo sapi | METTL26 | 22.579 | 2 | 35.13183 | yes | 8.81E-07 | 817.914 | 40.136288 |
| DVIVDFTQEEWK | 12 | Q99676 | Zinc finger protein 184 OS=Homo sapien | ZNF184 | 86.175 | 2 | 43.529083 | yes | 2.552E-07 | 754.867 | 47.541809 |
| LSTPAGVQVILR | 12 | Q9BQ48 | Large ribosomal subunit protein bL34m ( | MRPL34 | 10.165 | 2 | 49.52239 | yes | 1.022E-07 | 627.38245 | 919.48608 |
| GNEYQPSNIK | 10 | Q9BQ48 | Large ribosomal subunit protein bL34m ( | MRPL34 | 10.165 | 2 | 49.52239 | yes | 3.231E-07 | 575.2804 | 56.001991 |

| | | | | | | | | | | | |
|---|---|---|---|---|---|---|---|---|---|---|---|
| VVHLGVGTPGR | 11 | Q9BQ75 | Protein CMSS1 OS=Homo sapiens OX=9 | CMSS1 | 31.885 | 2 | 52.32419 | yes | 1.301E-07 | 546.31964 | 3455.6616 |
| MMDIPEIR | 8 | Q9BQ75 | Protein CMSS1 OS=Homo sapiens OX=9 | CMSS1 | 31.885 | 2 | 52.32419 | yes | 7.74E-07 | 502.74881 | 3395.2231 |
| ECFLIQPK | 8 | Q9BQ75 | Protein CMSS1 OS=Homo sapiens OX=9 | CMSS1 | 31.885 | 2 | 52.32419 | yes | 2.284E-07 | 517.77063 | 1640.3835 |
| SEPKPGLPEDLQK | 13 | Q9BQ75 | Protein CMSS1 OS=Homo sapiens OX=9 | CMSS1 | 31.885 | 2 | 52.32419 | yes | 5.057E-07 | 719.38281 | 1306.6553 |
| ANDLTHSLSSYLK | 13 | Q9BQ75 | Protein CMSS1 OS=Homo sapiens OX=9 | CMSS1 | 31.885 | 2 | 52.32419 | yes | 1.059E-07 | 724.87262 | 1169.7554 |
| QGGLNLSPLK | 10 | Q9BQ75 | Protein CMSS1 OS=Homo sapiens OX=9 | CMSS1 | 31.885 | 2 | 52.32419 | yes | 1.32E-07 | 513.80072 | 1068.4927 |
| KITDVLAK | 8 | Q9BQ75 | Protein CMSS1 OS=Homo sapiens OX=9 | CMSS1 | 31.885 | 2 | 52.32419 | yes | 9.722E-08 | 444.28165 | 279.25397 |
| PGLPEDLQK | 9 | Q9BQ75 | Protein CMSS1 OS=Homo sapiens OX=9 | CMSS1 | 31.885 | 2 | 52.32419 | yes | 7.018E-07 | 498.77167 | 223.349 |
| RVVHLGVGTPGR | 12 | Q9BQ75 | Protein CMSS1 OS=Homo sapiens OX=9 | CMSS1 | 31.885 | 3 | 52.32419 | yes | 2.578E-06 | 416.58255 | 17.850649 |
| NSPGSSQHPESPR | 13 | Q9BQE6 | LBH domain-containing protein 1 OS=Ho | LBHD1 | 31.566 | 3 | 48.289284 | yes | 1.167E-07 | 460.54813 | 101.84477 |
| EQIEGLIEDR | 10 | Q9BR77 | Coiled-coil domain-containing protein 77 | CCDC77 | 57.487 | 2 | 50.10399 | yes | 1.34E-07 | 601.30658 | 446.97583 |
| VFGLENELR | 9 | Q9BR77 | Coiled-coil domain-containing protein 77 | CCDC77 | 57.487 | 2 | 50.10399 | yes | 6.252E-07 | 538.79041 | 244.26486 |
| ALSDMQVCLFQEI | 13 | Q9BR77 | Coiled-coil domain-containing protein 77 | CCDC77 | 57.487 | 2 | 50.10399 | yes | 9.927E-08 | 798.87909 | 196.84874 |
| ILEVEGFK | 8 | Q9BR77 | Coiled-coil domain-containing protein 77 | CCDC77 | 57.487 | 2 | 50.10399 | yes | 3.013E-07 | 467.76584 | 174.21774 |
| LECDLQQR | 8 | Q9BR77 | Coiled-coil domain-containing protein 77 | CCDC77 | 57.487 | 2 | 50.10399 | yes | 2.689E-07 | 531.25586 | 103.76443 |
| EEEIAELQK | 9 | Q9BR77 | Coiled-coil domain-containing protein 77 | CCDC77 | 57.487 | 2 | 50.10399 | yes | 6.89E-06 | 544.77716 | 65.260506 |
| MAECEAENEDLLI | 13 | Q9BR77 | Coiled-coil domain-containing protein 77 | CCDC77 | 57.487 | 2 | 50.10399 | yes | 9.902E-08 | 776.33691 | 56.999947 |
| ANQDLALLCEVR | 12 | Q9BR77 | Coiled-coil domain-containing protein 77 | CCDC77 | 57.487 | 2 | 50.10399 | yes | 3.747E-07 | 701.36139 | 55.100533 |
| IVFLHNDR | 8 | Q9BSC4 | Nucleolar protein 10 OS=Homo sapiens ( | NOL10 | 80.302 | 2 | 49.608166 | yes | 2.83E-07 | 507.27997 | 153.84543 |
| SLPNILTDDR | 10 | Q9BSC4 | Nucleolar protein 10 OS=Homo sapiens ( | NOL10 | 80.302 | 2 | 49.608166 | yes | 1.018E-07 | 572.30383 | 153.40393 |
| LLEQQELR | 8 | Q9BSC4 | Nucleolar protein 10 OS=Homo sapiens ( | NOL10 | 80.302 | 2 | 49.608166 | yes | 3.824E-07 | 514.79041 | 132.35326 |
| IYSLSCGK | 8 | Q9BSC4 | Nucleolar protein 10 OS=Homo sapiens ( | NOL10 | 80.302 | 2 | 49.608166 | yes | 1.468E-07 | 464.23386 | 82.19709 |
| LMVNPFAYEEYR | 12 | Q9BSC4 | Nucleolar protein 10 OS=Homo sapiens ( | NOL10 | 80.302 | 2 | 49.608166 | yes | 1.23E-07 | 766.36615 | 62.103043 |
| SDKPLLVK | 8 | Q9BSC4 | Nucleolar protein 10 OS=Homo sapiens ( | NOL10 | 80.302 | 2 | 49.608166 | yes | 2.935E-07 | 450.28165 | 60.352028 |
| MGIYYIPVLGPAPI | 14 | Q9BSC4 | Nucleolar protein 10 OS=Homo sapiens ( | NOL10 | 80.302 | 2 | 49.608166 | yes | 3.258E-07 | 773.92615 | 45.483669 |
| LIEEEEEK | 8 | Q9BSC4 | Nucleolar protein 10 OS=Homo sapiens ( | NOL10 | 80.302 | 2 | 49.608166 | yes | 1.641E-07 | 509.75058 | 33.949883 |
| SILQEMER | 8 | Q9BTF0 | THUMP domain-containing protein 2 OS | THUMPD2 | 56.434 | 2 | 37.70119 | yes | 5.772E-07 | 503.25531 | 21.55411 |
| MSPFGSLVPVECY | 14 | Q9BTF0 | THUMP domain-containing protein 2 OS | THUMPD2 | 56.434 | 2 | 37.70119 | yes | 8.985E-05 | 807.38873 | 8.5126009 |
| LLPLTQYPDSSVR | 13 | Q9BTY7 | Protein HGH1 homolog OS=Homo sapie | HGH1 | 42.13 | 2 | 30.249231 | yes | 2.964E-06 | 744.90649 | 48.279419 |
| ALVNLAADPGLHI | 25 | Q9BTY7 | Protein HGH1 homolog OS=Homo sapie | HGH1 | 42.13 | 3 | 30.249231 | yes | 2.24E-06 | 832.45874 | 38.037369 |
| DQGAYLILR | 9 | Q9BTY7 | Protein HGH1 homolog OS=Homo sapie | HGH1 | 42.13 | 2 | 30.249231 | yes | 2.715E-06 | 524.79291 | 12.885743 |
| SAQTFLSEIR | 10 | Q9BWG4 | Single-stranded DNA-binding protein 4 C | SSBP4 | 39.389 | 2 | 51.749645 | no | 9.722E-08 | 576.3064 | 409.51349 |
| SSPGAVAGLSNAP | 17 | Q9BWG4 | Single-stranded DNA-binding protein 4 C | SSBP4 | 39.389 | 2 | 51.749645 | yes | 1.696E-07 | 769.89972 | 174.24789 |
| IEGVLLLGEPVR | 12 | Q9BXW7 | Haloacid dehalogenase-like hydrolase do | HDHD5 | 46.322 | 2 | 49.596436 | yes | 1.019E-07 | 647.89008 | 74.65815 |
| ALIAAQLDNAIEK | 13 | Q9BXY0 | Protein MAK16 homolog OS=Homo sapi | MAK16 | 35.369 | 2 | 51.37414 | yes | 9.728E-08 | 685.38794 | 232.20656 |
| NEYSLTGLCNR | 11 | Q9BXY0 | Protein MAK16 homolog OS=Homo sapi | MAK16 | 35.369 | 2 | 51.37414 | yes | 1.043E-07 | 663.80896 | 173.59511 |
| SSCPLANSQYATII | 14 | Q9BXY0 | Protein MAK16 homolog OS=Homo sapi | MAK16 | 35.369 | 2 | 51.37414 | yes | 9.722E-08 | 770.3772 | 108.93664 |
| MQSDDVIWDTLGI | 14 | Q9BXY0 | Protein MAK16 homolog OS=Homo sapi | MAK16 | 35.369 | 2 | 51.37414 | yes | 3.031E-07 | 832.38525 | 33.332787 |
| ITQYLIR | 7 | Q9BXY0 | Protein MAK16 homolog OS=Homo sapi | MAK16 | 35.369 | 2 | 51.37414 | yes | 4.147E-07 | 453.77402 | 21.348652 |

| | | | | | | | | | | | |
|---|---|---|---|---|---|---|---|---|---|---|---|
| ILATAGWDHR | 10 | Q9BYB4 | Guanine nucleotide-binding protein subu | GNB1L | 35.618 | 2 | 53.587593 | yes | 9.722E-08 | 570.30145 | 190.30922 |
| AVTTLDGHGGQC | 23 | Q9BYB4 | Guanine nucleotide-binding protein subu | GNB1L | 35.618 | 3 | 53.587593 | yes | 9.722E-08 | 832.42017 | 144.44806 |
| LCLWDLAEGR | 10 | Q9BYB4 | Guanine nucleotide-binding protein subu | GNB1L | 35.618 | 2 | 53.587593 | yes | 9.81E-08 | 616.80823 | 104.6618 |
| GSDEVQILEMPSK | 13 | Q9BYB4 | Guanine nucleotide-binding protein subu | GNB1L | 35.618 | 2 | 53.587593 | yes | 1.022E-07 | 716.85303 | 97.53804 |
| TSVCALKPK | 9 | Q9BYB4 | Guanine nucleotide-binding protein subu | GNB1L | 35.618 | 2 | 53.587593 | yes | 2.489E-07 | 502.28387 | 71.210022 |
| GTHELTNPGIAEV | 19 | Q9BYB4 | Guanine nucleotide-binding protein subu | GNB1L | 35.618 | 3 | 53.587593 | yes | 1.486E-07 | 692.69928 | 57.247059 |
| LWQADCSSR | 9 | Q9BYB4 | Guanine nucleotide-binding protein subu | GNB1L | 35.618 | 2 | 53.587593 | yes | 2.955E-07 | 561.75348 | 32.638016 |
| WTLAVPGR | 8 | Q9BYB4 | Guanine nucleotide-binding protein subu | GNB1L | 35.618 | 2 | 53.587593 | yes | 3.651E-05 | 450.25851 | 5.7554483 |
| VAVFTENASEVK | 12 | Q9BYD6 | Large ribosomal subunit protein uL1m O | MRPL1 | 36.909 | 2 | 53.85196 | yes | 9.996E-08 | 647.33789 | 730.17584 |
| FQILDFTSPK | 10 | Q9BYD6 | Large ribosomal subunit protein uL1m O | MRPL1 | 36.909 | 2 | 53.85196 | yes | 3.404E-07 | 598.32153 | 149.75798 |
| SSTSEGLLLK | 10 | Q9BYD6 | Large ribosomal subunit protein uL1m O | MRPL1 | 36.909 | 2 | 53.85196 | yes | 1.313E-07 | 517.79004 | 126.42082 |
| KFQILDFTSPK | 11 | Q9BYD6 | Large ribosomal subunit protein uL1m O | MRPL1 | 36.909 | 2 | 53.85196 | yes | 9.722E-08 | 662.36902 | 120.19321 |
| HRPLNLGPFVVR | 12 | Q9BYD6 | Large ribosomal subunit protein uL1m O | MRPL1 | 36.909 | 3 | 53.85196 | yes | 1.127E-07 | 468.94562 | 99.204872 |
| FLGSCLEDLAR | 11 | Q9C0B9 | Zinc finger CCHC domain-containing prc | ZCCHC2 | 125.936 | 2 | 52.715843 | yes | 9.722E-08 | 640.81879 | 766.20203 |
| VNWSDLSVTTVT | 13 | Q9C0B9 | Zinc finger CCHC domain-containing prc | ZCCHC2 | 125.936 | 2 | 52.715843 | yes | 1.138E-07 | 725.38281 | 215.88602 |
| LLPQVDSVLK | 10 | Q9C0B9 | Zinc finger CCHC domain-containing prc | ZCCHC2 | 125.936 | 2 | 52.715843 | yes | 1.125E-07 | 556.33972 | 155.21045 |
| TIGMLVPSPVAISA | 16 | Q9C0B9 | Zinc finger CCHC domain-containing prc | ZCCHC2 | 125.936 | 2 | 52.715843 | yes | 1.921E-07 | 812.97638 | 111.38457 |
| THQELQEFLLK | 11 | Q9C0B9 | Zinc finger CCHC domain-containing prc | ZCCHC2 | 125.936 | 2 | 52.715843 | yes | 1.15E-07 | 693.37482 | 75.750641 |
| LPLKPTHPAEPPPE | 27 | Q9C0B9 | Zinc finger CCHC domain-containing prc | ZCCHC2 | 125.936 | 4 | 52.715843 | yes | 2.037E-07 | 711.86859 | 53.397163 |
| VHSFFQSISSDSLH | 24 | Q9C0B9 | Zinc finger CCHC domain-containing prc | ZCCHC2 | 125.936 | 4 | 52.715843 | yes | 2.964E-07 | 669.59106 | 23.126665 |
| AVGHYNVAIPSDV | 17 | Q9GZU3 | Transmembrane protein 39B OS=Homo s | TMEM39B | 56.274 | 3 | 52.73117 | yes | 9.722E-08 | 623.65161 | 98.15789 |
| CYVPQNLGEALDI | 14 | Q9H568 | Actin-like protein 8 OS=Homo sapiens O | ACTL8 | 41.361 | 2 | 51.097305 | yes | 2.243E-06 | 841.40375 | 101.0305 |
| TVIIDHGSGFLK | 12 | Q9H568 | Actin-like protein 8 OS=Homo sapiens O | ACTL8 | 41.361 | 2 | 51.097305 | yes | 9.722E-08 | 643.85876 | 52.9296 |
| VELTPMQR | 8 | Q9H568 | Actin-like protein 8 OS=Homo sapiens O | ACTL8 | 41.361 | 2 | 51.097305 | yes | 3.776E-06 | 487.26041 | 52.310501 |
| ELMGDHVSSTK | 11 | Q9H568 | Actin-like protein 8 OS=Homo sapiens O | ACTL8 | 41.361 | 2 | 51.097305 | yes | 4.922E-07 | 602.28735 | 31.052353 |
| FLASSGHDQR | 10 | Q9H6Y2 | WD repeat-containing protein 55 OS=Ho | WDR55 | 42.071 | 2 | 55.707127 | yes | 9.722E-08 | 559.27289 | 333.75339 |
| VVGSVGQHTGEP | 23 | Q9H6Y2 | WD repeat-containing protein 55 OS=Ho | WDR55 | 42.071 | 4 | 55.707127 | yes | 9.728E-08 | 605.30121 | 84.136414 |
| VVGSVGQHTGEP | 23 | Q9H6Y2 | WD repeat-containing protein 55 OS=Ho | WDR55 | 42.071 | 4 | 55.707127 | yes | 1.09E-07 | 806.73254 | 84.136414 |
| VVGSVGQHTGEP | 23 | Q9H6Y2 | WD repeat-containing protein 55 OS=Ho | WDR55 | 42.071 | 3 | 55.707127 | yes | 9.728E-08 | 605.30121 | 84.136414 |
| VVGSVGQHTGEP | 23 | Q9H6Y2 | WD repeat-containing protein 55 OS=Ho | WDR55 | 42.071 | 3 | 55.707127 | yes | 1.09E-07 | 806.73254 | 84.136414 |
| FWDMAQLR | 8 | Q9H6Y2 | WD repeat-containing protein 55 OS=Ho | WDR55 | 42.071 | 2 | 55.707127 | yes | 9.733E-08 | 533.76056 | 146.73697 |
| IRDTPEDIVLEAPA | 23 | Q9H6Y2 | WD repeat-containing protein 55 OS=Ho | WDR55 | 42.071 | 3 | 55.707127 | yes | 9.722E-08 | 825.77118 | 131.77248 |
| AVVVDDYR | 8 | Q9H6Y2 | WD repeat-containing protein 55 OS=Ho | WDR55 | 42.071 | 2 | 55.707127 | yes | 2.69E-07 | 468.74289 | 105.46703 |
| AIHVLDVEQGQLE | 14 | Q9H6Y2 | WD repeat-containing protein 55 OS=Ho | WDR55 | 42.071 | 2 | 55.707127 | yes | 9.722E-08 | 803.93103 | 98.084412 |
| ELWSSGHHLK | 10 | Q9H6Y2 | WD repeat-containing protein 55 OS=Ho | WDR55 | 42.071 | 2 | 55.707127 | yes | 9.722E-08 | 597.3067 | 73.025063 |
| TWSTDDFFAGLR | 12 | Q9H6Y2 | WD repeat-containing protein 55 OS=Ho | WDR55 | 42.071 | 2 | 55.707127 | yes | 2.916E-07 | 708.33313 | 67.057915 |
| DILIEHYK | 8 | Q9H857 | 5'-nucleotidase domain-containing protei | NT5DC2 | 60.719 | 2 | 50.30636 | yes | 4.156E-07 | 515.78204 | 271.60843 |
| TGAIIPELER | 10 | Q9H857 | 5'-nucleotidase domain-containing protei | NT5DC2 | 60.719 | 2 | 50.30636 | yes | 9.838E-08 | 549.81134 | 175.884 |
| VDFTFYPR | 8 | Q9H857 | 5'-nucleotidase domain-containing protei | NT5DC2 | 60.719 | 2 | 50.30636 | yes | 7.3E-07 | 522.76111 | 70.230667 |

| | | | | | | | | | | | |
|---|---|---|---|---|---|---|---|---|---|---|---|
| GLMYQWIEQDME | 13 | Q9H857 | 5'-nucleotidase domain-containing protei | NT5DC2 | 60.719 | 2 | 50.30636 | yes | 9.819E-08 | 835.88129 | 60.058907 |
| KYDYNPSFAIR | 11 | Q9H857 | 5'-nucleotidase domain-containing protei | NT5DC2 | 60.719 | 2 | 50.30636 | yes | 3.189E-06 | 687.34607 | 28.302746 |
| IIHTGEKPYK | 10 | Q9H8G1 | Zinc finger protein 430 OS=Homo sapien | ZNF430 | 66.319 | 2 | 55.724957 | no | 9.722E-08 | 593.33496 | 245.72586 |
| DSFQEVILR | 9 | Q9H8G1 | Zinc finger protein 430 OS=Homo sapien | ZNF430 | 66.319 | 2 | 55.724957 | no | 2.046E-07 | 553.79565 | 80.377411 |
| GFNWSSTLTK | 10 | Q9H8G1 | Zinc finger protein 430 OS=Homo sapien | ZNF430 | 66.319 | 2 | 55.724957 | no | 4.235E-07 | 570.78784 | 77.165215 |
| VMLENYR | 7 | Q9H8G1 | Zinc finger protein 430 OS=Homo sapien | ZNF430 | 66.319 | 2 | 55.724957 | no | 2.284E-06 | 462.73401 | 23.750397 |
| LCQEQLELLSEK | 12 | Q9H8G2 | Caspase activity and apoptosis inhibitor 1 | CAAP1 | 38.368 | 2 | 50.749554 | yes | 9.723E-08 | 745.38196 | 71.718719 |
| NCSIEEIKK | 9 | Q9H8G2 | Caspase activity and apoptosis inhibitor 1 | CAAP1 | 38.368 | 2 | 50.749554 | yes | 1.013E-07 | 560.78699 | 34.353565 |
| STDSSSVSGSLQQI | 16 | Q9H8G2 | Caspase activity and apoptosis inhibitor 1 | CAAP1 | 38.368 | 2 | 50.749554 | yes | 1.065E-07 | 820.88409 | 25.835999 |
| AAVSWLTPAPSK | 12 | Q9H9L3 | Interferon-stimulated 20 kDa exonuclease | ISG20L2 | 39.154 | 2 | 50.872936 | yes | 6.828E-07 | 614.34021 | 757.17731 |
| ALQYFHPK | 8 | Q9H9L3 | Interferon-stimulated 20 kDa exonuclease | ISG20L2 | 39.154 | 2 | 50.872936 | yes | 2.232E-07 | 502.27164 | 402.48444 |
| GHVSSLAR | 8 | Q9H9L3 | Interferon-stimulated 20 kDa exonuclease | ISG20L2 | 39.154 | 2 | 50.872936 | yes | 1.454E-07 | 413.73013 | 309.68433 |
| MVAIDCEMVGTGI | 14 | Q9H9L3 | Interferon-stimulated 20 kDa exonuclease | ISG20L2 | 39.154 | 2 | 50.872936 | yes | 9.722E-08 | 754.35132 | 217.00072 |
| DTSHIPPLNR | 10 | Q9H9L3 | Interferon-stimulated 20 kDa exonuclease | ISG20L2 | 39.154 | 2 | 50.872936 | yes | 6.257E-07 | 575.3042 | 197.11359 |
| ADCPENATMSLK | 12 | Q9H9L3 | Interferon-stimulated 20 kDa exonuclease | ISG20L2 | 39.154 | 2 | 50.872936 | yes | 1.3E-07 | 668.79724 | 145.19775 |
| QHMVNATPFK | 10 | Q9H9L3 | Interferon-stimulated 20 kDa exonuclease | ISG20L2 | 39.154 | 2 | 50.872936 | yes | 1.328E-07 | 586.79767 | 115.10062 |
| KGETPTVDGTWK | 12 | Q9H9L3 | Interferon-stimulated 20 kDa exonuclease | ISG20L2 | 39.154 | 2 | 50.872936 | yes | 1.166E-07 | 659.83551 | 61.845345 |
| PSYLQTHEK | 9 | Q9HAH1 | Zinc finger protein 556 OS=Homo sapien | ZNF556 | 51.582 | 2 | 38.07047 | yes | 5.482E-07 | 551.78003 | 42.791862 |
| ASGSISQQDTSGEI | 14 | Q9HAH1 | Zinc finger protein 556 OS=Homo sapien | ZNF556 | 51.582 | 2 | 38.07047 | yes | 7.902E-07 | 697.82336 | 35.611214 |
| ESPSSTTPPIEISSSI | 16 | Q9HC44 | Vasculin-like protein 1 OS=Homo sapien | GPBP1L1 | 52.302 | 2 | 52.869293 | yes | 1.901E-07 | 837.91266 | 1208.3595 |
| PVPPPSKPNAWK | 12 | Q9HC44 | Vasculin-like protein 1 OS=Homo sapien | GPBP1L1 | 52.302 | 2 | 52.869293 | yes | 2.048E-07 | 659.36932 | 216.68883 |
| PVPPPSKPNAWK | 12 | Q9HC44 | Vasculin-like protein 1 OS=Homo sapien | GPBP1L1 | 52.302 | 2 | 52.869293 | yes | 2.165E-07 | 439.91531 | 216.68883 |
| PVPPPSKPNAWK | 12 | Q9HC44 | Vasculin-like protein 1 OS=Homo sapien | GPBP1L1 | 52.302 | 3 | 52.869293 | yes | 2.048E-07 | 659.36932 | 216.68883 |
| PVPPPSKPNAWK | 12 | Q9HC44 | Vasculin-like protein 1 OS=Homo sapien | GPBP1L1 | 52.302 | 3 | 52.869293 | yes | 2.165E-07 | 439.91531 | 216.68883 |
| LSSVVPSVYK | 10 | Q9HC44 | Vasculin-like protein 1 OS=Homo sapien | GPBP1L1 | 52.302 | 2 | 52.869293 | yes | 1.841E-07 | 539.81079 | 308.31592 |
| ESAFTSPISVTKPV | 26 | Q9HC44 | Vasculin-like protein 1 OS=Homo sapien | GPBP1L1 | 52.302 | 3 | 52.869293 | yes | 9.607E-07 | 848.80219 | 297.26807 |
| KGCAFQEKPPMEI | 14 | Q9HC44 | Vasculin-like protein 1 OS=Homo sapien | GPBP1L1 | 52.302 | 3 | 52.869293 | yes | 1.185E-07 | 564.28497 | 220.78261 |
| ESAFTSPISVTK | 12 | Q9HC44 | Vasculin-like protein 1 OS=Homo sapien | GPBP1L1 | 52.302 | 2 | 52.869293 | yes | 1.084E-07 | 633.83246 | 178.77565 |
| PVVLASGAALSSPI | 14 | Q9HC44 | Vasculin-like protein 1 OS=Homo sapien | GPBP1L1 | 52.302 | 2 | 52.869293 | yes | 9.722E-08 | 648.8797 | 73.726151 |
| GAYAGITGNPSGW | 18 | Q9HC44 | Vasculin-like protein 1 OS=Homo sapien | GPBP1L1 | 52.302 | 3 | 52.869293 | yes | 1.275E-07 | 602.28357 | 68.548096 |
| NLVPKPVPPPSKPI | 17 | Q9HC44 | Vasculin-like protein 1 OS=Homo sapien | GPBP1L1 | 52.302 | 3 | 52.869293 | yes | 1.782E-07 | 623.69635 | 28.404324 |
| PIGTPSGVWENPPS | 16 | Q9HC44 | Vasculin-like protein 1 OS=Homo sapien | GPBP1L1 | 52.302 | 2 | 52.869293 | yes | 6.413E-07 | 818.9201 | 27.903839 |
| HGEHLPR | 7 | Q9HC44 | Vasculin-like protein 1 OS=Homo sapien | GPBP1L1 | 52.302 | 2 | 52.869293 | yes | 7.917E-05 | 423.22247 | 19.907331 |
| STLAALIAAQR | 11 | Q9HCJ3 | Ribonucleoprotein PTB-binding 2 OS=H | RAVER2 | 74.339 | 2 | 51.481007 | yes | 1.07E-07 | 557.83258 | 172.60762 |
| GLLPEPNPVQIMK | 13 | Q9HCJ3 | Ribonucleoprotein PTB-binding 2 OS=H | RAVER2 | 74.339 | 2 | 51.481007 | yes | 2.202E-06 | 718.40253 | 162.12247 |
| VQVSFCAPGAPGR | 13 | Q9HCJ3 | Ribonucleoprotein PTB-binding 2 OS=H | RAVER2 | 74.339 | 2 | 51.481007 | yes | 9.722E-08 | 673.33771 | 87.904129 |
| GQGPSAEAHEGAI | 28 | Q9HCJ3 | Ribonucleoprotein PTB-binding 2 OS=H | RAVER2 | 74.339 | 4 | 51.481007 | yes | 1.908E-07 | 698.83185 | 84.004028 |
| GYGFVEYMK | 9 | Q9HCJ3 | Ribonucleoprotein PTB-binding 2 OS=H | RAVER2 | 74.339 | 2 | 51.481007 | yes | 3.065E-07 | 547.25476 | 81.085495 |
| TGIASSILDAISQGS | 23 | Q9HCJ3 | Ribonucleoprotein PTB-binding 2 OS=H | RAVER2 | 74.339 | 3 | 51.481007 | yes | 2.274E-07 | 781.39929 | 38.911083 |

| | | | | | | | | | | | |
|---|---|---|---|---|---|---|---|---|---|---|---|
| NLSGQPNFPCR | 11 | Q9HD45 | Transmembrane 9 superfamily member 3 | TM9SF3 | 67.889 | 2 | 37.36444 | yes | 6.058E-07 | 645.3064 | 26.04467 |
| SCNIPQTVENR | 11 | Q9NPI7 | Lysine-rich coiled-coil protein 1 OS=Hon | KRCC1 | 30.98 | 2 | 51.010506 | yes | 4.766E-06 | 659.31445 | 171.75066 |
| LPSETIQTYPR | 11 | Q9NPI7 | Lysine-rich coiled-coil protein 1 OS=Hon | KRCC1 | 30.98 | 2 | 51.010506 | yes | 9.722E-08 | 652.84589 | 63.613499 |
| LDSLSYCQFTR | 11 | Q9NPI7 | Lysine-rich coiled-coil protein 1 OS=Hon | KRCC1 | 30.98 | 2 | 51.010506 | yes | 1.194E-07 | 695.32703 | 50.55864 |
| LPQWLPAHDSR | 11 | Q9NPI7 | Lysine-rich coiled-coil protein 1 OS=Hon | KRCC1 | 30.98 | 2 | 51.010506 | yes | 2.299E-07 | 660.34637 | 42.890495 |
| YVNQMIINYVK | 11 | Q9NQX6 | Zinc finger protein 331 OS=Homo sapien | ZNF331 | 53.739 | 2 | 18.840052 | yes | 5.129E-06 | 692.86829 | 37.239258 |
| TDEAEPVGALLLE | 14 | Q9NW61 | Pleckstrin homology domain-containing f | PLEKHJ1 | 17.552 | 2 | 47.20726 | yes | 1.377E-07 | 756.89886 | 214.57098 |
| VYEIDITDASVVR | 13 | Q9NW82 | WD repeat-containing protein 70 OS=Ho | WDR70 | 73.202 | 2 | 49.281956 | yes | 1.049E-07 | 740.38812 | 81.41996 |
| FWDFAGMDASFK | 12 | Q9NW82 | WD repeat-containing protein 70 OS=Ho | WDR70 | 73.202 | 2 | 49.281956 | yes | 1.401E-06 | 711.31335 | 41.841022 |
| GEFMTCSNDATVI | 13 | Q9NW82 | WD repeat-containing protein 70 OS=Ho | WDR70 | 73.202 | 2 | 49.281956 | yes | 1.133E-07 | 744.31628 | 36.400982 |
| VLVDAGSALPR | 11 | Q9NX05 | Constitutive coactivator of PPAR-gamma | FAM120C | 120.577 | 2 | 52.598465 | yes | 1.061E-07 | 549.31934 | 818.81885 |
| SPQTPELVSALTFI | 14 | Q9NX05 | Constitutive coactivator of PPAR-gamma | FAM120C | 120.577 | 2 | 52.598465 | yes | 1.514E-07 | 773.41718 | 725.01947 |
| NLTTNQFLMQEV/ | 14 | Q9NX05 | Constitutive coactivator of PPAR-gamma | FAM120C | 120.577 | 2 | 52.598465 | yes | 9.742E-08 | 818.92181 | 517.77417 |
| LEIAGMVVGQWA | 15 | Q9NX05 | Constitutive coactivator of PPAR-gamma | FAM120C | 120.577 | 2 | 52.598465 | yes | 1.377E-07 | 787.41138 | 499.60605 |
| LPAPSQCALSR | 11 | Q9NX05 | Constitutive coactivator of PPAR-gamma | FAM120C | 120.577 | 2 | 52.598465 | yes | 9.722E-08 | 600.31372 | 467.29779 |
| CPGAVVPVDLLK | 12 | Q9NX05 | Constitutive coactivator of PPAR-gamma | FAM120C | 120.577 | 2 | 52.598465 | yes | 9.722E-08 | 634.35754 | 426.06671 |
| GVQGFQEFLEK | 11 | Q9NX05 | Constitutive coactivator of PPAR-gamma | FAM120C | 120.577 | 2 | 52.598465 | yes | 9.722E-08 | 641.32733 | 399.46414 |
| QTAQLIVGHVGNI | 13 | Q9NX05 | Constitutive coactivator of PPAR-gamma | FAM120C | 120.577 | 2 | 52.598465 | yes | 1.043E-07 | 682.88586 | 380.87933 |
| AVEYYSVTTK | 10 | Q9NX05 | Constitutive coactivator of PPAR-gamma | FAM120C | 120.577 | 2 | 52.598465 | yes | 1.294E-07 | 580.79535 | 361.82574 |
| LSSLPVGPSFLGFR | 14 | Q9NX05 | Constitutive coactivator of PPAR-gamma | FAM120C | 120.577 | 2 | 52.598465 | yes | 4.245E-07 | 738.91412 | 353.88727 |
| SLELHQGR | 8 | Q9NX05 | Constitutive coactivator of PPAR-gamma | FAM120C | 120.577 | 2 | 52.598465 | yes | 1.364E-07 | 470.25397 | 307.08356 |
| RLPVEVPSVILK | 12 | Q9NX05 | Constitutive coactivator of PPAR-gamma | FAM120C | 120.577 | 2 | 52.598465 | yes | 9.991E-08 | 675.42938 | 285.6543 |
| YMVQWPGGR | 9 | Q9NX05 | Constitutive coactivator of PPAR-gamma | FAM120C | 120.577 | 2 | 52.598465 | yes | 9.786E-08 | 547.26599 | 247.7829 |
| HELDTFLAQAVST | 20 | Q9NX05 | Constitutive coactivator of PPAR-gamma | FAM120C | 120.577 | 3 | 52.598465 | yes | 3.521E-07 | 778.38513 | 156.74022 |
| GFAGLHPIPPQGGI | 14 | Q9NX05 | Constitutive coactivator of PPAR-gamma | FAM120C | 120.577 | 2 | 52.598465 | yes | 4.337E-06 | 688.37769 | 135.0929 |
| AVSEYVSSIK | 10 | Q9NX05 | Constitutive coactivator of PPAR-gamma | FAM120C | 120.577 | 2 | 52.598465 | yes | 1.726E-07 | 541.79004 | 127.62154 |
| QLPPTAALAPGAP | 14 | Q9NX05 | Constitutive coactivator of PPAR-gamma | FAM120C | 120.577 | 2 | 52.598465 | yes | 1.464E-06 | 680.39081 | 118.25581 |
| PMFSHQVPQK | 10 | Q9NX05 | Constitutive coactivator of PPAR-gamma | FAM120C | 120.577 | 2 | 52.598465 | yes | 3.712E-07 | 599.80548 | 106.58685 |
| GSFGMQVVSVGGI | 15 | Q9NX05 | Constitutive coactivator of PPAR-gamma | FAM120C | 120.577 | 2 | 52.598465 | yes | 9.722E-08 | 703.85846 | 106.38069 |
| AMGSMPLPPQGR | 12 | Q9NX05 | Constitutive coactivator of PPAR-gamma | FAM120C | 120.577 | 2 | 52.598465 | yes | 4.369E-07 | 621.31012 | 68.747238 |
| DPSNLDVVGK | 10 | Q9NX05 | Constitutive coactivator of PPAR-gamma | FAM120C | 120.577 | 2 | 52.598465 | yes | 3.425E-07 | 522.27203 | 65.044571 |
| NQVGTISAGKPMF | 20 | Q9NX05 | Constitutive coactivator of PPAR-gamma | FAM120C | 120.577 | 3 | 52.598465 | yes | 3.422E-07 | 718.70898 | 61.018013 |
| KAVEYYSVTTK | 11 | Q9NX05 | Constitutive coactivator of PPAR-gamma | FAM120C | 120.577 | 2 | 52.598465 | yes | 9.982E-08 | 644.84283 | 23.734888 |
| TITGFQSLIQK | 11 | Q9NXD2 | Myotubularin-related protein 10 OS=Hon | MTMR10 | 88.274 | 2 | 48.425995 | yes | 2.333E-07 | 618.35333 | 96.042374 |
| TLPNIQEVQAAFV | 14 | Q9NXD2 | Myotubularin-related protein 10 OS=Hon | MTMR10 | 88.274 | 2 | 48.425995 | yes | 1.15E-07 | 779.43542 | 88.594696 |
| ISFITDDPMPLQK | 13 | Q9NXD2 | Myotubularin-related protein 10 OS=Hon | MTMR10 | 88.274 | 2 | 48.425995 | yes | 9.09E-07 | 752.8894 | 69.596481 |
| SGPLEACYGELGQ | 15 | Q9NXD2 | Myotubularin-related protein 10 OS=Hon | MTMR10 | 88.274 | 2 | 48.425995 | yes | 2.835E-07 | 812.37518 | 33.972595 |
| LLQPGGGPDVGTC | 16 | Q9NXH8 | Torsin-4A OS=Homo sapiens OX=9606 ( | TOR4A | 46.915 | 2 | 47.965744 | yes | 1.223E-07 | 746.39935 | 106.1228 |
| SAEAAAAQAEEDI | 14 | Q9NXH8 | Torsin-4A OS=Homo sapiens OX=9606 ( | TOR4A | 46.915 | 2 | 47.965744 | yes | 5.35E-07 | 716.33917 | 44.906078 |

| | | | | | | | | | | | |
|---|---|---|---|---|---|---|---|---|---|---|---|
| SFAENSSLIK | 10 | Q9NXT0 | Zinc finger protein 586 OS=Homo sapien | ZNF586 | 46.414 | 2 | 37.495842 | yes | 5.942E-07 | 548.28766 | 60.97506 |
| WSMVDVQFVR | 10 | Q9P016 | Thymocyte nuclear protein 1 OS=Homo s | THYN1 | 25.698 | 2 | 53.76923 | yes | 9.722E-08 | 633.8186 | 658.26984 |
| EPGIAGLMK | 9 | Q9P016 | Thymocyte nuclear protein 1 OS=Homo s | THYN1 | 25.698 | 2 | 53.76923 | yes | 1.3E-07 | 458.25204 | 297.24597 |
| EAYPDHTQFEK | 11 | Q9P016 | Thymocyte nuclear protein 1 OS=Homo s | THYN1 | 25.698 | 2 | 53.76923 | yes | 1.142E-07 | 682.80951 | 263.72192 |
| NLSSHWLMK | 9 | Q9P016 | Thymocyte nuclear protein 1 OS=Homo s | THYN1 | 25.698 | 2 | 53.76923 | yes | 9.722E-08 | 558.28693 | 258.64075 |
| TENSGEALAK | 10 | Q9P016 | Thymocyte nuclear protein 1 OS=Homo s | THYN1 | 25.698 | 2 | 53.76923 | yes | 2.303E-07 | 510.25381 | 150.59781 |
| LGEEAFFYHSNCK | 13 | Q9P016 | Thymocyte nuclear protein 1 OS=Homo s | THYN1 | 25.698 | 2 | 53.76923 | yes | 9.722E-08 | 801.35626 | 101.0488 |
| NNPHYDPSSK | 10 | Q9P016 | Thymocyte nuclear protein 1 OS=Homo s | THYN1 | 25.698 | 2 | 53.76923 | yes | 2.536E-07 | 579.76233 | 98.766098 |
| QTTCWDGVR | 9 | Q9P016 | Thymocyte nuclear protein 1 OS=Homo s | THYN1 | 25.698 | 2 | 53.76923 | yes | 3.284E-07 | 561.75348 | 78.245499 |
| SSYLLAITTER | 11 | Q9P227 | Rho GTPase-activating protein 23 OS=H( | ARHGAP2. | 162.192 | 2 | 52.22288 | yes | 9.722E-08 | 627.34045 | 106.42375 |
| TGLLHALSFR | 10 | Q9P227 | Rho GTPase-activating protein 23 OS=H( | ARHGAP2. | 162.192 | 2 | 52.22288 | yes | 6.796E-07 | 557.82202 | 57.685532 |
| TPWGINIIK | 9 | Q9P227 | Rho GTPase-activating protein 23 OS=H( | ARHGAP2. | 162.192 | 2 | 52.22288 | yes | 2.551E-07 | 521.30823 | 54.779068 |
| GPGDINLQDER | 11 | Q9P227 | Rho GTPase-activating protein 23 OS=H( | ARHGAP2. | 162.192 | 2 | 52.22288 | yes | 8.424E-07 | 607.29401 | 47.282421 |
| SAEALGPGALVSP. | 14 | Q9P227 | Rho GTPase-activating protein 23 OS=H( | ARHGAP2. | 162.192 | 2 | 52.22288 | yes | 3.478E-06 | 662.86462 | 41.510654 |
| AEGEDPGCANQAI | 16 | Q9P227 | Rho GTPase-activating protein 23 OS=H( | ARHGAP2. | 162.192 | 2 | 52.22288 | yes | 9.819E-08 | 830.3858 | 40.107853 |
| SIVSGYSTLSTMDI | 14 | Q9P227 | Rho GTPase-activating protein 23 OS=H( | ARHGAP2. | 162.192 | 2 | 52.22288 | yes | 1.337E-06 | 758.8692 | 21.549625 |
| SSSVLDSSLDHR | 12 | Q9P2B7 | Cilia- and flagella-associated protein 97 C | CFAP97 | 59.476 | 2 | 51.794655 | yes | 1.018E-07 | 651.81787 | 218.06462 |
| STLGQYSPLR | 10 | Q9P2B7 | Cilia- and flagella-associated protein 97 C | CFAP97 | 59.476 | 2 | 51.794655 | yes | 2.081E-07 | 561.30109 | 205.45195 |
| SAVDPSSGHPR | 11 | Q9P2B7 | Cilia- and flagella-associated protein 97 C | CFAP97 | 59.476 | 2 | 51.794655 | yes | 9.722E-08 | 555.27032 | 192.68311 |
| RLEAVKPTVGMK | 12 | Q9P2B7 | Cilia- and flagella-associated protein 97 C | CFAP97 | 59.476 | 3 | 51.794655 | yes | 1.128E-07 | 443.59546 | 187.56586 |
| SEQLMDYHR | 9 | Q9P2B7 | Cilia- and flagella-associated protein 97 C | CFAP97 | 59.476 | 2 | 51.794655 | yes | 1.395E-07 | 589.7666 | 143.92471 |
| IHVSIPNR | 8 | Q9P2B7 | Cilia- and flagella-associated protein 97 C | CFAP97 | 59.476 | 2 | 51.794655 | yes | 2.761E-07 | 468.27469 | 125.79228 |
| LYHSALNR | 8 | Q9P2B7 | Cilia- and flagella-associated protein 97 C | CFAP97 | 59.476 | 2 | 51.794655 | yes | 9.962E-07 | 487.26434 | 118.22163 |
| NMGYLNSSPLSR | 12 | Q9P2B7 | Cilia- and flagella-associated protein 97 C | CFAP97 | 59.476 | 2 | 51.794655 | yes | 1.222E-07 | 669.82715 | 113.6279 |
| EVCIPEIVSGSK | 12 | Q9P2F5 | Storkhead-box protein 2 OS=Homo sapie | STOX2 | 102.672 | 2 | 49.857273 | yes | 5.52E-07 | 659.3396 | 124.72176 |
| EYDFCDPLTR | 10 | Q9P2F5 | Storkhead-box protein 2 OS=Homo sapie | STOX2 | 102.672 | 2 | 49.857273 | yes | 1.001E-07 | 658.28479 | 67.588264 |
| EDVGTMQWLLER | 12 | Q9P2F5 | Storkhead-box protein 2 OS=Homo sapie | STOX2 | 102.672 | 2 | 49.857273 | yes | 4.965E-07 | 738.86121 | 63.918884 |
| INPDLTVENVMR | 12 | Q9P2F5 | Storkhead-box protein 2 OS=Homo sapie | STOX2 | 102.672 | 2 | 49.857273 | yes | 2.406E-06 | 700.86377 | 27.923153 |
| ELLDTELDLHK | 11 | Q9P2N5 | RNA-binding protein 27 OS=Homo sapie | RBM27 | 118.718 | 2 | 55.345356 | yes | 9.794E-08 | 663.35101 | 988.00836 |
| KIPQELNNITK | 11 | Q9P2N5 | RNA-binding protein 27 OS=Homo sapie | RBM27 | 118.718 | 2 | 55.345356 | yes | 1.869E-07 | 649.37738 | 927.36267 |
| AMSGLEGPLTK | 11 | Q9P2N5 | RNA-binding protein 27 OS=Homo sapie | RBM27 | 118.718 | 2 | 55.345356 | yes | 1.284E-07 | 552.29187 | 919.46783 |
| VVLEPDSR | 8 | Q9P2N5 | RNA-binding protein 27 OS=Homo sapie | RBM27 | 118.718 | 2 | 55.345356 | yes | 1.942E-07 | 457.75073 | 867.41034 |
| LSSGEDTTELR | 11 | Q9P2N5 | RNA-binding protein 27 OS=Homo sapie | RBM27 | 118.718 | 2 | 55.345356 | yes | 3.931E-07 | 604.2937 | 746.00592 |
| LSQLQVEAAR | 10 | Q9P2N5 | RNA-binding protein 27 OS=Homo sapie | RBM27 | 118.718 | 2 | 55.345356 | yes | 9.863E-08 | 557.81439 | 703.72858 |
| SHDVQEVLK | 9 | Q9P2N5 | RNA-binding protein 27 OS=Homo sapie | RBM27 | 118.718 | 2 | 55.345356 | yes | 2.628E-07 | 527.78003 | 613.29626 |
| LQLGTPPPLLAAR | 13 | Q9P2N5 | RNA-binding protein 27 OS=Homo sapie | RBM27 | 118.718 | 2 | 55.345356 | yes | 1.825E-07 | 673.91138 | 531.81049 |
| FGTIVNIQVAFK | 12 | Q9P2N5 | RNA-binding protein 27 OS=Homo sapie | RBM27 | 118.718 | 2 | 55.345356 | yes | 9.722E-08 | 668.88483 | 446.76801 |
| AFCADQLDVFLQK | 13 | Q9P2N5 | RNA-binding protein 27 OS=Homo sapie | RBM27 | 118.718 | 2 | 55.345356 | yes | 1.022E-07 | 777.8847 | 429.76642 |
| LNEHFSK | 7 | Q9P2N5 | RNA-binding protein 27 OS=Homo sapie | RBM27 | 118.718 | 2 | 55.345356 | yes | 2.814E-07 | 437.72452 | 418.02103 |

| | | | | | | | | | | |
|---|---|---|---|---|---|---|---|---|---|---|
| MIYSSSNLK | 9 | Q9P2N5 | RNA-binding protein 27 OS=Homo sapie | RBM27 | 118.718 | 2 | 55.345356 | yes | 4.528E-07 | 521.7655 | 307.95615 |
| NYLPLLEPVKPEPI | 14 | Q9P2N5 | RNA-binding protein 27 OS=Homo sapie | RBM27 | 118.718 | 2 | 55.345356 | yes | 1.4E-07 | 818.96906 | 137.2941 |
| NYLPLLEPVKPEPI | 14 | Q9P2N5 | RNA-binding protein 27 OS=Homo sapie | RBM27 | 118.718 | 2 | 55.345356 | yes | 0.0001773 | 546.31512 | 137.2941 |
| NYLPLLEPVKPEPI | 14 | Q9P2N5 | RNA-binding protein 27 OS=Homo sapie | RBM27 | 118.718 | 3 | 55.345356 | yes | 1.4E-07 | 818.96906 | 137.2941 |
| NYLPLLEPVKPEPI | 14 | Q9P2N5 | RNA-binding protein 27 OS=Homo sapie | RBM27 | 118.718 | 3 | 55.345356 | yes | 0.0001773 | 546.31512 | 137.2941 |
| LFESLYTK | 8 | Q9P2N5 | RNA-binding protein 27 OS=Homo sapie | RBM27 | 118.718 | 2 | 55.345356 | yes | 9.722E-08 | 500.77112 | 193.62645 |
| PQTSGAYVLNK | 11 | Q9P2N5 | RNA-binding protein 27 OS=Homo sapie | RBM27 | 118.718 | 2 | 55.345356 | yes | 1.166E-07 | 589.31421 | 175.74338 |
| MMSKPQTSGAYV | 15 | Q9P2N5 | RNA-binding protein 27 OS=Homo sapie | RBM27 | 118.718 | 3 | 55.345356 | yes | 1.5E-07 | 552.28125 | 57.187576 |
| MMSKPQTSGAYV | 15 | Q9P2N5 | RNA-binding protein 27 OS=Homo sapie | RBM27 | 118.718 | 3 | 55.345356 | yes | 2.48E-05 | 827.91821 | 57.187576 |
| MMSKPQTSGAYV | 15 | Q9P2N5 | RNA-binding protein 27 OS=Homo sapie | RBM27 | 118.718 | 2 | 55.345356 | yes | 1.5E-07 | 552.28125 | 57.187576 |
| MMSKPQTSGAYV | 15 | Q9P2N5 | RNA-binding protein 27 OS=Homo sapie | RBM27 | 118.718 | 2 | 55.345356 | yes | 2.48E-05 | 827.91821 | 57.187576 |
| DQPGTSAVPNLAS | 17 | Q9P2N5 | RNA-binding protein 27 OS=Homo sapie | RBM27 | 118.718 | 2 | 55.345356 | yes | 1.59E-07 | 835.42883 | 112.42033 |
| MLIEDVDALK | 10 | Q9P2N5 | RNA-binding protein 27 OS=Homo sapie | RBM27 | 118.718 | 2 | 55.345356 | yes | 2.189E-07 | 573.80737 | 107.9809 |
| LQISWHKPK | 9 | Q9P2N5 | RNA-binding protein 27 OS=Homo sapie | RBM27 | 118.718 | 2 | 55.345356 | yes | 1.262E-07 | 568.8324 | 107.09802 |
| QGNNNQNKPGFLI | 13 | Q9P2N5 | RNA-binding protein 27 OS=Homo sapie | RBM27 | 118.718 | 2 | 55.345356 | yes | 1.934E-07 | 743.87909 | 15.830462 |
| NAHFDYSPAGLAV | 15 | Q9UBI9 | Headcase protein homolog OS=Homo sap | HECA | 58.837 | 3 | 51.047623 | yes | 1.456E-07 | 552.27435 | 375.73453 |
| LNTFHVR | 7 | Q9UBI9 | Headcase protein homolog OS=Homo sap | HECA | 58.837 | 2 | 51.047623 | yes | 4.658E-07 | 443.74832 | 190.40182 |
| RLDLSELLTHIPR | 13 | Q9UBI9 | Headcase protein homolog OS=Homo sap | HECA | 58.837 | 3 | 51.047623 | yes | 2.886E-07 | 521.63861 | 187.17348 |
| NVVNCALCHR | 10 | Q9UBI9 | Headcase protein homolog OS=Homo sap | HECA | 58.837 | 2 | 51.047623 | yes | 9.722E-08 | 621.79517 | 174.29761 |
| NEAPCATPLICSFG | 15 | Q9UBI9 | Headcase protein homolog OS=Homo sap | HECA | 58.837 | 2 | 51.047623 | yes | 9.997E-08 | 846.89526 | 148.34483 |
| GPSHDLPR | 8 | Q9UBI9 | Headcase protein homolog OS=Homo sap | HECA | 58.837 | 2 | 51.047623 | yes | 1.175E-07 | 439.72757 | 142.98689 |
| LMHLYAVCVDCL | 17 | Q9UBI9 | Headcase protein homolog OS=Homo sap | HECA | 58.837 | 3 | 51.047623 | yes | 1.063E-07 | 681.9986 | 118.88002 |
| MEDDAQVGQGED | 14 | Q9UBI9 | Headcase protein homolog OS=Homo sap | HECA | 58.837 | 2 | 51.047623 | yes | 2.257E-06 | 781.84119 | 74.214767 |
| KGYDLAFR | 8 | Q9UBI9 | Headcase protein homolog OS=Homo sap | HECA | 58.837 | 2 | 51.047623 | yes | 2.959E-07 | 485.26126 | 59.29858 |
| GGHFDTPVQFLR | 12 | Q9UBI9 | Headcase protein homolog OS=Homo sap | HECA | 58.837 | 2 | 51.047623 | yes | 2.553E-07 | 687.35168 | 48.910393 |
| MEDDAQVGQGED | 15 | Q9UBI9 | Headcase protein homolog OS=Homo sap | HECA | 58.837 | 2 | 51.047623 | yes | 1.026E-07 | 845.88867 | 28.765078 |
| LVDEEPQLTK | 10 | Q9UG63 | ATP-binding cassette sub-family F memb | ABCF2 | 71.291 | 2 | 52.521976 | yes | 9.81E-08 | 586.3139 | 3072.6274 |
| MMASGLTER | 9 | Q9UG63 | ATP-binding cassette sub-family F memb | ABCF2 | 71.291 | 2 | 52.521976 | yes | 1.539E-07 | 498.23608 | 2730.8845 |
| ETTEVDLLTK | 10 | Q9UG63 | ATP-binding cassette sub-family F memb | ABCF2 | 71.291 | 2 | 52.521976 | yes | 1.027E-07 | 574.80591 | 2498.0867 |
| LAHEDAECEK | 10 | Q9UG63 | ATP-binding cassette sub-family F memb | ABCF2 | 71.291 | 2 | 52.521976 | yes | 3.461E-07 | 601.26135 | 2292.908 |
| LLTGELLPTDGMII | 14 | Q9UG63 | ATP-binding cassette sub-family F memb | ABCF2 | 71.291 | 2 | 52.521976 | yes | 1.15E-07 | 764.92383 | 1011.6622 |
| LLTGELLPTDGMII | 14 | Q9UG63 | ATP-binding cassette sub-family F memb | ABCF2 | 71.291 | 2 | 52.521976 | yes | 2.965E-07 | 772.92126 | 1011.6622 |
| WPGDILAYK | 9 | Q9UG63 | ATP-binding cassette sub-family F memb | ABCF2 | 71.291 | 2 | 52.521976 | yes | 2.321E-07 | 531.78455 | 1665.1483 |
| YYTGNYDQYVK | 11 | Q9UG63 | ATP-binding cassette sub-family F memb | ABCF2 | 71.291 | 2 | 52.521976 | yes | 1.114E-07 | 707.3197 | 1554.6368 |
| ILHGLGFTPAMQR | 13 | Q9UG63 | ATP-binding cassette sub-family F memb | ABCF2 | 71.291 | 2 | 52.521976 | yes | 1.429E-07 | 720.89264 | 947.71997 |
| LMELYER | 7 | Q9UG63 | ATP-binding cassette sub-family F memb | ABCF2 | 71.291 | 2 | 52.521976 | yes | 9.722E-08 | 477.24167 | 833.15179 |
| TPLHCVMEVDTEI | 13 | Q9UG63 | ATP-binding cassette sub-family F memb | ABCF2 | 71.291 | 2 | 52.521976 | yes | 1.164E-07 | 793.86871 | 517.88257 |
| LELEENQMK | 9 | Q9UG63 | ATP-binding cassette sub-family F memb | ABCF2 | 71.291 | 2 | 52.521976 | yes | 1.04E-07 | 567.27899 | 489.54355 |
| TLSFYFPPCGK | 11 | Q9UG63 | ATP-binding cassette sub-family F memb | ABCF2 | 71.291 | 2 | 52.521976 | yes | 1.507E-07 | 658.8208 | 468.58414 |

| | | | | | | | | | | |
|---|---|---|---|---|---|---|---|---|---|---|
| IPPPVIMVQNVSFK | 14 | Q9UG63 | ATP-binding cassette sub-family F memb | ABCF2 | 71.291 | 2 | 52.521976 | yes | 2.32E-07 | 784.94708 | 453.59125 |
| YGLIGLNGIGK | 11 | Q9UG63 | ATP-binding cassette sub-family F memb | ABCF2 | 71.291 | 2 | 52.521976 | yes | 9.722E-08 | 552.82422 | 391.7197 |
| ELEDFEMK | 8 | Q9UG63 | ATP-binding cassette sub-family F memb | ABCF2 | 71.291 | 2 | 52.521976 | yes | 1.241E-07 | 520.73389 | 362.87103 |
| LEELDADKAEMR | 12 | Q9UG63 | ATP-binding cassette sub-family F memb | ABCF2 | 71.291 | 2 | 52.521976 | yes | 4.86E-07 | 710.34283 | 329.61755 |
| TRLELEENQMK | 11 | Q9UG63 | ATP-binding cassette sub-family F memb | ABCF2 | 71.291 | 2 | 52.521976 | yes | 9.722E-08 | 695.85339 | 242.16599 |
| SMLLSAIGK | 9 | Q9UG63 | ATP-binding cassette sub-family F memb | ABCF2 | 71.291 | 2 | 52.521976 | yes | 9.722E-08 | 460.2677 | 240.74393 |
| LEELDADK | 8 | Q9UG63 | ATP-binding cassette sub-family F memb | ABCF2 | 71.291 | 2 | 52.521976 | yes | 2.616E-07 | 466.73221 | 201.19133 |
| FHWEQDQIAHMK | 12 | Q9UG63 | ATP-binding cassette sub-family F memb | ABCF2 | 71.291 | 2 | 52.521976 | yes | 9.722E-08 | 785.367 | 106.82305 |
| VALVGPNGAGK | 11 | Q9UG63 | ATP-binding cassette sub-family F memb | ABCF2 | 71.291 | 2 | 52.521976 | yes | 2.233E-07 | 491.78763 | 66.363892 |
| EVPIPEHIDIYHLTI | 15 | Q9UG63 | ATP-binding cassette sub-family F memb | ABCF2 | 71.291 | 4 | 52.521976 | yes | 2.564E-07 | 458.74799 | 38.796654 |
| VTISHAIGR | 9 | Q9UJJ7 | RNA pseudouridylate synthase domain-cc | RPUSD1 | 34.756 | 2 | 41.20331 | yes | 3.625E-07 | 477.27997 | 95.231499 |
| AFCQSSTLIR | 10 | Q9UJN7 | Zinc finger protein 391 OS=Homo sapien | ZNF391 | 40.66 | 2 | 49.763092 | yes | 1.004E-07 | 591.80042 | 111.84299 |
| STHLSLHQR | 9 | Q9UJN7 | Zinc finger protein 391 OS=Homo sapien | ZNF391 | 40.66 | 2 | 49.763092 | yes | 1.104E-07 | 539.79126 | 95.704979 |
| SSSLIIHQR | 9 | Q9UJN7 | Zinc finger protein 391 OS=Homo sapien | ZNF391 | 40.66 | 2 | 49.763092 | no | 2.113E-07 | 520.79602 | 89.080711 |
| NMPVQDLPR | 9 | Q9UJT0 | Tubulin epsilon chain OS=Homo sapiens | TUBE1 | 52.932 | 2 | 44.75801 | yes | 2.08E-07 | 535.27661 | 61.805126 |
| VLEDEFPEVYR | 11 | Q9UJT0 | Tubulin epsilon chain OS=Homo sapiens | TUBE1 | 52.932 | 2 | 44.75801 | yes | 6.194E-07 | 698.3432 | 31.374365 |
| VFGSLYQDQILEK | 13 | Q9UJT0 | Tubulin epsilon chain OS=Homo sapiens | TUBE1 | 52.932 | 2 | 44.75801 | yes | 1.886E-06 | 770.40631 | 29.465899 |
| LHIGEKPFK | 9 | Q9UL59 | Zinc finger protein 214 OS=Homo sapien | ZNF214 | 70.992 | 2 | 44.23539 | yes | 2.271E-07 | 534.81366 | 87.82222 |
| YKEELIEK | 8 | Q9UNX3 | Ribosomal protein uL24-like OS=Homo s | RPL26L1 | 17.257 | 2 | 46.372635 | yes | 1.577E-07 | 526.28711 | 1773.569 |
| HFNAPSHVR | 9 | Q9UNX3 | Ribosomal protein uL24-like OS=Homo s | RPL26L1 | 17.257 | 2 | 46.372635 | yes | 2.864E-07 | 532.77283 | 1040.3081 |
| LFSELPLAK | 9 | Q9Y291 | Small ribosomal subunit protein mS33 OS | MRPS33 | 12.629 | 2 | 51.141968 | yes | 1.007E-07 | 509.30258 | 186.55365 |
| SSLSEYAFR | 9 | Q9Y291 | Small ribosomal subunit protein mS33 OS | MRPS33 | 12.629 | 2 | 51.141968 | yes | 9.722E-08 | 551.26422 | 80.164696 |
| LFGEVTRPTNSK | 12 | Q9Y291 | Small ribosomal subunit protein mS33 OS | MRPS33 | 12.629 | 2 | 51.141968 | yes | 2.042E-07 | 674.86462 | 77.244194 |
| DEHQDFMDEQK | 11 | Q9Y291 | Small ribosomal subunit protein mS33 OS | MRPS33 | 12.629 | 2 | 51.141968 | yes | 3.546E-07 | 711.28552 | 23.062616 |
| GVVQLFNAVQK | 11 | Q9Y3B9 | RRP15-like protein OS=Homo sapiens O | RRP15 | 31.485 | 2 | 54.91815 | yes | 9.722E-08 | 601.84821 | 509.74164 |
| SEEGPGWTILR | 11 | Q9Y3B9 | RRP15-like protein OS=Homo sapiens O | RRP15 | 31.485 | 2 | 54.91815 | yes | 1.091E-07 | 622.81714 | 489.12384 |
| VSEEENLKK | 9 | Q9Y3B9 | RRP15-like protein OS=Homo sapiens O | RRP15 | 31.485 | 2 | 54.91815 | yes | 1.576E-07 | 538.28516 | 440.28171 |
| VKPDVVQDK | 9 | Q9Y3B9 | RRP15-like protein OS=Homo sapiens O | RRP15 | 31.485 | 2 | 54.91815 | yes | 3.898E-07 | 514.29279 | 336.021 |
| KDFISVLR | 8 | Q9Y3B9 | RRP15-like protein OS=Homo sapiens O | RRP15 | 31.485 | 2 | 54.91815 | yes | 9.722E-08 | 489.29257 | 214.65639 |
| GMDGSTNETASSR | 13 | Q9Y3B9 | RRP15-like protein OS=Homo sapiens O | RRP15 | 31.485 | 2 | 54.91815 | yes | 9.934E-08 | 656.77533 | 151.89282 |
| DDFMMGASMK | 10 | Q9Y3B9 | RRP15-like protein OS=Homo sapiens O | RRP15 | 31.485 | 2 | 54.91815 | yes | 3.354E-07 | 566.72723 | 63.303764 |
| KTPESKPTILVK | 12 | Q9Y3B9 | RRP15-like protein OS=Homo sapiens O | RRP15 | 31.485 | 2 | 54.91815 | yes | 1.518E-07 | 670.91101 | 34.372009 |
| LEWEMMCR | 8 | Q9Y3B9 | RRP15-like protein OS=Homo sapiens O | RRP15 | 31.485 | 2 | 54.91815 | yes | 9.722E-08 | 577.74323 | 19.102737 |
| AQIMGFMPVTYK | 12 | Q9Y3D5 | Small ribosomal subunit protein bS18m C | MRPS18C | 15.85 | 2 | 47.6767 | yes | 1.28E-07 | 693.35144 | 672.35419 |
| HITGLCGK | 8 | Q9Y3D5 | Small ribosomal subunit protein bS18m C | MRPS18C | 15.85 | 2 | 47.6767 | yes | 1.678E-06 | 443.23419 | 199.70349 |
| VLLFPPLSSR | 10 | Q9Y3T6 | R3H and coiled-coil domain-containing p | R3HCC1 | 49.092 | 2 | 53.594875 | yes | 9.722E-08 | 564.84241 | 138.32124 |
| FGSTLQLDLEK | 11 | Q9Y3T6 | R3H and coiled-coil domain-containing p | R3HCC1 | 49.092 | 2 | 53.594875 | yes | 2.613E-07 | 625.83496 | 108.54543 |
| TEDLLATFSEFQEK | 14 | Q9Y3T6 | R3H and coiled-coil domain-containing p | R3HCC1 | 49.092 | 2 | 53.594875 | yes | 9.722E-08 | 829.40143 | 70.289452 |
| VPSSDGLSGPCR | 12 | Q9Y3T6 | R3H and coiled-coil domain-containing p | R3HCC1 | 49.092 | 2 | 53.594875 | yes | 3.773E-07 | 616.29041 | 52.849022 |

| | | | | | | | | | | | |
|---|---|---|---|---|---|---|---|---|---|---|---|
| KPDQPLYVPR | 10 | Q9Y3T6 | R3H and coiled-coil domain-containing p | R3HCC1 | 49.092 | 2 | 53.594875 | yes | 0.0003266 | 606.84039 | 31.302505 |
| TVMPCGTVVTTV[illegible] | 16 | Q9Y426 | C2 domain-containing protein 2 OS=Hon | C2CD2 | 75.534 | 2 | 45.444614 | yes | 1.847E-07 | 832.44153 | 216.59863 |
| HLAGSASPSVVLI[illegible] | 15 | Q9Y426 | C2 domain-containing protein 2 OS=Hon | C2CD2 | 75.534 | 2 | 45.444614 | yes | 3.101E-06 | 740.43011 | 21.497227 |
| PFLPSSQSR | 9 | Q9Y485 | DmX-like protein 1 OS=Homo sapiens O | DMXL1 | 337.839 | 2 | 47.026638 | yes | 1.441E-07 | 509.76944 | 96.470589 |
| PGCIIALDPITK | 12 | Q9Y485 | DmX-like protein 1 OS=Homo sapiens O | DMXL1 | 337.839 | 2 | 47.026638 | yes | 1.418E-07 | 649.36285 | 21.628693 |
| LQTLATNLLASEA[illegible] | 16 | Q9Y4E6 | WD repeat-containing protein 7 OS=Hon | WDR7 | 163.811 | 2 | 34.008476 | yes | 1.1E-06 | 837.9491 | 29.274349 |
| SAFYVLEQQR | 10 | A0A087WUL8 | Neuroblastoma breakpoint family membe | NBPF19 | 440.409 | 2 | 47.532887 | yes | 1.31E-07 | 620.81964 | 80.557526 |
| STYYWPR | 7 | A1L429 | G antigen 12B/C/D/E OS=Homo sapiens | GAGE12B | 12.925 | 2 | 45.314674 | no | 2.652E-06 | 486.73233 | 1479.5133 |
| YVQPPEMIGPMR | 12 | A1L429 | G antigen 12B/C/D/E OS=Homo sapiens | GAGE12B | 12.925 | 2 | 45.314674 | yes | 1.887E-07 | 709.35199 | 134.12453 |
| GFLAGYVVAK | 10 | L0R6Q1 | SLC35A4 upstream open reading frame p | SLC35A4 | 11.133 | 2 | 39.188713 | yes | 4.735E-07 | 512.79492 | 88.833687 |
| IFEMLEGVQGPTA[illegible] | 15 | P0DMU7 | Cancer/testis antigen family 45 member A | CT45A6 | 21.232 | 2 | 53.17133 | yes | 9.722E-08 | 823.93219 | 113.94914 |
| LSAASGYSDVTDS | 14 | Q13670 | Putative postmeiotic segregation increase | PMS2P11 | 28.555 | 2 | 46.85033 | yes | 1.456E-07 | 700.83063 | 72.353851 |
| MLYAEFQR | 8 | Q2TB18 | Protein asteroid homolog 1 OS=Homo sa | ASTE1 | 77.094 | 2 | 54.11081 | yes | 1.259E-07 | 529.26038 | 172.14377 |
| QTILEPIPTSLK | 12 | Q2TB18 | Protein asteroid homolog 1 OS=Homo sa | ASTE1 | 77.094 | 2 | 54.11081 | yes | 9.722E-08 | 670.3952 | 146.82739 |
| SYAPAEIFLPK | 11 | Q2TB18 | Protein asteroid homolog 1 OS=Homo sa | ASTE1 | 77.094 | 2 | 54.11081 | yes | 1.453E-06 | 618.33716 | 93.422371 |
| QMLLLETLK | 9 | Q2TB18 | Protein asteroid homolog 1 OS=Homo sa | ASTE1 | 77.094 | 2 | 54.11081 | yes | 5.338E-07 | 544.82281 | 68.672104 |
| QLYEYLFNATR | 11 | Q2TB18 | Protein asteroid homolog 1 OS=Homo sa | ASTE1 | 77.094 | 2 | 54.11081 | yes | 2.124E-07 | 709.35913 | 54.332642 |
| TPTSGQSVSTCSS[illegible] | 14 | Q32NC0 | UPF0711 protein C18orf21 OS=Homo sa | C18orf21 | 24.827 | 2 | 43.68706 | yes | 2.487E-07 | 713.82758 | 20.989891 |
| GAEGLGTYCGLR | 12 | Q49AM3 | Tetratricopeptide repeat protein 31 OS=H | TTC31 | 57.105 | 2 | 50.17888 | yes | 1.889E-07 | 627.30078 | 180.00334 |
| LLVTEEEANR | 10 | Q49AM3 | Tetratricopeptide repeat protein 31 OS=H | TTC31 | 57.105 | 2 | 50.17888 | yes | 1.142E-07 | 587.30914 | 169.02654 |
| LAEELVAEEER | 11 | Q49AM3 | Tetratricopeptide repeat protein 31 OS=H | TTC31 | 57.105 | 2 | 50.17888 | yes | 9.862E-08 | 644.32501 | 136.79654 |
| EAAAVFQETLR | 11 | Q49AM3 | Tetratricopeptide repeat protein 31 OS=H | TTC31 | 57.105 | 2 | 50.17888 | yes | 3.989E-07 | 617.82495 | 133.86748 |
| VGDWPLSAR | 9 | Q49AM3 | Tetratricopeptide repeat protein 31 OS=H | TTC31 | 57.105 | 2 | 50.17888 | yes | 1.8E-06 | 500.76416 | 84.09491 |
| LVESDPQGLHR | 11 | Q49AM3 | Tetratricopeptide repeat protein 31 OS=H | TTC31 | 57.105 | 2 | 50.17888 | yes | 1.393E-07 | 625.828 | 65.335686 |
| SCLLHLTLQGQR | 12 | Q49AM3 | Tetratricopeptide repeat protein 31 OS=H | TTC31 | 57.105 | 2 | 50.17888 | yes | 3.06E-06 | 713.38519 | 8.6453066 |
| LQLWHHDYLLGH[illegible] | 18 | Q49AM3 | Tetratricopeptide repeat protein 31 OS=H | TTC31 | 57.105 | 4 | 50.17888 | yes | 2.307E-06 | 548.02484 | 7.4889836 |
| VAYMNPIAMAR | 11 | Q5BKY9 | Protein FAM133B OS=Homo sapiens O[illegible] | FAM133B | 28.385 | 2 | 46.673714 | yes | 9.096E-05 | 618.815 | 158.62769 |
| PTWEEVK | 7 | Q5BKY9 | Protein FAM133B OS=Homo sapiens O[illegible] | FAM133B | 28.385 | 2 | 46.673714 | yes | 1.499E-07 | 444.72672 | 140.36531 |
| LLDAVCLVPNK | 11 | Q5JUR7 | Testis-expressed protein 30 OS=Homo sa | TEX30 | 25.585 | 2 | 52.348995 | yes | 9.722E-08 | 621.34973 | 158.57596 |
| LRDEDLFR | 8 | Q5JUR7 | Testis-expressed protein 30 OS=Homo sa | TEX30 | 25.585 | 2 | 52.348995 | yes | 3.74E-07 | 532.28015 | 90.818298 |
| LQMEGVSVGQR | 11 | Q5R3I4 | Tetratricopeptide repeat protein 38 OS=H | TTC38 | 52.788 | 2 | 51.552242 | yes | 9.722E-08 | 602.31116 | 150.49422 |
| IVQLGGSNAQR | 11 | Q5R3I4 | Tetratricopeptide repeat protein 38 OS=H | TTC38 | 52.788 | 2 | 51.552242 | yes | 1.11E-07 | 571.81744 | 134.54962 |
| LPLSTTSNEACK | 12 | Q5R3I4 | Tetratricopeptide repeat protein 38 OS=H | TTC38 | 52.788 | 2 | 51.552242 | yes | 5.225E-07 | 660.82684 | 111.88899 |
| LFDATLTQYVK | 11 | Q5R3I4 | Tetratricopeptide repeat protein 38 OS=H | TTC38 | 52.788 | 2 | 51.552242 | yes | 9.722E-08 | 649.85315 | 90.409515 |
| SLGGIEGCLSK | 11 | Q5R3I4 | Tetratricopeptide repeat protein 38 OS=H | TTC38 | 52.788 | 2 | 51.552242 | yes | 2.983E-06 | 560.78699 | 88.680054 |
| DALKPNSPLTER | 12 | Q5R3I4 | Tetratricopeptide repeat protein 38 OS=H | TTC38 | 52.788 | 2 | 51.552242 | yes | 2.174E-06 | 670.86206 | 51.016689 |
| NSETVPEK | 8 | Q5RGS3 | Protein FAM74A1 OS=Homo sapiens O[illegible] | FAM74A1 | 14.779 | 2 | 23.043573 | yes | 3.73E-06 | 452.22455 | 175.85281 |
| EQLSICVR | 8 | Q5SVZ6 | Zinc finger MYM-type protein 1 OS=Hor | ZMYM1 | 128.717 | 2 | 49.254044 | yes | 1.052E-07 | 502.76331 | 73.757156 |
| VLSVTGILSK | 10 | Q5SVZ6 | Zinc finger MYM-type protein 1 OS=Hor | ZMYM1 | 128.717 | 2 | 49.254044 | yes | 3.553E-07 | 508.82114 | 35.661987 |

| | | | | | | | | | | | |
|---|---|---|---|---|---|---|---|---|---|---|---|
| LDEDIIPELR | 10 | Q5SVZ6 | Zinc finger MYM-type protein 1 OS=Hor | ZMYM1 | 128.717 | 2 | 49.254044 | yes | 2.719E-07 | 606.82715 | 30.275938 |
| DVISVQLEDTTSCI | 14 | Q5SVZ6 | Zinc finger MYM-type protein 1 OS=Hor | ZMYM1 | 128.717 | 2 | 49.254044 | yes | 7.81E-07 | 797.88507 | 29.101805 |
| WNEPLNETTAK | 11 | Q5SVZ6 | Zinc finger MYM-type protein 1 OS=Hor | ZMYM1 | 128.717 | 2 | 49.254044 | yes | 5.518E-05 | 651.81989 | 26.543846 |
| KEQVLER | 7 | Q5T6C5 | Ataxin-7-like protein 2 OS=Homo sapien | ATXN7L2 | 77.181 | 2 | 36.54462 | yes | 3.616E-06 | 451.25873 | 269.85074 |
| ADLPAADGAELEE | 16 | Q5T6C5 | Ataxin-7-like protein 2 OS=Homo sapien | ATXN7L2 | 77.181 | 2 | 36.54462 | yes | 6.87E-07 | 801.8783 | 121.63336 |
| APPPPPAPASSQK | 13 | Q5T6C5 | Ataxin-7-like protein 2 OS=Homo sapien | ATXN7L2 | 77.181 | 2 | 36.54462 | yes | 4.661E-06 | 622.83533 | 26.809584 |
| LLDGGSFTAESSR | 13 | Q5T8I3 | EEIG family member 2 OS=Homo sapien | EEIG2 | 39.309 | 2 | 52.2304 | yes | 9.722E-08 | 670.32806 | 111.01307 |
| VHLGIADLSAK | 11 | Q5T8I3 | EEIG family member 2 OS=Homo sapien | EEIG2 | 39.309 | 2 | 52.2304 | yes | 9.824E-08 | 562.32715 | 107.94229 |
| VDADDIVEK | 9 | Q5T8I3 | EEIG family member 2 OS=Homo sapien | EEIG2 | 39.309 | 2 | 52.2304 | yes | 4.301E-07 | 502.25076 | 60.88884 |
| VGSGAYEQVVIK | 12 | Q5T8I3 | EEIG family member 2 OS=Homo sapien | EEIG2 | 39.309 | 2 | 52.2304 | yes | 1.146E-05 | 625.34296 | 39.04863 |
| EVVQANCVR | 9 | Q5T8I3 | EEIG family member 2 OS=Homo sapien | EEIG2 | 39.309 | 2 | 52.2304 | yes | 1.499E-07 | 537.77167 | 20.359102 |
| VSGYSTCHSR | 10 | Q5T8I3 | EEIG family member 2 OS=Homo sapien | EEIG2 | 39.309 | 2 | 52.2304 | yes | 1.81E-07 | 577.25641 | 12.218544 |
| LQDLAAGPVGSLC | 14 | Q69YL0 | Protein NCBP2AS2 OS=Homo sapiens O | NCBP2AS2 | 10.891 | 2 | 50.295025 | yes | 9.819E-08 | 728.88245 | 139.46498 |
| LLAALLHSPQLVE | 14 | Q69YL0 | Protein NCBP2AS2 OS=Homo sapiens O | NCBP2AS2 | 10.891 | 3 | 50.295025 | yes | 7.991E-07 | 520.64709 | 18.10277 |
| LLAALLHSPQLVE | 14 | Q69YL0 | Protein NCBP2AS2 OS=Homo sapiens O | NCBP2AS2 | 10.891 | 3 | 50.295025 | yes | 0.0002091 | 780.46704 | 18.10277 |
| LLAALLHSPQLVE | 14 | Q69YL0 | Protein NCBP2AS2 OS=Homo sapiens O | NCBP2AS2 | 10.891 | 2 | 50.295025 | yes | 7.991E-07 | 520.64709 | 18.10277 |
| LLAALLHSPQLVE | 14 | Q69YL0 | Protein NCBP2AS2 OS=Homo sapiens O | NCBP2AS2 | 10.891 | 2 | 50.295025 | yes | 0.0002091 | 780.46704 | 18.10277 |
| TATYLPEVAK | 10 | Q6DCA0 | AMMECR1-like protein OS=Homo sapie | AMMECR1 | 34.5 | 2 | 52.90127 | no | 5.365E-07 | 546.80042 | 230.56297 |
| APITSEFR | 8 | Q6DCA0 | AMMECR1-like protein OS=Homo sapie | AMMECR1 | 34.5 | 2 | 52.90127 | yes | 3.08E-06 | 460.74545 | 121.32346 |
| VTISYAEYIASR | 12 | Q6DCA0 | AMMECR1-like protein OS=Homo sapie | AMMECR1 | 34.5 | 2 | 52.90127 | yes | 1.799E-07 | 686.85901 | 108.46899 |
| MNPASGALSPLPR | 13 | Q6DCA0 | AMMECR1-like protein OS=Homo sapie | AMMECR1 | 34.5 | 2 | 52.90127 | yes | 2.303E-06 | 655.8479 | 66.564789 |
| FTNDPYPLFVTWK | 13 | Q6DCA0 | AMMECR1-like protein OS=Homo sapie | AMMECR1 | 34.5 | 2 | 52.90127 | yes | 9.722E-08 | 814.41138 | 52.275364 |
| SELISIFQEAK | 11 | Q6P1M9 | Armadillo repeat-containing X-linked pro | ARMCX5 | 62.346 | 2 | 43.93132 | yes | 2.389E-07 | 632.84283 | 73.991112 |
| AEAGLQDGISGPA | 16 | Q6P1M9 | Armadillo repeat-containing X-linked pro | ARMCX5 | 62.346 | 2 | 43.93132 | yes | 5.175E-06 | 757.38391 | 38.581024 |
| QTQALADQFEDK | 12 | Q6P3X3 | Tetratricopeptide repeat protein 27 OS=H | TTC27 | 96.633 | 2 | 50.36941 | yes | 3.389E-07 | 697.33331 | 89.343689 |
| VTNDGEIWR | 9 | Q6P3X3 | Tetratricopeptide repeat protein 27 OS=H | TTC27 | 96.633 | 2 | 50.36941 | yes | 2.178E-07 | 545.26984 | 89.083649 |
| EFQECVECFER | 11 | Q6P3X3 | Tetratricopeptide repeat protein 27 OS=H | TTC27 | 96.633 | 2 | 50.36941 | yes | 9.81E-08 | 766.81085 | 86.750763 |
| AVIDGMTDR | 9 | Q6P3X3 | Tetratricopeptide repeat protein 27 OS=H | TTC27 | 96.633 | 2 | 50.36941 | yes | 2.212E-07 | 489.23965 | 68.885109 |
| QLFTDVATGEMSF | 13 | Q6P3X3 | Tetratricopeptide repeat protein 27 OS=H | TTC27 | 96.633 | 2 | 50.36941 | yes | 6.689E-06 | 727.85083 | 40.599724 |
| SSSQEAVQMLSSV | 14 | Q6P3X3 | Tetratricopeptide repeat protein 27 OS=H | TTC27 | 96.633 | 2 | 50.36941 | yes | 1.002E-07 | 754.87231 | 39.771297 |
| ALGLAHVAIK | 10 | Q6P3X3 | Tetratricopeptide repeat protein 27 OS=H | TTC27 | 96.633 | 2 | 50.36941 | yes | 2.617E-07 | 496.81619 | 38.935005 |
| NLELNDDTILNDIK | 14 | Q6P3X3 | Tetratricopeptide repeat protein 27 OS=H | TTC27 | 96.633 | 2 | 50.36941 | yes | 2.229E-07 | 815.42017 | 34.652885 |
| VRPLIIR | 7 | Q6P3X8 | PiggyBac transposable element-derived p | PGBD2 | 68.012 | 2 | 44.29153 | yes | 2.249E-07 | 433.80035 | 19.251831 |
| VVASDKETYELR | 12 | Q6P5R6 | Ribosomal protein eL22-like OS=Homo s | RPL22L1 | 14.607 | 2 | 51.447834 | yes | 9.722E-08 | 705.36719 | 187.79436 |
| TGNLGNVVHIER | 12 | Q6P5R6 | Ribosomal protein eL22-like OS=Homo s | RPL22L1 | 14.607 | 2 | 51.447834 | yes | 1.031E-07 | 654.85455 | 16.411037 |
| AQQLLDAVEQR | 11 | Q6ZSJ8 | Uncharacterized protein C1orf122 OS=H | C1orf122 | 11.471 | 2 | 53.712624 | yes | 9.934E-08 | 635.84113 | 473.52798 |
| QLLDTIAACEEML | 14 | Q6ZSJ8 | Uncharacterized protein C1orf122 OS=H | C1orf122 | 11.471 | 2 | 53.712624 | yes | 9.722E-08 | 831.91309 | 250.95348 |
| PGAPPQPAVSAR | 12 | Q6ZSJ8 | Uncharacterized protein C1orf122 OS=H | C1orf122 | 11.471 | 2 | 53.712624 | yes | 1E-07 | 574.31458 | 249.94408 |
| GLPLYPDPSR | 10 | Q6ZSR9 | Uncharacterized protein FLJ45252 OS=H | -- | 37.977 | 2 | 44.077007 | yes | 2.329E-07 | 557.79822 | 171.94524 |

| | | | | | | | | | | | |
|---|---|---|---|---|---|---|---|---|---|---|---|
| ITLWDANSEVEK | 12 | Q7Z5U6 | WD repeat-containing protein 53 OS=Ho | WDR53 | 38.99 | 2 | 46.624985 | yes | 1.511E-07 | 702.85388 | 63.561935 |
| AADQFDIYSSQQSI | 14 | Q7Z6I8 | UPF0461 protein C5orf24 OS=Homo sap | C5orf24 | 20.132 | 2 | 50.66953 | yes | 9.732E-08 | 794.36792 | 87.708031 |
| PSCLNEDAMR | 10 | Q7Z6I8 | UPF0461 protein C5orf24 OS=Homo sap | C5orf24 | 20.132 | 2 | 50.66953 | yes | 9.728E-08 | 596.75787 | 45.612656 |
| SVPMPLSNISVPK | 13 | Q86VQ1 | Glucocorticoid-induced transcript 1 prote | GLCCI1 | 58.024 | 2 | 44.06465 | yes | 2.335E-07 | 684.88141 | 56.700047 |
| SIDTQTPSVQER | 12 | Q86VQ1 | Glucocorticoid-induced transcript 1 prote | GLCCI1 | 58.024 | 2 | 44.06465 | yes | 2.72E-06 | 680.83881 | 28.209225 |
| ATQTPSCWAEEGA | 15 | Q86VQ1 | Glucocorticoid-induced transcript 1 prote | GLCCI1 | 58.024 | 2 | 44.06465 | yes | 1.568E-06 | 832.86469 | 26.857252 |
| VPCNVEGISPELEK | 14 | Q86VQ1 | Glucocorticoid-induced transcript 1 prote | GLCCI1 | 58.024 | 2 | 44.06465 | yes | 2.598E-06 | 785.8927 | 14.832313 |
| TLLLLQDTER | 10 | Q86X53 | Glutamate-rich protein 1 OS=Homo sapie | ERICH1 | 48.985 | 2 | 47.377632 | yes | 1.343E-07 | 601.34296 | 183.46201 |
| EEDGVDTIEEDLTI | 14 | Q86X53 | Glutamate-rich protein 1 OS=Homo sapie | ERICH1 | 48.985 | 2 | 47.377632 | yes | 1.587E-07 | 810.86542 | 88.015915 |
| NPNNVLIEQAELEI | 14 | Q86X53 | Glutamate-rich protein 1 OS=Homo sapie | ERICH1 | 48.985 | 2 | 47.377632 | yes | 1.527E-07 | 805.92285 | 22.380856 |
| EEADNPNTGIGAFI | 14 | Q8IVB5 | LIX1-like protein OS=Homo sapiens OX | LIX1L | 36.563 | 2 | 46.539795 | yes | 1.533E-07 | 745.84717 | 18.203917 |
| KAEAAVVAVAEK | 12 | Q8IZQ5 | Selenoprotein H OS=Homo sapiens OX= | SELENOH | 13.454 | 2 | 56.342667 | yes | 9.722E-08 | 593.34552 | 38.639462 |
| GEGLTPYQGK | 10 | Q8N2G6 | Zinc finger CCHC domain-containing prc | ZCCHC24 | 26.955 | 2 | 43.117493 | yes | 3.426E-07 | 525.26672 | 256.90002 |
| VLGYYCR | 7 | Q8N2G6 | Zinc finger CCHC domain-containing prc | ZCCHC24 | 26.955 | 2 | 43.117493 | yes | 2.724E-07 | 465.72873 | 239.82928 |
| GRPEQLGSPLHSSY | 23 | Q8N2G6 | Zinc finger CCHC domain-containing prc | ZCCHC24 | 26.955 | 4 | 43.117493 | yes | 3.102E-06 | 666.09375 | 22.639717 |
| TLPSVSETFPTLR | 13 | Q8NBP0 | Tetratricopeptide repeat protein 13 OS=H | TTC13 | 96.814 | 2 | 46.067295 | yes | 1.893E-07 | 724.39319 | 160.23711 |
| VMLNDPLPGQK | 11 | Q8NBP0 | Tetratricopeptide repeat protein 13 OS=H | TTC13 | 96.814 | 2 | 46.067295 | yes | 5.674E-07 | 606.32629 | 87.189911 |
| INEAVNDLTK | 10 | Q8NBP0 | Tetratricopeptide repeat protein 13 OS=H | TTC13 | 96.814 | 2 | 46.067295 | yes | 2.256E-07 | 558.7984 | 78.939804 |
| NQPIAMLYK | 9 | Q8NBP0 | Tetratricopeptide repeat protein 13 OS=H | TTC13 | 96.814 | 2 | 46.067295 | yes | 9.165E-07 | 539.29169 | 75.531357 |
| EYDGFTITITGDK | 13 | Q8NBP0 | Tetratricopeptide repeat protein 13 OS=H | TTC13 | 96.814 | 2 | 46.067295 | yes | 1.81E-07 | 730.3512 | 71.996895 |
| SMIEVLNTDSSPR | 13 | Q8NBP0 | Tetratricopeptide repeat protein 13 OS=H | TTC13 | 96.814 | 2 | 46.067295 | yes | 1.662E-07 | 724.85614 | 66.996429 |
| AEILSPLGR | 9 | Q8NBP0 | Tetratricopeptide repeat protein 13 OS=H | TTC13 | 96.814 | 2 | 46.067295 | yes | 6.393E-07 | 478.2822 | 58.349018 |
| VEDLLPIMK | 9 | Q8NBP0 | Tetratricopeptide repeat protein 13 OS=H | TTC13 | 96.814 | 2 | 46.067295 | yes | 2.018E-07 | 529.30176 | 57.072956 |
| GSSPCDSLLSLNTE | 15 | Q8NBP0 | Tetratricopeptide repeat protein 13 OS=H | TTC13 | 96.814 | 2 | 46.067295 | yes | 6.904E-07 | 804.38269 | 51.556229 |
| AMGLAALEVMQA | 15 | Q8NBP0 | Tetratricopeptide repeat protein 13 OS=H | TTC13 | 96.814 | 2 | 46.067295 | yes | 1.667E-05 | 794.4209 | 47.481853 |
| SIVDVGLAQAR | 11 | Q8NCJ5 | SPRY domain-containing protein 3 OS=I | SPRYD3 | 49.694 | 2 | 53.410698 | yes | 1.05E-07 | 564.8222 | 225.64236 |
| GDIMGCGIMFPR | 12 | Q8NCJ5 | SPRY domain-containing protein 3 OS=I | SPRYD3 | 49.694 | 2 | 53.410698 | yes | 1.525E-07 | 677.30927 | 107.72772 |
| VCGTLLEYLGK | 11 | Q8NCJ5 | SPRY domain-containing protein 3 OS=I | SPRYD3 | 49.694 | 2 | 53.410698 | yes | 1.011E-07 | 626.83392 | 99.747719 |
| DSNYFEVSIVDSGY | 15 | Q8NCJ5 | SPRY domain-containing protein 3 OS=I | SPRYD3 | 49.694 | 2 | 53.410698 | yes | 9.722E-08 | 843.9021 | 90.753059 |
| SHYFEVEIVDPGEI | 14 | Q8NCJ5 | SPRY domain-containing protein 3 OS=I | SPRYD3 | 49.694 | 2 | 53.410698 | yes | 2.123E-07 | 824.8963 | 45.795807 |
| MDDLNLHYR | 9 | Q8NCJ5 | SPRY domain-containing protein 3 OS=I | SPRYD3 | 49.694 | 2 | 53.410698 | yes | 2.505E-06 | 588.77698 | 29.745516 |
| CYIALGLAR | 9 | Q8NCJ5 | SPRY domain-containing protein 3 OS=I | SPRYD3 | 49.694 | 2 | 53.410698 | yes | 1.883E-06 | 518.78406 | 19.09318 |
| SDEGLPDGLSTK | 12 | Q8ND24 | RING finger protein 214 OS=Homo sapie | RNF214 | 77.668 | 2 | 51.538464 | yes | 1.171E-06 | 609.79608 | 253.1342 |
| AEILSLESR | 9 | Q8ND24 | RING finger protein 214 OS=Homo sapie | RNF214 | 77.668 | 2 | 51.538464 | yes | 2.76E-07 | 509.28238 | 178.08351 |
| AQMTNILQQIK | 11 | Q8ND24 | RING finger protein 214 OS=Homo sapie | RNF214 | 77.668 | 2 | 51.538464 | yes | 9.722E-08 | 644.35809 | 132.70049 |
| LSQNIAVQTDFK | 12 | Q8ND24 | RING finger protein 214 OS=Homo sapie | RNF214 | 77.668 | 2 | 51.538464 | yes | 1.018E-07 | 682.36444 | 88.226616 |
| NSPLLSVSSQTITK | 14 | Q8ND24 | RING finger protein 214 OS=Homo sapie | RNF214 | 77.668 | 2 | 51.538464 | yes | 9.819E-08 | 737.90924 | 87.073013 |
| LCLMCQK | 7 | Q8ND24 | RING finger protein 214 OS=Homo sapie | RNF214 | 77.668 | 2 | 51.538464 | yes | 5.598E-07 | 476.72427 | 81.608368 |
| VAASTQPLGR | 10 | Q8ND24 | RING finger protein 214 OS=Homo sapie | RNF214 | 77.668 | 2 | 51.538464 | yes | 1.424E-07 | 500.28271 | 75.813263 |

| | | | | | | | | | | | |
|---|---|---|---|---|---|---|---|---|---|---|---|
| EAELHLTYLK | 10 | Q8ND24 | RING finger protein 214 OS=Homo sapie | RNF214 | 77.668 | 2 | 51.538464 | yes | 2.464E-06 | 608.83228 | 75.482285 |
| QVENQLQVQLK | 11 | Q8ND24 | RING finger protein 214 OS=Homo sapie | RNF214 | 77.668 | 2 | 51.538464 | yes | 2.492E-07 | 663.87244 | 44.653473 |
| ASAETPRPQPVDK | 13 | Q8ND24 | RING finger protein 214 OS=Homo sapie | RNF214 | 77.668 | 2 | 51.538464 | yes | 2.04E-07 | 698.36499 | 40.690002 |
| DVASTAGEEGDTS | 15 | Q8ND24 | RING finger protein 214 OS=Homo sapie | RNF214 | 77.668 | 2 | 51.538464 | yes | 4.589E-07 | 754.34717 | 34.21595 |
| VDQDDDQDSSSLK | 13 | Q8ND24 | RING finger protein 214 OS=Homo sapie | RNF214 | 77.668 | 2 | 51.538464 | yes | 1.018E-07 | 726.31024 | 21.883749 |
| LLEPFEVR | 8 | Q8WUY1 | Protein THEM6 OS=Homo sapiens OX=9 | THEM6 | 23.865 | 2 | 46.949883 | yes | 5.124E-07 | 501.78458 | 311.83246 |
| AHTVLAASCAR | 11 | Q8WUY1 | Protein THEM6 OS=Homo sapiens OX=9 | THEM6 | 23.865 | 2 | 46.949883 | yes | 1.438E-07 | 578.79822 | 70.021637 |
| CGVLGALR | 8 | Q8WUY1 | Protein THEM6 OS=Homo sapiens OX=9 | THEM6 | 23.865 | 2 | 46.949883 | yes | 1.434E-07 | 423.23672 | 68.146622 |
| IGVADVDMSR | 10 | Q8WW59 | SPRY domain-containing protein 4 OS=H | SPRYD4 | 23.099 | 2 | 37.64663 | yes | 5.816E-07 | 531.76605 | 16.671032 |
| LQAENAAEWGR | 11 | Q96A19 | Coiled-coil domain-containing protein 10 | CCDC102A | 62.596 | 2 | 51.505505 | yes | 9.722E-08 | 622.80457 | 240.95659 |
| GSLLTILGSPSPER | 14 | Q96A19 | Coiled-coil domain-containing protein 10 | CCDC102A | 62.596 | 2 | 51.505505 | yes | 9.722E-08 | 713.89862 | 209.01704 |
| AQVGDLEEALAR | 12 | Q96A19 | Coiled-coil domain-containing protein 10 | CCDC102A | 62.596 | 2 | 51.505505 | yes | 6.904E-07 | 636.33313 | 183.87299 |
| LEGELSQWK | 9 | Q96A19 | Coiled-coil domain-containing protein 10 | CCDC102A | 62.596 | 2 | 51.505505 | yes | 1.873E-07 | 545.28241 | 177.76903 |
| MSEDLEDELGAR | 12 | Q96A19 | Coiled-coil domain-containing protein 10 | CCDC102A | 62.596 | 2 | 51.505505 | yes | 9.903E-08 | 682.80353 | 135.27956 |
| LAESPQLSK | 9 | Q96A19 | Coiled-coil domain-containing protein 10 | CCDC102A | 62.596 | 2 | 51.505505 | yes | 1.277E-07 | 486.77167 | 56.304897 |
| SMPEESEDCWEAR | 13 | Q96A19 | Coiled-coil domain-containing protein 10 | CCDC102A | 62.596 | 2 | 51.505505 | yes | 2.143E-06 | 813.31396 | 10.219779 |
| AAQPPAQPCQLCC | 14 | Q96B54 | Zinc finger protein 428 OS=Homo sapien | ZNF428 | 20.481 | 2 | 47.052464 | yes | 5.802E-07 | 777.3692 | 641.86108 |
| SPLGEAPPGTPPCR | 14 | Q96B54 | Zinc finger protein 428 OS=Homo sapien | ZNF428 | 20.481 | 2 | 47.052464 | yes | 2.519E-07 | 718.35358 | 166.97774 |
| ALGEEEEEPPR | 11 | Q96B54 | Zinc finger protein 428 OS=Homo sapien | ZNF428 | 20.481 | 2 | 47.052464 | yes | 1.413E-07 | 628.2937 | 156.01157 |
| LDPADFSVGYVDR | 13 | Q96B70 | Leukocyte receptor cluster member 9 OS= | LENG9 | 53.167 | 2 | 53.628746 | yes | 1.099E-07 | 727.3515 | 165.13956 |
| LAGVTEEALKPTA | 16 | Q96B70 | Leukocyte receptor cluster member 9 OS= | LENG9 | 53.167 | 3 | 53.628746 | yes | 1.301E-07 | 543.3053 | 69.638382 |
| LAGVTEEALKPTA | 16 | Q96B70 | Leukocyte receptor cluster member 9 OS= | LENG9 | 53.167 | 3 | 53.628746 | yes | 9.722E-08 | 814.45435 | 69.638382 |
| LAGVTEEALKPTA | 16 | Q96B70 | Leukocyte receptor cluster member 9 OS= | LENG9 | 53.167 | 2 | 53.628746 | yes | 1.301E-07 | 543.3053 | 69.638382 |
| LAGVTEEALKPTA | 16 | Q96B70 | Leukocyte receptor cluster member 9 OS= | LENG9 | 53.167 | 2 | 53.628746 | yes | 9.722E-08 | 814.45435 | 69.638382 |
| LAGAGEEAAAIGA | 15 | Q96B70 | Leukocyte receptor cluster member 9 OS= | LENG9 | 53.167 | 2 | 53.628746 | yes | 1.54E-07 | 685.37537 | 123.26168 |
| TDLVFGSGSAAGR | 13 | Q96B70 | Leukocyte receptor cluster member 9 OS= | LENG9 | 53.167 | 2 | 53.628746 | yes | 2.488E-07 | 619.31219 | 122.45389 |
| ALGTAADLGTLAP | 14 | Q96B70 | Leukocyte receptor cluster member 9 OS= | LENG9 | 53.167 | 2 | 53.628746 | yes | 9.722E-08 | 663.87244 | 66.117844 |
| VPHGSQVHLPK | 11 | Q96B70 | Leukocyte receptor cluster member 9 OS= | LENG9 | 53.167 | 2 | 53.628746 | yes | 1.265E-07 | 599.8382 | 46.70525 |
| LEAEGLSTLQSPGC | 23 | Q96B70 | Leukocyte receptor cluster member 9 OS= | LENG9 | 53.167 | 3 | 53.628746 | yes | 2.384E-07 | 809.43945 | 27.138151 |
| PLPEPSEQEGESVK | 14 | Q96CP2 | FLYWCH family member 2 OS=Homo sa | FLYWCH2 | 14.564 | 2 | 52.689075 | no | 6.47E-07 | 763.37268 | 116.88514 |
| AIEAAPQEPEQK | 12 | Q96CP2 | FLYWCH family member 2 OS=Homo sa | FLYWCH2 | 14.564 | 2 | 52.689075 | yes | 9.722E-08 | 655.83295 | 40.912476 |
| PGTEVIPAAPR | 11 | Q96CP2 | FLYWCH family member 2 OS=Homo sa | FLYWCH2 | 14.564 | 2 | 52.689075 | yes | 9.722E-08 | 554.31146 | 37.055962 |
| IPMETLETWEER | 12 | Q96D70 | R3H domain-containing protein 4 OS=Ho | R3HDM4 | 30.35 | 2 | 31.298412 | yes | 1.888E-06 | 767.36633 | 94.744278 |
| TLPLENASILSEGS | 20 | Q96H35 | Probable RNA-binding protein 18 OS=Ho | RBM18 | 21.649 | 3 | 47.52516 | yes | 1.311E-07 | 717.70618 | 128.36571 |
| LWIGNLDPK | 9 | Q96H35 | Probable RNA-binding protein 18 OS=Ho | RBM18 | 21.649 | 2 | 47.52516 | yes | 3.279E-07 | 528.29785 | 92.266083 |
| GYCFVNFETK | 10 | Q96H35 | Probable RNA-binding protein 18 OS=Ho | RBM18 | 21.649 | 2 | 47.52516 | yes | 1.328E-07 | 632.78699 | 44.401707 |
| SGAELALDYLCR | 12 | Q9BRJ6 | Uncharacterized protein C7orf50 OS=Ho | C7orf50 | 22.084 | 2 | 49.98083 | yes | 1.033E-07 | 684.33484 | 4138.9023 |
| ELDEEGSDPPLPGI | 14 | Q9BRJ6 | Uncharacterized protein C7orf50 OS=Ho | C7orf50 | 22.084 | 2 | 49.98083 | yes | 1.093E-07 | 755.85461 | 1191.8926 |
| EAGLVAQHPPAR | 12 | Q9BRJ6 | Uncharacterized protein C7orf50 OS=Ho | C7orf50 | 22.084 | 2 | 49.98083 | yes | 1.77E-06 | 623.33856 | 972.37885 |

| | | | | | | | | | | | |
|---|---|---|---|---|---|---|---|---|---|---|---|
| PGLDAEPELSPEE( | 15 | Q9BRJ6 | Uncharacterized protein C7orf50 OS=Ho | C7orf50 | 22.084 | 2 | 49.98083 | yes | 9.939E-08 | 833.8996 | 23.292107 |
| LGGIPDALPTVAAI | 15 | Q9BRP1 | Programmed cell death protein 2-like OS= | PDCD2L | 39.417 | 2 | 48.61843 | yes | 1.985E-07 | 724.41699 | 666.33704 |
| SQCLQVPER | 9 | Q9BRP1 | Programmed cell death protein 2-like OS= | PDCD2L | 39.417 | 2 | 48.61843 | yes | 1.122E-07 | 558.77698 | 408.91284 |
| IAACQEQILR | 10 | Q9BRP1 | Programmed cell death protein 2-like OS= | PDCD2L | 39.417 | 2 | 48.61843 | yes | 7.315E-07 | 601.32153 | 49.664154 |
| LISSVDPQFLK | 11 | Q9BVG4 | Protein PBDC1 OS=Homo sapiens OX=9 | PBDC1 | 26.057 | 2 | 24.689445 | yes | 3.409E-06 | 623.85571 | 47.162056 |
| IDVLDPEELK | 10 | Q9BVG4 | Protein PBDC1 OS=Homo sapiens OX=9 | PBDC1 | 26.057 | 2 | 24.689445 | yes | 3.862E-06 | 585.81628 | 28.922014 |
| AISYLATVPK | 10 | Q9H1X3 | DnaJ homolog subfamily C member 25 O | DNAJC25 | 42.404 | 2 | 49.662003 | yes | 1.012E-07 | 531.81335 | 100.32556 |
| SQFDSLEDHQK | 11 | Q9H1X3 | DnaJ homolog subfamily C member 25 O | DNAJC25 | 42.404 | 2 | 49.662003 | yes | 2.493E-07 | 667.30457 | 84.359291 |
| DCYEVLGVSR | 10 | Q9H1X3 | DnaJ homolog subfamily C member 25 O | DNAJC25 | 42.404 | 2 | 49.662003 | yes | 1.073E-07 | 599.28204 | 49.777977 |
| MDESTLLR | 8 | Q9H4I3 | TraB domain-containing protein OS=Hon | TRABD | 42.322 | 2 | 29.856926 | yes | 2.363E-06 | 482.74203 | 37.429432 |
| ELTQYFGVNDR | 11 | Q9H8W3 | Protein FAM204A OS=Homo sapiens OX | FAM204A | 27.022 | 2 | 55.972084 | yes | 2.874E-07 | 671.32532 | 250.392 |
| AEELSNQLATR | 11 | Q9H8W3 | Protein FAM204A OS=Homo sapiens OX | FAM204A | 27.022 | 2 | 55.972084 | yes | 2.486E-07 | 616.3175 | 192.29219 |
| IDQAVEEWNIEK | 12 | Q9H8W3 | Protein FAM204A OS=Homo sapiens OX | FAM204A | 27.022 | 2 | 55.972084 | yes | 9.722E-08 | 737.36462 | 188.83473 |
| AVACHNFVK | 9 | Q9H8W3 | Protein FAM204A OS=Homo sapiens OX | FAM204A | 27.022 | 2 | 55.972084 | yes | 3.981E-07 | 523.26599 | 107.03605 |
| VEVESIAETYGR | 12 | Q9NWW7 | Uncharacterized protein C2orf42 OS=Ho | C2orf42 | 64.077 | 2 | 52.83012 | yes | 2.89E-07 | 676.83826 | 45.015305 |
| QILDTPEMPLEITR | 14 | Q9NWW7 | Uncharacterized protein C2orf42 OS=Ho | C2orf42 | 64.077 | 2 | 52.83012 | yes | 9.722E-08 | 828.43726 | 42.914604 |
| LAVNCQAEATPLT | 15 | Q9NWW7 | Uncharacterized protein C2orf42 OS=Ho | C2orf42 | 64.077 | 2 | 52.83012 | yes | 9.722E-08 | 814.93744 | 41.989937 |
| KPVVASSLK | 9 | Q9NWW7 | Uncharacterized protein C2orf42 OS=Ho | C2orf42 | 64.077 | 2 | 52.83012 | yes | 1.233E-07 | 464.79492 | 37.981102 |
| GSFAPGHGPR | 10 | Q9NX00 | Transmembrane protein 160 OS=Homo sa | TMEM160 | 19.658 | 2 | 35.820663 | yes | 3.942E-06 | 491.74631 | 240.44949 |
| AHETAFLSWFR | 11 | Q9NX00 | Transmembrane protein 160 OS=Homo sa | TMEM160 | 19.658 | 2 | 35.820663 | yes | 7.773E-07 | 682.84094 | 84.942497 |
| ASDLSDFQSVSK | 12 | Q9NX31 | Oxidative stress-responsive serine-rich pr | OSER1 | 31.78 | 2 | 51.221737 | yes | 1.202E-07 | 642.30933 | 183.10072 |
| DGEEESLQTAFK | 12 | Q9NX31 | Oxidative stress-responsive serine-rich pr | OSER1 | 31.78 | 2 | 51.221737 | yes | 2.464E-07 | 677.31207 | 155.95546 |
| EDSSDATQVPQAS | 15 | Q9NX31 | Oxidative stress-responsive serine-rich pr | OSER1 | 31.78 | 2 | 51.221737 | yes | 9.733E-08 | 788.37848 | 126.55352 |
| FIHCSTIASSSSQL | 16 | Q9NX31 | Oxidative stress-responsive serine-rich pr | OSER1 | 31.78 | 3 | 51.221737 | yes | 9.722E-08 | 584.9577 | 102.73808 |
| MSHMAEMMYT | 10 | Q9NX31 | Oxidative stress-responsive serine-rich pr | OSER1 | 31.78 | 2 | 51.221737 | yes | 9.285E-07 | 616.23438 | 70.889313 |
| EFSAGESSTSLDAN | 24 | Q9NX31 | Oxidative stress-responsive serine-rich pr | OSER1 | 31.78 | 3 | 51.221737 | yes | 4.078E-06 | 825.39801 | 15.810413 |
| KIEHDVVMK | 9 | Q9UNZ5 | Leydig cell tumor 10 kDa protein homolo | C19orf53 | 10.577 | 2 | 50.58387 | yes | 1.046E-07 | 549.80243 | 1497.4771 |
| KIEHDVVMK | 9 | Q9UNZ5 | Leydig cell tumor 10 kDa protein homolo | C19orf53 | 10.577 | 2 | 50.58387 | yes | 1.366E-06 | 557.79987 | 1497.4771 |
| KNLEVGIR | 8 | Q9UNZ5 | Leydig cell tumor 10 kDa protein homolo | C19orf53 | 10.577 | 2 | 50.58387 | yes | 9.758E-08 | 464.78235 | 1075.1042 |
| IEHDVVMK | 8 | Q9UNZ5 | Leydig cell tumor 10 kDa protein homolo | C19orf53 | 10.577 | 2 | 50.58387 | yes | 2.359E-07 | 485.75494 | 661.30188 |
| AALIYTCTVCR | 11 | Q9Y5V0 | Zinc finger protein 706 OS=Homo sapien | ZNF706 | 8.498 | 2 | 47.782192 | yes | 1.26E-07 | 664.32849 | 2009.8657 |
| TPLPPELADVQA | 12 | Q9Y5V0 | Zinc finger protein 706 OS=Homo sapien | ZNF706 | 8.498 | 2 | 47.782192 | yes | 4.238E-07 | 625.83496 | 455.26035 |
| STPGVASTPQTPV! | 17 | A0A1W2PQ72 | Myb/SANT-like DNA-binding domain-cc | MSANTD7 | 39.761 | 2 | 48.90543 | yes | 1.088E-07 | 829.92084 | 25.592131 |
| VVGAVIDQGLITR | 13 | E9PRG8 | Uncharacterized protein C11orf98 OS=H | C11orf98 | 14.234 | 2 | 52.207066 | yes | 9.722E-08 | 670.89844 | 4474.7036 |
| TAMEVEAPSKPAF | 13 | E9PRG8 | Uncharacterized protein C11orf98 OS=H | C11orf98 | 14.234 | 2 | 52.207066 | yes | 9.722E-08 | 693.8559 | 609.07263 |
| TAMEVEAPSKPAF | 13 | E9PRG8 | Uncharacterized protein C11orf98 OS=H | C11orf98 | 14.234 | 2 | 52.207066 | yes | 2.493E-07 | 701.85339 | 609.07263 |
| APQDVEMK | 8 | E9PRG8 | Uncharacterized protein C11orf98 OS=H | C11orf98 | 14.234 | 2 | 52.207066 | yes | 3.278E-07 | 459.22348 | 382.12488 |
| TKAPQDVEMK | 10 | E9PRG8 | Uncharacterized protein C11orf98 OS=H | C11orf98 | 14.234 | 2 | 52.207066 | yes | 3.423E-07 | 573.7948 | 175.81578 |
| EFEAVLTER | 9 | Q96BQ5 | Coiled-coil domain-containing protein 12 | CCDC127 | 30.835 | 2 | 42.204685 | yes | 3.143E-07 | 547.27985 | 72.44017 |

| | | | | | | | | | | |
|---|---|---|---|---|---|---|---|---|---|---|
| AVAQLSLELEK | 11 | Q96BQ5 | Coiled-coil domain-containing protein 12 CCDC127 | 30.835 | 2 | 42.204685 | yes | 2.256E-06 | 600.84534 | 41.264935 |
| QNIYCSLFLPR | 11 | Q96BQ5 | Coiled-coil domain-containing protein 12 CCDC127 | 30.835 | 2 | 42.204685 | yes | 3.924E-06 | 705.86353 | 36.288898 |
| ASLTGTSSTASLTR | 14 | Q9UF83 | Uncharacterized protein DKFZp434B061 -- | 59.412 | 2 | 52.2605 | yes | 1.008E-07 | 676.85443 | 245.67776 |
| TPSMASLTR | 9 | Q9UF83 | Uncharacterized protein DKFZp434B061 -- | 59.412 | 2 | 52.2605 | yes | 1.167E-07 | 482.25003 | 111.92233 |
| RPSTASLTR | 9 | Q9UF83 | Uncharacterized protein DKFZp434B061 -- | 59.412 | 2 | 52.2605 | yes | 1.164E-07 | 494.78033 | 99.301048 |
| TPSWASPTTTPSR | 13 | Q9UF83 | Uncharacterized protein DKFZp434B061 -- | 59.412 | 2 | 52.2605 | yes | 9.722E-08 | 694.84387 | 75.174545 |
